# A functionally conserved *STORR* gene fusion in *Papaver* species that diverged 16.8 million years ago

**DOI:** 10.1101/2021.10.11.463683

**Authors:** Theresa Catania, Yi Li, Thilo Winzer, David Harvey, Fergus Meade, Anna Caridi, Tony R. Larson, Zemin Ning, Ian A Graham

## Abstract

The *STORR* gene fusion event is considered a key step in the evolution of benzylisoquinoline alkaloid (BIA) metabolism in opium poppy as the resulting bi-modular protein performs the isomerization of (*S*)- to (*R*)-reticuline which is required for morphinan biosynthesis. Our previous analysis of the opium poppy genome suggested the *STORR* gene fusion event occurred before a whole genome duplication event 7.2 million years ago. Here we use a combination of phylogenetic, transcriptomic, metabolomic, biochemical and genomic analysis to investigate the origin of the STORR gene fusion across the Papaveraceae family. The pro-morphinan/morphinan subclass of BIAs was present in a subset of 10 *Papaver* species including *P. somniferum* (opium poppy) and this correlated with the presence of the STORR gene fusion with one important exception. *P. californicum* does not produce morphinans but it does contain a STORR gene fusion that epimerizes (*S*)- to (*R*)-reticuline when heterologously expressed in yeast. The high similarity of the amino acid sequence linking the two modules of STORR along with phylogenetic gene tree analysis strongly suggests the gene fusion occurred only once and between 17-25 million years ago before the separation of *P. californicum* from the other *Papaver* species. We discovered that the most abundant BIA in *P. californicum* is (*R*)-glaucine, a member of the aporphine subclass of BIAs. Only the (*S*) isomer of this compound has previously been reported from nature. These results lead us to conclude that the function of the STORR gene fusion is not exclusive to morphinan production in the Papaveraceae.

---

The benzylisoquinoline alkaloids or BIA’s represent a structurally diverse group predominantly identified in the order Ranunculales [1; 2]. The morphinans are the best known class of BIAs, with powerful analgesic properties. They are synthesised in the genus *Papaver* and are currently commercially produced in opium poppy - *Papaver somniferum* from the order Papaveraceae. The commercial importance of opium poppy has led to its use as a model species for research into the biosynthetic pathway for morphinan production [2; 3; 4; 5]. The common precursor and central branch point in the pathway for production of the many structurally distinct subclasses of BIAs in the Ranunculales including morphinan, protoberberine, phthalideisoquinoline and benzophenanthridine is the 1-benzylisoquinoline alkaloid, (*S*)-reticuline [6](Fig. 1a). The gateway reaction leading to morphinan biosynthesis is catalysed by the STORR protein. Composed of P450 and oxidoreductase modules, this fused protein completes the epimerization of (*S*) - to (*R*)-reticuline, the first step in the morphinan pathway [2; 4; 7]. *STORR* is clustered with four other genes involved in synthesis of the first morphinan, thebaine and ten genes involved in synthesis of the phthalideisoquinoline, noscapine, which together make up the BIA gene cluster in opium poppy [5].

**Figure 1.**
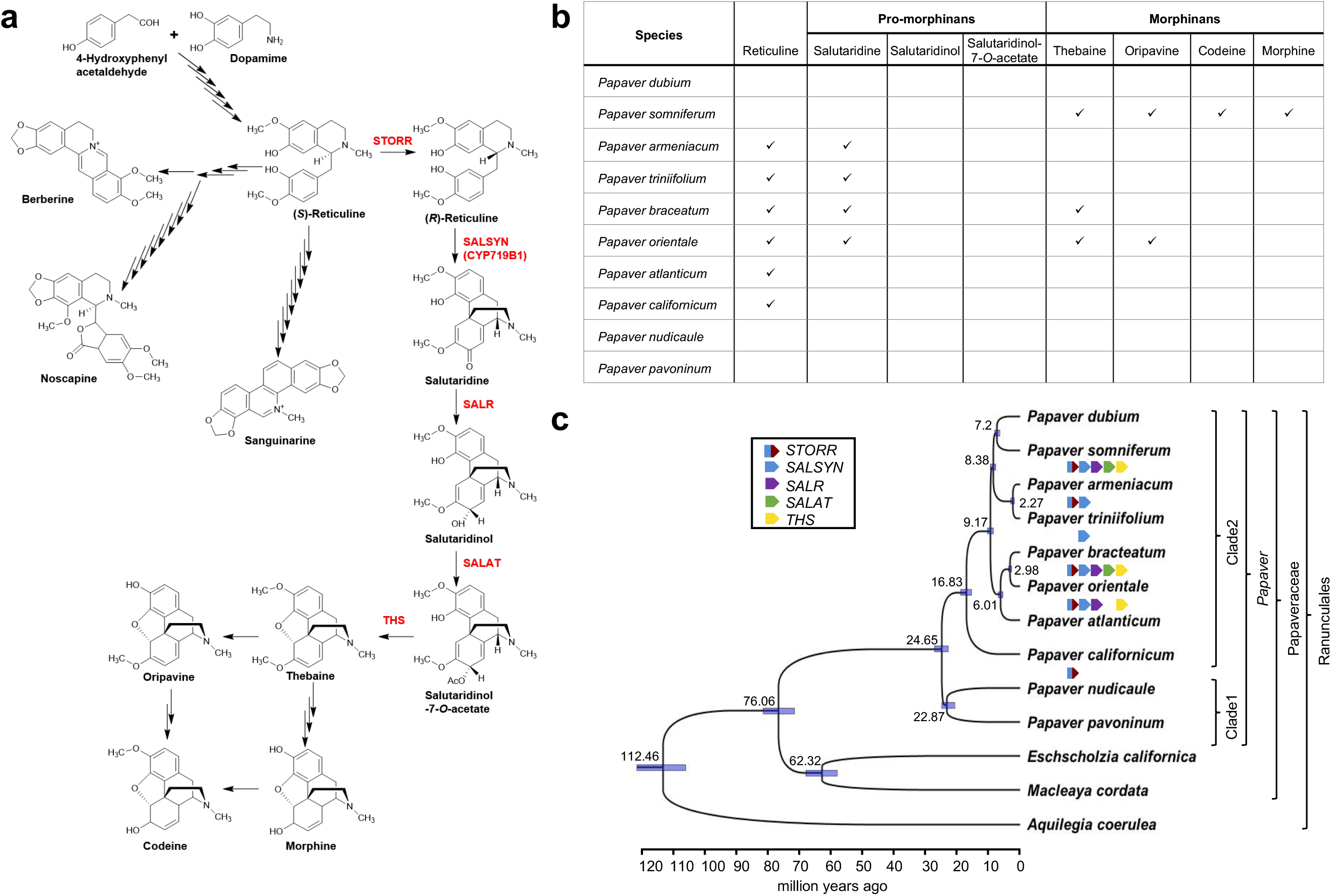
Metabolite and transcriptomic analysis to investigate STORR and its distribution in 10 Papaver species. **a) Schematic of the benzylisoquinoline pathway and enzymes in Opium Poppy with a focus on morphinan production.** (S)-reticuline is the central branch point of BIA metabolism in opium poppy for the production of structurally distinct compounds including the morphinans plus noscapine, berberine and sanguinarine. Conversion of (S) - to (R)-reticuline by STORR represents the first committed step in biosynthesis of the promorphinans (salutaridine, Salutaridinol and salutaridinol-7-*O-*acetate) and morphinans (thebaine, oripavine, codeine and morphine). Compound names are in black and enzymes specific to the promorphinan and morphinan pathway are in red. **(b) Metabolite analysis for promorphinan and morphinan compounds in 10 Papaver species.** The presence of the promorphinan and morphinans was determined by UPLC-MS analysis of latex and capsules material (supplemental table S2). Promorphinan compounds were identified in 4 species indicative of a functional STORR, with morphinan production identified in 2 species plus *P. somniferum*. **(c) Species phylogeny inferred by Bayesian inference species tree using eight single copy conserved ortholog sequences.** Phylogeny and divergence dates are estimated using BEAST2, which are shown on the nodes. Light-blue bars at the nodes indicate the range with 95% highest posterior density. Taxonomy groupings of the species are indicated and the ten Papaver species are placed into two different clades (Clade 1 and Clade 2) as described by Carolan *et al*. (2006) [20]. Presence of the *STORR* and the four pro-morphinan genes in the transcriptomic datasets are shown under species names (supplemental table S5).

Advances in genome sequencing technology and assembly offers the opportunity to compare the genome organisation of related species and provide insight into the role of events such as gene fusion and gene clustering in the evolution of specialized metabolites [8; 9; 10; 11]. Such an approach in combination with metabolomic, transcriptomic and gene function analysis has allowed us to determine the evolutionary sequence of events leading to the clustering of the genes encoding the *STORR* modules, the fusion of these genes to form a functional STORR and the clustering of the other four genes involved in morphinan production across a number of *Papaver* species.

The epimerization of (*S*) - to (*R*)-reticuline is followed by a sequence of conversions from (*R*)-reticuline to salutaridine, salutaridinol, salutaridinol-7-*O* acetate and thebaine catalysed by the products of *SALSYN*/*SALAT*/*SALR* and *THS* respectively (Fig. 1a) [14; 13; 15; 16]. To investigate the presence of the *STORR* gene fusion across the *Papaver* genus we selected nine other *Papaver* species together with opium poppy that provided good taxonomic coverage (Fig. 1b&1c; supplemental table S1). To determine the presence of promorphinan and morphinan compounds and related genes in these species metabolite profiling of latex from juvenile plants and capsule material post-harvest was carried out (Fig. 1b; supplemental table S2). As previously reported we found thebaine to be the most prominent metabolite in *P. bracteatum* and oripavine the most prominent in *P. orientale* [17; 18; 19]. These along with *P. somniferum* were the only species found to produce morphinans. However, promorphinans were identified as minor compounds in *P. armeniacum* and P*. triniifolium* (Fig. 1b; supplemental table S2). Identification of these morphinan and promorphinan BIAs is suggestive of the presence of a *STORR* ortholog in these species.

Transcriptomic analysis revealed expression of *STORR* and either none, some or all of the promorphinan genes in a subset of the nine other *Papaver* species, with the full complement in opium poppy as reported previously [5]. These are shown on the branches of a species tree generated using conserved orthologs of 8 low-copy nuclear genes identified from the same transcriptomic datasets (Fig. 1c; supplemental tables S3-6; supplementary datasets 1&2). The topology of the tree generated is congruent with other trees available for the genus *Papaver* and the wider family and order [20; 21; 22].

Similar to the opium poppy *STORR*, the homologs present in the other four *Papaver* species (Fig. 1c) encode full length P450-oxidoreductase fusion proteins (Supplemental Fig. 1). The direction of both modules and the 9-13 amino acid linker sequences are also conserved (Fig. 2a; Supplemental Fig.1) with overall sequence identity ranging from 90.1-99.7% at the nucleotide level and 87.7-99.4% with the predicted amino acid sequences (Supplementary tableS7; supplementary datasets 3&4).

**Figure 2.**
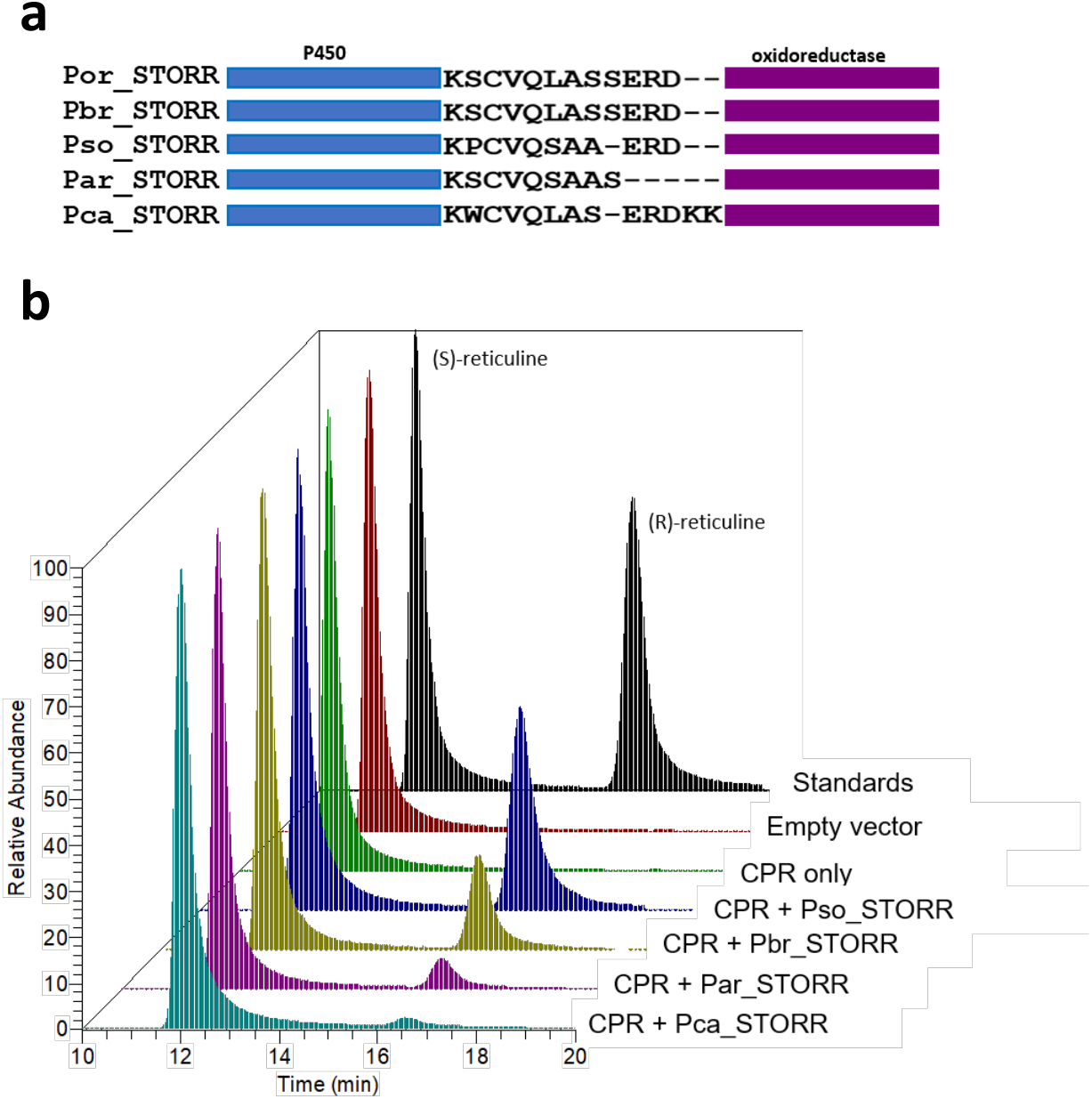
Functional analysis of STORR proteins in *Papaver* species. **(a) Alignment of the amino acid linker connecting the STORR P450 and oxidoreductase modules.** **(b) Functional characterisation of the STORR proteins by heterologous expression in *S. cerevisiae*.** STORR protein from *P. californicum, P. armeniacum, P. bracteatum* and P*. somniferum* were separately produced with a *P. somniferum* cytochrome P450 reductase (CPR) redox partner in *S. cerevisiae*. Microsomal proteins were assayed with (S)-reticuline assubstrate. Relative abundance is used to show the reticuline epimers present in the sample. Reticuline standards (black), pESC-trp empty vector (red), pESC-trp vector + cytochrome P450 reductase (CPR)(green), pESC-TRP+ CPR+ Pso STORR (blue), pESC-TRP+ CPR+ Pbr STORR (gold), pESC-TRP+ CPR+ Pam (pink) and pESC-TRP+ CPR+ Pca STORR (cyan).

The identification of *STORR* in conjunction with the metabolite data indicate that the fusion proteins are functional in *P. armeniacum* and *P. orientale* and *STORR* genes from *P. somniferum* and *P. bracteatum* have been functionally characterised previously [3, 4, 7]. In *P. californicum*, however, a notable inconsistency is the presence of STORR gene expression but lack of promorphinan gene expression and absence of any promorphinans or morphinans. The main BIA identified from our extraction of *P. californicum* was a member of the aporphine subclass, glaucine in the (R) configuration (Supplemental table 1 and supplemental fig. 2).

In order to investigate if the *P. californicum* STORR is functional, we expressed this gene in Saccharomyces cerevisiae, performed epimerization assays on microsomal fractions and found the same activity as the *P. somniferum*, *P. bracteatum* and P. *armeniacum* STORR proteins (Fig. 2b; supplementary dataset 5). Activity of *P. californicum* STORR microsomal fractions on alternative substrates (*S*)-glaucine and (*S*)-laudanosine were also investigated with no activity found (Supplemental Fig 3 and 4).

To understand the evolutionary relationship we performed gene tree analysis of the *STORR*s and the closest paralogues to the two *STORR* modules within the genomes of a representative subset of *Papaver* species. The presence of a closely linked gene pair of closest paralogs to the opium poppy *STORR* cytochrome P450 CYP82Y2 and oxidoreductase modules was previously discovered through analysis of a whole genome assembly [5]. The segmental duplication resulting in these paralogs was suggested to have occurred 20.0–27.8 MYA by the Ks estimation of the paralogous pairs (6). In order to establish if equivalent paralogous pairs are present in related *Papaver* species we used whole genome sequencing approaches to assemble draft genomes for *P. nudicaule* from clade 1 and *P. californicum*, *P. bracteatum*, *P. atlanticum*, and *P. armeniacum* from clade 2 (supplementary table S8). We compiled and annotated all homologous sequences that contain full length genes corresponding to either of the STORR modules in these draft assemblies (supplementary datasets 6&7). Together with the sequences retrieved from the searches in the annotated opium poppy genome, the transcriptomic data from 10 species from the present study, as well as the GenBank database searches, we constructed gene trees for the two gene subfamilies containing the coding sequences closely related to the STORR modules CYP82Y2 (Fig. 3a) and oxidoreductase (Fig. 3b) respectively. Consistent with previous work [6], both trees revealed robustly supported cytochrome P450 CYP82Y2 like (CYP82Y2-L) and oxidoreductase like (COR-L) groups. All P450 modules and oxidoreductase modules of STORRs are from clade 2 *Papaver* species and they form monophyletic orthologous subgroups highlighted in orange (Fig. 3a&3b).

**Figure 3.**
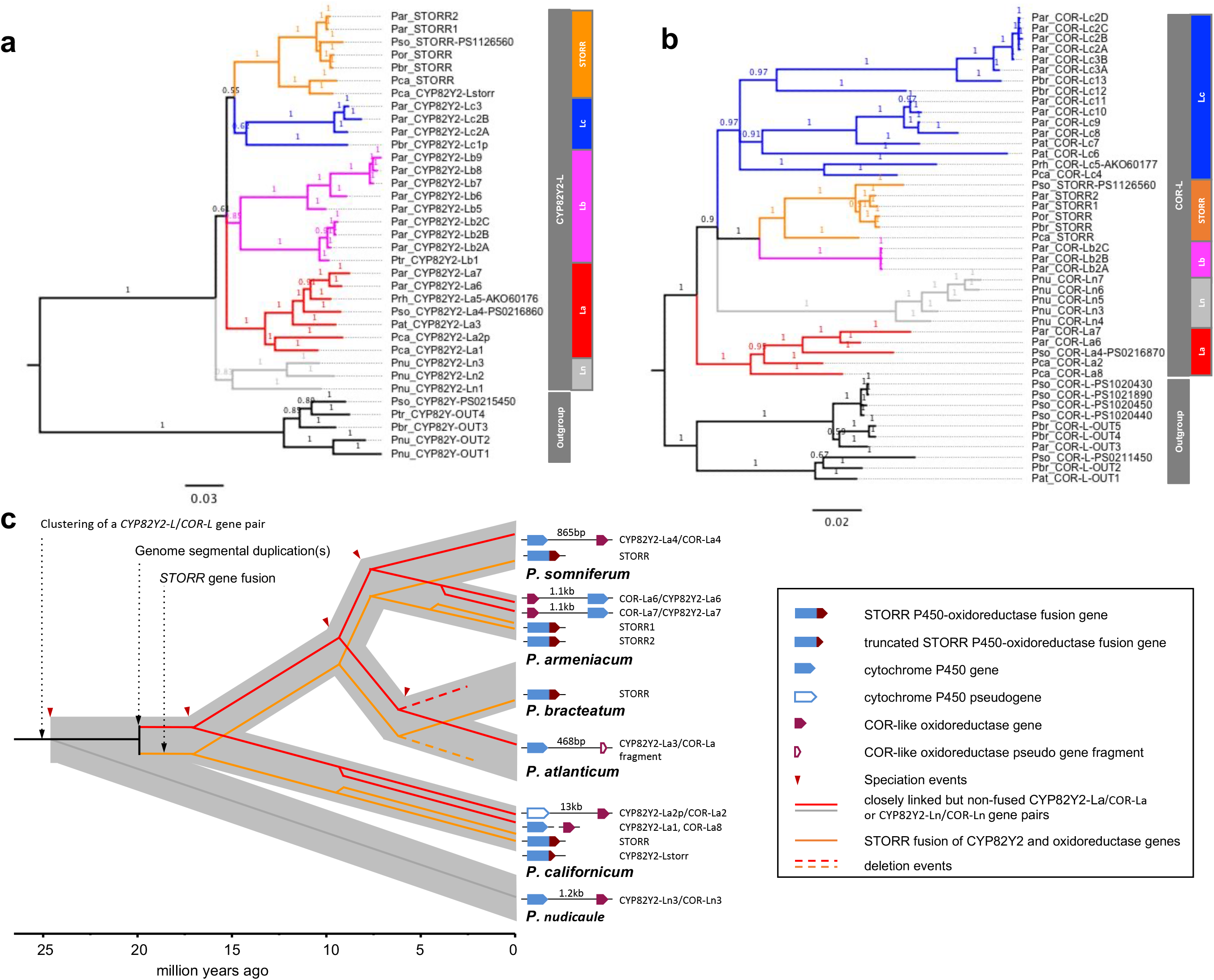
Evolutionary history of the formation of *STORR* fusion gene inferred from gene tree analyses of its P450 CYP82Y2 and oxidoreductase modules. **(a) Phylogenetic gene tree of***CYP82Y2-L* **P450 subfamily constructed using Bayesian Inference. (b) Phylogenetic gene tree of***COR-L* **oxidoreductase subfamily constructed using Bayesian Inference.** The posterior probabilities are shown on the branches and scale bar represents substitutions per nucleotide site. Both trees were rooted based on branch position of orthologous outgroup containing *P. somniferum* sequences CYP82Y-PS0215450 for *CYP82Y2-L* tree and COR_L-PS0211450/PS1020430/PS1020440/PS1020450/PS1021890 for the *COR-L* tree as described in Li *et al*. (2020)[6]. STORR, La, Lb, Lc and Ln are defined as orthologous subgroups and their members are highlighted by coloured branches. A three letter prefix followed by an underscore is used as a species identifier for each gene; including Par_ (*P. armeniacum*), Pat_ (*P. atlanticum*), Pbr_ (*P. bracteatum*), Pca_ (*P. californicum*), Pnu_ (*P. nudicaule*), Por_ (*P. orientale*), Prh_ (*P. rhoeas*), Ptr_ (*P. triniifolium*) and Pso_ (*P. somniferum*). **(c) Schematic representation of the evolutionary history of***STORR* **reconciling the species tree with gene trees.** The fusion event was preceded by the clustering of a gene pair of *CYP82Y2-L* and *COR-L* genes and subsequent segmental duplications. The grey background branches denote species divergence with speciation time points indicated by red arrows. The orange lines denote *STORR* and the red lines a gene pair of *CYP82Y2-La* and *COR-La*. The grey line indicates an ancestral *CYP82Y2-L* and *COR-L* gene pair prior to the divergence of *P. nudicaule* from the clade 2 *Papaver* species. The exclusive presence of clade 2 species in La and STORR subgroups and a single *P. nudicaule* Ln subgroup in both *CYP82Y2-L* and *COR-L* trees is consistent with the segmental duplication occurring after the divergence of *P. nudicaule* from clade 2 species but before *STORR* formation between 25 and 17 mya.

The STORR subgroups contain no other complete coding sequences for P450 or oxidoreductase apart from Pca_CYP82Y2_Lstorr, which actually contains the conserved linker sequence and the first 13 codons of an oxidoreductase before a stop codon, indicating deletion after the fusion. Therefore these two monophyletic STORR subgroups and the presence of functional STORR in *P. californicum* (Fig. 2b) lead us to conclude that all STORR genes from these *Papaver* species have derived from a single fusion event in the common ancestor of clade 2 species after their divergence from clade 1 between 17 to 25MYA (Fig. 3c). Our analysis of *STORR* gene evolution also reveals the importance of lineage specific deletion (*P. atlanticum*), duplication (*P. californicum* and *P. armeniacum*) and rearrangement after duplication (*P. californicum*) (Fig. 3c). In addition, we note that in *P. somniferum* one copy of STORR has been lost after a whole genome duplication event [5].

We also observed clade 2 orthologous subgroups (CYP82Y2_La and COR-La, highlighted in red), which contains the closest paralogue pair (Pso_CYP82Y2_La4-PS0216860 and Pso_COR_La4-PS0216870) from the opium poppy genome in the respective gene trees (Fig. 3a&3b; supplementary dataset 8). This and the presence of the corresponding copies in *P. californicum* indicates all sequences within the subgroups are derived from a single copy clade 2 ancestor as found to be the case for *STORR*. We found the same pairing arrangement of closest paralogue pairs in all our assembled genomes except *P. bracteatum* which appears to have lost this paralogous gene pair (Fig. 3c). In *P. californicum* and *P. atlanticum* we found evidence for the ancestral arrangement of the paralogous gene pairs but in both cases one member of the pair now is present as a pseudogene (Fig. 3c). The order of the P450 and oxidoreductase genes for both paralogous pairs in *P. armeniacum* have switched compared to the other species (Fig. 3c). Taken into consideration gene deletion, duplication, rearrangement and erosion it is still apparent from these findings that paralogous pairing of a CYP82Y2_La and COR_La existed in the clade 2 common ancestor (Fig. 3c), supporting the hypothesis that a segmental duplication would have occurred prior to STORR fusion/neofunctionalization.

Within both CYP82Y2-L and COR-L groups, the gene trees have identified single orthologous groups (CYP82Y2-Ln and COR-Ln, highlighted in grey) containing exclusive clade 1 *P. nudicaule* sequences, which are absent from all other subgroups (Figure 3a and Figure 3b). Among the members, there is at least one paralogue pair (Pnu_CYP82Y2_Ln3/Pnu_COR_Ln3) just over 1 kb apart (Fig. 3c), which was confirmed by direct sequencing of a PCR amplified genome fragment (supplementary dataset 8). This finding implies that the origin of the CYP82Y2_Ls and COR_Ls pairing event occurred before the separation of *P. nudicaule* from clade 2 *Papaver* species 25 Mya. This is consistent with the presence of additional paralogous pairs identified in the clade 2 Lb and Lc subgroups which will have arisen by multiple segmental duplication events (Fig. 3a&3b). One such duplication event will have led to the formation of *STORR* 17-25MYA.

As part of the 800kb BIA cluster, *STORR* is clustered with four promorphinan genes in the opium poppy genome, which also contains a syntenic region containing paralogs of the promorphinan genes but not *STORR* (Fig. 4a; 5). We found that *P. bracteatum* shows very good synteny with the opium poppy regions containing the pro-morphinan genes with a notable difference being an extra copy of *SALR* in the former (Fig. 4a; supplementary table S9). *P. armeniacum* shows synteny with *STORR* and *SALSYN* with gene order conserved in two smaller contigs, one of which extends to reveal five additional genes in the flanking region. Only *STORR* was found in the draft assembly of *P. californicum* the *P. atlanticum* draft assembly lacks *STORR* and the promorphinan genes (supplementary table S5). These findings are consistent with our metabolite profiling and transcriptomic analyses across these five species (Fig. 1b&1c), highlighting the role played by gene deletion events in formation of the diversity of BIA composition within the *Papaver* lineage.

**Figure 4.**
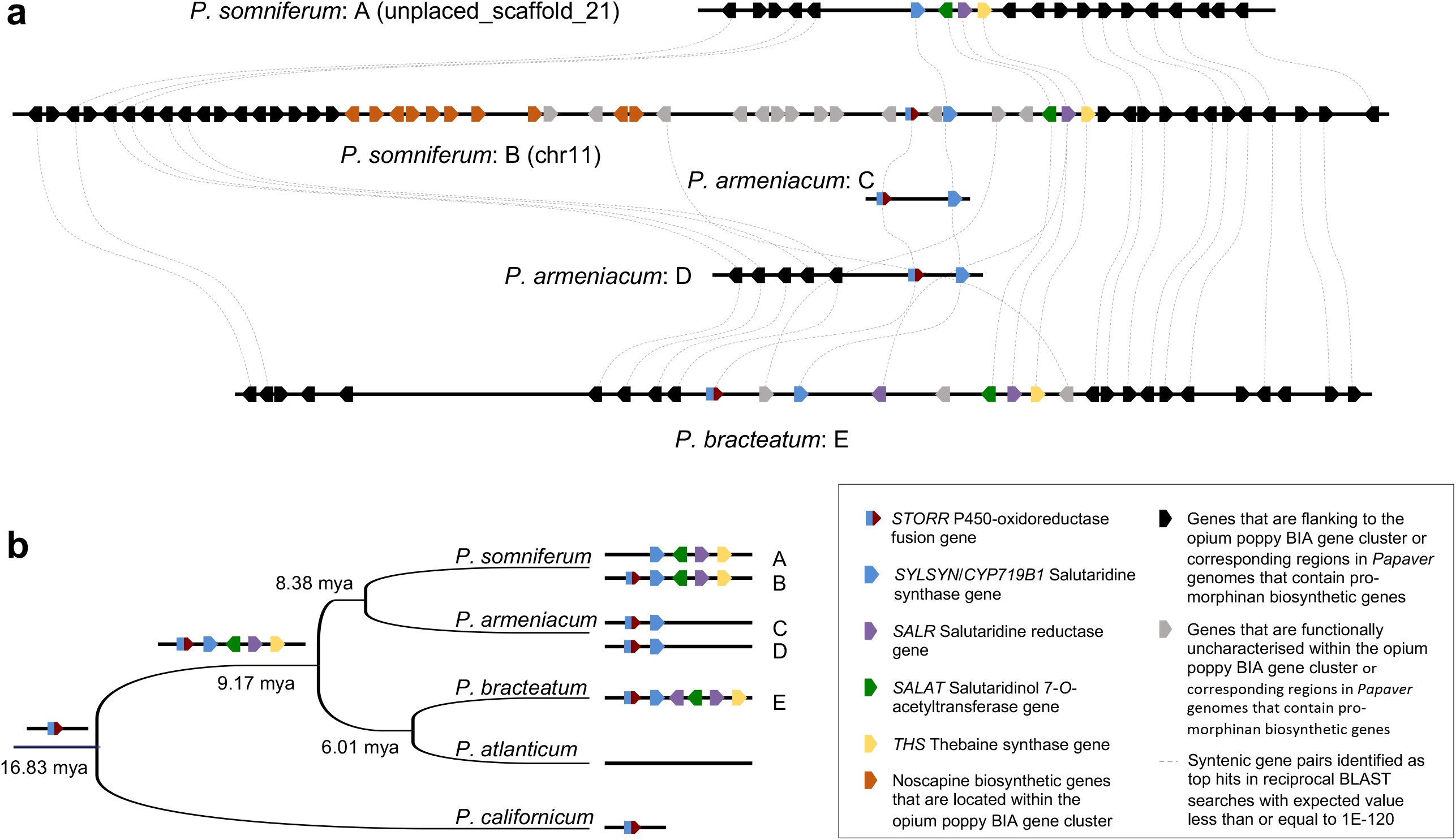
Evolutionary history of the STORR plus pro-morphinan component of the BIA gene cluster in *Papaver*. **(a) Graphic representation of synteny between contigs that contain *STORR* and pro-morphinan genes in *Papaver* genomes.** Arrangement of the genes and their orientations of the syntenic regions of Papaver species are shown as indicated in the inset (supplementary tableS9); including the corresponding regions in the opium poppy genome, one of which contains the BIA gene cluster and its flanking regions (*P. somniferum*: A&B) [5], the two contigs of 94kb and 294kb from the *P. armeniacum* genome (*P. armeniacum*: C&D) and the single 1.2Mb contig of the *P. bracteatum* assembly (*P. bracteatum*: E). Dashed lines denote syntenic gene pairs. **(b) Summary of the evolutionary history of the *STORR* plus pro-morphinan genes of the BIA gene cluster in *Papaver* as inferred from the species phylogeny and synteny in the genomes.** The phylogeny for the selected subset of Papaver species and divergent times at the branching points were extracted from the species tree in Fig. 1c. The emergence of *STORR* and its clustering with all four pro-morphinan genes are indicated on the ancestral nodes. These are derived from the presence and organisation of these genes in the genomes of these extant species, which are also shown.

Together these findings lead us to conclude that the STORR plus pro-morphinan component of the BIA gene cluster formed prior to the divergence of P*. bracteatum* and P*. somniferum* 9.2MYA (Fig. 4b), but after the formation of *STORR* at least 16.8 Mya. We demonstrate a functional STORR protein in *P. californicum*, which has persisted since its divergence from the common ancestor 16.8 mya. Genome rearrangements result in considerable structural variation even within a single species as evidenced by the loss of the noscapine component of the BIA gene cluster [3] and significant variation in copy number of genes associated with morphinan biosynthesis in *P. somniferum* [6]. That STORR persists in *P. californicum* suggests it is providing some selective advantage.

We found the most abundant BIA in *P. californicum* to be (*R*)-glaucine, which to our knowledge has not previously been reported in nature. Glaucine in the (*S*) configuration isolated from *Glaucium* species in the Papaveraceae is associated with bronchodilator, anti-inflammatory and neuroleptic effects [3; 24]. A chemically synthesised form of (*R*)-glaucine has been shown to increase the efficacy of serotonin [25]. Evidence for (*S*)-glaucine biosynthesis via (*S*)-reticuline has previously been proposed [26]. It is therefore interesting to speculate, based on the emergence of STORR at 16.8MYA in the *Papaver* lineage, that the *P. californicum* STORR could be involved in (*R*)-glaucine formation. Epimerization assays using *P. californicum* STORR with (*S*)-glaucine and (*S*)-laudanosine as substrate showed no activity implying that neither of these are intermediates in the biosynthesis of (*R*)-glaucine but rather the pathway is dependent on epimerization of reticuline by STORR followed by formation of the tetracylic aporphine structure.

Our findings clearly show that the STORR gene fusion was a single event that occurred between 17-25 million years ago in the *Papaver* lineage and was preceded by clustering and segmental duplication of the P450 oxidase and oxidoreductase genes that fused to form it. While this gene fusion is regarded as a key event enabling the evolution of the morphinan subclass of BIAs our findings show that it may also have enabled the production of compounds in other BIA subclasses such as the aporphine (*R*) -glaucine in *P. californicum,* which 16.8 mya branched from the common ancestor of those other *Papaver* species that now produce morphinans.

## Supporting information

Supplementary sequence datasets organised in relation to main text figures

## Acknowledgements

I.A.G. received support from the Biotechnology and Biological Sciences Research Council, United Kingdom (grant BB/K018809/1) and the Garfield Weston Foundation, United Kingdom. We are grateful to York Technology Facility, COEMS for support with metabolite profiling and RNA-seq library preparation as well as the University of York High Performance Computing service, Viking and the Research Computing team for support with bioinformatics.

## Author Contributions

T.C. conducted all RNA and genomic DNA preparation, molecular biology, biochemical and enzyme analysis, T.C, F.M, DH and Y.R.L conducted all metabolite analysis, T.C and Y.L. all transcriptomic analysis, Y.L. and A.C. species phylogeny analyses, Y.L. all gene tree analyses, Y.L. and Z.N. all draft genome assemblies and Y.L all synteny analysis. T.C., Y.L., T.W. and I.A.G. analyzed and interpreted results and were major contributors in writing the manuscript. All authors read and approved the final manuscript.

## Materials and Methods

### Plant material

The plant material used in the current study were voucher specimens sourced primarily from botanic institutions as detailed in supplementary table 1. Plants were grown under controlled long day conditions in the glasshouse facilities and in the experimental gardens located in the University of York. Samples for DNA cDNA and RNA were collected from young leaves of juvenile plants and flash frozen in liquid nitrogen. Metabolite analysis was conducted on latex harvested from 3 individual glasshouse grown plants at 70 days post germination and from capsules pre dehiscence. Metabolite extraction and UPLC analysis were carried out following the protocol as described in [3].

### Genomic DNA and cDNA isolation for PCR

Genomic DNA was extracted from frozen ground young leaf using the BioSprint 15 Plant Kit on the BioSprint 15 Workstation (Qiagen, Crawley, UK. The DNA was quantified on the nanodrop 1000 (thermofisher). cDNA was synthesised from RNA isolated from young leaf tissue using superscript V (Thermofisher) and used for gene specific amplification.

### Metabolite Profiling

Latex and capsule samples were collected and analysed as described in Winzer *et al.* (2012)[27]. Authentic standards for salutaridine, salutaridinol and salutaridinol 7 O acetate* (Toronto research chemicals, Canada), and also a MCONT standard mix (morphine, codeine, oripavine noscapine and thebaine) were included to confirm the presence or absence of promorphinan and morphinan compounds in the 10 species.

### Isolation of glaucine by preparative HPLC

20g of dried capsule material from *P.californicum* or *G. flavum* plants was ground to a fine powder and extracted with 25mls of 10% acetic acid in water. The plant material was removed by centrifugation at 4000g for 10 minutes and the extract dried down using an EZ-2 elite genevac. The dried residue was taken up in 2 ml of ethyl acetate and further spun to remove debris prior to purification. Isolation and purification of glaucine was performed using the interchim puriflash 4500 prep HPLC system with Advion Expression, compact mass spec (CMS). The ethyl acetate extracts were applied to a 12g BUCHI FlashPure EcoFlex silica 50μm irregular column and all fractions collected using a 0-100 % ethyl acetate in hexane gradient, followed by isocratic 100% ethyl acetate and 100% methanol. Samples from fractions thought to contain the glaucine peak (356 ion) were confirmed by running on waters acquity UPLC linked to a Thermo LTQ orbitrap mass spec on an acquity BEH c18 1.7μm 2.1×100mm column with the mass spec using APCI ionization in positive polarity. The glaucine containing fractions were dried down and resuspended in 10% acetic acid for further purification on a 4g TELOS Flash C18 column with fractions collected using a 2-80% gradient of Solvent B in Solvent A (where Solvent A is 10mMAmonium bicarb pH10.2; Solvent B is 100% methanol). This method yielded 3mg of glaucine from *P. californium* and 2mgs from *G. flavum* this was resuspended in 100% ethanol to a 2mg/ml final concentration for circular dichroism analysis.

### RNA-Seq sequencing and transcriptomic data analyses

Transcriptomic analysis was performed on RNA extracted using the Direct-zol RNA Miniprep Kit (Zymo Research, USA) according to the manufacturer’s instructions from young frozen leaf material. RNA was quantified on the Qubit 3.0 Fluorometer (Thermo Fisher Scientific) according to the manufacturer’s protocol and quality assessed by running 1ul on the agilent tapestation.

RNA sequencing was performed on the nine Papaver species (supplementary tableS1). RNA-Seq libraries were prepared from 1μg high quality RNA using the NEBNext Poly(A) mRNA Magnetic Isolation Module and NEBNext Ultra II Directional RNA Library Prep Kit for Illumina (New England Biolabs), according to the manufacturer’s guidelines. Libraries were subject to 150 base paired end sequencing on one lane of a HiSeq 3000 system at the University of Leeds Next Generation Sequencing Facility (Leeds, UK) except for P. californicum which was sequenced on NovoSeq 6000 at Novogene Co, Cambridge, UK. An average of 7.5Gb PE reads sequencing data were generated for each species (supplementary tableS3). These raw RNAseq datasets have been deposited at the National Center for Biotechnology Information (accession nos. xxx– xxx).

Transcriptomes were assembled with the Trinity (v2.2.0) RNA-Seq De novo Assembly software pipeline (28) after filtering out any of the remaining 1-5% ribosomal RNA in the raw reads for each species with mapping to rRNA_115_tax_silva_v1.0 downloaded from SILVA database (https://www.arb-silva.de/). These transcriptome datasets were used for identification of orthologous genes of selected BIA biosynthetic genes (supplementary tableS4) and the eight conserved ortholog sets (COS) genes (29; 30; supplementary tableS6; supplemental dataset2) by local BLAST searches. Sequences of the top matches were retrieved. Orthologous gene sequences belonging to gene families were identified and verified through subsequent gene tree analysis along with the related gene family’s datasets that were previously reported (6; supplementary tableS5; supplemental dataset1).

### Species phylogeny and estimation of divergence times

Estimations of divergence dates among the nine *Papaver* species together with opium poppy were based on the 8 COS gene sequences from the transcriptomic datasets (supplement table S6 and supplemental dataset2) along with other Papaveraceae species *E. californica*, *M. cordata* and one Ranunculaceae species *A. coerulea* with BEAST2 v2.5.1 (31).

Orthologous sequences of the eight COS genes were identified and retrieved from the annotated gene datasets of *P. somniferum*, *E. californica*, *M. cordata* and *A. coerulea* as described previously (6) after conducting BLAST searches. Multiple sequence alignments of each gene set were obtained with MUSCLE v3.2 (32). Conservative alignment blocks were generated with Gblocks v0.91 (33) to remove highly polymorphic regions and the alignments of all eight genes were subsequently concatenated in MegaX (34).

Species divergence times were estimated under the strict clock model implemented in BEAST v2.5.1 to generate a Bayesian posterior sample of time-calibrated phylogenies and the associated maximum clade credibility tree using two prior calibration points (Ranunculales 110 +/− 5MYA and Papaveraceae 77+/−4 MYA [6]). Priors were treated as fitting a Yule speciation process and lognormal distribution; the nucleotide substitution model used was GTR with 4 Gamma categories. The chain-length of Markov chain Monte Carlo (MCMC) was set to 30,000,000 runs, performed per 10,000 generations and collected every 1000th generation; with 10% of the total trees discarded as burn-in samples, the remaining trees were used for generating the consensus tree. Convergence of the runs performed by BEAST was assessed by effective sample sizes (ESS) using Tracer v1.5 [31]: ESS exceeded 1000 for all summary statistics, greatly above the threshold of 200 that is considered to indicate good sample quality. The species tree and divergence times were visualised using Figtree v1.4.3 [35].

### Heterologous expression of Papaver STORR for functional analysis

Full length cDNAs for STORR were cloned and sequenced after PCR amplification from *P. californicum* P. armeniacum and *P. bracteatum*. using the following primers Pbra_STORR Fwd-Pbra_STORR Rev, Pcal_STORR Fwd--Pcal_STORR Rev and Parm_STORR Fwd Parm_STORR Rev(supplemental dataset4).

Percentage identity matrix between pairs of STORR homologs in Papaver species were calculated with pairwise sequence comparison tool EMBOSS Matcher version 2.0u4 [36] (supplementary tableS7). Multiple sequence alignments of the amino acid sequences were obtained with MUSCLE v3.2 [32].

Geneart gene synthesis service (Thermofisher) created synthetic DNA sequences for the STORR gene of *P. californicum*, *P .armeniacum* and *P. bracteatum* identified from transcriptome sequence data. The sequences were codon optimised for expression in Saccharomyces cerevisiae. The codon optimised sequences are provided in supplemental dataset 5. The codon optimised genes were cloned from the Geneart pMK vector by digesting with NotI and PacI and inserting the cleaned DNA fragments behind the Gal10 promoter of the pESC-TRP expression vector. Vectors were sequenced for errors before being transformed into the S. cerevisiae G175 using lithium acetate protocol [37]. Yeast cultures were grown, and microsomal preparations were performed; the resulting crude microsomal preparations (12 mg mL-1 protein) were used for enzyme assays using 100um reticuine for the chiral analysis of the reticuline epimers. Reticuline was extracted from the assay reactions with dichloromethane, and epimers analysed by chiral LCMS [3], with the following modifications: a Lux 250 x 4.6mm 5u Cellulose-3 column (Phenomenex) was used, with LC performed using a Waters Acquity I-Class UPLC system interfaced to a Thermo Orbitrap Fusion Tribrid mass spectrometer under Xcalibur 4.0 control. MS1 and data dependent MS2 spectra were collected at 60000 resolution (FWHM), with the precursor m/z 330.1700 added to the acquisition method inclusion list. S-reticuline eluted at ~12min and R-reticuline at ~16 min. Data was processed using Xcalibur software.

### Whole genome sequencing and assemblies

10X Genomics whole genome sequencing and assemblies were performed on five Papaver species, including *P. nudicaule*, *P. californicum*, *P. atlanticum*, *P. bracteatum* and *P. armeniacum*. Further Oxford Nanopore Technology (ONT) sequencing and genome assemblies were carried out on two of these species *P. bracteatum* and *P. armeniacum*.

High molecular weight (HMW) genomic DNA was prepared for 10X Genomics sequencing for five species. Young seedling material was grown and sent to Amplicon Express (Pullman, WA, USA), where HMW genomic DNA was prepared by using their proprietary protocol for HMW grade (megabase size) DNA preparation. This protocol involves isolation of plant nuclei and yields pure HMW DNA with >100kb fragment length. The DNA samples were then used to construct the 10X Chromium libraries, which were subsequently sequenced on Illumina NovaSeq platform to produce 2 x 150bp reads at HudsonAlpha Institute for Biotechnology, Huntsville, Alabama. This produced a total of over 2 billion x2 reads or 600Gb of 10X Chromium library sequencing data for each species.

We used GenomeScope v1.0 [38] to estimate the genome size and heterozygosity level for each species. Firstly, a k-mer distribution was generated with Jellyfish [39] from the 10X Genomics short reads, and then used as input in the subsequent GenomeScope analysis (supplementary tableS8). De novo draft 10X assemblies were produced with the linked reads at approximately 50-60 times coverage of the estimated genome size, as required by the Supernova assembly software package (supplementary table S8) [40] for optimal performance.

Further sequencing of *P. bracteatum* and *P. armeniacum* was carried out using ONT long read sequencing platform by the Technology Facility at the University of York. HMW DNA samples were prepared as described in Vaillancourt *et al*. (2019) [41]. Long read sequencing was performed by the University of York Biosceince Technology Facility Genomics lab, using the Oxford Nanopore Technologies MinION and PromethION platforms. Sequencing libraries were prepared using ONT’s ligation sequencing kit SQK-LSK106, using a minimum of 5 μg high quality DNA. The resulting library was split into 4 to allow loading onto MinION (FLO-MIN106) and PromethION (FLO_PRO002) R9.4.1. flow cells, and sequencing for 48 hours (MinION) or 72 hours (PromethION), with a flow cell wash (using ONT wash kit EXP-WSH003) and library reloading step approximately 24 hours into the runs. Basecalling was performed using ONT’s Guppy toolkit, version 4.0.11. Just over 7 million reads were generated with average read length of 13,395bp and 95Gb in total for *P. bracteatum*, whereas 4.5 million reads with average length of 22,216bp covering 99G nucleotide bases for *P. armeniacum*.

We ran the FLYE version2.8-b1674 [42] de novo assembly pipeline with the ONT datasets, producing initial assemblies. This was followed by one round base polishing with the starting ONT raw reads using RACON (https://github.com/isovic/racon). Purge_dups [43] was then used to remove haplotigs and contig overlaps to produce a haploid representation of the genome. Final draft genome assemblies (supplementary tableS8) were achieved after two more rounds of base polishing using FREEBAYES software tool [44] after the Illumina short reads from the 10X Chromium libraries were mapped to the working assembly with Longranger software package (https://support.10xgenomics.com/genome-exome/software/pipelines/latest/what-is-long-ranger).

The presence of orthologous genes of selected BIA biosynthetic genes (supplementary tableS4) were conducted firstly by local BLAST searches in the 10X assemblies of *P. nudicaule*, *P. californicum*, and *P. atlanticum*; but the final draft ONT assemblies of *P. bracteatum* and *P.armeniacum*. Sequences of the top matches were retrieved and verified through subsequent gene tree analysis along with the related gene families datasets that were previously reported [6] (supplementary tableS5).

### Gene tree analyses of *CYP82Y2-L* and *COR-L* gene subfamilies

Gene tree analyses were used to understand the evolutionary events leading to *STORR*s in *Papaver species* and to reconstruct a detailed evolutionary history by analysing the entire set of homologous representatives to STORRs and their closest paralogous *CYP82Y2-L&COR-L* gene pairs within the five representative *Papaver* genomes, by reconciling the gene duplication events implied in gene trees with the Papaver speciation tree with the parsimony principles.

We searched the above five draft genome assemblies with BLASTN to identify regions corresponding to either of the STORR modules. The sequences of these regions were then retrieved and manually annotated with their sequence homology to other closely related genes. All *CYP82Y2-L* and *COR-L* homologous sequences that contain full length coding regions were retained for the further gene tree analysis.

As the draft *P.nudicaule* 10X assembly is highly fragmented, likely caused by high heterozygosity level in its genome, we obtained full length coding regions for all potential candidates by PCR amplification from both cDNA libraries and genomic DNA using gene specific primers. Fragments to confirm the P. nudicaule CYP82y2 and cor modules was carried out by RT-PCR and sequencing of fragments using the following primers PN_CYP+COR_1717256_Fwd - ATGGATTACTCTAATCTTCAG and PN_CYP+COR_1717256_Fwd - CTC TAA TCT TCA GTT TTT CGG. The amplified fragments were cloned and sequenced for verification.

We also retrieved *CYP82Y2-L* and *COR-L* homologous sequences after BLASTN searches in the annotated opium poppy genome, the transcriptomic data from the nine species from the present study, as well as the GenBank database as described in [6] Duplicated and partial sequences were removed together with other synthetic sequence entries in the Genbank database. These were combined with the above set from the five Papaver genomes (supplemental dataset6&7) for the gene tree analyses.

Multiple sequence alignments of the coding sequence sets of the *CYP82Y2-L* and *COR-L* were built with MUSCLE v3.2 [32] and conservative alignment blocks were generated with Gblocks v0.91 [33] by removing highly polymorphic regions. The gene trees were constructed using Bayesian analyses with MrBayes v3.2.6 [45]. The trees were then visualised using Figtree v1.4.3 [35]. Posterior probabilities were reported as supporting values for nodes in the trees and scale bar represents substitutions per nucleotide site (Figure3a&b).

The sequence regions containing all paralogous CYP82Y2-L&COR-L gene pairs were retrieved from the draft genome assemblies and manually annotated (supplemental dataset8, Fig. 3c).

### Synteny analysis of *STORR* plus pro-morphinan component of the BIA gene cluster among *Papaver* genomes

To investigate the synteny in the genomic regions containing *STORR* or the four pro-morphinan biosynthetic genes, we compared the gene order and orientation in the opium poppy 800kb BIA cluster and flanking regions to the contigs/scaffolds in the five representative *Papaver* genomes. The contigs/scaffolds were identified after BLAST searches for containing *STORR* or *SALSYN*/*SALAT*/*SALR*/*THS* genes in the draft genome assemblies and their sequences were retrieved (supplemental dataset9). A single contig was identified in P.bracteatum genome and its length is 1.2Mb, whereas the single scaffold in P.californicum is the shortest with 44kb in length. Two contigs were found in P.armeniacum to contain STORR/SALSYN genes, one is 294kb long and the other 94kb in length.

*Ab initio* gene prediction of these contigs and scaffolds was carried out using FGENESH, a web-based gene annotation tool [46] http://www.softberry.com/berry.phtml?topic=fgenesh&group=programs&subgroup=gfind) with Dicot plants option as training set. The predicted genes were functionally annotated by homologous BLAST searches in the swissprot database. Reciprocal BLASTN searches were performed between the coding sequence sets of predicted genes in these contigs/scaffolds and the annotated opium poppy genome [5] http://bigd.big.ac.cn/search?dbId=gwh&q=opium%20poppy&page=1). The top match records in both sets with an expected value less than 1E-120 with its corresponding queries were summarised in supplementary tableS5 and Figure 4a.

## ● Supplementary figures

**Supplementary Figure 1.**
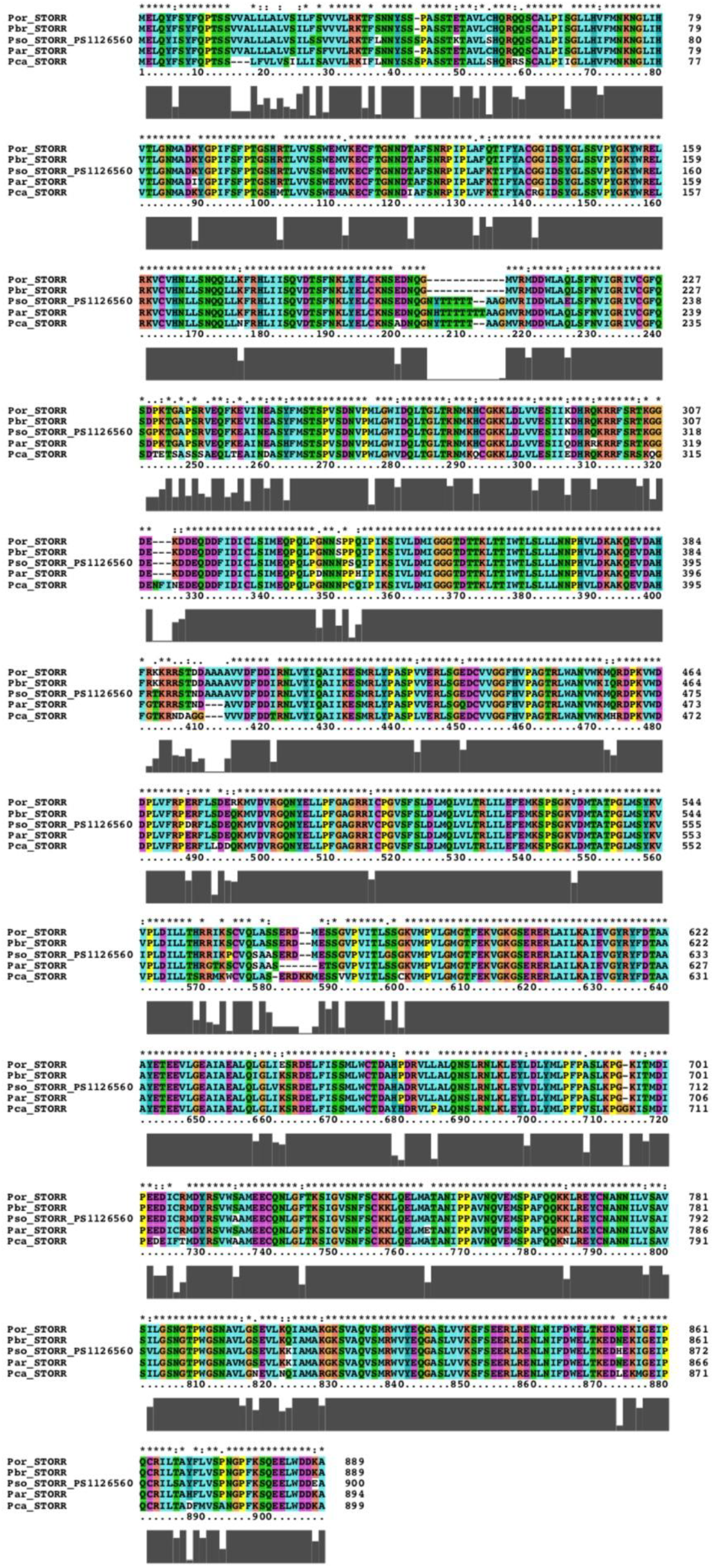
Multiple sequence alignment of STORR proteins from five Clade 2 *Papaver* species (Supplementary dataset 3&4)

**Supplementary Fig 2.**
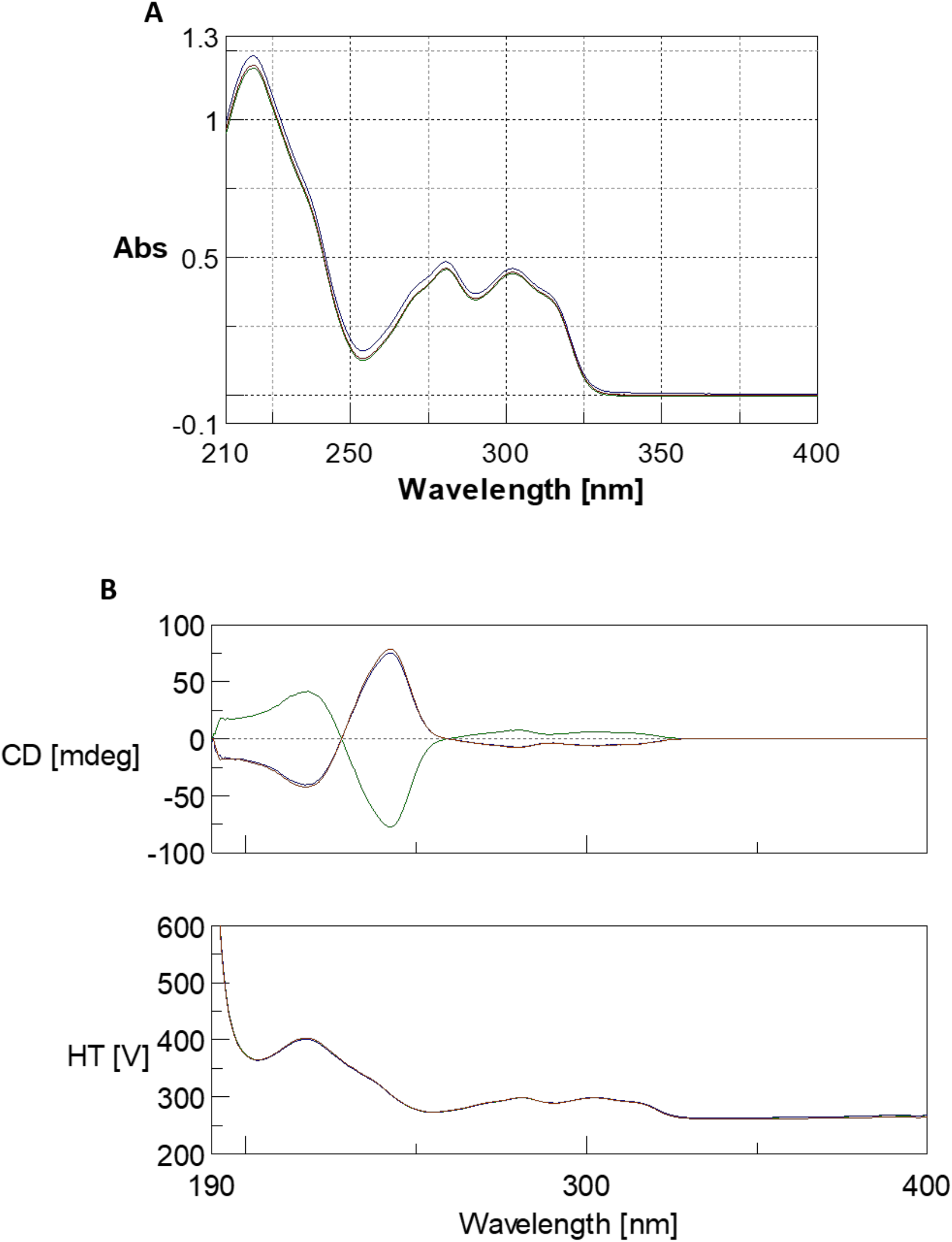
**A** - UV spectra for Authentic (S)-glaucine standard (blue), Glacium flavum glaucine preparative HPLC extract (green) and Papaver californicum preparative HPLC extract (brown). **B** - Overlay of the CD spectra for Authentic (S)-glaucine standard (brown), Glacium flavum glaucine preparative HPLC extract (blue) and Papaver californicum preparative HPLC extract (green). The concentration of samples run 0.125mg/ml

**Supplementary figure 3.**
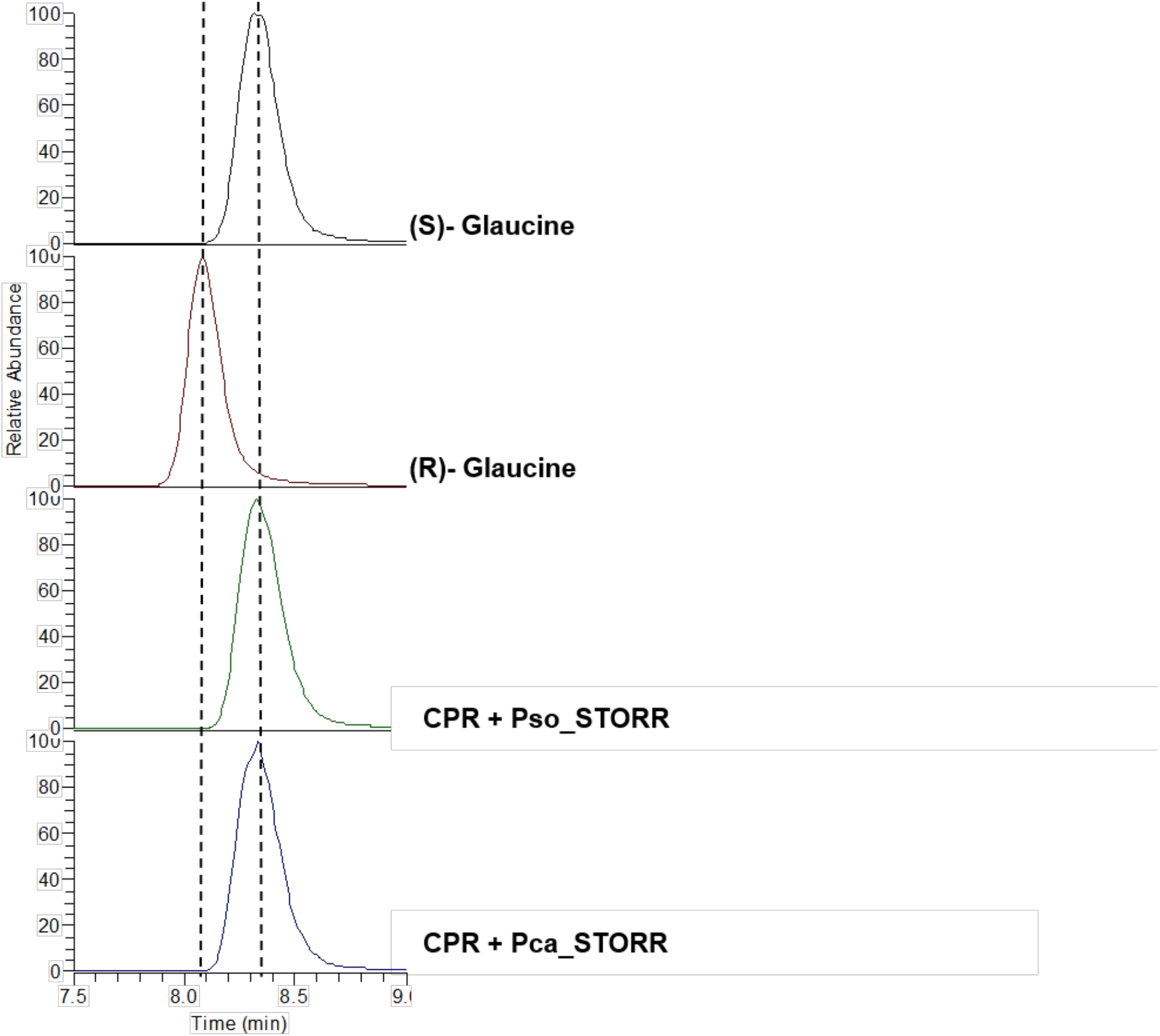
Enzyme assay with (S)-Glaucine substrate. Substrate specificity of the microsomal extracts from S. cerevisiae expressing the STORR protein from *P. californicum* alongside *P. somniferum* (control) were assayed with (S)-Glaucine as a substrate. Relative abundance is used to show the glaucine epimers present in the sample. Glaucine standards-(S)-glaucine (black) (R)-Glaucine (red), pESC-TRP+ CPR+ Pso STORR (green) and pESC-TRP+ CPR+ Pca STORR (blue).

**Supplementary figure 4.**
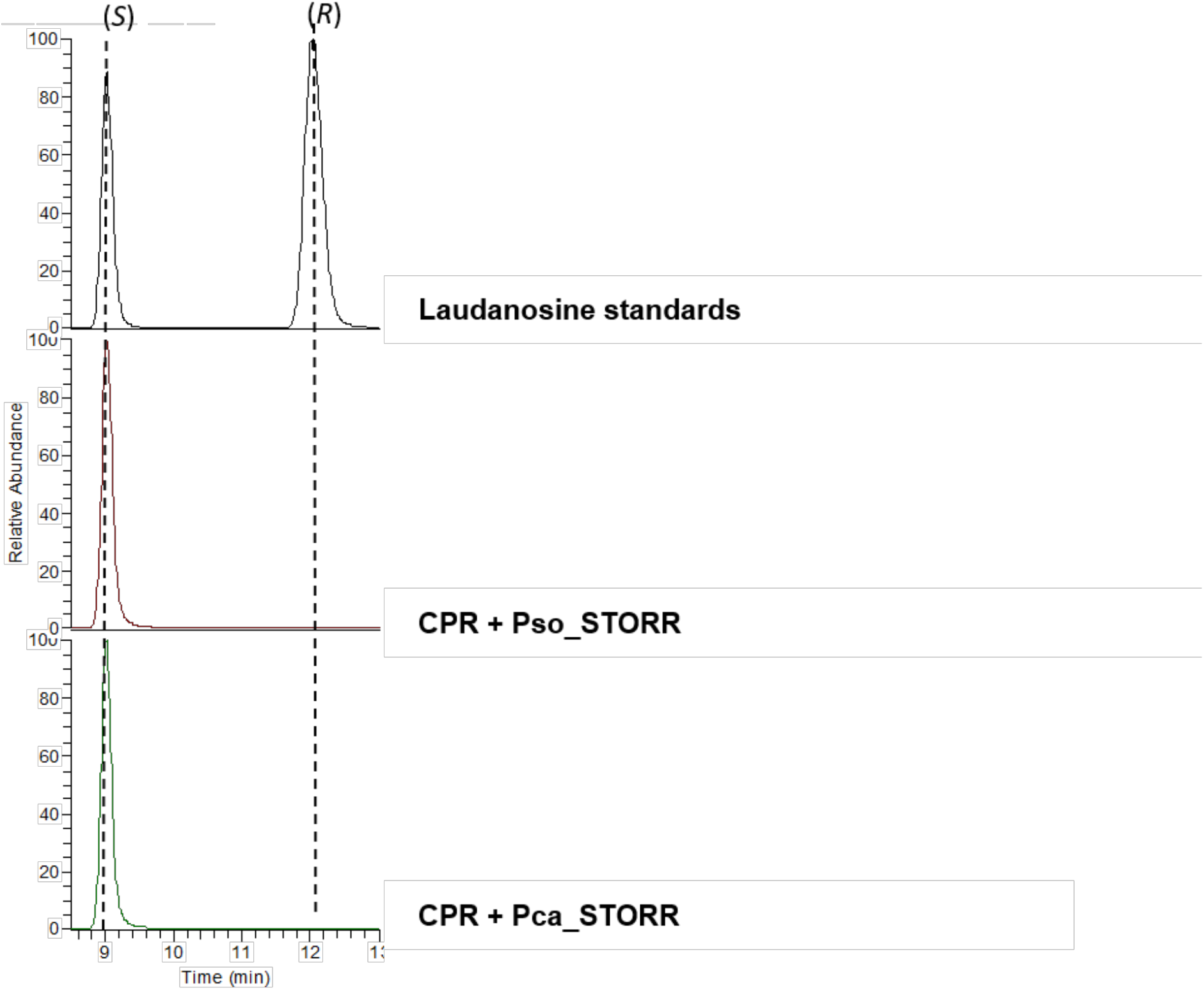
Enzyme assay with (S)-laudanosine substrate. Substrate specificity of the microsomal extracts from S. cerevisiae expressing the STORR protein from *P. californicum* alongside *P. somniferum* (control) were assayed with (S)-laudanosine as a substrate. Relative abundance is used to show the laudanosine epimers present in the sample. Laudanosine standards (black), pESC-TRP+ CPR+ Pso STORR (red) and pESC- TRP+ CPR+ Pca STORR (green).

## References

1. Ziegler, J. et al. Evolution of morphine biosynthesis in opium poppy. Phytochemistry. 70, 1696–1707 (2009).

2. Hagel, J. M. & Facchini, P. J. Benzylisoquinoline alkaloid metabolism: a century of discovery and a brave new world. Plant and Cell Physiol. 54, 647–672 (2013).

3. Winzer, T, et al. Morphinan biosynthesis in opium poppy requires a P450-oxidoreductase fusion protein. Science. 349, 309–312 (2015).

4. Farrow, S. C., Hagel, J. M., Beaudoin, G. A., Burns, D. C., & Facchini, P. J. Stereochemical inversion of (*S)*-reticuline by a cytochrome P450 fusion in opium poppy. Nature Chem. Biol. 11, 728–732 (2015).

5. Guo, L. et al. The opium poppy genome and morphinan production. Science. 362, 343–347 (2018).

6. Li, Y., Winzer, T., He, Z. & Graham, I.A. Over 100 million years of enzyme evolution underpinning the production of morphine in the Papaveraceae family of flowering plants. Plant Commun. 1(2), (2020).

7. Galanie, S., Thodey, K., Trenchard, I. J., Interrante, M. F., & Smolke, C. D. Complete biosynthesis of opioids in yeast. Science 349, 1095–1100 (2015).

8. Osbourn, A. Gene clusters for secondary metabolic pathways: an emerging theme in plant biology. Plant Physiol. 154, 531–535 (2010).

9. Nützmann, H.W. & Osbourn, A. Gene clustering in plant specialized metabolism. Curr. Opin. Biotech. 26, 91–99 (2014).

10. Hagel, J M. & Facchini, P.J. Tying the knot: occurrence and possible significance of gene fusions in plant metabolism and beyond. J. Exp. Bot. 68, 4029–4043 (2017).

11. Liu, Z., et al. Formation and diversification of a paradigm biosynthetic gene cluster in plants. Nature Commun. 11, 1–11 (2020).

12. Li, Q. et al. Gene clustering and copy number variation in alkaloid metabolic pathways of opium poppy. Nature Commun. 11, 1–13 (2020).

13. Gesell, A., et al. CYP719B1 is salutaridine synthase, the CC phenol-coupling enzyme of morphine biosynthesis in opium poppy. Journal of Biological Chemistry 284(36), 24432–24442 (2009).

14. Ziegler, J., Diaz-Chávez, M.L., Kramell, R., Ammer, C. and Kutchan, T.M. Comparative macroarray analysis of morphine containing *Papaver* somniferum and eight morphine free *Papaver* species identifies an O-methyltransferase involved in benzylisoquinoline biosynthesis. Planta 222(3), 458–471 (2005).

15. Grothe, T., Lenz, R. and Kutchan, T.M. Molecular characterization of the salutaridinol 7-O-acetyltransferase involved in morphine biosynthesis in opium poppy *Papaver* somniferum. Journal of Biological Chemistry 276(33), 30717–30723 (2001).

16. Chen, X., et al. A pathogenesis-related 10 protein catalyzes the final step in thebaine biosynthesis. Nature chemical biology 14(7), 738–743 (2018).

17. Nyman, U. & Bruhn, J. G. *Papaver bracteatum* –a summary of current knowledge. Planta Med. 35, 97–117 (1979).

18. Sariyar, G. Biodiversity in the alkaloids of Turkish *Papaver* species. Pure Appl. Chem. 74, 557–574 (2002).

19. Alagoz, Y., Gurkok, T., Parmaksiz, I. & Ünver, T. Identification and sequence analysis of alkaloid biosynthesis genes in *Papaver* section Oxytona. Turk. J. Biol. 40, 174–183 (2016).

20. Carolan, J. C. et al. Phylogenetics of *Papaver* and related genera based on DNA sequences from ITS nuclear ribosomal DNA and plastid trnL intron and trnL–F intergenic spacers. Ann. Bot - London. 98, 141–155 (2006).

21. Lane, A. K. et al. Phylogenomic analysis of Ranunculales resolves branching events across the order. Bot. J. Linn. Soc. 187, 157–166 (2018).

22. Hoot, S. B., Wefferling, K.M & Wulff. J.A. Phylogeny and character evolution of Papaveraceae s.l. (Ranunculales). Syst. Bot. 40, 474–488 (2015).

23. Cortijo, J. V. et al. Bronchodilator and anti-inflammatory activities of glaucine: In vitro studies in human airway smooth muscle and polymorphonuclear leukocytes. British journal of pharmacology. 127 (7), 1641–1651 (1999).

24. Zetler, G. Neuroleptic-like, anticonvulsant and antinociceptive effects of aporphine alkaloids: bulbocapnine, corytuberine, boldine and glaucine. Archives internationales de pharmacodynamie et de therapie. 296, 255–281 (1988).

25. Heng, H. L. et al. In vitro functional evaluation of isolaureline, dicentrine and glaucine enantiomers at 5-HT2 and α1 receptors. Chemical biology & drug design. 93 (2). 132–138 (2019).

26. Bhakuni, D. S., & Jain, S. The biosynthesis of glaucine in Litsea glutinosa. J. Chem. Soc., Perk. T. 1. 6, 1447–1449 (1988).

## References (materials and methods specific)

27. Winzer, T. et al. 2012. A Papaver somniferum 10-gene cluster for synthesis of the anticancer alkaloid noscapine. Science. 336(6089). (2012)

28. Grabherr, M.G. et al. Full-length transcriptome assembly from RNA-seq data without a reference genome. Nat Biotechnol. 15, 644–52 (2011).

29. Li, M.et al. Development of COS genes as universally amplifiable markers for phylogenetic reconstructions of closely related plant species. Cladistics. 24(5). (2008)

30. Levin, R.A., Blanton, J. and Miller, J.S. Phylogenetic utility of nuclear nitrate reductase: A multi-locus comparison of nuclear and chloroplast sequence data for inference of relationships among American Lycieae (Solanaceae). Molecular Phylogenetics and Evolution, 50(3). (2009).

31. Bouckaert, R., Vaughan, T.G., Barido-Sottani, J., Duchêne, S., Fourment, M., Gavryushkina, A., Heled, J., Jones, G., Kühnert, D., De Maio, N. and Matschiner, M., 2019. BEAST 2.5: An advanced software platform for Bayesian evolutionary analysis. PLoS computational biology, 15(4), p.e1006650.

32. Edgar, R.C., 2004. MUSCLE: multiple sequence alignment with high accuracy and high throughput. Nucleic acids research, 32(5), pp.1792–1797.

33. Talavera, G. and Castresana, J., 2007. Improvement of phylogenies after removing divergent and ambiguously aligned blocks from protein sequence alignments. Systematic biology, 56(4), pp.564–577.

34. Kumar, S., Stecher, G., Li, M., Knyaz, C. and Tamura, K., 2018. MEGA X: molecular evolutionary genetics analysis across computing platforms. Molecular biology and evolution, 35(6), p.1547.

35. Rambaut, A., 2014. Tree figure drawing tool. FigTree v1, 4.

36. Madeira, F., Park, Y.M., Lee, J., Buso, N., Gur, T., Madhusoodanan, N., Basutkar, P., Tivey, A.R., Potter, S.C., Finn, R.D. and Lopez, R., 2019. The EMBL-EBI search and sequence analysis tools APIs in 2019. Nucleic acids research, 47(W1), pp.W636–W641.

37. Gietz, R.D. and Schiestl, R.H., 2007. High-efficiency yeast transformation using the LiAc/SS carrier DNA/PEG method. Nature protocols, 2(1), pp.31–34.

38. Vurture, G.W., Sedlazeck, F.J., Nattestad, M., Underwood, C.J., Fang, H., Gurtowski, J. and Schatz, M.C., 2017. GenomeScope: fast reference-free genome profiling from short reads. Bioinformatics, 33(14), pp.2202–2204.

39. Marçais, G. and Kingsford, C., 2011. A fast, lock–free approach for efficient parallel counting of occurrences of k-mers. Bioinformatics, 27(6), pp.764–770.

40. Weisenfeld, N.I., Kumar, V., Shah, P., Church, D.M. &, Jaffe, D.B. Direct determination of diploid genome sequences. Genome Research. 27, 757–767 (2017).

41. Vaillancourt, B. & Buell, C. R. High molecular weight DNA isolation method from diverse plant species for use with Oxford Nanopore sequencing. BioRxiv, 783159 (2019).

42. Kolmogorov, M., Yuan, J., Lin, Y. and Pevzner, P.A., 2019. Assembly of long, error–prone reads using repeat graphs. Nature biotechnology, 37(5), pp.540–546.

43. Guan, D McCarthy, Wood J, Howe K, Wang y, and Durbin R, 2020. Identifying and removing haplotypic duplication in primary genome assemblies. Bioinformatics, 36(9), pp. 2896–2898.

44. Garrison, E. and Marth, G., 2012. Haplotype-based variant detection from short-read sequencing. arXiv preprint arXiv:1207.3907.

45. Ronquist, F. & J. P. Huelsenbeck. MRBAYES 3: Bayesian phylogenetic inference under mixed models. Bioinformatics. 19, 1572–1574 (2003).

46. Solovyev, V., Kosarev, P., Seledsov, I. & Vorobyev, D. Automatic annotation of eukaryotic genes, pseudogenes and promoters. Genome Biol. 7, 10.1–10.12 (2006).

