## Supplementary sequence datasets organised in relation to main text figures for "A functionally conserved *STORR* gene fusion in *Papaver* species that diverged 16.8 million years ago"

Supplementary Dataset 1 (associated with Figure 1c)

- BIA genes identified from the transcriptomic dataset of nine *Papaver* species.

Supplementary dataset 2 (associated with Figure 1c)

- Eight conserved orthologous gene set from the transcriptomic dataset of nine *Papaver* species and annotated genome datasets of *P. somniferum*, *Macleaya cordata*, *Eschscholzia californica* and *Aquilegia coerulea* species.

Supplementary dataset 3 (associated with Figure 2a)

- Protein sequences of STORR proteins from five Clade 2 *Papaver* species

Supplementary dataset 4 (associated with Figure 2a)

- Nucleotide sequences of full length STORR cDNAs from five Clade 2 *Papaver* species

Supplementary dataset 5 (associated with Figure 2b)

- Nucleotide sequences of yeast codon optimised full length STORR cDNAs from *Papaver* species

Supplementary dataset 6 (associated with Figure 3a)

- Nucleotide sequences of *Papaver* P450 cytochrome *STORR* and *CYP82Y2*-*L* genes

Supplementary dataset 7 (associated with Figure 3b)

- Nucleotide sequences of *Papaver* P450 cytochrome *STORR* and *COR*-*L* genes

Supplementary dataset 8 (associated with Figure 3c)

- Nucleotide sequences of genomic segments for *CYP82Y2*-*L*&*COR*-*L* gene pairs. Nucleotides of introns and intergenic regions are represented in lower case, exons in capital letters.

Supplementary dataset 9 (associated with Figure 4a)

- Nucleotide sequences of STORR-containing contigs/scaffolds in *Papaver* species

Supplementary Dataset 1 (associated with Figure 1c)

- BIA genes identified from the transcriptomic dataset of nine *Papaver* species.

>Pca_TRINITY_STORR Pca_TRINITY_DN1111_c0_g1_i3 len=3286

TTTGGATCAAAAGATTTTTCTATTATTTTGGAAAGGCTTATCAGTAGTACTAGTGCTTGCAAAAAGTAAGGATTAAAGATCTGTAGCTGGCTAGCTATATATACATGTGATGTGCAGGGTTCACCATGCATATCATTAAAATCCATTATCTATCCTACCTACTGGGTTGAATCATGGAGCTCCAGTACTTTTCTTATTTTCAGCCAACTTCCTCCCTTTTTGTTCTTGTATCCATATTACTGATCAGTGCTGTTGTTTTGAGGAAGATATTTTTAAATAACTACTCTTCATCACCTGCATCATCAACAGAAACAGCGCTGCTTTCTCATCAGCGGCGGTCGTCATGTGCATTGCCAATCATCGGTCTTCTTCATGTTTTCATGAATAAAAACGGCTTAATTCATGTAACTCTTGGAAATATGGCTGATAAATACGGACCGATTTTCAGTTTCCCAACAGGCAGCCATATGACTCTAGTTGTGAGCAGTTGGGAGATGGCAAAAGAGTGTTTTACAGGCAACAATGACATTGCTTTCTCAAACCGTCCTATCCCGTTAGTTTTTAAGACTATATTCTATGCATGCCGTGGTATTGACTCATACGGTCTTTCAAGTGTACCTTATGGAAAATATTGGAGGGAGTTACGAAAGGTGTGTGTGCATAACCTCCTCTCTAATCAACAACTACTCAACTTCAGACACTTGATAATTTCTCAAGTTGATACCTCTTTCAATAAGCTTTATGAGTTATGCAAAAACTCTGCAGACAACCAGGGAAACTACACTACTACTACTACCGCAGCTGGCATGGTGAGGATGGATGATTGGCTAGCGCAATTATCGTTCAACGTGATAGGAAGAATAGTCTGTGGATTCCAATCAGACACCGAGACAAGTGCTTCAAGCAGTGCGGAACAACTTACAGAAGCAATTAATGACGCATCGTATTTTATGTCGACATCTCCGGTGTCAGATAACGTTCCATGGTTAGGGTGGGTTGATCAACTAACTGGTCTTACTAGAAATATGAAGCAATGCGGAAAGAAATTAGACTTGGTGCTCGAGAGCATAATTGAAGATCATCGTCAAAAGAGACGGTTTTCTAGAAGTAAACAAGGAGATGAGAATTTTATCAACGAGGATGAACAAGATGACTTCATCGACATTTGCCTGTCAATTATGGAGCAACCACAGCTTCCTGGCAACAATAACCCTTGTCAAATACCTATCAAATCTATTGTCCTGGACATGATAGGTGGGGGGACCGACACCACAAAACTGACCACCATCTGGACCCTTTCATTGCTACTGAACAACCCCCATGTGTTGGACAAGGCAAAACAAGAAGTGGATGCACACTTCGGGACGAAAAGGAACGATGCAGGAGGAGTCGTGGTGGATTTTGATGATACTCGTAACCTTGTCTACATCCAGGCAATTATCAAAGAATCAATGCGGTTGTATCCAGCCAGCCCCTTGGTGGAGCGACTGAGCGGCGAAGATTGTGTGGTCGGTGGGTTTCATGTACCAGCAGGGACGCGGTTATGGGCTAACGTGTGGAAGATGCATCGAGACCCAAAAGTATGGGATGATCCATTGGTGTTTCGACCAGAGAGATTTTTGCTCGATGACCAGAAGATGGTTGATGTTAGGGGTCAAAACTATGAGCTGTTACCATTCGGAGCCGGTCGACGGATATGTCCAGGTGTATCGTTCTCTTTGGATCTAATGCAACTGGTACTGACTCGTCTTATTCTTGAGTTTGAAATGAAGTCTCCTAGCGGGAAACTAGACATGACAGCGACACCAGGTTTAATGAGTTACAAGGTAGTCCCCCTTGACATTCTGCTCACCAGTCGTCGCATGAAGTGGTGTGTGCAGTTAGCCTCTGAGAGAGATAAAAAGATGGAGAGTAGTGTTGTACCTGTAATCACTCTGAGCTCGTGCAAGGTGATGCCTGTTTTAGGCATGGGAACATTTGAAAAAGTTGGTAAAGGGTCTGAAAGAGAGAGGTTGGCGATTTTAAAAGCGATAGAGGTAGGTTACAGATACTTCGATACAGCTGCTGCATACGAAACTGAAGAGGTTCTTGGTGAAGCTATTGCTGAAGCACTTCAACTTGGCCTAATCAAATCTCGAGATGAACTCTTCATCAGTTCCATGCTCTGGTGCACTGATGCCTACCATGATCGAGTCCTCCCCGCTCTTCAGAATTCTCTGAGGAATCTTAAATTGGAGTATCTGGATCTATACATGTTACCCTTCCCGGTAAGCTTGAAGCCAGGAGGGAAGATAAGTATGGACATACCAGAGGATGAAATTTTTACGATGGACTATAGGTCTGTATGGGCAGCCATGGAAGAATGTCAGAACCTTGGCTTGACAAAATCAATCGGTGTCAGCAATTTCTCCTGCAAAAAGCTTCAGGAATTGATGGCGACCGCCAACATCCCTCCAGCTGTGAATCAAGTGGAGATGAGCCCGGCTTTCCAACAAAAGAATTTGAGAGAGTATTGCAATGCCAATAATATATTAATCAGTGCAGTTTCTATTTTGGGATCAAACGGAACCCCATGGGGCTCCAATGCAGTTCTGGGTAATGAGGTGCTAAACCAGATTGCTATGGCCAGAGGAAAATCTGTTGCTCAGGTTAGTATGAGGTGGGTTTACGAGCAAGGTGCGAGTCTTGTGGTAAAAAGTTTTAGTGAAGAGAGATTGAGGGAAAACTTGAACATATTCGACTGGGAACTCACTAAGGAAGACTTGGAAAAGATGGGTGAGATTCCACAATGCAGAATCTTGACTGCTGATTTTATGGTATCAGCTAATGGACCTTTCAAATCTCAAGAAGAGTTGTGGGATGACAAAGCTTGAAACATCGATCTTTAACTCTAGACGTTTATAAGAGAAGCTTCCCGGCAGTCACAGATCTCATTTCCCTTCCTATTGGTTTTGTACTCGCCTCTCTTGTTTTCTAAATAGAATCTGGTTTAGTTCCATCGTCTATAGTAAAGAAATAAATTTAGTACACTAAGCTTGTTTAATTTCTGTTGGATTGTAATTTGTAATGTCCAGAATTCTCTAGAGTAGTCTACATGTGTCTAGTGTGAACTCTAGATAGTAAAGTCGGCCAAAACAAACTACATAGCTAAGTTTTTTATACCAACTATAAGATTAGTCCTATGTCGAGAATTGTCTAGATAGTCTGTTGTAAAGGCTATAAATGGCCAAGAGTCAAGTATCATTTGTAAGTATCTATATCTATAATAACAAGGTAAGTTTACAAG

>Por_TRINITY_STORR Por_c53289_g1_i8 len=3106

TGTGCTCTTCCGATCTCCTACATACTGTAGTGGGTTGAATCATGGAGGTCCAATATTTTTCTTATTTTCAACCAACTTCCTCCGTTGTTGCTCTTCTACTTGCTCTTGTATCCATCTTATTCAGTGTCGTTGTTTTGAGGAAGACATTTTCGAATAACTACTCATCACCTGCATCATCAACAGAAACAGCGGTACTTTGTCATCAGTCGTGTGCATTGCCAATTTCCGGTCTTCTTCATGTTTTCATGAATAAAAACGGCTTAATTCATGTAACTCTTGGAAATATGGCGGATAAATACGGACCAATTTTCAGTTTCCCAACAGGTAGCCATAGAACTCTGGTTGTGAGCAGTTGGGAGATGGTAAAAGAGTGTTTTACAGGCAACAATGACACTGCTTTCTCAAACCGTCCTATCCCGTTAGCTTTTCAGACTATATTCTATGCATGCGGTGGCATAGACTCATACGGTCTTTCGAGTGTACCTTATGGAAAATATTGGAGGGAGCTACGAAAGGTCTGTGTGCATAACCTCCTCTCTAATCAACAACTACTCAAGTTCAGACACTTGATAATTTCTCAAGTCGATACGTCTTTCAATAAGCTGTATGAGTTATGCAAAAACTCTGAAGACAACCAGGGAATGGTGAGAATGGATGATTGGCTCGCCCAATTATCGTTCAATGTGATAGGAAGAATAGTCTGCGGATTCCAATCAGACCCAAAGACAGGTGCTCCTAGCAGGGTGGAACAATTTAAAGAAGTAATTAATGAAGCATCTTATTTTATGTCGACATCTCCAGTGTCAGATAACGTTCCAATGCTAGGGTGGATTGACCAATTGACTGGTCTTACGAGAAATATGAAGCACTGCGGAAAGAAATTAGACTTGGTGGTAGAGAGCATAATTAAAGATCATCGTCAAAAGAGACGATTCTCTAGAACTAAAGGAGGAGATGAGAAGGACGATGAACAAGATGACTTCATCGACATTTGCTTGTCAATTATGGAGCAACCACAGCTTCCTGGCAACAATAGCCCTCCTCAGATACCTATCAAATCTATTGTCCTGGACATGATAGGTGGGGGCACTGACACCACAAAACTGACCACCATCTGGACCCTTTCCTTGCTGCTGAACAACCCCCATGTGTTGGACAAGGCAAAACAAGAAGTGGATGCACACTTTCGAAAGAAAAGGAGATCAACAGATGATGCAGCAGCAGCCGTGGTGGATTTTGATGATATTCGTAACCTTGTCTACATCCAGGCAATCATCAAAGAATCAATGCGGTTGTATCCAGCCAGCCCCGTGGTGGAGCGACTGAGCGGCGAGGATTGTGTGGTCGGTGGGTTTCATGTACCAGCAGGGACGAGATTATGGGCTAACGTATGGAAGATGCAACGAGACCCTAAAGTATGGGATGATCCATTGGTGTTTCGACCAGAGAGATTTTTGAGCGATGAACGGAAGATGGTCGATGTAAGGGGTCAAAATTATGAGCTGTTACCATTTGGAGCCGGTCGACGTATATGTCCAGGTGTATCCTTCTCTTTGGATCTAATGCAACTGGTACTGACTCGTCTTATTCTTGAGTTTGAAATGAAGTCCCCTAGCGGGAAAGTGGACATGACAGCAACACCAGGATTAATGAGTTACAAGGTGGTCCCTCTTGACATTCTGCTCACCCATCGGCGCATAAAGTCGTGTGTGCAGTTAGCATCCTCTGAGAGAGACATGGAGAGTAGTGGTGTACCAGTAATCACTCTGAGCTCGGGCAAGGTGATGCCTGTTTTAGGCATGGGAACATTTGAGAAAGTTGGTAAAGGGTCTGAAAGAGAGAGGTTGGCGATTTTAAAAGCCATAGAGGTGGGTTACAGATACTTCGACACAGCTGCTGCATACGAAACTGAAGAGGTTCTTGGAGAAGCTATTGCTGAAGCACTTCAACTTGGCCTAATCGAATCTCGAGATGAACTTTTCATCAGTTCCATGCTCTGGTGCACTGATGCTCACCCTGATCGTGTCCTCCTCGCTCTTCAGAATTCTCTGAGGAATCTTAAATTGGAGTATCTGGATCTATATATGTTACCCTTCCCGGCAAGCTTGAAGCCAGGGAAGATAACGATGGACATACCAGAGGAAGATATTTGTCGAATGGACTACAGGTCTGTATGGTCAGCCATGGAAGAGTGTCAGAACCTTGGCTTCACTAAATCAATCGGTGTTAGCAATTTCTCCTGCAAAAAGCTTCAGGAATTGATGGCGACCGCCAACATCCCTCCAGCCGTGAATCAAGTGGAGATGAGCCCGGCTTTCCAACAAAAGAAGCTGAGAGAGTATTGCAACGCAAATAATATATTAGTCAGTGCAGTCTCTATACTGGGATCAAACGGAACCCCATGGGGCTCCAATGCAGTTTTGGGTTCTGAGGTGCTTAAGCAAATTGCTATGGCCAAAGGAAAATCTGTTGCTCAGGTTAGTATGAGATGGGTTTACGAGCAAGGCGCGAGTCTTGTGGTAAAAAGTTTCAGTGAAGAGAGATTGAGGGAAAACTTGAACATATTCGACTGGGAACTCACTAAGGAAGACAATGAAAAGATCGGTGAGATTCCACAGTGCAGAATCTTGACTGCTTATTTTTTAGTCTCGCCTAATGGACCTTTCAAATCTCAAGAAGAGTTGTGGGATGACAAAGCTTGAATATCGATCATTTAACTCTCTACATGTGCATTTATAAGAGAAGCTTCCCCTGCAGTGTGCTCAATCTTGATTTATTTTATCAAATTACATCTTACTATATGGGAGTCACAGATCTCTTTCCCTTCCTATATGTTTGAATACATTTGTATTGAGTGATGTAGCTAAAGAAATAAATGTAGTAAGTTTCTTTCCTAAGCTTGTTTTCCTTTACAAAAACTAGATGAGAGTCGGAGGTAAATAATGGTGAGCTCGTGTTCATCAGCAACAATAGTGCCATTAGGCACAATGTATATACCTTAGCAGAAAACACTAATCCAGACACAAAGAAATGTACGAATGCGTAAAGAGTATTCTGCCATGTATATACAACAGCTGATGCGAAAAAAAAAACAATACAAAACCTT

>Por_TRINITY_SALSYN Por_c54856_g1_i3 len=1864

TGTGCTCTTCCGATCTTTTTGATTCAATCTCTATTAATTAATAGTCAAATTCAATAGAGTAATTTCCTAATCTTTGCAAACTATCTTATATATTTCTGTAATTCATGGCTCCGATTAATATAGAGGGGAATGATTGTTGGATGATAGCATGCACTGTAATAATAGTATTTGCATTGGTGAAGTTCATGTTTTCTTTTTATCAATCTGCAAATACAACGGAATGGCCAGAAGGTCCAAAAACACTACCCATAATTGGAAACCTTCATCAGTTGGGAGGAGGTGTGCCCTTACAGGTTGCTTTGGCAAACTTAGCCAAAGTTTATGGAGGTGCGTTTACAATTTGGATTGGAAGCTGGGTTCCAATGATCGTCATAAGCGATATCGACAACGCTCGGGAAGTTCTTGTTAATAAATCTGCTGATTATTCCGCTAGAGATGTACCTGATATTCTTAAAATCATCACAGCAAATGGGAAGAATATTGCCGATTGTGATTCTGGTCCATTTTGGCATCATTTAAAGAAAGGTCTTCAAAGTTGTATAAATCCATCAAATGTTATGTCTCTATCTCGTTCACAGGAAAAAGACATGCAAAATGTCATCAAATCCATGCAAGAAAGAGCGTCAGCGCAGAATGGAATTTTAAAACCTCTTGATCATGCCAAAGAAGCGTCTATTCGATTACTGAGTAGAGTTATATTTGGTCAAGACTTTTCAAACGAGGATCTCGTTATTGGTGTGAAAGACGCCCTCGATGAGATGGTACGCATAAGTGGGTTGGCAAGTTTAGCTGATGCTTTTAAAATTGCTAAATATTTACCAAGCCAGAGAAAAAATATTCGGGATATGTACGCCACAAGAGACAGAGTATATAATTTGATTCAGCCACATATCGTCTCCAATCTTCCTGCAAATTCTTTCTTACATTTTCTTACGTCTCAAGATTACAGTGATGAAATTATTTACTCAATGGTACTTGAAATTTTTGGTTTGGGAGTAGATAGTACTGCAGCAACGGCAGTTTGGGCACTCTCCTTTTTAGTCGGCGAGCAGGAAATTCAAGAAAAACTTTACCGCGAAATCAACAACCTGACGGGTGGGCAAAGACCAGTGAAAGTTGTAGATTTGAAAGAGCTGCCATATCTACAAGCCGTGATGAAAGAAACATTGAGGATGAAACCCATCGCACCACTAGCGGTCCCTCATGTAGCAGCAAAAGATACTACATTCAAGGGGCGGAGAATCGTTAAAGGTACAAAAGTAATGGTGAATCTGTATGCTATTCATCACGACCCTAACGTTTTCCCTGCACCGTATAAATTCATGCCAGAGAGATTCTTAAAGGATGTTAACAGTGATGGACGTTATGGTGATATCAACACAATGGAAAGTTCATTGATACCATTTGGTGCTGGTATGAGAATTTGTGGAGGTGTAGAATTAGCCAAGCAGATGGTAGGTTTTGCTCTTGCAAGTATGGTCAACGAATTCAAATGGGATTGTGTTTCTGAGGGGAATTTGCCTGATCTTAGTGAAGCTATTAGCTTCATTCTCTACATGAAAAACCCACTTGAAGCCAAAGTTACCCCTCGTACAAAACCGTTCGACAGTAGATAAAATCTTATTTTTATCGAACTGCTAGTGTATGTTTATGTAAAATCAAGGGTGAAGATGCTTGATCGTAGTACACCAAAATGGTGGTGGCTATGTAGGAGTTAGATTAGCTCTTGATTTAGTAAATGTTTTTCTTTGCTAGAGACTGTCTCAAGTGTGTGTGTTTTAAGCGTGTGTTGTGTGTGTAATTACTTCCACCTGACAATTAAAGTGTACTTTAGTCGTCGGCTGTATCTGTCAGATCGGAAGAG

>Por_TRINITY_SALR Por_c56693_g3_i1 len=897

TGATATCTTGGTAAACAATGCTGGGGTTGCAGGTTTTTCAGTTGATGCTGATCGTTTTAAGGCAATGATAAGTGACATTGGAGAGGATTCAGAGGAGGTCGTGAAGATCTACGAAAAACCAGAAGCCCAAGAATTAATGTCAGAGACATATGAATTAGCAGAAGAATGTCTCAAAATAAATTACTACGGTGTTAAATCAGTAACCGAAGTTCTACTTCCTTTACTTCAACTATCTGATTCACCAAGAATTGTTAATGTTTCATCATCCACTGGAAGCCTCAAGTATGTATCCAATGAAACAGCTCTAGAGATACTTGGAGATGGTGATGCATTAACAGAAGAGAGAATTGACATGGTAGTGAATATGCTTCTTAAGGATTTTAAGGAAAATTTGATCGAAACAAATGGGTGGCCTAGTTTCGGAGCTGCATACACAACATCAAAAGCATGTTTGAATGCTTACACAAGGGTGTTAGCAAAGAAAATTCCCAAATTTCAGGTGAATTGTGTTTGTCCTGGTTTGGTTAAAACAGAAATGAACTACGGCATTGGAAATTATACTGCCGACGAAGGTGCTAAACATGTAGTCAGAATAGCTCTTTTCCCCGACGATGGACCTTCTGGTTTTTTCTATGATTGTTCTGAACTATCTGCATTTTGAGTTATATCGATTCAGCGTATCGTGCAGCAGCCAAATAAATAAAAATGGCATCAGTTGTTTGCTAGCACTGTCCTATCTACTGATAACTTTATCCCTTTGCTTCTGTTTTTAGCTTGTGTTGTGTGAATGTCCAGAGAGGATCGATCCATTTCTAATGTATTGAATCTTTATTTGGATGACAAATAATAAATCAGCTTGAGATTTTTCTTCCGATTTTAAAAAAAAAAGATCGGAAG

>Por_TRINITY_THS2 Por_c42204_g1_i1 len=877

GTGTGCTCTTCCGATCTCTTCTGTGCATATTTTAGTTTAGGGAACTAATCATCAAATTGTTGTTGATGGCTCCTCGTGGTGTTTCAGGTCTAGTCGGGAAACTTTCAACTGAATTAGATGTCAATTGCGATGCTGAAAAATATTATAACATGTATAAGAACGGAGAAGATGTACAAAAGGCAGTTCCTCATCTTTGCATGGACGTCAAAGTTATCAGTGGAGATGCAACCCGTTCAGGTTGTATCAAGGAATGGAATGTTAACATTGATGGTAAGACGATTCGCTCGGTAGAGGAAACAACACACAATGATGAAACGAAAACGTTACGTCACCGTGTATTTGAAGGAGACATGATGAAGGATTACAAGAAGTTTGATACGATAATGGAAGTCAATCCAAAGCCGGATGGAAATGGATGCGTTGTGACTCGGTCAATTGAGTATGAGAAAGTCAACGAGAATTCTCCAACTCCCTTTGATTATCTACAATTCGGCCATCAGGCCATGGAAGACATGAACAAGTACTTACGCGATTCAGAATAACTGATCGATGCCATTTATACATGAACCCATTCGGTGTTGGTGTACGTGACCTATGCACGTACATATTTTATATTTTATCAGTTGTGACTTGTGTGTATTTTATTGTTATAGATCATGGTAATACTCGTCTTAGTGACTCCATATTATGTCAGTGACCCAGTGTATGGGTGTGGTTTGATTTGTGATGTATGTTTCTAGTAGAATAAATAAATTTGGGATAAGTTGTTTCGGAAATGATTTATCCACCGAGGGAGAGATATATCAATAATTCTTCATTCGGCTCTGGCTTTTGGAACATTGACAAGTTCGCCAGATAATCTTTTTTTGCTCTGGAG

>Pbr_TRINITY_STORR Pbr_c40038_g1_i1 len=1888

TTTGAGTTCACCATAGATATATCATATCATTAAAATCTATCATCCTACATACTGTACTGGGTTGAATCATGGAGCTCCAATATTTTTCTTATTTTCAACCAACTTCCTCCGTTGTTGCTCTTCTACTTGCTCTTGTATCCATCTTATTCAGTGTCGTTGTTTTGAGGAAGACATTTTCGAATAACTACTCATCACCTGCATCATCAACAGAAACAGCGGTACTTTGTCATCAGCGGCAGCAGTCGTGTGCATTGCCAATTTCCGGTCTTCTTCATGTTTTCATGAATAAAAACGGCTTAATTCATGTAACTCTTGGAAATATGGCGGATAAATACGGACCGATTTTCAGTTTCCCAACAGGTAGCCATAGAACTCTCGTTGTGAGCAGTTGGGAGATGGTAAAAGAGTGTTTTACAGGCAACAATGACACTGCTTTCTCAAACCGTCCTATCCCGTTAGCTTTTCAGACTATATTCTATGCATGCGGTGGCATAGACTCATACGGTCTTTCGAGTGTACCTTATGGAAAATATTGGAGGGAGCTACGAAAGGTCTGTGTGCATAACCTCCTCTCTAATCAACAACTACTCAAGTTCAGACACTTGATAATTTCTCAAGTCGATACGTCTTTCAATAAGCTGTATGAGTTATGCAAAAACTCTGAAGACAACCAGGGAATGGTGAGAATGGATGATTGGCTCGCCCAATTATCGTTCAATGTGATAGGAAGAATAGTCTGCGGATTCCAATCAGACCCAAAGACAGGTGCTCCTAGCAGGGTGGAACAATTTAAAGAAGTAATTAATGAAGCATCTTATTTTATGTCGACATCTCCAGTGTCAGATAACGTTCCAATGCTAGGGTGGATTGACCAATTGACTGGTCTTACGAGAAATATGAAGCACTGCGGAAAGAAATTAGACTTGGTGGTAGAGAGCATAATTAAAGATCATCGTCAAAAGAGACGATTCTCTAGAACTAAAGGAGAAGATGAGAAGGACGATGAACAAGATGACTTCATCGACATTTGCTTGTCAATTATGGAGCAACCACAGCTTCCTGGCAACAATAGCCCTCCTCAGATACCTATCAAATCTATTGTCCTGGACATGATAGGTGGGGGCACTGACACCACAAAACTGACCACCATCTGGACCCTTTCCTTGCTGCTGAACAACCCCCATGTGTTGGACAAGGCAAAACAAGAAGTGGATGCACACTTTCGAAAGAAAAGGAGATCAACAGATGATGCAGCAGCAGCCGTGGTGGATTTTGATGATATTCGTAACCTTGTCTACATCCAGGCAATCATCAAAGAATCAATGCGGTTGTATCCAGCCAGCCCCGTGGTGGAGCGACTGAGCGGCGAGGATTGTGTGGTCGGTGGGTTTCATGTACCAGCAGGGACGAGATTATGGGCTAACGTATGGAAGATGCAACGAGACCCTAAAGTATGGGATGATCCATTGGTGTTTCGACCAGAGAGATTTTTGAGCGATGAACAGAAGATGGTCGATGTAAGGGGTCAAAATTATGAGCTGTTACCATTTGGAGCCGGTCGACGTATATGTCCAGGTGTATCCTTCTCTTTGGATCTAATGCAACTGGTACTGACTCGTCTTATTCTTGAGTTTGAAATGAAGTCCCCTAGCGGGAAAGTGGACATGACAGCAACACCAGGATTAATGAGTTACAAGGTGGTCCCTCTTGACATTCTGCTCACCCATCGGCGCATAAAGTCGTGTGTGCAGTTAGCATCCTCTGAGAGAGACATGGAGAGTAGTGGTGTACCAGTAATCACTCTGAGCTCGGGAAAGGTGATGCCTGTTTTAGGCATGGGAACATTTGAGAAAGTTGGTAAAGGGTCTGAAAGAGAGATCGGAAGAGC

>Pbr_TRINITY_SALSYN Pbr_c34114_g1_i1 len=1918

CTCTTCCGATCTTTTTGATTCAATCTCTATTAATTAATAGTCAAATTCAATAGAGTAATTCCCTAATCTTTGCAAACTGTCTTATATTTCTGTAATTCATGGCTCCGATTAATATAGAGGAGAATGATTTTTGGATGATAGCATGCACTGTAATAATAGTATTTGCATTGATGAAGTTCATGGTTTCTTTTTATCAATCTGCAAATACAACGGAATGGCCAGAAGGTCCAAAAACACTACCCATAATTGGAAACCTTCATCAGTTGGGAGGAGGTGTTCCCTTACAGGTTGCTTTGGCAAACTTAGCCAAAGTTTATGGAGGTGCATTTACAATTTGGATTGGAAGCTGGGTTCCAATGATCGTCATAAGCGATATCGACAACGCTCGGGAAGTTCTTGTTAATAAATCTGCTGATTATTCCGCTAGAGATGTACCTGATATTCTTAAAATCATCACAGCAAATGGGAAGAATATTGCTGATTGTGATTCTGGTCCATTTTGGCATCATTTAAAGAAAGGTCTTCAAAGTTGTATAAATCCATCAAATGTTATGTCTCTATCTCGTTTACAGGAAAAAGACATGCAAAATCTCATCAAATCCATGCAAGAAAGAGCGTCACAGCAGAATGGAATTTTAAAACCTCTTGATCATGCCAAAGAAGCGTCTATTCGATTACTGAGTAGAGTTATATTTGGTCAAGACTTTTCAAATGAGGATCTCGTTATTGGTGTGAAAGACGCCCTCGATGAGATGGTACGCATAAGTGGGTTGGCAAGTTTAGCTGATGCTTTTAAAATTGCTAAATATTTACCAAGCCAGAAAAAAAATATTCGGGATATGTACGCCACAAGAGACAGAGTATATAATTTGATTCAACCACATATCGTCTCTAATCTTCCTGCAAATTCTTTCTTACATTTTCTTACATCTCAAGATTACAGTGATGAAATTATTTACTCAATGGTACTTGAAATTTTTGGTTTGGGAGTAGATAGTACTGCAGCAACGGCAGTTTGGGCACTCTCCTTTTTAGTCGGCGAGCAGGAAATTCAAGAAAAACTCTACCGCGAAATCAACAACCTGACGGGTGGGCAAAGACCAGTGAAAGTGGTAGATTTGAAAGAGCTGCCATATCTACAAGCCGTGATGAAAGAAACATTGAGGATGAAACCCATCGCACCACTAGCGGTCCCTCATGTAGCAGCAAAAGATACTACATTCAAGGGGCGGAGAATCGTTAAAGGTACAAAAGTAATGGTGAATCTGTATGCTATTCATCACGACCCTAACGTTTTCCCTGCACCGTATAAATTCATGCCAGAGAGATTCTTAAAGGGTGTTAACAGTGATGGACGTTATGGTGATATCAACACAATGGAAAGTTCATTGATACCATTTGGTGCTGGTATGAGAATTTGTGGAGGTGTAGAATTAGCCAAGCAGATGGTAGGTTTTGCTCTTGCAAGTATGGTCAACGAATTCAAATGGGATTGTGTTTCTGAGGGGAATTTGCCTGATCTTAGTGAAGCTATTAGCTTCATTCTCTACATGAAAAACCCACTTGAAGCCAAAGTTACCCCTCGTACAAAACCGTTCGACAGTAGATAAAATCTTATTTTTATCGAACTGCTAGTGTATGTTTATGTAAAATCAAGGGTGAAGATGCTTGATCGTAGTACACCAAAATGGTGGTGGCTATGTAGGAGTTAGATTAGCTCTTGATTTAGTAAATGTTTTTCTTTGCTAGAGACTGTCTCAAGTGTGTGTGTTTTAAGCGTGTGTTGTGTGTGTAATTACTTCCACCTGACAATTAAAGTGTACTTTAGTCGTCGGCTGTATCTGTCACATCATTCTTGTGTTTTTAAATATCTTTATAAATGGTTTTCCCAATTTACTACCCTACATTTCCTATTTCC

>Pbr_TRINITY_SALAT Pbr_c40153_g1_i3 len=1733

TGCTCTTCCGATCTCGAGCTTCTCCAGCACTTGCGAACTAACTATCTTATTCAACTACCATTATCAATCCTGTTAAACAGGCAACAATGTATAGTGCTGCTGTTGAAGTGATCTCTAAGGAAACCATTAAACCCACAACTCCAACCCCATCTCAACTTAAAAACTTCAATTTGTCACTTCTCGATCAATGTTTTCCTTTATATTATTATGTTCCAATCATTCTTTTCTACCCGGACACCGTCGCTAATAGTACCGGTAGCAGTAGCCATCATGATGATCTTGACTTGCTTAAGAGTTCTCTTTCCAAAACACTAGTTCACTTTTATCCAATGGCTGGTAGAATGATAGACAATATCCTGGTCGACTGTCATGACCAAGGGATTGACTTTTACAAAGTAAAAATTAGAGGTAAAATGTGTGACTTCATGTCGCAACCGGATGTGCCTCTAAGCCAGCTTCTTCCATCTGAAGTTGTTTCCGCGAATGTCCCTAAGGAAGCACTGGTGATCGTTCAAGTGAACATGTTTGACTGTGGTGGAACAGCCATTTGTTCGAGTATATCACATAAGATTTCCGATGTAGCTACAATGGGTACGTTCATTCGTAGTTGGGCAAGCACCACCAAAACATCTCGTAGCGGGGGTGCAACTGCTGCCGTTATCACAGATCAGAAATTGATTCCTTCTTTCGACTCGGCATCTCTATTCCCACCTAGTGAACGATTGGCATCTCCATCAGGGATGGCAGAGATACCATTTTCCAGTATCCCGGAGGATACAGAAGATGATAAAACTGTCAGCAAGAGATTTGTGTTCGATTTTGCAAAGATAACATCTGTTAGCGAAAAGCTGCAAGTATTGATGCATGATAACTACAAAAGCCGCAGGCCAACAAGGGTTGAGGTGGTTACTTCTCTAATATGGAAGTCCGTGATGAAATCCACTCCAGCCGGTTTTTTACCAGTCGTAGATCATGCCGTGAACCTTAGAAAGAAAATGGACCCACCATTACAAGATGCGTCATTCGGAAATCTATCTGTAACTGTTTCGGCGTTCTTACCAGCAACAACAACGACAACGACGAATGCGGTCAACAAGACAATCAATAGTACGAGTAGTGAATCGCAAGTGGTACTTCATGAGTTACATGATTTTATAGCTCAGCTAAGGAGTGAAATAGATAAGGTCAAGGGTGATAAAGGTAGCTTGGAGAAAGTCATTCAAAATTTTGTTTCCGGTTATGATGCTTCAATAAAGAAAATCAATGATGTTGAAGATAAGGTGATAAACTTTTGGATAAGTAGCTGGTGCAGGATGGGATTATACGAGATTGATTTTGGTTGGGGAAAGCCAATTTGGGTTACAATTGATCCAAATATCAAGCCGAATAAGAATTGTCTTTTCATGAATGATACGAAATGTGGTCAAGGAATAGAAGTTTGGGCGAGCTTTCTTGAGGATGATATGGCTAAGTTCGAGCTTCACCTAAGTGAAATCCTTGAATTGATTTGATATTGCATTATCTACATGTGTTCCGTAATCATGATTTTCTCCATTTCCCTTTCCGTAGTTGGTTACAAAGAACGAAATAAAGGAAAATTAAAAAACTTGTACTGCTCGATGCTTTGACATTTCCATATTCATCCATAAATTCCCATCAGAAAAGAGTTTCAAATATTAGGGGATGAAGTGAAATCCTGGAATTGATTTGATGCATTTGCTAATCGAAGACAG

>Pbr_TRINITY_SALR Pbr_c20565_g1_i1 len=1330

TGCTCTTCCGATCTTTATAATCAATAGAAACCTCTCTTTACAAAATCAGTCAGAAAGAATCTTATGTTGATTTTCATTGCTTGAGTCTTCGCTTTTTCTTTTTATAGTTGTTAATCCAATCCAGTAATCATCAACAAGTATGGCTGAAACATGTCCAAATACTGTTACAAAGATGAGGTGTGCAGTTGTTACTGGCGGAAACAAGGGTATTGGATTGGAGATTTGTAAGCAATTATCTTCTAGTGGAATCATGGTTGTTTTAACTTGTAGAGATGTAACTAGAGGTCTTGAAGCTGTTGAAAAACTCAAAAATTCTAATCATGAGAATGTGGTTTTTCATCAACTTGATGTTACGGATCCAATTACTACTATGTCTTCTTTAGCGGATTTCATCAAAGCACGCTTCGGAAAGCTTGATATCTTGGTAAACAATGCTGGGGTTGCAGGTTTTTCAGTTGATGCTGATCGTTTTAAGGCAATGATAAGTGACATTGGAGAGGATTCAGAGGAGGTCGTGAAGATCTACGAAAAACCAGAAGCCCAAGAATTAATGTCAGAGACATATGAATTAGCAGAAGAATGTCTCAAAATAAATTACTACGGTGTTAAATCAGTAACCGAAGTTCTACTTCCTTTACTTCAACTATCTGATTCACCAAGAATTGTTAATGTTTCATCATCCACTGGAAGCCTCAAGTATGTATCCAATGAAACAGCTCTAGAGATACTTGGAGATGGTGATGCATTAACAGAAGAGAGAATTGACATGGTAGTGAATATGCTTCTTAAGGATTTTAAGGAAAATTTGATCGAAACAAATGGGTGGCCTAGTTTCGGAGCTGCATACACAACATCAAAAGCATGTTTGAATGCTTACACAAGGGTGTTAGCAAAGAAAATTCCCAAATTTCAGGTGAATTGTGTTTGTCCTGGTTTGGTTAAAACAGAAATGAACTACGGCATTGGAAATTATACTGCCGACGAAGGTGCTAAACATGTAGTCAGAATAGCTCTTTTCCCCGACGATGGACCTTCTGGTTTTTTCTATGATTGTTCTGAACTATCTGCATTTTGAGTTATATCGATTCAGCGTATCGTGCAGCAGCCAAATAAATAAAAATGGCATCAGTTGTTTGCTAGCACTGTCCTATCTACTGATAACTTTATCCCTTTGCTTCTGTTTTTAGCTTGTGTTGTGTGAATGTCCAGAGAGGATCGATCCATTTCTAATGTATTGAATCTTTATTTGGATGACAAATAATAAATCAGCTTGAGATTTTTCTTCCGATTTTAAAAAAAAAGATCGTAAGAGCGTCGTGTAGGGAAAGAG

>Pbr_TRINITY_THS2 Pbr_c30828_g1_i1 len=803

GTGCTCTTCCGATCTCTTCTGTGCATATTTTAGGGAACTAATCATCAAATTGTTGATGGCTCCTCGTGGTGTTTCAGGTCTAGTCGGGAAACTTTCAACTGAATTAGATGTCAATTGCGATGCTGAAAAATATTATAACATGTATAAGAACGGAGAAGATGTACAAAAGGCAGTTCCTCATCTTTGCATGGACGTCAAAGTTATCAGTGGAGATGCAACCCGTTCAGGTTGTATCAAGGAATGGAATGTTAACATTGATGGTAAGACGATTCGCTCAGTAGAGGAAACAACACACAATGATGAAACGAAAACGTTACGTCACCGTGTATTTGAAGGAGACATGATGAAGGATTACAAGAAGTTTGATACGATAATGGAAGTCAATCCAAAGCCGGATGGAAATGGATGTGTTGTGACTCGGTCAATTGAGTATGAGAAAGTCAACGAGAATTCTCCAACTCCCTTTGATTATCTACAATTTGGCCATCAGGCCATGGAAGACATGAACAAGTACTTACGCGATTCTGAATAACTGATCGATGCCATTTATACATGAACCCATTCGGTGTTGGTGTACGTGTACCTATGCACGTACATATTTTATATTTATCAGTTGTGACTTGTGTGTATTTTATTGTTATAGTACATGGTAATACTCGTCTCAGTGACTCCATATTATGTCAGTGACCCAGTGTATGGGTGTGGTTTGATTTGTCATGTATGTTTCTACTAGAATAAATAAATTTGGGATAAGTTGTTTCGGAAATGATTTATCCACCAAGGGAGAGATCGGAAGAGCGTCG

>Par_TRINITY_STORR Par_c33669_g1_i1 len=2037

ATGGAGCTCCAATATTTTTCTTATTTTCAACCAACTTCCTCCGTTGTTGCTCTTCTACTTGCTCTTGTATCCATCTTATTCAGTTTCGTTGTTTTGAGGAAGACATTTTCGAATAACTACTCATCACCTGCATCATCAACAGAAACAGCGGTACTTTGTCATCAGCGGCAGCAGTCGTGTGCATTGCCAATTTCCGGTCTTCTTCATGTTTTCATGAATAAAAACGGCTTAATTCATGTAACTCTTGGAAATATGGCCGATATATACGGACCGATTTTCAGTTTCCCAACAGGTAGCCATAGAACTCTCGTTGTGAGCAGTTGGGAGATGGTAAAAGAGTGTTTTACAGGCAACAATGACACTGCTTTCTCAAACCGTCCTATCCCGTTAGCTTTTAAGACTATATTCTATGCATGCGGTGGCATAGACTCATACGGTCTTTCGAGTGTACCTTATGGAAAATATTGGAGGGAGCTACGAAAAGTCTGTGTGCATAACCTCCTCTCTAATCAACAACTACTCAAGTTCAGACACTTGATAATTTCTCAAGTTGATACGTCTTTCAATAAGCTGTATGAGTTATGCAAAAACTCTGAAGACAACCAGGGAAACCATACTACTACTACTACTACTACCGCAGCTGGCATGGTGAGAATGGATGATTGGCTCGCCCAATTATCGTTCAACGTGATAGGAAGAATAGTCTGCGGATTCCAATCAGACCCTAAGACAGGTGCTCCAAGCAGGGTGGAACAATTTAAAGAAGCAATTAATGAAGCATCGCATTTTATGTCGACATCTCCAGTGTCAGATAACGTTCCAATGCTAGGGTGGATTGACCAATTGACTGGTCTTACAAGAAATATGAAGCACTGCGGAAAGAAATTAGACTTGGTGGTCGAGAGCATAATTCAAGATCATCGTCAAAAGAGACGATTCTCTAGAACTAAAGGAGGAGATGAGAAGGACGATGAACAAGATGACTTCATCGACATTTGCTTGTCAATTATGGAGCAACCACAGCTTCCTGACAACAATAACCCTCCTCATATACCTATCAAATCTATTGTCCTGGACATGATAGGTGGGGGAACTGACACAACAAAACTGACCACCATCTGGACCCTTTCCTTGCTGCTGAACAACCCCCATGTGTTGGACAAGGCAAAACAAGAAGTGGATGCACACTTTGGAACGAAAAGGAGATCAACAAATGATGCCGTGGTGGATTTTGATGATATTCGTAACCTTGTATACATCCAGGCAATCATCAAAGAATCAATGCGGTTGTATCCAGCCAGCCCCGTGGTGGAGCGACTGAGCGGCCAAGATTGTGTGGTCGGTGGGTTTCATGTACCAGCAGGGACGAGATTATGGGCTAACGTATGGAAGATGCAACGAGACCCTAAAGTATGGGATGATCCATTGGTGTTTCGACCAGAGAGATTTTTGAGCGATGAACAGAAGATGGTTGATGTAAGGGGTCAAAATTATGAGCTGTTACCATTTGGAGCCGGTCGACGTATATGTCCAGGTGTATCCTTCTCTTTGGATCTAATGCAACTGGTACTGACTCGTCTTATTCTTGAGTTTGAAATGAAGTCTCCTAGCGGGAAAGTGGACATGACAGCAACACCAGGATTAATGAGTTACAAGGTGGTCCCCCTTGACATTCTGCTCACCCATCGTGGCACAAAGTCGTGTGTGCAGTCAGCAGCCTCTGAGACTAGTGGTGTACCAGTAATCACTCTGAGCTCGGGCAAGGTGATGCCTGTTTTAGGCATGGGTACATTTGAGAAAGTTGGTAAAGGGTCTGAAAGAGAGAGGTTGGCGATTTTAAAAGCGATAGAGGTGGGTTACAGATACTTCGATACAGCTGCTGCATACGAAACTGAAGAGGTTCTTGGAGAAGCTATTGCTGAAGCACTCCAACTTGGCCTAATCAAATCTCGAGATGAACTTTTCATCAGTTCCATGCTCTGGTGCACTGATGCTCACCCTGATCGTGTCCTCCTCGCTCTTCAGAATTCTCTGAGGAATCTTAAATTGGAGTATCTGGATCTATATATGTTACCCTTCCCGGCAAGCTTGAAGCCAGGGAAGATAACGATGGACATACCAGAGGAAGATATTTGTCGAATGGACTACAGGTCTGTATGGTCAGCCATGGAAGAGTGTCAGAACCTTGGCTTCACTAAATCAATCGGTGTTAGCAATTTCTCCTGCAAAAAGCTTCAGGAATTGATGGAGACCGCCAACATCCCTCCAGCTGTGAATCAAGTGGAGATGAGCCCGGCTTTCCAACAAAAGAAGCTGAGAGAGTATTGCAACGCAAATAATATATTAGTCAGTGCAGTCTCTATACTGGGATCAAACGGAACCCCATGGGGCTCCAATGCAGTTTTGGGTTCTGAGGTGCTTAACAAAATTGCTATGGCCAAAGGAAAATCTGTTGCTCAGGTTAGTATGAGATGGGTTTACGAGCAAGGCGCGAGTCTTGTGGTAAAAAGTTTCAGTGAAGAGAGATTGAGGGAAAACTTGAACATATTCGACTGGGAACTCACTAAGGAAGACAATGAAAAGATCGGTGAGATTCCACAGTGCAGAATCTTGACTGCTCATTTTTTGGTCTCACCTAATGGACCTTTCAAATCTCAAGAAGAGTTGTGGGATGACAAAGCTTGA

>Par_TRINITY_SALSYN Par_c33168_g1_i1 len=1890

CGTGTGCTCTTCCGATCTGTTTTGATTCAATCTCTATTAATTAATAGTCAAATTCAATAGAGTAATTCCCTAATCTTTGTAAACCGTCTTATATTTCTGTTTCATGGCTCCGATTAATATAGAGGAGAATGATTTTTGGATGATAGCATGCACTGTAATAATAATATTTGCATTGGTGAAGTTCATGTTTTCTTTTTATCAATCTTCAAATACAACGGAATGGCCAGAAGGTCCAAAAACACTACCCATAATTGGAAACCTTCATCAGTTGGGAGGAGGTGTGCCCTTACAGGTTGCTCTGGCAAACTTAGCCAAAGTATATGGAGGTGCATTTACAATTTGGATTGGAAGCTGGGTTCCAATGATCATCATAAGCGATATCGATAACGCTCGGGAAGTTCTTGTTAATAAATCTGCTGATTATTCCGCTAGAGATGTACCTGATATTCTTAAAATCATCACAGCAAATGGGAAGAATATTGCCGATTGTGATTCTGGTCCATTTTGGCATAATTTAAAGAAAGGTCTTCAAAGTTGTATAAATCCATCAAATGTTACGTCTCTATCTCGTTTACAGGAAAAAGACATGCAAAATCTCATCAAATCCATGCAAGAAAGAGCGTCACAGCAGAATGGAATTTTAAAACCTCTTGATCATGCCAAAGAAGCGTCTATTCGATTACTGAGTAGAGTTATATTTGGTCAAGACTTTTCAAATGAGGATCTCGTTATTGGTGTGAAAGACGCCCTCGATGAGATGGTACGCATAAGTGGGTTGGCAAGTTTAGCTGATGCTTTTAAAATTGCTAAATATTTACCAAGCCAGAGAAAAAATATTCGGGATATGTACGCCACAAGAGACAGAGTATATAATTTGATTCAACCACATATCGTCTCTAATCTTCCTGCAAATTCTTTCTTACATTTTCTTACATCTCAAGATTACAGTGATGAAATTATTTACTCAATGGTACTTGAAATTTTTGGTTTGGGAGTAGATAGTACTGCAGCAACGGCAGTTTGGGCACTCTCCTTTTTAGTCGGCGAGCAGGAAATTCAAGAAAAACTTTACCGCGAAATCAACACCCTGACGGGTGGGCAAAGACCAGTGAAAGTTGTAGATTTGAAAGAGCTGCCATATCTACAAGCCGTGATGAAAGAAACATTGAGGATGAAACCCATCGCACCACTAGCGGTCCCTCATGTAGCAGCAAAAGATACTACATTCAAGGGGCGGAGAATCGTTAAAGGTACAAAAGTAATGGTGAATCTGTATGCTATTCATCACGACCCTAACGTTTTTCCTGCACCGTATAAATTCATGCCAGAGAGATTCTTAAAGGATGTTAATAGTGATGGACGTTTTGGTGATATCAACACAATGGAAAGTTCATTGATACCATTTGGTGCTGGTATGAGAATTTGTGGAGGTGTAGAATTAGCCAAGCAGATGGTAGCTTTTGCTCTTGCAAGTATGGTCAACGAATTCAAATGGGATTGTGTTTCTGAGGGGAAATTGCCTGATCTTAGTGAAGCAATTAGCTTCATTCTCTACATGAAAAACCCACTTGAAGCCAAAATAACCCCTCGTACAAAACCTTTCGACAGTAGATAAAATCTTATTTTTATCAAACTGCAAGTGTATGTTTATGTAAAATCAAGGGTGAAGATGCTTGATCTTAGTGCACCAAAATGGTGGTGGCTATGTAGGAGTTAGATTAGCTCTTGATTTAGTAAATGTTTTTCTTTGCTAGTGACTGTGTCTCAACTCAAGTGTGTGTATTGTGTGTGTTTCAAGTGTGTATTGTGTGTGTTTCAAGCGTGTGTTGTGTGTGTAATTACTGCCACCTGAAAATTACGTTAGTCGCCGGCTGTAACTGTCAAAAAAAA

>Ppa_TRINITY_4OMT1 Ppa_c27690_g1_i1 len=1557

CCGATCTGCACAACCCGAAACAAACATATTCATTTTAACAAGTATCAAACATTTCATATCATGGGTAGTTTAGATGCCAAACCAGCTGAAACACAAGAAGTTGGCAATGGCGGCATCAAAGAACAAGCTCAACTATGGAATATAATCTACGGTTTTGGCGATTCTCTTGTTCTTCGCTGCGCGGTAGAGATTGGTATTGCGGATATTATCAAAAAGAATAATGGGTCAATCACACTTGCACAACTCGCATCACAACTCTCAATCCCAAATGTCAACTCCGGTTACTTGTACAGAATACTAAGATACTTAGTACACCTGAATATCTTGCAACAAGAAACTTGTAATGATGGAGTCAACAAGGTTTACTCACTTAAACCAGTGGGTGCTTTACTTTTAAGAGATGCAGAAACAAGTATGGTACCAGTAATATTAGGCCTGACTCACAAAGATTTCATGGTTCCATGGCATTTTATGAAACAAGGCTTAGAAAGTGAAAGCGCAACAGCATTCGAGATAGGAACGGGAATGACAGTATGGAAGTTTCTAGAAGGTCATCCTGATCAAAGTCAGCTTTTTAATGAAGAAATGGCTGGAGAGTCAAGACTTGTAACTAAAACTCTAATTGAAGATTGTAGAGATATTTTTCAAGGTTTGGATTCTCTGGTCGACGTTGGTGGAGGAAATGGCACAATGATTATGGCAATCTATGAAGCTTTTCCTCATATTAAATGTACGCTTTATGATCTTCCTCATGTTATTGCTGATTCTCGTGATCACCCTAACATCGAAAAAGTTTCGGGTGATATGTTTAAGTCTGTTCCTAGTGCTCAAGCCGTGTTTCTCAAGATGATCTTGCATGACTGGACCGACGACGAGTGTGTCAGCATTCTGAAGAAATGTAAAGAAGCAATACCAAAAGAGACGGGGAAAGTGATCATAGTGGATGCAGCACTTGATGAAGAATCCACTCATGAACTCACAAAAACAAGATTGCTTCTTGATATTGATATGTTGGTTACTACTGGTGGTAGAGAGCGAACAGTAGATGACTGGAAAAATTTGCTCGAACGTGCAGGCTTTCGGTCTCATAAGATCAGACCCATTCGAGCTATTCAGTCTGTCATTGAAGCCTTTCCATAGTAAAAGCTCCAAGAAGTACGTTGCTTCTTTTTCAGAGAAGCATATGCGATTCCCTTCTCAACAATAAGCGCACACACAGCAAGGTGGCATTAGTTATCTTCTCATCAACAATATGGCATTAGTTCCTCCCCTACACCGGTGTGCTAGACTAGCTTGGCCGCTTCTTTTCTTGTTTTTTATGTTTGTTTTTGTGCTATCTAAGCACCCTTGATCTAACCGTCCGTTTTCCCTCAGTGAAAAAAAAAAACACAATAAGTCATCTTCAATTGAGGAGATGTTTTCGTATTTCGTATTTCCTACTATTGTAATTGAAATTTATATTGCTCGGTTGAATTCCATTGCGCACCCTTGATCTTCCCTCAATCTTCTTCACTAATAGAACTGCGTAATAAGTCTGAACAACATTGGAGGATGGAG

>Pnu_TRINITY_NCS Pnu_c45079_g2_i1 len=944

AACACTCCTAAGAGGCAATCTCTAGTGAAGAAGGAAATTCACTATGACTTGGAGGTACCAACCTCAGCTGATTCGATTTGGGACGTCTACAGTTCTCCTGAAATTCCTCGACTTCTTAGAGATGTTCTGCTTCCTGGTGTTTTTGAAAAGCTAGATGTCATTGAAGGCAATGGTGGCGTTGGCACTGTACTTGACATTGCGTTCCCTCTAGGTGCGGTGCCACGGAGTTACAAGGAGAAATTTGTGAAGATCAACCATGAGAAGCGATTAAAGGAAGTGATTATGATTGAAGGGGGATACTTAGACATGGGATGCACATATTACATGGATAGGATCCATGTCTTAGAGAAAACACCTAGCTCATGTGTTATTGAATCCTCTATTGTTTACGAAGTGAAAGAAGAGTATGCCGATAAAATGTCTAAGCTAATCACAACAGAACCATTGAAGTCCATGGCAGAAGTTATCTCCAGTTACGTTCTTAAGAATCAATTCCGCGTATTGGGGTTCGAGATTAAACCAAAATTAGGCTTGAGTATGTTGCTATGTATGATTATCTGTTTGGTTGTACTTGGTGGTATGTTGATCGGAGGTGTTCGACTCTAAGCTTCAGTGTTCTGTTAAGACGTACTACTACTTTTCAATTCACACGAGTCAAGAGAAAAACCGTGCCATGGGTGGAGTTCCGATTATGCTAGGGTGACTAGTGTCATTTGTCATTAGCAACTTATGGAGTCTACAAGATGAGAAATAAATGTAATAACAGTTTTAAGTAATGTACCTTATTTGTGTTTGCTTTAAGCATTTTCTGAAAATAAATAATGCTTTATCTGTTCAGTGACTACCTGCAATACTGCAAATACGGATGGGTGAATTTCGGAATATAAATGCACAGACGTAGATGTACAAGTCTGTTTGTCTGTATGTCCTTGTTTGAACCATGC

>Pnu_TRINITY_6OMT Pnu_c50693_g1_i1 len=1331

GTGTGCTCTTCCGATCTCAAAGCCCACGTTTTCTCTTGTGCTCCTAGAGAAATAATAATAAGCTAAGCTTATTATATAACAAGGTGGTTTAATGGGACACTGAAGAATATCTCAATCATTCTCATCAATCAACCACTAGTCAGCTACAATTTCCAGAGAGAAATGGAAGTAGTAAACAAGATCGATCAAGAAAACCAAGCAAAAATCTGGAAACAAATCTATGGTTTCGCAGAATCATTAGTTCTCAAATGTGCAGTTCAATTGGAGATTGCTGAAACACTTCACAACAATGTAAAACCCATGTCTCTGTCCGAATTAGCATCAAAACTTCCGGTTCAGCCCGTCAACGAAGACCGTCTGTACCGAATCATGCGCTACTTGGTGCACATGGGACTCTTCAACAAAGATGCTAGCACACAGAAATACTCAATGGCTCCACCATCAAAGTATTTGTTAAGAGGCTGGGAGAAATCAATGGTTGCTTCGATTTTATGCCTACAGGATAAGGATTTCTTGGCTCCTTGGCATCATTTGAGCGACGGGTTGACCGGTAACTGTGACGCGTTTGAGAAAGCGTTGGGGAAGAGTATTTGGGATTATATGAGTGAAAATCCTGGAAAGAATCAGTTGTTTAACTCAGCAATGGCTTGTGATACTAGATTGGTTACTTCGGCATTGGTCACCGAGGGCAAAAGTATTTTCAGTCATGGAATCAATACACTGGTTGATGTTGGTGGTGGCACTGGTACTGCAGTGAAAGCCATTTCCAAAGCTTTTCCGGATATCAAATGCACAATCTATGACCTTCCTCATGTCATAGCTGATTCTCCCGACATTCCCAACATTACCAAAATTTCTGGAGATATGTTCAAGTCTATTCCCAGTGCTGATGCCATCTTCATGAAGTGCATTCTTCATGACTGGAATGATGAGGAATGTGTTCAAATCTTAAAGCGATGCAAAGAAGCATTACCAAAAGAAGGGAAAGTGATTATCGTGGACATCGTCCTAGACATGGACTCGACTCATCCATATGCAAAAATTAGACTCACATTGGATTTGGATATGATGCTCAACACCGGTGGAAAAGAGAGAACTGAAGAAGAATGGAAGAAACTTATTGATGCGGCAGGTTTTGCTAGCTACAAAATTACCCAGCTATCTGCTGTCCAGTCTGTAATTGAGGCTTACCCTTATTAAGCATAACTCTATCCTGTTTTCCCTTTGATTAATTGTTGCCTTCTCTATGTGTTATGGACCTTATGGTCGTTTAAGTGTTATTTGCAAAGGAAATATGAGTGAAATTAATACTTGCATTGGATCAAAAAAAA

>Pnu_TRINITY_CNMT Pnu_c49398_g1_i1 len=1267

GACGTGTGCTCTTCCGATCTAGTTAGCATCATATCTCTACCAAACACTCTCTATAGACTTTTGTGTGTGTGTGTGTGTGTTTTTACAAGGAAGATGAAGCTAATGGCAAAGGAAGAGCTCTTGGAAAACATGGAACTTGGGTTGATACCAGACGAAGATATCAGAAAATTGATTAGAATCCAATTGGAGAAACGTCTGAAATGGGGTTACAAAGCAACTCATGAAGAACAACTCTCTCAACTTCTTGAACTAGTTCATTCTTTGAGAAGTATGAAGATGGCAACTGAGATGGAGAATTTGGATTTGAAACTATATGAAGCACCTATGTCATTCTTGAAGATACAACATGGAAGTAACATGAAACAAAGTGCGGGGTACTATACCGACGAATCAACGACGTTGGATGAAGCTGAGATAGCGATGCTAGATTTGTATATGGAGAGGGCACAAATCAAAGATGGTCAGAGTGTACTTGATTTAGGGTGCGGATTGGGAGCTGTAGCTCTTTTTGGTGCAAACAAATTTAAGAACTGTCAATTTACTGGAGTTACTAGTTCTGTGGAGCAAAAAGATTACATTGAAGGGAAATGCAAGGAGCTTAAGTTGTCAAATGTCAAGGTTCTGTTAGCTGATATAACAACATATGAGACCGAAGAAAGATTTGATCGTATTTTCGCAGTTGAACTGATTGAGCATATGAAGAACTATCAATTGCTTCTTAAGAAAATATCAGAATGGATGAAAGATGATGGACTTCTTTTCATTGAACATGTTTGCCATAAAACTCTAGCTTACCACTATGAGCCTATTGATGCAGAAGATTGGTACACTAATTACATATTCCCTGCTGGAACTCTGACTCTATCATCCGCTTCAATGCTTCTTTACTTTCAAGATGATGTTGCTGTTGTGAATCAATGGACTCTAAGTGGAAAGCATTACTCTCGATCCCATGAAGAATGGTTGAAAAACATGGACAAAAACATTGTTGAATTCAAAGAAATAATGAAGTCAATAACTAAAACAGAGGAAGAAGCAACCAAATTGCTCAATTTTTGGAGAATTTTCTGCATGTGTGGAGCAGAGTTGTTTGGGTATAAGAATGGTGAAGAATGGATGCTCACCCATCTTCTATTCAAGAAAAAATAAGAATTTTCCTTTCATGCTTTAATGCTTTCATTGCTCTAAAAAACTGTTTAAGAATGATGTGTAATTGTAACTTTTATGTAAAGAAGTCCACTTCTACAAATATTTTTGTATCACTAAG

>Pnu_TRINITY_CYP80B1 Pnu_c53666_g2_i4 len=1869

CTATGGTCATAAACACTTGCTCCAGCACACTTGGATTCTTCCTATATACATTAAAAGGAGAGCCAAATGAAAATCTAAAGAGAGAACAAAATGGAGATTGTCACAGTAGCTTTAATAGCCATTGTGATCACCACTTTGTTATACTTGATTTTTAGAGATTCAAGTGCTAAAGGCTTGCCACCAGGTCCAAAACCATGGCCAATAGTCGGAAACCTTCTTCAACTCGGTGAAAAACCCCATTCACAATTTGCACAACTTGCTGAAACCTATGGAGATCTCTTTACACTTAAACTAGGATCTGAAACAGTTGTTGTGGCTTCAACACCACTAGCAGCTAGTGAAGTTCTCAAGACGCATGATCGTGTTCTCTCCGGCCGGTACGTGTTTCAAAGTTTCCGTGTGAAAGAACATGTAAAGAACTCCGTTGTTTGGTCCGACTGTAATGAAACATGGAAGAAATTGCGCAAAGTTTGCAGAACAGAACTTTTTACACAAAAGATGATTGAAAGTCAAGCTGAAGTCAGAGAAAGAACAGTGTTGGAAATGGTGGGATATCTGAAGGAAAACGAAGGAAAGGAAGTGAAAATTGTCCAAGTTGTATTTGGCACATTGGTGAATATTTTTGGGAACTTGATATTTTCGCAAAATATTTTTAAGCTGGGTGATGAAAGTAGCGGCAGTGTCGAAATGAAAGAACGTCTATGGAGAATGCTGGAACTAGGTAATTCAACAAATCTAGCTGATTATTTTCCATTTTTAGGTAATTTGGATTTGCTTGGACAAAGAAAAGATGTTGCCGATTGTCTTCAAGGGGTATATGATATTTGGGGTGCTATTCTTAAAGAAAGAAAAATAGCCAAGCAACGTAGTAGCGATACGAAGAATGATTTTGTTGATGTTTTGCTTGATGCTGGACTTGACGATCAACAGATCAATTCCTTGCTCATGGAATTATTCAGCGCGGGAACAGAGACGAGTGCATCTACAATAGAATGGGCATTAACTGAGCTCACAAAAAACCCTGAAATAATAGCCAGTTTGCGATCAGAGTTGCTGGCTGTGGTAGGGCAGAGGCCGATCAAGGAATCCGACATACCTAACATGCCCTATCTTCAAGCATTTGTTAAGGAAACTCTACGGCTTCACCCAGCAACTCCATTGCTAATCCCACGTCGCGCACTGGAGACATGTCAAGTCTTGAATTACACGATCCCAAAAGAGTGTCAGATAATGGTGAATGCCTGGGGCATAGGTCGGGATCCAAAAACATGGACTGATCCACTGAAGTTTTCGCCAGAGAGGTTCTTGAATTCTAGCATTGATTTCAAAGGGAACGATTTCGAGTTGATACCATTTGGTGCAGGGAGAAGGATATGCCCTGGTGTGCCCTTGGTAACTCAGTTCATTAGTCTTATTGTGCCCACTTTCGTACAGAATTTTGATTGGGTACTACCAAAGGGAATGGATCCTAGCCAACTGATCATGGAAGAGAAATTTGGGTTGACACTGCAAAAGGAACCACCTCTATATATTGTTCCTAAAACTAGGGATTGATCTAGGAAATGGTTAAAGAACTACTTGGCCAGCCACGACGCAAATGTTAAAGGTGGAAATTTTCTTTTTCTTATTATAATTCCAAAAAGAGTTGGTGCTCAACTTGCCATCGGACTACCAACCATTTTTTTGTAATGAAAAATGTAAAGAAAAAAAAGAAAAAAAAAGTCCGACATTATTGATCCCCTTATCTTTCAAATATAAGTAATAACAAAAAAAAATTAAAATTCATATGTAGTAGAAACCAAGAAACAAGGTAGAAAATATAGAGACAAAGAAGGAGAAATTTCATTAAAGTGTGTAGATCGGAAGAGC

>Pnu_TRINITY_4OMT1 Pnu_c54622_g1_i7 len=1404

GTGTGCTCTTCCGATCCCAAAACGAACATAGTCTTCTTGAGAAATACTACACAAACTTCATAGCATGGGTAGTTTAGATGTGAATCTAGCTGAAACACAAGAAGTTGGCATCAAAGATCAAGCTCAACTATGGAATATAATCTATGGTTTTGCTGATTCACTTGTTCTTCGTTGCACGGTAGAGATCGGAATTGCAGATATTATCAAAAACAATAATGGGTCAATCACACTTGCACAACTTGCGTCGAAACTCCCGATTGCGAATGTCAATTCTGATTACTTGTACAGGATACTAAGGTATTTGGTACATTTAAATATCTTAACTTGCAGTGGTGCTGGAGTTGACAAGGTTTACTCACTTAAACCAGCCGGTACTTTACTTTTAAAAGATGCAGAAAGAAGTATGGTACCGATGATATTAGGCATGACTCAGAAGGATTTCATGGTACCATGGCATTTCATGAAAGACGGGTTGGCGGCCGGAAGTACAACCGCATTTGAAGAAGGAATGGGAATGAATATATGGACGTATTTAGAAGGCAATCCTGATCAAAGTCAGTTGTTCAACGAAGGAATGGCTGGAGAAACAAGACTTTTGACCAAGACTCTAATCGAAGATTGCAGAGACACATTCGAAGGTTTGGATTCACTTGTGGATATTGGTGGAGGAAATGGCACAACAATTAAGGCTATCTATGAAGCTTTTCCTCATATTAAATGCACACTTTATGATCTTCCTCATGTCATTGCTAACTCTCATGATCACCCGAACATCGAAAAAGTTCCCGGTGATATGTTCAAGTCTGTTCCCAGTGCTCAAGCTATCTTACTCAAGTTGATCTTGCATGACTGGACTGATGAAGAGTGTGTCAGCATACTAAAGAAGTGCAAAGAAGCAATACCCGAAGAGACAGGGAAAGTGATCATAGTGGATGTAGCACTTGAAGAAAAATCTCAACATGAACTTACTAAGACAAGATTGATTCTTGATATTGACATGTTGGTGAATACTGGTGGTAGAGAGAGAACTGCAGATGATTGGGAAAATTTGCTCAAACGTGCAGGCTTTCGGTCTCATAAGATCAGACCCATCCGCTCTATTCAATCTGTCATTGAAGCCTTCCCATAGTAAAAGCTCAATGGAGTACGTTGCTTCTTTATCTGAGAAGCGTATGCGATTCCCTTTATAACAATAAGCGTATGCAATATGGCATTAGTTCCATCCCACCAGTGCTGCACTAGCTTGTGCCTCTTTCTTTTCTCGTTTTTTCCGTTTCTTTTTGTGCTCGCTAAGCACCTTCAATAAAGAAAACATGAAGCCGTCTTCAAGTGAGGAGATGTTTTCCTATTTCCTGTTGTTCGTCCTTTCTGATTCCTCTTATTCTCAAATTGCATAAGTACGAG

>Pnu_TRINITY_BBE Pnu_c55138_g2_i5 len=1705

TGTGTCTAAATAGGCGAATACTCCTTATAAATGCTAAAATCAGCACTACTAGCAACACATCTTCCTAGTTATTACATACTGTATTTCCCAAACATGATCTGCAGAAGTCTAATATCATGTCTCCTGTTGTTATTTCTTGTACAAACATGCATACTAGGTGATGATGGTCTCCTTTTGTCATGTTTAAACTCCCATAGAGTTCACAATTTTACCGCAAGATCAACTGGTACAAATTCGGAGTACTTCAAACTACTCCATGTGTCTATTCAAAACCCATTGTTCGCGAAGTCGACGGTGTCGAAGCCGTCGTTGATCGTAATGCCCGGTAGCAAAGAAGAATTATCGAGCACCATTCATTGTTGTACAAAGGAATCATGGACAATCCGACTGAGAAGTGGTGGTCACAGCTATGAAGGGTTATCTTACACTGCTGATACACCGTTTGTAATTGTAGATATGATGAACTTGAATCGAGTTTCCATTGATGTTGAGTCAGAAAGTGCTTGGGTTGAATCGGGTGCAACACTTGGAGAACTCTACTATTCGATTGCGCAGTCAACAGATACACTGGGATTTACTGCAGGTTGGTGTCCGACTGTTGGTAGCGGTGGCCATATAAGCGGTGGTGGTTTTGGTATGATGTCGAGAAAGTACGGGTTGGCTGCTGATAATGTTGTGGATGCGATTCTCATAGATAGCAACGGAGCAATTCTCGACCGTGAGAAAATGGGTGAGGATGTTTTCTGGGCGATTCGTGGTGGCGGCGGAGGTGTTTGGGGTGCAATTTACGCGTGGAAAATCAAACTACTGCCAGTTCCGGAGAAGCTGACAGTTTTTCGCGTAACGAAAAATGTAGCAATCGAAGACGCTTCATCTTTGCTTCACAAATGGCAGTATGTTGCAGACGAATTAGACGAAGATTTTACGGTATCTGTGCTCGGAGGAGCACACGAAAACGAAGCCTGGTTAATGTTCTTAGGTCTGCACTTGGGACCAAAAGGTGCTGCGAAAACTATAATCGACGAAAAATTCCCTGAGCTGGGATTAACAGCGGAAGAGTTTCAAGAAATGAGTTGGGGTGAATCCATGGGTTTCTTAGCAGGATTAGAAACAGTCTCTGACTTAAACAACAGGTTTCTGAAATTTGATGATAGAGCTTTTAAGACAAAAGTTGATTTTACTAGAGAACCAATACCATTAGACGTGTTTAGACGTGCATTAGAGATGTTGTCAACACAGCCCGGTGGCTTTATAGCTCTAAATGGTTTAGGAGGGAAAATGAGCACAATTAGCACTGATTTTACGCCGTTTCCTCATCGTAAAGGTACTAGATTGATGTTCGAATACATAATCGCTTGGAACAAAGATGAAGAAACAAGAATCGGTGAATTTAGCGAATGGTTAGCGGAATTCTACCGTTATTTGGAGCCGTTTGTATCGAAAGAACCAAGGGTTGGTTATGTTAATCACATTGATCTTGATCTTGGTGGGATAGATTGGAGAACTAGTACTAGTAATGCAGTTGAGATTGCAAGAAATTGGGGCGAAAGATATTTCTTATCGAATTATGAACGTTTGGTTAAGGCTAAAACATTGATTGATCCTAATAATGTGTTTAATCATCCACAAAGTATTCCTCCAATGACGAAATTTGATCATTTTGAGGAAAATTACAAGGAAATGTAGTACTTTTTTTTTTTTTTTT

>Pnu_TRINITY_TNMT Pnu_c51404_g1_i2 len=1467

TGTGCTCTTCCGATCTTGCTGATACCACACAACTCTCATTCTCTCTAAAATTTCATTCACTTTCTCTTCCTACAATCACTCGAGTCTTCTTTACTAACCTGGACTCTGTCAGCATGGGTTCAATAGATGAGGTGAAGAAGGAAACAGCAGGGGAAACACTAGGGAGACTGTTGAAAGGAGAGATTAGAGATGAAGAACTGAAGAAACTAATTAGGCTTCAGTTCGAGAAACGTCTGCAATGGGGTTACAAGTCATCTCATCAAGAACAACTCTCCTTCAACCTTGACTTCATTAAATCCTTAAAGAAGCTGGAAATGTCAGGAGAGATTGAGACAATGAACAAGGAAACCTATGAATTACCAACAGAATTCTTAGAAGCAGTTTTTGGAAAAACTATCAAGCAAAGTGGATGCTACTTCAAACATGAATCAGCAACAATAGACGAAGCAGAAGAAGCTGCACATGAATTATACTGTGAGCGAGCACAAATCAAAGATGGACAAACTGTTCTTGACATCGGTTGCGGTCAAGGTGGTCTAGTTCTATACATTGCTCAGAAATATAAGAACTGCCATGTTACAGGGCTCACTAATTCCAAGGCACAAGTCAATTACTTACTCAAGCAAGCAGAAAAGCTTGGGTTGACAAACGTCGACGCCATTCTGGCAGATGTTACTCAATATGAATCTGACAAGACATACGATCGACTACTTATGATTGAAGCCATAGAGCACATGAAAAATCTACAACTGTTTATGAAGAAACTGTCAACATGGATGACCAAAGATAGTCTTCTCTTTGTGGACCATGTCTGCCATAAAACATTCAATCACTTTTTTGAGGCAGTCGACGAGGATGACTGGTACTCAGGTTTCATCTTCCCCCCAGGTTGTGCCACAATACTAGCTGCTAATAGTCTCCTCTATTTCCAGGATGATGTTTCAGTCGTGGATCATTGGGTTGTCAACGGAATGCACATGGCTCGCTCAGTAGACATATGGAGAAAAACACTAGACAAAAATATGGAAGCTGCAAAAGAAATTCTATTACCTGGACTTGGAGGAAGCCATGAAGCAGTGAACGGAGTTGTTACTCACATTAGGACATTCTGTATGGGAGGGTATGAACAATTCTCAATGAATAATGGAGATGAATGGATGATCGCACAGGTGCTTTTCAAGAAAAAGTAATGACCCAATTTCCACGTCAACTGCATTTCCAGAGTTCTGCAATAATGTGATCCTCTAGTAACAATGATCACGGATTGCACCTTGTGGGTTTAATCTTGTTTTTCGTTGCCTGCATTTGTACTGTGTTTTTTTATTTTACATCAGATTCAAGGCTTGAAGAATTTGATTGTTATTGTGATCGTGTCTCTTATTTATCAACTTAGCAGGCGTGGGAAAATTGTACCAACATTTTTGGCTTCTTTTGTCAAGTTATTTCTCCGAGATCGGAAGAGCGTCG

>Pnu_TRINITY_PSSDR1 Pnu_c36761_g1_i3 len=1017

GTGCTCTTCCGATCTCACCGTTCGGTTTGACCCCAAATTAAAGGAAGATGTAAGCCACTTGAAAGCCCTTCCTGGGGCCGCTGAGAAGCTCCAAATTTTTGAAGCAGATCTTGAAAACCCAGAAAGTTTTGATGATGCAGTTGACGGATGTATCGGTGTCTTTCATGTTGCTCAAGGAATTGGTTTTGAAGAAGCATACACTCAAGAAAAACTACTCAAAACGTCTGTTGAAGGAACTCTTGGAGTTCTCAAGTCATGCTTGAAGTCCAAAACTGTGAAAAGGGTTGTGTACACATCTTCGTCAGCTGCAGCAATGATGATAAGTAATCTCAAAGTTGTTAAAGAAATCGACGAGACAATGTGGTCAGAAGTCGACCATTTCATTAGCACACCCGAACATGTTATTCCTGCTGGTCTCTCATATGTGGTTTCAAAGACACTGGCAGAAAGAGCTGCCCTCAAATTTTCTGAAGAACATGGATTGGATCTTGTTACTATACTTCCTTCTATGATTGTTGGACCATTTATCACTCCCAATCTCCCCGGGAGTGTTGCTCTGGCTCTTTCTGTTATACTGGGTGATAGGACAATGATGATCCGTCTTAAACCTACAAGTGCAGTACACATAGATGATGTTGCATCAGCCCAGATATTTCTCTTTGAATGCCAAAATGCAAAAGGAAGGCATATTTGTTCTTCAGTTGATTTTACAATACATGACGTGGCTAAATTTATAGCTGAGAAATATCCGGAATTCCAATTGCCAACAGACTTACTAAAGGAGATTGAGGAAGAAAAACCAGTTCATCTTTCCCCACATAAGCTGTTGAGTATGGGATTTCAGTTCAAGTATGGTTTTGAGGATATGTTTGGTGATGCAATACGAAGTTGCAAAGAGAAAGGTTTCCTTTAGAATCTGATTAATAGTATGGTTCCCTGCGTGAGTTAGCTATCTAGCTTGAATTGCTGTTGCTTTGTGAAATGAGTGAATAAATGGGCAGGTCTTCTTTTGCGAAC

>Pnu_TRINITY_CYP82X1 Pnu_c37280_g1_i2 len=913

TGCTCTTCCGATCTCTGGAGATGACCCCGATTTGGTCATCAAGTCTACGATTTGGGTGAGGCTATTGCTACACTGTTCCAGTTTGCGTATGTACTTTCTTGTTCTAACTTTCACCTATTGCATATGGATGTGCAGGAAGCCATATTAGGTGGGACTGACACCATAACAACCACCTTAACTTGGATCCTCTCTCTGTTATTGAACCACCCCCAAGTGTTAAAGACGGCGAAGGAGGAGTTAGATACGCAAGTGGGAAACGATAGGCACGTAGAAGAGTCAGATACTCCTAAGCTCGTTTACATTAATGCAGTTATCAAAGAATCAATGCGATTGTATCCAAATGGGGCATTGCTTGATCGGTTGACAATGGAAGAGTGTGAAGTTGGTGGAATCCATGTACCAGCCGGAGGACGCTTAATCGTGAATGTTTGGAAGATCCAAAGAGATCCGAGTGTGTGGGAGAATCCTCTGGAGTTTAAACCAGAGAGGTTTTTGATTGATGGTGAAAAGATGCATGTGGATTTCAAAGGTCAAAATTATGAGTTCATACCATTCGGGATAGGCCGGAGGATATGTCCTGGTATGCTTTCGGCATTACAGGTGATGAATCTTGTGATTGCCCGCCTTATTCATGGGTTTGATATGAAAGCAGCGAGTGCCAATGAGAAAGTGGATATGGCAGAAAAGGCAGGCATGGCTTGCTATAAGATGGCGCCTCTTGAAGTCATAGTCACGCCTCGACAGTAGATGTTCTTAACTTCTTGTACTCGAAATGTAACGACAAATAAAAAGATGGGGCACTAAAAATACCATCGCTTTTTAAATCTTTTGATGGTATTTTTATTAGGGCCCACCATGAGATTTGTTTTGTTTTCTTTTAATAAGAAAACATTGGGTGTTTTTTTAAAAAAAA

>Pnu_TRINITY_CYP82X2 Pnu_c54954_g2_i2 len=1894

CGTCTATAAATATGAAGCTTACTGATGGAGTTGTTATCCTATCAGTTCCTTTTTGTTGGCAATGAAGTTGATTCTTTTTCAACCGATTACTGTTTCTTTTTCAACATTTCCTTTTACTGCTACAATAATTTCCATTATTTTTCTCTACAATGTCTTCTTGATAGTGACTAAAGAGAAGAAGAAGAAAGCACCGAATGCATCAGGAGCATGGCCATTGTTAGGTCATCTCGGACTCTTCATGAAGGATAAACCAATTTACAAAACTCTAGGAACCATGGCTGATACATACGGACCAGCATTCAACATTCAATTAGGCAGCCAAGAATACCTTGTTGTGAGTAATAAGGAAATGGTGAAAGAGTGTTTCACATCGAACGACAAGTTATTTTCGAACCGTTCAATCACGTTAGGCGCTAAATACATGTTGAATCACACCAGTACTGTCGTGTTTTCGCCTTACGGAACGTATTGGAGAGAAGTGCGGAAGATAATGATGCAACAAGTACTCTCTAACCGACGATTACAGGAACTGCAACATTCGAAAATAGACGAAATCGACAAGTCGTTTGCGAAGCTTAATGATTTATGCAGCAAAGATGCTAATGGCACAAAGGTTAGCATGGAGGAATGGTTAGAGGACATGACTTTCAACATCAGTGCCAGAATTGTATTTGGTTACCAGAGCGGAGGTGACACACCTGGAGCTACAACTTTGAATGTCGAGAGATACAAGAAAACGTTGAAAAAGGTGCTTCATCTTCTAGCATCCAGCTTTGCAGTTTCAGATTTCTTCCCATATCTGGAGTTTGTTGACCGATTGAGAGGCCTTGTAAAGAATATGAAAAACTTGGGAGATGAATTAAATTCAATTGCTGGATGTTTTATTGAAGAACATCGCCAAAAGAGACGAGCAGAATCATTATCCTCACCGTCATCTTTATCGAATGAAGTAGACGGTGATGTACAAGACTTCTTTGATGTGCTCATGCCATTAATGGAACAATCACGGCTTCACCTAGATGACCCAGATTTGCTCATCAAAACCACGATCTTGGAACCTTTTGCATTTTGGACAGAAAGTTCAACGGCAAGCCTAACCTGGGTCATCTCTCTACTGCTCAACCACCCCAACGTGTTAAAGCAGGCAAGGGAGGAAATAGATAGGCATGTGGAGAATGGTAAACAAGTGGAAGCGTTAGATATCCCAAAGCTGGCATACATGAACGCAATAATCAAAGAGTCGATGCGACTGTACCCATTCGGAGCTGTCCTCGAACGATACACGACCGAAGACTGCGAGGTTAGCGGGTTCAACGTACCCGCTGGCACACACGTAATGGTGAATGTATGGAAGATTCACAGAGACCCAAGTGTTTGGGAGAATCCTTCAGAGTTTCGACCAGAGAGGTTTTTGAGCAGCGACAAGGTGGGTGTGGACTTGTATGGTCAGAGTAATGAGTTCCTGCCATTTGGAACCGGCAGGAGGATGTGTCCGGGGATAAATTTAGCATTACAGATGATGCATTATGTGTTGGCACGTCTTGTTCAAGGCTATGAACTGAAACCGGCTAGCCCCGATGGGAAGGTGGATATGGAAGAAAAGATAGTCATTGCAAACATCAAGATGTGCCCTCTCGAAGCTATTATCAGCCTTCGGAGATAAATATTACCTTCCAATTTTACAGCAGCATCTGTGCTTGTAAATCTTTTCTGCTACGTGTGTTTTCTTTTTTCCTACTTTTTTTCTAAGTTCCTGGTGTTGGAAGTTATGTAACTATTATCACGCATTTAGTCTGCTTGAAGAGCCTGGTGTTAACACGCTCTGCATTCAACTAGGAATAACCGATTTGGCTGCAACTGGAATCCAATCGATGTGTTATTACTCTGTCTTGCA

>Pnu_TRINITY_CYP82Y1 Pnu_c52195_g1_i5 len=1680

TTTTGCACCTTACGGGAAATACTGGAGAGAGTTGCAAAAGATATCTACACATAAACTCCTTTCTAATCAGCAGTTAGAGAAGTTCAAACACTTGCGAATTTGTGAAGTCGATAACTCCTTTAAAAAGCTTCATGAGTTATGCAGCAATACACACATTGGGAACAAACAGGGAGGAGGTGATACTACTTCTGCAGCTAGTCTTGTGAGAATGGATGATTGGTTCGCGTACTTGACATTTAACGTAATAGGTCGGATTGTCAGCGGATTCCAGTCAAATTCAGTGACAGCAGGTACCACAAGCAACCAGGAAAAATACAAGCTTGCAATCGATGAAGTGTCGCATCTTATGGCAATGTTTGCCGTTTCAGACGTGGTTCCATGCTTAGGGTGGATAGATAGATTGACTGGCCTTGCAGGAAGGATGAAGAATTGTGGAAAGAAATTAGATGCAGTAGTTGGGAGTGCAGTTGAGGATCATCGCAAAAAGAAACGAACTTGTGGAAACAGCACAGAAGAAGCAGTGGTGGAGCACGAACAACAAGACTTCATCGATGTTTGCTTGTCGATTATGGAGCAATCACAGATTCCTGGAAACAACCCCGAAATAACTGTCAAATCTATCGTCCTGGACATGTTATCGGGTGGGAGTGACACCACTAAGTTGATCATGACATGGACCCTTTCTCTGTTGTTGAACCATCCAAATGTGTTGGACAAAGCTAAAGAAGAAGTGGATACGCACTTTCGGAAGAAGAAGAAGAAGATATCGAGTGATGCACTTGTGGTTGATGCTGCTGATGTTCCTAGCCTCGTCTACATCCAAGCAATCATCAAAGAATCATTGCGGTTATACCCCGCCAGCACATTAATGGAGCGAATGACGAGTGATGATTGTGAGGTTGGTGGCTTCCACGTACCTGCTGGTACACGACTATGGGTTAACGTATGGAAGATGCAACGAGATCCAAGAGTCTGGAACGATCCACTGGTATTCCGACCTGAGAGATTCTTGAGCAATGACAAGGTGATGGTGGACGTGAAGGGTCAGAATTATGAACTGATACCATTTGGAACAGGCAGACGGATATGTCCAGGTGCATCTTTTGCATTAGAAGTCTTGCATTTGGTTCTTACTCGTCTTATTCTTGAGTTCGAGATGAAGGCACCAAACGGGGAAATTGACATGAGGGCAAGACCAGGGTTTTTCAACAACAAGGTAGTGCCACTGGATGTTCTGCTCACCCCACGCACACTAGAACAAGATTCCGGTTAGCTGTCTGCTGGTTGTTGCACCCACTGTTATAATGCGCCTGTTAGCAATGGAAGCATGATGCTGGTTGCCACATTTTTCATCGTAATATGTCAATTGTGCGAAGATAAAGCTAGAACATAATTTCTGAACCAGCCGAACTGCCTGTTGATCAGTAGTCTACACGAAGCTGATAAAGATTTAGGTCCTTGTAACGAGAAATTGGGTCGTGGAAAAAGGACTCATGGTAAGAAGAAGATTTAGGATCATTTCCTGTTGTATTGTCTCTATTATTGTTTAATTATGCTCTTAGTATTGTGGTTGTAGTTGAAGATGGTTTGTAATCTTCCTGTGTTGCGCCTCCCTTTCTTGTACTCTCTTTTTTTTCTTTTATCAATAAAGCCGAACCCTGTTAAAAAGAGATCGGAAGA

>Pnu_TRINITY_CYP719A21 Pnu_c51429_g1_i3 len=1214

GTGCTCTTCCGATCTCTTTACTAGTGTGATATACAGTGACATAGGCTACTAATTGTGTATTTAAGTTGAGCGTGTGTGTTCTGCCTAGAAACATGCAAAAGAGTTGCTCAAACTCAAGCTAGCTAAAACACCAACAATGAAAGAAATTATGAGTGGTTCCTGGGTTCCTTCTGTTACTTCCACCGTACTAGCAGCTTTTATTGTGTTCATCATGATTTTCCGGAGAAGAAAATCAGCATCTGCAATGGAATGGCCAGTTGGCCCAAAAACATTGCCCGTCATAGGTAACTTGCACATTCTTGGAGGCACTGCTCTACATGTCGTCTTACACAAACTAGCTCAAGTTTACGGCAGTGTAATGACGATATGGATTGGTAGTTGGAAACCTGTCATCATTGTTTCTGATTTTGATCGAGCTTGGGAAGTTCTTGTTAACAAATCCTCGGATTATTCAGCTCGCGAAATGCCCGAGATCACTAAAATTGGTACTGCGAATTGGAGAACAATCTCAAGTTCTGATTCTGGTCCTTTTTGGTCCACTCTTCGGAAAGGTCTTCAGAATGTAGCATTGTCGCCTCACAATTTAGCATCCCAAGCTGCACTGCAAGAGAGAGATATAAGAAAATTGATTGAGAATCTGAAAGAAGAAGCACTGCTCAACAATGGAATCGTCAAACCACTTGATCATCTCAAGAAAGCAACTGTCAGAATAATCAGTCGGTTAATCTATGGGCAGGATTTCGATGACGATACGTATGTCGAAGGTATGCATGAGGTGATCGAGTTTTTGATTCGCATTAGTGGTTATGCTCAACTGGCTGAGGTATTCTATTACGCTAAATACTTACCAAGTCATAAGAGAGCTTTAAAAGGTGCCGAAGAAGCAAAAAGAAGAGTAATAGCATTGGTGCATCCTTTTCTTGCGTCGAACCCTACTCCTAATAGTTACTTGCATTTTCTCAAGGCACAGAATTATCCTGAAGAAGTAATCATATTTGCTATATTCGAAGCTTATCTTTTGGGTGTGGACAGTACTTCAACAACAACTGCATGGGCACTGGCATTCTTGATACGCGAGCCGTCTGTTCAACAGAAACTTTATCAAGAGCTTAAGAATTTCACAGCCGACAACAACCGCACAATGGTGAAAGTCGAAGACGCAAACAAATTACAGTACTTGCAAGCAGTTGTGAAAGAAACAATGAGGATGAAAC

>Pnu_TRINITY_9OMT Pnu_c50372_g1_i3 len=1380

GTGTGCTCTTCCGATCTGAAGTAGCAAATGAGTAGAAACATAGCTAGGATTGTCTATAAATATCCATGTATTTCCTTTAAGAGAAACATAAACACACCACACCACCACCACCACTTGACCATTGCCATCAAACCAGAGTACTAAAAATGGCTACCAATGGTCATAACCATCAAACATCCACCACCACATTCACTACCATTGCTGCTGCTAAAGAAAGCAACAATGATATCTCTTATCTTTCAGAAACGGCAAACTTAGGGAAGTTGATATGTATTCCAATGGCATTAAGAGCTTCCATGGAGCTAAATGTCTTCCAACTTATCTCAAAGTTCGGACCGGATGCAAAGGTTTCGGCTTCTGAAATAGTCTCCAAAATGCCGAAAGCTGCAAATAATCCCAAAGCGGCTGTCTATCTAGATAGGATTCTTCGATTACTCGGTGCAAGTTCCATTCTTTCTGTTTCCACTAGAAAGAATGTTGTTAATGGCAATGTCAATGGAGGAGGAGATCATTCGGATGAAGGATCAGTAGTGGTACATGATGAGATGGCGTATGGGTTAACAAGCTCGTCGCGTTGTTTGGTGCCCAGAGAAGAAGACGGGGTGTCATTAGTCGAAGAGTTGTTATTCACATCCGACAAGGTTGTTGTGGATAGTTTTTTCAAATTGAAAGATGTTGTTCTAGAAAAAGATATTGTGCCATTTGATGTTGCTCATGGTGCTAGCATCTTTGAGTATGCTGGTAAAGAACCGAGGTTGAATCAAGTTTTCAATGATGCAATGGGAGTTTTCTCTGTTGTGGTTTTCGAAGCTGTTTTCAGAGTTTATGATGGATTTCTTGGCATGAAAGAATTGATTGATGTGGGTGGTGGTATTGGTACTTCGGTTAGTAAGGTTGTTGCTAAATATCCTTTCATTCGTGGTATCAACTTTGACTTGCCTCATGTTATTTCCGTTGCTCCTCCATATCCAGGTGTAGAGCATGTTCAAGGAGATATGTTTGAGGAAGTACCCAAGGGTCAAAACATGTTGCTAAAGTGGGTACTTCACGATTGGGGTGACGAACGGTGCGTGAATTTGTTAAAGAATTGCTGGAATTCATTACCGGTGGGTGGAAAAGTTTTGATAATCGAGTTTGTTCTTCCTAATGAACTTGGTAATAATGCTGAATCATTCAACTCATTGATTCCCGATTTGCTTATGATGGCTCTGAATCCCGGTGGTAAAGAACGAACCATTTCCGAGTACGACGGTTTAGCCAAAAAAGCTGGGTTTATCAAAACTATCCCTGTCCCAATCTCCCAAGGTCTTCATGTCATTGAGTTTCATAAATGATTCAAAATGGGTTATTCCCATTGCATGCATTAAGTTACTATGTTTA

>Pnu_TRINITY_CFS Pnu_c45817_g1_i1 len=1876

CTCTTCCGATCTCTCAACTTCAATTGTCTTCTCTCTTCATTCATATCGTTTTGATCTTTCTTCATAAAGATAACAAAAAAATAGAGTGCTTCGTCAGATGGAGGAGACCTTCTGGTTGATTACTTGTGGTGTGGTAGTAGCATTTGCATTAGCAAAGCTACTATTCGGCAAGAAATCATCTCTGTCTACAATGGAATGGCCAGCAGGTCCAAAGAAGTTGCCCATAATTGGGAACCTTCATCAGTTAGGAGGTGCAGCGTTTCATGTCTGTCTGGCAAACCTGGCCAAAGCTTATGGAGGTGTGTTTACCATATGGGTTGGGAGTTGGCGTCCGTTTATAGTCATAAGCGATGTCGACAGAGCTTGGGAAGTTCTTGTCAACAAGTCCTCAGATTATTCAGCTAGAGACATGCCTGATATTACTAAAATCATATCGGCAAGCTGGAAGAATATTTCTCACTGTGATTCTGGTCCGTTCTGGCATAATCTAAGAAAAGGCCTTCAAAGTGTTGCCTTGACTCCTTTTAATGTTGCGTCTCAGTGTCATCTGCAAGAAAGAGACATGCAAAATCTGATCAAATCCATGAGAGAAAAAGCTTCGCGGACGAACGGGATTTTGAAACCACTTGATTATGTGAAAGAAGAGACTGTTCGGTTACTAAGTCGACTTATATTCGGTCAAGAGTTCGTAGACGAGGATTTCGTCGTTGGTATGCATCAAGCCCTCGACGAGTTGGTGAGTATCAGTGGGTATGCAAGTCTAGCTGATGCTTTTAAATTTTGCGAGAATCTACCAAGCCATAAGAGAACTGTCCAAGCAGTTCACGCCATCAAAAAGCGGTTCGACAGTTTGATTCGTCCGCATATTGCTTCAAAGCCTCCTACAAATACTTACTTGCATTTCCTTCTATCTCAAGATTTTAGTGAAGATGTTATAATCTCTGCCATTCTTGAAGTTTATGACTTGGGTGTTGATAGCACTGCATCGACAACGGTTTGGGCTCTTACGTTTCTAGTTCGCGAACAGAAGATTCAGGAAAAACTGTACCGCGAAATCAATAGCGTAACCGGAGGGAAAAGACCCGTGAAAGTTGAAGATCTGAACAAATTGCCGTATTTGCAAGCAGTGATGAAAGAAACAATGAGGATGAAACCCATTGCACCAATGGCAATTCCTCACAAAGCCTCGAAAGATACTTCACTAGCGGGCAAAAAGATCAATAAAGGTACAGTGGTAATGGTAAATCTTTACGCAATCCATCATAACCCTGACGTTTTCCCGGAACCGTATAAGTTCATGCCAGAGAGATTCTTAAAAGACGTTAACAGTGATGGAAGTTTGGGTGATATCAAGAAGATGGAAAGTTCATTGTTAGCATTCAGTGCTGGTATGCGAATTTGTGCAGGGATGGAGCTTGGGAAACTCCAATTGGCTTTCGGACTTGCAAGTTTGGTCAATGAGTTCAAATGGGATTGTGTTGCTGAAGGAAAATTGCCTGATCTCAGTGAAGAACATTGTTTCATTCTCTTGATGAAAAACCCACTTGAAGCCAGAGTTACTCCTCGAATCCATTAGTCATACTTCGCGGTAGGGTCAAAATTTGGAGTGCTGGCGTCAACCTCTCTAGCGTACGTAAAAGGCTGTAATGCATCCAAAGGTGTGATGGCTGGTGTTTATGCATGCATGATAAGTCGGTAATCTTTTTTGAGTTTGTTGGTTTGAATAAACAAGAATATCAACTGCATTTTATGTTTATCCTTTAGGACTTTTGTTTTGTATTTTGTGCGTGTACATATATCTCAGTCTTGTATCAGTCAACGTATCTTGGACTGGTCAATCACGATCGATACCTTAGTTAGATCGGAAGAGCGTCGT

>Pnu_TRINITY_SPS Pnu_c48423_g1_i8 len=1676

TTCCGATCTTTGTAACTAATACCAGAAATGGAGGAAACCATCTTTAGCAACTCATGGATTCTAACTACTTCAAGTCTAGTAGTAGTAACAATCATTACCATGCTAATGGTATTCAAAAGGAAAAAATCATCGACTATGGTGTGGCCAAAAGGGCCAAAAACGTTGCCAATCATAGGCAACATGCATATTCTTGGCGGCACAGCTCTGCAAGTTGTCTTACATAACCTTGCCAAAATCCATGGCAATGTTATGACGATATGGATCGGTAGTTGGAGACCGGTTATCGTCGTTTCGGATATCGATCGAGCATGGGAAGTACTTGTTAACAAATCGTCGGATTATTCGGCTCGTGATATGCCCGAAATCACTAAGATTATTTCTGCTGATTGGAAAACAATTTCGACTTCTGATTCTGGTCCTGCTTGGTCTACTCTTCGCAAAGGTCTTCAAAATGTTGCACTTTCGCCTCATAACTTGGCCGCACAAAGTCAATTCCAAGAGAAAGATATTGCCAAAATGATTCAAAGTTTGAAAAAGGAAGCTGCGAGCAACAACGGAATTGTCAAACCAATTGATCATCTTAAGAAGGCAACATTAAGATTAATCAGTCGATTGATTTTCGGCCAAGATTTCGACAACGATAAGTACGTTGATGATATGCATCATGCTATTGAAGAATTGATTCGTGTCAGTGGTTATGCTCAGCTCGCCGAGGCATTTTACTACGCGAAATACTTACCGAGTCATAAGAAAGCTGTTAGAGACGTCGAAGAAGCGAATCAGAGAGTCAAAAAGTTGGTGCGCCCATTTTTATCATTGAATCCTCCTACTAACACTTATTTGCACTTTCTCAATTCACAGAATTACGACGAAGAAGTAATCATATTCGCGATATTTGAAGCTTATCTTTTGGGTGTTGATAGTACTTCTCTGACTACGGCATGGGCACTTGCATTCTTAATACGCGAGCCAGTTGTGCAGGACAAACTTTACCAAGAACTTAAAAGTTTTGCTGTCAAGAACGATCGCGAGATTGTGAAAGTCGAAGATTTGAACAAGTTGGGATATCTACAAGCTGTTCTTAAAGAGACAATGAGAATGAAACCAATTGCACCATTAGCCATACCTCATAAAGCTTGTAGAGACACTTCGCTAAAGGGTAACAAAATTGATAAAGGCACCCGGGTTATGGTCAACATATATGCTTTACACCATAATGAGAAAGTCTGGAAAGAACCTTACAAGTTTATGCCAGAGAGATTTTTGCAAGCTCAAGATGTTGTCAATGGAAAAGCAATGGAGCAGTCGCTACTGCCGTTCAGTGCAGGTATGAGGATTTGCGCAGGCATGGAATTGGGTAAACTTCAGTTTAGTTTCTCCCTGGCTAATCTTGTCAATGCGTTTGAATGGTCTTGTGTTTCTGATGGAGTACTTCCTGATATGAGTGATCAACTAGGCTTTGTTTTGTTGATGAAGACACCGCTTCAAGCACGTATAGTCCCTCGTGCGTAATCGTCGTACATTGTTGATGGATCCGGTTGTAATGGTTTTTAATCAAGTCAATGCTTCTGTTAGAGGTCTTGTTTTAACTGCCCGTATGCGGTAGATGGTGGTCATTTCCCTTTGATGTACTGTATTAAGCTCCTAGTTTGGGGCATTTCAATGTTGTGTTCATG

>Pnu_TRINITY_MSH Pnu_c51942_g1_i3 len=1158

TCCGATCTCAATAATGTCGATAATCTTTATTTTTTTGTCGTGTCACTTTTTGTCACAAATGAGGCAGATAAATTCTCAACTTATGAGTTCACTACTTCACTTCCATCCATAGGGAAAAAAAAAATGGATTTTCAGTTCAAACCAATCTCCATAGCACTTGTTGTAATTGCAGTAATTTTTATCTTCTGGAACCATGGAAGAAGAACAAAACCCACCAAAAAACCAGCACCAGAAGCATCAGGGGGAAGACCTATAATGGGTCACCTACACCTTTTCAACGACGGTGAGTTAACTCATCGGAAACTCGGTCTCATGGCCGATACTTACGGTCCGGTTTTCAGTATCCGGTTCGGTTCGCATAGAACGTTAGTTGTAAGTGACTGGGAAATCGTCAGAGAGTGTTTTACTACAAATGATAAACTCTTCTCGAATCGTCCCGGTACGTTGGCTATCAAACTCATGTTTTACGATACCGATTCGGTTGGTTACGCACCGTATGGTGCGTACTGGAGGGATTTGCGAAAGATCACGACGTTGAAACTTCTTTCGAATCATCGCATCGATGCAATAAAACACCTAAGGACTTCTGAGGTGGAGTCGTGTTTTGAATCGCTTTATAACCAGTGGAAAATCGGGGAAAAGCGTGGCGAAAATGAAGGGTTCGCTCCGGTGAGAATGGACAGTTGGTTAGGCGAGTTGACGTTTAATGTTGTGGCGAGAATCGTTGCGGGGAAGAAGAATTTTTCGGCGAATGGCGATGCTGGGGCTCAGAGGTATAAAGCAGCAATGGATGAAGCTATGAGACTAATGAGATTTTTCGCATTCTCAGATGTCATTCCATCGCTAAGTTGGTTGGACCGTTTAAGAGGTTTGGTTGGAGATATGAAGAAATGTGCCTCTGAGATTGATGCAATTATGGCTACTTGGGTTGAAGAGCATCGTGTCAAAAGAATTTCAGGAAACGATTCGGAGCTCGAACATGATTTTATCGATGTTTGCTTGGATATTATGGAGCATTCTTCGTTGCCCGGTACTGATCCTGACCTCGTCGTCAAATCTACGTGTCTGGATATGATATTGGGTGGGAGCGACACTACAACGGTCACACTAACATGGGCGATGTCTTCACTCTTGAACCATCCCCAAGTGCTACAAAAA

>Pnu_TRINITY_P6H Pnu_c19781_g1_i2 len=1131

TGCTCTTCCGATCTGTTCGGGAAGCACACAACGCTGGTTGTGAGCGATACCCGAGTCGTAAAAGAATGTTTCACTACTAATGATACCCTCTTCTCCAACCGTCCATCCACCAAAGCTTTCGATCTCATGACTTATGCAGACGACTCCATAGCTTTCACTCCTTATAGTCCTTACTGGCGTGAGCTTCGAAAGATATCTACTCTCAAACTTCTCTCTAACAACCGTCTTGAATCCATCAAAGAACTTCGGACCTCAGAGGTGAGCGTATGTTTCAAGGAATTATACGAGCTAAGCAAGAAGAAAAACGAAAATGGAGCTCCTGTTCCGACTGACTTGAAGAGATGGTTCGATGAGGTCTCAAACAATGTGATTATGAGAGTCATATTTGGGAAACAAAACTTTGGATCTAAGGTTGTACGCGGGGAAGATCAAGAGGCAGTCAAATATAAAAACATCATGGACGAACTTTCGCGTCTTTCGAGTCTGACTATGTTATCGGACATGGCTCCTTTACTCGGTTGGTTGGATTACTTCAAAGGAGATCTTAGAGCAATGAAACGAAATGGCAGAGAACTGGACTCGATGCTCCAGAAATGGTTGGAGGAACACAAAAGCAAGAAAAATTCAGATGATCGACAGGATTTCATGGATGTCATGTTGTCAATTTCCAAGGAGACACAACTCTATGGCCACGACCAAGATACTTTCATTAAAGCTACTTGTCTAGCCATGATCATGGGTGGAACAAACAGTACGGAAGTGACTCTAACATGGATCTTATCGTTACTTATGAACAACCGATACGCACTGCAAAAGGCTCGAGAAGAAATAGATTTACTCGTCGGAAAGGATAAACTGGTGGAAGATTCGGATCTCAAGAATTTGACATACATGAATGCCATCATCAAAGAAACGATGCGATTGTACCCATTAGGTTTTCTTCTGGAACGTGATACCAAGGAAGACTGTGAAGTTGGTGGGTTCAACATCGAAGGTGGTACACGATTACTAATTAACGTGTGGAAGTTGCAACGAGATCCGAATGTATGGACAGACCCCATGGAATTCAAACCAGAGAGATTTCTAACAGAGAATGCAGATATAGACGTTAGATCGGAAGAGAGTCGTGTA

>Pca_TRINITY_NCS Pca_TRINITY_DN18392_c0_g1_i1 len=2188

TGGACTTTAATACCTTCTTGAGTTTTGTTTGCAGCATATAAAATATGATCTGGGATTTGAGTCTCAAACATCATCTCAAAACACTCATAGACTCGCAAGTAGAACTAGCTTAAGTGTTTTACAGAGAATCATCATGAGGAAAGAAATCAAATATGATATGGAGGTAGCTACCTCAGCTGATTCAGTATGGGCAGTTTACAGTTCACCGGATATTCCTAGACTTCTTCGGGATGTTCTACTTCCCGGTGTCTTCGAGAAATTGGATGTGATTGAAGGGAATGGTGGTGTCGGTACTGTTCTTGATATTGTTTTCCCTCCAGGTGCTGTGCCTCGACGTTACAAGGAGAAATTCGTCAACATCAACCTCGAAAAGCGATTGAAAGAAGTGATTATGATTGAAGGCGGATACCTGGACATGGGATGCACATTTTACTTGGACAGGATCCATGTAGTTGAAAAAACCCCGAGCTCATGTGTTATTGAATCGTCTATTGTTTACGAAGTGAAAGAAGAGTATGCTGATGCCATGGAAAAATTAATCACAACTGAACCATTGAAGTCGATGGCGGAAGTCATTTCTAATTACGTTATACAGAAAGAATCCACAAGAAACATTCTCAACAAGCAATCTTTAGTGAAGAAGGAGATTCGATACGATCTGGAGGTACCGACTTCAGCTGATTCAATCTGGTCAGTTTACAGCTCGCCCGATATTCCTCGGCTTCTTAGAGATGTTCTACTTCCTGGTGTCTTCGAGAAATTGGATGTCATTGAAGGGAATGGTGGTGTTGGTACTGTTCTTGACATTGTTTTTCCTCCAGGTGCAGTGCCTCGTCGATACAAGGAGAAATTTGTGAACATCAACCGCGAGAAGCGATTAAAAGAAGTGATTATGATCGAGGGAGGGTACTTAGACATGGGGTGCACATTTTACTTGGATAGGATCCATGTTATAGAAAAAACCTCTAAATCTTGTGTCATTAAATCGTCTGTTGTTTACGAAGTGACACAAGAGTGTGCTGATGCAATGTCTAAGTTAATCACAACCGAACCGTTGAAGTCGATGGCAGAAGTCATTTCTAATTACGCTATTAAGCAACAATCTGTTTCTGACAGAAACATTCCTAAGAAGCAATCTCTCATAAGGAAAGAAATTACTTATGAAACGGAGGTGCAAACCTCGGCAGATTCAATTTGGGACGTCTACAGTTCTCCTGACATCCCTCGACTGCTTAGAGATGTTCTGCTTCCGGGTGTCTTTGAAAAGCTAGATGTCATAGCAGGCAATGGTGGCGTTGGTACCGTGCTTGACATCGCTTTCCCTCTAGGTGCAACGCCGCGCAGTTACAAGGAGAAATTTGTGAAGATTAACCACGAGAAGCGATTAAAAGAGGTGGTTATGATTGAAGGGGGATACTTAGACATGGGGTGCACATTTTACATGGACAGGATCCATGTCTTTGAGAAAACCCCGAATTCATGTGTCATCGAATCCTCCATCGTTTACGAAGTGAAAGAAGAGTATGCTGGTAAAATGTCTAAGCTAATCACAACAGAACCATTGGAATCCATGGCAGAAGTCATCTCTAGCTATGTTCTTAAGAAACGATTCCAAGTATTTGGACTCGAGATTACACCAAAACTAAGATCAAATCTTTTCCTATGTTTGATAATCTGTTTGGCTATAGTTGGAGGTATGTTTGTTGGAGGTGTTCTACTCTAAGCTTAAGGGTTTTTCTTAAATGTGATTTAGGAAGTAAAGAATAGCTTTGATTCTTAAAACCATGGATTGCATTGCAGAGATTGAGTTCCGATTTACGCAATTTGTCATAAGTAATTTCACAATAAGTCTCGACAAGATGATAAAAGAATGTATGCTTCAAAAAAAAGATAAATGTAATAAAAGCTTCGTGCATTGTTTGAAAAACGGGTTTAGATATGAGTCACATGGAGCTTCAAGTATCGAACTATCGTAGGACTTCAAATGCTATCTTTAAAATGAAAACAAGTTGCAGTATTTCTATTAGCAAAGTTATGAAATGCATATTGGATGCATAAATATTGTGACCGCATTACATAAGTCATTACACAACTATTTTTTCGTGTAAAGAAATTAAGTCACTGGTATCATCACCCAATGACTTTACTAATATAAAAAGGTCGTTAATATCATCACCCAACAACCGGG

>Pca_TRINITY_6OMT Pca_TRINITY_DN182_c1_g2_i4 len=1258

CTAGAGAATAGCTCAAATCATACAATTGAATCAATTTTCATCAACAACTAATTTTTTTTCTAGAGAGACAGAGATGGAAGTAGTAAGCAAGATGGATCAAGAAGTCCAAGCAAAAATTTGGAAACAAATTTATGGTTTTGCAGAATCACTAGTTCTGAAATGTGCAGTTCAATTAGAGATTGCTGAAACACTTCATAACAATGTCAAACCCATGTCTTTATCCGAATTAGCATCGAAACTTCCCGTTATTCAACCCGTAAATGAAGATCGTCTGTACCGAATTATGCGTTACTTAGTTCACATGGAACTCTTCAACAAAGATGCTACCACACAAAAATACTCATCAGCTCCACCAGCTAAGTATTTGTTAAGAGGCTGGGAAAAATCAATGGTTGATTCAATCTTATGCATAAATGATAAGGATTTCTTAGCTCCATGGCATCATTTAGGTGATGGTTTAACCGGTAACTGTGACGCTTTTGAGAAAGCGTTAGGGAAAAGCATTTGGGCTTATATGAGTGAGAATCCTGAAAAGAATCAGCTGTTCAATGCAGCAATGGCTTGTGATACTAGATTAGTTACTTCAGCATTGGTTAATGAGTGCAAAAGTGTTTTCAGTAATGGAATCAGTACACTAGTTGATGTCGGCGGTGGTACCGGTACTGCTGTAAAAGCCATATCTAAAGCTTTTCCGGATATTAAATGCACTATCTATGATCTTCCTCATGTCATAGCTGATTCTCCTGAAATTCCCAATATTACTAAAATTTCTGGAGATATGTTCAAGTCTGTTCCTGGTGCTGATGCCATCTTCATGAAGTGCATACTTCACGACTGGAACGATGAGGAATGCATTCAAATCTTGAAGCGATGCAAAGAAGCACTACCAAAAGGAGGGAAGGTTATTATAGTGGATATTGTCCTCGACATGGCGGAATCGACTCATCCATATGCAAAAATTAGACTCACATTGGATTTGGATATGATGCTTAACACCGGTGGAAAGGAAAGAACTGAAGAAGAATGGAAGAAACTTATTGATGCCGCTGGTTTCACTAGCTATAAAATTACTCAGCTATCTGCTGTTCAATCTGTAATTGAGGCTTACTCTTATTAAGGATAATTTTATCCTGTTTTTTCCCTTTGAGTAATTGTTGCCCTCTATTTGAATTATGTTTGTGTTTAAAATAACTGCAAGATGAGCGAATTTCCATCTTGCATTGGAATCAATGTTTTTCGTTTACTTAGCAATTTTGAG

>Pca_TRINITY_CNMT Pca_TRINITY_DN14532_c0_g1_i3 len=1648

CCATGATTGAATTTGTCCCATAATTACTAACCTTTTCTTATCAAATTCCATATTATTATATAAACCACATCAGGTGGTAGTAGTTTAGTAAAATATACCACTATTACACTGTCTCTGTCTCTCTTAATCTATATAGATTTTGCAGTTTTTTTGTTTGCTAGTTGAATAAAAGAAGAAGATGCAGCTAATTGCAAAAGAAGAGCTCTATCAAAAAATGGAACTTGGGTTAATACCAGATGAACAGATTAGAGAAATGATTAGAATCGAGTTACAAAAACGTCTCAAATGGGGTTACAAAGAAACCCATGAGGAACAGCTTTCACAACTTCTTGAATTAGTTCATTCTTTAAGAAGTATGAAAATGGCGACTGAGATGGAGAATTTGGATTTAAAGCTATATGAAGCTCCTATGGAGTTCTTAAAAATCCAACATGGAAGCAACATGAAACAAAGTGCGGGTTACTACACCGATGAATCGACGACGTTGGACGAAGCCGAGATAGCGATGCTAGATTTGTATATGGAGAGGGCGCAGATCAAAGATGGTCAGAGTGTACTTGATTTAGGGTGTGGATTGGGAGCTGTAGCTCTTTTTGGTGCTAACAAGTTTAAAAAATGTCAATTTACTGGAGTTACAAGTTCTATGGAACAAAAAGATTACATTGAAGGAAAATGCAAGGAACTTAAGTTGTCAAATGTCAAGGTTGTATTAGCTGATATAACAACTTATGAGACTGAAGAAAGATTTGATCGTATCTTCGCAGTTGAACTGATTGAGCATATGAAGAACTACCAATTGCTTCTTAAGAAAATATCAGAGTGGATGAAAGATGATGGGCTTCTTTTCATTGAACATGTTTGCCATAAAACTCTAGCTTACCACTATGAGCCTGTTGATGCAGAAGATTGGTACACTAACTACATATTCCCTGCTGGAACTCTGACTTTATCATCTGCTTCAATGCTTCTGTACTTTCAAGATGATGTTGCTGTCGTAAATCAATGGACTCTAAGCGGAAAACATTACTCTCGATCCCATGAAGAGTGGTTGAAAAACATGGATAAAAACATTGTTAAATTCAAAGAGATAATGAAGTCTATAACTAAAACCGAGGAAGAAGCAACCAAATTGCTTAACTTTTGGAGAATTTTTTGCATGTGTGGAGGAGAGTTGTTTGGATATAAAAATGGTGAAGAATGGATGCTCACCCATCTTCTCTTCAAGAAAAAATGAGCGAAGAAAAAATGAAAAGACTTTTTTTTTATACTACCTCCGTTTCACCAAAAGTGTTATTATCATTTTTTCATTTTGACCTATTTTTAGATTAAAATGAAAAAAATAATAGTAACACTTTTACTGAAACGAAGGTAGTATTATTTTTTTGATTTTTTTATCTTAGACGGTAAACAGTTTGAAGTAGTTCCCATAGAGGGATTTCACCACCATTATTGCATACTAATTCGTTTCAAAGGAGATTGAACACTCCATCCTATAATCCTTATTCTCATGAAGGCTACTTTTGATGCCCTGTTATAGTAGTGTGGATAGATTGCACTACTATCCTGGCCACCTGAGTTATCATGTCTTCAGTTTGTTTCGCCTTTATTTGTTCATACATATGTTGTTACTTTTCTTGCAGACAATTTA

>Pca_TRINITY_CYP80B1 Pca_TRINITY_DN36732_c0_g1_i2 len=1573

GAGAGAAAGAAAAGATGGAGATTGTCACAGTAGCACTAATAGCAATTGTGTTCACTAGTTTTTTATACTTGATTTTTTTTAGAGATTCAAGTCCTAAAGGTGTGCCACCAGGTCCAAAACCATGGCCAATAGTTGGAAACCTTCTTCAACTTGGTGAGAAGCCTCATTCGCAATTTGCACAACTTGCTGAAACCTATGGTGATCTCTTTACACTCAAATTAGGATCTGAAACAGTTGTTGTAGCTTCAACTCCATTAGCAGCTAGCGAAATTCTAAAGACGCATGATCGTGTTCTCTCCGGTCGGTACGTGTTTCAAAGTTTCCGAGTAAAACACCATGTGGAGAACTCTATTGTTTGGTCTGAATGTAATGAGACGTGGAAGAAGTTGCGGAAAGTTTGTAGAACAGAACTTTTCACACAGAAGATGATTGAAAGTCAAGCTGATATTAGAGAAAGTAAAGCTACGGAAATGGTGGAATATTTGAAGAAAAATGAAGGAGATGAAGTGAAAATTGTTGAAGTTGTATTTGGGACGTTGGTGAATATATTTGGGAACTTGATATTTTCGCAAAATATTTTTAAGTTGGGTGATGAAAGTAGTGGAAGTGTAGAAATGAAAGAACATCTATGGAGAATGCTGGAGTTAGGTAATTCGACAAATCCAGCTGATTATTTTCCGTTCTTAGGTAGTTTGGATTTGTTTGGACAAAGAAAAGATGTTGCTGATTGTCTTCAAGGGATTTATAGTGTTTGGGGTGCTATGCTGAAAGAAAGAAAAATAGCCAAGCAACAGAACAGCAATAAGAAGTATGATTTTGTTGAGATTTTGCTCGATTCCGGACTCGGCGACCAACAGATCAATGCCTTGCTCATGGAAATATTTGGTGCGGGAACAGAGACAAGTGCATCTACAATAGAATGGGCGTTGTCTGAGCTCACAAAAAACCCTGAAGTAACGGCCAATATGCGATCGGAATTGCTATCTGTGGTAGGGAAGAGGCCGGTTAAGGAATCAGACATACCGAACATGCCATATCTTCAAGCTTTTGTTAAAGAAACTTTACGGCTTCATCCAGCAACTCCGCTGCTACTCCCACGTCGAGCACTTGAGACGTGCCAAGTTTTGAACTATACGATCCCAAAAGAGTGTCAGATAATGGTAAACGCCTGGGGCATTGGTCGGGATCCAAAAACGTGGACTGATCCATTGAAGTTTTCACCAGAGAGGTTCTTGAATTCGAGCATCGACTTCAAAGGGAACGATTTCGAGTTGATACCATTTGGTGGAGGGAGAAGGATATGCCCTGGTGTGCCCTTGGCAACTCAATTTATTAGTCTAATTGTGCCTACTTTGGTACAGAATTTTGATTGGGGACTATCAAAGGGATTGGATCCTAGCCAATTGATCATGGAAGAGAAATTTGGGTTGACATTGCAAAAGGAACCACCTTTGTGTATTGTTCCTAAAGCTCGAAATTAATCTTAATCAAGAAATTTAACTATGGATTCACACTCTTATTTGGTATGTGTTGTGTCTCAAGTTGATAAAGCTAACATTTCTTGAATCCCAGG

>Pca_TRINITY_4OMT1 Pca_TRINITY_DN20440_c0_g4_i1 len=908

ACACCCCAAAAAAACATAGCCTTCTTAAAGAAATATGTTATAGAAACTTCATATCATGGGTAGTAGTATAATATCTGAAACACAAGAAGTTGGCATCAAAGATCAAGCTCAATTATGGAATATAATCTTTGGTTTTGCTGATACTCTTGTTCTTCGTTGCGCAGTAGAGATTGGAATTGCAGATATCGTCAAAAACAATAATGGGTCAATCACACTTGAAGAACTTGCATCAAAACTCCCGATTCCAAGTGTCAATTCTGATTACTTGTACAGATTCTTAAGATACTTGGTACACCTTAATATCTTAAAACAAGAAACTTGTAATGCTGGAGTTACTAAGGTGTACTCACTTAAACCAATTGGTACTTTACTTTTAAGAGATGCAGAAAGAAGTATGGTACCAATGGTATTAGGTATGACTAAAAAAGATTTCCTGGTTCCATGGCATTTCTTGAAAGAAGGTTTGGGTAATGGAAGTACAACGGCATTTCAGAAAGGAATGGGAATGGAATTATGGGAGTATTTTGAAGGTAATCCTGATCAAAGTCAGCTGTATAATGAAATAATGGCTGGAGAAACAAGATATCTAACTAAAACTCTAATCGAAGATTGTAGGGATACTTTTCAAGGTCTGGATTCTCTTGTTCATATTGGTGGAGGAAATAGTACAACAATTATGGAAATCTATAAAGCTTTTCCTCAGATAAAATGTACACTTTATGATCTTCCTCATGTTATTGCTAACTCTCATGATCACCCTAATATCGAGAAAGTTTCCGGTGATATGTTCGAGTCTGTTCCCGGTGCTCAAGCTGTATTACTCAACTTGATCTTGCATGACTGGACTGATGAACAATGTATCGATATCCTAGCGAAGTGTAAAGAAGCAATACCGAAAGAGACGGGGA

>Pca_TRINITY_BBE Pca_TRINITY_DN70634_c0_g1_i1 len=1672

CTGTTTCTTTTTATGCTTTGTTGCACTAACATGGGTACTAGGTGACGATGTTAATGATAACCTCCTTCTGTCATGTTTAAACTCTCATGGTATTCGCAATTTCACCACACTATCAACTGATACAAATTCCGACTACTTCAAACTACTCCATGCATCGATACAAAACCCGTTGTTTGCGAAACCCACGGTATCGAAACCATCTTACATTGTAATTCCCGGTAGCAAAGAAGAATTATCGAGCACCATCCATTGTTTTACAAGAGAATCATGGAATATCCGACTACGAAGTGGTGGTCATAGTTATGAAGGGTTATCTTACACTGCTCATACACCTTTTGTTGTTGTAGATATGATGAACTTGAATCGAATTTCCATTGATGTTGAGTCAGAAACAGCTTGGGTTGAATCGGGTGCAACACTTGGAGAACTCTATTATTCGATTGCGCAGTCAACAGATATGCTGGGATTTACTGCTGGTTGGTGTCCAACTGTTGGTAGCGGAGGCCATATAAGCGGTGGTGGTTTTGGTATGATGTCAAGAAAGTACGGGTTGGCTGCGGATAATGTCGTGGACGCGATTCTTATAGATAGCAATGGAGCAATTCTCGACCGTAGAAAAATGGGTGAAGATGTTTTTTGGGCTATACGTGGTGGTGGCGGAGGTGTTTGGGGTGCAATTTACGCGTGGAAAATCAAACTACTGCCAGTTCCGGAGAAGCTGACCGTTTTCCGTGTGACAAAAAATGTAGGAATTGAAGACGCTTCATCTTTACTTCACAGATGGCAATATGTTGCAGACGAATTAGACGAGGATTTCACGGTATCAGTGCTCGGGGGAGCAAACGAAAATGAAGCCTGGTTAGTGTTCTTAGGACTACACTTGGGCCCAAAAAATGCTGCCAAAACTATAATCGATGAAAAATTCCCCGAACTAGGATTAATAGCGGAAGAGTTTCAAGAAATGAGTTGGGGTGAATCCATGGCTTTCTTAGCAGGAATAGAAACAATCTCTGAATTGAATAACAGGTTCCTGAAATTTGATGATAGAGCTTTTAAGACAAAAGTTGATTTTACTAAAGAACCAGTACCGCTGAATGTGTTTAGACGTGCATTAGAGATGTTATCAGAGCAGCCCGGTGGATTTATAGCTCTAAATGGTTTTGGAGGGAAAATGAGTAAACTTAGCACTGATTTCACCCCGTTTCCTCATCGTAAAGGCACTAAATTGATGTTCGAATATATAATCGCTTGGAACAAAGATGAAGAAATAAAAATCGGTGAATTTAGCGAATGGTTAGCGAAATTTTACGATTATTTGGAACCGTTCGTATCGAAAGAACCAAGGGTTGGTTATGTTAATCATATTGATCTTGATATTGGAGGAATGGATTGGAGAAATAAAAGTAGTACTAGCAATGCAGTTGAGATTGCTAGGAGTTGGGGTGAAAGATATTTTTCGTCGAATTATGAACGTTTAGTTAAGGCTAAAACATTAATTGATCCAAATAATGTGTTTAATCATCCACAAAGTATTCCTCCAATGATGAAATTTGAGGGGAATTACATGCTAAAGGAAATGTAGTACTAGGTTGATTTAGAAATTTACTTTTTTTTGTTGCAGGCATATTTGTCTATTTTAGCAACTAGTGAGCACAAAGCATGGCTGCCCATT

>Pca_TRINITY_TNMT Pca_TRINITY_DN69128_c0_g1_i1 len=1454

CTCGGGGGATCTCTCAACATTCTCTCTAGAGTTTCATTTTAGTTTGTCTTCCTAGAGAATTCCACTTAAGTTCCCTCATTACTAGCTTAGGTTGTGTCAGTATGGGTTCAGTAGATGAGGTAAAGAAGGAATCAGCAGGGGAAACACTAGGGAAATTGTTGAAAGGAGAGATTAAAGATGAGGAATTGAAGAAACTAATCAAGTTTCAATTCGAGAAACGTCTTCAATGGGGTTACAAAACATCTCATCAAGAACAACTCTCCTTCAACCTTGACTTCATTAAATCCTTGAAAAAGATGGAAATGTCGGGAGAGATTGAGACGATGAACAAGGAAACCTATGAATTACCGACAGAATTCTTAGAGGCAGTTTTTGGGAAAAGTATTAAACAAAGTGGATGTTATTTCAAACATGAATCAGCAACAACAGACGAAGCAGAAGAAGCTTCGCATGAATTATACTGCGAGCGAGCACAAATCAAAGATGGACAAACTGTTCTTGACATTGGCTGCGGTCAAGGTGGTCTAGTTCTATACATTGCTCAGAAATATAAGAACTGTCATGTTACAGGGCTCACTAATTCAAAAGCACAAGCCAATTACTTACTCAAACAAGCAGAGAAGCTTGGGTTGACAAACGTCGATGCCATACTAGCAGATGTTACTCAATATGATTCTGACAAGACATACGATCGACTACTTATGATTGAAGCCATAGAGCACATGAAAAACATACAACTGTTTATGAAGAAACTATCAACTTGGATGACAGAAGACAGTCTTCTCTTTATAGACCATATCTGCCATAAGACATTTAATCACTTTTTTGAGGCAATCGACGAGGATGACTGGTACTCAGGTTTCATCTTCCCCCCAGGTTGTGTAACAATACTAGCTGCTAATAGTCTCCTCTATTTCCAGGATGATGTTTCAGTTGTGGATCATTGGGTTGTCAACGGAATGCACATGGCTCGTACAGTGGACCTGTGGAGAAAAAAGCTAGACAAAAACATGGAAGCTGCAAAAGAAATTCTATTACCTGGACTTGGAGGAAGCCACGAAGCAGTGAATGGAGTCGTTACTCATATTAGAACATTCTGTATGGGAGGGTACGAACAATTCTCAATGAATAACGGAGATGAATGGATGGTGGCACAGCTGCTTTTCAAGAAGAAGTAATGAAGCAATGTCCACCACAACTGCACTTTTCCAGAGTTCTTTAATAATGCGATCCTCTGCTAACAATATTTGTTGGTTGTATCGATCTAGCTCAGGGGTTTAGCTTGTTTTTTGTTGTGTGCATTTCTACTTTTTTTCTTGTACTTTGGATTCAAAGTTTGGAGAATCTGAATCTTAATGTTATCGTGTAATGGGTTTATCAAAGCAACAGGGGTGAGAAAATTGTGCCAACATTTGCTATTCCTTTTATCTAGCGTTGGTTGCTACACATTGGCC

>Pca_TRINITY_9OMT Pca_TRINITY_DN29873_c0_g1_i1 len=364

TTTATAGAAACAGATATCGTGCCATTTGATGTTGCTCTTGGTGCTAGTATCTTTGAGTATTCTGGTAAAGAACCAAGACTGAATCAAGTTTTTAATGATGCAATGGGGGTTTTCTCTGTTGTTGTTTTTGAAGCTGTTTTTAGAGTTTACGATGGATTTCTTGGTATGAAAGAATTGTTAGATGTGGGTGGTGGTATCAGCACTTCCGTTAGTAAGATTGTTGCGAAATACCCTTTAATTCGCGGTGTCAACTTCGACTTGCCTCATGTTATTTCTGTTGCCCCTCAATACCCAGGTGTAGAGCATGTTGAAGGAGATATGTTTGAGGAAGTCCCAAAGGGTCAAAACATGTTGCTAAAATGGG

>Pat_TRINITY_NCS Pat_c33181_g1_i3 len=868

TGCACATCTTACCTGGACAGGATCCATGTTATAGAAAAAACCTCTAAATCTTGCATCATTAAATCTTCTGTTGTCTACGAAGTGAAGCAAGAGTGTGCTGATGCAATGTCTAAGTTAATCACGACGGAGCCATTGAAGTCGATGGCAGAAGTCATCTCTAATTACGCTATTAAGCAACAATCTGTTTCTGAGAGAAACATTCCTAAGAAGCAATCTGTGTTGAGGAAAGAAATTACTTACGAAACGGAGGTGCAAACCTCAGCTGATTCGATTTGGAACGTCTACAGTTCTCCTGACATCCCTCGACTACTTAGAGATGTTCTGCTTCCTGGTGTTTTTGAGAAGCTAGATGTCATTGCAGGCAATGGTGGCGTTGGTACTGTACTGGACATTGCTTTCCCTCTAGGTGCAGTGCCGCGGAGTTACAAGGAGAAATTTGTGAAAATCAACCATGAGAAGCGATTGAAAGAAGTGATTATGATCGAAGGAGGATACTTAGACATGGGGTGCACATTTTACATGGACAGGATTCATGTATTTGAGAAAACCCCCAACTCATGTGTCATTGAATCCTCTATCATTTACGAAGTTAAAGAAGAGTATGCTGATAAAATGGCCAAGTTAATCACAACGGAACCATTGCAGTCGATGGCAGAAGTCATCTCTGGTTATGTTCTTAAGAAACGATTCGAAGTATTTGGACTAGAGATTAAACCAAAAGTAAGGTCAAGTCTTTTGCTATGTTTGATCATCTGTTTGGTAATAGTGGGTGGTATGTTGATTGGAGGTGTTCCCCTCTGAGCTTGATAAAGATTTGCTTAAATGTAATTTTGTTTTAATAAATAATGGTTTTGAGCCTTAACACTAC

>Pat_TRINITY_6OMT Pat_c19346_g1_i2 len=1326

CTTCCGATCTATAGAGGATTATTTTATATAAAAGGGTCAAAAGGAGGCACTAGAGAATAGCTCAAATCATTCATTCATTCTCATCAACAACTAAAGTTTCTAAAAAAGAGAGAAATGGAAACAGTAAGCAAGATTGATCAACAAAACCAAGCAAAAATCTGGAAACAAATTTACGGTTTTGCAGAATCACTAGTTCTGAAATGTGCAGTTCAACTAGAGATTGCTGAAACACTTCACAACAATGTCAAACCCATGTCTTTATCCGAATTAGCATCGAAACTTCCCGTTGCTCAGCCCGTGAACGAAGACCGTCTGTTCCGAATCATGCGTTACTTGGTTCACATGGAACTCTTCAAGATAGACGCTACCACGCAGAAATACTCATTAGCTCCACCAGCTAAGTATTTGTTGAGAGGCTGGGAAAAATCAATGGTTGATTCAATTTTATGCATAAATGATAAGGATTTCTTAGCTCCGTGGAATCATTTAGGTGACGGTTTAACCGGGAACTGTGACGCTTTTGAGAAAGCGTTGGGGAAGAGTATTTGGGTTTATATGAGTGAAAATCCTGAAAAGAATCAATTGTTTAATGCAGCAATGGCTTGTGATACTAGATTGGTTACTTCAGCATTGGCTAATGAGTGCAAAAGTATTTTCAGTGACGGAATCAGTACACTGGTTGATGTCGGCGGTGGTACAGGTACTGCCGTGAAAGCCATATCTAAGGCTTTTCCAGATATTAAGTGCACCATCTATGATCTTCCTCATGTCATAGCTGATTCTCCTGAAATTCCCAATATTACTAAAATTTCTGGAGATATGTTCAAGTCTATTCCTAGTGCTGATGCCATCTTCATGAAGTGCATACTTCACGACTGGAACGATGAGGAATGCATTCAAATCTTGAAGCGATGCAAAGAGGCATTACAGAAAGATGGCAAAGTTATTATCGTGGACGTCGTAATAGACATGGATTCGACTCATCCATATGCAAAAATTAGACTCACATTGGATTTGGATATGATGCTTAACACTGGTGGAAAAGAGAGAACCAAGGAGGAATGGAAGATACTTTTTGATGCCGCTGGTTTTGCTAGCCACAAAGTCACTCAGATATCTGCTGTCCAGTCTGTAATTGAGGCTTACCCTTATTAAGGAGTAAGGATGATTGTATCCTGTTTTCCCTTTGATTGTTGCCCTCCATTTGAATTATGTCTGTGTTTAAAATATCTGCAAGATGGATGAATTTCCAGCTTGCATAGAAATCAATATTTTACTTTCTTGCTATTCAGTGGGGACTTCGCTTAAAAAAAAAAGATCGGAAGA

>Pat_TRINITY_CNMT Pat_c23107_g1_i2 len=1338

GATCTCTTTATCAAATTATAAAAGCCAGTGGTAGTAGTAGGTACAACCACTACCACCACCTCTGTCTGAACTTCTCTGGAAAATCTATAGACTTTGTGTGTGTTTTTCTTTGCAAGTTGAAGAAGATGCAGCTAATGGCAAAAGAAGAGCTGTTGAGAAACATGGAGCTTGGGTTGATTCCGGACAAAGAGATTAGAGAAAGGATTAGAATCGAATTAGAAAAGCGTCTCCAATGGGGTTACAAAGAAACACACGAAGAACAGCTTTCTCAGCTTCTTGAATTAGTTCATTCTTTGAGAGGTATGAAAATGGCAACTGAGATGGAGAATTTGGATTTGAAACTATACGAGGCACCTATGGAATTTTTAAAGATTCAACTTGGAAGCAACATGAAACAAAGTGCCGGTTACTATACCGATGAATCAACGACGTTGGACGAAGCTGAGATAGCGATGATGGATTTGTACATGGAGAGGGCACAGATCAAAGATGGTCAGAGTGTACTTGATTTAGGGTGTGGATTGGGTGCTGTAGCTCTTTTTGGTGCCAACAAATTCAGAAATTGTCAATTTACTGGAGTTACAAGTTCTATGGAACAAAAAGATTACATTGAAAGAAAATGCAAGGAACTTAAGTTGACCAACGTCAAGGTTTTATTAGCTGATATAACAACTTACGAAACTGAAGAAAGATTTGATCGGATTTTTGCTGTTGAATTGATTGAGCATATGAAGAACTATCAATTGCTTCTTAAGAAAATATCAGAATGGATGAAAGATGATGGACTTCTTTTCATTGAACATGTTTGCCATAAAACTCTAGCTTACCACTATGAGCCTGTTGATGCAGAAGATTGGTACACTAACTACGTATTCCCTGCTGGAACTCTGACTCTATCATCTGCTTCAATGCTTCTTTACTTTCAAGATGATGTTGCTGTTGTGGATCAATGGACTCTAAGTGGAAAACATTACTCTCGATCCCATGAAGAATGGTTGAAAAAAATGGATAAAAACATTGTTGAATTCAAAGAGATAATGAAGTCTATAACTAAAACTGAGGAAGAAGCAAACAAATTGCTCAACTTTTGGAGGATTTTTTGCATGTGTGGAGGAGAGTTGTTTGGTTATAAGAATGGTGAAGAATGGATGCTCACCCATCTTCTCTTCAAGAAAAAATGAGAAAATTTCTTTTATGCTTTCGTTTTGTGTGTGTTATTTAGCATCATAAGTATACCAATTCATTGTAATAAGAACTGTTTAGAATGATTGGTTGTAACTTTTATTTTATGTTTTTTTAATAAAAACTTCATCATCAAGCGCTTTATGGTTTCGTTAC

>Pat_TRINITY_CYP80B1 Pat_c45070_g1_i1 len=362

GTTGGTGAATATATTTGGTAACTTGATATTTTCGCAAAATATTTTCAAGTTGGGTGATGAAAGTAGTGGAAGTGTAGAAATGAAAGAACATCTATGGAGAATGCTGGAATTGGGGAATTCGACGAATCCAGCTGATTATTTTCCATTTTTGGGTAGATTCGATTTGTTTGGACAAAGAAAAGATGTTGCTGATTGTCTGCAAGGGATTTATAGTGTTTGGGGTGCCATGCTCAAAGAAAGAAAAATAGCCAAGCAGCATAACAACAGCAAGAAGAATGATTTCGTCGAGATTTTGCTCGATTCCGGACTCGATGACCAACAAATCAATGCCTTGCTCATGGAAATATTTGGTGCGGGAACAG

>Pat_TRINITY_4OMT1 Pat_c12815_g1_i1 len=1260

CTCTTCCGATCTCACAATTCCAAAGCCAAAAAAAAAAAAATAGTCTTCTCTTCTTGACAAATATACCGCAAAAATCTCATATCTAATGGGTAGTTTAGATGCAAAACCAGCTGCTGAAACACAAGAAGTTTCCATCAAAGATCAAGCTCAACTATGGAATATAATCTATGGTTTTGCTGATTCTCTTGTTCTTCGTTGCGCTGTAGAGATTGGAATCGCGGATATTATCAAAAACAATGATGGGGCAATCACACTTGCACAACTTGCATCAAAACTCCCAGTTCCAAATGTCAGTTCTGATTACTTGTACAGAATGGTAAGATACTTGGTACACCTGAACATCTTAGAACAAGAAACTTGTAACGGCGGAGTGGAGAAGGTTTACTCGCTTAAACCGGTCGGTACGCTACTTTTAAGAGATGCAGAAAGAAGTATGGTGCCGATGATATTAGGCATGACTCAAAAAGATTTCATGGTTTCATGGCATTTCATGAAAGAAGGTTTAGGAAGTGGAAGTACAACTGCATTTGAGAAAGGAATGGGAATTGATATATGGAAGTATTTAGAAGGTAATCCTGATCAAAGCCAGTTGTTTAATGAAGGAATGGCTGGAGAAACAAGACTTTTAACTAAATCTCTGATCGAAGATTGTAGAGATACTTTTCAAGGTTTGGATTCTCTGGTTGATATTGGTGGAGGAAATGGTACAACAATCAAGGCAATCCATGAAGCTTTTCCTCATATCAAATGTACACTTTATGATCTTCCTCATGTTGTTGCCAACTCTCATGATCTCCCTAACATCGAAAAAGTTCCCGGTGATATGTTCAAGTCTGTTCCCAGTGCTCAGGCTATCTTACTCAAGTTGATCTTGCACGACTGGACTGATGACGAGTGTGTCAACATTCTAAAGAAGTGCAGAGAAGCAATACCGAAAGAGACAGGGAAAGTAATTATAGTGGATGTAGCACTTGAAGAAGAATCCAACCATGAACTTACAAAGACGAGATTGATTCTTGATATTGACATGTTGGTGAATACTGGTGGTAGAGAGAGGACTGCAGATGATTGGGAAAATTTGCTCAAACGCGCAGGCTTTCGGTCTCATAGGATCAGACCCATTCGCGCTATTCAGTCCGTCATTGAAGCCTTCCCTTAGTAAAAGCTCTAAGGAGTACTTTGCTTCTTTCTCGGAGAAGTTTATGCGATGCACTTATTCACAATAAGCCATTTTCCTGTTTTTTGTTTTGTTTTGTTTTT

>Pat_TRINITY_BBE Pat_c18536_g1_i1 len=1611

GCAGAAGTTTAACAGTACGTTTCTTCTTATTCTTTGTTTTATTACAAACATGCGTACGCGGTGGTGATGTTAATGATAATCTCCTCTTGTCATGTTTAAACTCCCATGGTGTTCACAATTTCACCACACTAACAACTGATACAAATTCGGAGTACTTCAAACTACTCCATGCATCTATACAGAATCCGTTGTTCGCGAAGCCTACGGTATCGAAACCGTCGTTTATTGTAATTCCCGGCACCAAAGAAGAATTATCGAGCACCATTCATTGCTGTACAAGAGAGTCATGGACTATTCGACTGCGGAGTGGCGGTCATAGTTATGAAGGGTTGTCTTACACTGCTGATACACCTTTTGTGATTGTAGATATGATGAACTTGAATCGAGTTTCCATTGATGTCGAGTCGGAAACAGCTTGGGTTGAATCCGGTGCAACACTTGGAGAACTCTATTATGCGATTGCACAGTCAACGGAAACACTGGGATTTACTGCTGGTTGGTGTCCAACTGTTGGAAGTGGAGGTCATATAAGTGGTGGTGGTTTTGGTATGATGTCGAGGAAGTACGGGTTAGCTGCCGATAACGTCGTGGACGCGATTCTTATAGATAGCAAAGGAGCAATTCTTGACCGTGAAAAGATGGGGGAAGATGTTTTTTGGGCTATTCGTGGTGGCGGCGGAGGTGTTTGGGGTGCAGTTTACGCGTGGAAAATCAAACTATTACCAGTTCCGGAGAAGCTGACCGTTTTCCGTGTGACAAAGAATGTAAAAATTGAAGACGCTTCATCTTTACTTCACAAATGGCAATATGTTGCAGATGAATTAGACGAGGATTTCACAGTGTCTGTGCTCGGAGGAGTAAACGGTAATGAAGCCTGGTTAATGTTCTTAGGCTTACACTTGGGACCTAAAAGTGCTGCAAAAACTATAATCGATGAAAAATTCCCTGAACTGGGTTTGGTAGAGGAAGAGTTTCAAGAAATGAGTTGGGGTGCATCCATGGCTTTCTTAGCAGGATCAGAAACAATCTCTGAGTTAAACAACAGGTTCTTGAAATTTGATGATAGAGCTTTTAAGACAAAAGTTGATTTTACTAAAGAACCAGTACCGCTAAACGTGTTTAGACGTGCACTAGAGATGTTATCAGAACAGCCTGGTGGGTTTATAGCTCTAAATGGTTTTGGAGGGAAAATGAGCGAAATTAGCACTGATTTTACCCCGTTTCCTCATCGTGAAGGCACTAAATTGATGTTCGAGTATATAATCGCTTGGAATCAAGATGAAGAAACGAAAATCGTTGAATTTAGCGAATGGTTAGAGAAATTTTACGATTACTTGGAGCCGTTCGTATCGAAAGAACCAAGGGTTGGTTATGTTAATCATATTGATCTTGATATTGGAGGAATAGATTGGAGAAATAAAAGTAGTACCAGCAATGCAGTTGAGATTGCTAGAAATTGGGGTGAAAGATACTTTTCGTCCAATTATGAACGTCTAGTTAAGGCGAAGACATTGATTGATCCAAATAATGTGTTTAACCATCCACAGAGTATTCCTCCAGTGATGAAACTTGAGGAAAATTACATGTTAAATGAACTGTAATACTAGGTTG

>Pat_TRINITY_TNMT Pat_c35024_g5_i1 len=1269

GTGCTCTTCCGATCTATTCTCTCTTGCATTTCATTCTAGTCTTGTCTTCCGAGAAATCCACTTGGTTTCCTCACTATCCTAGATTGTATCAGCATGGGTTCAATAGATGAAGTGAAGAAGGAATCAGCAGGGGAAACACTAGGGAGATTGTTGAAAGGAGAGATTAAAGACGAGGAACTGAAGAAGCTGATTAAGTTTCAATTCGAGAAACGTCTGCAATGGGGTTATAAATCATCTCATCAAGAACAACTCTCCTTCAACCTTGACTTCATTAAATCCTTGAAAAAGATGGAAATGTCAGGAGAGATTGAGACAATGAACAAGGAAACTTATGAATTACCCTCAGAATTCTTAGAGGCAGTTTTTGGAAAAATGGTCAAGCAAAGTATGTGTTACTTCAAACATGAATCAGCAACAATAGACGAAGCAGAAGAAGCTGCACATGAATTATACTGTGAACGAGCACAAATCAAAGATGGACAAACTGTTCTCGACATTGGTTGTGGTCAAGGTGGTCTAGTTCTATACATTGCTCAGAAATATAAGAAATGTCATGTTACAGGGCTCACTAATTCAAAGGCACAAGTCAATTACTTACTCAAGCAAGCAGAGAAGCTTGGGCTGAAAAACGTCGACGCCATACTAGCAGATGTCACTCAATATGAATCTGACAAGACATACGATCGACTACTTATGATTGAAGCCATAGAGCACATGAAAAACTTACAAATGTTCATGAAGAAACTATCAACTTGGATGACAAAAGAAAGTCTTCTCTTTGTGGACCATGTCTGCCATAAAACCTTTGCTCACTTTTTTGAGGCAGTCGACGAGGATGATTGGTACTCAGGTTTCATCTTCCCCCCAGGTTGTGCAACCATACTAGCTGCTAATAGTCTCCTATATTTCCAGGATGATGTTTCAGTTGTGGATCATTGGGTTGTCAACGGAATGCACATGGCTCGTTCAGTAGACATTTGGAGAAAGGCACTAGACAAAAATATGGAAGCTGCAAAAGAAATTCTATTACCTGGACTTGGAGGAAGCCACGAAGCAGTGAATGGAGTCGTTACTCACATTAGAACATTCTGTATGGGAGGCTATGAACAATTCTCAATGAACAATGGAGATGAGTGGATGGTTGCGCAGCTGCTTTTCAAGAAGAAGTAGTGAGGCAATTTCCACGGCAACTGCATTTTCCAGAGCTGTGTAATAATGTGATCCTCAAGTAACAACAATTGTGGGTTGTATCACAGAGATCGGAAGAGC

>Pat_TRINITY_CFS Pat_c27072_g1_i1 len=1963

TGTGCTCTTCCGATCTCTCAGTTCTCTTCTTCTCTCTTCGTTCATCTTGTTTTGATCTTAATTTCTTCATAAACATAAAAAATATATTAAGAGTACTGTTTTCATAAAATGGAGGAGACCATCTGGTTGATAACTTGTGGTGTGGTAGTAGCATTGGCATTCGCAAAAATGCTTTTTGGCAAGAAATCATCTCTGTCTACAATGGAATGGCCAACAGGTCCAAAAAAGTTGCCCATAATTGGGAATCTTCATCAATTAGGAGGAGGAGAAGCATTTCATGTTTGTTTGACAAACTTGGCCAAAGTTCATGGAAGTGTTTTTACCATATGGGTTGGAAGTTGGCGTCCGTTTATAGTCATAAGCGATGTCGATAAAGCTTGGGAAATTCTTGTTAATAAATCTTCGGATTATTCAGCTAGAGATATGCCTGATATTACTAAAATCATATCAGCTAATTGGAAGAATATTTCTCATTGTGATTCTGGTCCATTTTGGCATAATTTAAGGAAAGGTCTTCAAAGTGTTGCTCTAACTCCTTTCAATGTCGCGTCTCAATACCATTTGCAAGAAAGGGACATGCAAAAATTGATCAAATCCATGCAAGAAAAAGCTTCTCTCCAGAATGGGATTCTGAAACCACTTGATTATGTAAAAGAAGAGACTGTTCGACTACTAAGTCGACTCATATTCGGTCAAGATTTCGTAGACGAGGATTTCGTTGTTGGTATGCATCAAGCCCTCGACGAGTTAGTAAGTATAAGTGGGTATGCAAGTTTAGCTGATGCTTTTAAATTCTGTGAGAATCTACCAAGTCATAAGAAAACTATTCAAGCAGTTCACGCTATCAAAAAGCGATTCGATACTTTCATTCGTCCACACATCGTTTCTAAGCCTCCTACAAATACGTACTTACATTTTCTTCTATCTCAAGATTTTAGTGAAGATGTTATAATCTCTGCTATACTTGAAGTATATGATTTAGGTGTTGATAGCACTGCATCAACAACAGTTTGGGCTCTTACATTTTTAGTTCGCGATCAAAAGATTCAGGAAAAGCTTTACCGCGAAATCAAAAGTGTAACCGAAGGGAAAAGACCCGTGAAAGTTGAAGATTTAAACAAATTGCCATATTTGCAAGCAGTCATGAAAGAAACAATGAGGATGAAACCCATTGCACCCATGGCAATTCCTCACAAAGCCTCAAAAGATACTTCATTAGTGGGTAAGAAAATCAATAAAGGTACAGTGGTAATGGTAAATCTTTACGCTATTCATCATAACCCTGACGTTTTTCCGGAACCATATAAGTTCATGCCAGAGAGATTCCTAAAGGAAGTTAATAGCGATGGAAGTTTGGGTGATATTAAGAAAATGGAAAGTTCTTTGTTGGCATTTAGTGCTGGTATGAGAATTTGTGCAGGAATGGAGCTAGGTAAATTGCAGTTAGCTTTTGGTCTTGCAAGTTTGGTTAATGAGTTCAAATGGGATTGCGTTGTTGAAGGGAAATTGCCTGATCTTAGTGAAGAACATTGTTTCATTCTCTTGATGAAAAACCCACTTGAAGCCAGAGTTACTCGACGTATCCATTAGTCACGGGTATACAAGTGATCAGAATCGGTCCCTGACTAGAGAGGGCTGTTAGTATAAAGCCAGTGATACCTCCCGAAAGAACAAAGTGTGATATTTATCCGGTTGATTGACAACTGGAAGTGCCTGACAAATTTGTGTTGTTTTTCTTCCATAAATGTTGTTGCTTTTTGTTGCTTTCGTGTTTATGCGTGCTTTACTTTAGCTTTTTTTATGTGTGTGCCTGTAATACCACCCGCCAGCAATTATTAAATCTTACCTTCGTTCCCCAATATGAAAACTTACTTTCCTTCCGCAATATATGTTGTTAAAGTCTGCGAGCAATAAGTGGAGCAGTCCTTTTATATAGTTGAGATTATAGATCGGAAGAGCGTCG

>Pat_TRINITY_SPS Pat_c34412_g1_i3 len=1977

TGTGCTCTTCCGATCTGAACTTTTCAATTTTGGGGGGACAATTATGCAAAATCCATAATTCCCAAGCCTAATTTGGCAACTTCAATTTTTTTAGGGAGGACAACTGTCCATCCTAGTCCCCATTAAGCTCCGCCATTGGTTTAAATGATACACCATCTTTTATTCGTGCTGCAACAGGATATGATAGATTATAGGTTTCGATATTTTAATTATAAATAGAAGTTATGCTTCTCCATATTCTTCACAGAAGCATAAGTACTTTGTAACCAATACATTAGGAAATGAAGGAAATGTTTAGCAGTTCATTGGTTCTGATTACTTCCACCACTATGGTAGTAATAACAACCATCACAATGCTACTGATTTTCGCAAGAAAAAAATCATCAACAATGGAGTGGCCAACTGGGCCAAAAACGTTGCCAATCATAGGTAACATGCATATTCTTGGAGGCACAGCTCTTCAAGTTGTCTTACATAACCTTGCTAAAATTCACGGCAACGTCATGACGATATGGATCGGTAGTTGGAGACCAGTTATTGTCGTTTCGGATATCGAACGAGCATGGGAAGTTCTTGTTAACAAGTCGTCAGATTATTCGGCTCGTGATATGCCCGAAATCACAAAGATTATTTCTGCTGATTGGAAAACAATCTCTACTTCTGATTCTGGTCCTGCTTGGTCTAATCTCCGGAAAGGTCTTCAAAATGTTGCGTTATCGCCTCATAACTTAGCTGCACAAAGTCAATTTCAAGAAAAAGATATCACCAAAATGATTCAAAGTTTGAAAAAAGAAGCTTCGAGCAATAATGGAATTGTTAAACCAATTGATCATCTTAAGAAGGCGACACTGAGATTAATTAGTCGATTAATTTTTGGCCAAGATTTCGATAATGATAAATATGTTGATGATATGCATCATGCTATTGAAGAATTGATTCGTGTTAGTGGTTATGCACAGCTTGCCGAGGCATTTTACTATGCGAAATACTTACCCAGTCATAAAAAAGCTGTTAGAGACGTCGAAGAAGCAAACCAAAGAGTAAAAAAGTTGGTACGCCCATTTTTATCATTGAATCCTCCTACTAATACTTATTTGCATTTTCTTAATTCACAGAATTATGCCGAAGAAGTTATCGTATTCGCAATATTTGAAGCTTATCTTCTGGGTGTTGATAGTACTTCTTTGACTACTGCATGGGCACTCGCATTTTTAATCCGCGAACCAGTTGTGCAAGACAAACTTTACCAGGAGCTTAAGACTTTTGCAGCCAAAAATGATCGCGAAATTGTGAAAGTTGAAGATTTAAACAAGTTTCAATATTTGCAAGCTGTTCTTAAAGAAACAATGAGAATGAAACCAATTGCGCCATTAGCAATACCTCATAAAGCTTGTAGAGACACTTCGCTAAAGGGGAACAAAATTGATCAAGGCACCAGAGTTATGGTTAACATTTATGCTTTACACCATAATGAGAAAGTTTGGAGAGAACCTTACAAATTTATGCCAGAAAGATTTTTACAGACTCAAGATGTTGTTAATGGAAAAGCAATGGAACAGTCACTACTACCATTTAGCGCAGGCATGAGAATTTGCGCGGGAATGGAATTGGGTAAACTGCAATTTAGTTTTTCTCTTGCTAACCTTGTTAATGCTTTTCAATGGTCTTGTGTTTCTGATGGAGTACTTCCTGATATGAGTGATCAACTTGGTTTTGTCTTGTTGATGAAGACACCACTTCAAGCACGTATAGTTCCTCGTGCGTGATCATCATATATCGACCATGGATCGGTTATAATGAATGGTTTTTAATCAAGTCAATGCTACCGTACTCTATATGCCTAGTTGTTGTTGGTGTTCCTTTGATGTACTTAATTTAGCTCCTAGTCTGGGGCATTTCAGTAAAGTATTAAATTTAATAAATTTAATAAAAGAAGTTACGAATTTCTTTTCAATGTTTTAAAAAAAAAAGATCGGA

>Pat_TRINITY_MSH Pat_c23964_g1_i1 len=1766

AAGAATGTTTTACTACAAATGATAAACTCTTTTCGAATCGGCCGGGTACGTTAGGTATTAAACTCATGTTTTACGATGCCGATTCGGTTGGTTATGCGCCGTACGGTGCTTACTGGAGAGATTTACGAAAGATTTCCACGTTAAAACTTCTTTCGAATCATCGTATCGACACCATAAAACATCTCAGGAGTCATGAAGTTGAGTCGTGTTTCGAGTCACTTTATAACCAGTGGAAAAACGGAGAAAAGCGTGGAGAATTCGCTCCGGTGAGAATGGATAGTTGGTTAGGCGATTTGACGTTTAATGTTGTGGCCAGAATTGTTGCGGGAAAGAAGAATTTCTCAGCAAACGGAGACGTCGGAGCTCAGAGATATAAAGCAGCAATGGATGAAGCTATGAGATTAATGAGATTTTTCGCATTCTCGGATGTCATTCCATCACTAAGTTGGTTAGACAATTTAAGAGGTTTGGTTGGAGATATGAAAAAATGTGCTTCTGAAATTGATGCAGTTATGGCGACTTGGGTTGAAGAACATCGTGTCAAAAGAATTTCAGGAAATAATTCTGAACTTGAACATGATTTTATTGATGTTTGCTTGGACATTTTGGAGCATTTACCCGGCGACGATCCTGACCTCGTCGTCAAATCTACGTGTCTGGATATGATATTAGGTGGGAGTGACACAACAACAGTTACCTTAACATGGGCAATGTCTCTACTCTTGAACCATCCCCAAGTGCTAAGAAAGGCAAGAGCGGAACTAGACACACAAGTAGGAAAGAACCGACAGGTCGACGACTCGGATATCCCGAATCTTCCATACATCCAAGCGATCATTAAAGAGACGATGAGATTGTACCCGGCCGGACCACTAATTGAACGGAGGACAATGGAGGATTGCGAAGTAGGTGGGTACCAAGTTCCGGCTGGAACGCGCTTATTGATTAACATCTGGAAAATGCAAAGAGATGGCAACGTATACAGAGATGACCCACTAGAGTTTAGACCCGACAGGTTTTTAACTAGTAATGCGGATGTTGATTTAAAGGGTCAGCATTACGAACTGATACCATTTGGAACGGGTCGAAGGATATGCCCGGGGGTATCGTTTGCGGTTCAGTTGATGCATTTGGCTCTCGCTCGGTTACTTCATGAGTTCGAAATCACAACGATGGAGCCAGAAGCGAAGGTGGATATGGCTGAAAGTGGTGGCTTAATTTGTTATAAGATAATGCCACTGGAAGCACTGATCAAACCTCGACTCCAGTTATGAAATGTTATAAGCATGCCCAGGTGAAGTCTCTGATCGGATTACATTATCCTATCTTAGAGCTTAGTTTTTGGTCATCTGCACAGTAAACACACATCCATAACTACTGCTGCCAAATCCAACTTAACTTTCACTTTTGATGCCAGCAAGCACAATCTAGAGCTCTCTTTAAAGCTGTTAAGTGGATCCACCAGAATCAATATCGGAATGTAATCGTCCAAACTGATAATCGTTCTCTTAGTCTTGCTATAAATAATCCTCATCTTCAGGTTTCCTGGTTAAATAAAGAATCTTTGTTAAGTTATTCTAAAACTATGTTTTTTAAATAATCATTGCTGGTTTTGTGTATTTGTTCCTAGATTTTGTTATGTAATAGGTCTACTGATAAGCTTGGCATGTGGTCTGGCTCATGATGTAACTGTTATCTCCCCTCTTTTAAAATGCTAAGTCATACAAGCTCCTACGTTGCACGGTGAGTGTGACACCCCCCTCAAA

>Por_TRINITY_NCS Por_c43817_g1_i1 len=716

ACTTAGAGATGTTCTGCTTCCTGGTGTTTTTGAGAAGCTAGATGTCATTGCAGGCAATGGTGGCGTTGGGACTGTACTGGATATCGCTTTCCCTCTAGGTGCAGTGCCGCGGAGGTACAAGGAGAGATTTGTGAAAATCAATCACGAGAAGCGGTTGAAAGAAGTGGTTATGATCGAAGGAGGGTACTTGGACATGGGCTGCACATTTTACATGGACAGGATTCATGTCTTTGACAAAACCCCAAACTCATGTGTCATTGAATCCTCTATTATCTACGAAGTTAAAGAAGAGTATGCTGATAAAATGGCTAAGCTAATCACAACAGAACCATTGGAATCCATGGCAGAAGTCATCTCTGGTTATGTTCTTAAGAAACGACTCCAAGTATTTGGATTCGAGATTCAGCCAACATTAAGATTCAATCTTTTGCTATGTTTGATTATCTGCTTGGTTATAGCTGGAGGTATGTTGATTGGACGTGTTCCACTCTAATGTTATGGGTTTTTCTTAAATGTAATTTTGGAAATAAAGAATAGGTTTGATCCTTAAAACCATGCATTGCATTGCAGAGGTTTAGTTCAGATTCGTGCAAGTTTTCATATGTAATTCTCCAATCTACAAGATCATATGCGAATTTAATAACAGTTTCAAGCAATGTTTGAAAAACGGGTTCGGAATGTAATAGTAATTTCAGTCATAAAAAGATCGGAAGAGC

>Por_TRINITY_6OMT Por_c35702_g1_i1 len=730

CCTGAAAAGAATCAATTGTTTAATGCAGCAATGGCTTGTGATACTAGATTGGTTACTTCTGCATTGGCTAATGAGTGCAAAAGTATTTTCAGTGACGGAATCAGAACACTGGTGGATGTCGGCGGCGGTACCGGTACTGCTGTGAAAACCATATCCAAAGCTTTTCCGGATATTAAGTGCACTATCTATGATCTCCCTCATGTCATAGCTGATTCTCCTGAAATTCCCAATATTACTAAAATTTCTGGAGATATGTTCAAGTCTATTCCTAGTGCTGATGCCATCTTCATGAAGTGCATACTTCACGACTGGAACGATGAGGAATGCATTCAAATCTTGAAGCGATGCAAAGAAGCACTGCCAAAAGGTGGCAAAGTTATTATTGTGGATGTCGTAATAGACATGGATTCGACTCATCCATATGCAAAAATTAGACTCACATTGGATTTGGATATGATGCTTAACACTGGTGGAAAAGAGAGAACCAAGGAGGAATGGAAGAAACTTTTTGATGCCGCTGGTTTTGCTAGCCACAAAGTCACTCAGATATCTGCTGTCCAGTCTGTAATTGAGGCTTACCCTTATTAAGGAACATTTTACCCAGTTTTCCCTTTGATTAATTGTTGCTTTCGATTTGGATTATGTTTGTGTTTATAATAACTGCAAGATGAATGAATTTCCAACTTGCATTGAATTTGAATCAATGTTCTTCGTTTAAAAAAAAAAGATC

>Por_TRINITY_CNMT Por_c52699_g1_i2 len=1286

GACGTGTGCTCTTCCGATCTCTCTGTCTCAACTTCTCCGGAAAATCTGTATAGACATTGTGTGTGTTTTTCTTTGCAAGTTGAACAAGATGCAGCTAATGATGGCAAAAGAAGAGCTGTTGAGAAACATGGAGCTTGGGTTGATTCCGGACCAAGAGATTAGACAAAGGATTAGAATCGAATTAGAAAAGCGTCTCCAATGGGGTTACAAAGAAACACACGAAGAACAGCTTTCTCAGCTTCTTGAATTAGTTCATTCTTTGAGAGGTATGAAAATGGCAACTGAGATGGAGAATTTGGATTTGAAGCTATACGAGGCACCTATGGAATTCTTAAAGATTCAACATGGAAGAAACATGAAACAAAGTGCCGGTTACTATACCGATGAATCAACGACGTTGGATGAAGCTGAGATAGCGATGCTGGATTTGTACATGGAGAGGGCACAGATCAAAGATGGTCAGAGTGTACTTGATTTAGGGTGTGGATTGGGTGCTGTAGCTCTTTTTGGTGCAAACAAATTTAAAAATTGTCAGTTTACCGGAGTTACAAGTTCTATGGAACAAAAAGATTACATTGAAAGAAAATGCAAGGAACTTAAGTTGACCAACGTCAAGGTTTTATTAGCTGATATAACAACTTACGAAACTGAAGAAAGATTTGATCGGATTTTTGCTGTTGAATTGATTGAGCATATGAAGAACTATCAATTGCTTCTTAAGAAAATATCAGAATGGATGAAAGATGATGGACTTCTTTTCGTTGAACATGTTTGCCATAAAACTCTAGCTTACCACTATGAGCCTGTTGATGCAGAAGATTGGTACACTAACTACGTATTCCCTGCTGGAACTCTGACTCTATCATCTGCTTCAATGCTTCTTTACTTCCAAGATGATGTTTCTGTTGTGAATCAATGGACTCTAAGTGGAAAGCATTACTCTCGATCCCATGAAGAATGGTTGAAAAACATGGATAAAAACATTGTTAAATTCAAAGAGATAATGAAGTCTATAACTAAAACCGAGGAAGAAGCAACCAAATTGCTCAATTTTTGGAGGATTTTTTGCATGTGTGGAGCAGAGTTGTTTGGTTACAAGAATGGTGAAGAATGGATGCTCACCCATCTTCTCTTCAAGAAAAAATGAGAAAATTTCTTTTATGCTTTCCTTTTGTGTGTGTTATTTAGCATCATAAGTATACCAATTCATTGTAATAAGAACTGTTTAGAATGATTGGTTGTAACTTTTATTTTATGTTTTTTTACTCCCTCGGTCTCAAAAAAAA

>Por_TRINITY_CYP80B1 Por_c44144_g2_i1 len=1822

GTGCTCTTCCGATCTCAAAAATCCAGGGAGAAAAATAAAGAGAAAGAAGATGGAGATCGTCACAGTAACACTTATAACAGTTGTCTTCACTACTTTGTTATACTTAATTTTCAGAGATTCAAGTCTTAAAGGCTTGCCACCAGGTCCAAAACCGTGGCCAATAGTCGGAAACCTTCTTCAACTTAGTGACAAACCCCATTCACAATTCGCCCAACTCGCCAAAACCTACGGTGATATCTTTACACTCAAATTAGGAACCGAAACAGTTGTTGTAGCTTCAACCCCGTTAGCAGCTAGTGAAATTCTCAAAACGCATGATCGTATTCTCTCGGGTCGATACGTGTATCAAAGTTTCCGCGTAGAAGGACATGTTAAGGTGTCTATGGCTTGGTCTGATTGTAATGAGACATGGCATAAATTACGGAAAGTTTGCAGAACAGAACTTTTTTCACAAAAGATGATCCAAAGTCAAGCTGAAGCACGAGAATGTAAGGCTATGGAAATGGTGGAATATCTGAAAAAAAATGAAGGAAACGAAGTGAAAATTATTGAAGTTATATTTGGAACACTAGTGAATATATTCGGAAACTTGATATTTTCACAAAATATTTTTGAGTTAGGTGATGAAAATAGTGGCAGTGTGGACATGAAAGAACGTATATGGAGAATGTTGGAACTTGGTAATACAACAAATCCAGCTGATTATTTTCCATTTTTAGGTAGTTTGGATTTGCTTGGACAAAGAAAAGAAGTTGCTGATTGTCTTCAAGGTATTTATGATGTTTGGGGTGTTGTGCTTAAAGAAAGAAGAATAGCCAATAAGCAAGATAAAAATATTAAGAAGAATGATTTTGTTGATGTTTTGCTTGATACTGGACTAGGTGATCGACAGATCAATGCCTTGCTTATGGAAATATTTGGTGGAACTGAGACTAGTGGATCGACAATAGAATGGGCATTCGCTGAGCTCACAAAAAACCCTCAGGTAGCAGCCAATATGCGAGCAGAATTGCTATCCGTGGTAGGGAAGAGGCCGATCAAGGAATCCGATATACCTAACATGCCCTATCTTCAAGCTTTTGTTAAAGAAACTCTACGGCTTCACCCACCAACTCCATTGTTAGTCCCACGTCGAGCACTTGAGACGTGTCAAGTCTTAAACTTTACAATTCCAAAAGAGTGTCAAATAATGGTGAACGCCTGGGGCATAGGTAGAGATCCAAAAAATTGGACTGATCCATTGAAGTTTTCACCAGAGAGGTTCTTGAATTCGAGCGTTGATTTTAAAGGGAACAGTTTCGAGTTTATACCATTTGGTGCAGGGAGAAGGATATGCCCTGGTGTGCCCTTGGCAACTCAGTTTCTTGGTCTTATTGTGCCTACTTTAGTACAGAATTTTGATTGGGGACTACCAAATGGAATGGATCCTACCCAACTGATTATGGAAGAGAAATTTGGGTTGACACTGCAGAAGGAGCCACCTCTGTATATTGTTCCTAAAGTCGTGAATTTTCAACTCTAAAGTTCACATGATTATGATCTCGTAAGTTTCTGTAATTTCCAGTAAAAAAATAAAGTGTTGTTTTTAATAGTATGAGTATGTACGCGTTTGAGTCAGTGTGTGTGCGAACTATATGAGCTTAGTATGTGAGTGTGTGTGCCGCACGTAAGAAATGTAAGAGAGATTTGTGTCGAGAATGATAGTAGCTATATATCCAGAGTCTGTCCTGTAATTCAGATATCTTTTCTCTTAATCAAATCTTTGATTCAAATCGTCTCCAATACAATGTAGACATTCTATTGAATGTTATCATGGTATCAG

>Por_TRINITY_4OMT1 Por_c50376_g1_i1 len=1355

TGTGCTCTTCCGATCTTACACCGTTAAGTTTGAACAGATTTCAAAACAACCACACACAACCCAAAGCCAAAACAAAAAAATAGTCTTCTCTTCTTGACAAATATACAGCAAAAATCTCATATCTAATGGGTAGCTTAGCAAAGTCAGCTGAAACACAAGAAGTTTCCATCAAAGATCAGGCTCAACTATGGAATATAATCTATGGTTTTGCTGATTCACTTGTTCTGCGTTGCGCTGTAGAGATTGGAATCGCGGATATTATCAAAAACAATGATGGGGCAATCACACTTGCACAACTTGCTTCAAAACTACCAGTTCCGAATGTCAGTTCTGATTACTTGTACAGAATGGTGAGATACTTGGTACACCTGAACATCTTAGAACAAGAAACTTGTAATGGCGGAGTGGAGAAGGTTTACTCGCTTAAACCGGTCGGTACGCTACTTTTAAGAGATGCAGAACGAAGTATGGTGCCGATGATATTAGGCATGACTCAAAGAGATTTCATGGTTTCATGGCATTTCATGAAAGAAGGTTTAGGAAATGGAAGTACTACGGCATTTGAGAAAGGAATGGGAATGGATATATGGAAGTATTTAGAAGGTAATCCTGATCAGAGCCAGTTGTTTAATGAAGGAATGGCTGGAGAAACAAGACTGTTAACTAAAACTCTGATCGAAGATTGCAGAGATACTTTTCAAGGTTTGGATTCTCTGGTTGATATTGGTGGAGGAAATGGTACAACAATCAAGGCAATCCATGAAGCTTTTCCTCATATCAAATGCACACTTTATGATCTTCCTCATGTTGTTGCCAACTCTCATGATCTCCCTAACATCGAAAAAGTTCCCGGTGATATGTTCAAGTCTGTTCCCAGTGCTCAGGCTATCTTACTCAAGTTGATCTTGCACGACTGGACTGATGAGGAGTGTGTCAACATCCTAAAGAAGTGCAAAGAAGCAATACCGAAAGAGACAGGGAAAGTGATTATAGTGGATGTAGCACTTGAAGAAGAATCCAACCATGAACTTACAAAGACAAGATTGATTCTTGATATTGACATGTTGGTGAATACTGGTGGTAGAGAGAGGACTGCAGATGATTGGGAAAATTTGCTCAAACGTGCAGGCTTTCGGTCTCATAGGATCAGACCCATTCGCGCTATTCAGTCTGTCATTGAAGCCTTCCCTTAGTGAAAGCTCTAAGGAGTACTTTACTTCTTTCTCAGAGAAGCTTATGCGATGCACTTATTAACAATAAGCCATTTTCCTGTTTTTTCTTTTCTTTTTCTGGCCATACTAGGCCATTGTTTCTGTTGTTCTTGCTCCTATTCAATGTAATTGAAGACTGATTGC

>Por_TRINITY_BBE Por_c50342_g1_i1 len=1183

TGTGCTCTTCCGATCTGGAGAACTCTATTATGCGATTGCGCAGTCAACGGAAACACTGGGGTTTACTGCTGGTTGGTGTCCGACTGTTGGCAGCGGAGGCCATATAAGTGGTGGGGGTTTCGGTATGATGTCGAGAAAGTACGGGTTAGCCGCGGATAACGTCGTGGACGCGATTCTTATAGATAGCAAAGGAGCGATTCTTGACCGTGAAAAAATGGGTGAAGATGTTTTTTGGGCTATTCGTGGTGGTGGCGGAGGTGTTTGGGGTGCAGTTTACGCGTGGAAAATCAAACTATTGCCAGTTCCGGAGAAGCTGACCGTTTTTCGTGTGACTAAGAATGTAAAAATTGAAGCCGCTTCATCTTTACTTCACAAATGGCAATATGTTGCAGATGAATTAGACAATGACTTCACAGTGTCTGTGCTCGGGGGAGTGAACGGAAATGAAGCCTGGTTAATGTTCTTAGGCTTACACTTGGGACGTAAAGATGCCGCGAAAAGCATAATCAATGAAAAATTCCCTGAACTGGGGTTAGTAGAGAAAGAGTTTCAAGAAATGAGTTGGGGTGCATCCATGGCTTTCTTAGCAGGATTAGAAACAATCTCTGAATTAAACAACAGGTTCTTGAAGTTTGATGATAGAGCTTTTAAGACAAAAGTTGATTTTACTAAAGAAGCAGTACCGCTAGACGTGTTTAGACGTGCACTAGAGATGTTATCAGAACAGCCCGGTGGATTTATAGCTCTAAATGGTTTTGGAGGGAAAATGAGCGAAATAAGCACTGATTTTACCCCGTTTCCTCATCGGAAAGGCACTAAATTGATGTTCGAATACATAATCGCTTGGAATCAAGATGAAGAAACGAAAATCGGCGAATTTAGCGAATGGTTAGAGAAATTTTACGATTATTTGGAGCCGTTCGTATCGAAAGAACCAAGGGTTGGTTATGTTAATCATATTGATCTTGATATTGGAGGGATAGATTGGAAAAATAAAAGTAGTACTAGCAATGCAGTTGAGATTGCTAGAAATTGGGGTGAAAGATATTTTTCATCGAATTATGAACGTTTAGTTAAGGCGAAGACATTGATTGATCCAAATAATGTGTTCAACCATCCACAGAGTATTCCTCCAATGATGAAATTTGAGGAAAATTACATATTAAATGAACTGTAATACTAG

>Por_TRINITY_TNMT Por_c52629_g3_i2 len=1194

CCCACTTTTAGAAAACTTTATGACAATGAGACGGATGATAATCTCTATAAATACTATGTACCAGGTTCTGCTCATACCACATCTATCAACATTCTCTCTTGCATTTCATTCTAGTCTTGTCTTCCGAGAAATCCACTTAGTTTCCTCACTAGCCTGGATTGTATCAGCATGGGTTCAATAGATGAAGTGAAGAAGGAATCAGCAGGGGAAACACTAGGGAGATTGTTGAAAGGAGAGATTAAAGATGAGGAACTGAAGAAGCTGATCAAGTTTCAGTTCGAGAAGCGTCTGCAATGGGGTTATAAATCATCTCATGAAGAACAACTCTCCTTCAACCTTGACTTCATTAAATCCTTAAAAAAGATGGAAATGTCAGGAGAGATTGAGACAATGAACAAGGAAACTTATGAATTACCCTCAGAATTCTTAGAGGCAGTTTTTGGAAAAACAGTCAAGCAAAGTATGTGTTACTTCAAACATGAATCAGCAACAATAGACGAAGCAGAAGAAGCTGCACATGAATTATACTGTGAACGAGCACAGATCAAAGATGGACAAACTGTTCTCGACATTGGTTGTGGTCAAGGTGGTCTAGTTCTATACATTGCTCGGAAATATAAGAAATGTCATGTTACAGGGCTCACTAATTCAAAGGCACAAGTCAATTACTTACTCAAGCAAGCAGAGAAGCTTGGGCTGACAAATGTCGACGCCATACTAGCAGATGTCACTCAATATGAATCTGACAAGACATACGATCGACTACTTATGATTGAAGCCATAGAGCACATGAAAAACTTACAACTGTTCATGAAGAAACTATCAACTTGGATGACAGAAGAAAGTCTTCTCTTTGTGGACCATGTCTGCCATAAAACCTTTGCTCACTTTTTTGAGGCAGTCGACGAGGATGACTGGTACTCAGGTTTCATCTTCCCCCCAGGTTGTGCAACCATACTAGCTGCTAATAGTCTCCTCTATTTCCAGGATGATGTTTCAGTTGTGGATCATTGGGTTGTCAACGGAATGCACATGGCTCGTTCAGTAGACATTTGGAGAAAGGCACTAGACAAAAATATGGAAGCTGCAAAAGAAATTCTATTACCTGGACTTGGAGGAAGCCACGAAGCAGTAAATGGAGTCGTTACTCACATTAGAACATTCTGTATGGGAGGCTATGAACAATTCTCAATG

>Por_TRINITY_CFS Por_c46926_g2_i1 len=700

GCAAGTTTAGCTGATGCTTTTAAATTTTGCGAGAATCTACCAAGCCATAAGAAAACTGTCCGAGCAGTTCACGCTATCAAAATTCGGTTCGATAATTTGATTCGTCCGCATATCGTCTCAAATCCTCCGACAAACACTTACTTGCATTTCCTTTTATCTCAAGATTTTAGTGAAGATGTTATAATTTCCGCCATTCTTGAAGTTTATGACTTAGGTGTTGACAGCACTGCATCAACAACAGTTTGGGCTCTTACGTTTCTAGTCCGCGAACAAAAGATTCAGGAAAAACTTTACCGCGAGATCAATAACGTAACCGGAGGGAAAAGACCCGTAAAAGTTGAAGATTTGAACAAATTGCCATATTTGCAAGCAGTGATGAAAGAAACAATGAGAATGAAACCCATTGCACCTACGGCGATTCCTCACAAAGCCTTAAAAGATACTTCACTGATGGGTAAGAAAATCAATAAAGGTACAGTCGTAATGGTAAATCTTTACGCTATTCATCACAACCCTGACGTTTTTCCGGAACCGTACAAGTTCATGCCACAGAGATTCCTAAAGGACGTTAATATTGATGGAAGCTTGGGTGATATCAAGAAAATGGAAAGCTCTTTGTTAGCATTTAGTGCTGGTATGAGAATGTGTGCAGGAATGGAGCTAGGTAAATTGCAGTTAGCTTTTGGTCTTGCAAGTTTGG

>Por_TRINITY_SPS Por_c43303_g1_i2 len=705

GCTCTTCCGATCTCAAGATTTCGATGATGATAAATATGTTGCTGATATGCATCATGCTATTGAAGAATTGATCCGTGTTAGCGGTTATGCTCAGCTTGCGGAGGCATTTTACTATGCAAAATACTTACCGAGTCATAAAAAAGCTGTTAGAGACGTCGAAGAAGCAAATAAACGAGTTAAAAAGTTGGTACGGCCATTTTTATCACTGAATCCTCCTACTAATACTTATTTGCATTTTCTCAATTCACAGAAATACGACGAAGAAGTGATCATATTCGCGATATTTGAAGCTTATCTTTTGGGTGTTGATAGTACTTCTTTGACTACTGCATGGGCACTCGCATTTTTAATACGCGAACCAGTTGTGCAAGAGAAACTTTACCAGGAAATTAAGGACTTCAGGGTCAAGAATGATCGCGAAATCGTGAAAGTTGAAGATTTGAACAAGTTGCAATACTTACAAGCTGTTCTTAAAGAAACAATGAGAATGAAACCAATTTCACCATTAGCTATACCACATAAAGCTTGTAGAGACACATCGCTAATGGGGAAGAAAATTGATCAAGGCACCAGGGTTATGGTTAACATATATGCTTTACACCATAATGAGAAAGTCTGGAAAGAACCATTCAAATTCATGCCAGAAAGATTTTTACAAACTCAAGATGTTGTTAATGGAAAAGCAATGGACAGATCGGAAGAGCG

>Por_TRINITY_MSH Por_c49980_g2_i3 len=1484

CTGGTCGATCAAAAAGAAACATGGATTTTCTTGATCTAATTCTTCTTCAGTCCAAACCGATTTCTATAGCCCTTCTTGTGATAACACTAGCTTTGGCGTTTTGGAACTATGGAAGAAGAAAACCGGCGAAAAAACTAGCACCTGAAGCGTCAGGTGGGAGACCTATAATGGGTCATCTCCATCTTTTCAACGACGGTGAGTTAACTCATCGGAAACTCGGTGTCATGGCGGATACTTACGGTCCTGTTTTCAATATCCGATTCGGTTCACATAAAACGTTAGTTGTTAGTGATTGGGAAATCGTTAAAGAATGTTTTACGACGAACGACAAGCTTTTCTCGAATCGGCCGGGTACGTTAGGTATTAAACTCATGTTTTACGACGCCGATTCGGTGGGTTATGCGCCGTACGGTGCATATTGGAGAGATTTACGAAAGATTTCTACGTTAAAACTTCTTTCCAATCATCGTATCGATACGATAAAACATCTGAGGAGTTCTGAAGTTGAGTCGTGTTTCGAATCGCTTTACAATCAATGGAAAAACGGAGAAAAGCGTGGCGAATTCGTTCCGGCAAGAATGGATAGCTGGTTAGGCGATTTGACGTTTAACGTCGTGGCGAGAATCGTTGCCGGGAAAAAGAATTTCTCGGCGAACGGTGACGTTGGGGCACAGAGATATAAAGCAGCAATGGATGAAGCTATGAGATTAATGAGATTTTTCGCATTTTCCGATGTAATTCCGTCGCTAAGCTGGTTAGACAATTTAAGAGGTTTGGTTCGAGATATGAAAAAATGTGCATCTGAAATCGATTCAATTATGGCTTCTTGGGTTGAAGAACATCGTGTTAAAAGAATTTCAGGAGATAACTCTCAGCTTGAACATGATTTTATTGACGTTTGCTTGGATATTATGGAGCATTCTTCGTTACCTGGGGACGATCCTGATCTCGTCGTCAAATCTACGTGTCTGGACATGATATTGGGTGGGAGTGACACAACAACAGTTACCTTAACATGGGCAATGTCTTTACTCTTGAACCATCCCCAAGTGCTACAAAAAGCAAAAGAAGAACTAGACACACAAGTAGGAAAGAACCGGCAGGTCGAAGACTCAGACATCCCGAATCTTCATTTCATCCAAGCGATCATCAAAGAGACGATGAGGTTGTACCCTGCCGGACCGTTAATCGAACGGAGGACGATGGAGGATTGCGAAGTAGCGGGGTACCAAATTCCGGCCGGCACGCGCTTAATAGTTAACGTCTGGAAGATGCAAAGAGATGGGAACGTATACAAAGGCGATCCGTTGGAGTTTAGACCCGACAGGTTTTTAACTAGTAATGCGGATGTGGATCTTAAGGGTCAGCATTACGAATTGATACCGTTTGGAGCGGGTCGACGGATATGCCCGGGAGTTTCGTTTGCGGTTCAGTTGATGCATTTGGTTCTTGCTCGATTACTACACGAGTTTGAGATCACGACG

>Por_TRINITY_P6H Por_c35030_g1_i3 len=1423

GTGTGCTCTTCCGATCTGCTGGCAGAAAAATATGGACCTGCCTTTACAATGAAGTTCGGGAATCACACAACACTAGTTGTGAGCGATACCCGAGTCGTCAAAGAATGTTTTACTACTAATGATACCCTCTTCTCCAACCGTCCCTCCACCATAGCTTTCGATCTTATGACTTATGCAACCGACTCGATAGCTTTCACTCCTTATAGTCCTTACTGGCGTGAACTCAGAAAGATATCTACTCTCAAACTTCTCTCTAACAACCGTCTTGAATCCATCAAACAACTTCGGACCTCGGAGGTAAGCGTATGCTTCAAGGAACTATACGACCTAACCAGCAAAAAAAATGATAACGGAGCTCCAGTTCCAATCGATTTGAAGAGATGGTTTGATGAGGTTTCAAACAATGTAATTATGAGAGTAATCTTTGGGAAACAAAATTTTGGATCCAAGATTGTAAGCGGGGAAGATCAAGAGGCAGTCCACTATAAAAAAATCACGGACGAACTTTCACGTCTTTCTAGTTTGACTATGTTGTCGGACATGGTTCCTTCACTTGGTTGGTTGGATTACTTCAAAGGCGATTTGAGGGCAATGAAACGAAATGGCAAAGAACTGAATTCGATACTCCAGAAATGGTTGGAGGAACATAAAAGCAAGAAAAGTTCAGATGATCGACAAGATTTCATGGATGTTATGTTGTCGATTTCCAAGGAGACCCAACTGTATGGCCACGACCAAGATACTTTTATTAAAGCTACTTGTCTTGCCATGATCATGGGTGGAACAAACAGTACGGAAGTGGCTCTAACATGGATATTGTCCTTACTTATGAACAACAGATACGCATTGCATAAGGCTCGAGAAGAAATAGACTTACTTGTTGGGAAGGATAGACAAGTGGAAGATTCAGATGTCAAGAATTTGACATACATGAATGCCATCATTAAAGAAACAATGCGATTATACCCATTAGGTTTTCTTCTAGAACGTGATACCAAGGAAGACTGTGAAGTTGGCGGGTTCAACATCGAAGGTGGTACAAGATTACTGATAAACGTATGGAAGTTACAACGAGATCCGAACGTGTGGACAGATCCAATGGAATTCAAACCAGAGAGATTTCTAACAGAGAATGCAGACATAGATGTTGGTGGTCAACATTTCGAACTACTGTCATTTGGTGCTGGTAGAAGGGTGTGTCCTGGTGTGTCCTTCGCACTACAGTTCATGCATTTGGTTCTCGCCCGTCTTATCCATGGCTATGATATGGAGACCCTAAATGGTGAAGTTGATTTAAGTGTTAGTAGCGGGGGACATGTTAACATCAAATCAACCCCGCTTGAGCTCGTCCTTACTCCTCGCCTTCACCCAGAGCTTTACGATTGTTAAACATTAAGCTTTCAAGTTGTTTATTATGATCGCT

>Pbr_TRINITY_NCS Pbr_c28236_g2_i2 len=728

ACTTAGAGATGTTCTGCTTCCTGGTGTTTTTGAGAAGCTAGATGTCATTGCAGGCAATGGTGGCGTTGGGACTGTACTGGATATCGCTTTCCCTCTAGGTGCAGTGCCGCGGAGGTACAAGGAGAAATTTGTGAAGATCAACCATGAGAAGCGATTGAAAGAAGTGATTATGATTGAAGGAGGATACTTGGACATGGGCTGCACATTTTACATGGACAGGATTCATGTACTTGAGAAAACCCCAAACTCATGTGTCATTGAATCCTCTATTATCTACGAAGTTAAAGAAGAGTTTGCTGATAAAATGGCTAAGCTAATCACAACGGAACCATTGCAGTCGATGGCAGAAGTCATCTCTGGTTATGTTCTTAAGAAACGACTCCAAGTATTTGGATTCGAGATTAAGCCAACATTAAGATTCAATCTTTTGCTATGTTTGATTATCTGCTTGGTTATAGCTGGAGGTATGTTGATTGGACGTGTTCCACTCTAATGTTATGGGTTTTTCTTAAATGTAATTTTGGAAATAAAGAATAGGTTTGATCCTTAAAACCATGCATTGCATTGCAGAGGTTTAGTTCAGATTCGTGCAAGTTTTCATATGTAATTCTCCAATCTACAAGATCATATGCGAATTTAATAACAGTTTCAAGCAATGTTTGAAAAACGGGTTCGGAATGTAATAGTAATTTCAGTCATAAAAAAATAATATTATCTTTTAAAAAAAA

>Pbr_TRINITY_6OMT Pbr_c21065_g1_i2 len=1222

TGCTCTTCCGATCTCTCATCAACAGCTAAAGTTTCTAAAAAGTGAGAAAAATGGAAACAGTAAGCAAGATTGATCAACAAAACCAAGCAAAAATCTGGAAACAAATTTACGGCTTTGCAGAATCATTAGTTCTGAAATGTGCAGTTCAATTAGAGATTGCTGAAACACTTCACAACAATGTCAAACCCATGTCTTTATCCGAATTAGCATCGAAACTTCCCGTTGCTCAACCCGTTAACGAAGACCGTCTGTTCCGAATCATGCGTTACTTGGTTCACATGGAACTCTTCAAGATAGACGCTACCACGCAGAAATACTCATTAGCTCCACCAGCTAAGTATTTGTTGAGAGGGTGGGAGAAATCAATGGTTGATTCAATTTTATGCATAAATGATAAGGATTTCTTAGCTCCGTGGCATCATTTAGGTGACGGTTTAACCGGTAACTGTGACGCTTTTGAGAAAGCGTTGGGGAAGAGTATTTGGGTTTATATGAGTGAAAATCCTGAAAAGAATCAATTGTTTAATGCAGCAATGGCTTGTGATACTAGATTGGTTACTTCTGCATTGGCTAATGAGTGCAAAAGTATTTTCAGTGACGGAATCAGAACACTGGTGGATGTCGGCGGCGGTACCGGTACTGCTGTGAAAACCATATCTAAAGCTTTTCCGGATATTAAGTGCACTATCTATGATCTTCCTCATGTCATAGCTGATTCTCCTGAAATTCCCAATATTACTAAAATTTCTGGAGATATGTTCAAGTCTATTCCTAGTGCTGATGCCATCTTCATGAAGTGCATACTTCACGACTGGAATGATGAGGAATGCATTCAAATCTTGAAGCGATGCAAAGAAGCATTACCAAAAGATGGCAAAGTTATTATCGTGGATGTCGTGATAGACATGGATTCGACTCATCCATATGCAAAAATTAGACTCACATTGGATTTGGATATGATGCTCAACACTGGTGGAAAAGAGAGAACCAAAGAAGAATGGAAGATACTTTTTGATGCCGCTGGTTTTGCAAGCCACAAAGTCACTCAGATATCTGCTGTCCAATCTGTAATTGAGGCTTACCCTTATTAAGAAATTTTTTACCCGGTTTTCCCTTTGATTAGTTGTTGCTTTCTCTTTGGATTGTGTTTCTGTTTATAATAATTGCAAGATGAATGAATTTCCAACTTGCATTGGATCAATATTTTTCATTTAAAAAAAAA

>Pbr_TRINITY_CNMT Pbr_c28136_g1_i1 len=1336

ATAAAAGCTAACACGTCAGTGGTAGTAGTTAGTACAACAACCACCACCACCTGTCTCAACTTCTCCGGAAAATCAAATTATACCTCACCTTGTGTGTGTTTTTCTTTGCAAGTTGAACAAGATGCAGCTAATGGCAAAAGAAGAGCTTTTGAGAAACATGGAGCTTGGGTTGATTCCGGACCAAGAGATTAGACAAAGGATTAGAATCGAATTAGAAAAGCGTCTCCAATGGGGTTACAAAGAAACACACGAAGAACAGCTTTCTCAGCTTCTTGAATTAGTTCATTCTTTGAGAGGTATGAAAATGGCAACTGAGATGGAGAATTTGGATTTGAAGCTATACGAGGCACCTATGGAATTCTTAAAGATTCAACATGGAAGAAACATGAAACAAAGTGCCGGTTACTATACCGATGAATCAACGACGTTGGATGAAGCTGAGATAGCGATGCTGGATTTGTACATGGAGAGGGCACAGATCAAAGATGGTCAGAGTGTACTTGATTTAGGGTGTGGATTGGGTGCTGTAGCTCTTTTTGGTGCAAACAAATTTAAAAATTGTCAGTTTACCGGAGTTACAAGTTCTATGGAACAAAAAGATTACATTGAAAGAAAATGCAAGGAACTTAAGTTGACCAACGTCAAGGTTTTATTAGCTGATATAACAACTTACGAAACTGAAGAAAGATTTGATCGGATTTTTGCTGTTGAATTGATTGAGCATATGAAGAACTATCAATTGCTTCTTAAGAAGATATCAGAATGGATGAAAGATGATGGACTTCTTTTCGTTGAACATGTTTGCCATAAAACTCTAGCTTACCACTATGAGCCTGTTGATGCAGAAGATTGGTACACTAACTACGTATTCCCTGCTGGAACTCTGACTCTATCATCTGCTTCAATGCTTCTTTACTTCCAAGATGATGTTTCTGTTGTGAATCAATGGACTCTAAGTGGAAAGCATTACTCTCGATCCCATGAAGAATGGTTGAAAAACATGGACAAAAACATTGTTAAATTCAAAGAGATAATGAAGTCTATAACTAAAACCGAGGAAGAAGCAACCAAATTGCTCAACTTTTGGAGGATGTTTTGCATGTGTGGAGCAGAGTTGTTTGGTTATAAGAATGGTGAAGAATGGATGCTCACCCATCTTCTCTTCAAGAAAAAATGAGAAATTTTCTTTTATGCTTTCGTTTTGGGTGTGTTATTTAGCATCATAAGTATACCAATTCATTGTAATAAGAACTGTTTAGAATGATTGGGTTGTAACTTTTATTTTATGTTTTTTTAATAAAAACTTCATCATCAAGCGCTTTATGGTTTCGAGATC

>Pbr_TRINITY_CYP80B1 Pbr_c33168_g1_i1 len=1726

GTGTGCTCTTCCGATCTGAAGCCAAACAAAAATATTATTATTGAGAGAGAGAAAAGATGGAGATAGTCACAGTATCACTTGTAGCAGTTGTGATCACTACTTTCTTATACTTAATTTTCAGAGATTCAAGTCCTAAAGGTTTGCCACCAGGTCCAAAACCCTGGCCAATAGTTGGAAACCTTCTTCAACTTGGTGAGAAACCCCATTCTCAGTTTGCAGAACTTGCTAAAACCTATGGTGATCTCTTTACACTGAAATTAGGGAGTGAAACAGTTGTTGTAGCTTCAACTCCATTAGCAGCTAGCGAGATTCTGAAGACGCACGACCGTGTTCTTTCTGGTCGTTACGTGTTTCAGAGTTTCCGCGTGAAAGAACATGTGGAGAATTCTATTGTGTGGTCTGAATGTAATGATACATGGAAGAAGTTGCGGAAAGTTTGCAGAGCGGAACTTTTTACACAGAAGATGATTGAAAGTCAAGCTGAAATTAGAGAAAGTAAGGCTATGGAAATGGTGGAGTTCTTGAAGAGAAATCAAGGGAGTGAAGTGAAAATTGTTGAAGTTGTATTTGGGACATTGGTGAATATATTTGGTAACTTGATATTTTCGCAAAATATTTTCAAGTTGGGTGATGAAAGCAGTGGAAGTGTAGAAATGAAAGAACATCTATGGAGAATGCTGGAATTGGGTAATTCGACGAATCCAGCTGATTATTTTCCATTTTTGGGTAGATTCGATTTGTTTGGACAAAGAAAAGATGTTGTCGATTGTCTGCAAGGGATTTATAGTGTTTGGGGTGCCATGCTCAAAGAAAGAAAAATAGCCAAGCTGCATAACAACAGCAAGAAGAATGATTTTGTTGAGATTTTGCTCGATTCCGGACTCGATGACCAACAAATCAATGCCTTGCTCATGGAAATATTTGGTGCGGGGACAGAGACAAGTGCATCTACAATAGAATGGGCGTTATCTGAGCTCACAAAAAACCCTGAAGTAACAGCCAATATGCGGTCGGAATTGCTATCTGTGGTAGGGAAGAGACCGGTTAAGGAATCCGACATACCGAACATGCCTTATCTTCAAGCTTTTGTTAAAGAAACTCTACGGCTTCATCCAGCAACTCCGCTGCTGCTCCCACGTCGAGCACTCGAGACGTGCAAAGTTTTGAACTATACTATCCCAAAAGAGTGTCAAATTATGGTGAACGCCTGGGGCATTGGTCGGGACCCAAAAAGGTGGACTGATCCATTGAAGTTTTCACCACAGAGGTTCTTGAATTCGAGCATTGATTTCAAAGGGAACGACTTCGAGTTGATACCATTTGGTGCTGGGAGACGGATATGTCCTGGTGTGCCCTTGGCAACTCAGTTTATTAGTCTTATTGTGCCTACATTAGTACAAAATTTTGATTGGGGACTACCAAAAGGAATGGATCCTAGCCAAATGATCATGGAAGAGAAATTTGGGTTGACACTGCAGAAGGAACCACCTCTGTACATTGTTCCTAAAACTCGGGATTTATCTTAATCGATAAATTTCAAAATTCACAAGGTTGTGGATTTCCCTCCGAAGTTTATTTTCAAATAAAAAAAAAGTGTTGTTTTTAAATATGACGCGATTGAAATATTCTGCTGTATAAGAATTTTCCACTATTATCTATTCATAGATATCAAACTATGCCAATGTTCCAGTTGTATCTCATATCTGGTTGCTTTAAATGATACACT

>Pbr_TRINITY_4OMT1 Pbr_c7980_g1_i1 len=1323

GTGTGCTCTTCCGATCTCCAAAACAAAAAATAGTCTTCTCTTCTTGACAAATATACAGCAAAAATCTCATATCTAATGGGTAGCTTAGCAAAGTCAGCTGAAACACAAGAAGTTTCCATCAAAGATCAAGCTCAACTATGGAATATTATCTATGGTTTTGCTGATTCTCTTGTTCTGCGTTGCGCTGTAGAGATCGGAATCGCGGATATTATCAAAAACAATGATGGGGCAATCACACTTGCACAACTTGCTTCAAAACTACCAGTTCCGAATGTCAGTTCTGATTACTTGTACAGAATGGTGAGATACTTGGTACACCTGAACATCTTAGCACAAGAAACTTGTAATGGCGGAGTTGAGAAGGTTTACTCGCTTAAACCGGTTGGTACGCTACTTTTAAGAGATGCAGAAAGAAGTATGGTGCCGATGATATTAGGCATGACTCAAAAAGATTTCATGGTTTCATGGCATTTCATGAAAGAAGGTTTAGGAAATGGAAGTACTACGGCATTTGAGAAAGGAATGGGAATGGATATATGGAAGTATTTAGAAGGTAATCCTGATCAGAGCCAGTTGTTTAATGAAGGAATGGCTGGAGAAACAAGACTGTTAACTAAAACTCTGATCGTAGATTGTAGAGATACTTTTCAAGGTTTGGATTCTTTGGTTGATATTGGTGGAGGAAATGGTACAACAATCAAGGCAATCCATGAAGCTTTTCCTCATATCAAATGTACACTTTATGATCTTCCTCATGTTGTTGCCAACTCTCATGATCTCCCTAACATCGAAAAAGTTCCCGGTGATATGTTCAAGTCTGTTCCAAGTGCTCAGGCTATCTTACTCAAGTTGATCTTGCACGACTGGACTGATGAGGAGTGTGTCAACATCCTAAAGAAGTGCAAAGAAGCAATACCGAAAGAGACAGGAAAAGTGATTATAGTGGATGTAGCACTTGAAGAAGAATCCAACCATGAACTTACAAAGACAAGATTGATTCTTGATATTGACATGTTGGTGAATACTGGTGGTAGAGAGAGGACTGCAGATGATTGGGAAAATTTGCTCAAACGGGCAGGCTTTCGGTCTCATAGGATCAGACCCATTCGCGCTATTCAGTCTGTCATTGAAGCCTTCCCTTAGTAAAAGCTCCAAGGAGTACTTTGCTTCTTTCAAAGAGAAGCTTATGCTATGCACTTATTAACAATAAGCCATTTTCCTGTTTTTTGTTTTGTTTTGTTTTGATTTTTTTTCTTTTTCAGGCCATACTAGGCCATTGTTTCTGTTGTTCTTGCTCCTATTCAATGTAATTGAAGATTGTAT

>Pbr_TRINITY_BBE Pbr_c45010_g1_i3 len=1637

TCTCCAACATGATGTGCAGAAGTTTAACATTACGTTTCTTCTTATTCTTTGTTTTATTACAAACATGCGTACGAGGCGGTGATGTTAATGATAATCTCCTCTTGTCATGTTTAAACTCCCATGGTGTTCACAATTTCACCACACTATCAACTGATACAAATTCGGAGTACTTCAAACTACTCCATGCATCTATACAGAACCCGTTGTTCGCGAAGCCTACGGTATCGAAACCGTCGTTTATTGTAATTCCAGGCAGCAAAGAAGAATTATCGAGCACCATTCATTGCTGTACAAGAGAGTCATGGACTATTCGACTGCGGAGTGGCGGTCATAGTTATGAAGGGTTGTCTTACACTGCTGATACACCTTTTGTGATTGTAGATATGATGAACTTGCACGGAATTTCCATTGATTTTGAGTCGGAAACAGCTTGGGTGGAATCCGGTGCAACACTTGGAGAACTCTATTATGCGATTGCGCAGTCAACGGAAACACTGGGGTTTACTGCTGGTTGGTGTCCGACTGTTGGCAGCGGAGGCCATATAAGTGGTGGGGGTTTCGGTATGATGTCGAGAAAGTACGGGTTAGCTGCGGATAACGTCGTGGATGCGATTCTTATAGATAGCAAAGGAGCGATTCTTGACCGTGAAAAAATGGGTGAAGATGTTTTTTGGGCTATTCGTGGTGGTGGCGGAGGTGTTTGGGGTGCAGTTTACGCGTGGAAAATCAAACTATTGCCAGTTCCGGAGAAGCTGACCGTTTTTCGTGTGACTAAGAATGTAGAAATTGAAGCCGCTTCATCTTTACTTCACAAATGGCAATATGTTGCAGATGAATTAGACAATGACTTCACAGTGTCTGTGCTCGGGGGAGTGAACGGAAATGAAGCCTGGTTAATGTTCTTAGGCTTACACTTGGGACGTAAAGATGCCGCGAAAAGCATAATCAATGAAAAATTCCCTGAACTGGGGTTAGTAGAGAAAGAGTTTCAAGAAATGAGTTGGGGTGCATCCATGGCATTGTTAGCAGGATTAGAAACAATCTCTGAATTAAACAACAGGTTCTTGAAGTTTGATGATAGAGCTTTTAAGACAAAAGTTGATTTTACTAAAGAAGCAGTACCGCTAGACGTGTTTAGACGTGCACTAGAGATGTTATCAGAACAGCCCGGTGGATTTATAGCTCTAAATGGTTTTGGAGGGAAAATGAGCGAAATAAGCACTGATTTTACCCCGTTTCCTCATCGGAAAGGCACTAAATTGATGTTCGAATACATAATCGCTTGGAATCAAGATGAAGAAACGAAAATCGGTGAATTTAGCGAATGGTTAGAGAAATTTTACGATTATTTGGAGCCGTTCGTATCGAAAGAACCAAGGGTTGGTTATGTTAATCATATTGATCTTGATATTGGAGGGATAGATTGGAAAAATAAAAGTAGTACTAGCAATGCAGTTGAGATTGCTAGAAATTGGGGTGAAAGATATTTTTCATCGAATTATGAACGTTTAGTTAAGGCGAAGACATTGATTGATCCAAATAATGTGTTTAACCATCCACAGAGTATTCCTCCAATGATGAAATTTGAGGAAAATTACATATTAAATGAACTGTGATACTAGAGATCGGAAGAGCGT

>Pbr_TRINITY_TNMT Pbr_c32804_g1_i1 len=1797

CCACAAGATGTTAATAATAATGATTAAAAGAACACCCACTTTCAGAAAACTTTATGACAATGAGACGGATGATAATCTCTATAAATACTATGTACCAGGTTCTGCTCATACCACATCTATCAACATTCTCTCTTGCATTTCATTCTAGTCTTGTCTTCCGAGAAATCCACTTTGTTTCCTCACAATCCTAGATTGTTTCAGCATGGGTTCAATAGATGAAGTGAAGAAGGAATCAGCAGGGGAAACACTAGGGAGATTGTTGAAAGGAGAGATTAAAGATGAGGAACTGAAGAAGCTGATCAAGTTTCAGTTCGAGAAGCGTCTGCAATGGGGTTATAAATCATCTCATCAAGAACAACTCTCCTTCAACCTTGACTTCATTAAATCCTTAAAAAAGATGGAAATGTCAGGAGAGATTGAGACAATGAACAAGGAAACTTATGAATTACCCTCAGAATTCTTAGAGGCAGTTTTTGGAAAAACAGTCAAGCAAAGTATGTGTTACTTCAAACATGAATCAGCAACAATAGACGAAGCAGAAGAAGCTGCACATGAATTATACTGTGAACGAGCACAGATCAAAGATGGACAAACTGTTCTCGACATTGGTTGTGGTCAAGGTGGTCTAGTTCTATACATTGCTCGGAAATATAAGAAATGTCATGTTACAGGGCTCACTAATTCAAAGGCACAAGTCAATTACTTACTCAAGCAAGCAGAGAAGCTTGGGCTGACAAATGTCGACGCCATACTAGCAGATGTCACTCAATATGAATCTGACAAGACATACGATCGACTACTTATGATTGAAGCCATAGAGCACATGAAAAACTTACAACTGTTCATGAAGAAACTATCAACTTGGATGACAGAAGAAAGTCTTCTCTTTGTGGACCATGTCTGCCATAAAACCTTTGCTCACTTTTTTGAGGCAGTCGACGAGGATGACTGGTACTCAGGTTTCATCTTCCCCCCAGGTTGTGCAACCATACTAGCTGCTAATAGTCTCCTCTATTTCCAGGATGATGTTTCAGTTGTGGATCATTGGGTTGTCAACGGAATGCACATGGCTCGTTCAGTAGACATTTGGAGAAAGGCACTAGACAAAAATATGGAAGCTGCAAAAGAAATTCTATTACCTGGACTTGGAGGAAGCCACGAAGCAGTAAATGGAGTCGTTACTCACATTAGAACATTCTGTATGGGAGGCTATGAACAATTCTCAATGAACGATGGAGATGAGTGGATGGTTGCGCAGCTGCTTTTCAAGAAGAAGTAGCGAGGCAATTTCCACGGCAACTGCATTTTCCAGAGCTGTGTAACAATGTGATCCTCAAGTAACAATAATTGTGGGTTGTATCACAGGGGTTTGGCTTGTTTTTGTTGTCTGCATTTCTACTTTGTTTTTTCTTTTAATTTGGATTCAAGATTTGAAGAACTCCATTCTTAATTATACCATGTAATGGATTTATCGAATCAATAGGTGTGATAAAATTGTACCAACTGCACTTTCCTGTTGTCTAGTTATTTGTCTATAATGGAAGTATGGAACTGAAGAAACTCTAGCACCTTTACTTTTCATTCACCTCTTACGTTTCTCATTTCAACCCGTTGCTGCTTAATTTTCCAGTGTTAATTCACACTGGCCATCTAATTCTTGATGAGCAACCTCCTCATTAAGTAAGCTACAATTCAGCTAAGTAATAATGAGACATGATATCTAGAGCTAGCTTTTGTAATCTTAATACTCAAAGAACCAGTGAGTCGGCAACACATTTAGATCGGAAGAGCACACGT

>Pbr_TRINITY_CYP719A21 Pbr_c51795_g1_i1 len=1694

TTTCATCTCCATTGCAGCCCAGTATTTTCCATCAGTGCCTACAAGCATGAAATTCTTTATGCAAAGAGTTAATCTGACTCAAGCTAGCTAGAATATATACCAACCATACAAGAAATATATGATCATGAGTAACTCTTGGATTCTTACGCTCATTTCTACCATATTAGCAGTCTTTGCTGCTGGGTTAATCATTTTCAGGAGAAGAATATCAGCATCCACAACGGAATGGCCTATTGGCCCAAAAACATTACCAATCATAGGTAACTTGCACATTATTGGAGGCACTGCTCTCCATGTCGTCTTACATAAACTGGCTGAAGTTTACGGCAGTGTGATGACGATATGGATTGGTAGTTGGAAACGTTTTATTATTGTCTCCGACTTTGATCGAGCCTGGGAAGTTCTTGTTAACAAATCGTCAGATTATTCAGCTCGTGAAATGCCTGAGATCACTAAAATCGGCACTACAAATTGGAGAACAATTTCAAGTTCTGATTCTGGTCCATTTTGGGCCACTCTTCGAAAAGGTCTTCAGAGTGTGGCATTATCCCCTCAGCATTTAGCATCGCAAACTGCACACCAAGAGAGAGATATAATAAAGTTGATCAAAAATTTGAAAGACGAAGCAGCTCTTAATTCTGGAACCGTTAAACCACTTGATCATCTCAAGAAAGCAACTGTTAGATTAATCAGTCGGTTAATCTATGGCCATAATTTTGATGACGACAAGTATGTTGAAGATATGCATGACGTGATCGTGTTCTTAATTCGTATTAGTGGTTATGCCCAACTTGCTGAGGTTTTCTACTATGCTAAATATCTACCAAGTCATAAGAGAGCTGTAAAAGGTGCCGAAGAAGCAAAACGAAGAGTAATAGCTCTGGTGCGTCCTTTTCTTCTTTCAAGCCCTCCTGCCAATACTTACTTACATTTTCTTAAATCGCAAAACTACCCAGAAGAAGTTATCATATTCGCTATATTCGAAGCTTATCTTTTAGGTGTTGATAGCACTTCTTCAACCACTGCATGGGCTCTAGCATTCTTAATACGCGAACCATCTGTTCAAGAAAAACTTTATCAAGAGCTTAAGAATTTCACAGAAAATAACAATCGCACAATGCTGAAAGTTGAAGACGTAAACAAATTACCATATTTACAAGCTGTTGTTAAAGAAACAATGAGGATGAAACCAATTGCACCACTGGCGATTCCTCATAAAGCTTGTAAAGACACTTCATTGATGGGCAAGAAAGTTGATAAGGGAACTAAAGTTATGGTTAACATTCATGCTTTACATCATATCGAAAAAGTCTGGAAAGAACCCTACAAATTCATGCCAGCGAGGTTTCTGCAGAAACAAGATAAGGCGATGGAACAATCACTATTACCATTTAGTGCAGGTATGAGAATTTGTGCCGGAATGGAATTAGGAAAACTTCAATTTAGTTTTTCTCTTGCTAATCTTGTTAATGCCTTCAAATGGTCTTGTGTTTCTGATGGAGTACTTCCTGATATGAGTGATCTACTAGGGTTTGTTCTGTTCATGAAAACCCCACTCGAAGCACGTATATTTCCTCGTCCATAGTGATGGAACTTTCTATTTCATGTTGTTTTTCTTTCTCTTTACAGCTAGGTACTGGTTTGATTTTTAGTGTAAGAAATAAGATCTAAATTCTCGGTTATCAGTTGAAAAA

>Pbr_TRINITY_9OMT Pbr_c36525_g1_i1 len=1382

CGTGTGCTCTTCCGATCTTGTCATCAAAAACAGTCCTAAGATGGGTACCAGTGGCGAAATTTTCAATACCTATGGTCATAATCATCAAACTGCCACAGTCACTAAAATCTGTGCTTCTAATGAAAGCAACAATGATATCTGTTATCTTTCAGAAACGGCTAACTTAGGGAAGTTAATATGCATTCCAATGGCATTAAGAGCTGCCATGGAGCTGAATGTGTTTCAACTTATCTCAAAGTTCGGAACTGACGCAAAAGTTTCAGCTTCTGAAATTGCCTCTAAAATGCCAAACACCACGAATAATCCTAAAGCGCCTATCTATTTAGATAGGATTCTTCGACTGCTCGGTGCCAGTTCTATTCTTTCTGTTTCTACCAGAAAAAAATTAAACGGAGGAGGAAATGATGGAGTAGTACATGAGAAGGTTTATGGGTTAACAAATTCGTCGTGCTGTTTGGTCCCTCGAGAAGAAGACGGGGTGTCATTAGTCGAAGAATTGCTATTCACATCTGACAAGGTTGTTGTGGATAGTTTTTTCAAACTGAAATGTGTGGTGGATGAAAAAGACAGTGTGCCATTTGAGGTTGCTCATGGTGCTAAGATCTTTGAGTATGCTGCTACAGAACCAAGAATGAATAAGGTTTTTAACGATGGAATGGCAGTTTTCTCTATTGTTGTTTTTGAAGCTGTTTTTAGAGTTTACGATGGATTTCTTGATATGAAAGAATTGTTAGATGTTGGTGGTGGTATTGGTACTTCAGTTAGTAAAATTGTTGCTAAATACCCTTTCATTCGCGGTGTCAACTTCGACTTGCCTCATGTTATTTCGGTTGCCACTCAATACCCAGGTGTAGAGCATATTGCAGGAGACATGTTCGAGGAAGTTCCAAAGGGTCAAAACATGTTGCTAAAATGGGTACTGCACGATTGGGGTGACGAACGATGTGTGAAGCTGTTAAAGAATTGTTGGACCTCATTACCTGCGGGTGGAAAAGTTTTGATAATCGAGTTTGTTCTTCCGAATGAACTTGGTAATAATGCCGAATCATTCAATGCGTTGATTCCCGATTTACTCCTCATGGCTCTGAATCCAGGCGGTAAAGAACGAACAATTTCCGAATATGATGATTTAGCCAAAGCAGCTGGATTTATAAAAACTATACCTATCCCTATCTCCAATGGTCTTCATGTCATTGAGTTTCACAAATGAATGGCTATTGAGTGCTTTCGTAATTAAACTATCAAGATAACTACATCCGTTTCATGCATTTTCTTTTCTTTTGTTTTTCTTTTGTATTCAAGGTGTGAACTAGTGTGTTAAGTGGACAAAACTAAGTAATCGTATTTTGTGTTTCTTAATGAGATAATGAAGTGTTCATGTG

>Pbr_TRINITY_SPS Pbr_c120288_g1_i1 len=531

ACCAATACCAAAAATGGAGAAAACCATCTTCAGCAATTCATGGATTCTTACCACTTCCACTTTAGTAGTAATAACAATCATTACCATGCTAATGGTATTCAAAAGAAAAAAATCATCGACTATGGTGTGGCCAACAGGGCCAAAAACATTACCAATTATAGGGAACATGCATGTTCTTGGCGGCACAGCTCTTCAAGTTGTCTTACATAACCTAGCCAAAATTCATGGCAACGTTATGACGATATGGATCGGTAGTTGGAGACCAGTTATTGTCGTTTCTGATATCGACCGAGCATGGGAAGTACTTGTTAACAAGTCGTCGGATTATTCAGCTCGTGATATGCCCGAAATCACTAAGATTATATCTGCTGATTGGAAAACCATTTCTACTTCTGATTCCGGTCCTGGTTGGTCTAATCTTCGCAAAGGTCTTCAAAATGTGGCGCTATCTCCTCATAACTTAGCCGCACAAAGTCAATTTCAAGAAAAAGATATTACCAAAATGATTCAAAGTTTGAAAAAAGAAGCTGC

>Pbr_TRINITY_MSH Pbr_c38445_g1_i2 len=1824

AATCAGAAAAAAAAATGAACTTTCTTGATCTGCTTTTTCTTCAATTCCAACCAATCTCTATTGCACTTGTTGTGATTGCACTAATTTTTATCTTATGGATTAATGGAAGAAAACCCGCTCAAAAGCTAGCTCCAGAAGCATCGGGGGGGAGACCTATAATGGGTCATTTACATCTTTTTAAAGACGGCGAGTTAACTCATCGAAAACTGAGTATCATGGCCGATGAGTACGGTCCTGTTTTCAACATCCGGTTCGGTTCGCATAAAACACTAGTTGTAAGTGATTGGGAAATTGTTAAAGAGTGTTTTACTACAAATGATAAACTCTTCTCAAATCGTCCGGGTTCGTTAGCTGTTAAGATGATGTTTTACAATGCCGATTCGGTTGGTTACGCTCCATATGGTGCGTATTGGAGAGATTTACGAAAGCTTTCAACATTGAAACTTCTATCCAACCATCGTATCGACACAATAAAACACCTAAGAACTTGTGAAGTTGAGTCGTGCTTCGAGTCGCTTTATAATCAGTGGAGAACAGGGAAAAAATGCGGCGAAGGATCGATGTTTACTCCAGTGAGAATGGACAATTGGTTAGGCGATTTGACGTTTAATGTTGTGGCAAGAATTGTTGCAGGAAAGAAGAATTTCTCAGCAAACGGCGACGTTGGAGCTCGGAGATATAAATCAGCAATGGATGAAGCTGTAAGGTTGATGAGTTTTTTTGCATTTTCGGATAAGATTCCATCACTAAGTTGGTTAGACAATTTAAGAGGTTTGGTTAGAGATATGAAAAAATGTGCATCTGAAATTGATACAGTTATGGCTACTTGGGTTGAAGAACATCGTGTTAAAAGAAGACTTTCAGGAAATAATTCTCAACTTGAACATGATTTTATCGACGTTTGCTTGGATATAATGGAGAATTCTTCATTACCCGGTGATGATCCTGATCTCGTCGTCAAATCTACCTGTCTGGACATGATACTGGGTGGGAGTGACACGACAACAGTTACCCTAACATGGGCAGTGTCTTTACTCTTGAATCATCCCCAGGTACTACATAAAGCAAAACAAGAACTAGACACACAAGTTGGAAAGAATCGGCAAGTAGAAGATACAGATATCCCTAATCTTCCCTACATCCAAGCGATCATCAAAGAGACGATGAGGTTGTACCCTGCCGGACCGTTAATCGAACGGAGGACGATGGAGGATTGCGAAGTAGCGGGGTACCAAATTCCGGCCGGCACGCGCTTAATAGTTAACGTCTGGAAGATGCAAAGAGATGGGAACGTATACAAAGGCGATCCGTTGGAGTTTAGACCCGACAGGTTTTTAACTAGTAATGCGGATGTGGATCTAAAGGGTCAGCATTACGAATTGATACCATTTGGAGCGGGTCGACGGATATGCCCGGGAGTTTCGTTTGCGGTTCAGTTGATGCATTTGGTTCTTGCTCGATTACTTCATGAGTTCGAGATCACGACGGTGGAGCAAGAAGCCAAGGTGGATATGGCTGAAAGTGGTGGTTTGCTTTGTTATAAGATAATGCCGTTGGAAGTACTGATCAAACCTCGACTCGAGATATGATCAAAGGAGGAGTCCAACATATAAACAAGAATGCGAATCGCCGTATCTCAGGTCCAACATAGTATAGTTATGTATGATACTTGTTTTCTCTTTTGTCTTGATTTTATGGGCTTTCTAAAGTTGTAAGAGCAGTACAATAAAATTAAAATTACGGGGACTTTAGCCCTACATATATATATATATATACTGGCCTAGCATCAGCTGCAACAGTACTTGCAGAAAGGTGCTCCAGTTA

>Ptr_TRINITY_NCS Ptr_c34808_g1_i1 len=1137

AGAAGTGATTATGATCGAAGGAGGGTACTTAGACATGGGATGCACATCTTACCTGGATAGGATCCATGTTGTTGAGAAAACCTCAAAATCTTGCATCATTAAATCTTCTGTTGTCTACGAAGTGAAGCAAGAGTGTGCTGATGCCATGTCTAAGTTGATCACGACCGAACCATTGAAGTCGATGGCAGAAGTCATCTCTAATTATGCTATTAAGCAACAATCTGCTTCTAAGAGAAACATTTCCAAGAAGCAGTCTCTACTGAGGAAAGAAATTACTTGTGAAACGGAGGTGCAAACTTCGGCTGATTCGATTTGGAACGTCTACAGTTCTCCTGACATCCCTCGGCTACTTAGAGATGTTCTGCTTCCTGGTGTTTTTGAAAAGCTAGATGTCATTGCAGGCAATGGTGGCGTTGGTACTGTACTGGACATTGCTTTCCCTCTAGGTGCAGTGCCGCGGAGGTACAAGGAGAAATTTGTGAAGATCAACCATGAGAAGCGATTGAAAGAAGTGGTTATGATCGAAGGAGGATACTTGGACATGGGATGCACATTTTACATGGACAGGATTCATGTATTAGAGAAAACCCCAAACTCATGTGTTATTGAATCCTCTATCATTTACGAAGTTAAAGAAGAGTATGCTGATAAAATGGCTAAGCTAATCACAACCGAACCATTGGAATCCATGGCAGAAGTCATCTCTGGTTATGTTCTTAAGAAACGACTCCAAGTATTCGGATTCGAGATTAAGCCAAAATTAAGATTCGATCTTTTGTTATGTTTGATCATCTGCTTGGTTATAGCTGGAGGTATGTTTGTTGCTCGTGTTCCACTCTAAGGTTAAGGGCTTTTCTTAAATGTAATTTTGGAAATAAAGTATTGGTTTGATCCTTCAAACCATGCATTGCATTGCAGAGGTTTAGCTTCGATTCGTGGAATTTCTCATATGTAATTCTCAAATCTACAAGATGAAAAATCAAATTTAATAATAGTTTCAAGCAATGTTTGAAAAACGGGTTTAGAATGTAATGTTAATTTTGGGCAATGTTTGATAAATGGGTATGTTTCTTGCTGGTGTTCCAGTCACTCAGAAAGAGAAATTATACTAGACAAAAAAATAAACATACTTTTTTG

>Ptr_TRINITY_6OMT Ptr_c35419_g2_i1 len=1355

GTGCTCTTCCGATCTGTCAAAAGATAGCACTAGAGAAATAGCTCAAATCATTCATTCATTCTCAATCTCATCAACAGCTAAAGTTTCTAAAAAGAGAGAAATGGAAACAGTAAGCAAGATCGATCAACAAAACCAAGCAAAAATCTGGAAACAAATTTACGGTTTTGCAGAATCACTAGTTCTGAAATGTGCAGTCCAATTGGAAATTGCCGAAACACTTCACAACAATGTCAAACCCATGTCTTTATCCGAATTAGCATCGAAACTCCCCGTTGCTCAACCCGTTAATGAAGACCGTCTGTTCCGAATCATGCGTTACTTGGTTCACATGGAACTCTTCAAAATAGATGCCACCACACAGAAATACTCATTAGCTCCACCAGCTAAGTATTTGTTGAGAGGCTGGGAGAAATCAATGGTTGATTCAATTTTATGCATAAATGATAAGGATTTCTTAGCTCCATGGCATCATTTAGGTGACTGTTTAACCGGTAACTGTGACGCTTTTGAGAAAGCGTTGGGGAAGAATATTTGGGTTTATATGAGTGAAAATCCTGAAAAGAATCAATTGTTTAATGCAGCAATGGCTTGTGATACTAGATTGGTTACTTCTGCATTGGCTAATGAGTGCAAAAGTATTTTTAATGATGGAATCAGTACACTAGTCGATGTCGGTGGCGGTACCGGTACTGCTGTGAAAGCCATATCTAAAGCATTTCCGGATATTAAATGCACTATCTATGATCTTCCTCATGTCATAGCTGATTCTCCTGAAATTCCCAATATCACTAAAATTTCTGGAGATATGTTCAAGTCTATTCCTAGTGCTGATGCCATCTTCATGAAGTGCATACTTCACGACTGGAACGACGAGGAATGCATTCAAATCTTGAAGAGATGCAAAGAAGCATTGCCAAAAGATGGCAAAGTTATTATCGTGGATGTCGTAATAGACATGGATTCGACTCATCCGTACGCCAAAATTAGACTCACATTGGATTTGGATATGATGCTTAACACTGGTGGAAAAGAGAGAACCAAAGAAGAATGGAAGAAACTTTTTGATGCCGCTGGTTTTGCTAGCCACAAAGTCACTCAGATATCTGCTGTCCAGTCTGTAATTGAGGCTTACCCTTATTAAGGAACATTTTACCCGGTTTTCCCTTTGATTAATTGTTATTGCTTTATCTTTGGATTATGTTTGTGCTTATGATAACTGCAAGATGAATGAATTTCCAACTTGCACTGAATTTGAATCAATGTTTTTCGGTTACTTAACTTTGTGCTATTTTGGGAGGATTTTTCTTAAATTTGCTTTTGGCAATTGAGATGTGCTTGTCATGCTTGTTAGTTCC

>Ptr_TRINITY_CNMT Ptr_c5952_g1_i1 len=1306

GCTCTTCCGATCTCACCACCTCTGTCTGAATTTCTCTGGAAAATCTATAGACTTTGTGTGTGTTTTGTCCTTTGGAAGTTGAACAAGATGAAGCTAATGGCAAAGCAAGAGCTGTTGAGAAACATGGAGCTTGGGTTGATACCCGATGAAGAGATTAGAGAAAGGATTAGAATCGAATTAGAAAAGCGTCTTCAATGGGGTTACAAAGAAACCCATGAAGAACAGCTTTCACAGCTTCTTGAATTAGTTCACTCTTTGAGAGGTATGAAAATCGCAACTGAGATGGAGAATTTGGATTTAAAACTATACGAAGCACCTATGGAATTTTTAAAGATCCAACATGGGAGCAATATGAAACAGAGTGCCGGTTACTACACTGATGAATCAACGACGTTGGACGAAGCTGAGATTGCGATGCTGGATTTGTACATGGAGAGGGCGCAGATCAAAGATGGTCAGAGTGTACTTGATTTAGGCTGTGGATTGGGTGCTGTAGCTCTTTATGGCGCAACCAAGTTTAAAAATTGTCAATTTACTGGAGTTACAAGTTCTATGGAACAAAAGGATTACATTGAAGGGAAATGCAAGGAACTTAAGTTGACCAATGTCAAGGTTTTATTAGCTGATATAACAACTTATGAAACCGAAGAAAGATTTGATCGGATTTTTGCTGTTGAATTGATTGAGCATATGAAGAACTATCAATTGCTTCTTAAGAAAATATCAGAATGGATGAAAGATGATGGACTACTTTTCGTTGAACATGTTTGCCATAAAACTCTAGCTTACCACTATGAGCCTGTTGATGCAGAAGATTGGTATACTAACTACATATTCCCTGCTGGAACTCTGACTCTATCATCTGCTTCAATGCTTCTTTACTTTCAAGATGATGTTGCTGTTGTGAATCAATGGACTCTAAGTGGAAAACATTACTCACGATCCCATGAAGAATGGTTGAAAAACATGGACAAAAACATTGTTGAAGCCAAAGAAATACTGAAGTCTGTAACCAAAACTGAGGAAGATGCAACCAAATTGCTCAATTTTTGGAGGATTTTTTGCATGTGCGGAGCAGAGATTTTTGGCTATAAGAATGGTGAAGAATGGATGCTCACCCATCTTCTCTTCAAGAAAAAATGAGAGAACTTCTTTTATGTGTGCTTTCATTCAGTGCGTCTTACTTTACATCGTACATATGCCAAATCATTGTAATAAGAACTGTTTAGAATGATTGGTTGTAACTTTTGTTTAAGCTTAATTTAATATATGTTTCATCATGAAGTGTTTTATGGAAAAAAAAAAA

>Ptr_TRINITY_CYP80B1 Ptr_c32430_g1_i1 len=1630

CTTGAATTCTTTACATAAACAGGAAGCCAAACAAAAATATACTTATACTTAGAGAAAGAAAAGATGGAGATTGTCACAGTATCATTTGTAGCAGTTGTGATCACTACTTTCTTATACTTAATTTTCAGAGATACAAGTCCTAAAGGTTTGCCACCAGGTCCAAAACCATGGCCAATAGTTGGAAACCTTCTTCAACTTGGCGAGAAACCCCATTCTCAGTTTGCTCAGCTTGCTGAAACCTATGGGGATCTTTTTACACTCAAATTAGGAAGCGAAACAGTTGTTGTGGCTTCAACTCCATTAGCAGCTAGCGAGATTCTAAAGACGCATGATCGCGTTCTTTCTGGTCGATACGTGTTTCAGAGTTTCCGGGTAAAGGAACATGTGGAGAACTCTATTGTTTGGTCTGAATGTAATGAAACATGGAAGAAACTGCGGAAAGTTTGTAGAACGGAACTTTTTACGCAGAAGATGATTGACAGTCAAGCTGAAATTAGAGAAAGTAAGGCAATGGAAATGGTGGAATATTTGAAGAAAAATGAAGGAAATGAAGTGAAAATTGTTGAAGTTGTATTTGGGACGTTGGTGAATATATTCGGAAACTTGATATTTTCGCAAAATATTTTCAAGTTGGGTGATGAAAGTAGCGGAAGTGTAGAAATGAAAGAACATCTATGGAGAATGCTGGAATTGGGGAACTCGACAAATCCAGCTGATTATTTTCCATTTTTGGGTAGATTTGATTTGTTTGGACAAAGAAAAGATGTTGCTGATTGTCTGCAAGGGATTTATGGTGTTTGGGGTGCTATGCTCAAAGAAAGAAAAATAGCCAAGCTGCTTAACAACAGCAAGAGGAATGATTTTGTTGAGATTTTGCTCGATTCCGGACTCGATGACCAACAGATCAATGCCTTGCTCATGGAAATATTTGGTGCGGGGACAGAGACAAGTGCATCTACAATAGAATGGGCCTTATCTGAGCTCACGAAGAACCCTGAAGTGACAGCCAATTTGCGATCGGAATTGCTATCTGTGGTAGGGCAGAGGCCGGTTAAGGAATCCGACATACCAAACATGCCTTATCTTCAAGCTTTTGTTAAAGAAACTCTACGGCTTCATCCGGCAACTCCGCTGCTGCTCCCTCGCCGAGCACTTGAGACGTGCAAAGTTTTGAACTATACAATCCCAAAAGAGTGTCAAATTATGGTGAACGCCTGGGGCATTGGTCGGGATCCAAAAACGTGGGCTGATCCATTGAAGTTTTCACCTGAGAGGTTCTTGAATTCGAGAATTGATTTCAAAGGGAACGACTTCGAGTTGATACCATTTGGTGCAGGGAGAAGGATATGTCCTGGTGTGCCCTTGGCAACTCAGTTTATTAGTCTTATTGTGCCTACTTTGGTACAGAATTTTGATTGGGGATTACCAAATGGAATGGATCCTAGCCAACTGATCATGGAAGAGAAATTTGGGTTGACACTCCAGAAAGAACCACCTTTGTATATTGTTCCTAAAACTCGGGATTAAGCTCAATCAAGAATTCTTGAATTCGCATGATTATGGATTTCTCGCTATTTTATTTGAAAATAAAATTAAAAGAAAAAAGTACTCCCTCCGTTACTTTTTAATA

>Ptr_TRINITY_4OMT1 Ptr_c32723_g1_i2 len=1218

GTGTGCTCTTCCGATCTCCTAAACAAAAATAGTCTTCTCTGCTTGACAGCAAAAAAATTCATATCTAATGGGTAGTTTAGATGTCAACCCAGCTTCTGAAACACAAGAAGTTTCCATCAAGGATCAAGCTCAACTGTGGAATATTATCTATGGTTTCGCTGATTCTCTTGTTCTTCGTTGCGCCGTAGAGATTGGAATCGCGGATATTATCAAAAACAATGGTGGGGCAATCACACTTGCACAACTTGCGTCGAAACTACCAGTTCCAAATGTCAGTTCTGATTACTTGTACAGAATGGTAAGATACTTGGTACACCTGAACATCTTAGAACAAGAAACTTGTAACGGCGGAGTTGAGAAGGTTTACTCGCTTAAACCGGTCGGTACTCTACTTTTAAGAGATGCAGAGAGAAGTATGGTGCCGATGATATTAGGCATGACTCAAAAGGATTTCATGGTTTCATGGCATTTCATGAAAGAAGGTTTAGGAAATGGAAGTACGACCGCATTTGAGAAAGGAATGGGGATGGATATATGGAAGTATTTAGAAGGTAATCCTGATCAAAGCCAGTTGTTTAATGAAGGAATGGCTGGAGAAACAAGACTTTTAACTAAAACTCTGATCGAAGATTGTAGAGATATTTTTCAAGGTTTGGATTCTCTGGTTGATATTGGTGGGGGAAATGGTACAACAATAAAGGCAATCTATGAAGCTTTTCCTCATATTAAGTGTACACTCTATGATCTTCCTCATGTTGTTGCCAACTCTCATGATCTCCCTAACATCGAAAAAGTTCCCGGTGATATGTTCAAGTCTGTTCCCAGTGCTCAGGCTATCTTACTCAAGTTGATCTTGCACGACTGGACTGATGAGGAGTGTGTCAACATTCTAAAGAAGTGCAAAGAAGCAATACCGAAAGAGACAGGAAAAGTGATTATAGTGGATGTAGCACTTGAAGAAGAATCCAACCATGAACTTACAAAGACAAGATTGATTCTTGATATTGACATGTTGGTTAATACTGGTGGTAGAGAGAGGACTGCAGATGACTGGGAAAATTTGCTCAAACGTGCAGGCTTTCGGTCTCATAGGATCAGACCCATTCGCGCTATTCAGTCTGTCATTGAAGCCTTCCCTTAGTAAGCTATGCAATGCACTTGTTATTGACAAATAAGCCATTTTCCTGTTGTTTGTTTGTTTGTTTTTTTTTTTTTTTT

>Ptr_TRINITY_BBE Ptr_c36782_g1_i2 len=1658

GTGCTCTTCCGATCTGTGCAGAAGCTTAACATTACGTTTCTTCTTATTCTTTGTTTTATTACAAACATGCGTACGAGGTGGTGATGTTAATGATAATCTCCTCTCGTCGTGTTTAAACTCCCATGGTGTTCACAATTTCACCGCACTATCAACCGATACAAATTCCGACTACTTCAAACTACTCCATGCATCGATACAGAACCCGTTGTTCGCGAAGCCTACGGTATCGAAACCGTCGTTTATTGTAATTCCCGGCAGCAAAGAAGAACTTTCGAGCACGATTCATTGTTGTACAAGAGAATCATGGACTATTCGACTGCGGAGTGGTGGTCATAGTTATGAAGGGTTGTCTTACACTGCTGATACACCTTTTGTGATTGTAGATATGATGAACTTGAATCGAGTTTCCATTGATGTTGAGTCGGAAACAGCTTGGGTTGAATCCGGTGCAACACTTGGAGAATTATACTATTCTATTGCGCAGTCAACGGAAACACTGGGGTTTACTGCTGGTTGGTGTCCGACTGTTGGTAGCGGAGGACATATAAGCGGTGGTGGTTTTGGTATGATGTCGAGAAAGTATGGGTTAGCCGCGGATAACGTTGTGGATGCGATTCTTATAGATAGCAATGGAGCGATTCTTGACCGTGAAAAAATGGGGGAAGATGTTTTTTGGGCTATTCGTGGTGGCGGCGGAGGTGTCTGGGGTGCAATTTACGCGTGGAAAATCAAACTTTTGCTAGTTCCGGAGAAGCTGACCGTTTTTCGTGTGACAAAGAATGTAAAAATTGAAGCCGCTTCTTCTTTGCTTCACAAATGGCAATATGTTGCAGATGAATTAGATGAGGATTTCACGGTGTCTGTGCTCGGGGGAGTAAACGAAAATGAAGCCTGGTTAATGTTCTTAGGCTTACACTTGGGACCTAAAAGTGCTGCAAAAACTATAATCGATGAAAAATTCCCTGAACTGGGGTTAGTAGAGGAAGAGTTTCAAGAAATGAGTTGGGGTGCATCCATGGCTTTCTTAGCAGGATTAGAAACAATCTCTGAGTTAAATAACAGGTTCTTGAAATTTGATGATAGAGCTTTCAAGACAAAAGTTGATTTCACTAAAGAACCAGTACCGCTTAACGTGTTTAGACGTGCATTAGAGATGTTATCAGAACAGCCCGGTGGGTTTATAGCTCTAAATGGTTTTGGAGGGAAAATGAGCGAAATTAGCACTGATTTTACCCCGTTTCCTCACCGGAAAGGCACTAAATTGATGTTCGAGTATATAATTGCATGGAACGAAGATGAAGAAACGAAAATCGTTGAGTTTAGCGAATGGTTGGCTAAGTTTTACGATTATTTGGAACCGTTCGTATCGAAAGAACCAAGGGTTGGTTATGTTAATCATATTGATCTTGATATTGGAGGGATACATTGGAGAAATGGAAGTAGTACTAGCAATGCAGTTGAGATTGCTAGAAATTGGGGTGAAAGGTATTTTTCATCGAATTATGAACGTTTGGTTAAGGCCAAGACATTGATTGATCCAAATAATGTGTTTAATCATCCACAGAGTATTCCTCCAATGATGAAATTTGAGGAAAATTACATTTTAAATGAACTGTAATACCAGGTTGATTTAGAAATTTACTTTTTGTACTAAATTG

>Ptr_TRINITY_TNMT Ptr_c36888_g1_i1 len=1370

TGTCGGGTGCCGACCTTTTAGTGCGTAATGTTGTGCCATAAACCCGTTTGTTGATCTCAATAATTTCGCAAGTATTTCATTTTGTTTCTCTCCCTAGTTTAGCCTAGATTGTCTCATCATGGGTTCAATAGAAGAAATGAAGAAGGAATCAGCAGAGGAAACTCTAGGGAGATTGTTAAGAGGAGAGATTGATGATGAGGAATTGAAGAAACTGATTAAGTTTCAATTCGAGAAACGTCTTCAATGGGGTTATAAATCATCGCATCAAGAACAACTCTCCTTCAACCTTGACTTCATTAAATCCTTAAAAAAGATGGATATGTCAGGAGAGGTCGAGGCATTCAAAAACGAAGTCTATGAATTACCAACAGAATTTTTAAAAGCAGCTTATGGAAAAAGTATGAAACTAAGTGGATGTTACTTTAAACATGAATCATCAACAATAGACGAAGCAGAAGACGCTTCACATGAATTATACTGCGAGCGAGCACAAATCAAAGATGGACAAACTGTTCTTGATATCGGTTGCGGTCATGGTGGTCTAGTTCTATACGTTGCTCAGAAATATAAGAACTGTCATGTTACAGGGCTCACTAATTCGAAGGAACAAGTTAATTACATACTCAAACAAGCAGAGAAGCTTGGGCTGAGAAACGTGGACGTCATACTGGCAGATGTTACTCAATACGAATCCGACAAGACATACGATCGAATACTTGTAGTTGGAGTTGTAGAGCACATGAAAAACCTACAACTGTTTATGAAGAAACTATCAACTTGGATGGCAGAAGATAGCCTTCTTTTTGTGGATCATAGCTGCCATAAAACATTTAATCACTTTTTTGAGGCACTCGACGAGGATGACTGGTACTCGGGTTACATCTTTCCCCCGGGTTGTGCTACATTTCTATCTGCTGATAGTCTCCTCTATTTCCAGGACGATGTTTCCGTTGTGGATCATTGGGTTGTGAACGGGATGCACATGGCTCGTACAGTGGATGCGTGGAGAAAGAAGCTGGATAAAAATATGGAAGCGGTAAAAGAAATTCTATTACCAGGACTTGGTGGGAACCATGAAGCAGTAAATGGCGTCATTACTCACATCAGGACATTGTGTTTAGGAGGGTATGTACAATTCTCATTCAACGATGGAGATGAATGGATGAACGCACAACTGCTTTCCAAGAAGAAGTCATGAATGAGTGGTCTAGGCACATATTCAACCCGCATTTCCGCGCCTTATTATATCAAATAATGGATTAGATGACAAATGTTATTTCTGATTAAGATAATGCCTTAAATGTACTGAATTTATGGCTACCTGTAATTTACAGTACTATAAATTTATGGCTACCTGAGATCGGAAGAGC

>Ptr_TRINITY_PSSDR1 Ptr_c32559_g1_i2 len=1183

CATAAATTCAAAGAGATGGAAGGAACAGGAAAGATAGTATGTGTAACAGGTGGAGCTGGATACTTGGCATCTTGGCTGATCATGAGATTGCTTGAACGTGGTTACTCTGTTCGGACCACCGTCCGGTCTGACCCAAAATTTAGGGAAGATGTGAGCCACCTGAAAGCTCTTCCTGAAGCTACAGAGAAGCTTCAAATTTTTGAAGCAGATCTTGAAAACCCAGAAAGTTTCGACGATGCAATCAACGGGTGTGTCGGTGTCTTTCTCGTTGCTCAAGGAATGAATTTTGCCGAAGAATATACTCTTGAAAAAATAATCAAAACATGCGTGGAAGGAACTCTTAGAATTCTAGAGTCATGCTTGAAATCTAAAACAGTGAAAAAGGTTGTGTACACATCTTCTGCTGATGCAGCAATGATGATAAGTAATCTCAAAGCTTTAAAAGAAATTAACGAGACAATATGGTCAGAAGTTGACAATTTCATTAGCAAACCGGAACAAGTTATTCCTGGATTGCCCTCATATGTGGTCTCAAAGGTACTGACAGAAAGAGCTTGCCTAAAATTTTCTGAAGAACATGGTTTGGATGTTGTTACTATACTTCCTCCGTTGATTGTTGGACCATTTATCACTCCCCATCCCCCTCCCAGTGTATCTATAGCTCTTTCGATAATTTCAGGTGATGTGTCGATGATGCTTGGTGTTAGACTTGAAAATGCGGTGCATATAGATGATGTTGCTTTAGCACACATATTTCTTTTTGAATGTGAAAAAGCAAAAGGAAGACATATTTGTTCTTCAGCTGATTTTCCAATGCATGATCTGCCTAAATTTATATCTGAGAATTATCCGGAATTTAACGTACCGACTGATTTACTAAAGGATATTGAGGAACAAGAACCAGTTCATCTTTCCTCAGATAAGCTGTTGAGTATGGGATTTCAGTTCAAATATGCTTTTGCAGAGATTTTCGGTGATGCAATACGATGTGCCAAAGAGAAGGGTTTCCTTTAGAGGCCAATTATAGTATGGTTCCGAGGAGATGTGGGAGTAGCTAGCCCAAGATGCCCTGCTTGCACTAGCTTATATTATTGTTGTTGTTTTTCAAATGAATAAATGGGCAGTTTTTTTTTGTTTTTTTTTTATGAACTTGATTTTACAACGAAGAGCAGATCGGAAGAGC

>Ptr_TRINITY_PSCXE1 Ptr_c37813_g1_i3 len=828

GCTTCCTATCATAATTTACTTCCATGGTGGAGGTTTCATTCTTTTCAATGCAGATTCAACTATAAACCACGACTTTTGTCAATCGATTGCTACACATTTACCCGCGCTGGTCGTTTCGGTAGACTACCGTCTTGCTCCTGAAAACAGACTTCCCGCTGCCTATGATGATGCTGTTGATGCTTTAAACTGGGTCAAAGACCAAGGTTCTGGCAAACTAAATAATAGTGAAGTATGGTTAAAAGAGTATGGCGACTTCTCAAAGTGTTTCATTATGGGGTGCAGCTCAGGTGCTAATGTTGCATATCATGCCAGTTTAAGAGCAATAGAAATGGATCTTGAACCAGTTATAATCAATGGATTAATATTACACAGCCCTTTTTTTGGTAGTCTTGAGAGAACTGAATCAGATTTAAAAGTGATCAACGATCTAGACTTGCCTCTTGCGGTAAGGGATGTCATGTGGGAACTGGCTTTGCCGCTTGGGTCTACTCGTGATCACATTTATTGTAATCCGAATATTGATAATGATGGATCATCATCCGGAAATATGGTGGGGTTAATCGAGAGGTGTTTTGTGGTAGGATTTTATGGGGATCCACTTATTGATCGACAAATTCAACTGGTGAAGATGCTAGAGGTAAAAGGTGTGAAGGTTGAAACTTGGATTGAACAAGGAGGGTATCATGGGGTGCTATGCTTTGACCCTACGATACGTGAAACCTTTTTGGAAAAACTAAAATATTTTATTTTAAACGAGACGAATTTATATACAACATGGAAAAACACAGCACCTAGCTCAGCAGGTCTAACACAGATCGGAAGAGCGTC

>Ptr_TRINITY_CYP82X1 Ptr_c36328_g1_i1 len=2019

CTCTTCCGATCTCTCATCATCATTCAAGAGAGACACAAATACCTTGATATTCTTTTATCATCAATGGAGTTATTCATAAAGTTACCGTTTATCCAACCAATTCCTTTCAGTATTATTCTTGTTACTACAGTTTCGATTGTTCTATTATACAGTGTCTTCTTCTTGGTTACTGATAAGAAAAAGAAGAAGAAGAAGAAAGCACCAAATGCTTCAGGGGCATGGCCGTTAATAGGTCATCTCCGTCTATTGATGAACGACAAGGAACCGTTGTATAGAGCACTAGGGAGCATGGCTGATAAGTACGGACCTGCATTCAACATCCGATTAGGTAACCAAGAAGTTCTTGTTGTGAGTAACTGGGAGATGGTAAAACAGTGTTTTGGTAATCAAAATGATAAGCTATTTTCGAATCGTCAAACTACATTAGCTGCAAAATACATGCTTAATCAAACAACTTCTAGCGGATTCGCACCATATGGACCATATTGGAGAGAGCTACGAAAGATAATGGTGCAGCAATTACTCTCTAAACAATCTTTAGAATCGTGGAAACATATGAAAATCAAAGAGATGGATGCTTCATTTAGTAAGCTTAACGAGTTATGCAACAACAACGGTACTGGAACAGCTACCCTAATTAGGATGGACGAATGGTTTGCTGAGTTGACGTTCAACGTGATCGCAAGAAATGTCTTTGGCTACCAAAGTGGCGGAAGGTCGACAGCGCTTACAAACGGTGATACGGAATCAAAGGGCGAGAGGTACAAGAAAACATTGGAAGAAGCACTTCATCTTATGTCAATCTTTGCGGTTTCAGACATATTTCCAAGTCTAGAGTGGGTAGATCGATTAAGAGGCCTTGTAAGGGATATGAAACGCTTTGGAGATGAGCTAAATTCGATTGCAGGGTGTCTTATTGAAGAGCACCGCCAAAAGAGATTACAATCCGTATCTAAAAGTGATAAAGGAGTTGGTGATGAACAAGACTTCGTTGATGTTCTCTTATCGGTTGCTGAACAATCGCAACTTCCTGGAGATGACCCTGATTTGGTCATCAAGTCTATGATTCTGGAAATCGTATCAGGTGGGAGTGAGACCACATCGTCAACCTTAACTTGGGCCCTCTGTCTGTTACTGAACCATCCGCATGCGTTAAAGAAGGCAAAAGAGGAATTAGATACGCACGTAGGAAAAGATAGGCATGTAGAAGAGTCAGATACCCCTAAGCTCGTGTATATTAATGCAATTATCAAAGAATCAATGCGATTGTATCCAAACGGGGCAATGCTTGATCGGTTGGCGTTAGAAGAGTGCGAAGTTGGTGGATTTCATGTACCGGCCGGGGGACGCTTATTTGTCAATGTTTGGAAGATTCAGAGAGATCCGAGTGTTTGGGAGAATCCTCTGGAGTTTAAACCAGAGAGGTGGTTTTTGAGTAATGGTGAAAAGATGGATGTGGATTACAAAGGTCACAATCATGAATTCATACCATTTGGGATAGGTCGGAGGATGTGCGCTGGTATGCTTTGGGCATCGGAGGTGATTCATTTGGTGCTGCCCCGTCTTATTCATGGGTTTGATATGAAAGCAGCAAGTGCCAATGGGAAAGTAGATATGGCGGAAATGGCAGGCATGGTGATTTGTTTTAAGAAGACACCTCTTGAAGTTATGGTCAATCCTCGAGAGTAGATGTTTTTATCCTTTTGCATATATAGTTTTTATATTGAAAATGTAGGCCAAGCGAGCAACATGTCGTACATTTGTTTCTTATCATACGAGTACTTTGATGCCTTTTTGCTGGTAACACATCAGAGTGACGAAGTTAGATGCTGATACAGACGGAGCCAGGGATTTTTGTCTTTGCGGACAAAATGCAAGGGTAAAGCAGTAATGCCGTAAAAATTGCGGCCGCAGGCTGCAAATTTTTTAAAGCCTTTAATTTTGTTTTGTTTTGCTTCAAATGTGTCGTTGTAAAACGTTTTAATTAATTGTTTTGATACTTTTCATTTGAGATCGGAA

>Ptr_TRINITY_CYP82X2 Ptr_c41184_g1_i2 len=2034

AACAACCTCAATCTATTCTAGCTAGAGTCGTGAGACTGTTATTTAGTTAGTTAAAAAAATTTCAATGAACAAGTTATTATTTCTCCAACGGGTTACTGTTTCTCCTTCGACCACCATTATCAGTACTTTTATTGTTACAATAATATCCATTGTTTTTCTCTACATTGTCTTGTTGATAAAGACGACTAAGAATAAGCAGAAGACAGGAGCACCAAAAGCATCGGGGGCGTGGCCGTTCATAGGTCATCTCAAACTATTCATGAAACAAGATACTCAGTTTTACAAAACTCTAGGAACCATGTCTGATAAATACGGGTCGGTGTTCACACTTCTATTAGGCAACCAAGAAATCCTAGTTGTGAGCAACTGGGAGATGGTAAAAGAATGCTTCACAACAAATGACAAGTCATTTTCGAATCGTCCAAGTACGTTAAGCACTAAATACATGCTGAATGACACCAATTCTGTTGTGTTTTCACCTTATGGAACGTATTGGAGAGAAATGCGGAAGATACTGGTGCAAAAACTACTGATCTCTAACCAAAGATCAGAGGCATTGAAAAATTTGAAAACGAAAGAAATCGACAACTCGTTTGTGAAGCTCAATGATTTATGCAGCAACGATGTTAGCGGAGGCACAAAAGTTAGGATGGACCAATGGTTGGCTGACATGATGTTCAACATTATTGCTAGGATTACATTTGGTTACCAAGGCGGAGGAGGCGATGCACCTGGCGCTTCTACAACATCCAAGAATGTCGAGAGATACAAGAAAACGTTGGACGAGATGTTTGTTGTTTTAGCGACGAGGTTTGCAGTTTCAGATATATTTCCATCTCTGGAGTTTATAGACCGATTGAGAGGTCTTGTAAAGGATATGAAAATCTTGGGAGACGAATTAAACTCAATTGCTGGATGTTTTATTGAAGAACATCGTCAAAAGAGACGAGAATCATTATCCTCATTGTCATCGTTATCAAATGAATCCGTTGGTGATGAACAAGATTTCATTGATGTTCTGTTGTCAATAATGGATCAGTCACGGCTTCCCGGAGATGACCCAGATTTTATTATCAAAATTATGATCCTGGAAGCTTTTGCAGGTGGGACGGACAGTTTAAGTGCAACCTTAACTTGGGCCCTCTCTCTACTGCTGAACCACCCAAACGTGTTAAAGAGGGCAAGGGAGGAAATCGATAGGCATGTGGAAAACGGTAAGCAAGTGGAAGCCTCTGATATTGCTAAGCTCGTATACATTTATGCAATAATCAAAGAGACGATGAGATTGTATCCAGTCGGAGCATTAAGCGAACGCTATACGACTGAAGAATGCGAGGTTGGTGGGTTTAATGTACCCGCCGGCACACGCTTACTGGTGAATATATGGAAGATCCATAGAGACCCAAGTGTATGGGAGAATCCATCAGATTTTCAACCAGAGAGGTTTTTGTGCAGCGATAAGGTGGGTGTGGATTTATATGGCCAGAATTATGAGTTGATACCATTTGGGGCCGGTAGGAGGGTATGTCCGGCTATAGCTTCATCACTGCAGACGATGCATTATGCGTTGGCGCGTCTTATTCAAGGATATGAAATGAAATCAGCCAGCCTCGATGGGAAGGTGAATATGGAAGAAATGATAGCCATGTCGTGCCACAAGATGAGCCCTCTCGAAGTTATTATCAGTCCTCGGGAGCTGAGGCGGAGTTAAATCTTATGTTCCAAATTTACATTAGCATCTTTGATTATGAAATGTATTGCTCTTGACTTTCTTTTTTGTTTTTTATAGTTTTAAGCTTGTATGTGATGATCAGAGAAAATGATGATGACTGAATCATCAAAAGAATCAATCGAAAAAGCTATGTTGTATTAGGCGCGCCGAGGAAAAATTAAAAAACTGCAGGAATAGGTATGATTTGGCGCCTTGGGCGCCTTTTTTTCTAGGCGTTCCCTAAGCGTCTCGCTTGCCTAGAACACTTAGGCGGGCGCCTAGGGCGTCTTTTACAACATAGAGAACAAGCAGATCGGAAGAGC

>Ptr_TRINITY_PSAT1 Ptr_c37012_g1_i4 len=1691

TATCGACCTTGTGGAACAATTAGGGGATATGCCAACAATGTCTAACGCTGCTGTACAAGTGATCTCGAAAGAAACGATTAAACCAACAAATCCAACACCATATCAACTTAGAAACTACAATATGTCACTTCTCGACCAATATTCTTCTCTAGTTTATGTTCCGATCATTCTTTTCTACCCTGCTGCCTCCGACGCTAATAGTACCGGAAGTAAGCACCGTGATGATCTTCACTTGCTTAAGAGGTCTCTTTCTGAAACGCTAGTTCACTTTTATCCAATGGCTGGTAGGATGAAAGACAACATGACTGTTGACTGTAACGACGAAGGTATCGACTTTTTCGAAGTAAGAATCAAAGGTAAAATGTGTGACTTCATGATGAAATCAGATGTACACTTGAGTCTGCTTCTTCCGTCTGAAGTCGCTTCCATGAATTTCGTGAAGGAAGCCCAGGTGATTGTTCAAGTGAACATGTTTGATTGCGGTGGAACTGCCATTTGTTTCTGTATATCAAACAAGATTGCAGATGCATGCACCATGATTACCTTCATTCGTAATTTGGCAGGCACCACCAACAAAGCTCGTCGTGGGAGCTCTACTGCCGCACCAACCACAAATCAGAAGTTGCTTCCTTCTTTCGATTCGGCATCACTCTTCCCACCTAGTGAACAATTGAAAACTCAAGTTTCCTATCCTACACAGGATAGTACCAGCGTAGATAAACTTGTCAGCAAAAGATTTGTGTTTGATGCGGCAAAGATTACATCTGCACGTGAAAAATTGCAATCCTTGATGCATGATAAATACAAATGCCATAGGCCGACAAGGGTTGAGGTAGTTTCCGCTTTGATATGGAAGTCAGCAGTGAAATCCGCTCCGCCCGGTTCTATATCCACTGTAACCCATGCCATGAACTTTAGAAAGAAAATGGATCCACCGTTACAAGATGCGTCATTCGGGAATCTTTGTGTGGTTGTTACAGCAGTATTACCAGCAACAACGGAAACAACAACAAATCCGGCAACCAAAACAGTTAGTAGTACGAGTAATGAAGAGCAAGTGGCACTTGATGAGTTAAGTGATTTTGTAGCCCTATTGAGGCGCGAAATAGATAAGGTAAAGGGTGATAAAGGTTGTATGGAGAAAATCATTCAAAATTTCATCCATGGTCATGATGCTTCCGTAGGGCAAGACAGTGATGTTGAAGATAAGGTGATAGCTTTGTTTATGACTAGCTGGTGCAAGTTTGGAATCTACGAAGCTGATTTTGGTTGGGGAACGCCAATTTGGGTAACTACTGTTCCATTAATTGAGCCAAAGTACAAGAACATGGTTTTCATGAACGATACGAAATGTGGTCAAGGAATTGAAGTGTGGGTGAATTTTCTGGAGGATGATATGACCAAGTTCGAGCATCACCTAAGCGAGATCCTCCAACTGTTATGATTTTCAACCATTTCCTTAATAGAGGTCAATTGTCGTGTTTGTCCTTCCCCTTAACTTTACCATCTTTATTTTTACGTGGTCATTGGGTTTACAAACGGAAGACTTTGTGTTGGCTTTTTTCCTCCGAAAATGCTATATGAACAGCTTTGTAACAGCTGATTAACAGCAAAATCAGACGTGGCATAAAACATAGCTATCCACTGTGGCTTTTCCTTTTACCACGTCGTCACAAAAATGAAAAGATCGG

>Ptr_TRINITY_PSMT2 Ptr_c2782_g1_i1 len=1408

GTGCTCTTCCGATCTATTCAATCAGAAACTTTCTTCTAAAATCTTTCAATACCAGTACTGTTAGTTTCCGAGAAGATTCTGCCACACTAATCCATTATGGAAATACATTTAGAAAGCCAAGAACAAGAAATGAAATATCAAGCTCAAATTTGGAACCAAATATGTGGCACCGTTGATACCTCTGTTCTGAGATGTGCAATTCAATTAGGTATATTTGATGCCATTCATAACTCAGGCAAAACAATGATTACCTTAACGGAATTATCAAGCATTATTTCATCATCCTCTTCATCTTCAATTGAACCCTGCAACTTATATAGATTAATGAGGTACTTATCCCATATGGATCTCATCGATATCGGAGAATGTTTGAATGAAGCAACTGTTTCACTAACAGGCACATCTAAGTTACTACTCAGAAACCAAGAAAAGAGTTTAATTGATTGGGTATTGGCAATTTCTTGCGAGATGATGGTTGTTGTTTGGCACGAACTAAGCAGCTCTGTTTCAACTCCTGCGGGTGAGCCTCCAATCTTCCAGAAGGTTCATGGCAAAAACGCTTTAGAATTAGCAGGGGAATTTCCAGAATGGAATGATCTGATCAACAATGCTATGACTAGTGATACTAGTGTAACTAAGCCAGCGCTAATACAAGGATGTGGCAAAATCCTGAACGGAGTTACATCGTTGATTGATGTCGGTGGTGGTCACGGTGCCACTATGGCCTACATAGTTGAAGCTTTTCCTCACATAAAAGGTGCGGTAATCGATTTACCACATGTTGTTGAAGCCGCTCCGGAGCGTCCAGGTGTTGAGTTCATCAGCGGTGATATATTCAAGTCTATTTCTAACGCTGATGCTGTGTTGTTGAAGTATGTCCTGCACAATTGGGAAGATACGGAATGTGTGAATTTACTGAAGAGATGCAAGGAAGCAGTTCCGGCAGACAACGGAAAAGTGATCATAATGGATTTAGTAATAGACGACGATGATGACAGTATTTTAACGCAGGCGAAATTGAGCCTTGATCTCACTGTGATGAACCATGGAGGAGGTAGAGAAAGGACTAAAGAAGATTGGAGAAATCTAATTGAGATGTCTGGATTCAGTAGGCATGAAATAATTCCAATATCTGCCATGCCATCAATTATTGTTGCTTATCCCTCCTAGTTAAGTCACCCACATGATCGGGAATAAGTTAGGCGCGTGTTGAATCTGTTGACATCGCAATTGTGCCTTTACTTTATGCATTTTCATTCCAGTAAAAACTGTTTTTGGGCATTCGGATTCTTCTGAGCCCTTGTATGTTTGTTTGTTTGTTGGTTGGTTTTCAAGTAACTGAAGATTCTTCTCTGTTTTCATGGCATGTGCATGTGACTAATTTGAGTATGCATATTGATGTTTCGTT

>Ptr_TRINITY_CYP82Y1 Ptr_c34673_g1_i1 len=2103

ATAATTTCAGTGCATCCATGGCGTCTCTGATGATCAAGAAATCTATCCATCTGTTTTTTGATCAGCCAACTTCAGTTGCCACTCTTATACTTGCTTTGCTGCTAACACTTTCGCCTGTTATTATTTACTATGAACAGAAGAAGAGGGATTTGAGGCGAAATCGCACCGCAATTACAACGACTCCATTACCAGAGGCATCAGGTGCATGGCCAGTGATTGGTCATCTTCTTCTTTTCATGAACGAAAACGATCTAAATCATGTAACTCTTGGTCACATGGCTGATAAATATGGACCTATTTTCAGCTTAAGATTCGGTAGCCATAGAACTCTAGTTGTTAGTAGTTGGGAGATGGTAAAGGAGTGTTTTACAGGTACCAATGACAAGTTGTTCTCAAATCGTCCTTCCTCCTTGGCGGTTAAACTTATGTTTTATGACACTGAGTCTTATGGTTTTGCACCTTATGGGAAATACTGGAGAGAGTTGCGAAAGATATCTACACATAAACTCCTCTCTAATCAGCAATTAGAGAAGTTCAAGCACTTGCGGATTTCTGAAGTTGACAACTCCTTTAAAAAGCTTCATGAGTTATGCAGCAACAACAAACATGGAGGTGATACTACATCTGCGGCTAGCCTTGTGAGAATGGATGATTGGTTCGCGTACTTGACATTTAACGTAATAGGACGGATCGTTAGCGGATTCCAATCAAATGCAGTGGCAGGTGCCACAAACAGCCAGGAAAAATACAAGCTTGCAATCGATGAAGTGTCAAATCTTATGGCAATGTTTGCCGTTTCAGATGTGGTCCCATGGCTTGGGTGGATTGATCGATTGACTGGTCTTACAGGAAAGATGAAGAATTGTGGGAAAAAATTAGATGCAGTAGTTGGGGATGCAGTGGAGGATCATCGCCAAAAGAAACTCAAAATTTCTAGAAATACCGCAGGAGCACTTATGGAGCATGAAGAAGAAGACTTCATCGATGTTTGCTTGTCAATTATGGAGCAATCACAGATTCCGGGAAACAACCCCGAAATCTCTGTCAAGTCTATTGCCTTGGTAATTCGTCTCATAAGCATGTTAGCAGATTTTACCTCTATATATACTTACATATTATTTTTTATCAATCACATATGTGCAGGACATGTTATCGGGTGGGAGTGACACTACAAAATTGATAATGACATGGACCCTTTCTTTGCTGTTGAACCATCCGGACATATTGGACAAAGCTAAAGAAGAAGTACTTACATACTTCAGGAAGAAAAGGATAGCGGATAACACACCCGTGGTTGATGCTGCCGATGTTCCTAACCTCCTCTACATCCAAGCAATCATCAAAGAATCAATGCGGTTATACCCTGCCAGCACATTGATGGAGCGAATGACAAGTGACGATTGTGAGGTTGGTGGCTTCCACGTTCCAGCTGGGACGCGATTATGGGTTAACGTATGGAAGATGCAACGGGACCCAAGGGTGTGGAATGATCCATTGGTATTTCGACCTGAGAGATTCTTGAGCAATGATAAAGAGATGGTAGATGTGAAGGGTCAGAATTATGAACTGATACCATTCGGAACAGGCAGGCGGATATGTCCTGGTGCATCTTTTGCCTTGGAAGTCTTGCATTTGGTTCTCACTCGTCTTATTCTTGAGTTCGAGATGAAGGCACCAGAGGGGGGAATTGACATGAGGGCAAGACCAGGTTTTTTCAACAACAAGGTAGTGCCACTAGATGTTCTACTCACCCCACGCACACTAGGTTAAGATTTCTGTATATGCTAATTAAATAAGATAAATAATGTTTGTTGTCGAGTAATTTTAGTTAATGTTAATTTGTGAGTAGTTAAGTAGTACTATTATAGTTGTCACTGTCAGCATCAGTAATTTGCTTTCTTCTTCTTTGGTGATTCAACAATAGCCTGTTTAGGAGTTGCGCACTCCCTTTGATGCTGGGTCTTTCTGTTGTTGATTGGTTTAGGAGTTGTACACTCCCTTTGTTGTATTGGTTGTCTCTGGTTAGGCTTTTGCCTTCCGCTTTGTTTCGCTTAGGCGCCTCTTTCCCTGTTTTCTTTTTTCTTTTTTTTTTTTCCTAACAGT

>Ptr_TRINITY_PSMT3 Ptr_c98941_g1_i1 len=494

GTGCTCTTCCGATCTAAAAGAAGAGAAAATTTTCTAAAGAGAGCTAAAAAGAAGAGAAAAAAGATGGAAGTAGTAAATAAGATTGATCAAGAAAACCAAGCAAAAATTTGGAAACAAATTTTTGGTTTTGCAGAATCATTAGTTCTGAAATGTGCAGTTCAGTTAGAGATTGCTGAAACACTTCATAATAATGTAAAACCCATGTCTTTATCCGAGTTAGCATCTAAACTTCCGTCTCAACCCGTTAATGAAGACCGTTTGTACCGAATTCTGCATTTCTTAGTTCACATGAAACTCTTCAACAAAGATGCTACCACACAGAAATATTCATTAGCTCCACCAGCGAAGTATTTGCTAAAAGGCTGGGAAAAATCAATGGTTCCTTCAATATTAAGCGTGACCGATAAAGATTTTACAGCTCCATGGAATCATCTTGGTGACGGTTTAACAGGTAACTGTAACGCTTTTGAGAAAGCGTTAGGAAAGAGCATTGG

>Ptr_TRINITY_CYP719A21 Ptr_c40355_g1_i2 len=1625

CGATCTAAAGAGTTAATCTGACTCAAGCTAGCTAGAATATATACCAACAATACAAGAAATGATCATGAGTAACTTATGGATTCTTACGCTCATTTCTACCATATTAGCAGTCTTTGCTGCTGTGTTAATCATTTTCAGGAGAAGAACGTCAGCATCCACAACGCAATGGCCTGTTGGCCCAAAAACATTACCAATCATAGGTAACTTGCACATTCTTGGAGGCACTGCTCTCCATGTCGTCTTACATAAACTAGCTGAAGTTTACGGCAGTGTAATGACGATATGGATTGGTAGTTGGAAACCTGTTATTATTGTTTCCGACTTTGATCGAGCTTGGGAAGTTCTTGTTAACAAATCGTCAGATTATTCAGCACGTGAAATGCCTGAGATCACTAAAATCGGCACTGCAAATTGGAGAACAATTTCAAGTTCTGATTCTGGTCCATTTTGGGCCACTCTTCGGAAAGGTCTTCAGGGTGTAGCATTATCGCCTCAGCATTTAGCATCGCAAACTGCACACCAAGAGAGAGATATAATAAAGTTGATCAAAAATTTGAAAGACGAAGCAGCTCTTAATTCTGGAATGGTTAAACCACTTGATCATCTCAAGAAAGCAACTGTAAGATTAATCAGTCGGTTAATCTATGGTCAGGATTTTGATGACGATAAGTATGTTGAAGATATGCATGACGTGATCGAGTTTTTGATTCGTATTAGTGGTTATGCTCAACTTGCTGAGGTATTCTATTATGCTAAATATCTACCAAGTCATAAGAGAGCTGTAACTGGCGCTGAAGAAGCAAAAAGAAGAGTAATAGCTCTGGTGCGTCCTTTTCTTCAGTCAAACCCTGCTACTAACACTTACTTGCATTTTCTCAAATCGCAACAGTACCCTGAAGAGGTTATCATATTCGCTATTTTCGAAGCTTATCTTTTAGGTGTTGATAGTACTTCGTCAACCACTGCATGGGCACTCGCATTCTTAATACGTGAACCATCTGTTCAGGAGAAACTTTATCAAGAGCTTAAGAATTTCACAGCCAATAATGATCGCACAATGCTGAAAGTCGAAGACGTCAACAAATTACCATATTTACAAGCTGTTGTTAAAGAAACAATGAGGATGAAACCAATTGCACCACTGGCGATTCCTCATAAAGCTTGTAAAGACACTTCATTGATGGGCAAGAAAGTTGATAAGGGAACTAAAGTTATGGTTAACATTCATGCTTTACATCATACTGAAAAGGTTTGGAAAGAACCTTACAAATTCATGCCAGAGAGGTTTCTGCAAAAACAAGATAAGGCGATGGAACAATCACTATTACCATTTAGTGCAGGTATGAGAATTTGTGCAGGAATGGAATTAGGAAAACTTCAGTTTAGTTTTTCTCTTGCTAATCTTGTTAATGCTTTTAAATGGTCTTGTGTTTCTGATGGAGTGCTTCCTGACATGAGTGATTTACTGGGGTTTGTTCTGTTCATGAAGACCCCACTCGAAGCACGTATAGTTCCTCGTTCGTAGTGATGGAAATTTCATCCCATGTTTTTGTTTTTCTTCATGTTTATTACTTTGTACTGGTTTGATTTGGTGTGAAGATAAGATCTAAATTCCCAGATCGGAA

>Ptr_TRINITY_9OMT Ptr_c36508_g1_i1 len=1495

GCTCTTCCGATCTCTTGATCATTGTCATCAAAAACAGTCCTAAGATGGCTACCAATGGCGAAATTCTCAATACCTATGGTCATAATCATCAAACAGCCACAGTCACTAAAATTGCTGCTTCTAATGAAAGCAACAACGATATCTGTTATCTTTCAGAAACGGCTAACTTAGGGAAGTTAATATGCATTCCAATGGCATTAAGAGCTGCTATGGAGCTAAATGTGTTCCAACTTATCTCCAAGTTTGGAACTGACGCAAAAGTTTCGGCTTCTGAAATTGCCTCTAAAATGCCAAACGCTACGAATAATCCTAAAGCAACTATCTATTTGGAAAGAATTCTTCGACTGCTCGGTGCCAGTTCTATTCTTTCGGTTTCTACCAGAAAAAAATTAAACGGAGGAGGAAATGATGGAGTAGTACATGAGAAGGTGTATGGGTTAACAAATTCGTCATGTTGTTTGGTCCCTCGAGAAGAAGACGGGGTGTCATTAGTTGAAGAATTGCTATTCACATCTGACAAGGTTGTTGTGGATAGTTTTTTTTAAACTGAAATGTGTGGTGGAAGAAAAAGACATTGTGCCATTTGAGGTTGCTCATGGTGCTAAGATCTTTGAGTATGCTGCTACAGAACCAAGAATGAACCAAGTTTTTAACGATGGAATGGCAGTTTTCTCTATTGTTGTTTTTGAAGCTGTCTTTAGAGTTTACGATGGATTTCTTGGTATGAAAGAATTGTTAGACGTTGGTGGTGGTATTGGTACATCGGTTAGTAAGATTGTTGCTAAATACCCTTTCATTCGAGGTGTCAATTTCGACTTGCCTCATGTTATTTCTGTTGCCCCTCAATACCCAGGTGTAGAGCATGTTCAAGGAGATATGTTTGAGGAAGTTCCAAAGGGGGAAAACATGTTGCTAAAATGGGTACTGCACGATTGGGGCGACGAACGATGCGTGAAGTTGTTAAAGAACTGCTGGAATTCATTACCTGTGGGTGGAAAAGTTTTGATAATCGAGTTTGTTCTTCCGAATGAACTTGGTAACAATGCCGAATCATTCAATGCGTTGATTCCCGATCTACTTATGATGGCTCTGAATCCAGGCGGTAAAGAACGAACAATTTCCGAGTATGATGCTTTAGCCAAAGAAGCTGGATTTATGAAAACTATACCTATCCCAATCTCCAATGGTCTTCATGTCATTGAGTTTCATAAATGATCAAGATGGTTATCTGAGTGTCATCCGTAATTTAACTAGTTACTTGGTTTTGTAATGAAATTACCAAGATAACTGTTTCCATTTCATGCATTTGGTGTGTTCTTCTTCTTCTTTTTTCTTTAAAAAAAATTGTATTTTGAAGGTTTGAATCAGTGTTTTAATAGGGGAAATTAAGTATGTTGTTTCGTGTTTCTTGATGAGATGATGGACCAATGATGTGCTTAATGTTTGTGAATCCCAGCTAGTATTAATAGCTTCTGGAATGGAATGTGTGTGAAGA

>Ptr_TRINITY_SALSYN Ptr_c34457_g1_i1 len=461

GAGCGTCACAGCAGAATGGAATTTTAAAACCTCTTGATCATGCCAAAGAAGCGTCTATTCGATTACTGAGTAGAGTTATATTTGGTCAAGACTTTTCAAATGAGGATCTCGTTATTGGTGTGAAAGACGCCCTCGATGAGATGGTACGCATAAGTGGGTTGGCAAGTTTAGCTGATGCTTTTAAAATTGCTAAATATTTACCAAGCCAGAGAAAAAATATTCGGGATATGTACGCCACAAGAGACAGAGTATATAGTTTGATTAAACCACATATCGTCTCCAATCTTCCTGCAAATTCTTTCTTACATTTTCTTACATCTCAAGATTACAGTGATGAAATTATTTACTCAATGGTACTTGAAATTTTTGGTTTGGGAGTAGATAGTACTGCAGCAACGGCAGTTTGGGCACTCTCCTTTTTAGTCGGCGAGCAGGAAATTCAAGAAAAACTTTACCGCGAA

>Ptr_TRINITY_CFS Ptr_c13501_g1_i2 len=926

TGATTATAAATTATGTTATCTAAAGAAGAAGAAGCAAGCAACCCACACCTTCAATTTTCTTCTCTCTCCATTCATCTTGTTTTGATATTTGTTCACAAAGAAAAATAAAAATTAGAGTACTGTTTTTGTAAAATGGAGGAGACCTTTTGGCTGATAACTTGTGGTGTGTTGGTAGCTTTTGCTTCAGCAAAATTACTATTTGGCAAGAAATCATCTCTGTCTACAATGGAATGGCCAGCAGGTCCAAAGAAGTTGCCCATTATTGGGAACCTTCATCAATTAGGAGGTGAAGCGTTTCATGTTTGTATGGCAAACTTGGCCAAAGTTCATGGAAGTGTTTTTACCATATGGCTTGGAAGTTGGCGTCCATTTATAGTCATAAACGATGTCGATAAAGCATGGGAAGTTCTTGTTAATAAATCTTCAGATTATTCAGCGAGAGATATGCCCGATATTACTAAAATTATTTCAGCAAATTGGAAGAATATTTCTCATTGTGATTCTGGTCCATTTTGGCATAATTTAAGAAAAGGTCTTCAAAGTGTTGCTCTAACTCCTTTTAATGTTGCGTCTCAATATCATTTGCAAGAAAGGGACATGCAAAATTTGATCAAATCTATTCAAGAAAAAACGTCGCACCAAAATGGGACCTTGAAACCACTTGATTATGTAAAGGAAGAGACTGTTCGGTTACTAAGTCGACTTATATTCGGTCAAGATCTCGTAGACGAGGATTTCGTTGTTGGTATGCATCAAGCCCTCGACGAGTACATCGAAACTATAAGTAAGGCAAGTCTAGCTGATGTTTTCAAATTTTGTGAGAATCTATCAAGCCATAAGAAAACTATTCGAGCAGTTCATGCTATCAAAAAGCGAATCGACAGTTTGATTCGCCCACACATTGTTTCTAAGAGATCGGAAGAGCG

>Ptr_TRINITY_MSH Ptr_c39855_g1_i3 len=1447

TCTCGAGACTATAAAAAACTTAAGGAGTTCTGAAGTCGAGTCATGTTTTGAGTCGCTTTATAATCAGTGGAGAAACAGAGAAAATAGTGTCGAAAAGTCAACTACGGTGAGAATGGATAGTTGGTTAGACGATTTGACGTTTAACGTTGTGGCGAGAATCGTTGCGGGGAAGAGGAAATTTTCGGCTAATGGAGATGTTGGAGCTCAGAGATATAAAGCAGCAATGGATGAAGCCACGAGATTAACGAGATTTTTCGCATTTTCCAATGTAATTCCGTCGCTAAGCTGGTTGGACAATTTAAGAGGTTTGGTGCGAGATATGAAAAAATGTGCATCTGAAATTGATACAATTATGGCTTCTTGGGTTGAAGAACATCGTGTGAAAAGAAATTCCGGAGGTAATTCTCAACTCGAACATGATTTTATTGATGTTTGCTTGGATATTATGGAGCATTCTTCGTTACCCGGCGATGATCCTGACCTCGTCGTCAAATCCACGTGTCTGGACATGATATTGGGTGGGAGTGACACCACAACAGTTACCTTAACATGGGCAATGTCTTTACTCTTGAACCATCCCCAAGTACTACAAAAGGCCAAAGAAGAACTAGACACACAAGTTGGAAAGAATCGGCAAGTCGACGACTCGGATATCCCGAATCTTCCGTACATCCAAGCGATCATCAAAGAGACGATGCGGTTGTACCCGGCCGGACCACTAATTGAACGGAGGACGATGGAGGACTGCAAAGTAGGTGGGTACCAAGTTCCGGCCGGCACGCGCTTATTGGTTAACGTCTGGAAGATGCAAAGAGATGGGAACGTATACAAAGGCGATCCGTTAGAGTTCAGACCCGACAGGTTTTTAACTAGTAATGCGGATGTGGATCTAACGGGTCACCATTACGAATTGATACCATTTGGGGCGGGTAGACGGATATGCCCGGGAGTATCGTTTGCGGTTCAGTTGATGCATTTGGTTCTTGCTCGATTACTTCACGAGTTCGAGATCACGACGGTGGAGCCAGAAGCGAAGGTGGATATGTCTGAAAGTGGTGGCTTGCTTTGTTATAAGATTATGCCGTTGGAAGTACTGATTAAACCTCGACTCGAGATATGATCGAACGAAAACCTCAACATAATAAAAGCAAGGCCGTCTATAACCCGTTTATAGTACTGAATGGTCCAATGTATGGGATTGTGAGACTAGTAGGTCTTGCAGAGTATTAGTAATGTGATGTACTTGTTTTCCTTTTGAGTCTTGATTTTGGGTCCAAATGTGCTGTAATCAACCCCATTTCTCAAGTTCCGTGCTAAATACCGGGGACTCAATTTTCATGTAATCTGTTGCTTTCTTTTCTTCTACCTATGGTAGAAATTCTGTACATTCTCTTACTTCTTAATACAACTTCTCTTTTGATCAAAAAAAAAGTTCGGATGAGCTCCT

>Par_TRINITY_NCS Par_c27622_g3_i1 len=1717

TGGTGTTGGTACAGTTCTTGATATTGTCTTCCCTCCAGGTGCGGTGCCTCGAAGTTACAAAGAGAAATTCGTAACCATCGATCGAGAAAAGCGATTGAAAGAGGTTATTATGATCGAAGGAGGATACCTGGACATGGGATGCACATTTTACTTGGACAGGATCCATGTAGTGGAGAAAACCCCGAGCTCATGTGTTATCGAATCGTCTATTGTGTATGAAGTGAAAGAAGAGTATGCTGATGCCATGTCAAAATTGATCACAACTGAACCGTTGAAATCTATGGCCGAGGTCATCTCTAATTATGTTATCCAGAAAGAATCATTTTCCGCAGGAAACATTTTCAACAAAATTTCTTTGGTGAAGAAGGAGATTCGATACGATTTGGAGGTACCAACCTCAGCCGATTCTATCTGGTCAGTTTACAGCTGCCCAGATATCCCTCGGCTTCTTAGAGACGTTCTGCTTCCCGGTGTCTTTGAGAAATTGGATGTTATCAAAGGGAATGGCGGTGTTGGTACTGTTCTAGACATTGTTTTCCCACCAGGTGCGGTGCCTCGTCGTTACAAGGAGAAATTTGTGAACATCAACCATGAGAAGCGACTAAAAGAGGTGATTATGATCGAAGGAGGGTACTTAGACATGGGATGCACATCTTACCTGGATAGGATCCATGTGGTTGAAAAAACCTCTAAATCTTGCATCATTAAATCGTCTGTTGTCTACGAAGTGAAGCAAGAGTGTGCTGCTGCCATGTCTAAGTTGATCACCACCGAACCATTGAAGTCGATGGCAGAAGTCATCTCTAATTATGCTATTAAGCAACAATCTGCTTCTGAGAGAAACATTTCTAAGAAGCAATCTCTACTAAGGAAAGAAATTACTTATGAAACGGAGGTGCAAACTTCGGCTGATTCGATTTGGAACGTCTACAGTTCTCCTGACATCCCTAGGCTACTCAGAGATGTTCTGCTTCCTGGTGTTTTTGAAAAGCTAGATGTCATTGCAGGCAATGGTGGCGTTGGTACTGTACTGGACATTGCTTTCCCTCTAGGTGCAGTGCCGCGGAGGTACAAGGAGAAATTTGTGAAGATCAACCATGAGAAGCGATTGAAAGAAGTGGTTATGATCGAAGGAGGATACTTAGACATGGGGTGCACATTTTACATGGACAGGATCCATGTATTAGAGAAAACCCCTAACTCATGTGTTATTGAATCCTCTATCATTTACGAAGTTAAAGAAGAGTATGCTGATAAAATGGCTAAGCTAATCACAACCGAACCATTGGAATCCATGGCAGAAGTCATCTCTGGTTATGTTCTTAAGAAACGACTCCAAGTATTCGGATTCGAGATTAAGCCAAAACTAAGATTCAATCTTTTGCTATGTTTGATTATCTGTTTGGTTATAGCTGGAGGTATGTTTGTTGCTCGTGTTCCACTCTAAGGTTAAGGGCTTTTCTTAAATGTAATTTTGGAAATAAAGTATTGGTTTGATCCTTAAAACCATGCGTTGCATTGAAGAGGTTTCTCATATGTAATTCTCAAATCTACAAAATAAAAAAAACAAATTTAATAATGTTTCAAGCAATGTTTGAAAAACGGGTTTAGAATGTAATAGTAATTTCCGGCAATGTTTGATAAACTTTTCTCAGAAAGAGAAATTGTACTTGGACAAAAAATAAACATACTTTTTTGATTGTTTATCTAAAAAAAA

>Par_TRINITY_6OMT Par_c22934_g1_i1 len=1357

GTACAAAGCCCGTGTTTTATAGTGCTCATAGAGAATATAGAGGATTACTAATTATTATATAAAAGGGTCAAAAGATAGCACTAGAGAATAGCTCAAATCATTCATTCATTCTCAATCTCATCAACAGCTAAAGTTTCTAAAAAGAGAGAAATGGAAACAGTAAGCAAGATCGATCAACAAAACCAAGCAAAAATCTGGAAACAAATTTACGGTTTTGCAGAATCACTAGTTCTGAAATGTGCAGTCCAATTGGAAATTGCTGAAACACTTCACAACAATGTCAAACCCATGTCTTTATCCGAATTAGCATCAAAACTCCCCGTTGCTCAACCCGTTAATGAAGACCGTCTGTTCCGAATCATGCGTTACTTGGTTCACATGGAACTCTTCAAAATAGATGCCACCACACAGAAATACTCATTAGCTCCACCAGCTAAGTATTTGTTGAGAGGCTGGGAGAAATCAATGGTTGATTCAATTTTATGCATAAATGACAAGGATTTCTTAGCTCCATGGCATCATTTAGGTGACGGTTTAACCGGTAACTGTGACGCTTTTGAGAAAGCGTTGGGGAAGAATATTTGGGTTTATATGAGTGAAAATCCTGAAAAGAATCAATTGTTTAATGCAGCAATGGCTTGTGATACTAGATTGGTTACTTCTGCATTGGCTAATGAGTGCAAAAGTATTTTTAATGATGGAATCAGTACACTAGTCGATGTCGGTGGCGGTACCGGTACTGCTGTGAAAGCCATATCTAAAGCATTTCCGGATATTAAATGCACTATCTATGATCTTCCTCATGTCATAGCTGATTCTCCTGAAATTCCCAATATCACTAAAATTTCTGGAGATATGTTCAAGTCTATTCCTAGTGCTGATGCCATCTTCATGAAGTGCATACTTCACGACTGGAACGACGAGGAATGCATTCAAATCTTGAAGAGATGCAAAGAAGCATTGCCAAAAGATGGCAAAGTTATTATCGTGGATGTCGTAATAGACATGGATTCGACTCATCCGTACGCCAAAATTAGACTCACATTGGATTTGGATATGATGCTTAACACTGGTGGAAAAGAGAGAACCAAAGAAGAATGGAAGAAACTTTTTGATGCCGCTGGTTTTGCTAGCCACAAAGTCACTCAGATATCTGCTGTCCAGTCTGTAATTGAGGCTTACCCTTATTAAGGAACATTTTACCCGGTTTTCCCTTTGATTAATTGTTATTGCTTTATCTTTGGATTATGTTTGTGCTTATGATAACTGCAAGATGAATGAATTTCCAACTTGCACTGAATTTGAATCAATGTTTTTCGTTTACTTAACTTTGTGCTATTTTGGGAGAGATCGGAAG

>Par_TRINITY_CNMT Par_c37271_g1_i1 len=1352

GCTCTTCCGATCTATCAACTTATAAAAAACCCACATCAGTGGTAGTAGTCAGTCATCAGTACAACCACTACCACCACCTCTGTCTGAACTTCTCTGGAAAATCTATAGACTTTGTGTGTGTTTTCTCCTTTGGAAGTTAAATAAGATGCAGCTAATGGCAAAACAAGAGCTGTTGAAAAACATGGAGCTTGGGTTGATACCCGATGAAGAGATTAGAGAAAGGATTAGAATCGAATTGGAAAAGCGTCTCCGGTGGGGTTACAAAGAAACCCATGAAGAACAGCTTTCACAGCTTCTTGAATTAGTTCACTCTTTGAGAGGTATGAAAATGGCAACTGAGATGGAGAATTTGGATTTAAAACTATACGAAGCACCTATGGAATTCTTAAAGATCCAACATGGGAGCAATATGAAACAGAGTGCCGGTTACTACACCGATGAATCAACCACGTTGGACGAAGCTGAGATAGCGATGCTGGATTTGTACATGGAGAGGGCGCAGATCAAAGATGGTCAGAGTGTACTTGATTTAGGCTGTGGATTGGGTGCTGTAGCTCTTTATGGTGCAAACAAGTTTAAGAAATGTCAATTTACTGGAGTTACAAGTTCTATGGAACAAAAAGATTACATTGAGGGAAAATGCAAGGAACTTAAGTTGACCAATGTCAAGGTTTTATTAGCTGATATAACAACTTACGAAACCGAAGAAAGATTTGATCGGATTTTTGCTGTTGAATTGATTGAGCATATGAAGAACTATCAATTGCTTCTTAAGAAAATATCAGAATGGATGAAAGATGATGGACTACTTTTCGTTGAACATGTTTGCCATAAAACTCTAGCGTACCACTATGAGCCTGTTGATGCAGAAGATTGGTACACTAACTACATATTCCCTGCTGGAACTATGACTCTATCATCTGCTTCAATGCTTCTTTACTTTCAAGATGATGTTGCTGTTGTGAATCAATGGACTCTAAGTGGAAAACATTACTCACGATCCCATGAAGAATGGTTGAAAAACATGGATAAGAACATTGTTGAAGCCAAAGAGATAATGAAGTCTATAACTAAAACTGAGGAAGAAGCAACCAAATTGCTCAACTTTTGGAGGATTTTTTGCATGTGCGGAGCAGAGTTGTTTGGCTATAAGAATGGTGAAGAATGGATGCTCACCCATCTTCTCTTCAAGAAAAAAATGAGAGAACTTCTTTTATGTGTGCTTTCATTTAGTGCGTGTTATTTTACATCATACATATGCCAAATCATTGTAATAAGAACTGTTTAGAATGATTGATTGGTTGTAACTTTTATTTTATGTTTTTTTAAAAATACTTCATCATGAAGTGCTT

>Par_TRINITY_CYP80B1 Par_c39418_g2_i1 len=1618

TGTGCTCTTCCGATCTCTTCATATAAACAAGAGGCCAAACAAAACTACATTTAGAGAAAGAAAATATGGCGATCATCACAGTAGCAATTGTAGCAGTTGTGATCACTACTTTGTTATACTTGATTTTCAGAGATTCGAGTCCTAAAGGTTTGCCACCAGGTCCAAAACCCTGGCCAATAGTTGGAAACCTTCTTCAACTTGGTGAGAAACCCCATTCTCAGTTTGCTCAGCTCGCTGAAACCTATGGTGATCTCTTTACACTGAAACTAGGAAGCGAAACAGTTGTTGTAGCTTCAACTCCATTAGCAGCTAGCGAGATTCTAAAGACACACGATCGCGTTCTCTCTGGTCGGTACGTGTTTCAGAGTTTCCGGATAAAAGAGCATGTAGAGAACTCTATTGTTTGGTCTGATTGCAATGAAACATGGAAGAAACTGCGGAAAGTTTGTAGAACGGAACTTTTTACACAGAAGATGATTGAAAGTCAAGCTGAAATTAGAGAATGTAAGGCTATGGAAATGGTGGAATATTTGAAGAAAAATGAAGGAAATGAAGTGAAAATTGTTGAAGTTGTATTTGGGACGTTGGTGAATATATTCGGTAACTTGATATTTTCGCAAAATATTTTCAAGTTGGGTGATGAAAGTAGCGGAAGTGTAGAAATGAAAGGACATCTATGGAGAATGCTGGAATTGGGGAACTCGACAAATCCAGCTGATTATTTTCCATTTTTGGGTAGATTTGATTTGTTTGGACAAAGAAAAGATGTTGCTGATTGTCTGCAAGGGATTTATGGTGTTTGGGGTGCTATGCTCAAAGAAAGAAAAATAGCCAAGCTGCTTAACAACAGCAAGAGGAATGATTTTGTTGAGATTTTGCTCGATTCCGGACTCGATGACCAACAGATCAATGCCTTGCTCATGGAAATATTTGGTGCGGGGACAGAGACAAGTGCATCTACAATAGAATGGGCGTTATCTGAGCTCACGAAGAACCCTGAAGTGACAGCCAATTTGCGATCGGAATTGCTATCTGTGGTAGGGCAGAGGCCGGTTAAGGAATCCGACATACCAAACATGCCTTATCTTCAAGCTTTTGTTAAAGAAACTCTACGGCTTCATCCGGCAACTCCGCTGCTGCTCCCTCGCCGAGCACTTGAGACGTGCAAAGTTTTGAACTATACAATCCCAAAAGAGTGTCAAATTATGGTGAACGCCTGGGGCATTGGTCGGGATCCAAAAACGTGGGCTGATCCATTGAAGTTTTCACCTGAGAGGTTCTTGAATTCGAGCATTGATTTCAAAGGGAATGACTTCGAGTTGATACCATTTGGTGCAGGGAGAAGGATATGTCCTGGTGTGCCCTTGGCAACTCAGTTTATTAGTCTTATTGTGCCTACTTTGGTACAGAATTTTGATTGGGGATTACCAAATGGAATGGATCCTAGCCAACTGATCATGGAAGAGAAATTTGGGTTGACACTCCAAAAAGAACCACCTTTGTATATTGTTCCTAAAACTCGGGATTAAGCTCAATCAAGAATTCTTAAATTCACATGATTCGGAGATCTCGCTAGATTATTCGAAAATAAAATTTTAAAAAAAAGTATTTAAATATG

>Par_TRINITY_4OMT1 Par_c27586_g1_i1 len=1264

GTGTGCTCTTCCGATCTCCTAAACAAAAATAGTCTTCTCTGCTTGACAGCAAAAAAATTCATATCTAATGGGTAGTTTAGATGTCAAGCCAGCTTCTGAAACACAAGAAGTTTCCATCAAGGATCAAGCTCAACTGTGGAATATTATCTATGGTTTCGCTGATTCTCTCGTTCTTCGTTGCGCTGTAGAGATTGGAATCGCGGATATTATCAAAAACAATGGTGGGGCAATCACACTTGCACAACTTGCGTCCAAACTACCAGTTCCAAATGTCAGTTCTGATTACTTGTACAGAATGGTAAGATACTTGGTACACCTGAATATCTTAGAACAAGAAACTTGCAACGGCGGAGTTGAGAAGGTTTACTCGCTTAAACCGGTCGGTACTCTACTTTTAAGAGACGCAGAAAGAAGTATGGTGCCGATGATATTAGGCATGACTCAAAAGGATTTCATGGTTTCATGGCATTTCATGAAAGAAGGTTTAGGAAATGGAAGTACGACGGCATTTGAGAAAGGAATGGGGATGGATATATGGAAGTATTTAGAAGTTAATCCTGATCAGAGCCAGTTGTTTAATGAAGGAATGGCTGGAGAAACAAGACTTTTAACCAAAACTCTAATCGAAGATTGTAGAGATACCTTTCAGGGTTTGGAATCTCTGGTTGATATTGGTGGAGGAAATGGTACAACAATCAAGGCAATCTATGAAGCTTTTCCTCATATCAAATGTACACTCTATGATCTTCCTCATGTTGTTGCCAACTCTCATGATCTCCCTAACATCGAAAAAGTTCCCGGTGATATGTTCAAGTCTGTTCCCAGTGCTCAGGCTATCTTACTCAAGTTGATCTTGCACGACTGGACTGATGAGGAGTGTGTCAACATTCTAAAGAAGTGCAAAGAAGCAATACCGAAAGAGACAGGAAAAGTGATTATAGTGGATGTAGCACTTGAAGAAGAATCCAACCATGAACTTACAAAGACAAGATTGATTCTTGATATTGATATGTTGGTGAATACTGGCGGTAGAGAGAGAACTGCAGATGATTGGGAAAATTTGCTCAAACGCGCAGGCTTTCGGTCTCATAGGATCAGACCCATTCGCGCTATTCAGTCTGTCATCGAAGCCTTCCCTTAGTAAAAGCTCTGACTATTCTGCTTTCTTTCTCAGAGAAGCGTACGTATGCGATGCACTTGTCAACAATAAGCCATCTTCAATTGAGGAGACGGTTTCCTATTTTCTGTTGTATTTTTTTTTTAT

>Par_TRINITY_BBE Par_c39416_g1_i2 len=1613

TGTGCTCTTCCGATCTCATCATGATGTGCAGAAGCTTAACATTACGTTTCTTCTTATTCTTTGTTTTATTACAAACATCCGTACGAGGTGGTGATGTTAATGATAATCTCCTCTCGTCGTGTTTAAACTCCCATGGTGTTCACAATTTCACCGCACTATCAACCGATACAAATTCCGACTACTTCAAACTACTCCATGCATCGATACAGAACCCGTTGTTCGCGAAGCCTACGGTATCGAAACCGTCGTTTATTGTAATTCCCGGCAGCAAAGAAGAACTTTCGAGCACGATTCATTGTTGTACAAGAGAATCATGGACTATTCGACTGCGGAGTGGTGGTCATAGTTATGAAGGGTTGTCTTACACTGCTGATACACCTTTTGTGATTGTAGATATGATGAACTTGAATCGAGTTTCCATTGATGTTGAGTCGGAAACAGCTTGGGTTGAATCCGGTGCAACACTTGGAGAATTATACTATTCTATTGCGCAGTCAACGGAAACACTGGGGTTTACTGCTGGTTGGTGTCCGACTGTTGGTAGCGGAGGACATATAAGCGGTGGTGGTTTTGGTATGATGTCGAGAAAGTACGGGTTAGCCGCGGATAACGTTGTGGATGCGATTCTTATAGATAGCAATGGAGCGATTCTTGACCGTGAAAAAATGGGGGAAGATGTTTTTTGGGCTATTCGTGGTGGCGGCGGAGGTGTCTGGGGTGCAATTTACGCGTGGAAAATCAAACTTTTGCTAGTTCCGGAGAAGCTGACCGTTTTTCGTGTGACAAAGAATGTAAAAATTGAAGCCGCTTCTTCTTTGCTTCACAAATGGCAATATGTTGCAGATGAATTAGATGAGGATTTCACGGTGTCTGTGCTCGGGGGAGTAAACGAAAATGAAGCCTGGTTAATGTTCTTAGGCTTACACTTGGGACCTAAAAGTGCTGCAAAAACTATAATCGATGAAAAATTCCCTGAACTGGGGTTAGTAGAGGAAGAGTTTCAAGAAATGAGTTGGGGTGCATCCATGGCTTTCTTAGCAGGATTAGAAACAATCTCTGAGTTAAATAACAGGTTCTTGAAATTTGATGATAGAGCTTTCAAGACAAAAGTTGATTTCACTAAAGAACCAGTACCGCTTAACGTGTTTAGACGTGCATTAGAGATGTTATCAGAACAGCCTGGTGGATTTATAGCTCTAAATGGTTTTGGAGGGAAGATGAGCGAAATTAGCACTGACTTTACCCCGTTTCCTCATCGTAAAGGAACTAAATTGATGTTCGAGTATATAATCGCTTGGAACGAAGATGAAGAAACGAAAATCGGTGAGTTTAGCGAATGGTTAGCCAAGTTTTACGATTATTTGGAACCGTTCGTATCGAAAGAACCAAGGGTTGGTTATGTTAATCATATTGATCTTGATATTGGAGGAATAGATTGGAGAAATAAAAGCAGTACTAGCGGTGCGGTTGAGATTGCTAGGAATTGGGGTGAAAGATACTTTTCGTCCAATTATGAACGTCTAGTTAAGGCCAAGACACTGATTGATCCAATTAATGTGTTTAATCATCCACAGAGTATTCCTCCAATGATGAAATTTGAGGAAAATTACATT

>Par_TRINITY_TNMT Par_c34579_g1_i3 len=1404

GTGTGCTCTTCCGATCTAAACTCTTTGTGGTTAGACGTCCACATGTTACACGCGAATTCATGTACTAAAACTCTTTATCTTACGTCGACATTTTAGTTTGTTGTGTTGTTTTGTGCCAGAAAACCGATTGATCTCCTATATAAACAATAGTACGTACTAAGTCTGGTTCACAACACACCACACCACACCTCTCTCATTCTCTCTAGAATTTTCATTTCATTAAGTTTCTCCCGAGCTAGCAGCCTAGATTGTGTCATCATGAGTTCAATAGAAGAGGTGAAGATAAAAGAATCGAAAGAAGAAACCCTAGGGAGATTGTTGAAAGGAGAGATTAAAGATGAGGAACTGAAGAAGCTGATTAAGTTTCAGTTCGAGAAACGTCTGCAATGGGGTTATAAATCATCTCATCAAGAACAACTCTCCTTCAACCTTGACTTCATTAAATCGTTGAAAAAGATGGAAATGTCAGGAGAGATTGAGACAATGAACAAGGAAACCTATGAATTGCCATCAGAATTCTTAGAGGCAGTTTTTGGGAAAACTGCCAAGCAAAGTATGTGTTACTTTAAACATGAATCAGCAACAATAGACGAAGCAGAAGAAGCTGCACATGAATTATACTGCGAACGAGCACAAATCAAAGATGGACAAACTGTTCTTGACATCGGTTGTGGTCAAGGTGGTCTAGTTCTATACATTGCTCAGAAATATAAGAACTGTCATGTTACAGGGCTCACTAATTCAAAGGCACAAGTCAATTACTTACTCAAGCAAGCAGAGAAGCTTGGGTTGACAAACGTCGACGCCATACTAGCAGATGTTACTCAGTATGAATCTGACAAGACATACGATCGTCTACTTATGATTGAAGCCATAGAGCACATGAAAAACTTACAGCTGTTCATGAAGAAACTATCAACTTGGATGACAGAAGAAAGTCTTCTCTTTGTGGACCATGTCTGCCACAAAACATTTGCTCACTTTTTTGAGGCCGTCGACGAGGATGACTGGTATTCCGGTTTCATCTTCCCCCCTGGTTGTGCAACAATACTAGCTGCTAATAGTCTCCTCTATTTCCAGGATGATGTTTCAGTTGTGGATCATTGGGTTGTCAACGGAATGCACATGGCTCGTTCAGTAGACATTTGGAGAAAGTCGCTAGACAAAAATATGGAAGCTGCAAAAGAAATTCTATTACCTGGACTTGGAGGAAGCCACGAAGCAGTGAATGGAGTCGTTACTCACATTAGAACTTTCTGTATGGGAGGGTATGAACAATTCTCAATGAACAATGGAGATGAGTGGATGATTGCACAGCTGCTTTTCAAGAAGAAGTAATGAGACACATTCAACCCGCATTTCCAGAGTTCAATAATAATGTGACCTTATTGTGTCAATAACTGC

>Par_TRINITY_PSSDR1 Par_c91731_g1_i1 len=1389

GTGTGCTCTTCCGATCTAAATAGGCGAACTATAACAGTTTGCACCTACAACATTCAATTTTTATTTTGGTAAACGAAAAAATAAGCTACAACTCCATGTTTTGGCTGAGTGTAGTCTGACTGTAGGTAAGCTACAACTGCATGTTACAGATAATCACTAACTGATTATTCAGGCATACCTAACGGTCCATATTGTTATAGTTCGCAAAACATTTTTGCGAACTATAAAGGCATGCATGGACAGAAAAAATACATCAGAGAGATATCATAAATTCAAAGAGATGGAAGGAACAGGAAAGATAGTATGTGTAACAGGTGGAGCTGGATACTTGGCATCTTGGCTGATCATGAGATTGCTTGAACGTGGTTACTCTGTTCGGACCACCGTCCGGTCTGACCCAAAATTTAGGGAAGATGTGAGCCACCTGAAAGCTCTTCCTGAAGCTACAGAGAAGCTTCAAATTTTTGAAGCAGATCTTGAAAACCCAGAAAGTTTCGACGATGCAATCAACGGGTGTGTCGGTGTCTTTCTCGTTGCTCAAGGAATGAATTTTGCCGAAGAATATACTCTTGAAAAAATAATCAAAACATGCGTGGAAGGAACTCTTAGAATTCTAGAGTCATGCTTGAAATCTAAAACAGTGAAAAAGGTTGTGTACACATCTTCTGCTGATGCAGCAATGATGATAAGTAATCTCAAAGCTTTAAAAGAAATTAACGAGACAATATGGTCAGAAGTTGACAATTTCATTAGCAAACCGGAACAAGTTATTCCTGGATTGCCCTCATATGTGGTCTCAAAGGTACTGACAGAAAGAGCTTGCCTAAAATTTTCTGAAGAACATGGTTTGGATGTTGTTACTATACTTCCTCCGTTGATTGTTGGACCATTTATCACTCCCCATCCCCCTCCCAGTGTATCTATAGCTCTTTCGATAATTTCAGGTGATGTGTCGATGATGCTTGGTGTTAGACTTGAAAATGCGGTACATATAGATGATGTTGCTTTAGCACACATATTTCTTTTTGAATGTGAAAAAGCAAAAGGAAGACATATTTGTTCTTCAGCTGATTTTCCAATGCATGATCTGCCTAAATTTATATCTGAGAATTATCCGGAATTTAACGTACCGACTGATTTACTAAAGGATATTGAGGAACAAGAACCAGTTCATCTTTCCTCAGATAAGCTGTTGAGTATGGGATTTCAGTTCAAATATGCTTTTGCAGAGATTTTCGGTGATGCAATACGATGTGCCAAAGAGAAGGGTTTCCTTTAGAGGCCAATTATAGTATGGTTACGAGGAGATGTGGGAGTAGCTAGCCCAAGATGCCCTGCTTGCACTAGCTTATATTATTGTTGTTGTTTTTCAAATGAATAAATGGGCAG

>Par_TRINITY_PSCXE1 Par_c36757_g1_i4 len=1082

CCAAAGTTCAAACCTCATCCAATAAAAATCCAACAATGGCAGATCCTTATGAATTCCTAATGTGCATTCACAATCCTGAAGAAGATACCCTAACAAGAAATTTCCCGATTCCTGCTACTCCCTTAGATCAAAACACCAAAGACATTTCTTTAAATCCCGATAGGAAAACCTCACTTCGAATCTTTCGACCCCCAACCGAAGAACTTTTTGCAACAACAAATAAGTTGCTTCCTATCATAATTTATTTCCATGGTGGAGGTTTCATTCTTTTCAATGCAGATTCAACTATAAACCACGACTTTTGTCAATCGATTGCTACACATTTACCTGCGCTGGTCGTTTCGGTAGACTACCGTCTTGCTCCTGAAAACAGACTTCCCGCTGCCTATGATGATGCTGTTGATGCTTTAAACTGGGTCAAAGACCAAGGTTCTGGCAAACTAAATAATAGTGAAGTATGGTTAAAAGAGTATGGCGACTTCTCAAAGTGTTTCATTATGGGGTGCAGCTCGGGTGGTAATATTGCATATCATGCCAGTTTGAGGGCAATAGAAATGGATCTTGAACCAGTTATAATCAACGGATTAATATTACACAGCCCTTTTTTTGGTAGTCTTGAGAGAACTGAATCAGATTTAAAAGTGATCAACGATCTAGACTTGCCTCTTGCGGTAAGGGATGTCATGTGGGAACTGGCTTTGCCGCTTGGGTCTAGTCGTGACCACGTTTATTGTAATCCAAATATTGACAATGATGGATCATCATCCGGAAATATGGTGGGGTTAATCAAAAGATGTTTGGTGATAGGATTTTATGGGGATCCACTTATTGATCGACAAATTCAGCTCGTGAAGATGTTGGAGGAAAAGGGTGTGAAGGTTGAAACTTGGATTGAACAAGGAGGGTATCATGGAGTACCATGCTTTGACTCTAAGATACGTGAAACGTTGCTGGGAAAAATAAAATATTTTATTTGATTCAGTTTATCTTATATTATTACTAATAAACAACGAATTACATGAAAGCTTTCGAAAAACATTTTCAGATGTGGCGTTGTACGTGAGCTTGTTGACGTAATTGGC

>Par_TRINITY_CYP82X1 Par_c39178_g1_i2 len=1874

GAACCATTAAACACTTAAGTCATGCCTTGATATGCTCAAATTTTAGTGTGTCATATTCACTATAACTATAAATTTCAATACAATTTCTAAACCTCATCATCATTCAAGAGAGACGCAAATACCTTGATATTCTTTTGTCATCAATGGAGTTATTCATAAAGTTACCGTTTATCCAACCAATTCCTTTCAGTATTATTCTTGTTACTACAGTTTCGATTGTTCTATTATACAGTGTCTTCTTCTTGGTTACTGATACGAAAAAGAAGAAGAAGAAAGCACCAAATGCTTCCGGGGCATGGCCGTTAATAGGTCATCTCCGTCTATTGATGAACGACAAGGAACCGTTGTATAGAGCACTAGGGAGCATGGCTGATAAGTACGGACCTGCATTCAACATCCGATTAGGTAACCAAGAAGTTCTTGTTGTGAGTAACTGGGAGATGGTAAAACAGTGTTTTGGTAATCAAAATGATAAGCTATTTTCGAATCGTCAAACTACATTAGCTGCAAAATACATGCTTAATCAAACAACTTCTAGCGGATTCGCACCATATGGACCATATTGGAGAGAGCTACGAAAGATAATGGTGCAGCAATTACTCTCTAAACAATCTTTAGAATCGTGGAAACATCTGAAAATCAAAGAGATGGATGCTTCATTTAGTAAACTTAACGAGTTATGCAACAACAACGGTACTGGAACAGCTACCCTAATTAGGATGGACGAATGGTTTGCTGAGTTGACGTTCAACGTGATCGCAAGAAATATCTTTGGCTACAAAAGTGGCGGAAGGTCGACAGCGCTTACAAACGGTGATACGGAATCAAAGGGCGAGAGGTACAAGAAAACATTGGAAGAAGCACTTCATCTAATGTCAATCTTTGCAGTTTCAGACATATTTCCAAGTCTAGAGTGGGTAGATCGATTAAGAGGCCTTGTAAGGGATATGAAACGCTTTGGAGATGAGCTAAATTCAATTGCAGGGTGTCTTATTGAAGAGCACCGCCAAAAGAGATTACAATCCGTATCTAAAAGTGATAAAGGAGTTGGTGATGAACAAGACTTCGTTGATGTTCTCTTATCGGTTGCTGAACAATCGCAACTTCCTGGAGATGACCCTGATTTGGTCATCAAGTCTATGATTCTGGAAATCGTATCAGGTGGGAGTGAGACCACATCGTCAACCTTAACTTGGGCCCTCTGTCTGTTACTGAACCATCCGCATGTGTTAAAGAAGGCAAAAGAGGAATTAGATACGCACGTAGGAAAACATAGGCATGTAGAAGAGTCAGATACCCCTAAGCTCGTGTACATTAATGCAATTATCAAAGAATCAATGCGATTGTATCCAAACGGGGCAATGCTTGATCGGTTGGCGTTAGAGGAGTGCGAAGTTGGTGGATTTCATGTACCGGCCGGGGGACGCTTATTTGTCAATGTTTGGAAGATTCAGAGAGATCCGAGTGTTTGGGAGAATCCTCTAGAGTTTAAACCAGAGAGGTGGTTTTTGAGTAATGGTGAAAAGATGGATGTGGATTACAAAGGTCACAATCATGAATTCATACCATTTGGGATAGGTCGGAGGATGTGCGCTGGTATGCTTTGGGCATCGGAGGTGATTCATTTGGTGCTGCCCCGTCTTATTCATGGGTTTGATATGAAAGCAGCAAGTGCCAATGGGAAAGTAGACATGACGGAAATGGCAGGCATGGTGATTTGTTTTAAGAAGACACCTCTTGAAGTTATGGTCAATCCTCGAGAGTAGATGTTTTTATCCTTTTGCATAGTTTCTATATTAAAAATGTAGGCCAAGCGAGCAACATGTCGTACATTTGATACAAGATTTTTTAAGTTTGTCAGATCGGAAGAGCGT

>Par_TRINITY_CYP82X2 Par_c28721_g1_i2 len=1639

TGTGCTCTTCCGATCTAGAAATCCTAGTTGTGAGCAACTGGGAGATGGTGAAAGAATGTTTCACAAGAAATGACAAGTCATTTTCGAATCGTCCAAGTACATTAAGCACTAAATACATGCTGAATGACACCAATTCTGTTGTGTTTTCACCTTATGGAACGTATTGGAGAGAAATGCGGAAGATACTGGTGCAAAAACTACTGATCTCTAACCAAAGATCAGAGGCATTGAAAAATTTGAAAACGAAAGAAATCGACAACTCGTTTGTGAAGCTCAATGATTTATGCAGCAACGAGGTTAGCGGAGGCACAAAAGTTAGGATGGACGAATGGTTGGCTGACATGATGTTCAACATTATTGCTAGGATTACATTTGGTTACGAAGGCGGAGGAGGCCATGCACCTGGCGCTTCTACAACATCCAAGAATGTCGAGAGATACAAGAAAACGTTGGACGAGATGTTTGTTGTTTTAGCGACGAGGTTTGCAGTTTCAGATATATTTCCATCTCTGGAGTTTATAGACCGATTGAGAGGTCTTGTAAAGGATATGAAAATCTTGGGAGACGAATTAAACTCAATTGCTGGATGTTTTATTGAAGAACATCGTCAAAAGAGACGAGAATCATTATCCTCATTGTCATCGTTATCAAATGAATCCGTTGGTGATGAACAAGATTTCATTGATGTTCTCTTGTCAATAATGGATCAGTCACGGCTTCCCGGAGATGACCCAGATTTTATTATCAAAATTATGATCCTGGAAGCTTTTGCAGGTGGGACGGACAGTTTAAGTGCAACCTTAACTTGGGCTCTCTCTCTACTGCTGAACCACCCAAACGTGTTAAAGAGGGCAAGGGAGGAAATCGATAGGCATGTGGAAAACGGTAAGCAAGTGGAAGCCTCTGATATTCCTAAGCTCGTATACATTTATGCAATAATCAAAGAGACGATGAGATTGTATCCAGTCGGAGCACTAAGCGAACGCTATACGACTGAAGAATGCGAGGTTGGTGGGTTTAATGTACCCGCCGGCACACGCTTACTGGTGAATATATGGAAGATCCATAGAGACCCAAGTGTATGGGAGAATCCATCAGATTTTCAACCAGAGAGGTTTTTGTGCAGCGATAAGGTGGGTGTGGATTTATATGGCCAGAATTATGAGTTGATACCATTTGGGGCCGGTAGGAGGGTATGTCCGGCTATAGCTTCATCACTGCAGACGATGCATTATGCGTTGGCGCGTCTTATTCAAGGATATGAAATGAAATCAGCCAGCCTCGATGGGAAGGTGAATATGGAAGAAATGATAGCCATGTCGTGCCACAAGATGAGCCCTCTTGAAGTTATTATCAGTCCTCGGGAGCTGAGGCGGACTTAAATCTTATGTTCCAAATTTACATTAGCATCTTTGATTATGAAATGTATTGCTCTTGACTTTCTTTTTTGTTTTTTATATTTTTAAGCTTGTATGTGATGATCAGAGAAAATGATAATGACAGAATCATCAAAAGAATCAATCGAACAAGCTATGTTGTATTAGGCGCGCCGAGGAAAAATTAAAAAACTGCAGGAATATGTATGATTTGGCGCCTTGGGCGCCTTTTTTTCTAGGCGTTCCCTGAGCGTCTCGCGTGC

>Par_TRINITY_PSAT1 Par_c36454_g1_i5 len=1651

GTGTGCTCTTCCGATCTTCCAATTTGCATTGTTTATCGACCTTGTGGAACAATTAGGGGATATGCCAACAATGTCTAACGCTGCTGTACAAGTGATCTCGAAAGAAACGATTAAACCAACAAATCCAACACCATATCAACTTAGAAACTACAATATGTCACTTCTCGACCAATATTCTTCTCTAGTTTATGTTCCGATCATTCTTTTCTACCCTGCTGCCTCCGACGCTAATAGTACCGGAAGTAAGCACCGTGATGATCTTCACTTGCTTAAGAGGTCTCTTTCTGAAACGCTAGTTCACTTTTATCCAATGGCTGGTAGGATGAAAGACAACATGACTGTTGACTGCAACGACCAAGGTATCGACTTTTTCGAAGTAAGAATCAAAGGTAAAATGTGTGACTTCATGATGAAATCAGATGTACACTTGAGTCTGCTTCTTCCGTCTGAAGTCGCTTCCATGAATTTCGTGAAGGAAGCCCAGGTGATTGTTCAAGTGAACATGTTTGATTGCGGTGGAACTGCCATTTGTTTCTGTATATCAAACAAGATTGCAGATGCATGCACCATGATTACCTTCATTCGTAATTTGGCAGGCACCACCAACAAAGCTCGTCGTGGGAGCTCTACGGCTGCACCAACCACAAATCAGAAGTTGTTTCCTTCTTTCGATTCGGCATCACTCTTCCCACCTAGTGAACAATTGAAAACTCAAGTTTCCTATCCTACACAGGATAGTACCAGCGTAGATAAACTTGTCAGCAAAAGATTTGTGTTTGATGCGGCAAAGATTACATCTGCACGTGAGAAATTGCAATCCTTGATGCATGATAAATACAAATGCCATAGGCCGACAAGGGTTGAGGTAGTTTCCGCTTTGATATGGAAGTCAGCAGTGAAATCCGCTCCGCCCGGTTCTATATCCACTGTAACCCATGCCATGAACTTTAGAAAGAAAATGGATCCACCATTACAAGATGCGTCATTCGGGAATCTTTGTGTGGTTGTTACAGCAGTATTACCAGCAACAACGGCGACAACAACAAATCCAGCAACCAAAACAGTTAGTAGTACGAGTAATGAAGAGCAAGTGGCACTTGATGAGTTAAGTGATTTTGTAGCCCTATTGAGGCGCGAAATAGATAAGGTAAAGGGTGATAAAGGTTGCATGGAGAAAATCATTCAAAATTTCATCTATGGTCATGATGCTTCCGTAGGGCAAGACAGTGATGTTGAAGATAAGGTGATAGCTTTGTTTATGACTAGCTGGTGCAAGTTTGGATTCTACGAAGCTGATTTTGGTTGGGGAACTCCAATTTGGGTAACTACTGTTCCATTAATTGAGCCAAAGTACAAGAACATGGTTTTCATGAACGATACGAAATGTGGTCAAGGAATTGAAGTGTGGGTGAATTTTCTGGAGGATGATATGACCAAGTTCGAGCATCACCTAAGCGAGATCCTCCAACTGTTTTGATTTTCAACCTTTTCCTTAATAGAGGTCAATTGTCGTGTTTGTTCTTCATCTTAACTTACCATCTTTATTTTTACGTGGTCATTGGGTTTACAAACGGAAGACTTTGTGTTGGCTTTTTTCCTCTCAATGGACTTTTTTTTTATTTTCTCTTTATTATTATTTAAACAGATCGGA

>Par_TRINITY_PSMT2 Par_c30871_g1_i2 len=1403

GTGCTCTTCCGATCTATTCAATCAGAAACTTTCTTCTAAAATCTTTCAATACCAGTACTGTTAGTTTCCGAGAAGATTCTGCCACACTAATCCATTATGGAAATACATTTAGAAAGCCAAGAACAAGAAATGAAATATCAAGCTCAAATCTGGAACCAAATGTGTGGCACTGTTGATACCTCTGTTCTGAGATGTGCAATTCAATTAGGTATATTTGATGCCATTCATAACTCAGGCAAACCAATGATTACCTTAACGGAATTATCAAGCATTATTTCATCACCCTCTTCATCATCAATTGAACCCTGCAACTTATATAGATTAATGAGATACTTATCCCATATGGATCTCATCGATATCGGAGAATGTTTGAATGAAGAAACTGTTTCACTAACATGCACATCTAAGTTACTACTCAGAAACCAAGAAAAGAGTTTAATTGATTGGGTATTGGCAATTTCTTGCGAGATGATGGTTGTTGTTTGGCACGAACTAAGCAGCTCTGTTTCAACTCCTGCGGGTGAGCCTCCAATCTTCCAGAAGGTTCATGGTAAAAATGCTTTAGAATTAGCAGGGGAATTCCCAGAATGGAATGATCTGATCAACAATGCTATGACTAGTGATACTAGTGTAACTAAGCCAGCGCTAATACAAGGATGTGCCAAAATCCTGAACGGAGTCACATCGTTGATTGATGTTGGTGGTGGTCACGGTGCCACTATGGCCTACATAGTTGAAGCTTTTCCTCACATAAAAGGTGCGGTAATCGATTTGCCACATGTTGTTGAAGCCGCTCCGGAGCGTCCAGGTGTGGAGTTCATCAGCGGTGATATATTCAAGTCTATTTCTAACGCTGATGCTGTGTTGTTGAAGTATGTCCTGCACAACTGGGAAGATACGGAATGTGTGAATTTGCTGAAGAGATGCAAGGAAGCAGTTCCGGCAGACAAAGGAAAAGTGATCATAATGGATTTAGTAATAGACGACGATGATGACAGTATTTTAACGCAGGCAAAATTGAGCCTTGATCTCACTGTGATGAACCATGGAGGAGGTAGAGAAAGGACTAAAGAAGATTGGAGAAATCTAATTGAGATGTCTGGATTCAGTAGGCATGAAATAATTCCAATATCTGCCATGCCATCAATTATTGTTGCTTATCCCTAGTTAAGTCACCCACATGATCGGGAATAAGTTAGGCGCGTGTTGAATCTGTTGACATCGCAATTGTGCCTTTTACTTTATGCATTTTCATTCCAGTAAAGACTGTTTTTGGGCATTCGGATTCTTCTGAGCCCTTGTATGTTTGTCTTCTCTGTTTTCAAGGCATGTGCATGTGACTAGTTTGAGTATGCATATTGATGTTTCGTTACAATTTTCTGAAATCATGGCTTATTATATCA

>Par_TRINITY_CYP82Y1 Par_c34116_g1_i1 len=1292

GTGTGCTCTTCCGATCTCGAATTTGAGTGCATTCATGGCGTCTCTGATGATCAAGAAATGTATCCATCTGTTTTTTGATCAGCCAACTTCAGTTGCCACTCTTATACTTGCTTTTCTGCTAACACTTTCGCCTGTTATTATTTACTATGAACAGAAGAAGAGGGGTTTGAGGCGAAATCGCACCGCAATTACAACGACTCCATTACCAGAGGCATCAGGTGCATGGCCAGTGATTGGTCATCTTCTTCTTTTCATGAACGAAAACGATCTAAATCATGTAACTCTTGGTCACATGGCTGATAAATATGGACCTATTTTCAGCTTAAGATTCGGTAGCCATAGAACTCTAGTTGTTAGTAGTTGGGAGATGGTAAAGGAGTGTTTTACAGGTACCAATGACAAGTTGTTCTCAAATCGTCCTTCCTCCTTGGCGGTTAAACTTATGTTTTATGACACTGAGTCTTATGGTTTTGCACCTTATGGGAAATACTGGAGAGAGTTGCGAAAGATATCTACACATAAACTCCTCTCTAATCAGCAATTAGAGAAGTTCAAGCACTTGCGGATTTCTGAAGTTGACAACTCCTTTAAAAAGCTTCATGAGTTATGCAGCAACAACAATCATGGAGGTGATACTACATCTGTGGCTAGCCTTGTGAGAATGGATGATTGGTTCGCGTACTTGACATTTAACGTAATAGGACGGATCGTTAGCGGATTCCAATCAAATGCAGTGGCAGGTGCCACAAACAGCCAGGAAAAATACAAGCTTGCAATCGATGAAGTGTCAAATCTTATGGCAATGTTTGCCGTTTCAGATGTGGTTCCATGGCTTGGGTGGATTGATCGATTGACTGGTCTTACAGGAAAAATGAAGAATTGTGGGAAAAAATTAGATGCAGTAGTTGGGGATGCAGTTGAGGATCATCGCCAAAAGAAACTCAAAATTTCTAGAAATACCGCAGGAGCACTTATGGAGCATGAAGAAGAAGACTTCATCGATGTTTGCTTGTCAATTATGGAGCAATCACAGATTCCGGGAAACAACCCTTTAATCTCTGTCAAGTCTATTGCCTTGGACATGTTATCGGGTGGGAGTGACACTACAAAATTGATAATGACATGGACCCTTTCTTTGCTGTTGAACCATCCGGACATATTGGACAAGGCTAAAGAAGAAGTACTTACATACTTCAGGAAGAAAAGGATAGCGGATAACACACCCGTGGTTGATGCCGCCGATGTTCCTAACCTCCTCTACATCCAAGCAATCATCAAAGAATCAATGCGG

>Par_TRINITY_PSMT3 Par_c19997_g1_i1 len=549

GCTCTTCCGATCTCTCCCATAAAAGCTAGCATTAGAAATTAATAAGCTCAAAAACTGTAAAATTTCTAAAGAGAGCTAAAAAGAAGAGAAAAAAGATGGAAGTAGTAAATAAGATTGATCAAGAAAACCAAGCAAAAATTTGGAAACAAATTTTTGGTTTTGCAGAATCATTAGTTCTGAAATGTGCAGTTCAGTTAGAGATTGCTGAAACACTTCATAATAATGTAAAACCCATGTCTTTATCCGAGTTAGCATCTAAACTTCCGGCTCAACCCGTTAATGAAGACCGTTTGTACCGAATTCTGCATTTCTTAGTTCACATGAAACTCTTCAACAAAGATGCTACCGCACAGAAATATTCATTAGCTCCACCAGCAAAGTATTTGCTAAAAGGCTGGGAAAAATCAATGGTTCCTTCAATATTAAGCGTGACCGATAAAGATTTTACAGCTCCATGGAATCATCTTGGTGACGGTTTAACAGGTAACTGTAACGCTTTTGAGAAAGCGTTAGGAAAGAGCATTCGGGTTTATATGAGTGAAAATCCTG

>Par_TRINITY_CYP719A21 Par_c31271_g1_i1 len=1662

TTCCGATCTGTTAATCTGACTCAAGCTAGCTAGAATATATACCAACAATACAAGAAATGATCATGAGTAACTTATGGATTCTTACGCTCATTTCTACCATATTAGCAGTCTTTGCTGCTGTGTTAATCATTTTCAGGAGAAGAACGTCAGCATCCACAACGCAATGGCCTGTTGGCCCAAAAACATTACCAATCATAGGTAACTTGCACATTCTTGGAGGCACTGCACTCCATGTCGTCTTACATAAACTAGCTGAAGTTTACGGCAGTGTAATGACGATATGGATTGGTAGTTGGAAACCTGTTATTATTGTTTCCGACTTTGATCGAGCTTGGGAAGTTCTTGTTAACAAATCGTCAGATTATTCAGCACGCGAAATGCCTGAGATCACTAAAATCGGCACTGCAAATTGGAGAACAATTTCAAGTTCTGATTCTGGTCCATTTTGGGCCACTCTTCGGAAAGGTCTTCAGGGTGTAGCATTATCGCCTCAGCATTTAGCATCGCAAACTGCACACCAAGAGAGAGATATAATAAAGTTGATCAAAAATTTGAAAGACGAAGCAGCTCTTAATTCTGGAATGGTTAAACCACTTGATCATCTCAAGAAAGCAACTGTAAGATTAATCAGTCGGTTAATCTATGGTCAGGATTTTGATGACGATAAGTATGTTGAAGATATGCATGACGTGATCGAGTTCTTGATTCGTATTAGTGGTTATGCTCAACTTGCTGAGGTATTCTATTATGCTAAATATCTACCAAGTCATAAGCGAGCTGTAACTGGCGCTGAAGAAGCAAAAAGAAGAGTAATAGCTCTGGTGCGTCCTTTTCTTCAGTCAAACCCTGCTACTAACACTTACTTGCATTTTCTCAAATCGCAACAGTACCCTGAAGAGGTTATCATATTCGCTATTTTCGAAGCTTATCTTTTAGGTGTTGATAGTACTTCGTCAACCACTGCATGGGCACTCGCATTCTTAATACGTGAACCATCTGTTCAGGAGAAACTTTATCAAGAGCTTAAGAATTTCACAGCCAATAATGATCGCACAATGCTGAAAGTCGAAGACGTCAACAAATTACCATATTTACAAGCTGTTGTTAAAGAAACAATGAGGATGAAACCAATTGCACCACTGGCGATTCCTCATAAAGCTTGTAAAGACACTTCATTGATGGGCAAGAAAGTTGATAAGGGAACTAAAGTTATGGTTAACATTCATGCTTTACATCATACTGAAAAGGTTTGGAAAGAACCTTACAAATTCATGCCAGAGAGGTTTCTGCAAAAACAAGATAAGGCGATGGAACAATCACTATTACCATTTAGTGCAGGTATGAGAATTTGTGCAGGAATGGAATTAGGAAAACTTCAGTTTAGTTTTTCTCTTGCTAATCTTGTTAATGCTTTTAAATGGTCTTGTGTTTCTGATGGAGTGCTTCCTGACATGAGTGATTTACTGGGGTTTGTTCTGTTCATGAAGACCCCGCTCGAAGCACGTATAGTTCCTCGTTCGTAGTGATGGAAATTGCATACCATGTTGTTGTTTTTCTTCATGTTTATTACTTTGTACTGGTTTGATTTGGTGTGAAGATAAGATCTAAATTCCCTAATATCATTTGAAAATGTTTAAGCGAATCGAAACAGATCGGAAGAGC

>Par_TRINITY_9OMT Par_c33130_g1_i1 len=1676

GTCATCAAAAACAGTCCTAAGATGGCTACCAATGGCGAAATGTTCGGTACCTATGGTCATAATCATCAAACAGCCACAGTCACTAAAATCGCTGCTTCTAAGGAAAGCAACAATGATATTTGTTATCTTTCAGAAACGGCTAACTTAGGGAAGTTAATATGCATTCCAATGGCATTAAGAGCTGCTATGGAGCTAAATGTGTTCCAACTTATCTCCAAGTTTGGAACTGACGCAAAAGTTTCGGCTTCTGAAATTGCCTCTAAAATGCCAAACGCTACGAATAATCCTAAAGCAACTATCTATTTGGAAAGAATTCTTCGACTGCTCGGTGCCAGTTCTATTCTTTCGGTTTCTACCAGAAAAAAATTAAACGGAGGAGGAAATGATGGAGTAGCACATGAGAAGGTTTATGGGTTAACAAATTCGTCACGTTGTTTGGTCCCTCGAGAAGAAGACGGGGTGTCATTAGTCGAAGAATTGCTATTCACATCTGACAAGGTTGTTGTGGATAGTTTTTTCAAACTGAAATGTGTGGTGGAAGAAAAAGACATTGTGCCATTTGAGGTTGCTCATGGTGCTAAGATCTTTGAGTATGCTGCTACAGAACCAAGAATGAATCAAGTTTTTAACGATGGAATGGCAGTTTTCTCTATTGTTGTGTTTGAAGCTGTTTTTAGAGTTTACGATGGATTTCTTCATATGAAAGAATTGTTAGATGTTGGTGGTGGTATTGGTACTTCGGTTAGTAAGATTGTTGCTAAATACCCTTTCATTCGCGGTGTCAACTTCGACTTGCCTCATGTTATTTCTGTTGCCCCTCAATACCCAGGTGTAGAGCATGTTGCAGGAGATATGTTTGAGGAAGTCCCAAAGGGTCAAAACATGTTGCTAAAGTGGGTACTGCATGATTGGGGTGACGAACGATGTGTAAAGCTGTTAAAGAATTGTTGGAACTCATTACCTGTGGGTGGAAAAGTTTTGATAATCGAGTTTGTTCTCCCGAATGAACTTGGTAACAATGCCGAATCATTCAATGCGTTGATTCCCGATTTACTGTTGATGGCTCTGAATCCAGGCGGTAAAGAACGAACAATTTCCGAATATGATGATTTAGCCAAAGCAGCTGGATTTATAAAAACAATACCTATCCCTATCTCCAATGGTCTTCATGTCATTGAGTTTCACAAATGAATGGTTATCGAGTCCTTTGGTAATTAATTAAACTACCAAGATAACTGCATCCAAGTCATGCATTTGCTTTTTTTCCTTTTATTTTGTTTTGTTTGTTTTTGTTTCAAGGTGTGAACTAGTGTGTTAAGTGGACAAAACTAAGTAATCGTATTTTGTGTTTCTTAATGAAGTCTTCATGTGATTGATGTTTGTGAATCCCATGGGTAGATCCTGGCTTATCATGTGATTCATGTCCTTGTTTTCACCACCTATTTTTAGTTTACTTATAAAACATGCTGAAAATACAGATCTAACCGTTTAATCCTAGTGCAAAAGAATTATGGATCCCCAGCCCTATCAGGAGTAAATAATCTTGATTCGGTATAGAATTGGTGAAGACAAGTTTCAAAACTCAGATCCATGTACTACAACGTTGACGACGGTACAAAAAGATTAAGTATGATGGTTTTTGTTACTTCTTTCTTTTGAATGATTTAGATCGGAAG

>Par_TRINITY_MSH Par_c19507_g1_i1 len=1116

GTGTGCTCTTCCGATCTGGTGCCTACTGGAGAGATCTACGAAAGATTTCAACGTTAAAACTTCTTTCAAACCATCGCATCGAGACGATAAAACATCTAAGGAGTTCTGAAGTTGAGTCGTGTTTCGAGTCGCTTTATAACCAGTGGAAGAACGGAGAAAAGCATGGCGAATTCGCTCCGGTGAGGATGGATAGTTGGTTAGGCGATTTGACGTTTAACGTGGTGGCGAGAATCGTTGCCGGGAAAAAGAATTTCTCGGCCAACGGTGACGTTGGGGCTCAGAGATATAAAGCAGCAATGGATGAAGCTATGAGATTAATGAGATTTTTCGCATTTTCGGATGTGATTCCGTCGCTAAGCTGGTTAGACAATTTAAGAGGTTTGGTGCGAGATATGAAAAAATGTGCATCTGAGATTGATACAATTATGGCTTCTTGGGTTGAAGAACATCGCGTGAAAAGAAATTCCGGAGGCAATTCTCAACTCGAACATGATTTTATTGATGTTTGCTTGGATATTATGGAGCATTCTTCGTTACCCGGCAATGATCCTAACCTCGTCGTCAAATCTACGTGTCTGGATATGATACTGGGTGGGAGTGACACAACAACAGTTACCCTAACATGGGCAATGTCTTTACTCTTGAACCATCCCCAAGTACTACAAAAAGCAAAACAAGAACTAGACACACAAGTAGGAAAGAACCGGCAAGTAGACGACTCGGATATCCCGAATCTTCCGTTCATCCAAGCGATCATCAAAGAGACGATGAGGCTGTACCCTGCCGGACCACTAATCGAACGAAGGACGATGGAGGATTGCAAAGTAGGTGGGTACCAAGTTCCGGCCGGCACGCGCTTATTGGTTAACGTCTGGAAGATGCAAAGAGATGGGAACGTATACAAAGGCGATCCGTTGGAGTTTAGACCCGACAGGTTTTTAACTAGTAATGCGGATGTGGATCTTAAGGGTCAGCATTACGAACTGATACCATTTGGAGCGGGTCGACGGATATGCCCGGGAGTATCGTTTGCGGTTCAGTTGATGCATTTGGTTCTTGCCCGATTACTTCATGAATTCGACATCACGACGGTGGAGCCAGAAGCGAAGGTCGATA

>Pdu_TRINITY_NCS Pdu_c29677_g2_i2 len=576

CTTCCGATCTGAAAAAACCTCTAAATCTTGTATCATTAAATCTTCTGTTGTCTATGAAGTGAAGCAGGAGTGTGTTGAGGCAATGTCTAAGTTGATCACCACTGAACCATTGAAGTCGATGGCAGAAGTTATCTCTAATTACGCTATCAAGCAACAATCTGTTTCTGAGAAGAACATTCCTAAGAAGCAAATTCTACTGAGGAAGGAAATTACTTATGAAACGGAGGTGCAAACTTCCGCTGATTCTATTTGGAACGTCTACAGTTCTCCTGACATCCCTCAACTACTTAGAGATGTCCTGCTTCCTGATGCTTTCGAAAAGCTACATGTCATTCAAGGCAATGGTGGCGTTGGTACTGTACTTGACATTGCTCTCCCTCTAGTGCCGCGAAGGTACAAGGATAAATTTGTGAAGATCAACCATGAGAAGCGATTAAAAGAAGTGGTTATGATTGAAGGCGGATACTTAGACATGGGATGCACATTTTACATGCATAGGATCCATGTCTTAGAGAAAACACCAAAGTCGTGTGTCATTGAATCCTCCATCGTTTATGAAGTGAAAGAAGAGTATGC

>Pdu_TRINITY_6OMT Pdu_c32757_g1_i1 len=1146

GCTCTTCCGATCTTTCTCATCAACAGCTAAAGTTTCTAAAAAGAGAGAGAAATGGAAACAGTTAGCAAGATTGATCAACAAAACCAAGCAAAAATCTGGAAACAAATTTACGGTTTTGCAGAATCACTAGTTCTGAAATGTGCAGTTCAACTAGAGATTGCTGAAACGCTTCACAACAATGTCAAACCCATGTCTTTATCCGAATTAGCATCGAAACTTCCCGTTGCTCAGCCCGTTAATGAAGACCGTCTGTTCCGAATCATGCGTTACTTGGTTCACATGGAGCTCTTCAAGATAGATGCTACCACGAAGAAATACTCATTAGCTCCACCAGCTAAGTATCTGTTGAGAGGGTGGGAGAAATCAATGGTTGATTCAATTTTATGCATAAATGATAAGGATTTCTTAGCTCCGTGGCATCATTTAGGCGACGGTTTAACCGGTAACTGTGACGCTTTTGAGAAAGCGTTGGGGAAGAGTATTTGGGTTTATATGAGTGAAAATCCTGAAAAGAATCAATTGTTTAATGCAGCAATGGCTTGTGATACTAGATTGGTTACTTCAGCATTGGCTAATGAGTGCAAAAATACTTTCAGTGATGGAATTAGCACACTGGTTGATGTCGGCGGTGGTACTGGTACTGCTGTGAAAGCCATATCCAAGGCTTTTCCAGATATTAAGTGCACAATCTATGACCTTCCTCATGTTATAGCTGATTCTCCTGAAATTCCCAATATCACTAAAATTTCTGGAGATATGTTCAAGTCTATTCCTAGTGCTGATGCCATCTTCATGAAGTGCATACTACACGACTGGAACGATGATGAATGCATTCAAATCTTGAAGCGATGCAAAGAAGCACTGCCAAAAGAAGGCAAAGTTATTATCGTGGATGTCGTGATAGACATGGATTCGGATCATCCATATGCAAAAATGAGACTGACATTGGATTTGGATATGATGCTTAACACTGGTGGAAAAGAGAGAACCAAAGAAGAATGGAAGATACTTTTTGATGCCGCCGGTTTCGCTAGCCACAAAATCACTCAGATATCTGCTGTCCAATCTGTAATTGAGGCTTACCCTTATTAAGGAACATTTTACCCAGTTTTTTCCCTTTGATTAATTGTTGCTTTGGATTTGGAT

>Pdu_TRINITY_CYP80B1 Pdu_c48941_g1_i1 len=991

GCTCTTCCGATCTTTTTAAGTTGAGTGATGAAAATAGTGGAAATGAAAGAACAGATATGGAGATTGCTGGAATTAGGTAATTCAACAAATCCAGCTGATTATTTTCCATTTTTGGGTAGCTTGGATTTGTTGGGACAAAGAAAAGATGTCGCTGATAGTCTGCAAGAGATTTATGGAGTTTGGGGTCCTATACTGACAGAAAAAAAACTAGCCAAGCAACAAAACACCCGTAGTACGACGAATGATTTTGTTGATCTTTTGGTTGATTCTGGACTTAATGATCAACAAATCAACGCCTTTCTCATGGTATGCTCAATTTACAGAAATGATCATTGTTAATATCGATCTTGGTGTTTACAGGAAATATTTAGTGCGGGAACAGTGACAAGTGCAACTACCATAGAATGGGCCTTAGCTGAGCTCATAAAAAACCCTGAGTTAACAACCAATATTAGATCAGAATTGCGGTCTGTGGTAGGGAAGAGGTCGGTTAAGGAATCCGACATACCTAACCTTCCCTACCTTGAAGCTTTTGTTAAAGAAACTCTGCGGCTTCACCCAGCAACTCCATTACTACTCCCACGACGAGCACTTGAGACGTGTCAAGTTTTCAACTATACGATTCCAAAAGGGTGTCAAATAATGGTCAACGCCTGGGGCATCGGCAGAGATCCGAAAACATGGACCGATCCGTTGAAGTTTGCACCAGAGAGGTTCTTGAATTCGAGCATCGACTTCAAAGGGAACGATTTCGAGTTCATACCATATGGCGCAGGGAGAAGGATATGCCCTGGTGTGCCTTTGGCAACTCAGTTTATCAGTCTTATTGTGCCTACTTTGATTCAGAATTTTGATTTGGGGCTACCAAAGGGAATGGTTCCTAGCGAACTCATCATGGAGGAGAAATTTGGGCTGATGCTGGAGAAGGAACCACCTCTGTATATTGTTCCTAAAACTCGGGAATAATTGATTACCTTTCTTTTCCTATCCC

>Pdu_TRINITY_CYP719A21 Pdu_c36265_g1_i5 len=1893

TCTTTAAATCAAATCATCGATGATCCTTTTACCTTTCTGAAACAGGATAAAGAAAAAGAGCACTGAGGCAATAATTCAATAGAAATTAGATAAACTACAAGAATAGTGGACACCCACATGAATGATTTGATTCTCAAATACAGCATTAACGGAGTATTAGCTAGCCGGGGGATCAGACCAAGTAATTGTATATAAGGTGAATGGTTGTTGTGCCTACAACCATGATATATTATTCTTCATGCAAAGAGTTAGTCAAACTCAAGCTAGCTAGAATATACACCAACAACAAAAGAAATGATCATGAGTAACTTATGGATTCTTACGCTCATCACTACCATATTAGCAGTCTTTGCTGCTGTGTTAATGATTTTCAGGAGAAGAACGTCAGCATCTACAATGAAGTGGCCAGTTGGCCCAAAATCATTACCAATCATAGGTAACTTGCACATTCTTGGAGGCACTGCTCTCCATGTCGTCTTACATAAACTAGCTGAAGTTTACGGCAGTGTAATGACGATATGGATTGGTAGTTGGAAACCTGTTATTATTGTTTCTGACTTCGATCGAGCTTGGGAAGTCCTTGTCAACAAATCTTCAGATTATTCAGCTCGTGAAATGCCTGAAATCACTAAAATCGGTACTGCAAATTGGAGAACAATTTCAAGTTCTGATTCTGGTCCATTTTGGGCCACTCTTCGTAAAGGTCTTCAGGGTGTAGCATTATCGCCTCAGAATTTAGCTTCCCAAACTGCATTACAAGAGAGAGATGTCAGAAAATTGATCAAAAGTTTGGATGAAGAATCTGCTCTTAATGATGGAATCGTTAAACCACTTGATCATCTCAAGAAAGTAACTGTTAGATTAATCAGTCGGTTAATCTATGGTCAGGAGTTTGACGATGATAAGTATGTTGAAGATATGCATGACGTGATCGAGTTTTTGATTCGTATTAGTGGTTACGCTCAACTTGCTGAAGTTTTCTATTATGCTAAATATCTACCAAGTCATAAGAGAGCTTTAAACGGTGCGGAAGAAGCAAAACTAAGAGTAATAGCTCTGGTGCGTCCTTTACTTGTTTCAAACCCTCCTCCTACCACTTACTTACATTTTCTTAAATCCCAAAACTACCCTGAAGAAGTTATCATATTTGCTATCTTTGAAGCTTATCTTCTAGGTGTTGATAGTACTTCTTCAACCACTGCATGGGCACTTGCATTCTTAATACGCGAACCATCTGTTCAAGAGAAACTCTATCAAGAGCTTAAGAATTTCACAGCCAATAACAATCGCACAATGCTGAAAGTCGAAGACGTAAACAAATTACCATACTTACAAGCTGTTGTAAAAGAAACAATGAGGATGAAACCAATTGCACCACTGGCGATTCCTCATAAAGCTTGTAAAGACACTTCACTGATGGGCAAGAAAGTTGATATGGGAACTAAAGTTATGGTTAACATTCATGCTTTACATCATACCGAAAAAGTCTGGAAAGAACCCTACAAATTCATGCCAGAGAGGTTTCTGCAGAAACAAGATAAGGCGATGGAACAATCACTATTACCATTTAGTGCAGGTATGAGAATTTGTGCGGGAATGGAATTAGGAAAACTTCAATTTAGTTTTTCTCTTGCTAATCTTGTTAATGCCTTCAAATGGTCTTGTGTTTCTGATGGAGTACTTCCTGATATGAGTGATCTACTAGGGTTTGTTCTGTTCATGAAAACCCCACTCGAAGCACGTATAGTTCCTCGTGCATAGTGATGGAACTTTCGTAGTGATCGATGAAACTTTCTTTTTCATGTTGTTTTTCTTTTTCTTTACAACTAGGTACTGGTTTGATTTTGGTGTAAAAAATAAGATCTGAATTCTCGATTATCAGTTGAAAAATGA

>Pdu_TRINITY_9OMT Pdu_c13499_g1_i2 len=1028

AAGTTTCGGCTACTGAAATTGCTTCTAAAATGCCAAACGCTGCGAATAATCCTAAAGCAGCTATATATTTAGATAGAATTCTTCGATTGCTCGGAGCAAGTTCTATTCTTTCTGTTTCTACTAGAAAAAAATTAATCAACGGAGGAGTAGAAGATGGAGTAGTACATGAGAAGGTGTACGGGTTAACAAATTCGTCGTGTTGTTTGGTTCCTCGAGAAGAAGACGGGGTATCATTAGTCGAAGAATTGTTATTCACATCTGACAAGGTTGTCGTGGATAGTTTTTTCAAACTGAAATGTGTGGTGGAAGAAAAAGACATTGTGCCATTTGTGGTCGCTCATGGTGCTAAGATCTTTGAGTATGCTGCTACAGAACCAAGAATGAATCAAGTTTTTAACGATGGAATGGCAGTTTTCTCTATTGTTGTTTTTGAAGCCGTCTTTAGAGTTTACGACGGATTTCTTGATATGAAAGAATTGTTAGATGTGGGTGGTGGTATTGGTACTTCGGTTAGTAAGATTGTTGCTAAATACCCTTTCATTCGCGGTGTCAACTTCGACTTGCCTCATGTTATTTCTGTTTCCCCTCAATACCCAGGTGTAGAGCATATTGAAGGAGATATGTTTGAGGAAGTTCCAAAGGGTCAAAACATGTTGCTAAAATGGGTGCTTCACGATTGGGGTGACGAACGATGCGTGAGGTTGTTAAAGAATTGCTGGAATTCATTGCCTGTGGGTGGAAAAGTTTTGATTATCGAATTTGTTCTTCCTGATGAACTCGGAAACAATGCCGAATCATTCAATGCGTTGATTCCCGATTTACTCCTGATGGCTCTGAACCCAGGTGGTAAAGAACGGACAATTTCAGAATACGATAATTTAGCTAAAGAAGCTGGATTTATGAAAACTATACCTATCCCAATCTCCAATGGTCTTCATGTCATTGAGTTTCATAAATGATTTCGAAATGGTAAATCGAGTTTACTAAGTTACTTGATTTGGGGTACACACCGACAGATCGGAAGAGCG

>Pdu_TRINITY_MSH Pdu_c96614_g1_i1 len=531

TTCTACGTTAAAACTTCTTTCGAATCATCGTATCGACACGATAAAACATCTGAGGAGTTCTGAAGTTGAGTCGTGTTTCGAGTCGCTTTATAACCAGTGGAAGAACGGAGAAAAGCGTGGCGAATTCGCTCCGGTGAGAATGGATAGTTGGTTGGGCGATTTGACGTTTAATGTTGTGGCGAGAATCGTTGCCGGGAAAAAGAATTTCTCGGCCAACGGTGACGTTGGGGCACAGAGATATAAAGCAGCAATGGATGAAGCCATGAGATTAATGAGATTTTTCGCATTTTCGGATGTGATTCCGTCGCTAAGTTGGTTAGACAATTTAAGAGGTTTGGTTCGAGATATGAAAAAATGTGCATCTGAAATTGATACAATTATGGCTTCTTGGGTTGAAGAACATCGCGTGAAAAGAAATGCAGGAGGTAATTCTCAACTCGAACATGATTTCATTGACGTTTGCTTGGATATTATGGAACATTCTTCATTACCCGGCGATGATCCTGACCTCGTCGTCAAATCTACGTGTCG

Supplementary dataset 2 (associated with Figure 1c)

- Eight conserved orthologous gene set from the transcriptomic dataset of nine *Papaver* species and annotated genome datasets of *P. somniferum*, *Macleaya cordata*, *Eschscholzia californica* and *Aquilegia coerulea* species.

>Pnu_TRINITY_Agt1 Pnu_c42321_g1_i1 len=1740

ACGTGTGCTCTTCCGATCTCATGCCGCATGATTTAGATGATTTAGACTTTGGTACATTGTAATCTTCCACGTGGAAGCTCTGTGATCTATCAGCATATACGTTTCATCTTGATCCTCATCTTCCACTCATTTTGGGTCTTTATGTATAAATATATGTATTTTTTCCTTAATTATTACACATATTTCCTTCCTCCCATTGTTTATCCCTCAAAGTTGGACTGGGAGAGTGAGTGAGAGAGCTGCTCTGAAGTTGTTCTCCGGAAACCGTTCTTGCAACAAAAAGCTTCAAAGAGGTACAATATGGATTACGTCAATGCACCTGGGAGGAACCATCTCTTTGTTCCTGGACCTGTCAATGTTCCAGAATCAGTCCTTCGTGCCATGAACAGAAACAATGAAGATTACCGTTCCCCTGCTATTCCTGCTATGACTAAAACTCTTCTTGAAGATGTGAAGAAGATTTTCAAGACAACTTCTGGAACCCCATTTTTGATCCCCACAACAGGTACTGGTGCATGGGAAAGTGCACTTACAAATACTTTGTCTCCTGGTGATCGGACAATTTCTTTCGTCATTGGACAATTCAGTCTTCTCTGGGTTGATCAGCAGCAACGCCTCGGATTCAACGTTGATGTTGTAGAAAGTGAGTGGGGCCGTGGTGCTGATCTTGACGTTCTAGCATCCAAGTTAGCTGAAGACACTGCTCATACCATTAAAGCTGTGTGCATCGTTCACAATGAGACTGCAACTGGAGTTACCAATAATTTGGCTGCTGTAAGAAAATTACTCGATCACTACAACCATCCTGCATTGTTTCTCGTTGATGGTGTGTCTTCAATATGTGCTCTTGATTTCCGTATGGATGAATGGGGAGTCGATGTAGCTTTAACTGGTTCACAGAAAGCTCTTTCTCTCCCTACTGGAATGGGTATTGTGTGTGCAAGCCCTAAAGCACTGGAAGCATCCAAGACCGCCAAATCAGTGAGAGTATTCTTTGACTGGAAAGACTACTTGAAGTTCTACAAAATGGGAACTTATTGGCCATACACTCCTTCAATCCAGCTTTTATACGGCCTCAGAGCTGCTCTTGATCTCCTATTTGAGGAAGGACTTGAAAATGTCTTTGCTAGGCACAACCGCTTGGCCACAGCAACTAGGCTTGCTGTTGAAGCCTGGGGTCTGAAAAATTGTACGCAGAGTGAAGAGTGGTTGAGCGACACTGTGACTGCAGTTGTTGTTCCACCTTACATTGACAGTGCTGAAATAGTTAGAAGGGGATGGCAGAGATACAATCTAAGCTTAGGTCTGGGTCTGAATAAAGTGGCTGGGAAAGTTTTCAGGATAGGCCATCTTGGAAACCTGAATGAGCTCCAACTGCTGGGTTGTTTAGCTGGAGTGGAGATGATTCTGAAAGATGTTGGGTACCCAGTTAAGCTAGGAAGTGGAGTGGCTGCGGCTTGTGCGTATTTCCAGAACACCATTCCAATGATCCAATCTAGGATTTAATGAGTCATCATCCATTACCCATGTGTAACTACTGTTCTTCCTCTTCTGATTTGCTTGGAGTACTTGTATAAAAGAAGCCATGAGAAACTCCTGTATTTGCACTCTTCTGAGGCTTACTCAGTAGTACAAGAACTCATTACTCTTTCATCCATTAATCCTGTATCTAAAATCTTTTTAAATTTTATCTAATCGCTGTTGCATATATCCAAGTACAAGATCGGAAGAGCGTCGTGT

>Pnu_TRINITY_At103 Pnu_c41539_g1_i1 len=1852

TATGTAAACAGTAATGGGGGGGACAAAAATCGTCCCCCCCATTAGCAGCGATGCAAATTTCCTTTCAGCAACTAAATCCTACAAAAACCTCCCATGTCTTATCTAACCATTTCCTTACTCAAACTGTGAAACTCCCTCTGCTTCTTCTTGTCACTAGAAACCACTACACACAATGGCTGCAGAGATGGCATTGATAAAACCCATCTCTAATTTCACTAGAACCAGCCCAAAATTTATCACCTCCACAAGAAAATCATTCTCCATTGTCAAAATGGCATCCCAATCCACTGCATCAACAACAAAGTCAGGGAAACCCAAGAAAGCAATCAAGGAGACCTTACTTGCCCCAAGGTTTTACACTACTGATTTTGATGAAATGGAAACTCTTTTTAACACTGAGATGAACAAAAACTTAAATGAGGCTGAGTTTGAAGCTTTATTACAAGAGTTCAAAACTGATTACAATCAAACTCATTTCGTTAGAAACAAGGAATTCAAAGAAGCTGCTGATAAATTGCAAGGTCCTCTTAGACAGATTTTTGTTGAGTTCCTGGAGAGATCTTGTACTGCTGAGTTCTCTGGGTTTCTTCTTTACAAAGAACTTGGAAGAAGACTTAAGAAAACCAATCCAGTTGTTGCTGAGATTTTCTCACTCATGTCCAGGGATGAAGCTAGGCATGCTGGATTCTTAAACAAGGGACTGTCAGATTTCAACTTGGCACTGGATCTTGGATTTCTTACGAAGGCTAGGAAATATACATTCTTTAAACCAAAGTTCATTTTCTATGCTACATATTTATCTGAGAAAATTGGCTACTGGAGGTACATTACCATATATAGGCACCTAAAAACCAATCCTGAATATCAATGTTATCCAATTTTCAAGTATTTTGAGAACTGGTGCCAAGACGAGAACAGACATGGAGATTTCTTCTCCGCGCTGATGAAAGCTCAACCCCAGTTTCTTAATGATTGGAAAGCAAAGTTGTGGTCTAGATTCTTCTGCCTATCGGTGTACGTGACAATGTACTTGAATGACTGCCAGAGAACAGCTTTTTACGAAGGGATCGGACTTAACACCAAAGAGTTTGATATGCACGTCATAATTGAGACAAACCGTACAGCCGCAAGGATTTTCCCTGCAGTTTTAGATGTTGAGAACCCGGAGTTCAAGAGGAAGTTGGATCGAATGGTGCAAATCAACGAGAAAATTATTGCAATCGGTGAAACTGAGGATACACCATTGGTGAAGAACTTAAAAAGGATCCCACACGTTGCAGCATTGGCGAATGAGCTCTTGGCTGCGTATCTCATGCCTCCTATTGATTCTGGGTCTGTTGATCTCGCAGAGTTTGAGCCACAAGTTGTCTATTAATAAGTTTTCCGTGTATTGAGTAAACAAAGCCGTTTCAATTAAATTCAGTGCAAAGTTTAATTGCTACAGTTAACCATTACTATGGACCTGCTGTGTAAAGAACTAAACATTTGTGACATAATGTATCACGATTCATGTACTCGCATCTTGCAGACATCACTCACAATGCTTTTTTTCCGTACATACTCATATTTAGGTTGTCACTTGTCCCAACTGGCAAGGCAAGTTCAAGAGCTAGACAATGGCCTTACTATCCGGTCAGTGGGATGACACATTTTGGTCACAAAACCTGTTGATTGGTCATTATACAACCCAGAACCCTATTGAGATGACCTGATATCACGGGCAGCAAATTCAAGGGCCAAAAAGTTTCGCCAGCAGCCACTCGAAACAAGGGAGATCTGACACATGTTACCAGTTTCTACACATGGGCGGACCCAGCCAGTATACACTGTATACTTACTCAGGCAACTAAAG

>Pnu_TRINITY_ChlP Pnu_c53611_g1_i1 len=1234

GTGTGCTCTTCCGATCTCTTGAATCCCCATCGCACCACCAAAAATGGCCTCCATTGCCGTCAATGCACGTTTCTCAGCCACAACATGCGACCGTAAAACCACCGAAATCAGCCGTGAGACACTAAAATCCTTCATGGGTCTTAGAAAAACCAATTTGTCACACCAGAATTCCATCTCACTTTCGCAATCTCTAACAACAACTACACATTCCAACTCTGCAAGAAGAAACTTTTCTATTAAAGCAGGGAAATCAAGTCCCAAAGTAACAGGAAGAAATCTCCGAGTCGCAGTCATCGGTGGTGGCCCTGCTGGTGGTGCAGCTGCTGAAACCTTAGCCAAAGGCGGCATTGAAACGTTCTTGATAGAGAGGAAAATGGATAACTGTAAACCATGTGGTGGTGCGATTCCACTCTGCATGGTTGGTGAGTTCGATCTTCCGTTAGATATCATCGACAGACGTGTTACCAAGATGAAGATGATTTCGCCCTCAAACGTTGCAGTCGACATTGGCCAGACTCTCAAACAGGACGAGTATATTGGTATGGTCCGTCGAGAAGTGCTAGACCAGTACCTTCGCGAGAGAGCAGAGAAAGCCGGAACGACTGTGATCAACGGTTTGTTCTTGAAGATGGATTTACCAAAAGAGCCAAATGCGCCATATGTTCTTCACTACAGTGCTTACGATGGCAAAACGGGCGGAGTCGGCACGAAAACCACGATGGAGGTCGACGCGGTCATCGGTGCAGACGGTGCGAATTCTCGCGTAGCCAAAGCAATTGGAGCCGGGGATTACGAGTACGCTATCGCATTCCAGGAGAGGATCAAAATCCCAGACGACAAGATGGTTTACTACGAGAACCTTGCGGAGATGTACGTCGGCGACGACGTTTCGCCAGACTTCTACGGATGGGTGTTTCCGAAATGTGATCACGTCGCTGTGGGTACGGGCACGGTCACGCACAAGGGAGACATCAAGAAATTCCAGGTGGCGACGCGGTTGCGCGCCAAGGAGAAGATTCTGGGTGGGAAGATCATTAGAGTGGAGGCACATCCTATTCCAGAACATCCAAGACCCCGCAGGTTGGCAGACAGGGTAGCCCTCGTGGGAGACGCGGCTGGGTACGTGACGAAATGCTCGGGTGAAGGTATATACTTTGCTGCTAAGAGTGGCAGGATGTGTGCCGAAGCCATTGTCGAAGGGTCCGAGAGAGGGACGAGGATGGTGAACGAAGCG

>Pnu_TRINITY_GbssI Pnu_c46969_g1_i1 len=2058

CAAGATTACTAATACTTACTTGACTTCTGTGCCGCTGTGACCACATCCATCATATAAGTGAACTCTACTGTCAGTCTAGAGAGAGACAGAGAAAGAGTGTGTGGATTTTGAGATTATCGAGAGTGACATATCTCTTGATTCTCCATCCTTGTGAACTCATCAGCTCTCGAGTCTGAACTGACAAGAATATAACCGATGGCAACCACAACTGCATCCCAGTTTGTATCACGAACCTCGCATGTCAATTGTGGAGGAAGTGAACCAAAAGACATACTGAAACAGATCAGTACGAAGAACCAAAGCATGACACATTCAGGGTTAAGATCTTTGAGCAAAGTAGATTTCTTACAAATGAAGGCTACTGTGAAAGCCGCCGCTTCTAGGGAATTGAGGAGGAACGCATTCCAGACAGACAAAACTACTGTGTCAGGAGACATAAAATGTTCTAGCGGAATGAACTTGGTCTTTGTTGCTGCAGAAGTAGGCCCCTGGAGTAAAACCGGTGGCCTTGGTGATGTTCTTGGAGGCTTACCACCAGCCATGGCGGCGAATGGACATCGTGTTATGACAGTCTCTCCTCGTTATGATCAGTACAAAGATGCTTGGGATACCAATGTGTCCGTTGAGATGAAAATTGGAGATAAAACTGAAACCATTCGTTTCTTCCACTGTTATAAACGTGGGGTCGACCGTGTATTTGTGGATCATCCACTGTTTCTTGAGAAGGTATGGGGTAAAACTGGATCTAAAATCTATGGTCCGAATGCTGGATCAGATTACCAAGATAACCAGCTTAGATTCTGTTTGCTTTCTCTGGCAGCTCTAGAAGCACCGAGGATTCTGAATCTTAACAGCAACAAATATTTCTCTGGAACATACGGAGAGGAGGTCGTGTTCATTGCTAATGATTGGCATACGGCCCTTCTACCATGCTACCTTAAAAGTATGTATCAATCGAAAGGGATGTATGAAACTGCAAAGGTTGCTTATTGCATCCACAATATTGCCTACCAAGGAAGATTTTCTTTTTCGGATTTTGTGCTTCTCAATCTTCCGGATTCATATAGGAGTTCTTTCGACTTCATTGATGGGTACGAGAAGCCTGTTAAAGGAAGGAAGATTAACTGGATGAAGGCTGGAATATTAGAGTCAGACAAGGTTGTAACTGTGAGTCCATACTATGCTGAAGAACTTGCCTCAGGCCCGGATAAAGGTGTGGAACTGGATAGCATTATCCGTAGAGCTGGCTGTTTGGGGATTATAAATGGAATGGATGTTCAAGAGTGGAACCCATTAACTGATAAACATATCGACGTCAAATATGATGCCTCAAATGTGACCGAAGCAAAGGCTCTTCTGAAAGAAGCACTGCAGGCAGATGTAGGATTACCTGTTGACCGTAATATCCCTGTCATAGGCTTCATTGGTAGACTTGAAGAGCAAAAGGGTTCAGATATTTTGGCAGCTGCAATCCCTGAATTTATTGAAGAAGAAGTTCAGATAATTGTCCTTGGTACTGGCAAAAAGCACATGGAAAAACAGTTAGAGCAGTTAGAGACTTTGTATCCTGAAAAAGCTAGAGGAGTGGCAAAATTCAATGTTCCCTTGGCTCATATGATCATAGCAGGGGCTGATTTTATGATGATCCCAAGCAGATTTGAACCATGTGGACTCATCCAACTGCATGCTATGCGTTATGGAACGGTGCCTATCGTTGCCTCGACGGGTGGACTAGTTGACACTGTGAAAGAAGGTTTCACTGGATTTCAAATGGGAGCTTTCAATGTCGAATGCGAGGCAATTGATCCATTAGACGTGACAGCAATCGCCACAACCACCAAAAAGGCCCTTGCAACCTATGGAACTCCAGCTTTAACAGAAATGATACAGAACTGCATGGCTCAAGATCTCTCTTGGAAGGGACCTGCCAAGAATTGGGAGGACATGTTGCTGAGCTTGGAGGTTGCGGGAAGTGAACCAGGAATCGAAGGCGATGAAATCGCTCCTCTTGCAAAAGAAAATGTTGCCACTCCCTGAGTCACAGAACCGCTCCAGTGC

>Pnu_TRINITY_Hmgs Pnu_c48920_g2_i1 len=1716

CCTTCTCTCCCTCTCTCTCAAAACAGCTCGATCGATCTAGGGTTCATTTGGAAGAAATTATCAAAAATGGGTTCAAATCAAAAGGATGTTGGAATTCTTGCTATGGAAATGTATTTCCCTCCTACTTGTGTTCAACAGGAAGCATTGGAGGCCCATGATGGTGCCAGTAAAGGAAAATACACCATTGGGTTGGGACAAGATTGCATGGCGTTTTGTTCGGAGTTGGAAGATGTCATTTCAATGAGTTTGACAGCCGTTAGCTCTTTACTCGAAAAGTATAAAATTGACCCTAAACAAATTGGACGCTTGGAAGTCGGCAGTGAGACCGTGATTGACAAGAGCAAGTCCATAAAGACTTTCCTGATGACAATCTTTGAGGAATGTGATAACACTGACATTGAGGGCGTTGACTCAACAAATGCGTGCTATGGAGGAACTGCAGCTTTGTTCAACGTTGTGAATTGGGTGGAGAGTAGTTCCTGGGATGGACGTTATGGTCTTGTTGTGTGCACTGATAGTGCGGTCTACGCAGAAGGGCCTGCCCGTCCCACTGGAGGAGCAGCTGCTATTGCTATGTTGATTGGCCCAGACGCACCAATTGCGTTTGAAAGCAAATATAGAGGATCCCACATGTCCCATGTCTATGATTTTTACAAGCCCAACCTTGCCAGTGAATACCCAGTTGTTGACGGAAAACTTTCACAGACTTGTTACCTTATGGCACTTGATTCCTGCTACAAGCGCCTATGCGCCAAGTATGAGAAATTTGAGGGTAAGCAATTCTCTATGGATGATGCAGACTACTTTGTATTTCATTCTCCTTACAACAAGCTTGTGCAGAAAAGTTTTGCTCGATTGTACTTCATGGACTTCTTAAGAAATGCCAGTTCTGTTGACGAGGCTTCCAAAGAAAAGCTTGCACCATTTTCAGCTCTATCTGGTGAAGAGAGCTACGCAAGTCGGGATCTTGAGAAGACATCCCAAACTGTGGCAAAGAATCTTTATGCTGAGAAGGTTGGACCTACCACTTTATTACCCAAACAAGTTGGTAATATGTATACTGCATCACTCTATGCGGCGTTTGCTTCTCTTCTTCACAACAAAAATAGCTCACTGGATGGTAAACGGGTAATAATGTTCTCTTACGGAAGTGGTCTAAGTTCCACAATGTTCTCATTCAGAATGAGGGAAGGTTCACATCCATTCAGCTTGTCAAACATTGCAACTGTAATGGATGTTGCAAACAAGTTGGAGTCGAGACATGAGGTTACTCCTGAAAAATTCGTCGAAACCATGCAACTCATGGAGCACAGGTATGGAGGGAAAGACTTTGTGACGAGCAAGGACACAAGCCTTTTACCTTCAGGCACATACTATCTTACCGAAGTCGACTCCATGTACCGCAGATTCTACTCCAAGAAGTCAAGTGCTGCAGTGCAAAATGGTCACTGAAAATGGTGAACCAAGGAAATGGCATCAGAAGAGCTGCCGAAGATTTGCTTATGCACCCCCATTGTTGTTGTTAGTGATCTAGAATAATTCTTATTATGATTTGTACAAGTTAATCTTTTCGAACTCGCCTATTAGTTATCAACTCATTGTAACAGGATTCATCTCCTCTCAGTATATCCATTCAAGATTCAAGTGTCAAAAAAAAAGAACGGAAGAGAGTAGTGCAGGGAGGGAGAGAGGAACAAAGAGAAGGAGCCGACTGGG

>Pnu_TRINITY_Nia Pnu_c54090_g1_i2 len=3028

GTGTGCTCTTCCGATCTTGCAAAAGCAAATCGAAAATATAGAAAACACAGAAGCAGCAGAGAGAGATAAAGTGAAACGAGAGAGAGTGATATATAAAAAGAGAGATCTTTCTTTCTCTTCTTTCTTTTGATTAGTCTCTAATGGCGGCTTCTGTTGAGAATCGACAGTACAACAGTCGACTTGAATCAGGTCTAGCAGTTCGTGGATTTAAACCCAACTCCACCCATCGATCTGATTCTCCTCCTATCCGTAGCTGTGCGTTCCCTATGTCGACCACCGAGCGTCTTTCTACACAGACAAAAACGGTCATCAGAGAAGATTTATCTAGCGACGACGAAGAGGATCAAAACGAGCAGTGGAAAGATTTCGGCAAAATCAACAATGAAATAGAGCCGTCTGTTTTCGACCAGAAAGATGAAGGAACCGCTGATAATTGGATTGAGCGCAACCCTTCCATGATCCGGTTAACAGGGAAGCACCCATTCAATTCAGAAGCGCCATTAACTCGGCTCATGCATCACGGGTTCATTACACCTGTTCCTCTTCATTACGTCCGTAATCACGGAGCTGTGCCAAAAGCCAAATGGGAGGATTGGACGGTTGAGATTTGTGGATTAGTAAAGCGTCCGTTGCGCATCACCATGGATATGCTAGTGAATGACTTCCCAAGCCGTGAGTTTCCAGCTACACTTGTCTGTGCAGGTAACAGACGTAAGGAACAGAATATGATAAAACAAACAATTGGGTTCAACTGGGGAGGTGCAGGAGTATCGACTTCAGTGTGGCGTGGTGCACGTTTATGTGACGTATTAAAGCGTTGTGGCATCATGAGCCGTAAGAAAGGTGCTCTTAATGTCTGCTTCGAAGGAGCTGAGGATTTGCCCGGTGGCGGTGGATCTAAATATGGAACTAGTATTAAGAAAGAGTTTGCAATGGATCCAGCACGTGATATCATCTTAGCTTACATGCAGAACGGTGAGAATTTGTCTCCTGACCATGGATTTCCAGTCCGTATGATAATTCCGGGTTTCATCGGAGGAAGAATGGTGAAATGGTTGAAGAGAATTATTGTCACAACACCAGAATCTCAAAATCATTATCATTTCCATGATAATAGAGTCCTTCCTTCTCATGTGGACGCTGAACTGGCTAACTCAGAAGCATGGTGGTATAAGCCAGAGTACATCATCAATGAGTTGAACATTAACTCAGTGATTACAACTCCTTGTCATGAAGAGATTCTTCCCATTAATGCTGCTACAACACAGAGAGCTTACACATTGAGAGGCTACTCATATTCCGGTGGCGGTAAAAAGGTAACTCGTGTTGAAGTAACTATGGATGGTGGAGAGTCGTGGTTAGTATGTACTTTGGATCATCCGGAGAAACCAAACAAGTATGGCAAATATTGGTGCTGGTGTTTCTGGTCACTAGAAGTTGAAGTTCTTGACTTGCTTGGTGCCAAAGAGATTGCGGTTAGAGCTTGGGATGAAACTCTAAATACTCAGCCAGAGAAGCTTATCTGGAATGTCATGGGTATGATGAACAATTGTTGGTTCAAAGTCAAAACCAATGTGTGCAAACCACACAAAGGAGAAATCGGTCTGGTGTTCGAGCATCCGACTCAGGCTGGAAATCAACCAGGTGGATGGATGGTTAGACAGAAACATCTTGAGAACTCTGAAGGTCCACACACCCTCAAGAAGAGTTTGTCCACTCCTTTCATGAACACCACGTCAAAAATGTATTCAATTTCCGAGGTGAAAAAACACAACTCGGCTGAATCGGCTTGGATCATTGTCCATGGACACGTTTATGACTGTACCCGGTTCTTGAAAGACCATCCAGGTGGTGCCGATAGTATTCTCATCAATGCAGGTACCGACTGTACCGAAGAATTTGATGCTATTCACTCTGACAAAGCCAAGAATTTACTAGAGGACTTCAGAATCGGTGAGCTAATTGCAACTGGTTACACTTCTGACTCGTCATCTTCACCAAACAACTCAGTTCATGGTTCGGGCAGCTTGAGTCACCTTGCTCCTATCAAAGAAATCGGCATAGCAAAACCAGTGGCTCTAGTTGGTAGAGACAAAATACAATGCAAGCTTATTGCCAAAACCTCAATTTCCCATGACGTTCGCGTCTTTAGGTTTGCATTACCTTCAGCTGATCAAGTCCTCGGGTTGCCTGTTGGGAAGCACATATTTGTATGTGCCACCGTTAACGACAAGCTTTGCATGCGTGCTTACACTCCATCAAGTACAGTTGATGAAGTTGGTTACTTTGAGTTGGTCATTAAAGTCTACTTCTGCGGAGTTCACCCCAAATTCCCTAATGGAGGTATCATGTCTCAGCACTTGGATTCTCTTCCTCTAGGTGGAACTATTGATGTTAAGGGTCCGTTGGGACACATTGAATACACGGGTCGTGGTAACTTCACCGTTCATGGCAAACACAAGTTTGCCAAACGACTCGCAATGCTTGCTGGTGGTACAGGAATTACACCAATATATCAAATTGTACAAGCAATACTGAAAGATCCAGAAGATGAGACGGAGATGCACATCGTGTTCGCGAATAGGACTGAGGATGACATCTTGTTGATGGATGAACTCGATTCTTGGGCAGCGAAGCACGAGAGGTTGAAGGTTTGGTACGTCGTTGGTGAGACATTGAGGGAAGGATGGAAGTACAGTGTAGGGTTCATAACCGAAGAGGTTCTACGGGAACATATCCCTCCGGCATCGGATGATACATTGGCATTGGCTTGTGGACCACCACCTATGATTCAATTTGCAGTGACTCCAAATTTGGAGAAGTTGAAGTATGATGTGAAGAATTCATTGTTGGTGTTTTGATCAATCTGTTCAAGCACTTGAAAGTTGAAACTACATTAAGAATATCATAAGGGGAGGACAAATTTATAAGGAAGTTCTCATTAGTTATTGTATATAATTAGTGTTGAAGGATGTAATGGTTTAGTTTGTTCTTAATAGTAAGGGGATTGTTAAATGTTAGTCTGTTT

>Pnu_TRINITY_Eif3E Pnu_c51577_g1_i2 len=1706

TGCTCTTCCGATCTGTTTTATTTAGGGTTTCTCTCTTTCTTCCTCCACTCCACCACCACCACCGCATCTCCTCAGAGGCGGTATCAACAACCAATCAATCATGGCGAAGTATGATCTAACATCAAGAGTAGCACTAAACCTAGATCGGCATCTTGTTCTTCCACTTCTTGAATTCCTTCAGGAACGTCAAATCCATTCCAATGTTCAAATCCTCAAAGCTAAAGACGCTCTTCTCAGCCAAACCAATATGGTTGATTACGCTATGGATATCTTCAAATCTCTTCATCAAACTGATGATGTTCCTCAAGAGATGATTAATAGACGCGCTGAAGTGGTTGGGAGGTTGAAATCACTAGAAGATGCAGCTGAATCTATTATTACGTTCCTTCAGAGCCCGAATATGATGAATGAACCAAGGCCTGACAAACAATACGTTCTTCAGTTGCTTCACGCAAACCAGATTGGTGCGGAGCAAATTGAGGCGTTGTATCAGTATGCAAAGTTTCAATTTGAATGTGGGAATTACTCGGGTGCAGCTGATTATTTGCATCAGTTCAGAGCGATCTCTAACAATCCTGATAGGATTTTGAATGCGTGCTGGGGAAAGCTTGCGTCTGAGATATTGATGCAGAATTGGGATGAGGCTCTTGGTGAGCTTCATGGCGTCAAAGATATTATTGAGTCAAAGAATTTCAAGAACCCATTGGCTTTGCTCCAGAACAGGATCTGGTTGATGCATTGGAGTCTCTTCATCTTTTTCAACCACGAGAATGGTAGAAATGGTATCATTGATTTGTTTACGAATGACAGGTACTTGAATGCCATCCAAACAAATGCACCCCATCTTTTACGTTATTTGGCAACTGCATTGATTGTTAACAAAAAGAGAAGACCTCAACTCAAAGAGTTGATCAAAGTCATCCAGCAAGAGCAATCTGCATACAAGGATCCCATCACTGAGTTCTTATACTGTTTGTATGTGAACTACGACTTTGATGGTGCACAGCAGAAGCTGCGTGAATGCGAGGATGTGATTTTGAATGATCCATTCCTTGGCAAACGAGCTGAAGAAGGTAACTTCAGTACCGTTCCTTTGAGGGATGAATTTCTTGAGAATGCCCGTCTCTTCATCTTTGAGGCTTACTGCAGAATCCACCAGCGCATTGATATTGGAACTCTTGCTGAGAAGTTGAATATGAATTTCGAAGAGGCAGAAAGATGGATAGTGAATCTTGTTCAGAATGCAAAGCTTGATGCAAAGATCGATTCTAAGCTAGGAACTGTTGTTATGGAGCCCACTTCTCTCAATGTGTATGAGCAGATCATTGAGAATACCAAATCACTCTCTCTGCGTACCTACAAATATGTAAATCTTCTTATAGAATCACAAACAGCAGCGGTACCGGCTGCCCGTTAGAGAAGAGATAGTATAGAGAGAAAACTTGTGTTGGAGCTCTTGCAATTTTCAAGCCTTTCGTGGGAAAGGAAGAGAATTTGGTCCTAGTAAGTATTTTGTTTTTGTGTATTTGTTTTTGAGTTACACCTAAAGCATAAAAAACCAAGAGTTGCAGGGGAACTTTGTTAGCTAATTATACTTTTTTGGGTTCTTTTAACGGTTAAAAGATTCTATCTTGTTGATTAATTTGACATCAAACCATGAAATTACAATGTTCTTTAATTCGGACATCGGAAGAGCGTCGTGTAGG

>Pnu_TRINITY_AroB Pnu_c44458_g1_i1 len=1658

GCTCTTCCGATCTCTCACCAACCCCCAATGGCTGCTGCTACTACAACTACTTCTTCTTCTGGTTCACAAATCAGAAGAAACCAGAATTCCCCAATTCCCAATGGCTTCTTCTTCTGCAACTTCATCTTCTCTCAATCTGATTGTTAGAAACTCTTGTGTAAGAGAGAATTCTGATTTATTTGTTAAAGGGAGGAATTTATCAGTGAAGTTGTCTTCTTCATTTGCACCTACTACAAATTCATTAAGATTGAATAGAATTGGAGGATCTTCTTTGCTTGGAATTGAATCGAAGAAGAGTAGAATAATATCTGCTGCAAATTCAGTAATGGAACAATCTTCAATTAGGGCACCAGTTACTGTTGATGTTGATTTGGGTAATAGAAGTTATCCGATCTATATTGGTTCTGGTCTTCTTGACCAGCCTGATCTTCTCCAAAGGCACGTACATGGGAAGAGAGTACTCGTGGTAACTAATACAACTGTTGCACCATTATACCTAGATAAAGTTATTAGCGCGTTAACTCATGGGAATTCAAGTATATCGGTTGAGAGTGTGATTTTGCCAGATGGAGAGAAGTATAAGAATATGGATACTCTCATGAAAGTTTTTGACAAGGCCATAGGATCTCGTTTTGATCGTCGTTCTACATTTGTCGCTCTTGGAGGTGGAGTTATTGGTGACATGTGTGGTTATGCTGCAGCTTCTTTCCTTCGTGGTGTAAATTTCATTCAAATTCCAACTACTGTTATGGCACAGGTGGATTCCTCTGTTGGGGGAAAAACTGGAATCAACCATACACTGGGGAAAAACTTGATTGGTGCTTTCTACCAGCCCCAATGTGTACTTGTAGATACAGACACATTACATACACTACCTGAGAGGGAATTAGCCTCTGGTTTTGCTGAAGTTGTAAAATATGGACTTATTAGGGATGCTGAGTTCTTCAAATGGCAAGAAAAAAATATGGATTTGTTATTGGCGAGGGACCCCAGTGCACTAGCATATGCAATCAAGCGTTCATGCGAAAACAAGGCAGAGGTTGTATCTTTGGACGAGAAGGAAGGTGGAGTGAGAGCAACGTTGAACTTGGGACATACCTTTGGACATGCAATTGAAACTGGATTTGGCTATGGTCAATGGCTCCATGGAGAAGCTGTAGCAGCTGGAACGGTCATGGCCGTCGATATGTCTTACCGTCTAGGTTGGATTGATGAATCAATTGTGAAAAGGACATACGATATTTTACAAAAGGCCAAGCTGCCAATTACCCCTCCAGATGTTATGACTGTGGAGATGTTCAAGTCTTTCATGGCTGTTGATAAAAAGGTGGCTGATGGACTACTAAGGCTCATCCTCCTCAAAGGTCCTCTGGGCAACTGTGTTTTTACAGGTGACTATGACAGAAAAGCCCTAGATGAGACCCTTCGTGCATTCTGTAAATCTTAATTAAATGGCCTCTTTCAACAATTATGTGCAGTTTTGTGTATTTTCTATTATAGACCCATTTGTTGAGGTCAAACCAAATTTTTGTATTTGTATCAATGTGACTTCTAGTTTTGCCATCAGGGCTCATGTTACTATACTTTAGAGTCTCTCTTATCCTCATTATAATAAAGAAAACAAAGTTTAGTTTTCTAAAAAAAGATCGGAAGAGCGT

>Ppa_TRINITY_Agt1 Ppa_c14011_g1_i1 len=1779

TGTATTATAGTATTAAACTACTACTGCCTATTTTAATGTTTTCCAGGATAATCCAAATGATTGTATGCCGCATGATTTAGACTTTGGTACATTCTGATCTTCCACGTGGAAGCTCTGTGATCTATCAGCATATAGGTTTCATCTTGATCCTCATCTTCCACTTATTTTGGGTGTCTCTATATATAAATATGTGTATTTTGTCATAATTATTACACATATTTCCTTCCTCCCATTGTTTATCCCTCAAAGTTGGACTAGGATAAAGAGAGAGTGAGAGAGCTGCTCTGAAGTTATTCTCTGGAGATAATTCTTGCAACCAGAAGCTTCAAAGAGGTACAATATGGATTACGTCAATGCACCTGGGAGGAACCATCTCTTTGTTCCTGGACCTGTCAATATCCCAGAATCGGTCCTTCGTGCCATGAGCAGAAACAATGAAGATTACCGTTCCCCTGCAATTCCTGCTATGACTAAAGTTCTTCTTGAAGATGTCAAGAAGATTTTCAAGACAACTTCTGGAACTCCATTTTTGTTCCCAACAACAGGTACTGGTGCATGGGAAAGTGCACTCACAAACACCTTGTCTCCTGGTGATCGTACAATTTCTTTCCTCATCGGTCAATTCAGTCTTCTCTGGATTGATCAGCAGCAACGCCTCGGATTCAACGTTGATGTTGTTGAGAGTGAGTGGGGTCGTGGTGCTGATCTTGAGATTCTAGCATCCAAGTTGGCTGAAGACACTGCTCATACCATTAAAGCTGTTTGCATGGTTCACAACGAGACTGCAACTGGAGTTACCAATAATTTGGCTGCTGTAAGGAAATTACTCGATCACTACAACCATCCTGCATTGTTTCTCGTTGACGGAGTGTCTTCAATATGTGCTCTTGATTTCCGTATGGATGAATGGGGAATCGATGTAGCTTTAACTGGTTCACAGAAAGCCCTCTCTCTCCCTACTGGAATGGGTATTGTGTGTGCAAGCCCTAAAGCAATAGAAGCATCCAAGACTGCCAAATCAGTGAGAGTGTTCTTTGACTGGAAAGACTACTTGAAGTTCTACAAATTGGGAACTTATTGGCCATACACTCCTTCGATCCAGCTTTTATATGGTCTCAGAGCTGCTCTTGACCTCCTATTTGAGGAAGGACTTGACAATGTGATTGCTAGGCACAACCGCTTGGCTACAGCAACTAGGCTTGCTGTTGAAGCCTGGGGTCTGAAAAACTGTACACAGAAAGAGGAGTGGTTGAGCGACACTGTGACTGCAGTTGTTGTTCCACCTTACATTGACAGTGCTGAAATCGTTAGAAGGGGATGGCAGAGATACAATCTAAGCTTAGGTTTGGGTTTGAATAAAGTGGCTGGAAAAGTTTTCAGGATAGGTCATCTTGGGAACCTGAACGAGCTCCAATTGCTTGGTTGTTTAGCTGGAGTGGAGATGATTCTGAAAGATGTGGGGTACCCAGTTAAGCTAGGAAGTGGAGTGGCTGCGGCATGTGCGTATTTCCAGAACACCATTCCGATGATTCAATCTAGGATTTAATGAATCATCATTCCACCACCCATGTGTAGCTGCTACTGTTCTTCCTCTTCTGATCATTTGCTTGGAGTACTTGTATAAAAGAAGGCATGATAAGCTCCTGTATTTGCACTCTTGTGAGGCTTACTTAGTAGTACAAGAACTCATTATTACTCTTTCATCCATTAATCTTGTATTTAAAATCTCTTTAAATTTAATCGAATCACTGTTGCATAAAAAAAAAAGATCGGAAGAGC

>Ppa_TRINITY_At103 Ppa_c18053_g1_i1 len=2076

ACGTGTGCTCTTCCGATCTCTCCCCTCCTCTTCTTATTCTTGTCACTGAGAAATTATTACACTCTCTGCAACAATGGCTGCAGAGATGGCATTGATAAAACCCATCTCTAATTTCACCAGAACCAGCCCTAAATTTATCACCTCCACAAGAAAATCATTTTCCATTGTCAAAATGGCATCCCAATCCACTGCTTCAACAACAAAGTCAGGGAAACCCAAGAAAGCAATCAAAGAAACGTTGCTTGCCCCAAGATTTTACACAACTGATTTTGATGAAATGGAAACTCTTTTCAACACTGAGATGAACAAAAACTTAAATGAGGCTGAGTTTGAAGCTTTGTTACAAGAGTTCAAAACTGATTACAATCAAACTCATTTCGTTAGAAACAAGGAATTCAAAGAAGCTGCTGATAAATTGCAAGGTCCTCTTAGACAGATTTTTGTTGAGTTCTTGGAGAGATCTTGTACTGCTGAGTTCTCTGGGTTTCTTCTTTACAAAGAACTTGGAAGAAGACTTAAGAAAACCAATCCAGTTGTTGCTGAGATTTTCTCACTCATGTCCAGGGATGAAGCTAGGCATGCTGGATTCTTAAACAAGGGACTGTCAGACTTTAACTTGGCACTGGATCTTGGATTTCTTACGAAGGCTAGAAAATATACATTCTTCAAACCCAAGTTCATTTTCTATGCTACATATTTATCTGAGAAAATTGGGTACTGGAGGTACATTACCATATATAGGCACTTAAAGACCAATCCTGAATACCAATGTTATCCAATTTTCAAGTATTTTGAGAACTGGTGCCAAGACGAGAACAGACATGGAGATTTTTTCTCTGCGCTGATGAAAGCTCAACCCCAGTTTCTTAATGATTGGAAAGCAAAGTTGTGGTCCAGATTCTTCTGCCTATCGGTCTACGTGACCATGTACTTGAATGACTGCCAGAGAACAGCTTTTTACGAAGGGATCGGACTTAACACCAAAGAGTTTGATATGCATGTCATAATTGAGACAAACCGTACAGCCGCAAGGATCTTCCCTGCAGTTCTAGATGTTGAGAACCCGGAGTTCAAGAAAAAGTTGGATCGGATGGTGCAAATTAACGAGAAAATTATTGCCATCGGCGAAACTGAGGATACACCACTGGTAAAGAACTTAAAAAGGATCCCACACGTTGCAGCATTGGCGAATGAGCTCTTGGCGGCGTATCTCATGCCTCCTATAGACTCTGGGTCTGTTGATCTCGCAGAGTTTGAGACACAAGTTGTGTATTAGTACGTTTTTCTGTGTGTTGAGTAAACAAAGCCGCTTCAATTCAGTGCAAAGTTTAATTGCTACAGTTAACCATTATTACTATGGACATGCTGTATAAAGAACTAAACATTTGTGACGTAATGTATCATGATTCGTGTACATATAGTTTAGGCTGCCATAACTGGTTAACTCTGTCATGACAGCGCTATTCCACATACGTCTCAGATGTATTCTGCCTAAACTTTCATGGGCTGCAAATTCCTATGGGCCTAAAAGTTCGGCAGCTAGCACTCAAAACTAATAGCTGGCAGGGAAAAGGTGTTCAAGACTTTTGTGAACACAATCCATTCTTTGATGGGTGCAATTTCTATTTTCTTCTTACATTTTCTACCACAGACATATATTCATGTGTAACTCATTTGTAGATAACACGTACTGAAGGATACACTAAAGACCTGCGTTAGTTACAAAAAGCTGGCTTCAGAGTGCAAGGATAGCCAAACTCAACTAAACTAAAGAAGGGCTATGCCATTTAAACTTGCTGGCCTACACTGACGACCTACTTGGGGAATTTCACTATGCTCTAACTAGCAAGAACAATAATAGAGGGGTTAAAATCCGTTGGTGAACACCCGCAATTATCTTCTCATTTGTGTAAACAGATTGGATTGCGGAGTTCCACTTGCTGGTGTGCTGCTCCCAAATGCAACACTTCCTTGCTGGTTGCCAAATGCTGCGAACCCACCACCACCTGTAGCAGCAGTAGCCCCAGCAAATCCACCACCACCAGAAGCTCCACCAAACAGATCGGAAGAGCACAC

>Ppa_TRINITY_ChlP Ppa_c17955_g1_i1 len=1667

GTGTGCTCTTCCGATCTCACAACCAAAAATGGCCTCCATTGCCGTCAATGCTCGTTTCTCAGCCACAACATACGACCGTAAAACAACCGAAATCAGCCGTGAAACACTAAAATCCTTTCTGGGTCTCCGGAAAACCAACCAAAACTCTATCTCACTTACGAAATCTCTAACAACAACAACCGCAACTCAACATTTAAACCCAAGAAGAAACTTCTCTTCCATTAAAGCAGGGAAATCAAGTCCCAAAGTAACAGGAAGAAACCTCCGAGTCGCAGTCATTGGTGGTGGTCCTGCTGGTGGTGCAGCTGCTGAAACACTAGCTAAAGGTGGAATCGAGACGTTCTTGATAGAAAGGAAAATGGATAACTGCAAACCATGCGGTGGTGCAATTCCACTTTGCATGGTTGGTGAGTTTGATCTTCCTTTAGACATCATCGACCGACGAGTTACGAAGATGAAGATGATCTCACCCTCAAATGTTGCAGTAGACATTGGGCAAACTCTTAAACCTGATGAGTATATTGGTATGGTACGTCGAGAAGTACTAGACCAGTACCTTCGTGAAAGAGCCGAGAAAGCTGGAACAACTGTAATTAACGGTTTATTCTTGAAGATGGATTTACCAAAAGAACCAAATGCGCCTTATGTGCTTCACTACAGTGCTTATGATGGCAAACCTGGAGGAGTTGGCACGAAAACCACGCTAGAGGTAGACGCGGTAATCGGTGCAGATGGTGCCAACTCTCGCGTAGCTAAAGCGATTGGAGCTGGTGATTACGATTACGCTATTGCATTCCAAGAGAGAATCAGAATCCCAGATGACAAGATGGTTTACTATGAAAATCTTGCCGAGATGTATGTTGGTGACGACGTCTCGCCTGATTTTTACGGATGGGTGTTTCCGAAGTGTGATCACGTAGCAGTGGGTACGGGTACGGTTACGCACAAAGGAGATATCAAGAAGTTTCAAGTGGCAACACGAAACCGTGCTAAGGATAAGATTCTAGGTGGGAAGATTATCAGAGTTGAGGCACATCCTATTCCAGAACACCCAAGACCCCGTAGGTTGGCAGATAGAGTAGCCCTTGTAGGAGATGCGGCTGGGTATGTAACAAAATGTTCAGGAGAAGGTATATATTTTGCTGCAAAGAGTGGCAGGATGTGTGCTGAAGCGATTGTCCAAGGGTCCGAGAGAGGCACAAGAATGGTCAATGAAGCGGATTTGAGAAACTATTTGGAGAAATGGGATAAAACTTACTGGCCAACTTACAAGGTTTTGGATATACTGCAGAAGGTGTTTTACCGATCGAATCCAGCTCGAGAAGCTTTTGTCGAGATGTGCGCAGATGATTATGTTCAGAGGATGACATTTGATAGTTACTTGTATAAGAGAGTTGTGCCTGGCAACCCGATTGAAGATTTGAAGTTGGCTTTTAACACTATTGGGAGTTTGGTTAGAGCTAATGCACTTAGAGGAGAGATGAACAAGATTACTAGTCTATGATGCATGAATGCAGTGAGTTTCTTTGTGAATGATGGGAATCTTAGCGGTTTTTGTTGATCTCTTTTTTCTTATAATTGTTAACTTCTCTCGATTAATTACATCAATAAATTGTATTAAAAACCTTTGAAGTCATTTCGATTTTGTTGATTACTTTAAAAAAAAA

>Ppa_TRINITY_GbssI Ppa_c19393_g1_i1 len=2583

TCGGACTTCTTCCTCTTGTAACAAGTTAAAAACCACATACTTGCCTTCTGTGCCGCATTCATCATATCAGTGTCTCACTCTACTGTCAGTCTCTAGAGAGAGAGAGAGAAGAGAAGAGAAGAGTGTTTTGTATTTTGAGATTATCGAGAGTGACATATCTCTTGATTCTCCAATAATCTACATCTTTGTGAACTCATCAGCACAATCCCTCAAGCTCTCAACTGACAAGAGTATAACTGATGGCGACCATAACTGCGTCCCAGTTTGTATCACGAACCTCACATGTCAATTGTGGAGGAGCTGAACCAAAAGACATGCTGAAACAAATCAGTATGAAGAACCAAAGCATGACACATTCAGGGTTAAGATCACTGAGCAAAGTAGATTTCTTACAAATGAAAAGCACTGTGAAAGCAGCTTCAAGGGAATTGAGGAGGAACGCATTCAAGACAGACAAAGGTACTCTGTCAGGAACCATTAAGTGTTCTAGTGGGATGAACTTGGTCTTTGTTGCTGCAGAAGTAGGCCCCTGGAGTAAAACCGGTGGCCTTGGTGATGTTCTCGGAGGCTTACCACCAGCCATGGCGGCGAATGGACATCGTGTTATGACAGTCTCTCCTCGTTATGATCAGTACAAAGATGCTTGGGATACCAATGTGTCAGTTGAGATGAAAATCGGGGACAGGATTGAAACTGTTCGTTTCTTCCACTGTTACAAACGTGGGGTTGACCGTGTATTTGTGGATCATCCACTGTTTCTTGAAAAGGTATGGGGTAAAACTGGATCCAAAATCTATGGTCCCAGTGCTGGATCAGATTACCAAGATAACCAGCTTAGATTCTGTTTGCTTTCTCTGGCAGCTCTAGAAGCACCAAGGATTCTGAACCTCAACAGCAACAAATATTTCTCTGGAACATACGGAGAGGATGTCGTGTTCATTGCTAATGATTGGCATACGGCCCTTCTACCATGCTACCTTAAAAGTATGTACAAATCTAAAGGGATGTATGAAACTGCAAAGGTTGCTTATTGCATCCACAATATTGCTTACCAAGGAAGATTTTCTTTTTCGGATTTTATGCTTCTCAATCTTCCGGATTCATTCAGGAGTTCTTTTGACTTCATTGATGGGTATGAGAAGCCTGTTAAAGGAAGGAAGATTAACTGGATGAAGGCTGGAATATTAGAGTCAGACAAGGTCGTAACTGTGAGTCCATACTATGCTGAAGAACTTGCTTCAGGCCCCGATAAAGGAGTGGAACTGGACAGCATTATCCGTAGAGCTGGCTGTTTGGGTATAATAAACGGAATGGATGTTCAAGAGTGGAACCCCTTGACTGATAAACATATCGACGTCAAATATGATGCCTCAACTGTGACCGAAGCGAAGGCTCTTCTAAAAGAAGCACTGCAGGCAGAAGTAGGGTTACCTGTTGACCGTAATATCCCTGTCATAGGCTTCATTGGTAGACTTGAAGAGCAAAAGGGTTCAGATATTTTGGCAGCTGCAATCCCTGAATTTATTGACGAAGAAGTTCAGATAATTGTCCTTGGTACCGGCAAAAAGCCCATGGAAAAACAGTTAGAGCAGTTAGAGACATTGTATCCTGAAAAAGCTAGAGGAGTTGCAAAGTTCAATGTTCCCTTGGCTCATATGATTATAGCTGGGGCTGATTTTATGATGATCCCCAGCAGATTTGAACCATGTGGACTCATCCAACTGCATGCGATGCGTTATGGAACGGTGCCTATCGTTGCCTCGACGGGTGGATTAGTTGACACTGTCAAAGAAGGTTTCACTGGATTTCAAATGGGAGCTTTCAATGTCGAATGCGAGGCAATTGATCCATTAGACGTGACAGCAATCGCCACAACCACCAAGAAGGCCCTTGCAACCTATGCAACTCCAGCTTTAACAGAAATGATACAGAACTGCATGGCTCAAGACCTCTCTTGGAAGGGACCTGCCAAAAATTGGGAGGACATGTTGCTGGGCTTGGAGGTTGCAGGTAGTGAACCAGGAATCGATGGCGATGAAATAGCTCCTCTTGCTAAAGAAAATGTTGCCACCCCTTGATTTGTGAACCGCTCCAGTGCAACTTAGAAGAGCTATCTCACAATCTAGAAAAGACGACGCCAATGCTGCATTAATAATGGTCTGACAATTTATCCAGTGTTAAGTGGCCTTACTGGGAAAGAGATACTGCTATTCTTTTTCTTCTTTTTTCTTGGTTAAGCAATCTACAGCACTCTATGCCATAAACGTAGGATAGCTAGCTCCAGTTTATTCATTTTCCTTTTCTTTTTTAGCCAGCTCCAGCTCTTGTATCTAGTCAAGAACTTCTACATAAATAACAACGGCTTTTTCAACTTCTGCTTCAGTTAGTTTCCATGTAAGGTTGTGTAAATATGCATGTAGCTGTACAAAAGATTTTCATGAGCGTATTTTTAAAGAGGTTTTCTTTTATATATAAACCGAAAGAAAGAGAAACCGTTGCAGACAACCTGCTATGAGGTTTTGTGCCTCCCACAGTTGAGGACTTTAGTTCCTTTCTGGCCAGGGGCTTAAGTTTAGGATGAAT

>Ppa_TRINITY_Hmgs Ppa_c31681_g1_i1 len=1839

ACGTGTGCTCTTCCGATCCAACAATCCCACCCCAACACAACTCACGAGTCACGAAGTTTGCATTGGTTCCGGAATCCGGTCTAACCGGTGCTCATGTATTCCCTCTGTCTATAAATACCGATTTATTCGTTGCCTTAATCTCTCACCACTTGCTTCTTCTTCCTTTCCTTCTTCTCTCCAAACACCTCGATCGATCGATCTAGGGTTCTCTTTTGAAAAGAATCCCAAACAAATTCTCAGAATTTAAAAAAAAAAATGGCTTCAAATCAAAAGGATGTTGGAATTCTTGCCATGGAAATGTATTTCCCTCCTACTTGTGTTCAACAGGAAGCATTGGAGGCTCATGATGGTGCCAGTAAAGGAAAATACACAATTGGGTTGGGACAAGATTGCATGGCGTTTTGTTCAGAGTTGGAAGATGTCATTTCAATGAGTTTGACAGCCGTTAGCTCTCTACTTGAGAAGTATAACATTGACCCTAAACAAATTGGACGTTTGGAAGTCGGGAGTGAGACCGTGATTGACAAGAGCAAGTCCATAAAGACCTTCCTGATGACAATCTTTGAGGAATGTGACAACACTGACATTGAGGGTGTTGACTCAACAAATGCGTGCTATGGAGGAACTGCAGCTTTGTTCAACGTTGTGAATTGGGTGGAGAGTAGTTCTTGGGACGGACGTTATGGTCTTGTTGTGTGTACTGATAGTGCGGTCTACGCAGAAGGACCCGCTCGTCCCACTGGAGGAGCTGCTGCTATTGCTATGTTGATTGGTCCAGATGCACCAATTGCCTTTGAAAGCAAATACAGAGGATCCCACATGTCCCATGTGTACGATTTTTACAAGCCCAACCTAGCCAGTGAATACCCGGTTGTTGATGGGAAACTTTCACAGACTTGTTACCTTATGGCCCTCGATTCCTGCTACAAGCGCTTGTGTGCCAAGTACGAGAAATTTGAGGGTAAGCAATTTTCAATGGATGATGCAGATTACTTTGTGTTTCATTCTCCATACAATAAGCTTGTGCAGAAAAGTTTTGCTCGATTGTACTTCATGGACTTCTTAAGAAATGCCAGCTCTGTTGATGAGGCTTCAAAAGAAAAGCTTGCACCATTTTCAGCTCTATCTGGTGAAGAGAGCTATGCAAGTCGGGATCTTGAGAAGACATCCCAAACTGTTGCAAAGGATCTTTATGCTGAGAAGGTTGGACCTACTACTTTGTTACCCAAACAAGTTGGTAATATGTACACTGCATCACTTTATGCGGCATTTGCTTCTCTTCTTCACAACAAAAATAGCTCACTGGATGGTAAACGTGTAATAATGTTCTCATATGGAAGTGGTTTAAGTTCCACAATGTTCTCATTCCGAATGAGGGAAGGTTCACATCCATTCAGCTTATCAAACATTGCAACTGTTTTGGATGTCGCAAACAAGTTGGAGTCGAGACATGAGGTGACTCCTGAAAAATTCGTCGAAACCATGCAACTGATGGAGCATAGGTATGGAGGGAAAGACTTTGTAACGAGCAAGGACACAAGCCTTTTACCTTCAGGCACATACTACCTTACCGAAGTTGACTCCATGTACCGCAGATTCTACTCAAAGAAGTCAAGTGCTACAGCCGCAGTGCAAAATGGTCACTGAAAACGGTGAAAGAAGAAAACGGCCGCAAAAGAGCAGCAGAAGATTTGCTTATGTATTTTCATTGTTGTGGTTAGTGTTTTAGAATAATTCTAAATGATTATTTGTACAAGTTAATCCTTCCAAACCCGTGGTTAGCTTATTAGCCATTGTAATAGGATTCATGTCCTCTCAGTATATCCATTCAAGTATTCA

>Ppa_TRINITY_Nia Ppa_c36353_g1_i3 len=3129

CTCGAGCATGGCAACCCTTTTCCAATGCACCCTTATATAAAGACCCAACCATGCTCCTCATTTTTCTTGCAACCATCAAATGCAAAAGCAAATCGAAAAAAGAGAAAACATAGAAGAAGAAGAAGAAGCAGAGATAGAAAAAGTGAGTGATAGAGCGGAGATCTTCATACAAAGAGAAAAGAGATCTTTTCTATCTTCTTTCTTTTGATTAGTCTCTAATGGCGGCTTCTGTTGAGAATCGACAGTACAACAGTCGACTGGAAAATGGTCTAGCAGTTCGTGGATTTAAAACCAATTCCACTCATCGATCTGATTCACCTATCCGTGGCTGTAATTTTCCCATGTCGACCACCGAGCGTCTTTCATTATCAAAACCAGTGATAGAAGATTTATCCAGCGACGACGAGGATCAAAACGAGCAGTGGAAAGATTTCGGCAAGATAAACAATGAAATAGAGCCGTCTGTTTTTGACTCAAAAGATGAAGGAACTGCCGATAATTGGATTGAACGAAACCCTTCCATGATTCGGTTAACAGGGAAGCACCCATTCAACTCGGAAGCACCATTAACTCGTCTCATGCATCACGGGTTCATCACACCTGTACCTCTTCATTACGTTCGTAATCACGGAGCGGTTCCAAAAGCCAAATGGGAGGATTGGACGGTTGAGATTTGTGGATTAGTAAAGCGTCCGTTACGCATCACCATGGATATGCTCATTAATGATTTCCCGAGCCGTGAGTTTCCAGCGACACTTGTTTGTGCAGGCAACAGACGTAAAGAGCAGAATATGATAAAACAAACAATTGGGTTTAACTGGGGAGGTGCAGGAGTATCGACTTCAGTATGGCGCGGTGCACGTTTGTGCGACGTATTGAAGCGATGTGGTATCATGAGCCGTAAGAAAGGTGCTCTGAATGTTTGTTTCGAAGGAGCTGAAGATTTACCCGGTGGCGGTGGATCTAAATACGGAACTAGTATCAAGAAAGAATTCGCAATGGATCCTGCTCGTGATATCATCCTCGCTTACATGCAGAATGGTGAGAATTTATCACCGGACCATGGATTTCCAGTCAGGATGATAATTCCAGGTTTCATTGGAGGAAGAATGGTGAAGTGGTTGAAGAGAATTATCGTCACAACTCCAGAGTCTCAAAATCATTATCATTTCATGGACAACAGAGTTCTTCCTTCTCATGTCGACGCTGAATTGGCTAACTCCGAAGCATGGTGGTATAAGCCAGAGTACATCATCAATGAATTGAACATAAACTCGGTGATTACAACTCCTTGTCATGAAGAGATTCTTCCCATTAATGCTGCAACAACACAAAGAGCTTACACGTTAAGGGGCTACTCATATTCCGGCGGCGGTAAGAAGGTAACTCGTGTCGAAGTAACTATGGATGGCGGAGAGACCTGGTTGGTGTGCACTTTGGATCATCCAGAGAAACCAAACAAGTATGGGAAATACTGGTGCTGGTGCTTTTGGTCACTTGAAGTTGAAGTTCTTGACTTACTTGGTGCCAAGGAGATTGCAGTTAGAGCTTGGGATGAAACTTTGAACACTCAACCTGAAAAGCTCATTTGGAATGTCATGGGTATGATGAACAACTGTTGGTTCAGGGTTAAGACAAACGTGTGCAAGCCACACAAAGGAGAAATTGGTCTAGTTTTTGAGCATCCCACTCAAGCTGGAAATCAACCAGGTGGATGGATGGTTAGACAAAAACATCTTGAAACCTCTGAAGGTCCACACACACTCAAGAAGAGTTTATCCACTCCTTTCATGAACACCACGTCAAAGATGTACTCAATTTCTGAAGTGAAAAAACACAACTCGCCCGACTCGGCTTGGATCATTGTTCATGGACACGTTTATGACTGTACTCGCTTCTTGAAAGACCATCCAGGGGGTTCCGATAGTATTCTTATCAATGCAGGTACCGATTGTACCGAAGAATTTGATGCGATCCACTCTGACAAAGCTAAGAATTTACTTGAGGATTTCAGAATCGGTGAACTCATTGCAACTGGTTATACTTCTGACTCATCATCTTCACCAAACAACTCAGTTCATGGTTCGGGAAGCTTGAGTCACCTTGCTCCTATCAAAGAAATCGGCATCGCAAAGCCGGTAGCTCTAGTTGGTCGAGATAAAATACAGTGCAAACTTATCGCCAAAACCTCGATTTCCCATGACGTTCGTGTCTTCAGGTTTGCATTACCTTCAGCTGATCAAGTCCTTGGGTTGCCAGTTGGGAAACACATATTTGTGTGTGCCACCGTTAACGATAAGCTGTGTATGCGCGCTTACACTCCATCGAGTACAGTTGACGAAGTTGGTTACTTCGAGTTGGTCATTAAGGTCTACTTTGGCGGAGTTCACCCCAAATTCCCCAATGGAGGTATCATGTCTCAGCACTTGGATTCTCTTGCTCTAGGTGGAACTATTGACGTCAAGGGTCCTTTAGGACACATTGAATACGCGGGTCGTGGTAACTTCACCGTTCATGGCAAACACAAGTTTGCCAAACGACTCGCGATGCTTGCTGGTGGCACAGGAATTACACCAATATATCAAATTGTACAAGCAATATTAAAAGATCCAGAAGATGAAACAGAGATGCACATCGTATTCGCAAACAGGACTGAAGATGACATCTTGTTGATGGATGAACTAGATTCATGGGCAGAGAAGCACGAAAGGTTGAAGGTTTGGTACGTCGTCGGTGAAACATTGAGGGAAGGATGGAAATACAGTGTAGGGTTCATCACAGAAGAGGTTCTACGGGAACATATCCCTCCAGCATCGGATGATACACTGGCATTGGCTTGTGGACCACCACCAATGATTCAATTTGCAGTAACTCCAAATTTGGAGAAGATGAATTATGACGTGAAGAATTCATTATTGGTGTTCTGATTGATGAGTTTAAAGTACTTGGAAAATAAAGTTATTATAGTTAAGAAAAAAGATAAGGGAGGACAAATTATAAGGAAGTTCTCATTAGTTATTGTATATAATTAGTGTTGAAGGATGTAATGGTTCAGTTCGTTCTTAATAGTAAGGGGATTGTTAAATGTTAGTCCGTTTATATTTCATGTAGAATTGAAAACAAAATGTG

>Ppa_TRINITY_Eif3E Ppa_c36589_g1_i1 len=1538

TGTGCTCTTCCGATCTCTCTATTTTCATCAAATCGTTTTTTAGGGTTTCTCTCTTTCTTCCTCCATATCCACCACCACAGACAGCATCTCCTTCTTCTCAGAGGTAGCAGCAGCCTCCTCCACATCACCCAAATCATGGCGAATCATGATATAACATCACGAGTTGCACTAAACCTAGATCGACATCTGGTTCTACCACTCCTTGAGTTTCTCCAGGAACGCGGTATTCATCCCGATGTTCAAATCCTCAAAGCTAAAGACGCACTTCTCAGTCAAACCAACATGGTTGATTACGCTATGGATATTTACAAGTCTCTTCATCAGACTGACGATGTTCCTCAAGCAATGATTAACAGACGTGCTGAAGTAGTTGGGAGATTGAAATCGCTTGAAGATGCTGCTGAATCGATTATTACTTTCCTTCAGAGTCCGAACATGGTTAATGAACCACGCGCTGATAAGCAATATATTCATCAGTTGCTACAGGGATTTCAGATTGGTGCTGAGCAAATTGAGGCACTATATCAGTATGCCAAATTTCAATTTGAATGTGGGAATTACTCGGGTGCATCTGATTACTTGCATCAGTACAGAACAATTGGTCAAAATCCTGACAGAATTCTGAATGCATGTTGGGGAAAGCTTGCGTCTGAGATATTGATGCAGAACTGGGATGAGGCTCTTAGTGAGCTTAATCACTTGAGAGAACTTATCGAAAATAAGAATTTCAAAAACCCATTGACTTTACTCCAGAACAGAATCTGGTTAATGCATTGGAGCCTCTTCATCTTTTTCAACCACGAGAATGGAAGAAATGCTATCATTGATCTATTCCTCAATGAAAAGTACTTGAATGCCATTCAAACAAATGCTCCCCATCTTTTACGATATTTGGCTACTGCATTGATTGTCAACAAAAAGAGAAGACCTCAACTCAAAGAGTTGATCAAAGTCATCCAGCAAGAGCAGTCTGCATACAAGGATCCCATCACTGAGTTCTTGTACTGTTTGTATGTCAACTATGACTTTGATGGTGCACAACAGAAGCTCCGTGAATGTGAGGATGTGATTTTGAATGATCCATTCCTTGGTAAACGAGCTGAAGAAGGAAACTTCAGTATCGTTCCTTTGAGAGATGAATTTCTTGAGAATGCTCGCCTCTTCATCTTTGAGGCTTACTGCAGAATCCACCAGCGCATCAATATTGGAACTCTTGCCGAGAAGTTGAATATGAATTTTGAAGACGCGGAGAGATGGATAGTGAATCTTGTTCAGAATGCAAAGCTTGATGCTAAGATCGATTCTAAGCTAGGAACTGTTGTAATGGAGCCCACTTCTCTCAATGTGTATGAGCAGATCATTGAGAATACCAAATCACTTTCTCTGCGTACCTACAAATACGTAAATCTTCTGTTAGAATCAGCACCAGCACAGGTGCAGGCTGCTAGATAAAAGTAAAGGGGGTAGTACAGTCAGAAAATTTTCAAGTTTGAAGTTTTCAAGTCATTGTTGCTGTTGAAGTTTTCAAGAATTTGGTC

>Ppa_TRINITY_AroB Ppa_c29084_g1_i1 len=1606

TGTGCTCTTCCGATCTCGGAATCAGAAACTGTTCTTCTCTGTAAACAAAATCCCTAATTCTCCGAATTGATGGCTTCTTCTTCTGCAACTTCACCTTCTCTTAATCTAACTCTGAAAAACTCTTGTTTACGAGAAAATTCTGAGGTTTTTCTTAGAAAGAGGAATTTATCATTGAAGTCTTCATTTGTATCTTCAAGTTCATTGCGATTGAATAGAATTGGTTCTCCTTTGGCTCGAATTGAATCGAAGAGTAGAGTAGTATCTGCGAATTCAGCTCAAGTAATGGAACAATCTTCAATAAAGGCACCAGTTATTGTTGATGTTGATTTGGGTAATCGAAGTTATCCGATCTATATTGGTTCTGGACTTCTTGAACAACCTGATCTTCTTCAAAGGCACGTACATGGGAAGAGGGTGCTCGTGGTAACCAATACAACAGTTGCGCCATTGTACCTAGATAAAGTTATCAGCGCGTTAACTCATGGGAATCCGAATATTTCGGTTGAGAGTGTGATTTTGCCTGATGGGGAAAAGTATAAAGACATGGATACTTTGATGAAAATTTTTGACAAGGCCATAGAATCTCGGTTTGATCGACGTTCTACATTTGTCGCTCTTGGAGGTGGTGTTATCGGTGACATGTGTGGATATGCTGCAGCCTCTTTTCTTCGTGGTGTAAACTTCATTCAAATTCCAACCACTGTCATGGCACAGGTGGATTCTTCTGTTGGGGGGAAAACTGGAATCAACCATCCATTGGGAAAAAACTTAATTGGTGCTTTCTACCAGCCCCAGTGTGTCCTTGTAGATACAGACACATTACATACACTACCTGAAAGGGAATTGGCTTCAGGTTTTGCTGAAGTTATAAAATATGGACTCATTAGGGATGCTGACTTTTTCGAGTGGCAAGAAAAAAACATGGACGCTCTATTGGCAAGGGATCCCACTGCACTAGCATATGCAATCAAGCGTTCATGCGAAAACAAGGCAGAGGTTGTATCCTTGGACGAGAAAGAAGGTGGAGTGAGAGCGACTTTGAACTTGGGGCATACCTTTGGACATGCAATTGAAACTGGATATGGCTATGGTCAGTGGCTGCATGGAGAAGCCGTTGCAGCTGGAACGGTCATGGCAGTCGATATGTCTTACCGCCTAGGTTGGATTGATGAATCAATTGTGAAGCGAACGTACGATATTTTACAAAAAGCCAAGCTGCCGATTAGCCCACCTGATGGTGTTATGACTGTGGAGATGTTTAAGTCTTCCATGGCTGTTGATAAAAAGGTGGCTGATGGACTACTAAGGCTCATCCTCCTCAAAGGTCCTCTAGGCAACTGTGTCTTCACAGGTGATTACGACAGAAAAGCCCTAGACGAAACCCTTCATGCATTCTGTAAATCTTAATCAAATGCCGACCTTGGTTTGGCTCAGTCTTGTGTTATTCTTTTTTATGAGCCCATCTGTTGAGGTCAAACCAAATTTTGTATTTGTACGTAATCAGTATTGACTTCTAGATTTGCCATCGGAGCAGAGGTACTATAGAGTCTCTTGCCCTCAATGTATAATGATAAATAATAATTGAAAAATTCTCTGTCAATCGGAC

>Pca_TRINITY_Agt1 Pca_TRINITY_DN805_c0_g1_i13 len=2004

TTTTTTTGGAGATAAAGATGTAATTGGACAACTGCAGTACATCATTGATTAATGCAAGTAATTACTGAGAAAACCAGACGTAATTGGGTATATGCAACAGTGATCAGATTAAATTTAAAGAGATTTTAAATACAAGATTAATGGATGACAGAGTAATGAGTTCTTGTACTACTAAGGAAGCCACAGAAGAGTGCAAATACAGGAGGAGATTATCATGTCTTCTTTTATACAAGTACTCCAAGCAAATCAGAAGAGGAAGAACAGTAGTTACACATGGATACTGAATGATTCATTAAATCCTAGATTCAATCATCGGAATGGTGTTCTGGAAAAATGCACTAGCCGCAGCCACTCCACTTCCTAGCTTAACTGGGTACCCAACATCTTTCAGAATCATCTCCACTCCAGCTAAACAACCCAGCAGTTGGAGCTCATTCAGGTTCCCAAGATGACCTATCCTGAAAACTTTTCCAGCCACTTTATTCAACCCCAGACCTAAGCTTAGATTGTATCTCTGCCATCCCCTTCTAACTATTTCAGCACTGTCTATGTAAGGTGGAACAACAACTGCAGTCACAGTGTCGCTCAACCACTCCTCTTTCTGTGTACAATTTTTCAAACCCCAGGCTTCAACAGCAAGCCTAGTTGCTTTGGCCAAGCGGCTGTGCCTAGCAATGACATTGTCAAGTCCTTCCTCAAATAGGAGATCAAGAGCAGCCCTGAGGCCGTATAAAAGCTGGATTGAAGGAGTGTATGGCCAATAAGTTCCCATTTTGTAGAACTTCAAGTAGTCTTTCCAGTCAAAGAACACTCTCACTGATTTGGCGGTCTTGGATGCTTCCAGTGCTTTAGGGCTTGCACACACAATACCCATCCCAGTAGGTAGAGAAAGAGCTTTCTGCGAACCAGTTAAAGCTACATCGATTCCCCATTCATCCATACGAAAATCAAGAGCACATATTGAAGACACTCCATCCACGAGAAACAATGCAGGATGGTTGTAGTGATCGAGTAATTTCCTTACGGCAGCCAAATTATTGGTAACTCCAGTTGCAGTTTCGTTATGAACCATGCAAACAGCTTTGATGGTATGAGCAGTGTCTTCAGCTAACTTGGATGCTAGGACATCAAGATCAGCACCACGGCCCCACTCACTCTCAACAACGTCAACGTTGAATCCGAGGCGTTGCTGCTGATCAATCCAGAGAAGACTAAATTGTCCAATGAGGAAAGAAATTGTCCGATCACCAGGAGACAAAGTATTTGTAAGTGCACTTTCCCATGCACCAGTACCTGTCGTGGGGATCAAAAATGGGGTTCCAGAAGTTGTCTTGAAAATCTTCTTAACATCTTCGAGAAGAGTTTTAGTCATAGCAGGAATTGCAGGGGAGCGGTAATCTTCATTGTTTCTGCTCATGGCACGAAGGACTGATTCTGGAATATTGACAGGTCCAGGAACAAAGAGATGGTTCCTCCCAGGTGCATTGACGTAATCCATATTGTACCTATAGACACTTGTGTTGTATTCAGACAAAATAGTCACCCCAATGGGATCCACTCATCGAGTACTTAATCAGAACCGAGTATATCTCTTAAATGAGTGGTGTCAATATTCGAATCTGTGAGACTCTTTGAAGCTTTTCGTTGCAAGAATGGTTCCCGGAGATTCACTTCAGAGCAGCTCTCTCACTATCACTATCTCCCAGTCCAACTTTAGGGGATAAACAATGGGAGGGAGGAAATACGTGTAATAAGCAGCATTAGGAAAAAAACATATATTTATATATATAGACCCTAAAATAAGTGGAAGATGAGAGGATCGAGATGAAACGTATATGCTGATAGATCACAGAGCTTCCACGTGGAAAATTAGAATCTACCAAACTCTAGATCATGCGTCATAAAATCATTTGGATTATTCTGGAAAATATTAAATAGCTAGTAACAATACTAAAACATATACTGCAAAAAATACGTTTGAGATGGAGATGAGGATGAGCAAAG

>Pca_TRINITY_At103 Pca_TRINITY_DN3782_c0_g2_i3 len=1750

TTTTTTTTTTTTTTTTTTTTTTTTGAACCGATAATTCAGATGCTTTTATTTTTCTTAATACTTTAACCGTACAGGAGACTCTCTTATATAACGAGAGTAGTTTTTATATAAGGTAGGAAATCTAAACAAATCCTAAACATAACAGGAAATCCTAAACATAACAAGAAATCTACTCAAATCCTAAACAGAATATATTGTTAACAAATCCTAAAGACTATGAGTTGGCATTTGTATCAGGATTGGTTTTCTGACTGGATTCAGAATTGGTCTTCTGGCTGTATGGTTCAACAGTGGGTACAGAAATCATGATACATTATGTCACTAACGTTTAGTTCTTTATACAGCAGGTCCATAGTAATGGTTAACTGTAGCAATTAAACTTTGCACTGAACTGGATTGAATTGAAGGGCCTTTGTTTACTCAATACACAACAAAACTTATTAATACACAACTCGTGGCTCAAACTCAGCGAAATCAACAGACCCAGACTCAATAGGAGGCATGAGATACGCAGCCAAGAGCTCATTCGCCAATGCTGCAACATGTGGGATCCTTTTTAGGTTCTTCACCAAAGATGATTCCTCAGTTTCACCAATTGCAAGAATTTTCTCGTTGATTTGCACCATCCGATCCAACTTCCTCCTGAACTCCGGATTCTCAACATCTAGAACTGCAGGGAAAATCCTTGCGGCTGTACGGTTTGTCTCAATTATGACGTGCATATCAAACTCTTTGGTGTTAAGTCCGATCCCTTCGTAAAAAGCTGTTCTCTGGCAGTCATTCAAGTACATTGTCACGTACACTGATAGGCAGAAGAATCGAGACCACAACTTTGCTTTCCAATCGTTAAGAAACTGGGGTTGAGCTTTCATCAGTGCTGAGAAGAAATCTCCATGTCTGTTCTCGTCTTGGCACCAGTTCTCAAAATACTTGAAAATTGGATAAACTTGGTATTCAGGGTTGGTCTTTAAGTGCCTATATATGGTAATGTACCTCCAGTACCCAATTTTCTCAGATAGATATGTAGCATAGAAAATGAACTTGGGTTTGAAGAATGTATATTTTCTAGCCTTTGTAAGAAATCCAAGATCCAGTGCCAAGTTGAAATCTGACAGTCCCTTGTTTAAGAATCCAGCATGCCTAGCTTCATCCCTGGACATGAGTGAGAAAATCTCAGCAACAACTGGATTGGTTTTCTTAAGTCTTCTTCCAAGTTCTTTGTAAAGAAGAAACCCCGAGAACTCAGCAGTACAAGATCTCTCCAAGAACTCAACAAAAATTTGTCTAAGAGGACCTTGCAATTTATCAGCAGCTTCTTTGAACTCTTTGTTTCTAACGAAATGAGTTTGATTGTAATCAGTTTTGAACTCTTGTAATAAAGCTTCAAACTCAGCCTCATTCAAGTTTTTGTTCATCTCAGTGTTAAAAAGAGTTTCCATTTCATCAAAATCAGTTGTGTAAAATCTAGGAGCAAGCAAGGTTTCTTTGATTGCTTTTTTGGGTTTCCCTGACTTTGTTGTTGATGTAGTGGACTGGGATGCCATTTTGACAATGGAGAATGATTTTCTTATTGAGGTGATAAATTTAGGGCTGGTACTAGTGAAATTAGATATGGGTTTTATCAATGCCATCTCTGCAGCCATTTTTGCAGTTTCTAGTGAATAGAAGAGTGTGAGCAAGGAGAAATGGTAAGATAAGACATGGGTTTTTGTAGGATTTTAGTCGATGAAAATTTAGAGTGGAATCTAAT

>Pca_TRINITY_ChlP Pca_TRINITY_DN882_c0_g3_i2 len=2298

AGCGATTCTTCGTACACACAGCTTTGTTTCATTGAAAGATTTAGGAGAAACAACAATGTTAGAACGAAAAGAAAGAAAAGATATCTTAGCTACTGAATTCATTATTAATCTGTTGCATCATTTATGAATCTATGCTATTCTGTTGGGAATTTGCCCTCTTGTTATGTCTGTTACTGTAGAATCTGTATCCATAAAAGTATCCTGGGTTGTAGTGTTGGTGCCCAATATATAGACTGCTTCCACTACAGCAATGCACCCAGCTTCCTCTGCAATTACACAACCATGCTTGTATATGCTCGACTATTAGAGCCCGAAGATTGCAACAGATATATATTCCAATCCTTAGGCAATCCTTAGGATAACAAATTGCATTGGTCAGCATCAAGAAAATTGCACTTTCTGAATCTACACGTTAAGATTGGGAAGTATCCTCACGTTAAATAACATATGGAGAAATGGCCGATGTGTTTCTATCCGTAATCCATTAGTTCCACGAAATGTGCGCTCGGTGCATCCATATGTGCAGCATTTTTCAAAGGGCAAAAGATGTTGCCACAGAAGAAATTGATACAATTTTATTGATGTGATTAATGAAGAAGTTAAATACACAACTATAACAAAAAGAGATCAACAAAAATCGCCAAGATTCGCATCACCCCCATTTATGTATCTTAATTCACCAGAAGAACAAACCTATTTTCACAAAAGAAACTCAACACATTCATCATCATAGACTAGTGATCTTGTTCATCTCGCCTCTAAGTGCATTAGCTCTAACCAAACTCCCAATAGTATTGAAAGCCAATTTGATATCCTCAATTGGATTTCCAGGCACAACTTTCTTGTATAGGTAACTATCAAATGTCATCCTTTGAACATAATCATCGGCGCACATCTCAACAAAAGCTTCGCGAGCTGGATTTGATCGGTAAAACACCTTCTGTAATATATCCAAAACCTTGTAAGTTGGCCAGTAAGTTTTATCCCATTTCTCCAGATAATTTCTCAAATCCCCTTCATTAATCATTCTTGTCCCTCTCTCGGATCCTTCAACAATCGACTCGGCACACATCCTGCCACTCTTGGCAGCAAAATATATACCTTCTCCTGAACATTTTGTTACGTACCCAGCCGCATCTCCTACAAGCGCTACTCTATCTGCCAATCTACGGGGTCTTGGATGCTCTGGAATAGGATGTGCCTCTACTCTAATAATCTTTCCGCCTAGAATCTTATCCTTGGCACGTAACCGTGTCGCCACTTGAAACTTCTTGATATCTCCTTTGTGCGTAACTGTACCTGTACCCACTGCAACATGATCACATTTAGGAAACACCCATCCGTAAAAATCGGGCGACACATCATCTCCTACATACATCTCGGCAAGATTTTCATAGTAAACCATCTTATCATCTGGGATTCTGATTCTCTCTTGGAATGCAATAGCATAATCATAATCACCAGCTCCAATTGCTTTAGCGACGCGAGAGTTCGCACCATCTGCACCAATAACTGCATCTACCTCTAACGTGGTTTTCGTGCCAACTCCACCAGTTTTGCCATCATAAGCACTGTAGTGAAGAACATAAGGCGCATTTGGTTCTTTTGGTAAATCCATTTTCAAGAATAGACCGTTAATTACAGTTGTCCCAGCTTTCTCTGCTCTTTCACGAAGATACTGGTCTAGTACTTCTCTACGTACCATACCGATATATTCATCTGGTTTAAGAGTTTGGCCAATGTCGACTGCAACATTTGAAGGTGAGATCATCTTCATCTTCGTAACACGCCTGTCGATGATGTCTAACGGAAGATCAAACTCACCGACCATGCAGAGTGGAATTGCACCACCACATGGTTTGCAGTTATCCATTTTCCTTTCGATCAGAAACGTTTCAATACCACCTTGTGCTAAGGTTTCAGCAGCTGCACCACCAGCAGGACCGCCACCAATGACCGCAACTCGGAGATTTCTTCCTGTTACTTTAGGACTTGATTTACCTGCTTTAATGGAAGTGAAATTTCTTCTAGCAGGGTTAAAATGTGGACTACTTGTTGTTAGAGATTGTGTAAGTGAGATAGAAGCATTTTGGTTGGTTTTTCTAAGACCCATGAAGGATTTTAGTGTTTCACGGCTGATTTCGGTTGTTTTACGGTCGCATGTTGTGGCTGAGAAACGAGCATTGACGGCAATGGAGGCCATTTTTGGTTGTGAAGGGGTTCTGCAAGTATGGAAGTTTTTTAAGATTTTTGTTATCCTTTGGTTTTTGATTTAATGTTTATTGTGATATGTTTGTG

>Pca_TRINITY_GbssI Pca_TRINITY_DN3397_c0_g1_i11 len=2821

CTTCTCTTTCTCTTATAACAAACTTACTTGTGATCTGTGCTGCTGTGTGTCCCTCTACCGTCAGTCTAGACTCTAGAGACAAAAAAGAGCGTGTATTTTGAGATTATCGAGAGTGACATATCTCTCCATTCTCCATCCATGTGAATTCATCAGCTGCATATATAATCTCTCAAGTCTCAACTGACAAGAATATTACCAATGGCGACCATAACTGCGTCCCAGTTTGTATCACGAACCTCGCATCTCAACTGTGGAGGAACTGCATCAAAAGACAAACTTAAACAAATCAGTATGAAGAACCAAAGCATGACTCATTCAGGGTTAAGATCTCTGAGCAAAGTAGATTTCTTACAAATGAAAAGCAATAAGAAAACAGCTTCTAGGGAATTGGGTAGGAACGCATTCAAGACAGACAAAGGTACTGTGTCAGGAGAAGTTAAATGTTCTAGTGGGATGAACTTGGTCTTTGTTGCTGCAGAAGTAGGCCCCTGGAGTAAAACCGGTGGTCTTGGTGATGTTCTTGGAGGCTTACCACCAGCCATGGCGGCGAATGGACATCGTGTGATGACAGTCTCTCCTCGTTATGATCAGTACAAAGATGCTTGGGATACCAATGTGTCAGTTGAGGTGAAAATTGGGGATAGAATTGAAACTGTTCGTTTCTTCCACTGTTGTAAGCGTGGGGTTGACCGTGTATTTGTGGATCATCCACTGTTTCTTGAGAAGGTATGGGGTAAAACTGGATCTAAAATCTATGGTCCGAATGCTGGATCAGATTACCAAGATAATCAGCTTAGATTCTGTTTGCTTTCTCTGGCAGCTCTAGAAGCACCAAGGATTCTGAATCTCAACAGCAACAAATATTTCTCTGGAACATACGGAGAGGAGGTCGTGTTCATCGCTAATGATTGGCATACTGCCCTTCTACCATGCTACCTCAAAAGTATGTACCAATCTAAAGGGTTGTATGAAACTGCTAAGGTTGCTTATTGCATCCACAACATTGCTTACCAAGGAAGATTTTCTTTTTCCGATTTTGCGCTTCTCAATCTTCCGGATTCATTTAGGAGTTCTTTTGACTTCATTGATGGGTATGAGAAGCCTGTTAAAGGAAGGAAGATTAACTGGATGAAGGCTGGAATACTAGAGGCAGACAAGGTCGTGACTGTGAGTCCATACTATGCTGAAGAACTTGCTTCAGGCCCTGATAAAGGTGTGGAACTGGATAGTATTATACGCAGAGCTGGCTGTTTGGGTATTATAAATGGAATGGATGTTCAAGAGTGGAATCCATTAACTGATAAACATATCGACGTCAAATATGATGCCTCAACTGTGACCGATGCGAAGGCTCTTCTGAAAGAAGCACTGCAGGCAGAAGTAGGGTTACCTGTTGACCGTAATATTCCTGTCATAGGTTTTATTGGTAGACTTGAAGAGCAAAAGGGTTCAGATATTTTGGCAGCTGCAATCCCTGAATTTATTAACGAAGATGTGCAGATAATTGTCCTCGGTACTGGCAAAAAGCCCATGGAAAAACAGTTAGAGCAGTTGGAGACTTTGTATCCTGAAAAAGCTAGAGGAGTGGCAAAGTTCAATGTTCCCTTGGCTCATATGATCATAGCTGGGGCTGATTTTATGATGATCCCGAGCAGATTCGAACCATGTGGACTCATCCAGTTACATGCTATGCGTTATGGAACGGTGCCTATTGTTGCCTCAACAGGTGGATTAGTTGACACTGTCAAAGAAGGTTTCACGGGATTTCAAATGGGAGCTTTCAATGTCAAATGCGAGGCAGTTGACCCATTAGACGTGACAGCAATCGCCACAACCACCAAAAAGGCCATTGCAACCTGTGGAACTCCAGCTTTAACAGAAATGATCCAGAACTGCATGGCTCAAGACCTCTCTTGGAAGGGACCTGCCAAGAATTGGGAGGACATGTTGCTGGGCTTGGAAGTTGCGGGTAGTGAACCAGGAATCGAAGGCGATGAAATCGCTCCACTTGCTAAAGAAAATGTTGCCACTCCTTGAGTGACAGAACCGCTCCAGTGCAACTTAGAGGAGCTATTTAACAATCTGACCCAGGCGATTCCAGTGCTGCAACGATTCCAAAATGGCTAGTCGTTTTTATTACTTCGATCATATCTAAGATTTGATAAAATTTATCCACTGCGAGCTGCATAGTCAGTGGAAGAGTGCAGGACGTGTATACGAAGTGGCTTTACTCTGGAAAAGAGATACTGGACCTGGTGGCTTTAACTACTTGTCAAATAAACTTCGTTATCTACCATGGAGGTGTCGTGCTGACCTTGTTGAACAATCTCATAATCTCGAGTTTGTGTTGCATAAATCCAACTTCAAAAGTCGTTTTATAATGTGATTATGATTAATATGTTCTAACGGGTTCCTTGCACATCTGATTAAGACTCTTTCTATTATGTGCTTGTATAAAGCTTATTACAAGTTCGGTCGGTTTTGTAATTTTTTTAGTTGATCTATAATAAACTTGTTTATCAAGTGGTTGGTTTATCCTTTTGTATTATATTTTTGTTTCTCCATTTTATTTCTGCAACTCTTGATAAAAACTAAGGCACTCGCCCCTTTCTTATAAGTTCTCCAAACAAATTGACGTCCAGTAGTTCTGCAGATGAAACCCCACCTACTACACACATGTGGTTGAGGTCAAGTATATAGTATGCGTTGAGTTTTGAGTGTTCATGTCATGTTGTTGCTTAAAAAAATGGTCATGTTGTAAGTGTGTGATATACACGTGAGTGGGTTAGAGAGTTGAGTGAGTCTGGTGATGACAATCGAG

>Pca_TRINITY_Hmgs Pca_TRINITY_DN5888_c0_g1_i2 len=1740

ACACTCTTCCCTTCTCTCCCTCTCCCTCTCTCTCCAAAACAGCTCGATCGATCTAGGGTTATTTTCAAGGAATTCAAACTAATTATCAGAATTCTAAAAATGGCTTCAAATCAAAAGGATGTTGGAATTCTTGCTATGGAAATGTATTTCCCTCCTACTTGTGTTCAACAGGAAGCATTGGAGGCTCATGATGGTGCAAGTAAAGGAAAATACACCATTGGGTTGGGACAAGATTGCATGGCATTTTGTTCGGAGTTAGAAGATGTCATTTCAATGAGTTTGACAGCCGTTAGCTCTTTACTTGAAAAGTATGGAATTGACCCTAAGCAAATTGGACGCTTGGAAGTCGGGAGTGAGACCGTGATTGACAAGAGCAAGTCCATAAAGACCTTCCTGATGACAATATTTGAGGAATGCGATAACACTGACATTGAGGGTGTTGACTCAACAAATGCGTGCTATGGAGGAACTGCAGCTTTGTTCAACGTTGTGAATTGGGTGGAGAGTAGTTCTTGGGACGGACGTTATGGTCTTGTTGTGTGCACTGATAGTGCGGTCTACGCTGAAGGACCCGCCCGCCCCACCGGAGGGGCAGCTGCTATTGCTATGTTGATTGGTCCAGATGCACCAATTGCCTTTGAAAGCAAATACAGAGGATCCCACATGTCCCATGTCTATGATTTTTACAAACCCAACCTAGCCAGTGAATACCCTGTTGTTGATGGAAAACTTTCACAGACTTGTTACCTTATGGCACTTGATTCCTGCTACAAGCGCTTATGCGCCAAGTATGAGAAATTTCAAGGTAAGCAATTCTCGATGGATGATGCAGACTACTTTGTATTTCATTCACCATACAACAAGCTTGTGCAGAAAAGTTTTGCTCGATTGTACTTCATGGACTTCTTAAGAAATGCCAGCTCTGTTGATGAGGCTTCAAAAGAAAAGCTTGCACCATTTTCAGCTCTATCTGGGGAAGAGAGCTATGCAAGTCGGGATCTTGAGAAGACATCCCAAACTGTTGCCAAGAATCTTTATGCTGAGAAGGTTGGACCTACCACTTTGTTACCCAAACAAGTTGGTAATATGTACACTGCATCACTCTATGCGGCATTTGCTTCTCTTCTTCACAACAAAAATAGCTCACTGGATGGTAAACGTATAATAATGTTCTCTTATGGAAGTGGTTTGAGTTCCACAATGTTCTCATTCCGAATGAGGGAAGGTTCACATCCATTCAGCTTGTCAAACATTGCAACTGTAATGGATGTCGCAAACAAGTTGGAGTCGAGACATGAGGTTACTCCTGAAAAATTCGTCGAAACCATGCAACTCATGGAGCATAGGTACGGAGGGAAAGACTTCGTAACGAGCAAGGAAACAAGCCTTTTAACTTCAGGCACATACTATCTTACCGAAGTTGACTCCATGTATCGCAGATTCTACTCCAAGAAGTCAAGTGTGTCAGTGCAAAATGGTCACTGAAAATGGTGAAAGAAGTAAATGGTAACAGATTAGCTGCCGAAGATTTGCTTTGTATTTTCATTGTTGTTGTTAGTGTTTTAGAATAATTCTTATCATGATTTGTATAAGTTAAATTATTCCAAACCTGTGGTTCGCCTATTAGCCATTGTAATGGATTCATCTCCTCTCAGCATATCCATCAGGTATTCAAGTGTTATGCTTCAAAAATAAAAAAATAAAAAGTGAAAAATTTCAAGTGTTAAAAAAAAAAAAAAA

>Pca_TRINITY_Nia Pca_TRINITY_DN1830_c0_g1_i1 len=3127

TTTTTTTGGAGTAAAATATACGATGATAATTAATATTGAGGAAAGAATGTATCAACTTCCACATTTTGTTTTCAGTTCTACATAAAATTTAAACAGACTAACATTTAACAATCACCTTACTTTAGGAGCAAACTAAACCATTACATACATCAACACTAATTATATACAATTACTAATGAGAACTTCCTTATAATTTGTTTATCCCCATATCATGTGTTCCTTTTTGTAGTTTCAAATCAGAACACCAACAATGAATTCTTCACATCGTAATTCATCTTCTCCAAATTTGGAATCACTGCAAATTGAATCATAGGTGGTGGCCCACAAGCCAATGCCAATGTATCATCCGATGCCGGAGGGATATGTTCCCGCAGAACCTCTTCAGTTATGAACCCCACACTGTACTTCCATCCTTCCCTCAATGTCTCACCAACAACATACCAAACTTTCAGCCTCTCGTGTTTCTCTGCCCAACAATCAAGTTCATCCATCAACAAGATGTCATCTTCAGTCCTATTTGCGAATACGATGTACATCTCTGTTTCATCTTCTGGATCTTTCAAAATTGCTTGTACAATTTGATATATTGGTGTAATTCCTGTGCCACCAGCAAGCATTGCGAGTCGTTTGGCAAACTTATGTTTTCCATGAACAGTGAAATTACCACGACCCGTGTATTCAATGTGTCCTAAAGGACCCTTAACATCAATAGTTCCACCAAGAGGAAGAGAATCCAAGTGCTGAGACATAATACCTCCATTAGGGAATTTGGGGTGAACTCCGCCAAAGTAGACTTTAATGACCAGCTCAAAGTAACCAACTTCATCAACAGTACTGGATGGAGTGTAAGCACGCATACATAACTTATCATTAACGTGGGCACATACAAATATGTGCTTCCCAACTGGCAACCCAAGAACTTGATCAGCTGAAGGTAATGCAAACCTGAAGACACGAACATCATGGGAAATCGAGGTTTTGGCTATAAGTTTGCATTGTATTTTATCTCGGCCAACTAGAGCTACGGGTTTCGCAATGCCGATTTCTTTGATAGGAGCAAGGTGACTCAAGCTGCCCGAACCATGAACTGAGTTGTTTGGTGAAGATGATGAGTCAGAAGTGTAACCAGTTGCAATGAGTTCACCGGTTCTGAAATCTTCAAGTAAATTCTTGGCTTTGTCAGAGTGAATTGCATCAAATTCTTCAGTACAATCGGTACCTGCATTAATGAGAATACTATCGGAGCCACCTGGATGGTCTTTCAAGAACCGGGTACAGTCATAAACATGTCCGTGAACAATGATCCAAGCCGAGTCAGGCGAGTTATGTTTTTTCACCTCAGAAATTGAATACATTTTTGACGTGGTGTTCATGAAAGGAGTGGACAAACTCTTTTTGAGTGTCTGTGGACCTTCAGAATTTTCAAGATGTTTTTGTCTAACCATCCATCCACCTGGTTGGTTTCCAGCTTGAGTCGGATGCTCGAATACTAGACCGATTTCTCCTTTGTGTGGTTTGCACACATTTGTTTTGACTTTGAACCAACAATTGTTCATCATACCCATGACATTCCAGATGAGCTTTTCAGGTTGAGTGTTTAGAGTTTCATCCCAAGCTCTGACCGCAATCTCTTTAGCACCAAGCAAGTCGAGAACTTCAACTTCTAGTGACCAGAAACACCAGCACCAATATTTGCCATACTTGTTTGGTTTCTCTGGATGATCCAAAGTACATACTAACCATGACTCTCCGCCGTCCATTGTTACTTCTACACGAGTTACCTTCTTACCGCCGCCGGAATATGCGTAGCCTCTTAACGTGTAAGCTCTTTGTGTTGTTGCAGCATTAATAGGAAGAATCTCTTCATGACAAGGAGTTGTAATCACCGAGTTTATGTTCAACTCATTGATGATGTACTCTGGCTTACACCACCATGCTTCAGAGTTAGCCAATTCAGCGTCGACATGAGAAGGAAGAACTCGATTATCCATAAAATGATAATGATTTTGTGATTCGGGAGTTGTAACAATAATTCTCTTTAACCATTTCACCATTCTTCCTCCGATGAAACCTGGAATTATCATTCGGACTGGAAATCCATGGTCAGGAGATAAATGTTCACCATTCTGCATGTAAGCTAAAATGATATCACGAGCTGGATCCATTGCAAATTCTTTCTTTATACTAGTTCCATATTTAGATCCACCGCCACCGGGTAAATCTTCGGCTCCTTCGAAACAAACATTAAGAGCACCTTTCTTACGGCTCATGATACCACAACGCTTTAGTACGTCACATAAACGTGCACCACGCCACACTGAAGTCGATACTCCTGCACCTCCCCAGTTAAACCCAATTGTTTGTTTTATCATATTCTGTTCTTTACGTCTGTTACCTGCACAGACAAGTGTAGCTGGAAACTCACGGCTTGCGAAATCATTAATGAGCTGATCCATAGTGATACGTAACGGACGCTTTACTAATCCACAAATCTCAACCGTCCAATCCTCCCATTTAAGTTTTGGAACAGCTCCGTGATTACGAACGTAATGAAGAGGAACAGGTGTGATAAACCCGTGATGCATTAGTCGAGTTAATGGAGCTTCTGAATTGAATGGGTGCTTCCCTGTCAATCGAATCATGGAAGGATTGCGTTCAATCCAATTATCAGCGGTTCCTTCATCTTTCGAGTCGAAAACAGAAGGCTCTATTTCATTGTTTATTTTGCCGAAATCTTTCCACTGCTCGTTTTGATCCTCGTCGTCGTCGCTAGATAAATCTTCTATCACCGCTTTTGTTTGTGTAGAAAGACGCTCGGTAATCGACATAGGAAAATTACAGCCACGGATAGGCGAATCAGATCGATGAGTGGAATTGGTTTTAAGTCCACGAACTGCTAAACCTGATTCAAGTCGACTGTTGTACTGTCGATTTTCAACAGAGGCGGCCATTAGAGACTAATCAAAAGAAAGAAGAAAGAAAAATCTGTCTTTGAATGCTCTCTATCACTCTCTTTTCGCTTTCTACTTCTATCTATGTTTTCTCTTTTTTCGATTTGCTTTTGCAAGCATTTGATAGTTGCAAGAAAAATGAGTAGCATGGTTGGGAAAAAAGAAGGAACCCGTGG

>Pca_TRINITY_Eif3E Pca_TRINITY_DN17491_c0_g1_i2 len=1701

CAGAAAACAAAAATATTACCATCCAAATTAAAGGTAATTTCATAATTTAATTTTAGATAAATCAACCGAAACAGTATTTTATCATTCATAGAACCCTAAGAAGAAGTATATTAAACTTACAAATTCTCCTGCAACTCTTGGTTTTTTATGCTTAAAGTGAAATTTTCAATAAACAAATAAACAAAAACAAAATACTGAGACCAAATTTTCTCCCTTACCCGCAAAAGGCTTGAAAACTTCAAGAGCTCCAAGACAAGTTTTCTCCCTATACTACACCTCTTCTCTATCGAGCAGCCGTTACCGGTGCTACTCCTGATTCTAAAACAAGATTTACGTATTTGTAGGTACGCAAAGAGAGTGATTTGGTATTCTCAATGATCTGCTCATACACATTGAGAGAAGTGGGCTCCATTACAACAGTTCCTAGCTTCGAATCGATCTTAGCATCAAGCTTTGCATTCTGAACAAGATTCACTATCCATCTCTCTGCTTCGTCAAAACTCATGTTCAACTTCTCGGCAAGAGTTCCTATATCAATGCGCTGGTGGATTCTGCAGTAAGCCTCAAAGATGAAGAGACGGGCATTTTCAAGAAATTCATCCCTCAAAGGAACGGTGCTAAAGCTTCCTTCCTCAGCTCGTTTACCAAGGAATGGATCATTCAAAATCACATCCTCACATTCACGGAGCTTCTGCTGTGCGCCGTCGAAGTCATAGTTCACATACAAACAGTATAAGAACTCAGTGATGGGATCCTTGTATTCAGATTGCTCTTGCTGGATGACTTTGATCAACTCTTTGAGTTGAGGTCTTCTCTTTTTGTTGACAATCAATGCAGTTGCCAAATAACGTAAAAGATGGGGTGCATTGGTTTGGATGGCATTCATGTACCTGTCATTGAGAAAGAGATCAATGATACCATTTCTTCCATTCTCATGGTTGAAAAATATGAAGAGGCTCCAATGCATCAACCAGATTCTATTCTGGAGTAAAGTCAATGGGTTCTTGAAATTCTTTGATTCAATAATCTCTCTCAACCGATTAAGCTCGTCAAGAGCTTCATCCCAATTCTGCATTAATATCTCAGATGCAAGCTTTCCCCAACATGCATTCAAAATTCTATCAGGATTAGTACAGATTGCTCTGTACTGATGCAAATAATCAGCTGCACCCGAGTAATTCCCACATTCAAATTGAAATTTCGCATACTGATATAACGCCTCAATTTGCTCAGCACCAATCTGGTATCCCTGTAGCAACTGAAGAACATATTGTTTATCAGGGCGTGGTTCGTTGATCATATTCGGGCTTTGAAGGAACGTAATAATCGATTCAGCTCCATCTTCAAGTAATTTCAATCTCCTAACTACTTCAGCACGTCTATTAATCATTTCTTGAGGAACATCATCAGTTTGATGAAGAGACTTGTAAATATCCATAGCGTAATCAACCATATTGGTCTGACTGAGAAGTGCGTCTTTAGCTTTGAGGATTTGAACATCGGGATGGATTTGCTTTTCCTGAAGAAATTCAAGGAGTGGAAGCACAAGATGTCGATCTAAGTTTAGTGCTATTCTTGATGTTAGATCATGATTCGCCATGATTGGTTGTAGAGGCTGAAAATCGCCTCTGAGGAAGGAGGCAGAAAGAGAGAAACCCTAAAAACTGATGAAAACAGAGAAAAGGGGAGATGAGACATGTG

>Pca_TRINITY_AroB Pca_TRINITY_DN2121_c0_g1_i1 len=1866

CTGTAAAGAGAGCAAGTTCCCCATGTAACTTTCTCCTTGGGGGAACACAATTTATATGATCATCCACCGTGCAAGGTTGAAATAAAGACTACCAAGATTTATTTTATTTTTATTTTTATTTTTGCTCAGTAAAAAGAATTATATACATTAAGAAAATAAGAGAACGGAACAAGGGTAAGCCACGCGAGGAGGCAAAATACACCCCCAACACAAGATAAAACAGAAAACTAAGAACCCTTCACGAACTACCACGATTCAAAATGAAAATTAAACCTTTCTTTTTTATTATCAATGAGGATACAACAAGACTCTATAGTAATAAGTGCCACGATGGAAAATCTAGAAGTCATTGAGGTACATAAATAACACAAAACTGCACATGGTTGATGAAGAAATGAGCCAAGATTGGCATTAAATCAAGACTTACAGAATGCACGAAGGGTCTCATCTAAGGCTTTTCTATCGTAATCACCTGTAAAAACACAATTTCCTAGAGGACCTTTGAGGAGGATGAGCCTTAGTAGTCCATCAGCCACCTTCTTATCAACAGCCATGAAAGACTTGAACATCTCCACAGTCATAACATCTGGTGGGGTAATCGGTAGCTTGGCCTTTTGTAGAATATCGTATGTCCGCTTCACAATGGACTCATCAATCCAACCTAGACGGTAAGACATATCAACGGCCATGACCGTTCCAGCTGCTACAGCTTCTCCATGGAGCCATTGACCATATCCAAATCCAGTTTCAATCGCATGTCCAAAGGTATGCCCCAAGTTCAGCGTCGCTCTCACTCCACCTTCCTTCTCGTCCAAGGATACAACCTCTGCCTTGTTTTCACATGAACGCTTGATTGCATATGCTAGTGTACTAGGATCCCTCGCCAACAAAGAGTCCATATTTTTTTCTTGCCACTCGAAGAACTCAGCATCTCTAATGAGTCCATATTTTATAACTTCAGCAAAACCAGAAGCTAATTCCCTCTCAGGCAGTGTATGTAATGTGTCTGTATCTACAAGTACACACTGGGGCTGGTAGAAAGCACCAATCAAGTTTTTTCCCAGTGTATGGTTGATTCCAGTTTTCCCCCCAACAGAGGAATCCACCTGTGCCATGACAGTAGTTGGAATTTGAATGAAGTTTACACCACGAAGGAAAGAAGCTGCAGCGTAACCACACATGTCACCAATAACACCACCTCCAAGAGCGACAAATGTAGAACGGCGATCAAACCGAGATCCTATGGCCTTGTCAAAAACTTTCATCAGAGTATCCATATTTTTATATTTCTCTCCATCTGGCAAAATCACGCTCTCAACCGAAATACTCGGATTTCCATGAGTTAATGCGCTGATAACTTTATCAAGGTATAATGGCGCAACTGTTGTGTTAGTTACCACGAGTACTCTCTTCCCATGTACGTGCCTTTGAAGAAGATCAGGTTGTTCAAGAAGTCCAGAACCAATATAGATCGGATAACTTCTATTACCCAAATCAACATCAACAATAGTTGGTGCTCTAATTGAAGATTGTTCCATTACTTGAGCTGAATTTGCAGATATTATTCTACTCTTCGATTGAATTCGAACCAAAGGAGATCCAATTTTATCCAATCGTAATGAACTTGAATGTATAAACGAAGACTTCAGCGATAAATTCCTCCCTCGAAGAAATAGATCAGAATTTTCCCGTACACAAGAATTCTTAAGAGATAAAGTAAGAGAAGGTGATGAAGAAGAAGCCATCAAATTGGGGAATTAGGGATTAGAGAGGGAAACAAAACAAAATTATTGTTCTTCTGATTTTTTTGAAACCAGAAGAGAAGAAGTAGGTTTCTTTTTTGAAGCATAGAAGTAGTAGTTGAAG

>Pat_TRINITY_Agt1 Pat_c8785_g1_i1 len=1581

ACGTGTGCTCTTCCGATCTCTGCTTATTACACACATATTTCCTTCCTCCCATTGTTTATCTCTTAAAGTTGGACTGGGAGAGAGAGAGAGAGTGAAAGTCTGAAGTAGTTCTCCAGAAACCATTCTTGCAACGAAAAGCTTCAAAGAGGTACAATATGGATTACGTTAATGCACCTGGGAGGAACCATCTCTTTGTTCCTGGACCTGTCAATATTCCAGAATCAGTCCTTCGTGCCATGAGCAGAAACAATGAAGATTACCGTTCCCCTGCAATTCCTGCTATGACTAAAACTCTTCTTGAAGATGTTAAGAAGATTTTCAAGACAACTTCTGGTACCCCATTTTTGATCCCCACAACAGGTACTGGTGCATGGGAAAGTGCACTTACAAATACTTTGTCTCCTGGTGATCGAACAATTTCTTTCCTCATCGGGCAATTTAGTCTTCTCTGGATTGATCAGCAGCAGCGCCTAGGATTCAACGTTGATGTTGTAGAGAGTGAGTGGGGCCGTGGTGCTGATCTTGAAGTCCTAGCATCCAAGTTAGCTGAAGACACTGCTCATACCATCAAAGCTGTTTGCATGGTTCATAACGAAACTGCAACTGGAGTTACTAATAATTTGGCTGCTGTAAGGAAATTACTTGATCACTACAACCATCCTGCATTGTTTCTTGTGGATGGAGTGTCTTCGATATGTGCTCTTGATTTTCGTATGGATGAATGGGGAATCGATGTAGCTTTAACTGGTTCACAAAAAGCTCTTTCTCTACCTACTGGGATGGGTATTGTGTGTGCAAGCCCTAAAGCACTGGAAGCATCCAAGACTGCCAAATCAGTGAGAGTGTTCTTTGACTGGAAAGACTACTTGAAGTTCTACAAAATGGGAACTTATTGGCCTTACACTCCTTCAATCCAGCTTTTATACGGCCTCAGAGCTGCTCTTGATCTCCTATTTGAGGAAGGACTTGACAATGTCATTGCTAGGCACAGCCGCTTGGCCAAAGCAACTAGACTTGCTGTTGAAGCCTGGGGTTTGAAAAACTGTACACAGAAAGAGGAGTGGTTGAGCGACACTGTGACTGCAGTTGTTGTTCCACCTTACATCGACAGTGCTGAAATAGTTAGAAGGGGATGGCAGAGATACAATCTAAGCTTAGGTCTGGGTTTGAATAAAGTGGCTGGAAAAGTTTTCAGGATAGGTCATCTTGGGAACCTGAATGAGCTCCAACTGCTGGGTTGTTTAGCTGGAGTGGAGATGATTCTGAAAGATGTCGGATACCCAGTTAAGCTAGGAAGTGGAGTGGCTGCGGCTTGTGCGTTTTTCCAGAACACCATTCCTATGATTGAATCTAGGATTTAATGAATCATTCATTATCCATGTGTAACTACTGTTCTCCCTCTTCTGATTTGCTTGGAGTACTTGTATAAAAGAAGACATGATAATCTCCTGTATTTGCACTCTTTTGAGGCTTACTTAGTAGTACAAGAACTCATTACTCTTTCATCCATTAATCCTGTATTTAAAATCTCTTTAAATTTAATCTCATCACTGTTGCATATATCCAATAGATCGGAAGAGC

>Pat_TRINITY_At103 Pat_c12751_g1_i1 len=1467

GTGTGCTCTTCCGATCTCTCACTCAAGAAGCTACACACTCTGCAAAAAATGGCTGCAGAGATGGCATTGATAAAACCCATCTCTAATTTCACTAGAACCAGCCCTAAATTTATCACCTCAACAAGAAAATCATTCTCCATTGTCAAAATGGCATCCCAATCCACTGCATCAACAACAAAGTCGGGGAAACCAAAAAAAGCAATCAAAGAAACCTTACTTGCTCCTAGGTTTTACACAACTGATTTTGATGAAATGGAAACTCTTTTTAACACCGAGATGAACAAAAACTTGAATGAGGCTGAGTTTGAAGCTCTATTACAAGAGTTCAAAACTGATTACAATCAAACTCATTTCGTTAGAAACAAAGAGTTCAAAGAAGCTGCTGATAAAATGCAAGGTCCTCTTAGACAGATTTTTGTTGAGTTCTTGGAGAGATCTTGTACTGCTGAGTTCTCAGGGTTTCTTCTTTACAAAGAACTTGGAAGAAGACTTAAGAAAACCAATCCAGTTGTTGCTGAAATTTTCTCACTCATGTCCAGGGATGAAGCTAGGCATGCTGGATTCTTAAACAAGGGACTGTCAGATTTCAACTTGGCACTGGATCTTGGATTTCTTACAAAGGCTAGAAAATATACATTCTTCAAACCCAAGTTCATTTTCTATGCCACGTATCTATCTGAGAAAATTGGGTACTGGAGGTACATAACCATTTATAGGCACTTAAAGACCAACCCTGAATACCAAGTTTATCCAATTTTCAAGTATTTTGAGAACTGGTGCCAAGACGAGAACAGACATGGAGATTTCTTCTCCGCACTGATGAAAGCTCAACCCCAGTTTCTTAACGATTGGAAAGCAAAGTTGTGGTCTCGGTTCTTCTGCCTATCGGTGTACGTGACCATGTACTTGAATGACTGCCAGAGAACAGCTTTTTACGAAGGGATCGGACTTAACACTAAAGAGTTTGATATGCACGTCATAATTGAGACAAACCGTACAGCCGCAAGGATTTTCCCTGCAGTTTTAGATGTTGAGAACCCGGAGTTCAAGAGGAAGTTGGACAGGATGGTGCAGATCAACGAGAAACTTCTTGCAATTGGTGAAACTGAGGAATCATCTTTGGTGAAGAACTTAAAAAGGATCCCACATGTTGCAGCGTTGGCGAATGAGCTCTTGGCTGCGTATCTCATGCCTCCTATTGATTCTGGGTCTGTTGATCTTGCTGAGTTTGAGCCACAAGTTGTGTATTAATAAGTTTTTTGTGTATTGAGTAAACAAAGGCCCTTCAATTCAATTCAGTGCAAAGTTTAATTGCTACAGTTAACCATTACTATGGACCTGCTGTATAAAGAACTAAACATTTGTGACATAATGTATCATGATTTCTGGACGCACCTCATGTGGACATAACTCGTTATGCATATTTTTTTCCGCACACATATAGCCCAGGTTGTCACAACTGGCAAG

>Pat_TRINITY_ChlP Pat_c64005_g1_i1 len=1768

TGTGCTCTTCCGATCTCAAAATCTTAAAACTTTCATTCATACTTGCAGAAACTTGAAGAATCCATCACCACCAAAAATGGCCTCCATTGCCGTCAATGCTCGTTTCTCAGCCACAACATGCGACCGTAAAACAACCGAAATCAGCCGTGAAACACTAAAATCCTTCATGGGTCTTAGAAAAACCAACCAGAATGCTTCTATCTCACTTACACAATCTCTGACAACCAGTACTCCACATTTCAACCCTGCAAGAAGAAATTTCACTTCCATCAAAGCAGGTAAATCAAGTCCTAAAGTAACAGGAAGAAATCTCCGAGTTGCGGTCATTGGTGGTGGTCCTGCTGGTGGTGCAGCTGCTGAAACCTTAGCTAAAGGCGGTATCGAAACGTTCTTGATCGAAAGGAAAATGGATAACTGTAAACCATGCGGTGGTGCAATTCCACTATGCATGGTTGGTGAGTTTGATCTTCCGTTAGACATCATCGACAGACGTGTTACTAAAATGAAGATGATTTCGCCTTCAAATGTTGCAGTCGACATTGGCCAAACTCTTAAGCCAGATGAGTATATCGGTATGGTGCGTCGAGAAGTGCTCGACCAATATCTTCGTGAAAGAGCAGAGAAAGCTGGAACAAAGGTAATTAACGGTTTATTCTTGAAAATGGATTTACCAAAAGAACCAAATGCGCCTTATGTTCTTCACTACAGTGCTTATGATGGCAAAACTGGTGGCGTTGGCACAAAAACCACGCTAGAGGTAGATGCAGTCATCGGTGCAGATGGCGCGAATTCTCGCGTGGCTAAAGCAATTGGAGCTGGTGATTATGATTATGCCATTGCATTCCAAGAGAGAATCAGAATCCCAGATGAGAAGATGGTTTACTATGAAAATCTTGCCGAGATGTACGTCGGTGATGATGTGTCGCCTGATTTTTACGGATGGGTGTTTCCGAAATGTGATCACGTCGCAGTGGGTACTGGTACAGTTACGCACAAAGGCGATATCAAGAAATTTCAAGTGGCTACACGACTGCGCGCCAAGGATAAGATTCTAGGCGGGAAGATTATTAGAGTAGAGGCACATCCTATTCCAGAACATCCAAGACCCAAGAGGTTGGCTGACAGAGTAGCGCTTGTTGGAGATGCAGCTGGGTACGTAACAAAATGTTCAGGAGAAGGTATATATTTTGCTGCTAAGAGTGGCAGGATGTGCGCTGAAGCGATTGTTGAAGGATCCGAGAGAGGGACAAGAATGGTTAACGAAACGGATTTGAGAAATTATCTGGAGAAATGGGATAAAACTTACTGGCCAACTTACAAGGTTCTGGATATATTGCAGAAGGTGTTTTACCGATCGAATCCAGCCCGAGAAGCTTTCGTTGAGATGTGCGCGGATGATTATGTCCAGAGGATGACATTTGATAGTTATCTGTATAAGAGAGTTGTGCCTGGAAATCCAATTGAGGATATAAAATTGGCTTTCAATACTATTGGGAGTTTGGTCAGAGCTAATGCACTTAGAGGCGAGATGAACAAGATTACTAGTCTATGATGAAGAATGCGTTGAGTTTTACTTTTGTGAAAATAGGTCTGTTTTTCTGGTGAATTAAGATAGATAAAATGTGAAGCGATCGGAATCTTGGCGATTTTTGTTGATCTCTTTTTGTTATAATTGTGTATTTAACTTCTTGATTAATCACATCAATAAAGTTGTATCAATTTCTTCACTCTACGGCAATAAAATGATTTATCACCTGTAAGGTACTGC

>Pat_TRINITY_GbssI Pat_c12579_g1_i1 len=2097

TTCTCTCGTAACAAAGTTAACAACTCCTACTACTTACTTGTGATCTGTGCTGCTGTGTCTCACTCTTCTGTCAGTCAGTCTGGAGACAAAAGAGTGACATATTCTCCATCCTTGTGAAACTCATCAGCATATAATAATCTCTCAAGTCTCCACTGACAAGAATATTACGGATGGCGACCATAAGTGCGTCCCAGTTTGTATCACGAATCAACTGCGGAGGAACTGCACCAAAAGACATACTTAAAAAAATCAATATGAAGAACCAAAGCATGACTCATTCAGGGTTAAGATCTTTGAGCAAAGTAGATTTCTTACAAATAAAAAACCCTCAGAAAGTAGCTTCTAGGGAACTGAGGAGGAACGCATTCCAGACGGACAAAGGTACTGCGTCAGGAGAGGTTAAATGTTCTAGTGGGATGAACTTGGTCTTTGTTGCTGCTGAAGTAGGCCCCTGGAGTAAAACCGGTGGTCTTGGTGATGTTCTCGGGGGCTTACCACCAGCCATGGCGGCAAATGGACATCGTGTTATGACAGTCTCTCCTCGTTATGATCAGTACAAAGATGCTTGGGATACCAATGTGTCAGTTGAGATGAAAATTGGGGACAGAATTGAAACCGTTCGTTTCTTCCACTGTTATAAGCGTGGGGTGGACCGTGTATTTGTGGATCATCCACTGTTTCTTGAGAAGGTATGGGGTAAAACTGGATCTAAAATCTATGGTCCAAATGCTGGATCAGATTACCAAGATAACCAGCTTAGATTCTGTTTGCTTTCTCTGGCAGCTCTAGAAGCACCAAGAATTCTGAACCTCAACAGCAACAAATATTTCTCTGGAACATACGGAGAGGAGGTCGTGTTCATCGCTAATGATTGGCATACGGCCCTTCTACCATGCTACCTTAAAAGTATGTACCAATCTAAAGGGATGTACGAAACTTCCAAGGTTGCTTATTGCATCCACAACATTGCTTACCAAGGAAGATTTTCTTTTTCCGATTTTGCGCTTCTCAATCTTCCGGATTCATTTAGGAGTTCTTTTGACTTCATTGATGGGTATGAGAAGCCTGTTAAAGGAAGGAAGATTAACTGGATGAAGGCTGGAATATTAGAGGCAGACAAGGTCGTAACTGTGAGTCCATACTATGCTGAAGAACTTGCTTCAGGCCCGGATAAAGGTGTGGAACTGGATAGTATTATACGGAAAGCTGGCTGTCTGGGTATTATAAATGGAATGGATGTTCAGGAGTGGAACCCATTAACTGATAAATATATCGACGTCAAATATGATGCCTCAACTGTGACCGAGGCGAAGGCTCTTCTGAAAGAAGCACTGCAGGCAGAAGTAGGGTTACCCGTTGATCGTAATATTCCTGTCATAGGTTTCATTGGTAGACTTGAGGAGCAAAAGGGTTCAGATATTTTGGCAGCTGCAATCCCTGAATTTATTGACGAAGAAGTGCAAATAATTGTCCTCGGTACTGGCAAAAAGCCCATGGAAAAACAGTTGGAGCAGTTAGAGACTTTGTATCCTGAAAAAGCTAGAGGAGTGGCGAAGTTTAACGTTCCCTTGGCTCATATGATCATAGCTGGAGCTGATTTTATGATGATCCCAAGCAGATTTGAACCATGTGGACTCATCCAGTTACATGCTATGCGTTACGGAACGGTGCCTATTGTTGCCTCGACGGGTGGATTAGTCGACACTGTCAAAGAAGGTTTCACGGGGTTTCAAATGGGAGCTTTCAATGTCGAATGCGAGGCAATTGACCCATTAGACGTGACAGCAATCGCCACAACCACCAAAAAGGCCCTTGAAACCTATGGAACTCCAGCTTTAACAGAAATGATCCATAATTGTATGGCTCAAGATCTCTCTTGGAAGGGACCTGCCAAGAATTGGGAGGACATGTTGCTGGGCTTGGAAGTTGCAGGTAGTGAACCAGGAATCGAAGGCGATGAAATCGCTCCTCTTGCTAAAGAAAATGTTGCCACTCCTTGAGTGACAGAACCGCTCCAGTGCAACTTAGAGGAGCTATTTCACAATCTAACCCAGGCGATTCCGTTGCTGCACCGACAAGGCAGATCGGAAGAGCGTC

>Pat_TRINITY_Hmgs Pat_c1543_g1_i1 len=1835

CAGACGTGTGCTCTTCCGACCTGTCGGACCGGTCATGAATTTCCTCCGTCTATAAATACCGATTTATTCGTTGCCTAAATCTCTCACCACTTCTTCTTCATTCTCCCTTCCTTCTCTTCCCCTCTCTCTCCAAAAACAGCTCGATCGATCTAGGGTTCTCTGAAAGAATTCAAACTAATTATCTAAAGATGGGTTCGAATCAGAAGGATGTTGGAATTCTTGCTATGGAAATGTATTTCCCTCCTACTTGTGTCCAACAGGAAGCATTGGAGGCTCATGATGGTGCCAGTAAAGGAAAATACACCATTGGGTTGGGACAAGATTGCATGGCGTTTTGTTCCGAGTTGGAAGATGTCATTTCAATGAGTTTGACAGCCGTTAGCTCTTTACTCGAAAAGTATAAAATTGACCCTAAACAAATTGGACGCCTGGAAGTCGGCAGTGAAACCGTGATTGACAAGAGCAAGTCCATAAAGACCTTCCTGATGACTATCTTTGAGGAATGTGATAACACTGACATTGAGGGTGTTGACTCCACAAATGCATGCTACGGAGGAACTGCAGCTTTGTTCAACGTTGTAAATTGGGTGGAGAGTAATTCCTGGGATGGACGATATGGTCTTGTTGTGTGCACTGATAGTGCGGTCTATGCAGAAGGACCTGCCCGCCCCACTGGAGGAGCAGCTGCTATTGCTATGTTGATTGGTCCAGACGCACCAATTGCCTTTGAAAGCAAATATAGAGGATCCCATATGTCCCATGTCTATGATTTTTACAAGCCCAACCTAGCCAGTGAATACCCGGTCGTTGACGGAAAACTTTCACAAACTTGTTACCTCATGGCACTCGATTCCTGCTACAAACGCTTCTGCGCTAAGTACGAGAAATGCCAGGGTAAGCAATTCTCGATGGATGATGCAGACTACTTTGTATTTCATTCTCCTTACAACAAGCTCGTGCAGAAAAGTTTTGCTCGATTGTACTTCATGGATTTCTTAAGAAACGCCAGCTCTGTTGACGAGGCTTCAAAAGAAAAGCTCGCACCATTTGCAGCTCTATCTGGTGAAGAGAGCTACGCCAGTCGGGATCTTGAGAAGACATCCCAAACTGTTGCAAAGAATCTTTATGCTGAGAAGGTTGGACCTACCACTTTGTTGCCCAAACAAGTCGGTAATATGTACACTGCATCACTCTATGCGGCATTTGCTTCTCTTCTTCACAACAAAAATAGCTCACTGGATGGCAAATGTGTAATAATGTTCTCTTATGGAAGTGGTTTGAGTTCTACAATGTTCTCATTCCGAATGAGGGAAGGTTCACATCCATTCAGCTTATCAAACATTGCTACTGTAATGGATGTTGCACACAAGTTGGAGTCGAGACATGAGGTTACTCCTGAAAAATTCGTCGAAACCATGCAACTCATGGAACATAGGTACGGAGGGAAAGACTTCGTAACGAGCAAAGACATGAGCTTGTTAACTCCAGGCACATACTACCTTACCGAAGTTGACTCCATGTACCGCAGATTCTACTCCAAGAAGTCAAGTGCCGCAGTGCAAAATGGTCACTAAAAAAATAGTGGAGGATGGAAATATCTAGCAGAAACGCTGCCGAAGAATTGCTTTTGTATTCTCATTGTTGTTGTTAGTGTTTTAGAATAATTCTTATTTATGATTTGTACAAGTTAAATTATTCCAAACCTCTATGGTTCGCCCATTAGCCATTGTAACAGGATTCTTCTCCTCTCAGTATATCCATTAAGTATTCAAGTGTTCCTCTCCATCTCTCCCCCTGATATAAAAGTTTCCTTTCAGATCGGAAGAGCGTCGTGT

>Pat_TRINITY_Nia Pat_c18416_g1_i1 len=3113

CAAATCGGAAAAAAAAAAAAGAAAACATAGATAGAAGTAGAAAGCAGAGAGAAAGTGAAACGAGAGGAGGAGCATTCAAAGAGAGATTTTTCTTTCTTCTTTCTTTTGATTAGTCTCTAATGGCGGCTTCTGTTGAAAATCGACAGTACAACAGTCGACTTGAATCAGGTCTAGCAGTTCGTGGATTTAAAACCAATCCCACTCATCGATCTGATTCGCCTATCCGTGGCTGTAATTTTCCCATGTCGACTACCGAGCGTCTTACTATACAAGCAAAACCGGTCATTGAAAATTTATCTAGCGACGACGATGAAGATCAAAACGAGCAGTGGAAAGATTTCGGCAAAATAAACAATGAAATAGAGCCGTCTGTTTTTGACTCGAAAGATGAAGGAACCGCTGATAATTGGATTGAACGCAACCCTTCCATGATTCGGCTAACAGGGAAACACCCATTCAATTCAGAAGCACCATTAACTCGTCTCATGCATCACGGGTTTATCACACCTGTTCCTCTTCATTACGTTCGTAATCACGGAGCTGTTCCAAAACTTAAATGGGAGGATTGGACAGTTGAGATTTGTGGATTAGTAAAACGTCCGTTACGTATCACCATGGATCAGCTCATTAATGATTTCACAAGCCGTGAGTTTCCAGCTACACTTGTCTGTGCAGGTAACAGACGTAAAGAACAGAATATGATAAAACAAACAATTGGGTTCAACTGGGGAGGCGCAGGTGTATCGACTTCAGTATGGCGTGGTGCACGTCTATGTGACGTATTAAAGCGTTGTGGAATCATGAGTCGTAAGAAAGGTGCTCTTAATGTTTGTTTCGAAGGAGCTGAAGATTTACCCGGTGGCGGTGGATCTAAATATGGAACTAGTATAAAGAAGGAATTCGCAATGGATCCTGCTCGTGATATAATTCTATCTTACATGCAAAATGGTGAACGCTTGTCTCCTGACCACGGATTTCCAGTCCGTATGATAATTCCAGGTTTCATCGGAGGAAGAATGGTGAAGTGGTTAAAGAGAATTATCGTCACAACTCCAGAATCCCAAAACCACTACCATTTCATGGATAATCGTGTTCTTCCTTCTCATGTCGACTCTGAACTGGCCAACTCAGAAGCATGGTGGTATAAGCCAGAGTACATCATCAATGAGTTGAACATAAACTCTGTGATTACAACTCCTTGTCATGAAGAGATTCTTCCCATAAATGCTGCAACAACACAAAGAGCTTACACATTAAGGGGATACGCATATTCCGGCGGCGGTAAGAAAGTAACTCGTGTCGAAGTAACAATGGACGGCGGAGAGTCATGGTTAGTGTGTACTTTGGATCATCCAGAGAAACCAAACAAGTATGGCAAATATTGGTGCTGGTGTTTTTGGTCACTAGAAGTTGAAGTTCTCGACTTGCTTGGTGCCAAGGAGATTGCAGTCAGGGCTTGGGATGAAACTCTAAACACTCAGCCTGAAAAGCTCATCTGGAATGTCATGGGTATGATGAACAACTGCTGGTTCAAAGTCAAAACAAATGTGTGCAAACCACACAAAGGAGAAATCGGTCTAGTTTTCGAGCATCCAACTCAGGCTGGAAACCAACCAGGTGGATGGATGGTTAGACAAAAACATCTTGAAAACTCTGAAGGTCCACATACCCTCAAAAAGAGTTTGTCCACTCCTTTCATGAACACCACGTCAAAAATGTATTCAATATCCGAGGTGAAAAAACACAACTCGCCTGACTCGGCTTGGATCATTGTTCACGGACATGTCTATGACTGCACCCGGTTCTTGAAAGACCATCCAGGTGGTTCCGATAGTATTCTCATCAATGCAGGTACCGATTGTACCGAAGAATTTGATGCAATTCACTCTGACAAAGCCAAGAATTTACTTGAAGATTTCAGAATCGGTGAACTCATTGCAACTGGTTACACTTCTGACTCGTCATCTTCACCAAACAACTCAGTTCATGGTTCGGGCAGCTTGAGTCACCTTGCTCCTATCAAAGAAATCGGCATTGCGAAACCCGTAGCTCTAGTTGGCCGAGATAAAATACAATGCAAACTTATTGCCAAGACCTCGATTTCCCATGACGTTCGTGTCTTCAGGTTTGCATTGCCTTCAGCTGATCAAGTCCTTGGATTGCCAGTTGGGAAGCACATATTTGTGTGTGCCCACGTTAACGATAAGCTATGCATGCGTGCTTACACTCCATCGAGCACAGTTGACGAAGTTGGTTACTTTGAGTTGGTCATTAAAGTCTACTTCGGCGGAGTTCACCCCAAGTTCCCCAATGGAGGTATTATGTCTCAGCACTTGGATTCTCTTCCTCTTGGTGGAACTATTGATGTTAAGGGTCCGTTGGGACACATTGAATACACTGGTCGTGGTAATTTCACCGTTCATGGCAAACACAAGTTTGCCAAACGACTCGCAATGCTTGCTGGTGGCACAGGAATTACACCAATATATCAAATTGTGCAAGCAATATTGAAAGACCCAGAGGATGAAACGGAGATGCACATCGTATTCGCAAACAGGACTGAAGATGACATCTTGTTGATGGATGAACTTGATTCTTGGGCAGAGAAGCACGAAAGGTTGAAAGTATGGTACGTTGTTGGTGAAACATTGAGGGAAGGATGGAAGTACAGTGTGGGGTTCATAACTGAAGAGGTCTTGCGGGAACATATTCCTCCGGCATCGGAAGATACATTGGCGTTGGCTTGTGGACCACCACCTATGATTCAATTTGCAGTGACTCCAAATTTGGAGAAGATGAATTATGATGTGAAGAATTCATTGTTGGTCTTCTGATCACGAGTCAGTTGAAGTATTTGAAAGGTGAACCTACAAAGAGGAACACATATGATCAAATTTAAGAAAATGATAAGGGGAGGACAAATTATAAGGAAGTTCTCATTAGTTATTGTATATAATTAGTGTTGAAGGATGTAATGGTTTAGTATGTTCTTAAAAGTAAGGGGATTGTTAAATGTTAGTCTGTTTAAATTTTATGTAGAATTGAAAGCAAAATGTGGGAAATGATACTTTCTTTCCTCAATATTTATGATCATTATATATTTTACTCCCTCC

>Pat_TRINITY_Eif3E Pat_c8804_g1_i1 len=1968

CAGACGTGTGCTCTTCCGTTCTCTCCGCTTTCCTTTCTCAGTTTTTCTCATCAGTTTTTAGGGTTTCTCTCTTTCTTCCTCCCACCACCATTGCGCCGCTCCTCATCCTTCTCAGCAGAGGTTGTTCTTCAGCCTCTCCAACAACTAACAGCAACCAATCATGGCGAATCTTGATCTAACATCAAGAATAGCACTAAACTTAGATCGGCATCTCGTTCTCCCACTCCTTGAATTTCTTCAAGAGAAACAAATCCATCCCGATGTTCAAATCCTCAAAGCTAAAGACGCACTTCTCAGCCAGACCAATATGGTCGATTACGCTATGGATATCTACAAATCTCTTCATCAAACTGATGATGTTCCGCAAGAGATGATTAACAGACGCGCTGAAGTAGTTAGGAGGTTGAAATTACTTGAAGATGCAGCTGAATCGATTATTACGTTCCTGCAAAGCCCGAATATGATTAGCGAACCACGCCCTGATAAGCAGTATGTTCTTCAGTTTTTACAGGGATACCAGATTGGTGCTGAGCAAATTGAGGCATTGTATCAGTATGCCAAATTTCAATTTGAATGTGGGAATTACTCCGGTGCAGCTGATTATTTGCATCAGTACAGAGCAATTTGTACCAATCCTGATAGAATTTTGAATGCATGTTGGGGAAAGCTTGCATCTGAGATATTGATGCAGAACTGGGATGAGGCTCTTGATGAGCTTAATCGGTTGAGGGAGATTATCGAATCAAAGAATTTCAAGAACCCATTGACTTTACTCCAGAATAGAATCTGGTTGATGCATTGGAGCCTCTTCGTCTTTTTCAACCACGAGAATGGAAGAAATGGCATCATTGATCTATTTCTCAATGACAGGTACATGAATGCGCTCCAAACAAACGCACCCCATCTTTTTCGTTATTTGGCAACTGCATTGATTGTCAACAAAAAGAGAAGACCTCAACTCAAAGAGTTGATCAAAGTCATCCAGCAAGAGCAATCTGCATACAAGGATCCCATCACTGAGTTCTTGTACTGTTTGTATGTCAACTATGACTTCGATGGTGCACAACAGAAGCTCCGTGAATGTGAGAATGTGATTTTGAATGATCCATTCCTTGGTAAACGAGCTGAAGAAGGAAACTTTAGTACCGTTCCTTTGAGGGATGAATTTCTTGAAAATGCCCGTCTCTTCATCTTTGAGGCTTACTGCAGAATCCACCAGCGCATTGATATTGGAACTCTAGCCGAGAAGTTGAATATGAAATTTGACGAGGCAGAGAGATGGATAGTGAATCTTGTTCAGAATGCAAAGCTTGATGCTAAGATTGATTCTAAGCTAGGAACTGTTATAATGGAGCCTACTTCTCTCAATGTGTATGAGCAGATCATTGAGAATACAAAATCACTTTCTCTGCGTACCTACAAATACGTTAATCTTGTTTTAGAATCAGCAACACCACCAGTTACAGCTGCTCGTTAGAGAAGAGGTAGTATTATAGCGAGAAAATTTGTGTTGGAGCTCTTGAAGTTTCCTTGTGGGGAAGGAAGAAAATTTGGTCTCAGGTATTTTGTTTTTGTTTACTTGTTTTTCAGTTTCGCCTAAAGCATAGAAAACCAAGAGTTGCAGGAGAACTTGTAAGTTTAATATACTTCTTTTTAGGGTTCTTTGAACGATAAAATACAGTTCCTGTTGATTTATCTAACATTAAATTATGAAATTACATTTAATTCGGATGGTAATGTTGTTTTCTATTATTTCCCATTCTCAAGAGAGATACACATTCTTTATGGATAGCAACCCTGTAACGACCAGTTCCATTTCCATCCGTTCTTTGGGCGACTCGGTAGAACATGTAACACCAAGTTTGATAATCTTTTTCAAAGCCTCGCATAACTTGGTCCTTTTCTCAACTTTGATTGTAACCTCTTCATTGTTATTTAGTAGATATGTAGGAATAAGGTTAGGATCCA

>Pat_TRINITY_AroB Pat_c23141_g1_i1 len=2182

ATTTTCTCACCAACCCCCAGAGCCCAGAGGCTGCTGCTTCAACTACTTCTTCTTCTTCTTCTCGTCTGGTTTGAAAAATAATCAGAAGAAGTTTTGTTTTGTTTCCCTCTCTAATTCTTCCCCAATTTTTTGATGGCTTCTTCTACTACTTCTTCACCTTCTCTTAATCTAACTCTGAAGAACTCTTGTGTACGAGAAAATTTTGATCTATTTATTAGAGGGAGGAATTTATCACTGAAGTCTTCATTTATAAATTCAAGTCCATTACGATTGAATAGAATTGGATCTTTAAAAATTCAATCGAAGAGTAGAATAATATCTGCAAATTCAGCTCAAGTAATGGAACAATCTTCAATTAGGGCACCAACTATTGTTGATGTTGATTTGGGTAATAGAAGTTATCCGATCTATATTGGTTCTGGACTTCTTGAACAACCTGATCTTCTTCAGAGGCACGTACATGGGAAGAGAGTACTCGTGGTAACTAATACAACAGTTGCGCCATTATACCTAGATAAAGTTGTCAGCGCTTTAACTCATGGAAATCCGAGTATTTCGGTTGAGAGTGTGATTTTGCCAGATGGAGAGAAATATAAAAATATGGATACTCTGATGAAAGTTTTTGACAAGGCCATAGGATCTCGATTTGATCGCCGTTCCACATTTGTCGCTCTTGGAGGTGGTGTTATAGGTGACATGTGTGGTTACGCTGCAGCTTCTTTCCTTCGTGGTGTAAACTTCATTCAAATTCCAACTACTGTCATGGCACAGGTGGATTCCTCTGTTGGGGGAAAAACTGGAATCAACCACACACTGGGGAAAAACTTGATTGGTGCTTTCTACCAGCCCCAATGTGTACTTGTAGATACAGACACATTACATACACTGCCTGAGAGGGAATTAGCTTCTGGTTTTGCTGAAGTTATAAAATATGGACTCATTAGGGATGCTGAGTTCTTCGAGTGGCAAGAAAAAAATATGGACTCTCTGTTGGCGAGGGATCCCAGTGCACTAGCATATGCAATCAAGCGTTCATGCGAAAACAAGGCAGAGGTTGTATCATTGGACGAGAAGGAAGGTGGAGTGAGAGCGACGCTGAACTTGGGTCATACCTTTGGACATGCAATTGAAACAGGATTTGGCTATGGACAATGGCTCCATGGAGAAGCTGTAGCAGCTGGAACGGTCATGGCAGTCGATATGTCTTACCGTCTAGGTTGGATTGATGAATCAATTGTGAAGCGGACATACGATATTTTACAAAAAGCCAAACTGCCGATTACCCCACCAGATGTTATGACTGTAGAGATGTTCAAGTCTTTCATGGCTGTTGATAAGAAGGTGGCTGATGGACTACTAAGGCTCATCCTTCTCAAAGGTCCTCTCGGCAATTGTGTTTTTACAGGTGATTACGATAGAAAAGCCCTAGACGAGACCCTTCGTGCATTCTGTAAGTCTTAATTTAATGCCGGCCTTGGTTGGGCTCATTTCTTCATCAACCATGTGCAGTTTTGTGTTATTTTTGTTGACGTGAAACCACATTGTCTTATGTATATGTATAATTCAATAACTTCCTAGATTTGCCATCAGCGCATGTATTACTATAGAGATTTTTTCCAGCGTTTTTTGGGTTACATATTCACTTCTATCTTAGATCAAAGGTTTCTTTCTAATCGATCCAACAAAAAAAAAAACATAGTACAGAAATAGGAGTGCTACTGGCCCTCCATTTGGAACTAGAACTGTTGCAGGTCGTATTTGGAAAGAAAGCTATGGTTCTGCTAAGCAGCAACACACATTTACGATTGAAGTATTAAAGACCAAAGAAGAGGAACCACTTCCTCCACTCCTCCCTCTTCTAATTAAGGGTAGAAACCTTTATAATCCAAACACAATGAGATAGGCGGATGAGGAGATGAGCAGGCAGGTCTAATAAAGAAGCAAGAATGAACAAAAGGAAATGTGGAAGAAGAGTACTAGACAAAGGTGAACTTCTCAAGTCTGTCCTGCTTGTCAACAATATCGTCTTGGGTTTTGGTTGAAAGTCCCAACTCCCAAAATATGAATTCAGTTAATTTTTTTTTGTTTTTGCCATTTCTTGATTTATTTGAATCTATTTAGTATTTAGCCTTTAGCAGTACACATATGTAGTTGCTGCTATCACAAAAGAGATCGGAAGAGC

>Por_TRINITY_Agt1 Por_c44419_g1_i1 len=1576

ATAAATATATGTTTTTTCTTCCTAATGCTGCTTATTACACACATATTTCCTTCCTCCCATTGTTTATCCCTTAAAGTTGGACTGGGAGAGAGAGAGAGTGAGAGTCTGAAGTAGTTCTCCAGAAACCATTCTTGCACCGAAAAGCTTCAAAGAGGTACAATATGGATTACGTTAATGCACCTGGGAGGAACCATCTCTTTGTTCCTGGACCTGTCAACATTCCAGAATCAGTCCTTCGTGCCATGAGCAGAAACAATGAAGATTACCGTTCCCCTGCAATTCCTGCCATGACTAAAACTCTTCTTGAAGATGTTAAGAAGATTTTCAAGACAACTTCTGGAACCCCATTTTTGATCCCCACAACAGGTACTGGTGCATGGGAAAGTGCACTTACAAATACTTTGTCTCCTGGTGATCGGACAATTTCTTTCCTCATCGGGCAATTTAGTCTTCTCTGGATTGATCAGCAGCAACGCCTTGGATTCAACGTTGATGTTGTAGAGAGTGAGTGGGGCCGTGGTGCTGATCTTGAAGTCCTAGCATCCAAGTTAGCTGAAGACACTGCTCATACCATCAAAGCTGTTTGCATGGTTCATAATGAAACTGCAACTGGAGTTACTAATAATTTGGCTGCTGTAAGGAAATTACTTGATCACTACAACCATCCTGCACTGTTTCTTGTGGATGGAGTGTCTTCGATATGTGCTCTTGATTTTCGTATGGATGAATGGGGAATCGATGTAGCTTTAACTGGTTCACAAAAAGCTCTTTCTCTACCTACTGGGATGGGTATTGTGTGTGCAAGCCCTAAAGCACTGGAAGCATCCAAGACTGCCAAATCAGTGAGAGTGTTCTTTGACTGGAAAGACTACTTGAAGTTCTACAAAATGGGAACTTATTGGCCATACACTCCTTCAATCCAGCTTTTATACGGCCTCAGAGCTGCTCTTGATCTCCTATTTGAGGAAGGACTTGACAATGTCATTGCTAGGCACAGCCGCCTGGCCAAAGCAACTAGACTTGCTGTTGAAGCCTGGGGTTTGAAAAACTGTACACAGAAAGAGGAGTGGTTGAGCGACACCGTGACTGCAGTTGTTGTTCCACCTTACATCGACAGTGCTGAAATAGTTAGAAGGGGATGGCAGAGATACAATCTAAGCTTAGGTCTGGGTTTGAATAAAGTGGCTGGAAAAGTTTTCAGGATAGGTCATCTTGGGAACCTGAATGAGCTCCAACTGCTGGGTTGTTTAGCTGGAGTGGAGATGATTCTGAAAGATGTCGGATACCCAGTTAAGCTAGGAAGTGGAGTGGCCGCGGCGTGTGCATTTTTCCAGAACACCATTCCTATGATTGAATCTAGGATTTAATGAATCATTCATTATCCATGTGTACCTACTGTTCTCCCTCTTCTGATTTGCTTGGAGTACTTGTATAAAAGAAGACATGATAATCTCCTGTATTTGCACTCTTTTGAGGCTTACTTAGTAGTACAAGAACTCATTACTCTTTCATCCATTAATCCTGTATTTAAAATCTCTTTAAATTTAATCTCATCACTGTTGCATGTAAAAAAAAA

>Por_TRINITY_At103 Por_c49245_g1_i1 len=1436

GACGTGTGCTCTTCCGATCTCACTCTAGAAGCTACACAGACTCTGCAAAAATGGCTGCAGAGATGGCATTGATAAAACCCATCTCTAATTTCACTAGAACCAGCCCTAAATTTATCACCTCAACAAGAAAATCATTTTCCGTTGTAAAAATGGCATCTCAGTCCACTGCATCAACAACAAAGTCAGGGAAACCCAAAAAAGCAATCAAAGAAACCTTACTTGCTCCTAGGTTTTACACAACTGATTTTGATGAAATGGAAACTCTTTTTAACACTGAGATGAACAAAAACTTGAATGAAGCTGAGTTTGAAGCTTTATTACAAGAGTTCAAAACTGATTACAATCAAACTCATTTCGTTAGAAACAAAGAGTTCAAAGAAGCTGCTGATAAAATGCAAGGTCCTCTTAGACAGATTTTTGTTGAGTTCTTGGAGAGATCTTGTACTGCTGAGTTCTCAGGGTTTCTTCTTTACAAAGAACTTGGAAGAAGACTTAAGAAAACCAATCCAGTTGTTGCTGAAATTTTCTCACTCATGTCCAGGGATGAAGCTAGGCATGCTGGATTCTTAAACAAGGGACTGTCAGATTTCAACTTGGCACTGGATCTTGGATTTCTTACAAAGGCTAGAAAATATACATTCTTCAAACCCAAGTTCATTTTCTATGCCACATATCTATCTGAGAAAATTGGGTACTGGAGGTACATAACCATTTATAGGCACTTAAAGACCAACCCTGAATACCAAGTTTATCCAATTTTCAAGTATTTTGAGAACTGGTGCCAAGACGAGAATAGACATGGAGATTTCTTCTCCGCACTGATGAAAGCTCAACCACAGTTTCTTAACGATTGGAAAGCAAAGTTGTGGTCTCGGTTCTTCTGCCTATCGGTGTACGTGACCATGTACTTGAATGACTGCCAGAGAACAGCTTTTTACGAAGGGATCGGACTTAACACTAAAGAGTTTGATATGCACGTCATAATTGAGACAAACCGTACAGCCGCAAGGATTTTCCCTGCAGTTCTAGATGTTGAGAACCCGGAGTTCAAGAGGAAGTTGGACCGGATGGTGCAAATCAACGAGAAACTTCTTGCAATTGGTGAAACTGAGGAATCATCTCTGGTGAAGAACTTAAAAAGGATCCCACATGTTGCAGCATTGGCGAATGAGCTCTTGGCTGCGTATCTCATGCCTCCTATTGATTCTGGTTCTGTTGATCTTGCTGAGTTTGAGCCACAAGTTGTGTATTAATAAGTTTTTTGTGTATTGAGTAAACTAAGGCCCTTCAATTCAATTCAGTGCAAAGTTTAATTGCTACAGTTTACCATTACTATGGACCTGCTGTATAAAGAACTAAACATTTGTGACATAATGTATCATGATTTCTGGACGCACCTCATGTAGTTGTTACTTGTTATGCATATTTTTTTTCTT

>Por_TRINITY_ChlP Por_c54793_g1_i1 len=1619

TGTGCTCTTCCGATCTCAGAAACTTGAAGAATCCATCACCACCAAAAATGGCCTCCATTGCCGTCAATGCTCGTTTCTCAGCCACAACATGCGACCGTAAAACAACCGAAGTCAGCCGTGAAACACTAAAATCCTTCATGGGTCTTAGAAAAACCAACCAAAATGCTTCTATCTCACTTACACAATCTCTAACAACAAGTACTCAACATTTCAACCCTGCTGCTAGAAGAAATTTCACTTCCATCAAAGCAGGTAAATCAAGTCCTAAAGTAACAGGAAGAAATCTCCGAGTTGCGGTCATTGGTGGCGGTCCTGCTGGCGGTGCAGCTGCTGAAACCTTAGCTAAAGGCGGTATCGAAACGTTCTTGATCGAAAGGAAAATGGATAACTGTAAGCCATGTGGTGGTGCAATTCCACTCTGCATGGTTGGTGAGTTTGATCTTCCTTTGGATATCATCGACAGACGCGTTACGAAAATGAAGATGATTTCGCCATCAAATGTTGCAGTGGACATTGGCCAGACTCTTAAACCAGATGAGTATATCGGTATGGTACGTCGAGAAGTGCTCGACCAATATCTCCGTGAAAGAGCAGAGAAAGCTGGAACAAAGGTAATCAACGGTTTATTCTTGAAAATGGATTTACCAAAAGAACCAAATGCGCCATATGTTCTTCACTACAGTGCTTATGATGGCAAAACTGGTGGAGTTGGCACAAAAACCACGCTAGAGGTAGATGCAGTCATCGGTGCAGATGGTGCGAATTCTCGCGTAGCCAAAGCAATCGGAGCTGGTGATTATGATTATGCTATTGCATTCCAAGAGAGAATCAGAATCCCAGATGAGAAGATGGTTTACTATGAAAACCTTGCAGAGATGTACGTTGGTGATGATGTGTCGCCTGATTTTTACGGATGGGTGTTTCCAAAATGTGATCACGTTGCAGTGGGTACCGGTACAGTTACTCACAAAGGCGATATCAAGAAATTTCAAGTGGCTACGCGGCTGCGCGCCAAGGATAAGATTCTAGGCGGGAAGATTATCAGAGTAGAGGCACATCCTATTCCAGAACATCCAAGACCCAAGAGGTTGGCAGACAGAGTAGCCCTTGTAGGAGATGCAGCTGGGTACGTTACAAAATGTTCTGGAGAAGGTATATACTTTGCTGCTAAGAGTGGCAGGATGTGCGCCGAAGCGATTGTTGAAGGATCCGAGAGAGGGACAAGAATGGTTAATGAAACGGATTTGAGAAATTATCTGGAGAAATGGGATAAAACTTACTGGCCAACTTACAAGGTTCTGGATATATTGCAGAAGGTGTTTTACCGATCGAATCCAGCTCGAGAAGCTTTTGTTGAGATGTGCGCGGATGATTATGTTCAGAGGATGACATTTGATAGTTATCTGTATAAGAGAGTTGTGCCTGGAAATCCAATTGAGGATATTAAATTGGCTTTCAATACTATTGGGAGTTTGGTCAGAGCTAATGCACTTAGAGGCGAGATGAACAAGATTACTAGTCTATGATGATGAGTTTGTGAAAATAGGTTTGTTTTTCTGGTGAAATAAGATAGATAAAACGGGGGTGATGCGAATCTTGGCAGATCGGAAGAGCGTCGT

>Por_TRINITY_GbssI Por_c59766_g2_i1 len=2325

TACGGACTTCTTCTTCTTCTTCTCTCGTAACAAAGTTAACAACTCCTGCTACTTGTGATCTGTGCTGCTGCTGTGTCTCACTCTATTCTCCATCCTTGTGAACTCATCAGCATAAACTGATAAGAATATTACCATGGCGACCATAAGTGCGTCCCAGTTTGTATCACGAATTAATTGCGGAGGAACTACACCAAAAGACATACTTAAACAAATCAATATGAAGAACCAGAGCATGACTCATTCAGGGTTAAGATCTTTGAGCAAAGTAGATTTCTTACAAATAAAAAACACTCAGAAAGCAGCTTCTATGGAACTGAGGAGGAACGCATTCCAGACAGACAAAGGTACTGCGGCAGGAGAGGTTAAATGTTCTGGTGGAATGAACTTGGTATTTGTAGCTGCTGAAGTAGGCCCCTGGAGTAAAACCGGTGGTCTTGGTGATGTTCTTGGAGGATTACCACCAGCCATGGCGGCAAATGGACATCGTGTTATGACAGTCTCTCCTCGTTATGATCAGTACAAAGATGCTTGGGATACCAATGTGTCAGTTGAGATGAAAATTGGGGACAGAATTGAAACCGTTCGTTTCTTCCACTGTTATAAGCGTGGGGTTGACCGTGTATTTGTGGATCATCCACTGTTTCTTGAGAAGGTATGGGGTAAAACTGGATCTAAAATCTATGGTCCGAATACTGGTTCTGATTACCAAGATAACCAGCTTAGATTCTGCTTGCTTTCTCTGGCAGCTCTCGAAGCACCAAGAGTTCTGAACCTCAATAGCAACAAATATTTCTCTGGAACATACGGAGAGGAGGTCGTGTTCATCGCTAATGATTGGCATACGGCCCTTCTACCATGCTACCTTAAAAGTATGTACCAATCTAAAGGGATGTACGAAACTGCCAAGGTTGCTTATTGCATCCACAACATTGCTTACCAAGGAAGATTTTCTTTTTCCGATTTTGCACTTCTCAATCTTCCGGATTCATTTAGGAGTTCTTTTGACTTCATTGATGGGTATGAGAAGCCTATTAAAGGAAGGAAAATTAACTGGATGAAGGCTGGAATATTAGAGGCAGACAAGGTCGTAACTGTGAGTCCATACTATGCTGAAGAACTTGCTTCAGGCCCGGATAAAGGTGTGGAACTGGATAGTATTATACGGAAAGTTGGCTGTCTGGGTATTATAAATGGAATGGATGTTCAGGAGTGGAACCCGTTAACTGATAAATATATCGACGTCAAATATGATGCCTCAACTGTGACCGAGGCGAAGGCTCTTCTGAAAGAAGCACTGCAGGCAGAAGTAGGGTTACCTGTTGACCGTAATATTCCTGTCATAGGTTTCATTGGTAGACTTGAGGAGCAAAAGGGTTCAGATATTTTGGCAGCTGCAATCCCTGAATTTATTGACGAAGAAGTGCAAATAATTGTTCTCGGTACTGGAAAAAAGCCCATGGAAAAACAGTTGGAGCAGTTAGAGACTTTGTATCCTGAAAAAGCTAGAGGAGTCGCGAAGTTCAACGTTCCCTTGGCTCATATGATCATAGCTGGGGCTGATTTTATGATGATCCCAAGCAGATTTGAACCATGTGGACTCATCCAGTTACATGCTATGCGTTACGGAACGGTGCCTATTGTTGCCTCGACGGGTGGATTAGTCGACACTGTCAAAGAAGGTTTCACGGGATTTCAAATGGGAGCTTTCAATGTCGAATGCGAGGAAACTGACCCATTAGACGCGACAGCAATCGCCACAACCACCAAAAAGGCCCTTGAAACCTATGGAACTCCAGCTTTAACAGAAATGATCCAGAACTGCATGGCTCAAGACTTCTCTTGGAAGGGACCTGCCAAGAATTGGGAGGACATGTTGCTGGGCTTGGAAGTTGCAGGTAGTGAACCAGGAATCGAAGGCGATGAAATCGCTCCTCTTGATAAAGAAAATGTTGCGACTCCTTGAGTGACAGAACCGCTCCAGTGCAACTTAGAAGAGCTATTTCACAATCAAACCCAGGAGATTCCATTGCTGCACCAACAAGGCTCTGATTACTTCCATCATATCTAAGATTTGATAAAATTTATCCACTGCGAGTTACATAGTCAGTGAAAGAGTTCATGATGTGTATATGAAGTGGCTTTACTGGAAAAGAGATACTTCTAGTCATTTTCTTTTTCTTCCTTTTCTTCTTTTGATTTAAGCAATCTACGCCACTCAGTGTCCAATGCGGGATGCCAAAAGAATATGCTAGCTAGCTCTAGATCTTGTATCTAGTCAAGAAGATCTTTATATAAATAAAAACGACTTTTTCACCTAAAAAAAAA

>Por_TRINITY_Hmgs Por_c52524_g1_i1 len=1897

AGACGTGTGCTCGTCCGATCTCAAAAGTCCGCCTTTGAATTTCACAAACTCACGTTGGCTGCATTTTTTTTTCTGGTTCCGGTCGGACCGGTCATGAATTTCCTCCGTCTATAAATACCGATTTATTCGTTGCCTAAATCTCTCACCACTTCTTCTTCATTCTCCCTTCCTTCCTTCTCTTCCCCTCTCTCTCTCCAAAAACAGCTCGATCGATCTAGGGTTCTCTGAAAGAATTTAAACTAATTATCTAAAGATGGGTTCGAATCAGAAGAATGTTGGAATTCTTGCTATGGAAATGTATTTCCCTCCTACTTGTGTCCAACAGGAAGCATTGGAGGCTCATGATGGTGCCAGTAAAGGAAAATACACCATTGGGTTGGGACAAGATTGCATGGCGTTTTGTTCCGAGTTGGAAGATGTCATATCAATGAGTTTGACAGCCGTTAGCTCTTTACTCGAAAAGTATAAAATTGACCCTAAACAAATTGGACGCCTGGAAGTCGGCAGTGAAACCGTGATTGACAAGAGCAAGTCCATAAAGACCTTCCTGATGACTATCTTTGAGGAATGTGATAACACTGACATTGAGGGTGTTGACTCCACAAATGCATGCTACGGAGGAACTGCAGCTTTGTTCAACGTTGTGAATTGGGTGGAGAGTAATTCCTGGGATGGACGATATGGTCTTGTTGTGTGCACTGATAGTGCGGTCTATGCAGAAGGACCTGCCCGTCCCACTGGAGGAGCAGCTGCTATTGCTATGCTGATTGGTCCAGACGCACCAATTGCCTTTGAAAGCAAATATAGAGGATCCCATATGTCCCATGTCTATGATTTTTACAAGCCCAACCTAGCCAGTGAATACCCGGTCGTTGACGGAAAACTTTCACAAACTTGTTACCTCATGGCACTCGATTCCTGCTACAAACGCCTCTGCGCTAAGTACGAGAAATGCCAGGGTAAGCAATTCTCGATGGATGATGCAGACTACTTTGTATTTCATTCTCCTTACAACAAGCTCGTGCAGAAAAGTTTTGCTCGATTGTACTTCATGGATTTCTTAAGAAACGCCAGCTCTGTTGACGAGGCTTCAAAAGAAAAGCTCGCACCATTTGCAGCTCTATCTGGTGAAGAGAGCTACGCCAGTCGGGATCTTGAGAAGACATCCCAAACTGTTGCAAAGAATCTTTATGCTGAGAAGGTTGGACCTACCACTTTGTTGCCCAAACAAGTCGGTAATATGTACACTGCATCACTCTATGCGGCATTTGCTTCTCTTCTTCACAACAAAAATAGCTCACTGGATGGCAAACGTGTAATAATGTTCTCTTATGGAAGTGGTTTGAGTTCTACAATGTTCTCATTCCGAATGAGGGAAGGTTCACATCCATTCAGCTTATCAAACATTGCCACTGTAATGGATGTTGCACACAAGTTGGAGTCGAGACATGAGGTTACTCCTGAAAAATTCGTCGAAACCATGCAACTCATGGAGCATAGGTACGGAGGGAAAGACTTCGTAACGAGCAAAGACACAAGCTTGTTAACTCCAGGCACATACTACCTTACCGAAGTTGACTCCATGTACCGCAGATTCTACTCCAAGAAGTCAAGTGCCGCAGTGCAAAATGGTCACTAAAAAAATAGTGAAGGATGGAAATATCAAGCAGAAAAGCTGCCGAAGAATTGCTTTTGTATTCTCATTGTTGTTGTTAGTGTTTTAGAATAATTCTTATTTATGATTTGTACAAGTTAAATTATTCCAAACCTCTATGGTTCGCCCATTAGCCATTGTAACAGGATTCATCTCCTCTCAGTATATCCATTCAAGTGTTCCTCTCCATCTCTCCCCCCCTGATATAAATAAAAGTTTCCTTTCTCGGAGGCCTGAGCAGGG

>Por_TRINITY_Nia Por_c55953_g1_i1 len=3111

TGCTCTTCCGATCTGCTTGCAAAAGCAAATCGGAAAAAAAGAAAGAAAACATAGATAGAAGTAGAAAGCAGAGAGGAGGAGCATTCAAAGAGAGATTTTTTTCTTTCTTCTTTCTTTTGATTAGTCTCTAATGGCGGCTTCTGTTGAAAATCGACAGTACAACAGTCGACTTGAATCAGGTCTAGCAGTTCGTGGATTTAAAACCAATCCCACTCATCGATCTGATTCGCCTATCCGTGGCTGTAATTTTCCCATGTCGACTACCGAGCGTCTTTCTATACAAGCAAAACCGGTCATTGAAGATTTATCTAGCGACGACGATGAAGATCAAAACGAGCAGTGGAAAGATTTTGGCAAGATAAACAATGAAATAGAGCCGTCTGTTTTTGACTCGAAAGATGAAGGAACCGCTGATAATTGGATTGAACGCAACCCTTCCATGATTCGGCTAACAGGGAAACACCCATTCAATTCAGAAGCACCATTAACTCGTCTCATGCATCACGGGTTTATCACACCTGTTCCTCTTCATTACGTTCGTAATCACGGAGCTGTTCCAAAACTCAAATGGGAGGATTGGACGGTTGAGATTTGTGGATTAGTAAAGCGTCCGTTACGTATCACTATGGATCAGCTCATTAATGATTTCACAAGCCGGGAGTTTCCAGCTACACTTGTTTGTGCAGGTAACAGACGTAAAGAGCAGAATATGATAAAACAAACAATTGGGTTTAACTGGGGAGGCGCAGGAGTATCTACTTCAGTATGGCGTGGTGCACGTTTATGTGACGTATTAAAGCGTTGTGGAATCATGAGTCGTAAGAAAGGTGCTCTTAATGTTTGTTTCGAAGGAGCTGAAGATTTACCCGGTGGCGGTGGATCAAAATATGGAACTAGTATAAAGAAGGAATTTGCAATGGATCCTGCTCGTGATATAATTCTAGCTTACATGCAAAATGGCGAACGTTTGTCTCCTGACCACGGATTTCCAGTCCGTATGATAATTCCGGGTTTCATCGGAGGAAGAATGGTGAAGTGGTTAAAGAGAATTATCGTCACATCTCCAGAATCCCAAAACCACTATCATTTCATGGATAATCGTGTTCTCCCTTCTCATGTCGACGCTGAACTGGCCAACTCAGAAGCATGGTGGTATAAGCCAGAGTACATCATCAATGAGTTGAACATAAACTCAGTGATTACAACTCCTTGTCATGAAGAGATTCTTCCTATAAATGCTGCAACAACACAAAGAGCTTACACATTAAGGGGATACGCATATTCCGGCGGCGGTAAGAAAGTAACTCGTGTCGAAGTAACAATGGACGGCGGAGAGTCATGGTTAGTGTGTACTTTGGATCATCCAGAGAAACCAAACAAGTATGGCAAATATTGGTGCTGGTGTTTTTGGTCACTAGAAGTTGAAGTTCTCGACTTGCTTGGTGCCAAGGAGATTGCGGTCAGGGCTTGGGATGAAACTCTAAACACTCAGCCTGAAAAGCTCATCTGGAATGTCATGGGTATGATGAACAACTGCTGGTTCAAAGTCAAAACAAATGTGTGCAAACCACACAAAGGAGAAATCGGTCTAGTTTTCGAGCATCCAACTCAGGCTGGAAACCAACCAGGTGGATGGATGGTTAGACAGAAACATCTTGAAAACTCTGAAGGTCCACATACCCTCAAAAAGAGTTTGTCCACTCCTTTCATGAACACCACGTCAAAAATGTATTCAATATCCGAGGTGAAAAAACACAACTCTCCTGACTCGGCTTGGATCATTGTTCACGGACATGTCTATGACTGCACCCGGTTCTTGAAAGACCATCCAGGTGGTTCCGATAGTATTCTCATCAATGCAGGTACCGATTGTACCGAAGAATTTGATGCAATTCACTCTGACAAAGCCAAGAATTTACTTGAAGATTTCAGAATCGGTGAACTCATTGCAACTGGTTATACTTCTGACTCGTCATCTTCACCAAACAACTCAGTTCATGGTTCGGGCAGCTTGAGTCACCTTGCTCCTATCAAAGAAATCGGCATTGCGAAACCCGTAGCTCTAGTTGGCCGAGATAAAATACAATGCAAACTTATTGCCAAGACCTCGATTTCCCATGACGTTCGTGTCTTCAGGTTTGCATTGCCTTCAGCTGATCAAGTCCTTGGATTGCCAGTTGGGAAGCACATATTTGTATGTGCCCACGTTAACGATAAGCTATGCATGCGTGCTTACACTCCATCGAGCACAGTTGACGAAGTTGGTTACTTTGAGTTGGTCATTAAAGTCTACTTCGGCGGAGTTCACCCCAAGTTCCCCAATGGAGGTATTATGTCTCAGCACTTGGATTCTCTTCCTCTTGGTGGAACTATTGATGTTAAGGGTCCGTTGGGACACATTGAATACACCGGTCGTGGTAATTTCACCGTTCATGGCAAACACAAGTTTGCCAAACGACTCGCAATGCTTGCTGGTGGCACAGGAATTACACCAATATATCAAATTGTGCAAGCAATATTGAAAGACCCAGAGGATGAAACGGAGATGCACATCGTATTCGCAAACAGGACTGAAGATGACATCTTGTTGATGGATGAACTTGATTCTTGGGCAGAGAAGCACGAAAGGTTGAAAGTATGGTACGTTGTTGGTGAAACATTGAGGGAAGGATGGAAGTACAGTGTGGGGTTCATAACTGAAGAGGTTTTGCGGGAACATATCCCTCCGGCATCGGAAGATACATTGGCATTGGCTTGTGGACCACCACCTATGATACAATTTGCAGTGACTCCAAATTTGGAGAAGATGAATTATGATGTGAAGAATTCATTGTTGGTCTTCTGATCACGAGTCAGTTGAAGTATTTGAAAGGTGAAACTACAAAGAGGAACACATATAATCGAATTTAAGAAAATGATAAGGGGAGGACAAACTATAAGGAAGTTCTCATTAGTTATTGTATATAATTAGTGTTGAAGGATGTAATGGGTTTAGTATGTTCTTAAAAGTAAGGGGATTGTTAAATGTTAGTCTGTTTAAATTTCATGTAGAATTGAAAGCAAAATGTGGGAAATGATACTTTCTTTCCTCAAAAAAAAAAGATCGGAAGA

>Por_TRINITY_Eif3E Por_c50471_g1_i2 len=1661

GACGTGTGCTCTTCCGATCTCAATTTTTAGGGTTTCTCTCTTTCTTCCTCCCACAACCACTGCGCCGCTCCTCCTTCTGCTCAGAGGTTGTTCTTCAGCCTCTACCACAACAACAGCAACCAATCATGGCGAATCATGATCTGACATCAAGAATAGCACTAAACTTAGATCGACATCTCGTTCTACCACTCCTTGAATTTCTTCAAGAGAAACAAATCCATCCCGATGTTCAAATCCTCAAAGCTAAAGATGCACTTCTCAGCCAGACCAATATGGTCGATTACGCTATGGATATCTACAAATCTCTTCATCAAACTGATGATGTTCCTCAAGAGATGATTAACAGGCGTGCTGAAGTAGTTAGGAGGTTGAAATTACTTGAAGATGCAGCTGAATCGATTATTACGTTCCTGCAAAGCCCGAATATGATTAACGAACCACGCCCTGATAAGCAGTATGTTCTTCAGTTGTTACAGGGATACCAGATTGGTGCTGAGCAAATTGAGGCATTGTATCAGTATGCCAAATTTCAATTTGAATGTGGGAATTACTCGGGTGCAGCTGATTATTTGCATCAGTACAGAGCAATTTGTACCAATCCTGATAGAATTTTGAATGCATGTTGGGGAAAGCTTGCATCTGAGATATTGATGCAGAACTGGGATGAGGCTCTTGATGAGCTTAATCGGTTGAGGGAGATTATCGAATCAAAGAATTTCAAGAACCCATTGACTTTACTCCAGAATAGAATCTGGTTGATGCATTGGAGCCTCTTCATCTTTTTCAACCACGAGAATGGAAGAAATGGCATCATTGATCTATTTCTCAATGACAGGTACATGAATGCGCTCCAAACAAATGCGCCCCATCTTTTACGTTATTTGGCAACTGCATTGATTGTCAACAAAAAGAGAAGACCTCAACTCAAAGAGTTGATCAAAGTCATCCAGCAAGAGCAATCTGCATACAAGGATCCCATCACTGAGTTCTTATACTGTTTGTATGTCAACTATGACTTCGATGGTGCACAACAGAAGCTCCGTGAATGTGAGAATGTGATTTTGAATGATCCATTCCTTGGTAAACGAGCTGAAGAAGGAAACTTTAGTACTGTTCCCTTGAGGGATGAATTTCTTGAAAATGCCCGTCTCTTCATCTTTGAGGCTTACTGCAGAATCCACCAGCGCATTGATATTGCAACTCTTGCCGAGAAGTTGAATATGAATTTTGACGAGGCAGAAAGATGGATAGTGAATCTTGTTCAGAATGCAAAGCTTGATGCTAAGATTGATTCTAAGCTAGGAACTGTTGTAATGGAGCCCACTTCTCTCAATGTGTATGAGCAGATCATTGAGAATACCAAATCACTTTCTCTGCGTACCTACAAATATGTAACTCTTGTTCTTGAATCAGCAGCACCACCGGTTACGGCTGCTCGTTAGAGAAGAGGTAGTAGTATAGCGAGAAAACTTGTGTTGGAGCTCTTGAAGCTTACTTGTGGGGAAGGAAGAAAATTTGGTCTCGGTATTTTGTTTTTGGTACTTGTTTTTCAGTTTCATCTAAAGCATTAAAAACCAAGAGTTGCAGGAGAACTTGTAAGTTTAAAAATATACTTGTTTTTAGGGTTCTTTGAAGGATAAAATACAGTTCCTGAGATCGGA

>Por_TRINITY_AroB Por_c48498_g1_i2 len=1632

CTCTTCCGATCTCTTTCACCAACCCCCAGCCAGACCCCAGTGGCTGCTGCTTCAACTACTTCTTCTTCTTCATCTCGTCTGGTTTCAAAAAAATCAGACGAAGAAATAATTTTGTTCTGTTTTATTTCCCTCTCTAATCCCTGATTCCTCAATTTTTGATGGCTTCTTCTACTACTTCTTCACCTTCTCTTAATCTAACTCTGAAGAACTCTTGTGTACGAGAAAATTTTGATCTATTTCTTAGAGGGAGGAATTTATCACTGAAGTCTTCATTTATAAATTCAAGTCCATTACGATTGAATAGAATTGGATCCTTAAAAATTCAATCGAAGAGTAGAATAATATCTGCAAATTCAGCTCAAGTAATGGAACAATCTTCAATTAGGGCACCAACTATTGTTGATGTTGATTTGGGTAATAGAAGTTATCCGATCTATATTGGTTCTGGACTTCTTGAACAACCTGATCTTCTTCAGAGGCACGTACATGGGAAGAGAGTACTCGTGGTAACTAATACAACAGTTGCGCCATTATACCTAGATAAAGTTGTCAGCGCTTTAACTCATGGAAATCCGAGTATTTCGGTTGAGAGTGTGATTTTGCCAGATGGAGAGAAATATAAAAATATGGATACTCTGATGAAAGTTTTTGACAAGGCCATAGGATCTCGATTTGATCGCCGTTCCACATTTGTCGCTCTTGGAGGTGGTGTTATAGGTGACATGTGTGGTTATGCTGCAGCTTCTTTCCTTCGTGGTGTAAACTTCATTCAAATTCCAACTACTGTCATGGCACAGGTGGATTCCTCTGTTGGGGGAAAAACTGGAATCAACCACACACTGGGGAAAAACTTGATTGGTGCTTTCTACCAGCCCCAATGTGTACTTGTAGATACAGACACATTACATACACTGCCTGAGAGGGAATTAGCTTCTGGTTTTGCTGAAGTTATAAAATATGGACTCATTAGGGATGCTGAGTTCTTCGAGTGGCAAGAAAAAAATATGGACTCTCTGTTGGCGAGGGATCCCAGTGCACTAGCATATGCAATCAAGCGTTCATGCGAAAACAAGGCAGAGGTTGTATCCTTGGACGAGAAGGAAGGTGGAGTGAGAGCGACACTGAACTTGGGTCATACCTTTGGACATGCAATTGAAACAGGATTTGGCTATGGACAATGGCTCCATGGAGAAGCTGTAGCAGCTGGAACGGTCATGGCAGTCGATATGTCTTACCGTCTAGGTTGGATTGATGAATCAATTGTGAAACGGACGTACGATATTTTACAAAAAGCCAAACTGCCGATAACCCCACCAGATGTTATGACTGTAGAGATGTTCAAGTCTTTCATGGCTGTTGATAAGAAGGTGGCTGATGGACTACTAAGGCTCATCCTTCTCAAAGGTCCTCTCGGCAATTGTGTTTTTACAGGTGATTACGATAGAAAAGCCCTAGACGAGACCCTTCGTGCATTCTGTAAGTCTTAATTTAATGCCGACCTTGGTTGGGCTCATTTCTTCATCAACCATGTGCAGTTTTGTGTTATTTTTCTTTTGTTGACATCAAACCACATTGTTTTATGTATATGTATGTAATTCAGTAACTTCCTAGATTATAGAGATTACTCTGAAC

>Pbr_TRINITY_Agt1 Pbr_c5449_g1_i1 len=1732

TGTGCTCTTCCGATCTGATTTCTTGCCGCATGATCTAGAGTTCAGTACATTCTAATCTTCCACGTGGAAGCTCTGTGATCTATCAGCAGCATATACGTTTCATCTTGATCCTCTCATCTTCCACTTATTTTGGGCCTGTCTATATATATAAATATATGTTTTTTCTTCCTAATGCTGCTTATTACACACATATTTCCTTCCTCCCATTGTTTATCATTAAAGTTGGACTGGGAGAGAGAGGGAGCGAGAGTCTGAAGTAGTTCTCCAGAAACCATTCTTGCACCGAAAAGCTTCAAAGAGGTACAATATGGATTACGTTAATGCACCTGGGAGGAACCATCTCTTTGTTCCTGGACCTGTCAACATTCCAGAATCAGTCCTTCGTGCCATGAGCAGAAACAATGAAGATTACCGTTCCCCTGCAATTCCTGCCATGACTAAAACTCTTCTTGAAGATGTTAAGAAGATTTTCAAGACAACTTCTGGAACCCCATTTTTGATCCCCACAACAGGTACTGGTGCTTGGGAAAGTGCACTTACCAATACTTTGTCTCCTGGTGATCGAACAATTTCTTTCCTCATCGGGCAATTTAGTCTTCTCTGGATTGATCAGCAGCAACGCCTCGGATTCAACGTTGATGTTGTAGAGAGTGAGTGGGGCCGTGGTGCTGATCTTGAAGTCCTAGCATCCAAGTTAGCTGAAGACACTGCTCATACCATCAAAGCTGTTTGCATGGTTCATAATGAAACTGCAACTGGAGTTACTAATAATTTGGCCGCTGTAAGGAAATTACTTGATCACTACAACCATCCTGCACTGTTTCTCGTGGATGGAGTGTCTTCGATATGTGCTCTTGATTTTCGTATGGATGAATGGGGAATCGATGTAGCTTTAACTGGTTCACAAAAAGCTCTTTCTCTACCTACTGGGATGGGTATTGTGTGTGCAAGCCCTAAAGCACTGGAAGCATCCAAGACTGCCAAATCAGTGAGAGTGTTCTTTGACTGGAAAGACTACTTGAAGTTCTACAAAATGGGAACTTATTGGCCATACACTCCTTCAATCCAGCTTTTATACGGCCTCAGAGCTGCTCTTGATCTCCTATTTGAGGAAGGACTTGACAATGTCATTGCTAGGCACAGCCGCTTGGCCAAAGCAACTAGACTTGCTGTTGAAGCCTGGGGTTTGAAAAACTGTACACAGAAAGAGGAGTGGTTGAGCGACACTGTGACTGCAGTTGTTGTTCCACCTTACATCGACAGTGCTGAAATAGTTAGAAGGGGATGGCAGAGATACAATCTAAGCTTAGGTCTGGGTTTGAATAAAGTGGCTGGAAAAGTTTTCAGGATAGGTCATCTTGGGAACCTGAATGAGCTCCAACTGCTGGGTTGTTTAGCTGGAGTGGAGATGATTCTGAAAGATGTCGGATACCCAGTTAAGCTAGGAAGTGGAGTGGCTGCGGCTTGTGCGTTTTTCCAGAACACCATTCCGATGATTGAATCTAGGATTTAATGAATCATTCATTATCCATGTGTAACTACCGTTCTCCCTCTTCTGATTTGCTTGGAGTACTTGTATAAAAGAAGACATGATAATCTCCTGTATTTGCACTCTTTTGAGGCTTACTTAGTAGTACAAGAACTCATTACTCTTTCATCCATTAATCCTGTATTTAAAATCTCTTTAAATTTAATCCATCACTGTTGCATATATCCAATAGATCGGAAGAGC

>Pbr_TRINITY_At103 Pbr_c49641_g1_i1 len=6426

TCTTCTTTACAAAGAACTTGGAAGAAGACTTAAGAAAACCAATCCAGTTGTTGCTGAGATTTTCTCACTCATGTCCAGGGATGAAGCTAGGCATGCTGGATTCTTAAACAAGGGACTGTCAGATTTCAACCTGGCACTGGATCTTGGATTTCTTACAAAGGCTAGAAAATATACATTCTTCAAACCCAAGTTCATTTTCTATGCTACATATTTATCTGAGAAAATTGGGTACTGGAGGTACATAACCATTTATAGGCACTTAAAGACCAACCCTGAATACCAAGTTTATCCAATTTTCAAGTATTTTGAGAACTGGTGCCAAGACGAGAACAGACATGGAGATTTCTTCTCCGCACTGATGAAAGCTCAACCCCAGTTTCTTAACGATTGGAAAGCAAAGTTGTGGTCTCGGTTCTTCTGCCTATCGGTGTACGTGACCATGTACTTGAATGACTGCCAGAGAACAGCTTTTTACGAAGGGATCGGACTTAACACTAAAGAGTTTGATATGCACGTCATAATTGAGACAAACCGTACAGCCGCTAGGATTTTCCCTGCAGTTTTAGATGTTGAGAACCCGGAGTTCAAGAGGAAGTTGGACAGGATGGTGCAGATCAACGAGAAACTTCTTGCAATTGGTGAAACTGAGGAATCATCTTTGGTGAAGAACTTAAAAAGGATCCCACATGTTGCGGCGTTGGCGAATGAGCTCTTGGCTGCGTATCTCATGCCTCCTATTGATTCTGGTTCTGTTGATCTTGCTGAGTTTGAGCCACAAGTTGTGTATTAATAAGTTTTTTGTGTATTGAGTACACAAAGGCCCTTCAATTCAATTCAGTGCAAAGTTTAATTGCTACAGTTAATCATTACTATGGGCCTGCTGTATAAAGAACTGAACAT

>Pbr_TRINITY_ChlP Pbr_c20508_g1_i1 len=1789

TGCTCTTCCGATCTATTAAATCAAAAAAAACAAGGATAACAAAATCTTAAAACTTTCATTTATCCTTGCAGAAACTTGAAGAATCCATCACCACCAAAAATGGCCTCCATTGCCGTCAATGCTCGTTTCTCAGCCACAACATGCGACCGTAAAACAACCGAAATCAGCCGTGAAACACTAAAATCCTTCATGGGTCTTAGAAAAACCAACCAAAATGCTTCTATCTCACTTACACAATCTCTAACAACAAGTACTCAACATTTCAACCCTGCTGCTAGAAGAAATTTCACTTCCATCAAAGCAGGTAAATCAAGTCCTAAAGTAACAGGAAGAAATCTCCGAGTTGCGGTCATTGGTGGCGGTCCTGCTGGCGGTGCAGCTGCTGAAACCTTAGCTAAAGGCGGTATCGAAACGTTCTTGATCGAAAGGAAAATGGATAACTGTAAGCCATGTGGTGGTGCAATTCCACTCTGCATGGTTGGTGAGTTTGATCTTCCTTTGGATATCATCGACAGACGCGTTACGAAAATGAAGATGATTTCGCCATCAAATGTTGCAGTGGACATTGGCCAGACTCTTAAACCAGATGAGTATATCGGTATGGTACGTCGAGAAGTGCTCGACCAATATCTCCGTGAAAGAGCAGAGAAAGCTGGAACAAAGGTAATCAACGGTTTATTCTTGAAAATGGATTTACCAAAAGAACCAAATGCGCCATATGTTCTTCACTACAGTGCTTATGATGGCAAAACTGGTGGAGTTGGCACAAAAACCACGCTAGAGGTAGATGCAGTCATCGGTGCAGATGGTGCGAATTCTCGCGTAGCCAAAGCAATCGGAGCTGGTGATTATGATTATGCTATTGCATTCCAAGAGAGAATCAGAATCCCAGATGAGAAGATGGTTTACTATGAAAACCTTGCAGAGATGTACGTTGGTGATGATGTGTCGCCTGATTTTTACGGATGGGTGTTTCCAAAATGTGATCACGTTGCAGTGGGTACCGGTACAGTTACTCACAAAGGCGATATCAAGAAATTTCAAGTGGCTACGCGGCTGCGCGCCAAGGATAAGATTCTAGGCGGGAAGATTATCAGAGTAGAGGCACATCCTATTCCAGAACATCCAAGACCCAAGAGGTTGGCAGACAGAGTAGCCCTTGTAGGAGATGCAGCTGGGTACGTTACAAAATGTTCTGGAGAAGGTATATATTTTGCTGCTAAGAGTGGCAGGATGTGCGCCGAAGCGATTGTTGAAGGATCCGAGAGAGGGACAAGAATGGTTAACGAAACGGATTTGAGAAATTATCTGGAGAAATGGGATAAAACTTACTGGCCAACTTACAAGGTTCTGGATATATTGCAGAAGGTGTTTTACCGATCGAATCCAGCTCGAGAAGCTTTTGTTGAGATGTGCGCGGATGATTATGTTCAGAGGATGACATTTGATAGTTATCTGTATAAGAGAGTTGTGCCCGGAAATCCAATTGAGGATATTAAATTGGCTTTCAATACTATTGGGAGTTTGGTCAGAGCTAATGCACTTAGAGGCGAGATGAACAAGATTACTAGTCTATGATGATGAGTTTGTGAAAATAGGTTTGTTTTTCTGGTGAAATAAGATAGATAAAACGGGGGTGATGCGAATCTTGGCGATTTTTGTTGATCTCTTCTTGTTATAATTGTGTATTTTAACTTCTTGATTAATCACATTAATAAAATTGTATCAATTTCTCTACTCTACCGCAATAAAAATGTTTAATCACCTGCTGAGAGATCGGAAGAGCGTC

>Pbr_TRINITY_GbssI Pbr_c52348_g1_i1 len=2589

ATCCTTGTGAACTCATCAGCATAAACTGATAAGAATATTACCATGGCGACCATAAGTGCGTCCCAGTTTGTATCACGAATTAATTGCGGAGGAACTACACCAAAAGACATACTTAAACAAATCAATATGAAGAACCAGAGCATGACTCATTCAGGGTTAAGATCTTTGAGCAAAATAGATTTCTTACAAAGAAAAAACACTCAGAAAGCAGCTTCTATGGAACTGAGGAGGAACGCATTCCAGACAGACAAAGGTACTGCGGCAGGAGAGGTTAAATGTTCTGGTGGAATGAACTTGGTATTTGTTGCTGCTGAAGTAGGCCCCTGGAGTAAAACCGGTGGTCTTGGTGATGTTCTTGGAGGATTACCACCAGCCATGGCGGCAAATGGACATCGTGTTATGACAGTCTCTCCTCGTTATGATCAGTACAAAGATGCTTGGGATACCAATGTGTCAGTTGAGATGAAAATTGGGGACAAAATTGAAACCGTTCGTTTTTTTCACTGTTATAAGCGTGGGGTTGACCGTGTATTTGTGGATCATCCACTGTTTCTTGAGAAGGTATGGGGTAAAACTGGATCTAAAATCTATGGTCCGAATACTGGTTCAGATTACCAAGATAACCAGCTTAGATTCTGTTTGCTTTCTCTGGCAGCTCTAGAAGCACCAAGAGTTCTGAACCTCAATAGCAACAAATATTTCTCTGGAAAATACGGAGAGGAGGTCGTGTTCATCGCTAATGATTGGCATACGGCCCTTCTACCATGCTACCTTAAAAGTATGTACCAATCTAAAGGGATGTACGAAACTGCCAAGGTTGCTTATTGCATCCACAACATTGCTTACCAAGGAAGATTTCCTTTTTCGGATTTTGCACTTCTCAATCTTCCGGATTCATTTAGGAGTTCTTTTGACTTCATTGATGGGTATGAGAAGCCTGTTAAAGGAAGGAAAATTAACTGGATGAAGGCTGGAGTATTAGAGGCAGACAAGGTCGTAACTGTGAGTCCATACTATGCTGAAGAACTTGTTTCAGGCCCGGATAAAGGTGTGGAACTGGATAGTATTATACGGAAAGCTGGCTGTCTGGGTATTACAAATGGAATGGATGTTCAGGAGTGGAACCCGTTAACTGATAAATATATCGACGTCAAATATGATGCCTCAACTGTGACCGAGGCGAAGGCTCTTCTGAAAGAAGCACTGCAGGCAGAAGTAGGGTTACCTGTTGACCGTAATATTCCTGTCATAGGGTTCATTGGTAGACTTGAGGAGCAAAAGGGTTCAGATATTTTGGCAGCTGCAATCCCTGAATTTATTGACGAAGAAGTGCAAATAATTGTCCTCGGTACTGGCAAAAAGCCCATGGAAAAACAGTTGGAGCAGTTAGAGACTTTGTATCCTGAAAAAGCTAGAGGAGTGGCGAAGTTCAACGTTCCCTTGTCTCATATGATCATAGCTGGGGTTGATTTTATGATGATCCCAAGCAGATTTGAACCATGTGGACTCATCCAGCTGCATGCTATGCGTTACGGAACGGTGCCTATTGTTGCCTCGACGGGTGGATTAGTCGACACTGTCAAAGAAGGTTTCACGGGATTTCAAATGGGAGCTTTCAATGTCGAATGCGAGGCAATTGACCCATTAGACGTTACAGCAATCGCCACAACCACCAAAAGGGCCCTTGAAACCTATGGAACTCCAGCTTTAACAGAAATGATCCAGAACTGCATGGCTCAAGACTTCTCTTGGAAGGGACCTGCCAAGAATTGGGAGGACATGTTGCTGGGCTTGGAAGTTGCAGGTAGTGAACCAGGAATCGAAGGCGATGAAATCGCTCCTCTTGCTAAAGAAAATGTTGCCACTCCTTGAGTGACAGAACCGCTCCAGTGCAACTTTGACGAGCTATTTCACAATCAAACCCAGACGATTCCATTGCTGCACCAACAAGGCTCTGATTACTTCCATCATATCTAAGATTTGATTAAATTTATCCACTGCGAGTTACATAGTTAGTGGAAAAATTCATGATGTGTATATGAAGTGGCTTTACTGGAAAAGAGATACTGCTAGTCATTTTCTTTTTCTTCTTTTGATTTAAGCAATCTACGCCGCTCAGTGTCCAATGCGGATGCCAAAAGCGTGGCTATCTAGCTCTCTAGATCTTTTTATAAATGAAAACGACTTTTTCACCTTCTGCTTCTTCGGCTGCTTCAGTTAGTGTGAGCCTTGGATTCGGAAACGCCCTACATAAGAAACTACAAATTTGTTGTGAACTGCTTCAAGGCGGATGATGACCCACCGAAAATGCCAATGAAATAAGAGATAATCTCACCGTAGAGTATCTTGGCATTGGCGAAGGAGGCACTATCGATATACTGCATGAAAGGATGTTAATTAGGTAATCAGTATATCCACTGCTGCACTTGAAATAATGTGAATTGGTATTTATATTCTGTACTGCTTCATTTGAAGGCATGTTGGTTGGTTCCTAGATTTTTTGTTGCTTTGCTTAAACGGATGTCAATTAGTACTTGGAATTTGTATGCATGATTAATGAGACTGCGGACCAGAAGAAGGTTAAATGAA

>Pbr_TRINITY_Hmgs Pbr_c37327_g1_i1 len=1913

TTTTAATTTCTCTCACTCACAAAAGTCCGCCTTTGAATTTCACTAACTCACGTTGGCTGCATTTTTGTTTCTGGTTCCGGTCGGACCGGTCATGAATTTCCTCCGTCTATAAATACCGATTTATTCGTTGCCTAAATCTCTCACCACTTCTTCTTCATTCTCCCTTCCTTCCTTCTCTTCCCCTCTCTCTCTCCAAAAACAGCTCGATCGATCTAGGGTTCTCTGAAAGAATTTAAACTAATTATCTAAAGATGGGTTCGAATCAGAAGAATGTTGGAATTCTTGCTATGGAAATGTATTTCCCTCCTACTTGTGTCCAACAGGAAGCATTGGAGGCTCATGATGGTGCCAGTAAAGGAAAATACACCATTGGGTTGGGACAAGATTGCATGGCGTTTTGTTCCGAGTTGGAAGATGTCATATCAATGAGTTTGACAGCCGTTAGCTCTTTACTCGAAAAGTATAAAATTGACCCTAAACAAATTGGACGCCTGGAAGTCGGCAGTGAAACCGTGATTGACAAGAGCAAGTCCATAAAGACCTTCCTGATGACTATCTTTGAGGAATGTGATAACACTGACATTGAGGGTGTTGACTCCACAAATGCATGCTACGGAGGAACTGCAGCTTTGTTCAACGTTGTGAATTGGGTGGAGAGTAATTCCTGGGATGGACGATATGGTCTTGTTGTGTGCACTGATAGTGCAGTCTATGCAGAAGGACCTGCCCGTCCCACTGGAGGAGCAGCTGCTATTGCTATGCTGATTGGTCCAGACGCACCAATTGCCTTCGAAAGCAAATATAGAGGATCCCATATGTCCCATGTCTATGATTTTTACAAGCCCAACCTAGCCAGTGAATACCCGGTCGTTGACGGAAAACTTTCACAAACTTGTTACCTCATGGCACTCGATTCCTGCTACAAACGCCTCTGCGCTAAGTACGAGAAATGCCAGGGTAAGCAATTCTCGATGGATGATGCAGACTACTTTGTATTTCATTCTCCTTACAACAAGCTCGTGCAGAAAAGTTTTGCTAGATTGTACTTCATGGATTTCTTAAGAAACGCCAGCTCTGTTGACGAGGCTTCAAAAGAAAAGCTCGCACCATTTGCAGCTCTATCTGGTGAAGAGAGCTACGCCAGTCGGGATCTTGAGAAGACATCCCAAACTGTTGCAAAGAATCTTTATGCTGAGAAGGTTGGACCTACCACTTTGTTGCCCAAACAAGTCGGTAATATGTACACTGCATCACTCTATGCGGCATTTGCTTCTCTTCTTCACAACAAAAATAGCTCACTGGATGGCAAACGTGTAATAATGTTCTCTTATGGAAGTGGTTTGAGTTCTACAATGTTCTCATTCCGAATGAGGGAAGGTTCACATCCATTCAGCTTATCAAACATAGCAACCGTAATGGATGTTGCACACAAGTTGGAGTCAAGACATGAGGTTACTCCTGAAAAATTCGTCGAAACCATGCAACTCATGGAGCATAGGTACGGAGGGAAAGACTTCGTAACGAGCAAAGACACAAGCTTGTTAACTCCAGGCACATACTACCTTACCGAAGTTGACTCCATGTACCGCAGATTCTACTCCAAGAAGTCAAGTGCCGCAGTGCAAAATGGTCACTAAAAAAAATAGTGAAGGATGGAAATATCAAGCAGAAAAGCTGCCGAAGAATTGCTTTTGTATTCTCATTGTTGTTGTTAGTGTTTTAGAATAATTCTTATTTATGATTTGTACAAGTTAAATTATTCCAAACCTCTATGGTTCGCCCATTAGCCATTGTAACAGGATTCATCTCCTCTCAGTATATCCATTCAAGCGTTCCTCTCCATCTCCCCCCCCTGATATAAATAAAAGTTTGCTTTCTCGGAGGCCTGAGCAGGGATTAGCCTCTAAAAAAAA

>Pbr_TRINITY_Nia Pbr_c39067_g2_i1 len=3079

AAAAAAAAAAAAAAAAAAAACATAGAAAGAAGTAGAAAGCAGAGAGAAACTGAAACGAGAGGAGGAGCATTCAAAGAGAGATTTTTTTCTTTCTTCTTTCTTTTGATTAGTCTCTAATGGCGGCTTCTGTTGAAAATCGACAGTACAGCAGTCGACTTGAATCAGGTCTAGCAGTTCGTGGATTTAAAACCAATCCCACTCATCGAGCTGATTCGCCTATCCGTGGCTGTAATTTTCCCATGTCGACTACCGAGCGTCTTACTATACAAGCAAAACCGGTCATTGAAGATTTATCTAGCGACGACGATGAAGATCAAAACGAGCAGTGGAAAGATTTCGGCAAAATAAACAATGAAATAGAGCCGTCTGTTTTTGACTCGAAAGATGAAGGAACCGCTGATAATTGGATTGAACGCAACCCTTCCATGATTCGGCTAACAGGGAAACACCCATTCAATTCCGAAGCACCATTAACTCGTCTCATGCATCACGGGTTCATCACACCTGTTCCTCTTCATTACGTTCGGAATCACGGAGCTGTTCCAAAACTTAAATGGGAGGATTGGACAGTTGAGATTTGTGGATTAGTAAAGCGTCCGTTACGTATCACTATGGATCAGCTCATTAATGATTTCACAAGCCGTGAGTTTCCAGCTACACTTGTCTGTGCAGGTAACAGACGTAAAGAACAGAATATGATAAAACAAACAATTGGGTTCAACTGGGGAGGCGCAGGTGTATCGACTTCAGTATGGCGTGGTACACGTCTATGTGACGTATTAAAGCGTTGTGGAATCATGAGTCGAAAGAAAGGTGCTCTTAATGTTTGTTTCGAAGGAGCTGAAGATTTACCAGGTGGCGGCGGATCTAAATACGGAACTAGTATAAAGAAGGAATTCGCAATGGATCCTGCTCGTGATATAATTCTAGCTTACATGCAAAATGGTGAACGTTTGTCTCCTGACCACGGATTTCCAGTCCGTATGATAATTCCGGGTTTCATCGGAGGAAGAATGGTGAAGTGGTTAAAGAGAATTATCATCACAACTCCAGAATCCCAAAACCACTACCATTTCATGGATAATCGTGTTCTTCCTTCTCATGTCGACGCTGAACTGGCCAACTCAGAAGCATGGTGGTATAAGCCAGAGTACATCATCAATGAGTTGAACATAAACTCTGTGATTACAACTCCTTGTCATGAAGAGATTCTTCCCATAAATGCTGCAACAACACAAAGAGCTTACACATTAAGGGGATACGCATATTCCGGCGGCGGTAAGAAAGTAACTCGCGTCGAAGTAACAATGGACGGCGGAGAGTCATGGTTAGTGTGTACTTTGGATCATCCAGAGAAACCAAACAAGTATGGCAAATATTGGTGCTGGTGTTTTTGGTCACTAGAAGTTGAAGTTCTCGACTTGCTTGGTGCCAAGGAGATTGCGGTCAGGGCTTGGGATGAAACTCTAAACACTCAGCCTGAAAAGCTCATCTGGAATGTCATGGGTATGATGAACAACTGCTGGTTCAAAGTCAAAACAAATGTGTGCAAACCACACAAAGGAGAAATCGGTCTAGTTTTCGAGCATCCAACTCAGGCTGGAAACCAACCAGGTGGATGGATGGTTAGACAGAAACATCTTGAAAACTCTGAAGGTCCACATACCCTCAAAAAGAGTTTGTCCACTCCTTTCATGAACACCACGTCAAAAATGTATTCAATATCCGAGGTGAAAAAACACAACTCGCCTGACTCGGCTTGGATCATTGTTCACGGACATGTCTATGACTGCACCCGGTTCTTGAAAGACCATCCAGGTGGTTCCGATAGTATTCTCATCAATTCAGGTACCGACTGTACGGAAGAATTTGATGCAATTCACTCTGACAAAGCCAAGAATTTACTTGAAGATTTCAGAATCGGTGAACTCATTGCAACTGGTTACACTTCTGACTCGTCATCTTCGCCAAACAACTCTGTTCATGGTTCGGGCAGCTTGAGTCACCTTGCTCCTATCAAAGAAATCGGCATTGCGAAACCCGTAGCTCTAGTTGGCCGAGATAAAATACAATGCAAACTTATTGCCAAGACCTCGATTTCCCATGACGTTCGTGTCTTCAGGTTTGCATTACCTTCAGCTGATCAAGTCCTTGGGTTGCCAGTTGGGAAGCACATATTTGTATGTGCCCATGTTAACGATAAGCTATGCATGCGTGCTTACACTCCATCGAGCACAGTTGATGAAGTTGGTTACTTTGAGCTGGTCATTAAAGTCTACTTTGGCGGAGTTCACCCCAAGTTCCCCAATGGAGGTATTATGTCTCAGCACTTGGATTCTCTTCCTCTTGGTGGAACTATTGATGTTAAGGGTCCGTTGGGACACATTGAATACACTGGTCGTGGTAATTTCACCGTTCATGGCAAACACAAGTTTGCCAAACGACTCGCAATGCTTGCTGGTGGCACGGGAATTACACCAATATATCAAATTGTGCAAGCAATATTGAAAGACCCAGAGGATGAAACGGAGATGCACATCGTATTCGCAAACAGGACTGAAGATGACATCTTGTTGATGGATGAACTTGATTCTTGGGCAGAGAAGCACGAAAGGTTGAAAGTATGGTACGTTGTTGGTGAAACATTGAGGGAAGGATGGAAGTACAGTGTGGGGTTCATAACTGAAGATGTCTTGAGGGAACATATTCCTCCGGCATCGGAAGATACATTGGCATTGGCTTGTGGACCACCACCTATGATACAATTTGCAGTGACTCCAAATTTGGAGAAGATGAATTATGATGTGAAGAATTCATTGTTGGTCTTCTGATCACGAGTCAGTTGAAGTATTTGAAAGGTGAAACTACAAAGAGGAACACATATGATCGAATTTAAGAAAATGATAAGGGGAGGACAAATTATAAGGAAGTTCTCATTAGTTAATTGTATATAATTAGTGTTGAAGGATGTAATGGTTTAGTATGTTCTTAAAAGTAAGGGGATTGTTAAATGTTAGTCTGTTTAAATTTTATGTAGAATTGAAAGCAAAATGTGGCTGATTGGTTCCATAACACATGC

>Pbr_TRINITY_Eif3E Pbr_c12083_g1_i1 len=1661

TGTGCTCTTCCGATCTCAATTTTTAGGGTTTCTCTCTTTCTTCCTCCCACAACCACTGCGCCGCTCCTCCTTCTGCTCAGAGGTTGTTCTTCAGCCTCTACCACAACAACAGCAACCAATCATGGCGAATCATGATCTGACATCAAGAATAGCACTAAACTTAGATCGACATCTCGTTCTACCACTCCTTGAATTTCTTCAAGAGAAACAAATCCATCCCGATGTTCAAATCCTCAAAGCTAAAGACGCACTTCTCAGCCAGACCAATATGGTCGATTACGCTATGGATATCTACAAATCTCTTCATCAAACTGATGATGTTCCTCAAGAGATGATTAACAGGCGTGCTGAAGTAGTTAGGAGGTTGAAATTACTTGAAGATGCAGCTGAATCGATTATTACGTTCCTGCAAAGCCCGAATATGATTAACGAACCACGCCCTGATAAGCAGTATGTTCTTCAGTTGTTACAGGGATACCAGATTGGTGCTGAGCAAATTGAGGCATTGTATCAGTATGCCAAATTTCAATTTGAATGTGGGAATTACTCGGGTGCAGCTGATTATTTGCATCAGTACAGAGCAATTTGTACTAATCCTGATAGAATTTTGAATGCATGTTGGGGAAAGCTTGCATCTGAGATATTGATGCAGAACTGGGATGAGGCTCTTGATGAGCTTAATCGGTTGAGGGAGATTATTGAATCGAAGAATTTCAAGAACCCATTGACTTTACTCCAGAATAGAATCTGGTTGATGCATTGGAGTCTCTTCATCTTTTTCAACCACGAAAATGGAAGAAATGGCATCATTGATCTATTTCTCAATGACAGGTACATGAATGCACTCCAAACAAATGCACCCCATCTTTTACGTTACTTGGCAACTGCATTGATTGTCAACAAAAAGAGAAGACCTCAACTCAAAGAGTTGATCAAAGTCATCCAGCAAGAGCAATCTGCATACAAGGATCCCATCACTGAGTTCTTATACTGTTTGTATGTCAACTATGACTTCGATGGTGCACAACAGAAGCTCCGTGAATGTGAGAATGTGATTTTGAATGATCCATTCCTTGGTAAACGAGCTGAAGAAGGAAACTTTAGTACTGTTCCCTTGAGGGATGAATTTCTTGAAAATGCCCGTCTCTTCATCTTTGAGGCTTACTGCAGAATCCACCAGCGCATTGATATTGCAACTCTTGCCGAGAAGTTGAATATGAATTTTGACGAGGCAGAAAGATGGATAGTGAATCTTGTTCAGAATGCAAAGCTTGATGCTAAGATCGATTCTAAGCTAGGAACTGTTGTAATGGAGCCCACTTCTCTCAATGTGTATGAGCAGATCATTGAGAATACCAAATCACTTTCTCTGCGTACCTACAAATATGTAACTCTTGTTCTTGAATCAGCAGCACCACCGGTTACGGCTGCTCGTTAGAGAAGAGGTAGTAGTATAGCGAGAAAACTTGTGTTGGAGCTCTTGAAGCTTACTTGTGGGGAAGGAAGAAAATTTGGTCTCGGTATTTTGTTTTTGGTACTGGTTTTTCAGTTTCATCTAAAGCATTAAAAACCAAGAGTTGCAGGAGAACTTGTAAGTTTAAAAATATACTTGTTTTTAGGGTTCTTTGAAGGATAAAATACAGTTCCTGAGATCGGAAGAG

>Pbr_TRINITY_AroB Pbr_c21965_g1_i1 len=1662

TGCTCTTCCGATCTCTTTCACCAACCCCCGGTGGCTGCTGCTTCAACTACTCCTCCTTCATCTCGTCTGGTTTCAAAAAAATCAGACGAAGAAATAATTTTGTTCTGTTTTATTTCCCTCTCTAATCCCTGATTCCTCAATTTTTGATGGCTTCTTCTACTACTTCTTCACCTTCTCTTAATCTAACTCTGAAGAACTCTTGTGTACGAGAAAATTTTGATCTATTTCTTAGAGGGAGGAATTTATCACTGAAGTCTTCATTTATAAATTCAAGTCCATTACGATTGAATAGAATTGGATCCTTAAAAATTCAATCGAAGAGTAGAATAATATCTGCAAATTCAGCTCAAGTAATGGAACAATCTTCAATTAGGGCACCAACTATTGTTGATGTTGATTTGGGTAATAGAAGTTATCCGATCTATATTGGTTCTGGACTTCTTGAACAGCCTGATCTTCTTCAGAGGCACATACATGGGAAGAGAGTACTCGTGGTAACTAATACAACAGTTGCGCCATTATACCTAGATAAAGTTGTCAGCGCTTTAACTCATGGAAATCCGAGTATTTCGGTTGAGAGTGTGATTTTGCCAGATGGAGAGAAATATAAAAATATGGATACTCTGATGAAAGTTTTTGACAAGGCCATAGGATCTCGATTTGATCGCCGTTCCACATTTGTCGCTCTTGGAGGTGGTGTTATAGGTGACATGTGTGGTTATGCTGCAGCTTCTTTCCTTCGTGGTGTAAACTTCATTCAAATTCCAACTACTGTCATGGCACAGGTGGATTCCTCTGTTGGGGGAAAAACTGGAATCAACCACACACTGGGGAAAAACTTGATTGGTGCTTTCTACCAGCCCCAATGTGTACTTGTAGATACAGACACATTACATACACTGCCTGAGAGGGAATTAGCTTCTGGTTTTGCTGAAGTTATAAAATATGGACTCATTAGGGATGCTGAGTTCTTCGAGTGGCAAGAAAAAAATATGGACTCTCTGTTGGCGAGGGATCCCAGTGCACTAGCATATGCAATCAAGCGTTCATGCGAAAACAAGGCAGAGGTTGTATCCTTGGACGAGAAGGAAGGTGGAGTGAGAGCGACACTGAACTTGGGTCATACCTTTGGACATGCAATTGAAACAGGATTTGGCTATGGACAATGGCTCCATGGAGAAGCTGTAGCAGCTGGAACGGTCATGGCAGTCGATATGTCTTACCGTCTAGGTTGGATTGATGAATCAATTGTGAAACGGACGTACGATATTTTACAAAAAGCCAAACTGCCGATAACCCCACCAGATGTTATGACTGTAGAGATGTTCAAGTCTTTCATGGCTGTTGATAAGAAGGTGGCTGATGGACTACTAAGGCTCATCCTTCTCAAAGGTCCTCTCGGCAATTGTGTTTTTACTGGTGATTACGATAGAAAAGCCCTAGACGAGACCCTTCGTGCATTCTGTAAGTCTTAATTTAATGCCGACCTTGGTTGGGCTCATTTCTTCATCAACCATGTGCAGTTTTGTGTTATTTTTCTTTTGTTGACATCAAACCACATTGTTTTATGTATATGTATGTAATTCAGTAACTTCCTAGATTATAGAGATTACTCTGAACAAATCCAAATAAATACATATTACCTTCTATCTTAAAAAAAAA

>Ptr_TRINITY_Agt1 Ptr_c4523_g1_i1 len=1570

GTGTGCTCTTCCGATCTATTTCCTTCCTCCCATTGTTTATCCCTTAAAGTTGGGCTGGGAGAGAGAGAGAGAGTGAGAGTCTGAAGTAGTTCTCCAGAAACCATTCTTGCAACGAAAAGCTTCAAAGAGGTACAATATGGATTACGTTAATGCACCTGGGAGGAACCATCTCTTTGTTCCTGGACCTGTCAATATTCCAGAATCAGTCCTCCGTGCCATGAGCAGAAACAACGAAGATTACCGTTCCCCTGCAATTCCTGCTATGACTAAAACTCTTCTTGAAGATGTCAAGAAGATTTTCAAGACAACTTCTGGAACCCCATTTTTGATCCCCACAACAGGTACTGGTGCATGGGAAAGTGCACTTACAAATACTTTGTCTCCTGGTGATCGAACAATTTCTTTCCTCATTGGGCAATTTAGTCTTCTCTGGATTGATCAGCAGCAACGCCTTGGATTCAACGTTGATGTTGTAGAGAGTGAGTGGGGCCGTGGTGCTGATCTTGAAGTCCTAGCATCCAAGTTAGCTGAAGACACTGCTCATACCATCAAAGCTGTTTGCATGGTTCATAATGAAACTGCAACTGGAGTTACTAATAATTTGGCTGCTGTAAGGAAATTACTTGATCACTACAACCATCCTGCATTGTTTCTCGTGGATGGAGTGTCTTCGATATGTGCTCTTGATTTCCGTATGGATGAATGGGGAATCGATGTAGCTTTAACTGGTTCACAAAAAGCTCTTTCTCTACCTACTGGGATGGGTATTGTGTGTGCAAGCCCTAAAGCACTGGAAGCATCCAAGACTGCCAAATCAGTGAGAGTGTTCTTTGACTGGAAAGACTACTTGAAGTTCTACAAAATGGGAACTTATTGGCCATACACTCCTTCAATCCAGCTTTTATACGGCCTCAGAGCTGCTCTTGATCTCCTATTTGAGGAAGGACTTGACAATGTCATTGCTAGGCACAGCCGCTTGGCCAAGGCAACCAGACTTGCTGTTGAAGCCTGGGGTTTGAAAAACTGTACACAGAAAGAGGAGTGGTTGAGCGACACTGTGACTGCAGTTGTTGTTCCACCTTACATTGACAGTGCTGAAATAGTTAGAAGGGGATGGCAGAGATACAATCTAAGCTTAGGTCTGGGTTTGAATAAAGTGGCTGGAAAAGTTTTCAGGATAGGTCATCTTGGGAACCTGAATGAGCTCCAACTGCTGGGTTGTTTAGCTGGAGTGGAGATGATTCTGAAAGATGTCGGATACCCAGTAAAGCTAGGAAGTGGAGTGGCTGCGGCTTGTGCGTTTTTCCAGAACACCATTCCGATGATTGAATCTAGGATTTAATGAATCATTCATTATCCATGTGTAACTACTGTTCTCCCTCTTCTGATTTGCTTGGATTACTTGTATAAAAGAAGACATGATAATCTCCTGTATTTGCACTCTTTTGAGGCTTACTTAGTGGTACAAGAACTCATTACTCTTTCATCTTAATCCTGTATTTAAAATCTCTTTAAATTTAATCTCATCACTGTTGCATATATATCCAATTACGTCTGGTTTTCCCAGATC

>Ptr_TRINITY_At103 Ptr_c20176_g1_i1 len=1783

CGTGTGCTCTTCCGATCTCTCTGCAAAAATGGCTGCAGAGATGGCATTGATAAAACCCATCTCTAATTTCACTAGAACCAGCCCTAAATTTATCACCTCACCAAGAAAATCATTCTCCATTGTCAAAATGGCATCTCAGTCCACTGCATCAACAACAAAGTCAGGGAAACCAAAGAAAGCAATCAAAGAAACCTTGCTTGCTCCAAGATTTTACACAACTGATTTTGATGAAATGGAAACTCTTTTTAACACTGAGATGAACAAAAACTTGAATGAGGCTGAGTTTGAAGCTCTATTACAAGAGTTCAAAACTGATTACAATCAAACTCATTTCGTTAGAAACAAAGAGTTCAAAGAAGCTGCTGATAAATTGCAAGGTCCTCTTAGACAGATTTTTGTTGAGTTCTTGGAGAGATCTTGTACTGCTGAGTTCTCAGGGTTTCTTCTTTACAAAGAACTTGGAAGAAGACTTAAGAAAATCAATCCAGTTGTTGCTGAAATTTTCTCACTCATGTCCAGGGATGAAGCTAGGCATGCTGGATTCTTAAACAAGGGACTGTCAGATTTCAACTTGGCACTGGATCTTGGATTTCTTACAAAGGCTAGAAAATATACATTCTTCAAACCCAAGTTCATTTTCTATGCTACATATTTATCTGAGAAAATTGGGTACTGGAGGTACATAACCATTTATAGGCACTTAAAGACCAACCCTGAATACCAAGTTTATCCAATTTTCAAGTATTTTGAGAACTGGTGCCAAGATGAGAACAGACATGGAGATTTCTTCTCCGCACTGATGAAAGCTCAACCCCAGTTCCTTAACGATTGGAAAGCAAAGTTGTGGTCTCGGTTCTTCTGCCTATCGGTGTACGTGACCATGTACTTGAATGACTGCCAGAGAACAGCTTTTTACGAAGGGATAGGACTTAACACTAAAGAGTTTGATATGCACGTCATAATTGAGACAAATCGTACAGCCGCAAGGATTTTCCCTGCAGTTCTAGATGTTGAGAACCCGGAGTTCAAGAGGAAGTTGGATCGGATGGTGCAAATCAACGAGAAACTTCTTGCAATTGGTGAAACTGAGGAATCATCTTTGGTAAAGAACTTAAAAAGGATCCCACATGTTGCAGCGTTGGCGAATGAGCTCTTGGCTGCGTATCTCATGCCTCCTATTGATTCTGGATCTGTTGATCTTGCTGAGTTTGAGCCACAAGTTGTGTATTAATAAGTTTTTTGTTTATTGAGGAAACAAAGGCCCTTCAATTCAATTCAGTTCAGTGCAAAGTTTAATTGCTACAGTTAACCATTACTATGGACCTGCTGTATAAAGAACTAAACATTTGTGACGTATAATGTATAATGATTTCTGGACGCACCTCATGTGGAGGTTACTCGTTATGCATATTTTTTTCTGCACACATATAGGCCAGGACTTGTCACAACTGGCAAGGCAAGTACAAGAGCTAGACAATGGTCCTACTATCTAGGCGATGTATGAGCATTTCACCTGAAGAGACGTGACATTTTAAAACTGGCAGAACGGATTAAATACCCATGTTGGGTCCTAAAGCTGTTAATTGTCATCATGACAGTGGTTTCATTGTTCTCGGATGTTATTCTCGATCATATGAGATCCAAGTTTGAAGTATTTTGCTACCGAGTATTTAGACGATCCCATAAAACGGGCTCCAAGGCCAAAAAGTTTGAAACAATGGAGATTTAACAGTTGGCGATCAAGATAAAACAGTTTAAAGGACACAAAAGGCGCACATGTTCC

>Ptr_TRINITY_ChlP Ptr_c23889_g1_i1 len=1830

CATAATTCTAACGGCTAAAAATCACTGAAGATCACAAGCACCATATCCACAAGCATATCACAATAAACATTAAATCAAAAAAACAAGGATATCAAAATCTTAAAACTTTCATTCATCCTTGCAGAAACTTGAATCCCATCACCACCAAAAATGGCCTCCATTGCCGTCAATGCTCGTTTCTCAGCCACAACATGCGACCGTAAAACAACCGAAATCAGCCGTGAAACACTAAAATCCTTCATGGGTCTTAGAAAAACAAGCCAAAACGCTTCCATCTCACTTACACAATCTCTATCAACAAGTACTCCACATTTCAACCCTGCTAGAAGAAATTTTACTTCCATCAAAGCAGGTAAATCAAGTCCTAAAGTAACAGGAAGAAATCTCCGAGTTGCGGTCATTGGTGGTGGTCCTGCTGGTGGTGCAGCTGCTGAAACCTTAGCTAAAGGTGGCATTGAAACGTTCTTGATCGAAAGGAAAATGGATAACTGTAAGCCATGTGGTGGTGCAATTCCATTGTGCATGGTTGGTGAGTTTGATCTTCCCTTAGACATCATCGACAGACGTGTTACGAAGATGAAGATGATTTCGCCTTCAAATGTTGCAGTCGACATTGGCCAGACTCTTAAGCCAGATGAGTATATCGGTATGGTACGTCGAGAAGTGCTCGACCAATATCTTCGTGAAAGAGCAGAGAAAGCTGGAACAAAGGTAATTAACGGTTTATTCTTGAAAATGGATTTGCCAAAAGAACCGAATGCTCCTTATGTTCTTCACTACAGTGCTTATGATGGCAAAACTGGTGGTGTTGGTACAAAAACCACGCTAGAGGTGGACGCTGTCATCGGTGCAGATGGTGCGAATTCTCGTGTAGCGAAAGCAATTGGAGCTGGTGATTATGATTATGCTATTGCATTCCAAGAGAGAATCAGAATCCCAGATGAGAAGATGGTTTACTATGAAAATCTTGCCGAGATGTACGTCGGAGATGACGTTTCGCCTGATTTTTACGGATGGGTGTTTCCAAAATGTGATCATGTCGCAGTGGGTACGGGTACAGTTACGCACAAGGGAGATATCAAGAAATTCCAAGTGGCTACACGTCTACGTGCCAAGGATAAGATTCTAGGCGGGAAAATTATCAGAGTAGAGGCACATCCTATTCCAGAACATCCAAGACCCAAGAGGTTGGCAGATAGAGTAGCGCTTGTAGGAGATGCAGCTGGGTACGTTACAAAATGTTCTGGAGAAGGTATATACTTTGCTGCCAAGAGTGGCAGGATGTGTGCAGAAGCGATTGTTGAAGGATCCGAAAGAGGGACAAGAATGGTTAATGAAACAGATTTGAGAAATTATCTGGAGAAATGGGATAAAACTTACTGGCCAACTTACAAGGTTCTGGATATATTGCAGAAGGTGTTTTACCGATCGAATCCAGCTCGAGAAGCTTTTGTTGAGATGTGCGCGGATGATTATGTTCAGAGGATGACATTTGATAGTTATCTGTATAAGAGAGTTGTGCCTGGAAATCCAATTGAGGATATTAAATTGGCTTTCAATACTATTGGGAGTTTGGTCAGAGCTAATGCACTTAGAGGAGAAATGAACAAGATTACCAGTCTATGATGAAGAATGCGTTGAGGTTTTCTTTTGTGAAAATAGGTTTGTTTTTCTGGTGAATTAAGATAGATAAAATGGGAGGTGATCGGAATCTTGGTGATTTTTGTTGATCTCTTTTTGTTATAATTGTGTATTTAACTTCTTGATTAATCACATCAATAAAACTGTATCAATTTCTTTACTCTACGGCAATAAAAATGAAAAATTCTG

>Ptr_TRINITY_GbssI Ptr_c19836_g1_i2 len=2476

GTGCTCTTCCGATCTGTTTCATCCTCTACGGACTTCTTCTTCTCTCGTAACAACTCCTACTACTTACTTGTGATCTGTGCTGATGTGTCTCACTCTACTGTCAGTCAGTCTAGAGACAAAAAAGAGTGCGTATTTTGAGATTATCGAGAGTGACATATCTCTCTATTCTCCATCCTTGTGAACTCATCAGCATATAATAATCTCTCAATTCTCAACTGACAAGAATATTACCATGGCGACCTTAAGCGCGTCCCAGTTTGTATCACGAATCAACTGCGGCGGAACTGCACCAAAAGACATACTTAAACAAATCAATATGAAGAACCAAAGCATGACTCATTCAGGGTTAAGATTTTTGAGCAAAGTAGATTTCTTACAAATAAAAAACACTCAGAAAGCAGCTTCTAGGGAACTGAGGAGGAACGCATTCCAGACAGACAAAGGTACTGCGTCAGGAGAGGTTAAATGTTCTAGTGGGATGAACTTGGTCTTTGTTGCTGCTGAAGTAGGCCCCTGGAGTAAAACCGGTGGTCTTGGTGATGTTCTCGGAGGCTTACCACCAGCCATGGCGGCAAATGGACATCGTGTTATGACAGTCTCTCCTCGTTATGATCAGTACAAAGATGCTTGGGATACCAATGTGTCAGTTGAGATGAAAATTGGGGACAGAATTGAAACCGTTCGTTTCTTCCACTGTTATAAGCGTGGGGTCGACCGTGTATTTGTGGATCATCCACTGTTTCTTGAGAAGGTATGGGGCAAAACTGGATCTAAAATCTATGGTCCGAATGCTGGATCAGATTACCAAGATAACCAGCTTAGATTCTGTTTGCTTTCTCTGGCAGCTCTAGAAGCACCAAGAATTCTGAATCTCAACAGCAACGAATATTTCTCTGGAACATACGGAGAGGAGGTCGTGTTCATCGCTAATGATTGGCATACGGCCCTTCTACCATGCTACCTTAAAAGTATATACCAATCTAAAGGGATGTACGAAACTGCCAAGGTTGCTTATTGCATCCACAACATTGCTTACCAAGGAAGATTTTCTTTTTCCGATTTTGCGCTTCTCAATCTTCCGGATTCATTTAGGAGTTCTTTTGACTTCATTGACGGGTATGAGAAGCCTGTTAAAGGAAGGAAGATTAACTGGATGAAGGCTGGAATATTAGAGGCAGACAAGGTCGTAACTGTGAGTCCATACTATGCTGAAGAACTTGCTTCAGGCCCGGATAAAGGTGTGGAACTGGATAGTATTATACGCAAAGCTGGCTGTCTGGGTATTATAAATGGAATGGATGTTCAGGAGTGGAATCCATTAACTGATAAATATATCGACGTCAAATATGATGCCTCAACTGTGACTGAGGCGAAGGCTCTTCTGAAAGAAGCACTGCAGGCAGAAGTAGGGTTACCTGTTGACCGTAATATTCCTGTCATAGGTTTCATTGGTAGACTTGAGGAGCAAAAGGGTTCAGATATTTTGGCAGCTGCAATCCCTGAATTTATTGACGAAGAAGTGCAGATAATTGTCCTCGGTACTGGCAAAAAGCCCATGGAAAAACAGTTGGAGCAGTTAGAGACTTTGTATCCTGAAAAGGCTCGAGGAGTGGCGAAGTTCAACGTTCCCTTGGCTCATATGATCATAGCTGGGGCTGATTTTATGATGATCCCAAGCAGATTTGAACCATGTGGACTCATCCAGTTACATGCTATGCGTTATGGAACGGTGCCTATTGTTGCCTCGACGGGTGGATTAGTCGACACTGTCAAAGAAGGTTTCACGGGATTTCAAATGGGAGCTTTCAATGTCGAATGCGAGGCAATTGACCCATTAGACGTGACAGCAATCGCCACAACCACCAAAAAGGCCCTTGAAACCTATGGAACTCCAGCTTTAACAGAAATGATCCAGAACTGCATGGCTCAAGACCTCTCTTGGAAGGGACCTGCCAAGAATTGGGAGGACATGTTGCTGGGCTTGGAAGTTGCAGGTAGTGAACCAGGAATCGAAGGCGATGAAATCGCTCCTCTTGCTAAAGAAAATGTTGCCACTCCTTGAGTGACAGAACCGCTCCAGTGCAACTTAGAAGAGCTATTTCACAATCGAACCCAGGCGATTCCATTGCTACACCAACAAGGCTTTGATTACTTCCATCATATCTAAGATTTGATAAAATTTATCCACTGCGAGTTACATAGTCAGTGGAAGAGTTCATGATGTGTATATGAAGTGGCTTTACTGGAAAAGAGATACTGCTAGTCATTTTATTCTTTTTCTTTTTCTTCTTTTCTTCTTTTGATTTAAGCAATCTACGCCACTCAGTGTCCAATGCGGATGCCAAAAGCGTAGACTAGCTAGCTCTAGATCTTGTATCTAGTCAAGAAGATCTTTATATAAATAAAAACGACTTTTTCGCCTTCTGCTTCTTCAGTTGCTTCAGTTAGTGTGAGCCGTAGGTTCGGTAACGCCCTA

>Ptr_TRINITY_Hmgs Ptr_c14855_g1_i1 len=1831

GTGCTCTTCCGATCTTTTGAATTTCACGAACTCACGAGTCACGAAGGCTGCATTTTTTTTGGTTCCGGTCCGACCGGTCATGAATTTCCTCTGTCTATAAATACCGATTTATTCGTTGCCTAAATCTCTCACCACTTCTTCTTCATTCTTCCTTCATTCTCTTCCCCTCTCTCTCTCCAAAAACAGCTCGATCGATCTAGGGTTCTCTGAAGGAATTCAAAGTAATTTTCTAAAGATGGGTTCAAATCAAAAGGATGTTGGAATTCTTGCCATGGAAATGTATTTCCCTCCTACTTGTGTTCAACAGGAAGCATTGGAGGCTCATGATGGTGCCAGTAAAGGAAAATACACCATTGGGTTGGGACAAGATTGCATGGCGTTTTGTTCCGAGTTGGAAGATGTCATTTCAATGAGTTTGACAGCCGTTAGCTCTTTACTCGAAAAGTATAAAATTGACCCCAAACAAATTGGACGCCTGGAAGTCGGCAGTGAAACCGTGATTGACAAGAGCAAGTCCATAAAGACCTTCCTGATGACTATCTTTGAGGAATGTGATAACACTGACATTGAGGGTGTTGACTCCACAAATGCATGCTACGGAGGAACTGCAGCTTTGTTCAACGTTGTAAATTGGGTGGAGAGTAATTCCTGGGATGGACGATATGGTCTTGTTGTGTGCACTGATAGTGCGGTATATGCAGAAGGACCTGCCCGCCCCACTGGAGGAGCAGCTGCTATTGCTATGTTGATTGGTCCAGACGCACCAATTGCCTTTGAAAGCAAATATAGAGGATCCCATATGTCCCATGTCTATGATTTTTACAAGCCCAACCTAGCCAGTGAATACCCGGTCGTTGACGGAAAACTTTCACAAACTTGTTACCTCATGGCACTCGATTCCTGCTACAAACGCTTATGCGCTAAGTATGAGAAATGCCAGGGTAAGCAATTCTCGATGGATGATGCAGACTACTTTGTATTTCATTCTCCTTACAACAAGCTCGTGCAGAAAAGTTTTGCTCGATTGTACTTCATGGATTTCTTAAGAAACGCCAGCTCTGTTGACGAGGCTTCAAAAGAAAAGCTTGCACCATTTGCAGCTCTATCTGGTGAAGAAAGCTACGCCAGTCGGGATCTTGAGAAGACATCCCAAACTGTTGCAAAGAATCTTTATGCTGAGAAGGTTGGGCCTACCACTTTGTTGCCCAAACAAGTCGGTAATATGTACACTGCATCACTCTATGCAGCATTTGCTTCTCTTCTTCACAACAAAAATAGCTCACTGGATGGTAAACGTGTAATAATGTTCTCTTATGGAAGTGGTTTAAGTTCCACAATGTTCTCATTCCGAATGAGGGAAGGTTCACATCCATTCAGCTTATCAAACATTGCAACTGTAATGGATGTCGCGCACAAGTTGGAGTCCAGACATGAGGTTACTCCTGAAAAATTCGTCGAAACCATGCAACTCATGGAGCACAGGTACGGAGGGAAAGACTTCGTAACGAGCAAAGACACAAGCTTGTTAACTTCAGGCACATACTATCTTACCGAAGTTGACTCCATGTACCGCAGATTCTACTCCAAGAAGTCAAGTGCTGCAGTTCAAAATGGTCACTAAAAAAATAGTGAAAGGATGGAAATATCAAGCCGAAAAGCTGCCTAAGAATTGCTTTTGTATTCTCATTGTTGTTGTTAGTGTTTTAGAATAATTCTTATTTATGATTTGTACAAGTTAAATTATTTCAAAACCTGCAGTTCGCCTATTAGCCATTGTTACAGGATTCTTCTCCTCTCAGTATATCCGTTCAAGTATTCAAGTGTTCAAAAA

>Ptr_TRINITY_Nia Ptr_c26044_g1_i1 len=3096

CTCTTCCGATCTCAACTATCAAATGCTTGCAAAAGCAAATCCGAAAAAAGAGAAAACATAGATAGAAGTAGAAAGCAGAGAGAAAGTGAAACAAGAGGAAGAGCATTCAAAGAGAGATTTTTTCCTTTCTTCTTTCTTTTGATTAGTCTCTAATGGCGGCTTCTGTTGAAAATCGACAGTACAACAGTCGACTTGAATCAGGTCTAGCAGTTCGTGGATTTAAAACCAATCCCACTCACCGATCTGATTCGCCTATCCGTGGCTGTAATTTTCCCATGTCGACTACCGAGCGTCTTTCTATACAAGCAAAACCGGTCATTGAGGATTTATCTAGCGACGACGATGAAGATCAAAACGAGCAGTGGAAAGATTTCGGCAAGATAAACAATGAAATAGAGCCGTCTGTTTTTGACTCGAAAGATGAAGGAACCGCTGATAATTGGATTGAACGCAACCCTTCCATGATTCGGTTAACAGGGAAGCACCCATTCAATTCAGAAGCACCATTAACTCGTCTCATGCATCACGGGTTTATCACACCTGTTCCTCTTCATTACGTTCGTAATCACGGAGCTGTTCCAAAACTTAAATGGGAGGATTGGACAGTTGAGATTTGTGGATTAGTAAAGCGTCCGTTACGTATCACTATGGATCAGCTCATTAATGATTTCGCAAGCCGTGAGTTTCCAGCTACACTTGTCTGTGCAGGTAACAGACGCAAAGAACAGAATATGATAAAGCAAACAATTGGGTTTAACTGGGGAGGTGCAGGAGTATCTACTTCAGTATGGCGTGGTGCACGTTTATGTGATGTACTAAAGGGTTGTGGTATCATGAGTCGTAAGAAAGGTGCTCTTAATGTTTGTTTCGAAGGAGCTGAAGATTTACCCGGTGGCGGTGGATCTAAATACGGAACTAGTATAAAGAAGGAATTCGCAATGGATCCTGCTCGTGATATAATTCTAGCTTACATGCAAAATGGTGAACGTTTGTCTCCTGACCACGGATTTCCAGTCCGTATGATAATTCCGGGTTTCATCGGAGGAAGAATGGTGAAGTGGTTAAAGAGAATTATTGTCACAACTCCAGAATCCCAGAACCACTATCATTTCATGGATAATCGGGTTCTTCCTTCTCATGTCGACGCTGAACTGGCCAACTCTGAAGCATGGTGGTATAAGCCAGAGTACATCATCAATGAGTTGAACATAAACTCAGTGATTACAACTCCTTGTCATGAAGAGATTCTTCCTATAAATGCTGCAACAACACAAAGAGCTTACACATTAAGGGGATACGCATATTCCGGTGGCGGTAAAAAGGTAACTCGTGTCGAAGTGACAATGGACGGCGGAGAGTCATGGTTAGTGTGTACTTTGGATCATCCAGAGAAACCAAACAAGTATGGGAAATATTGGTGCTGGTGTTTTTGGTCACTAGAAGTTGAAGTTCTCGACTTGCTTGGTGCCAAAGAGATTGCAGTCAGGGCTTGGGATGAAACTCTAAACACTCAACCTGAAAAGCTCATCTGGAACGTCATGGGTATGATGAACAACTGTTGGTTCAAAGTGAAAACAAATGTGTGCAAACCACACAAAGGAGAAATCGGTCTAGTTTTCGAGCATCCGACTCAGGCTGGAAACCAACCAGGTGGATGGATGGTTAGACAAAAACATCTTGAAAACTCTGAAGGTCCACATACCCTCAAAAAGAGTTTGTCCACTCCTTTCATGAACACCACATCAAAAATGTATTCAATTTCCGAGGTGAAGAAACACAACTCGCCTGACTCGGCTTGGATCATTGTTCACGGACATGTTTATGACTGTACCCGGTTCTTGAAAGACCATCCAGGTGGTTCCGATAGTATTCTCATCAATGCAGGTACCGATTGTACCGAAGAATTTGATGCAATTCACTCTGACAAGGCCAAGAACATGCTTGAAGATTTCAGAATCGGTGAACTCATTGCAACTGGTTACACTTCTGACTCGTCATCTTCACCAAACAACTCAGTTCATGGTTCGGGCAGCTTGAGTCACCTTGCTCCTATCAAAGAAATAGGCATTGCAAAACCCGTAGCTCTAGTTGGCCGGGATAAAATACAATGCAAACTTATTGCCAAGACCTCAATTTCTCATGACGTTCGTGTCTTCAGGTTTGCATTACCTTCAGCTGATCAAGTCCTTGGGTTGCCAGTTGGGAAGCACATATTTGTATGTGCTCACGTTAACGATAAGCTATGCATGCGTGCTTACACTCCATCAAGCACAGTTGATGAAGTTGGTTACTTTGAGCTGGTCATTAAAGTCTACTTCGGCGGAGTTCACCCCAAGTTCCCCAATGGAGGTATTATGTCTCAGCACTTGGATTCTCTTCCTCTTGGTGGAACTATTGATGTGAAGGGTCCGTTGGGACACATTGAATACACTGGTCGTGGTAATTTCACCGTTCATGGCAAACACAAGTTTGCTAAACGACTCGCAATGCTTGCTGGTGGCACAGGAATTACACCAATATATCAAATTGTACAAGCAATATTGAAAGATCCAGAGGATGAAACGGAGATGCACATCGTATTCGCAAACAGGACTGAAGATGACATCTTGTTGATGGAGGAACTTGATTCTTGGGCAGAGAAGCACGAAAGGTTGAAAGTATGGTACGTTGTTGGTGAAACATTGAGGGAAGGATGGAAGTACAGTGTAGGGTTCATAACTGAAGAAGTCCTGCGGGAACATATTCCTCCGGCATCGGAAGATACATTGGCGTTGGCTTGTGGACCACCACCTATGATCCAATTTGCAGTGACTCCAAATTTGGAGAAGATGAACTATGATGTGAAGAATTCATTGTTGGTGTTCTGATGAATCAGCTGAAACTACAAAGAGGAACAGATATGATCAAATTTAAGAAATTGATAAGGGGAGGACAAATTATAAGGAAGTTCTCATTAGTTATTGTATATAATTAGTGTTGAAGGATGTAATGGGTTTAGTATGTTCTTAAAAGTAAGGGGATTGTTAAAATGTTAGTCTGTTTAAATTTTATGTAGAATTGAAAGCAAAATGTGGGAATTGAAACTTTCTTTCCTCG

>Ptr_TRINITY_Eif3E Ptr_c28012_g1_i1 len=1880

GTGTGCTCTTCCGATCTCTTTCCTTTCTCAGTTTCCACCAGTTTTATTTAGGGTTTCTCTCTTTCTTCCTCCCACCACCAGTGCGCCGCTTTTCCTTCTCCGCAGAGGTTGTTCTTCAGCCTCTCCAACAACAACAGCAACCAATCATGGCGAATCATGATCTGACATCAAGAATAGCACTAAACTTAGATCGACATCTCGTTCTCCCACTCCTTGAATTTCTTCAGGAGAAGCAAATCCATCCCGATGTTCAAATCCTCAAAGCTAAAGACGCTCTTCTCAGCCAGACCAATATGGTTGATTACGCTATGGATATCTACAAGTCTCTTCATCAAACTGATGATGTTCCTCAAGAGATGATTAATAGACGTGCTGAAGTGGTTAGGAGATTGAAATTGCTTGAAGATGCAGCTGAATCGATTATTACTTTCCTTCAGAGCCCGAATATGATTAATGAACCACGTCCTGATAAGCAATATGTTCTTCAGTTGCTACAGGGATACCAGATTGGTGCTGAGCAAATTGAGGCATTGTATCAGTATGCGAAATTTCAATTTGAATGTGGAAATTACTCGGGTGCAGCTGATTATTTGCATCAGTACAGAGCAATTTGTACCAATCCTGATAGAATTTTGAATGCATGTTGGGGAAAGCTTGCATCTGAGATATTGATGCAGAACTGGGATGAGGCTCTTGATGAGCTTAATCGGTTGAGGGAGATTATTGAATCGAAGAATTTCAAGAACCCATTGACTTTACTCCAGAATAGAATTTGGTTGATGCATTGGAGCCTCTTCATCTTTTTCAACCACGAAAACGGAAGAAATGGCATCATTGATCTATTTCTCAATGACAGGTACATGAATGCACTCCAAACAAATGCACCTCATCTTTTACGTTACTTGGCAACTGCATTGATTGTCAACAAAAAGAGAAGACCTCAACTCAAAGAGCTGATCAAAGTCATCCAGCAAGAGCAATCTGCATACAAGGATCCAATCACTGAGTTTTTATACTGCTTGTATGTCAACTATGACTTCGATGGTGCACAACAGAAGCTCCGTGAATGTGAGGATGTGATTTTGAATGATCCATTCCTTGGTAAACGAGCTGAAGAAGGAAACTTTAGCACCGTCCCTTTGAGGGATGAATTTCTTGAAAATGCCCGTCTCTTCATCTTTGAGGCTTACTGCAGAATCCACCAGCGCATTGATATTGGAACTCTTGCCGAGAAGTTGAATATGAATTTTGATGAGGCAGAGAGATGGATAGTGAATCTTGTTCAGAATGCAAAGCTTGATGCTAAGATTGATTCTAAGCTAGGAACTGTTGTAATGGAGCCCACTTCTCTCAATGTGTATGAGCAGATCATTGAGAATACCAAATCACTTTCTCTGCGCACCTACAAATACGTAAATCTTGTTCTAGAATCAGCAGCACCACCGGTTACCGCTGCTCGTTAGACAACAGGTGTAGTACAGCGAGAAAATTTGTGTTGGATCTCTTGAAGTTTACTTTTGGGGAAGGAAGAAAATCTGGTCTTGGTATTTTGTTTTTGTTTACTTGTTTTTCAGTTTCATCTAAAGCATTAAAAACCAAGAGTTGCAGGATAACTTGTAAGTTTAATATACTTCTTTTTAGGGTTCTTTGAACGATAAAATACAGTCCGATTTATCTAACATTAAACTTTGATGAAATTACCTTTAGTAATTTTTTTGTTTTCTATTATTTTCCCATTCTCAAGAGAGATACAAATTCTTTATGGATTGCAACTCTTTTACAACCTGTACCATTTCCATCCGTTCCTTTGGAGCTTCCACAGAACATATAACACCAAGTTTGAAAATCTCTGTTAAAGCCTCGCATAATTTGGCCCTTTT

>Ptr_TRINITY_AroB Ptr_c26222_g1_i1 len=1590

GCTGCTTCAACTACTTCTTCTTCTTCTCGTCTGGTTTGAAAAATAATCAGAAGAAGTTTTGTTTTGTTTCCCTCTCTAATTCTTCCAATTTTTGGATGGCTTCTTCTACTACTTCTTCACCTTCTCTTAATTTGGCTATTAAGAATTCTTGTATACGAGAAAATGCTGATCTATTTCTTAGAGGGAGGAATTTATCACTGAAGTCTTCATTTATACATTCAAGTTCATTACGATTGAATAGAATTGGATCTTTAAAAATTCAATCGAAGAGTAGAATAATATCTGCAAATTCAGCTCAAGTAATGGAACAATCTTCAATTAGGGCACCAACTATTGTTGATGTTGATTTGGGTAATAGGAGTTATCCGATCTATATTGGTTCTGGACTTCTTGAACAACCTGATCTTCTTCAGAGGCACGTACATGGGAAGAGAGTACTCGTGGTAACTAATACGAAAGTTGCGCCATTATACCTAGATAAAGTTGTCAGCGCATTAACTCATGGCAATCCGAGTATTTCGGTTGAGAGTGTGATTTTGCCAGATGGAGAGAAATATAAAAATATGGATACTCTGATGAAAGTTTTTGACAAGGCCATAGGATCTCGATTTGATCGGCGTTCCACATTTGTCGCTCTTGGAGGTGGTGTTATTGGTGACATGTGTGGTTACGCAGCAGCTTCTTTCCTTCGTGGTGTAAACTTCATTCAAATTCCAACTACTGTCATGGCACAGGTGGATTCCTCTGTTGGGGGGAAAACTGGAATCAACCACACACTGGGGAAAAACTTGATTGGTGCTTTCTACCAGCCCCAATGTGTACTTGTAGATACAGACACATTACATACACTGCCTGAGAGGGAATTAGCTTCTGGTTTTGCTGAAGTTATAAAATATGGACTCATTAGGGATGCTGAGTTCTTCGAGTGGCAAGAAAAAAATATGGACTCTCTGTTGGCGAGGGATCCCAGTGCACTAGCATATGCAATCAAGCGTTCATGCGAAAACAAGGCAGAGGTTGTATCCTTGGACGAGAAGGAAGGTGGAGTGAGAGCGACACTGAACTTGGGGCATACCTTTGGACATGCAATTGAAACAGGATTTGGCTATGGTCAATGGCTCCATGGAGAAGCTGTAGCAGCTGGAACGGTCATGGCAGTAGATATGTCTTACCGTCTAGGTTGGATTGATGAATCAATCGTGAAGCGGACATATGATATTTTACAGAAGGCCAAGCTGCCGATTACCCCACCAGACGTGATGACTGTAGAGATGTTCAAGTCTTTCATGGCTGTTGATAAGAAAGTGGCTGATGGACTACTAAGGCTCATCCTTCTCAAAGGTCCTCTCGGCAATTGTGTTTTTACAGGTGATTACGATAGAAAAGCCCTAGACGAGACACTTCGTGCGTTCTGTAAGTCTTAATTTAATGCCGACCTTGGTTGGGCCAATTTCTTCATCAACCATGTGCAGTTTTGTGTTATTATCTTTTGTTGACGTGAAACCACATTGTTTTATGTATATGTAATTCAATGACTTCTTAGATTTGCCATCAGCGCATGTATTACTATAAAGTCTCATATCATCATCT

>Par_TRINITY_Agt1 Par_c18822_g1_i1 len=1579

GTGTGCTCTTCCGATCTCACACATATATTTCCTTCCTCCCATTGTTTATCCCTTAAAGTTGGGCTGGGAGAGAGAGAGAGAGTGAGAGTCTGAAGTAGTTCTCCAGAAACCATTCTTGCAACGAAAAGCTTCAAAGAGGTACAATATGGATTACGTTAATGCACCTGGGAGGAACCATCTCTTTGTTCCTGGACCTGTCAATATTCCAGAATCAGTCCTCCGTGCCATGAGCAGAAACAACGAAGATTACCGTTCCCCTGCAATTCCTGCTATGACTAAAACTCTTCTTGAAGATGTCAAGAAGATTTTCAAGACAACTTCTGGAACCCCATTTTTGATCCCCACAACAGGTACTGGTGCATGGGAAAGTGCACTTACAAATACTTTGTCTCCTGGTGATCGAACAATTTCTTTCCTCATTGGGCAATTTAGTCTTCTCTGGATTGATCAGCAGCAACGCCTTGGATTCAACGTTGATGTTGTAGAGAGTGAGTGGGGCCGTGGTGCTGATCTTGAAGTCCTAGCATCCAAGTTAGCTGAAGACACTGCTCATACCATCAAAGCTGTTTGCATGGTTCATAATGAAACTGCAACTGGAGTTACTAATAATTTGGCTGCTGTAAGGAAATTACTTGATCACTACAACCATCCTGCATTGTTTCTCGTGGATGGAGTGTCTTCGATATGTGCTCTTGATTTCCGTATGGATGAATGGGGAATCGATGTAGCTTTAACTGGTTCACAAAAAGCTCTTTCTCTACCTACTGGGATGGGTATTGTGTGTGCAAGCCCTAAAGCACTGGAAGCATCCAAGACTGCCAAATCAGTGAGAGTGTTCTTTGACTGGAAAGACTACTTGAAGTTCTACAAAATGGGAACTTATTGGCCATACACTCCTTCAATCCAGCTTTTATACGGCCTCAGAGCTGCTCTTGATCTCCTATTTGAGGAAGGACTTGACAATGTCATTGCTAGGCACAGCCGCTTGGCCAAGGCAACCAGACTTGCTGTTGAAGCCTGGGGTTTGAAAAACTGTACACAGAAAGAGGAGTGGTTGAGCGACACTGTGACTGCAGTTGTTGTTCCACCTTACATTGACAGTGCTGAAATAGTTAGAAGGGGATGGCAGAGATACAATCTAAGCTTAGGTCTGGGTTTGAATAAAGTGGCTGGAAAAGTTTTCAGGATAGGTCATCTTGGGAACCTGAATGAGCTCCAACTGCTGGGTTGTTTAGCTGGAGTGGAGATGATTCTGAAAGATGTCGGATACCCAGTAAAGCTAGGAAGTGGAGTGGCTGCGGCTTGTGCGTTTTTCCAGAACACCATTCCGATGATTGAATCTAGGATTTAATGAATCATTCATTATCCATGTGTAACTACTGTTCTCCCTCTTCTGATTTGCTTGGATTACTTGTATAAAAGAAGACATGATAATCTCCTGTATTTGCACTCTTTTGAGGCTTACTTAGTGGTACAAGAACTCATTACTCTTTCATCTTAATCCTGTATTTAAAATCTCTTTAAATTTAATCTCATCACTGTTGCATATATATCCAATTACGTCTGGTTTTCCAGATCG

>Par_TRINITY_At103 Par_c28985_g3_i1 len=1931

GTGTGCTCTTCCGATCTCTAGAAGCTACACATACTCTGCAAAAATGGCTGCAGAGATGGCATTGATAAAACCCATCTCTAATTTCACTAGAACCAGCCCTAAATTTATCACCTCACCAAGAAAATCATTCTCCATTGTCAAAATGGCATCTCAGTCCACTGCATCAACAACAAAGTCAGGGAAACCAAAGAAAGCAATCAAAGAAACCTTGCTTGCTCCAAGATTTTACACAACTGATTTTGATGAAATGGAAACTCTTTTTAACACTGAGATGAACAAAAACTTGAATGAGGCTGAGTTTGAAGCTCTATTACAAGAGTTCAAAACTGATTACAATCAAACTCATTTCGTTAGAAACAAAGAGTTCAAAGAAGCTGCTGATAAATTGCAAGGTCCTCTTAGACAGATTTTTGTTGAGTTCTTGGAGAGATCTTGTACTGCTGAGTTCTCAGGGTTTCTTCTTTACAAAGAACTTGGAAGAAGACTTAAGAAAACCAATCCAGTTGTTGCTGAAATTTTCTCACTCATGTCCAGGGATGAAGCTAGGCATGCTGGATTCTTAAACAAGGGACTGTCAGATTTCAACTTGGCACTGGATCTTGGATTTCTTACAAAGGCTAGAAAATATACATTCTTCAAACCCAAGTTCATTTTCTATGCTACATATTTATCTGAGAAAATTGGGTACTGGAGGTACATAACCATTTATAGGCACTTAAAGACCAACCCTGAATACCAAGTTTATCCAATTTTCAAGTATTTTGAGAACTGGTGCCAAGATGAGAACAGACATGGAGATTTCTTCTCCGCACTGATGAAAGCTCAACCCCAGTTCCTTAACGATTGGAAAGCAAAGTTGTGGTCTCGGTTCTTCTGCCTATCGGTGTACGTGACCATGTACTTGAATGACTGCCAGAGAACAGCTTTTTACGAAGGGATAGGACTTAACACTAAAGAGTTTGATATGCACGTCATAATTGAGACAAATCGTACAGCCGCAAGGATTTTCCCTGCAGTTCTAGATGTTGAGAACCCGGAGTTCAAGAGGAAGTTGGATCGGATGGTGCAAATCAACGAGAAACTTCTTGCAATTGGTGAAACTGAGGAATCATCTTTGGTAAAGAACTTAAAAAGGATCCCACATGTTGCAGCGTTGGCGAATGAGCTCTTGGCTGCGTATCTCATGCCTCCTATTGATTCTGGATCTGTTGATCTTGCTGAGTTTGAGCCACAAGTTGTGTATTAATAAGTTTTTTGTTTATTGAGGAAACAAAGGCCCTTCAATTCAATTCAGTTCAGTGCAAAGTTTAATTGCTACAGTTAACCATTACTATGGACCTGCTGTATAAAGAACTAAACATTTGTGACGTATAATGTATAATGATTTCTGGACGCACCTCATGTGGAGGTTACTCGTTATGCATATTTTTTTCTGCACACATATAGGCCAGGACTTGTCACAACTGGCAAGGCAAGTACAAGAGCTAGACAATGGTCCTACTATCTAGGCGATGTATGAGCATTTCACCTGAAGAGACGTGACATTTTAAAACTGGCAGAACGGATTAAATACCCATGTTGGGTCCTAAAGCTGTTAATTGTCATCATGACAGTGGTTTCATTGTTCTCGGATGTTATTCTCGATCATATGAGATCCAAGTTTGAAGTATTTTGCTACCGAGTATTTAGACGATCCCATAAAACGGGCTCCAAGGCCAAAAAGTTTGAAACAATGGAGATTTAACAGTTGGCGATCAAGATAAAACAGTTTAAAGGACACAAAAGGCGCACATGTTCCCTGTTTCTACACATGAAGCAAGCTTCAACTCGTTGTTTCTTAGTTTGGGAAGCTTCAATTTGTTTCGAAAATGATAATTACCTTCATAGAGATAGGTGCCCGACACTTTTGAGTAGCCATAAAAATCTTCCTT

>Par_TRINITY_ChlP Par_c33794_g1_i1 len=1759

CCATCACCACCAAAAATGGCCTCCATTGCCGTCAATGCTCGTTTCTCAGCCACAACATGCGACCGTAAAACAACCGAAATCAGCCGTGAAACACTAAAATCCTTCTTGGGTCTTAGAAAAACCAACCAAAACGCTTCCATCTCACTTACACAATCTCTATCAACAAGTACTCCACATTTCAACCCTGCTAGAAGAAATTTTACTTCCATCAAAGCAGGTAAATCAAGTCCTAAAGTAACAGGAAGAAATCTCCGAGTTGCGGTCATTGGTGGTGGCCCTGCTGGTGGTGCAGCTGCTGAAACCTTAGCTAAAGGTGGTATCGAAACGTTCTTGATCGAAAGGAAAATGGATAACTGTAAACCATGTGGTGGTGCAATTCCACTCTGCATGGTTGGTGAGTTTGATCTTCCGTTAGACATCATCGACAGACGTGTTACGAAGATGAAGATGATTTCACCTTCAAATGTTGCAGTCGACATTGGCCAGACTCTTAAGCCAGATGAGTATATCGGTATGGTACGTCGAGAAGTGCTCGATCAATATCTTCGTGAAAGAGCAGAGAAAGCTGGAACAAAGGTAATTAACGGTTTATTCTTGAAAATGGATTTGCCGAAAGAACCAAATGCGCCTTATGTTCTTCACTACAGTGCTTATGACGGCAAAACTGGTGGTGTTGGCACAAAAACCACGCTAGAGGTAGATGCAGTCATCGGAGCAGACGGTGCGAATTCTCGTGTAGCTAAAGCAATTGGAGCTGGTGATTATGATTATGCTATTGCATTCCAAGAGAGAATCAGAATCCCAGATGAGAAGATGGTTTACTATGAAAATCTTGCAGAAATGTACGTCGGTGATGATGTGTCGCCTGATTTTTACGGATGGGTGTTTCCAAAATGTGATCACGTCGCAGTGGGTACCGGTACAGTTACTCACAAAGGAGATATCAAGAAATTTCAAGTGGCTACTCGGCTGCGTGCCAAGGATAAGATTCTAGGCGGAAAGATTATCAGAGTAGAGGCACATCCTATTCCAGAACATCCAAGACCCAAGAGGTTGGCAGACAGAGTAGCGCTTGTAGGAGATGCAGCTGGGTACGTTACAAAATGTTCTGGAGAAGGTATATACTTTGCTGCCAAGAGTGGCAGGATGTGCGCAGAAGCGATTGTTGAAGGATCCGAAAGAGGGACAAGAATGGTTAATGAAACAGATTTGAGAAATTATCTGGAGAAATGGGATAAAACTTACTGGCCAACTTACAAGGTTCTGGATATATTGCAGAAGGTGTTTTACCGATCGAATCCAGCTCGAGAAGCTTTTGTTGAGATGTGCGCGGATGATTACGTTCAGAGGATGACATTTGATAGTTATCTGTATAAGAGAGTTGTGCCTGGAAATCCAATTGAGGATATTAAATTGGCTTTCAATACTATTGGGAGTTTGGTCAGAGCTAATGCACTTAGAGGAGAAATGAACAAGATTACCAGTCTATGATGAAGAATGCGTTGAGGTTTTCTTTTGTGAAAATAGGGTTGTTTTTCTGGTGAATTAAGATAGATAAAATGGGAGGTGATCGGAATCTTGGTGATTTTTGTTGATCTCTTTTTGTTATAATTTGTGTATTTAACTTCTTGATTAATCACATCAATAAAACTGTATCAATAAAAATGAAAAAGCCATTTGCGCATCCAGGGAATCGAACCCTGGTCAGTACCGTGGGAGGGTACTATGATACCACTACACCAGATGCGCTGTTTGTGAC

>Par_TRINITY_GbssI Par_c91557_g1_i1 len=2413

TCCGATCTCTTCTCTCGTAACAACTCCTACTACTTACTTGTGATCTGTGCTGATGTGTCTCACTCTACTGTCAGTCAGTCTAGAGACAAAAAAGAGTGCGTATTTTGAGATTATCGAGAGTGACATATCTCTCTATTCTCCATCCTTGTGAACTCATCAGCATATAATAATCTCTCAATTCTCAACTGACAAGAATATTACCATGGCGACCTTAAGCGCGTCCCAGTTTGTATCACGAATCAACTGCGGCGGAACTGCACCAAAAGACATACTTAAACAAATCAATATGAAGAACCAAAGCATGACTCATTCAGGGTTAAGATTTTTGAGCAAAGTAGATTTCTTACAAATAAAAAACACTCAGAAAGCAGCTTCTAGGGAACTGAGGAGGAACGCATTCCAGACAGACAAAGGTACTGCGTCAGGAGAGGTTAAATGTTCTAGTGGGATGAACTTGGTCTTTGTTGCTGCTGAAGTAGGCCCCTGGAGTAAAACCGGTGGTCTTGGTGATGTTCTCGGAGGCTTACCACCAGCCATGGCGGCAAATGGACATCGTGTTATGACAGTCTCTCCTCGTTATGATCAGTACAAAGATGCTTGGGATACCAATGTGTCAGTTGAGATGAAAATTGGGGACAGAATTGAAACCGTTCGTTTCTTCCACTGTTATAAGCGTGGGGTCGACCGTGTATTTGTGGATCATCCACTGTTTCTTGAGAAGGTATGGGGCAAAACTGGATCTAAAATCTATGGTCCGAATGCTGGATCAGATTACCAAGATAACCAGCTTAGATTCTGTTTGCTTTCTCTGGCAGCTCTAGAAGCACCAAGAATTCTGAATCTCAACAGCAACGAATATTTCTCTGGAACATACGGAGAGGAGGTCGTGTTCATCGCTAATGATTGGCATACGGCCCTTCTACCATGCTACCTTAAAAGTATATACCAATCTAAAGGGATGTACGAAACTGCCAAGGTTGCTTATTGCATCCACAACATTGCTTACCAAGGAAGATTTTCTTTTTCCGATTTTGCGCTTCTCAATCTTCCGGATTCATTTAGGAGTTCTTTTGACTTCATTGACGGGTATGAGAAGCCTGTTAAAGGAAGGAAGATTAACTGGATGAAGGCTGGAATATTAGAGGCAGACAAGGTCGTAACTGTGAGTCCATACTATGCTGAAGAACTTGCTTCAGGCCCGGATAAAGGTGTGGAACTGGATAGTATTATACGCAAAGCTGGCTGTCTGGGTATTATAAATGGAATGGATGTTCAGGAGTGGAATCCATTAACTGATAAATATATCGACGTCAAATATGATGCCTCAACTGTGACTGAGGCGAAGGCTCTTCTGAAAGAAGCACTGCAGGCAGAAGTAGGGTTACCTGTTGACCGTAATATTCCTGTCATAGGTTTCATTGGTAGACTTGAGGAGCAAAAGGGTTCAGATATTTTGGCAGCTGCAATCCCTGAATTTATTGACGAAGAAGTGCAGATAATTGTCCTCGGTACTGGCAAAAAGCCCATGGAAAAACAGTTGGAGCAGTTAGAGACTTTGTATCCTGAAAAGGCTCGAGGAGTGGCGAAGTTCAACGTTCCCTTGGCTCATATGATCATAGCTGGGGCTGATTTTATGATGATCCCAAGCAGATTTGAACCATGTGGACTCATCCAGTTACATGCTATGCGTTATGGAACGGTGCCTATTGTTGCCTCGACGGGTGGATTAGTCGACACTGTCAAAGAAGGTTTCACGGGATTTCAAATGGGAGCTTTCAATGTCGAATGCGAGGCAATTGACCCATTAGACGTGACAGCAATCGCCACAACCACCAAAAAGGCCCTTGAAACCTATGGAACTCCAGCTTTAACAGAAATGATCCAGAACTGCATGGCTCAAGACCTCTCTTGGAAGGGACCTGCCAAGAATTGGGAGGACATGTTGCTGGGCTTGGAAGTTGCAGGTAGTGAACCAGGAATCGAAGGCGATGAAATCGCTCCTCTTGCTAAAGAAAATGTTGCCACTCCTTGAGTGACAGAACCGCTCCAGTGCAACTTAGAAGAGCTATTTCACAATCGAACCCAGGCGATTCCATTGCTACACCAACAAGGCTTTGATTACTTCCATCATATCTAAGATTTGATAAAATTTATCCACTGCGAGTTACATAGTCAGTGGAAGAGTTCATGATGTGTATATGAAGTGGCTTTACTGGAAAAGAGATACTGCTAGTCATTTTATTCTTTTTCTTTTTCTTCTTTTCTTCTTTTGATTTAAGCAATCTACGCCACTCAGTGTCCAATGCGGATGCCAAAAGCGTAGACTAGCTAGCTCTAGATCTTGTATCTAGTCAAGAAGATCTTTATATAAATAAAAACGACTTTTTCGCCTTCAAAAAAAAAGATCGGAAGA

>Par_TRINITY_Hmgs Par_c21336_g1_i1 len=1749

GTCATGAATTTCCTCTGTCTATAAATACCAATTTATTCGTTGCCTAAATCTCTCACCACTTCTTCTCCATTCTCCCTTCCTCCTCTTCCCCTCTCTCTCCAAAAACAGCTCGATCGATCTAGGGTTCTCTGAAGGAATTCAAAGTAATTATCTAAAGATGGGTTCAAATCAAAAGGATGTTGGAATTCTTGCCATGGAAATGTATTTCCCTCCTAGTTGTGTTCAACAGGAAGCATTGGAGGCTCATGATGGTGCCAGTAAAGGAAAATACACCATTGGGTTGGGACAAGATTGCATGGCGTTTTGTTCCGAGTTGGAAGATGTCATTTCAATGAGTTTGACAGCCGTTAGCTCTTTACTCGAAAAGTATAAAATTAACCCCAAACAAATTGGACGCCTGGAAGTCGGCAGTGAAACCGTGATTGACAAGAGCAAGTCCATAAAGACCTTCCTGATGACTATCTTTGAGGAATGTGATAACACTGACATTGAGGGTGTTGACTCCACAAATGCATGCTACGGAGGAACTGCAGCTTTGTTCAACGTTGTGAATTGGGTGGAGAGTAATTCCTGGGATGGACGATATGGTCTTGTTGTGTGCACTGATAGTGCGGTCTATGCAGAAGGACCTGCCCGCCCCACTGGAGGAGCAGCTGCTATTGCTATGTTGATTGGTCCAGACGCACCAATTGCCTTCGAAAGCAAATATAGAGGATCCCATATGTCCCATGTCTATGATTTTTACAAGCCCAACCTAGCCAGTGAATACCCGGTCGTTGACGGAAAACTTTCACAAACTTGTTACCTCATGGCACTCGATTCCTGCTACAAACGCTTATGCGCTAAGTATGAGAAATGCCAGGGTAAGCAATTCTCGATGGATGATGCAGACTACTTTGTATTTCATTCTCCTTACAACAAGCTCGTGCAGAAAAGTTTTGCTCGATTGTACTTCATGGATTTCTTAAGAAACGCCAGCTCTGTTGACGAGGCTTCAAAAGAAAAGCTTGCACCATTTGCAGCTCTATCTGGTGAAGACAGCTACGCTAGTCGGGATCTTGAGAAGACATCCCAAACTGTTGCAAAGAATCTTTATGCTGAGAAGGTTGGGCCTACCACTTTGTTGCCCAAACAAGTTGGTAATATGTACACTGCATCACTCTATGCAGCATTTGCTTCTCTTCTTCACAACAAAAATAGCTCACTGGATGGTAAACGTGTAATAATGTTCTCTTATGGAAGTGGTTTAAGTTCCACAATGTTCTCATTCCGAATGAGGGAAGGTTCACATCCATTCAGCTTATCAAACATTGCAACTGTAATGGATGTCGCGCACAAGTTGGAGTCCAGACATGAGGTTACTCCTGAAAAATTCGTCGAAACCATGCAACTCATGGAGCACAGGTACGGAGGGAAAGACTTCGTAACGAGCAAAGACACAAGCCTGTTAACTTCAGGCACATACTATCTTACCGAAGTTGACTCCATGTACCGCAGATTCTACTCCAAGAAGTCAAGTGCTGCAGTTCAAAATGGTCACTAAAAAAATAGTGAAAGGATGGAATTATCAAGCCGAAAAACTTCCGAAGAATTGCTTTTGTATTCTCATTGTTGTTGTTAGTGTTTTAGAATAATTCTTATTTATGATTTGTACAAGTTAAATTATTTCAAAACCTGCAGTTCGCCTATTAGCCATTGTAACAGGATTCTTCTCCTCTCAGTATATCCATTCAAGGATTCAAGTGTCCC

>Par_TRINITY_Nia Par_c14618_g1_i1 len=3146

GCTCTTCCGATCTACCATAACCATGCTCCTCATTTTTCTTGCAACTATCAAATGCTTGCAAAAGCAAATCCGAAAAAAGAGAAAACATAGATAGAAGTAGAAAGCAGAGAGAAAGTGAAACAAGAGGAAGAGCATTCAAAGAGAGATTTTTTCCTTTCTTCTTTCTTTTGATTAGTCTCTAATGGCGGCTTCTGTTGAAAATCGACAGTACAACAGTCGACTTGAATCAGGTCTAGCAGTTCGTGGATTTAAAACCAATCCCACTCACCGATCTGATTCGCCTATCCGTGGCTGTAATTTTCCCATGTCGACTACCGAGCGTCTTTCTATACAAGCAAAACCGGTCATTGAGGATTTATCTAGCGACGACGATGAAGATCAAAACGAGCAGTGGAAAGATTTCGGCAAGATAAACAATGAAATAGAGCCGTCTGTTTTTGACTCGAAAGATGAAGGAACCGCTGATAATTGGATTGAACGCAACCCTTCCATGATTCGGTTAACAGGGAAGCACCCATTCAATTCAGAAGCACCATTAACTCGTCTCATGCATCACGGGTTTATCACACCTGTTCCTCTTCATTACGTTCGTAATCACGGAGCTGTTCCAAAACTTAAATGGGAGGATTGGACAGTTGAGATTTGTGGATTAGTAAAGCGTCCGTTACGTATCACTATGGATCAGCTCATTAATGATTTCGCAAGCCGTGAGTTTCCAGCTACACTTGTCTGTGCAGGTAACAGACGCAAAGAACAGAATATGATAAAGCAAACAATTGGGTTTAACTGGGGAGGTGCAGGAGTATCTACTTCAGTATGGCGTGGTGCACGTTTATGTGATGTACTAAAGCGTTGTGGTATCATGAGTCGTAAGAAAGGTGCTCTTAATGTTTGTTTCGAAGGAGCTGAAGATTTACCCGGTGGCGGTGGATCTAAATACGGAACTAGTATAAAGAAGGAATTCGCAATGGATCCTGCTCGTGATATAATTCTAGCTTACATGCAAAATGGTGAACGTTTGTCTCCTGACCACGGATTTCCAGTCCGTATGATAATTCCGGGTTTCATCGGAGGAAGAATGGTGAAGTGGTTAAAGAGAATTATTGTCACAACTCCAGAATCCCAGAACCACTATCATTTCATGGATAATCGGGTTCTTCCTTCTCATGTCGACGCTGAACTGGCCAACTCTGAAGCATGGTGGTATAAGCCAGAGTACATCATCAATGAGTTGAACATAAACTCAGTGATTACAACTCCTTGTCATGAAGAGATTCTTCCTATAAATGCTGCAACAACACAAAGAGCTTACACATTAAGGGGATACGCATATTCCGGTGGCGGTAAAAAGGTAACTCGTGTCGAAGTGACAATGGACGGCGGAGAGTCATGGTTAGTGTGTACTTTGGATCATCCAGAGAAACCAAACAAGTATGGGAAATATTGGTGCTGGTGTTTTTGGTCACTAGAAGTTGAAGTTCTCGACTTGCTTGGTGCCAAAGAGATTGCAGTCAGGGCTTGGGATGAAACTCTAAACACTCAACCTGAAAAGCTCATCTGGAACGTCATGGGTATGATGAACAACTGTTGGTTCAAAGTGAAAACAAATGTGTGCAAACCACACAAAGGAGAAATCGGTCTAGTTTTCGAGCATCCGACTCAGGCTGGAAACCAACCAGGTGGATGGATGGTTAGACAAAAACATCTTGAAAACTCTGAAGGTCCACATACCCTCAAAAAGAGTTTGTCCACTCCTTTCATGAACACCACATCAAAAATGTATTCAATTTCCGAGGTGAAGAAACACAACTCGCCTGACTCGGCTTGGATCATTGTTCACGGACATGTTTATGACTGTACCCGGTTCTTGAAAGACCATCCAGGTGGTTCCGATAGTATTCTCATCAATGCAGGTACCGATTGTACCGAAGAATTTGATGCAATTCACTCTGACAAGGCCAAGAACATGCTTGAAGATTTCAGAATCGGTGAACTCATTGCAACTGGTTACACTTCTGACTCGTCATCTTCACCAAACAACTCAGTTCATGGTTCGGGCAGCTTGAGTCACCTTGCTCCTATCAAAGAAATAGGCATTGCAAAACCCGTAGCTCTAGTTGGCCGGGATAAAATACAATGCAAACTTATTGCCAAGACCTCAATTTCTCATGACGTTCGTGTCTTCAGGTTTGCATTACCTTCAGCTGATCAAGTCCTTGGGTTGCCAGTTGGGAAGCACATATTTGTATGTGCTCACGTTAACGATAAGCTATGCATGCGTGCTTACACTCCATCAAGCACAGTTGATGAAGTTGGTTACTTTGAGCTGGTCATTAAAGTCTACTTCGGCGGAGTTCACCCCAAGTTCCCCAATGGAGGTATTATGTCTCAGCACTTGGATTCTCTTCCTCTTGGTGGAACTATTGATGTGAAGGGTCCGTTGGGACACATTGAATACACTGGTCGTGGTAATTTCACCGTTCATGGCAAACACAAGTTTGCTAAACGACTCGCAATGCTTGCTGGTGGCACAGGAATTACACCAATATATCAAATTGTACAAGCAATATTGAAAGATCCAGAGGATGAAACGGAGATGCACATCGTATTCGCAAACAGGACTGAAGATGACATCTTGTTGATGGAGGAACTTGATTCTTGGGCAGAGAAGCACGAAAGGTTGAAAGTATGGTACGTTGTTGGTGAAACATTGAGGGAAGGATGGAAGTACAGTGTAGGGTTCATAACTGAAGAAGTCCTGCGGGAACATATTCCTCCGGCATCGGAAGATACATTGGCGTTGGCTTGTGGACCACCACCTATGATTCAATTTGCAGTGACTCCAAATTTGGAGAAGATGAACTATGATGTGAAGAATTCATTGTTGGTGTTCTGATGAATCAGCTGAAACTACAAAGAGGAACAGATATGATCAAATTTAAGAAATTGATAAGGGGAGGACAAATTATAAGGAAGTTCTCATTAGTTATTGTATATAATTAGTGTTGAAGGATGTAATGGGTTTAGTATGTTCTTAAAAGTAAGGGGATTGTTAAAATGTTAGTCTGTTTAAATTTTATGTAGAATTGAAAGCAAAATGTGGGAATTGAAACTTTCTTTCCTCGATATTTATGATCAAAAAAAAA

>Par_TRINITY_Eif3E Par_c31839_g1_i1 len=1665

GTGTGCTCTTCCGATCTTCTCAGTGTTTTTTCATCAGTTTTAGGGTTTCTCTCTTTCTTCCTCCCACCACCACTGCGCCGCTCCCCATCCTTCTCAGCAGAGGTTGTTCTTCAGCCTCTCCAACAACAACAGCAACCAATCATGGCGAATCATGATCTGACATCAAGGATAGCACTAAACTTAGATCGACATCTCGTTCTCCCACTCCTTGAATTTCTTCAGGAGAAACAAATCCATCCCGATGTTCAAATCCTCAAAGCTAAAGACGCTCTTCTCAGCCAGACCAATATGGTTGATTACGCTATGGATATCTACAAGTCTCTTCATCAAACTGATGATGTTCCTCAAGAAATGATTAATAGGCGTGCTGAAGTGGTTAGGAGATTGAAATTGCTTGAAGATGCAGCTGAATCGATTATTACTTTCCTTCAGAGCCCGAATATGATTAATGAACCACGTCCTGATAAGCAATATGTTCTTCAGTTGCTACAGGGATACCAGATTGGTGCTGAGCAAATTGAGGCATTGTATCAGTATGCCAAATTTCAATTTGAATGTGGGAATTACTCGGGTGCAGCTGATTATTTGCATCAGTACAGAGCGATTTGTACTAATCCTGATAGAATTTTGAATGCATGTTGGGGAAAGCTTGCATCTGAGATATTGATGCAGAATTGGGATGAGGCTCTTGATGAGCTTAACCGGTTGAGGGAGATTATTGAATCAAAGAATTTCAAGAACCCATTGACTTTACTCCAGAATAGAATCTGGTTGATGCATTGGAGCCTCTTCATCTTTTTCAACCACGAAAACGGAAGAAATGGCATCATTGATCTATTTCTCAATGACAGGTACATGAATGCACTCCAAACAAATGCACCTCATCTTTTACGTTACTTGGCAACTGCATTGATTGTCAACAAAAAGAGAAGACCTCAACTCAAAGAGCTGATCAAAGTCATCCAGCAAGAGCAATCTGCATACAAGGATCCAATCACTGAGTTTTTATACTGCTTGTATGTCAACTATGACTTCGATGGTGCACAACAGAAGCTCCGTGAATGTGAGGATGTGATTTTGAATGATCCATTCCTTGGTAAACGAGCTGAAGAAGGAAACTTTAGCACCGTCCCTTTGAGGGATGAATTTCTTGAAAATGCCCGTCTCTTCATCTTTGAGGCTTACTGCAGAATCCACCAGCGCATTGATATTGGAACTCTTGCCGAGAAGTTGAATATGAATTTTGATGAGGCAGAGAGATGGATAGTGAATCTTGTTCAGAATGCAAAGCTTGATGCTAAGATTGATTCTAAGCTAGGAACTGTTGTAATGGAGCCCACTTCTCTCAATGTGTATGAGCAGATCATTGAGAATACCAAATCACTTTCTCTGCGCACCTACAAATACGTAAATCTTGTTCTAGAATCAGCAGCACCACCGGTTACCGCTGCTCGTTAGACAACAGGTGTAGTATAGCGAGAAAATTTGTGTTGGATCTCTTGAAGTTTACTTTTGGGGAAGGAAGAAAATCTGGTCTTGGTATTTTGTTTTTGTTTACTTGTTTTTCAGTTTCATCTAAAGCATTAAAAACCAAGAGTTGCAGGATAACTTGTAAGTTTAATATACTTCTTTTTAGGGTTCTTTGAACGATAAAATACAGTCCAA

>Par_TRINITY_AroB Par_c5571_g1_i1 len=1617

CTCTTCCGATCTGCCCAGAGGCTGCTGCTTCAACTACTTCTTCTTCTTCTCGTCTGGTTTGAAAAATAATCAGAAGAAGTTTTGTTTTGTTTCCCTCTCTAATTCTTCCAATTTTTGGATGGCTTCTTCTACTACTTCTTCACCTTCTCTTAATTTGGCTATTAAGAATTCTTGTATACGAGAAAATTCTGATCTATTTCTTAGAGGGAGGAATTTATCACTGAAGTCTTCATTTATACATTCAAGTTCATTACGATTGAATAGAATTGGATCTTTAAAAATTCAATCGAAGAGTAGAATAATATCTGCAAATTCAGCTCAAGTAATGGAACAATCTTCAATTAGGGCACCAACTATTGTTGATGTTGATTTGGGTAATAGGAGTTATCCGATCTATATTGGTTCTGGACTTCTTGAACAACCTGATCTTCTTCAGAGGCATGTACATGGGAAGAGAGTACTCGTGGTAACTAATACGACAGTTGCGCCATTATACCTAGATAAAGTTGTCAGCGCATTAACTCATGGCAATCCGAGTATTTCGGTTGAGAGTGTGATTTTGCCAGATGGAGAGAAATATAAAAATATGGATACTCTGATGAAAGTTTTTGACAAGGCCATAGGATCTCGATTTGATCGGCGTTCCACATTTGTCGCTCTTGGAGGTGGTGTTATTGGTGACATGTGTGGTTACGCAGCAGCTTCTTTCCTTCGTGGTGTAAACTTCATTCAAATTCCAACTACTGTCATGGCACAGGTGGATTCCTCTGTTGGGGGGAAAACTGGAATCAACCACACACTGGGGAAAAACTTGATTGGTGCTTTCTACCAGCCCCAATGTGTACTTGTAGATACAGACACATTACATACACTGCCTGAGAGGGAATTAGCTTCTGGTTTTGCTGAAGTTATAAAATATGGACTCATTAGGGATGCTGAGTTCTTCGAGTGGCAAGAAAAAAATATGGACTCTCTGTTGGCGAGGGATCCCAGTGCACTAGCATATGCAATCAAGCGTTCATGCGAAAACAAGGCAGAGGTTGTATCCTTGGACGAGAAGGAAGGTGGAGTGAGAGCGACACTGAACTTGGGGCATACCTTTGGACATGCAATTGAAACAGGATTTGGCTATGGTCAATGGCTCCATGGAGAAGCTGTAGCAGCTGGAACGGTCATGGCAGTAGATATGTCTTACCGTCTAGGTTGGATTGATGAATCAATCGTGAAGCGGACATATGATATTTTACAGAAGGCCAAGCTGCCGATTACCCCACCAGACGTGATGACTGTAGAGATGTTCAAGTCTTTCATGGCTGTTGATAAGAAAGTGGCTGATGGACTACTAAGGCTCATCCTTCTCAAAGGTCCTCTCGGTAATTGTGTTTTTACAGGTGATTACGATAGAAAAGCCCTAGACGAGACACTTCGTGCGTTCTGTAAGTCTTAATTTAATGCCGACCTTGGTTGGGCCAATTTCTTCATCAACCATGTGCAGTTTTGTGTTATTATCTTTTGTTGACGTGAAACCACATTGTTTTATGTATATGTAATTCAATGACTTCTTAGATTTGCCATCAGCGCATGTATTACTATAAAGTCTAGATCGGAAGAGCGTCG

>Pso_Agt1 PS0919200.1 transcript offset:2050 AED:0.22 eAED:0.22 QI:2050|0.83|0.71|1|0.83|0.71|7|0|34

GTGCTCCAAAACACTAGGTGAGTCCATTATACTAGGATAGTAAAGTCGATAGACTGACGGTGATATCAAGTCGGAAAAAGAACACGGTGTCGCCCCATGCTCCAAAAACACACTAGTCATGTATAAACAAAGCAAGTGTGTGGTTATTGGCTCGCAACTACAAGGAGAGGCTGTGGTTGCAAACCCGTTATCCCTTTATCTCTCTCGCTTTCTCTCTTTGTCCTTGTCCTTTGCTCATCCTCATCTCCATCTCAAACGTATTTTTAAATAGAAATTATATGAATGCACTACGTTATAGTACGTATTTTAATATTTTCCACAATAATCCAAATGATTTTATGCCGCATGATCTAGAGTTCAGTACATCGTAATCTTCCACGTGGAAGCTCTGTGATCCATCAGCAGCATCTTGATCCTCTCATCTTCCACTTATTTTGGGTCTGTCTATATATAAACATATGTTTTTTTTCTTCTAATGCAGCTTATTACACACATATTTCCTTCCTCCCATTGTTTATCCCTTAAAGTTGGACTGGGAGAGAGAGAGAGAGAGAGAGAGAGAGAGAGAGTGTGTGAGAGTGTGAAGTAGTTCTCCAGAAACCATTCTTGCAACGAAAAGCTTCAAAGAGGTACACTATGGATTACGTTAATGCACCTGGGAGGAACCATCTCTTTGTTCCTGGACCTGTCAATATTCCAGAATCAGTCCTTCGTGCCATGAGCAGAAACAATGAAGATTATCGTTCCCCTGCAATTCCTGCTATGACTAAAACTCTTCTTGAAGATGTTAAGAAGATTTTCAAGTCAACTTCTGGAACCCCATTTTTGATCCCCACAACAGGTACGGGTGCATGGGAAAGTGCACTTACAAATACTTTGTCTCCTGGTGATCGGACAATTTCTTTCCTCATCGGGCAATTTAGTCTTCTCTGGATTGATCAGCAGCAGCGCCTCGGGTTCAACGTTGATGTTGTAGAGAGTGAGTGGGGCTGTGGTGCTGATCTTGAAGTCCTAGCAGCCAAGTTAGCTGAAGACACTGCTCATACCATCAAAGCTGTTTGCATGGTTCATAACGAAACTGCAACTGGAGTTACTAATAATTTGGCTGCTGTACGGAAATTACTTGATCACTACAACCATCCTGCATTGTTTCTCGTGGATGGAGTGTCTTCGATATGTGCTCTTGATTTCCGTATGGATGAATGGGGAATCGATGTAGCTTTAACTGGTTCACAAAAAGCTCTTTCTCTACCTACTGGGATGGGTATTGTGTGTGCAAGCCCTAAAGCACTGGAAGCATCCAAGACTGCCAAATCAGTGAGAGTGTTCTTTGACTGGAAAGACTACTTGAAGTTCTACAAAATGGGAACTTATTGGCCATACACTCCTTCAATCCAGCTTTTATACGGCCTCAGAGCTGCTCTTGATCTCCTATTTGAGGAAGGACTTGACAATGTCATTGCTAGGCACAGCCGCTTGGCCAAAGCAACTAGACTTGCTGTTGAAGCCTGGGGTTTGAAAAACTGCACACAGAAAGAGGAGTGGTTGAGCGACACTGTGACTGCAGTTGTTGTTCCACCTTACATTGACAGTGCTGAAATAGTTAGAAGGGGATGGCAGAGATACAATCTAAGCTTAGGTCTGGGTTTGAATAAAGTGGCTGGAAAAGTTTTCAGGATAGGTCATCTTGGGAACCTGAATGAGCTCCAACTGCTGGGTTGTTTAGCTGGAGTGGAGATGATTCTGAAAGATGTCGGGTACCCAGTTAAGCTAGGAAGTGGAGTGGCTGCGGCTTGTGCGTTTTTCCAGAACACCATTCCTATGATTGAATCTAGGATTTAATGAATCATTCATTATCCATGTGTAACTACTGTTCTCCCTCTTCTGATTTGCTTGGAGTACTTGTATAAAAGAAGACATGATAATCTCCTGTATTTGCACTCTTTTGAGGCTTACTTAGTAGTACAAGAACTCATTACTCTTTCATCCATTAATCCTGTATTTAAAATCTCTTTAAATTTGATCTCACCACTCTTGCATATATCCAACTATGTCTGGTGTCCCCACGCACCGGCATGCTGGGACCGTGGGTTTCGCAAATAAACAGGCCAAGTGCTCAGCCCGCGGTCAATTCTCCATTGACTTTGACCAGTAA

>Pso_At103 PS0934400.1 transcript offset:0 AED:0.38 eAED:0.38 QI:0|0.25|0|1|1|1|5|0|400

ATGGCTGCAGAGATGGCATTGATAAAACCCATCTCTAATTTCACTAGAACCAGCCCTAAATTTATCACCTCAACCAGAAAATCATTCTCCATTGTCAAAATGGCAACCCAATCCACTGCATCAACAACAAAGTCAGGGAAACCAAAGAAAGCAATCAAAGAAACCTTACTTGCTCCTAGGTTTTACACAACTGATTTCGATGAAATGGAAACTCTTTTTAACACTGAGATGAACAAAAACTTGAATGAGGCCGAGTTTGAAGCTTTATTACAAGAGTTCAAAACTGATTACAACCAAACTCATTTCGTTAGAAACAAAGAGTTCAAAGAAGCTGCTGATAAATTGCAAGGTCCTCTTAGACAGATTTTCGTTGAGTTCTTGGAGAGATCTTGTACTGCTGAGTTCTCAGGGTTTCTTCTTTACAAAGAACTTGGAAGAAGACTTAAGAAAACCAATCCAGTTGTTGCTGAAATTTTCTCACTCATGTCCAGGGATGAAGCTAGGCATGCTGGATTCTTAAACAAGGGACTGTCAGATTTCAACTTGGCACTGGATCTTGGATTTCTTACAAAGGCTAGACAATATACATTCTTCAAACCCAAGTTCATTTTCTATGCTACATATTTATCTGAGAAAATTGGGTACTGGAGGTACATAACCATTTATAGGCACTTAAAGACCAATCCTGAATACCAAGTTTATCCAATTTTCAAGTATTTTGAGAACTGGTGCCAAGACGAGAACAGACATGGAGATTTCTTCTCCGCGCTGATGAAAGCTCAACCCCAGTTTCTTAATGATTGGAAAGCAAAGTTGTGGTCTCGATTCTTCTGCCTATCGGTGTACGTGACCATGTACTTGAATGACTGCCAGAGAACAGCTTTTTACGAAGGGATCGGACTTAACACTAAAGAGTTTGATATGCACGTCATAATTGAGACAAACCGTACAGCCGCAAGGATTTTCCCTGCAGTTTTAGATGTTGAGAACCCGGAGTTCAAGAGGAAGTTGGACAGGATGGTGCAAATCAACGAGAAACTTCTTGCAATTGGTGAAACTGAGGAATCATCTTTGGTGAAGAACTTAAAAAGGATCCCACATGTTGCAGCATTGGCGAATGAGCTCTTGGCTGCGTATCTCATGCCTCCTATTGATTCTGGGTCTGTTGATCTTGCTGAGTTTGAGCCACAAGTTGTGTATTAA

>Pso_ChlP PSUN36800.1 transcript offset:0 AED:0.03 eAED:0.03 QI:0|-1|0|1|-1|1|1|0|491

ATGGCCTCCATTGCCGTCAATGCTCGTTTCACAGCCACAACATGCGACCGTAAAACAACCGAAATCAGCCGTGAAACACTAAAATCCTTCATGGGTCTTAGAAAAACAAGCCAAAACGCTTCCATCTCACTTGCACAATCTCTGACAACCAGTACTCCACATTTCAACCCTGCTAGAAGAAATTTCACTTCCATCAAAGCAGGTAAATCAAGTCCTAAAGTAACAGGAAGAAACCTCCGAGTTGCGGTCATTGGTGGCGGTCCTGCTGGTGGTGCAGCTGCTGAAACCTTAGCTAAAGGCGGTATCGAGACGTTCTTGATCGAAAGGAAAATGGATAACTGTAAGCCATGCGGTGGTGCAATTCCACTGTGCATGGTTGGTGAGTTTGATCTTCCTTTAGACATAATCGACAGACGTGTTACGAAGATGAAGATGATTTCACCTTCAAATGTTGCAGTCGACATTGGCCAGACTCTCAAACCAGATGAGTATATCGGTATGGTACGTAGAGAAGTGCTCGATCAATATCTTCGTGAAAGAGCAGAGAAAGCTGGAACAAAGGTAATTAACGGTTTATTCTTGAAAATGGATTTGCCAAAAGAGCCAAATGCGCCTTATGTTCTTCACTACAGTGCTTATGATGGCAAAACTGGTGGTGTTGGCACAAAAACCACGCTAGAGGTGGACGCAGTCATCGGTGCAGACGGTGCAAATTCTCGCGTAGCGAAAGCAATCGGAGCTGGTGATTATGATTACGCTATTGCATTCCAAGAGAGAATCAGAATCCCAGATGAGAAGATGGTTTACTATGAAAATCTTGCTGAGATGTACGTCGGTGATGACGTGTCGCCTGATTTTTACGGATGGGTGTTTCCTAAATGTGATCACGTCGCAGTGGGTACCGGTACAGTTACGCACAAAGGAGATATCAAGAAATTTCAAGTGGCTACACGTCTGCGCGCCAAGGATAAGATTCTAGGAGGAAAAATTATCAGAGTAGAGGCACATCCTATCCCAGAACATCCAAGACCCAAGAGGTTGGCAGACAGAGTAGCGCTTGTAGGAGATGCAGCTGGGTACGTAACAAAATGTTCAGGAGAAGGTATTTATTTTGCTGCCAAGAGTGGCAGGATGTGCGCGGAAGCCATTGTGGAAGGATCCGAGAGAGGGACAAGAATGGTGAATGAAACGGACTTGAGAAATTATCTGGAGAAATGGGATAAAACCTACTGGCCAACTTACAAGGTTTTGGATATATTGCAGAAGGTGTTTTACCGATCGAATCCAGCTCGAGAAGCTTTTGTTGAGATGTGCGCGGATGATTATGTTCAGAGGATGACATTTGATAGTTATCTGTACAAGAGAGTTGTGCCTGGAAATCCAATTGAGGATATTAAATTGGCTTTTAATACTATTGGGAGTTTGGTCAGAGCTAATGCGCTTAGAGGCGAGATGAACAAGATTACTAGTCTATGA

>Pso_GbssI PS0814490.1 transcript offset:244 AED:0.16 eAED:0.16 QI:244|1|1|1|0.92|0.92|14|550|609

GTAACTTTTAATAACTTGTTAGTTCTGAATTACTGAGGTTTTCTTGGCTAGATTTTGTTAATTTTATGTTAGATTTTTTTGTCTATGATGAAGTTGTTCTCTGGACTAGTGTTAGACTTCGGTGTATATTTCGACTGTCTGCATTGTAATCAACCGCAGGAGAAAAATATCCGAGATTGTATGCTTTGGTTCTGATTGTTTTGGAAAATTGTCAAAAAAAAAAAAAAGGACAAGAATATTACGGATGACGACCATAAGTGCGTCCCAGTTTGTGTCACGAATCAACTGCGGAGGAACTGCACCAAAAGACATACTTAAACAAATCAATATGAAGAACCAAAGCATGACTCATTCAGGGTTAAGATCTTTGAGCAAAGTAGATTTCTTACAAATAAAAAACACTCAGAAAGCAGCTTCTAGGGAATTGAGGAGGAACGCATTCCAGACAGACAAAGGTACTGCGTCAGGAGAGGTTAAATGTTCTAGTGGGATGAACTTGGTCTTTGTTGCTGCTGAAGTAGGCCCCTGGAGTAAAACCGGTGGTCTTGGTGATGTTCTTGGAGGCTTACCACCAGCCATGGCGGCAAATGGACATCGTGTTATGACAGTCTCTCCTCGTTATGATCAGTATAAAGATGCTTGGGATACCAATGTGTCAGTTGAGATGAAAATTGGGGACAGAATTGAAACCGTTCGTTTCTTCCACTGTTATAAGCGTGGGGTTGACCGTGTATTTGTGGATCACCCACTGTTTCTTGAGAAGGTATGGGGTAAAACTGGATCTAAAATCTATGGTCCGAATGCTGGATCAGATTACCAAGATAACCAGCTTAGATTCTGTTTGCTTTCTCTGGCAGCTCTAGAAGCACCAAGAATTCTGAATCTCAACAGCAACAAATATTTCTCTGGAACATACGGAGAGGAGGTCGTGTTCATCGCTAATGATTGGCATACAGCCCTCCTACCATGCTACCTTAAAAGTATATACCAATCTAAAGGAATGTACGAAACTGCCAAGGTTGCTTATTGCATCCACAACATTGCTTACCAAGGAAGATTTGCTTTTTCCGATTTTGCGCTTCTCAATCTTCCGGATTCATTTAGGAGTTCTTTTGACTTCATTGACGGGTATGAGAAGCCTGTTAAAGGAAGGAAGATTAACTGGATGAAGGCTGGAATATTAGAGGCAGACAAGGTCGTAACTGTGAGTCCATACTATGCTGAAGAACTTGCTTCAGGCCCGGATAAAGGTGTGGAACTGGATAATATTATACGCAGAGCTGGCTGTCTGGGTATTATAAATGGAATGGATGTTCAGGAGTGGAACTCATTAACTGATAAATATATCGACGTCAAATATGATGCCTCAACTGTGACCGAGGCGAAGGCTCTTCTGAAAGAAGTACTGCAGGCAGAAGTAGGGTTACCTGTTGACCGTAATATTCCTGTCATAGGTTTCATTGGCAGACTTGAGGAGCAAAAGGGTTCAGATATTTTGGCAGCAGCAATCCCTGAATTTATTGACGAAGAAGTGCAGATAATTGTCCTTGGTACTGGCAAAAAGCCCATGGAAAAACAGTTGGAGCAGTTAGAGACTTTGTATCCTGAAAAAGCTAGAGGAGTGGCGAAGTTCAACGTTCCCTTGGCTCATATGATCATAGCTGGGGCTGATTTTATGATGATCTCAAGCAGATTTGAACCATGTGGACTCATCCAGTTACATGCTATGCGTTATGGAACGGTGCCTATTGTTGCCTCGACGGGTGGATTAGTGGACACTGTCAAAGAAGGTTTCACGGGATTTCAAATGGGAGCTTTCAACGTCGAATGCGAGGCAATTGACCCATTAGACGTGACAGCAATCGCCACAACCACCAAAAAGGCACTTGAAACCTATGGAACTCCAGCTTTAACAGAAATGATCCAGAACTGCATGGCTCAAGACCTCTCTTGGAAGGGACCTGCCAAGAATTGGGAGGACATGTTGCTGAGCTTGGAAGTTGCAGGTAGTGAACCAGGAATCGAGGGCGATGAAATCGCTCCTCTTGCTAAAGAAAATGTTGCCACTCCTTGAGTGACAGAACCGCTCCAGTGCAACTTAGAAGAGCTATTTCACAATCTAACCCAGGCGATTCCATTGCTGCACTGACAAGGCTCTGATTACTTCGATCATATCTAAGATTTGATAAAATTTATCCACTGAGAGTTACATAGTCAGCGACTCAGTGGAAGAGTTCATGATGTGTATATGAAGTGGCTTTACTGGAAAAGAGATACTGCTACTCATTTTCCTTTTCTTCTTTTCTTTTGATTTATGCCATCTACGGCACTCTCGATGTCCAACGTGGATGCCAAAAGCGTAGGCTAGCTAGCTCTAGATCTTGTATCTGGTCAAGAAGATCTTTATATAAATAAAAACGACTTTTTCACCTTCTGCTTCAGTTGCTTCAGTTAAATTTAAAGCCTGAAGTGTTGTGAGCTGTGGGTGGTAGTTGAATAAAGTGAATGTTTAAAACAATTGTCTTGTTACAAGAAAGAAAGAAAGAAACACACAGTATTTAAAGACAACAGACATAACAGGTGACCAATCCTGTAAGACAGATAAAGGACCAATAAGAGAGAGC

>Pso_Hmgs PS0403850.1 transcript offset:352 AED:0.27 eAED:0.27 QI:352|0.81|0.73|1|1|1|23|0|1078

CTCCCGTAAATAGTGTCTTAAACAACGTGGTAGTTGAACTCGTAATCAGAAGATTAATTAATTTCCCGTCAATTAAATCTCTAGTAATTCCTCAAATACCTCTCTTTCTTTTCAATTTCTCACTCACAAAAGTCCGCCTTTGAATTTCACAAACTCACGAGTCGCGTTGGCTGCATATTTGTTTTTGGTTCCGGTCGGACCGGTCATGAAATTTCCTCCGTCTATAAATACCGATTTATTCGTTGCCTAAATCTCTCACCATTTCTTCTTCATTCTTCCTTCCTTCCTCTCTCTCCAAAACAGCTCGATCGATCTAGGGTTCTCTGAAGGAATTCAAAGAAATTGTCTAAAGATGGGTTCAAATCAGAAGGATGTTGGAATTCTTGCTATGGAAATGTATTTCCCTCCTACTTGTGTTCAACAGGAAGCATTGGAGGCTCATGACGGTGCCAGTAAAGGAAAATACACCATTGGGTTGGGACAAGATTGCATGGCGTTTTGTTCCGAATTGGAAGATGTCATTTCAATGAGTTTGACAGCCGTTAGCTCTTTACTCGAAAAGTATAAAATTGACCCTAAACAAATTGGACGCCTGGAAGTCGGCAGTGAAACCGTGATTGACAAGAGCAAGTCGATAAAGACCTTCCTGATGACAATCTTTGAGGAATGTGATAACACTGACATTGAGGGTGTTGACTCCACAAATGCATGCTACGGAGGAACTGCAGCTTTGTTCAACGTTGTGAATTGGGTGGAGAGTAATTCCTGGGATGGACGATATGGTCTTGTTGTGTGCACTGATAGTGCGGTCTATGCAGAAGGACCTGCCCGCCCCACTGGAGGAGCAGCTGCTATTGCTATGTTGATTGGTCCAGACGCACCAATTGCCTTTGAAAGCAAATATAGAGGATCCCATATGTCTCATGTCTATGATTTTTACAAGCCCAACCTAGCCAGTGAATACCCGGTCGTTGACGGAAAACTTTCACAAACTTGTTACCTTATGGCACTCGATTCCTGCTACAAACGCTTTTGTGCTAAGTACGAGAAATGCCAGGGTAAGCAATTCTCGATGGATGATGCAGACTACTTTGTATTTCATTCACCTTACAACAAGCTCGTGCAGAAAAGTTTTGCTCGATTGTACTTCATGGATTTCTTAAGAAACGCCAGCTCTGTTGACGAGGCTTCAAAAGAAAAGCTTGCACCATTTGCAGCTCTATCCGGTGAAGAAAGCTACGCGAGTCGGGATCTTGAGAAGACATCCCAAACTGTTGCAAAGAATCTTTATGCTGAGAAGGTTGGACCTACCACTTTGTTGCCCAAACAAGTCGGTAATATGTACACTGCATCACTCTATGCGGCATTTGCTTCTCTTCTTCACAACAAAAATAGCTCACTGGGTGGTAAACGTGTAATAATGTTCTCTTATGGAAGTGGTTTAAGTTCCACAATGTTCTCATTCCGAATGAGGGAAGGTTCACATCCATTCAGCTTATCAAACATTGCAACTGTAATGGATGTCGCGCACAAGTTGGAGTCCAGACATGAGGTTACTCCTGAAAAATTCGTCGAAACCATGCAACTCATGGAGCACAGTGTTAAGAGTGAAGGAGTAGTAGCTAAGATATTTAGGGTTTCATTCTTGAGTGGTGTTTTGATTTCTGAAGTTATGGAGCTTTATACGAAATGGGTTCGAAGGAATAAAGATTATGTTCATTCTATGGAGTCCCTCGCCAATGGATTGACATGGCTACTACCAGAACGCTTCTCTGAGACAGAGATAGGACCAGAAGCAGTTACTGCTTTTTTGGGTATAATCACTTCTATAAATGAACATATTATCGATTCGGATCCATTGCAACGGCGAATGGGCTCTACGGAAGAGTCGTCATCATCATCGTTTCCTTGGTCACTTGGGATATCAGTTCTCAAGGACTTGGAAACATTGGTGGAAGTCATGGCTCAGCATTTCTATGGAGATGATAAGAAATGGAATTTCCTAGCTGTCACTGAGGGTACCAAATTTTTAGTTAGGTTAGCTTTGTTTCGGGATAGTGGGTATAAAATGGTACTAAATGGAGGGGAGATAGCTAATGTTGATAAGAGACCTGAGGCGTCGAACTCTGAACCCAGAATGGGGAATTCTGTAAGACGAGATCGTTTTGGGCGTGATCCACGAAGTCTTGAAGGAAGGGCGTTGTCAGCCTTAAGCAAGTTTGGAGAAGACGCTAGAATGATTTCTGAACCTAAGTGGTTAAATAGATTTCAGAACCACCGTGATCCTGCCATGGAAACTCCACCACCTGTTCCCAAAAAACCTACACTTTCAAAAATTTGGTCTGAGAAGGGAGTTCCTGGGGTTTTGTTTCTCACTGCTGAGGTGTTGCATATGAGTAGACCACTAATTTATGTTTTGCTTATGAGGAAGTATGGAGTTCGGTCATGGTTACCTTGGTTTCTTGCTCTCTCTGTGGATCTGACTGGAATGAGCATACTTTCTCATGTTAGACATTCTGGACACAGTAGATATGGAGGAAGTTCTTCTCTTTCAGCATCTGAAAAGGATGAGGTTAAAAGGCGAAAGATGTTGTTGGCATTTTACCTCATGAGAGATCCATTTTTTAGCCGGTATACTCGGAAAAGACTTGACACAGTTGAGAGAGTCACAGACCCTATCCCAATAATCGGGTTTCTTACAGCTAAAATCATTGAGCTTGTTATTGGAGCTCAAACTCGCAAAGTTGTCGCCTTTTGTGCTCTTGAAGACTCTGGTAGTCTTCATTATGCCCATCTCGTCTTTACCCAAATCCCGAATCCTACTCCTTACATCTGCAACTCTATAATTCGAGAGAAGAGTGTGGTTTCATGGGCGACAATGGTTGATGCTTATACGCAATCGGATCAACCCGTAAAAGCCCTTGAGCTTTTCCAGAGAATGGAAGTGGAACAGGTTAATCCCAACGAGGTTACGTTGCTGAATGTTTTAACAGCTTGCGCCAGGGCGAGAAATTTGGAAATGGGGAAGTGGGTTCACAAGTATATAGATGACAATGGAATTGGGTTCAGTTTGGTTCTTACAACAGCTCTTATGGATGTTTATTGCAAATGTGGTTGTTTCTCTCTAGCTCGGGAATTGTTCGACCAGATACCTGAAAAGAATTTGTTTCACTGGAATATAATGATCAATGGACATGTTGAAGACAGTGACTATGAGGAAGCCTTATCACTCTTTCACGAAATGCAAATCAAGGGAATTAAAGCGGATAAGGTGACTATGGATATGATATCTAAGTTAAAGATAGCAGGTTATGTACCTGACAATTCTGAAGTGCTGTTTGAAATGGACGAAGAGGAAAAGGAGAATGCCCTTAGTATGCACAGTGAGAAGTTAGCAATCTCTTTTGGACTCATAAGTACTAGCCCTGGTACTCGCATTCGAGTAGTGAAGAATCTTCGTGTTTGCAATGACTGTCACTCTGCTGCAAAGCTGGTATCCAAAGTGTACAATAGGGAGATAGTCTTGCGGGATAGAAACCGGTTTCATCATTTCAAGGATGGGTTGTGTTCTTGTAAAGACTTCTGGTAA

>Pso_Nia PS0312620.1 transcript offset:413 AED:0.12 eAED:0.12 QI:413|1|1|1|1|1|4|415|904

TTTCTGTTATATAAAGTCTAGAGGATTGTATATTTATTTATGTGGCTATAATCGGCAATTACAAATCACAGACCATAGTTTGCATGCACTGTAACCACGGGTTCCTCATTTTTTCTTAGTCGGCTGGCTCGGCTATTAAAATTTGTGACATCACTACCCAAGATTAATTTGTCCCACCTTTAGAGCCATCTCTATCAGAAGAGCATGGCACAAGCACATCCCTTTTCCAATGCATCTTATATAAAGCCCTAACCATGCTCCTCATTTTTCTTGCAACTATCAAATGCTTGCAAAAGCAAATCGGAGAAAAGAGAAAACATAGATAGAAGTAGAAAGCAGAGAGAAAGTGAAACGAGAGGAGGAGCATTCAAAGAGAGATTTTTTTCTTTCTTCTTTCTTTTGATTAGTCTCTAATGGCGGCTTCTGTTGAAAATCGGCAGTACAACAGTCGACTTGAATCAGGTCTAGCAGTTCGTGGATTTAAAACCAATCCCAGTCATCGATCTGATTCGCCTATCCGTGGCTGTAATTTTCCCATGTCGACTACCGAGCGTCTTTCTATACAAGCAAAACCGGTCATTGAAGATTTATCTAGCGACGACGAGGAGGATCAAAACGAGCAGTGGAAAGATTTCGGCAAGATAAACAATGAAATAGAGCCGTCTGCTATTGACTCGAAAGATGAAGGAACTGCTGATAATTGGATTGAACGCAACCCTTCCATGATTCGGTTGACAGGAAAGCACCCATTCAATTCAGAAGCACCATTAACTCGTCTCATGCATCACGGGTTTATCACACCTGTTCCTCTTCATTACGTTCGTAATCACGGAGCTGTTCCAAAACTTAAATGGGAGGATTGGACAGTTGAGATTTGTGGATTAGTAAAGCGTCCGTTACGTATCACTATGGATCAGCTCATTAATGATTTCGCAAGCCGTGAGTTTCCAGCTACACTTGTCTGTGCAGGTAACAGACGTAAAGAGCAGAATATGATAAAACAAACTATTGGGTTTAACTGGGGAGGTGCAGGAGTATCAACTTCAGTATGGCGTGGTGCACGTTTATGTGACGTATTAAAGCGTTGTGGTATCATGAGCCGTAAGAAAGGTGCTCTGAATGTTTGTTTCGAAGGAGCTGAAGATTTACCCGGTGGCGGTGGATCTAAATATGGAACTAGTATAAAGAAGGAATTCGCAATGGATCCTGCTCGTGATATAATTCTAGCTTACATGCAAAATGGTGAACGTTTGTCTCCTGACCACGGATTTCCAGTCCGTATGATTATTCCAGGTTTCATCGGAGGAAGAATGGTGAAGTGGTTAAAGAGAATTATCGTCACAACTCCAGAATCCCAAAACCACTATCATTTCATGGATAATCGGGTTCTTCCTTCTCATGTCGATGCTGAACTCGCCAATTCTGAAGCATGGTGGTATAAGCCAGAGTACATCATCAATGAGTTGAACATAAACTCTGTGATTACAACTCCTTGTCATGAAGAGATTCTTCCCATAAATGCTGCAACAACACAAAGAGCTTACACATTAAGGGGATACGCATATTCCGGCGGCGGTAAGAAGGTAACTCGTGTCGAAGTAACAATGGACGGCGGAGAGTCATGGTTAGTGTGTACTTTGGATCATCCCGAGAAACCAAACAAGTATGGCAAATATTGGTGCTGGTGTTTTTGGTCACTGGAAGTTGAAGTTCTCGACTTGCTTGGTGCCAAAGAGATTGCAGTCAGGGCTTGGGATGAAACTCTAAACACTCAGCCTGAAAAGCTTATCTGGAACGTCATGGGTATGATGAACAACTGCTGGTTCAAAGTCAAAACAAATGTGTGCAAACCACACAAAGGAGAAATCGGTCTTGTTTTCGAGCATCCAACTCAGGCTGGAAACCAACCAGGGGGATGGATGGTTAGACAAAAACATCTTGAAAACTCTGAAGGTCCACAGACCCTCAAAAAGAGTTTGTCCACTCCTTTCATGAACACCACATCGAAAATGTATTCGATTTCCGAGGTGAAGAAACACAACTCGCCTGACTCGGCTTGGATCATTGTTCACGGACATGTTTATGACTGTACCCGGTTCTTGAAAGACCATCCAGGTGGTTCCGATAGTATTCTCATCAATGCAGGTACCGACTGTACCGAAGAATTTGATGCAATTCACTCTGACAAGGCCAAGAATTTACTTGAAGATTTCAGAATCGGTGAACTCATTGCCACTGGTTACACTTCTGACTCGTCATCTTCACCAAACAACTCAGTTCATGGTTCGGGCAGCTTGAGTCACCTAGCTCCTATCAAAGAAATCGGCATTGCGAAACCCGTAGCTCTAGTTGGCAGAGATAAAATACAATGCAAACTTATTGCCAAGACCTCGATTTCTCATGACGTTCGTGTCTTCAGGTTTGCATTACCTTCCGCTGATCAAGTCCTTGGGTTGCCAGTTGGGAAGCACATATTTGTATGTGCCCACGTTAACGATAAGCTGTGCATGCGTGCTTACACTCCATCAAGCACAGTTGATGAAGTTGGTTACTTTGAGCTGGTCATCAAAGTCTACTTCGGCGGAGTTCACCCCAAGTTCCCCAATGGAGGTATTATGTCTCAGCACTTGGATTCTCTTCCTCTTGGTGGAACTATTGATGTGAAGGGTCCTTTAGGACACATTGAATACACTGGTCGTGGTAATTTCACCGTTCATGGCAAACACAAGTTTGCCAAACGACTCGCAATGCTTGCTGGTGGCACAGGAATTACACCAATATATCAAATTGTACAAGCAATATTGAAAGACCCAGAGGATGAAACGGAGATGCACATCGTATTCGCAAACAGGACTGAAGATGACATCTTGTTGATGGATGAACTTGATTCGTGGGCTGAGAAGCACGAGAGGCTGAAAGTTTGGTACGTCGTTGGTGAAACATTGAGGGAAGGATGGAAGTACAGTGTAGGGTTCATAACTGAAGAGGTCCTACGGGAACATATCCCTCCGGCATCGGAAGATACATTGGCGTTGGCTTGTGGACCACCACCTATGATTCAATTTGCAGTGACTCCTAATTTGGAGAAGATGAATTATGATGTCAAGAATTCATTGTTGGTGTTCTAATCAGATTAAGTACTTAGAAGGTGAAACTACAAAGAGGAACACATGTGATCAAATTTAAGAAAATGATAAGGGGAGGACAAATTATAAGGAAGTTCTCATTAGTTTATTGTATATAATTAGTGTTGAAGGATGTAATGGGTTTAGTATGTTCTTAAAAGTAAGGGGATTGTTAAATGTTAGTCTGTTTAAATTTTATGTAGAATTGAAAGCAAAATGTGGGAATTGATACTTTCTTTCCTCGATATTTATGATCATTACATATTTTACTCCCTCCATTTTCATTAATAGTATGTCTGGGTATTTCTGCTTTTCCAGAAACAGAAGTCAAAAACAATATGCTTTTGTTTCACAAAAAAATAATTTTTTTTATAAGTCATTCTAAAATGTATATTCAAACTAATTTTTGATTTTTTTA

>Pso_Eif3E PS0149380.1 transcript offset:246 AED:0.14 eAED:0.14 QI:246|0.87|0.77|0.88|1|1|9|0|454

GACAAACTTCCAAAAATTGTCACCAACTTGTAGCCTTATAAGTAAAAAAGATCCGATCCAAACTCGACCTTGACCGACTGTATTGACTTTGACCTTCTCTAAATAAATCGAATCGTGTGTTAAACTTATCTCTTTTACTCGCCTTTCTCAGTTTCCATCAGTTTTAGGGTTTTTCTCTTTCTTCCTCCCACCACCACTGCGCCGCTAGAGGTTGTTCTTCAGCCTCTACAACAACAACAACCAATCATGGCGAATCATGATCTGACATCAAAAATAGCACTAAACTTAGATCGACATCTCGTTCTACCACTCCTTGAGTTTCTTCAGGAGAAAAAAATCCATCCCGATGTTCAAATCCTCAAAGCTAAAGACGCTCTTCTCAGCCAGACCAATATGGTTGATTATGCTATGGATATTTACAAGTCTCTTCATCAAACTGATGATGTTCCTCAAGAAATGATTAATAGGCGTGCTGAAGTGGTTAGGAGAATGAAATTGCTTGAAGATGCAGCTGAATCGATTATTACTTTCCTTCAAAGCCCAAATATGATTAACGAACCACGTCCTGATAAGCAGTATGTTCTTCAGTTGTTACAGGGATACCAGATTGGTGCTGAGCAAATTGAGGCATTGTATCAGTATGCGAAATTTCAATTTGAATGTGGAAATTACTCAGGTGCAGCTGATTATTTGCATCAGTACAGAGCAATTTGTACCAATCCTGATAGAATTTTGAATGCATGTTGGGGAAAGCTTGCATCTGAGATATTGATGCAGAACTGGGATGAGGCTCTTGATGAGCTTAATCGGTTGAGAGAGATTATTGAATCAAAGAATTTCAAGAACCCATTGACTTTACTCCAGAATAGAATCTGGTTGATGCATTGGAGCCTCTTCATCTTTTTCAACCATGAGAATGGAAGAAATGGCGTCATTGATCTATTTCTCAATGACAGGTACATGAATGCTCTCCAAACAAATGCACCCCATCTTTTACGTTATTTGGCAACTGCATTGATTGTTAACAAAAAGAGAAGGCCTCAACTCAAAGAGTTGATCAAAGTTATCCAGCAAGAGCAATCTGCATACAAGGATCCCATCACTGAGTTCTTATACTGTTTGTATGTCAACTATGACTTCGATGGTGCACAACAGAAGCTCCGTGAATGTGAGGATGTGATTTTGAATGATCCATTCCTTGGTAAACGAGCTGAAGAAGGAAACTTCAGTACTGTTCCTTTGAGGGATGACTTTCTTGAAAATGCCCGTCTCTTTATCTTTGAAGCTTACTGCAGAATTCACCAGCGCATTGACATTGGAACTCTTGCCCAGAAGTTGAATATGAATTTCGATGAGGCAGAGAGATGGATAGTGAATCTTGTTCAGAATGCAAAGCTTGATGCTAAGATTGACTCTAAGCTAGGAACTGTTGTAATGGAACCCACTTCTCTCAATGTGTATGAGCAGATCATTGAGAATACCAAATCACTCTCTTTGCGTACCTACAAATACGTCAATCTTGTTTTAGAATCAGCAGCACCACCGTTCACCAGTGTAGGATCGACGATCTGCATAACTCCAGTATTAAGAGCCATCTTTGCAAAACTATGA

>Pso_AroB PS0624870.1 transcript offset:275 AED:0.15 eAED:0.15 QI:275|1|1|1|1|1|8|305|442

GCCGTTTGTCAGTGTATTGAGAGGGTTTTACAGCAAGAAAAAAGCTTTTGGGTAGTTAAAAGGATATTTCAGTCTTTTCGACGTCAAAGAGGGTAGGCGGGTATTTTCGCACGTGCAACATATGTGCTTTTAACGATTTTCACGATTTTTCAATAATTCTCACCAACCCCCAAGAGGCTGCTACTTCAACTACTTCATCTCGTCTGGTTTCAAAAATAATCAGAAGAAGTTTTGTTTTGTTTCGTTTCCCTCTCTAATTCTTCCCCAATTTTTTGATGGCTTCTTCTACTACTTCTTCACCTTCTCTTAATTTGGCTATTAAGAATTCGTGTGTACGAGAAAATTCTGATCTATTTCTTAGAGGGAGGAATTTATCACTGAAATCTTCATTTATACGTTCAAGTTCATTACGATTGAATAGAATTGGATCTTTAAACATTCAATCGAAGAGTAGAATTATATCTGCAAATTCAGCTCAAGTAATGGAACAATCTTCAATTAGGGCACCAACTATTGTTGATGTTGATTTGGGTAATAGAAGTTATCCGATCTATATTGGTTCTGGACTTCTTGAACAACCTGATCTTCTTCAGAGGCACGTACATGGGAAGAGAGTACTCGTGGTAACTAATACGACAGTTGCACCATTATACCTAGATAAAGTTGTCAGCGCATTAACTCATGGAAATCCGAGTATTTCGGTTGAGAGTGTGGTTTTGCCAGATGGAGAGAAATACAAAAATATGGATACTCTGATGAAAGTTTTTGACAAGGCCATAGGATCTCGATTTGATCGTCGTTCCACATTTGTCGCTCTTGGAGGTGGTGTTATAGGTGACATGTGTGGTTATGCTGCAGCTTCTTTCCTTCGTGGTGTAAACTTCATTCAAATTCCAACTACAGTCATGGCACAGGTGGATTCCTCTGTTGGGGGAAAAACTGGAATCAACCACACACTGGGGAAAAACTTGATTGGTGCTTTCTACCAGCCCCAATGTGTACTTGTAGATACAGACACATTACATACACTGCCTGAGAGGGAATTAGCTTCTGGTTTTGCTGAAGTTATAAAATATGGACTCATTAGGGATGCTGAGTTCTTCGAGTGGCAAGAAAAAAATATGGACTCTCTGTTGGCGAGGGATCCCAGTGCACTAGCATATGCAATCAAGCGTTCATGTGAAAACAAGGCAGAGGTTGTATCCTTGGACGAGAAGGAAGGTGGAGTGAGAGCAACGCTGAACTTGGGTCATACCTTTGGACATGCAATTGAAACAGGATTTGGCTATGGTCAATGGCTCCATGGAGAAGCTGTAGCAGCTGGAACGGTCATGGCAGTAGATATGTCTTACCGTCTAGGTTGGATTGATGAATCAATTGTGAAGCGGACATACGATATTTTACAAAAGGCTAAGCTGCCGATTACCCCACCAGATGTGATGACTGTAGAGATGTTCAAGTCTTTCATGGCTGTTGATAAGAAGGTGGCTGATGGACTACTAAGGCTCATCCTTCTCAAAGGTCCTCTCGGCAATTGTGTTTTTACAGGTGATTACGATAGAAAAGCCCTAGACGAGACCCTTCGTGCATTCTGCAAGTCTTAATTTAATGCCGACCTTGGTTGGGCTCATTTCTTCATCAACCATGTGCAGTTTTGTGTTATTTTCTTTTATAGACTATAGACTCATCTGTTGACGTGAAACCACATTGTTTCTGCATATGTAATTTAATGACTTCCTAGATTTGCCATCAGCGAATGTATTACTATAGAGCCTCATACTTACATCTTACTATAATAAAAAACAAGTTTAGTTTTTCAAATTTGGGTGCATGTTACATGGGCCTGTACTCAAAAAAAAATTGTTTGATAACCAAAAGTTTTATTGAAAGTACAACACAATATTTTGAC

>Pdu_TRINITY_Agt1 Pdu_c37367_g1_i1 len=1530

TATTTCCTTCCTCCCATTGTTTATCCATTAAAGTTGGACTGGGAGAGAGAGAGTGAGAGTCTGAAGTAGTTCTCCAGAAACCATTCTTGCGACGAAAAGCTTCAAAGAGGTACAATATGGATTACGTTAATGCACCTGGGAGGAACCATCTCTTTGTTCCTGGACCTGTCAATATTCCAGAATCAGTCCTTCGTGCCATGAGCAGAAACAATGAAGATTACCGTTCCCCTGCAATTCCTGCCATGACTAAAACTCTTCTTGAAGATGTTAAGAAGATTTTCAAGACAACTTCTGGAACCCCATTTTTGATCCCCACAACAGGTACTGGTGCATGGGAAAGTGCACTTACAAATACTTTGTCTCCTGGTGATCGAACAATTTCTTTCCTCATCGGGCAATTTAGTCTCCTCTGGATTGATCAGCAGCAACGCCTCGGATTCAACGTTGATGTTGTGGAGAGTGAGTGGGGCCGTGGTGCTGATCTTGAAGTCCTAGCATCCAAGTTAGCTGAAGACACTGCTCATACCATCAAAGCTGTTTGCATGGTTCATAATGAAACTGCAACTGGAGTTACTAATAATTTGGCAGCTGTAAGGAAATTACTTGATCACTACAACCATCCTGCATTGTTTCTCGTGGATGGAGTGTCTTCGATATGTGCTCTTGATTTCCGTATGGATGAATGGGGAATCGATGTAGCTTTAACTGGTTCACAAAAAGCTCTTTCTCTACCTACTGGGATGGGTATTGTGTGTGCAAGTCCTAAAGCATTGGAAGCATCCAAGACTGCCAAATCAGTGAGAGTGTTCTTTGACTGGAAAGACTACTTGAAGTTCTACAAAATGGGAACTTATTGGCCATACACTCCTTCAATCCAGCTTTTATACGGCCTCAGAGCTGCTCTTGATCTCCTATTTGAGGAGGGACTTGACAATGTCATTGCTAGGCACAGCCGCTTGGCCAAGGCAACCAGACTTGCTGTTGAAGCCTGGGGTTTGAAAAACTGCACACAGAAAGAGGAGTGGTTGAGCGACACTGTGACTGCAGTTGTTGTTCCACCTTACATTGACAGTGCTGAAATAGTTAGAAGGGGATGGCAGAGATACAATCTAAGCTTAGGTCTGGGTTTGAATAAAGTGGCTGGAAAAGTTTTCAGGATAGGTCATCTTGGGAACCTGAATGAGCTCCAACTGTTGGGTTGTTTAGCTGGAGTGGAGATGATTCTGAAAGATGTTGGATACCCAGTTAAGCTAGGAAGTGGAGTGGCTGCGGCTTGTGCGTTTTTCCAGAACACCATTCCGATGATTGAATCTAGGATTTAATGAATCATTCATTATCCATGTGTAACTACTGTTCTCCCTCTTGTGATTTGCTTGGATTACTTGTATAAAAGAAGACATGATAATCTCCTGTATTTGCACTCTTCTGAGGCTTACTTAGTAGTACAAGAACTCATTACTCTTTCATCCATTAATCCTGTATTTAAAATCTCTTTAAATTTAATCTCATCACTGTTGCATAAAAAAAAAA

>Pdu_TRINITY_At103 Pdu_c33183_g1_i1 len=1611

GTGCTCTTCCGATCTCAAAAACTTGAATGAGGCTGAGTTTGAAGCTCTATTACAAGAGTTCAAAACTGATTACAACCAAACTCATTTCGTTAGAAACAAAGAGTTCAAAGAAGCTGCTGATAAATTGCAAGGTCCTCTTAGACAGATTTTCGTTGAGTTCTTGGAGAGATCTTGTACTGCTGAGTTCTCAGGGTTTCTTCTTTACAAAGAACTTGGAAGAAGACTTAAGAAAACCAATCCAGTTGTTGCTGAAATTTTCTCACTAATGTCCAGGGATGAAGCTAGGCATGCTGGATTCTTAAACAAGGGACTGTCAGATTTCAACTTGGCACTGGATCTTGGATTTCTTACAAAGGCTAGAAAATATACATTCTTCAAACCCAAGTTCATTTTCTATGCTACATATTTATCTGAGAAAATTGGGTACTGGCGGTACATAACCATTTATAGGCACTTAAAGACCAACCCTGAATACCAAGTTTATCCAATTTTCAAGTATTTTGAGAACTGGTGCCAAGACGAGAACAGACATGGAGATTTCTTCTCCGCACTGATGAAAGCTCAACCCCAGTTTCTTAACGATTGGAAAGCAAAGTTGTGGTCTCGGTTCTTCTGCCTATCGGTGTACGTGACCATGTACTTGAATGACTGCCAGAGAACAGCTTTTTACGAAGGGATCGGACTTAACACTAAAGAGTTTGATATGCACGTCATAATTGAGACAAATCGTACAGCCGCAAGGATTTTCCCTGCAGTTCTAGATGTTGAGAACCCGGAGTTCAAGAGGAAGTTGGATCGGATGGTGCAAATCAACGAGAAACTTCTTGCAATTGGTGAAACTGAGGAATCATCTTTGGTGAAGAACTTAAAAAGGATCCCACATGTTGCAGCGTTGGCGAATGAGCTCTTGGCTGCGTATCTCATGCCTCCTATTGATTCTGGATCTGTTGATCTTGCTGAGTTTGAGCCACAAGTTGTGTATTAATAAGTTTTTTGTGTATTGAGGAAACAAAGGCCCTTCAATTCAATTCAGTTCAGTGCAAAGTTTAATTGCTACAGTTAACCATCACTATGGACCTGCTGTATAAAGAACGAAACATTTGTGACATATAATGTATCATGATTTCTGGACGCACCTCATGTGGAGGTTACTCGTCATGCGTATTTTTTTCTGCACACATGTAGCCCAGGACTTGTCACAACTGGCAAGGCAAGTACAAGAGCTAGACAATGGTCCTACTATCTAGGCGATGTATGAGCATTTCACCTGAAGAGACATCGACATTTTAAACTGGCAGAACGGATTAAATACCCATGTTGGGTCCTAAAGCTGTTAATTGTCATCATGACAGTGGTTTCATTGTTCTCAAATGTTATTCTCGATCATATGAGATCCAAATTTGAAGTACTTTGCTACCGAGTATTTAAATGATCCGGGCTCCAAGGCCAAAAAGTTTGAAACAGGGGAGATTTAACAGTTGGCGATCAAGATAAAACAGTTTAAAGGACACAAAAGGCGCACATGTTCCCTGTTTTTACACATGAAGCTTCAACTCGTTGTTTCTTAGTTTGGGAAGCTTAAATTTGTTAGATCGGAAGAGCGTCGTGTAG

>Pdu_TRINITY_ChlP Pdu_c39955_g1_i1 len=1375

TCTTTAAATCAAAAAAACAAGGATAACAAAATCTTAAAACTTTCATTCATACTTGCAGAAACTTGAATCCACCACCACCAAAAATGGCCTCCATTGCCGTCAATGCTCGTTTCTCAGCCACAACATGCGACCGTAAAACAACCGAAATCAGCCGTGAAACACTAAAATCCTTCATGGGTCTTAGAAAAACCAACCAAAATGCTTCCATCTCACTTACACAATCTCTAACCACAAGTACTCAACACTTTAACCCTGCTAGAAGAAATTTCACTTCCATCAAAGCAGGTAAATCAAGTCCTAAAGTAACAGGAAGAAATCTCCGAGTTGCGGTTATTGGTGGTGGTCCTGCTGGTGGTGCAGCTGCTGAAACCTTAGCTAAAGGTGGTATCGAGACGTTCTTGATCGAAAGGAAAATGGATAACTGTAAGCCATGTGGTGGTGCAATTCCACTGTGCATGGTTGGTGAGTTTGATCTTCCTTTGGATATCATCGACAGACGTGTTACTAAAATGAAGATGATTTCGCCTTCAAATGTTGCAGTCGACATTGGCCAAACTCTTAAACCAGATGAGTATATCGGTATGGTACGTAGAGAAGTGCTCGACCAATATCTTCGTGAAAGAGCAGAGAAAGCTGGAACAAAGGTAATTAACGGTTTATTCTTGAAAATGGATATGCCAAAAGAGCCAAATGCGCCTTATGTTCTTCACTACAGTGCTTATGATGGCAAAACTGGTGGAGTTGGCACAAAAACCACGCTAGAGGTAGATGCAGTCATCGGTGCAGATGGTGCGAATTCTCGTGTGGCTAAAGCAATTGGAGCTGGTGATTATGATTACGCTATTGCATTCCAAGAGAGAATCAGAATCCCAGATGAGAAGATGGTTTACTATGAAAATCTTGCCGAGATGTACGTCGGAGATGACGTCTCGCCTGATTTTTACGGATGGGTGTTCCCAAAATGTGATCACGTCGCAGTGGGTACCGGTACAGTTACACACAAAGGAGATATCAAGAAATTTCAAGTGGCGACACGGCTACGTGCCAAGGATAAGATTCTAGGCGGAAAGATTATCAGAGTAGAGGCACATCCTATTCCAGAACATCCAAGACCCAAGAGGTTGGCAGACAGAGTAGCGCTTGTTGGAGATGCAGCTGGGTACGTGACAAAATGTTCCGGAGAAGGTATATATTTTGCTGCTAAGAGTGGCAGGATGTGCGCCGAAGCGATTGTTGAAGGATCCGAGAGAGGGACAAGAATGGTGAATGAAACGGATTTGAGAAATTATCTGGAGAAATGGGATAAAACTTACTGGCCAACTTACAAGGTTCTGGATATATTGCAGAAGGTGTTTTACCGATCGAATCCAGCCCG

>Pdu_TRINITY_GbssI Pdu_c27554_g1_i1 len=2450

CGTGTGCTCTTCCGATCTTACGGACTTCTTCTTCTCTCGTAACAACTCCTACTACTTGCTTGTGATCTGTGCTGCTGTGTCTCACTCTACTGTCAGTCAGTCTAGAGACAAAAAAGAGTGTATTTTGAGATTATCGAGAGTGACATATCTCTCTATTCTCCATCCTTGTGAACTCATCAGCATATAATAATCTCTCAAGTCTCAACTGACAAGAATATTACCATGGCGACCATAAGTGCGTCCCAGTTAGTATCACGAACCAATTGCGGAGGAACTGCACCAAAAGACATACTTAAACAAATCAATATGAAGAACCAAAGCATGACTCATTTAGGGTTAAGATCTTTGAGCAAAGTAGATTTCTTACAAATAAAAAACACTCAGAAAGCAGCTTCAAGGGAACTGAGGAGGAACGCATTCCAGACAGACAAAGGTACTGCCTCAGGAGAGGTTAAATGTTCTAGTGGAATGAACTTGGTCTTTGTTGCTGCTGAAGTAGGCCCCTGGAGTAAAACCGGTGGTCTTGGTGATGTTCTTGGAGGCTTACCACCAGCCATGGCGGCAAATGGACATCGTGTTATGACAGTCTCTCCTCGTTATGATCAGTACAAAGATGCTTGGGATACCAATGTGTCAGTTGAGATGAAAATTGGGGACAGAATTGAAACCGTTCGTTTCTTTCACTGTTATAAGCGTGGGGTTGACCGTGTATTTGTGGATCATCCACTGTTTCTTGAGAAGGTATGGGGTAAAACTGGATCAAAAATCTATGGTCCGAATGCTGGATCAGATTACCAAGATAACCAGCTTAGATTCTGTTTGCTTTCTCTGGCAGCTCTAGAAGCACCAAGAATTCTGAATCTCAACAGCAACAAATATTTCTCTGGAACATACGGAGAGGAGGTCGTGTTCATCGCTAATGATTGGCATACGGCCCTTCTACCATGCTACCTTAAAAGTATATACCAATCTAAAGGGATGTACGAAACTGCCAAGGTTGCTTATTGCATCCACAACATTGCTTACCAAGGAAGATTTTCTTTTTCCGATTTTGCGCTTCTCAATCTTCCGGATTCATTTAGGAGTTCTTTTGACTTCATTGATGGGTATGAGAAGCCTGTTAAAGGAAGGAAGATTAACTGGATGAAGGCTGGAATATTAGAGGCAGACAAGGTCGTAACTGTGAGTCCATACTATGCTGAAGAACTTGCTTCAGGCCCGGATAAAGGTGTGGAACTGGATAGTATTATACGCAAAGCTGGCTGTCTGGGTATTATAAATGGAATGGATGTTCAGGAGTGGAACCCATTAACTGATAAATATATCGACGTCAAATATGATGCCTCAACTGTGACCGAGGCGAAGGCTCTTCTGAAAGAAGCACTGCAGGCAGAAGTAGGGTTACCTGTTGACCGTAATATTCCTGTCATAGGTTTCATTGGTAGACTTGAGGAGCAAAAGGGTTCAGATATTTTGGCAGCTGCAATCCCTGAATTTATTGACGAAGAAGTGCAGATAATTGTCCTCGGTACTGGCAAAAAGCCCATGGAAAAACAGTTGGAGCAGTTAGAGACTATGTATCCTGAAAAAGCTAGAGGAGTGGCGAAGTTCAACGTTCCCTTGGCTCATATGATCATAGCTGGGGCTAATTTTATGATGATCCCAAGCAGATTTGAACCATGTGGACTCATCCAGTTACATGCTATGCGTTACGGAACGGTGCCTATTGTTGCCTCGACGGGTGGATTAGTCGACACTGTCAAAGAAGGTTTCACGGGATTTCAAATGGGAGCTTTTAATGTCGAATGCGAGGCGATAGACCCATTAGACGTGACAGCAATCGCCACAACCACCAAAAAGGCCCTTGAAACCTATGGAACTCCAGCTTTAACAGAAATGATCCAGAACTGCATGGCTCAAGACCTCTCTTGGAAGGGACCTGCCAAGAATTGGGAGGACATGTTGCTGGGCTTGGAAGTTGCAGGTAGTGAACCAGGAATCGAAGGCGATGAAATCGCTCCTCTTGCTAAAGAAAATGTTGCCACTCCTTGAGTGACAGAACCGCTCCAGTGCAACTTAGAGGAGCTATTTCACAATCTAACCCAGGTGATTCCGTTGCTGCACCGACAAGGCTCTGATTACTTCGATCATCTCTAAGATTTGATAAAATTTACCCACCGCGAGTTACATAGTCAGTGGAAGAGGTTATGATGTGTATATGAAGTGGCTTTGCTGGAAAAGAGATACCGCTACTCATTTTCCTTTTCTTCTTTTCTTTTGATTTAAGCAATCTACGGCACTCAATGTCCAATGCGGATGCCAAGAGCGTAGGCTAGCTAGCTCTAGATCTTGTTTTAGTCAAGAAGAACTTTATATAAATAAAAACGGCTTTTTCACCTTCTGCTTCAGTTAAATTTCCAAAGCCTGAAGTTCTAGGTAGATGGCCCACG

>Pdu_TRINITY_Hmgs Pdu_c38849_g1_i1 len=1379

GTGTGCTCTTCCGATCTATTCTCCCTTCCTTCTCTTCCCCTCTCTCTCCAAAAACAGCTCGATCGATCGATCGATCTAGGGTTCTCTGAAGGAATTCAAAGAAATTATCTAAAGATGGGTTCAAATCAAAAGGATGTTGGAATTCTTGCTATGGAAATGTATTTCCCTCCTACTTGTGTTCAACAGGAAGCATTGGAGGCTCATGATGGTGCCAGTAAAGGAAAATACACCATCGGGTTGGGACAAGATTGCATGGCGTTTTGTTCCGAGTTGGAAGATGTCATTTCAATGAGTTTGACAGCCGTTAGCTCTTTACTCGAAAAGTATAAAATTGACCCTAAACAAATTGGACGCCTGGAAGTCGGCAGTGAAACCGTGATTGACAAGAGCAAGTCCATAAAGACCTTCCTGATGACAATCTTTGAGGAATGTGATAACACTGACATTGAGGGTGTTGACTCCACCAATGCGTGCTATGGAGGAACTGCAGCTTTGTTCAACGTTGTCAATTGGGTGGAGAGTAATTCCTGGGATGGACGCTATGGTCTTGTTGTGTGCACTGATAGTGCGGTCTACGCAGAAGGACCTGCCCGCCCCACTGGTGGAGCAGCTGCTATTGCTATGTTGATTGGTCCAGACGCACCAATTGCCTTCGAAAGCAAATATAGAGGATCCCATATGTCCCATGTCTATGATTTTTACAAGCCCAACCTAGCCAGTGAATACCCGGTCGTTGACGGAAAACTTTCACAAACTTGTTACCTTATGGCACTCGATTCCTGCTACAAACGCTTTTGCGCTAAGTACGAAAAATGTCAGGGTAAGCAATTCTCGATGGATGATGCAGACTACTTTGTATTTCATTCTCCTTACAACAAGCTTGTGCAGAAAAGTTTTGCTCGATTGTACTTCATGGATTTCCTAAGAAATGCCAGCTCTGTTGACAAGGCTTCAAAAGAAAAGCTTGCACCATTTGCAGCTCTATCCGGTGAAGAGAGCTACGCGAGTCGGGATCTTGAGAAGACATCCCAAACTGTGGCAAAGAATCTTTACGCTGAGAAGGTTGGACCTACCACTTTGTTGCCCAAACAAGTCGGTAATATGTACACTGCATCACTCTATGCGGCATTTGCTTCTCTTCTTCACAACAAAAATAGCTCACTGGATGGTAAACGTGTAATAATGTTCTCTTACGGAAGTGGTTTAAGTTCCACAATGTTCTCATTCCGAATGAGGGAATGCTCACATCCATTCAGCTTATCAAACATTGCAACTGTTATGGATGTCGCTCACAAGTTGGAGTCCAGACATGAGGTTACTCCTGAAAAATTCGTCGAAACCATGCAACTCATGGAGCACAGGTACGGAGGGAAAGACTT

>Pdu_TRINITY_Nia Pdu_c45852_g1_i1 len=3006

TTGCTTGCAAAAGCAAATCGGAAAAAAGAGAAAACATAGGAGTAGAAAACAGAGAGAAAGTGAAACGAGAGGAGGAGCATTCAAAGAGAGATTTTTTTCTTTCTTCTTTCTTTTGATTAGTCTCTAATGGCGGCTTCTGTTGAAAATCGACAGTACAACAGTCGACTTGAATCAGGTTTAGCAGTTCGTGGATTTAAAACCAATCCCACTCATCGATCTGATTCGCCTATCCGTGGCTGTAATTTTCCCATGTCGACTACCGAGCGTCTTTCTATACAAGCAAAACCGGTCATTGAAGATTTATCTAGCGACGACGATGAAGATCAAAACGAGCAGTGGAAAGATTTCGGCAAAATAAACAATGAAATAGAGCCGTCTGCTATTGACTCGAAAGATGAAGGAACTGCTGATAATTGGATTGAACGCAACCCTTCCATGATTCGGTTAACAGGAAAGCACCCATTCAACTCAGAAGCACCATTAACTCGTCTCATGCATCACGGGTTCATTACACCTGTTCCTCTTCATTACGTTCGGAATCACGGAGCTGTTCCAAAACTTAAATGGGAGGATTGGACAGTTGAGATTTGTGGATTAGTAAAGCGTCCGTTACGTATCACCATGGATCAGCTCATTAATGATTTCGCAAGCCGTGAGTTTCCGGCTACACTTGTCTGTGCAGGTAACAGACGTAAAGAGCAGAATATGATAAAGCAAACAATCGGGTTTAATTGGGGAGGTGCGGGAGTATCTACTTCAGTATGGCGTGGTGCACGTTTATGTGACATATTAAAGCGTTGTGGAATCATGAGTCGTAAGAAAGGTGCTCTAAATATTTGTTTCGAAGGAGCTGAAGATTTACCCGGTGGCGGTGGATCTAAATATGGAACTAGCATAAAGAAGGAATTCGCAATGGATCCTGCTCGTGATATAATTCTAGCTTACATGCAAAATGGTGAACGTCTGTCTCCTGACCACGGATTTCCAGTCCGTATGATAATTCCGGGTTTCATCGGAGGAAGAATGGTGAAGTGGTTAAAGAGGATTATCGTCACAACTCCAGAATCCCAAAACCATTATCATTTCATGGATAATCGGGTTCTTCCTTCTCATGTCGACGCTGAACTGGCCAACTCCGAAGCATGGTGGTATAAGCCAGAGTACATCATCAATGAGTTGAACATAAACTCTGTGATTACAACTCCTTGTCACGAAGAGATTCTTCCTATAAATGCTGCAACAACACAAAGAGCTTACACATTAAGGGGATACGCATATTCCGGTGGCGGTAAAAAGGTAACTCGTGTCGAAGTAACAATGGATGGCGGAGAGTCATGGTTAGTGTGTACTTTGGATCATCCAGAGAAACCAAACAAGTATGGCAAATATTGGTGCTGGTGTTTCTGGTCACTAGAAGTTGAAGTTCTCGACTTGCTTGGTGCCAAAGAGATTGCAGTCAGGGCTTGGGATGAAACTCTAAACACTCAGCCTGAAAAGCTTATCTGGAACGTCATGGGTATGATGAACAACTGCTGGTTCCGAGTCAAAACAAATGTGTGCAAACCACACAAAGGAGAAATCGGTCTAGTTTTCGAGCATCCAACTCAGGCTGGAAACCAACCAGGTGGATGGATGGTTAGACAGAAACATCTTGAAAACTCTGAAGGTCCACATACCCTCAAAAAGAGTTTGTCCACTCCTTTCATGAACACCACATCAAAAATGTATTCGATTTCCGAGGTGAAGAAACACAACTCGCCTGACTCAGCTTGGATCATTGTTCACGGACACGTTTATGACTGTACCCGGTTCTTGAAAGACCATCCAGGTGGTTCCGATAGTATTCTCATCAATGCAGGTACCGATTGTACCGAAGAATTTGATGCAATTCACTCTGACAAGGCCAAAAATTTGCTTGAAGATTTCAGAATCGGTGAACTCATTGCAACTGGTTACACTTCTGACTCGTCATCTTCACCAAACAACTCAGTTCATGGTTCGGGCAGCTTGAGTCACCTTGCTCCTATCAAAGAAATAGGCATTGCGAAACCAGTAGCTCTTGTTGGCCGAGACAAAATACAATGCAAACTTATTGCCAAGACCTCGATTTCCCATGACGTTCGTGTCTTCAGGTTTGCATTGCCTTCAGCTGATCAAGTCCTTGGTTTGCCAGTTGGGAAGCACATATTTGTATGTGCCCATGTTAACGATAAGCTATGCATGCGTGCTTACACTCCATCAAGCACAGTTGATGAAGTTGGTTACTTTGAGCTGGTTATCAAAGTCTACTTCGGAGGAGTTCACCCCAAGTTCCCCAATGGAGGTATTATGTCTCAGCACTTGGATTCTCTTCCTCTTGGTGGAACTATTGACGTTAAGGGTCCGTTGGGACACATTGAATACACTGGCCGTGGTAATTTCACTGTTCATGGCAAACACAAGTTTGCCAAACGACTTGCAATGCTTGCTGGTGGTACAGGAATCACACCAATATATCAGATTGTGCAAGCAATATTGAAAGATCCAGAGGATGAAACGGAGATGCACATCGTATTCGCAAACAGGACTGAAGATGACATCTTGTTGATGGATGAACTTGATTCTTGGGCAGAGAAGCATGAAAGGCTGAAAGTATGGTACGTTGTTGGTGAAACATTGAGGGAAGGATGGAAGTACAGTGTAGGGTTCATAACTGAAGAGGTCCTACGGGAACATATTCCTCCGGCATCGGAAGATACATTGGCGTTGGCTTGTGGACCACCACCTATGATTCAATTTGCAGTGACTCCAAATTTGGAGAAGATGAATTATGATGTGAAGAATTCATTGTTGGTGTTCTGATGAATCTGCTGAAACTACAAAGAGGAGCACATATGATCAAATTTAAGAAAATGATAAGGGGAGGACAAATTATAAGGAAGTTCTCATTAGTTTATTGTATATAATTAGTGTTGAAGGATGTAATGGGTTTAGTATGTTCTTAAAAGTAAGAGATCGGAAGAGCGT

>Pdu_TRINITY_Eif3E Pdu_c36930_g2_i1 len=1643

GTGCTCTTCCGATCTGTTTTATTTAGGGTTTCTCTCTTTCTTCCTCCACCACCACTACCGCGCCGCTCCTCCGTCTTCCTCAGAGGTTGTTCTTCAGCCTCTACCTCAACCACAAGAACCAATCATGGCGAATGATGATCTGACATCAAGAATAGCACTAAACTTAGATCGACATCTCGTTCTACCACTCCTTGAATTTCTTCAGGAGAAACAAATCCATCCCGATGTTCAAATCCTCAAAGCTAAAGACGCTCTTCTCAGCCAGACCAATATGGTTGATTACGCTATGGATATCTACAAGTCTCTTCACCAAACTGATGATGTTCCTCAAGAGATGATTAATAGGCGTGCTGAAGTGGTTAGGAGGTTGAAATTACTTGAAGATGCAGCTGAATCGATTATTACGTTCCTGCAAAGCCCAAATATGATTAATGAACCACGTCCTGATAAGCAATATGTTCTTCAGTTGCTACAGGGATACCAGATTGGTGCTGAGCAAATTGAGGCATTGTATCAGTATGCCAAATTTCAATTTGAATGTGGGAATTACTCGGGTGCAGCTGATTATTTGCATCAGTACAGAGCAATTTGTACCAATCCTGATAGAATTTTGAATGCATGTTGGGGAAAGCTTGCATCTGAGATTTTGATGCAGAACTGGGATGAGGCTCTTGATGAGCTTAATCGGTTGAGGGAGATTATTGAATCAAAGAATTTCAAGAACCCATTAACTTTACTCCAGAATAGAATCTGGTTGATGCATTGGAGTCTCTTCATATTTTTCAACCATGAGAATGGAAGAAACGGTATCATTGATCTATTTCTCAATGATAGGTACATGAATGCACTCCAAACAAATGCACCCCATCTTTTACGTTATTTGGCAACTGCATTGATTGTCAACAAAAAGAGAAGACCTCAACTCAAAGAGCTGATCAAAGTCATCCAGCAAGAGCAATCTGCATACAAGGATCCAATCACTGAGTTCTTATACTGTTTGTATGTCAACTATGACTTCGATGGTGCACAACAGAAGCTCCGAGAATGTGAGGATGTGATTTTGAATGATCCATTCCTTGGTAAAAGAGCTGAAGAAGGAAACTTTAGCACCGTTCCTTTGAGGGATGAATTTCTTGAAAACGCTCGTCTCTTCATCTTTGAGGCTTACTGCAGAATCCACCAGCGCATTGATATAGGAACTCTTGCCGAGAAGTTGAATATGAATTTTGATGAGGCAGAGAGATGGATAGTGAATCTTGTTCAGAATGCCAAGCTTGATGCTAAGATTGATTCTAAGCTAGGAACTGTTGTAATGGAGCCCATTTCTCTCAATGTGTATGAGCAGATCATTGAGAATACCAAATCAATTTCTTTGCGTACCTACAAATACGTAAATCTTGTTTTAGAATCAGCAGCACCACCGGTAACGGCTGCTCGTTAAAGAAGAGGTAGTAGTATCGCGAGAAAATTTGTGTTGGAGCTCTTGAAGTTAACTTTTGGGGAAGGAAGAAAATTTGGTCTCAGTATTTTGTTTTTGTTCACTTGTTTTTCAGTTTCATCTAAAGCATTAGTTGCAGGAGAAGTTGTAAGATTAATATACTTCTTTTTAGGGTTCTTTGAACGATAAGATACAGTTTATGTTA

>Pdu_TRINITY_AroB Pdu_c34912_g2_i1 len=1479

GTGCTCTTCCGATCTGCTTCAACTACTACTACTTCTTCTTCTCGTCTGGTTTGAAAAATAATCAGAAGAAGTTTTGTTTCCCTCTCTAATTCTTCCCCAATTTTTTGATGGCTTCTTCTACTACTTCTTCACCTTCTCTTAATTTGGCTATTAAGAATTCTTGTCTACGAGAAAATTCTGATCTATTTCTCAGAGGGAGGAATTTATCACTGAAATCTTCATTTATACATTCAAGTTCATTACGATTGAATAGAATTGGATCTTTAAAAATTCAATCGAAGAGTAGAATGATATCTGCAAATTCAGCTCAAGTAATGGAACAATCTTCAATTAGGGCACCAACTATCGTTGATGTTGATTTGGGTAATAGAAGTTATCCGATCTATATTGGTTCTGGACTTCTTGAACAACCTGATCTTCTTCAGAGGCACGTACATGGGAAGAGAGTACTCGTGGTAACTAATACAACAGTTGCGCCATTATACCTAGATAAAGTTGTCAGCGCTTTAACTCATGGAAATCCGACTATTTCGGTTGAGAGTGTGATTTTGCCAGATGGAGAGAAATATAAAAATATGGATACTCTGATGAAAGTTTTTGACAAGGCCATAGGATCTCGATTTGATCGCCGTTCCACATTTGTCGCTCTTGGAGGTGGTGTTATAGGTGACATGTGTGGTTACGCTGCAGCTTCTTTCCTACGTGGTGTAAACTTCATTCAAATTCCAACTACTGTCATGGCACAGGTGGATTCCTCTGTTGGGGGAAAAACTGGAATCAACCACACACTGGGGAAAAACTTGATTGGTGCTTTTTACCAGCCCCAATGTGTACTTGTAGATACAGATACATTACATACACTGCCTGAGAGGGAATTAGCTTCTGGTTTTGCTGAAGTTATAAAATATGGACTCATTCGGGATGCTGAATTCTTCGAGTGGCAAGAAAAAAATATGGACTCTCTGTTGGCGAGGGATCCCAGTGCACTAGCATATGCAATCAAGCGTTCATGCGAAAACAAGGCAGAGGTTGTATCCTTGGACGAGAAGGAAGGTGGAGTGAGAGCGACGCTGAACTTGGGTCATACCTTTGGACATGCAATTGAAACAGGATTTGGCTATGGTCAATGGCTCCATGGAGAAGCTGTAGCAGCTGGAACGGTCATGGCAGTAGATATGTCTTACCGTCTAGGTTGGATTGATGAATCAATTGTGAAGCGGACATACGATATTTTACAGAAGGCTAAGCTACCGATTACCCCACCAGATGTGATGACTGTAGAGATGTTCAAGTCTTTCATGGCTGTTGATAAGAAGGTGGCTGATGGACTACTAAGGCTCATCCTTCTCAAAGGTCCTCTCGGCAATTGTGTTTTTACAGGTGATTACGATAGAAAAGCCCTAGACGAGACCCTTCGTGCATTCTGCAAGTCTTAATCTAATGCCAACCTCAGAGCCGTCTTCTGGACTAGGCCAGCTA

>Mco_Agt1 OVA09444.1

ATGGACTATGTTTATGGACCTGGGAGGAACCATCTCTTTGTTCCTGGACCTGTCAACATCCCGGAACCTGTCATTCGTGCCATGAACAGAAACAATGAGGATTATCGTTCCCCTGCAGTGCCAGCCATGACCAAAACCCTCCTAGAAGACATTAAGAAGATCTTCAAGACCACTTCCGGGACCCCGTTTATGATCCCCACCACAGGTACTGGTGCATGGGAAAGTGCACTTACAAATACGTTATCTCCTGGTGATAGAACTGTTTCTTTCCTCATCGGACAATTCAGTCTGCTCTGGATTGATCAGCAGCAGCGTCTCAAATTCAATGTCGATGTCGTAGAAAGTGAGTGGGGGCAAGGTGCTAATCTTGAAATTCTCGCATCGAAGTTAGCAGCGGACACTGCTCATACTATTAAAGCTGTTTGCATTGTTCATAACGAGACTGCAACTGGAGTTACCAATAACTTGGCTGCTGTAAGAAAAATACTTGATCACTACAGGCATCCTGCATTGTTTCTCGTTGATGGAGTATCGTCAATATGCGCTCTTGACTTCCGTATGGATGAATGGGGAGTAGATGTAGCTTTAACCGGCTCACAGAAAGCTCTTTCTCTGCCCACTGGGATGGGAATAGTGTGTGCAAGCCCAAAAGCACTAGAAGCATCCAAGTCGGCAAAATCAGTGAGAGTGTTCTTTGACTGGAACGATTACTTGAAGTTCTATAAATTAGGAACTTTTTGGCCATACACTCCTTCAATCCAGCTTTTATATGGCCTGAGAGCTGCTCTGGATCTCATATTTGAGGAAGGTCTCGATAATGTGATTGCTAGGCACAGCCGTCTGGCCAAAGCAACAAGGCTTGCTGTTGAGGCATGGGGTTTGAAAAATTGTACACAGAAGGAGGAATGGTTTAGTGATACTGTGACTGCAGTGGTTGTTCCACCTTACATTGACAGTTCAGAAATTGTTAAAAGGGCATGGCAGAGATACAACCTAAGCTTAGGTCTGGGTTTGAATAAAGTGGCTGGAAAGGTCTTCAGAATAGGGCATCTTGGCAACCTCAATGAGCTCCAATTGCTGGGTTGTTTAGCTGGAGTGGAGATGATTCTCAAAGATGTCGGATACCCGGTTAAGCTGGGAAGTGGAGTTGCAGCGGCCTGCGCATATCTGCAGAACAGCATTCCGATGATCACTTCTAGGATTTAA

>Mco_At103 OVA13800.1

ATGTCATCACAACCCACTGCAACAACAACAAAATCAACAACAGCAAAACCCAAAAAGGCAATCAAGGAAACCCTACTTGCACCAAGATTTTACACCACTGATTTTGATGAAATGGAAACCCTTTTCAACACTGAGATCAACAAAAACCTAAACCAGTCTGAGTTTGAAGCACTTCTACAGGAGTTCAAAACTGACTACAACCAGACCCATTTTGTCAGAAACAAAGAATTCAAAGAAGCTGCAGATAAGATGCAAGGCCCTCTTAGACAGATCTTTGTAGAGTTCTTGGAGAGATCTTGTACTGCTGAGTTTTCTGGGTTTCTTCTCTACAAAGAGCTGGGAAGAAGACTTAAGAAAACCAATCCAGTGGTGGCAGAGATTTTCTCTCTCATGTCCAGGGATGAAGCTAGGCATGCTGGATTCTTAAACAAGGGTCTATCTGATTTCAATTTAGCACTGGATCTGGGATTTCTTACAAAGGCTAGAAAATATACATTCTTCAAGCCCAAATTCATTTTTTATGCTACATATTTATCCGAAAAAATTGGATACTGGAGGTACATAACCATTTACAGGCATCTAAAGGCCAACCCAGAATTCCAAGTTTACCCAATTTTCAAGTATTTTGAGAACTGGTGCCAAGATGAGAACAGACATGGAGATTTCTTCTCTGCATTGATGAAGGCTCAACCTCAGTTTCTTAACGACTGGAAGGCGAAATTGTGGGCTCGATTCTTCTGCTTATCGGTTTACGTGACAATGTACTTAAACGACTGCCAGCGAACTGCTTTCTATGAAGGGATCGGACTTAACACCAAAGAATTCGACATGCACGTCATAATTGAGACAAACCGCACCACTGCGAGGATTTTCCCAGCTGTTCTAGACGTTGAGAACCCCGAGTTCAAGAGGAAGTTGGATCGCATGGTCGAAATCAACCAGAAAATTATCGCGATCGGAGAGAGTGATGATATACCATTGGTAAAGAACCTAAAGAGGATCCCACACGTCGCAGCATTGGCGAATGAGCTCTTGGCTGCCTATCTCATGCCTCCTATCGAGTCTGGATCTGTGGATTTTGCAGAGTTTGAGCCACAAGTTGTTTATTAA

>Mco_ChlP OVA09813.1

ATGGCGTCCATTGCCCTAAATGCCCGCTTCACCACCACCACATGTGACCGTAAAACAACCGAACACAGCCGCGAAACACTCAAAACCTTCATTGGCCTCCGCCAAACCACTCTAGCAAACCACAATTTTCACTCTCTAACCAAGAACCTACCATCTCAACAAATCAACCGTAGACTCAGTATTAAAGCCGCAAAATCTAGCCCTAAAATCACAGGCCGAAATCTACGAGTTGCAGTTATCGGTGGAGGTCCAGCTGGTGGCGCAGCCGCCGAAACCCTAGCCAAAGGCGGAATCGAAACTTACTTAATCGAACGGAAACTCGACAACTGTAAGCCATGCGGTGGTGCAATCCCACTCTGTATGGTCGGTGAATTCGATCTTCCACTTGATATCATCGATCGACGAGTCACCAAGATGAAGATGATCTCACCGTCCAATGTCGCAGTCGACATTGGCCAAACCCTGAAACCTGACGAGTACATCGGTATGGTACGCCGAGAAGTACTGGATGCATACCTCCGTGAACGAGCAGAAAAGTCAGGCGCAACTGTAATTAACGGCCTGTTCCTAAAAATGGATTTACCAAAAGAACCAAACGCGCCCTACGTTCTGCATTACAGTGCCTTCGACGGAAAAACTGGTGGAGTTGGCGAACGGAGCACGTTAGAGGTCGACGCCGTAATCGGCGCAGACGGCGCGAATTCTCGCGTCGCTAAAGCCATCAACGCCGGTGACTACGATTACGCCATCGCTTTTCAAGAGCGAATCAAAATCCCAGACGACAAAATGGTGTACTACGAAAATCTCGCCGAGATGTACGTAGGTGATGACGTGTCACCCGATTTCTACGGATGGGTCTTCCCGAAATGCGACCACGTGGCGGTGGGTACGGGTACGGTGACACACAAGGGGGACATCAAGAAGTTCCAAGTAGCGACACGGAACCGCGCGAAAGATAAGATTTTGGGCGGGAAAATCATAAGGGTAGAGGCGCATCCTATCCCGGAGCATCCAAGGCCGCGCAGGTTATCAGGGAGAGTAGCCCTAGTAGGGGACGCGGCCGGGTACGTGACCAAGTGTTCAGGGGAAGGGATCTATTTTGCTGCGAAGAGCGGGAGGATGTGCGGGGAAGCCATTGTTGAAGGGTCTGAGAATGGGAAAAGGATGATTGAAGAGGGAGATTTGAGGAATTATTTGGAGAAATGGGATAAGACGTACTGGCCAACTTACAAGGTGTTGGATGTGTTGCAGAAGGTGTTTTACAGGTCGAATCCTGCGAGGGAAGCGTTTGTGGAGATGTGTGCGGATGAGTATGTGCAGAAGATGACCTTTGATAGTTATTTGTATAAGAGGGTGGTGCCTGGGAATCCATTGGAGGATTTAAAGTTGGCGATTAATACGATTGGGAGTTTGGTTAGGGCAAATGCACTTAGAGGAGAGATGGATAAGATCAGTCTATGA

>Mco_GbssI OVA04705.1

ATGGCGACCGTAACTGCTTCCCACTTTGTATCACGAACCTCGCATGTCAACCGCGGCGTGACTGAGCCTAAAGACATACTGAAACAAATCAGCCTGAAGAACCAAAGTATGACTCATTCTGGGTTGAGGTCTCTGAGCAAAGTGGATTTCATTCACATGAAAAACACTGCAAAAGCAACTTCTAGGCAATTGAGGAGGAATGCTGTCAAAACTGACGAAGCCAGTGTATCAGGAGATATTAGATGTTCAAGTGGGATGAACTTGGTCTTTGTGGCAGCAGAAGTAGGTCCATGGAGCAAAACTGGTGGACTTGGTGATGTTCTTGGAGGTCTGCCACCAGCCATGGCGGCTAATGGCCACCGTGTTATGACGGTCTCTCCTCGCTACGATCAGTACAAAGATGCATGGGATACAAATGTCACAGTTGAGATGAAAATTGGGGATAGAATTGAAACAGTCCGCTTCTTCCACTGTCACAAACGGGGGGTTGATCGTGTTTTTGTGGATCATCCGTGGTTCCTTGAGAAGGTATGGGGAAAGACTGGATCAAAAATCTATGGTCCGAAAGCTGGATCAGATTACCAGGACAACGAACTTAGATTCTGTTTGTTATCTCTGGCTGCTCTGGAAGCACCAAGGATTCTGAATCTCAACAGCAATAAATATTTCTCTGGAACATCTGGGGAGGATGTCGTGTTCATCGCTAATGATTGGCACACTGCCGTTCTACCATGCTACCTTAAAAGTATGTACCAATCTAAAGGGATATACAAAACTGCAAAGGTTGCTTTCTGCATACACAACATTGCCTATCAAGGAAGATTTGCGTTTTCTGATTTTTCACTTCTCAATCTCCCCGATTCGTTTAGGAGTTCGTTTGACTTTATCGACGGGTATGAGAAGCCGGTGAAAGGGAGGAAAATTAACTGGATGAAGGCTGGAATATTAGAATCAGACAAGGTCTTAACTGTGAGCCCATACTATGCTGAAGAACTTGCTTCAGGCGAGGATAAAGGTGTGGAATTGGATAGCATCATCCGCAAAACTGGCATTACCGGTATTGTGAATGGCATGGACGTCCAGGAGTGGAACCCATTGACCGACAAATATGTTGATGTCAAATATGATGCTACAACTGTCATGGATGCAAAGCCTCTTCTGAAGGAAGCACTGCAAGCAGAAGTCGGGTTGCCTGTAGACCGAAATATCCCTGTCATAGGCTTCATTGGTAGACTTGAAGAGCAGAAGGGTTCAGATATTTTGGCCGCTGCCATTCCTGAATTTACCAACGAAAATGTGCAGATAATAGTCCTTGGAACCGGCAAAAAGCCCATGGAGAAGCAACTGGAGCAGTTAGAGGTTATGTATCCTGATAAGGCTAGAGGAGTGGCGAAGTTCAATGTTCCCTTGGCTCATAAGATCATAGCTGGAGCTGATTTTATGATGATCCCAAGTAGATTTGAACCATGTGGACTCATTCAATTACATGCTATGCGATATGGAACAGTCCCTATCGTTGCCTCGACTGGTGGACTAGTTGACACTGTCAAAGAAGGTTTCACCGGATTTCAAATGGGAGCTTTCAATGTTGAATGCGAGGAAGTCGATCCAGCAGACGTGACTGCAATTGCCACAACCACCAAAAGGGCACTTGCAACCTTTGGTACTCCAGCTTTAGCAGAGATGATTCAAAATGGCATGGCTCAAGACCTTTCCTGGAAGGGACCTGCCAAGAAGTGGGAGGAGTTGCTGCTAAGTTTGGAAGTTGCTGGTAGCAAACCAGGAATCGAAGGCGAGGAAATCGCTCCTCTTGCTAAAGAGAATGTTGCCACTCCTTGA

>Mco_Hmgs OVA12377.1

ATGGGTTCGAACCAGAAAGATGTCGGAATTCTCGCCATGGAAATCTACTTCCCTCCTACTTGTGTTCAACAGGAAGCATTGGAGGCTCATGTTGGTGTCAGTAAAGGAAAATACACCATTGGGCTGGGACAAGATTGCCTGGCATTTTGTTCAGAGTTGGAAGATGTCATTTCGATGAGTTTGACAGCTGTTACTTCCCTCCTTGAAAAGTATGGGGTTGACCCTAAACAAATTGGTCGCCTGGAAGTAGGCAGTGAGACTGTGATAGACAAGAGCAAGTCCATAAAGACCTTCCTGATGCAAATCTTTGAGGAATGTGATAATACCGACATTGAAGGAGTTGACTCAACAAATGCATGCTATGGAGGAACTGCAGCTTTGTTCAACGTTGTGAATTGGGTGGAGAGTAGTTCCTGGGATGGACGTTATGGACTTGTTGTGTGCACTGATAGTGCGGTCTATGCAGAAGGTCCAGCTCGGCCTACTGGAGGAGCAGCTGCTATTGCTATGTTGATAGGTCCAGATGCACCAATTGCGTTTGAAAGCAAATTTAGAGGGTCGCACATGTCTCATGTCTATGATTTTTACAAGCCCAATCTTGCAAGTGAATACCCGGTTGTTGATGGGAAACTTTCACAAACTTGCTATCTCATGGCGCTTGATTCTTGCTACAAACGTTTATGTGCTAAGTACGAGAAAATCCAGGGGAAGCAATTTTCACTTTCTGATGCAGACTACTTTGTGTTTCATTCTCCGTACAACAAGCTTGTGCAGAAAAGCTTTGCTCGATTATGCTTCATCGACTTCTTGAGGAATGCCAGCTCTGTTGACGAGGCTTCTAAAGCAAAACTTGCACCATTTTCTGCTCTATCCAGTGAGGAGAGCTACGCAAGCCGGGATCTTGAAAAGGCATCCCAGACTGTTGCAAAGAGTCTTTATGATGAGAAGGTGCAACCTACCACTTTGATACCTAAACAAGTCGGCAATATGTACACTGCATCACTTTATGCAGCATTTGCATCTCTTCTTCACAACAAGAATAGCTCACTGGATGGGAAACGTGTAATAATGTTTTCTTACGGGAGCGGTTTATCTTCAACAATGTTCTCATTCCGACTTAGGGAAGGTTCACATCCATTCAGCTTGTCAAACATTGCAACTGTAATGGATGTCGCCAGGAAGTTGGAATCAAGACACGAGGTTACTCCTGAGAAATTTGTCGAAACTATGCAATTAATGGAGCACCGTTATGGAGGCAAGGACTTTGTAACGAGCAAGGACAGGAGCCTTATACCATCAGGCACGTTCTATCTCACCCAGGTCGACTCCATGTACCGGAGATTCTACTCCAAGAAGGCGGGAAATGAAACCACCACCGCCGACAATGGTACATTGCTCAACGGTCATTGA

>Mco_Nia OVA08942.1

ATGGCGGCTTCTGTTGAGAACCGGCAATACGGTCGGCTCGAATCGGCCGGTCTAACATCGGTGGTTCGTGGTTTTAAAACAAACACGAACCACCGATCCGACTCTCCGGTTCGTGGCTACAATTTCCCTTCAAATTCATCAGATCATTTATCATCACCACGCACTCATAAAAAAACTTCTTTCATGGATGATTCTTCGAGCGAAGACGAAGAAGATCAGTACGAGTGGAAAGATTTCCGCAAAATAAACAACGAAATCGAGCCGTCCATTTTCGACCCGAAAGATGAAGGAACGGCGGATAATTGGATCGAACGTAACCCGTCCATGATCCGTTTAACAGGGAAACACCCATTCAACTCTGAAGCACCACTGACTCGGCTCATGCATCATGGGTTTATCACACCGGTTCCACTTCACTACGTTCGTAACCATGGAGCTGTTCCTAAAGGTAAATGGGAAGATTGGACGGTCGAGATTTGCGGGCTTGTGAAACGACCTTTACGACTCACTATGGATCAACTCGTTAACGAGTTCCCGAGTCGTGAGTTCGCTGCTACACTTGTTTGTGCTGGTAACCGACGTAAAGAACAAAACATGGTGAAACAAACGATAGGATTTAATTGGGGCGGTGCGGGTGTATCAACTTCTGTGTGGCGCGGAGCACGGTTACGCGACGTGCTTAAACGTTGTGGTATTATGAGTAAAAACAAGGGTGCCCTTAACATTTGTTTCGAAGGTGCTGAAGATCTGCCTGGTGGTGGTGGTTCCAAATATGGAACCAGTTTGAAGAGGGAGATGGCAATGGATCCTTCGCGAGACATCATATTAGCTTACATGCAGAACGGTGAACGTCTATCGCCTGACCATGGGTTCCCTGTACGGATGATCATCCCTGGATTCATTGGCGGAAGAATGGTTAAATGGTTGAAGAGAATCATTGTTACTACTCAAGAATCACAGAATCATTACCATTTCAAGGATAATCGTGTACTTCCTTCTCACGTGGATGCTGAACTGGCAAACGCTGAAGCTTGGTGGTACAAATCAGAGTACATCATCAATGAATTGAACATAAACTCTGTCATCACAACTCCTTGCCATGAAGAGATACTTCCGATCAATGCTGCAACTGCTCAAAGACCGTACACAGTCAAGGGTTACGCATATTCCGGGGGAGGTAAGAAGGTGACACGTGTGGAGGTGACAATGGATGGTGGAGAGACATGGCAAGTGTGCACCTTGGATCACCCAGAGAAGCCAAACAAGTATGGGAAATACTGGTGCTGGTGCTTCTGGTCACTGGAAGTTGAAGTCCTTGACTTACTTGGTGCCAAAGAGATTGCTGTTAGAGCCTGGGATGAAACCATAAACACTCAACCCGAAAAGCTCATCTGGAACGTCATGGGTATGATGAACAACTGCTGGTTCCGAGTTAAGACAAACGTGTGCAAGCCACACAGGGGAGAAATCGGTATAGTTTTCGAACACCCAACTCAGGCTGGAAACCAACCCGGTGGATGGATGGTTAGACAGAAGCATCTTGAATCCTCTGAAACTTCACAGACGTTGAAGAAGAGCTTGTCCACTCCTTTCATGAACACCTCTTCAAAGATGTACTCGATGTCCGAGGTGAAAAAACATAGCTCAGCTGACTCGGCTTGGATCGTTGTCCACGGTCACGTTTACGATTGCACTGGCTTCTTGAAAGATCATCCTGGTGGTACAGATAGTATTCTCATCAATGCAGGTACTGATTGCACCGAAGAATTCGATGCAATTCACTCCGATAAAGCCAAAAAATTGCTTGAAAGTTACAGAATCGGTGAACTCATTACAACTGGGTACACCTCTGACTCATCTTCACCAAACAACTCTGTCCACGGCTCTTCTACAGTTCTAACCTACCTTGCTCCCATTAAAGAAGTTGCGGTTTCTAAACCTGTAGCTTTGATACCACGTGAGAAAATCTCATGTAAGCTTATCTCCAAGACCTCAATTTCCCATGACGTTCGTCTCTTCAGGTTTGCATTACCTTCATCAGATCAAGTACTTGGGTTACCAGTTGGGAAACATATATTCGTCTGTGCCACCATTGACGATAAGCTCTGCATGCGTGCTTACACTCCATCCAGTTCTGTAGACGAAGTGGGTTATTTCGAACTCGTCATTAAAGTCTACTTCAAAGGTGTGCATCCGAAATTCCCTAACGGCGGCCTCATGTCGCAGTACTTGGATTCTCTTACTTTAGGTTCAACAATTGAGATCAAGGGTCCATTGGGACATATCGAATACACGGGTCGTGGTAACTTCACCGTTCACGGCAAACATAAGTTTGCTAAACGACTCGCAATGATTGCCGGTGGCACAGGTATTACGCCGATCTATCAAGTGATCCAAGCAATTTTGAAAGATCCAGAAGATGAAACAGAGATGTATGTTGTGTTTGCGAATCGAACAGAAGATGATATTCTGTTGTGGGAAGAACTTGATGCTTGGGCTGAGAAGTACTCAGACAGGTTGAAAGTTTGGTATGTTGTTGGAGAAACATTAAGAGAAGGATGGAAGTACAGTGTGGGGTTCATTACAGAACACATTCTAAGAGAACATATACCCGAAGCATCGGACGAGACGTTGGCGTTGGCTTGTGGACCACCACCAATGATTCAATTTGCAGTGACACCAAACTTGGAGAAGATGAACTATGATGTCAAGAATTCATTGTTGGTGTTTTGA

>Mco_Eif3E OVA12941.1

ATGGCGGACTACGACCTGACCCAACGAATCGCTCCTAACCTGGATCGGCATCTTGTTTTTCCTATCATGGAATTCTTGCAAGAGCGTCAGCTCCATCCCGATGACCAGATCCTCAAAGCCAAGATCGAACTTCTCAGCGAGACCAACATGGTTGATTACGCTATGGACATCTACAAGAGCCTTTACCACACTGAAGATGTCCCTCAAGAGATGATTGATAGGCGAGTCGAGGTAGTTGGGCGATTGAAATCTCTGGAAGAGTCCGCAGCTCCATTGATTTCGTTTCTTCAAAACCCTAATATGGTTCAGGAACTCCGACCAGATAAACAATACAATCTCCAGATGCTGCATGATCGATACCAGCAGATTGGTCCGGAGCAGATAGAGGCTTTATACCAGTACGCCAAATTTCAGTTTGAATGTGGAAATTACTCTGGTGCAGCTGACTTTCTGTATCAGTACAGAGCATTATGCACCAACACTGACAGGAGTCTGAGTGCATTGTGGGGAAAGCTTGCAGCTGAAATACTGATGCAAAACTGGGATATTGCTCTTGAAGAGCTTAATCGGCTGAGAGAAATAATTGATTCAAAGAATTTTAAATCGCCCTTGAATCAGCTGCAAAATAGAATCTGGCTGATGCATTGGAGCCTCTTCATATTTTTCAACCATGACAGTGGTAGAAATGGAATCATTGATCTGTTTCTTTATGATAGGTATCTAAATGCCATCCAAACAAATGCACCCCATCTTTTACGTTATTTGGCCACTGCATTGATTATTAACAAAAAGAGAAGACCTCAGCTCAAAGAGTTGATCAAAGTCATTCAGCAAGAGCAATCCTCTTACAAAGATCCTATCACAGAGTTCTTGCATTGTTTATATGTGAATTATGACTTTGATGGTGCACAACAGAAGCTTCGTGAGTGTGAGGATGTGATATTGAATGATCCCTTTCTTGGTAAACGCGTTGAAGAAGGAAACTTTAGTACTGTACCTTTGAGGGATGAATTTCTTGAAAATGCTCGTCTCTTCATCTTTGAGGCTTATTGCCGTATACATCAGTGCATTGATATAGGAACGCTTGCAGTGAAGTTGAATATGAATTATGACGAGGCTGAGAGATGGATAGTGAATCTTGTTCAGAATGCAAAGCTTGATGCTAAGATTGATTCCAAGCTGGGGACTGTTGTGATGGAACCCACCCATTTTAATGTATACGAGCAGATAATAGAGAATACCAAGGCACTCTCCTTGCGTACTTACAAATACGTAAATCTTGTTCTAGAATCTGCATCGGCACAGGCTGCACGATAA

>Mco_AroB OVA19598.1

ATGGCTTCTTCTGCAACTTCTCCTTCTCTTAGTTTCTCTATCGAGCTCTCTTCTACAGATCCCTGTCGCTCTCGCTCCGAGACCAGCGATCTCTTTCTCCGGTGCCAAAATCGCGATGTAAAATCGCTTAAATCCTCCTTCATTTCACCAGCTTCACTACGGTTGAATCGAATTGGATCTCCTTTATCTAGATGCGATCGAATTCGATCCAGTAAGATTTCCGCAAATTCAGCTCAAGTAATGGAGGCGCAATCTCCCTTTAAGACGAATTCTAGGGTTCCCACGATCGTTGACGTCGATTTGGGCAACCGAAGCTATCCGATCTACATCGGTTCCGGACTTCTGGAAGAACCTGATCTCCTTCAAAGGCATGTTCATGGGAAGAGAGTACTTGTTGTCACTAACACAACAGTTGCGCCATTATACCTTGATAAAGTTATTAACGCGTTAACTCATGGAAACTCGAGTATTTCAGTTGAGAGTGTGATTTTGCCAGATGGAGAAAAGTATAAGGATATGGATACCCTCATGAAAGTCTTCGACAAGGCCATAGAATCTCGATTTGATCGCCGCTCTACATTTGTGGCCCTTGGAGGTGGTGTGATTGGTGACATGTGTGGATTTGCTGCTTCTGCTTTTCTTCGTGGTGTAAATTTCATTCAAATCCCAACTACTGTTATGGCACAGGTGGATTCTTCCGTTGGGGGAAAAACTGGAATCAACCATCCACTAGGGAAAAACTTGATTGGTGCTTTCTACCAGCCCCAATGTGTACTTGTAGACACAGACACATTACATACGTTGCCTGAGAGGGAATTAGCTTCTGGTCTTGCTGAAGTTGTAAAATATGGGCTTATTAGGGATGCTGAGTTCTTCGAGTGGCAAGAAAAGAATATGCACCTGCTATTAGCGAGGGACCCCAGTGCACTGGCCTATGCTATCAAGCGTTCATGCGAAAACAAGGCAGAGGTTGTATCCTTGGATGAGAAGGAAGGTGGACTGAGAGCGACATTGAACTTGGGACATACATTTGGCCATGCAATTGAAACCGGGTTCGGCTATGGTCAGTGGCTCCATGGAGAAGCTGTCGCTGCTGGCACGGTCATGGCTGTTGATATGTCTTACCGTCTAGGTTGGATTGATGAGTCAATTGTGAAGCGGGTGTACCATATTTTACAACAGGCCAAGCTACCGATTGCCCCACCTGAGGGGGTTATGACTGTGGAGATGTTCAAGTCTTACATGGCTGTTGATAAGAAGGTGGCTGATGGACTGCTAAGGCTCATCCTCTTGAAAGGTCCTTTGGGTAATTGTGTTTTCACCGGTGATTATGATAGAAATGCCCTGGATGAGACCCTTCGTGCATTCTGTAAGTCTTGA

>Eca_Agt1 Eca_sc002690.1_g1690.1

ATGGATTACACATATGCACCAGGGAGGAACCAATTGTTTGTTCCAGGACCTGTCAATATCCCAGAACCAGTTCTTCGTGCTATGAATAGAAATAATGAGGATTATAGATCCCCTGCTATACCAGCTTTGACCAAAACCCTAATTGAGGATGTTAAGAAGATTTTCAAGACTACCTCTGGAACCCCTTTCTTGTTCCCCACTACAGGCACTGGTGCATGGGAAAGTGCACTTACAAACACATTATCTCCCGGAGATAGGACTGTTTCGTTCCTCATCGGACAATTCAGTCTGCTATGGATTGATCAACAGCAACGCCTCAAATTCAACGTCGATGTCGTAGAAAGTGAGTGGGGACAAGGTGCTAATCTTGAAGTTTTAGCTTCAAAGTTGGCAGAAGACAAAGCTCATACTATTAAAGCTGTTTGCATAGTTCACAATGAGACTGCAACTGGAGTTACCAATAACTTGGCTACTGTCAGGAAAATACTCGATCACTACAACCATCCTGCATTATTTCTCGTCGATGGAGTGTCCTCAATATGTGCACTCGACTTCCGTATGGATGAATGGGGAATAGATGTAGCTTTAACAGGATCACAGAAAGCTCTGTCTCTTCCGACTGGAATGGGTATAGTATGTGCAAGCCCTAAAGCACTAGAAGCCGCCAAGACCGCGAAATCAGTTAGAGTGTTCTTTGATTGGAATGACTACTTGAAGTTCTATAAACTAGGAACTTACTGGCCATACACCCCTTCAATCCAGCTTTTATATGGACTTAGAGCTGCTCTTGATCTCTTGTTCGAGGAAGGTCTCGATAATGTTATCGCTCGGCACACCCGTTTCGGCACAGCAACTAGGCTTGCTGTTGAGGCCTGGGGTTTGAAAAATTGTACACAAAAAGAAGAGTGGTTTAGTGACACTGTCACTGCAGTCATTGTTCCGAGTTACATCGACAGTTCTGAAATTGTAAGAAGGGGATGGAAAAGATACAATCTCAGCTTGGGTCTTGGTTTGAACAAAATAGCTGGAAAGGTCTTCAGAATAGGCCATCTAGGGAACCTGAACGAGGTCCAACTGCTCGGTTGTTTGGCCGGAGTGGAGATGATACTGAAAGATGTCGGGTACCCGGTGAAATTGGGAAGTGGAGTAGCAGCAGCTTGTGCTTATCTTCAGGACACAACTCCCATGATTCCTTCTAGGATTTGA

>Eca_At103 Eca_sc194615.1_g1740.1

ATGGCTGCAGAAATGGCATTGGTAAAACCCATCTCTAAATTTACTAGAACCAGCCCTAAATTCTTCAACCCAACCACAAGAAATTCTTCCTTCACAATCAAAATGTCATCACAACCCACTTCAACAACAACAACAACAAAATCAGCAAAACCAAAGAAAGCAATCAAAGAAACCCTACTTGCTCCAAGATTTTACACAACTGACTTTGATGAAATGGAAACCCTTTTCAACACTGAAATTAACAAAAACCTAAACCAAGCTGAATTTGAAGCACTTCTACAGGAATTCAAAACTGATTATAATCAAACCCATTTTGTTAGAAACAAAGAATTCAAAGAAGCTGCTGATAAAATGCAAGGTCCTCTTAGACAAATCTTTGTTGAGTTCTTAGAAAGATCTTGTACTGCTGAGTTCTCTGGTTTCCTTCTCTACAAAGAACTTGGTAGAAGACTTAAGAAAACTAATCCGGTGGTGGCTGAGATTTTCTCACTTATGTCTAGGGATGAAGCTAGGCATGCTGGATTCTTGAACAAGGGTCTCTCTGATTTCAATTTGGCATTGGATTTGGGGTTCCTTACGAAGGCTAGAAAATATACATTTTTCAAGCCCAAGTTCATCTTCTATGCTACGTATTTATCCGAAAAGATTGGGTATTGGAGGTACATAACCATTTACAGGCATCTCAAGGCAAACCCAGAATACCAAGTTTATCCAATTTTCAAGTATTTTGAGAACTGGTGCCAAGATGAGAACAGACATGGTGATTTCTTCTCTGCACTCATGAAAGCTCAACCTCAGTTTCTTAATGATTGGAAGGCGAAATTATGGGCTCGATTCTTCTGTTTATCGGTTTACGTGACGATGTATTTGAACGACTGTCAACGAACAGCCTTCTACGAAGGAATTGGACTTAACACCAAAGAGTTCGACATGCATGTTATAATTGAGACTAACCGAACAACAGCGAGGATTTTCCCCGCAGTCCTAGACGTGGAGAACCCAGAGTTCAAGAAGAAATTGGACCGAATGGTGGAGATCAACACCAAAATACTTGCCATCGGCGAAACTGAAGATATACCCTTTGTGAAGAACTTGAAGAGGGTTCCTCTTATTGTAGCGTTGGCGTCAGAAATCTTGGCTGCCTATCTCATGCCTCCTATCGAATCTGGATCTGTTGATTTTGCAGAGTTCGAGCCACAAGTTGTTTATTAA

>Eca_ChlP Eca_sc002150.1_g1300.1

ATGGCTTCCATTGCCGTAAACCCTAGTTTCTCCGCCACCTCATGCAACCGTAAAACCACCGATATCAGTCGTGATTCCTTCAACACCTTCACCGGCCTCCGCAAAACCACTCTTTCAAACCACAATTCTCTCTTTCTAACCAAATCCCTACCACAACATCAGACCCTAAACCCTAAATTTTCCATCAAAGCTGGAAAATTCAGCCCTAAAATCACCGGTCGGAACCTTCGAGTTGGGATAATTGGTGGGGGACCAGCTGGTGGCGCCGCCGCAGAAACCCTAGCTAAAGGCGGGATCGAAACTATCTTAATCGAACGAAAACTCGATAACTGTAAACCTTGTGGTGGTGCAATTCCATTATGTATGGTTGGTGAATTTGATCTTCCACTTGATATCATCGATCGACGAGTCACTAAGATGAAGATGATCTCACCGTCAAATGTAGCAGTTGATATCGGACGAACACTAAAACCTGATGAGTACATTGGTATGGTACGTCGTGAAGTACTAGATCAATACTTACGTGAACGAGCTGAAAAGGCAGGAGCAACTGTAATTAACGGTCTTTTCCTCAAAATGGATATTCCTCAAGAACCAAACGCGCCTTATGTTCTTCACTATAGCGCTTTCGACGGAAAAACCGGCAGTGTTGGTGCAAGAACCACGATAGAAGTTGACGCCGTTATCGGAGCTGACGGAGCAAATTCTCGCGTAGCCAAATCCATAAATGCTGGAGATTATGATTACGCTATTGCATTTCAAGAGAGAATCAAAATTCCTGACGAGAAAATGGTTTACTACGAAGATCTCGCCGAGATGTACGTAGGCGATGACGTGTCACCCGACTTTTACGGATGGGTCTTTCCAAAATGTGACCACGTGGCTGTCGGTACAGGTACCGTAACTCACAAAGGGGATATCAAGAAATTCCAGGTAGCCACAAGGAATAGGGCTAAAGATAAAATTTTGGGTGGAAAAATCATAAGAGTAGAGGCGCATCCTATCCCAGAACACCCGAGACCACGTAGATTATCGGGACGGGTAGCCCTAGTTGGAGATGCAGCCGGATACGTTACAAAATGTTCAGGAGAAGGTATATATTTTGCAGCTAAAAGTGGAAGAATGTGTGCCGAAGCAATGGTTGAAGGAAGCGTATTGACACCACTTCGTATGGGTGCCTATAGATGA

>Eca_GbssI Eca_sc002013.1_g0590.1

ATGGCGACGTTAACTACTTCGCAACTTGTATCACGAGCCTCAAATTTCAACTACGGTGGAAGTAGTGAGCCTAAAGACACATTAAGACAACTTAGCATGAAGAACCAAACCATGACTCGTAGTGGGTTAAGGTCTCTGAGCAAAGTGGATTTCATTAATATGAAAACAAGTAGTAAAACAATGATAAGAAAGGCAACCAAAACAAGGAAACCCACAGGAGAAATAACATGTTCAAGTGGGATGAACTTGGTTTTTGTTGCAGCAGAAGTAGGTCCATGGAGTAAAACTGGTGGACTTGGTGATGTTCTTGGTGGTCTTCCACCAGCCATGGCGGCAAACGGTCATCGCGTCATGACTGTCTCACCTCGGTACGATCAATACAAAGACGCTTGGGATACAGATGTTGTAATCGAGATCAAAGTCGGGGATAGAACTGAAACAGTTCGCTTCTTCCACTGTTATAAACGCGGGGTTGATCGTGTTTTCGTGGATCACCCATTGTTCTTAGAGAAGGTATGGGGTAAAACTGGATCTAAAGTTTATGGTCCTAATGCCGGATCAGATTACCAAGACAATCAACTTAGATTCTGTTTATTATCTCTGGCAGCTTTGGAAGCACCAAGGGTTCTAAATCTCAACAGCAATAAGTATTTCTCTGGACCATATGGAGAGGATGTTGTATTCGTCGCAAACGATTGGCATACTGCCCTTCTACCATGCTACCTTAAAAGTATGTACCAATCTAAAGGATTTTACACGTCTGCCAAGGTAGCCTACTGCATACACAATATTGCCTACCAAGGCAGATTTTCTTTCTCGGATTTTTCACTTCTCAATCTTCCTGACTCATTTAGGAGTTCGTTTGATTTTATCGATGGGTATTTGAAGCCCGTGAAAGGAAGGAAAATCAATTGGATGAAAGCTGGTATTTTAGAATCAGATAAGGTTATTACTGTCAGTCCATACTATGCCGAAGAACTTGCTTCTGGAGAGGATAAAGGTGTTGAATTGGATAACATCATTCGAAAGACTGGCATTGCCGGTATTGTGAACGGCATGGATGTCCAGGAGTGGAACCCATTGACCGATAAATACGTTGACATCAAATACGATGCTACAACTGTGATGGATGCAAAGCCTCTTCTGAAGGAAGCATTGCAAGCAGAAGTCGGGTTGCCTGTGGACCCAGATGTACCCGTTATAGGTTTCATTGGTAGACTTGAAGAACAAAAGGGCTCTGATATTTTGGCTGCTGCCATTCCTGAATTCATCGATGAAAATGTGCAGATAATAGTCCTTGGAACCGGCAAAAAGCCCATGGAGAAGCAGCTGGAACAACTTGAGATAAGCTACCCCGACAAGGCCCGAGGAGTTGCAAAATTCAATGTTCCATTGGCTCATATGATCATAGCAGGAGCTGATTTTATGATGATTCCTAGCAGATTTGAACCATGTGGACTCATTCAATTACATGCTATGCGATATGGAACGGTACCTATTGTTTCTTCAACAGGAGGACTAGTTGATACTGTCCAGGAAGTTGATCCAAAAGATGTAACTGCCATTGCTACAACCACCAAGAGAGCACTTGCAACCTATGGAACTCCAGCATTAACAGAGATGATTCAAAATGGCATGGCTCAAGATCTCTCTTGGAAGGGACCTGCCAAGAAGTGGGAGGAAATACTGCTGAGTTTGGAAGTTGCAGGGAGCGAACCAGGCATCGAAGGAGAGGAAATAGCTCCACTTGCAAAAGAGAATGTTGCTACTCCTTGA

>Eca_Hmgs Eca_sc194483.1_g1080.1

ATGGGTTCAAATCAGAAAGATGTTGGAATTCTCGCCATGGAAATCTATTTCCCTCCTACTTGTGTTCAACAGGAAGCTTTGGAGGCTCATGATGGTGCGAGTAAAGGAAAATACACTATTGGACTCGGACAAGATTGCATGGCATTTTGTTCAGAGTTGGAAGACGTCATTTCAATGAGTTTGACAGCTGTTGCTTCCCTCCTTGAAAAGTACAAAATTGACCCTAAGCAGATTGGACGTCTGGAAGTCGGCAGTGAGACCGTGATAGATAAGAGCAAGTCCATAAAGACCTTCCTGATGCAAATCTTTGAGGAATGCGATAATACCGATATTGAAGGAGTTGATTCAACAAATGCTTGCTACGGAGGAACTGCAGCTTTATTTAACGTTGTGAATTGGGTTGAGAGTAGTTGCTGGGATGGACGTTATGGACTTGTCGTGTGCACTGATAGTGCGGTTTACGCAGAAGGCCCAGCTCGGCCCACTGGAGGAGCAGCTGCTGTTGCTATGTTGATAGGTCCAGATGCTCCAATTGCATTTGAGAGCAAATACAGAGGCTCACACATGTCTCATGTCTATGACTTTTACAAGCCCAACCTTGCGAGTGAATACCCGGTCGTTGATGGAAAGCTTTCACAAACGTGCTATCTAATGGCGCTTGATTCCTGCTACAAACGTTTATGCGCCAAGTATGAGAAATTCGAGGGCAAGCAATTTTCAATTTCAGATGCAGATTACTTTGTATTTCATTCTCCTTATAATAAGCTTGTGCAGAAAAGCTTTGCTCGATTATGCTTCATGGACTACTTAAGAAATGCCAATTCAATTGAGGAATCTTCGAAAGAAAAACTTGCCCCATTTGCAGCTTTATCTGGCGAAGAGAGCTATGCTAGTCGAGATCTTGAAAAAGCATCCCAGACTGTTGCAAAAGGTCTATACGATGAAAAGGTACAACCAACCACTTTGTTACCGAAACAAGTCGGCAATATGTACACTGCTTCACTCTATGCCGCATTTGCATCTCTTTTGCACAACAAAAATAGCTCACTGGATGGTAAGCGGGTAGTAATGTTCTCGTATGGAAGTGGTTTAAGTTCAACAATGTTCTCATTCCGACTTAAGGAAGGTTCGCATCCATTCAGCTTGTCAAACATTGCAACTGTAATGAATGTCGCCAAGAAGTTGGAATCAAGACATGAGGTTACACCCGAAAAATTTGTCGAAACAATGCAACTGATGGAGCACAGATATGGAGGTAAGGACTTCGTAACAAGCAAGGACACAAGCCTTATACCATCGGGCACTTTCTATCTCACTGAAGTCGACTCCATGTATCGAAGATTCTATTCCAAGAAGAAAAGTGAAGCAACCACCACAACTGCTGATAATGGTGCAGTTCTCAATGGTCACTGA

>Eca_Nia Eca_sc194756.1_g0980.1

ATGGCGACTTCCGTCGAAAATAGAATGTATAATGGTATTGAATCGGGTCTAACATCGGTGGTTCGTGGTTATAAAACGAACCACCGATCCGACTCTCCCGTTCGTGGTTCTTTTAATAATTTCTCTTCTTCTGAATCACAACAACTTCCCTCCAAGAAAATTTATAAGGAAGATGACGATGACTCGAGTGACGATGAGTCAGGTGAGTGGAGCAAAGAAATTGCTAAAATTAACAATGAAATCGAGCCGTCGATTTTCGACCCAAAAGATGAAGGGACGGCTGATAATTGGATTGAAAGAAACCCATCAATGATTCGATTGACAGGTAAACACCCATTCAACTCGGAGGCACCACTGAATCGACTCATGCATCACGGGTTTATAACACCCGTTCCACTTCACTATGTTCGAAATCATGGAGCCGTACCTAAATGTAAATGGGAAGATTGGTCGGTTGAGATTTGTGGTCTAGTAAAACGTCCAACTCGACTCACTATGGATCAACTCGTTAACGAGTTTTCGTTTCGTGAGTTTCCCGCTTCACTCGTTTGTGCTGGGAATCGGCGTAAAGAACAAAATATGGTTAAACAAACCATCGGTTTTAATTGGGGTGGCGCAGGTATATCGACTTCTGTTTGGAGGGGTACACGTTTACGTGATGTGCTTAAACGATGTGGGGTAATGAGTAAAAATAAAGGTGCCCTTAACATTTGTTTCGAAGGAGTTGAAGACTTGCCTGGAGGTGGTGGTTCTAAGTATGGTACTAGTATTAGAAAAGAAGTAGCAATGGATCCTTCAAGAGACATCATATTAGCTTACATGCAAAACGGTGAGCGCTTACATCCAGATCACGGTTATCCTGTAAGAATGATCATCCCAGGGTTCATCGGAGGTCGAATGGTTAAATGGTTGAAAAGAATTATCGTTACTACTCAAGAATCACAGAATCATTATCATTATAAAGACAACAGGGTTTTGCCTTCTCATGTGGACGCTGAACTAGCTAATTCTGAAGCTTGGTGGTATAAACCAGAATACATCATTAACGAGTTGAACATAAACTCTGTGATAACAACACCTTCTCATGAAGAGATTCTACCAATCACTGCCTCAACAACTCAGAGGCCCTACACCTTGAAGGGATACGCATATTCCGGAGGAGGTAAGAAAGTCACACGTGTGGAAGTGACTTTAGATGGTGGAGAGAATTGGCAAGTGGGCACTTTGGATCACCAAGAGAAACCAAACAAGTATGGGAAATATTGGTGTTGGTGTTTTTGGTCACTTGAGGTTGAAGTCCTTGACTTGCTTGGTGCTAAAGAGATTGCTGTTAGGGCTTGGGATGAATCCCTCTGCACTCAACCTGAAAAGCTTATTTGGAATGTCATGGGTATGATGAACAACTGTTGGTTTCGAGTTAAGACGAACGTATGCAAACCACATAGAGGAGAAATCGGACTCGTTTTCGAACACCCAACTCAAGCAGGAAACCTACCAGGAGGCTGGATGGTAAAACAAAAGCATCTTGAATCTTCTGAACCTTCACATACTTTGAAGAAGAGTTTATCAACACCATTCATGAATACAACTTCGAAAATGTATTCCATGTCTGAAGTAAGAAAACATAGCACACCTGATTCAGCTTGGATTATTGTCCATGGTCATGTCTATGATTGTACAAAGTTCTTGAAAGATCATCCTGGTGGTACAGATAGTATCCTCATTAATGCAGGTACAGATTGTACCGAAGAATTCGATGCAATTCACTCCGATAAAGCTAAGAAATTACTCGAAGATTATCGAATTGGTGAACTTATCTCAACTGGGTATACCTCTGATTCTTCTTCTTCTTCACCAAACAACTCTGTTCATGGCTCTACAAATCTAAGTCATCTTGCTCCTATCAAAGAAGCTACTTCTAAACCAGTAGCTTTAGTCCCAAGAGAGAAAATCCAATGCAAACTTATCATGAAAACGTCCATTTCTCATGACGTTCGTTTATTCCGATTCGCTTTACCTTCTTCTGATCAAGTCCTTGGGTTACCAGTTGGAAAACATATCTTCATTTGTGCAACCATTGATGATAAGCTTTGTATGAGAGCTTATACCCCAAGTAGTACTATTGATGAAGTTGGGTATTTTGAATTAGTCATTAAAGTTTACTTCAAGAATGTTCATCCTAGGTTTCCAAATGGTGGTCTTATGTCTCAATACTTGGATTCACTCTCTATTGGTTCATCTCTTGATATCAAAGGTCCATTAGGACATATCGAATATACGGGCCGTGGAAATTTTACGGTCCATGGAAAGCCCAAGTTTGCTAAAAAGCTTGCTATGTTAGCCGGTGGAACGGGTATTACACCCATATATCAAATAATGCAAGCAATCTTGAAAGATCAAGAAGACGAGACGGAAATGTATGTTGTTTATGCTAATCGAACCGAAGATGATATTTTGTTGAGAGAAGAACTTGATTCTTGGGCTAAAAAGTATTCAGACAGATTGAAGGTTTGGTATGTGATCGAAAATGCTAAGGAAGGATGGGAGTATAGTACTGGTTTTATAACAGAAAAGATTCTTAGAGAACATATCCCTGAATCATCTGAAGAAAGTCTGGCTTTAGTTTGTGGACCTCCTCCTATGATTCAATTTGCAGTTAAACCACATTTGGAGAAGATGAACTATGATATCACTAATTCTTTATTGGTGTTTTAG

>Eca_Eif3E Eca_sc194760.1_g2160.1

ATGGCGAAATACGACTTGACCACACGAATTGCTCCTAACCTCGATCGACATCTTGTCTTCCCTCTTTTGGAATTCCTCCAAGAGCATGAGCTTCATCCTAGTGATGAAATACTCAAAGCCAAGATCGAACTACTCAGCCATACCAACATGGTTGATTACGCCATGGACATCCACAAAAGCCTCTACCACACTGAAGAAGTCCCACAAGTTGGGAGACTGAAAGCTCTGGAAGAATCTGCCGCCCCATTGATTTCCTTTCTTCAGAACCCTAATATGGTTCAGGAACTCCGTTCTGACAAGCAGTACAATCTTCAGATGCTGCAAGACAGATACCAGATTGGTCCCGAGCAGATAGAGGCATTATACCAATATGCCAAGTTTCAGTTCGAATGTGGAAATTACTCTGGCGCAGCTGACTTTCTGTATCAGTATAGAGCGTTATGCACAAACAGTGAGAGGAGTTTGAGTGCATTGTGGGGAAAGCTTGCTGCTGAGATACTAATGCAAAACTGGGAGGTTGCTCTTGAAGAACTTAATCGGCTGAGAGAAATAATTGAATCAAAGAACTTCAAGTCACCATTAAACCTTCTACAGAATAGAATCTGGCTAATGCATTGGAGCCTCTTTATATTTTTCAATCACGAAAGTGGAAGAAACGGAATCATTGATCTGTTTCTCAATGATAGGTATTTGAATGCCATCCAAACAAATGCACCCCATCTCTTACGCTACTTAGCAACTGCGTTGATCATTAACAAAAAGAGAAGACCTCAGCTTAAGGAGTTGATTAAGGTCATTCAGCAAGAGCAGTCCTCTTACAAAGATCCTGTCACAGATTTCTTATATTGTTTATATGTGAATTATGACTTTGATGGTGCACAACAGAAGCTACGTGAGTGTGAGGAAGTGATAATGAGTGATCCCTTTCTTGGTAAACAAGATGAAGAAGGAAACCCAAGTTCCGTATCTTTAAGGACCGAATTTCTTGACAATGCTCGTCTTTTCATCTTTGAAGCTTACTGCAGGATACATCAATGCATCGAGATAGCAACACTTGCAGAGAAGTTGAATATGAATTACGATGAGGCTGAGAGATGGATAGTGAATCTCGTTCAGAATGCAAAACTTGAAGCTAAGATTGATTCCAAGTTGGGAACTATTGTTATGGAACCCACCCATCTTAATGTGTACGGACAGATAATAGAGAACACCAAGGCGCTCTCGTTGCGTACTTACAAATATGTAAATCTCGTGCTAGAATCTGGACAGGCGCTAGCTACTCGTTGA

>Eca_AroB Eca_sc001139.1_g2710.1

ATGGCTTCTTCTGCAACTTCTTATACTCTTACTTTCACTCCAAAGTCATCATCATCTGTTGATCCTTGTCGTTCTAAGACGTGTATTTCATCTCTTCAATGTCCGAATCTCGATAAAAATCCCCTAAAATCATCTTTGATTTCATCAGGTTCACTATTATTGAATCCAATTAGATCTCCTCTCACTAGATCTAATCGGATTATTTCAAAGAAGATAGTAGCAAATTCAGCTCAAGTTATGGAGCAATCTCCTCGTGATACTGGTTCTAGGGTTTCGACAGTTGTGGATGTTGATTTGGGGGACCGAAGTTATCCAATTTATATCGGTGCTGGACTTCTTGAAGAACCTGATCTTCTCCAAAGGCATGTTCATGGGAAGAGAGTACTAGTGGTAACCAACACAACAGTTGCACCACTATACCTAGATAAAGTTACTGCTGCATTAACTCATGGAAACTCAAATGTTTCAGTTGAGAGTGTGATTTTGCCTGATGGGGAGAAGTATAAAGATATGGATACCCTTATGAAAGTATTCGACAAAGCTATAGGATCTCGTTTTGATCGTCGTTCTACATTTGTCGCTCTTGGAGGTGGTGTTATTGGTGATATGTGTGGATTTGCTGCTGCTTCTTTTCTCCGTGGTGTAAATTTCATTCAAATTCCAACTACTGTCATGGCACAGGTGGATTCCTCAGTTGGGGGCAAAACTGGAATCAACCATCCACTTGGGAAAAACTTGATCGGTGCTTTTTATCAACCACAGTGTGTCCTTATAGACACAGACACGTTAAACACACTACCTGATCGGGAATTAGCATCTGGTCTTGCTGAAGTTGTCAAATATGGGCTTATTAGGGATCCTGAGTTTTTCGAGTGGCAAGAAAAGAATATGCATTTGCTATTAGCAAGGGACCCTGATGCATTTGCTTATGCTATCAAACGTTCGTGTGAGAACAAGGCAGAGGTTGTATCTCTAGATGAGAAGGAAGGAGGATTGAGGGCGACACTAAACTTGGGGCATACATTTGGCCATGCAATTGAAACGGGATTTGGCTATGGGCAATGGCTCCATGGAGAAGCTGTAGCAGCTGGAACGGTAATGGCTGTTGATATGTCTTACCGTCTCGGCTGGATCGACGAATCAATTGTGAAGCGAGTTTATAGTATTTTAGAGCAAGCCAAGCTACCGATAACCCCACCTGATGTCATGACTGTCGAGAAATTCAAGTCATACATGGCTGTTGATAAAAAGGTGGCTGATGGGCTACTAAGACTCATCCTATTGAAAGGTCCTCTGGGCAATTGCGTTTTTACCGGTGATTATGACAGAAATGCACTCGACGAAACCCTTCGTGCATTCTGTAAGTCTTGA

>Aco_Agt1 PIA48213.1

ATGGACAATGTCTATGCACCCGGAAAGAACCATCTTTTTGTTCCAGGGCCGGTCAATATCCCGGATCCTGTCATTCGCGCAATGAGCAGAAACAATGAAGATTATCGCTCCCCTGCAGTGCCAGCCTTGACAAAGACCCTTCTTGAGGATGTTAAGAAGATCTTTAAGACAACAACTGGGACACCATTTATGATCCCCACCACAGGTACTGGTGCATGGGAAAGTGCATTAACAAACACGTTGTCTCCTGGTGATCGGACTGTTTCATTTCTGATTGGGCAATTCAGTCTTCTCTGGATTGATCAACAACAGCGCCTCAAGTTCAATGTTGATGTTGTTGAAAGTGACTGGGGCCAAGGTGCCAATCTTGATATTCTGGCATCGAAATTAGCTGAAGACACAGCACATACCATTAAGGCTGTCTGCATTGTTCACAATGAAACAGCTACTGGTGTTACCAATAATTTGGCCGCTGTAAGGAAGATTCTGGATCACTACGGCCATCCTGCCCTGTTTTTGGTTGATGGAGTGTCCTCTATATGTGCTCTTGATTTCCGTATGGACGAGTGGGGAATAGATGTTGCTTTAACAGGCTCTCAAAAAGCTCTTTCCCTCCCTACTGGAATGGGTATTGTGTGTGCTAGCCCCAAAGCACTTGAAGCATCCAAGACTGCCAAATCAGTAAGAGTATTCTTTGACTGGAATGACTACTTAAAGTTCTACAAAATAGGAACATATTGGCCATACACTCCCTCCATCCAACTTTTGTATGGCCTGAGGGCAGCTCTAGATCTCATATTTGAGGAAGGACTTGATAATGTGATTGCAAGGCACAATCGTTTAGGCACTGCGACTAGGCTTGCTGTTGATGCATGGGGTTTGAAAAATTGTACCCAGAAAGAGGAATGGTTTAGTGACACTGTCACTGCAGTGGTTGTTCCACCTTATCTTGACAGTGCAGAAATTGTTAGAAGGGCATGGCAGAGGTACAACTTAAGCTTGGGTCTGGGATTGAATAAAGTAGCTGGAAAGGTGTTCAGAATAGGGCATCTTGGTAACCTTAATGAGTTGCAATTGTTGGGTTGTCTGGCTGGCGTCGAAATGATACTCAAGGATGTAGGATACCCTGTAAAATTGGGAAGCGGAGTTGCAGCAGCTTGTGCATATCTGCAAAACAACATTCCTATGATCCCCTCGAGGATTTAA

>Aco_At103 PIA48687.1

ATGGCTGCTGAAATGGCACTCATCAAACCAATCTCCAAATTTAACAGAACCCTTCCTAAGTTCAGAAACCCCATTACACCAAAACCATGTTACTCCTATGTCAAGATGTCATCATCAGCACCAACAACTTCACCCAAGAAGAAATCCACTGCCAAACCCAAGAAAGCAATCAAAGAGACCCTTTTGGCTCCAAGGTTCTACACTACTGATTTTGATGAAATGGAGACTTTATTCAACACTGAAATCAACAAGAATCTTAATGAAGCTGAGTTTGAAGCATTGCTTCAAGAATTCAAAACTGATTACAATCAGACCCACTTTGTTAGGAACAAAGAATTCAAAGAAGCTGCTGATAAATTACAAGGTCCTCTTAGACAAATCTTTGTCGAGTTTCTGGAAAGGTCTTGCACTGCTGAGTTTTCTGGTTTTCTTCTCTATAAAGAGCTTGGAAGAAGACTCAAGAAAACAAACCCAGTGGTGGCCGAGATTTTTTCATTAATGTCCCGAGATGAAGCCAGGCATGCTGGGTTTTTGAACAAAGGTCTATCTGACTTTAATTTGGCATTGGATTTGGGATTTCTCACAAAGGCTAGGAAATATACATTCTTCAAACCCAAGTTCATTTTCTATGCTACATACCTATCTGAGAAAATTGGTTATTGGAGATACATTACAATTTACAGGCATCTAAAGACCAACCCAGAATACCAATGTTACCCAATTTTCAAGTATTTTGAGAACTGGTGCCAGGATGAGAACAGACATGGAGATTTCTTTTCTGCATTGATGAAAGCACAGCCCCAATTCCTTAATGATTGGAAGGCCAAGTTATGGTCACGATTTTTCTGCTTATCGGTCTACGTGACAATGTATCTCAATGACTGCCAGCGAACAGCTTTCTATGAAGGCATCGGACTGAACACCAAAGAATTCGATATGCATGTCATCATTGAGACTAACCGCACAACCGCCAGGATCTTCCCAGCAGTTTTAGATGTTGAAAACCCTGAGTTCAAGAGGAAGTTGGATCGAATGGTGGAGATTAACACAAAATTGATTGCTGTTGGGGAGAGCGATGACATACCTTTAGTGAAGAACTTAAAGAGGATTCCAATTATCGCAGCATTAGCATCTGAGCTCTTGGCAGCTTACCTCATGCCCCCTATCGAATCTGGATCTGTAGATTTTGCAGAATTTGAGCCACAAATTGTGTATTGA

>Aco_ChlP PIA62715.1

ATGGCGTCAATTGCCCTCAATGCCCATTTCACCACCACAACATGTGACCGGAAAACCACCGAAATCACCCGAGAATCACTCAAATCCTTTGTCGGTCTCCGTAAAACCACATTTGAAAACCACCATTCACTTACTCTAACAAAATCACTTCCAATCCGGTTAACTAACAGAAGACTACACAATATTGTAGCCACTAAGTCCAGCCCAAAACTCACAGGTCGAAATCTTCGTGTTGCGATAATTGGTGGTGGTCCAGCTGGTGGTGCAGCTGCTGAGACGCTAGCTAAAGGTGGTATTGAGACGTTTTTAATTGAGCGAAAGCTCGATAATTGCAAACCATGCGGCGGAGCAATTCCGCTATGCATGGTTGGCGAGTTTGATTTACCGTTAGATATTATCGATAGACGAGTAACAAAAATGAAGATGATTTCACCATCCAATGTAGCTGTCGACATTGGCCAAACATTAAAGCCACATGAGTACATTGGTATGGTGCGACGTGAGGTGCTTGATCAGTACCTACGTGAACGTGCAGAGAAAGCCGGTGCAACTATTTATAACGGTCTTTTTCTTAAGATGGACATGCCAAAAGAACCCAATGCACCTTATGTTCTGCACTATAGTCTGCATAATGGTAAATCTGGTTCTGTTGGAGAAAAATGTAAGTTAGAAGTAGACGCTGTGATTGGTGCAGATGGAGCAAATTCTCGTGTGGCGAAGTCTATTGGTGCTGGGGATTATGATTATGCTATTGCATTTCAAGAGAGGATTAAAATACCAGATGACAAGATGGTGTATTATGAAAATTTAGCTGAGATGTATGTTGGCGATGATGTGTCGCCAGACTTTTATGGTTGGGTTTTCCCAAAGTGCGATCATGTGGCAGTAGGTACTGGGACTGTGACACATAAGGGTGATATCAAGAAATTTCAGTTGGCTACACGAAATCGGGCCAAGGACAAGATTTTGGGTGGGAAAATCATTCGCGTTGAGGCACATCCTATTCCAGAGCACCCAAGGCCAAAGAGGTTATCAGGGAGAGTAGCGTTAGTAGGTGATGCGGCAGGGTACGTAACCAAATGCTCTGGCGAAGGAATCTACTTTGCAGCAAAGAGTGGAAGAATGTGTGCTGAAGCTATTGTTGAAGGATCAGAAAATGGAACTAGATGGATTGAAGAGTCTGATTTAAGGAAGTACTTAGAGAAATGGGACAAGACTTACTGGCCAACATACAAGGTGTTGGATGTATTGCAGAAGGTGTTTTATAGATCAAATCCAGCAAGAGAAGCTTTTGTCGAGATGTGCGCAGATGAGTATGTGCAGAAGATGACTTTTGACAGTTATTTGTACAAGACAGTTACGCCCGGGAATCCATTGGAAGATTTGAAGTTAGCTGTGAATACTATTGGGAGTTTGGTGAGAGCTAATGCACTTAGGAAGGAAATGGACAAGATTAGTGTATGA

>Aco_GbssI PIA34528.1

ATGGCGACTATTAACGGTTACCACTTTGTATCAAAAACCTTACATGGAATGTGTGGACGAACTTCTGCACAAGAGGGTAAAACCACATTGACACAAATTGGCCTGAAGAACCAAATTATAATTCACGATGGATTGAGATCTGTGAACAAAGTGGACATTATTGGCATGAGAAGCAAATCAAGAGCAGTTGTCACTAAAGCCAGTATAAATGGGTCTAGTACTGAGAATGCAAAGCCATCTGAGGAGATTGTATGTAGTGTAGGGATGAAGTTAGTATTTGTGTCATTTGAGATGGGTCCGTGGAGCAAAACTGGCGGACTTGGTGATGTTATTGGAGGTCTACCACCGGCCATGGCAGCCAATGGGCATCGTGTTATGACTATTGCCCCACGCCACGATCAGTACAAAGATGCATGGGATACGAATGTTGTAGTTGAGATTAAAGTTGGGGATAACATAGAGACTGTCCGCTTCTTCCACTCTTACAAACGGGGAGTGGATCGAGTTTTCGTTGATCACCCATGGTTCCTAGAGAAGGTATGGGGTAAAACTGGATCTAAACTTTATGGTCCTATAGCTGGAAATGATTACCAGGACAATCAACTAAGGTTCAGCTTGTTATGCCAGGCAGCTCTGGAAGCACCAAGGGTTCTAAACCTCAATAGCAGTAAATATTTTTCTGGGCCTTATGGTGAGGATGTTGTATTCATAGCTAATGATTGGCACACTGCTATTCTTCCATGCTACCTTAAAAGCATGTATCAGTCTCAAGGGATCTACAAAAGTGCCAAAGTTGCTTTCTGCATACACAACATAGCCTACCAAGGGCGATTCCCCTTTTCAGACTTCTCACTTCTTAATCTTCCCGATTCATTCAAGAGTTCCTTTGATTTCATGGATGGGTATGAAAAACCTGTAAAAGGAAGGAAAATAAATTGGATGAAGGCTGGTATACTGGAATCACATATAGTCATAACTGTGAGTCCATACTATGCTGAAGAACTGCAATCAGGAGAGGATAAAGGTGTGGAGTTAGATAACATCATTCGAAAGACTGGTATATTTGGTATTATCAACGGAATGGATGTCCAAGAGTGGAACCCATTGAGCGATAAATATATTGACATCAAATATGATGCTACAACTGTGATGGATGCAAAGCCTCTTATAAAGGAAGCACTGCAAGCAGAAGTTGGATTGCCTGTGGACCGAAATATCCCTGTCATAGGCTTCATTGGTAGACTTGAAGAGCAGAAAGGTTCAGATATTCTGGCAGCTGCTATTCCTCAATTCGTAGATGAGAATGTTCAAATTATTGTCCTTGGTACGGGCAAAAAGCACATGGAGAAGCAGATTCAGGAATTGGAGATTATGTATCCGGACAAGGCTAGAGGAGTGGCTAAATTCAATGTTCCTTTGGCCCATATGATCATAGCGGGGGCTGATTTTATGGTGATCCCAAGTAGATTTGAACCATGTGGCCTCATTCAATTACATGCTATGAGATATGGGACGCCATGTATTGTTGCTTCCACTGGTGGACTTGTTGACACAGTTAAAGAAGGTTTTACAGGATTCCATATGGGCAGCTTCAGTGTTGAATGTGATGAAGTTAACCCAGAGGATGTGACTGCCATTGCCACAACCGTCAAAAGAGCTCTTACAGTGTACGATACCCCTGCAATGACTGAGATGATTCAAAACTGCATGGCTCAAGACTTCTCTTGGAAGGGACCTGCTAAGAAGTGGGAAGAGGTTCTGTTGACTTTGGAGGTTTCTGGCAGCCAACCAGGAATTGATGGTGAGGAAATTGCTCCTCTTGCTAAGGAAAATGTTGCAACTCCTTGA

>Aco_Hmgs PIA49717.1

ATGGATTCAAATCAGAATAATGATGTCGGAATTCTTGCTATGGAAATCTACTTCCCACCTACATGTGTTCAGCAGGAAGCATTGGAGGTTCACGATGGTGCAAGTAAAGGGAAATACACCATAGGACTTGGGCAAGACTGCATGGCATTTTGTACAGATGTGGAAGATGTCATCTCAATGAGTTTGACAGCCGTTTCTTCCCTCCTTGAGAAGTATCAGATTGACCCAAAACAAATTGGTCGCTTGGAAGTAGGCAGTGAGACTGTAATAGACAAGAGCAAGTCTATTAAGACCTTCTTGATGCAAATCTTTGAGGAACATGGTAATACTGACATCGAAGGAGTTGATTCAACAAATGCCTGTTATGGTGGAACCGCAGCCTTGTTCAATGTCGTGAACTGGGTAGAGAGTAGCTCTTGGGATGGGCGCTATGGACTTGTTGTTTGCACCGATAGTGCGGTCTATGCAGAAGGTCCAGCACGCCCTACTGGAGGAGCAGCTGCTATTGCTATGTTGATAGGTCCAAATGCTCCAATAGCTTTTGAAAGCAAATTCAGAGGGTCACACATGTCTCATGCCTATGATTTTTACAAGCCTAACCTTGCAAGTGAATACCCGGTTGTTGATGGGAAGCTTTCACAAACTTGTTATCTCATGGCACTTGATTCTTGCTACAAACGCTTCTGTGCTAAGTTTGAGAAATTCCAGGGAAATCCATTTTCCTTGTCTGATGCTGACTATGTTGTATTTCATTCTCCATATAACAAGCTTGTGCAGAAAAGCTTTGCTCGGTTGTACTTCATGGACTTCTTGAGAAATGCCAGCTCTGTTGATGAAGCTGCAAAAGAAAAACTCGCACCATATTCATCTTTGTCGGGTGATGAGAGCTATGCTAGTAGGGATCTTGAGAAGACATCTCAACAAGTTGCAAAACAGCTATACGATGCCAAGGTGCAACCGACCACTTTCGTACCTAAGCAAGTTGGCAACATGTACACTGCATCACTCTATGCAGCATTCGCTTCACTTATTCATAACAAGCATAGTGAACTGGATGGTAAACGGGTAGTTATGTTCTCATATGGAAGTGGTTTAACTTCCACAATGTTTTCATTTCGACTTCGGGAATGTCAAAATCCTTTTAGTTTGTCAAACATTGCAACTGTGATGGATATCTCGGGGAAATTGAAATCAAGACATGAGGTTACTCCAGAAAAATTCATCGATAATTTGCAACTAATGGAGCATCGCTATGGGGGCAAGGACTTTGTGACCAGCAAAGACGTGAGCCTTTTATCTCCTGGAACATTCTATCTGACTCAGGTTGATTCCATGTACCGAAGATTCTATTCCAAGAAGGCTAAGAAAGAAACCAACGATGCCATCTGTGAAAACGGTTCAGTACTCAATGGTCACTAG

>Aco_Nia PIA47304.1

ATGGCGGCTTCAGTTCAAACTAGACAGTTCAATGGAACTGACTCACAATTTAGAGGTGGTGGTTGTAACTTTATTTCCACACTTGAAAAAAAACCAGTTATTCAACAAAAAGATTATTACTATTCTTCAAGTGAAGATGAAGAAGATGATGAATGGAGTAACTTTACTGATCAACACAAAGATGTAGAAAAAGAGTCTATTCTAGACCCGAAAGATGAATCAACAGCGGATAATTGGATTGTTCGTAATCCATCTATGATTCGTCTTACAGGCAAACATCCATTCAATTCAGAACCACCATTGAATCGACTTATGCAACATGGATTTATTACACCAGTTTCTTTACATTATGTAAGGAATCATGGTTATGTTCCAAAAGCTAAATGGGAAGATTGGAGTATTGAGGTATGTGGACTTGTTAAACGTCCTATGAAACTTAGTATGGATCAACTTGTTAAAGAGTTTCCTAGTTGTGAATTTCCAGTTACTCTTGTTTGTGCTGGAAATCGTCGTAAAGAACAGAATCAGGTCAATCAAACTATAGGATTTAATTGGGGTGCTGCTGGTACCTCAACTTCTGTATGGCGTGGGGTTAGATTGCGTGATGTACTTAAACGTTGTGGTATTTATCATCGTAAAAATGGTGCACTTAATGTGTGCTTTGAAGGTATAGAAGATCTACCAGGTGGTGGCGGATCCAAGTATGGTACCAGTTTAAGAAAAGAGATTGCTATGGATCCATCAAGAGATATTATACTAGCTTATATGCAGAATGGTGAGCTTTTATCACCAGATCATGGTTTTCCGGTAAGAATGATAATTCCAGGATTTATTGGTGGTAGAATGGTTAAGTGGTTGTCAAGAATCATAGTTACCACACAAGAATCACAAAATCATTACCATTTTAAGGATAATCGCGTTTTACCTAGTCATGTGGATGCCGAACTGGCGAACACAGAAGCTTGGTGGTACAAGCCGGAATATATCATCAATGAACTGAACATAAACTCGGTGATTACTACACCATCTCATGAAGAGATCTTATTGATAAATTCTGGAACAACACAAAGACCCTACACAATAAAGGGATATGCATATTCAGGTGGAGGAAAGAAGGTGACCCGTGTCGAGGTGACGTTGGATAATGGAGAATCCTGGCTTCTTTGCACATTGGACCACAAAGAGAAGCCAAATAGGTATGGCAAGCATTGGTGCTGGTGCTTCTGGTCTCTTGAAGTGGAGGTTTTGGACCTGATTGGTGCCAAAGAAATTGCTGTTCGTGCTTGGGACGAGACTCTCAACACGCAGCCCGAGAAGCTGATCTGGAATGTGATGGGCATGATGAACAATTGTTGGTTCCGGGTCAAAACAAACATGTGCAAGCCCTACAAGGGAGAGATAGGGCTTGTTTTCGAGCATCCAACTCAGGCAGGTAACCAGCCTGGTGGTTGGATGGTTAGAATGAAGAACCTTCAATCGGCTGAGACCCAAAACACAATGAAAAAGAGTGTGTCTTCGCCGTTCATGAATACTTCCACAAGAATGTATTCAATGTACGATGTGAAGAAACACAATTCTGCTGAGTCAGCATGGATTGTTGTCCACGGCTATGTTTACGATTGTACTCCATTCTTGAAAGATCATCCTGGAGGTATCGATAGTATCCTCATCAATGCCGGGTCAGATTCTACTGAAGAGTTTGACGCCATTCATTCCGATAAAGCAAAAAAATTGCTTGAGAAGTACCGTATTGGAGAGCTGATCGATTCTGTTTCTGACTCTTCCGCTTCAACAACACCTAACAACTCAGTCCATGGTGCCACCAACTTGATCCACCTTGCCTCTATAAAAGAGGTTGTACTACCAAAGTCGAATCCAGTGGCCCTGATACCACGTGAGAAAATCCAATGCAAACTGATATCTAAGACCTCACTCTCTCATGACGTGCGCCTATTCAGGTTCGCATTACCATCAGCAGATCAAGTTCTTGGGTTACCAGTAGGGAAGCACATATTTCTTTGCGCAACAGTAGACAATAAGCTATGTATGCGAGCATACACACCAACCAGCACCATTGATGAAGTGGGCTACTTTGAACTAGTCATCAAGATATACTACAAGGGCGTACACCCTACATTTCCAAATGGGGGGATAATGTCGCAGTATTTGGAGTCGCTTCCTTTGGGCTCGTCGATTGATGTTAAAGGCCCTTTGGGCCACATCGAGTATACCGGGCGTGGTAACTTCACAGTCAATGGGAAAAACAAGTTTGGTAAGAAACTTGCAATGCTTGCAGGTGGAACTGGAATCACACCTATATATCAAGTGATGCAAGCTATCTTGAAAGACCCTGAGGATGAAACAGAAATGTATGTGGTGTATGCCAATCGAACAGAAGATGATATATTGCTGAAAGAGGAGTTGGACAGTTGGGCAGAGCAATATAGTGATAGGGTTAAAGTGTGGTATGTGGTGCAAGTTGCGCCTCCAGAAAGGGAAGAAGCATGGAAGTACAGTGTTGGATTTATTACAGAAGAGATCTTAAGAGAGCATATTCCTCTTGGATCTAAAGATACACTAGCATTAGCTTGTGGGCCACCGCCAATGATACAGTTTGCGGTTAATCCCAACTTGGAGAAGCTGAGGTATGATGTGAAGGACACCTTGTTGGTGTTCTAA

>Aco_Eif3E PIA48328.1

ATGGCGAACTACGATTTGACTCGGTATATCGCGCCGAATCTAGATCGGCATCTTACGTTCCCACTTCTTGAATTCGTTCAAGATCGAAGACTTTATCCTGATGATGAAACTCTTAAAGCAAAGATCGAACTTCTTAACAAAACAAACATGGTTGATTATGCTATGGATATTCACAAGAGTCTCTATCAGACTGAAGAAGTTCCTCAAGATATGATTGATAGAAGAGTTAAGGTTGTTGCGAGATTGAAGGCTTTGGAAGAATCAGCTGCTCCATTGATTACTTTCCTTCAGAATCCTATTATGTTGCAGGAGTATCGTGCTGATAAACAATATAATCTTCAGATGTTGCAGGAACGATATCAGATTGGTCCAGATCAGATAGAAGCATTGTATCAGTATGCAAAATTTCAGTTTGAATGTGGTAACTACTCTGGTGCAGCAGATTTTCTCTATCAGTACCGTCTTTTATGCACAAACAGTGAGAGGAACTTGAGTGCATTATGGGGAAAGCTTGCAGCTGAGATACTGATGCAGAACTGGGATGTTGCACTTGAGGAGCTTAATCGTTTGAGAGAGATAATTGATGCAAAGAACTTTCAATCTCCTCTTAATCAGTTGCAAAGTAGAATTTGGCTAATGCACTGGAGCCTCTTTATATTTTTCAATCATGATAGTGGCAGAAGTGGGATAATAGACCTTTTTCTCTATGACAGATATCTAAATGCTATTCAAACAAATGCACCCCATCTTCTTCGCTATTTGGCCACTGCATTAATTATCAACAAGAAGAGAAGACCTCAGCTTAAAGAGTTGATTAAAGTAATTCAGCAAGAACAGTGCTCTTATAAAGATCCCATCACAGAGTTCTTACAGTGTTTGTTCGTGAACTATGATTTTGATGGTGCACAACAAAAGCTGCAAGAATGCGAGGAGGTAATTTTGAATGACCCTTTTCTTGGAAAGCGAGTTGAAGAAGGCAACTTCACTACTGTACCTTTGAGGGACGAGTTTCTTGAAAATGCCCGTCTCTTCATATTCGAGACCTACTGCAGAATACATCAATGCATTGATATTGGAATGCTTGCTGAAAAGTTAAATATGAGTTATGATGAGGCTGAGAGATGGATAGTGAACCTTGTGCGAAATGCAAAACTTGATGCAAAAATTGATTCGAAATCAGGAACTGTTGTGATGGAACCCAATCATGTCAATGTGTATGAACAGATCATTGAAAGCACCAAGGCACTCTCTTTGCGTACTTATAAATTCGCAAATCTTGTCCTAGAATCTGCACAGGCACAGACTGCTGCATAA

>Aco_AroB PIA31892

ATGGCGTCTTCTGCAACCTCGACTTCATTTCAATTCTCTCTAAAACCTTCTATAAATCTAAATCCAATATCTCGATCTGAATCTTCTTCTAATCTATCTTTTCGATTTCAGAATCAGAATAATCACGAATCATCACTACTTTCTTCTTCTTCTTCTTCACTTCGATTGAATCGTAGTAGTAGTTGTAATAGTATTATTAGTTCTGTTTCATTGAAATTAAACAGAATTGTTGCTTCAGCAGCATCAACATCAGCAGTAATGGAAGAGCCATCAACTAACTCTTCTTCTTCTACTACTACTGATCCGACAATTGTTAATGTCGATTTAGGTAATAGAAGTTATCCGATCTATATTGGATCTGGACTTCTTGATCAATCTCATCTTCTTCAAAGGCATGTTCATGGGAAGAAAGTTCTAGTGGTAACTAATACAACTGTTGCACCTCTTTACCTGGAAAAAGTTACTAGTGCGTTGACTCACGGGAGGTCGGATATTGTGGTGGAGAGTGTTATTTTGCCTGATGGTGAAAAGTACAAGGACATGGAAACCCTCATGAAAATCTTGACCAAGGCTGCAGAATCTCGATTGGATCGTCGCTCTACATTTGTTGCTCTTGGGGGTGGTGTAATTGGTGACATGTGTGGGTATGCTGCTGCTTCTTATCTTCGTGGTGTGAATTTCATTCAGATTCCAACTACTGTAATGGCACAGGTGGATTCTTCGGTTGGGGGAAAAACTGGGATAAACTTGCCCCAAGGGAAGAATTTGGTTGGTGCTTTCTACCAGCCGCAATGTGTTCTAGTAGATACAGACACATTAAACACCTTGCCTGATAGAGAACTAGCATCCGGACTTGCAGAAGTGATAAAGTATGGACTTATTAGGGATGCCGAATTTTTTGAGTGGCAAGAAAAAAATATGTCGGCACTATTAGCAAGGGACCCAAATGCAATGGCCTATGCCATTAAACGTTCTTGTGAAAATAAGGCTGAAGTTGTTTCACTAGATGAGAAGGAAGGAGGATTGAGGGCTACATTAAATCTGGGTCACACTTTTGGTCATGCAATAGAAAGTGGGTATGGCTATGGTCAGTGGCTTCACGGAGAAGCTGTTGCTGCTGGAACAGTCATGGCTGTGGACATGTCATATCGTCTCGGCTGGATAGATGAGTCAGTCATGAAGCGTGCATTTAATATTCTAGAACAAGCGAAGTTACCCACTACCCCACCCGAAATTATGACTGTGGAGATGTTCAGGTCTTACATGGCGATTGACAAGAAGGTGGCTGATGGCCTATTGAGGCTAATTCTTCTGAAAGGTCCTCTTGGTAATTGTGTTTTTACTGGTGATTACGATAGAAAGGCCCTAGAAGAAACCCTTCATGCATTCTGTAAGTCTTGA

Supplementary dataset 3 (associated with Figure 2a)

- Protein sequences of STORR proteins from five Clade 2 *Papaver* species

>Pca_STORR

MELQYFSYFQPTSSLFVLVSILLISAVVLRKIFLNNYSSSPASSTETALLSHQRRSSCALPIIGLLHVFMNKNGLIHVTLGNMADKYGPIFSFPTGSHMTLVVSSWEMAKECFTGNNDIAFSNRPIPLVFKTIFYACRGIDSYGLSSVPYGKYWRELRKVCVHNLLSNQQLLNFRHLIISQVDTSFNKLYELCKNSADNQGNYTTTTTAAGMVRMDDWLAQLSFNVIGRIVCGFQSDTETSASSSAEQLTEAINDASYFMSTSPVSDNVPWLGWVDQLTGLTRNMKQCGKKLDLVLESIIEDHRQKRRFSRSKQGDENFINEDEQDDFIDICLSIMEQPQLPGNNNPCQIPIKSIVLDMIGGGTDTTKLTTIWTLSLLLNNPHVLDKAKQEVDAHFGTKRNDAGGVVVDFDDTRNLVYIQAIIKESMRLYPASPLVERLSGEDCVVGGFHVPAGTRLWANVWKMHRDPKVWDDPLVFRPERFLLDDQKMVDVRGQNYELLPFGAGRRICPGVSFSLDLMQLVLTRLILEFEMKSPSGKLDMTATPGLMSYKVVPLDILLTSRRMKWCVQLASERDKKMESSVVPVITLSSCKVMPVLGMGTFEKVGKGSERERLAILKAIEVGYRYFDTAAAYETEEVLGEAIAEALQLGLIKSRDELFISSMLWCTDAYHDRVLPALQNSLRNLKLEYLDLYMLPFPVSLKPGGKISMDIPEDEIFTMDYRSVWAAMEECQNLGLTKSIGVSNFSCKKLQELMATANIPPAVNQVEMSPAFQQKNLREYCNANNILISAVSILGSNGTPWGSNAVLGNEVLNQIAMARGKSVAQVSMRWVYEQGASLVVKSFSEERLRENLNIFDWELTKEDLEKMGEIPQCRILTADFMVSANGPFKSQEELWDDKA*

>Pso_STORR_PS1126560

MELQYISYFQPTSSVVALLLALVSILSSVVVLRKTFLNNYSSSPASSTKTAVLSHQRQQSCALPISGLLHIFMNKNGLIHVTLGNMADKYGPIFSFPTGSHRTLVVSSWEMVKECFTGNNDTAFSNRPIPLAFKTIFYACGGIDSYGLSSVPYGKYWRELRKVCVHNLLSNQQLLKFRHLIISQVDTSFNKLYELCKNSEDNQGNYTTTTTAAGMVRIDDWLAELSFNVIGRIVCGFQSGPKTGAPSRVEQFKEAINEASYFMSTSPVSDNVPMLGWIDQLTGLTRNMKHCGKKLDLVVESIINDHRQKRRFSRTKGGDEKDDEQDDFIDICLSIMEQPQLPGNNNPSQIPIKSIVLDMIGGGTDTTKLTTIWTLSLLLNNPHVLDKAKQEVDAHFRTKRRSTNDAAAAVVDFDDIRNLVYIQAIIKESMRLYPASPVVERLSGEDCVVGGFHVPAGTRLWANVWKMQRDPKVWDDPLVFRPDRFLSDEQKMVDVRGQNYELLPFGAGRRVCPGVSFSLDLMQLVLTRLILEFEMKSPSGKVDMTATPGLMSYKVIPLDILLTHRRIKPCVQSAASERDMESSGVPVITLGSGKVMPVLGMGTFEKVGKGSERERLAILKAIEVGYRYFDTAAAYETEEVLGEAIAEALQLGLVKSRDELFISSMLWCTDAHADRVLLALQNSLRNLKLEYVDLYMLPFPASLKPGKITMDIPEEDICRMDYRSVWAAMEECQNLGFTKSIGVSNFSCKKLQELMATANIPPAVNQVEMSPAFQQKKLREYCNANNILVSAISVLGSNGTPWGSNAVLGSEVLKKIAMAKGKSVAQVSMRWVYEQGASLVVKSFSEERLRENLNIFDWELTKEDHEKIGEIPQCRILSAYFLVSPNGPFKSQEELWDDEA*

>Par_STORR

MELQYFSYFQPTSSVVALLLALVSILFSFVVLRKTFSNNYSSPASSTETAVLCHQRQQSCALPISGLLHVFMNKNGLIHVTLGNMADIYGPIFSFPTGSHRTLVVSSWEMVKECFTGNNDTAFSNRPIPLAFKTIFYACGGIDSYGLSSVPYGKYWRELRKVCVHNLLSNQQLLKFRHLIISQVDTSFNKLYELCKNSEDNQGNHTTTTTTTAAGMVRMDDWLAQLSFNVIGRIVCGFQSDPKTGAPSRVEQFKEAINEASHFMSTSPVSDNVPMLGWIDQLTGLTRNMKHCGKKLDLVVESIIQDHRRKRRFSRTKGGDEKDDEQDDFIDICLSIMEQPQLPDNNNPPHIPIKSIVLDMIGGGTDTTKLTTIWTLSLLLNNPHVLDKAKQEVDAHFGTKRRSTNDAVVDFDDIRNLVYIQAIIKESMRLYPASPVVERLSGQDCVVGGFHVPAGTRLWANVWKMQRDPKVWDDPLVFRPERFLSDEQKMVDVRGQNYELLPFGAGRRICPGVSFSLDLMQLVLTRLILEFEMKSPSGKVDMTATPGLMSYKVVPLDILLTHRGTKSCVQSAASETSGVPVITLSSGKVMPVLGMGTFEKVGKGSERERLAILKAIEVGYRYFDTAAAYETEEVLGEAIAEALQLGLIKSRDELFISSMLWCTDAHPDRVLLALQNSLRNLKLEYLDLYMLPFPASLKPGKITMDIPEEDICRMDYRSVWSAMEECQNLGFTKSIGVSNFSCKKLQELMETANIPPAVNQVEMSPAFQQKKLREYCNANNILVSAVSILGSNGTPWGSNAVMGSEVLKKIAMAKGKSVAQVSMRWVYEQGASLVVKSFSEERLRENLNIFDWELTKEDNEKIGEIPQCRILTAHFLVSPNGPFKSQEELWDDKA*

>Por_STORR

MELQYFSYFQPTSSVVALLLALVSILFSVVVLRKTFSNNYSSPASSTETAVLCHQRQQSCALPISGLLHVFMNKNGLIHVTLGNMADKYGPIFSFPTGSHRTLVVSSWEMVKECFTGNNDTAFSNRPIPLAFQTIFYACGGIDSYGLSSVPYGKYWRELRKVCVHNLLSNQQLLKFRHLIISQVDTSFNKLYELCKNSEDNQGMVRMDDWLAQLSFNVIGRIVCGFQSDPKTGAPSRVEQFKEVINEASYFMSTSPVSDNVPMLGWIDQLTGLTRNMKHCGKKLDLVVESIIKDHRQKRRFSRTKGGDEKDDEQDDFIDICLSIMEQPQLPGNNSPPQIPIKSIVLDMIGGGTDTTKLTTIWTLSLLLNNPHVLDKAKQEVDAHFRKKRRSTDDAAAAVVDFDDIRNLVYIQAIIKESMRLYPASPVVERLSGEDCVVGGFHVPAGTRLWANVWKMQRDPKVWDDPLVFRPERFLSDERKMVDVRGQNYELLPFGAGRRICPGVSFSLDLMQLVLTRLILEFEMKSPSGKVDMTATPGLMSYKVVPLDILLTHRRIKSCVQLASSERDMESSGVPVITLSSGKVMPVLGMGTFEKVGKGSERERLAILKAIEVGYRYFDTAAAYETEEVLGEAIAEALQLGLIESRDELFISSMLWCTDAHPDRVLLALQNSLRNLKLEYLDLYMLPFPASLKPGKITMDIPEEDICRMDYRSVWSAMEECQNLGFTKSIGVSNFSCKKLQELMATANIPPAVNQVEMSPAFQQKKLREYCNANNILVSAVSILGSNGTPWGSNAVLGSEVLKQIAMAKGKSVAQVSMRWVYEQGASLVVKSFSEERLRENLNIFDWELTKEDNEKIGEIPQCRILTAYFLVSPNGPFKSQEELWDDKA*

>Pbr_STORR

MELQYFSYFQPTSSVVALLLALVSILFSVVVLRKTFSNNYSSPASSTETAVLCHQRQQSCALPISGLLHVFMNKNGLIHVTLGNMADKYGPIFSFPTGSHRTLVVSSWEMVKECFTGNNDTAFSNRPIPLAFQTIFYACGGIDSYGLSSVPYGKYWRELRKVCVHNLLSNQQLLKFRHLIISQVDTSFNKLYELCKNSEDNQGMVRMDDWLAQLSFNVIGRIVCGFQSDPKTGAPSRVEQFKEVINEASYFMSTSPVSDNVPMLGWIDQLTGLTKNMKHCGKKLDLVVESIIKDHRQKRRFSRTKGGDEKDDEQDDFIDICLSIMEQPQLPGNNSPPQIPIKSIVLDMIGGGTDTTKLTTIWTLSLLLNNPHVLDKAKQEVDAHFRKKRRSTDDAAAAVVDFDDIRNLVYIQAIIKESMRLYPASPVVERLSGEDCVVGGFHVPAGTRLWANVWKIQRDPKVWDDPLVFRPERFLSDEQKMVDVRGQNYELLPFGAGRRICPGVSFSLDLMQLVLTRLILEFEMKSPSGKVDMTATPGLMSYKVVPLDILLTHRRIKSCVQLASSERDMESSGVPVITLSSGKVMPVLGMGTFEKVGKGSERERLAILKAIEVGYRYFDTAAAYETEEVLGEAIAEALQIGLIKSRDELFISSMLWCTDAHPDRVLLALQNSLRNLKLEYLDLYMLPFPASLKPGKITMDIPEEDICRMDYRSVWSAMEECQNLGFTKSIGVSNFSCKKLQELMATANIPPAVNQVEMSPAFQQKKLREYCNANNILVSAVSILGSNGTPWGSNAVLGSEVLKQIAMAKGKSVAQVSMRWVYEQGASLVVKSFSEERLRENLNIFDWELTKEDNEKIGEIPQCRILTAYFLVSPNGPFKSQEELWDDKA*

Supplementary dataset 4 (associated with Figure 2a)

- Nucleotide sequences of full length STORR cDNAs from five Clade 2 *Papaver* species

>Pca_STORR

ATGGAGCTCCAGTACTTTTCTTATTTTCAGCCAACTTCCTCCCTTTTTGTTCTTGTATCCATATTACTGATCAGTGCTGTTGTTTTGAGGAAGATATTTTTAAATAACTACTCTTCATCACCTGCATCATCAACAGAAACAGCGCTGCTTTCTCATCAGCGGCGGTCGTCATGTGCATTGCCAATCATCGGTCTTCTTCATGTTTTCATGAATAAAAACGGCTTAATTCATGTAACTCTTGGAAATATGGCTGATAAATACGGACCGATTTTCAGTTTCCCAACAGGCAGCCATATGACTCTAGTTGTGAGCAGTTGGGAGATGGCAAAAGAGTGTTTTACAGGCAACAATGACATTGCTTTCTCAAACCGTCCTATCCCGTTAGTTTTTAAGACTATATTCTATGCATGCCGTGGTATTGACTCATACGGTCTTTCAAGTGTACCTTATGGAAAATATTGGAGGGAGTTACGAAAGGTGTGTGTGCATAACCTCCTCTCTAATCAACAACTACTCAACTTCAGACACTTGATAATTTCTCAAGTTGATACCTCTTTCAATAAGCTTTATGAGTTATGCAAAAACTCTGCAGACAACCAGGGAAACTACACTACTACTACTACCGCAGCTGGCATGGTGAGGATGGATGATTGGCTAGCGCAATTATCGTTCAACGTGATAGGAAGAATAGTCTGTGGATTCCAATCAGACACCGAGACAAGTGCTTCAAGCAGTGCGGAACAACTTACAGAAGCAATTAATGACGCATCGTATTTTATGTCGACATCTCCGGTGTCAGATAACGTTCCATGGTTAGGGTGGGTTGATCAACTAACTGGTCTTACTAGAAATATGAAGCAATGCGGAAAGAAATTAGACTTGGTGCTCGAGAGCATAATTGAAGATCATCGTCAAAAGAGACGGTTTTCTAGAAGTAAACAAGGAGATGAGAATTTTATCAACGAGGATGAACAAGATGACTTCATCGACATTTGCCTGTCAATTATGGAGCAACCACAGCTTCCTGGCAACAATAACCCTTGTCAAATACCTATCAAATCTATTGTCCTGGACATGATAGGTGGGGGGACCGACACCACAAAACTGACCACCATCTGGACCCTTTCATTGCTACTGAACAACCCCCATGTGTTGGACAAGGCAAAACAAGAAGTGGATGCACACTTCGGGACGAAAAGGAACGATGCAGGAGGAGTCGTGGTGGATTTTGATGATACTCGTAACCTTGTCTACATCCAGGCAATTATCAAAGAATCAATGCGGTTGTATCCAGCCAGCCCCTTGGTGGAGCGACTGAGCGGCGAAGATTGTGTGGTCGGTGGGTTTCATGTACCAGCAGGGACGCGGTTATGGGCTAACGTGTGGAAGATGCATCGAGACCCAAAAGTATGGGATGATCCATTGGTGTTTCGACCAGAGAGATTTTTGCTCGATGACCAGAAGATGGTTGATGTTAGGGGTCAAAACTATGAGCTGTTACCATTCGGAGCCGGTCGACGGATATGTCCAGGTGTATCGTTCTCTTTGGATCTAATGCAACTGGTACTGACTCGTCTTATTCTTGAGTTTGAAATGAAGTCTCCTAGCGGGAAACTAGACATGACAGCGACACCAGGTTTAATGAGTTACAAGGTAGTCCCCCTTGACATTCTGCTCACCAGTCGTCGCATGAAGTGGTGTGTGCAGTTAGCCTCTGAGAGAGATAAAAAGATGGAGAGTAGTGTTGTACCTGTAATCACTCTGAGCTCGTGCAAGGTGATGCCTGTTTTAGGCATGGGAACATTTGAAAAAGTTGGTAAAGGGTCTGAAAGAGAGAGGTTGGCGATTTTAAAAGCGATAGAGGTAGGTTACAGATACTTCGATACAGCTGCTGCATACGAAACTGAAGAGGTTCTTGGTGAAGCTATTGCTGAAGCACTTCAACTTGGCCTAATCAAATCTCGAGATGAACTCTTCATCAGTTCCATGCTCTGGTGCACTGATGCCTACCATGATCGAGTCCTCCCCGCTCTTCAGAATTCTCTGAGGAATCTTAAATTGGAGTATCTGGATCTATACATGTTACCCTTCCCGGTAAGCTTGAAGCCAGGAGGGAAGATAAGTATGGACATACCAGAGGATGAAATTTTTACGATGGACTATAGGTCTGTATGGGCAGCCATGGAAGAATGTCAGAACCTTGGCTTGACAAAATCAATCGGTGTCAGCAATTTCTCCTGCAAAAAGCTTCAGGAATTGATGGCGACCGCCAACATCCCTCCAGCTGTGAATCAAGTGGAGATGAGCCCGGCTTTCCAACAAAAGAATTTGAGAGAGTATTGCAATGCCAATAATATATTAATCAGTGCAGTTTCTATTTTGGGATCAAACGGAACCCCATGGGGCTCCAATGCAGTTCTGGGTAATGAGGTGCTAAACCAGATTGCTATGGCCAGAGGAAAATCTGTTGCTCAGGTTAGTATGAGGTGGGTTTACGAGCAAGGTGCGAGTCTTGTGGTAAAAAGTTTTAGTGAAGAGAGATTGAGGGAAAACTTGAACATATTCGACTGGGAACTCACTAAGGAAGACTTGGAAAAGATGGGTGAGATTCCACAATGCAGAATCTTGACTGCTGATTTTATGGTATCAGCTAATGGACCTTTCAAATCTCAAGAAGAGTTGTGGGATGACAAAGCTTGA

>Pso_STORR_PS1126560

ATGGAGCTCCAATATATTTCTTATTTTCAACCAACTTCCTCCGTTGTTGCTCTTCTACTTGCTCTTGTATCCATCTTATCCAGTGTCGTTGTTTTGAGGAAGACATTTTTGAATAACTACTCATCATCACCTGCATCATCCACAAAAACAGCGGTACTTTCTCATCAGCGGCAGCAGTCGTGTGCATTGCCAATTTCCGGTCTCCTCCATATTTTCATGAATAAAAACGGCTTAATTCATGTAACTCTTGGAAATATGGCTGATAAATACGGTCCGATTTTCAGTTTCCCAACAGGTAGCCATAGAACTCTCGTTGTGAGCAGTTGGGAGATGGTAAAAGAGTGTTTTACTGGCAACAATGACACTGCTTTCTCAAACCGTCCTATCCCGTTAGCTTTTAAGACTATATTTTATGCATGCGGTGGCATAGACTCATACGGTCTTTCGAGTGTACCTTATGGAAAATATTGGAGGGAGCTTCGAAAGGTCTGTGTGCATAACCTCCTGTCTAATCAACAACTACTCAAGTTCAGACACTTGATAATTTCTCAAGTCGATACGTCTTTCAATAAGCTGTATGAGTTATGCAAAAACTCTGAAGACAACCAGGGAAACTATACTACTACTACTACCGCAGCTGGCATGGTGAGAATCGATGATTGGCTCGCCGAATTATCGTTCAATGTGATAGGAAGAATAGTCTGCGGATTCCAATCAGGCCCTAAGACAGGTGCTCCAAGCAGGGTGGAACAATTTAAAGAAGCAATTAATGAAGCATCTTATTTTATGTCGACATCTCCAGTGTCAGATAATGTTCCAATGCTAGGGTGGATTGACCAATTGACAGGTCTTACGAGAAATATGAAGCACTGCGGAAAGAAATTAGACTTGGTGGTCGAGAGCATAATTAATGATCATCGTCAAAAGAGACGATTCTCTAGAACTAAAGGAGGAGATGAGAAGGACGATGAACAAGATGACTTCATCGACATTTGTTTGTCAATAATGGAGCAACCACAGCTTCCTGGCAACAATAATCCTTCTCAGATACCTATCAAATCTATTGTCCTGGACATGATAGGTGGGGGCACTGACACCACAAAACTGACCACCATCTGGACCCTTTCCTTGCTGCTGAACAACCCCCATGTGTTGGACAAGGCAAAACAAGAAGTGGATGCACACTTTCGAACCAAAAGGAGATCAACAAATGATGCAGCAGCAGCCGTGGTGGATTTTGATGATATTCGTAACCTTGTCTACATCCAGGCAATCATCAAAGAATCAATGCGGTTGTATCCAGCCAGCCCCGTGGTGGAGCGACTGAGCGGCGAAGATTGTGTGGTCGGTGGGTTTCATGTACCAGCAGGGACGAGATTATGGGCTAACGTATGGAAGATGCAACGAGACCCTAAAGTATGGGATGATCCATTGGTGTTTCGACCAGACAGATTTTTGAGCGATGAACAGAAGATGGTTGATGTAAGGGGTCAAAATTATGAGCTGTTACCATTTGGAGCCGGTCGACGTGTATGTCCAGGTGTATCCTTCTCTTTGGATCTAATGCAACTGGTACTGACTCGTCTTATTCTCGAGTTTGAAATGAAGTCTCCTAGCGGGAAAGTGGACATGACAGCAACACCAGGATTAATGAGTTACAAGGTGATCCCCCTTGACATTCTGCTCACCCATCGTCGCATAAAGCCGTGTGTGCAGTCAGCAGCCTCTGAGAGAGACATGGAGAGTAGTGGTGTACCAGTAATCACTCTGGGCTCGGGCAAGGTGATGCCTGTTCTGGGCATGGGAACATTTGAGAAAGTTGGTAAAGGGTCCGAAAGAGAGAGGTTGGCGATTTTAAAAGCGATAGAGGTGGGTTACAGATACTTCGATACAGCTGCTGCATACGAAACTGAAGAGGTTCTTGGAGAAGCTATTGCTGAAGCACTTCAACTTGGCCTAGTCAAATCTCGAGATGAACTTTTCATCAGTTCCATGCTCTGGTGCACTGATGCTCACGCTGATCGTGTCCTCCTCGCTCTTCAGAATTCGCTGAGGAATCTTAAATTGGAGTATGTGGATCTATATATGTTACCCTTCCCGGCAAGCTTGAAGCCTGGGAAGATAACGATGGACATACCAGAGGAAGATATTTGTCGCATGGACTACAGGTCTGTATGGGCAGCCATGGAAGAGTGTCAAAACCTTGGCTTCACTAAATCAATCGGTGTTAGCAATTTCTCCTGCAAAAAGCTTCAGGAATTGATGGCGACCGCCAACATCCCTCCAGCTGTGAATCAAGTGGAGATGAGCCCGGCTTTCCAACAAAAGAAGCTGAGAGAGTATTGCAACGCAAATAATATATTAGTCAGTGCAATCTCTGTACTGGGATCAAACGGAACCCCATGGGGCTCCAATGCAGTTTTGGGTTCTGAGGTGCTTAAGAAAATTGCTATGGCCAAAGGAAAATCTGTTGCTCAGGTTAGTATGAGATGGGTTTACGAGCAAGGCGCGAGTCTTGTGGTAAAAAGTTTCAGTGAAGAGAGATTGAGGGAAAACTTGAACATATTTGACTGGGAACTCACTAAGGAAGACCATGAAAAGATCGGTGAGATTCCACAGTGCAGAATCTTGAGTGCTTATTTTTTGGTCTCACCTAATGGACCTTTCAAATCTCAAGAAGAGTTGTGGGATGATGAAGCTTGA

>Par_STORR

ATGGAGCTCCAATATTTTTCTTATTTTCAACCAACTTCCTCCGTTGTTGCTCTTCTACTTGCTCTTGTATCCATCTTATTCAGTTTCGTTGTTTTGAGGAAGACATTTTCGAATAACTACTCATCACCTGCATCATCAACAGAAACAGCGGTACTTTGTCATCAGCGGCAGCAGTCGTGTGCATTGCCAATTTCCGGTCTTCTTCATGTTTTCATGAATAAAAACGGCTTAATTCATGTAACTCTTGGAAATATGGCCGATATATACGGACCGATTTTCAGTTTCCCAACAGGTAGCCATAGAACTCTCGTTGTGAGCAGTTGGGAGATGGTAAAAGAGTGTTTTACAGGCAACAATGACACTGCTTTCTCAAACCGTCCTATCCCGTTAGCTTTTAAGACTATATTCTATGCATGCGGTGGCATAGACTCATACGGTCTTTCGAGTGTACCTTATGGAAAATATTGGAGGGAGCTACGAAAAGTCTGTGTGCATAACCTCCTCTCTAATCAACAACTACTCAAGTTCAGACACTTGATAATTTCTCAAGTTGATACGTCTTTCAATAAGCTGTATGAGTTATGCAAAAACTCTGAAGACAACCAGGGAAACCATACTACTACTACTACTACTACCGCAGCTGGCATGGTGAGAATGGATGATTGGCTCGCCCAATTATCGTTCAACGTGATAGGAAGAATAGTCTGCGGATTCCAATCAGACCCTAAGACAGGTGCTCCAAGCAGGGTGGAACAATTTAAAGAAGCAATTAATGAAGCATCGCATTTTATGTCGACATCTCCAGTGTCAGATAACGTTCCAATGCTAGGGTGGATTGACCAATTGACTGGTCTTACAAGAAATATGAAGCACTGCGGAAAGAAATTAGACTTGGTGGTCGAGAGCATAATTCAAGATCATCGTCGAAAGAGACGATTCTCTAGAACTAAAGGAGGAGATGAGAAGGACGATGAACAAGATGACTTCATCGACATTTGCTTGTCAATTATGGAGCAACCACAGCTTCCTGACAACAATAACCCTCCTCATATACCTATCAAATCTATTGTCCTGGACATGATAGGTGGGGGAACTGACACAACAAAACTGACCACCATCTGGACCCTTTCCTTGCTGCTGAACAACCCCCATGTGTTGGACAAGGCAAAACAAGAAGTGGATGCACACTTTGGAACGAAAAGGAGATCAACAAATGATGCCGTGGTGGATTTTGATGATATTCGTAACCTTGTATACATCCAGGCAATCATCAAAGAATCAATGCGGTTGTATCCAGCCAGCCCCGTGGTGGAGCGACTGAGCGGCCAAGATTGTGTGGTCGGTGGGTTTCATGTACCAGCAGGGACGAGATTATGGGCTAACGTATGGAAGATGCAACGAGACCCTAAAGTATGGGATGATCCATTGGTGTTTCGACCAGAGAGATTTTTGAGCGATGAACAGAAGATGGTTGATGTAAGGGGTCAAAATTATGAGCTGTTACCATTTGGAGCCGGTCGACGTATATGTCCAGGTGTATCCTTCTCTTTGGATCTAATGCAACTGGTACTGACTCGTCTTATTCTTGAGTTTGAAATGAAGTCTCCTAGCGGGAAAGTGGACATGACAGCAACACCAGGATTAATGAGTTACAAGGTGGTCCCCCTTGACATTCTGCTCACCCATCGTGGCACAAAGTCGTGTGTGCAGTCAGCAGCCTCTGAGACTAGTGGTGTACCAGTAATCACTCTGAGCTCGGGCAAGGTGATGCCTGTTTTAGGCATGGGTACATTTGAGAAAGTTGGTAAAGGGTCTGAAAGAGAGAGGTTGGCGATTTTAAAAGCGATAGAGGTGGGTTACAGATACTTCGATACAGCTGCTGCATACGAAACTGAAGAGGTTCTTGGAGAAGCTATTGCTGAAGCACTCCAACTTGGCCTAATCAAATCTCGAGATGAACTTTTCATCAGTTCCATGCTCTGGTGCACTGATGCTCACCCTGATCGTGTCCTCCTCGCTCTTCAGAATTCTCTGAGGAATCTTAAATTGGAGTATCTGGATCTATATATGTTACCCTTCCCGGCAAGCTTGAAGCCAGGGAAGATAACGATGGACATACCAGAGGAAGATATTTGTCGAATGGACTACAGGTCTGTATGGTCAGCCATGGAAGAGTGTCAGAACCTTGGCTTCACTAAATCAATCGGTGTTAGCAATTTCTCCTGCAAAAAGCTTCAGGAATTGATGGAGACCGCCAACATCCCTCCAGCTGTGAATCAAGTGGAGATGAGCCCGGCTTTCCAACAAAAGAAGCTGAGAGAGTATTGCAACGCAAATAATATATTAGTCAGTGCAGTCTCTATACTGGGATCAAACGGAACCCCATGGGGCTCCAATGCAGTTATGGGTTCTGAGGTGCTTAAGAAAATTGCTATGGCCAAAGGAAAATCTGTTGCTCAGGTTAGTATGAGATGGGTTTACGAGCAAGGCGCGAGTCTTGTGGTAAAAAGTTTCAGTGAAGAGAGATTGAGGGAAAACTTGAACATATTCGACTGGGAACTCACTAAGGAAGACAATGAAAAGATCGGTGAGATTCCACAGTGCAGAATCTTGACTGCTCATTTTTTGGTCTCACCTAATGGACCTTTCAAATCTCAAGAAGAGTTGTGGGATGACAAAGCTTGA

>Por_STORR

ATGGAGCTCCAATATTTTTCTTATTTTCAACCAACTTCCTCCGTTGTTGCTCTTCTACTTGCTCTTGTATCCATCTTATTCAGTGTCGTTGTTTTGAGGAAGACATTTTCGAATAACTACTCATCACCTGCATCATCAACAGAAACAGCGGTACTTTGTCATCAGCGGCAGCAGTCGTGTGCATTGCCAATTTCCGGTCTTCTTCATGTTTTCATGAATAAAAACGGCTTAATTCATGTAACTCTTGGAAATATGGCGGATAAATACGGACCAATTTTCAGTTTCCCAACAGGTAGCCATAGAACTCTCGTTGTGAGCAGTTGGGAAATGGTAAAAGAGTGTTTTACAGGCAACAATGACACTGCTTTCTCAAACCGTCCTATCCCGTTAGCTTTTCAGACTATATTCTATGCATGCGGTGGCATAGACTCATACGGTCTTTCGAGTGTACCTTATGGAAAATATTGGAGGGAGCTACGAAAGGTCTGTGTGCATAACCTCCTCTCTAATCAACAACTACTCAAGTTCAGACACTTGATAATTTCTCAAGTCGATACGTCTTTCAATAAGCTGTATGAGTTATGCAAAAACTCTGAAGACAACCAGGGAATGGTGAGAATGGATGATTGGCTCGCCCAATTATCGTTCAATGTGATAGGAAGAATAGTCTGCGGATTCCAATCAGACCCAAAGACAGGTGCTCCTAGCAGGGTGGAACAATTTAAAGAAGTAATTAATGAAGCATCTTATTTTATGTCGACATCTCCAGTGTCAGATAACGTTCCAATGCTAGGGTGGATTGACCAATTGACTGGTCTTACGAGAAATATGAAGCACTGCGGAAAGAAATTAGACTTGGTGGTAGAGAGCATAATTAAAGATCATCGTCAAAAGAGACGATTCTCTAGAACTAAAGGAGGAGATGAGAAGGACGATGAACAAGATGACTTCATCGACATTTGCTTGTCAATTATGGAGCAACCACAGCTTCCTGGCAACAATAGCCCTCCTCAGATACCTATCAAATCTATTGTCCTGGACATGATAGGTGGGGGCACTGACACCACAAAACTGACCACCATCTGGACCCTTTCCTTGCTGCTGAACAACCCCCATGTGTTGGACAAGGCAAAACAAGAAGTGGATGCACACTTTCGAAAGAAAAGGAGATCAACAGATGATGCAGCAGCAGCCGTGGTGGATTTTGATGATATTCGTAACCTTGTCTACATCCAGGCAATCATCAAAGAATCAATGCGGTTGTATCCAGCCAGCCCCGTGGTGGAGCGACTGAGCGGCGAGGATTGTGTGGTCGGTGGGTTTCATGTACCAGCAGGGACGAGATTATGGGCTAACGTATGGAAGATGCAACGAGACCCTAAAGTATGGGATGATCCATTGGTGTTTCGACCAGAGAGATTTTTGAGCGATGAACGGAAGATGGTCGATGTAAGGGGTCAAAATTATGAGCTGTTACCATTTGGAGCCGGTCGACGTATATGTCCAGGTGTATCCTTCTCTTTGGATCTAATGCAACTGGTACTGACTCGTCTTATTCTTGAGTTTGAAATGAAGTCCCCTAGCGGGAAAGTGGACATGACAGCAACACCAGGATTAATGAGTTACAAGGTGGTCCCTCTTGACATTCTGCTCACCCATCGGCGCATAAAGTCGTGTGTGCAGTTAGCATCCTCTGAGAGAGACATGGAGAGTAGTGGTGTACCAGTAATCACTCTGAGCTCGGGCAAGGTGATGCCTGTTTTAGGCATGGGAACATTTGAGAAAGTTGGTAAAGGGTCTGAAAGAGAGAGGTTGGCGATTTTAAAAGCCATAGAGGTGGGTTACAGATACTTCGACACAGCTGCTGCATACGAAACTGAAGAGGTTCTTGGAGAAGCTATTGCTGAAGCACTTCAACTTGGCCTAATCGAATCTCGAGATGAACTTTTCATCAGTTCCATGCTCTGGTGCACTGATGCTCACCCTGATCGTGTCCTCCTCGCTCTTCAGAATTCTCTGAGGAATCTTAAATTGGAGTATCTGGATCTATATATGTTACCCTTCCCGGCAAGCTTGAAGCCAGGGAAGATAACGATGGACATACCAGAGGAAGATATTTGTCGAATGGACTACAGGTCTGTATGGTCAGCCATGGAAGAGTGTCAGAACCTTGGCTTCACTAAATCAATCGGTGTTAGCAATTTCTCCTGCAAAAAGCTTCAGGAATTGATGGCGACCGCCAACATCCCTCCAGCCGTGAATCAAGTGGAGATGAGCCCGGCTTTCCAACAAAAGAAGCTGAGAGAGTATTGCAACGCAAATAATATATTAGTCAGTGCAGTCTCTATACTGGGATCAAACGGAACCCCATGGGGCTCCAATGCAGTTTTGGGTTCTGAGGTGCTTAAGCAAATTGCTATGGCCAAAGGAAAATCTGTTGCTCAGGTTAGTATGAGATGGGTTTACGAGCAAGGCGCGAGTCTTGTGGTAAAAAGTTTCAGTGAAGAGAGATTGAGGGAAAACTTGAACATATTCGACTGGGAACTCACTAAGGAAGACAATGAAAAGATCGGTGAGATTCCACAGTGCAGAATCTTGACTGCTTATTTTTTAGTCTCGCCTAATGGACCTTTCAAATCTCAAGAAGAGTTGTGGGATGACAAAGCTTGA

>Pbr_STORR

ATGGAGCTCCAATATTTTTCTTATTTTCAACCAACTTCCTCCGTTGTTGCTCTTCTACTTGCTCTTGTATCCATCTTATTCAGTGTCGTTGTTTTGAGGAAGACATTTTCGAATAACTACTCATCACCTGCATCATCAACAGAAACAGCGGTACTTTGTCATCAGCGGCAGCAGTCGTGTGCATTGCCAATTTCCGGTCTTCTTCATGTTTTCATGAATAAAAACGGCTTAATTCATGTAACTCTTGGAAATATGGCGGATAAATACGGACCGATTTTCAGTTTCCCAACAGGTAGCCATAGAACTCTCGTTGTGAGCAGTTGGGAGATGGTAAAAGAGTGTTTTACAGGCAACAATGACACTGCTTTCTCAAACCGTCCTATCCCGTTAGCTTTTCAGACTATATTCTATGCATGCGGTGGCATAGACTCATACGGTCTTTCGAGTGTACCTTATGGAAAATATTGGAGGGAGCTACGAAAGGTCTGTGTGCATAACCTCCTCTCTAATCAACAACTACTCAAGTTCAGACACTTGATAATTTCTCAAGTCGATACGTCTTTCAATAAGCTGTATGAGTTATGCAAAAACTCTGAAGACAACCAGGGAATGGTGAGAATGGATGATTGGCTCGCCCAATTATCGTTCAATGTGATAGGAAGAATAGTCTGCGGATTCCAATCAGACCCAAAGACAGGTGCTCCTAGCAGGGTGGAACAATTTAAAGAAGTAATTAATGAAGCATCTTATTTTATGTCGACATCTCCAGTGTCAGATAACGTTCCAATGCTAGGGTGGATTGACCAATTGACTGGTCTTACGAAAAATATGAAGCACTGCGGAAAGAAATTAGACTTGGTGGTAGAGAGCATAATTAAAGATCATCGTCAAAAGAGACGATTCTCTAGAACTAAAGGAGGAGATGAGAAGGACGATGAACAAGATGACTTCATCGACATTTGCTTGTCAATTATGGAGCAACCACAGCTTCCTGGCAACAATAGCCCTCCTCAGATACCTATCAAATCTATTGTCCTGGACATGATAGGTGGGGGCACTGACACCACAAAACTGACCACCATCTGGACCCTTTCCTTGCTGCTGAACAACCCCCATGTGTTGGACAAGGCAAAACAAGAAGTGGATGCACACTTTCGAAAGAAAAGGAGATCAACAGATGATGCAGCAGCAGCCGTGGTGGATTTTGATGATATTCGTAACCTTGTCTACATCCAGGCAATCATCAAAGAATCAATGCGGTTGTATCCAGCCAGCCCCGTGGTGGAGCGACTGAGCGGCGAGGATTGTGTGGTCGGTGGGTTTCATGTACCAGCAGGGACGAGATTATGGGCTAACGTATGGAAGATTCAACGAGACCCTAAAGTATGGGATGATCCATTGGTGTTTCGACCAGAGAGATTTTTGAGCGATGAACAGAAGATGGTTGATGTAAGGGGTCAAAATTATGAGCTGTTACCATTTGGAGCCGGTCGACGTATATGTCCAGGTGTTTCCTTCTCTTTGGATCTAATGCAACTGGTACTGACTCGTCTTATTCTTGAGTTTGAAATGAAGTCCCCTAGCGGGAAAGTGGACATGACAGCAACACCAGGATTAATGAGTTACAAGGTGGTCCCTCTTGACATTCTGCTCACCCATCGGCGCATAAAGTCGTGTGTGCAGTTAGCATCCTCTGAGAGAGACATGGAGAGTAGTGGTGTACCAGTAATCACTCTGAGCTCGGGCAAGGTGATGCCTGTTTTAGGCATGGGAACATTTGAGAAAGTTGGTAAAGGGTCTGAAAGAGAGAGGTTGGCGATTTTAAAAGCCATAGAGGTGGGTTACAGATACTTCGACACAGCTGCTGCATACGAAACTGAAGAGGTTCTTGGAGAAGCTATTGCTGAAGCACTTCAAATTGGCCTAATCAAATCTCGAGATGAACTTTTCATCAGTTCCATGCTCTGGTGCACTGATGCTCACCCTGATCGTGTCCTCCTCGCTCTTCAGAATTCTCTGAGGAATCTTAAATTGGAGTATCTGGATCTATATATGTTACCCTTCCCGGCAAGCTTGAAGCCAGGGAAGATAACGATGGACATACCAGAGGAAGATATTTGTCGAATGGACTACAGGTCTGTATGGTCAGCCATGGAAGAGTGTCAGAACCTTGGCTTCACTAAATCAATCGGTGTTAGCAATTTCTCCTGCAAAAAGCTTCAGGAATTGATGGCGACCGCCAACATCCCTCCAGCCGTGAATCAAGTGGAGATGAGCCCGGCTTTCCAACAAAAGAAGCTGAGAGAGTATTGCAACGCAAATAATATATTAGTCAGTGCAGTCTCTATACTGGGATCAAACGGAACCCCATGGGGCTCCAATGCAGTTTTGGGTTCTGAGGTGCTTAAGCAAATTGCTATGGCCAAAGGAAAATCTGTTGCTCAGGTTAGTATGAGATGGGTTTACGAGCAAGGCGCGAGTCTTGTGGTAAAAAGTTTCAGTGAAGAGAGATTGAGGGAAAACTTGAACATATTCGACTGGGAACTCACTAAGGAAGACAATGAAAAGATCGGTGAGATTCCACAGTGCAGAATCTTGACTGCTTATTTTTTAGTCTCGCCTAATGGACCTTTCAAATCTCAAGAAGAGTTGTGGGATGACAAAGCTTGA

Supplementary dataset 5 (associated with Figure 2b)

- Nucleotide sequences of yeast codon optimised full length STORR cDNAs from *Papaver* species

>Pca_STORR_yeast_coden_optimised Papaver californicum

ATGGAATTGCAATACTTTTCTTACTTCCAGCCAACCTCCTCTTTGTTCGTTTTGGTTTCCATTTTGTTGATCTCCGCCGTTGTTTTGAGAAAGATCTTCTTGAACAACTACTCTTCATCTCCAGCCTCTTCTACTGAAACTGCTTTGTTGTCTCATCAGAGAAGATCATCTTGTGCCTTGCCAATTATTGGTTTGTTGCACGTTTTCATGAACAAGAACGGTTTGATCCATGTTACCTTGGGTAATATGGCTGATAAGTACGGTCCAATTTTCTCTTTTCCAACCGGTTCTCATATGACCTTGGTTGTTTCTTCTTGGGAAATGGCTAAAGAATGTTTCACCGGTAACAACGATATTGCCTTTTCTAACAGACCAATTCCTCTGGTTTTCAAGACCATTTTCTATGCCTGTAGAGGTATCGACTCTTACGGTTTATCTTCTGTTCCATACGGTAAATACTGGCGTGAATTGAGAAAGGTTTGCGTTCATAACTTGTTGTCTAACCAGCAGTTGTTGAACTTCAGACACTTGATCATCTCTCAAGTTGACACCTCTTTCAACAAGTTGTACGAATTGTGTAAGAACTCCGCCGATAATCAAGGTAACTACACTACTACAACTACTGCTGCTGGTATGGTTAGAATGGATGATTGGTTGGCTCAATTGTCCTTCAACGTTATTGGTAGAATTGTCTGCGGTTTCCAATCTGATACAGAAACTTCTGCTTCTTCCTCTGCTGAACAATTGACTGAAGCTATTAACGATGCCTCCTACTTCATGTCTACTTCTCCAGTTTCTGATAACGTTCCATGGTTAGGTTGGGTTGATCAATTGACAGGTTTGACCAGAAACATGAAGCAATGTGGTAAGAAGTTGGACTTGGTCTTGGAATCCATTATCGAAGATCACAGACAGAAGAGAAGATTCTCCAGATCTAAACAAGGTGACGAGAACTTCATCAACGAAGATGAACAGGATGATTTCATCGACATCTGCTTGTCCATTATGGAACAACCACAATTGCCAGGTAACAACAACCCATGTCAAATTCCAATCAAGTCCATCGTTTTGGATATGATTGGTGGTGGTACTGATACCACTAAGTTGACTACTATTTGGACCCTGTCCTTGTTATTGAACAACCCACATGTTTTGGACAAGGCTAAACAAGAAGTTGACGCTCATTTTGGTACTAAGAGAAATGATGCTGGTGGTGTTGTTGTCGATTTTGATGATACACGTAACTTGGTTTACATCCAGGCCATCATCAAAGAATCTATGAGATTATATCCAGCCTCGCCATTGGTTGAAAGATTGTCTGGTGAAGATTGTGTTGTTGGTGGTTTTCATGTTCCAGCTGGTACTAGATTGTGGGCTAATGTTTGGAAAATGCACAGAGATCCAAAGGTTTGGGATGATCCATTGGTTTTTAGACCAGAAAGATTCTTGTTGGACGACCAAAAGATGGTTGATGTCAGAGGTCAAAATTACGAGTTGTTGCCATTTGGTGCTGGTAGAAGAATTTGTCCAGGTGTTTCTTTCTCCTTGGATTTGATGCAATTGGTCTTGACCAGATTGATCTTGGAATTCGAAATGAAGTCTCCATCCGGTAAGTTAGATATGACTGCTACTCCAGGTTTGATGTCCTACAAAGTTGTTCCTTTGGATATCCTGTTGACCTCTAGAAGAATGAAGTGGTGTGTTCAATTGGCTTCTGAAAGGGATAAGAAGATGGAATCTTCAGTTGTCCCAGTTATCACCTTGTCCTCTTGTAAAGTTATGCCAGTTTTAGGTATGGGCACCTTTGAAAAAGTTGGTAAGGGTTCTGAAAGAGAGAGATTGGCTATTTTGAAGGCCATTGAAGTTGGTTACAGATACTTTGATACAGCTGCTGCTTACGAAACTGAAGAGGTTTTGGGTGAAGCCATTGCTGAAGCTTTACAATTGGGTTTGATTAAGTCCAGGGATGAGTTGTTCATCTCTTCTATGTTATGGTGTACCGATGCCTACCATGATAGAGTTTTGCCAGCATTGCAAAACAGCTTGAGAAACTTGAAATTGGAGTACCTGGACTTGTACATGCTACCATTTCCAGTTTCATTGAAACCAGGTGGTAAGATCTCTATGGATATTCCAGAAGATGAGATTTTCACCATGGATTACAGATCAGTTTGGGCTGCTATGGAAGAATGTCAAAACTTGGGTCTGACTAAGTCTATCGGTGTTTCTAACTTCTCCTGCAAAAAGTTGCAAGAATTGATGGCTACTGCTAACATTCCACCAGCTGTTAATCAAGTTGAAATGTCTCCAGCTTTCCAGCAGAAGAACTTAAGAGAATACTGTAACGCCAACAACATCTTGATTTCCGCTGTTTCTATTTTGGGTTCTAATGGTACTCCATGGGGTTCAAATGCTGTTTTGGGTAACGAAGTTTTGAACCAGATTGCTATGGCTAGAGGTAAATCTGTTGCTCAAGTTAGTATGAGATGGGTCTATGAACAAGGTGCTTCTTTGGTTGTTAAGTCCTTCAGTGAAGAAAGGTTGAGGGAAAACTTGAACATCTTCGATTGGGAATTGACCAAAGAGGATTTGGAAAAAATGGGTGAAATCCCACAATGCAGAATTTTGACTGCTGACTTTATGGTTTCTGCCAATGGTCCATTCAAATCCCAAGAGGAATTATGGGATGATAAGGCCTAG

>Pbr_STORR_yeast_coden_optimised Papaver bracteatum

ATGGAATTGCAATACTTTTCTTACTTCCAGCCAACCTCTTCTGTTGTTGCTTTGTTGTTGGCATTGGTGTCCATTTTGTTTTCCGTTGTTGTTCTGAGAAAGACCTTCTCCAACAATTATTCTTCTCCAGCTTCTTCTACTGAAACCGCTGTTTTGTGTCATCAAAGACAACAATCTTGCGCCTTGCCAATTTCTGGTTTGTTGCATGTTTTCATGAACAAGAACGGTTTGATCCATGTTACCTTGGGTAATATGGCTGATAAGTACGGTCCAATTTTCTCTTTTCCAACCGGTTCTCATAGAACCTTGGTTGTTTCTTCTTGGGAAATGGTCAAAGAATGTTTCACCGGTAACAACGATACTGCCTTTTCTAATAGACCAATTCCTTTGGCTTTCCAGACCATTTTTTACGCTTGTGGTGGTATTGACTCTTACGGTTTATCTTCTGTTCCATACGGTAAATACTGGCGTGAATTGAGAAAGGTTTGCGTTCATAACTTGTTGTCTAACCAGCAGTTGTTGAAGTTCAGACACTTGATCATCTCTCAAGTTGACACCTCTTTCAACAAGTTGTACGAATTGTGCAAGAACTCCGAGGATAATCAAGGTATGGTTAGAATGGATGATTGGTTGGCTCAATTGTCCTTCAACGTTATTGGTAGAATCGTTTGCGGTTTCCAATCTGATCCAAAAACTGGTGCTCCATCTAGAGTCGAACAATTCAAAGAAGTTATCAACGAGGCCTCCTACTTCATGTCTACATCTCCAGTTTCTGATAACGTCCCAATGTTAGGTTGGATCGATCAATTGACTGGTTTGACCAAGAATATGAAGCACTGTGGTAAGAAGTTGGATTTGGTTGTCGAATCCATCATCAAGGATCACAGACAAAAGAGAAGATTCTCTAGGACAAAAGGTGGTGACGAAAAGGATGACGAACAAGATGATTTCATCGACATCTGCTTGTCCATTATGGAACAACCACAATTGCCAGGTAACAATTCTCCACCACAAATTCCAATCAAGTCCATCGTTTTGGATATGATTGGTGGTGGTACTGATACCACTAAGTTGACTACTATTTGGACCCTGTCCTTGTTGTTGAACAATCCACATGTTTTGGACAAGGCTAAGCAAGAAGTTGACGCTCATTTTAGAAAGAAAAGAAGATCTACTGACGACGCTGCTGCAGCTGTTGTTGATTTTGATGATATTCGTAACCTGGTCTACATCCAGGCCATTATCAAAGAATCCATGAGATTATACCCAGCCTCTCCAGTTGTAGAAAGATTGTCTGGTGAAGATTGTGTTGTTGGTGGTTTTCATGTTCCAGCTGGTACTAGATTGTGGGCTAATGTTTGGAAGATTCAGAGAGATCCAAAGGTTTGGGATGATCCATTGGTTTTTAGACCAGAAAGATTCTTGTCCGACGAACAAAAGATGGTTGATGTCAGAGGTCAAAATTACGAGTTGTTGCCATTTGGTGCAGGTAGAAGAATTTGTCCAGGTGTTTCTTTCTCCTTGGATTTGATGCAATTGGTCTTGACCAGATTGATCTTGGAATTCGAAATGAAGTCTCCATCCGGTAAGGTTGATATGACTGCTACTCCAGGTTTGATGTCTTACAAAGTTGTCCCATTGGATATCTTGTTGACCCATAGAAGAATCAAGTCTTGCGTTCAATTGGCCTCTTCTGAAAGAGATATGGAATCTTCTGGTGTCCCAGTTATTACTTTGTCCTCTGGTAAAGTTATGCCAGTCTTAGGTATGGGTACTTTCGAAAAAGTTGGTAAGGGTTCCGAAAGAGAGAGATTGGCTATTTTGAAGGCTATCGAAGTTGGTTACAGATACTTTGATACTGCTGCTGCTTACGAAACTGAAGAAGTTTTGGGAGAAGCTATTGCTGAAGCCTTGCAAATTGGTTTGATCAAGTCTAGAGATGAGCTGTTCATCTCTTCTATGTTGTGGTGTACAGATGCTCATCCAGATAGAGTTTTGTTGGCTCTGCAAAACTCTTTGAGGAACTTGAAGTTGGAATACCTGGACTTGTACATGTTGCCTTTTCCAGCTTCATTGAAGCCAGGTAAGATTACCATGGATATTCCAGAAGAGGATATCTGCAGAATGGATTACAGATCAGTTTGGTCTGCTATGGAAGAATGTCAAAATTTGGGTTTCACCAAGTCTATCGGTGTCTCTAATTTCTCCTGCAAAAAGTTGCAAGAATTGATGGCTACTGCTAACATTCCACCAGCTGTTAATCAAGTTGAAATGTCCCCAGCTTTCCAGCAGAAAAAATTGAGAGAATACTGCAACGCTAACAACATCTTGGTTTCCGCTGTTTCTATTTTGGGTTCTAATGGTACTCCATGGGGTTCAAATGCTGTTTTAGGTTCTGAAGTCTTGAAGCAAATTGCTATGGCTAAGGGTAAATCCGTTGCTCAAGTTTCAATGAGATGGGTTTATGAACAAGGTGCTTCTTTGGTCGTTAAGTCCTTTAGCGAAGAAAGGTTGAGAGAGAACTTGAACATTTTCGATTGGGAGTTGACCAAAGAGGACAACGAAAAGATTGGTGAAATTCCACAATGCAGAATCTTGACCGCTTACTTTTTGGTTTCTCCAAATGGTCCATTCAAGTCCCAAGAGGAATTGTGGGATGATAAGGCTTAG

>Par_STORR_yeast_coden_optimised Papaver armeniacum

ATGGAATTGCAATACTTTTCTTACTTCCAGCCAACCTCTTCTGTTGTTGCTTTGTTGTTGGCATTGGTGTCTATCTTGTTCTCATTCGTTGTTCTGAGAAAGACCTTCTCCAACAATTATTCTTCTCCAGCTTCTTCTACTGAAACCGCTGTTTTGTGTCATCAAAGACAACAATCTTGCGCCTTGCCAATTTCTGGTTTGTTGCATGTTTTCATGAACAAGAACGGTTTGATCCATGTTACCTTGGGTAATATGGCTGATATCTACGGTCCAATTTTCTCTTTTCCAACCGGTTCTCATAGAACCTTGGTTGTTTCTTCTTGGGAAATGGTCAAAGAATGTTTCACCGGTAACAACGATACTGCCTTTTCTAATAGACCAATTCCTTTGGCTTTCAAGACCATTTTTTACGCTTGTGGTGGTATCGACTCTTACGGTTTATCTTCTGTTCCATACGGTAAATACTGGCGTGAATTGAGAAAGGTTTGCGTTCATAACTTGTTGTCTAACCAGCAGTTGTTGAAGTTCAGACACTTGATCATCTCTCAAGTTGACACCTCTTTCAACAAGTTGTACGAATTGTGCAAGAACTCCGAAGATAATCAAGGTAACCATACCACTACTACTGCTGCTGGTATGGTAAGAATGGATGATTGGTTGGCTCAATTGTCCTTCAACGTTATTGGTAGAATCGTTTGCGGTTTCCAATCTGATCCAAAAACTGGTGCTCCATCTAGAGTCGAACAATTCAAAGAAGCTATTAACGAAGCCAGCCATTTCATGTCTACATCTCCAGTTTCTGATAACGTCCCAATGTTAGGTTGGATCGATCAATTGACTGGTTTGACCAGAAATATGAAGCACTGTGGTAAGAAGTTGGATTTGGTTGTCGAATCCATCATCCAAGATCACAGAAGAAAGAGAAGATTCTCTAGGACAAAAGGTGGTGACGAAAAGGATGATGAACAGGATGATTTCATCGACATCTGCTTGTCCATTATGGAACAACCACAATTGCCAGATAACAACAACCCACCACATATTCCAATCAAGTCCATCGTTTTGGATATGATTGGTGGTGGTACTGATACCACTAAGTTGACTACTATTTGGACCCTGTCCTTGTTGTTGAACAATCCACATGTTTTGGACAAGGCTAAGCAAGAAGTTGACGCTCATTTTGGTACAAAAAGAAGATCTACAAACGACGCCGTTGTTGATTTCGATGACATTAGAAACTTGGTCTACATCCAGGCCATTATCAAAGAATCTATGAGATTATACCCAGCCTCTCCAGTTGTAGAAAGATTGTCTGGTCAAGATTGTGTTGTTGGTGGTTTTCATGTTCCAGCTGGTACTAGATTGTGGGCTAATGTTTGGAAAATGCAGAGAGATCCAAAGGTTTGGGATGATCCATTGGTTTTTAGACCAGAAAGATTCTTGTCCGACGAACAAAAGATGGTTGATGTCAGAGGTCAAAATTACGAGTTGTTGCCATTTGGTGCAGGTAGAAGAATTTGTCCAGGTGTTTCTTTCTCCTTGGATTTGATGCAATTGGTTTTGACCAGGCTGATTTTGGAATTCGAAATGAAGTCTCCATCCGGTAAGGTTGATATGACTGCTACTCCAGGTTTGATGTCTTACAAAGTTGTCCCATTGGATATCTTGTTGACCCATAGAGGTACTAAGTCTTGTGTTCAATCTGCTGCTTCTGAAACTTCTGGTGTTCCAGTTATTACTCTGTCCTCTGGTAAAGTTATGCCAGTTTTAGGTATGGGTACTTTCGAAAAAGTTGGTAAGGGTTCCGAAAGAGAACGTTTGGCTATTTTGAAGGCTATCGAAGTTGGTTACAGATACTTTGATACAGCTGCTGCTTACGAAACCGAAGAAGTTTTGGGCGAAGCTATTGCTGAAGCTTTACAATTGGGTTTGATCAAGTCTAGGGATGAGTTGTTCATCTCTTCTATGTTGTGGTGTACAGATGCTCATCCAGATAGAGTTTTGTTGGCCTTGCAAAACTCTTTGAGGAACTTGAAGTTGGAGTACTTGGACTTGTACATGTTGCCTTTTCCAGCTTCATTGAAGCCAGGTAAGATTACCATGGATATTCCAGAAGAGGATATCTGCAGAATGGATTACAGATCAGTTTGGTCTGCTATGGAAGAATGTCAAAATTTGGGTTTCACCAAGTCAATCGGTGTCTCTAATTTCTCCTGCAAAAAGTTGCAAGAGTTGATGGAAACTGCTAACATTCCACCAGCTGTTAATCAAGTTGAAATGTCCCCAGCTTTCCAGCAGAAAAAATTGAGAGAATACTGCAACGCTAACAACATCTTGGTTTCCGCTGTTTCTATTTTGGGTTCTAATGGTACTCCATGGGGTTCAAATGCTGTTATGGGTTCTGAAGTCTTGAAAAAGATTGCTATGGCCAAGGGTAAATCCGTTGCTTTGGTTTCAATGAGATGGGTTTATGAACAAGGTGCTTCTTTGGTCGTTAAGTCCTTTAGCGAAGAAAGGTTGAGAGAGAACTTGAACATTTTCGATTGGGAGTTGACCAAAGAGGACAACGAAAAGATTGGTGAAATTCCACAATGCAGAATCTTGACTGCTTACTTCTTAGTTTCTCCAAACGGTCCATTCAAGTCTCAAGAGGAATTGTGGGATGATAAGGCTTAG

Supplementary dataset 6 (associated with Figure 3a)

- Nucleotide sequences of *Papaver* P450 cytochrome *STORR* and *CYP82Y2*-*L* genes

>Pbr_CYP82Y-OUT3 Pbr_CYP82Y_contig_5568_CDS

ATGGAGTTCTTTAAGAAATATTTCCGTTCGTCTTCACATCTTCAAATGTACAATGCTGCTCTTCTGCTTGCTCCTTTACTCATATTTCTCACTATCATCTACAATGTACAGAAGCTGAGTGTTGAGAGAAATAACAAGGGTCAGAAGAAGAAGCATTACTCATCGTCATCATGTACAACACCAGTTACCAAAACTCCGGTGCCACAAGCTTCGGGTGCATGGCCTGTGATAGGTCATCTTCGTCTTATCACGAACGCCAAAGAGTCAACTCATATAACTCTTGCAAAAATGGCTGATAAGTATGGACCCATTTTTAGCATACGATTCGGTAGTCATAAGACCTTAATAGTGAGTAGTTGGGAAATGGTAAAAGAGTGTTTTACTGGCACCAACGACAAGTTATTATCAAATCGACCTCCCCATTCAGCGTTTGAGCATATGTTCTATCAAAATGGATATGGAGTTGGTTTGGTACCTTATGGAGAATATTGGAGGGAGTTACGCAAGATAACTACTCATAACCTCCTCGCTAATCAAACATTTTTGAAGTACAAACACTTTGTAACTCATGAAGTGGATACCTCGTTTAGAAAGCTTTATGAGTTATGCAACAATAAGAATCATACCACTGACAGCAAGGGAGTTACTACTCCTGCAACAGCTCTAATAAGAATGGATGATTGGCTCACACACTTGACGCACGACGTGGTTGGTATGATTGTTGGGATAGGCGGATCCCAATCTAATGCTGAGACAGGTGCCGTGAGCAACCGCGAACGATTCATACTTGCACACCATGAAGCATCACATTTTGTGACCACATCTTCAATTTCAGATATAGTTCCATGGCTAGGGTGGATTGATAGATTAACTGGTCTTACGAGAAAAATGAAAATTTGTGGCAAGAAATTAGACATGACACTTGGGAGTATAATTGAAGAACGTCGGCAGAAGAGACAATTTTGTAAAACTAAAGGAAAAGAATACAATTCAACGAACATAGCTGTCGAGGATAAGCAAGAAGACTTCATTGACATTTGCTTTTCGCTTATGGAGCAGTCACAACTTCCTGGGGAGAATCCTGAAATATCCATCAAATCTTTTATCCTGGATATGATGTCGGCTGGGACGGACACAACCAAATCGGTCTTGATATGGACCATTTCTTTGTTGTTGAACCATCCTCATGTGCTAAACAAGGCAAAAGAAGAAGTAGATGCACACTTTGAAATGAAAAGGAGATCAACTGGTGATATAATTGTCGTGGATGTTGATGATGTCCGTAACCTTGTTTACATCCACGCTATAGTTAAAGAATCAATGAGGTTATATCCAGTGAGCCCAGTCATGGAGCGAATGACTAGTGCCGATTGTGCTGTTGGAGGCTTCTGTGTTCCTGCTGGGACACGAGTATGGGTTAACGTATGGAAAATGCAACGTGATCCAACAGTGTGGGTAGATCCAATGGTGTTTCAACCTGAGAGATTCTTGAGCGACAGTAATAGCAAGATAGATATTTCAAAGGGTCACAATTATGAACTCATACCATTTGGAACCGGTAGGCGAAAATGCCCTGGTTCATCTTTCGCATTGGAACTGATGCATTTGGTACTTACACGCCTTATTCTTGAGTTCGAAATGAAGGCACCGGGAGGGAAAGTGGACATGACAGAAACATCGAGCGTCATTCTTAGTGCCAAGGTAGCACCAGTGGACGTTCTAATCAGTCCAAGAACACCCTAG

>Pso_CYP82Y-PS0215450 Pso_CYP82Y_PS0215450

ATGGAGTTCTTTAAAAAATATTTACATTTTTCTTCACATCTTCAAATTTCCAGTGCTGCTCTTTTACTCATATTGCTTACCATTATCTATAAGCTGATCAGCATTGAGAGAAATAAGAAGGGTCTCAAGAAGCAGTACTCATCATCATCATGTACAACATCAGTTACCAAAACTCCGGTGCCACAAGCTTCGGGCGCATGGCCTGTGATAGGTCATCTTCGTCTTATCACGAACGCCAAAGAGTCAACACATATAACTCTTGCAAAAATGGCTGATAAGTACGGACCCATTTTTAGCATACGATTCGGTAGTCATAAGACCTTAATAGTGAGTAGTTGGGAAATGGTAAAAGAGTGTTTTACAGGAACCAATGACAAGTTATTGTCGAATCGTCCTCCCCATTCAGCGTTTGAGCATATGTTCTACCAAAATGGATATGGAGTTGGTTTGGTACCATATGGAGAATATTGGAGAGAGTTACGGAAGATAACTACACATAACCTCCTCGCTAATCAAACATTTTTGAAGTACAAACACTTTGTAACTTCTGAAGTGGATACTTCGTTTAGAAAGCTTTATGAGTTATGCAACAACAACAAAAAGCATAACAGTGACAGAAAGGGTGCTACTACTCCTGCAATAGCTCTAATAAGAATGGATGATTGGCTCACACACTTGACGCACGACGTGGTTGGTAGGATAGTTGGCATAGGCGGATCTGAATCAAATGCTGCGACAGGTACCATGAGCAGCCGCGAAAGATTCATACTTGCACACCATGAAGCATCCCATTTTGTGACCACATCTTCAATTTCAGATATAGTTCCATGGCTAGGGTGGATTGACAGATTAACTGGTCTTACGAGAAAAATGAAAAATTGTGGCAAGAAATTAGACATGACACTTGGGAGTATAATTGAAGAACGTCGGCAGAAGAGACAATTTTGTAGGACTAAAGGAAAAGAATACAATTCAACCAACATAGCTGTTGAGGATAAGCAAGAAGACTTCATTGACATTTGCTTTTCACTTATGGAGCAGTCACAACTTCCTGGGGAGAATCCTGAAATATCCATCAAATCTTTTATCCTGGATATAATGTCGGCTGGGACGGACACAACCAAATCGGTCTTGATATGGACCATTTCTTTGTTGTTGAACCATCCCCATGTGTTAAGCAAAGCGAAACAAGAAGTAGATGCACACTTTGGAATGAAAAGGAGATCAACTGATGATATAGTTGTGGTGGACGTTGATGATGTTCATAACCTGGTTTACATCCACGCTATATTTAAAGAATCCATGCGGTTGTATCCAGTGAGTCCAGTCATGGAGCGAATGACTAGTGATGATTGTGTTGTTGGTGGCTTCTGTGTTCCTGCTGGGACACGAGTATGGGTTAACGTATGGAAAATGCAACGTGATCCAACAGTGTGGGAAGATCCAACGGTGTTTCAACCTGAGAGATTCTTGAGCGACAACAAAAGAAATATTGATATTGCAAAGGGTCATAATTATGAACTCATACCATTTGGAACCGGTAGGCGAAAATGCCCTGGTTCATCTTTCGCATTGGAACTGATGCATTTGGTACTTACACGCCTTATTCTTGAGTTCGAAATGAAGGCACCAGGAGGGAAAGTGGACATGACAGAAACATCAAGCGTCATTCTTAGTGCCAAGGTAGCGCCAGTGGACGTTCTAATCAGTCCAAGAACACCCTAG

>Ptr_CYP82Y-OUT4 Ptr_CYP82Y_gc30928_g2_i1

ATGGAGTTCTTTAAGAAATATTTCCATTCGTCTTCACATCTTCAAATTTCCAATACTGCTCTTCTGCTTGCTCCTTTACTCATATTTCTCACTATCATCTACAATGTACAGAAGCTGATCAGTGTTGAGAGAAATAACGACGGTCGGAAGAAGAAGCAGTACTCATATACAACATCAGTTACCAAAACTCCAGTGCCACAAGCTGCGGGTGCCTGGCCTGTGATAGGTCATCTTCGTCTTGTCACGAACGCCAAAGAGTCAACTCATATAACTCTGGCAAAAATGGCTGATAAGTACGGACCCATTTTTAGCATACGATTCGGTAGTCATAAGACCTTAATAGTGAGTAGCTGGGAAATGGTAAAAGAGTGTTTTACTGGCACCAATGACAAGTTATTGTCGAATCGTCCTCCCCATTCAGCGTTTGAGCATATGTTCTATCAAAATGGATATGGAGTTGGTTTGGTACCTTATGGAGAATATTGGAGGGAGTTACGGAAGATAACTACACATAACCTCCTCGCTAATCAAACATTTTTGAAGTACAAACACTTTGTAACTTCTGAAGTGGATACCTCGTTTAGAAAGCTTTATGAGTTATGCAACAACAACAACAGCAAGGGAGCTACTACTCCTGCGATAGCTCTAATAAGAATGGATGATTGGCTCACACACTTGACGCACGACGTGGTTGGTAGGATAGTTGGCATCGGTGGATCTGAATCAAATGCTGAGACAGGTACCATGAGCAGCCGCGAAAGATTCATACTCGCACACCATGAAGCATCCCATTTTGTGACCACATCTTCAATTTCAGATATAGTTCCTTGGCTAGGGTGGATTGATGGATTAACTGGTCTTACGAGAAAAATGAAAAACTGTGGCAAGAAATTAGACATGACACTTGGGAGTATAATTGAAGAACGTTGGAAGAAGAGACAATTTTGTAGAACTAAAGGAGAAGAATACAGTTCAACGAACATAGCTGTTGAGGATAAGCAAGAAGACTTCATTGACATTTGCTTTTCGCTTATGGAGCAGTCACAACTTCCTGGGGAGAATCCTGAAATATCCATCAAATCTTTTATCCTGGATATGATGTCGGCTGGGACGGACACAACTAAATCGGTCTTGATATGGACCATTTCTTTGTTGTTGAACCATCCTCATGTGCTAAACAAAGCAAAAGAAGAAGTAGATGCACACTTTGAAATGAAAAGGAGATCAACTGGTGATACAATTGTCGTGGACGTTGATGATGTCCGTAACCTTGTTTACATCCACGCTATAATCAAAGAATCCATGCGGTTGTATCCGGTGAGTCCAGTCATGGAGCGAATGACTAGTGATGATTGTGTTGTTGGTGGCTTCTGTGTTCCTGCTGGGACACGAGTATGGGTTAACGTATGGAAAATGCAACGTGATCCATCAGTGTGGGAAGATCCAACGGTGTTTCAACCTGAGAGATTCTTGAGCGACAACAAAAGAAAGATTGATATTGCAAAGGGTCATGATTATGAACTCATACCATTTGGAACTGGTAGGCGAAAATGCCCTGGTTCATCTTTCGCATTGGAACTGATGCATTTGGTACTTACACGCCTTATTCTTGAGTTCGAAATGAAGGCACCAGGAGGACAAGTGGACATGACAGAAACATCGAGCGTCATTCTTAGTGCCAAGGTTTCACCAGTGGACGTTCTAATCAGTCCAAGAACACCCTAG

>Pnu_CYP82Y-OUT2 Pnu_CYP82Y_7065

ATGGAATTCTTTAAGAAATATTTCCATTCTTCTTCCATTAATGCTGCTCTTCTCCTTGGTCTTTTGCTCATATTTCTTAATATTATTTACAATGTATACAAGCTGGTCAGTATTGAGAGAAATAAGAAGAGTCAGAAGAAGAAGCAGTACTCAGTTACCACAACTCCAGTGCCAGAAGCTTCAGGTGCATGGCCTGTGATAGGTCATCTTCGTCTTATCACGAACGCCAAAGAGTCGACTCATATAACTCTAGCAAAAATGGCTGATGAATATGGACCAATCTTTAGCATAAGATTTGGTAGTCATAAGACTTTAATAGTGAGTAGTTGGGAAATGGTAAAAGAGTGTTTTACAGGGACCAATGACAAATTATTGTCCAATCGTCCTCCGCATTCAGCATTTGAGCATATGTTCTATCAGAATGGATATGGAGTTGGTTTGGTACCTTATGGAGAGTATTGGATGGAGTTACGAAAGATAACTACACATAACCTCCTCGCTAATCAAACATTTACGAAGTACAAGCACTTTGTGACTTCTGAAGTGGATACCTCGTTTAAAAAGCTTCATGAGCTATGCAACAACAACAACAAGCATAACAGTGACAGCAAGGGAGCTACTACTCCGGCAATAGCTCTAGTCAGAATGGATGATTGGCTCACACACTTGACGCACGACGTGGTTGGTCGGATGGTTGGCATAGGCGGATCCGAATCAAATGCTGAGACAGGTGCCATGAGCAACCGTGAAAAATTCATACATGCACACCATGAAGCATCAAATTTTGTGACCACATCTTCAATTTCAGATATAGTTCCATGGCTAGGGTGGATTGATCGATTAACTGGTCTCACGAGAAAAATGAAAAATTGTGTAAAGAAATTGGACAAGACACTCGGGGATATAATTGAAGAACGTCGTCAACAGAGACGATTTCGTAGAACTAAAGGAAGAGAATACAATACTACAGACATGGCTGTTGCGGACGAGCAAGAAGACTTCATTGACATTTGCTTTTCGCTTATGGAGCAGTCACAGCTTCCCGGGGAGAACCCTGATATATCCATCAAATCTTTTATCCTGGATATGATGTCGGCCGGGACGGACACAACCAAATCGGTCTTGATATGGACCATTTCTTTGTTGTTGAACCATCCTCATGTGCTCAACAAGGCAAAAGACGAAGTAGATGCACATTTTGGAATGAAAAGGAAATCATCAACCGGTGATAAAGTTGTCGTGGACGTTCACGATGTCCATAACCTTGTTTACATCCATGCTATAATCAAAGAATCAATGCGGTTGTATCCAGTGAGCCCAGTCATGGAGCGAATGACTAGTGACGATTGCGCTGTTGGTGGCTTCCGTGTTCCTGCTGGGACACGAGTATGGGTTAATGTATGGAAAATGCAACGTGATCCAACAGTGTGGGAAGATCCAATGGTGTTTCAACCTGAGAGATTTTTGAGCGGCAACAAAAGCAAGATCGACATTGCCAAGGGTCATAATCATGAGCTCATACCATTTGGAACCGAGAGGCGAAAATGCCCTGGTTCATCTTTCGCATTGGAACTCATGCATTTGGCACTTACACGTCTAATTCT

>Pnu_CYP82Y-OUT1 Pnu_CYP82Y_1315731

ATGGAGTTGCTTAAGAAAGATTTTCCTTTTTCTTCACATCTTCAATTTTCAAATGCTGCCCTTCTACTTGCTCTTTTAATCATATTTGTTACTATCATCTACAATGTACAGAAGCTGATCAGTATTGAGGGAAATGAGAAGAGTCCGAAGAAGAAGAAGCAGTATTCAGTTACCACAACTCCAGTGCCAGAAGCTTCGGGTGCATGGCCTGTGATAGGTCATCTTCGTCTTGTCACGAACGCCAAAGAGTCAACTCATATAACTCTTGCAAAAATGGCTGAGGAATATGGACCAATTTTCAGCATCAGATTTGGTAGCCACAAGACCTTAATAGTGAGCAGTTGGGAAATGGTAAAAGAGTGTTTTACAGGCACCAATGACAAATTACTGTCGAATCGTCCTCCCCATTCAGCGTTCGAGCATATGTTCTATCAGAACGGATATGGAGTTGGTTTGGTACCTTATGGAGAATATTGGATGGAGTTACGAAAGATAACTACGCATAACCTGCTGGCGAATCAAACGTTTTTGAAGTACAAGCACTTTGTGACTTCTGAAGTGGATACCTCGTTTAAAAAGCTTCATGAGCTATGCAGCAACAACAAGCATAACAGTGACGGCAAGGGAGCTACTACTACTCAGGCAATGGCTCTAGTAAGAATGGATGATTGGCTCACACACTTGACGCACGACGTGGTTGGTAGGATCGTTGGCATAGGCGGATCCGAATCAAATGCTGAAACAGGTGCCATGAGCAACCGCGAAAGATTCATACATGCACACCATGACGCATCAAATTTCGTGACCACATCTTCAATTTCAGATATAGTTCCATGGCTAGGGTGGCTTGATCGATTAACTGGTCTTACGAGAAAAATGAAAAATTGTGGAAAGAAATTGGACAAGACACTAGGGGATATAATTGAAGAACGTCGTCAACAGAGACGACTTCGTAGAACTGTTGCGGACGAGCAAGAAGACTTCATTGACATTTGCTTTTCGCTTATGGAGCAGTCACAGCTTCCTGGGGAGAACCCTGATATATCCATCAAATCTTTTATCCTGGATATGATGTCGGCCGGGACGGACACAACCAAATCGGTCTTGATATGGACCATTTCTTTGTTGTTGAACCATCCTCATGTGCTCAACAAGGCAAAAGACGAAGTAGATGCACATTTCCGAATGAAAAGGAAATCAACCGGTGATGATAAAGTTGTCGTGGACGTTGATGATGTACACAACCTTGTTTACATCCATGCTATATTCAAAGAATCAATGCGGTTGTATCCAGTGAGCCCTGTCATGGAGCGAATGACTAGTGATGACTGCGCTGTTGGTGGCTACCGTGTTCCTGCTGGGACACGAGTGTGGGTTAATGTATGGAAAATGCAACGTGATTCAACAGTGTGGGAAGATCCAATGGTGTTTCAACCCGAGAGGTTTCTGAGCGGCAACAAAATCAAGATTGACCTTTCGAAGGGGCATAACCATGAACTCATACCATTTGGAACCGGCAGGCGAAAATGCCCGGGTTCGTCTTTCGCATTGGAGCTGATGCATTTGGTACTTACACGTCTAATTCTTGAGTTCGAAATGAAGGCACCGGGAGGGAAAGTGGACATGAGCGAAACATCAAGTGTCATTCTTAGTGCCAAGGTAGCACCTGTAGACGTTCTGATCAGTCCAAGAACACCATAG

>Pso_STORR-PS1126560 Pso_STORR_PS1126560

ATGGAGCTCCAATATATTTCTTATTTTCAACCAACTTCCTCCGTTGTTGCTCTTCTACTTGCTCTTGTATCCATCTTATCCAGTGTCGTTGTTTTGAGGAAGACATTTTTGAATAACTACTCATCATCACCTGCATCATCCACAAAAACAGCGGTACTTTCTCATCAGCGGCAGCAGTCGTGTGCATTGCCAATTTCCGGTCTCCTCCATATTTTCATGAATAAAAACGGCTTAATTCATGTAACTCTTGGAAATATGGCTGATAAATACGGTCCGATTTTCAGTTTCCCAACAGGTAGCCATAGAACTCTCGTTGTGAGCAGTTGGGAGATGGTAAAAGAGTGTTTTACTGGCAACAATGACACTGCTTTCTCAAACCGTCCTATCCCGTTAGCTTTTAAGACTATATTTTATGCATGCGGTGGCATAGACTCATACGGTCTTTCGAGTGTACCTTATGGAAAATATTGGAGGGAGCTTCGAAAGGTCTGTGTGCATAACCTCCTGTCTAATCAACAACTACTCAAGTTCAGACACTTGATAATTTCTCAAGTCGATACGTCTTTCAATAAGCTGTATGAGTTATGCAAAAACTCTGAAGACAACCAGGGAAACTATACTACTACTACTACCGCAGCTGGCATGGTGAGAATCGATGATTGGCTCGCCGAATTATCGTTCAATGTGATAGGAAGAATAGTCTGCGGATTCCAATCAGGCCCTAAGACAGGTGCTCCAAGCAGGGTGGAACAATTTAAAGAAGCAATTAATGAAGCATCTTATTTTATGTCGACATCTCCAGTGTCAGATAATGTTCCAATGCTAGGGTGGATTGACCAATTGACAGGTCTTACGAGAAATATGAAGCACTGCGGAAAGAAATTAGACTTGGTGGTCGAGAGCATAATTAATGATCATCGTCAAAAGAGACGATTCTCTAGAACTAAAGGAGGAGATGAGAAGGACGATGAACAAGATGACTTCATCGACATTTGTTTGTCAATAATGGAGCAACCACAGCTTCCTGGCAACAATAATCCTTCTCAGATACCTATCAAATCTATTGTCCTGGACATGATAGGTGGGGGCACTGACACCACAAAACTGACCACCATCTGGACCCTTTCCTTGCTGCTGAACAACCCCCATGTGTTGGACAAGGCAAAACAAGAAGTGGATGCACACTTTCGAACCAAAAGGAGATCAACAAATGATGCAGCAGCAGCCGTGGTGGATTTTGATGATATTCGTAACCTTGTCTACATCCAGGCAATCATCAAAGAATCAATGCGGTTGTATCCAGCCAGCCCCGTGGTGGAGCGACTGAGCGGCGAAGATTGTGTGGTCGGTGGGTTTCATGTACCAGCAGGGACGAGATTATGGGCTAACGTATGGAAGATGCAACGAGACCCTAAAGTATGGGATGATCCATTGGTGTTTCGACCAGACAGATTTTTGAGCGATGAACAGAAGATGGTTGATGTAAGGGGTCAAAATTATGAGCTGTTACCATTTGGAGCCGGTCGACGTGTATGTCCAGGTGTATCCTTCTCTTTGGATCTAATGCAACTGGTACTGACTCGTCTTATTCTCGAGTTTGAAATGAAGTCTCCTAGCGGGAAAGTGGACATGACAGCAACACCAGGATTAATGAGTTACAAGGTGATCCCCCTTGACATTCTGCTCACCCATCGTCGCATAAAGCCGTGTGTGCAGTCAGCAGCCTCTGAGAGAGACATGGAGAGTAGTGGTGTACCAGTAATCACTCTGGGCTCGGGCAAGGTGATGCCTGTTCTGGGCATGGGAACATTTGAGAAAGTTGGTAAAGGGTCCGAAAGAGAGAGGTTGGCGATTTTAAAAGCGATAGAGGTGGGTTACAGATACTTCGATACAGCTGCTGCATACGAAACTGAAGAGGTTCTTGGAGAAGCTATTGCTGAAGCACTTCAACTTGGCCTAGTCAAATCTCGAGATGAACTTTTCATCAGTTCCATGCTCTGGTGCACTGATGCTCACGCTGATCGTGTCCTCCTCGCTCTTCAGAATTCGCTGAGGAATCTTAAATTGGAGTATGTGGATCTATATATGTTACCCTTCCCGGCAAGCTTGAAGCCTGGGAAGATAACGATGGACATACCAGAGGAAGATATTTGTCGCATGGACTACAGGTCTGTATGGGCAGCCATGGAAGAGTGTCAAAACCTTGGCTTCACTAAATCAATCGGTGTTAGCAATTTCTCCTGCAAAAAGCTTCAGGAATTGATGGCGACCGCCAACATCCCTCCAGCTGTGAATCAAGTGGAGATGAGCCCGGCTTTCCAACAAAAGAAGCTGAGAGAGTATTGCAACGCAAATAATATATTAGTCAGTGCAATCTCTGTACTGGGATCAAACGGAACCCCATGGGGCTCCAATGCAGTTTTGGGTTCTGAGGTGCTTAAGAAAATTGCTATGGCCAAAGGAAAATCTGTTGCTCAGGTTAGTATGAGATGGGTTTACGAGCAAGGCGCGAGTCTTGTGGTAAAAAGTTTCAGTGAAGAGAGATTGAGGGAAAACTTGAACATATTTGACTGGGAACTCACTAAGGAAGACCATGAAAAGATCGGTGAGATTCCACAGTGCAGAATCTTGAGTGCTTATTTTTTGGTCTCACCTAATGGACCTTTCAAATCTCAAGAAGAGTTGTGGGATGATGAAGCTTGA

>Par_STORR1 Par_STORR_contig_11585_CDS

ATGGAGCTCCAATATTTTTCTTATTTTCAACCAACTTCCTCCGTTGTTGCTCTTCTACTTGCTCTTGTATCCATCTTATTCAGTTTCGTTGTTTTGAGGAAGACATTTTCGAATAACTACTCATCACCTGCATCATCAACAGAAACAGCGGTACTTTGTCATCAGCGGCAGCAGTCGTGTGCATTGCCAATTTCCGGTCTTCTTCATGTTTTCATGAATAAAAACGGCTTAATTCATGTAACTCTTGGAAATATGGCCGATATATACGGACCGATTTTCAGTTTCCCAACAGGTAGCCATAGAACTCTCGTTGTGAGCAGTTGGGAGATGGTAAAAGAGTGTTTTACAGGCAACAATGACACTGCTTTCTCAAACCGTCCTATCCCGTTAGCTTTTAAGACTATATTCTATGCATGCGGTGGCATAGACTCATACGGTCTTTCGAGTGTACCTTATGGAAAATATTGGAGGGAGCTACGAAAAGTCTGTGTGCATAACCTCCTCTCTAATCAACAACTACTCAAGTTCAGACACTTGATAATTTCTCAAGTTGATACGTCTTTCAATAAGCTGTATGAGTTATGCAAAAACTCTGAAGACAACCAGGGAAACCATACTACTACTACTACTACTACCGCAGCTGGCATGGTGAGAATGGATGATTGGCTCGCCCAATTATCGTTCAACGTGATAGGAAGAATAGTCTGCGGATTCCAATCAGACCCTAAGACAGGTGCTCCAAGCAGGGTGGAACAATTTAAAGAAGCAATTAATGAAGCATCGCATTTTATGTCGACATCTCCAGTGTCAGATAACGTTCCAATGCTAGGGTGGATTGACCAATTGACTGGTCTTACAAGAAATATGAAGCACTGCGGAAAGAAATTAGACTTGGTGGTCGAGAGCATAATTCAAGATCATCGTCGAAAGAGACGATTCTCTAGAACTAAAGGAGGAGATGAGAAGGACGATGAACAAGATGACTTCATCGACATTTGCTTGTCAATTATGGAGCAACCACAGCTTCCTGACAACAATAACCCTCCTCATATACCTATCAAATCTATTGTCCTGGACATGATAGGTGGGGGAACTGACACAACAAAACTGACCACCATCTGGACCCTTTCCTTGCTGCTGAACAACCCCCATGTGTTGGACAAGGCAAAACAAGAAGTGGATGCACACTTTGGAACGAAAAGGAGATCAACAAATGATGCCGTGGTGGATTTTGATGATATTCGTAACCTTGTATACATCCAGGCAATCATCAAAGAATCAATGCGGTTGTATCCAGCCAGCCCCGTGGTGGAGCGACTGAGCGGCCAAGATTGTGTGGTCGGTGGGTTTCATGTACCAGCAGGGACGAGATTATGGGCTAACGTATGGAAGATGCAACGAGACCCTAAAGTATGGGATGATCCATTGGTGTTTCGACCAGAGAGATTTTTGAGCGATGAACAGAAGATGGTTGATGTAAGGGGTCAAAATTATGAGCTGTTACCATTTGGAGCCGGTCGACGTATATGTCCAGGTGTATCCTTCTCTTTGGATCTAATGCAACTGGTACTGACTCGTCTTATTCTTGAGTTTGAAATGAAGTCTCCTAGCGGGAAAGTGGACATGACAGCAACACCAGGATTAATGAGTTACAAGGTGGTCCCCCTTGACATTCTGCTCACCCATCGTGGCACAAAGTCGTGTGTGCAGTCAGCAGCCTCTGAGACTAGTGGTGTACCAGTAATCACTCTGAGCTCGGGCAAGGTGATGCCTGTTTTAGGCATGGGTACATTTGAGAAAGTTGGTAAAGGGTCTGAAAGAGAGAGGTTGGCGATTTTAAAAGCGATAGAGGTGGGTTACAGATACTTCGATACAGCTGCTGCATACGAAACTGAAGAGGTTCTTGGAGAAGCTATTGCTGAAGCACTCCAACTTGGCCTAATCAAATCTCGAGATGAACTTTTCATCAGTTCCATGCTCTGGTGCACTGATGCTCACCCTGATCGTGTCCTCCTCGCTCTTCAGAATTCTCTGAGGAATCTTAAATTGGAGTATCTGGATCTATATATGTTACCCTTCCCGGCAAGCTTGAAGCCAGGGAAGATAACGATGGACATACCAGAGGAAGATATTTGTCGAATGGACTACAGGTCTGTATGGTCAGCCATGGAAGAGTGTCAGAACCTTGGCTTCACTAAATCAATCGGTGTTAGCAATTTCTCCTGCAAAAAGCTTCAGGAATTGATGGAGACCGCCAACATCCCTCCAGCTGTGAATCAAGTGGAGATGAGCCCGGCTTTCCAACAAAAGAAGCTGAGAGAGTATTGCAACGCAAATAATATATTAGTCAGTGCAGTCTCTATACTGGGATCAAACGGAACCCCATGGGGCTCCAATGCAGTTATGGGTTCTGAGGTGCTTAAGAAAATTGCTATGGCCAAAGGAAAATCTGTTGCTCAGGTTAGTATGAGATGGGTTTACGAGCAAGGCGCGAGTCTTGTGGTAAAAAGTTTCAGTGAAGAGAGATTGAGGGAAAACTTGAACATATTCGACTGGGAACTCACTAAGGAAGACAATGAAAAGATCGGTGAGATTCCACAGTGCAGAATCTTGACTGCTCATTTTTTGGTCTCACCTAATGGACCTTTCAAATCTCAAGAAGAGTTGTGGGATGACAAAGCTTGA

>Par_STORR2 Par_STORR_contig_40428_CDS

ATGGAGCTCCAATATTTTTCTTATTTTCAACCAACTTCCTCCGTTGTTGCTCTTCTACTTGCTCTTGTATCCATCTTATTCAGTTTCGTTGTTTTGAGGAAGACATTTTCGAATAACTACTCATCACCTGCATCATCAACAGAAACAGCGGTACTTTGTCATCAGCGGCAGCAGTCGTGTGCATTGCCAATTTCCGGTCTTCTTCATGTTTTCATGAATACAAACGGCTTAATTCATGTAACTCTTGGAAATATGGCTGATATATACGGACCGATTTTCAGTTTCCCAACAGGTAGCCATAGAACTCTCGTTGTGAGCAGTTGGGAGATGGTAAAAGAGTGTTTTACAGGCAACAATGACACTGCTTTCTCAAACCGTCCTATCCCGTTAGCTTTTAAGACTATATTCTATGCATGCGGTGGCATAGACTCATACGGTCTTTCGAGTGTACCTTATGGAAAATATTGGAGGGAGCTACGAAAAGTCTGTGTGCATAACCTCCTCTCTAATCAACAACTACTCAAGTTCAGACACTTGATAATTTCTCAAGTTGATACGTCTTTCAATAAGCTGTATGAGTTATGCAAAAACTCTGAAGACAACCAGGGAAACCATACTACTACTACCGCAGCTGGCATGGTGAGAATGGATGATTGGCTCGCCCAATTATCGTTCAACGTGATAGGAAGAATAGTCTGCGGATTCCAATCAGACCCTAAGACAGGTGCTCCAAGCAGGGTGGAACAATTTAAAGAAGCAATTAATGAAGCATCGCATTTTATGTCGACATCTCCAGTGTCAGATAACGTTCCAATGCTAGGGTGGATTGACCAATTGACTGGTCTTACAAGAAATATGAAGCACTGCGGAAAGAAATTAGACTTGGTGGTCGAGAGTATAATTCAAGATCATCGTCAAAAGAGACGATTCTCTAGAACTAAAGGAGGAGATGAGAAGGACGATGAACAAGATGACTTCATCGACATTTGCTTGTCAATTATGGTGCAACCACAGCTTCCTGACAACAATAACCCTCCTCATATACCTATCAAATCTATTGTCCTGGACATGATAGGTGGGGGAACTGACACAACAAAACTGACCACCATCTGGACCCTTTCCTTGCTGCTGAACAACCCCCATGTGTTGGACAAGGCAAAACAAGAAGTGGATGCACACTTTGGAACGAAAAGGAGATCAACGAATGATGCCGTGGTGGATTTTGATGATATTCGTAACCTTGTATACATCCAGGCAATCATCAAAGAATCAATGCGGTTGTATCCAGCCAGCCCCGTGGTGGAGCGACTGAGCGGCCAAGATTGTGTGGTCGGTGGGTTTCATGTACCAGCAGGGACGAGATTATGGGCTAACGTATGGAAGATGCAACGAGACCCTAAAGTATGGGATGATCCATTGGTGTTTCGACCAGAGAGATTTTTGAGCGATGAACAGAAGATGGTTGATGTAAGGGGTCAAAATTATGAGCTGTTACCATTTGGAGCCGGTCGACGTATATGTCCAGGTGTATCCTTCTCTTTGGATCTAATGCAACTGGTACTGACTCGTCTTATTCTTGAGTTTGAAATGAAGTCTCCTAGCGGGAAAGTAGACATGACAGCAACACCAGGATTAATGAGTTACAAGGTGGTCCCCCTTGACATTCTGCTCACCCATCGTGGCACAAAGTCGTGTGTGCAGTCAGCAGCCTCTGAGAGTAGTGGTGTACCAGTAATCACTCTGAGCTCGGGCAAGGTGATGCCTGTTTTAGGCATGGGAACATTTGAGAAAGTTGGTAAAGGGTCTGAAAGAGAGAGGTTGGCGATTTTAAAAGCGATAGAGGTGGGTTACAGATACTTCGATACAGCTGCTGCATACGAAACTGAAGAGGTTCTTGGAGAAGCTATTGCTGAAGCACTCCAACTTGGCCTAATCAAATCTCGAGATGAACTTTTCATCAGTTCCATGCTCTGGTGCACTGATGCTCACCCTGATCGTGTCCTCCTCGCTCTTCAGAATTCTCTGAGGAATCTTAAATTGGAGTATCTGGATCTATATATGTTACCCTTCCCGGCAAGCTTGAAGCCAGGGAAGATAACGATGGACATACCAGAGGAAGATATTTGTCGAATGGACTACAGGTCTGTATGGTCAGCCATGGAAGAGTGTCAGAACCTTGGCTTCACTAAATCAATCGGTGTTAGCAATTTCTCCTGCAAAAAGCTTCAGGAATTGATGGAGACCGCCAACATCCCTCCAGCTGTGAATCAAGTGGAGATGAGCCCGGCTTTCCAACAAAAGAAGCTGAGAGAGTATTGCAACGCAAATAATATATTAGTCAGTGCAGTCTCTATACTGGGATCAAACGGAACCCCATGGGGCTCCAATGCAGTTTTGGGTTCTGAGGTGCTTAACAAAATTGCTATGGCCAAAGGAAAATCTGTTGCTCAGGTTAGTATGAGATGGGTTTACGAGCAAGGCGCGAGTCTTGTGGTAAAAAGTTTCAGTGAAGAGAGATTGAGGGAAAACTTGAACATATTCGACTGGGAACTCACTAAGGAAGACAATGAAAAGATCGGTGAGATTCCACAGTGCAGAATCTTGGCTGCTCATTTTTTGGTCTCACCTAATGGACCTTTCAAATCTCAAGAAGAGTTGTGGGATGACAAAGCTTGA

>Pbr_STORR Pbr_STORR_contig_2915_CDS

ATGGAGCTCCAATATTTTTCTTATTTTCAACCAACTTCCTCCGTTGTTGCTCTTCTACTTGCTCTTGTATCCATCTTATTCAGTGTCGTTGTTTTGAGGAAGACATTTTCGAATAACTACTCATCACCTGCATCATCAACAGAAACAGCGGTACTTTGTCATCAGCGGCAGCAGTCGTGTGCATTGCCAATTTCCGGTCTTCTTCATGTTTTCATGAATAAAAACGGCTTAATTCATGTAACTCTTGGAAATATGGCGGATAAATACGGACCGATTTTCAGTTTCCCAACAGGTAGCCATAGAACTCTCGTTGTGAGCAGTTGGGAGATGGTAAAAGAGTGTTTTACAGGCAACAATGACACTGCTTTCTCAAACCGTCCTATCCCGTTAGCTTTTCAGACTATATTCTATGCATGCGGTGGCATAGACTCATACGGTCTTTCGAGTGTACCTTATGGAAAATATTGGAGGGAGCTACGAAAGGTCTGTGTGCATAACCTCCTCTCTAATCAACAACTACTCAAGTTCAGACACTTGATAATTTCTCAAGTCGATACGTCTTTCAATAAGCTGTATGAGTTATGCAAAAACTCTGAAGACAACCAGGGAATGGTGAGAATGGATGATTGGCTCGCCCAATTATCGTTCAATGTGATAGGAAGAATAGTCTGCGGATTCCAATCAGACCCAAAGACAGGTGCTCCTAGCAGGGTGGAACAATTTAAAGAAGTAATTAATGAAGCATCTTATTTTATGTCGACATCTCCAGTGTCAGATAACGTTCCAATGCTAGGGTGGATTGACCAATTGACTGGTCTTACGAAAAATATGAAGCACTGCGGAAAGAAATTAGACTTGGTGGTAGAGAGCATAATTAAAGATCATCGTCAAAAGAGACGATTCTCTAGAACTAAAGGAGGAGATGAGAAGGACGATGAACAAGATGACTTCATCGACATTTGCTTGTCAATTATGGAGCAACCACAGCTTCCTGGCAACAATAGCCCTCCTCAGATACCTATCAAATCTATTGTCCTGGACATGATAGGTGGGGGCACTGACACCACAAAACTGACCACCATCTGGACCCTTTCCTTGCTGCTGAACAACCCCCATGTGTTGGACAAGGCAAAACAAGAAGTGGATGCACACTTTCGAAAGAAAAGGAGATCAACAGATGATGCAGCAGCAGCCGTGGTGGATTTTGATGATATTCGTAACCTTGTCTACATCCAGGCAATCATCAAAGAATCAATGCGGTTGTATCCAGCCAGCCCCGTGGTGGAGCGACTGAGCGGCGAGGATTGTGTGGTCGGTGGGTTTCATGTACCAGCAGGGACGAGATTATGGGCTAACGTATGGAAGATTCAACGAGACCCTAAAGTATGGGATGATCCATTGGTGTTTCGACCAGAGAGATTTTTGAGCGATGAACAGAAGATGGTTGATGTAAGGGGTCAAAATTATGAGCTGTTACCATTTGGAGCCGGTCGACGTATATGTCCAGGTGTTTCCTTCTCTTTGGATCTAATGCAACTGGTACTGACTCGTCTTATTCTTGAGTTTGAAATGAAGTCCCCTAGCGGGAAAGTGGACATGACAGCAACACCAGGATTAATGAGTTACAAGGTGGTCCCTCTTGACATTCTGCTCACCCATCGGCGCATAAAGTCGTGTGTGCAGTTAGCATCCTCTGAGAGAGACATGGAGAGTAGTGGTGTACCAGTAATCACTCTGAGCTCGGGCAAGGTGATGCCTGTTTTAGGCATGGGAACATTTGAGAAAGTTGGTAAAGGGTCTGAAAGAGAGAGGTTGGCGATTTTAAAAGCCATAGAGGTGGGTTACAGATACTTCGACACAGCTGCTGCATACGAAACTGAAGAGGTTCTTGGAGAAGCTATTGCTGAAGCACTTCAAATTGGCCTAATCAAATCTCGAGATGAACTTTTCATCAGTTCCATGCTCTGGTGCACTGATGCTCACCCTGATCGTGTCCTCCTCGCTCTTCAGAATTCTCTGAGGAATCTTAAATTGGAGTATCTGGATCTATATATGTTACCCTTCCCGGCAAGCTTGAAGCCAGGGAAGATAACGATGGACATACCAGAGGAAGATATTTGTCGAATGGACTACAGGTCTGTATGGTCAGCCATGGAAGAGTGTCAGAACCTTGGCTTCACTAAATCAATCGGTGTTAGCAATTTCTCCTGCAAAAAGCTTCAGGAATTGATGGCGACCGCCAACATCCCTCCAGCCGTGAATCAAGTGGAGATGAGCCCGGCTTTCCAACAAAAGAAGCTGAGAGAGTATTGCAACGCAAATAATATATTAGTCAGTGCAGTCTCTATACTGGGATCAAACGGAACCCCATGGGGCTCCAATGCAGTTTTGGGTTCTGAGGTGCTTAAGCAAATTGCTATGGCCAAAGGAAAATCTGTTGCTCAGGTTAGTATGAGATGGGTTTACGAGCAAGGCGCGAGTCTTGTGGTAAAAAGTTTCAGTGAAGAGAGATTGAGGGAAAACTTGAACATATTCGACTGGGAACTCACTAAGGAAGACAATGAAAAGATCGGTGAGATTCCACAGTGCAGAATCTTGACTGCTTATTTTTTAGTCTCGCCTAATGGACCTTTCAAATCTCAAGAAGAGTTGTGGGATGACAAAGCTTGA

>Por_STORR Por_STORR_cc53289_g1_i4

ATGGAGCTCCAATATTTTTCTTATTTTCAACCAACTTCCTCCGTTGTTGCTCTTCTACTTGCTCTTGTATCCATCTTATTCAGTGTCGTTGTTTTGAGGAAGACATTTTCGAATAACTACTCATCACCTGCATCATCAACAGAAACAGCGGTACTTTGTCATCAGCGGCAGCAGTCGTGTGCATTGCCAATTTCCGGTCTTCTTCATGTTTTCATGAATAAAAACGGCTTAATTCATGTAACTCTTGGAAATATGGCGGATAAATACGGACCAATTTTCAGTTTCCCAACAGGTAGCCATAGAACTCTCGTTGTGAGCAGTTGGGAAATGGTAAAAGAGTGTTTTACAGGCAACAATGACACTGCTTTCTCAAACCGTCCTATCCCGTTAGCTTTTCAGACTATATTCTATGCATGCGGTGGCATAGACTCATACGGTCTTTCGAGTGTACCTTATGGAAAATATTGGAGGGAGCTACGAAAGGTCTGTGTGCATAACCTCCTCTCTAATCAACAACTACTCAAGTTCAGACACTTGATAATTTCTCAAGTCGATACGTCTTTCAATAAGCTGTATGAGTTATGCAAAAACTCTGAAGACAACCAGGGAATGGTGAGAATGGATGATTGGCTCGCCCAATTATCGTTCAATGTGATAGGAAGAATAGTCTGCGGATTCCAATCAGACCCAAAGACAGGTGCTCCTAGCAGGGTGGAACAATTTAAAGAAGTAATTAATGAAGCATCTTATTTTATGTCGACATCTCCAGTGTCAGATAACGTTCCAATGCTAGGGTGGATTGACCAATTGACTGGTCTTACGAGAAATATGAAGCACTGCGGAAAGAAATTAGACTTGGTGGTAGAGAGCATAATTAAAGATCATCGTCAAAAGAGACGATTCTCTAGAACTAAAGGAGGAGATGAGAAGGACGATGAACAAGATGACTTCATCGACATTTGCTTGTCAATTATGGAGCAACCACAGCTTCCTGGCAACAATAGCCCTCCTCAGATACCTATCAAATCTATTGTCCTGGACATGATAGGTGGGGGCACTGACACCACAAAACTGACCACCATCTGGACCCTTTCCTTGCTGCTGAACAACCCCCATGTGTTGGACAAGGCAAAACAAGAAGTGGATGCACACTTTCGAAAGAAAAGGAGATCAACAGATGATGCAGCAGCAGCCGTGGTGGATTTTGATGATATTCGTAACCTTGTCTACATCCAGGCAATCATCAAAGAATCAATGCGGTTGTATCCAGCCAGCCCCGTGGTGGAGCGACTGAGCGGCGAGGATTGTGTGGTCGGTGGGTTTCATGTACCAGCAGGGACGAGATTATGGGCTAACGTATGGAAGATGCAACGAGACCCTAAAGTATGGGATGATCCATTGGTGTTTCGACCAGAGAGATTTTTGAGCGATGAACGGAAGATGGTCGATGTAAGGGGTCAAAATTATGAGCTGTTACCATTTGGAGCCGGTCGACGTATATGTCCAGGTGTATCCTTCTCTTTGGATCTAATGCAACTGGTACTGACTCGTCTTATTCTTGAGTTTGAAATGAAGTCCCCTAGCGGGAAAGTGGACATGACAGCAACACCAGGATTAATGAGTTACAAGGTGGTCCCTCTTGACATTCTGCTCACCCATCGGCGCATAAAGTCGTGTGTGCAGTTAGCATCCTCTGAGAGAGACATGGAGAGTAGTGGTGTACCAGTAATCACTCTGAGCTCGGGCAAGGTGATGCCTGTTTTAGGCATGGGAACATTTGAGAAAGTTGGTAAAGGGTCTGAAAGAGAGAGGTTGGCGATTTTAAAAGCCATAGAGGTGGGTTACAGATACTTCGACACAGCTGCTGCATACGAAACTGAAGAGGTTCTTGGAGAAGCTATTGCTGAAGCACTTCAACTTGGCCTAATCGAATCTCGAGATGAACTTTTCATCAGTTCCATGCTCTGGTGCACTGATGCTCACCCTGATCGTGTCCTCCTCGCTCTTCAGAATTCTCTGAGGAATCTTAAATTGGAGTATCTGGATCTATATATGTTACCCTTCCCGGCAAGCTTGAAGCCAGGGAAGATAACGATGGACATACCAGAGGAAGATATTTGTCGAATGGACTACAGGTCTGTATGGTCAGCCATGGAAGAGTGTCAGAACCTTGGCTTCACTAAATCAATCGGTGTTAGCAATTTCTCCTGCAAAAAGCTTCAGGAATTGATGGCGACCGCCAACATCCCTCCAGCCGTGAATCAAGTGGAGATGAGCCCGGCTTTCCAACAAAAGAAGCTGAGAGAGTATTGCAACGCAAATAATATATTAGTCAGTGCAGTCTCTATACTGGGATCAAACGGAACCCCATGGGGCTCCAATGCAGTTTTGGGTTCTGAGGTGCTTAAGCAAATTGCTATGGCCAAAGGAAAATCTGTTGCTCAGGTTAGTATGAGATGGGTTTACGAGCAAGGCGCGAGTCTTGTGGTAAAAAGTTTCAGTGAAGAGAGATTGAGGGAAAACTTGAACATATTCGACTGGGAACTCACTAAGGAAGACAATGAAAAGATCGGTGAGATTCCACAGTGCAGAATCTTGACTGCTTATTTTTTAGTCTCGCCTAATGGACCTTTCAAATCTCAAGAAGAGTTGTGGGATGACAAAGCTTGA

>Pca_STORR Pca_STORR_141675

ATGGAGCTCCAGTACTTTTCTTATTTTCAGCCAACTTCCTCCCTTTTTGTTCTTGTATCCATATTACTGATCAGTGCTGTTGTTTTGAGGAAGATATTTTTAAATAACTACTCTTCATCACCTGCATCATCAACAGAAACAGCGCTGCTTTCTCATCAGCGGCGGTCGTCATGTGCATTGCCAATCATCGGTCTTCTTCATGTTTTCATGAATAAAAACGGCTTAATTCATGTAACTCTTGGAAATATGGCTGATAAATACGGACCGATTTTCAGTTTCCCAACAGGCAGCCATATGACTCTAGTTGTGAGCAGTTGGGAGATGGCAAAAGAGTGTTTTACAGGCAACAATGACATTGCTTTCTCAAACCGTCCTATCCCGTTAGTTTTTAAGACTATATTCTATGCATGCCGTGGTATTGACTCATACGGTCTTTCAAGTGTACCTTATGGAAAATATTGGAGGGAGTTACGAAAGGTGTGTGTGCATAACCTCCTCTCTAATCAACAACTACTCAACTTCAGACACTTGATAATTTCTCAAGTTGATACCTCTTTCAATAAGCTTTATGAGTTATGCAAAAACTCTGCAGACAACCAGGGAAACTACACTACTACTACTACCGCAGCTGGCATGGTGAGGATGGATGATTGGCTAGCGCAATTATCGTTCAACGTGATAGGAAGAATAGTCTGTGGATTCCAATCAGACACCGAGACAAGTGCTTCAAGCAGTGCGGAACAACTTACAGAAGCAATTAATGACGCATCGTATTTTATGTCGACATCTCCGGTGTCAGATAACGTTCCATGGTTAGGGTGGGTTGATCAACTAACTGGTCTTACTAGAAATATGAAGCAATGCGGAAAGAAATTAGACTTGGTGCTCGAGAGCATAATTGAAGATCATCGTCAAAAGAGACGGTTTTCTAGAAGTAAACAAGGAGATGAGAATTTTATCAACGAGGATGAACAAGATGACTTCATCGACATTTGCCTGTCAATTATGGAGCAACCACAGCTTCCTGGCAACAATAACCCTTGTCAAATACCTATCAAATCTATTGTCCTGGACATGATAGGTGGGGGGACCGACACCACAAAACTGACCACCATCTGGACCCTTTCATTGCTACTGAACAACCCCCATGTGTTGGACAAGGCAAAACAAGAAGTGGATGCACACTTCGGGACGAAAAGGAACGATGCAGGAGGAGTCGTGGTGGATTTTGATGATACTCGTAACCTTGTCTACATCCAGGCAATTATCAAAGAATCAATGCGGTTGTATCCAGCCAGCCCCTTGGTGGAGCGACTGAGCGGCGAAGATTGTGTGGTCGGTGGGTTTCATGTACCAGCAGGGACGCGGTTATGGGCTAACGTGTGGAAGATGCATCGAGACCCAAAAGTATGGGATGATCCATTGGTGTTTCGACCAGAGAGATTTTTGCTCGATGACCAGAAGATGGTTGATGTTAGGGGTCAAAACTATGAGCTGTTACCATTCGGAGCCGGTCGACGGATATGTCCAGGTGTATCGTTCTCTTTGGATCTAATGCAACTGGTACTGACTCGTCTTATTCTTGAGTTTGAAATGAAGTCTCCTAGCGGGAAACTAGACATGACAGCGACACCAGGTTTAATGAGTTACAAGGTAGTCCCCCTTGACATTCTGCTCACCAGTCGTCGCATGAAGTGGTGTGTGCAGTTAGCCTCTGAGAGAGATAAAAAGATGGAGAGTAGTGTTGTACCTGTAATCACTCTGAGCTCGTGCAAGGTGATGCCTGTTTTAGGCATGGGAACATTTGAAAAAGTTGGTAAAGGGTCTGAAAGAGAGAGGTTGGCGATTTTAAAAGCGATAGAGGTAGGTTACAGATACTTCGATACAGCTGCTGCATACGAAACTGAAGAGGTTCTTGGTGAAGCTATTGCTGAAGCACTTCAACTTGGCCTAATCAAATCTCGAGATGAACTCTTCATCAGTTCCATGCTCTGGTGCACTGATGCCTACCATGATCGAGTCCTCCCCGCTCTTCAGAATTCTCTGAGGAATCTTAAATTGGAGTATCTGGATCTATACATGTTACCCTTCCCGGTAAGCTTGAAGCCAGGAGGGAAGATAAGTATGGACATACCAGAGGATGAAATTTTTACGATGGACTATAGGTCTGTATGGGCAGCCATGGAAGAATGTCAGAACCTTGGCTTGACAAAATCAATCGGTGTCAGCAATTTCTCCTGCAAAAAGCTTCAGGAATTGATGGCGACCGCCAACATCCCTCCAGCTGTGAATCAAGTGGAGATGAGCCCGGCTTTCCAACAAAAGAATTTGAGAGAGTATTGCAATGCCAATAATATATTAATCAGTGCAGTTTCTATTTTGGGATCAAACGGAACCCCATGGGGCTCCAATGCAGTTCTGGGTAATGAGGTGCTAAACCAGATTGCTATGGCCAGAGGAAAATCTGTTGCTCAGGTTAGTATGAGGTGGGTTTACGAGCAAGGTGCGAGTCTTGTGGTAAAAAGTTTTAGTGAAGAGAGATTGAGGGAAAACTTGAACATATTCGACTGGGAACTCACTAAGGAAGACTTGGAAAAGATGGGTGAGATTCCACAATGCAGAATCTTGACTGCTGATTTTATGGTATCAGCTAATGGACCTTTCAAATCTCAAGAAGAGTTGTGGGATGACAAAGCTTGA

>Pca_CYP82Y2-Lstorr Pca_CYP82Y2_141914

ATGGAGCTCCAGTACTTTTCTTATTTTCAACCAACTTCCTCCCTTTTTGCTCTTGTATCCATATTACTGATCAGTGCTGTTGTTTTGAGGAAGAAATTTTTGAATAACTACTCATCACCTGCATCATCAACAGAAACAGCGCTACTTTCTCATCAGCGGCGGTCGTCATGTGCATTGCCAATCATCGGTCTTTTTCATGTTTTCATGAATAAAAACGGCTTAATTCATGTAACTCTTGGAAATATGGCTGATAAATACGGACCGATTTTCAGTTTCCCAACAGGTAGCCATATGACTCTAGTTGTGAGCAGTTGGGAGATGGCAAAAGAGTGTTTTACAGGCAACAATGACACTGCTTTCTCAAACCGTCCTGCCCCGTTATCTTTTAAGACTATATTCTATGCATGCGGTGGTATTGAGTCATACGGTCCTTCGAGTGTACCTTATGGAAAATATTGGAGGGAGTTACGAAAGGTGTGTGTGCATAACCTCCTCTCTAATCAACAACTACTCAAGTTCAGACACTTGATAATTTCTCAAGTTGATACCTCTTTCAATAAGCTGTATGAGTTATGCAAAAACTCTGCAGACAACCAGGGAAACTACACTACTACTACCGCAGCTGGCATGGTGAGAATGGATGATTGGCTCGCGCAATTATCGTTCAATGTGATAGGAAGAATAGTCTGCGGATTCCAATCAGACACCGAGACAAGTGCTTCAAGCAGTTTGGAACGATTTACAGAAGCAATTAATGAAGCATCGTATTTTATGACGACATCTCCAGTGTCAGATAACGTTCCATGGCTAGGGTGGGTTGATCAATTAACTGGTCTTACGAGAAATATGAAGCAATGCGGAAAGAAATTAGACTTGGTGGTCGAGAGCATAATTGAAGATCATCGTCAAAAGAGACGGTTTTCTAGAACTAAACAAGGAGATGAGAATATTATCAACGAGGATGAACAAGATGACTTCATCGACATTTGCATGTCAATTATGGAGCAACCACATCTTCCTGGCAACAATAACCCTTGTCAAATACCTATCAAATCTATTGTCCTGGACATGATAGGTGCGGGGACCGACACCACAAAACTGACCACCATCTGGACCCTTTCGTTGCTACTGAACAACCCCCACGTGTTGGACAAGGCAAAACAAGAAGTGGATGCACACTTCAAGGACGAAAAGGAACGATGCAGGAGGATTGGTGGTGGATTTTGATGATACTCGTAACCTTGTCTACATCCAGGCAATCATCAAAGAATCAATGCGGTTGTATCCAGCCAGCCCTGTGGTGGAGCGACTGAGCGGCGAAGATTGTGTGGTCGGTGGGTTTCATGTACCAGCAGGGACGCGGTTATGGGCTAACGTGTGGAAAATGCATCGAGACCCAAAAGTGTGGGATGATCCGTTGGTGTTTAGACCAGAGAGATTTTTGTGCGATGACCAGAAGATGGTTGATGTAAGGGGTCAAAACTATGAGCTGTTACCATTCGGAGCCGGTCGACGGATATGTCCAGGTATATCGTTCTCTTTGGATCTAATGCAACTGGTACTGACTCGTCTTATTCTTGAGTTTGAAATGCAGTCTCCTAGTGGGAAAGTAGACATGACAGCGACACCAGGTTTAATGAGTTACAAGGTAGTCCCCCTTGACATTCTGCTCACCAGTCGTCGTGTGAAGTGGTGTGTGCAGTTAGCCTCAGAGAGAGATAAAAAGATGGAGAGTAGTGTTGAAACGCAAAATTCAACATATTATTAA

>Pbr_CYP82Y2-Lc1p Pbr_CYP82Y2_contig_533p_CDS

AGTTTTTTAATTTTCAACTTGCAACTTCTGTTGTTGCTCTTCTACTTGCTCTTGTATCAATATTACTTTGTGTTCTTTGTTTGAGGAAGAAATTTCTGAAAAACTACTACTCTTCAACAGCCATTTCTCATCAGCGGTCGTGTGGGTTGCCAATCATAGGTCTTTTTCATGTTTTCATGAATAGAACCGACTTAATACATGTGACTCTTGGAAATATGGCCGATAAATACGGTCCAATTTTCAGTTTCCATTCAGGTAGCCACAGGACTCTGGTTGTGAGCAGTTGGGAGCTGGTAAAGGAGTGTTTTACAGGCAACAACGATACCATTTTTTCGAATCGTCCCATCCGGCTAGCGTTTAAGATTATATTCTATGCAGGTGGTATAGACTCATACGGTATTCCGATTCTACCTTACGGAAAATATTGGAGGGAGTTACGAAAGATCTGTGTTCATAACCTCCTCTCCAATCAACAATTACTCAAGTTCAGACACTTGATAATTTCTCAAGTTGGCACCTCATTCAATAAGCTTTATGAGTCATGCAACAACGATGGAGACTCTACTACTACTACCTACTACTGCGGAAGCTGTTATGGTGAGAATGGATGATTGGCTGGGGAAATTATCCTTTAACGTGATTTGAAGACTAGTCTGCGGATTCCAATCAGACACCGAGACAAGTGCTACAAGTGAAAGCAGTGTGGAACGATTTACAGTAGCAAATGAGGGAGCATCGCGTTTTTTGTCGGTAACTGCAGTGTCAGATACCTTTCCATGGTTAGGGTGGATTGACCAATTAACTGGTCTTACGAGAAAAATGAAGCACTACGGAAAGAAATTAGACCTAGTGGTTGAGAGCATAATTGAAGATCATCGTCAAAACAGACGATTTTCTGAAACTAAAGAAGATCACGAGAATACCGCTGAGGATGAACAACTAGACCTCATCGACATTTGCTTATCAATTATGGAGCAACCACAGCTTCCTGACGACAGTATCCCTCGTCAAGAAATACCTATCAAATCTATTGTCATGGACATGATAGGTGGGGGGACTGACACCCCAAAAATGACCATGATCTGGACCCTTTCCTTGCTGTTGAACAACCCCAATGTTTTGGAGAAGGCGAAACAAGAAGTGGATGCGCTCTTTGGAACGTAAAGTAGATCAACAAATGATACAGTAGTCGTGGTGGATTTTGATGATATTCGTAACCTTGTCTACATCCAGGCAATCATCAAAGAATCAATGAGGTTGTATCCAGTCAGTCCACTGGTGGAGCGACTGAGCAGCGAAGATTGTGTGGTCGGTGGGTTTCATGTACCAGCAGGGACTCGACTATGGGCTAACGTATGGAAAATGCAGAGAGACCCAAAAGTATGGGATGATCCGTTGGTGTTTCGACCAGAGAGATTTTTGGGTAATGACCAGAGCATGGTCGATGTAAGGGGTCAAAACTTTGAGCTGTTACAATTCGGAGCCGGTCGACGGATATGTCCAGGTATATCGTTCTCTTTGGATCTAATGCACCTGGTACTTACTCGTCTTATTCTTGAGTTTGAAATGAAGTCTCCTAGTGGCGAAGTGGATATGACAGCAACGCCAGGTTTACTGAGTTACAAGGTAGTCCCCCTTGACATTCTGCTCACCCGTCGTCGAATGCTTTAG

>Par_CYP82Y2-Lc2B Par_CYP82Y2_contig_30250b_CDS

ATGGAGGACTATTCTCATGATCTCCCTTTTTCTTACTTTCAACCTCTTCTACTTGCGTTTTTATCTATATTACTTAGTGTTGTTTGTCTGAGGAAGAATTTTTTGAAAACTACAGTACCATCATCATCAACACCAACGGCAACAGCCATTTCTCATCAGCGGCAAAAAAGGTCTTGTGCGTTGCCAGTCACTGGTCTTCTTCATGTTTTCATGAATAGAAACGACTTAATTCATGTAACTCTTGGAAATTTGGCCGATAAATACGGACCGATTTTCAGTTTCCCAACAGGCAGTCATAGGACTATGGTTGTGAGCAATTGGGAGGTGGTAAAAGAGTGTTTTACAGGCAACAATGACACTGTTTTCTCAAACCGTCCTGTCCTCTTAGTGTTTAAGATTATGTTCAATGCAGGTGGGATTGACTCAGAAGGTCTTTCGATTGTACCGTATGGAAAATATTGGAGGGAGTTACGCAAGATCTGCGTGCATAACCTCCTCTCTAATCAACAATTACTCAAGTTCAGACACTTGATAATTTCTCAAGTTGATGCCTCTTTCAATGAGCTTTATGAGTTATGCAAAATTAATTCTGAAAACCAGCAGCGAGACTCTGGCATGGTAATCAGAATGGATGATTGGCTAGGGAAATTATCCTTCAACGTCATAGGAAGACTAGTCTGCGGATTCCAATCACAAACCGAGACAAGTGCTACAAGTAGAAGCAGTATGGAACGATTTACAGCAGCAAATGATGAAGCATCGAGTTTTATGTCGATATCTGCAGTGTCAGATACCTTTCCGTGGCTAGGGTGGATTGATAAATTAACTGGTCTTACAAGAAAGATGGAGCACTGCGGAAACAAGTTAGAGTTGGTGGTTGAGAGCATAATTGAAGATCATCGTCAAAAGAGAAGATTTTCTGGGACTGAACAAGCTGATGACTTCATCGACATCTGCTTGTCAATTATGGAGCAACCACAGCTTCCTGGCGACAACAAACCTTGTCAAATACCTATCAAATCTATTGTCCTGGACATGGTAGGTGGGGGAACTGATACCACAAAACTAACCACCATCTGGACCCTTTCCTTGCTGTTGAACAACCCCATGTGTTGGACAAGGCAAAACAAGAAGTGGATGCGCACTTTGGAACGAAAAGAAGATCAATCATGGTGAATTTTGATGATATTCGTAGCCTTGTCTACATCCAGGCAATAATCAAAGAATCAATGCGGTTGTATCCAGCCAGCCCACTGGTGGAGCGACTTAGCAGCGAAGATTGTGTAGTTGGTGGGTATCATGTACCAGCAGGGACGCGATTATGGGTTAACGTATGGAAGATGCAACGAGACCCAAAAGTATGGGATGATCCATTGGTGTTTCGACCAGAGAGATTTTTGAGCGATGATAAGAAGACGGTTGATTTACGGGGTCAAAATTATGAGTTGTTACCATTTGGAGCGGGTAGTCGTATATGTCCAGGTATATCGTTCTCTTTGGATTTAATACACATGGTACTTACTCGTCTTATTCTTGAGTTTGAAATGAAGTCTCCTAGTGGGGAAGTGGACATGGCAGCAACACCAGGTATAATGAGTTACAAGGTAGTCCCGCTTGACATTCTGCTTACCCGACGTCGCATTCTTTAG

>Par_CYP82Y2-Lc2A Par_CYP82Y2_contig_30250a_CDS

ATGGAGGATGATCTCCCTTTTTCTTACTTTCAACCAACTTCCGCTGTTGCTCTTCTACTTGCGTTTTTATCTATATTGCTTAGTGTTGTTTGTCTGAGGAAGAATTTTTTGAAAACTACAGTACCATCATCATCAACACCAACGGCAACAGCCATTTCTCATCAGCGGCAAAAAACGTCTTGTGCGTTGCCAGTCACTGGTCTTCTTCATGTTTTCATGAATAGAAACGACTTAATTCATGTAACTCTTGGAAATTTGGCCGATAAATACGGACCGATTTTCAGTTTCCCAACAGGCAGTCATAGGACTATGGTTGTGAGCAGTTGGGAGGTGGTAAAAGAGTGTTTCACAGGCGACAATGACACTGTTTTCTCAAACCGTCCTGTCCTCTTAGTGTTTAAGATTATGTTCAATGCAGGTGGGATAGACTCACAAGGTCTTTCGATTGTACCGTATGGAAAATATTGGAGGGAGTTACGGAAGATCTGCGTGCATAACCTCCTCTCTAATCAACAATTACTCAAGTTCAGACACTTGATAATTTCTCAAGTTGATACCTCTTTCAATGAGCTTTATGAGTTGTGCAAAATTAATTCCGAAAACCAGCAGGGAGACTCTACTACTACTAACGCAGAAGCTGGCATGGTGATCAGAATGGATGATTGGCTTGGGAAATTATCGTTCAACGTCATAGGAAGACTAGTCTGCGGATTCCAATCACAAACCGAGACAAGTGCTACAAGTAGAAGCAGTATGGAACGATTTACAGCAGCAAATGATGAGGCATCCAGCTTTATGTCGATATCTGCAGTGTCAGATACCTTTCCGTGGCTAGGGTGGATTGATAAATTAACTGGTCTTATAAGAAAGATGGAGCACTGCGGAAAGAAGTTAGAGCTAGTGGTTGAGAGCATAATTGAAGATCATCGTCAAAAGAGAAGATTTTCTGGGATTAAACAAGGAGATGACAACAGAACCAAGGATGAACAAGCTGATGACTTCATCGACATTTGCTTGTCAATTATGGAGCAACCACAGCTTCCTGGCGACACCAAACCTTGTCAAATACCTATCAAATCTATTGTCCTGGACATGGTAGGTGGGGGAACTGATACCACAAAACTAACCACCATCTGGACCCTTTCCTTGCTGTTGAACAACCCCCATGTGTTGGACAAGGCAAAACAAGAAGTGGATGCGCACTTTGGAACGAAAAGAAGATCAACAACAAATGATGCAATCATGGTGAATTTTGATGATATTCGTAGCCTTGTCTACATCCAGGCAATAATCAAAGAATCAATGCGGTTGTATCCAGCCAGCCCACTGGTGGAGCGACTTAGCAGCGAAGATTGTGTAGTTGGTGGGTATCATGTACCAGCAGGGACGCGATTATGGGTTAACGTATGGAAGATGCAACGAGACCCAAAAGTATGGGATGATCCATTGGTGTTTCGACCAGAGAGATTTTTGAGCGATGATAAGAAGATGGTTGATTTAAGGGGTCAAAATTATGAACTGTTACCATTCGGAGTGGGCCGGCGGATATGCCCCGGCATATCGTTCTCTTTGGATTTAATGCACCTGGTACTTACTCGTCTTATCCTTGAGTTTGAAATGAAGTCTCCTACTGGGGGGAAAGTGGACATGACAGCAACACCAGGTATAATGAGTTACAAGGTAGTCCCGCTTGACATTCTGCTTACCCGTCGCGTTCTTTAG

>Par_CYP82Y2-Lc3 Par_CYP82Y2_contig_26129_CDS

ATGGAGGATGATCTCCCTTTTTCTTACTTTCAACCAACTTCCGCTGTTGCTCTTCTACTTGCGTTTTTATCTATATTGCTTAGTGTTGTTTGTCTGAGGAAGAATATTTTGAAAACTACAGTACCATCATCATCAACACCAACGGCAACAGCCATTTCTCATCAGCGGCAAAAAACGTCTTGTGCGTTGCCAGTCACTGGTCTTCTTCATGTTTTCATGAATAGAAACGACTTAATTCATGTAACTCTTGGAAATTTGGCCGATAAATACGGACCGATTTTCAGTTTCCCAACAGGCAGTCATAGGACTATGGTTGTGAGCAATTGGGAGGTGGTAAAAGAGTGTTTCACAGGCGACAATGACACTGTTTTCTCAAACCGTCCTGTCCTCTTAGTGTTTAAGATTATGTTCAATGCAGGTGGGATAGACTCACAAGGTCTTTCGATTGTACCGTATGGAAAATATTGGAGGGAGTTACGCAAGATCTGCGTGCATAACCTCCTCTCTAATCAACAATTACTCAAGTTCAGACACTTGATAATTTCTCAAGTTGATGCCTCTTTCAATGAGCTTTATGAGTTATGCAAAATTAATTCTGAAAACCAGCAGCGAGACTCTGGCATGGTAATCAGAATGGATGATTGGCTAGGGAAATTATCCTTCAACGTCATAGGAAGACTAGTCTGCGGATTCCAATCACAAACCGAGACAAGTGCTACAAGTAGAAGCAGTATGGAACGGTTTACAGCAGCAAATGATGAGGCATCCAGCTTTATGTCGATATCTGCAGTGTCAGATACCGTTCCATGGCTAGGGTGGATTGATAAATTAACTGGTCTTACAAGAAAGATGGAGCACTGCGGAAAGAAGTTAGAGCTAGTGGTTGAGAGCATAATTGAAGATCATCGTCAAAAGAGAAGATTTTCTGGGATTAAACAAGGAGATGACAAGAGAATTGAGGATGAACAAGCTGATGACTTCATCGACATTTGCTTGTCAATTATGGAGCAACCACAGCTTCCTGGCGACAACAAACCTTGTCAAATACCTATCAAATCTATTGTCCTGGACATGGTAGGTGGGGGAACTGATACCACAAAACTAACCACCATCTGGACCCTTTCCTTGCTGTTGAACAACCCCCATGTGTTGGACAAGGCAAAACAAGAAGTGGATGCGCACTTTGGAACGAAAAGAAGATCAACAACAAATGATGCAATCATGGTGAATTTTGATGATATTCGTAGCCTTGTCTACATCCAGGCAATAATCAAAGAATCAATGCGGTTGTATCCAGCCAGCCCACTGGTGGAGCGACTTAGCAGCGAAGATTGTGTAGTTGGTGGGTATCATGTACCAGCAGGGACGCGATTATGGGTTAACGTATGGAAGATGCAACGAGACCCAAAAGTATGGGATGATCCATTGGTGTTTCGACCAGAGAGATTTTTGAGCGATGATAAGAAGACGGTTGATGTAAGGGGTCAAAATTATGAGTTGTTACCATTTGGAGCGGGTAGACGTATATGTCCAGGTATATCGTTCTCTTTGGATTTAATACACATGGTACTTACTCGTCTTATTCTTGAGTTTGACATGAAGTCTCCTAGTGGGGAAGTGGACATGACAGCAACACCAGGTATAATGAGTTACAAGGTAGTCCCGCTTGACATTCTGCTTACCCGTCGCATTCTTTAG

>Pca_CYP82Y2-La2p Pca_CYP82Y2_143837p

ATGGTAATTATCAGTCGACTTGGACTGATTATCCAGTGCACATTTCTCACTAGAAGAAGCAACACTTCTGTTGTTGCTCTTCTACTTGCTGTTGTATCTATATTAATTGGTGTTGTTGCTTTCAGTAGCAGAGTTCTGAAAAACTACTCATCACCATCATCAACAGCAACAGCCGTTTCTCATCATCGGCCAAAAGCATCCTGAGCACTGCCAATGATCGGTCTGTATCATGTTTTCATGAATAAAACCGGCTTAATTCATGTAACTCTTGGAAATATGGCTGATAAATATGGACCCATTTTCAGTTTCTCAACAGGTGGCCATAGGGCTCTAGTTGTGAGCAGTTGGGAGATGGCAAAAGAGTGTTTTACAGGCAACATTGACATTGTTTTTCAAATCGCCCTATGCCGTTATCGTTTTAGCTTATATTCAATGCAGGTGGTATAGACTCGTACGGTCTTTCACAAGTACCTCATGGAAAATATTGGAAGGAGTTACGAAAGATCTGTGTGCATAACCTCCTCTCTAATCAGCAATTACTCAAGTTCAGACACTTGATATATTCTGAAGTTGATAAATGTTTCAATAAACTGTATGATCATGCAACAACAATAAAAACTCTGAAAGTGAGGGAGACGACTCTACTACTACTGCAGCTGGGATGGTGAGAATGGATAATTGGCTTGCACAATTATCGTTCAACGTGATAGGAAGAATAGTTTGCGGATTCCAATCACACACCGAGACAATGCTACAAGCAGTGTGGAACGATTTACAGTAGCAATTGATGAAGCATCGCGTTTTATGTCGATACCTGCAGGGTCAGATACCTCATGGAAAATATTGATCAGTTAACTGGTCTTAGGAGAAATATGAAGCACCACGGAAAGAAATTGAAGATGATCGTCAGAAGAGACGACTTTCTAGAACTAAAAAAGGAGATGAAAATACTACCGATGAGATGACTTCATCGACATTTGCTTGTCAATTATGGAGCAACCAGAGCTTTCTGGAAACAATTACGCTCGTCAAATGCCTATCAAATCTATTGTCGTGGACATGATAGGCGGAGCGAATGACACTACAAAACTGACCACAATATGGACCCTTTCCTTGCTGCTGAACAACCCCCATGTGTCGGACGAGGCAAAAAAAGAAGTGGATGCACACCTTTGGATCTAAAAGGAGATCAACAAATGGTGAAGTCACGGTGGATTTTGATGATACTCGTAACCTTGTCTACATCCAGGCAATAACCAAAGAATCAATGCGCTTGTATCCGGCCAGCCCATTGGTGGAGCGACTAAGCAGCGAAGATTGTGTGGTTGGTGGGTTTTATGTACCAGCAGGGACACGACTATGGGTTAACATAGGAAGATGCAATGAGACCCAAAAGTATGGGATGATCCGTTGGTGTTTCGACCAGAGAGATTTTTGAGCAATGCGCAGAAGATGGTTGATGTAATGGGTCAAAATTATGAGTTGTTACCATTTGGAGTCAGTCGACAGGTATGTCCAGGTATATCGTTCTTTTTGGATCTAATGCAACTGATACTTACTCGTCTTATTCTTGAGTTTGAAATAAAGTCTCCTAGTGGGAAAATGGACATGACAGCAACACAAGGTTTAATGAGTTACAAGGTAGTCCCCCTTGGAATTCTGCTCACCCGTCGCATGCTTTAG

>Par_CYP82Y2-Lb2B Par_CYP82Y2_contig_48062b_CDS

ATGGAGTACTCTAGTCTCCAGTTTTTTTACTTTCAACCAACTTCCTTTGTTGCTCTGCTACTTGCTCTTGTATCTTGTGTACTTAGTGTCGTTGCTTTGAGAAAGGAATTTTTTAAAAACTACTACTCATTATCATCATCAACAGCAACGGCCGTTTCTCATCAGTGGCCAAAAGCATCGTGTGCATTGCCATTGATCGATCTTCTTCATGTTTTTTTCAATAAAACCGACTTGATGCACGTAACTCTCGGAAATATGGCTGATAAGTTCGGACCGATTTTCAGTTTCCCAACAGGTAGCCATAGGACTCTAGTTGTAAGCAGTTGGGAGATGGCAAAAGAGTGTTTTACAGGCAACAATGACACTGTTTTTTCAAACCGTCCTATCCCGTTAGCGTTTAAGATTATATTTTATGCAGGTGGTGTGGACTCATACGGGCTTGCACTTGTACCTTATGGAAAATATTGGAGGGAGTTACGAAAGATATGTGTGCATAACCTCCTCTCTAATCAACAATTACTCAACTTCAGACACTTAATAATTTCTCAAGTTGATACCTCTTTCAATAAGCTTTATGACTTGAGCAACAAGAATAAAAATACTACTACTGAAGCTGGCACGGTGAGAATGGATGATTGGCTTGCACAATTATCGTTCAATGTGATCGGAAGAATAGTCTGCGGATTCCAAACACACACCGAGACAAGTGCTACAAGCAGTGTGGAACGATTTACAGAAGCAATTGAAGAAGCATCACGTTTTATGTCGATAGCTACAGTATCAGATACCTTTCCATGGCTAGGGTGGATTGATCAGTTAACTGGTCTTACGAGAAAAATGAAGCACTATGGAAAGAAACTAGACTTAGTGGTTGAGAGCATAATTGAAGATCATCGTCAAAACAGGAGATTTTCTGGGACCAAACAAGGAGATGACTTCATCGACATTTGCTTGTCAATTATGGAGCAACCACAGATTATTCCCGGAAACAATGATCCTCCTCGTCAAATACCTATCAAATCTATTATCCTGGACATGATAGGTGGGGGGACTGACACCACAAAACTGACCACCACCTGGACCCTTTCATTGCTCCTGAACAACCCCCATGTGTTGGACAGAGCAAGAGAAGAAGTGGATGCACACTTTGGAACAAAAAGGAGAGCAACAAATGATGCAGTAATGGTGGAGTTTGATGACTTTCGTAACCTTGTTTACATCCAGGCAATCATCAAAGAATCAATGAGGTTGTATCCAGTCAGCCCAGTTGTGGAGCGACTGAGCAGCGAAGACTGTGTGGTCGGTGAGTTTCATGTGCCAGCAGGGACGAGATTATGGGCAAACGTATGGAAGATGCAACGAGACCCAAATGTATGGGATGATCCATTGGTGTTTCGACCAGAGAGATTTCTGAGCGATGACAAGAAGATGGTTGATGTAAAGGGTCAAAATTATGAGTTGTTACCCTTCGGAGCCGGTCGGCGAATATGTCCAGGCATATCGTTTTCTTTGGATCTAATGCAACTGGTACTTACTCGTCTAGTTCTTGAGTTTGACATGAAGTCTCCTACCGGAGGGGAAGTGGACATGACAGCAACACCAGGTTTAATGAGTTACAAGGTAGTCCCCCTTGACATGCTGCTCACTCGTCGTCTTCGCATGCTTTAG

>Par_CYP82Y2-Lb2A Par_CYP82Y2_contig_48062c_CDS

ATGGAGTACTCTAGTCTCCAGTTTTTTTACTTTCAACCAACTTCCTTTGTTGCTCTGCTACTTGCTCTTGTATCTTGTGTACTTAGTGTCGTTGCTTTGAGAAAGGAATTTTTTAAAAACTACTACTCATTATCATCATCAACAGCAACGGCCGTTTCTCATCAGTGGCCAAAAGCATCGTGTGCATTGCCATTGATCGATCTTCTTCATGTTTTTTTCAATAAAACCGACTTGATGCACGTAACTCTCGGAAATATGGCTGATAAGTTCGGACCGATTTTCAGTTTCCCAACAGGTAGCCATAGGACTCTAGTTGTAAGCAGTTGGGAGATGGCAAAAGAGTGTTTTACAGGCAACAATGACACTGTTTTTTCAAACCGTCCTATCCCGTTAGCGTTTAAGATTATATTTTATGCAGGTGGTGTGGACTCATACGGGCTTGCACTTGTACCTTATGGAAAATATTGGAGGGAGTTACGAAAGATATGTGTGCATAACCTCCTCTCTAATCAACAATTACTCAACTTCAGACACTTAATAATTTCTCTAGTTGATACCTCTTTCAATAAGCTTTATGACTTGAGCAACAAGAATAAAAATACTACTACTGAAGCTGGCACGGTGAGAATGGATGATTGGCTTGCAGAATTATCGTTCAATGTGATCGGAAGAATAGTCTGCGGATTCCAAACACTCACCGAGACAAGTGCTACAAGCAGTGTGGAACGATTTACAGAAGCAATTGAAGAAGCATCACGTTTTATGTCGATAGCTACAGTATCAGATACCTTTCCATGGCTAGGGTGGATTGATCAGTTAACTGGTCTTACGAGAAAAATGAAGCACTATGGAAAGAAACTAGACTTAGTGGTTGAGAGCATAATTGAAGATCATCGTCAAAACAGGAGATTTTCTGGGACCAAACAAGGAGATGACTTCATCGACATTTGCTTGTCAATTATGGAGCAACCACAGATTATTCCCGGAAACAATGATCCTCCTCGTCAAATACCTATCAAATCTATTATCCTGGACATGATAGGTGGGGGGACTGACACCACAAAACTGACCACCACCTGGACCCTTTCATTGCTCCTGAACAACCCCCATGTGTTGGACAGAGCAAGAGAAGAAGTGGATGCACACTTTGGAACAAAAAGGAGAGCAACAAATGATGCAGTAATGGTGGAGTTTGATGACATTCGTAACCTTGTTTACATCCAGGCAATCATCAAAGAATCAATGAGGTTGTATCCAGTCAGCCCAGTTGTGGAGCGACTGAGCAGCGAAGACTGTGTGGTCGGTGGGTTTCATGTGCCAGCAGGGACGAGATTATGGGCAAACGTATGGAAGATGCAACGAGACCCAAATGTATGGGATGATCCATTGGTTTTTCGACCAGAGAGATTTCTGAGCGATGACAAGAAGATGGTTGATGTAAGGGGCCGTAATTATGAGTTATTACCATTCGGAGCCGGTCGGCGAATATGTCCAGGCATATCGTTTTCTTTGGATCTAATGCAACTGGTACTTACTCGTCTAGTTCTTGAGTTTGACATGAAGTCTCCTACCGGAGGGGAAGTGGACATGACAGCAACACCAGGTTTAATGAGTTACAAGGTAGTCCCCCTTGACATGCTGCTCACTCGTCGCATGCTTTAG

>Par_CYP82Y2-Lb2C Par_CYP82Y2_contig_48062a_CDS

ATGGAGTACTCTAGTCTCCAGTTTTTTTACTTTCAACCAACTTCCTTTGTTGCTCTGCTACTTGCTCTTGTATCTTGTGTACTCAGTGTCGTTGCTTTGAGAAAGAAATTTTTGAAAAACTACTACTCATCAACAGCAACGGCCGTTTCTCATCAGTGGCCAAAAGCATCGTGTGCATTGCCATTGATCGATCTTCTTCATGTTTTTTTCAATAAAACCGACTTGATGCACGTAACTCTCGGAAATATGGCTGATAAGTTCGGACCGATTTTCAGTTTCCCAACAGGTAGCCATAGGACTCTAGTTGTAAGCAGTTGGGAGATGGCAAAAGAGTGTTTTACAGGCAACAATGACACTGTTTTTTCAAACCGTCCTATCCCGTTAGCGTTTAAGATTATATTTTATGCAGGTGGTGTGGACTCATACGGGCTTGCACTTGTACCTTATGGAAAATATTGGAGGGAGTTACGAAAGATATGTGTGCATAACCTCCTCTCTAATCAACAATTACTCAACTTCAGACACTTAATAATTTCTCAAGTTGATACCTCTTTCAATAAGCTTTATGACTTGAGCAACAAGAATAAAAATACTACTACTGAAGCTGGCACGGTGAGAATGGATGATTGGCTTGCACAATTATCGTTCAATGTGATCGGAAGAATAGTCTGCGGATTCCAAACACACACCGAGACAAGTGCTACAAGCAGTGTGGAACGATTTACAGAAGCAATTGAAGAAGCATCACGTTTTATGTCGATAGCTACAGTATCAGATACCTTTCCATGGCTAGGGTGGATTGATCAGTTAACTGGTCTTACGAGAAAAATGAAGCACTATGGAAAGAAACTAGACTTAGTGGTTGAGAGCATAATTGAAGATCATCGTCAAAACAGGAGATTTTCTGGGACCAAACAAGGAGATGACTTCATCGACATTTGCTTGTCAATTATGGAGCAACCACAGATTATTCCCGGAAACAATGATCCTCCTCGTCAAATACCTATCAAATCTATTATCCTGGACATGATAGGTGGGGGGACTGACACCACAAAACTGACCACCACCTGGACCCTTTCATTGCTCCTGAACAACCCCCATGTGTTGGACAGAGCAAGAGAAGAAGTGGATGCACACTTTGGAACAAAAAGGAGAGCAACAAATGATGCAGTAATGGTGGAGTTTGATGACTTTCGTAACCTTGTTTACATCCAGGCAATCATCAAAGAATCAATGAGGTTGTATCCAGTCAGCCCAGTTGTGGAGCGACTGAGCAGCGAAGACTGTGTGGTCGGTGGGTTTCATGTGCCAGCAGGGACGAGATTATGGGCAAACGTATGGAAGATGCAACGAGACCCAAATGTATGGGATGATCCATTGGTGTTTCGACCAGAGAGATTTCTGAGCGATGACAAGAAGATGGTTGATGTAAAGGGTCAAAATTATGAGTTGTTACCCTTCGGAGCCGGTCGGCGAATATGTCCAGGCATATCGTTTTCTTTGGATCTAATGCAACTGGTACTTACTCGTCTAGTTCTTGAGTTTGACATGAAGTCTCCTACCGGAGGGGAAGTGGACATGACAGCAACACCAGGTTTAATGAGTTACAAGGTAGTCCCCCTTGACATGCTGCTCACTCGTCGTCTTCGCATGCTTTAG

>Par_CYP82Y2-La7 Par_CYP82Y2_contig_26608_CDS

ATGGAGTACTCTGATGATCTCCAGCTTTTTGGTTTTCAGCCAACTTCCGTTGTTGCTCTTCTACTTGCTATTGTATCCATATTACTTGGTGTTATTGCAGTTTCTCATCATCGGGCAAAACCATCGTGTGCATTGCCAATGATCGGTCTTTATCATGTTTTCATGAATAAAACCGGCTTAATTCATGTAACTCTAGGAAATCTGGCTGATAAATACGGACCTATTTTCAGTTTCCCAACAGGTGGCCATAGGGCTCTAGTTGTGAGCAGTTGGGAGATGGCAAAAGAGTGTTTTACAGGCGACAATGACATTGTTTTTTCAAATCGTCCTATGCCGTTATCGTTTAAGATTATATTCAATGCAGGTGGTATAGACTCAGCCGGTCTTACACAAGTACCTTATGGAAAATATTGGAGGGAGTTACGAAAGATCTGTGTGCACAACCTTCTCTCAAATCAACAATTACTCAAGTTCAGGCACTTGATAAATTCTCAAGTTGATACCTCTTTCAATAAACTGTACGAGTCATACAACAAAAATAAAAACTCCGAAAACGAGGGAGATTCTGGCATGGTGAGAATGGATGACTGGCTTGGAGAATTATCATTCAATGTGATAGGAAGAATAGTCTGTGGATTCCGATCAGACACTGAGACAAGTGGTACAAGCAGTGTGGAACGATTTACAGTGGCGATTGATGAAGCATCACGTTTTATGTCGATACCTGCCGTATCAGATACCTTTCCATGGCTAGGGTGGATTGATCAGTTAACTGGCCTTACAAAAGATATGAAGAACCATGCAAAAAATTAGACTTAGTGGTCGAGAGCATAATTGAAGATCATCGTCAAAAGAGACGGTTTTCGAGATCTAAAAAAGGAGATGAGAATACCACCGAGGATGAACAAGATGACTTCATCGACATTTGTTTGTCAATTATGGAGCAACCAGAGCTTCCTAGCAACAACTACGCTCGTCAAATGCCTATCAAATCTATTATTGTGGACATGATAGGCGGGGCGACTGACACCACAAAACTGACCACAATCTGGACCCTTTCCTTGCTGCTGAATAACCCCCATGTGCTAGACAAGGCAAAAGAAGAAGTGGATGCACACTTTGGAACGAGAAGAAGTTCAACCAATGGGGAAGTCATGGTGGATTCTGATGATATTCGTAGCCTTGTCTACATCCAGGCAATCATCAAAGAATCGATGCGGTTGTATCCAGCCAGTCCAGTGGTGGAGCGACTGAGCAGCGAAGATTGTGTGGTTGGTGGGTTTCATGTACCAGCAGGGACACGATTATGGGTTAACGTATGGAAGGTTCAACGAGACCCAGAAGTATGGGATGATCCGTTGGTGTTTCGACCAGAGAGGTTTTTGAGCAATGAGCAGAAGATGGTTGATGTAAGGGGTCAACATTATGAGCTTCTACCATTTGGAGCCGGTCGTCGGATATGTCCAGGTGTATCGTTCTCTTTGGATCTAATGCACCTGGTACTTACTCGTCTTATTCTTGAGTTTGAAATGAAGTCTCCTGGTGGGGAAGTGGACATGACAGCAACACAAGGTTTAATGAGTTACAAGGTAGTCCCCCTCGATATTATGCTCACCCGTCGCATGCTTTAG

>Par_CYP82Y2-La6 Par_CYP82Y2_contig_38806_CDS

ATGGAGTACTCTGATCTCCAGATTTTTGGTTTTCAGCCAACTTCCGTTGTTGCTCTTCTACTTGCTCTTGTATCCATATTACTTGGTATTATTGCAGTTTCTCATCATCGGGCAAAACCATCGTGTGCATTGCCAATGATCGGTCTTTATCATGTTTTCGTGAATAAAACCGGCTTAATTCATGTAAATCTTGGAAACATGGCTGATAAATACGGACCCATTTTCAGTTTCCCAACAGGTGGCCATAGGGCTCTAGTTGTGAGCAGTTGGGAGATGGCGAAAGAGTGTTTTACAGGCGACAATGACATTGTTTTTTCAAATCGTCCAATGCCGTTATCGTTTAAGATTATATTCAATGCAGGTGGTATAGACTCGGCCGGTCTTTCACAAGTACCTTATGGAAAATATTGGAGGGAGTTACGAAAGATCTGTGTGCACAACCTTCTCTCAAATCAACAATTACTCAAGTTCAGGCACTTGATAAATTCTCAAGTTAATACCTCTTTCAATAAACTGTACGAGTCATGCAACAAAAATAAAAACTCTGAAAACGAGGGAGATTCTGGCATGGTGAGAATGGATGATTGGCTTGGAGAATTATCATTCAACGTGATAGGAAGAATAGTCTGTGGATTCCGATCAGAAACTGAGACAAGTGGTACAAGCAGTGTGGAACGATTTACAGTAGCAATTGATGAAGCATCGCGTTTTATGTCCATACCTGCAGTGTCAGATACCTTTCCATGGCTAGGATGGATTGATCAGTTAACTGGTCTTATAAAAGATATGAAGCACCACGCAAAAAAAATTAGACTTAGTGGTTGAAAGCATAATTGAAGATCATCGTCAAAACAGACGGTTTTCGAGATCTAAAAAAGGAGATGAGAATACCACCGAGGATGAACAAGATGACTTCATCGACATTTGTTTGTCAATTATGGAGCAACCAGAGCTTCCTGGCAACAACTACGCTCGTCAAATGCCTATCAAATCTATTATTGTGGACATGATAGGCGGGGCGACTGACACCACAAAACTGACCACAATCTGGACCCTTTCCTTGCTGCTGAATAACCCCCATGTGCTAGACAAGGCAAAAGAAGAAGTGGATGCACACTTTGGAACGAGAAGAAGTTCAACAAATGGGGAAGTCATGGTGGATTTTGATGATATTCGTAACCTTGTCTACATCCAAGCGATCATCAAAGAATCGATGCGGTTGTATCCAGCCAGTCCAGTGGTGGAGCGACTGAGCAGCGAAGATTGTGTGGTTGGTGGGTTTCATGTACCAGCAGGGACACGATTATGGGTTAACGTATGGAAGGTTCAACGAGACCCAGAAGTATGGGATGATCCGTTGGTGTTTCGACCAGAGAGGTTTTTGAGCAATGAGCAGAAGATGGTTGATGTAAGGGGTCAACATTATGAGCTTCTACCATTTGGAGCCGGTCGTCGGATATGTCCAGGTGTATCGTTCTCTTTGGATCTAATGCACCTGGTACTTACTCGTCTTATTCTTGAGTTTGAAATGAAGTCTCCTGGTGGGGAAGTGGACATGACAGCAACACAAGGTTTAATGAGTTACAAGGTAGTCCCCCTCGATATTATGCTCACCCGTCGCATGCTTTAG

>Ptr_CYP82Y2-Lb1 Ptr_CYP82Y2_gc30928_g1_i2

ATGGAGTACTCTAATCTCCAGTTTTTTTACTTTCAACCAACTTCCTTTGTTGCTCTGCTACTTGCTCTTGTATCTTGTGTACTTAGTGTCGTTGCTTTGAGAAAGAAATTTTTGAAAAACTACTACTCATCAACAGCAACGGCCGTTTCTCATCAGTGTCCAAAAGCATCGTGTGCATTGCCATTGATCGATCTTCTTCATGTTTTTTTCAATAAAACCGACTTGATGCACGTAACTCTCGGAAATATGGCTGATAAGTTCGGACCTATTTTCAGTTTCCCAACAGGTAGCCATAGGACTCTAGTTGTAAGCAGTTGGGAGATGGCAAAAGAGTGTTTTACAGGCAACAATGACACTGTTTTTTCAAACCGTCCTATCCCGTTAGCGTTTAAGATTATATTTTATGCAGGTGGTGTGGACTCATACGGGCTTGCACTTGTACCTTATGGAAAATATTGGAGGGAGTTACGAAAGATATGTGTGCATAACCTCCTCTCTAATCAACAATTACTCAACTTCAGACACTTAATAATTTCTCAAGTTGATACCTCTTTCAATAAGCTTTATGACTTGAGCAACAAGAATAAAAATACTACTACTGAAGCTGGCACGGTGAGAATGGATGATTGGCTTGCACAATTATCGTTCAATGTGATCGGAAGAATAGTCTGCGGATTCCAAACACACACCGAGACAAGTGCTACAAGCAGTGTGGAACGATTTACAGAAGCAATTGAAGAAGCATCACGTTTTATGTCGATAGCTACAGTATCAGATACCTTTCCATGGCTAGGGTGGATTGATCAGTTAACTGGTCTTACGAGAAAAATGAAGCACTATGGAAAGAAACTAGACTTAGTGGTTGAGAGCATAATTGAAGATCATCGTCAAAACAGGAGATTTTCTGGGACCAAACAAGGAGATGACTTCATCGACATTTGCTTGTCAATTATGGAGCAACCACAGATTATTCCCGGAAACAATGATCCTCCTCGTCAAATACCTATCAAATCTATTATCCTGGACATGATAGGTGGGGGGACTGACACCACAAAACTGACCACCACCTGGACCCTTTCATTGCTCCTGAACAACCCCCATGTGTTGGACAGAGCAAGAGAAGAAGTGGATGCACACTTTGGAACAAAAAGGAGAGCAACAAATGATGCAGTAATGGTGGAGTTTGATGACATTCGTAACCTTGTTTACATCCAGGCAATCATCAAAGAATCAATGAGGTTGTATCCAGTCAGCCCAGTTGTGGAGCGACTGAGCAGCGAAGACTGTGTGGTCGGTGAGTTTCATGTGCCAGCAGGGACGAGATTATGGGCAAACGTATGGAAGATGCAACGAGACCCAAATGTATGGGATGATCCATTGGTTTTTCGACCAGAGAGATTTCTGAGCGATGACAAGAAGATGGTTGATGTAAGGGGCCGTAATTATGAGTTATTACCATTCGGAGCCGGTCGACGAATATGTCCTGGTATATCCTTCTCTCTGGATCTAATGCAGCTGGTACTTACTCGTCTTATTCTTGAGTTTGAAATGAAGACTCCTACCGAGGGGGAAGTGGACATGACAGCAACACCAGGTTTAATGAGTTACAAGGTAGTCCCCCTTGACATGCTGCTCACTCGTCGCATGCTTTAG

>Pnu_CYP82Y2-Ln3 Pnu_CYP82Y2_1340598

ATGGATTACTCTAATCTTCAGTTTTTCGGTTTTCAACCAACTTCCATTGTTGCTCTTCTACTTGCTCTTGTATCTGTTTTACTTGGTTTTCTTGGTTTGAGGAAGAAATTTTCGCACTACCGGCCAAAAGCATCGTGTGCACTGCCACTGACCGGTCTTCTTCATGTTTTCATGAATAGAGATGACTTAATTCATGTAACTCTTGGAAACATGGCTGACGAATACGGACCGATTTTCAGTTTCCCAACAGGCAGCCATAGGACTCTGGTTGTGAGCAGTTGGGAGATTGTAAAAGAGTGTTTTACGGGCAGCAATGACACTGTTTTTTCCAATCGTCCCGTTCCGTTGGCTTTTAAGATTATATTCTATGCAGGTGGTATAGATTCGTATGGTCTTGCACTTGTACCTTATGGAAAACAATGGAGGGAGTTACGAAAGATCTGTGTGCATAACCTCCTTTCTAATCAACAGTTGCTCAAGTTCAGACAGTCGATAATCTCTCAAGTTGATGCCTCTTTCAGTAAGCTTTATGAGTCGTGCAACAGCAATAGAAACTCTGACGACCAGGGAGTCTCTAGTACTACTGCGTCTGGATCAGTCTCTAGTACCAGTGCAGCTAGCATGGTGAGAATGGATGATTGGCTTGCACAGTTATCGTTCAATGTGATAGGGAGAATAGTCTGCGGATTCCAATCACACACAGAGTCTGTGGAGCAATTTACAGTAGCAATGGATGAAGCATCGCGTTTTATGTCAATCACTGCAGTGTCGGATACCGTTCCATGGCTAGGGTGGATTGATCAACTAACTGGTCTTACCAGAAAAATGAAGCACTCTGGAAAGAAATTAGACTTGGTGGTTGAGAGCATAATCGAAGATCATCGTCAAAAGAGACAATTTGCTAGAACAAAAGGAGATGAAAATACTACCGAAGACGAACAAGATGACTTCATCGACATTTGCTTGTCAATTATGGAGCAGCCACAGCTTCCTGGCGACAACAACCCTCGTCAAATACCTATCAAATCGATTGTCCTGGACATGATAGGCGCGGGGGTTGACGCCACAAAACTGACCACGATCTGGACCCTTTCCCTGCTGCTGAACAACCCCCATGTGTTGGACAAGGCAAAAGAAGAAGTGGATGCACACTTTGGAACGAAAAGGAGGAGATCAGCAAATCATGCAGTCATGGTGGATTTTGATGATATTCGTAGCCTTGTCTACATCCAGGCAATCGTCAAAGAATCAATGCGGTTGTATCCAGCCAGCCCGATTGTTGAGCGACTGAGCAGCGAAGATTGTGTTGTCGGTGGGTTCCATGTACCAGCAGGGACGCGGTTATGGGCAAACGTATGGAAGATGCAACGAGACCCAAAAGTATGGGGTGATCCGTTGGTGTTTCGACCAGAGAGATTTTTGAGCAATGACCAGAAGAAGATGGTTGATGTAAGGGGTCAAAATTACGAGTTGATACCGTTTGGAGCTGGTCGGCGGATATGTCCAGGGATATCGTTCTCTTTGGATCTAATGCACCTGGTACTTACTCGTCTTATTCTTGAGTTTGAAATGAAGTCTCCTGGCGGCGGGGAAGTAGACATGACGGCGACTCCAGGCTTGATGAGTTACAAAGTAGTCCCGCTTGACATTCTGCTCACCGCTCGCATGCTTTAG

>Pnu_CYP82Y2-Ln2 Pnu_CYP82Y2_1286804

ATGGATTGCTCTAATCTTCAGTTTTTCGGTTTTCAACCAACTTCCCTTGTTGCTCTTCTCCTTGCTCTTGTATCCATATTACTTGGTTTTCTTGGTTTGAGGAAGAAATTTTCTGAAAACTGCTCATCGCCATCGTCAACAGCATCGTGTGCACTGCCACTGGCCGGTCTTCTTCATGTTTTCATGAATAGAGCCGACTTAATTCACGTAACTCTTGGAAACATGGCTGACAAATACGGACCGATTTTCAGTTTCCCAACAGGCAGCCATAGGACTCTGGTTGTGAGCAGTTGGGAGATGGTAAAAGAGTGTTTTACAGGCAGCAATGACATTGCTTTTTCCAATCGTCCCGTTCCGTTAGCGTTCAAGATTATATTCTATGCAGGTGGTATAGATTCGTATGGTCTTCCACTTGTACCTTACGGAAAATATTGGAGGGAGTTACGAAAGATCTGCGTGCATAACCTCCTTTCTAACCAACAATTGGTCAAGTTCAGGCACTTGATCACCTCTGAAGTTGATGCCTCATTCAGTAAGCTTTATGAGTCATGCAACAGCAATAGAAACTCTGACGGCCAGGGATCAGTCTCTAGTACTACTGCATCTGCCATGGTGAGAATGGATGATTGGCTTGGACAACAGTCGTTCAATGTTATCGGGAGAATAGTCTGCGGATTCCAATCACACACTGAGATAAGTGCTTCAAGCAGTGTGGAGCGATTTACAGTAGCAATGGATGAAGCATCGCGTTTTATGTCGATAACTGCTGTGTCGGATAACATTCCATGGCTAGGGTGGATTGATCAACTAACTGGTCTTACGAGAAAAATGAAGCACTCTGGAAAGAAATTAGACTTAGTGGTTGAGAGCATAATCGAAGATCATCGTCAAAAGAGACAATTTTCTAGAACAAAAGGAGATGAAGAGGACGAACAAGATGACTTCATTGACATTTGCTTGTCAATTATGGAGCAGCCACAGCTCCCTGGCGACAACAACCCTCATCAAATACCTATCAAATCGATTGTCCTGGACATGATAGGCGCGGGGGTTGACACCACAAAACT

>Pnu_CYP82Y2-Ln1 Pnu_CYP82Y2_1388589

ATGGAGTACTCTGATCTCAAGTTTTTTAGTTTTCAACCGACTTCCATTGTTGCTCTTCTACTTGCTCTTGTATCTATATTACTTAGTGTTGTTTGTTTGAAGAAGAAATATTCCAACACATCAACAGCAGCAGCAGCCGTTTCTCATCAGCGGTCGTGTAGATTGCCAATCATCGGTCTACTTCATGTTTTCACGAATAGAACTGACTTGAGCCATGTAACTCTTGGAAACATGGCTGATAAATACGGACCGATTTTCAGTTTCCCAACAGGTAGCCATAGGACTCTGGTGGTGAGCAGTTGGGAGTTGGTAAAAGAGTGTTTTACAGGCAGCAATGACAAAGCCTTTTCGAATCGTCCGATCCCATTGGCGTTTAAGCTTATGTTTTATACAGGTGGTATAGATTCATGCGGTCTTGCACTAGTGCCGTATGGAAATCAATGGAGGGAGTTACGAAAGATCTGTGTGCATAACCTCCTTTCTAATCAACAATTACTCAAGTTCAGACACTTGATAATTTCTGAAGTTGATACCTCTTTCAGTAAGCTTTATGAGTTATGCAACAAAAATAAAATCTCCGACAACGAGGGAGACTCTACTACTAATGTAGTTGGCATGGTGAGAATGGATGATTGGCATGCAAAATTATCGTTCAATGTGCTAGGACGAATAGTGTGCGGATTCCAATCACACACCGAGACAAGTGCTGCTAGCAGTGTGGAACGATTTACAGCAGCAATGGATGAAGCATCGCGTTTTATGTCGATATCTGCAGTGTCGGATACCGTTCCATGGCTAGGGTGGATTGATAACCTAACTGGTCTCAAGAGAAAAATGATGCACTGCGGAAAGAAATTAGACTTGGTGGTTGAGAGCATAATTGAAGATCATCGTCAAAAGAGACAACTTTATAGAACTAAAGAAAGGGATGATGAACAAACCACCGACGAGGAACAAGATGACTTCATCGACATTTGCTTGTCAATTATGGAGCAACAACAGCTTCCTGGCATTAATAACCCTCCTCAATTGCCCATCAAATCCATTGTCCTGGACATGATAGGTGGGGGGACTGACACCACAAAACTGACCACGATCTGGACCCTTTCCTTGCTGTTGAACAACCCCCGTGCATTGGACAAGGCAAAAGAAGAAGTGGATGCGCACTTTGGAATGAAAAGCAGACCAGCAAATCATGCAGTCATTGTGGATTTTGATGATATTCGTAACCTTGTCTACATCCAGGCAATCGTCAAAGAATCAATGCGGTTGTATCCAGCCAGCCCACTGGGGGAGCGACTGAGCAGCGAAGATTGTGTGGTCGGCGGGGTTTATGTACCAGCAGGGACGCGATTATGGGTTAACGTATGGAAGATGCAACGAGATCCAAGAGTATGGGATGATCCGTTGGTGTTTCGACCAGAGAGATTTTTGAGCAATGACCAGAAGATGGTTGATCTAAAGGGTCAACATTATGAGCTGATACCGTTCGGAACCGGCCGGCGGATATGTCCAGGATTGTCGTTTTCTTTGGATCTAATGCACCTCGTACTTACTCGTCTTATTCTTGAGTTTGAATTGAAGTCCCCCGGTGGCGGAGAAGTGGACATGACAGCAAC

>Pca_CYP82Y2-La1 Pca_CYP82Y2_179126

ATGGAGTACTCTGATCTCCAGTTTCTTAATTTACAGCCAACTTCCGTTGTTACTCTTCTACTTGCTCTTGTATTTATATTACTTGTTGGTTTCAGTAACAAAGTTCTGAGAAACTATTCAACGCCATCATCAACAGCAACAGCCGTTTCTCATCATCGGCCAAAAGCATCCTGTGCACTGCCAATGATCGGTCTGTTTCATGTTTTCATGAATAAAACCGGCTTAATTCATGTAACTCTTGGAAATATGGCTGATAAATATGGACCCATTTTCAGTTTCCCAACAGGTGGACACAGGGCTCTAGTTGTGAGCAGTTGGGAGATGGCAAAAGAGTGTTTTACAGGCAACAATGACATTGTTTTTTCAAATCGCCCTATGCCGTTATCGTTTAAGCTTATATTAAATGCAGGTGGTATAGACTCACACGGTCTTTCACAAGTACCTAATGGAAAATATTGGAGGGAGTTACGAAAGATCTGTGTGCATAACCTCCTCTCTAATCAACAATTACTCAAAGTTCAGACACTTGATATATTCTGAAGTTTGTAACTCTTTCAATAAACTGTATGCGTCATGCAACAGCAATAAAAACTCTGAAAACGTAGGAGTCGACTCTACTACTACTACTTCAGCTGGCATGGTGAGAATGGACGATTGGCTTGGACAATTATCGTTCAACGTGATCGGAAGAATAGTCTGCGGATTCCAATCACACGCCGAGACAAGTGCTACAAGCAGTGTGGAACGATTTGCAATTGCAATTGATGAAGCATCGCGTTTTATGTCGATACCTGCAGTGTCAGATACCTTTCCATGGCTAGGGTGGATTGATCAGTTAACTGGTCTTACGAGAGATATGAAGCACTACGGAAAGAAATTAGACTTGGTGGTTGAGAGCATAATTGAAGATCATCGTCAAAAGAGACGATTTTCTAGAACTAAAGGAGATGAGAATACCACCGAAGGTGACTTCATCGACATTTGCTTGTCAATTATGGAGCAACCACAGCCTCCGGGAAACAACTACGCTCGTCAGATGCCTATCAAATCTATTGTCGTGGACATGATAGGCGGGGCGACAGACACCACAAAACTAACCACGATCTGGACACTTTCCTTGCTGCTGAACAACCCCCATGTGTTGGATAAGGCAAAAGAAGAAGTGGATGCACACTTTGGAATGAAAAGGAGATCAACAAATGGTGCAGTCATGGTGGATTTTGATGATATTCGTAACCTTTTCTACATCCAGGCAATCATCAAAGAATCAATGCGGTTGTATCCAGCTAGCCCAGTGGTGGAGCGACTGAGCAGCGAAGATTGTGTGGTTGGTGGATTTCATGTACCAGCAGGGACACGATTATGGGTTAACGTATGGAAGATGCAACGAGACCCAAAAGTATGGGATGATCCGTTGGTGTTTCGACCAGAGAGATTTCTGAGCAATGAGAAGAAGATGGTTGATGTCATAGGTCAAAATTATGAGTTGATACCATTTGGAGCCGGTCGACGGATATGTCCAGGTATATCGTTCTCTTTCGATCTAATGCACCTGGTACTTACTCGTTTAATTCTTGAGTTTGAAATGAAGTCTCCCTCTGGGAAAGTGGACATGACAGCAACACAAGGTTTAATGAGTTACAAGGTAGTCCCCCTTGACATTCTACTCACCCGTCGCAAGCTTTAG

>Pat_CYP82Y2-La3 Pat_CYP82Y2_9004p

ATGGAGTACTCTAATCTCCAGTTTTTTAATTTTCAGCCAACTCCCATTGCTGCTTTTCTACTTGCTCTGGTATCTATATTACTTGGTGTTATTGCAGTTTCTCATCATCGGGCAAAACCATCGTGTGCATTGCCAATGATCGGTCTTTATCATGTTTTCATGAATAAAACCGGCTTAATTCATGTAACTCTAGGAAATATGGCTGATAAATATGGACCCATCTTCAGTTTCCCAACAGGTGGCCATAGGGCTCTAGTTGTCAGCAGTTGGGAGATGGCAAAAGAGTGTTTTACAGGAAACAATGACGTTGTTTTTTCGAATCGTCCTATGCCGTTATCGTTTAAGATTATATTCAATGCAGGTGGTATAGACTCAGCCGGTCTTTCACAATTACCTTATGGAAAGTATTGGAGGGAGTTGCGAAAGATCTGTGTGCACAACCTTCTCTCAAATCAACAATTACTCAAGTTCAGACACTTGATAAATTCTCAAGTTGATTCGTCTTTCAATAAACTGTACGAGTCATGCAACAAAAATAAAAACTCTGAAAACGAGGGAGACTCTGCTACTGCAGCTGGCATGGTGAGAATGGATGATTGGCTTGGAGACTTATCGTTCAACGTGATAGGAAGAATAGTTTGCGGGTTCCGATCGGACACCAAGACAAGTGCTACAAGCAGTATGGAACGGCTTACAGTAGCAATTGATGAAGCGTCACGTTTTATGTCAATACCTGCAGTGTCAGATACCTTTCCATGGCTAGAATGGATTGATCAGTTAACTGGTCTTACGAAAGATATGAAGCACCACGCAAAAAAATTAGACTTGGTGGTTGAGAGCATAATTGAAGATCATCGTCAAAAGAGACGATTTTCCAGAACTAAAAAAGGAGATGATAATACTACCGAGGATGAACAAGACTTCATCGACATTTGTTTGTCAATTATGGAGCAACCACAGCTTCCTGGAAATAATTACGCTCGTCAAATGCCTATCAAATCTATTGTCCTGGACATGATAGGCGGGGGGACTGACACCACAAAACTGACCACAATCTGGACCCTTTCCTTGTTGTTGAACAACCCCCATGTGCTAGACAAGGCAAAAGAAGAAGTGGATGCACACTTTGGAACGAAAACGAAATCAACAAATGGGGAAATCATGGTGGATTTTGATGATATTCGTAACCTAGTCTACATCCAGGCAATCATCAAAGAATCAATGCGGTTGTATCCAGCTAGTCCAGTGGTGGAGCGACTGAGCAGCGAAGATTGTGTGGTTGGTGGGTTTCATGTACTAGCAGGGACACGATTATGGGTTAACGTATGGAAGGTTCAACGAGACCCGGAAGTATGGGATGATCCGTCAGTGTTTCGACCAGAGAGATTTTTAAGCAATGAGCAGAAGATGGTTGATGTAAGGGGTCAACATTATGAGTTGTTACCATTTGGAGCCGGTCGACGGATATGTCCAGGTTTATCGTTCTCTATGGATCTAATGCACCTGGTACTTACTCGTCTTATTCTTGAGTTTGAAATGAAGTCTCCTCGTGGGGAAGTGGATATGACGGCAACACAAGGTTTGATGAGTTACAAGGTAGTCCCCCTTGACATTCTGCTCACCCGTCGCATTCTTTAG

>Pso_CYP82Y2-La4-PS0216860 Pso_CYP82Y2_PS0216860

ATGGAGTACTCTGATCTCCAGTTTTTTAATTTTCAGCCAACTTTTGTTGTTGCTCTTCTAGTTGCTCTTCCATCCATATTACTTGGTTTTATAGCAGTTTCTCATCATCGGGCAAAACCATCGTGTGCATTGCCAATGATCGGTCTTTATCATGTTTTCATGAATAAAACCGGCTTAATTCATGTCACTCTTGGAAATATGGCTGATAAATACGGACCTATTTTCAGTTTCCCAACAGGTGGCCACAGGGCTCTAGTTGTGAGCAGTTGGGAGATGGCAAAAGAGTGTTTTACAGGAAACAATGACATTGTTTTTTCAAATCGTCCTATGCCGTTATCGTTTAAGATTATATTCAATGCAGGTGGTATCGACTCAGCCGGTCTTTCACAAGTACCGTATGGAAAATATTGGAGGGAGTTACGAAAGATCTGTGTGCACAACCTTCTCTCAAATCAACAACTACTCAAGTTCAGGCACTTGATAAATTCTCAAGTTGATACCTCTTTCAATAAACTATACGAGTCATGCAACAAAAGTAAAAACTCTGAAAATGAGGGAGACTCTACTACTACCGCAGCTGGCATGGTGAGAATGGATGATTGGCTTGGAAAATTATCATTCAACGTGATAGGAAGAATAGTCTGCGGATTCCAATCAGACACCGACACAAGTGCTACAAGCAGTGCGGAACGATTTACAGTAGCAATTGATGAAGCATCACGTTTTATGTCCATACCTGCAGTCTCAGATACCTTTCCATGGCTAGGGTGGATTGATCAGTTAAGTGGTCTTATAAAAGATATGAAGCACCACGCAAAAAAATTAGACTTAGTGGTTGAAAGCATAATTGAAGATCATCGTCAAAAGAGACGATTTTCTAGAACTAAAAAAGGAGATGAGAATACCACCGAGGATGAACAAGATGACTTCATTGACATTTGTTTGTCAATAATGGAGCAACCAGAGCTTCCTGGAAACAATTACGCTCGTCAAATGCCTATCAAATCTATCATCGTGGACATGATAGGCGGGGCGACTGACACCACAAAACTGACCACAATCTGGACCCTTTCCTTGCTGCTGAATAACCGCCATGTGCTAGACAAGGCAAAAGAAGAAGTGGATGCGCACTTTGGAACAAAAAGGAAATCAACAAATGGGGAAGTCATGGTGGATTTTGATGATATTCGTAACCTTGTCTACGTCCAAGCGATCATCAAAGAATCGATGCGGTTGTATCCAGCCAGCCCAGTGGTGGAGCGACTGAGCAGCGAAGATTGTGTGGTTGGTGGGTTTCATGTACCAGCAGGGACACGGTTATGGGTTAACGTATGGAAGGTTCAACGAGAGCCAGAAGTATGGGATGATCCGTCGGTGTTTCGACCAGAGAGATTTTTGAGCAATGAGCAGAAGATGGTTGATGTACGGGGTCAAGATTATGAGTTGTTACCATTTGGAGCAGGTCGACGGATATGTCCAGGTGTATCGTTCTCTATGGATCTAATGCACCTGGTACTTACTCGTCTCATTCTTGAGTTTGAAATGAAGTCGCCTGTTGGGGAAGTGGACATGACGGCGACACAAGGTTTAATGAGTTACAAGGTAGTCCCCCTTGACATTCTGCTGACCCGTCGCATGCTTTAG

>Prh_CYP82Y2-La5-AKO60176 Prh_CYP82Y2_AKO60176

ATGGAGTACTCTGATCTCCAGCTTTTTGGTTTTCAGCCAACTTCCGTTGTTGCTCTTCTACTTGCTCTTGTATCCATATTACTTGGTGTTATTGCAGTTTCTCATCATCGGGCAAAACCATCGTGTGCATTGCCAATGATCGGTCTTTATCATGTTTTCATGAATAAAACCGGCTTAATTCATGTAACTCTTGGAAACATGGCTGATAAATACGGACCCATTTTCAGTTTCCCAACAGGTGGCCATAGGGCTCTAGTTGTGAGTAGTTGGGAGATGGCAAAAGAGTGTTTTACAGGCGACAATGACATTGTTTTTTCAAATCGTCCTATGCCGTTATCGTTTAAGATTATATTCAATGCAGGTGGTATAGACTCAGCCGGTCTTACACAAGTACCGTATGGAAAATATTGGAGGGAGTTACGAAAGATCTGTGTGCACAACCTTCTCTCAAATCAACAATTACTCAAGTTCAGGCACTTGATAAATTCTCAAGTTGATACCTCTTTCACTAAACTGTACGAGTCATGCAACAAGGGAAATTCTGGCATGGTGAGAATGGATGATTGGCTTGGAGAATTATCATTCAACGTGATAGGAAGAATAGTCTGCGGATTCCGATCAGACACTGAGACAAGTGCTACAAGCAGTGCGGAACGATTTACAGTAGCAATTGATGAAGCATCGCGTTTTATGTCCATACCTGCAGTGTCAGATACCTTTCCGTGGCTAGGATGGATTGATCGATTAACTGGTCTTATAAAAGATATGAAGCACCATGCAGAAAAATTAGACTTAGTGGTTGAAAGCATAATTGAAGATCATCGTCAAAAGAGACGATTTTCTAGAACTAAAAAAGGAGATGAGAATAACACCGAGGATGAACAAGATGACTTCATTGACATTTGTTTGTCAATTATGGAGCAACCAGAGCTTCCTGGAAACAACTACGCTCGTCAAATGCCTATCAAATCTATCATCGTGGACATGATAGGCGGGGCGACTGACACCACAAAACTGACCACAATCTGGACCCTTTCCTTGCTGCTGAATAACCCACATGTGTTAGGCAAGGCAAAAGAAGAAGTGGATGCACACTTTGGAAAGAAAAGGAGTTCAACCAATGGGGAAGTCATGGTGGATTTTGATGATATTCGTAGCCTTGTCTACATCCAGGCAATCATCAAAGAATCAATGCGGTTGTACCCAGCCAGTCCAGTGGTGGAGCGACTGAGCAGTGAAGATTGTGTGGTTGGTGGGTTTCATGTATCAGCAGGGACGCGATTATGGGTTAACGTATGGAAGGTTCAACGAGACCCAGAAGTATGGGATGATCCGTCGGTGTTTCGACCAGAGAGATTTTTGAGTAATGAGCAGAAGATGGTTGATGTAAGGGGTCAAGATTATGAGTTGTTACCATTTGGAGCCGGTCGACGGATATGTCCAGGTGTATCGTTCTCTTTGGATCTAATGCACCTGGTACTTACTCGTCTTATTCTTGAGTTTGAAATTAAGTCTCCTGGTGGGGAAGTGGACATGACAGCAACACAAGGTTTAATGAGTTACAAGGTAGTCCCCCTTGACATTCTGCTCACCCGTCGCATGCTTTAG

>Par_CYP82Y2-Lb5 Par_CYP82Y2_scaffold_75401_CDS

ATGGAGTACTCATCCGATCTCCAGAACTACTGTTCTTATTATTTTCAACCAACTTCCCTTGTTGCTCTTCTACTTGCTCTTGCATCCACTGTTGTTGTTTTGAGGAAGAAGAAATTTTTGAGAAACTACTCAGAAACAGCGGTTTCTCATCAGCGGTCGTCGTGTGCATTGCCAATCATCGATCTTCTTCGTATTTTCATGAACAAGAATGACTTAATTCATGTAACTCTTGGAAATATGGCGGATAAACACGGACCGGTTTTCAGTTTCCCAACAGGCAGCAGCCATAGGACTCTTGTTGTAAGCAGTTGGGAGATGGCAAAAGAGTGTTTTACAGGCAACAATGACACTGTTTTTTCAAACCGTCCTATCCCGTTAGCGTTTAAGATTATATTTTATGCAGGTGGTGTGGACTCATACGGGCTTGCACTTGTACCTTATGGAAAATATTGGAGGGAGTTACGAAAGATCTGTGTGTATAACCTCCTCTCTAATCAACAGTTACTCAAGTTCAGACACTTGATAATTTCTCAAGTTGATGCATCTTTCAACAAGCTTTATGAGTTATGCAACAAGAATAAAAACGAGGGAGACTCTTCAGTTGGCATGGTGAGAATGGACGATTGGCTTGGGAAATTATCGTTCGATGTCATAGGAAGAATAGTCTGCGGATTCCAATCACACACCGAGACGAGTGCTACTAGCAGTGTGGAACGATTTACAGAAGCAATTGATGAAGCATCGCGTTTTATGTCGATATCTGCTGTGTCAGATACCATTCCATGGCTAGGGTGGATTGACAAATTAACTGGTCTTACCAGAAACATGAACAACTGTGGGAAGAAACTAGACTTGGTTGTTGAGAGCATAATTGAAGATCATCGTCAAAAGAGACTATTTTTTAGAGCTAAAGAAGGAGAGACCACCGAGGATGAACAAGATGATTTCATCGACATTTTCTTGTCAATAATGGAGCAACCACAGCTTCCTGGAAACAGTGACCCTTATCAAATACCTATCAAATCTATTGTCCTGGACATGATAGGTGCGGGGACTGACACCACAAAATTGACCACCATCTGGACACTTTCCTTGCTACTGAACAACCCCCATGTGTTGGACAAGGCAAAGGAAGAAGTGGATGCACACTTTGGAAAGAAAAGGAGATCAACAAATGATGCAGCAGCAGCAGTCGTGGTGGACTTTGATGATATTAGTAATCTTGTCTACATCCAGGCAATAATCAAAGAATCAATGCGGTTGTACCCAGCCAGCCCACTGGTGGAGCGATTGAGCAGCCAAGATTGTTTGGTAGGCGGGTTTCATGTACCAGCAGGGACTCGACTATGGGCAAATGTATGGAAGATACAACGAGACCCTAACGTGTGGGATGATCCATTGGTGTTTCGACCAGAGAGATTTCTGAGCGATGGCCAGAACAAGATGGTTGATGTAAGGGGCCGAAATTATGAGTTATTACCATTCGGAGCCGGTCGACGGATATGTCCTGGTATATCGTTCTCTCTGGATCTAATGCAGCTGGTACTTGCTCGTCTTATTCTTGAGTTTGAAATGAAGACTCCTAGTGAGAAAGTGGACATGACAGCAACACCAGGTTTAATGAGTTACAAGCTAGTCCCCCTTGACATTCTGCTCACCGGTCGTCGCATGACAAAGCTTGAAACATCATTCATCTCCAGTTTATAA

>Par_CYP82Y2-Lb6 Par_CYP82Y2_contig_27277_CDS

ATGGAGTACTCATCCGATCTCCAGAACTACTTTTCTTATTTTCAACCAACTTCCCTTGTTGCTCTTCTACTTGCTCTTGCATCCATATTACTCACTGTTGTTGCTTTGAGGAAGAAGAAATTCTTGAAAAACTACTCATCACCACCATTGTCAACAGAAACAGCGGTTTGTTATCAGCGGTCGTCGTGTGCATTGCCAATCATCGATCTTCTTCGTATTTTCATGAATAAGAATGACTTAATTCATGTAACTCTTGGAAACATGGCTGATAAACACGGACCGGTTTTCAGTTTCCCAACAGGTAGCCATAGGACTCTTGTTGTAAGCAGTTGGGAGATGGCGAAAGAGTGTTTTACCGGCAACAATGACATTGCCTTTTCAGATCGTCCTCTCCCGTTAGCATTCAAGACTATATTCTATGCAGGTGGTATAGACTCATACGGTATTTCGCAAGTACCGTACGGAAAAAAATGGAGGGAGTTACGGAAAATATGTGTGCATAACCTCCTCTCTAATCAACAGTTACTCAAGTTCAGACACTTGATAATTTCTCAAGTTGACGACTCTTTCAACAAGCTTTATGAGTTATGCAACAACAATAAAAACGAGGGAGACTCTTCAAGTGTTCAGTTGGCATGGTGAGAATGGACGATTGGCTTGGGAAATTATCGTTCGATGTCATTGGAAGAATAGTCTGCGGATTCCAATCACACACCGAGACGAGTGCTACTAGCAGTGTGGAACGATTTACAGAAGCAATTGATGAAGCATCGCGTTTTATGTCGATATCTGCTGTGTCAGATACCATTCCATGGCTAGGGTGGATTGACAAATTAACTGGTCTTACCAGAAACATGAACAACTGTGGGAAGAAACTAGACTTGGTTGTTGAGAGCATAATCGAAGATCATCGTCACAAGAGACGATTCTTTAGAGCTAAAGAAGGAGAGACCGAGGATGAACAAGATGATTTCATCGACATTTGCTTGTCAATAATGGAGCAACCACAGCTTCCAGGAAACAGTGACCCTTATCAAATACCTATCAAATCTATTGTCCTGGACATGATAGGTGCGGGGACTGACACCACGAAACTGACCACCATCTGGACACTTTCCTTGCTGCTGAACAACCCCATGTGTTGGAAAAGGCAAAGGAAGAAGTGGATGCACACTTTGGAAAGAAAAGGAGATCAACAAATGATGCAGCAGCAGCAGTCGTGGTGGACTTTGATGATATTTGTAATCTTGTCTACATCCAGGCAATAATCAAAGAATCAATGCGGTTGTATCCAGCCAGCCCACTGGTGGAGCGATTGAGCAGCGAAGACTGTCTGGTAGGCGGGTTTCATGTACCAGCAGGGACTCGACTATGGGCAAACGTATGGAAGATACAACGAGACCCTAACGTGTGGGATGATCCATTGGTGTTTCGACCAGAGAGATTTCTGAGCGATGGCCAGAACAAGATGGTTGATGTAAGGGGCCGAAATTATGAGTTATTACCATTCGGAGCCGGTCGACGGATATGTCCTGGTATATCCTTCTCTCTGGATCTAATGCAGCTGGTACTTACTCGTCTTATTCTTGAGTTTGAAATGAAGACTCCTAGTGAGAAAGTGGACATGACAGCAACACCAGGTTTAATGAGTTACAAGGTAGTCCCCCTTGACATTCTGCTCACCGGTCGTCGCATGACAAAGCTTGAAACATCATTCATCTCCAGTTTATAA

>Par_CYP82Y2-Lb7 Par_CYP82Y2_contig_28363_CDS

ATGGAGTACTCATCCGATCTCCAGAACTACTTTTCTTATTATTTTCAGCCAACTTCCATTGTTGCTCTTCTTCTACTTGCTCTTGCATCCACTGTTGTTGTTTTGAGGAGGAAGAAAAACCACTCATCACCACCACCATTGTCGACAACAGAAACAGCGGTTTGGTCGTCGTGTGCACTGCCAATCATCGATCTTCTTCGTATTTTCATGAATAAGAAGAATGACTTAATTCATGTAACACTTGGAAATATGGCTGATAAACACGGACCGGTTTTCAGTTTCCCAACAGGTAGCAGCCATAGGACTCTGGTTGTAAGCAGTTGGGAGATGGCAAAAGAGTGTTTTACCGGCGAGAATGACATTGCCTTTTCAGATCGTCCTCTCCCGTTAGCATTCAAGACTATATTCTATGCAGGTGGTGTAGACTCGTACGGTATTTCGCAAGTACCGTACGGGAAAAATTGGAGGGAGTTGCGGAAAATCTGTGTGCATAACCTCCTCTCTAATCAGCAGTTGCTCAAGTTCAGACACTTGATAATTTCTCAAGTTGACGACTCTTTCAACAAGCTTTATGAGCTATGCAACAAGAATAAAGACGAGGGAGACTCTTCAGTTGGCATGGTGAGAATGGACGATTGGCTTGGGAAATTATCGTTCGATGTCATTGGAAGAATAGTCTGCGGATTCCAGTCACACACCGAGACGAGTGCTACTAGCAGTGTGGAACGATTTACAGAAGCAATTGATCAAGCATCGCGTTTTATGTCGATATCTGCTGTGTCAGATACCATTCCATGGCTAGGGTGGATTGACAAATTAACCGGTCTTACCAGAAACATGAACAACTGTGGGAAGAAACTAGACTCGGTTGTTGAGAGCATAATTGAAGATCATCGTCAAAAGAGACGATTTTTTAGAGCTAGAGAAGGAGAGACGAATACCACCGAGGATGATTTCATCGACATTTGCTTGTCAATAATGGAGCAGCAACCACAGCTTCCTGGAAACAGTGACCCTTATCAAATACCTATCAAATCTATTGTCCTGGACATGATAGGTGCGGGGACTGACACCACAAAACTGACCACCATCTGGACACTTTCCTTGCTGCTGAACAACCCCCATGTGTTGGACAAGGCAAAGGAAGAAGTGGATGCACACTTTGGAAAGAAAAGGAGATCAACAAATGATGCAGCAGCAGCAGCAGTCGTGGTGGACTTTGATGATATCTGCAATCTTGTCTACATCCAGGCAATAATCAAAGAATCAATGCGGTTGTATCCAGCCAGCCCACTGGTGGAGCGATTGAGCAGCCAAGATTGTCTGGTAGGCGGGTTTCATGTACCAGCAGGGACTCGATTATGGGCAAACGTATGGAAGATACAACGAGACCCTAACGTGTGGAATGATCCATTGGTGTTTCGACCAGAGAGATTTCTGAGCGATGGCCAGAACAAGATGGTTGATGTAAGGGGCCGAAATTATGAGTTATTACCATTCGGAGCCGGTCGACGGATATGTCCTGGTATATCGTTCTCTCTGGATCTAATGCAGCTGCTACTTACTCGTCTTATTCTTGAGTTTGAAATGAAGACTCCTAGTGAGAAAGTGGACATGACAGCAACACCAGGTTTAATGAGTTACAAGCTAGTCCCCCTTGACATTCTGCTCACCGCTCGTCGCATGACAAACCTTCAAACATCATTCATCTCCAGTTCATAA

>Par_CYP82Y2-Lb9 Par_CYP82Y2_contig_30186_CDS

ATGGAGTACTCATCCGATCTCCAGAACTACTTTCTTATTATTTTCAACCAACTTCCCTTGTTGCTCTTCTTCTACTTGCTCTTGCATCCACTGTTGTTGTTTTGAGGAGGAAGAAAAACCACTCATCACCACCACCATTGTCGACAACAGAAACAGCGGTTTGGTCGTCGTGTGCACTGCCAATCATCGATCTTCTTCGTATTTTCATGAATAAGAAGAATGACTTAATTCATGTAACACTTGGAAATATGGCTGATAAACACGGACCGGTTTTCAGTTTCCCAACAGGTAGCAGCCGTAGGACTCTTGTTGTAAGCAGTTGGGAGATGGCAAAAGAGTGTTTTACCGGCGAGAATGACACTGCCTTTTCAGATCGTCCTCTCCCGTTAGCATTCAAGACCATATTCTATGCAGGTGGTGTAGACTCGTACGGTATTTCCCAAGTACCGTACGGGAAAAATTGGAGGGTGTTGCGGAAAATCTGTGTGCATAACCTCCTCTCTAATCAGCAGTTGCTCAAGTTCAGACACTTGATAATTTCTCAAGTTGACGACTCTTTCAACAAGCTTTATGAGCTATGCAACAAGAATAAAGACGAGGGAGACTCTTCAGTTGGCATGGTGAGAATGGACGATTGGCTTGGGAAATTATCGTTCGATGTCATTGGAAGAATAGTCTGCGGATTCCAGTCACACACCGAGACGAGTGCTACTAGCAGTGTGGAACGATTTACAGAAGCAATTGATGAAGCATCGCGTTTTATGTCGATATCTGCTGTGTCAGATACCATTCCATGGCTAGGGTGGATTGACAAATTAACCGGTCTTACCAGAAACATGAACAACTGTGGGAAGAAACTAGACTCGGTTGTTGAGAGCATAATTGAAGATCATCGTCAAAAGAGACGATTTGCTAGATCTAAAGAAGGAGAGACGAATACCACCGAGGATGATTTCATCGACATTTGCTTGTCAATAATGGAGCAGCAACCACAGCTTCCTGGAAACAGTGACCCTTATCAAATACCTATCAAATCTATTGTCCTGGACATGATAGGTGCGGGGACTGACACCACAAAACTGACCACCATCTGGACACTTTCCTTGCTGCTGAACAACCCCCATGTGTTGGACAAGGCAAAGGAAGAAGTGGATGCACACTTTGGAAAGAAAAGGAGATCAACAAATGATGCAGCAGCAGCAGCAGTCGTGGTGGACTTTGATGATATCTGCAATCTTGTCTACATCCAGGCAATAATCAAAGAATCAATGCGGTTGTATCCAGCCAGCCCACTGGTGGAGCGATTGAGCAGCCAAGATTGTCTGGTAGGCGGGTTCCATGTACCAGCAGGGACTCGATTATGGGCAAACGTATGGAAGATACAACGAGACCCTAACGTGTGGAATGATCCATTGGTGTTTCGACCAGAGAGATTTCTGAGCGATGGCCAGAACAAGATGGTTGATGTAAGGGGCCGAAATTATGAGTTATTACCATTCGGAGCCGGTCGACGGATATGTCCTGGTATATCGTTCTCTGGATCTAATGCAGCTGCTACTTACTCGTGTTATTCTAGAGTTTGAAATGAAGACTCCTAGTGAGAAAGTGGACATGACAGCAACACCAGGTTTAATGAGTTACAAGCTAGTCCCCCTTGACATTCTGCTCACCGCTCGTCGCATGACAAAGCTTGAAACATCATTCATCTCCAGTTCATAA

>Par_CYP82Y2-Lb8 Par_CYP82Y2_contig_30192_CDS

ATGGAGTACTCATCCGATCTCCGGAACTACTACTTTTCTTATTATTTTCAACCAACTTCCATTGTTGCTCTTCTTCTACTTGCTCTTGCATCCACTGTTGTTGTTTTGAGGAGGAAGAAAAACCACCCATCACCACCACCATTGTCGACAACAGAAACAGCGGTTTGGTCGTCGTGTGCACTGCCAATCATCGATCTTCTTCGTATTTTCATGAATAAGAAGAATGACTTAATTCATGTAACACTTGGAAATATGGCTGATAAACACGGGCCGGTTTTCAGTTTCCCAACAGGTAGCAGCCATAGGACTCTGGTTGTAAGCAGTTAGGAGATGGCAAAGAGTGTTTTACCGGCGAGAATGACATTGCCTTTCAGATCGTCCTCTCCCGTTAGCATTCAAGACCATATTCTATGCAGGTGGTGTAGACTCGTACGGTATTTCCCATACCGTATGGGAAAAATTGAGGGAGTTGCGGAAAATCTGTGTGCATAACCTCCTCTCTAATCAGCAGTTGCTCAAGTTCAGACACTTGATAATTTCTCAAGTTGACGACTCTTTCAACAAGCTTTTATGAGCTATGCAGCAAGAATAAAGACGAGGAGACTCTTCAGTTGGCATGGTGAGAATGGACGATTGGCTTGGGAAATTATCGTTCGATGTCATTGGAAGAATAGTCTGCGGATTCCAGTCACACACCGAGACGAGTGCTACTAGCAGTGTGGAACGATTTACAGAAGCAATTGATGAAGCATCGCGTTTTATGTCGATATCTGCTGTCAGATACCATTCCATGGCTAGGGTGGATTGACAAATTAACTGGTCTTACCAGAAACATGAACAACTGTGGGAAGAACTAGACTCGGTTGTTGAGAGCATAATTGAAGATCATCGTCAAAAAGAGACGATTTTTTAGAGCTAAAGAAGGAGAGACGAAGCACCACTGAGGATGATTTCATCGACATTTTGCTTGTCAATAATGGAGCAACCACAGCTTCCTGGAACAGTGACCCTTATCAAATACCTATCAAATCTATTGTCCTGGACATGATAGGTGCGGGACTGACACCACAAAACTGACCACCATCTGGACACTTTCCTTGCTGCTGAACAACCCCATGTGTTGACAAGGCAAAGGAAGAAGTGGATGCACACTTTGGAAAGAAAAGGAGATCAACAAATGATGCAGCAGCAGCAGCAGTCGTGGTGGGACTTTGATGATATCTGCAATCTTGTCTACATCCAGGCAATAATCAAAGAATCAATGCGGTTGTATCCAGCCAGCCCACTGGTGGAGCGATTGAGCAGCCAAGATTGTCTGGTAGGCGGGTTTCATGTACCAGCAGGGACTCGATTATGGGCAAACGATGGAAGATACAACCAGACCCTAACGTGGTGGAATGATCCATTGGTGTTTCGACCAGAGAGATTTCTGAGCGATGGCCAGAACAAGATGGTTGATGTAAGGGGCCGAAATTATGAGTTATTACCATTCGGAGCCGGTCGACGGATATGTCCTGGTATATCGTTCTCTCTGGATCTAATGCAGCTGCTACTTACTCCGTGTTATTCTTGAGTTTGAAATGAAGACTCCTAGTGAGAAAGTGGACATGACAGCAACACCAGGTTTAATGAGTTACAAGCTAGTCCCCTTGACATTCTGCTCACCGCTCGTCGCATGACAAAGCTTGAAACATCATTCATCTCAGTTCATAA

Supplementary dataset 7 (associated with Figure 3b)

- Nucleotide sequences of *Papaver* P450 cytochrome *STORR* and *COR*-*L* genes

>Par_STORR1 Par_STORR_contig_11585_CDS

ATGGAGCTCCAATATTTTTCTTATTTTCAACCAACTTCCTCCGTTGTTGCTCTTCTACTTGCTCTTGTATCCATCTTATTCAGTTTCGTTGTTTTGAGGAAGACATTTTCGAATAACTACTCATCACCTGCATCATCAACAGAAACAGCGGTACTTTGTCATCAGCGGCAGCAGTCGTGTGCATTGCCAATTTCCGGTCTTCTTCATGTTTTCATGAATAAAAACGGCTTAATTCATGTAACTCTTGGAAATATGGCCGATATATACGGACCGATTTTCAGTTTCCCAACAGGTAGCCATAGAACTCTCGTTGTGAGCAGTTGGGAGATGGTAAAAGAGTGTTTTACAGGCAACAATGACACTGCTTTCTCAAACCGTCCTATCCCGTTAGCTTTTAAGACTATATTCTATGCATGCGGTGGCATAGACTCATACGGTCTTTCGAGTGTACCTTATGGAAAATATTGGAGGGAGCTACGAAAAGTCTGTGTGCATAACCTCCTCTCTAATCAACAACTACTCAAGTTCAGACACTTGATAATTTCTCAAGTTGATACGTCTTTCAATAAGCTGTATGAGTTATGCAAAAACTCTGAAGACAACCAGGGAAACCATACTACTACTACTACTACTACCGCAGCTGGCATGGTGAGAATGGATGATTGGCTCGCCCAATTATCGTTCAACGTGATAGGAAGAATAGTCTGCGGATTCCAATCAGACCCTAAGACAGGTGCTCCAAGCAGGGTGGAACAATTTAAAGAAGCAATTAATGAAGCATCGCATTTTATGTCGACATCTCCAGTGTCAGATAACGTTCCAATGCTAGGGTGGATTGACCAATTGACTGGTCTTACAAGAAATATGAAGCACTGCGGAAAGAAATTAGACTTGGTGGTCGAGAGCATAATTCAAGATCATCGTCGAAAGAGACGATTCTCTAGAACTAAAGGAGGAGATGAGAAGGACGATGAACAAGATGACTTCATCGACATTTGCTTGTCAATTATGGAGCAACCACAGCTTCCTGACAACAATAACCCTCCTCATATACCTATCAAATCTATTGTCCTGGACATGATAGGTGGGGGAACTGACACAACAAAACTGACCACCATCTGGACCCTTTCCTTGCTGCTGAACAACCCCCATGTGTTGGACAAGGCAAAACAAGAAGTGGATGCACACTTTGGAACGAAAAGGAGATCAACAAATGATGCCGTGGTGGATTTTGATGATATTCGTAACCTTGTATACATCCAGGCAATCATCAAAGAATCAATGCGGTTGTATCCAGCCAGCCCCGTGGTGGAGCGACTGAGCGGCCAAGATTGTGTGGTCGGTGGGTTTCATGTACCAGCAGGGACGAGATTATGGGCTAACGTATGGAAGATGCAACGAGACCCTAAAGTATGGGATGATCCATTGGTGTTTCGACCAGAGAGATTTTTGAGCGATGAACAGAAGATGGTTGATGTAAGGGGTCAAAATTATGAGCTGTTACCATTTGGAGCCGGTCGACGTATATGTCCAGGTGTATCCTTCTCTTTGGATCTAATGCAACTGGTACTGACTCGTCTTATTCTTGAGTTTGAAATGAAGTCTCCTAGCGGGAAAGTGGACATGACAGCAACACCAGGATTAATGAGTTACAAGGTGGTCCCCCTTGACATTCTGCTCACCCATCGTGGCACAAAGTCGTGTGTGCAGTCAGCAGCCTCTGAGACTAGTGGTGTACCAGTAATCACTCTGAGCTCGGGCAAGGTGATGCCTGTTTTAGGCATGGGTACATTTGAGAAAGTTGGTAAAGGGTCTGAAAGAGAGAGGTTGGCGATTTTAAAAGCGATAGAGGTGGGTTACAGATACTTCGATACAGCTGCTGCATACGAAACTGAAGAGGTTCTTGGAGAAGCTATTGCTGAAGCACTCCAACTTGGCCTAATCAAATCTCGAGATGAACTTTTCATCAGTTCCATGCTCTGGTGCACTGATGCTCACCCTGATCGTGTCCTCCTCGCTCTTCAGAATTCTCTGAGGAATCTTAAATTGGAGTATCTGGATCTATATATGTTACCCTTCCCGGCAAGCTTGAAGCCAGGGAAGATAACGATGGACATACCAGAGGAAGATATTTGTCGAATGGACTACAGGTCTGTATGGTCAGCCATGGAAGAGTGTCAGAACCTTGGCTTCACTAAATCAATCGGTGTTAGCAATTTCTCCTGCAAAAAGCTTCAGGAATTGATGGAGACCGCCAACATCCCTCCAGCTGTGAATCAAGTGGAGATGAGCCCGGCTTTCCAACAAAAGAAGCTGAGAGAGTATTGCAACGCAAATAATATATTAGTCAGTGCAGTCTCTATACTGGGATCAAACGGAACCCCATGGGGCTCCAATGCAGTTATGGGTTCTGAGGTGCTTAAGAAAATTGCTATGGCCAAAGGAAAATCTGTTGCTCAGGTTAGTATGAGATGGGTTTACGAGCAAGGCGCGAGTCTTGTGGTAAAAAGTTTCAGTGAAGAGAGATTGAGGGAAAACTTGAACATATTCGACTGGGAACTCACTAAGGAAGACAATGAAAAGATCGGTGAGATTCCACAGTGCAGAATCTTGACTGCTCATTTTTTGGTCTCACCTAATGGACCTTTCAAATCTCAAGAAGAGTTGTGGGATGACAAAGCTTGA

>Par_STORR2 Par_STORR_contig_40428_CDS

ATGGAGCTCCAATATTTTTCTTATTTTCAACCAACTTCCTCCGTTGTTGCTCTTCTACTTGCTCTTGTATCCATCTTATTCAGTTTCGTTGTTTTGAGGAAGACATTTTCGAATAACTACTCATCACCTGCATCATCAACAGAAACAGCGGTACTTTGTCATCAGCGGCAGCAGTCGTGTGCATTGCCAATTTCCGGTCTTCTTCATGTTTTCATGAATACAAACGGCTTAATTCATGTAACTCTTGGAAATATGGCTGATATATACGGACCGATTTTCAGTTTCCCAACAGGTAGCCATAGAACTCTCGTTGTGAGCAGTTGGGAGATGGTAAAAGAGTGTTTTACAGGCAACAATGACACTGCTTTCTCAAACCGTCCTATCCCGTTAGCTTTTAAGACTATATTCTATGCATGCGGTGGCATAGACTCATACGGTCTTTCGAGTGTACCTTATGGAAAATATTGGAGGGAGCTACGAAAAGTCTGTGTGCATAACCTCCTCTCTAATCAACAACTACTCAAGTTCAGACACTTGATAATTTCTCAAGTTGATACGTCTTTCAATAAGCTGTATGAGTTATGCAAAAACTCTGAAGACAACCAGGGAAACCATACTACTACTACCGCAGCTGGCATGGTGAGAATGGATGATTGGCTCGCCCAATTATCGTTCAACGTGATAGGAAGAATAGTCTGCGGATTCCAATCAGACCCTAAGACAGGTGCTCCAAGCAGGGTGGAACAATTTAAAGAAGCAATTAATGAAGCATCGCATTTTATGTCGACATCTCCAGTGTCAGATAACGTTCCAATGCTAGGGTGGATTGACCAATTGACTGGTCTTACAAGAAATATGAAGCACTGCGGAAAGAAATTAGACTTGGTGGTCGAGAGTATAATTCAAGATCATCGTCAAAAGAGACGATTCTCTAGAACTAAAGGAGGAGATGAGAAGGACGATGAACAAGATGACTTCATCGACATTTGCTTGTCAATTATGGTGCAACCACAGCTTCCTGACAACAATAACCCTCCTCATATACCTATCAAATCTATTGTCCTGGACATGATAGGTGGGGGAACTGACACAACAAAACTGACCACCATCTGGACCCTTTCCTTGCTGCTGAACAACCCCCATGTGTTGGACAAGGCAAAACAAGAAGTGGATGCACACTTTGGAACGAAAAGGAGATCAACGAATGATGCCGTGGTGGATTTTGATGATATTCGTAACCTTGTATACATCCAGGCAATCATCAAAGAATCAATGCGGTTGTATCCAGCCAGCCCCGTGGTGGAGCGACTGAGCGGCCAAGATTGTGTGGTCGGTGGGTTTCATGTACCAGCAGGGACGAGATTATGGGCTAACGTATGGAAGATGCAACGAGACCCTAAAGTATGGGATGATCCATTGGTGTTTCGACCAGAGAGATTTTTGAGCGATGAACAGAAGATGGTTGATGTAAGGGGTCAAAATTATGAGCTGTTACCATTTGGAGCCGGTCGACGTATATGTCCAGGTGTATCCTTCTCTTTGGATCTAATGCAACTGGTACTGACTCGTCTTATTCTTGAGTTTGAAATGAAGTCTCCTAGCGGGAAAGTAGACATGACAGCAACACCAGGATTAATGAGTTACAAGGTGGTCCCCCTTGACATTCTGCTCACCCATCGTGGCACAAAGTCGTGTGTGCAGTCAGCAGCCTCTGAGAGTAGTGGTGTACCAGTAATCACTCTGAGCTCGGGCAAGGTGATGCCTGTTTTAGGCATGGGAACATTTGAGAAAGTTGGTAAAGGGTCTGAAAGAGAGAGGTTGGCGATTTTAAAAGCGATAGAGGTGGGTTACAGATACTTCGATACAGCTGCTGCATACGAAACTGAAGAGGTTCTTGGAGAAGCTATTGCTGAAGCACTCCAACTTGGCCTAATCAAATCTCGAGATGAACTTTTCATCAGTTCCATGCTCTGGTGCACTGATGCTCACCCTGATCGTGTCCTCCTCGCTCTTCAGAATTCTCTGAGGAATCTTAAATTGGAGTATCTGGATCTATATATGTTACCCTTCCCGGCAAGCTTGAAGCCAGGGAAGATAACGATGGACATACCAGAGGAAGATATTTGTCGAATGGACTACAGGTCTGTATGGTCAGCCATGGAAGAGTGTCAGAACCTTGGCTTCACTAAATCAATCGGTGTTAGCAATTTCTCCTGCAAAAAGCTTCAGGAATTGATGGAGACCGCCAACATCCCTCCAGCTGTGAATCAAGTGGAGATGAGCCCGGCTTTCCAACAAAAGAAGCTGAGAGAGTATTGCAACGCAAATAATATATTAGTCAGTGCAGTCTCTATACTGGGATCAAACGGAACCCCATGGGGCTCCAATGCAGTTTTGGGTTCTGAGGTGCTTAACAAAATTGCTATGGCCAAAGGAAAATCTGTTGCTCAGGTTAGTATGAGATGGGTTTACGAGCAAGGCGCGAGTCTTGTGGTAAAAAGTTTCAGTGAAGAGAGATTGAGGGAAAACTTGAACATATTCGACTGGGAACTCACTAAGGAAGACAATGAAAAGATCGGTGAGATTCCACAGTGCAGAATCTTGGCTGCTCATTTTTTGGTCTCACCTAATGGACCTTTCAAATCTCAAGAAGAGTTGTGGGATGACAAAGCTTGA

>Pbr_STORR Pbr_STORR_contig_2915_CDS

ATGGAGCTCCAATATTTTTCTTATTTTCAACCAACTTCCTCCGTTGTTGCTCTTCTACTTGCTCTTGTATCCATCTTATTCAGTGTCGTTGTTTTGAGGAAGACATTTTCGAATAACTACTCATCACCTGCATCATCAACAGAAACAGCGGTACTTTGTCATCAGCGGCAGCAGTCGTGTGCATTGCCAATTTCCGGTCTTCTTCATGTTTTCATGAATAAAAACGGCTTAATTCATGTAACTCTTGGAAATATGGCGGATAAATACGGACCGATTTTCAGTTTCCCAACAGGTAGCCATAGAACTCTCGTTGTGAGCAGTTGGGAGATGGTAAAAGAGTGTTTTACAGGCAACAATGACACTGCTTTCTCAAACCGTCCTATCCCGTTAGCTTTTCAGACTATATTCTATGCATGCGGTGGCATAGACTCATACGGTCTTTCGAGTGTACCTTATGGAAAATATTGGAGGGAGCTACGAAAGGTCTGTGTGCATAACCTCCTCTCTAATCAACAACTACTCAAGTTCAGACACTTGATAATTTCTCAAGTCGATACGTCTTTCAATAAGCTGTATGAGTTATGCAAAAACTCTGAAGACAACCAGGGAATGGTGAGAATGGATGATTGGCTCGCCCAATTATCGTTCAATGTGATAGGAAGAATAGTCTGCGGATTCCAATCAGACCCAAAGACAGGTGCTCCTAGCAGGGTGGAACAATTTAAAGAAGTAATTAATGAAGCATCTTATTTTATGTCGACATCTCCAGTGTCAGATAACGTTCCAATGCTAGGGTGGATTGACCAATTGACTGGTCTTACGAAAAATATGAAGCACTGCGGAAAGAAATTAGACTTGGTGGTAGAGAGCATAATTAAAGATCATCGTCAAAAGAGACGATTCTCTAGAACTAAAGGAGGAGATGAGAAGGACGATGAACAAGATGACTTCATCGACATTTGCTTGTCAATTATGGAGCAACCACAGCTTCCTGGCAACAATAGCCCTCCTCAGATACCTATCAAATCTATTGTCCTGGACATGATAGGTGGGGGCACTGACACCACAAAACTGACCACCATCTGGACCCTTTCCTTGCTGCTGAACAACCCCCATGTGTTGGACAAGGCAAAACAAGAAGTGGATGCACACTTTCGAAAGAAAAGGAGATCAACAGATGATGCAGCAGCAGCCGTGGTGGATTTTGATGATATTCGTAACCTTGTCTACATCCAGGCAATCATCAAAGAATCAATGCGGTTGTATCCAGCCAGCCCCGTGGTGGAGCGACTGAGCGGCGAGGATTGTGTGGTCGGTGGGTTTCATGTACCAGCAGGGACGAGATTATGGGCTAACGTATGGAAGATTCAACGAGACCCTAAAGTATGGGATGATCCATTGGTGTTTCGACCAGAGAGATTTTTGAGCGATGAACAGAAGATGGTTGATGTAAGGGGTCAAAATTATGAGCTGTTACCATTTGGAGCCGGTCGACGTATATGTCCAGGTGTTTCCTTCTCTTTGGATCTAATGCAACTGGTACTGACTCGTCTTATTCTTGAGTTTGAAATGAAGTCCCCTAGCGGGAAAGTGGACATGACAGCAACACCAGGATTAATGAGTTACAAGGTGGTCCCTCTTGACATTCTGCTCACCCATCGGCGCATAAAGTCGTGTGTGCAGTTAGCATCCTCTGAGAGAGACATGGAGAGTAGTGGTGTACCAGTAATCACTCTGAGCTCGGGCAAGGTGATGCCTGTTTTAGGCATGGGAACATTTGAGAAAGTTGGTAAAGGGTCTGAAAGAGAGAGGTTGGCGATTTTAAAAGCCATAGAGGTGGGTTACAGATACTTCGACACAGCTGCTGCATACGAAACTGAAGAGGTTCTTGGAGAAGCTATTGCTGAAGCACTTCAAATTGGCCTAATCAAATCTCGAGATGAACTTTTCATCAGTTCCATGCTCTGGTGCACTGATGCTCACCCTGATCGTGTCCTCCTCGCTCTTCAGAATTCTCTGAGGAATCTTAAATTGGAGTATCTGGATCTATATATGTTACCCTTCCCGGCAAGCTTGAAGCCAGGGAAGATAACGATGGACATACCAGAGGAAGATATTTGTCGAATGGACTACAGGTCTGTATGGTCAGCCATGGAAGAGTGTCAGAACCTTGGCTTCACTAAATCAATCGGTGTTAGCAATTTCTCCTGCAAAAAGCTTCAGGAATTGATGGCGACCGCCAACATCCCTCCAGCCGTGAATCAAGTGGAGATGAGCCCGGCTTTCCAACAAAAGAAGCTGAGAGAGTATTGCAACGCAAATAATATATTAGTCAGTGCAGTCTCTATACTGGGATCAAACGGAACCCCATGGGGCTCCAATGCAGTTTTGGGTTCTGAGGTGCTTAAGCAAATTGCTATGGCCAAAGGAAAATCTGTTGCTCAGGTTAGTATGAGATGGGTTTACGAGCAAGGCGCGAGTCTTGTGGTAAAAAGTTTCAGTGAAGAGAGATTGAGGGAAAACTTGAACATATTCGACTGGGAACTCACTAAGGAAGACAATGAAAAGATCGGTGAGATTCCACAGTGCAGAATCTTGACTGCTTATTTTTTAGTCTCGCCTAATGGACCTTTCAAATCTCAAGAAGAGTTGTGGGATGACAAAGCTTGA

>Por_STORR Por_STORR_cc53289_g1_i4

ATGGAGCTCCAATATTTTTCTTATTTTCAACCAACTTCCTCCGTTGTTGCTCTTCTACTTGCTCTTGTATCCATCTTATTCAGTGTCGTTGTTTTGAGGAAGACATTTTCGAATAACTACTCATCACCTGCATCATCAACAGAAACAGCGGTACTTTGTCATCAGCGGCAGCAGTCGTGTGCATTGCCAATTTCCGGTCTTCTTCATGTTTTCATGAATAAAAACGGCTTAATTCATGTAACTCTTGGAAATATGGCGGATAAATACGGACCAATTTTCAGTTTCCCAACAGGTAGCCATAGAACTCTCGTTGTGAGCAGTTGGGAAATGGTAAAAGAGTGTTTTACAGGCAACAATGACACTGCTTTCTCAAACCGTCCTATCCCGTTAGCTTTTCAGACTATATTCTATGCATGCGGTGGCATAGACTCATACGGTCTTTCGAGTGTACCTTATGGAAAATATTGGAGGGAGCTACGAAAGGTCTGTGTGCATAACCTCCTCTCTAATCAACAACTACTCAAGTTCAGACACTTGATAATTTCTCAAGTCGATACGTCTTTCAATAAGCTGTATGAGTTATGCAAAAACTCTGAAGACAACCAGGGAATGGTGAGAATGGATGATTGGCTCGCCCAATTATCGTTCAATGTGATAGGAAGAATAGTCTGCGGATTCCAATCAGACCCAAAGACAGGTGCTCCTAGCAGGGTGGAACAATTTAAAGAAGTAATTAATGAAGCATCTTATTTTATGTCGACATCTCCAGTGTCAGATAACGTTCCAATGCTAGGGTGGATTGACCAATTGACTGGTCTTACGAGAAATATGAAGCACTGCGGAAAGAAATTAGACTTGGTGGTAGAGAGCATAATTAAAGATCATCGTCAAAAGAGACGATTCTCTAGAACTAAAGGAGGAGATGAGAAGGACGATGAACAAGATGACTTCATCGACATTTGCTTGTCAATTATGGAGCAACCACAGCTTCCTGGCAACAATAGCCCTCCTCAGATACCTATCAAATCTATTGTCCTGGACATGATAGGTGGGGGCACTGACACCACAAAACTGACCACCATCTGGACCCTTTCCTTGCTGCTGAACAACCCCCATGTGTTGGACAAGGCAAAACAAGAAGTGGATGCACACTTTCGAAAGAAAAGGAGATCAACAGATGATGCAGCAGCAGCCGTGGTGGATTTTGATGATATTCGTAACCTTGTCTACATCCAGGCAATCATCAAAGAATCAATGCGGTTGTATCCAGCCAGCCCCGTGGTGGAGCGACTGAGCGGCGAGGATTGTGTGGTCGGTGGGTTTCATGTACCAGCAGGGACGAGATTATGGGCTAACGTATGGAAGATGCAACGAGACCCTAAAGTATGGGATGATCCATTGGTGTTTCGACCAGAGAGATTTTTGAGCGATGAACGGAAGATGGTCGATGTAAGGGGTCAAAATTATGAGCTGTTACCATTTGGAGCCGGTCGACGTATATGTCCAGGTGTATCCTTCTCTTTGGATCTAATGCAACTGGTACTGACTCGTCTTATTCTTGAGTTTGAAATGAAGTCCCCTAGCGGGAAAGTGGACATGACAGCAACACCAGGATTAATGAGTTACAAGGTGGTCCCTCTTGACATTCTGCTCACCCATCGGCGCATAAAGTCGTGTGTGCAGTTAGCATCCTCTGAGAGAGACATGGAGAGTAGTGGTGTACCAGTAATCACTCTGAGCTCGGGCAAGGTGATGCCTGTTTTAGGCATGGGAACATTTGAGAAAGTTGGTAAAGGGTCTGAAAGAGAGAGGTTGGCGATTTTAAAAGCCATAGAGGTGGGTTACAGATACTTCGACACAGCTGCTGCATACGAAACTGAAGAGGTTCTTGGAGAAGCTATTGCTGAAGCACTTCAACTTGGCCTAATCGAATCTCGAGATGAACTTTTCATCAGTTCCATGCTCTGGTGCACTGATGCTCACCCTGATCGTGTCCTCCTCGCTCTTCAGAATTCTCTGAGGAATCTTAAATTGGAGTATCTGGATCTATATATGTTACCCTTCCCGGCAAGCTTGAAGCCAGGGAAGATAACGATGGACATACCAGAGGAAGATATTTGTCGAATGGACTACAGGTCTGTATGGTCAGCCATGGAAGAGTGTCAGAACCTTGGCTTCACTAAATCAATCGGTGTTAGCAATTTCTCCTGCAAAAAGCTTCAGGAATTGATGGCGACCGCCAACATCCCTCCAGCCGTGAATCAAGTGGAGATGAGCCCGGCTTTCCAACAAAAGAAGCTGAGAGAGTATTGCAACGCAAATAATATATTAGTCAGTGCAGTCTCTATACTGGGATCAAACGGAACCCCATGGGGCTCCAATGCAGTTTTGGGTTCTGAGGTGCTTAAGCAAATTGCTATGGCCAAAGGAAAATCTGTTGCTCAGGTTAGTATGAGATGGGTTTACGAGCAAGGCGCGAGTCTTGTGGTAAAAAGTTTCAGTGAAGAGAGATTGAGGGAAAACTTGAACATATTCGACTGGGAACTCACTAAGGAAGACAATGAAAAGATCGGTGAGATTCCACAGTGCAGAATCTTGACTGCTTATTTTTTAGTCTCGCCTAATGGACCTTTCAAATCTCAAGAAGAGTTGTGGGATGACAAAGCTTGA

>Pso_STORR-PS1126560 Pso_STORR_PS1126560

ATGGAGCTCCAATATATTTCTTATTTTCAACCAACTTCCTCCGTTGTTGCTCTTCTACTTGCTCTTGTATCCATCTTATCCAGTGTCGTTGTTTTGAGGAAGACATTTTTGAATAACTACTCATCATCACCTGCATCATCCACAAAAACAGCGGTACTTTCTCATCAGCGGCAGCAGTCGTGTGCATTGCCAATTTCCGGTCTCCTCCATATTTTCATGAATAAAAACGGCTTAATTCATGTAACTCTTGGAAATATGGCTGATAAATACGGTCCGATTTTCAGTTTCCCAACAGGTAGCCATAGAACTCTCGTTGTGAGCAGTTGGGAGATGGTAAAAGAGTGTTTTACTGGCAACAATGACACTGCTTTCTCAAACCGTCCTATCCCGTTAGCTTTTAAGACTATATTTTATGCATGCGGTGGCATAGACTCATACGGTCTTTCGAGTGTACCTTATGGAAAATATTGGAGGGAGCTTCGAAAGGTCTGTGTGCATAACCTCCTGTCTAATCAACAACTACTCAAGTTCAGACACTTGATAATTTCTCAAGTCGATACGTCTTTCAATAAGCTGTATGAGTTATGCAAAAACTCTGAAGACAACCAGGGAAACTATACTACTACTACTACCGCAGCTGGCATGGTGAGAATCGATGATTGGCTCGCCGAATTATCGTTCAATGTGATAGGAAGAATAGTCTGCGGATTCCAATCAGGCCCTAAGACAGGTGCTCCAAGCAGGGTGGAACAATTTAAAGAAGCAATTAATGAAGCATCTTATTTTATGTCGACATCTCCAGTGTCAGATAATGTTCCAATGCTAGGGTGGATTGACCAATTGACAGGTCTTACGAGAAATATGAAGCACTGCGGAAAGAAATTAGACTTGGTGGTCGAGAGCATAATTAATGATCATCGTCAAAAGAGACGATTCTCTAGAACTAAAGGAGGAGATGAGAAGGACGATGAACAAGATGACTTCATCGACATTTGTTTGTCAATAATGGAGCAACCACAGCTTCCTGGCAACAATAATCCTTCTCAGATACCTATCAAATCTATTGTCCTGGACATGATAGGTGGGGGCACTGACACCACAAAACTGACCACCATCTGGACCCTTTCCTTGCTGCTGAACAACCCCCATGTGTTGGACAAGGCAAAACAAGAAGTGGATGCACACTTTCGAACCAAAAGGAGATCAACAAATGATGCAGCAGCAGCCGTGGTGGATTTTGATGATATTCGTAACCTTGTCTACATCCAGGCAATCATCAAAGAATCAATGCGGTTGTATCCAGCCAGCCCCGTGGTGGAGCGACTGAGCGGCGAAGATTGTGTGGTCGGTGGGTTTCATGTACCAGCAGGGACGAGATTATGGGCTAACGTATGGAAGATGCAACGAGACCCTAAAGTATGGGATGATCCATTGGTGTTTCGACCAGACAGATTTTTGAGCGATGAACAGAAGATGGTTGATGTAAGGGGTCAAAATTATGAGCTGTTACCATTTGGAGCCGGTCGACGTGTATGTCCAGGTGTATCCTTCTCTTTGGATCTAATGCAACTGGTACTGACTCGTCTTATTCTCGAGTTTGAAATGAAGTCTCCTAGCGGGAAAGTGGACATGACAGCAACACCAGGATTAATGAGTTACAAGGTGATCCCCCTTGACATTCTGCTCACCCATCGTCGCATAAAGCCGTGTGTGCAGTCAGCAGCCTCTGAGAGAGACATGGAGAGTAGTGGTGTACCAGTAATCACTCTGGGCTCGGGCAAGGTGATGCCTGTTCTGGGCATGGGAACATTTGAGAAAGTTGGTAAAGGGTCCGAAAGAGAGAGGTTGGCGATTTTAAAAGCGATAGAGGTGGGTTACAGATACTTCGATACAGCTGCTGCATACGAAACTGAAGAGGTTCTTGGAGAAGCTATTGCTGAAGCACTTCAACTTGGCCTAGTCAAATCTCGAGATGAACTTTTCATCAGTTCCATGCTCTGGTGCACTGATGCTCACGCTGATCGTGTCCTCCTCGCTCTTCAGAATTCGCTGAGGAATCTTAAATTGGAGTATGTGGATCTATATATGTTACCCTTCCCGGCAAGCTTGAAGCCTGGGAAGATAACGATGGACATACCAGAGGAAGATATTTGTCGCATGGACTACAGGTCTGTATGGGCAGCCATGGAAGAGTGTCAAAACCTTGGCTTCACTAAATCAATCGGTGTTAGCAATTTCTCCTGCAAAAAGCTTCAGGAATTGATGGCGACCGCCAACATCCCTCCAGCTGTGAATCAAGTGGAGATGAGCCCGGCTTTCCAACAAAAGAAGCTGAGAGAGTATTGCAACGCAAATAATATATTAGTCAGTGCAATCTCTGTACTGGGATCAAACGGAACCCCATGGGGCTCCAATGCAGTTTTGGGTTCTGAGGTGCTTAAGAAAATTGCTATGGCCAAAGGAAAATCTGTTGCTCAGGTTAGTATGAGATGGGTTTACGAGCAAGGCGCGAGTCTTGTGGTAAAAAGTTTCAGTGAAGAGAGATTGAGGGAAAACTTGAACATATTTGACTGGGAACTCACTAAGGAAGACCATGAAAAGATCGGTGAGATTCCACAGTGCAGAATCTTGAGTGCTTATTTTTTGGTCTCACCTAATGGACCTTTCAAATCTCAAGAAGAGTTGTGGGATGATGAAGCTTGA

>Pca_STORR Pca_STORR_141675

ATGGAGCTCCAGTACTTTTCTTATTTTCAGCCAACTTCCTCCCTTTTTGTTCTTGTATCCATATTACTGATCAGTGCTGTTGTTTTGAGGAAGATATTTTTAAATAACTACTCTTCATCACCTGCATCATCAACAGAAACAGCGCTGCTTTCTCATCAGCGGCGGTCGTCATGTGCATTGCCAATCATCGGTCTTCTTCATGTTTTCATGAATAAAAACGGCTTAATTCATGTAACTCTTGGAAATATGGCTGATAAATACGGACCGATTTTCAGTTTCCCAACAGGCAGCCATATGACTCTAGTTGTGAGCAGTTGGGAGATGGCAAAAGAGTGTTTTACAGGCAACAATGACATTGCTTTCTCAAACCGTCCTATCCCGTTAGTTTTTAAGACTATATTCTATGCATGCCGTGGTATTGACTCATACGGTCTTTCAAGTGTACCTTATGGAAAATATTGGAGGGAGTTACGAAAGGTGTGTGTGCATAACCTCCTCTCTAATCAACAACTACTCAACTTCAGACACTTGATAATTTCTCAAGTTGATACCTCTTTCAATAAGCTTTATGAGTTATGCAAAAACTCTGCAGACAACCAGGGAAACTACACTACTACTACTACCGCAGCTGGCATGGTGAGGATGGATGATTGGCTAGCGCAATTATCGTTCAACGTGATAGGAAGAATAGTCTGTGGATTCCAATCAGACACCGAGACAAGTGCTTCAAGCAGTGCGGAACAACTTACAGAAGCAATTAATGACGCATCGTATTTTATGTCGACATCTCCGGTGTCAGATAACGTTCCATGGTTAGGGTGGGTTGATCAACTAACTGGTCTTACTAGAAATATGAAGCAATGCGGAAAGAAATTAGACTTGGTGCTCGAGAGCATAATTGAAGATCATCGTCAAAAGAGACGGTTTTCTAGAAGTAAACAAGGAGATGAGAATTTTATCAACGAGGATGAACAAGATGACTTCATCGACATTTGCCTGTCAATTATGGAGCAACCACAGCTTCCTGGCAACAATAACCCTTGTCAAATACCTATCAAATCTATTGTCCTGGACATGATAGGTGGGGGGACCGACACCACAAAACTGACCACCATCTGGACCCTTTCATTGCTACTGAACAACCCCCATGTGTTGGACAAGGCAAAACAAGAAGTGGATGCACACTTCGGGACGAAAAGGAACGATGCAGGAGGAGTCGTGGTGGATTTTGATGATACTCGTAACCTTGTCTACATCCAGGCAATTATCAAAGAATCAATGCGGTTGTATCCAGCCAGCCCCTTGGTGGAGCGACTGAGCGGCGAAGATTGTGTGGTCGGTGGGTTTCATGTACCAGCAGGGACGCGGTTATGGGCTAACGTGTGGAAGATGCATCGAGACCCAAAAGTATGGGATGATCCATTGGTGTTTCGACCAGAGAGATTTTTGCTCGATGACCAGAAGATGGTTGATGTTAGGGGTCAAAACTATGAGCTGTTACCATTCGGAGCCGGTCGACGGATATGTCCAGGTGTATCGTTCTCTTTGGATCTAATGCAACTGGTACTGACTCGTCTTATTCTTGAGTTTGAAATGAAGTCTCCTAGCGGGAAACTAGACATGACAGCGACACCAGGTTTAATGAGTTACAAGGTAGTCCCCCTTGACATTCTGCTCACCAGTCGTCGCATGAAGTGGTGTGTGCAGTTAGCCTCTGAGAGAGATAAAAAGATGGAGAGTAGTGTTGTACCTGTAATCACTCTGAGCTCGTGCAAGGTGATGCCTGTTTTAGGCATGGGAACATTTGAAAAAGTTGGTAAAGGGTCTGAAAGAGAGAGGTTGGCGATTTTAAAAGCGATAGAGGTAGGTTACAGATACTTCGATACAGCTGCTGCATACGAAACTGAAGAGGTTCTTGGTGAAGCTATTGCTGAAGCACTTCAACTTGGCCTAATCAAATCTCGAGATGAACTCTTCATCAGTTCCATGCTCTGGTGCACTGATGCCTACCATGATCGAGTCCTCCCCGCTCTTCAGAATTCTCTGAGGAATCTTAAATTGGAGTATCTGGATCTATACATGTTACCCTTCCCGGTAAGCTTGAAGCCAGGAGGGAAGATAAGTATGGACATACCAGAGGATGAAATTTTTACGATGGACTATAGGTCTGTATGGGCAGCCATGGAAGAATGTCAGAACCTTGGCTTGACAAAATCAATCGGTGTCAGCAATTTCTCCTGCAAAAAGCTTCAGGAATTGATGGCGACCGCCAACATCCCTCCAGCTGTGAATCAAGTGGAGATGAGCCCGGCTTTCCAACAAAAGAATTTGAGAGAGTATTGCAATGCCAATAATATATTAATCAGTGCAGTTTCTATTTTGGGATCAAACGGAACCCCATGGGGCTCCAATGCAGTTCTGGGTAATGAGGTGCTAAACCAGATTGCTATGGCCAGAGGAAAATCTGTTGCTCAGGTTAGTATGAGGTGGGTTTACGAGCAAGGTGCGAGTCTTGTGGTAAAAAGTTTTAGTGAAGAGAGATTGAGGGAAAACTTGAACATATTCGACTGGGAACTCACTAAGGAAGACTTGGAAAAGATGGGTGAGATTCCACAATGCAGAATCTTGACTGCTGATTTTATGGTATCAGCTAATGGACCTTTCAAATCTCAAGAAGAGTTGTGGGATGACAAAGCTTGA

>Pnu_COR-Ln5 Pnu_COR_PCR_genomic_CDS

ATGGAGAGTAGATATGGTGTACCAGTCATCACTCTGAGCTCCGGCAAGGAGATGCCTGTTCTAGGCATGGGAACATTTGAAACAGAAGGTAAAGGATCTGAAAGAGAGAGGACGGCGTTTCTGAAAGCCATAGAGGTGGGTTACAGATACTTCGATACAGCTGCTGCATACGAAACCGAAGAGGTTCTTGGTGAAGCTATAGCTGAAGCACTTCAACTTGGCTTCATCAAGTCTCGAGATGAACTCTTCATCAGTACCATGCTCTGGTTCGCTGATGCTCACGCTGATCGTGTCCTCCCCGCTCTTCAGAATTCTCTCAGGAATCTTAAACTGGAGTATGTGGATCTGTACATGTTACTCTTTCCGGTAAGCTTGAAGCCTGGGTCGAAGATAACCATGGACATAGAGAAAGATGATTTACTTCCACTGGATTACAAGTCTGTATGGGCAGCCATGGAAGAGTGTCAGAACCTTGGCTTCACTAAGTCAATCGGTGTCAGCAATTTCTCCAGCAAAAAGCTTCAGGAATTGATGGCACACGCCAAAATCCCTCCGGCTGTGAACCAAGTGGAGATGAGCCCCGCTTTCCAACAAAAGAATTTGAGAGTGTATTGCAAGGCGCACAACATATTGATCATTGCGATCTCTGTTTTGGGATCGAATGGAACCCCATGGGGCTCCAATGCAGTTATGGGTTCTGAAGTGCTTAACCAGATTGCTACGGCCAGAGGAAAATCCGTTGCTCAGGTTAGTATGAGATGGGTTTACGAGCAAGGTGCGGCTCTTGTGGTAAAGAGTTTCAGTGAGAAGAACATGAGGGAAAACTTGAAGATATTCGACTGGGAACTGACTAAGGAGGACTTGGAAAAGATCGGTCATGAGATTCCACAGCGCAGGGTATTAGACGCTCATTTTCTGGTGTCAGCCCATGGACCTTTCAAATCTCTAGAAGAGGTGTGGGATGACAATTAA

>Pnu_COR-Ln3 Pnu_COR_938mRNA_col1

ATGGAGAGTAGATATGGTGTACCTGTGATCACTCTGAGCTCCGGCAAGGAGATGCCTGTTCTAGGCATGGGCACATTTGAAACAGAAGGTAAAGCATCTGAAAGAGAGAGGACGGCGTTTTTGAAAGCCATAGAGGTGGGTTACAGATACTTCGATACAGCTGCAGCATACGAAACCGAAGAGGTTCTCGGTGAAGCTATAGCTGAAGCACTTCAACTTGGCTTCATCAAGTCTCGAGATGAACTCTTCATCAGTACCATGCTCTGGTTCGCTGATGCTCACCCTGATCGTGTCCTCCCCGCTCTTCAGAATTCTCTCAGGAATCTTAAACTGGAGTATGTGGATCTATACATGTTACTCTTTCCGGTAAGCTTGAAGCCTGGGTCGAAGATAACCATGGACATAGAGAAAGATGATTTACTTCCACTGGACTACAAGTCTGTATGGGCAGCCATGGAAGAGTGTCAGAACCTTGGCTTCACTAAGTCAATCGGTGTCAGCAATTTCTCCAGCAAGAAGCTTCAGGAATTGATGGCACACGCCAAAATCCCTCCGGCTGTGAACCAAGTGGAGATGAGCCCCGCTTTCCAACAAAAGAAATTGAGAGAGTATTGCAAGGCCAACAACATATTGATCATTGCAATCTCTGTCTTGGGATCCAATGGAACCCCATGGGGCTCCAATGCAGTTATGGGTTCTGAAGTGCTTAACCAGATTGCTATGGCCAGAGGAAAATCTGTTGCTCAGGTTAGTATGAGATGGGTTTACGAGCAAGGTGCGGCTCTTGTGGTAAAGAGTTTCAGTGAGAAGAACATGAGGGAAAACTTGAAGATATTCGACTGGGAACTGACTAAGGAGGACTTGGAAAAGATCGGTCATGAGATTCCACAGCGCAGGGTATTAGACGCTCATTTTCTGGTGTCAGCCAATGGACCTTTCAAATCTCTAGAAGAGGTGTGGGATGACAATTAA

>Pnu_COR-Ln4 Pnu_COR_qc49714mRNA_con2_1

ATGGAGAATAATGGTGTACCTGTGATCACTCTGAGCTCCGGCAAGGAGATGCCTGTTCTAGGCATGGGAACATTTGAAACAGAAGGTAAAGGGTCTGAAAGAGAGAGGCTGGCGTTTTTGAAAGCCATAGAGGTGGGTTACAGATACTTCGATACAGCTGCAGCATACGAAACCGAAGAGGTTCTTGGTGAAGCTATAGCTGAAGCGCTTCAACTCGGCCTAATCAAGTCGCGAGATGAACTCTTCATCAGTAGCATGCTCTGGTTCGCTGATGCTCACGCTGACCGTGTCCTCCCCGCTCTTCAGAATTCTCTCAGGAATCTTAAACTGGAGTATGTGGATCTATACATGTTACTCTTTCCGGTAAGCTTGAAGCCTGGGTCGAAGATAACCATGGACATAGAGAAAGATGATTTACTTCCACTGGACTACAAGTCTGTATGGGCAGCCATGGAAGAGTGTCAGAACCTTGGCTTCACTAAGTCAATCGGTGTCAGCAATTTCTCCAGCAAAAAGCTTCAGGAATTGATGGCAAACGCCAAAATCCCTCCGGCTGTGAACCAAGTGGAGATGAGCCCCGCCTTCCAACAAAAGAATTTGAGAGAGTATTGCAAGGCGCACAACATATTGATCATTGCGATCTCTGTCTTGGGATCCAATGGAACCCCATGGGGCTCCAATGCAGTTATGGGTTCTGAAGTGCTTAACCAGATTGCTATGGCCAGAGGAAAATTTGTTGCTCAGGTTAGTATGAGATGGGTGTACGAGCAAGGTGCGGCTCTTGTGGTAAAGAGTTTCAGTGAGAAGAACATGAGGGAAAACTTGAAGATATTCGACTGGGAACTGACTAAGGAGGACTTGGAAAAGATCGGTCATGAGATTCCACAGCGCAGGGTATTAGAAGGTCATTTTCTGGTATCAGCTAATGGACCTTTCAAATCTCTAGAAGAGGTGTGGGATGACAAATCTTTCTAG

>Pnu_COR-Ln7 Pnu_COR_598mRNA_col1

ATGGAGAGTAGATATGGTGTACCAGTCATCACTCTGAGCTCCGGCAAGGAGATGCCTGTTCTAGGCATGGGAACATTTGAAACAGAAGGTAAAGGGTCTGAAAGAGAGAGGACGGCGTTTTTGAAAGCCATAGAGGTGGGTTACAGATACTTCGACACAGCTGCTGCATACGAAACCGAAGAGGTTCTTGGTGAAGCTATAGCTGAAGCACTTCAACTCGGCCTAATCAAGTCGCGAGATGAACTCTTCATCAGTACCATGCTCTGGTTCGCTGATGCTCACGCTGACCGTGTCCTCCCCGCTCTTCAGAATTCTCTCAGGAATCTTAAACTGGAGTATGTGGATCTGTACATGTTACTCTTTCCGGTAAGCTTGAAGCCTGGGTCGAAGATAACCATGGACATAGAGAAAGATGATTTACTTCCACTGGATTACAAGTCTGTATGGGGAGCCATGGAAGAGTGTCAGAACCTTGGCTTCACTAAGTCAATCGGTGTCAGCAATTTCTCCAGCAAAAAGCTTCAGGAATTGATGGCACACGCCAAAATCCCTCCGGCTGTGAACCAAGTGGAGATGAGCCCCGCTTTCCAACAAAAGAATTTGAGAGAGTATTGCAAGGCCAACAACATAGTGATCATTGCGATCTCTGTTTTGGGATCGAATGGAACCCCATGGGGCTCCAATGCAGTTATGGGTTCTGAAGTGCTTAATCAGATTGCTACGGCCAGAGGAAAATCCGTTGCTCAGGTTAGTATGAGATGGGTTTACGAGCAAGGTGCGGCTCTTGTGGTAAAGAGTTTCAGTGAGAAGAACATGAGGGAAAACTTGAAGATATTCGACTGGGAACTGACTAAGGAGGACTTGGAAAAGATCGGTCATGAGATTCCACAGCGCAGGGTATTAGACGCTCATTTTCTGGTGTCAGCCAATGGACCTTTCAAATCTCTAGAAGAGTTGTGGGATGGCAATTAA

>Pnu_COR-Ln6 Pnu_COR_598mRNA_col3

ATGGAGAGTAGATATGGTGTACCAGTCATCACTCTGAGCTCCGGCAAGGAGATGCCTGTTCTAGGCATGGGAACATTTGAAACAGAAGGTAAAGGGTCTGAAAGAGAGAGGACGGCGTTTCTGAAAGCCATAGAGGTGGGTTACAGATACTTCGACACAGCTGCTGCATACGAAACCGAAGAGGTTCTTGGTGAAGCTATAGCTGAAGCACTTCAACTTGGCTTCATCAAGTCTCGAGATGAACTCTTCATCAGTACCATGCTCTGGTTTGCTGATGCTCACGCTGACCGTGTCCTCCCCGCTCTTCAGAATTCTCTCAGGAATCTTAAACTGGAGTATGTGGATCTGTACATGTTACTCTTTCCGGTAAGCTTGAAGCCTGGGTCGAAGATAACCATGGACATAGAGAAAGATGATTTACTTCCACTGGATTACAAGTCTGTATGGGCAGCCATGGAAGAGTGTCAGAACCTTGGCTTCACTAAGTCGATCGGTGTCAGCAATTTTTCGAGCAAGAAGCTTCAGGAATTGATGGCACACGCCAAAATCCCTCCGGCTGTGAACCAAGTGGAGATGAGCCCCGCTTTCCAACAAAAGAATTTGAGAGAGTATTGCAAGGCCAACAACATAGTGATCATTGCGATCTCTGTTTTGGGATCGAATGGAACCCCATGGGGCTCCAATGCAGTTATGGGTTCTGAAGTGCTTAATCAGATTGCTACGGCCAGAGGAAAATCCGTTGCTCAGGTTAGTATGAGATGGGTTTACGAGCAAGGTGCGGCTCTTGTGGTAAAGAGTTTCAGTGAGAAGAACATGAGGGAAAACTTGAAGATATTCGACTGGGAACTGACTAAGGAGGACTTGGAAAAGATCGGTCATGAGATTCCACAGCGCAGGGTATTAGACGCTCATTTTCTGGTGTCAGCCAATGGACCTTTCAAATCTCTAGAAGAGTTGTGGGATGGCAATTAA

>Prh_COR-Lc5-AKO60177 Prh_COR_AKO60177

ATGGATAGTAGTGGTGTACCTGTAATCCCTCTGAGCTCCGGTAAAGGGATGCCTGCTTTAGCTCTAGGAACATTTGAAACATTTGGTAAAGGATCGTCCGAAAGACAGAGATCGGCGTTTTTGAAAGCGATAGAGGTGGGTTACAGATACTTTGACACAGGTGCTTCATACGGAACCGAAGAGGTCCTTGGCGAAGCTATAGATGAAGCACTTCAACTTGGCCTTATCGAATCTCGAGAGGAACTGTTTATCAGTTCCAATCTCTGGTGTACTGATGCTCACCCTGATCGTGTCCTCCTTGCTCTTCAGAATTCTCTAAGGAATCTTAAATTGGAGTATCTGGATCTATACATGATACCCTTTCCGATAAGCTTGAAGCCTGGGAAGATAATGGAGGAGATAACTATGGACATACCAGAGGATGAAATGTTACCTATGGACTACAAGTCTGTATGGGCAGCCATGGAAGAGTGTCAGGACCTTGGCTTCACTAAGTCGATCGGTGTCAGCAATTTCTCCTGCAAAAAGCTTCAGGAGTTGATGGCGACCGCCAATATCCCACCAGCTGTGAATCAAGTGGAGATGAGCCCCGTTTTCCAACAAAAGAAACTGAGAGAGTATTGCAAGGCCAATAATATATTAGTCAGCGCAGTCTCGGTTTTGGGATCGAACGGAACCGCATGGGGCTCCAATCAAGTTATGGGTTCTGAGGTGTTGAAACAAATTGCTACTGACAGAGGAAAATCTGTTGCTCAGATTAGTATGAGATGGGTTTATGAGCAAGGTGCGAGTCTTGTGGTAAAAAGTTTCAATGAAGTGAGGATGAGGGAAAATTTGAACATATTCAACTGGCAACTTACTAAGGAAGATTTGGAAAAGATCAGCGAGATTCCACAATGCAGGATATTAACTGGTGATTTTTTAGTTTCAGCTAATGGACCTTTCAAATCTCTACAAGAGTTATGGGATGACCAAGTTTGA

>Pca_COR-Lc4 Pca_COR_141851a

ATGGAGAGTAGTGGTGTGCCTGTAATCACTCTGAGCTCGGGCAAGAGGATGCCTGCTTTAGCTCTGGGAACATTTGAAACATTTGGTAAGGGATCGTCTGAAAGACAGAGGTAGGCGTTTTTGAAAGCGATAGAGGTGGGTTATAGATATTTCGATACAGGTGCTTCATACGGATCCGAAGAGGTTCTTGGCGAAGCTGTAGCTGAAGCACTTCAACTTGGCTTAATCAAATCTCGAGATGAACTCTTCATCAGTTCCAATCTCTGGTTCACTGATGCTCACCCTGATCGTGTCCTCCTTGCTCTTCAAAATTCTCTAAGGAATCTGAAATTGGAGTATCTGGATTTATACATGATACACTTTCCGATAAGCTTGAAGCCTGGGAAGATAATGGAGGTGATAACTATGGACATACCAGAGGATGAACTGTTACCAATGGACTACAAGTCTGTATGGGCAGCCATGGAAGAGTGTCAGAACCTTGGCTTCACTAAGTCGATCGGTGTGGGCAATTTCTCCTGCAAAAAGCTTCAGGAACTGATGGCGACCGCCAGTATCCCACCAGCTGTGAATCAGGTGGAGATGAGCCCCGTTTTCCAACAAAAGAAACCGAGAGAGTATTGCAAGGACAATAATATATTAGTCAGTGCAGTGTCAGTTTTGGGATCCAACGGAACCGCATGGGGCTCCAATGCAGTTATGGGTTCTGAGGTGCTTAAGCAAATTGCTATGGACAGAGGAAAATCTGTTGCTCAGGTTAGTATGAGATGGGTTTACGAGCAAGGTGCGAGTCTTGTAGTAAAAAGTTTCAGTGAAGTGAGGATGATGGAAAACTTGAACATATTCGACTGGGAACTGACCAAGGAAGACTTGGAAAAGATCAGCGAGATTCCACAACGCAGAATATTAACTGCTGATGTTTTGGTATCAGCTAATGGACCTTTCAAATCTCTACAAGAGTTATGGGATGACCAAATTTGA

>Pat_COR-Lc6 Pat_COR_8802

ATGGGGAGTGGTTGTATTCCTGTTATCACTTTGAACTCGGGCAATGAGATGCCTGTTTTAGGTATGGGAACATTTGAAACATTTGATAAAGGGGGTGAAAGAGAAAGATTGGCGTATTTGAAAGCGATTAAAGTGGGTTACAGATACTTCGATACAGGCGCTGCATACGGAACGGAAGAGGTTCTTGGCCAAGCTATAGCTGAAGCACTTCAACTTGGCCTAATAAAATCTCGAGATGAACTCTTCATCAGTACCATGATCTGGGCCACTGATGCTCACCCTGATCGTGTTCTCCTCGCTGTTCAGAGATCTTTAAGGAATCTTAAATTGGACTATGTGGATCTCTACCTGATACCATTTCCGGCAAGCTTGAACCCAGAAGGGGAGATACCTAATTACATACCAGAGAACGAAACTTTTTTAAAAATGGACTGCAAGTCTGTATGGGCAGCTATGGAAGAGTGTCAAACCCTTGGCTTTACCAAGTCTATCGGTGTCAGCAATTTCTCCTGCAAAAAGCTTCAGGAAGTGATGGAGACCGCAAATATCCCTCCAGCTGTGAATATGGTGGAAATGAACCCTACTTTCCAACAAAAATATCTGAGAGAATATTGCAAGGCCAATAATGTATTAGTCAGTGCGTACTCTGTTTTGGGATCCACGGGAACTTCATTGGGCTCCAATGCAGTTATGGGTTCTGAGGTGCTTAAGCAAATTGCTACGGACAGAGGAAAATCTATTGCTCAGGTTAGTATGAGATGGGTTTATGAGCAAGGTGCGGGTTTTGTGGTAAAAAGTTTCAGTGAAGAGAGGATGAGGGAAAACTTGAACATATTCGACTGGGAACTGACTAAGGAAGACTTGGAGAAGATCGGTGAGATTCCACAATGCAGAGTACTAACTATGGATTTTTTAGTATCACCTGATGGAGCTTTCAAATCTTTAGAAGACTTGTGGGATGGCGAAGCATGA

>Pat_COR-Lc7 Pat_COR_8998

ACGGAGAGTAGTGATGTACCTGTAATCACTTTGAGCTCAGGCGAGAGGATGCCTGTTTTAGGTATGGGAACATTTGAAACATTTGGTAAACCGTCTGAAAGAGAGAGGTTGGCGTATTTGAAAGCGATAGAGGTGGGTTACCGGTACTTCGATACAGGTACTGCATACGGAACTGAAGAGGTTCTTGGCGAAGCTATAGCTGAAGCACTTCAACTTGGCCTAGTCAAATCTAGAGATGAACTCTTTATCAGTTCCATGCTTTGGAGCACTGATGCTCACCCTGATCGTGTCCGACCCGCTCTTCAGAATTCTCTGAGGAATCTCAAATTGGAGTTTGTGGATCTATACCTGATACCCTGGCCAGTAAGCTTGAAGCCAGAAGGGAAGATAACTATGGACATACCAGAGGACGAAATTTTTGCAATGGACTACAGGTCTGTATGGGCACCCATGGAAGAGTGTCAGAACCTTGGCTTCACTAAGTCTATCGGTGTCAGCAATTTCTCCTCCAAAAAGCTGCAGGAATTGATGGCGGTCGCCAACATCCCTCCAGCTGTGAATCAAGTGGAGATGAACCCCGCTTTCCAACAAAAGAATCTGATAGACTATTGCAAGGCAAAAAATATATTTGTCATTGCATTCTCGGTTTTGGGATCCAACGGAACGTCATGGGGCTCCAATGCAGTTATGGGTTCTGAGGTGCTTAAGAAAATTGCTATGGCCGGAGGAAAATCTGTTGCCCAGGTTAGTATGAGATGGGTTTACGAACAGGGTGCGAATTCTTGTGGTGAAAAGTTTCAATGAAGATAGAATGAGGGAAAATTTGAACATATTCGAGTGGGAACTGACCAAGGAAGACTTGGAAAAGATCGGTGAGATTTCACAGCGCAGAGTAATAATTGCAGATTTTTTGGTATCGGCTAATGGACCTTTCAAATCTCTGGAAGACTTATGGGATGAGTAA

>Par_COR-Lc11 Par_COR_contig_31561_CDS

ATGGAGAGTTGCGGTGTACCTGTAATCACTCTGAGCTCAGGCAAGAGGATGCCTGTTTTAGGTATGGGAACATTTGAAACATTTGGTAAACCGTCTGAAAGAGAGAGGTTGGCGTATTTGAAAGCGATAGAAGTGGGTTACAGATACTTCGATACAGGTGCTGCATACGGAACTGAAGAGGTTCTTGGCGAAGCTATAGCTGAAGCACTTCAACTTGGCCTAATCAAATCTAGAGATGAACTCTTCATCAGTTCCATGCTTTGGAGCACTGATGCTCACCCTGATCGTGTCCGACTCGCTCTTGAGAATTCCTTGAGGAATCTCAAATTGGAGTTTGTGGATCTATACCTGATACCATGGCCGGTAAGCTTGAAGCCAGAAGGGAAGGTAACTATGGACATACCAGAGGACGAAATTTTTGCAATGGACTACAGGGGTGTATGGGCAGCCATGGAAGAGTGTCAGAACCTTGGCCTCACTAAGTCTATCGGTGTCAGCAATTTCTCCTCCAAAAAGCTTCAGGAATTGATGGCGGTGGCCAACATCCCTCCAGCTGTGAATCAAGTGGAGATGAACCCCGCTTTCCAACAAAAGAATCTGATAGAGTATTGCAAGGCCAACAATATATTTGTCATTGCATTCTCGGTTTTGGGATCCAACGGAACGTCATGGGGCTCCAATGCAGTTATGGGCTCTGAGGTGCTTAAGCAAATTGCTTTGGCCAGAGGAAAATCTGTTGCCCAGGTTAGTATGAGATGGGTTTACGAACAAGGTGCGATTCTTGTAGTGAAAAGTTTTAGTGAAGATAGAATGAGGGAAAATTTGAACATATTCGACTGGGAACTGACTAAGGAAGACTTGGTAAAGATCCGTGAGATTACACAGCGCAGAGTAATAACTGCAGATTTTTTGGTATCAGCTAATGGACCTTTCAAATCTCTGGAAGACTTATGGGATGAGTAA

>Par_COR-Lc9 Par_COR_contig_56378_CDS

ATGGAGAGTTGCGGTGTACCTGTAATCACTCTGAGCTCAGGCAGGAGGATGCCTGTTTTAGGTATGGGAACATTTGAAACATTTGGCAAACCGTCTGAAAGAGAGAGGTTGGCGTATTTGAAAGCGATAGAAGTGGGTTACAGATACTTCGATACAGGTACTGCATACGGAACTGAAGAGGTTCTTGGCGAAGCTATAGCTGAAGCACTTCAACTTGGCCTAATCAAATCTAGAGATGAACTCTTCATCAGTTCCATGCTCTGGAGCGCTGATGCTCACCCTGATCGTGTCCGACTCGCTCTTGAGAATTCCTTGAGGAATCTCAAATTGGAGTTTGTGGATCTATACCTGATACCATGGCCGGTAAGCTTGAAGCCAGAAGGGAAGGTAACTATGGACATACCAGAGGACGAAATTTTTGCAATGGACTACAGGTGTGTATGGGCAGCCATGGAAGAGTGTCAGAACCTTGGCCTCACTAAGTCTATCGGTGTCAGCAATTTCTCCTCCAAAAAACTTCAGGAATTGATGGCGGTGGCCAACATCCCTCCAGCTGTGAATCAAGTGGAGATGAACCCCGCTTTCCAACAAAAGAATCTGATAGAGTATTGCAAGGCCAACAATATATTTGTCATTGCATTCTCGGTTTTGGGATCCAACGGAACGTCATGGGGCTCCAATGCAGTTATGGGATCTGAGGTGCTTAAGCAAATTGCTTTGGCCAGAGGAAAATCTGTTGCCCAGGTTAGTATGAGATGGGTTTACGAACAAGGTGCGATTCTTGTAGTGAAAAGTTTTAGTGAAGATAGAATGAGGGAAAATTTGAACATATTCGACTGGGAACTGACTAAGGAAGACTTGGTAAAGATCCGTGAGATTACACAGCGCAGAGTAATAACTGCAGATTTTTTGGTATCAGCTAATGGACCTTTCAAATCTCTGGAAGACTTATGGGATGAGTAA

>Par_COR-Lc10 Par_COR_contig_46708_CDS

ATGGAGAGTTGCGGTGTACCTGTAATCACTCTGAGCTCAGGCAAGAGGATGCCTGTTTTAGGTATGGGAACATTTGAAACATTTGGGAAACCGTCTGAAAGAGAGAGGTTGGCGTATTTGAAAGCCATAGAAGTGGGTTACAGATACTTCGATACAGGTGCTGCATACGGAACTGAAGAGGTTCTTGGCGAAGCTATAGCTGAAGCACTTCAACTTGGCCTAATCAAATCTAGAGATGAACTCTTCATCAGTTCCATGCTTTGGAGCACTGATGCTCACCCTGATCGTGTCAGACTCGCTCTTGAGAATTCGTTGAGGAATCTCAAATTGGAGTTTGTGGATCTATACCTGATACCATGGCCGGTAAGCTTGAAGCCAGAAGGGAAGGTAACTATGGACATACCAGAGGACGAAATTTTTGCAATGGACTACAGGGGTGTATGGGCAGCCATGGAAGAGTGTCAGAACCTTGGCCTCACTAAGTCTATCGGTGTCAGCAATTTCTCCTCCAAAAACTTCAGGAATTGATGGCGGTGGCCAACATCCCTCCAGCTGTGAATCAAGTGGAGATGAACCCCGCTTTCCAACAAAAGAATCTGATAGAGTATTGCAAGGCCAACAATATATTTGTCATTGCATTCTCGGTTTTGGGATCCAACGGAACGTCATGGGGCTCCAATGCAGTTATGGGCTCTGAGGTGCTTAAGCAAATTGCTTTGGCCAGAGGAAAATCTGTTGCCCAGGTTAGTATGAGATGGGTTTACGAACAAGGTGCGATTCTTGTAGTGAAAAGTTTTAGTGAAGATAGAATGAGGGAAAATTTGAACATATTCGACTGGGAACTGACAAAGGAAGACTTGGTAAAGATCCGTGAGATTACACAGCGCAGAGTAATAACTGCAGATTTTTTGGTATCAGCTAATGGACCTTTCAAATCTCTGGAAGACTTATGGGATGAGTAA

>Par_COR-Lc8 Par_COR_contig_81375_CDS

ATGGAGAGCGGTGTGCCTGTAATCACTCTGAGCTCAGGCAGGAGGATGCCTGTTTTAGGTATGGGAACATTTGAAACATTTGGTAAACCGTCTGAAAGAGAGGTTGGCGTATTGAAAGCGATAGAAGTGGGTTACAGATACTTCGATACAGGTACTGCATACGGAACTGAAGAGGTTCTTGGCGAAGCTATAGCTGAAGCACTTCAACTTGGCCTAATCAAATCTAGAGATGAACTCTTCATCAGTTCCATGCTCTGGAGCGCTGATGCTCACTCAATCGTGTCCGACTCGCTCTTGAGAATTCCTTGAGGAATCTCAAATTGAGTTTATTGGATCTATACCTGATACCATGGCCGTGTAGCTGAAGCCAGAAGGTAACTATGGACATACCAGAGGATGAAATTTTTGCAATGGACTACAGGTGTGTATAGGCAGCCATGGAAGAGTGTCTGAACCTTGGCCTCACTAAGTCTATCGGGTGTCAGCAATTTCTCCTCCAAAAACTTCAGGAATTGATGGCGGTGGCCAACATCCCTCCAGCTGTGAATCAAGTGGAGATGAACCCGCTTTCCAACAAAGAATCTGATAGAGTATTGCAAGGCCAACAATATATTTGTCATTGCATTCTCGGTTTTGGGATCCAACGGAACGTCATGGGCCAATGCAGTTATGGGATCTGAGGTGCTTAAGCAAATTGCTTTGGCCAGAGGAAAATCTGTTGCCAGGTTAGTATGAGATGGGTTTACGAACAAGGTGCGATTCTTGTATGAAAAGTTTTAGTGAAGATAGAATGAGGGAAAAATTTGAACATATTCACGACTGGGAACTGACTAAGGAAGACTTGGCAAAGATCCGTGAGATTACACAGCGCAGAGTAATAACTGCAGATTTTGGTATCAGCTAATGGACCCTTTCAAATCTCTGGAAGACTTATGATGAGTAA

>Pca_COR-La8 Pca_COR_143993

ATGGAGAGTAGTGGTATGCCTGTAATCACTCTAAGCTCCGGCAAGGGAATGCCTGTTTTAGGTATGGGAACATGTGAAACACTTCGTGAAGGGTCTGAAAGAGAGAGGTTGGCGATTTTAAAAGCGATAGAGGTGGGTTACCGATACTTTGATACAGCTGCTGGATACCAAACTGAAGAGGTTCTTGGTGAAGCTATTGCTGAAGCACTTCAACTTGGCTTAATCAAATCTCGAGATGAACTCTTTATATCTTCCATGCTCTGGTGCACCGATGCTTACCCAGATCGTGTCCTCCTCGCTCTCCAGAGTTCTTTGAGGAAACTTAAACTGGAGTATCTGGATCTATACCTGATACCATTTCCGGCAGCGGTAAGCTTTAAGCCAGGGAAGATAACCATGGCCTCACTGGAGGACGTAATTTCTCTTTTGGACTACAAGTCTGTATGGGCAGCCATGGAAGAGTGTCAGAACCTTGGCTTCACTAAGTCGATCGGTGTCAGCAACTTCTCCGTCAAAAAGCTTCAGGAATTGATGAGGACCGCCAACATCCCCCCAGCTGTGAATCAAGTGGAGATGAGCCCCATTTTCCAACAAAAGAATCTGAGAGAGTATTGCAAGGCCAATAATATATTGATCACTGCAATCTCGGTTTTGGGATCCAAAGGAACCCCATGGGGCTCCAACGCAGTTTTGGATTCTGAGGTGCTTAACCAGATTGCTATGGCCAGAGGAAAATCTGTTGCTCAGGTTAGTATGAGATGGGTTTACGAGCAAGGTGTGAGTCTTGTGGTAAAAAGTTTCACTGAAGAGAGGATGAGGGAAAACTTGAACATATTCGACTGGGAATTGACTAAGGAAGACTTGGAAAAGATCAGTGAGATTCCACAGTGCAGAGTATTAACTGCTGATTTTTTGGTATCAGCTAATGGGCCTTTCAAATCTCAAGAAGAGCTATGGGATGACAAAGCTTGA

>Pca_COR-La2 Pca_COR_143837b

ATGGAGAGTAGTGGTGTGCCTGTAATCAGTCTAAGCTCGGGCAAGGGGATGCCTGTTTTAGGTATCGGAACGTGTGAAACACTTGGTAAAGGGTCTGAAAGAGAGAGGTTGGCGATTTTGAAAGCGATAGAGGTGGGTTACAGATACATTGATACAGCTGCTGCATACGGAACTGAAGAGGTTGTTGGTGAAGCTATTGCTGAAGAACTTCAACTTGGCCTAATCAAATCTCGAGATGAACTCTTCATCTCTTCCCAACTCTGGGGCACCGATGCTCACCCTGATCGTGTCCTCCTCGCTCTCCAGAATTCTCTGAGGAATCTTAAATTGGAGTATCTAGATCTATACCTGATACCATTTCCGGTAAGCTTTAAGCCAGGGAAGATAACTATGGATGTACCAGAGGACGAAATTTTTCTTATGGACTATAAGTCTGTCTGGGCAGCCATGGAAGAGTGTCAGAACCTTGGCTTCACTAAGTCAATCGGTGTCAGCAATTTCTCCATCAAAAAGCTTCAGGAATTGATGGCGACCGCCAACATCCCTCCAGTTGTGAATCAAGTGGAGACGAACCCCATTTTTCAACAAAAGAATCTGAGAGAGTATTGCAAGGCCAATAATATATTGGTCACTGCAATCTCGGTTTTGGGATCCAAAGGAACCCCATGGGGCTCCAATGCAGTTTTTGATTCTGAGGTGCTTAACCAGATTGCTATGTCCCGAGGAAAATCTGTTGCTCAGGTTAGTATGAGATGGGTTTACGAGCAAGGTGTGAGTCTTGTGGTAAAAAGTTTCAATGAAAAGAGGATGAGGGAAAACTTGAACATATTCGACTGGGAACTGACTAAGGAAGACTTAGAAAAGATCAATGAGATTCCACAGTGCAGAGTATTAACTGCTGATTTTTTGGTATCAGCTAATGGGCCTTTCGAATCTCTAGAAGAGCTATGGGATGACAAAGCTTGA

>Pso_COR-La4-PS0216870 Pso_PS0216870

ATGGAGAGCAATGGTGTGCCCATAATCACTCTAAGCTCGGGCAGGGGGATGCCTGTTTTAGGTATGGGAACATGTGAAACACTTGGTAAAGGGTCTCATGAAAGAGAGAGGCTGGCGATTTTGAAAGCGATAGAGGTCGGTTACAGATTCATCGACACAGCTGCTGCATACGGAACTGAAGAGGTTGTTGGTGAAGTTATTTCTGAAGCACTTCAACTTGGCCTGATCAAATCTCGAGATGAACTCTTCATCTCTTCCCAGCTCTGGTGCACTGATGCTCACCCTGACCGTGTCCTCCTCGCTCTCCAGAATTCTCTGAGGAATCTTAGATTGGAGTATCTGGATCTATACCTGATATCATTTCCGGTAAGCTTGAAGCCAGGGAAGATAACTGCGGACGTATCAGAGGACGAAATTTTTCTGATGGACTACAAGTCTGTATGGGCAGCCATGGAAGAGTGTCAGAGTCTTGGCTTCACTAAGTCTATCGGTGTCAGCAATTTTTCCATCAAAAAGCTTCAAGAATTGATGTCAAACGCCAAGATCCCTCCGGTTGTGAATCAAGTGGAGATGAGCCCAACTTTCCAACAAAAGAATCTGAGAGAGTATTGCAAGGCCAATAATATACTAGTCACTGCAATCTCGGTTTTGGGATCCAAAGGAACCCCATGGGGCTCAAATGCAGTTTTGGATTCTGAGGTGCTTAACCAGATTGCTATGGCCAGAGGAAAATCTGTTGCTCAGATTAGCATGAGATGGGTTTACGAGCAAGGTGTGAGTCTTGTGGTAAAAAGTTTCAGTGAAGAGAGGATGAAGGAAAATCTGAACATATTCGGCTGGGAACTGACTACGGAAGACTTGGAAAAGATTGGTGAGATTCCACAGTGCAGAATATTAACTGCTGATTTTTTGGTATCAGCTAATGGACCTTTCAAATCCCAAGAAGAGCTATGGGATGACTGAGCTTGA

>Pat_COR-L-OUT1 Pat_COR_9059

ATGGAGAGTAGTGGTGTACCTGTAATCGCTCTGAGCTCGGGAAAAGGGATGCCGGTTTTAGGTATGGGAACATTTGAATCAGTTGGTAAAGGGTCTGAAAGAGAGAGGTTGGCGTTTTTAAAAGCGATAGAGGTGGGTTACAGACACTTCGATACAGCTGCTATATATGAAACTGAAGAGTGTCTTGGTGAAGCTATAGCTGAAGCACTTCAAATTGGCCTAATCAAATCTCGAGATGAACTTTTCATCACTTCCAAACTTTGGTGCACTGATGCTCACCCTGATCGTGTTCTCCCCGCTCTTCAGAATTCTCTGAGGAACCTTAAATTGGAGTATCTGGATCTATACCTGATACACTTTCCGGTAAGCTTGAAGCCAGGGGAGCTTGTTATGGACGTACCTGGGGACGCAATTTCTCTAATGGACTACAGGTCTGTATGGGCAGCCATGGAAGAGTGTCAGAACCTTGGCTTCAGTAAGTCAATAGGTGTCTGCAATTTCTCCAGCAAAAAGCTTCAGGAATTGATGGCTGTAGCCAACATCCCTCCAGTTGTAAATCAAGTGGAGATGAGCCCCGTTTTCCAACAAATTTTTTTGAGAGAGTATTGCAAGGCCAACAATATATTAATCACTGCATACTCAGTTTTAGGAGCTAAGGGAACCGCATGGGGTTCCAATGCAGTTCTGGGTTCTGAGGTGCTTAACCAGATTGCTACGGCCAGAGGAAAATCTATTGCTCAGGTTAGTTTGAGATGGGTTTACGAACAAGGTGCGAGTCTTGTGGTGAAAAGTTTCGATGAAGACAGAATGAGGGAAAACTTGAACATATTCGACTGGGAACTGACTAAAGAAGACTTGGAAAAGATTGGTGTCATTCCACAGTGTAAAATAATAATGGAGGATTTTATGGTATCAGCTAACGGACCTTTCAAATCTATAGAAGAGTTCTGGGATGATGATATATAA

>Pso_COR-L-PS0211450 Pso_PS0211450

ATGGAGAGTAGCGGTGTACCTGTAATCGCTCTGAGCTCGGGAAAAGGGATGCCGGTTTTAGGTATGGGAACATTTGAAACAGTTGGGAAAGGGTCTGAGAGAGAGAGGTTGGCGTTTTTAAAAGCGATAGAGGTGGGTTACAGACACTTCGATACAGCTGCTATATATGAAACTGAAGAGTGTTTTGGTGAAGCTATATCTGAAGCACTCCAAAATGGCCTAATCAAATCTCGAGATGAACTTTTCATCACTTCCAAGCTCTGGTGCACTGATGCTCACCCTGATCGTGTTCTCCCAGCTCTTCAGAATTCTCTGAGGAACCTTAAATTGGAGTATCTGGATCTATACCTGATACACTTTCCGGTAAGCTTGAATCCAGGGGAGCTTGTTATGGACGTACCACGCAACGAAATTTTTCTAATGGACTACAGATCTGTATGGGCAGCCATGGAAGAGTGTCAGAACCTTGGCTTCACTAAGTCAATAGGTGTCAGAAATGTCTCCATCAAAAAGCTTAAAGAATTGATGGCGATAGCCAACATCCCTCCAGTTGTAAATCAAGTGGAGATGAGCCCCGTTTTCCAACAAAAAAAATTGAGAGAGTATTGCAAGGACAACAACGTATTGATCACTGCATACTCAGTTTTAGGAGCTAAGGGAACGGCATGGGGTTCCAATGCAGTTCTGGGATCTGAGGTGCTTAACCAGATTGCTACGGCCAGAGGAAAATCTATTGCTCAGGTTAGTTTGAGATGGGTTTACGAACAAGGTGCAAGTCTTGTGGTGAAAAGTTTCGATGAAGACAGGATGAGAGAAAACTTGAACATATTCGACTGGGAACTGACTAAAGAAGACTCGGAAAAGATTATTGTCATTCCAGAGCGTAAAATAATAATTGAAAATTTTATGGTATCAGTTAATGGACCTTTCAAATCTGTGGAAGAGTTCTGGGAT

>Pso_COR-L-PS1020450 Pso_PS1020450

ATGGTGAACACTGGTGTACCTGTTATCACTCTGAGCTCGGGGAAAGGGATACCTGTTTTAGGTATGGGAACATTTGAAACAGTTAGTAAAGGGGGTGAACGAGAGAGGTTGGCGTTTTTGAAAGCGATAGAGGTGGGTTACAGACACTTCGATACTGCTGCGTGCTACCAAACTGAAGAGTGTCTTGGTGAAGCTATAGGGGAAGCACTTCAACTTGGCCTAATCAAATCTCGAGATGAACTCTTTATCACTTCTAAGCTCTGGTGCACCGATGCTCACCCTGATCGTGTTCTTTTGGCTCTTCAGAGTTCTCTGAGGAATCTTAAGTTGGAGTATCTGGATCTATACCTGATACACTTTCCGGTAAGCTTGAAGCCAGGGAATGAGGTCACTATGGACGCAGCAGGGGGCAAAATTTTTCTAATGGACTACAAGTCTGTATGGGCAGCCATGGAAGAGTGTCAGAACCTTGGCTTCACTAAGTCAATCGGTGTCAGCAATTTCTCCTGCAAAAAGCTTCAGGAATTATTGGCGATAGCGAACATCCCTCCAGTTGTAAATCAAGTGGAGATGAGCCCAGTTTTCCACCAAACGAATCTGAGAGAGTATTGCAAGGCCAATAATATACTGGTGACTGCATACTCGATTTTGGGTGGCAAAGGAACCGCATGGGGCTCCAATTCAGTTTTGGGTTCCGAGGTGCTTAACCAGATTGCTATCGCCAGAGGAAAGTCTATTGCTCAGGTTAGTATGAGATGGGTATACGAGCAAGGCGCGATTCTTGTGGTGAAAAGTTTCAGTGAAAAGAATATGAGGGAAAACTTGAACATATTCGATTGGGAACTGACTAAGGAAGACTTGGAAAGGATTGGTGAGATTCCACAGCGCAGATTAATCATTCAGGAATTTATGATATCATCTAATGGACCTTTCAAATCTCTAGAAGAGTTCTGGGATGAAAAAGCAGCTTGA

>Par_COR-L-OUT3 Par_COR_contig_31510_CDS

ATGGTGAATACTGGTGTACCTGTAATCACTCTGAGCTCGGGGAAGGGGATACCTGTTTTAGGTATGGGAACATTTGAAACAATTGGTAAAGGGGGTGAACGAGAGAGGTTGGCGTTTTTGAAAGCGATAGAGGTGGGTTACAGACACTTCGATACTGCTGCGTGTTACCAAACTGAAGAGTGTCTTGGTGAAGCTATATCGGAAGCACTTCAACTTGGACTAATCAAATCTCGAGATGAACTCTTTATCACTTCTAAGCTCTGGTGCACCGATGCTCATCCTGATCGTGTCCTTTTGGCTCTTCAGAATTCTTTGAGGAATCTTAAGTTGGAGTATCTGGATCTATACCTGATACACTTTCCGGTAAGCTTGAAGCCAGGGAATGAGGTTACTATGGACGCAGCAGGGGGCGAAATTTTTCTAATGGACTACAAGTCTGTATGGGCAGCCATGGAAGAGTGTCAGAACCTTGGCTTCACTAAGTCAATCGGTGTCAGCAATTTCTCCTGCAAAAAGCTTCAGGAATTATTGGCGACAGCGAACATCCCTCCAGTTGTAAATCAAGTGGAGATGAGCCCAGTTTTCCACCAAAAGAATCTGAGAGAGTATTGCAAGGCCAATAATATACTGGTGGCTGCATATTCGATTTTGGGAGGCAAAGGAACCGCATGGGGCTCCAATTCAGTTTTGGGTTCCGAGGTGCTTAACCAGATTGCCATGGCCAGAGGAAAGTCTATTGCTCAGGTTAGCATGAGATGGGTGTACGAGCAAGGCGCGATTCTTGTGGTGAAAAGTTTCAGTGAAAAGAATATGAGGGAAAACTTGAACATATTCGATTGGGAACTGACTAAGGAAGACTTGGAAAGGATTGGTGAGATTCCACAGCGCAGATTAATAATTCAGGAATTTATGATATCATCTAAAGGACCTTTCAAATCTCTAGAAGAGTTCTGGGATGAAAAAGCTGCTTGA

>Pso_COR-L-PS1020440 Pso_PS1020440

ATGGTGAACACTGGTGTACCTGTTATCACTCTGAGCTCGGGGAAAGGGATACCTGTTTTAGGTATGGGAACATTTGAAACAGTTGGTAAAGGGGGTGAACGAGAGAGGTTGGCGTTTTTGAAAGCGATAGAGGTGGGTTACAGACACTTCGATACTGCTGCGTGCTACCAAACTGAAGAGTGTCTTGGTGAAGCTATAGGGGAAGCACTTCAACTTGGCCTAATCAAATCTCGAGATGAACTCTTTATCACTTCTAAGCTCTGGTGCACCGATGCTCACCCTGATCGTGTTCTTTTGGCTCTTCAGAGTTCTCTGAGGAATCTTAAGTTGGAGTATCTGGATCTATACCTGATACACTTTCCGGTAAGCTTGAAGCCAGGGAATGAGGTCACTATGGACGCAGCAGGGGGCAAAATTTTTCTAATGGACTACAAGTCTGTATGGGCAGCCATGGAAGAGTGTCAGAACCTTGGCTTCACTAAGTCAATCGGTGTCAGCAATTTCTCCTGCAAAAAGCTTCAGGAATTATTGGCGATAGCGAACATCCCTCCAGTTGTAAATCAAGTGGAGATGAGCCCAGTTTTCCACCAAACGAATCTGAGAGAGTATTGCAAGGCCAATAATATACTGGTGACTGCATACTCGATTTTGGGTGGCAAAGGAACCGCATGGGGCTCCAATTCAGTTTTGGGTTCCGAGGTGCTTAACCAGATTGCTATCGCCAGAGGAAAGTCTATTGCTCAGGTTAGTATGAGATGGGTATACGAGCAAGGGGCGATTCTTGTGGTGAAAAGTTTCAGTGAAAAGAATATGAGGGAAAACTTGAACATATTCGATTGGGAACTGACTAAGGAAGACTTGGAAAGGATTGGTGAGATTCCACAGCGCAGATTAATAATTCAGGAATTTATGATATCATCTAATGGACCTTTCAAATCTCTAGAAGAGTTCTGGGATGAAAAAGCTGCTTGA

>Pso_COR-L-PS1020430 Pso_PS1020430

ATGGTGAACACTGGTGTACCTGTTATCACTCTGAGCTCGGGGAAAGGGATACCTGTTTTAGGTATGGGAACATTTGAAACAGTTGGTAAAGGGGGTGAACGAGAGAGGTTGGCGTTTTTGAAAGCGATAGAGGTGGGTTACAGACACTTCGATACTGCTGCGTGCTACCAAACTGAAGAGTGTCTTGGTGAAGCTATAGGGGAAGCACTTCAACTTGGCCTAGTCAAATCTCGAGATGAACTCTTTATCACTTCTAAGCTCTGGTGCACCGATGCTCACCCTGATCGTGTTCTTTTGGCTCTTCAGAGTTCTCTGAGGAATCTTAAGTTGGAGTATCTGGATCTATACCTGATACACTTTCCGGTAAGCTTGAAGCCAGGGAATGAGGTTACTATGGACGCAGCAGGGGGCAAAGTTTTTCTAATGGACTACAAGTCTGTATGGGCAGCCATGGAAGAGTGTCAGAACCTTGGCTTCACTAAGTCAATCGGTGTCAGCAATTTCTCCTGCAAAAAGCTTCAGGAATTATTGGCGATAGCGAACATCCCTCCAGTTGTAAATCAAGTGGAGATGAGCCCAGTTTTCCACCAAACGAATCTGAGAGAGTATTGCAAGGCCAATAATATACTGGTGACTGCATACTCGATTTTGGGTGGCAAAGGAACCGCATGGGGCTCCAATTCAGTTTTGGGTTCCGAGGTGCTTAACCAGATTGCTATCGCCAGAGGAAAGTCTATTGCTCAGGTTAGTATGAGATGGGTATACGAGCAAGGCGCGATTCTTGTGGTGAAAAGTTTCAGTGAAAAGAATATGAGGGAAAACTTGAACATATTCGATTGGGAACTGACTAAGGAAGACTTGGAAAGGATTGGTGAGATTCCACAGCGCAGATTAATAATTCAGGAATTTATGATATCATCTAATGGACCTTTCAAATCTCTAGAAGAGTTCTGGGATGAAAAAGCTGCTTGA

>Pso_COR-L-PS1021890 Pso_PS1021890

ATGGTGAACACTGGTGTACCTGTTATCACTCTGAGCTCGGGGAAAGGGATACCTGTTTTAGGTATGGGAACATTTGAAACAGTTGGTAAAGGGGGTGAACGAGAGAGGTTGGCGTTTTTGAAAGCGATAGAGGTGGGTTACAGACACTTCGATACTGCTGCGTGCTACCAAACTGAAGAGTGTCTTGGTGAAGCTATAGGGGAAGCACTTCAACTTGGCCTAGTCAAATCTCGAGATGAACTCTTTATCACTTCTAAGCTCTGGTGCACCGATGCTCACCCTGATCGTGTTCTTTTGGCTCTTCAGAGTTCTCTGAGGAATCTTAAGTTGGAGTATCTGGATCTATACCTGATACACTTTCCGGTAAGCTTGAAGCCAGGGAATGAGGTTACTATGGACGCAGCAGGGGGCAAAGTTTTTCTAATGGACTACAAGTCTGTATGGGCAGCCATGGAAGAGTGTCAGAACCTTGGCTTCACTAAGTCAATCGGTGTCAGCAATTTCTCCTGCAAAAAGCTTCAGGAATTATTGGCGATAGCGAACATCCCTCCAGTTGTAAATCAAGTGGAGATGAGCCCAGTTTTCCACCAAACGAATCTGAGAGAGTATTGCAAGGCCAATAATATACTGGTGACTGCATACTCGATTTTGGGTGGCAAAGGAACCGCATGGGGCTCCAATTCAGTTTTGGGTTCCGAGGTGCTTAACCAGATTGCTATCGCCAGAGGAAAGTCTATTGCTCAGAATATGAGGGAAAACTTGAACATATTCGATTGGGAACTGACTAAGGAAGACTTGGAAAGGATTGGTGAGATTCCACAGCGCAGATTAATAATTCAGGAATTTATGATATCATCTAATGGACCTTTCAAATCTCTAGAAGAGTTCTGGGATGAAAAAGCTGCTTGA

>Par_COR-Lb2C Par_COR_contig_48062a_CDS

ATGGAGAATATTAATGGTGTACCTGTAATTACTCTGAGCTCAGGCAAGGGTATGCCTGTTTTAGGTATGGGAACATTTGAAATAGTTGGTAAAGGGTCTGAAAGAGCGAGGTTGGCAATTTTGAAAGCGATAGAGGTGGGTTACCGATACTTTGATACAGCGGCTGCATACAAAACTGAAGAGGTTCTTGGTGAAGCTATTGTTGAAGCACTTCAACTTGGCTTAATAAAATCTCGAGATGAACTCTTCATCAGTTCCATGCTCTGGTGCTCTGATGCTCACCCTGATCGTGTCCTCCTCGCTCTGCAGAATTCTCTGAGGAGTCTTAAATTGGAGTACGTGGATCTATACATGATACCCTTTCCGTTAAGCTTGAAGCCAGGAGGGAAGATAGGTATGGACACACCAGAGGACGAGATTTTAGTAATGGACTACAGGTCTGTATGGGCAGCCATGGAAGAGTGTCAGAACCTTGGCTTCACTAAGTCAATCGGTGTCAGCAATTTCTCCTGCAAAAATCCCTCCTGTTGTGAATCAAGTGGAGATGAGCCCGGCTTTCCAACAAAAGAATCTGAGGGAGTATTGCAAGGCAAATAATATATTAGTCAGTGCAGTCTCTATTTTGGGATCAAACGGAACCCCATGGGGCTCCAATGCAGTTTTGGGTTCGGAGGTGCTTAGCCAGATTGCTATGGCTAGAGGAAAATCTATTGCTCAGGTTAGTATGAGATGGGTTTACGAGCAAGGTGCAAGTCTTGTGGTAAAAAGTTTTAGTGAAGAGAGGATGAGGGAAAACTTGAACATATTCGACTGGGAACTCACTAAGGAAGACTTGGAAAAGATACGTAATGAGATTCCACAGTGCAGAATCTTGACTGCGTATTTTATGGTATCACCTAATGGACCTTTCAAATCTCAACAAGAGCTCTGGGATGACGAAGCTTGA

>Par_COR-Lb2B Par_COR_contig_48062b_CDS

ATGGAGAATATTAATGGTGTACCTGTAATTACTCTGAGCTCAGGCAAGGGTATGCCTGTTTTAGGTATGGGAACATTTGAAATAGTTGGTAAAGGGTCTGAAAGAGCGAGGTTGGCAATTTTGAAAGCGATAGAGGTGGGTTACCGATACTTTGATACAGCGGCTGCATACAAAACTGAAGAGGTTCTTGGTGAAGCTATTGTTGAAGCACTTCAACTTGGCTTAATAAAATCTCGAGATGAACTCTTCATCAGTTCCATGCTCTGGTGCTCTGATGCTCACCCTGATCGTGTCCTCCTCGCTCTGCAGAATTCTCTGAGGAGTCTTAAATTGGAGTACGTGGATCTATACATGATACCCTTTCCGTTAAGCTTGAAGCCAGGAGGGAAGATAGGTATGGACACACCAGAGGACGAGATTTTAGTAATGGACTACAGGTCTGTATGGGCAGCCATGGAAGAGTGTCAGAACCTTGGCTTCACTAAGTCAATCGGTGTCAGCAATTTCTCCTGCAAAAATCCCTCCTGTTGTGAATCAAGTGGAGATGAGCCCGGCTTTCCAACAAAAGAATCTGAGGGAGTATTGCAAGGCAAATAATATATTAGTCAGTGCAGTCTCTATTTTGGGATCAAACGGAACCCCATGGGGCTCCAATGCAGTTTTGGGTTCGGAGGTGCTTAGCCAGATTGCTATGGCTAGAGGAAAATCTATTGCTCAGGTTAGTATGAGATGGGTTTACGAGCAAGGTGCAAGTCTTGTGGTAAAAAGTTTTAGTGAAGAGAGGATGAGGGAAAACTTGAACATATTCGACTGGGAACTCACTAAGGAAGACTTGGAAAAGATACGTAATGAGATTCCACAGTGCAGAATCTTGACTGCGTATTTTATGGTATCACCTAATGGACCTTTCAAATCTCAACAAGAGCTCTGGGATGACGAAGCTTGA

>Par_COR-Lb2A Par_COR_contig_48062c_CDS

ATGGAGAATATTAATGGTGTACCTGTAATTACTCTGAGCTCAGGCAAGGGTATGCCTGTTTTAGGTATGGGAACATTTGAAATAGTTGGTAAAGGGTCTGAAAGAGCGAGGTTGGCAATTTTGAAAGCGATAGAGGTGGGTTACCGATACTTTGATACAGCGGCTGCATACAAAACTGAAGAGGTTCTTGGTGAAGCTATTGTTGAAGCACTTCAACTTGGCTTAATAAAATCTCGAGATGAACTCTTCATCAGTTCCATGCTCTGGTGCTCTGATGCTCACCCTGATCGTGTCCTCCTCGCTCTGCAGAATTCTCTGAGGAGTCTTAAATTGGAGTACGTGGATCTATACATGATACCCTTTCCGTTAAGCTTGAAGCCAGGAGGGAAGATAGGTATGGACACACCAGAGGACGAGATTTTAGTAATGGACTACAGGTCTGTATGGGCAGCCATGGAAGAGTGTCAGAACCTTGGCTTCACTAAGTCAATCGGTGTCAGCAATTTCTCCTGCAAAAATCCCTCCTGTTGTGAATCAAGTGGAGATGAGCCCGGCTTTCCAACAAAAGAATCTGAGGGAGTATTGCAAGGCAAATAATATATTAGTCAGTGCAGTCTCTATTTTGGGATCAAACGGAACCCCATGGGGCTCCAATGCAGTTTTGGGTTCGGAGGTGCTTAGCCAGATTGCTATGGCTAGAGGAAAATCTATTGCTCAGGTTAGTATGAGATGGGTTTACGAGCAAGGTGCAAGTCTTGTGGTAAAAAGTTTTAGTGAAGAGAGGATGAGGGAAAACTTGAACATATTCGACTGGGAACTCACTAAGGAAGACTTGGAAAAGATACGTAATGAGATTCCACAGTGCAGAATCTTGACTGCGTATTTTATGGTATCACCTAATGGACCTTTCAAATCTCAACAAGAGCTCTGGGATGACGAAGCTTGA

>Par_COR-Lc2D Par_COR_contig_30250d_CDS

ATGGAGACTAATGGCGTACCGGTAATCACTCTGAGCTCAGGCAAGGAGATGCCTGTTTTAGGCATGGGAACATTTGAATCATTTGGAAGTAAAGGGTCTGAAAGAGAGAAGTTGGCATATTTGAAAGCCATAGAGGTGGGTTACAGATACTTTGATACAGGTGCCGCATATGGAACCGAAGAGGTTCTTGGCGAAGCTATAGCTGAAGCACTTCAACTCGGCCTAATCAAATCTCGAAATGAACTCTTCATCAGTTCCATGCTCTGGTGCACCGATGCTCACGCTGATCGCGTCCTACTCGCTCTTCAGAATTCTCTAAGGAATCTTAAGTTGGAATATCTGGACCTATACATGTTACCTTTTCCGGCAAGCATGAAGCCAGCAGGAGAGAAAATAACAATGGTGGACTTCCATGACATAACAGAAGAAGACGAGATTTTTGCAATGGACTACAAGTCTGTATGGGCAGCCATGGAAGAGTGTCAGAACCTCGGCTTGACTAAGTCAATCGGTGTCAGTAATTTCTCCAGCAAAAAGCTCCAGGAATTGATGGTCAACGCCAAAATCCCTCCGGCTGTGAATTCTGTGGAGATGAGCCCGGCTTTTCAACAAAAAAATCTGAGAGAGTATTGCAAGGCCAATAATATATTGATCATTGCGATCTCCATTTTGGGATCCACCGGAACCGAATGGGGCTCCAATGCAGTTATGGGTTCTGAGGTACTTAAACAAATTGCTATGGACAGAGGAAAATCTACTGCACAGGTTAGTATGAGATGGGTTTACGAGCAAGGCGCAATTCTTGTAGTAAAAAGTTTCAGTGAAAAGAATATGAGGGCCAACTTGAAGATATTTGACCGGGAACTCACCAAGGAAGACTGCGAAAAGATCGGTACTAGGATTCCACAGTGCAGAACCTTAAAAGGTGATTTTATGGTATCACCAAATGGTCCTTTCAAAACTCTACAAGACTTATGGGATGACTGA

>Par_COR-Lc2C Par_COR_contig_30250c_CDS

ATGGAGACTAATGGCGTACCGGTAATCACTCTGAGCTCAGGCAAGGAGATGCCTGTTTTAGGCATGGGAACATTTGAATCATTTGGAAGTAAAGGGTCTGAAAGAGAGAAGTTGGCATATTTGAAAGCCATAGAGGTGGGTTACAGATACTTTGATACAGGTGCCGCATATGGAACCGAAGAGGTTCTTGGCGAAGCTATAGCTGAAGCACTTCAACTCGGCCTAATCAAATCTCGAAATGAACTCTTCATCAGTTCCATGCTCTGGTGCACCGATGCTCACGCTGATCGCGTCCTACTCGCTCTTCAGAATTCTCTAAGGAATCTTAAGTTGGAATATCTGGACCTATACATGTTACCTTTTCCGGCAAGCATGAAGCCAGCAGGAGAGAAAATAACAATGGTGGACTTCCATGACATAACAGAAGAAGACGAGATTTTTGCAATGGACTACAAGTCTGTATGGGCAGCCATGGAAGAGTGTCAGAACCTCGGCTTGACTAAGTCAATCGGTGTCAGTAATTTCTCCAGCAAAAAGCTCCAGGAATTGATGGTCAACGCCAAAATCCCTCCGGCTGTGAATTCTGTGGAGATGAGCCCGGCTTTTCAACAAAAAAATCTGAGAGAGTATTGCAAGGCCAATAATATATTGATCATTGCGATCTCCATTTTGGGATCCACCGGAACCGAATGGGGCTCCAATGCAGTTATGGGTTCTGAGGTACTTAAACAAATTGCTATGGACAGAGGAAAATCTACTGCACAGGTTAGTATGAGATGGGTTTACGAGCAAGGCGCAATTCTTGTAGTAAAAAGTTTCAGTGAAAAGAATATGAGGGCCAACTTGAAGATATTTGACCGGGAACTCACCAAGGAAGACTGCGAAAAGATCGGTACTAGGATTCCACAGTGCAGAACCTTAAAAGGTGATTTTATGGTATCACCAAATGGTCCTTTCAAAACTCTACAAGACTTATGGGATGACTGA

>Par_COR-Lc2B Par_COR_contig_30250b_CDS

ATGGAGACTAATGGCGTACCGGTAATCACTCTGAGCTCAGGCAAGGAGATGCCTGTTTTAGGCATGGGAACATTTGAATCATTTGGAAGTAAAGGGTCTGAAAGAGAGAAGTTGGCATATTTGAAAGCCATAGAGGTGGGTTACAGATACTTTGATACAGGTGCCGCATATGGAACCGAAGAGGTTCTTGGCGAAGCTATAGCTGAAGCACTTCAACTCGGCCTAATCAAATCTCGAAATGAACTCTTCATCAGTTCCATGCTCTGGTGCACCGATGCTCACGCTGATCGCGTCCTACTCGCTCTTCAGAATTCTCTAAGGAATCTTAAGTTGGAATATCTGGACCTATACATGTTACCTTTTCCGGCAAGCATGAAGCCAGCAGGAGAGAAAATAACAATGGTGGACTTCCATTACATAACAGAAGAAGACGAGATTTTTGCAATGGACTACAAGTCTGTATGGGCAGCCATGGAAGAGTGTCAGAACCTCGGCTTGACTAAGTCAATCGGTGTCAGTAATTTCTCCAGCCAAAAGCTCCAGGAATTGATGGTCAACGCCAAAATCCCTCCGGCTGTGAATTCTGTGGAGATGAGCCCGGCTTTTCAACAAAAAAATCTGAGAGAGTATTGCAAGGCCAATAATATATTGATCATTGCGATCTCCATTTTGGGATCCACCGGAACCGAATGGGGCTCCAATGCAGTTATGGGTTCTGAGGTACTTAAACAAATTGCTATGGACAGAGGAAAATCTACTGCACAGGTTAGTATGAGATGGGTTTACGAGCAAGGCGCAATTCTTGTAGTAAAAAGTTTCAGTGAAAAGAATATGAGGGCCAACTTGAAGATATTTGACCGGGAACTCACCAAGGAAGACTGCGAAAAGATCGGTACTAGGATTCCACAGTGCAGAACCTTAAAAGGTGATTTTATGGTATCACCAAATGGTCCTTTCAAAACTCTACAAGACTTATGGGATGACTGA

>Par_COR-Lc2A Par_COR_contig_30250a_CDS

ATGGAGACTAATGGCGTACCGGTAATCACTCTGAGCTCAGGCAAGGAGATGCCTGTTTTAGGCATGGGAACATTTGAATCATTTGGAAGTAAAGGGTCTGAAAGAGAGAAGTTGGCATATTTGAAAGCCATAGAGGTGGGTTACAGATACTTTGATACAGGTGCCGCATATGGAACCGAAGAGGTTCTTGGCGAAGCTATAGCTGAAGCACTTCAACTCGGCCTAATCAAATCTCGAAATGAACTCTTCATCAGTTCCATGCTCTGGTGCACCGATGCTCACGCTGATCGCGTCCTACTCGCTCTTCAGAATTCTCTAAGGAATCTTAAGTTGGAATATCTGGACCTATACATGTTACCTTTTCCGGCAAGCATGAAGCCAGCAGGAGAGAAAATAACAATGGTGGACTTCCATTACATAACAGAAGAAGACGAGATTTTTGCAATGGACTACAAGTCTGTATGGGCAGCCATGGAAGAGTGTCAGAACCTCGGCTTGACTAAGTCAATCGGTGTCAGTAATTTCTCCAGCAAAAAGCTCCAGGAATTGATGGTCAACGCCAAAATCCCTCCGGCTGTGAATTCTGTGGAGATGAGCCCGGCTTTTCAACAAAAAAATCTGAGAGAGTATTGCAAGGCCAATAATATATTGATCATTGCGATCTCCATTTTGGGATCCACCGGAACCGAATGGGGCTCCAATGCAGTTATGGGTTCTGAGGTACTTAAACAAATTGCTATGGACAGAGGAAAATCTACTGCACAGGTTAGTATGAGATGGGTTTACGAGCAAGGCGCAATTCTTGTAGTAAAAAGTTTCAGTGAAAAGAATATGAGGGCCAACTTGAAGATATTTGACCGGGAACTCACCAAGGAAGACTGCGAAAAGATCGGTACTAGGATTCCACAGTGCAGAACCTTAAAAGGTGATTTTATGGTATCACCAAATGGTCCTTTCAAAACTCTACAAGACTTATGGGATGACTGA

>Par_COR-Lc3B Par_COR_contig_26129b_CDS

ATGGAGACTAATGGCGTACCGGTAATCACTCTGAGCTCAGGCAAGGAGATGCCTGTTTTAGGCATGGGAACATTTGAATCATTTGGAAGTAAAGGGTCTGAAAGAGAGAAGTTGGCATATTTGAAAGCCATAGAGGTGGGTTACAGATACTTTGATACAGGTGCCGCATATGGAACCGAAGAGGTTCTTGGCGAAGCTATAGCTGAAGCACTTCAACTCGGCCTAATCAAATCTCGAAATGAACTCTTCATCAGTTCCATGCTCTGGTGCACCGATGCTCACGCTGATCGCGTCCTACTCGCTCTTCAGAATTCTCTAAGGAATCTTAAGTTGGAATATCTGGACCTATACATGTTACCTTTTCCGGCAAGCATGAAGCCAGCAGGAGAGAAAATAACAATGGTGGACTTCCATGACATAACAGAAGAAGACGAGATTTTTGCAATGGACTACAAGTCTGTATGGGCAGCCATGGAAGAGTGTCAGAACCTCGGCTTGACTAAGTCAATCGGTGTCAGTAATTTCTCCAGCAAAAAGCTCCAGGAATTGATGGTCAACGCCAAAATCCCTCCGGCTGTGAATTCTGTGGAGATGAGCCCGGCTTTTCAACAAAAAAATCTGAGAGAGTATTGCAAGGCCAATAATATATTGATCATTGCGATCTCCATTTTGGGATCCACCGGAACCGAATGGGGCTCCAATGCAGTTATGGGTTCTGAGGTACTTAAACAAATTGCTATGGACAGAGGAAAATCTACTGCACAGGTTAGTATGAGATGGGTTTACGAGCAAGGCGCAATTCTTGTAGTAAAAAGTTTCAGTGAAAAGAATATGAGGGCCAACTTGAAGATATTTGACTGGGAACTCACCAAGGAAGACTGCCAAAAGATCGGTACTAGGATTCCACAGTGCAGAACCTTAAAAGGTGATTTTATGGTATCACCAAATGGTCCTTTCAAAACTCTACAAGACTTATGGGATGACTGA

>Par_COR-Lc3A Par_COR_contig_26129a_CDS

ATGGAGACTAATGGCGTACCGGTAATCACTCTGAGCTCAGGCAAGGAGATGCATGTTTTAGGTATGGGAACATTTGAATCATTTGGTAGTGAAGGGTCTGAAAGAGAGAAGTTGGCGTATTTGAAAGCCATAGAGGTGGGTTACAGATACTTTGATACAGGTGCCGCATACGGAACTGAAGAGGTTCTTGGCGAAGCTATAACTGAAGCACTTCAACTCGGCCTAATCAAATCTCGAGATGAACTCTTCGTCAGTTCCATGCTCTGGTGCACCGATGCTCACGCTGATCGTGTCCTACTCGCTCTTCAGAATTCTCTAAGGAATCTTAAGTTGGAATATCTGGACCTATACATGTTACCTTTTCCGGCAAGCATGAAGCCAGCAGGAGAGAAAATAACAATGGTGGACTTCCATGACATAACAGAAGAAGACGAGATTTTTGCAATGGACTACAAGTCTGTATGGGCAGCCATGGAAGAGTGTCAGAACCTCGGCTTGACTAAGTCAATCGGTGTCAGTAATTTCTCCAGCAAAAAGCTCCAGGAATTGATGGTCAACGCCAAAATCCCTCCGGCTGTGAATTCTGTGGAGATGAGCCCGGCTTTTCAACAAAAAAATCTGAGAGAGTATTGCAAGGCCAATAATATATTGATCATTGCGATCTCCATTTTGGGATCCACCGGAACCGAATGGGGCTCCAATGCAGTTATGGGTTCTGAGGTACTTAAACAAATTGCTATGGACAGAGGAAAATCTACTGCACAGGTTAGTATGAGATGGGTTTACGAGCAAGGCGCAATTCTTGTAGTAAAAAGTTTCAGTGAAAAGAATATGAGGGCCAACTTGAAGATATTTGACTGGGAACTCACCAAGGAAGACTGCCAAAAGATCGGTACTAGGATTCCACAGTGCAGAACCTTAAAAGGTGATTTTATGGTATCACCAAATGGTCCTTTCAAAACTCTACAAGACTTATGGGATGACTGA

>Par_COR-La6 Par_COR_contig_38806_CDS

ATGGAGAGTAATGGTGTGCCCATAATCACTCTTAGCTCGGGCAAGGGGATGCCTGTTTTAGGTATGGGAACATGTGAAACCCTTGGTAAAAGATCTCATGAAAGAGAGGGGTTGGCGATTTTGAAAGCGATAAAGGTGGGTTACAGATGCATCGATACAGCTGCTGCATACGGAACTGAAGAGGTTGTTGGTGAAGCTATTTCTGAAGCACTACAACTTGGCCTGATCAAATCTCGAGATGAACTCTTCATCTCTTCCCAGCTCTGGTGCACTGATGCTCACCCTGATCGTGTCCTCCTCGCTCTTCAGAATTCTCTGAGGAATCTTAAATTGGAGTATCTGGATCTATACCTGATATCATTTCCGGTAAGCTTCAAGCCAGGGAAGATAACTACGGACGTACAAGAGGACGAAATTTTTCTTATGGACTACAAGTCTGTATGGGCATCCATGGAAGAGTGTCAGAACCTTGGCTTCACTAAATCTGTCGGTGTCAGCAATTTTTCCATCAAAAAGCTTCAGGAATTGATGGCGGCTGCTAAAATCCCTCCAGTTGTGAATCAAGTGGAGATGAGCCCCACTTTCCAACAAAAGAATCTGAGAGAGTATTGCAAGGCCAATAATATACTGGTCACTGCAATCTCGGTTTTGGGATCCAAAGGAACCCCATGGGGCTCTAATGCAGTTTTAGATTCTGAGGTGCTTAACCAGATTGCTATGGCTAGAGGAAAATCTGTTGCTCAGGTTAGTATGAGATGGGTTTACGAGCAAGGTGTGAGTCTTGTGGTAAAAAGTTTCAATGAAGAGAGGATGAGGGCAAACTTGAACATATTCGACTGGGAACTGACCAAGGAAGACTTGGAAAAGATCGGCGAGATTCCACAGCGCAGAGTATTAACTGCTGATTTTTTGGTATCAGCTAATGGACCTTTCAAATCTCTAGAAGAGCTATGGGATGACTAA

>Par_COR-La7 Par_COR_contig_26608_CDS

ATGGAGAGTAATGGTGTGCCCATAATCACTCTTAGCTCGGGCAAGGAGATGCCTGTTTTAGGTATGGGAACATGTGAAACCCTTGGTAAAGGGTCTCATGAAAGAGAGGGGTTGGCGATATTGAAAGCGATAAAGGTGGGTTCTGAGATACATCGATCACTGCTGCATACCGGCCCTGAAGAGGTTGTTGGTGAAGCTATTTCGAAGCACTACAACTTGGCACGATCAAATCTCGAGATGAACTCTTCATCTCTTCCCAGCTCAGTGCACCGATGCTCACCCCGATCGTGTCCTCCTCGCTCTTGAGAATTCTACGAGGAATCTTAAACTGGAGTATCTGGATCTATACATGATATCATTTCCGGTAAGCTTCAAGCCAGGGAAGATAACTATGGACGTACGAGAGGATGAAATTTTTCTTATGGACTACAAGTCTGTATGGGCAGCCATGGAAGAGTGTCAGAACCTTGGCTTCACTAAATCTATCGGTGTCAGCAATTTTTCCATCAAAAAGCTTCAGGAATTAATGTCGAACGCTAACATCCCTCCGGTGGTGAATCAAGTGGAGATGAACCCCACTTTCCAACAAAAGAATCTGAGAGAGTATTGCAAGGCCAATAATATACTGGTCACTGCAATCTCGGTTTTGGGATCCAAAGGAACCCCATGGGGCTCTAATGCAGTTTTAGATTCTGAGGTGCTTAACCAGATTGCTATGGCTAGAGGAAAATCTGTTGCTCAGGTTAGTATGAGATGGGTTTACGAGCAAGATGTCAGTCTTGTAGTAAAAAGTTTCAATGAAGAGAGGATGAGGGAAAACTTGAACATATTCGACTGGGAACTGACCAAGGAAGACTTGGAAAAGATCGGCGAGATTCCACAGCGCAGAGTATTAACTGCTGATTTTTTGGTATCAGCTAATGGACCTTTCAGATCTCTAGAAGAGCTATGGGATGACTAA

>Pbr_COR-L-OUT5 Pbr_COR_contig_972_CDS

ATGGGGAATACTGGTGTACCTGTAATCACTCTGAGCTCGGGGAAGGGGATACCTGTTTTAGGTATGGGAACATTTGAAACAGTTGGTAAAGGGGGTGAACGAGAGAGGTTGGCGTTTTTGAAAGCGATAGAGGTGGGTTACAGACACTTCGATACTGCTGCGTGCTACCAAACTGAAGAGCGTCTTGGTGAAGCTATAGAGGATGCACTTCAACTTGGCCTAATAAAATATCGAGATGAACTCTTTATCACTTCTAAGCTCTGGTGCACCGACGCTCACCCTGATCGTGTCCTTTTGGCTCTTCAGAATTCTCTGAGGAATCTTAAGTTGGAGTATCTGGATCTATACCTGATACACTTTCCAGTAAGCTTGAAGCCAGGGAATGAGGTTACTATGGATGCAGCAGGGGGCGAAATTTTTCTAATGGACTACAAGTCTGTATGGGCAGCCATGGAAGAGTGTCAGAACCTTGGCTTCACTAAGTCAATCGGTGTCAGCAATTTCTCCTGCAAAAAGCTTCAGGAATTATTGGCGACAGCAAACATCCCTCCAGTTGTAAACCAAGTGGAGATGAGCCCAGTTTTCCACCAAAAGAATCTGAGAGAGTATTGCAAGGCCAATAATATACTGGTGGCTGCATACTCGATTTTGGGAGGCAAAGGAACCGCATGGGGCTCCAATTCAGTTTTGGGTTCCGAGATGCTTAACCAGATTGCTATCGCCAGAGGAAAGTCTATTGCTCAGGTTAGCATGAGATGGGTATACGAGCAAGGCGCGATTCTTGTGGTGAAAAGTTTTAGTGAAAAGAATATGAGGGAAAACTTGAACATATTCGATTGGGAACTGACTAAGGAAGACCTGGAAAGGATTGGTGAGATTCCTCAGCGCAGATTAATAATTCAGGAATTTATGATATCATCTAATGGACCTTTCAAATCTCTAGAAGAGTTCTGGGATGAAAAAGCTGCTTGA

>Pbr_COR-L-OUT4 Pbr_COR_contig_16531_CDS

ATGGGGAATACTGGTGTACCTGTAATCACTCTGAGCTCGGGGAAGGGGATACCTGTTTTAGGTATGGGAACATTTGAAACAGTTGGTAAAGGGGGTGAACGAGAGAGGTTGGCGTTTTTGAAAGCGATAGAGGTGGGTTACAGACACTTCGATACTGCTGCGTGCTACCAAACTGAAGAGCGTCTTGGTGAAGCTATAGAGGATGCACTTCAACTTGGCCTAATCAAATCTCGAGATGAACTCTTTATCACTTCTAAGCTCTGGTGCACCGACGCTCACCCTGATCGTGTCCTTTTGGCTCTTCAGAATTCTCTGAGGAATCTTAAGTTGGAGTATCTGGATCTATACCTGATACACTTTCCAGTAAGCTTGAAGCCAGGGAATGAGGTTACTATGGATGCAGCAGGGGGCGAAATTTTACTAATGGACTACAAGTCTGTATGGGCAGCCATGGAAGAGTGTCAGAACCTTGGCTTCACTAAGTCAATCGGTGTCAGCAATTTCTCCTGCAAAAAGCTTCAGGAATTATTGGCGACAGCAAACATCCCTCCAGTTGTAAACCAAGTGGAGATGAGCTCAGTTTTCCACCAAAAGAATCTGAGAGAGTATTGCAAGGCCAATAATATACTGGTGGCTGCATACTCGATTTTGGGAGGCAAAGGAACCGCATGGGGCTCCAATTCAGTTTTGGGTTCCGAGATGCTTAACCAGATTGCTATCGCCAGAGGAAAGTCTATTGCTCAGGTTAGCATGAGATGGGTATACGAGCAAGGCGCGATTCTTGTGGTGAAAAGTTTTAGTGAAAAGAATATGAGGGAAAACTTGAACATATTCGATTGGGAACTGACTAAGGAAGACCTGGAAAGGATTGGTGAGATTCCTCAGCGCAGATTAATAATTCAGGAATTTATGATATCATCTAATGGACCTTTCAAATCTCTAGAAGAGTTCTGGGATGAAAAAGCTGCTTGA

>Pbr_COR-L-OUT2 Pbr_COR_scaffold_17762_CDS

ATGGAGAGTAGTGGTGTACCTGTAATCGCTCTGAGCTCTGGAAATGGGATGCCAGTTTTAGGTATGGGAACATTTGAAACAGTTGGGAAAGGGTCTGAGAGAGAGAGGTTGGCTTTTTTAAAAGCGATAGACGTGGGTTACAGACACTTCGATACAGCTGCTATATATGAAACTGAAGAGTGTCTTGGTGAAGCTATAGCTGAAGCACTTGAAATTGGCCTAATCAAATCTCGAGATGAACTTTTCATCACTTCCAAGCTTTGGTGCACTGATGCTCACCCTGATCGTGTTCTCCCCGCTCTTCAGAATTCACTGAGGAACCTTAAATTGGAGTATTTGGATCTATACCTGATACACTTTCCGGTAAGCTTGAAGCCAGGGGAGCTTGTTATGGACGTACCAGGGGACGCAATTTCTCTAATGGACTACAGGTCTGTATGGGCAGCCATGGAAGAGTGTCAGAACCTTGGCTTCACTAAGTCAATAGGTGTCAGCAATTTCTCCAGTAAAAAGCTTCAGGAATTGATGGCGATAGCCAACATCCCTCCAGTTGTAAATCAAGTGGAGATGAGCCCCGTTTTCCAACAAAAAAAATTGAGAAACTATTGCAAGGCCAACAATATATTGATCACTGCATACTCAGTTTTAGGAGCTAAGGGAACCGCATGGGGTTCCAATGCAGTTCTGGGTTCTGAGGTGCTTAACCAGATTGCTACGGCCAGAGGAAAATCTATTGCTCAGGTAGTTTGAGATGGGTATACGAACAAGGTGCGAGTCTTGTGGTGAAAAGTTTCGATGAAGACAGGATGAGGGAAAACTTGAACATATTCGACTGGGAACTGACTAAAGAAGACTCGGAAAAGATTGGTGTCATTCCACAGCGTAAAATAATAACTGAGGATTTTATGGTATCAGCTAATGGACCTTTCAAATCTATAGAAGAGTTCTGGGGTGATGATATATAA

>Pbr_COR-Lc13 Pbr_COR_contig_13678_CDS

ATGGAGATTAGTGGTGTACCGGTAATCACTCTGAGCTCAGGCAAGGAGATGCCTGTTTTAGGTATGGGAACATTTGAATCATTTGGTAGTAAAGGGTCTGAAAGAGAGAAGTTGGCGTATTTGAAAGCCATAGAGGTGGGTTACAGATACTTTGATACAGGTGCCGCATACGGAACTGAAGAGGTTCTTGGCGAAGCTATAGCTGAAGCACTTCAACTCGGCCTAATCAAATCTCGAGATGAACTCTTCGTCAGTTCCATGCTCTGGTGCACCGATGCTCACGCTGATCGTGTCCTACTCGCTCTTCAGAATTCTTTAAGGAATCTTAAGTTGGAATACCTGGACCTATACATGTTACCTTTTCCGGCAAGTATGAAACCAGCAGGAGAGAAGATAACTATGGTCGACTTCCACGACATAACAGAAGAGGATGAGATTTTTGCAATGGACTACAAGTCTGTATGGGCAGCCATGGAAGAGTGTCAGAACCTCGGCTTGACTAAGTCGATCGGTGTCAGTAATTTCTCCAGCAAAGAGCTCCAGGAATTGATGGTCAACGCCAAAATCCCTCCGTCTGTGAATTCTGTGGAGATGAGCCCGGCTTTCCAACAAAAGAATCTGAGAGAGTATTGCAAGGCCAATAATATATTGATCATTGCCATCTCTATTTTGGGATCCACCGGAACCGAATGGGGCTCCAATGCAGTTATGGGTTCCGAGGTGCTTAAACAAATTGCTATGGCCAGAGGAAAATCAACTGCACAGGTTAGTATGAGATGGGTTTATGAGCAAGGTGCAATTCTTGTAGTAAAAAGTTTCAGTGAAAAGAATATGAGGGCAAACTTGAAGATATTTGACTGGGAACTCACCAAGGAAGACGGAGAAAAGATCGGCACCGGGATTCCACAGTGCAGAACCCTAAAAGGTGATTTTATGGTATCACCTAATGGTCCTTTCAAAACTCTACAAGACTTATGGGATGAATGA

>Pbr_COR-Lc12 Pbr_COR_contig_5136_CDS

ATGGAGAGTAGTGGTGTACCTGAAATCACTCTGAGCTCGGGCAAGAAGATGCCTGTTTTAGGTATGGGTACATTTGAAACATTTGGTAAAGGGTCTGAAAGAGAGAGGTCGGCGTTTTTGAAAGCCATAGCGGTGGGATACAGATACTTCGATACAGGTGCTATGTACGGAACTGAGGAGGTTCTTGGCGAAGCTGTAGCTGAAGCACTTCAGATTGGTCTAATCAAATCTCGAGATGAACTCTTTATCAGTTCCATGCTCTGGGGCACTGATGCTTACTCTGATCGAGTCCTCCTCGCTCTTCAGAATTCTCTGAGGAATCTTAAATTGGAGTATCTGGATCTATACTTGATACCCTTTCCGGCAAGCTTGAAGCCAGGAGCTAGGGATATAACTACGGACGTACGTCACTTACCAGCGGATGAAATCCTTCCGATGGACTACAGGTCTGTATGGGCAGCCATGGAAGAGTGTCAGAACCTTGGCTTAACTAAGTCAATCGGTGTCAGCAATTTTTCCTGCAAAAAGCTTCAGGTATTGATGGCCACCGCCAAAATCCCTCCAGCTGTGAATCAAGTGGAGATGAGCCCGGTTTTCCAACAAAAGGATCTGAAAGAGTATTGCAAGGCCAATAATATATTGATCATTGCATACTCCGTTTTGGGATCCAACCGAACCGAATGGGGCTCCAACGCAGTTATGGGTTCTGAGGTGCTTAAACAAATTGCTATGGCCAGAGGAAAATCTATTGCTCAGGTTAGTATGAGATGGGTTTACGAGCAAGGTGCGATGCTTGTGGTAAAAAGTTTCAGTGAAGAGAGAATGAGAGAAAACTTGAACATATTCGACTGGAAACTAACTGAGGAAGACTTGGAAAAGATCGGTCATGCGATTCCACAGTGCAGAGTCTTACCTGCTGATTTTATGATAACAGCTAATGGACCTTTCAAATCTCTGGAAGAGTTTTGGGATGACAAAGCTTGA

Supplementary dataset 8 (associated with Figure 3c)

- Nucleotide sequences of genomic segments for *CYP82Y2*-*L*&*COR*-*L* gene pairs. Nucleotides of introns and intergenic regions are represented in lower case, exons in capital letters.

>Pca_STORR_genomic

ATGGAGCTCCAGTACTTTTCTTATTTTCAGCCAACTTCCTCCCTTTTTGTTCTTGTATCCATATTACTGATCAGTGCTGTTGTTTTGAGGAAGATATTTTTAAATAACTACTCTTCATCACCTGCATCATCAACAGAAACAGCGCTGCTTTCTCATCAGCGGCGGTCGTCATGTGCATTGCCAATCATCGGTCTTCTTCATGTTTTCATGAATAAAAACGGCTTAATTCATGTAACTCTTGGAAATATGGCTGATAAATACGGACCGATTTTCAGTTTCCCAACAGGCAGCCATATGACTCTAGTTGTGAGCAGTTGGGAGATGGCAAAAGAGTGTTTTACAGGCAACAATGACATTGCTTTCTCAAACCGTCCTATCCCGTTAGTTTTTAAGACTATATTCTATGCATGCCGTGGTATTGACTCATACGGTCTTTCAAGTGTACCTTATGGAAAATATTGGAGGGAGTTACGAAAGGTGTGTGTGCATAACCTCCTCTCTAATCAACAACTACTCAACTTCAGACACTTGATAATTTCTCAAGTTGATACCTCTTTCAATAAGCTTTATGAGTTATGCAAAAACTCTGCAGACAACCAGGGAAACTACACTACTACTACTACCGCAGCTGGCATGGTGAGGATGGATGATTGGCTAGCGCAATTATCGTTCAACGTGATAGGAAGAATAGTCTGTGGATTCCAATCAGACACCGAGACAA

gtgagcattttcctacagtttatgttaaaaatgattgattttataggttgagatcaaagggtatacatatatatcttattgcgtgcacatgcataaacag

GTGCTTCAAGCAGTGCGGAACAACTTACAGAAGCAATTAATGACGCATCGTATTTTATGTCGACATCTCCGGTGTCAGATAACGTTCCATGGTTAGGGTGGGTTGATCAACTAACTGGTCTTACTAGAAATATGAAGCAATGCGGAAAGAAATTAGACTTGGTGCTCGAGAGCATAATTGAAGATCATCGTCAAAAGAGACGGTTTTCTAGAAGTAAACAAGGAGATGAGAATTTTATCAACGAGGATGAACAAGATGACTTCATCGACATTTGCCTGTCAATTATGGAGCAACCACAGCTTCCTGGCAACAATAACCCTTGTCAAATACCTATCAAATCTATTGTCCTG

gtatgcaatcgtagtaggatgacaaaaccaaaggctagacaccgattagtagaatcttatccattgcattcctgcattattaattggttgaagtggtagatcaaagagtgtacaaacatgatctgtttttgatgggttttcttgactattgttgtgttctgactctaatcatgtatatatgcag

GACATGATAGGTGGGGGGACCGACACCACAAAACTGACCACCATCTGGACCCTTTCATTGCTACTGAACAACCCCCATGTGTTGGACAAGGCAAAACAAGAAGTGGATGCACACTTCGGGACGAAAAGGAACGATGCAGGAGGAGTCGTGGTGGATTTTGATGATACTCGTAACCTTGTCTACATCCAGGCAATTATCAAAGAATCAATGCGGTTGTATCCAGCCAGCCCCTTGGTGGAGCGACTGAGCGGCGAAGATTGTGTGGTCGGTGGGTTTCATGTACCAGCAGGGACGCGGTTATGGGCTAACGTGTGGAAGATGCATCGAGACCCAAAAGTATGGGATGATCCATTGGTGTTTCGACCAGAGAGATTTTTGCTCGATGACCAGAAGATGGTTGATGTTAGGGGTCAAAACTATGAGCTGTTACCATTCGGAGCCGGTCGACGGATATGTCCAGGTGTATCGTTCTCTTTGGATCTAATGCAACTGGTACTGACTCGTCTTATTCTTGAGTTTGAAATGAAGTCTCCTAGCGGGAAACTAGACATGACAGCGACACCAGGTTTAATGAGTTACAAGGTAGTCCCCCTTGACATTCTGCTCACCAGTCGTCGCATGAAGTGGTGTGTGCAGTTAGCCTCTGAGAGAGATAAAAAGATGGAGAGTAGTGTTGTACCTGTAATCACTCTGAGCTCGTGCAAGGTGATGCCTGTTTTAGGCATGGGAACATTTGAAAAAGTTGGTAAAGGGTCTGAAAGAGAGAGGTTGGCGATTTTAAAAGCGATAGAGGTAGGTTACAGATACTTCGATACAGCTGCTGCATACGAAACTGAAGAGGTTCTTGGTGAAGCTATTGCTGAAGCACTTCAACTTGGCCTAATCAAATCTCGAGATGAACTCTTCATCAGTTCCATGCTCTGGTGCACTGATGCCTACCATGATCGAGTCCTCCCCGCTCTTCAGAATTCTCTGAG

gtaaaatttttgatgaaaattttgaacccaattaggcttatatattcttaaacatctatcaaattttagataaatcaaattgttttgtgtgatatag

GAATCTTAAATTGGAGTATCTGGATCTATACATGTTACCCTTCCCGGTAAGCTTGAAGCCAGGAGGGAAGATAAGTATGGACATACCAGAGGATGAAATTTTTACGATGGACTATAGGTCTGTATGGGCAGCCATGGAAGAATGTCAGAACCTTGGCTTGACAAAATCAATCGGTGTCAGCAATTTCTCCTGCAAAAAGCTTCAGGAATTGATGGCGACCGCCAACATCCCTCCAGCTGTGAATCAA

gtgagtacttttacttgaaagttttgataaaaattagtgttttccttttccggtgctaattaaatttccaacatttgacaccttcacatgcataaaaatactacgtcttatag

GTGGAGATGAGCCCGGCTTTCCAACAAAAGAATTTGAGAGAGTATTGCAATGCCAATAATATATTAATCAGTGCAGTTTCTATTTTGGGATCAAACGGAACCCCATGGGGCTCCAATGCAGTTCTGGGTAATGAGGTGCTAAACCAGATTGCTATGGCCAGAGGAAAATCTGTTGCTCAG

gttggtcaaattctctgcacatgccgtagagcagtctctattctttggccatcgtatttgcatcacattataccgaatttaacttgcatttatattacgtacgtactgatcag

GTTAGTATGAGGTGGGTTTACGAGCAAGGTGCGAGTCTTGTGGTAAAAAGTTTTAGTGAAGAGAGATTGAGGGAAAACTTGAACATATTCGACTGGGAACTCACTAAGGAAGACTTGGAAAAGATGGGTGAGATTCCACAATGCAGAATCTTGACTGCTGATTTTATGGTATCAGCTAATGGACCTTTCAAATCTCAAGAAGAGTTGTGGGATGACAAAGCTTGA

>Par_STORR1_genomic

ATGGAGCTCCAATATTTTTCTTATTTTCAACCAACTTCCTCCGTTGTTGCTCTTCTACTTGCTCTTGTATCCATCTTATTCAGTTTCGTTGTTTTGAGGAAGACATTTTCGAATAACTACTCATCACCTGCATCATCAACAGAAACAGCGGTACTTTGTCATCAGCGGCAGCAGTCGTGTGCATTGCCAATTTCCGGTCTTCTTCATGTTTTCATGAATAAAAACGGCTTAATTCATGTAACTCTTGGAAATATGGCCGATATATACGGACCGATTTTCAGTTTCCCAACAGGTAGCCATAGAACTCTCGTTGTGAGCAGTTGGGAGATGGTAAAAGAGTGTTTTACAGGCAACAATGACACTGCTTTCTCAAACCGTCCTATCCCGTTAGCTTTTAAGACTATATTCTATGCATGCGGTGGCATAGACTCATACGGTCTTTCGAGTGTACCTTATGGAAAATATTGGAGGGAGCTACGAAAAGTCTGTGTGCATAACCTCCTCTCTAATCAACAACTACTCAAGTTCAGACACTTGATAATTTCTCAAGTTGATACGTCTTTCAATAAGCTGTATGAGTTATGCAAAAACTCTGAAGACAACCAGGGAAACCATACTACTACTACTACTACTACCGCAGCTGGCATGGTGAGAATGGATGATTGGCTCGCCCAATTATCGTTCAACGTGATAGGAAGAATAGTCTGCGGATTCCAATCAGACCCTAAGACAG

gtgagcattttcctagagtttctgtagaaaatgttggattttacagagccgatcctgagtgtagaagggccctgtgcgaacgtgaccgtagggccttcatagtgtcaaattatttttttcatgataattttttttcaactaataaatcagtatttttaagttaaaacttgacttatgcagcacatgagtccatctctagtccttaacgtatatgcactcaacgagtatgcgaaaactgtttttgaacattattatatatggtttgcattcataagataactagttcttttttaaattctagatgctaaattgcattctgaatagtgaattaaagtttagatgtcagttttcgtacctaaatataaaaattaagtataacgttaatctcacatttcgtaactaaaaaggttttcaaatttaaactaagttctatattatattctaaaaagagttcaaatttaaactatgtattatactgactaaaattataatcaggagcatttgaattatatacaccctaagagcaactgcagtggtgcgatcatataccaaagatcaaagagcaaaaaaatttgaaaatttgggtttagtccgtgttgcaacgcagtgggggagactaaaatttggtcaggcgtaagaggaagttacgcctgaatcaggcgtaagaggaggttccgccccattgtcatcaggcgtaactttgaggttacgccttaaattttttttttttttttttttttttttttaatggggcgtaagtttgagctacgccccatttcaggcgcaactttaagttgcgccccatttttttttgttttgttttttagtgtgaggcggagttttttgttacgcctgttcgggcgtaacgcaccaattgcgcccgactaaagaccagactaaagactaaatatggtctgggttttggttatagtctggtctttggtctatactccttccccactacgtctgctatttgatctttagtatgggttttagtcgcccactacggatgctctaagcctttggtctctcgaggttgaatctctctctctgggtttgttttgtcaaaattttaattttgcaaatgaaatgggctttggtattaaggtgtccactagtaggaaaatgtccctaaccagagctaggccctgtgcggtcgcaccccctgcacaccctcaggcccggctctgtcagcatcgattttatactatgagatcatcaaagactattcttctaactgtgcacatgtataaacag

GTGCTCCAAGCAGGGTGGAACAATTTAAAGAAGCAATTAATGAAGCATCGCATTTTATGTCGACATCTCCAGTGTCAGATAACGTTCCAATGCTAGGGTGGATTGACCAATTGACTGGTCTTACAAGAAATATGAAGCACTGCGGAAAGAAATTAGACTTGGTGGTCGAGAGCATAATTCAAGATCATCGTCGAAAGAGACGATTCTCTAGAACTAAAGGAGGAGATGAGAAGGACGATGAACAAGATGACTTCATCGACATTTGCTTGTCAATTATGGAGCAACCACAGCTTCCTGACAACAATAACCCTCCTCATATACCTATCAAATCTATTGTCCTG

gtatgcaaatattgatccgtggtatggctggagcattaattgtattccagcggttaattttggttgtggtagctcatagagtgcagaaacatgatttggttttgattggtatcattgtataaatgcag

GACATGATAGGTGGGGGAACTGACACAACAAAACTGACCACCATCTGGACCCTTTCCTTGCTGCTGAACAACCCCCATGTGTTGGACAAGGCAAAACAAGAAGTGGATGCACACTTTGGAACGAAAAGGAGATCAACAAATGATGCCGTGGTGGATTTTGATGATATTCGTAACCTTGTATACATCCAGGCAATCATCAAAGAATCAATGCGGTTGTATCCAGCCAGCCCCGTGGTGGAGCGACTGAGCGGCCAAGATTGTGTGGTCGGTGGGTTTCATGTACCAGCAGGGACGAGATTATGGGCTAACGTATGGAAGATGCAACGAGACCCTAAAGTATGGGATGATCCATTGGTGTTTCGACCAGAGAGATTTTTGAGCGATGAACAGAAGATGGTTGATGTAAGGGGTCAAAATTATGAGCTGTTACCATTTGGAGCCGGTCGACGTATATGTCCAGGTGTATCCTTCTCTTTGGATCTAATGCAACTGGTACTGACTCGTCTTATTCTTGAGTTTGAAATGAAGTCTCCTAGCGGGAAAGTGGACATGACAGCAACACCAGGATTAATGAGTTACAAGGTGGTCCCCCTTGACATTCTGCTCACCCATCGTGGCACAAAGTCGTGTGTGCAGTCAGCAGCCTCTGAGACTAGTGGTGTACCAGTAATCACTCTGAGCTCGGGCAAGGTGATGCCTGTTTTAGGCATGGGTACATTTGAGAAAGTTGGTAAAGGGTCTGAAAGAGAGAGGTTGGCGATTTTAAAAGCGATAGAGGTGGGTTACAGATACTTCGATACAGCTGCTGCATACGAAACTGAAGAGGTTCTTGGAGAAGCTATTGCTGAAGCACTCCAACTTGGCCTAATCAAATCTCGAGATGAACTTTTCATCAGTTCCATGCTCTGGTGCACTGATGCTCACCCTGATCGTGTCCTCCTCGCTCTTCAGAATTCTCTGAG

gtaaatttttttgatgcaaattaggcttacatgtatttgcacataccagcaggtacacttaaacatctatttttagataaatgaaattgtgttgtgtgatatgtacatag

GAATCTTAAATTGGAGTATCTGGATCTATATATGTTACCCTTCCCGGCAAGCTTGAAGCCAGGGAAGATAACGATGGACATACCAGAGGAAGATATTTGTCGAATGGACTACAGGTCTGTATGGTCAGCCATGGAAGAGTGTCAGAACCTTGGCTTCACTAAATCAATCGGTGTTAGCAATTTCTCCTGCAAAAAGCTTCAGGAATTGATGGAGACCGCCAACATCCCTCCAGCTGTGAATCAA

gtgagtactcttacttgaaagttttgataagacatcggaagatttctaatgcttattaaatttccaacattatgcataaaactactactctttataatgttgattaag

GTGGAGATGAGCCCGGCTTTCCAACAAAAGAAGCTGAGAGAGTATTGCAACGCAAATAATATATTAGTCAGTGCAGTCTCTATACTGGGATCAAACGGAACCCCATGGGGCTCCAATGCAGTTATGGGTTCTGAGGTGCTTAAGAAAATTGCTATGGCCAAAGGAAAATCTGTTGCTCAG

gttggtcaaattctctgcatatgccatagtgtggtctctgttctttggccatcgtatttgcatcacattgtattccttccttatacggaatgtaacttacatttgtattacgtactgatcag

GTTAGTATGAGATGGGTTTACGAGCAAGGCGCGAGTCTTGTGGTAAAAAGTTTCAGTGAAGAGAGATTGAGGGAAAACTTGAACATATTCGACTGGGAACTCACTAAGGAAGACAATGAAAAGATCGGTGAGATTCCACAGTGCAGAATCTTGACTGCTCATTTTTTGGTCTCACCTAATGGACCTTTCAAATCTCAAGAAGAGTTGTGGGATGACAAAGCTTGA

>Par_STORR2_genomic

ATGGAGCTCCAATATTTTTCTTATTTTCAACCAACTTCCTCCGTTGTTGCTCTTCTACTTGCTCTTGTATCCATCTTATTCAGTTTCGTTGTTTTGAGGAAGACATTTTCGAATAACTACTCATCACCTGCATCATCAACAGAAACAGCGGTACTTTGTCATCAGCGGCAGCAGTCGTGTGCATTGCCAATTTCCGGTCTTCTTCATGTTTTCATGAATACAAACGGCTTAATTCATGTAACTCTTGGAAATATGGCTGATATATACGGACCGATTTTCAGTTTCCCAACAGGTAGCCATAGAACTCTCGTTGTGAGCAGTTGGGAGATGGTAAAAGAGTGTTTTACAGGCAACAATGACACTGCTTTCTCAAACCGTCCTATCCCGTTAGCTTTTAAGACTATATTCTATGCATGCGGTGGCATAGACTCATACGGTCTTTCGAGTGTACCTTATGGAAAATATTGGAGGGAGCTACGAAAAGTCTGTGTGCATAACCTCCTCTCTAATCAACAACTACTCAAGTTCAGACACTTGATAATTTCTCAAGTTGATACGTCTTTCAATAAGCTGTATGAGTTATGCAAAAACTCTGAAGACAACCAGGGAAACCATACTACTACTACCGCAGCTGGCATGGTGAGAATGGATGATTGGCTCGCCCAATTATCGTTCAACGTGATAGGAAGAATAGTCTGCGGATTCCAATCAGACCCTAAGACAG

gtgagcattttcctagagtttctgtagaaaatgttggattttatactatgagatcatcaaagtctaactgtgcacatgtataaacag

GTGCTCCAAGCAGGGTGGAACAATTTAAAGAAGCAATTAATGAAGCATCGCATTTTATGTCGACATCTCCAGTGTCAGATAACGTTCCAATGCTAGGGTGGATTGACCAATTGACTGGTCTTACAAGAAATATGAAGCACTGCGGAAAGAAATTAGACTTGGTGGTCGAGAGTATAATTCAAGATCATCGTCAAAAGAGACGATTCTCTAGAACTAAAGGAGGAGATGAGAAGGACGATGAACAAGATGACTTCATCGACATTTGCTTGTCAATTATGGTGCAACCACAGCTTCCTGACAACAATAACCCTCCTCATATACCTATCAAATCTATTGTCCTG

gtatgcaaatattgatccgtggtatggctggagcattaattgtattccagcggttaattttggttgtggtagctcatagagtgcagaaacatgatttggttttgattggtatcattgtataaatgcag

GACATGATAGGTGGGGGAACTGACACAACAAAACTGACCACCATCTGGACCCTTTCCTTGCTGCTGAACAACCCCCATGTGTTGGACAAGGCAAAACAAGAAGTGGATGCACACTTTGGAACGAAAAGGAGATCAACGAATGATGCCGTGGTGGATTTTGATGATATTCGTAACCTTGTATACATCCAGGCAATCATCAAAGAATCAATGCGGTTGTATCCAGCCAGCCCCGTGGTGGAGCGACTGAGCGGCCAAGATTGTGTGGTCGGTGGGTTTCATGTACCAGCAGGGACGAGATTATGGGCTAACGTATGGAAGATGCAACGAGACCCTAAAGTATGGGATGATCCATTGGTGTTTCGACCAGAGAGATTTTTGAGCGATGAACAGAAGATGGTTGATGTAAGGGGTCAAAATTATGAGCTGTTACCATTTGGAGCCGGTCGACGTATATGTCCAGGTGTATCCTTCTCTTTGGATCTAATGCAACTGGTACTGACTCGTCTTATTCTTGAGTTTGAAATGAAGTCTCCTAGCGGGAAAGTAGACATGACAGCAACACCAGGATTAATGAGTTACAAGGTGGTCCCCCTTGACATTCTGCTCACCCATCGTGGCACAAAGTCGTGTGTGCAGTCAGCAGCCTCTGAGAGTAGTGGTGTACCAGTAATCACTCTGAGCTCGGGCAAGGTGATGCCTGTTTTAGGCATGGGAACATTTGAGAAAGTTGGTAAAGGGTCTGAAAGAGAGAGGTTGGCGATTTTAAAAGCGATAGAGGTGGGTTACAGATACTTCGATACAGCTGCTGCATACGAAACTGAAGAGGTTCTTGGAGAAGCTATTGCTGAAGCACTCCAACTTGGCCTAATCAAATCTCGAGATGAACTTTTCATCAGTTCCATGCTCTGGTGCACTGATGCTCACCCTGATCGTGTCCTCCTCGCTCTTCAGAATTCTCTGAG

gtaaatttttttgatggaaattaggcttacatatatttgcacataccagcaggtacacttaaacatctatttttagataaatgaaattgtgttgtgtgatatgtacatag

GAATCTTAAATTGGAGTATCTGGATCTATATATGTTACCCTTCCCGGCAAGCTTGAAGCCAGGGAAGATAACGATGGACATACCAGAGGAAGATATTTGTCGAATGGACTACAGGTCTGTATGGTCAGCCATGGAAGAGTGTCAGAACCTTGGCTTCACTAAATCAATCGGTGTTAGCAATTTCTCCTGCAAAAAGCTTCAGGAATTGATGGAGACCGCCAACATCCCTCCAGCTGTGAATCAA

gtgagtactcttacttgaaagttttgataagacatcggaagatttctaatgcttattaaatttccaacattatgcataaaactactactctttataatgttgattaag

GTGGAGATGAGCCCGGCTTTCCAACAAAAGAAGCTGAGAGAGTATTGCAACGCAAATAATATATTAGTCAGTGCAGTCTCTATACTGGGATCAAACGGAACCCCATGGGGCTCCAATGCAGTTTTGGGTTCTGAGGTGCTTAACAAAATTGCTATGGCCAAAGGAAAATCTGTTGCTCAG

gttggtcaaattctctgcatatgccatagtgtggtctctgttctttggccatcgtatttgcatcacattgtattccttccttatacggaatgtaacttacatttgtattacgtattgatcag

GTTAGTATGAGATGGGTTTACGAGCAAGGCGCGAGTCTTGTGGTAAAAAGTTTCAGTGAAGAGAGATTGAGGGAAAACTTGAACATATTCGACTGGGAACTCACTAAGGAAGACAATGAAAAGATCGGTGAGATTCCACAGTGCAGAATCTTGGCTGCTCATTTTTTGGTCTCACCTAATGGACCTTTCAAATCTCAAGAAGAGTTGTGGGATGACAAAGCTTGA

>Pbr_STORR_genomic

ATGGAGCTCCAATATTTTTCTTATTTTCAACCAACTTCCTCCGTTGTTGCTCTTCTACTTGCTCTTGTATCCATCTTATTCAGTGTCGTTGTTTTGAGGAAGACATTTTCGAATAACTACTCATCACCTGCATCATCAACAGAAACAGCGGTACTTTGTCATCAGCGGCAGCAGTCGTGTGCATTGCCAATTTCCGGTCTTCTTCATGTTTTCATGAATAAAAACGGCTTAATTCATGTAACTCTTGGAAATATGGCGGATAAATACGGACCGATTTTCAGTTTCCCAACAGGTAGCCATAGAACTCTCGTTGTGAGCAGTTGGGAGATGGTAAAAGAGTGTTTTACAGGCAACAATGACACTGCTTTCTCAAACCGTCCTATCCCGTTAGCTTTTCAGACTATATTCTATGCATGCGGTGGCATAGACTCATACGGTCTTTCGAGTGTACCTTATGGAAAATATTGGAGGGAGCTACGAAAGGTCTGTGTGCATAACCTCCTCTCTAATCAACAACTACTCAAGTTCAGACACTTGATAATTTCTCAAGTCGATACGTCTTTCAATAAGCTGTATGAGTTATGCAAAAACTCTGAAGACAACCAGGGAATGGTGAGAATGGATGATTGGCTCGCCCAATTATCGTTCAATGTGATAGGAAGAATAGTCTGCGGATTCCAATCAGACCCAAAGACAG

gtgagcatttgcctagagtttctgtagaaaatgatggattttatactattagatcatcaaagactattctgtatatcttaatcaagcatacatatcttctaacagtgcacatgtataaacag

GTGCTCCTAGCAGGGTGGAACAATTTAAAGAAGTAATTAATGAAGCATCTTATTTTATGTCGACATCTCCAGTGTCAGATAACGTTCCAATGCTAGGGTGGATTGACCAATTGACTGGTCTTACGAAAAATATGAAGCACTGCGGAAAGAAATTAGACTTGGTGGTAGAGAGCATAATTAAAGATCATCGTCAAAAGAGACGATTCTCTAGAACTAAAGGAGGAGATGAGAAGGACGATGAACAAGATGACTTCATCGACATTTGCTTGTCAATTATGGAGCAACCACAGCTTCCTGGCAACAATAGCCCTCCTCAGATACCTATCAAATCTATTGTCCTG

gtatgcaatattgatccgtagtatgatgacaaaacaaaaggctggagcattaattgtattccagcgattaattggttgcggtagctcatagagtgcagaagcatgatttggttttgattggtattattgtataaatgcag

GACATGATAGGTGGGGGCACTGACACCACAAAACTGACCACCATCTGGACCCTTTCCTTGCTGCTGAACAACCCCCATGTGTTGGACAAGGCAAAACAAGAAGTGGATGCACACTTTCGAAAGAAAAGGAGATCAACAGATGATGCAGCAGCAGCCGTGGTGGATTTTGATGATATTCGTAACCTTGTCTACATCCAGGCAATCATCAAAGAATCAATGCGGTTGTATCCAGCCAGCCCCGTGGTGGAGCGACTGAGCGGCGAGGATTGTGTGGTCGGTGGGTTTCATGTACCAGCAGGGACGAGATTATGGGCTAACGTATGGAAGATTCAACGAGACCCTAAAGTATGGGATGATCCATTGGTGTTTCGACCAGAGAGATTTTTGAGCGATGAACAGAAGATGGTTGATGTAAGGGGTCAAAATTATGAGCTGTTACCATTTGGAGCCGGTCGACGTATATGTCCAGGTGTTTCCTTCTCTTTGGATCTAATGCAACTGGTACTGACTCGTCTTATTCTTGAGTTTGAAATGAAGTCCCCTAGCGGGAAAGTGGACATGACAGCAACACCAGGATTAATGAGTTACAAGGTGGTCCCTCTTGACATTCTGCTCACCCATCGGCGCATAAAGTCGTGTGTGCAGTTAGCATCCTCTGAGAGAGACATGGAGAGTAGTGGTGTACCAGTAATCACTCTGAGCTCGGGCAAGGTGATGCCTGTTTTAGGCATGGGAACATTTGAGAAAGTTGGTAAAGGGTCTGAAAGAGAGAGGTTGGCGATTTTAAAAGCCATAGAGGTGGGTTACAGATACTTCGACACAGCTGCTGCATACGAAACTGAAGAGGTTCTTGGAGAAGCTATTGCTGAAGCACTTCAAATTGGCCTAATCAAATCTCGAGATGAACTTTTCATCAGTTCCATGCTCTGGTGCACTGATGCTCACCCTGATCGTGTCCTCCTCGCTCTTCAGAATTCTCTGAG

gcaagtctttttgatgcaaattaggcttatatatatttgcacataccagcaggtacacttgaacatctattttcagataaatgaatttgttttgtgtgatatgtatag

GAATCTTAAATTGGAGTATCTGGATCTATATATGTTACCCTTCCCGGCAAGCTTGAAGCCAGGGAAGATAACGATGGACATACCAGAGGAAGATATTTGTCGAATGGACTACAGGTCTGTATGGTCAGCCATGGAAGAGTGTCAGAACCTTGGCTTCACTAAATCAATCGGTGTTAGCAATTTCTCCTGCAAAAAGCTTCAGGAATTGATGGCGACCGCCAACATCCCTCCAGCCGTGAATCAA

gtgagtactcttacttgaaagttttgataacacatcggaagatttctaatgcttattaaatttccaacattatgcataaaactactactctttataatgttgattaag

GTGGAGATGAGCCCGGCTTTCCAACAAAAGAAGCTGAGAGAGTATTGCAACGCAAATAATATATTAGTCAGTGCAGTCTCTATACTGGGATCAAACGGAACCCCATGGGGCTCCAATGCAGTTTTGGGTTCTGAGGTGCTTAAGCAAATTGCTATGGCCAAAGGAAAATCTGTTGCTCAG

gttggtcaaattctctgcatatgccatagtgtggtctctgttctttggccatcgtatttgcatcacattgtattccttccttatacggaatgtagcttacatttgtattacgtactgatcag

GTTAGTATGAGATGGGTTTACGAGCAAGGCGCGAGTCTTGTGGTAAAAAGTTTCAGTGAAGAGAGATTGAGGGAAAACTTGAACATATTCGACTGGGAACTCACTAAGGAAGACAATGAAAAGATCGGTGAGATTCCACAGTGCAGAATCTTGACTGCTTATTTTTTAGTCTCGCCTAATGGACCTTTCAAATCTCAAGAAGAGTTGTGGGATGACAAAGCTTGA

>Pso_STORR-PS1126560_genomic HN1 Chr11 PS1126560.1

ATGGAGCTCCAATATATTTCTTATTTTCAACCAACTTCCTCCGTTGTTGCTCTTCTACTTGCTCTTGTATCCATCTTATCCAGTGTCGTTGTTTTGAGGAAGACATTTTTGAATAACTACTCATCATCACCTGCATCATCCACAAAAACAGCGGTACTTTCTCATCAGCGGCAGCAGTCGTGTGCATTGCCAATTTCCGGTCTCCTCCATATTTTCATGAATAAAAACGGCTTAATTCATGTAACTCTTGGAAATATGGCTGATAAATACGGTCCGATTTTCAGTTTCCCAACAGGTAGCCATAGAACTCTCGTTGTGAGCAGTTGGGAGATGGTAAAAGAGTGTTTTACTGGCAACAATGACACTGCTTTCTCAAACCGTCCTATCCCGTTAGCTTTTAAGACTATATTTTATGCATGCGGTGGCATAGACTCATACGGTCTTTCGAGTGTACCTTATGGAAAATATTGGAGGGAGCTTCGAAAGGTCTGTGTGCATAACCTCCTGTCTAATCAACAACTACTCAAGTTCAGACACTTGATAATTTCTCAAGTCGATACGTCTTTCAATAAGCTGTATGAGTTATGCAAAAACTCTGAAGACAACCAGGGAAACTATACTACTACTACTACCGCAGCTGGCATGGTGAGAATCGATGATTGGCTCGCCGAATTATCGTTCAATGTGATAGGAAGAATAGTCTGCGGATTCCAATCAGGCCCTAAGACAG

gtgagcatttgcctagagtttctgtagaaaatgatggattttatattatgagatcatcaaagactattctgtatatcttaatcaagcatacatatcttctaacagtgcacatgtataaacag

GTGCTCCAAGCAGGGTGGAACAATTTAAAGAAGCAATTAATGAAGCATCTTATTTTATGTCGACATCTCCAGTGTCAGATAATGTTCCAATGCTAGGGTGGATTGACCAATTGACAGGTCTTACGAGAAATATGAAGCACTGCGGAAAGAAATTAGACTTGGTGGTCGAGAGCATAATTAATGATCATCGTCAAAAGAGACGATTCTCTAGAACTAAAGGAGGAGATGAGAAGGACGATGAACAAGATGACTTCATCGACATTTGTTTGTCAATAATGGAGCAACCACAGCTTCCTGGCAACAATAATCCTTCTCAGATACCTATCAAATCTATTGTCCTG

gtatgtaatattgatccgtagtatgatgaccaaacaaaaggctggagcattaattgtattccagcgattaattggttgcggtagctcatagagtgcagaaacatgattttgttttgattggtattattgtataaatgcag

GACATGATAGGTGGGGGCACTGACACCACAAAACTGACCACCATCTGGACCCTTTCCTTGCTGCTGAACAACCCCCATGTGTTGGACAAGGCAAAACAAGAAGTGGATGCACACTTTCGAACCAAAAGGAGATCAACAAATGATGCAGCAGCAGCCGTGGTGGATTTTGATGATATTCGTAACCTTGTCTACATCCAGGCAATCATCAAAGAATCAATGCGGTTGTATCCAGCCAGCCCCGTGGTGGAGCGACTGAGCGGCGAAGATTGTGTGGTCGGTGGGTTTCATGTACCAGCAGGGACGAGATTATGGGCTAACGTATGGAAGATGCAACGAGACCCTAAAGTATGGGATGATCCATTGGTGTTTCGACCAGACAGATTTTTGAGCGATGAACAGAAGATGGTTGATGTAAGGGGTCAAAATTATGAGCTGTTACCATTTGGAGCCGGTCGACGTGTATGTCCAGGTGTATCCTTCTCTTTGGATCTAATGCAACTGGTACTGACTCGTCTTATTCTCGAGTTTGAAATGAAGTCTCCTAGCGGGAAAGTGGACATGACAGCAACACCAGGATTAATGAGTTACAAGGTGATCCCCCTTGACATTCTGCTCACCCATCGTCGCATAAAGCCGTGTGTGCAGTCAGCAGCCTCTGAGAGAGACATGGAGAGTAGTGGTGTACCAGTAATCACTCTGGGCTCGGGCAAGGTGATGCCTGTTCTGGGCATGGGAACATTTGAGAAAGTTGGTAAAGGGTCCGAAAGAGAGAGGTTGGCGATTTTAAAAGCGATAGAGGTGGGTTACAGATACTTCGATACAGCTGCTGCATACGAAACTGAAGAGGTTCTTGGAGAAGCTATTGCTGAAGCACTTCAACTTGGCCTAGTCAAATCTCGAGATGAACTTTTCATCAGTTCCATGCTCTGGTGCACTGATGCTCACGCTGATCGTGTCCTCCTCGCTCTTCAGAATTCGCTGAG

gtaaattttttgatgcaaattaagcttatatatatatatatgtgcaaagcctttgcacataccagtaggtacacttaaaacatctatcatattttagataaatgaaattgttttgtgtgatgatatag

GAATCTTAAATTGGAGTATGTGGATCTATATATGTTACCCTTCCCGGCAAGCTTGAAGCCTGGGAAGATAACGATGGACATACCAGAGGAAGATATTTGTCGCATGGACTACAGGTCTGTATGGGCAGCCATGGAAGAGTGTCAAAACCTTGGCTTCACTAAATCAATCGGTGTTAGCAATTTCTCCTGCAAAAAGCTTCAGGAATTGATGGCGACCGCCAACATCCCTCCAGCTGTGAATCAA

gtgagtactcttacttgaaagttttgataaaacatcggaagacttctaatgcttattaaatttctaacattatgcataaaactactgctctttataatgttgattaag

GTGGAGATGAGCCCGGCTTTCCAACAAAAGAAGCTGAGAGAGTATTGCAACGCAAATAATATATTAGTCAGTGCAATCTCTGTACTGGGATCAAACGGAACCCCATGGGGCTCCAATGCAGTTTTGGGTTCTGAGGTGCTTAAGAAAATTGCTATGGCCAAAGGAAAATCTGTTGCTCAG

gttggtcaaattctctgcatatgccatagtgtagtctttgttctttggcatcgtatttgcatcgcattgtattccttccttgtacggaatgtaacttacatttgtattacttactgatcag

GTTAGTATGAGATGGGTTTACGAGCAAGGCGCGAGTCTTGTGGTAAAAAGTTTCAGTGAAGAGAGATTGAGGGAAAACTTGAACATATTTGACTGGGAACTCACTAAGGAAGACCATGAAAAGATCGGTGAGATTCCACAGTGCAGAATCTTGAGTGCTTATTTTTTGGTCTCACCTAATGGACCTTTCAAATCTCAAGAAGAGTTGTGGGATGATGAAGCTTGA

>Pnu_CYP82Y2-Ln3-Pnu_COR-Ln3-genomic 1181 bp intergenic region one intron less in CYP82Y2, experimentally validated

ATGGATTACTCTAATCTTCAGTTTTTCGGTTTTCAACCAACTTCCATTGTTGCTCTTCTACTTGCTCTTGTATCTGTTTTACTTGGTTTTCTTGGTTTGAGGAAGAAATTTTCGCACTACCGGCCAAAAGCATCGTGTGCACTGCCACTGACCGGTCTTCTTCATGTTTTCATGAATAGAGATGACTTAATTCATGTAACTCTTGGAAACATGGCTGACGAATACGGACCGATTTTCAGTTTCCCAACAGGCAGCCATAGGACTCTGGTTGTGAGCAGTTGGGAGATTGTAAAAGAGTGTTTTACGGGCAGCAATGACACTGTTTTTTCCAATCGTCCCGTTCCGTTGGCTTTTAAGATTATATTCTATGCAGGTGGTATAGATTCGTATGGTCTTGCACTTGTACCTTATGGAAAACAATGGAGGGAGTTACGAAAGATCTGTGTGCATAACCTCCTTTCTAATCAACAGTTGCTCAAGTTCAGACAGTCGATAATCTCTCAAGTTGATGCCTCTTTCAGTAAGCTTTATGAGTCGTGCAACAGCAATAGAAACTCTGACGACCAGGGAGTCTCTAGTACTACTGCGTCTGGATCAGTCTCTAGTACCAGTGCAGCTAGCATGGTGAGAATGGATGATTGGCTTGCACAGTTATCGTTCAATGTGATAGGGAGAATAGTCTGCGGATTCCAATCACACACAGAGTCTGTGGAGCAATTTACAGTAGCAATGGATGAAGCATCGCGTTTTATGTCAATCACTGCAGTGTCGGATACCGTTCCATGGCTAGGGTGGATTGATCAACTAACTGGTCTTACCAGAAAAATGAAGCACTCTGGAAAGAAATTAGACTTGGTGGTTGAGAGCATAATCGAAGATCATCGTCAAAAGAGACAATTTGCTAGAACAAAAGGAGATGAAAATACTACCGAAGACGAACAAGATGACTTCATCGACATTTGCTTGTCAATTATGGAGCAGCCACAGCTTCCTGGCGACAACAACCCTCGTCAAATACCTATCAAATCGATTGTCCTG

gtaaagaatcaacttaatccgtaggataataaaacatggattatggaaatcgtaggtatgcttagtgttatggtttgattatatactcatatatgcag

GACATGATAGGCGCGGGGGTTGACGCCACAAAACTGACCACGATCTGGACCCTTTCCCTGCTGCTGAACAACCCCCATGTGTTGGACAAGGCAAAAGAAGAAGTGGATGCACACTTTGGAACGAAAAGGAGGAGATCAGCAAATCATGCAGTCATGGTGGATTTTGATGATATTCGTAGCCTTGTCTACATCCAGGCAATCGTCAAAGAATCAATGCGGTTGTATCCAGCCAGCCCGATTGTTGAGCGACTGAGCAGCGAAGATTGTGTTGTCGGTGGGTTCCATGTACCAGCAGGGACGCGGTTATGGGCAAACGTATGGAAGATGCAACGAGACCCAAAAGTATGGGGTGATCCGTTGGTGTTTCGACCAGAGAGATTTTTGAGCAATGACCAGAAGAAGATGGTTGATGTAAGGGGTCAAAATTACGAGTTGATACCGTTTGGAGCTGGTCGGCGGATATGTCCAGGGATATCGTTCTCTTTGGATCTAATGCACCTGGTACTTACTCGTCTTATTCTTGAGTTTGAAATGAAGTCTCCTGGCGGCGGGGAAGTAGACATGACGGCGACTCCAGGCTTGATGAGTTACAAAGTAGTCCCGCTTGACATTCTGCTCACCGCTCGCATGCTTTAG

tccttctgcttgatcgatcaccttcagtatgtgtgcgtgtgtgtccttgtttacccttgcttgatgtttgttttcggtcgattgcctatgtactctaatgccctgcccttgcacattagtatggtcgatctcaggtattgcctaattctttttatccatgtttactattagagtatatcacgtataaaaggagtcgaggtgaccgtgaagacaagcaaccgagacgaccgtgcagtgtgtgcacatgaaggtaagatacatggacagaagatgtgcaaaatattgtgtaatctttatagtttcctagaataagtattgtgattaaattctaggatatttgcattatattctcaactctataaataagagtttcttgtaaatgtaatgtatccaattatcttattatggatcaataatactatcaaccctatggggtttctccgtggagtaggtcgacatcaacagtcgaccgaaccacgttaaacattgtctccttctttactttggtatgtagattgttcttattaatttactttcttttatcattggttaatttattattgtctcatatcaatgatatccatcttagttttgatctacatatcaaaatttaagatttacttcttgcgtttgggtaactagctgcttcttgctctcaatgatcaaatattataactgttgagattattagtcccacattgtatgggcaccctaagatcctgcagttaataatcctatgggccactccactcattgccaattggttttggatgcccacagttcaataataacatagctaagagcatctcgaacgagagtattcctaaatttaaggctttttgacaataattctatgatctatctatcctttagattcaaaaataattctgctccttgccagtgggtaacactctagccctacctaaataaaaatcacggatccgtccctgacttttaaaaatcaaaataaagtatccaactttctaacttacttcaaaaatcaaattattgtagtatctaacagtcaattactgttgaccacgagcttaaacatggtgaccacgtgtttcccgtgcattaccttctttttaggtgagaagaggtggatcctgatttctctctttatactgtgaactgtcttcttaccgaaagaatcaataattgtgtgagaaggaacaaag

ATGGAGAGTAGATATGGTGTACCTGTGATCACTCTGAGCTCCGGCAAGGAGATGCCTGTTCTAGGCATGGGCACATTTGAAACAGAAGGTAAAGCATCTGAAAGAGAGAGGACGGCGTTTTTGAAAGCCATAGAGGTGGGTTACAGATACTTCGATACAGCTGCAGCATACGAAACCGAAGAGGTTCTCGGTGAAGCTATAGCTGAAGCACTTCAACTTGGCTTCATCAAGTCTCGAGATGAACTCTTCATCAGTACCATGCTCTGGTTCGCTGATGCTCACCCTGATCGTGTCCTCCCCGCTCTTCAGAATTCTCTCAG

gtgataaagtttgttgcaaattgaacccatattaaggacaaattaatatagttttgtgtaatatgtag

GAATCTTAAACTGGAGTATGTGGATCTATACATGTTACTCTTTCCGGTAAGCTTGAAGCCTGGGTCGAAGATAACCATGGACATAGAGAAAGATGATTTACTTCCACTGGACTACAAGTCTGTATGGGCAGCCATGGAAGAGTGTCAGAACCTTGGCTTCACTAAGTCAATCGGTGTCAGCAATTTCTCCAGCAAGAAGCTTCAGGAATTGATGGCACACGCCAAAATCCCTCCGGCTGTGAACCAA

gtgagtgaaaacattataagtcactttctcttgaagttttgatttcaaagacatctaattataatgatgttgattattaag

GTGGAGATGAGCCCCGCTTTCCAACAAAAGAAATTGAGAGAGTATTGCAAGGCCAACAACATATTGATCATTGCAATCTCTGTCTTGGGATCCAATGGAACCCCATGGGGCTCCAATGCAGTTATGGGTTCTGAAGTGCTTAACCAGATTGCTATGGCCAGAGGAAAATCTGTTGCTCAG

gttcgtttgattctccgtgtcgtatatatcacatgccatagtatgtatttgcacagaatttaacatttttagcttacatttgtactattactgatccaacag

GTTAGTATGAGATGGGTTTACGAGCAAGGTGCGGCTCTTGTGGTAAAGAGTTTCAGTGAGAAGAACATGAGGGAAAACTTGAAGATATTCGACTGGGAACTGACTAAGGAGGACTTGGAAAAGATCGGTCATGAGATTCCACAGCGCAGGGTATTAGACGCTCATTTTCTGGTGTCAGCCAATGGACCTTTCAAATCTCTAGAAGAGGTGTGGGATGACAATTAA

>Pca_CYP82Y2-La2p-Pca_COR-La2-genomic 13406 bp intergenic region, Pca_CYP82Y2-La2p pseudogene

ATGGTAATTATCAGTCGACTTGGACTGATTATCCAGTGCACATTTCTCACTAGAAGAAGCAACACTTCTGTTGTTGCTCTTCTACTTGCTGTTGTATCTATATTAATTGGTGTTGTTGCTTTCAGTAGCAGAGTTCTGAAAAACTACTCATCACCATCATCAACAGCAACAGCCGTTTCTCATCATCGGCCAAAAGCATCCTGAGCACTGCCAATGATCGGTCTGTATCATGTTTTCATGAATAAAACCGGCTTAATTCATGTAACTCTTGGAAATATGGCTGATAAATATGGACCCATTTTCAGTTTCTCAACAGGTGGCCATAGGGCTCTAGTTGTGAGCAGTTGGGAGATGGCAAAAGAGTGTTTTACAGGCAACATTGACATTGTTTTTCAAATCGCCCTATGCCGTTATCGTTTTAGCTTATATTCAATGCAGGTGGTATAGACTCGTACGGTCTTTCACAAGTACCTCATGGAAAATATTGGAAGGAGTTACGAAAGATCTGTGTGCATAACCTCCTCTCTAATCAGCAATTACTCAAGTTCAGACACTTGATATATTCTGAAGTTGATAAATGTTTCAATAAACTGTATGATCATGCAACAACAATAAAAACTCTGAAAGTGAGGGAGACGACTCTACTACTACTGCAGCTGGGATGGTGAGAATGGATAATTGGCTTGCACAATTATCGTTCAACGTGATAGGAAGAATAGTTTGCGGATTCCAATCACACACCGAGACAA

gtgagcatttccatacaggatgttaagaacatttggatactttgagatcattactattcagcatatctaaccgtgtgcgcacatgtataaacaa

GTGCTACAAGCAGTGTGGAACGATTTACAGTAGCAATTGATGAAGCATCGCGTTTTATGTCGATACCTGCAGGGTCAGATACCTCATGGAAAATATTGATCAGTTAACTGGTCTTAGGAGAAATATGAAGCACCACGGAAAGAAATTGAAGATGATCGTCAGAAGAGACGACTTTCTAGAACTAAAAAAGGAGATGAAAATACTACCGATGAGATGACTTCATCGACATTTGCTTGTCAATTATGGAGCAACCAGAGCTTTCTGGAAACAATTACGCTCGTCAAATGCCTATCAAATCTATTGTCGTG

gtatgcagtcaacttattgatctgtaggataataaaacaaggcttctattaaatatagcctatagtccaaacagtttaagttaaaattataaatcaagtaaatcttaattctttaataataatccacttaaatcatattagtagaaaagtatgtgctatgtacatatttatattggctcctttaacaatataattttttttaaaaacaaataattatttaaagttaaaaaagtttatatttttatttatactcattgtgtagatgatgcaaagagctttccaaatatacaaattgttgtaaaaatttaatttgtattttaaaagatataaaagtttgaagttttataatttatttttaaaacaaaactacctccgtttcaggaaaagagttactatcacttttttaattggcctatttttaggctaaaatgaaaaaaatgataataacacttttccagaaacagagggagtagtatttttataaatatgtaaagataggttttatatacaattttacacttcaatacatatatatagtggtattattgatgtcagaggttattattgtaatatcaggtttagaccaaccaaaaataagtatttccctatattcttaaaatttcttttcctaaattaaaatataaaccctagttttttatctttaaatccaaactaaattatatttcatattatttataactaatctctcacgattctcacaaattgaccaattataaatctcaattcttatttttcccccaaaacaaactaaacttttctctaaaatgatgttatttttctcaatcatgattacattagcatcgattagaatatatattccaattaaccccttaaaaaatctctacaataataccaattgaaatggaatcaacaaaggttttgattcttttcataaaatcaatcttaactccataaacatattctacacttattatttttttaagttgatgttttttttttaatcatgattatattaaaatatactattatttaaaattatcaattatatcacagatcttatggtaaatttaattatgaaagttatgtagttaactcattttttgtcgattaagggatttgataatttattaaattatattgagctatatataataattgtattgtatataataatgagttatgttgaataattataaaaatctatttggttatcttaatcaattattttatatgaattatagttataaaagtataattacatttaagagaagcctaaaacaaagggtgcccgacaagtcgaaattgtagaatcttatccgttgtattccaacattggttgaagttgtagcgcaaagggagttcaaacatctgtttatgatgggtatgtttattgtttatttatggtctgactctaatcatacgtgcag

GACATGATAGGCGGAGCGAATGACACTACAAAACTGACCACAATATGGACCCTTTCCTTGCTGCTGAACAACCCCCATGTGTCGGACGAGGCAAAAAAAGAAGTGGATGCACACCTTTGGATCTAAAAGGAGATCAACAAATGGTGAAGTCACGGTGGATTTTGATGATACTCGTAACCTTGTCTACATCCAGGCAATAACCAAAGAATCAATGCGCTTGTATCCGGCCAGCCCATTGGTGGAGCGACTAAGCAGCGAAGATTGTGTGGTTGGTGGGTTTTATGTACCAGCAGGGACACGACTATGGGTTAACATAGGAAGATGCAATGAGACCCAAAAGTATGGGATGATCCGTTGGTGTTTCGACCAGAGAGATTTTTGAGCAATGCGCAGAAGATGGTTGATGTAATGGGTCAAAATTATGAGTTGTTACCATTTGGAGTCAGTCGACAGGTATGTCCAGGTATATCGTTCTTTTTGGATCTAATGCAACTGATACTTACTCGTCTTATTCTTGAGTTTGAAATAAAGTCTCCTAGTGGGAAAATGGACATGACAGCAACACAAGGTTTAATGAGTTACAAGGTAGTCCCCCTTGGAATTCTGCTCACCCGTCGCATGCTTTAG

tactcctacttgctcgaccatcaaacatatataagttgtttgtaatcatcaattgtggcataaaccacccgatgttatcgagtctgtcctttatggctctatctatatatgtaatatctgttttgtgttcaatgaagtttttgtttgttttacaaaaagaaaaaaaaagagttatattttttgtttgattcaagacgccttgtaacccaagaaaacgttgtcaaaaaggcagggttatcaaagctccttagaaaattatcctgtttttattcgacttaatctaattcagatagttctaagagatttgttaaaactataactagttgccttttggaagaataaaagttggtcgagaaacaatctaagtggagccatcatgctttttgtttgttgatttggttgagtcagccttgcttgtgtatgtttagaacccccttcctgttaggatttatttttagagtttatttttgtaaatatattaggattttttggttagaaaatgtcaagatggtattgtgaggaccagtacaatgaatatccgactgaatttcaaaatacattatccgaactattaacatggcgtggatagtagtcgggaccaccaccaaccttttggaggttatttctcataccaaggtgatctcaattactattcacacatacacccgtcttatgaacatgaaaatgagcaaaattcgagttctttgtcttggagggattccctaaagaagatgcaatctagtacttataatgaacctattatgatcacccgagcaattgcgatctagtactccttatgatcaaattgtttctctagagaaatccatcaaggagttgaatgaggatacaagaaggttcatattaagtatagggggaacaattagtagtccaacttaattgcgatctagtactccttatgatccaattgtttctctacagaattccatcaaggagttgactgaggatacaagaaggatcggattaagtgtgagaggaacaaatagtagtccgactctttctctagaagaaccacatgagaagttgcaatctagtaatccttatgatcatttgtttttctagaagaatccctcaaggagttgactgagaatacaaaaagatttgcgttaagtctggatgaaagtcttgcaaagaataatattgctcaaaatgatcttaatttccaatatagtgctacctttgataattattcacctagtccagaggacgagtatagaattggtaacactacttatttacacgaggttcaaacatcttcatattattattatgatgcggataatattggtgaagaatttgaaatatacaggcagagtgatcaagaaattatcactccaattgtgtgcatgatgatagtgctatttctatttcaaatccaaataattttaatgattattcacctattcaaaaggaagaggatttgactagagataccactattttagacggtgtagaattttcttgttttgaatatgataatagtttagagaaacgagtttgtcctgagactaatgatttagaatctaacgattcgaagacaatagtctctgacgaagatgatagactcttagagaattttaatgtgatagaggaagatttgattcaggttgatataacagtttccttgtggattttgttgtcctagacacccaacttttcaggtcccagagcctcaaattccagttatcttaggtcgtccctttttagctacatacaatgtcgtcatcaattacggaaatggaataattaatttttcttttggtaacatgactatttaactgaatctatttaatatgagtagacaaccttcttgtgagatctatgatttagaagaagtgaacgtaactagtactttagttcagaaatatttaccaaacactttgttagaagatccattagacaagtgcctagcccaatttgggtttgatgttgacgataatagtgttattaatgaggtgaataatttgttagattttgcacctttgatggacacttataatatatgtaaatctaagtttgaacttctaccagtttattagtctaccttatttccttctatagaggaagctgaatatagttcttaccttaaggaccctagaatccaacaaacagtttttaatccatctgagagtaatatccaccctatgtttgggggtaccaatgattcttgtgactgtcaagtatccctgcttaaaatccctcacttgggacttgacatatgcacctctaccatttttcaagatcatcttcaaactagattccctgaacctgttgatgtccagaaagaagttcagttgttagagacccatcctatggttgatgtggtttacctaggaaataataccaagattgactttgttttcccaccaaaatatttttttccatatgtgggaacatttgaatttcaaatgtgtcagttattaagttttgagactaagcctaattactttaggaaattagaatcgaaaaatttgcttaagaatgaccaccttcattgtagtcaactgtgtggttcaaatctgtttgactttgaggatccacaattattcaggttattattatgtgcttcaaatttattagttgagttactccagactttacaacatgatcccgttgatcccgcctgtaaggaaatctaacccattaaaatattttatctagactttttcatagagcccgaaactgaacctcaattagatgtagttgtcttaaacgagacactagatagggattagggtgtgttcggtttgttcactttctgggcctgctgcagtttccttctatttttgataatattattttattgagatgacccacggttgttccgactgttgttgtatgattctataaggtgactaattctttcttagtctggatgaagactttaaacttggcacttcttgggaggtaacccatgctcatgcgacatggtaatacttttccttaactcctttccttcgatccttaacagtttttccttgtcattcttttaataattcatctttagaacattgaggataatgttagatttaagtttgggggtaagggagaaacttttattgaaatatgaaactccaaaatctagaaacttatgaacaattcaggatgacactaaccattctaggtagatggaaacttagttctttctctataggaattggctttcccagctaaatggtctattcataaaaggatgcctagctcaagtgttagaaataacatgatagttgtcaccatatctcggagtctatctcgaccatatcaccttctatttttatttttatgatttttgtgtttatttaagtgatttggtggggcaccaaaatcaaattgttatcgctgtcaggatggagtagattgtttgagataaagtcatattattgtattatttagaccattatttaaaaaaataaaataaaataaaatgagtattgacactttgatagcaatatttgtgtgttctccctgtctgtgttgcctaagcaagcgtgaaacgtaggatccatgagaattatcatgtaagatgctgataatcagcatgcgcttggatttatttgacccattcatgttgccagtacatttaaaaataaacagttgttttatttttaataaattgatcgttgaagccttgtatataccagctgagtcgacattagtataaggagtaagactagtgggagcgtatctcaatccattaggattcatcctggcactaaccttcagaccgatatgggaaacaccattcaccttaatcaacatgcgcgaacttcatttttccacattcatcttctaaatcttttcatgtgattgtactttgatagattctgagtatcgtggttgaacgactatctaagtagagttttgaattcttatatgaattttagtacgccgaatgaaaactcgtgtacaataattggactcacatagggtaatgatttccatggccgaaaaattaaatatgccaactaaatagcttgtttagtgtttaccctcaatcgttcacatgtagctagggtttggagtaaaggttttttgggtatatctcttgtaaaccctaacgagactataactcgtccgaagggagacctatgggtttaaaggcttgttgcacacgctaagtgcaatcgcgatgcctgcgaaagtgagttaggatttttaaattagatttgctcggtgactagcagataataagcttggggttatttgatagacacatttttgtgtctgatttgtctcaattctaaatattgttagttctcatttttgtactaattttggcgttttattatgtttgtaggtattttaggaaataaataattatggaaaaatcggctgaaaaggttataaagacacccaagaaattactaaaggcacccaccagttggataaaggcacccaccatctcttccatatcaaatatagattttggcggcaaagttcttctctcaaaacaacaaggaataaaatcaagttttcaggaaattcttatgggctttaagataaaatattttcttgccatacttgtaagagcttagacttgatagggaaactcgaaaagacacggaaaatttatcctccccaaaaacagttcgcgtcagaatcttaaattactttcgcaacaacatatccttttcaaggtgtctgagagtatttggataactaaggaaaagaaaatcaacaaaatatagctataaagagaatgaaaaatcaaaacaaaatcaagttatgatgatcgtgagaaagaaaataatctcgtgagaattctcattgaaagaaaaatattcgtgaagatttgagagaggatctgggttttaattgtataaggaagagttctggggggagggggtgctggggggggggcttctggagaggccagaagcgcaaaatcaaaatccagaaaaactcatgtgatgagaaagatgaagaaaacgttgcgaaaaggttaaagtttccgtggctaaaaggttgaagattatgttgctaaatcttttagctttgcaacatttatctttatttggaacgtttatctttagtgaaacacttttccctctgtaacagtgagtctgtaatgcttataaacgttgcaaactcgttgttttcattactttcactctttttcatcatttgtaaacactttgagctatgtaccaaatcttttgagggtgtttttaacatgaggagctaaacccctttgctgaggcgatggaggaagccattgttccaagaaaagtggtatatttctattttaattaatttttgcaatttcttttatgaatatttgctttgataaaaaaaattgatttaaggattttattaaaaaattatgatttctttcgatagactaggttttttgatgttctatgctttcgatttacaattgttattttgagagtctacttttgcaacatattagaatcactttaaatgtaagaaattacatgaatattgattaggttgagtcactttaattggtcaatggtgaaacccataacctcagtatttgtgaatattatttttcatattttgtgtatatttttagtttaaaattagtttattaaatctaaaatcaatctccacaagtttgagagtttgaaacccttttcttaccactcttacaaaactatcatctaggtattcatatgtacatacttttctaattccaatccccttactagttccactatgctgacatagcatgggtatggactagtattaggaaatgtatggaaatccttaagcataaaaacatatgagtccgtgcccaaattctaggcttgatatttgggatgaaaagtggattcctggaattgaatttcctattcatcctttggttatagaataacaaactccaagaaaaaatgctcaggtaatcaataaaaatttctctcctgagataggtagtaaaaacatttctctaccaactcataatgttgattctgacgagctaaagtggtcgtaggtaaaaaaaatggtatcccttttgtaaagtatttatataaggtgttgactgcatgtgttgaggttccaacccgatttggagcaagatctggagcttaaaatgccagcataaaatccagttgtttgtatgaaaaattcattcaaatactctttcacttaccaatatattagctaagatgacacctaaaaatcttcccttttgttgtctatgcaatgctgagaagattctttacacttgctttttcatttttcatttttctttgcagtttagtttgatccgggtattaataacatgttggtttgtaattgtggtttggtctattaggaaacaaaaatgaaatgttatctttaagaatcataaggttgatccattgaaactcattaaagatatacaatttcgtttgtcttactattgtttgagtagagaaagaaattcccaatgatctgtaaacaataacttccctgatcttgtgagtatgcagagaaattagttataattgatgcttcttttgactaattttcaggaaggtatggattggcatccattatgttgaatgaaaatagggataatgtccaggaaaaaacttctgggatgactattgtacctaagaaaacagaaagtttggcactttagaaagctagttattctaagacactgaacctaaaggaaagttctaatatattgtttatgggagactgcaagactgtgaaggaaaattttaaaggcgatactaactcttttcctaagacggagggagtattaactagtaatagaatttagtgggataggctaaatgggaccaatctactctcgcactataatggtcttcaaaggttactcattggttcatcctaacacataggttcggtttgcttaccagtgatatctttaaaaatattcatctaatagaatccctcagtctgttttcgactggtactaattgttcaggggatttcaagttacaagaaaaaaaaataaaaaagtaagaaaattaattaggtttcaagcaatgaggagaaatatgatcaaagaatccttcgtcgttattaaaaattcactacatccgcatatttgtttatctccttttaaacatattttcaacaattgtaggacattaatgctttttttttctatcctagaaattccttttatcactaagtagaataggttctctcctaaattatatccgtctgaaccaccaagaatatgttctcaaggtgtaacctagttgaatatgttatggtttttgaattagattgatccgaggatcaaacacataggtttgctattcttactagtgttatgtttacaccgatcaatttaacaaaatctctatcaattcccacgagttactacttgtgcaaggggtttcaggtgataagcaagctaaattatatataataaaaaaaaattcaatcaatgagcaataatatgatcaaggaatccttcgacgttgcattcactaaaccaacatacttgtttatctgttttgatcacgcattgttaacaaccgtaggataacgatatgtttttctatcctcaaaatttattttataaccatttataatgagttctctactgaattatatcggtttgagaccaccaagaatacgttctcaagttgtaacccagtcaaatatgttagggtttgtgaattatgttgatcctatcccactaattataaattgatttaacgtatattaacatacttgttagtaaacattcactaaatcaacatacatgttatctcttgtgatcacacattattagcaaccgtaggataacaatacgtttttatatcctaaaaatttattttatcaccaagtggaataggttatctactgaattctttctgtttgatgctaccaagaatatgtgtcaaggtatcacccaatagagttctttagggtttataaattatgttgatgctagttaacaatttgagtaacaaattgagttacaaattgagtaatgctttagggttactcaaattcttttaggttgatcctttaatccatgttgtttgcatccatgacaccgaactgatgttccatatatacattaatttaattttagaatagttaaaagggttccaataagagtttagaattctatcattagggtttaatgggaaggaaaccaatcggtttacattgttattggggtgtggttgaatactttttttattggactgtgtggtcggcaaaagtgtttcacatacatgttgtcgaatatatcacaaccattgctggataattaatttgttggtgtgggtttgatttattagacaaaaaatagtatatggggttgacctactgatgtataaagggagaccgtagtgaagggagttaataaaatagaatcagcagtttaattgtgcacatattttaacccgtaagtgtttacgtatgccggctcgatttgattttttgatttgattctagcagttattcaatgagagacgttttcttaaaaactaggtggtgaaagtgaaaatgtcaaattaacatcctttgaagaagtgtgcacaaaaattgatggtaagaaggttgacgtttttacgcataaataacttgttaattaagtttcacgtgtcaaggaaatactagttaaatttcaacaaaactttttattacccttgataagtatttttataatggaaataatattattagtttatttactcaatttgttgttagtgggatatgatcaatctacaaacacatcgaaggaaggttggaacttttatgcttaaataacttgttagttaatttcacgcattaaagaaatactagtttagtttcaagaaaatataatattatcttttatgactactttgttgtgacccacactacgttctaatttaatataatgtattcttggtggttcagaatatagaatatattaaagaacctattatattttttaataaaagaaatttcgaggataggaaaaattattggtatcctacaatagttgataatgtgtaattaacaagagacaaaaaatatgttaatttagtgaatgcttagtaatatttaaggattccttgatcattttttcctcatcgcttgaaacctaattaattttattttactttgcttatgacctaaaatcccctatataatcagtaacttatggagaactaatggattctgttaagtgaatatttgtaaaaaataacactaataaaccgactaaatgtttgtaaaaataatactaataaaccgactgaacctatgtgtttaggaaaaccaaaaaataacttttgaaaaacattacaataaaaaataatactattcctaattagcgtagcctattaaattttattactaattaatatgaatctagcctattaattaaataaatatataaaatttgttagccttacttttatttgacttttaactttttttagttggtctttattaataatttggtacgtaaaaaatttatatttactcaacttattactcaaattttatacattttctaatgaaatctacagtcaggtccattttctctacttttgttccccacaaacccacgagctgaaaacttcacccgcacgcccatttactataagaaaaattctctcacaaacccataatttataaaattatgtttttttttcttcttcattttcttttcttcttcctctgaatggatgtctattctcatggaacagtatgttaactatcgtcgatttggtcttcaatctagttaattagagaaatcaaaagagatcatttctaatggatgtaaagctatggaatgacaagagagagcgagagattattataattttgttgatctgtatgaaattattaaagcaaccagaaaagatacaaaaaccttatgtgcgtgatttaatcatatcatcggtgtatgaacccgagagtttaaagttgattgctcagttcaaaaccctaacttcatatcttgtgggtgcactccctagtattatcgacctgtgaaagtgaaatacttttcttttatagatgtcgacgctcatactcaactggagtacattcgttttgttagaaattatggatttgattgatatccaacaaaaggattgcattctttggcctccactaagcatcttcatttgaattcgtatttggaatcccattccttcatttacagtgttagggttccttccgacctccattggtattcatctttacaatttctggtggattgacgagtaagagttacaattcgaatcattaggaggagtcttgtttcatctccctggaaagcgcgagcactaattgaggtgtgtatttgtatttatccccgtgacggtgtatatttgtatttatttccaatcagtgaagtagatagtttataattgtactatccattgtaattgtagaatttttgtaagcttattttcttgttttttgtgtctatgatgttcgtagtattgcatgtgcatttttcttgtaaccacttttttgttagcattgaaataaaaatttgcatgcataatctgttatgcagctacaaaaaggactgcataacctgttatgcagtcaagaaaatggttgcatgatctgttatgcatttacaaaaaaagttgcataatctgttatgcagtcgagaaaatggatgcatattgcattatgcatccattttttggcaccattatcaaccagaaaatgactttataatttgttttacaactgcaaaaattcccagataactgatagacacatttttgtgtctgatttgatctcaattgtatgtattgttggtactcgattttgtatttattatggttttttatttctttgtaggtgtttttggagaaataaactcttacggaaaaattggctcgaaaagtgatgtttgaacctccctagaaagaaatgttaaaggcacccactctttggttaagggcacccattccaacattttgaaattagattttggcgggaaaactaatcttcacgtgaacaggaaacaacatagctttggaaactagtttggtttatggaatgttctctatttaggaaaccgaattgtttgggaaagaactaaaaagagaaagagatcaatttgattctcttattagcatccagaaacgaagaaatcaggaggaaaagaattctgtttttggaataatattctcggtgatctgccaagaatcaaaggaaagatttgagaggatttgaggttagttttgagatgatagtatatatagaagagttctgggacgtaataaaggggagggaaggaagggtgaagcttagaaagaatcaagagctcgaagaagaagtgtacaggaacaagaaaatttcagctgttgatcaacgaagaagtagaacaaactacaaggaagaactgtcgttcattagcgaggagtccgtccttctttccattctttttaaataccttctgtaacagactgaagtactcacagtaacgcttataagcgtatcaaaattcctatgagcgtttctattcccattgtaagcattatttgagttataaaaacgatttttgagggttttgccattacaatgaggagctaaacttcatctttgaggctatggaggaagctacttttccaacaattatgtggtaagtatttaattaatttatgcaatttatttatgattaattgcctggttaaaaattgatttaaaagatttttgttaagcaattgtgattccgttagatggtgcatgcctcggcttaggtccctagatgctccatgcttttgttatacagttgttgttttgaaaatctatttttgcaatgatttagaatcaaattaaaatgttagaattgcatgattaaaattattatcactttgaaattggaaaatagtggaatcctagcctcagttttctctcaactttattaaagaaatatttgagtttttcatatttttagctttagaaattagatttaaaaacaactcttcacaagtctcaacgaactttatttaccactttaaaattacatcaataaccttttatgcagtcgagagaatggatgtataatctgttatgcaactacaaaaccggctgcataatctgttatgcagtcgagaaaatggcagcatattgcattatgcatccattttgttgttaatattgttatcgaccaaaaaatggcttcataatcgattatgcagctacaaaaatgactgcataatattgcattatgcatcgttttttttcatggaatgttaattttgaggcatatagaccgttttaatcaatggttgcataaccacactagtggatgcatgaatcaaaatacggctgcacaatgtgttatgcatcttttttttgttggaatgatttttttaggcatatacatcgttttagtggttgcaaaggcaaattagtgggttcatagaccaaaatatggctacaaaatgtgttatgtgtcctgtatggtgtttgcataatgtattattcatcttttgtgtggctgcataatgactacgtatccaatttttgaaatttgtgcctgaaaagacaaacaccttcgattttttcgtaaaaaatctaaatttaatattgttgtttatactcgttgtgtaaatctctttaaaagctttccaactagataaaatttgtaaaattccaaggcacgaatttttagatatgttatattctagtttggttactaattatactcctgaaaaaatatgatgcataacgagttattcagatattttttttaacaattatgggtgtcacaaaaactggaaaatattttagacttcacaataaaaataataatttttttagacgtgaacctaaaattccattttccaattattagttgggctaatcttgggctaatcgagtaaagtaattagtgggctaggcttatacctaatattaaaactactaataagtttttaactaaggcatgtataccggttaatgtgccttagccaaaaaagtaaacattgaattgtacacatttgtacacaattgataacttggtcaacgatttggtaaacagtccattcttgagtaaaggagtaaccttatagaaaattataaatttgttgtagtaattaaataatattgatgtatgtatttagaagaacaaaaatttaacaaaaattcttagtgtaccatccatgtgaataaaataaaaattgcatcagctcctataaatttttaaaaatagaattttttttttgtgagtgaacagtttttaagttgttaaaatgaacgatacatctgacactcttagaagaatgaaaacatgcaaaatctattaaaatccttgcaagaagatacaagcagcgcgtcaatcccaaggtccccagcagttaggtgagaaaagttggatccattttcacttgttatctaatattatcttctcgtcggattgtacatacgtacatgtgcagacttgcagttaggctctgagagagagagacagtaag

ATGGAGAGTAGTGGTGTGCCTGTAATCAGTCTAAGCTCGGGCAAGGGGATGCCTGTTTTAGGTATCGGAACGTGTGAAACACTTGGTAAAGGGTCTGAAAGAGAGAGGTTGGCGATTTTGAAAGCGATAGAGGTGGGTTACAGATACATTGATACAGCTGCTGCATACGGAACTGAAGAGGTTGTTGGTGAAGCTATTGCTGAAGAACTTCAACTTGGCCTAATCAAATCTCGAGATGAACTCTTCATCTCTTCCCAACTCTGGGGCACCGATGCTCACCCTGATCGTGTCCTCCTCGCTCTCCAGAATTCTCTGAG

gtaaaaaatatttggtgcaaattgaatccaagttttatgtgcaaagcctttgtacatacatatatataaagagttcttaaacatctattaaattttataaaaatcaattttttttgtgtaatatag

GAATCTTAAATTGGAGTATCTAGATCTATACCTGATACCATTTCCGGTAAGCTTTAAGCCAGGGAAGATAACTATGGATGTACCAGAGGACGAAATTTTTCTTATGGACTATAAGTCTGTCTGGGCAGCCATGGAAGAGTGTCAGAACCTTGGCTTCACTAAGTCAATCGGTGTCAGCAATTTCTCCATCAAAAAGCTTCAGGAATTGATGGCGACCGCCAACATCCCTCCAGTTGTGAATCAA

gtgagtaaaacgtaataaaaattagtgtttgcctttttttacctcgaagatacctaatgctaatcaaaatttccatcatttgacaagcatataacccttcgcataaaactctcataatgttgattaag

GTGGAGACGAACCCCATTTTTCAACAAAAGAATCTGAGAGAGTATTGCAAGGCCAATAATATATTGGTCACTGCAATCTCGGTTTTGGGATCCAAAGGAACCCCATGGGGCTCCAATGCAGTTTTTGATTCTGAGGTGCTTAACCAGATTGCTATGTCCCGAGGAAAATCTGTTGCTCAG

gttggtcaaattctacacatatatatttacagtggttctggaccgcccatatgaactatttgaaaacgtctcctcatatagagtaggcttttggttaatatttaatattcattttattccggtattaaaaatatatcaaaagatccttttttttagttaggtgttttgtttttagcttagttgtttgaaaaaggttgaatcttttatgattgatcgttttgaacaatttattctgctgaagatgcaattaaaaaaaaacattttttttcttgtttttctttataaattcttttgctccgtccacttgaaattaacaggggttatttttatgcccaactgtttaaaaagatgtgaaggaacaagttttgcatagtctaattaaacggtccataaagaatttttaatatattcatcacatgacatagtatttgcgtcacatgtattccttattgtacatgcacaatttaacatctttaaacttacatttgtattactatttatag

GTTAGTATGAGATGGGTTTACGAGCAAGGTGTGAGTCTTGTGGTAAAAAGTTTCAATGAAAAGAGGATGAGGGAAAACTTGAACATATTCGACTGGGAACTGACTAAGGAAGACTTAGAAAAGATCAATGAGATTCCACAGTGCAGAGTATTAACTGCTGATTTTTTGGTATCAGCTAATGGGCCTTTCGAATCTCTAGAAGAGCTATGGGATGACAAAGCTTGA

>Pat_CYP82Y2-La3-Pat_COR-Lap-genomic 468 bp intergenic region, Pat_COR-Lap pseudogene with 5’ fragment including first exon

ATGGAGTACTCTAATCTCCAGTTTTTTAATTTTCAGCCAACTCCCATTGCTGCTTTTCTACTTGCTCTGGTATCTATATTACTTGGTGTTATTGCAGTTTCTCATCATCGGGCAAAACCATCGTGTGCATTGCCAATGATCGGTCTTTATCATGTTTTCATGAATAAAACCGGCTTAATTCATGTAACTCTAGGAAATATGGCTGATAAATATGGACCCATCTTCAGTTTCCCAACAGGTGGCCATAGGGCTCTAGTTGTCAGCAGTTGGGAGATGGCAAAAGAGTGTTTTACAGGAAACAATGACGTTGTTTTTTCGAATCGTCCTATGCCGTTATCGTTTAAGATTATATTCAATGCAGGTGGTATAGACTCAGCCGGTCTTTCACAATTACCTTATGGAAAGTATTGGAGGGAGTTGCGAAAGATCTGTGTGCACAACCTTCTCTCAAATCAACAATTACTCAAGTTCAGACACTTGATAAATTCTCAAGTTGATTCGTCTTTCAATAAACTGTACGAGTCATGCAACAAAAATAAAAACTCTGAAAACGAGGGAGACTCTGCTACTGCAGCTGGCATGGTGAGAATGGATGATTGGCTTGGAGACTTATCGTTCAACGTGATAGGAAGAATAGTTTGCGGGTTCCGATCGGACACCAAGACAA

gtgagcatttgtttcttctccacagtctatgttaaaaagaattggatacttttgagatcattactattcactatatcttaattagtagtcatatatatatctaaccgtgtgcacatgtataaacag

GTGCTACAAGCAGTATGGAACGGCTTACAGTAGCAATTGATGAAGCGTCACGTTTTATGTCAATACCTGCAGTGTCAGATACCTTTCCATGGCTAGAATGGATTGATCAGTTAACTGGTCTTACGAAAGATATGAAGCACCACGCAAAAAAATTAGACTTGGTGGTTGAGAGCATAATTGAAGATCATCGTCAAAAGAGACGATTTTCCAGAACTAAAAAAGGAGATGATAATACTACCGAGGATGAACAAGACTTCATCGACATTTGTTTGTCAATTATGGAGCAACCACAGCTTCCTGGAAATAATTACGCTCGTCAAATGCCTATCAAATCTATTGTCCTG

gtatgctatcaacttattgatagtaggatagtgaagggatacgaaatctttttttgatgggtatgtttattggtatggtctgactctagtcatatatgcag

GACATGATAGGCGGGGGGACTGACACCACAAAACTGACCACAATCTGGACCCTTTCCTTGTTGTTGAACAACCCCCATGTGCTAGACAAGGCAAAAGAAGAAGTGGATGCACACTTTGGAACGAAAACGAAATCAACAAATGGGGAAATCATGGTGGATTTTGATGATATTCGTAACCTAGTCTACATCCAGGCAATCATCAAAGAATCAATGCGGTTGTATCCAGCTAGTCCAGTGGTGGAGCGACTGAGCAGCGAAGATTGTGTGGTTGGTGGGTTTCATGTACTAGCAGGGACACGATTATGGGTTAACGTATGGAAGGTTCAACGAGACCCGGAAGTATGGGATGATCCGTCAGTGTTTCGACCAGAGAGATTTTTAAGCAATGAGCAGAAGATGGTTGATGTAAGGGGTCAACATTATGAGTTGTTACCATTTGGAGCCGGTCGACGGATATGTCCAGGTTTATCGTTCTCTATGGATCTAATGCACCTGGTACTTACTCGTCTTATTCTTGAGTTTGAAATGAAGTCTCCTCGTGGGGAAGTGGATATGACGGCAACACAAGGTTTGATGAGTTACAAGGTAGTCCCCCTTGACATTCTGCTCACCCGTCGCATTCTTTAG

tactcctgcttgctcgatcacccagtatgtgtatgtccttgttaaataatccaccggttatcgggtctgcccttcatattgtgtttgtgttgagtgaagtttttgtttgtttttgaaaagaaaaaaagtaataataacaaaaaaaaaaaaaaaaaaaaaaaaagataatgggggcaatctaaaaaacggtaataattttacgattttcctgatatccttgtactcattaagctacagtgaaattttggtttgatcaacaaaaaaaatgtacatgtaacatgcatgcacatgcaaaatcttcttaaatccttgtaagaaaatgcaagcaggtcaacccaacatgccgttagctgagaaaaggtggatccttttacgttagttatctaatacgtatcatcttttcttcttatcagaagaaaaataaatgtgatgtacatgttagcttctgagagagagaaagagagtaag

ATGGAGAGTAATGGTGTACCCATAATCACTCTTAGCTCGGGCAAAGGGATGCCTGTTTTAGGTATGGGAATTGATGTCGAACCCCAACATCCCTCCAGTTATGAATCAAgtgagtaaaacatatgttactctaacttcaaagttttggtaaacattagtgtttaccttcttttacctcgaagatatctaatgcgaatctccaacatttgacaagcatataacccttcacataaaacttctataatgatgattaagGTGGAGATGAACCCCACTTTCCAGAAGAAAGCAAGGTTCGTataaacaggaaaggaagatgaagaaaaaaacacaactgaatgttataaaataaaacgaatcacacagttgacttactagtaggtggttgaatttcaagggagcagtcatcttccactggtatcttcttcctccttttggagacactagtttctccttttgaaattcaaccacctactagtaagtaattattattcttagattccctaacatttgtttgagaacactatgcataggggtgaggttgaagtatttgtgctagtaaaaagtttcatgtaaaacccaggtgcgcaactttctgaggagccataaacatttctgacaagcaagttaagcaacaagtttatacatgatacttccagtggctgcagatttgaattaatagacctgatttgagacaaactatatttgtaagtctaaatgttgcattaaaatgagtaaaaactgaaaaaagaacaggaacatttataagattactcagaacccaaaaattggtgcaatggagcaaggttatggaggagcttacaattgcaatcttcggtggctgatataaagagtgggaagattgaatcaggaccgagcttcacttggcacacagttggatgaaggtgtcacatgcaaatcttacatccatctgaagttcataaagtagagagataactgtaaatcaaaaggagaagagacgtgataccacaacgtgtcaacaataaccatgaaccaacaggttctggtgttggttcatacctgggtaactctattaactatagaatcatcagcataatttccttagtgaatacatggcagtctctcaaatcgaatatcctatgcagtactatcaaacgataaatggcatgtcagacagtttatcttaagaacaatccccagagttaagaataaaacatgctagctgatttgagaacaatgaaacttctcaaattaacatctaacctcaaagctactcg

>Par_COR-La6-Par_CYP82Y2-La6-genomic 1046 bp intergenic region, Par_flye_v1.0_contig_38806_28735_CORL-CYP82Y2L_pair contig_38806:c33061-28735

ATGGAGAGTAATGGTGTGCCCATAATCACTCTTAGCTCGGGCAAGGGGATGCCTGTTTTAGGTATGGGAACATGTGAAACCCTTGGTAAAAGATCTCATGAAAGAGAGGGGTTGGCGATTTTGAAAGCGATAAAGGTGGGTTACAGATGCATCGATACAGCTGCTGCATACGGAACTGAAGAGGTTGTTGGTGAAGCTATTTCTGAAGCACTACAACTTGGCCTGATCAAATCTCGAGATGAACTCTTCATCTCTTCCCAGCTCTGGTGCACTGATGCTCACCCTGATCGTGTCCTCCTCGCTCTTCAGAATTCTCTGAG

gtaagaaaattttctgcaaattgaactcattgtttatgtgcagagcgttgtacacacatctacacctagagttcattataaacatctattaaattttagaaaactcatttttttttgtgtgatatatatatatatatag

GAATCTTAAATTGGAGTATCTGGATCTATACCTGATATCATTTCCGGTAAGCTTCAAGCCAGGGAAGATAACTACGGACGTACAAGAGGACGAAATTTTTCTTATGGACTACAAGTCTGTATGGGCATCCATGGAAGAGTGTCAGAACCTTGGCTTCACTAAATCTGTCGGTGTCAGCAATTTTTCCATCAAAAAGCTTCAGGAATTGATGGCGGCTGCTAAAATCCCTCCAGTTGTGAATCAA

gtgagtgaaacaaaagttgaacttcaaagttttgataaaaattagtgttttccgttttctttacctcgaagatacctaatgctaatcaaatttccaacatctgacaagcatatcttcacataaaactcttataatgttgattaag

GTGGAGATGAGCCCCACTTTCCAACAAAAGAATCTGAGAGAGTATTGCAAGGCCAATAATATACTGGTCACTGCAATCTCGGTTTTGGGATCCAAAGGAACCCCATGGGGCTCTAATGCAGTTTTAGATTCTGAGGTGCTTAACCAGATTGCTATGGCTAGAGGAAAATCTGTTGCTCAG

gttggtcaaattctctgcatatatatcacatgccatagtattttccatcacatgtattaatccttactataatttaacacctttaacttacatttgtattactatttag

GTTAGTATGAGATGGGTTTACGAGCAAGGTGTGAGTCTTGTGGTAAAAAGTTTCAATGAAGAGAGGATGAGGGCAAACTTGAACATATTCGACTGGGAACTGACCAAGGAAGACTTGGAAAAGATCGGCGAGATTCCACAGCGCAGAGTATTAACTGCTGATTTTTTGGTATCAGCTAATGGACCTTTCAAATCTCTAGAAGAGCTATGGGATGACTAA

gcttgaaacatcatttaattgtagttgctcacagaagcttccctggcaatgcgctcaatgactatttattttcaacttacaaatatcaaggaaacatttaagaagcaatgcttatgttttttgaatcgttttattaatttttcgatattgaatagtatattaaatcaaactaaaaatggactaagttagcccttcttttggcaacaaacagaacctaatcaagttatcaaatatcaaaccagaagaggactaagttaacccttcttttagcaacaaactcaacctaattcaagaggcaggcagacacgtcaatcaaattcttttagcaaggaacacaacctaattaagttcaactgaacaacaagcagacagagatgccaatcaaatatcaacatagtttttagggatcacaaaaggaagaaattaaggaacgatggatccttactaaacacccaaaaatcaaatgttatatcaatgtggtggaatactttaaagtttaaaccacaaaaaaattccatccgtctacaggatatttggtaattatccatacagtatggatatgcagcgtgcaacagaagtacttataaacttgtggtgtaccatctttaagtatagttttggggaagacaccacacgaagaagaagatgacattttacgagctaccagtcattgttatcagctagtaattcggtcttctaattgctggttagtatacgtaaacgaatgggatcccatgacaatgggaattgacagaatattttaacaaatctcttagaccattagatgagagatatatagaatggcccatcgttatgctgacacagcaatgtaaaacacgactgtcagttgaaacgacatttctcacccgaagaaacaacactcgctctagctagctatacatgtgcaggcagggttcagccatttcgacacccggggttttctttgttgtcacatagaattgaatcggtaaatagggaaaatatacagtgcgttggataagaattctccatttttttttaagttgatcatccttttgttttgaatt

ATGGAGTACTCTGATCTCCAGATTTTTGGTTTTCAGCCAACTTCCGTTGTTGCTCTTCTACTTGCTCTTGTATCCATATTACTTGGTATTATTGCAGTTTCTCATCATCGGGCAAAACCATCGTGTGCATTGCCAATGATCGGTCTTTATCATGTTTTCGTGAATAAAACCGGCTTAATTCATGTAAATCTTGGAAACATGGCTGATAAATACGGACCCATTTTCAGTTTCCCAACAGGTGGCCATAGGGCTCTAGTTGTGAGCAGTTGGGAGATGGCGAAAGAGTGTTTTACAGGCGACAATGACATTGTTTTTTCAAATCGTCCAATGCCGTTATCGTTTAAGATTATATTCAATGCAGGTGGTATAGACTCGGCCGGTCTTTCACAAGTACCTTATGGAAAATATTGGAGGGAGTTACGAAAGATCTGTGTGCACAACCTTCTCTCAAATCAACAATTACTCAAGTTCAGGCACTTGATAAATTCTCAAGTTAATACCTCTTTCAATAAACTGTACGAGTCATGCAACAAAAATAAAAACTCTGAAAACGAGGGAGATTCTGGCATGGTGAGAATGGATGATTGGCTTGGAGAATTATCATTCAACGTGATAGGAAGAATAGTCTGTGGATTCCGATCAGAAACTGAGACAA

gtgagcatttctctatgttcaaaaaaattggctacttttgagatctttactattcaggatatcttaattagtagtcataaatatctaacagtgtgcacatgtataaccag

GTGGTACAAGCAGTGTGGAACGATTTACAGTAGCAATTGATGAAGCATCGCGTTTTATGTCCATACCTGCAGTGTCAGATACCTTTCCATGGCTAGGATGGATTGATCAGTTAACTGGTCTTATAAAAGATATGAAGCACCACGCAAAAAAAATTAGACTTAGTGGTTGAAAGCATAATTGAAGATCATCGTCAAAACAGACGGTTTTCGAGATCTAAAAAAGGAGATGAGAATACCACCGAGGATGAACAAGATGACTTCATCGACATTTGTTTGTCAATTATGGAGCAACCAGAGCTTCCTGGCAACAACTACGCTCGTCAAATGCCTATCAAATCTATTATTGTG

gtatgctatcaaattattcatcagtaggataataaataaaagggtgactgatgagtagaatcttgtccgctgtattccaacagcattggttcaagtggtagtgcaaacggagtacaaacatctgtttttgatgggtatgtttatttgtttatatatagtctgattctactcatatatatatgcag

GACATGATAGGCGGGGCGACTGACACCACAAAACTGACCACAATCTGGACCCTTTCCTTGCTGCTGAATAACCCCCATGTGCTAGACAAGGCAAAAGAAGAAGTGGATGCACACTTTGGAACGAGAAGAAGTTCAACAAATGGGGAAGTCATGGTGGATTTTGATGATATTCGTAACCTTGTCTACATCCAAGCGATCATCAAAGAATCGATGCGGTTGTATCCAGCCAGTCCAGTGGTGGAGCGACTGAGCAGCGAAGATTGTGTGGTTGGTGGGTTTCATGTACCAGCAGGGACACGATTATGGGTTAACGTATGGAAGGTTCAACGAGACCCAGAAGTATGGGATGATCCGTTGGTGTTTCGACCAGAGAGGTTTTTGAGCAATGAGCAGAAGATGGTTGATGTAAGGGGTCAACATTATGAGCTTCTACCATTTGGAGCCGGTCGTCGGATATGTCCAGGTGTATCGTTCTCTTTGGATCTAATGCACCTGGTACTTACTCGTCTTATTCTTGAGTTTGAAATGAAGTCTCCTGGTGGGGAAGTGGACATGACAGCAACACAAGGTTTAATGAGTTACAAGGTAGTCCCCCTCGATATTATGCTCACCCGTCGCATGCTTTAG

>Par_COR-La7-Par_CYP82Y2-La7-genomic 1094 bp intergenic region, Par_flye_v1.0_contig_26608_84691_CORL-CYP82Y2L_pair contig_26608:c107061-84691

ATGGAGAGTAATGGTGTGCCCATAATCACTCTTAGCTCGGGCAAGGAGATGCCTGTTTTAGGTATGGGAACATGTGAAACCCTTGGTAAAGGGTCTCATGAAAGAGAGGGGTTGGCGATATTGAAAGCGATAAAGGTGGGTTCTGAGATACATCGATCACTGCTGCATACCGGCCCTGAAGAGGTTGTTGGTGAAGCTATTTCGAAGCACTACAACTTGGCACGATCAAATCTCGAGATGAACTCTTCATCTCTTCCCAGCTCAGTGCACCGATGCTCACCCCGATCGTGTCCTCCTCGCTCTTGAGAATTCTACGAG

gtaaaagaattttacgcaaattgaactcattgtttatgtgcaaagcgttgtacatacatctacatctagagttcataaacatctattaaatttcagaaaactcaaattgttttgtgtgatatatatgtag

GAATCTTAAACTGGAGTATCTGGATCTATACATGATATCATTTCCGGTAAGCTTCAAGCCAGGGAAGATAACTATGGACGTACGAGAGGATGAAATTTTTCTTATGGACTACAAGTCTGTATGGGCAGCCATGGAAGAGTGTCAGAACCTTGGCTTCACTAAATCTATCGGTGTCAGCAATTTTTCCATCAAAAAGCTTCAGGAATTAATGTCGAACGCTAACATCCCTCCGGTGGTGAATCAA

gtgagtaaaacatatcttactcgaacttcaaagttttggtaaaaattagtggttaccttcttttaccttgaagatacctaatgctaatcgagtttccaatatttgacaagcatgtaacccttcacatataacttttataatgatgactaggtatcaccagggccttacggccctggctcaacctctcttgatagtcttcagtttacatatattaaaacagactacatcttagttaatgtatccaaatacttttaaaacataagtaagcaatttagttacttagaccccatgatgttcacatgtcgatataaaaaattattgtaatactgtctcattagacacctaaaatacatctacattaaagaaaaaaaagtatttcatgaaatacacggcgataggaaaaaaaatgtaatattgtttttgcttatgaaaaaaactttggattaaatgtcattacaatagtattgcatatgaaaaaaaaattcaacccttataagtaaatcagtaatctcaaaaaaaaaaaaaaaaaaaaaaaagaatagccctacatagatgcgtacatattcgtacctgatgtacatatgtcatgttcaaccaaaacagaaaaatagtaacacatgttgatgaaaaaacgacatggaaacaaccataaaggttaacgtaaaacgtagcataacattacgttggaatccaattcatagtaagaacacggagtggattctgctcgaatgaactcgacatcttttgaggaaattcatagtcttttcgcagatacacctcaatatcaaacttacaataacatagatgatgatagaaaatgaaaatcaaataagaatcaccaataaaagctccaaatagtggcaaaagagttgccacaatacgggaatctaaaaaaaaaaaagggttgatccttaaaacagcaaatctttcgaactgttaccttaggttcattttgaactgtttccataagttaatttatttaggttacacctttggcacgacattggttagttggttacaatttatgttaaccaagcccaagaatgttgtttcacacaaagatcttgtatccttagaaataagttctgctttgcatgcatggataaattagattaaatacatgcttaccttgttatcatcgtattgcagcatatcaatcacttcccgatagtctttccattcttcgtgggatcattttgcagggccaggtcgatctttgaaatcttgtttcttctggagcaagcctaaataaattaattgttcattttcagtgtgcattaattaataattacattgtccacttttgatgatgcatgattacatttttctgaattcattgacctccggtatgtcagtatttgtggctgaaccaaaacaattgtatctacatttaaatcagttgtcagttgagcagatcactaactatctctttttgttgcgttgtggctgtataaaatcagatcactaaccttaaaactcagttatgaacgttgcaatgattataacaattattgacacatttccatgtttgtcacctcccaatccgctgatctgcaccattcttcccagaaagttaccttcatgtcgatccctttacctcggcaaaaagtaaaagcagttagtcatagatcaacaacgtgattatctagtacattggattgctatttccatcgttgacacttcgttagtcctgtactggtatcgcaggataccatggtagcatctgagacggactccagttttccaaaagcatctgcacgaatgcacatcttgtcttacattacataagcaattaaaaggtaatgcaatctatacattaaatagacaatgtctacacatgcaagtgcttatacatataatgaatgatcacaaacctaaaggactaatatggacagtattctaagaaatatttagccacaaaagtgtttgtctctgcttattttggtgtttgccacacatcaagaaaaatagtcccaaatgctttaacatcaaacccaaagataccatgataactctccattatcaattacaatttcatcacctcatctcactgagtactaactggtattgttcaactcagtcgtaagttaaatactctaaatgaatcttatagctagaacaaaaaaaatcaacaaaatgaatgaaaagatataagcagatgatcgattatcattatcataacaataagatctggaactcaggatttcataacctcatctattatcatttagcttggacaactttcctagtgaaagagttattagtcgtgaaattgttgacattaacacctacgatctttcaaatcttaaacagacaacaaaccaattaagcaaatagtgtatggtttaattccaaataacaaaccaattaagcaaatagtgtatggtttaattccaaatattaatatagaattgaaactatcttagtacaacacatccaaaaactattctgatcaagtgaaattcaatcaatacatacttaaaaaccctagctaataagatccagaataatgtaatgtacccttggaaagacaggaaaaacaattcaacaaaagaaaaagtgtataagggggaaaaatacctcttgttgagggatattacaaaggttaacatcaccgtgctcttgggaacctacttgctgggcaacacgttgattctgattaaaagaccatcaaaacaatgcaattaatttaaaattccacaataataatactgataattatgtattaatccatgtatcctagcaagaactgataattatgtattaatccattaaccctagaaagaagactgcagatttaataacacatctatagaaaaattccttaagtttttacactgaattgcggagaggaggcatgtaaaaattggattatcaacttctaccgagattaaatttgtctgttgttctgaataagctgttttagttttttttcctctcatcgtttgggctattcttccctctccttttctctacgcttccgctgtgacatcgacccactatttgctgcaggaactgtggacaataatgtggatggaaaactgatagaacagattacaccttcgtttgattttcgtttatttgtgtatgattgggagctgatatatatatatggtaaaactcttgcgcgagatgatatatatgccaaaatgccaatatatatatatatatatatggtaaaactcttgagtttaataagtgtggccctggtttcgtgaacttgggaatttgaactgaacacaaccaatgttataaaaatatgtggtgtttgtcttcgtgatcaaagagtcaatcagtggacaggatctatccgggagggttaccaagcggtttttgaaaagtttttctaagcttatatagcttaaaatataatatagctaaaaatataacataatccaatgaataaaaatataatttggagatttgcttaaagaaccaatatctaattaaagatttaatacctaaacaataatagaattgaacattaaccaaaaatactaagaacttgaaatagtaaaaattgtcttttaaagttattaaaaaaaattattccctaaagaaattaatgcaaaaatgttgtaacatagatagtaagttgttgaaacatattgatgcttctatgtttaggtaatgattggggcggtgtttaaataatcgtatcattgtagccggggcctgtggattagaatttttaaagttaaatatagtccccactttaaaattagccgtgattaggttgaaatctagagggtggaacctataagaagatcccgatgtcttgctgccctgatcaaatttagtcagccgatcagtatcctcgcacaaaatttttgtcgtccggtaatgattgaggaggtgtctaggtaattgtgtgatagtatcctggtattacgaccccgaccatggattaaaatttgtagagtaaaatatagtctccactttaaaattagccgtgattaggttgaaatttacagggtcgaacctataaggaagatcacgatgtcttgctgtcatgatcaaatttggtcagctaggttagtatcctcacacaaaattttagtcgtctggtaatgattgggaaagtgtttagataatcgtgtaatagaagccggggcgtacggccctggccgtggattagaatttctagagttaaatatagtccatactttaaaattagtcgtgattaggttgaaatctagagagtggaatctttaaggaagatcccgatgtcttgctggcctgatcaaatttggtcagccggtcagtatactcacacaaaattttagtcgccggtaatgattggggcggtgctcaggtaatcatgtcatagtagccgaggcctacggcccctgccctgaattagaatttcttgagttaaatatagtcccccctttaaaattagttgtgattaagttgaaacctggtcatgaataacaataatccttgcccatatttttaaatgttcctacctctaccgaaactatatacatcctataacgtaatcacaaatatatacatccccgtgattatagaaggctccaatttgatatcaatagcaactttagattgcaaaataatgtggatatataggtaaacaatcgatacagagaaaaaaaatcaaaagtttcatttaattcttagcaggggatatattacctgtgaaatcacttcgaattgtgagaggaaatcagctcccaaataaaaaaataaaaataaaaatccagcaagcaaaagagcatatggcagtaaaaaaaaatccagcggaaaaataactgaattaagtaaaataattgaatgaagaaaggagttgaatgctatccgtaggccccgggctcaacctctatcaacctttttatgttgtttggtcgtccatctgtatccttccacaatattttagtcaacatgtaagttaatcaatattagtcgtgattaggttgaaatctaaaggatgaaacctataaggaatatctcgatgtcttgctggcctgatcaaattttgtcacccggtcagtatcctcgcacaaaattttagtcgcccgataatgagtgaggcggtgtttaggtaatcatgtcatagtagccgaggcctacgactcctaccctgaattagaatttctagatttaaatatagtccccactttaaaattagttgtgattaagttgaaactcggtcatgaataacaataatccttgcccatatttttaaatgttcctacctctactgaaactatatagatcctataacgtaatcacaaatatatacatctccgtgattatagaaggctcaaatttgatatcaatagcaattttagattgcaaaataatgtggatatataggtaaacaatcgatacagagaaaaaaaatcaaaagtttcatttaatgcttagcaggggatatattacctgtgaaatcacttcgaattgcgagaggaaatcagctcccaaatatcaaaattaaaaaaaataataaaaaattcagcaagcagaagagcatatggcagtaaaaaaaatccaacagaaaaataactgaattaagtaaaataattgaataaagaaaggagttgaatgctaaccgtagggcggggctcaacctctatcaacctttttatgttgtttggtcgtccatctgtatcctcccataatattttagtcaacatgtaagttaatcgatattagtcgtgattaggttgaaatctaaaggatgcaacctataaggaagatctcgatgtcttgttggcctgatcaaatttggtcacccggtcagtatcctcgcacaaaatagcaatcgtccgataatcagtgaggcggtgtttaggtaatcatgtcatagtatccgaggcctacggccccttccctgaattagaatttctagagttaaatatagtctccactttaaaattagttgtgattaagttgaaacccggtcatgaataacaataatccttgcccatatttttaaatgtacctacctctactgaaactatatacatcctataacgtaatcacaaatatatacatccccgtgattatagaaggatccaatttgatatcaatagcaactgtagattgcaaaataatgtggatatataggtaaacaattgatacagagaaaaaaatcaaaagtttcattaaattcttagcaggggatatattacctgtgaaatcacttcgaattgcgagaggaaatcagctcccaaatatcaaaaaaaaaaattaaaaaaaaaaaaaatccagcaagcagaagagcatatggcaataaaaaaaattcagcagaaagataattgaattaagtaaaataattgaatgaagaaaggagttgaatgctaaccgtaggccggggctcaacctctatcaacctttttatgttgtttggtcgtccatctgtatcctcccacaatattttagtcaacatgtaagttaatcggggataacagaacctgtatttaaatagcccggttattctaatatcgtttttagttagccgggacacaataataagggcctacaaccccggcctagtgctttgctggagcctgcggtctggtgttttgtcgacggccgggtcaaatcaatggtgagccaagttttggagtttagaagtctacacttattgaaattgacaattttccggtggtgagccatgtcatgcaaattactatgcgttggttaatttattgcatcaacatttttttttagtagtagatgtgtctttcgtggtccaataatttataaacacagacctgcctattatttaggaacataggaagtataataggttgagtttataggttaggcagggccaacggccccgatgcttagtcgggtcatattggtcgagggtcatacccattaggaattaaagtttagaagccaacaattactgcttaactttcttatccaggtgagaagcttttttgtccgttggtcggggtcgtacccgttaggaactttcactttattatccatggtcgggatcgtacccgttaggaattatacttttcgagtttagaagccaacaattaccggcttaactttcttatctaggttagaagcttttttgcccaacggatgaccgattcctatcagaggacaatttcgagggtctgatattgtggtagcgacctttaaggttaatggatacgaatattgtgccatcctaacttctgtatgaaaatgacccattgaatgaattatatctttgggtacttctgtatacaattaaggttcaactccaaacatctaggaaggaataataatctggaacctccaccaggtaaattgaacttccgtcatattcttttaatttcgatgtataatttcacccactgcaatttttatggagataataaaattgcaatttctgttatgtgcaatttttcttcattattcttttaaggaaaactgcattagaaacacccactgctatggccaaggagctaaatggagatcctcatcttcattgcacttgaacattatcatagtagatgatacaatttctcccgagtgtgagatgggggtaattgttccacctataagtttcggggattatccacccgatatgacctatattcctaaaaaacaactataaacttgagaatagaaaagaaaagagtcagatgtttaaccatgaactttacaatgcacagtgatttacatgctataaaaatccatttacaactgcaccacaaagaagaatccatttaaaatagtgaagctcagttaagctttgtaatgccctttataaaaaaaaaaaaaaaaaaaaaactccgcttgcaaacaataaaatgaattatagttttggttgcagcctactgctacgaacataagaaccgtaaatgaatcttataagattccaatctggcaacatgtacaaactacctcaaaagaaacatctgtgaacttgccattggtaagggtcttcctcagatttcttagtttctgcagccaaagaagcagtcaacctctatattgcaaatctagcttctttgtcaatcttgtgagaatctacaacaacttatatgcaaaatacccagataagaaacgaaacagttagaatcaaccaaaacaggaaataccatttcaattattagtagaacacgcatctaaatacccgtgattggacctgatgtgaatcgaacacgtaaccttctgacatggagtcagatgcgctaccattgcgccacagatccgtggtctaaaagacttgtctaagtatgttacagaagcaggatttactcacttatcttgcagctgcattcttgtgtttcattcataagattatggccatcattctttccgccatatgcaatctgcagccaattgataaaatcataatcaacaatctgtacgaataaggaaatctaaaaagaaaccatgcacattatccaagcacagagaaaagtaaaagatagattaaagttaccacacaactacatgtcaaacaaatttaaactcgaaaaccatcttcacagatcaagtcaatcactttgtcagggaggatgcaacaatatcatagcattaactataacaaattagtatataaatttattagcgtaactcaagcacgcttcgtgtgctagtgactttacggatacaaataaaatgcatgcaatcgtgtatcccataacatcattagactactacatcaacccaacccacacactttcaattttttacccaccaaaagtaataggttcaaatatcatttgtattaattaacaccgaattacagataagagaaagcaaaaaagaataataccaagattgctgaagtttttttatgggtttcctacaaacactaatacttgataccgagtctccggaaatgggtgtactttagcctcaaatgtaacaaaattcttggacttcgatatccaagagtttcctttcattatgtgtttgacctcagatcgcctaatgtcattttcatatcttgctgaagtaatatactaaatatggaaaagaaacatcattacatagttcatatcttacctgcaacaacccattaaaaaataggaattcaaaatgacgaagaacggatgaagttattgatgtgaactgtggtctcaaatccggacaaagcaaaaagagagaattcagaactgagggagatatgtatcctttcttccactgatgtaatcaacaatcccctgtaagatttttgattaaatatccatcttggtaaaggtccaatataaacgacacttcacatgccgaaagaagaagcctacaaaaataatcctacaatcatgaattccacatatagatgttcaaacaacatcaaattatgtacagaaaactaagtactactgagtttacaactacaatctagtttctttctatgtggaattcatgatctagtttctttctatacctttcgcaagagcaggaatgatattcgacaacccttgatgaaattttaacctgccaataccatttaaaatagattatttaaacagttgcaaacacgaaactcaacgaatttagaaacaacaaatactggtttccgcgggcgttctgttgatatattttataaaaatcattagaacatgaaccttatcatctaaactacaatacctacaatacccatattttgaaatttgaaaatcaaatttaaaagattaaagaaaacacataaagataataggtgataggaattcaaattaagaagaagtgatggatggagttgttgttgactgtagtttcaaatcgggacaaaggaaaaagagagaactgacaacttaattgatgagaaaaaccctagatattttgttaggtaaaaaaattaaactataacatgctctaatttcataagaaacaccaccgaccaagatatcctattgaagaaacaccgataaaatattgaaaaaaactgataataagtagaaattgaacagttttattaattaatgtagagaagatgagaaacacaccagcaatgttcaattgactgagaaggcgctgaaatttgttgttgatcccagaactggacatgatttctagatgttagggttgctgtgccaaaaaaaacctagaagtagagttttgatcgaattgaaaccattaaatagttttttcaaacaaagaaaatccggctaaattaaaaaataaataaaacatattcaatgaacaaattaagcaattgagagatcgaatttctttttaggaataaaaaacaatagggttacctgaaacatcaacttctctttagctttgcttcaatggaattcctcagagaaagtttattaacacttctctttaactgaatcaggatcaccggggaaaaaaatctcttggttgcaccatctctccgattgcgtacccaacctttaagacctaattctttagcattctcgccttccgctttcataacttgataaaccaaaccgaaacttcacaaaaatcttaaacatactcttgactcgcaatcaaccgagaaaacccatacatcaattaaaaaaaaaaattgaacaaaatttcatctatgttagtgtatggagagaaaattagggtttttttaggattccatgatattctaatgtaatgtaagaccgtaattttaaacacataattaaattcatacctcatatttgttatagaaaggacgatacgttcacaaaactatggatgaaatcaatatctccaatgaccaagcgagaatagcttaattcataaaactaaaccgtgttaacgaaataagggtgggtttgaggatgagtgaggccgtccagatattgttatatgtttttctctcttcctaggaaacgatttttttcagcgagagatggcttcaccttcacatccttcgctgagttacgtaagtagcagcggctgttcttgcttagaagataaacaaagcaggtgtatggtcttctctgtgaagattagagatgtgatgaaaacttggttagtaaatatgaattctagggtttgttctttaatttttagcttttctcctctttgagaaaagtcaccatttgttagtctttttgatcggaattatgaattgactcattaatttgattttgattatgttttctggagttttctggagttcgaactaaaattattcaaatttatttcgattagaattagttgtattagggtttttatgatgtgaactccagctagatcaataggaatgatattcgacaacccttgatgaaattttaactgccaataccatttaaaatagattatttaaacagatttgcaaacacgaaactcaacgaatttagaaacaacaaacactttggtttcgcgggtgttctgttgatatattttataaaaatcattagaacatgaaccttatcatctaaactacaatacctacaatacccatattttgaaagttgaaaatcaaatttaaaagattaaagaaaacacataaagataataggtgataggaattcaaattaagaagaagtgatggatggagttgttgttgttgtagtttcaaatcgggacaaaggaaaaagagagaatcgacaacttaattgatgagaaaaaccctagatattttgttaggtaaaaaaattaaactataacatgctctaatttcataagaaacaccaccgaccaagatatcctattgaagaaacaccgataaaatattgaaaaaaactgataataagtagaaattgaacagttttattaattaatgtagagaagatgagaaacacaccagcaatgttcaattgactgagaaggcgctgaaatttgttgttgatcccagaactggacatgatttctagatgttagggttgttgtgccaaaaaaaacctagaagtagagttttgatcgaattgaaaccactaaatagttttttcgaaacagcaaaatccggctaaattaaaaaataaataaaacatattcaatgaacaaattaagcaattgagagatcgaatttctttttaggaataaaaaacaatagggttacctgaaacatcaacttctctttagctttgcttcaatggaattcctcagagaaagtttattaacacttctctttaactgaatcaggatcaccgggggaaaaaaatctcttcagttgcaccatctctccgattgcgtacccaacctttaagacctaattctttagcattctcgccttccgctttcataacttgataaatcaaaccgaaacttcacaaaaatcttaaacatactcttgatcgcaatcaacagaaaacccatacatcaattaaaaaaaaattgaacaaaatttcatctatgttagtgtatggagagaaaattagggttttttttaggattccatgatattctaatgtaatgtaagaccgtaattttaaacacataattaaattcatacctcatattggttatagaaaggacgatacgttcacaaaactatggatgaaatcaatatctccaatgaccaagcgagaatagcttaattcataaaactaaaccgtgttaacgaaataagggtgggtttgaggatgagtgaggccgttggagatattgttatatgtttttctctctttaggaaacgattttttcaaatgagagatggcttcaccttcacatccttcgctgagttacgtaagtataatggctgttcttgcttagaagataaacaaagcaggtgtatggtcttctctgtgaagattagagatgtgatgaaaacttggttagtaaatatgaattctagggtttgttctttaatttttagcttttctcctctttgagaaaagtcaccatttgttagtctttttgatcggaattatgaattgactcattaatttgattttgattatgttttctggagttttctggagttcgaactaaaattattcaaatttatttcgattagaattagttgtattagggtttttatgatgtgaactccagctagatcaataggaatgatattcgacaacccttgatgaaattttaacctgccaataccatttaaaatagattatttaaacagttgcaaacacgaaactcaacgaatttagaaacaacaaatactggtttccgcgggtgttctgttgatatattttataaaaatcattagaacatgaaccttatcatctaaactacaatacctacaatacccatattttgaaagttgaaaatcaaatttaaaagattaaagaaaacacataaagataataggtgataggaattcaaattaagaagaagtgatggatggagttgttgttgactgtagtttcaaatcgggacaaaggaaaaagagagaactgacaacttaattgatgagaaaaaccctagatattttgttaggtaaaaaaattaaactataacatgctctaatttcataagaaacaccaccgaccaagatatcctattgaagaaacaccgataaaatattgaaaaaaactgataataagtagaaattgaacagttttattaattaatgtagagaagatgagaaacacaccagcaatgttcaattgactgagaaggcgctgaaatttgttgttgatcccagaactggacatgatttctagatgttagggttgctgtgccaaaaaaaacctagaagtagagttttgatcgaattgaaaccactaaatagttttttcgaaacagcaaaatccggctaaattaaaaaataaataaaacatattcaatgaacaaattaagcaattgagagatcgaatttctttttaggaataaaaaacaatagggttacctgaaacatcaacttctctttagctttgcttcaatggaattcctcagagaaagtttattaacacttctctttaactgaatcaggatcaccaggggaaaaaaatctcttcagttgcaccatctctccagttgcgtacccaacctttaagacctaattctttagcattctcagcttccgctttcataacttgataaatctgaaaccgaaacttcacaaaaatcttaaacatactcttgactgcaatcaaccagaaaacccatacatcaattaaaaaaaaaattgaacaaaatttcatctatgttagtgtatggagagaaaattagggttttttaaggattccatgatattctaatgtaatgtaagaccgtaattttaaacacataattaaattcatacctcatattggttatagaaaggacgatacgttcacaaaactatggatgaaatcaatatctccaatgaccaagcgagaatagcttaattcataaaactaaaccgtgttaacgaaataagggtgggtttgaggatgagtgaggccgtccagatattgttatatgtttttctctctttccggaaacgattttttcagcgagagatggcttcaccttcacatccttgctgagttacgtaagtagcagcggctgttcttgcttagaagataaacaaagcaggtgtatggtcttctctgtgaagattagagatgtgatgaaaacttggttagtaaatatgaattctagggtttgttctttaatttttagcttttctcctctttgagaaaagtcaccatttgttagtctttttgatcggaattatgaattgactcattaatttgattttgattatgtttttgagtttttacggagttcgaactaaaattattcaaatttatttcgattagaattagttgtattagggtttttatgatgtgaactccagctagatcaataggaatgatattcgacaacccttgatgaaattttaacctgccaataccatttaaaatagattatttaaacagttgcaaacacgaaactcaacgaatttagaaacaacaaatactggtttccgcgggtgttctgttgatatattttataaaaatcattagaacatgaaccttatcatctaaactacaatacctacaatacccatattttgaaagttgaaaatcaaatttaaaagattaaagaaaacacataaagataataggtgataggaattcaaattaagaagaagtgatggatggagttgttgttgatttgtagtttcaaatcgggacaaaggaaaaagagagaatcgacaacttaattgatgagaaaaaccctagatattttgttaggtaaaaaaattaaactataacatgctctaatttcataagaaacaccaccgaccaagatatcctattgaagaaacaccgataaaatattgaaaaaaactgataataagtagaaattgaacaggttttattaattaatgtagagaagatgagaaacacaccaacaatgttcaattgagcgagaaggcgctgaaatttgttgttgatcccagtaaccggacatgatttctagatgttagggttcttgtgccaaaaaaacctagaagtagagttttgatcgaattgaaaccactaaatagttttttcaaaacaaccaaaatccatttaaattaaaaaataaataaaacatattcaatgaacaaattaagcaattgagagatcgaatttctttttaggaataaaaaacaatagggttacctgaaacatcaacttctctttagctttgcttcaatggaattcctcagagaaagtttattaacacttctctttaactgaatcaggatcaccaggggaaaaaaatctcttcagttgcaccatctctccagttgcgtacccaacctttaagacctaattctttagcattctcagcttccgctttcataacttgataaatctgaaaccgaaacttcacaaaaatcttaaacatactcttgactgcaatcaaccagaaaacccatacatcaattaaaaaaaaaaattgaacaaaatttcatctatgttagtgtatggagagaaaattagggtttttttaggattccatgatattctaatgtaatgtaagaccgtaattttaaacacataattaaattcatacctcatattggttatagaaaggacgatacgttcacaaaactatggatgaaatcaatatctccaatgaccaagcgagaatagcttaattcataaaactaaaccgtgttaacgaaataagggtgggtttgaggatgagtgaggccgtccagatattgttatatgtttttctctctttccggaaacgattttttcagcgagagatggcttcaccttcacatccttcgctgagttacgtaagtagcagcggctgttcttgcttagaagataaacaaagcaggtgtatggtcttctctgtgaagattagagatgtgatgaaaacttggttagtaaatatgaattctagggtttgttctttaatttttagcttttctcctctttgagaaaagtcaccatttgttagtctttttgatcggaattatgaattgactcattaatttgattttgattatgttttctggagttttctggagttcgaactaaaattattcaaatttatttcgattagaattagttgtattagggtttttatgatgtgaactccagctagatcaataggaatgatattcgacaacccttgatgaaattttaacctgccaataccatttaaaatagattatttaaacagttgcaaacacgaaactcaacgaatttagaaacaacaaatactggtttccgcgggcgttctgttgatatattttataaaaatcattagaacatgaaccttatcatctaaactacaatacctacaatacccatattttgaaagttgaaaatcaaatttaaaagattaaagaaaacacataaagataataggtgataggaattcaaattaagaagaagtgatggatggagttgttgttgactgtagtttcaaatcgggacaaaggaaaaagagagaactgacaacttaattgatgagaaaaaccctagatattttgttaggtaaaaaaattaaactataacatgctctaatttcataagaaacaccaccgaccaagatatcctattgaagaaacaccgataaaatattgaaaaaaactgataataagtagaaattgaacagttttattaattaatgtagagaagatgagaaacacaccagcaatgttcaattgactgagaaggcgctgaaatttgttgttgatcccagaactggacatgatttctagatgttagggttgctgtgccaaaaaaaaacctagaagtagagttttgatcgaattgaaaccactaaatagttttttcagaaacaaagaaaatccggctaaattaaaaaataaataaaacatattcaatgaacaaattaagcaattgagagatcgaatttctttttaggaataaaaaacaatagggttacctgaaacatcaacttctctttagctttgcttcaatggaattcctcagagaaagtttattaacacttctctttaactgaatcaggatcaccaggggaaaaaaatctcttcagttgcaccatctctccagttgcgtacccaacctttaagacctaattctttagcattctcagcttccgctttcataacttgataaatctgaaaccgaaacttcacaaaaatcttaaacatactcttgactgcaatcaaccagaaaacccatacatcaattaaaaaaaaattgaacaaaatttcatctatgttagtgtatggagagaaaattagggtttttttaggattccatgatattctaatgtaatgtaagaccgtaattttaaacacataattaaattcatacctcatattggttatagaaaggacgatacgttcacaaaactatggatgaaatcaatatctccaatgaccaagcgagaatagcttaattcataaaactaaaccatgtagcaatgtttacgaagttatataaattgataagatgaaacaatcaaaaaaaaaaaaattaaacaaaagaaaccgagattgatatgaacccttgcagtgaataagatgaaacaatcaacaccaaacaaaaataggacggatttctttaggaatcgtaccgagagctacgagagaacaaactggattaatcatggaaacaaccgattttgtgatacaatacaacttttcagaaacaataagaattttgttaggttaaacaataagatgaaacaatcagaattgctcgcagtttcctggacggaaaaggaattccccgtgacctagcctatcattcctggccgtccaagtcataactcgatcaggatgctagtggccgccagatcagggtccccgtcaaatcctgtccgtctaatgactctcaaatctcagagtcaaacaccacctttttttataacagtgattaag

GTGGAGATGAACCCCACTTTCCAACAAAAGAATCTGAGAGAGTATTGCAAGGCCAATAATATACTGGTCACTGCAATCTCGGTTTTGGGATCCAAAGGAACCCCATGGGGCTCTAATGCAGTTTTAGATTCTGAGGTGCTTAACCAGATTGCTATGGCTAGAGGAAAATCTGTTGCTCAG

gttggtcaaattctctgcatatatatcacatgccatagtattttccatcacatgtattaatccttgctataatttgacaactttaacttacatttgtattactatttag

GTTAGTATGAGATGGGTTTACGAGCAAGATGTCAGTCTTGTAGTAAAAAGTTTCAATGAAGAGAGGATGAGGGAAAACTTGAACATATTCGACTGGGAACTGACCAAGGAAGACTTGGAAAAGATCGGCGAGATTCCACAGCGCAGAGTATTAACTGCTGATTTTTTGGTATCAGCTAATGGACCTTTCAGATCTCTAGAAGAGCTATGGGATGACTAA

gcttgaaacatcatttaattgtagttgctcacagaagcttccctggcaatgcgctcaatgactatttattttcaacttacaaatatcaaggaaacatttaagaagcaatgcttatgttttttgaatcgttttattaatttttcgatattgaatagtatattaaatcaaactaaaaatggactaagttaacccttcttttagcaacaaacagaacctaatcaagttaccaaatatcaaaccagaacaggactaacttaacccttcaacaaactcaacctaattctatacaagaagcaggcacacaaccaaattaagttcaactgaacaacaagcaggcagagatgccaatcaaatatcaacatagtttttagggatcacaaaaggaagaaattaacttctattggagtaacagatgaacaatggatcctaactaaacacccaaaaatcaaatgttataccaatgtggtggaatatactttaaagtttaaaccacaaaaaaaaaaaaatccatccgtctacagaacatttggtaattatcaatatataatatggatatgcagcgcgcaacaaaagtacttataaacatgtggtgtaccatcttcaagtatagttttggggaagacaccacacgaagaagaggatgacattttacaagctaccagtcgctaccagtcattgttatcagctagcaactcttcggtcttataaatttatcgttttatgtatttttaaatgcatataattgctggttagtatacgtaaacgaatgggatcccatgacagtgggaattgacaaaatattttaacaaatctcttagaccattagatgagagatatataattgctcatcgttatgctggtacagcagtgaaaaacacgactgtcagttgaaacgacatttctcacccgaagaaacaacactcgctctcgctagctatacatgtgcaggagggttcggccatttcgacacccgagattttctttgttgtcacatagaattgaatcggtaaatagggaaaatagtgcgttggataagaattctccacttttgtttttttgttttgtttgagttgatcatccttttgtgttgaatt

ATGGAGTACTCTGATGATCTCCAGCTTTTTGGTTTTCAGCCAACTTCCGTTGTTGCTCTTCTACTTGCTATTGTATCCATATTACTTGGTGTTATTGCAGTTTCTCATCATCGGGCAAAACCATCGTGTGCATTGCCAATGATCGGTCTTTATCATGTTTTCATGAATAAAACCGGCTTAATTCATGTAACTCTAGGAAATCTGGCTGATAAATACGGACCTATTTTCAGTTTCCCAACAGGTGGCCATAGGGCTCTAGTTGTGAGCAGTTGGGAGATGGCAAAAGAGTGTTTTACAGGCGACAATGACATTGTTTTTTCAAATCGTCCTATGCCGTTATCGTTTAAGATTATATTCAATGCAGGTGGTATAGACTCAGCCGGTCTTACACAAGTACCTTATGGAAAATATTGGAGGGAGTTACGAAAGATCTGTGTGCACAACCTTCTCTCAAATCAACAATTACTCAAGTTCAGGCACTTGATAAATTCTCAAGTTGATACCTCTTTCAATAAACTGTACGAGTCATACAACAAAAATAAAAACTCCGAAAACGAGGGAGATTCTGGCATGGTGAGAATGGATGACTGGCTTGGAGAATTATCATTCAATGTGATAGGAAGAATAGTCTGTGGATTCCGATCAGACACTGAGACAA

gtgagcatttctctatgttcaaaaaaattggctacttttgagatctttactattcaggatatcttaattagtagtcataaatatctaacagtgtgcacatgtataccag

GTGGTACAAGCAGTGTGGAACGATTTACAGTGGCGATTGATGAAGCATCACGTTTTATGTCGATACCTGCCGTATCAGATACCTTTCCATGGCTAGGGTGGATTGATCAGTTAACTGGCCTTACAAAAGATATGAAGAACCATGCAAAAAATTAGACTTAGTGGTCGAGAGCATAATTGAAGATCATCGTCAAAAGAGACGGTTTTCGAGATCTAAAAAAGGAGATGAGAATACCACCGAGGATGAACAAGATGACTTCATCGACATTTGTTTGTCAATTATGGAGCAACCAGAGCTTCCTAGCAACAACTACGCTCGTCAAATGCCTATCAAATCTATTATTGTG

gtatgctatcaaattattgatcagaaggataataaataaaagggtgactgatgagtagaatcttgtccgttgtgttccaacagcattggttcaagtggtagtgcaaacggagtacaaacatctgtttttgatgggtacgtttatttgtttatatatagtctgattctactcatatatatatgcag

GACATGATAGGCGGGGCGACTGACACCACAAAACTGACCACAATCTGGACCCTTTCCTTGCTGCTGAATAACCCCCATGTGCTAGACAAGGCAAAAGAAGAAGTGGATGCACACTTTGGAACGAGAAGAAGTTCAACCAATGGGGAAGTCATGGTGGATTCTGATGATATTCGTAGCCTTGTCTACATCCAGGCAATCATCAAAGAATCGATGCGGTTGTATCCAGCCAGTCCAGTGGTGGAGCGACTGAGCAGCGAAGATTGTGTGGTTGGTGGGTTTCATGTACCAGCAGGGACACGATTATGGGTTAACGTATGGAAGGTTCAACGAGACCCAGAAGTATGGGATGATCCGTTGGTGTTTCGACCAGAGAGGTTTTTGAGCAATGAGCAGAAGATGGTTGATGTAAGGGGTCAACATTATGAGCTTCTACCATTTGGAGCCGGTCGTCGGATATGTCCAGGTGTATCGTTCTCTTTGGATCTAATGCACCTGGTACTTACTCGTCTTATTCTTGAGTTTGAAATGAAGTCTCCTGGTGGGGAAGTGGACATGACAGCAACACAAGGTTTAATGAGTTACAAGGTAGTCCCCCTCGATATTATGCTCACCCGTCGCATGCTTTAG

>Pso_CYP82Y2-La4-Pso_COR-La4-genomic 865 bp intergenic region, Pso_PS0216860_PS0216870_pair PS_v6.0_Chr2_PS0216860_PS0216870

ATGGAGTACTCTGATCTCCAGTTTTTTAATTTTCAGCCAACTTTTGTTGTTGCTCTTCTAGTTGCTCTTCCATCCATATTACTTGGTTTTATAGCAGTTTCTCATCATCGGGCAAAACCATCGTGTGCATTGCCAATGATCGGTCTTTATCATGTTTTCATGAATAAAACCGGCTTAATTCATGTCACTCTTGGAAATATGGCTGATAAATACGGACCTATTTTCAGTTTCCCAACAGGTGGCCACAGGGCTCTAGTTGTGAGCAGTTGGGAGATGGCAAAAGAGTGTTTTACAGGAAACAATGACATTGTTTTTTCAAATCGTCCTATGCCGTTATCGTTTAAGATTATATTCAATGCAGGTGGTATCGACTCAGCCGGTCTTTCACAAGTACCGTATGGAAAATATTGGAGGGAGTTACGAAAGATCTGTGTGCACAACCTTCTCTCAAATCAACAACTACTCAAGTTCAGGCACTTGATAAATTCTCAAGTTGATACCTCTTTCAATAAACTATACGAGTCATGCAACAAAAGTAAAAACTCTGAAAATGAGGGAGACTCTACTACTACCGCAGCTGGCATGGTGAGAATGGATGATTGGCTTGGAAAATTATCATTCAACGTGATAGGAAGAATAGTCTGCGGATTCCAATCAGACACCGACACAA

gtgagtatttgtctatgttaaaataaaaattgggtactacttgatactaccttcttcgtttttttagaaaaatatttaattatttttcattttaacttaaaaatagggcaaattaaaaaaaaaaaaaaaaactaatagcatctccttttaaaaacagagggagttgtatatgttaattagtagtcatacatatttaactgcgtgcatatgtataaacag

GTGCTACAAGCAGTGCGGAACGATTTACAGTAGCAATTGATGAAGCATCACGTTTTATGTCCATACCTGCAGTCTCAGATACCTTTCCATGGCTAGGGTGGATTGATCAGTTAAGTGGTCTTATAAAAGATATGAAGCACCACGCAAAAAAATTAGACTTAGTGGTTGAAAGCATAATTGAAGATCATCGTCAAAAGAGACGATTTTCTAGAACTAAAAAAGGAGATGAGAATACCACCGAGGATGAACAAGATGACTTCATTGACATTTGTTTGTCAATAATGGAGCAACCAGAGCTTCCTGGAAACAATTACGCTCGTCAAATGCCTATCAAATCTATCATCGTG

gtatgctatcaacttaatgatcagtaggatagcaacatttgttgaagtggtagcgcgaagggatacaaacatctatttttgatgtgtatgtttattgtttatgttagaactcgggcatccaactcaaaaccaattggcaatgagtggagaggccctaagaaattataaaccgcaggatcttaattgtccatacaatatgggactaataatcccaacagtttatatatgtatgatctgactcttaatgatcatatatgcag

GACATGATAGGCGGGGCGACTGACACCACAAAACTGACCACAATCTGGACCCTTTCCTTGCTGCTGAATAACCGCCATGTGCTAGACAAGGCAAAAGAAGAAGTGGATGCGCACTTTGGAACAAAAAGGAAATCAACAAATGGGGAAGTCATGGTGGATTTTGATGATATTCGTAACCTTGTCTACGTCCAAGCGATCATCAAAGAATCGATGCGGTTGTATCCAGCCAGCCCAGTGGTGGAGCGACTGAGCAGCGAAGATTGTGTGGTTGGTGGGTTTCATGTACCAGCAGGGACACGGTTATGGGTTAACGTATGGAAGGTTCAACGAGAGCCAGAAGTATGGGATGATCCGTCGGTGTTTCGACCAGAGAGATTTTTGAGCAATGAGCAGAAGATGGTTGATGTACGGGGTCAAGATTATGAGTTGTTACCATTTGGAGCAGGTCGACGGATATGTCCAGGTGTATCGTTCTCTATGGATCTAATGCACCTGGTACTTACTCGTCTCATTCTTGAGTTTGAAATGAAGTCGCCTGTTGGGGAAGTGGACATGACGGCGACACAAGGTTTAATGAGTTACAAGGTAGTCCCCCTTGACATTCTGCTGACCCGTCGCATGCTTTAG

tactcctgcttgctcgatcacccaatatgtgtccttgttaaataatccaccggttattgagtctgcccttcatgggtctgtatatgtaatctgtgtttgtgttaagtgaattttttgttcgattttggagaaaacaaaataataataataagctttaggcaaattgttgctttaggccagtagaaatgatcaagtaggacttcctcacgatccgctggtgaaataaataattttcagtaacgaaaagtttcctccgttttttgaaaaaagatatttttacttttttattttaacttatttttagatcaaattgaaaaataaaaatatatcttttccagaaacggaggaagtaattcttttcttatttacctcacttttatatttactcaaaactaaaaaatagaaccgataactaaaatgtgggcctattcgttttttacgtggttcacgaggacggtattccgataaccattgttcagactattatttagtaaagaaatgataatgattatgaggggtcatttagaaaaaaaaggataataattttatgaatttcctttatagttgttgatgcaaactttgtacacataataaatgtatatctttatatcctcgatcgtactcattgctcattagagctacagttagatttttgtttgataatcattttttttataggtaacaacaacatgcacatgcaaaatctactcaaatccttgcaagaaaatacaagcaggtcaaccccaacatgccgttaggtgagaaaggtggatcaatttacatttgttatcttctcttcttatcaaaagaaaaatatatgctgcggacatgttagcctccgagagagagagagaaatagagcaag

ATGGAGAGCAATGGTGTGCCCATAATCACTCTAAGCTCGGGCAGGGGGATGCCTGTTTTAGGTATGGGAACATGTGAAACACTTGGTAAAGGGTCTCATGAAAGAGAGAGGCTGGCGATTTTGAAAGCGATAGAGGTCGGTTACAGATTCATCGACACAGCTGCTGCATACGGAACTGAAGAGGTTGTTGGTGAAGTTATTTCTGAAGCACTTCAACTTGGCCTGATCAAATCTCGAGATGAACTCTTCATCTCTTCCCAGCTCTGGTGCACTGATGCTCACCCTGACCGTGTCCTCCTCGCTCTCCAGAATTCTCTGAG

gtaaccttttctggtgcaaattgaacccaaggtttatgtgtaaataagcagctacacgtatagagtttttaaacatctattaaattttagaaaacttaaattgttttgtgtaatatatgcatttag

GAATCTTAGATTGGAGTATCTGGATCTATACCTGATATCATTTCCGGTAAGCTTGAAGCCAGGGAAGATAACTGCGGACGTATCAGAGGACGAAATTTTTCTGATGGACTACAAGTCTGTATGGGCAGCCATGGAAGAGTGTCAGAGTCTTGGCTTCACTAAGTCTATCGGTGTCAGCAATTTTTCCATCAAAAAGCTTCAAGAATTGATGTCAAACGCCAAGATCCCTCCGGTTGTGAATCAA

gtgagtaaaacatatgtcactctaacttcaaagttttgataaaaattagtgtttgccttcttttacctcgaagatatatatatacatctaatgcgaatccaatttccaacattcgacaacctgatgattaag

GTGGAGATGAGCCCAACTTTCCAACAAAAGAATCTGAGAGAGTATTGCAAGGCCAATAATATACTAGTCACTGCAATCTCGGTTTTGGGATCCAAAGGAACCCCATGGGGCTCAAATGCAGTTTTGGATTCTGAGGTGCTTAACCAGATTGCTATGGCCAGAGGAAAATCTGTTGCTCAG

gttggtcaaattcttatcaccagccatagtacgtattttgcatcacatgttaattttttttttttaataacttacatttgtattgctatttag

ATTAGCATGAGATGGGTTTACGAGCAAGGTGTGAGTCTTGTGGTAAAAAGTTTCAGTGAAGAGAGGATGAAGGAAAATCTGAACATATTCGGCTGGGAACTGACTACGGAAGACTTGGAAAAGATTGGTGAGATTCCACAGTGCAGAATATTAACTGCTGATTTTTTGGTATCAGCTAATGGACCTTTCAAATCCCAAGAAGAGCTATGGGATGACTGA

>Par_CYP82Y2-Lb2C-Par_COR-Lb2C-genomic 1773 bp intergenic region, Par_flye_v1.0_contig_48062_pair1 contig_48062:c119099-114200

ATGGAGTACTCTAGTCTCCAGTTTTTTTACTTTCAACCAACTTCCTTTGTTGCTCTGCTACTTGCTCTTGTATCTTGTGTACTCAGTGTCGTTGCTTTGAGAAAGAAATTTTTGAAAAACTACTACTCATCAACAGCAACGGCCGTTTCTCATCAGTGGCCAAAAGCATCGTGTGCATTGCCATTGATCGATCTTCTTCATGTTTTTTTCAATAAAACCGACTTGATGCACGTAACTCTCGGAAATATGGCTGATAAGTTCGGACCGATTTTCAGTTTCCCAACAGGTAGCCATAGGACTCTAGTTGTAAGCAGTTGGGAGATGGCAAAAGAGTGTTTTACAGGCAACAATGACACTGTTTTTTCAAACCGTCCTATCCCGTTAGCGTTTAAGATTATATTTTATGCAGGTGGTGTGGACTCATACGGGCTTGCACTTGTACCTTATGGAAAATATTGGAGGGAGTTACGAAAGATATGTGTGCATAACCTCCTCTCTAATCAACAATTACTCAACTTCAGACACTTAATAATTTCTCAAGTTGATACCTCTTTCAATAAGCTTTATGACTTGAGCAACAAGAATAAAAATACTACTACTGAAGCTGGCACGGTGAGAATGGATGATTGGCTTGCACAATTATCGTTCAATGTGATCGGAAGAATAGTCTGCGGATTCCAAACACACACCGAGACAA

gtactaagcatttgaattgcccacattttgtttttatatatgtttaattcttaattagtcatatatatctaattgtgtgcaataaacag

GTGCTACAAGCAGTGTGGAACGATTTACAGAAGCAATTGAAGAAGCATCACGTTTTATGTCGATAGCTACAGTATCAGATACCTTTCCATGGCTAGGGTGGATTGATCAGTTAACTGGTCTTACGAGAAAAATGAAGCACTATGGAAAGAAACTAGACTTAGTGGTTGAGAGCATAATTGAAGATCATCGTCAAAACAGGAGATTTTCTGGGACCAAACAAGGAGATGACTTCATCGACATTTGCTTGTCAATTATGGAGCAACCACAGATTATTCCCGGAAACAATGATCCTCCTCGTCAAATACCTATCAAATCTATTATCCTG

gtataggataacaaaacaaacatcctttttttggatgggtatgttatattaattgttatggaatgactgactctaatcatatatatttgcag

GACATGATAGGTGGGGGGACTGACACCACAAAACTGACCACCACCTGGACCCTTTCATTGCTCCTGAACAACCCCCATGTGTTGGACAGAGCAAGAGAAGAAGTGGATGCACACTTTGGAACAAAAAGGAGAGCAACAAATGATGCAGTAATGGTGGAGTTTGATGACTTTCGTAACCTTGTTTACATCCAGGCAATCATCAAAGAATCAATGAGGTTGTATCCAGTCAGCCCAGTTGTGGAGCGACTGAGCAGCGAAGACTGTGTGGTCGGTGGGTTTCATGTGCCAGCAGGGACGAGATTATGGGCAAACGTATGGAAGATGCAACGAGACCCAAATGTATGGGATGATCCATTGGTGTTTCGACCAGAGAGATTTCTGAGCGATGACAAGAAGATGGTTGATGTAAAGGGTCAAAATTATGAGTTGTTACCCTTCGGAGCCGGTCGGCGAATATGTCCAGGCATATCGTTTTCTTTGGATCTAATGCAACTGGTACTTACTCGTCTAGTTCTTGAGTTTGACATGAAGTCTCCTACCGGAGGGGAAGTGGACATGACAGCAACACCAGGTTTAATGAGTTACAAGGTAGTCCCCCTTGACATGCTGCTCACTCGTCGTCTTCGCATGCTTTAG

tcctgcttgcttcatcaattaattagcaactccgaggagcgatcaaggggattcatttgtgtgatttaaagttgatgtaaaaaatggttgattattctaatgagccttaaagaagttgtttgtttatgcgggtctctaatcagtttctgttgagtgtaaattgtcttatttctctacttagtatcattttcaatcttaatataatcttattgtccgttttctgttttctccaatattctttatttacctataaattagaattagaatactgtgaagagacactagtcccacatcgctagattccgcatacaatataatatggaagggccgctccactcattgccaattagttttgaatcggatgcccgtagactttaacaaattataagacacgaaagagcaccatcctagatgaaaaggattataaaaaataaaataaaataaataaaaccttgaccagccagcgaccaaattaattgtttggaaggatgataaaaagtgataagaacgatgataaaaagtgacagaccctttaaacatgttaaaaagtgtttgcaattttcgagcagcaatggcattgacacatcttgcacaacctgtctgtcgtgtgaaacacttttgtcgactacactcccataataataataataacaagcttggtaaaggggcggcaagaaatggtagaggggcggcaagtgtgatagcaatgtccttcttattgattagggggtgtcctatagggggtgtagattgactagattacccttttcaccttaaattaaccaaaatcaaatcaatttttttttaaacctaataaatcaaaaaaaatcaaatcagttttttttttcatcttctcttctcctccatcaaccgtaacctccattaaaactgaaaatttcatcgtcttcgattaatcggcgatttgaaaaatgtttggaaaaacccagttagggtttagtttatcatgttggatttaagttaattttgtatcaaaaattagggttttgtatgaaaattgaaattgaaatttctgggtttatgtgagttacggatggtaataactcaattacaaaccgtaaatggagattttgaagaacatacggttggtaatagttcaattaccaaccgtatgttcttatatctgagaattttcttcagttacggttggtaactattttagtttaccaaccgtaactcagttaccgttggtatcgagcatttggttaccaaccgtaacttagttacggttggtatcgagcatttagttaccaaccgtaactggttttacgattggatattccccaaatttaaccagttacggttggtattcgataacttacgaaccgtaactaagttggtaacaagcaaaattggaaccgtaagttcttctgaaaattcaaaagttttgattttgttatgtactttagataattttgagcgagaaaaaactaatgggaactaatttagaagtataccgggtcactggaatcactcattttggttgagtgaatcaaattctagtggattctcttttcttttctattgtggaacaagctaggccacactaatagaattaaaataggtacagagagcagaggaaaatcttcagttccatgaatgatcacgagcttcccgtgcattacttaatttcgatggaaaaatacagaaaagacatgcatgagcatgggcaagagtcatagacacggcataacgtgcaattgttcaatctataaatatagatgttgtgcgagcgagtgagcctctgagagaaag

ATGGAGAATATTAATGGTGTACCTGTAATTACTCTGAGCTCAGGCAAGGGTATGCCTGTTTTAGGTATGGGAACATTTGAAATAGTTGGTAAAGGGTCTGAAAGAGCGAGGTTGGCAATTTTGAAAGCGATAGAGGTGGGTTACCGATACTTTGATACAGCGGCTGCATACAAAACTGAAGAGGTTCTTGGTGAAGCTATTGTTGAAGCACTTCAACTTGGCTTAATAAAATCTCGAGATGAACTCTTCATCAGTTCCATGCTCTGGTGCTCTGATGCTCACCCTGATCGTGTCCTCCTCGCTCTGCAGAATTCTCTGAG

gtaaattaagcctttgaacacctacatttctttcttaaacatgtatcatcatcagattgtagaatactcgaaatctgttaccatcatcaagttatagaattctcaaaattgttttgtttaataatatatgtatttgtag

GAGTCTTAAATTGGAGTACGTGGATCTATACATGATACCCTTTCCGTTAAGCTTGAAGCCAGGAGGGAAGATAGGTATGGACACACCAGAGGACGAGATTTTAGTAATGGACTACAGGTCTGTATGGGCAGCCATGGAAGAGTGTCAGAACCTTGGCTTCACTAAGTCAATCGGTGTCAGCAATTTCTCCTGCAAAAATCCCTCCTGTTGTGAATCAA

gtgagtaaacacttatcttcaggaattaaagtttttgataattaaattccctataataattaattaatgttgattaag

GTGGAGATGAGCCCGGCTTTCCAACAAAAGAATCTGAGGGAGTATTGCAAGGCAAATAATATATTAGTCAGTGCAGTCTCTATTTTGGGATCAAACGGAACCCCATGGGGCTCCAATGCAGTTTTGGGTTCGGAGGTGCTTAGCCAGATTGCTATGGCTAGAGGAAAATCTATTGCTCAGGTTG

gtcagattctcgagacgcatatatcgatcatcacgtacatcccatagctagcatctgcattattccttgtacagaatttatttaacatcttaaatgcattttgtgttactgatggatcag

GTTAGTATGAGATGGGTTTACGAGCAAGGTGCAAGTCTTGTGGTAAAAAGTTTTAGTGAAGAGAGGATGAGGGAAAACTTGAACATATTCGACTGGGAACTCACTAAGGAAGACTTGGAAAAGATACGTAATGAGATTCCACAGTGCAGAATCTTGACTGCGTATTTTATGGTATCACCTAATGGACCTTTCAAATCTCAACAAGAGCTCTGGGATGACGAAGCTTGA

>Par_CYP82Y2-Lb2B-Par_COR-Lb2B-genomic 1761 bp intergenic region, Par_flye_v1.0_contig_48062_pair2 contig_48062:c193882-188986

ATGGAGTACTCTAGTCTCCAGTTTTTTTACTTTCAACCAACTTCCTTTGTTGCTCTGCTACTTGCTCTTGTATCTTGTGTACTTAGTGTCGTTGCTTTGAGAAAGGAATTTTTTAAAAACTACTACTCATTATCATCATCAACAGCAACGGCCGTTTCTCATCAGTGGCCAAAAGCATCGTGTGCATTGCCATTGATCGATCTTCTTCATGTTTTTTTCAATAAAACCGACTTGATGCACGTAACTCTCGGAAATATGGCTGATAAGTTCGGACCGATTTTCAGTTTCCCAACAGGTAGCCATAGGACTCTAGTTGTAAGCAGTTGGGAGATGGCAAAAGAGTGTTTTACAGGCAACAATGACACTGTTTTTTCAAACCGTCCTATCCCGTTAGCGTTTAAGATTATATTTTATGCAGGTGGTGTGGACTCATACGGGCTTGCACTTGTACCTTATGGAAAATATTGGAGGGAGTTACGAAAGATATGTGTGCATAACCTCCTCTCTAATCAACAATTACTCAACTTCAGACACTTAATAATTTCTCAAGTTGATACCTCTTTCAATAAGCTTTATGACTTGAGCAACAAGAATAAAAATACTACTACTGAAGCTGGCACGGTGAGAATGGATGATTGGCTTGCACAATTATCGTTCAATGTGATCGGAAGAATAGTCTGCGGATTCCAAACACACACCGAGACAA

gtactaagcatttgaattgcccacattttgtttttatatatgtttaattcttaattagtcatatatatctaattgtgtgcaataaacag

GTGCTACAAGCAGTGTGGAACGATTTACAGAAGCAATTGAAGAAGCATCACGTTTTATGTCGATAGCTACAGTATCAGATACCTTTCCATGGCTAGGGTGGATTGATCAGTTAACTGGTCTTACGAGAAAAATGAAGCACTATGGAAAGAAACTAGACTTAGTGGTTGAGAGCATAATTGAAGATCATCGTCAAAACAGGAGATTTTCTGGGACCAAACAAGGAGATGACTTCATCGACATTTGCTTGTCAATTATGGAGCAACCACAGATTATTCCCGGAAACAATGATCCTCCTCGTCAAATACCTATCAAATCTATTATCCTG

gtataggataacaaaacaaacatcctttttttggatgggtatgttatattaattgttatggaatgactgactctaatcatatatatttgcag

GACATGATAGGTGGGGGGACTGACACCACAAAACTGACCACCACCTGGACCCTTTCATTGCTCCTGAACAACCCCCATGTGTTGGACAGAGCAAGAGAAGAAGTGGATGCACACTTTGGAACAAAAAGGAGAGCAACAAATGATGCAGTAATGGTGGAGTTTGATGACTTTCGTAACCTTGTTTACATCCAGGCAATCATCAAAGAATCAATGAGGTTGTATCCAGTCAGCCCAGTTGTGGAGCGACTGAGCAGCGAAGACTGTGTGGTCGGTGAGTTTCATGTGCCAGCAGGGACGAGATTATGGGCAAACGTATGGAAGATGCAACGAGACCCAAATGTATGGGATGATCCATTGGTGTTTCGACCAGAGAGATTTCTGAGCGATGACAAGAAGATGGTTGATGTAAAGGGTCAAAATTATGAGTTGTTACCCTTCGGAGCCGGTCGGCGAATATGTCCAGGCATATCGTTTTCTTTGGATCTAATGCAACTGGTACTTACTCGTCTAGTTCTTGAGTTTGACATGAAGTCTCCTACCGGAGGGGAAGTGGACATGACAGCAACACCAGGTTTAATGAGTTACAAGGTAGTCCCCCTTGACATGCTGCTCACTCGTCGTCTTCGCATGCTTTAG

tcctgcttgcttcatcaattaattagcaactccgaggagcgatcaaggggattcatttgtgtgatttaaagttgatgtaaaaaatggttgattattctaatgagccttaaagaagttgtttgtttatgcgggtctctaatcagtttctgttgagtgtaaattgtcttatttctctacttagtatcattttcaatcttaatataatcttattgtccgttttctgttttctccaatattctttatttacctataaattagaattagaatactgtgaagagacactagtcccacatcgctaggccgctccactcattgccaattggttttgaatcggatgcccgtagactttaacaaattataagacacgaaagagcaccatcctagatgaaaaggattataaaaaataaaataaaataaataaaaccttgaccagccagcgaccaaattaattgtttggaaggatgataaaaagtgataagaacgatgataaaaagtgacagaccctttaaacatgttaaaaagtgtttgcaattttcgagcagcaatggcattgacacaccttgcacaacctgtctgtcgtgtgaaacacttttgtcgactacactcccataataataataataataataacaagcttggtaaaggggcggcaagaaatggtagaggggcggcaagtgtgatagcaatgtcctttttattgattagggggtgtcctatagggggtgtagattgactagattacccttttcaccttaaattaaccaaaatcaaatcaatttttttttaaacctaataaatcaaaaaaaatcaaatcaatttttttttttcatcttctcttctcctccatcaaccgtaacctccattaaaactgaaaatttcatcgtcttcgattaatcggcgatttgaaaaatgtttggaaaaacccagttagggtttagtttatcatgttggatttaagttaattttgtatcaaaaattagggttttgtatgaaaattgaaattgaaatttctgggtttatgtgagttacggatggtaataactcaattacaaaccgtaaatggagattttgaagaacatacggttggtaatagttcaattaccaaccgtatgttcttatatctgagaattttcttcagttacggttggtaactattttagtttaccaaccgtatctcagttaccgttggtatcgatcatttggttaccaaccgtaacttagttacggttggtatcgagcatttagttaccaaccgtaactggttttacgattggatattccccaaatttaaccagttacggttggtattcgataacttacgaaccgtaactaagagttacggttggtaacaagcaaaattggaaccgtaagttcttctgaaaattcaaaagttttgattttgttatgtactttagataattttgagcgagaaaaagctaatgggaactaatttagaagtataccgggtcactggaatcactcattttggttgagtgaatcaaattctagtggattctcttttcttttctattgtggaacaagctaggccacactaatagaattaaaataggtacagagagcagaggaaaatcttcagttccatgaatgatcacgagcttcccgtgcattacttaatttcgatggaaaaatacagaaaagacatgcatgagcatgggcaagagtcatagacacggcataacgtgcaattgttcaatctataaatatagatgttgtgcgagcgagtgagcctctgagagaaag

ATGGAGAATATTAATGGTGTACCTGTAATTACTCTGAGCTCAGGCAAGGGTATGCCTGTTTTAGGTATGGGAACATTTGAAATAGTTGGTAAAGGGTCTGAAAGAGCGAGGTTGGCAATTTTGAAAGCGATAGAGGTGGGTTACCGATACTTTGATACAGCGGCTGCATACAAAACTGAAGAGGTTCTTGGTGAAGCTATTGTTGAAGCACTTCAACTTGGCTTAATAAAATCTCGAGATGAACTCTTCATCAGTTCCATGCTCTGGTGCTCTGATGCTCACCCTGATCGTGTCCTCCTCGCTCTGCAGAATTCTCTGAG

gtaaattaagcctttgaacacctacatttctttcttaaacatgtatcatcatcagattgtagaatactcgaaatctgttaccatcatcaagttatagaattctcaaaattgttttgtttaataatatatgtatttgtag

GAGTCTTAAATTGGAGTACGTGGATCTATACATGATACCCTTTCCGTTAAGCTTGAAGCCAGGAGGGAAGATAGGTATGGACACACCAGAGGACGAGATTTTAGTAATGGACTACAGGTCTGTATGGGCAGCCATGGAAGAGTGTCAGAACCTTGGCTTCACTAAGTCAATCGGTGTCAGCAATTTCTCCTGCAAAAATCCCTCCTGTTGTGAATCAA

gtgagtaaacacttatcttcaggaattaaagtttttgataattaaattccctataataattaattaatgttgattaag

GTGGAGATGAGCCCGGCTTTCCAACAAAAGAATCTGAGGGAGTATTGCAAGGCAAATAATATATTAGTCAGTGCAGTCTCTATTTTGGGATCAAACGGAACCCCATGGGGCTCCAATGCAGTTTTGGGTTCGGAGGTGCTTAGCCAGATTGCTATGGCTAGAGGAAAATCTATTGCTCAGGTTG

gtcagattctcgagacgcatatatcgatcatcacgtacatcccatagctagcatctgcattattccttgtacagaatttatttaacatcttaaatgcattttgtgttactgatggatcag

GTTAGTATGAGATGGGTTTACGAGCAAGGTGCAAGTCTTGTGGTAAAAAGTTTTAGTGAAGAGAGGATGAGGGAAAACTTGAACATATTCGACTGGGAACTCACTAAGGAAGACTTGGAAAAGATACGTAATGAGATTCCACAGTGCAGAATCTTGACTGCGTATTTTATGGTATCACCTAATGGACCTTTCAAATCTCAACAAGAGCTCTGGGATGACGAAGCTTGA

>Par_CYP82Y2-Lb2A-Par_COR-Lb2A-genomic 1772 bp intergenic region, Par_flye_v1.0_contig_48062_pair3 contig_48062:c261486-256589

ATGGAGTACTCTAGTCTCCAGTTTTTTTACTTTCAACCAACTTCCTTTGTTGCTCTGCTACTTGCTCTTGTATCTTGTGTACTTAGTGTCGTTGCTTTGAGAAAGGAATTTTTTAAAAACTACTACTCATTATCATCATCAACAGCAACGGCCGTTTCTCATCAGTGGCCAAAAGCATCGTGTGCATTGCCATTGATCGATCTTCTTCATGTTTTTTTCAATAAAACCGACTTGATGCACGTAACTCTCGGAAATATGGCTGATAAGTTCGGACCGATTTTCAGTTTCCCAACAGGTAGCCATAGGACTCTAGTTGTAAGCAGTTGGGAGATGGCAAAAGAGTGTTTTACAGGCAACAATGACACTGTTTTTTCAAACCGTCCTATCCCGTTAGCGTTTAAGATTATATTTTATGCAGGTGGTGTGGACTCATACGGGCTTGCACTTGTACCTTATGGAAAATATTGGAGGGAGTTACGAAAGATATGTGTGCATAACCTCCTCTCTAATCAACAATTACTCAACTTCAGACACTTAATAATTTCTCTAGTTGATACCTCTTTCAATAAGCTTTATGACTTGAGCAACAAGAATAAAAATACTACTACTGAAGCTGGCACGGTGAGAATGGATGATTGGCTTGCAGAATTATCGTTCAATGTGATCGGAAGAATAGTCTGCGGATTCCAAACACTCACCGAGACAA

gtactaagcatttgaattgcccacattttgtttttatatatatatatcttaattagtcatatatatctaattgtgtgcaataaacag

GTGCTACAAGCAGTGTGGAACGATTTACAGAAGCAATTGAAGAAGCATCACGTTTTATGTCGATAGCTACAGTATCAGATACCTTTCCATGGCTAGGGTGGATTGATCAGTTAACTGGTCTTACGAGAAAAATGAAGCACTATGGAAAGAAACTAGACTTAGTGGTTGAGAGCATAATTGAAGATCATCGTCAAAACAGGAGATTTTCTGGGACCAAACAAGGAGATGACTTCATCGACATTTGCTTGTCAATTATGGAGCAACCACAGATTATTCCCGGAAACAATGATCCTCCTCGTCAAATACCTATCAAATCTATTATCCTG

gtataggataacaaaacaaacatcctttttttggatgggtattttatattaattgttatggaatgactgactctaatcatatatatttgcag

GACATGATAGGTGGGGGGACTGACACCACAAAACTGACCACCACCTGGACCCTTTCATTGCTCCTGAACAACCCCCATGTGTTGGACAGAGCAAGAGAAGAAGTGGATGCACACTTTGGAACAAAAAGGAGAGCAACAAATGATGCAGTAATGGTGGAGTTTGATGACATTCGTAACCTTGTTTACATCCAGGCAATCATCAAAGAATCAATGAGGTTGTATCCAGTCAGCCCAGTTGTGGAGCGACTGAGCAGCGAAGACTGTGTGGTCGGTGGGTTTCATGTGCCAGCAGGGACGAGATTATGGGCAAACGTATGGAAGATGCAACGAGACCCAAATGTATGGGATGATCCATTGGTTTTTCGACCAGAGAGATTTCTGAGCGATGACAAGAAGATGGTTGATGTAAGGGGCCGTAATTATGAGTTATTACCATTCGGAGCCGGTCGGCGAATATGTCCAGGCATATCGTTTTCTTTGGATCTAATGCAACTGGTACTTACTCGTCTAGTTCTTGAGTTTGACATGAAGTCTCCTACCGGAGGGGAAGTGGACATGACAGCAACACCAGGTTTAATGAGTTACAAGGTAGTCCCCCTTGACATGCTGCTCACTCGTCGCATGCTTTAG

tcctgcttgcttcatcaattaattagcaactccgaggagcgatcaaggggattcatttgtgtgatttaaagttgatgtaaaaatggttgattattctaatgagccttaaagaagttgtttgtttatgcgggtctctaatcagtttctgttgagtgtaaattgtcttatttctctacttagtatcattttcaatcttaatataatcttattgtccgttttctgttttctccaatattctttatttacctataaattagaattagaatactgtgaagagacactagtcccacatcgctagattccgcatacaatataatatggaagggccgctccactcatcgctaattagttttgaatcggatgcccgtagactttaaaaaattataagacacgaaaagagcaccatcctagatgaaaaggattataaaaaataaaataaaataaataaaaccttgaccagccagctgaccaaattaattgtttggaaggatgataaaaagtgataagaacgatgataaaaagtgacagaccctttaaacatgttaaaaagtgtttgcaattttcgagcagcaatggcattgacacatcttgcacaaacctgtctgtcggtgaaacactttttgtcgactacactcccataataataataataacaagcttggtaaagggcggcaagaaatggtagaggggcggcaagtgtgatagcaatgtccttcttattgattaggggtgtcctataggggtgtagattgactagattacccttttcaccttaaattaaccaaaatcaaatcaatttttttttaaacctaataaatcaaaaaaaatcaaatggagttttttttttcatcttctcttctcctccatcaaccgtaacctccattaaaatcaaaaatttcatcgtcttgattaatcggcgatttgaaaaatgtttggaaaaaacccggttagggtttagtttatcatgttgatttaagttaattttgtatcaaaaattagggttttgtatgaaaattgaaattgaaatttacgggtttatgtgagttacggatggtaataactcaattacaaacaaatggagattttgaagaacatacggttggtaatagttcaattaccaaccgtatgttcttatatctcgagaattttcttcgattacggttggtaactattttagtttaccaactgtatatctgctttggattggtatcgatcatttggttacaaccgtaacttagttacggttggtatcgagcatttagttaccaatcgtaatcgggttttacgattggatattccccaaatttaaccgattcacagttggtattcgataacttacgaaccgtaaattaagttggtaacaagcaaaaaattggaaccgtaagttcttcctcaaattcaaaagttttgattttgttatgtactttagataattttgagcgagaaaaaactaatgggaactaatttaagtataccgggtcaccggaatcactcattttggttgagtgaatcaaattctagtggattctcttttcttttctattgtggaacaagctaggccacactaatagaattaaaataggtacagagagcagaggaaaatcttcagttccatgaatgatcacgagcttcccgtgcattacttaatttcgatggaaaaatacagaaaagacatgcatgagcatgggcaagagtcatagacacggcataacgtgcaattgttcaatctataaatatagatgttgtgcgagcgagtgagcctctgagaaaag

ATGGAGAATATTAATGGTGTACCTGTAATTACTCTGAGCTCAGGCAAGGGTATGCCTGTTTTAGGTATGGGAACATTTGAAATAGTTGGTAAAGGGTCTGAAAGAGCGAGGTTGGCAATTTTGAAAGCGATAGAGGTGGGTTACCGATACTTTGATACAGCGGCTGCATACAAAACTGAAGAGGTTCTTGGTGAAGCTATTGTTGAAGCACTTCAACTTGGCTTAATAAAATCTCGAGATGAACTCTTCATCAGTTCCATGCTCTGGTGCTCTGATGCTCACCCTGATCGTGTCCTCCTCGCTCTGCAGAATTCTCTGAG

gtaaattaagccattgaacacctacatttctttcttaaacatgtatcatcatcagattgtagaatactcgaaatctgttaccatcatcaagttatagaattctcaaaattgttttgtttaataatatatgtatttgtag

GAGTCTTAAATTGGAGTACGTGGATCTATACATGATACCCTTTCCGTTAAGCTTGAAGCCAGGAGGGAAGATAGGTATGGACACACCAGAGGACGAGATTTTAGTAATGGACTACAGGTCTGTATGGGCAGCCATGGAAGAGTGTCAGAACCTTGGCTTCACTAAGTCAATCGGTGTCAGCAATTTCTCCTGCAAAATCCCTCCTGTTGTGAATCAA

gtgagtaaacacttatcttcaggaattaaagtttttgataattaaattccctataataattaattaatgttgattaag

GTGGAGATGAGCCCGGCTTTCCAACAAAAGAATCTGAGGGAGTATTGCAAGGCAAATAATATATTAGTCAGTGCAGTCTCTATTTTGGGATCAAACGGAACCCCATGGGGCTCCAATGCAGTTTTGGGTTCGGAGGTGCTTAGCCAGATTGCTATGGCTAGAGGAAAATCTATTGCTCAGGTTG

gtcagattctcgagacgatatatcgatcatcacgtacatcccatagctagcatctgcattattccttgtacagaatttatttaacatcttaaatgcattttgtgttactgatggatcag

GTTAGTATGAGATGGGTTTACGAGCAAGGTGCAAGTCTTGTGGTAAAAAGTTTTAGTGAAGAGAGGATGAGGGAAAACTTGAACATATTCGACTGGGAACTCACTAAGGAAGACTTGGAAAAGATACGTAATGAGATTCCACAGTGCAGAATCTTGACTGCGTATTTTATGGTATCACCTAATGGACCTTTCAAATCTCAACAAGAGCTCTGGGATGACGAAGCTTGA

Supplementary dataset 9 (associated with Figure 4a)

- Nucleotide sequences of STORR-containing contigs/scaffolds in *Papaver* species

>Pca_10x_v1-0_scaffold_141675_minus P.californicum-10x_141675:c44375-1

CTCACTAGAAATTAAGGTGTAATTTAGACACTAAAATGTGTCTATCAGACTGGCGCTGAGAAGAAGAGTTATAGTGTGAA

TCTGGCGAAGGCCTTGGGGATCACACTATTCTGAGGTTTGAATGTTGTTTCCCTGAGACTCATTGGGTGAACTCAAAAAG

TTGTATCATAAGGGTTTGGGTTGGATTCACATCTCAGATCAGAAGGTATACATTGCTTCTAGCGGCCTAGTGATAATCAT

TCTCTATGGAACAACCAATCTTTTTTTCACTGTCAAGAATTAGGCCTACCTAGACAATATCACTTCTCATTATTGTAGAG

TTCTGATACAACATTCAAACGGTGACTGCAAGATTAAAATTGGCTTGGAATTATGACCCAATGAAAGCTTTAAAGCTGAC

ATATATGCAACCTAAGAGGAGTTAAAGGAAAGGAAATTCAGATAGAGAAGGGTTTAACGCTGCAGTGTTAAGGCTTCACA

AACATCATCCTAAAACCCTGGCCTGCAATGTTAAAGTAATTGCAGATTTCGGATATTTCAGAGATTTTCTTGAAGTACTT

TATAGATGTTGGTTCTACAACAATGGAAAAAGACTAAACAGAGAAAAATTTGTTAACGTTAGGAAATTGATATACACTCT

GAAGTTGATGCACACACAAGGACAGAGCCACCTGCTCAACAAATTGTCCTTAACACAATAGTTCTCAGGTACTCCTCAAT

AATGAGAGATGCTATACCCCAACAGAGAAGATGTGTCCAGGATAGAACTTCCCGTTATGATTTGAATTAACACAATACAA

ATCACCCTCTAGGACTTAACAACAAGATAGATAGAAGGGAAATAAACAATCAATTGCGCAAAACAAAGCAAGAAGAGAGG

AAAAAAGCTAGGGATAGTAGCCTGCTCTGGAGTAAGTTGAAATAGAGAAATCATTTTTCTTTTATAGACAAAGCATGAGG

TGAAATAAGTGGCGCAATTAATGAGAATTAATTTTATAATTAATCACAAAATTAATTGTGAAATTTAATTACTAAACGGT

AATAAAACGGTCAAAAGATATTTTCTTTTATGAAAAGAGTTATTCTCGCCGGGTCATACGCCCCCACATTTAAAATAAAA

TTTAATGAAAACTTTATTTTCAAAAACGTAAATCTTGGTTTGCCCGCACCCTCAAGCATACTCAATCCCGCAGCACATAA

GCACCGCAAGCTCGCAAGCGTGCGTGCTTCGGTGCGAAGCACCGCACATGTGTGAAAGGTATATGTGCACCAACTATCTC

ATTGGCTTAGTCCTCCCCCAACCACAAAAACTTCTAAACCCTAATGGCCCTCCCTTAGTTGCATTTTTTGTGGTAATAAA

GGGAGAACTCATCTCTAGTCTCTCCAATGTGGGACTAAAGGTTCTTCACAAAAAAGTGAAAATGACTTAGTGACTCTTAG

TGACCCATTGTTCCTAAGATCCATTCCTACCAAGACCAATAAATAATTTTAAGTTCATTTTCTCCAACAATCTCCCACAT

GAATGAAAATTAGAAAGCGAGAAATACCTCTTAGGCAATAGGTGTCCATAAGGTCTTGAACCTTTTCTTAGTGAGAAATT

AGCGGAACTAGTTGATCGACAGTGAACGCGATGTCTTGAACTTCCATTGTTTGGTGTAGTTCATCAATAACTTCCACGCG

TGATATCTCTCTAGGTGTTGTTAAGTTCGTGCCATTGTGTCCATTTTGGCTCTGGACGTATCCCGTTTCTCAGAATGCTC

TAGAGAATCATCTCAGTTCTCATAGGAAGCGGCCCACTTCCTCATTCAGATAGGTGAGTTCTATCAAGAGTGTTTACTGC

TACACCCCACTTCAATCTTAAATTGAAACTACAGAACTCATTAAGACTTAATAAAAAGTCAACCTCACATGCAGTCACAC

TATCACGTCTACACCATAGGGGAAGGACAAAGAATGAAATTCTATGATAGTAGTTCCTTTAACCCACCACAAATTAGTTG

CCTCATTCGAAACCTTGATCTTTGGATCTCCAGTCAGCAAGGTTGAGTATCCTTCGTGGCAAGTTTATTTGATGAGCTTA

AGCCCCATCCCCCTTGATGCATTACTAACTATCTTCCGGACAAACCTTTCGACAAAGGGTCCGCGATATTATCTTTAGAC

CTCACAAAGTCAACAGAAATAACGGCAGTTGAGATTAGTTGTTTCAGGGAATTGTGACTTCCACGAATGTGACTTCCCAT

TATAGTTGTTGTTTTTAGCATTAGCTATTGCAGCTTGACTATCGCAATGTATAGATATAGCTACATCTCTAATCGATGCA

ACAACCTAGAATACCCCCCTGAGTTTTCTGAACAAGCTCAAACATTATTTACTTAAATGAAGTTCAACTGAAAATGCAAA

ATATGAAACCCATAAACATGTAGCTTCACAAACATCCCATAATAGTTTTCTAAACTTCCCACTTTTATCATAAGGGAAAA

TAAATGAAATAGAAAAAAAAGCTTTTGTCAATTAAAAGAAATAGAAATAAATATCATGTTATTAAATAAATCTAAAATAA

ATGTATTGTAGGTGGAAATATTTCACAATCCAAGTTTTTGGAGGAAAATTGTTTGGTAGGTGAAATAAAGAGTGGTAGAA

ATTGGGGGCAAATCATTTTCAAGATTTTTTAATTTTTTTTGAATTAAAACTAAAATCAAATAAGCAAATAAGATATTCAG

CTCTAAGAATCCTTGAAAATGAGACAGTTTACAATAATGCCACAATTTGCAACTTATAAGATGTTGCAAAAACGTTCCAG

GTGTGGTGGTTGTGTTGAGAAGGGACGGAAAATATTTTACAGAGAGACGTCAAGACAAGTGTGTTTACCTTGCTGCATAT

GCGGACACAAATATTTTACAGAGACATATCTAGATAGGTATGTTAACCTTGCAGCATATGCCGACGAAGACCATATCGTT

CGAGTTGGAGTTCAGTTATGTGGTGATTAATCATGGCTAGGCACTAATTATCAATTTCTATATACATAAACACTAAATAA

AATGTCATCTTCTGGATTGCGAATGGTAAATAATAAATTGAATGACCAGACATTGAAAAAAAAACAAACAAACTAAACTC

CCTAAATAGTGAAACAATTTTGAAATTTAAGTGCCATTTAGACTAATTATGTGTGTGATTTCGACACTAAGAGTGTCATT

TTAACTAATTAGGTCAAGTTCTATATATCTCTCAAACAATTTAATTTGTCCATTTTAAGCTACTATATATGGACCTCTTA

AATAGATAAATTCACCTCTTAAACGCTCAATTTTGAAGGTTTGAAGATCACGTCAGAGTTTTTGTCGAGTGATAAACACT

TTATCTCTCTCTGGCGACAAACACTCTGTCTCTAGCTAATTATTTTAAGTATTTTATATTCTTCTCTCATATTTTTTTAG

TCTTTTAGGGCGTCCACAGTGGATCGACCATCCCAAAAGGCCAAAGACTAAATTTTTTAGTCTTTGGTCGGCCCAATAAC

AACCATATATAGTCGGACGGTGTTCTAGTGGAACGCTTGGTGTTGGGCTGTGTCCCAACATAGCGCTTGGAAAGACGGGC

AAAATTCTAGCATAGTGTTTGAAACAGGCAGAACCCTAGCATAACGCTTGATTTGGTTCGGGAGGGGAAGATAACGCCTG

GTAGTGGACGGTATCCTAAGACCACGCCTAAGAAACGGGCTGGGACATACCGGAATTTTTTTGCTTCCAATTGGTCGGGT

TTCGGTCCATTAGACTAGCTCCAACTTCCCGATACGCCACTCGAAAAACTGAACTTTTAAATTTATCGGACCAATTCGAT

TTTTACCAAGCAAATGTTTATCAAATTATCAAGTCGGTTACCAGCTTCGCTAAATATTTTGTCTTACAAAGAAACTGTTC

ATTTGAACTGCTTTATCAGCAGACAAGCACAATACTGACATCGAGGGTCATCGAATCTGTCAAAGCTCCATCGTTTTAGA

TATAATTTTTGATATTTATATAACCAACCAAATGATATGAGCCACCTAGTGTAGCTGAAAATCATCTCCAAGCGAACATA

AAGTTTTGATACTGGGATTCTGCTGAACAAAAACCTTGTTTTAAGGGTCATCTCATTGTCAGGGTAGTCGGATACCGGCC

ACTGTTTCCAAAACAGTCTTCCGGGTAGCCACCGAGGTGGTTGGATGCTAGCCGGCCTGATAACACACCGTTTTAAACAG

GTGAGGGATGATCAGTCCCAGTGTCATGCCACCGTTTCCAAAAGGGCCTTCCGGGTAACCACCGATATGGTGGGTTCCCA

ATTAAGGCCTGACAATCTCGTCGTCCCAGACGCTTCATCCACATGTGGTGGGGGTGTCAACTATGGCCTAGTAACTCACA

GTCGGTCCAATCTGATAAGCTTCATTTCTCAGATGATATGAACTACTTTATAGAGTTAGTAATCATCTCTAGATCGATGA

TATGAACTATCACCATCGGTAGCAACCATCATTCTCAAATGGTATGAGTTGCCTTCAAAGGCAACAACCATCTCTAAGTC

TCACTAGAAGTATGAATTGTCTTCTTCACTTTTCACTTAGATGATATGAACAACCTCGAAGAAGTTGTAATCATCTCTCA

AGGTAGTATGAACAGCCAGTACTGCAACTACCTGGTCGTGTCGTAATCAACCAAAATAATCGAGTTCTTACTCTTCAAAA

ATAGATTTAGGTATAGCAAATAAGAACAAGTTCATTCCCACAGAGAGGCAATTAAATTGTTATTATTCTTAGTTACGCAA

GTAAACAGGGGGGTTTTATGTTGTGAATTAAGCTAAATTAAACGAAAGCAAATTAATAGAAACGTGTATCAGGGAGAGAA

ACATTGGTTAGAGAATTCATCGTCCGTTTTCCAACATGTTATCAATCAAAGCATACTCAACATTCATGCATTAGTAATAC

CAAAAACCCTAAAGTAGCCAGTTATCTCGTCTACTTCGGCAAAACTCCTAATATCATCAAAGAATGAGTACCTCCACCCT

TTGAATTCTTAGCACCGGATAACCTGAGTGAGTACCTCCACTCGAGTTCAATCCAATGATAGCTATGGTCTTTGTGAGTT

GAGATTAATCTAAGTAACCCGGCGCAATTCGCCACTCGGATTCAACCTAGGTCAAGCTTCTACACATACTTGCATGGCAA

TTTTAAGTATCGTTAACAAATATGCTATACTACTTGTAACCGGAGGCCCTCGACAGAATCGACTCTCAGAACTCCAGTTT

AGTTGTAGATATCAATACTTATTTACCTAATTTGCAACTCAAATAAATCAGCAATCAATCATATATAACACTATATTCGG

TAGAACATGAACGATATTAAATAACATGAAATCATGAGATATGCGCCATTCAATTAATAAACATGTTTTCCAACTTCAGA

TTACATCCCTAACCAATATGAAAGGTTTAGCTCAACATGGGTTTCTCAAAACCCATGGAAATTAATCAAAATATACTGTA

CTTAAACCCGGAAAGAGTACGAAGAACGGATACAGCTTCGAAGGATGGTTGCAATGGTGGACAGCAGCCTTTTACGGAAG

TGCTGTTTGTTTTTACGCCTCTTCTTCTTACCTCCTCTCCTGTCTTGTATGGTATGGTAGATGTCCGTGATATGATAGAT

AGCCAGCAGCAAAAACTCCCTTTTCTTTTATATATCAAGTCCTTCTTTTTAGCTCAGCAGCAGCACCATGTATCCAGCCC

TTTTACGCCAATAGGATTCTTGTTTGTACGTGAGTGGCTAAGTCCAAAATATTGTTGTTTCACCACCTTTCCTTACTTGC

CTCAAATTCTCCTTTTCCTACAAGAACTCCTATATTCTTTATCTCAATTTTCACCAAAAGAATCCATGAGAAACCGATTT

TTCTTCTTATAGCTAGAGTTTGCCACACTTCCCTTTTGCTAGTCCTCGAGCAAACTATCAATCTAAACTAATAAAAAGAA

ACAAATACAACTCCATTTCGTCGAGTAAACGGTTACACTTAGCATGTGCAACAAGTCTTTAAACCCCTAGGTGTCCCTAG

TGGACGAGTATAGTCTCGTGAAGGTTTACAAGAGGTATACCCACAAAACCTGCTTCAACTTCCAAAAATTCCAATGCACC

TGCATCCAAAAAGCTACGAAGACATATAGTTTCTACATACCAACAATAAGGACTCAGAAATATATAAAAAAAAAATTATT

CTCATGCTTAATCCTTGAGTTTGGAGAAAACCACCAAATAAAAGAGTGCCGGTTCGAGAGTGACCGAATATTTCTGCCTT

ACCAATAGGATTTTATGAATTTGACACTCAAAAGACGCACTTTATGGTTCTCACATATGTGCAGGCAAGAATAAAGGTAG

AAGGCCTACACATGTTGACCTTCCGAATCATATTAAGATGTTATCCCATAAAAAGTGGGACAAGAGAGAACCCCTTTTTA

TTTTTCTAGTGACCTCTACGAAAGGAAATCAACCATTATGCGAAGATGGGAATTGTTACTTACCTTCTCATCCGGTCGAC

AATCAGCACCACTCTCCCAACATCCCATAGAAACGTCTTCGGTATCAACTGTAGATAATGAACTCTTGAACTTCGTAGAA

CTTGACGATTTTTAGCTCTTTTTCTCTTCACGGCCTGTACCACTTCAAGCACTGTAACTCAGAATGAATATCTCCGGGTA

CTGTAACTCAGAATGAATAGCTCTAGGTCAGCAAAAAATAAATTTTCTTCTTTTTTTTCTCTTTATTCCATATTCAACAA

CTCTTTTTCACGTTTTTTCACGTTTTTTTTTTTTTTTTTTTTTTTTTTTTTTTTTTTTTTTTTTTTTTTGTACGTGAGTA

TGCTTATTCTCCTGGATACTTTTGCTATTTTTCTCAATTTCTTGATTCGTGAGTTTTTTCTCTTCCGGTGAGTTCTATTT

CGACTCTAAATCTGTATAGAATCCACACCTTTCGATTTCTTTCGCTTCTGTAAGTCTTCAACATGTTTAATTGTATTGTT

GCTTTACTCAAATATTAATTTTTCTCTTTGAATTGTGGCTTGCACCTTGACGGTCTTCAACTTTCCTTCAAATTTTGATG

TTCCGCTTGTTGATGATGCTATGTTTCTATGGACGTTGTCGAGAGAACGATGCCAACTGCAAACCAAACTACAAGAATAA

GTTAGCAATTCTGATGACACCCTCATGATTGATAAAAATAGCACTCAAGAGATCAAAAATAGTGTTGAAAATTGGTGCGA

GGTTAGGTAGTTCATCATTTTACCCGGAATTAACGCAGATGAAACACTCTAAACGTATAATTGAAGGTCTGGTGCCTCAC

TCATGGGTATTGAATGATCACAAGTCTAACCTGTACTCTCCTACCGGCAATTTAGGTCCATTTTGGTCGGCTCTAGCCTC

TCAAAAACTAAGCACTCCGAATGTAGTTCTCTAATATTTCTAAGTTCAGTTGTTTCGAGTAGAAGTCCGTATGTAAACGT

AGCTAGCCGTATGCTAAGTGACTCTAAAATCTCTAAAAGTTGAAGGTTTTCATGCAAATGTGTGTATGCTTCAAAATTCT

ACCTAAAAATCTATATGCCTAAATACTCCCCACACTTAAACTTTACATTGTCCTCAATGTGAATTGAATCGTCCAATGTA

ACGTCAAGATGGGAAATTCTACATGCTCTAAAAATAAAAATCAGCAACAAAAGATAGAAATACAAAACCAATGGGTTGCC

TCCCATTTAGCGCTTAGTTTAACGTCTATAGCCCGACTCTCTGATCGCCCGTAGACCAATAATCTGAAAATTTGGGGATC

CACCCATAAAACAATCTGCATTGCAAACGTTAACTTCACAAAAATATTAATACAAGAAAATAATATTGAAGTTATCGCTC

TATAGAAGTTTCCCCCACACTTAGTCTTTTCTACACTCGGAAGTGGATAAAGAACATAGAACTCGGGAACTTCTATAGAC

TCCTCTTTGCTAACCTGCACAGAATGAGAATCAAGTGCTTCCAAAAGAAGCAAAGTAATTTATCAGTACTTCTGAGGCAC

GTAAATCCAAGCCAAAAAGAGGTAAAGGGAAAGGATTAAGAAACAAAGGTTTAGGAAAACCTTCTAAATTATCGTGAATT

GTGGAATCCTTCTCTTTTTCAAAAATATGTTATGAATTACTTACTTCTTGTATGTTGTGGTCCTCTATCAAACCTTCTGA

CTGCTCTGCCTCATTAAAAATCTCGGCACTACCATTATTAATATCATCCTCGGAAAGTTCAATTGGGTCTTCCACAGTAT

TAATCAATTCGGTTTCAATCTCAACACACTCCTCTTCATTTTCATACTCACTGTAATAAGAATCTGCATCGGTTTCCCAA

CCTTTGTCAGAACGTCGATTAATTTCCATATGGATGGCCTTACTAATTTCTTTGAGAAGCTCTTCCGACGCTCCGTCATC

CTCCAACTCGCGGAATAATTGAAAAAGTTTATCAATATTCAAAAGTTTTCCACCATTAACTACAGTGGAATCTATTGAGC

ACGACTCTTCCCTTTGAATGGGTGAATGATCATTATTAAGATCAAGACTCGAGTCAACTACACAATCATCATCAACCAGA

ACTGGAGTTGGAGTAACGTCATCACTGTTGTCGGGATTTTCGAAAGGTTCAAAATCATAATCGTATCCCATAGAATAACG

TCGTTCACGTTCTATTTCCAAAGCTCTTATAGCTTCTGCAATTTCTTCATCTGATGCCCCGTCATCTATCAATTGCTCAA

ACCTTCTCTCAAGTTCTCTCAAACTTGGTAGTCTTACACCATTACTTGTGGTGGAGTTTCTAAACCTGATCTCATCCCGT

TGAATAGGTGAATGATCATAATTACGATCAAGAGTAGATTTAGAAAAACCATCATAAGCATAATTAAAAGGGTACGTGGA

ATATTGGTCATAACCAAAGGATTGGTTGTGATTATACTCCCCGTATGGTTGGAAATTGTCGTAGGGTTGAGACTAAAACT

AAATTGATTACCATCATAAGCTTGATTTTGTGCACTGTACATGGAGGAGTACCTAAAAAAAATTCTCAAAATTCGGTTAG

AAAAGAAAAAGAAAAAAAAAAAATCAGCAAGAAAAATAAGCAGAAAAATAAAAATAAAAAGAAACTAAAAATACATAACA

ATTCGCTATGCTTCCCCGGCAGCGGCGCCAAAATTTGGTCGTGTCGTAATCAACCAAAATAATCGAGTTCTTACTCTTCC

AAAATAGATTTAGGTATAGCAACTAAGAACAAGTTCGTTCCCACAGAGAGGCAATTAAATTGTTATTATTCTTAGTTACG

CAAGTAAACAGGGGGGTTTTATGTTGTGAATTAAGCTAAATTAAACTAAAGCAAATTAATAGAAACGTGTATCAGGGAGA

GAAGCATTGGTTAGAGAATTCATCGTCCATTTTCCAACATGTTATCAATCAAAGCATACTCAACATTCATGCATTAGTAA

TACCAAAAACCCTAAAGTAGCCAGTTATCTCGTCTACTTCGGCAAAACTCCTAATATCATCAAAGAATGAGTACCTCCAC

CCTTTGAATTCTTAGCACCGGATAACCTGAGTGAGTACCTCCACTCGAGTTCAATCCAATGATAGCTATGGTCTTTGTGA

GTTGAGGTTAATCTAAGTAACCCGGCGCAATTCGCCACTCGGATTCAACCTAGGTCAAGCTTCTACACATACTTACATGG

CAATTTTAAGTATCGTTAACAAATATGCTATACTACTTGTAACCGGAGTCCCTCGACAGAATCGACTCTCAGAACTGAAG

TTTAGTTGTAGATATCAATACTTATTTACCTAATTTGCAACTCAAATAAATCAGAAATCAATCATATATAACACTATATT

CGGTAGAACATGAACGATATTAAAAATAACATGAAATCATGAGATATGCGCCATTCAATTAATAAACATGTTTTCCAACT

TCAGATTACATCCCTAACCAATATGAAAGGTTTAGCTCAACATGGGTTTCTCAAAACCCATGTAAATTAATCAAAATATA

CTGTACTTAAACCCGGAAAGAGTACGAAGAACGGATACAGCTTCGAAGGATATTTGCAATGGTGGACAACAGCCTTTTAC

GCAAGTGCTGTCTGTTTTTACGCCTCTTCTTCTTACCTCCTCTCCTGTCTTGTATGGTATGGTAGATGTCTGTGATATGA

TAGATAGCCAGCAGCAAAAACTCCCTTTTCTTTTATATATCAAGTCCTTCTTTTTAGCTCAGCAGCAGCACCATGTATCC

AGCCCTTTTACGCCAATAGGATTCCTGTTTATACGTGAGTGGCTAAGTCCAAAATATTGTTGTTTCACCACCTTTCCTTA

CTTTCCTCAAACTCTCCTTTTCCTACAAGAACTCCTATATTCTTTGTCTCAATTTTCACCAAAAGAATCCATGAGAAAAC

CGATTTTTCTTCTTATAGCTAGAGTTTGCCACTTTTAGGAAACCGACTAGGAATAGGAAATGTGCTAAGTTTCCTAATTC

CATCCCCACAAGTTATGGCAGCTCCAAAGCTTTCCTAATTCCATAGAACCTCCACAAGCTAGCCATGAGTCCACCAAGGA

TAGCCACCTTCCACTCGGACCACTCTAGCTCAACAACACCTCCACATGACCACTTAGCACAAGCGCACTTTTGCTTCAGC

CAACTTAGCGCCTGGGCGCTGGCAGCACCAACTAGTTAGCCACCATATGGCTTTGCCCATATCTTCCTCATACGAGATCA

GAATGACCTCGTTCTTTTGCCGTTGGATTCGTAATTCAAATTCCTACATTATGGAAGGTGAAAGGACTGAATTCAGCAGA

GTTTTACTGGCTCCTCGGTAATTCTTCCCTTTAAGCTCCATTCTGCGGATTTCCGGACTGTAACCTGAAAAGATGACAAA

ACTTCGAAAGTAATCGAAATACATAATAATTCACTCAAATATAGCAAATAAGGATCAGGAAAAACTACTTAGAATGTATG

ATTTATGCACCTATCACTACCTCTTACTGCCAAGGTTAAAATCATCTCCGAATGGATGGTATGAACTGCAAAGCAGCAAC

CAACCTTTTCCCGACGATATGAACAACCTTAAGAAGTAGTGCTGAACCGCATAGCGGCTGGCTACGTACCCTCTCCGTAG

CTCGAAGGGATCAAGCACGACCATAGTTCGTCTTTTTCGTTAAGATAGTATGAACTACCCTCAATGGTAGTAACTATTTC

TTATTAGATGATCTAAACTATCATAATTTGATGGCAATCATCTCTTACATAGATTATATGAACTGTTTGTCATAGCAGCA

ACAATCTCTGAGATAGTATGAACTGCTTACCCAAGGAAGTAACTATCTCTAAATATAGCAATAAGTTAGATAATATAGAC

AGTATGAGTTGCGTAGCAACAACTGTCCTTCACACATCCCATCACCAACAGTCACTTCAGAAACCACCCATTGACAAGAT

GTCAACATATATTTGTACTTGTACCTGCAACTTTTAATTAACAGGTTAATAAAATATAAAATATACACAAGGATACTTTA

TATAAATCATCCATTACATTAATCATAACGAGGTTAAAGAGCATACATTATAGTTCTAAGCTGTAGAATAGAGATTTAGT

GCTAAAGTGTCATTATAAAAAAATCTCCACAGTTAATTCGAACAAAATATATTAAACAACACACTTAAACAATTTATAAA

CATTTATATACTGTGTTATCATTCAAAGGATTCATGGGATTGAATAGCAACCACACTTTTAATTATAAAGCTCACTACTT

TATTGTAGATGTTTTCATGCAAAAATACCTCACTCTCTGTACATATCTTATGTACATTAGCCAACTAGTTATGGAACACA

AGTGTATCAGTTTTTCACCGTATCGTTCTGTATAGAAAACAGAGGGGATAATCATTGATATCTACTTAGGTTGTGCCAAA

TGTTGTAAATTAATTCTTTGGTGGATTCCACCACTCTTACACCCCTCTTACACCACCACCTTATCATCTCCTCTTCCACC

TCTTCACATCTTTCACTTCTTGGTAACCCCCACATTTAGTGTAATAACTTAGGTTTAATACCTTAGTTATGAGAATGCAT

TCCATTACTTACTAATGTTAATGCTATCTTTATCATCCTCTCTTGTATTAGCCTATAAATTGAGGCTCATTCCTTTGTAA

TGTATACTACTATTTAGAATTGAGAAATTCTCTCTTCTTCTCTCTATATTTCTTTTGTGCAATAGTTCATCTTTTGATTT

AGTTTATAACACGTAATCAACATACTAGCTCTCCTTAGCGGGTCGTCAATTGACCACGATGATGATCAAATTTCTATTGT

GATGAATCTTACACAATAAGGTAAAATACTATATATCTTTTTTCTTGTTTTTAAGAAAAACTAAAATCAAAATCTTTTAC

TTTATATTCATGAGTTGAAAAAAAAAAAGACTGTTCCACCTGAAGTTGGAACTCAATGTGACACAATATGGACAAAAGAA

GTTCACTCGGTTAGTCGGTCGGTCCCTGAAGTGACCCGTAACAAAGTGACATCATGTCTCTATCCGCGGTTCCTGAAGTA

ACATGACCCTGATGTTAAGACAATTTTATTTTACAAATAATGTGTCCACGTGTACACTGGCAATGAATATTTCGTCAAAC

CCGAAGATGACGAAATTGATATTTAAATTATGTTGGTACTGTATCTGTATATGTTCTTTATTTATTTTGATTTTTGTTTA

TTTTAATTATTTGTCTTGAAAAAAAAAAAAAAAAAAAAAAAAAGGATTCCAGAAACAATCCATGGGATGTTATTAACTCC

AGAAGTAGTCTAACATGCAAATTATAAATTTTTGTAGCAATCCAAATTATTGTGGGTTCCCGCAGCAATCCATAAATATA

GACTCCAGAAGTGGTCTATGAATTTTTTTGGATTTCAGTTATCCATAAAATTCTAGGTTCCCGAAGTAGCCCATAGTTAC

TAAATACTTTTGTTAATGGTTTTATTATTTTTCTCTCGTCAATCTATACTTTTGTTGATGGTTTTATTATCAAAACTCAA

AATATATGTCATATTCTTTGCTACAAGAGCACAATTTTTACAGCATTAAACGTTTCCGACATATTGATTGATTAGTACCA

ATCAAACGGAATTTCCCGCTCAATTTTTGGTAGTACCAATACAAGAGCATATGACTATCGTTTATAATAAATGCTAATCA

GGGTGCACACTGATAAAGTATACGATGTCAAATTTTAATTACATATATGAGCATGGGTGATCTAAAATTATTAATTGTAA

TATTATGATGAACAATCCGAAGAAAAAAAAAAGTTTTTCGGATGACCTACTTGGATCCAAGGTCATTCACCATAATATAT

TTTGCATGAATAATACTAGATGTGATGAGACTTCTCGCCATTAGAGGGAGGAAAGCAGTCGTTATAAGAACCTCATGAAT

TGGTCTCATCATAATTCTTGCACCAATTCGGTATAAGGTGGAGAAGTTATTCATTCATTATGATATGCTTGAAGCATTTT

ATTCATACAAGAAGAATAATTAAAACTCATTTGTTGTTGCACGTATTTTGGCATTAGTTTATGTCCCTGCAGGGCATGCC

ATGAGTTAACAATAATTGAGTCGTCTACAACTCATATGAATTGTTGATGAAAAAAATGAAATAAAATAATATATATTCCT

TGATAGAGGAAGCCAATCACGCTGGATCAAAATAGTGCTCCCGGGGAGGCTATTGATAATGTTCTCAATGAGACTATTTA

ACTACGATCTTTAATTACCGAGCCATATTTATCTCCTAAAGAGGAATTCATTCCTAAGTGGACAAGTGGTGTCGATATTA

AAGAATGATTTTTATACGCGTATATAATCGGAGTTATTGTCGAACATGTATTCAGAGTAGCGACAACCATACATGTATTA

TAGACAAAATCATTTTTCGCAGAGAAACTGCCGTGAAGTATCGACAATATGTATGGTTGACTAATATGGGTGAAAGGCAT

CCATATTTGGTTGTCCTCTCACTAATAAACATCTTTTGACATAAATAAAAATTATGAACCCGGAAGGTTACAAGTTTGAT

GCTTAATTAGTTTGCATGAAAGTGAGACGAGAGGAACAAGGACGCTAATTGCACAGCACCAATCGTGGCATCTAGTTTCT

ACAGACACTTGAGATCTGTGAAATTACATCATCTACTCTTAAATGATTTGAAAATACCCAATACTTTGGTAAGTATTGTG

TACCCCGTTGATAGATGAATTTTAAGGGAAACTATTATTTGCCAGTTTTACATGTTATAGTCTGACATAATTGTTTGTTA

TCGAAGATAACGTTCGTGTAAACTTCTAAAAGATTACCACAGGTTATACTTCCTTATTGAAAGAATTCAAGATAAAAGAT

CTTGGAATCATGTTCAAAATTATTAAGCATTATAATGGTCCATGATTTTGTCAACTTTAGAAAGTTTTGACATGTTTACA

TGGATAAGTACGTAACCTCCTAATAAAACTTGTGTAATAGCAATCTACTTTCATCTCGGTAACGGGTATGTAACACTCTC

TCTGGTTCGTATTACGTACTAATTACAAGTTGTTTTTGTGCTTGATTTTATGTATAACTTTATATGTTATACTTGAAATT

GTTCGGTAGTGAATTTGTTTGCCAAACTTAATTACCACTAATATGGAGGCATTGTGTATATTGGTTTTTGTAGTACCTCT

AACATTTGATCTTTGCTTTGACGGTTTATTTAAAGTCTCGTAAGTAATGAACTACAGATAATCAGTTTCCATTTGTCAAA

TTGTTAGACGATACAACATCCTTTTTGCAGCAAAAACGATGATCATCTATTATTGTTTGAAAGGAATTTACGAAGAATAA

AAGATATATCAGATCCTACAAATCAACCATTGGGAATTTGACAAAGACAATTCAACTTGAGATTTGAAATTAAGTTATGG

AAACATAAAGGTCTCCATTACCAACAAATTTTTTTTGAGCTGAAGATGGTTGCTGATTAGCTACTTGTGGTAAATCAACT

GAGATATCTTGTTTCTGCTGGAGGATATATTGGACTTTACGAGTTGTACTCTTTTTCCTTAGTCAAGGTTTTTGTCGAAC

TGAGTTTTCCTTCTCAAGGTTTTAACAAGGCAATATATATATGTCCAACTCCTTTTCTTAATCTATCTACAGTTGTACTC

TTTTTCCTTAGTCAAGGTTTTTTCCTTATGGGTTTTCCCGACAAGGTTTTGATGAGGCAACGAACTTGAAGAAATTTTCT

TAAAGAGATATTCGAAGACATTATTGAAAGACAAACTTATATCTTGTTTTGAACAAAGATGGTCAGTGATTCTACTTGAA

GATCAACATTATGTAATCAGTCTTTCTATACTTCTTTGGGTGCACATCATCAGGGGGAGAACATCAGTATTGTGTGATAG

AAATTTATGCGTGTTGTACTCTTTTTTCTTCGCTTGGTTTTGTCCCAATGGGTTTTCCTAGCAGGGTTTTTAATGAGGCA

ATATTTGACGCGTCCAACACCAAGATATGATTTGTACTCTTTTTCCTTCGTCTAGGTTTTATCACTTTTGGGTTTTCCTG

GCAAGGTTTTAACGAGGCAATATCATAATAGAGTGGTGTCATCCAAGGGGGGTGTTGTAAATTAATTTGTTGGTGGATTC

CACCACTCTTACACCCCTCTTACACCACCACCTTATCATCTCTCATTCCACCTCTTCACATCTTTCACTTCTTGGTAACC

CCCACATTTAGTGTAATAACTTAGGTTTAATACCTTAGTTATGAGAATGCATTCCATTACTTCCTAATGTTAATGCTATC

TTTATCATCCTCTCTTGTATTAGCCTATAAATTGAGGCTCATTCCTTTGTAATGTATACTAATATCTAGTAATGAGAAAT

TCTCTCTTCTTCTCTCTATATTTCTTTTGTGCAATAGTTCATCTTTTGATTTAGTTTATAACACCAAATTGATTTTTGTC

TCATCCAGACATAGTACTAATAACAACTAAATAACGTACCACAATATTCCTATCAATTAGATTTAGTCCAGCCCACTGGT

ACATAAACTGATGGAATAATTTATACTAAAAATTACACCAAACTCTACATCTGTATATCTCGTTACCGAGAAAATAAAAT

GCAAGCAAGAACCTCATGAATCAAACATCAAACCGCTTGAAGTAGTGGTCAGTGGGAAAACCTATTATTATACAGCTAAA

CCACATACTTTTCTTTAAGAAAAATGCAGAACACTAGAATTTGTTATATAGCCAAAAAAACCAAAACGAATTCCATTTTA

TACATCAAAAATAGGTCTAACTTGCAAGGTCTTACTTTATAAAAACCCAGCTTAAATGAGTAAGTACTTAAGCTTCATAT

ATAGAGCGGAATAACCCATCGTACCGGACTACCCCAAATTAAAAACTGATCATACTGAACTCCCAAGTAAATATATTACT

GCGGTTCGTTCCTTTGAACCAAAACAGTAAAGTACAAAGCACAACCAAGGTATATTAAGCATCCAATGTTATCAGGCCAT

ACACAAATTTTGTAAAAAGCCAACAGCTACAAATGCAGACTATAAAGTGAATTAAATAGACAATGGTGATTTCAAACTAT

ACCTCATAAAACCATTCTGGAACATCGACAAGATATTAGGTCGTCAACAAGACCACTCCGCAGTACCTCCAAGTTAAAAT

GTTAATTTAAAACCACTTCCTCAAGCTAACCAGAAAAAGAAATTAAGTTTACATTCATTCATGCCGCTGTTTAGTATGCC

TACATATTACCACACAGATCATGCATTTAACTACCGACCATGCCAATGTAAATTTAAAAGATATAATTCAAGCCTTCCCA

CTTCACCACATATACTCCATCTGAGATCCTAAAGTGTAAGATCAATATTGTAGCATTATTTAACCACCACAACTTAATTA

CAACCATCAGTCCGTAGACATGAACTAGAAATCTATACACCACACAATAAGCAACATATTAATCCAAGAGAATTAGTGCA

CCATGTATTATATGAACTCAAGGTTATACATGCATGTGGAAGTCAACAGTAGGAACATATAGCATATATCTTCAATTTCT

TAAATTTTGTCTTGTCAAAATGGGTGTCACCATTTAAGAAAGGCATAGAAACAAGGTGAGTGCAAAAATTAGCCATATAC

AGAGGATCAAAATATGTACTTGCTTGTTGTAAATAAACCCAAGAAAACTAAAACATGAGGAATAAGAGATCATCATCGCC

TAGTAATAGCTGCAGTGGTAGAATAAACTTGTGCACAACTCTCTTTAATCCAGGCCTTTGTAGCATAACAGTCAAATAAA

CTTCATCATTCTTAGTTCCACTACCATCAGTCTCACTGGCAAGTAGCCCATTTCACCATCTTATGACCTAATTTCTTAAT

CCATGGACTATAAATCTCTTTAAGATCCAACTTCATATCTTTCTTCAGGTGACACAAACCCACCAATTAATAAGAACTAC

CTTGGTCAATTGATTGAAAATCCATTTCCTCCCGATAAATCCTCAATTTCAAGAATAGAAAGAAAAGAAATCCCCTAACC

AGTTTGTTGGGGATAAAATGATGTTCGCCCGAAGTGACATCATCACGTCTCGAAATGACGTCATCAAATCTGATGCAACT

TGGACCGAAGCGACGTCATCATTTCCCGAAGTTATGTCATCAATTCAAAACTCCAACTTTGACCGAAGCGACGTCATCAT

TTCCCAGAGTGACGTCATCAATTCCGAACTCCAACTTGGACCGAAGCGACGTCATCATTTTCCCAAAGTGACGTCATCAA

TCTCGATGTACCAAGTCAATTCCGATAGACCAAGTCAATTCCGCACCAACTTTTGGATTGAGGTCATTTTGAAGATAAGG

ATAATAAGAACACGTGGCGGATCTCGTTGACACATGGCGAAACGTGGTGAATCCTCTACGACACGTGGCGAGCTTAGAAG

AATTTTCATCTACACGTGACGAACTTTGATGACATGTGGCGAAGGGCTTTTTATTAATTATCATTTATTTTATCCCTATA

AATACAATTTTAGAGTGAAGAAGAAGGGGATTTTTTGGCCCGATGGAGGCTTCCTCTACTTCATTCTCAACTGTAACCCA

GATATGAATAAAAACTCCAACTTCATTACCCATGGATGTAGGCTAATCATGGCCGAACCACGTTAAATCTCATGTGTTAT

CTTCATTTTTGTCTATCTTTATTATAATTTTTGGTTGTTTTTTGCCCATAAGAAATTCGGCAGAAACACAGTTTAAGAAA

CTAATTCTCATGAACGACATCAACAATTCTATGTAACCATAGCTCAAAAGAAAACTTCCTGATTTCCATTTTCAAACCAA

GTTCTTAATTTTTCAAAGGAAGGGCGGAGATTGCCAGAGAAGAAAATACGAAGAAGGGGAAAAGAAGACGGTGAAAAGAA

GAGAATTTTCTTTGTTTTTATCACTCTGGTACGGAAATGAAAGAAGTGTCCTTTGTTGTTCTTGTCTAGCCATGTGTCAG

ACATGTTTAAATTTGGTTTCAAACACATCTACTAGTTATGCATATTGTTTCTACCAAGGGATAACTCATTTAGGACACAC

ATCAGATTTAGTTAAACCCGAGAAAAGTACGGGTATGACACGGGCGGTGTCCCAGCATCAGACCTGATGGGGCGGTATCC

AAAAATCACGCCTGGTGGAGTATCACATGTGTAACAGAGCGTGTGAGATGAATGTGACGGGCGATGTCCTAGCGCCCCGC

CTGGGATTCCAAGCGAGTTGCTAGCAGTCCGCCCTATTCTCTCTTCCTTCTTCTACAGCCGCAACACCACTTTAGATCAT

TTCTTCTCCTGTGAAGCTCCCACTTATAAACTCCGCCACCATAGCTCCACCAGCAGCACCCAGCCTGGTTTTGTGAGTAA

CCCATCGTCAATTTCGAAGGTAATAACACTAAACTTTATTTTTATAAGTGATACCTTAATTTATAATCCGAAGAATACAT

TTAGTTTTGTTTAACATTGTATTTTGATTTTGAATATGAGAATTGAAGTTATAGTGTTTTAACTCAATTGAATTTAGGGG

TATTGAATTTATAATTGATAATGAGAATTTTAGGATTTAATTGAATAGAAGAAATTGTTGAATTTAAGTTAAAATTCGGA

TTAATTTTTTGTTGGTATTTTAGTTGAATTGAAGTTATAATTGATAATTATTTGATTAACTGAATAATATTATTCTTTAT

GAGATTTGGGTACTCTATTAAGTTGAATTTTATTTAGGCCTAAAATTTACGCAGATTCTGAAGATGAACGTAAATGATCT

GTATGTTGTTGAAAGAGCGCGATGACCTGGGGGTTGGATCGACTTATCAGGATTGAACCTGAAGGGGGTGGGGTTGTTGA

TCCACCTAGGGATGTTGATTTGGTGAAAGAGGTAGTTCAACCACCAATCATTGTTGATTTAGAGAATGAGGAAGTTCAAC

CACCAGTGGTTGATAACCGAATGTCTGTTATAAGGGATCGAACAGATGTGTTGGTTAGGACACTGACCCGTTCGCTTGAA

CTTTCTACTTAGATGTTATAATCTTGTTCTTTATGTTGTTGAACCTTATTAGTAATTTTATAGCAAATATGATTCTGTTA

ATTTACCGTGAAAGAGTTGCTTGTAGTTTGTGAACAACATATGCAGTTGCAAATTTATTTTCTTTATTACCTAGTTATAA

TAGTATGACAATATCTTAGTTCATACATATTCCAACAGTTGTTTACATATTTATTGAAACTTAACCTTCTTACATGTTTG

ACATAAGTAGTATTTGTTTCCAATGGATTCAGGTTTTTTAGAAGTGAGCCGCTTCATCTTACCATCACAACATGGACCTG

TGCAGGTTTTTTAGAAGTGATCATCTTCCCCACATGAGTAGCAGTAAAATTGATTCTCTTGTAACCACTTTTCATCATCT

GGTAAAGTTGGTGGAATAAGTGGTGGAATACCTTGGTGTGAAGCTACATAAGCATTTAGCCAGATGAAAACTCCACAATA

GTTGCACTTCACATAAGGTTGAGATCATGTATTTCCTTTTTTTGCTTTTCTGAGAAACAGATACCCCTTGCAGTTGGTTA

GTGTGCATTTTCTACCTTCCCAAGGATAATCTCTAGCAACATGACCAGCAGAAATACAAAGAAAACACGTGGTAATTGGT

GATGAGCTTTAACCCTATATTATGTATGTTTTGTTTGAAATTTTTTCAAGAATTTGTTGTTGTAGCAGAAGATGAGACTT

TGCTGGTGATGGTGGTGGAATAGGGTATAACAATGGTCATTTTATAACTGGAATACCAACGGTCATTTTTTTTGAAAAAA

AAGAAAATCAGACCGTTGGATCTGAATTAATCTTAGATACTGTTAAAGGCATGCGAGTTTTTAAAAAATAAAAAATTTAA

AATTTAAAATAAAATTATGATACATTAAAAGATAACAATATATCGGTTAAAAATAAACTGAAAATAACTAAATTAGTTCA

TAAAACTGCCCTTCACAAAGGTTTAAGGTATTCCAAGAATCCTTTATGGTTAGTTTAATCTTTTCTTCAAAATCTCTCCC

TTCATAAGTTCGATGTACTGTCTTGCTACCGGATCCATCCCCGAGAGATCCATCGTCACTATACGAAACTCCTCGTCTGC

TTGTTTCCTAGCTTGTTTTGCCGCAAACTTTGCCGAAGAGTCAGCTAGTTTTGCTGTAAACTCGGCTTCTCTCTTCGCCA

TCTTCATTTATGTTTGTTCTGTATATTTGGTGTTGAAGTTTTGTTGCTCTTTAATAATAGTTTCCATCATTTCCTGTTGG

CCGCTACATTTTTCTTTCCCTTTTGTTTTCTGTCGTTTCGCCTCTTTTTTTCCCATTTGACAAATATAGTGTTAGGGATA

TCTCCTATGTCTTCATATCCACTGGAAGCGTCTAGGCTTAAGTCCTGTGTTTCCCGGTTGTCCTCACTGCTTTCATAATC

TGACTCAGCTTGTGATGAATCGAAAGCAGTTATCGATCTTTTGGAATATTTTATATTATCTAACATAATTGTGTAGCACT

CTTCATGCTTAAAACTTTTCCCACCATTAGCTTCCTTGAAAAGCCTCCTTACTTCTGCAATCTATTTTTTGAATTCAGTT

TGTGAAGGATATACAATTCAAACAATGTAATCGTCAAAATTAAATGAGTAGTTACCTTATTTTTGGGGGCTCATCCAGTA

TGGTGTTCACGTTCCACTTCCGCCTCCTTTGCAGATAAAAGTGCCATATCTTTACTTATAATTTGAAATCGTTTCTGAAG

GGAAGTCCAACTGCGTTGTGGCATCGAAGGTATGGCTTCTTTCATCTGATATAATATGTGAGACCATAATCTTTCTTCTG

ATTTAGATGTACCAATACTACAATCTTGAGATATTGACAAATGAATTTTACACATCGTTACATCTTCAAATGTAGTAAAA

TTTGGACCTCGTGGTCTTGGTGGTGCCATAATTCTAATCGATTCGAAAAGTAAACTGAAATGGAAAAGAACTTAATTATT

TTCGAATACTAACTAGATAGCAATGAGTGATTTTAGTAAAATGAAATTGGATTAGTATTCCATATATAATGTCATAATAT

AGTCGTTGCCGTTGCAACGGTAAACTCAACGGTAACATCACTCAACGATAACAACACAGGCGTTACTCCACAGTCAGGCG

GGCTTCTAGCAACATACCATTATCAGTTATATTGCAAGCGGAGTTATAGGAGACCGCCTGGTGTTGGGCGTTGCTTTAGA

ACACCATTTCGATAGGGCGGAGATTTTGAGTCCGCCTGGTGGTGGCCGAGATGCTAGAAGCACGCCTCATCAGACGTTAT

CTTAGTATTACACTTGATCAGGCGGACCACTTCAATACCGTCCGTGTTTTCAAGCGGTATCCTACGAAGCCGATTACCAT

CAAGCGGTATCCTACGAAGCCTCTTACAATCAAAAGTTATCCTAAGACACCGCTCGACTAAACTATGTTTAGTCCAGAAA

AATGTCTAAATCTAGTCTGAAACAATGCCAAAACCAGACCATCGACTGCTTTTAGTCTTACCGTTGCGCTTCAATTTTAT

GGATTATCGACTACATATAATCTTTAGTTCGTTGTTTGATCGTCCCATTGTGGACGCTTTTAGCATACTTTTATCATACA

AAGGGTTCACTAAATATATTATAATATATAAAACCCCACCTAACATATATAAATAAATTTTTTATTGTTAAGAGTCTCAC

AAAACATGATTAAACATTATTCTACTTAAATTTACCAGCTTTACCATAGTGGAGAAACACATTTATCCGTCCATCAGCTG

GCTGGGTAATTAACCCATTTGATAGGGGTGAATAGGAACTGGCTATATGGATGGATATTGGCACTTCCCAAGAAAATAAG

GAATCTAACCAATCACAAAAGGCAGGGGAGGTGGTGTTTAAGCCAGCCCCGCATAGCTCAAGCCACAAATTCACGTGCTT

CATGTGGTTCATACGGTGCTGTGTTGTGCTAAAGATTCAAATTTGATATGTTGTGCCGTGCTAGAGGGCGTGCTTTACTA

TATTGTACCAGAAAAATAATCACTTTCCAAAATCTCCTTGTAAGTACCAGGGGTAACAATTTCACGAATTTTTATTATGT

TATAATATATTCAGACGCGTAATAACACGGATTTTATATATCATCATAAAAAGAAAAGAATTTTTTTTCTCATTGTTTGA

ACCTTTATCCTTTAAATTGATAATAATGTTATATAATGTGTAACTATAAAATTAATAATAAAATAAGAATAATAATTTAT

TTTATTTGGATATGTTTGGCGAACCTGAAAAATCATATTACAGAAAACATAAATAAAGAAAACGAATATCAGAACAATAA

AAATTAAAGAAGAAAAAAATTCAAAAACTAGCTCGTCGTATGGTGCATATCAATCAGATAAGCAGATATAATTGATTTGC

ACCCACCACATCCACAACACTTATCATCTCTGTACCCATACAATATCAAAAGCAAATCAATCAGAAAAGCATTTACAATT

GATTTTGTCATGCTTTTTGCTGAAAAATATTTTATGCCATTGTGATTACTGATGCACGACATTTTCCACTAAAAACATAA

AAGGTAACTAATGCACGTGAAACCCGTGATAATCAACATTGAAAATGGCTCTCCAAATTGTTCTTGAGATTTAAAAAAGT

ATGAAAGTGAGTGGATGTCAGTCTAGTGCAGTGGTAAATGTCAGGATTTTTACCTACTCTAATGTCAGTTCCGTTCTGGT

ATTAGTTATCCGGACCACGTAATACATGAGTTTAGAATGATATAATGAATAATAAATAACCAAGTATAATTAATAATAAA

GAACGAGAAATTAAATTAAATTCACATGATATATAATATTTATAACGGTTCAACAATATATATAACGTGTATGTATTGTC

TATGTCCACTAGAACCGCCAACTCCAATCCATTATAAAATAATCAGACACAAACATAAAACACCTTATATTAGTTCAGGG

ACCAGATAAAGGGATGCTTATATTGGGGCTCGCATCCCTAAAACAAATTTCGACTTAGTTTTAGAAATTTCCAACCTAAA

ACAAATTATATACGGATTTATGAATTACTGGATTATACGGCCTTACAATTTTTGTCATGTAATCACCAAATTATCATTCT

CTATCATATTTTTTGACTTTTAAAATCCTACATACACCAATCTAAATTATACCACCAACATACATGTTATTTACTCCCTC

CGTTTCTAAATGATAAACCTTTTGGTTTAATAAAAAAAAAAACAATCATCTGAGCAATATTTTTATATTATTTTTTTTGC

TTATTTATCTTTAGGACAAAGTGGTCTTTTTGTGTTTAAATAATAAAAATAGTAGAGAGAAATCATAGAAAAAGTGGTCC

TCACGTGGATAAAATAATAATCTTAAAATATTTGATTTTTTATTTATTGTAACAAACTTTTTTTTAACATATCCAAATAT

AATAATCTTACCTATCATTTTAAAATGGATGGAGTATTGATTATATTACACATATAATACATATAGTTCAATAAAATCAT

AAAATAATAACACTTTATAAATAAATTTTTTTACAGACTAATATATAGCAAATTTTATATATCAAATTTGTATCTCTAAA

ACGAATATACGTATATTCTGAATTTTCCCGGACTTCCTCCTCGTACAACTTTATAACCCACGAGACCTCGTACATTCCTT

GTACTTTGCAACCCACGAGAGGTCCGTGTATATAGATTAATGAAATGTGCGTCCCAAATTCCTGGGACCTGCCAATAAAG

TGGCCAGTTAATCATTGAGAAAAAGTATATATATTAACTTCTCAACTACTGACGGCAAGTCCACCTCTTGTCTGAATTAA

TAGCCACAAATATTTGATGTGGCCGACAGATGAAAATGATTCCAAGTTACTTTGAGTAAAAAGACATATTATTATATATC

TTATCATTTGCTATTTTAGTTTATATTCAAGATGCAAATATACACAAACTATACAAAGTGAAGCAGTAATTGAAGGTAGT

CTAAGATATGCTAGAGAAATAATTTAACTGACAACTTTATTTAATTTTACTTTCAAAAGTTACAGCTGATGGTATTATGG

AGGGGCATTCATATTAAATAGTTAGAATCCTATGTTTCATAAAAATAATATTTTTATAAAAAATTAACTTTTTATAATTT

AAAATATTCTTATTTAATCTTTGTAATTGAATTCGACTTCTGGAGAAGTATAAATAGGATCACTTCCACTTCCACTTCTG

GTTTTCAAAAACTCTATGTACATATGCATTGCATGACACATGTTATACTTTTTTATCCGAATGCAAAATAGATTTCTCTT

TTATAAACTTATGAGTTAAACGAGTTAGAACATCATTTATGCGACTAATCAATATATAAGTGTATGTTTTTTTTGGATTT

GTCCTTAAAATATGTTTAATTTTTGGGCGGAGAAATGTATTAAAATAGGGAGTGACATAAGTATGTTGAAAACAAAATTT

CATATTTTTATCTATGTTTGTTATGTAAATAATAATTGACTGGAGCATTTCAGATATCTACTTCTTTAAAAGACTAACTT

AATGGTATCTCCATATTAAACGGTAACCAAAACTCCCTATTTAGTTAAAAATGAATAATATCCACCTACTAATATTGTTT

AATATTGCTCGAATATACAAATACTAAAATGGTAGGGATTACCCATCTTACCCGTAAAAATTAAGAAAAAGGAGGACATT

TTTTTTATATTTATTCCAATAAATAGTATACATATTTAGAAAACTATGTAAAAAAAAATAATATGCTAAAATATTATTCA

TATGCATCACGACGGGACGGGGCCACATGAAATCAAAAGTTCTAATTGAACCTGGTGCATAACATGGGTTACATACTGAT

GTAGATGAGCAGGCTACAACCTATTTGGAGGCTAAAGACCCGAGTGTCGTAATTTATCTCAAGAGTGGTCCATTTTGTTA

CGATAAATAATGTTATTTTATGAGTACTTTCATTATTTTAGGTCGACCAGAATTTTCCAAAAATAGTAAATTAAGGTTTG

AGTTTGTTTAGTGGTTTTTTTTATCCTTTCTTGACTAGCAAGTTAAAGTTTTTTCTCAAGTCTATAAAAGAGTCCCCCAT

ATATTGTTTTAAGCGGATGAATTTGATTAATAAGAATTATTTTCTTAAACTTATTCAAGTTTAATTGATTATTATTTAGA

ATCGGAAAATCGAAGTCTAAATAATTATCGATGTTCATAGTCGATATTCCTAGGAATATTTGAAGTCTGAACAAAACCTC

TCATTTTTTTTATTTTCTTTATTATCCGCTGCATCAGTTTGTATCAGAGCAAAGTCGATTATGAAGGTTTCATTCCATGG

AGATCTAGAGAATAGTTCATTGAAGTCGTCCAAGATTTACTTGGTCGATATAACCTTTGATAGGAAGAAAAAGATGATAA

ATACCAACCCTTTTGCTAAAGGGAATTGCGAAGGTAGAACAAATCCACGAGAGGTTTCTGATCGTTTGAAGGCAGGAATA

AAAGTAGAGATCCTATAATTTCATGGTGGTTTGGAACCGGAAGAGTTTCTCTCCTTTGTGAATACTGTTGAAGAAGTATT

GGAATTCAAAGAAGTACCAGATAAAAAACATGTTTCGTTGGTAGCGACTATGTTTCGTGGAAGGGCTAACTCGTGGTGGT

AACAACACAAGTTGCTACTCAGTCGTAAGGGAAAACATAAGTTAATATCATGGGAGAAGTTAAAGAAACACATGAGAACA

AAGTTTTTACCACATAACTACATTGGATTGATGTACCAGCAGCTGCAAAACCTATGCAAGGGAGTACGTTCTACAGACGA

TTACACCAAGGAATTTTACCGACTACTTTCTCTTAACGACTTGTCAGAGCCAGATGAGAAAAAATTCGCTCGCTATATAG

GTGGTCTCCCTCAACCATATCAGGATACACTCAACATATTTTATTGCTGCTCTATTTTGGATGCACGTCATAGAGAAAGG

CATATAGAGAAAAAAATAGGGCGTAAAAGCTTCAATTGGGGTATCAAAGTGGCATCATCTCCACCAAGTAACTGTAGTAC

TTCAAAATATAGTATTGCTAGTAAACTTAATTCTTCTTCTACACCAAATAAAACCCAAGTTAACATTTTTGGAGGTAGGT

GTAATAAGTAAGGAGAATCTGGGTACAAAGCCAGTGACTTTCACAAAGCAGAACGAGGGAACATGGTATTTTTCAATAAA

AACGACCGTGAAGAAAATGTCTGGGTAACTGATTATGAAGAAAATCATGAGGATGATTCTGAACCTGTCGATGAAGTAGT

TGCTGGAGATAAAGGAGTTAACTTTGTAGTAAAGAAGTTATCTCACTCCACGTAAAGATCAGGGCGACGGCTGGTTAGTA

ACAATATATTCCATTTGACATGTACGATCCAAGGAAAAATTTGACGTCTAATTATTGATCCTGGAAGTTGCGAGAATATT

GTTGATGAAAAGGTCGTGAAACGATTCAAGATGGAAACCAAGGCACATCTATATCCATACAAGCTTTCTTGGCTTAAGAA

AGGGAATGAGGTGAAAGTAAACAAACATTGGTTGATTTCTTTTCTATTGGTAACAAATATAAAAATAAAATATGGTGTGA

CGTTACCAGTATGGATGCTTGTCACTTGTTATTGGGTCGTCCATGGGAGTTTGACAGGAAAAGTAGTCACGACGGATATA

AGAATACCTACAGTTTTTTATGGAACGACGTACGTATAATACTTGTACCCAGTAAAGAGATTGCACCTACACTATCTACT

GGAAAAACTACCAATCTATTGCCGTTTAAGCAGTTTGAAGATGAGATAGAAAATGCAGGTGAATTGTATGTGTTGATTGG

CAAAGAAGAAAAACAGGATGAAAGTAGTATCCCCATTGCATCTCAACCTTTAATTAAGGAATTTATGGATGTTTTTCCTG

ACGATTTTCCGGACGAGCTACCACCTCTCCAAGATATACAACATCATATTGATCTCGTTCCTTGATCAAGCTTTCCAAAT

AAAGCGCATTACAGAATGAGTCCTAAGGAGCATGAAGAACTTCGTCGTCAAGTGGAAGATTTATTACAGAAAGGATTGAT

CCGACATAGTCTAAGTCCTTTTGCAGTACCAGCCTTGTTGATTCTAAAGAAATATGGAACATGGAGAATGTGTGTTGAGC

GTCGAGCCATCAACAAAATCACAGTCAAATACCGTTTTCCAATTCCCAGATTGGATGACTTGTTAGATCAGTTGTGTGGA

GGAAGGTGTTTAGCAAACTCGACTTGAAGAGTGGGTACCATCAGATCAGAATTCGTGAAGGTGATGAGTGGAAGACTGCT

TTCAAAACTCGAGAAGGATTATAACAGTGGTTGGTTATGCCATTCGGTTTGACCAATGCTCCAAGTACATTCATGTGTAT

CATGAATCAAGTACTCAGACCCTTTCTCGGTACTTTTCTGGTTGTTTATTTTGATGACATTCTTCTTTATAGTACTAACA

TGGACTTTCATGTTCAACATCTTCGAGAAGTGCTCTCAGTTCTACGCAGGGAAAAATTATTTGCGGTCATCAAGAAGTGT

CCATTCATGACAGATCGAGTTTTATTTCTCGGTTATGCCGTAACCAAGGTAGATGAGTCCAATGTTGAGGTTGTGAAAAA

TTGGACTACACCTACTACATCATACGAGGTAAGAAGTTTTCATGGTTTCGCTTCTTTTTACCGGAACTTTATTCCCAACT

GCAGCACTATTATGGCACCCATTATTGAATGCATGAAAGGAAAAAAGTTTTCATGGTCTGAGGAAGCTGACGAAGTATTT

GAAGTTGTCAAGACAAAGCTGATAACTGCACCAATATTGGTGTTACCTGATTTCAGCAAGCCGTTTGAACTATACTGTGA

TGCATCAAAGGTGGGAATTGGAGCTGTTTTGAGTCAGGAAAGACGACCTGTAGCGTATTTCAGTGAAGTTGAATGGTGTT

AAGCTCAACTATAGTCCTTATGATATGGAGTTTTATGCTATAGTTTAAGCCTTGAAGCGTTGACGTCACTCTCTAATTCA

TAACGAGTTCATTATGTTTTTTGATCATGATTCTATAAAGCACATCAATAGTCAAGATAAGATTTCTTCTAGACATGGTA

AGTGGGATGCTTATTTGCAAAAGTTCACTTTTGTACTGAAGCACAAAGATGGGTCGCAGAATCAAGTTGATGATGCCTTA

AGCCGAAGGAGTATGACACTCAACACTATGCGAACCGAGGTAATAAAATTTGATTCATATTTTGACCATATTCTAACCGA

TCCAAACTTTGAGGAGATTGTGGATGATATCAGTTCAGGCAAACGAAGTGATTACCTTATGCATGAAGGATTTCTCTTTA

AAGGTAACCAATTGTTTATTCCTCATTCTAGTCTAAGATTGAAGATTATTAAGGAGTTACATGACGAGGGGAACATGAGA

AGGGATAAGAATTTCCAACTTGTTTCGAAGTCATATTACTGGACAACAATGAGGCGTGAAGTGGTTAAATTCGTGGATAG

ATGTCAAGTTTCAAAAGGAAAAGCAACTAATGCACGGTTGTATATCCCATTGAAAGTTCAAAGTCAACCCTAGGCAGATT

TAAGCATGGATTTCGTATTGGGTTTACCTCGAACTCATAGGGGAAATGATTCAAGCTTTGTAGTTGTGGACCGCTTTTCA

AAAATGTCCCATTTTATTGCTTGCAAGAAGACTACTAATGCATTGAAGGTAGCTGAATTGTTCTTCCGTAAAGTGTATCT

TCTATATGGATTCCCATGTTCAATCTCTAGTGATCGAGACAAAAGATTCCTTGGTTATTTCTGTAAAACTTTGTGGAAAT

TAGCTAATACAAGGCTTAACTTCAGTAGTGCGTTCGCTGGGTGATATACTACGGGATATGTTTGGAGACAATCCGAAGAG

TTGGGATTAGAAACTTTGCCAAGATGAATTTTCATTTAATCATTCAGTAAACCGGAGTCGGGGTACGAGTCCGTTCGAGG

TTGTTTATGAATACAATCCACGCTATTAGATTTAGCAACAATTCCTGATCTGAAGAGAATTGAAGTTCGTGTTGAAGAAT

GTATCCAACAGTTGCAACAGGTTCACAAGTTGGATGAAGAAAAGTTTCTCCGAGATAATAAAAGAGTCCTTTATATATTG

TTTTAAGCGGTAGAATTTGATTAATATCTTTTTAAACTTGTTCAAGTTTATTTGATTATTATTCAAAGTTGGGAAACCAA

AGTCTAAATAATTATCGTTGCTCATAGTCTATATTCCTAGAAATATTTGAAGTATGAACAAAACCTCTCTTTTCTTTCTT

TTTTTTATTATCTGCTGCATCACATACTAATATTGTTATGAGAAGTTAAATATGCCCGGTGGGTAGCATGGACACAAATC

TAGTATAAAGTTAAAAAAAAGTAGGAATTACTGATTAATCCATACATTGATATGTATTGTTTAAAAACATGAATTATGCT

TTTAGTCAGTAGATAATGATATGTACTGTATATTAATGTGAACTAGGTTTTTAGTCAGTACACTGAGTAATATGAGATAT

GTATTGTCGGGTATGTACTGAGTAATAGGAGATATATACTGTAGGATCACCGTGTGAGAAATAAAATTGGAAAACATTGG

TGAAAAAAAAACCAGTTGTGGCTAATTCTTTAAAATAATTTTCTTCTAATGATTGGGACTGATCTCTTCTTCCATTGTTT

TACTTGTCCGGTTTTTTGTTCTTTTCTTCATGTTGTCAACATTTATTTTGTTTAGATATGGGTTTCTCAGTAGATTTTAT

GATAGTTTCAACGGATTTTTGTAATGATTTCAGTGAATCTAGGTTTTTCTGGTTAATTTTCTTCGATCTTGTTGACGATT

GATTAGGAAGATCACTTTAATATTAGGAGAAATAAGGTACAAAAGGATGAGAAGGATTTAGGTGTATTGTAGAGAGAAAA

CTCTTTTCTGGAAGAAGAAGAAGAAGAAATAACGAAGAGTAAGGGAAAACATGATTTATTCACACGTGAGTTGTAACGGG

TAGGAAGGTAATTTTGTGATAAAATATTTTTGGACCTCCGGTTAAATGTTTTTTCTGGATGGACCATAGATTAAACGGAT

TCAGATATTTTGGACTAACCAACAAATTTCCTAAAAAATTATATGATATGAGCGACAAGGTGTTGCACCTCAATGCATTG

CATTGTGGCTGCCTACTACCCAACCCGTACCTTGAAGGGATCAAGCATGACCGTAGTTCATCCGAAGGGTCAATTCGAAA

GGAACAAGCAACATAAGCGTATTAATGCAAAGCCAACATCGATTCCAATTTCGTTGCAAGGCCAAAGGCCAAGCTAAGCT

TAGTCCGTACAGGCAAAGGACTTTCGCAAAATATATGTACCAGCTTATGCATAACTATCACTACTTAGCATGCATAATGC

ATTCTCGAGTACGCAACACGAATGAATGATGCTTTCGTACAATTCAAGCTTGCCCGGGTATGCTACAAGCATATTGACTC

TTATGCATCAAGTATGCACCAACGCATTGGCCATATTGCATGTATGTGCATTACCGCTCATGCACATTCTTGCAAATCTC

CCTCCATATATGTCCTAATCGTATTTTCATATATTTTGAATAGAAGGGTAAAGGACATCCATCGATTGCACTTTTTAATA

GTGCATTACTCGGGTACACATCACGGATTAGTGAAGCTTCCTTGGGTACGTTTCCTCGGGTACGCAACATAGAGCCAGTG

ATACATCTCAAGCATGCATCACGGTATAATAGTACCTTAACACTATAATGCATTCTCGGGTATGCTTCATGGCCATTTGT

GCTTTTCCTGGGTAAGGGTCATTTTGACTTGTGATGCATTCTCGAGTATGCATAAGGAATAATAGTTCTTAAGCACTATA

ATTCAAGGCCAATGCATCCACAACTTGATGCATTCCTAAGTAAGCTTTATAAGCTTATGGTGAAACTTCATCACACGCCA

TGACGCATCCATGAGTAAGCCTTATATGAAAATTACGGGTGTTACATTCTTACCATCTTCAAAGAATTTCGTCCCCGAAA

TTCAACTCAAAAACTATTGTGGACAAGCTTCATCAATTTGGATTTTCTGCGCATATGTCCGATACCATGATGCACAGAAA

TTCTCTAAGGGTCCAAATTTTGTTAATAAAACATATATTAGAAACTTCTGAATATGCGTCCCATACCGCATATCCGGGAA

TCCTGACAGGGTCTGTAATTGTCGTACACGCCGATTAAGGCCTTCCAAACATACATCCCATAACGTATGCTCGAGGATCA

TAAAAGGTATTACAACTCTACTAAGAAAATCAATTTGAATCCCTAAACATGTGTCCCATACCATATGCTCAAAAATGCAA

AAGGATTTACTCAAACAGGATCTTAAGTGCACGTTCCATACCATACATTTAAGATTCCCCAAAGTTCACGAAGATTGAAA

GAATCATCACAAATTAATAATTAGTTCCGAACATAGTACTGTGAGGACCAAACTCTAATGACAGTGCAGTCAGATGAAAG

CCATCGTCCCTGACGATCTTCCACTCGTAAGTGGGAGCTATCCAACCTGACAACGCACTCCGTCCCTAACGGATGTGGGG

TGATCATTCCCAATGCCAAGGGTAGTGGAATACCATCCGTCGTTCCAAATGGGTTTTCTGGTAAACTCACAGAACTAGGT

CAAGCTCATTCAAATCTTAGCGGAGTAACTGGCAAGTTACAATTTCTTTTTGTATAGGCACTTGGCCTGATTTATCCGCT

CTATTTTCGTACGAAAGTTGGCATACTCTTCCAACTTTTTCTATTTCCTTTGAAGGAAAATTTTCTAGATACAATATAAA

CACTCTCTTAAGAGTCAAAAACGCAAAGTCCCTAGTGATTTCTCAGAACTTGCTCTGATACCATTTGTCACGGAACGAAA

AATCCCACCTCCGGCGCTATGCACTGGTCAGGCCAGAACTCTCCGATGCGCGGCTTGAAAGACTGTAACCAGACCTCAAA

CCTAACTTCTCGCTGGCAAATGCAGGTAGATTTTGCTTCCCAAGGATGCTTTGCAAGCATGATACTCGAAACGTAATTCC

CACACTACTCCAATTCACCCCTTGTTCTGCAAAGGGTGCAAGGCCATAAAATCCAATGGCGATGCCTTTGGCATGCATCA

CTTGCTTACGAGCATTGTCCAGGCCTAAGTCACTCCTTGGACTAGCACCTAGGCCATTTTAAGCCTAATTTCCAATTTTT

TAACTCTACTTCGACCATGTGCTGGAGCCTTCACTGATCTTGCTCTTGCAAAGGGTGCAACATCCAGGCATATGCCATTC

CATGCCTTAACGGTTGATAGGAAGTATGAACAATCCATTGAATGGAATTGCAACTACCTCATCAAGTGTGGCAACAATCA

TCCCTTGACTTGGAATTGTCGAGCCAGATGATATGAGTTACCATAATATCGCAATCACCTCTAAATTAGATGATATGAGC

GACAAAGTGCTGCACCTCAATGCATTGCATTGCGGCTGCCTACGTACCCAACCCGTATCTTGAAGGAATCACGTATGACC

GTAATTCATCCGAAGGGTCAATTCGAAAGGAACAAGAAACATAAGCCTATGAAGGAAAATCCAACATCGATTCCAACTTT

GTTGCAAGGCCAATCTAAGCCTAGTCCCTATAGGCCAAAGGACTTTTGCAAAGTATATACCAGCTTATGCACAACTATCG

CTACTTAGCATGAATAATGCATTCTCGAGTACGCAACATGGATGAATGTTGCTTTCGTATAATTCAAGCTTACCCGGTAT

GCTACAAGCATATTGACGCTTATGCATCAAGTATGTACCAACGTATAGGCCACATTGCATGTATATGCATTACCACACAT

GCACATCCAAACAAAGCTCCCTCCCTATATATGTCCTAATCGCATTTTCATATATTTTGAGTAGAAAGATAAGGGACATC

CATCGACTGTACTTTTTAATAGTGCATACTCGGGTACACATCACGGATTAGTGAAGCTTCCTCGGGTACGCGTCCTCGGG

TACGTAACATAGAGCCAATGAAACATCTCAAGCATGCATCACGGCATAATAGTACCTTAACACTATAATGCGTTCTCGGG

TATGCTTCATGGCCATTTGTGCTTTTCCTGGGTAAGGGTCATTTTTGACTTGTGATGCATTCTCGAGTATGCATCACGCA

TAATTGTGCTTAAGCACTATAATTCAAGGCCAATGCATCCACACCCTGGTGCATTCCTAAGTAAGCTTTATAGGCTTAGG

ATGAAGCTTAATCCCACGTCATGACGCATCCCTGAGTAAGCGTCATGCGAAAATTACGGGTGTTACAAAAGCACTAGGAT

ACTAGCAATGAGCAATATTCTATTTATCAAATGAACATTAACTAACAAATAAAAGGGTGAGAACTGAGAAAAAATGGGTG

CATGGAGGAAACTAAAAATACTAATTTCTATTTTCCGTTGCTTTCGATCAGCCTCAAGAAATTGGAAGATTTCAAATAGG

TAGCTAGCGCAAGGTTTTTTTTTTTTTTTTCTTATATACGAAACACCACATAAGCATAGCGTGCCAACCCTTTAAAACGT

GTTAATAAATCAATTAGATAGAAGGTGTCATATAGTATGATACTGCGAAATATACCGTCCATATTTATTGTTGCATAAAT

AACACATTACACAGATTTAAACTTTATGAAATAGCCTATACCGTTTCAGTACGGATAATACCATAGTATATATATACAAC

TTAGTTGATGTAAAGTTCAATTAAATTTAAACCACACATCTTGTTATATATCTTGATAAATAAAAGAAGATAATTTATAT

AAGTAAAAGACTACACTTATAAATTACATTTAGTAAAAAAAGAAAAAATCGAGAAAAAAATATATTTATGTGTAGATGTC

ATTGGTAGCTTAATGTAGTATAGCTCCAGTTTAATCAACCGTGAGTCAATTTGTTAGGGTTGAACAACTGATAAATACAT

ATTTATATTTTTATAAGTGTCGATAATATATATTGTTAGTGTTTATTTTTATTTAGTATCTATTGTATTTTTGTGGGTAC

ACTAATAAAACCGGATTAAGTGAGAAAAACGGTTAAAACGTTCACGGACAGTGTGAGAAAACGTTTTTTTTTTAAGCATT

AGCAAACTTTTGTTGAGAAGAAATAAAAATTTGTTTGAGCCAATTTCATTTAATCAGACAGGTGATAACGGTTTGTAATA

TCTGACAATAAATAATACGTTTAGTGGTGGGGCCGGGTCAACGTAGAATCACTGCTACAAATGTAAGCCCTGGTGACATT

AATGTCTAAAATGAGGAGAAGCACCGAATCTTTTTTATCTTTATCTTTATCTTTTTCTGTTTCTTTTTCTTTTCCTTCCC

ACTTCGGGGTTAAAGAGAAAGCCATTTTTATTTATTGTGTAAGAAATGAGGATTTTTGGATCAAAAGATTTTTCTATTAT

TTTGGAAAGGCTTATCAGTAGTACTAGTGCTTGCAAAAAGTAAGGATTAAAGATCTGTAGCTGGCTAGCTATATATACAT

GTGATGTGCAGGGTTCACCATGCATATCATTAAAATCCATTATCTATCCTACCTACTGGGTTGAATCATGGAGCTCCAGT

ACTTTTCTTATTTTCAGCCAACTTCCTCCCTTTTTGTTCTTGTATCCATATTACTGATCAGTGCTGTTGTTTTGAGGAAG

ATATTTTTAAATAACTACTCTTCATCACCTGCATCATCAACAGAAACAGCGCTGCTTTCTCATCAGCGGCGGTCGTCATG

TGCATTGCCAATCATCGGTCTTCTTCATGTTTTCATGAATAAAAACGGCTTAATTCATGTAACTCTTGGAAATATGGCTG

ATAAATACGGACCGATTTTCAGTTTCCCAACAGGCAGCCATATGACTCTAGTTGTGAGCAGTTGGGAGATGGCAAAAGAG

TGTTTTACAGGCAACAATGACATTGCTTTCTCAAACCGTCCTATCCCGTTAGTTTTTAAGACTATATTCTATGCATGCCG

TGGTATTGACTCATACGGTCTTTCAAGTGTACCTTATGGAAAATATTGGAGGGAGTTACGAAAGGTGTGTGTGCATAACC

TCCTCTCTAATCAACAACTACTCAACTTCAGACACTTGATAATTTCTCAAGTTGATACCTCTTTCAATAAGCTTTATGAG

TTATGCAAAAACTCTGCAGACAACCAGGGAAACTACACTACTACTACTACCGCAGCTGGCATGGTGAGGATGGATGATTG

GCTAGCGCAATTATCGTTCAACGTGATAGGAAGAATAGTCTGTGGATTCCAATCAGACACCGAGACAAGTGAGCATTTTC

CTACAGTTTATGTTAAAAATGATTGATTTTATAGGTTGAGATCAAAGGGTATACATATATATCTTATTGCGTGCACATGC

ATAAACAGGTGCTTCAAGCAGTGCGGAACAACTTACAGAAGCAATTAATGACGCATCGTATTTTATGTCGACATCTCCGG

TGTCAGATAACGTTCCATGGTTAGGGTGGGTTGATCAACTAACTGGTCTTACTAGAAATATGAAGCAATGCGGAAAGAAA

TTAGACTTGGTGCTCGAGAGCATAATTGAAGATCATCGTCAAAAGAGACGGTTTTCTAGAAGTAAACAAGGAGATGAGAA

TTTTATCAACGAGGATGAACAAGATGACTTCATCGACATTTGCCTGTCAATTATGGAGCAACCACAGCTTCCTGGCAACA

ATAACCCTTGTCAAATACCTATCAAATCTATTGTCCTGGTATGCAATCGTAGTAGGATGACAAAACCAAAGGCTAGACAC

CGATTAGTAGAATCTTATCCATTGCATTCCTGCATTATTAATTGGTTGAAGTGGTAGATCAAAGAGTGTACAAACATGAT

CTGTTTTTGATGGGTTTTCTTGACTATTGTTGTGTTCTGACTCTAATCATGTATATATGCAGGACATGATAGGTGGGGGG

ACCGACACCACAAAACTGACCACCATCTGGACCCTTTCATTGCTACTGAACAACCCCCATGTGTTGGACAAGGCAAAACA

AGAAGTGGATGCACACTTCGGGACGAAAAGGAACGATGCAGGAGGAGTCGTGGTGGATTTTGATGATACTCGTAACCTTG

TCTACATCCAGGCAATTATCAAAGAATCAATGCGGTTGTATCCAGCCAGCCCCTTGGTGGAGCGACTGAGCGGCGAAGAT

TGTGTGGTCGGTGGGTTTCATGTACCAGCAGGGACGCGGTTATGGGCTAACGTGTGGAAGATGCATCGAGACCCAAAAGT

ATGGGATGATCCATTGGTGTTTCGACCAGAGAGATTTTTGCTCGATGACCAGAAGATGGTTGATGTTAGGGGTCAAAACT

ATGAGCTGTTACCATTCGGAGCCGGTCGACGGATATGTCCAGGTGTATCGTTCTCTTTGGATCTAATGCAACTGGTACTG

ACTCGTCTTATTCTTGAGTTTGAAATGAAGTCTCCTAGCGGGAAACTAGACATGACAGCGACACCAGGTTTAATGAGTTA

CAAGGTAGTCCCCCTTGACATTCTGCTCACCAGTCGTCGCATGAAGTGGTGTGTGCAGTTAGCCTCTGAGAGAGATAAAA

AGATGGAGAGTAGTGTTGTACCTGTAATCACTCTGAGCTCGTGCAAGGTGATGCCTGTTTTAGGCATGGGAACATTTGAA

AAAGTTGGTAAAGGGTCTGAAAGAGAGAGGTTGGCGATTTTAAAAGCGATAGAGGTAGGTTACAGATACTTCGATACAGC

TGCTGCATACGAAACTGAAGAGGTTCTTGGTGAAGCTATTGCTGAAGCACTTCAACTTGGCCTAATCAAATCTCGAGATG

AACTCTTCATCAGTTCCATGCTCTGGTGCACTGATGCCTACCATGATCGAGTCCTCCCCGCTCTTCAGAATTCTCTGAGG

TAAAATTTTTGATGAAAATTTTGAACCCAATTAGGCTTATATATTCTTAAACATCTATCAAATTTTAGATAAATCAAATT

GTTTTGTGTGATATAGGAATCTTAAATTGGAGTATCTGGATCTATACATGTTACCCTTCCCGGTAAGCTTGAAGCCAGGA

GGGAAGATAAGTATGGACATACCAGAGGATGAAATTTTTACGATGGACTATAGGTCTGTATGGGCAGCCATGGAAGAATG

TCAGAACCTTGGCTTGACAAAATCAATCGGTGTCAGCAATTTCTCCTGCAAAAAGCTTCAGGAATTGATGGCGACCGCCA

ACATCCCTCCAGCTGTGAATCAAGTGAGTACTTTTACTTGAAAGTTTTGATAAAAATTAGTGTTTTCCTTTTCCGGTGCT

AATTAAATTTCCAACATTTGACACCTTCACATGCATAAAAATACTACGTCTTATAGGTGGAGATGAGCCCGGCTTTCCAA

CAAAAGAATTTGAGAGAGTATTGCAATGCCAATAATATATTAATCAGTGCAGTTTCTATTTTGGGATCAAACGGAACCCC

ATGGGGCTCCAATGCAGTTCTGGGTAATGAGGTGCTAAACCAGATTGCTATGGCCAGAGGAAAATCTGTTGCTCAGGTTG

GTCAAATTCTCTGCACATGCCGTAGAGCAGTCTCTATTCTTTGGCCATCGTATTTGCATCACATTATACCGAATTTAACT

TGCATTTATATTACGTACGTACTGATCAGGTTAGTATGAGGTGGGTTTACGAGCAAGGTGCGAGTCTTGTGGTAAAAAGT

TTTAGTGAAGAGAGATTGAGGGAAAACTTGAACATATTCGACTGGGAACTCACTAAGGAAGACTTGGAAAAGATGGGTGA

GATTCCACAATGCAGAATCTTGACTGCTGATTTTATGGTATCAGCTAATGGACCTTTCAAATCTCAAGAAGAGTTGTGGG

ATGACAAAGCTTGAAACATCGATCTTTAACTCTAGACGTTTATAAGAGAAGCTTCCCGGCAGTCACAGATCTCATTTCCC

TTCCTATTGGTTTTGTACTCGCCTCTCTTGTTTTCTAAATAGAATCTGGTTTAGTTCCATCGTCTATAGTAAAGAAATAA

ATTTAGTACACTAAGCTTGTTTAATTTCTGTTGGATTGTAATTTGTAATGTCCAGAATTCTCTAGAGTAGTCTACATGTG

TCTAGTGTGAACTCTAGATAGTAAAGTCGGCCAAAACAAACTACATAGCTAAGTTTTTTATACCAACTATAAGATTAGTC

CTATGTCGAGAATTGTCTAGATAGTCTGTTGTAAAGGCTATAAATGGCCAAGAGTCAAGTATCATTTGTAAGTATCTATA

TCTATAATAACAAGGTAAGTTTACATTGAACTGATCTTGCTCCACCACTCTCTCTATCTACTAGGAGCTCTGTATAGCAA

TTATCTCCACACTAGTGGGTAGAAATTGCAATACGTGGACATTTTAGGAACGGAAAAACCAAACATAACCGGTTACTATT

CCTATATACCTGTCAAAATTAGGTTTTATTGGTTGGGTGTGTGTGACAATAATATGACTCTTTTGAGCTTAATTTGATCA

TGAATTCATGTGTATTCAATAGTACCAGTTGATTTACTGGTTAATGATCGTCGATTGGGAATAGGGTTTGGTTGTGTTTT

TCGTTTTATATGCATCTGTGAATGAAGATATTAGTTATGTTGAGTTTATTATACCTTTGTGTTAAAATATATCAAAGGTT

TATTAAGAATAAAACCTCCAGGGGGACAAATAAGATGATGCACACATCTAACCTTTTATCTCAGTCCCGTAATTGTTTTT

TTTTTTTTTTTTTTTTTTTTTTGTGTGCAGGAAAGATGAATAGATTGATATTGATTAAACCATTAAATTGTTATAGGCCT

TCTCCAAATCTTCTACCAGGAGACTTGGAGTGTAGACCCAGCGGCAGAAATAGGAATTTACATGTGGGGGCTAAAAAAAA

AATTCAATGAACACTCAATTTTAGTAATCCCAAATACGATAATAATAAGTGTGATAAAGCCATAATGGTAAATCAGTTAC

GGAGCTAGAATTCATCTTAGCGGTGGAAATTTTCAATAGCAGAAGTTTTATATAAGGTATATTTATTAATTTTTTAGGCC

TATTTTTTAGAACGAAATTGTAACCACATTTAAAATACAGTATTTTATACGTTCATTCTCATAATATAGTTGATTTGATC

TAGAATTATTCTCATATTATAGTTGAAATGTCTAATTCATCTAAGACGATTTCAATAATTATCAGAATGAAATGACAACA

CCCCTTAGCCTCCCCTGATTCCGTCGCAGTTGTACTGCAAAGTTTTTGCGAGTTCTTTTTATTCTTTATTTAGCAAAGTG

GGCAACAATCACGATAACATCACTCGATTTAGTATTAAGTTATATTGTGATTGGAGAACTGCAACAAACAGATGGAATAG

TACATATGTAATCAAAAATGAAAGTGATTAGTATTTCACATTTTGTATCAACTAACCTAATTAGTAACAAAAAAATTCTT

CATTTAGGACATCTCACCATGCTTTTTTTATAACTAATGGGCTAACTCAAATCTTACCTGGGCTAAATGATATTTTATCA

CCAATTATAAAGTCTTAAATAGGTTTTCCAAAATTTTGGGGTGGGCTACAACCTACATAAGCCACCCCCTACCTCCGTCA

GTGCGTAGACCATTCATCATTTGATAAGTTTGATCTTGCATACTTTACTAAAGTATATGTTCACAAGAGTTTGATGTTCT

CTTGACTGATGTACATAGAAACTATCAATTGGTCATAATATAAAATATATTGTGTTAGTTTGGTCCTGTTTATTTGGTCC

AATTGCTCTAAGATCTTTTTTTTTAAAACAGAGTCAAAAATATCATTTCTCTCTCTTCTAAAAATAAATAAAAAATCTTA

AAAGTTAAATATCTTTCAAACTACAAGTCGAAATTTAACAAATTTCATATATTCGGAAAGCTCTCGACGAGATCTACAAA

ATAAGTACCACTTTCACTATTTTTTGATGTGTTAACTTTTATATTATCGTTTAGCTAAAGATAAACTATGAAATGAACTA

TCATTACGAGTTCATTCTACAAAATGAACTGCTAGAATTGACTACCATTGTAATGAACTAACTTTGTGAGTTCATTTTAC

ATAATGAACTATCATTACATGCAAGAATGAACCACCATTACGAGTTTATTTTACAAAATGAACTGCTCGATGAACTCCCA

TCACGGGTTCATTCTACAAAATGAACTGCTAGAATGAACTACAATTGTAATGAACTAACTTTCCGAGTTCACTTTACAGA

ATGAACTACCATTACATGCTACAAAATAAATTGTTAAAATGAACTAGCATTTTGAGTTCATTCTACAAAATGAACTGCTA

GATAAACTACCAACCGCTAGAATGAACAACCATTACGAGTTCATTCTACAAGATGAACTACTAGAATGAACTACCATTGT

AATGAACTAACTTTGCGAGTTCATTTAACATAATGAACTACCTTTATATACTACAATGAACCACCATTACTAGTTTATTC

TATAAAATGAACTGTTAGAATGAACTCTCATTACGAGTTCATTCTATAAAATGAATTGTTAGAAAAACTAGCATTTTGAG

TTCATTCTACAAAATGAACTACTAAAATCAACAATCATACGAGTTCATTCTACAAAACGAACTGCTAGAATAAACTACTA

TTACGAGTTCATTTCTACAAAATGAAATGCTAGAATGAACTATTATTAAGAATTCAATTTACAAAATAAACTCTTTAATA

GTGTTTATTTTTTAGTATGAATTAATATTACCGATATCATGCAGATTAAAATTTCGAAAAATAGTGAAATGGGTACTCGT

TTTGTATATCTCGTCGAAAACTTTCCAAATATATAAAATTGGTTAATTTTCAACTTGTGGTTCGAGAGATATTTAATTTT

TAAGTTGTTTTTCCTCGTTTAAAAAAAATAAGAGAGAAAAAAGATAATTTAGTTATTTTAATATTTTTAAATAAATAAGG

ACCAAACTAAACTTCCTACTTTATCGGTGGACCAAATTAACTTAGGATTATGTAAAATAGGACCAAATTAAACATTTCCC

TTATTAATAAGCTTAATCCTAACTAAACAATCCTGGACAACACTATTTATAGCCTTTTTGTTACTAAAAGAGCTATTACT

GGCTGACACAATTTATTATTGTCTGTCTCCAAGTGCAATTGTTGAATTCCCTTTGGCACAACCCATTCCACTGCTTCCTA

AAACAGCTAGCGCTTCAGCTTGTTCTTCATCTATAGTTCTACTTGTACTCCCTTTTGCTGCTCTAAATTCACCTACACAA

TTCCTCAAGATTAGTCCAATGCATGCAATACTGGTGTCAAACTTCTTATCAAAAGAAGCATCTTCATCTATAGTTCTACT

TGTACTCCCTTTTGCTGCTCTAAATTCACCTACACAATTCCTCAAGATTAGTCCAATGCATGCAATACTGGTGTCAGACT

TCTTATCAAAAGAAGCATCTGTATTTATTTTTTTCCGTCCAACTTCAGGACATTTCCAACTTTTTGTTGAAACAGTTGGA

TGATTTTGATCTGTAGTGGAGATATTCACAAAATTTTCCCATTCAGAGAAATTTTTATGAGTCAGCATGACTATGTCTTG

AGGATTAACCTTCACATTATCAAACACAACTGAACATCTGTATCTCTAGACTGTCCACATGCTAAAACATATAAATCTTA

AGGATGTATTGTATGTCAGCAAAATAATATGTATAAACAGTTTTGCCTATAAATACTGAGGATGATTTCAGCGTTATGGC

AAGGTTTTTAGGTTAGCAGAAGTGTGTTAGAGAATATTGTGAGCGTAGAGAGTCTCACCCCAAGTTTGTAACAAGTGTTT

CACCGTTAATGAAGTCTTCTTAGTTCCTCAAAGTGGTTAGGAAATGAATCTTTACTAGGAGATACTATTACGTTTATTTA

CATTTTTTCATGTCTTATACCTAGATAGTAAGCTTTAATATGTGATTTAATCTAACGATAGAGATCAACGTCACTGTTTA

GTTAGAAAAATGGTTACTACAATAATGTATTATAACAAGATCAATTATCACCTTTTCAATGAATCTTTCCTATTGACCGA

TCAATTTAAATTATCACCTTTTAAGAAATGAATACAGGGCCTAAAAATAATTAAGAACTACATATTAAAATTTGGAATGG

TTGTTATAGGGGTTATGATACAGGAAGATTTATGGAAAATAAATTTATTATTTTACTACTATTATTTATATTTTATGAAT

ATTATTTTTAATAAAAAATAATTTCCGGATTTATTATTTAGGGTAATTTTAAAAGAGTCTAGAAGTTTATTTTAAGAGTT

TTAATAAGTTAGGAAAGTTTCCTAAAGGTAGTAGATACCTTGGGAGACAAGTTTATAGGTTTTTGATAAGTTTAGAGTTT

GTTTAGTAGGAGTTATCTATACAAGGAGCTCCAAAAACGAAAATTAGAGTTTGAGTCGGTTTAGGGGTTTTTTCATCCTT

TCTTGACTAGCAAGTTTGGGTTTTTCCCCAAGTCTATAAAAGAATCCTCCATGTATTATTTTAGGGAGTTGAATTTGATT

AATAAGAATTATTTTCTTAAACTTGTTCAAGTTTAATCGATTATTGTTTATAGTCGGGAAACCGAAGTCTAAATAATTAT

CGTTGTTCAAGTCGATATTCCTAGGAATATTTGAAGTCTGAACAAAACCTCTCTTTTCTTTCTTTTCTTTATTATCCGTT

GCATCAGTTGGTATCAGAGCCAAGTTTGATTCTAGGGTTATTTGATGTTATTAAGACGAGTAGTCAAGGTTAATAAATTG

TCACCAAGACAAGATCCGAGAAGATGTGAATATCTGATCAAATCAAAGAAGGAGATTGTGGATGTGATAGAATATTATTT

GAAGAAATCACTGAAATCTCGATATAAACCATGCTTGCTTATAGAAGAAAACAATATTACAGATAACGACTTTGAAGACT

TGTTTTGCCGTCATGACAATGATGCACAAAATCAGGTTATTGTTGAGAGATATACAACTCCAATCTTTGATAAATATCCT

GATGATGAGGAGGATATGAATTTTTCTTCTCAAAAAGGTGAAGAACATCACGTTTCTGACAAAGAAAAACAACCCCAAGA

AGTTTCACAGGATTTAATGGAGTTTCATGGTGAGTCATCTAATTTTGATAAACTTATTGTTTTAGAAATTCCTGATATTG

ATAAATTGTCTTTGCCCGAGAATAAGCTAGTTGTTGATTATTGTGATGCACAAATGGTCTCTCATATCTACCCTAACAAG

AGAGAATTTGTTAAATCAACTTATGAAGATCCGTTTTGGGCTTCCATCATATCTAATGCTTCTATGCGGTGCATAAATTA

TTTTTTACAACATCAATTCATGTGTACGAATATTGTCATCAACGAGTTACACATGCAGAACATGGTACCGGCAAGTGATA

GACACATTTTAGTGTCTAAATTGCACCTTAATTTCTAGTGAGTTAGTATTCGATT

>Par_flye_v1-0_contig_11585_plus contig_11585

AAGGCACTAAAAAATCTCCTTGGTTGGCATACTTATTGACTACGGGAGGAAGTACCACGATGCGAAATTCGATTCTTTGT

ACACGAGTTTGCACTCAAGCATACTAAAATTCATATATAAGTCACGGAGCTCTACTATGAGATAGTCTCACTATGGACAT

CAATACCCGGAGTCAAAACTAATCACATGGATAGATCAAGAAGATGGATATGAGAAAAACATAGATGGTTTTGATGTTTA

CTAAGTGATCGGTGTTTCCCATATCAATACAAGAAGGCCAACGCCAAAATGAACCTATCTAATGGACTGAGATACTGAGT

CGACTAATATCATTAACCGGCATATACAAAGGGAACCGGTGGTCGACAATCCTAATTCTAGATCAACACGTGCGGGCATA

TACAAGGGAAGGCGGTCAATTTATTCTTTGAACATTAATTTTTTTTTTTTTTTTTTTGCATGATTAGATCTTTTGGATCC

AAGCGCATGCTTCTTGTCACCCAGTTACATGATAGTTCCCGTGGATCCCTGCATTCCACGCTTGTTTAGCGACGGAGACG

GGGAAAAACACACATATTGCTATTCCCAAAGTGTTAATTTTATTTCCGGTTGGTCTAATTGGTCTAGTCTTTTTTTTTTT

TTTTTTTTTTTTTTTTTAAAGTAACTCAATCACTCTATTTCACCTAGCAGTGGTAACAACTTGAATCGTGAGCCCACCTA

ATCACTTAGAGAAACATAGTTTAAAAACAAAACAAAATAAAAACGAAGTGAAAAGGACTCAACGAGATTGGTGAAACTAT

CATGTTATTTCTAACACCTCGAGCTCAGTGCTTTTATGAATAGACTCTTTAGATGTTTCCATCTAATCAGATTGGTTCCT

CAACTCCTACAACCAAAATGCTTCCATCCACTTAGATTGGTTAGTGCCATCCTTAATAAGCATAAATTTCTAGGCTCACG

GAGTTTATTTATTGTAACAAAAAGGTAACAAAAGTTTCTACCCCACCCCCAAACTTAAATCTAACATTGTCCTCAATGTT

CTAAAGATAAAATTAAAAGCATGAACAAAGAGAAACTGTTACCATTTGAAGTAAAAGAGTTAAGGAAAGATATTACCGTG

TCGCATGAATATTGGGTTACCTCCCAAGAAGTGCTAAGTTTAACGTCTTCAGCCAGACATTACGTACCCATAAAATGGCT

AATTACCCTTTCAAATCGTATATCAATAGTCGAAACAATTGTGGATCATCAAAATCAATTAAAACTGCCACAATTATGAG

ACAACCGCACCAAATCGAGAAAAATGAGCAGACGGGAGCACATCCTCATCCAAGGTTTCTGCCCAGACAACTGGTTTTTT

CTGTGGTGCCCACTTGGGCTTCATATGAGTATCTTGATAAACGGATTTACCCAAATAACGAGTCTCAAATAGCTGGATTC

CCCCTGGATAGTATCAGGAGGATCGGGTTGTGGAGTCAGAAGAGACTCGGGTAATACCTCAGACACACTAGGCTCGAGTC

CCGAGTTAGGGACTTCAAATAGGGATACTTGACGATTTACCCCCAAACTTAGAATTTTTGGTGTCTCTAGATAGACTAGT

CACAATATTCCTAATTTCTAGACCATTAGGTTCTGAAAAGGTCAATTGCTTTCTCTAATTTATAATCGGGTGGTTCATCC

TCTAAATGCATCTGAGGCACACAAAGCGTAGGAATTATATGTTTCATATCCTCTATGGGCAACCTAAAGTTTCCTTATTG

TCTTGAAGTACGGTCATTTGGCCTGCTCTCATGATCACTATCCAAATCGGAAGCTATAGGCAAAATCTCAGATGGGCCTA

AAAATGCATAATTAGGGTCCAACTTAGGAGGATCTCTAGGCTCTTCGAAACAAGGGCTTAAGGTAGATTCGGTAGCTAGT

AATGGTTCGAACCTAGCTTTCCATTTATCGATATCTAACATAGGAATAGAATCCAAAAGAGCATTTACTTGTTCAATGTT

GCTATCATCGTATAATCCATGCTAAAATGAGTTATACAACTCTCTAATGGATCTTCCAATATGATGTTTGGTAATGACTC

CTGAACTAAGGTTCCTATCATGTTCACCTCTTCAATGCATGTGTCATCTAGCTCAGAATGCAGCTTATTGACATTAAAAA

TATTCAACTCAATAGTCATATTACGCAGATAGATTCATAATACCAGGTTAAATGAGTTTATGATCGCATTAGACGTAGCT

AAAAATGGGCGACCTAAAATCACAGGTATCTGGTTCTCTGGGTCAGGGACAGGTTGGGTATCTAAGACCACGAAATCCAC

AGGATAAATAAACTTGTCGACCTCAATAAGAACATCTTCTATCACACCACGAGGAATTTTAACAGACCTATCACCAATCT

TTAGTGTTATCTTGGTAGGTTTCAATTCACCAAGTCCTAGCTGTAAGTACATGGTACGGCAGTAAGTTCACACTAGCTCC

TAAGTCAAGTAAAGCTTTATCTACCCGATGTTTTCCTATCGTGCAAGCAATGGTAGGGGCACACAGGTCTTTATATTTAG

GAGTTGTGTTGTTACGAATGATCGAACTTACATGACTAGCTAAAAAAGGCTTTTTAGGGACACTAAGCTTGCGCTTTCGC

GTACACATATCCTTAAGGAATTTGGCATAAGCCGGAAGTTGCCTAATTGCTTCTAGTAACGGGAAGGTTTATGGTAACTT

TCTTAAAAACCTCCATTATGTCATTAAAGTTGGATTCACTCTTTGTTGGTACTAATAGCTGTGGAAATGGGGCTCTAGGC

ATAAAATAGGGCCTCTCAGGAACAGAATCGGCATCATCAGAGATTTTATCAGTCCCCTCAGCTATGGATGGTCCTGAGGG

GTGAACTACAGTATGCTCACTATCGGGCATGGTTACCGATTGTCTACTACTCTACCACTCTAAGGGTTCTAATTGAATTC

AACGATGAGATGGTTTTTCACCTACTTCATGAACACCTCTAGGGTTTGGTTGTGTTTGACTAGGAAACCTACCTTTTTCT

CTCAAGGACTCATTTATCAGACTAACTGCGTTTTTAACTCGGCAATAGCTTGACTATTTTCTTGCCCTATCCTGTTACTA

GCTTCTACACTTTGTGAAAGTAATTGGTGCATCTTTTCAGTGTTTTGAATAAATGAGGCAAGAGTTTCTTCTAGACTTAA

AATTTTCTTATCGGACTGATTTTGAAACTGAGCTATTCCTGAAGGATTCTTAGTATAGCCAAAACCTGGGGGAGCATTCG

AATTACTAAACTGACCTTGGTCCTTAGACCATGAGAGGTTCGGATGGTTTCTCCAACCAGATTATAGGTTTTACCGAATA

TGGGTCAAACTTTTGACGGTTATCAAATCTAGTGTTATTATTATAAAGAGCATTGGCATGCTCTTCAATAGTCTGGCCTT

CCCAAAAAGGCTCCACCCTACCACTAGCGTGACCCAATTCTAAAGCTTCTAACCTTTTAGCTATTCGCAATTTTGGCATC

GATTCAAAGCCTCCTTCTACCCTATTAACGTTTCCTCTTCCTAGAAGAATTGTTTTCTTGGGTTCCCTACTATTTTCCCA

TTCTTTGGGTCTTTTCGGCGATTTCATGCAAATATGTCATCGCTCGATCAACTGTTTGGTTTTCAAACCCACCCACACAT

AGACTCTAACGTGGTCGTAGTGGAATAATCTAAACCCTCATAAAAGGATCCTCGAACTAACCTAACCTTTTCAAGACCAT

GATGAGGACATTGGGCTAATAGATCATTGAACCTTTCCAAATACCTATACAAAGATTCTCCCTCCTGTTGAGAAAACGTG

CAAATTTGCGTCATAATAGACGATGTTTTATGCCTAGGGAAAAACTTGTTCAAAAGGCGGATGTAAGTTGTTCATATGTT

TCAATTGAACCAGAAGCCAAACTATATAGCCACGATTTGGCTTTATCTTTCAAGGAAAAGGGGAATAACTTTGAGTTTCA

AAGCATCATCATCAAGGTTTCTAATCTTGGTGGTACTACAAATTTCCTCAAAGTCCCTAACATGAAAATAGGGGTTTTCA

TTTTCTTTCCCTAAGAAGATTGGGAGCAGTTGTAAGGTCCCAGGTTTAAGTTCATAAGTTGCTTCCGTTCCAAATTAACT

TTGATACAAGAGGGACGAGTAGCCCTAGTCGGGTTCAACAAGGCTTTCAAAGTTGCCATTTCTGGCACTATCGGAATATT

AGGGATTGTCTCCCCAGACGGACGTTCAAAAACGGACTCTTCAAAAATTGGACTTTCTAAATTGAGGTAATCGAGACGCT

TAAAACTACTAGGTTTCTCTTTAACAAATCTACCTAGGGCGTCTCTTTTACGTTCAGGCATGCAAAAGAATAAGGAAGGG

GATTCTATGCAATCACAAATCAAGGCCGACTCAACCAGAGCAAACCTAGAGATTTCTAGCAAACAAAAAGCATGATGGCT

CCACTTAGATTGTTTCTAGACCAGCTTCTAATCCTTCGAAAGGGAATTCGTTACAATTTAAGCAAACCCCTATGGAATCA

ATCTGAAGTCAAAGTGAGTTGAATAGAGGCGAGGGAAGCTCGGTAGAGCTTTGATACCCAAGGCCTCACCGGAATTACAA

TGCGGCGCAGTCACGCATTCAACTCAGAAACCTTCCGAACTTCGAAGTGTGCTTAAAAGAGTAACCAATATTTTTTGAAT

AACTTTCCTATTAAGCTCGTTACCTATCGGTCTTGTTCTAGTCAAAATTTTAAGGCTTAGGTTCGCGTAGGTTACGTGTT

CCTAAAGGCGGGCAAGAAGAGAACGGTGATGAAATCCGAACCCTTATCTTGTATGGCCGGGCCTTGCCCTTTACTAGAAA

ATTAAATCAGTAGTATTCAAGTCCTCAAAGCATATATGCAAACGAAGGAGTCCAAGAACTCGCTCCGACGGGGGATTCGC

GAGTGTTCAGAATCTTACCTCCCGTTCCAGGACGGGGGATGAATCGGTTGTAGTCGACTCGGGCCACCGACTCCTATGTC

GATTATGAAGACAAGGTGCAGGGACAATATCGTAATTGTCCTCCTTTCAAAACGATTTTATATTTAAAGTACCCTTAGTA

GGGTAAAAAAAATAATGTCAAAGTCCCAAAAATGTCCAAAAAGTCCAAAAATAAAAAGAAAAATTACAAAAATAATTATA

AACCCTAATACGAGTTTTCCGAAAAACCCAAAAAATAAATAAAAATAAAAATAAAATTCTTTGTCTCTTCCTCGTTTCTT

TTCGCTTTGTCTTTGACTCCAAGTCTTTAATTATTCACCAAATCTTTGGCTCAGCTTTCTTTTTCGATCGAAATCAATAC

CTGTAACCACAAGACAAATACCCAAGAAAACGTAAAATAGAACAAAAGAATAAAAAAACTAAAAAAAATAAATCTAAAAC

CCTAAAAACAAGTCCGCGTCGGCGGCGCCAAAAATTGATGTGGTTTTTATGTTGTTGTAGAAAGTGGTAAGAAAGATGTC

GTTACTCTCAGACTTGTGAAGGATTTTTTTAGATTCAAATAAAACTAGAAAACTCAAAGAAGGTTTTTATCAAGATTTTA

AAAAGCACCGAGACTCAGGATTCCACCAAACTCCAATTTTTATGTGATTTAATCTAATCAATATTCGTGCAACTCTTTAT

GTTTTAAAGTGATTCTAATATTTTGCAAAGGTAGATTTTTAGAATAACAATTGTATTTCGAAAGCATAGAATATCAAAAA

CCTAGGTCTAAGCATAGTCTATCAAGAGAAATCACAACTGCTTAGGAAAAATCATTCAATCAATTTTCAATCAAGGCAAA

TAATCTTAAAAGAGATTGCAAAAATTAATTAAAATAGAAATTACCACACACATCATGGAAAATTGGCTTCCTCCACGCCT

CGGCTAAGGGGTTTAGCTCCTCATATTAATCACTTGCTCAAAATATCGGTTCATAGCTCAAAAGTTGATTAAAAGGATGA

AAAACAATAAAGCAGTGTTTGCAACGGCTGTAATGGCGTTACAAAAAAGTTGTTACAATGGACCGTTACGAGAGGCAGAC

TTATGCTGCTTTGTTGAATTGTCGTTGAATCTATGACCCCTTAAACGATTTGTCCTTAATAGTCCTTTGTTCTTCGTGTT

CTTCAGCTACACCAGCAGAACCGATTTTCTGCAACTCTGTTGTTGATCTCTACTGTGCTCTAAAGCTCCCCAAAGTATCC

GTCACCCTTCCTTGACTTGAACCCAACCTATATATATCAATGACTCCCAAAAACGCGATTTAATTCCTTCTTTTCTTGGC

GAAAGTCTCGGACTTTTTCTTTCCTGCCAAGCTTCTCTACGCGTTTCGAGTTGTGAGACTTCAAAAGCTGGAAGATACGA

ATCCCAGTTAGTATATCCTTCCTTCACGAAGGCGCCTAACTTCCAAAAATAAACATAAAACTTGCCGAGAATAACTCGTC

CCTGTTTTGTTTCTCACGCATCCACGAGTTATCGAGAACGCATATCCAGCCCATTTCAATCGATCTAACCTCACATAATA

AGCCTGTCTGCTCTTGGTAAGTCAGCCCATCAAAAATCGGCCATTGAATCACGTTATTTCACGTCAAGCTGATGCTCCCA

AATTCACGATAATAATATATTTTCCCTTAAAATTCGGTTCAAACGGGGAAGAACAGGATGCCCCCTATCCTACGATGGGT

TCTTTGATAGTAAATGCCCTAGGGGTTACCCCTTATCAACTGGATACCCTTATCCATTTCAGGGTCCGAATAGCAAGTGT

CCTCCGGGCGCAACTTTTCGAGCCGATTTTTCCAAAAATGTTTATTGGCCAAAAATACCTACACACGCATAAAACACCAT

AATAAGTACAAAAATGAGCACTATCAATATATATAATCGAGACAATTTAGACACAAAAATGTGTCTATCACTAGGCAGAG

ACGAGAAAGAGATGGGTTTGAGTTTGAAGAAGGCAGGGAAACAGAACGAGGTTGAGAATAAGTTCACGTAAGAAACAGAA

AAGAAGTTCAGGAAACTTACGGTGGAAAACTTCACCGTAGGAGATTACAGTTGAGGCTACGACGATATTTTCAAACCGTA

CATAATCTCAGGTGAATATACGGATGGGAAATTCCACCGTATGCAAACACAAAGCGAATCTACGGAGGGTTTTCCCACCG

TACGTAAACTCGGGGAAGAGAATAAAGTTTTCGTGAAAACTTTAAGGAAAAAGGCTATTATTGGGCGTCACATTCCATTA

GAGGGGCGTCACCTAATAAGACAAAAACGGGTCACCCGTAAGTAATTTCAAAAATCCCTTATCTCAAATAATTGTGTAAT

GACCAAACAACCCTTATTTATTTAGTTAATTTTAATTAAATTTAATTATTATCTAATTAATTTAAGAAGATGAATTTTGA

TATATATTTGTAAATTCTGGGACAAAGTTTAAAACCAATGGTTGAAAATCAACAGGTACACCTCTAAACTATCTCCCATG

GGTTCAATTTCGTTGAAAACTAGCTTAAGAACGATAATAAATTTGATATTTTTAGACTTTCATGGCAAATTACGGTTGGG

TTGGGCTTTGTTCCAACCGTAATTCAGTTTTACGTCTGGGTTTGGAAGAGGGTTTTGCTTTAGTCCAACCGTAATTCAGT

TTACGCCTGGGAAATGTAGTGCATGAACCTAGGCGTAAACCCAAAGTTGGAATTCTTCAAAGCATCCATGATAAATTACG

CCTAGGTTCGACATATAGAAAACCTGGACTTCCAACATTACGCCTGGGAAGTGTAGTGCAAGAACCTAGGCGTAAACCCA

AAGTTGGAATTCTTCAAAACATCCAGGAAGAATTACGCCTAGGTTCTACATATAGCAAACCTAGACGTCCTAAATTACGC

CTGGAAAATCAACTAAACCTAGGCGTAAAATTGGAATTACGCCTGGAAATCAACTGAACCTAGGCGTAAAATTGGAATTA

TGCCTGGTAAACAACTAAACCTAGGCGTAAAATTGAAATTACGATTGAAAAATCTAGTACCGTACCCAGGTGTAACACTA

GCAGAAAGTAAATAAAATCCTAATTTTACTTCGATTTCATTCGATTTAAGCACAAAAACGGGTACAAAAAAGATGGGTTT

GTTTGATTATTACCTTACATACACCATGGGTTGTTGTTGAGGAGTTTGAAAACATGATTCTTCAATTGATTGAATCGGTG

GAATGAGTTGAGGATGAAGTTGAGGTTGATTGAGTTGTTGATTACCATCACTAGGATCAGTTTGCTTCTTCCTCATCTAT

TTTTAGTAGTTTTAATTGAAGGGTAATTTTGAAAAATTGCTATGGAATCAAAATATCCCTTAACCAGATTTTTGGTCACC

AAATATCCAAGTGAAGCCCCTCTAATGGAGTGTGACGCCCCAATAATAGCCTTCTAAGGAAATGGACTGTCAGTCCTCTG

TATCCGATAGGCCATGAAGTATTGGTGAGTAATTCAAATGGGCTGGATTTTCAAATTTCGTGCGAACCCAATATGCGGAG

AATTTGAGTGTTGTTGAGGCCGAGACTTTGGGAAGTGACTAGTTTTAGGGAACTTGCGACTTTTGGCAACTTTTTTTCGA

CCATAGGGATTTGAAGCACGGCGTAGCTACAACTCTGGGTACTATATAAGAGGCTCATCTAATATTTTTAGAGATATCAT

ATAATCTTTTATCAAGTTTTGGAGAGGAGAACTCAAGAGAAGAAAAGCTAGGGTTTGGAGCTATTTCCATCATAGTAACT

ATTTCTTTACTATTTTATCATATGAAACTCATGGGTTTTATTAGTACCATGAGTAGCTAAACTTCTTAGTGATTGGGGAT

GAATTCTAAGTTCTAGACATGATTTGAGTTATTATATAAATATATTTTGAAGTTCTTCACATGATTATTGATTGTTTATA

CTAATTAGAATATTTATGATTGATTGATAGATTGTTTAGGTGGCATTGAATTAATTTGTTGATTCAATCAAATGCCAATA

TTGAGTTAGGAGATAAGTAATAGTCGAATAATTCATACAAAAGTAGAGAACACAAAACCTTGCAGAGGGATTCTGTATAG

TGATTGTGTGTATAAACATCACTAGAAAGTGGACCTTGAGCTAAGAGTCTAACTACTAGGATCAATCTATAATTCGCAAA

AGATAAAGCATTCGGAAAAATTACACCTTGAGTGAGCTACTACTAGGTGGTTTGAAGAATAGAATCAGTATTGGAGAGCT

TTCGTACTTAAAGTGATTTAAGGAATTTAGGGGATAGCAGAGCTAGTTTGTTATTCCTTGTTGATAATAATTTGTGATTA

ATTGTATTAATGAATGAATATAATAACTTGATGATTGTTTAGTAACGAAGAAGGAATCCCTAATCATATTTCTCTCCATT

GCTTCCAAATTTTATTACTTTCAATTGTTTTGTTATTTACTTTGTTCTAAAATCAAAAACAAACCCCCATTGTGACACTT

TTTGACAACTAAATCTCCCTGCTCTTCGTGGGAACGAACTTGACTTCCATTATATTATTTTTAATTGGGTGGAAATAAGA

TATTAATTTGATTGCACCTACGACACGCATCACCATACCTACTTCGGATGAAATTCTTGAAACCATTAGAAACATGGCTC

CGCGGAAATCGCCAGGTCCGGATGGTTTCCCTGGTGGGTGTTTTAGAGACAACTGAATCTTTAGAAGTTATAAATCATGT

GAAATCGTTTTTCCGAAACAAGTTTCTTCTTAAGCAGCTGAAACATGCCTTTATTACGCTGATCCCAAAGGTTAATAACC

CTACATCTCCCAATGACTTCAGACCTATTAGTCTAACAAATACTATCTATAAAATCATCTCACAAATACTTACATATAGG

TTAAAACCTTTACTTGATTCTCTTATTTCGCCTTTTCAGTCTGCGTTTATAGCAAATAGGCAAATACAAGATAATATATT

AATCATTCAAGAAATTTTACACCCCTTTAAAATCAGGAAAATGAAAAATAATAAAAATGGGTTTATGGCTATCAAGTTAG

ATTTATCAAAAGCTTTTGATCGTCTTGAGTGGCCTTTTATTCTTTCTATTTTTAAAAAGTTGGGTTTCTCTGAAGACTAG

TGTCATCTAATTTTTGAATGCATTAGTACAGTTTCTTACTGAGTCTTTGGTTAATGGTGCTCTGGTGAGATCTTTTTTCC

ATCTAGGGGGATAAATCATGGGGATTGCCTTTCTCCTTACATTTTTATTATATGCATGGAAACATTATACGAACTCTTAT

TAAAGGGAGAAAGAGATAATCGATCCAGGGCTTTAAATTAAGGAAACATAGTCCTTCGATTTCTCACTTGTTCTTCGTTG

ATGAGCGCATCTTTGTTTTAGAAGGCCTCCTTAACTTATGCTAGAAATCTTATGAATATAATTGACTCCTTTGCTCAAGC

TTTCAGTCGGGCTATAAATTTGAAAAATCTGGATTATTTACTAGCAATAAAATGCATCATAATAGCATGATAAATTATTA

GCTAGAACCCTCGGAATTAAATTTCTCAGTACTTCTGAGAAGTATCTTGGTACCTATTTGTTTGTTAATAGGGATAAGAC

TAAGACCTTTCAATTTTCGTTGATAAATTCTATAGCGAGGTTAAGTAATTGTAAGAAAACAAACTTAAATATTGCTGGTA

GAACTATTGTCACAAAACATGTCCTTTCTAGCCTATTTGTTTATCATATGTCTTGCTTCCCTTTCCCTAAAAAGATCTTT

GCTAAGATAGATTTAATTCAGAGAAACTTTTGGTGGTCAAAGAAAGACCCTAGGAGAGCTGCTTATTATCGCTCTTGGAG

TGATATTGGTAAATCAAAATTGAATGGTGATCTTGGTATTAAAATTTCTTATGCTACTAATAAAGTATTTATAGCAAAAA

TTGGTTAGAGAATGTATAAGAAGCCGATCTTTTAGTAACTAGGTTCTTCAAGGATAAGTACTTTCTAAATCAAAATTTGT

TAGAGATTGATAAAGCTACAAATACTTCATCTTGGATTTGGAAGGGTATTGTAAAAGGTTTGACATTTATTAAAGATAAT

ATTGTTTACAAAATAAATGATGGTAAGGCTACAAGTATTGGTCCTCTAACTGGTTACCTTTCGCAGCTCTCCTCCGACTT

CGATTAACCCAAATTTTGCAAACTTCAACTTTAATACGTGAACTTATTGATGTGCAAAATATCTCTGGAATATTGGTCTT

ATTACTTCGTTGTTTTCTCGGGATGATGTCAACAAAATTCTTTCTCTTAGAATAAATACAACCAAACAAGACAATATCAT

GTGGGCTCACACAAGAAATGGAAATTTTACACCTTATAAAGTGTATATGAATGAATCTATAACTCTTTGAAGATGCTTCT

TTCTGGAAGAAAGTTTGGTCTTTAGATTGTCTTCCTAAGATTAATTATTTCATATGGAGTTTTTTTGCTTATATACTACC

TGTTAACTCTCTTCTTATGCTTATAATCCTGCGGTGGATGACTGTTGTTGTCTATGTAAGAATGCTTCTGAGACAGTGAT

GCACTTGTTTTTTCAGTGTCCTATAGTGTCCCATATATGGTTTGCCCTCTCTTTGCAGCATATTGTAAGTGGGGATTTTG

ATTGGGTGGATAATAATCTAGGTATCTCTCCTTATGCAGTTGATTGGCCTAGTATTGGTGCTACAGTTATGTGGTCTGTT

TGGAAATTTAGATGTGATGTGATCTTTAAAAAATTCCACTATTAACTTGGACAAGGTTATTATGGATACTAGAAGGTTGT

TCAATACATTTATTTATCCTCCTTCTTTGAATACTTTGAGTAGAGTGAATACTACAACTCCACTGCAGGATGTTGACTTT

GTTATTTTTGCTGATGGTTCTTATAAAGATCTTAATATGGGTATTGGAATTGTCTTATGTGATAATGCAGGAAGAGTAAT

TCAGTCAAGAGCAGACTTTGGACTGATTACAGTGTTGTTGGTCTTTGAGGCAGCAACCCTAATTTTGGCTATCTCTTGGC

GGAAGAGATGAATCATCTAAAGTTACGTTTTGTCGGCGATTTGCCTTCAATTAGTGAAATTCGTCAAAGAAGGCAAAGGT

GAAGTGGATTGGCGTAGTAGTGATCTTTTAGAAGAATGTCGGATTTCCCTTTTAAATCGTTTTAGTTTTAATGTTGTGTA

TATTAAGAGAATCAAAAATAAGGTAGCGGACCGGCTTGCACGTCGGGATAGAAAGTTTCTTTTGAAAACTTCATGGGTTT

CTATTCCTCCCTTTTTGGACTCTTTGTTGAGGAAAGAACCTTTAACTGATGTTTGTAATTTTATTCTTAGTTAATGTTAT

GGTGTGCTTCTCAGCAAAAAAAAAAAAAGAAACGACTTGTATTATTTTTACACGTTTAAAGGGACGATATGTGTTTACAC

TCTAGTAAAATCGAAATCAAAATTAAAGAAGCATTTTTGTTCTTATTGACCGAGTAATACAGTCACACTACGCCCTTAGA

TGAATATTTACTCATACATAGTAATCGTTAATATTTTTTATTTCAAAATAACTTTTCTTGATTTTTAGAGAAAATAATTA

GCATTTTAATCACGTGTTATTTTATATAATATATACATTCAAATGGTATGGAAACAAGAAAAAATAAATGAAAAAAAAAC

AAATTCATATCTTGAGACATACTGTGTTTTTAATACACATTCCATAATTAAATGATTGTTAATGAATAATTGTGAGATAG

TGTCTTCCGGCATAAGTAAGTTATGGACATTAGCATAGAAATCTATTTTCCACATCCGATCAGTATTGCATTAAAAATAT

TTACCCATTCTCGTTCTTTAACAAATTCACATATTATGCAACAGAAACTTCTAGAAATATCTTTTCAGCAATGAAACCCA

TTCCTTTGGGATTTTGATATTTTTTTAAAATATGAATTATCCATAAAGAAATTGTTTATCCTTGCTGACCTTGATCGCGT

TGGGAAAATGAATTTTCTAATATCAAGTTGACCATTCTTCTTAAAAGATGAAGGATTCAAGAGGATTTGAAATGCAGCAT

TTGTATGTGGTTATTACATATCCATTATTACTTTATATTTCGATCATCAATTATCCTTGCAAGGGTGATTATCATATATC

CATAACTCTATATTGTGATTATCCACACGGTCTATTATAAAGAAATATGATCATCGACTTGTAACAACTTTTCGCCACGA

CCAGGAGAGCTTCCCGGAGTAGTATCGACAGTAGACCGCTGGTCATCAAAGTACTCACCGTCACAGGTGAGACAGAATTT

CTTAATCTCTAACATATCACGAAGAACCAATACTCTGTCCACAAAATTCATGAAGCTCCTGGTAGAAGATCGTGTAGGTG

GGGTATTAGGAGATTTCCATTCGCGAAAATCAAGAACATGAACAGATATCCAGAGATTTTTCCATCCTTTAGCCAAAATT

GTTGTTGAAACAGCACATTTGGTAGGAACTAAAGAGAGAATGTGACGGATTATGGGTTTAGGCAACCTGCTGATTCTATC

CTCTCCTCCCACACCCTCCTCCTCCATGTTGTTTCTAAAAAGCAACCAAGATTGGTTCGTTTAAAGTCAGACTACAGGTA

TTTAATCATGTTCAAACACATGCTTCATAAACATTAGAGAAATCAATAGAATTATCATACAAGGGTTTGTTTGTATTTCC

TGTCATCACCAAAAATCATGGGGGTGGTGTTATCGTCCTACGAAGAAACCAAGAATAACTTAACAGTAAAAAGGATATGC

CCAACAAGTGATTCTGGTAAACTAGCAAAGCACAAAGCAAATTGATTGAAAATGCACGTGCGAGATAAATTTGGGTATTT

CCAGTTTACTCCAAACTGCTAAACCTAGACAAACACAACAAATTGAAATACTTGGGATAGTTTTAACCAAAATTCAGCTT

TGAAACCCCTAAGATATACGATTATTATCAATCATTTTCAACAACACACATATTTATGAACATCTATAAACATAAAATCA

AGGCAAACCCTAAAATACCAAATTGAAATCTAAGCGAAATATTTTATCCCCCTTAAATAAAATTAGGGGAAGAAATTAAT

GCGAAGTTCCTACTAAATTGAAATTTACCTCTTGAAGCAGTCGAATCAAATTTGCAGCTTGAGTTCATAAAGAGGTTTTC

TCACAGGGTTTCAACAAATCTGATGAAGAAGCCGGGAAGTGTGTATCTGGGGTGTTGGAAGGAGCATATTAGTAGAAAGA

TGTAGAAATGAGTTAGAGAGGGATTTGATCAACTGAGAGACGAGACGAGACAACAGAGTTCTAGGGTTCTCCCATTTCTG

CTCTCGATTCGAAGATGGGATACGGGGAATTAAAATGATTTTCCTTTTTTTTTAGTAGGAAATACCATATGGTCCTAGGA

ATAATATATTAAAGAGAGTCTAGTCCAGTTGACTCAGTCATATGTCTGGTTGGTCCTATTTGGGAAAATATATTATAGTT

TGGTCCTAAAAATTATGGTTTGGTCCTAGGGTGACACATATATACCACAATTACTATAAAATATCTCATTCTGAGATTTA

CTTTCTCTGTCATCTTCTTTTCTTTTTCAGATCTACTTTTCTCTCTCTCTTTTTTTTTTTTTTTCTTGATGATTTGATCA

GATTTGATGTTAAAGAAGATGATGAAGACGACGATTATCTCATGTTCTTTGCTGCTTTGCTTGTGTTGAAGATGAAGATG

TACTCCGTTTTCAAGGTTTTTTTTTGTTGTCTTGATAAAGAAGAAGATGAAGACGATGATGTTGTTGTTTCTGTTGAAGA

CGACGATGAAGATGGTGTTGAAGACGACGATGAATATGATGTTGTTGAAATCGATGAAGAAGATGAAGATTTGGGTGTTT

TTCGTCGTTTTTTGTTGTTGTTGTTGTTGAATATGAATGATTGACGATGAATTTTGATGAAGTTCTTTTTCTTTGTTGTT

GTTGCTGATGTTTTTATATGGAGTTCATTCCAGAAATGAAGTCCGGTTTGTATAGGTTTTTATACGGACTTCATTTCTGG

AATGAAGTCTGTTTTTTTAGGTTTTTATCTGGAATGAACTCTGGTTTGTATAGGTTTTTATATGGACTTCATTTCTGGAA

TGAAGTCTGTTTTTTATATTTTTATATGGAGTTCATTCCAGAAATGAAGACCGATTTGTATAGATTTTTATATGGACTTC

ATTTACGGAATGAAATCTGGTTTTAGGTTTTTATATGGACTTCATTTCTGGAATGAACTTTGGTTTGTATAAGTTTTTAT

ATGGACTTCATTTCTGGAATGAAGTCTGTTTTTTTAGGTTTTTATATGGACTTCATTTCTGAAATGAACTCTGGCTTGTA

TAGGATTTTATATAGACTTCATTTCTAGAATGAACTGTTACAAGAAAATAAGTAAAACTTAACACGGAAGGGTACTTTTG

TCATTTTAATATTTTTAAATAATTATGGACCAAGCAAAGAAAGGCATTTTTTCCTAAAAGACCAAGCAGACATGGGCCCA

CCTAAAAGGAACCAAGCGATATTTTTTTTTTTTATTATATAAAAAGATAGAGGGCATTATAGTCTTTCTGTTCGACTCGG

GAAGTGATGTCCGGCGGCTCGGAGTGTATCATAGATATAGAAACAAGAGGATGGTTTCCGTTTTTATATCTGATTAGTGA

GGGGTAGATTAGGATTTACGTAAAATACATAGATAAGGGGTGATCCCGAATTGATATACGGATCAGTTTTGTCCTTTTCT

TATATTCCCAATTCCTTTTTATATACCCAATTTCGTGTTCCTAAGAGCAACTGCAATCATACGACTAAAACCAAAGTCCA

AAAACTAAAAAAAAAAAAAAAAAAAATGGTTTACTCTGTGCTGCGACGCTACGGGGCAGGACTAAAAACCAAAAAAATTT

CATCAGGCGCAACTTTAAAGTTGCGCCCCAATCAGGCGAACTTTAAAGTTGCACCCCATTTTTTTTTTTTAAATTCGGGC

GTAACTTTAAGTTACGCCCCACCATCAGGCGCAGTATAAACTTCCGCCCCACCATCAGGCGTAACTATAAATTACGCCCG

TTCAAACGCAACTTTAAATTACGCCCGACTAAAACCAAACTAAAGACCAAATATGGTTTGGTTTTTGGTTTTACTCTGGT

CTTTGATCTTTACTCTTTCTCCACTGCGTCTGGTTTGTGGACCAAAATTTTAGTTTTGATCTCCCGCCATGGATGCTCTA

ATAAACCATGCCGCCCTTATAACAAGATTGAATTAAAATGATTTTCCTTCTTTCTTTTATATAAAATAAAAAGATAGAGG

GCATTATAGTCTTTCTGTTCGACTCGGGAAGCTGATGTCTGACTGACTCGGAGGATACCTTCAAAAAAAAAAAAATAGAA

ACAAGAGGATGGTTTCCATATCCGAGGCCGGACCGATTTTTGGAAAGAACGGAATTCGGAAGCCAATCGGCATATAAAAG

GGGAGTCTTTGGTGAAGTTTTTGCATAGGTATGTTTGATTTCTCGTGAGTTAATAATTCGAGCATCGATCTAAAATAACA

TCTCTCGAGTTTTTGAATTGAATCAGTAAGATTTTGATTTCATCTGAATTAGTTGAATTGAATCAGTAAGATTTTGATTT

CATCTGAATTAGTTGAATTGAATCAGTAAGATTTTGATTTCTAAGATGAACAAAAGAAGAAGAGAGAAATCACTTTCAAA

CCGTCTTGATGACAAGTCAGTAGAAGATATACTTGTCAACCTTCCTGCCAAGTCTCTTGTGCGTTTTAAATCTGTATCCA

AAGCTTGGAAGTCATTCATCGAGGAGGATTCGTCTTTCATCCAGTCACATCTAAGTCAATCCGAAGCTCAACTTGTCATC

ACCACCACATGCGGCGGTTATTCGCTAAAAGTCTTTTCACCAAAGGATGGGTTTCAAGGCGGGGAGGCTATTCATCAAGT

CACACTTCCATGGCCACTGGGCGCAACCGTACCACGGCCTATCCATGGATTATTCTGTTTTATTGATCACACTGATAATT

ATGCTACTCGGATCTACAACCTCGCGACTCGGCAAGCTACTCCGTGGGCACAAACATGCATACCACCACAGAGGGGCGCT

TCGGTTACTCAGACGCCAACTTATGGTTTCGGTTTCGATCCTTTGACCAAAACCTACAAACTAGTATGTGTCTGGGAGAT

AGCCAGGGACTCCATCTGTAGTTATGAAGTCGACCACATTTGCGAAGTTCTTACTGTAGGAGCGGGGGAAGGAGGAAATC

AATGGAGGAAAATAGATGAGGAACCACCAGTTCGGTTATGCGAACTGCCCGGCGTCTCGCAAACGGTTCGATTCTTTGGA

GGAATCTTTGGTAAGAGTCTTTCCTTGGATCCCTGATGAGGAATTGATCGTGGCATTTGATGTTGGAGCAGAAAAGTTTA

GAGTCATTCGGATCCCGAGACTTCATCGTTGGTCCTCGTGAGATTTTAGAAGAAAATCGGGCAGGGAAGTTTACAATAAC

ACATATAGATGGGCGCATGGCTTTATTTGACAGAGCAGACGCCTGCATCGTAAAATTATGGATATCGGATGACGACACTT

CTTCTTCTAACAAGGAGACGGCAACAAATTGGACCGAGGAAGCCACAATATTTCTTCCTCCTTCCTTTGAAGAAGAGGGA

TATCTTTACTTTAATCCAGTCCAAGGAACAGATGTGATAGTCATACAATCACCTATTTCGGACGACGAGAGAAGTCCGTT

AGATTTTATAAGTCTGTATATTTTTAACCGGACTACCAAGACTCTTCGAATGATTGACATCACTGGTATAGCACCGGCAT

TGGAATTTCCCTACCTTTATGATGTGCACATTGTTCACGAAAGTCTTCTACCTATTCAGCAAGATGCAGAAAAGAAGAAA

CTTGATCAACCCAGACCACCACCACTGGTTCAAAGATGGAAGCAGACACCTCACATGCTTTTCTGATCACCCCTTTTGTT

TGATTAGATTTTCCCTTGTAATTTGGATTATGTTTAATCTAATACAGTATGTTTTATCTTCTTCGTGATAGTTCCTGGGT

CTTTTCCTTTTCATATCAGTTCATTCAGTATGGGAATTCCTAACCTTAAGCTGAACATAATATGTAATGTCAGCCCTACC

AAGATCACTTTTCTATAACTTGAAGCAATATCAAATTCGCTAATAATATAATTGAAGTGTATGTTGTTGTTGTTGTGAAA

TCAAGCAAGCAACCATGGTAATTTTAATTTCACCGAACTTATACTTCATAAAAAAAAAAAAAAGGCTTGGGATTTAATAA

GCTTTAAATTCACCAACAACAATAATGAACAACCAAGCCTTTATTCCCAAACAAGTCGGGATATGCTAAGCTTTTAATAC

ACCATCACTCAACATTCTGATAATCCCTAGTCATGGTACTACTGCAAGTCTCATTTCCTTTAATGTGTTATCTTCCCCAA

ATTTAAGTAACCACTCACTTAAAGGCTAGGTTTTACTTGTATTCTATGTCAAGGGTATTGATTTTACTCTTGCAAAGTGG

CAACACATAGCTACTATTACACGGAAGAAATGGCATTTAAGAATCATAGATGCAGATCTTCACAATGATATATTTGTTCT

CGACCTAAAATAAACACTAGGTATACATGGCATACAAAGTTAGAGAAAACAATCATTATCACGGTTCAAATGAGTTCTTC

TACCAACCTAACAATGAACTATACATGGAAGAGAACATGAAAAATCATATGGAAAGTAAATCAAATGGGCAGCGACAAAT

AAATAAGGGTGGATATGAAGTTTCACCTCTGTTAGGAACAAGCTAGCAATGAAAGAGAGCAAAGATACTTGTGAGCAGTA

ACTAACTGGAGTTGCTCCTCCTTCCTGTCCTGCTGTAAGAAAAGTTGAAAAGCTTAGAAGATTAAGCAACTGCCTTGCGC

CATGGTACTTTTAGTTTCACTGAACTAACAATTAACAAAAAGGCTTGGGATTTGAAAAGTTTTTATTCACCAACAACAAC

AAAGAACAACCAAGCCTTTATCCCCAAACAAGTTGGGATATGCTAAGCGTTTTTAATACACCATCACTTCTGATACTCAC

TAGTCATGGTACTACTGCAAGTCTCATTTCCTTTAATGTATTATCTTCCCCAGATTTAACTAACCTCTCACTTAACTGTT

AAGAGTATAACAAGAATTCTAAATGATCGTAAAGATCAAGCGGATAGGATTGAGGCGACGTCCCAAGGCGGCGGTGACGT

CATCGCCTCGATCGTGGACGATAAATTAATGTGCAACCTTAGACACTCAACTGAAATAAAATACAAACAAAATGGCAACA

GAAAAAATCTAGCAAATAACTGACCCAACTCACTATGCAAAGCAATTGAATTAAACACAAGTGTTGTGAGCAAACCAGCT

AACCCATTTGCTCAATTGTATTGGACTCAAGATATCTATATCTTTATCCCAAAGAAGGTCAACTGCTGCAACACAAATGA

TGCGTCGGAAAACTGACTTTGTTTTAGGCTATGGGAATCCGTTGTAGGCTAGATTTTACTTGTATTCCATGTCAAGGGTA

TTGATTTTACTCTTGCAAAGTGGCAACACATAGCTGCTATTACAAGGAAGATATGGCATTTAAGAATCATAGCTGCAAAT

CATAGCAAAGATATATATATATTTGTTCTCAACCTAAAATAAACACTAGGTATACATGGCATACAAAGTTAATAGAAAAC

AATTATTATCAAGGTTCAACAGAGTTCTTCTACCAACCTATACATGGAAGAGAACATGGAAAATCATATGGAAAGTAAAT

CATTAAGCTCAATTGATAAAAATATTAATTAGAATCATTAGCTAATCAAAAACGAATATGGAACTTTCAAATCTATACCC

ACCCAGTTGACGGCCTCCCCTTCAAGTCTGTCTCTCTTCATATTTATAGGTGCTGAGAAACTGCCCGGAGGCCTAATCCA

ATACCCACCAATATGTTGAAACTCTTCCGAATTCAGGCAATTACCGAAGTCCGGTCACTGCCCATTCAAGTCTCCTCACC

TATTTGAGTTCACTGCGACAATTTCTCAAACACCCACTAGTCTCTTCAGCTCCATTTAGTCGAACCCCATTGCAGATTCA

GTTCAGCCATCTGCTGATTATCCCCAACTGATTGTAAATCAAATCAGTTAGCCTACGAATTAGAAACAACTCCTCTCGAT

CAAACAGACCCATACAAAATCAATTCGAGACAATAGTTCCTTTTCTCTTTCATGTCTACGAGAATCCAAACCCATCTTGA

TATCTTCTTAATCTTCGTAAACCTTCTCAACAAAGACTCTGTTGCTGTTTCTTTTGACAAAGAACAGACGATGATGTTCC

TTGGTTGTTTTGATGCTTAAGAAATACTAGTATATCACCAAGAAATGTGGTCCCGAAACTGCCACGTCCCACTTTTTTTC

ATGTCATTTGCCTTTAATTGAAAAATGCTCTCTTCGATGCACCCATAGTCATGGTCCACTAGACTGGGTTAGGGCAGCAA

GTACTATATGGCGCACCATTTGCCATTTTCATTTGCCTTTGTCGCAGGTTAACTGTATTATTTTTGGGTAGTATTTATTT

TTAAGAATGATACACCACAGTTTATCAGGATTTTTAATCAATCGCCAGGCAAGTCTGGTGAGCAAAGCAAGATTAAATTT

ATGAGGATTTTTTATACCTAAACCTCCATTAGCTTTACTAGTACAAATGAAATCCCAAGCCTTAAGATACAAACCTCTAT

TGTTTTTATTTTTCTGCCACCAGAAGTCTCTTTGAAGAGCATCCAGTTTGTCTAAGGTTTTTTTAGGTAAGAGAAAACAC

ATCATTTGGTACATAGGCATAGCACTAGTGATAGAGCAAATCAAAGTGGTTCTACCTGCTTGGGAAAGAAAATTGGAAGA

ATATGAAATTGTATCCAAAATACTCAATTGAATTAACCACAAGTGAACCCATGCTTAATTCAACCGGAATACTCAACTAA

ATTAACCACAAGAGATCCCATGATTAATCCAATTGGAAATGCTCAACATAAGAGAACTTACGGAGCCAAACAGTATATAC

ATAAAAGATCAATACTGCAGAATAGACAATCTATCTTCCATCACTATTTACACAATGACATACAATAGACTTAATCTGTA

AATAAAAGCTCATCCTTTCTTCCATCAATATTTGCATAATGACATAAAAGGCTTAAATTTTGTTTTTGTCAAAAGTAAAT

TCATTCTTTTATCAATACTTGCATATCGACATACGAAGGACTTAAATTTTGACAACATATGGGACAATAGTTCACAGACG

CAAACACACATATCCCATAACAAATTGCAATATAAAACCATAAAGATTAATACTGCAAAATTATCTTCCAAACAAACTTT

AGAATTTAAACAAATAAATCCAAAAATATTTGAAGATGAAAATCGTTGGACATAGCTATGTGTAATCACAATAATGGTTA

TTCCAAACTCTAGTTATTCTTCTTAAAAATGCAAGAATAAAATTCTCATAAGAAGCTTTCCTAGACATAAAAGAGATTAC

TCATCATCTTTATCTTCTGAAACCATCCAAGAGCCTTGAATAATATCCATGTCTTCCTTGATTTCAGGAAACTTGGTTGC

AATCAACCACAATTGTTCTTGCACACACTTTACCCTTGAATTGATCTTAACCATACCTTTGTGAATTTGTTGAAGAAGAC

TCAAAGTGGTAGGATCACAAGTGCTGTCAAGGGAAACACATCTTTGTCTTTTACACACATTCTCCTTCATACGCTTAAAG

GTACGTGGGCGAACAAGTTTGGGAGTTCCAAGAGAGTTGCCAATTGGAGCAAATGAGAGTGATTGACAAATCTATCCAAT

TAATCAAGGGAAGACAACTTCTTGTTGGGCGAAATCATCGAGATCATTTGAAAAATGATAAACCCACAAGTGTCGAGACT

TGGGGTACCTAGTTCAAGGAAATAGACCAATTCCGCAAACTCGCGACTCCATCAAGAATTTTCAATTGTGGAGGGACAAA

GATTAGAGATACCAAGCTTTCCGAAGGACATCAAAGCAAGACTATGATAATTTGTAGGAAACTTTCCTTCATTACAATTA

ACATTCTTTCCACAAAGACCATGAGAGATGGATTCATATGATGGTCTTTCATCACTTGGTCTAGGGAGGCAAAGGAATTT

TGGTAATCCTTGATATCAACTCTCTATCTACAAGAAACTTTTCTCTTTTTATCATGGTTTTGAATTCCATGTTCTCAAGA

TCAACATCATGAATGTTGGCATATAAAATCCTTGTGAGATTGTCAAAGCCTTCACCAAGACCATTAAAGATGTTTCCAAG

ATTAAACTTTTCAAAGCAAATCAACTCTTCTTTATGACCACTAGACAAATCCAACTTCTTTTCAAGAATGAAACCACTAG

AAGCAATCCTTTAAAAGATCATACTATTAGAATCATTGACAAATCTAATTCTAATAGAGTCTTGAGATTGAGGAAGAGTC

ATGCTAGAAGAAGATGCGTCCAAAGTTTTTGAAACCTTGAAATTTCCCAAAAAACCTAGAAAATGAGGGTACAAGGTGAA

GGAAGAACTTAATTGATTTGTCTCTTGAAAACCCTTTCATTTTTTTCTCTCCAAAGATGCTTTAATGGAGGAAATACATG

AACCTAGAAAGGTTCACATACCCGAACGTTACAAAGGAGAAGAAAGGTACGCGTACCAGCTTCTTATGTAGCCTCAAATG

GGCTTGGACCCGTTCACAAATTGGACCCACAAAATAAAAGTAGGATTTTTAAATCGTTTACCATCCAAGGTACGCACACT

GGAAGAGATCAGTATAGTGCAATAAAAGAATAGTAGGGAGCCTACAAAACTTTTAGTTGACCTTGGAAGGAAGAAATAAA

AAAAGATTTACAAAATAAATTAACCCAGACAACACTGTCAACCGTTACTTACCCCACATCAAGTTATATACAGATGGGGT

ATAGCCAGTCTTGTTATCAAGAGACTAAATGTGCCGAGAAATTCTCATGCACTTTTTCAGGTTGATGTTTGTGAATATTA

GTATAATCATAGTAATGATTGTATACCTGGTCAACGGAATTTTTTGAATCAATTTTCTTATACCCAAGAAAATACTTTTT

TCAAAGATTAGAACCTATACCGAACTCACAAGAAATCTTTTTATCAATTTCACATGAGTTGGAAGTAGGTCTAGTTCGTA

CATCATGAATACATGATTTGACAGGTACAGAACTTTTATGGGAACAATTCTTATTCTCAAAGGAATTAATCTTTTCTAAG

AATTAAAAAATGCTTTCAAACGCTCTACATAGATCTTTGTCAATTGATCCATTGTGATGGTATTCCTCATAAAGATTAAG

GTTTAATCTAGTGAGATTCCATATCTTTGAGCGTCGACTACATTTTTTAATAGACTCTTCTTAGAAGAACTTTTAACTGT

TCCTTCATATTGTTCCTCATCATCTAGACTTTACAAGGACTTGGATGAAGCGAGATTATCACAAGAGACCAGAGGTCTAA

TAAATAATATTTGAACACTCTTTACAGAATTCTGGCATGCGTTACAGATGCCAAGAGCGAGTTTCCTAAAGGAGTTTCCC

ATAGGGACATACTTTTAGGAAACAATCAAACACAAACTTGTTAGGTTTATGGTGTTTGCCTGCTCTGATACCAATTGAAA

AAGTGGGGGTAAAACAACCTCACCCAATAATTCTCTTAGCAATCTGTATGGACTAAATCCAATATACTTTTAAGAGAATC

AACTGGATAGTCAGACTCAATCTTAATAAAAAGTATATCAAAGAGTTATAACTCAATTTCTCGATTCAATCCTCACTCAA

GCAAATAGAAATCTGCGAGTCTAATTGAATACAAGAGAAATTAACTTGAACTGTACCAAAGACCAATGTTCAAGTATCAA

TCAATTTCAATAAACAACCAAAGGTCGGATTTCCCAATTGACTAATTTAACGCACAACTTGTGATATTTTATTTATATAA

TAACAAAATATAATGCGAAAAAGAAATAATATAGACACCGAAGTTTTGTTAACGAGGAAACCACAAATGCAGAAAAACCC

CGGGACCCAGTCCAGATTGAAGACACATTGTATTAAGTCGCTACAGACACTAGCCTACTACAAACTAACTTCGGTCTGGA

CTGTAGTTGAACCCCAATCAATATCACACTGATCCAAGGTACAGTTGTGCTTCTTACGTCTCTGATCCTAGCATGATACT

ATGCACTTGATTCCCTTAGCTGATCTTACCCACAACTAAGAGTTGCTATGACCCAAAGTCGAAGACTTTAATAAACAATT

CTGTCTCACATAGAAAAGTCTACAAGAATAGATAAATCTGTCTCCCACAGAAATACCTACAAGTTTTTGTTCCGTATTTT

GATAAATCAAGGTGAACAGGAACCAATCGACAACCCGGACTTATATTCCCGAAGAACAGCCTAGAATTATCAATCACCTC

ACAATAATCTAAATCGTATGATGGAGAAACTAGATATCGTGGAATCACAAACGATGAGACGAAGATGTTTATGATTACTT

TTTATCTTGCCTATCGGAGAAATTAATCTCGAGCCAATTTTATAATTGTACTCAATACGATAGAAACATCAAGATCAGAT

CATGCAACTACAGAGAAAATAGTTGGGTCATGCTTCACAATCCCAATGAAGTCTTCAAGTCTTTAACCTACATGGTCTCG

GTAGAAACCTAAGGTTAAAGGAGAATCGACTCTAGCTTATACAACTAGTATCACACATGAGGTGTGGGGATTAACTTTCC

TAGTTGCTAGAGTTCTCCTTTATATAGTTTCCAAATCAGGTTTTGCAATCTAAGTTACCTTGGTAACAAAGCATTCAATA

TTCACCGTTAGATGAAAACATGATTAGATTCAATCTAATATCTTTCAAACTTTAGATCGAACTTAAATTTTTATACACAA

ATGAAATATACGTCATTTAGGTTTAAGTAACCGTACCTAAACATGTACAAGTTGTTGTTTAACAGTAGTTAACCAAATGG

TTAGCCATATGAGCACTTTCATATCAACCTTATTCATCTTCACAATAACTAGTTTAAATGACTCAAATGAACTAGTTCAA

GAGTTTTTCAATTGCTTAGATCTCATAAAAGTATACAAGACACAATCGAAGCAAAATCGGTTTTGATTCACTCGAATCGA

TTCATGAACTTCATAGCCATGGTTTGCAAATATGCATTCCTTAGTTTATATAAATTTAAGTTCACGAATAAACCAATTTT

AGAAAGTAACACACTTAAGTATGCCTACGTGTATGCGTACTTACATAACCGGATTTGAGTTTGTTTTAGTTTCCAAACTC

AGCAAAAATTCACGGATTTGAACTTTTACGCATACTTAGGTAACCGGGTTGAGTTGGTTTTGATTTCCAAACTCAGCAGA

AATTCACGGACGTGAACTTCCGCCAGTATACCCAATCTCCATCAACTATTTAGTACATACACAAGTATGCATACATTTGG

TTCCTGGTTTAGGACTTATACACTAATGCGCGAACACATTAAATGCTTAAATCCAAAGATGGTTACATGTTCTGAACTCT

TTATTTCAATCATTGAAACATTCTTATAGGATGTTAAATAGAGGTTATTCTCACACTATTGAAATAATCAACATGACTTT

CGTCACTTGTAAAGATGAACTTGGCCAAAGAAAAAGCTTACCAACACATATTTCGAGAGCGAGATAAACTCGGCTTGAAA

TAGCAAATGTGTATAATCGAAGTCTATATAGCAATACGACTATTGTCTCAAGATAGGAGATAGAGTAGATAGACTTTTTG

AGTGATATATAAGTTCAAGTCTCCACATACCTTTTAGTCGATGAAGTTCCACCAGTTCCTTGAGTAGTTCTTCGTATTTG

TATGATGAACGCCGTGGAGTCTAGAGCTCAACTACACTTTCTATCCTAGTCCGAGACTTAGCTATAAGTAGACTAGAAAT

CAAGACTTATAGTTCTGTAAGGTCATCCTATTATATACTTGAACCTTGAAAATCTGAATTCAATCAAAACAAAGTCTTTT

ACTAATAATGATGAGACATGCTAGATCGTTTAAAGTTTATGCAATCATTATTTGCATTCAAAGCAAGTTCATAAACTTGT

TTCAATGTTTAAGAATTTATTTCATCCCAAGCAAATTTAAACTTCTCAATATAGATTACAAAAGTTTGAAAACTATATGT

AAGCATGTGAGAAGTCTTCTTTGTAATACAAAGAAGAATATGGACCAACGTATTTTTGGATTAAGGACCGTGCATAAAAC

GTTAGCTCATAAGTGACTAATATGTTCCGTGAACCATCAAGCAAAGTTGGAGTTCGGTAACACACAAAAAGTGTCTGGTT

TGCGAACTAAAAAATTCGGTTTCCCAACTTGAGGTATTTGGAATATTGAGGAACTTATCCAAAAGTCCATAGTTAGCGAA

CTAACATTCAATTCGTGAATAAAGAGCAAATTCAAAAAGTTCCTGGAAGATATTGAAAATTGCATAGTTTGTGAACTCCT

TTATGCACATTGGAATTCAGTAACATAAATTTTGTGCGTAGTTCACGAACTAGAAAAACAATTCTTGAACATTGATGTAA

CTTAATATTTTTTAGGATCTGCTTTCTGCAATTCTTGAACATTGATGTAACATAATTTTTTTAACAATGATACTTTTTCT

CAGTGTCATAATCCATAAGTCCTATGGACATTCGATATGTGAGTTCATGTTTGCAGATTAATCAATAAAAAAATCTATCA

AGTATTTTTGTTATCTCTTCGTAGACATGATATGTAACATTAGCTAATTGATCAACCATAGCATCTCATATATTATGTAT

ATACTACATCAAGCACTACAACCCAAAGATATAACAAATTATCCAATATTAAATTCGACAAGATATTGAACTACACGACA

CCCTACAATTGTACATTAGTTATTCACGAGAATTAAGGAGATATGGCACCCCATAAACAAGTATACATAAACAAAGTGAA

ACTGCGACTAGTTCATCAGTTTCTACAAATTATAACTAGTTCATCAATATTAGTTTAATTTAATTTTCACTACCAACTAA

TAATTTACAATTTAAATCTCCGAAATTCCAACCTAAGACTATTTTAGATCTAAACATAAAAAGACGGGCGAATAACAATC

AATCCCTTACTTAAATTATATACATTGTTTCTGGAACTATATTTAGCTTTGAAAAAATGAGGCAGACGTAAAGAACTATT

TTTTATTGAGTTTCATCATCGGTAAAAAATAAAAAAAGAAAAACATACCAATTTGTCAAGAACATACCTCTTTGAACTCA

ATTATGCCTACATCATTTCTCAGTCCACGTACTTGTAGACATATTATCGAATTATTCCCCCTTGAATATATGTTATTAAT

GGATGTGGTTTCATCCAATATACTGTATGTTAAATAAAAAATGAATAGGAAGAGGAAGATAAATAGTGTGGATGAAATTA

GCAAGGAAAGAAGACGAAAAACTATATTTTGAGTGACTGGTATTCTTTTTACACGTGTAAAGAGACAATATGTATTTACA

CTCGAGTAGAATCCAAATACAAATTTGAGAAGAATTTTTGTTCTTATTGGCCGAGTAACAATTAAATTTTTTCCTTCCGA

TGAATATTATCCATACATAATAATCGTTAATTTTTTATTTCAAAATAAGTTTTCTTGATTTTTAGAGAAAATAATTAGCA

TTTTAATCACTTGTTATTTTGTATAATATATATATATATATATATATTAAAATGGTATGGAAACAAGAAAAAATAATATG

CAAAAAATAAATTCAAATCTTGAGATATCATGTGTTTTTAATACACATTCCATAATTAAATGGTTGTTAATGCACAATTG

TGAGATATTGTCTTCATCATAGAAATCTCTTAATCACCATATCCGGTGAATATGCTTCAGTATCATAAAGATTTTCCCAT

TTTTGTTCTTTAACAACTTCACATATTTTGCAACCAAAACTTCTGGAAATATCTTTTCAACAATAAACCCATTCCTTTGG

GATTTTGGTATTTTTTAAAACATGAATTATCCATAAAGAAGTTTTTTTTTAATTCTTGCTGACCTTTATCGCGTTGGGAA

ACTGATTATTCTCAATATCAAGTTGACCATTCTTCTTAGAAGATGAACGATTCATGAGGACTCCAAATGCAGCATTTGTA

TGTGGTTATTACATATTCATTACTTTATAAACGATCATCGGTAATCCTTGGAAGGGTGATTATCAAATTTTCTTCTCCTC

TTCTTTCTTTGTGGCTAGATGTGCAAGAGATTAGAGTCACAAACACTAGAAAACCTTGTAGATAAGGACATGTGTCAATT

TTGTGGTGGTTGGTTTTTGTTTCTTACTTTTATACCCTCTATGCTTAACCATATTTACCTTTCCATCAAAGCAGTAAGTA

GATTTTTTTTCTTTCACTTTTACATCCATCTGTCACTTTAAACAGTTTTTTCGATTTGCTTAGACAGATTCCATGATGAA

AATCAGTGGAGTTCAATTACTGATAATTTATACCCATTGTGTTGTTTTAAAGAAGAAAGAACTCGAGTCATTTTTCAACA

ATTCAATTCCTAACATAAAAATGCAGTCTTACAAGATTAAATCCAAAATTCACATCTGTATGCAGTAATTTTTTCATAAA

CCCTTACACAATTTTCTCAACAAAAAAAAAAAAAGAAAAAAAAAAACCATATGTGGCATAGACTTTTGGATCATCACATA

TGCAAGCATATCCTTCTAATCCTCGATATAAACGTTGCAAGACACAAGCTTACTTAGTTAAGTTTGCGGGGCCGGGGCAA

AATGAAAAAGATAATAAAATGGTAATAGCTAGAAATTTCAATTAATATGATTCATATCGATCAAAAAAATAAATCATTCT

TCAAAGCAAGAAGAATTTAGTGACACTGTAAAAAAATAATAACCAACCAAATTTACATCTACAACCAACTTTGAAATCCC

CACGACGATAATCAACACTATCTATTGAAATCTCTCCCACAAAAAATTTCCGAAAGTTTAGGAGGTGTAAAATATTTGGC

GACAGATAGATACTCTTGCAGTAATTTTCTTCTTTCACTTCTTGTTTGTTCTTCTCTATGATTACAACTTCAAGATCCAA

ACACAAACCAACAAAAGAGAGAGGAACAAACAAGAAGTTGCACTACCATCTTCTAACAAGAAGTGCTAGTGTATGGTAAC

AGAGGCATGAGATACATTTAGGATTCACTTCACATAACCACCTCGTATTTAAGGATCTCGTCTCTCTGTTTGATAACACA

TGATATAGGTTGTTAAGTTTTGATAGAGGTAAAAGTCTTGGGTTTCTTCAATTGGAGCGGGATTTATCTTTGATAAGACG

TTTAAATTCATTAGATTCCTCTTCTTTTCCTTCTACTTCTTCTGCTGGTATTTGAGGAGCCACTGAAGAGACAGAAGAAC

CTCAATGTCCACCTGGTTTGAGACAGTATGAAACTATGGCTGTTTTAAGACCTGATATGACTCAAGATGAACGTCTTACT

TTTACCCATAAGTATGTATGAGGAGGTAAATTAGATGAAGGAATTTGTTCTTTAATCTTAACCAGTTTTCAAAGTTTCCA

CTGAAGATCTTAGATAACTTTTTGGTGTTGCATTCTTTTTTTCCTTATTACTTGCAGCTTCATTGTATCATTTGTCCTGT

GTCTTGCGAAATCTGAAAGGAAAATCGCTTCTAGTTTAGGAATGTAGCTTCAACCTATGAGCAGTCATGCAAACACATTG

GAAGTATTGCATACACTACTTGGTTGCACTATCAAAAGACTTAAAAAATTCTACTCTTTGTCACCCTGCTTGTTCATATT

GAAGCTGTTTGCTTTTTAGTTCTATTTTTGAACAAGTATGAATTGGCCTTTGTTTGCAGTTAATTGTTGCTGGTGGTGGA

ATGTACGTCGAGGTGTTTGACAGAGGAGTGGTTCCGCTGGGCAGGAGAAACGCATCTATCTACTTTCTACCTATCTCACC

ACACCAGAGTCCGTAAATGATCTAGAGTAGAGTCCCGCGACGTTAGAGGATCATCCAACTTGAAGATCCGGAAGTGGGAG

TAGTAGAATCTGACTTTTTTGACTGTACCTATTTTCGATTTTAGTCCTTGTTTTGGGTCTTTTTTTTTGTTGTAGTTGCT

GTGTGTGGGAATATTTGGCTTGATAAAATGGGTACCAGGGGAAACTTCACCTCTCACAATTGATTACTTGTATTTTTGAT

TAACCATTTTTTAACATCAAACTCTTCTGCCTGTCATTCTAAAATGCTGGATATTATCAATTAATGATCCTCAAAACATC

CAGATAAACGCTTTATAGAATAAGAAATTCTGGAAAATCGATTAAAAATCTTGTTACGGAAATATTAAATTCAGATGTGA

TAATTATAACTAATTAACCGTTAACCTGTAAATTTAAGCTTTTAACAGAAAAATGGTAAAAAAAATAAATGGCCCCCACC

AAAAAAAATAAAAATGAAAAAGAACCACTAGGAACACGCCACGTGGCATCTTATCCACTGGTGTGTTTAGAATTTATATT

CTTGTCTAGAGCAGGTACTAAGAAGAGGAGGAAGAAGAAGGAGACAAGTAAAACAGTAGATCCGAAGAAGTGAGAAATCA

ATACCTAAACACGATAATGGAGGTAAAAAAAATAAAAACACTACTTTCATTTACTAGCATGGTCAAAGCATCAGTTTTTT

GCGTTAAATTTTGTATTACTCTCCTTTATTTTGTCTTATCATAGTCAATTTTGTTTTGGAGTTCTTAAGGTTCGACCCAT

AATTATCGTGTGGTGGGAGTTTTAAGTTCCTTGATTTCCGTTTTCAATTAAAAACACAAAAAATTTGTTCTAGGGAAGGA

GTCGACTCAATTCAGTGGCATTGGTTTTTTTTTTCTTCTATGTATATAGTATTAAAATCAATTTTACTTATGCATTTATG

GTGTTTCAATATGTAGTTATGGTATTACTTGGTTCCTGTTTTACTAATTTTGGTTTCAGATCTCTACTAAGAAGATGAAG

GAGGTAACAAGCTTGGTAGATGTTTTGAGACACCAAAATATAGTTGATGTCATGTTTAACAAATATTCAATACCCAAATA

TAGTTGATGTTTTGAGACACCAAAATATAGTTGATGTCATGTTGGACTTTGGAGCCGGAGAGGGTAAAGACTTAAAAGAG

ATTCAAGTAAAACAGTAGATCTGAAGAAGTGAGAAATCAATACCTAAACAAGATAATGGAGGTAAAAAATACACTACTTT

CCTTTACTAGCATGGTCAAAGCAGCAGTTTTATGCGTTAAATTTTGTATTCCTCTCCTTTATTTTGTCTTATCATAGTGA

ATTTTTTTTTGGAGTTCTTAAGGTTCTGACCATAATTATCTGTGTGGTGGGAGTTTTTAAGTTCCTTGATTTCCGTTTTC

AATTAAAAAACACAGACAATTTGTTCCCAGGGAAGGAGTTGACTCAATTCAGTGGCATTGGTTTTTTTTCTTCTATGTAT

ATAGTATGAAAATCAATTTTACTTATGCATTTATGGTGTTTCAATATGTAGTTATGGTATTACTTGGTTCCTCTTTTACT

AATTTTGGTTTCAGATCTCTACTAAGAAGACGAAGGAGGTAACAAGCTTGGTAGATGTTTTGAGACACCCAAATATAGTT

GATGTCATGTTGTACTTTGGAGCCGGAGAGTGTTAAGACTTAAAAGAGAGAGACAAGTAAAACAGTAGATCTGAAGAAGT

GAGAAATCAATACCTAAACACGATAATGGAGGTAAAAAAAACACTACTTTCCTTTACTATAATGGTCAAAGCAGCAGTTT

TTTGCGTTAAATTTTGTATTACTCTCCTTTATTTTGTCTTATCATAGTCAATTTTGTTTTGGAGTTCTTAAGGTTCTGAC

CATAATTATCTGTGTGGTGGGAGTTTTTAAGTTCCTTGATTTCCGTTTTCAATTAAAAAATACAAAAAATGTGTTCCTAG

GGAAGGAGTCGACTCAATTCAGTGGCATTGGTTTTTTTCTTCTTCTATGTATATAGTATTAAATTCAATTTTACTTATGC

ATTTATGGTGTTTCAATATGTAGTTATGGTATTACTTGGTTCCTGTTTTACTAATTTTGGTTTCAGATCTCTACTAAGAA

GATGAAGGAGGTAACAAGCTTGGTAGATGTTTTGAGACACCCAAATATAGTTGATGTCATGTTTAACAAATATTCAATAC

CAAAATATAGTTGATGTTTTGAGACACCAAAATATAGTTGATGTCATGTTGGACTTTGGAGCCGGAGAGGGTAAAGACTT

AAAAGAGAGATTCAAGTAAAACAGTAGATCTGAAGAAGTGAGAAATTAATACCTAAACAAGATAATGGAGGTAAAAAACT

ACACTACTTTCCTTTACTAGCATGGTCAAAGCAGCAGTTTTATGTGTTAAATTTTGTATTCCTCTCCTTTATTTTGTCTT

ATTATAGTGAAATTTTTTTTTGGAGTTCTTAAGGTTCTGACCATAATTATTTGTGTGGTGGGAGTTTTTAAGTTCCTTGA

TTTCAGTTTTCAATTAAAAAACACAGACAATTTGTTCCCAGGGAAGGAGTTGACTCAATTCAGTGGCATTGGTTTTTTTT

TTCTTCTATGTATATAGTATGAAAATCAATTTTACTTATGCATTTATGGTGTTTCAATATGTAGTTATGGTATTACTTGG

TTCCTCTTTTACTAATTTTGGTTTCAGATCTCTACTAAGAAGATGAAGGAGGTAACAATCTTTGTAGATGTTTTGAGACA

CCCAAATATAGTTGATGTCATGTTTAACAAACTTCCACAGCCAACAATGATAAAATAATGTTAACATCTAAACAATTCTG

GTGGAAAACTAGTATTCAGTCTAAATTACATGACATTAAGTCATTGAGAAAAGTTTCAAATCTCATAGATTTATTCGATG

TAGAGATTGTTTCTAAACTTATGCCAGCTTCGAAACCTTTCATCTGAAACTATTGTGAAACTTGCATTAGTGTCTTCTTT

ACTGTCTAAGAATATAAATATTCAAAGGTATTTAAAAGGCAAATCTTCTAGAAATTTCCCCAACATCATGGATGATGCTT

CGATGAAAGAACTACTTAGTCGTCTTCCTGCAGAAGAATTGAAGAGATTTGCAACCGTCTCCAAGAGTTGGAATTCACTT

CTTTTTAATTCATCTTTAATAGAATTGCATTTGGATAATGCTAAATTAAAGTCTCGAATACTCATGGTTACTTCTGAAGT

CAATTCATTGTTTTTTCACAGTATGGTTTTCAAGATAAGTGTCAAGTTCTTCTTAAAGTGTCCATCCCCTGGTGTTCATG

CTCTGCGATGAAGCCTGTAAATGGCTTGATGTGTTTATATGATAGAAGAACACATATAACAAGGATTTTTAACCCTAGTA

CAGGACAGAAAACTCCATGGGCCAAAACATCAATAGGGATAGAAGATTCAGGGACTTTGGTGATTGCAATTAGGGGTTTT

GGTTTTGACCCTTTGACTAAAAAACACAAAGTTTTAGCGATTTGGGAAATTTCAAAAGAACCATCATTTGGGGCATACCC

AAAGATTTTGGGGCTTCTTGAATAAGACAAAAACACAAGTGGAAATAGTGTTTTTAGGGTGCCCCTTATCTAGCCATTTT

TGTTAATGGCAAAAATGCCCTTATTATTTATATTTTCAATTTTTTTTTAAATTAAAATAAAAAAAATATTTTCTTAAAAA

AATATTATATTTTTTTAATTTAATTTTAAACGTTTTAAGATTTTTTTTTTTTGTTAATTTTTTTTATTTACGAATCTAAG

TTCGGCGGACAAGTGATATGCCGACCTAAACATAGGTAGGTTCGGCTGATAATATGTCAGCCGAACTTGTGTTTTGTGAA

AACATACCTAGGTTCGGCAGATTCGATAACAAAAATAAGACATCTGAACTTAATATTATTCTTCATAAAGTGAAGTTCGG

CTAACATCTTTCTGTTTAATTATCAACCGAATTGTTCATCGACACTTTCAGGATGAAAAAAGAGAAGTTCGGCTGATAAT

TTTGATTTTATTGTTAGCCGAACTTTTTTCTGTAATTGTCGTAAAAAAAATTTCCATGCAATAAAAAAAAAAATTCAAAA

TTTTCAAAAAAAACACATTCAAAAAAAAGTTAAAATTAACAAAAAAAAAAAAACTAAAAATGTTAATATTAAAAATTAAT

TAATTAATCAAGGGTATTTAAGTAACTTAACTACTCATTAGACACCCCTTATCACCCACACTAGTTAGGGTCATCCATGT

GTATACCCCAAATGAGGGTTCTTTCAAAAAATCTAGCTGAGAATGGAATGCTAATTGGAAATGTTGGGTATTATTGTGAG

GCCTTCACTGTTGGAGAAACCAAGTGGAGAAGAATATTTCATCTACCCCCAAATTAATGTTAAGTGGACCCTGTGTTTAT

GCAAATGGTTCCATCTATTGGAGAAATCGTGGTATTCACTTTGCAAAAGCTCCTGAAAATGAACTTATTGTGGCATTTGA

CCTCCACGATGAAACATTCAGGGTAATTGAAATTCCTAAAAATGTTATGAATACTGTTCCTGGAACTTCTCCAGGCGTGG

TGAAGAGTTATTACAAATGGATAATCATATTTCTGTAATAGATCGAGTGGATGATCGCAATATCAAGTTGTGGATATCTG

ATGATGAACCCTCGAAGGAAAGTTCATCTATAAATTGGAAATTAGAATATATTAAACTTCTTTTTTCGTGAAAACTAGGC

CATAACATTTATGTCACTCAAGTTGAAGGTACAAAAGAATTAGTCATGAGATCGACAATTTCATCCATGGAATCGAGTCC

CAAAGAATTGGTATCCTTTTTCCTTCATGATCGGGTTAGAAAGAGTTGTCGGCAAATTGACATCCCTGGTTTGTCACTCA

AGTATCCATTTCGATTTGGCATGTTCATTTATAGTCCACAAAATTGGTTAAAAAAGACCAAAATCAACAATTCCTAGGTG

AAAATGACATTTAGATTTTGATACTGTTTAAATGAATAAAAATGTAAAAATAGCCAGGATGTAAACAATTTCATCCTGCC

CATTTTCAAATATTTTTTCTTATTTTTAATTTACATCACGATGCATCCAGTTTCATCCTTACTATTTTTTAAGTTTAAGT

AAGGATGAATCCAGTTTCATCCTTGTTATTTTTTTAGTGTCCATTTCACCCATACAAATTTTTACTCGTCCATTTGAACC

ATGTTTTAAAAATATTTGGACAAATGACCCATTTTCCGTTTATAGTCAAAGTCTACTACCGTGACAAGTTTATGAAGACA

TGGAAAGCGAAGATCCTGAACCACTACTAGTAGAGATGGAAGACTGTGGGCATGAGGCCTTTTGATTTGGATAGATGATG

AAAAATATGGAGTTTTATTAATTTCTCTTTTTATTTGGTGCTTACAGTTTACTCCCTCCGTCCCACTATTAGACGACCTA

GTTGAAGTTTGCACAATTCTTAAGGCAAGGAATGGGAAAGAGTATTTTTAAGTATTTTTTACAATTATGCCCTCATGGAT

AGTAATTTGTAAAATTTAGAAATGATTTATCTCTCAAACTATACCATGGATTTTCGTAAACTTTGTATCATTGGAAAGCA

TTTTAAAACACCTACATAACGAGTATAAACATGGATATCAAATTATATACATTTCTTATACTAACGATAATCAATCAAAA

GGATAATTTTAGAAATATTTCCTTGTTTAATGAAATAGGTCAACTAATGGTGGGACAAAATTTTAAACTAAGTGGGTCAT

CTAATAGTGGGACGAAGGGAGTATTTATTTTCCTCAAATCAGTACTGACTTTAGGTACCACTGATCGATCACGTATCAGT

ATGTAATCAGTTACTATATAACCAATTGAAAAGGCGTACATAAATAGCCAGCCATATCTAAATCCCAATTAGTATATAGA

TGTATCACAAATGTATTTTGTCAAAAGATGCACGCAGAGACGAAAGCAAGTGCGACTGTTATGTCCCGCTTCAAGGATAA

GTCACTATGGACGTATTTGCCCAATACATAGTCGCTCGTCGTATATAATATTAATTTGCATTGATAAATATTTTATTCGT

TATTTGTATGATTAACACATTAATCTAATTACCATTATTTATTCAACAAATTCGATTGATTGATACGAATTGATTTTTTG

TTACGATAAATGGACAAAACAATGGCTGGCCCAATAGCTGCTTCAGCGGCTGCAATAGCTAAAACAAAAATGGAAAAAAT

GTCTCCTTTTAATTGACGACTATCAAATAAATCAGAAAATGTTACGAAATTCATATTAACTGCATTCAATATAAGCTCAA

GACATATAAGTGCTCTAACCATATTTCGACTTGTGATCATTCCATAAATCCCAATAGAAAATAAATAGGCACTCAAAACA

AGTTCATGTTCAAGCAGCATTAACCAAACCCTCCTCAATCTGAATTTCTTTAATTTCAATATGAACAACAATTCAACCTT

AACTAATTTGGTTGACTCGAATCTAACAAGTACAGAACTAAGGAAGATATTGGTAATAGATTGAACTAAAAGAAAAAATT

TACAATTTATACCTGAAAAATATGACCATGGAATTGAAGGAAAATTGATAGGCTAAGCCAGAACAAAATAATTCTTTTTT

TTTCGGTATTTATTACTGACGAGCCATAGCAATTGCACCTATCAAAGAAACTAAAAGAATTATAGAAATAAGCTCAAAAG

GAAGAAAAAAATCAGTTGATAAATGAATCCCGATTTGTTGACCTTTATTTATCAAATCTTGCTCCAAAATCTGGTTTGAT

CTTGTAGTCCAAATAACCCCGTACCATGACGTATCTGGAATATTAGTTATTAGTGAAACAAAAATACTTGTACAAACCGG

TGAAGTTACCCCATCCCCAACAGTCAAAGATGAAATCATTGTAATATTCTCGAGTCGTTCATGAACATAACGTGAATATA

ATTAAGACATTTATGGCTCCCACATAAATAAGGAGTTGTGCAGCAGCTACAAAATGGGAATTCGATGGAATATGAAATAA

TGATATACAAACAAGAACCAATCCCAATGAAAAGGCAGAAAAAATAGGATTAGTAAGTAATACCACTCATAGACCTCCTA

CTTTAAGACCCAATCCCAAAAAAACTAAAAGAAAATCATGTATTGGTCCAGTTAAATCCATTATATGTAAAATAAGAGAT

AAATAAGTCGAAATGTTTCATAATCTTATTGACTAGACCATAAAAAGAAATTTACCCTATTTTTATGATAAGTTACTTAT

TTAATTTAATCCTATACGGGGTGTGGGTTGATGTAGATAGATTAATTTATTCCATTTCTCTATCCGATTGCTTATATATC

GAAATTCTCGCGAATATAGTTCATTCTTTTTTATCCAAAGTAAGCCGTGATTCTCGCGAATCACCAATCAATAAGAGCGC

TTTTTTTTTAGTTATTAACCGAACAAAATAAAAGAAGCTCCGCTCCTTTAGATGAAAACGGAATCTTAGTAATCGGTAAT

AGTTCTTGAACCAAGGAGTTTACCCGTGGCTTTTTTATTTGAGTAGAATTAATAATGGTTCGAATCGTGTAATCTCCAAT

TACTGTCATTGGTAACCGACCCAAAGCAATTTGATTATAATTCAATTCGTGACGATCATATGTAGAAAGTTCATATTCTT

CAGTCATTGATAAACAGTTTGTTGGACAATATTCGACGCAGTTCCCACAAAAATACAGATTCCAAAATCAATACTATAGT

TAAGCAATCGCTTTTTTCGAATATCCGTTTCCAATTTCCAATCAACTACGGGCAGATCTATAGGGCATACACGAACACAT

ACTTCACAAGCAATGCATTTATCAAATTCAAAGTGGATTCGACCGCGGAAACGCTCTAATGTGATCAATTTTTCATAAGG

ATATTGAATAGTTACGGGTAAACGATTCGTGTGGGATAAGGTAATCATGAAACTTTGACCAATATATCTTGCAGCTCTTA

CTGTTTGTTGACCATAATTAATGAACCCAGTTACCATAGGGAGAATATCGTGAATATCTATGAATAATTTTAAGTTTCTT

TCTCTTGTTTGAGAGAAGTTCTAAATCGAGAATAGAATATCGTATGCCCTTAGAGTGAAAAGAGTTGGAAGGAAGTTGTC

AATAATAGATTCCCTAGAGAAATAGGCAAAAGAAACTTCCACCCAAGATTTAATAGTTGATCCATTCTCATCCTAGGTAA

AGTCCATCTTGTTGTAATAGGAATGAACATGAACAAATAAGCTTTAGCTAATGTAATAAAAATACCAATTGTCATTCCAA

AAACTCCACCCACTTCATAAATTTCGAAATGAGGAAGAAATATGTATGGAAGAGAAAGATTCCAACCCCCTAAATAAAGA

ACTGTTACAAATAATGAAGAAACTAGTAGATTTAGATAGGAAGCAACATAAAATAAACCAAATTTTATACCTGAATATTC

GGTTTGATAACCTGCTACTAATTCTTCTTCGGCTTCTGGTAAATCAAAGGGTAACCTCTCACATTCTGCTAGGGAAGAAA

TTACAAAAACTGCAAAACCTATAGGTTGACGCCACAGATTCCACCCCCAAAAACCATATTTTGACTGCGCCTCAACTATA

TCAACTGTACTTGAACTGTTAGATAATTATAGTAGGTGATAACATCACTGCTTCCCATGATACTTCAGCCTCATACGGCT

CCTCGATGGCCTTAACGAAATCTAAGGACTGGTTCGGTATTATCTCGATATGTGTTAGATAGAATCAAGATGTATCGAAG

AGTCACAAATTAAACCAATGCAATTCCGTCTGCTATAATAGAAAAAAGGGTGCTTCCGAATTGATCTTATCCTTTATAAT

TTAAGTTTGTCTTTATTCAGTAATAACTTAATCCTTGAATAAAATACTCGTTGTCTAAGTAGGTATTTCAACCTATTTAT

TTCGTTCCTATTCTTCTTTCTAAGAAAAGTAAAAGTGGGCTCAAAAAAAAAGGATTACTTCGTTCCTGATAATTATTTAT

TTCACCTGTGGATAGGACCATACTCGGGATCGGGATCGTGGGGAAGTACTACTTTATCGTTTCTACCAACTTAAAGCCCC

ATTATGATTCCGTTTATGCAGAAATACCTTTTTTGGTAACTTACATTATCTCTGTTACTAATCCTTTGTGTATTTTGGTG

TTCCTAACCGTCCACTCATTTTTGCTCGATCCCCGGTATGGTAATACATACAATATAATAGAATAGTTAAAACAAAACTC

TAGCCAGTTGATCTTTTCTACCCGCTTCAAACCGTGATGACTAATCAACCAACCTTGGGGTAATTTCTTTTTTTTTGCTT

AACTCTTGTACATAGGGAATGAGTCTCAACCTTTTTACTGCTAATTTAGAAGCTGTTTTGTTTCACTCATATAACTATCT

GGTTTAGGTCATCACCCCGAATGACGAATAAAAAAACGAGGATCTCTATTCACTCCATTCAACTTCTTTCCTAGAAAAAA

ATTAGGTAAAAGTTCATGCCCCAACGAATCGCACGTAGAGATATTGATAACACACATGGAGTTAATGGTATTTCATAACT

AATCGATTGAGCAGCAGCTCGTAGACCACCCAAAAAAGAATATTTATTATTTGACCCATATCCTGACATAAGAAGTCCAA

TAGGAGCAATACTTGAAATAGAAATCCATAAAAAAACACCTATACTGAGATCAGCTAAAACAAGACGATATCCAAAAGGA

ATTACTGAATAACTTAGTAAAATGGATATGACTGCTATAGAGGGTCCGAGACTGAATAAAAGAATATCCCCTCTAGATGG

GAGAAGATCCTCTTTAAAAAGTAATTTTGTCCCGTCTGCTAGAGCTTGAAGAATTCCCAAAGGACCCGCGTATTCGGGTC

CAATACGTTGTTGTACCCCCGCAGATATTTTTCTTTCTAACCACACAATAACTAGTATTCCTATCGTGATTCCCAATACA

GGAGTAAAAATAGGGACAAGCAACCATATAAGACCATAGACCCCCTTTAAGGATTCCAATCTGAAAAAAGAATTGATAGC

CTGTAATTCTGTCGTATCAATTATCATATTAACGATCAACTTCTCCCATAATGATATCTATACTACCTAGTATCGTCATA

ATATCCGCCAATTTCATTCTTTTAACTAACTGAGGAAGAATTTGCAAATTGATAAAACCCGGAGGACGAATTTTCCATCT

CCAAGGAAAAGCACTATGATCCCCTATCAGAAAAATCCCCAATTCTCCCTTTGGGGCTTCTACTCTCACATAAAGTTCTT

GTTTTGACAATTCAAAAGTAGGAGAGTTTTTTTACTAATAAATCTATAGTCAAAATCATTCCAGTCAGAATCCCCTGCTT

TATCAAAGTGTTGGGCTTCTAAATTTTAGCAGCACTATGTACGGGAGAGGGCATCAAGGGGTCCATAAAACAGAACAACC

CATCTATAGACGATGGTGTTTTTTTTAATTCCATGCACCATGGAACTGTCACTTTATGAATGGCCACTCCACCTTTAAAG

CCATCCTTTGGTGAATAGACTGTTATGGAATGACCGCACTGCTGCTGGGAGATGAGAAGTTTAGGAGGATGTGCTTTTGA

TCGATACATCTCTATGCATACACCATACTGCTCTTTTATATAATGGTTGTCCAAATTCCCTTTTTATCTAGATGTCCAAG

ATATAGAAGGAAAGCATGCATGCACCCATAGATGTGCATGCATGCATGACCGGTCACAGGTGAGTGATTCCTAATTAGCC

TAAGTTTTTAAAAGGTTGGCAGATTTCCCAATCTGAGTTGCTCAGTTTCACCCATCCGCATATAAATAAGGACTTGACAA

AGTTACAACTCTGGTAGATGTTTGAGATTTCAAACGCAGGAGGTGAGTTTGGTTCCATATTAGTAATTGGTTTCTCTTTT

TCAGACCAGTAGAAAGAAGATAATGAATATGGAGGAAACAAACCTTTCAAATAGCGACACCATGCATCAAATACTTAGCA

ACCTTCCCGTTAAGTCCCTTATGAAGTTTAAATGTGTATATCCAAAAGCTGGGAATCACTTATTCATAATCCATCTTTCA

TTAAGCTACATCGATCAAAAGCACAACTTCAGCTTCTCATCATGATTTACAAGGTTGATTCCATAATAGTTTATACACCG

AAAAATGGGTTCGAAGGCGGTGAAGCTCTTTATAAAGCGACGATCCCATGGAGTAGAGGTGTTATACTGAAGCATGTTAA

CGGGTTGTTCTGTTTTATTGATCCCAGACGTTCCGCATCCTGTGTTTACAACCTTGCCACTCAACAGACTACTCCTTGGA

CTTGTATACCAGTTCCATCGGTGATAAAGCCTGGATATGAAATTAATAAAACTCCTACGTATGGATTCGGTTTCGATCCT

TCAACTGGACAACACAAGGTACCGTGCACTTGGGTTATATCTCAGCATGGCGCAATGACATTTGGGAAAGCTGACCATAT

TTGTCATGTTTTTACCCTGGGAGATACTCAATGGAGGAAAATAGATGAAGTGCCACCGGTTAGGCCAGATGGAATTTCGG

TTTATGCAAACGGTTCGATTTACAGAAGAAACCGCGATGATTACTTTTTCAAAGAGCCAGAGGTCGAAGTCATCGTGGCA

TTTGACGTCGGAACCGAAAAGTTCAGAGTGATTCCAATCCCAGACTTTGTCATTGCTTCTTCTGAGACTCCGGAAGACTC

GTGTCGTCACCTTGACCAACTCTTACTGGAAGTAGATGGACACATAGCTCTGATTAACAGGATAAGCGAGAGTATGGTGA

AGTTGTGGGTTTCTGATGATGACTGCATACAGAAGAACACCGATGTGAAATGGGTTGAAGAAACCATCATACTTCCTTTT

CAGTGGCCAAGGGGAGAACGTCTGCATTTCCGTGCGGTTCAAGGAACAAAACAAATGATGATACACAAAGCACCACTAGC

GAGAGGCTTTATAGCTATATACATCTATGACCGGATTACGAAAAGCTCCAGAGAAATCAAAATCGTTGGTTTACCACGAC

TCTCCAGCCCATTTCGTTACGGCATGTTCGCAACCCATGAGACTCTTCTGCCCCTTCAAAACAAGGAAAATAATCAACAC

TAAAATCACGGTCAAATGATTTGAGTTCGAAACTTGTAAGCACCAAATTCTTTAATCACGGTCAATTATAGCAGTAATGT

TCATTATTTTATGCTTCTACTTGGGATCCTGGCTCTGCTTCTGCTGTCATTCGTGACACAAAAACAACTTGGATAAAAAT

CATGATATGGTAAAAGGGTGTCACTTCATGCACTCATGAGTGATAGTAAAAAGGTGATTGTGATTTCTTAACCAAAATAA

GATAAGATAAAAGATCATTCATGGCCTACTTTATGATCTTTCTCCTTGTGATGTGCCTTAATTGTTTATATCTTGGAAGA

TCAATAAGTAGGGACTAAAAAGATTGAGCACTCACTTTGTCAAGTGTGCATCCTTAATCATTTGAAGAGAAGTGAGAGTG

AAAGATCCAAGTATATCCAACACCAACATCATATGCTGCATTCTTCCATGTTGCATTTTGATAATTCCATATTGACTAAG

CCACGAAAGTTGATGATGGGCACTATGATTTTATATCTAGGATTCTTCTAAAATCTTGCATTTTTTGCTTTTATCAAAAA

TAGTTGTTATTGGTGAACCTCTTGTGAATGTGCAAGGATATAAACATTAAATGGAGGCGTTGCCACTGTTTTGCAAATGA

AGACATATATACATCATCTTGCAAATACCAGAAGATCGGTTAAAACGGTAATAAATGCATGGTTATATTGTGGTAGTATA

TTGCTAATCCCACATCACCCAATACCTCCATCACCTTGTATTTGTAAGCATGCTTATATTGCATAATGAACAAAAGTAAA

TCATGATACGATTTTTTATTCAATAGATAACTTGCACTACAAACTCGTTAGAATTTGGTATTGAGTTTGTAGTTAGCTTA

TAACTGCTAATTAATTAAGAAAAGTTGAGTTTGAGTTGTCTACAAAAACCAAAGGATATCTATGTTTAATATTGAGATGG

ACATAACAATGGAAAGTAAAATAGAAAACTCTAAAAAAGATGATAGAAGTTGGGTACAAAATCAAAGATCATCATCATCT

TATATTAAAACTATAAATATACTTCCAATATTCTCATTCATCTTCGTAACTTCTACTCATGGATATGCCGCCTACCGACA

ATTGCCATTAAATGTTGACACTATCGTTTATCTTATTACTTCGCAGATGGGTTATGAATGCAGATGGGTTATAAATATAT

TAATTTTGAGAGATATTAACAAACTTTGAGATTAAATTCTACTAGTATTATAAATTATCATTTATACAAGAACTGTGTTG

GCCTCTTTAATGGATTAGAAGATTTTTTAAAGCTTATGGCCAACTCACCTTTGTAATGGGTTAAAGATTTTATTAAAATT

TATGGGGCTAATTCACCTTGAAAAACCAGTCTTACAAGGTTACAGTTGATCAAAAACGGATGCACTTCACAAATTAAGCT

TAATACTCCCTCCGTCTCTAAAAGATAGGCAGGTTTGTGTGTAAAATTTACACACAAACCTGCCTATCTTTTAGAGACGG

AGGGAGTAGTATTACCATGAATGGAAGGTTTAAGTGTGTTAAACAATCCCTTAGAGGGTTAAACAATGTCCACACATCAC

CTACCCAAGCTCAAACTCCATGCTAAACTCCATGAACCTAATTCATGTTGGAAATCGACTTACAAGGTTAAGATTTTCCT

AGAGATACAAACTTCACAAATCCATGAGATTGTGCACTATTTAATCCAGATTTATACTCTAGCTTCAACGTAAGTTCATT

GGTTGCTAACAGATAATATTAGTTTACTGGGAGGGTGCTAAAATTCATATAAGAGTATCAGATATGGATTTTGGCACCCT

CCCTATTAACAGTGTTGATAAGTTTTGGTAATAGTAGTGGAGCACATGGAACCAATGAGGCAATGAGTTTAAAAGAGGGG

TTAAGATGAATGGGTTAAAAAGTTTAAGCATTTTGAGGGCCATATATCCAAGGTTGCTAATAGGGGGGACTGGTTTTTCT

TGGGGTATACCAAATGCTAATAAGACAAATCATCATTTGCCTTGGGGTTGGTACACCCCCAAGCAAATTGGTACCCCCAT

TAGCAGCCTTGCATATATCATAGTGCTTTAAGAAGTGATGGTGGGCTATTGTCCCATCATGCAAAAAAGAAAAATGTCTA

TCAACACACTTAACAATAATTTGTCTTGTTGATGAATTTTTTGACACATAAAAAAAGAAGTATATCTATTTTGTATCATT

ATTAAAGGTGCATTATACTTATCAAAAACAATGTTTGCACATTTCTTTTCATGGAGCCAGGGGGGTCGACAAACACCTTT

CTCTTGATAAGATATTTCTCATGTGCCTATACCCTATTCCTGCATCTTGTTCAACTTAAACATGAAACATTACATTATCA

AAAAAAATCCCAACTTTTTCTTCTCTTTAAAGCTCGCTCACTCCTTAAGAATCCCACTTACTGAGTCTTGGGAGTTGGAT

GCATATATATAAGAGTGATGTACCCTCACAATCACAGAGTACATAGTTCAAACTTCAAAGATTAAGTTGCCTAATTTTTT

AGAAGAAAAGGAAGAGATTGTTTATAGTAGTAAAAGGAGATTAATGGGTGTTCCTAATCAAAATCAAAATAATCATTTCT

ATCCTCAAGAACTTCAATTAAAACTATACCAAGCATTCATATTTTCAGTACCAATTCTTTTCTCTATAATTCTTCTCCTC

TTGTTTTATCTCTTTTACCTCAAAAGAACTAGAGATCAAGCAAGCAATCACCCCACTTCTCTTCCAACAAGTGCTCCAAG

AAGTACTAGTTTGAATAATCAACCCACTTCGATCCTTCCATTCATGGTACGTACTTACCATAATCTCTTAATTTCTCTTC

AACTATATATTGAGTGTTTTTTTTTTTATATATATAACTTTTTCTTGTACTAAATCTACTTTTTTGTTGTTTGTAGTCGA

TTGATGTTGGTTTGATCAAGGAAGAACTAAAGGAAAAGCTTCCAATAGTCACATTTGATGAAAGCTTAAGAGACAAAGAT

TCACTGTGAGTACATACACATCCATCTTAACTTGGTGAAATTTGTTACTTTTTTTATATCTGAGTTTCGAGTTCTGGTAA

CTGGTATCATCGTCCAGTGACTTTACCACTGTGATAATCTTTGGAGATTTGTGGGCATTGTTTTCTTGCATCCAACTTTG

CAAACATGCATTACAATGTTGCATATGGCACTATTGTGCCCAAAATTGGGCACTTGAGTGAAAAAATTGAGTAACTTTTG

TCAAGTAACTGAAGGCATGAATCTTGCAGTAAATTTTTTATAGGACATGGTAGGTCCATTTACATTATTATTTTTTTATA

TGGTAAAAAGTGTTGGTAGGTCAAATAGTTGGATGGTTGAGATCTATGATACTAAAGTGATGGTGCACATAAGTGGCAGT

GTCATCATTCCAAGTTCTTTGTGATTTTCAATGTCACTAACTTTTCATGTAGAATCTCTTATACCATCCATTTTTTTTTT

TTTTTTTTTTGTCAAGAATGAAAGAAAAAATAAGTAGCATACAAACTCAATGGTTCAGAGAAATTCAGCATCTTCAACAT

ATTAAAACTACTAGTCATATTGTGAAAGGCATCCTATGGGTCAAAGATGCAAAAACTTGCATATTCTCCTAGTGAATTTA

GCTTAAGATAAAAAAAAAATCAACAACAAACAAAAATGTTTATTGAAAAAGAAAAAAAGAACATAAATTAATCAGTAAAT

AAGGCTTACATATTCTCATTCGCATTCTTCTAGAATTTGGTAACTCTTCTGTAGCAGCCTTGATATCAAGGAAAGACAGT

TATGAAAAACTTAATTATAAGATAATTCGTGGAGGAAAAACACCCTCAAAAACCGTTTGATACGGAAAAGGTAATGAGTC

TTACCTGAATGTCATGAAACCACCAAAAAAAATATTTTACCAGTGATTTCAAAATAGTAAACTTTTCACGGTTTGTTTAC

AATTTCTCAAAAAAAATTAACGATCTAAAAGAGAATCAAACACTTTCCTCTATCTCACAGTGAGAAAGTGGAGATAAGAA

TCTCCATTGTACAAAATCAAATAAATAATTACTGGTAAGATGGTACCATCCATGGCCATGGTCTCAAAAGCCTCCAATAT

GCGCCAACCTATGTGCGGAGTAGATAGATCAGGTGCAACACGTACCCATGCATTCAAAAATCTCCATCAATCAGAGAGAT

ATACATTGTCGAGCTTAGTGCTCAAAGATTCTCTTCTCTTCTTGGTGGTACCAAAATCACAGTACGGTTTTTGTTTTTTT

TTAAAGAAAAAGGGAGTTCATGATATTTACTTGCACCGAATTCCCTAGTGGTATGTTTGGTTTGTGATGCTGATATTTCA

TCTATTTGCAATTCCTGATGCAATCATGTATTTTGCAGATGTTGCGTATGTCTGGGTGATTTTGAGCTCAGGGAGCAATT

GCAACAGATTCCGTCGTGCAAACATGTGTTTCACGTTGAATGCATTCATCACTGGCTCCGAACTAACTCGACGTGTCCGC

TGTGTCGATGCTTCATTCTTCCGGTCAAGAAGGCGGATCCTGATCCTGCAGTACACCCTCCACTTCCACCGGTTTGATCC

GCGGAGCTCAATTTTCATGGTTAAAGAAAACAGACAAAAACAAAGCAAATTCCGATCACCATTGCACCCTCGGAACCAGA

AGCAATTGCTGGGTTTTACAAGGTGGCCGGAAAAGTGCGCCAGAGGAAGGGCAACCAAATCTTTATTCAAACGAGCAGTC

TATACACACATGCAGCATATACATAGATGAAGCCTAGTATTTCATTTCTCATTTTAAACTTGCCATGTATCACCCAATTT

CTCAATGTTATAAGTTTTTCTGCTACCAGAAATCTCAATTAAGACCGGCAACCAGTGGCGGAGCTAGGACTGTAACATAT

TTGCCTGTTTGGCGGAGCTAGAACTGTAACATATTTGCCTGGTTGCCGGAGTTAGGACTGTAATAAATTTGTCAGCTCAA

TTATTTCTGCCAACCATCTAGTTAAATTCAGGCAACTCTACCACCACCACCACTAAAGTGATCAATCATATGTCTGTCTA

ACGTCTCATTCAAATAATGAGAGATAGAGATTGAGAGAGATAGGTCATACTGTCTCTCTTAAATGCACATCTCTATCTCT

CTTTTACACTTTAAGGTCCTTTTCCTTGTCCATTTCCAATGGATTCTTGTGTTTCAACTGGAAATTAATCTTCTTCTCTA

CCTAAACACACCGGAGATAATTGAATTCATAGACCTCGTACAACACTTTGGCCCACTGATTCAACCAACATTCTACCCCC

ATAATCACAGTTGAGAAGATCAAACATCTTGATCTTGATTGACCGCTGCGTGTGGAACGCGCACGTAACCGTGCACACCC

AACATGTTTAGGCGGCGACTCAGAAAAAAAAAGGAAAACCAAAATTTAGATGCTTCAAGACTAATCGAGAAGTTCAGTTC

CTTTTAATGAGACAAGATTCGATTAAAACACTCAAAATATCGAAAATGAAATAAACTGGCTTTGGCGAAACCAAACTTGC

ATATATACAGGAAGTCGTTTACAAATATTAAGAAAAGCAAGTACAGATTCATCTTCCTCCCATGGCTACGGTGACGACTT

ATATTCACACCCTCAAAAGGTTAACCCTGCTCTATCATTACTACTAATACTACTGCGACCACTCGAGTCTACTACTACGT

ACGGACAGTGCTAATATATATACCAAACAAAGAGTTATCATCGCTCGTTACTACTTTGTTTACAGACCCATCATTTCATT

CTCTTGAATTCCAACAACCATATCAGGAAGCTCTAAAAAAACCGAGCTGCGCAAATTAGTTATACTTTTTGTTCTCTCCT

CAAAATATGTAATAGATTGTTGTTGTTGCTTTCAAAAGAGAACCAACCTCCCAAGAAAAGATAATGGAACAAACTACGGC

CTTTTCATTTATCTGTTTTTAAATTTGAGTCTAAGCTTACAAGAGTACCTTCGCCACCACTTAAAGCCGATAATAACTTG

CATCCTGGACTCATCCTTCTGACTTCTTGGCTGGAATAGATGAAAATCTTCTTCACCATCTTGCAGAATTCCCTGCAAGT

ATAAAAAGATAGCGACTGTCGGTGAGAATTGAAATGTTGCAGTAGAAAAAACTGCAATGAAATGCAACTGTAGAAAGACG

AAGTGATAGATTATGAGGTGTTAGGAAGACGCTTACAGCCATGGATCATCACCAACGAGCATCATATCCCCTTCATCATC

GGTAAATACAACTTCCCATTTGTCACGAGGGGATAACTCTCCTTTAATCTCAAACATCTCCTCCAACTCATTTATAAGCT

GGTCATACCCATCCAAAGTGGTCAAATCCACTGCTCGACCAACTGCTATCCCTTGCATGTGTACCTATATAAGCAAATGA

AGAATCAAAGTCAATTCTCTAGTGGATTGAAAACAAAATCAATTGAGGACATAGAAAGTTTATTTACTAGACAGCCCAGG

TTCAATACCTTAGTACGACTTCTGGTAGATGTCGAGAAGCTCTTTGGCTGTGTTTGCAATAGTACTTGTTGCTGTTCTTT

GGATGCATTCGAAAGAGCAGATTGTGGATCTGACTCAGACCCAGATAAGGCTGAGGGGCCAGTTATTTCAGCAACGACAC

TGGAACAAGTGATGTCATTTACTGGTTCCTTAACTATATTAGCTGGGGTATTTGTGTTATTAATCAGCTCAATCCCAAAC

AAACGGTAACCACTTGGAGGAGTCTCAGATTTCTTTACCGATTCTACTTGGTCCTGCATCTGATGGTCAGCGTTTGAAAA

GCCCAAGTGCCCTGACCACATAGAAGCTGTTCTATCATCTTGAAACAAATCGAGGGAATTATTTCCAAGGGACTCTCTCT

GGATCCTCGTGATACTGTTGCTGTTGACAAAATTTCCCATGTTTTCATTTTGTTTGGGAAGCCAGATGCTTTGACTTTCA

CTACTTTGAATGTCCATGCCACTCGACGCAGTTACTTCATTGGATTGCTTTGCTCCAGGATACCAGAGTGGTGAAGCTAC

AGAACTTGTAGTAGTTCCTACGGATAAATAAAACAAAAGGATTAGCACATATTTCTATAACCAAATGAAATCTTATGCAC

GAACGGATATATCAATATCAGATTCCGTACCATGAATTGGGAGATCTGTGGAGATTCGAAGCCTTTTATTCTTTGATACC

AAAGGTTGAATCGGCTTTGGAGGAGCTGGAGAAGCAACAAAAGGCTCAATCTCCCATGGAGAAACTCTGTCAGGTCTTGG

TGTGGAGCACGGTTCATCCCATTGAACCTAGAAAATAAACGTATAGCCTGAGATGGGTGACAAGATAACAAATCATTGAA

CTACATAAATGTTATATCGAAGATCTACCTTCAATGATCTCCACTTGGAATCTGCCCACTGGGGAGAAACATCATTAGTA

CCAATTATCGTGCCAGAAAACCTGACCAATCATAAACATCATTCAGGTTAGATTAATAATCCCGGAAAACAAATAACTCA

AGAAACCACCAATAGAATCAATCTTATATGTTAATATACGTAGTCACATACCTTCTCTCAGGAGAATCTTCCCCTTCAAA

TCTCATCTTAAAACGCATTCCCACAGAGAATCCATTTTTCAAAGATTCTAAATACTTGTTCAGGCCAATTATAAATTGAC

TTGTCCTGATTACCAAAACAGAAGGACGAACAAAGTTAGGCAAATGAAGGGAATCCAAGAAACTTTCAGATTTAATTAGA

GGCGGAATATCTCAAACCTTGGCTTGTTGTAAACTATGAAGAGGGTTTTCGTCGATACAGCATGAGATGCTGTTGCAAGC

ACTCCAAGATGCATGCTCTGGCTGGATATCACAGATGAGGGCATATTGCTTTGTTGCCTAGCATGACGCCTTACTCCAAC

ACGTAAATTTCCATTCTCCCCTCTATACATGAAGTAAACAAGAGATCGATCAGAAGCAATTCTAAAAAGAGATATCGAAA

AGAGTTGGAAATTTAAAGCCCTTGTTACCTGAGAAAAACAAAAGTATCCCCAGCAACCAATCTCTTAGATGTGACAAAGG

TACTCCAACCAGTTGTAAGTAAATGTCTTCGAGGTTGACCTGTCCATGTAATAATCAGCAGTAGTCAATTATAAGGACAT

GGATAAAGAAGTATGGGCTTCCATCATACATGAATAATTGCTAGATCTATATTTCGTTCTTGGTTTTCAAACTTATCAGC

TTATGTTGTGATCATTAAAATTTTGAATAAACTACTACTGTTACATACCTCTAAAAATATGCTTAAATCGCCACTCATAG

CCATGAAGATCCCTTGCAATCAATTCTTGAGTTGGTGTTGCTTGGGTCATATCCTACAGAACCCAAAGACAAAGCTTACA

GAATTAGATCCAGACTACACAAAAAAGAGAAACATAACGAGTGAAATGAAATGAAACAAGTCTTACCAACGGAGGTAAAC

TTTCATTAGCATGTTTCCGAAGAACGGAGAACCCTCCATGAGTACTAGTATCAGAAGCAGTCAAAATCTTGCAAAAAGAT

TGAACACCTTCTCTTGGTGGTTCTTGTATACATGGATCAGGACTATTAGGTTCATTCGGCTAAATACAAAAACAAAAGCT

CATAAGTATAAATCCGACAATCAAATTCGACAAAATCAGTGAAAGAAAACACTTCAATGAGAGATTGATCGACTCCTTAC

ATCATCTTCTGGTTGCAAAGTAACTCTTGCATAAACCTCATCAGTTTCTTGTTCAGCCTAAAGACGAAACAAAACATTCA

AAAACTCCAATTAGCATCATACTAGACATCATAACATAAAGATCTAACCATAGAGAACAAGAACAAAGATCAATACATAC

CAGTAATTCGATATGAACAACTCGACAAAGTATCTTTGAAGGTAGATTAAACAAAGGAATTCTCTGAACTAATTCTTCAT

TTGTTGATGCTTCTAACTACAAAAACAAAACAACCCCCAAATAATGCACCAAATTAGCTCAAAATTTGAAAAAAAAAAAC

AGAAATGGATGAACAAGGGAGAATGAAAATCAGTTTAGGGATGATGATTCTTACTTGCTCCATATGACCTTGAGGGAAGT

AGTAAACCCTTTCACCAATTTTTGGAACATCAACAAGAGGACCAGCACAAGCTCTCCATAACTCTGAATATAGATCATCA

CATCCATAACCTACACAAAATCAAAGTCAACCAAAGTCAAAATCAAAATCAAAACCCAGAAAAATTAAGAGAAACCCATT

ATCAAACTCAGAAAAGGGAGCTTCAATTTCTAATTTCTGAATCAAAACCCAGAAAAATTAAGAGAACCCCATCATCAAAC

TCAGAGCTACATTTCTAATTTCTGAATTAAAACCCAGAAAAATTAAGAGAACCCCATTATCAGACACAGAAAAGGGAGTT

TCATTTCCAATTTCTGAATCAAAGCCCAGAAAAATTAACCCCCATTATCATCAAAACCATTAAAATAGAAAGAATCTCAC

CTGTATTATTTTGCACTTCCCCAGGAAAACCTGGGTTGTTATCAAACATAAACGCCATTAAAACCAGTAAACCACCAGCT

TCTCCTCCCTCTGTCTTCTTCTCTCAAACACCAAAATCAAACAAAACCCAAACCTCAAAAATCCACTATAAAAACCCCCC

CAAACTTCAAGCAATCTCTCTTTCTACATATAAAATCCCAAAACCGTTTCTCCACAATCAATAAACCCACCACCACACCA

CACTATAAATGCTCTTTAATCACCAGCAGCAGCAACAGTTTTCTTCCGTTACAAAACTCTCAAAAACGTTTCTTCTTCTT

CTTCCTCTGAGAAAAAAAAGGAAACTGCTCCACAAAGCTTGAGCTTTAATAATGAACTTTGAACAAACAAAAAGAAAACA

AACCCACTTTTTAAGTAGACTCTCTCAGACTGCAGACCTGCAAGAACCTCGTTTTAACATGAAAGAGAGCTCTCTTTATC

ACTTTTTTTTTTTTTTTTTTTTACTAACCAGAAAACGACGTTACCAAATCCCCAAACCAAACTTACTTCGGTTTTTCTCT

CTCTCTCTCTCGCGCTCTGATTATTGATTAAGTAGGGAGAAGAAGAAGGCAAGTTTCAGATCACAGTTTTCTATTAAGAA

AAAAACAGAGCAAACTGGTAAACGGAAAAGTTGTGTTTCTGATGCCGTCTGATAACGGCGGCGCCTTTTTAATGTCTCTT

TAGCTTTTTGTGCCTTTACTCACTGACAATTTTTCTGTGCACATTTATGTGTTAGGGTAACGGTAACGGTTTTCTTTATT

ATTTTTATTAATGCCTTAATGATGTTTGCATAAATAGATATAAATTTCGGTTATAAGATGCTAAGTCAGGTGATCGGGTC

CTCGAGTGTATAATGATGAATGGTTGTGCGCTTCTCACCCGTTTAATTTATGTGCATTTCTGTGATAGCAAATCAGGTAT

TCATGTGGCATCCGGATGTGTGATATAACAATATGACCATCTCCGATTTCTACAAAAAAAGATACTATCATTTTCTTCTT

TTTTACCTTAAAAATAAGCAAAATTAAAATTAAATAGAAATTAATAATATATATCTCTTTACCAGAAACGTATGGAGTAT

TATTTTTTAGGTGTGGGGCTATAATAGAGTCACCCACACAAATGTATATAACTACATAATGAATATGGAAGCTGATGCTA

ATTATCTGGCATATTCATTATGCCAATTTGCTATGCATGATAATGTGAATTTTAGGTGAAATGCCAGTTATCTATCAAGT

TTTATCTCTTCAACCCTCAGTTTCCATGAGGTAGTTCAATAAAGTGTTGGGCTTCTTTAACATAGCCGCTAATTACGGGT

TCCAGTCTACGGTAGGGAGGGCATAAAAATTCATATAAAATAAACTAGAAATTTCTTGTTTTATATGAATTTGGGTACTC

TCACAAGTAATCTGGCACACTCATTAGTAATGTTGGCCATCTTTCCTTCCGAAATAAAAATAAGAAATGAATATGCACAT

GGTTTTTTATTAATGTACATCTTTGTATTAGCAATTTGGATAGTTCAGAGAGGTGAGCATGTTGTGCACTCATCTGCGTG

ATAGCACTAGTCTTACTATAATTGGAGACAAAATTAATTAGTCTTGATTTTTCTTCATATGTCTGGTTTAAGCAAAATTA

TTTGAAACATATGAAAATAAACAAACACTTACAATTATTAACCATCAATGGGAGTAGAAATTGACTTATACTAAACATAC

TAGGAGATCTGATCCTCTAAAACTGAACTTATTTGTTCGTAAATGAATTCTTGCTTTTAATCACAACAAAGATTACCTGT

CCAAAAATTAGTCCATTTTTTTAAAAAAATCATCGATTTATTTTTCTTATCGATAAAAATTTGGTGATAACAAGTTCCGG

ACTCAGATTCTGGATATCATCACTGTCAACCAGTAACTTAACCACTGTACCGGGAGTGAGTCATTTTTTTAGTAGCTCCT

CTATCGAGATATCATTCTTGCTTTTAATCACAACAAAGGTTACTTGTCCAAAAATTAGTCCATTTTTTTAATAAAATCAT

CGATTTATTTTTCTTATCGATAAAAATTTGGTGATAACAAGTTAGGACTCGGATTATGGATATCATCACTGTCACCCAGT

AACTTAACCACTGTACCGGGAGTGAGTCATTTTTTTAGTAGCTCCTCTATCGAGATATCATTCTTGCTTTTAATCACAAC

AAAGATTACTTGTCCAAAAATTAGTCCATTTTTTAAAAAAAAATTATCGATTTATTTTTCTTATCGATAAAAATTTGGTG

ATAACAAGTTCCGGACCCAGATTATGGATATCATCACTGTCACCCAGTAACTTAACCACTGTACAGGGAGTGAGTCATTT

TTTTAGTAGCTCCTCTATCGAGATATCATTCTTGCTTTTAATCACAACAAAGGTTACTTGTCCAAAAATTAGTCCATTTT

TTAAAAAAAAATCATCGATTTATTTTTCTTATCGATAAAAATTTGGTGATAACAAGTTAGGACCCAGTTCGGATATCATC

AACATCACTTGATAACTTAACCAGCATCATTAGGAGTGAGTCATTTTTTTAGTAGCTCCTCTATCGAGATATCATTCTTG

CTTTTAATCACAACAAAGGTTACTTGTCCAAAAATTAGTCCATTTTAAAAAAAAAAAAATCATCGATTTATTTTTCTTAT

CGATAAAAATTTGGTGATAACAAGTTCCGGACTCAGATTCTGGATATCATCACTTTGTCATGAGAATTTAACCTTGTGTC

ACGAGAGTGAGTCATTTTTAGTAGCTCCTCTATCGAGATATCATTCTTGCTTTTAATCACAACAAAGATTACTTGTCCAA

AAATTAGTCCATTTTTTTAAAAAAATTATCGATATTTTTCTTATCCATAAAAATTTGGTGATAACAAGTTTCGGACTCGG

ATATTCTGGATATCAACACTGTCACCCAGTAACTTAACCATTGTACGAGCAGCGAGTCATTGGCCCTAAAAAAATTGACG

TGCACCGTGAATCTGAACATAACAAGTTCCGGACTCAGATTCTGGATATCATCACTGTCACTCAGTAACTTAACCACTGT

ACCGGGAGTGAGTCATTTTTTTAGTAGCTCCTCTATCGAGATATCATTCTTGCTTTTAATCACAACAAAGGTTACTTGTC

CAAAAATTAGTCCATTTTTTTAAAAAAAATCATCGATTTATTTTTCTTATCGATAAAAATTTGGTGATAACAAGTTCCGG

ACCCAGATTCTGGATATCATCACTGTCACCCAGTAACTTAGCCACTGTACCGGGAGTGAGTCATTTTTTTAGTAGCTCCT

CTATCGAGATATCATTCTTGCTTTTAATCACAACAAAGATTACTTGTCCAAAAATTAGTCCATTTTTTAAAAAAAAATTA

TCGATTTATTTTTCTTATCGATAAAAATTTGGTGATAACAAGTTCGGACTCGGATTCCGGATATCATCATTTGTCACCCA

AGAACTTAACCTTTGTACTGGGGGAGTGAGTCATTTTTTTAATAGCTCCTCTATCGAGATATCATTCTTGCTTTTAATCA

CAACAAAGATTACTTGTCCAAAAATTAGTCCATTTTTTTAAAAAAATTATCGATATTTTTCTTATCCATAAAAATTTGGT

GATAACAAGTTTCGGACTCGGATATTCTGGATATCAACACTGTCACCCAGTAACTTAACCATTGTACGAGCAGCGAGTCA

TTGGCCCTAAAAAAATTGACGTGCACCGTGAATCTGAACATAACATGGCGGTGTTGTATACTGTGGGTACATTTTGGTTG

CATGTCAAGTTTTCAACGGACCCGACTGACAAATTCAGGTGGGGTGTGAGAGGAGTACATAGTATCACGAGTTTACATAA

TTTCCCGTGAAACTTTCCGTCACCGTCACCGTTAGAAAAAGTTTTCTCTCACCCGTCGATTTTTGTCTGCTTCAAAGCAG

GCAGTCCTGCTCCTGCAAACAGAACTCCTCTTTCTTTCCCATCAAAATCGACAATGAAAATCTGTGGGACCATCTCCACA

CGTGGCGTTATCATAGTGGTTTCAACTTTGTTCGGTTACACGTCCGTTAATACCGAGGTCTTCTAAGTTATCCGGTTACG

GATGAGATTAAAATGTCACGTAAACAATCGTTAGAGCAGGCATCCATTATCGGAGGATCGGGATCGATAAGAACCATTCT

ATACTTATGTCGATTTTTCTTGGGATTTATCCCGAATAGAGATCTAATGGCAGAAGGACAACCCTTCTAACATTGAGATA

GACGATGGTTAAAGTGGGTACTGGAGATTTTTTTCGGGATAATTCTCAAGATATCCAGCATTTTCGTATCAATAAATACA

ATCAAAATGATAAGTCTTATCGGATCGTTGGTGCTCACAGAGATACTTTATTTCGCCAATAATCTGCAACGATTAATATG

TGATCAGTCTGATGATATTTCTATATCAGAATTGAGTGACATGTGAATATGCAATAAAAACTGAATTAGATAGCATGTTC

TGGGAGTAGACCAAGATGGTTTGTGTTGTGGACCATTTTGCACTCGTATTGTGCAGGTGTGGTGAAAGCTATTTGAGAGT

CTTATGACATCATCACTTTTGTTTATTTTATTACTGAAAATGACTTAAAAGTAAAATAAATTAGAATTCCAATAATATGC

AATTTAAGGCTTTAAATATGAGAAAGACCGATTCAATTGACATATATCTAGCGATTCTAGCCCATTTGGTTGTCACTTTT

CACCATTTATGTCCAAGGGGAACGATAGACATCCCGCGGGTTGTCCCGACAGGAAGAGTGGGTGTGGTCTGGGAGTGACC

CGCTAATTTTATAGGGGATGGGTTTAGGGTGGACGATGCCCACTTCCTATCGGAACAACCCGCAGGATATAAATCATTTC

CACATGTCAAGATGGATTCCTAGCTACGCAATTAATTACTTAAAATAAGAATAAAAAAAAAATCATTTGAGACCCATTTG

TGTTCACTTTTCACTATTTATGTCAAAATGGATTCGTAGTTACATAGCCTCCTATAATTGATGTCGTATTACCCAAGTTA

AATAGTTTCATGAATATGTAAATATCATGTCAAAATGGATTCATAGTTACATAGTCATAGCCTCCTATAATTGATGTCGT

ATTACACAAGTTACACAGTTTTATGGATATGTAAATATCTGGAACATGAAAATAACTTCGATCAAGACGTGTTACAACCG

TACGAGCAAATCTAGTTTAAAAACAAGTTGATTCCCTTTACTGTAGAATCATGGCAACAGGGTTGATGGCATGATTGTTT

ATTTCGGTAAATCTTGATTACAAAGAAAAGGTTAATTTTAAGTATTAGATTAAACTCTCTGTCAAACTCCATAGGTTTAG

GACCACCCTTACCTGTTAACACTCAAACTAATCACCAATTATTAACCCTAACATTAACATTGTATCCCTTTTCTCCTTTT

CTCATGATGTCATTTTCAAACCTATAATTAGTGCATAAAATTAAAAAAACATCTAATCAAATATTAATTTAGTCGAATTA

GTCGGAACTTACGACATCGATCGACAAATTATCGTACGCTTATTGGTTAACTTGGATAAGAATCTTATTTAGTGTGTGTG

TGTGATTAATATTTAATCAAATAATTTTACAACTTTTGCTTAAAAATATTTTTGTTAATTAAGAACATGGATTACAAACA

AGGACAAACCAATCAAGTTGTTAATACGTGTGTGGGATTGTCCTTATAATCAGGTTTAGGGTCTGCCTTTAAAAAAAAAT

ACAATTATACTTAAATTAATACATTTTTTTTTATTAAGGGAAGTGAAGTACATCAATTTTAATTTTGCCTCATGAATGGA

ACTAAGAGGGAGATAATGACTCATTCTTATTATCAAGGGTCATTGAATTTGGTTAAGGGTCAAAATACTTTAATTTCCTC

GTCACAACTTTAAACCCTAAAATATTGTTTTGGTATATAAATGGGAGAAGCACATTATGGATTCTTCTTCTTCTTCTCTA

GCTAGCTAGCATTCAACTAGCAATTACAAAGTGGTTTTGTTTTCTTTTACAATCTTGCAACCAAATTGGATGCAAGGCTC

ATAAAACTGTCAAGAATTTGTACATTTAAGGTTGGGTGAATATTTTTCTGATGCGCTGGTTGGAGGGAATGGAATCAGTT

TGCCAGGAATACAAATTCATTTTGATATTCAGTAAATAGGAACGGAGGGGAACATGAATAAGATTTCCGAATAATATAGG

GAAATCCAAACCTATAGGGGCGATGGGTAGAGTGTAAACGTGCGGGCCCGGCACAGCCCAACCCACGAAGTCACGTGCTT

AGCGTCTTTGGGCCGGGTCAGTCTGCAGGAGATTCTGAGCGTCTTGTGCTTAAATGGCGTGCCGTCTTGTCTTCATGTGA

TAAAATAAACGTGTTGTGCCTTGCTAGCACACTTATTTTATGTTCTCCAAAATAAATATTTTTGAGATATTTTAGTTAGT

ACATGTATTATTATAGAAATAAATTAGCATATCGAAGATGAAATGTCGAAATTGGCTAAAACAATGATTTTTTGTGAAAT

ATGCATCAAATGTGATGTATTGTGTAGTATTTTACGCATGACGTGGCGTGCTTTCTTCGCGTCCGGGTGGCTATGCTGGG

TGTGCTAACACGTGCCGCGTGATGTCTTGGGCCGTGTGCCTATTTTGTCCGGCACATGACCACGGAACATTAAACAAACG

TGTTGTGCCGTCTTTTGGGTAGGTCACGTCTTTGTCTTAGTGATGTACTCGAATTTTAACGTCTTTGTCTTTGTGCCATA

AACGTCTTTGGCGAGTCCCACTTTACACCTACAGGTGTAGGATTATCTGGAAATTGATTGTTGGAAATGGGATTCCTTAG

TCCATCCAAACACGTGAATCGGAAAATTTAGCTTGTGCTTCCTTCATAAGCAAATTATCAAGATTTTTAGCTTTTTCATT

TTCTCATCTGAGTCTATTGATCGGATACAAGTACAGTAACAATTACCATGGTGATATAAAAAAAAATTTAATTAATTAAA

ACTATCGCGTAATTATAATGATAATCTGAATTTAAGACTCATTGACATCACACTTTTGACATATCAAGAAGAAAACGCCA

TGATTTTCTCTTTCACCGATAATAATACACGAATTAATCTCATAAATTTTTGGTCGTACATTGCTAGAACATTTTTTATC

TAGTAGCATCATAACAACGCTAAGAAAGAAGCACTTCAGTGCGAGGAAAGACCTGCCCATCACATCTTTGAATTCTTAGG

TGACGATATATTTAGACGAGGAACAATATGTAAAATATAGGGAATGTCAAAAAAAAAAAAAAAAAGTTGCATACAATCAA

AAACTTTGAATTATACTAAGAATCTTGGAACTTTCATCCTTTTTTTACCTTCCTTGGATAGTTTTCACAAGAACTTTCTC

AAATGGAACGTGCTAAAGTTTGAAAAGGGGTTGTGATGAGGAAATCCTTACATAAAATAATAATAATGGATCAATAAGAA

AAGAGTTTTGAAAAGAGAGTTTTCCCCTATCTCAATAGAAAATGACATCCATCTACAACAATGAGGGCATCCTTCCTTTT

TTTTTTTTTTTGATGATAGGGAGTGACCTATTGCTGGCTAGGAGGAGAAGAAGTGAGTGATTGTGCAATGTACACCCCAT

GTGGAGACCAATTTGGTTGGTGTTCCTTGGTTGCCCTTAATCTTGACCAATCTCATTCTTCTCTTTATACCCACCTACAA

CATTTTTTTTAAGTACTAACTTTATTTTTTAATTGATGTCATTGGGGGTTCCACAAAGTCACCCCATAAGTTTGAACACT

ATTTTGTGATTAGTCCAATAGTTATATGATTTCATTTTCTTAATTAGCTGGACAATTAGCATTAACAAATTATTAATAGA

TTTTGAAGAAGCATTTAGGCCCCTTTTTTTAATTCATGTCTGCTCCCACAAGAACTCAAATGGGGTTTGGAATCTTATAT

AGTCATTAGAACATGTGGAGAGATCATTATAGGCAAATTAATGAAGTTTAGTAAACCTTCGATTTATTGGAATCCTCTTG

TATATACCATTAGAAAAGATAATGGAATCAAATTGTGTGTTTAGTGTAGCTAGCTACCAAGGTAGGTCAGGATCTTGACT

TTGGTCAAACATCTTTGACCTAGATGATAGGCTTTGGTCATCATCAAATAGTTGGAATCAAATCCAACCAATTAGGGTTG

TTCATTTGACTAACACTAATTGGTTTAACAAATCTTTTCTGCCCTTCCAAACATTTAATTTCCCTTTTTATTCAATTTAT

TAAGGTTGGTAAGATATAGTAGTGTGACCTAATTTTTGCTATTTCTATGCCAATAAAACTCTCAATTGGTCCTTCCTATT

ATGCAGTGTGCTGATTGGTTTTAATATTGTACAAATCATGTTTAGGTCATGTTTTAATGCTATACAGTACAGATCATGTG

TTCCAAAATGTGGTTACTTTTTAGTAGAATGATGTCATAAAAACATATATTTACTTTTTTTTTTTATTATTTTTGAGGAG

GATTTGTGTATATTTTTAATTTATTATTAATTATATAAAAAATAAAAGACCAAGCACTATGTGCCCCTTTTTCCTACCAT

ATCCCTCGTATGTACGAAAAAAAACCCAAAAATCATTTTTAATTGTCTTCCATATCCCTACGTGTGTAGGGATATCCCTC

CCTTTCCCTACACGGAGGATCACATCTGATCTGATCCGTATAGTAGAGAAGGGAAGTCGCACGACTACTCAGTTCAATTG

CCGTATAATTTTACGCTTTAGGGTGTTTTTTTATCAAACTTTAATCAAATTTGGAGAGCAATGAGGAAATCAGTCGGGGA

GAGAGGTTAATGTCCAAACACTCCACTCCATGCGAGTTGCATTTTACCTCCAAAATTGACTCACTTCTTCGATTTAATGG

TAGTTTAATGTCAAAACATTCCACTCCCTAGGAGTTGCTTTTAGATCAACTATAACGACACCCTCATTCTACCATACTAT

GACCGTGTAAAAAACGCTCAAAATGTGGAATGTCCTAACATTTTTCACAACTTTATCAAACAAGCAAGGAAGAAGAAAAA

ATCTCTACCAAATACGAATTCTCGTGAAAACCATTTTTGGTCCACCTAAAATTAAAGAAAAAAAATAACTTTTGCATATT

CGAATTGGTTTAGTATTCGCCGAAAATCTAGCCGGTTACTAAATATCCAGAACATATTAAAAACACCTCATAAAATTACT

TTTCACTCCCAAAACTTCTCGAACATTTTAGGTTTTTGGGGATAAGTTTCTCTTTCCATAGTGTTTGTTTTTGGGGAGCC

TCGGTGCTAAAACTTCTCAGGCAAGGGCAGAGCCAAGGGTTGACAAACCCTAGCTCTAGCCACCTCTAAAATCCCGAAAG

TGATGTAAAAGCCCTTAATTTGATCCCATAATCTATGTTTAGCCCATTAATGTAAAATATTTAGTCCACTAAAATCTATA

TACTTATTAGAATTTGTTTAGCCCGTCTTTATTGTACTAATTCCTGGTTTCGCCACTGTCCTGAAGCCTCGTAAAACGGG

ACATGTAGTAAAAAAAAATGGTTATTCTTCCTTGACAATGCAGCAACAAACTTACCCATTGTTTTCAAACAAGATACTTG

ACCAAACTCAAAAATCAAATCCTAAACTAATTAGACATTTAGTGACCCAATGCAATGCAAGGAACTTTAAACAACTAAGT

TACACACAAACTTAGTTACTAGCTGCTGCTAGTAGGAATATGTTCCAATCATGGAGTTAACTTGTTTTGATGGGATTTGG

GACCACTGTACCACCTGTCTAAACAAGCTGTTTGTTTTAGTTTTTAGGTCATTTTGTTTTTGCTTTTGGCTTTGGGAATA

GTGGGTAGTGAAACTAGGGTTACTATCATTGGAGATCAAGTCTATTAGGCCCTTCCCTTAATTCATTTATTCAAAGACAA

GGTACCAATAATATAAACACTGATTTTCCTTTCTTAATTAATTAATCAAATAATTAGATGTGGTACCACCATTGTTGTAA

GGCTCAAAGTATATTCTTTAGTATCCACCATTATGAAATGTTTAATCAACAAGATAATAACAATAATGTAGCATATAATC

ATACTCGGGAAATCCGGTTTATTAATGTTTTATATTTTCGGTGTCAACATATACAGATAGTAGCCTGATTTTGATTCTTC

TGCTCGAAAAAATAAAATAATAAAATCCAAATATTTTGGAAATGTGATATTTTTGTATCGAAATTGTGATTTTGCTTGAA

GTAGTAAAAGACTTCGGGTTGGAAGTTTGGAAGAAATTGAAACAGTCATTGTATGGTATTGAGCAAATGACAAACAATAA

AACCTAATTTTTTTTGTTATATTATGTTTAACATTTCTTTACCTAATTAATTTATTTTTTGTTATATTATGTTTTAACAT

TTCTTTACCTAATTAATTTAACTTTCTTCTTCTTCTTCTTCCAATTGGGTAAGTGGATGAAAACAATAGGCAACTTCCAA

CTTTGAATTTTGGTCCCAAATTATATGTTTGTAAATAATCAGACAATGACCAAATTTTGAGTACTAGACTTGTTCAATGT

GAGTTGTTTTTTCTTAACAATTTGGAAAAAGCAGGCACTATAAATTACAATAGTAGTACTCATTCCTTTTTGGGCATTGC

CCATTTGATGATGCAATCCCACCAAGTGTTTGAAAATTCTCAATTGGAATTGGGCTTATCATATACATGTATAGAGGTAA

GTTTGCTTCTTGTCTAGCTAACCTCTGTCATCAAATCTTTTTATCTATCTAACCTACTCTACAACAATTTTAGTTTGATA

ATTTTTGAAGAACCAAAGAAAGCTCCAATTTGAGGGTTGTTGAGTTGATGATAATGTCATGGTCGAAAAGCAGTTAGGCA

CGATACTAAGTAAGCTAAGTATAAGCATTTCCGTGGTAAAGTTGTTAGGTAATGATGTTTAGGATTCGAGTTCAATAACA

TGTCTGCATGATAGACTCCCGATAATATTTTTTCTTTGCTGCGGTGTGTGAATAGCGAACAGGGCAGTTCTAACCAACCC

CCACTGCCCTTTTCTAGTAGTAAAATTGGCTTTCATGACCAACAAATGGTTCTATGGTCTAGCGGTTAGGACATCAGACT

CTGAATCTGGTAACCCGAGTTCAATTCTCGGTAGAACCTGTTTTTTTCTGTTTTTTTTTTCTTGATTATACCCGTTCATT

TTTTTTATGGAAGTCAAAACCTAAGTTTTAAATCCAAAATCAAAATGAAAATCACAAAGGATATTACACGAGGAACTTTC

ATCCAATATGCAGGGATGAACATCTAGCCAGCTAGGTCAGTTCAAAGCCCACAAGCAAATCTTAGCAGACTAGTTGGGTG

TTTGCAAGTCACAATCTAACCAGCCCCAAAGAAGGGAAAGAAAGGTTGTTGCGACTAGTAATGAAAAATCAAAAAAGATG

AAATGACAACTACTAAAAAAACAAACAGCATAGAAACTAAGAAGATGTATGAATGATTTTCCATATGAAGTGCCCAGTAC

TATATGAGGTGAGACTCCGACAACTTTTCCCCAAAGAAAAGTGCCTCAAAAACTGACCGACTCATCAACAACACTATCTA

TACTCTCGACGACAACTTTTCCCTGAAGAAAAAGTGACTCAATTTAACTTACTCATCAACAACACTATACTTTGGGAGGG

ATACAGTTTTTTCCTCTTTTCTCCCTTTTTCCCATTACAAAAGACACCAAGTAAAAAAAGACAGTGCTATCCATATTTGA

ACCCCTCCAAATATGAAACTGAGAAAAAGTTGAGTAGGAAAAAGACAACAAAATAAATGCAAAATGACCTGACGACGACC

CGATTTGCAGTAATCTTCAATATCTATCAGCATTGGAGACACAAATGACACGAACCAGTTCACTGCAATCTCCAGCGTAT

AGAAGCAACCGACCCCACCTCAAAAAAATTGAATAGCCTTTTTTTCTTTTTCTTGTAACCTACTATTCCTGCAAAATAAA

CATAGAACAAAAAGAATGGCTACAAATTAGCTTCTTTCGCTTAATGCAGATATTAAAGAAGAACTAAGAAACCCTCACAA

CAATAACCAATAAATGAACAGATGAACAAGACTACAGGCTTAAATTACATGTTAAAGCTAACAAAGCGACAATGTCTTTA

TTTTTGAAATTTTAGTATACACCCATTGAATTGGTTATACTGCTTCAATATGCAACAGCACAGAGGTTCACAGATTACGC

GAAATTGTAAAGTGTCAATGTTTTGGAGACTATGCAAGTGAGTCCTGAGGATACTACTCCCAAACCCAAATCAGCCACAA

AAGTCTGAAGGTCAATGTTTTCAAGCACCTTAATCCTATACAAACTTGAACAGGAAGGTTACCTTTCGGTCCATCACCAC

AATTAACAATGCATGTTTACAATAGATATATCCAAGTATTAGTTCTATCGATCATTCGATGCCAAGATTGCTAAGGAAGT

GTTGCAAGGTTTGCACAATTCATTTTCTGCCATTGGGAGCCAGATCCTAGTAATGGAGCCATTTCCATCCACCACAAAGA

CTTATTCCTTGATTCGCCAAGAGGAGAAACAACAAAAGATCAATACAACTCCTTATTTCTAAAAAGTTAGTATTGGTCTC

TTGTTGTTTCTCTTCCGACACATAAATCTTTGTGGTATATGAGTAAATGGTTCCATTAGTATGATCAGACTCCTAATGAC

ATAAAAGTGACTCCGTAGACCCTGCAAGAATTTCATAGCACGATCTTGTTGCAAACACTCCTTCATAGATTTTCCGTGTC

CACATGAAACTAAGTTAGCTGACAAAAAACCTTCACATGTTCAAGAAGTTTTGCTATCTATTATAGTTTGGACTTTAGCA

TGCACCGGGAAAAAAGAAAAAAGAAAATCCCGTGAAATGCCTGAGTAGTGGTCTGCGAGCAGATTCACAAGCAGAGAAAA

TTACATACTTATATGCAGATACTGTTGAGATATGTGACATACCCTTTTTAAGAGGGAAAATTGAATATCTAAACAACAAA

ATCAAGGAGCAAAAGGCACTAAAAAAATGATAGATGATACCATTGGTCACTATGGTTAACCTAAATAGAGATAAAATGAT

GATATATGGTAAAAAGGTACATTCAAGTTTAGAAAACTTGTTTCGGACAGATTGATTTCTACTCATAACTGCACCTAATT

CTAACAAACCCACAAGAGAAACCTAACAAGCAAGGCTACACTAGGCCCTCGATAATAGATGTAAGAGGTAAGGGATAAGT

AAACCATAAAAAGTACAGATCACAAAAGGAAAAAAAGAAGAAGCAAAAAGCGCACAACAACACTACGAACAAACAGCTTA

CGTTAAAACAACTTACGTTAAAACAACTTACTCAGACTATGGTGCAATTGTATGTGCTGCAGAGCTGGAGCAATGCTGTT

TGATTGGCTGGATCCAAAGACTGTTGAATAATTGTTGGATACCTTCCAGGACTCATAGCTGAGAGCCTTGCCAATGACCT

TACCAAGAACTCCTTCGGGTCCCTTATATCACTAACAGGATCTTCATCTTTCTTTCCAGCATTGTGGAGACGCACAAAGC

TAGCACTGTAACCCACATTCTCCCCAATTTCTGGCACTTCTGGGTCTTCATCAACCCTTTCTTCTTCTGGTCGAGAGAGA

AGGGTAACAATGCTGTTCAACATTTTCCCCCAAGTAGCAGCAGCTGCAGGATCCAAGAGGACTGTAGACTCACATATTAA

TCTGGTCGAAGCAACAGCAGTTAACTTCAACTCAATTGTCCCAATTATGGATTTCAGATGAGGTATCCAAAACTGCTCGA

TTATACTACCAAGTAATTTTCCATCATTCTGAATGGCGTTGATTGAATCAGTGAGTGCTGGAGGTCCATACTTGACCAAA

AATAGTGACATATATATCACCATACACTTGATGAATTTAACCTTGGGACTCTGCATGAGCTTTGTAAAGAGCACAGTCCA

GATGTGAGTAACAAAAGGTGCGATCACATCATACCTGAGGTTCTCTATAATGGTGTTCAGCACATAAAACCCTAGCTCTT

CTGTGCTTGGTACGTTGACGAGCTTTTGAAATATCCCGAGCACTTGGCCAAGCCTACCCCCTTGGCTCAGTTCATCAGGA

GCCTTTTTCAGATAGGCCTGAAGAAGCTGAACAAGAGCAGGTGTGTTAGCGTGTCCCTTCCATGTCTCGGTTGCAAGGAG

AAGCTCAAAGATCGACATATAACTAGCAGGAAGAGGTGGTTTATTCAGATTGACTAGCTGCGCTAATAGCTGGAAAGCAT

AAGGCCAAAATTCAGTGACATTCTGGCCCAAAATATATTGGAGTATTGGAAATATGCTAGCTTCAAAGGTTGGTATCAAA

GAAGGATCCTTCTCACACACTCTCCCAACCAAACCCGCTACTGCCTCAAACAGATAGTGATTGTACACAGGGTTCTTCGG

ATTTTTGCAAACCTCGCTGAGAAGATGTGTCAAACCAGTAATACAATGTCCAGCCACTTCTTGAGAAAAATCAGCAACTC

CCAGAACCCGCATTATACACTTCATTACGTATGAATTCTCCTCTGATTCGGGAAACTTCAGGGCATTGAAAAGGTTAGTC

ATGAGACTGGGAAGGAAAGGGTTGATGTCAGTGGAATTAAACCTCCTCTGCCCTCCATCCTTGATTAACAAGAGCTTTTC

AACACAATTTGCAGCGTAAGAGTGAACGACATTTGACTCTGATGCGAGGAACCGAACAACATCACCCATCAATCCAACGG

CAGTCTGCTTCGGTATCTCATTCCTGAAAACTGTAAAGAACTTCAAGGCACCGGCTTTCAACATTGCAGAACCGGACTGG

CTTTGAATCTCTGGGATAATAACTGATCCAAAAAAATTCACCGACATCGACAAGATCAGTTGAGGCAGAAGTTCCTCCAG

CCTTCCTTGTTGCAAGAGAAACAACCAAGTAAATAGCACAATCCTTATCCCGCCAATTCTCAGCTGGATTTGCATTAAAA

ACAGCCAACATATTTTTAATTTGTTCTACAACAATGGATTGAACTTCTGTTTTATAGTTGGTGGCAATTCCTTTGAGTAG

CTCACATGCTATCCTCCTCCTAGTGTCAACATCACTCCCCTCAATATCCCTCCGAACAAACTCCACGTAATTGAATTCAA

ACAACTCCTCATCCTCTTCCCTTAATCTTGCATTCGGAATCACAATAGACTGACAAATCTGCTGCAAGACATCAGCTCCG

GCAAACAAGTTGTGGTGTACACTCGTACTGATAGTAGTTAAAAACTTAATAGCAGTAATAGTTAACCTGTCACGACCTGT

CGCCGATGAAACCCCAACCAAAAGTGTCCAAACTGCAAGTGCAAAATCCTTCAAGTAACCTGCAAACTCCTCCTCATACC

TCTCCAGATATAAACTAATATTCTCACACACAGCCGCTCTAAGATCATCAACAAGTGCCAAACCATCACCACCACCACCT

TCCAGAACCGGATAAGTACTAGTCAAATACTTCTTAAACTCCCCATCCACTCCTTCATATGATCTCTTGAAAAACTCGGG

TATTTCCATAAAATTCAAACTAAAGAAAATCCTACAACAAAGCCTCTGACTCTCCATAAGTGGTTTCAGATTGGCAGCCG

TTACAGCACCACCACCAACCATACTATCAATCACATTGGCAGTCCTCATGAAAACCTCAAGCAACGGCGCAGCAAAACCA

TCAAGACAATATTTCAAATCAAGCAACAAATCATTCGTCTTATACTGATACCTAAACTTCTTAAAAATCGAATTGGCAGT

ACCTAATATCCCATTAACAATAACATAATCATTATTCTCCATACCTTGTCTCAAACTCTGAACAATCTGAGGTAACAACG

CAGGCCAAGCTTTAGGAAAATCATGTTTCCCAATCACAACCAACGCTTCACTCAATTGACTCTGTATCCTCGGGGAAGCA

GACAACATCAATGGCAAAATCAACTGTTTAATCTGTTCCTTCTCAGTATCATCAATTGGTGACAAAACAGGAGCATCAGG

TTGATTATTATTACCACCACGATGATTCTTGTTCGTTTCTGCTGAAATCCATCTCGATTTAAGATGATTTTTGAAATTAA

CAGCAGCAGAAATACGGATCTGATCATCGATGTTGGTCTCAGCAACTAAACGGAGAACGGCGAGACCGAAATTAGGTTTA

TCAGATGCTTCTGATAAGTATGATTCAGCTTGTCTACGAGGTTCATGATTTGGTGATAGTGTGTGTAAGAAACATTCTGA

TAAGATTTTTAGGGTTTCTGCATTCCACTCCATGACGATGGATTTCTAGGGTTTAACGATGAAGAAAGAAATCAAAATCT

GTAAACAAGGGTTTAAGATGATGATGAAGAGATTAGGGTTTTAGGGTTTTACCTTTGAAGAAGATGAAGGACTGAGCTAA

GGAGAGAGAAACTTCTTCGGCTTCGTCTCTCTCTGATTTGAAAAATGAGGAACTGAGTGATATGAAAGGAAAATCAGGGT

TGAATCGGGGTTTATATAGAGGAGGATGTTTTGGTTTAGGTCCTTTGATTTGTTGCTTTATTTGTGATACCCCTTTGAGA

AATGTTTTTTTTTTGGTAATTACTTGACGGTAAATGCTCTTGTGGAAAGGATATTCTGTTTGGTTGCTTTGTTGTTGAGA

CTTGTCGAGGCTAACCTTACCGACTTTGAATTCAAGGAAATTTTTCTTGGAGGATGATTTTGTGAATTAATTTGGGATGG

ATCGGTTGAGTTGGCTGATTAGGCCATTCGGCTTGATGAAACGCAGTGTCTAGACGCTTCTGCCAGTTTGATTTTTCACA

TGTAAGTGTTCATGTTATCATTTGAATTCAGTAACTGAGCCAGGAATGGAATAACTCGGGGTCTAGTCTCCTCCTAAATC

TTCATAAACCACATATCCACTCTTCATTTTTTCTGGTATCTAGTTTTTAGTCCACCAGTTGATGTACATAGTGCCAAGAC

GTTTACATATTCCTAATTCCAGCAGTAACGCAAGGACAAAACGTATTACATGGTTTGTTCAGAACTTGGGTGCTTCATAG

AATTTGAAGAGCAGACTAGTAACAAATCAGTGTTTGGTCATAAACAATTTTATTCAAAGGAAGTTCAGTACAGGACTCCT

AAGCCTTGGTAAAGAGATTAATGCAAATAAGAAAAATTGCAGGACGACCAAGCGTTACTGCTGCGGCAATATGTGTACTG

CAGCATAGCTAGTTGGTAAGAAACAAAATATCTCATATTGACTGCATCAAGGGATTAAAAACACATGTACCACCGCTGGG

ATTCTATCTGACATTCTGGAGTGAAGAGGAAAAAAGAAGAATGTTTCTTTAGTAGTTTAGGGATTGAAAAGAGCAAAGAC

GACACCTGAATTGTCCCCGCTGAAGCCAAATGAATTAGATATACCTGTAGGTACTGCACACAAACAAGTTGGTAACCAGT

GTAAGCTATACGAATGGGAAATATGAGCAAAGGAAAGCTAACTGCTTAGACATTTTTTTGCAGGATTAAGAATCATGTAA

TAGGCATTTTGATGTCGCTGTGACAGATCTTAATGTATTTGGAAGTGAATACATACCAACATGAACTTCATGCTGCTTCT

TCACATTTGGGACGGTGTCAATTGTAACGTCAGACTCCAAATTCTGCGTCAAAATCAAAACAACGAATATACACAGTCAG

AAATGAAGATAAACATAGTTTAGTGAAAGAATATTAGTCATAATGGAGCTTTACATACATCTTAGTTAATGGTTGGATGT

AACCATCCAGTGGTGATAGCTTTGATTGTAGCTATGGCTTCTAATCCACCAGCTGCACCGAGCCAGAAAGTAGAAACGGT

GTGTCAGAGGATGAGTAATCTTGCCTTAGTTCCATTCATCTTGATCTCGGACGTGTTCTTAAACACCTTCTTGATGGCAT

TAACCTCTGCTAAGTCTCCCGCAAGTGTCGACGTTGCATGAGCATTGATATAGTTCACCTAAAAATTTCAGACATTAACT

CAAAATTTCAAAGAAACGAGGGTAAATTTACTAAGACTGGAATGTGAATTCGACGGGCATCATTTATTTATTTTCACCTC

TTCTGGAGAACCTCCAGCCTCTTCCAAACTCTTGTTTATGCATGAAGAAACACCAAGACTGTCTCTGTCATATGATGAGT

ATCACAGGTAACGGTGCCCCCCCAAATACTCCGCTATTATAGAGAAACTTCATGACACAGCCTAAAGGGAACCAAAAGCG

ATATACAGAAACTGCATGACACAGCTCAGAGCTTAGTCACCAGGAGCGGGAGCGAGGTCGGGAACGTGAGCGAGATCGAA

ATCGAAATTCGTGTAATTTTCAAAAATGAAATTCGAGGAGCGTGTCGGGAACGTAAATTAGGTTTCATAGATATATTAGT

TATACTGCTAATTTTCCTAAAACAACATGGTAAAACATTAGAAAATCTTAATAGAGTGCCTACTTTCTAGTTTCCTACAA

ACATATCTCAATCATTCACTTATTGTTTCGAGCTTTCTGAAGTCGATTCTGTTCTTATGTATCCTGTGAACTTGGGGTTG

GTAGTGGAGGTGTAGACGGTGTAGGGGAGGATGGAATATTTCCTTATATCCATTTAAATAGGAGTCTGAGATATACACTG

ACCCTAACTTCTTGACAAGAAAAATACTAAAACTTAGCTAGGTAAGTAATACAGTTGTTACATGTTTATATGAAAGTTAA

CCTATAAAATAAAGGAGCAAGATACATGATTTACTAGATAAAGATAGGAAGATATGAGAATATACGGTTAGTAGGATATG

AAGCCAGATATTAGATATGCTTCCACATAATTATTACGGTTGGATAGAGGGAAACGTTTCTCACCCAATAGGAGCGCATT

TCTCCCTTGTTAGAAGCACGCTCCCATCACGATTTTCACTGAGAAGCGCAAATTCGCGAATCTCAGGGAAATTCGTGAGG

ACTTTCACGAATCTGGTGACTAAGGCTCAGAGCTATGTTCTGTGAGGCGCTAGGTGAGGCTCCCCCTGCACTACTCGAGT

ACCTAGGCAAGCGCCTAGCTCGCCTTTAACAACATAGGCTCAGAGAATTAAAATGGGGTAATTACCAGCACACCAGAACC

TTCACCCATAACAAATCCATCACGATCTCTATCCCACAGCCTCGAAGCTTTTTGGGGTTCACCGTTTTTCTGAGACAAAG

CTCTGCAAGCTATGAAACCACCAACACCAATTGGGATAATAGCAGCTTCAATCACACCAACTACCATGATATCAGCTTCA

CCTCTTCTGATATGGAGTATGGAATAAAGAACGGAGTGATCTTCTTGGATCCTTTCTGGAGAAGAGATTCAAATCCACTG

CTGAATGCAGTTAATCCTCCCATACCTGTCTCTACTAGAACTCCAATCTTCGACCGGTCCATCTGTTGCATGCGCAAAAT

CCAAGTACATTGCAATAAGCCAAACAGTATATGCAAACACGAACTACGTTACAGATTGTACTGCAATAAGACTTACAGAT

TCAAGAATTTCGGTTCCAAGGTTGGCATCTTCAAGAGCTCTTTTACCAGCAACTAAGCAAGTACCTCCAACAGTCATCAA

GACGGCGGTCATGCTTTCCGTCAATGTAACCTTCGGATGAAAAGTTACATATCTGACCAGCAAATCTTACAGAAAAAGGG

TGAAGCATCAAACCTATGTATAAGACTACTACCACTCTCTCTTTCAAGTAATGTATTATAGAAGGTATCAACATCATTTC

CAAACACCGAAACAAGTCCCATTCCTGTTATAACAATTCTTTTCTTCGGATCCGTCTCTCTTTTCGGAGCTGATGGGATG

CCATAGCTCTGACTTGCCTACTCTTCAAAACTGAAGGCAGGCAGGAAATGTACAAAATCATATTCAATTCAAAGATGCAT

ACATTAAAAGTACTGCGTAGTTGGAAGAAGCTGAAATAAGCCTAAATCCCGGCCAGTGCATTGACTCAACAACCTAATGT

GCAACATAACTTAAATTATCCGCAATGGATTCTACCTAAGAACATGCTAGCAAATTTGCAACAAACAATTACAATGTGCA

TCAAAATAAAACAACACGTACCTGATTTCGCCCACAATTTGCTTGGTTTCAAAAGATCTAAAAAGACTATTATAATGATT

ACCTGAAACTAATTCATTCTTAACAAATTCTTTGTTTCGAAAGAGAAGACTACTAGTAGAACAAGCACTACCAGCAATGG

CTACAATTGTCACGAACTAATCAACAATTGAAATTGAGGAAAAACAAATTATAGGGCAATCAAGAAAAAGATAAAAGAAA

GTTTTTATATAAACAGATTAAGATGCCTAGATATTTTTCTAATAGGATATTGTTGATCAAGTTACAGATAACATAACAGA

CAACAGACAACAAAAAACACAAACTAAATTTTGTAGTTCAGTCAGTCAGACATCGCTGAAAACTTATATTCAGCTTCTAT

TTCTATGCTTTCACTTACAAACTCTCTGGCTCTTCCTTTTCCTTAGGTTAAGCTCCTTGCCCTTGTCAGCATCAACCCCT

TCCATTAGATCTTTGGAGGACTTGTCATCAAGCTGACTCAATGCATTCTTAAGTTGCTTTTGAATTTCACTCTTCTCAGC

TTCAACACTCTTGAGATTATCTTTCAAAGACCTAACCTCGGATTTCAAATGCTTGTGAGCATCAATTAGAACTTTGGAAT

GCTTCTCAATTGACGAAATGGTAGTTTCAATATCATCAAATGCATCTTCCTCTTCACTAACTTCCTTGAATTTCTTTCTT

CTTCCGGAATCCAATTCCAGTTGGTGTTTTTCAAAATGAGCTAGTTTCATTAGCAGATCGTCCCTATTCTTCTCAATCTT

ATTCTTCTCCACAGCAATTTGTTGCTTCTCTCTTTTCAGTTCCTGGACCATCTTCCTTGAATCAACCAATTTCTGCTCAA

TCAAACCTTTCTCAACCAATAAACCCTCAATTGCTTCGTCCTTTTCCTTGTAATTTTCTGCAACTTTCTTTCTCAACATC

TTAAACTTCTCCTCGAAATCAATGATTAAATTCTGCTCATAAGCTCTCTTATTCTCGGATTGTATAGCCTGAACAACGAG

TTTCATTTTCATATCAGAAACAATTTGTCTTTCACGAACAAGATCTCCATTAATAAGATCCACAGATTTCTTCAACTTGT

TGATTTCGCCTTGTTTCTCTTGCAAAACACCATCATACTTGGATTTCAACTCATCCAGATTTTGTTTCTCTTCCACTTCA

ACTTTAACCTTCTGATCTATTTCTCTCTTCATTTCTTGTTGCTGCTGCTGCTGCTGCGATTCTTTCTTCTTCTTCGCCAT

TGATTTTAGTCTAGGGTTTGTGACTGGGATTTCAAAATGAGGAACGAGTGATATGAAGAAAGAACAGGGACAGGGTTGAA

TCGGGGTTTATATAGAGGAGAATGTTTTGGTTTAGGTCCTTTTACTTATGGTTTTATTTTGTGATACACCTTTGAGAAAA

GTTTTTTCTTTTTTTTTGATAATAACTTGACGGTAAATGCTCCTATGGGAAAGGATATTCTGTTTGAATTTTTGGGGTGC

TCGAGGCTAACCTAACCGACTTTCAATTTAAAAAAGTTTTTTCTTTAAGGATGATTATGTAAATTAACTTGGGGATGTAT

CAGTTTGAGTTGGCTGATTAGATCATTTGGCTTGCAGTGTCAAGACTAGAGCTGGCAATTAGGGTCAAAACCCACGGGTT

GGACCTAACCCGGACCGTGTAAACCGGACCTAACCTAGCTTGTTAGTTAATAGGTCCGGGCTAGAGTTAAGGTTTTAGGA

CCCGTTGGTAAGTAGGGTTGACCCGTCTAACCCGCGGGTTAATCCGCCAACCCGTGTAATAAACTGATTACACAGTAAAC

TAAGATATTTGTTAACCTGTTACCCGAGTTGAAGGTAATTTTTGACGTATTACACCAGGTGACCGGTCAAAGGTCAGGGC

TTTTTCTTATCTTCTCACATATTCACATAACCTTATCTCTTCAGTTCCAATTTCCCAAGTTAGGAAACATAACTTTTGTT

TCCTTTCCTTCCACTTTCCTTTCCTCAACTAAACTAGATCTGTAACTACAATGGGTTCGATGATCAATCAGTTCAAGTCA

GTTTCATCATCCCTCAACAACTAAATTGATCTGTAACTATTCTTCACCTTTTAATCTCTTCTTAGTGGTGGTTTTTTTTT

GATTAATTTCTCTAAATAAATCTTTTTTGTTGTTGTTTGTTTGTTGATTCAAGTTTCTGTTTCTAATGGTGTGTTTTTTA

ATCTGTGATTTTTTTTTTAGGTAAGTGATCGCTGTCGTATGCGTCAAAAATAATTTCACTTTCCTAAAGTATATGATAAG

AAAGAGTTCGTATCCACAGAGAGGCGATTGTAGTTATTTCGTAATCAATTTCTTAAAGTAACCAATTGGGGGGATTTCTT

ATTTCTATTTAAGCAATAAAAACAAATTAAAAGCAAAGTAAATTGTAGAAAGTAATCAATGGAGAGAGATATTGGTCAAG

GAATTCATCTTCAATCTTAGACATGTGCTTGCATATATGAATAGAATTACGATTTAATCTCTTGTGTTATCAAAAGCCTA

GAGAATCAAGAAAGCAAGATAATCTATTAATTTCTCAAGTTCATCAACACACACATTAGGGCTGCGTATGTGAATTCTAC

CTAAAGAATAACCTAACGCCTTGCGCTAATCAAGTTCAATTCCTAGGAGCATTAAGTTTCAGAAATAGGCTAATCAATGC

AATTGTCATGCGTTGCGCAAAACAATTCGGACAATGGTCAAACTCTACATATGATTACGAGAGTGTATCACTACAAACCG

TAAACATATCATGCTACTCGTATTCAAGGTTATCGCTTGATGAAAACCCTAAAATAGCTATGGACTAAGATGCAAGTTCA

ATTAATTGGCCACTTAACTAATCAAACACCATAATCCTATCACAATACTTCAATCATGTGATCAAAGAAACCGCGAGAGA

ATTGAACAATAAACAAGAGAGAAAACTAATACATAAAGCACACACAAGTTTTAGAAATCGGACTACATCCTTAACCAATA

ATAGAGAATTTAGCTACTCATAGCTATGGAGTTCATCATCATCAATATTCAATAATAAAACAAAGAAGAAGAACAAAAAC

CAAACCCTAGAAGCGTAAACAAAAACGGCTCTCCAAAGTCCAAGAATAAGATATCATATATGATAAGTTGTAGAAGACCT

TCCCTTTTATAGCTTTTCTCCAACTTCTAAAATTAGGTCTAGTCTTCTAAGTCCATTATGTGAAAATCCACCAAGCCCAC

AAGAGGGAGAAACCCGTGTAGAAAATATGCCCCAAAACGGTCGCCGGAAGTCTGGGATTGTGGCCGGAAATCGGCCGGAA

AACAGCCCCGGTGACCGGCAAAGTCCGCCCGGTCACCGGTCAACGGGTCGGATCGGTCGGGTCAACGGGTCAACTCAGTC

GGATCGGTCGGGTCAACGAGTCACCTCGGGTCAAAGCGAATCGGTCGAGTCAACTCAAGTCATACCGAGTTGTGTGAGTC

AGGGCTGAGTCACTGTCTGAGTTAAACCCAAGTCAGCTAAAAACTGAATCAGCTGTGTCAGGTGTAGCTGGTCTGACTGA

ACCTGACATGACCCAGGCTATTGCACTGGCCTGACCGTGTGTAGCACAATCCTTGTGCATTACTACCAGCCCTGATGGCT

ACTCCTTCCTTTGCATAGCAAACTCAAACCCATTCCAGCTCCAGCCATTGTGACCATGGCAGCATCACAAAACATTCCCA

TTTCCATCTCAACTTGCACTTTCTCCATTTGCAGAAACATCAAGCTATGGCAATCAGCTCCAATACTCAATTCCAGTTCA

GCTCATAACATCCTGCAAAACCACCTGAGTTGCAACTCCAATCCACCTCTACCACAGTTCTTTCATATTAACCTGCACAA

TATACACCTCCCTGTAAACTTCCAACCGCCAACTCCAATTCCACAGCTCAAGCCACACTGCAGCATCTGCTATCATCAGC

CTCAAATCCATTTGCATTTCATTAGAATCACAGCAACACCACCAGAGACCTCCTCCATTCATTCATCTCAGACCCATATT

TTCAGCTCAGTTCCATATGCTCATTTATTCACAATACAACTCTTCCATGATGTGCACAGTTGCACTAATTCGACACAGCT

CGAGCCAATCCAGCCTTGCAGTCTCTCTCGCTTCCACTGTCATCACTAGCACTTCCTGTTTCAGTTCAACTTCACATTGA

GCCCAAACCTGTCCTAAACATGCATCTAAGTCAGTTAGCTCAATTAGCTCAGGCCAAACATTCATCTTGTCGCCAGCACC

AACTATGTATTCTGCCGTGCCACTGCCTCATTTCATAGCTCTCCAATTGCAACATTCAGTTCCTGCAGCTCTTGCATTCC

ATGTAACTCATTCACAGGCCCCTGTAACAAAATCACCAATTGTAGCTACAACTTCTTCAGCACACAGAACCCATTCAACA

CAGTTCCTTGCACCAGTCCATCCAAGTGTCATGATCAGCTGCAACATTATCACCACATCCTTCCAGCTTCAATTTGCCTC

ACTGACTCATCTGAGCTGCAACCTTCAATTCTAACCCATTAGTGCAACCTAATATCCTCCATCTGCTAGCCACACACACC

AATTCTATCTATTGCCAGCTCCACCAGAATCCATCACCATCTGCATCAAGCCTGCAATCTCCTAACCATTGCCACAGTCT

TCATTCAACTCAGGGTCACCTGCCAACTTCATTCTGATGCCCCTGCAATAACAACCCATCTTCATCTGATATTCAATTCT

TAGCTCAATAGCAAAGGCTTCATGTCTTCCCTGGCAAATACACAACAACTTCAACATCAGGGCTTCCAACTGCAGCTCAT

TGCCTCAGTCCATTCTCAACAACTCCCTTATCCTGCTCACTCAATCCTCAGAGCACAAACCACCATTGCAGCACTAACAT

CACCAGGTTAACTTCAACCACTAGAGACTCAACCATTTCAGACCCCTTTCAGTTCATCCACACTCACTGCCATCACTAGG

ACAACAGATAATCTCCATCTCAATCTCAGAATCCCATCTGATTTCAATAGCTGAATTCATCTCTGTTTCAATTTGAATAC

AATTTCAGCTCCAAAACCCCTTCATTCACCGATTCGATCCAAACCCCATCTCTGGTTCGTCGATCTGAACTCATTGTTAG

CAGTCCAACAGATCCCATTTCATCTCAGTTCAGCTCTGGATCTCGAACCTCACAACAACAACCAAACCGTAGCTTCTTCC

ATTCAAATCGACCATCGGAATCATCTCTTGCTTCTGAGTTCATCTCGTTCTGAATCTCAACATCTTCAGCACTCAATTTC

TCCTCTCTGCATTTCTCCCTTTCTCAGACAAACAATAGCCGTTTCCTCTCGGCTCTAGCTTTGAGATGATAATGCTTGAA

AGAAGAAATAATACCCACAGAATTCCAGGGTCTTTGGTAATGAGTAAAACTGATATAGGCACCAGTTTTCGGATGAGGGC

TACATGTTGATATTTGGCTTGTCGCCTCAAAGCAAACCGATCGCCTAATCTGTTTCCTTGCTCAATTGTCGGTTGCGAGG

AGGCGACAGCTCATTCCTCGCCTCTTTTGCTGACAGCTAGCTCTACGTTTCGCCCGCCTGTGAATTGACATGTTTTCTCC

TTCTTTTGCAAAAAAAACATAATGACTCCATTGCACCTAATAGCTCAAAAACACAAATAAAGTACATAATTCACAAGAAA

AATAGCAAAGAAAGCATAAACTCTATATAGTAAATACGGTGTATTCTACACCTATCAGATTCCCCCACACTTAGACTTTG

CTAGTCCTCGAGCAAATCAAACAAAACAATAAAATACAACAACTCCGTGTCGCCGAGGCTACGGTCACTTTTAGCATAAA

TAACAAGCCTTTAAACCCCTAGGTGTCCCTAGTGGACGAGTTATAGTCTCGTGAGGGTTTACTAGAGATATACCCACAAA

ACCTATTTTTCCAACTTACAAGTTCTCCGTACTGATAGTATCCAACGCGCTAAGAAGACATAGAGATTCTGCACACTAGT

CATAAGAAGTTAGACACATAAATGAACACAATCTCATTCTTAAAAGTTGAGAGAGAGATAACCATCCCTTCTCGACTAAA

ATTCGGGTTCGGAGTGATCAAAAGTCTCGACTCTGCCTCACAAGAAAGGTTTTTATGATATTGCTCTCAATAATAATGAC

TTTTTATGGGTCACACATGTGTCCAGGTAGGAATGAAGAGAGAATCCTGCGACACGTTGAATGGTAGGAAGGCATAACCC

TCCGAAGTATTTTGAAAACCCACATCTTTTATATATTTTTGAAAACCCATTTTTCGACGATCTCTTGAAAGGAAATCAAC

CATTTAACGAAGGTGGGAATATGCTTACTTACCTCGTTATCCGTTCAACGTGCGATTAGCACCACTCTCAAAGATCCATA

GAAACGTCTTCGGTCTTCTATCAGTGTCTTCATCTTCGTCTTCGGTCTTCAACTCTTGATGATAACTCTCGAACTTCGTA

GAATCTCGACAATATGATTATTTCCTCTAATTCCTTGTCGCTTGATACTTCATTATAGATATTCCTTCTTTCCATAGGCT

TTATTGAAAGAATACAAAAATAAAAACAAACTATACAAACCCAATTTTTTTTTTTTTTTTTTTCTTTTCTTCTTTTTTTT

TTTTTTTTTTTTTTTTTTCTTTTTTTTTTTTTTTTCGTTTTTCTTTTTTTTTTTCAATTTTTTTTTTTTTTTTTTTTTTT

TTTTTTTTTTTTTTTTTTTTTTTTTTTTTTTTTCATTTTTTTTTTTTTTTATTGAGACAAGAGAAAGCAAAATACAAATA

AAATAAAAATGTAAGTACAATGCCATGGTATAAGTACTTTACCGCTTGAATCTTCACAACTTATATGTCTTGATATCATA

GACTTCTCGAATTCTCATCCTTAATAATTTCCGCTCTTGTATTATAGTCTTCAACTTATAAAAATTCCGCCCTGTTTTGT

TGCTTTACTCAAATATGAAATCTGCCCATTCCTGTAGTGTTGCTTGTAAACCTCGAACGTATTCAACTTTCCTTCGAATT

TGTGATGCTCCTCTTGTTGATGATGATGGTGTTTCTATGGACCTGTTGGTGTAGAGAGATAAGGTGGATGGTGCTAACTG

CGAACAAGATCACAAGTGGTATGTAAGTTAGCAATTCGATGTCACCAACATGGTTGATAAAAATAGCACTCAAGAGATCA

AAATAGTGCTTCTATGGTGGGAAATTGGACAGCTCACCATTCGGCCCGGAATTTACGTACGGTGACACACACTCTAAAAG

CACAATTTAGACAGGTACAAAGAATTGAAGGTTGGTGCCTCACATTATGTAACATGGGTAGTCTCCACTATAGGGGATCA

CTGTTCTAATACCGAATTCAGTCTGCACCTTATTTGCCACTGGCATGGTTAAATCCAACTTTAACTCTCATCCAATTAAG

AAACCCGATCATGTTCTCTGTCATCTCTAAGTTCAGTCACTATTGTGAGATTCTCGATGTAATCTTAGCGACATGTATAC

TATGTGACTCTAAACTAACTACAAGTTGGAGTTTTCTTATTCACTAGTTCACAAAAAAGTTGAAAAATTGCAAAAATTTA

CAACATAAACCACTCCCCCACACTTAAACTTTACATTGTCCTTAATGTAATTGAATCATCCAAAGTAAGTCAAGGTGGGA

AATCCTAATAAGAAATCAGCAAAAACAAAGAAAACAAAACAAGAAAAACTAAAAACAAGAAAATAAATAAATACAAGACA

AGAAATACACATCCTTTGGGTTGCCTCCCAATTAGCGCTTGGTTTAAAGTCGTCAGCCCGACTAACTCCTTGCTTTCTAT

GACTCCTTCAGAGGGAGTACTTCCACAAGATTAACTCCTTCCACGACATCCGAAGAGAAGTTCTCATAATATGGTTTCAA

CCGGTGCCCATTCACTTTAAGTTCATTACCCGTAGCTAAACTACGAATTTCCTTTGCACCATGAGAAAACACATTAGTCA

CAATAAAAGGTCCAATCCAACGTGACCTTAATTTACCGGAAATAATTGCAAACGAGAATTAAACAAAAGCACTTTCTCGC

CCAATGCAAAATTGTTTCTTAGAAATCATATTATCATGAAAAGCTTTCGTTTTTCCTTGTAAATCCGTGAGCTTTCAAAT

GCATCATTGCGAATCTCCTCTAGCTCTTGAAGTTGTAACTTCCTTTGCTTTCCACCCCATCCAACTCCACGTTGCATTGC

TTAATAGCCCAAAACGCCTTATGCTCAATTTCCACGGGTAAGTGACAAGGTTTACCATAAACAAGCCTAAAAGGAGACAT

CCCAATAGGTGTTTTATATGCGGTTCTATACGCCCACAAAGCATCATTAAGTCGCGAGCTCCAATCTTTCCTTGTGGGAT

TCACCGTCTTTTCAAGAATGGATTTCACCTCTCGGTTGGAAACTTCGGCTTGTCCATTTGTTTGCGGATGGTAGGGAGTT

GCAATCCTATGCGACACATTGTACTTCCTTAACAAGCTTTGCAAGAACCTGTTTTTGAAGTGTGAACCTCCATCACTAAT

TATAGCCCTTGGAATTCCAAACCTTGCAAATATATTAGCTTGCACAAAATCTGAAACAACTTTAGAATCATTAGTTGGGG

TGGCTTTAGCTTCCACCCATTTCGAGACATAATCCACAGCAAGCAAAATATAATAATTCCCATTAGAATTGACAAAAGGG

CCCATAAAATCAATTCCCCATACATCAAAAACCTCCACAAATTGAATAATAGTTTGCGGCATTTGATTTCGAGCACCTAG

ATTGCCGGTTCTTTGACACCTATCACATGTGATACAAAATGCATGTGCATCCTTAAACAAAGTTGGCCAAAAGAAACCAC

ATTCTAATACCTTGAGAGCGGTTCTCTTGCTCCCAAAATGTCCTCCACAAGCATATGAGTGGCAAAAAGACAAGATAGAA

TTAAATTCAATTTCGGGAACACACCTACGTATCACTTGGTCACTACAATGCTTCCATAAGTATGGATCATCCCATATGTA

TTGGTTGCCTAACTTCTTAAGTTTGTCCCTCTCAAAGACGAGACAAGTTCTTTGGGATTTGTTTAGACACTAAGTAATTA

ACAATATCAGCGTACCATGGTTCTCCAAAGGCAATTGAGAATAGTTGCTCATCCGGAAACTTTCACGCAACGGATAGTCT

CCTTGTCCAGCAAGTCGGCTCAAGTGATCCGCAACGGTGTTCTCAATTCCTTTCTTGTCTTTGATTTCAAGATCAAACTC

TTGCAAGAGTAAAATCCATCGTATGAGCCTCGGTTTCGCCTCTTTCTTTGTAAGCAAGTATCTAAGTGCTGCATGATCAG

AAAAGACAACAACTTTTGTACCAATTAGATAAGAGCGAAACTTTTCTAAAGCAAAAACAATAGCTAATAACTCTTTTCAG

TGGTTGTGTAGTTTTGTTGAGCCTCATTTAACGTTCTAGAAGCATAATAAATGGCATGAGGAAGCTTCCCCACACGTTGC

CCAAGCACCGCACCAACCGCATAGTCACTTGCATCACACATGAGTTCAAACGGCTTACTCCAATCCGGTGGTTTAATGAT

AGGGGCTGAAATCAAAAGTTCTTTCAAACGATTGAATGCCGCCACACACTTCTCATCAAAGTTAAAAACCACATCTTTTT

GCAATAGGTTACAAAGTGGTGATGCTATCTTGGAAAAGTCCTTGATGAACCTACGATAAAAACCTGCATGACCAAGAAAA

GATCGTATCTCCCTCACCGTAGTGGGAGGGGTTAGAGCACGAATAAGGTCTATCTTAGATTTATCAACTTCCAAACCTTT

AGAAGATATTATATGGCCTAAAACTATGCCATGAGTTACCATGAAATGGCACTTCTCCCAATTTAGCACAAGATTTGTTT

CAACACATCGTTCAAGGATTAGAGCCAAGTTGTTCAAGCAAAGGTCAAATGAATCACCAAAGACACTAAAGTCATCCATG

AACACTTCGATTATGTTTTCAACATATTCCGAAAAGATACTTACCATACACCTTTGAAAAGTAGCCGGAGCATTGCATAA

GCCAAAAGGCATCCTTCTATAAGCAAAAGTACCAAAAGGACAAGTGAAAGTGGTTTTCTCTTGGTCCTCCGGCGCTATCA

CAATCTGATTATAACTCGAATAGCCATCAAGAAAGCAATAGTAAGAGTGTCCGGCTAAACGTTCAAGCATTTGATCAATA

AAAGGTAAAGGAAAGTGATCTTTTCTAGTGGTGGCATTCAATTTTCTATAATCAATACAAACCCTCCAACCGGTTTGCAC

ACGAGTTGGGACAAGTTCATTCTTATCATTCTCCATCACCGTTACTCCCGATTTCTTTGGTACCACTTGTACCGGGCTTA

CCCACTTGCTATCGGATATGGGGTAGATAACTCCCACGTCCAAAAGCTTGAGTATTTCCTTTTTCACCACTTCCATCATG

GGAGGATTCAAACGGCGTTGCGCTTCCCTTGTAGGCTTTGAATCATCCTCCACACGAATTTTATGCATACAAACGGCAGG

GCTAATACCCTTGATATCGGTGATTGTCCATCCTATGGCCGTCTTGTACGTCCTTAGCACCCTTAGAAGTTTTTGCTCCT

GTAATTCATTAAGCTCGTTAGAAACAATCACGGGTAAATCTTCATTGTCACCCAAGTACAAGTACTTTAAGTGACTTGGA

AGAGGTTTCAATTCTAAACTTGGAGATTTAACAATAGATGGCACAAGTTTCTCATTATTGCAAGGTAAAGAAGCATATGA

AATATTCTTAGTTGCTGATTTTTCCGGTAATGAGTCAAGAGAACCAATGGTTTCTTTCATCTCATCACTATACGCATAAC

CGTTGTCAATAGGTGTAGTGAGCACGGTTTCCAAAGCATCAACACCATTCAACTCAAACATTTGTTGCGCCAATGTGTCC

ATCATATCAATGGAGAAACAAGAGTGGACATCGCTAGGATATCTCATCGTCTCGAAAATGTTGAAACGTATTATTTCTCC

ATCAAACTCCATAGTTAGCGTTCCCTTATCAACATCAATTATGGTTTTCGCCGTTTTCATGAAAGGACGCCCAAGTAGCA

ATGGAAGAGACGAGTCCGGTGAACCTTCATCCATCTCTAACACGCAAAAGTCCGCTGGAAATACAAGTTGATTAACCTGC

ACAAGAACATCCTCAACAATACCCATTGGGTAGACATTAGAACGATCGGCTAATTGCAACACAATTCCAGCCTTTTTAAG

AGGACCAAGATCAAGTGAATTGTACATAGACGCAGGCATGACATTAACAGAAGCACCTAAATCCAACATAGCTTTGTTAA

ATTTGGTGTTACCAATTGTGACGGGAATGTCAAAACTACCCGGGTCCTTGCACTTAAGTGGGAGTTTGTGTTGTAAGATT

GACGAAGCATTCTCCCCCACACTAAGCACTTCATTACCTTTGAGCCTTTGCTTATTGGTATACAAGTCCTTCAACACCTT

TGCATACTTTGGAACTTGCTTGATTGCATCTATGAGTGGTATGTTCACATGGATCTTCTTTAGAACGTCTAAAATCTCCT

TTTCTAGTTCCGCCTTCTTAGAACTTGCAAATCTGCGAGGGAAAGGTAGAGGAGGAACATAAGTAGAAACCGGAGTTTTA

GAGTTAGGAATAGGTACCTCAGAAGTTTGTGGAGAATCCTCGGTTTTCTCCTCCAAAACAGGTTCCTTTGGTGTTTCCAC

CGTGCCCGAATCAAAGAACCGGGTTGCACAAGTTGTGTACCACTTCTCAATGTAATTGCATTAACACCCTCTTTTGGGTT

GATAGGTTGAGAAGGAGTTTCCCACCATTTTGTGCCTTCAATTGATTCAACTCCATAGCCATAGAACTAACTTGAGTTTG

CAACTCCTTAACCGAACTCTCCGTTCGTTGCATAAAAGATTTCAACCCAACTCCTCCATATTGGACCCCATTTGAGGTTG

TTGTTGTTGTTGAAATTGTTGTCTCGGTTGGAATTGTTGTTGTTGAGGTTGAAATCCACTCGGGTTGTTGAAACTAGGTT

GTTGAATCGCTCCTTGCTTATTGGCATAGCTAAAGTTGGGGTGATCTCTCCATCCCGGATTGTAAGTGTTGGAGTAGGGA

TCATACCGTCTTTGTTGATTTGGAAAGATCGCATTAGCTTGCTCGTCCTCATAAGATTGAGAGGAACTTGGTATCACCGC

GGCTGCAACTTGTTGAAGCATTGCGTTCATGTTACCCATTTGTTGACGAAGCTCTTCAACCTCGCTCACCTCACTCACCC

TTCTTAGAAGCACATCCGAACCACGACTCAAGAATTGTTGTGAGTTCGCTGCCATGTTTTCAATCAACTCCCTAGCTTGT

GCCACCGTCTTGTTAACTAGTGCACCACCACCCGCAACGTCGAGAAGATTCCTATCACTTGATTGGAGTCCTTCATAGAA

GTATTGGATGATCATAGCGTCACTGAATTGGTGGTGAGACTGACTTGCCACCAATCTCTTGTAGCGTTCCCAACATTCAT

ATAAGGATTCTCCCACATTTTGCTTCATCCCACAAATCTCCTTGCGAATTTGAGTAGCCTTTGATGCAGGAAAATATCTC

TCTAAAAAGAGTTTCTTCAACCCATTCCATGTTATGACACTTAGGGAGGTAAATAGAACAACCAATCCTTAGCACTATCC

ATTAAAGAGAAACGGAAAACTTTCAACATTGCACCATCCGCTGGAAGCATGGCCGTGACAATGGTGTGGAAGTCCATAAG

ATGTTTGTTTGGATCTTCATTCACCATGCCATGGAACTTTGGTAATTCTCTAAGCAACCCCGGTTTAATTTCAAAAGTAG

AGTCCGTTTGAATACACCCCTGTTGTATATCAAGCCTATGAGAAGCTAAGTCCTTGAGTGAACGATTTGCATCACCCATT

GCTCCTTGTTCTTCCACGTGTGTTTCCTCTACGATTGGAACCTCTTTTCCTTCTTCTTGTTGTTGGAAAATCTCTTCTCC

TACTTCCTCCTCGTGCTCGTCCGTTGTGTCTTGACTTGGTTGGAGTAATGTGCCGCGTCGAGTAGCTATCCCGCACCTTG

GATAATTCCAATCCTTAGAAGCCAGAGATTAGGTACCTGCAAACAATTCTCGCAAACAAGTAAGAAACAAGACAAGAAAC

AAAAAGCAAAGATGAAAGCGAGAAATTGTAAAAAGAAACTAAAAACTTAATAACTAGCCGTGTGCCTCTCCGTGCAATGG

GCGCTAAAATTTGATCATTTGTCGTATGCGTCAAAAATAATTTCACTTTCCTAAAGTATATGATAAGAAAGAGTTCGTAT

CCTGAGAGGCGATTGTAGTTATTTCTTAATCAATTTCTTAAAGTAACCACTTGGGGGATTTCTTATTTCTATTTAAGCAA

TAAAAACAAATTAAAAGCAAAGTAAATTGTAGAAAGTAATCAATGGAGAGAGATATTGGTCAAGGAATTCATCTTCAATC

TTAGACATGTGCTTGCATATATGAATAGAATTACGATTTAATCTCTTGTGTTATCAAAAGCCTAGAGAATCAAGAAAGCA

AGATAATCTATTAATTTCTCAAGTTCATCAACACACACATTAGGGCTGCGTATGTGAATTCTACCTAAAGAATAACATAA

CGCCTTGCGCTAATCAAGTTCAATTCCTAGGAGCATTAAGTTTCAGAAATAGGCTAATCAATGCAATTGTCATGCGTTGC

GCAAAACAATTCGGACAATGGTCAAACTCTACATATGATTACGAGAGTGTATCACTACAAACCGTAAACATATCATGCTA

CTCGTATTCAAGGTTATCGCTTGATGAAAAACCCTAAAATAGCTATGGACTAAGATGCAAGTTCAATTAATTGGCCACTT

AACTAATCAAACACCATAATCCTATCACAATACTTCAATCATGTGATCAAAGAAACCGCAGAGAATTGAACAATAAACAA

GAGAGAAAACTAATACATAAAGCACACACAAGTTTTAGAAATCGGACTACATCCTTAACCAATAATAGAGAATTTAGCTA

CTCATAGCTATGGAGTTCATCATCATCAATATTCAATAATAAAACAAAGAAGAAGAACAAAAACCAAACCCTAGAAGCGT

AAACAAAAACGGCTCTCCAAAGTCCAAGAATAAGATATCATATATGATAAGTTGTAGAAGACCTTCCTTTTATAGCTTTT

CTCCAACTTCTAAAATTAGGTCTAGTCTTCTAAGTCCATTATGTGAAAATCCACCAAGCCCACAAGAGGGAGAAACCCGT

GTAGAAAATATGCCCCAAAAACGGTCGCCGGAAGTCTGGGATTGTGGCCGGAACTCGACCGGAAAACAGCCCCGGTGACC

GGCAAAGTCCGCCCGGTCACCGGTCAACGGGTCGGATCGGTCGGGTCAACGGGTCAACTCGGTCGGATCGGTCGGGTCAA

CGAGTCACCTCGGGTCAAAGCGAATCGGTCGAGTCAACTCAAGTCGACCAGAGTTGTGTGAGTCGAAGTCGAGTGTCTCG

AGTTAAACCCAAGTCGACCTAAAACCGAATCGGTGTGTCGAGTGTGTAGCTGGTCCGACCGAACCCGACATGACCCGGGC

TATTACCGGCTCGGCCAGTGTGTAGCACAATCCTTGTGCATTACTACCATACGATGGCTACTCCTTCCTTTGCATAGCAA

ACTCAAACCCATTCCAGCTCCAGCCATTGTGACCATGGCAGCATCACAAAACATTCCCATTTCCATCTCAACTTGCACTT

TCTCCATTTGCAGAAACATCAAGCTATGGCAATCAGCTCCAATACTCAATTCCAGTTCAGCTCATAACATCCTGCAAAAC

CACCTGAGTTGCAACTCCAATCCACCTCTACCACAGTTCTTTCATATTAACCTGCACAATATACACCTCCCTGTAAACTT

CCAACCGCCTAACTCCAATTCCACGGCTCAAGCCACACTGCAAAGATCTCGCTATCATCAGCCTCAAATCCATTTGTATT

TCATTAGAATCACAGCAACACCACCAGAGACCTCCTCCATTCATTCATCTCAGACCCATATTTCCAGCTCAGTTCCATAT

GCTCATTTATTCACAATACAACTCTTCCATGATGTGCACGGTTGCACTAATTCGACACGCTCGAGCCAATCCACCTTGCG

GTCTCTCTCGCTTCCATTTGTCATCACTAGCACTTCTGTTTCGATTCAACTTCACATTGAGCCCAAACCTGTCCTAAACA

TACATCTAAGTCAGTTAGCTCAATTAGCTCAAAGCCAAACATTCATCTTGTCGCCAAGCACCAACTATGTATTCGTAGTA

CCCTTTGCCTCATTTCATAGCTCTCCAATTGCAACATTCAGTTCCTGCAGCTCTTGCATTCCATGTAACTCATTCACAGG

CCCCTGTGACAAAATCACCAATTGTAACTACAACTTCTTCAGCACACAGAACCCATTCAACACAGTTCCTTGCACCAGTC

CATCCAAGTGTCATGATCAGCTGCATCATTATCACCACATCCTTCCAGCTTCAATTTGCCTCACTGACTCATCTGAGCTG

CAACCTTCAATTCCAACCCATTAGTGCAACCTAATATCCTCCATCTGCTAGCCACACACACCAATTCTATCTATTGCCAG

CTCCACCAGAATCCATCACCATCTGCATCAAGCCTGCAATCTCCTAACCATTGCCACAGTCTTCATTCAACTCATGGTCA

CCTGCCAACTTCATTCTGATGCCCCTGCAACAACAACCCATCTTCATCTGATATTCAATTCTTAGCTCAATAGCAAAGGC

TTCATGTCTTCCCTGGCAAATACACAACAACTTCAACATCAGGGCTTCCAACTGCAGCTCATTGCCTCAGTCCATTCTCA

ACAACTCCCTTATCCCGCTCACTCAATCCTCGAGCACAAACCACCATTGCAGGACACTAACATCACCGGGTTAACTTCAA

CCACTAGAGACTCAACCATTTCAGACCCCTTTCAGTTCATCCACACTCACTGCCATCACTAGGACAACAGATAATCTCCA

TCTCAATCTCAGAATCCCATCTGATTTCAATAGCTGAATTCATCTCTGTTTCAATTTGAATACAATTTCAGCTCCAAAAC

CCCTTCATTCACCGATTCGATCCAAACCCCATCTCTGGTTCGTCGATCTGAACTCATTGTTAACGAGTCCAATGATCCCA

TTTCATCTCGGTTCAGCTCACGGATCTCGAACCTCACAACAACAACCAAACCCTAGCTTCTTCCATTCAAATCGACCATC

GAGAATCATCTCTTGCTTCCGAGTTCATCTCGCTCAATCTCAACATCTTGACACTCAATTTCTCCTCTCTCGCATTTCTC

CCTTTTCTCAGACAAACAACAGCCGTTTCCTCTCGGCTCTAGCTTTGAGATGATAATGCTTGAAAGAAGAAATAATACCC

ACAGAATTCCAGGGTCTTTGGTAATGAGTAAAACTGATATAGGCACCAGTTTTCGGATGAGGGCTACATGTTGATATTTG

GCTTGTCAGTCTCAGCAAACTGACTGCCTAATCTGTTTCCTTGCTCAATTGTCGGTTGCAGGGAACTGACAGCTCATTCC

TTTCCTCTTTTGCTGACAGCTAGCTCTACGTTTCGCCCGCCTGTGAATTGACATGTTTTCTCCTTCTTTTGCAAAAAAAA

CATAATGACTCCATTGCACCTAATAGCTCAAAAACACAAATAAAGTACATAATTCACAAGAAAAATAGCAAAGAAAGCAT

AAACTCTATATAGTAAATACGGTGTATTCTACACCTATCAGTAAGTTGGGTTCGTAACTTCGTTTTGTGCTCATACTCGG

CGGTGGATCATCTTTTACCGGTGGGTTTATTGTGGCTTTATTTTTGGAATTTGATTCATTGTTTTTGATTGTTTTTTTAT

GATGGGTACTCTTGTAGACTTGTACTCTTTTTCAATTTTGGATTATGTTAACAGGTTTTTCTTGATTTTATTTTTTTCTT

GATTTTATTTTTTCTGTTGCAGCGGCTACTTCCCCTGGTGGTGCTGCTGGAACTTGGTAATTGACTAATTGGTATGTAAT

TTAAGAGACGCGATACCTTTATCTTTCTTAGCCACCAACAAGACAAAGTGGGAATAATCAATCAATATGAATATATGATT

AACTAGCTCCCCTCTTTTACGTCAGTATGTACACTTCTTAATTATCTCATGTGCGCCTCCAAAATTCTTAAGAGAAGAAT

ATAGTTTTCACTTTTCACTGCCACCTCAAGGACATCTTTCATATAATTGTCAGATGCAATGGCATTGGTACTCTGAACTA

GAAGTTATTTAAGCACTTGAGAAAAGATACATCTGACTGATGGTGTGTTGGGTATTTTTAATGTGATAATTAAAATCTAT

TAGTTATTGTGCAAAAATTAGATGCCATGTGAACGTAGTAACGTCTTTGTTTGCATCCATATTGGTCATTCTGTGTTGTG

AAATTTATGAATAGTTAGTGACCCAATTTTGATTGGTGATATGCGATCTCGACCAACTATATATACAATCCAAATTTTGT

ATCTATCTATGATTCAACCAATAGTGAGTTTTTATGTTAGCTTGGAATTATGAGCGAGACACTATTTCATTTCATGTTTC

CAAAGGGGGACGAACCCACCCATGTGACAATTTTAACTAGAGGTTTCATTTGCACCCACTCCGTCCAAAGGTTTTGAACT

CGCACCTGCTTAATTACTTAATCAAATTTAATTCCTCAAAGACCATATGGTGTTGCGATCTAACTACTGCTTTACTAGTT

AGAAATAAGATGTTCTGATTTTTTTTTTGTATGATTAACCTGAAAAACGCTTTGCCTTAATCTTTCTGTTCACTGCTAGT

GTTCAAGATATGTTATTGTTGTATTATATTTATATGCCATTGTCCAATGGATAGGATTAAATCTGATTTTGACCTTTTTG

ATTTTCTTATAGCTTCAAAAAATGTCCAATATTTGGGCTCTCTTTGATTTGGTAGAACAAGCTGATGGTTCAGTAAAGGC

GATGTGCAATGTTTGTGGCCAGAAATATTCTTATGATAGTCACAGAGGTGGAACTTCAACTTTGTCAAGGCATAAATGCC

CAAGGCGTGAAACACAAAGGGTGGGACAAATGTTAACTGGAAAGAAAGGACAAGTGTCTACTCGTGCACGCAAAATATAT

TAGATGAAGTTCCAGGAGCTTGCAAGAAATCTTCCATCAAGTTTAGTGGAATTGCCAGAATTTAGGGACTTGTGTATTTG

TGTGCCTATCTAAATGATGATGTTAAAATCCTCTTCACTGATGAAATACTGCAATTCCAATTCTATTTATAGTCTTTTTA

GAGGTTTAGTTTTCTTTGTTAAATAGTTGAATGCCAGTGATGGAGGCTGCTGGACTTTTGTTATGCTTATTGTTGTAAAA

GTAAACCATTGTAAGTGTATTTGTTTAGAGTGGTTTCAAGACTTGCAGCTGTAACTTACCAGTAAGCAGTATTTCAATTT

TGGAAACATAAGAATTTGTGCATACGATAAAACCCAGAGCCAACCCGTCAGCCCGCAGAGCCAACCCGTCTAGGGTTAGG

GTTGAGGTTTCTGGACCCGTTCTATAAATTTATAGGATTAGGGTTCAACATTATCAACCCGTAACCCGTATAACCCGCCA

CGGGTTGACCCTAACCTGGATCGTTACCACCTCTAGTCAAGACGTTTCTGTGAGTTTGATTTTTCACAGTACTTTCTGAA

GAGTGTTCATGTTATCATTTGAGATATTTAACAAGGAACAATTTTCTTAACACATTTGAAGACGAGACGTCTTTCAAACC

CTCTATTTTCTTTATCGAATCGCATTCATCTAAAGTCCGAACATATTTTCAGTGATGGACCGAAAAATTATGCCTGGGGA

GCCTGGCCGCGCAGCCATAACTTTCCCGATGCGCCATGATGATGCTACTTACCGGGTCTTAAGACTAGAAAACTGCACGT

TCATTTTCCCTTGCAAGCATGCTCATGGAGCATGATGCGGCTAACTTACTGTGACATACCAGAAAATTATATATGCCTGT

GGCACCTGGGGGCGCAGGCCATAACTTCCCCGCTGCGCCATGATGATGCTACTTACCGGGCCTGAAGACTAGGTAAATGC

ACGTTCATTCCTTGCCTCAACGTAGGGAATTCATTACTGAATTTCCTATCTAGATGAGTAACCGTCATACTGCCGAGTTT

TGTCCAGGCGTAAGGCATGCCTTGGACTTACACACAGGTCATTGCAGGCCTAATACCCCCTTTTCACTTAGATTAAGGAA

TTTCCAAAAGCATGCACTTGTCTTGCTCGTGCCGAGGGTGCATCAATCAGGCTTGTGCCGTACTTTCCTTAGGCTTATGC

AAGCTCCACCAACTTGCATATCAATGTAGATTACTTTTAAGGCTATGCCAATGCGCCAATGGTACATGCCTAAGTCGAAC

GCCTTGATAGGAAGTATGAGCATCCAAGGAATGTAATGCAACTAGCCTCATCAAGTTTGGCCACAATCATCTCTTGCGTC

GAACTTCCGAGCCAGATGATATGAGAGGCCAATATGCCGCAATCATCTCTAAATTAGATGATATAAGCAACAAAGTGTTG

GACCGGCAATGCTGCGTTGCGGCTGCCTATGTACCTTCCCTACTCTAAAGGGATCAAGCACAGACCGTAGTTCATCCTCG

AAGGGATAGCAACTTAAACCAACTCATTTGCAAGGTCGCGGCCTTGCAATAATGATAATTGCCTAGTCCAAATATGCCAT

TTGTTCAAAATACACAAAGGTTGCACATTATCAGGTCCGCGACATGGCATAGTGATGCCTAAGCATCAAATGCCCTATTT

TCAAGGCTACGCAGCTATAAATGAGGCTTACGCATCATTTATACTCTTTTGGTTAAGCTATGAAGCTTACTTGGCACATA

GTCCGCATATCCACCTTCAACCAACTACCCCTCCCTATATATGTTTATTTGACATTTCTATAACTTTGGAAAGGAAAGGG

GATTAACATGCCTGCTTCACTATTCATGGCCGTTGCCATGTATTTATGAATAACCGGCAATGATTCCTGGGTTTACTTCA

GTTTTGCACAATGATTGCGCAACAATTACTCTGGCATACTGCACAATGATTATGCAACACTCGTTCTGGCTATTGTACAA

TGATTGCGTAACATTTACTCCGGCGTATTGCACAATAATTGTGCAATATTCACTTGGCCTGCCCGCTCTAGCATGATGCA

CAATAGTTGTGCAATATTGGCTCTAGGCCAATAGCCTTCATAATGCGCAAAGGATGCGCTGTAAGGCTCTAGCCATCTAA

CCAACTGGCCTAATGCATCATTTGATGCGAGATTTGGCCCTTTGCCAAATGCCAAACATAATGCGTCATTTTGGCGCGAT

ATTGGCCTTAAGCCAATCTACCTCTTACCAGGCAACGAAAGCTTTGGATGAAGCTTAATTCCAGGCCATGGCACATCTTT

TAGCATGCGCCATGCTAAAAACACGGAGTGTTACATTTTTCGTGTACAGGACAAATCTATTATCTTATTTATAAAATTTC

CCTTAGGAAAATAGTATTTTGGTGTCGAGTCTCATAAATGCTTTATGAAGAAAGAATATTCTGTTTGATTTCAGCTTGCT

TGTTGTTGAGGCTAACCTAACCGATTTTAATTGTAAGGTATGTTTCTTTTGGAGGACAATTTTGAGAAAGAGGTTGGATC

GGTTGTGAGTGGGGCTGATCAGGTAATCTGACTCACCCATTTGGTCCAATAAACTCGCTGTAAAACTTCTATCGGTCCAG

TGAGGCCGATTTTGTGCACACTCCTTAGCATGTTGTGTTCATCCTTTGATTTAATTCTGGGATGTTTCAAAACCCTCTAT

TTTCCTAGCAATTTGCATTAATCCTTGGGACAAAATGTATTCCATGGCTTGTTCTCAACTTGGGAGCTTTATAAAAATTT

TATTCAAAGAAAGTTCAATACAAGACTTAAGCTAACCTCATCAGGTGGGTAAAGAGATTAATGCTAGAAAACATAAAAGA

AAAATTGCAGGAGCCTGCTGTGACAATATGTGTGATGCAGGATAGCTAGTTGCAAAGAAACGAAATATCCCATATTGACT

GCATTAGGGGATTAAAAACACATCTACCACCGCTGGGATTCTATCTGACATTATGAAGGAGTGAAGAGGAAAAGAAGAAG

AATGTTTCTTTAGTTTAGGGCTTGAAAGGAGCAAAGACGACAACTGAATTGTGGCCGCCGAACCCAAATGAGTTAGATAT

ACCTGCAGGTAATGCACTCAAATAAGTTGATAACCAGTGTAAGCTATTCGAACAGGAAATGAGCAAAGGAAAGCTATCAT

GTAATAGGCATTTTGATGCCTCTGTGGCAAATCTTAATGTATTTAGAAGTGAATACATACCAACGTGAACTTCATGCTGC

TTCTTCACATTTGGAACGGTGTCAATTGTAACGTCAGACTCCAAATTCTGCATCAAAATCAAAACAGTGCATATGGTGAA

ATGCAGTCAGAAAAGCAAAATCAAATTAAAAATAAGAGTTAAAGAAAGTTAGTCTTAATGGAGTTATACATCTTGGTTAA

TGGTTGGATGCAACCATCCAGTTGTGATAGCTTTGATTGTAGCTATGGCTTCTAATCCACCAGCTGCACCGAGCCCATGT

CCAATCATCGACTGAAACCAGAAAGACATACAAAAACCTCAAATCCATTATCGTTGTTCTTTTAGCATGCTAGAATGTTG

CTGGACTGACAAAAGAATGGCTAGAGCTTATCATTGTCAGAAAGTAGAAACAGTGTGTCAGAGAATGAGCAATCTTGCCT

TAGTTCCGTTCATCTTGATCTCGGATGTGTTCTTAAAGACCTTCTTGATGGCATTAACCTCAGCTAAGTCTCCCGCAAGT

GTTGATGTTGCATGAGCATTGATATAATTCACCTAAAAATGTCAGATATTCATTAGAAATTTGATCAGTTGCCAAAAAAA

TGCAGGAAAGTGTACTAAGACTAGTATGTGAATATATAACGGGCATCATTTATTCCCACCTCTTCTGGAGAAACTCCAGC

ATCCTCCAAGCTCTTGTTTATGCATGAAGAAACTCCAAGACCGTCTGAACGAGGATCTGTCATATGATGAGCATCACAGG

TAACCGCGCCTCCCAAATACTCCGCTATTATGTTAGCTCCTCGTTTCATTGCACTCTCTAGGCTCTCCATAACCTACTCA

ACAGAAAGAAAAAACAACAGCCCAACCAATTCAATTATAACAGCATTCACTGCCTACAGGGAACCAAATGCAATATACAG

AAACTGCACGACACAGCTCAGAGCTATGTTCTGTGAGGGCAGGGGGCGCCTCACAGAACATAGCTCTGCGCCTAGTGCTC

GCCTTTCACATCATAGGCTCAAGGGGTAAAATTGGGGTAATTACCAGCACACCAGAACCTTCACCCATAACAAACCCATC

ACGATCTCTATCCCACGGCCTTGAAGCTTTTTTGGGTTCACCGTTTCTCTGAGACAAAGCTCTGCAAGCAATGAAACCAC

CAACACCAATTGGGATGATAGCAGCCTCAGTCCCACCAACTACCATGATATCAGCTTCACCTCTTCTGATATGATTTGCG

GCAGCATAGAAGCAGTAATTTGCAGTGGCGCACGCAGTGGAAATGGAGTAGTTTGGACCCATTAGACCCGTATCAATAGC

AAGTAATGCCGATCCCATGTTCGTAATGGAGTACGGAATGAAGAATGGAGTAATCTTCTTGTATCCTTTCTGGAGAAGAG

ATTCAACTCCACTGCTGAATGCAGTTAATCCTCCCATTCCTGTCCCTACTAGAACTCCAATTTTCGACCGGTCCATCTGT

TGCATACGCAAAATCCATGCGCATTGCAGTAAGCCAAAGCGTATATGCAAAAACGAACTATTTTACAGATTGCAAACAAG

GACTTACTGATTCGAGAGTTTCGGTTCCAAGGTTGGCATCTTCAAGAGCTCTTTTACCAGCAACTAAGCAGTACCTCCAA

CAGTCATCCAGACGGCGGTCATTTTTCCCGTCAATGTAACCTTCAGACGAAAAGTTGCGAATCTGACCAGCAAATCTGAC

AGAAAAGGGTGAAGCATCAAACCTATCTATAAGACTAATACCACTCTCTCCATCAAGTAATCTGTTATAGAAGGTATCAA

CATCGTTCCCAAACACCGAAACAAGTCCCATTCCTGTTATAACAATTCTCTTCTTCGGATCGCTCTCTCGTTTCGGAGCT

GATGGGATTCGCGAAGCCATAGCTCTGACTTGCCTACTCTTGAAAACTGAACACATACAGGAAATGCACAAAATCATATT

CAATTCAAAGATGCATATACATTTGAACTACCGCACAATCGGAACAAGCCTAAATCCCGTCCTGGGCATCCACTCGAACA

CCTAATGTGCAAATGCGCATCTAAAAAAACACAAAAACCGTACCTGATTTCGCCCGCACAGAACCAATCTGCTTGGTTTC

AAAAGGTCTTATAAGACCATTAGAATGAACCCCGGATGATCCTAATTCCTTTCTGAGTAAATTTTTGTTCCCGAAGAGAT

GAGTACCAGTAGAACAAGCACTACCAGTCATGGCTGCCATTGTTACACGAACTGATCAAGAATCCGCACAATCGAAATTG

TGGAAAAACAAAATTATGGGGCGATCAAGAAAAAGATTAAAAATGGGGAGTATTTTAAAAGAGAAATTTTCTAACGAAGA

GATGATGATTTAGGTTTGGAATTCATTGGCAAGTCCGTCACCTAACTTGTTATCACCTCCATTTGACGATGTTTACAATT

TTCATGGTTTTTATTATTTACATCACAATGACACCATTTGTATAACACACAAATTTCATTTTGTTCAACAGCTTTAACTA

CCACTTGTATGGGATAACTATCGTATACTTTTTCCTACTACTTTGACAAATAGTGTCCCGATAAATCTTAGCCGTTAATC

GGAATTATTTTTATTTAAAGGCTTTTGTTAACTTGTATAACGGTATACATGTTAAGGTTTAATAAAGAATTATTTTATTT

CTTTGGAAATTGTGTTTTTTGTCTTATTCTACTAGAAATTGGTTTTTCTTGTACTAAAAACAACAAATTTTGTTCCCAAT

GCAAAGAATGGTTGTTTTCAATGCAATAAAGCAAACAAATCTTGTTTCCAATGCCAGAATTTGTTATTTTAGTGAACCTT

AACATGTATACGGGTATACAAGTTAACAAGACCCGTATAATATACTCGAATTAATGAGATTATGAGGAGACATGTGGAAA

TAATTAGGAAAAGGCTGGTGGCAACTGGCAATGAAGCCATTGTCTTGAAAGATTGATTGGCGAACCTAACTCCATTAAAT

TTGGTTATTACTCCTACTAATAAGACCAGACAAGATTTTTAGTAGATGGGAGATGTGGTATGTCCATGATGCATCCTTTG

CAACTGGCCTAGTAAAAGCATTTGAGTGTCTCTTTAGTTTTGGGACTAGTTAGGGGCAAGTGAGGTTAGTCAAGCCAAAC

GAGGTGTTAGTAGTTATGGAATGAGAAATGAGATTCTAAGAATGCACGGATGTGTGGCATGTTTCAAACCCCAATGATTC

GCATTCACCCCAACAGATTTACTGTTCAAGGATGATTAGGGTGTTCAAGCTAAGTTTTGGTCCCTTGGCAATTTTTGAGT

TACAAACAAATTCTGAGCCACACTGCAGATTTTGATTCAAAGAAAAATGTGCTTGTGCATTGATATTTTCCGACAACTGA

AAGAAAAATAAATGTTTTCAGGTAGCAACACATTTGAATCCTTGGCAAATGAACAACCATGCCAAGAACAAAATACAACA

AATAACCAAATCTACCAACAAAACTATTCTAACATTTTCTCTCCATCTCTTCCGAGCTTCTTGCTGCAATCTTACTTCAT

CTACGTAGCTACATTCTCTGCTCGACGTGCCTCGGTTCCATGCATTCACTCCTTGTAATACGCCATGGACGAATTCCTCT

TCTTTATGATCATCAATACTTTTCTTATTCTTCTTTTTGTTCTTTATACCTATATTAATCTTATTCTCTGATCTTCCTAG

CAACGGTGTCGAACAAATTTTACTTCCTGTAGACGATCTCGAGTCAAAGCTTTCGTTTCGACTAGCATTTGATAAAGCAT

CCGGTGACGATACGACCGATAATTCATCATTAAAAGATACTAATAGCTCTCGGAAAATCGGACCAAATTCTTCGTTACAG

CAACCGGGAGCCATGTCTATTGGCCCTTGTTCATTGTTGCAGAAGTTTGCAACGAACGCGTCTTTAACAAGCGCAAGAAA

GTCAAGGCCGCGAGACAGTTCCCGTTCGCTGCCATCCACATTCGGAATAGTGTCTTTCTTAAAGATTGCAAAATATGTAA

AATGACCGTCTATCAAGAAACTATACGTCCTGGTTGAAGTTGTCTGGGAGAATTTTTGATGAAGTTTTGGAGCTTTTTCA

AGATATTGCGAAGCTAGTTTTTCGATATCGGTATCACCGGCCATGTTGAGTTCGGCTAAGATCATCGTTCCTTTTGAAAT

ACAAGCGTAAACAAATTGATTGTTTGGCTCCGCCGGAATCATTTGATATTTTGTAATGCAAAAAATAGAAAATTAACGGA

CAAGCAGAAGAAGACTTATCTCATGCATGCATGGTTTCTCCGGCAAGGAAAAGCTTCCGAGAAGAATGAACGAACATTGA

AGAGATTCAAGAGAACACATGAAATGAATTGGGGCGAACGGGGTTGTACTATTTATAGAATTTTGTTAAGATCTGAAAAT

TTTGATCAACTGGTATTTAAAAAATGCGAATTTTTCCTTCTGATTTTCTTAAAAATCTTCAGATTTGTGGAAGTTTGGGA

GGGGTAATTTAGTTGACAAGAAATCCCATGAAAATTGAAAAAACCAATGAAAATTGGCGAGTCTTTTCACAAATGTTGCT

TGGGTTATGAAATCCAAGGACAATTTTCAAGAAAAAGAAAACAAATGAGATGAGAGATAATTTTGTGAGTTGGGAAATAT

TTTGCTAAAACTTGAAGGCTTGAGACCAAGACATCTTGTAGCGCGCTAGACTTGGGATGCACCAGGACTAGCCGAGAAGG

CTTCCATGAATTCAGCATCATTCTCCATAGCCTTAAAAACCTCAATTGGAAGGCAAATTTCAATTTCAACACTTGCTTCA

ACTAGTCCTGGATTCACAGTAGTTACTCCATAAGATTTACCGTTTGCACCAGTTTTTACAGCAACCACCCTAAATCGTTA

CCATACATGTTGAACCACTTGGAACTCCTCGCCTATGTTGTAAATGGCGAGCGCCTTTGGCGCTTAAGCTCCAAAGGCGC

ACTAACCAGCGCCCGAGGAATCAAATTCTACACTACATTCCACTCTGGCACCTTAGGCGAGCGAGTATGGCGCTTTAGGC

GCCTCGAGGAAGTCTTAAGCTCTAGATTAAGCTCCAAAGGCGCCATGGTTGGTGGTCCTCCGTCTTCTTTTCGTTGTTAT

TTCGTCCTTACTTCTTCAGCATCTCTGCCTCCATTCTTCTTCATCATCTCGCCAATATCAAAGTATAGAAACTTTATATA

CGGAGAAATATCAACATCAATCAACATTGAGTCATTTACAATCAATGATAAAAATATGAAAATAAATAGTAATCAAGTTG

GAATAACGTGATTAATAATCAAGTTTTAAGGAAATAACGTGATTGAATATATTTTCTTTCTTTGGCAGAAAGTCTCGGCT

TTTTCTCATTCCTTTTTCTTCTCAAGTGGTTTCCTATTTCTATACAAACTCTCCAATTATTCCTCCGCCAGCAAATCCTG

GATTATAACTTCCATAATTAGCCAAACAGGACCGAGATTCTTCCATGTGTCCGTGAATATCTTCTTCTTCCCTTTTTAGA

GCATCTCCAATAGAGTTTGGGTCTCTAAAAAAATTGATGCCACATAGGATTTTAGATACTCTAACATAAAAAATTCTCTC

AAATGGTGGTTGAATAGCAAAAAATTAGCATGAGGTGGCACTTTGAGTTTGGATGAGAGTATCTAAATTTAGATACTCCA

ATCCATACCCACGTCATATATTTTTCTGTTCAAAATTATTTAGGCCCACATTTTAAATTAAGTAACCTATAACTAAATGA

CATTTGTATTAAAATATATATAGATACATCTACCATTGGAGATGCTTGTTAAAATAGGATTCTATTTACCTCTTTTTACA

GCTGTTTCACATCTCTTCCGATCCTCATTAAATGCACTGCTTTCTTTTGTAGACTTGATCGTCAGTGTGATTTGACTAGG

GCTTACACGCGTCCTTCAAGAGGAGAGGCGGTAAATATTCTTGGTTTTGATGTAATCACCTTTTACATCTTTATTCGGTG

TATCTATTCTATCATCTTATCTTATGATCCCCTCCTGTGATGCTTACACGGGGCGATTATATATTTTCTCTCACATCCCG

GGTATCCTAAGTCATGACTATACCAAAGTCATGCATTTCAATCATGGTGTTGCGATAATGCATGCAAAGAACAACAAGGT

GCAGGTTCTTTGGCATACAAATAATGCGAGTCCGAGAAAGCGAAGTTTTCAATATGATTTGGAGGCCAAATCTAGTGAAG

ATGCTGATGTTTGCAAAGTGCAGCAAAGGTTATATGTTTTGACGCATACCAAGAGTGGTAGGCGCATATGATTGAAGAGT

AAGGCCAAGTCTCGTATAAGTCCTGTTATGTTGCGTGGTGCAACAATAGTCGGAAAATGAAGGAAAAATACTATAACAAT

GATCAGTGTGAGGAATCATTGCTTGCGTGTACTTAGTCACAAATGATTCAAGTGAAATTCAAGCCTAATTCATGGAATTG

GCAAGAGGAATTTCTAGTTTGCATTTGGGCGTTGATATGCGATTGTAGTGGGAGTTGGATCGTAGGACAACGTGACAACA

ACAATACAAGTCAGTAGGATCAATGACAAGGCAGTAATGCTAAGTGATGGCATATGGTACTAAGCATAAATTTTTGGTAA

GACAAAAGGCCAAACAAGTTTGCTGATATCTGAGATAGGCTTGGAAGCATACAAGGCCAATGAACAAGTATAGCGAGTAT

ACTTTGTTGTGTTCAATGAGGAAATTGAACAGATTGGGTGGGGGAGGTCTATTACAGTCCTCAATAGTGACACGCAAAGA

TTGCGCAAACATGAGTTATTGGCCTAGCCAATGAAGCTGCATATCTCGACGATGTAGCGCCAACATGGATGAGACAAGAA

AGGATACTTAGCACATGCAATACTATGCGATAGTGTTGCCTATCCTTGATTCATAGTTGGCCCAGTTCAAGGAAGGAATT

AGGGATGCAATACAAGTCATGGGATGACTTTGGCATTGGTCCAAGCATAGCCAAGGCTTGGCCAATGCATATACAATAAT

GGCATTGCCAAATACTCCATTGATGACTATTGCTTGGCAAAAAGAATGTATGTGTTGCACTTAACATATGAAGCTGAGCT

AAGCAACAAAACAAGACATGATTGGCATGACTTTACTTGGATGTTGGCATCCAAATGAGCTAAGTGCATGGCCAAGAGAA

ATGTAATGCTAAGTTGGATGCATATTGGATTAGGCCAAGTTGAGTGCAGCGTGATGATGGCCAAGGCCAAGTTGAGTGCA

GTGCGACATTGGCTGGATGCTGGCCAAGGACAAGTTGAGCGTAGCGCATGTTGGATGGATGTTGGCCAAAGCTGTCAGTT

GCAAGTTGTGTGTGTATCACTCGCATGCCAGAGCAAAGAGTGGCTTGAGACATTTATATAGTGGCTTTATAACCGAGTTT

GGCAAAGCCTAACTCTGAAGAACAAATGTGGCATCATTTGGCTTGACAACCCAAAAGTTGACAGTTAAAAAGCATATTAG

CGGTTAGGAAAACGACCTTATGCGATACTTGGCTGTTCCATGCGTGTGCAAGTGTTGTGCACGGTTTTTGGCTATTTGTT

TACTATATAAATAATGTAGGAGGCATTAGAAATCAGCAGGAGATAGAAGGCTACAGAATTTGTATAGTTAATTAGTATCA

TGTTTGAGAGAATAAAGCATCATCAGGTGGGAGATGGCCTGTTTTGGTCCATGTGATGGCCAAGTTTTGTAATCTTCCTT

TAGTGCAATAAAATGGGTTTGCTGCATTATATATTTTTGTTCTTATGTTGCTGATTTGTGTGAGATTTTCAAATGGAATT

GATGCATGGCAAACCTCCTATGCTAATGCGGGCATAAAGTGTTATGGAGAAAGAAGAAAAGACTTCTGTTGCGTAAATCC

TTAAGAGTCAAGGAAGATGCATACTTTGATGCAAGTGTGCAAGGCTGAGTACTTGTGGGTGAAGTTGATTCACTGAACCG

CAGAATATTGCCGGTCTTGTCCCATAAAAATTTTAGGCCGTTAAATGAGCATGTGTCAGCTAGCTAGATAAGTGTATAGC

GCATGCATACGATATTATATTTCTATAACATCTAAATTTAGAAATTAAACAATAATACTATAAATTACAGGTAGCCATAA

ATTATGTACTCTAAAGAGCGACTGCAGTGGTGCGACTAAACCCAAAGATCAAAGACCAAAAAAAAAAAGACCAAATTTTA

GTTTGGTCCGTCGTGTGACGCAGTGGGTGGCGAGTAAATTTAGTCGAGCGGGGAGTATAAGTTCGCCGGGTGTGGGGCGT

AAACTTTATCCCCGCCCGACAAAAAAAAAAAAAAAAAAAAAAAAAAAAAAAAAAAATGGGGCGAACATATAATGTCCGCC

TGGTGTGGGGATCACATATAATGTTCGCCCCACACCAGGATCACATATAATGTTCGCCTGGAACAAAAAAAAAAAAAAAA

AAAATGTGGGGCGTAAACTATATCACCGCCTGGGTGTGGGGCGTAAACTATATCCACGCAAAAGGGGGCGTAAACTATAT

CACCGCTTTGGTTTTGGGGCGTAAACTATATCACCGCTCGACTATATTTGGGTTTAGTCTTTGGTCCCGACCAAATATAA

TCGCCCACTGTTCATTTAGTCTGGTATTTGGTCGATGGTCCGTCCCGCTACGCCTGGTTCAGAGACCAAATAAATGGGTT

TAGTCGCCCACTGCAGTCGCTCTAAGGCGCTATAAAATCGAAAATGCTTTAGATACCGACCGAAAATCTCGACAATTATC

CAGCAAGACAGAGGGATTGGGGGTTCCACTTTACATTTTATATCTTAGATGCTAGAACGGCTGCCTAACTCCACACGATC

CCCTGTCGGGATGATTCAACGGGATGTTTTGTCGGTCAGTGTAGCATCCCTCTCTATAAAATCAATCAATTAAAGCCACA

AAAAGTGGTACAACCGATTACCCATCACAACCACCTTCTAGCTAGTATTGGGGTTTCTTTGGAGAAAATTAGACTACCCC

TGTTTCACCTTTTGGTCTTTAGGTTTAACTGTTAGACTTTTACATTTTTACTGCTTTATCTTTTAACTTTTGTGAAAGGA

TTCACTATTTAATGTTGAGATTATATTAGTACCATATTGTCTGGGCAATTTAAGATCCTGCGGTTTATAATCCTTTGGGC

CTCTCCACTCATTGCCAATTGGTTTTGAGTTGGATGCCCACATTCTAACTTGGTATCAGAGCGGGCTATTTGTGACGAGT

CATTAGACCTCGCCTTTACTCTGACCCAACGAGGCTACACGTGAGGAGGAGTGTTGAGATTATTAGTCCCACATTGTCTG

GGAAATCTAAGATCCTACGGTTAATAATCCTCGCCTTTACTCTGACCCAACGAGGCTACACGTGAGGAGGAGTGTTGAGA

TTATTAGTCCCACATTGTCTGGGAAATCTAAGATCCTACGGTTAATAATCCTTAGGGCCTCTCCACTCATTGCCAATTGG

TTTTGAGTTGGATGCCCACATTCTTACATGGTATCATAGCAGGCTATTTTTGTCGAGTCATTTGACAACAACTTTACTCT

GCGTCACCCGATTTATTGTCCACGTGTTAGACCTAACGAGGCTACACGTGAGGGGGTGTGTTGAGATTATATTAGTCCCA

CATTATCTAGGCAAATCCTGCGGTTTATAATCCTTTGGGCCTCTCCACTCATTGACAATTGGTTTTGAGTTGGATGCCCA

CATTCTAACATTTAACTCGTGTTTCACACTATCTCCTCTATTTAAGTCTCTCATGTGACTCTTGTAAGGTCACCGATGAT

AATCTCAATACGTTAGGGGACGTGTTGAGATTATATCAGTCCCACATTGTCAGGGCAATCTAAGATCCTGCGGTTTATAA

TCCTTTAAGCCTCTCCACTCATTTCACTTATTGCCAATTGGTTTTGAGTCACTATCTCCTCTATTTTAAGTCTCTCTCAG

GTGACTCTTGTAAGGTCACTGATGATAATCTCAATTATATGAAGTACTTTTTCTTATCAACAACTTCTAACACATCAACA

CTGACAATTTCATATATTTAAGTATTTCATGTGACTCGTGTAAGGTCACTGATGATAATATGAAGTACTTTCTCTTATCA

ACAGCTTCTAACACATCAACACTGACAATTTCATCAATCTTAATCAGGAAACCTCTGCCATTTAATCCATAACTTAAAAT

TTGGAACAGTATGTTGCCAAATTCCAAAAATTATGAATTTCCTAGCATGCTGTGTTCGGCTTATTACAACTTCTACTAAT

GTGCTGGAAAATCCGGCTATTAAAATTACTAGATAAAATCTAATGCATTCAATAATCTATTGATTTTTCCAAGATAAGTT

AGCAAGAGATCCATTAACAGTAGCAGCTACATTCTTTGAGTTTGTCTCCAGTTTAGGTATTTTATCTTTGTTGTTACTGC

CTAGCACATTGCTTTCAGCTTCCTCGGCCGTCCGTCAGATTTCCATGCTTGAGATGATATTCAATGGGGTTGTGTACGTA

TTTATAGAATTCCATTTTTTTTTTATAAAGAGAAAGAAAGATTACTGACTGCTAACAATCATAAAGAGGTTTTGTGCCTC

CCAAGTTTGAGGACTTTGGTTTCTTCCACCAATTAAAGTAAGTTGCCGAAGATCACTGAAATTTCTGGACTGAATGCCCT

TACAAAAGACCTTTTCAAATAAAAGTAAAGAAGACAATAGATTGTCGACTATCTTATTTGGCTGCCAAGCTGTTGTGCGC

TGGACACCTTTCAGAGTGCTATTTAGATTTTTGTTGGTACCTAATAGATTAAAAGTTCTGCATTGCAAAAGAGAAGCCCA

CTGGAGAGCATTCTGAGCCGCTAGGCACACTAGTGGAGAATGCAAAAAAAAAGTCTGGCAGAAAAAGCAGTAAAACCTCT

AAATAAAACTGTTTTTGGGAGCCCCAATCCTGGAGTCTTATTTCCCAAAAGGACTGCTTCTAAGATTCTTTTAATTTTTT

AAATTAGGACTTCCTTCATCAGCCTGACAATAAATGCTGACCGACGCAGTCCTTTCATTAATTTTATATTGAAACCTTTA

AAAGTCTTTCATCTGGGCTGAATAATTTCAGAAAAGCTGATTCGTCTGAACCCCAGTCTAACTGGTATTTCTGACTAAAC

CTATTTTCAAAAAATCTGAATCATCTGAATCGCCTGACTTATTTCTGACTGAACCTAAAGACTACAATCTCATTAAACTG

AAATTACAAATTCATTTCATAAGCTGATCAAAACAATCATCATTCAATTCAAAATTTGAAATTTATTGTACAATATAAAC

TGAATTGACATGTGAATTGAGATGACAAGTTCAAAACCCCATAACACTCTAGACAACTGCTGAATTCCCGCCATTTTAAG

TTCTCAAAAACCTACAGACGATGAAATAAACTAAAATCAGCAAAAAGATAAACTGAGCTGCCCAACTGATCGGAATAGTA

TACCTGCAGGAAAATAAGTTTGCGCATTAGAAGTTTGACAAAAAAGTTTCCGGCAGCATAGTATGAATGATTCTCAAAAA

CAAGCAGTTCTCTAATATATGACTTATAGTTAATCTTCCTCGCATAACTGAAACTTGAGTGTTGTGCGAATAATTTCAGA

TACTAAAACTTGTACATTCACGCCAATAATAGCTGGCAATATCCAGGCTACACAAAGACAAGAAAATTATGGCAAACCTG

CCAATATAATTGAACATCTGTTCATTGCTCATTAACTAAAACACACACGTAAATTCATACAGATGCTACAAACATCAAGT

AACCTGATATTCAAAGCATTCTAGGAAGTTTGTTCTTGTGAAGATGCTCATTAAATTTTATGCATACCAAAAACTCAATC

AAAATTTATGCCAACACATGCAAAAAGAAAAAGGAGAAAGCATATATACCTCGACCAATACACAGATGGTTACATAATCC

TCAACTGCAGTCTATGGCTTTGACACTCGATCCTTGCCTTGTTTATCAAAACTTGTGCATCTGCCTTATCCAGATTAATA

ATCTTCCTCACTGATCTGCAACAACACCAAAGAGACAAAAAAAATTTATCAGAGCAAGCTACGCAAAGACATCACAAGTC

ATAATTACAAGCCTTGAAAAGAAAACAAAAAGTTTTAAGTGATTCAGTCTTCATATGGGATGTATAAGCCTGACGACAAA

AAGTTTATAGAAATATGCAGTCTGACACTAAACAAGTTGCTCGGACTGCAAACTTTAAGAACATATAAGCAGAAACTATA

AGCAACAACCAACAACATTCAACTAACATAATTAGACACTTAATTGTATTGCACCAATTACGAGCAAAATATAACACATG

AAACTTAAAGAAGAATGTACAAAAGAGGAACCAATTACCTCGTTTGCTACTTGAACATCCAAATACACATCGCTGAATGG

AGGAAAATGCATCTTACAACAGTTACACCAAAAAGAAAATGAATACTCGAAACTGGAAAACCATCAGATACAAAGATCAA

TATCTTCAAAAATCTTTAGATTAATCACACAATTCTCAAAATGAGCACTCTACTCCTTTTGATTGTCAGTTATAAAAACC

AAGGTACCCAATATAGCTATATGCATCAGTATATATTAAAGTTACATAATCCTGCAAGATCATCTGCATGACTCAAACAA

CAATGCATACGTGAGTAACTCCAAAATGCTCACCATCATTGCCATTAACAGTCTTCACAACCCATAGTACAAGAGCAGCA

ACTCAGCAACCATATAACAAAAATCATATACCAAATTTCAGACAAATCATATCCACCACGTAGCAGAGCCAGTAATCAAC

ATTAGGTAAATGTGGGGAAAGGAGATAAGATAAGATTATTACCTATTAGCTTGATCATTAGAATGATGTAACGTAGGGGT

GGAAGCAATATCATAAGATCATGCTATTTCTTTAAGTACTTGCATAAACCACTCTTCTTACCATCGAGAGAAGTCCCTGC

AAACATCAAACACAACAAAAAATATCCATCATTATTCTTATCAAACCCCATTTCATAACTAACATATAGGGATAAAACTA

GAAAGATTTAGTCAAACAAACAATCGAAAAAACTAAAAAAAATTAACAACTCAAGAAACAATTCTAAAACTCAAAATCCC

ATACTTCTTAGCTACATTTACTGTCAGTTCCAAACCTAAATCACCCATAACAGAAACACAATAAACATTTCAGAAATCAA

ATAAACATAAAAGAAATGATCCAAAACAAAACATTGCAGAGACTAAAGCACAAAGGTATACAAAATAGCTACAACCTTTG

AGATTTTGATACTAATTAGATGATGTAGGGTTTTCTCTGCCTCAATGAAGTTGTTGATGAACTCTGCTTCTTCCAATATA

TTTAAATAAAGTGCTTGTTTAGAGATTAATGGCCGCCAAACATTAAAAAACAACAAAAACATAGTTAGGGTTCTGTTTCT

TTTACAAAAAATCTAAATTTATTTTAGGATTTGGAGACATACCAAGAAATCAGAAAATTCGGGTTTGTGAGTAGAAACCG

CCAAAGCTGATGTTTCAATTTCGAGGGAAACCCATGCTTTAAAACTCCTTCAAATACAATAATCGCAAACAAAAATCGCA

GAATCAAGCAAAAATGAGAAAATCAAGCAAAAATGGAAAAGAACAAGTAACAACAGGTCCATACCTGATGAAATCTTCTT

ATTGGAGTCTTCTTCGCATCCAGAAATATAAGCAATAAAGGTAGCTGATGTCGTTGATGATGAATCTATATCTTCAAATC

GAATTAAACAATACCAATCAAATCAAATCTAATCAAATAACAATAATCACAAGTAAAAACCAAATCACAGTTAAACATGA

GTTTAATCTCTATTTCGATATCTTCTTCTTAACCGTTAATCAAACTAAATCTAGGGTTACAAAAAAAGGTTTGCTTTTTT

ATTTCTGTTGATACATGAGATTTGATTTTGATATGGGGTTATGATTAATCTTCTATTTGTAAGTGTGAGGTTCGATCTGT

GGGAGAGGGTTTTTTTTTCGAAAACTGAGAGAGAAGTGAGGTTGTGAAATTTTGAGATGGAAGAGGAACGAGATAGATAA

GATTTGTTTCTAGATGACTATAGAAACAATTTTCCATCGCACTGTTAAATTGAAGTGGGACCATAAACCAATTTTCGCTC

GCAGTCCTAATATAAATCCACGTGGAAATATCTTTATTTCCTAAATTGCCCCTTTATGTTAAGACTGTTATGGTAGCTAG

ATTTTCTTTTGAGTGAAAAAAATCTTCCCTCGGTAGCCCAATATATAAAGAAATCCCTAGGAAGACTTATTTATAAATCC

TAAACAGACTTAAAAAATTCATTTTCCACTAGTGGCAGTCTCCTTCTTCTCTGCTGTGAGCTCTGACCCCTCTAGTTTTC

CTTCCAATTGTTTTCCCCGTGGGATCTGAAATAGTAACACTTACACCTGTTAGGCCAGTATGAGAGTTAAAGCTAGACAT

ATAAGCAAAATAGATGTGCATGCAGTTATTATCATCATTATTCCTAACATTATTATAGGCAGTAATAGTAGGAACCCTAT

TGGCAACTTTCAGACAAGCATTGAGATGGTTTAAAATGCGGGTCGCAGTGGTTTTAGGATTGGTTTTTATTCCCTGGAAA

GCCACATCACATCTTGCTTTCCAGGTAAACCAACAAAGAGTCGCAACTATATTCTCCCAAGTACCCCAATGGGTCTGTGG

GTGGTTTCTAGAGAACCAGGAGGAGCACCAGTTGGTAAGGGTATTGTAGTTTAAAGTAGACGCAGTCAAATTTAGCGAGC

AATGTGCCCACACTGCTCTAGTAAAATTGCATTGCATTAGTAGATGTTCTCCTGTCTCAATAAGTGCCCTACACAAAACA

CAAATATGTAGGGAAGAACCCCTAACATTGATTGACATCACCTGTACAAAACCCTCATTCTACCGCGCCAAGGCATGCCA

ACCATGCCAGCCGCATCACTGTGCAAATGGCAAACCGTTCCAACCACATCTACCGCACCAAGGCATGCCAACCATGGCAG

CCGCACCACTATGGCAATGACAAACCGTGCCAGCCACATCTACCATGCCAAGGCATGCCAACCATGCCAGCCGCACCACT

GTGCCAATGGAAAACCGTGCCAGCCACATCTCCGCGCCAAGGCATGCCAACCATGCTAGCCGCACTACTGCGCCAATAGA

AAACCACGCAAAGCCAAACCGCACCTTGCCTTGGCCCTAGCCATGCTAGCCGCACCATATGCCAAACGGCATGCCAAGCC

CTTGGCCCCACATGCCAAGCCAAACCGCACCTTGTCTTGGCCCCTAGCCATGCTAGCCGCACGATGTGCCAAACGACATG

CCAAGCCATTGGCCCCACCTCGCCAAGCCAAACACATATTGCCTTGGACCTAGCCATGCTATCCGCACCATGAGCCAAAC

GGCATGCCAAACCATTGGCCCCACCTCGCTAAGCCAAACGCACCTTGCCTTGGCCCTAGCCATGCTAGCCGCACCATGTG

CCAACGGCATGCCAAACCATTGGCCCCACCTCGCCAAGCCAAATGCACCTTGCCTTGGCCCTAGCCATGCTAGCCGCACC

ATGTGCCAAACGGCATGCCAAACCATTGGCCCCAGCCATGCTAGCCGCACCATGTGCCGCGGCCCCACTATACCATGCCA

GCCTTCCCTATGCGGCTACTAAAACGAAGGCGTGCGACAATCAACGGTCGCCATTCAATCTCGTGGAGCAAATCCTAGCC

GTCCAAGGTTACTGTGACAACGCAAGGTAACCTTTGGCTCGAAAACCCTAGTTTGGCCACGCCAAAACGCGCCAGGGCAT

GCCAACCTAGTCGCGCCACCATGCTGGCCTTCCCTATGCGGCTACTAAAATGAAGGCGTGCGACAATCAATGGTCGCCCT

TCAATCCTAGGATCAAATCCTATCCGACTAAGGTTACGAGACAACGCAAGGTAACCTTTGGCCCGAAACCCTAGTTTGGC

CACGCCAAACCGCGCCAATGGCATGCCAACCTAGCCGCGCCACCATGTTGTCCTTCCCTATGCGGTTACTAAAACGAAGG

CGTGCGATAATCAACGGTCGCCCTTCAATCCCAGGATCAAATCCTAGCCGTCCAAGGTTACCAGACAACGCATGGTAAGC

TTTAGCCCGAAACCCTAGTTTGGCCACGCCAAACCGCGCCAAGGCATGCCATGCCAACCTTCCCTATGCGAATACTAAAA

CGAAGGCGTGCGACAATCAACGGTCACCCTTCAATCCTAGGATCAAATCCTAGCCGTACAAGGTTACCAGACAACGCATG

GTAACCTTTAGACCGAAACCCTAGTTTGGCCACGCCAAAACGCGCCAAGGCATGCCATGCCAACCTAGCCGCGCTACCAT

CCGGACTTCTCTACCCGGCTATCAAAAACGGAAGCGTGCAACGATCAACGGCTACCTTTCCTCTTAAGATGTAAAATTTC

AACCGTCGAAGGTCACCAAGCTAGCAAGTCTCCCAAGCTCGACTTGACAGGCAACAACCATGCTACATGTTTTCGACGAA

AACATTCGAGACATCAATACATATCACAAACTGGGGGATCCACTTTGGGTATTGGTTTGGCGGTTTACAGCGTGCGGCGT

ACACTACGCCCGTCATAAGGAAGTGTAATAAGGATGAGGCGGTTAGTAAATAAAAGGAGTAATGGTGAAACACTCTTTTT

TATGGAACATCAATTCTATGCGCTAACGGTTACCACCTCCTCCCATTTACTCACCCATTTTTCCATTCCTTAATGAGACC

CGCAGTACGTTTCACTTGACTTGTTGTAGATGGTGGATTTTCGACAAAGGCTAAAATCGAACCCATAATTATAAGTATTC

CACGATCCAAAGAATCGACGAATATTGAGGCTCATGTCTCTTATCGAAAGCATGAAAGCTCATGCCACAAGAATCTCTCG

ATCGAGAATACTGTCTCTCGGAGTATATATGGATATGTAGTCTCTCACCGAGGCAACGACATATCTCATGAATACCGAGC

AATTTCAAGTAACTACTTCTACATAAAATCGAAAGATTAGATGGCTCCTAGACGCTTGCCTTCTAGTATAGAGCAATAAC

GTCTTCAGGATGTAACAACGTTTTCCCTGACTATATGAAAGAAATATGACGCTTCAACAAGCTCATAACACTAGATTCCT

GAATGATTAATGCTCCTAGACGATCGTACTAGTATAGAGCCATTAATTAATTATTCTCAAAATCAATAGATTCATTAACC

GAATCCCTGGAAAATATTCATTATTTTCATAATATTTCGTTACTTCAATTTATGGTTCTTCCCAGCAGAATTCCATGGGT

TAGTAATAAATATTCAATACACGGGCATCTGAGCTACCTTTGAGTAAGCCCCGATCAAAACGGTGCAAGTGTATTTAACA

ATAATGTACTAACGAGCACCTGAACACACGAATGAGTATTTCAGAATACGACGAGACGATGAACCTATGCAAAATAGGAA

GAAAATAATAAATAATAAAATATTAATCAAGGCGCTAGGGACTGGGCTCACCGGATGGCCGGTCATGCCGTGACCGATCC

CACGCCCAAATTCTATATTTTATCATATTTTTATTATTTTCCTTGTTCTCATGAAAATCCTTTCATTTGAACAAATATCC

TTTGTTTTAAGGAAACTTCTCAGTTTCATGGTGTTTCCTTCAAATCATGAAATAATATAAAATTAGGAAAGGTGTCACGG

GACTGTTATCTGTCCGGCTATGCCCTAGTCATGGCCGGTCCCACACCTCATGAATCCTCAATATTTTATTATTATTTCCC

TAATTTCATGAAAACTCCTTAATCTCATGATATTTCCTTGATCCCATGAAAAAGAATATAAAATTAGGGAAACCCGTGGG

ACCGGCCATAGCCGGCCGATCATGCCCTAGCTGGCCCCACACGTCTTATTTTATTATTATAATATTTTTCATGTTTCCTT

AATCCCATGAAATTCCTTGATTTCATGATGTTTCCTTCAAATTATGAAAATTCCTTCAAATCATGAAATAATACATAAAA

TTAAGGAAACTCACGGGACCGGGCCCTAGACGGCCAACCATGACCAAGCCGGTCCCACACGTCCCTTATTTTATTATTAT

TATTTTTTATGTTTTCTTCATCTCATGAAAACTCCTTGGTTTTATGATATTTCCCTAAAATCATGAAAACATAATAAAAT

TAGGAAAAATACCATGGGACCGGCCCCTTGGCCGTCCGGCCATGCCTTGGCCATAGCCGGTCCCACACTTTATGATTCCT

TCATTATTTTATTATTTTTCCTTCATCTCATGAAGTTTTCATGGTCTCAGCAAAAATCCCTGATTTCTCCATATTTTCTC

AAAATGTTGCTCAAACGCACACCAGAAAATATCAAAATTCTCATAACTGAGATACGAACACTTTGGTGATGTGAGCATGT

TACCTTGACCGACCAGGGTTGGCTCTTTGGCTTGAAAGAGGCCGGTCCCATGTATTTTGACAATTTTGACCTAATTTGCG

CAATTGCACATATTAGGTCCAAAACTCTTCCAAACAACTTGGAACTTCATAAGACGATCAATATTCGATCTCGTGACTAC

CCAGGACCGGTTTCATGACCCCTTGGTCGGTCCCTCACATCTTCATAATTAATTAGGTTTTATTACCTAAGGTTCAGACG

AGCATTATTTTTTAATAAATGATTAAACCAGCATTTAATCATCCTTTCATTAAATCAGCTGGTCTTCAGGAGACCTTGAT

ATTTGCTCACGTGAGCATCTGGGCACTACATGGTTGATCCATAATCCCATGTAGCCGGTCCCTCCTTCGCTCCGTGGTTA

ATTTTCAATAAACCATCAGTTAATCATCAATTGATCAAATTAGGGTTTTGAGTCCAAGGTTCATAATTCCAGATTCAAAT

GCTAATAATTTTACGTTGACCCCATGATCAACATTTTTTTTTTATCATACTCGGTTCAATTGCCAGTATCTTATTTGGTC

ATGCTACCAATAATTCGTCGAATGAGCAACATTTGCTCAGATGAGCAATATTTGCTCAAACCGAGGAATACTGGTTCAAC

TATCAATATTCAACAATTCATCGAATGAGCAATATTTGTTCATGTTAGCTTATTCACGCATTATATCTCTGTCTCAACAG

ACATGTTCAATTCATGAGTCTCAGAACATTTCCAAATCACGTTCGACTCAACAAAACAAGGACTCATCGTTCCATGAAGT

CATGAAGTCAGCAACTGACTAACTAAATACACGTCTACCTTGCAGACTCCACAATCACACGAGACATCAATCGTGTTACA

TGGGGGGATATTAACTAGGGTTTTGGTCTAGCGGTCTACGGCACGTGTGTTCATACATACGATGAGAATGTGAGCAAGTC

GTGTAATCAGTTGAAGGAGTTAACAAAGTAGTGGATGGAAAATCAACCAAGTATCCGCACGTTGAGCAACTGGTTTTAAC

ACGATTTCCACTTCCCCATTCTTTGATTCCATCAACTGTCACACTTCATGAGATCATGGTGTTTTCAATTCCAGCAATAT

AAATAAATCTCTGAACCATGATTGAAACAAGGATAATCTCACGTCAAACAGACAACACGAGATTAACAATCCAACTCTCA

ACACAACAAATTCAATCATTCAGAATTTATCTTATTTTCAGAATACACAACACACCTACAATCTTCGATCACCATTGATC

TCACACATTTCTCCGCTTCCCTCCTACAGACCGACCCATCCTCTCTTGTGACCGAATTTACTCTGGAACGGCCATTGTCT

TGGTTTAGGCCGGAGTACTACAGAGTGATATCTCGAATTCAAAGCACTCCCTTCTTGCAGTGCATCTGTGTGAAGTTAAG

AAATTTGATCGGTTCAAAGAGTCTCCTCCGCGCGATCATCTCCTCAATTCCGTAAAAACCAGCGAATCGTTTTTCCCCAT

CTACAGATTGGCGACCACAGTGGGAGATCATTCTCTTGGTTGCAATCTAAATTCCCACAAAGATGGTCGGTCTTAGGTCT

GGTTCAGATTACCAACCCGGATTCAACACCCGGTTCAAATATTGCAAGCTCTAGCAGCATCAAGTTAACACTACTCCTCC

AAAGGCAACCGCCACTACTACTCCTCCAAGCACTGGTGCTCCAGTGATCACCCCTGCAAGCGACAACACCGACGCCATCA

ATAATCCACTCGGTCGGTCTCCCGAAGAAGTCAGAGAAAATCCACCTACCATAGGTGATCTCATGAAGGTACAAGAAACT

CTTCCCAAAAATCAGACTGACATGGCTGCTACTCAGAAGGAGCTATTCAACTATCTCAAGACTCTTACAGACAAGAGAAA

ACTCAAGAGAAAGGAAAAGCTAAAGAAAAATCTTCAGACATTGATCCCGAAGTCACTCCAATCCATGTAGTAGATGATGA

GGAAGTTCCCAAGGCTACAGATGATCCACCAATGAAAGAACCATCCAGTTTCATCACTCGTGAGGACCTGGAACGCCTCT

TGGAAGATCATGGAAAAGACAAGACGCCTTCTGTCCACCATCACCAACCTCCATACCCGACTTCTACGCAAATGATTCCT

CTACCAAAAGGTTATACCTCTCCAACATTTACTCTTTATGACGGAACAGGCAATGCTCGAGAACATGTTTCTCGGTTTCT

GGAAGCGTTGGGAGAACATGAGCACAACCATACTGTCTGTCTCAAGGAATTTTCAAAGTCTCTGAAAGGCAGGGCATATA

CCTGGTATAATAACATCGCACCAGGAAGTATCAAAAGTTGGGGAGAAATGGTTAATGCTTTCTATAGAAAATATTTCTTC

GTTTCAAAGCAAATCACTCTGTCCGATCTCGGAAGAATGTTTCAAAAGAGTAATGAACACCCTAACGATTATGTGAAAAG

GTTCAGAGTTTAAGCCTTGGATTGTCATGATCCAAACGTCACAGAGCAACAACTCGTGGAATTGTGCATCAATGGAATGA

TCCCAGTCTATAGAGCTTTACTAGAAAATCTCAGGTTCCAGACTTTCTCAGAACTCCATGAAGCGGCCAAGAGATCAACA

ACTACTGCACCTGCCCTATTAGAGAGAGCTAAGACTATAAAAGCTGAATAGCCATGGGAGACTCGAGGTAGCAGGCGTCT

AATTAACAAATAGTAAAACCTACAACCTTCCACAAATGGTGTTGCAGAAGGGAGTAAACGAAAAGCTAAGCCACATCGCA

AACATACTTCCTCAACCCCTTCAAAGGCGCAAAGGAAGGATAATATGCAAGTTCCTACTCAACACACAGAAGCTGATCCA

GAGGCACCTGACTTCCCCTGTTCCATTGAGGAAGTTATTGAACTTCTAGATGCCTGGGTTCAAGACGGCACGATCAAGCT

CCCATATGTCAAGAAATAAAACCAACTGAAGAAGCCATGGAAAATCCTCGTTACTGTCGCTTTCATAGGTTTGTCAACCA

TCCCACAAGTGATTGCAGAACTTTGAAGCGCATATTCAAAGAAAATATCGAATCAGGAGAACTCAACCTGGGATCGAAGG

AGTGCACGAGAGATCCTCTCCCCGTCGTAACCTTCACTCCTTCTCAACCACCTGTCAAAGAAGCGTTCAATCCTTAATTG

AGCATATCCGTGAATTGTTATACTTATCAAAAGCTCGAGAGGGTAGACATGTTCGTTGCACTCGAACCACATCGTGTCGG

AAAGCGTTTGTTGATTCAAGAAGCATTCCCCGAGCCGCCCTCAACAAAGATGAAGAAATTTACGACCGGGACTCCTCACC

ACGGTCCATCTCAAGGAAACGAGTTCAAACGAGCATTGATTGATGTTGGCACATTGTTAATATTATCCTCTCGAAAACTC

TCGAGCTTCGGTATCACATGACAAGAAGCGATTCATGCTCCAAGTAATAATCACAAATCATGAAGGAACACCCAGAGATG

CTTATGGCCACACCACTCTTAAGGTGAACATAAACTCAGCCTGGACAGAAACAAAATTTTTCATAATTCGAGAAGATCTA

GGATATGATATGATTCTTGGATGAACACGGATTCACACTGGAAAGGTAACACCAACACCAGTTGCACATTTCTCTATTGA

AGAAACCTCTGATTCGGAAGAAGAAAAACAAGATACTTCACCATTCGAGTATTTGAAGGAGGTCAGAGCCCTAGCAGAGC

TAACACAAAAACCTGAAGAAGATGAAAATGCTTTCATGAGAAGGTTTCGCACTCAGGCAAGTCTTATTCCTCCTCATGTG

TCTATATCATCAGTTTTCAAATATGCTTATTTGGTCCGAGGACTTTACTCCACCAACAACTTAGCAATTGCAAAATGCTA

TGAATGGATAACCTCGCCCCAACAATATTTCACCAAATGGTTAAGTTCAGAACTTCTTCAAACCGAGCATATCATCAAAA

AGTTTATTGCTGAAGACATATCCAAGCATGATAAACAGTACCTCAAATACTCTCCTCTGCATGAGTGTCCATACATACCA

CAGCCATGGAACTCCACCGCACGAAGAATTCCAAACCAGCGAGAAAATATTCAGGGCACGAATGATCAAGTCTAACAAAT

TAGGGAAGGAGATTTGGTTCTCAAAGCAACTAAGCGTAACTCCCATTCTCGCAAGGAATTCTAACTGGGAAGGACCCTAC

ATGGTCTCTAAGTCTCATTGCTTAGGTGGATACTACAAACCGATCAACATGGATAACAAGGAAAGCGTGGCCGTGATACA

TGAGAATTGGCTCAAGGCCTTTGATGTCTAAGGTTTTGGATATTCGAGGTATTGGATGTTCATGGCACGAACACCTAAAA

TATTTAGCATTTAGAGTTTTCTTTCATTATATTTCAATTCCTCCAAAATGTTTGCAATACTTGCATTCAGTAATTTAAAT

TCAACAATTTATCAAAATTCAGTTTACTTAAAGAAAATCTCCATCAAACCCAAAGAAAATCCTCAATCATCTACCGATCC

AAAACCAATCAAAACCAAAAATAAAACCTCACATCTCTAAAAAGTTCAGAAGTTCAAGAGATTATGAAAGGAAATCATAG

AAAATCAGCAAAGAAAGATTTTTGTTCCTCTGCATCCTTCAAGACTTTTAAGGCCGCTTTCTCCATGTTCAGCTTTTCAG

TCAGCCTGGCAATCTTGTCTTCCTTTTCCATCACAGCATCATTGGGAAAGGGTATGATATTCTCACATGAGTCCTTAATA

GCATTTATCTTCTTACGAAGCCACTTCACATTGAAGCCAAGTCTCTCAGAATTCTCCAAGTTGAACTCCCAATCAAAGAG

TACATCTGGAGTCACAGTATTTGTAGCTCTAGTGCATATATCGCTTACAACACGCACGACATCACTTACTTGCATGGTTA

AGGAATAAGCACGCTTTGGGTCTCTAGTAACGATGATGTGACCAAATTTCTTCCATATTTTCTCGTACAAGTTGGCATAT

CCAATGGGAATGCCGAATCCACCAACTAGTTCGTGGTATGGGAGTGATGCATCAAATGGAGGATCAAAAGCAACTCCTGT

ACTTGGGTATTCATGCACCACTATATGGTTTTCAATCACTGGAGCAATCTCTACAACAACAGTTTCTTCAGTCTCTTTTA

CTTGCGGCTCAGTTTCTTTCATCACTGAAGCAACAACAATCGAGGTCTCGATCCCCTTAATTGCTTCAAGCAACAACACT

CTCATGGCCTTGAAGTGCAGAGATAGTTGCTTCTTTATCAATAACCCAGGGTTGCTCATTTCATCATTATTGATTCCTTC

ATTCTCAACTTCAGTATTTTGTACCTAATACGAAGATTACTTGAGATTAGGATCATCCAAGCATGAAACCAAAAGAAATC

GTAAACTACAGCATATAAATGGTCTAGCATCATCAAGGTTCTTCATTCTACATGTTATATGTCAACAACTAACGTGAAAG

TCACAAAGACACGTTTTACAACAAGCCCGCCTGAGAAGATTTTACACTCCCCTGTACAAAACGGTGAACTAGGAGCAATT

GGTGAAGAATCAATGGTTGGGTTATATACCATTCTCTTCGTATGGATGTTATCTACAACATATCAAAATTTCATAGCAAT

CGGATAAACGAGCAGGAAGATGTGGTTGAAACTTCAGACTACATAACAGCTGAAAATTTTCTGAAAATCTCCTGGATGTT

TACTGTTTTGTCGATTTCGACCCATAAACATGCTCAAAACTTAAAAATCTTCTTTGAAAAAGCTTGGAAATTTCCTGGGC

ATAAGGATACATATGCGGATCACGAAAATCTGATATCGGGCTAAGATTTTATGCACTTTTGCTAAGAGGCTGAGTTCTGA

ATTTTCTGGTGACGTGTTTTCGGCAAAACTTTTCATAAATTCATATACATTCTCATTTATTGACTCGTGGATCAACAAGC

TTATGCTTCTATGAAAAAAATACAAGAGAAACACCTACTTGGGTAACCTCTGAAGCGTTCAAAGAACCAGAATTGCTCGC

GTCAGCATCATCAACACCATTATCCGGTTTAGAGCTTTGGGAATTTCCTTTATCACTATTCTCCGAACGGGCATCATGAG

GGATTTCCACCTCCATGGCGACATCCTCCTGGAAATACCCTTAAATAATTATCATTTTATTTACAAAACATCATAATTGA

ATGTACGAGGAAATACTTACGTCCTCTTCTTCTTCAATCAAGTCGGAAGTCTTCTCCCCAGCAGAAGAGAACTGAAGACC

GTCAATACTTGATGGAGTGAAATTCTGCAACGGAATAACAAAAGTCGTTTTCAAGATCCTAATTACACGGATGGGAAGAA

GTCATGATGAAAGAGATATCGATAAACTCACCAAGCAAGTCTTGGGGACTTTATGGTGTTAGGTAACATCTTATAATTCT

TACTCTTGCCTTCTCCGCCATGAACAATTAAACCTTCTCCGAGAGGTTTACATCAATTCGTGGTCTTACATTACGACCAT

ACGTGGAAAAGAAAATTTTGATGCAATAATCAAAATACAATCATCATAATCTTATAAAGACTTTGTAAAATACTTACTTG

TGAACATCTTGGAGATATTTTCCGTTTGTCACTACCAGGGAAATGCTTCAATCTTGGTCGAGTAGCAATCATCTCAGGAG

AAGGGGGCTCCTTCACTAAGGTTTGATTAACTCTTACAAAATCAGATAAACGCTTGAGTTGTTAATCCCAAAAGTTTATG

TAACCCGGAGTTACATATGAAACTCTTTCAGAGCTGGGGAACCAGAATTTGCTTCCTTCCACATGTGTGAGGTCCAGACG

GGTCATAAGCTGATCAACTGAGGGTTTTCTTCTTCGGAAAGTCGGAACACACTGATCCATACCCATTTGCCGAGCCGCCC

TATCAATATTGTATTCTTCCACTGAGAATTCACCATGCATTATGGATATTAGATAACCGAGAGTACAACTTGGCATAAAA

GCTCTCTCACCGTCGTCCAAATCAGCACCAGATTTCAAAGTCATGCTCTCTTCATCATAAAAGTTCTCAACTGAGAGATG

TAAGGGGGAACTAATACCCAAGGGCGGAAGTTGAATTCAGATTCACAATCCAGAACATCAACAAAGCAAGTCTTCAGTCG

AGATTGCTTTGTAGACCAACGCATGATCCTGACATTAATTTTCTCAGAGAAAACATGACGAACATCAGTATTCAATCCTT

GCAAAATGGTACGTGGAATAGGTGCATAATTCTTGAAGTTCTCCCACATCAAAGCTTGAAGAAACATTGTATTAGTGTAA

CATTCAATCTTCATAAAACCGTTCGAGACACTAGAGTCCATGATCAAGGAGTCCAAGTTAGAATACAAAGAACCCAAGAA

CAAGCTACCTATTGGGAGTTGCTCGCCCTGAGCCATATTAATCGCAAAAGGAATCACGAATGACTTCAACAAGTTTTCTG

CATCTTCAAATACGTCTCTGGACAGCCATAAACACAAGAAAGCAGCAATAGATAACGTACCGTCAACAGGCCACTCCAGA

GGAGTATCTCTTGGCCACTATTGTATTAACCAATGAGCATAACAACCTTTGCTGCCAGATGTTTTATTAATCTCCTCCAT

CGTGGTTGTCAAGATGTTATAGATTGCAACTTCCTCTAGTGAAGAAAGATTTTCAGGAAGATTTCCCATGATTGGAAGAT

GCAACAACGCAACTACATCTTCTAAAGTAATGGTAAACTCTTCCCATCTGCATATGAAAGAGTGAGTTGGAATACACCAT

CGAGCAAGTAAAGCTACAATACCTCGCGTATCATGAAAAATGAATAAATCTCCAGAAGCTTGAATAGCTCGTATTACTCG

TGCGTCATGCAACTTAGCTCGAACACGAGGAAAACCCAGCATGTACCTCGCCCATCCAGAAAAATTTTCCAAGGGCCGTC

GAGAAAGCTTGAATTCAATGATTGATGCGAAAAATATCCTCGTTCGAGACGTAACCGAATAATTACTCCGAGGAGTCACC

AATTTTTTCGAGTAACCTCTCTTGGATTTATCAAGACGAAATCTTGGAGTTTTGAGACGCTTTCCGTGCCCTTGATCTAT

GAAGAATGCATCCTCTATGTTCGGAACGTTCTTGATTCCAGGAGTTTCATCTTCGCCAATAGAAGTCTCATCCGTGCATG

AATCATCCCTAGAGGCATCATTGTCTATGTGTGAATCTTCCCTGGAAGCATCATTATCCACACACATATCATCTCTGGAA

GCATCATTGACTTGAGAATCAGACATCCTTGCAAAGTAGCTGTAACGTGCACACAATACTTCGCTTCAGTATACCATTCA

TACAAAATGCAAACGTCTACAAAATGGTAATTCTCAACTCTTACTTTTTTACGATTCTACTAGCGATAGTGATTGTAATG

CAAAAGTTAACACTTCAAAACTTGCTCGTTTCTTCAAATCTACAAAAAACGAGCATAAACTCAGATTTTCATAAAATCAT

GAATATTTCTACTGCGTGAGCATTCACCATGGAATTATGAAAACTAAAATATTATTTCATCATAAATAATTGTTTACTTA

CAAAGAATACACAAAACTCAAGTTTTGATTTATATTTATATATATATATATATAACGTGATTAATTCACGTAAATAAAAA

AAATGTTTTGAGATGAGTAAAGCTCATGAAAATAAAATCCGTGGGATATCCCACGGTTTCATAAAAAAATACATATCCAT

GGTTTTATAAATTTTTTTTTCCTTTGTACGATCGTCTTTCTTTGAAACGAAGGTTTATTTCGGTTCTGTGAAAGCTCACA

TCTTTTCAAGGTAAAGTTTTGGTCTTCATGAAAGCTCATAAACAAAAATTTTATCACTAGACACAAGTCTATACATAAAA

AAATCTCTACTAGGTCAGGGATTATTAATTGGTTCTCAAACTAACGATGAAACGAACGTACGTTTTCTAAAAATAAGGGT

TTTCAACAAAACTCACACTCAAATATACACATATATATAATTTCAACATGATCAAAGAATGAACTCAGCTTACTTCTGTA

AAAAAAAAAAAAAAAATTCACGTACCTATCGGTGCCGGCCGGAATATGGCCGGACGGTGGTTGGACGGTGCCGGCGATCA

GCAACAGCTCCGGCAAAAAAATTATCCTTCTCTCTCCTGTGCTCGAACGTGAGGATGATGGGAGTATGAAGGTGGTGGTC

GGTCGTGAAGGTGAAGTAAAAAAAAAAGGAATTACTCCTTTTATATCAAGTTTTATTTCCAAAACCTAATTCTATAAGGA

TTTCCCTATTGGGCCCGGCCCGTGAATCCTAAAACGACCAAGGTCCCGTCATCAAAATCAGCGGTTCAACACCAATTTAT

GCTCATTAAACGATGCTCTATCATGCCACTAGAATCTTAATTCAAGTCGTGGTTTGCTTGTACTATCAATAATCCGGTAA

GGTATCTAATGACTAAGTTCAAATTGCTTCCCGGATGTCAAGACAGTAGAATCAAAAAATCCGCTCAGGAAACGACGAAT

CGATTACCAGGCGGCTCAATTACCAGTCATTACACGGAAAAGGACGAATGGGGACTTCCGAGGGGTTTAGAAGCGAACGA

TCCGCCCAGGGATCGGCAACACGACAAGGAATGACTGGGGACTTTCCACAGTCATGGCGACGTTTCCCATAAAACTCGGT

CTAACGCTCGGGAGATTTCTTTTTCAACAGGAAGAGATGAAAAACCGAGTAAGTCTTACCTAGACAAAATACATGGTTTG

TTATTTCAAGTTTTCACGAGAATGATAAAACCGATAAGTCATGTTATGTTTCATCCCATGTCATCTACCAACACGCGGTT

CCATTGTTTGCCTGGAACGTTTGCGCCACTCGCAATCATAACCAAAACAATGCATTCCAAAAGAACAATCCAAAACCATC

AAGGTAGCATTTGTCTATCAACTCAAAAATCCAGAAAATCAAAACCATAAACTAGGCGTTTCATTTGCCTTTCAAAAAAA

AAAAGAAACCTAAACTAAGTAGATCAGAAAAAAAAAGAAAAAAAAAAAAGAAAAAATGTGTTCAAGGAAAGTTTTCCAAC

AATGGCTTGCCCTTCTTGGCGAGAGCCTCCTTTGCGCTTTGAAGATCCGCCTTTTTCAATTTTAGTTTCCTGGACAGCTT

TTAGACTTCCACTTCCTGCTACTTCACCATATCCGCAGAAGGGCCTTCGACTTTTTCGGCCAGCATCTTTTCAACCTCAG

AGAAACCTTTGACGAACCAAGGAGCATTGAACTCAAACTCCATGCACATGTCACGATGGAACCTCCCGACTTGTGACCAC

ATCCCCGTAAACAAGATGACCAAATCCACGTTGCACATGTCGTCAGTCGAAGACAGAATTTCCTCCACACGCTTAACCAA

CGCGTACCGACTAGTAATTTGTTTGGTCGTGGCAATATGACCGTAGTTTTCCCAGATCTTAGTGTATAGTACCGCATGTT

TGGCTGGCACGCTAAACCCTTCAATCAGCATGTGATCCCGATGAGGAACCAAGTACGGAGGATTAAAAGGGGCGCCTGCG

ACTACATCAACAACCGGCAAGGGACTCCTAACAACAATCGCACCTGCGGCCACTGGAGTACTTTCCTTTGTATGAGCAGC

AACGACGTTTTCATCTTGATCAACATCCATTTCAAGATCGCCCGGTACGGTTGCCACAGTTGGAGACAAAGTTTTATCTT

CCTCAGTCTCTATCTCAGAGCCATCTTCAGAAACGACTTCCTCCTGTAATACCGCGAATTAGACGTTTTTTTTTTTTAAA

AAAAAAAAAGAGGGGGAACATGCGGAAAAAACTCACTGATTCACTTTCAGACTGACTAGAGGAAGATTCAGTACTGCTGT

CAGAAGAACTACCCTCGCCATCACTGGACTCCGAACCGTCATCCTTCTCGGGTACCCCATCACCATGGATAGGTTCAACA

CCGACATCCTCTGTCTCGGGAAGCAATGTTTTCTGTCTACTTGGAAGTTTGATAAAGGCGTTCACACTGTTGAGACCCTG

GACAAAATATGAAGATGGATACTCAGAAAATAAACGAGAATATACACGGCCCAACTGAAGTCAAGAGGAAAATCATACAT

ACCTGTATGTTGGACGAACTAAAAGTCGGTAGAGCCGTGGAATCTCGACGGAAACCACTGCGCGATTTTTATACCTTCCT

CCTTCGCTTGGAGACGGGTCCGCGCCCGGGACCGAAGGATTTTCTTTTGGTTGAATAAGTATCCCTCAACAACGGATGCT

TTGCTAAAATCGCAGCATAAGACTACCACGGATGTTCTCCACTACAACTTTCACAAATTCCTGGATGACAAGGAATTCAC

TCTGCCAGAACGTGTTATACTTGGAGGTAGTCGAAGGATTTCGTTCCGAAAGCAAGAAAGGGGGCTTTTACCCGTGCACG

AGAACACTAGCATCAAGAACGGTCGTGTAATGAGTTCTCTGAAGCTACCAGCTGACGAACCAGAGGGACCCCTTGGTCGA

AACCCATATGCCGAGCCTCCAGATCAATATTGTAAGAGATTGCCCGACACGATCCTCGAAAGAAAGATGGTATGTAACCC

GGCGTGCAACTTCGCATGAATACAGCCTCGCCGACATCGAGGTTCCCTTCGTCAGAAAGCAAAGACATGTTTGGAGCGGA

AGAAAAAGTATTGACCTGGACTATGGATGTACACACTGGGGCCCATCGACGAAAATTAACGGCGCACGAATTATCGAGGA

AATCAGCCAGGTTTGAGCCAACTCTTGGGCGCCTATACGACCAGCGAAGTATTCTCGAACCACCCCAAGACGCGGGAAGT

GAAACCAATGGTTTAGGAGCGTGTTTTTCAAGATGCTCCCATAGCCACGCCTGGAGAAAAGCAACGAGAACATACGAGTC

CACTTTCAAGTAACCATTTGAAACGCGCATATCCACAAACTGGATGATCCAAGTGCGTGTACAGAGAACCGAGAAACAAC

CCTCCAATGGGGAGAATGACTCCTGTTGCCAATATTATAGCAAATTTGAGGAGTTCCGGCCTTATCTCCTTCTTACCAGA

TCCGTCATCGAAAATATCCCGGGATAACCACAGATCCAAGAATGCCGCGACATGGAGTGTACTGTTCTTTTGATCCGGTG

CCAGCTCATCAGGAAACCATTGACACACCCACCAGCTGTAGAAACACTTCGTCTCGTTTTATTTTCGAACAAACCCTTTC

GACTTCGCAACTGTGGTGGTATGCATTTTTTCTTCATCATCTCGATGACTTAATGTTGAGGTTTCTGTTATGGGAAGATT

GCAGAATGGCTACACTTTCTAAGGATATAGTCATTTCTCCGCACCTGCATATAGTTTGTGTGGGTATCTAGACACCACCT

GGAAATGAAAGCAACCAAACCCGGGATGTCTTTCATAATTTGAAGTGATGCAGAAGCCGCGACAGCATCAATGATTTGCG

CTTTGGTTAAACTAGCTCTTACACAATCTTGACTTATCATAAATCGCATCCATTTATCAAAGGAACGCAACAGACTCACG

GAGATGTTGAAATCAATAAGAGAGGCAGAATACTCTTTCCTTTGAAGACAAAACCGTATTACGTTCCCAGGCGGAACCGA

CCGATCCCCTCCCTCACCTTCCGGACCGGTCAGAAGAATCGACACATGCTTAAGATGACTTCTCATGGCAGAAACTATCG

TGAAGAACTCATCAGCCGCGTTTGGCTTGGAACTGCCAGCATCCTTCGGTACAACAGAAATTGATGACATCCTAAAGAAA

ATTCCCTTACGCTGCTTTCACCTCTGAGATAACTACAATGGAACAATACGAAGAGAGATTATTACGGAAAGTTCCTGGAA

ATTATTTTATCAAACGCAAGAGGGACATCCTGGACGCAAGCCAATTGGTTTCAAAGTCATTTTTGCGCATATGCCAAGAA

ATGGGGACTGACAGACTCTGCATCACCACTTCATGCAACGGGAAATTGCGCCTATCCGAATAGGCGCGCCCCACAACGGG

AACCATGCACACGACCACACCAGGAAACGAAGGAAACCCTAAAAGCTAGATTTTTACGGGAAGGGAATGACAATGCTCGT

GCACACCTACTTCGCATAGTAATTGAGGCGGCCAACATCACTCACGCCTAAATTTTACTACAAGGTCGTGCAAAGCATAC

CTATGGCTGGGATTCATACCAACTCAGAAAAATGATATTTCTACAAGTTCATGAGAAGGTTCACCCACAGCTAAGGTTCA

CATTAAAAAAACAAAAAGAAAAAAGAGAGGAAAGAAAGCAAGGTAAGGAGGAAGTATGGGAAGACATACCAGGGATTCGC

TTGCCCGCTTTGCTTTACTGTTACCACACCTAAATACAATAAATAGGTATGAAACAAAAGGAGAGGCCGGCCCTCTTTTT

ATGTGCGTGATATTTGGGAATCCGAACAAGGAAAGGATTTCCTATTTTGAGCCAAGTAAGGAAAAGAGGCAACCGGTCCC

ACGAAGACAAAATCACCATGCCAAGACTGTCCGCGCCATTGCCTTGGCCCCACCTGCCAAGCCGTCCGCCATGCCTTGCC

TGTCCAATACCGCGCCATTGGCTTGGCCCCACCTGCCAAGCCGTCCATCATGCTTTGTCTCTCCAAGACCGCGTCATTTC

CCTGGCCCACATGCCAAGCTTGTCCGCGCCATGCCTTGCCACGCCGAGTCCAAGCCAGCGCAACGCCTGCGCCCCCTTTC

CAAACCATACTGACACCAACAACTCGCCCGGCCAGGCCGGCACGATGCTCCCTAACCCGGCCCAAGGTGATGCATAGCCG

ACTAAGCCGACAATTTTCATCTCACCACATCACGGTCCGCACCACCAGTACAATTTATGCAAGGCCGCCAACACGGGAAG

CGCAAACTCTCTTTACCTCTCGAAGAGGTTAGTCGGGCTTTCCTGACCATCTTATCACGACTTGTTTCAATTTCTTATCC

TAGCCTGGCACCATCTTATGCTTAGCACGCAACGATGCTCCCGTTCCGAGCTAGCAGGGGACTTAATGTTGATGGTGACT

TTTAGCCTAGGGTTAAAATCGTAAAACCGTACATATGGCATGACGTAATTATGAACATTAGCACATTTATCGAATCGATC

AGCATGCTTACGTAAGAATTACTAGGGTACCTTTTTTTTTCTCTATGTATCATGTTCCACGGCAGAACCCCTAACATTGA

TTGACATCACCTGTACAAAACCCTCATGCTACCGCGCCAAGGCATGCCAACCATTCCAGCCGCACCACTGTGCTAATGGC

AAACCGTGCAAACTACATCTACCGCGCCAAGGCATGCAAACCATGTCAGCCGCACCACTGTGCCAATGGAAAACCGTGCC

AGCCACATCTACCGCGCCAAGGCATGCCAACCATGCCAGCCGCACTACTGCGCCAATAGCAAACCATGCCAAGCTAAACC

GCACCTTGCTTTGGCCATAGCCATGCTAGTAGCACCATATGCCAAACGGCATGCCAAGCCTTTGGCCCCACATGCCAAGC

CAAACCGCACCTTGCCTTGGCCCCTAGCCATGCTAGCCACACCATGTGCCAAACGACATGCCAAACTATTGGCCCCACCT

CGCCAAGCCAAACGCACCTTGCCTTGGCCCTAGCCATGCTAGCCGCACCATGTGCCAAACGGCATGCCAAACCATTGGCC

CCACCTCGCCAAGCCAAACGCACCTTGCCTTGGCCCTAGCCATGCTAGCCGCACCATGTGCCAACGACATGCCAAACCCG

TAGCCCCACTATACCATGCCGGCCTTCCCTATGCGGCTACTAAAACGAAGGCGTGCGACAATCAATGGTCGCCCTTCAAT

CTCAGGAGCAAATCCTAGATGTCCAAGGTTACCAGACAACGTAAGGTAACCTTTGGCCCGAAACCCTAGTTTGGCCACGC

CAAACCGCGCCAAGGCATGCCATGCCAATGCCATGCCAACCTAGCCGCGCCACCATGATGGACTTCCTTATGCGGGTACT

AAAACAAAGGCGTGCGACAATCAACGGTCGCCCTTCAATCCTAGGATCAAATCCTAGCCGTCCAAGGTTACCAGACAACG

CATGGTAACCTTTAGCCCGAAACCCTAGTTTGGCCACGCCAAACCGCGCCAAGGCATGCCATGCCAACCTAGCCGCGCCA

CCATGCTGTCCTTCCCTATGCGGCTACTAAAACGAAGGCGTGCGACCATCAACGGTCGCCCTTAAATCCCAAGATCAAAT

CCTAGCCGTCCAAGGTTACCAGACAACGCATGAACCTTTAGCCCGAAACCCTAGTTTGGCCACGCCAAACCGCGCCAAGG

AATGCCATGCCAACCTAGCTGCGCCACCATGCTGGCCTTCCCTTCACGGTTATTAAAAACGGAAGCGTGCAACGATCAAC

GACTACCTTTCGTCTTAAGATGCAAAATCTCGACCGTCGAAGGTCACCAAGCTAGCAAGTCTCCCAAGCTCGACTAGCCA

AGCAATAACCATGCTACATGTTTTCCACGAAAACATTCGAGACATTAATACATATCACAAACTGGGGGATCCACTTTGGG

TATTGGTTTGGCGGTTTACAGCGTGCGGCGTATACTACGCCCGTCATAAGGAAGTGTAATAAGGATGAGGCGGTTAGTAA

ATACAAGGAGTAATGGTGAAACACTCTTTTTTATGGAACATCAATTCCATGCGATAACGGTTACCACCTCCTCCCATTTA

CTCACCCATTTTTCCATTCCTTAATGAGACCAGATTACGTTTTACTTCGACTTGTATAAATAGGTCTTACCTATTTCCAC

CAAACAACAAGTTCAGTCAGGACCAATACAACACTCAGAAACATTTGCTATCTTTCCATCTGTTAGATTCCCATTTTCTG

ATACAAGTCGTGAAACCACTACTCTTCCAGAATCAACCTGGTCTGAAAACTTTCTTCGCTTCCCTCCCCAAAACCAACCC

ATTCTCCTCCTTTTTGTGACCGAAGCAAGTCTGGAATGGCCATTTCTTGGTTTAGGCCGGATTTGTACAGATTGATATCT

CGAATCTAAAGTACTCCCTTGTAGTACATTTTTTAGGGTTTAAATTCGTTTCTCACCCACACACCCGAAATTACCGAAAT

CAGCAGAAACCATTTTAACCCTCAAATAATTGGCGACCACAGTGGGAGGTTGATCTTTCAGTCATGATGTCAATTTTCAA

TCCTCGATCTCGTACGTCGACCCGGACGTCAAGACAGCAGAATCAAACAATCCGCTTAGGAAACAACGAATCAGTTACCA

GGCAGCTCAATTACCAGTCGTTACACGGCAAAGGACGACTGGGGACTTCCGAGGGGTTTAGAAGCGAACGATCCGCCCGT

GTATCAGCAACACGACAAGGAATGACTGGGGACTTTCCACAGTCATGGCTTCCAACAAAACTCGGTCTAACGCCCGGGAG

ATTTCCTTTTCAACAGGAAGAGATGAAAAACCGAGTAAGTCTTACCTAGACAAAATACATGGTTTGCTATTTCAAGTTTC

CACGAGAATGATAAAACCGATAATTCATGTTATGTTTCATCCCATGTCATCTACCAACATGTGGTTCCATGGTTTGCCTG

GAACGTTTGCGCCACTTGCAATCATAACCAAAACAATGCATTCCAAAAGAACAATCCAAAACCATCAAGGTAGCATTTGT

CTATCAACTAAAAATCCAGAAAATCAAAACCATAAACTAGGCGTTTCATTTGCCTTTCAAAAAAAAAAAAAAACGTAAGC

TAAGAAGATCGAAAAAAAAAAAAAAAAAAAAAAAAACTGAAATATGTTCAAGGAAAGTTTTCCAACAATGGATTGCTCTT

CTTGGCGAGAGCCTCCTTTGCGCTTTGAAAAGCCGCCTTCTTCAATTTTAGTTTCCTGGACAGCTTTTCGACTTCCACTT

CCTGCTACTTCACCATATCCGCAGAAGGGCCTTCGAATTTTTCAGCCAACATCTTTTCAACCTCAGAGAAACCTTTGACG

AACCAAGGAGCATTGAACTCAAACTCCATGCACATGTCACGATGGAACCTCCCGGCTTGTGACCACATCCCCATAAACAA

GTATGACCAAATCCACGTTGCACATGTCGTCAATCGAAGCAATTTCCTCCACACGCTTATCCAACGCGTACCGACTAGTA

ATTTGTTTGGTCGTGGCAATATGACCGTAGTTTTCCCAGATCTTAGTGTATAGTACCGCATGTTTGGATGGCACGCTAAA

CCCTCCAATCAGCATGTGATCCCGATGAGGAACCAAGTACGGAGGATTAAAAGGGGCGCCAGCGACTACATCAGGAATCG

GCAAGGGACTCCTAACAACAATCACACCCGCGGCCACCGGAGTACTTTCCTTTGTCTCGAGCAAAGAACGACGTTTTCAT

CTTGATCAACATCCATTTCAAGATCGCCCGGTACGGTTGCCACAGTTGGAGACAAAGTTGTATCTTCCTCAGTCTCTATC

TCAGAGCCATCTTCAGAAACGACTTCCTCCTGTAATACCGCGAATTAGACGTTTTTTTTTTTTAAAAAAAAAAAAAAAGG

GGGAACATGCGGAAAAACTCACTGATTCACTTTCAGACCGACTAGAGGAAGATTCGATGCATCGTTGAGAAGAACTACCC

TCGCCATCACTGGACTCCGAACCGTCATCCTTCTCGGGTTCCCCATCACCATGGATAGGTTCAACACCGACATCCTCTGT

CTCGAGAAGCAATGTTTTCTGTCTACTTGCAAGTTTGATAAAGGCGTTCACTTTGTTGAGACCCTGGACAAAATATGAAG

ACGGATACTCAGAAAATAAACGAGAATATACACGGCCCAACTGAAGTCAAGAGGAAAATCATACATACCTGCATGTTGGA

CGAACTAAAAGTCGGTAGAGCTGTGGAATCTGACTGGAAACCACCTGCGCGATTTTTATACCTTCGCTCCTTCGCTTGGA

GACTGTCCGCGCCTGGACCGAAGGATTTTCTTTTGGTTGAAGAAGTATCCCTCAACAACGGATGCTTTGCTAAAATCGCA

GCAGAAGACCTACCACGGATGTTCTCCACTACAACTTTCACAAATTCCTGGATGACAAGGAATTCACTCTGCCAGAACGT

GTTATACTTGGAGGTAGTCGAAGGATTTCGTTCCGAAAGCAAGAAAGGGAGGATTTTGTCCGGTACGAGAACACTAGCAT

CAAGAACGGTCGGTAATGAGTTCTATGAAGCTACCAGCTGACGAACAAGAGAGACCCCTTGGTCGAAACCCATATGACGA

GCCGCCAGATCAATATTGTAAGAGACTGCCCGACACGATCCTCGAAAGAAAGATGATATGTAAACCGACGTGCAACTTCG

CATGAATACCACCTCGCCGACATCGAGGTTCCCTTCGTCAGAAAGAAAAGACATGTTTGGAGCGGAAGCAAAAGTATTGA

CCTGGACTATGGATGTACACACTGGGGCCCATGGACGAAAATTAACGACGCACGAATTATCGAGGAAATCAGCCAGGTTT

GAGCCAACTCTTGGGCGCCTATTCGACCAGCGAAGTATTCTCGAACCACCCCAAGACGCGGGAAGTGAAACCAATGGTTT

AGGTGCGTGTTTTTCAAGATGCTCCCATAGCCACGCCTGGAGAAAATCAACGAGAACATACGAGTCCACTTTCAAGTAAC

CATTTGAAACGCGCATATCCACAGCCAGATGATCCAAGTGCGTGTACAGAGAACCGAGAAACAACCCTCCAATGGGGATA

ACGACTCCTGTCGCCAATCTTATTGCAAATTTGAGGAGTTCCGGCCTTATCTCCTTCTTACCAGAGCCGTCATCGAAAAT

ATCCCGGTATAACCACAGAGCCAAGAATGCCGCGACATGGAGTGTATTATTCTTTTGATCCGGTGCCAGCTCACCAGGAA

ACCATTGACACACCCACCAGCTGTAGAAACACTTCGTCTCGTTTTATTTTCGAACAAACCCTTTCGACTTCGCAACCAGG

GTGGTACGCATTTTTTCTTCATCATCTGACAGCTTAATGTTGAGGTTTCTGTTATGGGAAGATTGCCAGAATGGCTACAC

TTTCTAAGGATATAGTCATTTCTCCCCACCTGCATATAGCTGTGTGGGTATCTAGACACCACCTGGAGATAAAAGCAACC

AAACCCGGGATGTCTTTCCTAATTTGAAGTGATGCAGAAGCCGCGATAGCATCAATGATTTGCGCTTTGGTTAAACTAGC

TCTTACACAATCTTGACTTATCATAAAGCGCATCCATTTATCAAAGGAACGCAACGGACTCACGGAGATGTTGAAATCAA

TAAGAGAGGCAGAATACTCTTTCCTCTGAAGACAAAACCGTATTACGTTCCCAGGCGGAACCGACCGATCCCCTCCCTCA

CCTTCCGGACCAGTCAGAAGAATCGACACATGCTTAAGATGACTTCTCCTGGCATAAACTATCGTGAAGAACTCATCAGC

CGCGTTTGGCTTGGAACTGCCAGCATCCTTCGGTACAACAGAAATTGATGACATCCTAAAGAAAATTCCCTTACGCTGCT

TTCACCTCCGAGATAACTAAAATGGAACAATACGAAGAGAGATTATTACGGAAAGTTCCTGGAAATTATTTTATCAAACG

CAAGAGGGACATCCTGGACGCAAGCCAATTGGTTTCAAAGTCATTTTTGCGCATATGCCAAGAAATGGGGACTGACAGAC

TCTGCATCACCACTTCATGCAACGGGAAATTGCGCCTATCCGAATAGGCGCGCCCCACAACGGGAACCATGCACACGACC

ACACCAGAAACGAAGGAAACCCTAAAAGCTAGATTTTTACGGGAAGGGAATGACAATGCTCGTGCACACTACTTCGCACA

GTAATTGAGGCGGCCAACATCACTCACGCCTAAATTTTACTACAAGGTCGTGCAAAGCATACCTATGGCTGGGATTCATA

CCAACTCAGAAAAATGATATTTCTACAAGTTCATGAGAAGGTACACCCACAGCTAGGGTTCACATTAAAAAAACAAAAAG

AAAAAAGAGAGGAAAGAAAGCAAGGTAAGGAGGAAGTATGGGAAGACATACCAGGGATTCGCTTGCCCGCTTTGCTTTAC

TGTTACCACACCTAAATACAATAAATAGGTATGAAACAAAAGGAGAGGCCGGCCCTCTTTTTATGTGCGTGATATTTGGG

AATCCGAACAAGGAAAGGATTTCCTATTTTGAGCCAAGTAAGGAAAAGAGGCAACCGGTCCCGCGAAGACAAAATCACCA

TGCCAAGATCTGTCCAGCGCCATTGCCTTGGCCCCACCTCGCCAAGCCGTCCGCCATGCCTTGCCTGTCCAATACCGTGC

CATTGGCTTGGCCCCACCTGCCAAGCCGTCCATCATGCTTTGTCTCTCCAAGACCGAGCCATTTCCCTGGCCCACATGCC

AAGCTTGTCCGCGCCATGCCTTGCCACGCCGAATCCAAGCCAGCGCAACGCCTGCGCCCCCTTTCCAAACCATACTGACA

CCAACAACTCGCCGACCCAGCGGCACGATGCTCCCTAACCCAGCCAAGGTGATGCATAGCCAGACTAAGCCGACAATTTT

CATCTCACCACATCACGGTCCGCACCACCAGTACAATTTATGCAAGGCCGCCAACACGGGAAGCGCAAACTCTCTTTACC

TCTCGAAGAGGTTAGTCGGGCTTTCCTGACCATCTTATCACGACTTGTTTCAATTTCTTATCCTAGCCTGGCACCATCTT

ATGCTTACCACGCAACGATGCTCCCGTTCCGAGCTAAGCAGGGGACTTAATGTTGATGGTGGCTTTTAGCATAGGGTTAA

AATCGTAAAACCGTACATCTGGCATGACGTCATTATTAACATTAGCACATTTATCGAATCGATCAGCATGTTTACGTAAG

AATTACCAGGGTACCTTTTTTTTCTCTATGTATCATGTTCCACGGCAGAACCCCTAACATTGATTGACATCACCTGTACA

AAACCCTCATGCTACCACGCCAAGGCATGCCAACCATTCCAGCCGCACCACTGCGCCAATGGCAAACCATGCAAACTACA

TCTACCGCGCCAAGGCATGCAAACCATGTCAGCAACACCACTGTGCCAATGGCAAACCGTGCCAGCCACATCTACTGCGC

CAAGGCATGCCAACCATGCCAGCCACACCACTGTGCCAATGGCAAACCGTGCCAGCTACATCTCCGCGTCAAGGCATGCC

AACCATGCCAGCCGCACTACTGCGCCAATAGAAAACCATGCCAAGCTAAACCGCACCTTACCTTGGCCCTAGCCATGCTA

GCGGCACCATATGCCAAACGGCATGCCAAGCCCTTGGCCCCACATGCCAAGCCAAACCGCACCTTGCCTTGACCCCTAGC

CATGCTAGCCACACCATGTGCCAAACGGCATTCCAAACTATTGGCCCCACCTCGCCAAGCCAAACGCACCTTGCCTTGGC

CCTAGCCATGCTAGCCGCACCATGTGCCAAACGGCATGCCAAACCATTGGCCCCACCTCGCCAAGCCAAACGCACCTTGC

CTTGGCCCTAGCCATGCTAGCTTCACCATGTGCCAACGACATGCCAAACCCGCCGCCCCACTATACCATGCCGACCTTCC

CTATGCGGCTACTAAAACGAAGGCGTGCGATAATCAATGGTCGCCCTTCAATCTCAGGAGCAAATCCTAGCTGTCCAAGG

TTACCAGACAACGTAAGGTAACCTTTGGCCCGAAACCCTAGTTTGGCCACGCCAAACTGCGCCAAGGCAAGCCATGCCAA

TGCCATGCCAACCTAGACGCGCCACCATGCTGTCCTTCCCTATGCGGCTATTAAACGAAGGTGTGCGACAATCAACGGTC

GCCCTTCAATCCCAAGATCAAATCCTAGCCGTCCAAGGTTACCAGACAACGCATGAACCTTTAGCCCGAAACCCTGGTTT

GGCCACGCCAAACCGCGCCAAGGAATGCCATGCCAACCTAGCTGTGCCACCATGCTGGCCTTCCCTACACGGCTATTAAA

AACGGAAGCGTGCAACGATCAACGGCTACCTTTCGTCTTAAGATGCAAAATCTCGACCGTCGAAGGTCACCAAGTTAGCA

AGTCTCCCAAGCTCGACTTGCCAAGCAACAACCATGCTACATGTTTTCCACGAAAACATTCGAGACATTAATACATATCA

CAAACTGGGGGATCCACTTTGGGTATTGGTTTGGCGGTTTACAGCGTGCGGCGTATACTATGCCCGTCATAAGGAAGTGT

AATAAGGATGAGGCGGTTAGTAAATACAAGTAGTAATGGTGAAACGCTCTTTTTTATGGAACATCAATTCCATGCGTTAA

CGGTTACCACCTCCTCCCATTTACTCACCCATTTTTCCATTCCTTAATGAAACCAGATTATGTTTCACTTCGACTTGTAT

AAATAGGTCTTACCTATTTCCACCAAACAACAAGTTCAGTCAGGAGCAATAAAACACTCAGAAACATTTGCTATCTTTCC

ATCTGTTAGCTTCCCATTTTCTGATACAAGTCGTGAAACCACTACTCTTCCAGAATCAACCTGGTCTCAAAACTTTCTTC

GCTTCCCTCCACAAAACCAACCCATTCTCCTCCTTTTTGTGACCGAAGCAAGTCTGGAATGTCCATTTCTTGGTTTAGGC

CGGATTTATACAGATTGATCTCTCGAATCTAAAATACTCCCTTGTAGTACGTTGTTTAGGGTTTAAATTCGTTTCTCACC

CACACACCCGAAATTACCGAAATCAGCAGAAACCATTTTCACCCTCAAATAATTGGCGACCAAAGTGGGAGGTTGATCTT

TCAGTCATGATGTAAATTTTCAATCCTCGATCTCGTACGTCGACCCGGACGTCAAGACAGTAGAATCAAACAATCCGCTC

AGGAAACGACGAATCAGTTACCAGGCGGCTCAATTACCAGTCGTTACACGGCAAAGGACGACTGGGGACTTCCGAGGGGT

TTAGAAGCGAACGATCCGCCCAGGTATCAGCAACACGACAAGGAATGACTGGGGACTTTCCACAGTCATGGCTTCCCACA

AAACTCGGTCTAACGCCCGGGAGATTCCCTTTTCAACAGGAAGAGATGAAAAACCGAGCAAGTATTACCTAGACAAAATA

CATGGTTTGCTATTTCAAGTTTTCACGAGAATGATAAAACCGATAAGTCATGTTATGTTTCATCCCATGTCATCTACCAA

CACGTGGTTCCATGGTTTGCCTGGAACGTTTGCGCCACTTATAATCATAACCAAAACAATGCATTCCAAAAGAATAATCC

AAAACCATCAAGGTAGCATTTGTCTATCAACTCAAAAATCCAGAAAATCAAAACCATAAACTAGGCGTTTCATTTGCCTT

TCAAAAAACAAAGAAACCTAAACTAAGAAGATCGAAAAAAAAAAAAAAAAAAAAAAACAGAAATGTGTTCAAGGAAAGTT

TTCCAACAATGGCTTGCTCTTCTTGGCGAGAGCCTCCTTTGTGCTTTGAAGAGCCGCCTTCTTCAATTTTAGTTTCCTGG

ACAGCTTTTCGACTTCCACTTCCTCGCTACTTCACCATATCCATGAAGGGCTTTCGATTTTTTTTCACCAAAGATCTTTT

CAACCTCGGAGAAACCTTTGACGAACCAAGGAGCATTGAACTCAAACTCCATGCACATGTCACGATGAAACCTCCAGCTT

GTGAACACATCCCCAGAAACAGTATGACCAGTCACGTTGCACATGTCGTCAATCGAAGACAGAATTTTCTCCACACGCTT

AACCAACGCGTACCGACTAGTAATTTGTTTGGTCGTGGCAATATGACCGTAGTTTTCCCAGATCTTAGTGTATAGTACCG

CATGTTTGGCTGGCACGCTAAACCCTCCAATCAGCATGTGATCCCGATGAGGAACCAAGTACGGAGGATTAAAAGGGGCG

CCTGCGACTACATCAGCAACCGGCAAGGGACTCCTAACAACAATCGCACCAGCGGCCACTGGAGTACTTTCCTTTTCTCG

AGCAAACAACGACGTTTTCATCTTGATCAACATCCATTTCAAGATCGCCCGGTACGGTTTCCACAGTTGGAGACAAAGTT

GTATCTTCCTCGGTCTCTATCTCAGAGCCATCTTTAGAAACGACTTCCTCCTGTAATACCGCGAATTAGACGTTTTTTTT

TTTAAAAAAAAAGAAGAATAGGGGGAACATGCGGAAAAACTCACTGATTCACTTTCAGACTGACTAGAGGAAGATTCAGT

ACTGCTGTCAGAAGAACTACCCTCGCCATCGAAAATATCCCGGGATAACCACAGAGCCAACAATGCCGCGACGTGGAGTG

TACTATTCTTTTGATCCGATGCCAGCTCATCAGGAAACCATTGACACACCCACCAGCTGTAGAAACACTTCATCTCGTTT

TCTTTTCGAACAAACCCTTTCGACTTCGCAACCAGGGTGGTACGCATTTTTTCTTCATCATCTGACAACTTAATGTTGAG

GTTTCCTGTTATGGGAAGATTCAGCAGAATGGCTACACTTTCTAAGGATATAGTCAGTTCTCCCCACCTGCATATAGCTG

TGTGGGTATCTGGACTCCACCTGGAGATGAAAGCAACCAAACCCGGGATGTCTTTCCTAATTTGAAGTGATTCAGAAGCC

GCAACAGGATCAATGATTTGCGCTTTGGTTAAACTAGCTCTTACACAATCTTGACTTATCATAAATCGCATCCATTTATC

AAAGGAACGCAACGGACTCACGGAGATCTTGAAATCAATAAGAGAGGCGAAATACTCTTTCCTCTGAATACAAAAACGTA

TTACGTTCCTAGGTGGAACCGACCGATCCCCTCCCTCACCTTCCGGACCGGTCAGAAAAATTGGCACATGCTTAAGATGT

CTTCTCCTGGCAGAAACTATCGTGAAGAACTCATCAGCCGCGTTTGGCTTGGAACTGCCAGCATCCTTCGGTACAACAGA

AATTGCTGACATCCTAACGAAAATTCCCTTACGCTACTTTCAACTCCGAGATAACTACAATGGAACAATACGAAGAGAGA

TTATTACGGAAAGTTCCTGGAAATTATTTTATCTGACGCAAGAGGGCCATCCTGGACGCAAGCCAATTGGTTTCAAAGTC

ATTTTTGCGCATATGCGAAGAAATGGGGACTGACAGACTCTGCATCGCCACTTCATGAAACGGGAAATTGGCGCCCAGCC

GAATAGGCGCGCCCCATAACGGTAACCATGCACACGACAACACCAGGAAACGGAGGAAACCCTAAAAGCTAGATTTTCGC

GGGAAGGGAATGACAATGCTTGTGCACACCTACTTCGCACAGTAATTTAGGCGGCCAACATCACTACGGCACTCACGCCT

AAATTTTACTACAAGGTCGTGCAAAGCATACCTATGGCTGGGATTCATACCAACTCAGAAAAACGATATTTCTACAAGTT

CATGAGAAGGTACACCCACAGCTAGGGTTCGCATTAAAAAAAAAAAAAAAAAAGAGGAAAGAAAGCAAGGTAAGGAGGAA

GTATGGGAAGACATACCAGGTATTCGCTTGCCCGCTTTGCTTTACTATTACCACACCGAAATCCAACAAATAGGTATGGA

ACAAAAGGAGAGGCCAGCCATCTTTTTATGTGCATGATATTTGGGAATCCGAACAAGGAAAGGATTTCCTATTTTGATCC

AAGTAAGGAAAAGAGGCAACCGGTCCCGCGAAGACAAAATCACCACGCCAAGACTTTCCGCGCCATTGCCTTGGCCCCAC

CTGCCAAGCCGTCCGCCATGCCTTGCCTGTCCAATACCGCGCCATTGCCTTGGCCCCACCTGCCAAGCCGTCCACCATGC

TTTGCTTCTCCAAGACCGCGCCATTTCCCTGGCCCCACATGCCAAGCTTGTCCGCGTCATGCCTTGCCATGCCGAGTCCA

AGCGCCGCAACGCCCGCGCCCCTTTCCAAACCATACCGACACCAACAACTTGCCGAGGCCAATGCAAGCACGATGCTCCC

CGAACCGACCAAGGCGATGCATAGCCAGACTAAGCCGACAATTTTCATCTCACCACATCACGGTCCGCACCACCAGTACA

ATTTATGCAAGGCCGCCAACACGGGAAGCGCAAACTCTCTTTACCTCTCGAAAAGGTTAGTCGGGCTTTCCTGACCATCT

TATCATGACTTGTTTCAATTTATTATCCTATCCTGGCACCATCTTATGCTTACCTCGCAAAGATGCTCCCGTTCCGAGCT

AAGCAGGGGACTTAATGTTGATGGTGGCTTTTAGCCTATGGTTAAAATCGTAAAACCGCACATCTGGCATGACGTCATTA

TGACCATTAGCACATTTATCGAATCGATCAGCATGCTTACGTAAGAATTACCAGGGTACCTTTTTTATTTCTATGTATCA

TGTTCCACGGCAGAACCCGTAACATTGATTGACATCACCTGTACAAAACCCTCATGCTACCGCGCCAAGGCATGCCAACC

ATGCCAGCCGCACCACTGTGCCAATGGCAAACCGTGTGACCACATCTCCGCGCCAAGGCATGCCAACCATGCCAGCCACA

CTACTGCGCCAATAGCAAACCATGTCAAGCCAAACCGCACATTTCCTTGGCCCTAGCCATGCTAGCCGCACCATATGCCA

AACGGCATGCCAAGCCCTTGGCCCCACATGCCAATCCAAACCGCACCTTGCCTTGGCCCCTAGCCATGCTAGCCGCACCA

TGTGCCAAACTGCATGCCAAACCATTGGCCCCACCTCTCCAAGCCAAACACACCTTGCCTTGGCCCTAGCCATACTAGCC

GCACCATGTGCCAAACGGCATGCCAAACCATTGGCCCCACCTCGCCAAGCGAAACACACCTTGCCTTGGCCCTAGCCATG

CTAGCCGCACCATGTGCCAACGTCATGCCAAACCCGCGGCCCCACTATACCATGCCGGCCTTCCCTAAGCGGCTACTAAA

ACGAAGGCGTGCGACAATCAACGGTCGCCCTTCAATCTCAGGAGCAAATCCTAGCCGTCCAAGGTTACCAGACAACGCAA

GGTAACCTTTGGCTCGAAACCCTAATTTGGCCACGCTAAACCACGCCAAGGCATGCCAACCTAGCCGCGCCACCATGCTG

GCCTTCCCTATGCGGCTACTAAAACGAAGGCGTGCGACAATCAACGGTCGCCCTTAAATCCCAGGATCAAATCCTAGCCG

ACCAAGGTTACCAGACAACACAAGGTAACCTTTGACCCGAAACCCTAGTTTGGCCACGCCAAGGCATGCCAACCTAGCCA

TGCCACCATGCTGGCCTTCCCTATGCGGCTACTAAAACGAAGGCGTGCGACAATCAACGGTCGCCCTTCAATCCCAAGAT

CAAATCCTATCCGTCCAAGGTTACCAGACAACGCATGGTAACCTTTAGCCTGAAACCCTAGTTTGGCCACGCCAAACCGT

GCCAAGGCATGCCATGCCAACCTAGCCGCGCCACCATGCTAGCCTTCCCTATGCGGATACTAAAACGAATGAGTGCGACA

ATCAACGGTCTCCCTTCAATCCCAAGATCAAATCCTAGCCGTCTAAGGTTACCAGACAATGCATGGTAACCTTTAGCCCG

AAACCCTAGTTTGGCCACGCCAAACCGCGCCAAGGCATGCCATGCCAACCTAGCCGCGCCACCATGCTGGCCTTCCCTAC

ACGGCTATCAAAAACGGAAGCGTGCAACGATCAACGGATACCTTTCCTCTTAAGATGCAAAATCTCGACCGTCGAAGGTC

ACCAAGCTAGCAAATCTCCCAAGCTCGACTTGCCAAGCAACAACCATGCTACATGTTTTCCACGAAAACATTCGAGACAT

CAATACATATCACAAACTGGGGGATCCACTTTGATTATTGGTTTGACGGTTTATAGCGTGCGGCGTACACTACGCCCGTC

ATAAGGAAGTGTAATAAGGATGAGGCGATTAGTAAATACAAGGAGTAATGGTGAAACGCTCTTTTTTATGGAACATCAAT

TCCATGCGTTAACGGTTACCACCTCCTCCCATTTACTCACCCATTTTTCCATTCCTTAATGAGACCGCAGTACGTTTCAC

TTCGACTTGTATAAATAGGTCTTACCTATTTCCACCGAACAACAAGTTCGATCGGGAGCAATACAACACTCGAAACATTT

GCTAGCTTTCTATCTGTTAGCTTCCCATTTTCTGATACAAGTCGTGAAACCACTACTCTTCCAGAATCAACCTGGTCTCA

ACACTTTCTTCGCTTCCCTCCCCAAAACCAACCCATTTTCCTCCTTTTTTTTGACCGAAGCAAGTCTGGAACGGCTATTT

CTTGGTTTAGGCCGGATTTGTACAGATTGATCTCTCGAATCTAAAGTACTCCCTTGTAGTACATTGTTTAGGGTTTAAAT

TCGTTTCTCGCCCACACACCCGAAATTACCGAAATCAGCAGAAACCATTTTCACCCTCAAACAACTGCCAGAAATAAGAA

ATAGTAGGATAACAGAAAGTAGGTAAAACTAGGGTTTTAGCATAAAATTTCAAATCCAAATCCTTATGCACAATAGAATT

GAATTCAGGGAGATACCACAAATAAGAAAATAAGAGGGCTTTTGAAAAAACCTGGTCAGCTCCCTCACAATTCTCCATAG

CTGGTGATTCCAAGGTCTTATCTTGCTGAGCCGTTGCTATTGCATCCTTAACTATGGTAGCAGCAGATTTCCTTTGCCTA

GAGCTTTTGAGAGATTTTGACTTAGCCAGAGATACAGTTTTAGATCCTCCATTAAATTCAGCCAGCTTAGGGGAAAATTG

TTTCAAAATTCCTTCAGATTGAGGAGCAATTTCTAAAGACTTAGCTTTTTTTATCTTTCAGAATAGAATCTTCAGTGATG

AATTGACGTTTCCCCTTCAAAGTGCTCACCTGATTCAGGTTAGGGATTCTACTGATGTTGTCTGAAGGTTTGCGAATCCA

GGCTCGAGAATTAGGTTTGAGATTGTTCTGGGAAGGGGGATTAGAATTCCGTTGATATATGAAAATTTGTATCAATTGGG

ATTTAGAAAATGCATTATAATTTGAGGGAGAAAGGATTTAGAAAATGCATTATAATTTCGGGTCACGCTTTGGCTCAATA

ATTTTGCATTTTTGGGGTCGAATAATTCATTAGATTGTTGTCAAAGTGATGTTGTATTAATTTAGCCTTCCTTGTATCAT

TCCAGCGACTTTTTTGAGTCTTGCAATTATATTTTTTTGGCTTATGGGGAATTATTTTATTCTTTGTTAGAGACCAAGAT

AAGACCAAAAACAAAACAAACATCAAAGTTTATATAATCCGAGATACATTGCATAAAGAGAAAAATATATAGAGGTGATG

AAATGATCGCCCGGCCGGGTGCAAGAATTTCTTCTGTTCTACTAGGAAATTAAGCAGAGACAAGTACATAAAATAAAAAT

TCTTCAAATCCAATATAGGGATGTGAACAACGGATGACAAGCAAAACCCCTGAGAAGCAACCCGCAATTATTGAATAAGG

CGCGGAAATGCTGGTTGAATATGTGCCTGGACCACTCATTACTTCTTCTTGAAAAGCAGTTGTGCGTTCATCCATTCATC

TCCATCGTTGAATGAGAATTGTACATACCCTCCTAAACACAATGTCCTGATGTGAGTAATGACGCCATTTACTGCTCCAC

CAAGTCCTGGTAATAGAATTTCTTTTACCGCTTCCATATTTTTATCTAGCTTCTTTCTCCACGCATCCCTGCAGTCAAGA

GGAATTTTAATCACTCAAAAAAGAGTTTGGTCTGTTGTTCCATTAAGTAGGATTTATTTTTTGTGTCGAGATTAATTATA

AAATCCTTACAATGTACGAGCCATGTGCATCCCGTTCACAACCCAATGATCCACAACCGAAACATCATCCTGCAAATAGA

TAACACTATTACAAACTCCACAAATATATTATGAAGATTGAGTATAATAGGTAATGACTTACTCGAGCTTTACCTGGAAA

TAGAGGAGACTATCAGCAGATAGAACTGTAACACAACCCGGGGAAAAGATGTAACCAGAGTACCAGTCATCCTCGTCGAT

TGCCTGCACAAGCCAAGCCAAGCTTTAAATTAAATACACATACGTAAATCAAAAATGCCCAAACTACAACTCAGTGAAAT

TCTGACCTCAAAAAAGTGATTAAATGTTTTATGGCAGCTATGATCCACAAAAAGAAGGCTATCTTCTGCCATCCAAGTTG

ATAGTTTCTTCATAAACAGTTGTAGGTTTTTCATGTGCTACAAAACATGAAAAAAGAAAGGTATTCAGCTTTTGCTTGCG

CCATTCTACTTTTCTCTCTATATATTAATAATCACAAAATTGGTTAAAAAGACCAAAATCAATAATTCCTGGGTGAAAAA

GACATGTAAAATTTGATACTGTTTAAATGGACGAGAATGTAAAAATAGCGGGGATGTAAGCAGGTTTTCATCCTACCCAT

TTTCAAGTACTTTTTTCTTATTTTTAATTTACATCAGGATGCATCCAGTTTCATCCTCACTCTTTTTTAAGTTTAAGCCA

GGATGAATCCAGTTTCATTCTTGTTATTTTTTTATGTCCATTTCATCCATACTAATTTTTACTCGTCTATTAGAACCATG

TTTTAAAAATATTTGGACAAATGACCCATTTTTCGATTAATAATTGGATACAAGTGCTTATGAGAATGAGTGTACCTCTA

CAACTCCAATTACAATTATTCGATCGTATGTCTTGTCAGATTCGTATTGAGTAACATCTGCCAGTATGGCGTCCACGTTT

CTCAGCCCAAGCTTCCTATATATGAAAAGATATCGAATAGTTAAGTGCAATACCAATAGTCGTCAGAATATTGTCGTTTA

CTCATAAAGTTAACTTACTCTGCTTGTTTGAGTATGTAATTTACTTGTTCCTTTGAATTAGTGAGCCCCGTAACATGACA

ATTCTTATATTTCTGAGCAACGTATAGAACTAGACCACCATGACCGCAACCGATATCAAGAACAGTTTGTCCATCTTTGA

TTTGTGCTCGCTCGCAGTATAATTTATGTGAAGCTTCTTCTGCTTCGTCTATTGTTGATGATTCATGTTCGAAGTAACAC

CCACTGCGTTGATTGTTTTAAGATAATGATCAGAAACTATATGATACAACACATATGGCATTGCTGATTTTGTTGCTCGA

GTTGTACCTTAGTTTCATACTTTTTCCATAAGCTGCTTTTAAAAATTCTGTTGGTAATTCATAGACTTCGTTTGTGAATG

CCTCGACCTCTCCTGACATATCCATCTTTTTTATAGCTACACAGGAAATGAGTTTGTCAGATTATTATTATGTCAATAGT

ATCGAACTTAAAACAAATAAGTTGTATTACAATGACCTGATTTCGTCGGCACCAAAATCTTTACAGATGAAAAGAAAATA

GAAATTGATTTGTACTATAATTTAAGGGAAGAGAGAAATGTAGATACATTTAATGAAGTCAAGGTTGAAGGAGAGTTGTT

CTTGATGCGATGATTTATAACCCCATTGAAGACGTTTCTCGAATTGAAACTTAATCAGTTTCTTCAATTCCTCATCTTTA

ATCTCTCCTCTTAACAATCTCCCTAGAGTTGCCTCTGCTGATTCCTTCTTCATTTCTTCTATTGAACCCATGATGAGACT

ATCTAGGCTAAACTAGGGAGAGAAACAAAATGAAATACTTGCGAAATATTGAGAGAGGTGGTGCGTATGAGCCAGGCTTA

TTACACAGTTTATATAGAGACCAACATCGGGTTTATGGCACAACATTATGCACTAAAAGGTCGGCACAAGAAAACATATC

TCAAAATTAGCATGCCAACATGCTTCTAGAAGTACATGAATTCCGCGTGTTATGTTATCTAAGTGCAGCAGATCTGTTAT

AAACAATCTTGGTAAAGGGGCGGCATGAATCGGTAGAGGGGCGGCAAGTGTGATAGCAATGTCACTCTTATTGGTTAGGG

GGTGTGCTATAGGGGGTGTAGATTGACTAGATTACCCTTTTCACCTTAAATCAACCAAAATCAAATCATTTTTTTTTAAA

CCTAATGAATCAAAAAAAAATCAAATCAGTTTTTTTTTTTTCATCTTCTTCTTCATCTTCTCTTCTCCTCCATCAACCGT

AACCTCCATTAAAACTGAAAATTTCATCGTCTTCGATTGATCGGCGATTTGAAAAATGTTTGGAAAAACCCAGTTAGAGT

TTAGTTTATCGTGTTGGATTTAAGTTAATTTTGTATCAAAAATTAGGGTTTTGTATGAAAATTGAAATTGAAATTTCTGG

GTTTATGTGAGTTACGGTTGGTAATAACTCAATTACTGAACGTAATAGCGGAGATTTTGAAGAACATACGGTTGGTAATA

GTTCAATTACCAACCGTATGTTCTTAATCTGAGAATTTTGTTCAGTTACTATTGGTAACTATTTTAGTTTACCAACCGTA

ACTGAGTTACGGTTGGTTTCGAGCATTTAATTACCAACCGTAAATCGAGTTACGGTTGGTATCAGCATTTAGTTACCAAC

CGTAATCTTGGTTTTACGATTGGATCTTCCCCAAATTTAACCGATTCACGGTTGGTATTCGATAACTTACGAATCAGTAA

CTAAGAGTTACTGGTTGGTAACAAGCAAAATTGCAACCGTAAGTTCTTGAAAATTCAAAAGTTTTGATTTTGTTATGTAC

TTTAGATAATTTTGCGCGATAAAAAAACTAATGGGAACTAATTTAGAAGTATAACGGGTCACCTTTGGAATCACTCATTT

TGGTTGAGTGAATCAAAATCTAGGGTTTCACAAATATCACATAAGTTTTCTTTTCTTCTCCTCTAATTTTTTTTTTTTAT

AACTCTAAAAAATCGATGTTAGAGTTATAATAAGTGAAAATTAATTATCAACACAAATCTTGATTATCATTACTAATCTT

GATTATTAACAATAAGGGCTAATTAGTCATTTCCAGATTTTTATTGATAAGGGACACTTGAATTTACTTATAATGACCTT

TTTTGTCTTATTAGGTTGCCTCCCCTCTACCAATTCTTGCCGCCCCTTTACCAAGATTGGTTATAAAGCCAACTCCCCTT

GCTGTAAAAATTTTGGACCATTTTGCCCTTGTATCATACATTCACAAGAAATGCATAAAAATAAAACTGAGGCTATATTT

GGTTTATTAACATACCTTGAAACCAATTCTTTTTACACGTGCATTAGCATGTGCGGAACATTCTAGATTTAGAACAAAGT

GGACAGGGAAAAAAAAAAAAAAACTCTAATACTTTAACCCTAAAAAAGGACAAGCAGATGTGAAATTATCATACATAATT

CAGCAAAACGTGTGGACCAGTCTGAACCCTAGAAACTAGATATGAATTCCTCTAGATAGCTTCCACAACTAGTTATCATT

TTGATTTAGTATACCGCAGAAGATGATAAAGGTTTCCAACCCTACCACAAGTGTGCGTTAAAAAAACTTATTGCTGCTAT

AAGAGTTAGTAATTGTACAATTTTTGGCTCTATGAATGGACTATTATTGCCTTAAATATCTCAATTATTGAAGCTGCTCA

AGAAAATAATTTTTGTTTCTTTGCTTATATCATTTCAAGCAGCCCTTTCACAAGTGCAGTCGATAAAGGCAGTATATAAT

TTTTGGCTCACTCGATGGTCTCTATTATTCCATTGTTATCTAGCTAGCTAGCTAAAAGATAAGCAATTCAATTTGCACCG

GCAAAATCTATTCAATTGGACCAGACTAGTGTGAGTCTAGCATCTATCCAAGAAGAATTCCATATATGCTATGATGATGA

GGCCTCCATGAATTCAGTATCGGTGTCCATGGCCTTAAAAACTTCAACTTGGAAGGAAAATCTCAAAATCAATACTTCCG

TCCAACTGGTCCTTAAGACTCACAGCGGGTTCCACAAAGAACGATTTTTCGGGTTGTTAATTGGGCCGACCGAAGGGCCC

GTAGGGAGGGAGTCCCTTTTATGTCCACTTTTTTGTTAAATGAAACATAACGAAAATTTTCAATTTCTTAAAAAGATTGA

ATTTTCTCCCGTTTCCAGTGAAGTGACTAAAAGACCCTTATACTTAAAAACCACTTCCTTCATTATCCTCACCTTTATTT

TCACCGCTACCATGCCACTCCCGTTCAAGACACCACCTCTAATGAACAACACCATATTTCTTCGCCTATCATCACCATCA

TCGTCACAACCTCCGCCACAACCACCACCTCCGTCAATATTTCCACCACCATCACCAATAGCACCTCCACAACCACCAGA

AACTCCGTCGCCACCTCTAGCGACATCAATTCAACAAGTCACCACCAGATCCATTCTCAAACCCATCTTGTAATTCCATC

AACTATCTTCGCTGCATCACCACTCCAAAATACAGTAAACCCTAAATTAAGCCCCAGCAACCACCATCTCTGCAAGGATA

GGAGACTCCCAAGGATAACACTTGTCTTGCTGATGATAATGCTTCTTATCTTCTGTAATTGTCTTTGCTTGTGTGCTTTA

AACATCTCTACCTCTTTTGATTTTTGATGGATATCTGGGCGTTCCCAAGCTTGGGGGACATAACATTCCTTGTTACAGTT

ATTTCCTTGTTTAATTGCATTTTCATTTCATTCTGCTTTCTGCATTCAAAAAATTAAAAATAGGGCAAAAAACAGGTGCT

GTCTGTAGTCGAATATTAGCAATTTATATCAAAGTATTAGGCATGCCTCAGTCATGCGAAAGAATGCTCTAAAGTCTGAC

TGAGAAGGTGAAAAGCATGCTTTTTAGTATTCCCATACTTATCTCCGCTTTTCTCATTCGAAGGAAATTATGCGTTATAA

CAATCTTGTACGAAATTTTGATTTAATATCAAGGTAAGGTATGTGTATTAAAAATAGATATGCAAATTTTTGTTCTGTAA

GAGGGTGCCGTCTCATTAAAAATAGATATGCAAATTCATCTCCCTAGTAGAGGATGTCGTCCCTTTACTTTCCCCCTGAA

TGCCCTTTCAAGGAGGTCACCCTAACCATTTAGGTTGAGCTCCTCCAATCCAATAATCGACAATCAGTTTTTTTCGTGAC

CTCATATGCTCCACCGGTATGGCTTTTGGTTATGAGATCACACCCTAGGTGGGTATAAATCGGGACCGGGTGTATCCAAG

CCAGGAACCATTCTATCAACTGGTCATGTTTCTACACCACCTGCCGGAGATCTTAATTCGCTGATTCGTTAATTCAGTGG

CCTCCATCGAATAGGCTTTTATTTCGCCCCTAACTCATGACCCAAGGTTGATATCCCCTGAGCCATGTCACCAGGGTTTT

CCCAATATGACGTGTGTTAGGTCTTAAATCGCCTCTTGCACAGTTGTGTGAACTAAGTTGAACGCCAAAATTGGGTACCT

AGCCTCTAGTAGAGGTTTTAATATGTATCTTTGCTCCCTCCTAATGAGGTCTTACTTGCTTGCTTGAGAGATCTTATTTC

GCCCTTTTGCTCACTTACTTATAGAATAAACTTTGTTTTTTCATTAATATTTCTTTGATCGGTACATAACTTGATTTCTT

CATTATTTTACTTGAATACATACTCCCTCCGTCCCTAATTAGATGAGCTACTTGTAGTTTGCAAAAATTTTAAGGCAAGG

AGGAAAGTGGAGTATATTTAAGTATTTTTTACAATTATATCCTTATGGGCAATAATTAGTAAAATTTAGAAACGATTTAT

CTCTTAAACTATACCACGGTTTTTTATGAACTTTATACCATTGAAAAGCATTTTAAAACACCTACGCAACGAATATAAAC

ACGACTATCAAATTATACATATTTCTTATAGAAAAAATAATCAATTAAAAGGGTAGTTTTAGAAATATCCCCTTGGTTAG

TGAAATAGCTCATCTATTTGTGGACAAAACTAAAAACCAAATAGCTCATCTAATTAGGGACGGAGGAGTATTATACTACT

CCTTCGCCTTCATATTACTTTCCTTCGCATGATAAATCAGGGGCGATACGCCGATATATCATTTCGTTTACCTAATTATT

TTTGTCTTCCTCCAGTATTTCGCATGCTGAGCCTCTACCTTCTTTTGCTCTTATCGTGTTAATCATCATCTCTTCTGATC

TCCAGTCTTGCCTTACGTTTTTCTTCTGTTTCTCCTTCTGCTTTGCTTTTCCTTCATAATGCTTTACATCTAGCTCATTG

CAAACTTTCGCAATAGTAGCCCCAAATTCAAAATTTTCAAAACCCAATAGAATTTGAATTTCTATTTTCACAATCGGTGG

ACACGGGTGGATTTTATCTACCGATTATGCTTAAATTTTGTCACAGAAAATTGCAGTTTTTGATGCAAAAATAATCGGTA

GATACAATCAACTTTATACCTACCGATTATGTTTTTGCAACTGTAAATCTTAGAAAAATTTCCGATCAATTTTTTGATTT

CAAACAATCAAATCCTACTTTTGGATCACAACTACTACTCCCTCTGTCCCACTAATAAGTGACCTAGTTCAAGTATGCAC

AATTCTTAAGGCAAGCAAGGGGGAAAGTATTTTTAAGCACTCTTTACAATTATACCCTTAAATAGTGAAATTTTGAAATG

ATTTATCTCTCAAACTACACCACGGATGTTCGCAAACTTCGTACCATTGAAAAGCATTTTAAAACACCTACGTAATGAAT

ATAAACATGACTATCAAATAATGCATATTTCTTATAATCAATGAAAAGGATAATTTTAGAAATATCTCCTTGCTTAGTGA

TATAGGTCACTTAATGGTGGGACAAAATCTAAAAGTAACTATGTCACTTATTAGTGGGACGGAGGGAGTATCATTTATCC

ATTTTGTTTGTTTGACCTTTTTTTTTTTTTTTTGATGTTCTAAGTTTCAATTTTAAGGTTAATGAATTTTGGGTTTGACT

TTAAACAATTAATCATAAAGTTTGATCGATGATTGTTGCAATCAAAATTAACATGATGTAACAAAATTTCAATCGGTAGA

ATAAAAAAGAAGAATGTTATGATTATGAAAAATAAAAGACATATGTTTAGATTTAATATGAAGGTGGAGGACAACATAGA

CAATTTCTATTTTAAGACACCCCATAGCCATCTTCCTTGGATGGGTATAAAAAAAATCGGCCCTTGTTATTTCGGAATCC

CTAAGGGTGCACACGGTACGGTTCGGCTCGGTTCTTGCCGAGACCGAGTACCTGTACCTCCCTAGGCTCGGTTCCAAAAT

TTTGGACCGGTGCACTGCACTGCACCCTGCCCGCACTATTCGGGATCGATGCGGTACTAGGTACCGGTTCCCAGTACTAG

ACTCGGTTCCAAACAAAAATTTCTGTTAATAAGAAAATTTCCGTTGTTCGTGCATTTTCTCAAGCAATCAATGTGCGTAT

TGCGAGCCGTGATGAAGGTTATGCACTAGTTTTATTTTAAGGACCGCAAATCAACTCCTTCCCCGCAAACTAATTCGATT

CCAACAACAATAACATTCACAAATTCTCGAGAGCACACAATATTCTCAATGATTCATTCTAAAACCATCAAAGACACTTT

CATTCAAATCAACGGTAGCTCATCCTCGTTCTTCTTCACCGAACTTTTAGCTTCCAAACTCGAGCTTCTCGATTTGTAAG

ATTGAATCGGACACGAAGGTTTTCTTGGAGGGCAAAAGCATTTGACGAAATTGATAGGCTTAAAAGGGACCGCAGTGTGT

TAATGTAGGAAGTTGTGAGATTACTAAAGCATCAAAGAATTACGAGAGTGAAATTCTAGTCATTGAAAAAGATTGCAAGT

AATTGAAAGCAAGCAGGAAAACTTGATGGGTTTCCTTGCGAAAGCGCTAAAGAACTCGATTTTGTTCATCAAATCGTTTA

GCCGGAATGAGCGGACCAAAGAATTGGGACATGGAATTGGAATTGGAAAGAAAAGAAGATTGATTGCTAGTACAAGCGGA

AGTCTTGCACCTGGAAGGGATATTGGAGGATATCCAGATTGAGGATTACCTTGGCCATCTTTGTAAAAAGAAGAGACCGT

GATTGAATCGGGACGTGGAGACATTATACTCGGCGACACTCGGTCGTGCAGTGCAAGACGGATTTGGGCGGGACCGTGTA

CCTATACCTCTCGGCCTCGGTTCCTATTTTTTGACCGGCACTGCATTCCCCTTCCTCGGTTCGGTACAAAAACAGGTACG

GTTCCACCGGTTCCGGGACGGTCCGTCGGTCCGGCTCGGTTCCTGTGCACCCTTAGGAATCCCCCTAAAAATGCCAAAAC

TAATGAAATCTTCCGATTATTTTAGATCCTTGATGAAAGTGATCCTACGGCCAAATCCAAAGACCTCGAGTCGAGACTAA

AACAGATTTAAGAAATAACTGCCTCCATTTTCTCCTCCGTCATGGGTTTATCAAAGATGAAATCTTCTTCTCATATCATT

AAAGTGAAGAAATCGTGGACAGATCGAGACTCGAAATCAGTAAGGATTTTTGTGATTTTGATTGATATAATCATCTGATT

CATATTACGTTCTTATGAACTTCTAATTAGATTGAGGTTTTCTTATATTGAGATTGTTATTCTTCATTTGATGGAATCAT

CTGTTGGTGGTATGAGATTTTGGTTTTCAGTATTTTGGATTTGATATAAATTCGAATTCAAATTTGATTTAATCATCTGA

TTCACCAAAATTAGGACTTTGGTGTGATAATTGGAATCATCTGTTGATGGTATGCGATTAGGGTATATCAGTATTTTGGA

ATTTAATATAAATTCGAATTCGAATTTGAAATCAAGACTTTGGGAATTACTATGTTGAAAGGTAACACGGTCTAAAGATT

TTGTCATGTTTTCGTCTTGAATAGTAGTCATCGACCTGATGTTTGGTAGGTGACATAACCTTGCTGTGTGCTCCCTTTTA

AGTTTTTTGGATTTGGTAGTGGTTGGGCACATTTTTGTTTCATAGTCATTGGTGTTACTTGTCAAACTACTGGAATCTAG

CGTGATGTACTGGCTAATTAAGTTTTCTTAGAATGGGTGTCGATAAATGCTTGATGTGCGTTTAAACCGTTGCACTCAAA

CAACCTAGCTAGTCAAATCTAGCCTTAGTTTGGAACTGCGTACTTAAATTATTATGAGTACTTAACTATTAGGTGAATGA

TCTTACCATTTTAGTATATTACTATACGAGCACTTGCTTCTAAGAATTTTATCTGACCACTTTTAAGTGCTCTAGACTAG

CATCCTTAGGTGATGATACTAATATTTATTTATTTTATTTTAGCCTGACATCAGTAGTATTCGTGTTAACTGTAGGAGAG

ATCTTTCTGTAGGCTTAGCAGAAGCGGCGGTCTGTGCACTTTGCTTGCAAGCATTAGGCCGGGTACTAAATTGAGCATCA

AAGTTATTTGGTTTATCTGTATATCTGTGTTTATTTTTATTTGTATTCCATCCTTCATGCTTGAAGTAATCTTTTCTTTA

ATCATGGATAGGATATCCACTTTGCATAAATCCACCTCGATCAAAGTAAATGACTGTTGTTTTTTACGTAACTAGTTGTG

AATTGGTGTAATGCTTGTTGCGAATTTTTGCTTCCTTCAGCAAGCACTTGTCTGTAAGAGGTCAAGAATTTTTCGTTTAG

ATCAACAAGCACATGTTTGTTTCTAATTCCTGCAATTTTTAACATGTCTTGACCACTTATGGACCAATAAGAATTAGTGT

AGTACTTGTATCTGCTAGCAACACAGATACAGTGCGTTGATATTGAAGTGGTCAAGAAATTTTTAAATTAGCCCAGGCTC

TCCTGGCCCCTGAAGTTGTCCTTATTAGGCCAGGGGTTTACTGGGGTTGGGTTGTCATTCCATTTGTTGTTTTTAAGTAT

TTACATGCGTCGTTCACTTTGACATTTGTAGGTATAGATTCTTGATATAAAATTGATGTCTATGTGAACTTATTAAGGGC

AATGCTTTGTTGCTATTGATGTGTGCATAAACTTGTTTATGATAACTAAAGATTCATATGAAATTTCATAATGTGAATTT

CTACTCGTATAAGAACATTTTCTGGAGTTTGTGGACTCCTTTATATCTTGTTGGAATTTTAATTCAAGTTATGTCCATAT

TGGATCCAATAAACCTTGTTTGTAGTGTCTCTTTGGTTTATAACCTTACATTTCTTGAGGAAGAGGAAGTTTTTAATGTG

AGTGGGAAATACTTATAGGGTTTAAGTAGTGGATCTTGCTGTGTCACTTGCCTTGTCAAGGAGAAGGGCATCACTATCTC

AAACATGGGGGACTGCAGAATGGTTCTGCGTAGATGTGGTGTTGGTTGATGCTCTCATGAGGGAGCACACAGCTCGACAG

GAAGATGAGCCTCTTTTTTTTTCTTCTAGGTTGTGAAGTTTTTGGTTGATAAAAACAATAACAGGAACGATTTGAACACT

CATTTTTCAGGTAATTGGAGATCTTTGCTTGAGTGGTATCATTTATACCTAGTATTGTGATCGAAATGTCCAGATAGACC

TAGTATTGGAGATCTTTGCTTGAGTGGTATCATTTATACCTAGTATTGGAGATCTTTGCTTGAGTGGTATCATTTTTCAG

GTAAGCACTTCAGATATGCATGTTGATATTGTTAAGTTCAAGCTTATGGATGTAGTCTTAAACCTTTCTTGTTAGGTAAT

ACGGGCAATGATCAGGGCGAGAAATGAACGGCAACAGTTCAGCTGTACATACAAGTCTTGTAGTGCATATACTAAAGGTA

TGCATTAGTTGACTGCAAATTACAGAATCAGAATTGTAGTACCTGCAAGTTACAGAATCAGAATTTTAATAACTGTATCG

TAACTGCAAGTTACATAATCAGAATTGCTTAGGAACATGACAGTTCAATTGAATTCATGAAATGTTAAACCCTTGACATT

TTTAACCTTGGGAAATTTGTTTTGAGTTGTCGTTTGTGTAAATTGAATGATTGTTTGTGTAAATTGAATGATTGGTTAAG

TAGTTTGAAAGATTTCAGTAAATCTGAGGGAATTAGGATGAATCGGGGTACATGTAGGAAAATTTTGACTAATTCAACAG

AGACTGACCAGTTTTTTTTTTTTTGTTGCAAAAAATTGGCAGTGGCAGAAAGACGTTACACAATTTAGTACCAGAACCAG

GATATCCTATCCGGTACTTAATAATCTGTATTGAGAAAAAAACTGCCATTGCGAAAAAATTGCACAGGATGTTTCGCAAC

GGCTGGCTGACGTTGTAATACTAAAACCGAAATTTGGTGTAGTGCAATTTCCTAACTCCCAATAACAACATTGCTTAATT

TGGTTAATACCCGTTAACATCCGTTTCCAAATCCAGCTATCTTTCCTATTTGGTTTGCTATTTATATGAAGAATGTCGCA

TGTTCGATAATATTTGGCTTTCATAATAATTGCACAAAGAGATTTAGGATTCTGACTTGGATTCCATCCGATTCAAGCTA

CCATTGCTATATTAATTTTTAAAAATTTTTTTATCCCTAAACCTCCTCCTTCCATACTTTTACATAATGAAAGTCCAGTT

TTTCAAATTCACTTTTTTTCCCATTTTCATTTTCTCCCCACCAATAATCCCTTTGAAGTTTTGTTATCCTTTTTTAAAAC

GCGTGGGTATAACAACCACACCCAATATTTCGTTCGGCAATATGCATGGACTAAACTCCAATATAATTCCTAGAGAACCG

ACTTAAAAGTGAGACTCAATCAAGAAGGATATCAAAGAGCTTTATCTCTTTCTCAATAAAATCAGTAAATCAACTAGATA

AAAATTCGTGAGACTGATTGATACGAGAATAACTTGGATGGTATCAAATATCAATGTCCGAGAGTCAATCTAATTAAGTT

CTATCCAACAAACACGGTCGGATTTATTAATTGTTATTTAACTACGCACAACCTGTGATATTTCAATTATATAAACAAAT

ATAATGTAGAAAAGAAATAATACAAACACGGAAATTTTGTTAACGAGGAAACCGGAAATGCAGAAAAACCTCGGTGCTTA

GTCCAGATTTGAACACCACATTGTATTAAGCCGCTACAGACTCTAGCCTACTACAAGTTAACTTCAGAATGGAATGTAGT

TGAGCCCTAACCAAGTCTCACACCGATTAAGGTACAGTCGCGTTCCTTACGCCTCTTGAATCCCAGCAGAACTCTGCGCA

CTTGATTCCTTGTTAGCTGATCTCACCCACAACTAAGAGTTGCTACGACCCAAAGTCGAAGACTTATAAACAAATTTGTC

TCCCACAGAAAAATCTATTCTGGTTTTGTTTCCTTCTTTAGATAAATCAAGGTGAATAGGAAACTCAACTAATAATCCGG

TCTTATACTCCCGAAGAGCAGCCTAGAAATATTAGTTACCTCACAGTAACCTAACTGATTAACAGAAGAAGTTATTGCGG

AATCACAAGAGTCTGAGACAAAGAACTGTTGTGATTACTTTTTATATCTTATTTATCGGAGATAGATCTCGAGTAAATCT

TAAAGATGATAAAACTCAATACGATAGAAAAAGTAAGATCAAAATATGCAACTACAGAGAAAATAGTTGGATCTGGCTTC

ACGAATCCCAAATGAAGTCTTCAAGACGTTAACCTAATAATGGTTTTGGAAAACCTAGTTTAAAAGAGAATCGACTCTAG

TTTGCAACTATAGGACACACGAGAGTGTCGGGATTAGGTTTCCAGATGCTACAGTTCTCCCTCATATAGTCTTTCAAATC

AGGATTGCTTATAATCAAAGCTAAGATAGCTTGGTAACAAAACAATTGATATTCACCGCTAGATGAACTCCTAATTTAAT

ATTCAAGTTAAGATAGCTTGACTTATACACAAATGAAATATACACTCATTTAGATGAGAACAAACCGTACCTAAACCGTG

TACAATGAGTTGGTTCGATAATAGTTAATCGAAGTTAGTCATACAAAAAAGAAAAAGATAAGGGAAGATTGGGTATCCCT

GTCTCAAACCCCTTGTATGTTTAAAGAACTTCGTGGGAGACCCATCTAGCAAAACAGAGATAAGGTTGTTGTCGAAATAC

ATTGGTGTACTAAATTACACCATTTGTTATTAAATCCCATTTTAGTCATGACTTGAATTAAGAATTCCCATTCGACTCTG

TCGAATGCTTTAGACATGTCGATTATAAGACCCATAAAACCCTTCTTAGGTTTTGAGTTTTTCATCTTATGAATATATTT

ATGAGCTAAGGTGATATTCTCTCGAAATATTCCTTTCAAGAATGAAGCTGATTGATATGGTGCAATTATTTTTCCAAGAA

GAATTTTCATTCTTTCGCCATCGTTTGGAAATAACTTTATAGCAAGTATTGGCCGAGTCTTATTGGTTTGATATCCACAA

GAATAACAGATTTAGGGTGTTGGGAATTAAAGAGATAAAAGAGGCATTCATCTCTTTTGATAAATGACCGATTTCAAAGA

AAGTTTGGACCATGGTTATTAGGTCAGTTTCGATAGTTTCCCATTTCCGTTGAAACAAATTGGGAGGGAAACCATCCGGT

CACGTGCTTTTATCAGGTTCCATGGAGAACACAATCTTCTTAATTTCACAATCTAGGGGTATGACACATAGCATCTCATT

ATGAACATGATTTAAGCTGGTAGGGATAAGAGTTATCAAATTTGGATCAATTTCAGGCTTTAAAGTAATATCAATATCAT

GGAAATGTTTAGTTATACAATATGCAATTTCCTCTTTCCTATTTACCAAACATTATCGTCATTTTTTGTTGTCTCTATTT

TATTTCTTTTATACCTTTTTTAGCTTTACTATGAAAAAACCTTGTGTTCCTATCTCCTTGGGCAATATTCTGGTCTCTAC

TCTTGTTTTTCCAAAATTTTCCTTTAATAGCATACCAATTAGAGAGGTCATTTCTCAGGTTAATTAAATCATGCATCTTG

TTTGGGTAATTGACCAAAGAATTGATTTTTTTTTCCGGTTTTTTCTTTATATTTTCTATGTTACTATTTATATTGCCAAA

ACTAGTTCTATTCCATAATTTCAGGATGTGTTTCATTCCGCCTAGTTTCTTGTTTACTTTATAGGCTGAGGACCCTTTGA

CATTTGTGTTCCAATTGTCCTGAATTAATAATTTATAGTTCTCATGATTTGTCCACTCTCCAAAGAATTGAAAAGGAGTT

GGACCATTATTTTACAGTCCTCATGCTTCGTCCACTCTCCAAAAAATTTAAAAGGAGTTGGACCATTATTCCAATGGTTA

TACATAGTCACTTTTATAGGCATATGATCAGAAGAGTTGGGGTACTTCCTATTCCATTTTGAAAAGACGAGGGTACCCAA

ATACACCACAATCTTTTATTTATCAACTTATAACTCCTCTACCGAATGTGATCGTCTATGGAAAGATTCGAGACAATATG

ACAATTTAGTTCACACTTCGTGTGATCGTCTACGGATACGAGATCGAGAGAATACAACAACAAGATCACTTGTGTGATTG

ACAATGGATACAAGATCAAGACGATACGAAAACAAAGTGTGTACTTGATAATAGGTTCGGTAATAACCGAAACACTATAG

GATCAATATCAAGTATTATGGAGTTAACATACTTGTGTATTTTACTTAATTATAATAACAATTATAATGCAGAAAAAACA

AAGTAAAAGGCATAACAAGATTTTGTTAACGAGGAAACCGCAAATGCAGAAAAACCGCGGGACCTAATCCATTTTTGAAT

ACTCTCAGAATTAAGCCGCTATACAAAATCAAATACCAACTGCGTATAGTTGAGACCAAGCATACTACTCCTAGCTACTT

ATTTCCCTTAGAATCCCTGCGCCTTCGACTTCTAGAGTCACACACGTGAATAGAAATTCCTTTGGATCGTATTACAAACA

ACAAAGGAAGAATATTTTTTGTAACTACTCTAATCAATCTTGCTAGAAGATAAGATGAGTCGTTTACAAAGGTTCTTATG

TTTAATCTAATAAAATCCTTTGTCTGGTTAGATCAATCTATCCAATAACTACCGAAATATCAAGTTTAGATTCGCAGTCA

AACGATATACATCTCAAATAGAAACTATAGAAAATGCCGATCTCACGCAACTAATCAATCATATCTATCAAACATAAACT

GATTCTAGTTGGATACCAACCGATCAAGGTTTGTGCACTAAATCCACAAGATAGGAAAACTAATACGAATTCTTGTTCGT

CTTCAAAACGATATTGCCTTCAAATACCTACACAACAACACTTGAATCCTCTTGTAGAGTCTGTTAACAATGGATTATCA

CAAGATGTCTTTAGATCTAACAACAGTTCTAAATATCCAGTCGATACTTTGATCTAGTCTGAGTAAATCTTATATGAGAA

GAGAAGATTCTCAAGAATAAACAAACTAGGTGCAATTAAAGTTTCAACAACTGTTAGTCAATCAAATAAATAATAAAAAA

CTAATAACACTATAATTATCTAGTTTCTCACCAATGGTACTCATAGAGCTTCTTGATCCCACAAATGTCTTTAAACGAGC

GGTCTTAAGAGATATCACATAATTAAGATACTTTCCTCTCTTAATAGACGGCTCCACTAGAAACAATAAGAAATGAAGTT

TTCCTGGATCTTAGAATAGTTTGCTAGAAATGAAAAATTAGGTATTTATAAACCAAGGTTGTTTGGACAAGGCTTCGAGT

CTCCCTCAAGGTTTCGAGTCTCCCTCAAGGTTTCGAGTTTCTCTCTAGCATATTTTCTGCAAATGAGAGACATAACAGTT

GGTTAACAGTTGTGAATCCGAGCAACAGTTCTCAGGGTCGAATCTCCCTTTGCGCAATATTTTTTTGACTTATCAAAAAA

AAAAAAAAAGTTTGGACACCAAGGAATTTCCAAGACCGAAAATATTCTTAAGATATGCAATGAATGCCCAAATTCGGTTT

TTCTAATTCCAATAAATGCAGTCCAATATTTTCCGAAATCTCTCATAGAAAATCTCTAATTAGTAAATGCACATTGTTAG

AGCATTGCTCGGTTGAACCCACTAGCGTTGGTATGCCAAGTTAGTTGTCAATTTTAGTTTCCAAAATTCATTCTTGATTT

TGCATACTAAAGATAGTTTCGGACTAGATTAGGTCTAAGAAGTTGAATCGAAGCTATCCTTAAAGGATGAAGATTGAAGA

CTACAAGAAGACATCTATAAGTACTTCATAAACAAAGGTATGTGATTGATTTCATTTGTATTCTTGTTCAATTCTACCTT

TCTAATCTATCGAGATAAATGCTCACTCTATGGAACAATTCCTATGACGGTAAATGTACATTTATGTATATATACTTGAG

GTTATTTGATTCATGACTTTAGTGACAATAATCTTTATCCCTATAGAGGATCTCATGTTATAGGGAATGAGTTTATGAAT

TCAACGAGCCAAGAATCACTCAATTCCGAGTTATAACGATGAAGTTAGGAGTGATTTATTAAAGTAACAATTATAAGTGT

TGTTGTATTTGTTACAGTTCGTTAGTAGGAAACAATCCATGGACTGTTTGTAACGGTTCACGGACTTGGGCCCAATTACG

TGACTAGAAGTCTTTTTTAGTCAAGTTGGTGATGTTTCTTTATCTAGGCAAGTTAGCTATTACTCCAAACATGTTTGATT

ACCATATTAATTAAAGTGTTTGTGATTCCTTCCTAATTTAATTTGTGATTTATTCACATAAATACAATTTGCTAAATTAA

TTATTTATTTAATTATTTTGGCAAGAGTTGTAACTTAGGAAAGACTCCCACAATTAGGAGAGTCTTGAAAAAAAGGAAAA

TAGCTTTGCTTAGTTTTCAAGCTACCTTTCACTATAAAGAAAGGGTTCGTGAGTGCTACACGTTTTCATCCCCTTTGGAA

AATTGGTGTAAGCTCATAAGGTTGTTTTCCATGTCTTATGTTCTTTATGAACAAGAGTGAGATAAAAGTCTTTCTATTGG

GGATCACAAAGCCGAGTTGGGTTTAATTTTCGTGATTGATTATACAACTTGTAATCTAGTTTTTTCACCTCTTGTATATT

CTAAGGTTTTCTCTTCTGATATAAAACCATTCAAGAGTTGTTGATCATAAATCCTAGGTTTTGATAGATTTCAGTTTCCG

ATCGTATACCAACACTTGTTAGTCGAAATCGGAAGCTAGAAAGAAAACTTCGATTAAGGTATCTCGTAAAAATCCTTAAG

AAGTTTTAAACAAGATATCTCTAGATTTATCTAGTTGTTCTTGTTGTAGAATTATTAGGGTTTTCTTAACTTGGAAATTG

ATCTTTGGAATCACCCAAGTTCACTTACGAAAATCAGGTTCACGGACTTCTGATGAGTACGCATACTGATTGTAAGTTAA

AACCCTAATCTACAAGTTGGTTTTGGTTTCACTACTTTTTATTTGGTAGTTGTTCAAAACAAACACAAGATTTGCAAAAC

CTTGATTCACATCTAAAGGGAAATCAAACCGGTGTTTTGTGCAAACAAAATATCAAATCATATCAATCGTGTTGTTGTAT

CTTCTAAAGATAACCTTGAATTCTGCTAGAAGATTTGCGTGAAGAATTTCTAAAGAGAATCGGTATTCTGCTAGGAGTTC

TACAAGTGATGAAGTAGTTATTACATCTAGTCCGAATAGGTAGTAGGAATTTGGTGTAACAGCTTATAATCATTGTGTGT

TTATTCACGGACTAGGTCCAGGGTTTTTCTCCATTTGCGGTTCCTCGTTAACAAAATTTCTGGTGTCACGTGTTATTTCT

TTTCCACATTATATTGTTTATCTTTATAATTGAAATATCACGGAGTTGTGCATAAGTTCAATCAATTGGGAATCCAACCT

TTGGTTCTTGATTAAATTGATTGACACTTGGATATTGGTTTTTGATACGAGTCAAGTTATTTCTTATATTCAATCGGTCT

CGCAAATTCCTATTTGTTTGATTGCAGATTGAATTGAGAAATAGAGATATAACTCTTTGATATACATTTCTTAAGATTGA

GTCATTGTCTAGTTGATTATCTTGAAAGTATATTGGAGTTAGTCCATACAGATTGCAAAACGAATTATTGGGTGTGGTTG

TTAGACCCCCCGCTTTTTCACACATTACTAAGTTTTATTCTCTAGAGATATGCATTAATTGCAGGTAATTAAAGCATAAA

AACCAAAAACCTTAATTAAAAGATTCTTAATTATTTGATTCTGAGGATCGCCTTGAGTACCAAGGAATATCTTTGAACAA

TAAATGATAAGAGTTAATGCTCATGTTAAAATTATGTTGACATCTTTACACAACAAACCTAAATCCTAAAGTACACAAGT

TCATGAAGAAATGTAATTCATGGAAAATCGGATTTGCATCTTGGAATCGGTTTTACCACAATTAGAACAAGTTTAGAATA

GGTTCTACCGAATTCTAAGACTACTATGTGATCAGTCATACCAAGTCATAATGGTTAGTTGTGATCGGTCCTACCAAGTC

ACATACATGGTTCGGATCGGTTATACCAATCACAACCTGTGTATTGTTAGGTTAATTAGGGTATTAAGATTCATGTGTAT

TGTTTGATGTTTTTCAGGTAAGTATGATTATTGATTTATGCTAAGATACACCTAATTTTTTTTTACATTGATGCATGTTT

GATGTTTGCTTCAATTTGAACCTCAACCTCTACTGAATGGGTTATATGTAAAACCGATGTTGTATTCTTACATATTTCAA

ACTCTATGAAATGCAAAAAATGATCTTGTATTTTTTTCTTTGTTGATAGATTTTAATTCATGCTGATGAGAGTTCCTTAG

CCTTATTAGAATTATGTCTCATCTGAACTTGTACTTCATACTGATAATTAGAGATTTATGTGGCGGATGAGATTTTTTTC

TTTGCCTGATTTGATCTATTTATAAGACAATTCTTTCTAGGTTGCAGTTGTTATTCTTTATTTTGAGGTGTAATTTTTCT

TCCTGTTTGTGTTATAGTCTATATGAGTCTACAAAAATAACATATATAGATTTGTTAACAGCGTACCCTGTGAATTTTGA

CAAAATGCACATTAAGTGTTTGTGTGATTTACTTTAAAGATATGGGTTTTCTTAAATACTTGTGTTGTTATTGTCTTCCA

GTTGCATAATACTTAAAGTGTTTTCAGAACTGAAATTTATTAGGTCTGTTTGAACCCTGTATGACATGGTTTTCATCCAA

TATTGAAGCAAATCAAATCATATTCAAGTAACCCAATTTACTTTTACATATGTGCAGGTACTATTAAACATTCTACGGAA

AATGTAGCACCAATTGGTAAGAAACGTCAAGTTAGACCTACAGTGTGGGATCAATTTGATATGGAAGAGGCGCCTTCAAG

ATATGCAAGGTGTCGTTATTGTAATGCAAGATTTGCTCTTTGATCTTATGAAGAATGGTACCTCTGGGTTGATTGCTCAT

CTAAAGAGATATAGACTAATAGTTAATCAGATAAGAAATTGATTGAATGGGATCAATTTAATATGGAAGAGGCGCCTTCA

AGATATGCAAGGTGTCGTTATTGTAATGCAAGATTTTCTACTGATCTTATGAAGAATGGTACCTATGGGTTGATTGCCCA

TCTAAAGAGATATAGACTAATAGTTAATCAGATATGAAATTGATTGAAGTCTACTTTGTACTAGTATTAGTACATATTTG

AAATCTTAATGCCTTAAAAATTATGTTTGTAGCGAGTTAAGTTAAATCGGTTGCATGTTACTAGAAATTTTGGTGAAAAT

TTACTTGTCGGATATGCGAAACCGACTAAAATGAAAGTCTGTTTATAAAATGTGGTTAACCAAAACCTACAAATTTAAAA

CCGAAGCCGAGAGGTTCAAAACCGAGTGGAACCGAATAGGCTAAATGGTTTCATTGCTTTTCATTATTGGAAAACCGAGT

ACTTTGGTTTCGGTTTTGGTTTTGCCAATTACCAGGAAAACCGAACCATGTGCAACGCTAATCACAGGATCGGTTCTACC

TAAATATGGTACCTACTTGTGATCGGTCATACCAGTCACTAGGGTCGGTCATACCAATTACAAGGATCGGTCACACCAAT

TACGGAGGTGGTCTTACAGATTACAAGGATCGATCATACCATCTCATGGTGATTACTTAGGATCGTTTACACCCAGCAGG

TCATACCAAATCATAAGTCTAGGTATTGTGATTGGTTATACCAAGATACATAACCCGAGTTAAGATTGGTTATACCAACA

CACACATATTGGTCATCCAAAGAATATGCAATGAATATTTGGACCAAAATTATACCTATTGATTTCCCTTTCAATTCATG

AAACAAGTTCACAAACTTACTTCCTTTAAAAGATGTAAAACATTGTTTCTAGGATGAAATCTTCACCATACACCCATACA

CATAATCATAAAACTATGTACAAGATTATGTCGATGTCATATCTACGAAAAAGATAAGTGTTATACTTCGTAATCTAATT

CCTTACTACTATGATCATGTCTGTCCACTACAGTATTATATACAATATACACAGTATATACAACATAGCATGTTATGTTT

TCAATATAGCACGGCTTGACAGATACGTTAGGAATTGAAACAATTTCAAGTCAATATTACTAACCTCAAGTGGAAGGATG

ATGTCGTCGTTTTAGTTGTTACTTCTTACGATTCTTGGTCTTGGAGTAATAATTCTATGTCTCAACATTCCTAGACTTTC

TAGTCTAACCTAAACGAAGTTGACTATAGTATTTAATCAAGCGACTCTAGATGAGTTTCGATACTAAAATACGACAACCA

AAATTGACATACCAACTCTTGGTGAGTTCAACCGAGCTATGCTCTAACACATTTATCGTTCACCAAAGCTCTGTCTAATC

TTTATTCAATATTTTCATCATTTTGCCTATGGTTATTCCAGGTGAAAGTACAACCTGTAAATCCTAGATCCATAAGATTG

CACTTGTTAATTAACTTATTAAAAGTATTGGCCACATTTTTCTTAAAAGGATAACGACTTTGATTTTCTTCTTCATGAAG

AACAACATTTAAATCCCCTATGATTATCCAAGGTTTGTTTATTGTAGCAAACCATTTATTTTATGGTTTTCCAAGATTTG

GTTTCAAATTTATGTATGGACTCCCATAAAATAGGTGAGAGTCCAACTACTGTCATTTATATGATTCGTAATAGTAGCAT

TTAGTTAAAAGCATGGTCCTCTATTTTTATGTTAACATTATTTTTCCAAATGAGACAAAGACCTCTAACACTACGTCTAG

GAGAAACTAAAAATAGTTATGAACATTCACAATCTCATGATATTTCCCATCTCCTTATTTTCTTTGTTTAGTTTCGATAG

GAATATGATATACGGGTTTTCCTTATTTAAAGTATCTAGGAGACTGTTCCTGGTGTTTAATTTCCCTAGTCCTTGACAGT

TCAAAGCAATCATACTAATAGACTGCTAGAAGTATGATATTTCAGGATAGCTAAATTATACTAGCACAATACAATTTACT

ATGTCATGAGTACTACTTAAATATTTTAAAATTTTCTAAACTCCAATAGCTTGAGCAGTAGTTTACTGCTCTTGAAATAT

CAATTGGTATACATAGAAAACCACAAAAAAAAGATATGCAATTATGCGCGCACGAGAGATGGAATCATTAAACAAGAAAA

AATCCATTTTACAATAAGTAAAGCCTCAATACTTGTACGTAGGCTACAATATTGAATAACATATTGGATTAATGCTGGTG

AAAAGCGAATATATTGATTATTAATGATCTTAATAAGCGATGACATTTTTTTTTTGTTATAATCGATTTTTATCAGCTAG

GTTTTCTTTTTACATGAATATTAACGACAGGGTGGTTTTCGAGAATATTTACTTGGTATTTGTTGTATTTTACTTTGACT

GCATGTTAACTAGAATTCGATGATAATTCTTCCCTCCTCATATCCATTATATTATTTCCTACCTAATAACAGGACTCGTG

ATTTTGTTTACTCATTATAAATTGAACTTAATAATTAAATTTTGAGTTCTAGTTAAAAAAATCAAAGTTGTAATGAAAAC

CCGAAAAAAACCAAAAAAACGGTGGGACGGTTTTGGTTTCTCATATTATAAACCGAGCATAGTCTAGTTGGTTTTTGGTT

TTTCAGAAGAGACATTGGTTTGTGCCAAACCCAAACCCAAGCAAAACCAAACCACGAGCAACCCTTGTCAAGACACTTGA

AATGTTGCTTGGGTAGTCTTCTTTGGTGATTGTACACTTACGTATGTATATTTATAACCAAAGTCTGATTATTGAAATTT

AATCATTATAAGCTAAATTTTCTTATCCCATATCAGGACTCGAAAGAGCAATTTTTTTTTTTTTGTCACATATCAGGATA

TGGAAGAGCAAAAATTGTGCAGAAATATAGATATGCAAAGACATTAAATGTAGCAGTAGTTGCCAAGGCAAGTTGATTCA

GCAGCCAGAACGAGTTCCGGACAAGGTTGCGAGATATATACCAGATATGCCCATCTTAGTTCAATGATGCATCTTAGAAG

TTCTGGAAATGAACTTGGCATGAATGTTTTTTTAGAACTTGTCAAGTCAAACAACTCATTTTACTGGTTGCTTGCAGTAA

GACATCCAATTTAGGCGCAAGGTTTTGAAGGAAATGGTGGCGAAACACATTGCTCCAAGGGACAGATCAATCTGTTTTGA

GATTAATATCATGCCAATGCAAATTGGGAAATATGGATTCTGCAGCAGACGTCCCTTGAAGGGATAAATCAATTGAGTAT

TCCTACTGAGACAACTCATTACGGGGTTTTATAATTTTTGCAAGACCGGGCAACATGATACAACCATTGTGTTTGTGGAA

AACGTATTAAGAGAGCGATCCTCTTGAACCCAATAACCCAAAAAATGATTCGAGTATCTTTAGAACTACGGGCTCATTGG

AAAGCAGAAACATTTATCCTACAGTTACTGAAAAAAAGATATCTAGTTTCCTATTGTTGTTTTGCTCAATGGCTGTCCTG

GTAGCTATTTCAAGCCTACTAGAGGTCTTACGCAAGTGGATCCTCTTTCCCCTTACTTATTTATAATATGTATGGATATT

TTCTCCAGAATGCTTATAAATAGTAAGTATGAGTTTATCTACCTTTAACTATCATCGTCCCCTACTCAGAGTCACTTGAC

TATCAATTTTAACCTTATGATCTATAGTGAAGAAACCCAATCCCCCGTGGACATAGGCATTAGTGCTTAGAACCACGTAA

ATCATTGTCTTATTTACATTCCACACTTATGATTTTGGTTTATCTCTTTTAAACATCGTTTTAGTCATCATAGGTGCATC

TGCAACAATAAGCAGAGTCGAGACCATGACGAGCCTCAGGCTCAGAGTTTCCAGTGAGAGTTGGGAGCCAATTCATAGTC

ATTTCGTTCATTATTGTATTTCATCATTGCTGCGAACATTGATTGTGGACACGATGCTACCATCGTTACTTCTGTGTTCA

CAATTTGGCGCTAGAAAGCGGGACTCTTACGTCCCGTGTAAAGATTTATCTTTCCTTGTGAACTCTTTTTATCGTTCAAT

ACTGTTTTTTAGTACATATTTTGGGTGAGAAACCAGCAAAAAAATATCAGATTTAAAAGGATTTTGGGTGAAACAGAGGC

TTTTCATTATAAAAACCAACGGATCTGAGAGATCGACTCAAAATACTAGACAAAAAATCGGATTTGTTACCAAGAAAAAA

AATGTTCGGATCCATATTTTTGTTTATAAAAACCCTTGGATCTATTTCTCTTGCAAAAGGTCCTTAAATCTACCTCTAGA

TCTGCATTCTCCCTTCGGCAGATCTGTAATATGGTTTTGTTTTGGCTCGCGCCTAACAAAAAAGTACCTTCGATTTTTCT

TACATTACCGCTATCCTTGTGACTAACAAAACACACGTGGTTATGCTTTCTCTGCCAACTGTTTATTCTTGCAGAACATG

GATAAGACAAAGGATATGAGAAAGGCCAAGGAAGCGTCAAAGGAAACACCAAGGACCACACCAGTCGTAACACGCAGCAA

GCAAAAGGGTGAAAAGCTCAAAGTAGCAAGCAAAAGGATGGATGGAGCCAAAAGTACCACACCCATAGATGCCAATGCAA

TAGTAGCGGCGACCCTCAAAGCTGCAGCAGCCGAAGCCGAAACCTCCAAAGCAGCAGCAGCAACAACAACCAGAATGAAT

GAACAACGAGAAGGGGTTGCGAAATCAACGATGCAGCAACCTGTCTATGGAATGCCACTCACTGACGAGAAAAAGCAACC

ACTACTGACTACCCATCCCCTGTTAATGGCAAACCAGCTGGCACCACAACCTATGGAGCTTCAGGTCTTGCCAACCGATC

CACCGATCCCTCTGATCCAGACTGTATATGAAGATCATATTATAGCCGCTGGTGGACAAGCACAGATAGGAACTCCGAAC

CAAGGATCAAACCATTCCATCCAGCTAGCAGCAGAGCTCGAAGAGTTGAAAAGGAATCAGAAGATATATGCAGATGTTGT

AGCGCTACTTGCATGAGAGAACCAAGAGCTCAAGGATAAGATCGCCCACAATGCCGAAGTAGAGAACATACATGATGAGG

CGAATTCCAAGGAACCACAGCCGAACCATGACCGAAGGATGGTAGTGGCCAACATCAACAACCCCGAACCACTGAACCGA

CGTATGGCAGAAAGGAACAGATTGTCCGATCCCGATTACGCGCCCGAAGATTCAGACTACTTCGATAGTGATAACCGAAG

AGAGGGACGATCATCCATGAGAGACGAGGCAAGGAGGAGGAAGATTAGAGGACCACTGATCGCACGTCTGGCAAGAGCCC

TCATTCCACAAAAGTGTTCCATCCCAGCGTTTGAATGTTACGACGGATCTAGCGACACCGTAGCTCACCTTCGGTATTAC

AACCGAACTGTGGCCCGATGGGACCGAGACGATGTGGTCCTCTGCAGATACTTCCCCTAAAGTTTGAAAGGATCGACATT

ATCTTGCTTTGACAACCTGATGCCTAATTCCATTGATTCCTACAGTCGACTCACAAAGATATTCTTGGAAACTTACATGT

ATAACAAGACCGTCAACGCAGGAATGTACAAGCTCTTCTCACTAGCGGTCGCCTTCAAAGAAACAATCAGAGAGTACACA

GATAGGTGGCACAAGATCTGTCAAGCGATAGGGAATGTAGACCCGGTAATTAGCAACAATTGTTACAAGCGGGGATTGGA

TAGGATGAGCCCATTGTTCTTTGAGATTCATGGAAGTATTCCCACTACTGAAAGAGATCTCCGGGTAATCATCGAAAAGC

ACGCCAGGCTTGAGGAGATCCAGCGGGACAACCCTAGAGCGCAGATGCAGAAATCTCATCATACAAATTTGGTAGAGCAA

GATAACGGCTCCAAAAGGACAAATTCAACCGAGCGCCTCAACGAAGATAAGAGAGGACGAAGAGATGATTATTGACGCAA

CGATCAAAAGTTCGAAGATCAAGTTTAGGCGAAGCTCAACACAAGATATGTTCGTATTTTGATAGAAATCAAGGATCACA

CCAATCTCGAATGGCCATGGTCAAAGGGAAAACAACCACCGCGATCGGAGAAATCAAGGGGTTACTGTGAATACCATTGC

TTCCACGGTCACCAGACAGAGAAGTGCAAAAACCTCAAAATAATGATCCGAAACTAATAGATGTTGGAGATCTCAAACAA

TATGTTCTAAAGCCACATGACAAGAAGAAAAAACACATCGGAACAAGTAAGTCCACCTACCAGAAATGATCGATTGATCA

ACACCATTTCTTGCTCGGAGCCTTGCGGCCCATCACTAAATGACCAGATCTGGAAAAGGTTGAGAAAAAAATTCGAGTAC

TATTGCGAGCTATACAAGATCGATGGAGAGGAGGTAGATGGACACGAATATTGGATGAATACACCGCTAACTTTTTCTGC

CGATGACGTTGAAGAAGATATGGTAGACCATAACGACCCTCTGGTGCTCACACTACCTGTCTCAGGGTGGAATATAAATA

AGATTTTGATCGACGGAGGGAGTTCAGTCAACGTTTTGTTCTATGATACCTTCAAGCAAATGAAATTAAATGACGAGCAA

GTGATGTCTTCTTATTACACTATTTATGGATTCAACGGGGTAGCCACGAAGCCCTTAGGGGATATTGTCTTGGAAGTTAA

AGACGGACCAATAAAGGTCAGCACCAGGTTCAGCGTAGTAGATGCACCGTCCCCCTATAATTCCATCATCGGACGAAGAT

GGGTGCACAAACTCAAAGGAGTAGCTGCCACGTACCACCAATACCTTAGATTTCCAACACCCGAAGGGGAAATGAAGATC

AAAGGTGATCACGTTACAACACGGGAATGTCAGGCTCAGCAAAATAAACTAAACGACGAGCAGGAAGAGCGGCGCAAGGC

CCGAATGGGCGAAAACAAAGCAGCGGCCAAAGACAATGCGATCGACCTGTATCTTGAAAAAATCTCAGGGAAAAGCCTAA

CTAAGAGTAGTACCGTTCTAACTACCGAAGCAGGCACTTCAGGGGTAAAGGAGGTTAAAGAGTCCACCAAATAGAAATTA

AAGAACGTTTCTCTCCTAGGGAACCGAAGCCTACATTCACGCCGGTAGAGCCCACCAAGAAGATCAACATAGGAACAGAA

GAAGATCCTAAGATGATTAAGATAGGGACTCTGATGGATAAAGGCGAGAAGAAGCACTAATCAAACTTCTCAAGGAGTAC

ACGGATATTTTCGTATGGAAGTTAGGAGATATGCCAGGAATCGATCCCAAGATCATACATCACGAACTTCGGATAAAGCC

AGGCACAAGCCCCTTCAGGCAGAAATTACGCAAAATAGCACCAGAATATCATCAAGCGGTCGAAAAAGAGCTTAACAAAC

TTATGGAAGCAGGATTCATGAGAGAAGTTAGGTACCCCACATGGATTTCGAACATGGTCATTGTTCCAAAGAAGAATGGT

GGGGTAAGGATATGCATCAACTTCACAAATCTCAACAAAGCATTCCCGAAGGATAGCTACCCGCTCCCAAGCATCGATCA

ATTGGTGGACGCCATAGAAGGACGGGGAAGAGCTATCATTCATGGATGGATACTGGTTACAATCAAATAGCCCTAGCAGA

AGAAGATCAACAACACACAATATTCTTTACTCCACACGGACTCCATGCTACACTCGCATGCCCTTCGGACGAGAGGAATG

CGGGAAACATACCATAAGATAGTTGATGCAATCTTCAAGCCATGAATTGGGAAAACCTTAGAGGTCTATGTTGATGATAT

GCTCGTAAAAAGTAAGCTGCGCAAGGATCACCTCCAGGACTTGAAAGATATCTTCAGTGAAATGAAGAAATATATCATGA

AAGTAAACCCGGAAAAGTGCACCTTCGGCGTCACATCAGGAAAGTTCCTCAGATATTTGGTGACGAATAGGGGCATCGAA

GTTAATCCCGCTAAAATCCAAGCCATCGTAGAGATGACATCCCCAAAGAATTTGAAGGAAGTCCAAAAACTCAACGGATC

CATAGCGGCACTCGGGAGATTCATCGCTAGATCATCGGACAAATAAAAACACTTCTTCAACATCCTCAAGAAAGAGAACA

AGTTTGCATGGACCGCTGAGTGTGAAGAAGCTTTCCAGAAGATCAAAGAATACTTGGCAACGATCCCGATCCTCCAAAAA

TAGGAACCCGATGAGTTGCTGGAATTATATATAGCTGCAACCGAAAATGCTGTTAGTGTTGTATGTCATGTGCCTCATGT

GAAGAGACTGCCCCATATCCTGTAAGTGTCAGTAATTAATATTAGTTGTTGGTGGTAGTGTTTAGCTTAATAATAGGTGT

ATTTATCAGTTGTATGGGATGGGTTAGATTTAGCCATCTGTGATGGTAGTAGCCGTTAGATGCAGGGCAGGGAAGCTTAG

TTTTATAAGCTAATTGTTCTGTAATGAAAACTCATGAAAATTATTAATCAAAATATCGATTTATCGATTTAGGCTGTGTT

CTTCGGAACCATAGAAACTTGGTTCTTTGAATCAAGTGGTTACTATCCTCATCTTGTCATTGTACATCTCGATACATGAC

TGTATTATTTGTGAATCTATGATCTAGCGATGGGTTATACGAAACTTGGTGCATAATTGTTAGGATCGAAAATATATTTG

TCTACTTATTTGATATTTCCATGCTTGGGTTAAATCCTTTGATGGGTTATGCTTAGAATAGCCAGATATTATTTGTGAAT

CTATATTTAGTTTTGCATAACTTTCTCGCTAATGCAATTTGATGGTTCATGCTTTGGTTTAATCTTTTGATGTGCCATAC

TTAGGGTGTCCTAGCGATATTTGCGACTCTAATACATATAATAGGTGATGTCGATAGAACGAACGATAAATAAATATTCA

TGAATTATGTTTCATTTCAGTAACTGAATAGAAATAATGGTGGATACTAAAAACCCTGGTTCGCGACTTTGTATTGGATT

CATTCTATTAGTTTAGTTCATTTTCTTCGATCTTTTTATTATTTTCTAAAAATCCAAAAACATCATTTGTGGTTAGTTAA

TTAGAGATATTGGTTACATTATTTTCACCGCTCCCTGTGGGTTCGACTCGTACTTTCCTCTCGTCTACTAGTTAGACACC

GTGCGATTGCGATTATTATATATAGGGTTTCCCGAAATCCTATCAATTTTTGACGTGGTTGTGGTTAGACTTTTGAGGTG

TCCCAATGATATTTGCGAATCTAATAAATATAATAGGTGATGTCGATAGAACAAGGGAGAATGATTGGCAGCAACAAGAC

ATGGTCTTTGTGGATTGATGGAAGTTTGAGTCGGGCAGTGAGACGAAGAATTGAGATGCTGGTGTTTGGCCTGACATGAA

AGACCAATTGACGGACTGAAAACCGAGAAGGATTGAATCAGTTGCAGTTGCAGGAAGTGGTGATTTCATGGACAAAATGA

ATCAATTAGGTTCGAGTTATTGAGATAATGAATGCATGGATGATTGCAGATGGTGGCTGTATACGTAAAGCTTCCAAGTT

TCAAGGAAATGACAGTGAAAGCTTCCATGCAATCTGATTATGAGTGATAAAATAGTGAAGAACTGGTGATCAAACTGCCT

TGGTTTTGGGTTGTGTACGAGAAGATGGAAATATAATGGAAAATGAGTGATGGAGCTGTTGATCTCAATGGAACGACAGT

TGAAGGAAATGCTAGAAGATAACGAAAGGAAGATGTTGTTTGGAGTTAAGTACGGATGGCAATTGGTCATGGTGGAGCTT

GAACTCGGATGGATTTCGGAAGCTGGAGCTACTAGACCCGAATGGACTGGGCCTTGGCAGATGTTGCTAAGGCTTAGTTG

AACTCTTGGCAGCATATATATTACGAAAAGAGAAAAATAAGGAAGGGGAGACGAAAAGAAAAAGCTGCGGCTACAAAGCC

GAGAGCTGTGGAGAAGAAGACGATTTGAAGTTCCATTATCTCTATTCCGTTTCGATTTATGAGTGATTTTATTTTACATG

TTTATGGCTTTTGAGAAAGCCATGTTTTGCTAAATTTATATAACTAAGGTTTTTATGTTGAAACTACATGGGTTTAGGCT

TATTTGGATTGATTATATAGCAATAGACCATTATGATATGAATGAATTAATCTTTGAATATTGCTTATCGATTTATTGAA

AATGAGTTGTTGCTTCACCGGTTTCTTATAACCTTTGTTCGTAATTGTTAGGTTCAAGAATATATTTGTCTCATTATTTG

ATATTCCATGCTTGGTAATTTCTTTGATGGATTATGCTTAGAAGAGCCATATTATATTTGTGAATCAATATTTTAGTTTC

GTATAACTTTATCGCTGATGCAATTTGATGGTGCATGCTTGGTTTTATTCTTTGATGCCCATGCTTAGAGTATACCAGTG

ACATTTGTGACTCTAATACATATAATAGGTGATTCCAATTAAGCAAACAATAAATATATATTCGAATTATGTTTCATTTG

GTGATTAAATAAAAGTAGTAGTGGATATCATAAAACCCTGGTTACCGACTCTCGCATCTTGGTTACATCTTTATTATTTT

AGTTTTGTAATTTTATTTACCAATTCTTAAAACTTCTGAAATATTGATTTGATTGGTTACTCTTTGGTATTAGTTAGTAG

TAGTATTTTCACACTTCTCGTGGGAACGACCTATACTTTCCATTGTTCTACTAGTTAGACACTGTGCACTTGCAGTATAT

TTTATTGTAGGTTTCCGAGTGTTGATTTTGGTTTTCAGTTTAGGGCTAAAATCGTGAAACCATGTATTTAATACGCTGTC

ATCCTACAAAGGAACAAACGCCATTTGAAAGTGATGAGTGCATAAACCTCCTCCCTTACGTATGTATTGAGAAATTCAAC

ATATTTATGAAATTTATCCAGAATGGTGCATGAAAGCAACAATCCCGGATATTATTCACTCCACACGGACTCTACTACTA

CACTCGCATGCCCTTCGGACAGAGGAATGCAGGGAAACATACCATAAGATGGTTGATGCAATCTTCAACCCATGGATTGG

GAAAACCTTAGAGGTCTATGTTGATGATATGCTCGTAAAAAAAGTAAGCTGCGCAAGGATCACCTCCAACACTTGAAAGA

TATCTTGGCGGAATGAAGAAATATATCATGAAAGTAAACAGGAAAAGTGCACCTTCGGCGTCACATCGGGGAAAGTTCCT

CGGATATTTGGTGACGAATAGGGGCATCGAAGTTAATCCCGCTAAAATCCAAGCCATCGTAGAGATGCCATCCCCAAAGA

ATTTGAAGGAAGTCCAAAAACTCAACGGATCCATAGCGGCACTCGGGAGATTCATCGCTAGATCATCGGACAAATAAAAA

CACTTCTTCAACATCCTCAAGAAAGAGAACAAGTTTGCATGGACGGCGAGTGTGAAGAAGCTTTCCGTAAGATCAAAGAA

TACTTGGCAACGATCCCGATCCTCCAAAACCGGAACCCGATGAGTTCTTTGGCATTATATATAGCTGCAACCGAAAATGC

TGTTAGTGTTGTATGTCATGTGCCTCATGTGAAGAGACTGCCCCATATCCTGTAAGTGTCAGTAATTAATATTAGTTGTT

GGTGGTAGTGTTTAGCTTAATAATAGGTGTATTTATCAGTTGTATGGGATGGGTTAGATTTAGCCATCCTGTGATTGTAG

TAGCCGTTAGATGCAAAGGCAGTGGAAGCTTAGTTTTATAAGCTAATTGTTCTCAGTAATGAAAACTCATGAAAATTATT

AATCAAAATATCGATTTATCGATTTAGGTCGTGTTCTTCGGAACCATAGAAACTTGGTTCTTTGAATCAAGTGGTTACTA

TCCTCATCTTGTCATTGTACATCTCGATACATGACTGTATTATTTGTGAATCTATGATCTAGCGATGGGTTATACGAAAC

TTGGTGCATAATTGTTAGGATCGAAAATATATTTGTCTACTTATTTGATATTTCCATGCTTGGGTTAAATCCTTTGATGG

GTTATGCTTAGAATAGCCAGGTATTATTTGTGAATCTATATTTAGTTTTGCATAACTTTCTCGCTAATGCAATTTGATGG

TTCATGCTTTGGTTTAATCTTTTGATGTGCCATACTTAGGGTGTCCTAGCGATATTTGCGACTCTAATACATATAATAGG

TGATGTCGATAGAACGAACGATAAATAAATATTCATGAATTATGTTTCATTTCAGTAACTGAATAGAAATAATGGTGGAT

ACTAAAAACCCTGGTTCGCGACTTTGTATTGGATTCATTCTATTAGTTTAGTTCATTTTCTTCGATCTTTTTATTATTTT

CTAAAAATCCAAAAACATCATTTGTGGTTAGTTAATTAGAGATATTGGTTACATTATTTTCACCGCTCCCTGTGGGTTCG

ACTCGTACTTTCCTCTGTCTACTAGTTAGACACCGTGCGATTGCAGTTATTATATATAGGGTTTCCCGAAATCCTATCAA

TTTTTGACGTGGTTGTGGTTAGACTTTTGAGGTGTCCCAATGATATTTGCGAATCTAATAAATATAATAGGTGATGTCGA

TAGAACAAGGGAGAATGATTGGCAGCAACAAGACATGGTGCTGTGGATTGATGGAAGTTTGAGCTGGGCAGCTGAGACGA

AGAATTGAGATGCTGGTGTTTGGCCTGACATGAAAGACCAATTGACGGACTGAAAACCGAGAAGGATTGAATCAGTTGCA

GTTGCAGGAAGTGGTGATTTCATGGACAAAATGAATCAATTAGGTTCGAGTTATTGAGATAATGAATGCATGGATGATTG

CAGATGGTGGCTGTATACGTAAAGCTTCCAAGTTTCAAGGAAATGACAGTGAAAGCTTCCATGCAATCTGATTATGAGTG

ATAAAATAGTGAAGAACTGGTGATCAAACTGCCTTGGTTTTGGGTTGTGTACGAGAAGATGGAAATATAATGGAAAATGA

GTGATGGAGCTGTTGATCTCAATGGAACGACAGTTGAAGGAAATGCTAGAAGATAACGAAAGGAAGATGTTGTTTGGAGT

TAAGTACGGATGGCAATTGGTCATGGTGGAGCTTGAACTCGGATGGATTTGAAGTTTGGAGCTACTAGACCCGAATGGAC

CGGGCCTTGGCAGATGTTGCTAAGGCTTAGTTGAACTCTTGGCAGCATATATATTACGAAAAGAGAAAAATAAGGAAGGG

GAGACGAAAAGAAAAAGCTGCGGCTACAAAGCCGAGAGCTGTGGAGAAGAAGACGGTTTGAAGTTCCATTATCTCTATTC

CGTTTCGATTTATGAGTGATTTTATTTTACATGTTTATGGCTTTTGAGAAAGCCATGTTTTGCTAAATTTATATAACTAA

GGTTTTTATGTTGAAACTACATGGGTTTAGGCTTATTTGGATTGATTATATAGCAATAGACCATTATGATATGAATGAAT

TAATCTTTGAATATTGCTTATCGATTTATTGAAAATGAGTTGTTGCTTCACCGGTTTCTTATAACCTTTGTTCGTAATTG

TTAGGTTCAAGAATATATTTGTCTCATTATTTGATATTCCATGCTTGGTAATTTCTTTGATGGATTATGCTTAGAAGAGC

CAGATTATATTTGTGAATCAATATTTTAGTTTCGTATAACTTTATCATCGATGCAATTTGATGGTGCATGCTTGGTTTTA

TTCTTTGATGCCCATGCTTAGAGTATACCAGTGACATTTGTGACTCTAATACATATAATAGGTGATTCCAATTAAGCAAA

CAATAAATATATATTCAGAATTATGTTTCATTTCAGTGATTAAATAAAAGTAGTAGTGGATATCATAAAACCCTGGTTAC

CGACTCTCGCATCTTGGTTACATCTTTATTATTTTAGTTTTGTAATTTTATTTACCAATTCTTAAAACTTCTGAAATATT

GATTTGATTGGTTACTCTTTGGTATTAGTTAGTAGTAGTATTTTCACACTTCTCGTGGGAACGACCTATACTTTCCATTG

TTCTACTAGTTAGACACCGTGCACTTGCAAAGATATTTTATTGTAGGTTTCCGAGTGTTGATTTTGGTTTTCAGTTTAGG

GCTAAAATCGTGAAACCATGTATTTAATACGCTGTCATCCTACAAAGGAACAAACGCCATTTGAAAGTGATGAGTGCATA

AACCTCCTCCCTTACGTATGTATTGAGAAATTCAACATATTTATGAAATTTATCCAGAATGGTGCATGAAAGCAACAATC

CCAGATATTATTCACTCCACACGGACTCTACTACTACACTCGCATGCCCTTCGGACAGAGGAATGCAAGGGAAACATACC

ATAAGATGGTTGATGCAATCTTCAACCCATGGATTGGGAAAACCTTAGAGGTCTATGTTGATGATATGCTCGTAAAAAGT

AAGCTGCGCAAGGATCACCTCCAACACTTGAAAGATATCTTCAGCGGAATGAAGAAATATAGCATGAAAGTAAACCCGGA

AAAGTGCACCTTCGGTGTCACACCAGGAAAGTTCCTCGGATATTTGCTGACGAATAGGGGCATCGAAGTTGATCCCGCTA

AAGTCCAAGCCATCGTAGAGATGCCATCCCCAAAGAATTTGAAGGAAGTCCAAAAACTCAACGGATCCATAGCGGCACTC

GGGAGATTCATCGCTAGATCATCGGACAAATGAAAACACTTCTTCAACATCCTCAAGAAAGAGAACAAGTTTGCATGGAC

CACTGAGTGTGAAGAAGCTTTCCAGAAGATCAAAGAATACTTGGCAACGATCCCGATCCTCCAAAAACCGGAACCCGATG

AGGTGCTGGCATTATATATAGTTGCAACCGAAGATGCAATTAGTGTTGTATGTCATGTGCCTCATGTGAAGAGAGTGCCC

CCATATTCTGTTAGTGTCAGTAATTAGTATTTGTTGGTGGTAGTGTTTAACTTAATAATAAGTGTATTTATCAGTTGTAT

GGGATGGGTTAGATTTAGCCATCTGTGCTGGTAGTAGCCGTTAGATGCAGGGCAGGGAAGCTTAGTTTTATAAGCTAATT

GTTCTGTAATGAAAACTCATGAAAATTATTAATCAAAATATCGATTTAGGTTGTGTTCTTCGGAACCATAGAAACTTGGT

TCTCTGAATCAAGTGGTTACTATCCTCATCTTGTCATTGTACATCTCGGTACATGACTGTAAGGACCCTCAAGCTCGTCA

ACGCTATTGGACCGGTCAAGTGAAGAATTAGCCGATAATGACTCAATTAGTTTAACACAACCGTTAATTAATAAGAATTA

ATCTAACAACGATCTAGATATGAAAATAGTACTATTGGATATATATCTCGAAAAGTTATGCGAAGTGGACTACTCGAACG

TGTCATTCAGACATCGGACGAAGAAGATATTTGTTATTTTAGGAGAAGGGTAAAATGGTCATTTTAAATAAAGTATTATC

AAGGAATCCAGCTTTTCTTCCCAGGGCACCCTCTGTGGATAAACTGCAATTCGATAAGGTCGAGGATTTTCTATTGATTT

TTGTTAAAGGGAGAGCAAAAATGGATCTGAGGAGAGCAAAAATCGTAGTTAACTCGCGAGTTAAAACCTTTTCGATATAT

TTTTCTAACTCGGACAATAATATCCTCCTAAATACATCCACTAACCGTACATAATCCACAAGCACCCACGACTTCCACCA

GCAACCCCACCGCCAACACCTAAACAGTTGAATCTAAGGGAAACCATTAAAACTATGGTAGGGTGTTAAAAATCAAGATA

ATCTTGACGGTAAAACAATTAACAAAGTCGTGATGAAATCAACGAGAAACTGAACAGATTTCATAAGAAAAATTGGGGGA

AATTTTGTGACAGTAGCGATCCATTGTTCAACAAAAAAAATTGAGGTGATCATCAAGATATACAAAAAAAAAAATCACAC

GACCAGCTAAGATTGCTGCTTGATCTTTTAGTATGAAATTACATACTATGATTACCATGATGACTAATAATACTAGTAAA

AGTTTGTGCCTATATATTTTTCTATTTCTCATTCATTTTCTTTTCTTCTTCTTTCTTCTCTGTTCTTCATCTCTTTCCTC

TGTTCTTCACGCTCGTGATTCTGTGAGAGGGTTTGGGTGTTTTGAGATCGGAAACTTAAGGGAAAGCTTTAAGAGTTGAT

CGTTGGTGTGATGTTGTGGTTGTTCGCGGGATGAAGATGGTAGAGTTGTTTCTATATAGCGAAGATAAGATTTCTGAAGT

TAGGGTTTCTGTCAAATTGGGAAGAGAATCGAGATGTTGATGGATGAAGAAGAAAAGGGTTGTTGTTAAATAAACAAAGT

AAAGTTTCGGTAACAGTGAATCGAAGAACTAGGGTTTCTCAGTGGATTGTTTAGAGGTTCGATTAATGATGAATAAGAAA

CTATTGATGGTGTTGAGGAAAAGACATTTGAAGAGAACTGGATGGTGCTTCATTAAATTGATTTGAGGTAATTGTTCTTT

TCTGTGAATTTTGTGAAGTTATGCAGTTCTGTTATTGAATTCATGGATTTGTATATAGACGGTGAATTGAGATGGAATTA

CGAATGTTTAGGGTGTATCTTGAACTACAAAAAAATAAAGATGATGCAGGGATTGAATTGTGTTTTAGAATTGTTATCGA

GATGAGTGGAAGGGAAGAAGATTTGGTGATCTGAAGTTAAGATCATGTATGTGTTGTTAATGTCAATGCTTAGAAATGGA

ATTGTTAGTGTTAGTATTCATAGGTGTAATACTAGATGGAGTTGCAGTTGAGTGGAGAAGATGAGGTTGTGCTATCGTTT

GTGTTTGAGAATTTGTGGAAATGGAAGTGGAAATCGAGTATCTGGTGGAGGTATAATTGCAGGTAGTGGTTCTGAAATAG

TGAAGCTGAGTTATGAAGGTGGGATTGGGACTCTCAAGTGAAATATGGTGGTGTTGTTGTGTTGCTGATGGCATAAAGTT

AGTGGTGCAAAGGATTGGTAATTTAGAGTTGAAGTTGCAGGGGATATGAGTGAATTTAGTTTAAATGAATGAGTTATGAA

GGTGTATGAGTTTGTTTTGCCAGTCACAGGACTTGGAAAAAGTATATGATGTGGTATGAATGTAATGGGATTATGGGAAC

AGATTGTAATTAGGGTAATTGCAGGCCTGAAGTCATTGAGCTTTGACGTTGAGAGTTAGGCGAGTGTAAGGCTTTGCAGA

GAGGATGAATTAGTATTAGTTGTTGTTAGTGTTGCTGTAAAATCTTGGAACTGCATTCTCATGTAGGCTCTATTGTGATT

ATGAAATAATCTTGTAGTTATTTGGTAGAATTGAATTTGTAATAGAGTTTTGGATTTGTGTTATATGTTGTTGAGACACC

ATTATGGTGCGTCTCAAGTTTTGGCCTGTGCAATGGCCTTGGCCAGATTGCATTCAGTTTAGCTGATTGAATCTGTTTTA

GCCAAGTTGTAACCCCAGTCAGTCGTCTGACTCGTCTGACTCGGTTTGTCTGACTGAGTTGACTCTAAATCAGCCGACTT

TGACCGATTCATGTTTGACTCAGTGGACCGTTTGTCTTGACCATTTACCCAGGACTTTGACCTTGGACTTGACCAGACTT

TGACCTGGACTTAGATGTTGACTGTTAGTCGACTCCTCTGACCAAACATTGACCGTTAGTTGACCAGTTAGACCGGACCT

TAGATTAATGGGCCAGAATAGAGGACTTGGGCTTAGGACTAGTTTTATTAATTAGATGTTTTGGACCGTTTATGGTCTGA

ACTAGCATTTGACTCCTTCTACAAAGTTGTAGATTTTCATAAGACTTATTAATGGCCTTAGAACCAGTAGATAGACTTGT

ATGATTCTTGTGTAAACCATTTTGACTAGATTGTTAGAGTTCTTGTGTGATTAATAGTTAGTTATTAACAACGATCGAAT

CAAAGGATGAACCAGTGGATCGTGGATCTCGAGGTAGGCGAGGCTTGTCATCAAAAGAGGTGGGAATACTTTTAACTCTT

TTGAAACCATTGTTGTTTCTTTTACAAAAGTTTATGTTTAACAAACTCATGTATTGTCATGCATGTGCTTTTCATGCATT

GTTTAGTTTGTATTATGATTGTTCTTTTAATTCTTTGAATCTTTCCATGAATTGGAAAGGACTTGTAATGTATGTCATTA

TGAATTGTTATGGGAAATAAGAATTTCCACTTGTGGTATATAGGATACACTCCTTCACATTTATACCTACATGAATTCAG

CTTTTATGCATATTACGTGTGGAACAACCTGGCATGAATATATTATTAGGTGTGGAACAACCCGGCATCACTATTATACA

TTGCTTGACGAAGGAGTTTGTCCATATACATGTTTGTGCATTTCCAGTGACTTAGTCACTTTGATGTTTGGCAAAACATG

TATTGTTTTTCCAAATCTTGCTCATGATTTCTTTAAAGTTAAATGTTATTCCTGCTGGGCACCCCTTTTAGGGATGATGT

GCTCACCCATTCCCACTTTCAGTTTTCAGAGACAGATCAGAGTTGTGCAGCGAAAGATTCGAGGAGTTGACTCTGTTAAT

TAGTTAACTTCTGCTATGTTTATTATGTTATTCTGTTTGTAAACCAAATGACTATTATATATGTATCATTTGAGAGGAGT

TTTGTATATATATATTCCCGAGCGGGGATAGTTTTGGGTTAGGTTTTGTAGCAGTTTTCTTAATTGAATGTTTAAGTAAA

TGATTTAGATTCTTAGTGCCTCTTTAATTAGTTAATCTCTTGGATTAGTCAGCTGCTAGCTTAGGGGGCTCTATAATGAC

AATTGGTATCAGATTCATTCGATCCATGGAGGTATTCACTGAAACTCATCAACATCAACAGATATCTGAAATTCAGACTT

TATCAAGCGAAAAATGGGTACAAATGAATGGAAAATTGGTACCAATATTGAAATTGGAAAGGTTAGCAAGTGATGAATTT

GACACTTGCTTTTCGTGGTATCTGTTTTATTTGATGAACAAGAACTGTGTAAATCCAGAGGACGACTTTCGAAGGTGGGG

CAATTTTTTAGACATAGCGGAGAAGAAATTGGATCAGATCATGTGTTTATCTCATCAACCTCCTCGCAGAGTGAAAACTG

CTTCTATAGTTAGTGATGAATTTTCAGAATCAGAGTTGATTCCTTCTTATTTCTTCGAAGAAGTTTTAACTGTGGATAAC

TCAATTCCTTCCATTTGTCAATAAAGCAGATGGTTCTCTCAATTTAGGGTCAAATGAATTCCAAATCGGAAAAGTCCAGC

AGTTCCAAAATAATGTCGTGAATGGGTTCGACAGGAAAAACCAGATCAAATTGGGTGAGCACTCTTCCATTATTCTCAAT

TCTTCTCGTAGCATATTTAATCCAGGAAAGGATTCCAAACTCATTAAACGTTTAACACATGGAATTTATGTTCCATAAAA

AGAACGTTTCACCATAAACTCCCTTTATTTACTAGCGGCTTCATCCTTATGACACTTCCTTATGACGGACGTAGTGTACG

TCGCGCGCTGTAAAACGCCAAACCAATACCCAAAGAGGATCCCCCAGTTTGTGATATGTGTTGATGTCTTGAATGTTTTT

GTGGAAAACATGTAGCATGTTGTTGTTGCCTGGATGGGGAGACTTGCTAGCTTGGTGACCTTCGACGGTCGAGATTTTGC

ATCTCAAGAGGAAAGATAGCCGTTGATCGTTGCACGCTTTTGTTTTGGTAGCCGCGTAGGGAAGGCCAGCATGGTGGCGC

GGCTAGGTTGGCATGATACTGGCATGGTATGGTTTGGCGTGGCCAAACTTAGGTTTTCGGGCCATAGGTTACCATGCGTC

GTTTGGTAACCTTGGATGGCTAGGATTTGCTCCTGGGATTGAAGGGGGACCGTGGATCGTCGCACGCCTTCGTTTTTGGT

AGCCGCGTAGGGAAGACCAGCATGGTGGCGCGACTAGGTTGGCATGGCATGGTTTGGCGTGGCCAAACTTAGGTTTTCGG

GCCAAAGGTTACCATGCGTCGTCTGGTAACCTTGGATGGCTAGGATTTGCTTCTGGGATTGAAAGGGGGCCGTGGATCGT

CGCACGCCTTCGTTTTGGTAGCCGCGTAGGGAAGGCCAGCATGGTGGCGCATCTAGGTTGGCCTGCCATTGGCATGACAT

GGTTTGGTGTGGCCAAACTTAGGTTTTCGGGCCAAAGGTTACCATGCGTCGTCTGGTAACCTTGGATGGCTAGGATTCGC

TCCTGGGATTGAAGGGGGGCCGTGGATCGTCGCACGCCTTCATTTTTGGTAGCCGTGTAGGGAAGGCCAACATGGTGTCG

CGGCTAGGTTGGCATGGCCAAACATAGGCTTTCGGGCCAAAGGTTACCATGCGTCGTCTGGTAACCTTGGATGGCTAGGA

TTTGCTCCTGGGATTGAAGGGGGACCGTGGATCGTTGCACGCCTTCGTTTTTGGTAGCCGCGTAAGGAAGGCCAGCATGG

TGGCGCGGCTAGGTTGGCATGCCGTTGGCATGGCATGGTTTGGCGTGGCCAAACTTAGGTTTTCGGGCCAAAGGTTACCA

TGCGTCGTTTGGTAACCTTGGATGGCTAGGATTTGCTCCTGGGATTGAAGGGGGATCGTTGATCGTCGCGCGCCTTCGTT

TTTGGTAGCCGCGTAGGGAATGCCAGCATGGTAGCGCGGCTAGGTGTGTTTGGCATGTGGGGCCAAGGGCTTAACATGTC

ATTGTCACATGGCGCGGCTAGCACGGATAAGGCCAAGGCAAGGTGCGGTTTGGCTTGGCATGTGGGGTCCAAGGGCTGGG

CATGCCATGGGCGCATGGTGCGGCTAGCATGGCTAGGGCCAAGGCAAGGTGCGGTTTGGCTTGGCATGGTGGGTCCAAGG

GTTTGGCATGCCGTTGGCACATGGCGCGGCTAGCATGGCTAGGGCCAAGGCAAGGTGAGGTTTGGCTTGGCATGTGGGCC

CAAGGGTTTGGTAGGTGCGAGTAGCATGGCTAGGCATGCCTTGGCGTGGTGGGTGTGGCTGGCACAGTTTGCCCTTGGCT

CAAGAGTAAGGCTAGCATGGTTGGCATGCCTTGGCGCGGTGGGTGTGGCTGGCACGGTTTGCCATTGGCACAAGGGTGCG

GCTAGCATGGTTGGCATGCCTTGGAGCGGTGGGTGTGGCTTACACGGTTTGCATTGGCACAAGGGTGCGGCTAGCATGGT

TGGCATGCCTTAGCGCGATGGGTGTGGATGGCACGGTTTGCCATTGGCACAAGGGTGCGGCTAGCATGGTTGGCATGCCT

TGGCGCGGTGGGTGTGGCTGGCACGGTTTGCCATTGGCACAAGGGTGCGACTAGCATGGTTGGCATGCCTTAGCGCGGTG

GGTGTGGCTGGCGCGGTTTGCCATTGGCACAAGGGTGCGGCTAGCATGGTTGACATGCCTTGGCGCGGTGGGTGTGGCTG

GCACGGTCTTCCATTGGCGTGGGAATCATGGTTTTGGCCTATCAAAACCCTAATTAGTAATTAATTAGAAAAAAGGTACC

TTGGTAATTTTCTCGCAGACATATTAAAGGGTTCAATGATGCGTTCAAGTCCTGTGACGTCACTGTTTGGCTAGGCAAAG

GCCAAATGTTTCTGAATACAGTCTAATTGTCTCCCAAGTGGAACATGAGATCTCCATTGCTATAGATTTGATCATATGAG

ATACTAGATCCCTTTTTAGTTCCACTCATCCTCTTGATAATTCCTTGAGAGACAGTTTTAGTATCTTTTTCATCTCCAAA

GCTGTTCCCCATTGCGTTTTCACATAACTTGGTGATGAGACAAGGATATCCAAGATTTCTATGGAATGCTATGGTATTGA

TAGATTCAATGATTTTAATCATTTGATTGATAATTAACCCACAAATGTCAATTTGCTTTTTGCCTTCTAGAAGGTAGTAT

ATAAATTCAGCAAATTCTTTACTCCATTGTGAAGAATCTCTTGTACAAGAAAAGAGGGTTGCTACAACAAGTTTACCGAA

AATTCTTAAGTAGAAGGGTACATCTTTGGTTAGCATCTTTCCCTTTTTCCATTCAACAGTTTTACCAGCGATAAGTTTTT

AAATGTTATCGTGCTCAATTTCAGAACCGGTAATCATATGATTCCCTTTCACTTTGGAATTGATGATTTTTGATATCATC

TTTATGTCGACATGATGAAAAATGCTTCCAATCAAAGTACTAAAAGTAATATTTCCATGATTAATATTGTGAATATTATC

ATAAAAGATTCTTACTATATCAGGTGAATATTTTTCAAACACGAACATATTTCCCCAGATTCGATCTTGAACAAATTTCA

CATAATTGGTTCCTGAAACATCTGGATCGAATCTCTTGTCAGCAGTGAAACTTACTCCCTTTAAAGCTTGATATCTTTCA

AATCATGTATTATCCACGAAGATTGTACGATCATGTTTGTCTTTGATGAAGTGAGGATACTTGATTTTCTCAATATCTCC

AGGAGGAATAAATGGGTTTAAATCCAGTTTGTCTGTTTCAGAGTTTTCTTCCCTTGATTTCCTTTTTCTCATATTTGCGA

AAATATGAAAAACCCTAGTTACAGTTTCGATCAATCTTCCCCAATATGTTTGTGAACATAGCAAGAGGTTATCTTTTATA

GGAAGAATTGATATCAAAATCCGGAAATAACTAGTTTTAGGAAATTCTATCAAAATCGGAAATCGAGGCTTGTGGGGAGA

CAATTTTATATCTAGGTTTTTTTTTTTGGAAAACCAGTTTCTCATGGTCTGGATCTTTGTCTCGGATTGATTCTATGTTT

CTGTCCAAAAATCTGTTTACTCTTTTATTACCAGAATTTGATAACATCATTTGAAATTCACTCAAGAATTTTCTAATAGA

GTATGCAGAGATAAATTTGAACATAATATGATATTTTTAAAAATACCATCAGAATCATAAATCTTGTAATATTTCAAAAG

ATCACTTGCTTCATTAAAACAGTTGTGAGGTGTATGAAGCATTCCATTGTTAATAGAATTATTAACTGAAAAAACATTCC

TTGCAAGGTTACAAAGTTTCCCAAATTGCATCCTAGGTAGCGAAACATTGGCAGTCTTAGATCTGAAAATGTACATATAT

ATACTTTGATAGGATATCCAAAATATATTTAAGATTCCAGATTTTTCAAGAACACGATAGCCATGGTTCATATAATAAAA

AATTCTGAGAGGAATCTTATGCATTTTCGAGATACAACCTGTCTCATAGAGCATTGTATCATTGTTACTAAACATGTTAG

TATCCAAGGCAATGATATACAATCCCAGCTAATTATGTACGTCCCTCATGACATACATAGACATAATTCAAATTCCTAAT

ACATTTAGGAATAAGAGTATTACATACTTCATTTGTGTCTTCCACCACGAGTTTTTGAAGTAGTCAAATCCTGCCGTTAA

GTTCACTGAATCAAACATAGATCCAATTCTTATAGGTTGGATCGCACCAAACACATATTATTAGATCTTTTCGTGTTTGC

CTTCTCTGATACCAATTGAAAAGACGAGGGTACCCAAATATACCTCAATCTAAAACTTTTCCACCTATAAGTCCTTTTTC

CGAAAGTGATTGTCTATGGATTGAGTCGAGACAATACAACTAATCGGTTCACACTTCGTGTGATCATCTATGGATACGAG

ATCGAGACAATACGACAACAAGATAACTTGTGTGATTGACTATGGATACAAGATCGAGAAAATACAACAACTAAGTATGA

TTACTTGATAAAAGGTTCGGACTTAACCAAACACAATAGGATTGCAATCAAGTAATTAAGAATTAACGTTTGTGTATTTT

ACTTCTAATTATAATAAAACAATTATAATTGCGGAAATAGAAAAGTAAAAGACACAGCAAGATTTTGTTAACGAGGAAAC

CGCAAATGCAAAAAAACCCCGGGACCTTGTCCAAAATTGAATACTCTCAGAATTAATCCGCTATACAAAATCTAAACAAA

CTTCGTATAGTTGAGACCAAGCAACTAAACCTATAGTTCACCTAGTTCCCTCAGTATCCCTGCGCCTCCAACTTGTAATA

AGTCACGCACTTGGAATAATTCCTTTGGTTCGTATTCCAAACAGTAAAGGAACAACAAATCTGTTTGGTAACAACTCGTT

TCAAACAAAGTGATATGAGTTTGACAAAAGGCTCTTCCGTTTATCCCAATAAACTCCTTTGTCGGGTTCTTAGATCTATC

TATTCAACAACTACCATAGTAATTGTTTAGACTATGCAATCAATACTATTAATCACAAAAAAATATATTGATGCCGATCT

ACTTAACTAATCAATCAAGGTTATCACAAAGATAAACCGATTATAGTTGGATCCCTCCAGCCGATCAAGTTTTGTGCACA

CCAAAAATTTTGAACTCAAATAAGAAATCTTCTTTGACTTCAAATCTTCTTAGATCTTCAATAAACACCTGCGCACAAAT

AACTTGAATCTCTTGTGATCAATCACACACATAATGGAGTCTGTTAACAATGGATTATCACAAGATGTCTTTAGATCTAT

AAACAGTTCTAAAGATCCCCGTCGATACTTCGATCTAGTTTGAGTGAATCTTATATCAGAAGAGAAGATTCTCAAGCATA

AACAAACTAGGTGCAATCAAAGTTCAACAACCGTTATTCAATCAAATCAATCAAATACTAATAATAAACTACAATTATCT

AGTTTCCCACCAACGGTATTCGTAGAGCTTCCCAACCCTAAAAAAGTCTTTAAAACGAGCGGTCGTAAGAGATTTCGCCT

AATTAGGGTACTTTCCTATCCGAATAGACGGCTCCACCAGTAACAACACAGCTAGGTATTTTTGTTGGCTCTGAGGATTA

GTTTGCTTGAAATGAAAACTTCAATATTTATAAACCAAGGAAGTTTGGACACCAAGGAATTTCCAAAACCGAATATTCTC

AAAGATGTGCAATAAAGTCCAAATACGATTTTCAAAATTTCTGGAAATGCTCTTTCCAAATATTGACCGAAATCTCAGTA

GAAAATCTCCAATTAGTAAATGCACATTACCAATTTTTATTTTCTAAAGATATGCATTTAATTGATGGAAATTAAAAGCA

TATAAAACTAAAAACTTTAATTAAAAGATTCTCAATTTATGTCGATCCGGGATTCTCCTTTAGTTATTAAGGAATATCTT

TGAACAATAAAAGATAAGAATTACTGCACATGTTCAAATTATGTCGACATCTTTACTTTGTAAATCATTTTCCATATTTA

CAATCTTGGAACCGATTTGCCACACTTCCAAATAAGTTTAGAATTGGTTCATCTGACTTTCAAGAACTATGTGATTGATT

ATCCTATCAAATCACCAATCATGGGTTTAATGGTTTACCAAAAAAAGTTTCGGTTCTACCTCCATGTGGGTACTAGGATC

AGTCACACTAGGTTTCCAAAAATTGGTTGACTAGGTAATAGGATAGGTTACCACATATTTATGGTATTTAACTTGTAATC

GGTGGCACAAGTCATAGGATCGGTTACCAATTACTAAGACTTGTTGCACCTCTTACACGTATCGATTCCACATGCATGTG

ATCGGTTGCGCCTCTTACTAGGATCGGTTCCCCAATGTCTAGAGTTGGTCATACCAATTACTGATACGTATTGATCTCTG

TACCTTGGTAACTATTACTATAAAGGCGGAATTCAGAATAATAAGAGACGAGAAACTTAGAAAACAGAACACAAGTAGTA

ATTCGAAGAAAATATTTTCTCTTAGATTATGAACAAGAAAACAATTACAATTCTCTCTTTGTTACGCTCACAATCTTCTC

TAAACAGTGGATTAACCTTAAACTGTTTGCTAAGCTTGCACAAGTATTGTCGTAAGCTTTCTCTAGGTTTCCTGTGCCTA

GTATCTATCTTAGCTATTAGCCAAAGCCCTCTATTTATAGCCTTTATCGTACTCAGCCGACCAAGGACTTCTATTAGGAA

TCTATTTCCTAAACTAAGAGTATTGCCTAAAACAAAACTATTCTCTAACTAGGAATTATGTCTAAACAAGGATTCATGCC

TAGAATAGGTTTCTTCCCTCAACTTAGCTTGAGGTTAACTAAGTTCCCTAAAGGAGTACGGATACACCTGTATCATGGGT

CCCCCTTTTAAGAACTTGAACTCCTGTGAGAGTTAAAAATGAAGGTTAGGGAACTCGTGAGTCCCTTTTTCTTTGTTAAA

CCACTTAGCCTTGCTAGGAACTTGACCTTTACGTCCAATCAGAAAAGCTTCTTTCTCTCCTTGTCGTGTTTTAGTCACTC

TTCGCTCCAATATGCGGTCTTCCTTGAGTGGTTCCTCTCTATCAAACCATAAGTCTTCTACTCTTGGTAAGATCATAGTG

TCGTCGCCGTCATCATCAAGTAATGGTGGTTCAAACAACTTTAAGTATTCGGCATTTATTACCGAGTACATCCCTAGATA

AGGTGGCAGATCTAGACGAAAGGCATTGTCACCAATCTGATCGAGAATACGAAATGGTCCATACCTCAATGGTTTCAACT

TCTTACCTTCCCCCTTCAGTCGTTCCTTACATAATCTTAACCATACCAAGTCACCAACACCGAAATTACAAGGTACAAGA

TGCTTGTCATTCTTTGCTTTGTATTTGGCTTGTGACTTCTTTATTTGCGCTTCAACAGTCGCGTGAATCTGAGATATCTT

CTCCAAGAACTTTTGTGCCTTTTGCCGTTCACTTCCTTCCTTTCCCCCCATTGAGGTGTTACTAGAAAATACTAAATCGA

AAGGACTAGGTGGTAAGTAACCATAGTACGTCTCGAAAGGAGATTTCCCCGAAGAACTGTGAACTGCTCTATTGAAAGCA

AATTGTAGATACGGTAAACTCTCATCCCAAGTTTTAGGATGCTTAGAATTATATCCCCTTATCATGTGAACCATAGTCCT

ATTGACCACTTCTGTTTGTCCGTCTGTTTATGGATGAAAAGCTGTACTCTTCTTCAACCTAGTGTACATCATTTTCCATA

AAGAATCCCAAAAGTGACTAAGGAATCGACTATCCCTATCAGAAATAATCGAAGTAGGTAAACCAAAATGCTTCCACACG

TGCTCAAAGAAGAGCTTTGCTGCATCGGCTCCCGTTACTATCTTTGTACATGGAATTAATGCAATCATCTTGCTGAATCG

GTCGACCACAACGAACAAGTAATCATGACCAGACTTGGTCTTTGGTAACCCACCAAGAAAATCCATAGAAATACTCTCCC

AAGGCCTTGATGGTACTGGGAATGGTAGATATAACCCACGTTTTCTGTTGGAAGGTTTAGATGTGCTGCACAACTTACAC

CCTCTAACAAACTTAGCCACATCATCTTGCATCTTCGGCCAAAAGACATAACGCCGGAGGTTAAGTAGAGTTTTATTCAT

CCCAAAGTGTCCAGCGACTCGGGAAGTGTGTGCCTCTCTCAACCACTGTAAGCGATCTCCATCTTATGGTATGCAAAGTA

ATTGACCTTTGTATAACAATCCATCTCGGAGTTCATACTCGCTTTTCAAACCACTCTTTACACCTTCTAGCACCTTCTTG

AAGTCTTTGCTGGATGCGTATAACTCCGCATACTCTTGAGGGACAATTGGCTGAATTCTCATGGCTACCATTAATGCTCG

AACTGGAGGCCGAGATAACATATTTGCTAACTTGTTTGTTACTCCCTTCTTGTACCTAATGAGAATATTGAAAGTCATCA

AGTAAGACATCCACTTCATGTGCCGGGCTTGTTGTAGTTTCGTCTGTGCATATAGATACTCCAACGGTTTGTGATCTGAA

TGTACCACTACTTCTTTACCTAAGAGATACACTCGCCAATGTTTGATACATTGATATAAAGCATAAAATTCTTTGTCATA

AGGTGCGTAGTTCTTAATAGCACCTGAGAACAATTCCGAATGATACTCTACTTTGGTTTACCATCCCGTAACAACACTCC

TCCCAATGCATAGTTAGAAGCGTCGGTCTCAACTTCAAAAGTACGTTGTAGGTTAGGCAATGCCAACACTAGTGCTTCTG

AAATCTTTCTTTTTAGTAACTGAAAAGCGTCTTCATGGGTTTGCTGCCATTCAAACTTTGCCTTAGCTCCCGTCAAACCA

TGTAATGGTCCTGCAATATTTGAAAAATGCCGGATAAACTTACGCAAGTAGGTGCAAGCCCCTAAGAAACTCCTGATTTC

AGTTACAGTACTAGGTCTAGGACACTTAGTGATCACTTCAACCTTCCTTGGATCAATCTTCCGTTGTCCACCACCAACAA

TAAATCCGAGATACACTAACTACTCCTTCACAAACTCCCACTTCTTTACGTTCAACTTCAGTTAGCCTTCATGTAACACT

TTAAACACCTTCTCAACGTGAACGAGATGCTCTTCCCAAGTTTTGTTGTATATTAGGATATCATCCAAGTAAACAATCAC

AAAATCTTCTATAAAAGGTCGTAACAAATCATTCATCAAACGCATAAACATTGCTGGAGCATTACACAAACCAAAAGGCA

TCACCAACCATTCGAATAAGCCTTGTTTGTTTTGAAAGCAGTTTTCCATGTATCATCTTCTCGAACTCTCATCTAATGAT

ATCCGAATTTGAAATCTAACTTTGTAAAGACTCGAGCTTTTTCCAACTGATCCATCATGTCGTTTATCCTAGGTAAAGGA

TAACGATTCTTGACTGTGATCTTGTTCAATGCTCTATAATCAATACACATGCGCCAGCCTCCATCTTTCTTTAGAACCAA

CAACACTGCTGATCCACAAGGTGAAGCACTTGGCACTATCATTCCTAATTCTAGAAGTTCCTAGACTTGTTTCTTGATCT

CATTAGATTCCTATATGGTCGTGCGATAAAGACCTACATTAGGAAGAGAACTCTCCCCGGTCAACATGATATCATTCTCC

ATACTCCTCTTTAAAGGTAATCCTATGAAATCTGAAAATAAATCCTTGTACTTCAACTTCAACCTTTCTAGGTCATCTTC

TTGTTGTGCAGTCAAGGCTGTCAAACAAACTCTACTCGGCTTCTCCTCAAGAGAACGGATCACAAGGAGAATGAACTTCG

TGCTAGCATTCACCAACCGCTTAGCCTGAGTGGATGTGATCAGACTTTCTCCCTCAAGAGGAGAATCAATAGCCTGAATC

ACAAAATTTTCTCCATCTTTGGTAAAGGTGTACTTTTGTTCCCTCCTGTGTAGAGTTGCATCTCTATCCCATAGGTAAGG

ACTACCTAGTACCACCTGACAAATATCCAAAGGAACTACGTCGCATGTAACTTCGTCAATATATGACTCATCTAGAGCAA

ATTTGAACGTGCATTGTTGTGAAACTTGTAAGCCCCCTTCCTTTTGAATCCATCCTAAAGGGTATGGTTTTGGATGTTTG

AACGTCTTCAAACCTAACTTCTGCACCAAAGAATTTGAAAGTAAATTCTTCTGACTTCCCGTGTTAATAACAACATCAAC

TAGTGAAGTTTTGACTTGCATCTTCACTGTGAAGAGTTGCTCCCTAAGATCATCTTTTGAAGATCTTTCTTTTGACCCTG

TCATAAGAGAAAGCTTTGAATTGGGTTCCGTTAGTCCGTGGACTTCTTTTGGAGTTTGAGAAACTAGTGATGCCTTCTTG

TCTTCTTTCCTCTGCAACCCTTTAGGCTTTAGATGAGGGTTCTTAACCCAACACTTGTCGACAATGTGACCTGTTTGCTT

GCAATGGGTGCAAGACAGACCTTCCTTGTCACTAGATTTCCCTTCTTCTTTGCTCCTTTCACCTCCTTTCACAGTGGTAC

CTCCAGATTTCATCTTAGTATTTCCCTTTGAATCAGATTTCTTAAGCCTGCCCTCAATGGCATTAGATTTTATACTTGCT

TCGGAGATAGTATCCACCGAAAACATCTTCAACTCCTTCCGTATATACTCATGGAACCCGGAAATATATATTTCATGTAG

ATAACATAATCTCTCAAGTCTAAGTCCAAAATCATAGCTTCATTCTGAAATTCCGAAGTATATTCTTGTACGGTTTGGTT

GTAAGTCTGCTTCAAGTTGTACCACTTGAACCATCTCTCCTCAAGGTAGCCAACCAGATAAAACTGTTTCCTTACAACTG

CCTTGAATCTTTTCCACGACAATACTGCTTTACCTAGGTTTTGTCTTTGATAAGCTTTCCACCAGGTTAGAGCATGATTT

TGTAGCTTCAATGCCGCGAAAGCAATCTTTTCCTTTGAACCATATTCAAAGAAATAAAAATACGTCTCTAGACGTTCAAT

CCAATCATCCAATTTTTCTACATCGACAATCCCATCGTAAAGTGGAATATCAACACGGAAATCAATTTTCAGATTCTTAT

CATGAGGTTTAGTTGATGTAGGACTTTTAAGAAACTCTTTCTTCTTTTCTTCTTCCTCTTCTTCTCCTTCTTCCGAATCG

TCTTCTTCCGAATCATCTTTTTCCGAAGCTTCAGGTGAAGGTATAATCTTTTTCTTCGTCTTCAGCTTGGTTGTATGAGA

ACGAATACATTCACTCAGCTCCTGAAGGGTAACTTGAATTGACTTCATGTCTGTTTGTAACGTAGCCACCTTTTCTTCCA

TAGTTTGTGGTTCGCCTTCCTTAGGATCACCATCTGACTTTGTCATCCTTGATCCCCTTGTTTTCTTAACGTCTTCCAAC

TTCTCGAGTACCTCACCAAGCTTTATGTAAAGAGATCTAGTGAAAACCATAAACCACAGGGATTTACCCTAAAAAAACTA

ACCTTGCTCGTGATGCCAACTTGATACGTATTGATCTTTGTACCTTGGTAACTATTACTATAAAGGTGGAATTCAGAATA

ATAATAGACGAGAAACTTAGAAAACAGAACACAAGTAGTAATTCGAAGAAAATATCTTCTCTTAGATTATGAACAAGAAA

ACAATTACAATTCTCTCTTTGTTACGCTCATAATCTTCTCTAAACAGTGGATTAACCTTAAACTGTTTGCTAAGCTTTCT

CTAGGTTTTCTGTGCCTAGTATCTGTCTTAGCTATTAGCCAAAGCCCTCTATTTATAGCCTTCATCGTACTCAGCCGACC

AAGGACTTCTATTAGGAATCTATTTCCTAAACTAAGAGTATTGCCTAAAACAAAACTCTTCCCTAACTAGGAATTATGCC

TAAACAAGGATTCATGCCTAGAATAGGTTTCTTCCCTCAACTTAGATTGAGGTTAACTAAGTTCCCTAAAGGAGTACAGA

TACACATGTATCAATTACAATATATCGATCATACCATCTCAGGTGATTACTTAAGATCGGTTTCACTAATAAAAGTCATA

CTAATACAAAAGTCAGGCATTGTGAATAGTTTTACCAAGATACATAAACAATTTATGATCAATTATACTAAACACACATT

TTGGTAATCCAAAGATTTGCAATGAATAACAATACCAATAAGCCTAGCGATTTCCCTTTCGATTCACAAAACAAGTTTAT

GAATTGTACTTCCTTTAAACGAATGTAAAACATTGTTTCCTAGGACGAAATCTTCACCCATACCCATACATAATCACAAT

AACATTCATACGATTATTTCGATGTATTATATACGAAGTTCAAAAGATAGACGTTATACTTCGTATTGTAATTCCTTAAT

ACTATGTCTAACTAGAGTATAATCATTCACAGCTTCGCAATTATGTTTTCAATATGCACAACTTGAAAGATACGTTAGGA

ATGAAACAGTTCAAGTCAAATATTACTAACTTCAAGAGGAAGGATGATGTCGTCGTCATAGTTCTTTACTTATTCACATT

CTTCAAAGTTTTCATGTAATACTTGTAAATCTCATATCCTAGTAATTTTCTAGATAACCTATACGAAGTTGACTCTAGTA

AATAATCAAGCGACTCTTTAAATGAGTTTTGATTCACTAAAATATGACAACCAAACTTGACATACCAACGCTTGGTGGGT

TCAACCGAGCTATGCTCTAACAAGCTGTCTCTCGAGATCTGAAACAAAAGCAAACAAGTAATCAGGAGAAAGGAATAAAC

ATCTACCCACGACTAAACAAACATATACCTTCGACATCAGCCGAAGCTATTTCAAATTCATTTACTACAGCATCCCGAGC

ATCGTCAAGAAAATCGTATTCGTCGGAAAGTTTTTTATAATCGAGCCGAAGGTTTTCGAGGTCAAATTAGCACTGATCTA

ACTCTACCTTCTTAATGCGATCTATGATGAAGACGATTGACTTTGTTAGTCATCTCCTCGATCCCTTTGGTGAGTCTATA

CATGTTAATGAACGAGCAACATCGGCACGAGAAGCAGTTAAGGATTTTCTAAGAGCATGCGCCTCCGCCATGGGATTGTC

TCTTTCTCGAACAATCCACTGTATATAATCCTGAATTTCAGGGTCAGGGGACGAACGGGCTTGATGAAGCTCCACTTCTA

GATCCGTTACTTTGGCTTCCAATCGATAAGCTTTATCCCGAGCCTCATAAAGAGATTTGCGTGTCCATTCCATAATACTA

TTAAATTGATCCCGAGCGGCGTACATGGAATTCTGCATTTCAAACATTTGATCTTGTTTCTTCAGGTAACCCGCCAACAA

TTTATTACGCTGATCGGCATGAAAATCCACTTATTAGCTTATCACTTCTGAATCGGACAATCATTGTTAGAAAGAACAGC

AAGGTGCTTTCATCAAAGGAAGGAATATTCATGAAAAAATTGTTTTAGCATCTGAGCTTGTAAATGAGATGAGCATTAAG

AGAAGGGGTGGAAATGTGGGCTTGAAAATTGATATTAATGCTTTGGATTCAGGAGGATTTTATGTCAGCAAGAATTTCGC

TTGTAAAATGGGGACCATGTGGATTCTTTGGTTGGGAGAGGACTCGAGGCAAGGTGATCCAGCCACCAATTCTTTTTGTC

CTAGTCGAGGAAGTGTTGAGTAGGAACATTACAAAGTTGACTCAAGAGGGCAAAATTCAACCAAAGTTGGAAGTCAACCA

ACTCATTTGATGTTTGCAGATGATATCTTCATATTCTGTAATGGTCATAAGAAGACTGGATTGCTTAACCGAGTTACTAC

ACAAGTATCAAGTTTCCTCCGGTCGAAGTGAATAGAGCAAAAAGCAAATGTTTTGTAGGAGGAGTTTCATGAAGACATTT

AACAATCGAGATGGAACTATCTTTCTTCCTGATAAATACTTAGGTGTCATCCTTAACCAAGGAAGAGTCAAGACTCATCA

AGTTTGGGGGATAGTGGAAATGATGCGAGAAAATGTTAGTTCGGATGGATGGGGAAGTTATTAGCTTTTTCAAAGAAGAC

TAACTTTGGTAAAATTTGTGTTATGTAGCATGCCCATCTACAATATGTGGGTGTACAAATGGCCTAAGAGTGTAATTCAA

GCATGTGAAAGAATCATTAGAAAAAAAGAGGGTGGTTTGGGGTTGAGGAGGCGGAAGTGATAAATAAAGCTCTACTTTTA

AAATTGCGAAAATTGAAAACGAGAGAAGATGTTGAATGGACTGTGTTTATGAGAGCAAAGTATAAAGATAAAAAAGGTGA

ATGGATAAAGTATCATAAACAGTCTACTATTTGGCCTGGGATTAAATGGGTGATCTGCGATTTTCAAGAAGGAAGTAGAT

GGATTGTGGGTGATGGAAAAAGATATATCTGTATGGAATGATAAATGGATTAAAGATTATGCCATAATTGAGAGGTAAGA

GATACTTACATTCTGCAAAATAAATTAATGAATGTACAAGATCTAATCCCATTCTGATGCTATGTGGCATTATTTTGAGG

TTTGCCGATCGGTGGAGGACATGATAAAAAGATTTGGTCAAAAGATTTAGTTGGCAACTTTACAAGATCAAATGCAACTC

AAATGATTAGAAAAAAATTTCTAGAAAGCTTGGACTCATCTGAAGAAGCGAGTAAGAAAAAAGGTTTCTCAACAACTCAA

AGTGTTATTTGTGTGGTAATGCTCAAGATACAATGGAGCACATTCTTTTGGCAATGTGATTTTAGTGAAGATATATGGCA

TCGGGCTGTGTGGAGTGTTTCAATTTTCTTATCCACAAAAACTTTGATGAAGTTTTAAAAGCTTTATTAGAAATTTGGTT

CATTTGTGCTTTTTAATACAATGGTCGTGGTTTGCCGTAAATGTCGTGATTTTTGAAGGTGAGTATCCATCTCTGTGGAG

AAAATGAAAAGAAGAATACAGGATGTGCAAAGGAAGGAAGTTTTAGAATGAGAGGCAAAATGTGGGGAGTGCTCTGATCT

CAATATCATGTTACATTTTGGCATTTCGAGGATACAAGTCAAACCGGTGGAGAGTAAAACAATGTTTCTTCAATTTTCCC

TCTCAAAATCAAGATGGAGCTTAAAGGCAATCCGGGGTTCAAATTTTGGATTTGGTTATTGGTAGAGATCACTATGGAAA

ACACATGGGTTCCGGTGGACTTGGAATTGCCACAAACTATTTGGCGAGGTAATGGCTCTAATTATAGCATGTGAGTGGGC

AATGTCTAAAGAATTATTGAATGTAGTGTTCGGATTAGATTCTAAAGTCACAATCATAATGGCAAGAAACATACCTTGGA

TTGAGGGATGGATTAGAATCACTAAGCCATTCCAATCATCAAGTCTGGGAGGTCAATTTCTCACCCGATAAAATGGCAAA

AAGAGGTGTGCTTTTAAACAAGGAGAGTACATTATTTTGAAGAGAAACCCGTATTCCCGAGCAATCTGAAAATGAGACTG

ATATATACTTTAGATTCTAACTATAATGTATCCTGGTAAGATAATTGGGCTTTGGAAGAATGGGTTGATTGCCTTACGTT

TTTTGTTCTTTCTTTTGTATTTTTTGGCTTTAAAGGAAAAACTTTCCTTTTGTAATTTGTAAATATTTATTTTAAGAATA

AAATTTGCGATTAAGCAAAAAAACTTCATTTCTATTAGTCAATAAGAGATACTGATTATTGATATTTTGCTCCTTGTGGG

AACTAATCTGCACTTGCCTCTCGTCTACTTGTTAGACACCGTGCGATTGCGGTTATTATAATATAGGCATCTCCGATCCT

ATCAATTTTTTGGCTCCGCTTTTGCCGGGGATATGCCAATGCCTTACGTGCCTTAGTTGTTTATGCTCCCGATAATCAGA

ATTACTTATACTTTCTGTAAAAGCACTAAAACAATGTTGTTAGCCAAGATATTGAGTCCTAGCTAATATAAATATGGGTA

TAAAATGTAGCACTAACGTGCTTATCAACACCCCCATACTTATATTTTGCTAGTCCTCGAGCAAAACAAAGAATTGACCC

GCTCAATAGCTGGTCATCATTTTTTTTTAGACCCGCTCAATCTGGTCATAAGAAAATGATTGATCATTTTTTTTTCTTAA

ATTTGAGACCCGCTACACAAATGGTGATAGAATAAGAGATGAGAGAGAACTGGTCATAACATGCAAACAAAATAAAAATC

TACACACCGTCATCACCCTAACAATCATTTTTTTTTTTTTTTTTTTTAGACCCGCTACACCGGGTCATAAAAATACCTAG

GAACCTATAATAGACCCGCCTCAAGGGTCATAAATTATAAGAAAATAAAACAAAAAGTAAAAAGTTAACCACTCCCCCAC

ACACTCAAACATTACATTGTCCTCAATGTAATTGAGTCATCCAACGTAAAAATTAAGATGGGAAATTCATAATATGCGAA

AATAAACAATTAAAAATGAAATAAGAACATAGATGTACCTACCGGAATATACAAATAAGACTCACCATAAATCAACAACC

TAAATATTTGGGGATGCAGCACAATATATACAACCGATGGTAACCTCAAATCCTCGCTGAAGCAACGTAAAGATATAGAG

ATTATCAAAAGGAAAGAGCTACCAAACCTAATTTTTGCAAAGGATATATCGTCAATAACAAGAATTACCGACTCAAGTGG

TTAAGAGCATGCGTTCCGAATGATCATGCCTATGAGAATAATTCCAATCATTATTTACACACTCGTGAGTTCTTGATAAT

GATAAGGTGCATACGCATACTCCTCATGTGAAATATCGATGACACCTATGAAAAAATCAAATAAGCTACGTAAAAAGAAC

TAAAAGAAAATATAAAAATAAAAGCAAAAATAAGCTATGTACAGAAAATAAAATAAAAATGTACAAAATACCAATTACCC

ATAACTCATCAAAGAAATCCCAAGCATAAATTATCAAACAAGAAACAAATAATTAGGTGAAGCTAATTAAACGAATTAAA

ATTAAGGTGAAACAACAACATATATTCTAGGTAAATCGATGGACAAAATTCTAGGAATGAAAATTGCATTCATAAAAGCA

TGTTGCCAAATATTCAATTTATTATTATAGCCTATTGCAAAATCAATGCGCTCCATGGGATTGTCTCTTTCGAACAAGCC

ACTGTATATAATCCTGAATTTCAGGGTCAGGGGACGAACGGGCTTGTCGAAGCTCCACTTCTAGATCCGTTAATTCCAAT

CGAGAAGCTTTATCACGAGCCTCATAAAGAGATTTGCGTGTCCATTCCATAATACCATTAAATTGATCCGGCATCATCTT

CGACACCAACTCTTTAATCCGAGAAGCTTGGCGATTCCTTTCACCCCGAAGCCATACTAATTCTTCGGCATCACCCGCTG

GAGACATGGCCGATTAACTCAGAAAGAACACTAAGAAGGCAAAGGTGGAAGAAAACAAAAAGGGACAAACCTTTAAACGA

CGAAGACTCAGCACGAGCCTTCTCCAAATCTCCACGAAGAGAAGCCACCTCTTGACGAAGTTGTCCTTCAGTGGCACCAG

CCTGCTGCTTCCCCTTCAGCTGTTACTTCAGCTCCAATATCTTATTGTCCTTGGCCAAGAGAGATTCCTAATAAGTAAAT

AGATCATCATACCTCTGAAGTAGCTTCGCCTCCAGCTTGAGTGCCTTAGCTATGAATAATTGATATAGAGCATGGTTGCA

GTGCTCGCTCCTTTCCAACTGCAACAACCAGGGGAAAATAAGAATACTAAAAAAGAAAAAGCTGTGCTACTTCGACATAG

AACAAGAACAACATCTTACCTCGAGGATACGCTGCTGGGTAAAGCCATATTGATGACCATCAGCTATGGCCATAATCTCG

GAGATCGAAGAAGGAGGAGCGGTCAAGAGATCAGAGGGGACTGCCACACCTAGGTCTCATTTCCAGGCTTCGTTTACCAT

GTCAAAAGGTGACAACTCCATGTTCCGATGCGTGAAAGCATCAGAATCCCTCTCACCAGGAATAATAGGGGCATGGAAAT

GTACATTCATAAGAACCTTCTTCTTTAACCAGTCGAAGGTCTCATCGTCCCCACCACGCATCACAGCGTTGACAAAACCA

TCGCAAGATACACCAACAGCTGTTTTTCCATCAGCGGAGGTATCATTACCTGCAGAGGTACCTCCGCCAGCAGCGGTCTC

CTTTCCTTTTGCTTCTTTAAAAGCGCGAAGTTCGGCATCCTCGCCATCAGAACTCTCATCTTCGACTACTTTCCCACCAA

CGTCCTTGTCCATTACCTCCCAATCATCGCTAGGAGACAACAAGAAAGAATCTGAATCTATCCCCATCACAACACCAATA

TCTACCTGACTGCCAGCGGAGTAGATCTTACCGAAGGAGAATTCTTGGTTCGGTTCCGAGCTGTTCATCAGCTCGGGAAT

TGCTGATTGAAGATCAGCGTCTTCATTTCTTTGACGAGGAGGACTGCCCTCGGGTTCAGCCATAAGATCATCTTTGTGCC

ATTCAGGAGCAATCTCTTCTTCTGATTCTTCTTTATTATCAAGAGTAGTCTCAGTACGTTCTTGTTCATCGGTCTCCTCC

TCCTCTTTATCGTCAAATTCTTCTTCGTCACCCACTGGACTGGTAACTTCAGGAATCCCCGAATCTTCAACCTCCACAAT

AGAAGATTCAAGGGGACAAATCTTCTTCTTCCTCGAAGACTACAGTAAGCACACATCAATCTATGGTTCAAGTGGAAAAA

CACATACATATACAACACATTTTGTTAAATCATACCTTGATCACAATAGTGGCATCCCTCGAAGGAAGAAGAGTCTCGGA

ACCTTCTTCGTCACCAACATCATCGGATTCAACAGCTTCAAAGGTATAATTGAAGTTCATACCGGTAAAATTTAATTCCC

AAGGACAAAAATCACCGTAGCGGGATGGGGCACCATCACGAGGTTTACTTTGAGGAGTAGGCTTCCACCCATGCTGACCA

GGGATCCAACCATGGGCCCAAGGACCAATAACTTCAATAGGCGTAGCATGCCACTCATAGTCATGATCACGACGAAAACG

CTCCTTGAGGGGGAAGAGCTCTGGCTTATCAGTTTTCTCAGTACCAGGAACGAAGCGTAGCTTTGAGTCACTCACTTCAC

TCAGATGACGAACCTCGCCATGAGGAGCACCCATCGCGCGAAGACCAACACTCCACCGTTTTCGATTCATGCTATTTACA

TAGTATCTAAAAGAAGCATTGAAATTATCAAGGGTATACAAACTCTTATCCTTAGGATCAGGAGTATAACGTGTCATCAC

CGTCTCACCCTTGCTCCGAAGGTAACATTCACGAAGAGTTCGTAGGTAGTTCCCACTCCATTGAACTACTGAGCGGCAGT

GGGTATGATGAGTAGAGCCCTCTCGATGCTCCAAAACATCATAGTAGAATGAATCTCCCCAATTATACAAAGGAAGCATT

AAACCCGCTTCAAAGGCACCAACGGAATAGATGCTGGGAATCATACTTATAATTCTTTATCATGTCATAAGTAAGATCAT

CACCCTGATTATAAAAAAGGACCTCAAAATGCTCAAGACCATGCTTTACGTTAAACTCTTTGAGGTCAATATGCTTGAAG

GACACAGGAGCCCTCTTTTTCTTCTTCGGAGCTGGAGCAATTCGTTCAGAAGGGATAACCCCAGAAGATGGTGTTCTACC

CGATGATGTCTTCTTAGGAAGACTCATTTCCGAAGCCTTTCTCTTTGTCGACGGATTCCTCGAAAAATCTACATCCTTTG

GCGAAGACCTCTTCGACGAAGCAGTTTTTGGTAGAGCTTTAGATGTATGTGAAGACGACGGGGCATTAGGAGCGATACGT

AGAGGTTGAACTTCCAATGGATCCGAGTGTTGTGAAGAAGTTCTAACTGTACGACGATCAACCGCATAGCGAGATCCTCT

TGTCGGATCTCCCCTCTTTTGAGAAGAAGCCTTCAGATCATATGACGAAGAAGTTCTATGAGGACGAAGATCCGACTGGG

AAGATCTGGACGACCCACCCTTTGACGGAGATATATCAGGACTTCTAGTTGGAGGAAATCTATAAGGACTCGTCGGCAGA

GATTTGTAAGGGACACGTCTACGATCATTGCCTGACATAGATACCTGAAATGGGCATAAAAGGGTAATGCAGGTCAAAAA

CAGATTTGTGATCAACTTTTTGCATCAAGAACACCATTAGAACATGAGAGAAAAACCCAGAAGTAATCAAAACACATGAA

TCCAACTTTCGTGATGAAGAATAGGGGCAAGTATATATACTTTCATATATTATGTTCTCCATCACGAAACTCATGAGCAC

AACAACAACGGAAGAATCGAAGATAATCAAAATCATGGATATAACAATGATGAACAACACACCTGAAACAATGGCAGAAC

CCAGAAAACAGCCGGACGAAATCAATGATCACCGATCGAAGCTACTTAAATAGGAAAGGTACTTGTAGGATATGATTAGT

ACGAGGAAGGAATATCACTAGCAGAATTGAAGAACAAGAAGACAAATACGAAATCAAAATTAAAGTCTCTCCAGAAGACA

GAGACAGAGAAGAATGATAGTGAGAATGAGAATGAGGAGAATAAAATTTATTTCACTCATCCCCTCTCTCATATTTATAG

CATCCATAAAAAGAAGAAAACTTCCCTGGAATTCGAGGGGAGTTATTGCAAGGAGTGGGAAACGTGCCAAAAATTCTAGG

AGAAGTTATTCCGGGAAGAATGTGGAGAGTTTTCCTCAAAATTTGACCTAATTCAAACTTAGAAGAAAAGGGGCAAATTG

TATACACATAATCCCGCTATTAGCCATGTGGCATAAGTATTACACGTGCCGCACATAAGGCGAGTAAACATCAATCCTCG

ATGTCATGTTACCACACCCAAGGGGCCGTATCATCACCAATTGCCACGTACTCCTCACCGAACGGATCCAACGATATAAA

GATCTGGGAGCACCAGATGTATAAGGAGCTGCGAAGCACCATAGAGGATCCGAAATATACTTTATCCCATCCCAAGATAG

ATCCAAAATGAACGGAGAGGATAGGTATAACTGACGAGGATGTGACAGCAACAGTAGCAGCTATCGGGCGCGTGCAGTCT

GCCCCAACCACCCGCCTTAAATGCCATAAACATAGGATACGTGTCAAGCATCCTATGGAAGAAGGAGAGGCATGCCATCG

TTATGACATATATCCGTTAGAGGCATCGGGCGAGCGCGAAGATGGTTATCGGCATCGACACGTCTACCAAGAAGATAACC

ACTAGTAAAGACAGTGAGGAGAGGCCCACACAAGTACTCTATAAATACCCAGCTCGACTAAGAGAGAAGGGGTCGACCAA

ATTTTAGGAGAGAGAACTATTGGTAGTGAAAAGATAGATTGTATACAGTAAGTATGAGTTTATCTACTTTTAACTATCAT

CGTCCCATACTCAGAGTCACTTGACTATCAATTGTAACCTTATGATCTATAGTGAAGAAACCCAATCCCCCGTGGACGTA

GGCCTTAGTGCCGAACCACGTAAATCATTGTCTTATTTACATTCCGCACTTATGATTTTGGTTTATCTCTTTTAAACATC

GTCTTGGTCATCATAGGTGCATCTGTGACAATAAGCAGAGTGGAGACCATTACAAGCCTCAGGCTCAGAGTTTCCAATGA

GAGTTGGGAGCCAATTCATAGTCATTTCGTTCATTATTGTATTTCATCATTGCTGCGAACATTGATTGTGGACACGACGC

TACCATCGTTACTTCTATGTTCACACCTTAATAATTGTGCAAACTACAAATAACTCAATTAATTTGAGACGCAGGAGTAG

TATACTGGCGGACATGTGAGGAGCTAACCTGGGGTTTAGAAGATTACATTGACCAAAAACTTAGTCCTTTAGAAATGATG

TTGTTCAAAATGAATATTGGCATCAAAAAAGAAATCAACTAATTAAAGTGTAAAGAATCACATGTGAGTACGAACTAAAG

CACAAAAGTTCTCCAAATAGCTGATGAGCCTCTACATTAATGCGATATGGACTCATGGTGAGTGTTGCACTCACCTGAGG

GTGCACTTGGGTGAGAAGCCATTGATATCTTCCCATGGTACATGGTAGCTATGTCGTACCATTTCATTTTCGTAGCAATT

ACTTATCTAAGAATTCAAACAGATACCCCCCTCCACCATTTGCTTATCTAATAATCACATGGTTCCAATTTAAGCACCTT

AAAGCTTATGTATGTGTTTCTCACCCCTTATCTGTAAGAGCAGAATCTCGCAACAGTTATACCTCAGCGAAGCTTTAACT

AAAGGAAGGTTGCGAAGATCATCGCAGAGTGTTAAAGTAGTGATGTAGCGAGTGACATCAGCTTAGCCATAAGGAAAGTG

TGAAGATTCTGCGAAGATAGAGATTACCTTGCGAAGAAAACCCTTATAATCTTGCGAGAACAATGCAGATCAGATCCGAA

AATAAAGGCTGGCTTAGATGTCATCCACTATGTAATATCCTATATAAAGGACCAAGCAAAGACGAGAGAGGGGAGATCTT

TTTGAGAGTGAGAGGAGAACAAATAGGATAGAGAAAGTTCAGAGCTATTTTGATTTCCTTGTATTCTCATTACTTTTGTA

AAAATCATTAAAAAGATAAATAAAACTTTGTAAATTTATCATGGAATTTGGCTAGAATAGCATATAATCTTTGAGGGGTG

TGGTTGTAGGAAATCCTGCAACCACATATTGGCGCTAGAAACAGGGAGTGAAAATTTGAAGATTATCTGAAAACAAGATT

GAATGAGGATATTATGATTCGACAGAATTTGAAGTATTTGATGAAGAATTTGTGAGGTGTATAACAAGTTATTGAAATCA

AAAGAGATGATGAAGAAAATCAGTAAATTGATTGAGGAAAAGATCATCAAAATCAAAGAAGACGGCAAGAAATTGGAATG

TAACTTAATTACAAAGACAATGATTAATGAGTTTCATGAAAATCTCAAAGGAAGAATACATAAACGAGATTTTGAGGGAA

AGAAAATTATGACACAATAAAAGGAGCCACCATTGAAGGAGTGGGAAAAACAGAAAATTACATTCACAGCAGCAGAAATA

CCGGTAGAAAATCTAAACCACACATCTCCGTTAGTAATTACAGTAACTGCAGCAAGAAAGGATAAAGATGGACCAGCGAA

ATCTATAGAAGAATGAGCGGTGAACAGAACTTTAATAGATACAAGAAGTTCAATCGATATAATATTTTACCATGCATTCA

AAGGGATGGGGTACAAAGATGAAGAGATGACTTGTTCAACATATCATATCCATGGTTTTGGTAAATTTACAACAATACCT

AAAGGAGAAATAGTGGTGCGAATATTATTAGGAGAAATTGAAACACACATGACATTATGTGTGGTGGATATAGAGTCACC

TTACAACATGCTATTGGGAAGACCATGGATACATGCAATTAAAGCGGTAGCATCCACATTGCATCAGTGCATCTAATTTC

GTATGCCTAGCGGAGTAGGTGAAATCAGAGGAGATAACAATGAGGCGAAAATGTGTCATCAGGTGGAAGTAGAAAATTAT

GAAGGACGTGCAAAGAAGCGAAAGCTTCGCTGGAAAATGGCGAAAGAAGCAAGGAAAGAAGAAGAATTTATAATCTACAT

GATAAAAGCAAAATACCAAGCGAAATTTCGCAAGAAGGAGACAAACCGATAAAAACGATAAAAGAGCAAACACCAATAGG

AGAGTCAAAGGCGGCCTACACATCAGAAGAACCAACAAAAGAAGTGAATATTGGAACTGAAAAAGAGCCAAAAATGTTAA

AGATCAGAACTAAAATGGATGCAGATGAAGAAGAAAGAACAATCAATCTATTGCGAGAATTTAAAGATGTCTTCGCATGG

AACATGGATGAAATACCTGGGATAGATCCTTCAGTTGCGTGTCATCATCTGGAAATCAAAAAAAATATAAAACCATACAA

CAGAGAATATGAAAAATAACAACAACATATCACCATCAAATAGAACAAGAGTTACAGAAGATGTTGGATGCAAGAATTAT

ACGACAAGCAAAATACCCAAAATGGATTGCAAATATGGTAGTGGTACCTAAGAAAAATAATGGGATAAGGATTTGCATAG

ATTTCACTGATCTAAATAAAGCTTGTCCAAAGGACAATTTTCCTTTACCAGACATCCCATAATTGCTTGAATCAGCGGCA

GGGAATGATAGAATATCTTCTTTAGATGGATATAAGGGTTACAATCAAATTCTTCTCGCTGAAGAAGATCAGGAGCATAC

CGCCTTTTTTGCACCAAGAGGATTATATTGTTACACGAAAATGCCGTTTGGTTTGCGAAATGCTGGAGCGACATATCAGA

GAATAGTAGAGAAGGTGTTTGCAAAATGGATTCATAAGACATTAGAAGTGTACGTGGATGACATGTTGGTGAAAAGCAAA

GAAGATAAAGATCATGTGCAAGATTTAGAAGAAATCTTTGAACAAATGCGGCAGTTCAATATCAAAGTAAATCCTGAAAA

ATGTATCTTCGGTGTTGCGTCAGGAAAAATTTTAGGATACATTGTGTCAAAAGAAGGAATACAAGTTGATCCGAAGAAGG

TGCGAGCAGTTCGCGACATGCCAACACCAGCGATGGTGAAAGATGTACAGAAATTAAATGGACTCTTGGCTTCGCTGGGA

AGATTCATTTCGCGATCATCAGACAAGTGCAAGCATTTCTTCAATATACTCAAGAAGGGAACAAAATTTAAGTGGACAGA

TGAATGCGAGAAAGCACTGTAAGGGATAAAGGATTACTTGATGAATATATCCATCTTACAAAAAGCAGAGCCAGGGGAAG

AATTGCTGATTTATCTTGCGACCACTTCGCATGCTTTAAGTGCCGTTTTGTTGCGAGTAGATGCCGGAGTAGAAAAGCCA

ATTTATTACATTAGCAAGACGTTTAATGCAGCAGAGAAGAACTATTCAAAGATTGAAAAACTAATACTGGCATTAGTTTA

TGCAGCCTTTAAGCTTCGCATATATTTTCAAGCCCATAGAATAAGGGTATTAACAGAGGTTCCCATTGAATCAGTAATGA

CAAATTCAAAGAGGTCAGGTAGGATTGAGAGGTGGAATGCTCAAGTGGGGCATTTTAAGATTAAATATGAAATATTATCT

TCGCCAAAGTCACAAGTTGTTGCGAATTTTTTAGCCGAATTTCCATTGGAAAATGATGAAGCAGTGGAAGAGATGATGGA

TATATAAGAAGAACATCGGAATCCAAGAGACCTGTTAACAGAGCCTGATAGATGGGAAATATTGGTGGATAGATCATCAA

ATGGAGAAGGTAAGGGAATTGGAATTATCTTGATTTAACCAAAATGAACAAAGATGGCTTTTTCTTTTAGATTGGAGTTC

GCATCTACTAACAATGAAACGGAGTATGAAGCAGTGGTGCATGCTTTACGACTCGCAATAGAAATGAAATTAGAAAACGT

AAGGATCACTAGCTATTTGCAATTAGTCATTTGACAAATAAAAGGAGAATACACTACAAATGAACCCTCATTGAAGAAGT

ACAAAAGGCTTGTTGGGGAGTTGTCAGCGAAAATCCCGAAAATAAAATGGAGACATATATCAAGGAGGGATAATAGGTTA

GCGGATGCTTTTGCTTTTATCTCAAGCATGATGGTAGATCCGACTGCAAGATACATAAAAATTCATACGTTTCATTTGTT

GTCAATCAACAAAGAAGAAGAAGAAGAAGAAGTAGATGTGATGATAGTAGAAGGTGTGGAAAATGAAGAGGCAAATAAAA

TCACAGATTGGAAAATTGAACTTCATGCATACTTGGAGAGAGGAGAAGTACCAAGGAATAGATTAGAGGCACATAAATTA

AAGAGTCGTGCGACAAATTATGAACTCAGAGACGGGGTATTGTATCGAAGGTCATTTAATGGACCTTCGCTGAGATGCTT

GACACGAAAAGAAGGAGAGGAAATACTGAAAGCGTTACATTATGGAGATGTTGGTAATCATAGTGGAGGAAGATCTTTGG

CATACAAAGCGAAAACACAAGGATATTATTGGCCGTACATGCATGAGGATGCAAAAGAAGTATCAAAACGTTGCGAAGAG

TGCCAGCGACACGGTAAGAGAATACATGCACCAGGAGCACTTTTATCATCTTCAACAAGTCTTTGGCCTTTTGGAAAATG

GGGTCTAGACATTGTGGGACCATTTTTACCAGGATCAGGGCAAAGGAGATATTTTATTGTCGTAACAGACTATTTCACAA

AGTGGGCAGAAGTAAAGGCAGTACAGCATATATGCAATAAAGACGTTTTCACTTTTATATTCGAAAACATCATTTGTAGA

TTTGGAATTCCGGCACAACTGGTGTCTGATAATGGAAAACAGTTCGAAGGGCAGAATATAGAAATGTTGCTTAATGCATT

TAAGATAAAGTGTGGGAAATCCACACCATTATATCCACAAAGTAACGGACAAGCGGAGGAAACAAATAAGACAATAGCAG

ATACATTGAAGAAGAAGTTGGAGGGGCATAATAAAGCATGGTGTGAACAGGTACCTAATGCAGTTTGGGCTTATAACACA

ACTAGGAGAGAAGCTACGTGAATGTCACCTTTCTGTTAAACATATGGAGTTGAAGCGATGTTACCAACTGAAGTCATTAT

ACCCACTACAAAAAGAGAAGCATGGGAGAAACATCTAAGTGCGGATATGATTTTATCAAAGCTTGATGATTTAGAAGAAA

AGAAGGAAAAAGCCATACAACATATGGAGAATTATCAGAGAAGATTAGCTCGAGAATACAATAAGTGAGTCAAGATAAGA

GAGTTTCAGCCAGGTGATTTGGTTCTACGAGAAATACCGCCGTATCAGCGAGGAAATGATGGTAAACTAGAGAAAAAATG

GGATGGACCTTATATCATTAAAGAAATAGTTGGAACAGGGGCATATAGACTGATGGATCCTGAGGGAATAAACACTGGTC

ATAAGGTTGATAGGCCATGGAATAGATTATATCTAAAGAAATATTATCCATAAAATATTTGGCGAAAATCACCAGGAAGT

AGAGAAAAAGAAGAAATTTCATAGTTGCGAAATTATATTTACAGAATGAAGAATGCGATGATTTCGCATAAATTATTTTT

ATCGAAACAAGTATGCGATAATTTCGCATAAAATGTATCAGATAGAACAGGTGAAAGATATGCGATAATTTCGCCAGAAT

TATGAATATTACAAAGAAAATAAAAGAAAATATGACAATAAACAAAAAAGATAGAGGCACCCAACTTTGGTGTACACCCT

AGTTCGCGAATAAAATTTCGCAAGAGGCAAATGTTAGACCTAAAGTGCGTCATTGAAGGGGAGACCTATTACATGGCTCA

GGGAAAGGTTACACCGCACCGGAACCAGGGTCATGGAGACTTGGGGTTTTTAAACCCATCCAGTAGAGGCGCCTTGGATT

CTCAGCTTAAGCTTATGACTTAGGTTGGGCGACTAAGGTAAGAAGGTCTCTTCTAAGGAGTGCGGAACTCGATCAAGGAG

CCGAGGTACACGGTTGAGCCCAGAGTGCCTGGGGCGTTTGAAGCGTCTTGGCCATTCTTGGCAAGTCTTGGCTATGATGC

ACTCGGCCCGAAGAAAACTCCACTTAGGGTGCGGTCTTGTAACCATGAGCCATATCGACGAAGGGATAAAAGGACGCGAA

AACAACTGGTTGTTGTAAAGAAATCGGATGAGAGGAGCCAACCTTGTATGGTAGAGGTACGCCTCCTTGAAGGGGCAACC

AGGGGGATGATAAGGGCGGCACCCTCCATTAGGGAGCTGATAAGTGTCTAAAGACGCAAATGTCATGAAACTTTTTACTC

ATGGGAGTATTAAGGCTTGTCATATTTGTGCAACGATCCTGTAGAGGTAAAAATACTAGAGAAAGATAAGACGAGATCAA

TGGGTTGATGCACGAAAGTGTAAAGACGTCCTGCGATTTCGTCGCAAGATGACAATATCATGGGGCTTTACGTTCGCAAG

AATATTTAGGCCTGTCATATTGTCGCATGCAAAATATCCTGTTGAGGCAAAAATACAAGAAAATGATAAGACAAGATCAA

TGGGTCAATACATGAAAATGTAAAGACGTCCTGCAATTTTGTCGCAAATATAAAAAGTGGCAAGATAAGACCTTTTCTTA

GGAAGGCAAAATAAGACCACATAACAAAATGAAATTTTGCAAAATTCAAATATATATTTGATAGCAAAATTTCGCAGGAA

AATAAGAACATGTCTGCGATATTGACGTCCAAATGAACTTATACAAACTAACCATTTAAGTATTGTGCAGGAGACAAATT

CAGTTTTCGCAAGATAGATAATGAGAGGCAAGTATGGGAATTAATAAACAAGATGCATTATCTATCTCGCTGATAAGGAG

TTATTAAAAAGATAAGTGAATTTCAAGATTCATGAAAGTACACTGCTTGGCAGAAAAGTAAGGCGAAGATAAAATAACTT

AAATAACTAGAAAAATTTCAGATTTCAACATGTGATTCGCCAAGAGGGGCTGGTGTATGGCCTTCTAAAATCATTTCATC

TTCACAAGCCTTTTTATTTTTGTTGGTCCATCCTAGCTTGTTCTCCATCTTGATTATCCTCAACATTAACTTCCTTGATG

GGCTCATTTTCTTCATTTTCTTCAAGAATAGCGTCATCATCACTTTCGAAGTCAAAATCACTATCAGGATCACGAGGAAC

TTCATCAGCATCTATGTCAAGAGGCATAATTTTTGTAGGAGGAAGTGATTTATCTAGTAAAATTCCATTCACAAGAACTT

CATCCTTGAGATGAACTTGGCGAAGAAGTGCCTTCTGGTGATTTCTACTTTCAATATCTTTGAAATGTTGAAGGTAGTCA

AATCTTCTTTCTGCACGATCACGAGAATGAGCAAGGGAAGTAGCAAGTGTAGCATTTTCCTTGCGAATCTTGGATAATTT

CGATTGCAAGGAAGATATAGAAGAATGAAGACTTCTCTCAGAATCTACGAACAATAAGAACACAAAAGATTGTTATAGCG

AAATTCACAAAAGAAGAGTATTGCGAAGATAATTAAGCGTAAAAAGTTGGATGAGGCAGTCAGATTCTTATGACTACCTT

CCTTCTGTGAAATAAGATATTGGGCTAAAGCCAAATTATTATTCTCCAAGTTGTTCATTGCATCATCAAAATTATCCCCA

ATTAAACCTTTCAGGCGAGATCGAAATTTCTTATCATTAGCGATAAGTAAAGCATTTTCCTTCGCTAAAGAAGCATAGGA

ATCTTCTAGCTGAGTATGTTGCTCTTTAAGTTGGTTTGCCTTTACGCTCAAAGCTATTCTACCTTGGATGAGTTTATTCT

TCTCAGCAATAGACGATTCAGTTACTTCTTTAAGTTCACTTCTTAACGAGCGAAGAGATCACTCTTGTTCATCACATATT

TTTTGACTATATTAAGTTGTTGCGAAGATATCGCTACTTTGTTTAGGAGGATTCTCTTCTCTTAAGACAAGGTTCCATTT

TCATCTGACAAAGTTTCAATCTTCAAGTTAGCTTTAATTAATGAATAAGAGAGTTGGGATAGCTTGCTACTTGATACTTC

TTGTTCTTGGATTAGTTGGGAATTCTCATTAGAAAGGTTGTTAATTTGATCAACAGAGTACGAGTAAAGATTATCTAATT

GGTTATATTGGTTTAACAAAGCTTCATTGTCAGCACGAGCTTCTTCAGCGGCATATTGAAAGTCGTATCTCTCCATTATT

CGTTTCTGAATTAGAACTTAACGTGCGAACCAGAAAAGTGAATACAAAAGAAGTAAAACTAACAAAGAAGATAAGTTATA

AATGAAAACAAGAAGATCATACCTTTCAGTTTATCATTTTTTCTGCGAAGATTCTCGTAATCTAGCGAAACATTCTCAAG

CTTTTGATTTTTGAGACGAAGATCATCACATTTCAAAGTCATAGCTTGGAGATTCTCCTTCAAAGATTCAATTTCTAAAG

ATGCTTGCGAAGTTGTTCTTTCAGCATTAAAATATTTACCTAGGGTTTCGCAGACGTTAAGCTTGAGAGGGTCAACAGAA

GATCTTCGAGCATCATCAAGAACATCAGACAAAACTCTAAAGGCAATGTCTACTCCTTCCAACAACTATATCTTAACAGG

AAGGGTTTCAGATGAAGAATTAGTGATAATTGAAGGTATGGGAAGTGAATCATCTTTCGCAATGGATATCTTTTTTCCAG

GCGAAGAAGAAAAAGGTTCTGTAGGCGAGGTTGCGACAGAATCAATCTTTGCGGCATTCGCAAGAGGAGAAGAAGGGGTT

TCATTCGAATAAGGGTCAACTGATAGGACGACATCTGTAATGATATCCTCATGGGAAGGAGCAATAAGTAATAAAGATTC

AACTGGCGAAGCCGTAGGAGAAGAAGCTAAGGGTTTGAAATCAAGAAATGAAACGGAAGAAATAACAGTCTTTAGTTTCT

GAGATTTCTTGTAGGGTTTGTTAACAGCTTCATCATCCTGCGAATATCAACTCAAATAAGAAACAAAGGCAACAATATTG

TTATTAAAGAAGATATGTTTCTTATTACCGTGGAATTTGCAGGCTTTCTTTTGACCACAACAGTGAATTTGGTAGAAGAG

GTGCTTTTGCTGAAACAAAATCAATATGGGAATTAGACAAGATAATAAAATTAGTTAAAAAAGTCATTAAAGTAAGATCA

GTTCCACAAAAATGAAGAAGAAAAGCAAAACTAACCTGGGTGACTCCATGAAAACAGTAGCAGACAGATAAGCTTGTAAG

AACTAAAGATGAAGAAAATGGCAGAGAATGAAGATTTACAGAGAGCTCGGAGACGAGGAAGATGAACTAAAGGAAGAAAA

TAACATTCCCATCAATCTTCTCCTTAAATAGGCAAAAAGGATGACCGGTTGAAATTGAATGTGCCGGTAAAAAAGGGGAA

CAATAAACACGTGTAGACGGTTACGCAAAATTCACGGACGGTTGTCAGTTGAAAAAGATAGGGAAAAATGAGCGGTTGCG

GCATTTAGAGATAGAAATTCTCATATGATTCTCTTCTTCGCAGAGAAGACATGAGAAAAGGCAAGATGTAGGAGCAGAAT

CGCGCAACAGTTATACCTCAGCGAAGCTTTAACTAAAGGAAGGTTGCGAAGATCATCGCAGAGTGTTAAAGAAGTGATTA

GCGAGTGATGTCAGCTTAGCCATAAGGAAAGTGTGAAGATTCTGCGAAGATAGAGATTACCTCGCGAAAAAAAACCCTTA

TAAGCTTGCGAGAACAATGCAGGTCAGATCCGAAAATAAAGGATGGCTTAGCTGTCATCCACTATGTAATATCCTATATA

AAGGACCAAGCAAAGACAAGAAAGAGGAGATCTTTTTGAGAGTGAGAGGAGAACAAATAGGAGAGAGAAAGTCCAGAGCT

ATTTTGATTTCCTTGTATTCTCATTACTTTTGTAAAAATCATTAAAAAGATAAATAAAACTTTGTAAATTTATCATGGAA

TTTGGCTAGAATAGCATATAATCTTTGAGGGGTGTGGTTGTAGGAAATCCTGCAACCACATTATCCCGTCTATTTATATA

TGCTTGTTTAAATTTGAGTAGCACTTAGCTTTTTTAGCCGTATTCTTCATAAGCTGTGTTTGTTTTTTCTTTACTTTTAG

TGTCCTTTTGTCAATCGAGAGGGAAAGTCTCTGCCGTTATGTGAGTTTGGCACCTATACTATTCCACCTCCAAGTTGCTC

TTGGACATTTCACGAGCTGGAGTGTTCCAAAGAACTTGGGTCATGTCAGAAACTATATAAAACTTGTGCAAAAATCGTGT

TGGCGTCTATGTATACTTGTTGTTAGTCCCGTTTCTACGTATGCCTATTGTCTTTAACCAATTCCCCTGTTCTTTTAATG

CAATTTGAATGCACATCGCTGTAGTCAGTACATTTCTGATAAGCATGAAGCAAATATCTCCACTTTTAGTACTGCTTTAG

AACTTTATTGAGCAAATATCTCCACTTTTAGTACATTTCTTGGATGGTTTAGCCCCACCAAACAATAACTCCTGACACAG

TACAAACGCCTTCTATAAAAGCCACATCTAGCGTGGGTGACCTTAATTATCCCCACGAGGTTCTTGCATTACTCTCGTGG

GCTGATATGGGACACTGCTCCACCCCCACAATCAGACACCTTTATAGAAAGCACCGGCATATTATTACATCAATCAATCC

ATTATAAAAGATAATTGATATGTATAAGGGATGATATGGGACACCGCTCCAACACACAATCAAACACCATTATAGAAAGC

ATTAACATATTAACATCAATTCTTTATAAAAGATCATCTTTATACATGTATGAGTAGCAATAGGGGTGTGCATGGGATGA

TCTTTATAAAAGATCATCTTTATAGGTTTTCATCTCATCCGCATCCAATCCAATGCACTACGGATTTTAAGTTTGGCATC

CGCATCCAATCCAGATCCAACGGATTTGCATCCAATGGATTAAAGGTGTAAAACGTGCCGGCCCGGCACATCCCGGCACA

CGAAGTCGCGTGCTTCACATGCCATGTGCCGTGCTGTACTGTGCCAACGATTTAAACGTGTTGTGCCGTGCTGTGCTTTA

CTAGTCAAAAGCTAGGCACAAAGACGACCACGGGCACAACACACGAATAAACGTGTCGTGCTGTGCCGTGTTAGCACACG

TGCTATAAACAATTTGAAATCACTTAATGTATACATTGAGTCGGACATTATCACAATAATTTAATTAGACAACATATCAT

TTGAAATTATTTCAAGTAAAATTAACAAGTTTATTCAATAAAAAGAAATAGCCCATCAAATATAAAATTGGATTATTGTC

ATGTAAATTAATGTTCTAGTTAAATATAGGTGACAACATTATAACAACGATATTGAAATATATTTTCCAAATATTAAGGT

ATTATATGTGTTGTTATAAATAAGCCAACATAAAATGTTAAAAACTAGCTAGAATAGTGACTCTTTTGTGAAATACACAA

AAAATGTGATGTATTATGCCTTGTTATATGGTGTATTTGTACATGTTATGACGTGCTCTCTTAACGTGCTGTGCTGCACC

GGCATATTATTACATCAATCCATTATAAAAGATAATTGATATATATATATATATATATATATATATATATATATATATAA

GGGATGATATCGGACACCGCTCCAAAACACAATCAAACACCATTATAGAAAGCATTAACATGTACGTGGAGGTGTAAATT

GGGTTGTGCCAACGCGAGCATAACCCAGCTCAGTACGCTAAAATTTAAGCACAGCCCAGTCCAGGACGTGACTTAGCCCA

GCACGGCACTGCACGTTAGTTTTTTGGGACGGGCTGGGTTTGACTAATCGACACAGTAGCCCAGCACAACACGCGGCACG

GATAAACACCTGGCACGACACAACACCGCACACTAAGAAAGCACGCCGTAATATGTATAAGATACTACACAATGTGGCAC

AATACATCACATTTTGTGTATATTTCACAAAAGAGCCAATATTCTAGCCAATTATTGACATTTTATTATCGGTATGTTGA

CTTATTTTCATAATAACACATATAATACCTAAATATTTGGAAAACATTTATATTAGAAAATATAAAATAAATGTGCCCGA

CACAGCACAACACGACACAACACGTTTATTTTTCTGGCACAGTCCAACACAGCACGCCTTTAAGCACGTCACAACACGAC

ACGTCATAATCTCCGGCACAGCACAACACTACACATAACACGCTAAGCACGTGACTTCGTGGGCTGGGCAGTGCTGGGCC

GGCACGTTTCACACCTCTACATGTATGGGCTGATACGTGAAACTGCTCCACCACATAATCAATCCTTTATAAAACATTGA

ACTTTATATACAGTATATTTATATAAGTTATGATAGGTTGACCATGGATTCTGTTCCGCATATTTTGTAGATTGCTTTTT

TCAACTAAACCACTTTATAATAAACTTTTTACCAAAAATTGTGTTTGGTAATTTTTTTTTTTAAAGTACTTTTGAAAAGA

AGCACTTTCATTTCAGACTTGCTTTTAAAAGCTACTCGCACTAACTTTTAAAAGCAGAAGCCATAATTTGATTTAAGGGA

TTCTCAATTTTGGAAACTGCTCAAATGGGCCCAACTACAACTGTAAGATATGCTGGGATTAAAATTCGAGCTTAAGTTAC

CATGAGACGCACCGAAATAAAATAATCCTCCTATTCATGTGCTTCATGTATGGAATAACTTGATTGCTAAAGACATGTAC

TGGTATCAGCGGGGCGGGTACCATTTCAAATAAGACACAAAACTTACAACTAGTGGCTCAACCACTTCAAATACGATTAG

AGTTATCAGATATCAAGTAATTTTTCAATAATTTTCGTTATGTAACAGAAATCATAAAAACCTTTTTTGTCCAAAATTCA

GTTTAAAAAGTTCAAAATATCAAATTCTGTCCAGTCGTTTTGTAAAAAATCTGTTGGGTATGAAGGTCCAGCCGTTTTAT

AAAAGAAAATATCAGGTAAAAAGATCCAGCCAATTTATATAAAAGAAAGTGGCGGGTAAGAAGGTCCGCCCATTTTATAA

TAGATACGACCGAGGAAGAAAAAGAAACCTTAATTCTCTATATTGATTCACTGATTCATAAGTGATAAAGAGAAGAGACT

GGGGAGAGTCTGACCAAGACAAATTAGATTGAACAAGAAGAAGATTTAGAAGAGAATAGATGGAGGTGTAAATTGGGCTG

TGCCAGCACGAGCATAACATGCTAAAATCTGAGCCCAGCCCAGCCCAGCGCGTTAGTTTCATGGGTTGTGTTGGGTTAGC

CTAATCGGCGCAGTAGCATGGCACAGCATGCGGCACGGATAAGCACGTGGTCCAGCCCAGCACAACACGTTAAGAAAGCA

CGTCATAGCATGCACAAGTATACAATATAACAAGGCATAATACATCACATTTTGTGTATATTTCACAAAAGAGTCATTAT

TCCAGCGCGACTTCGTGTGCCGGGCCGACACGTTTTACGCCTCTAAGAATAGGTAAAAAAAAAAAAAAGTTTGATTTTTC

TTAGTTTACGTCGAAGAAGAAGATATTATACGCGGGTATAAAACGATAACAGATCAATATATACTATCGGTGTTACCGTA

ACGGAAACTGTCTCAGCTGGTTCAGTTACATTAAGCCAAAAAGTTCTAATAACTTCCCGATCCCAAGATACGCTAACCGT

TATGTCGGAAAAAATTCTATCGTATCCAGGATTATCTGTAACGTCTATCGGATTTCGTCTGCTTGCAGGACCGTCGGATC

ATTCAAATGATACCAGAAACGACGTTAATGCTAGTAGTGCCTCTAACAATCACGTGGAACGTGTGTTTTTTTTTTTTCTT

CTAGTGGTGCTTATATATATTTGAGTTCACCATAAATATCATATCATCAAAATCTATCATCCTACATACTGTACTGGGTT

GAATCATGGAGCTCCAATATTTTTCTTATTTTCAACCAACTTCCTCCGTTGTTGCTCTTCTACTTGCTCTTGTATCCATC

TTATTCAGTTTCGTTGTTTTGAGGAAGACATTTTCGAATAACTACTCATCACCTGCATCATCAACAGAAACAGCGGTACT

TTGTCATCAGCGGCAGCAGTCGTGTGCATTGCCAATTTCCGGTCTTCTTCATGTTTTCATGAATAAAAACGGCTTAATTC

ATGTAACTCTTGGAAATATGGCCGATATATACGGACCGATTTTCAGTTTCCCAACAGGTAGCCATAGAACTCTCGTTGTG

AGCAGTTGGGAGATGGTAAAAGAGTGTTTTACAGGCAACAATGACACTGCTTTCTCAAACCGTCCTATCCCGTTAGCTTT

TAAGACTATATTCTATGCATGCGGTGGCATAGACTCATACGGTCTTTCGAGTGTACCTTATGGAAAATATTGGAGGGAGC

TACGAAAAGTCTGTGTGCATAACCTCCTCTCTAATCAACAACTACTCAAGTTCAGACACTTGATAATTTCTCAAGTTGAT

ACGTCTTTCAATAAGCTGTATGAGTTATGCAAAAACTCTGAAGACAACCAGGGAAACCATACTACTACTACTACTACTAC

CGCAGCTGGCATGGTGAGAATGGATGATTGGCTCGCCCAATTATCGTTCAACGTGATAGGAAGAATAGTCTGCGGATTCC

AATCAGACCCTAAGACAGGTGAGCATTTTCCTAGAGTTTCTGTAGAAAATGTTGGATTTTACAGAGCCGATCCTGAGTGT

AGAAGGGCCCTGTGCGAACGTGACCGTAGGGCCTTCATAGTGTCAAATTATTTTTTTCATGATAATTTTTTTTCAACTAA

TAAATCAGTATTTTTAAGTTAAAACTTGACTTATGCAGCACATGAGTCCATCTCTAGTCCTTAACGTATATGCACTCAAC

GAGTATGCGAAAACTGTTTTTGAACATTATTATATATGGTTTGCATTCATAAGATAACTAGTTCTTTTTTAAATTCTAGA

TGCTAAATTGCATTCTGAATAGTGAATTAAAGTTTAGATGTCAGTTTTCGTACCTAAATATAAAAATTAAGTATAACGTT

AATCTCACATTTCGTAACTAAAAAGGTTTTCAAATTTAAACTAAGTTCTATATTATATTCTAAAAAGAGTTCAAATTTAA

ACTATGTATTATACTGACTAAAATTATAATCAGGAGCATTTGAATTATATACACCCTAAGAGCATCTGCAGTGGTGCGAT

CATATACCAAAGATCAAAGAGCAAAAAAATTTGAAAATTTGGGTTTAGTCCGTGTTGCAACGCAGTGGGGGAGACTAAAA

TTTGGTCAGGCGTAAGAGGAAGTTACGTAATCGAGGTAAGAGGAGGTTCCGCCCCATTGTCATCAGGCGTAACTTTGAGG

TTACGCCTTAAATTTTTTTTTTTTTTTTTTTTTTTTTTTTAATGGGGCGTAAGTTTGAGCTACGCCCCATTCAGGCGCAA

CTTTAAGTTGCGCCCCATTTTTTTTGTTTTGTTTTTTAGTGTGAGGCGGAGTTTTCTGTTACGCCTGTTCGGGCGTAACG

CACCAATTGCGCCCGACTAAAGACCAGACTAAAGACTAAATATGGTCTGGGTTTTGGTTATAGTCTGGTCTTTGGTCTAT

ACTCCTTCCCCACTGCGTCTGCTATTTGATCTTTAGTATGGGTTTTAGTCGCCCACTACGGATGCTCTAAGCCTTTGGTC

TCTCGAGGTTGAATCTCTCTCTCTGGGTTTGTTTTGTCAAAATTTTAATTTTGCAAATGAAATGGGCTTTGGTATTAAGG

TGTCCACTAGTAGGAAAATGTCCCTAACCAGAGCTAGGCCCTGTGCGGTCGCACCCCCTGCACACCCTCAGGCCCGGCTC

TGTCAGCATCGATTTTATACTATGAGATCATCAAAGACTATTCTTCTAACTGTGCACATGTATAAACAGGTGCTCCAAGC

AGGGTGGAACAATTTAAAGAAGCAATTAATGAAGCATCGCATTTTATGTCGACATCTCCAGTGTCAGATAACGTTCCAAT

GCTAGGGTGGATTGACCAATTGACTGGTCTTACAAGAAATATGAAGCACTGCGGAAAGAAATTAGACTTGGTGGTCGAGA

GCATAATTCAAGATCATCGTCGAAAGAGACGATTCTCTAGAACTAAAGGAGGAGATGAGAAGGACGATGAACAAGATGAC

TTCATCGACATTTGCTTGTCAATTATGGAGCAACCACAGCTTCCTGACAACAATAACCCTCCTCATATACCTATCAAATC

TATTGTCCTGGTATGCAAATATTGATCCGTGGTATGGCTGGAGCATTAATTGTATTCCAGCGGTTAATTTTGGTTGTGGT

AGCTCATAGAGTGCAGAAACATGATTTGGTTTTGATTGGTATCATTGTATAAATGCAGGACATGATAGGTGGGGGAACTG

ACACAACAAAACTGACCACCATCTGGACCCTTTCCTTGCTGCTGAACAACCCCCATGTGTTGGACAAGGCAAAACAAGAA

GTGGATGCACACTTTGGAACGAAAAGGAGATCAACAAATGATGCCGTGGTGGATTTTGATGATATTCGTAACCTTGTATA

CATCCAGGCAATCATCAAAGAATCAATGCGGTTGTATCCAGCCAGCCCCGTGGTGGAGCGACTGAGCGGCCAAGATTGTG

TGGTCGGTGGGTTTCATGTACCAGCAGGGACGAGATTATGGGCTAACGTATGGAAGATGCAACGAGACCCTAAAGTATGG

GATGATCCATTGGTGTTTCGACCAGAGAGATTTTTGAGCGATGAACAGAAGATGGTTGATGTAAGGGGTCAAAATTATGA

GCTGTTACCATTTGGAGCCGGTCGACGTATATGTCCAGGTGTATCCTTCTCTTTGGATCTAATGCAACTGGTACTGACTC

GTCTTATTCTTGAGTTTGAAATGAAGTCTCCTAGCGGGAAAGTGGACATGACAGCAACACCAGGATTAATGAGTTACAAG

GTGGTCCCCCTTGACATTCTGCTCACCCATCGTGGCACAAAGTCGTGTGTGCAGTCAGCAGCCTCTGAGACTAGTGGTGT

ACCAGTAATCACTCTGAGCTCGGGCAAGGTGATGCCTGTTTTAGGCATGGGTACATTTGAGAAAGTTGGTAAAGGGTCTG

AAAGAGAGAGGTTGGCGATTTTAAAAGCGATAGAGGTGGGTTACAGATACTTCGATACAGCTGCTGCATACGAAACTGAA

GAGGTTCTTGGAGAAGCTATTGCTGAAGCACTCCAACTTGGCCTAATCAAATCTCGAGATGAACTTTTCATCAGTTCCAT

GCTCTGGTGCACTGATGCTCACCCTGATCGTGTCCTCCTCGCTCTTCAGAATTCTCTGAGGTAAATTTTTTTGATGCAAA

TTAGGCTTACATGTATTTGCACATACCAGCAGGTACACTTAAACATCTATTTTTAGATAAATGAAATTGTGTTGTGTGAT

ATGTACATAGGAATCTTAAATTGGAGTATCTGGATCTATATATGTTACCCTTCCCGGCAAGCTTGAAGCCAGGGAAGATA

ACGATGGACATACCAGAGGAAGATATTTGTCGAATGGACTACAGGTCTGTATGGTCAGCCATGGAAGAGTGTCAGAACCT

TGGCTTCACTAAATCAATCGGTGTTAGCAATTTCTCCTGCAAAAAGCTTCAGGAATTGATGGAGACCGCCAACATCCCTC

CAGCTGTGAATCAAGTGAGTACTCTTACTTGAAAGTTTTGATAAGACATCGGAAGATTTCTAATGCTTATTAAATTTCCA

ACATTATGCATAAAACTACTACTCTTTATAATGTTGATTAAGGTGGAGATGAGCCCGGCTTTCCAACAAAAGAAGCTGAG

AGAGTATTGCAACGCAAATAATATATTAGTCAGTGCAGTCTCTATACTGGGATCAAACGGAACCCCATGGGGCTCCAATG

CAGTTATGGGTTCTGAGGTGCTTAAGAAAATTGCTATGGCCAAAGGAAAATCTGTTGCTCAGGTTGGTCAAATTCTCTGC

ATATGCCATAGTGTGGTCTCTGTTCTTTGGCCATCGTATTTGCATCACATTGTATTCCTTCCTTATACGGAATGTAACTT

ACATTTGTATTACGTACTGATCAGGTTAGTATGAGATGGGTTTACGAGCAAGGCGCGAGTCTTGTGGTAAAAAGTTTCAG

TGAAGAGAGATTGAGGGAAAACTTGAACATATTCGACTGGGAACTCACTAAGGAAGACAATGAAAAGATCGGTGAGATTC

CACAGTGCAGAATCTTGACTGCTCATTTTTTGGTCTCACCTAATGGACCTTTCAAATCTCAAGAAGAGTTGTGGGATGAC

AAAGCTTGAACATCGATCATTTAACTCTACACGTGCATTTATAAGAGAAGCTTCCCCTGCAGTGTGCTCAATCTTGATTT

ATTTTATCAAATTACATCTTACTATATGGGAGTCACAGATCTCTTTCCCTTCCTATAAGTTTGAATAAATTTGTAGCGAG

TGATGTAGCTAAAGAAATAAATGTAGTAAGTTTCTTTCCTAAGCTTGTTTTCCTTTACAAAAACTAGATGAGAGTCGGAG

GTAAATAATGGTGAACTCGTGTTCATCAGCAACAATAGTGTGATTAGGCACAATGTATATACCTTAGCAGAAAACACTAT

TCCAGACACAAAGAAATGTACGAATGCGTAAAGAGTATTCTGCCATGCATATACAGCAGCTGATGCGGAAAAAAAACAAT

ACAAAACCTTCTGTGAAACCAGACGTATTATTGTCCATGTGAGAGTAAAATATCTAATTGCCACGTGTAAGCATCACAGG

AGGTGATCATAAGATAAGATGGTAAAACAGATACACTGAATAAAGATGTAAAAGGTGATTACAACAAAACCAAGAATATT

CACTGCTTCTTTTCAAGTCTAAAAGAAAGCAGTGCATTTAATGAGAATCAGAAGAGGTGCGATACAGCTGTAGAAGGAGG

TACATCCCTCTCGTGAATTCGGACGAAGGGCGAGGTACTTCTACCCGAAGAAGGCGCCACACGATAAGACCCAAAGGGCG

ATATATAGTATCATCCGAAGAAGGCATCACACGAGGTAAGTCGAAGGGAGCAATGGCGAGCCAGTCTGAAGCGAAGAGGC

GTATATATTGTATATATCGTCATACACCCAGGAGCACCGTATCGGACCTGAATGAGTTTACTTTGCCTATTGTTGATGGT

GGTTTTTAAACAAAGGGTGAAATCGTAAAACCCAAAATAACCTAATGTCACCGGTGACACATTTATTAGGACATTTAATG

CATGTAGTCAGTAATTAATAGGGGGTATTTTTTCAAATTAGGGTTACAACGAACAAAACCCTAATTCCCCTCCGTGCCAA

TGGCAAATCATGCCAGCCACACCTAGGCCAAGCTAGCCGCACCTCTATGCCAATGGCAAAGCATGCCAGCCATACCAAAC

GCGCCGAAGCATGCCAACCATGCTAACCGCACCTCTATGCCAATGGCAAATCATGCCAGCCATACCAAACGCGCCAAAGC

ATGCCAACCATGCTAGCCGCACCTCTATGCCAATGGCAAATCATGCCAGCCATACCAAACGCGCCGAAGCATGCCAACCA

TGCTAGCCGCACCTCTATGCCAATGGCAAATCATGCCAGCCATACCAAACGCGCCAAAGCATGCCAACCATGCTAGCCGC

ACCTCTATGCCAATGGCAAATCATGTCAGCCATACCAAACGCGCCGAAGCATGCCAACCATGCTAGCCGCACCTCCGTGC

CAATGGCAAATCATGCCAGCCATACCAAAACGCGCTAAAGCATGCCAACCATGCTAGCCACACCTCCGTGCCAATGGAAA

GTCATGCCAGCCACACCTAGCCCAAGCTAGCCGCACCTCTGTGCCAATGGCAAATCATGCCAGCCACACCAAACGCGCCA

AAGCATGCCAACCAAGCTAGCCGCACCTCCGTGCCAAAGGCAAATCATGCCAGCCACACTTAGCCCGAGCTAGCCGCGCC

ATGCCAAACACACTCTGCCTTGTCCCTAGCTACGCTGTTCATGCCATGCCAAGCTAAACACCCTTACCTTGGCCCTAACC

ATGCCATGCCAAAATGGCGCCCACGCCTAGGCTTACAAATCATGCAACCTAGGCCAAATTGGCCACGCATGCCATTGGCA

CATGCCAACACGCTCGCTGGTAGCTTTTCTTACCAAGGCACGACCCAAATTGGCCACGCATGCCATTGGCACATGCCAAC

ACGCTCGCTGGTAGGTTTTCCTACCAGGGCGCGACCCAAATTGGCCACGCATGACAATGGCACATGCCAACACGCTCTCG

GTAGCTTTTCCTACCGGGCGCGACCCAAATTGGCCACGCACGCCATTGGCACATGCCAACACGCTCGCCGGTAGCTTTTC

CTACCAGGGCGCGACCCAAATTGGCCACGCATGCCATTGGCACATGCCAACACGCTCGCTGGTAGCTTTTCCTACCAGGG

CGCGACCCAAATTGGCCACACATGCCATTGGCACATGCCAACACGCTCGTTGGTAGATTTTCCTACCAAGGCACGACCCA

AATTGGCCACGCATGCCATTGGCACATGCCAACGCGCTCGCTAGTAGCTTTTCCTACCAGGGCGAGACCCAAATTGGCCA

CGCATGCCATTGGCACATGCCAACACGCTCGCTGGTAGCTTTTCTTACCAGGGCGCGACCCAAATTGGCCACGCACGCCA

TTGGCACATGCCAACACGCTCGCTAGTAGCTTTTCCTACCAGGGCGCGACCCAAATTGGTCGCGCATGCCATTGGCACAT

GCCAACACGCTTGCTGGTAGCTTTTCTAAACACGGGCCATGATTCAACTTGGCCACGCATGCCATTGGCGCATGCAAAAC

ACGCATGTTGGTAGCCCAACTTGCAAGTCGTGCAACCTAGGCCAACTTGGCCGTACGCGCCATTGGCACATGCCAAAACC

CACGCTGGTAGCCTAACTTGCAAACCGTGCAACCTACCGCCAAGGCATTCCGCACATGCCATTGGCACATGCCACGCACG

CCACTAGCAGCTTTATGCATTGCCACTTACACTCAACGTACAACCTACCACCAAGGCATTCCACACATGCCATTGGCACA

TGCCACACACGCCACTTGCGCTCGAGGTGCAACTCGCAACAAAGGCATTCTGCACATGCCATTGGCACATGCCACATGCG

CCCCTAGCAGCACCACTTGCACTCAAGATGCAACCAACACTCGAGACATCAACACAAACTGGGGGATGCTTTCCAGGGTC

TTGGTTTGGCGGTCACTGTAGGATCATAATATCGCAATGCATAGGGCTCGCGAGATCAATAAACGACAGTACCGCGAAGG

AGTCGCAGGAAACTAAAATAGACGACGCAATGGAAAACGACACACAAAACAACTGAAAGGCATGGAACCTGGCGAGGGTT

AAGAAGTGTCGCGAGATTGAATAGTTGTACACTGGCGACACTTCGCGAGACATTCCCGAAAATAAAGGATAGCTTAGCTG

TCATCCACTATGTATTACCCTATATAAAGGGTCGAGTGAGTCTGTTGTAAGGGGGGGAAGGTTTTTTGGAATTGTTGCGC

AAGTCATTAGGAGAGAGAAAGTTTAAGAGTGTTTTATTTGCTAAGTGTAATTCCATTTTCCCTGTGTAGATAACTCATGA

AATTAATCAATAAACATTCTTTATTATGCATCTTTCAATGGACTAATTAACCAAAATATAGTGAGGGGTGTAGTGTGGGA

AAACCTGCTACTACATATGCGCTAGAAACAGGGACGAGCGGTGATTGACGAAGAAATTGGAGAGAATTTGGATTAACAGA

GAAGAGATTTGGAAAAGATACTCAAGAATTCATTGAAAGGAAAGATGGCTAGAAGAAGAAATGTACAAGAACAACCAACG

ATTGTAAGAAGAAGTGGGAGACTTGCTGGAGAAAGGAGGAATGAAACAGGAGAATCGTCAAGGATAAATAGGAGAAATGA

CGGAGAAGATCAAAATCAGCCGCCGATCTCCCCAGTAGCAAACGCAGCTATGGATTATGATAGAGTAAGTGTACATACTT

GGAGATCTGATTCTGAAGACGAGAGAGAGAGGAAGAACAGCAGTTGGTGAACAATGGACAAGGGGAACAAGGGATGAATG

GAGAAATGAATCCAGGAAATGAAGAAATGGCAGCGTTAGAAAGATTGAGGCAAACAATACACGAGGAGAGACGACTAGAG

GCAGAGGAGCGAGCAAACTTGACAAGACAAAATCATGAATTGAGGATGGAAAATATGAGATTGCAAAGTAGGAGATCTCG

AAGTAGCGTGAGATCAAGTTCTAGATCAAACAGAGAAGAGAGAAGAAGAGTTTTACCAACAGAGAATACCGAGGAAGAGA

CACAAGATCCAGACACAGAGCACGGCGAGAGAAGAGGAGAAATAGAAGATCGTTATATTCCTCAGCCCGAAAATCTTCGA

AATGAGCGAAATAGGGAAAATCAGGAAAACATGCAACTCCAAGGGCCAAATGACCCATTTCTTTGAAGGAAGAATAATCC

TACGCGGGAGAGAAGAGTTGAGAAGAAACCGAGAGATCGAGGATGACATTGAGAGACACTACATTGAACAAGCGCGACTA

GCTCGCGAGAGAGAAAGGCTAAGAGAAGAAATGGAAAATCAAGAGATGGAGGAAGCAATCCGGTGAGAATAACCATGACA

ACCGCGAGAGAAGGAGGGAACAAAATCGTGCTAATAGAAATATTAATGAAGAGTATGAGAGGGTGCAAATGATGAGAGCT

GATAGGCAGCCAAGAAGAGAGAACAACCGATATAATGGCGTGCGAATTAACGAAAGATGCCAAGAAGAGCGTCGCGAGGA

AAGAAGAGGAGAAGATAATGAGCAGGAAGGTCAAAATAGGCACGAGCGAAGAAGGAGAGAAGCAGAGCTAAAAAGACCAG

TAGAATCAAATTCTGAGGCAAATGAACAAATCTTAAGAGAGCTAGCTGAAATGAGAGGATTACTGAACAATAGAAGGGAA

GGAGGAAGGAGACAATTGGATGAAGCCATCGCAGAAGCCGCGAAGAACCCTTTTACGCTAGAAATACAGATGGGAGGAAT

ACCTCCTAAGTGCAATCTACCTGCATTAACCAACATCTTTGATGGAACAACCTGTGCGATACAACATATTAAGGCTTACA

TGAGATGTATGTTACAATGGGAGAACCATGATACCATATTGTGCAAATATTTCGCGTCAAGCTTAATAGGAGAAGCATTA

AAGTGGTTTGAAAGCCTACCAAGGAATACAATTACATCATTTCACCATTTACAAGCCACATTCCTGGGAGCATACATAAG

TAACAACTCTTCGCGACCAGGTATTGAGGATGTATTCGGGCTGAAACAAAGGATTGGCGAGAGCTTAAGACAGTTGACAG

TAAGATGGAGAACCATGTGCAGCGAAATGGCAGGACGGGTTGAGGAGAGATATCTCATACTCTCATTTGTGAATGCTCTC

TTCGCGACTAACTTGTTATACATTCAAATTTTTAGAATCAAAAATACAATCACGATGGCAGAATCATGAGAATTACAGGA

AGAATATATTGCCTTAGAAGAAAGACAAAATGAAATGAATTCTTATCCCGTCGCGAACACGAACTCCAACGCCGCGAACG

CAAGTTTGTTACCTCAATTCATCAATACGATGGCGAACACTTCGCAAGGACAAGCGAGCAAGGAGGGCTGATGAGGGACA

TCAGAAGGTAGTAGCCATGGGAAGCCATGATCAAGAAGAGTTTGAACGCGAAAGGAATTACTACAATCGTAACAGCAATA

ATAAGATTCAGAGACTAGATCATGTGCAAGAAACTTATGGCGGACGAAGGCAAAATAACCGTGGAGGGCAGAGAAGTCAT

AGAACAGTATGGGAACAAATTAAGATGCCTCCACTGAATGCTAGCGTTGAAAAGATCTGGGAAGCTATACTTGTGATGGA

AAATGTACCAGCACCTTGGAATGTTGGGATGGAACCACCGCCAGGAGATAGAAGCCATGAGTTTTGTTCTTACCATCGCT

TCCATGGGCATCAAACAAACGATTGTAGAAACATAAAGAGAATTATTCTCAGGATGATTGATCAGGGGAAGCTGAATCAC

TTTTTAAGGGGAGAAGAAACAGGTCAGACATTACCACCACCACCAGAGATGCACAGAACAGTGTCAGCGAAGGGAAAGGA

AACATTTGTGATAGAGGTAGGAACAAAAGCCAAGAATTTGAACTGCAACTCCATTGTCCACTCCTATCGCACCATCGAAG

ATTTTCATGACAACGTGCTTAGTAGAGTCTTTGCGAGAGATGCTAATGATAAAGAAATTATGAGTCTGGCGAGATCAACG

CAGTTAGAACAGTGGCAGTTGAAGACTATTTCTTTTTCTGCAGATGAAGTGCCAAAAGGAAGAGAAACACATGATAGCCC

GTTGGTTGTGAAACTAGAGATTAACCCGAAACCAGTCGATGGAGAAGATGATGAGGCAGAAACTTGCTGGGCGATCAACC

GAATCTTAATCGACACCGGAAGTTCGGTGAATATCCTCTTCTACCATACTTACAAGACGATGGGAGGAAAAGATGAGGAT

CTGATCCCCTCTACATACAAGATATATGGCTTTAACGGCACTGCAAACAAACCAAAGGGAGAGGTTCACATGAGAATACC

TCTACAAGGATTATCAACTGGTATTTTATTCTGTGTGGTTGACGTGGAATCTCCATACAATGCGCTGATTGGGAGGCCGT

GGCTTCATGAAATCTTGGGAGTAGCGTCAACATATCATCAATGTATTAAATTCCCACATCCAAATGGCATTGGTATTATA

CAGGGAGATTGGAAGGAGTCGCAACAGTGTCATGAAGTAGATGTTGAAAGCTGCGAAGAAAGAAATGCCAAAAGAAAGGG

TTGGAAGCAGCATGTGAGAGATGCGCAGAGAAAGGAAAAACTGATGGTGGACGCCATAGAGGTAAGGACTGAACATACCT

CGCGAGATGAATGCCTAGAGACAGGCGAATATTATGGTAATAAGGAAGCAAAGGATATGAGGAGCAAAAGGCATAAGAAA

TGAATAAACAACAAAAAGCAGTTGAAGTAATGATATGCGAGAATCTCGCAGAAGAGTAAAAATGAGAGTTCGACTAAGAA

AGAAAAGTGAAATACAACAAGTGAAATAGAACTCTGCGAATACTTCGCAAACAAATATCAGCAAATAATGAAACATGAAT

GATTGAATGATTTAACAAATATAAATGCGCAAATGAATGGGACGAACCTTTATGGCGTACACCCTAGTTCGCGAATAATA

TTTCGTAAGAGGCAAATGTAAGACCTAAAGTGCGTCATATAAGGGGGTATCTCTCGCCGTGGTTCGGGGGAAGGTTACAC

CGCTCCCGAAGCCCGAACCATGAAAATTTGGGGCCCATGAAAGCCTATCCGGAAGAGGTACCTTGGATTCCCAACTATAT

ATGTAGGTTATACCCTGGTTGGGCGACCGAGGGTCTCTTCCTAGGAGTCGATGTATGCTCAAGGCGCGAGGTACACGGTT

GAGCCGAGTGCCGGGATGGATGAACCTACCCCTGGCCATTCTTGGCAAGTCTTGGCATGCGCTGCACTCGCCCGAAGAAA

ACTCCACTTAGGGTGCAGTCTCGTAACCATAAACCATATAGGTATAAGGGATAAGAGGCTGCGAAAACAAGCGGTTGTTG

TTAAGACCGGATGGGAGGAGCTAGTCTTGTATGGCCGATGTTTCGCCTCCTTGAAGGGGCAAGGGGAGATGATAAGGGCG

ACGCTCTCCATTAGGGGGGGTGTCAAAACAGACGCAAATGTCATGAGGCTTTGTTTTCGCAAGGAGACTTAGGCTTGTCA

TATTTGTGCAATGATCCTGCAGAGGCAATAATACTAACGCGAGGGTAAGGCGAGATCAATGGGTCTTCACACGAAAGTGT

GCGGACGTCCTGCGATTTCGTCGCAAGACGGCAATGTCATGGGGCTATGTGTTCGCAAGGATATATAGGCCTGTCATATT

GTTGCAACCAAAACATCCTGCAGAGGCAAAAATACTTGAAAAAATGATAAGGCAAGATCAATGGGTCTTTACATACAAGT

GTAAAGACGTCCTGCGATTTTGTCGTAATGAACGGTGGCAGGGGAGGACCTCTAACTAGGGAGGTCTGACTTGGAAGACC

ACACAAAATGATAAGGCAAGGTCAATGGGCCTTTGCATGAAAGTGTAGAGGCGGCCTGCGACCTTGTCGCAGCACAAAGA

GTTAGCATTCTCAGCTCATAAAGATATCCGCGACCTCGTCGCCGCCATACATTTCAACAACAATAAAAATTATTTACATG

AGGATAACTAACTTCCTCATCAGAAGATAAATATCAGAGGCAAGTATGGGAATGGAGAAGTTGGAATATTGTTATTCATC

TTCTGATGAAAGAATTATACATTAATAATGTTAGCCAATTATATTACAAGGAAGAACTACCAAAAAAGGGGAGTAGAGAT

TGAACTCATGAAAAAAGAGTTCACTATAATAACAAAGATACAGCAATGTACAGATATTACACGCAGCAAAAGAAAACAAA

AAGAATTATATCTTTGGTGATTCTTCTTCTTCATCTGATTCTTCAGCTTCATTATTCTCATCGCCAGAATTTTCTTCTTC

GCCAGCTTCACTGTCAGAAACATCAGAGACAGGTAGATTTTCAGGAACCTTTTCAAGAAGCCAGGGATATGTATGCAACG

GGATGTTCTGGTGTTTACAAACATACTCCATGCCTCCGTTGCGAAAACGAACAGCATCATTGTTAAAGTTCTCCACTGTT

GTTAGAAATTGCTTCTTCAACTTGGAATATCTATTCTCGCGAAGGGAGAGATTTTTTGCGAGCTGTTCTTTTTCGGCTTC

GAGTTCTTCGATTCTCGCTGGAAAATTCCCGGCGAGTTGTTCTTTCTCAGATTCAAGTTCTCTAATTCTAGTTTGAAGCT

TCTTCTCAGATTCTGCAAATAAGTAACAACAAATTAATAATTTACAAGCAATATTACAGAAGCAAGAAGTAGTAAGGTGT

AATAAATAACCTTCTCTATCAGATATCATATCTTTAATCAAGGAAGTATGCTCGGCGTGAAGACTAACATCAACACCTAG

GTTATTCCTGGCATTATCTAAAACTTTCGCAGCCCAGGCGAATTCATCGTCATTATTTATAACGAGAAGAGAACGAACTT

GCTCTTCTCGCTGCTTAAGTATGGTATTCTGACGAAGGAGTTCATCTCGCTCAATTTGCACACCAAGGAGAGATTTATGG

AATTCTGCTAACGCTTGTGCACCTCTGTTAGTAATACGATCATTCTCATCTATGAGGTCACTAATTTTCTTCCTCGATTT

TTTGATTTCTTTATTGGACTCGAGAAACATTCGTTTAAATTGGTTGGCAGCACTTAAACCAGATCTATGTTCGATTTCTA

AAAGACGATATTTGGACTGAAGTTCATTTAATGAAAGGGAGTTAATTTTGTTTGATGAACTACTTAAAGAGTTGTTGAGG

TGAGCGAGAAGAAGGTTATTATCCATAGGATCAGGAAATGTATGGAGAAGAATAGCCTCATCGGAGAGATTATATACATC

TGCGAAAGTAATAAGTCAGGTAGTAATAGGCGACCATTAAAATAAAATTAATAAAAGGAGGTAAAATTTTATACCATAAA

GTTGCTGATTGGCATTTTGAAGAGAGGACACCTTATCCTTTTGGGAAGTAAGTTCGGTGTTTAACTTTTGAATTCTCTCG

TCTTTCTCACGAGATGACGCAGCTTCGGTCTCTAATTCCTGAACCTTCTCGCTCAACCGCAAATTCTTCTTTTCAAGATT

ATTGCGCCTTCTCTCAAGGTCTGAGGCAACTACAAAGGAAGATCTTCCAACCTATGAAAGGAGATAGAAAAGCTGTTAAT

ACATATACAACAATATAAAGTCAATGGGGAGACATGATTGTATACATACCAAGGATCCAAGCGCATTCAAGAAGTCAGGA

GACATAGTTCTAGAGACTCCATTGAAAGAATCATTGCCTTCCAGTAGAGTGGCATTGCACACGTTTGCGAGAGTCCCGCA

AAGAGAAGCAAGGGGAGCATCTCCCATTCCTTGGAAAGAATCAGAGAAGATACTGGACAACTGACCCATAGAAGAACTAG

TTAACGGGGTCCCTGTAGCGAGAGAGCGATTCGAATCATGGTCATCCTTGTCAGAAGATTCAGAGCTTGAGCTAAGGCGA

GAAGAGGATTGGATCTTTCGCTTCTTAGTTAAGGGGGCAGAGGATTTACTCTTGCGAGCGACAGTTTTACCCTTATTTGT

AGTCTTCGCAGCAATAATTTTTTCAACTTCAACCTTCGCATACGTGTAGAGATAAGAAACAGAGGCAACAATATTGTTTA

AAATAAGAAGAAAGGAAAGGATTCTTACTTCACTCGCAAGCGATCGAAGCAATGTCAGAGTATCCTCCGACGGATCATTT

TTGTAGCTTCTCTTCAATCTAGCTATCTGAAAGATTTCACAAACTCAGCAATTATAGAAAGAAAATTACGAAAACCAAAT

TTGAAGTCGGAAAAATACCTCTTTCTTCTTCTCAGGCCATTTGAATTCCCAAGGATGATATCCTGCGAGATTCGCAGGAA

GAATATGCGAACCCGTGATGTAAGGACCATTTAGGATCAAGGGTACGACCCACCATCTCTCATCTTTGGAGTGGCGAGGA

GTAGTGTTGTTACCCGGATGCCAATCAATATCCTGCATGACCAATTTATCCTCAGAGACATTATCCTTTTTCTTCAGGCG

AACGCCCCAGCGAGACTTCGCGTTCACCATAGTTATCACTGAATAATTTTCAAAAAAGTTTTCCACAGTGTATTTCTCAG

CATTTATTTCCAAGTCTTTGAATTTGGGATTTCTGACAGAGTTAGAATATAAAGATCCTTTGCCTGCACCGCGGTTCGCG

AACTCAAGCATCAGGCGAATACAGTCGCCACTTAATTGATAAACAGCCCGAGAGAATTTTTCATGGGCTAAAATTTCATA

AAATAAAGGCAATTCAGGATTGTACAAGGGAATAGGAAGTCCCGCGAGAAGCTGACCGAGCGAGACTATAATCTTCTGAT

CATTACATTGATGATCAGAGAAAAGATTTACAGAAAGAATGGATGAGGGTTCTTGATCAGGAATGGTTGACAGAGTAAAA

CCCTTCTCGATCAATTTCTGTTGAACTTCTTCAAAAGAAGTTTTGTATCCACGACTGATGGAAGCCATCTATCAGTATGG

GAATGGATGAAAGATCGGAGATTAAAAACAGCGCTAGGAAGGATGAAGAAAAGCTTTGCAGCAGCAGGAGCGCAAATTGC

GGAAGGAGAGAGTCGAAGAAAGATTGCAGAGAGAAACCAAAGAATGAAAAGGAGGGGAAAAGAAAACTATATATAGGGAA

AAAGATCCTCGAGAAAAACGAGGTCATTTAATGCGAAGAGACATGAACGGTCAAGAAAAAGTGGTTAGAAGAAACGTGTC

AAGAGAAGGCATTAATAATCAAAATCCGTGTAAAAGTACAAAACTGAAGATAAGCAGTCGCGGCAACAACATTGTAAAAA

TCTCATGGTATTCTTCCCCTCGCAGGAGAGATAAAGAGACGATACTTCATATATACCTCAAAGAAGCTCCGGAACTTCGC

AAAAGAATAGAGGCAAGTACGTACTTTTCAAGTACAATATTCTTCTGCTCGTTCCTTCATCAACAAAGATCAAAGATTAC

TACAACAACAAATTTTTCCTTCAAGATCGCCCTTCAAGCAATATTCAAGGAAAAAGAGGCAAGATGTAGGATCATAATAT

CGCAATGCATAGGGCTCGCGAGATCAATAAACGACAGTACCGCGAAGGAGTCGCAGGAAACTAAAATAGACGACGCAATG

GAAAACGACACACAAAACAACTGAAAGGCATGGAACCTGGCGAGGGTTAAGAAGTGTCGCGAGATTGAATAGTTGTACAC

TGGCGACACTTCGCGAGACATTCCCGAAAATAAAGGATAGCTTAGCTGTCATCCACTATGTATTACCCTATATAAAGGGT

CGAGTGAGTCTGTTGTAAGAGGGGGAAGGTTTTTTGGAATTGTTGCGCAAGTCATTAGGAGAGAGAAAGTTTAAGAGTGT

TTTATTTGCTAAGTGTAATTCCATTTTCCCTGTGTAGATAACTCATGAAATTAATCAATAAACATTCTTTATTATGCATC

TTTCAATGGACTAATTAACCAAAATATAGTGAGGGGTGTAGTGTGGGAAAACCTGCTACTACAGTCACAACTTGCAGCGT

ACACTACGCTCACATTAAGAAAGCGTGAAAAGGAATAAAGCGGTCAAATAAATGAGGAAATAATGGTGAAAACGTTTTCA

CCCATTAACACCATACGGTTACCACCTCCTCCCATTATTTCACCCTTTACTCTCTTCCTTCATGAGGCCAGGATACGTTT

CAATCTCGACTTGTATAAATAGGTCTTACCTATTTCCACCAGACAACAAGTCTCGGTCAGGAGAAAAATACAACACTCTC

AGAAAAGGCAACTTTGCAAGCTCTCCAAAAAGGCTTTCATCTCTTGCAAGCTTCCGAAGCTTTCACTTTCTGATACAAGT

CGAGACACTACTCTCCCAGAATTATTCTGGTCTCAACACCTTTCTTTGCTTCCCTCCCCAAATCAACCCTTCTCCTCCAC

TTTGTGACCGAAGCAAGTCCGGAACGACCATTTCTTGGTTTAGGTAGGACTTGTATGCGGATCGATCTCTCGAATCTATA

AAGTGCTCACGTGCAAAGCACATTTGTTTAGGGTTTGAATCAGTTTCTCACTCACCTACCGAATTTACCGAAATCGGCAG

AAACGGTTTTTACCCATAAACAATTGGCGACCACAGCGGGAGGATTGATCTATCGGCTACAATGTCAGACTGTCAATCTC

GTTCTTCGACCCAGAAGTCAAGATGGTAGGGGAAAGCGACTCAATCTGACTCAACTGCCAGACAACCCGGTTACCAGTTG

TAAAAGCTACACCTAAAAACTGGGGACTCGTTGAAGATGCTACCAGCGATTGGGGACTTGCGACTCAATCCGACTCAACT

GCCAGACAACCCGGTTACCAGTTGTTAAAACTACACCAGAAAGCTGGGGACTCGCTGAAGATGCTACCAGCGATTGGAGA

CTTGCTGGCGATGCTACCAGCAAAGCCGCCTTTTACTTAGAGAGAGGCGTGTTTGCAAAAACCTATTAAGTGAGTAAATT

TAGATTGGGCAAGATATACGATATGTTTTTGGGTTCTTAATAAAACCGTCATACTACTGCGTCTTGTGCCCATTGCTAGC

ACACACTTTCATAAAAAGCAATAATCCTTTCCCAAAACATAACCAGAGTTATATTATCAAAACCATTCATCCCAGAAATA

ATCCAATATCCTCAAAAGATACCAGCTGTCTATCAAAACCGGTCGAACTTTGAACCAGACAACCAGTATGTTTTTCAAAA

AAAAAAAAGAAGAAGAACCTAAATGAAAGTTTAACAGAGAAACATTCAAGGAAAATCATCCAACAGAGGTCTGCTTTTCT

TGGAGAAAGTCAACTTTGTGTCTTCTCTTCATAGTTAACATCTTAGAAAGCTTTTCGAGTTCTACCTCTCCTGTCGTGCA

ATCTCCTCGGCGGAAAGGCTTTCGATTGCTGCATCTTTCAACTTTTGGATCTCAGCAAGGCCTTCGACAAACCACGAAGT

GTTGAACTCCTTTTCTACGCACACGTCGCGATGGTGCTCCCCGATCGGCAATAACGTCTCGTAAACGGTGTGACCCGGTC

ACCTCACGCATATCCTCGATGGAAGATTAAGCCTCTTCCACACGCTTAACCAGCGCAAATCGACTAGTGACTTTCTTGGT

TGTGGCCATATGTCCGTACTTCTCCCAGATTCTAGTGTACAACGCCTCATACTTAGCTTGACGCTGAAGCCGCCAACCAA

AGACATGACTGGATAAGGAACCAAGTACGGGGATCATAAATGGCGCCTGCAACTGGCCTCACGACTCACAAAGGGTCTTC

TTCCATGTCTACGGCAGCAACCATCAGAGAATCATCCACCGTTTGAACCACATCTGTGCCCGGCGATTGGCTAACTCCTA

TCTCCAAGTCACCGGCGGTGGTCACCAAAGGCAAGTTTTGTGTGGCTCCACCCCAGAATCGTCCTCCGAACCGGATTCCT

CCTGCGAACTTGAGTTAGACATTTTTCAAAAGAAGAAACATAAGAAGAAAGAAAAGAAGCGCGAAACTCACCGAACCACT

TCCCGAAAAAGAACCAGTCTTTGCCAGAGGAATCACTCTCACTTGTCGAACTTTCTTATTTCTCTGACTCATCACCGCTA

TGAGTAGGCTCGGTGCCGCCGGAATTCTTCAAAGAAATTGCCGTCTCCTGGCAACGTGCAAGGGCAGCGAAAGCATTAAC

ACTACCGAAGCCCTGGAAGAAACAAGACACATGTTTTCAGCAGAAATTCACGTGTTTTGAGACAAAACAACAAGGAGGCG

ACACATACCTGGTTATTGGAAGAACTGAAAGACACGAGAGCTGCGGGAGGGTCTTTTGAAATCGCCCGGAACCCGCCAAC

ACGATTCTTCGTTCTCTGTTCCTTGGCCTGGGGACTTTCCGAGTTGCTAGGCAACTTTCGCTTGAGATCTCTCAACATTG

GATACTTCTTCAAAACTGCTGGGGAAGGTTTACAACGAACATTCTCCACCACGCTGTTTACAAAATCTTGGATCATAATA

AACTCATCGCACCAGAATTCATTATAGCCAAAGGTCGTCGAAGTTACTCTTCCCGTTTGAAAGAAGGGAAGGCGGATACC

CGGCTCGAGGGCACTGGCATTGAGGGCGGCACGCAATGAATCCTTAGGAATCACCGGCTGGTAAGCAAGAGGCACCCCTT

GATCGAAACACATGTGACGAGCCGCACGGTCAATATTGTAAGCAGCTATCTGGAAAGACCCCATGAAGAAAGACGGCATA

TGGACCGACAAATTCAAACCGGACATGCAGCTTTGAATGAACAAGTTTTCACCAACGCTCAAATTGCCATCAGAAGACAA

AGACATGATCGGAGCAGAAACAAAGGTGTCAATTTGAGCCACCGACCAATGGACCGGCTCCCATGGGCGAAAACTGATTT

CATACGACTAATCGAGAAGACTAACCAAGCTCATGCCAACCCTTGGACGCCTGTTCACCCATCGAAGAATTCGGGAACCC

CCCCAACTCGCGGGAATCGAAGACAACGGTTTTGGGGCATACTTCTCAAAGTGTTCCCATAACCAAGCTTGCAGGAAAGC

AACGTGCACATATGAATCCACTTTCATAAAACCATTCGAAACTCGCATGTTCGCAACTAGGTGATCCAGATGCACATACA

TGGAACCAAGAAATAGCTTGCCAATGGGAAGAACTTCGCCTTTTGCCAACTTCACAGCAATCACTAGCAACTCTGGTCTT

ATCTCCTTCTTACCTTAATCAAGTCGTCAAAAATGTCTCTAGACAACCAGAGAGCCAAAATCTTTGCAACATGAAGATCA

TCGTTTCTGTTTAGGTTCGAGCTCGTCAGGAAACCATTGAGATATCCACCACTGTAAAGGCACCTTTCATCCTCTTAGGG

ACAGAATCCTTCGACTTAGCAGTGGAGTCTTTCCGAGCGAAATCTCCCAACATAGCGGCAAGGTCATCCCGCATCTTCCC

CTCACTTTCGAGATAATTGGATGTTAAGATTCCCGATTCTGGGAAGATTCAAAGAAGAGAGCCACACTCTCCAAGGAGAA

AGTTATCTCACCCCATCTACATATGATCGTATGGGTATCCGAACACCATCTGGAGATGAAAGCGACCAAGCCCGGATATC

CTTCCTAACTGGAGATTTATTGAAGCTCGACAAAGCATCAATAATCTGCGCTCGAGTCAAGTTGGTATTTATGTGCTCCT

TACTCACCATGAACTCAACCCATCTATCAAAAGAACGCAATGGGCTTTGACACATCCTGAAATCAATGTTGGAAGAGGAA

TATACCCCTTTATGAAGATAGAACCGTATGGTGTTCTCTGGGCGAACCGAACGTGTCTTCTTTTCATCTTCACCACCGGT

TAACAAAATTGGCACGAAAACACAGGTGATTTTTCCGAGCAGAGGCAAGTTTTTTGGCGAAGAATTCATCATCCGTGTTT

GGTTTGGAACTATGTACATCCTTCGATGAGGCAAAATTTTGTTCAGGTAACATCTTGCCGGATTTCTTCATAACAACTTA

ATAACAAGATATCTGCACACGGAGAAAAACAAAATTATCTCCCTGACACATTACCACGCCCAACAAAAGGAAAACCTGAA

ATACAAAAAAAAGAGAGGGCGTATTCTTTACTTGCTCAAAACTGCACACGCAGTATTTTCTTCAGGGTTCAAAATTATAT

ACGCAGTATTTTATACGGGCTCAAGTGCGTCTGCAGTATCATTGTTCGGGTTCAAGATTATATATGGAACATTTTTCTAC

GGGTTCAAAACTGCACGCGCGGTATTTTCTTCAAGTCCGAAATTCATATATGTAATATCTCCTATGGGTTCGAAGTCATA

TGAAGCACATATACAATATTCTCTACGGGTTCAAAAATACACACGCAGTATCTTCTTCAGGGTTCAAAAAATTATATATG

CAGTATTTTATACGGGCTCAAGTGCATCTGCAGTATCCTTCTTCGAGTTCAAGACTATATACGCAGTATTATACGGGCTC

AAGTGCATCTGCAGTATCCTTCTTCGGGTTCAAAATTATATGTGGAGCATTTTTCTACGGGTTCAAAACTGCACGCGCAG

TATTTCTTTCAGGACCGAGAATCATATATGCAATATTTCCTACGGGTTCGAAGTCATATGAAGCGCATATACAGTATTCT

TCTTCGAGATTCATATACAGTATTCTCTACGGGTTCGAAACTGCACACGCAGTATTTTCTTCAGGGTTCAAAATTATATA

CGCAGTATTTTATACGGGCTCAAGTGCGTCTGCAGTATCCTTCTTCGGGTTCAAAATTATATATGGAGCACTTTCCTACG

GGTTCGAATAATATATGCAGTATTTTCCTTGGTTGCATACATGCAATATTTTCTACAAGAGACACGCCAAGAAATTATAT

GCGCAATATCATTGCAAGGATTATAAAGAAAAACGCCTACTCTTAGGGTTTGCACCGACATAGGGAAACAAGAGAAGTAC

TGGAAGACATACCTGGAAGCCGGTTTTTCTTTGTAACTCTGCCGCTAGTAGGAAAGGATGAAAGGAATAAGGTGCAAGAT

GAGAGAAAGGGATTTTATATATATATGGGCGTGAAGATTCTGAAATAGGGAAGGATTTCCCTTTACCAAAAATCGTGCGG

GCTCACAAGAACGCCCAACTTCCCAATGCCGCCCAAGTCAAGTTGGCCGGTCCAAAGCAAGCACGCTCGCTTCTTTCCCT

ACCTTGCGCCGGTCCCACCTGCTAGCACGCTCAGTTTCCAAACTCGCAACAACTTCCCAATGCCGCCCAAGTTGCGCTGG

TCCCACCAGATAGCACGCTCAGTTTCCCAACTCGCAACAACTTCTCAATGCCGCCCAGCTTCCTGTCTCTCACGACGCTA

ACTTCCCAATTGTGCCAGCCTCCCGATTTGCGCAAAACCGCCAGAGGCAAGGATTCCAAATCCTATTTATAAAAAGGTCA

GACAAGCCTCCTCAATTCTCTTTACGATTTGCTGCTGATAATTTGTCACCATCTTATGCTTACCTCACAACAAATCTCCT

TTTCCGAGCTAAGCAGGGGACTTAATGTTGATGGTGGTTTTTAAACAAAGGGTGAAATCGTAAAACCCCAAATAACCTAA

TGTCACCAGTGACACATTTCTTAGGACATTTAATGCATGTAGTCAGTAATTAATAGGGGGTATTTTTTCAAATTAGGGTT

ACAACGAACAAAACCCTAATTCCCCTCCGTGCCAATGGCAAATCATGCCAGCCACACCTAGCCCAAGCTAGTCGCACCTC

TATGCCAATGGCAAAGCATTTCAGCCATACCAAACGCGCCGAAGCATGCCAACCATGCTAACCGCACCTCTATGCCAATG

GCAAATCATGCCAGCCATACCAAACGCGTCAAAGCATGCCAACCATGCTATCCGCACCTCTATGCCAATGGCAAAGCATG

CCAGACATACCAGACGCGCCGAAGCATGCCAACCATGCTAACCGCACCTCTATGCCAATGGCAAATCATGCCAGCCATAC

CAAACGCGTCAAAGCATGCCAACCATGATAGCCGCACCTCTATGCCAATGGCAAATCATGCCAGCCATACCAAAGGCGCC

GAAGCATACCAATGATGCGTGTCGTAATCACAACAAATTAATTAATACTTTACTACCTAATTAACAAGTAATATAGGGTA

GTCAAGGTCGATCCCACGAGGAGCACGAGATTCTAGTTGTAAAAATTGTCACGAAAGGGGGGTGTAAAAAGCAAAAGAAA

ATAAACAAAGCAATTAAAAGATAAAGTTGAAAGCAATAAGAGAGACAAGATCAAGGAATCCTTCTTCGTTGCAGAACAAT

GATTAAAGTTATGTTCTTTTCTACTTGTTATTAGTCACAAATTATCACCAACCGTAGGTAAAGCAGCTAGCTCAGTGCTA

AACCCCTAAATCCCTTGTATCACCGGACACGGAAGTTCTCGACTACCGGATTCTATTTACCAAACCACCTAGTAGTAGCT

CACTCTAGATGTAATTTAAATAAACGCATTAAGATTTATGAACTTATTAGGTTAATATTAGTAGTTAGACTCTTAGATCA

AGGTCTACTTGCTAACGTTGTTTCTACACACAATTGCTCCACGGAATCACTAGCTTTAGATCAAATTAACAAATCAATTT

AGTTAGCCACCCAAACAATCTATCAATCAATCATGAATGTCTTTAAGATCATTCCTATTAGTAAAAACAATCGATAATTA

TGTGAAAAACTTCAAGATAGAATTGAAAACAAAAACTCAAATCATATTAAGAACTAGGAATTCATCCTCAATCAATAAAA

TTAGCTACTCATGTTAAGGAGTTCACACAAAGAATAAGAAAAGAAAAGTGTTGTGTGTGTAGAGATGTGTAGAAGTTGTA

GAAAACTTCTCTATTTATAGGCTTCAATATTCTCCAAATCTTCCAAGGAATATAATCCTTTTTCTTTTGCAATTCAATTT

CCAAATCCAAGCTTTTCTCAACTCTTCCACCTTCTCTTTCCAAATTTGAGTCCAATTCTCCATCCAATTCCAAGCCCAAG

TCTTCACACCCATGAGAGAAGTCCATAGAAAATTCTGGAATTTTTGTTCACGGAAAATCGGGATTGTGGCGGAATTTGGC

CGGAAAACGTCACCGTAGGAACTTTGAAATTCACTCTGCGGTCACTTTGGAATCTTTGGCGGAATATTCCGTCGGTCACA

GTTAGGACTAACGGTCAAACAGGTCAACCGTCAGTCATTTGTTAGTCAAAGGTCAAGGAGAAGGAGTGAGTCGGACTCGA

GTTAGCTTCGACTCGCCTAATCTGGGCTTACTCGGATGAGTCGATTCTTGAATCACCGCTTACTCGGTGACTTAATCTAA

TTGACTCACCTGGAATCAGTTGAACTCGGTTGGGATCGGTCATCGGGATCGTCTAGGATCGGTCCGAGTCGACTCGCCGG

GAGGATGATTTGCCGAGTTTCGTTTATTTCAGGATTTCGGCCAATTTCGAGGCCTCTACTTCATCGGTTCTGTGAAATTC

CTAAGCATTCATCTAGTTTCAACCATTTCAAGCTTCACTCATACCCAAGCTTGCTCGAGGATGGCTTCAAATCATTTCAT

ACTTGCGGTCAACACCATCTTCGTTATCGATTCAATCCCCCGCAATGCAAACACCACCATCTTCTCGTGGCGTCGGACCA

TCTCCAATTTCGTATTCTCGACATCACAAAGAGCTCAATCAACCTGTTTGATTCATCCGAGCCACAAACCCATCTCTAAT

CTCGCTGCTAGTTTCTTCCGATTCAACCCAAAATCTCTTTTAGTTCATGGCGGCTATCATCTTATCTTCCTGTGCAATTC

TTGCTACGGCAACTAAATCCGATTATACCCATCTCTCAACAACATCGATTTCTTCTCATAAATATCATCTTGACTCATGG

CCATCTTCGATTCAATCTCTCGGCCTCGAGTGCAGCCTCATCTCCACTTCACCGATTCAATCTCACGGATACCCTTCTCA

TACTCGATTCCATTCAATAGCAGCAGTGAGGTCTCAATATCGACAAATCCCATCACCGAACTCAAACCAACAAATTCAGA

GCAAGCCATTCACAGAAACTCATTCAATTTCATCATCAGCTTCTCTCCCAGATCTTCTTCATTTCGCCTCGATTTCACTC

AGCAACAACGTCTCATCACCAACAATTACAGCTTCTCGATTTTAAACTCAAACTCAAATTCCTCCCAAGTCTCTGAGCTC

GATCTCTCTGTGTTAAATTCCAGAATGAAATGAAAAATGTGAGAGACGAGTTCTCTCGATTAGGGTTGGGGCGATAAGGA

TATGTACACCCCACAGATAGGACCTGATAATTACACCCACTAGTAGGTGATAAGGGACACCCTGATAACAACCCCCCTTA

GCCTTGACTGGGCCTGGTTTTAGTACTCCTCCAATTGGGTTGCCCCTTAACCTCATCTGGGCTCCAAATATCAATCGCCT

GCAAATTGACCTTTTTGCCCTTTTGGCTCAAATTGGGTAATTTCTTCTTCTTTTCCTCCGTGTGACTCCAAAGTAGCTAC

AAAACTCAAAACAAGCGTAAAATACATAATTTACAAGAAAAATAGCAAAGAAAGCATAAACTACTAGATAGTAAATTAGG

TGTATTCTACACCTATCAACCAACCATGCTAGCCGCACCTCTATGCCAATGGCAAATCATGCCATCCATACCAAACGCGC

CAAAGCATGCCAACCATGCTAGTCGCACCTCTATGCCAATGGCAAATCATGCCAGCCATACCAAACGCGCCGAAGCATGC

CAACCATGCTATCCGCACCTCCGTGCCAATGGAAAATCATGCCAGCCATACCAAAACGCGCCGAAGCATGCCAACCATGC

TAGCCACACCTCCGTGCCAATGGCAAGTCATGCCAGCCACACCTAGCCCAAGCTAGCCGCACCTCTGTGCCAATGGCAAA

TCATGCCAGCCACACCAAACGCGCCAAAGCATGCCAACCATGCTAGCCGCACCTCCGTGCCAATGGCAAATCATGCCAGC

CACACTTAGCCCGAGCTAGCCGCGCCATTCCAAACACACTCTGCCTTGGCCCTAGCTACGCTAGTCATGCCACGCCAAGC

CAAACACCCTTACCTTGGCCCTAACCATGCCATGCCAAAATGGCGCCCACGCCTAGGCTTGAAAATCATGCAACCTAGGC

CAAATTGGCCACGCATGCCATTGGCACATGCCAACATGCTCGCTGGTAGCTTTTCCTACCAAGGCACGACCCAAATTGGC

CACGCATGCCATTGGAACATGCCAACACGCTCGCTGGCAGATTTTCCTACCAGGGCGCGACCCAAATTGGCCACGCATGC

CATTGGCACATGCCAACACGCTCGCTGGCAGCTTTTCCTACCAGGGCGCGACCCAAATTGGCCACGCATGCCATTGGCAC

ATGCCAACACGCTCGCTGGTAGCTTTTCCTACCAGGGCGCGACCCAAATTGGCACGCACGCCATTGGCACATGCCAACAC

GCTCGCTGGTAGCGTTTCCTACCAGGGCGCGACCCAAATTGGCCACGCTCTCCATTGGCACATGCCAACACGCTCGCTGG

TATCTTTTCCTACCAGGGCGCGACCCAAATTGGCCACGCATGCCATTGGCACATGCCAACACGCTCGTTAGTAACTTTTC

CTACCAAGGCACGACCCAAATTGGCCACGCATGCCATTGGCACATGCCAACGCGCTCGCTTGTAGCTTTTCCTACCAGGG

CGCGACCCAAATTGGCCACGCATGCCATTGGCACATGCCAACACGCTCGCTGGTAGCTTTTCCTACCAGGGCGCGACCCA

AATTGGCCACGCACGCCATTGGAACATGCCAACACGCTCGCTGGTAGCTTTTCCTACCAGAGCGCGACCCAAATTGGCCG

CGCATGCCATTGGCACATGCCAACACGCTCGCTGGTAGCTTTTCTAAACACGGGCCATGATTCAACTTGGCCACGCATGC

CATTGGCGCATGCAAAACACACATGCTAGTAGCCCAACTTGCAAGCCATGCAACCTAGGCCAACTTGGCCGTACGCGCCA

TTGGCACATGCCAAAACCCACGCTGGTAGCCTAACTTGCAAACCGTGCAACCTACCGCCAAGGCATTCCGCATATGCCAC

GCACGCCACTAGCAGCTCTATGCATTGCCACTTACACTCAACGTACAACCTACCACCAAGGCATTCCACACATGCCATTG

GCACATGCCACACACGCCACTAGCAGCTCTGGCATTGCCACTTGCGCTCGAGGTTCAACTCGCAACAACGGCATTCCGCA

CATGCCATTGGCACATGCCACATGTGCCCCTAGCAGCACCACTTGCACTCAAGATGCAACCAACACTCGAGACATCAACA

CAAACTGGGGGATGCTTTCCAGGGTCTTAGTTTGGCGGTCACATCTTGCAGCGTACACTACGCTCACATTAAAAAAGCGT

GAAAAGGAATAAAGCGGTCAAATAAATGAGGAAATAATGGTGAAAACGTTTTCACCCATTAACACCATACGGTTACCACC

TCCTCCCATTATTTCACCCTTTACTCTCTTCCTTCATGAGGCCAGGATACGTTTCAATCTCGACTTGTATAAAAAGGTCT

TACCTATTTCCACCAGACAACAAGTCTCGGTCAGGAGAAAAATACAACACTCTCAGAAAAGGCAACTTTGCAAGCTCTCC

AAAAAGGCTTTCATCTCTTGCAAGCTTCCGAAGCTTTCACTTTCTGATACAAGTCGAGACACTACTCTCCCAGAATTATT

CTGGTCTCAACACCTTTCTTTGCTTCCCTCCCCAAATCAACCCTTCTCCTCCACTTTGTGACCGAAGCAAGTGCGGAACG

ACCATTTCTTGGTTTAGGCCGGACTTGTACAGATTGATCTCTCGAATCTATAAAGTGCTCACGTGCAGCACATTTGTTTA

GGGTTTGAACCAGTTTCTCACTCACCTACCGAATTTACCGAAATCGGCAGAAACGATTTTTACCCATAATTACAGGAAAA

ATATACATCTACAACTGGATATTTACAACGCTTCGCTAAGTATTGTAGAAAGTCCTATGTTTTACAACTGGTTTGAAAGG

AAGAGTTGTCGTATATCATCACTGCCAACCCTTAGGAAACAAGGGGTTGCGCTAAGAAAACAAACGACAACTCCTTAAGC

AAAACAGTTGCGACGACATTTAGATATAACAACTTTGTTCCATATATGTAGTGAATTAGCCTATCACAATTCTCACAAGT

GTTGTTGAAGAACACAAAAAATATGATAATGAAAAATATGTTAGAGGGTAGATTCGAACATGAAACCTAGTGCATGGAAG

TCTAGCTCATCAACCACCCTTTCAGTTAACAAGTTTGGACTTTTTTTTCTTTTTTTTAAATAGAAACCTACAACAACACT

TATAAGTGTTGTTGAAGTAAAAATATAAATGTTGGAAAAAATGAGGTTATAGGGGAGATTCGAACATGAATCTACTGCAT

GGACGTCTAGCTCATCAACCATCCTTACCATTCACAAGTTTTGGCTTTTTTTTTCTTTTTTAAATAAAAATGTATAACAA

CACTTATAAGTGTTGTTGAAGTAAAAATATAGAACATTAGAAGAAATGAGGATGGGGGGATTTGATCCTCAACCTCCTAT

ATGGAAGTCTACACCACTAACCATACCTACACTATTACATGTTCTGTCAAGTTTGTGGTTCTTTAATACACTAACTTTAT

GACAACCCTCATAAGAGTTGTAAAACAAAATGTAATGCTCAAAAATAATGCTCAGGTGACTGAGCCGCAAACAGCACGAA

GCCTTCTCTGTAAACAATCACATACACGCCTTCTATGTAAATAATTACATACATTTTTCACTATCTGTAATATTTACTTT

ACTCACTTCAGATAAGGGTACTCTGGTAATTTTTACAGTCTCCATAATATTTATTACCTTTATTTCTAGTAAGGGTAATA

TGGTGATTTTAACAAATTATAATATTTAGTTTACTTATTTTCCGATAAGCATGGTCTGGTCATTTTCATAGTCTTCATAA

TATTTATTGCATTAATTTCCAAGAAGGGTTATATGGTCATTTTCACAAGCAGTAATATTTACTTTAATTTTTTTTGATAA

GGGTAGTCAGGTCACTTTCACAATCTCCGTAATATTTATGGCACAAATTTCCATGAAGGGTAATATGATCATTTTCAAAA

TCGTGATATTATTTACTTTACTTATTTTCAAACGTGATAAAGGATAGTTTAGTAATTTTCACAATCTCCGATAAATATTT

TCTGTTTTATTACCTTTATTTAGCCAGTATAATGCCAAGTTTTTAAAAAGTCGTAAGGGTAATATGGTCATTTTCTAGGG

TTTCAATAATGGGAAGAAGAAGCTATCGATCTGAAGATTCAAGATATGGATTCATCATCAACAAAATCAGCTACGTGAAA

ATCAGGTATAAACCTTATTCTTGTTAATCTTCGAATTTGAAAGCAAGTCAGTTATATTTTTTTGTATTCAGTTATATGAA

AATTTGATGGTTATGGATTTAAATTGGGATGTTCATTTTAACGTCATCAGAAGGATAAATTTTCCCTATTCGTAAATATA

GTAAATATGTTTCTTTGGATTATTATTTCTTATCCATTTTGCAAATATAATATATTGATGAGTGGACCAAGTTACAACTC

AAGCACAGTGCAGGGGGACTAAAGTTTCTCAAATAGATTTTAGGTGAGAGCATCTGTATATTTCTAATGAATAAAGTTTT

ACATTGAAAATTTGTCTACTGTATTTTCTACATTTCTTAAAACAAGGGACACAGTTGTTTTAATGTTTCCAGCTGCATAA

ACTGTAATTCAAAAATGGATGGATCTGTTTTGCACCATATCGAAAGTATTTGAGTATTATGGAATTGTTGGTGCTTTCTT

ACCAAGTTTACTTTTGACTTAGTATATTGTCTTTTAGTCAACTAAAACCCGTTTTCTAGGGGGAAAAGATCTATCAAAAT

AACTTTGTTAGTGCCAGTTTCGATCAAATGCTTTCCCAGGGGAGCAAGGTATGCATAAAATATGCATAAATAAGGTTTGT

TTAAGGTTTCAAGCCGAAAAGGATGAAAGGTGAAATCTACACTGTTAAATATCAATCACACAATTCTATGTGAAAAGGAA

TCTTTATTTCAGATTCAGCCAGCAATGGAAAAGGTAAAATGGATGGTGGAAGTGGATTATGGTTTACAGCTAGTGGACAT

AGTTCTCTCAGTTCGGATTTCTCTGCGCTCCTTCTTGAATTAGATTCTGAACCATGATAATGTGTTACCTGCAGTCAATT

TTTACTGTCTTGGTTTGTTGTATGGAGTCTAAAGACAAGTTTCACTATCAAAATTTTTGTACAGTTAATTACCAGATATT

TTGCCAGTATCTTTGGAACTATTTTCCTAGGAGCTGAAGTTCAAGGTTTTACTTTTGAAGCCTCACCCCTTCTAAGTTTT

TATTTATATTGTTGTAGGTTTGGCTGAAACTGATCCGAATGAGCATTATTGGAAGGGTTTTCTGAAGTAGTGCCTATTGA

ACTGGATATTCCAGAAGGGTAAGTTTCATAAAATGCTAAGCTTGCCATCTAGGTAATAATATGTGGAACTGTATATGTGA

ATCCCATTTCATTAGAATGTTAAGTTAATTACACCTCTCCTCAGTTCTATGTCTCATGTTACTTGGGAACCAACTAACGG

AATAGCATCTAGTCTTTCTAAGGAAGATTCTCAACGATGAACCAATGGCAGAACCAATCAATCTCATTTTTATTATTAGT

ATTGGATTTAGTTTCAGTATGTGATCTTGGATTGCTAGAATTTATTTCTAAAATTGTATCCAGAGATTCTATTTTTTTGA

ATATTATTTTGGTAAAAATATTATTTTCAATTCGTTCTTTTAGAATTCTATTTCAATTTTAGGTTTTCAAATTAAATTTT

TGTTTCACGTCCAGAATAAGTTTTAGATAGACATTCAGATTCACAAGATCTGTATCATTAGCCATAAATTAAGGAATATT

CTCGAGATTTTTATTTTTATGATAGCGCCTCCCTTATGACAGATATTAAACGTTATTTTATTTAACGTTTCTATATAACA

TACTATAACCGTGGTACTAAATAACAGCTTCTATTTGTGACTTAAAGTTGCATATTTTATTCGTCATATTTAATAACGTC

TTTTTTCTTGAAAACCGTGATAAACCATGGATCGTTATTTATAGCCCCTTTTGGACTAGTGCTTATTTCGGATAAGGATA

GTTTAGTAATTTTCACAAGATGCTTCGAACTGGGATGTTCACAATGCATAAAATATAGGTTTCGCAGAATATCTCGAAGG

AATCTTACCTGTAAATGGATGGATTCGTCGTTTTTACCAAGTTTCTGACTTAAAGTAGTCAACTCGCGGCCAGGTTTCGA

TCTTGGATCCCGAGATGCATACAATATACAGAAATAAGGTTTGTTTAAGGTTTCGTATAATATTTCGAAAGAATCTTACC

TGTAAATGGATGGTTTCTTCGTTTTTACCAAGTTTCTGACCTAGAGTAGTCAACTCGCGGCCAGGTTTCGATTTAAGATC

CTGGTATGCATACATTTATGCAGAAATAGGGTTTGTTTAAGGTTTCGTAGAATATCTAGAAGAAATCTTACCTATAAATG

GATGGATTCGTCGTTCTTACCGAATTTCTTAATTAGAGTAGTCAACTCGCGGCCTGGTTTCTATCTCGGATCCTAGTACG

CATACAATATGCAGAAATAAGGTTTTTTAAGGTTTCGTAGAATATCTCGAAGGAATCTTACCTGTAAATGGATGAATTCG

TCGTTCTTACTAAGTTTCTGACTTAGATTAGTCAACTCACGGCCAGGTTTCGATCTCGGATCCTGGTACGCATACAATAT

GCAGAAATAGGGTATTTTTAAGGTTTCGTTGAATATCTCGAAGAAATCTTACCTGTAAATGGATGGTTTCGTAGTTCTTA

CCAAGTTTCTGAGTTAGAGTAGTCAAATCGCGGCCAGGTTTAGCTTTAAGAGCCTGATACGCATACAATATGTAGAAATA

TGGTTTGTTTAAGGTTTCGTAGAATATCTAGAAGAATCTTACCTGTAAATGGATGGTTTCATCATTCTTACCAAGTTTCT

GACTTAGAGTAGTCAACTCGCGACCAGGTTTCGATCTCGGATTCTGGTACCCATACAATATGCAGAAATATGGTTTGTTT

AAGGTTTCGTAGAATATCTAGAAGAAATCTTACCTGTAAATGGATGGTTTCGTCGTTCTTACCAAGTTTCTGACTTAGAG

TAGTCAACTCGCGGCCAGGTTTCAATTTAAGAGCCTGATACGCATACAATATGCAGAAATATGGTTTGTTTAAGGTTTCG

TAGAATATCTAGAAGAAATCTTACCTGTAAATGGATGGTTTCGTCGTTCTTACCAAGTTTATGACTTAGAGTAGTCAACT

CGCGGCCATGTTTCGATTTAAGAGCCTGGTATGCATACATTTATGCAGAAATAGGTTTTTTTTTAAGGTTTCGTAGAATA

TCTTGAAGGAATCTTTCGTCGTTCTTACCAAGTTTCTGACTTAGAGTAGTCAACTCACGGCCAATTTTCGATCTCATATC

CTGGTACGCATACAATATGCAGAAATAGGGTATTTTTAAGGGTTCATACAATATCTCGAAGAAATCTTACCTGTAAATGG

ATGGTTTCTTCGTTCTTACCAAGTTTCTGACTTAGAGTAGTCAACTCGCGGCCAGGTTTCGATCTCGGAGCCTGGTGTGC

ATACATATATGCATAAATAGGGTTTATTTAAGGTTTCGTAGAATATCTCGAAGGAATCTTACCTGTAAATGGATGGTTTC

GTAGTTCTTACCAAGTTTCTGACTTAGAGTAGTCAAATCGTGGCCAGGTTTCGCTTTAAGATCCTGATACGCATACAATA

TGTAGAAATATGGCTTTTTTAAGGTTTCGTAGAATATCTAGAAGAAATCTTACCTGTAAATGGATGGTTTCGTCGTTCTT

ACCAAGTTTCTGACTTAGAGTAGTCAACTCGCGGCCAGGTTTCGATTTAAGAGCCTGGTATGCATACATTTATGCAGAAA

TAGTGTTTGTTTAAGGTTTCGTAGAATATCTAGAAGAAATATTACATGTAAATGGATGGATTCGTCGTTCTTACCAAATT

TCTGAATTAGAGTAGTCAACTCGCGGCCAGGTTTCGATCTCGGATCCTAGTACGCATACAATATGCAGAAATAGGGTTTT

TCTAAGGTTTCGTAGAATATATCGAAGGAATATTACCTGTAAATGGATGGATTCGTCATTCTTACCATGTTTCTGACTAA

GAGTAGTCAACTCGCGGCCAGGTTTCGATCTCGGATCCTGGTACGCATACAATATGCAGAAATAGGGTTTGTTTAAGGTT

TCGTAAAATATTCTGAAGAATCTTACCTGTAAATGGATGGATTCATCATTACTACCAAGTTTCTGACTTAGAGTAGTCAA

CTCGAGGCCAGGTTTCGATCTCAGAGCTTGGTTTGCATATAATATGTAGAAATAAGGTTTGTTTACCTGAGAAGGTCCGA

AACTCTTGTTTGCATACCTAGTATGCGAACACAGTAGTTAAGTTCTAGAATCAGTATTGTATGAATCTCATACTCATGAA

CTAAAACATTTAAGATCTAAGGAATGCAAAGTTTGCAAACAGTGGATTAATGCAAAGTTCGAAATACAATTAGATTTATT

TGATTATCTTTCATGATTGACTGATCATCATATTTGATTAAGTGTTAGATAAATATATGGTTAAGTTAAAACTGTTCATA

TGGCTAACTTCGGTTAACTATATGGTTGGTTCGATCTCGGAGCCACTTTCTTTTTTCGAAGTTATTATAAAAGGGTTTGT

TTAAGGTTTTAGAGAATATTCTGATATATCCTGCTAAGGTTGAAGTCGAAACCCTCCCAGAGCTTCTTGGTTCATAGTTT

GTATGACGGTATACCACATTTGAAGTTCGAATCGACATTGAGTTTCATATTTATTGATTTAGATGATGTATCATGTTAAG

TTTTAAGTCTGAACCCTCCTGGAGCTTCTCAATTCATAGTTTGTATGCCGTTATGCCACATTTGAAGTTCGAATCGACAC

TGAGCTTCATATTTATTGAGTTAGATGATGTATCATGTTAAGTTTTAAGTCTGAACCCTCCTGGAGCTTCTCGGTTCATA

GTTTGTATGCCGTTATACCACATTTGAAGTTTGAATCGACAATGAGCTTCATATTTATCTATTTCGATGATGTATCATGC

TATGTTTCAAGTCAGAACCCTCCCGGATCTTCGTGTCTTCATAGTTTGTATGTATTTATATCACATTTGAAGTTTGAATC

CATCCGGAATTTCGTGGTCCATCTCTAGTAATCTTATAACCTGAGTCTACCGGTATGAACTAAAGCTACTTCATGACATG

TATGACTAGTCAAAATTACTTCATGACCTCTGTGACTAGTCGAAATTCAAACCGGAAGCACAAAAAAAAGTATAAAAGCA

CAAAGAGAAAGCACAAAGAAACACACCAATTATCGAATTCATTTTTATTTTCCCGGTTCATTCTTATACATTTTTTGGAT

TTTTTTTATGTCAGAATATCGATAAAAATGCCATAAATTTATTAATATTTTTTTTAAATTTTTTTTTCGTGTTGAAGGTT

GATAATATAACCGAATAATGAGTTCGTATTAACACAAAGAGAAACACATGCTTCTCAATCCGTATGAGCATCCCAAGTAC

AATCTCAACCGTCAAGATCTACCCTTAATCCGTATGAGTGGATCAAATTAAATCTTGTTCGTACATTCTTTTACACCTAA

CCCTCCTCAACACTAAGAACTCCAATTCCTATCAATTTTCTCTTTCTTCCTCTCCGTCTGTTTTCTTCCTCAGCCTGCGC

CTCCGACTCTCCCTCTCTTTTTTTTTTCTTCATCTATATCTAATGGGTGTTGTTGATATTTACTTCTTCCATATGGATCT

GCATCATACCCATCTCTATTCAACATCATAGAACCCTAATTTCATCTTCAATCGAATCTCGAGACCAGATTCAGAAACCC

TAGCTCAGTTCATCTCCAATACAAGTTTAGAATTTCAATTGAGACTTAACCCATGGTTCTTCATCAAACCACAACTCTAA

TTTCATCTCTTAAAACTTCAGAGCATCAATCGGTTCTATAAATCCTAATTCTGAATATAACAGCATCGACTTCTTCCTTT

CCACAAATTAATAAACATCAATCACCACCACCAACCGTTATCCCTAACTTGTCTAATATATCGACCTCACAACCTCAGAG

TTCTCACAAAATCGCCTGAGGCGAAGAACAGAAGAAGAAGAAGAAGATGAGAGAAGAAGAAAGAAGATTAGGGTTCTGTT

CTTCCCCAATTTAAGAATTAAGGTTTTGAGAATTGAAGATGGTGATGAAGGTTGCAGGGTTGTTGATGATTGTAGAAGCA

ACAGATGAGATTTTTGGTGTTGATTGTTGTTGAACTGGAAAAATTGAGAAGATGGATTAATCTTAAGGTATTGGGCTAGT

ATTTGAGGTAATAGTCCTAATCTATGATGTCTGTTAAGTTATGCAGTTATGTTAGATCAATATTATTATCTGTGTTTGAA

TCTGTATGATTTGGTGAAGCTGTTGATTTTATGATTTTGGTATTGAGTAGGATTTTGTAATGTTGGTGGTTGTGGTTATG

AGCTGTTAGTATGAAATGGAAATGGAGTTTGCCTGTTGAACTGAAGTGCAGGTGGTGGCTTTGTAAAATGAATGTAATTG

TGTGTATTGGTGATGATACTGAGTGAGTGTTATCAGAGATGTGATGTTGAGATTCACAGATGCGGGGGAAGATTGACAAG

GTTTGTTTGCGGACTCTACAAACAAAATCTTGTCTCTGTATTTTCCCTGGATGTTGTTATTCGTGTTCTCTAATTGGTTC

TAATGTGCAGTTTTCAGGATGTGGGATTGATGAGAGGGAGAACTATGAGTACATAATTGTTGGTGGGATTATTCTTCACA

AGTTAAGTGGAGAACTTATAAATACAAGTATAGGATCTGAAGGAGCTAAATGGATATTTGTACATCCAAGAAACTTTATG

CAGGAGAGGTAATAAATTTTAAGATGATCATGTAAACTACAATCTATGTTGCCGCACCATTCTGCTACTTTGGATGTTGT

TGAGTTGCTCATACTGTTGCTTACTGTGTGCAGAAAAAGAAAGGGTGTTTTCATCATTCCAGTTTTCTTGCTGGAGGAAC

TACTTTAGCAGCTGGTAGGCTTGTTGTCCAGAATGGAAAGCTCAAGGTGAATTATTGGCGTTATAATCAATCCTAAGTAA

CTTGGTCTACTTACTAAACCGAACTTAGTTCAACGAAATGTCATTTTTTTTCATCCGTTTTGGGCTCTAATTCTTTATCC

TATTGCAGACAATTTCTGCATATAGTGGACATTATCGCCCAACTGAAGAAAACCTTGACAGCTGTAAATATTTTCTCAGT

GAAAATGGGGTCAACCTTGATCAAGTTCAGGTATTATGCATTGTGTTTATTGGTGTTCCAATCTCACTACCTCGCAATCC

AGGTAGTCAAGGGACAATAGTTAGGGGAAATGGTGAACACTTACTGTTTTTGTTTATGTATCTTCCGAGTGAATGAGCCT

TATCTTCAATTTTTGGTATTGTAGATTCGCTCTTCCACTGATGACTACGAGAGCAATGGAGACAACAAATTAGTTCATGG

GAGTGCAGCATTAGAATCCTTTGCAATTAAAACTGAAGCTTCGGCATCTTCTGATGATTGTCTTCATCTTGAAATTTCGA

CTTGTAAGGTTCAATCCAGTTCCTTCATGTAATGCAATTCTGAAAATCCACTTGTTGGTTGTTGTGTAGCCTTTCTAGAT

TACATAAGTTTGCTTCTTTTGTTTCCCATATTTGTTTTGGACTTAAGAGTTATGATGTAGTTATTTGTTAGGCTTAAAAA

ATCTAAGTTTTGGGAATTATTTGGTGACAGTTATTTAGTAAGGGTTGGCTTTGGATATTAACTTGTATTCTGAATAATCT

GCTTGGATGCAATTACTGGGAATGTTGTTGTAATGGTATTGACACCAGTAATAGAATGTTCTTATATGCAGTGCTTCCAC

TTTTAGTTAGTTTTTTTTTCTCTTCGTTTATATTATTGGTAAAAAGAAAAGAAGATATTTTTGAAAATATAATTTCTTTG

TTTGTTGGTTTGGTGCCAATTTCAGGTAATTGGAGATCCAGTTCTGGGTTTGAGGATGACTCTCGTTTGAGCTTTCATGA

AAGAGTCCATGGTGCTCGTAATAATCGGCCTCGAAGTCTAACCAAAGGGGGAAGTGGTAGTGGGTGGGGCACAAGAAGCA

GGAGCCGCAGCCCTAATAGATTTAAATAGATTTGAATGAGGGTATAACAAACAAAAGGATGAAGTAAAGGAAGCTTTTGA

CTAGTGATTTGTAAGAAGGAATTTGGAGCCAATGTTTACTCTGACTGATGAAGAGAAGTGGGATTGATCCCAGGTTCTGT

ACTGCGAGCAATGAACTTGTCCCTGATCTCATAGGTTATAGTTTCTCTTTTTCTTTTCGTGATATGATTTTTTGCTGGTG

TTGTAGTAGAAAGAAGAAATAGTGGTAAAAGTCTTAACATTGTAAGGTTGTCATTTTTTGCAACTTAGAATGAATGTAAA

TGTGAATTGAAATACTTTGGCGGATTAATGTGAATGTGAAATACGGAAATTTGCGCTATAATTTTTTTTTAATGTAAATG

TGAAATACTTTACACAAACCAAAGGTGTGAAAAATTTGATTCCAAGATATTTCTACAATAACAAATTTTTTTTCAAAATA

GTTTTTATGCACACAACTTCTAATAAGAGTTGTGTTCTATAATTCAACTACACAAACCAAAGGTGTGAAAGAATGTCTTT

ACAACATAACTAGCTATTGTCCAAGATATTTCTACAACCCTAATACAGTGTTGTAAATGATGATTTTTCCCGTAGTATAA

ACACCTATAAATACCAGTCCATGTAAAGGGAGATGCACATGTTATGTAATATTAGATGGTCTTTCTACAGAGAATTCTAG

AGAGATATTTAGAGAAGGAAAGAGTGGGAAGTCTAAGTGATATTTGTAATTCATTCAAGGGTAAGACCTTAATCAATAAA

GAAATACTTTCTTTCCCATGGATTTACCTACGATAAAAAGCTTGTAACCATGTTATACAGTGGAATTCGTGCTGATATAT

ATCTAGTCCAAAAGTCAATTGAACTAAGTCGGAGCAAAGTGGTGGAGGCTAGTTGAAGAGTTTTTAGGTTAAATCGTTTT

ATGGATATGCAACAATAACTGATTGAATGGCAAAAATAGGAAAGAATTCAATCTTGCTGAAACCAGCCGCTTCCACTAAA

GTTCTCAATTCATCTTTAGTCCTTTCTCTTCCTCCCGAACATACCATCATATCTAAATCAAGGCTCAATCTTCTAGATGG

TTACTCGAAATACACAAACACAGGTGTTGATGGTGGTTTTTAAACAAAGGGTAAAACCGTAAAACCCCAAATAACATAAT

GTAACCGGTGACACATTTATTAGAACATTTAATGCATATATTCAGTAATTAATAGGGGGTATTTTTTCAAATTAGGGTTA

CAACGAACAAAAACCTAATTCCCCTCCGTGCCAATGGCAAATCATGCCAGCCACACCAAAGCGCGCCAAAGCATGCCAAC

CATGCTAGCCGCACCTCCGTGCCAATGGCAAATCATGCCAGCCACATTTAGCCCGAGCTAGCCGCGCCATGCCAAACGCA

CTCTGCCTTGGCCCTAGCTACGCTAGTCACGTCATGCCAAGCCAAGCACTCTGCCTTGGCCCCAGCCACACCTAGCCCAA

GCTAGCCGCGCCATGCCAAATGCACTCTGCCTTGGCCCTAGCTATGCTAATCACACCACGCCAAGCCAAACACACTTACC

TTGGCCCTAGCCGCACCTCTGTGCCAATGGCAAATCATGCCAGCCACACCAAAACGCGCCAAAGCATGCCAACCATGTTA

GCCGCACCTCCGTGCTAATGGAAAATCATGCCAGCCACACCTAGACCAAGCTAGCCGCACCTCTGTGCCAATGGAAAATC

ATTCCAGCCACACCAAACGCGCCAAAGCATGCCAACCATGCTAGCCGCACCTCCGTGCCAATGGCAAATCATGCCAGACA

CATTTAGCTCGAGCTAGCCGCGTCATGCCAAACGCACTCTGCATTGGCCCTAGCTACGCTAGTCACGTCATGCCAAGCCA

AGCACTCTGCCTTGGCCCAAGCCATGCCATGCCAAAATGGAGACCGCGCCTAGGCTTCCAAACCATGCAACCTAGGCCAA

ATTGGCCACGCATGCCATTGGCACATGCCAACGCGCTCGCTGGTAGCTTTCCCTACCAGGACGCGACCCAAATTGGCCAC

GCATGCCATTGGCACATGCCAACGCGCTCGCTGGTAGCTTTTCCCACCAGGGCGCGACCCAAATTGGCCACGCATGCCAT

TGGCACATGCCAACGCGCTCGCTGGTAGCTTTCCCTACTAGGACGCTACCCAAGTTGGCCACGCATGCCATTGGCACATG

CCAACACGCTCGTTGGTAGCTTTTCCCACTAGGGCGCGACCCAAGTTGGACACGCACGCCATTGGCACATGCCAACACGC

TCGCTGGTAGCTTTTCCTACCAGGACGCGACCCAAATTGGCCACGCATGCCATTGGCACATGCCAACACGCTCGCTGGTA

GCTTTTATTACCAGGCCACGACCCTAATGGACGAGCATGCTATTGGCACATGCCAACATGCATCGCTGGTAGCTTTTCGC

ACCAGGGCGTGACCCAAGTTGGCCACGCACGCCATTGGCACATGCCAACACGCTCGCTGGTAGCCTTTCCTACCAGGACG

CGACCCAAATTGGCCACGCATGCCATTGGCACATGCCAACACGCTCGCTGGTAGCTTTTCTTACCAGGCCACGGCCAAAA

TGGCCGAGCATGCCATTGGCACATGCCAACACGCATCGCTGGTAGCTTTTCCACCAGGCCGTGACCCAAATTGGCCGAGC

ATGCCATTGGCACATGCTAACACGCATCGCTGGTAGCTTTTATTACCAGGCCATGGCCCTAATGGCCGAGCATGCCATTG

GCACATGCCAACACGCACCGCTGGTATCTTTTCTTACCAGGCCATGGCCCTAATGGCCGAGCATGCCATTGGCACATGCC

AACACGCATCGATGGTAGCTTTCTTACCAGGACATAGCCCTAATGGCCATGCGTGCCATTGACACATGCCAAACACACAC

GCTGGTAGCCTAACTTGCAAACCGTGCAACCTACCACCAAGGCATTCTACACACGCCATTGGCACTTGCCACACACGTCA

CTAGCGGCTCTGGCATTGCCACTTACACTCAACGTACAACCTACCACCAAGGCATTCCGCACATGCCATTGGCACATGCC

ACACACGCCACTAGCAGCTCTGGCATTGCCACTTGCACTCAAGGTGCAACCCGCAACAAAGGCATTCCGCACATGCCACA

CGCGCCACTATCAGCGCCACTTGCACTCGAGGTGCAACCAACACTCGAGACATCAACACAAACCGGGGATGCTTTCCGGG

GTCTTGGTTTGGCGGTCACAGCTTGCAGCGTACACTACGCTCACATTAAAAAGCGTGAATAGGAATAAAGCGGTCAAATA

AATGAGGAAATAATGGTGAAAACGTTTTCACCCATAAACACCATGCGGTTAATTACCACCTCCTCCCATTATTTCACCCT

TTACTCTCTTCCTTCATGAGGCCAGGGTACGTTTCAATCTCGACTTGTATAAATAGGTCTTACCTATTTCCACCAGACAA

CAAGTCTCGGTCAGGAGAAAATTACAACACTCTCGGAAAAGGCAACTTTGCAAGCTCTCCAAAAGGCTTTCATCTCTTGC

AAGCTTCCTAAGCTTTCACTTTCTGATACAAGTCGAGACACTACTCTCTCAGAATTATTCTGGTCTCAACACCTTTCTTT

GCTTCCCTCCCCAAATCAACCCTTCTCCTCCACTTTGTGACTAAGCAAGTCCGGGAACGGCCATTTCTTGGTTTAGGCCG

GACTTGTACAGATTGATCTCTCGAATCTATAAAGTGCTCACGTGCAGCACATTTGTTTAGGGTTTAAACTAGTTTCTCAC

TCACCTATCGAATTTACCAAAATCGGCGGAAACGATTTTTACCCATAAACAATTGGCGACCACAGCGGGAGGATTGATCT

ATCAGTTACAATACCAACTCCGACTCCTGATTTTGTTTCCCGAACGTCAAATCCCAAAATGATGGAACTTAGGTCTGAAG

CGGCTGCGAGAACACCAAAGATAGAATGCATCCACGATGTTAGCGTCTCTCCCAATAACGTTGTTGCATCACTGTGACTA

CGGTAACATCGAATGCACTCCCTCAAGTGTAACACCTCCAATCACTGTGGGACAGAGCGCTGCCTCCTCTAGTATCTCCG

CACAACCCTCTAATCATGGCAACGAAAATATCTCTTCGGGTGTCACACCACCGATCACGGGGCTAGAGCCGCCGCTACGA

CCCCAAATGCTTCGTCTAGCATGGCGTCTCCGACCACAACTAAGGTTCCAACTACCCCGCCGACGGGACGAGAGACCGGA

GGGACCAGAGGAAGTCGTCCTTACGTCACCATCGCTGATTTGATGGAACGACAAGAGATGCTCGCCAGAGCCCAGACGGA

TATGGTTTCAACTCAGAAGGAGGTACTCATCTTTCTCAAGACCCTGACAGAGCGACTACCACGTGAGTCGCGTCCTCCAG

AGCCGCTTAGAAACATCTCCAACAACCCCGACACCAGCACCTCGGCGGAGCACGCCTCCACGGATGAGGAAACTCGCGCT

AGAATCTCGTTGGAAAGAAATCAGCCGCCCAACTTCGTCACGAGAGAATATTTGGAACAACTTCTCCGCAACCGAATCAA

GAATGATGAGTTGGACGTGCATCATCACCAACCCCCTTACCCATCCGAAATGCAGAGAATCCCTCTCCCAATGGGGTACT

CTTCTCCTCAATTTACCTTGTACGATGGTACCGGGAATGCACATGAGCATGTCTCTCGTTTCCTGGAATCCCTAGGTGAA

CATGAATATAATCACATTCTTCGTTTAAGGGAATTCTCAAAATCACTTACTGGAAGGGCGTATACATGGTACAACAGCAT

TTCACCGAACAACATTCCCAACTGGAGTAGCATGGTCACTGCGTTTTACAAGAAATACTTCTTCGTGTCTGAACAAGTTG

CTTTCTCAGACCTCGGAAGAATGTACCAGTGGGATAACGAACATCCGAATGATTTCGTCAAAAGGTTCAGGGTTCAAGCA

CTCGACTGTCTAGACCCAAACGTGACCGAGCAACAACTGGTGAACGCATGCATCAATGGTATGATTCCAATTTACCGCAC

CCTGCTAGAAAACTTGCGCTTCCAAACGTTCTCCGAACTCCATGAAGCCGCCAGACGATCTGCAACAACAGCACCAGCAT

TACTGGAAAGAGCTAGGCCAAATTGAACCGAAGACACACGTGAAACTTGCGGGAGTAGACGCCTCATCAGCAAACAATTT

AATGCCGGCTCATCCTCCGTGAACATAGTAAACGAAAGGTGAAAAAGGAAAGCCCCAACTACGAGCCACCAGCCAAAACG

TGAAGCGCCGGCGCAACGGGAACCCCCGCCATGAAAGGCCGTCACTAAAACTCCTATGCAACATCATGAGACCGAAGGAA

ATACCTTTGATTTCCCGTTTCCCATCAAGAAGGTGATTGACCTCCTAGAAGCCTGGGTTCAAGATGATATTATCAGACTG

CCACCTACCATACCAACTGAGGCTGAGATGGCGAACCCAGAGTACTGCCACTACCACAGGTTCGTTCACCACCCGACCAG

CGATTACAACAGACTAAAGCATATTTTCAAAGAAAATATAGAAGCGGGAGGACTCCAACTAGGAGATGAAAGAGTGCATC

GAGACCCACTCCCAATAAGAAACTGCGGGTCCTCTGGGGACTCCGTGCACGAGACTGTACAATCCATGATGCAACATGTA

TGCGAAATTCTCTATCTTTCCAAGGTGCAGCGCCAAGATATTTTCGCAGCTCTCAATCGTGTTGTATCCGGCAGACGACT

TTTTCTCTCCTCGGAAACCATGCTAGGGGCCCAAGCTTTCCAAACAGGCGGAACCAAAAAGGAGAACTGGGGACTGCTGA

CCACGTCCCATTTAAGAGGTAACGAATTTAACGACACGCTCATCGACACGGCCTCTGACCACAACATCATTACTTTCAAG

ACACTAAAGGCAGCAGGATTTACAAAACAAGACGCTACCTATTCCCCGTCTGTGATTAAGAGATTGAGAGGAGAAACTTT

GAACGCGCATGGCTACGTCAACCTAGAGATCAAGGTGGGAGACGCTATAACGCATAGCAAGTTTCACATTGTCAAAGAAC

TAGATCATGACATGATATTGGGACGCTTATGGGTGCTCGACAACAAAGCAAAATTGGATGTACACCCTCTGCCGACGTCC

ACACCCGAAGGAATTCCCACGGCGGCGCCAAAAATTGATGGTTATTTTCAATGATGTTGTAGAAAGTGGTAGTAAGGTTG

TCGTTCACTCGGACTTGACGAGATTGATTTTAAGAATTAATAAAAGAAAATAAAAGAAAAAAAATATATACAAAAATTGT

CACAGGATGAAGAGTGTTACTGGGACTAGGATTTCGTCGAATTCATAACATAGGGTCCAATTTATTTATTCTCAACAATT

AAAGCTCAATAAATAAAATTAACACGGACTCTGATTTTTCCAAGGTAGATTCTTAAAAGATTGGTTGTAAATCCTAAGCA

TGATGTATCAAAACATTTAAGCTAAGCATACCTCATCAAATTAAATGACAACCAATTAATTCAAATCATATTTCAATTAA

AATTAATGCAAAAGTCTTAAAAAGAATTAAATAATTTTACCCATGTATGAAATTCGTCTTCCTCCATCGTCCCAGTGTTG

GAGTCTAGCTCATCATGTTGGAAACACGCTCAAAATATGATTTTATTGCTCAAAGGTGCTTACAAATGATGAAATGGATA

AAATGATGAAAATCGGGTGTTTGCAACGTTTATAATTGTTGCAAACACCATTCTGGCTTGTTGCTTTAAGACTGCCTCTT

AACTACGACCCTCAGGAACGACTGTCGCCTGTGGGTCTTATGTTCTTCGTGTTCTTCCTTAATGGCAGCAGCAACAGATT

TTCAAACTCTCGATGCTCTGTAATGCTCTAAACTCTCCCAAACTCTCCATCTCCGTTCTATGACCTCCCAACACCTATTT

ATAACCTTTTTGGAGCAAGAATAACTCCTCATTAATCCCAGAAATCTTCCCATTACTCGGCAGCAATAATATTGTTCTCG

GAATATATTCTTTCCATTTTTTGTAGCAGCACACGCCTGCAGCTGTTATTTTCTTCCTCATTTACCTTCTACACTCGTCT

GGCAACAACCCACACACTCCAAACACGAGCAGTGCACGTCTCAGAGCTCACAGCGATTCACGAAATACCTATGCAAACCC

GTGCTTCTCTGCTTTCTCCTAGCGAGTTATCTAGCCAAATTTAGGTCACTAATCGACCCATACCAACCCTGTTAGGCTCT

AACAGGTTATCCCAAGCAATCTCAGCCATTGAGTCTCTCCAAAATGCCTTCAATTCTCGAACCAAAATTTTGCAGGTGAG

AAAATAATTTTTCCCGCCAAAACTAGATTTTCAAACGATGAAGAAGGTCAGCCCTTATCCAACAGCTGGGGGTGCGAATA

TCAATTGGGTGCCCCTTAGTAATTGGGTTATCCCTTATCCAAAGTGGGAGTCCGAATATCAAATGTCCTCCGGGGGTGTT

CCGAGCAACTTTTCGAGCCGAATTTTCCAAAAAATATTTTTTCCAAAAATACCTAAAAATACATAAAAACACAATATTAG

TACAAAAATCGAGTTCCAACAATAGCAACAATGAGGACAATTCAGACACAAAAAATGTGTCTATCATCAGCAGCCATTAT

AAATTCTCCTGTTGCAAAAAGCCGATGCAACCTAATTTTTACCCTTACAAATTCTCATGTTACAAATTCTCTTTTGCCCT

TCAGGCAAGAGCCGTTACAAATTTTTTTCGTTATCCATAACAAGGTTCGGCTGTTACGAATCGTAATAGGATAATTTATT

ACGTGTGCTAATCACACGCTTGGTCACGTTCGGATGTTATTAATCCATTTTCACATTATTTGTAATGCATACATTCTCTT

ACTTGCCCTTGGTAAATAGGTATGACCTAATTCTAACATACATGGCCATCTTCATTTTAATTCAGAAAAGAGAAAGGCGC

AGAGGACTATGTATGAAAATGAAAGGTTGATGAGAGAGTTAGAAACTCAAAGAAAAGAAAATCAAGCAGCAGGGTAAGGA

AATAGATAATGCGATGGCTGAATTGGGGAGGTTACTTATCTATGAAGCGTCAAGGACCTAGTACGTTGGCTTATCTATAG

GTTACTTAATAGGCCAGAGCCTGTGTGACTACTAAACACTGATGATGAGCAATATGCTTTGGAGTACCAAGCTGGTGGAT

CATGCATCGTTACCAGAGATGTAGATGGTTAGCAGTTGAGTAGAGGAATCAGAATCACTCTCTTTCTCAAAGAAGGTAAG

GTAATCAAAATAACTTCGGTCATTATTGACTGAGTCGGACCAGGTCAATGTTGACTGGCAGTAATTTTTTGATGTTACAA

AAAAGTTCGTAATGGCTTAGGGCTGTTACGATGTGTTACAAAATAATTTCGTAACGGTTGTTACGAAATTGCCACATAGT

AACAAATAGCGGCCGTTACAACAGCCGTTACAAAATAAATTCGTAACGGCCATTACAAAAAATTTGTAGTACCCCTTTTC

ATTACAACAGAAAAGCTTTACAACCCAGATTCGTAACGCGTACTTGCGTAAGGTACAGGTCCGGGAATTTTTGCTGAAGT

TTGGAAGTGTTAAGGTATGCATACACGTTCGCATAATAGCGAACCTAAACACAGTCTGGTCACTAAGGTATGCATACCCG

TTTGCAGACCAAAGTGAATCATGTTCTAAAATCGTCTTGCTCATGAACTAATACATTTATATAATAAGGAATGCATTCTT

TGCAAACCGTGGCTATAATGTTCATGAATTGATTCGAGTGAATCAAACCGATTTTGCTTCAATTGTGTTCTTGTATAATT

CTATGAGAATATAACAATTGAACAACTCTGTAACTAGTTTCATTTGAATCATTTGAACTAGTTATGGTTAAGATGAATAT

GGCTAAATACCACATGTATGATTCTTAATTACCTAACTAATTTGTGATCGAAATTCTTAAGTATAACAGGTTAGAAGCAA

TAATCAAAGCCATTGGATGTTGGTTTTTATTGTAAAAAAAACCTTCTTCATACGCTTATGGTAACTATTCTTGGTGTAAA

TCCTTGTAGAAAAGGTTATTCTTCCTTCTAAGCATATCAATTCTTGTTTATACAAGTTTGTATCTTAGAAAAGATGATCA

CAATATTTGATTTTCATGATTACATATTGTTTATTGTTTCCATGAGATAATTCTAATTTTTTAGTAACACGATCTCATGA

GATACTTTCAATGATTGGTTCTTAAATCATATCTCCTTGTGGGATTTCTAAAATCCAATATATAATCCTTCTCCCTGGAG

TAAGGTCACTCTTGTTGTTCTGTCGGAAATAACATTTTATGGGAGAGAGTTCTTAAATGAACTTTTTAGGGAGTTCTTAA

TTGTCATATATTTGTAGGGAGAAGCGGGTTTGGAACATGAAGGGGTTATCTTGTATCTTTATAAACTCCTTGATGAATGC

ATTTAGTTTCGGCTATATGATTGCATCTAAACAAGTTGATATGCTGTTATCTTTTGGTCATGAAGTATCTCTATGGAAAT

TTCATTAGGATCCCACTAGTTTTCGTACCTTTGCCAATTTATATTGACAAAAAGGGGGAGAATTAATGTGTAGTTTACAC

TACAGATACATATGATTTACAGATCATTATGTAAGGGGGAGTGGTTTTCATTGTGAGATGAAGTATTGACTAAGGGGGAC

TGATACATATCACCGTAGTATTATTGTCAAAGTTGTGATACAATTGAACTTTGATAATGTGTATAATACTATGATGGATC

TTCAACAAGTATAGGATGAACACATGCATAGTTGAAGAACCAAGTAAATCAAGCATGTCGGATTTTGGTTCTCTAGTTCA

GTTAGAACCTTGCAGGTTAACACGACTTCCTAGGCGCTCCCATACAGAGCGACCTCTCCGATTTTGATTAAAATCTACCA

ATCGATCTTGATTGTTCAGATCAATAGTCCTTGATCGAGACTGGCTCACACGTTCTAGTCTCAGGAGAGCATTTTCTTTC

TCCACCTCATTCAGCTGCTGGCGCAGTAGATCTTCCACTGTACTAGGAGTCGCGGTTCGTCCATCTTGCTGATCAAGTTG

ATTATCGTTTCTCCTTGATGGGCTGCCGAACACCAGTTCTGTGTTTACCCGAGCTCGCGGGTTTCCTTCACTGGGTACGG

GGTCAACGTCCATTCCTGCGCTATCGATTAATGGTTGTTTTTCGCTTCTCCGAGTATTTGGATCTGGAGTTCTGTCATGA

ACTCGATCCCGATCTCTTATGTTTCCAGTAGGAACAGCCGGTGCAGATTGAACCTCGTTTGGTTCAGTCATTGCGCGTCG

CGGGTAGGTAGCAGAAACGATCCTTCGAGCAGTTAGCTTTACTTTTGCCATTATCTTGATCCGAGAAAAAACACGATCTA

TTTTCTCGAATCCCGTTAGTCGGCGCCAATGTTTTTAAGATAAAAGATTTTTACCTCAAACACGAATCTGATGTTGATGT

TTATCGATCTGCTGCCTTTATCTTCACGTGAACAGTGCTGGGGGTACCTGCAAAAAACCCTCCGATGCTTAAGTTAGCTA

GAGTTTTTCATAGGAGTTTTGTGTAGAGAAGATTGCGTACCTTTTAGGGTTGTGAACCTATGTATTTATATGTTGCAGAT

TAAGGATTAACCCTAATTTCGAGGTAAGCATAACTGTGTCTACCTGCCCTGCAAGTCTCCTGGAATACTTGGCTAACAAT

CCTTCCAAATATTTCCAGAAACTTCCTCTAGGGTTAAAGGTGCAAAGGGTTGGCCCAAACAAATGTGTAAAATGCCAAGC

CCATGAAGAGTAATTCACTTAAGCCCATAGTGGGGCGTATAAATCCGCAAGCCTAACAGCATGGGGCTGAAAATTCCATC

AAGCTATTGGCTTGACTCGACTAATCAACTCAGCTGGTTGAGTTGGCTCAGTGGGGCGCAAATATCCTTCAGTGTAAAGA

AATGGGGCGCAAATATCCAGAGGGTACGAGAATAAATTACTTCGTGATGCACTGGAATTTTCTCGCCACTGCCGGACAAA

ATTCGGTCCAGCCGTCGGACACAGACGCATGACCTCGTCGCCTGACGGAGCCGCCTGATACTACCGCCGGGAGCCGCCGG

GCAAGGCGGGCACAACCTGCTGCTCATCGTTCTTTGGAGAAGACCGCCGTCGCGGCACGCTGCTTTATTCTTGGCATTGA

CGGAATATTCCTTTTTCAGACGGAATATGCTAACGGCGTTAGTCTCTGTTGGCATCATGATGACGTCATCGACCTTGAAC

TGCTGACTGTGCTTGTGCTGCTAATGTGGCAGTCTCTGATTGGCCCACACTGGTGACGTGACAATGGTGACATGGTTCCT

TCTTGCCTTGACTTGGATATTTGCATATGGGCTTCTATGTATAGTGCTTATGAGGCCTAGTTAGCCATAGTTGGCTAACA

AGAGGAGCATATTAGGCCCAATGAATCATACTGAGCTAATAAGAGGAGCTTTTTAGACCTAAGTAAGCCCATTTTTGGTA

TAAAATACTAGGGTACAAACAAAGTTAATAAAAATATTAATTTATCGATTAATTAATATCGCGCTTAATTAATAAATTTT

GGTGGTCCCAATAGCATTAATTTATAGAGTTTCTACTATACTTAGCAAACAGATGCCCTCTCCACCATTTGCTTATCAAA

TATACACATGGTTCTAATTTAAGCACCTTAAAGCTTATGTATATGTTTCTCACCCCTTATACCCTCTATTTATATATGCT

TGTTTAAATTTGAGTAGCACTTATATTTTTTAGCCGTATTCTTCGTAAGCTGTGTTTGTTTTTTCTTTACTTTTAGTGTC

CTTTTTTCAATCGAGAGGGAAAGTCCAGAGCTATTTTGATTTCCTTGTATTCTCATTACTTTTGTAAAAATCATTAAAAA

GATAAATAAAACTTTGTAAATTTATCATGGAATTTGGCTAGAATAGCATATAATCTTTGAGGGGTGTGGTTGTAGGAAAT

CCTGCAACCACATTATCCCGTCTATTTATATATGCTTGTTTAAATTTGAGTAGCACTTAGCTTTTTTAGCCGTATTCTTC

ATAAGCTGTGTTTGTTTTTTCTTTACTTTGAGTGTCCTTTTGTCAATCGAGAGGGAAAGTCTCTGCCGTTATGTGAGTTT

GGCACCTATACTATTCCACCTCCAAGTTGCTCTTGGACATTTCACGAGCTGGAGTGTTCCAAAGAACTTGGGTCATGTCA

GAAACTACATAAAACTTGTGCAAAAATCGTGTTGGCGTCTATGTATACTTGTTGTTAGTCCCGTTTCTACGTATGCCTAT

TGTCTTTAACCAATTCCCCTGTTCTTTTAATGCAATTTGAATGCACATCGCTGTAGTCAGTACATTTCTGATAAGCATGA

AGCAAATATCTCCACTTTTAGTACTGCTTTAGAACTTTATTGAGCAAATATCTCCACTTTTAGTACATTTCTTGGATGGT

TTAGCCCCACCAAACAATAACTCCTGACACAGTACAAACGCCTTCTATAAAAGCCACATCTAGCGTGGGTGACCTTAATT

ATCCCCACGAGGTTCTTGCATTACTCTCGTGGGCTGATATGGGACACTGCTCCACCCCCACAATCAGACACCTTTATAGA

AAGCACCGGCATATTATTACATCAATCAATCCATTATAAAAGATAATTGATATGTATAAGGGATGATATGGGACACCGCT

CCAACACACAATCAAACACCATTATAGAAAGCATTAACATATTAACATCAATTCTTTATAAAAGATCATCTTTATACATG

TATGAGTAGCAATAGGGGTGTGCATGGGATGATCTTTATAAAAGATCATCTTTATAGGTTTTCATCTCATCCGCATCCAA

TCCAATGTACTACGGATTTTAAGTTTGGCATCCGCATCCAATCCAGATCCAACGGATTTGCATCCAATGGATTAGAGGTG

TAAAACGTGCCGGCCCGGCACATCCCGGCACACGAAGTCGCGTGCTTCACATGCCATGTCTTGCGTCTTTGTCTTTGTAG

GCCAACGATTTAAACGTGTTGTGCCGTGATGTGCTTTACTAGTCAAAAGCTAGGCACAACACAGCCCACGGGCACAACAC

ACGAATAAACGTGTCGTGCTGTGCCATGTTAGCACACATGTTATAAACAATTTGAAATCACTTAATGTATACATTGAGTC

GGACATTATCACAATAATTTAATTAGACAACATATCATTTGAAATTATTTCAAGTAAAATTAACAAGTTTATTCAATAAA

AAGAAATAGCCCATCAAATATAAAATTGGATTATTGTCATGTAAATTAATGTTCTAGTTAAATATAGGTGACAACATTAT

AACAACGATATTGAAATATATTTTCCAAATATTAAGGTATTATATGTGTTTTTATAAATAAGCCAACATAAAATGTTAAA

AACTAGCTAGAATAGTGACTCTTTTGTGAAATACACAAAAAAATGTGATGTATTATGCCTTGTTATATGGTGTATTTGCA

CATGTTATGACGTGCTCTCTTAACGTGCTGTGCTGCACCGGCATATTATTACATCAATCCATTATAAAAGATAATTGATA

TATATATATATATATATATATATATATAAGGGATGATATCGGACACCGCTCCAAAACACAATCAAACACCATTATAGAAA

GCATTAACATATTACATCAAATCTTTATAAAAGATCATCTTTATACATGTACGTGGAGGTGTAAATTGGGTTGTGCCAGC

GCGAGCATAACCCAGCTCAGTACGCTAAAATTTAAGCACAGCCCAGTCCAGTACGTGACTTAGCCCAGCACGGCACTGCA

CGTTAGTTTTTTGGGACGGGCTGGGTTTGACTAATCGACACAGTAGCCCAGCACAACACGCGGCACGGATAAACACCTGG

CACGACACAACACCGCACACTAAGAAAGCACGCCGTAATATGTATAAGATACTACACAATGTGGCACAATACATCACATT

TTGTGTATATTTCACAAAAGAGCCAATATTCTAGCCAATTATTGACATTTTATTATCGGTATGTTGACTTATTTTCATAA

TAACACATATAATACCTAAATATTTGGAAAACATTTATATTAGAAAATATAAAATAAATGTGCCCGACACAGCACAACAC

GACACAACACGTTTATTTTTCTGGCACAGTCCAACACAGCACGCCTTTCAGCACGTCACAACACGGCACGTCATAATCTC

CGGCACAGCACAACACTACACATAACACGCTAAGCATGTGACTTCGTGGGTGGGCGAGTCTTTGGCCGTAAACACGTTTC

ACACCTCTACATGTATGGGCTGATACGTGACACTGCTCCACCACATAATCAATCCTTTATAAAACATTGAACTTTATATA

CAGTATATTTATATAAGTTATGATAGGTTGACCATGGATTCTGTTCCGCATATTTTTAGTATAGCCCCGTTGTAGATTGC

TTTTTTCAACTAAACCACTTTATAATAAACTTTTTACCAAAAATTGTATTTGGTAAAAAAAAATTTTAAAGTACTTTTGA

AAAGAAGCACTTTCATTTCAGACTTGCTTTTAAAAGCTACTCGCACTAACTTTTAAAAGCAGAAGCTGCGGATCCACATA

ATTTGATTTAAGTGATTCTCAATTTTGGAAACTGCTCAAATGGGCCCAACTACAACTGTAAGATATGCTGGGATTAAAAT

TCGAGCTTAAGTTACTATGAGACGCACCGAAATAAAATAATCCTCCTATTCATGTGCTACATGTATGGAATAACTTGATT

GCTAAAGACATGTACTAGTATCAGCGGGGCGGGTACCATTTCAAATAAGACACATAACTTACAACTAGTGGCTCAACCAC

TTCAAATACGATTAGAGTTATCAGATATCAAGTAATTTTTCAATAATTTTCATTATGTAACAGAAATCATAAAAACCTTT

TTTTGTCCAAAATTCAGTTTAAAAAGTTCAAAATATCAAATTCGGTCCAGTCGTTTTGTAAAAATTTGTTGGGTAAGAAG

GTCCAGCCGTTTTATAAAAGAAAATATCAGGTAAAAAGATCCATCCAATTTATAAAAGAAAGTGGCGGGTAAGAAGGTCC

GACCATTTTATAATAGATACGACCGAGGAAGAAAAAGAAACCTTAATTCTCTATATTTGATTCACTGATTCATAAGTGAT

AAAGAGAAGAGACGGGGGAGAGTCTGGCCAAGAAAAATTAGATTGAACAGGAAGAAGATTTAGAAGAGAATAGGTAGAGG

TGTAAATTGGGTTGTGCCAGCACGAGCACAACCCAGCACAACATGTTAAAATATGAGCCCAACCCAGCACAGCACGTGAT

TCAGCCCATCACGGCCCAGCACGTTAGTTAAATGGGACGTGCTGGGTTAGACCAATTGGCACATGATAGCACTCAAGACA

AAGACGCTAAGACGGATAAGCACGTGGTACTGACCCAGCACACCACGTTAAGAAAGCACGTCATAGCATGCACAAGTATA

GAATATAACAAGGCATAATACATCACATTTTGTGTATATTTCACAAAAGAGTCACTATTCCAGCTAGTTTTTGACATTTT

ATGTTGGCTTATTTTTATGACAATACATATAATACCTAAATATTTGGAAAATATATTCCACTATTGTTGTTATAATGGCA

TTACCTATATTTGACTAGAACATTAATTTACATGACAATGATCCAATTTTATATTTGATAGGCAATTTCTTTCTATTGAA

TAAACTTGTTAATTTTACTTGAAATAATTTCAAATAATATGTTGTCTATTTAAATTCTCGTGATAATGTCCGACTTAATG

TATACATTAAGTGATTTCAAATTTTTTATAACACGTGTGCCAACACGACACAGCACGACACGTTTATTTGTGTGCTGTGC

CCGTGGGCTGTATTGTGCTTAGCTTTTGACTAGTAAAGCACAGCACGACACACACGTTTAAATCGTTGGCACGTGAAGCA

CGCGACTTCGTGTGCCGGGCCGACACGTTTTACACCTCTAAGAATAGGTTTAAAAAAAAAAGTTTCATTTTTCTTAGTTT

ACGTCGAAGAAGAAGATATTATGCACGGGTATAAAACGATAACAAATTATATACTATCGGTGTTACCGTAACGGAAACTG

TCTCAGCTGGTTCCGTTACATTAAGCCAAAAAGTTCTAATAACTTCCCGATACGCTAACCATTATGTCGGAAAAAATTCT

ATCGGATCCAGGATTATCTGTAACGACTATCGGATTTCGTCTGCTTGCAACTGATTGGGACCGTCGGATCACTCAAATGA

TACCAGAAACGACGTTAATGCTAGCTAGTAGTGCCTCTAACAATCATGTGGACCGTGTGTTTTATTTTCTACTATTAACT

TAGTGGGACTCACTATACCTGATAGTGGAAGGTGACAAGGTGTGGGGAAATGTGCGTGGGAGGAAACTCACGCTCATGGC

CAGAGGTGTATTCTCTTTCTTTTGTTTGTAATGGATAGCAGCCAGTTCCGGCATGCTCGAAAATCAGATAACTGTGCCGG

AAGTAGACTGATTAAAACCAGAAAAAATAGGTTTTGAATACCACTTATGAAAATTTCCAGCAAATGTCGGCATGGTTTTT

TGGTTGAATACCATGCCGGCACCAAGTTTTTTCAGCTACAACAATGGTGGATTCAAAGAAAAAAAAAATCAAATTTTCAA

CCTCTTAGCAGCAATTCTCTCATCACAATCATCTTCCCATTTTAAAAAAAAAATTCCACTCAAAAACCATTTTCCCTCCT

TCTACTCTCAACAAACCCACACAAAATTCAAATAAAAATAAAATACACACTAATTTATCCCCAAATACTTAACATTTTAC

TAATTATAATTAATCATTAATCCTGATTAGTGGGGGCTAGATTAGGAATTAAAAAAATAATTACATAGGGGTCGCCCAGA

ATTGCTATTGGGTTGTCCTGTAGCTATATCCCATTAATTCCCATATCCCCCAATTAATTAATACAAAACAAGCGGGTCGG

CCGTTTTTAGTTGTGATTGATTTTAGGACAAGGTGTCCCCCCTCTAAACTATCGTTTTGGATTCGCCATTGCACGAGCTT

CTCGTGCATGACCGTTTGCTGTCTTCTTGTGCATGACGGTTTGTTATCCCGATGTCCTCTTTCCCTTATAAATGTAGTTC

TCAGTGCTTGAATACCTAAGACATTTGGGTTTTGATTCAATCTCTATTAATTAATAGTCAAATTCAATAGAGTAATTCCC

TAATCTTTGTAAACCGTCTTATATTTCTGTTTCATGGCTCCGATTAATATAGAGGAGAATGATGTTTGGATGATAGCATG

CACTGTAATAATAATATTTGCATTGGTGAAGTTCATGTTTTCTTTTTATCAATCTGCAAATACAACGGAATGGCCAGAAG

GTCCAAAAACACTACCCATAATTGGAAACCTTCATCAGTTGGGAGGAGGTGTGCCCTTACAGGTTGCTCTGGCAAACTTA

GCCAAAGTATATGGAGGTGCATTTACAATTTGGATTGGAAGCTGGGTTCCAATGATCATCATAAGCGATATCGATAACGC

TCGGGAAGTTCTTGTTAATAAATCTGCTGATTATTCCGCTAGAGATGTACCTGATATTCTTAAAATCATCACAGCAAATG

GGAAGAATATTGCCGATTGTGATTCTGGTCCATTTTGGCATAATTTAAAGAAAGGTCTTCAAAGTTGTATAAATCCATCA

AATGTTACGTCTCTATCTCGTTTACAGGAAAAAGACATGCAAAATCTCATCAAATCCATGCAAGAAAGAGCGTCACAGCA

GAATGGAATTTTAAAACCTCTTGATCATGCCAAAGAAGCGTCTATTCGATTACTGAGTAGAGTTATATTTGGTCAAGACT

TTTCAAATGAGGATCTCGTTATTGGTGTGAAAGACGCCCTCGATGAGATGGTACGCATAAGTGGGTTGGCAAGTTTAGCT

GATGCTTTTAAAATTGCTAAATATTTACCAAGCCAGAGAAAAAATATTCGGGATATGTACGCCACAAGAGACAGAGTATA

TAATTTGATTCAACCACATATCGTCTCTAATCTTCCTGCAAATTCTTTCTTACATTTTCTTACATCTCAAGATTACAGTG

ATGAAATTATTTACTCAATGGTACTTGAAATTTTTGGTTTGGGAGTAGATAGTACTGCAGCAACGGCAGTTTGGGCACTC

TCCTTTTTAGTCGGCGAGCAGGAAATTCAAGAAAAACTTTACCGCGAAATCAACACCCTGACGGGTGGGCAAAGACCAGT

GAAAGTTGTAGATTTGAAAGAGCTGCCATATCTACAAGCCGTGATGAAAGAAACATTGAGGATGAAACCCATCGCACCAC

TAGCGGTCCCTCATGTAGCAGCAAAAGATACTACATTCAAGGGGCGGAGAATCGTTAAAGGTACAAAAGTAATGGTGAAT

CTGTATGCTATTCATCACGACCCTAACGTTTTTCCTGCACCGTATAAATTCATGCCAGAGAGATTCTTAAAGGATGTTAA

TAGTGATGGACGTTTTGGTGATATCAACACAATGGAAAGTTCATTGATACCATTTGGTGCTGGTATGAGAATTTGTGGAG

GTGTAGAATTAGCCAAGCAGATGGTAGCTTTTGCTCTTGCAAGTATGGTCAACGAATTCAAATGGGATTGTGTTTCTGAG

GGGAAATTGCCTGATCTTAGTGAAGCAATTAGCTTCATTCTCTACATGAAAAACCCACTTGAAGCCAAAATTACCCCTCG

TACAAAACCTTTCGACAGTAGATAAAATCTTATTTTTATCAAACTGCAAGTGTATGTTTATGTAAAATCAAGGGTGAAGA

TGCTTGATCTTAGTGCGCCAAAATGGTGGTGGCTATGTAGGAGTTAGATTAGCTCTTGATTTAGTAAATGTTTTTCTTTG

CTAGAGACTGTGTCTCAACTCAAGTGTGTGTATTGTGTGTGTTTCAAGTGTGTATTGTGTGTGTTTCAAGCGTGTGTTGT

GTGTGTAATTACTGCCACCTGAAAATTACATTAGTCGCCGGCTGTAACTGTCACATCATTCTTGTGTTTTTAAATAACGC

GTTATTACACTACGTGTCTATACCTCCTTTGGTACATGTTGTCTAATAATACGCAACCGTTCATCCAAACATAGTTGTGT

AAACATAAGCAAAGCTTTCTCTTCTTCATCACTATCACTAGAAGTATCCAAATAGATTCCATGTTTGAATCTCACATAAG

AACGCATTAGACTCATTGTGCACCTTAGAAAAAATAAAAACCCAATATGAATTTGTGTATAGCATACATAAATACAAGTC

TATCAAGGTTCTACATTGATATGCATTTAGATGAAACATTTCATTCAACATAAATCAAGACACACAACAAATAATCATGC

ATTTTTCACTACATATATGTTTTAATAACGTTATTCAATTGCATACTACATATCCAACCAACCTAGTATAAACAATTTCA

TCAAAACTAGAATCAAATTAATTTAAAAACCCTAACATTAGAATCACTCACCTTAGATTATTTTTTTGATGAAATTCAAA

ATTGAAGAATGGATTTGTTTCACTAGTTGATTGTATGCAATGGCCATGGAATAATTCGAAATTCGAAATCGGGATGGGAG

GTCACCATAGCAAGCTTGAGAGGATGAAAGTTAGAGAAGAAGAAGAAATGGCCGTCTGCTGCGAGTTCTGTGTGTTTAAA

CCTACGAAACGCGTACTGAGTACCCGTATTTTGGTCAAGTCAACGAGTTGACTAATATGCGTACTGAGTACCCGTTTGGA

ACTGTACATACGCGTACTGAGTACCCGTTTGTGTAGGGAAGGGATGGGAGGGCACCATAGCAAGCTTTCCTTCCATGTGT

CATCCCTTCCCTTCAACGCTCTTATTATTTTCCTTTTGTGAGATTTACCTTCTCTCCCCTGTTCTGAATATTATTTCACT

TTACATTGAAGATTGTTGTTAGGTTTACTAGATCGTATAAAACCTTTCGACAGTAGAAAAAATCTTATGTTTATTTATTT

TTTGATGGGAAGAGAGAATTGTATTAGATAAAGAGAAAAGCAAGAACACAGAACTAAGAAAATTAAGAAAAAACAGTTTC

CCAGTTGTCGAGTGTGTAATAGAAGTTGATGTGCACTCTCTTCCCAAACACAGCTGCCCAAGACAGAACAAGACTTTTTG

CGTCATTTGCCAAATCTTTGTCAGTCTTAGTCAATTTTCGGTGTCAAAGTTGAGTTTATATTTTTTAAGATATATTTTTG

TGTTTCCAATATGTATATGTATATTTTCTCAAGTGCATGTATGAAAGTTTTTCTTTCTGTCTATTCAAGATAGATCTCCA

TATAAGAATAACCCCCATATAAGAATATTTTTTATCGATCCCTAGAATATTCTTATACAAAAGTTTTACTGTATCTGAAA

TTTTCACAGTCAGCTGGAAGCTTGGAACTCACTAGAGGTGGCCAAAATAAATTCCTAGATACCCACATATCACATATGAC

ATAATAAGATTAAAATAGCCCGGAGGTACCAAAAATAAGCTAGGGGATAGACGTCCACTACCCTATACATATCTCAATTT

ACTACTGTTCATCATGTTCGCATGTAGTGTATGTAATCTTTCATCACATTCTCCGTCCAATTAATAGGAAAAAGCAGACA

AATAACTAAATGTTGTTTATCTAAAGTAGGTAAAAATATATGTTTTTTTTTTTGTTTATTAGCGGAGATGATACCATTTA

ACGATCTGACTATCAGGATCCGTTATTGGTTTGTTTTATATGTAACAATACCGCTTATATTGGTAATTTGGTATACCTTT

ATAAATCGTGATTTTTGATGGGGTAAAAAGATCTGAAAATGGGTTATTTGTCCAAATAAATTTAAAACATGGTTCAAATG

GATAAGTAAAAATTATCATGGGTGAAATGGACACCAAAAAAATAGCAAGGATGAAACTGGATTCATCCTGGCTTAAACTT

AAAAAATAGCGAGGATGAAACTGGATGCATCCTGATGTAAATTAAAAATAAGAAAAAGAATTTGAAAATGGGTAGGATGA

AACTGGTTACATCCTGGCTATTTTTATATTTTTGTCCATTTAAACAGTATCAAAATCTAAATGTCTTTTTCACCCAGAAA

TTATTGATTTTGATCTTTTTAACCAATTTTGTGTAAAAATATTAGCTTATCAAGTACGCATAGTGTTTCTAGGGTACATA

TACGAACACCCGTTAGGCTGAGTCTAGTGGAGAAATTGGGTATGATGATGGTGACCCAAAATCGTGGTCTAAAACCAAAC

AAACGAAGTTACCGTTCTCTCTGCTCGAATGCAGGTAAGATGGGATAGGATTTAAATCCTCGCCAAGAAGAAGAGAAAAG

CGAACATAATCTAATAGTGGACCCGCTTTATATATTTAAAAATAATAAAACTTATAAAAAAAGAGTATCTTATCAATTTT

TACATTATATTTGAGGATTATGATTCATACACAACTTTGGTTGGTAACCGAGCAACGAGCTGGACGCCAGCTCCCTTTTG

AATAGCAACACCCCAGGGACGGAGCCAGGATTTTTTCACAAGGGGGGCGAAGATAAAGAGGACAAACACAAACCATTACA

AAATAGAAAAATAAACCACAGCTATTTAAGTTAAAAATGATCAAAATGTTTACCAATGCTATTTAAGTTAAATTGCTAAA

AAATTTAGCAATTTCTTTTTCGATATGCTCAATCAAAGAATCTCCAAGGAATTCATCACCCATTTTGCTGCGAAGTCTGG

TCTTGATAATTTTCATTGCAGAAAAAACTCATTCTACCGTTGCAGTAGAAACTGGTAGGGTTAGCACAAGTTCAATCACT

CGATGTATAAGAGAATATGTAACCGACCCAAGATTTTAATTTTAAATATGGAACAAAAAAAATCTAGTGACTAGTCCAAA

ATTTATAAATTGAAATTTACATTCAATTCAGTCATATCTAAAACCCAAATCAAACAAGCCTAAATGTTAGATATGATATG

ATTCATGAAAGACAAAAATTTAGTCTACAATTTAAAATTTACACTCAATTATTTAGAAACTAAATTGATAAAAGAAGTAG

AAAAATACCTTCAATGCAATCACTTTCTTATCTCCTCGTATAAAAATGAAATATTTGCAGAGATACGAAGTTTTTATGTT

GGTTTCTTGTTCACTTTCTTTCTCAATATTCTTTATGTAAAAAACAATATTGAAAATATCAAATCGGATCTCATATTCTC

AAGCAGATAAACTATAAAAGGAAACACGTTTTAGTAAGCCATAAATACGTTTTAGACTAAGAAAAGTCATACTCAAAAAT

ACAAAAACAGATCCCACACCCCACGTGTTTATACGAAACCTTAACAAATTTAAATTAGTATTTCTAAATCATAATTTGGG

GCTGATATTAGACTTACAATTTATTTTAGGGGGGCTGTATTAGATTTTTTCCCCAAAACTAAGAAATTTGACGACAACTA

ATGAAGATTTTCCGCCAACCCTGATGTATACACAGCTCCGTCCCTGCAACACCCAATTTATTAAAAAAAAAACTGCATAG

AACACTAGCAGCATCATTACTAACTAAATAATTTTTCAAACGAAAAAAAAAAGAAAGATTTATTTAAAAGTAATAACATT

CGTTAAAATGGCATATAATATATTTTGGGCGAACATAATTTTTTTGTCGATATCATAATTACCTCACTTACATTAGACTA

AAACCTCCCTGCAATGTAACTTGGCGCCACTATTCAGACACCACCTTACCTCCTCCACTACCCTTAACATTTAATAGTAA

AACGCGAAAAACGCTGAAGTTTTAACACTTTTCAAAGTGGAAACTACTTTTGTGAGTGTGATTTGTGATCATAATCGACA

AAAAAAAAATTAAAAGAAAATGTTGTGTTCACAAAAATGAAAAGTAGACAAATAATTGATAGTACGGACTAGATCATAAT

CGACAAAAAAAAATAATTAAAAGAAAATGTTGTGTTCACAAAAATGAAAAGTATGACAAATAATTGATATCATAATCGAC

AAAAAAATAATTAAAAAGAAAATGTTGTGTTCACAAAAATGAAAAGTATGACAAATAATTGATAGTACGGACTACCAACC

ACTAGATATCTATAAAGGGTAAATTTATTAAATACCCTTAGTTTTGACACTCAATAAATATATCCTTATATAACAATATA

AATACCTTTATCATTTAAATACTTATCTTTTAAAAAACAGATTACCTTAATACCCTCACTTACAATATTATTTTCCATTT

AAAAATCTTATCCATTAAAAAATAAATTTTCCTATTATTTCACATCCACCAGCATGAGCAATGGCTGCACCATTTCACCA

CCACCAATAGATCACGGATCCCGGATGGTGATAATCAAAACAAATCTTCAAAGAAACCAAACCGCCACCACCACTCCACC

ACCACCGCCACTCCGCCACCACCACCGCCACTCCGCCACCGCCACTCCGCCACCACCACCACTTACTAAATACAACCACC

AATAATATTCCACCACCGTCACACACCACCACCACTGTTGTTGCCGCCGCCATTACTATTAGGTGTCAACAACTGAAGTT

GATCCAAAAAATCAAAAGTTCAAAAATATTTGTCAAATTTGGTTGTTGATTCCATTTTTCCATTGTTGATACACAAACAT

TGGTTGTTGACTCCCTAAAAACTAAGGTCAATTTCTATTGTGGATACACAAATATTTGTTCACTAAGATCTATTTCTATT

GTTGATATACAAATATTGGTTGTTGATTCCATTTTTCTATTGTTGATACACAAACATTGGTTGTTGACTCCCTAAACTAA

GGTCAATTTCTATTGTTGATACACAAATATTTGTTGTTGACTTCTTACACTAAGATCTATTTCTATTGTTGATACACAAA

TACTGGTTGTTGACCACCATTTCTATTGTTGATACAAAAACATTGGTTGTTGACTCCCTAAACTAAAGTCAATTTCTATT

GTTGATACACAAATATTTGTTGTTGACTCCTTAAACTAAGATCAATTTCTATTGTTGACACACAAATATAGGTTGTTGAT

TCCATTTTTCCATTGTTGATACACAAACATTCGTTGTTGACTCCCTAAATAGTTGATGGAGTAGTATTCAACAACTTGTT

GTCCAACCTCACCACCACCACCGCCAGCAGCGTACAACATTTCCAACACCACAAACAACCAGAATCACCATTATCTTCAT

GACCAACTTTAATTAATCATTCCAACCACAACCAACACAATCTCCATCATCAAAATTCATCGTTCTAACCTCACCAACAT

CACCAACACCACCTGCACACAGAATATCCACCAACACCACAACAACCATAACCACCACTATCTTCAAGATTTAACGAACC

TAAATCTCAATTGGAATCACTATAAATAAAAAAGTTTCTGATGTAACCCCAAATTTCAAAACAAAAACAGATTCCAACAC

ATACCTCTTCCACCACCGCCCCACTAGAACCATCACCACCACCATCATCATTGCTTTTTGAGCTTCAATAATTCGAAGAA

CTTCAACTCTAAAATCATACACACCGCAACATGATGTCGGCTTCAATGAATCTAACATGCAAAGATCTCGTTCCATTTTC

AACTTGAAAACCAAAAAAATCGAATCTATCGATCCGAACGGAAAATCATACACACCCAATCTCCACCACCACAACCACCA

CAACAACAGCAATAACCACCACTATCTGTATCACCAATTTTATCATTCCGACCCAACCACCACCAAAAACAACAAAAACT

AGTTATACCACCACCGCCGCCATCACCAGGCACCAGCAGAAACTTAACTAAAGAAGAGAAAGAAGATAATCGATGCAATC

TCATTTTATCTCCATCACCAATTTAGAAAACGAAGGAATTTGAATTTGGATCTGTAATCCAATAAATAAATAAATAATAA

TCAATCACACTTCTCAATTTGTTTGTTCATAATCTGGCTGCGGCGGCTTCGACGATGATGGTGAAATTTTGGAAAACAAT

GATGTATTGTTGATGGTGGCGATGAAGATGGTGGTGGTGGGTTTAGTTTGGTTTAATTTTCGATCGAAGATCTCTCTCCC

AATCTGGATATAAATCTCTTCTATCTCACTCAGAAATGGTAGTGGTGCTGGTTATGGTGGCGCAGGACGACGTGGTGGTG

GCAGCGACGGAGCATGTAGTGGTAGTTGCAGAGGTGCGACGATGAAAACGATGATGGTGCTGATAAAAACGGAAAGAGAG

AGAGAATTGAATAAAAGAGGAGATGATGTAAGGGTATATTTGTAATGTTGTATAATCCTTAATGGATCCCTGATGGGCTT

TGGATGGAGAGGGGTATTTTTATTGTAGGATAAGGGTAGTTGTGTAGCCCGTCAGAATTGAGGGCAATTATATAATTCCC

ACATATATAAATGTATCGTAGCATTACCGGTTCCCAACTCTATGGTTGGTCCCAGAAAAATAGCATTACCGGTTCCCAAC

TCTCTCTCTATATATATATATAAATGTAGGGTTCTATTAACTTATATACCAGGGCATACGTAAGGGACATTAAATACTCT

ACTTTTTACCTTGGATTTAGGGATCATTTATAGAAAATGCAACAATCTCGGTTAATCTGATATTAAATACCAATGAAACG

TAAATACCTACTCTACCCTTATTAATTAATGATTATTTTATCCTGATAGGAAATAAGAGGCGGATTGAATCGTTTATTAC

AGTGGGACCTAGTCTTATGTTTAGCAAATAACTCCTAATTAATGTTGAATAAATTGTGTCCGCACATCTATGTAGGATGT

GTCTTTTTTTTTTTTTTTTTTTTTTGAAGCATGCATATATTATAAAATCAACTAAACCGAAATTACACGAGTGAATGTGA

TTCCCCTAGAATCGCTTAACAGAATATGGTTAAGAAAAGCCGGGTTCGCCCGGTCCCATATTGAGGTTGAGAAGTTTCCA

TCTTGGCTACCGTCTGCCGCATAGTTCGCTAGACCATCTGCTGCTTGGTTTCCTTCTCTGTAATTGTGCTGAATAGCACA

ACATGGAATTAGGCCCATCCGATGTTTAATTTCTGCAACCATTTCTGATATATACCAAGGGGCTTCTGTTGGAGTTTTAA

TGAAGTGTATGAGACTTTCAGAATCTGCCTCAAATAACACTTTAGACCATCCTCTTTACGTGGCAGTTCTAGTTGCTAAT

ACTGATTGCCCATGTTTCTGTTTGTGAGCGATGATGTTAATCCTAGAGGTTGAGCAAGAGCCATTATCGTTCGCGCATTC

GAGTCCCCGCAAATGAAACCTACTCCCGCCATACCCGGGTGGCCTCTAGCAGCTCCATCGGTGTTGACTTTGATCCATTA

CAGTTGGGTTACACCAACCTACTTGAATTGTTATGGTTGTTTTGGTTATGTTTGTAGCAGTTCGAGACGGCGGTTCTCCC

GGGAGAACTGGAGGGGACATCGTTAGAGCCCATTCAAATTCCTGTTGTTGTGCCAAGGATTTTTGCATAATATCTGTGAT

AGTGGATGGTGCTTGATTGAACACGAGTTCATTTCTTGTCGTCCAAAGTGACCATAACAGAAAACAACATGCCGTGATAG

AAGATGTATTGCCTCGAGTTTGGAGTATTTGTGATATTGTTGTTTCAGGGGTTATAGTAATTGATTGGTTGTTGCCTTGA

CGTTGGTTTGTTGATATTTGGTTTGAAAGGATGGTCCAAGTGGACATTGCTATCGGGCAGTGAATAAGTATGTGTTCAGA

TGTTTCTGGTGCGGAGTTGCAGCGGGAGCAAAGGTTGGTATTGGAGAGGTTGATGTGATGTAGTTTAGATAAGGTTGGTA

AACCTTCATTTATAGCTTTCCACAAGAAAAGTTTAATTTTTGGTGGACATGGTAAAGTCCATAAAAATTTGGTAGTTGGT

AGGAGGTTAGGGTGTGAGAGGTTCTCTTGTTGGATTAGACAACTATACCCAGATGCAGTGGTGTATTTTCCTGACTTAGT

GTGGGGGCAGATTATTTTATCCGGGGTGCTGACATTAGGAAGGTGGATGTTTATTATGTGTTGGATGGCATTAGGTGGGA

GGTTGTTTGGCGATTATGGCTCCAATTATGAGAAAATGGGTCAATTAACTCACTTCTGATTGTATAAGGGAACACTTTGG

GTGGGATCGAGATTTTGAAAGTGTGGTATCCAATTAGCTTGGATAGGGGTATCTAAACCATTTCCTATTTGGTGAAATAA

ATGTTGTTTTATCGACGGTACGAGAGACGCCATTTGTTTCCATTGTGGGCTGGCATTAGAAGGAACTGTGAGCTCACCTT

GGAATATTGGTTGATGTTTGAGGTACTTCTCATTGAAAAGTCCGGTGTAGATGGAATGTGGCTGGGCATGAATGTTCCAT

ATCCGTTTCATGAAAAGAGCTTTGTTGTGGTGGCTACTCTTTCTGATTCCGAGTCCTCCCTCTGCTATCGGCTTGTGAAG

TTTGTTCCATCCTATTGGGTGAAACTTTCTCACAGAAGAATCATGTCCCCAAAAAAAATTCCTAGCAATCCGAACAATTT

GTTTGTGTACCTGGGTGGGGAAAAGCTGTGTTTGCATAAGATGATTGGATGTTGGTGTTAAAGAGGTTTTAATAAGAACT

AGCCTCCCTGCCTGGTTAAGGCATTTTGTCATCCATCCTTGAGCTTTATGAGCCAGACGTTGGAGTAGAGGTTCGAATAT

GTGTCGAGAGTTTCTTCCGTGCTTGAATTGGACTCCTAAATATATAGGTGGAATTGATGTGGTTGACATGTCGTAAAATT

GTTTCAGAGAATCAATGTTATTCTGCTGTAGCCTCGGGTGGTGGATGATTGTTGATTTCGTTGTGTTAATTATTTGTCCT

GCTGAAGTGTTGTAAGATTGAAGTATCCTCTTTAGGTGATTGCTACTTTCCTGTGTAGCTTTGTACGTGATTAGTAGATC

GTCTGCATACATTAGATGTAAAATTGGTGGTGCCTTTTTTGATATTCGGAGACCTCCGACCATATTCCTCGCTTCAAGAA

TAACAAGATTTGTTGATAGAATTTCTGCACAAATAATGAAGATGTAGGAAGATATAGGATCACCTTGTCTGATACCCCTG

GTAGGAGTGATAAATCCATGAGGTGTGCCATTGATGTTGATCGAGTAAGTGACTGATTTGATACATAACATGATATACTC

CACAAATATTGATTGTAGGATTCAAATATCGCACAACATATTCTCTTAGCGTGCGAGATTCAATTATTTTCTTAACTCGC

GAGAGTGTCTCGAGAGAAGGTAGCAAGACTCGAGAGAATGTCTCGAGAGAAGGTAGCAAGAGCCTGCGAAATCATTAAAA

GGGCTCGCGAGAATATGAGCTGTATGTGAGCTGTCATCCATTATGTAAACACCTATATAAAGAGGAGAGCAGGAGAGAGA

AAAGGGAGGAATAATTTTGAGGAGACACATTTTGGGGAGGACACTTTTAGGAGGAGAGAGAGATTTTTAGAGAGAGATTT

ATTTTCTTTATCATCTTCTTCCCCAAATAATTAGATCTAGAGCTCTACAAAATATCCATTAATGGAGATTGTAAATCTAA

TTCTTATGTAACCTCAAAATCATAGTGAAGAATCGTGGATGTAGATCACATTGATCGAACCACGTAAAAAATCTTATGTG

TCTATTTAAATTTCTAGCACTTATATTTTGAGATTTAGGTGATTAATAAGGTTTAGATCTACGAAATTGAGTAAGGGGTG

TAGTGTGGGATTTCCTGCAACTACATATGCGCTAGAAACTGGGACCTAATCTTGTTGGTTCAAGAAAGTTTAGAAAAAGC

GACTCTTTTGTTGGTGATGAATTCAATTGGATCTATAAACTTTAAGAGAAATTTCCATGAAAAGTAGTTCTTGTTCATGC

TTTTTATTAGATCTATAATATTTGGGTTCAAAAATGGAAGTTTTATGGAAGAAATCACATTTTCGGGAGAAAACAACTGT

GATGAAATTCAAGGAAGAAAGCTAGTTTTTTAATGGAAGGTTTCATATTCAGGAGAAAAGAAAAAGGCAAATTGTTGAAG

GAAATAAAGGAAGAATGAAGAAAAGGATAAGAACAAAATCTTTGTGAATTTTATCACAAAAAGAAGCTCGCATGAGATGG

TTTGAGTTGGAGCGAAGAAGCTTGATGTCGCAAGATCAAATCTCGCCACAGTCATATTTTTTCAAATATTTTTGAACTTT

CAAGAGGGAAAAAATGGAATAAAACTATAGCATTGTGTGTTTCCTTCGCAAGCGACAAGTAGCCCGGATGGTCATTCGCG

AAGAGAATAAGCTTGGGGTTGCGGTTCGAATCCCACCTTTTACATATTTTTGAACTTCCGAGAGCAAAGAAGTGAATAAA

GTTATAGTATTGTGTGTTTCTTTCACAAACAGACACTAGCACAATTGGTCACCTGCGAAGAGTATAAACATAAGGTCGCG

AGTTCGAGTCTTACCCTTCGCCAAGAATTTTTGTCATCTTTTTCCTTGAATATCAAAGGGTGAATATAAGTCATGTATTA

GCATTCTGCGATCCTTTCGCACAGGAGGAGTAGCACAATTGGTCACTGGCGAAGAGTATAAACATAAGGTCGCGGGTTCG

AATCCCCCACCTCGCAAAGAATTTTTTGCTTCGCAAAGAAGGGAAAAAATGGATAATTTCGCAAGGTTGTTTCATTCTTT

CGCAAACAAGAGGCAGCACAGTTGGTCACCTGCGAGGTTAATGAGTTCATGGTCGCGAGTTCAAGTCCCATTGAACGTCC

ATTTTTGCATGGATTTTTCGCAATGCAGTATCTCGCAAGAAGAAAGTAAGTGATTATTTCGCGAGAAACATTTCGCAGAG

GCAGGGCTAGCACAGTTGGTCAGGAGGCATGAATGCAAGCAGGAGGTCGCAGGATCAATTCTTGTCTTTCGCACATTTAT

TTTTGTTATGCGAAATTTCGCATGACTTTATTATCTCGCAAAGAGGAAAGTTGATGATTATCTCGCAACTTAATCTCGCA

AATGAGGGTTAGAGCAGTTGGTATGGTGCGAGAATATACAAGAGGCGGGATGCTAGCTTCGCAAGCAAATCATATACTTA

GCAAGAAACAACAACAACAAACCACAAAAGCTAAGTACCACTTTCCTTAAATATTTTACTTACAAAAAATAAAAGATTTT

GATTTGATTTACAAAAGGCGAGATAATCGCAGAACTACAAAGATTTCACGATTACAATAATTCGAGAAAAATCTCGCAGA

TATTACAAGAGAAACCGAGAACAATATCTCATAAAGATCAATTTATATATTTCCCTTAAATACTCGCTTTGTTTCAAATG

AGAATGATATCCAACATGGAGAGTAGTAGCGGGCAAACCTTTATGGCGTACACCCTAGTTCGCGAATTTACTCTAGGATT

TCGCAAGAGGCAATGTACCTAAAGTGCGTCATATAAGGGGTACCTATTAGGATCATTTCCATGAAGGATGTTTTATCGGA

TGAAATGAAATGAGTTATACAAAGGAAATCAAAACAAAGAAAAATGATTTGAACTTGAAATTAAATAAATTTCTTTGATA

AAAGAAATGTGGAATACTAATCTAAATTTTTATTTGCATTACAGCATAGCATGAGGCAAAGAACATGGAGCGGGAGTTTC

GCATGATGGATTCACAATACTTCTTATATTTAGAGGATTAAGTACTTCTTATACTTCTCAAAGGATACTTCATACTTCTT

ATATATTTCGAGGATTGAGTACTTCTTATACTTCTTCAAGGATACTTCATACTTCGCATACTTCAAGAGATAATACTTCT

TATTCACTTCGCAAAGCTTCGGAACTTCGCAAAAGAATAGAGGCAAGTACGTACTTTTCAAGTCCAGTATTCTTCCGCTC

GTTCCTTCATCAACAAAGATCAAAGATTACTACGACAACGGATTTTTCCTTAAAGATCGCTCTTCAAGCAATATTCAAGG

AAAAAGAGGCAAGATGTAGGATTCAAATATCGCACAACATATTCTCTTAGCGTGCGAGATTCAATTATTTTCTTAACTCG

CGAGAGTGTCTCGAGAGAAGGTAGCAAGACTCGAGAGAATGTCTCGAGAGAAGGTAGCAAGAGCCTGCGAAATCATTAAA

AGGGCTCGCGAGAATATGAGCTGTATGTGAGCTGTCATCCATTATGTAAACACCTATATAAAGAGGAGAGCAGGAGAGAG

AAAAGGGAGGAATAATTTTGAGGAGACACATTTTGGGGAGGACACTTTTAGGAGGAGAGAGAGATTTATTTTCTGTATCA

TCTTCTTCCCCAAATAATTAGATCTAGAGCTCTACAAAATATCCATTAATGGAGATTGTAAATCTAATTCTTATGTAACC

TCAAAAATCATAGTGAAGAATCGTGGATGTAGATCACATTGATCGAACCACGTAAAAAAATCTTATGTGTCTATTTAAAT

TTCTAGCACTTATATTTTGAGATTTAGGTGATTAATAAGATTTAGATCTACGAATTTGAGTAAGGGGTGTAGTGTGGGAT

TTCTGCAACTACATATGCGCTAGAGCGCAGGACCTAATCTTGTTGGTTCAAGAAAGTTTAGAAAAAACGACTCTTGTTGG

TGATGAATTCAATTTGATCTATAAACTTTAAGAGAAATTTCCATGAAAAGTAGTTCTTGTTCATGCTTTTATTGAGATCT

ATAATATTTGGGTTCAAAAATGGAAGTTTTATGGAAGAAATCACATTTTCGGGAGAAAACACTTTGTGATGAAATTCAAG

GAAGAAAGCTAGTTTTTTAATGGAAGGTTTCATATTCGTGGAGAAAAAGAAAAAGGCAAATTGTTGAAGGAAATAAAGGA

AGAATGAAGAAAAGGATAAGAACAAAACGTTGTGAATTTTATCACAAAAAGAAGCTCTCACAGATGGTTTGAGTTGGAGC

GAAGAAGCTTGATGTCGCAAGATCAAATCTCGCCACAGTCATATTTTTTCAAATATTTTTGAACTTTCAAGAGGGGAAAA

AATGGAATAAAACTATAGCATTGTGTGTTTCCTTCGCAAGCAGCAAGTAGCCCAGATGGTCATTCGCGAAGAGAATAAGC

TTGGGGTTGCGAGTTCGAATCCCACCTTTTACATATTTTTGAACTTCCGAGAGCAAAGAAGTGAATAAAGTTATAGTATT

GTGTGTTTCTTTCACAAACAGCAGCTAGCACAATTGGTCACTGGCGAAGAGTATAAACATAAGGTCGCGAGTTCGAGTCT

TACCCTTCGCCAAGAATTTTTGTCATCTTTTTCCTTGAATATCAAAGGGTGAATATAAGTCATGTATTAGCATTCTGCGA

TCCTTTCGCACAGGAGGAGTAGCACAATTGGTCACTGGCGAAGAGTATAAACATAAGGTCGCGGGTTCGAATCCCCCACC

TCGCAAAGAATTTTTTGCTTCGCAAAGAAGGGAAAAAATGGATAATTTCGCAAGGTTGTTTCATTCTTTCGCAAACAAGA

GGCAAGACACAGATTGGTCACTGCGAGGTTAATGAGTTCATGGTCGCGAGTTCAAGTCCCATTGAACGTCCATTTTTGCA

TGGATTTTTCGCAACAGTATCTCGCAAGAAGAAAGTAAGTGATTATTTCGCAGAAACATTTCGCAGAGGCAGGGCTAGCA

CAGTTGGTCAGGAGGCATGAATGCAAGCAGGAGGTCGCAGGATCAATTCTTGTCTTTCGCACATTTATTTTTGTTATGCG

AAATTTCGCACAGACTTTGCTTATCTCGCAAAGAGGAAAGTTGATGATTATCTCGCAACTTAATCTCGCAAATGAGGGTT

AGAACAGTTGGTATGGTGCGAGAATATACAAGAGGCAGGATGCTAGCTTCGTAAGCAAATCATATACTTCGCAAGAAACA

ACAACAACAAACCACAAAAGCTAAGTACCACTTTCCTTAAATATTTTACTTACAAAAAATAAAAGATTTTTGATTTGATT

TACAAAAGGCGAGATAATCGCAGAACTACAAAGATTTCACGATTACAATAATTCGAGAAAAATCTCGCAGATATTACAAG

AGAAACCGAGAACAATATCTCGCAAAGATCAATTTATATATTTCCCTTAAATACTCGCTTTGTTTCAAATGAGAATGATA

TCCAACATGGAGAGTAGTAGCGGGCAAACCTTTATGGCGTACACCCTAGTTCGCGAATTTACTCTAGGATTTCGCAAGAG

GCAATGTACCTAAAGTGCGTCATATAAGGGGGTACCTATTAGGATCATTTCCATGAAGGATGTTTTATCGGATGAAATGA

AATGAGTTATACAAAGGAAATCAAAACAAAGAAAAATGATTTGAACTTGAAATTAAATAAATTTCTTTGATAAAAGAAAT

GTGGAATACTAATCTAAATTTTTATTTGCATTACAACATAGCATGAGGCAAAGAACATGGAGCGGGAGTTTCGCATGATG

GATTCACAATACTTCTTATATTTAGAGGATTAAGTACTTCTTATACTTCTCAAAGGATACTTCATACTTCTTATATATTT

CGAGGATTGAGTACTTCTTATACTTCTTCAAGGATACTTCATACTTCGCATACTTCAAGAGATAATACTTCTTATTCACT

TCGCAAAGCTTGAACTTCGCAAAAGAATAGAGGCAAGTACGTACTTTTCAAGTCCGGTATTCTTCCGCTCGTTCCTTCAT

CAACAAAGATCAAAGATTACTACGATTAACGGATTTTTTCCTTAAAGATCGCTCTTCAAGCAATATTCAAGGAAAAGAGG

CAAGATGTAGGATTCAAATATCGCACAACATATTCTCTTAGCGTGCGAGATTCAATTATTTTCTTAACTCGCGAGAGTGT

CTCGAGAGAAGGTAGCAAGACTCGAGAGAATGTCTCGAGAGAAGGTAGCAAGAGCCTGCGAAATCATTAAAACGGCTCGC

GAGAATATGAGCTGTATGTGAGCTGTCATCCATTATGTAAACACCTATATAAAGAGGAGAGCAGGAGAGAGAAAAGGGAG

GAATAATTTTGAGGAGACGCATTTTGGGGAGGACACTTTTAGGAGGAGAGAGAGATTTTTAGAGAGAGATTTATTTTCTT

TATCATCTTCTTCCCCAAATAATTAGATCTAGAGCTCTACAAAATATCCATTAATGGAGATTGTAAATCTAATTCTTATG

TAACCTCAAAAATCATAGTGAAGAATCGTGGATGTAGATCACATTGATCGAACCACGTAAAAAATCTTATGTGTCTATTT

AAATTTCTAGCACTTATATTTTGAGATTTAGGTGATTAATAAGGTTTAGATCTACGAAATTGAGTAAGGGGTGTAGTGTG

GGATTTCCTGCAAGAGATAATACTTCTTATTCACTTCGCAAAGCTTCGGAACTTCGCAAAAGAGTAGAGGCAAGTACGTA

CTTTTCAAGTCCAGTATTCTTCCGCTCGTTCCTTCATCAACAAAGATCAAAGATTACTACGATTAACGGATTTTTCCTTA

AAGATCGCTCTTCAAGCAATATTCAAGGAAAAAGAGGCAAGATGTAGGATTCAAATATCGCACAACATATTCTCTTAGCG

TGCGAGATTCAATTATTTTCTTAACTCGCGAGAGTGTCTCGAGAGAAGGTAGCAAGACTCGCGAGAGTGTCTCGAGAGAA

GGTAGCAAGACTCGCGAGAGTGTCTCGAGAGAAGGTAGCAAGGCTCGAGAGAGTGTCTCGAGAGAAGGTAGCAAGACTCG

AGAGAATGTCTCGAGAGAAGGTAGCAAGAGCTGCGAAATCATTAAAAGGGCTCGCGAGAATATGAGCCGTATGTGAGTGT

CATCCATTATGTAAACACCTATATAAAGAGGAGAGAGGGGAGAGAGAAAAGGGAGGAATAATTTTGGGAGACACTTTTAG

GAGGAGAGAGATATTTTTAGAGAGAGATTTATTTTCTTTATCATCTTCTTCCCCAAGTAATTAGATCTAGAGCTCTACAA

ATTACCCATTAATGGAGATTGTAAATGTAATTCTTATGTAACCTCAACAATCATAGTGAGAACCGTGGATGTAGATCACA

TTGATCGAACCACGTAAAAAATCTTATGTGTCTATTTAAATTTCTAGCACTTATATTTTGAAATTTAGGTGATTAATATG

ATTTAGATCTACAAAATATAGTGAGGGGTGTAGTGTGGGATTTCCTGCAACTACATTGATGGAAATCCCAGTTGTGTAAA

AGCTTTTTTGATGAATCCCCAATCAAGGCTGTCATAAGCTTTCTGAATATCTAGCTTTAGAGATATCTTGGGGTCTTTAG

TTATTGGAGACGTTTTAATGTGGTGGAAAAGCTCTCTGGCAATTGCTATGTTATCATGTATTGATCTTTGAGGTACGAAG

GCACTTTGGTATGGGCTTATTAAGAATGGGAGAAGGGGGCGTAATCGATTGACAAGGATTTTGGATATGATTTTGTAACT

GACATTACATAGGCTAATAGGTCTGAATTGGGATGGTTTTGTTGGATTGTCGACTTTTGGAATGAGTGTTATGTTTGTGT

GATTCATGAGAGGGTGTATGTTTAAAGAATTGAAAAAACTTCGTACCATAGCTATAAGTTGTAGGTGAGTGGTGTCCCAG

AAAACTTTAAAAAACTTACTGGTGTATCCGTCCGGTCCCGGAGCGCTTTCGAAATTAATGGAGAACAGAGCTTGCTTGAT

TTCTTTCACAGTAACTTCCGCAGAAAGACTGTCACATTGTGCCGATGTCAATCTTGGAGTTATTTCCTCAAAGAGTGTTT

CATCTATCGTTGTATTTGGGGCAGTGTATTGTGAAGTGAAATTGTCCAGGATTATCTGGATGACCTGGTCTTTATCACTG

ACCCAGTTGTTTTGATGATCTTGTAGTTGTTTTATGTTATTACGGCTTCGGTGGATGGTTGCCTTTGTGTGGAAGAAAGT

CGTATTATTGTCCCCCGATTGAAGCCATTGAATTCTGGATCTTTGCTGTCAATATGTTGCATGGCATTGGAGTAATGTGA

TGTGTTGACCGCGTAGCTCTTTGTCGCGTTGTATCCAGAAATTTAGTTCCTCGCCGGTGTGAATAGCGCATTGATGTTGA

GCCCTTTGAATTTCCGCTGTTGATTTTTTAATTAATGTTGGTAGGTGACCAAAAGTGTTCTTGTTCCATTCCATCAGAGT

TTGTCCCGTTGTGATTAATCTTAGTTGAAGGTCTAGAGCAGAGTCTGAGTCACTTTGTTGCGTCCATGCTGATTCGATAA

TCTGCTTGAGTTGTGGATGTGTTAGCCACAAATGCTCAAACTTGTAATTTTTTTGTCCTTGCCAATCGATTGCTTTAGTG

TGCAGCAGGATGGGAGCATGATTTGAAGCAATCCTCGGGAGGTGAGTAACTCCTGCATGAGGATTTGTTGTTCGCCAATG

TTGGGTTGCAATTGCCCTATCAAGCCGTTCCATTATATTGTCTTGACCCTGTTGGTTGTTTGTCCAGGTGAATGGGTCCC

CTCGAAAACCTAAATCTATGAAGCCCATTTGATCAATCATCGTTAGAAATGGCTGGGTTTGACTAAATGTGATCTGTCTT

CCTCCCTTTTTTTCTGATTGATCCATTATGTCATTAAAATCTCCTATCAGAAGCTATTCCGTTGATATGTTGATATTGGG

GAAGATTGTTGGGAAATCTTGCCAAAAATCCGTTCTTGCATCTGGGTGAGGAGGACCGTATATACCTGTGCAGAGCCATG

GTTGTGCAGGGGGTGGAGGTTTGATAAGAATGTGGATGTAACTTTGGGACTGAAGTAACACTTGAATGTTTAGGTTGTTC

CACATGAGACAGAGTCCTCCTCGTCGACCCACCTTTGGAACAGAGGTTAGGTTTTGAAACTGAAGAGATTGAGCAAGATT

TGACATATGACGATCATTGGCTAAAGTTTCAGAGAGAAATAAGATATCGGGCTTATGTAATGTTACCAGTCCTTTTAGAT

GTTGAATCGTTGGTTGATTGTTGATGCCCTGGCAATTCCAGGCTAGGATATTCATGATGTGGTTTAAATAGTTTTGGGGA

ATTGGAGAAGGTTTAGTTCATATAATTTAAGTGATTGTTTTAGGTGAGCGAATTAAAAAGGAACAAGGATATTCATGATG

TGGTTTAAATAGTTTTGGGGAATTGGAGAAGGTTTAGTTCATATAATTTAAGTGATTGTTTTAGGTGAGCGAATTAAAAA

GGAACAAATGAGGGGTTGAAGAGTATTTTGAAATGGGGTAGAACTAAAACTTGAGGAGAAAAATCAAATACAGAGAGAAT

TAGAAATTTGAGGGTTAACTGGTAGACTAATAAGGCTAAGGCTCAAAGCATGATTAAAAAAAACAAATCTAGTTTAATTG

GGAAAGGAATTTTGGAACGCTGGAGTGCCATACTTGCACCGAAAATGAAACTCGCATGAACATTGTTTTTCATAAACCAA

ATATTTTATTCTTCTCCATTTTATCTTTCGAGTTTGATGCGATTTTTTCCCATTTATTTCGTTCCGCATATTAGGTTTTA

AATTGTTAACTATCAAAATTCGGGATATCTAAAAATATTACAGTAAAGATTTCTAGAGAGTATTTTAGGAGCCCTTAACT

TTAGTTGTTTATATGTCTCGTGGATTTTTCTTGGACAAAAACAATTATCAAAACAATAGGATTTTGTATCCAGTTAAGTT

AATAATTCAAAGAAAGCCCATAATCTTCACTAATTTATTTAGCCAAAGACCTTAGTCATCATGTGGGGCATACGTCAAAA

CGCATATTCACCAAATTTTCTTTCTGATAAATCATATAAAAAGTAAATAAGGGTAAAAACGGAATAGATGTTTTCTAATT

AACATGAAAACATTTCCCCACGATAACTATCTTTCCAATAAAAGTTATCAAATTCTAACATAAAAAAAAAAGTTATTAAT

GAGATTTTACCAAATTTCAAGACAAAAAAAAGTAATGATTGAATTCTTAGCACGTAGTATTATACTCCCTCTGTCCATTT

TTATCTGTCCTAATTTCTACTTTGGTATGTCCATTTTTATCTGTCCGCTCTGTTTATAGACATACTTTTCCCTTAAACTA

TCAAATATACCCTTTCTTTTTTATCATTATAAAATCATTTTTAATATAAACTTAATTCAAAATCATAATATCATCACCAT

ATATAAGGAAAAAATCCATTCAATATTTTTTTATACTAATTGCCATTCCTTTTTTAAATAAAAACATTTATGGGTAGTTA

CAAAATTCCTAATACTTCTATTTATTCTTACATGGCAAAGAATCAAAAAAATTTATAACAAAATCTCAAATTCATGGTAG

AGTATATCTAAAGAGGAAATAATCTACAATTTCAATTTTTTTTCCCTCTCAAATAAAAATTCCAAAGGCTCCAAAGTTTT

AGTACGGGAAACCAAAATCAAATGGAGGAAATAAAAAATTCTTAAATTTTGGAGTATAAGTTTGCTTTTATTTCTTTATA

ATATTTCCTAAAACAATAATACTATCCAAATTAGAATTTTTAACACTAATAATTTTAATTTTGGTAATAACGTAATATTA

AATAGGCAAGTCAAGGGATAGTCATGACATTTTATAGGATTTCCTTAATATCTTGACAAAGTCAAAGCGGACTCTTAAAA

ATGGACGGAGGGAGTACAAGTTAATAAGACCCATAAATATATCGTTAGCATTCCTATCGCCAAAACACCAACTTGAGTAA

TTATTTAGTGGAAACTACTGTATTACCGGTTCCCAACTCTCTCTCTCCAATTTCATTACCGGTTCCCAACTCTATCTCTC

CAATTTCCAACTTGAGTAATTATTTAGTGGAAACTACTGTACGGTTGGTCCCAGAAAATAGCATTACCGGTTCCCAACTC

GGATAGACTAAGAAAAGTCAAACTCAAAAATACAAAAACAGATCCCACACCCCACGTAAACACGAAACCTTAACAAATTT

AAATTAGTATTTCTAAATCATACTCAAAAATACAAAAACAGATCCCACACCCCACGTAAACACGAAACCTTAACAAATTT

AAATTAGTATTTCTAAATTATACCCTCCGTTTCTTTTTAATAGACAAGTTTGGTTTTTAAAAAAATTAAGGAAATTAAGA

GAGCCAATCATTGAAAGTGGTCTCTCATGATCTTGTCGATAAAAAAGTGAAGTGAAAATGGTCCCATGCACTTGTCGTGA

AAATAAGTAAAGTGAAATGGTCATGACACTTGTCATCAAAAGAAGTTAAGAGAAAAATGGTCCCACAAAAATAAAGTAAC

ATTTGACTTTCCCAACATATTTTTTTTGTAACATTTATTTATAGAAACCAGCCTATTATTATTTGACGGAGGGAGTAATA

AATTTATTATTTAGGGGGCTGATATTAGACTTACAATT

>Par_flye_v1-0_contig_40428_minus contig_40428

ATGGAGAAGGTAAGGGAATTGGAATTATCTTGATTTAACCAAATGAACAAAGATGGCTTTTTCTTTTAGATCGAGTTCGG

GATTTTACTAACAATGAAACGGAGTACGAAGCAAAGGTGCATGCTTTACGACTCGCAATAGAAATGAAATTAGAAAACGT

AAGGATCACTAGCTATTTGCAATTAGTCATTTGACAAATAAAGGAGAATACACTACAAATGAACCCTCATTGAAGAAGTA

CAAAAGGCTTGTTGGGGAGTTGTCAAATAAAAATCCCAAAATAAAATGGAGACATATATCAAGGAGGGATAATAGGTTAG

CTGATGCTTTTGCTTTTATCTCACATGATGGTAGATCCGACCGCAAGATACATAAAATTCATATTTCATTTGTTTCAATC

AATCAACAAAAGAAGAAGAAGAAGAAGAAGTAGATGTGATGATAGTAGAAGGTGTGGAAAAATGAAGAGGCAAATAAAAT

CACACATTGGAAAATTGAACTTCATGCATACTTGGAGAGAGGAGAAGTACCAAGGAATAGATTAGAGGCACATAAATTAA

AGAGTCGTGCGACAAATTATGAACTCGTAGACGGGTATTGTATCGAAGGTCATTTAATGGACCTTCTACGAGATGCTTGA

CACAAAAAGAAGGAGAGGAAATACTGAAAGCGTTACATTATTGAGATGCTGGTAATCATAGTGGAGGAAGATCTTTGGTA

TACAAAGCGAAAACACAAGGATATTATTGGCCAGTACATGCATGAGGATGCAAAAGAAGTATCAAAACGTCATAAGAGTG

CCAAATACACGAAGAGAATACATGCACCAGAGCACTTTTATCATCTTCAACAAGTCTTCGGCCTGAAAATGGGGTCTAGA

CATTGTGGGACCATTTTTACCAAAATCAGGCAAAGGAGATATTTTATTGTCGTAATGCACTATTTCACAAAGTGCGTAAG

TAAAGGCAAGACAAAGATATACGTTAATAAAGACGTTTTCACTTTTATATTCGAAACATCATTTGTAGATTTGGAATTCC

TATTAACCGGTGTCCGAGTAATGGAAAATAGTCAAGGGCGTAATATAGAAATGTTTTAATGCATTTAAGATAAAGTGTGG

GAAATCCACACCATTATATCCACAAAGTAACTGATAAGCTGAGGAAATAAATAAGACAACAGCGTATACATTGAAGAAGA

AGTTGAGGGGGGCATAATAAAGCATGGTGTGAATGTGTACCTAATGCATTTGGGCTTATAACACAACTAGAGAGAAGCTA

CGTGAATGTCACCTTTCTCAAAAACATATGGAGTCAAGCGATGTTACCAACCCAAGTCATTATACCCACTACAAAAAAGA

GAAGCATGGGAGAAACATCTATGGGATATGATTTTATCAAAGCTTGATGATTTAGAAGAAAAGAAGGAAAAAGCCATACA

ACATATGGAGAATTATCGTAGAAGATTAGCTCGAGAATACAATAAGTGAGTCAAGATAAGAGAGTTTCATCCAGTGATTT

GGTTCTACGAGAAATACCGCCGTATCGAGGAAATGATGGTAAACTAGAGAAGAAATGGGATGGACCTTATATCATTAAAG

AAATAGTTAGAACAGGGGCATTTAGACAATGGATCCCGAGGAGGGAATAGACACTCGGTCATAAGGTTGATAGGCCATGG

AATAGATTATATCTAAAGAAATATTATCCATAAAATATTTGGCTAAAATCACCAGAAGTAGAGAAAAGAAGAAATTTCAT

AGTTATGAAATTATATTTACAAATGAAGAATGCATGATTTCGCGTAAATTATTTTTATCTAAACAAGTATGCTATAATTT

CGCATAAAATGTATCGATAGAACAGTGAAAGATATGCGATAATTTCGCTAATTATGAATATTACAAAGAAAATAAAAGAA

AATATGACAATGAACAAAAATAGAGGCACCAACTTTGGTGTAGACCCTAGTTCGCGAATAAAATTTCGCAAGAGGCAAAT

GTTAGACCTAAAGTGCGGTCATTGAAGGGGAGACCTATTACATGGCCTGTGGAAAGGTTACACCGCCCCGAACTGGGTCA

TGGAGACTTGGGGTTTTTAAACCCATCCAAGAGAGGCGCCTTGGATTCTCAGCTAAGCTTATGACTTAGGTCAAGGACTA

AGGTAAGAAGGTCTCTTCTAAGGAGTGCTAGAACTCGATCAAGGAGCCGAGGCACACGGGCTGAGCCTTTGAAAGTGCCT

GGGGCGTTGAAGCGCCTCGCCATTCTTGGCAAGTCTTGGCTATGATGCACTCGGCCCAAGAAAACTCCACTTAGGTGCGG

GCTCAGTAACCATGAGCCATACGACGAAGGATAAAGGACGCAAAACAACTCGGGTTGTTGTAAAGAAATCGGATGAGAGG

AGCCAACCTTGTATGGTAGAGGTACGCCTCCTCAAGGGCAAACTGTGATGGATGATAAGGGCGGCACTCTCCATTAGGAG

AGCCGATAAGTGTCTAAAGACGCAAATGTCATGAAACTTTTGACTCATGGGAGTATTAAGGCTTGTCATATTTGTGCAAC

GATCCTAGAGGTAAAATACTAGAGAAAGATAAGACGAGATCAATGTGGGTTGATGCACGAAAGTGTAAAGACGCCCGCGA

TTTCGCTACGCAAGATGACAATATCATGGGGCTTTACGCTCGCAAGAATATTTAGGCCTCATATTGTCGCATGCAAAATA

TCCCGCTGAGGCAAAAATACAAGAAATGATAAGACAAGATCAAACGGGTCAATACATGAAAATGTAAAGACGTCCCGCAA

TTTTGTCGAAATATAAAAAGTGGCAAGATAAGACCTTTCTGTAGAAGGCAAAATAAGACCACATAACAAAACGAAATTTT

GCAAAATTCAAATATATATTTGATAGCTAAATTTCTCAGAAAATAAGAACATGTCTCATATATTGACGTCCAAATGAACT

TATACAAACTAACCATTTAAGTATTGTAGAGACAAATTCGGCCAAGATAGATAATGAGAGGCAAGTATGGGAATTAATAA

ACAAGATGCATTATCTATCTCATTGATAAGGAGTTATTAAAAAGATAAGTGAATTTCAAGATTCATGAAAGTACACCGCT

TACGTAAAAGTAAGGCGAAGATAAAATAACTTAAATAACTAGAAAAATTTCGTGATTTCAACATGGATTCGCTAAAAGGG

GCAAAATGGCCTTCTAAAATCATTTCATCTTCACAAGCCTTTTTTCTTTTTTTGTTGGTCCATCCTAGCTTGTTCTCCAT

CTTGATTATCCTCAGCATTAACTTCCTTGATGGGCCATTTTCTTCATTTTCTTCAAGAATAGCGCCATCATCACTTTCAA

GTCAAAATCACTATCATGATCACGAGGAACTTCATCAAAGATCTATGTCAAGAGGCATAATTTTTGTAGGAGGAAGTGAT

TTATCTAGTAAAATTCCATTCACAAGAACTTCATCCTTGAGATGAACTTGGCGAAGAAGTGCTCTCTCGGGGTGATTTCT

ACTTTCAATACCTTTGAAATGTTGAAGGTAGTCAAATCTTCTTTTCCGCACGATCACGAGAATGAGCAAGGGAAGTAGCA

GTGTAGCATTTTCCTTGCATGAATCTTGGATAATTTTGATTGCAAGGAAGATATAGAAGAATGAAGACTTCTCCTTTAAT

CACGAACAATAAGAACACAAAAGATTGTTATAGCGAAATTCACAAAAGAAGAGTATTGCAAGATAATTAAGCGTAAAAGT

TGGATGAGGCAAATAGATTCTTATGACTACCTTCCTTCATAAATAAGATATTGGCCAAAGCCAAATTATTATTCTCCAAG

TTGTTCATCAGCATCATCAAAATTATCCCCAATTAAACCTTTCAAAGTAGATCGAAATTTCTTATCATTAGCGATAAAAG

CATTTTCCTTCGCTAAAGAAGCATAGGAATCTTCTAAACAGTATGTTGCTCTTTAAGTTGGTTTGCCTTTTACGCCCAAA

GCTATTCTACCTTGATGAGTTTATTCTTCAAAAGAATGGACGATCTGGCTTACTTAAGTTCACTTTTAACGAGCGAAGAG

ATCGCTCTTGTTCATCACATATTTTTTTGACGTATATTAAGTTGTTGCGTAAGATATCGCTACTTTGTTTTAGGATTCGG

ACAAGGTTCCATTTTCATCCGACAAAGTTTCAATCTTCAAGTTAGCTTAATTGAACAATAAGAGATTGGGATAGCTTGCT

ACTTGATACTTCTTGTTCTTGGATTAGTTGGGAATTCTCATTAGAAAGGTTGTTAATTGATCAACAGAGTAATGAGTAAA

GATTATCTAATTGGTTGTATTGGTTTAACAAAGCTTCATTGTCGCACGAGCTTCTTCAGCGGCATATTGGAAAGTCGTAT

CTCTCCATTATTCGTTTCAATTGTAACTTAACGCGAAACGTAAAGTGAATACAAAAGAAGTAAAACTAACAAAGAAGATA

AGTTATAAATGAAACAAGAAGATCATACCTTTCAAGCCATCATTTTTCTCAAGATTCTCGTAATCTCAGCGAAACATTCT

TAAGAGTTTGATTTTTGAGACGAAGATCATCACATTTCAAAGTCATAGCTTGAGATTCTCCTTCAAAGATTCAATTTCTA

AAGATGCTTCATGGGTTGTTCTTTCAAAGATTAAAATATTTACCTAGGGTTTCGTACGTTGCTTGGAGTGAGGGTCAAGA

AGATCTTCGATCTTCATCAAGAACATCGACAAAACTCTAAAAGGCAATGTCTACTCCTTCCAACAACTATATCTTAACAG

AAGGGTTTTATGAAGAATTAGTGATAATTGAAGGTATGGGGAAGTGAATCATCTTTCGCAATGGATATCTTTTTTTCCAT

GCGAAGAAGAAAAGGTTCTCAGAGGCTTAGGCTGGACAGAATCAATCTGCGGCATTCGGCAAGAGGAGAAGAAGGGGTTT

CATTCGAATAAGGGTCAACCTGATAGGACGACATCTATAATGATATCCTCATGGAAGGATGCAAGTGCAAGTGGAAGGAA

GATTCAATCGGGCGGCTTGCAGGAGAAGCAGGCCAAGGGTTTGAAATCAAGAATGAAACTAAGAAAAACAAAGCTTTAGT

TTCGAGATTTCTTGTAGGGGTTTGTTAATTAAATTTCATCATCCTCATAATATCAACTCAAATAAGAAAAACAAAGGCAA

CAATATTGTTATTAAAGAAGATATGTTTCTTATTACCGTGAATTCGCAGTTTTCTTTGACCACACACAAATAATTTGCTA

AGAGGTGCTTTTGCTGAAACAAAATCAATATGGGAATTAGACAAGATAATAAAATTAGTTAAAAACCTGGCGAGATTGGG

GTATACCCAAACTAATTGGGGTATACCCAATGAGAAGGGGACATGGTGTGTCATTATTAAAAGTGTGATTTTACTTTTAA

TACCCTTGAAATAATTAAATCTTAAAATAAATTCATTTCTTAATTTCATTAACCATAAAATTAAAAACTAACCAACTATT

AATCTTCTTCTTCTTCATCTTCATCTTCCATCAACCAAACCCATATAAAAATCTCAACTTTGATTTTCCAAAAAATGTTG

ACTCACAAATGGAAAGAAGACTATAAAATGATGTTTAAACTTAAACCCATCATTGTGTTGTGTGATTTGATTGTTATATC

AAAGTTAGAAATCGAAATTTATGATTTCCTCTCGGCCGGCAAGGTCGTGACATGCATATCTTGCCGACTTTTCATACTAG

CTTCATGGCGCACCTGATCAGAAAAATTTCAATGTTCTAAGCAGTGGTATTTTGAAGACCTTGTTGTTAACAATACAAAT

TAAGGGTATTTTGAATACCTTGCCTACTAAAAACAGCCTAAGGGTATTTTTGAATACCTTGCCTGGCCAATAAATTTTTT

TTATGTCATCTCCGTATGTCAAAAAACAAAAAGTCTGCAAGGTCGACGATGAAGATCATGCCGCCACTTTTGGCTGGGCG

AGGCCTGGGCGATGAAGACCTTGCCGACTTTTCATACTTTAACTAAAAGTGTTGATTTATTCTTTGATTTTGAGTATGAT

TTCATATAAATGTATTTGCAATCCCCTTTTTTAAGTATTTTGGTCAGGTGCTTCATTTTCGCTGCAAGAGATATTCTCAA

AAAAAAAAAAAAAAAATCATCCCACAAAAGTCAATCAAAAACAAAATTTCATAACTTAAAATTCATAAGTTTTAATCTAT

TCAATCAATTTAAACTTAATTAATTAATCTAATCAAACTTTTTAGTGTTAATTAAACGAGGGCAATTTTGCCATTTTGAA

AATATTTGGTTAAGGGGTGACCTCTTTTACTTCAAAATGACCCTATTTTGTCTCATTGGGTATACCCCAATTAGTTTGAG

TATACCCCAATCTCGCCAGGTTAAAAAAGTCATTAAAGTAAGATCAGTTCCACAAAAATGAAGAAGAAAAGCAAAACTAA

CTCCGGGTGACTCAATGAAAACAAGAGGGCAGATAAGCTTGTAAGAACTAAAGATGAAGAAAATGGCAGAGAATGAAGAT

TTACAGAGAGCTCGGAGACGAGGAAGATGAACTAAAGGAAGAAAATAACATTCCCATCAATCTTCTCCTTAAATAGGCAA

AAAGGATGACCGGTTGAAATTGAATGTGCCGGTAAAAAAGGGGAACAATAAACACGTGTAGATTTGTTACGCAAAATTCA

CTTTTAGTTGTCACTGAAAAGATAAAGGAAAATGAGGGGTCGGCATTTAGAGATAGAAATTCTCATATCATTCTCTTCTT

CGCAGAGAAGACATGAGAAAAGGCAAGATGTAGGAGCAGAATCGCGCAACAGTTATACCTCAGCGAAGCTTTAACTAAAG

GAAGGTTGCGAAGATCATCGCAGTGTTAAAGAAGTGATTAGCTTAGTGATGTCACTTAGCCATAAGGAAAGTGTGAAGAT

TCCCGCGAGATAGAGATTACCTCAAAAAAAACCCTTATAAGCTTGCGAGAACAATGCAAGTGCATCCGAAATAAAGGATG

GCTTAGCTGTCATCCACTACATGTAATATCCTATATAAAGGACCAAGCAAAGACGAGAGAGGGGAGATCTTTTTGAGAGT

GAGAGGAGAACAAAATAGGAGAGAGAAAGTCCAGAGCTATTTTGATTTCTCAGATTCTCATTACTTTTGTAAAATCATTA

AAAAGATAAATAAAACTTTGTAAATTTATCATGGAATTTGGCTAGAATAGCATATAATCTTTGAGGGTGTGGTTGTAGGA

AATCTTGCAACCACATTATCCCGTCTATTTATATATGCTTGTTTAAATTTGAGTAGCACTTAGCTTTTTTAGCCGTATTC

TTCATAAGCTGTGTTTGTTTTTTCTTTACTTTTAGTGTCCTTTTGTCAATCGAGAGGGAAAGTCTCTGCCGTTATGTGAG

TTTGGCACCTATACTATTTCCACCTCCAAGTTGCTCTTGGACATTTCACGAGCTGGGTGTTCCAAAGAACTTGGGTCATG

TCAGAAACTATATAAAACTTGTGCAAAAATCGTGTTGGCGTCTATGTATACTTGTTGTTAGTCCCGTTTCTACGTATGCC

TATTGTCTTTAACCAATTCCCTGTTCTTTTAATGCAATTTGAATGCACATCGCTGTAGTCAGTACATTTCTGATAAGCAT

GAAGCAAATATCTCCACTTTTAGTACTGCTTTAGAACTTTATTGAGCAAATATCTCCACTTTTAGTACATTTCTTGGATG

GTTTAGCCCCACCAAACAATAACCCTGACACAGTACAAACGCCTTCTATAAAAGCCACATCTAGCGTGGGTGACCTTAAT

TATCCCCACGAGGTTCTTTTGCATTACTCTCGTGGGCTGATATGGGGACACTGCCCACCCCACAATCAGACACCTTTATA

GAAAGCACCGGCATATTATTACATCAATCAATCCATTATAAAAGATAATTGATATGTATAAGGGATGATATGGGACACCG

CCCAACACACAATCAAACACCATTATAGAAAGCATTAACATATTAACATCAATTCTTTATAAAAGATCATCTTTATACAT

GTATGAGTAGCAATAGGGTGTATGGGATGATCTTTATAAAAGATCATCTTTATAGGTTTTCATCTCATCCGCATCCAATC

CAATGCACTACTTGATTTTAAGTTTGCATCCGCATCCAATCCGATCCAACTGATTTGCATCCAAATAGATTAAAGGTGTA

AACGTAGCCTGGCACATCCTTACACGAAGTCGCGTGCTTCACATGCCGTGTGTCGTCACATCTTGCTAACGATTTAAACG

TGTTGTGCCGTAATCGTGCTTTACTAGTCAAAAGCTAGGCACAAAGATGTTTTACGGCACAAATACACGAATAAACGGTC

GTGTTTGTGCGTGTTAGCACACGTGCTATAAACAATTTGAAATCACTTAAATGTATACATTGAGTGGACATTATCACAAT

AATTTAATTAGACAACATATCATTTGAAATTATTTCAAGTAAAATTAACAAGTTTATTCAATAAAAAGAAATAAGCCATC

AAATATAAAATTGGATTATTGTCATGTAAATTAATGTTCTAGTTAAATATAGGTACAACATTATAACAACGATATTGAAA

TATATTTTCCAAATATTAAGGTATTATATGTGTTGTTATAAATAAGCCAACATAAATGTTAAAAAACTAGCTAGAAAAGT

GACTCTTTTGTGAAATACACAAAAATGTGATGTATTATGCCTTGTTATATGGTGTATTTGTACATGTTATGACGTGCTCT

CTTAACGTGTCATGGCCGCACCTAAGATATTATTACATCAATCCATTATAAAAGATAATTGATATATATATATATATATA

AGGGATGATATCGGACACCGCTCCAAAACACAATCAAACACCATTATAGAAAGCATTAACATGTACGTGGAGGTGTAAAT

GGGTTGTGCTAACGCGAGCATAACCAAAAGATCGCTAAAATTTAAGCACATACAAAGACGTGACTTAGCCCAAAGACGGC

AATCGCACGTTAGTTTTTTTGGGACGGCCGGCTTGACTAATCGACAAAGAGCCCAGCACAACACGCGGCACGGATAAACA

CCTAGCACGACACAACACCGCACACTAAGAAAGCACGCCGTAATATGTATAAGATACTACACAATGTGGCACAATACATC

ACATTTTGTGTATATTTCACAAAAGAGCCAATATTCTAGCCAATTATTGACATTTTATTATTCCAGTATGTTGACTTATT

TTCATAATAACACATATAATACCTAAATATTTGGAAAACATTTATATTGGAAAATATAAAATAAATGTGCCCGACACAGC

ACAACACGTTTATTTTTCCGCACAGTCCAACACAAAGACGCCTTTCAAAGACGTCATAATCTCCGGCACAGCACAACACT

ACACATAACACGCTAAGCACGTGACTTCGTGGGCTGGGCAGTGCTGGGCCGGCACGTTTCACACCTCTACATGTATGGGC

TGATACGTGAAACTGCTCCACCACATAATCAATCCTTTATAAAATATTGAACTTTATATACAGTATATTTATATAAGTTA

TGATAGGTTGACCATGGATTGTGTTCCGCATATTTTTAGTATAGCCCGTTGTAGATTGCTTTTTTCAACTAAACCACTTT

ATAATAAACTTTTTACCAAAAATTGTGTTTGGTAAATTTTTTTTTTAAAGTACTTTTGAAAAGAAGACTTTCATTTTCAA

ACTCTTTTAAAAGCCACTCAAGACTAACTTTTAAAAGCAGAAGCCATAATTTGATTTAAGGGATTCTCAATTTTGGAAAC

TGCTCAAATGGGCCCAACTACAACTGTAAGATATGCTGGGATTAAAATTCGAGCTTAAGTTACTATGAGACGCACCGAAA

TAAAATAATCCTCCTAATCATGTGCTTCATGTATGGAATAACTTGATTGCTAAAGACATGTACTGGTATCAGCGGGGCGG

GTACCATTTCAAATAAGACACAAAACTTACAACTAGTGGCTCAACCACTTGAAATACGATTAGAGTTATCAGATATCAAG

TAATTTTTCAATAATTTTCGTTATGTAACAGAAATCATAAAAACCTTTTTTGTCCAAAATTCAGTTTAAAAAGTTCAAAA

TATCAAATTCTGTCCAGTCGTTTTGTAAAAAATCTGTTGGGTATGAAGGTCCAGCCGTTTTATAAAAGAAAATATCAGGT

AAAAAGATCCAGCCAATTTATATAAAAGAAAGTGGCGGGTAAGAAGGTCCGCCCATTTTATAATAGATACGACCGAGGAA

GAAAAAGAAACCTTAATTCTCTATATTGATTCACTGATTCATAAGTGATAAAGAGAAGAGACTGGGGAGAGTCTGACCAA

GACAAATTAGATTGAACAAGAAGAAGATTTAGAAGAGAATAGATAGAGGTGTAAATTGGGCTGTGCCAGCACGAGCACAA

CATGCTAAAATGAGCCGAGCCTGCAATTGACCAAATACAAGTTAGTTTCATGGGTTGTGTTGGGTTAGCCTAATCGGCAA

AGAGCATGGCACAAAGATGCGGACGGATAAGCACGTGGTCCAAATTTCAAAGACAACACGCGTTAAGAAAGACGCTATAA

TTTGCGCAAGTATACAATATAATAAGGCATAATACATCACATTTGTGTATATTTCACAAAGAGTCATTATTCCAAATAGC

GGCTTCTTAGTGTCTTGCGACACGTTTTACGCCTCTAAGAATATGTAAAAAAAAAAAAAAAAAAGTTTGATTTTTCTTAG

TTTACGTCGAAGAAGAAGATATTATGCGCGGGTATAAAACGATAACAGATCAATATATACTATCGGTGTTACCGTAACGG

AAACTGTCTCAGCTGGTTCCGTTACATTAAGCAAAAAGTTCTAATAACTTCCCGATCCCAAGATACGCTAACCGTTATGT

CGGAAAAATTCTATCGTATCCAGGATTATCTGTAACGTCTATCGGATTTCGTCTGCTTGCAGGACCGTCGGATCATTCAA

ATGATACCAGAAACGACGTTAATGCTAGTAGTGCCTCTAACAATCACGTGGAACGTGTGTTTTTTTTCTTCTATTAACTT

AGTGGGACAAACCAGGAGATTGGACCCACCAAAGCCTTATGACTAGTAGTGCTTATATATATATATATATATATATATAT

ATATATTTGAGTTCACCATAAATATCATATCATTAAAATCTATCATCCTACATACTGTACTGGGTTGAATCATGGAGCTC

CAATATTTTTCTTATTTTCAACCAACTTCCTCCGTTGTTGCTCTTCTACTTGCTCTTGTATCCATCTTATTCAGTTTCGT

TGTTTTGAGGAAGACATTTTCGAATAACTACTCATCACCTGCATCATCAACAGAAACAGCGGTACTTTGTCATCAGCGGC

AGCAGTCGTGTGCATTGCCAATTTCCGGTCTTCTTCATGTTTTCATGAATACAAACGGCTTAATTCATGTAACTCTTGGA

AATATGGCTGATATATACGGACCGATTTTCAGTTTCCCAACAGGTAGCCATAGAACTCTCGTTGTGAGCAGTTGGGAGAT

GGTAAAAGAGTGTTTTACAGGCAACAATGACACTGCTTTCTCAAACCGTCCTATCCCGTTAGCTTTTAAGACTATATTCT

ATGCATGCGGTGGCATAGACTCATACGGTCTTTCGAGTGTACCTTATGGAAAATATTGGAGGGAGCTACGAAAAGTCTGT

GTGCATAACCTCCTCTCTAATCAACAACTACTCAAGTTCAGACACTTGATAATTTCTCAAGTTGATACGTCTTTCAATAA

GCTGTATGAGTTATGCAAAAACTCTGAAGACAACCAGGGAAACCATACTACTACTACCGCAGCTGGCATGGTGAGAATGG

ATGATTGGCTCGCCCAATTATCGTTCAACGTGATAGGAAGAATAGTCTGCGGATTCCAATCAGACCCTAAGACAGGTGAG

CATTTTCCTAGAGTTTCTGTAGAAAATGTTGGATTTTATACTATGAGATCATCAAAGTCTAACTGTGCACATGTATAAAC

AGGTGCTCCAAGCAGGGTGGAACAATTTAAAGAAGCAATTAATGAAGCATCGCATTTTATGTCGACATCTCCAGTGTCAG

ATAACGTTCCAATGCTAGGGTGGATTGACCAATTGACTGGTCTTACAAGAAATATGAAGCACTGCGGAAAGAAATTAGAC

TTGGTGGTCGAGAGTATAATTCAAGATCATCGTCAAAAGAGACGATTCTCTAGAACTAAAGGAGGAGATGAGAAGGACGA

TGAACAAGATGACTTCATCGACATTTGCTTGTCAATTATGGTGCAACCACAGCTTCCTGACAACAATAACCCTCCTCATA

TACCTATCAAATCTATTGTCCTGGTATGCAAATATTGATCCGTGGTATGGCTGGAGCATTAATTGTATTCCAGCGGTTAA

TTTTGGTTGTGGTAGCTCATAGAGTGCAGAAACATGATTTGGTTTTGATTGGTATCATTGTATAAATGCAGGACATGATA

GGTGGGGGAACTGACACAACAAAACTGACCACCATCTGGACCCTTTCCTTGCTGCTGAACAACCCCCATGTGTTGGACAA

GGCAAAACAAGAAGTGGATGCACACTTTGGAACGAAAAGGAGATCAACGAATGATGCCGTGGTGGATTTTGATGATATTC

GTAACCTTGTATACATCCAGGCAATCATCAAAGAATCAATGCGGTTGTATCCAGCCAGCCCCGTGGTGGAGCGACTGAGC

GGCCAAGATTGTGTGTCGGTGGGTTTCATGTACCAGCAGGGACGAGATTATGGGCTAACGTATGGAAGATGCAACGAGAC

CCTAAAGTATGGGATGATCCATTGGTGTTTCGACCAGAGAGATTTTTGAGCGATGAACAGAAGATGGTTGATGTAAGGGG

TCAAAATTATGAGCTGTTACCATTTGGAGCCGGTCGACGTATATGTCCAGGTGTATCCTTCTCTTTGGATCTAATGCAAC

TGGTACTGACTCGTCTTATTCTTGAGTTTGAAATGAAGTCTCCTAGCGGGAAAGTAGACATGACAGCAACACCAGGATTA

ATGAGTTACAAGGTGGTCCCCCTTGACATTCTGCTCACCCATCGTGGCACAAAGTCGTGTGTGCAGTCAGCAGCCTCTGA

GAGTAGTGGTGTACCAGTAATCACTCTGAGCTCGGGCAAGGTGATGCCTGTTTTAGGCATGGGAACATTTGAGAAAGTTG

GTAAAGGGTCTGAAAGAGAGAGGTTGGCGATTTTAAAAGCGATAGAGGTGGGTTACAGATACTTCGATACAGCTGCTGCA

TACGAAACTGAAGAGGTTCTTGGAGAAGCTATTGCTGAAGCACTCCAACTTGGCCTAATCAAATCTCGAGATGAACTTTT

CATCAGTTCCATGCTCTGGTGCACTGATGCTCACCCTGATCGTGTCCTCCTCGCTCTTCAGAATTCTCTGAGGTAAATTT

TTTTGATGGAAATTAGGCTTACATATATTTGCACATACCAGCAGGTACACTTAAACATCTATTTTTAGATAAATGAAATT

GTGTTGTGTGATATGTACATAGGAATCTTAAATTGGAGTATCTGGATCTATATATGTTACCCTTCCCGGCAAGCTTGAAG

CCAGGGAAGATAACGATGGACATACCAGAGGAAGATATTTGTCGAATGGACTACAGGTCTGTATGGTCAGCCATGGAAGA

GTGTCAGAACCTTGGCTTCACTAAATCAATCGGTGTTAGCAATTTCTCCTGCAAAAAGCTTCAGGAATTGATGGAGACCG

CCAACATCCCTCCAGCTGTGAATCAAGTGAGTACTCTTACTTGAAAGTTTTGATAAGACATCGGAAGATTTCTAATGCTT

ATTAAATTTCCAACATTATGCATAAAACTACTACTCTTTATAATGTTGATTAAGGTGGAGATGAGCCCGGCTTTCCAACA

AAAGAAGCTGAGAGAGTATTGCAACGCAAATAATATATTAGTCAGTGCAGTCTCTATACTGGGATCAAACGGAACCCCAT

GGGGCTCCAATGCAGTTTTGGGTTCTGAGGTGCTTAACAAAATTGCTATGGCCAAAGGAAAATCTGTTGCTCAGGTTGGT

CAAATTCTCTGCATATGCCATAGTGTGGTCTCTGTTCTTTGGCCATCGTATTTGCATCACATTGTATTCCTTCCTTATAC

GGAATGTAACTTACATTTGTATTACGTATTGATCAGGTTAGTATGAGATGGGTTTACGAGCAAGGCGCGAGTCTTGTGGT

AAAAAGTTTCAGTGAAGAGAGATTGAGGGAAAACTTGAACATATTCGACTGGGAACTCACTAAGGAAGACAATGAAAAGA

TCGGTGAGATTCCACAGTGCAGAATCTTGGCTGCTCATTTTTTGGTCTCACCTAATGGACCTTTCAAATCTCAAGAAGAG

TTGTGGGATGACAAAGCTTGAACATCGATCATTTAACTCTACACGTGCATTTATAAGAGAAGCTTCCCCTGCAGTGTGCT

CAATCTTGATTTATTTTATCAAATTACATCTTACTATATGGGAGTCACAGATCTCTTTCCCTTCCTATATGTTTGAATAA

ATTTGTAGCGAGTGATGTAGCTAAAGAAATAAATGTAGTAAGTTTCTTTCCTAAGCTTGTTTTCCTTTACAAAAACTAGA

TGAGAGTCGGAGGTAAATAATGGTGAACTCGTGTTCATCAGCAACAATAGTGCGATTAGGCACAATGTATATACCTTAGC

AGAAAACACTATTCCAGACACAAAGAAATGTACGAATGCGTAAAGAGTATTCTGCCGTGCATATACAGCAGCTGATGCGG

AAAAAAAACAATACAAAACCTTCTGTGAAACCAGACGTATTATTGTCCATGTGAGAGTAAAATATCTAATTGCCACGTGT

AAGCATCACAGGAGGTGATCATAAGATAAGATGGTAAAACAGATACACTGAATAAAGATGTAAAAGGTGATTACAACAAA

ACCAAGAATATCCACTGCTTCTTTTCAAGTCTAAAAGAAAGCAGTGCATTTAATGAGAATCGGAAGAGGTGCGATACAGC

TGTAGAAGGAGGTACATCCCTCTCGTGAATTCGGACGAAGGGCGAGGTACTTCTACCCGAAGAAGGCGCCACACGATAAG

GCCCAAAGGGCGATATATAGTATCATCCGAAGAAGGCGTCACACGAGGTAAGTCGAAGGGAGCAATGGCGAGCCAGTCTG

AAGCGAAGAGGCGTATATATTGTATATATCGTCATACACCCAGGAGCACCGTATCGGACCTGAATGAGTTTACTTTGCCT

ATTGTTGATGGTGGTTTTTAAACAAAGGGTGAAATCGTAAAACCCCGAATAACCTAATGTCACCAAGTGACACATTTATT

AGGACATTTATGCATGTAGTCGGTAATTAATAGGGGGTATTTTTTCAAATTAGGGTTACAACGAACAAAACCCTAATTCC

TCCGTGCCAATGGCAAATCATGCCCAGCCACACCTAGCCCAAGCTAGCCGCACCTCTATGCCAATGGCAAAGCATGCCGG

CCATACCAAACGCGCCGAAGCATTCCAACCATGCTAGCCGCACCTCTATGCCAATGGCAAATCATGCCAGCCATACCAAA

CGCGCCAAAGCATGCCAACCATGCTAGCCGCACCTCTATGCCAATGGCAAATCATGCCAGACCATACCAAACGCGCCAAG

CATGCCAACCATGCTAGCCGCACCTCTATGTCAATGGCAAATCATGCCAGCCATACCAAACGCGCCAAAGCATGACAACC

ATGCTAGCCGCACCTCTATGCCAATGGCAAATCATGACAGCCATACCAAACACGCCGAAGCATGCCAACCATGCTAGCCG

CACCTCCGTGCCAATGGAAAATCATGCCAGCCATACCAAAACGCGCTGAAGCATGCCAACCATGCTAGCCGCACCTCCGT

GCCAATGGCAAGTCATGCCAGCCATACCTAGCCCAAGCAAGCCGCACCTCTGTGCCAATGGCAAATCATGCCAGCCACAC

CAAACGCGCCAAAGCATGCCAACCATGCTAGCCGCACCTCCGTGCCAAAGGCAAATCATGCCAGCTACACTAAGCCCGAG

CTAGCCGCGCCATGCCAAACACACTCTGCCTTGTCCCTAGCTACGCTGGTCATGCCATGCCAAGCCAAACACCCTTACCT

TGGCCCTAACCATGCCATGCCATGCCAAAATGGCGCCCACGCCTAGGCTTGCAAATCATGCAACCTAGGCCAAATTGGCC

ACGCAAGCCATTGGCACATGCCAACACGCTCGCTGGTAGCTTTTCCTACCAAGGCACGACCCAAATTGGCCACGCATGCC

ATTGGCACATGCCAACACGCTCGCCGGAAGGTTTTCCTACCCGTAGCGACCCAAATTGGCCACGCATGCCATTGGCACAT

GCCAACACGCTCGGCTGGTAGCTTTTCCTACCAGTGCGCGACCCAAATTGGCCACGCATGCCATTGGCACATGCCAACAC

GCTCGTTGGTAGCTTTCCCTACTGTGGGCGCGACCCAAATTGGCCACGCACGCCATTGGCACATGCCAACACGCTCATTT

GGTAGCTTTTCCTACCGTATACGACCCAATTGGCCACGCATGCCATTGGCACATGCCAACACACATCGGGGTAGCTTTTC

CTACCACGGCGCGACCCACCTGGGCCACGCATGCCATTGGCACATGCCAACCCGCTCGCTGGTAGCTTTTCCTACCAAGG

GACGACCCAAATTGTCACGCATGCCATTGGCACATGCCAACGCGCTCGCTAGTAGCTTTTCCTACCAGGCGCGACCCAAA

TTGGCCACGCATGCCATTGGCACATGCCAACACGCTTTGCCGGTAGCTTTTCTTACTGTGGCGCGACCAAATTGGCCACG

CACGCCATTGCCGTGCCAACACGCCGTAGCTTTTCCTACCTTGGCGCGACCCAAATTGGCCGCGCATGCCATTGGCACAT

GCCAACACGCTCGCTGGTAGCATTTCTAAACACGGGCCATGATTCAACTTGGCCACGCATGCCATTGGCGCATGCAAAAC

ACGCATGCTGGTAGCCCAACTTGCAAGCCAGGCAACCTAGGCCAACTTGGCCGTGACGCGCCAATGGCACATGCCAAAAC

CAACGCTGGTAGCCTAACTTGCAAACCGTGCCACCTACCGCCAAGGCATTCCGCACATGGCATTGGCACATGCCACGCAC

GCCACTAGCAGCTCTATGCGTTGCCACTTACACTCAACGTACAACCTACCACCACGGCATTCCACACATGCCATTGGCAC

ATGCCACACACGCCACTAGCAGCTCTGGCATTGCCACTTGCGCTCGAGGTGCAACTCGCAACAAAGGCATTCCGCACATG

CCATTGGCACATGCCACATGCGCCCTAGCAGCACCACTTGCACTCAAGATGCAACCAACACTCGAGACATCAACATAAAC

CGGGGATGCTTTCAGTGGTCTTGGTTTGGAGGTCACAGCTTAGAGGCGTACACTACGCTCGCATTAAGAAAGCGTGAAAA

GGAATAAAGCGGTCAAATAAATGAGGAAATAATGGTGAAAACGTTTTCACCCATTAACACCATACGGTTACCACCTCCTC

CCATTATTTCACCCTTTACTCTCTTCCTTCATGAGGCCATGGTGTACGTTTCAATCTCGACTTGTATAAATAGGTCTTAC

CTATTTCCACCGAACAACAAGTCTCGGTCGGGAGAAAAATACAACACTCTCGGAAAAGGCAACTTTGCAAGCTCTCCAAA

AAGGCTTTCATCTCTTGAAAGGTTCCAAGCTTTCACTTTACGATACAAGTCGAGACACTACTCTCCCGCACTATTCTCGG

TCTCAACACCTTTCTTTGCTTCCCTCCCCAAATCAACCCTTCTCCTCCACTTTGTGACCGAAGCAAGTCCTGAACGGCCA

TTTCTTGGTTTAGGCCGACTTGTGCATCGATCTCTCGAATCTATAAAGTGCTCACGTGCGGACATTTGTTTAGGGTTTGA

ACCAGTTTCTTACTCACCTACCGAATTTACCGAAATCGCGCGAAACGGTTTTTACCCATAAACAATTGGCGACCACAATG

GAGGATTGATCTATCGGCTACAATGTCGACTATCAATCTCGTTCTTCGACCCGAAGTCAAGATGGTAGGGAAAGCGACTC

AATCGACTCAACCGCCAGTACAACCCTAAGTTACCGGTTGTAAAAGCTACACCCTAAAAACCTGGGACTCGTTGAAGATG

CTACCAGCGATCGGGGACTTGCGACTCAATCCGACTCAACCGCGGTACAACCCTTTGTTACCGGTTGTTAAAACTACACC

GTAAAGTGGGACTCGGCAAGATGCTACCGACGATTGGAGACTTGTTTGGCGATGCTACCAAAGAAAGCCGCCTTTTACTT

AGAGAGAGGCGTGTTTGCAAAAAACCTATGAAGTGAGTAAATTTAGATTGGGCAAGATATACGATATGTTTTGGGTTCTT

AATAAAACCGTCATACTATTTTGCGTCTTGTGCCCATTGCTAGCACACACTTTCATAAAAAGCAATAATCCTTTCCCAAA

ACATAACCAGAGTTATATTATCAAAACCATTCATCCCGAAATAATCCAATATCCTCAAAAGCTACCAAAATTGCCTATCA

AAACCTATTCGAACTTTGAACTGGGACATCTAGTATGTTTTTCAAAAAGAAAAAAAAAAAGAACCTAAATGAAAGTTTAA

CACAGAAACATTCAAGGAAAATCATCCAACATAGGTCCGCTTTTCTTGGAGAAAGTCAACTTTGCGTCTTCAAGAGCCGC

TCTCTTCATAGTTAACATCTTAGAAAGCTTTTCGAGTTCTACCTCCTCGTCGTGCAATCTCCTCGGCGGAAAGGCTTTCG

ATTCTTTGCATCTTTCAACTTTTGGATCTCAAAGAAGGCCTTCGACAAGCCACGAAGTGTTGAACTCCTTTTCTACGCAC

ACGTCGCGATGGAGCTCCCGTCGGCAATAACGTCTTCGAAATGGTGTGACCAGTCACCTCACGCATATCCTCGATGGAAG

ATTAAGCCTCTTCCACACGCTTAACCAGCCGCAAATCGACTAGTGACTTTCTTGGTTGTGGCCATATGTCCGCACTTCTC

CCAGCATTCTAGTGTACAACGCCTCATACTTAGCCACGCTGAAGCCGCCAACCAAAGATATGACCTGGATAAGGAACCAA

GTACGGGGATCAAAATGGCGCCTCGTAATCGGCCTCACGACTCACAAAGGGTCTTCTTCCATGTCTACTGCAGCAACCAT

CGAGAATCATCCACCGTTTGAACCACATCTGCATTCGGCGATTGGCTAACTCCTATCTCCAAGTCACCTAGCGGTGGTCA

CTTAAGGCAAGTTTTTGTGTGGCTCCACCCCAATCGTCCTCCGAACCGGATTCCTCCTGCGAACTTGAGTTAGACATTTT

TCAAAAGAAGAAACAAGAAGAAGAAAGAAAGAAGCGCGAAACTCACCGAACCACTTCCGAAAAAGAACCAAAGGTCTTTG

CAGAGGAATCACTCTCACTTGTCGAACTTTCTTCTTTCTCGACTCATCACCGCTATGAGTAGGCTCGGTCCGCCGAAATT

CTTCGAATAAATTGCCGTCTCCCCGGCAACGTGCAAGGGCAGTAAAGTATTAACACTACCGAAGCCCCTGGAAGAAACAA

GACACATGTTTTCAGCGAGAATTCACGTGTTTTGAGACAAAACAACAAGGAGGCGACACATACCGGTTATTAGAAGAACC

GAAAGACACGAGAGCTGCGGGAGGGTCTTTTGAAATCGCCGAACCCGCCAACACGATTCTTCGTTCTCTGTTCCTTGGCA

CGGGGACTTTCCGAGTTGCTAGGCAACTTTCGCTTGAGATCTCTCAACATTGGATACTTCTTCAAAACTGGGGAAGGTTT

ACAACGAACATTCTCCACCACGCCGTTTACAAAATCTTGGATCATAATAAACTCATCGCACCGAACTCATTATAGCCAAA

GGTCGTCAAGTTACTCTTCCCGTTTGAAAGAAGGGAAGGCGGATACCCTACTCGAGGGCACTCGTCATTGAGGGCGGCAC

GCAATGAATCCTTAGGAATTACCTACTGGGTAAGCAAGAGGCACCCCTTGATCGAAACATATGTGACGAGCCGCACGGTT

AATATTGTAAGCATTATCCGGAAAGACCCCATGAAGAAAGACGGCATATGGACCGACAAATTCAAACCTGGACATGCAGC

TTTGAATAACAAGTTTTCACCAACGCTCAAATTTCCATCGAAGACAAAGACATGATCGGAGCGTAAACAAAGGTGTCAAT

TTGAGTCACCGACCTATGGACCGGCTCCCATGGGCGAAACGATTTCATACGACTTATCGAGAAGACTAACCAAGATCATG

CCAACCCTTGGGCGCTGTTCACCCATCGAAGAATTCGGGAACCCCCAACTCGCGGGAATCGAAGACAACGGTTTGGGGCA

TACTTCTCAAAGTGTTCCCATAACCAAGATTGCAGGAAAGCAACGTGCACATATGAATCCACTTTCATAAAACCATTCGA

AACTTGCATGTTCGCAACTAGGTGATCCAGATGCACATACATGGAACCAAGAAATAGCTTGCCAATGGGAAGAACTTCGC

CTTTTGCCAACTTCACAGCAATCACTAGCAACTCTGGTCTTATCTCCTTCTTACTGGCAACCGTCGTCAAAAATGTCTCT

AGACAAACCGAGAGCCAAAATGCTGCAACATGAAGATCACTGCTTTCCGTTTAGGTTCGAGCTCGTCCAGAAAACCATTG

AGATATCCACCAGTCGTAAAGGCACCTTTCATCCTCTTTCCGGACGAATCCTTCGACTTCGCGGTGGAGTCTTTCCGAGC

GAAATCTCCCAACTTAGCGGCAAGGTCATCCCGCATCTTCCCCTCACTTTCGGATAATTGGATGTTAAGATTCCTGATTC

ATCGGAAGATTCAAAGAAGAGAGCCACAGGTCTCCAAGGAGAAAGTTATCTCACCCCATCTACATATGATCGTATGGGTA

TCCGAACACCATCTGGAGATGAAAGCGACCAAGCCCGGGATATCCTTCCTAATCGGAGATTTCTTGAAGCTCGACATCAA

TAATCTGCGCTCGAGTCAGTTGGTATTTATGTGCTCCTTACTCACCATGAACTCAACCCATCTATCAAAAAGAATGCAAT

GGGCTTTGACATATCCTGAAATCAATGTTGGAAGAGGAATATACCCCTTTATGAAGATAGAACCGTATGGTGTTCTCTGA

GCGAACCGAACGTGTCTTCTTTTCATCTTCACCACCGGTTAACAAAATTGGCACGAAACACAGGTGATTTTTCCGAGCAG

AGGCAAGTTTTTGGCGAAGAATTCATCATCCGTGTTTGGTTTGGAACTATGTACATCCTTCGATGAGGCGGAATTTTGTT

CGGTAACATCTTGTCGGATTTCTTCATAACAACTTAATAACAAGATATCTCGCACACGGAGAAAAACAAAATTACCTCCA

CCGACCGATAGACGCATTTATGTGTCTATTTTGTTCTCGATTTTCTATATTGTTAGTACCCATTTTGTATTTATTTTGGC

ATTTTATGTTTTTGTAGGTGTTTTGGAGAAATAAACTTTTGCGAAGAAATTGGCTCGAAAAGTGGCATTTGGACCGCCGG

GAGGAATTACTAAATGCACCTCGAAGAAGTGTTATTGGCCCCCTGAGAAAGTTATAAAGGCGCCCCAAGTTTTGGTTATT

TCCACCCAAAAATTACTATTTCCGCCCCAGTTGCTAAAGGGACTCCACTGTCGGATAGGGGGAATCCATCATCTTCATTT

AAACTTTGAGTTTTGGCGGGAAAGTTCACAGTTCACGGCATTTTAGGGTTCGAGTTTTGGAAGATTTACTCAAGAGATTC

GATCGCTGTATTTAGTTGATATGGGCCTAGTAAGCCTAAACAGGGAAGTTATGGGTGATAGTCTCGATTAAATTGGGCTG

GATAGTTGTCTTGGAGCAAAACAGAGTGTGTGGAATACATCGGGTTCTGTTGGTTTATTTACGGAAATCCAGTGAGACTA

AAACGCACATAAACTTTCCTAAGATTCACATATACCTTATAGAGAGTTTGGAAAGGATTGATACTTGGAAACGCGTAAAC

GCTTCGGCGAGAGATTGTTGCCTGACTTGCACGGAAAGGAAAGGAAGAAATTTAATCGGGTTTTGGAGGGTTTGGAGGTG

TATAAAAGGTATGCTCGAGTCCCAGAGGTGGGTATTGAAGTTTAGAAGAAGTTAGGGGAAGTTTAGAACACCACATCGCC

GAGAAATCAAAGTTGCAGGAATTGGAGTTCTGCTGGTATAGGAGGAAGAACACGAAGAACAAAGGCCATCTAGGACAGTC

GTTTAAGGCGTCTTAAATACAACGACAATTCAACAAAGGCATAAGTCGGCCCTTGTAATGTTTCATTGTAAATAAATGCT

TTGTAACTCCGATACACCGTTACAAATCGGGTTTGTGTTATTATTTTCTCATCCTTTTAATCAACTTTTGGGCTATAAAC

ATGTATTCGAGCAAGTGATTAATATGAGGAGCTAAACCCCTTTGCTTTGGCGACGGAGGAAGCTATTTTTCCAACAAATA

TGTGGTAATATTTATTTAATTAATTTCTGCCAATTCTTTTTATGATTATTTGCATTGAATCGAAATTGAAATAATGATTT

TTGCTAACTTGTTGTGTTTTCACTTGATGGTGCATGCTTAGCTTAGGGCTCTTGATGTTCCATGCTTGAGTTTAACAACT

AGTGTTTTGAAAATCTACTATTGGCAATATTTAAGAATCAATTGAATGTATGAGTTGCATGATAATTATTAATATTGAAT

CACACAAAAGTTGGAAAATAGTGGAATCCCGAGTCTCAAATGCTTTCTAAAATCTTGATACAACCATCTTTGAGTTTTCT

AGTTTTATTTGAGTCTAAAAATCCTTCACAAGTCTGAGAGGAACGACATCTTTCTTACCACTTTCTACAACAACATAAAA

ACCACATCAATTTTTGGGCGCCGACGCTTGGACTTGTTTTTAGGGTTTTAGATTTATTTTTAGTTTTTTTTATTCTTTTG

TTCTATTTTACGTTTTCTTGGGTATTTGTCTTGTGGTTACAGGTGTATTGGATTTGGATCGAAAAGAAAGCCGAGCCAAA

GATTTGGTGAATAATTAAAGACTTGGAGTCAAAGACAAAGCGAAAAGAAACGAGGAAGAGACAAAGAATTTTATTTTTAT

ATTTATTTATTTTTAGGGTTTTTCGGAAAAACCGTATTAGGGTTTATAATTATTTTTGTAATTTTTCTTTTTATTTTTGG

ACTCTTTGGACATTTTCGGGACTTTGGACATTATTTTTAAACCCTAAAGAAGGGTAGTCTTAAATATAAACTGTTTGCAG

AGAAGGAGGACAATTACGATATTTTCTTCGCACCTTGGTTCTGCATACGACATAGGAGTCGATGGCCCGAGTCGACTACA

ACCGATTCATCCCCGTCTCGGAACGGGAGGTAAGACTCTACACACTCGCTCGAATCTCCCATCGGCCGAGTTACTGGACT

CCTTCGTTTGCATATATGCGAGGACTTGAATACTGCCGCTTTAATTTTCTAGTAAAGGGCAAGGCCTGGCCATATAAGAT

AAGGGTTCGGATTTCATCACCGTTCTCTTCTTGCCCGCCTTAGGAACACGTAACCTACGCGAACCTAATCTTTAAAATTT

TGACTAGAACAAGACCGATAGGGTAACGAGCTTAATAGGACAAGTCGTTCGAAAAATATTGGTTACTCTTTTAAGCACAC

TTGAAGTTCATGAAGGTTTCGTGAGTTGAATGCGTGATCGCTAGCCGCATTGTAATTAGGTGAGGCCTTGGGTATCAAAG

CTCCACCGAGCTTCCCCTCGCCTCGATTCAACTTACTTTGACTGGATTGATTCGGAGGGGTTTGCTCAAATTGTAACGAA

TTCCCTTTCGAAGGGTTAGAAGCTGGTCTAGAAACAATCTAAGTGGAGCCATCATGCTTTTTGTTTGCTAGAAATCTCTA

GGTTTGCTTTGGTTGAGTCGGCCTTGATTTGTGATTGCATAGAATCCCTTCCTTATTCTTTTGCATGCACAACGTAAAGA

GACGCCCTAGGTAGATTTGTTAAAGAGAAACCTAGTAGTTCTAAGCGTCTCGATTACCTCAATTTAGAAAGTCCGATTTT

TGAAGAGTCTGTTTTGAACGTCCATTTGGGGAGACAATCCTAATATTAGATAGTGCCGAAATGGCAACTTTGAAAGCCTT

GTTGAACCCGACTAGGACTACTCGTCCCTCGTGTATCGGGTTAGTGAAATGAAGCAACTTATGAACTTAAACTGGGACCT

TACAACCTGCTCCCAATCTTCTTAGGGAAAGAAAATGAAAACCCTATTTTCATGTTAGGGACTTTGAGGAAATTTGTAGT

ACCCTAAAGATTAGAAACCTTGATGATGATGCTTTGAAACTCAAAGTTATTCCCCTTTCCTTGAAAGATAAAGCCAAATC

GTGGCTATATAGTTTGGCTTCCGGTTCAATTGAAACATATGAACAACTTACATCTCGCCTTTTTGAACAAGTTTTCCTAG

GCATAAAACATCGTCTATTAGGACGCAAATCTTGCACATTTTCTCAACGGGGAGGGAGAATCTTTGTATAGGTATTTGGA

AAGGTTCAATGATCTATTAGCCCAATGTCCTCATCATGGTCTAGAAAAGGTTAGGTTAGTTCGGATCCTTTATGAGGGTT

TAGATTATTCCACTACGACCACGTGAGAATCTATGTGTACGCAATGGGTTTGAAAACCAAACAGTTGATGAAGCGATGAC

ATATTTGCACGAAATCGCCGAAAAGACCCAACAATGGGAAAATAGTAGGGAACCCAATAAAATAATTCTTCTAGGCGGAG

GAAACGTTAATAGGGTAGAAGGAGGCTTTGAATCGGATGCCAAATTGCAAAGAATAGCTAAAAGGTTAGAAGCTTTAGAA

TTGGGTCACACTAGTGGTAGAGTGGAGCCTTTTTGGGAAGGCCATACTATTGAAGAGCAAGCCAATGCTCTTTATAATAA

TAACACTAGATTTGATAACCGTCAGAAATTTGACCCATATTCAGAAACCTATAATCATGGTTGGAGAAACCATCCGAACC

TCTCATGGTCTAAGGACCAGAACCAAGGTCAGTTTAGTAATTCGAATGCTCCGCACTCAAGCATACTAAAATTCATATAT

AAGTCACAGAGCTCTACTCAGATAGTCGCACTATGGACATCAATACCCGGAGTCAAAACTAATCACATGGATAAATCAAG

AAGATGGATATATAGAAAAACATAGATGATTTTGATGTTTATTAAGTGAACGGTGTTTCCCATATCTCGTCCAAGGCCTC

CGCCAAAATGAACCTATCCTAATGGACTGAGATACTAGTCGACTAATATCAACACACTGGCATATACAAGGGAACCAAAA

CAATATAATCCTAATTCTAGGTCAACACAACCGGGCATATACAAGGGTTCCGCAGGTCGACTTTATTGAATTTATTCTTG

GTCGATGGTCGTGTTCTCACAATTTTTTTTTTTTTTTTTTTTTTTTTTTTAGTATCTCAATCACTCTATTTCACCCTAGC

ATTGGTAACAACTTGATTCGTGGGCCCCACCTAATCACTTAGAGAAACATAGTTTAAAAACAAAATAAAATAAAAACAGA

AGTGAAAAGGACTCAACGAGATATGGTGAAACTATCATGTTATTTCTAACACCTGAGCTCTGTGCTTTTATGAATAGACT

CTTTAGATGTTGCCATCTAATCAGATTGGTTCCTCAACTCCTACAACCAAAATGCTTCCATCTACTTAGATTGGTTAGTG

CCATCCTTAATAAGCATAAATTTCTAGGCTCTGGAGTTTATTAATGCAATTAAAAGTTTCTCTCATACCCCCAAACTTAA

ATCTAACATTGTCCTCAATGTTCTAAAGATGAAATTTAAAGCATGAACAAGGAGAAACTGTTACCATTTGAAGTAAAAGA

GTTAAGGAAAGATATTACCGTGTCGCATGAGTATTGGGTTACCTCCCAAGAAGTGCTAAGTTTAACGTCTTCATCCAGGA

CTAACAAAGGATTAATCACCTCATAGAATCATAAAGTAATAGCCGTAATTTAAATGGGTCATCAAAACCAAATAAAGCTA

CCACAAATAAAAGGAATCACGCAACATGCCAAGAAAATTAACAAATAATGCACACCCTTGTCTAGTTTCTGTTTAAGAAA

ACTACATCTAGTTGTGGTTCAGGTTCAGGTTCTATAACTGGGTCGAGATAAAATATTTTCATGGACTGGAATTCCTCAAA

GCTAAGATCTGATTCAGGTATTAGAGTCTGGAAAAACTTAAATAAAAACTTAGAAGCACATAATAATAACCTGAATAATT

GCGGATCCTTAAAGTCAATCAGTTTTGACTCACACAATTGACCACGATGGCGGCGGTCAATCATAAGAAAATGTGTCGAT

ACTAATCTCTTAAAGTACATTTGAAACTCATATACGTTCCCACAATTGGTTGAAAAATTTTTGGTGGGAAAACAAAGTCA

ATCTGGGTATCATAGCCTGGATAAACCACATCAACCGGAGGATGGGTTTCTAACAACTGAACCTTTCTATGAAGATACTC

TTGTGAAATGGTAGAGGCATATATTCCAAGTCCCCCCCAAACTTAGAATTTTGGGTGTCTCTAAATAGACTAGTCACAAT

ATTCCTAATTTCTAGACCATCAGGTTCCTGAAAAAGGTCAATTGCTTTATCTAAGTTATAATCAGGTGGTTCATCCTCTA

AATGCATCTGATGCATACACAACGTAGGACTTCTGTCCTTAATATCCTCTATGGTCCACCCTAAAGCTTCCTTATTGTCT

TGAAGCACGGTCACATCGGCCTACTCTCCCGATCACTATCCAAATGGAAGCTATAATAACGGGGAAAGTGTCGGATGGGC

CTAAAAATGCATAGTTAGGGTCCAACTTAGGAGGCTCTTCTAAACAAGGGATTAAGGTAGACTCGAAAGCTAATAATGTC

TCAAACCTAGATTTCCATGTATCAGTATCTAACATAGGAATAGAATCCAACAGAGCATTTACTTGTTCAATGTTGCTATC

ATCGTCGAAATCCATGATAAAATGGGCTAGACAACTCTCTAATGGATCTTCCAATATGATGTTTGGTAATGACTCTGAAC

TAAGGTTCCTATCATGTTCACCTCTTCAATGCATGTGTCATCTAGCTCGAATGCGACTTATTGACATTAAAAATATTCAA

CTCAATAGTCATATTACCAAAGATAGATTCATAATACCAGTTCGGCGCTTTATGATCGCATTAGACGTAGCTAAAAATGG

GCGACCTAAAATCACGGGTATCTCGGTTCTCTGGGTCGGACAGGTTGGGTATCTAAGACCACGAAATCCACAGGATAAAT

AAACTTGTCGACCTCAATAAGAACATCTTCTATCACACCACGAGGACTTTTAACGGACCTATCATTAAGCTGCAGGTGTT

ATCTTGGTAGGTTTCAATTCACCAAGTCCTAGTCAGAAGTACACATGGTACGGCAAAGTAAGTTCACACTAGCTCCTAAG

TCAAGTAAAGCTTTATCTACCCGATGTTTTCCTATCGTGCAAGCAATGGTAGGGGCACAGGGTCTTTATATTTAGGAGTT

GTGTTGTTCAATGATCGAACTTACATGACTAGCTAAAAAGGCTTTTTAGGGACACTAAGCTTGCGCTTTCGCGTACACAT

ATCTTTAAGGAATTTGGCATAAAGCGGAAGTTGCCTAATTGCTTCTAGTAACTAAGGTTTATGGTAACTTTCTTAAAAAC

CTCCATTATGTCATTAAAGTTGGATTCACTCTTTGTTGGTACTAATAACTCATGGAAATGGGGCTCTAGGCATAAAATAG

GGCCTCTCGGGAACCGAATTGGCATCATCGGAGATTTTATCGGTCCCCTCGGCTATGGATGGTCCCGAGGGGTGAACTAC

AGTATGCTCACTATCGGGCATGGTTACCCTATTGTCTACTACTCTGCCACTCCTAAGGGTTCTAATTGAATTCAACTGAT

GTGATGGTTTTTCACCTACTTCATGAACTCCTCTAGGGTTGAGTTGTGTTTGACTAGGGAACCTACCTTTTTCTCTCAAG

GACTCATTTATCAGACTAACCTGCGTTTTTAACTCGGTAATAGCTTGACTGTTTTCTTGCCCTATCCTGTTACTAGCTTC

TACACTTTGTGAAAGTAATTGGTGCATCTTTTCAGTGTTTTGAATAAATGAGGCAAGAGTTTCTTCTAGACTCAAAATTT

TCTTATCTGACTGATTTTGAAACCGAGCTACTCCTCTGAAGGATTCTTAGTATAGCCAAAACCCGATTCTTTAGTGCCTT

CCAAGGTTACGCGTAGATAGTTAGGGTCTGGAGTATAAAGGTTTTGTGGGTACACCTCTTGTAAACCCTCCCGAGACTAT

AACTCGGCCACTAGGGCCACCTAGGGGTTTAAAGGCTTATTGCATACGCTAAATGCAATCGACGATGCCTGCGACAGTGA

GTTAGGAATTTATTTTCTAGATTTGATTTGCTCGGGACTAGCAAATAATAAGTTTGGGGGTATTTGATAGACGCATTTAT

GTGTCTATTTTGTTCTCGATTTTCTATATTGTTAGTACCCATTTTGTATTTATTTTGGCATTTTATGTTTTTGTAGGTGT

TTTTGGAGAAATAAACTTTTGCGAAGAAATTGGCTCGAAAAGTGGCATTTGGACCGCCGGGAGGAATTACTAAATGCAGC

TCGAAGAAGTGTTATTGGCCCCCTGAGAAAGTTATAAAGGCACCCCAAGTTTTGGTTATTTCCACCCAAAAATTACTATT

TCCGCCCCAGTTGCTAAAGGGACTCCACTGTCGGATAGGGGGAATCCATCATCTTCATTTAAACTTTGAGTTTTGGCGGG

AAAGTTCACAGTTCACGGCAGATTTTAGGGTTCGAGTTTTGGAAGATTTCTGAAGAGATTCGATCGCTGTATTTAGTTGA

TATGGGCCTAGTAAGCCTAAACAGGGAAGTTATGGGTGATAGTCTCGATTAAATTGGGCTGGATAGTTGTCTTGGAGCAA

AACAGAGTGTGTGGAATACATCGGGTTCTGTGGGTTTATTTACGGAAATCCAGTGAGACTAAAACGCACATAAACTTTCC

TAAGATTCACATATACCTTATAGAGAGTTTGGAAAGGATTGGACAGCTTGGAAACGCGTAAAACGCTTGGCGGAGAGAGA

TTATTGCCGTGACTTGCACGGAAAGGAAAGGAAGAAATTTAATCGGGTTTTGGAGGGTTTGGAGGTGTATAAAAGGTATG

CTCGAGTCCCAGAAGGGGGTATTGAAGTTTAGAAGAAGTTAGGGGAAGTTTAGAACACCACAGTTGCAGAAATCAAAGTT

GCGGGAATGGAGTTCTGGGTGTAGGAGGAAGAACACGAAGAACAAAGGCCATCTAGGACGATCGTTTAAGGCGTCTTAAA

TACAACGACAATTCAACGGTGCATAAGTCGGCCCTTGTATATGATTCATTGTAACATTTGCTTTGTAACTCCGATACACG

TTACAAATCGGTTTGTGTTATTATTTTCTCATCCTTTTAATCAACTTTTGGGCTATAAACATGTATTCGAGCAAGTGATT

AATATGAGGAGCTAAACCCCTTTCCGAGGCGACGGAGGAAGCTATTTTTCCAACAAATATGTGGTAATATTTATTTAATT

AATTTCGCAATTCTTTTTATGATTATTTGCATTGAATCGAAATTGAAATAATGATTTTTGCTAACTTGTTGTGTTTTCAC

TTGATGGTGCATGCTTAGCTTAGGGCTCTTGATGTTCCATGCTTGAGTTTAACAACTAGTGTTTTGAAAATCTACTATTG

ACAATATTTAAGAATCAATTGAATGTATGAGTTGCATGATAATTATTAATATTGAATCACACAAAAGTTGGAAAATAGTG

GAATCCCGAGTCTCGGTGCTTTCTAAAATCTTGATACAACCATCTTTGAGTTTTCTAGTTTTATTTGAGTCTAAAAATCC

TTCACAAGTCTGAGAGGAACGACATCTTTCTTACCACTTTCTACAACAACATAAAAACAACATCACTGACACATTACCAC

GCCCAACAAAAGGAAAACCTGAAATACAAAAAAAAAAGAGGGCGTATTCTTTACTTGCTCAAAACTGCACACTCAGTATT

TTCTTCAGGGTTCAAAATTATATACGCAGTATTTTATACGGGATCAAGTGCGTCTGCAGTATCATTCTTCGGGTTCAAAA

TTATATATGGAGCATTTTTCTACGGGTTCAAAACTGCACGCGCGGTATTTTCTTCAAGTCCGAAATTCATATATGTAATA

TCTCCTATGGGTTCGAAGTCATATGAAGCACATATACGGTATTCTCTACGGGTTCAAAAATACACACCCAGTATCTTCTT

CAGAGTTCAAAAATTATATACGCAGTATTTTATACGGGCTCAAGTGCGTCTGCCACGGTATCCTTCTTCGAGTTCAAGAC

TATATACGCAGTATTATACGGGCTCAACTGCGTCTGCAGTATCCTTCTTCGGGTTCAAAATTATATGTGGAGTATTTTTC

TACGGGTTCAAAACTGCACGCGCAGTATTTCTTTCAGGACCGAGATTCATATATGCAATATTTCCTACGGGTTCGAAGTC

ATATGAAGCGCATATACAGTATTCTTCTTCGGGATTTATATACAATATTCTCTACGGGTTCGAAACTGCACACGCAGTAT

TTTCTTCAGGGTTCAAAATTATATACGCAGTATTTTATAAGGGCTCAAGTGCATCTGCAGTATCCTTCTTCGGGTTCAAA

ATTATATATGGAGCACTTTCCTACGGGTTCGAATAATATATGCAGTATTTCCCTTGGTTGCATACATGTAATATTTTGTA

CAAGAGACACGCCAAGAAATTATATGCGCAATATCATTGCAAGGATTATAAAGAAAAACGCCTACTCTTAGGGTTTGCAC

CGACATAGGGAAACAAGAGAAGTATTGGAAGACATACACAGAAGCCGGTTTTTCTTTGTGACTCTGCCGCTAGTAGGAAA

GGATGAAAGGAATAAGGTGCAAGATGAGAGAAAGGGATTTTATATATATATGAGCATGAAGATTCTGAAATAGGGAAGGA

TTTCCCTTTACCAAAAATCGTGCGGGCTCACAAGAACGCCCAACTTCCCAATGCCGCCCAAGTTGCGCCGGTCCCGCCAG

CAAGCACGCTCAGCTTCCCTACCTTGCGCCGGTCCCACCAGCTAGCACGCTCAGTTTCCCAACTCGCAACAACTTCCCAA

TGCCTCCCAAGTTGCGCCGGTCCCACCAGCTAGCACGCTCAGTTTCCCAACTCGCAACAACTTCTCAATGCCGCCCACCT

TCCTGTCTCTCACGACGCTAACTTCCCAATTGTGCCAGCCTCCCAATTTGCGCAAAACAGCCAGAGGCAAGGATTCCAAA

TCCTATTTATAAAAAGGTCGACAAGCCTCCTCAATTCTCTTTACGATTTGCTGCTGATAATTTGTCACCATCTTATGCTT

ACCTCACAACAAATCTCCTTTTCCGAGCTAAGCAGAGGACTTAATGTTGATGGTGGTTTTTAAACAAAGGGTGAAATCGT

AAAACCCCAAATAACCTAATGTCACCAAAGTGACACATTTATTAGGACATTTAATGCATGTAGTCAGTAATTAATAGGGG

GTATTTTTTTCAAATTAGGGTTACAACGAACAAAACCCTAATTCCCCTCCGTGCCAATGGAAAATCATGCCACCACACCC

TAGCCCAAGCTAGCCGCACCTCTATGCCAATGGCAAAGCATGCCCGACCATACCAAACGCGCCGAAGCATGCCAACCATG

CTAACCGCACTCTATGCCAATGGCAAATCATGCCAGCCATACCAAACGCGTCAAAGCATGCCAACCATGCTATCCGCACC

TCTATGCCAATGGCAAAGCATGCCAGACATACCAGACGCGCCGAAGCATGCCAACCATGCTAATCGCACCTCTATGCCAA

TGGTAAATCATGCCAGCCATACCAAACGCGTCAAAGCATGCCAACCATGCTAGCCGCACCTCTATGCCAATGACAAATCA

TGCACCATACCAAACGCTACCAAGCATACCAACCATGCTAGACGCACCTCTATGGCAATGGAAAATCATGCCCGACCATA

CCAAACGCGCCAAAGCATGCCAACCATGCTAGCCGCACCTCTATGCCAATGGCAAATCATGCCACCATACCAAACGCGCC

AAGCATGCCAACCATCCTAGCCGCACCTCCGTGCCAATGGCAAACCATGCCGGCCATACCAAAACGCGCCGAAGCATGCC

AACCATGCTAGCCGCACCTCCGTGCCAATGGCAAGTCATGCCAGCCACACCTAACCCAAGCTAGCCGCACCTCTGTGCCA

ATGGCAAATCATGCCAGCCACACCAAACGCGCCAAAGCATGCCAACCATGCTAGCCGCACCTCCGTGCCAATAGCAAATC

ATGCCAGCCACACTTAGGCCGAGCTAGCCGCGCCATGCCAAACACACTCAGCCTTGGCCCTAGCTACGATGGTCATGCCA

CGCCAAGCCAAACACCCTTACCTTGGCCCTAACCATGCCATGCCAAAATGGCGCCCACGCCTAGGCTTGCAAATCATGCA

ACCTATTCCAAATTGGCCACGCATGCCATTGGCACATGCCAACACGCTCGTTGGTAGCTTTTCCTACCAAGGCACGACCC

AAATTGGCCACGCATGCCATTGGCACATGCCAACACGCTTCGTTGGTAGATTTTCCTACCAGTGTGCGACCCAAATTGGC

CACGCATGCCATTGGCACATGCCAACACGCTCGGGTAGCTTTTCCTACCGGGCGACCCAAATTGGCCGCGCATGCCATTG

GCACATGCCAACACGCCGTCGGGTAGCTTTTCTAAACACGGGACATGATTCAACTTGGCCACGCATGCCATTGGCGCATG

CAAAACAAATATGCTAGTAGCCCAACTTGCAAGCCGTGCAACCTAGGCCAACTTGGCCGTACGCGCCGATGGCACATGCC

AAAACCCACGCCGGTAGCCTAACTTGCAAACCGTGCAACCTACCGCCAAGGCATTCCGCACATGCCATTGGCATATGCCA

CGCACGCCACTAGCAGCTCTATGCATTGCCACTTACACTCAACGTACAACCTACCACCAAGGCATTCCACACATGCCATT

GGCACATGCCACACACACCACTAGCGGCTGGCATTGCCACTTGCGCTCGAGGTGCAACTCGCAACAACGGCATTCCGCAC

ATGCCATTGGCACATGCCACATGCGCCCTAGCGGCACCACTTGCACTCAAGATGCAACCAACACTCGAGACATCAACACA

AACCGGGATGCTTTCCGGGGTCTTGGTTTGGCGGTCACATCTTGCGGCGTACACTACGCTCACATTAAAAAAGCGTGAAA

AGGAATAAAGCGGTCAAATAAATGAGGAAATAATGGTGAAAACGTTTTCACCCATTAACACCATACGGTTTACCACCTCC

TCCCATTATTTCACCCTTTACTCTCTTCCTTCATGAGGCCATGGTACGTTTCAATCTCGACTTGTATAAATAGGTCTTAC

CTTTTTCCACCAGACAACAAGTCTCGATTAGGAGAAAAATACAACACTCTCGAAAAAGGCAACTTTGCAAGCTCTCCAAA

AGGCTTTCATCTCTTGCAAGCTTCCTTTCACTTTCTGATACAAGTCGCGACACTACTCTCCCGAACTATTCCTGGTCTCA

ACACCTTTCTTTGCTTCCCTCCCCAAATCAACCCTTCTCCTCCACTTTGTGATCAAAGCAAGTCCTAACGACCATTTCTT

GGTTTAGGCCGGACTTGTACAGATTGATCTCTCGAATCTATAAAGTGCTCACGTGCAGCACATTTGTTTAGGGTTTGAAC

CAGTTTCTCACTCACCTACCGAATTTACCGAAATCGGCAGAAACGATTTTTACCCATAAACACCTATAAATACCAGTCCA

TGTAAAGGGAGATGCACATGTTATGTAATATTAGATGGTCTTTCTACAGAGAATTCTAGAGAGATATTTAGAGAAGGAAA

GAGTGGGAAGTCTAAGTGATATTTGTAATTCATTCAAGGGTAAGACCTTAATCAATAAAGAAATACTTTCTTTCCCATGG

ATTTACCTACGATAAAAAGCTTGTAACCATGTTATACAAAGGAATTCGTGCTGATATATATCTAGTCCAAAAGTCAATTG

AACTAAGTCGGAGCAAAGTGGTGGAGGCTAGTTGAAGAGTTTTTAGGTTAAATCGTTTTATGTATATGCAACAATAACGA

TTGAATGACAAAAATAGGAAAGAATTCAATCTTCTTTGAAACCGGTAAACTTCCACTAAAGTTCTCCATTCATCTTTAGC

CCTTTCTCTTCCTCCCGAACATACCATCATATCTAAATCAAGGCTCAATCTTCTAGATGGTTACTCGAAATACACAAACA

CAGGTGTTGATGGTGGTTTTTAAACAAAGGGTAAAACCATAAAACCCCAAATAACCTAATGTCATAGGTGACACATTTAT

TAGAACATTTAATGCATATAGTCGGTAATTAATAGGGGGTATTTTTTCAAATTAGGGTTACAACGAACAAAACCCTAATT

CCCCTCCGTGCCAATGGCAAATCATGCCAGCCACACCAAAACGCGCCAAAGCATGCCAACCATGCTAGCCGCACCTCCGT

GCCAATGGCAAATCATGCCCCACATTTAGCCCGAGCTGGCCGCGCCATGCCAAACGCACTCTGCCTTGGCCCTAGCTACG

CTAGTCACGTCATGCCAAGCCAAGCACTCGCCTTGGCCCCGGCCACACCTAGCCCAAGCTAGCCGCGCCATGCCAAATGC

ACTCTGCCTTGGCCCTAGCTACGTTAATCACACCACGCCAAGCCAAACACACTTACCTTGGCCCTAGCCCGAACCTCTCG

TGCCAATGGCAAATCATGCCCGCCACACCAAAACGCGCCAAAGCATGCCAACCATGTTAGCCGCACCTCCGTGCCAATGG

CAAATCATGCCCGACCACACCTAGACCAAGCTAGCCGCACTCTGTTCCAATGGAAAATCATGCCGACCACACCAAACGCG

CCAAAGCATGCCAACCATGCTAGACGCACCTCCCGTGCCAATGGCAAATCATGCCCGACCACATTTAGCTCGAGCTAGCC

GCGCCATGACAAACGCACTCGCCTTGGCCCTAGCTACGCTAGTCACGTCACGCCAAGCCAAGCACTCGCCTTGGCCCTAG

CCATGCCATGCCAAAATGGCGCCCGCGCCTAGGCTTCCAAACCATGCAACCTAGGCCAAATTGGCCACGCATGACATTGG

CACATGCCAACGCGCTCACGGGTAGCTTTCCCTACCAGGACGCGACCCAAATTGGCCACGCATGCCATTGGCACATGCCA

ACGCGCTCATTGGTAGATTTTCCCACCAGGGCGCGACCCAAATTGGCCACGCATGCCATTGGCACATGCCAAAAGCGCTG

TTGGTAGCTTTCCCTACCCGAGGACGCTTTACCCAAGTTGGCCACGCATGCCATTAGCACATGCCAACGCGCTCGTCGGT

AGCTTTTCCCACTAGGGCGCGACCCAGGAGTTGGCCACGCACGCCATTGGCACATGCCAACACGCTCGCCGGTAGCTTTT

CCTACCCGGGACGCGACCCAAATTGGCCACGCATGCCATTGGCACATGCCAACACGCTCGGCGGGTAGCTTTTATTACTG

TGCCAGGACCTAATGGACGAGCATGCCATTGGCACATGCCAACATGCATCTCTGGTAGCTTTTCCCGCTGACGCGACCCA

AGTTGGCCACGCACACCATTGGCACATGCCAACACGCTCGTTGGTAGCCTTTCCTACCCGATGCGCGACCCAAATTGGCC

ATGCATGCCATTGGCACATGCCAACACGCTCGGCTAGTAGCTTTTCTTACCAGGCCACGGCCTAAATGGCCGAGCATGCC

ATTGGCACATGCCAACACGCATCGCTGGTAGCTTTTCCACCAGGCCGTGACCCAAATTGGCCGAGCATGCCATTGGCACA

TGCTAACACGCATCGCTGGTAGCTTTTATTACCAGGCCATGGCCCTAATGGCCGAGCATGCCATTGGCACATGCCGACAC

GCATCGCTGGTAGCTTTTCTTACCAGGCCATGGCCCTAATGGCCGAGCATGCCATTGGCACATGCCAACACGCATCACTG

GTAGCTTTCTTACCAGGACATAGCCCTAATGGCCATGCGTGCCATTGACACATGCCAAACACACACACTGGTATCCTAAC

TTGCAAACCGTGCAACCTACCACCAAGGCATTCCATACACGCCATTGGCTCTTGCCACACACGTCACTAGCGGCTCACAG

CATTGCCACTTACACTCAACGTACAACCTACCACCAAGGCATTCCGCACATGCCATTGGCACATGCCACACACGCCACTA

GCAGCTCTGGCATTGCCACTTGCACTCAAGGTGCAACCCGCAACAAAGGCATTCCGCACATGCCACACGGGCCACTATCA

GCGCCACTTGCACTCGAGGTGCAACCAACACTCGAGACATCAACACAAACCCCGGGATGCTTTCCAGGTCTTGGTTTGGC

GGTCCACGGCTTACGGCGTACACTACGCTCACATTAAAAAAGCGTGAATAGGAATAAAGCGGTCAAATAAATGAGGAAAT

AATGGTGAAAACGTTTTCACCCATAAACACCATGCGGTTACCACCTCCTACCATTATTTCACCCTTTACTCTCTTCCTTC

ATGAGGCCAGGGTACGTTTCAATCTCGACTTGTATAAATAGGTCTTACCTATTTCCACCCCGGACAACAAGTCTCAGTCG

GGAGAAAAATACAACACTCTCGAAAAGGCAACTTTGCAAGCTCTCCAAAAGGCTTTCATCTCTTGCAAGCTTCCTAAGCT

TTCACTTTTTGATACAAGTCGAGACACTACTCTCCCGCACTGTTCCGGTCTCAACACCTTTCTTTGCTTCCCTCCCCAAA

TCAACCCTTCTCCTCCACTTTGTGACCGAAGCAAGTCCTGAACGGCCATTTCTTGGTTTAGGCCGGACTTGTGCAGATTG

ATCTCTCTAATCTATGAAGTGCTCACGTGCAAAGCACATTTGTTTAGGGTTTAAACTAGTTTCTCACTCACCTCACCGAA

TTTACCAAAATCGGCGGAAACGATTTTTACCCATAAACAATTGGCGACCACCGCGGGAGGATTGATCTATAAGTTACAAT

ACCAACTCCGACTCCTGATTTTGTTTCCCGAACGTCAAATCCCAACATGGTGGAACTTAGGTCCGGAGCGGCTGCCGAGA

ACACCAGCATAGAATGCATCCCTGATGTTAGCGTCTCTCCCAATAACGTTGTTGCATCACCTGCTGACTACAGTAACATC

GAATGCACTCCCTCAAGTGTAACACCGCCAATCACCAAGGACAGAGCCGCTGCCTCCTCTAGTATCTCTGCACAACCCTC

TAATCATGGCAACGAAAATATCTCTTCGGGTGTCACACCACCAGTCACCAGGGCTAGAGTCGCCGCTACGACCCCAAATG

TTTCGTCTAGCATGGCGCCTCCGACCACAACTAAGGTTCCAACTACCCGCCGACGGGACGAGAGGCCTAGAGGGATCGGA

GGAAGTCGTCCTTACGTCACCATCGCTGATTTGATGGAACGACAATAGATGCTCGCCCAGATCCCGAGACGGATATGGTT

TCAACGAGAAGAAGGTACTCATCTTTCTCAAGACCACGAGAGAGCGACTACCACGTGAGTCGCGTCCTCCCGAGCCGCTT

AGAAACATCTCCAACAACCCGACACCAGCACCTCGGCGTAGCACGCCTCCACGGATGAGGAAACTCGCGCTAGAATCCGT

TGGAAAGAAATCAGCCGCCCAACCGTCACGAGAGAATATTTGGAACAACTTCTCCGCAACCGAATCAAGAATGATGAGTT

GGACGTGCATCATCACCAACCCCCTTACCCATCCGAAATGCGAGAGAATCCCTCTCCGAGGGGGTACTCTTCTCCTCAAT

TTACCCCCGCGATGGGTACGGAATGCACATGAGCATGTCTCTCGTTTCTGGAATCCCTAGGTGAACATGAATATAATCAC

GTTCTTCGTTTAAGGGAATTCTAAAAATCACTTACTTTGGAAGGGCGTATACATGGTACAATGACATCCCACCGAAAAGC

ATTCCCAAGCTGGAGTAGCATGGTCACTGCGTTTTACAAGAAATACTTCTTCGTGTCCGAACAAGTTGCTTTCTCGGACC

TCGGAAGAATGTACCGGTGGGATAACGAACATCCGAATGATTTCGTCAAAAGGTTCGGGGTTCAAGCACTTTCACATGTC

ACGACCCAAACATGACCGAGCAACAACGGTGAACGCATGCATCAATGGTATGATTCCAATTTACCGCACCCCTTTAGAAA

ACTTGCGCTTCCAAACGTTCTCCGAACTCCATGAAGCCGCGGACGATCTGCAACAACAAGACAGCACCAAAGACATTCTT

TGGAAAGAGCTAGGCCAAATTGAACCGAAGACACTCGTGAAACTCGCGGGAGTAGATGCCTCATCAAAGAAACAATTTAA

TGCCGGCTCATCCTCCGTGAACATAGTAAACGAAAGGGAAAAAGGAAAGCCCCAACTACGAGCCACCGGCCAAAAACGTG

AAGCGGCGGGCGCAGCGGGAACCCCGCCACGAAAGGCCGTCACTAAAACTCCTATGCGACATCATGAGACCGAAGGAAAT

ACCTATGATTTCCCGTTTCCCATCAAGGAGGTGATTGACCTCCTAGAAGCCAGGTTCAAGATGATATTATCAGGCCCGCC

ACCTACCCGGACGTCTACCAACCGAGGCCGAGATGGTGAACCCCAAACGCCACTACCACGGGTTCGTTCACCACCCGACC

AGCGATTACAACAGACTAAAGCATATTTTCAAAGAAAATATAGAAGTGGGAGGACTCCAACCGAGGAGATGAAAGAGTGC

ATCGAGACCCACTCCTAATAAGAAACTGCGGGGTCCTCAGGGGACTCGTGCACGAGACTAGTACAATCCATGATGCAACA

TGTATGCGAAATTCTCTATCTATCTAAGGTGCAGCGCCAAGATATTTTCGCAGCTCTCAATCGTGTTGTATCCGACAGAC

GACTTTTTCTCTCCTCGGAAACCATGTCGGGGGCCCAAGCTTTCCAAGCAAAGGAACCAAAAAGGAGAATCGGGACTCGA

CCACGTCCGTTTAAGAGGTAACGAATTTAACGACACGCTCATCGACACGACCTATGACCACAACATCATTACTTTCAAGA

CACTAAAGGCATGATTTACAAAACAAGACGCTACCTATTCCCCGTCGTGATTAAGAGCTTGAGAGGAGAAACTTTGAACG

CGCATGGCTACGTCAACCTAGAGATCAAGGTGGGAGGCGCTATAACGCATAGCAAGTTTCACATTGTCAAAGAACTAGAT

CATGACATGATATTGGGACGCTTATGGGTGCTCGACAACAAAGCAAAATTGGATGTACACCCTCTGCCGACGTCCACACC

CGAAGGAATTCCCACGGCGGCGCCAAAAATTGATGGTTATTTTCAAGGATGTTGTAGAAAGTGGTAGTAAGGTTGTCGTT

CACTCGGACTTGACGAGATTGATTTTAAGAATTAATAAAAGAAAATAAAAGAAAAAATATATACAAAAATTATCACAGGA

TGAAGAGTGTTACTGGGACTAGGATTTCGTCGAATTCATAACATAGGGTCCAATTTATTTATTCTCAACAATTAAAGCTC

AATAAATAAAATTAACACGGACTCTGATTTTGCCAAGGTAGATTCTTAAAAGATTGGTTGTAAATCCTAAGCATGATGTA

TCAAAATATTTAAGCTAAGCATACCTCATCAAATTAAATGACAACCAATTAATTCAAATCATATTTCAATTAAAATTAAT

GCAAAAGTCTTAAAAAGAATTAAATAATTTTACCCATGTATGAAATTCGTCTTCCTCCATCGTCCAGAGTGTTGGAGTCT

AGCTCATCATGTTGGAAACACGCTCAAAATATGATTTTATTGCTCAAAGGTGCTTACAAATGATGAAATGGAGAAAATGA

TGAAAATCGGGTGTTTGCAACGTTTATAATTGTTGCAAACACTGTTACAGAACTGTTGCTTTAAGACTGCCTCTTAACTA

CGACCCTCAGGAAGGACTGTCGCCTGTGGGTCTTATGTTCTTCGTGTTCTTCCTTAATGGCAGCAGCAACAGATTTTCGA

ACTCTCGATTATTTGCTCTGTAATGCTCTAAACTCTCCCAAACTTTCCATCTCCGTTCTATGACCTCCCAGCACCTATTT

ATAACCTTTTTGGAGCAAGAATAACTCTTCATTAATCCCAGAAATCTTCCCATTACTCGGCGGAATAATATTGTTCTCGG

AATATATTCTTTCCATTTTTTGTAGCAGCACGCGCCTGCAGCTGTTATTTTCTTCCTCATTTACCTTCTACACTCGTCTG

GCAACAACCCACACACTCCAAACACGAGCAGTGCACGTCTCAGAGCTCACAGCGATTCACGAAATACCCATGCAAACCCG

TTCTTCTCTCGCTTTCTCCTAGCGAGTTATCTAGCCAAATTTAGGTCACTAATCGACCCATACCAACCTGTTAGGCTCTA

ACAAAGTTATCCCAAGCAATCTCGGCCATTGAGTCTCTCCAAAATGCCTTCAATTCTCGAACCAAAATTTTGGGTAAGAA

AATAAGTTTTCCAGCAAAACCGGATTTTCAAACGATGAAGAAGGTCAGCCCTTATCCAACAGCTGGGGGTGCGAATATCA

ATTGGTTGCCCCTTAGTAATTGGGTTATCCCTTATCCAAAGTGGGAGTCCGAATATCAAATGTCCTCAGGGGTGTTAGAG

CAACTTTTCGAGCCGAATTTTCCAAAAAATATTTATTTCCAAAAATACCTAAAAATACATAAAAACACAATATTAGTACA

AAAATCGAGTTCCAAATGATACCAACAATGAGGACAATTCGGACACAAAAATGTGTCTATCATCAGCAGCCATTATAAAT

TCTCCTGTTGCAAAAAGCCGATGCAACCTAATTTTTACCCTTACAAATTCTCATGTTACAAATTCTCTTTTGCCCTTCAG

GCAAGAGCCATTACAAATTTTTTTCGTGATCCGTAACAAGGTTCGGCTGTTACTAATCGTAATAGGATAATTTATTACGT

GTGCTAATCACACGCTTGGTCACGTTCGGATGTAATTAATCCATTTTCACATTATTTGTAATGCATACATTCTCTTACTT

GCCCTTGGTAAATAGGTATGACCTAATTCTAACATACATGGCCATCTTCATTTTAATTCATAAAAGAGAAAGGCGCAGAG

GAATATGTATGAAAATGAAAGGTTGATGAGAGAGTTAGAAACTCAAAGAAAAGAAAATCAAGCAGTAGGGTAAGGAAATA

GATAATGCGATGGCTGAATTGGGGAGGTTACTTATCTATGAAGCGTCAAGGACCTAGTACGTTGGCTTATCTATAGGTTA

CTTAACAGGCCAGAGCCTGTGTGACTACTAAACACTGATGATGAGCAATATGCTTTGGAGTACCAAGATGGTGGATCATG

CATCGTTACCAGAGATGTAGATGGTTAGCATTTGAGTAGAGGAATCAGAATCACTCTCTTCCTCAAAGAAGGTAAGGTAA

TCAAAATAACTTCGGTCATTATTGACTGAGTCGGACCAGGTCAATGTTGACTGTCAGTAATTTTTTGATGTTACAAAAAA

GTTTGTAATGGCTTAGGGCTGTTACGATTGATAGGTGTAGAATACACCGTATTTAATATCTAGTATTTATGCTTTCTTTG

CGAGTTTTCTTGTAAATTATGTATCTTATGTGTGCTTTTGAGTTTATTAGGTGCAATGGAGTCATTTGGAGCAAAAAGAC

CAATTGGAACTCGTTCAAATACAAGATTTCTACTTTTCATTGTGTAGAGGACGAGAAGACGAAGCCAACGGCAAAGAACG

AAGTCATTCCGAGTTCGTGTGGAGAAGTTACAAGCAAAACGAGAATTTGAAGGCTAAAAAGAGCAAAATGGTCGGACAAG

TGCTATTGGTAACCCCACTGTACGTTAAGGGGTAGCCATTCTGTTAAGGGGTGAACTGTTCCAAAGGGACTACTACTGGG

AGCCCATCCATCTGTTAAGGGGCAACTCTCTTTAGGAGGAGTGTTAAAGACAACCAGCTAAGGTTATGGGGCTATTCCTC

GGGTTCACATCATTAGACCGTGGGTGCGAATAGCCATTCCCACATAAAACTCAATTGGGGTCGACTGTGTTTATACTCAT

TCTTCCCATTTGATGAATCGAGAGAGGATGACTCGAGTTTTGAGAGGATTAGGGTTTGAATTTGTGAGATAAGAAATCGA

GAGATGGAAACAATGAGAGCTGACGATGGTGGTGACGAAGGATGGAGGCTGCTCAATCTGAGATGTGAGTATTTTTTGCT

TTGAGATCGAGAAATTGGATCGAAGTAAGAATTGAGACTTGGGTTTCTCTCAAGATCGAGTCTCCGGAAGAAGATGGTGG

AATTGATGGATTCGATGAGGATGATAAAGCTCTCGGATAAAGAAGAGGATGTTGCCGTGAGTTAATGAGTATAGTCCGGT

TGAGATGAAGAATTCAAAGTCCGAGTTAATGGAAGGTCGGAAATGAAGATGATTATGGTGGTCACGATCCGGGGCTCGAG

CTTTTAAAGGGATTGTGATATGGGCTTTGAATAATGTTCATAGCCAAGAATCAATTGTTGCCATGGGTCTTTGTGGAGGA

ATGGTTATTCCTTGGCAAGTGTAGCCGGAATTGAAGCTCAAGTTATGCGATTCGGTATTGAAGAAGTGATGTGTCCGTGG

TACTTGTGGTTGTTTTTATTGTTATGGAGATTTGCAATGGAATATTTGCGGTGGTGGTTGAGTGCAGCAGTGACGGTGAG

GGATGTTTGCTGAGTTAGAATGGATTTGAAGGTGGTGCAGAGAGTAATGAATGCTCCTGGCGATTGAAGCATCAGAGAGG

ATTGAAGCTTGGGTTTCTCTGGAATTGATTCAAGATAGACTGGTGGTACCTGCAATGGAGTGAGTTACTATTGTTGCAGC

TGATTTGTGGCAGTGAGCTGCTGCCAAGGAAGAAGGATATTAGGATGATTCTGTAGGCATTGAGTCGATAGAGACCAGGG

AATGGAATGGAAAGCTGCAGAGTGTTAATTATGAAGCTGGTGGTTTTTACAGGTCAGGTGCAGTCCAAGGATTGGTGTTG

CCATGACCTGAACTGCATTGCAGGGCTGAGCAATGCTGGTGTGTTGTTTGAACTGAGCAGAAGTGGCTGAGGCTCATGGT

ACTGTGGTACTAGGCCTGGGATAGTGATGGAAAGGTTTCATGGCTGAAAAGGAGCAAAGGGTCTGTGGTTTGTGGTGGTA

CTGAGCTGGAGGTGATTAAGATGCTACTTGAGAAGTTGATGGAGTTGATGGGTCTTGAATCTGTGAAATTAAGGCTGGAT

TGAATGACAAGGTTGCAGGTCTGAAGGCATGTGTGCTGAGGCTTTAGGGTGTTGGCTAGAGAAACAATGTTGCAGGGCAA

AGCACAAGCAAATGAAGCCCTGAGCTTAGGTGAAGCTGGGAGTGGATTTTTGTTGCGGGGGGATTTGAATCGATGAGGAG

CTAAGGAGTTGAAACAATGTTTGGATGTTATGAGCCCCGAGCTTAGGTGTTTGTGTTGAAATTGAAGCTACAAAAGGTAT

AGCCATCATGGTGATGAAGCACAATCATAGTGCATGACCAATCATAGACACACACTGTGCAATGGCCTAGTCCCGGCCAA

GGCAACTCAGACCACTCGGCCTTGACTAAGAAGCTGACTTGGCTACCTGACTCACCTGACTGGACCGGTTTGACCGATTT

GACCCGACTGGTTTGACTGATTTGACCCGACTGAGTTGGACCGTTGACCGGTGACCGGGTTAACTTTGCCGGTCACCTGA

GATAGTTTTCCGGCGACTTTCCGGCGGGATTCCGGCCATAATCCCAGTTTTCCGGCGAGCGTTTTTGGGACATAATTTTT

ACACGGGTTTCTCTCCATGGTGGGCCGGGTGAATTTCCAACTAATGGACTTTGAAGACAAAACCTAGTTTTGGAAGGAGT

GGAGCTATATAAAGAGAAGACTTTTATTATCTATGGGATATCTTTTATATACAACGTTTTATTATCTCTTCTACTTTGTT

CTAGGGTTTAGAACCATGAAGTCTTTTCTCTTTCTTCTTGTGATGAACTCCTTGAATATGAGTACCTAAGTTTATTATTG

GTTATGGAATAGTTGGATTTCACAAAGCATGATTCTTTGATTCAATATATTAGTTTTGTCTCATAATTATCGTTTGCAAT

TCTCTGAAAATATTGTTTTATGATTGGTTGATTGTTTTGTCAAGTTGCAAATTGGTAGAACTTCTAATCACATCTATTGC

TAGTTTAGGGTTTATTATACGCAAAAGTGATATTCCTTGGATACAAGTAGCATAATATGATTATAGTTTGTAGTGATATA

TTCTTATGATCATATGTGAATCGTAACCTTTGTTAAATTGAATTAGTGCGCTTTCCACGTTCAATTGCTTTGATTGGCCT

GATTTCATAGAACTTAACGCATCTAGGGAATTAAACTTGATTAGTGCTTTTACACGTTAGGTTGTTCCTTTAGGTAGAAT

TCATATTCACATCTTTTAATGTGTTTGTTGATGACATGAGGAGATTTGCGGGATATTCTTGCGATCAAGGTATTCTAGGG

TCTTTGATAAAAGTTGAGATTAAAATATCAATTCTAGTTATTGTGTTGAAAACATGTGCGTGAATGAAAACGAAATCCAT

GACCAATATCTTCTTATCCTTGATTTTATTCGTTTACTTTTCTTTATTTGTTTTATTTAGTTTATTATAAAATCACAAAT

CCACCCGGTGTTTTATATAGGAAACCATAACAATTTGACAACCTAGACCTCTCTCTCTGGGATCGATCCTTTCTTGCCCT

TGCTATATTTATATTTTGAGTAGTAAGAAAGTGTAATTATATTTGTGTCGCTACGGACAGCGATCAAATTTTAGCGCCAA

GTGTTTGGGAAGGCAACGGTTGTTAATTGTTTTTGGTTTCTTATTTTTGTATTTAGTTTTTTTTTGTTTTATTTTATTTT

TCTTATTTTCTGGGTTCTAACTTGGTTTGAGAATCTTATTTCAGGTGCTAGTTTCTCATGGAAGGAGCATATTGGAGCAA

TGGATATGATTTTCAACCTCCTCAACACTATGAAAAAATTACCCACCATCTATGGGATGCAATCAAAACCAATTTATTGG

CTATGATGCATATCCTAGAGAACCTTATGGTGATTTTCATACATACCAACAACAACCTTGTAGTGGGTATGTAAGTCAAG

CACCTAGGTGGGATAATTCTACATCTAATTGGCTTTATTCGAGTTGTATGCAGATTATGAATGGGTTACAAAACATGCAA

CAAAGGCTAGACCAACTTGACCGTGATAGAGAGAACGTCGCACAAACCAATTTTTGGAATATTCCTTCTTACCCGACTCC

GGATCGTAGTTATGATCATTCACCCATTCAACGGGAAGAGTCATGGGAAAGAGAACATATTGTAGTCAATGGTGGTAAGC

GTGTGAACGTCAAGAGACTTGCGCGAAATTATACAAGTTGGAAGATGATGGAGCCTCGGAAGAGCTAGTCAATGAAATTA

GTAGGACCATTCATATGGAACTTAAACGACGTCGTGATAAGGGTTGGGAAACCGACGACGATTCTTATTATGGTGAGTCC

GAAGATGAAGACGAAGATGAATGTGTTAAAAAGGACCAACCTTTGGAGATATTTGATGACATTAGTGTAGTTGACTCAAC

TATTGATCCTTATAATGATCAACCACCTATTCAAAAGGTAATTGAGTATGATGAAACTAGTATTTTACAAGATTTTGCAA

AAGAGTATGTGCCTTACAACAAAGAGGAGTATGTTGAGAATGAGACCGGTTTGACCAATATTGTGGAAGACCCAATTGAG

CTTGCGTATGATTGTAATGATGATGTTTATGTCGATACTTGATTACGGCGAAACGGACGAAGAATGGTTGCAAGGTTTGA

TAGAGGACCATGATATACAAGAGGTGAATAATTCACTTGAGTTTTTCAAGGATGAAGAGGATTCCATAATACAAGAGATT

GTGAAAGGTTTATCTAACCCTTTGCCTATTATTTCTTCTCCTTTACCTCTTATTGGTTTGAATGTATGTGCTTCGAAAAC

ATTGTTGAATGACTTTGTTTCTCGTTTTCCGCCATTAGAGAAACTTGATTCTCATACTATGCAGGTTATGGAACCCGAAA

CCGTGGAAGTACCCGATGTCTATATTCTTTATACACTTCCAAGTGTGGACAAGAATAAGTGTGGGGGAAGCTTTTATCGG

TTCATAGCTTCACTTTTGCTTTATGTTACTAATATTTGTGTGAAGCTAGTGTTTGAAAAACAGGTTCTATGGAAGGATCC

CCAACTTTTCCGATTATTGGTGTATGGGGAATGAATCTTACAAGAAGCTAAGTCGGGCCGTCGACTTTAAAACAAAAGCG

CTTAGTGGGAGGCAACCCACCGGTTTTGTATCTCTTATCCTTTTTCTTTTTATTTTTGTTTCGTTTATTTATTTTTTACT

GTTTTTATATCATATTTGCATCCCTATGTTTAATTTCATTGAGTCCGTAATCTATCTTAGAAGAGAATTATGACGAATAC

CTTTTCGTCAGAAAATTTCAGACTGCGCGTAGTTCAGTCCAGAGACCATAACTGTCAGACCGTTTATCAAAACACTGTCA

AATTTTGACACTGTGTAGAAAACACGTAGGTGAACAACTTTTGTGAAATGGAGTTTTCCAAATTCCTTACCAATTTGCAC

AGTTTTTGCGTTTTATCTGCTGTTACGTCAGAATTCAGAACTTTCGGTTTTACACATTGAGGACAATGTGAAGTTTAAGT

GTGGGGAGAATTTATATGTTTGTATAGTTTGTTTTTGACACCATTAGTGGATGGAACCATTTTGAATGCAGGAGTTGAGG

AACCAATTAACTAAAAGGGAAGAATATAGCTCAGAAATCACAAATTTATATGATAGTTGTTACCATATCTCGGAATATCA

CCATATCGCATTCTATTTTTATTTTTCCATATTTATCTATTGTTCTCTAAGTGCTTTGGTGGGTCACTAAAAATCGACGT

GTTATTACTGCTAGGATGAAATAGAGTGGTTGAGTTACATGAAAAAAAAAAAAAAAAAAAGAAGAAAAAAAAAAAAAAAA

AAAAAAAAAAAAAAAAAGACCGATCATTTAGACCAAAACGGAATAAAAAGACACTTGGGAATAGCGGTATGTGTGAAACG

TAGTCCCCGTCTCAAAATCGCCTAAACAAGCGTGGAATGCGGGATCCAAGGCAACTATCATGTACTTGCGACAAGAAGCA

TGCGCTTGGATCCAAAAGATCTAATCATGCCGTCTAAAAAAAAAAAGAGAAAATTGGATCCAGTTTGATCCAATCATATA

AAAAAAAAAAAAGAGAAAAAAATTATTATTGTGTTGAATAAAGTCACTTTTGGTTCCCTTGTATATGCCAATTGATTTGA

TGTAGCATTCAAGCACTAAGTAGCTCAATCCAATAGGATTAGGTTCATCTTAGCAATGACCTTCGAGACAGAGATATGGG

AAACATCGAACACTTGTGAATATTCGCCAACATCTACATCTATATCCATCTTCTTAATCTTTTCATTGTGATTGTGGTTT

GTTAGTTTCCGAGTATTGTGGCTTGTTCGACTATCTAATTAGAGTTTATTTACTAATATATCAATTGAATTTGCAAGGTA

CCTCCTCCTGTAGTTATTTTGGATTCGTTCATTGATTTGTCACACATAGACGGAGTTTGAAATATAGGTTTTGTAGGTAT

ATCTCTAGTAAACCTTCACGAGACTATAACTCGTCCACTAGGGACACCTAGGGGTTTAAAGGCTTGTTATTTATGCTAAA

ATTAACCGTAGCCTCGACGACACGGAGTTGTTGTATTTTAGTTTTGTTTTGTTTGATTTGCTCGAGGACTAGCAAAGTCT

AAGTGTGGGGGAATTTGATAGGTGTAGAATACACCGTATTTAATATCTAGTATTTATGCTTTCTTTGCGAGTTTTCTTGT

AAATTATGTATCTTATGTGTGCTTTTGAGTTTATTAGGTGCAATGGAGTCATTTGGAGCAAAAAGACCAATTGGAACTCG

TTCAAATACAAGATTTCTACTTTTCATTGTGTAGAGGACGAGAAGACGAAGCCAACGGCGAAAGAACGAAGTCATTCTGA

GTTCGTGTGGAGAAGTTACAAGCAAAACGAGAATTTGAAGGCTAAAAGAGCAAAATGGTCGGACAAGTGCTATTGGTAAC

CCCACTGTACGTTAAGGGGTAGCCATTCTGTTAAGGGTGAACTGTTCCAAAGGGACTACTACTTTGGGAGCCCATCCATA

TGTTAAGGGGCAACTCTCTTTAGGAGGAGTGTTAAAGACAACCAGCTAAGGTTATGGGGCTATTCTGGGGTTCACATCAT

TAGCACTGTGGGTGCGAATAGCCATTCCCACATAAAACTCAATTGGGGTCATTGTGTTTATACTCATTCTTCCCATTTGA

TGAATCGAGAGAGGATGACTCTGAGTTTTGAGAGGATTAGGGTTTGAATTTGTGAGATAAGAAATCGAGAGATGGAAACA

ATGAGAGCTGACGATGGTGGTGACGAAGGATGGAGGCTGCTCAATCTGAGATGTGAGTATTTGCTGCTTTGAGATCGAGA

AATTGGATCTGAAGTAAGAATTGAGACTTGGGTTTCTCTCAAGATCGAGTCTCTGGAAGAAGATGGTGGAATTGATGGAT

TCGATGAGGATGATAAAGCTCAGATAAAGAAGAGGATGTTGCCGTGAGTTAATGAGTATAGTCTGGTTGAGATGAAGAAT

TCAAGGGTCTGAGTTAATGGAAGCTGGGAAATGAAGATGATTATGGTGGTCACGATCTGGGGCTCGAGCTGTTTAAAGGG

ATTGTGATATGGGCTTTGAATAATGTTCATAGCCAAGAATCAATTGTTGCCATGGGTGCTGTGGAGGAATGGTTATTCCT

TGGCAAGTGTAATGGAATTGAAGCTCAAGTTATGCGATTCGGGTATTGAAGAAGTGATGTGTCTGGTGGTACTTGTGGTT

GTTTCTGTTATGGAGATTTGCAATGGAATATTTGCAGTGGTGGTTGAGTGCAGGCAGAAAGCAGCTGATGGATGTTTGCT

GAGTTAGAATGGATTTGAAGGTGGTGCAGAGAGTAATGAATGCTCCTGGCGATTGAAGCATCAGAGGCGATTGAAGCTTG

GGTTTCTCTGGAATTGATTCAAGATAGACTGGTGGTACCTGCAATGGAGTGAGTTACTATTGTTGCAGCTGATTTGTGGC

AGTGAGCTGCTGCCAAGGAAGAAGGATATTAGGATGATTCTGTAGGCATTGAGTCGATAGAGACCAGGGAATGGAATGGA

AAGCTGCAGAGTGTTAATTATGAAGCTGGTGGTTTTTACAGGTCAGGTGCAGTCCAAGGATTGGTGTTGCCATGGCCTGC

ATTGCGGGCCGCAATCTTTGGTGTGTTGTTTGAACGGTGAGCAATCTTTGGTGTGTTGTTTGAAAGCATGGCTGAAAAGT

GGTGAGGCTCATGGTACTGTGGTACTAGGCCTGGGATAGTGATGGAAAGGTTTCATGGGTGAAAAGGAGCAAAGGGTCTG

TGGTTTGTGGTGGTACTGTGCTGGAGGTGATTAAGATGCTACTTGAGAAGTTGATGGAGTTGATGGGTCTTGAATGCACG

TGAAATTAAGGGTGGATTGAATGACAAGGTTGCGGGTGTTCTGAAGGCATGTGTCTTTGAGGCTTTAGGGTGTTGGCTAG

AGAAACAATGTTGCGGGGGCAAAGCACAAGCAAATGAAGCCACAGAGCTTAGGTGAAGCTGGGAGTGGATTTTTGTTGCA

GGGGGATTTGAATCTGATGAGGAGCTAAGGAGTTGAGCTGGATGTTATGAGCTGTTTGTGTTGAAATTGAAGCTACAAAA

GGTATAGCCATCATGGCTGATGAAGCACAATCATAGTGCATGACTGACACGGCTGTGTCTGTGGCCTAGTCTGGCCAAGG

CAACTCAGACCACTCAGCCTTGACTAAGAAGCTGACTTGGCTACCTGACTCACCTGACTGGACCGGTTTGACCGATTTGA

CCCGACTGGTTTGACTGATTTGACCCGACTGAGTTGGACCGTTGACCGGTGACCGGGTTAACTTTGCCGGTCACACCGAG

ATAGTTTCCGACTTTCCTATGGGATTCCGCCATAATCCCGAGTTTTCCGGCGAGCGTTTTGGGACATAATTTTTACACGG

GTTTCTCTCCATGGTGGGCCAGGTGAATTTCCAACTAATGGACTTTGAAGACAAAACCTAGTTTTGGAAGGAGTGGAGCT

ATATAAAGAGAAGACTTTTATTATCTATGGGATATCTTTTATACACCACGTTTTATTATCTCTTCTACTTTGTTCTAGGG

TTTAGAACCATGAAGTCTTTTCTCTTTCTTCTTGTGATGAACTCCTTGAATATGAGTACCTAAGTTTATTATTGGTTATG

GAATAGTTGGATTTCACAAAGCATGATTCTTTGATTCAATATATTAGTTTTGTCTCATAATTATCGTTTGCAATTCTCTG

AAAATATTGTTTTATGATTGGTTGATTGTTTTGTCAAGTTGCAAATTGGTAGAACTTCTAATCACATCTATTGCTAGTTT

AGGGTTTATTATACGCAAAAGTGATATTCCTTGGATACAAGTAGCATAATATGATTATAGTTTGTAGTGATATATTCTTA

TGATCATATGTGAATCGTAACCTTTGTTAAATTGAATTAGTGCGCTTTCCACGTTCAATTGCTTTGATTGGCCTGATTTC

ATAGAACTTAACGCATCTAGGGAATTAAACTTGATTAGTGCTTTTACACGTTAGGTTGTTCCTTTAGGTAGAATTCATAT

TCGCATCTTTTAATGTGTTTGTTGATGACATGAGGAGATTTGCGGGATATTCTTGCGATCAAGGTATTCTAGGGTCTTTG

ATAAAAGTTGAGATTAAAATATCAATTCTAGTTATTGTGTTGAAAACATGTGCGTGAATGAAAACGAAATCCATGACCAA

TATCTTCTTATCCTTGATTTTATTCGTTTACTTTTCTTTATTTGTTTTATTTAGTTTATTATAAAATCACAAATCCACCC

TGTTTTATATAGGAAACCATAACAATTTGACAACCTAGACCTCTCTCACAGGATCGATCCTTTCTTGCCCTTGCTATATT

TATATTTTGAGTAGTAAGAAAGTGTAATTATATTTGTGTCGCTTACGACAGCGATCAACGATGTGTTACAAAATAATTTC

GTAACGGTTGTTACGAAATTGCCACATAATAACAAATAGCGGCCGTTACAACAGCCGTTACAAAATAAATTCGTAACGGC

CATTACAAAAAATTTGTAGTACCCCTTTTCATTACAACAGAAAAGCTTTACAACCCAGATTCGTAACGCGTACTTGCGTA

AGGTACAGGTCCGGGAATTTTTGCTGAAGTTTGGAAGTGTTAAGGTATGCGTACACGTTCGCATAATGGCGAACCTAAAC

ACAGTCTGGTCACTAAGGTATGCATACCCGTTTGCAGACCTAAGTGAATCATGTTCTAAAATCGCCTTGCTCATGAACTA

ATACATTTATATAATAAGGAATGCATTCTTTGCAAACCGTGGCTATAATGTTCATGAATTGATTCGAGTGAATCAAACCG

ATTTTGCTTCAATTGTGTTCTTGTATAATTCTATGAGAATATAACAATTGAACAACTCTGTAACTAGTTTCATTTGAATC

ATTTGAACTAGTTATGGTTAAGATGAATATGGCTAAATACCACGTGTATGATTATTAATTACCTAACTAATTTGTGATCG

AAATTCTTAAGTATAACAGGTTAGAAGCAATAATCAAATCCATTGGATGTTGGTTTTTATTGTAAAAAAACCTTCTTCAT

ACGCTTATGGTAACTATTCTTGGTGTAAATCCTTGTGGAAAAGGTTATTCTTCCTTCTAAGCATATCAATTCTTGTTTAT

ACAAGTTTGTATCTTAGAAAAGATGATCACAATATTTGATTTTCATGATTACATATTGTGTATTGTTTCCATGAGATAAT

TCTAATTTTTTTAGTAACACGATCTCATGAGATACTTTCAATGATTGGTTCTTAAATCATATCTCCTTGTGGGATTTCTA

AAATCCAATATATAAATCCTTCTCCCTGGAGTAAGGTCACTCTTGTTGTTCTGTCGGAAATAACATTTTATGGGAGAGAG

TTCTTAAATGAACTTTTTAGGGAGTTCTTAATTGTCATATATTTGTAGGGAAAAGCGGGTTTGGAACATGAAGGGGTTAT

CTTGTATCTTTATAAACTCCTTGATGAATGCATTTAGTTTCGGCTATATGATTGCATCTAAACAAGTTGATATGCTGTTA

TCTTTTGGTCATGAAATATCTCTATGGAAATTTCATTAGGATCCCACTAGTTTTCGTACCTTTGCCAATTTATATTGAAA

AAAAGGGGGAGAATTAATGTGTAGTTTACACTACAGATACATATGATTTACATATCATTATGTAAGTGGGAGTGGTTTTC

ATTGTGAGATGAAGTATTGACTAAGGGGGACTGATACATATCACCGTAGTATTATTGTCAAAGTTGTGATACAATTGAAC

TTTGATAATGTGTATAATACTATGATGGATCTTCAACAAGTATAGGATGAACACATGCATAGTTGAAGAACTAAGGAAAT

CAAGCATGTCGGATTTTGGTTCTCTAGTTCAGTTAGAACCTTGCAGGTTAACACGGCTTCCTAGGCGCTCCCATACAGAG

CGACCTCTCCGATTTTGATTAAAATCTACCAATCGATCTTGATTTTTCAGATCAATAGTCCTTGATCGAGACTGGCTCAC

ACGTTCTAGTCTCGGAGGCATTTTATTTCTCCACCTCATTAATTCGCCGGCGCAATAGATCTTCCTCTCGTACTGGAGTC

GCGGTTCGTCCATCTTCTTTGATCAAGTTGATTATCGTTTCTCCTTGATGGGCTGCCGAACACCAGTTCTGTGTTTACCC

GAGCTCGCGGGTTTCCTCCACTGGGTACGGGGTCAACGTCCATTCCTGCGCTATCGATCAATGGTTGTTTTTCGTTTCTC

CGAGTATTTGGATCTGGAGTTCTGTCATGAACTCGATCCCGATCTCTTTCATTTCCAGGAAGGAAATGTAGGTGCGAATT

GAACCTCGTTTGGTTCAGTCATTGCGCGTCGCGGGTAGGTAGCAGAAACGATCCTTCGAGCAGTTAGCTTTACTTTTGCC

ATTATCTTGATCCGAGAAAAAACACGATCTCTTTTCTCGAATCCCGTTAGTCGGCACCAATGTTTGTAAGATAAAAGATT

TTTACCTCAAACACGAATCGATGTTGATGTTTATCGATCTCTGCCTTTATCTTCACGTGAGCAGTCTTTGAAAATTACCT

GCAAAAACCCTCCGATGCTTAAGTTAGCTAGAGTTTTTCATAGGAGTTTTGTGTAGAGAAGATTGCGTACCTTTTAGGGT

TGTGAACCTATGTATTTATATGTTGCAGATTAAGGATTAACCCTAATTTCGAGGTAAGCATAACTGTGTCTACTTGCCCT

GCAAGTCTCCTGGAATACTTGGCTAACATTCCTTCCAAATATTTCCAGAAACTTCCTCTAGGGTTAAAGGTGCAAAGGGT

TGGCCCAAACAAATGTGTAAAATGCCAAGCCCATGAAGAGTAATTCACTTAAGCCCATAGTGGGGCGTATAAATCCGCAA

GCCTAACAGCATGGGGCTGAAAATTCCATCAAGATATTGGCTTGACTCGTACTAATCAACTCAGCTGGTTGAGTTGGCTC

AGTGGGGCGCAAATATCCTTCATGTAAAGAAAATGGGCGCAAATATCCATAGGGTACGAGAATAAATTACTTCGTGATGC

ACTGGAATTTTCTACTCGCCATCAGGACAAAAATTCTCCCATCCGTCGGACACGGAGCCATGACCTCGCCGACTGAGTTG

CGTGGCTTTGATCTCCGGGAGCCGCCGGAGCAAGCCGCCAGCCAAGCTAGCCGCTCTACTTGCTTTGGAGAAGCCACCGG

CCGTCGCCGACCCTTTCTTTATTCGGCATTGACGGAATATTCCTTTTTCAGACGGAATATGCTAACGGCGTTAGTCTCTG

TTGGCATCATGATGACGTCATCGACCTTGAACTGACTTGGTGTGCTTGTCTGCTAATGTGGCGATCTCGATTGGCCCACA

TTGGTGACGTGACAATGGTGACATAGTTCCTTCTTGCCTTGACTTGGATATTTGCATATGGGCTTCTATGTATAGTGCTT

ATGAGGCCTAGTTAGCCATAGTTGGCTAACAAGAGGAGCATATTAGGCCCAATGAATCATACTGAGCTAATAAGAGGAGC

TTTTTAGACCTAAGTAAGCCCATTTTTGGTATAAAATACTAGGGGTACAAACAAAGTTAATAAAAATATTAATTTATCGA

TTAATTAATAAATTTTGGTGGTCCCAATAGCATTAATTTATAGAGTTTCTACTATACTTAGCAAGAAGATGCCCCTCCAC

CATTTGCTTATCAAATATACACATGGTTCTAATTTAAGCACCTTAAAGCTTATGTATATGTTTCTCACCCCTTATACCCT

CTATTTATATATGCTTGTTTAAATTTGAGTAGCACTTATATTTTTTTAGCCGTATTCTTCGTAAAGTTGTGTTTGTTTTT

CTTTACTTTTAGTGTCCTTTTGTCAATCGAGAGGAAAGTCCAGAGCTATTTTGATTTCCTTGTATTCTCATTACTTTTGT

AAAATCATTAAAAAGATAAATAAAACTTTGTAAATTTATCATGGAATTTGGCTAGAATAGCATATAATCTTTGAGGGTGT

GGTTGTAGGAAATCCATGCAACCACATTATCCGTCTATTTATATATGCTTGTTTAAATTTGAGTAGCACTTAGCTTTTTT

AGCCGTATTCTTCATAAGCTGTGTTTGTTTTCTTTACTTTGAGTGTCCTTTTGTCAATCGAGAGGAAAGTCTCTGCCGTT

ATGTGCGTTTGCCACCTATACTATTCCACCTCAAGTTGCTCTTAGACATTTCACGAGCTGGGTGTTCCAAAGAACTTGGG

TCATGTCAGAAACTATATAAAACTTTTGCAAAAATCGTGTTTGGCGTCTATGTATACTTGTTGTTAGTCCCGTTTCTACG

TATGCCTATTGTCTTTAACCAATTCTCCTGTTCTTTTAATGCAATTTGAATGCACATCGCTGTAGTCAGTACATTTCTGA

TAAGCATGAAGCAAATATCTCCACTTTTAGTACTGCTTTAGAACTTTATTGAGCAAATTTTTAATATATACCTCTTACAA

CATTTTCATAAACAGTTTCCTTTTCACCAGGACCAGCTTTCTTGGATGGTTTAGCCCCACCAAACAATAACTCCTGACAC

AGTACAAACGCGTTCTATAAAAGCCACATCTAGCGTGGGTGACCTTAGTTATCCCCACGAGGTTCTTGCATTACTCTCGT

GGGCTGATATGGGACACTGCTCCACCCCCACAATCAGACACCTTTATAGAAAGCACCGGCATATTATTACATCAATCAAT

CTGTTGGAAACAATGAAGAAATGATGAAATCGGATCTCAAACAGCTGAGTCGCGGACTAGGTCGCTATCTTTAAGGTATT

TCGCGACTCTGCCTCTTGATTGTGCTAGCAGTCAAAAAATTGTCGAACACCTATGGGATAAAGCAGTGATAACTCTCTTA

TTGATTTCTCTGCACTGAGAAATCGAACCAACAACTACTCTTTCTTCACTTACTCGGTACTCAAAAATAGATGAAAAACA

AGCTGTATTTCATCTTCTCCAAACTGTCTTTTATAACTCAAAAAGAAATCAATACGGTTTTAATGAATATAAATTCGTAA

TTAGTACTCACTTGAAAATCCTCCATAACGAATGAAATCGCCAAAAATTAATAGAGTTAATTAGAGAATATTAAGTACAA

AAACCGTTTTGGAAATCAAGAGTTAAAACACCAAAAAATCGGTTTTTACGAATCACTTAGTTTTGGAAATTAAGGCCCCC

AAGGCCCCGCTCTCCAGGATTTTTAATAAATATTATAGATATATAAATATACCTATTCAAATGCATGGGGTGAGACTTGA

ACCCAAGTCTATAGTGTGGGAGCCCAAAATTATACCACCACACTATTTCTCTAAGTTTGATATAATATTACAAAACTATG

TTTTTAAATATATATATCTCCCAACACAATCCATTATAAAAGATAATTGATATATATAAGGGATGATATGGGACACCGCT

CCAGCACACAATCAAACACCATTATAGAAAGCATTAACATATTAACATCAATTCTTTATAAAAGATCTCTAACTGTGTCG

TGCTTTCGACGTATATGTCGATTCTTATTGTTATTATAACAATTATCTACTACTCCGATAGCCGCGGTACTATCGCAATA

GATCAAAGTGGCGGTATTGGTTTTCCCATCTTGGAATGTCGACAACACACATTTCACCATCCCGCCTCTTCACTAGATGC

TTCTCTGCTATCAATTCGGCCTCCATCGTAGATTGGGCTATAATTGTCTGTTTCTTTGATCTCAAGATACTGGCGCCCCA

AAGAATAGTAAAAACATATCCATTTGGTGGACTTGGAATCATCTGAAAGAATATTCCAATTAGCATCGCTAAATCCTTCA

AGGACCGCGGGATACCTTTGATAGTGTAATCCTAGGTTTGTGGTTCTTTTTAGATACGCATAACCCTCTCAACGACATCC

AGGTGTTTCAAACTAGGATTACTTGGTAAACACCACATTTAACTCCCCTTGCATACGCAATGTCCTGGTCTAGTACAATC

AGTGCGCATAACATAGGACCTTTATTATACTAGCATATTCGGATTGTCTAACACTTTCTCCGGTGTTCCTTAACAATTTC

ACGGTAGGATCAAAAGGAGTACAAACGGGTTTATAATCAAAATATTCATACTTCTTTAGGATTTTTTCAATGTAGTGAGA

TTGATCCAAACAAATACCACTATTAGTTCTAGTTATTTTGATTCCCAAGATTACACTAGCTTCACCCAAATCTTTCATGT

CAAAATTCGAACTAAGCATGCTTTTAGTTTCATTGACAACGAGATAAACTGGAACCAAATATTAGCAAATCATCCACATA

CAAGCAAACAATCACACATACATCATTCTCGATTTATAGTAGATGCATTTGTCACCTCATTTATTACTCGGTTACTTGAA

ATCATTAAATTATCAAATTTCTCATGACATTGTTTAGGTGCTTGTTTCAAACCATAAAGAGATTTTCTAGCTTACATACT

TTATGTTCTTGTCCAGAATTACAAAACCTTCGGGTTGTTCCATGTAAATTTCCTCTTCTAGTTCACCATTCAAAAAAGCG

GTTTTAACATCCATTTGATGAATAACAAGATTATGAGTAGCAGCAACAGCAATCAAAACACGTATAGATGTTAATCTAGT

AACGGAGAATAAGTATCAAAAAAGTCTACGTTCTCTCGCTGCCTATAACCTTTGGCAAACCGTCTAGCCTTATGCTTATC

TATTGTTCCATCGGGTTTGAGTTTCCTTTTCGGGACCCATTTACAACCTATTGGCTTACAACCGGGAGGTAAATCTACAT

AACTCAAGTTCCATTAGACATGTGAGAATCCCATTCATTATCTACAGCTTCTTGCCGGCTAGACTCTGCGGAACTAAACG

CCTCTTGTAAATTAGAAGGTTCTTCTTCTATAGCGTAAAATTCAAAATCGGGATTTTTATCTTTTGTCTCGATCCTAGCT

CTTTTCTGCATCTAGGTTCAAATTCGGTGATCCTAGTTTCTTCACTTCTAGGTTCTTCTAATCTAGACTCGGAATCAACA

CTAGGTAAAGTGCTAGGTGTATTACCCCACTAATTCTAGATTTAAAAGGAAATCTATCTTCAAAGAAGTCAACATCAACG

ATTTCAATAATAACCTTATTATTAATATCAAAGAACCTATAAGCTTTCTTGTTTTGAGCATAACCAATAAAAACACATTC

ATAAGCTCTACTTTCTAACTTATGTCTTTTAGGATCGGGTTTCTAACAAAAGCCAAAAACAGCCCCAAACTCTAAAATAA

GATACATTAGGTTTTCTATTTCTCCAAAGTTCATAAGGAGATATCATGGTTTTATTATTGGGAATGCGATTCAAAACATG

GCAAACAGTCAACAAACATTCACCCCACCAATGAGAAGCGGCACCGAAACTAAGCAAACAAGCAACGGTAAGTTCGGTCG

GTGTACGATTCTTTCTTTCATTTTCCATTCATTTGAGGATTATATGGAGCAGTTCTTTCATGTATTATGCCATTAGTTTG

ATAAATTTCGGTAAAGGAGCGAGAATCATATTCGATTGCCTCTATCACTACGAAACCTCTTAATTTTTCTACCAAATTGA

TTTTCAATTTCATTTGATAAAAATTTTAAACTTTTCAATAGCTTCATTCTTATGGCTCACAGCAAATAAACATACGTATA

ATTAGAACAATCATCAATAAACGTTATTATATATCTCTTCCCATTTCTAGTTAGGATACCTTCATATTCACACAAATCAG

AATGTATTAAGTCTAGTGGTTGGGATTCTCTTACAACAGATTTATGTGAAGTCTTAGTTATTTTTGCTTGACTGCAGTAT

TCACATTTTCAAATTCATTTTTCGAAAATTCGGGAAGGACACCTAACTTGCTCATGTTATTTACTAACTTTTTACCCACA

TGACAAAGCCTACCATGCCAAACATTAAAATTGCAAACCATATAAGCGGAAAGAATCAATTTTATTCATAGCATCAACAT

TAAGCTTAAACATTCCATCGGTAGCATAACCCTTTCCTACAAAATTCTTATTCTTAGTTATAGTGTAAACATCGGACCCA

ATAGTTTGATACAACCCGACCTTGTTGAGAAGATAGCCGGAAACAAGGTTCTTTCTCATTTCTGGAGTGTGAAGGACATC

TTTCAACATAAGTGTCTTCCCAAGTAAACTTCAACTCTCTTAGTACAGAACCTTTCACCACGGTCTTTGTGAGAGTCTCC

CAACAACACTTTCTTATCCTTTGTTTCAAAGATAAGTCTTAAACATGGACCTATCAAAACAACAATGACGCAAGCACCAC

TGTCTATCCACCACCCATCGGATCCACCAACCATGAAGAACTCGGATCGAGACCATGGCAATTATAACGTCACCCACGAT

GTTGGCTTGTGCGTTATTCTTTTCCTTGTTAAACCTACAATTCCTAGCCATGTGGTTTGGTTTACCACTGGTATAACAAA

GAAACTTCAGTGGAATTCTGTCCATTTTTCGATGGGTGTTGGTTCTTGTTTTGATTAGGCCTTCCTCCTTTCTTTTGGTT

CGGGTTAGGTTTCATATCCCTTTTCTTAGGTTTCAAACTTGGATGAGTCTTATTGTTAGAAACAATGAGAATCTCTTCCT

TCTTATCTTGTTTCCGAGCTTCTTCCTCAACCACCGAGCTTAACAATCAAACTTTCAAGAGAAAATTCTTTAGTCTTGTG

ATGCAAAGTATTACGAAAATCTTTCCGGCTAGGTGGTAACTTGTCAATCATTACGAAAACTTGAAATTGTTCATCGGCTT

TCATACCTTCGGCCACAATTTCGTGGTAAATTTTTTTGAAGTTCATGAGCACATGCAACAACGGACCTATCATCCACCAT

TTGGTATCTCAAATACCGGCTCACGACATACTTCTTTGAGCCGGCTTCCTCGGTGTCGTATTTTTTACGTAATGCATCCC

ACATATCTTTAGCGATTTCAAAAGTTGAATAATACTCATATAAATCATCGAGATAATGCGTTAAGAATGTAGTTTTTGCA

ATCAAAGTCGTCATCGATCCATTTGTCAATGGCCTTTTGTTTCTCCTCGGCGGTTTGAGAAACAACATCTTTGGCACCAT

CGGAAGGAGATGGATCGATGGCAGCGACACCATCATCGGCATCGGCAACGAGTCTTATCGGCGGTAGAAGTTTTATCGGC

GGCGGGTTCCATGGTTCCAGGATGAACACTCGACACGATCATATGCGGCTTCATGAGTTTGAGATAAAAGAACATCTTGA

CTTCCATCTTACGAAATGCGGCCCTTCAAATTTGAATGGCTTGTTGGTATCATCAGCCCTTCCTTTCAACCTCCATAGCA

AAATCAACTACCTTAAAATTGTTGGAAACAATGAAGAAATGATGAAATCGGATCTCAAGCAGTGAGTCGCGGACTAGGTC

GCTATCTTTAAGGTATTTGTGACTACGCCTCTTGATTGTGCTAGCAGTCAAAAAATTGTCGAACACCTATGGGATAAAAC

AGCTGATAACTCTCTTATTCGATTTCTCTGCACTATGAGAAATCGAACCAACAACTACTCTTTCTTCACTTGCTCGGTAC

TCAAAATAGATGAAAAACAAGCTGTATTTCATCTTCTCCAACCTGTTCTTTTATAACTAAAAAGAAATCAATACGGTTTT

AATGAATATAAATTAAGTAATTAGTGCTCATGAAAATCCTCCATAACATAAGAAAACGGTAACAATATAATTACCAAAAA

TTAATAGAGTTAATTAGAGAATATTAAGTACAAAAACCGTTTTGGAAATCAAGAGTTAAAACACAAAAAAAATCGGTTTT

ACAATCACCGTTTTGGAAATTAAGGCCCCAAGGCCCCGCTCTCCCGGATTTTTAATAAATATTATAGATATATAAATATA

CCTATTCAAATGCATGGGGTGAGACTTGAACCCAAGTCTATAGTGTGGGAGCCCAAAATTATACCACCACACTATTTCTC

TAAGTTTGATATAATATTACAAAACTATGTTTTTAAATATATATATCTCCCAACACAATCCATTATAAAAGATAATTGAT

ATATATAAGGGATGATATGGGACACCGCTCCAGCACACAATCAAACACCATTATAGAAAGCATTAACATATTAACATCAA

TTCTTTATAAAAGATCATCTTTATACATGTATGAGTAGCAATAGGGGTGCATGGGCCGGTATGGACCGGTTTTCATCTCA

TCCGCATCCAATCCAATGCACTACGGATTTTAAGTTTGGCATCCGCATCCAATCCAGATCCAACGGATTTGCATCCAATG

GATGCACGGATGAACGGATTGGATGCGGATAATCCAATGGATTTGTTTTAGTTAAAATTTATAGTAAAGAAAATAAAGAC

AATACAAATGACTCCTTATAAAACCTACATATCATATATATGTATACAAATATAAAAATGTCATGTACAACACCCCAAGC

AAACACCTTAAAAATTCATAAATGATTCAGAGTATTTTCTAGAACTATATAAATGCGCTTAATTTAACGGATATGGATGT

CCAACGGATTTTCAAGGCTGCATCCGGGCCCGATCCGTAATCCGTTGGATTTCAAAAATTCCATCCACATCCAACCCATT

AACGAACGGTCCGGGCATCCATCCGCAAAAGAACAGTTGGTCCCGGTTAAATCCTGGGATTACGGGCCAAATGCTCACCC

CTAAGTAGCAATAGGACAATGGAAGAACAAATGAGTAACAGTTTCTATATCAGCACTGCACAGAGGGCAAGATTCATCAA

TAGCAGGGTTATAATTTTTCAACAAAGCATAGATTGGAAGCATATGAGCAAATATCTTCCACGTGAAAAATTTAATTTTC

GGCAGACAATTCAGGTTCCAAACCTTTTTCCAAAACAAAGCTTCTTCACCATTACTAAATTCATTTTTATGAACTCTATA

AGCAGATTTTATAGTAAAAGACCCATCTTTCGTATGAGCCCACATAATTTGATCCTTAAGAAGCAAATTAATCTTAATAG

TTCTAATCCTAACAACATCTTCAGGGATCAAACAGAGAATTTAACAGATCAATATTCCAAGTACCAGATTGCTTATCAAT

AAGTTCACTAACTAGATTAAATTCATCATAATTAGGATTAAGAGAAACGGGGAGGAGAAGAAGTACCGGGCAACCAAAAC

GAGAATCCCATATTCTTGTATCATTACCATCATTTATTTTAAAAACTATATTAGACTTTAAGAAAATTAGACTATTCACT

ATACCCTTCCAAATCCAAGAATTTGAATCCATGACTTTATCAATTTCCAACAAATTATGATTAATAAAGTACTTGTCTTT

TAAAAACTTTGAAATTAAGAGATCAGGATTTTGAATAATTGTCCAACCAAGTTTTGCTATAAACACCCTGTTAGTAGCAT

AAGTAGATCTAATACCAAGCCCACCACATCTTTTATATTTCCCAATACACCCCAAGAACGAAAATAAGCCGCACGTCTAG

GGCCTTTCTTAGACCACCAGAAAATTCTTTGAATGGAATCAATTTTAGAAGTGATCTTCTTAGGAAAAGGAAAACATGAC

ATATGATAAAGCTGTAGACCGGATAACACATGCTTTGTTACAACAGTTCTACCAAAGAACATTTAGACTAGATTTCTTTG

TGTTTCCTAATTTAGAGTAAAATTTATCTATTAAGAATTGAAAAGATTTGGTTTTATCTCTATTAACAAAAAGAGGGGTA

CCTAAGTATTTTTCGAAGAAGAACTCATAAATTTCATCCTAAAGCTTTTGAAAGAAGTTTCTTATGCTTATGGTGCATTT

TAGAACTAGTAAAGAAACCATACTTTTCAAAATTGATAGCCTGACCAGAGGCCTTAGCGAAAACATTGATAATTTTCATC

AATTTCTAGCATACATTAAAGAAGCTTTTGAAAACAGCATACAATCATCAGCGAAGAATAAATGAGATATAGAAGGACCA

TCTTTTCGAATTTTGAAACCTTGAATAAGATTAAGAGCTTCACCTTTCAAGACGAGTTGAGATAAAACTTCCATGCAGAT

AATGAAAATGTATGGTGAAAGACAATCTCCTTGTCTAATACCTCTAGAAGGGAAAAAGACCTCACCAGGAGAACCATCAA

CCAGAGTTGAATAGGACACCGTACTAATGCATTGTTCAATAATCTGACACCAATCATCGGAGAAACCTAGTTTTTTAAAG

ACTGAAAGAATAAAAGACCACTCCAGCCTATCAAAATCTTTGGAAAGGTCAAGTTTTATAGCCAGAAAACCATTTTTAGC

ATTTTTGTTTTTTCTTTTAAAAGAGTGTAAAATTTCATGACTAATAACAATATTATCATGAATTTATCTATTAGCAATAA

AAGCGGCAATCGAAAGGGAGAAATAAGGGAATCAAGTAAAGGTTTGAGACGATTTGTAAGAATTTTAGAAATGATTTTTT

GGATTGTGTTTGACAAACTAATAGGTCTGAAATCCTGAGGAGAAAAAGGGTTATTCACCTTAGGTATTAAAGCTATGAAA

GAATGGTTTAATTGCTTGAGGAGAAATCTTGTTCGAAATTTTTTTTGAACATGATCAATAACTTCCTTCGAAATTTCTTG

CCAATTGTCTTTGAAGAAGCCAGCAGGAAAACCATCAGGACCCGGTGATTTCCATGGAGCCATTTCATTAATAGTAGCAC

TGATTTCCTCAATAGTAGGAATAGCTACTAAACGAGAATTTAAATCATCTGAAATGATAGGCTTTACATCTTTTAGAGCC

TCGTCAATATCAGAGTTATTAGGGTTCGAAGAATTATATATGTTCTTGAAATGATGAGAAAGGAGAGTCAGAACTTGTTC

CCTTTTTTCTAACCAATTACCTTTCTCATCTTTGATTGTGTGTATAGTGTTATACATATTTCTGAGTTTGACAGTATTAA

GAAAGAATTGAGTATTCTTATCATAATGAGTAAAATAGTCAATCCTAGATTTTTGTTTATAAAAAGAGTTTTCAATCTCA

TACCAATAATCGAGCTGTTTAGAGTAGTTTAAGATGTAATCCCCAATAGCAGAAGTATAAGGTAAAATCTGTAACTTCTC

AATTTCAGCATTTAATTTACCAATTGTATTTTTAACATGACCAAACGACTCAATGTTCCATTTAGTGAGGTCGGTCTTAA

GATTTTTAAGCTTTCCTGCCACTATGAAACTAGGAGACCCACTAACTTTCATATCCCAAGAATTTGTAAGAACATTTTTG

AATTCAGGACTTAACTACCAAAACTTAAAAAATTTATAAGGAATATACAGATTTTGTTCATTGTGAAAACACTTGAGAAG

AATAGGGTTGTGATCAGAGTAGACTCTAGCAAGATTAGTGATAAGAGTTTGAGGAAGAACATCTACCCAAGAGTCATTCA

TAAAACCTCTATCTAATTTCTCAAAAATCAGCTCAGAAGGATTTTTAAATCTTCTATTACACCAAGTAAAAGGATTACCA

AAAGCAAAAACCTTATTCAATCTTAACTCCACTAATTTATTTTTAACGAAAGATGGCGCCAAAGAAGTATCAGGATTTCC

TCTCTGTTTTTCTGAACTATGAAGAATAAAATTAAAATCACCCAGAAGCAACCAAGGTCTAATATCATCTTCATTCATGT

GACTTAGAAAATTCCACTGATTTTTCATTCCAGAAACATTTAAAGCTCCATACACAAAAGATACCATACAATTTTTAATA

CTAGGCGATATATCACATAAAATATGAATCATGTTAAGAGAGCTACTGATGACTTCTATATTGGATTTAGCAAAGTAACC

TAGTGCTAAGCTTCCAGATTTTCCAACTGCCGGCACTACGAACCAAGAGTCAAAAGGAAGATGATGACTGAAATTTTTGA

TAGTGTCATTATCTACCATAGTTTCCATAATACAAACAACATGCGGTCAAAATTTTCTATATAAATTAAGAAAATGTATT

TTTTATCTGAGCAACATCCATTCATATTCCATCAAATGAAATTCATGGTTCAGATAAAGAAAAACTAAACAAGAAACAAA

GTAGAGGTAAAATTAAAATATCAAGACAAAGCAAAAACTATGATTATGAAAAGCAGAGAGAAACAACGATTTTAGAGTTT

AAAGTACTTACAAGGTCATGAGCTAAAGAAGGCTCTGCATCCATAGAATCCACAGAATCCACCATCATGAGATTAACCAC

CTTTTAATATTCTTCATCACTAAAAATAGGTTCTGGGGGAAGAGTAGCAAAAGAGGTTGGTTCCAAAATAACAACTCTAT

CTGATAATTCTTCTATAATTGGATTGCTCAACATAGCATTGTCAGTAAACCAAATAACACAAATTGGGAGTAACAAAAAG

CTTATAACATCCCACAAACAAAAACAACTTTAAAATCTGTTCCACAAGTAACCATCACTTGCCAAGAACTCAAAGATTAG

ACAATGAACTAGTAGATTAGGCGATCATGGAATCGGTTCCACAAATAACCCCAACTTGCCAAGAAACATAGACAGAAACA

AAGAGACCAGAGAGTAGGCAATAATGATTCCAATCCTAAGCATTTCTAATATCAAGAGGCCCTTTATAGTGAAATAATAC

TATGGCCCTTTAAAATATCTTTCAATCACAGGCCCATAATTAGTTAAAACCCATCTTGACATCTTCACTTGACTCCACAT

GAACAGACGTAATGAAACCCAAGTAAGTTTAGCAAGATGATTACAGTTGTCTTGAAGATAAGTTTCGAAGCATATAGAAT

CAAAGATTGTTCTGGTCAAAAGCTACACAAAGCTTAAATAAACTTTCTTATTGGGGCCTCACATCATGTTACAAAAAACA

AAAAGTAAACGAATAAAGACGAAGTAATTTACAAACTCATGCAAAACTCACAGTAGTCCCTTTGCAGTACCATTACCTTT

TCAATAAAAGAATTTGATTTCTTCAGGCAAATTCCAATAGAAATTCCTAGTTCTTCCAACGTAAACCACCTGATTCCTAT

GTTGAAGAAGCCAAGCTAATAAGACCATCCGCATCCCTACAAAGAATAAGAAGCTTCGAGTAATTCAAATGAAGTACTTC

AAAAAAATTGCCGTAAAAAAGCATAATCTTTCAGCCATTCACACAATAATAATAGACACTGAAGCACTTGTAAAGATCAA

AACATCATGTCACTGTCCTTAAAACATTTTATTCACCGAGCTAGGATCAGATTTGCTCCTTCTTGAAGACAAAGCAAACA

GAGGTGTTTGCTTTTAAGGAAGGAAGACAGTGCTCAAGGAGGTGGGTTGTGCCATTAGGTGCTTTGCATCATACTTCCCC

TATCTCAAAAACCACATTAGCGTGTCTATCTGGCTGGACATTCAATAATGTAGCCGTATTGTTAATTTATTGACGACCTC

CAATAATTTCTTGTCGTGATTCGGTTCACACCATTGTCGAAGGAGTTGGAACATGATATAATAATATAAACATAATAAGT

AGTACTCATAGAAAAGGGCATCAGGCATCCATATTGTCATGAACAAATTTGTGGTGCAAGCTCAGTCCTTTTACATTGAC

GTGAAAAATTTAAACTGTAAAAAAGTCCAACATATAATGCACAATGCAAACTAACAACTTATATAAAACCTTCCACATCA

AAACAATAACAAAACCTAAGTGAACCATGAAGCACTTCATGGATCCATGCCATTTCAATATCAATAAGAGCAAAGATATA

TCACTGACCCTTGAGTTACTTCTTAAAAAGAACAGAAATCTCAATATTATAACAAATCCCTACAAAAGTAGTATCTCCTA

ATGCTAGAGAGCATATCCAATAAAGACTATTCTTCACTGATTAACCTGACTAGATGAGCAATATTGACGAAATCCAAAAA

TCCAGATATCAAGAAAGAAACCAAAAATTAACCCCACTGTATACTAACTAACTGTAAATTCCTCTTCCTCTTCCACATAT

TAATCTCAAGATGGTGAAAAAACCACGACCCTATGGATGCTACCTCCTTCACTCATGACCTAATATGAAAAACAGAAAGG

GAAAAGCTTAATCAGATAACTTAGAGATTCACAACAGATAAATCAAAATAGAAATAAGCTAAACAAACAGAAATTTGTAC

ATGATATAAAAAGAAAATTTGTTTTAAAAGCTTTAAAAACACGAACCCTAAATGATTAATCCATGAAAATTGTAAAATCA

AAAATTGGGGTTTTTAGACAAGGGAAAATTTAATTCATTAAGAATAAGCAAACAAGAAGAGAAAAATAAAGATTAGGGTT

AAACTATCATGAGATTCAAAAGAAACTTAGACAAAACATCTTATCTTATATCTTATCTTATACTAATATAAAGAAAAGAG

GTATTTTTGGTGAAGCCTTTTCACATGACAATAATACCCTCATTAACTAAGCTTTTAGTTAATATTTGTATATTTACATG

AATGCCATTGTTTTAAAAATAGATCAAAAAATTTAAAACTTAGGTATCTTTAGAAATACATATCGGATTTTTGCAAAATT

AATATATTTGGAAAGCTCTTTAAATTATCTACGTAATGAGTACAAACAAGGATATTAAATTTTTATTTTAAGATTTTTTT

TTCCATTATTATCAATTGGTATTGTTAAAAGAATCCAATTCAAGCATGTGCATCGCACATGTGGCCTTACTAGTATGAAT

TAATTGAAGAAAACAAAGTAAGAATTCATCAGAGAAGATCGAAGAAACTTGAGGTTGACTAAGTCAACGTCGCACGAAAA

ATCGCCTCTTTGTCTCGCCTCTTTCTCTTATAGACAAAATAAAGCATCAGCGTATTACATAAATCCTTCGTAAAATATAT

AACTGATATATATATATAAGGGATGATATGGGACACTTTATAAAAGATAATTGATACATATATAACCTAGCTAAATTTGG

GCCTATAGCTATTAATTGGGGCCTATAGCTAGATGACAAAAAAGGTCATTAGAAATCACCTGCACACCACCCCTTATCAA

AATAATTACCTAACTACTTATTTTACCCCTGAATAATTAGCACTAATTAATCGTATTAGGGGTTAATTTTGTTTTGGATG

TGAGATTAACTAATGTTAGAGAGTTCATCAAAACATCAAACATTTTTTTTTTTCGTACTCAAAATCGGTTTACGTTCATT

CACTTGTACACCGATTCTGGATTGGTGTTTACGAATTAACAGATGACATCTATAATCGGTTAATGTTGACATATTTATAA

TCTAATTTTGGTAATCGGATTTTACTTATACATCAATTCTTTATAATCTTTATATATATTCAGTGGCGGAGCCAGCCTTA

GTGAATGAAGAGGATTAAAAAAATTTTTATAAAATCTGTACACTAGGTGGGCTAAATTTTTAAAACTAAGTGGACTAAAT

GCAAAATATGCTACAAATTCAAGAGCTTTTTAATAGTTTTGGTGATTTTAGGGGGGCTAGAGCCAGGGTTAGTCAACCCT

TGGCTCCGCCTAATATATATATATAGGCTGATATGTGACACCGCTCCACCACATAATCAACAAATCAAGGCTGATCAGAT

CTGATCACTCGTGTAGGGAAGGGAAATCACACGGGTGCTCAGTACCCGTATTATTTTACGTTTTACGCGTACTCAGTACG

CTTATTGCCCTTGATACGGTTACTCAGTACGCGTAGCTTTTTGCACGGGTACTGAGTACGCGTTTCGTAAATAATATTTA

TTTGACCTTTTTTGGTTAACACTTCTCTCACACCCGATCCTAACCATCCATCACTAATAACGGCTTTATTTTCTACACTG

TCGGATCCTAACGGCCATTTTTTCAAAATCCTCTATAAATACCACTCTAATTCTCTACTCAAACCACACAAAATCAATTG

ATTTCTTATTTTCTCCTTCTTACATCAATTGATCTCTTCTTTCTCTTCATCTTACTAAACGTCTTTGGTATTTTGATTTA

TTGTACCACCTTTATTTATTTATTTTTATTTATTTTTATATCTTAGTTTCATTCAAAATGGTACTTAATTTTGTCACACA

AGAAGAGGTTGATCTTACTACTTGCTTTTATTTATGTGAGTATGGACGAAATTGTTGGTGCGGGCGCAAAAACGCTCATT

TTGGATACGTATTCATGAAATTTTCATACAACGCAATGGTAATCCATATGGAAGAGATTGGGATGCTTTAAAAAGCAAGA

TGAGACATATTAAAAAAGATGTCAAAGAATTCGTGTCCATTCTTAATGCTATATATCGGCATACCAAAAGCGGTGTGAAC

CACGAAGATTTGGTAAGTTTTATTAAATTTTTAAGTTTTTCCATATAGTTGGTGATAATAGCAATAAACCTAATGTTTCT

TATATAATTTTTTAGACAAAAGATGGTCACCTTCAATTTCAGGAACAAAGAGGACAACCATTCAAGTATGATGCATGTTA

TGAATTGTTTAGAGCAAAGGTCCCTGGATATAATTATGAACTGACTGTGAGCCAAAACAGTGAGTTGTTTGATAATCATC

GTAACTTTGTATCAGTCGAAGATGGTACTGAAGTTCTTCCCAATGGAGAAACTCGTTGGAAATACAAGGAAAATGCACGT

CCTATAGGAACAAAGAAAGCCAAAAACTAGCCGTGACGAAGAAGGATCGTGAGCAAGGTATATATAGAGAAGATGATGAG

GGAACATATTCTTTTAAGATGGAAGAAGATAAGGTCTATCAAAAAAGTTCTAGCATATTGGAGTATCTTAAGGAAACACG

ATCAAGGAAGGAAAAAATGAGAATGGAACAAGAATGCCAATACCAAACTTTTTGGGGTATACTCTACACCAAATTCTGGG

ATTAGAAACAGAACTGAGCCGTTACTAAAAGGGGTTGTAACGGCTCTTGCCTGTTTCAAAAAGGGGTGTTACAATTTTTT

TGTAACGGCAATGGGGCCGTTACGAATTTTGGGTTGTAACAGCGTTTGTAACAGGCATGAACCGTTACTACGTGGCGAGA

CGTAACAGGCAAGAGCCGTTACGAATTATTTGTAACCAGACAGTCGTAACACCTCCAAGCCGTTACGAATGTTTTTGTAA

CAGCAAAAAAATTACTGCAGTCAATATTGACCTAGTCAAAATTAGTCAATACTGACCAATTTTGACCTGGGCAATATTGA

CCGGCAGTGCATTTTCTGCCGGAAATACAAATAACCAGCATTCCTCCCGTTCTTCCACATTGAACAATAGTTTCTTCATT

CAATTCATATCAATATTCATTCTTCTTTCAATTCATATCAATATTCAGTCGATTCATACCCCATTCAAATTCTCAATCAA

TTCATATAAAAAAAAGAACATCAATTCTTCCAAGTACAACTTTTTTTTTTTGTTACCAAATCTCAAGGGAATACATAGAA

ACTTCCTAAGTTCAGCTCCTTCTCAGTTAGGTCAGTATAAGTTCAACTCAAAAACTACATACAACTTACTCAAAACTTGC

AGATAGTTGCCAAGCATTCTTCATCAAGTTCTTCGTAATCCCTGACTTATTCATAAGGATAATAACCCGGCCGCTTGAAG

TTCCTGCATAAAGAATCTGGCCTGAAAAAAGAAGGTAGAAAACAAAATCAAAACCACTAAGAATCCAGGATTTAAATTGC

AGGGTAACCAACTGAAAGCAGATAGAATTCTTCAAATTTGATCATAAATTAACACCTCTTCGTCCTTGAGAATAATTGAA

GCTAATAGGATCAAAAATAAGTATGAGATAAAGCAGAACTTACAAAAAGCCTTACCTGTAAGCTAGTCTATTGCTCCAGC

AACATTTCGCGAAACCTACCAACGTCATTATTCTTCAATGCCTCGATCACCTGTAATGCTGGTCTTCCCATCTCAAAGAC

AATATTGCTCTCGCGCTTGATGTCCTCCAGCTTTCTAAAATGTGTCAAAGTAACAAAAATTTCAGAAACACAACTGAAAC

TGACATATATAGGTACCAAATTAAATTCATAGAGAAAACACGTTAAGAGGGAAGTCATAGAAACTCAAACAAATACACAT

TTATTTATACATGCAATTTTGACGTAATTTGCCCAGCTTAGTGCTTTTAATGCAATTATCCTAAATGATACTCTGCAAAC

GATATGTAAAGGTTCCTGAAGTTCTTTAAACTGATCCAAAACTTAGCACTAGATCCAAAACTACTTATGTAAACTGCAAA

AATCAGAATATCATTTAATTGGCACGAGCCAACACACCTTCCTCACATATTATTTCTTTAACTCCTCCTGAAATTTTCCT

TCAGGGGAAAAAATTTAGGCTCTACCACCCATCATAGAGGAGATACCTATGAAGAAAGGATGGTTATGAGTATCATTTGA

TACAAATACAGATATTAGATATACTATAAACTACCTAGTTCACTTTAAGAATCTCTAACTGAATGAGAAAAAACAGTTAT

TTAGATGATTCAAGAATCCAAATGATACCCTGTCTTTGTGAGATGCTTCCTAGTAAAATGCACATCAAATGTGTTGATGT

GTGGAATATACCCTTTGTTCCAAAATGCATACTACGGACCTTTCAAAGCTTGTACTGCAACAGTGAAATATTGTTTCATG

CATTACGTATTCTTAGTAAAATGTAAGTCTACGAACTCCAAGAAGACAACTTCTTTTAAACTATTACACATCTAGAGAGT

GTTGATTAGACAACAAAAAGAAAAGGCTTAATTATATCCTGTACAACTGTGAAAAACAACAAAATAGAAATACTGACCAT

GGACTCTTCATCCTTGATTAAGTGGAATCTGAAACGGCCCTTGGTATCGTAAAGAAGACAGGCGTTTTCATTTGTCTTGG

GACTTGACACAACATCTGAAAAACAAAGCCAAAAACATCACAAAGGTAAGTACCAGAATAAATGGTAGGAATCTTCCACA

GGATTCACGTTTTGTTAAACTGAGAATACAAAATTGGCCACGCGGTGTCTTGGGAAATCTAAACATTGGCATACAACGAA

ACATGAACTAGTTATTGATTCCATCAATAATGATCATGGGTTCTTAATCATTTCTGAGAATACAGATAAAGAATGAAAAT

CTAGCATCCAGTCGGTAATGATTTTAGGCCACCCAGTTCACTTTGGATATAAATAGAATGGAAAAGAAATATTATGCACC

AAAATAGCTACACGTACAATGTCCAAATTCATAACTTGCTCAGAATTTCGAGTAGTCTTTTAAGTATGAACGAACTCAAT

GAAATCCAATTTCTGTATCCTATGCTTTACATGATCAGTCTAACCACATTATCAAAATCACTAACTATACCGCCAATATT

ATTGAACCAGTTTATTGTCAACTATTACTGATTTACTGCTACATGCCTAAATAAAGCAAATCGGTGCAAAACATGGACAT

TCCTTCTACAAAAATCCCGAATATAGAAATTTTAACTTAAGAAGATTAAAATGCAAACAAACCACAATCGATTAAAGAAC

TGGAGATAACCTATTGAATGTGTCCTTTGGAGGATTTTCTGACAGAAGTATAATTGAATGGGAAAGTAGTTGTCCGATTT

TGCAATCAAAGACCTATCTTGAACTAACCACATATTAGGTCATATTGCAAGATACACCAAATGATAGATGCAAGTCAAAA

TTTACCATCAAAAACATATGTTAGATAGCAACACAGATTTGTTTAATTGTGTTCAACGGAAGACTGAACAGTAGCAATGC

CATGCTCAAAAATATCCCTAGAGTAATTACTCAAGTTTTGCTTCAGGATATTATGATAATACACCAAAAATAATTAGAAA

CAAGATGGTCACTACAACACGCTTTCATCAATTCGAACCGAATTAATATCTAACTACATAAAAGATAATAACCCATAGAC

ACATTTTTTAGAAACCCAATGAGTTAGACCACATACCTTAGTGTCACGAGCCCCATGAGGTACTAGCAATCTTTGGCCGA

AATGCATTCAGTTTACGTTTTAAAGTAGCTTCCACATCCGCTTTCCTAGAGGTCTTGTTCTCAACATTGTTAGAAAGATT

TGTGCAGTTGCATCCATCATGGTAACTCCTGAAGCAAAGCACTCATTAAATCAAATTAGATAACTTAGGAAACCACCAAA

GAACAAAAAAAAAAATCAAACTAATACTGCCCCTAAATCAAAGTGTGAATTACATGGAACAAGCACCACACTTTAGAAGC

AGTGCGAAACTGAACCCAGATGCATTAGTAGAATAATGTGTGAAATGCTGACCCTAGTATGCACGATGTTGAAATTCCAT

TTAAAGGTACCCATCATCATGTAGCATACCGTCCAAAACCAAACCAGTTACCCTTAACCATTAACATACTTTTCCATTCA

TAATCCTTCTTCATCAGTACGTGACATATATATAGACTTACAACCCATAACCGTACTGCCCAATACACAACCTAATCTAG

ATAAACTAAGAACACTTATCTGAACAACCATGCGAACTACCCTATTGCCTGACCAACCATGTGAGCGTATCATTCCGCCA

TCCTCGAAAACAACCTTGTCCTCAAGGTTGTGCACAATGAAAATTAGTCTTTAGCAACATACATTACATTAACAAGACAC

CAAATTAGTACATATACTAAAAACATCCTAACAGACCCATAAACTTGCACATATCATTAACAGCCTTGTCCTCAAGGTTG

TGCATAAGGAAGGCCAAGTGAATGCGCATAAGAGATTGCAAGAGTTGATTCTTTTCCACTGTCACCTGTAGGCAGTTCTG

TAGGTTGATTCTGCTCTTGTACAATCTGTGGCCAAGTGCCTTCCACCATGAATAGTTGTGGGTTCTTGCAAGTATGACCC

TGGAACTACTTTTCATTACAGTGAATATTGTCTTACCGTTGCTAATTCCAGTTACCCTTCCTGCTACGGTTAAAAGATCT

ATTATTACGAGCATGAGCTGTTCTAGCTTTCTTTGCAACCTCATTAGCTAAAGGAAAATGGATATTCCCTTGGAAACTCG

ACCTGTGCTGAGCAAGTGCTCCCTGTTTACAATAACAAATTTCAGCAACCCCTTACAAAACCCAAACCGAACATACACTT

CCCATAAATATAAAAACACAAGGTGAATAAGCCTCATCTTCCATTATACCTGTCGAAGAGATGACCTAGAATCCATGAAC

TGCCTCACTTTCAAAGGGTTCCCTTTAACACCATGCCCTTGGGCATTACCGAGACCAATTTATGGTTTTTATTCTGCAAA

CCATATATGACTCCACATAAGGATAACACAAAAACAAGCCTAAAATCCAAATTTCTGGTCTCTTATAAACCTATACGAAC

TATAAATCAAGAAATCACTTACAAAAATTAGGTGATTTTTAAGTCCTCCTCTTTGTGTTGGTTTCTTGGACATCTTGATG

ATAATGTCATATCCATCTTCTTCTCTGTAAAAGCAATTGCTTTGTAAGTGGTTTTAAAGTCTCCATCTACAAAAAGCAAA

CAAAATCATCGAAAATCAAAAAAATTCAACATCAACGAAAAAACCCTAGAAAAGAACGACAACGAAGAAATTAAGAAACT

GATGATGATGACACTGAAATCGAAACCCTAAACTGATGATGATGATGATTAACAGATGATTGAAGAACTGAAATCTTCAG

TTGAGAGTGAAGATGAGAACAACCAGACGATTGAAAAATCTAGAAACCCTAAAAATCGAAAAACTTATTGAAGTGATCGA

TGAACTCTAACATAGATTGATGATTAAAAAGATTTCTGTGAGATTTCTGTGGTGATTTCATCAACTCATCGGTTAAAAGA

GAAGAGGCAGGAGCAGAAAAAAAAAAAAAAAAAAGAAATGAAAAATGAAAACTCGATCTCGATTTCCATAGGGTATATCC

TCAAAATCACGCATGGGAGGAAATATCCCTAGTGTTACTCATCTTGGGTGGAAACACATTTGTACGCGTGTGAGTAACAC

ACGCACCAACTCCAGTGTTACGACCCAGTGTGACGACGTGTGGCTGGCACGCCTAAAAAAAAATTAAGTAGAATCACTGG

TCTTTGTTACGAATTATTAAAAAATTTGTAACGGCTAGCGCCTGTTACAACCGTTTTCGTAACGGCCATTTGTAACGGGA

ATTTGTAACGGCCCATAAGCCGTTACGAATTTCTGTTACTATTCCCTTAATTTGGTGTAGTGAATGGAAGATGAAATCAA

TTTGCAAACGCTTCAACAACAGACTAATCAAATGCATGGACAACAAATGCATTTTGATCAAGACAATGCAATAATGTTGA

TGGACCTTAGTCGTTTAGCTCCCAACGAAAAACAATATTATCAACGCAAGCAAAAGGAGATTTTGAAGGGGAAGGGTATA

ATCAGCGTTGAAGATTCAGAAGATGAAGATGTAATCCAACTCTAATTGTTTTATCTTTCATTGAGTTAAGTTTTTAATTT

ATAGTACATGTTTAAATTAAAAGTGTTGTCTTAAATTTCTAGTTAGTATTGTTGTTAACTTAATTTTAAATGTGTTGGAA

CCACTCGATATTCAAATCCAAATTCAAATAAATAACTTCATTTTATCACTTCATTAAACTAATGTTCATAAAAAACTTTA

AAAAGATAGTACAACAGATGATTCGATAGGTCTGAACACATTAATTTTCCGTGGTGTCAGATTCAACAATCAAGGCCGCT

TCGAAATGATAAAACGGTTCTTTTCTCCAAGTTATATATGCATCTTGTGCAGCGAACCGGTCACCTCTGTACCTATTGAG

AAATTCATTGTACCTGATGCATAATTTCCTAACATGATTTGTCCTTGAACAAATGTAGGATGGTTGATAATTAAAGCGTG

GTTTCTTTCCCGTCTCTTTAACGAAAGAACCCCAACCAACATTAGTCCAGGAAGATTGTATGATCTTTAGGATCTTCTGG

AAAATTATCCTTGAGAAGGTAAGGGTTTCTAATCCATTCACTCTCTTCTTCAACATCCTTTAACCTTTCCCTCGGTTGGC

ATTGTTGTTGACCTCGTTCTTTTGCCTTGCGTTTTTCTTCGTATTCCTTGAGTGTATGTTGGTATTCAGTCGAACAACCA

GATGAATATTTGGTTTTTTTGTGAAATTTCCTTTGCTTAACACTTCATCCATCACATTAGTTAATTTGGTAGGTGATGAA

GACGTCGTTGCGTTCGCAGTAGCTTGATGACCTCTTTTGCATATCTCCTTTTCCATTACAGCAATTGATGGAGTGGTTCG

CTTGCATCTTCTAATTATGTCTTCGACTGAGTTGGTAGTTGCTGATTGATTTTCCGTTGAATAGATGTTTAGCTTTGTTT

TTCTCCTCCAAGATTAATGTTGCAAAGAATATGTAGATTGATGTTGCAAAGAAAAATAGGTAGACTGATGTTGCAAAGAA

GAATGGGTTATCTTTATTATATAGGATTCCTTCCCAATGGCTCTATTTATAACAACCCGTGCCCAACGACTCTTTTTCCC

AACGACTATATTTTCCAACGGGTGTTTTTCGCAACGACTAGATTTCCCAACGACTCTATTTCTTTATATAAATACTGAAC

ATTTTTTTGTTCTTACCATCAACAATCTCAATACATTCTTACATATAAACAACCAAAAATATGTATTCTAACAGTTCGAG

TAGAAATGGAAAACAAATAGTATGTGTGGAAGCAAAAGAAGTTTTTCCAAGTTGTTTCATGGATACACATAGTTTGCTTC

AATGCCCATGGATATATACAAAATGTCCACAACGAGGTTGCACTGGCACTAGGAAGGTAAAAGAGTCGATACCAGAGAAG

ATAAGGTATTTGGAATGTGAGTGTGTTAAATTTTCGGGAGAAGCAAATTTTCGGGAGAAGCACAATGTCGATACCGACGC

TCGAGAATGTCCAATGTCTGAGAAAAAGCTTGTACAATTGTAGTAAACTTTGGATAGATACCATCACTAAGAAATAACCC

ATATTGTATTGGTGACCGTTGATGACATAATTGCATGGAGGTGCTACACCCTTTAGCATGTTATCAAAAAGGGGTGAGTG

TGCCAACACATTCAAATCGTTGTTTGATCCAGGCATTCCAAAAAGGCATGCTAAAACCACAAATCGTATGATGCCACCGC

CTCTAATACCAAACTACTCTCTTTATCTTTACCCCGGTAAGTTCCTTGCCAAGCTACCGGACAATTATTCCATGGCCATT

GCATACAATCAACACTACCAAGCATTCCTGGAAAACCTCGGTCGGCATTCTCAGCTAGTAACCTATGAACATCTTTAGGA

GTGGGTTCACGCAAATATCTTGCTCCAAAAGTCATGCAAATGGTGTGGCAAAATCTTTTTAGATACATGTATATTGTAGT

AGCACCCATACGAGTGTAATCATCAACACTATCAGCTGGCGTACTTTTGCATAAACATTTCATCACGATAACTAATTTCA

TATAAGTCTTCCCAATAGTACCAGTTGCATCCGGACGTTGATGCCATTCCGGATCCACTTCAAGCAATTTTCCTAACAAT

TTCTCGAATAAGTCTTAGACGTTGTTTGAATTGTCTAGAGGTATAACCGGTTCCAGGCGTAAAATAATCATTCATCATTT

TCACATCATGCCACACTCGATAACACGTTATTACACTACGTGTCTATACCTCCTTTGGTACAGGTTGTCTAATAATACGC

AACCGTTCATCCAAACATAGTTGTGTAAACATAAGCAAAGCTTTCTCTTCTTCATCACTATCACTAGAAGTATCCAAATA

GATTCCATGTTTTAATCTCACATAAGAACGCATTAGACTCATTGTGCACCTTAGAAAAAATAAAAACCCAATATGAATTT

GTGTATAGCATACATAAATACAAGTTTATCAAGGTTCTACATTGATATGCATTTAGATGAAACATTTCATTCAACATAAA

TCAAGACAAACAACAAATAATCATGCATTTTTCACTACATATATGTTTTAATAACGTTATTCAATTGCATACTACATATC

CAACCAACCTAGTATAAACAAATTCATCAAAACTAGAATCAAATTAATTTAAAAACCTTAACATTAGAATCACTCACCTT

AGATTATTTTTTTGATGAAATTCGAAATTGAAGAATGGATTTGTTTCACTAGTTGATTGTATGCAATGGCCATGGAATAA

TTCGAAATTCGAAATCGGGATGGGAGGTCACCATAGCAAGCTTGAGAGGATGAAAGTTAGAGAAGAAGAAGAAATGGCCG

TCTGCTGCGAGTTCTGTGTGTTTAAACCTACGAAACGCGTACTGAGTACCCCGTATTTTGGTCAAGTCAACGAGTTGACT

AATACGCGTACTGAGTACCCGTTTGGCACTATTCATACGCGTACGGAGTACCCGTTTCGTGTAGGGAAGGGATGGGAGGG

CACCATAGCAAGCTTGCCTTCCATGTGCCATCCCTTCCCTACATTTGAGGGATAGGGAAGGGATAGCTAGCACCATAGTG

CTAGCTCTAACAACAATTGAAATCTAATTTCTTTTTGGGTTTTGAATAAAAAAACCCCGAAAACAAACAATTCGAGTGAT

TGAGTTAAATCCCCTTATTCATTCCTTCTAAGAGGAACTCTACAATGATTTACCTATAAATCTTTGAAATTCACTCACCA

AATTTTTTTAAAAATCCACCCAGAATCGGTTTATGTGTGCACTGTGTTAGTCCGATTGTAGTTGATGTTACAAGCAATCA

GTTTATATTGTACACTTACATAAACTGATGAACATCAACCACAATCGGTTTACGCTAATGTACACATAAATCGATCCGGG

TTCGAGTCGGATATGTAGAGAAATCAAAAACCGGATTATTCGTGCATCTATATAATCCGATTATAACTTTTAACGGCTAT

TTATTTTTTTGGAATTATAGCCGTTGGTCCTTTGCGTATATAAAGAGGATAACCTATTCTCTATCATACACTACCTAATT

CAAAGAAATCTTTCTCCGTTGTTAATTTATTTCATTTGAAGATGTCAAGTCCATCATCATCGACTACCAATTCAATCGAA

CACATACTTAGAATATGCAAGCGAACAAACAAATCAATTGAGGCGATGGAAGAAGAGATACACAAAAGAATTAAAAAAAT

GATGATACTAATGGCCTGTGTTCGAAACTCGCCAACATAATATTCTTTACCAATAAAAAAATAAAAAAATAAATATTAAT

AATTAATGAGTGATTCTTAATGGCCTGCTCAAAAAACCGCTCACTCACAAAACAAGCGGGTCCGCCGTTTTTTAGTTGTG

ATTGATTTTAGGACAAGGTGTCCCCCCTCTCAACTATCGTTTTGGATTCGCCATTGCACGAGCTTCTCGTGCATGACCGC

TTCTCGTGCATGACGGTTTGTTATCCCGATGTCCTCTTTCCCTTATAAATGTAGTTCTCAGTGCTTGAATACCTAAGACA

TTTGGGTTTTGATTCAATCTCTATTAATTAATAGTCAAATTCAATAGAGTAATTCCCTAATCTTTGTAAACCGTCTTATA

TTTCTGTTTCATGACTCCGATTAATATAGAGGAGAATGATTTTTGGATGATAGCATGCACTGTAATAATAATATTTGCAT

TGGTGAAGTTCATGTTTTCTTTTTATCAATCTTCAAATACAACGGAATGGCCAGAAGGTCCAAAAACACTACCCATAATT

GGAAACCTTCATCAGTTGGGAGGAGGTGTGCCCTTACAGGTTGCTCTGGCAAACTTAGCCAAAGTATATGGAGGTGCATT

TACAATTTGGATTGGAAGCTGGGTTCCAATGATCATCATAAGCGATATCGATAACGCTCGGGAAGTTCTTGTTAATAAAT

CTGCTGATTATTCCGCTAGAGATGTACCTGATATTCTTAAAATCATCACAGCAAATGGGAAGAATATTGCCGATTGTGAT

TCTGGTCCATTTTGGCATAATTTAAAGAAAGGTCTTCAAAGTTGTATAAATCCATCAAATGTTACGTCTCTATCTCGTTT

ACAGGAAAAAGACATGCAAAATCTCATCAAATCCATGCAAGAAAGAGCGTCACAGCAGAATGGAATTTTAAAACCTCTTG

ATCATGCCAAAGAAGCGTCTATTCGATTACTGAGTAGAGTTATATTTGGTCAAGACTTTTCAAATGAGGATCTCGTTATT

GGTGTGAAAGACGCCCTCTATGAGATGGTACGCATAAGTGGGTTGGCAAGTTTAGCTGATGCTTTTAAAATTGCTAAATA

TTTACCAAGCCAGAGAAAAAATATTCGGGATATGTACGCCACAAGAGACAGAGTATATAATTTGATTCAACCACATATCG

TCTCTAATCTTCCTGCAAATTCTTTCTTACATTTTCTTACATCTCAAGATTACAGTGATGAAATTATTTACTCAATGGTA

CTTGAAATTTTTGGTTTGGGAGTAGATAGTACTGCAGCAACGGCAGTTTGGGCACTCTCCTTTTTAGTCGGCGAGCAGGA

AATTCAAGAAAAACTTTACCGCGAAATCAACACCCTGACGGGTGGGCAAAGACCAGTGAAAGTTGTAGATTTGAAAGAGC

TGCCATATCTACAAGCCGTGATGAAAGAAACATTGAGGATGAAACCCATCGCACCACTAGCGGTCCCTCATGTAGCAGCA

AAAGATACTACATTCAAGGGGCGGAGAATCGTTAAAGGTACAAAAGTAATGGTGAATCTGTATGCTATTCATCACGACCT

AACGTTTTTCCTGCACCGTATAAATTCATGCCAGAGAGATTCTTAAAGGATGTTAATAGTGATGGACGTTTTGGTGATAT

CAACACAATGGAAAGTTCATTGATACCATTTGGTGCTGGTATGAGAATTTGTGGAGGTGTAGAATTAGCCAAGCAGATGG

TAGCTTTTGCTCTTGCAAGTATGGTCAACGAATTCAAATGGGATTGTGTTTCTGAGGGGAAATTGCCTGATCTTAGTGAA

GCAATTAGCTTCATTCTCTACATGAAAAACCCACTTGAAGCCAAAATTACCCCTCGTACAAAACCTTTCGACAGTAGATA

AAATCTTATTTTTATCAAACTGCAAGTGTATGTTTATGTAAAATCAAGGGTGAAGATGCTTGATCTTAGTGCACCAAAAT

GGTGGTGGCTATGTAGGAGTTAGATTAGCTCTTGATTTAGTAAATGTTTTTCTTTGCTAGTGACTGTGTCTCAACTCAAG

TGTGTGTATTGTGTGTGTTTCAAGTGTGTATTGTGTGTGTTTCAAGCGTGTGTTGTGTGTGTAATTACTGCCACCTGAAA

ATTACATTAGTCGCCGGCTGTAACTGTCACATCATTCTTGTGTTTTTAAATAACGCGTTATTACACTGCGTGTCTATACC

TCCTTTGGTACATGTTGTCTAATAATACGCAACCGTTCATCCAAACATAGTTGTGTAAACATAAGCAAAGCTTTCTCTTC

TTCATCACTATCACTAGAAGTATCCAAATAGATTCCATGTTTGAATCTCACATAAGAACGCATTAGACTCATTGTGCACC

TTAGAAAAAATAAAAACCCAATATGAATTTGTGTATAGCATACATAAATACAAGTCTATCAAGGTTCTACATTGATATGC

ATTTAGATGAAACATTTCATTCAACATAAATCAAGACACACAACAAATAATCATGCATTTTCACTACATATATGTTTTAA

TAACGTTATTCAATTGCATACTACATATCCAACCAACCTAGTATAAACAATTTCATCAAAACTAGAATCAAATTAATTTA

AAAACCTAACATTAGAATCACTCACCTTAGATTATTTTTTTGATGAAATTCAAAATTGAAGAATGGATTTGTTTCACTAG

TTGATTGTATGCAATGGCCGGGGAATAATTCGAAATTGAAATCGGGATGGGAGGTCACCATAGCAAGCTTGAGAGGATGA

AAGTTAGAGAAGAAGAAGAAATGGCCGTCTGCTGCGAGTTATGTGTGTTTAAACCTACGAAACGCGTACTGAGTACCCGT

ATTTTGGTCAAGTCAACGAGTTGACTAATATGCGTACTGAGTACCCATTTGGAACTGTACATACGCGTACTGAGTACCCG

TTTGTGTAGGGAAGGGATGGGAGGGCACCATAGCAAGATTTCCTTCCATGTGCCATCCCTTCCCTTCAACGCTCTTATTA

TTTTCCTTTTGTGAGATTTACCTTCTCTCCCACATCGAATATTATTTCACTTTACATTGAAGATTGTTGTTAGGTTTACT

AGATCGTATAAAACCTTTCGACAGTAGAAAAAATCTTATGTTTATTTATTTTTTGATGGGAAGAGAGAATTGTATTAGAT

AAAGAGAAAAGCAAGAACACAGAACTAAGAAAATTAAGAAAAAACAGTTTCCCAGTTGTCGAGTGTGTAATAGAAGTTGA

TGTGCACTCTCTTCCCAAACACAGCTGCCCAAGACAGAACAAGACTTTTTGCGTCATTTGCCAAATCTTTGTCAGTCTTA

GTCAATTTTCGGTGTCAAAGTTGAGTTTATATTTTTTTAGATACATTTTTGTGTTTCCAATATGTATATGTATATTTTCT

CAAGTGCATGTATGAAAGTTTTTCTTTCTGTCTATTCAAGATAGATCTCCATATAAGAATAACCCCATATAAGAATATTT

TTTATCGATCCCTAGGATATTCTTATACAAAAGTTTTACTGTATCTGAAATTTTCACAGTCAGCTGGAAGCTTGGAACTC

ACTAGAGGTGGCCAAAATAAATTCCTAGATACCCACATATCACATATGACATAATAAGATTAAAATAGCCCGGAGGTACC

AAAAATAAGCTAGGGGATAGACGTCCACTACCTATACATATCTCAATTTACTACTGTTCATCATGTTCGCATGTAGTGTA

TGTAATCTTTCATCACATTCTATATGTAACAATACCGCTTATATTGGTAATTTGGTATACCTTTATAAATCGTGATTTTT

GATGGGGTAAAAAGATCTGAAAATGGGTTATTTGTCCAAATAAATTTAAAACATGGTTCAAATGGATAAGTAAAAATTAT

CATGGGTGAAATGGACACCAAAAAAATAGCAAGGATGAAACTGGATTCATCCTGGCTTAAACTTAAAAAATAGCGAGGAT

GAAACTGGATGCATCCTGATGTAAATTAAAAATAAGAAAAAGAATTTGAAAATGGGTAGGATGAAACTCAAGTTACATCC

TCTGGCTATTTTTATTTTTTGTCCATTTAAACAAAATGTCAAAATCTAAATGTCTTTTTCACCCAGGAATTATTGATTTT

GGTCTTTTTAACCAATTTTGTGTAAAAATATTAGCTTATCAAGTACGCATAGTGTTTCTAGGGTACATATACGAACACCC

GTTAGGCTGAGTCTAGTGGAGAAATTGGGTATGATGATGGTGACCCAAAATCGTGGTCTAAAACCAAACAAACGAAGTTA

CCGTTCTCTCTGCTCGAAAGCAGGTAAGATGGGATAGGATTTAAATCCTCGCCAAGAAGAAGAGAAAAGCGAACATAATC

TAATAGTGGACCCGCTTTATATATTTAAAAATAATAAAACTTATAAAAAAAGAATATCTTATCAATTTTTACATTATATT

TGAGGATTATGATTCATACACAACTTTGGTTGGTAACCGAAACAACGAGTGGGCCTTGACTCCCTTTTGAATAGCAACAC

CCGGGGACTTGAGCCGGGATTTTTTCACACAAGGGGGCGAAGATAAAGAGGACAAACACAAACCATTACAAAATAGAAAA

ATAAACCCATGACTATTTAAGTTAAAAATGATCAAAATGTTTACCAATGCTATTTAAGTTAAATTGCTAAAAAATTTAGC

AATTTCTTTTTCGATATGCTCAATCAAAGAGTCTCCAAGGAATTCATCACCCATTTTGCTGCGAAGTCTGGTCTTGATAA

TTTTTATTGCAGAAAAAACTCATTCTACCGTTGCAGTAGAAACTGGTAGGGTTAGCACAAGTTCAATCACTCGATGTATA

AGAGAATATGTAACCGACCCTAGATTTTAATTTTAAATATGGAACAAAAAAAATCTAGTGACTAGTCCAAAATTTATAAA

TTGAAATTTACATTCAATTCAGTCATATCTAAAACCCAAATCAAACAAACCTAAATGTTAGATATGATATGATTCATGAA

AGACAAAAATTTAGTCTACAATTTAAAATTTACACTCAATTATTTAGAAACTAAATTGATAAAAGAAGTAGAAAAATACC

TTCAATGCAATCACTTTCTTATCTCCTCGTATAAAAATGAAATATTTGCAGAGATAAGAAGTTTTTATGTTGCTTTCTTG

TTCACTTTCTTTCTCAATATTCTTTATGTAAAAAACAATATTGAAAATATCAAATCCGATCTCATATTCTCAAGCAGATA

AACTATAAAAGGAAACACGTTTTAGTAAGCCATAAATACGTTTTAGACTAAGAAAAGTCATACTCAAAAATACAAAAACA

GATCCCACACCCACGTCTTTATACGAAACCTTAACAAATTTAAATTAGTATTTCTAAATCATAATTTGGGGCTTGAATAT

TAGACTTACAATTTATTTTAGGGGGGCTGTATTAGACTTTTTCCCCAAAACTAAGAAATTCGACGACAACTAATGAAGAT

TTTCCGCCACCCCTGATGTATACACAGCTCCGTCCCTGCAACACCCAATTTATTAAAAAAAAAAACTGCATAGAACACCA

GCAGCATCATTACTAACTAAACAATTTTTCAAACGAAAAAAAAAAGAAAGATTTATTTAAAAGTAATAACATTAGTTAAA

ATGGAATATAATATATTTTGGGCGAACATAATTTTTTTGTCGATATCATAATTACCTCACTTACATTAGACTAAAACCTC

CCTGCAATGTAACTTGGCGCCACTATTCAGACACCACCTTACCTCCTCCACTACCCTTAACATTTAAAAGCAAAACGCGA

AAAACGCTGAAGTTTTAACACTTTTCAAAGTGGAAACTATTTGTGAGTGTGATTTGTGATCATAATCGACAAAAAAAAAA

AAAAAAAAATTAAAAGAAAATGTTGTGTTCACAAAAATGAAAAGTAGACAAATAATTGATAGTACGGACTACCAACCAAA

GCTTGTAATTTTGGAAATTATTTTATTATGTGTCCTATAAATAGTAGTAGTAATTATTTAGTATGGTTGGTCCCAGAAAA

ATAGCATTACCGGTTCCTAACTCTATCTCTATATATATATAAATGTATCGTACGGACTACCAAATGAAAAGTAGACAAAT

AATTGATAGTACGGACTACCAACCAAAGCTATAAATAGTAGTAGTAATTATTTAGTGGAAACTACTTTTGTGAGTGTGAT

AATGATTTGTGATCATAATCGACAAAAAAAAAAAATTAAAAGAAAATGTTGTGTTCACAAAAATGAAAAGTAGACAAATA

ATTGATAGTACGGACTAGATCATAATCAACAAAAAAAAATAATTATTAAAAGAAAATGTTGTGTTCGCAAAAATGAAAAG

TAGACAAATAATTGATAGTACGGACTACCAACCACTAGATATCTATAAAGGGTAAATATATTAAATATCCTTAGTTTTGA

CACTCAATAAATATACCCTTATACAACAATATAAATACCTCTATCATTTAAATACTTATCCATTAAAAAACAGATTACCT

TAATACCCTCACTTACAATATTATTTTCCATTTAAAAATCTTATCCATTAAAAATAAATTTTCCTATTATTTCACATCCA

CCAGCAGCACCATTTCACCACCACCAACAGCACTGTCGCCACCACCACTCCACCACCACCACCACTCCACCACCACCACC

GCCAATCCGCCACCGCCACTCCGCCACCACCACCACTTACTAAATACAACCACCAATAATATTCCACCACCGTCACACAC

CACCACCACTGCTGTTGCCGCCGCCATTACTATTAGGTGTCAACAACTGAAGTTGATCCAAAAAATCAAAAGTTCAAAAA

TATTTGTCAAATTTGGTTGTTGATTCCATTTTTCCATTGTTGATACACAAACATTGGTTGTTGACTCCCTAAAAACTAAG

GTCAATTTCTATTGTGGATACACAAATATTTGTTGTTGATTCCTTACACTAAGATCTATTTCTATTGTTGATATACAAAT

ATTGGTTGTTGATTCCATTTTTCTATGGGATATTTAATGTTTGGTACCCAACAAATGACCAACACATTAGTTTGGTACCC

AACATTTTAAAAATTAAGGGATGGTACCTAGACCCGTCATTAACTGTCAGGTCAATACGTTGACCACTGAAAGACGTAAC

TTCTGTGACTGAACTTCAAGTATTACCTTATTAGCCTTGAGAATGGTATAAATGACTAAGATTTCCCTAACGTCCAGTTA

GTGATCATGTCATCATCTGTACGTGTCACACAATTAAACTGCCACGATTAATCTAGTGTAACCATTAAAACTCATTCAAA

TCTTATATCTAGCCTAAATCAGTTGTTATCGCCGGCAAACTAGACAGAGGAGGCTGTTTTAGGTTCGTAAACATGAGTTG

GGGATTCAAGTGATTTTTGTCAGATTTATCTATTTAAATCTTAGGGTTTCGCTAGATCTACCAGTGTTTAAGTTGGGTAT

CATCATATTTGGGATTCGAAGACAAAATCAAGACAGTTTCAAGGCCAACCGAGAAGCAAAATTTGAAGCAGATCCGTGTA

TTTTTGTAGTTTAAGATTATCTGTCTTTCTTTCCTACATCATTGCCATTTTATCTGTTTAAGTTCTTGATTTTTTGTTTT

TTTATTTTTTTTTATTCAAGTGAATGGGATCTGGATGTAGGAAGAAGATCGGAAGAGATTAGGAGAAGACATGGGTGAAT

TTTATGTTGGAAATGTTGACCCAAATAGGAGAATGAAAACCACAGATTAGGTTGTTGCTAACTGGTTATGTTGACACAAA

GTAATTTTGAAAAAGTGACGGTGAAATGGCTTCTGCCCTTTTATGCATGGTTAGGCAATGGTGTTTGTTGATTCCAGATG

TGTTAGATTGAAGAAAGAAATTCAGAGAAATTCAGAGCTATCATCTTAATCAAAGAAACTGTACTACAATTCCATTGTTA

ATTGTGCAAATAATTTTGATGAATCAAAAGTTACGATGGTGATATGTTTCGATGATGAAATCAATTGATCTATAACGGGA

TGAATCGGCTGTAATGGGTCAAATATTGGTTTTCAACTCAGTGCTAACGTTTAGTTAACGACGTCGAAAAATTAAAAAAA

AATTGAAATTTATTTTTAATTTAATTATCCAGGTAACGGTCTGTTAGGTGCGGGGTTTGACTTGACAGTCGATAACAATT

CCAGGTACCATCCCTTAATTTTTTAAATGCTGGGTACCAAAGCAATGTGTTGGTCATTTGCTGGATACCAAACATTAAAT

ACCCCTTTTTCTATTGTTGATACACAAACATTGGTTGTTGACTCCCCTAACCAAGTCAATTTCTATTGTTGATACACAAA

TATTTGTTGTTGACTTCTTACACTAAGATTTATTTATTGTTGATACTGCAAAATCTTTGGTTGTTGACCACCATTTCTAT

TGTTGATACAAAAACATTGGTTGTTGACTCCCTAAACTAAAGTCAATTTCTATTGTTGATACACAAATATTTGTTGTTGA

CTCCTTAAACTAAGATCAATTTCTATTGTTGACACACAAATATAGGTTGTTGATTCCATTTTTCCATTGTTGATACACAA

GCATTCGTTGTTGACTCCTAAATAGTTGATGGAGTAGTATTCAACAACTTGTTGTCCAACCTCACCACCACCACCACCAT

GTCTACAACATTTCCAACACCACAAACAACCAGAATCACCATTATCTTCATGACCAACTTTAATTAATCATTCCAACCAC

AACCAACACAATCTCCATCATCAAAATTCATCGTTCTAACATCACCAACACCACCTGCACACAGAATATCCACCAACACC

ACAACAACCATAACCACCACTATCTTCAAGATTTAACGAACCTAAATCTCAATTGGAATCACTATAAATAAAAAAGTTTC

TGATGTAACCCCAAATTTCAAAACAAAAACAGATTCCAACACATACCTCTTCCACCACCGCCACCACTAGAACCATCACC

ACCACCATCACCATTGCTTTTTGAGCTTCAATAATTCGAAGAACTTCAACTCTAAAATCATACACACTTAAAGATGTCCG

GCTTCAATGAATCTAACATGCAAAGATCTCGTTCCATTTTCAACTTGAAAACCAAAAAATCGAATCTATCGATCTGAATC

TGGAAAATCATACACACCCAATCTCCACCACCACAACCACCACAACAACAGCAATAACCACCACTATCTTTATCACCAAT

TTTATCATTCCGACCCAACCACCACCAAAAACAACAAAAACTAGTTATACCACCACCGCCGCCATCACCAGGCACCAGCA

GAAACTTAACTAAAGAAGAGAAAGAAGATAATCGATGCAATCTCATTTTATCTCCATCACCAATTTAGAAAACGAAGGAA

TTTGAATTTGGATCTGTAATCCAATAAATAAATAAATAATAATCAATCACACTTCTCAATTTGTTTGTTCATAATCTGGG

CATTTGCGCTGGCTTGACGATGATGGTGAAATTTTGGAAAACAATGATGTATTGTTGATGGTGGCGATGAAGATGGTGGT

GGTGGGTTTAGTTTGGTTTAATTTTCGATCGAAGATCTCTCTCCCAATCTGGATCTAAATCTCTTCTCTCTCACTCAGAA

ATGGTAGTGGTGCTGGTTATGGTGGCGGAGGACGACGTGGTGGTGGCAGCGACGGAGCATGTAGTGGTAGTTGCAGAGGT

GCGACGATGAAAACGATGATGGTGCTGATAAAAACGGAAAGAGAGAGAGAATTGAATAAAAGAGGAGATGATGTAAGGGT

ATATTTGTAATGTTGTATAATCCTTAATGGATCCCTGATGGGCTTTGGATGGAGAGGGGTATTTTTATTGTAGGATAAGG

GTAGTTGTGTAGCCCGTCAGAATTGAGGGCAATTATATAATTCCCACATATATAAATGTATCGTAGCATTACCGGTTCCC

AACTCTAGGGTTGGTCCCAGAAAAATAGCATTACCGGTTCCCAACTCTCTCTCTCTATATATATATATATAAATGTAGGG

TTCTATTAACTTATATACCAGGGGCATACGTTAAGGGACATTAAATACTCTACTTTTTACCTTGGATTTAGGGATCATTT

ATAGAAAATGCAACAATCTCGGTTAATCTGATATTAAATACCAATGAAACGTAAATACCTACTCTACCCTTATTAATTAA

TGATTATTTTATCCTGATAGGAAATGAAAGGCGGATTGAATCGTTTATTACAGTGGGACCTAGTCTTATGTTTAGCAAAT

AACTCCTAATTAATGCTGAATAAATTGTGTCCGCACATCTATGTAGGATGTGTCTTTTTTTTTTTTTTTTTTTTTGAAGC

ATGCATATATTATAAAATCAACTAAACCGAAATTACACGAGTGAATGTGATTCCCCTAGAATCGCTTAACAGAATATGGT

TAAGAAAAGCCGGGTTCGCCCGGTCCCATATTGAGGTTGAGAAGTTTCCATTTTGGCTACCGTCTGCCGCATAGTTCGCT

AGACCATCTGCGCTTGGTTTCCTTCTCTCAGAATTGTCTTTGAATAGCACAACATGGAATTAGGCCCATCCAGATGTTTA

ATTTCGCAACCATTTCATATATACCAAGGGGCTTCATTTGGAGTTTTAATGAAGTGTATGAGACTTTGAATCTGCCTCAA

AATAACACTTTAGACCATCCTCTTTCGTGGCAGTTCTAGTTGCTAATATCTGATTGCCCATGTTTCGCTTTGTGGAGCGA

TGATGTTAATCCACAGAGGTTGAGCAAGAGCCATTATCGTTCGCGCATTCGAGTCCACACAAATGAAACCCTCCCGCCAT

ACTTGGGTGGCCTCTAGCAGATCCATCGGTGTTGACTTTGATCCATTCTCAGTTGCAGGTTACACCAACCTACTTGAATT

GTTATGCTTGTTTTGGTTATGTTTGTAGCGATTCGAGACAGGTTCTCCCGGAGAACCGGAGGGGACATCGTTAGAGCCCA

TTCAAATTCATTGTTGTGCCAAGAATTTTTGCATAATATCTGTGATAGTGGATGGTGCTTGATTGAACACGAGTTCATTT

CTTGTCGTCCAAAGTGACCATAACAGAAAACAACATGCCGTGATAGAAGATGTATTGCCTCGAGTTTGGAGTATTTGTGA

TATTGTTGTTTGAGGGGTTATAGTAATTGATTGGTTGTTGCCTTGACGTTGGTTTGTTGATATTTGGTTTGAAAGGATGG

TCCAAGTGGACATTGCTATCGGGCAGTGAATAAGTATGTGTTCAGATGTTTCTGGTGCGGAGTTGCAGCGGGGGCAAAGG

TTGGTATTGGAGAGGTTGATGTGATGTAGTTTAGATAAGGTTGGTAAACCTTCATTTATAGCTTTCCACAAGAAAAGCTG

AATTTTTGGTGGACATGGTAAAGTCCATAAAAATTTGGTAGTTGGTAGGAGGTTAGGGTGTGAGAGGTTCTCTTGTTGGA

TTAGACAACTATACCCAGATGCAGTGGTGTATTTTCCAGATTTAGTGTGGGGCCAGATTATTTTATCCGGGGTGCTGAAA

TTAGGAAGGTGGATGTTTATTATGTGTTGGATGGCATTAGGTGGGAGGTTGTTTAGCTGGCTCCAATTATGAGAAAATGG

GTCAATTAACTCATTCTGATTGTATAGGGGAACACCGGGTGGGATCGAGATTTTGAAAGTGTGGTATCCAATTAGCTTGG

ATAGGGGTATCTAAACCATTTCCTATTTGGTGAAATAAATGTTGTTTTATCGGCGGTAAGAGAGACGCCATTTGTTTCCA

TTGTGGGCTGGCATTAGAAGGAACTGTGAGCTCACCTTGGAATATTGGTTGATGTTTGAGGTACTTCTCATTGAAAAGTC

CGGTGCAGATGGAATGTGGCTGGGCATGAATGTCCCATATCCGTTTCATGAAAAGAGCTTTATTGTGGTGGCTACTCTTT

CTGATTCCGAGTCCTCCCTCTGCTATCGGCTTGTGAAGTTTGTTCCATCCTATTTGGTGAAACTTTCTCACAGAAGAATC

ATGTCCCCAAAAAAAATTCCTAGCAATCCGATCAATTTGTTTGTGTACCTGGGTGGGGAAAAGCTGTGTTTGCATAAGAT

GATTGGATGTTGGTGTTAAAGAGGTTTTAATAAGAACTAGCCTCCTGCCTGGTTAAGGCATTTTGTCATCCATCCACGAG

CTTTACGAGCCGGCGTTGGAGTAGAGGTTCGAATATGTGTCGAGAGTTTCTTCCGTGCTTGAATTGGACTCCTAAATATA

TAGGTGGAATTGATGTGGTTGACATGTCGTAAAATTGTTTAGAAGAATCAATGTTATTACTGTCGTAGCCTCGGGTGGTG

GATGATTGTTGATTTTGTTGTGTTGATTATTTGTCCTGTTTGAAGTCTTTGTAAGATTGAAGTATCCTCTTTAGGTGATT

GCTACTTTCCTGTGTAGCTTTGTACGTGATTAGTAGATCGTCTGCATACATTAGATGTAAAATTGGTGGTGCCTTTTTTG

ATATTCGGAGACCTCCGACCATATTCATCGCTTCAAGAATAGCAAGATTTGTTGATAGAATTTCTGCACAAATAATGAAG

ATGTAGGAAGATATAGGATCACCTTGTCTGATACCCCTGGTAGGAGTGATAAATCCATGAGGTGTGCCATTGATGTTGAT

CGAGTAAGTGACTGATTTGATACATAACATGATATACTCCACAAATATTGATGGAAATCCCAGTTGTGTAAAAGCTTTTT

GATGAATCCCAATCAAAAATTTGTCATAAGCTTTAAATATCTAGCTTTAGAGCTATCTTGGGGTCTTTAGTTATTGGAGA

CGTTTTAATGTGGTGGAAAAGCTCTCGGCAATTGCTATGTTATCATGTATTGATCTTTGAGGTCTGAAGGCACTTTGGTA

TGGGCTTATTAAGAATGGGAGAAGGGGGCGAATCGATTGAAAGGATTTTGGATATGATTTTGTAACTGACATTACATAGG

CTAATAGGTCTGAATTGGGATGGTTTTGTTGGATTGTCGACTTTTGGAATGAGTGTTATGTTTGTGTGATTCATGAGAGG

GTGTATGTTTAAAGAATTGAAAAAACTTCGTACCATAGCTATAAGTTGTAGGTGAGTGGTGTCCCAGAAAACTTTAAAAA

ACTTACTGGTGTATCCGTCCGGTCCCGGAGCGCTTTCGGAATTAATGGAGAACAAAGCTTGCTTGATTTCTTTCACAGTA

ACTTCCGCAGAAAAATGTCACATTGTGCCGATGTCAATCTTGGAGTTATTTCCTCAAAGAGTATTTCATCTATCGTTGTA

TTTGGGGCAGTGTATTGTGAAGTGAAATTGTCCGGGATTATCTGGATGACACAAGTCTTTATCAACACCCGAGTTGTTTT

GATGATCTTGTAGTTGCTTTATGTTATTACGGCTTCGGTGGATGGTTGCCTTTGTGTGGAAGAAAGTCGTATTATTGTCC

CCCGATTGAAGCCATTGAATTCTGGATCTTTGATGCCAATATGTTGCATGGCATTGGAGTAATGTGATGTGTTGACTGCG

TAGCTCTTTGTCGTGTTGTATCCAGAAATTTAGTTCCTCGCCGGTGTGAATAGCGCATTGATGTTGAGCCCTTTGAATTT

CCCATTTGTTGATTTTTTTAATTAATGTTGGTAGGTGACCAAAAGTGTTCTTGTTCCATTCCATCGAGTTTGTCCGTTGT

GATTAATCTTAGTTGAAGGTCTAGAGCCCAGTTAAGAGTCACTTTGTTGCGTCCATGCGATTCGATAATGCTTGAGTTGT

GGATGTGTTAGCCACAAATGCTCAAACTTGTAATTTTTGTCCTTGCCAATCGATTGCTTTAGTGTGCAGCAGGATGGGAG

CATGATCTGAAGCAATCCTCAGGAGGTGAGTAACTCCTGCATGAGGATTTGTTGTTCGCCAATGTTGGGTTGCAATTGCC

CTATCAAGTCGTTCCATTATATTGTCTTGACCCTGTTGGTTGTTTGTCAGGTGAATGGGTCCCCTCGAAAACCTAAATCT

ATGAAGCCCATTTGATCAATCATCGTTAGAAATGGTCAGGTTTGACTAAATGTGATCTGTCTTCCTCCCTTTTTTTTCTG

ATTGATCCATTATGTCATTAAAATCTCCTATCAGAAGCTATTCCGTTGATATGTTGATATTGGGGAAGATTGTTGGGAAA

TCTTGCCAAAAATCCGTTCTTGCATCTGGGTGAGGAGGACCGTATATACCTGTGCAGAGCCATGGTTGTGCAGGGGGTGG

AGGTTTGATAAGAATGTGGATGTAACTTTGGGACTGAAGTAACACTTGAATGTTTAGGTTGTTCCACATGAGACAGAGTC

CTCCTCGTCGACCCACCTTGGGAACAGAGGTTAAGTTTTGAAAATGACGATCTTTGGCTAAAGTTTCAGAGAGAAATAAG

ATATCGGGCTTATGTAATGATACTAGTCCTTTTAGATGTTGTATCGTTGGTTGATTGTTGATGCCACAGCAATTCAGCTA

GGATATTCATGATGTGGTTTAAATAGTTTTGGGGAATTGGAGAAGGTTTAGTTCATATAATTTAAGTGATTGTTTTTAGG

TGAGCGAATTAAAAGGAACAAATGAGGGGTTGAGGAGTATTTTGAAATGGGGTAGAAC

>Pbr_flye_v1-2_contig_2915_minus contig_2915

GAGTAAAAACAAGGGAACATTATTTAAATACCTTTACTATAGGGCCCCTTCCAAAATACCTTTATCAACTTATACAAATA

CCCCTAAGCATAATACATTACTAAAATACCCTCTTTTGTGTAATGTATATATAAAACCATTACCACTAACCACTATTCAA

CCACAAAAAAAAAACATCTAATGTCCGCCATCACCACTGTCCACCACCAACCACCACCACCGCCGACCACCACCACCACT

GTCCACCACCGCCAACCGCCATCGCCGCCAACCACCACCACTGTCCACCAACACCGCCGCCACCCGCCGCCATCCACCAC

TGCCACCAATCAACACCGCCATCGTCAACTACCACCACGGACAACCAGCGCCATTGTCCATCCACCACCTCCACCAACCA

CCACCACTTTCATCCACCACCACCGTCGCCACCAATCACCACCAGCAACCACCAATTCCACCACCACCATGTGGAGTAAA

CTTGATGAAGTTGTTTCCAAATCTGAAAGCAACTAGCTCAAAGTTGTCTCCAAATTTGAAAGCAACTTTCTCAAGTTGCC

TCCAAATATGAAATCAACTAGCTCAAGTTGTCTCCAAATTTGAAAGCAACTTTCACAAGTTGCCCCCAAGTCTAGAGTCC

AAACTTTCTCAAGTTGCTCTGTCTCGAATGAATAGGCTGGCAAACTCAGTTTTGACACATCCTAATGCAACTAAGCTTCC

CCTCTTAGATTATGGACATCAACAACACCTACACTTCATGCCTCAGAATTTGAGCTAAAAAGTCATGAATCGCACTAGTA

TCATCACCTCACACTTCATAACTCGGCCATAATCATAACCTCGCAGCTCCATGCCTTCCACTTATGCCATATATAACCAC

AACTACGCCAAGACATCTTCAAGCATAACCGCTAAACCAAATCCATCTCTCGTGTCTTTCCACAAGACAAAACACTTCAA

CCACGCTTAGCTTCCAGCATGACTTCAATCCTTCTCTTGCAACCACCATCATTGCAATCATGCAGACCTACTTGTACCTT

CTTTGATAGCAACACCAACATCATTCAGTCCACAATCTAAGTCACTGATCACCATTACAACTGCATCCCTTGTCAGTTCA

GCTCACAAACTCAACTTCAACTCTACCAGAACCACCGTCTTAGCATTAATCACCTGCAATATCATCTCCCATTCTCTGCA

ATTTCATCATTGCCTGCAATTCATTCCTAATCATACATCTAGCATTTGCATCTCCCCAACCAATTGACCACTGCATTTCA

GTTACTACCAGCAATATATTTGGCATTACTTCCTGGTTCATCCCAGCACATCTTCTAAGCTTCACTTGCACCTAAATCAA

CCCTGCAACAGTCATTGAACCAGTTGTTGCACATTTCAACCAACAACAACATCAAACAACACTTCATTCCCATTCTTTGT

CTAGTTCCAGCTCATAAGCATTCATCTAATATTATTTTTGTAGGTTAACTCGCTTGTCGTGCTAATTTACTTGCAACTCC

ATTCAAACACATACCAACACACATCCAACCCCCTCTGCATTTCATTTAAAAGCACAGTAACACTATAAACTGCAATCTCA

ACCTGCACCCTGATATAATATACCAACCACCACCCTGTAGTAAATTATCTTCTCCATTCATAAACTCCACCTTCATCACT

AAACCGCCATTTCCATCGAAGAAACCAAATCCATCTCTCTCTGCAACTACAACTTCAAACACCAACTCAAATCTTCAGAA

TTCTACCAAAACACCAACTAAACTCGCCAAAAGGCAATATCTCACCTAGCTCTTTCATCTGCTTCTTTCTTTACTTGTCA

CAATAATGGCGACGACCACCATCTTCTCTCAAAAATCAACAAAGCAGCAAACCCAACCAAAACAACAACACAGTTGGGAT

TATAGTCAAAACACAAAATCAAAGCACAAAATAATAATTATCCCTATAATTATTGAAAATGACATCCACCACCACCGTTG

CAACAACACCAACACCACCACCACCAGTCCACAACCAAAACACCAACCTTCACCATTATTCACCCTACGAGATAAATCAT

AAAGTTTTTGAAATCCTAAAATTACTTTTTAGAAATTTCAAATCTGAAATCGACCAATTGAAACTCCCGATTGAGTTAAA

AGGATAATGATTAACTTACCTTTTCAATCGTTTTTGAAGATGAATCCTTCAAATTAAACTGATCTTGAATCTCGTCAATT

CTTCCCTTCCATTTACAGTAAACCTTCGAGAAAAATGATGTTGACATCGATCTACGTTTTTCAAACTAGCTCGAATTGAA

TAAAATTAAACCTAATACATATTTTCAGAATACGTTTGTAGTTCAATCTAAATTAATGTACCAATAACACCATCAGTAGC

TTCAAACTCCTAACCCTAGCTTCACCGATTTAACCACCACCACCAGAAACTAATTGAACCTTACCAGTTTCAAAATCAAC

AAAAACTCTAGCTCAAACCATTGAATCCGAATCTTTAACTAAAAACCCAACAAAAATCTGCATCTCAACTCGAAATTACT

GAATCGAAAATAAAAAAATCAAATCGATCTCTGAACTGTTTTGAAGTCGAACACAAATCATGGAAAATCTGTTGATACTA

GTGGCGGAGCTAACGGAGGTGGTGGTGCTAGTTGTGGTGGCGGTGATAGATCTTGGTAATGGTAGGGGTGACGGAGATGA

TGGAGGTGGGAGCGATGATCTTCTGTTATTAGAAATGAAGAAGAACAAAGGAAGAAAGGTTAAAACAGGGATTATGGTTA

TTATTATTTTAAAGAAAATTTCAATTATGCCATCAAGGGTGTTTGTAAAGACTCATAAGGGTGTTTGTGCAAGGCCCATG

GATAGTAGGGGCATAAATAACATTCCCACTAAAAACAAATAAGGTTTCTCTCTCATATAGAATCTTAGATGGATAAGATT

TTATCGGGATATCCTTCCAAGGAAAGGGTACTATCCAAACGTCGTTGGATTGGTCAACTAAAGTCCTAACACAAGGACAT

CTTGACTTTAGTCAGGGATTAACTAGTGAGTGTCCTTGTATAAGTCAACAAAAGTCTTGAAATTACGGCACTTTGACTTT

AGTCAACGACATTTTCGGTGTATAGTTAATTCACGATTTCTAGTATGTGTCGTTCCATTATGTCTCAGATAGCCCATTTT

CCACTAGTGTAACAAAATACAAGCATGTGTATTTCTGCAGCAGGGCAGGTTTGGTGGTGACGGGTACCTTACCATAAATC

GTGAGGGGGAGATTGTTGGGCTATCCCTTCTTATGGCCCAAACATGTGGAGTCCCCTTAACCAATGGGAGGAGGAAGATA

GAAGGTGCCTTGTTTTTTGGTTATAAGGCAAATAGGCATCAGGAGACGGGTTTTATGTTTTCTTCTTCATTTTGTGAAGA

AAGAGAAAGAGAGAAAGAAGAGAAAGAGAGAAAGAGTGTAGCCACCATTAGAGTTGATCTTTGGTGATGATTTTGGTGTT

ATCTTGAAGAGAAGAAGAAAGGGTTCTTGTCTAAGGATCTCTTGGTTGTTGTGGTGAAGATTGAGCTATCAATTTCCACA

TCTCTTCAAAGGGGGAAAATTCTAGGGTTTGTTCAAACCCTAGTAGTAAGTTTCTAAATCACTTCCTATTGTCAAATTGA

TAGTGTCTATTATCTATTTTGTGAATGAATTGTTGATCCATCCTTAGGGATGAGTGTAATTTAGGCAAATTGGTATCCAA

GTGCTATAGGCACCTTGAGGTTGTTGTTATCCCCAAATTGGGAGAATTGTGTGTAACCCATTATTGTTATAGTGGAAGAT

TTGCCCGTGGTTTTTTACCCTTCACCTTGGAGGGGTTTTTCCACGTAATTACCGGTGTTGTGTGATTGCTTCTATATTGC

GTAGTTGTGTTATTTCCGCATTGGTATGTCATTGCTTGTGTAGGTTTCGTATTGCGCATAGCGGCTCGGATTTCGTAAGC

CTCCCCCAACAGAACAAACCCTTGAAGAGAAGAACAAACCCTTGAAAAATTAAGCCCGTTCAAAACTGACCCAAAGTGAA

AATCATCCTGGTAGATGACGACGTCTCAGTTTCAATCCAGCGGCTGTTGAGTTCCATCCGTTTGGTTGCGGTTACCGGAG

GCGTAACTGATAAATCCACCGCCGGCGACAGAGTTAAACCTAGTTTTTTGAAATCATCCTGGAAATCTTCAAGCTGCTTT

GATGTAGTGGCAAGGGGAAATTTTCTGGGTGCCATTGTTGAAAGAAGAGAAGATGATGTGGAAATATTGAGGAAGAATTA

AAGAAAGCAGAAAGCGAACAGAGTTTACAGAAGAATGGAGTCGGGGAAGAAGGAACAGGAGACTGTAAAAAGATGAAAAG

GTAAAGAAGAGAGACTATTTATAAAGAGCCAAGGCATGACCGGGAGAGGTGGTTACACGAGGAGAGGAACCGTCTAGTGA

ATGTTGACAAGAATCGTGGGAGACAAGATGGCGGAAAATCCGGGTACGTGGCGGTTACGAATTTAATGCGAGGTTCTCTT

TACTTTCTTGGTACCCGCAAAAGGAATGAGAAAAGGAAAAATGTATGTGCAGAAAACCACTCTGCCACGTGTCCGAGGAA

GATTGAGAGAGATAATAAGGAGAATCTTCGAGGAAGATATTATACACCAATACCGAAGGAGTGTAAGTGAGTAGTCTAAT

CAACTGTCAGATCTGCCTTAACGAGGCTCCGTCAGCAAAGAAGAACTAACTTGCTCGGAGAGACATCTAGCGAAGAAGAA

AAAACTTGGTTATGGGCAATAGAAAGGATCTTGGTGGACACAGGGAACTCGGTGGATATTCTATTTTACCGCGCATTTAA

AATAATGGGCTACCGAGATTCGGATCTAGTTCCATCAACCTATAATATCTATGGGTTTAACGGAGTAGCATCAAAGCCGA

AAGGGGAACTAGTCATGAAAATCTTCGCAGGAGAACTTGAAACCGAGGTAACAGTATGCGTGGTGGATGTAGAATCTCCC

TACAACGCTCTGATTGGGCGGTCATGGATACATGGGATCAAGGGAATATCATCTACATATCACCAAGCTATCCGATTTAC

TATGCCAAGCGGTATTGGTGAGATAAAGGGAGATCAGAAAGAAGCCAGAGAGTGCAATGATGAAGATGTGCATAACTATG

ACGAGAAAATAAAGCGAAAGAAGGAACAAAAGAAGAGAGTGCGAGAAGCAAAGAGGGAAGAAGAATTAATGGTGTATATG

GTCAAAGCAAAAGAGGGATGGGCTATTCCAAGTGAAATACCCGAAGATAAGGATGAGCCTTTGTAGCAATTAAAGGGTCC

GTCTCCGCTCGGAGACCCAATCATGAACTTCGCAGAAGTAGAACCCACGCAAGAGATAAACTTGGGCAGCGAAGATGAAA

CGAAAATGGTGAAGATCGGAACAACGATGGAGGCGGACGAAGAGGAAAGGTTGATAAAGATGTTGAAAGAATACAGTGAT

GTATTTGCATGGTGCATGAACGAGATGCCTGGGATAGATCCTGAAGTTGCTTGCCACAGACTGGATCTAAACCCAGCAAT

CAATCCAATAAGGCAACGAATACGAAAGATATCAACTGCTTATCACGAGCAAATAGAGCAAGAATTACAGAAAATGTTAA

ACTCGGGAATAATAAGACCCGCGAAGTATCCGACATGGATAGCCAACATGGTGATAATACCAAAAAATACTACGGGAATA

AGAATATGCATAGATTTCAGCGATTTAAATACCGCTTCTCCAAAGGATAGTTTCCCTCTACCCAATATACCGCTGATGGT

GGAGTCAGCATCAGGGTACAAGAGGTTAACTTTGGTGGAAGGTTATTGTGGATACAACCAGATACCACTCGCGGAGGAAG

ACTAGGAGCACACGGCTTTCTTCGATCCTAGAGGGTTGTATTGCTACACGAGAATGTCGTTCGTCTTGAAGAATGCGGGA

GCAACTAATCAACGGATGGTCGAGAAGATTTTCGCAAAATGGGTGCACCACAAGCTAGAGGTTTACGTCGACGACATGCT

GGTAAAAACTAAGGAATCAGGGGACCATGTTGAAGACTTGAAAGAAATCTTCGAGCGGATGAGGGCATACAAAATGAAGA

TAAATCCTGCGAAGTGTGTATTCGGTTTGGAATCAGCGAAGTTCCTGGGACACATCATGTCCGAGAAAGGGATTGAAGTA

GATCCATCCAAGGTACAAGCCATATTGGACATGCCATCGCCAGCAATGGTAAAAGACGACGCAAAAAACTGAATGGGCAA

CAAGCGGCTTTGGGAAGATTCATTTCTAAATCCTCGGATAGGTGCAAACACTTCTTCAACATTTTGAAGAAAGGAGCAAA

ATTTTCGTGGACACAAGAATGCGAACAGGCCTTACAAAATATAAAAGAGTATCTTATCAACTTATCAATTCTCCAAAATC

CCGAGCCAAATGAAGAACTACTGATTTGTCTCGCAGCTACACCAAATGTCATCAGTTCAGTGTTGCTTCAGACAGATGAA

GGCGTTGAAAAGCCTATATTTTACGTCAGTAAAACACTCAATGCCACTGAAAAGAACTACACAAAGATAGAAAAGTTTAT

TCTAGCACTGGTGTATTCATCATAAAAACTAAGAACCTATTTTCAGACTCACAAGATAAAAGTTCTAACCAGAATCCCTA

TAGAGTCATATTTAAAGAATACAAGGAGAGTTGGGTGAGTTGAAATATGGAATACCCGAATAGATCAATTCGGGTTAAAA

TACGAAGTCACACATAGTTGCGGAATTCCTGGCCGAGTTCCCAATAGAAGAATACGAAGGAGTGGAAGAGATGTTGGATA

TAGATGAGGAGCGTCCGGATCCTACAGATCTACTGAAAGAAAATCACCCAAGGAGATGGGAAATATTTGTAGATGGATCA

TCCAACAGCGCTGGAGGAGGAATGGTAATAGTATTTACTTCGCCTAAAGGAGCAAGGACAGTATTTTGCTTCCGACTGGA

ATATAAAGCTACTAACAACGAGACAGAATATCAGGCAGTCATACAAGCCTTGAGGCTGGCTATCGAGATGAAGTTGGAAG

AAATAAGAATAACTAGCGATTCTCAGCTGGTAGTCAGGAAGATAGAAGGAAGGTATCAAGCGAATGATGCAGTGATGAAT

CGATATTTGCAACTCGTGAAAGAGTATTCAAAGAGAATACCCAATATAATTTAGAGGCAAGTAAGCAGAGATAATAACAG

ACATGCGGACGCTTTAGCCTTTATTGCATCAATGATAGAAGACACCTCGGAGAAATATATCCGAATTGAAAGGTTAATGC

ACCCCTCAGTAACTAACGAGTTGGAAGATGAGAAGTCCATGATGATAATAGAGGGTAGCACATCCGACGAATTGGGAGAA

GATGACTGGAGGAAACCTATTCACAAGTACCTCGAAAAGGGAGAACTACCTCGGGACAGATTTGAGGCACACAAGATCAA

GGGGAAGTCTACCAACTATGAATTAAGAGAAGGAATCCTATATCGGAGATCATTCCTAGGACCATTGTTGCGATGCTTGT

CACAAAAATAGGGGGTACAAATCATGAAGCCAATACATTACGGTGACGCTGGAAATCACAGCGGAACTAGATCATTGGCT

TACAAAACTAAAACGCAAGGATACTTCTGGCCAAGCATGCATAAGGACGCAAAAGGTGTAGCAACGAGGTGTGAAGCTTG

TCAAAGATTCGGAAGACGCATACACGCACCCTCGACGAACTTGAATTCTGTGCTAAGTGTGTGGCTATTTGCCATATGGG

GAATAGACATCGTGGGACCATTTGTTGCTGTAACCTTGCAGCGAAAGTACCTAATAGTGGCAACAGACTACTTCATGAAG

TGGGTTGAATCCAAACCAGTAAAGAATACGCGAGATGGAGATATATACGACTTTATACTTGAGAACATCATATGCAGATT

TGGAATTCCACTGCAGATAGTCTCGGATAACGGAATGTAATTCGAGGGAGAGAACGTGAAAATTCTATTTAGCACTTTTA

AAATACAAGCCGCAAAGTCGACTCCGCTATACCCGCAGAGCAATGGACAAGATGAAGCAACAAACAAGACAATTACATCC

ACTTTAAAAAAGAAACTAGACGGGCATCTCCATAATTGGTGCGAAGAAGTCCGAAGGTGCTATGGTCATACAGAACCACC

AGACGAGAGGCAACAGGGATGTCACCTTTCTGCTTGGCATATGGAGTAGAAGCGGTTCTGTCTACTGAGATCATGTTCCC

CACTACAAAACAAGAATCCTGGAAAAAATGTCTAAACTAGGACATGATCCTAGCAAAATTAGATGACTTGGAGGAAAACA

GGGAGATTGCTCTACAACACATGACGAACTATCATCGGAGGCTGTCTCGCGGGTACAATATGAAAGTGAAAGAAAGGAGT

TTCGTTCCGGGGGAACTTGTTCTGCGAGAAATACCACCATATCAAAGGGGATCCGTAGGAAAGCTGGAACCTACGTGGGA

AGGACCTTTTATCTTGAAAAGGAGAGTAGGAAATGGAGCTTATGAACTCGCGGATTGCGAAGGAAAGGACACAGATAGCC

ACGGGGACATCATATATGGCGAAGCGGAAAGGCATGGTCTAAGGCTAGACCACCCATGGAATCCAATTTACATCAATAAA

TATTATCCTTAAAAAGCAGTAACTCGTAGAGAGTTGTTCAAAAAAAATAAAATAAAATAAAATAAAATGTATGTGTTATT

TTGTGAACAAACAGAAAATGGAATGAAACATCTATCGAGCGTAAATAATAGATCAGCAACAATAAAGAAAATAACAAGGC

GTGAATTCAAAATATACATGCATACAAGGGCAAAGGTTGGACCTCACTAGGAGGTAGCTTAAGACCCACATAAGATGAAG

AAAAGGAAATCATAAGTCCTTAATAAAACCTCTACATAGGGAGGCCGAGACCTCGTGGCGTGAAACCTAGTTCGCAAAAA

GTGCAAGAGGAAAATATAAGACCTAACGCGTGTCACTAGATAACCTCGGGATATGGCTAATAGCCATCAGCTAGGGGCAA

GAAGTAAGACCCATACGCCGAGGGCCCCAATTGGGTAACCACTCGGTAAGGGGTGTAAGAACAAGCCAGTGGAGAAGAAG

ATGGCCTGGTGTAGTTCTTCCGCTCGGTCCCAAAACCACCTCAAATAGGGTGTGACTTCGTAACCAAGAGCCATATCGGC

GAAAGGATGCGAGGTCACGAAATAACTGATGCCTAAGCCCAAAAGCACGGGGGTATTCAGAGGGGAAGGGACGGCACCCT

CCTTTAGGGAGCTGAGTGACCACAAAGTCGGCAATGTTACGGGGCTGGTACTCACGGGAGTAGGTAGGCCTGTGATATTG

TTGCACATAAGACCTTAATGAGCGAGATAAGGTGAGGTTGATGGATCTTTACTTCGCAAGAGTAAAGAGCGCCTGAGACT

TCATCGAAAAGCCCTGACCAACGGGGTGGGGCACAATAAGACCTTAACTAGGAGGCAAAATAAGAACCCGAACTGATGAT

AGAGATGTGAGGAAAAAATATGAGACCTGAGGCAAAACAAAAAGGCAAGTGTATACTTGCAAAGAGGCATGTATGTACTA

CAAAGAAAAAATCAAGCACATTACAAAGCTTGATTACACAAGCCATACAATTCTAAGAAGTACTTCTTACATTGCATAAG

AAGCACTCGCAACTGAAATCTTTTACATGATGATATATGCACTAACCCAATTATGTTATGAATTGAAAGATAATAATCGC

ATCGAAATGATTATAATGCGAGATCGACTCGCATGAGAAATTAAGGCAGAGAAAGTAAGGTCAAATGGAGCTAAAAGCAC

TAGGTATGCTACATAAATCCACCAAAGAATAAGAGGCAAGTATATACTTTAAATACGATTTTATTCTTTCTAGCATTTAT

TTCGCATTAAGGAAAAGGAGCAATTACTTCGGAAACACCCAAAGAATAAGGGGCAAGTATATACTTAAGCAACACTAATT

ATTCTTTGATTATTTCCTTCGTCAAGTATATACTTGGTATACCGGAGCGATTCCGAAATACCAGGGCAGGGAAACCATTA

ACAAACATGCAGGTTGTATATAAGTGCAACAAATGAAGCAACTAAAGGGAAAAGCTAATATATTAAGGCAAAATAAGAAC

AAAAATTGTCATGCCCCCCAAGCTTGGGAGCACCCAGATAAAATGTTAGAAGGACAAAGAACCAAAGTATCACAAACATC

AAAGAAAATAATCAGGGATTGTTAACTCCCTCTTCATCAAGATTGCCATCAAATCCCGCAGCCCCATCATTGTTACAAGC

TCCAGCATTGGCACCATCCCCAGCATCGTTATCTCATGTGCTTTGAGGAACCTCGTGATGAGGTCTCTCAATGTCTTCAT

CAATCTCTGTATCTTCGTTCTCATCTTCGGCGAAGACAGTAGGTTCATCATCGGAGAAGATCTCGAGATCCATATCAGGG

ACGTCAAGACCACGCGAGTTGCAAAACTCGGTGAAGAATTTCACCTTCTCTTCCTGGGAATTTTGGCGAGTCCACTTCGC

TACCTTATTAGCGTCTTCCTCCATCTTATCATAGTCCTCGTTCAAGGATGCAAAGTCTTCCTGTAATGTGGACAGCTTCT

TGGCCAAGTCGTACTTTTGGCGAGACAAGTGCAAAACCTGATTCCTCAAACTCTCCTCGGATTCTGCAAGATAACAGTGA

AATGAGAAACAAGAATTACCAGGCAAAACACGAAGCAACAACAACATCGAGATTAAAGTGATCAAGTCTTGATATTTCAG

ATTCGAGAAAGCTATGATGATCTAGGGATGATTTTTCCATATCCTCTCAAGGGCCGGACATGGGTATATCATCATCAGGG

GATCCTATAACAAGTTTACCGCTTCTGGGAGGCATAACTAGCTTCTTCTTCGTAGGAGGAACGGGGGCAGATTTCTTTCC

TTGAAGCCTCTCCAAGCGAACCAGGATGGAGGACATGTCACTATTCAATTCGTTGTTCTCATTCCTTAGCCTAGTGTTCT

CAGTCTTCAGCTCCTCATGCCCATATTTTAACTCGGAAAGCTCACTCCGGAGCTTGCGAAGTTCAGCTTCGGATTTATCT

TTCCTCTTAATTATTATATCTTTGTCCTCCTCAAGACGCCTAACTTGACCCATGGATAAGGAGTGAAGGTTACGGGAAGT

AGTGAGAGAATTACTGATCCGTTCAAGCTCAGTTATCAACCGTTCATTCTCCTCATGAAGATGGAAATTCAACTCTTCAT

TATTAGTGGATTCATTAGCTAGACGCTCAACCAGAGAATATGATGTCATTAGCTTTTCATTCAAGATCTCATTTTCGGAT

AAATAACAATTAATCTGAGACAAAAGGTTTGCTTTTTCTGCTTGGTAATCGTCTGCGGTCGACTTGGCTTCTTCAAAATT

CATATAATACTTCGCAGAATGGGCTTGACGAAAAACAACAAGATAGTAAAAATGAGAACGAGATAGAAATAAAGACCGGG

AAAAGATTAAACTTGCTGCTAAGCTCCGAAATCCTTTCATCTTTCTTGACTGAACGCTCCCTCTCATCAGCGAGATCATT

CCTAAGATCCTGAATTCGTTCATTAGCGGTCTTCAGATTCTCGCGGGCAACCTTCATATCCACAAGTTGAGCCATATGAC

GGTTGAACTCCTGTGAAGTATTTCCAGTAAGATGGGGATTTGTACGTAAAATTCATTTAAGATTGATAATACTAATCAAG

GCCATAATAGCAGCGTACTTGTTGGTGGAGAGATACAACGAAGCGTTCTGCATGAAAGATTGATCTTGAGCTAGAAAAGT

ATCGGCGCATAGCGAGATTACAGATTCCATAGTTGCAGACCTGGTGGAAGGGTTTTTCCGAGTTAAAATAAGGGCGTCCC

TTAAGAAATCCATGTCGGGGGAATAGGCCGGATCCTCTACTTCAAAAACCTTCCGAGTCTTTTCACTAGAAGGACCTGTT

GATGGTTTAACCGGGGACTCTTCCACCGTGATACGAGGAGTTGGCATGGGGACCTTCAAAGAATGAGTGGAACCCTTTAG

CATCTCGGGCTTGTTCTTTCCAGGAGGAGAAGGAAACACTGCAGGCTTCTAGGCTGAAGGTATACGGGTGAGATTCCCCA

TATTCAAGACCCTGCGACGTTTTTGGGGTTCAGCGGCAGGAACGGATTGAGAAGGAACCTGAAAATATGGATCAGTAACT

CGGAAGTGAATAAAGGCTAAGGCAACAAGGAGAGAGATACAATCTCGGAAGTAACTATAACAGAGAAATACCTGAGTAAC

AGCAGCAGCTGGGCTTTCCTGATGATCTCGCTTCCTATGAATACGAGGGTTTACAACGGGAGGTTCCACGAATTCTTCCT

CCTGTCGAAATCAAAAGTTAGATTCCTAAAGAAAAATTAAAATACCAAAAGCGAAGTAGAAAGCGGGCTACCTGGTCAGT

AGCAGGAGATCCAACAGTTAGTTTCTTGGTAAAGGAGGATCTATCATTAATCAATCCGGGACCAAGCAATACGTACTACC

AAAAAAAAAGAAAAAAAAAAGATAAGAAACTCAGATAAAAGGAAACGAAATGAAACGAGGGGGAAAGATTAGCGAACTTA

CATCGGAGGGCTTCTCGGACCAGCAGAACTTCCAAGGCTCGAAGGAAGTGAGATGAACGTGACTTGGGAGGGGGCCCTCC

CGAGCATTTCCGGATTCATCATAACCCCAGACAAGGGGGCCCTCAATAACGACTGGGAACATCATCGAGTCCTCATCATT

ACTCAGACGTAAAGAGGGGCCTCTGGGAGCCTCCCGAAGAACTAACGGTCTCGACATCCAAAAGAAGAACTTTGGAAGGA

TCCTTCAGGGACTTCCGATGTAGAGCAACACCCCATTCTTGATACTTCCTTATCTGACCTTGAGTCTTATCGAAGTAATT

CTCCGAGAAAGAAGTAATGTTGTAATCCGAAGGGTTGAAATTCTCTGCCGCAGCCGGGATCTGGGACAAACCACCAGAGG

AACGCCTCACAAAATCGTTACAAATATGAATTGAATTACCGTTCAGCTGGAAGGCTCCCCTGTTGAGATGAGAAAGAATA

TCACAGAGAAGAGGATTCCGGGCGTCATACAGAGGAAAAATTAAGCCCGTTCAAAACTGACCCAAAGTGACAATCATCCT

GGTAGATGACGACGTCTCAGTCTCAATCCAGCGGCTGTTGAGTTCCATCCGTTTGGTTGCGGTTACCGGAGGCGTGACTG

ATAAATCCACCACCGGCGACAGAGTTAAACCTAGTTTTTTGAAATCAGCCTGGAAATCTTCAAGCTGCTTTGATGTAGTG

GCAAGGGGAAATTTTCTGGGTGCCATTGTTGAAAGAAGAGAAGATGATGTGGAAATATTGAGGAAGAATTAAAGAAAGCG

AAGCGAACGAGTTTACGAGAAGAATGGAGTCGGGGAAGAAGGAACAGGAGACTGTAAAAAGATGAAAAGGTAAAGAAGAG

AGACTATTTATAAAGAGCCAAGGCATGACCGGGAGAGGCGGTTACACGAGGAGAGGAACCGGCTAGTGAATGTTGAAAAG

AATCGTGGGAGACAAGATGGCGGAAAATCCGGGTACGTGGCGGTTACGAATTTAATGCGAGGTTCTCTTTACTTTCTTGG

TACCCGCAAAAGGAATGAGAAAAGGAAAAATGTATGTGCAGAAAACCACTCTGCCACGTGTCCGAGGAAGATTGAGCGAG

ATAATAAGGAGAATCTTCGAGGAAGATATTATACACCACTCCCGAAGGAGTGTAAGTGAGTAGTCTAATCAACTGTCAGA

TCTGCCTTAACGAGGCTCCGTCATCAAAGAAGAACTAACTTGCTCGGAGAGACATCTATCCCCCTTGGTTTATGTTTTCT

CCCGTACCGGCTTCATCTCGCACATCTCGAACAGTCCTTTCTTCATTAAGTTGGTTATTTTGCTGCACCAAGGTCGCGTT

CTGCCTCATGAGTATGGTTCTTTGCTCCATTAGAGCTCTCTGTCTATCTCTTTCTAGACGCAGCACCTCCCTGATCTTCT

CTAATGTCATGTTTTCTTCACCGGCGTCCTCCTCGCCGTTTCCTTGAACAGGTTCTTCTTCCTCCGTTTCGGTGTTATCG

GTATGAATCGATCCCTCTCGGTAATGTTCATCCTCTTCCTGTTGGATTTCGCCTTGTCTCCCTGCTCTATTTGACGTTTC

ACCCCTCTCCATCCTTCCTCCTTCTCTTCCTTCTAGACGCAAGCTTCTCCTTGACGCGATGAGGTGTCGTGCTCTTTCCG

GTACTCTTCCCATGGATAACCTGCTAAACTTTCTTTCTCCGGTTTACTGGCTTTTGACCAGATCTAACTTGCTCACCTTC

TGGGTAGTAGATCTCAGATCTACTTTCTTTCGATGCTACTATGTTCACTGCTCTCCAATCTTTGTGAAATAATAACTTTC

TTAACGATTTATATAACAACGGATCTCTCAAACTATCTCAAGTGAGACTAAAATCTCAACTTGAGATCCCTGTTTCTTGG

CGCCAAAATGTAACTACGGAAAATCCATAGTTACACCCACTTACTAATTTGATAAACACGAAGTTAAACAAGAATGAACT

ATTAACAGATCTTTTGTTATTGAAATCTTATCGTACCTAAGTTACAAACTACACAAGAACCCACTCAAGGATTAAATCGT

TCTCCGCACAAAATATTCTCTCTCTCCTAATTTTCTCCCACCCCTTTCAAAGATGGTTAAGTCCTCTTTTTATAGGGGAA

AGTTACATAGTGGGATACAACCTGATTTATCTCGGAAATACACTTTTTACTTCGCTAAGGACAACTTTATGCTTCCTTTT

CGCCTTAAAGATCGGCTCTGGTCGGGGACAACTGTCTAACTTCTTGCACAACAATCCAAACTTCGCACAATATACTTCGG

GAATTTTTTTATTTCTTGGACAACAAGTCGAACTCCTCATAATTTACTTCGGGGATCTTTCTATTTCTTGGACAACAAGT

CGAACTCCACGTAATTTACTTCGGGGATCTTTCTATTTGTTGGACAACAATTCGAACTTCATACTCCGTATTTACTTCGC

TTAGACGTCTTTCCGAGCAAGTTAATACTATCTTCTGGTAGATCTGACACTTGATTTGACTATTCGCTTACACCTCTCTG

GGAGTGGTAAACCATATATTTCTCGAATATTCTCCTTATTGTCTCGCTCAATCTACTTCGGACACGTGGCAGAGTATTTT

TCTGTACCTACACCAAGTGAGATAGAATTTCTGAATATTATGCTTCTTATGACAACATAACACTGTATCCAAATAATCCA

TAAATTGACGGGTTTTCAAACGATTTTCTTTAGCAGTACGATTGACATCCCGACACCCACGAAGATCAAGAGTGGGTATA

CAGTATGATACACATACTATGGGAAAAATATACATCTACAACTGGAGATTTACAACACTTTCCTAAACATTGTAGAAAGT

CCTATGTTTTACAATTGATTTCAAAGGAAGCGTTGTTGTATATCATCACTACCAACACTAGGGAAACAAGGGGTTGGGCG

AAGAAAACAAACGACAACTCCTTAAGCAAAACTGTTGCGACAACATTTAGCTATAAAAACTTTGTTACTTAAACATAGTG

ACTTATCCTATCACAATTCTCATAAGTGTTGTTGAAGAAGACAAAAATATGATAATGAAAAATACGTTAGAGGAGAGATT

CGAACATGAACCTAATGCATGGATGTCTAACTCATCAACCATCCTTACGGTTCACAAGATTTGGTGTTTTTTTTTTTTAA

CAGAAATGTATAACAACACTTATAAGTGCTGTTGAAGTTAAAATATAGAATGTTGGGAAAAAAAGGTTGGGCGGATTCGA

ACATGAACCTAATGCATGGATGTCTAGATCATCAACCATCCTTATGGTTCAAAAAAATTGGTGATTTTTTTTTCCTTTTA

TTAGAAACGTATTACAACACTTATAAGTGTTGTTGAAGTTAAAATATAGAATGTATTACAACACTTATAGGTGTTGTTGA

AGGTAAAATATAGAATTTATTACAACACTTATAAGTGTTGTTAAAGTTAAAATATAGAATGTTGGGAAAAAAGAGGTTGG

GGGATTCGATCCTTAGCCTCTTGCATGGAAATCTACACCACTAACCATTCCTACACTATCACAAGCGCTATCAAGTTTGT

GGTTCTTTAATATACTAACTTTATGACAACCCTTTATAAGAGTTGTAAAACAAAATATAATGTTAAAAAAAAAAGGCTCA

TTTGACTGAGCCGCAAACATCGTGAATCCTTCTCAGAAAATTACTTATTTCTGATAAGGGTAGTCCTGTCATTTTTCACA

ATCTCCATAATATTTATTGCACTTATTTCCGTTAAGGGTAATATGGTCATTTTCACATTTTATAATATTTACTTTACTTA

TTTCCGATAAGGGTAGTCTGGTCATTTTTCACAGTCTCCATAATGTTTATTGCACCTATTTCCATTAAGGGTAATATGAT

CATTTTCACATTCTGTAATATTTACTTTACCTTTTTCCGATAAGGGTAGTCTGGTCATTTCACAGTCTCCATAATATTTA

TTGCATCAAATATGATATATTTCGTCTTCAGCTCATCCGAACGAGAAACTCTGGTCTAACTATAGTAATATTTGAACCTG

TGATGTTCAATTCTGATAAATTTTGTTTTATGTTCATCCGAACGAGAAACTAAAGTCTAACTCTAGTAATATTTGAAATT

GTGATGTTCAATGATAAATTTCTTATTCAGTTCATCCGAACGAGAAACTCAGGTCTAACTCAAGTAATATTTGAACCTGT

GATGTTCAATTCTAATATATTTCGTCTTCAGTTCATCAGAACGAGAAACTCTAGGCTAACTCTAGTAATCTTTTAAATGG

ATGGATTCATCATTCTTACCAAGCTTTGGGGAATATTCCGAAGAAAGCTTACATGTAAATGGATGGATTCATCGTTCTTA

CCAAGCTTTCCAACTTATAGTAGTCCAATCCTGGACAAGTTTCAACCTCGGAGCCAGGGTATGCATACAATATGCCAGAA

TAGGCATTGATTAAGGTTTCAGGAAATATTCCGAAGGAATCTTACATGTAAATGGATGGATTCATCGTTCTTACCAAGTT

TCCGACTTACAATAGTCCACTCATGGCCAGGTTTCAACCTCGAAGCCATGGTATGCATACAATATGCCAGAATAGGGCTT

GTTTAAGGTTTCAGGAAATATTCTGAAGGAATCTTACGTATAAATGGATGAATTCATCGTTCTTACCAACTTTCGGCATA

TAATGGTCCACTCAGGGCCAAGTTTTAATCTCAGAGTCAGGTTATGCATACATTATACAAAAATAGGTTTTGTTTAAAGT

TTCGGGGAATATTTGGAAGGAATCTTACATGTAAATGGATGGATTCATCATTCTTACCAAGTTTCCGACTTCTAATAGTT

CATTCACAGTCAGGATTCGATCTCGGAGCCAGGGTACGCATATAATATGTCAGAATATGGTTTGTTTAAGGTTTCAGAGA

AAATTCCGAAGGAATCTTACTTGTAAAAGGATGGATTCATCGTTCTTACCAAGTTTCCGACTTATAGTAGTCCACTCCCG

GCCAGGATCAGATCTCAAAGCCAGGTTATGCAAACAATATTCCAAAACAGGGGTTGTTTAAAGTTTCGGGAAATATTCCA

GAGGAATCTTACCTGTAAATGGATGTATTCATCGTTCTTACCAAGTTTTCGGCTTATAGTAGTCCAGTCACGACCATGTT

TCAACCTTGGTGCAAGGGTATGCATACAATATGCTAGAATAGGGTTTGTTTAAGGTTTCGAAGAATATTCCGAAGGAATC

TTACATGTAAATGGATGGATTCATCGTTCTTACCAAGTTTCTGACTTCTAGTAGTCCACTTACTGCCATGTTTCGATTTC

GGAGCCCGGGTATGCATATAATATACCAGAATATGGTTTTTTTAAGGTTTCAGAGAATATTCCGAAAGAATCTTACTTGT

AAAAGGATGGATTCATCGTTCTTACCAAGTTTCCGACTTATAGTAGTCCACTCCCGGCCAGGATTCAATCTCAGAGCCAG

GTTATGCATATAATATGCCAAAACAGGGTTTGTTTAAAGTTTTGGGGAATATTCCAGAGGAATCTTACCTGTAAATGGAT

GGATTCATCGTTCTTACCAAGTTTTCGGCTTATAGTAGTCCAGTCACGGCCATGTTTCGACCTTGGTGCCAGGGTATGCA

TACAATATGCTAGAATAGGGTTTGTTTAAGGTTTCGAAGAATATTCTGAAAGAATCTTACATGTAAATGGATGGATTTAT

CGTTCTTACCCAATTTCTGACTTCTAGTAGTCCACTTACTGCCATGTTTCGATTTCGGAGCCCGGGTATGCATATAATAT

ACCAGAATATGGTTTGTTTAAGGTTTCAGAGAATATTCCGAAGGAATATTACTTGTAAAATGATGGATTCATCGTTCTTA

CCAAGTTTCCGACTTATAGTAGTCCACTCCCGTCCAGGATTCGATCTCAGAGCCAGGTTATGCATGATATGCCGGAACGG

GGTTTGTTTAAAGTTTTGGGAATATTCCGGAGGAATCTTACCTGTAAATGGATGGATTCATCGTTCTTACCAAGTTTTCG

GCTTATAGTAGTCCAGTCACGGCCATGTTTCGACCTTGGTGCCAGGGTATGCATACAATATGCTAGAATAGGGTTTGTTT

AAGGTTTCGAAGAATATTCCGAAGGAATCTTACATGTAAATGGATGGATTCATCGTTCTTACCAAGTTTCTGACTTCTAG

TAGTCCACTTACTGCCATGTTTCGATTTCGGAGCCCGGGTATGCATATCATATACCAGAATATGGTTTGTTTAAGGTTTC

AGGGAATATTTCGAAGGAATCTTACTTGTAAAAGGATGGATACATTGTTCTAACTAAGTTTCTGACTTATAGTTGTCCAC

TCCTCGCCATGTTTCAACCTCGGAGCCAGTTTCTTTCTCGAAGTTGACAAATTCGTGTGTGTTTAAGGTTTTGAAGAATA

TTCCAATGGAAACTTACTTGTAAAAGGATATATTCATCGTTCTTACGATTGAAGTTAGAACCCTCCCAAAGCTTCATGAT

TGATAGTTTTTTGTCGGTATACCATATTTGAAGTTCAAATCCCCTCTAAGCTTCATATTTATCGATTTCGATGATGTATC

ATGTTACAATTGAAGTCAAAACCCTCCTAGAGCTTCTTGGTTCATATTTTGTATGCCTTTATACCACATTTGAAGTTCTG

ACTTATACCGAATATGAAGTTCGAATTGATGTATATTGCTAAGTTTGAAGTCAGAACCCACCAAAAGCATCATGATTAAA

AGTTTGTTTGTCGTTATACCGAATATGAAGTTCGAATCGACACTGAGCTTTATATTTATTGACTTCAATGATGTGTCATG

CTAAGTTTGAAGTCTGAACCCTCCCAGAACTTATTGGTTCATAGTTTGTATGTCGTTATACCATATTTGAAGTTCGAATC

AACACCTAAGTTTCATATATATCGATTTCAATGATGTATCATGCTAAGTTTGAACCCGTAGCTTCTTGGTTCATAGTTTG

TATGCCATTATACCATATTTGAAGTTCAAATCGACACTGAGATTCATATCTATCGATTTCGATGATTATCATGCTAAGTT

TGAAGTCTGAACCCTCCTGGAGCTTCTTGGTTCATAGTTTTATGCTTCTTTAGTCTACAAGTGTGAACTAAAGCTACTTC

ATGACCTGTATAACTAGTCGAAATTACTTCGTAACCTATGTGACTAGTCAAAATTCAAACCGGAAGCACAAAGAGAAAGC

ACAAAGAAACACACCAATTATCGAATTCATTTCTTATTTTTAAGCTTCATTCTTATATTGATCTCTAGCTCCCTGTTGGA

TTTGACTGATTTTTTGATATTGATCTCTATCTCCCTGTTGCTGGGGTCTTCCTCTTAGTTGTGCAGACTGTCGAATTGAG

TGGAAAGTTTCAGGACAGCCAGGGATAAGATCTCCAGACATACCCCTACCTGCAATGCTGAAGGTGTTACTAATGCTACT

AGTGGAAATTTCCCCAACTAAGTTTGATTTTTCTTTTAATTAAGTATAATTTTGGTACTTTTGGTTCTTCTTCAGATTAT

TGGAGAGATTACTTTGAGCTTAAAGATCTTGAAGTAAGTAATAGAAAACAATTGTTGATTTTGATCGGCTTTGAGAACAA

AATTGATTTTCTCTGTGCTTTTCCCTTTCCCTAATTTTGATTATGATTTTGTATTTGCAGAATGAGTTTACTAAGGAGGA

ATTAGAGATGTTAGTAATACTGTCTGGAGGAGTGAAATTTTGGTTGAGGTTTTATTACAGGGTATGAAAGATGTTATGCA

AGGTATGAATAACAATGGACTTCACCTTTGATGTCTGCTACATATTCAGCAAAGTTTAGGGATTTTCCAAATAATTTTTT

GCAACTAATTTTAATCTTGTTTCTTTGGTCATCATTAATCCCTGAGCATACTTTTATTTTATCAACATATTTTGGAATTG

AAGTAGCATACCAGTTGATATCTAAGTTCAAGCATGTATGAGAGTTAAAAATCTTTTATCAATACTGCACGGACAATGTT

TTTGAGGTTAATTTATGGTGCGTATTGGTGAAGATTTAAACAAGGATTTTTTCACGGAAAGTGTTTGCATTAGTCTCCTC

TATACCTCTGAAAAAGGGCCTGGAGTTTAGACTTGTCGATTGAAAGTTATAGTATCAAACATACAGTCAGCCGCAACTGA

AATTTCTGAATGGCAGCACTTGTTGGGGTTTTGCTGGATGATTTTGGGGCAATTGGTCTCTGATTTAGTCCCCATAGTGT

ACTTGAAAGTTTTTAAGGGTTGTTAGACCTTTCCAAAATGATATCACATGCCAGATTTGGGTAGGTCCTTTAGAAAATAT

ATCTTGCCAAATATAGCATGTCAGAGCCGAAAACTGACCTTATAATGTTTGTGTAAATGCTTTAAAGTGAATATTGAACT

TTTTTTCGTGATATTTTATTATGAGTACTTGTACTGTGGTATGAATTATATTAGTTCAACTGTGATGTGGTGTGACTTTA

GTGACACTTGATCTATAAAATGTGACTTCTGTCCTGTTTGGTAAAGGCGTATCTTGTTTGTCTAATTCATTATTTTTGTT

CTTTATAGGTGCGTGAATGTGCTGTTGACTAACCCTATATGGGTGGTTGTTACCCATATGAAGGTAAAATAATCTCCACT

TGTGCTCTTAGGCTATTTGATAATATTAAACATAGTCTTCGTCTATCCGTGTTGAAAGAAAATGCTTGAAAATAATCTTC

ACTTTTGCTCTTAGGCTATAAATGATACCTGAGAGCAGAAATGTAATGTGTACCTTCTACAAGAATGGGTTTTTGCTAAG

CTCTTGTTGCAGGTGAACATGAGAACAAAGAGAATTGCAGATGGGTTTGTAGTGGAGCCTTGGTTGTCTGGTTGACAAGA

TGATCTGTTTCCGTGAGCTGATGAGGATCTGAATTGAAGAAGAAGATGATTTGGTTAGAGTGGATTTTGGTTAAGTGGTT

TATACTTACTGCTGGAGAAAGTGAGGTGCTGTTGGTGAGATGAATTGGGTGGTTTGCTGGGAAGTTAGTGAAGGATACTA

GAAGATACTGTGGTGTTAAGTATTGCAGCAAGAGTGTTCAAATTGAAGTCCTGGTAATGAAGAGATTGAGGCCAAGAAAT

CAGTTGTGGTGGTTGCTGTTGAGAGTGTGAGATTGAAGGGTTTGATATGAGAATTAGATTGGAGCTGCAGACTAAGTGCC

GTGTTGACGAGCTATATTTCTTAAGGAAGATATTTGTAGGGAGGCCAATATCTGCTGCGTTCTGATTTATATGATATGAT

ATTTTGAGTCATGTTCAAAGAATTTATCATTTTATTTTCTTTCGTTGCGCAACACACTAGGTGATTTGAAGTTAACTAGG

AAATGCAATTGGAGTGGTAGGTTAACTTGAATGAATGTAAATGTGTTACGCTGACAGATTAATGTGAATGTGAAATACTT

TCGGCAATACGGTTTCAGCTATAATTTTTTTTTTCAAAATTATTTTATGCACACAACTTCTAATAAGAATTGTATTCCAT

ATTTCAACTACACAAACCAATGGTTGTGAAAGAATGTGTCCAAAAAGTTTCTGACAATGAATTAGTGTGCTGTCCAACAT

AGTTCTACAACACTAACAAGTGTCAAGTTAATCCACAACTTGTAAGCAAACAATTTTTGCGGTTTCATTCATATTAGTAA

TGCAAAAGCATGTCTTTAGTGTTGATATTTCCACAATACACATCCTAAGAATTGTCTAAGATAGTCTCAGTTCAATTCTT

ATACAAATACACATCTAAGGCGACTTGTTTCCAAATAATTTTCAAGTTGTGTAATGCCATCCTGGCTCCGGCTGCCCAAG

AGAATCACAACTTTGGAAAAATATTGTCATCTCTTGAACCAGAATCACAGTTACAACACTTTCCAACCATTGTAGAAAAG

CTTGGACTATTGAAGTACCCAAAAATTACTACATTACCCCTTCCATTGATGACCAAAACCCATTATTCTTAACATGTTTT

TCAAAAAAAAACCAACTGAAATAGGCTTGTCGTAGGGATGTACTACAACTCTCAATAGCTAGAACTTTGGATGTAAGTTT

GTAAACACTAGTGAGCAAACAAATAGGCCACAGTCTTTTATTGTTTCATCAATCTCTTTTAGGAACCAAGGCAGCTGAAG

GTGTTATGTCTTGTATCTATTCTTCCGCTAGACAGAACTAGTTAAGTCATGTTTCAAGAACTTCGGCAACACTGGTCCTG

GTGCCTTGTCAAGCTGAATAAACTTCATCTTCGAGTAGTTAATTTGATCCAAAATTAATGTCCTCTGACCTATAAAAGGT

AAGTTCCTTGTGAAAGAATGTCTTTTCTACATTAAAGACCCCTTTCAAATTTTGAAAGTTCCCCATAGATTGCCTTGGCC

TCTTAGATTGTGTTGTAAACGTTGTTTTTTCCGTGGTGACACTTCCATCTTTTGGATAAAATGCGGGTAGACAGAGCATA

TTTGGCTGAAAGGAAAGAAAGAATGTGATGAAGTAGTCCTTCAGGTAGATTACTTATCTTATCCTCTCCCTCAACCACAT

CGTTGTTGTCTCTTTTGTGTGAATTTTCAGGGTTCTTCTGTTTTTTAACACCATGACTCCTTCCTGCACAGAGTAAATGG

AGTGATTATAGAAGAATCTGCGCTCCAATATATACTTTCACTAAATGAAACCTTGTTATCAGTAGAGGATGACAATTTAT

CCAATATAATTGTAACCGCACTAATATGCAGCAAAATTTTGATCAATTTTTTTTTTTTTTTTGAAGGAAAATTCAAAATT

TTGTACTTATTGTTAAGAATATCAATGAAGGCACGCAGCATATGGAGATGGAAATTACAATCATTTTTTGTTACTACCAG

AAAAAATACATGGACAAATCCCTTTTTATTTGATAACCAAAAACAAATATCAATCAACAGATAAACACACAATAATGCAA

AATAAACTCTTAACAAAAGAAATCCGAGACCCAAAATTAAATTAAACATCAAGAAACATATATACTCAGGCAATCGAACA

TACTCACAAATTAAAACCTAGGAAATGGTGACTCCCAGATACAAAATCATTCCAGATCATGAACTAGAGAAGAACAGGAA

TTACCTTGATCACAAAAATGGATAAGATTTAAGATTTCCCTACACCCTAGTAAAGCTTTTGGGAATGGAGAATGAACAGT

AAAGATACCTCGTGTCGTATAAAGATTGTTCCGAGTGGCTGACACATCGTTTGACACATTTTGTTGTGGGACCCGTAATA

TCACCATGTTGATGGTCTCCATTAATTTAGCACAAATCAATGGACTAATTGAACTAAACGGTTTTAATTTTCAAAAACGG

AAAAAAAATAGTTAAAAGGTGAGGAATTCAAGGCCAAAGAATTTCAGCCTTGAAAGAAATGCATATAATATGGGCATAAG

AAGCCTAATATTTTGATTTTCGTGGAAAGAAATGCATATTAAGTAGTAGAATTGTCATTTTATTCCTCACTGTTGATAGT

ATACAACCAACATTTTATTTGATCACCGTTATATTTGTGATCCATCAGCACTCAGCTAGACAATGCAGATAATAACAGTA

AACAATTAAACTTGCATAGTTTGTATTCACTAATCTTGGCGCTCCAAAACACAGCCCAACCGAGTCTAAGTTCTTCGAAG

ATGAGCCTCTTCATCAGCGTCAAAGTTTCCCGAATGTTCAGGGCATTGCGGAGCTCTTCTCCGCTCTTACCCATAACTAG

ACCGCTAATCGTCTGGCACATCAAGCTATGCAGACTTGCAATGTTCAAATAATCAGCTGCCAAGAGAGAGCAATGCTTTA

TTGGACTTTGCATATTCAGCAGGCCAGTTCTGCTGTTGAACACATCTGTCACGGTACTCGATAACCTTTTCGAGCACTCG

ATAATTAGGCAGTGGGATGCCATCACCAGACCCCATGCCTTCAATCGCGTGCTTAATTGTATGGGATATGCCGATGGCAT

TCTGACTTACAAAAAACTTGGCTCCTTCAGAGCTTTTCAACAAAACCATCTTATCAATGGATGATGATGACATTGTCATA

TACACAGAGTAGTTATGGCTACGCATATATATATGCTCTTGATATTTAACTATATTTTTTATTTTCCAAAAATTTCTCTA

TTAAATGTGATGTTCTTTAGTATTTTATTTCAGAAATACGCTCCAAATTAAAGAAGATGTGATTCCATGTCACATATTAA

TTTTGGGCATACTTTTGTAATGAAATAGAATACTTTTTCATGTCATACATCAAAGAACATCACATGGGCTAGTAAAGATT

TTGTATACCCAAAGAGTGGTGAAATCTTCAAGTAAATTATTGTATAACAGATTGCCATCGACATATCCCATACCATTAAA

CATGTAATTAGAATCTTCAAGAAATCTTTTCTTAAATTCCACTCGACGTATCAAATCTGCAAAAATCTAAGATGATATAA

CAATAATGTAAAATAAGATCAGTAATTAATCAAAATGAATAGCGTAATAAGTTATATTTTCAGGAAATTATACTATGAGA

TTTTTTTCTTAATGGCTAAAGATATTTGAGTTAGAGATGGAGAAGGAGAGGTTAATACATACCGGAAAATACAATCTAAT

TAGAATTAAATTTCATTCAATACTCTCCATTAATTTAGCACTAATCAACGGCTAATTAAATGAAATGGTTTTAATTTTCA

GAAAACGGCAAAAATAATTTAAGAGAGGGGGAATTGAAGGCTGAAGAGTTTCAAACTTTCAGCCTGGATGAGGACACTTC

CAAAATTTCAAAGAGAGGGAGAAATTCATGGATGCTTGATCTGCATGATGAAACTGATCATACTGCTAAGATTCCTCATG

CAATTACCCACCAAGTATTTATACAGTAGATGGGTCGCCCTAAGTGTTCGTGTATTACGAGTCTTTTCTCTAGTAATATG

GAAAATGCCAGTATTGACGTCCATATCACTTTGATAATCCAAAACATTTGTAGGGTAAAAGCAAGCGGTATTCTTTTGAC

TTAATAACAACCAAAAGTCTAAAGGTCATAAGTTAATTTGGGAACCAAATTTTGAAACGATTTTTGAAGGAAAAAAAATA

TAATGTCCGTTTTGTTTCAGAAAATTCCTTTGTTTCTGGCTTTTGATTTCGCCTTGTTCAATTGGCTAGTGCAAGGGGCT

TGATTTTCATTCATAAAGTTAGGGGTCCGGACTCCGGATTATGGTGTTGCCACTGTCACAGAAGGAACCAAAACTGAAAG

TTCTAGGAGTGCTAAAACAGTAGGACGGGTTACATATTAAATTTCTAGGTGGCTAATTAATTCCTGTTCCAGAAAATGCC

TTTGGAAAATAGTTTTTTTATATATAAATTTTAGAGTTTTGGGGATAGCTAGAGCACCCAGAAACCACCACCAGTTTAAG

TCAGCGGTTGTTGCTGTGTAAGTATCCGTTCAGAAAATAAGTAAATTAGTATCCATGTGATGATACATATGGTTCTACTG

AGGCGCTGGTTCAACTTTTCTTTATATTTGTACATGTACATTTTCCGACTAGTAACAGATAAAACTTTGAAGAATAAAAT

ATGAACATTAGCAAAAGAAACTGGAAATAACAAAACTTTCAAATCAATAGAACTCCTTGTAGACAATGGTGGTAATTGGT

AAATCAACGACTAGCCATCCGATCTCCTGTCTCGAAAACAAAGCCCAATCTGCTGGTTAGTGTTCTTTGCAAATGAGCCT

TGTCTTCTGAATCCAAATTAATTATTTCCCTTAAATTCATGGCAATACGAATATCTTCTCAGCTGTTACCTGCAAACGCA

CAATAAATCTTGCACGCTCAAAATATACATTCAAATAATCTGCTATACGAAAAAAGATCTAAGAGTAAAATGGTTCATTG

ACCTTGGCATATTCGGAGGAAATTCTGTTGCTCATGATATTTCTGACAGTATTGAGCACAAGAATCCTCTATTAAGGTAT

CAAAACCTCAATTGGAACTCCATCTACTAGGAAGCTCTTTCCGTCAGAGCTAGCTGTTCAATAGTGAAATTTTAAGAAAC

TAGATCGGAATTTTCGGATAACCGTAGATGGAGTTACAATTGCGGTATCCAATACCATGAAGATAGAACTCATTAGTTTA

GCACTAATCAATCACTAATGAAACGTTTTGAAAAAAGGAAAAAAAAAAAAAAAAAAAAGAGGGATTGAGAAATTCATGGG

CAGGGTTTTTAAAAAACAAGACCGACGGACAGGTGAAGAAGATCAGAAACACTTTTAACAAGTGCCAGAATGACTTTCCA

GCAGAAGCAGAATGCTAAACAAAGAAAACTCCATTCTTTATAGGCTACGTACACAATAAATACAGTAATAAAAATCTTGA

AAATAAAAATTTAAGATTGTGTAGATTCTTATTAAAACTTATTGATACGATTCCAACTCTACAAACGGCAACAGAATCCT

GTTTTCCTACAGAAACTACTACTAATATTGCCTAAAAATTTCGAAAAATATATTCGATATTTTCATATTGCATATTTCCA

TGCATGGGTATGTTAGATTTGATCTTATGAGAATATGAGATAGTTCCTTGGCACCAAATCTTCATACAAAAAGAATAGGG

AACAAAATAAAGATCATTCGGAAAAGGCATGACATAACTGACATGGGATTTTGTAGCCTAGAAAATTCTGAGATCGAGTT

TGGATAGTGATAGTATCACTTTTTCATATATAAGCTAAAAATATACTCCCTCTGTTTCAAAAGGCCTGTCCACTATTCCA

TTGTTTCAAACTTATGTCGAGTTTCGATTTATAGTCATTTTTTTTTTCCAATGAACTCTCTAATATGTCCTTAAGTTTTT

ATATTTATAAAATCTTTTATAATGGACTCAATCTAAAATATTAATGAAATTTGTGTAATTAAGGAAACAAATATATCCTT

CATTCCTTTTTATATTTCCTATCTACAATCCCATAACAATTTTTTGGTAATTTTTATTGAAATAAATTAGGCAAGTAATT

AAGGGTAGGCATGTATACTACTATCTTTATTATCTTGACAAAATCAACGCGGACTGTCTTTTCGAAACGGAGGAAGTATA

TACTCCCTATATTTTTGAAAAAGGACACATTACTTTCTAGGCTAAAATAAAAAATCTTATAAGGATAAAGCAAAAAAATT

GATAATTTTTTTTTTTGATGCTAAGGCATATGCCCGTCCTGCCCCCTAGCAGCGTAGCCACCGGCTTGCCCCCACTGAGC

AAACTCGGCCCTTCTAGAACCCATGTTAAAGTTAGTTTTTAAATTACACATGAGTGGAGACTCGAACCGCTGACCTCCTA

CTTGAGAGTCGGGCTCCTGGCCAACTAAGCTAACCCCTGATGGTTAAAAAATTGATAATATTGCTTTTCCACAATATTGG

GGCTTAAAAAGGCTTGGCTTTTGGGGGGCTAGATAGGGTCATGAAATAGTGAAAGTGAACACCCCTATCCAAATATTTAC

ATAATGACTAATATGACCCCATATGATTAGAGATAATAAAGTTGATTAAAATGACTAATCATGATTATCTAATTAATCAA

CACTACTTCATTTCTTTAAGTTAAGTGTTGAAGATGAAAATCAAAACAAAAAAAATTAGAGAGAAGAGAAGAAGAAGAAG

AGAAAAAAAAATTGGGAAAAAAAAATGAAAATCAGTGTTTCAAGCAAACCTTTTGATGTAGGTGAACCCTCATCTTCAAA

ATACGATAACATCTTAATACCCATCAACAACGATCAGTTCTTCGCTGATTTTTCAGAAAATCATGAAATATTTTCTCATA

CTCAAAATAACCCTTATGAACCTTACTATGATTTCGAAGGACCAACCCAAGAAGAAGAAGAAATCGAAGCAACAAATGCT

CAAGATTACCAGATATACCTCTATTCTAACTCCTATTTCAGTTTTTAAGCTTAGCGTTGCAAGAAGTTTCGATAATTTCT

TTCCGATAACCATAGGTTTCCAGCCGCCAGGTCCAATCATGGCTAGGATCCCATGACCTCCTGGTCGTTATCCTGTCATA

CAGCTCACAATTCATAAACTCCCAGCCATTATCAAAATATTACGGATGGAACGATGAATAATCCTGGCCGTAACTGAAAA

ATTACGGCCCAGTTTTTTCTAGTGCATGGAAGCATTTAATATTGTCGGCGGCGATTTGGTTTCTTGTCCGAAAATATTAA

ATGTTATTTAATGCTATCGGCGCCGGTTGGGTTACCCAGCCGTAACTAGTTTTACGTCTCTTTTTCTAGTACGTAAGGTG

ACTCATCCAGTTACAGGTCAATTTACGGCCCGGAATTCTTGGACGTAATTAATTTACGTCTCGAAATGTCATTTTTCGAC

CCAGCCGTATACGTTTTGACGGCTGGAATATCCCTGGTAAACCCAGCCGTGTCGTTTATTAAGGCTGAATATTTACTTTT

CGACCCATCCGTTTCTGTGTTTTCAAAAAATATTTTTTGACATATGTCATTCTATTATAGATTGTACCGTACATTCCGGT

GGAAGACCAAGAGGTGGTTATGGAAGATCAAGAGGTGGTTATTTAAGAGGAAGAGATGGTTATGGAAGATCTACCCCAAC

CTATGCAAGTTTCCGAGGACACAAGTGCTCACTATGCAAACAACTATGAGTCGAAATATAAGGAAGATGCAAAGAAGTGG

GGTCGATCACAAGGGTTAAAAAAGATGTGCATCATAGTCCAGAATAAACAAGTTATAGACACCACCTTTCAAATGGTTTA

CGAGTGTAGTGGAAAGTATGAGAGCTATTGGAAAAAGGGTAGTGAGTATAAATGAAAAACCACAAGTAAGAAGAGACGTT

ATAATACGAAGAAATGCGGATGCCCTTTTAAGATTCAATTTAAATATAAAAATGATAAGAAAGTGTGGTTTATGGTGAAA

GTCGACTCGGGTTATCATAATCATCCTATTCCGGAGAGTTTGATCAACCATCCTTATTTGGCAAGGCTCAACCCCTTAGA

AAAGGAGTTAGTACACGTGTTGAGTAAAAGATGCACAAGACCTATTGATATTCTTAGTGGCGTGAAGGTGTTAAACCCCC

AAAATTCATCCTCATTGGCAACTATCTATAACGAAAGAGAAAAATTTAGAAAATAGTCATGGCAAGAGAGAGTGCTTATG

TAACAATTATTGTTTTTGTTTGAAGAGGCAAAGTACTCTACCTCCTATGAAATGAATGAAGAAAAGGAAGTGGAATATCT

TTTCATCGCCAATACGGAATGTGTACACTTCGCAAGATGCTTTCATCAAATTCTCTTCATGGATTGCACGTAAAAAACTT

ACAAGTACAACATGCCGATATTGAACTTTGTTGGGAAAACATCTACCAAGACGATATTTACCGTTGTGTTTTGTTTCTTG

AAAGACGAGACGAAGGAAATTTATATGTGGGCATTGCAAAAGGTGAAGTTATTATATCAATAAGGTTATACTCCACAGGT

GTTGATTACGGATAAGGAGCAAGAATTGATGTGGGCCACACCACGTATTTTCCCCTACGCGCATCATCATATTTGCACGT

TTCATCTATGGAACAATGTGCAAACTAATTGCAAGAGGGTTATATGGCCCGCAAAGAAGCCCGAGATGGATAGGATTAAA

AATCTACCGTTGGAGAAACAACAAGAAGAGAAAGATAAGTTTGAGATCAATGAAAAAATAGCCGAGGTGCTTTCGGCGGG

TTTTCAATTTGATTGGGAAGAACTAATATGGTCGCTCACGGAACCATTATATATCCTAAGGGGGCGTTTTGTAATGGAAA

AATGGTCAACCTACCCGGATGTTATAAAATATTTTAAGAACGAGTTATTGGATCCCCACAAAGAAAAGTTTGTCAATGCA

TGGGTAAACCGTTATAAACATTATGACAATCAAGCCACAAGAACGGTGGAGTCGGATCACTATCAATTCAAGGGTAAGTT

TAATGGTTGTGATGGTGGATTCTTTGTTGTTCTATGGTCGAGTATTTCAAGCGTGATCTCGATAGAATTAAAGGGGATTT

TAAAAATATCCGATCTAAAGCGGTTGTGGACTACCGGGATAGCCCTTGGCTCCGGGGAATAAGTTTATATGTTTCTCAAT

GGGCAATCTTAAAAATTATAACGGAAGTGGAGGCTTGGGAGGAACACAATTTTCCACCGGGAATGAGATGTAATTGTCCC

ATTAAGTCGGCTTTAGGGCTCCCATGTAGGCATGTGGTAGCAAATTATCCCACAAGGATGATTCCGATCGAAGATATAGA

TCCATTTTGGAAACAACTAACATTTATGGATCCATTACCAAATCAAGGTCTTGGGACATGTTTTGCGACATGGAGGTACA

CAAGGAACTTTTTGAGAAGTATGCTTCTTTACTACCCCCACAAAGGAAACTTTGGTTGCATGAATTGAGGAAATTCAACC

GTCCATTCACTAGAGAAGATATTTTAGAACCTAACAAGGGGCGGGGCAAGGGTAGGCCTTCGACCAAAATAACAAAGAAA

CATCAAAGGGAAGAAGCTAAGAGAAAGGTTCAAGAGGGTCCACAATTGACTCCTTCATGCGAAGAGATCTTTCGGATTTA

GAGAAGTTGAACGACTTGTACAAACTTAAACGAAAGGAAATGGAAAAAGGAGGACCAAGTCGAGCTAGTGAAATGAAGGT

AAAAGGCCGCAATAAGAATGTGGTTATTGCATCCCAAGAAAGTTCAAAAAGAAAAGTTGATAGTGGAGTCAAAATTAAAG

ACCCGGTTGTTCCATCCTAACAAGGTTAAACAACAAAAGAAGCAAGTAGTGTCCAAAGAAAAGTTTTTGGTGCGGTTGGT

ATTAAGAAATTAGTGAGAAGCCGAGGTAATGATGTCAAAGGAAAAATACCGATGGTCGAACAGATAATGATAATTATTGC

TAGGCAAAAAATAATGCAAGTGATGCACCTAGAGTATGAAGTTGAACTAAGCAGCAAGGAAGACATGATTGGCATGACTT

TACTCGGGTATTGGCATCATATAAGGCGTGAAACATTGGGTTGGACTGTCAAGAGCTTAAGGTTCAACAATAGTGTTGGT

TCTCACTTAATTAATTCGTTTACAACAAGATGTAAATATATATATTTACAAAGGGAAGAAGAGAAATTATTCTACAAAGA

AAATACGGAAGGGAGGAAGAAAATAAGCCATCCTATAGCCACCCTACAACTAATGTTACAACTACTTCCACCTACTTATA

CTAATTCATTAGTCAATCTTGATGACTAATGTTACAACTACTTCCACCTACTTCTACTAATTCATTAGTCAATCTTGATG

ACTAATGTTGCAACTACTTCCACTATTTACAATTCATTCTTCAACACTCTCCCTCAAGGTGGTGCATATATATCACATAT

GCCCAACTTGACAAGTGAATCATTAAAGATTTTATTCTATACTGTCTTGGTTAGAACATCCTCCAATTGTTGTGTTGTAC

GTATATATGGTACTTCAATAATATTTTCTTTGAGTTTCTCATAGATGAAGTTCCTATCAATTTCCACGTGTTACGTTCGA

TCATGTTGCACTGGATTCTCTGCAATTTTGATTGCTGAATGATTATCACAAAACAACTTCATAAGTTTTTTCAAGGAAAC

CCAAGCTCTCCCATAAGCCTTTTCAACCATAAAAATTCACAAATTCCTTTTATCATTGCCCTATATTCAGCTTTAGCACT

TGACAGCGACACAACCTGTTGTTTCTTAGATTTCCAAGTTACTAAGTTACCTCCAACGAAGGTAACATACCCAAATGTTG

ATCTTCTCCTATCACCTTTGTCTCCCCAATCCGAGTCACTGTATCCTGATACATCGAGATGTCCATTTTATGAAAACATC

AACCCTCTTCCTGGTGCAGACTTCAAATATCTCAGTATCCTGATGACTGCATCCATGTGTTGTTGACCCGGATTATGCAT

AAATATACTTACAACACTTACATCATATATAAGCAATATCAGGTCTCGTATGAGAAAGATAAATCAAGAGGACAACTAAC

CTTTGGTATTTTCCTTTGTCAGCTGGTACTTGATCAGAAAACTCTTGAAAACCATGATTTTGTTCCATAGGAGACCCTAT

CGGCTCACAACCAAGCATACCGGTTTCCTTCAATAGATAAATCACATACTTTCGTTGAGACAAGAAAATTACCTTTTTAG

GTCGTGAAACTTCAATCCCCAAGAAGTATTTAACTCCACCAAAGAATTATTTAACTCCATTAACAGTGAAAATAAAAAAT

AGGTCCACATGCTCCTATTTCGCTCCATCAAAGAAGTATTTATTATTACAGGCAAGGGATACATTTACCACACTTGCGCA

AATAGCTCACTTCGCAAATGATTGGTGTTGCATGTACTGTTTAGAATGCAACGAGGAATTTGAGGAAACAATGCACCAGA

AGAAGTGTCCGGAGTATTTTGGATATAGTAACTTAGACCCGATGTGATACTTATGTTTTTTTTTTTCCATAATCTATTTA

TTTAAGAAAAATGAACATTTCAAATACATAACATCAACAAAAATTAGACAGATTGCGTTTGACAATAATGGCTACTGATA

GTCCGGAAATGACAAAAGAAATGAATTGTTGTAAAAGAATTATTGGGATATACAACGGATTACTTGATATGAATGAAAGA

GGTATAACTATTTTGATTCTGAAATCACAGAAAGTTCACAGTAGAAGATGACATCAAGTTTAAAGTTAGTAACAAACATA

ATATGGCTTACTAAAATGTTAAAAACAAACAAGATATATAAACTTAGAAAGCCGTGGGACTTGAAGAGGCCAATTTTGAA

AACTACAATCTTAGAAAGCAACGATCCACAATACCATAATTGTCTTTTATAAATATTCAACGTAGAATATGTCAAATTTT

GGTAAAGGAGTACAATATGTGTTCAAGGTTCTTTATGAGGTAATGTGTCGCCAGATTGTATTCCAAGTGAAGGTAAGAAC

ATATGGAAGTGACAAAATGTCCAAGACTTTTGAAGTTACCAAAGCCTTCTTTAAGACGAATTGCTGGGTACAATACTTCG

TTCAAGATTTAATGTCAGAGGTTCACATATATACAACCCTTCGAAAAGGTAATGGGTACACATTTTAAACATTGTCATCT

AATGGGTGTTGAAAAATTACAGGAAAATCATGTCACATTGGATAGTTTCTATCATTCGTTCGATGCAAAAAAAAAATACT

ATTCTCTGCCTATTTGGAGGATCGACAACATGACCATGATAGATGAGTCAGTTGGAACGAAATAAGAGCATGTGGAGACT

TTTATGTATGATATTGATTAGTTGATTTCACTGAGATGAGCGCCGCTGACGACATATTTCCAAAAAATATTTGAACTTCC

ATGTTCATGTAGAGCATTATCTCATTATAATCAATGAACCTAAATAAAAAGTGTCGGGAAGTTAAGATTATATTTCTTTC

GTTTGAGCAAATCTCCGGCGGTAAAATATTAAAGGAAAAAAAAAAGGAAAAAAGAGAGGAAAATCCCCAACATTTTTATG

ACAAACACATATTAATTCATAAGAAATTGATAAGTATTCGGGACACAAATTATACGATATATTAAAAATTAACCAACATC

ATGGTTCGTACCGTAACGGCACAGGTGCGCGTGGAAATCAAAATCCGTAAAGATATGGTTTTGCTACTTGCTTTTGTAAG

GAAACCTAGAACAACAAAACCCCACCGAACATGGTTACATAATATCTTGTATAATAAGGGTATTAGACAATCATGGCTTA

ATAATTTAGCCATCTAAACAATCAAGAATTCGAGTAAAAACTAATAACAAAATGAAACCAAAAACCATGACAGAAAAATC

CAAGCAAATAAATGCACATGCACTAATTAAAATCATGAATTAGAGTAAATTTCAACAATAATTCCATATTTTACCTTTCA

TCTTGGGAAATAAAAAGAGTGAGAAATTCATGAATATCTGATTGTTGTGATCCGAGATAAAATCGATCCAAAACATTATA

ATTTTAATAGTAAATAATTTCAGATAACGATTATCTCAAAACTGAATATTTTGGCACAAATGACAAACGTGTGCTTTCTA

GAGTGCTTATTGATCCTCGATAACACTATTACATTTTTTGTGTAGAGCCAAATAAGATATAAGATTGTATCACTAAACCA

TACATTATTATATCATACTCCCTCCGTCCAATTTTACTTGTCAAAATTGGATTTTTCAAAAACTTGAAAGATATTCAAAT

TACTTCCACATACACCGTTGATCTTCCAAATTGTTTGTTTAATATACAAATTTTGATATCCTAGTTTTTGTTTCTGTCTT

GATAATTATAAACTTTTAGTGGAAAATATTTCAATTTTTTGGTACAAATTTGCTAAAAAAAATACAATTTTAGCAAGTAA

AGTTGGACGGAGGGAGTATCATTTTTATGAATTGAATCGTACAATAATCATTCATCTTCACTGGAGAGTTTACGTTTGAA

TCACCAATGGTCATCAAATATCGCATAGATGTAAGTTCATATATCGGAGCTTTAATGCCTTTTGTTAACTTCTTACATCG

AAAATCTTAACAACATAACATATTTTCCTGTCCACTTGAAAGATGGCAATGTAAAATCGCAAAACCCTTAAAATTTTGCT

CTGAACCACTTTACATAGACATTCAATGTTGTGCTGAAGAAAATCATTAACTACCAATCAACAATTCGACTAATTGTCAT

TTGAGTTTTAGCATATAACTTTCATAGCCATTTTGTAACTTTACTTTTTGTGCATCGCTAATGCTAGTCAATGTCCTAAC

TACATCTGTGAATAACTTCAAAGAATCAATTTGTCAAAAGTGAGGGGAAAAATTATGCGCTAAACACATCCAACCTATTA

GTTGCAATGCCCACCAATTGGAATCCAGCTGAACCAATTATACAATATTTTTCTTAAAACATTACGTACACTCCAAACAC

AAAATTATCAATATACTTGTCTTGTGGCAAGCTTTTGTGCGACAAATTCTATTAAAAAAAAGAAGAAATTTTTTTATAGG

GGGCATCATAGCAAAAACCTCCATCAAAGATTTTTTATGTTTTACATGGGTAAGGTTTTACAAAAAAAAAAAAAAAAAAA

AAACGAATGCTTAATTTGATTTCTAATCTCAAACACAAATCATTAATAAAAGAAAAATGAAAGGGTAAATGATAAATGTG

TGAAAATTGAACCTTTTTACATATATTTGCGAACATCTCTCTTAATATACACAAACAAATAAATGGGCTATCTATCAAAT

ACTACAAAAACCAATTTAATATACAACATATTAAATAGAAATGCTGAAATCGTTGCAAATTGTATGTGGTAAGACAAAAA

GCTTAAAATTGAACTATGACATTGTATTGAGGTGATCCACATATATTTAAACCTCATATCAAATCAAATATAGGTATAAT

ATCATAACGTGATAAATCTAACAAAACTGTAACTTCGCAATTGACCAATTTATATAGGCCTTCCATCCTATATATACAAA

GAATACTTCATTTTAATACCCAAAATTTTAAAATCCAAATTAAAAACTTAACAAAATTTAAAGAATTGAAAACTTTAAAA

AATATACCTAAATACAGAAGAGGGTAGCATAGAGACAATTCATATGAACCTTGAAGCCTGACAATTTTATGAGAGTTAAG

TTTATTACAACAAAGAAAAACTGATTTATCCAAATAATACATAATCACTTTCCTTTTATTAATGAAAAATGTCAGACACG

AATTAATCTTTTGCAAGCACAAATATGGACCGCAACTTATATAAGCTAACTTAGCCAATACCATAACAACATGAGCTCGA

TAAAAATATGTATCATTCATTTTTTGTTACTACATAAAGTGACAACCCCGATACTTATTAAATTGTCATTTTATTTTATT

TTTGTATTGTAACCAAGGGTATTTTAACTAACACGCATTTAAAATTGTCCACTACACAACACCCACCAACATCATAAGAG

CCGCCCACCTGTACTTCTTTTTGTACCACCACCAGCCACTAACACCTCACAACCACCACTATTATTTTTTAGAAAATTTA

TTATTTATTAATGGAGAGATATTTATTAGGATCTATAAATAATCAGATATATATTTATTATAGGAAATCAATACAAACTA

ATCATAAAAAAATTAATGGGACATTAATGATATAATAATAAGATTAAGACTGTTAGATGTAATTGTAACTTTCAATCTCA

GACATATAAAAATAAATAACCACAACCGTTGATTTAGCTTTTGGGAGAACAAATTAGGACCATTCTTTATCTTGTCTAAC

TTGTTCAAGCTTTGTTTTATAGTGAAAAACCGTTGATTGCGTTGATTTATTTTGGCAACACCAATCTAATTAGTCCTATA

AAATGTTAATCTTAAATTTTCTAAATATATTTTTATTAAAATTACTTATTAATATCATCCCAGAAATATATGGTACCTTC

CTTTATCAAAAGTGTTAAAATCAACCAACCACATCTTCCACACTACGATGTTATCGATGTTGTAACGACAACATAATATT

AATATTGTCCATCACTACTCACCAGCACTCACCGTCACTACTTCTTCCGCCACAAACCAGGGGGAAGCCGGGATTTTGAG

GAAAGAGGGAGAAAAAAATCTTAGCAAGTTGGATACATGTGAAAAATGGACTTCGTATCCCATTTTATGATAAAAAGAAC

AAAAATAAATATAATTGTGCTTGGCGCCGCCCTTAACCACCACTAATCATTCGGCCTCCACCTCTTCTACTACTCATTGT

CGCCAAAAATGTCTATTGATATTATTTATCAAAATTACTTATTAATAGAAATACATATTATAATTAATTAAGAAAACATT

CATAATTAAAAAAATGATTAGTCATTTATTATGAGCAATTAGTGAAACACATATTAATAAAACACTTTATAATGAATAGA

TTATTTATTATGTGCAATTAGTGAAACACTAATTAATAAAGCAATTATGATATTTAAGTTGAACAAACCATCACCGTTGA

TTTATCTTTTCTATTAATTCCAACCAAGGTGGAACACTGGTTAGAGGCCACTCTTATGACACAATAGGTCATGAGTTTGA

ACCTCCCCGTTATAATTTTTGAAAATCCATATCTTGACTTGAGTATTCATTTTTCATTCATAATATTGGAATTAATGCCA

CGCGTGCGAGTTGTATGGGGTGGTCAGGAGAACAACAACCCTTGATTTATCTTTGTATGGATAAATCCTGACCACCACTA

ACACAGGCTAACTTATCCTAGCTTTGTTTTGTACTAATGATGTCAATTATTCTTACATGGAGGTAAATTCCTTAAGACGG

GAACGGAAAATAAAATGACTTATTTCAATATGCAGTTTATTCCTAAATATAATTGGCTCAAACCGGAACCAGAAAATTTT

ATTCTCATATGGAGGTTATTCCTATATAGAGGTTCTACTGTATTCGTGTTCCTGTTTGTTTGTCCATTTACTTGCTTAGT

CTAAGTAATTTGTGAGAATAGTCCTGCATTTTATTTTGACACATTTAATAGAATTAACTTATATTCAATGCAAAATAAGA

TACATATGGACTTCGTCATGGCATGAGAATAGATCACATATAAGAGAGACCCTGATCACATATAAGAGAGACCCAATGCC

ACCGAACACTGACGTCTAGTCTAACTGTCGACCAGTGACAACAATGAAATGTCATAATCAAAACCAGCAAGAAAATATAC

ATGATTTGTTGTATTTTTGTAGTATTTTATAGTGCATGGCCCTTAGAAGGTGTCATTACTGTTAGCCAGTCGTTACATGA

AGGTTAGGTTCATGTCCTTCTCAATGCCCTCACTGCACCTAGTTTAATATTATTCCTATGATTTTATCATGTGGGAAAAG

TATCCTAGTCGACGATTTTGCACAAAAACAAACATGTTTACACAGTTGTTGTGGTAAATATTATTCATATTTAAAACTAG

TCGTGCACAACCTATTATTTAAGGAAGCCTATTATTGGGCTGCAAAAGTCTTGGTCAATTTGGGGCTATTTTGGGTCACC

AAATAGCAATTGGATTACCCCTATCAAATACTTATATAATGACTAAAGTGCCCCATATGTTAATAAACTTGATTAACATT

TGTAATCATTATTAGGAGATGAATTAAGAGTAATTCATGTGTTTAAGCTAAGTGTAGATGAAATGAAAACAACAAAAAAA

AATCAGAGGGAAGAGGAGAAGAAGAAGGAGAGGAAAAAAAAAATTGGGGATTTTATTTTGAAAATGAGTGATTCAACCAA

GACTTTTGATGTAGGTGAGCCTTCATCTTCAAAATATGATAACATATTAATACCCATCAACAACGATCAGTTCTTGGCTG

ATTTTTCAGAAAATCAAGAAATTTTATCTCAAACTCAACCTAACCCTTATGAACCTTACAATGATTTCGAAGGACCAACC

CAAGGAGCAGAAGAAGAAGAAGATATCAATGCTGAAGATTACCAGGTATGCCTCTTTTCTAACTCTAATTTCAGTTTTTA

AGCTCAGCTTTTCGATATTTTCCTTCCGAAAACCCTAGGTTTCAAGACATCAGCAGTAATAACGGCTGGGAAACCAAGAA

CTCCTGGCCGTTGTGATGGTTTACGGCCAGGAAATAATGAACTCCCATTCGTTACCAAGTACTTACGGCCGGGACGATAA

AATTTCCTGGCCGTTAATAAAAAAAATACGGCCAGCTTTTTCTACGTGCATGGGAAGCATTTAATACTGTCGTTGGCGGT

TGGGTTACCCGGTCGTAACTTGTCATACGTCTCGTTTTCCATGCCGTAGGTTAACTGTTGCAGTTACAAGACAATTTACG

TCTTGGAATACCTAGATGTAACAACGTTACGGTTCGGTATCATACTTTTCGACCCAGTCGTTTTATTTCTTACGGTTGGG

ATATAAGTTATCGACCCAGCCGTTGACCCTATGATGGTTGGAATGTCACTGGTTTACCCAGACGTTTTTGAGTGTTTAAA

TTTCTTTTTGAATAATTATGTGACATATGTCATTCTATCATAGATTGTACCATACATCCCGGTGGAAGACCAAGAGGTGC

GTATGGAATATGAAGAGGTGCGTATGGAAGATGAAGAGGTGTTTATTGATCAACCCCAACTTGTGCAAGTTTCCGAGGAC

ACAAGTGCTCACTATGCAAACAACTATGAGTGGGAAGATAAGGAAGATGCAAAGGATTGGGCTCGAGCACAAGGGTTGAA

AAAGATGTGCATCATAGTCCAGAATAAACAAGTTAAATTCGACCGCTTTGAAATGGTTTGCGAGTGCAGTGGAAAGTAAG

AGATCCATTGGAAAAAGGGTAGTGAGTATGTACTAAAAACCACAAGGAAGAAGGGAGTATATAATACGAAGAAGTGCGGA

TGCCCTTTTAAGCTTCAATTTAAATGTAACGCCGATATGGACATGTGGTATATGGATAAAGTCGTCTCGGGTTATCATAC

TCATCATATTCCGGAGAGTTTTATCAACCATCCTTATCCGGCAAGGCTCAACACCTTAGATAAGGAGTTAGTACACGTGT

TGAGTAAAAAACGCACAAGACATATTGATATTCTTAGTACCGTTCAAGTGTTAAACCCCCAAAACTCATCCTCATTGTCA

ACAATCTATAACGAAAGAGAAAAATTTAGAAAAGAGTCATGGAAAGAGAGAGTGCTTATGCAACAATTATAGTTTTTGTT

TGATGAGGAAAAGTACTCTACCGCCTATGGTACGAATGAAGAAAAGGAAGTGGAATATCTTTTCATCGCCAATCCGAAAT

GTGTACAATTGGCAAGATGCTTTCATCAAATTCTGTTTATGGATTTCACGTATAAAACTAACAAGTACAACATGCCGATA

TTGAACTTTGTTGGGAAAACAGCTACCAAGTCGACATTTACCGTTGCGTTTTGTTTCTTGAAACACGAGACGAAGGAAAG

TTATTTGTGGGCATTGCAAAAGGTGAAGTTATTGTATCAAGAAGGTTATACTCCACATGTGTTGATTACGGATAAAGAGC

AAGCATTTATGTGGGCCATACCCCGTGTTTTCCCCTACGCGCGTCATCATCTTTGCACATTTCATATATGGAACAATGTG

CAAACTAATTGCAAGAGGGTTATATGGCCAGCAAAGAAGACCGAGATGGAGAGGATTAAAAATCTACCGTTGGAGAAACA

ACAAGAAGAACAAGAAAAGTTCGAGATGAATGACAAGATAGCCGAGGTGCTTTGGGCGGATTTTCAAAAAGATTGGGAAG

AACTAATATGGTCGCTCACGGAACCACTATATTTACAAAGGGAGAGTTTTGTAATGGAAAAATGGTCAACCTACCCGGAT

GTTATAACATATTTGAAGAACGAGTTATTGGATCCTCACAAGGAAATTTTTTTTAGTGCATGGGTAAACCGTTATAGACA

TTATGACAATCAAGCCACAAGCACGGTGGAGTCGGCTCACTATCGGTTCAAGAGTAAGTTAAATGGTTGGGATGGTGGAT

TCGTTGTTGTTTTGGAAGCTATGGTCGAGTATTTCAAGCGTGATCTCCATAGAATTAGAGGGTTATTTGAAAAGAGCCTA

TCTAGAACGGTTGTCGACTACTTGGAAATCCCTTGGTTCCGGGGATTAAGTTTATATGTTTCTCAATGGGCAATCCTAAA

AATTATAACGGAAGTGGAGGCTTGGGAGGAACACAATTATCCACCGGGAATGAGATGTAATTGTCCCATTAAGTCGGTTA

TGGGGCTCCCATGTAGGCATGCAGTAGCAAATTATCCCACAAGGATGATTCCGATCCAAGATATAGATTCATTTTGGAAA

CAACTAGATTTCAAAGATCCATCACCCGATCAAGGTTTTGGGGAAGTGTTTTGCGACACGGAGGTACACAAGGAACTTTT

TGAAAGGTATGCTTCTTTACTACCGCAGAAAAGATAACTTTGGTTGTATGAAATGCGGAAATTCACCCGTCCATCCACTA

CAGATAATATTTTTGAACCTAACAAGGGGAAAGGAAAGGGTAGGCCTTCTAAGAAGATAACAAAGAAACAGGAAAGGGCG

GAAGGCAGGAAAAAGGTTCAGGAGGGTCCACAATTGACTCCTTCAATGCGAAGAGATAATTAGGGTTTTGATAAATTGAA

CGAGCTCTACACCCTTAAAAGAAAGGAAATGGAAAAAGGCAGACCAAGCCGAGCTAGTGGAACAAAGGTAAAAAGGCGCA

ATAAGAATGTGGTTGTTGCATCCCAAGAGAATTCAAAAAGAAAAGTTGATAGTGGAGTCAAAATTAAATACCCGGTAGTT

CCATCCCAACAAGATTCAACAACAAAAGAAGCTAGTAGTGTCCAAAGAAAATTTGGTGGTGCGGTTGGTATTAAGAAATT

AGCGAGAAGCCGAGGTAACAAAGGAAAAGCACCGATGGTCGAACCACAAAGAGTTGTTAGTAACCTTGAGAAAGAGACGG

TTTACATGGATGTAGGTCCGCTGAAATCACCCCCTATAGATGAGGAAGGTAATTATATCCGAACTACTTACACAATATAC

AAAAATTACATGCACTTTGTACCACATTTTATTCATCACTATGTCAAGTCCACCGACGATGTTGAGGGCGACGGAAATTG

TGGTTACCATGTGTTCGTAGATCAACTTGGACCTTTTGAGGAGGAAAAAGAATGGGGTGATGACCAAATTACTTATGTAA

GGCAAAAGTGTTTGCTTGAACTACGTATTTTCCCCGATTTCTACAAGGAAATGATGAGAATCGAAATAAATGACACGGAG

GTGATGCTAAACGAGGCGTTCAAAGCATTTGAACAGCGTTTAATGGGAAACACTACATTGACAAGTGAGTATTGGATGAG

AATGCCAATATGTGTCCATCTACTCTCCAAAGCATTCCATTGCGTTATTCATTCCATCTCCTATGCACATAGTGTTACTT

ACGCACCAACAAGACGTATTTTTACCGAGGAAGAATCTCCAAAGATAGTCATCATGGGGTTCGTGATGATGACCCATTAT

ATCGGCCTCTTATTGAAAGATGGATGCCCATTACCTCCATTAAGGAGGTTAGGCGGATGTGACGGCGAAATACCAATGGG

TTGGTGCCATAGGTTCGAGGAAAGATTTCATTTGTGGGAATCATTACAGTTGTCGAGGGACGGACCGTTGTGTGTACTAA

TGAGTTCCGATGAGGATGACTATGAGTAAATGTCGTGTCAAGGTATAGGATAACTTTTTTTTTTGTAGGGTGGTGATTAA

AATGATAATATGATAACAATGTGGTTTGAACCTATGTATTTTGAATCACTATATATATATATATATATGAAGATTCTAGT

TTTACTCTTACTATGTGTTTGTAATGTTGATATGATAAAAATATGGTTATTTAGAGGTATTTACGGCCAGGTCGTACGCC

GACACGACCTAGCCGTAATTACTCAACTGGGATATTCTTCTGCAACTCGGCTGAGAAACCCAACCGTGTAACACAAAACG

TTTAGGAATCCAAGCCGATTTTATGGGTATTGTCAGAACTACGGCTGATCATCCTGGCCGTAACTCTTCCTGGAATTTCA

GTTTCATTTTTTACAAAGTTACGGCTGAGAAACCCAACCGTGAAACCGAAAACGTCCAGGAATCCTACCCGTATTTCTGT

GTATTGTCAATACTACGGATTTTACGGCTGAGAAACCAAACCGTGAAATAAACGTCGGCCAGGAATCCTAGCCGTAAAGA

TGGGTATTGTAACTTCTACGGATGTAAATCCTGGCCGTAATTAGTATCATATTAAACTTAAACAACAATCAGATACATTA

ACAAGCAAAACACTAGTATCCATTCATTCAAAGTTAAGTCTGCATAATGAAAAGTGAGTCTGAAAAGTAAATCACCCAAA

TCCATAACCTAAAACATGCTAAAAGTCTGACTTAAACAAGTACATAACCTACAATCATCCACTACGACCAACAATCGAAG

GAACTTCATCAGAAGATGATGAAGAACTGTCGGGAGTTTGGGAAAGACTGATCCAATCATCCTCGTCGTTAGTATAGAGA

TCATCCTTGGATGGATTATGTAACTCCCGAAGCCTCCTAATCAGTGTTCTTTGTTCGTCGCAATCCAAATATAGAACCTC

CTCAATGTTACGGATCAGTCCCCTGGAAATCCACTTAGCTCCCTTGACCTGTTTTCACACCATTCAATTAATAATGGTAC

ATAATTTGAGCAACGTATACAAAAACAAAAGAATATCACACATTGGTACCATCATCGTAGCAATATCACACATTGGTATT

TCCTGGGGTTCTTTCTCCTTTTTAGAGTTCCTCCTAAAAATTATGAAAAACATATCAGAAGATACGAACAATTCACAAGG

AGTGATACAATTACGGTTTGGAAATAACATATAACCCATCCGTAAAACAAGTTTCGGATAGAAATAAATAAAAATGGCTA

GGAAATACAATTTACGGTTAGATACTCCCAGCAGAAACACTCAGCACCAAATTTTACTCTGCAAATACAGGAGAACTTAC

GGCTTGGAAATACCCATCCGTTAACTACAGTAACAGTTGGGTTTTCAGATTTTTCGAACCTATTAAGACATTAACGGCTG

GGAGTTCTTGACGTCCTAACCGTTTTATACCATTACGGCCAGGACGATATGATTTCCCAGCCGTAATTGCTTCAGAAAAC

CTTTTCTGGAAACCCTAAAACTATCAATCGAACCTAGTTTTTCAAACAAAAGGCAAAATAACTAGTGGGTTTTTCGATTC

TTACCTAGTAGGATTCATTTGTGGAGGATTCGAAGGTGTTTGAAAAGATTGGTTCACCACATCTCCATTAAAAGGAACAT

CAATAACTTGCCCTGTTTCAGAATCAGGGTTCAATCGGCCGGATCTTGTTACTCTTGATCTTGATGCTATCTAGAGCTTC

TTCATGATGTTAACATGATGTTATATACAGACCCGTGACAAATTAACATCATGTTATTGTGCATGTTAGACTATAAATAG

TCTTGAGAAGCAGTGTAAAAGAAACGAAAATCAATAAAACTTTCTCTTATATCTACGACTATATACTAATCAATTCCCAT

GTTCTTCATGTATCTCTTATGTCCATCATTTATTTTACGATACTGATCAAACAATGGTTGTGAACCGATAGTTACGATTG

TTGACACCATTAATACACACACAAAAAAAATGATCAACAAAAGCGAGAATGAAAAATTAAAAGAGTTCAGCGAGAGTAGA

AATCTTAAAGATGTTCAAGCGAACTGGGTAGAGGTGTAAAACGTGCCGGCCCGGCACAGCCCGGCACACGAAGTCGCGTG

CTTCACGTGCCATGTGCCGTGCTGTGCTGTGCCAACGATTTAAACGTGTTGTGTCGTGCTGTGCTTTACTAGTCAAAAGC

TAGGCACAACACAGCCCACGGGCACATCACACGAATAAACGTGATGTGTCGTGCTGTGCCGTGCTGGCACACGTGTTATA

AACAATTTGAAATCACTTAATAGTATACATTAAGACGGACATTATTACTAGAATCTAAATAGACAACATATCATTTGAAA

AAAAATTCAAGTAAAATCAACAAGTTTATTATTCAATAAAAAGAAATAACCCATCAAATATAAAATTGGATCATTGTCAT

GTAAATTAATGTTCTAGTCAAATATAGGTAATGACATTATAAAAACGATATTGGAATATATTTTCCAAATATGTAGGTAT

TATATGTGTTGTTATAAAAATAAGTCAGCATAAAATGTCAAAAACTAGCTAGAATAGTGACTCTTTTGTGAAATATACAC

AAAATGTGATGTATTATGCCTTGTTATATTGTGTACTTGTGCATGCTATGACGTGTTTTTTTAACGTGCTGTGTTGTGTT

GTGCCACGTGCTAATCCGTGCCGCGTCTTTGTCTTTGGCTACTTTGTGCCAATTAGTCAAACCCAAAGATATCCCATTTA

ACTAACGTGTTGTGCCGTCTTTGAGCTAAGTCACGTCTTTGAAATTTGCTCGTCTTTGGCACATCCTAGTTTACACCTCT

AGAACCGGGTGCATGCGCTTCTCCCGATTCCCTATACAAACTTGTATAGGGTATTTTGGCCAGTTATTATTAAACATTTT

TAATATCTTGGGAAAAATACCCTAAATTGATTATTGAAAAAGTGTCTTTTTAATGTGTGGATATTTGAACAATATCAAAT

GTTACACATCTTTTACACCCAGAAAAAAAGTTTTAATGGTTATTTAACCAATTTTGTGTAAATAAAAAATATAGAAGAAA

CAGTGAAATAAAAGAAATTCTCATTTTATTTGAATAAGAGGATATCAATATTGTTTCCGTTGTTGTAGGAAACTTATATT

CTTGTTTCTGTTGTTGTTCATAAAGCCTTGTCAATCCAAATAAGACAAAATCATTTTTTACCACTAAATTGGTACACCCC

GGTACCTCTATTAGCAGTGTAATGGGTGGTTCATCAGTTTTTTCTAAAATCCTTATGAATTTCAGAATTTGTAATAGCTC

ATTCGAGAATGCAGATTGACCATTCGTATATAACATCCGATAGTTAAAAGAACATTCACTGGCCTTCCAAATTAACATTG

CATTTTAACACCATAAAAGTTTTACCAGCTCAACAACCAAAATTTACTAAATAAGTGAATGAAGTTATCTTTCACCACCA

TCATTATATATAGTGGGATAAACATCCACCAATTAAGAATCTCATCCTAATACTCTAATACAACTCATAAACATATAGAC

TTCCAAATTCTCTTATCTTCAAGAAACAAAGAGATTAGTACAATGGGGTTCTTCTGCAAATACTTGATCTTATCATTGAT

GATTAGCTTCGTCGTGCTTAACAGCAATGTCGCTGACGCTGCTGAGGGAGCCCCTGTGAAGTATATTGATCTCCCAGAAT

TGACTTCCTGCAAAGGTCCCGATTGCCCCAAGAAACCTCCTGTCCCCGCAAACGAGTATAGCCGTGGTTGCCACAAGCAT

CAAAGATGTCGCACTAATTAAAATGGTTAATCGACAGTTTATCCACACGATGATTTTTGAAATATTGTATTCATGCTAAT

ATAAAATAAATATCAAGTATGATTGTATTCTGTCTAGCAACCTATACGTGATGCTATTTCTTAATTTTCAATGTAACAAT

TTTTTGAAAAAAATAAAGATATACGGTAGAACCTCTATATAAAATTATCCTAGGGACTACATAAAAATATTCTTATGCAG

AGGTTATTCTTATGGAGGTCTACCTTGAATGCATCGATTTTTTATTTTTACATATGTACCTAGGAAAATTACATATATAT

GCATATGAAACATTGTAATATACCTAGGTTATAGAAATATTAATTAAATTTCGTTTACTTCTACAAATAAGGATAAATGA

AGCTTGCGATAAATCACAATCCACACAAAATTCAACCACGAATTGGAAAATTTTATTGAATTGTGCAAAACAAAATATCA

AACCAACTTGGAAGTAAAATGATCTTGCAAAATGGTTACAAGAATCTGTTCATCTTAAATTGTGATGTATGACAAGACCT

ATAATATTTATATTATAATGTCAATTATTCTTATATGGAGGTAAATTCCTTAAGACGGGACCAGAAAATAAAATGACTTA

TTTCAATATGGAGTTTATTCCTAAATATAACTGGCTCAAGCCGGGACCAGAAAATTTTATTCTGATACGGAGGTTATTCC

TATATTTTATTTTGACACATTTAATAGAATTAACTCCAGACTAAATAATAAACCCCCCACGGTGAAAGACTTTAGGCCGC

TGAGTTTAATAAGTAGCTTCTACAAAATACTGAGTAAGCTGTTAGCGGAAAGAATGAAGGTGGTAATGCCAGATTTGATT

TCAAATTCTCAAGGAGCGTTTATCAAGGATAGACAAATTCTAGACGGCATCTTGGTGGCGAACGAGTTAATTGACAGCAG

GTTAAGACAACAAATTCCAGGGATTATGTGCAAGATTGATATGCAAAAAGCCTTCGACAATGTGAGTTGTCCATCACTTT

TCTCTATTTTGGCAAAGAATGGTTTTGGCATGAGATGGATACAATGGATGAGATGGTGTATAACAGAAGCGCAGTTCTCG

ATCCTTGTTAATGGAGAAGCAACAAAGTTAATAAAACCATCGAAAGGAATTCGACAAGGTGATCCTTTGTCGCCATTTTT

ATTTCTTCTAGTGGTAGAAGTTTTGGCGTTGCTCATCAACAACTCGGTGGCTCAAGGAAGACTCAGTGGATTCAAGGTGA

AAGAAGGGGGCACTATTGTGTCACATATGCAATTTGCAGACGACACAATTATCTTCTTAGATGCAACATCTATAGAGGTT

AGAAGACTTCTGCTGATTCTAGTCATGTTTGAGATACTAACCGGCCTTCGACTGAATTTAGAGAAAAGCACGATGATAAG

CATAGGAGCAGACCATCTAGTCAATGATTTAGCAACGGAGTTGGGATGCAGAGTTGAATTCATTCCTGTAACATACTTGG

GAATGCCAATTGGCTCAAATAGAAGGAGTGAAATCATATGGGAGATAATCATTCACAAAATGCAAAAAAAACTGGCACCG

TGGAAAAGGAGGTACTTGAATAAAGCTGGTAGAGTGGTTTTGATTAGAAGTTCTCTCGAGAGTTTGGCGGTTTATTTTAT

GTCTTTACACCACATCCCTGTCTCAGTAGAAAAAAGATTGAATACGATAATGAGAAAAAAATTATGGGGTGAGGATGATG

AAAGGAGAAAGATGAGTTGGGTGTCTTTTAAACGGATATGCAAACCAAAGAAGAAAGGAGGATTGGGTATACGAAATTTG

CGGTTGGTAAATAAAGCCTTGTTATCCAAATGGCACTTCAGGTACTGTAAGGAGAAGACATCTCTTTGGAGGAGAATTGT

GTGTGAAAAATCAAAGGCAGAGGAGGATTGTTTGCTTCCTTTGCAGGTTAAGGAACCCATTTGGAGAAGCATGTGGTTCG

ACATCTTGAGACAGAATAAGGTCTTTATGGAATCTGTGAATGTGCAGGTTAAAGACGGATCAAGAAGTAGATTCTGGCAC

GACGGATGGTGCGAAAAGAACTCCTTCAGGTGGAAATTTCCAAGGCTTTACAGGATTAGTACACAAAAGATGGCGAGGGT

TAATTCAATGTTTCAGACAACTTGGAACTTCCAATTCAATAGAGACTTGAATCCGGCGGAGAGGATTGATCTTTTGGAAA

TGAAGCATGATTTAAGACTTGTCATACTAACTCAGGGAGAAGAAGATTGTTTGGAAGGAGAGATTTCGGCCAAAGAAATT

TATAAAAACCTGTCTGACACGGATCAAGAATGGGAGGGTGAGCGAATTTTATCAAGTAAATATGCACCTCCTAAAGTTTT

ATTCCTATTTTAGGCTTCTATGCTTGAGTCCATCCCAACTAGGCATATGTTACAGCATAGGGGAGTAGACATCCAGTCCA

ATCTGTTCCTCTTTTGCAACACAGAGGTCGAAACTTTAGAGAACTTATTTATGCACTGCAAGGAGGTTTTAAAACTATGG

GATTACTTCTTATCTTCGCTGAGTGTTAATTGGGTGATGCCACTAAAGTTCAAGGACATGATGCTAAGTTGGAGGTTGGA

AAGAGCATCAGTCAAGGTGAAGATGGCCTGGAACCTACTGCCATTTGCTATTTCCTGGGAACTATGGAATGAGCGCAACA

AGAGGGTGCACGGAGGAAGACCGAAGTCAGGATATGAGTTGGAAATTTCGGTAAAGCAATGGATTCTTCTTTGGAGTTCA

TCATCGAAAGTTTTTAGCCATTTATCATCAACTCAACTTTTCTTTCAATGGGAGACGATTTTGAATCTATAGTTGCAGTT

TCTTCTTTCTTTTGAAGGTCCCTTGAACTTGTAAATACCATGTACTTTTTGCTCTTTAATATATCCTTCTCTTTTCAAAA

AATAATAATAATAATAATAATAATAAACCCCCACTTGAAATAATTTACATTATCATCTATGCATGATTAGTTTCATTTTC

ATAGCAAATTCCAAACTTGGTGTTATACGCGATTTTACTGCGAAGGTAGGAAGATCAATTCCTAAGTTTTTTTTTTTTAA

GCATAATTTATTAAGCAAAATTAGTTTACAATTTGTCGTGGGTATGTGGTACCCACTGAGTCTGAAAATAAGATTTGAGA

AATGAAAGATGGGATTGCCTGGTCCCATATTTTTGTTGACGAGTTTCCTGTTCGACTTCCATCCGCTGCGTGATTGGCTA

ATCCGTCTGCTGCTTGGTTTCCTTCTCTGTAGTTGTGTTGGATGGCCGCATGCGGGATCCTATGGAATCTCATTTTTATT

TCGTCAATCATTTCTGCTATGTGCCAGGGTGGGGGGTAGACGAGGTGATGAAGCGCAAAATGTTTTCTGAGTCTGTTTCA

AATATAACTTTTGGCCAAAGACTTTATGTTGCTGTTCTTATGGCTAAGAGCATCGCCCAGGTTTCCGAAATTAAAGCAGT

CGTAATTACCAGTGGTTGAGCAAGCGCTAAAAGTGTTCGGGCATTTGAATCCCTGCAGATGAAACATGCTATTGCAAAAC

CTGGATGTCCTCTGGCTGTTCCATCCGTGTTAATTTTGATCCAGTTGAAATTTGGCATTGCCCATCTTACTGATATTGTG

GATATCCTTTTATCTCTTGTTGGGTGATTAAATTTTGGTGTATCTCCCGGAAACACTGGAGGTGTGGAGTGAATGGCTCG

AAAGTATTCTTGTTGCAGTTTTTGTGCCCGGGCTAGGATTTCTTCCGGAGTTGATTGCTTTTGTTAGTAAATTAGTTCAT

TCCTAGCTAGCCATAAGGTCCAAAAAAGAAAACATCAAGAGGAGATTATCGATGTCGAAGCATGTTGTTGAAGGATTTGA

GAAATGGTTGTATGTTGCGTTAAGGTGAAAGCTTGGAGAGGATTGGAGTTTGATGAATTTGTAGATTGATTAAATAAGGT

GGTCCATGAATTAGAAGATGTTGGACAGTGAATTAGAATGTGACTTGCTAGCTCTGGATTTGTGTTGCATCTGGGTCATA

TGTCCGAGTTGGTCAAATGAATGTGGTATAGGAGAGATAAGGTTGGTAATCCTTCGTTGATGGCTTTCCACAAGAAAAGC

CTAATTTTTGGTGGACATGGTACGGTCCATATAAACTTGGTAGGAGGTAATGGGATGTGTGTTGGTTGACCTTGGGTTAG

ACATTTGTATCCCGTCGAAGTAGTGTAGCTTCCAGATTTTGAGTGTGGCCAGATAGTTTTATCTTGTGGCCCATTTTGGG

GTAGATGAATGTTTATGATTTTTTGGATTATAGGGTTGGGGAGGTTATTTAATTTTTCTTGATCCCAAGTGTGGGTTATT

GGGTCAATGATATCTGCTACCAATTGGAATGGATAGCACATTGTTGGGTCTAGTTTTGGTGAGTGCGGGATCCAGTTGGC

TTCTATTGGAGTTGATAATCCATTTCCAATCTGGTGGAAGATTAGACTTTGCATGGTTGGGACAAGTGTTGCTAGTTGTC

TCCATTGAGGACTGGCAGAGGAAGGTATATGATCAATGCCTTCTAGTATTGGTTGTTGGTTAAGATATTTTGCACTGAAT

ATACTTCCACAGATGTAGTTAGGTTGTTCATGAAGATTCCATATTCTTTTCATGAGTAAAACTTTGTTATGATCGCTGCA

CTTCCTTATTCCCAGTCCTCCCTCAGATATTGTTTTTGTCACTCTATCGTATCCTATAGGGTGTAGTTTTTTGGTAGTTG

AGTCGTGTCCCCAGAAAAAATCTCTTGTGATACGATCTATTTGTTTGTGAATATCGCTCGGTAGGTTTTGTGTTTGCATG

ATGTGATTTTTTGTGGGTATTAAAGATGTTTTGATTAGTGTTAGTCTTCCGGCTGGAGTTAGGCATTTTGTCATCCATCC

TTTGGCTTTTCTGGCTAATCTTTGTAAGAGTGGTGCAAATGTTTGATTGGAGGCCCTTCCATGCTTGAATTGGACTCCTA

GATAGATCGTAGGTTTCGATGTGGTTGACATGTTGAAGATGATTTATCTTATTTAGGTCTAGTTTTGGGTGATGGATGAT

AACTGATTTGCTGGTATTTATTAATTGACCGGCAGAAGTGCTATAGGAGTGAAGTAGGTTTTTTAGGTGTTGATTACTCT

GATTTGAAGCTTTGTAAGTAATCAGTAAATCGTCCGCATACATTAAGTAATAATTGGTGGTGCTTTATTTGATATGCTAA

TGCCTTGAACTAAGTTTTGGTTTTGAAGGTTTCCTAGATTTGTAGATAAGATTTCAGCGCATATGATGAAAATATAGGAG

GAAAGTGGGTCTCCTTGTCTTATACCTCTATTTGGGCTTATGTATCCATGGGGAGTACCATTTATTTTTATTGAGTACGT

AACTGTTGTGACACATAACATAATGTAGTTTATAAATGCTTGTGGAAATCCCAGTTTTTTGAATGATATTTTGATGAACT

CCCAATCTAAGCTATCATAGGCTTTCTGGATGTCTACTTTGAGAGCTATTTTTGGATCTTTAGTTCAGTTTGATGTTTTG

ATATGATGGAAGAGTTCCCAGGCTATTGCAATATTGTCATGTATAGATCTTTGGGTACGAAAGCACTTTGAAAATGTCTT

ATTATAAAGGGTAAAATGGGTCTAAGCCGGTTGACTAAGATTTTGGATATGATTTTATATGTGACGTTACAGAGTCCTAT

GGGTCTGAATTGTGAGGGTTTTGAAGGATAATCGATTTTAGGTATTAGAGTGATGTTTGTGTGATTCATGAGAGGATGCA

TATTTAGATCATTAAAAAACCTTCTAACCATTTCTATCAGTTGGGTGTGTGTTGTTTCCCAGAAAACTTTAATCAATTCC

TAAGTTAAAATCTATGTAGTAATACAATGAGCATGAATTTTTAACGGCTGAAAACTGCTTGTTGATTGTGTTACTCAAGT

GCCTGATACCATCACTCTTCTACCAAAACTTTGACTTTGACTTTCATTTTTTTTTTTTTTTTTTAAATAAGACATTATAC

TGGGCCTCCGTTAATTTACGGATTCGGTCCCTTCAGAAACAAACTCACGATGAGAAGCTCTTCCAATGACTAAATTTCTA

TGCTCTTCCAATGACTAAATTTCTATGCTGGTTTTGGATTGTTCTCTGTTGATATATATGTATATATATACTGTCTTTTC

CAACAAAAATTAGAAAATAAACTATATATATATATATGAGGGAGGATCCCCTCACACTCTTAATTTTACATTATCACACA

TTTTGGACGTACTTTTTGCGGATAAAAACCAAACGATATCGTATTTGAAGCTAATATTTTGGGGATATATTCTTCGTATA

AAGATCTACGTACCCACCAAAAATGAGCACATTCCGAAACGTATAACTTTACCATCTAATACTTTGAAACTCAACCGTCG

AAAGTCGACAAATTCTCAACCGTTTAAATTAAGATTGATAGATGATGAGGTTTGGTATTTCGGAATGTGCTCATTTTTGG

TGGGTACGTAGATCTTTATACGAAGAATATATCCCCAAAATCTTAGCTTCAAATACGTTATCGTTTGGTTTTTATCCGCA

AAAAGGATTCCAAAATGTTTGATAGTGTAAAAAAAGGGGCGTAAAGGATCCTTCCCCTATATATATATAAGAAACTGTTT

TTATTTTTTATTTTTATTTTAAATATCAATTTATTCAATAATGACCAGTACATTTTTATTTGCATAAAATCAGTAATGCT

ATGCGATCAATCCCCAACATTTGCAACAAAAATTAATAACTTCAGCATCAGTAGATCAGGGACCAGTCATCCAATTTCCA

CTCTCAAAAACACAGCCCAATCTGGTCAGTGTTCTTTGCAAATGAGCCTGAGTCCAGATTTCCCTTAATGTTCAAGGAAG

CCAAAATCTCTTCTCGGGTCTTACCAGAAACCAAATTGCTGATCGTGAGGCACATCAACTCCTATTTTTGTTGTTCATAA

TACTTTTCACAGTATTGAATGACTTTTTCCAAAACCACTCTGCGGACATTTGGCACTGGGATGCCATCATCTGTGCACGT

ATCTTCTAGTACATTCCTTATGGTATCGAAAACTGCAATTATAACTTCCTCTACTAGAAAGCTCTTTCCATCTGAGATTG

TCAGTAACACCATTTTTTCATTAGCTGATGACATAGTTCGGATTAAGATACTCCTATGTGCATAGTAGCACTAATGGATA

TATTAATGACAATTCAACCAAAACTTTTTGTTAATAAATACATATTTTCTAATTTATGTAGAAAGGATTTGATTTGATTT

TAAAATCTCTATTCTATTAATGAACCTTCTCCGAAATATAAAGACAAGAACGAAATAAAGAGTACCAAATCTTCCGCATA

AATTACAAAACACGCATATATAATATAAAAAGCCTTTGAAGGTCAGATTACACGCATATATTAACAAAAAAATCTGGGGA

AAAAAATCGTTGAATTTAAAGAGTTATATTGGAATGGGTAGAAGAAGTTATAATTGAGGTATCCAATACTATGAAGATTT

TTATTCTGTTCACCAAATCTGTATACAACAAACAGGAGAATCCCTATGTAATTGAAGTATTGAACTACAGTAAACAAGTA

TGATAAGAAAAAGAATTGGCAAGAATGGAATTTAATAGGCTATATTTGTAGGAAAGATATATGAGATTTTTCTTAATGGC

TATATTTGTAGCAGAAGATATCTTCTCCCTTAAAGGCCAGACAAATAAAGCTCTCACACTTAACGGTGATGAAACCCTCG

CTTCCTATTGGCCTAGAAAGTTCTTTTCTAATGCAGAAGCTATGGTGGGGACATTTGGCATAACTTTATTGGTTGTAATA

ATTAGAATTTGGATACATTAATATACTTAACTGTCCGTTAGTAACGGTGTTAAAAAAAAGTGGGTAAACAATACTTATGT

TAGAATTTAAGATTGAATTATTTAGTTACACCGTGTTTACACATATACTTAATTGTACTTTAATAAAGGTGGACAAAACT

AACTTCCAAACCGTTAAATAAATTAAGAAAAAAAAAAGATTAGAAAAAAGCGTGAAACTCTTAAAAAGGAATTCCTATTA

AAAAGAAAATCCTTACCCACTAAAAAAGGAAATCGGTTGAAATCAGGAGATCGTGCTAAAAAAGGAATTATTCTTTTTCT

TTAATAAGAGTACTTAACGTCTGTAATATAAACAAACTGTAAGTACATAGATTAGGAACAACAATAGCGTACTTGTGCCT

TAGCATAAAGAATTTGACATAATAAAAGGTGCTCACGTATGTTTGAAGGAATATACAATTAGTTAAATTAGTTGATACAT

GTAGTAAAATGGTAATATGATCTTAGCCCATAATTAATTGATTTGTGGGCATAATAATATGATCATTAGCCCATAATTAA

TTGATTTGTGACCATTAGCCCATAATAAAATGGTAATATGATCTTGGTCTGACCATTAGCCCATAATTAATTGATTTGTG

GGCATAATAATATGATCTTAGCTACTTCATGCGAACAAGCTATGTTTTCATGGGATAGTAAACATGATGATCAAACATAT

GTACATATACGACACTATCCTTGGCTTTACTTTGCACATGCATATCATGGGATAGTAAACATGACGAGAAAACATATGTA

CATATGCGACACTATACTTGGCTTTACTTTGCACATGCACATCACAACGTGATTAAGTGATTCGTAGTTGGTCCGTTTTT

GATAGTTAAAAATAATATACTCCCTCCGTCCTAGTTTACTTGCCAAAATTGGGGATTTTTTTTGTCCTAAATTACTTGCT

AAGTTTCATATTTTTTCACTTTTTATCCTAATTTACCCCTCTGTTGGAATCATGTCATATTATTGAGAATTGTGTAATTC

TATTTCCTATACACAACTTTAAATTCTCATCTCTCTCTATATCCTTTATTTTTTATTTTTTTTGAGTTCTTCATGTAAAG

ACCAGAACACTAGACATGTAAAAACTATGTATCTTAAATGCTAGCTTGGGGTAATTTTGGAAAAAAATCCACTCTCTACA

TTTCTTAATCTGTGTGCAAAAACCAATTTTGGCAAGTAAACTAGGACAGAGAGAGTATATAGATAAATACGATGCCTCCC

CACACTAAATTTTTGTATATAGATTAAGGCGTTCTTTTATATACCCCTAGAAATACTAATGATACCCCTTTATTTATGTA

CCCCCAAAGTTTATTTACCTACCCCATCAGTTTTACTAAATTCATATACCCCTATAGATATAAATTTCTTCTTCTTCGTA

ACAATTTTAATCGCACTTCCAGATATAACCTTTTTATATCATTATGGGAACATACGAATACAATTGATAAATTACAATCC

CATTTTTCAAACTTTGGACTATTGTCTGTTTTTTTCGTTTCATAAAAAAAATAATTTGTTGATGAATTAACCAGCGGAAA

TTCTTTTTTTCCTGCTGCCATCTTATCAAAGAAAGAATGATAAAATTATGATCCTCGGCTTAGTCGGCTAAAGGATTACG

TGGGAAAGTTATCATATCTTTCGTTGTGATACAAGGTAGTCTTATGCCGAAACTACCAATCGACCGCAACTACCAATCGA

AAATTCTTCTTCTTCCTCCGCCATCAAAACGGTGTATTTGAAGCTCCATTTCTCTGACTGTAAAATATACTAATAAATAG

ATGAAGAAGGTGGTGGTGGGGAAGAAGATTTTAACAAGGCATCTAGACCATGAGATATACATGATAGATTGACTTTTTTC

TGCCATTGTTGAATTTGATAGATGAAGCGAGGTTTGGTGACTTGTGCTCCCCTTTAATAACTAGTGTGTAGTTGTGTACT

ACACAGAACGTTTTCAAGCTAACTCATGTACTGGAAAAAACGTATTCAGTTGGCCTATGAATCAACAAAGGCTAGCTCAT

GCTAGAACATGGAATTTGCATGTGATGCATGAGCACAAAAACCAATACTGTTAACATGTGAAATTTCATTTCTGAAATTT

TAATGGAAACATGGGTATAAATTATTGATTTCCCCGGATTTTAGCATAATTTGATATCTTTTGTTATCAGTATGAATAGT

CGAAGAAGATAACAAAAATGAAGAAGATAAAATTTATGATTTGGAGTTTACTGAGTAAGAAAGAAATGATTTTTTTTTTC

TGTTTTATTTTAGGAAGGGGTATCACCATCAAAATATCGAAATAATTTGATAACTTTTATATTAACCGAATTTTAACTTC

CATAACTAACTAAAGGGGGTACCTTTAGTATTTTCTAGGGGTATATAAAGGAACACCCATAGATTATTGTCTTTGTCCTC

CAGCCGGTTCATTCTTTGGCCGAAATAGGTTTTTTTTGAGTTTTGTGAAACGAACGTTTTGGTGAGGACACTTAGCAATG

TATGATTGGTTTAAAAAGAATATGAAAAAGATTAAAAAAGGAAACCAAATTCAATTTGGAAGTTCATGAAGTGACAATAT

TTATGTGAGGACACTTGGCAATGTATAATTGGTTCTACGAAAATAAAAAGAATAAATAAATAAAATCCGCACATATCCTC

CACCTATATAAAATCCAAAGATACATCTCCATGGGGAGGGGCAAAGTCCCGCGCACACCAAATCAAAAATGCATTGTCAC

GTGGTGGAGTGAAGCCCCGTGCACACTAAATTAAAAGGAATAACCAAATAAAATCTACACATATTTCCCACCTATTTAAT

TTAATTTCCCCGCCACACCAAATCAAAAATACACCTCCACGGGGCGGGGCGAAGCCCCGCGCTCGCCAAATCAAATACAT

CGTCACGGGGAGAGAAGAAACCCCGCACACACCAAGTTAAAAGAAAAAATATGCCCGGTGCGTAGCACGGGCACAAATCT

AGTGCAATTATATAGTAGAAAAATTGGTTTAATTAGCATTAAAACTGATTTAAGAGACTCCGTCAATATAGCACTAAGCA

ATCACAGAATCACTAATGAAACGTATGGTTATTAGCTTGCATGCATGATGAAACTGCATAAATTCCTCACAAAATTGCAT

GGGAACAAATGGTATTCATGGTACACAGTGGATGCGTTTGCCTACTTACTTTCCTAGCCTTGATGGCCAACCGAAGGAAA

CAATAGTCGCTCCTAGGCCATGAGTAATGGAGATGGCGAAACGTTACAAACTAGTTATGAATGCCATTCCACGGGAGCTG

TTGGCCGGGAAAGGATTTTAAAAAAGGTGATATTGTAGGAAGGTACTATTCCGGCTCAAAAAATTCAAAAAACGGCAATT

TAGACTTTTGGTAACTATATTATACCTTTTTCACCCTCCACAATAGATTTTTAGCTCGATTTTGGTCAAACGAGGACTAG

ATTGTGTAATATTGGCCAGTTTATTGAGAAACTTGAGTCCTTTGACTGATCCAAGTACGCCAGTGTTCTCTTACTGGACA

ATCGAAAGCGTATTGTTGACTGTTCACTGAGTACTGTAGACCAGTGAACTTGAGTCCAGTTGACCGGTCAATATTTTGAA

CGCACTATTAGGGCCCTCCAAAAATTGGTCTTACTATATTCATTATGAAGTCATGTAGGTCATCATTTCAGGGACATAGT

CGGTACTTGTATGGTCATTTTTTCACTTTTTAGTCATACTTACCTATTTCGGAATTTTTGTGAACGCTGGGGTGTTGCAC

GGAATTTCTTTGCCGCAAAAATCCGCTAGGCTAAGATGTGATCGGGTCATATACTTTAAGTATGTGGTCCCTCTTCTTTT

GCTTACACGTATTAAAAAATTGGTTTTTAATTAAAACCATATGCTACTTTTTTTCCATAAAAATATTAAAACTACCTTTG

TCCTAAAAACCACCCCTCTAAATCCCCTGAAAAACAAAAAAACTGAAACTTTCCAAGACGTGGCAAAAAAAGAAGAGGGG

CCACATACTTCTACGATGTGGCTCGACCACATCGTAGCTGCTCCGGCAAAAATCTCCATTCTGGGGTAGTAAAAATTTGA

CATCGGGGTTCGACTGTCTTTTGCAGTGGTAAAATCTAACCCCTTTCATCCATAGTACGGGCAAAATGTAGAGAGATCAA

TTTCCACTAGAGGAGTAATTGGTTTCCCTATATTTTAACTAACCTCCTCTATAGATAAGGCCGAAGTTGGACTTATTGTT

GAGAATGAAAACTTAGATCGTAGTGTGTCCAACTGGCAGAGAGATTAACTCAAACTTACAGCGTCTTTTTGAAGTTTATC

TAAAGAATCAATTCAAAATAAAGTGTGGCTCTGCAGATCGTACGTAAAACACTGAATTAAAAATATCACAACGACAAAAT

TAAGGGACTAGCTAAATTATTATCAGCTGGTTGATGTTGGTGGTGAGCCTGCAAAATACGCGACGGCACCAACGAATGGT

GCATTAAAGTAAGTAGCTGGTTCAGATTGCTCGTGGTTATTTCGATCATCACTGAAGCTGTCGCCTTGATCAGGACCACC

CATAATGGCTCCAACAAGTACGTTCGGGTTCGGTGATCCGGAATTGAGATGTTGAAATCCATCTTTACAAGCTATCTGAC

TAGGTTGTTGGCGAATAGATGGTATAGAAGAACCTCTATGATGAACTTTTTGTGGATATTTCTCTCCGAATCCTACCATG

TATGACATTTTCGCAGGATTATCACCTAAAATATAATCTACTTGTTTCTTTGCTAATGCAACGATGTCTTTCGCTGCGAA

ACCCTTTGATCCACACGACACGGATTCTCCTGATGCTTGTAAGTACTTTGCATATGTCACGAGGAGGAAAGCTGTGCTAG

TCACGTATTGCATATTACATTCACCTTGTTTGAAGAGAAGTCCTCCTGTCATTACGAATATTCAAATTGGTATGAGAGCT

GATAAATGAAATCTAACAACGGAATTCGGAAAATGTTCACATGCGTCAGACCTGGCGTATATTTGGCACTAAATGTAGGT

GATCCAGAGAGGAGAGAGCAGAAGTAGTTGTCAGCATGGTTTTTATAGGACTGTAAATCTTGAACTTTCTTTTGTAGGTA

GAACTGCAACGTAGAAACTTGACGTTATTAAGGGAAATGAAGATTTTGTGTACTCATAAAAACTTGTGACTATAATATGC

AAACCTTGGACAGAAGAATTTTTGTGCCAACTCGTTTATCGTCCCAACTGAAAGAGAAGTCGTCATCGTCTCCACCATCA

GTATGCCCGTTGGTTTTAATGTAATTCAGGTACTTATCGTCTTTTGTCGCTCTATATAGCCAGGCTGCTGCCCATAGAAG

TTCATCCTGACATGCATGGATATTATGAGAGATTGCTGTGTCTTATGTCTTCGTAGGAGAATGATGGAGATTTCTTGATG

GCTTACTTACATTGTATCCAGAATAAGAACAGTAAAATGGGCAAACAACTGAACTGAGAGAATCACTATATGATCCTCTA

TGCTGGTCCGCGAACTCAAATACCTGTATCATAACAAATGTATCTATCAACACCCTTAAAAAATTGCGGTTCATGATGGA

TATTTGATTGGTATGTGTGGCAAATTGATTTACTTTAATCGCAGTTTGAAGTAACTTGGTAGAGTATGGAGGATCGGATC

TTTTAAACACAACCGAAGCAGCAGCTAATGCGGCAGCAGTCTCTCCCGCGACATCAGAACCTGGATTTTGGGTAGACACC

TTGTACACATTCCGGGAGGAATCCAAATCTTCAGGCCTTTGCCAGCATTCGTGATCTGCGTTAGGATCTCCCACCTTCAT

AAACAACAACAACCAAGTTAGCAATCTACGACTAGAAAACCTTTTTTTTTTTTTTGAGATCTTAGTAATGTGATTCACTC

ACTTGGACATATAATGTATCAGGCTTAGCTGTAGCTGCCTTTAATAAATAATCAGTGCTCCAACGAATGGCGTCTCTGGC

ATGGGCAGCTTGGTGTTTCATATTTCGACCAAATTCAGCGACACTCCAAGACAACATTGTCGTTGTAAACGCCAAAGGAA

GACCAAATTTGACATTATCTCCTGCATCATAATATCCTCCAGAGAGATCCACCTGCATTTGTTGCAAAGCACAACACTCT

ATATTTACTACTTGAAAGACTAGTTAATTAAACTTGAAACTTGATTATATATATATATATATATATATATATAAGAAAAC

ACTAACATTTTCAGTAGACCCGTCCTTTAATCCAGAGTCGCCTCGCCAGGTAACTCGTTGATTAGGTGGTAGTTTTCCCG

ATCGTTGCCCTTCGAAGAACATAATGGATTTCTCGAGTGCTTCAGCATAATTGTTAGAATCAAAAGCACAGCAATAACAT

GAAAAAAGAACTACGACCAGAATGGATATAGAAATAAGCTGCATCGTGGGAATTCTAGCCAAACTGAAAATTTGTAAACA

CGGTTAATCGTTCTGATTGTAATTTATCGATTGCTGATGAATGAAAAAAGAAACAGGTTGATTGTTAGATGAGAGAGAGA

GTCAGATCCCAAGTTTTCTTGGCCATCCATTTTGCACTAAATAAACTCTCAACCAAGTTCTGAACATTAACTGAGGAGGT

GATAGCACCAAATTTAGGATATTTAGAAGTAGTTTAATACTAATTTGAGGTAGATAAACATAAAATTGGGATAACATACT

AGAATTCCTTCTTTTAATCGCTGAAGTCCTAAAACTGAGCTAAAGTTTGGTTTGAGTGCATGGGCAACCTCTCCACCACA

CATAATATGAGCTAAAGTTTGCTAAAAGAATTGCTTGTAGTCGTGGGGGAGTGATTTTTAGCTATATGTGCATCTTTTTT

TGCAGCTCTAGTTTGTTTCTTCTTTGCCATGAGAATCCAATATATAAATATCGATGTGTGCGTTTTCTACAGGCTTAATT

TCAGAGATCGATTATTTTACCGTCTTTGTTTGATGTCATGATGTGAGATGTGAAACAAGAACAAGGCCATACCTTCATTA

GAAGTTTGATTAAAATATATAGAACTAGCTAGAAATGTCAAGAAATCAAAATATATAGTAGTAGTAAAACAATTTATTAA

CAGACAACAAAATAAATAAGAAATACTAGGACAAAAGTAACACCCAACTCTGGCGTGATTTTAAGATTGAAAAATGAGGT

ACAGGCTTCTTCTCTAGTTCTTAAACAATTTATCATTTAGCACGTTAATCGCCCTCTCAGCATCTTCCTTGTTCACAAAG

TTCACAAAAGCAAATCCTCGGCTCACGCCAGTCCTCCGGTCGACAGGAATATAAACACGGCTGACAGTACCAAATCTTCG

GAACATCTCAAGCAAGTCAGGCTCACGAGTTTCTTCCGAGAGGTTGGTGACACGAACAGAGTTTTCTTCGTTCCTCCGTG

GTCTCATCTCTGATCCGAGTTGTCTCGCGCCATCTCTCTGGCGGGGAGATCATCTTTGCTACCAGGAGGTCGAGGTCGAG

GTTGAGGCCTTTCAAGACGAATCTCTTCAGTAGAAACCATGGTTAGTCTAGCACCAACATCTTCATGAACAGCATCACCA

AACTTATTCCAAGAACGACGTTACAGTAATTTTGTTAATACGATTGTCATGTAGTCCGCTTACCACTGGTGGTGGTAAAA

GGGAATGATTAAGATCTTGTCCATCATCTTCTAAATCTCCCCAACAAACTGAACCTGGTGTTTGCATAGTATCTGTGCTC

ATGAAAGCAAAATCAAACAGACACTTTCCTGTTTTGAGTTACAAAACGTAATGCGCGTAGGAAACTCTACGTAGTGAATC

CTTAACACATTTATCCATCGGTTTCGGTCCTACTCTATAATCACCCTTGACAGAATCTTTCGGGCTTATCTTTGGCATGT

CTGACACAAATGGGCTTCAAGCCTTTCACACTAACTAGCCAGGCCCATCTGTTTTTCTCCTTTCGGATTTTTTAATAAAT

GTGGCTCAAAATTTTGTTACAACAAATTTCACACTACCTCAGGTAGAGAGAGAGAGAGAGAGAGATTCAGAGAAAAGAAA

TCTACAGACAGAAAAATGGCAATAATGAAGAAATTGAAGATGTTTGTGGCACAGGAGCCAGTGGTTGCAGCTTCATGTCT

CCTTGCAGGAGTTGGTAACTTCATTCTCCATTGATTTTCTCATTGTTTTTAATCACATTTTCAGTGTTTTCAATTTAGAT

TTTGATATGGTTTGTTTGACCATGAGATCAAGTTTTCTGTCAATATCCAAATTTGATGTTTCTTCATTTCACCTAAATCT

GTGTTTAAGATCACCATGTTTAACCATACAGTAAAAATAACAGATTCAGTCATTCAAATGTCATCTAACCATCAGAAATC

GAAACCTTAAACGTCAGAGACATCGATGAATAATCTGCATATGATTGATTGTTCGACATCAGTATCAATACCATAACATG

GTTTCGTTTTATGATTATCAGGGCTTTTCCTTCCAGCTGTGGTCAGACCAATTCTGAACCCTCTGGAAACAACAACAAAC

CAATTCCCACCACCTGTTTTCGCTCACCATGTAAGTATCATTTTGCTATTTCACATCTAATTTGATCTCTGGGTCTGCCT

GCCAACATTTCTAGTGTATGCAGAAGAGAAAAGAGACGAAGACAAGATAAACACACAAGAACAACAGAGACATATATGTC

CCTGCAAGCCATGGAATTTCTAATGACAGTGATGTAAAAAACATGACCAAGGAGAAGACATCCAATGGCAAAAAAGATGT

TAATATTGATGTTATTAGATGCAGTCTTTGCTTCCATTAGTTTTCCAATGGTTGAGTTTCGTAATTCTATGTCTTTCATG

TTGCAGGTGGTTCCAGGTAAGAAGAAAATGGAGTAACCCATAAATCTGACAGCCGGTGATTTCTCCTTTCCGGCAGACTG

AGATTTTTTTTCTTTTAATCATTTGATGGCTCACAAAATAAAACCATTTTTCTGCACTACCCATCTGTACATATAATTTC

TAAACCCTTGTAGCATCGAGCCGATTGTTTTATTTTCGGTTTTGCAATTTTGTAAATGAATGATTAGGGAACATGAGCAT

AACTTTCATGTCCTTCGAACAATAAAAGTTTTGTTTCTGTAGAACTAAAATATCATGCTATTGGAAAGACACCTTCGACT

GTCATTACTCATGAAATTGGCTATATAAAAACAAAATACGATTTCACTAATCCGAAGGAAAACAAACAAACTTGATTAAC

GCTCATAAGTACATTAAAGTTACCTTTTCTTTCAAAGATGAAGGTGTAAAAGGAAAATCCATATTCTAGTACTTCCTCTT

CCTGATCTTGAAACTGGATGATCGGATAACGTCGTCATCCCTAATGAGGCGGGACTCCAGAGCAGTTATGGACTCTGGTT

TGGTGAAATAGGTAAAAAGTAGATAGATACCATCCAAATAGGTGTTGGTTTCTCCTGCCTTGTTCTTCTTTTTGATGCTG

TATGCAAGAGGAATCACTCCTCTGTTAAATACCTCGACGTACATTCCACCACCAGCAACAATCAACTGCAAACAAATGAG

ACATTTCAAATTTGTTCATAAAGAGAACTAAAAAGCACCGCTCCAGTATGAACAAGCAGAGTGACAAAGCGTAGCATTTT

AAATCTTTTGATAGTGTAACGAGTAATATATGCAATACTCCCAATGTGTTTGCATGACTGGTCATAGGTTGAAACTAAAT

TCCCAAGCTAGATATCGTTCTTCTTAATGATTTGCAGAACATAGAACAAATGCTATTCGATAATGAAGCTGCAAGTACTA

ATCAGAAATATCCAACACTGGAAAAGCTAACTTTCGATACATAGAACAAACACTACTCGAGAACGAAAAGATGCAAGTTC

ATTTACTACTTCGGAGTACTTCCAAATTCACAGAATTTCTACAGCATTATGGTAGCATCTGAACAGTGACAATTGCTAAT

CCAACAGCCCTTCATTAAGGAATACAAGTCATCTTAACACCAATACATTGAATTTTCAGTGACAATTGCTACTCCAGAGC

ATATAACAGTGTTGGCCTTGATTGTGAAAGCCTCAATCTTATTGCCCAGTGCAAGCAAATTGATAGCTCCCACCATTTTG

ACTTATTGCGAGCAGGAAAACACACATCTCTACCAAACCATTATACTTGTTCTTCTTATCAATTGCAAGTTATCTATCTC

TAAGGACATTCAACAAACACCTTAACTGCATTTCCATGGTGGATTCTTACAAACAAATTCAGGCGGAACTAAAACAAAAT

TAAGGGTTTCATTTAATTTACCTCTTCATACTTCTGGGTAAGTGTAAGACGTTCATCTTCAGTCATGTCAGGTCTTAGCA

CAGCCATAGTTTCATATTGTCTCAAACCAGGTGGACATTGAGGTTCTTCTTTCTCTTCAGTAGCTGAAATACCACCAGAC

GATGGTGGTTCATCATCTTCTCCAACACCTCCTTCAAAGAATGAACCTGAAAAATCTAAAGTTAAAGCTCTAACTGATAA

GCTTTCATGGGTTCTTCTCAGAATCTCAACTTTGTTGAAATTACTTTGAAAGAAAACTTGAGGAGTGAATGAAACAAGAG

ATGACGAAGAAGAACTGGAGAGAATTTTGGGTGATGAGAAGGAAGTGGAAGCTAACGAAAGTGATGATGTTGCCATTGTT

GAGAAGACGAGGAATGAATTGAAGGGTTGTCACTGGACTTTCTTTTAGAAGTGAGAAAATGAGGAAGAAGACTAGAAAGT

TGTAGCCATGGGTGGATGATTGATTGGATAAGGCTGTGGGATGTTTACACCCAACCTTTAATATTTCTTACACCTCTTTT

ACACCGGGTAAAAGAGTGAAACACTATACGTATTTTTTGTTGGTAGGATTGTGTTGGCGCGATCGATGGAACTCACATTA

GTGCTTCAGTTCCAAGTGGTGTTGCACAAATACCATATTGAGGCAGGAAGACCTTACCCACTCAAAATGTGATGTGTGTT

TGCGATTTTGATATGTGTTTCACTTATGTTTTAGCTGGTTGGGAAGGCACAGCTAATGATGCAAATATGTTGGGATACTT

AGCTCCATATAGAGGTCAACGGTACCATCTCAATGATTATCCCGGAACCAGGTTACCTCTTAGTGCAAGGGAGAAATTTA

ACCATGCACACTCGTCATTGCGTAATATAATTGAACGGTGTTTTGGACAACTGAAAGCTAGTTTTCCTATTTTGAAGATG

ATGCCATCTTATAGCTTGCGAAAGCAAAGAAACATTTTTAGAGCATCACGTGCAATACACAATTTTATTAGAAGACACAC

CATGGAAGACGATTTATTTTTAGAGTATGAAATTGATGACATGATTATTCCCGGTGAAGGGCCAAGTGATCCTTATCAAA

CTACAACTTACGCCAATTTGACTGCTAGTGAATCTCTAAAAAGAAGACGTGAAATGGATTCTTATCGAGACGGGATTCGC

ACGGGCATGTCCTTGCATTACCGCCTGCCTATGTGAAAATGTTGTATAAGTTCTTGTAACTTTAATTAGATTTCGGAAAA

CATTATGCTCAACATGCATGGATATAAGTTATATTTATTGTAGATGCATTTTACTACAAACTACATCCAAAGGATAGAAC

GCAATGCTCATCTCAAATGAAAGTAAACACTCCACAATGAACACAAGATTTACGTGGTTCGGTATGATTACCTACGTCCA

CGGGGGTGAAAGGATGAATCTTATTACTTTGAAGTGTTACAAAGGTGTTGCTCTCCGTGGGATTTCCTTACAAGATGCAC

TTACTACTCACTTTTCTTCTTCCTCGTGCTTACTCGAGTTTTTCTCCTTTTCTCTCCCTCCAAATACAACCTACCTCTCT

TGAATACAAGTCCCTCTATTTATAGGAGAAGTTCTATATAACGAAATTAAGGTGTTTGTCATGTTATATCTATTTATCAG

TTCTATGATCGGGCCGGTTCACGTCGTCTCATGTGGCCGATGTATCCTGACTCCTCGCCTATCCTCCGTTCGGGCCGATA

TTTTTAGGTGCATTATTAACGCAATGTCGCCTATCTTTTCTGTCGTCTTTATGGTTATCCGATGTGACCCGTTCTTCTTT

ATTCGTTTGACCATAGTTCCTCGTTTGACCCGCCATGCCTTATCCTGTCAGCTTTATCAGAAGTCTGGGTCATGTCTTGT

CCGACATTATTACTCCCGATCCACGTGGATCCTTAAATCTCGCGCCCGATAATCTTCGCTCTATGCTTCGCGTCTCTTCG

TATTTTGGTACACCGGTTCTAGGATGTTGCCACCTGGATCTGTAGATAATTTACACCTACATTTATGCCCCTTCTTTCGC

TCGTGTAGCAGACACGAGTGGAAGAAAGTAGTTTCAACCGCCACGTCCTCGTCCATGCCGCATATATCGCCATGCAGCGC

CCCACTATCCCCACTTTATTAACTGTACCCGTTGCCATCGGTTACAACTTGTTCTCTTTAAATAGCCCGCTCCTTTTTTT

CTTTTCCTTCTTCTTCTCCACTTCTTACTTCCGCCATTGCAACCTCTTCTTCAGGATTTCTTCATTACTTTGCGCTTAAA

AACCCTAGCTTTCTTATTTTTCTTCCTTCATCTCATCTTCCCCAATCGTTACAGTAGCTGTTGTCGCCTCAGCTTTTTTT

GTTGGAAACCGAGCTTTGCATTATCTCCATTAATGGCTTCTTCTCCTTCCATGAATGAGAAAAGGTCTGTTGATGATTTT

TTTTCCGAGTTTGATGCTCCTCCCCTTGCTTCTCTGTTTTTGCTTGAAAGTCCTTCTAGGTATGATGATCGCGAGATAAA

ATATATATTTGCTGATAGCTCTTCTTCTAGTTCGTGCAGGTCCGAGAAAAGGGTGAAGGATCTCTCTGATGAAGAGTTCA

TTGAGAAGCTTAAGGAGAAGTACGATCTTCAAATTTACGAAGTGACCCTTCTCTCAGGCCGAAACGGTATCTTGAAGAAA

ACTGATATGAAGAAGCACGAGCAGTCGGCTTATTGTGTGGTCATCACCTTGGGTCAACTGGAGCTGGGACTTCGTCTTCC

CTTATACAACCCTGATAGCCCATTTTATTACGAGATTTTGTCCTAGATTCGGCCTTCTCGTGCTTTAGCTCAATGGACTT

GCAATACTTTTCGAATTATGAACGAGTGGAGGGATCGCCGCGATTGTCATGGTTCTACCGCCGAGTGGTCGACCTACAAG

CCTGAGGACGCTCCTGATTATACCGTTGCCAATTTCCTTGTCAACTATCGAGCTGGTACTTCAACCTTAGGAGATCTTGC

CGGTTTTGCTTCTAGTCCTCATCGGGTCAAAACTATGCCCGAGGGTGTTGAAAGGATGATGCTGGATGCCGATCCCATCG

GCCAGGGTTCCAACAAGCGAGCCCGCCATTCTAATGACCCGATTTATCCCCAAGTTCTGCTTCTAGTCCAAGGTCCGATC

CTTCATGGTGACTTCCCGTCGCCTACTTTTCCCGTCGAGCCCTATTCTTTCTGGGTGGTTAATGATGACAAGGCTGATAA

GGTATGCACTTTTCTTGATTTTCATCTTGTTAGCTTCGGGCGTCAGTTAGTCATTTGTTGTTGTTTAAAAATCTCAGGTC

GGCTCCACCAAGACTATCGGGTCTAGGAAAAGAGGTGCTGGCACAAGGGTTGAGGAAGACCAGGGGAAGGTAACAAACAT

AATGATTTTATTATAAGTACATGTACTTTCCCCTGTTTTTTTCCTTCATATGTTCTGACGAGAAACTTGTGCAGAAACAC

CCAAGGCACGATGATGATGAAGCGGGCGATGTTGACCCCACTCCGCTGAGGAGTAATCCTGGTAGAGGTAAGAAAATTGC

TAATCCACCGCCGAAGACGGGCCCCATCATTGATATTTTCGATGAGGATGAGCTGTTTGGAGAGAATGTGTCGGGTGACA

AAGGCGATGAACAAGGTATGGGAGACCAACAGGATCCCTCGGAGACGCCCGTCGTAGAGAAGGTCGTTACTGATATACCG

GTAGAGCTTCCTTGTGTGCCCCAGATTACTGAGGGATCGTCTTCTAGTGCTGAAACTTCCTTTATTGTCTCTGGCCCGAT

CATGAACTCCTTGGGTGCGCTTACTTCCTCCAACTTTGGGAAGTATACTGATGAGGAAATATTGAGCGTTGTGCCCCGTT

ATAGAGACATCTCCCTGACACTTGACGCTGTGGTGAGCGACTCGTTTCTGTATGGTAATGCCCCGCTTCTGATTTCTTTT

TCTTAGTGTCGCTTATGATGCAAACTATTTTTTTTGTGTTATATTATGAACCGCGTACGCTTGGATGCCTCCAATCGCCT

TCGAGATAAGCAGTTGATAGAGTCTCTCAAACTTCAACTTCAGAATGAGAAGGGTCGCGCGAACGAACTGGAGATCAAAC

TTGCCGATGCTCAAGGTATATGATCATTCTTAACTTGTGTTGATTGAGTTTATCGTTCCATTATCCTAAGTCGGGTGTCC

TTCTTCCTAGACGAGGTATCTAGTATAGTTATAGACGCTACTTCGGAGTTTAGAATTATGAAGAGAAAATGGGATATTGA

GAAGGATCTAGTGGAAAATCTTCGCCAACGAATGGCAAACAAAGACGACAAAATAGACCGTCAAACTGAGGAAATTGCAA

GACTTGAAGACGAGTTAGAACGACGTCGCCGGTTTGATTATGATCCTGCCGAGATGGATAATTTACGGGCTCACTTGGGA

TATTTTCAGCGTCTTGCAGCTGAAAAAGCTACTTTATCCACAAGTTTGGAGCAAAGGGTAGACGCTCTACAATTACAGAA

TCGAGACCTTCACGAGGACCGTCGTCGAATACTGAGAGATAAAGATGTGGTGGAGCGAGCTAATAGCGATTTACTGGGCC

GAGTGAGTGAATTTGATAAATTTGTGGGCGATTTATTGTGGAGTAGATCGCAGCTTGCAGTGCTCCAAGCGTCGCGTAAT

CGGTTAATGGACGATTGGACCGAGCATAAGAAGTATTTCCCCAGTAACGAGTTCAATGAGAAACACTATAACTAATTAAC

CCTGAAGTTGTCCAAGGCCGAGGATGAGTTGTCCAAGTCGGTAGAGTCGCTTCAGTCGGTGCTACCCGCCCTGTCAGGTC

TCTTAGACGATGTGTTGAGGCCGTTCTTTTAGATCTAGATTGCGCTTTATTCTTGCAAAACTGTACCCGATGAATAACAT

TCTGTCTTTGTGATTTTGTAGCCGAGCAAAAACGCTCTCGGGAGCTAGAGCTGGAGGTTCGCCGATTGAAACAAGCCGAG

GTCGATTTGAGCACCAAGTTAGCTGAAGCTGAGGAAAACGCTTCCCAGATTACCCAACAACTCGAGGATGAATGCGTCGC

TCATGTGGAAGAGTTAGACCAAAAGGTTCAGGCGGGAATTATGGACTGGGTGGAGAAGAAGCGGCTCGAGGTCGCGCATA

GGAAGGACGGCGCGGCTCCTCCCAAGGATTAAACTTTATTTTTGTGGTGCTGCGTCACTATGTTGCCCCTTCCATCGGGT

TGTAAATCTCTCTGAACACTTCTCCCCTTTTAAATTTCCGCTACCTCGCTTGTTTTTTTTTAATGGGGGTGCTGCGTCAC

CGATGGTGTTTGTTTTATTTTCATTAGATGCTTCATTTCGCTTGGTCACATGATGCTTTTCGTATTTTCTTAATAGTCGC

ATGACGCAATCGAGTTCTTCTTTGTTTCATATGTTATTCCTGCAATCATTGTTCAAGTAAGTCAGTATTCTAACCATTTA

TGTCGAACACACCCAACAAGGAGGGTCATCGGGCCCTATGCCATGTCCTGATTGGGGTATCTCATCACTATCTCTTTGGA

TTCAGTGGAAGGGTACCAGTCATCCTAGTTCTAGGCCCGACAATCCCGAGTGGTTATCCACAAACCGATCTCGGGCAAGT

GGACACGCCCACTTGCGATGGTGGTGACCTTTCAGATGTAAGTAGGCCACCTAGATGAGTAGACCTCGCATCTAAACAGC

CTTTGGAATTGGTTTCCAACCACCAGAACCCTTAGGCTACCTCGGCACTTGCACACTTCCACTGCCCCGAGTTCGAATAT

CCAGGCGACTGTAAGGCACCTCGGCACTTCCACACTGGCGTTACCCCGAGTACGAATGCCCATTCGAGTGTAAGTCACCT

CGGCACTTCCACACTGGCTTTGCCCCGAGTACGAATGACCATTCGAGTGTAAGTCACCTCGGCACTTACACACTGGCATT

GCCCCGAGTACGAATGACCATTAGTCACTCATGCCCTATCCCATAACCCTCACGGTTTCACAGAGATAGAAGACACGTTT

TCTTACCGTTTTTCCGCATCACTTGTCAAGGATGGTAGAGTTTCAGGAATCGGATGTTCCATGGTTTGAGCTCTCTTGTT

CCGTCGAGTCACAGTAGTCGATATGATCCCTCTCCCGCCTTAGATTCGATAATATATGGCCCACCCCACTTAGGTGCTAG

TTTTCCTCCTTTTTTGTCGCGCTCCCATTCGGCCAAAAATTTCAATACCTGCTGCCCGGGATGAAATTCTCTTGGTCGAA

CGCGCTTGTTGTATTCCCGTTGTAACCTCCTTTGATAATTTTCCATTTTTTGTAACGCCATTTCTCTTGTTTCTTCCAAG

TCGTCGAGTATAGCCAAGATCATATCGGATGACACGTTTCGCCTCCAAGCTTCGGTTCTTGTGGTAGGAATCATCCCTTC

TGTTGGTAGAATCGCCTCGGTTCCATAGGTAAGCATGAATGGAGAGAATCCCGTAGCTTCCCTTCGTGTGGTCCTGTATC

CCATAGGACGTTATATAGCTCCTCGCACCAGTTGCCATATTCCCCGTCCAGTTTTTTCTTTATGTTATCGGTGATTGTCT

TATTAGTGATCTCGGCCTGCCCATTGCACTGGGGGTAAATCGGGGTTTCTTTGCTCTTCCTGATGTTGTAAGTGTTGAAT

AGAAGATCGATTTTCTTGCCCTGTAGTTGTTTTCCATTATCAGAGAAGATTGCAGTCGGGACCCCGAAGTGACAAATGAT

ATGTTCCCAAATAAACTTAAAGACATCTTTGTCCCGAATATGCCTTAGTGGTGCCGCCTCTACCCATTTGGTAAAGTAAT

CAGTGGCCACGATCAAATACTTCCTTTTCCCCGTTCCTACATGTAATGGCCCGACAATGTCCACCCCCCATTTATCAAAA

GGCCAAGGACTTATGATTGAGTTCAACGGTACGCCGGGAGCATGGATTTTCTTCCCAAAACGCTGACATTCTTCGCAAGC

TCTTGCCATTTGCTTTGCGTCCTCGTTCATATACATCCAAAAGTAGCCCATTGTTTTTGCCTTAATCGACAAATTTCTTC

CCGAGCTGTGATTTCCAGCCGCCCCATGGTATAGCTCGTGTAGGATTATATGGCCCTCTTCCCTGCTTAAGCATCGTAGG

AGTGGTCCTAGGTAAGATTTTCGATATAAGATCCCTTCCCTTAACTCGTAGGCGCCCGATTTGCTTTTGATCTTGTTCAT

CTCGGGACGACTCTTAGGTAGCTCACCCGTCTCTAAGTATTGATGTATTGGTTTTCTCCAGTCGCCTTCTTTGTATCTCT

CGACTGTGCTAACGGCGACCTCGATCCGTACGTCTGGTGTTTCTGGGATAGACGGGTTAAGAAGCCTCATAACGCGTATC

CCTTCGATGCTCGGGTCTACTAGTTGTGAGGAGATGTACGATAGAGCGTCCGAGTGACTGTTATCCTTTCGGCAGATATG

GCGGAACTTAATATCTGGTATTTTCTCGATGTAAAATTGGGCGAGCTCCCAATATTTTTGCAGCACCGGATCGTGTGTAG

CGTATTGTCCCGTAATTTGCCGGATCACTAGTTGTGAGTCACTAGTCAATCGAATCTCATGTAGGCCCATTTCCACAATT

ATTCGTAAGGCATGGATCACCGCCTCATACTCGGTGACATTGTTTGTCGCCTTGAATTCTAACCTGAACGAATGCACTAT

TTTCCTCCCCTTTGGCGTGGTAAACACCAACCCGAGACCCGATCCGTCTTTGTTTGATGCCCCATCAACAAATACTTCCC

ATCGTGTTGAACTACTGGCGGATAGTAGGTCCGCTGTCTCCTCTTGTTCTTCTGCCATTCCCAGTATATATTCTACCTCG

TCCTCATCACTGAGGGGGAAATCTTCCAGGAAATCGGCCACTACTTTCCCTTTGATCGCGGTTCTTGTCTCGTATATCAC

GTCATATTGTTTAATTATGGCTCCCCACTGGAGATTCTCCCCAATCGCCCAGCACTATCCAAGACCGCTTCGATATGCGA

TTTTGTGAGCACCCGTATTCGATAAGCTTGAAAATATGTTCATAATTTATAGGCAGCAAAGGCTAGTGCGAGCAACATTT

GCTCCATTCTGGTGTAGTTCTTCTCGGCCGGACTTAACATTTTGCTAGCGAAGTATACCGTCTTCTCCACGCCTATCTCT

GTTATGACCAATACCGCACTGATGGCATAGGGCGTCGCGCCTAAGTATAGAGTGAGAATTTCTGACGGCTCGGGCCTTTG

AAGAACCGGGAGGCTAGCTAGATGTAGCTTTATATAGTCCGATCTCACTTCTCCTCGGGGGCAGGTCCGATCTATTTTCT

GATCGCCTGATATGTCATTTATCTTCTCAACTTCCTTGAACACCTCTTTCAAAGAAGCGTTTAGAGGGGTATACGATCGG

GCTTCCTGCTTTGGCTTCTTCCATTTGGCCATCCATTCCCTCTTCTCACTGCTCTTCGGGGGCGGGTCTGATCTCTTTTC

TGATCGCCTGATATGGTCGTCCATGGCCGTTTTAGTTTCGGTACCAACTTCAGCACTGGCTTCTTGTACTCCTCTATCTC

TTGATTCGGCCTGAATTTCTTCCAAGGCGATGTAGTGTTCTTACATTCTTCTCATTTTCTTTAAGGTTTTAGGTTTTTCA

GAAAACATGGCAATCCAGATGGGGTCCAATCTCCCCAACGCGTTTTCAAACCCAAAGATGGCTTGGTCCACCGGTACCTT

CCCAATTTCAGTACACAGTTTCCTCCACCTGTCGGTCAGGGTTCTCAAGGGCTCTCTGAATCTTCGGGCGAGAGTAAACA

GCTTGTTCATTCTTGGCCGAGGTCGACTGTTATGCATGTAGGTCTCAAGGAAGGCCTCGACTAGCGCATCGTACCCGTTA

ATCGAACCAGGTGTGAGCGAATAAAACCAATCTCGCGCCTCGCTCTCCAAACTAGCTGGAAAGAATTTGCACATAACTAC

CGGGTTGCTTCTCCATTGGATTAGTGACATGGTATACATCTTAAGATGCTCGATCGCATCGCCCGTTCTATCAAATCTAG

AAGGGAATGAAGGTAAAGTACACTTAGGTGGGAACATCATTTGTTCCAGTTCTTGAGAGAAAGGCGTCTTCTCGGCTTCA

CTCATAATCTCGGCCAACTTATTCTCTTCGCTTCCGCCCGAGAATTCCTTGATTTTTTCTTCCAACTCCTTAAGTTTGTC

TTTAGTTGTCTGATCTTGTCGCTCGTCCCTTCCTCGCCTTTCTTCGTCGGTGGACGAGTCCTCGGGTAGTCCATATCTCT

GCATGGGGGGTACCGGAGGTAATGGTGGCTGACCCAACTGATTGGGATGATTAAGGTTACTTGGCCCCATCGTAGGTGGT

GAGTGTTCGGATCGGACACGATAACCCGATGTCCCTCGGCTGGACGACCCTCTTGATCTGGTCCGTTGGTTTCTCAGCTG

GTAGTTCTCATACTCCAAGGCTAAATTCCGCTGCTGGAGTTCCCTTAGTGCCGCTTCTTATCTCCTCTGACTCTCCAAAA

TGGCCAACAACGTCGGGTCTGTACTTTCATGACTGGAATGTCCATTGGGTTGGTGTGGCCGAGGCTGTAATGGTTCAGAT

GTGGGTGGTGGGGGTAGTATCGGGTTGGTTACACGATCGACCACGTGAACAGTTGTCGGGACGATGTTACTGGTTTCCGC

TCTCCCCAGCTGAATCTGCCTAAGACGCTCTTGTTCCCTTCCTAAGGATTTCTGGTACTCAGCTTGTTGCCTCATCCTTT

GCCTATGAGCTTGTGCTGCAAGTGCTTCCTCATCGTCATCGGTGTCTGAAGCTGTGAGTCCATCGATTTGGTCAACTCTT

CTCTGAGTCGGTATAACCTGGTCCAGAGGTAAGGTGTTCTCATCTCTCCCCAAATCTACTTCCTCGTCTACGACAGGCAT

TCCTCGCGCAGCTGCATGAGATTCCTCGGGCGGTGATTAGCGACTTTCTTCTTGCCTCTGCCCCGTTATCCCTGCCTGCA

TGCTGCTTCTAGTGAATGTACGATTCCTTAACATCCTAATCGTCCCTGAAGTGCTAGCCATCGGTCACTTCGCATGATAA

GCCGAGTATCGCCCGTCCCTACGTCAGCAAATACGAAAAGAACCCCTTACTTTGACCTAATTTTCAAAATAAAAAATTCA

GACATAAGGGCAAAAAGTTCGACTGTACGACTGATGTCACTGTGTAAGAACTACTTCAGTTGTTAGGTGGGTACTGTCCT

TCTACACTATTTTTTACCTATGTTGAATCTGTGTCTGACCTTATTTTCCCGTCGCTATCACTCCGTACGACATCGCTCTC

TGTACTCGACCTATGCATTTAAGTTAGTTTTGTCCTTTATTCTACATAAATCATTGATCTTATGCACATGATGCTTAGAA

AAAAGTACGGCAAGCAACCTCTAGAATCAAATCATACTTGTACCAGCTCCACTTTCTCGAGATCTAATTATGAACTCTCC

GACGACTTATGCTACACCGCCGGTTAAAAAATTCTATACGGTTTGACAGTTTATCTCACGATGAAAACAGCTACAAACAC

AAACTTTTGGTTATTCCAAGCACCCGATTTCATGTCATAGCTAAAAACAACTCGTTTAACGCTATCAATTATACAACTAT

CAAGAACAAACAGTCGTCCTTTCAACAGATTCTTCACATATCTTCTAACCAGAACTGTTAGTCGCTGGTGAGCTCGTGAA

GTCTCGCTATTGAACAAACTATTTAAGATGTTAAAAAGACGCTGCTACTGTTCTGGCTTTGAAGTAAATGACTCAAAACA

ACCGTTCTTTCGGCCAACAGAGTTCATCATAGCTTCAACTTCAACACGTTAATCGTGTATTGAGGAGCTGTTGATTGTCT

CTGAAGGTTTTTACGGCAAGGTATCCGGAGATCTTAGTCGTGTCCCCTTAGTCGGCGCCAGAATGTAGATGCATTTTACT

ACAAACTACATCCAAAGGATAGAACGCAATGCTCATCTCAAATAAAAGTAAACACTCCACAATGAACACAAGATTTACGT

GGTTCGGTATGATTACCTACGTCCACGGGGGTGAAAGGATGAATCTTATTACTTTGAAGTGTTACAAAGGTGTTGCTCTC

CGTGGGATTTCCTTACAAGATGCACTTACTACTCATTTTTCTTCTTCCTCGTGCTTACTCGAGTTTTTCTCCTTTTCTAT

CCCTCCAAATACAACCTACCTCTCTTGAATACAAGTCCCTCTATTTATAGGAGAAATTATATCCAACGAAATTAAGGTGT

TTGTCATGTTACATCTATTTATCAGTTATGTGATCGGGCCGGTTCACGTCGTCTCATGTGGCCGATGTATCCTGACTCCT

CGCCTATCCTCCGTTCAGGCCGATCTTTTTAGGTGCATTATTAACGCGATGTCGCCTATCTTTTTTGTCGTCTTTATGGT

TATCCGATGTGACCCGTTCTTCTTTATTCGTTTGACCATAGTTCCTCATTTGACCCGCCACGCCTTATCCTGTCAGCTTT

ATCAGAAGTCTGGGTCATGTCTTGTCCGACATTATTACTCCCGATCCACGTGGATCCTTAAATCTCGCGCCCGATAATCT

TCGCTCTATGCTTCGCGTCTCTTCGTATTTGGGTACACCGGTTCTAGGATGTTGCCACCTGGATCTGTAGATAATTTACA

CCTAAATTTATATATTATGAAGTACTGATCTCGTTTTCACAATTATGTTTAAGTTCATATCAAAATATTTCTTGGTTCAG

ATTAATCATTTTCAACACTAAAAAATATAAAAATAGTGATGGTATTTTGTCTTTTAACTTCTGATTTTGTAAATTTGCCA

AACTAATTTATTTGCTTTTCAACTTATATGTTAGCTAGTTGCTACTAACTCTTCGACTTCTCAAAAGTAACTTCTAGAAG

TTCAGAAGCAGAAGCACTTTTACTTCCGGTATCCAAACATCCCGACACTTTTAAAACGACTTTTGGAAATAAAAAAATAA

GGATGAAGACTTGAAAGTTGTTATCCTTGTAACATCAGTAGCAACTAATCCAACATATTCGGCCATTGTTGGGCGGCTTC

ATTCTGAGCTAGCCTCCAACAAGACTTTTTTTTTCTGAATCACAAACTTTTATTAACTTTCCAAAATTTAAAAAAAACCA

ATTACAAGGAGACAATAACTTTCAAACATAAAAGTAAAGACACAAGCCCACAAATCAGACACAAAGCCCAAAATGAAAGC

CCAAATAATATGAGAAAAATGAACTAAACTTGTCAACCAAATCTATAATATGCAACCTCTGGCATTTCAACTCTTGGAAA

TAAATGAGGCCTTCCCATATGAATCTGTCTTTCTCCTGCAGCTAATAAGGCACCTCTCTTGGATACACCATCAGCTGAGA

AATTAACCTCCATGGAACAATGTTGAAACTTTACTTGCGTTAGCTTCCTTATATCCATATACCATCTACTCTTAAGAAAC

CATAGAACTTTCCCCTGAGCAAATTCTTCAAGCACTGTTTTTGAATCAGAAACAATGATAACATTCTCCACACTCCACTG

AATTTTCAGTTCTAATGCACTCATAATTCCATAAGACTCAGCAAAATAATTTGAAGCTATTCCAATGCCTCCAGCAAGGG

TACCCATCACCTAACATACTGAATTTCTAACTACCACACCAAAACCTGCAATTCTAGGATTTCCTATTGAAGATCCATCA

CAACAAAACATGACAAAGTTTTGCTTAGGAGGAAACCTGAAATATGAAATGTATTAGCCCTAACCGAAACTTTTCTATCA

GCAAATTGTCATAATTCTAATTCCACTTGGTGCCAGTGATTCTTAAACCACCCTCATTCACTAATTTCATAATTATGCAT

TTGAATCTTTGCATATTTGGTTTAACTTCCGCAAAAAATTTCATGTTCTTCTGAAACCAAAGTTCTTTTATAATAATAAA

GGAAGCAGACTTCCAAACCTTCTTAATCAGAGGACTACAGTTTGAGGCAGATTTCCATATGTCTTCCCAATATCTTGTCT

TATTGAATTGAAATATATTACTTATCCAACTCTAAATTTCTGAACTGAATTTACATTCCCAGAGAAAGTAATTCATACAA

TCTTGATCAACTTCACAAATGCAACATCTTGAGACCATTTCATAGCCATTTTTCACCATAATTTTGTCATCAATGTATAT

TCCTTGTATCATCTTCCACACATTACTGGCTACAGAAGGATGAAGAGAGGGATACCATAATTTTTTATACCAGTTAACTG

GATGCTCCTTATGCCTTATCAAATTGACTTCTTCTGAGACAGAGAATTCCCCTTTTAAATTTCTTGTCCAAACTAAAATA

TCATCACCTCCAAGAATTCCAGGCATTTTAAGATTATGAAAGACCTGGTTGAGATCATCATTATAATTCCATACACCATC

TGCCAAGATATCAGAAACTTTCAAGATGGTGTTCTCTGCAACAATTTTATGATGTTTGAATCTATCAAATAAAGGCCCAT

ATTCTGACCAATAATCAAACCAGAGAGAAATGTGTTTACCATTGCCAATTAACCATCTTGTATCTTCTTTAAGAGCTTCC

CATGCCCACTTTAAACCTGGCCAAACCGTAGATTTATGCCACTTAGTGCTCCACTAACCATTTTTATCCTTGTATTTTTC

ATGAAAAAACAAGGCCCAGTCCTCCTTTGAATTACTGATTTTCCACATCATCTTCATAAGTAAAGCCCTGTTAATATCAG

AAAGCCTTCTAATGCCCAAACCTCCTTCACTAAAGGGAGTACAAACTCTTTTCCATGAAAAATTAATGTACTTTCTTGTT

TCACCATCCCCTGACCAGAGGAAATTCCTCATAATTCTTTCACAATTTTTAATGACTGATGAAGGCCATTTATAGACTGC

CATATTATAAACTGGAATACTACTAAGCACTAACTTTATAAGAACTAATATTTCTTGAAAAGACAGTAACTTACCTTTCC

AAGAGGCTAACTTCCTTTGCATTACTTCTACCATGGGCCAAACAGTTGATACTTTAACCCTTCCAGCTGTGAGAATTACA

CCAAGATACTTTTCTGGGAATGTGCTTCTTGGTAGGCTCGGAAAGATACCTACTATTGTGTATACTGCAAGTGCACAGTG

TCGACAAGCAACACAGGGCAAGCACAGGTCGAACCACAGAGACTTGGAGAGTGTAGAAGTTGATTCTAAATCCTGATTCA

TAGAGTAACCAAAAGGGGTTTGATTTTGTCGATACGACGATATGATTTTCAAGTGACTGAAATAATAAAAGTAAAAGATG

CAAATAACAAGATATGTAAACAATGATAATATGGATACTAGGGTATTCAAATCCACCAATAACCTATGATATGCATTCAA

CTTGTAATATTATTCTTGTCCTTATAGTAGATTCAAGTCTACCGTATAACTAAGGTGTTTCTCCAATGGATAATTTAATC

ACAACTAAAGTATTTCTCCAAAGTATTAATTGTCTAACTAAGCACAACTAATCTAAAGATCAATATTTGGTTAAGCATGA

GTTGTAGCATAATCAACATAGAGATATTCTAACTACAATGGATACATGGCATACATTGTGAGAGCAAGATGTTGACAAAC

TTGGATTAAATTATATCTTTAAACAATTCAAAACTTTCAAGAACAAGATACAAGATATGAATACATACCAATGAGATTAG

GGCTTCATCTAAACCCTAGTAAATGAAATTAGAACATGAAGAACATTACAACAAAATTGAAATTTGATAAACTAACATGA

ACAAGCTATTCTTGGATGAGTTTTCAACACAACAACCCTCTCTATTTATACACAAAAACTCTCTCAACACTTGGATTCGA

AATTTATTGTAGCAACCACTCAACAATTAAACAAAACATGTTTTCTTAATTGATGGATATAATGGAATACAATGGGGGTG

AGGTGGATGTTTTAGAAAGAATCCAATGGGGGTGAGGTGGATATTTTGGAAAGAATCCAATGGGGGTGAGGTGGAAATGG

ATTTGGGTCTTGGGTTTTGGATCATGGGTTTTTTAATCCCATGAATTTGTAAACAGGTTGCCACTGATCTTGTGAGCGGG

TTGCAACGCTTCCAACTTTCTTCTCTTTTGTCTTCTCAATTCTTCCAAACTTTATTTAGTACTTAAAAACACAAAATTAA

TTAATACAAATATTTATTCTTGAAGATGGTAAGAATGCAGAATTTGGGATAATATATGACTTTGAAGCGCGAAAGATGAG

TTAAGTGCCAATGAAAAGGTGTATGAATATGCAATATTTGGCACTCATCACTTCTTTCCATCTGAATGGAATTATTTATT

TGATTTTTCCTTAGCTTAGAAGTGCCATCAATGAAGAGTTTGATTTTACTTTTATTTATAATTTTCCCTGAACACTTCTG

ATAAGCCTCCAACAAGATTGGATTAAATGATTAGGGGCTTTTTTGTTTTAATATATCTATATCCTATTCTCAAAAAAAAC

TAGAAAGTTGTATCTATATGTGGAAGATTGGATTGGGTAAGACAATCAGATGTTTACACCCAATCTTGTTTTTTTTATCT

CACTAACACCGGGTCAAAAAAAGTGAAATCGTCGGAAATGTATACTTGTGATAAATTTTTCGTTTCGAGAAAGATTTGGA

CAAATAATGGATCTCCAAAAAGCATATGATCTATATTCCATATAGATTGAGTAGTTCCGAATAATTTTTTCCTCAATATT

AACTAAGTGCGAATAGAACTCAAGACGATATCTACCCGTCTTGTGATGGTATCTTGACACACATGAATTAGCCTCTTTTG

ACAGTATCTAGATGTCCTTTCTAACTACAAGAATCCATCTGGCATCCAAACAAAGCGTGAATAGAAATTAAGAAAGAAAA

AGGAGTCAAATGAACAGATGTCCTCGACAAAATGATAAAAGTGATCAGTTAGAGCATTTCCTATGGAATGAACAAACATG

AAAAATATATGTTCAAATCAACAAATCTGATGGTTTGAAAATTGAGTCGGAACATCCAGCACGCGTCTGTGGTAAGGACG

GGCGTCATTTCTACCAACGCTGGTATTTTTCCTTATGTCGTCAGCGTTCCTACACATGAAGTACGTCATCAGTTAGAACG

CTAGCGTATTTGTTAATGGCGCCGCGTTCTTTCTGAGAGAACCAACTTCTCTTCCCATCCCGTCTCCATCGTTCCTTTCC

TTTACCTATAAATTCACTTCGTCTACGGACTTTAGACTTACACCTTCCTCAACATTATAAAATTTCTATACCGTCTCTAA

TTTTTTTTTCCTTTTCAAACAAAACTATTAATATCAATTCAATCTTTCAGAATAATTTCAACATAAACAAATATGACATT

CTCACAACAACGATCACAACAACAAACACCACAAAAAAAAACAATCACCACAACAAACACAAGCACCACAACAAACACAA

CAATCACAATAAAGGAACACAAGAATTCGTGGTCGTAAGTATACGATGGATGAAGACGAGTAACTTTGCAGAAATTATGT

ATTATATACATTAGATGATATCAATGGTATTTATCAAAATAAAAGAATTTTTTATGGGAAGATCTTACAAACCTTTTGTG

AAGAAACCGGTAACATTCATGGTCGTGATGTTTATGGGTTGTCTCATCGTTTTTGCACAATTTCAGGAGCCGTTAGTGAA

TACTAGCTTTGATCACTCAAATGTGGAACAACAAAAAAAGTGGTGAAGGCGAATCCGATGTGGATCGAAGATGTCGGGAA

GAGTGGCAAAGAACCCACAAAGGTTCAAGCTTCCAACATGAAACTTGTTTCACTATTTTGAGGGTGCTTAACAAGTTTAA

TCCATATTTGCTGGAAGGTCATCAAGTGCCTATAAGATTGTTGTGTATTTATGAGATCAAATTCGAGCCGTATTTATAGA

GGTGGAAACTAGCCGTTCGGTGACCGTTACAAAAATTTCCGTTTCCGTTGGAAGTCGTGCCTCAAACACTGGCGTTCGTA

GCTCGGCCGCTGGAGTCATCAACAAAAGCGCCATAGATTGTACTTCGGGCGCCAGCGTCAGAGTTGATGTCGCGCGTTTA

ACTAAGAATCGCTCACTTCTTTTCCATAGTGGAGAATCCTGTTTGACCATCCACGTGGCTCAACAAACAAATACATTTGA

ACATTTCCATCAGAATTGCTTTTAGGCATACCTAGCTAGTTGATACTGACTCATATGAGAAAAAAAGATAAAACCATAAG

AATATATGAATTAGAATTATAACGCTTTTGGATACACATCCAGAAATTTCTTTTTGACTTCTGGAAGTAAAAAAGTCAGA

AATTAGTTTGGGTAACAATTTCAGAATGACTTCCAGAAGCAGAAAAGTTAACTTTTTGTGAAGCAAAATGTTTATGCTTC

TGACTTCTGACTTATGGAAAAGCAGAAGTAAGAAGCAGATTCGTATCCAAATATGCGCTTAACATTAGAACAAAATCAAA

TATGTTTTTCTTCTTTTTTTAGGATTAAAAAATTTGGTGCTAATGAGTTTCGAATTTAAGTCGAACGACCACCCGACAAC

TTTGCACTATACTACAGTTTCATAAGAGAAGTAAAAGTCTTTGTGGAGACAGTAAGATCAACCATGTACCTATTCGTACA

AAACATAAGATAGATATACAATACAATGAACTTTTATAAGTTGTTTTGATACCGAAATCTCTTCTGAATTTCTAAAAGTT

ACTTCCGAGAAGTGAAAAAGTAAATAAGTTAGTTTGGCAGATGCATAATGTAAAAGTTGAAAATTCAATAAGTAACATTG

GAGAATTAACAAACTCGAAAACAAAACAAAAAATCTTATAAGTCAAAATATAATTGTTATTGTTATTTCTGTTTTTTTTT

TGTATTTAAAATTGACAATTTTGGCACAAACATGGATTTATCGTGAGAATGATTATTATAGAACATCCACTTTAGCATGG

ATATAGATCAGTGGCAAACCAGAAGCCTGCTGAAATACTGTCCGTGATATATTTTGATCCAACGGATATGGGGATCCGAG

ATATTGAAAAGCTTTCTGAGCCCTATGGCGAAGGAACTTCTTTTGTACTACCAGATATATCAAATCTTACTTTGTACTAC

TAGATATATTAAATCCTACACACATATACATGACTTGGAGAAAATAAAAAAAAAAATAACCCACAGACACATACCGTACC

AGGGTGAAAACCTTGTTAAACAAATGACCTCATAAAAAAATAACCCACAGATATATTTATGCCTCCTCCACAACCACCCT

CATATCCAGAATAGTTTTTACATCTTCATGGCCTCTTCTTTAAGAACTAAAAATTCTGAAAATTTAGAAAACTTGTTAAA

CAAATGAATTCATCTCTAAACATTCCAGAAGATCTATTCCCAAAAATCTTTGTTAACAAGGCTAATATTCTACCAAAGAA

GAAGATAGACTGGAAATGGGTTTTTATTGGGAGATTGTGCAGTAGAATATCATATGAAACAAGAGAAATATCAAGATTCA

TCAGCAATAACTGGGAATTACAAAGAAATTCTGAAGTTTCCCTATGGAAAAAAAGAAGAAACTATTATCAAATTTGACTA

GTAATGAAGATTATAAGGTTTTCAGTAAAGGGGGAACAAGAGGAATATTTGACAAGCTCATTGTTCTAACTGAGGTGCCA

CAATCAGGACCAGACTTCATTGATTAAGCTAATTTCTACAAAGTCTGTATCTGGGCTTTGGTCAAAAGAATTCCAACCCA

CATCGTTCCTGACATAAGAAACTTGATTGGTCCAGCCGGAATCTACCAGGGATCTAGGAAAGCTTATGGAAGAGATGATG

ATAAGAAAAATCTTGAAGTCCAGATCACTATAGATGTAACTAAGGCTCTAAAATATGGCATTGATGCTTATGAAGATGAA

AACACTTCCCACTGGCTTGAATTCGCTTATGCTCTTAATGGTATAGTTTTTTTGTAAAATCTACAGAATTATTGACCATC

CTGGTCCTTTGTGCATTGAAGAAAAAGGAAATACTCAGAAACCTCCAATCAACCAATACATTTATACCTATGAAAATATT

TTCCCTTCAACTCCCATCACTCCATTATCATCCAACAAAACTATCACTCCGCCTACAAGCCCCAAAAAACCACCCACAAT

ACTACTACACTGATAAAGCTGGAGAGGAAGAATTTGCTTTCAAAATGAGGAAGGCTCAACATCATACTCTTCTTAGAAAC

CATTGTGAAGAATTGGATGGACCAACACAACACATGATTTTTCAAGCTAGTTTAGAAAAACCTATTATAGGAACCATTCC

AACCACCAATCTCGATGCCTTAATTAGCATGTCAGAAGAAGCACAAGAAGCTGATAGAGAATCACAGAAGCAAAAAATTT

CAGGGGAATAAGAAAAACAACATCAAAGCTGCAGCCACCTCACCATTGGCAAGAAAGAGTTAGGATTCTTATCTTGTAAT

TTTCTTTTTCTTTTGGGACTTTGGACTTGGAATTCTATTTTTATTTTTATATTTTAAACCTCACGGAAGGATTATTAAAT

ATCAACTGTTTGGAAGTGAACAATATCATTTGAAATCATCCCCGCCTAGAACGGGAGGTAAGTTTTCATTCGCTTTTCAA

TCCTCGTTAATGAACATACACCTCGGAAAGGGCAAGGACGGGATAAGGGTTGACATCACCCAAAAACGTGATTTTGAATA

TTAAGGTAACGACAAAAAATATTGGTTACTTGTCCGGTTTCTGTAAATTGAATGCGTGACTGCGCCGCCTTGTCGGAGCT

CTGTGACTTCCCTCGCCTCTATTCAATTTACTTAAACTCGGATTTGCTCAGGATGGATTAGAAGCAAAAACAATCAAGTG

AATGGACATCAATCATCCTTGTTTGTGATTGCATAGAACATGGTTAACATGCTTGATGTGCTAGCTAAGAAGCTATGATT

GTGTCTTTAGAGACCATTTGCCTAAATTTAAAGGGGTTGAAGTAAAATCCCTACCACCATGTGAAAGAGTTCGAGGAAAT

TTGTGGGACTCTTCGGATCAACCAAATGACGGATGAGCACCTCAAAATGAAACTTTTTCCCTTTTTCTTTGATAGACAAA

GCGTGACTTGGCTTTGGTGCTCAAACAGTCATTTCATACATGGGATGCTCTTCTGAAAGAGTTCTTCAAAAAGTTCTTTC

CCAAACCATAAGACTACGACACCGTCAAAGTCCGAATAGTTTTGTGCAATTAGAGGAGGAAACTTTGGCAAAATATTTGG

AGAGATTCAATGAATTATTGCTCCAATGTCCTCACCATGGTTTTGAAAAATGGAGACTTGTGCAAATTTTGTATGAAGGT

TTAGATGTCAAAAAACCGAACCTTGAGTCTTGGGATTTCTTAATCGAAGAAGTCGCTGAAAGACTGTGATGGGAATCCCC

GTGAACCAAATAGATTACCCTTTCATCCAATGTTCATGGAAGTGCAAAGATGGCATCCTTGGCGGAGAATAGAAAAGCTA

GAGTTAGAAAAGAATGGAAAAACTAATATCCCTACTCTCCGTAGTCAAATTGAAATGGCCACTTGTGCCGCATGCCATGG

TCCAAATCTAGTGGACACTTGTCCAGACCTCCATGCATTTCAAGAATCTAAAACCGAGGAAGCCAATGTATTGTACCAAA

GGCAAGAGAACAACCCATATTCTCGGACCTACAACCCAGTGGTGGAGAAATCACCTAACTTTTCATGGTCTAAGGGGCAT

GTGCAAGGGGAACATCAAATAATAACCAAGGATACCCTTATCCTGAAGTGCAACCACATTCTCAAGACTGGAGAGAAAAG

CTTCTAGCTTCAAGAAAGTTTCCGGTTGTTGACCTCAACTTCAAAAGACCACGGACCAACGTTTTCAAGTCTAGAACTTA

AAATGGGTCAAATTTGTGAAGCATTGAATGAAAGAGACAAAGGCAAACTTCCTAGTCAACCCCAACAAAATCCAAAAACT

GTATATCGTTGCAGAAAATTCAACTCAAAATGAGGCCTCCAATAATCAAGTAAAAATGCGTCACCACTCTTCGTAGTGGT

AAAGTTGTTGAAAACAATGTATGTACCACAAATCAGTGAGTCTACATCATCGACTCAAAGTGCATAAAATACCACAGAAA

ACACCTATTGTTGAAAAAATGGCAACATAAGAATTCCCATAATTCTAATGTCTCTCCACCATCTCATGTACCTGTTGCTC

CATTCCCACAAAGATTGGTTCAACAAAAAGAGAGAGTACAAACGAGTTAACATCAACATCCATTTCTTGATGCAATCAAG

CAAGTCCCTACTTATGCTAAATTCCTTAGCACGGTAACTAAGAAAGACCATTTTCCCGCTGCCCTTCATCTGCTAGAAAA

GTTGTCCGGACATAGGTACTATTGTTTCTTAGATGGCTAGTTTACTCGCAAAGATGTTGCATTGTGTTTGATGATGCTTG

TGTGCATGCTTGGGAGGAGCTCAAACTTCTTTTAACTTCTACCCCTATAGTACGACCTCCTGAAAAACCTTTCAACCATC

CATCTCGCGTAGTTAGAGGAAAATTTAGAACAGACAACCATTTTCCCAAAAACCAATATCTCTCTCCAACATCAAAATGC

TGATATTGAGATCCAAAACATCCCTACTTTATCTGATATACAAAAAAACATGATGGGAGAACATAACAAACAGAAAAAAC

TCTTATGGGAAAGGAGAAGAAGTGAACTGAATTTGAAATCCGTCAACGCTGAACAACACCCTCAGATGGTTGACCCCAAG

TGGAAAATTTACAACTTGTCCAGCCAGTCGGAGTATCATGATATGCAATGGAATGGACATCAATCAGCTAACATGAGAAA

CACTCAACATTCCCCAATGGAGGAGGAACAATTGACAAACAACACTACTAGTAATCTGGAGGGAGCTAACACATCTGCAA

CCAACCAGGTAATACACAACAATCTCCTAGTTAAATTTAACATTCATATTACTTTAGTTATGAAGATGAATTCTGGATTA

CTAAGGACTATAAACCTAGGAAACTATACTTAAACTTAAGATACATAATCACTATTTTCTCTTATCCTATCATATCCTAC

TTTTGGAAATTTTATTCGATAATAGCTTGGAACTGTCAAGTATTAGGACGTGCTGACATTATTAGATATCTAGACAGCAT

GCTAAATAAACATAACCCTGAGATCATTTTCCTCTCTGAGACTAAACAACAGTATAGAAATCTTAAGAATATCCTAAGAA

ACATTGGAGTCACTAATTTCTGGTTTGTTGAACCAAAAGGCAAAGCCAAAACTGCTGGAGGCCTTATTCTTGCTTGGAAA

ACTAACTTAGATGTCATTATTAAGGACTTCTCTGATTACCACATTAATGCTACCATTAAACCCCTGCATAGAGACCCTTG

GATTTTCACTGGCTACTATGGAAGCCCATTTACTATTAATGATAAGTTGCATTCCTGAAAACATGATTGCTATTAACTCT

CTACCTTGGTTGATCATGGAGGATTGTAACTTTGTTCTACATGAACATGAGAAATTTAGTCAACATCCGATTAATCAAGC

TGAAGCTTCTATTTTCCTAAGCAAGATTGAAGAAGCAAATCTATCTGATCTAAGCTATACTGAATGTCCTTTCACTTGGA

CAAATAAAAGAACTGGTCATCAACTCGCTGAACAAAGAATGGACAGAGGTCTAGCAAATGAAGCTTGGCTGGATAACCAT

CCCAATACCACCATCTCAAACCTCACTACCATTGGCTCTGATCAAAACCCAATCATCCTCAATACCAACCCCAATTGGAA

ACATGGTCAAATTCCCTTCATATTCTTTGGTCCATGGCTCAATCATGAAGATTGTAAAAACATTATAGTTGATTGCTGGA

AAACAACTCATAAAGGTTCTTTAGCCATAAAAATAGCAAGGAAACTAAGAGACATAAAAGTCAAGATGAAAAAATGGAAC

AAAGATGTTTATGGAAACATAAAGACTAATCTAGAAAAAAAGTATACAACATCTAGACTGGATAACTAAAAATCAGTTTA

AACTAACTAGGGGGCAGATATTAAAGAAGCAAGGCTTCAAGTTGAACACTGGCATCATGTTTAAGAAGCCTTCTGGAAAC

TAGAAGCAGAGATCAACATATCAAGCTGGGAGATATAAATACCGGTTATTTTCACATTGTTGCCAAGAAAAGATACAGAA

GGAACATGATTGATTCCATTCAAAAGGATGATGGCTCTTGGATACAAGACAATGCAGAAATCACCAAGACTTTCAGAAGT

CACTTTGCCAATATGGCCACAGCTGAAACCATTAACATCAACCCTTTCATTATACAGCTCATACCCACTACTATTACTGA

CCTAGAAAACCATACTCTTATCAAAACACCTGAACCTATGGAAATTAAAAATATTCTCTTTAGTATGGATGGAGATAAAG

CCCCGGGTCCCAATGGTTTTCCTACAAACTTCTTTCAAGCCAACTGGGACATTTTGGGGGAAGATATTATTCACATGGTT

CAGAATTTTTTCACAACTTGACATATCCTTAAAGAAATGAACTCCACTTTCATATCCCTCATCCCCAAAACTGAAAAACC

TACCACACCAGCTCAATTCAGACCCATCTCCCTTTGCAACACAACCTACAAAATCATTTCTAAACTCATTTCTCAGAGAT

TAAAACCTCTTCTACCCAAAATAATATCCCCATTTCAATATTCTTTCATTCATGGAAGGAAGATATCTGATAATATAGCC

ATATCCCATGAAATTATTCATCACATGAACACCAGAAGGGAAAAAAAGGAAGCGATGGTATAATAGGTATAAAGATAGAT

ATGGAAAAAGCCTTTGACAGAGTGAACTGGGAATTTCTCCTTCAAGTTCTATCTCAAATGGGCTTCAACTCTCATTGGTG

CAACCTAATCCATCAATACATCTCCACCACAAGCATGGCCGTTCTAGTTAATGGATCTCCAGGAAATTTTTTCAAACCGT

CTAGAGGCCTAAGGCAAGGGGACCCTCTATCCCCATATCTATTTCTCTTCTGCATGGAAGCTCTCTCCAGATATCTAACA

CATGCTGAGATCCAACAACAAATACATGGAGTCAAAATTTGTAAGGATGCTCCAGCTATTAATCATCTCCTCTTTGCAGA

TTGCTGCATGATATTCTGCAAAGCTAATATACAGGAATGTAACAATATCATACAAATCTTCAAGGAGTTTGGAAAATATT

CAGGACAGTTAATAAATTTCTCAAAATCCGGCATCTTCTTCAGTAAGAACACTCCTCCAGATATTGCTGATATTATAAGT

AATACTATGAAGGTCCAAAAGATTAACACAACAGACAAATACCTGGGATCTCCACTCTTCACTACTAGGAGTAAGGTTCA

AGCTTTCAAACCATGTGTTGAGAAACTAAAATTCAGATTTGCTGGATGGAAAACAAATCTCTCAACTGTTGGGAAGATGA

CAATGATACAATTAGTAACCTCCACATCTAGTATATACCAGATGAATTGTTTCAAAATCCCAAAGGCTACTTGCAAGGAG

ATCAATGCTATTCAAAGAGATTTCTTATGGAATAAAGAACAAGACAAATCCAAAGGAGTTTATTATTTTGCTTGGGATGC

AGTAAAAAAACCAAAAGAACTGGAGGTCTGGGGTTCAAGAGTATGGAACTTTTCAACATGGATATTCTTACTAAGATTGC

TTGGAGACTAATCTCTGAACCAGACTCACTATGGAGCAGCACTATGAAGGCTTCACATTACCCCCATACTGAAGTCATAA

GGTTAGATACAAAACCAAAAAACACTGACTCTTGGATCTGGAAAGGAATTCTGGATAGCATAGCCCTGATTCAGCAGCAT

TACATTTGGAAAGTGGGCAATGGATTGAAAATTCACATTTGGGATGACAGATGGATTCCAAACTACCCTAGAAGACTCAT

AAAACCTACCTTATGTCCAGAAAACCTCATAACTGTTAACCAACTATTCAATACCTATGGGGATTGGGATACCAACATTC

TAAATCAATGGTTCACACCTGAAATCCAAAAAATCATTCTTCAAACACCCCGCTACCAAAATGAGAAGGATAAAGTCATA

TGGCCCCTCACCCCCAACAACAAATTTTCTGTCCATTCCCTATATGACAAGCAGATCCAAGAAAAATATGCTATGGATGT

TCAAATTGCTCCTTGGAGTAAAATATGGGGTCTTGATACCAACCCTGCAATCCATCTATTCATCTGGAAATGTGCCCATG

GCATCCTCCCAACCAATGCCAAGACATCTAGCATCCTGAATTACATTGATCCCATATGCTCTCACTGCAAGAATGGTGTC

GAAGATATAACACATGTTCTATTAGCTTTCCCTGAGGCTTCTGAAGTTTGGAGAAAGCTATTTGGAGAACATCATAACCT

CTTCATCAACTTCAATTCTTTCCATGATTGGCTCAACAACTGTTTCCATGCAGGCAACATAAATGCTAGCTGGAATGAAA

CCTATGCTACCATATACTGGTTCATATGGAAAGCAATATGTGATTGGGTCTTCAGAAATATCACTCCAACAGCCTATGCC

ACTGCTCAAGCTATTACTAAACACTTATTCAGTCATTCCAGGATTCACCAGAAGCCATGGCATATTCTAAATAATCTCTA

TTTTTAGTCCAACAACAACAACTCTCCTCATGAGAACTACAGTCAACATAGACAAACATATGATAAATTTGGAATCACTA

TTAGCATGCACATTGATCCAGCTGATCTCTATCAGGCTACTAGTGATACTCAATCTTATACTAACTACTCTCTTGTTGCT

ATGTCCTATTCAAGTCAATGTATGGGAACCAGAAACTTCAAGGACTTTACAAATGGAATCAGCAGAGCATAACGGTCAAG

CAGAGGAGCTCAACTAGCCCTGACCTGGGAAAAACAATGCAGCAAGCTAACATCACCTTTGTTGCAGAGTCTGAGGAAAC

TCTAAAAGATATAAAAGCTCAATACCAATGGGAAGTTCAACAAAGATTCAAAGGTCGAAGCACACAACAGCATGGCAGCA

ACACCAAACGCCACACAATTGATTTTGTTCTAATGAAGCTCTCTAGAGTGACTACTCAACAAAACTATGCAGCCATCAAC

CTAGCCATACTGGCTCGACAATCGTCTCTTTTCACTGACTCTTCATGGAGAGGAACGAATGTATACACTATCATGCATGC

TTTACAAATTCCTCCAAAACGTTATGATATGAATTATTGTAATAACTATACTTATATTTTGGGAGATGATCCTGTTGCTA

GAGTTAGCACAAGCACCACAATGGTTGTAACTGGGAGTAGGCAATCTTCCTAGATGCCTTGTCTCTTGTTTTCGTATAAA

AAAAAAGATTGCTCCTTTTATTTTCTCTTTCTAGAATGTCCCCGAACACATACCATCGTGCGAAGTACCGACACACCGGT

ATTGGTGACATGTCGGTGTCCCACTGCACCGTGCATCGTGCATCAGACATATGACGCGCGTGGGACACTGACATGATACA

CGTGCAAAAATTGATACTCAAATTATGATGAGTTTTTAGTGTGTAATCTAGTGTAATCTAGTAGTATTTATGCTAAAAAA

TTATGACGAGTATTTATACTCAGAAATAGAAGTTTATTTTACTGGTTGGGTAATCTAGTGTGTCACGGAAATTAGTATAC

ACGTGAGCTAACTTAACACCATAAAATTAAGTTTTTTTAATATATTTGCATGATTAATATGTTTGATATACCATATCATT

ATATATGATGCCGGTCTCGATGTGCTTATCGTGTCTGTGTTAGTGCTTCCCGGCTCCTGATAAAACCTTTGTTTGTTTTC

CAATTTTTTTATTTTATTGAAAAGATCCAGCCAGCGTTTGCAATACACAAAACAAATATAAAGTTAATAAATAAACAAAA

TACATACTGTATATTATTCTAACTAAACATAACACAAGTGCAGTGAATAATTCCAACCGAAAGGGAAAAACTAATTAAAC

ATAATTCAAATAAGAAGGACCGCTATCCAACAGTTTTGTCTTCCATCTTTAAACCATTGGTGTGTGTTGATCGAGTTTCT

TTTCTGCATCTTCAACAGGTACAAGGCTTTCATGAACACAGTCCACATCATAACGATACGGATGACTCAATAACGGTGTT

ATACCAGTGATGTCAATCATCCGAAAACTCTTGTTTGTCCGATTATAGATATTGAGACTTAGGGATCCTCTCGGACTTCT

CGTGTCGTCCCGAATAAGTGTTTTTATGACCCATCTCATCTGTCCCTTGAACTGGATGAACACAGAGATTCTTCTCTTCT

TTGAAGGAAGGAGGAAGTATAATCGTTTCCTCGCTACCGGTGTTATTGTAAACTTTTCATCCCGCTTTAAACTATCTTCC

AATAAAATCTCAGGAGGACCAACGATGAAGTCTGGGATCTCAATGACTCTAAACCTTTCGGTCCCAATATCAAATGCCAC

AATCAACTCCTCATCTGGAGGTGTGATAGAATCTGCACCAGCATTCCTCCAGTAAATTGAACCGTTTGCATAAACACCAG

CAGATTCAAATAACTGAACCGGAGGCACCTCATCTATATTCCTCCATTGATTTCCTCGTCCTCCCACGGTAAGAACTTGG

CATATATGTTGGACCTTAGAACTTGTTGAGAAATAGGCCTTGGGTATCTCCAGAACATATACTAGTTTGTAGGTTTTAGT

CAAAGGATCAAATCCAAATCCATAAGTTGGCGTCTGAGCAGTATCGAAAGCACCCCTCCTCAGCTGCAATGGTACGGGTG

CCAACTGTGCCCACGGAGTAACTTGTCGAGTCCCGAGGTTGTAGATGCGAGGTTGTAGATGCGAGTAGCAGTATCTGATT

CGTCAATAAAACAGAACAAGCCATTGATAGGCTTTGGTACGTTTTCGTCAAGTACCCATGGAAGTGTGACTCTGTGAACA

GCCTCTCCACCTTCAAACCCTTCCTTGGGTGAAAAAGGTTGTACGGTTTTGCCGCAATACCTGGAGATGACAAGTTGTGG

TTTCGATCAACTTAGATGGGACTGGATGAAAGATGAATCCTCCTCAATGAATGACTTCCAACTCTTGGCTACACACTTGA

AACGCACAAGAGACTTGACAGGAAGGTTGACAAGTATATCTTCTACCGATTTGTTATCAAGAAGGTTTGAAAGTGTTTTC

CTTTGAGATTTGTCCATCTTCTTGTCCTCTGTTACTACTGATGCGAAACCGGCCAATCCCTTGGAAGACAAGACTGCTAT

TATATTCGGATCTGTTTCCCAATTCCTTTCAAACCAGAAGTCGGTTTGGGCCTGGATATGGAAGGCAGACATAAGAAGAA

CGGTTTTTTTTGGGACCATTGTTTTATTTTGGGAAAGACATTAGAAGTGAATATAGGTCACTCTTTATCTAAATATTTAT

ATTAATATCAAAATTACCCATTTCAATTAAGTTAGTGTAATGATTAGTTGTATCATTAGGTTAACATAATTAGTGATTAG

TGCAAGATTAAATCAAAAAAAAAAAAACTAAATAATTGAGAGAAGAAGGTTATCTTTTTGTGAAAAAAAACACTAAATAA

TTGAGTGATTAGTTTGGTTGGAAAATGTTTTTTGCGACGTAACTGAACAGATATTCGGTTGGGAAATATTTTGTGAAATA

ACCGAACTAAGGTTCGGTTGACAAATGTTTTTTGCAAAGTAACCGAACTATTAGGGTCAATTAGGAAATGTTTTCTGCAA

CTTAACCGAACTGATGTTCTGTTGGGAAATATTTTGCGAAATAACCGAACTAAGGTTCGGTTGGGAAATATTTTTTGCAA

CGTAATAGTTTGGTTAGGAAATGTTTTTTGCAACTTAACCGAACTGATGTTCTGTTGGGAAATATTTTGCGAAATAGCCG

AACTAAGGTCGGTTGGGAGATTTTTTTGCAACTTAACCGAACCCTAATGGTTCGGTTTGGAAATGTTTTTTGCGACGTAA

CCAAACTATTGTTCGGTTAGGAATTATATTTGCGAAATAACCAAACTGTTCTTCATGTTCATCATTTCCATTAGGTTCGG

CTAGTTCGACAAAGTTTTTCACATAATTCTTAGACGAACCTCTCTCTGCAACTCATTTTGATGGTTAATACTCATTTTTT

TCAATCGAATGAGGAACGTCAAGGAAAGATAGAAGAAGAAAATAATAATATTTTTTTTGAAAACTAGAAAATTTCGATTA

CCCCCTATTTTTGTTTTTGTTTTTGTTATTGTTTTTGATTTTGATTTTGGTTAGTAGATTTATTTAGATTAGATATATAT

GATTAGATTTATATAGTTATTAGGTTAATTTTAATTTAAATTAAAGTAAACGGTAAATGTGTCTAAGAAAAGATGCAATT

GAGAGTTCCCAAAATCCTCGTAAATTCCAAAACAAGAAAATATGTCTACAAGGGAAAAAAACTACAAAAAGATAACGTTT

ATATCCCGAAATGATGACGTACATGATGAACGCAATAAAAACAATTATTAGAGTGCACTAATAATGCGTTATAAATAGAC

GGTTCGTTAGCTGGACTGACATTAATGACCGGTCAACAACACTATATTCGAGTGTCGTGTAGGAGGGCACCCCGTCCTGG

GGACTCAGTCAATTAGTCATAGCGCATTAACTAGACTCAGGCAAACGTTAACTGGGAACTGGTCATAGCATGTTAGTCAA

GACGGAGGTTTCAGTCAACAGCCAGTTAACAAAACCGTTACGAAAAATCCATCGTGGGAGTAAAATCGACATAAAATCGT

TGCTAATTATATCGGCAATAAACAAGAGGTAGTGCTCAAGCCTCAAGAAATTAGCCGTCTATTCCGTGGGTGGATCGAGT

TTCTTCTTTTCTGAATCTTGTTGAATAGATAGAAGACTTTCATGAACAATGCGTACGTCGTAGCCATATGGATAGTTCAA

TAATGGTGATAAACCGGTGATGTCAATCACTCGAAATCTCTTCATAGTCCGGTTATAAATATACAAACTTATAGATTCTC

TTGGGGTCCTGTCACTAGTGGGAAGATTATGTGATTTCACGACAATCTCATCTGTTCCTTCCACCGGACTAAAGGAAAGA

TCATACCCGCTTTCGAACGAACAAGGAAGTGATATGGTTTCATCGGTCCAATTTGTTGTCTTCTTCTGCTCATTGTAACA

ATCATCATCAGAAATCCATAATTTTACGATATGCTGGTCTACTCGATCTATCAAAGCCATATGCCCATCTATCTGTGTTA

TACTGAACTTTTCTGCCCGTAAACCGCCATATTTTTTCAAAGTCTCCGAAAAACCAACGACGAAGTCTGGTATCCGGATT

ACTCTAAACTTTTCGGTTCCAACATCAAATGCCACAATTAATTCATCAGCAGGAGGGGTGCAAAAACTCGCGCCAGCATT

CCTCCAGTAAATCGAACCGTTTGCATAAACGCCATCAGGTGGTCCATATAACCTAACTGGTGGCACCTCATCTATTTTCC

TCCATTGGTTTTCTCCTCCCACTGTGAGAACTTCGCAAATACGGGTGATTTCTCTATTGAACACAGACCTGCCAGATATC

TCCCAAACACATAGTACTTTGTAGTTGTTAGTCAAAGGATCGAAACCAAATCCATACGTAGGCGTCTGCTCAACAAAAGG

ACCCCTCTGCAATGGTATGCAAGTTTGTGCCCATGGCGTAACTTGTCGAGTTCCGAGGTTGTAGATGCGAGTAGCACTAT

TGGAGACGTCAACAAAACAGAACAAACCATTGATAGGCAATGGTTTGTTATTAACAGCCCAAGGAATTGTCACTTTATGA

ACAGCCACTCCACCGTAATAAATCGCCGTGAGAAGTTGAGGAGGATGTTTTTTTGATCTACTTAGATGCGACTCTACGAA

AGATGGATCCTGAATGAATGACTGCCAACTTTTGGATACACATTTAAAACGCACAAGAGACTTGACAGGAAGATTGACAA

GAATATCTTCAACTGATTTGTCATCAAGACGGTCTGAAAGTGTTTTTCTTTGTAATTTCTCTCTTCTGTCCATCTTAGAA

AATTTACAAGGATCTGAAATTGAGATTGAGTGTGTTCGCTATCAGATTGCCCTTTTATATGCGTATGGGTTTCCTAATTC

CTTTCTAACCAGAAGTCGGTCCGGCCCCAGAAATGGAAAGCAGATTAAAGAACGGAAACTTTCTATATATGGTATAGTAT

TCACGCTAGTAAAATATGATAAAAAATTACTTCTAAAATATGGTAGAAAAATTGCACTGGAGAAGCTAAGATATAGTAAA

ATATGACAAATTGACCAGGTGAAACTTTGGTGGTCAACTCTAAGTATATGACATTCCAGGCAGGGGTGAGTAATTTTTTT

AGGAAATTACAAAATCTAAATTAAGTAGTTCATTAAAGACTGGTGTAGGAAGAAGATAAATACTGGAGTATGATAATTAA

TAGAGGGAGCTATTTTTACTAAAGGGAAAGCCGACTTTGGGGATCAGCAATTAATAATTCTTCACATTTTTGTGGCAAAT

GGATGACTCCTATAATCATCATCTTCCGAAGATCAAATCAAGGGTTTTAATTTCGCACATGTCATGAGTAACTACGGCAA

AATTAGATTGAAACAAACAAGGAGAAGAAGAGGTGCACCTCATTCGGTGTATCTAGAGTTTTACAAAATTAGTATCCGGA

AATCAGTGATCATTGAACATATCACACTAAGGGAAGGCATTAAGCATATATATGTGAAAACTGTCCCGTTTTTATCATAT

TGATGTTGCACATTGGATCTAGCTTTGGTGCATCTACTGTCCGACACAAACTTCGCTAAGATTTCATTTTGTTTTCAGAT

TTGTATCCATGTCTCTTTAACTAGTATAGGAGTTCCAAACACACCACAAGGAGACCTACTATTGTTAGGGGAAAAGCTTT

TTCTAAATCCACCCAAAAATCAGCTTTCCCTTTAGTAAAAGTGGCTTTCCCCCTAACAATTATCATAGCTAAGATGGTTA

TTGACGGAGAAGGATTGGATGGTGAGAAATTTCTCTTAAGAAGAAGTTTGGCACGTTATATCTCACATGGGCTGCAATAA

AGCTCAAAGAGGATTTATTGCAATGGGCCTTTGGAAAGAAGAAATTTCCAACAGCCTCTGGTTTCACAATGAAAGCTGTT

TTTGGGCATACAGGAAAATTTCTGGGCATACAAAATTATTTTTCCACCCAGGGTGGCTTTATAAGGGGTAGTGTAATTAT

TTATACATCCTCAAACAATTTAAATTACTAATATATCCTTACCCTTAAAAAAGAAAAACTAAAATCAAAATCAGTTTCCC

ATTATATACATCTCATCTTCTTCCTTTTCTAATCCATCAATCGAAGAACAAATAGAAGTTGAATGTGCAGAATCTGAAAT

TGAACCAAAAATTCAACTGATTGAAAAAACAACTCCTAACATGGAATTAGAAGGTATTTATCACAAACCCATCTCTTATT

TTATTGTTTATGATTGATTGAATTGGTTTAATTCGGAACAAAGTTGGTTTTTAACGATCACAGATTGATGTTCGGCTGAT

AATTTGTCTCCGTTATCAGCCGAACTGTTCATTATTTTTCACCCTGAAAGTGTTTATGAACAGTTCGACTTATATTAATT

TTTCACCTATCAGCCGAACTACTCTGTTTCATCAACACATTGAAAGTTACCTTAGTTCGGCACATATTTTGTGTTTCGGG

ACAACCGAACATTTTTAGATACTCCAAAGAAAAATTAATGGTTAGGTTCGGCTTGTACGATAATCATAATACGTGCCGAA

CCTAGTACGTTTATAAGGTTCGGCTTACAATTGAGTTGTGTTTTTAGCCGAACTTTTCTTCACAAAAGCTTCATGTAAGT

GTATATGAACAGTTCGACTTGAAATTTCATGCAATTTTAAGCCGAACCTATTCACAGAAAGAGATGCATAGGGGTTCGGC

GTATTCGATAATCATAATATGTGCCGAACCTGACACATTTACGAGGTTCGTCTATTACGATATTCACAATATGTGTCGAA

CCAATACAAAATTATTAACCCAAGAATGTACATGATCAGGTTCGGCTTATATTCTACTTATTCAAATAGCCGAACAATGT

GTTTCCAACAGGTTCGGCGCTTACGAATTTCTAAATATAAGCCGAACCCTAAACATTTTTTCTTCCAGACAGTTCAGTGT

AGGGTTCGGCTTGTTAGGACAATTCAAAACAAGCCGAACTCTGGTCTACAAGGTACGTTCGGCTCTTAATCAACACTTAC

AGCTAGCAGAACTGTTCATCCAAACTCGCCAAACCTCAGTGAAAAAACACAAACTTTTGATGATTTTAACCTAAAAAAGC

GAGATTAAACATAAGATATGTTGGAAAATTTCAAGGGTTTGTTAAAAGTCGAACTGTTAAACATAAAATAGGTCGAACCG

CGAAAGCTATCTAAATGATTACAATTCAAAGGTTTGTTGAAAGTAATCTTAAAACTAATTACTAATCATTAATCACTAAT

CAGAATTATTTTAACTAATCATTACTATTAACACTAATCCTAAAAGGGCATATTTTCCATTACCAAAAATATTTGGTTAA

AGGGCTTCTAATATTACTATTTACACTCCATTTTTGTTTTTACTTGGTATGCCCCGAAAGATTTGTGTATGCTCAAAAAC

AACTTTCTTCGGAATGGATTTCTTTTGAAACCTGTTTTAGGTAGGGGTGAGGGTGGGCAGCAAGGACCAACTCTTATTAT

ATTCCATACGGTCAAACGGATCTAAAAAAAAAAAAAATTGAGGTATGATTGACGTTTTATATTGGAGATTTTTTGTTGAA

ATATGACCACTGCAATTGAAGGAAACATACTAGAGATTCACTAAATAAAGATGCCAGACAATAAAATTCTGATTGATTAG

AAAAACTCACTGAACAAAAATAGCCTCGATACATTTGGGGGAAACAGAAGGGCAGATACATTTGGGAAACAGAAGGGAAG

ATACATTTGGTGCAACTCCTGTCATCAGATCTGGAGATGGGCAGATACATTTGGGGGAAACAGAAGGGAACTGGGAAGAC

ACACCAAACCGAAACAGACGTTGGACTGGGACTGAAAATTATGGCCTTCACTCAACGGAAAAGGCTAATTCAGGAACAAG

GTTTGGACCTCTACAGAACCAACAAAATTCTGCTGATGTGAATTCAGAGAACACCAACTCGGCAGGAAACCGCAATGACC

TGGAAGAGAAGGTGGTACAGAACTTGGAAGACCTGCAGCAAAATGGAGAAGAATATGAAGAGAGGCTGATTGAAACGACA

GTCCATTCCCTGGTTCAGGTGGCCTGTAGTTACGGAAGCATTAATCCGGCTCAACAGGAAACATTGAAAAATAATATGGC

TGCTCTCGTATTTAGTTACAGGCAGAGATTCAAAAAACAAGAAGAAGATTTTGAGGGGGACGACGGCGATGAGGAATACT

CGGACATGGAGGGCTCTTATAATGAGAACGAAGAGGATAATGGGAACGAAAGGACTGTGACGACAACGTTTAAACATGAC

TGTGAAGTTAATAATACTGCGGAAGAAATACGTATTGCGAGGGAAGAAATAATTTTAAATGGATTTTAAAATTCTGTCTT

TGAACATTAGAGGCATGAATAGCTTTGACAAGCAAATTTCAGTTAAGAATTTGGTGGCTTCCCAAAAGTGTGTGATTTGC

TTGTTACAGGAAACAAAAATTCAAAGTATGTCTTACTGGTTTGTGAGGCAGCTGTGGTACGATGACAACTTCGACTGGGC

TTACATTCCTTCGATTGGCAGTAGTGGAGGATTATTATCCATTTGGGATATCAGTAGACTGGAGAAACTGGACGAGAGAA

CGGGTTTCAATAACATCACCACGGTCTTATCAACAAGCATCAACGGGTTTCAATGGGCATTGACAAACGTTTATAGCCCT

TGCGAACAGAATCCTAGAGCGGAATTATGGGAGGATTTGGAAAATTTGAGAAGTTGGTGGGTTGGTCCTATTTGTATTGG

GGGAGATATGAATGCTGTCCGTTCTAATGAGTAAAGGAGTCGTGGAGAGGGAGATAGCCGTAATACGGCATTCCTAAATA

ATTACATCCTAGAGAATGAGTTAATTGATCAGCCTTTGATCGGTGGCGCTTTCACTTGGTCGAACAACCAAGCTAATCCT

CTTTTATGTATGCTGGATCGTTTCCTTTTTAGCCACGACTTCGAAGCGGCTTTTCCGAATGCTCTACAGGTTGTTTTGAC

AAGAACTTCGTCTGATCATAACCCGTTAATGGTCATTTTGGAACCGGTAATGACTGCGAAGCCGTATTTCAAGCTGGATA

GGATATGGACAAAACATAAAGACTTCGGCAGTAATGTGAGGCGTTGGTGGGGGATGATGAATTTTGTTGGTAGTGCTAGT

ACTCGTTTCTTTTTGAAGATCCAAAACTTGAAACATTTAATTAAGCCATGGAGAATTATGGAATTTGGAAATTTGGGAAA

GGAAAAGTCGGACCTGACTAGCAAAATTCACGCGCTTAACATTTTGGAGGAAACATATGCTCTTTAACATTCGTAGCTGG

AGGAGAGAATGAGGTGTAAGCTGAAGTTAAGAAATGTGGAACTAACGGAAGCTAGGAAGTGGCATTTGAGGTCAAAACAA

AACAACTTCAGATGGGGAGATTCGAATACGAGGTACTTCCATAGTATAGAAAGTGCAAGGAAAAAGAGAAATACAATTCT

AAAACTTCAAATAGGAGGGGTGGATTGCTTTGACCAACTTTTCATAAAGGAAGAACTGCGAAGTTACTACAGTAACCTGT

TTTCACAGGGCGACGAGGTGAATTCGAATATGGAAAATTTGCAATTTCCGAAAATTATAGAGGCGGACATATTATGGCTT

GAGCGTGATTTTTCGGAGGAAGAGGTTTGGGAGGTGGTTAGGAAAATGGGGTGTAACAAATCTCCGGGTCCGGACGGCTT

CACGGCGGAATTCTTTAAGAAGTGTTGGCCGACGATTAAAGAGGATTTCATGAAGTTGGTCAAAGATTTCCACAGGTATG

AATCAATCAATTGGCGCTTCAATTGCTCTTTCATCACACTTGTTCCCAAAAAATAGGACTCTTGTACTACTAAAGACTAC

AGACCTTTAAGTCTGATTGGTATAGTGTATAAGATTTTATCAAAGTTACTTGCGAATCGTCTCAAGAGGGTGATTCCAGC

TCTAATCTCGGACTTCCAAGGGGCATTTATTCACGGTAAACAGATCCTAGATGGTGTTTTAATTTCAAATGAGTGTGTGG

ACAGCAGATTGAAGTCAAGGAAACCGGGAATTCTGTGTAAGATAGACATGGAGAAAGCTTTCGATAATGTTAATTGGAAT

GCCTTATTCACAATTTTACAAAAGCACGACTTTGGAACCAAATGGATCTCTTGGATCAAATGGTGTGTATCTTCTACACA

TCTTTCTCTTCTTGTTAATGGTGGTTCGACAGAAAAATTTAAACCGGGCAAGGGTTTAAGACAGGGAGACGCGCTTTCGC

CGTATTTGTTTCTTTTGGTGCCGGAAATTCTTTCTAAGCTAATTAATGAGGCGGTGGAAAGAGGTAATCTCTCTGGTTTT

CAGGTGGTCAATCGGGGTACTATCATATCTCACCTTCAATTTGCGGACGACACTTTGATTTTTCTTGATGCTTCGGTGGA

GGAAGTTAGACGTCTTTTTATCATTTTGTCTGTTTTCGAAATCATTACGGGTCTGAAGCTGAATTTAAAGAAGAGCACTA

TGATTAGTGTGGGAGCCGATGAGGTGATTGATACTTTGGCTAGAGAGCTAAGCTGCAAGACAGAGAAATTGCCGTTTACT

TACCTCGGGATGCCGATTGGTGCTCATTGGCGTAGCACTTCAGTGTGGGATCATGTCTTGATTAGAATGGAGCAAAAGCT

GGCGTCGTGGAAGAAAAGGCAACTTAATAAAGCAGGTAGATTAATTCTTATTAAGAGTTGTCTTGCTAGCTTACCAATTT

ATTACTTATCTCTTTTTCACTTACCTGTGTCTGTGGAAAAAAGAATGATTAAGATCATGCGCAACTTTTTATGGGGTGCG

GTGGAAGGTAGAAAGAAATTGGTGTGGGTTTCTTGGAAAAAGCTAAGTGTTCCAAAGGTGAATGGGGGTTTAGGTGTGAA

AAATTTGAGGAGAATTAACCATTCTTTGTTGATAAAGTGGATTTGGAGATTTTCGAAGTCCAAAAATAGTCTGTGGAGAA

GGTTAGTTAACGAAGAAAATTTCCATAATAATGATTTGTTAATTCCGGATGTGGATAATAAGCCGCATGGTAGGAGTTTG

TGGAGGAACGTTACTAAAATGGTGACGATGGTTCAGAATATGGTGACTTTTACGGTAAGAAATGGAAAATGCATCCGCTT

TTGGAAGGACAGATGGTTGGATAGGGGGATATTGCAAGACTTGTTTCCTGTTATTTTCAAGGCGGCGAGGGCTAGGGATG

CTACAATTTATGACATGATAGAAAATGGAGTTTGGGTTTGGGACTTCAAGTGGCCGTTAAATGCAAACGAGCAACTGGAA

TGGGACCTTCTAAGGCGTGACTTAAATCAGGTTCCGATGCTTGTAGAAGAGGATGATGAAATGGAGATTTTGGAGAACTT

CAATGTTTCAAGATGTTATGAAAGACTATCAGGCGATCTGGAAGAGTGCGGATTTGATGAATTCCTTTGGAGAAGTAACA

TTCCAAACAAGGTGAGTTTCATCATTTGGGCTACTTTTCATAATTCGTTACCAACCAGGGATATGTTGCGTTACCGGGGT

GTAAATATGGATAGTGAGCAGTGTGTCTTGTGCAACGAAGTCAGGGAATCGGCAAATCATATGTTCCTTCATTGTGAGTA

CTCTTTTCAGATTTGGAATTATTTCATCAAATCATTTAGGATAGCTTGGCCATTACCAAATTCTTTGCTTCAATTGTTCA

AAGCGTGGTCTAACAACGTGTTGCAAGGGAGAGGTAAAGAGGTTTGGCAAATCGGTAGTTTTATCGGTAGGTTTGGCAAT

TGGTAGTTTTATGGTCTTGCGACAAAGAGACTTTTAAGTTTGTCGATCCTAGTTTAATTTGGAGTAATTGGGATACTCTT

ATGTGTATGTAGACTTCTTTGTATTAACTTGGGTCTAATCCCTGGTTAATTTGTCCTTTTTTCTTTTTAATATAATTTTT

CTCTTCAAAAAAAATAAAAATAGCCTCGATACTTACTAACAGAGTACATCATGCATTGATTGATACGGAAGGTATATTAG

TCAACGGATTTGGTAAATGGGCCGTGATCCATTTAGCTAACCCGCGTATTTTTGTGATGAAATCTCAGCCATTCAAAATC

ACCACATCCATTGACTATGGGACTTTGTGATGGACTGTCATTTCAACCGTATGAGCCGATCCTGCCTCATGACGATAATG

TGTGACACGTACTCTACCACACATCATGAATAATACCACCAACATCGGTAGTCTACAGGCTACCTAGGTGTACATTAGAG

TGCATTTTCCCTTCTCTTTTTATCACCTGAAAGTTCGATTAAGAATGTGTTTGTTACTTACAGATTCGATGTTTGAATCT

GACTCGGTGAGTCAGATGTCAGATCGTTTGTTTTTTATTTTGAATCAGATCGCGAGTCAGATTTGACTCAAACCCATTTG

AATTGAGGAATAAAATAATCCTGACTCACTGGACCAAGCCACTCACTCGTATATTTTTGAGTTAGATCCAGATGAGTCTG

GTGATATCAGTCAGACGAATCGGGTGATAGCATTCAGATCCAAACCAGACGAGTCAGATTGAGACTGAATTGAAAAAACA

AACGAGGTGTAAGGCAAGTGCTGTGTGGCCGTGTGGGTGAGTTGAAGGAAAATAAGTAGGACGTGGCAGCTTCGGGACCA

CATTTCTTGGTGATATTTCTTAAGCATCGAACAACAACGGAGGAACACATCGTCTGTTCTTTGTCAAAGAATTTCTCATT

TCCTTTGTTTACGAAGAATTAGGGGATGTTCTTGAATACCAAAATCAGTGGAAGGAATTAGGGTTGCTTCCATTGATTCA

TGTCTGTGTACGAGAAGATTAAGAAGGGGTTTAGTTGATGGTAGTTGTGTAGACATGAAAGATGGATTAAGTTCCATAAT

CTGCAGATGTTTCTAATTCATAGGCTCACTGATTTGCTTTACAATCAGAAGATGGCTGAACTGAATCTGCAATGGGGTTC

GACTAAAAAAGAGAAATGGAGCTGAATTTGGTTGGAGACTAGTGTGTGTTCGAGAAAATGTCGCAGTGGACTCAAATAGG

TGAAGAGATTTGAATGGACGGAAATGGGCAGTGACCGGACTTCGGCTTTAAGTTGTAGAAAAGTGATTTTGATATTGAGT

ATTAAATTAGTGCACACATAATTCGAATTACAAGGGAAAACCTAATTAAACAAAAGGGGTGATCAGAAAAGCATGTAAGG

TGTCTGCTTCCATCTTTGAACCAGTGGTGGTGGTCTGGGTTGATCAAGTTTCTTCTTCTTTTCTGCATCTTCCTGAATAG

GTAGAAGACTTTCGTGAACAATGTGCACATCATAAAAGTATGGAAATTCCAATGCCGGTGTTATACCAGTGATGTCAATC

ATTCGAAGAGCCTTGGTAGTCCGGTTAAAAATATACAGACTTATAAAATCCAACGGACTTCTCTCTCCGTTCGAAATAGG

TGATTGTATGACTATCTCGTCTGTTCCTTGGACTGGATTAAAGTAAAGACGTCGCACAAGAGACTTGACTGGAAGGTTGA

TTAGTATGTCTTCTACTGATTTTTCATCAAGACGTTCGTTCTCTCTTCTTGTTTTGTCCATCTTAGATCTGGTGAAATCA

AAATCTTACTGATTCAATTCAAAAACTCCAGAGATATTATTTTAGATTGATGCTGGAATTATTAACTCACGAGAAATCAC

TATGCAAAAACTTCAGCAGAGACTGCCCTTTTATATGCGGATTGGTTTCCCAATTCCGTTCTATCCACAAGTCGGTCCGG

CCTCGGATATGGAAAGCAGACATAAGAACGGAAACCATCCTCTTGTTTCTATATCTATGATGATATGATACACTCCGACC

AGTGTAGTTAATCTCGGCCGGTTTCGGAAGCTAAGCCGACGAAATGCCATATCTCGGCGCGACGAGATCCTTGCGATTTT

TCCGTTTGAGTACCATTTTACTCGTGGAACCTCATCATTTGAATTAAATGCAAAGAAATTTCGTTAATGACTTGTAGAAT

CTGTAATTGAAGCTTATCCTATTCACTGTACAACTGCTTCCACACGTTAGTATTCACTGTACATCGCGTCGTGCTATCAA

ACAAATGAGTGCAATATTTTTTAACTCATTTCACTTCACATAATAGCAAATCTGACGGTTTAAAGATTCGTGCTGATAAA

CACAAGTCTTTCATTCGTATTCAACATCTTATTTCTGGGACTAATTTTTGTCAATAATTGGAGTCTACAAGACTGCACAA

AGACCCGCTGTCTGACAGCAACAAACGCAAAAGAAACAAGGCCAGGGTTATAAAAAACCTCACATTCCATTTCAGCCTTC

CTAGCTCGCATTGAGACCTTTCTATTTTTCCGCTCCATGGTTAAATGGCTATTATACCTATCAAAATCCGCCTCTGAACT

GAATTCAATTCCAGAACAACCTTCATCCGTAACTCTTTCTTCAATGTTCCTTCCATCCACCTTGTCTGTATTCTTGCTAG

ATAAAATAGAGGGTACTCCCTTCTCTTTACCACCACAAGCTAGAGACTATCTCAGGGGAATACTTCTTGCCTTTTTCGAC

TATCCACTCGTCCTCATTCTTGTCGCTGGCAGTTTTCTCTGCCCGAGGCTTGGTAATAGCCTTCTGCTTACAGGTACATC

TGAATTCAACTCAACATCCCCTAAGATAGTCCCAACACCAGCTTTTTCCACAAGTTCATCAACTGACCTTTTAAAACCAC

CAGTTGCAGTCATACCTTGCTTTGCTGGTGTTGTTTTCCAAGTCACTGTCCCTTCTGGCTTCTTAATGCTTGCAGTCTTG

GTTGGACTTTTCGCCTTTGCATTGATCCCTTTGGACTATGTCCTGTGTTTGTAGCGGCTTGAATCGGAATAGCTGTCTCC

ACACTAATCCCCAAATCAACCATGCAGCTGAATTTTGCTCAGAAGCTCCATCTGTGCCTACCCCTTCAACTGCTGCATTG

GTGACTAGCGGTTCATTAACTTCCAGCATAGTTTCTGCAATATCCTCTCTGTCCATTTCTTCCCAGGTCAAGATCACTGT

CTGTTGAAATCTCACAACTACCCGCTGAAAGCTTGAGAGTGTCATCCTGATTTTTGTCAGCTAACTCACTCTCCATTGCT

CTGGACAAAGATTTCTCTGGAATCTTGCCATGTTTCTTACGAGTCCATCCATATTTCAGCTTCTTTGTGCAGTTCTTGGA

GGAATGTCCAAAGACCATGCACAACTCACACTTGGGTGGTTTCCATTGAACCATCCTGAATGAATTTCACGACCTCCAAC

TGAAACACTAAAGTTCTCTGTACGGATATCTTTTGCAGCTAGACCCTCATTGTTATCATGCTTTACAGATACAACTGGCA

TCCCTGTTTTGGCTGCTTCCATTTCCGCCAAATGCATCAGATAATGTGCCTGTGTGGGTGAAAAGGGACAAAACAAAATG

GAAATCATAACCAACTGATACATTGTTAAATTAATCGGTAGTTTGACTCAGATGCCGTCCTCTTACCCCTCTTTGCCGCT

CATCCTTACTTTTTTTTTTCTCTGAAGCTTTGCTGCATCACAATCAGTCAACGTAGGACCTTCCTGCAGCACCCCAGGTG

GTTACTTTGGAACTGCGTCCTGGTCCATTCCATCAAACATCCGCAAGGGTGTGGCAAGACCTATATTTGAGAACAAATCA

TTCAACAGTCTTCTAGTACAACAAAATAACATGCAAGTCAATCCAAAAACAGTACTAAACATAAAATAGTAAGCAAATTG

GTTGTACATATCATATTGGTTGACATCTAGAATGGTGGCAGTGGCAGGTAAAATTAAAATCGGTTTCCCCACATTAAAAT

GAATCAAAGGAATGTTTTCAAGGGATCAAATTTCTTCATAGAGCTAAGAAATTCACATAACACTCGTTTGAAGTTCAAAT

CCTCTCAAGAGCAATTCATATACTACTCAATTGCACAATTGCTTACAACAGCTCCAGAGCTTATAAAGGTGTTTAGTAAA

TTTTGAAACAGCCTGCTAAACTAGATATCCTTCTCCGTTAAGATTTGGCGGAACATAGAACAAACGCTACAAGTACTAAC

CAAAAATATGCAACACCGAAAACGTTATTTTGTCTCTTCCGCAAAAAATTATAAAACTGGTTAAGCTTAAGGATTAAACA

ACCTTGGCAGAATCACAAAGCTAAGGAGAATCAATCTCATATACTACTTCCAATCTCACAGAACTTGTAGAGCATTATGA

TGGTAGCATCTGGACAGAAAACAGGCTCAAAAACTAATTCGGGGAAAAATTACAAAGACACATTCTATCATTTTCAGAGA

CAAGCTTCCCTACAAACAAGAATCATTATCTGAAATTTTTGAAATGAATCCGGAGATCCAACAACCCTTTCATCTTAACG

ACCAATACATCGAATTTTCCGCCACAATTTCTACTCCAGAGCATGTAACAGTGTTGTCCCTGATTGGGAAAGCCTCAATC

TTATTGCCCAGTGCAAGCAAATTGATAGCTCTCACCATTTTGACTTGTTGCAAGCAGGACAACACACATCTCAACCTAAC

TATTCTACATGTTCTTATAAATGCAAGTTATAACTTTGAAAGCAATACATCTACTAGTAGTTCTTACACCCAATTGTTCA

ATTGACTACAATAGCTCCAGAGCTTATAACGGCGTAGCTACTTTCGTAACAGCCCCTATCTATTAGCTCACTGGAAACAC

ACAGAGAGCTCCCACCATTTTAACTTGTTGCAAGCAGGAAAACACACATCTCCACCTAATTATTCTACTTGTTCTTATCA

ATGCAAGTTATATGTCTCTAAGAACATTCAACAAACACCTGAATTGCATTTTCATGGTAGAATCTTACAAACAAATTCAG

GCGGAACTAAAAGAAAATTAAGGGCTTCATTTTCGGGTTTACCTCTTCATACTCCTGGGTAAGATTAATACGTTCATCCT

GAGTCATATCAGATCTTCAAACAGCCATAGTTTCATATTGTCTCAAACCAGGTGGACATTGAGGTTCTTCTTTCTCTTCA

GTAGCTCCTAAAATACCACCAGATGATGGTGGTTCATCATCTTCTCCAACGCCTTCTTCAAAGAATGAACCTAAAAAATC

TAAAGTCGAGGCTTTAACTGATAAACTTTCATGGGTTCTTCTCAGAATCTCAACTTTGTTTAAATTACTTTGAAAGGGAA

CTTGAGGAGTGAATGAAACAAGAGATGATGAAAAAGAAGTAGAGAGAAGTTTGGGTGATGAGAAGGAAGTTGAATCTAAT

GAAAATGATGATATTGCCATGGCTGAGAAAAGGAGGAATTAATCAAAGGGTTTAAACTTTAAAGCTTCACTGGATATTCT

TTTTTACAAGTGAAGTGAGAAAATGAAGAAGAAGAAGACTATAAAGTTGAAGCTAATGAAAATGATGATGTCGCCATTGT

TAACGAGGATTGAATCAAAGGTTTTCACTGGATTTTTTCAAGAAGTGAGAGGAAGAAAACTAGAAAGTTGTAGGCATGGA

TTTGTTGGATTTTTTTTTTTTTTTTTTTTTTTGAAAAATGGATTTGTTGGATGATAAACTTTCATGGGTTCTTCTCAGAA

TCTCAACTTTGTTGAAATTACTTTAAAAAGGGAACTTGAGGAGTGAATGAAACAAGAGATGATGAAAAAGAAGTAGAGAG

AAGTTTGGGTGATGAGAAGGAAGTTGAATCTAATGAAAATGATGATATTGCCATGGCTGAGAAAAGGAGGAATTAATCAA

AGGGTTTAAACTTTAAAGCTTCACTGGATATTCTTTTTTACAAGTGAAGTGAGAAAATGAAGAAGAAGAAGACTATAAAG

TTGAAGCTAATGAAAATGATGATGTCGCCATTGTTAACCAGGATTGAATCAAAGGTTTTCACTGGATTTTTTCAAGAAGT

GAGAGGAAGAAAACTAGAAAGTTGTAGGCATGGATTTGTTGGATTTTTTTTTTTTTTTTTTTTTTTGAAGCATGGATTTG

TTGGATGGATGGATGGATGATTCAACTGGATAACGCTTATACCGAACTTTTATTTATTCTTACACAGAGGTAAGAAGGTG

AAATTGTCATAAGTGTTATGCGGTCCAACTAGATTCCTAGGGTTATATATAAAGTAAACGATGTAATACTTGTACCCTAT

TATTCTAATAAGATATTCTTCTTCTTGCCTCTACATATTATTCATCCACCTCTGATTCTCCTCAAACACGTTATCAGCGC

GAGTTCTACCGTGAAGCTAAAGTTGAATCAATGGAATCGTCATCGTCAAATCAGATAAGTTGGATTTTTTTTTTCTTCTC

TTTTCCCTAAAATAAAAATTATAAGGAATTGATTTGATGTTTGAATCTGTTTTATACCATATTGATTTGATTGGTAGTCG

TGTTTTGCTTTGTCTCGCATACACTATGATACAAGAATTTTCTTCTTTTGACTATTTTCTCACATGATGGATTAACAAAT

CTGTGTGCCTGGTTTGTCTGCCGTGTTGTGATTTATTTTTTTTAATTTTTTTTTCAACCAGACATGATCTAGGTTCTTTA

TGTTATGTATGAGCCACCACTATTGATTCATGTTTCCATATAGCATGATGTTAATTTACCATGAACATAAAAACAATATG

AAGAGTACAAGAATTTTTATAAACTAATCAAGACTTTGTAGAGATTGATGTTTGCACTGTTTGTTTCTCATTTTATTTTT

GATTAATCTATGTAGAAAATAATAATAATAACAATAGTCAATAGATTCATAAATTAGATTTGATTGCAAAGTTGTTGACC

GTAACGTATGTACCGTATCGCCAAACAGCTATATTATTTATTTTAATTTCATTTTACAGAAATCGTACGTTATCTTGCTA

GATTTTTGTCTTTTTATTCTTTTTAACCATGTTATGATAATTATCCCGTGTACTTTAATTTGATAAATAGTACAAAAAAA

ATAATGTTTGTATGTGATCATGTATATTCTTTCAAATGTACTGTTTTTCTTTTTCTTTTTCTTCAAGCAATCATTTTCTC

CATATTCCTATAAGTCTGTACTACTATAGTGGTCAACTATTAATGGTGTTATGATCGTTGATTGGGATTTTGTTATAATG

GAGTGATTATGTTTTAATTTAATTTATTCCTATGTTTAAATTAACATAATATATTCATCCGTATGTTTGTTTGGGCACGA

AACTTGATTCGAATTTTTTCTCTAATCACAATTTTTATTTTTTTGGCGTCGGTTAATGACATTGATATGTTGACCGTCGT

TTGTAATTCTCTGGGAAGGAGAAGAAGGAAAGACTGACCCAATCTGGTCCATCCAACTGGTTCTGGTCCAACCCAGCCGA

CCCGGTCCATCTGGTCCGCTCTAACCCGGTCATTCCGGTTTAGCTGAATCCTATTATGTTCTGGTAATGGGGCATGCATG

TACAGGTGGGTTCTGGTTCTGTATGTGTGGATAAAACCAAATTTTGGCTTGAGGTGAATGGTGTCTTTTGTTCAATTGGT

TGTACCAACTCAATTTCAATATATTCATTTGGGGCTTAGAAGTACTTGAGCACCATTATTGGATACTTGATATTTTCCCA

AAAATTTGAATGTTCCTTGGAATTTTCAAAACATATGTTCAAATAAATCACATGTATTCTAATTATTATTATTTTTGTGT

GGAAGAACTCCCAATTATTTTTATTTGAGTCGGAAAATTTTCATTTCTCTACTTCATCCATAATACTAACGAAACGGTAT

ATGTTGAAGTATGACAACAAGAGAAATTTATTTTTTTAGTAATATATTCAACCTAAAGTAATTGGTTGACATGTGTAGTG

AGATTCTCTTGGCCTAATGAGATAAAGAAATTTCTCAAATTAAGATGAATATAGAAGAAAAAAAATTAAAATGGAAAGAA

CCCCAAAGTAATGAGGGTCAGACGCACCGACTCATATAAAGCATGTGAAAAGGGTTAGACGCACCGACTCATATTAAAAA

AAAAATTACCAGCAGTAATGATACCAAAGTATAGGTATCAATTGAACATGGGATCAGCTACAATGTAATTCTAGCTCCCA

TCATAAATATAAGAGTTGGAAATGCACCATGGTCATAGGTGTTAGTATATACAATATAATGTGTATGTGTATCATAGTAC

CAACCAATTTTCACATCAACTTTAATGGCAACAAGAAATTTGCCTAGCCAATTGATTATCTTATTCAGTTGAACTGGATC

CAATGTTTGTACTTATGGAATGAAAACATGAGACATAAATTCTCCAAAGAACATGAGATATATTGGGTGACACGTAATAT

ATATTCACTCTAAAGTACTTGAATTGAATCCAACTTTTATTATGTAATAAGGCCACACCACCTGCTAAAACTCTCAAGAC

CCAAGTGTCTAACTCATAATTTTTTTCTTCCATTATCTTAAGGAATTTAAACTAGTTGTTGAACTACTTCCTTATAATAT

GACTATGATGATTGGATTATTGAGCGAAGTATTGTTCAAAATTTGTTTCCAAGATGAAACAAAGTCATAAAAGCTCTATA

TGTACTGAGAGCTAATCACCTTGTTAAATTCCAAAGAAATCCATGAGAGTGGATATCATATGGAAACTCATAGTGATGAG

AATGTAATTGATTGCATCTTTTATGCCTCTAATATATTTGGAAGGAAATATATTTTAGAGAAACTAATGAGTCAATCTAG

TGCATTGTAATTTACTATAATATTTGGATTATTGAATCATATTGACTCCGATGATGAAATGTTGGAAATTGACTCACACA

GGCTTTGGGCATAACCATAAAATAACTTTGGTTGTGACATGATGATCGTTTTGAAAGGACTCATGCGAACATCACTTTAT

TTGAGTGAACAAAAATGGGAACAGTTTTGACAGAATGCGAAGTAATGACATATCTCTCAATAAGTGTTTTCTAAAGTACT

AGATGCACAACCCCCTCCTCCCTCACATTATGCTTTTAAGTAAGAGAGCATATCACTCATTTTTACAAAGTCTTCAGTAA

AACAGGATAGAGACCATCCTAAGATGCAAATGGTAAAAATTCCATATTACAAAGAAAACAAGGCGAATTTTAGAAAAATA

TTCATGCAAAATGCGGACCATTAAAGTGACTGATGGATCTGGTGAATGTTTTAACATTTTGGACTAGTTATAGTTCCCAA

GAAATTTGTGTTTGAAAACTCCTAGCACATATTACACAATGCAAAGTGCAAAGGCTAACCACCCTTGTTAATGCTAGAGA

ATTTACATCAAAAGGACTTGATGAATATTGCATGTTTAATAAGATCAATATAGAACATCTTATACCCAATGGTCTCACGG

AGGCTACCATCAAAGGTTACAGATGGTGTCTAAGGAATAGGTTATGCGTACCAACCTACTTTTTATTGCTTCGGGGATAT

GCAATGTTACACGCATCTTACTTAATTCGTTTTTATAAACCCAATATTAGTTAACCTTTTTTGCGTACCAGTTGGTAACT

GGATTTAATCCTGACATTCGTGTTCTTACGCATTTTTGGTTGCATTATATATGTGAAATTACGACTCCACATCGTACTAT

GATGGGTCTTCAAACCTGTTAAGTAGTTATGTTGGAAATGAGTTCCCAACAATTATCGGCATTTTTGAAACCTTTGGCAG

GAGATCTCTTACCGCTAGATTTGCGGGTTGTCATTTTAATGAGACATTCTTCTCGTCGTTAGGGGGAGATAAGAAAAATT

ATTTTCTAAAGGAACGGCTGGAATTGTCGTGGTGTGTTCCCACTATGTCTCATGTTGATTCTTGTACTCCACAAATTTAA

ATGTGAAGTGACAAAGAATAATCGATTTTCAGAATGTACACTGATGTCGCTAAAGTGACAAGATCACACATTCCAGCTGC

AAATATTCCTGTAAGGTTAGAAATCCTCAGTATGGGGTACACCATAGGCTAAGGTGTTGCAACTACACTTGGTGGAAGTG

TAGTTAAGGCCGTGGCTCCATAAAGGAATATGGAGATGCCACTAGGTTCGATCAATGCTCACCTAGAAAGGAAATAGATG

AGTAAGGCACAACCTAATTTGTATCATTACTGAAATCATCTCATTTGATTGTCTCTGACTATGTCCATGAATCAATACTG

GAGGATGCTCCGAAGTTTAACGATTCCAGAGAACAATGACATCCAAAGTGGATAATGAGAACACACACGAGTCAATGGAA

AGATCGTGTGACCATATTAATGATTATTATTTTCATATACAGTTGCTAAAGGAAATCAAGCACGATGAGATCGAAGCATG

CTTCTTATTGTTTGAATATCAACAATAAAGAGCATGTTTGGCCTACACAATCCAGGTAGAATTTGATTCTTTGAAAAATA

TATAGGTATTTGGTGTGGTAGTGCTAACCAACCAAGTGTAAAGCCTGTGGACATACATGATTATTTGTCACAAAGCACAG

TGAGAATAATAAAGTCGTAATGGTATAAATACTCTCCTTGTGGCGCGAGGTATCTCACAAAGCACTGGAATTGTTTATTT

TGTTGTAATGAGCATTATAGCATTTTGCTACTTAGTTAGCTTGGTAATTTCAAAAGGACTTGAAATGCAGCATATGTATG

TGGTTGTTACTTATCTCTGAAACAATCAAGATATAGAAGATATTTACAAAATTAATTAATAGATTTTTGTTGCCCAAACC

AAATGACTCTAAACCACAGAGTGCGTTTACAGTTAGATAGAACGCTTACTTATAGATTCAAGCAATCCGGCGGATGTGGT

ATACCCGTCTAAGTGGCTATTTGATTTGGAGGGGATAGACAAATAAGTTATTTTTCCTTGCGTATTCACAAGAAAGTTCC

TAATTTAGAATTGTAGCTATCTATGTCGATGGTACGGACATGATGGGTACTCTTGATGTAATAAGAGACCTTAAAAGCTA

TTTGAAATCCGAATTTGAGATGAAAAATCTAGGGAAAGCTCGATCGAATACCGAGCTTGTGGTATATTATTACACCGGTT

TGCATATATCTAAAATTGTCAGGCAATTTAACAAAGACATGCATCCTGATAGCACTCCCATGATTAGTCGAGGTTCAAAT

GTAAGTAAGTGACCATTTCGTCTAAAGGAAGATGACGAAGATGTGTTGGGAGATGAAATTCCCATATATAAGTACAATAG

ACGCATTGTTGTACTTAGTATAATAATGTACTCGATAAAATTTTGCATCCTCAAGGAACTTGTTAGCTAGTTATAGCTCA

GCACCAACGCAACGTCATTGGAATGGTATAGTGAATGTAATCATGTACTTAAAAGTAACCATTGACATAAATTTGTTTTA

TCCCTACAAAGACATTAAAAGGAATGCTAATGAAAGTGCAATCCAAAAGTTGTTGATTATGAAGGAACAATAATCTCTTC

TCCAACGAAAGTAATCAGGGGGAGATACAGTAGACTATTTTCTAGGTCATTACCGATATTCAGTTTCGAAAAGTACATGA

CTATAGGAATAGTCTGGAACAACTCTGAAAGTAATCAGGGGAGATGCCGACATCAGGGGGAGGATCCAAGGATATGATGT

CGACATATTTTACTTCGAAATTGAAGATGTGTTGTACTCTTTTTCCCTTCAATCGAGAATAGTTTTTCCCAAAGGGTTTT

GTTACTCGACAAGGTTTTTAGTGAGACAACACTAAAAACATAAGTATGTTGAATGTTGAAGACATAAATATCGCGTTGAT

ATTACTGAAAATATCTGAATCAAAGAAATGAAATGCGATAGTCTGTTAAGCAACGTAACTTCCAGGATCAACAACATGGT

CTTATAAGTTTTCAAGTCACTAAGGTACAGTGTGATAAACTCCTTTTGACTGCATTAGACTAATGAAAATTGTCTAACAT

CAGGGGGAGCATCTAATGGTGTGTTGAACTCTTTTCCTTCACCGAGGTTACTTTTTCCCACAGGGTTTTGTTGCTCGGCA

AGGTTTTTAATGAGACAACAAAAAATACCGGGAATGTAATTTCCTAGGTAAGGCTATTGTCTTTCCTACGTGGATTTTCT

TCCTTAAAATTTGTGAAGCAAATAGTCAACTTCAACAGAGCAAAGTGATCATCTGCAATGGATCACCTTTACCTGCATCT

GTCATGTTGTACTCTTTTCCCTTCGTCAAGGTTTTATCCCACTGGGTTTTCCTTGTCAAGGTTTTAATGAGGCAACATAT

TTGAGTCCAACTATATTCTAAGTTTGCTTAATATTGTACTCTTTCTATTTAGTTCAGGTTTTGTCCCTCTGGGTTTCCCT

GGCGAAGTTTTAACGAGGCAATTAACTTAGACTCATCGATCTTTGAAGATCGTATTACATGTGATGAACTACACGTAAAG

TACGGGACAACATGTGAAGTCCTACATGTGAAGAAGGTTTTATCCCACTGGGTTTTTCCTTATCAAAGTTTTAGCGAGGC

AATTGATTTTGGCACAAGTCATCAAGGAGACCGACATTTGAAGAACTACATGTGAAGCACTATGTATAAAGCTTTGTTAT

TGAACCGCACAAGGGGGAGTGTTACAGGGCCTACGTCCCGTTGGATGTGCGGTCCAACTAGATTCCTAGGGTTATATATA

AAGTAAACAATGTAATACTTGTACCCTATTATTCTAATAAGATATTCTTCTTCTTGCCTCTACATATTATTCATCCACCT

CTGATTCTCCTCAAACAAAAAGTAGTTACTTGTGGTGAATTTTTCATTTCGAGTCCGTTTCGAACAAATAATGGATCATT

GGAATACATATGATGTTATTCTAGAAAATATATATGCTTAGTTCTAAAGATATTTTTTTTCACCTAATTGAACTGAGAAC

AACTTTGAGAAAGAAAAATTCATTGTATTTACGAAGATTCAAATAAAGTTTGAATATAAATTAAGACGGAGGAAAAAAAA

GGAGTCAAATGAAAAGATGTCTTCAACATTAAAAATTGTTAAAAGTTAGATGTGAAAGGTAGTCAGGCTTATTTTTTTGT

GAATAGGACGGCTAAAATTACCACTTAAATAAAGAGAGTTAGCTGTACCTAATTGATCCGACTCTCATGAGAATAAAAAA

AAATTGCTTCTTTTTCTTCATTTTTCGGTTAGGGTTTTAAACGGTAAAATATTTTAAAAACAGTGGAACTAGCTAGGAGT

GGATTCTTGATACATAGATAAAAAGATACTCAATGGAATAAAGAAGGGAAAAAACGGGTGAAACACTGATATACGGCCGA

AAATAAATTGCATATCTTTCTACTACTGACTTAACTTGTAGATATGTTTGAATAGTTTTATTCATATCCCAACATAAACA

CGATAAATCAGTAATTGCGTGAATTAGAAGCTTGTCAATAATATTTAATCTTAAACAAATTGTCATAACTAAAAATAAGA

AAAAGGTTTGACCTTCTATCGGTCTTAATATTGTGTGTCTAAAAAGTAAAAGTATATAAAAAGCCAGTCATCTATCGTAA

AATAATTTTACACGCATTACACTTCCTAATAAATAATAATTATGTATTTTAAATGAATAAAGAAACTACTTTTACCTTTG

GATATCTTTCAAAAATTGGATTTTTATACAAATTCCAACATAAACATGATAAATACACAACTGAATGAATTAGATAAAGA

ATTAGAAAGAAGCGATAAAGATGCAATTGTTATCCAATGTCTTTCTCCCACCAAGAAAAAACATCGCAAATGTATTCGAC

ATCCCTAAGATGATAACCAACAATATCTCTTGAAATCTCCCTTATGCAGAGAACCTTGTGCTGACGTGTAACAGAGTCAC

GCCTAAATCGAAAAGTAATAAGCAGTGAAGTCCCACTCGGTTGTAACGGGAGTGAAGTAGAAATAGTTGGAGGTACTCTT

CGTGTCCCCATTTTGTAAATAAGAGTTATAATCTCTCGGGTTGAAAAGGTAGAATAAACATTTTAGATGGTCTAACATGC

CATCTTCATTCCATCTAATTGACTCTCTAAATATTGTTTCACCACCACCTTCGAAGCCAAAATTAAAAGAGTCTTGGCCG

GAAAAGTAAACATTTGAAGACATAAATAGAAATAAATAACTTGACTTTACCTCTTACTTTAGTCGTAGATACTTTTGGAT

AACTAGATTACTATTCAACTCTAGATTACTATTCAACTCCCAACATGCATGAACACGAAAAATACATATCTAAATAAAAC

AAGAAACTACCTTTACCTTTAGCAGATTTTAATAGGAGGCTTTCTCTTCGTGTTCCTACACAAAATTTGTAGGTTTGCAA

CAGTTTGAATTGGAACCTTGTTCGTGATATTTTTGACTACCACTCACTTCATATACATCCAAATCCTGACTCCCCGTCTG

CGCAAGCCATCCGATGAAATATGGAAACATTTTGGAACTTAAAACTGGGTATAATACTGAAAAATAAATAATAATGTACA

GTGGATTTTTAGACAGAAAAAATATTCTTTCAACAACAATTCAAAATACACAGAAGATTATAAAGACAACAACAAATTAG

TTCGTGTGATTTTTCAACTTGTTGACATTGTTAGAGAATAATTTAAGACCTTTTTAGGAGTGAGAGCATAAAGTGATTCG

AGGAGTGTGGAAATTGCATTTGACTCATCGATTAGGACCTATCCTATATAATTTGTATTTTTTTTATACAATTTTTTTAA

TAATTTGTAAACAGAAGTCAAAATTTAACCTGGTGAAAGTCATTTCCAAGTACCAAACCCTAAGTTGGACAAAAGTTAGA

CTAGACAATAATATTTGACAGAATGTAAGCACTAAACACCTATTAACCCCCAAAATAACCCTTAATACTTTCACCCACTA

GATAAAAAAAAACATGCATTTTCCAAACAACTTCACAACTCTGAACCCTAGTAAATACCACCACCACCATCATCCTCTTT

TCCTTCTCTTAGCTTACAAATGCATGCCATCAAAAAAACAAACATATACTAGATGAGACGAATTTTTGTGAAATATTTAT

TGATTTCTACTTTATAATCAATCTTTAATTTATTATAAAGACCCGAGAGAATGTTTTGTTTTGCAAATCAAATAAAAAAT

GGAATCCAGTGGATTCAATAGTTTGCGAGAAACCACAAGGAAAGTGTTCATCTTACTTAGTTGTTGAAAAATATGATCAT

TATACTGATTCTTATAATTAAATTTCAAATGCGCACGAAAACATTGAGAATCGCTTTTTCTTCTTTTGTTTATACTTAAT

TGTATGACTTTCTGCATTTTTGGGTTCAATAATAAAAATTTGATGTTCAAAAGATTTCTAATAGGTGTTTCTTTAACCAG

AAAATTCGTGTTGCTGGTGGAGTTGTGCGGGAAATGTATGTGAATATTTAATAGGGGCAGTTTAGACGGAGTTAGGAAAT

TATATTGGATAGGTTCTATTTGGCAAGTGAACTACGATTTCCGCTCTCCTCAAATTACTTTTTGCTCTCCTTAAAATAAA

AATCTAATTCAAATCCTAACATACACACCCGTTGAAGCCGCAACTGATTAAATTGAAAGTTTGTTAGCAATAATTTGAAC

CACCTCAGTGTATCTAGAAATTTGCAAAAAGGTTCAAACCTCTACAAGTCATGTCGTCTCTTTAAAGTTATTAAATATAA

GAGAATAAAATTGAGTCAATCATTATATCATACTAGAATAATTTTAGGTAGCAAAATCAATAAGATAACAAGAAATTGAG

TGACCGAATAAACACATAAAACACCGAAAGGTATGAGGAAATAGATGTTTTGTTTTAGATAAATAAAGTTTTACCTCCAT

TCTTAAATTCTCCGAAAAATTATGAGGTGGGAAATATTTGGTAATGGATAGGTATACTCTTGTGGTAAGTTCCTTCTTTC

TTTTCCTTTTTCTCTCTCTAATTACAATTACAAGATCCAATCACAAACTAATAAGATAAGAAAATTTAGATTAATAAATG

ACTTGTTGTAGGTAAATAAAGTTTTTACTTCCATTCTTAAATTTTACAAAATTTAAAAAGGTAGAAAATGTTTGGTGATG

GATAGATACAATAAATACTCTTGAATTCTTGTTTACAATCACAAGATCCAAACACAAACTAATAAAAGAGGAAATTATAG

ATTAGTAAACGACTTGTTGTAGGTAAAAAAAAAAAATTACGTCCATTCTTGAATTTTATAAAATTTAAAGAGGTAGAAAA

TATTTGGTGATAGATAGATACAATAGATACTCATGTATTCTTCTTTCTCTTCATATTCCTCTCTATAACTACAACTATAA

AACCCAAACACAAACTAATAAGAGAGTAAAACATATAGACTAATGATTTTACCCCCGTAAAAGAATAAATTGACATTATA

ATAAAGAAAATATTGTCATTGATTGATGATATAATACGCGTCAAACAAAACTTAGATTTATACATATGACAACAAAAATA

CAATTTTCATGAATACTTTGAACTCTTGAAAATAAACATTTGTTTCTCATTATTGTGTGTAAATGGTTGTTTTTTATACG

AATGTGTTAAAAATACATAGCTCTTAATAAATGTTTTAAGTTAATATAGTTACCCAAACTTGGTAAAAGAGTTTTTTAAT

TAAATATGTGTTAATTGAAGCAAAGAGACTTCAAGATTCCTCGGGTATCGCTGCAATGATTTGGCCACAGTATTCTGTAG

GAAGCATGAAAAACAAACATTCTGAAATTAAACAGAGTTGTACAGAGGGTCAATATCTTCCTCCATGTTACTATTCTTCT

TTCTTCTTTGATATACTATCCACCCCGTGCGTCCTTCACTCGTTTGTACCCTCTTAATAACCTGGACAATATGGGAATTT

TAGATTATTTTCATAGATATAAAGATACTTTTCCCCAAAACAAGGGAAGTTGTTTGATAGAAAAACAGAAATTTAAACAC

ACAGAAATTTCAACAGCAGAAATAATAAAGGTTACCTGAGAATCCTGTAATGGTGGTTTTCTTTGTAGTAGAGAGCTTCG

TAGATAATAATTCAAATGGAAAAACACACAAGATAAACTAGCGAGAGATTGAGTGGGAAGCTTCTTAGCAATGATGCTAA

CCACCATAGGGTCTTTAAAAACTTGCAAAAGGCCCTGAAAGTAATAGAGAATAAAAGTGATATGTAGTTAGACAAAATAA

AAGTTAGAATGAAGTAATATTCCCAACTTCTGGAAACACATTTGAACTAAGTATTAATTACTAGGAATAGGTTGTTTGAG

AATTCAAGCCACAAACAATAATAAAAGCAGGAGATTAAATTAACTGCAGAAAAGAAGGTAGTTGTTGAGTTTCTGTAAGA

AACTTTTACCTCTATATTTAATAAAACGCTCAAAAATTACAGAGAGGATTTTGATAATTGAGGATAATTTTCTTATTTTT

GTATCTATAGTCTCTTCTTGTTTCTCAACCTTCCTTTCTCCCACTCTTTAACCTCATAAAGAAGATAAGACGTAAGACAC

AACAATAGATAGACACCATATGTATGATTTCCAATGCTTCATTTTTCTTTATCATCTCGGTTGTATTTGTTTTTCATTCA

ACTCGTGCGGATAGATATCAACTAACTCACTTGATTTAACTCAGAACTATTGCATGAGCCGTGAATCGGAAACCGACAAC

TACAAATAAACAAAAAAATTGGGAGTTAGTGATTCCTATTTTTAGACTAAGTTTGGATTATGGCTCTGCTACTGTTGTTA

ATTGTGAAACAAAAACTACTTATGATGGATCGTGATACGGTGAAAAGGTGTCACTTCATGGTTTCAGAGTGAAAGTAAAA

AGCCGATATCATAGTAATATTATGGTAACCGGTATAGTTACTGGATGATGATATCCATAACCTGAGTTTAAAATGTGTCG

ATCAATTTTTATTTAATATCATAGAAATAAACTAAGATATAAAAAGTTTTCCCAGACGTAATACACCTAAAGATATCCAA

AAGGGATTTTCCTTTATCTTCATTGGTGCATTTTTACAAAAACTTAGAAAAGGGTAAAGTACAATGAAAAAGAAAAGAGG

GATTAATTAATTACTTCTAGTGGATTTTCATGAAGAAAACTTCCGATTCAAAGAGAAAAAATGAAACAAGTTTTATCAAA

GTAAGGCACGATAAATGAAGAAAATTTAGGTATTAATTACAAGAAACAAGTTGTTTGAGAATTGAAGGCACACAAACACT

ATAAGATTAAATTAACGGAAGAAGAAGAAAATAGTTGTTGAGTTTCAGTAAGGAAATTTTACCTCCATATTTATAATCAA

ACGCTTGAAAATTACAGGAAAAATATTGTAAACTGGTGATGATCTTGTTCTTTCTTTTTCACTAGTCTCTTCTTATTTCT

CAACTTTCCCTTTTCTCACTCTTTTCACCACATAAAGAATATAAGACCTAAGACACAACAATAAATAGACGCCATGTGTC

GGATTTCCAATGGTTTAAATTTTCTATTTTTTTATTATCTCGGTTGTGTTTGTTTTTCATTCAAATCATGCAAATTAGTA

TAAACTAACTCACTTAATTTAACTCAGAACAGTTTTATGATCCGGTGAATCGGAAACGAAAAGTTGCAAATTAACAAAAG

AATTGGAAATGGACAACTGCAAATATTTTTAACTAGTTCTTTTCTAAACACAAACCTACCCAAAGAACTTTCAACCAACA

ACATCAATATGTTGGGATTTAGATATTGTGTATACAACCACAAGAGGCTTGTCAATTTTGTCATACGATAATAACATTTT

GATGCAATTCCCTTGGATATATTCTAACTATCATAAAGAATATTTTCCTGACATCAAAATGGTGAATGAATTACAACTGG

AAAGATTAGGTATTTGGGATGTCAATATCCTGTAACTCTAATGAACAAACACAATTGCTAGATAATGATGTAAAAGAAAA

GAACATGAAAAGTTAGAAAAATTGTGCAAAATCTTCGAACTTAAAGAAAAATTGAAGAAATTTATGGGCAACCACACTAC

ACATATCCTGGTTTTAGCCACGCTTCATTTTTGGAGGTGCGCAACGGATATAAACGTTGCACAATTTACAACGTTAATAT

CCGTTGCACAAATTATTTATATTTCATTTACTAAAATAAACTAGTGGTGCAACGTATTGTTAACGGCTATAACCATTGCA

CAACATATGGTTTTTTCATGGAGTATTAGTTTTCTGATTTTGTTTTTCTTAAATTTTTTTAAACATATATCTCTCTTTCT

ATCTCTTTCTATTTCTCATCGAAGAACTTCTTTTTCAGTCCAGAAAAGAGAGAAACTTTTCTCTTTTTCTTCGACAAATC

TGCTTAGATTTTAGGTTTTTCATGAAATTACCCTCGTTTCAACAAATCTGCAATCTAATCAAAACCCATCGAAATCTGAA

ATCAAATCTATACTTTTTTCTAAACCTAATTTTTTCCCTGGAATCGAAAATCAAATCATTCAACCTCTGATTTTGGTGGA

TTATCATCAGGTATAAGCTAGTTTTTTGTTAATTTTATTTTTATCTGAAATCATTTGATAGAGTTAAGGTTTTCGATTTT

GAGCTAACTAGTATATGAAAAAGTTAGGGTTTTGATTGTGGTGTTCCTTTTTTTTTTTTTTTTTTTTTATATAGAAACTA

ATTATTTGATTTGAAGTCATCTGATGCTTAATTGGGTCATTTTTTTAATAGAGTTTTAATTTTTGTGTAGAGTCTTTGAG

TCATCTGATGTTTTCCAAGGGTTTGTAGTGATTTTACTTCGTTTGGTAGAGGTTCTGTAAACTATATTCTGATTCTAGAT

GTTGTTATTATTGTTGTTGTTGTTGTTCGAAACCAAGGTATAATTTTTCTATGCTCTTAAGTCTTTGCACTTGCACTCCA

TGAAGCTTTATTCTTATTTTGTATTCAATATGAATCTAACTCAAATGTAAATAAAGAGGAAGTTTTATTTTGTAAATAAT

GTTACTGATCCAAAAAAGATAAAATTGAAGTGGAGGTTCCGGTGGAGGTAAAATATATTCAAAAGGGGATGGGGAAATGG

ATGCATGGAAAGGAATCAAAGTAGCCTTGATCAGTAAATCACTTGTCCACTACATATTCTACTATACTTATCTTGGCTTC

TATGTTTTCATTTTTTTTCATTACTGATTCTCTTGCTCTTATACTTGTTGTCTCCTGTCTAAATTTTATCCAACTAAGAC

AAAACGTATTAAAAGATCAATTGAGAGAAATATGGCTCCTTACTCCGACCTTAATGGTGTGCATTAAAAATTGTTGCTGA

TGATAATATTAATTACTTTCACATCAGGTAATAATATACATTCTTCATTTGCTTTATTTTTCCATCTTTTGTTCGGTTGT

GTCTGTGCAGTGCGGGTATGTACGTATCTGTCGCATACTTTCACATGTAATTTTTACTACAATTTTTTTTTTTTGAGCTA

TTTTTATAGAATTAGGATGATAATTTTCCTAAGGTTCAATCCTTGAAACTGATCTAGTGCATGCATTTTTTTTATTGAGA

ACAAGTGGTAAATTGTTCATGTGTTTTTTCTTCCAGGAAGACCTTGGGAAATCTCTGTTAACAGCTATAAGCTAGAAAAC

CCAATGAATTTATGTTGTACAATGAGATGTTTTCAATGAAGCAATCATTGAAGAACAAGAAAAAGTTATCCAAGAGAGAC

AAAACCAAATGGTGAGGTAAATGGGATTTTTAAAGATCTTGCTGTGTTGGTCCATGGGCAAGGAGTTGTCATCGGTAGCA

CACCTCCCTCCTCGAGATCGAATTATCTCTAGAGCCACGCTTTTCTTTTTAGTTGACATTGATTCACATGTTGAGAACAA

AATCTCAACTTGCAGAAGCGGCAAAGACCCAAAAGTCAAAAATCATCCCTGGTGAGCTTTGGATAGTCCGTATTGGGAAT

GTAACTATCTTAATTAATCTTTTATCTCCACATACTGACTTGATGTTCCTGCCAATGACGCAAATGTGCTTGTTGTTGGT

GATTTTTGGGGCTGTCTTGCCCATTGTGACAATAGTCCTGACAGGTTAAGTAGTAGAAACTTGGTGTCTCAATCGAATTC

CTTGTCCAAGCCAAATCTGTTGGACAAAGTCTTATATTTAAACTATGCTCATCATGCTTGAAATATGGTTTTACAAATCT

GCTGACTTTAATCACAATGTCATTAGAGCTTTATCACCAAGTACTTTTTGTAAGTTTTATTTTTTTCTGATCTATACCTC

TACTAATCCCAGGAGTGATTTACCTTGAGGTAACCCATCTCCATCATATCCAGCTGCTCTACATACATGGGAACCTGAGA

TAAGCTAGCAAAGATTTGTTAACTGGCACTTCACTCTCTTGTGTATTAGATAATTCACAGGAAGCTCAGAACGCGATAAA

TGACTTCACTGGTGAGACCTTTCTTCCCCTATAGTTTCTGGTATATCATTTGTGGGTACAAAATATATTGTATCTAATTT

CCAACCGCATCTTTTTAGGTAAATGTCTCGGGAATAGGCAAATTAGACGCAACTGGGCTGCTAAATAAGCTACTTAAAGA

GAAACAAAATAATGAGAACCAGAATATGGTTGTATTTACAAACAGATCATCAGGATCTTGAATTTCTCCTAACATTTATT

GAATCTGGGTTTGGTTTACTCATAAATCTGCCAGTGTACAAAAGTGTTGCTTGATATTCAGTTTTTAGACTAGTCTAGGT

GTTGAGAAGTAACTCTGAAGAAGTTGGTTGTGTTTGAGTTTCCTTTAAATATCAATTGTTCTGCAGATGGAGGTCAAGAA

AATGAAAACGAGACTCCTGAAAGCAACCCATCATTTACTGCTGTTGATAAACTGTTGAGACATCATTAACCTTTTTAAAT

TGAAGACAATATGGATTCCTTTACAGGTTATGCAACCTGAACTTCATCGTCAGCTTCATATCCTCAGGGTCGGGTTTATT

GAGGAAGTACGCATCCAGATAGACAAAGGGTTTGGGTTCATTAGGTGCAATACCCACGAAGAAGCAGCCATGGATATTCA

GATGACCAATGGGAGGATAGTCTGTGTAAAATCCATGAAGGTATTTTTGCGCTCCTCTTTATTATTGATATTGTACATCT

CTTCGTTATTGGTATTCCTCATCATGGTATTGTCTCTTCTTTATTATGTGCGCTTTTGTAATTAATCCGGGAATCGAAGT

CATATTACTAAACTAAGAGACATAAAAGCATGTTCATTGTGTTCCTCCACCCATTATTTTTAGGTAAGGCAGTGTATTTG

GTCGGATGCTTATCAAACCATCTACTCTTAGTTTATGACTCTGGTAAATATCTCCTTGGTTGAGTTGAAAGACCCAATTG

AAAACATCAGTGCTAAGTTGGTGAGACAAGCTGCTGCTAAGTTTATGATTCTAGTAAGTAAGAGTCTGTCGCGAGACAGA

TGGTTGGTATTCTTCTATGTAGTGCAGGGAATTACATTAAAGCTGACTCCTGGTTCTGTAGTTGCTCTAGTTGGCCTAAG

TGGTGGTGTAAAAGTAAACATTTTATGTTTATGCAATATCTAACTATCATGACTTCCATTTCCAATTCTTTCCATAAATG

TCTTGTTGCAAATTACAATTCTATTGCATGTGAAAGTGAAGTTTGTGAAATGATTTTCGAAGGTTTCATCACTTGTAGTA

AATTTCTTTTGTTTTATGTGCTCATTAGGCGACATTGATGCTTATTATGAGAAGGTATTGGAAGTTTCAAGCAACAATAC

ATGACTCCAAAAAACTGAATTGGTGATTTCTTGGACTTCGTTTTTGTTTATGCCACAACAAAAGACTAAAATGCATTCTT

CTAAAAGAATTATTTGGATGAACCGCGGCAATGCATATATATGAAAGCTATTTAGTTGGAGTATTCATCATCTTATAATA

GGTTTCTATTCATTTTCTTATGCTTTGGAATGAAGATTTGATTGTTCCAAAAGTTGTTTAATATTGTATAGGTTTATGGT

AAGTTCATCTATCTATACTGATGATGTATGAATCGTTCTTATCAATTGTGATGATCAATAATTGTTCTTGATTGATTGTA

ATGATGCGAAGATCCATCCAAATGATTAATTGGGTCTTCTTGTTCATCTAGAATGGTATATTCAGGTGCAGAATATATAT

TCAGGTGCAGAATATATATTCAGGTGATAAATCTGCCGGAATTTAAGCCGGCGAAGAAAAAAATGATCTGAAAATAATTT

TGCAACGGATATAAACGTAGGAAATATATCCGTTGCACCAGTTTCCTACAATAAGCAACGGATACAATTGATACCAAACA

AATAAAACGTCGACAAAAATGGAAGTGGGACCCACAAAAGCGTTTTCTATTAATAATAAGCGTTGTACAAATTAGTATAT

ACGTTGCATAAATTAGAAAAATAAAACGTTGCACCACATATTGTGCAACGCTTATAAGCGTTGCTCAATATAATTTGTGC

AACGAATAATAAGCGTTGCACAATAATGCGCAACGCTGTTTTACCAACACTTATTAACGTTGCACAAATTATGCGCAACG

TTTATAAACGTCACTAAAAGGGTATTTTGGTGTAGTGCCTCTATAGACAAAATCACCATAAAAGGATCCAAAAGATCAAG

ATATGCATGGAAATCATTTATGGACTGGACTAGTTAAGTGGCTTTTTATGTTTTAGTTTATGTAGCACCCCAAAATCCAG

CTATTTAGAAATCTGAGAGACGATTCTACATTCTGAAGCATTAAAGATCTAAAACATTTACTTATGCATTAATTTGAAGT

TTTAAACAAAGATTAACCCAATTTTAAACCTTATTTAAATTTATTGAAAAGGCGAGGGTACCCAAATATACCTCAAGTTA

AAACTTTTCCTTCATATAAGTCCTTTCTCCAAAAGTGATTGTCTATGGACTGAGTCGAGACAATGCAACTAATCGGTTCA

CACTTCGTATGATCGTCTATGGATACAAGATCGAGACAATACAACAACGAAGTATGTTTACTTGATACAAAGGTTCGGAC

TTAACCAAACTCAGTAGGATTCATGTATCAAGTAAATAGGAATTAACGTTTGTGCAATTTACTTTAATTATAATAAAACA

ATTATAATGCGGAAATATAAAGTAAATGACACAGCAAGATTTTGTTAACGAGGAAACCGCAAATGCAGAAAAACCCCTGG

ACCTTGTCTAGAATTGAATACTCTCAGGATTAAGCCGCTATACAAAATAATACCAACTTCGTATAGTTGAGGCCAAGAAA

TTAAACCTATAGTTCACCTAGTTCCGTCTGTATTCCCACGCCTCCAACCTGTAAACAAGTCACGTACTTGGTGGTTCGTA

TTCCAAACAGTAAATGAACTACAAATCTGTTTGGTATCAACTCTTCAACCAAGTTATATGAGTTTCACAAAGGCTCTTAT

GTTTATCTTAACATAAACTCCTTTGCCAGGTTCCAAGTTCTATCTTATATATGTTCAATCACCAAAAGGTAATCCTTAAG

ATTAAGCCAACAACACTCTTAATCCAAAGAATTGTGTTGATGTCGATCTTCTCAATCAATCAATCCAATTTACCACAAGG

ATAAACCTATTATTAATTAGATCCTCTTTTACCAAAACAAGTATTGTGCATACCAAAGATTATAAAAACCAAGTCTGAAA

TCTTCTATCTCTTCATTGTCTTCAAATCTTCTTAGATATTCAATAAACACCCTCACACAACTACTTGAATCTCTTGTGAT

CAATCACACATAGAACGGAGTCTGTTGACAATGGATTATCACAAGATCGTCTTTAGATCTAACAACAATCTAAATATCCA

TGTAGAAACCTTGAACTAGTTTGAGTGAATCTTATATGAAAAGAGAAGATTCTCAAGCATAAACAAACTTGGTGCAATCA

AAGTTCAATGATAAGCACGTATGTGCTACATTTTGTACCCATATTTATATTATCTAGGCTTGATATTTTGACTAATGATA

TTATTTTAGTGCTTTTGTAGAAAATACAAGCAAATTGGATCGCCGGGAGAATAAATTGGCTAAATTGGAAGCTAAATGGT

ATTTGCGACGAAAACAACCAAAATGTGGCTGGCCTGGTATTTCTATATATCAGCAGCACTTATATATCACGGCACCAAAG

GACTCGGAGCTTAATCATTTAATTATGAGGTGCCTTGTAATTTCCTTGTGTTCGTCGCACCAACTGAACAATTTCTCGAA

CAGAAGATAAATGAAGCATCAGTCAACAAATAAATGCCTTTGATGTCAATGGAACTGTGAACTTTGTTAACATACTAGCA

TATATCACGGACCCAACCATTTCATGGCCAAGAGAGTATCGTATAGTGGATATCTTTATTAATTCAATCTCTTGAATTCC

ACTGTGGGTACAAATAAGTGGTGGACGTGGCCACTTACTCCACATTTATATCGGAGAATATTCTTTACTTCACCAAACAG

TAGAATAAAATCTCATTTCTCATAAGTAGGTGTTATTCATTTTATCAAGAGAAATGGTTTCGAGCAGCATCTTTGGCGAA

GCAGCTAATGCTCACTATCTTTTCAGATCCCTCATCCCTCTCAAATATCTTGGTCTGTATTGAGACGATATTCATGGGGT

TCTCAGATTTGAAGAAATCAAATCAGTGTTGGCTGCTATTTTGGTGAGATGGTTTACAAAGAATTAGGGCTTGTTCTTGA

ATACTAAAATAGTGGGTATGTATCAGGGGTCAGAAGTGGTTCAAGTGACTTTGATGACTGCATGGGTTTTGAATGGATTG

ATGGTTGCTTATAGCTATGGATGGTTTGATTGGTGTTGTGGTCGACATTGGAAGATGGATACAGATTGTTAACTGGTCGC

ACGAGCCGTGCTATTGCTGGTGGAGGTGTTTAAATGGTCACTGCAGCGGTGGTCGTTCGATGCAGCAAAGAAGTCGAGAA

ACTGAAGACCTAGAAATCGGCGGGGTGTTTATTTTCTTAAGTGGGGTTGATCACTGGTTGAAAGTCCTTTCAAATTCTGG

TACGTCAGGAAGGTTATGTTTGCATTCAATTTGATTCGACATCAACATCAGTTGATCAGCGGGGTTCGAATAAGAAGTTG

GTGTTGGTTATTAACAACTGTTTTCAGTTTTTGTACGAGAATGGAGTTGTATATGAAGCTGTGTATGCATGAAAGAAGAA

GAGTTTGTTGATTTCTTGTAGTTTCACCGCAAGACTTTGGAACTACTTATTACCACAGAGAGGTATCAATGGTGTTTGGC

CGAATACCGTGCTTTCCTTAATGCAAAATTGGGGTGATTTGAGGCATCGAGAGGTGAATAAGGTTGTTTGGTCTTTAATT

CCGGCAGCAGTGCTTTGGTCTATCTGGACCGAAAGAAATCAACAAACATTTGAGGAGAAGACCAGTGACTTTTGGGCGGT

TAGTAGTGGTGCAAGGTCGATGTTGCTCAGTTGGGTATCCATTATAAAACCGAACATTGACGTAGGTAGATGTCTTCATG

AGTGGGATCTTGTGTTTTTGTAATGTGCATGTAGTTTTTCTTTTTTTCTTCGTCTCCAGTCTTCTCTTTTACTTCTTGTA

AAAGAGATTGTTTTTGTTCTTCTCTTCTCTCTTAATAAATTCTTCTTTTATCAAAAAAAAAAAAAGTTATTTGAGTTTTC

GATTACAGCTTGGGTTTGTGGTGACTTTGAACCGTGAGAAAGAATTGGCAGCAACCAAACTAGATGCTGGTGTTGAAATA

TGGTGTTGTATGTGTTTGGCTGGAATTGTGGAGATGGGTGTGTTGGCTAGTTTAGTCGAGAATAGAATGGTCGATGGGTA

CCGAGATTTAGAACATGAATCAGTATGGCAGTCCATGCAGAGGTGTGCTATTGGTAGCTTGCTGGTGTTTGTTGGAGGAG

TTCTCAGTCGAGAAAACCAAACCAGAAAAGTGTTTGCAGGAGGGGTTGTGCTCGAGTGGCTATTGATTACGCCAAGTGAG

ATGACGAAATTGAGATGGTCAGAGAAGTTGCGAAGTCGCGGCTGATGGTTAAACAGATGTTGGACTTTGATAGTGGGCCT

GAATATAGTGAGTATCAGTCCGGGCCTTGGAAATTATGCTAGGTTTTTGGTGGAATTTCAACATCAACTTAATAACTAAA

TTTTTCCTTAAGTTTAGGAGGAAAAATAGTTCATCTCCAATGGTGTTGTTTGTGATGTTTTATGAATTAAATGTAAATTT

TATTTTTAGTAATTTTTAAATTTTATTAGAGGTAAAAATGGTTATTTAATATATTAAAATTAATTTAGGTAAGCAAAAGC

TTACTAGGATAGCTTGACATTGTTTAGGTGAATGTCTAAGAGCTTCTCCAATGGAATGTATGTAAAGATAAATGTTTAAG

TTCACATAGGATAGACATTCATTTTCCCTAAAATTCTTCTCCAACCCCAACTTCTTAAATCAATGTGAAGATGACATGGC

TTCATTTTTGTTAGACAATGTCAAGGTGAATGTCAAGTTTTAGACATTCATGTTGACAATGTCAAGCTATCCTAGTAAGC

TTTTTTCTTACCAAAATTAATTTCAATATATTAAATAACAATTTTTAGCTCTAATAAAATCTAAAAACTAATAAAGATAA

ATTTTACATTTAATCCATCAAACAAACAACACCATAAGGATCGAGAAGATACAGACAAGAGGTGAGATCCTGGAGGCCGG

TTCTGGGAGACGAAGAGGAGGGATTGGATGCGGATGCGCGGCAATGGATTGGATGCGATGAGGCACAGCCATTTTTAATA

AGGATCGAGTCTAGGCTTTGGGGTTTTACTTCTCCATTTTGATTTAGCTATTTTGATTTTAATCTTTTTATGGATTTTGA

GAAAGCCATGGCTAGCTAGAACCATGTTAGAATTTAGGTAGAACCCAATTTTATTATGAAAACTCTGCATTTATATGCTT

AATAATGAGTTGATTGATTAGTAGTTTTTCTTCATATAGCTTGGTATTTAATTTCTCATGTGATTTTCTTGATTATTTAT

GCTTTTGAATTGATATTTCGTGCTTCAGTTAAATCCCTAGACGTGGTATGCAAGGATAGATAGGTAATGTTTTAGGATCA

CTAGCTATGAAGCGAAGAAAACTATAGCGGAGAAACAGTATTTGAAAAATCATGGAATGTATTTTTAAAGATTGGTAGAG

TTTGCATAATTAATTGGTGGAAACCGAAAACCCTAATAACCCATTCTCATTTGTTTATCATTAGATTTATTATTATTTCA

TTTTGTTCCAAATTTCTGAAAAAATTGCTAAAACATCATCTTGTTGATTGGTTACTTTTGATATTAGTTAGTAGTAATAT

TTCACACTCCTCGTGGGAACGACTTGTACTTGCCATCGTTCTACTAGTTAGATATTGTGCACTTGCAGTATTTATTGTGG

GTTTTTCGAGCCTAATATTCAACCACCATTAGTCAATCAAATCAATAAGAAAACTAAAGATAAACCGCAATATCTAGTTT

CCCACCAACGGTACGCACTAGAGCTTCTCAATCCCAAAGAATACTTTAAACTGAGCGGTCGTAAGAGATTTCACCTAATT

AGATTACTCTCCTCTCTGATAGGCGGCTTCACTAGTAACACTACAATAGAGGAAGTTTACAATTACGAAGGATTAGTTTG

CTAGAAATTCAAACTCAAAGTATTTATAGACAAGGAAATTTGGACACCAATGAATTTCCAAAACCGAAAATATTCTCAAG

ATATTTCCATATTAACCCAAATTCGGTTTTCATAATCCCTGTGAATGCACTGTCTAAATATATGATCGAAATCTCCTTGA

AAAATCTTTAATTAGAAAATGCACATTACTAATTCTTATTTTCCATATATAATTAAGAACCTTAACTAAAAGATTCTTAA

CTTATTTAAATTCGATCTTGGGATTTACTTCCTTCTTAGTTGTTAAGGAATATCTTTGAACAATAAAAGAATACACATTA

ATGCGGGTGTTCAAAGTGTTTCGACATCTTTCTCTTGCAATTTCTTTTTCATACTTACTCACTTGAAACCGATTTACCAC

ACTTCCAAACAAGTTTAGAATTGGTTCATCTGACTTTCAAGAACTATGTGATTGATCAAGTAACACTTCTACCAAAAAAA

GTTTCAGTTATACCTCCATGTGAGTACTGTGCATAGTCACACTAGAAAACCAAAATTTGGCTGACTAGGTACTAGGATCG

GTTCCCCACATATATTTGGTATCTAACTTGTATGCACTGCACATGTCCATAGGATCGGTTCCCCATTGCCTTAAAACGTG

TTGCACATGTTCATAGGATCGGTTCCCCTTTCTTCTATAAACCTTGCTACACCTCATACAAGGATCGGTTCCCCTTGTGA

TGTATTACACCTCTTACTAGGATCTGTTCCCCTCTCCCAAATTTGGTCAGACATACACAAACCCGATCATACCCCAGGTG

ATTACTTAAGATCGGTTTCACTAATAAAAGTCATACCAATACATAAGTCGGGCCTCATGTGAATAGTTTTACCAAGAACA

CAAACAAGTCATGAGTGGTTGTACTAATCACACATATTGGTCGTTCATAAGATATGCAATGAAGAACGCCTGGCGATTTC

CTTTTCGGTTCACAACACATATCATGAACTTACTTCCTTAGAACACATGTAAACATTGTTCCCTAGGATGAAATCCTCAC

CTCATACCCATACATAATCACAATAGCATTCATACGATTATGTTGATGTATTATATACAAAGTTTAATGGTTAAGAAAGA

AACTTCATATTGTATTCCTTAATACTATGTCTATCTAGAGTGTTCATGCTTCGCAATTTTGTGTTTTCAATATGCACAAC

TTGAAATATACGTTAGGGAATGAAACAGTTCAAGTCAAATATCACTAACCTCAAGTGGAAGGATGATGTTGTGGTTGTAG

CTCCGTACTTCTTCACTCCTTCAAGTCTTCTCAATACTTGTAATGTCTCATCCTAATACTTTCAAGCTATCCAATACGAA

GTTGACTCTAGTACATTTAAATCAAGCGACTCTCAAATGAGTTTTGGTTCACTAAAATATGACAACCAAACTTGACATAC

CAATGCTTGGTGGGTTCAACCGAGCTATGCTCTAACATTTATGTACAAGAACAATTCTTAGATGATACATACACAATAGT

GGGTATTGATTTGAAAACAAAATATAAGCACATAATAAACATAACGAAAATCTAACCTACAAAGATACTCGGTTATGAGC

ACTACTATTCTGTTTTTATAATTGAAAGTTGGAACTGAGTGAGTACATCATCCTTAAAGAGGTATCCAATAGAAATAATA

AATAACTTTCAAGTAATCACTAGCATGATTCGAAGACAATGCAGGTTTTATTGAAAACCAATTAAACACGGAAAACAGAT

AAATCAAATGAGATAATAATTGTACTGAAAAAAAATTGTTACCAACCGATCTGGCACACACATTCAGAAAAATAATATAG

TTGACTGACATAATCCTTGGAAGTAATAAGGTTGATTGGATTCTTTAATGATCATTACAGCGTCCTTTTCCTAGTCATAT

TTAGATTTGTAGTCATACCAACACCTCATTCATACACACATCCTACTTGAAAATTACTCAAACAAAAAAATAGTGGTATC

ATTATATCCGATAAAAGAATCTATCTTGGAATTGCACACTCAATCCATGCGTCAATCTAACAATCATATATAAGATTCTC

ATCATCCTTCTTTCTAAAAATTAAAACATAATGTAATAGCTTTTTCAAAGTAATTTTAACGTTCAATAAACATCCATGGA

TGAGACATGAGTTTCCCTACAAAAATAGGGAATACTTTTGTGTTCCCTTCAGTACCTATATACTATTGTGTGCCCTTATG

TACCTATAGATTGAACAAAAAAAATAGATATAAATCGGGATACACATCTCATACCAATCAAAAGGAAGTCTCCTTCATTC

CATTCTAACAACAAGTAAAATTTGTGATTTCTTTCCATTCCATGAAAAGAAGTAAATTTTATAAATAAGTAAATACTCCA

AAAAATAGATAATAGTTTTTTTTGAAAATCCAATCCCATCCAAAAATGTAACCAAAAAAAAAAACTAATTTGTATTGTTA

GAGCACTACTCGGTTGAACTCGCAAGCGTTTTTATCTTAATCTTGTTTGTCAATGTCAGTGATCAAAACTATAAGTCTTG

ATTTCTATCTACTTAAAGTAATGTCTCGGACTAGGATAGATTGTGTATTTGAGTGATAGGTGTAGAATACACCGTATTTA

CAATCTAGAGTTTATGTTTTCTTTGCTATTTTTCTTGTGAATTGCATCTCTTATGTTTAGTTTTGAGTTATTAGGTGCAA

TGGAGTCGTTTGAAGCAAAAGAAGGAGAAATCAGCCATTTGAAGCGAAAAGGGCGAAAAGGTTAATTCACAGGCAATATA

TGGAGCATGCTAGGTTGTGCATGAGGAGTTATGCAGGAAATTGAGTTGTCAGTCTCCTAGAACTGACATTTGAACTAAGA

ATTAGTCTGTCGGTTTTCTGAAACCGACAAGCCATGTTTGAGAAGGTTGGCCCTTAACCACCAAAGTTGGGTGCATATAG

CAATTTATGTTATTACCAAAACCAAAACTCAAGAAGAGAAAATCATTTCTCTAATAATTTGTTTTATTTCTCCTTCTATG

CCTGTATGAATGGCGGCTTCATTGGAACAAGGTCGAGAAGAATCAGAAGTTTCTGCTGCTAAAATCAAAAGATACTAGAA

GGGTTTGTGGCTCTGATATGAATGATGGGTCTGCGAATATAATGGGTTCTGAGTTAATTTTAAGGGTTTGCTTTCAAATA

TCAGAAAGAACTGGAATTTGTATAAGGATTTGCTGCCAAGGGTTTGATGGATTGAGGAGATGGAGGTTAAGAATGGGTTT

GTGCTTTGTGCTAAAATTGTGTTTGCAGGGAATAGAAAATGATACAAAGAGAAGGAAGGTAACAGCTAGCTTTGATCTGC

AGTGTAGTTCGATATAAATCCTGAGCCACACAGTTCTTGCCAACCTCAACTGAGCCATCTACTTCTACAAGTCTGATGGG

TGTAGTTGAACTGGAGTTAGGTCAGTACATAGAATGAAGGAAGCTGGAGCTACGAGTCTGTGTGACATTGGAGTTAGGTC

AGTACATAGAATGAAGGAAGCTGGAGCTACGAGGCTGTGTGACATTGGTCCGTTTGGTGCGTCCAAGATAGCCAACACAG

GGGATCAGTTACAGTTTGTTGCCATGGTTTAATGGTCGTGTTTTGAGAAGTCAGGAGAAGGAAAAATGGTTCCAAGATTG

AGTTGTTATTGAGTTCTCAAGGTACGAACAGTGAACCTCTTGGTCTGAATTTCAGTAGCAAGGTGGGATTGCTGCTATGA

ATTGTGTATGGCTATGGTGCAGTTCAGGGAGGATAGTGCACAGCTGAAGTGGAAGCAACTGCAAGTGGTGGTTGTAATGC

ACTGGTTGCTGGTGCCTGTTGAGCCTTCATGTGTATTGATTTAAGTTACAAGAATGGATGTAAGAATGGGTTTATATATG

AATGTTAGGATCCAGGTACAAATTGCAGTTGCGGTTCAACTTACAAGGAAATAGTACCGGTGATGTTGTTGGTGGTGATG

GAATGAAGCCTAACAGAGATCAACGGAATGCAGTAGTGTTGTGTGCTGAGATATGGGGATCGTAAAGATTCAATAAATCG

TGAAGGGGATATATGTTGGATACAAGAGTTGGCGGCTGGAGATATGAGCTACGCGTCATACTTTTGGGTACGTTTTCGCA

AGAAACAAGGAATTGGCTAAAATGGTGTTAATTTGTCAATGTTTAAACTGCAGAGAAGTAGAGTTGAGCTCATGAGGTTA

TACATTTGGCTGGTGACATCTTCCAATGATTCAAGTGGCCTGAGACAATTTTCCGGTGCCGGATTCCCGCGACAGTGGCC

GGATTCATGTGATGACGCCAGCAACCCGGCGACCCAGTTTTGGACCCAAAATTTCCAGAGAATTATTTCCACTCATGGGC

TTAGTGGAATTCTTGGAGATGGGTTTTGAGAAGAGAGTATCCTTTTGGGAATTGGAGAAGCATATAAATAGAAAGTCTTT

CTTTAAACCTAGATATCTCCATTAGGGAGAATTTATTATCTTCTACTTTTGTTTTCTAGGGTTTAGAAACATGATAGGTT

TTCTCTTCCTTCTTGTGATGAACTTCTTTAACATGAGTACTTAATTTTATTATTGGTTATGGAATAATTATATTTCACAT

AATATGCTTTTTGATATGATAAATTAGTTTTGTCTCCATATTATCGATTATGATTCACTATTTATATTGTTACATGATTT

GAATGCTTGTTTGATTGAATCGCGAATCGGTTGAGTTTGTTTTCATCTCTATTGCTAGATTAGGGTTTATTATACGCAAA

AGTGATAGATTTCTAATTACAAGTAGCATAAATATGGTTATAGCTTGTAGTGATGCATTCTTATAATCATATGTGAACCA

TAACCTTTGTTAAATCGGATTAGTGCGATTTCCACATTCGATTGCTTTGATTGGCCCGATTTCATGGAACTTGATGCATC

TAGGGAATTAAACTTGATTAGTGCTTTTACACGTTAGGTCGTTCTCTAAGATAGAATTCATATTCGCATCTTTTAATGTG

TTTATTAATGATAAGAGGAGATTAGCGGGATATTCTTTCGATCAAGGTATTCTAGGGCCATTGATAAAAGTTGATATTAA

ATATCAATTCTAGTTATTAATAAAAGAACATGTTAGTGAATGAAAACGAAATCCTTAACCAATATCTTTCCTTGTTTGAT

TTGATTTGATTTGCTTTGTTTTTTTGTTTTGCTTTTATTTTATTTTAGTATATTAAAAATCAGAAAAACCCATATTAAAA

ATCAGAAAAATCCCCCCTTTGTTTATATAAGAAATCGAAACAAAAGACAACGTAGTGCTCTCTGTGGGAACGATCCTTTC

TTGCCCTTGCTATATTTTATATTTTGAGTAGTGAGAAAGTAAATTTATTTTTTGACGCCAACGATAGGGATCATTGATCT

TTAGACATCACGACGTTCATCATTGACGATGAAGAACTACTAAGGAGAGCTTGTGGAACTTCATCGATAAAAGGTATGTG

GAGATTAAAACTCATCTATCACTTGGAAAGTCTATTTCTACTCTATCTCGTATATTGAGACAAAAGTCGTATTACGATAG

TTTTCGACTATACACATTTGAGGTATCGAGCTGAGTTTAACTCGCTTACATATTTCTCGAAATATGTGTTGGTAAGATTT

CACTTCAACCAATTTCATCTTACATTCTTGACGAAAGTCAAAAGATGATCATGTAAAATTTGTTGAGTAACATCTTACAT

GGTTTATCTGATACAATCATTTGGTGTAGATTTGGAATGCTTCGTTATGATTATTTCAATAACTTGAAAATTGCTTTGAT

GCTAATAGTGTGAGAAAACGACTATTGTCATCCTCTAAGAAAGTTTCAATGATTGAAATAGAGTTTAGAACAATTGTATT

TGCACATAGTATGTGTACTTACATATATGTAATCCATGTCCGGGAACCATAGTATGCATGCCAGTATGCGTACATGTTGG

ATTGTAAAAGTCCGGGTACCTAAGTATACATACCGGTACGTGTATTTACGTAAAGTATAGGTCCGGGAATTTCTGTTGAA

GTTTGGAAATGTTAAGGTATCCATACACGTTCGTATACTGGCGAACCCAAATCCAGTCTGGTCACTAAGGTATGCGTACC

CGGTTGCATACCTAAGTGAATCATGTTTTAAAATCTGTTGGCTCATGAACTACATTTATCAAATAAGGAATGCAATCTTT

GCAAACCGTGGCTATAATGTTCATGAATTAATTCAAGTGAATCAAACCGATTTTGCTTCAATTGTGTTCTTATATACTTC

TATGAGAATATATCGATTGAACAACTCTTTAACTAGTTTCATTTGAGTCATTTGAACTAGTTGTGGTTAAGATGAATAAA

GTTGATATGAAGGTGTTCATATAACTAACTTTGGTTAACTACTGCTGATCCAACATGGAGTACATGTTTAGGTACGGTTA

CATAAATCTAAATGAAGGTACATTTCATTTGTGTATAACAAGCTAAGTTTGATCCAACGGTTGAAAGATATTAGCTTGAG

TTGAATCATGTTTTCATCTAACGGTGAATATTGAATGCTTGATTACCAAGGTAACTTAGATTGCAAACCCTGATTTGAAA

ACTATATACACTATAAAAACAAGTCATATTAGTGAGGGTTCATAAACCATCACAAATAGAAAAAACCGTCACTAATAGGA

AAAAACAGTGACTAATACTCATTTGTGACGGACGACCAACCGTCTAGACTATAGGAGTCACAAATGTTATTGGTGACGTT

TTTTCTATTTTTGTGGCGGCATCACAAATTTTTTCTATCTAGACGGTTTTTGACCAATTGTTTGCTAAATTTTTTCCGCA

ACGTCTTTTTTCCTATTTGTGAGTTTGTCATTTGTTTTTTAGAGTCTTCTTTAAATTTTTCAGAATATGAATCTTAGATA

GAAACTTGCATTCATATATAAAAGAGTTTTAGAAAGTTCTCCATTGAACTTGTATTCATATATAACAAACTTTTACAAAG

CTTTGAAACTTACATATATATTTGTGACAACCTACGAACACAACTGATGGTATGAAAACAAAAAAAGACATCATCCCATT

CCCTTATTATTCAAAAAAAAAAAAAAACGAGAACCTATGTATTAGTCACCCATCCATATACACCAACATCTTCCATTCAT

GGACTGTTCCTGATCTCTGCGTCAGTCGATGGGGTAACAGTCTTATATGTCCTGTAACACACAAACCATTTCAAGCATTA

TCGTCGCAGAAAATTTCATGTTCGTTTCAAGGATTTAAGGAGGAAATAACTTAGTGAAGCAACTCAGTTACCTCAATTGT

GCCTTCAGCTCTGCAATCATATCCAAGGCTTCTTGATAAGCTGTGTTGGTAGTTCCCGTCTCATCATGTGCAACAAGTAT

CCTCCACTGGCTCTCAACAACATTATCTGGTGGGTTAGCAAGACGTTCTTCTACAGAATTCTAACAACTTGTTTAGACAG

ATTCAAATGGATTGTGCTACCAAGAAACTCGAGACAATTAACGAAATCTAGTAAAAGTGCAATATAGTCAATAACAATGC

ATTCAATTCTTAACACCAAAAACTTACCAGGGCACTCATCGCCAACAACACCCACCAGCAGAACCACTCGGTTTCCCCCA

CCAACCACCACCACTCGCTATCCAACCAAATAAACAACATATATTCAGAGTCTTATTTCTACTCTTGTGATTACTGTAAT

TATAGACTTAAACAACTCCAATACTTGAATCTTATATGGCATTTGAATTCTGATACTAAACCATAATAAATGCCAAAATC

TTAAAAAGTACAACTAAATGTGCCACCCAGATTTGGCCACCATAGCCACCAAACAATTTCGAATTTCATTTTGGAATCCA

CAAGAAAAGCATTCCATCCACTAGTGCTAAGCATTAACAATGTCAGACGTATTTGTACATGGTATGGTGTAAACGGCAAT

CTTACGCGACCTATTGCACTGACATTGCATGTCGGAGCAGTACTTAAATAAACATAACATTAAATTATGATTACTCTCCA

AAAATTCCATTATTATAAGTAGAATCAATCACTTCGGAGGTTACTGATATTGACAAATGTTTGGAAATTTTTGGGGTCTA

CATTTAATTTTTGAAAGATGGCACAAGGAGTAGAGAATACCTGCTATGTAATGGATAATGCTATGGGAACCACTATTATT

CGCTGCAAAGAGAAAGTGCTTAGATGCCTTAGCTCTATCGAAATAAAAACCTCCAGATACAAGTTTCTCCTTGAGTTGCA

GCAGAACCAGAGGCCGCAGCTAGAGCGGCAATGGCAAACTTGCTTGGATCCTGAGATTCAAAAGAAAATCGTTAAAAAAA

TGTAAACCTACATACAAATCAATAAAAAAATCACCATAAACTAAAAACACGAACCTTCAAATATTCCTTCACTTGCTCAG

CCAAAAGGGAATGAGTACTCTGTGACAAGAGCAACGTCCAAAAGGTTCTTGTATGCATTGATGAAAATGTGAGGAGCAGC

AGCAAGAGTGGGGTATGCAACAGCCAGGGACAAGGAAGTAACCATGGAAACACTTCATCAGTAATATCAAGCACCTCAGG

ACTGAAAACTGACCCATTGTCATAGACAATAAGGACAACAAGACCATATGAAAATACCCAACTTAGCAAGCAAAGAATCC

TCAGAAGATACGACCTTTTCTCCCTTCTTAATAAGTTCAACAGGGGTGCATCAGGTGCAGCTTTCTATAATAATGGCACC

ATCAAATTTGTCAAGCAAAACATCCCTGCTACATTTACGGCAGAGTTTAACTCTAATCTTGGAAAAATAAAAAATAGACA

AAATCGAATCAGGGAACAAACAAAAATCATAATCAATTCATCCAATTTGGAAACTATCATTTCTGGAGATTAATCACTAA

AATCCAAAACAAATCACATCTTGAACCAAGTATTTGGAAAACATTCGAAACACGTACACCTACCTCTAATTTTGATTGTC

ACATGCTAAGCACAAACCAATTGGTAATATTCAAGATCAACCTAATTTCACAACTTAAGCATTTTCACAAACACATGTAT

ACACATCAAATTCCATCAACACAAACCAAAAATTTAAAAAATTAAAGAAATTGAAGAAATTGGAATTTAAGAACACTCTT

GTAGGTCTTTAAAATTTTCTCCTTTAGATCCAGCATAACAAAGAAAGAAGAAATTAAAAGTAATAAATAAAAAAACCCTA

GAAATTAAAGAAAATAAATCTTCTTTAAAGGATAAAAGTTTTCACCTTAACAATCTCATGTAGTTCTTTCGGTTCCTCTA

TTAGTTCCTGTGAAACAAAACAACAAGCAATTAACAGTAATATTTATCAAAATTAAAAACCCATTCATGGAACAACATAG

AAGAGATGGACGAGATGATAAAAGAGAAACAATCAAAAATTTCACCTTAACAATCTCTGGTAGTTCCTTAAAACTTCCTC

GTTTAGTTCCTGCAAAACAAAGCAACAACCAATTAAAAGTAATATTTAACAAAATTAAAAACCCATTGATGGAACAACAT

AGAAGAAATGACAATATGATAACAGATCGAGAAGAGATGTACGAAATGATAACAGAGAATGAGGGACGAGAGCCAACCTC

TCAGTTCTTCAGCGGAACAGAAGGTAAGTGAAAAACAAATGTAAACTAGCTAGATTTGTTTCTAGAGAAGATAAGGTTGA

TAAAGTGTATGTAAAAAGAAAATTTATTTTTTTAAACAAATGGTGGAAATTTGGCGAAATTTCAGCAGGGGAAGATGAAA

ATTAGAGCCAAATTCTGAAAGTATTTTTGACGGCTGAAGAAACCGCAGCAATTAAAGGCATTTACTCTATCGGAACTCTG

GAACCGACCAAAAAAAATAGCTCATTTGACTAATCCAACTATTTCCTACGATTGGTCAAATACCCTCACAAATAGTCACT

ATTTGTGACGCCCCTAAATAAAAACCCTCACTAATATGACCTATTTTTGTAGTGATAAAGGAGAACTCTAGCAACTGGGA

AATCTAATCCCTACACCTCATGTGTGTTACTAGTACTAGAGTCGATTCTCCTCTAACCTTAGGTTTTTCACGAAATCATG

TAGGTTAACGACTTAAAGACTTCATTAGGATTGTGAAGACATACCCAACTATTTTCTATGTAGTTGTGTGTTCTGATCTT

GCCCTGGTCTATCACATTGAGACTATCTTCTCTAAGATTTGTTAACGAGGAAATCGCAAATGCTGAAAAACCCCGGGACC

TAATCCAGATTGAACATTGTATTAAGCCTCTACAGACACTGGCCTAATACAAACTAACTTCAGTTTGGACTGTAGTTCAA

TCTCACATTGATCAAAGGTACAGTTGCGCTCCTTACGTCTATAATCCCAGAGGATACTAGGAACTTGATTCCCTTATCTG

ATCTCACCCACAACCAAGATTTTCTATGACCCAAAGTCGAAGACTTGAATAAACAAATCTGTCTAACACTGAAAAGTTTA

CAGGAATAGATAAATTTGTCTCCCACCTACGAGTTTTGTTTTGTCTTTTGATAAATCAAGGTGAACAGGAACCAATTAAT

ATACAAGACTTATATTCCTGAAGAAGATCCTAGAAATATCAATCACCTCACAATAATCTTAATCGACTAGCGAAACATGA

TATTGTGGAATCACAAACGGTGAGACGAAGGTGTTTGTGACTACTTTTTTATATTGCCTATCGGAGAAATAAATCTCAAG

CCAATCTTATGATTATACTCAATCACAATAGAAACAGCAAGATCAGATCACGCAACTACAAATAAAATAGTTGGGTATGG

CTTCACAATCCCAATGAAGTCTTTAAGTCATTAACCTACAGGGTCTCGAGAGAAACCTAAGGTTAAAGGAGAATTGACTC

TAGCTTATATAACTAGTATCACACAGGATGTGTGGGGATTAGGTTTCCCAGTTGCTAGAGTTCTCCTTTATATAGTCTCC

AAATCAAGGTTTGCAATCTAAGTTACCTTGGTAACAAAGCATTCAATATTTACCGTTAGATAAACCTGATTAGATTCAAG

CTAATATCTTTCAACCGTTAGATCAAATCTTAGCTTGTTATACACACATGAAATACTTCATTTTGGTTTGAGTAACGGTA

CCTAAACGTGTACATATAGTTGGTTCAACAATAGTTAACCAATGGTTAGCTATATGAGCACTTTCATATCAACCTTATTC

ATCTTCACCATACTAGTTCAAGTGACTCAAATGAACTAGTCAGATAGTTGTTCACTTGCTTAGATCTCATAGAAGTATAC

AAGACATAATCAAACCAAAAATGATTTTGATTCACTCGAATCGATTCATTAAAATTATATCCATGGTTTGCAAATATTGC

ATTCCTTAATATATATATAAATTTTTAGTTCATGAATAAACCGTTTTTAGAAAATAACCCACTCAAGTACGCATACTAAA

GTACCCGGATTGAGCTTGTTTTTAGTTCACAAACTCCAGCATAAATTCACGAGATGTGAACTTCCTGCAGTACGCGTACG

GGTACACGAACTTAGCTTCGGACACCCTAAACCAGTAAAATACGCATACTTTGGTTCAAGGATTTTGGACTTACACAAGT

ATGAGTTCACATACAATGTTTATATCCATTCAAGGTTATATATTCTAAACTCTCATTTCAATCATTGAAACATTCTTATA

GGATGTTATATCACAAACTATTTTTCATCAAAGCGATTTTCAAGAAATTAAAATGATCAACATGCTTTCTTCACGAGTAA

AGATGAACTTGGCCAAAGCAAAAGCTTACCAATACATATTTCGAGAAATAGATAAGAGAGATAAACTCGGCTAGAAATAT

CACTAGTCCAAAACGTGATATTTAGGACACTTGTTGTTCGTAAAAAAAGACAGATCATTGCTGTCGTTAAATATTTGACA

GTTGAAATTCGTCGAACATTTTTCCTATGGCATATAACATGAGTAATGAGTCGTTTTCTTGGTTTTTATTTGCCCCCCAT

TCCCTTAAAGTTCCACCCCTACCGATATAAGAATAATAATAGATAAATTCTAATCGTTCTGATAAAAAAATTAGCGCTTA

TTCACCCCCGAATTTTTTGATTTTTCAACCAGAAAAGATAATTCCCTCCACGATTAATATTTATGTGTCGTTGATAATTT

AAATCGGATGGCTGGTATTAGGGGTTTGATTGAAATATTTTTGGATCCGTGTCCATACAAGTAAGCCAATTAAAACCCTT

GAACTACATTAGTCATTTTTGACCTTTGATAAAGGTAATAATCGGATCTAGAAACTTCTCAAGACTGAAAATTGAAAGAA

ATAACCGACTCCGTATTCTCCTCCTTTGTGTTTATAAAAAATTATACAGTTTCCCATAAACTTCAATTCTTATTTCGTTT

TCTCATATCATTACGTGAAGAAATCCACGGCGGACGGGAGAAGATAAAAGACGTGAAATCAATAATAATGATTCTTGTGA

TTTTGATATTATTATCTTGATTTTTCTTTTTTACGAAAACCCAAAATCTCGTTTCTTCATTTAGCTTCTTCTTTTTATTT

ACTGATTCGAATTTTTGATCTTGGGTGCAATAGCAGAAGAGAAACCAAACAAGGAGAAGCTCCTATAATTTTCAAGGTAT

GCTTTTCTTTCTCTTTATTTTACCTAAATTGGGAATATTCGTTACATGTTTCTTTCTCATCTTTTTAATTTCATATACCT

ATGTTTATGTGAATTATGGGATCTATCCCTTTTGCTCGCTTGATTGATTTGCTTCACCAGATTAGTATTTTCCATGGGCT

CAGATTTTTGTTTTCTTATTACTTGTTGATTAAAATTCCGGTATGATTTTCTGATTTCAAAGAAGATTGGTTATTGTGCT

TGGATATCAGGATTGACTATTGTCCTTAGTTGTAGATATTGTTTTAAAGCTAAATATGGTGCCCGTATTTTTCATGTGGT

ATGAACCGTTGTTGGCATGTGAACTTATTGTTTTGTTTTAATGGTGGAAAATGTCTAGTTGATGGATGCGTCTTGAGGAT

TTCAAAGAAAAAAGCAGATGTGGCTAAGAGATATTAACTTTAGCGCACTGAAAATGATGAGTACATATCGAATCCTTAGA

AATGAAATTAGTAATAGCAATATGGAGTTATGTGCTGGAGATGGAACCGTTGGTGAATTTTTTTCTTGTGATACGATTGG

ATGGCCACCACAATATGTAATTGTGCACTTAAAATTAAATCAAGCAGTTTTTATTTTCTTCTTCTTCTTTGCTATTCTGA

ACATGAACGGGATATGTAGCTCTATTTTTGAACTAGATATGTTTGATTTGTCAAGTTATTACTCGAATTCAAGTTAATTT

TAATAAAATGATGAGTTGCATTTGTTAAGAAAAAATTACGGATGTTCCGGTCTCAAGTATTAATATAATAACAATAGTAT

TAAAAAAGAAAAGAGAACAAAAGATTGAAATAAATAAATAATAAGAAAGCAAAAATAATATTCAGATTGAGTCAGCTAAA

TCTGATTGGGCTGAATCTTAAAACCTGAATTTCGACAGGCATAGTTAGTCGTTATTTTCCTACAATTGGAATCCGTTGAT

ATTATTTAACGGAAATAATCTGTCCTTTTAGAAATAGTACACCAAAAATATGACGGTTTTCTGGACAGGTCGAGTTTGTC

CTGTATCCCAATTTTCGACAGATTCGGTATGTCCTAAATTTCGCATTTTGGACCATTGTAGCAAATGTGTATAATCGAAG

TCTATATAGCATTACGACTTTATCTGCAAGATAGGAGATAGAATAGATAGACTTTTGAGTGATAGATAAGTTCAAGTCTC

CACATACCTTTTTTCAATGAAGTTCCACCAGTTCCTTGAGTAGTTCTTCGTCATTACATGATGAACACCGTGGAGTCTAG

AGATCAACTACACTTAATATCCTAGTCCGAGACTTAGCTATAAGTAGACTAGAAATCAAGACTTATAGTTTTGGAAACTA

ATTGACAAACAAGCTTGAGATATAGCAACGCTTGCGAGTTCGACCGAGCCGTGATCTAACATACCTTGAATCAGTGTGAG

ATTCATTAGGGTTCAACTACAGTCCAGTCTGAAGTTCATTTGTAATAGGATAGTGTCTGTAGCGGCTTAATACAGTATGG

TGTACAAAACTGGACTAGGTCCCGGGGTTTTTCTGCATTTGCGGTTTACTTGTTAACAAAACTTCTGGTGTTTGTGTTAT

TTCTTTTCCGCATTATATTTTGTTATATAATTAAAATATCACAGGTTGTGCATTGAATCGATCATTTGGGAAATCCAACC

TTTGGTTGTTGATTGAAATTGATTGATCCTTGAACATTGGTCTTTGGTACCATTAAAGTTATTTCTCTTATATTCCATCG

GGCTCGAAAATTCCTATTTGCTAATTGCATATTGAATTGAGAGATAGAGATATAAAACTCTTTGATATACTTTTTTCTAG

ATTGAGTCTGACTGTCTAGTTGATTCTCTTGAAACTGTATTGGAGTTTGTCTATTTAGATAGCCAAACGAAATATTGGAT

GTGGTTGTTGTACCCCCCTCTTTTTCAATTGGTATCATAGCAGGAAAACACATTTAAGACGTTATTAGTCTATGTTTGTA

GCAATCTGAATATGGACAGTGTTATCTCAGATAAATGCATTACTAGAAATAAAAGGTTTGTTTCTATAAAATCTATTAAA

GAGAAAGATTTTTCAATCTCGAATGTATACGAACCTAGTGTTCCAACGAAATAGACTTCTATTGACAAGTTTTACCCGGA

TTACCTTACTGACGAGTCTGACTCTGAAGTTGAGAGGGAAGCATCCATAGAGAATGCAGTAATTCTAAAACTTGTTAGAA

TCCAAGATGATGTCAACCGGTTGAAACACAAAGTCAATACCCTTGTTGGTATGGTAAATGAGCATGACTCCCTTCTAAAG

GTCCTTCGAGAGGAAAAGCCTCAGTTTTTTCTTCACGAACTCTTCTTGACAAAAAACTCAAAAAAAATTCTTTGGAAATT

GAACTTCTTTTGAGTAAACTCTCTATTCAAGAATGTTCCAAAGAGGCTACTATACCAATAGCTTCTGATTCAACTGTAAA

AAATTCAGTTCAAGAAGCAAAACCGAAAGCCCTTCCTGTTGTTAAAGATGTGCTTTTGTCTTCTTCTAATCAAGGAATAT

CCACATCACATGGAAGGAAAACATCTCCCAACAATGGTGTTAAGACTGTTCTGTCTAATCACTCAAGTAATAAGAAAGTA

TGTCTCTTTTGTGATTCAGAAGGACATGTTGAACAAAAGAGGAAATCGAGGTTGAATGACAATTCTATTGTTCGTCTTCG

ACACACCATAGATTTAATTCTCAAAGGTGTAACTGATATTCGTACGTCTAGACCAATTTGTTTTAGTACTCACCATGTGT

ATCCTAACAGGAAATTCAGTTCGATGTGTTCCTCAAATGTTCAAATACGAAACAGTCTTTCTAACCAATTCGTTTAAGAA

GAACTCATGGGTCTTCTTTAATCAAAAAGGGAGATAATGTTGTTCATGATGTGAAAAAGGTACCAGAAGAAGGAAGCAAA

AATGAAGAGATGGAAGGATATAAAGAGATTATTAACAGGCTGTCAATTCCCTCTAGTCAAAAAATTTCAGGTAAGATTAC

ATACTATGCGTCGTTCTTTGATGATAGTAAATTTTATGATAACTTTTCGCATAATAAAGGTTTTCCTCCAAAGATCAGGA

GAATGAGGTTTAACCGAAACAACATTCATTTGATAAGATTAAGGCTCATACTTGTGCTAATTGATGACAAGGTTGAAATC

TCACTCACAAAAATCCTGATTGGGTGAGACATGATCAGGTATACTTGATAGCAAAGTGCTTTCTGATTCTTTAGCTATTT

TATTATCATTCTTAGCTGGGTTATCATCACAAACATGTTTGCATAAGTATTTGTTTGTTTGTTTTGTTAAGATTTTGATT

TTTTTGATTTTTTTTGAAATTTTCATTATGACATGTTTGCTACTCCTCTGCTCTAGATTTTTTTCTTGAGAGGTTATAAA

ATTAATCGAGCTAAGATTTGTGAAATTTTTCACTCAGAAAAATAATGATTTGCAAACAGTTTTTTATTTACCTTCTTCGT

AGGTTTATTTTTTTCTCCCATGGGCCATGTTTTCGTTCGTACGGTTACCCATGTTACGTGTCACGAACCTACTCTGAAAA

TCATAAAAATTGTGTTCCTGAGAAACCTTTTAAGTAACATCTTTTTAGAAGAGTACGTATACTCCAAGTTATTTTTCTTG

TCAAGAATGGAATCTCCTTCGGGAAATTGTAGCTTTGCAAAAGCTTTTCTTGATAAGGGTGTTGGCTTCGAAAAGGAGGA

GAAAATAACAATTGTGGAAGTTAATGTCATCTCTCCTCATTCTCTTGATAGTTTTTCAGACATTGAGGATGATGAAGAGA

TTTCTACTGAAGGTATCCAAGATTTTATCGGTCAGTTTAACGATGTAAAACTGTTTCTTGTTAATACCTTAAAAAATCTT

GATTTGTTAAGAACTGAGTTGAATAAGGTTAACTCTGACCTTGCTCTCATGAAAACACGTTTTGAGAATAGTCTCAAGAA

TAATATCGTGTCCGAAGATGCAGTCAATTCCATCTGTCACAAGCTTGAAGGTCTTGAAACTCATGAAGCTTCCGTATGTG

AATGCTGATTTCAGCTTGTCATCTATTTGGTTTTTAGAGTTTGTTTTTGTTTAGAATTTTTTCTTTTTGATCAATATCTA

GGAAACTTCTTATGAGAATTTATTTTTGTGTTTTTAAGAAGGATAACTAGGGTTTGGAATAGCCATTATTGTGAGTACAC

ATAGCTATTGCCGACATTTTTCATCTTGAATTTTTTTAGATTTATTTATTTAAATTCTAAAGTTTATTTGGAAGATGATT

TTTCCAGTATTAATCTGTATGATTTTATATATTGCAAATTGTTATGGGATATTTTGTTTACGTCCGTGAACCTTGACTGT

CCCATATTTATTAAAAAATTAACACTATTATATGTCGGTATGAAAGTATTGTTAAAAGATAGTATGAACTTTTGACTGCA

AAAGTTAAGCCTTTTATGTCATTATGCAAATATTGATAGAAGAAAGGATGAACTTTTGTTCTCAAAGATTAAGTCTATTA

TATGTCATTGTGCAAATAGTGATAAAAAATATAATGAATCTTTGATTATTCCGCAGTATTGGCCTTTCCCTGATCCACAT

CTTTTTGTATGTACTGTGTTGCTCCGTAAGTTTTCTTATGATGAGCATGGCTAATTGAATTGATCATTTTCCTTGTGGTT

AATTCAATTGTGTTTTTTTGATACAAATTCATGTTCTCATGTGATTTTATTATGTCCAAATAAATCATTCTTTTTGTGTA

TAAAAGTAAGGTCGCTCTTGTTTTCCTTTCAGGAATGAAATTTTATGGGGGAGAGTTCGTTTTGAACTTGTGATTAATTG

CCAAATCTTTGTGGGGAGTGTGGCTGTGGAATATTGTAGGAGTTTTCTTGTATCTTTATAAACTCTTTGATGAATGCATT

TAGCTTAGGCTATATGATTGCATCCAAATAAGTTGAAGTATCTCTATGAAAATCTCATGAGGATCCCACTAGTTTTCGTA

CCTTTGCCAATTTTTATTGACAAAAAGGGGGAGAATTAATGTGTAGTTCACACTAAAAATACATATGGTTTACGGATCAT

TATGTAAGGGGGAGTGGTTTTCATATTGAGATGAAGTATTGACTGGGCGGGGGGATACAAATCACCATAGTATTGTTGTC

AAAGTTGTGTTACAATTGGACTTTGATACTGTATAATGATACTATGACATTGTATAACAATAATTGAGAGAAACTATTTT

CTCATTGTTATTTCTACGGATCTTCAACAACTATGATGCTGAATTGAACATCTATGGAATCATGGGAGTACTTGGAAGTG

ACGAAGATTTCAAGTAAAGTTTAAGAACCAAGAAGATCAAACATTTGGATGAGAAGCTACAAAGTCTTATTTATTTTGTA

ATCCATATGTATTGATGGTTTTGTCACTCAAATTGACAAAGGGGGAGATTGTTAGAGCTGATCGGTCGAACTCGCAAGCG

TTTCTATCTCAAGCTTGTTTGTCGATCTTAGTGATCAAAACTATGAGTCTTGATTTCTAGTCTACTCAAAGTAATGTCTC

AGACTAGAATAGATTGTGTAGTTAAGCTTTAGACTTCACGACGTTTATCATTGAAGATGAAGAACTACTAAGGAGAGCTT

GTGAAACTTCATCAACAAAAGGTATGTGGACACTGAAACTCATCTATCACTTGGAAAGACTATTTCTACTATATCTCCTA

TATTGAGACAAAAGTTGTATTAAGATAGTTTTCGACTATACACATTTGAGATTTCGAGTTGAGTTTAACTCGCTTACATA

TTTCTCGAAATATGTGTTGGTAAGCTTTCGATTCAACCAAGTTCATCTTATATTCTTGACGAAAGTCAAAAGATGATCAT

GTGAAAATTGCCAAGTAACATGTTACATGGTTTGTGTGATACAATCATTTGGTGTAGACTTGGAATGTTTCGTTATGATT

ATTTCAATAACTTGAAAATTGCTTTGATGCTAATATTGTGTGAAAACAACTATTGTCATCCTCTAAGAAATTTTCAATGA

TTGAAATAGAGTTTAGAACAATTGGATTTACACATAGTATGCGTACTTGCATATATGTAATCCATGTCCGGGAACCATAG

TATCCGTACTTGCATATATGTAATCCATGTCCGGGAACCATAGTATGCATACCGGTATGGGTACTTGCGTACTTGTTGGT

TTGTGAAAGTACGAGTACCCGTATGCATACTTGTTGGTTTGTGAAAGTACGGGTACCTAAGTACACATATTGGTACGCGT

ACTTGCATAAGGTGTAGGTCCTGGAATTTCTGCTGAAGTTTGGAAATGTTAATGTATGCGTACCCATTCACATACTGGCG

AACCCAAACCCAGTTTGGTGACTAAGGTATGCGTACCCGTTTGCATACCTAAGTGAATCATGTTCTAAAATCGGTTGGCT

CATGAACTAATATATTTATATAATAAGGAAGGTAATCTTTGCAAACCGTGGCTATAATGTTCATGAATTGATTCGAGTGA

ATCAAACTGATTTTGCTTCAATTGTGTTCTTGTATACTTATATGAGAATATAGCAATTGAACAACTCTTTAACTAGTTTC

ATTTGGGCCATTTGAACTAGTTTTTGTTAATATAAATAAGTTTGATATGAAGGTGTTCACATAACTAACTTCAGTTAACT

ACTGTTGAGCCAACATGGTGTACATGTTTAGGTACAGTTACATAAACCTAAATGAAGGTACATTTCTTTGTGTATAACAA

GATAAGTTCGATCTAACGGTTGAAAGATATTAGCTTGAGTTGAATCAGGTTTTCATCTAACGGTGAATATTGAATGCTTG

GTTACCAAGGTAACTTAGATTGCAAACCTTGATTTGAAAACTATATAAATGAGAACTCTAGCAACTGAGAAATTTAATCC

CCACACCTCATGTGTGTTATTAGTTGTGACTAGAGTCGATTCTCCTTTAACATTAGGTTTTTCACGAACCTTGTAGGTTA

ACGACTTAAAGACTTCATTGGGATTGTGAATCCATACCTAACTATTTTATCTGTAGTTGTGTGTTCTGATCTTGCCCTGT

TCCATCGTATGGAGTACTATCTTCTCTAATATTTGCTCGAGATTAATTCTCTGATAGGAAATATAAAAGTAGTCACAAAC

ATCTTCATCTCATCGTTTGTGATTCCACAATATGTTTTTTCGTTACCATATGATTAAGATTATTTTGAGGTGATTGATAA

TATTATACTATTCTTCGGGAATATAACTCTGGGTTATCAATTGGTTCATGTGCACCTTGATTTATAAAAAGATGGAACAA

AACTCGTAGGTATTTCTGTGAGAGACGGATTTATCTATTCCAATAGACTTTTCTGTGTGAGACAGATTTGTTTATCAAGT

TGTGGATGAGATCAGCTAAGGGAATCAAGTGCGTAGTATCTTGTTTCAACTACAGTCCATTCCGAAGTTCATTGGTAGTA

GTCTAGTGTTTGTAGTTACTTAATACAGTGTGATGTTCTAAAATGGACTAGGTTTCGGGGGTTTTCTGCATTTTCTGGTG

TCTGTGTTATTTCTTTTCCACATTATATTGTGCGTTGAATCGATCAATTGGGAAATCCAACCTTTGGTTGTTGATTGAAA

TTGATTGATCCTTGAACATTAGTCTTTGGTACCATTCAAGTTATTTCTCTTATATTCAATCGGGATCGCAAATTCATATT

TGCTGATTGCAGATTGAATTGAGAGATAGAGATATAAAACTCTTTGATATACTTTTCTCTAGATTGAGTCTGACTGTCTA

GTTGATTCTATTGAAAGTATATTGGAGTTTGTCTGTTCAGATTGCCAAACGATATATTGGGTGGGGTTGTTGTACTCCTG

CTTTTTCATGTATTACATAGATACACATATATATCATGCTTAAAATAGATATGTATTATTTTATTAAAAAAAAAAGATAC

CGCATTTTTGACAAGCCACACGCCTACCACGAGACCCTCAATCCACTAGTTCATCTTCTGATCGATGTTAATATAGACTA

ACTGTTAATCACATAAGCATTCCAAAAAATCTAGTCAATATGCTCACACAATATCCATAACACAATATCAAACAAGTTCA

CAATTATTAGCTCGGTGAAATTACTAATTAATACTTCTAATTCTAATAAATGAATTCCATTTTTATTCTAATATTTTCTA

ATTTTATCATAAATAAAAGTGTAGTCATGAAATCAAAGAATTAGGTGTTATTGGAAAGCTCTCGCAAAACTCAATAATTT

TGCAAAAGGAACGGAAAGTTAATTCTGATTATAAGTAATAACTCGTGAATCTGTTTCAATATTAAATTTGATTATGAATA

GTTGAAAATCAAAAAAAGAATTTGGGACACCAAATTATAAGTGGTCGAAAGAGACAAGCCCAATAGGACCGCCTGATCAT

CCTAGCCCATAAAAATCCGATTCCAAAAAACCGGAGTAATAAGACTCCAAGTTCCAGATCCGATTCCAAATACGAGGCAC

TTTAGTTTTCTGCTCAGCAAACACAAATCGATTTCACTCACAATGGAGAAAATTAGGTTAATGGGTTCTTCTTCAGATTC

CACAATTAAAAGGGTTGATCGAAAGGATGGTGTAACAGATGAAGTTTGGTTTGAAATCTTTGTTTATCTTCCTCTTACTG

AAGTCCTCAAATTCAGATGCGTTTCGAAGGTATGGTTATCAATACTCTCAAACCCTTATTTCATTAATAAATGGTATAAA

CTCAACTTTTATCACTCCCATGGGGATTAATTCAACATGCTTATGGAGCAGAAAACCGATCCCGAGATTACAAATTCCAT

AAAAAATTTACACTTGCATATCCTGATTCACATTCTGAGTTTATAGCTCATCATCAAACTGGGTTTTCTTTTAGCTTTCT

AAATTCAATTTTCCCCGATCAAACAAGTACATACTCTTCCTACAATCTTTATGTAGTAGGTTCAAGCAATTGTTTGGTGT

GTACAGCGAGTACTGATTATCAGACTAATTTTCATGTTTGTAATCCACTCGGAAAGAAATGGGTTTCTCTACCTCTGGTA

CCGTTGAGAAGACAAGATAAAGATGTTGTGACTGATTTCATATCTAATGGATGTTTTTCTGAATCTATGAGTTATAAGAT

TATGCAATTACCAAAGTCTGTAGTTACAACAAAGGAATTTGAAGTTGAAATGTTTTGTTCTGAATTGGGAGAATGGGTTC

TCTATGATGTTTCATGTCCTCAAAATGTTACTTGGAATTTCACCGTTTATGACAATCTTTTTAGTCACAATGGTGTTTTC

TAATGGTGTCAAAAGCAAAACCGCAGGATTTTAGCTGTGGGTTTTAATAGGAGCGGAACTTATGGGAGTGAATGTAGGTT

GATTGGTATTCCTAATGGAGCAATATATGGGATAAAATCAACCACTTATGGGAATATCTCGGGGAGTCAGAAGGATTGAT

TTGTTTTGCTAGCTTCAAGGATGTAGATAGGAATTTGAGCGTTTGGGTGCTTGATGGGGGAGACTGGTATATGTTGCACA

AGGATATTCAACTTCATGACTGGTTTGCATAAATATAATCTCAGTTGTATGATGGATCAGTTACGGTTAATTTGATCAAA

GAATTTGAGGTTGTAGGTTTCAGTCCATTAGACAAAAATGTGGTTATACTTGGTTCCGAGAAGTATATATGGGCATATAA

CACCAAGACCAGGGTATATGAACAGTTATGTCATCCTATTGAGAGTACCTCACCAAATCCTTTTATATCAATGTTTCCAT

ACTCACACCACTTTCTCCGTGTGGCACATGAATCTTGGATGTTCAAATATGAATACGATTTGAAAGTAATTAAAGGTGTT

GCAACTGTTCCTATCTACTTGTGGGTACAGGTCCATTCATGTTATTGGTTCTTTCTTAATTTACTCCATCTAAGTAATGT

TTTTTTTAATTGTTATTTTTTTAATTTAACGATACATTCCTTGTTCAAAATTCAAACTAGAATGTTTCTATTTCCTGGAA

CAAAGGAATGAACATCATCACCAGTGAAGGGCGAGACACCAGTGACATCCATGAAAACATCTTGACCATTTTCCCAGTTA

ATCACAAAATATCAGCGTGCCTCAAATCTTTACCATCATTCGACACCATACCAAGAAGGACTTCCTTACGCGCAGGCACA

CCTGTTTTGTAACAAATATCATCCATAACATCACGAACAACGTCATGTCGAAACTTAATTCCAACATCTTTAGCACAATG

AAGCGCATGATCTCCGAAGATGTCCATGGATTTGTTACGACTTGAGCATAAACCATTCTCAACAAACAAATGGATACCAA

GGCGATAGCAGAGCACAGCTCGGAACTATCTAAAATTCTGCATGGAAGTGGGGGGCCGGGTGTGGGTGGGTGGGGGGGGG

GGGGGGGGGGTAGTATCATCGACACAATAAGAAGGAGGTGATCCACATACCTGGATAAAATTCTGAAGAGCCATCTGATA

AGCAGATCCTTTGTTTGTTGGGGATAAATTACCGAGAATGGTCTTCTGGGTAGAAGTTGTCTGACTCTGGGAAGCTAGGT

AACAATATGTATGGGTCTCAGTCGTGGTATAAATACCTAAACCACCGTCTTTGATAGTCAAGGTAGCCAATCTATGATGT

AGAGGACCAAAACCCGCACCATCACCAGTGATTAGTAATCTCAAATACTTAAGCAGATGGTCATCATAAATATTTGTGGC

TGATTGCAAGGCTGCTGGATTGGTAGTACGCATTGCGAAATATAATCTAGATACACCAGTGCAATTGCGAAGTAATAGCA

TCTCGCTTTGATGGTCCTTGAGTTTCTTGATTGCATTCATTAATTGAACAATTTTATTCACCTTGTTCAAGATCATGTCA

CTGTTAAAGTTCAAATCCAAACTCACTGGTCCACCTAAAAGTTTAACACCATTAGAGGGTCTACCAATGTCATGGGGAAA

TACCCCCTCAGCCGTGCTCCTTGGATCAAAAGAAAGCCAAAATACTTTGGTTTTCTTTATATTGAGATGCAACCCCCTAC

TTGGCCCTTCAGTTTCTATTATGTTCAAGGCCTTGGACACCTCCATAGTATCACCAATAATAGTACCATCATCAAGGTAC

CATGCATGTAAATCAATCTTGCACTGAGAAGCAATAGACTTCACTAGAGGGTGAAGTGTCAATGCAGATAGCAAGGGACC

GAGAGGGTCACCTTGCTGAACTCCAAGTGCGGACGATAAAATATATTGGTTATAATATAGCTTAGCAGGTCGTAAGTAAC

AAAATTCAACCCAGCGGGAAATACCTAGGCAATGTAGACGAACTTCCTGGATGAGTTAAGATCTGTTCACCATATTAAAC

TCATTATAAAAGTTAATGAGTAACATTGACTTTTTATCAGAATGCCTTTTCATCTCCAGTACGCAATTTACAGCATGTAA

AATGCTTTCCCCCAAATAGGCACTCCAACACCAAATTGGTAGTCACCTAAGTAAGAACTCATGTCATTTCCGACGGAGAA

ATCCGCTACCTTCGAGACCAATCTATGCCAAACCATACCTATAGAAATGGGTCGAATACCACCACCAGGTTTAAGTAATG

GAGTTAAGGGCACACTTGCAATGAACTCACCTAAAACCTCCGGACATTTTCCAGCCAACCAAAGGTTGAAAACCCCTGTA

ATAGAAGCTAGTAAATTATCCGCTACAGCTGCAGCCGCTCCACTCATTACATCAACTAAATGTTGAGCACAGAGGCCATC

ACAGAACGAGAGTCAACAATAATTGGGTAGACCAAGATAGCGTCGTCGAGAATGACAGGGGGAGGCACAGGTGGGTGTTT

GTCTTATAAGTCAAGATAAGTTGCATCATTGCATGGAGCCACCCAGACGAGGAAAGTACCCGTATTGCTGCAACAAAATG

TCCGCACGCAATCTTCTTCCGGAAAGCTACCATGTTGACAGCTTAATGCTTCTTCGAGTCTGTCTTCCTAGATCTCATGT

GAATAGGATGATGGAATAATTTATTCACCAGGACAACGCAGCCACCGGGCTCCCTCCAAACTGAAATTGCATGGTTGATT

GTCGCAAACTGCAATCTCTTCTTATTGCCGGACTTTTCTTCAGCTGAGTTCCTCGGTCTAAATAGATTCAAAGTGAAGAT

TGGAAGAAGCATCAGCTGCAACCATGCCGATAAGCTACCCAAACTACTGATAATCCTGTCTAGACAAGTCTTCAGCTCCC

TTGAAAATGCGAGTCTGGACTTGTGTGGGAAACAGGTTACAGTTGTAAACTGTTTTTGAAGAACATCATTTAGCAGGTCC

GTGTTGAGGGATTCTACAACATCACCACTATGAATCTCAACCCCATCATCACAGACTGACAACAGAAGAATTCAGAGCTA

CAATACTTAACTCAACAGGCTTATAGACACTGTGAATAAGGAAATCGGAAGCAATTCCATTGGATGGCCCTGTAACAACA

TCTCCATGAACAACCCCTTCACGGCCCTTACATGGCATCTTTCAGGAATGTAGGTGCATACATCTACAACATAGCCACAT

ATTTAACAAACCAAGAACCTTCTCCCAGGCGTGATATATATGCAAGTTCGATGAGATCATCTCCATGCAGTTGGCTTTGT

TCTCAACAAATGAGAAGTTCCGATCATGTAGATGTTTCAAAAAGTATCCTTTGGTATATCCCCTGCCATTAACGCCATCA

AAGCTCTTATAAGGGCAGTGACAAAGGCCATTACCAACAAGAGCTTCTTCTATAATGGACTGTGAAGAAGTTGAGGATAA

AGAAAACTGAGAGGAGTGGGTTATCATCGACAAACGCCGTGAAGACGAGGACCTGGTGGTTCTAGCCATGAAGGATGGAG

ACGAACCCTAAAGAAAAGTTAGAAAAAAAAAAGATACAGGAACCCTAAAAACAAAAACGATAAAACAAAATACTAAAAAA

GGGAAAAGATGAATAAAAATCCGTGGAAAAGAAATCAAAAACAAAGAGAAAGACCTGCGAGAAAACCCAAATCAATCATG

GTGTTCCAGGGAAACAAACCCGGTTTTCAGTCAAGACAATCAAGACAAAAATATTAATTATTTTATTATTGCTTTTCAAA

TCAATCAAGACAATCACACACATACAGTACACATGCCGAGTAGACAATCATGATCAGAAACTTTTCGAAAATTAAAATAA

ATCACTGGTTGAAAAAGGACATACAACTCTTCGACATAAATAATAACAAAAAAATCTTATGAAATAAAACATGTAAAAAG

AATAAACAGAAAAACGAAACAGTCCAAAAAAAATTATTTGATGATGGCATATGAAAACATTTAATAACATATGATGACAT

AACAATAGATGAAAATATTAAATTTATACAACATGAGGCATATTAAAATCAAACTTTATCACATAGAGATCATTAACATG

ATTTTAGTATGAATAAATAAAATGACATCAAACATGTATCACTCATATTTGTTAGATACACAAACAGATTTAAACATGTT

ATACAGTTGATTGAACAACAAATGATATTATAATAATCATAATAATCTCGATAAACGAATAGGAACAAACAGATTGTGTA

CAACAAACAAACATCTCACAACAAACATTAAAATAATATATATATACAATATACGCAATCACAAATTAAAAATTCATTCA

ATCAATCAGAACATGATCGACGCATGTGATACAAATATACATGTGAAGAAATAAAAAAAAATCACATACATACAACATGA

GAACATGGAAGTCAAGATTACATGTTAATTTATAAGAAAATTGATAAAAGGTTTAGGTATACCTCAACATATTGCTTGAT

TGAGAAAAAAATAATCAGAAATACGGATCTAACTAAAAGTAGCTCTTCCTTCAGTGATTTTATAATAACCCAAAAGAAGA

AGAAAAAAAAATTAGTGGTTGAGCTCGTGCAGATAACGTTTTGGAGTAACAATTAGGTTAAAGCAAGAGATAGAGAAGGA

GAAGAACTTGGTATTGATAACGAGGAATCAAACTGTTACATACATATAATTTTCTCTTATATACAGGAAAGGACCGCACA

TCCATGGGCCACATGCCCTATAACAAAAAACAAGTTTTAAGGCATGATAAATGCAAAAAATCTCGGTATTAATCACAGGT

TGTTCGAGAATTCGACAATATGATAAATGCAGAAAATCAAATTAACGGCAGAAGAAGAAACTATGCATAGTTTTAGTAAG

AATCTTTTACATACCTCCTTAATAAAATAACAGAGAGAATTGATTTTTGTTACTCTCCGGTCTGTTTTCTTCTTTCTATA

TCTCTCACTATTTCTTTTAACCACACAATGGTGGAGATAAGAACTAAACACACCAAAAATATAAGTGCCACGCGTTAGAT

TTTTAACAGATCATATAATTTTTATTTATGTTTGAGGTAATTGTTAAGCTGGTGCACCCTTGTTTTTTGTTAAGAGGAGC

CGGTCGCCGGCAATTTGACCTCTTTTTCACTCTGGCTAGATAATTATTTTTCTGAGTTCCTAGATAACCTATAATCAATT

AGTGCTTAATAATAAATTGGATGAGTTCAATTAGTGGTAATTAATACCCATTGTGTTGTATTTCTAAACAAGAAAGGAGA

AGCCATTGTTCACCACAGTCAAAAATTAAATATAATTTTAAAAAAAATCTTTTGAGAATTGTTTTATTTAAGTATGCAAC

TGTCTGAGATCAAATTACCAAGGTTCAAATTCTTAAACAAGATAGGTCCACTGGGATAACCAAATGACGTGTTGTTGTGA

AAGTTCAACGTTTTCAGAGACAAAGCTTAAGGTGAACAGAAGCAACATAGCTACAAGGATAACTCGGACACTTTTGTCAC

TCAACATTACAATATTATCTACCCTTCAGGGTTCTGTAATAATCAAAACATACACATTAAGTAATGAGATGAGGCTCAGA

TTAGCAGCATAGCTAATATTTAGGTATGTTTTAATGAATTTACCTAGGTTAAAGAATAAATTCACATTATAATAAGGAAA

ATATCGTTATGCATAATTATTATAATTAACGAAATATCAAGTTTCAAATAATTATTCTCAACTAACGAAATGTCAAGTTT

CCAATAATTATTATATGAAATGTTCAAATTAAACCTAGTTTTTATGCATATAACGAAAGAAATTTAATTTTTTAGAGTAC

CTTGAACATTTCTAAATAGATATAAATGTATTAAAAACATAGCAACCACTAGTAAATATTTATAATAAGGAAAATATATT

TACCTAAACTTGGAAAAGGGTTTAAAGATGAGATATTTGTAGCAAAAAGGTTTGAAAGTATGTTTTTGTTTGGATTCCTA

AAAAGTTGACACATGATCTCTTAGATTTTTCTGAACCGTAGTTAACTTTTCGGTTAACAAGATAGTCAAAAAAGGTGTAC

TTCTAGGAAGTTGTGGAGTGAAGATTGCCTAATATTTACCGGGAATATCTTATTTTGTCAATTATTAGTGGAAAATGATT

GTCAAGTAATGTTTTGATGATCTCATTGTTGTGTATGTGTTTGGGTAAATCAATCTCGAAGTATCATCTTCTTCGTTCTA

TCTACTACTTTGTCGATTAAGATTGCATACTTTTAATCAAATTGCGATATCTCCTCTATGCTTCCAACGGTAAGTTACAA

CGACCATTCCATACCTAGAAATGTTAGAGAGTTGTGGGCGGCATGGTATTCCAACTGATCTTCTTCTTAGGTCTTCATGC

TCATATTATTACTAAGAGATTAATGTTTGATGGAACCATGATTTTATGGTTGTTGGGACAAAGATTTTGTATACGAAGCA

AGAGAGGCTTGTCAATTATGTTATATGGATAATAATAGTCTAATGCAATGCCCTTGCATGTATTTTAAGTGTCCACAAGA

AGATTTTCCTAACATCAAAATGGTCGTTGAATTACAACTAGAAAGATAAGATATTTGCAATTAATGTCAATATCTTATAA

CTGTGCTGAAGAAACATAGTTGCTAGATAATGATGTAAAAGCTAAAGAACATGGAAATCTAGCAAAATCGTGCAAAAACT

TCAAACTCAAAGCCGAATTAAGTACAATCACTATGGAGAAAATCACCATAAAGACATTGTCGCACAATGATTGATTTCTC

AAAATTATGACCAATTGCTCAAACCAAGATTGTAATGCTTATATGCACATCTACTTCAACAAATCAATACATGCGTGGAA

ATCTTTTTGGACTAGACTAATTAAGTATCTTTTCATGTTTTAGTTTATTAATTGTGTTGTGTTATCAATGAAAATTGTCT

TTGTCTTTGCACCGTTAGTTAACTATCTTAGTTATTTTAAGCCAACCTTATTTTATAGCAAACAACATTGAACTAAAGAA

GACTAAAAATACGTAACCTACTCCATTATTTTTAATTAAAATTAACCAAAACTAATAAGAGCAATGCATTACATTCAATT

CAAACTAAATTAAAACTCCTCAATCAACATCTTCATTCAATTGGAGGCGGTAGAGCATAACAAGCTTTATAACTGAAAGG

TTTTCCTTTAGATAGATAACTGTCAAAACCAAATACCTAAGTATGCAATTTTAAGATTCTGAGGCTAATGCCAGATTTTA

CATAGTCTTTAAAACAAAACAAAAAAAACTCTATGAATCCTAAGAAATTTAATTGATTTACCGAATCTCGGGATTTATAA

TCCCATGAGATCATAAATTCCACGGTATTGTGAAGAATCCTATTTTAGGGCATACGTAAAAATTTCAGTGCATACTGAAT

AAAGACAAAAACTTATTATAAATAGTAAAATCAATATCCCCTTAACCAAATATTTTTAGTAATGGCAAATCTTCCCTTTT

AATGTTAGTGTTAATACTAATGATTATTTATTAATAATCCTAATTAGTGATTAGTATTAATAATAAGTGATTAAGATAAT

TTAGTTATAAATTTGTGTAGATGAAATTTTTTGAGGAAGAAAAAAAAAGTTAGTTTTTTTTGAAGAAGGAGAAGGGGAAA

GTGAGAGAAAAACTTGTATTTTGATTCAATGGATTAGGAGGGTCAATCTTCATCTAAAAAACCTAGGAAAATACACATCA

ATTGCAACAATGATCCGAAAATGGCTGATTTGTTAGATTATGAGAATGATTTGTTCAAACAAAAACTCTAACTCAAACTC

AAACTCAAACTCAAGATGATGATATTTATGAGCCAAATCCAAATGATGAATACATATATGAGGAACCCAATGCTTGCAGC

ACACTTCTATCTTATGTTTAATCTCACTTTTGTAGCTTGAAATCATCAAAAGTTTGTATTTTTTTCACTATGGTTCGACG

AACCAGGAATATGTTCGTCTTAAATTTATCAACCGAACCCACCTGGTTATTGAAAAATGATGAACAATTCGGTTAGCTGC

AAGTGTTATGGTTAGCCTCCGAACCTTTATTTTGTTATAGGTTCGACACATATTGTGAATATCGTAATAGCCGAACCTAA

CCATTAATTTTTCTTTGTGGTACAGAGAATAGTTCGGCTTATAATGGAAATATCATAACCAGCCGAACCTAGACTTTTCC

ATGTGCACTATAAACTAGATGCACATGGTTCAGCTCAAACTCAATACTATTGAATAAGGCGAACCCAACAACTGTCATTT

GATTCTCCAACACAGTGAAGTGTTCGAATTGTAATTCAAAATTATTTCAAGTCGAACCAACCGCGATCAACCAATAAATA

AGCACAGTGGGTTGTTCGGCTTAAAATAAGTATTATCGAATACGCCGAACCCCATCTTCGCCCCTATCTGTCACTAGTTT

CGGCTTAAAACTCATATGAAATTTCAAGATGAACTGTTCATATGCAGTTTCAGAAAGTTGTTATGAAGAATAGTTCGCTC

AAAGTTCAACTCAATTTTAAGCCGAATCCAATAGTGTAACCGTCAAAAACCCTAGTTTTTCAGAAATTAAACCTATTCAA

TCAATGAAAAACAATAAATAAAGAGATGGGTTTGTAGAACTTACTTGTGTATCCTCATTTCTGGAGTTGGTGCTTCAATT

GGTTGATTTTGTGGTTCAATTTAAGTTTCTACGCCTTCATATTCATTTGGTTCTTCTTCTTCAATTGATGGATTAGATAT

GGATTAGAAGACGATTTTGAATGCCTACTCATTGTTTTTTTCCTTTTTACGAACCATTATAGTTGAGTTTAATCGACATT

AAATTTTAGCCGCAATGAATTTTTTCTTCTTCTTTGGAAGAGTTCGGATAGAGTAGGGGAGGGAGGAAGAACAAGAGATT

ATGTGAGTGAAACTAATTTATTTATTTATTTTAAACTGATTTTAGGTTTTTATATTAAAGTAAAGGTATATTAGTAATTA

AAATCGGTTGAAGGGTATATAGGTAATTGAACTTCCCAAGGGATACCCTTTATAGGGCACCCTATATGGGATTATTGGTT

TGTATGCCCTGAAAGTTTTATGTATGCCCTAAAAGGGGATTCATTGTGAATCCCAAGACCAAACCTAGGGGTCATCATTT

TATCCGTAATCTGCGGATTTAATAGGGACCAACCGTATTTTTGCGGATGGATGTCCGGACCATTCACTAATGGGTTGGAC

ACGGATGAGATTTTTGAAATCCAACAGATTACGGATTGGGCCTGGATGCAACCTTGAAAATCCGTTGGACATCCATATCC

GTTAGATTAAGGGCATTTATATAGTTTCTAGAAAATATATGGATAGTTTTGGAAATTTTAAGGTGTTTGCTTGGAGTGTT

GTCCATGGAACTTCCGTATTTATATACATATATAAAATGTGTAGGTTATATAAGAATTCATCTATATTGGTTTTATAAAT

TTTTACTAACACAAATCCATTGGATTATCCGCACCCAATCCTTTCATCCGTGCATCCGTTGGATGCAAATATGTTGGATG

TTGGATGCAGATGCAGATGCCAAATTTAAAATCCGTAGTGGATTGGATTGGATTGGATGCATATGAGATAAAAACCGGTC

CATACCTAGAGCTGCACATGGGACGGTATAGTCCGGGTTTCTTATGAACCCGGTACCTGTACCTGGAGTTGACCGGGTCT

ACATTTTAAGGACCGGTACATGTACCTGGACTGGTACCTGTACGTGTACACCGGTCCGGTACAAGTCGGTACCGGGTTTT

TACCCACTGCAGATTATGTCTCCATCTAAACTCAAAAAAAAAAAATGTCCAAAATTCCAAATCCATTACAGTGATTAATC

AATTTAAGAAATAATCTTTATCAAGTACTGAATCACAATGAAATAAATTCATACTTTTAATCAATACAAGATCAAGAGAT

AAAAAAATCCATTACAGTAATTAATCAATCCATTAGAGCACAAACCCATTCATCCTCATCTCATGAACACCACAAATCCA

TCTCAACCTGAATCAAATTTCAATTTCATACACAACCCAACATGTATTTCCTTCTTGACTTCAAATCCGTAGCCAATCCC

TTGAATCATCGTCTTCTTACACACCTGTTGAGATTCAAATCTTCAAACCCTAACTTCAATTGTTCTTCACCAGAACATTA

ATACCGTGTTTAAACAAACTAAAATAAAATAGAAAAAATTAAGAACATAATCGAACAAAACCATCATCAACAGAATTGAT

TGAATAATCAACTGATCAATTGAAGAAACTAATTAGAACCAATATGAAATTTCAATCCATAAATCCATCTAAAAAACCTT

TCAATTCTCATTTAACTGATTCAACCCAAACATGCAATTAAACAAAACCCTAAATCGATTAAAGAAGTAACAAAAAAGAA

CTAAAGAAATAAAAGAAGAAGAAGATTAATAACTGAATGGAGATTATTTTGGATTCATGGTGTTGGTATCCCTCTCTGTG

ACGATGAAGAAGGTGGTGGTCTGGCAAGGTATGGCGAAGAAGAAGAGAGAAGAAAGATGGGCTTCCGTCTCTGTAAGGAG

AAGAGATAATGAAGGAGAGATTACAAAAATAGATTATAAAACCTAGCATCAATGGTCAAGATGCATTTAAATCAATGGAT

ATAATCACTCCGGGACATATGGTCCGGGACATGCCGGGAATCCCCCAAGAACCGGTTCCAGTACCGGCCTACACCGGTTC

TCTACTTTAGGAACCAGTACCTGTCCCGTCTACTCCGGTCCGGTACGGTTTTGATGCGGGATTTCCGATTCCTGGCCGGG

TCACCGGTTCCCAATCGGATCTTGTGCAGCTTTATCCATACCTGTCCATGCTCACCCCCTGGTCCAAACGCACCTTCATG

TGTTCGCTTGGAAAATGAAGAACATGCCTGACATAGGCCCATTACCCATAGGTTGATAGTGAAAGTAAAATACTGTATGA

CCCAATTTTTTTAAAACCGGAGTAATGAGAATCCAAGAACCGGAGTAAAATACAAGTCTCTTCGTGGGCAATGCCCTAGT

TTAGTTTTCTGCTCAGCAACCACAAATCGATTTCACCCGCAATGAACAAAATTAGGTTAATGGGTTCTTCTTCAGATTCC

ACCATTGAAAGAGTTGATGGAAAGGATGATGTAACAGACGATGTTTGGTTTGAAATCTTTGTTTATCTTCCTCTTACTGA

AATCTTCAAATTCAGATGCGTTTCGAAGGTATGGTTATCAATACTCTCAAACCCTTATTTCATTAAGAAATGGTATAAAC

TCAACTATTTATCACTCCCATGGGTATTAATTCAACATGCTTATGGATATGAAAACCGATCCCAGGATTACAAATTCAAT

AAAGACTTTTCACTTGCATATCCTGATTCATATTCTGAGTTTATATCTCATCATCAAACTGGGTTTTCTTTTAGCTTTCT

AAATTCAATTTTCCCCAATCTTTATGTAGTAGGTTCATTTACATACTCCTCCTACAATCTTTATGTAGTAGGTTCAGGCA

ATGGTTTGGTGTGCACAACGAGTACTGATAATCAGACTAATTTGCATGTTTGTAATCCACTCAGAAAGAAATGGGTTTCC

CTACCTCCGGTACTTTTGAGAAGACAAGATAAAGATGTTGTGACTGGTTTCATATCTAATGGATGTTTTTCAGAATCTAC

AAGTTATAAGATTATGCATTTACCAAAATCTGTATCTGCAACAAAGGAATTTGAAGTTGAAATGTTTTGTTCTGAATTGG

GAGAATGGATTCTCTATGATGTTTCATGTCCTCAAAATCTTACTTAGAAGTTCACCGTTTATGACAATCTTATTAGTCAC

AATAGTGTTTTCTACTGGTGTCAAAAGCAAAACCGCAGGATTTTGGCTGTGGGTTTTAATAGGAACGGAACTTATGGGAG

TGAATGTAGGTTGATTGGTATGCCTAACGGAACAATAGATGGTGATAAAATCAAGTTCTTATGGGAATGTCTCGGGGTGT

CAGAAGGATTGATCTGTTTCGCTAGGTTCAACGATGTGGACATGAGTTTGAGTGTTTGGGTGCTTGATGGGGGAGCACAA

GGATATTCAACTTCATGACTGGTTTGCAGAAATAGAATCTCAGTTGTGTGATGGATCAGTTGCGGTTAATTTGATCAAAG

AATTTGAGGTTGTAGGTTTCAGTCCATTCGACAAAAATGTGGTTATACTTGGTTCCCAGAAGTACATTTGGGCATATAAC

ACCAAGACCAGGGTATATGAACAGTTATGTCATCCTATTGAGAGTACCTCACCAAATCCTTTTATATCAAGGCTACCATA

CTTGCACCACTTTCTTCGATTGGCACATGAATCTTGGATGTTCAAATATGAATAAGATTTGAAGGTAATTAAAGGTGTCG

CAACTGTTCCTGTTTACTTGTGAGGACAGGTCTGCTTCTTTCTTAATTACTCCATCTGAATAATGTTTTTTCTTTGTAAT

TTGTTATTTTGACCGTACATTCCTTGTTCAAAATTCAATACTAGAATGTTTCATGATTTGCATTACCCTTATTTTCATTT

TTACTTTCTTTCTTTCTTCCTTGAATGGTTGTTCTATGTTAGCTTTTTTTCTTAGTCAATGAGGCAATAACCCTCTTATT

GTTGAATTTGGTGTCATTAAGGCTGTAGCTTAGAGGGTAGGTGTTTTCTCAGAACATCCATCATCAGAATTACAGTTTTT

GTCAGGTTTATTTGTAAATCTTGGAGATCTGTCGAATACATTTTGAAGAGAGTCCCAGCGGAACCTACTGTTGCTTGCAA

ATCAGTATGTAAATCTTGGAGATATGTCGTCGCTGAATCAATCTCTATGATGCATAGGTATCATCATATTTCTGCATCTG

TTGATCCTGGTACGCTGGGTTTTCTCTCTAAAACTGATAAGAATCAGGGATGTTGGATTTATATACTTTGAGTATAATGA

GATGTATGCACCACCCATTGATAAAATTTGAAGAATGAATATAAATCTTTATAGGATCTTTACTGAATGTCCTGTCCTCC

CTAGCATTGTAGGTTCGTGTAATGTCCTACTCTGTCTTTCTGGAAGCGAATCTGAATCTGTATATATTTTCAACATGATC

ACTAGAGAACAGGTCAAGCTTCCTCATGTGGAAAGAATAGAAAGAAATAAAAGGAAAAGCAGCAGAACAAAGGATCCATA

TTATAAGTGGACTAGTGGCCTTGGTTTCATATCATCAACTAATGAGTACAAAATTTTATCACTATATGTGTTCAGAAACA

CTATAAAAGTCTATATATATACATTAGGCAGTCACATCCGATGGAGATATGCGAGAACAATAAATTTTGGATTCATGTTA

AATCGTAATCGTTGGGAACACGGTGTGTTTGCCAACGAATCTTTTTATTTCTTCGACTGTGCTGTGAAAAAAAATTATTA

CCTTCGATTTGGCCAAGGAAAAGTTTTCTTTTATTTCGCCCCCTTCAAAACTTTCTAGTAGGGATGTTACACTAGATATA

AACCTAAATAATATAGGAGTTCTAGATACACACTTGGCAGACGGATTCAACATAGGAAGAAGAAGAAGGACAGCAATTGA

GACGAATAGAGAAGGACAGAAAAATAGAAGAAGAAATGATTCATGGAAAGGAGGAGAATGAAATGCTAGAGGAATTAAGA

TATAATAGAATGTAAACAAAGAAGATTAAGAAGAAGAAAACTTGTTATTACTGGTTAAACAAAATATACAAAGAATAATA

GAGTTTAGACATATTTATTACAAAGGAATTATGCTAGAGGAATCAAGATATAACAGAATGTAAACAAAGAAGATAAAGAA

GAAGAAAACTTGTTATTACTAGTTAAACAAAAGATACAAAGAATAATAGAGTTTAGACATATTTATTACATAGTAATTAT

GCTGTCTGAATCAGACAGATTACTCAGACAGCTAACTCAGAAAACCATACTAACTCAAATACTAAGTCAAGTCAAACATA

TTTGACTTGATTTCCTATTAGCCCCCCTCAAACTGATAATGAGAAGAAATCAGTTTAAAAACAGCAACGATTTTTAAGGA

ACAAGAAAAGAAACATGTTTAGGAACAAGCATACAACAATTTGCATCAAAATGACCAGTAACTGTTGAAGAAGATGAACC

ACTAATTGCTGAAGAAGTAGAACCACTAACTATTGAAGAAGCACCCTGATCAACAAATGAATCAATATGTGAAGATATAG

GAGAACCCAGTAATTGCTGAAGTAGTGAATGAAACACTGGAGCACATAAACCTTTAGTAAAATGATCTTCAAGCTGATTT

TCTGAAGAGATGTGATGAACACTAAGAATCCACTATGAATCAGCTCTCTAACCACATGATACTGAATTTTTATATGTTTT

GTCCTGTAATGAGAGACAGGATTAGCAGACAAGTAAATAGCACTTGTATTATCACAATAGAGTGAAATAGGAAATTTAAC

ATTAACATGTAACTCAGACAGCAAAGAAGAAAGCCACTCCAAATCAGCTGTATCCACATATAAACATTTATATTCTGCTT

CAGTTGTTGATCTTGCAACTGTAGGCTGCTTTTTACTAAACCAAGAAACTAAAGAATCACCAAGAAAAATAGCATATCCA

CAAGTTGATCTAGCAGTTTCAGGACAACTTCCCCAGTCTAAATCTGTATAAGCTGTTAATTTTCAATATCACCCTTTCTC

AAAGTAATGCCATATCCAACAGTGCCTTTTAAGTACCTAAGAATCCTCTTTACCAACAACAAATGAACATCTGTAGGAGC

ATGCATAAATTGGCTTACATAATTTACCCCAAAACAGATATCAGGTCTAGTTAAGAATAAGTACTGTAACATTCCAACTA

AAGTCCTGTAAGTTGTTGCATCTGATAATAACTGACCTTCATGAACTGAAACTCTAGGACCCTTAGCTACCGGTGTAGTG

CAGGTATTACACTCTGTCATGCCTTCTTTAACTAATAACTCAAGAGTATACTTTTTCTGTGTCAACAAAACTTCATTTTG

TGTTCTTGTAGCCTCTATGCCTAAGAAATAATGCAAATCTCCCAATTGCTTCATTGCAAACACGGTACTTAAATATTGTA

CAAGATGATCAAGCAAAGAATTTGAAGAACCAGTAAGAATTATATCATCTACATACAACAATAGTAACATCATTTCTGAA

CCAACAACAAAAACAAACATGGAGAAATCTGAAATTGCTCTAACAAAACCATATGAAATTAAGGCTGAACTGAACTTCTG

AAACCAAGCACGTGGAGCATGTTTTAAGCCATATAAACTTTTCTTAAGATGACACACTTTATTTGGAAATTCTGTACTGA

CAAAGCCTGGTGGCTGTTTCATGCAGACATTTTCTGTTAAATCACCATGCAAAAAGGCGTCTGAAACATCCAATTGCCTT

ATAGGCCAATTTCTTGTAACAGCTAAACAAAGAACAACTCTAATTGTAGTAGATTTAACTACACGACTAAAAGTTTCCCC

ATAGTCAATACCATCTTGTTTATTATATCCTTGAGCAACTAGTCTTGATTTTAGTTTATCAACAGTCCCATTTGATTTTA

GTTTAACCCTGTAAACCCATTTACATCCTAAGATGTTCATGTGAGGCTCAGGATCAACTAAGTCCCAAGTTCCATTAATC

TGCAAAGAAGTATTCTCTTCTGCCATCGAAACAATCCATTGTGGATCTTGCATTGCCATTTTGAAATTTTTAGCCTCAGT

AGGAGTTTGAAGAATAGAAGTAAGAGCAGAAGAATGTTCAGTAGAAAAATTCATGACATGATCTGGAAAAGTTTTTGTCT

TAGTAATACCCTTTTTATATCTAGTGATAATAGAGGAATAAGGAATAGAAGGAACAAAAATAAATGACTCATTGTCTAAA

ACAGGAGAAACAAAGGAAGAAGTAGAGAAATAAAACTCATTCTAAAAAAAAATCACATGTCTAGATATATAAAGCTTTTT

AGTCATAGGATTAAAACACTTGTACCCCTTATGTAAAGTACTATATCTAACAAAAATACATTTAACAGTTTTTGGAGACA

ATTTGTCAACTCTTGAATGCCCTAAGAAAGGATAACATATGCATCCAAAAACACGTAAAAAAGGAATAATCAGGCAGCTC

ATTGTAAAGAACTTGAAAAGGAGACTTATCATGAAGAATAGGAGTAGGAGTTCGATTAATAAGATAAGTAGCTGCTAAAA

AAGCATCATACCAGTACTCTTTTGGACAAAAGCATTAAAAAGCAATGTATTACCCATTTCAACAATGTGCATGTGCTTTC

TTTCAGCAAGGCCATTTTTCTCAGGTGTTTTAGGACAAGAAATTCGTAAAAGAATACCAGAAGTGTCTAAAAAAGTTTTG

AAAACCCCTTTCACTAGTTCACTAGCACCATCTGTTTGAAAAGCAAGAATTTTAGTATGAAATAAATTTTCAGTCAGATT

CTTAAAATGCTGAAAACATTGCAAAGCTTCTGTTTTTACTTGCATTGGATAAATCCAATGGAATCTACTAAAATCATCAA

TAAAAAGAATATAATATATATGCCCTGCTATGAAAGGAACAGGAGAAGGACCCCACTCATCACAATGAACAAGTGCAAGT

GGAGAGGAAGCATGAGAAGAAGAAAGATGAAAGGGAAGCCTTTTACTTTTTGCTAGTTGACAAGGATGACAAGGTGTCTC

TCTGCTTTGTGATGATATAACAAAAATATCATATTCACTACCCAAACATTCATAACAAAATTCATAAGATCTGTATCACT

AACCTTCTCACCAATATCAGCCAAAGAATCTTCTATACTATTAACCTTCTGAAGACACTACACCAAATCCTGGTTTTAGC

AACGCTTCATTTTCAGAGGTGCGCAACGGATATAGACATTGCGCAGTTAACAACGTTAATATCCGTTGCGCACTAGATTA

TCCGCTGCACAAATTATTTATATTTCATTTACTAAAATAAACTAGTGGTGCAACGTATGGTTAACACCTACAACCGTTGC

ACAGACTATGGTTTTTTTTTAATATATAGTATGCTTATGGGCTTTTAGTATTTATATTTTAATACCTATTTTTATTATTT

TCGGAAATAAAAATCCCCCTAATTTTTCCAGACAACTTTAATACACCTACCGCTCTCTCTCTCTCTCTTTCCATTTTTTT

TTTCAGTCGATGACTTCTATGTTCAGTTCAGTGCGAGAGAAGAGGAGAACTTTTCTCTCTTTCTTCACCAGATATTCTAT

GTTCAGTCGATAACTTCTTCCTTTCAAAAAATCAAAACCCATCATTCGATAATTTCTTCGTTTCAACAACTTGAAATCTT

ATAATCGATTCATCAGTTGAAATCTTCCTAAACTCTAATTATATAAGCTCCATTTTTTCCCTGAAATCTAAAATTAAATC

GGATTATCAGCAGCAATTTGTGGATTTTCATCAGGTATAATCGTTCAGCCACTTCTACTGCTTCTTAATTTTATTTTTAT

CGAAATCTTTTGAAGTTTTTATGATTTTGACCTAACACTGTGAATGAATTAGGAGTTTTGGTTGTTTTGGTCTTTATTTT

GGTAATATGTACTTTTGAAGAATTGTGTTTTTTTCTATTTAGTTGAAGTTGAAGATATTTTGAATGACTCGACTGCTTAT

CTGAAACACACCTTTGTGATTCCATGGAACCAATCATATGTACTTTTCTAGTAGTCATCTCTGGTGTCGATCTATACACC

ATAGTTGCTTCTAATCCTCCCATCATTACTGTGATTATTCTCTTCTTTCTCCATTTATCTTCTTCTTTTCTCGATCTATG

CTTTGGTGTGCTAAATTGGACATAATCCTGGATTTTTGAACCTTGGTTATTATTTTACTAAAATTGTGTCTGATTGTGAC

TATATATGAAGGGTAGTAGAAAGATCATGGTTCCCATGTTGCATATTGGTGCACGATGACTTGTTGATTTTCATTGTTTT

CTTTCTCAAAAAATTTCAGTATTGAAGTATCACTTATGCATCATAGATGTTGATTGATTTTGTTGGGTTTTTTGTGTTTG

TGTATTGTGTACAGGACTAAGTGTGATAGAGGTATGAATACTTTCATATTGAGTATGCTATTGAAGCAATTAAGGTAAAT

AACTAAACTAAGAAGGCCTAATGTTTGATTTCTGACTGAATTTTTTAAACCTAATTCCTGTGTTTATGGTCATGATTGGT

TTGGGTTTTTGTTAGGTTGTAGCCTAGATCAACTGCAATTGGGTTAAAGACTAAAGGAGGAGTAGTTGTTACAGTCGAGA

AGTGTATCACTTGCCATTGCTGGTTCGTTCGTTCTTTGTTTCATAGAATAGTCATGTTTAATAATTTCCTATTAATCGGT

CTGAAGCGTATCACTTCGCCATTGCTGGTTTGTTCTGTGTTTTATAATTTTGTTTGTGGTTAATAATTACTGGACGAATT

GGTCTTTGGACCTGATAATTGGTGAAATAGGCTTGTGGGGAAGTAGCCTATGGGAGAGGACATTTCCCTCTATGGCGTCT

GTTTTTATGATTGGCGATGATTGATTAGTCTTCTTTGGATACTTCATGTTAGAAATTTTAAGTAATATGTATACGCGAAA

CCGAGTAGAACTTGTAAGGAGTTATCCAACACTTCTTTGGGATTTGTAGGATTTGTACCATACTAGGTTAGTTGTTATCC

TAGATGATGTTCTCTAGTTAACTTCTTTGCCATATCTTTGCTGGAATGGGTTGTAGAAAATGTGTTGTCATTGTACTTGG

TGATAGTAAGCCAGTATTCATCATACACTTGCCAATGCCACTTGTATGGGTGTTTAAGGCTTAGTAGTTGTTTATCTGGA

AGAGTTATGGGTCATACCAAAGTTATGAAGCTGATGTTCATAGGGTTAGTAGTGATGTACTGCCCAGATAGAGTGAGGTA

AAGAAAATGTGAGGAGACAAATGACAATCAAGTTGTAATTAGTTAGCATTTAATGGTTTATTTAATGGTTTAAACCAGTG

AAGTTCTTTTTTATTGTGGAATCAAGATCTATATATCATCCTTTACTGATAGACGTCTACAACACCTAATTCTAGTATCG

ATTTCTTAGAATTTTATTAGCTAGTTTTCTTATAAATTTGTATATTACGCTTGTTTTTCATGTTTTTAGGTGCAATTGAA

TAATCTCGCATAAAGGAATAGAAACCAAGCTATTTGTGTTAGTTTGACTACATGGCATCAAATTTGTACGGGGAGTGGAG

TGCTAGCTGCATTTCCAGTTCTGCAAGGTGGAACCAAGTCTAGTGGCCCATTAAAACAACTCATTCCAGCAAAGGGGCCT

ATTCTACATGACCTTGTGTTGTTGGCATGTGTGTGCTAAAGGGTAGTAAAAATAGATGAAGTTAGGTATATTTTGAAACC

AGATTCACTTCACAAGCAGAGACAAGACTCAAGTATGCCGTTGTTCTAGTAGCCGTCGACCTAGTCTGCAGATTAGTTTT

CCATCAATGTAGTCATGTCCATGATGGGTGTTCACCAAGGAGCAGCAACTATACTTCAGACAACAAGGAGAAGATAAAGG

TGATTAATTAATGGAGGTTTTGTGGGTGAATTTGTGTGTGAAAATGAGATGAATGAAAAGGGGATATAGAGATGATTGAG

TAATGGGTTATAATGGATGAATGTGACGAATCAGGAGCTCGAGATGGATAGAAAATTTAAGCCGCAAACCAATGAAGAAC

TGAACTTGAAATTCAGTGACTAAATGTGGAGTTTCAAGGAAATATCTCAGCTGAGAATACAAGGGAGTTGTTGTTGTTGG

GGACTAGAGTGGTTATTGAATATTTTTACACTGAATGGAGTTATAGGTTTAGGTTATGCTTTGTAGGGAACTTGAATTTG

GGAAGAAATGGAGCCAAGAGAAGAGAACAAAGGCCAGCAGTTGAATACACACAGCAGGTGTTTGACAAAGCCAGGGAGAT

CATTTATGTACCAGCTTTTGTTGGTTGTTGATTGAAATAATATGTGCTGGTGATTATCGACAGTTCTGGAGTGCATGTAG

GAGCTGAGAGTTGTTGGTTACCGTGGACAGAATGGGTTGTCTGCAATGAGTAGTGGCTGTGAAGGTCAGTGAATGAAGGT

GATTGTGTAAGTTGGTTTGCAGTCGAGCTGAAGTGTTGATGTTGTCCATTGTGTTGATGTTGTAGTGGATTCGTGACTTG

CGGCTGTTTACAAGGGTTGTTATTGGTTACTATGGATGGGAATTGCGCTGGAGCTGGAAACTATGTCGAACTGATTCCAA

TGCCGAAAGATGAAGATATGATCTTGGCAGTGATGTTCTATCGGTGATGCCATGGTTACAGGGAACAAACTCAGGGAATT

GAAGGGAATTACAGGGAAAAAACCCTAGACTCCCTATGATTTTTTTTTTCAAACACCTCATGTTCATTGTTAATGACCTG

TTTAGTACTATACATTAATTGCAGAGATCTAAGCCTCAACTTGTTAAGTGGACCAATTCCTTCAATCTTGGGCATGGGCA

ATTTGACTTATACCGAGAAACTGTAAGTTTAGTATCTTAGTTTACCATATTGATTAAAAAATGCTACTAATATTTATGTA

TTCATACGTCTAGGAATTTCATACGTTATGCATGGTGCACACCTGCATATTATAATTGGTACGGCTTTGTTATTTTAGGT

ACTTGCACATTAACAAACTTATAGGAAGCATTCCACCCAAGTTTGGAAAGATGACTAGGATCCACTATCTGTGATTATAC

TTTTCGTTTGACATCTGATAGCTGCAAACTAATTGTTTCTCCAAAGTAACACTTCGATTTTGCACTTTAGGGAGTTGAAT

GATAATCAACTAAGTATAATCTCTGCAGTAACACTTCCATTTTGCACTTCAGAGAAGGTTGGTTTTGTCTATAACATAGT

GTTACTGTTAAAGCAAAGAAGGATCGTTTAAGAATCTGTACTAACTACATTCCACCAGTTACAAATTCAAAAGAAATTCA

AGACCAGAATTTCACTAGAAAGGTTAGTGTTCAACCTAATGCTTTCAGGAAGAGAACTACATGCGAATTATATTATCTCA

TACTTATGTGCCTTTGATTCTTTGGCATCTACAGGTGGTAGAATTTGTGAGCGGAGACTGTATCATAGTAGATGATAATG

CTGTATCGTTTGGCAGCCCATTATCAGAACGACGTGTCAATCTCTCTAGTATCAGATATGCAAAGCTGGGCAATCCCCTT

AGGGATGAAAACCAACTGCTAGTATCAACAGTCTGTACGCAAGGGTTGTTGTTGCAATGGTTTGTGTGGCTGCTTGCAAT

ATTCTGGTGTTTGCACTAGATTTAGCTGAGTCTTAGAAGTCGGTAAGAAGGAATAGGGTTATTGGAAGTCGATACATAGA

AGGACAACTGAGCTGAAATGGGTTTTTGAATCTGAAAAGTGCAGTTTGGTTTGAGATTTTTCTGTGAAGTCAAATCAACA

AAGGAAATGGTATGTGTGTTAGAGTTGATGTGTGATGCTGTCCCTAGTGGAGATGTGATTCAAACTTGTGACAGTATGAT

GGGGTTTACTAAAACTCAGCTAAGAAGAGAGCAACATAGACACAGATAAGGAGGTTTATGTCATTGATATCATGTGTTTG

TAAATGGCAACAAAGTTACTGGAATTTGCGGGAGTTTTGGTTGAGTTTTGAGCTAGTGTTGCTGCCATGAGACGGTGGAT

TGTTTTCCATGGTGCAGGTGGTCTATTTTATGGAGTTTGGAGCTGGCCTTTGAAGGCAAGGAGGGAACATAAGTTAACTC

TGGCTGGAATATGGAGTTGCTAGGAGGTCCATAAGCTGAGCACTAAACATGTGTTATCGTAACTGAAGCACTATGCCCTA

GTCTGTGCAGCTATCTATGGAGTTTGGAGCTGGCCTTTGAAGGCAATGAGGGAATAGTACTATCTAGCTACTTAATTTGT

TGATATATGAATCGTTCTTGATGGATTGTGATGATCAATAATTGTTCTTGTTCGATTGCAATTTAGGAGTAGAAATATGC

CGAAAAATGTGCCGGCGGTATCTTTTTTATAACCGGAAAATTAATTTGCAACGTATATAAACATTGTACAATTATACATT

GCACAAATTACCAGAAAAGAGCAACGGATGTAACTGTAACCAAATAAATAAAACGTTGAGAAAATCAAAATGGGACTCAC

AAAAGTGTTATTCTATAAATAATACCCGTTGCACAAATTATTTATACGTTGCAAAAATAAGAAAAGGTGCGACGGTTGTG

CAACGTTTATAAGTGTTGTACAAATTAATTTGTGCAACGGATAATATGTGTTGCACACCAATGCGCAACGCCGTTTTGCC

AATGTTTATTTACGTTGCACAAATTATGCGCAACGTTTATAAACGTCCTTAAATGGCCGTTTTGGTGTAGTGAGATATTG

CACAATAGACATATTACTATGAAATTGATTCCTTAACATACTCTTCCTAGCAAGAAACTGAGTTCTAAAAATCATTTGAA

GAAAATCCCAAATTTCCTTTGCAGTAGATAGACCAAGAACATCACCACTAACAGTTGGTGTAAAGGTAGCTTTAAGACAA

GTAGAGACGAATCTATCTAATCTTCTCCAAGAAACCCATTCTGGGTTAAAAAATTCATTACCATCAACCATTTTTTTCTG

AGACGGCTCAACTATTTCTCCACTAACATACCCATATAAATCACAAGAAAATAACACACCCTCAAATTGATCTTTCTAGA

GAAGATAATTTGAATGATCTAATTTCATGGAGACAAAATTTGATATATTTGTAAAAGGAAGAGATGTAGAATTACCAGCG

GAAGACATGATATTATGATGTTGAGTGTTCTGTAGAATTGCAAAATCTGAAATTAAAGTACCACCAAGACCACGAGTTGT

GTTGATTTTGCTATTGATTTGTTGATGATTGACGTTATTGCTGTAATCAGTTGTTCGTGTAGTTACCATAAAATTTTCCA

AGAACCCTAGAAATGTGTTTTGGGGAAAAATTTCTTGAATATGATGTTACCCAAATTGAAGCTTTTGTTTTAAAAATTGT

TTATAAAATTGTTATAAGTTATGTAAAAGAACCCTAAGACCGGAACCTATCTGATACCATGAGAAATCTAGGTTTTCATG

ATAATCAAGATATAACAGAATGTAAACAAAGAAGATGAAGAAGAAGAAAACTTGTTATTAATGGTTAAACAAAAGATACA

AAGAATAATAGAGTTTAGACATATTTATTACAAAGGAATTAAGCTGTCTGAATCAGACAGATTACTCAGACAGCTAACTC

AGAAAACCATACTAACTCAAATACTATAAGTCAAGTCAAACATATTTGACTTGATTTCCTATTAAACCGATAAACTTTCA

TGGGTTCTTCTCAAAATCTTTAACTAATTACAAGATTCGTTTTGAAAGAAAACTTGAGTGAATGAAACAAGGGATGAAGA

AGAAGTGGATAGAAGTTTGGGTGATGAGAAGGAAGTGAAAGCTAATGAAACTGATGATGTTACCGTTGTTTGGAAGAGGA

GGAATGAATCAAAGGGTTTTCACTGGATTTTTCTGTTTTTTTTTAGAAGTGAGAAAGTGAAGAAGAAGAAGACGAAGACT

AGAAATTTGTCGCCATGGAAAATGGATCGATGATTTTGATTGTGAGATGTTTACCCCCAACCTTTATTTTCCCATACACT

TCATTTACATTGGATAAAAGAGTGAAATTGTCCCGAATGTTTAGCGTGGTGAATAATTTGTGTAGTTGTGTGCATGTTTG

GCTCATAAAAACTCTTAACATACGACAACACTAAAACAACTCTTCACATTTATCTGACTGGTGGGTCCCAAAATTGGCAA

CCTGCTAATCCCCCAAAGGGCTAATGTGAGGAGGTTGTTTTAGAATTGTTTGGTTGCATGGTATGTGTAGAACCATTCCT

TTTTTAATTGGTATAAATTTTATTTTCTTATTAGAGGTAAAAATGGTTAGTTAAAAGTCAATTCAATTAAGTAAAAGCTT

AGTAGGATGGTAGACAATGTCATTTATACATTGGTTTAAGACATCAGGGTTGGAGAATGATTTTAGAGAAAATGGATGTG

TATCCTAGGTGGATTTTGACATTTATCTCAACACATGTGCCATTTGAGAAGCTCCAAGTTCTAGGACACTAAGGTGTTTG

AATTCTTTGAAAAGTGGAGGATTATTCAACAATATTCACCACCCATTACTCCACAAGACAATGGAGTTGCAGAAAGAAGA

ATAAGAATATTCAGGAAATGGCAAGGGTAATGCTCCATAACATAAACATATTGTTGAAGTGTAATACCCGGTATTTTTAT

TTACCGTTTCGGGTCGGGTTGTACACAACTTATCTAACTATGTGGTCCAGATGGGGTTGATTGAAACTAAAAAGGAGGGT

GCAACTTATCTAAAATTTGCTATACTTTTGGTGGGCTCCAGATTGGACTACTCAAGGGGTCAGAAAGTGATTGAAACTAA

AAGGATAGTTTTGCTAGGGTGCAAGTATAATCACCTAAATATGAGGAAAATCAAAAGACATTTAAAATTAAAATAAGTAA

TAATAAAAATGTGTATTATGTATTATTATTAGTAATGTTAGTATTTATATATAATGTTTTAGAAAATAAGAACTTGGAAA

GTTAGAAAAGAAAAGTCTAGGAGAAGAAGATGAGATGGAGCTAGAGATTTTTAGAGAAGAAGGATAAATAAAATCATAGG

AGGACCAATGGAATTGAGGTAGAATGTTTAACCCTTTTCTCTTTGTGTTTTTCTTTGATATACTTTGATTTCAATTTTGT

TTATTTTTATATCATTAGGGTTCCGAAATTTGTCTATTTAATTTTATGCACACTATTGATGATCAGACTTCTATTATTGC

TTTATAACATCATTGGGTATCTATGGTTAAAGTTTGAGAACAATATGCCAAAAATTCACCGTTGAATGTGTGTTTGAAAT

TGGATGACAGTTTCTGAAATTTCTGGACAGTTACGAGTTTATGTGGTTTTGATGTTATTAGGCAATTTTAGTGATGTTGA

AACAATTTGAAGTCTTAACAAAAGTTGTAGATAAATGTATTATCTTCCAATATGCTTTGGATTTGCTCAATTTTAGACTA

GTGTGAATTTATAGTGAATTCTTTGGTGATAACATGTAATCTGCCCAGTTTTTGGGATTCCATTGTAAATTCAATATATC

TTGTTTGAACCGCTGGTTTAGTGTGAATTTTAGATATGTTATGTATGAATGTGTCAAGAATGTTCTTGTATAATTTGAGG

ATGATCTGATAAAGGTATCAGCATGAAACATATAATGTAATGTGGTTGCAGCTAGTGTGTTTGATTAAATGCTTGAATGA

AATTCATGTTGTTAAGCCATGGTTGCTTGTGTTGTGTGTAAAAACGATTTATAAAAAGGAAATGAGTTCATGTTTTTCCG

TTAGATGAAATATGTTTTCATGCTAACGGCGGGTAATGATCCCCGAGAGTTAAATGGGTAATGATCCCCATGGGAACTTG

TGTTTCCCGATCATCGATGGCGGCTGTCCGTTCCGGTAAACGATAGGTGGTGTACGAACCAAGGTTTTCGTACTGTTAAT

GAAACGGGTAATGATACCCAAGAGCAAATGGGTATGATCCCCATAGGAACTTTTGTTCCTGACCATCGATGGAGGTTGTC

CGTGCCGATAAACGATAGGTGGGTGTACGAACCAAGGTTTTCGTACCGTGAACTCAAGATTTGTTGGTGAGATGTGTTGA

AATATAGATTAAATGATTTATTGGAAATATAATTGATTTTATAAACTCATGAGTTGAAACTTAAATGGATGAATTGTGAA

TTGATGTGTTAAATCCCAATTGGATGATTAGTGAACTCATGGTTTCTATGTGACGATATTAATACGTGAACAAATGTGTG

GAATCATTGATTAATGACTTCTTGACAATTATTTTGGTGTTGTAAAGTTATGGTTTGTTGTGACAATATAAATATTATTA

GTGTTATGAACTCATGTATTGAATCTGAATTTAGATGATTTGTTGGAAATATTATTGGCGCTGTGAATTCTGATTGGAAT

CTAAATAAGATAATTTATTAGAAACATTATTGGTAACGTGATTGTCGAATATAACAATGATTCGCCGAAGTCGTGATGTG

ATTCTTGGTATATGTTTTTGCTTGATAAAATAATTATGTTTGCGTATCTTTGGAGAATTGGGAATGCATTCTATTGAGCT

GTCATCTCACCCCAATTGACATCCTTGCAGATTTATCAGAGTGGCGCAGCGAAAGCTAGAAATGAGTAGCTTAGTGATTG

AATTGTTTTATTGTATTGCTTGAAATGTAATTTGAAATAATATGTTTTGTTGAATTCGTTGTTAAGATAATAAGTGGATT

TATATGCTTCATTTTATGTTATGTAAAGTTTCAACTATGTTTTGTGCAAAAATAATTTAGCAAAAATATGTATGCATTAG

TCTCCTTCCGACCCGTAACTCTTCGAGTTCGGATCTCCATATGGGTTTGACCCTGGTCCGGAACTCGGGGTGTTACATGA

AGTTTTGGGGAGAAAATGTTTATACAACATGCTATTTGATCAACCTTGTTTACTTACGGTCTAGAACCCTAGCTAAATAC

TCTGTAGAAGTTGTGGTACGGTAAGAAGTCCAACTTACATTATCTCCGAGTGTTTGGTAGTAAGTGCTACATTATGAAAG

ATCGAGAACAGAAAGGTAAATTTGATTACAAGAGCGATGAAGCTATTTTTATTGGCTATGAATCTGACAGTCGTGGTTTT

CGATGGAATCAGAAAATGTTATTATTGACGCTCTGCGCGATTTTTGTCATGCTGACTCTCCTGCTGAATTGCCTCCAATG

GAGACAATTGAGAAAGTCAAAGAAATCCCAGAATATGTAGAAGATGTACCAATGGTTGCAGATGCTGATATTTCAAGTGA

CGAGGAGAAAATCCCTAATCATGGTATTCCTGTTGAACGTGTCCCCTCATGACATCTTTGGGTTCAAAGGAATCATGATA

CTCACAGTATTATTGGATGAACATATTACACAGTAACACTACACCAAAATGGCCATTTAGGGACGTTTATAAACGTTGCG

CATAATTTTTGCAACGGAAATAAACGTTGGAAAAACGTCGTTGCGCATTGTTGTGCAACACATATAATCCATTGCACAAA

TTAATTTGTACAACGCTTATAAACGTTGCACAACCGTTGCACCTTTTCCTATTTGTGCAACGTATAATTAATTTGTGCAA

CGGATATTATTTATAAAATAACACTTTTGTGGGTCCCATTTCGATTTTTTCAACGTTTTAATTATTTGGTTACACTTACA

TCCGTTGCTCATTCCCGGTAATTTGTGCAACGTATAATTGTACAACGTTTATATCCGTTGCAAATTAATTTTCAGGTTAA

AAAAAGTTACCGCCGGCACATTTTCCGGCATATTTCTGCTCCTGAATTGCAATCGACCAAGAACAATTATTGATCATCAC

AATCCATCAAGAAAAATTCATATATCAACAAATTAAGTAGCTAGATCTCAAATGGGGTTCATCTTGAAATATCGACTAGA

ATTCTATGAATTCAGGCTTCAATAAGTCATCTGATAGTCTAGTTATTCATCATCACAACCGATCAAGAAGAATTCATGCA

TCAACATGTAATAGATGAACATATCATAAACCCATACAATTAGAGTAGCAGCTTCTTGATTCTCTGATAAGTGTAGACAC

AAACCCATACAATAAATTGGGTAGATTTACAAAAGAAACAACAATCTACTGGAAACCCATAGATTTGAGATTATCTAGCT

TAGTTAAAGAATCCTAGTTTCTGAATTTGGTTCACCTACAAGTACAAGTCATATCTCTAAGTACAAGTCCTTGAGAAGTA

TCGATTTAGAAACTTTTTGGTTTACCTACAAATCATAAAATCAGCTAAAAAGGAGAGTATGTAGTTCAAGGTCGAAAGAG

GGAATATATGTGCATATATTCGTGAATCATTGATCAATACAAACGTTTAGACCCATCAGTTTTATATACCTTTACTGTGC

CATCTTCAGAACCCATGTATATCCAGCTTCCATCACATTGTAACTCCAAAGACGTAACATTATTAGTATGCAAATCATAC

ACCATCATCTTCCATTGAGAAATAGAAAGACTGAAGTTAGTACACAAGTAAAAGATGAGAAAATGAAACCTACAGTGGTG

AAACAACCAACTCATTAAAACTTACAGGGTATGTGATACTTGAGGTGACAATTTGAACATAAGAAAACTTTCATGAATGT

TAAAACAATGATTATGTCTCTAACAATTAGAATAGAAGTTGAAGAGACAGTGTATATGTAAAATATGCAAAAGAGTTTGT

CCTAATGCAGGGTTTAACCATCCCCCAAACTTCTACTAAAGCAACACCTAAAAATAGGAGAGTACTACATATAGTCAGCA

TATATGAAGAGATACAAAAATACAAGTTCAAACTTACTCAAGGTCAAACTTCAGCTGTAGAAAATCAAAAGCAAGCTTTT

GCACTTACACAGCAAGCTCAGCCCCTGTGTTTGTATGAAAGATGTTAATCAAAATTGTTGCGGCAATCCCACTTGGGTAT

GCTAGCTTGTAATCAATGACCATTACCTGTATACGTCACATAAAAGAAAACATAAAAAAACCAATTTCACAATTTATACG

ATCCTTAGGTCCAACACAAGAACATAACGGACTACTCTCACAAATACAAAAACAAGCTAAATATACATATTACCATAATA

ATGTGCAGCATATAATCAACAACTTTGAAGTAAACTAAAACATATTCAAATGAACAAGCCAGATATACCTTTCAAAGAAT

AACTAAACTAAACAATCCAAGGAAACTGACAACAAACATGAAACCAATCATCTAACCCAAAGCTGGATTCTTAATAACTT

CACCTCGGTTACCACGGTAACCAGGACCAATGAGTTTATATGTTTTCCCATCCATTGTAGCAGAAGTTTCTTTGCCCAAG

AGATTCCTTCAATAAGAGCCGGTTCAAATCATTCACTGCAGCAACATCATCTTATTCCTCTGGTTTCAACAATATTCAAG

TTCCCTACAACAACACAAATTAGTTTTCTACTTACACAACACAAATCTTGTGTCAGGAACCCGATGATAGATGCAAGGAC

GGATCCAGAACTTATATCTCATGGAGTCAAAAAAAAAAAATGTTTTTGCGGCTCACAAAAAATTGTTTCTGATCCATATT

TTAGTCAAGTAACACGGAATCTTGTACGGTAGAAAATCTTACCACTTTCACGATAGTACCATTAATCCTATTCTAAATAC

ATCAATTTTGGCAAAGAAATTGTGCACATTGATGCGAAGAGCCAACAACAGTAACATGGCATACCAAAACAGTTCGAAGA

TTATGTTTTTGATTCCCAATGATTGCATCTACGAATCATCCTGTCATATGTGTGCAAACAATCATTTATGCAACAAATTA

TTTAAAGATAATCGGGGCCAGCCAATGACACACTTAATCCACCAGTTCTGCTGCGTACTCCACCCGAACTTGATAATAAA

TGAATCCATTAAAGAGAAGATTTGAAAACGTCCCTTCATAAACAAATAAAACTAAACCATCTTCTTAAATGAGAAAGAAA

TTCATGAAAGCCATTACAAAACCATGAAAAGCCAAAAGGCCATGAAATTTTACTGAGAAAAAACTGGACCCATCTAAGAC

TAAAAACAACTAAGCTAGACAATAGTAGAACCAGATATCCAACTTTAAGTAAACAGCAAAATGAAGTAAACACCATAAGA

ACCTGGTTGGGAAATAGTGACATAAGAAACAAAGCCATCAACAAAAAGAATGCACATAGCTCGTGGAACTTTACAAGAAA

ATGAACAATTAAGAACAAACTTTGAACCTATCGATGCTTGATCATTTCTAAGTGAAAATTGGAAAATAAAAATAAAATCT

GTATACCTATCTTGAAGTAAAGTCTTCTGGCTGTTGATGCCCACGAACTTTATTGCTAACTGGAAATATTTTTCTACTGC

CAACTGCTTCTGCAGACCAATTTGACATCGTCTCACGAGAGCCATGAGAAGATGGGTCCAGTTTTTTCTTCATGAATTTA

ACGAGAGCCATGAGAAGTTTCACTCAATGTTTTCCTGCAATGACAACGTTAACAGAAATTAAACAATGGCATCGTACACA

AATGTGAGATGTAACGTGACTGGTATTACATGTTAGGATGAAGAAAAAAAAGTATGACAATCCCAACTGATAAAAAATAT

AATTGGAAAGTCTAGGTTAAGTACTAAAAACTAATTTATCATTAAAAATTGAGTTGATCAAACTTTTGGTTACAACAGCT

AACTTTTGGTATTAAAAGCTTCTCTCCATCAGCTTGCACAAATTAAGTCATGGCTGACGGATCTTTCAACAAGGTAGATA

GGCGACTAATAACACTTGGGATACCACGAGAAGGAAGGGTAGGATACTTCAGTTGCAATACAGAACAACTTATCAGAATA

TTACATTGAAATAATGGAGAGAAAAAAAATCCCAAGATCTGGGAATCCTAGAACTACAATTTTACACGAGTTATGTTTAT

TTACAAACCTGAGCACAAATCCATTCTTATAACAGAGGTTTCATCACATACACCGAACAAATGATGCACCACAACCACCA

ACTAAACCCAAAAATATGTATCACTAACCACACGACCAACACCTGCTAGCACCACCAACAGAATCACCCACACCATCAAC

ACCACTCGCCAACACTTGAAACACCATAAAACTGAAATCAAAGGAAACCCATGCTTTAAAACTATATCAAATATAGTAAA

CAAAAAGAAAATTTGCAAATCCTTATAGAATCAAGCGATAATAACAAAAACAAAACCAAGAAAAAAGATTGGGGCTTATA

CCTGACGTCTTCTTCTTGTACCCTTCAGCTCTGTCAAATATGAGATGAAGGTTTTTCGATTGAAAGAGAAAAATTAACTA

GATCAAGAGTTACAAAATTAGGGATAATAAAGGAATATGGTTTTGTTTCTGAGATGGGTTTTGATTCTTAAATGAGGTTA

TGATTAGGTTTGAAATCAGGCGTTATTCAGTGATGAGAGTTGTTTAGATATGCAGGATTTGTAATCTCCCTCTCATACAG

TAAACGGTCGAGCTTAAATAAGATGAGGATTGGATATTCATTTTGGGGAGCAAATAAACTAAATCTAGGGTTACAAAAGA

GGGGTTCTCTTTTAGATATGATACATGGGATTTGATTTTGATATGGGGTTAGGATTCGTATTTGATTTGTGGAGTTTTCT

GTTAAATCTGGGTTTTGATTTTTGGTTCTCCAGTTGAAATCATACCAAATGGTTGTTCTTGGTTATGAAAATAGGACCGA

GAGAGAGAAGAAAACACGAGTTATGAGATTTAATGAAATACATTCAGAAACTTAAAAATAAAATAATTATTAAAAAGCCC

ATGAAAATATCCAATATTATAAAAATATCATGATGTACAACGTTTTTATATGATTCAAAACCGTTGCTCCATCATAAAAA

TATCATAATAGGAAATAAACCAATATTACCTATTTTGTGCAACGTATTTATTGGTGTGCAACGGATATAGACGTTTCAAA

GTGTGCAACGTCTATATACGTTGCACACCAGCGAAAATGAAGTGTTGCAAAATACAGGATATGGTGTAGTGTAAGACGAG

AAGACAACTTCAAAATATTTGCAATTTTGGATGTTATCTATCGCAAGTGGAACCAAGAAACGTTGACGAAGCTCTTAGCG

ATCCCTTCTGGGTGAATGCAATGCATGGAGAGTTAAATCAATTTCAAAGACAAGTTGTATGGGAACTTGTACCTTGTCCC

TCTAATATAAATATTGTTGGTACTAAATGGATATTCAAGACCAAGTCAGATGAATTTGGCACAATTGTCAGAAACAAGAC

TAGATTTGTCGCTCAAGGATACTCACAGATTAAAAGTATCTATTTTGACGAAACCCTTGCTCATGTGGCACGCCTTGAGT

CCATAATATTATTACTAGATCATGCTTATTTTCTCAAGATTAAGATATTTCAAATGAACGTAAAATCCGCATTCTTAAAT

ATATGGATGTTTAACGGAAGAGGTATATGTTTCCCAACCTAAAGGATATGAAAGTTTAGAATTTCCCGATCATGTCTTTA

GGCTTAGAAAGGCATTGTATGGATTAAAATAAGCACCCCACTTCTCTTCTTAGAAAAGATTTTTTGAGGGGTGGAGATGA

TAAAACACTTTTTACCAAATAGAGTGAGAATGAGGTTGTTATTGCTCAAGTCTATGTAGATGATATCATCTATGGATCGA

CCTCGGTAAAACTTGCAAAATATTTTCAAGTCTCTCTTGGAAGAGAATTCGAGATGAGCAACGTTGGTGAATTAAAATTC

TTTTTAGGATTGCAGACAACAACACAAGGATGGTATATACTTATCTTAAGAAAAATACTCTAGGGATCTTATAACAAGGT

TCGGTCTTGAGAATTCTACTCCAAAACTTACTCCGATGCCCACTTCCGGTAAACTGCACAGAGATGAGAAAAGAATCAGT

GTTTTGTATCTTACTGAAACTAGACCTAATATTGCTTTTACTTTTGGTTGTCGGGCTAATCCGAAGAAATCTCATCTTTC

TGCTACAAAACGGATCATGCGATATATTAAACATACTATTAATTGGTTATGATCTGTCTTATACTTTTGATATTAACACT

GAGCTTACAGCTTATTCAGATGCTGATTGAGAAAGATACGTGGAAGACAGAAAAAGTACATCTGGGGGATTTTACTATGT

AAGATTGAATCTTATGTCATGGCATAACAAGAAACAAAACTCACAATCCATTTCAACATGTGAATAAGAATAGATCGCTG

CAGGTTCGTGTTGTACTCAACTTTATGGATGAAACAAATTTTTTTTGACTATGGGATTGATACTGGAATAATAAAAATCT

TCTGTGATAACTCAAGTACGATTCACATCATTGAAAAATCTTGTTGAGCACTTATAACTAAGCACATTGACATAAGGTAC

CACTTTATAAGAGATCTCTATGAAAATGGTATTATTAATATGGATTTTGTGCCATATGAGAAACAATTGGATGATATTCT

CACCAAACCTCTAGACACCGCAACATTACAACACCTTCGGCAATCTATTGGAGTCGGTTTGGTTCATTAATGAAATCTAG

CGTTTTGACCCTGGCTCTATCTATATCTTTTGGGAAATAAAGCTTGAGTTCTAGTAAAAATGGCTATTTAAGTTCCGCAT

TTGTTTCTCTTAGTTGTGTTTTGCTTTTTATCTTGTCTTTGAAATATCAAAGTATACGTGAAATCCTTATTTTCTCGTAA

AAGTAAGGTCGCTCATGTTGTTCTCGTGGGAATGACATCTAATGGGGGAGAGTTCTTAAATGAACTTGTGTTTAATTGAG

GATAACATTAATTTAATTGAGGATATATTCGTAATTATTGAAAATAATATGACAGGGGGTTGTTGCATTTACTACTCCGC

GACCATCTTTTACCCTTTACTCAGCCCCAAAATATAAGTGGGTGCAGCCCAATATGCCTCAAAACAAGAAACTTGTTTTG

AGGCATACTGAGTTTTTTTAGGCATATTAAATGATGATACCATCTAGGGTATGCTTATGAGGGGTGTCCTAGCCGTGATG

AAGTTACTAGGTTACCCTTCAACTAAATTAAAAACATAAACTAAAACTAATCTGAATGAGTTCAATCTTTCTTCTTCTCA

TCTCTCGCAGCCGAATTCTTCCCTGCATAAACGTTATTTTATTTTTTTTTCGTCGCCGATTAACCATTCAATCAACCAAA

TTTTCAACAAATTGAAGAATAAAAGCTAAAAAAAAGAGATACTCCTTCATCATTTGGAAATCGCCCATTTTGAATCACAA

TAATCTCAAAAACCTAGTTTTTCTTTTCTCTTCTTCTTACTCTAAAAAAATATGATTTCTCTACTAATATAACATTAGCA

ACTTAACTTAATCAATAAATTAACGTTACTAATTAATCATATTCATTACTACTAATTAAATAAGGGTAGTTATGCCATTA

TAAAAAAAGAGTAGATAAGGGGTGATGGGTTTTTTTGACCTAGTGACCCTATTTTGACACCATTAGGTATGCCTCAAAAT

ATTTGAGTATGTCTCAAAACTGGTTTCTTCCCAATTCTCCATTTTGATTTCATTTTCCCCTTTTTAACAAACCCTAACCC

TCGTACTCGGCTCAAATCAACTATTTCTCCCGGATCACTTCCAAAATTAAGTCAACAATGAAGGTAAGTAAATCTTGATT

GAATTAGCCTTTAATTTATTCAGGTTTTCAGATTTTCTTTTTGCTGAAAATATAGAATCTTCGAGTGATAAATCACGTTA

GAGATTGTGTTGGGACCTTAGTCCCACATTGGTAGATATGGCTTAATTGGGTAGTTTATAAGTCAATGGTATCCCTCATC

TAGTCAACTGGTTTTCTAGTTGAGTTCTACCCATGGGCTTATGACGAGTTTATGGTCGGTACCCTAACATTTGGTATCAG

AGCCAGGTTCCACGAACCATACCCGCTCCACGGAAGGGGTGACCTATAGATTGGATTAAGGTGTAGAGGCACCTATGTAT

GGGCATAGTTGTCACCGCATTGAATACTGATAGTTGAGTGGAGCCTCCATAAGGAAAGGGCTACCGCCGGGTCAGTATCG

ATAGAGTGGGACAACGAGGACGTTGGGTCTGGAAGCGTGGTGGGATGTTGGGACTTTAGTCCCACATTGGTAGATAAGGC

TTAATTGGGTAGTTTATAAGTCAATGGGATCCCTCATCTAGTCAACCGGTTTTCTAGTTGAGTTCTACCCATGGGCTTAT

GACGAGTTTAGGGTCGGTACCCTAACATATTGAAAATTGATTATCTGCCGAGTTATGGAACGTCTTTCATCAAAAATCCT

ATTAGACATACTTTCTCGACTACCAGTTGAATCTGCGTTAGATTGCAAAGGTATTTATGTGGGAGACAAATTTATCTATT

CATGTAGACTTTTCTGTGTGAGACAGATTTGTTTATTAAAGTCTTCTACTTTGGGTCATAGCAACTCTTAGTTGTGGGTG

AGATCAGCTAAGGGAATCAAGTGTGTAGTATCCTGCTGGGATCAGAGACGTAAGGAGCGCAACTGTACCTTGGATCAGCG

TGAGATTGATTGGAGTCCAACTATAGTCCAGACCAAAGTTAGTTTGTAGTAGGCTAGTGTCTGTAGAGGCTTAATACAGT

GTGTGTTCAATCTGGACTAGGTCCCGAGGTTTTTCTGCATTTGCGGTTTCCTCGTTAACAAAACTTCTAGCGTCTGTGTT

ATTTATTTTCCGCTTTACATTTTGTTATATAATTGAAATATCACAAGTTGTGCGTTGAATCGATCAATTGTTAAATCCAA

CCTTTGGTTGTTGATTGAAATTGTTTGATCCTTGAACATTGGTTTTTGGTACCGTTCAAGTTATTTCTCTTGTATTCAAT

CAGGCTCGCAGATTTCTATTTATTTGGGTTAGGATTGAATCGATAAATTGAGATATAACTATTTGATATACTTTTTATTT

AGATTGATTCTGAATGTCTAGTTGATTCTCTTAAAAGTATATTGGAGTTAGTCCATACAGATTCTAAGCGAAATATTGGG

TGAGTTTGTTATACCCCCGCTTTTTCAGTTAACCTACAGGGTTTCGTGAAAAACCAAAGGTTAAATGAGAATCGATTATA

GTCGCAACTAGTATCACACAAGAGTTGTGGGGATTAGGTTTCCCAGTTTCTAGAGTTTTCTTTTTTTATAGTTTTCAAAT

CGGGGTTTGCAATATAAGTTACCTTGGTAACAAAGCATTCAATATTCACCGTTAGATGAAAACCTGATTAAACCCAAGAT

AATATCTTTCAACCGTTAGATCAAACTTAGCTTGTTATAAATAAATGAAATGTACCCTCATTTAGGTTTATGTAACCGTA

CCTAAACGTGTACACCATGTTGGCTCAATAGTAGGTAAATGAGGTTATCTATATGTACACTCTCATATCAACCTTATTCA

TCTTAACCATAACTAGTTCAAATGACTTAAATGAAACTAGTTAAAGAGTTGTTCAATTGATATATTCTCATAGAATTATA

CAAGAACAAAATTGAAGCAAAATCGGTTTTATTCACTAGAATCAATTCATAAACATTATAGCCACAGTTTTCAAAGAATG

CATTCCTTATTATATAAATGTATTTGTTCACAACAAAACCGATTTTAGAACTTTAACCATTCAAGTATGAAAACGGGTAC

GCATACTTGAAGTAGTCGGGCCAAGTTTGGTTTTCGCCAGTATGCGAATGGGTACGCATACCTCCAAACCCAACAGAAAT

TTCCGGTCACAAATCTACGCCAGTATGTGTACGGGTAAGTGCACTTAGGTTCCCGAACATGGATTACATATATGTAAGAG

TCCACAACATGTTTATAATCCAATGTTGGTTAAGTGTTCTAAAATCTTATTTCAATCATTGAAACTTTCTTAGAGGATGA

CAATAGCCGTTTTCACACACTATTAGCATCAAAGCAATTTTCAAGTTATTGAAATAATCATAACGAAACATTCTAAGTCT

ACACCAAATGACTGTATCACACAAACCATGTAAGATGTTACTCGGAGATTTACATGATCATCTTTTGACTTTCGTCAAGA

ATATAAGATGAACTTAGTTGAAGCGAAAGCTTACCAACAAATATTTCTAGAAATATGTAAGCGAGTTAAACTCAGCTCGA

AATCTCAAATGTGCATAATCAAAAACTATCGTAATACGACTTTCGTCTCAATATAAGAAATGGAGTAGAAATAGATTTTC

CAAGTGATAGATGAGTTTTAGTCTACTTAAATTGTGATGACGTTCCACAAGCTCTCCTTAGTAGTTCTTCGTCTTCATTT

GATGACCGCCATGAAGCCTAATGCTCAACTACACAATTTATCCTAGTCCGAGACATTACTTTAAGTAGACTAGAAATCAA

GACTTATAGTTTTGATCACTAACATTGAAAAACAAGCTTGAGATAGAAACGCTTGCGAGTTCGACCGAGCAGTGCTATAA

CACTGACAACTTTCTGGGACATATTTGGACAAGAAATTGACTTGCAACTCAAGGAAACACGGTCCCGGAAGGATTCTAAA

GCTTAGCTTTGAGAGTTCTATATAAGAAGACATCCAGAGCCTTAAGAAAACATCTCTGATTATATATTTTTGTTCTTCAT

AAAATATTCTAGGGTTTACCCCTTTAGGGGGAAACCGGGTGTAGATAAACCTTTGTTGATTAAGGATGAATTCAATGTTC

TAGCGTGAAGCTTTATCAGTTATATAAATCTATTCGAAGTTTTAATCATCTTCGATTATGTTTTCTATATCTAATGTTTA

TGAGTGATTCTAAGATTGATTTAGGATGCATACTAGATTATTCTATTGATCGTTCTATCGCTAGTAGAACTTATGGGATA

AGTAATCGTCGATTGACTCTCTACACAATTAGACATCGCGAGACCTTGCAGAGGGATTATGTGGCAAGTGGACCTTGACC

TAAGAGTTCAACTACTGGGATCAACCTAAATTCGCAAAGCTTAAAGCATTTGATTGGATTACACCTTGAGTGATCTACTA

CTTGGTGGTTCGTTGAATAAGATTTTGGTAGTGGAGAACTTCCGTATCCAGTGTCAGAACGAGTTTTGGGAATAACATTA

ATCTAGATGTTATTCTACGGTTGGTGATGAATGACTCGACCAGGAGTTCATACTTTTGAACCGTTGGTGATTGAGAGGTT

CTTAACAGCCGTGAATGAGTGTTACGGCTGGGTCGACACGATACCCAGCCAAAACTGCAAAAAAAAAACCCATATTTTTT

ATCCAATTTTTAACGGATTTGTATCAAGAAAATTATTATTGGGAGTTGAGTAAAGGATCTCTTGAGTATTTTAAGGATCA

TCATAGTTAACACTAATGGAGGATGAAAATAGTTTGATAAGGGGTTATTGATTTACTATTTCACGACCATCTTTCACCCT

TTCCTCAGCCCCAAAACATAAGGATTCCATGGTTTGTCTCTCAAACAACATGTTAGCCATGATACCCGGTCTGGTAACAT

GAATTCACCAGTTTCTATTCTCTGAGTTGGTATCAGCCAAACCGATCCTTTCATCGCTTCTGCAAAAAAAACCTAAACCA

CAAAAAAAAAATCAACAATCAATACCCAAATACAACTTCAATGGCACTCGATGTTGATGGAACCAAGGCAAAACCAAACG

AACTCTTCCGAGAAAGTGAAGAAATCAATCCAGATTTCACTGACTTTGAAGCTCGGGACTCTCTTGGAGTTCATACTTTT

GAATGAGGCTTGAACGTAAGACACCACCAAAACAAAGAACCTGCGGCTTATATTGCATTACCAGGAGGTACAAATTTGAA

ACACAATGGGTGACCGATACTGGTGCCACCCATCATTTGATTCCGAAACTTGAGTATACCTCATGAATATGATGGTACAG

ATCAAGTAAAAGTTGGTAATGGTAATTCTTTAGAGATAAAAAATACTGTTCTTTTACTCATAATTCTCGTTGCTATCGAT

TAAATAACATCTTTCATGTGCCACAAATCAAGACTAATCAAATCAGTTTTGCAAAGATAACAATGTTTATTTTGAGTTTC

ACACGGGTTTCTTTGTTGTGAAGGACAATTCCACGGGCGTGACGCTGCTGCAGGGGAGGAATAGGAATGGGTTATACATC

TTTGATGGAGTAGGTGGCATGAATGGTTCAATTTCTCCTAAGTCTTTCGTCTCTGTAGAAGTTCTTAAATTCCTACGTGT

TATTCCAAAGCATGTACCAATCCCAAGTGGAGAAAGGCAATGGACGAGCAAATAAATGCTCTACTAAGAAATGGAACTTG

GACTCTGGTAAAATACGAAGATGGTATGAATGTCGTAGGATCAAAATGGGTGTATCGCGTTAAGCAAAACCCGGATGGTA

GTATTGATCGTCATAAGGCACGATTGGTGGCGAAGGGGTTTCACCAACAAGAGGGGCTGGACTATGGAGAAACCTTTAGT

CCCGTGATAAAACCTTGTACGATTCGGCTGGTTCTCTCGATTGCAGTAATGAACAAGTGGTCTCTTCGGCAGTTGGATGT

AGAAAATGCGTTCTTACATGTGGTAACATTATTTTCAAAGTTTGAAGATCCGACTTTGTCTGAGAAGTGTTGTGGGTGCA

TTGCGGTATCTTCATATCACTCGACCAGATATTTCATTCGGATGAGAGGTTCTTAGCATTGGGATCAAACGGATGAATGA

GTGTTACCAAGTTCGTCGACTCGACGATAGCCTAGATGATCGTAGGTCTAAAACTTTGGTGGAAATCAAAAAAAAAAAGC

TGTCTCCAAGTCCCAGATTTTTTATCCAATTTTTAACGGATTTGTATCAAGAAAATTATTATTGGGAGTTGAGTAAAGGA

TCTCTTGAGTATTTTAAGGATCATCATAGTTAACACTAATGGAGGATGAAAATAGTTTGATAAGGGGTTATTGATTTACT

ATTTCACGACCATCTTTCACCCTTTCCTCAGCCCCAAAACATAAGGATTCCTCGATGACGGTATTTCCAGTTGTCCATTT

CACTTTCATTTTGGTTCCCCCTTTTAACAAACCCTAAAACTCGTAACTCGGCTCAAATCAACCATTTCTCCCTGTCGATC

ACTTCCAAAATCAAGTCAACAATGGAGGTAAGTAGATCTTGATTGAATTAGCCTTTAATTTATCCAGATTTTCTTTTTGC

TGAAAATATAGAATCTTCAATTTGATTATCTGCCCAGTTATGGAACGTCTTCCATCAGAAATCGTATTAGACATGCTTTC

TCGACTACCAGTTGAATCTGCGTTAGATTGCAAATTAGTTTCCAAAAGTTTTCTGATTCTGCTTACTACTAGAAAGCAAC

TGTTTGCTAAGATGCACCATCACCGTCAGTTACTGCAACTTGCTGATGGCCATGAGGCTGCCTCTTCTAGTGTGAAATTG

GGCATTCTATTCTCGTTTGTAGATTTAGGTGATGAGGATTCAGTAGGCCTCTTTTACGGGGACAAGTATAACGATGAGAT

TAATATCCATGAGAATTTATGTTACAAGGCACTCAAAAGAACCAACATCTGTTGTAATGGTGCATCAAGATATAACGGTG

AAAATGTCTTAGTTGGTTCCTGCAATGGTTTGGTTTGTCTCAACAGATATCACTTTTCACGTCGTGAATCTATCTTCATA

TGCAATCCCGTCACTGGTGAATATGTCAATCTTCCGAATCCTGACCCGCGTCCTGATTATACAGCCGTGTTTGGGTTTGG

ATTTGTTGTGCATTCCGTAACTAATTACGAATACAAGGTTGTTATGATCTACTATCGTATTATCCGCGGCATCTTTTCTC

CTCGTGGGCATCCTGGAAGGGTCATGGTATACACAATTGGTAGTGGCAGAGGGTGGCGAAACAAGGGTACGACCACTTAC

CAACTTGGTCGTCAAAATCTGATGGGAGTGTTTGCAAATGGAGCTCTTTACTGGTTACATGATCATGAGGATATAGTAGC

TTTCGGTTTGGCAGATGAGGAGTTCAGATTAGTCGAACCACCACCTTGTCTTAATCCTGTCCCATCTGAATGTTATTCTG

AACTGCGGAACTTGGGAGGGAAGTTGTGCCTTTTTCAAGACAAATACAAAGAAAATCGCCTAGAAATATGGTCATTAAAG

AGAAACTGTGTGGAGGAGAGTTGGACTAGAGACTTCGGTATAGCATACGGCTCGGCTAGCCAGTTTCATATGACGCCTTT

ATTGGTCACAAAGAAGAATCAAGTTCTCATGGTGGAGGATAAAGAATTAACACTTTACTGTTATGACCCGGAAACAAGTA

TTATAATTCAGCTGTGGGATGATGACTCGATGACTAATGTGACGTCTCTCCATGCAATCTCCCACATGAGTAGCTTCGTC

TCCCTGAAAGATCTAGGAGAAGAATCAATGATGAAAAAAAGAGATATAGGACAGTGTGATCAATCAGTTTAATAGAGCAG

ATTTCGAACAATCAACTTTGCACTTTGCTGTAGCTAATTTAAATCTTTGTAAGCTTTAGATATTCACACTGGCTTAACTA

AATAAAATCATCTCTTTTGGGGATTCAGGTGGACTTAAGCAGGTGTTGGCACTCGTCATAGCAAGATATTCACACTCGTT

CTTTTGGTATAGTGTTTGGTAAGTATAATGCATCTGGGTATTTGACAGGCAGGGGATCGATGACAATGTAGCTACACTTG

TGAATCTTTTGGCAGGGGAAAACACAAATACACATCTCTTTTTGCAATTTTCCGTTGAACGTTAGTATATTTAGTATTTG

GATTATTACTAGCATTACTATTTTGCAGTTAACTCTCAGCATATACAGCCAAGCCAACCTTCCTCAAGATTCTAAAGATG

GTCAGTAAAGATGTGGTTTATGATTTGACACCTTGAAATGCGACAGGGATGCCAAATGTTTATTTGGAGAATTAACTCAT

GATATGCTTCCAGTCTCAAGACTCTCAGCCAGATTCCAGAACAATTTGTGAGATGTCTTCTTCCACAAATTGTCAGATGT

TGGAAACTAAAGACTATTTGGCTGCTGTCACTATGTGGCATATTTGGAAAGCAGAATCATCCAGCAAATTCATAACTCAC

TGGCTGCTTTTTTATGATTAATTTCTTTCTCAATTCATTTGTAAACCAGAATAAATAAATAATATTTACAATCTAATTCC

TACAGGCAACCATTTTCATAAATAAATTAAACAATTTGTGAGATGTCAGCCTGCATCCAGTTTCTTTTGTGCATCTTGTT

GAATAGGTAACCAAACATAATTCTAACAAAAAGGCATCATCCTAACAGTTTGTCAGGTGTTTGCTTCCAGTGAAATCCCT

AAAGTGCACTTAGACGAGCTTCTCCTTTCTGGAGTTGAATCTCTCAGCATCCGCATTATGAACACATTGCAAAGATTCAA

CAGGCATCCCTTCTTTCGCTGCTTCCATTTCCGACAAATGCATCAGATATTGTGCCTGCGTGGGTGACAGAGGACAGAAC

AAAATGGAAATCATAACCAACTCCTATATTGTTAAATTAATCGGAAGCTTGATTTAGATGTCCTCGTCTCACCTTTCTTG

ACGGCTCATCCTTCCTCTTTTTTCTTTCAAGCTCTGCGGTATCACATGCAGTCAAAGTAGGACCTTCCTGCAGCACCTCA

GGTTGTTTCTTTGGAACTGTATCCTGGTCCACTCCATAAAACATCCGCAAGGGCACGGCAGACCTATATTTGGGAAAGGA

ACTAATCATTCAATAGCCTTCTAGGACGATGAAATAGCGTGCAAGTTAATCCGAAGAAGGTCCAGAGAAACATAACATAG

CAAGCAAATTGGGTGATATATCATATTGGTTGACATCTAGACCGGTGGCAGTGGCAATGGCATGTAAAATTATAATTGGG

GAGCAAAGGTATATTTTCAACGGATTGATCATGGTAACAAAAAAATATCCAAACCCGAGGATTCTGAGTTCTAACTTCTA

ACCCTATCAGGAAATCTGTGTTTTCAACAACAAAATGACAGTCTCCAAAGAACCCGAAGAAGTATGTCTAAACAGCCCCA

AAAATGAACCTGTAAAGAACCCTTACTACAATATCCCCTATGTTTCAGTTTCCAAAGAACATTCCTACAAGTAGTTACTT

AAAAATATATTGAATGATCACTTCAAATAGCATAACATAGTAGTAATACAGATTTCCTATTGAACCACCTCGCTTCCTAA

AATCCGGCAAAAAACAGAGAGATGAGATCCAAGAATCCTAGGTAGTAACTCCTAAATGTAGATTCTCATCCCTCCCTTGC

CTAACAATTGTAGATTTGTAGTACAAATAGCATTATCTCTAAAACAATTGTTTATGATACAGCTTACAATTCGTATGCTT

AAAAGTAATTTTACTTCAAAGATCAACTCATCTTGTTTTGAGCCTTTTTTATATAAAATGACATCAACTATGTTCCTCTC

GACTACAAGATAGTGGAAGGTTACAGTCTATATCTATTTTTTTATTGTACTTGTATAATTCATTCACCAGTTTTTCGACT

ACTTAAGCACCAGCTTCAAAATATGCTATCTAAACTTGCCCATCCACGTATACTAGATATGAATCTAACATTTTGCTAAC

CAAAGATGGGCCTTGAAAGAGACATATATACCTGACGAAGTGTACATAGGTTGAAGAGATCGCAGTTTCCTTTCATGGAC

TCCCTATAAAATAAAGGTAAAATTTCAGTGACTCTGATATGTGAAAGTGTATGGTTTTAACATGCTGGTGGTGGAAGAAT

ATGAAAAAATTCGTTAGTACAAAGCAGGTAAACGGCAGATCAGAAACTCACAGGATAACAGCGCATCACGTACAAAAGGG

GATTTGAAATTAAATCGAGCTCATGCTCGTGGCAGTGGCATGTAAAATTAAAATTGGTCTCCCACACATGACGTTAACAA

CAAAATTATGCAAACCCGAGGCTTCTAACCCTATCAAACAATCTGTGTTTTCAACATCAAAATGACGTCTAAGGTGTATA

GCAATCAATACAAATGTCCCATCAAAGATCAATCCCGCTCCAACAGAAGAAACCCAAGGCTTTTAACTCTATCAAACTTA

ACAACCTATATCATGTGTAAAGAACCATTTGCCACAATATTCCATATCTTTTAGTTTCCAAAGAACATTCCTACAAGTAG

TTACTTAAACCTCCAAATATCACTCCGATAGAAGATGAATAACACAAATTAAGAATAACAGAACATGGTAGACATATAAA

TCTCCTATATACTTAATTTGAAGCACCTTCCTTCAACCATATCCGGCAAAAAAGAGAAAGATGAGCTCCTAAATTCCGAG

GAAGTAACTCCTAAATGTAGAACTCATCCCTCATTTGCCCAACAGTTGTAGTACAAATAGCATATTCATCTAAAACAACT

GTATACGACACAAGATAATAACATGTAGACGTGAATCTGTAATATAATGAACACTTATTGCAACTTTAACAAAAATTCAG

TTTTCTAAAATGTTATCTATGCAAAAGCAACTCTGCTATTCTGACCACCTGGATATCATTTAAAACAAGATCACATCAGC

GTTGCTCTCGTCTACAAGATAGTTTAAACAGGCTCTATTTTCAATGCTACAGTGTATATCTATTTTTCTATTTTACTTGT

ATAATTATTTCACCAGACTACTTAAGCATTAGGTTCAAATATGCTATACTTAGATCGCTATTAAAAGAACATTTGAATGG

AATTACTTGGATCAGAAGTTATGCATCCAGATAAGTAGGTATAGCAAAATAATCATTTACCAACTCACATAGTGAACTAA

GAGCAAACTTTACTTAAGCTCAAATAGTTCGAGCAAGCATAAAGGTTTCATAAAAGCAGTAACAACGTCTACCCAAGAAT

TAATTTGAGAATTATTGACTGTCCTATTCCATTGGGACTAACCACTGCAAAGCAATTTAAGTTTTCAGAAAATCATGGGA

AACGAAGATTATAATGACAGGCTTGGTTCGGCAGCATGTCAAGAAATTTCCCCGTAATATTACAGAAAAGTGAAAGATAA

TATGTTGACCCTTACTTGCAATGCTGCTGTTAAGATCAATGCCGGAGTTCAAATTCTTAAATTTATCGGTAGTTAGATTT

AAATTTCGTCGTCTCACCTCCTCTTTTCCGCTCATTCTTACTCATTTTCCTTTCAAGCTTTGCTGCATCACGTTCAGTCA

ACGTAGGACTTTCCTGTAGCACTTCAGGTTGTTACTTTGGATCTGCCATTCCATCAAACATCCGCAAGAGCGTGCCAAGA

CATATATTTGAGAACAAATCATTCCATAGCATCCTTAACGTGCAAGTTAATCCAAAGACAGTCCTATACAATACATAGTA

AGCAAAAATTGGGTGTACATATCACACTGGGTGACATGTAGAATGGTGGCAGTGGCAGGTAGCATTAGAATCGGTTTCCC

CACATTAAAATGAATCAAAAGTATATTTTCAAGGGATCAAATTCCTTCAAAGACTCTCATTTGAAGTTCATATCCTCTTA

GAGAGCTTACAGTCAATTACACTATTGCCTACACCAGCTCCTGAGCTTATAATGGCGTTTAGCTACTTTCGGAACAGCCT

GTTATTGCATTCCTAAACTAGATATCATTCTCCTTTAAGATTTGACAGAACAAACGCTACAAGTACTAATTAAAACTGAA

AACGCTATTTTTGCTCTTCCAGAGAGAAAATAGGAAATTGGTTAAGCTTACAGATTAAATACTTTCACAAAGCTTACAGC

ATCTGAACATAAAACATGCTCAAAAAACTAATTTGGGGAAACATTGCAAACACAAATTTTATCATTCTCAGAGACAAACT

TAAATCAGCAAAGACCAATTGAGAGCTCAATCTTACTTCCCAGTGCAAGCAAATATAAAGCTCTCCCCATTCCGGATTTT

TGCAAGCAGTAAAACACACATCTTCACCCTTATCTATCTAAGAACATTCAACAAATACCTGAATTGCATTTTCATGGTGG

AACCTTACCCAAAAAAAAGACAGGAACAATTAAAAGAAAATGAAGGGTTTCATTTGGTTTACCTCTTCATATTTCTGAGT

CATATCAGACCTTAGAACAGCCATAGTTTCATATTTCCTTAAACCAGGTGGACATTGAGCCAGGTTCTTCTTTCTCTTCA

GTAGCTCCTAAAATACCACCAGATGATGATGGTGGTTCATCATCTTCTCCAACACCTTCTTCAAAGAATGAACCTGAAAA

ATCCTAAATCAAGGCTTTAACTGATAAGCTTTCATGGGTTCTTCTCGAAATCTCAACTTTGTTGAAATTACAGGATTTGA

TTTGGAAGAAAACTTGAGGTGTGAATGAAACGAGAGATGATGAAGAGAAGTTTTGGGCGATGAGTAGAAAATTGTAGATA

CATTTGGATAAGGCTGTGAGATGTTGACGGTTTACCCTAGAGCTGTCAACGGGGCAAGCTAGGGTCAACCCGGCCCTAGC

TTGTGAACCCGTTGACAAGCTAGGGTCAACCCTAGTGCTATTCATCAACAAGCCCAAAATCCCAACCCTAGCTTGTACGC

GGGTTGGCTTGTAGGGTTCACGGGTTGTCTTGTTAGTTATCATAACAACCAAAAAAAATGTTGAAAGTTAAAAAAAAAAA

AATGTTTAAAATTACGCTGTTAGGGAGGATTGAACCTAGCTCAGCTAGGTGTTCACTTAGCATAACAACCACTGGACTAC

AAATGAACACGTTTCTATTAGATAAAATCTTTAAAATAGGTAACTAACAAGCCAACCCGTGAACCCGCGGGTTGACCCTA

GCTTGGCTTGTTTTCTACTAGGGTTAGATATGCCAACCCTAACCCGGCTTGTTTAGACAACAAGCCAAGCCGGGCCGGGT

TGAAACAAGCCAAGCTAGGTCCAACCCGCGGGCCGGGTTAGGAATTGACAGCTCTAGTTTACCCCCACTTACACCGGGTG

AAAAAGTGTACTCTCGTAGATATTTGCTTGGGGTAAATTTTCACTTGCAGAGTCAGATTTGGACAAATAACCGATATTTG

GAATTAGCGTTGTCAATGGGTATTTATTACCCGGAACCGGACCCTTTAGATTCGGATCCGGTTCTGGGTCCTAAAGTTGG

ATCCATTAGCAACTTAGGACCCGACGGGTAATACCTACGGGTAATGGGTACCCGGTTATTTATCTTTTTAGCAAAATTAT

AGGTATTTTGTTAGTTTCGACTTTGATTTTTTTTTCCATTTTTCAATTTTTTGTTACATTTAATCTTTATTTAGTACATT

AATAAAAAAATAATAATAAATTTTTACAAAAATAATTTAGACTCAAGTTAAAAAGAGAAAGAAATTAAGATTTTAAGATT

TTCTTTACATTTTTTTTTTAAGGAGAACTTTATTTGTAGCCCAATTTAAGATGTGTTATCTTGCATTGGAGATTGCAAAA

CTTCACTCTTAATGAATGCACATTTGGTAAATCTTTGTTAATTTGCATTCTCCAATGCAAGTTTACACCATCTATCCTAA

TTTTTTTTGTTTTTTTGCACCTTCCCGATGCAATGATTGGGTGTTATCTTGTGCATTCATGCACAAGATAACAAAACCCT

TTATATATTACTTTCACATAATTTGTGTAGTTCTGAAGAATTATTTTACTTTTTACAAAAGTTATAGAAATAAATGCAAG

ATGTAACCAAAAAGAGGAAACAAAAGTATTTGATTCCCCAAATTTATTTCATTCTCCTTTTAAAGGTCAATTTGAAATCT

GGCTTTTACATTTTTGATTTATACTACGTTCGTTTGCATTAGAGATATGGATCTGAGCCGATCCCTAACTAATCGTGAGT

CAAATGTTATATCGTTTGTTTTCTATTTTGAGTCAGATATGATTTGAACTCTGACTCATACCTAATTTCTGATTTAGATC

CGTTTGAGTTAGGTAACGAAATATCCATGACTCATGAGACCAAACAACTTACTCACATATTTTTGAATCAATGGAGTCAG

ATCTGACTCAAAACAAACACATTGAGCCAGCTTTGACCCAAGTCTTCCTTTGACATTCTATTGGATTTTCTAATTGAGAA

AGTGGAGATTCAGCAGATCCACACGACTTCGATCCAGATGACTCAAGTGAGTCAAGAATAAACAAACATGGTGTTAGTTA

AATCGAGAAGCAAGTTATGGTATCTGAAAGCCCTCTTTAACTCACCAAAAAAAATCATCATATATAATTGCATCCAAACA

AAGTATGCGTAATCAAGAAAAAAAAAACCAAATAAATATATTAACTCAACAGGATATAATAGAATGATAAAATCGTTAAA

TATCGCGTCAAAGCTAGTCGACAACCTAAATCATAAGAGTAGAGCCAGTGGGAGCTCTCAAATATGTACATTTAATGCTT

TATGTGGATGGAAAAACTAAAATTCTCACAGTGCGAAGTAGTTTTTCACAGATGATTCAAACGGTTGGGTTAACTTTGGC

CGGTTTTTGCAATTTCATCTGAGCATTAGTTCAACTTTAGCTGATCGGCCAAGGTTTGTGGATAATGCCTACGTGTTTAA

CCATCGCCAACGACTATATCTCCATTTTCCCTATATAATATAAAAATCTCCCCCATCCAACAATTAACGCACAATTCATA

TATGTTTTTTATATTCTCAAAAAAAAAAAAAAAAACAACAACAAATTTCTCCAATATATTACTTGTTTTCAGTCTTTTCA

ATCACCTTTTTTTTTATGGATCATAATTTAACTGGTGGCAAGATTTTCCTTCAAAAATCATATTTTGAACTGGTGTTTAT

CCCCCTCTCCTCCTTCTTGTATTGTCATCGAGATGGTATCTAGACACATGTGCGTTAGCTTCTTTAGATGGTATCTAGAG

TCCTAGACGCCCTTTTTAACAAGAGAGAATTCATCGTATCCACTTGCATCCAAACAAAGTGTGAATAAAAGTTAAGAAAG

AAAAAGAAAGCCAAATGAAAAGATGTCCTCGCGACAAAATCATAAAAGTGAGTTAGTCATACCTAGTTAAGGTCGACTAA

TATGAGTCAAAAAATACAACCATAAACACACATGAGTTAAAATTAACAGTAGAACAAAATCAGCATAAAAAAATTATCGT

TAGAGAATTTGGCGCTAATGAGTTTCGAATCCGTTAACTTTACCATTATATTACAGTGACATCGACATTAAACCAACTAA

TTAAAAGTTAAGACTTACGAGTCTTCGACATTGAACAATAAGACATTTTCTCAGGATCGGGTAATTGCAGTTAACCATTG

AAATGGAGTAATTCCAGGTTAACCAGAAACAATAATGAAAATTTGAGGTTGTGTTTTTGCTTGACAAAGATGTAACTCTA

TGAATGAAGAATCAGAAATAAGAGAATTCCAAGCTCTAGAAACACATTGAATGTGAGAAGAGTTTTAGTAGGAATATGGA

TGGTTGAGTATGTGATGTATTAGTTCCAAAATTTTAACTTTGCTTTTCTGATCCTGGAAATTTTAGGGAATGTTTGCAAT

TTCAAAGAAAACAAACTTTTAAGGAAAAAACTATTCTCCCTCTTAATTAACAAGAATAAGTATTAAAGTGAAGTGAAGTG

AAAAAAATACCACCTACATATCTATTTTCAATTCATTGGGAATATTTTTAATTTCAAAGAAAATATACTTTTAAGGAAAA

ACAAATATTTTGTGGCTCTTAATTACCAAGAATAAGTATGGTGTTCAAGTGAAAAAAGTAATCTACTACAGTTCATAGAG

AAAATAAAGGGGAAAATATTTTCATTTCAGAGAGAAAATAAATGAAAAAAATAAATAAGTGATGGACTCAATCACTCCTA

ATCACAAGGAAAGTTGAATAACTAAACAGATAAAGGCACAAAATATTGCTAATTTAGAGAGAAAATAAAAGCAAACATTC

AATAGAAAATACATTGCTGCTAATAAAACAAGAAGGTTAGTACTTTTACCTCTTGGTATTTCTCTAGATAATTTAATATT

GTATACTTCCTCTCATGTGTATGCATAAAAGTTGTATGTTAGCAACAGTTTGAATTGGAACCTTTTTCATGATTGTTTTA

GCACCTCTTCGTACATGAAAATAGTCTCCAAAAAAATTTTGGATTTGCCTCCCCCATACTTGTCATTCCCTCCTCCTGTT

TATTCCTCATCTTCTTTCTTTTCTAAAAATAAAAAGAAAAACAAGGATGGTAAGAATTTGGTGCAATATGGTAAGCCACT

TCATGTATCTCAGAAATTGGAAAAGGTTTTTGCTATCACTACTAGTCATGTCGCCATTCTATAATAAGTAAATAAAATAC

AAGTGGGTCCATTAATCATGATCATATTAAATGTTTAAAGGAAACAATAAAATCAACAGGACAATAAAAAAATATTGAAA

ACAATAACATTAATGAATGCATAGAGATAATTGCAAGTAAGTTTCTATTTTCTCTATGTGCAATAAACATATGAATACTA

CTAAGCATATCCTAATGGATTGTGAGTTTATCAAATCAGTACTCACTGTTGTTCCAAGATATAGAAGATCTATGAAAAAT

CAAGGGTTTGTACAAAATTGGATATCTTCATGGTTTTATCTTTTTCAAAATGTATTCTGAAGTATGGGTTCAATGGAAAC

TATATTGATTGTTAGTTACAAAACAAAAAAAAAAACAAGAAGGTTACTAGTTTTACCTCTTAATATTTCTCTAGATAACT

TCAAAATATATCCTTTTTCTCCATGTTCGTGTATATAATTCTTAGATTTGCAACACTTTGAATGGGAATCCTTTCCGTGA

TTAATTTAGCCACTTCATCGCATAGGAAAATAGTCTGCAAAAACTTTGGTATTGCCATTCTCTTTACCTCAAGCTTTTAT

GACTGTTTTCTGTGAAAAGAGAAATTAATTAATAAATAAAAGGATTGAATTGTGAAATAAAGAGTAATAATAAATTGAAT

CACTGTTTCGCATTTCCAGATATGTACATTCAGGATTCCTCAAGAGCTTATTAACAAATTACATACAATGCAGAGGAAGT

TTTGGTGGGGATAAAATAAATAGGGGAGCTAATATGACTTCTTGGCATAATTTTAGTAAACCCAAATCTATGGTTGGTTT

TCCTTTAAGGAATCTAGGAAATCTGAATAATGCTTTACTTACTAAGATGGCTTAGAGGGTTTGTAATAATTCCAACCAGC

TGGTGGTACAAGTCCTCAAGTCGAAATATTTGAATAACAATGAGTGTGTTCATGTGACTGTTCCTAAACAAAATAGTTCA

TGGGTTTGGAAATGAATAGAGCATGGTATTTCTAATATTCAAGGGAATTATTGTGTGAAAGTTCATAATGACAAGAACTT

CCAACAATTACCTATCATCTTCCAACAACTTATAAAGATCTCTAATAAATAAGATTACTTTCAATTTAATTTAATTTAAT

TTCAATTCCAACAACTGTAATTATTCGAAATTTAATTTAATATAACTTATAGCTTCCAACAACTGTAATTGTTTGAAATT

TAATTTCAATTCCAGTTACTACATATGACTAATAAACAAAGATCTCTCATACGTAGCTAGGTTAGATACTTATACAATTA

CGGTTAATATTTTTTCATTAGGGTGGGAGAAAGTGAACTCCCATCCGACTTGGAAATGATAAAAGACTCATCAATGATAA

AAAAAGATTTTACGCTCTAAAAAAATGAACTCCCGTGCATGGTGAATCTGAAACTTTTCACCTATAACAAGGAAATAATA

AAGACTCAGTCGTAAATAGGATTTTACAGCTAAGTTACTTTAGTTACCCATCCATAGGTTCAACTGAAATTTCAACACAA

AAAATTCAAGTTTTATTTTATTTTCAAACACTAACTACAAATCAATCCATTAGCCTAATTTAACCACTATTTATCCGATT

TTAACCTAAAAACGAGTATTGATTATCCGTTTTCTATTGAGTTACACGATTTTTATGTAAGACCCCACTTTACCCCGTTA

AATTACATTGTTTATAAGTTAAAATGGATAATGAATGTCCATTTTCTACTTTCTAAAAACTCCATACGCTAAAACTCCAT

CTGCTACTTCTTTCCCGCAGCAATACATAATGTGTTTGGGATGAGGTACTTTTGGATGCGGTTATTGGGATGGGAAGAAG

CTGGGAACCTCCACTAGCCCAATATAATAAATGTCATTATCCGAAGCTGGGAAAAACCAATCTAATAAGCTTGGAAAGAA

ATTGATAAATTCAATTTAATTATTCTCTCTATATACATTAATAGAACTACACTATTGCTCCTATATATATTTTTTAGTGT

ATATTCTGAATCGACCATTGCGAGAGAAGTTACTTGAAAATACGAGGGTACCCAAATACACCAGAATCTTTAAATTATTC

ACCTATAGGTCCTCTTTCCGAAAGTGATTGTTTATCGAAAAGGTCGAGACAATAAAACAAATCGGTATTCACACTATGTG

TGATCGTCTATGGATACGAGATCGAGACAATATAACAATCAAAGTGTGATTAATTGATAATAGGTTCAGACTTAACCAAA

CTCTATAGGATCACTATCAAGTAAAGCGGAGTTAACGTTTGTGTATTTTACTTCTAATTATAATAAACAATTATAATTGC

GAAAATAGAAAAGTAAAAGACACATCAATATTTTGTTAACGAGGAAACCGTAAATACAGAAAAACCCCGGGACCTAGTCA

AGAATTGAATGCTGTCAGAATTAATCCATTATACAAAATCTAAACTAACTTCGTATAGTTGAGACCAAGCAACTAAACCT

AGTTCACTTAGTTTCTTCTGTATCCATGAGCTTTCGACATCAATAAGTGCACGCACTGGAATAGTTCCTTTGGATCATAT

TCCAAACAGTAAAGGAACAATAAACCTGTTTGGTAACAACTCTCTTGATTCTTTCCAGATAAAGATATCTCAAGGAATAT

GCAAAGGATCTTTCGTTTAAACTAATTGATAAACTCATTTGCCTGGTTAGATCAATCCAATCTCTATTACCTAAGTAATA

TGATTCGGATTTGTAACAATCAAAATAGAACTCAAGAGAACTATTAGGTGTTACCGATCTCACGCAACTAATCAATCGAA

TATATCTGATTCTAGTTGGATCCCAGCGGATCAAGGTTTGTGTTACACCAAGATATGATAAACTAATAATAAATCTTCTT

TAATCTTCAATTAAACATGCACAACACCACTTGAATCTCTTGTGATCAATCATGCACAGAACAAATTCTGTTAACAATGG

ATTATCATAAGACGTCTTTAGATATACAAACAGTTCTAAAGATTATGTCGACACTTCAATCTAGTTTGAGTGAATCTTAT

ATCAGAAGAGAAGATTTTCAAGAATAAACAAACTAGGTGCAATCAAAGTTCAACAACCGTTAGTCAATCAAATCAATCGA

AAACTAATAAACTGCAACTATCTAGTTTCCCACCAACGGTACTCGTAGAGCTTTTGATCCCATAAAATTCTTTAAACGAG

CGGTCGTAAGAGATTTTGCCTAATTAGGGTACTTTCCTCTCCGGATAGACGACTCCACCAGAAAAAACAAAAAGATGAAT

TTTGCCTGACTCTTAGGATAGTTTGTTATAAATTCAAACTTATGTATTTATAGACCAAGGATGTTTGGACACCAAGGAAT

TTCCAAAACCGAAAGTATTCTCAAGATATGCAATAAATGCCCAATAGGTTTTCATAATTTCTGGAAATGCTTTGTCCAAT

ATTTTCCGAAATCTATCAATAAAAAATCTCCAATTAGTAAATGCACATTACAAATTTTTATTCTCTAAAGATATGCATTT

AATTGATGGTAATTAAAGTACATAAAACTAACAACCTTAATTAAAAGATTCTTAATTTATTTCGGAATGGGATCTCCTTG

AGTAATTAAGGAATATGTTTGAAAAATAAAAGATAATAGTTATTGTTCGTGTTCAAGGTATGTTGACATCTTTTCTTTGT

AAATCATCTTTCATATTTACAATCTTGGAATCGATTATACCACACTTCCAAACAAGTTTAGAATTGGTTCATCTGTATTC

CAAAAAAACTATGTGATTGATCAAATATCAAATCACACTCATGGATTTAATAGGTTTTACCAATCACTGGGATCAGTTAT

ACCTTAATATGGAACACTTGTGATCCGTCACACAAGTTACCATGACCGATTACATCAGTTACCAGGATCGGTTATGTCAA

TTACCAGGATATGTTACCATATACTCATGGTATTGCTTGTGATCGGTCACACCAATTACTAGGGTCGGTCACACCAATTA

CAAATATCGATCACACTATCTCGGGTGATTACTTAGGATCAGTTACACTAATTAATCAAAACTAGTCATACCAAATCATA

AGTCAGGCACTGTGATTAGTTATACCAATATACATAATAAGTTATGATCGGTTCTACCAACTCACACATATTGGTAATCC

AAAGATATGTAATGAATAACAATACCAATAATCCTAGAGATTTCCCTTTCAATTTATAAAACAAGTTCATGAATGTACTT

CCTTTAAACAAATGTAAAACATTGTTTCCTAGGATGAAATCTTCACCTTTAGCCATACACATAAGCACAATAGCATTCAT

ACGATTATGTCGATGTTATATCTACGAAGTTCAAAAGATAAGCATTATACATCGTAGCCTAATTCCTTAATACTATGATC

ATATGATTATGTCTTACTACTAGAGTAAAATATTCATTCACAGCTTCGCAATTATGTTTTCAATATAGCACGACTTGAAA

GATATGTTAGGAATGAAATAGTTCAATTCAATATTACTAACCTCAAGAGCAAGGATGATGTCGTCGTTGTAGCTTGTTAC

TTCTTCACGTTCTTCAGGTATTCTAATAATACTTGTATGTCTCATAATCCTAGAATTTCTGGTCTAACCTATACGAAGTT

GTCTCTAGTACATAATCAAGCGACTCTTAGATGAGTTTTGATACACTATAAAATATGACAACCAAACTTGACATACCAAC

GCTTGGTGGGTTCAACCGAGCTATGTTCTGACATTACTCCATAAAGAAGAAAAAAAAACGTAAATATTCACTCTCCAGAA

GATTGTTATAATACCAGCGTACATAAACTCATGGCAAACCCTCTAAAAATTCCTCGCGGATATCAAACCTAGAAATAGAA

CAGTCCAGCTCGTTAGGAACGTGATATGTAGATGGGCTCCAAGAATCCTAGAAATACGTGGAAAACGGTCTCTGACATTT

GTCATATCATAAAGCGCCATTATCTAACCCAAAGACCCAACAAACAAAGCAGCTTCTACGACGATATTTCTGCCGAACAT

ATCAAAAGACGAAGATTGGGGTAAATGGTTGAAGTGGAAGTTACGATGAGGTTTCGCCTTTGATCTATGAAGATGTAAAC

TTATGAAAGAATAATTAGAATTAAGTGATCCCCAACTTTTAGATGCACATTTAAAAAATAATAGCAACCAAATTTTACAT

CCACAACTAACCCCGACATCCCCACGCCGATAATCAACACCATCTATTGAAATCTCTCCCATAAAAATTCTCTGAAAGTT

TAGGAGGTGTAAAATATTTGGCAATAGATAAATACTCTTGCAGTAATTTTCTTCTTTCACTTCTTGTTTGTTCGTCTATC

TAATTACAACTTCAAGATCCAAACACAAACCAACAAAAAGAGGGGAACATACAAGAAGTTGCACTACCATCTTCTAACAA

GAAGTTTATGAGATGTAAAATATATATTTACCGATATATCATAGACAGAACCTCTTCTAACAAGAAGTTGCACTACCATC

TGCTAGTGTACTTCTCCAATGCTTTCAATCTCATAGACAGAACCTCCATGATCTCTGAGGTACGTAGTTTATGTCCTTCG

ATATCTTGCTGGGTAGAATTTTTCTACGCTTCTCCGGCTCGTTTGTACTACATGGACCACGTACTTTTGTCATCTCAGGG

AGTGCATCAAGGCGACCCTTTGGGTCCTTTATTATTTGCACTTACTCTTCACCCACTGGTTATGAGTATTGCAGAGCGAT

GTGACCTAGAAATGCAAGCTTGGTACCTAGATGATGGTATTGTGATTGGTGATACTATGATGGTTTCAAAGGCACTCAAG

ATCATACAAGAATAAGGTGTTGCTAGGGGGCTGGTCTTGAACATCAAGAAAACAGAATTATTTTGGCCATCCCCGGACCC

TAGAGCTGATATTTTTGGGGTGTTTCCATCGGACATTGGTAGGCCGGAAAAAGGAGTTAAGCTACTAGACGCACCTGTTA

GTCTTGATGCTCAATTTAATGCGGAGATGGTGGCAAAAAGGGTGCTTAAAACAGTGGAACTCATTGATGCAGTCAAGAAC

CTTGAAGATCCACAGTGTGAACTTCTCTTATTTCGGAATTGTGCTGGAGTTTCTCGATTATACTTTGCCTTGCGTACCAC

CTTACCTGTCTATTTACTAGAGGATCAAACTATGTTCGATTCTCGGCTTTTCCAGTTCCTACGGCATGTGATCACAGTGG

ATGGGCCTGGTTTTGGTCCCTTACAACAAAGGATTTCCACTTTGCCCTTCAGACTAGGAGGTTTAGGGGTCTACTCGATG

CAAGATACTATGCATTATTGCTATTTAGTGTCGTGTTTGCAAACACTAGAACTTCAGTCTGTTATTTTAAGGCACACTGG

CATCAATGGTTTTAGCCCACATCTTACGCAAGCTGCTGCCTCTTTCAACTCATTGTGTAGTCCTTTTTCTCCTATATGCG

TCGATAGCACTGCCACCAATTCAATGGAAACTCTGGCAAATGTTTATTTTGATGCAGTAAAGAAGAACTTGACCTCCAAT

TTCGAGTTTACCACGCGTGATTTAATGCTATGGCAGAGTAACCAAATTCAACATGCTCTGGACTTCCTATTGGCTATCCC

TATTGCTAGTTTGAATCAACAAATAGGTGCAAGGCAATTCAGTGCGGTTCTCTGCTACAGACTTGGAATTCCTTTGTTTG

AGTTGGATAGTTTGTGCCCTTTCTGCAAAAGGGATATGGATGTGTTTGGTGACCATGCCTTGCATTGTGCGAATGAAGTT

GGCTTAAAGTTCCGACATGACTTGGTCCGAGATACTATCGCAGACATGTGCTATCGTAGCGGAGTGCTGGAAAGGAAAGA

AGTGGACTTGGGTCTCCTCGCAGACAATGACACTTCTCTGAGACCTGCTGACATTCTTGTTCTCAACTGGGATAATGGTC

GATATGTCTGCATGGATGTGACAATTGTTTCGCCCTTTGTTGGAGGTGGAGTTCGCTCATTGGAGCCCGGACAAGTCATC

AAGGATGTTGTTGCTCGTAAAAATTCCAAATACTTGGATAAGTGCACTTCTCAAGGTTACGTATTTGGTGCTTTGGCCTT

TACTACTCTTGGCGAACTTGGAGACGGAACTACTAACTTTATTAAACAACTACGCAATTACATGGCTAGCCATGATTCTA

ATGTTCGAGTTGGTGATTTTTTATTTCATAGGTTAAGTGTGGTAGTTCAAAAAGGGGTTGGAGCCCAGCTTGTTGCTCGG

CTGCCGACAAGTTTCTTGTCCGATAAGGATGTAATCGATTTTTATTAATAAAATAGAAAAAAGATAAATTAAAAAAAAAA

ATACAAGAAAGGATAGAATGCACTTTTTTCTTTTTTTTTTCATTATTTGCACTACCATCTCATTCATGGCATTTGATAAA

GCATTGTCTAGTTAATCCATGTTAGAGTTTAGTTTGAAACCAATTCAACTTTGTATGCTCTACAATGATGATTTGATTAA

CTAAAAAAAATACTTTACCCATAAATACAAGAAAGGATAGAATGCAGCGTATCCAGATTCTCTATAAGGCGGGGTTGTTT

TGATACAATGCTAAATCCATATCTAGGAAGTCTCTGATACCATGTTAACAGGGGGTTAGTCAGTTTGGATTAGTAGAGAC

ATTCAGAGACTTTTTATGCAAGAGATTTTGGGTGACTCTCCCTGAGTTTAGAGATTTTGAGATACTCTCTAAATGATTTT

CAAGGAATTATGGTGCATATATGAATTTAAGCTTACACACATGAGACAAACAATATAGTATTATTTATTTTACAAAATGA

AAAAAATGAATATTTCAATGAAAATTTATTAGTCCATGCATGGATAACAAAAATTGTTACCATTTTTCCAATAAAATCAA

AACGAGACGAACAGAAGGATTTGAATCTCTAGTAAACTTATTGGGTTTTGGAAGACAACAAAAAAATTTCTTCAATTTTT

ATGAATTTAATTGTATTGTTTTTAATAATTTGATATTTAATCCTTCAAGAAAAGAGAAAAAACAATTTCTCTCAAAATAT

CACCTACTTAGGTGAGACTTTTATTATCCATTTATGGTATGTTTTGTAATAAAAACCTCTACAAGTCTCTCAAAACCATA

AATCATTGATTTCAAATTCCAAATCCAATAACAGAGAATTATGATGACTATTAGAAAGTCACGGTCCAATAACAGAGATT

ATTTGAGATTTTAAAGAACTCTCTATTGACTAACATCTCCACAAATCTCACTCGACTAACCCCATATAAATTAAATCCCA

ATATCAACAAGTCTCTGATAAAGTCTATGATAGACTGATACCATGTTAATAATCCCAATATCAACAACATGATTTAACAT

ATGACAATAGGAATGGGCCTGTGAAGCTTCATGATCCACTTCTTCAAAAACGATTTAGCAAAAAATTATCGTCCATGCTA

ATCTCAAGAGTTAAGTTTCCGGGGCCGGGGAAAAATTAAAAAGATGATAAAATGGTAATAGCTAGAAATTTCAATTAATA

TCATTCATATCGATCAAAAAAAAAAATCGTTCTTCAAAGCAAGAAGAATTTAGTGACACTGGAGAGAATATACAAGATTT

CTATTCATACATTTTTTAAGAACGCGCAATTGGGGGATATCAAAACTAATTAGGGGATATAGCTACAGGACAAAAACAGT

GTCCCAATAGAAATTTTGGGTGACCCCCTATGTAATTATTTTTTATATTCCTAATCTAGCCCTCACTAATCCTAATTAGT

GGTTAATTTTAATTAGTAAAATGTTAAGTATTTGGGGGATAAATTAGTTTGTTTTTAGTTTTATTTTTAATTGTGTGGGT

TAGTTGAGAGTAGAAGGAGGGAAAAAAATTGTGTGAATTTTTTTTTTTTGAAAATGGAAGATGATTGTGATGAGAGAATT

GCAGCAGCTGAATCTTTAGCTCTACTACAACAACAATCACATGTTCAATCTTCGGTGTATGATAATAACTCACAATTAGA

GTTATATGATGAACCAACACCCTTGAATCATGAACACGATGAACATGAAGTGTTTTTTGAAGAACCAACCCAAGAAAGTA

AACATGTTCAGCATACAAGCAACCCCAACACACAGGTAATGCTTCAAAGAGGTCGAAATTTGATTTTTCTTTTTCAAACC

GTCATTATTTCTCTGAAAAAGCTTAGTACCGGCACACTCGTAGTATGAATTCTACGCCGGAACTAGCGCTACCGGCGTGG

TATTCATACAATACGCATGCCGGTACAATATATCCCCCATTTTAAACACGTTCCCGGTGTTGTGTTCCAACTCCGTTTAA

TGCCGGTACAATATATACCCCATTTCAAAATATTTCCCGGCGTGGTTTTCATAGTTTTTTCTATGCCGGTATAGCATATC

CCCCATTTCAAAAGCTTCCCGGCATGTTATTGAAACAATCTATGTATGCCGGTATTTAGTTATGCTTTTTACCTAACACC

GAATATTTCATAGAATATTCCAACCGGCATAGCTTACTCCTAACTGACGATGTCGGTACTTAAAGAAAAGCATCCAGGAG

CTTAAGATTTTTGAACTCCAATACCGGCGTAGTATATAAAATATCTGGAACGCCGGCACTCGGTTCCGGTTGGAAAACTC

ACATGTCGAGAATGCCGGTTATTGTTACCGGCATGTACCCGGTTTAGATAAAATGCCGGTATTGACTCTTCCGGCCTTTG

TATTGTAGAACATTCAATGCCGGAACTGTTGTCGAGCACGCCGGAAATGGTACCCGGCATGGTTATTTTTTTATTTTCTA

TGAACTAATTCAAAATTTTTTTTCATTTATTCCTTAGATTGTGACATACATTGATCCGACAAATTCCAAACCGCTTGGTC

CGGATACGGCCGAATACTATAAGATGCCCCCGGTATGTTACCAAACAATACTTTTCACCTAGTTTTTAGAACGTACTATG

CGGATTGCTTGTAATGAACCTTATAATCTTCCATTGTTGTCCTAATGTAGGGAATTAAATCTCCGGAGGAAGCAATTGCT

TGGGCAAAAGAGATGGCTCTTAAGAACAAGTGTGTGTTGGTGAGGAATACCCAAGGTTCAAATACGCGGTTTGATATGGT

TTGCAAGAGAAGTGGGAAATACACGACGAATAATAGCCATAAGAGAAAGGGTTATGTATATGAAAAGAAGACTAATAGGG

TATACAAGACTCGTACGAGGAAGTATGAATGCCCCTTTAAGTTAGTATTCTATTTAAGAGACAAAGAAAAGAATTGGGAT

TGTACGGTGTGGTTTGGCCAACATTACCACTTGGATCCGAAAGATTATAATGGTCACTCCATAGTTGCAAAGCTAAAACC

TCATGAATTCGAGGATGTAAAGATAATGACCAAGGCATGTATCAAACCAAGAGTGATTCTCAGAACCTTCAAGGAAATGG

ATGATCAGAACGTTTCTTCTCTAAGTACAATTTATAGCGCACAAGCAACCATTAGAAGGGTGGAATGGGAAGGGAGGACC

GTTATGCAAGAATTTGAGAAGCTAGTTTGGGATCACAACTACACGCGCATCATTAAAAGAGGACCGGATGAAAATCCCCT

TCAAATATTCATTGCGCATCCTTTGCTTTCACAATTGGCTCATACATGTTGTGGTGTTCTTTTGATGGATTGCACCTACA

AGACAAACAAATACAACATGCCGTTGTTCAACATCGTGGGGAAAACATCGGACAAGGTAACATTCACATTGGCTTGGTGT

TTAATGGAAAACGGGAGGGATTACAATTATCATTGGGCATTACGACAATTGAAATTATTGTTCCCGGAAAATCAATTTCC

GAGGGTCATCGTAACCGATCAAGATGAGGCACTAATGGGTGCTATATCTGATGTATTTCCGGATACACAAAATTTCCTAT

GTACGTATCATATAGGTAAAAATGTGACAAAAAATTGTCATGCTTTGTTTGAACCCTCTAAGGTGGATATTAAGAAGAGA

ATGATTTCTCAACTAGAGGAAGATGTTCGCAAGAAGAAACTATCACCCGAAATAGAAGAAGATGAAAGAGCAAAGATTAA

AGAGCGAGTGAACAAAGAACACGATGAAAATCATAAAAAATGGTTGGAATTTATAAAGGACTGGGAAAAAGTTTATTGGT

CACTTACCGAGGATATATATGAAAAGAGATTGAATATGTTTAGTGCTAAATATAAGGAAGTGTATCCAAGCGCCGTCTCA

TATTGTCGGGATAAATTGTTGGATAAGTACAAGGAAAAATTTGTGCGCGCATGGACAAACCGGTATAGGCACTTCAAAAC

TGAAGCAACTAGTGCAGCGGAATCCACTCATGGTAGATTTAAAGATCTCATACAGTCCGGGCAAGGCAACGTGGTTACGA

TCACCGAGGCCATGGAGCAAAACTTCAAGGCTGATATCGATCGGATCAAGAAGGCTTTTCAGAAAAGCTTAATGGAAAGG

ATGACTCAATTTCTTCAATATACTAAGTTTCTCTAAGGAATAGAGTTCAACGTCTCTCAATGGGCAATAAAACATATGAT

GAGAGAAAGGGCATATGAGAAAAATTATGGAAAACCAGGTGATGTGTGTATTTGTCCGGACATGTCTTCATTGGGGCTTC

CGTGTCCTCATATGCTTGTGAAGTATAAAGAAGTTATACCTATTGAGGTTATTGATCCGTTTTGGAGGCAACTATCTTTT

GACCCTCCCCCGACGGAAGCTCCCGGACAATCCCCATGGGATACAAGTGAAGGAAGGAATTATACGAAGCTTACTCACGT

GGAAACTCGGCTAGCCGACAAGTATTGTTGAGCCAACTAAGGCTTCTATCTCGCCCATGGATCAAACCAAGTGAAGAGCC

AGGAAAGGGAGATCCGATAGGTAGACCACAAACTAAAGCTTCAAGGAGAAAACAAAGGAAAGAATTTAAAGAAATGACTA

GACGTGAAGAAGAACTCGCGTCAACCAAGTTACGAGATCTATCCGCAAGTGAGATTTCGGAACTCTGATTCGCGGAAGAG

CCGGTTACAATTAAAAGGGGTAGACCGAGGAAGGAACCGTTAACAAGTCAACAATAAGCTTCAGTTTCTGTATCCACACC

AAAAACTGATGGATCGGAGGGTCCAAAGAAAAGGGGTAGGCCACCAAAGGTACCAACATTGAGCTTGGAAGAACAACAAG

TTGAGCATTCCGAAGCCACAACCGTAGTCGATGTATCGGATGTTCGAAAGAAAAGGGGTAGGCCGCCAAATGTACCAACA

TTGAGCTTGCAAGAACAACAAGCTTCGGTTTCTGTATCCACACCAAAAACTGATGGATTGGAGGGTCCAAAGAAAATGGG

TAGGCCACCGAAGGTACCAACATTGAGCTTGCAAGAACAACAAGTTGAGCATTCCGAAGCCACACCAGTAGTCGATGTAT

TGGAAGTTCCAAATAAAAGGGGTAGGCCGCCAAATGTACCAACATTGAGCTTGCAAGAACAACAAGGTGAGCATTCTGAA

GCCACACCAGTAGTCGATGTATTGGAAGTTCCAAAGAAAATGGGTAGGGGTAGGCCAAGGAAGAATATTCGTCCTCACTT

TCCTCTTCATCTTGCTCTTGTTGCTTATATTCCTCGTCTTCCTCCTCAACAGCTGCAGTAGGTTCACGGATTACAAGAAT

ACTACGATCACTCTCCCAAATATCCCCTCTTGTCTCAAAACCCAAACGCTTTTTGTTATTCCCTCGAGGAGGCAAAGACA

GAGGAGTATCAAGACCATCCTTCCTTTTTTTTCTTAGTCGCAACAGCTTTAGCTTTTCCTTTGTTTGCCGCCTTTGCTTT

GGCCTCCTTGGCTTTAGATCTATTTCAAAAGAAGCACGAAATTAATACCACTAACTTTCATTTTTAGTACCGGCATACAA

AACATGCCGTTAGCTAGTTCCGGCATGTACGTATAATAAAATTTCATGCCGGGTATAAATGCCGGCATTTAAAATTAAAC

GTCGAAAATGCCGGTATACTTCAGAAATTTTCCTGGATAAATTTACACCAATACCGGCATCGTTATATAATAAACGTCCA

TGCCGAATTAAAGTAACGGCATACAAAATTAAATGTCGAAAATGCCGGAATAAATTCTAAAAATCCTGGATACCAAAGAA

AGACTTCCGATATGACTTCTAAAATTCAATTTATGCTGGTAGAACTCACTAAACAATGGGTATCTGTATATTAGGAGTTC

AATACCGTCGTGGTATTCAACCTAGATTGAGTGCCGGTACCAATTTACGGTGTGGTATTCATACTAAATCTTCTGCCGGT

ATATCTACCGGCGTGGTATTTATCCTAGATTGCATGCCAGTTTCAACTACCGGTGCGGTATTCATACTAAATTAAATGCC

GGAACTGCTGAAACTGAAAATTTGCAGCTAGGTTTTTGCGATTTTGACCTAAACCCATAGCTACAAATCTAGTGAAACAC

TAATTAAACAGTTCAAATGACTAACCTTATCACAGAAGCCATCTTAGCTAGTAGTTCATCGTCTGATAACTCTGCTTCTT

TGTCTTCCTCCTCTTGAGCAATCAAATCAAGTTTTTTTTTCAATTTATGTTGAGCAATCGTTCTTGGGTTCGATTGAATT

GGATTCGCTTGACTTGGGTTCGATTTGGAAGTTGTTTTCTTCCGTGGAGGCATGGTTATGATTCGGGTAGGTTAATCGAC

GAAGAAATTTTTGATTTAACGATTAATCGTCGATGAAGAACGATGAAGAAGATGAAAAAAAAATCGGAACCCTCACACGA

CTATGCCGGTATAGAAATGGGGGATAACTGATTTTGTGTTTGGTTTTGATTTTAAGTTTTTAATTTTTTGGGGAGGGTAA

AACAGTCTTTTCATAAGTTTTTTTAATTAATCTAAGGGTGTTTTAGTATTTTCATACCCAAACAACACCCCTTAGCAGAC

ACCCAAGGTTGGGGGAAGTAATTCATATCCCCCAATTAGTATGGATATCCCCCAATTGCGCGTTCTTTTTTAACCATTTT

TCCAATAAAATCAAAACGAGACGAACAGAAGGGATTTAAATCTCTAGTAAACTTATTGGGTTTTGGAAGACAACAAAAAA

AATTCTTCATTTTTTATGAATTTAATTGTAGCTTATTTGGTTATTGTTTTTAATATTTTGATATTTAATCTTTAAAGAAA

AAAGAACAAAGTCTCTCAAAATATCACCTAGGTGAGACTTTTTTTAATTCATTTATGGTATGTTTTGTAATAAAAGTTTC

TACAAATCTCTCAAAATCATAAATCATTGATTTCAAATTCCAAATCCAATAAAAGAGAATTATGAAGACTTTTAGAAAGT

CACGTACCAATAACAGAGATTGTTTGAGATTTTAAAGAACTCTCTATTGACTAACAAGAGATTTTAGAAGTCTCTAAAAT

TCTTTCAACATCTCCACAAATCTCAATTGACTAACCCCATGTAAATTAAATCCCAATATCAACAAGTCTCTGATAGACTG

ATACCATGTTAATAGTGAACCATTTTTTGTGGACCATGGTTTTTTTGAGGGGACCATGATTTGATTAGGCCACCTTTCCT

ATAATTATAAGGGGTGTCCTAAAATGTGGAGATGACTAGTTTACCCTTTTACCTAATTAAGATTATAACTAATCTACAAC

TTTTTCTTCCTCTATTCTACAATTTTTCTTCTTCTCTTCTTCGTCAATTTATGATCATCTAAACCTCATCATATAAAAAT

CTAATCAAAATCATCAAATTTTCATCATCGTCGATTCATTGAATTTTCATCGTCGATTAATCGATCTTTTATAAACAAAC

CTATCCATAAGTATTTTCCCACAAACCCATTACTTCTAATTCGTTTTTGATTGAAATTTTGAGCCATAGTCATCAAAATA

TGTCGAATTTGGAGGTTACAGTAGTAAGTTCGGTTAGGATAATTTAGAAAATAAGCCTACTTTTACAACCGAACTTTTAA

AAATGAAGAACACGAAGAACACTTCGGCTAAGATTTTTTTTTTTTGGCCTAACCGAACACTTGAGTTCGGTTGAGTCGCA

AAAATAATTTCAAAACTGAACTCTTCTCTATATAGCCAAACGTTGAGTTCGGTTTAATAGCATAAAAAAATCATAACCGA

ACTTTACTCTGAAAACTTATGTAATGAGTTTAGTTAACTCGCAAAAATAATTTCAAAACCGAACTCTTCTCTATATAGCT

AAATGTTGAGTTCGGTTTAATTGTAAAAAAAAAAATCATAACCGAATTTTACTCTGAAAACCTATATAATGAGTTCGGAT

AACTCGCAATTTTTCTGCTAACCGAACGTAGAGTTCGGCAATGTTTTATAAACGTTGCGATCATGCCGAACCTATAGATT

TTAAGTGCATGGAAAAACTAAGGGTTCGGCTGAATGACTGGTTCGTAGTGATTTTGCGAACTTACCGAACTCTGTTGTTT

AAAGTTTGGTTACAACGAGGTTGAGTTGACTGAACCGATCAAAACTCTGTAAACTCAAGCCTAGTAGTTCGGTTACATTT

GATTCTTCATCATCTAGCCCAACAAACAAAAACCCCATTAAAGCTCTGGTTCGGTTGCATGTGATTTTGGATTTATTAAC

TGAACATCATTTGCAGAAAGTTCGGTTACATGTTCTTCATATAATATAACCGAACTTTGTTGACCTGGTTGAAATTTTTT

TGTTACAATTTTGATTTTGAGGATATTTTAGGCCATTTATAGCAACATTCACTAAGTTACAAGCATACTTCGGTACCCAG

AGGATGCTTTTTCTCCATATTCTCCCATCTCAAAATAATTTTTGTTCTCCATCTTCTTCTTCTTCTATCACTTTAATCAC

TCAATAATTCTACTTCTTTTAAAAAAAAACAGCTTGGTGTGAGGGAGCCCAAAGTTAACATGGGGTCCTTTGAAGGTATG

TGGACCCCACCTTTGTGTCTCACCCCGGTTTTGGGGGAGTTTATCTCTCGGTACGTTAGGCTTTTTTGTAAAAAAAAAAC

AATCTATTAATGATTAGAATATTAACTAATAAATTACAAAGTTAAAAATTACATACTAGGCAAGCGAGCGACAAGCTGTG

CCGCTAGCCCTTTTTTGAATAGCAAAACTTATCCTATTAAAAATAAAATCTCGTGAGCCATTACATTGCTGTGAGGGGTA

GCCCAGCCAAATAGGGCAAAGGGGACGCATGCCCATTGTCAACACATGCTTTCTCATGGATATCTTACCTGAGTTAACAT

GGGGTCCTTTAAACATGGTATGTCCAAAAATAAGGACCCCACCTTTGCTTCTTTCTTTGTCTCACCATCATACAAAATAT

CCCTCACATGATCATGTTTTTTATAACCGGGGGAGTTTATCTCTCTCGGCGGGATAAAGCGGGATCATTAGGCTTTTTCT

GCCCTATCCCCTAAAAAAAAAAAACAATCTATTAATTTAACCTTAATCAATAATCAATCACACTAACTAATCATTATACA

AAAATAATCAAGATACCAGGCATGTAGGGATAATATGCAGTGCTCATTTATCTTAGCTATAAGCTGATGAAGTACGAGCA

AATAACAAAGGCCCTAAAGGGTCACCTTGTTGTACTCCTGTGGATGAGTAGATAAAGGGAGTCCCACACATAGAGCCGTG

CGGGTTGACCATACAAGTATTCTACCCAAGGCGATATAGATGGACAAAGCTTACGTCGCTTTGGACACCGGTCGACGCCA

AATTGTCCCAACAGCAATGGGACGAAACAACCCTAGTTATGGCCGTTAATTTTGGTACCTTCTTCGGCCACTAGCGGTGT

CTCCTGCAACGTACAGGTTGGTTTCCTTGGAGGCGGCATCGTCGGATGCTTATCCTCAAGTGCTTTCCTTGTGTTCTCGT

TGTATGGCCACCACCGGACGACGATGTCTTTAACGTATCATAATATATAAATATTTAGATAAAAGATGACCTAGAATTAC

TTCTAATATCTTACCCAAAAAAAAACAATGGTCCCAGAAAAGACTAGTCCCCAAAAAAAAAAACGTTCTTAATAATCCCA

ATATCAACAACAAGATTTAACATATGACAATAGGCCCATTCTTAAATGGGCCTGTGAAGCTTCATGATCCATTTCTTCAA

AAACGATTTAGCAAAGAAATTCTCGTCAATGCTAGAAATTTAAATTAAAATATCATTTATATCGATCAAAAAAAAAAAAA

AAATCATTCTTCAAAGCAAGAAGAATTCACCGACCCTAGAGATTACAAGACTGTTGTTCTTCTGTAAGGAATTTCATGCA

TGGTGATTATTAAAATGATGAAACTTGCGTTCACTAGCCAAAAACCACACAAGCGACAATGACACATTCGGGTCACGAGG

TAGATTTTGAGTTGCATTGTAGATGTTAAGCTAGTTGAACCATGTTGGATCACTGAGCAGATTCTGATTCACGAGTTAGA

TTTAGGGTGACGCATCACTTGGTAGACTGAACCAGGTAGTACAGAGCTCTCGCGAAAATCCAAATTTATCCATCAAACTC

TTCAACACGGATGACTTAATAAAGAACCCATGAATCATGATTTAGGTAGAATCATCGTCAAAAGATGTTACTTTGATGTA

TAACCCTAGCACAATCAACAGTGTAGTACAGAGCTTTTCGGTTCTTCACGACAAAAAGCCCACTTCCAGGCGCCTGCAAG

ACCACATATGCAAAGGCTCAAACCAAACAGACAAGCTACACAATGCATAAAGAAGATGGACAAAGAAGGTAAAACATAAG

CCTCAGAACTGCAGACAACCGTCTGACAGTTCAAGGTTTTTGTACAACGTTTACAGTAGAAGCTACAAACACAACTAGTT

TGCTCCAATCTGTGAAAGGGAGAACTTTGGCCAAAACCTCCTAGTCAAGAAAATTATTTGATAACAGAAACAATATACAG

ACCACTATAAAAGAAACAATCACATAAACTACGAAAAACGCCTAGTCACATATTCCCTTTCCAAATTACACACACATCAG

AAGAAACGGAACAACCACCACCCACAATACCAGAACTCCAATGACTAGAAATCAAAAGAACTCTCTAGAAATTCTTGATA

AAAACCAGTCCAGTTTCAAAAAGGGTTTCGAAGTGTATTAGGAAGAAATGAATCAACTGCTGTGTTTAAGAGTGTCATGC

GGCACTTCCTAATAAACATAAGTACTAACAAAGATAATGAAAATCAAGCGAGAATTCATAATTAAATTAAAAAGAGGTAA

CGAAACTAGGTTTTTCTGAAACTTATCAGATTTTTAATATTCAAAACGGAACGCATTGGCAGTTTACTGAATCTAACACG

GCTGCTTAAGAGATTCTTCTAAAATGTGGCATGACGCTAGTTTCTGCATTGACAGACCTTTCCCCTATTAAGGTCTAATA

CCATCCAGAATCACCTTGATTATGTTTTCGTTGTTTAAAAACTAAAAACCAACGAACTACAAAAGATGGTACAATTATTC

CTCTTCTATAACTAAATGAATCGTTGATAGTAGAACTCTCAAACCTTAAAGCAATTACCTCTGAAGAAACATTCTTATGT

GGTGGGTACATGAATGAAAGAGAGATCGATTTTTGAGCCCGAGGATTCTTGTTTTGAGCCTTTTTTATAAATCACATCAA

CAACTTTGCTCTAGACTACAAAATAGTGCAAACCGGATCTATATCCAGCGCTACAGTGTATATCTATTTTTCTATTTTAC

TTGCATAATTCTTTATGACTAAGACATTTTGTTAACCAAAAATGGGCCTTGAGAAAAGCATATATACCTGACAATGTGTA

CATAGGCTGGAGTGCAAGATTCCAGGTAAACCAAAGAACCCAGTGAGATCGAAGATTCTGTTCATGGACTCCCTAGAAAG

AATAATAAAGATAAAAATCAGTGACTCTGATATGTGTATGTGTAAATCGCTAGCATAAAGCACTATCTTTTGTAGCATAA

AGCACGTAAACGGCAGATCAGAAACTCACGGGATAGCAGTGCATCACGTACAATAGGGATTTGAAGTTAAATCGAGCCAC

CAATGGGTGCTGATTAAAGAAGGTAATATGAACCTGGAGAATGCAAACCACTCATTAGATCAGAGTATACGTAGATCACT

ATTAACAGAACATTTGAGCTAACCCTTTTTAACCACATGAAAACACAGCTAATCAAGTTAATAAACTCAAATAGTTCGAG

CAAGCCTAAAGGTTTCGCAAAAGCAGTAACAACGTCTACTCAAGAAATTAATTTGAGAATTGTTGACTTTCCTATTCCAT

GGGGACTGACCACTGCAAAGCAATTTCAGTTTTCAGAAAATCATGACAAAGGAAAAACTACAATTGCAGACTTGGTTTGG

CAGCATGTCACGAAATTTCCCCATAATATTACAGAAAAGCAAAAGAAGAGGTTGACCCTTACTTGCAAAGCGACTGCTAA

GATCAATACCGAAGTTCAAATTCTTAAACAGTGCGACACAGGACAAAACAAAATATTAATCGGTAGTTTGATTTAAATGT

CGTCGTCTCACCTCCTCTTTGCCGCTCATCCTTACTTTTGGCTTTTGAAGCTTCGCCGCGTCATGCTCAGTCAACGTAGG

ACCTTCCTGCAGCACCCCAGGTTGTTACGTTGGAACTGCGTCCTGGTCCATTCCATCAAACATCCGCAAGGGTGTGGCAA

GACCTATATTTGAAAACAAATCATTCAATAGTCTTCTAGTACAATAAAATAATGTGCAAGTCAATCCAAAGACAGTCCTA

AACATAACATAGTAAGCAAATTATTTGTACATATCATATTGTTTGACATCTAGAATGGTGCCAGTGCCAGTGGCAGGTAA

AATTAAAATCAGTTTCCCCACATCAAAATGATCAAATTCCTTCACAGAGCTAAGAAATTTCCATAACACTCGTTTGAAGT

TCATATCCTCTTAGAGAGCAATACATCTAATAGTTCTTACATTCCATTGCACAATTTCCTACACCAGCTCCAGAGCTTAT

AACGGGGTTTAGCTTCTTTAGGAACAGCCTGTTAATTTGCAAATAGGTATTGAAGTTGAATTCCTAAACTAGATATCCTT

CTCCTTTAAGATTTGGCGGAACACAGAACAAACGCTACAAGTACCAATCAAAACTATGCAGCACTGAAAACGTTATACTG

CTCTTCCGAAGAAAACTAACAAAACTGGTTAAGCTTAAAGAATGAAAAAACCTTCCCAGAATCACAAAGCTAAGGAAAAT

CAATCTCATTTACTACTTCCGAATTCACAGAATTTGTAGACCATTATGATGGTAGGATCTGAACAGAAAACATGCTCAAA

GAACAAATTTGGGGAAACATTGCATGAACAAATTCTGTCATTTTCAGACACAAACTTCCACCCTAGAAACAAGAATCATT

ATCTAAAAGTTATGCAGTGAATTCAGAGATCCAACAACCCTTTCATCTTAAAGACCAATACATCAAATTTTCAGAGCATG

AAACATTCTTGCCCCCTGATTGTGACTTATTGCCCGATTGCAAGCAAAGAGAGAGCTCCCACCATTCTGATTGCAAGCAG

TAAAACACACATCTCCATCTAACTATTCTACTTGTTCTTATCAATGCAAGTTATAATAATCTAAGAACATTCAACAAACA

CCTGAATTGCATTTTCATGGTGGAACATTCATTATAGACAAAATCAGGAGATGAAGGGTTTTATTTTGGGGTTTACCTCT

TCATACTTCTGAATAATAGTAAGACGTTTATCTTCAGTCATATCAGGTCTGAGAACAGCCACAGTTTCTCAAACCAGGTG

GACATTGTCACATTGAGGTTCTTCTTTCTCTTCAGTAAATACCACCAGATGATGGTGATTCATCATCTTCTCCAACGGAC

CAACACCTCCTTCATAAAATGAACCTGAAAATTCTAAAGTTAAATCCTCAACTGATAAATTCTCATGGGTTCTTCTTCTC

AGAATCTCAACTTTGTTTAAATTCAAAGATTTGTTTTGAAAGAAAACTTGAGGTGTGAATGAAACGAGAGATGATGAAGA

AGTTGTGAGAAATTTAGTGGATGAGAAGGAGGTCGAAGCTAATGAAACTGATGATTTTTCCATGACTGAGAAGAATCAAA

GGGTTTAAAGCTTCACTGGACTTTTTTTTGGAAGTGAGAAAATGGAGAAGAAGACTAGCAAGTCATGGTCCCATGGATGG

ATGATTGATTGGATAAGGCTGTGAGATGTTTACACCCAAACCTTCATTTTTCTTTACACCGGGTAAAAGGGTGAAAATGT

CAACAACAAAAAAATACAGTGTGGTAAATTTTTCGTTTCTCAGATTTGGGCAAAATAATGGATGTGTTGGGGGACTTGGG

ATGCATACCAATAAAGTGAAAATATCATTTTTCTCTTGAACTGGAGGGAGTATTATATAGTTTTGGTATTTCTGAAACTG

TAGTGATTTTCAGAAATCAAAGTATTATAGTTTTGGTACTTCTGAAACTGTAGGGATTTTTAGAAATCAGAAAGTATGGG

ATGGAGACTCGAAAGTTGTATCCATGGATGGGATGATTGGGTTGGATAAAGCTATGAGATGTTTACCCAACCTTTTTTTC

TTCTTCTTACACTTCACTTACACCGGGGGAAAGAAGTTGAAGTTATCAAAAAATATTTACTTGTGGTAAATTTTTCGTTT

CGAGTCATATTTGGACAAATAATGCAACGGTGGAGTGCATATGGCTTATATTACTTTTCTAGAAAATTTGTGTGTATTTG

AAGAAAAATCATTATATCTACTAGCATCCGAACGAAGTCTGTGTAATAAAGAAAACAAGAGTCAAATAAATAGATGAGCT

CAGCGGAATTATAAAATCGCCAAAATTTATGCCAAAGATAGTCCACACCCTAACTCATAAGAGCAAAGCCTGCGGGAGTT

CTCAAATATGCACATTTGATGATTCAAATGGATGGAAAACTCAATTTGTCATAATGGGTAATAGTTTCTCGAAGATGATT

GAACCGCTCAGCCGAACCTTGGCCATTCGTTGCAAGATTAGCCGAGCGACTATTGATAGAAAGATAATAAACTATACAAA

AAAACATTCAATAATGGTAAAATAAATAAATATTGTTGTTCAAATCAGAAAAAAAAAGTAATTCGTTTTTACTATAATGC

TTGCTAGCTCTTATACTAGCCAAAAAATAAAATTCATGGCTTGAGCCTGAAATTCTAAACAGGCCACATATTAATACAAC

GGTGTAAGTATTGTTCTTTCTATAAACACAATAACAAAAATTTCAACGAAAGATAAGCCTTACATAGTACAGAGTATTCT

GGATAGGCATATGCAAGCTTAACCAGCGGGGAACGAACTTGGTTAAGTGGATCTCTCATAGAAGAAATTCAACATATGCT

TTGAAAAAGATAAATAAGGTTCAACGGGTACCGACAAGTTAAGGAGGTGATTATGGTGTGATAAAGTCAAAATTTCTCAT

TTTTTATTAATGTTGTGATTCTGCAAGAATAAGGATGGATATTAATCCTTAATGAGTCTATTATTGATTACAAGTGTCCC

AAAAAAACACCTTAAGAGACTTACCTATATGTGTACGGAAATCAAGTTGTTTCCTAAGAGAATAAAACTATTCTATAAAG

CAGGTTCATGACTCTATGATAGACGCACATGTCCATTAATGTGCTATAAGCTACATAACACACTATTCTCTGGAATCAAT

ATATATATATGCGTAGAGATCCTGATTTTATCGCGTACTTGGTTCAAGACTGGTTTCACCATAAAACCTCGATAAGGGTT

TGAATTTCTTACACTAACTCAAGGTTCAAAGGAAAAAGACACCTATCATTGATGTATTGGTTTTTACGATGCATGAAGAT

TATTCTCAACTTTTCTCGAACTACTTTGTCTTGATTCTACTCGGCTGCGTAGGCATTTCGGTTTTGTAAGTTTTTTTTTT

CTCTTTTCTTTTTATTTTATGGTTATAGATAAATGGGGAGGATTATTGGATATTTTTAATATCATAGAAAATAAAATAAT

TTTCGTAAAGGTTTGCCTCTACGATTGGGAAGCGTCTAATGTCGAATAGATGCAATGGATGTTTGGAGATGACGATTGAA

GTGGCATCTCCAAAGGGTGTGATGGTTCACATAAGACAAATCCATTCCTATTAGACATTATGGATGTCCATTTGAATTGA

AGAGACATTTCCAATAGGTAAGGGTCCCGTATAAATAATGATGATTGATTTTTAATCATGTATGAAAAACATCAAAATTT

CTATTTTCTCTCTTCTAGGCATCATCAGAATCCGATTATTGTGTGTTCTTGGTTGAATCAAATGGTACAAACCTTAAATC

CAATTTCGTTGTTATATGATTTCATTGAATCTTGATGACAATATTTCATATACCCTTTTATAATCGACACTTTGATTGAG

AGGTTCGAAATAATTGTCTAAAAGAATCGAAGGCCTTAAAGTCAGAAGTACAATAATTCTCTTTTGGAGTAGTAAAACAT

TTCTCTTAGTGTGTCATCCTCCCCATTAGCCTAAATTTTCCAACATAATTGTGCTGAAAAGTGTTGACAATGCCCCATTA

AAAAAATCAACACCTAAAAGTTTAACTTTTGAAATAATAATGAAATAATAAAAATGAGTGTAAAATCTAACCTTTGTTTA

ACGACCGTTTCCGCACAATTACACCGTAACCAATACATATAATATAACGAGGAACGCAATATGTGTTGGAAAAAACTGGG

TTGCAATAGAAGATGAAATAACACATGAAGAGATTGTTGTACTCCAAACTTTCAAGGCTCATAAGATTTCTTTTAGACAA

TTATTTCTACCCTCCCAATAAATTGGCGATTACAACATGTATGTGAAATAATGCCTTCAGGATACAATGAAATCCCTACA

ACAGAAATTGAATTCAAGGTTTGTACCCCTTGTTCCAATCAAGAACACATGTAAACAAAATTTCTGATGATGCTTTAGAA

AGAGAGAGAACAAAAAAGGTTTTTTCGATATGTTTGAAACGAGTTTTGGGAGATGCTATTTATAGAGGTTTTCGCATCTG

TTGGGAATAGACATATTTTCACCAACAGGTTTCTCTTCAAGAAACGATGAGTCTCTTCATCATCATACATCAATGGCGTC

TGTTGGATTCGAAGATTCTGGATATGTTTGTAACTGTCGAAACAGTTACGTACCCCTCTTAGATAGCGCAGTTGAAAATA

TTCCAACAATATGAACCATTTGCCTCCCACAAACTCGCAAATTAATACCTAAATCCAAAAACATCATATTAGCACATATG

TAATAAAATAATATTATTAATTAAAATGAATTACGATAACAAAATATAGCATTCTCATTATCGTATCATGCAAATAGAAA

GTTGATAACATTTTCATCTATATCGAAACCTTTAATCTCCCTTATTTATCTATTTAAAAACTTTGCAGTTTCTTTCACAG

AAATCAAACACTAGTTTACAATAGTCTCCTTTAGAAATTTCAAACAAAAATATTAAATTACGAATACGGTACCCTTGATT

TAAGATGTAGATAAAGAATATTACCTCAGTTAGTGCGAGATTCATCGAATACTCTTCATAATCTAAATTTCCTGATACTC

CACAACATATGTTATGGTAAATGACAAACATATTCTCGTATGTTTGTGATACTGTAATATTAAACAACCCATAATTTTTT

TTCAAATTTTTATGTAGGGTTTAGGGATTAAAATCATAATCTCCACCAGATCGGTTCAATTCAAAACCGTATAAACATAT

TCTTTATTAACAACTAATAATCTACTATTTCACATTCTTAACTTTCCAAAATCCCAAACAGACTATTTGAAATCGAAGCA

CGGAAAAGAAGCCTTCATGATCAAACCCTTAGTCAGATTTCACCAAAATAGCCAGTGCCACATGTGGGGATTTGTAGGAT

ATGCTTGCATATGGGCTGATCTAAAATAGCCAGTGCCACATGTGGCTTATTTCTTTTTCTTTTTTCTTGAGAATTGTTTT

TTTTTTCATGAGAAAATTGTGTAAATGTGTTTAGAAAAGATTACTTGTGACTATATTCATATATAGTAGGCAAGGAAGAT

TTATATCATAGATTGATCTATTTATTTTGAAATCCCCTTCTTCATACATATTTGAATTTTGGATTTAATCTTGTAAGACT

GCATTCTTAATCAAAGATGAATGAATTTGTAAATTTACTTTATTTTCTTCTTATGCTAAAAACATTTTTTTATTTTTTGT

TAGGAATTAAATTGTTGAAAAATTACTCAAATTCTTTCTTCTTTAAAACAACACAATGGGTATAAATTAGCACTATTGAA

CTCCACTTTCCATTATGGAATTTTTCTAAGTAAATCGAAAAAAATTGTTAAAAGGGACAGATAAATGTAAAACTAAAAAA

TCTAAGACAATGAGTAATTATTTCTTTAACATAAGGTAAGAAAGGTTAAGTACAGAGAGTATAAAAGTAAGAAATAAAAA

CCAACCACCATAAAACGGACACATGTCCTTATAAATATGGTTTTGTAGTGTTGGGAATCTTATCTCTTGGACTTATGATC

ACTTAGAAAGAAGGAGGAAGAGGAGAAAAATTGAGAAAGAAATGAGGGGTGTGCGAAACATTGAAATTTTTCCTGGTTTT

AAGAATGGAGGTACACCTTCTTCATTTATTGTTATCATGCCTTATAATTGGCCTTAAAAAATTTCATTGATGCATTTAAT

GTTTAATTCCAGTTTCATAGATAGATTTCTTAAAATTTGTTGATGTTCCTTAAAAGATTAGTTAGAAGAAGTGTTTAATA

AAATTACTTAGATTTAATGATTTTATTGAAGACCGACTCACTCTCACTTTTATTTCTTTAAGAGAGATAATGGTAGCATT

TTGCAACTTCTTAAAGAGAGTGAGGTGGTTCAAAGTATTGTTGACAAATTTCTTATTCGATTTGACATTGATTTATCCTT

TATGGTTATCAACTTGTAATAACTCTTCGCCACGCCGTGGAGAGCTTCCAGGAATAATATTCATAACATATCTAGGGATT

TCAATTAATCTGAATGTTTCATCGTGGAGGTCGAATGCTACAATAAGTTCATTTTCAGGAGCTTTTGAAAAGTGAATACC

ACGATTTCTCCTATAGATGGAACCATTTGCATAAACACAGGGTCCACTTAACTTTTCCAATTAGCATTCCATTCCCAGGA

TAATTTTGTGAAATTTCCCAAATAGCTAAAACTTTGTGTTTTCCACTCACAGGGTCAAAACCAAATCCACGAATGGCAGT

CACCAAAGTCCCTGCATCCTCTAACCATATTGATGTTTTGGCCCAAGGAGTTCTCTGCCCTGTGTTAGGGTTAAAAATCC

TTGTTATATGCTTTCTTCTATCATATAAACATATCAACCCATTTACAAGCTTCATCGCAGAGCATGAACACCAGGGGATC

GACACTCTAAGAAGAACTTCACACCTATCTCGAAAACCATACTGTGAAAAAACTATGAAATTGTCTTCAGAAGTAACCAT

GAGTATTCGAGACTTTAATTTAGCATTATCCAAATGTAATTCTATGAAAGATGAATTAAAAAGAAGTGAATTCCAACTCT

TGGAGACGGTTTCAAATCTCTTCAGTTCTTCTACTGGAAGATGACTAAGTAATTCTTTCATCGAAGCATCATCCATGATG

TTCGAGAAATTTCTAAAAGATTTCCATCTTAAATACCTTTGAATATTTATTTTCTTAGACAGTAAAGAAGATACTAATGC

AAGTGTCACAATAGTTTCCGATGAAAGGTTTCGAAGCATAAGTTTAGACACCATCTCTACATCGAATAAATCTATGAGAT

TTGAAACTTTTCTCAATGACTTAATCTCTTGTAATTTAGACTGAATATTAGTTTTCCACCAGAATTGTTTAGATGATAAT

ATTACATTTACCATTGTTGGCAGTGGAAGTTTGTTAAACATGACATCAACTATATTTGGGTGTCTCAAAACATCCACCAT

GCTTGTTACCTCCTTCATCTTCTTAGTATATCTCTGAAACCAAAATTAGTAAAAGTGGAATCAATTAATACCAAAGTTTC

CATAACTACATATTGAAACACCATAAATGCATAAGTAAAATTGATTACTGAAAAGAAAACCATCAAACCTAAAATTTTAT

CTATTTTCATACTATATACATAGAAAGAAAAAAACAATGCCACTGTATTGAGTCAACTCCTTCCCTAGAAACAAGGAACA

AATTGTTTGTGTTTTTTAATTGAAAACGGAAATCAAGTAATTTAAAAACTCCCACCACACAGATAATTATGGTAAAACAT

CAAAACCTTAAGAACACCAAAAATGGATATACAAAATTGACTATGGTAAGAGAAAATAATGGAGAGTAATACAAAATTTA

AAGCATAAAACCGTTGTTTGACCATGGTAACTGATGCTAGTAAAGGAAAGTACAATAGAAACTCTTTTAATTAACACTCA

ATTATTTAATAAACTCTCTTAAATAATAGTTTTATCCGGTCCCGAGTCGGGGACAACGTGCCAAATTAATAATTCGTTAA

AATTATAAGATAATACGTTTTTTATTACCTATGTAGACCCCATGTGAAATATAAAATAATAATGCACTATATATATTCAA

TACATTAATTATCTACGAAGATCTTTTGAAAAATGATTCAATTCTCTTCTGTTTTTTCCTAAAATCAATATCACATTGAA

TCTCATCTCTAATTTTTCTTATCATCTTAAGAATTTGTGGTGTTGTTTTCTCATAGCGCAACAAGAAATTATTCAGTGTG

ATTTATGCTTTAATAGCCTCGTTTCGTGAAGAGGGCTCTAGTGAAGAACTTTCATCATCATCTTCCACATCTTTATCAAT

TCCCATGACATTCTTAATTATTTCTTCATCAGTCAACAACTGCGTAACTTCATTTTCTTTCGGATAATTTAAGACATCTT

CGACATTCATTGAATTGCGATAGCCCAACTTTTCAATCAAATTTCGCAAATCTTGAGTGATTTCACTTTCTAACTGTTCA

TCCAAATTTTCCAAACTTGGTGTTATTTGTTGATCGAAGTTTGCAATGACGAAAGCAGTTTGCAAGAGTACTTGTACGAA

CATCGATAGTCCATGCTGAAATTGCAAGATTCATTGCATCTAACACATTTATTTTTTCCAGATTTGATGTCCTTTATTCA

TAGCCTTCCGGAAGATATTTATAAAATCGACGACGATAATGCATTTTATAAGAATATAATTTGATATTGATATTGTGATT

ATGTATGAGATAAAAAAGAACACACAATTAGGGGATATGAAATAGTAATGTTACTGTTCATACCGGCATGGTTTTTCAAC

GAGGGAAATAAAAACTTTCCATGCCGGTATGAACAGTAACATTACTATTCATATCCCACAATTAATCATTCATAAAAAAA

GATATGGTGTGGTATTAAGTTTATATAGAAACTAATTATTTGATAATAAACTTATCCATAAGTTAATAAGATATTAATTA

ATATATAAATTAATACTTATTAATTTATCAGTTAATTAATATCTCTCTTAATTAATAATGTTTGGTAGTCTCGATAGCAT

TAATTTCTAGAGTTTCCACTGTAGTGTTTTTCTTTTTACCTCCATTATCGTGTTTAGGTATTGGTTTCTCACTTCTTCGG

ATCTACTGTTTTACTTGTCTCCTTCTTCTTCCTCTTCTTTGCACCTGCTCTAGACAAGAATATGAGTCCTAAACACACCA

GTGGATAAGATGCCACATGACGTGTTCTTAGTGGTTCTTTTTACTTTTTTTTTGGGTGGGGGCCATTTCTTTTTGGCTTT

TCTTAAATATAGCCCACACATTTATTAACCATAGCCCAAAAAAGTTTAAGTTAAAATGAAAAATCAAACAAGTTCTAATT

TTCTACTAACAATTGATCTTAAATCTCCCTTTTACACAAAATTAAAATAAACCCGAACTTTTTTGTTAATCATACTTGAA

ATTAATTATTATGTTTTTTTTAGTAATCTCACAAATTGAACAATTATTATAAACTCTAATTCTTACTTTTTCTCCAAACA

AAATAAACTCTAATTCTTAATTTTTTCTAAAATAAAAGGCTTTTCTTAATCATGATTACACTAGCATTCTCCTAGAACTC

CGTAAACATAATTCATAAAACACAAAAAAATAATAATCTAAGATGATGTTTTTTTCTTCTTCTCAATCATGATTACATTA

AGATTGATTGAGGGTATTGATTATGAGTCCGACTTTGTGCACCCGCGAGTGCACATTCCTCCCCTCATTAGCTGAATTTA

ACGGACAGTTTGTTTTGTTTCTTTGTTTCTTTTTTATAATTTTACCGGGAAATAGAAAAGCAGAAAGTTATATTCTATTC

GATATTTTATCTCTTTCCACTCCTTCTTCTTCTTCCGTTTCGTTTTCCTTTCTTTCTGTATCTACCATGAAAAATCAAAA

GATCGTATTATTTGGTCACAACAGAATTTTTCAATTATTGGTCATAGTATTGATTAAGGATTAAGAATTGGTTACTTTTT

TCTTTTTTATAATTAAGGATTAAGGATTTGGTTTGGAACCATATTAAAAACCATTATAACTAAAATAAAATCATCTAATC

AACATCTAAAATGTGTATTCAACTCCTGGAAACAGTTGTTATAATGTTATTGGTGAAAAATGATGTTGCTGGAGCTGAAA

TCAAATTATATCTTTCCACTTCACTAATTCCAAATTTATCATTCATAATAATCATATTTATCAAAACCCAAAGTACAAAA

GTAAGGGTAATAAATCAATCAAATTTCCTATAATTAGTAATAACAACTTTCTTTATTGCGATTTCCAACTTGACAACTCC

CCTGTCTTTAACCATGGCAGTTTAGTCCCTTTGTGAATCTTCTTTTTGATCAAAAAAATAATTAATCCATGACAGTCTAT

AAAATCACTTCTCTGTACGAACTAATATCAAGCAATATACCCACATGATCAAGGGAATCACAACTCAGGTGCATATGATC

CAATTATCCAACCTATGTTGTTCCCATCCAAGTCTAATAATCACTTAACCCGCCACTCGGTACAAGCTTTTAAAGCAAAA

CAGAACGTAATTGGGTGAAAGAGGTAGAAATATGGGTTTCAAGATGAAATAGGAAGATAGCAATATGTTGAGTATAATCA

AAATCACCATTTTATGGTTTTATCAAGATGTAGCGAAGAGAGAGACAAAAAAAAAAGATAGAGACCAAAGAGAAAGAACA

ATATTAATGGAAAGATATATATTTTTAGTTTCCGATGTATATTTTTTCTTTAAACAAATTATTAATTGGATTTTGAGAAA

ACAGATAATCGTGGAAAAGATTCCGCAAATAAGGAGGGGAGGAACGTGCACCCCGGTTAAAGCCGGGGTGCACAAAGTCG

GACTCATTGATTATTCTTCAAAAATATTAATTATGTCAAATATCTTACATAAATTTAATTAGATAATTAGCTTTTTTTGT

TGTTGTTGATGTTGATTTAAGAATTTGATGATTGTCTTATTTTTTGGGCTAAATATAATAATTATATACTAATCTTGTGG

GTTATGTTAAATGAATATTAAGGTGTGTTTGGTTAGCTTAAGCACTTATTTTAAATGGGCTATGGTTAATAAATATATGG

GCTATATTTAAGAGAAGACTTCTTTTTTACCAATTTGCTGTTAAAAGCTTAAATTTACAGGTTAACGGTTAATTAGTTAT

AATTATCACATCCCGAATTTAATATTTCCATAACAAGATTTTTAATCACCGGGTATGACGATTTTCCAGAATTTCTTATT

CTACAAAGCGTTTATATGGATGTTTCGAGGATCATTAATTGATAAGAAGAATTAATTGGGGGATATGAATTACTTCCCCC

AACCCATGGTGTGTGCTAAGGGGTGACTTTTGGTTATGAATATACTAAAATACCCTTCCCTCAAAATAAAAAATAAAAAC

TTAAAATCAAAAAACAAAATCATATCCCCCATTTCCATCTCCACTTCTTCCCTCAAAATAAAAATTAAAAACTTAAAATC

AAAAATCTAAATCATATCCCTCATTTCCATCTCCACTTCTTATTAATCTTATTTAGGGGTAGTGTTTTGAAACCCAAATA

TTCTCCATTCCCTTTCAAATTCTAAACTCTCCCCACTCTATATCAAATTCTCTTCACCTCACTTTGGAAACGAATTTTTT

TTTTATTCGGAAGTGATGAAGATCATGCTGAAAGTGATGATGACCATGCCGAAATTGATGAAGACCATGCGGTAAAGATC

GGGTTTTCGGCATATGGTCTTTATCAATTCCGGCATGGTCTTCATCACTTCCATTTTGAATTTTTCCTGGCTTTAAAAAG

CTTGGTACCGGCATGGTCAGTTGATATGAAGACCATGCCGGTATTTTTTCCCAAGATAGTTTTCATATTTATTTCTTATG

CGTAAAAAACAATACCGGCATGGTCTTCATCAACTCGACCATGCCGGCACCAAGCTTTTTAAAGCCAGGAAAATTCAAAA

TGGAAGTGAGGAATACCATGCCGGAATTGATGAAGACCATATTCCGAACACGGATGACGGCATGCCCGATAATTTCCTAT

CAATGCCTGGTAGTCAAGTGAAAAAAATCAACATTCGGATACTTTTCATTACATTCCGGCATAGGTATTTAAGCAAAATA

CTATGCCGGAACAGGTACCGGCACGCTCGAAAAAACTATTACCATGCCGGTACCTGTTCAAAATCAGAAAAAAAATCAAC

ATTCGGAAGATCAAAACCAGGAAAAAAATCGTACTACGCCCCCCCCGAACCCCCCGGATAGTGGCAAACAATTTATGAAC

ATTTCCAACAATTGCCGGCATGAGTTTTTGGTTGAATACCATGCCGGCACCAAGCTTTTTCAGCTGCAACAATGGTGGAT

TCAAATAAAAAAATCAATTTTTCAACCTCTTATTATCAGTAATTCTCTCATCACAATCATCTTCCCATTTTCAAAAAAAA

AGAAATTCCTCTCAAATATTTTTTTCCCTCCTTCTACTCTCACCCAAACACAAATAAAAATACACACTAATTTATCCCCC

AAATACTTAACATTTTACTAATTATACTTAATCACTAATCCTAATAAAAAAAAAATTACATAGGGGGTCACCCAAAATTG

CTATTGGGACACCTATTTTGTCCCGTAGCTATATCCCCCAATTAATTCCCATATCCCCAATTAATTCTTCATTGATAATA

TTCAGCATTTTAAAATGACAAGCAGAAGAGTTTGATGTTAAAAAAAGCTTAATAAAAAATACAAGTAATCAATTGTGAGA

GGTGAAGTTGTAAGTTTTATTTATCCTTATTGTTTTTTCAAATAAAACTCTTGAATGGTCAATGTGCAAAACACACGTTT

GCGAAAACTTTGGGTCAAAAGTCACACGGGTTATTCAGTCGAGATAAGGATGAGAACCAGTTGAAACTGACTGGAAAACG

TTCTCAACTGCTTGGTGGTTGGAAGTTCATGAAGATGGTTGTATCCCCATTTTTCGCTCCACTTCCCGGTGATGGGCAGG

GATTTTGGAGATAATATATAAAGAGGGGAGTCCCTCTTCTTCTCAAAAACTACAACATTTTTAGTATCCCATTTTACTGC

TAACATTGAGAGAGAAAATAAGACTTCATTCTCGCACATTGGCTGGTTCCTCATAATTCAATTCGTAATCATAAGCTCTG

TTTCTAATTCTCTTATTTACTTTGAATCATGATTCTAGTTTTGGGTAAGATCATGTACTTGTATGCATTGGTTGTAATGA

ATGCATAATATATGAAATTCCTACATTGGTATCATGAGCATGGTTGTTCCCTTAATTGAGAATCTGATTTTGAATTAAGA

AATATAATGATTTGTTATTGGAGTATGTTGCCTAGTATGACTGACCTTTTACTCTCAGATTAAATTTACATAAAACCGAT

CATATGCATTCAAGGTTAATGATAAAATGTCCAAGAGATTTTCTTTTCTGAATAGCATATTTAAGCTTGTAAACATTGAA

ATTATACTTGGTTTTATCATCAGATACATACGTAGTTGAATTATATGTTGTTTGATTTGGTTTGTAACACCATAAATTGA

TATGTATGTACCATTTTACATGATGGATTGTTTCTGATGAAATAATAAAAAGGTTTATCCTATATTGGGTTAGGGAGAAC

CATTAAAGTGATCATTCAAGATTCAATAATTGGTGTAAGTTCTTAGTCTTGCCCAAAGGTGTCTGACTTGCAAAAATTAT

TCTTTCATGAATTTAGTTTATAAATATTTGGATCATAAGTGTGTACTTGTAAACAGATGAATTACTAGTACTGCCCAAAG

GTGTTTTAGTAAGTTTAAATGTGAACTAGGTAGAACATAATAGCTTGTAATATTTGACAAGAAGTGGATAACCCTTACCC

AAAGGAAAGGAAGTGTCCACTTCAAAATTGAATGCTAACAAGTTAGTTATGTAGGCGATTCTTATCGCAAATAGGGAGGT

GAATTGCATGTTGTTTGTACTTCCCAAAGGTGTACTTAGATGATGTGTCGCTGTCTCTCAAAGTGGAAGGCTTGTAAAGC

ATAATTAATACACTTTGAGTATTTCAGCAAAAACCCAATAAGTCTGACTTACATTTTTTTCCAGTAGAATTATTGTTGTT

TTTTCTTGCTGTGAATAGAAATTTAACCTATTATTATAATGGTTAAATTTTAATAGTTATGGATGGAACTATCATCCAAC

GACATCTGACTGAAATTGAACCACTTAGTGGTTCAAATTTCAAGAAATGGAAGTCACAACTGGATATTGTCTTGGGATGT

ATGGAATATGACATTGCTCTTACTGAAGAGATGCATGTGATTCCTAGAACAAATGCACCTGAACCAGTTAGGACAAAGTA

TAACAAATGGGTTAAGGCAAATAGGATGGCACTGATGATTATTAAGTCTTCAATAAGTGATGCTATTCGTGGTGGCATTC

CAACTAAGGATACTGCAAAAGAACTAATGAAGGAAATATCAAATCAGTTTACTGGGTCCTCTAAGGCACTTCAATTTACC

TATTTAACTCAGCTCATTTCTCTCAAGTATGATGGTCATGGAAATGTTCAAGTTGTCTAATCTTATCATTTCTCTTAAGG

AGTTGGAAATGGATTTAGGTAATGATTTTCTTGTGCATTTGGCTATTGCTTCTCTTCCTAGTTCCTTTGAGACCTTCAAA

GTGAACTACAATGCTCAAGAACATAAGTGGACTATGAGTCAGTTAATTCCTTATTGTGTAATGAAGGAAGAACAACAAAA

GCAAGGAAAGTTTGAATCTACAAACCTGACTTTTCATGGAGCTAATAAGAAAAGTTTCAAGAAGAAATGTTTCAAGAAAG

ACTTTAAAAAAAAAAATGGATCTGATCACGCAGTTACTGATAAGAAGAAAGGATCAGACCAAGACTCTACCTCTGCTGGA

AAGTCTAAGTAAGATAATCCAAAATCATTTGAGGAAATTGAATGTTTTTTCTTTCGTAAGAAAGGACATATGAAGAAAAG

TTGCTTTAAGTACAAGGAATGGTTAAAGAAGAAAGGAGGTAGTAACTTAGCTTTTGTTTGTTTTGAAACAAATTTAATAG

ATACTCCTAGTAATACATGGTGGATTGACTCTGGGTCCTCTGTGCATATCTCAAATTCTTTGCAGGGGTTCATAAGGAAG

CGTGAACCCATGCCTGTGGAGAAGTTGTTTCTTGCAAATTATCAACCAGCTAAAGTCGAGGTTGTTGGAACCTTTAGACT

GGTTTTAGATGGTGGATTTATCTTGGATTTAGAAGATACTTTCTTTGTTCCTAGTATTAGGCGTAATTTGGTTTCTGTTT

CTAGACTGGATTCTGTTGGTTATGGTTTTATGCTTGAAGGAGGTAGACTTTTGCTTAGATTTAATATGCAGAATGTTGGA

TTTGGTGTTTTAATTGATGAATTACATAAACTGTCTTTAGATACTTGTTATGCTTCATCGGTATTATATATGTCAATTGA

TTCTCTAGCTACTTCGTCTTCTGGTTTTAAAATATCTCTTCTGAAAGAAAACTCTTTTATGTTGTGGCACAAGCGACTAG

GTCATATATCTAAAGAACGGTTAAGAAAACTAGTCGTAGATGGAATACTAGAAAACTTGGATTTCAGTGATTTTAACACT

TGTGTAGATTGTATCAAAGGGAAGTTAACAAAAAGAAGAAAATTAACTTCCACTCGCAGTAGTGATCTTTTGGAACTGAT

TCACACAGACATATGTGGACCTTTTCCTACGCCTTTTCTTGGCGGTCAGCGTTATTTTGTATCCTTTATTGATGATTTTT

CAAGGTATGCCTATGTGTACCTTATTCATCATAAGTCCGAGGATTTAGATATGTTTAAAATTTTTCATCTTGAAGTTGAA

AATCAGTTAAATAATAGGATAAAAGTTGTTCGTTCTTACCGTGGAGGAGAATACTATGGTAGGGCAGATCAAGATGGTGT

TAATTTTGGGAAATTTCTTTAATATCTTCAAGAACATGGTATAGTTTCACAATATACAACTCTTGGTACTCCTGAGCAGA

ATGGGGTGGCTGAAAGACGCAATAGAACTTATCAAAACATGGTCCGTAGCATGATTTGTCATTCTGATTTACCTGAAAAC

TTATGGGGTGAAGCTTTGAAGACTGCAGTTTACCTACTAAATCGGGTTCCCAGTAAAGTTGTACCGAAAACCCCTTTCGA

ACTTTGGAAAGGAAGGAAACCAAGTTTACATCATCTTCATGTATGGGGTTGTCCCGCGGAGGCGAGATTCTATGATCCTG

ACATTAAAAAGCTGGATCCAAGAACTGTAACATGTCGTTTCATAGGATACCCGGAAAGATCTAAAGGTTTTCATTTCTAC

TGTCCGGCGAGGTCTATGCAAATTGTAGAAACAAATAATGCTAAGTTTATGGAGTTTGGTGATGGTCATTTAGACCATCG

AGAAATGGTGTTTGAGGAGGTTATAGATGATATATTTGTAATTCCAAGTGCATCAACTGAGTTTCCATCCACAGAAATCT

TTTCACCCATAGATTCTATAATTGATGAACATGTCCATTCCCAAACTGTTGAAGTTGTTCAACCTGTAGCTCCTCATACT

GTTGAGAGTCTCACACATATTCCTGAAGAAGTATTTCCCATTGAGCCTGTAGTACCAAGGAGATCCACTAGGGCAAGGAA

ACCTGCAATTTCTGACGTTTACCAAACCTATATTCTTGAAGGTGATGTTCAGGAAACTGATGATCCTTCTAGTTATTTGG

AAGCAGTAAGGCGAAACAATGGCAGAAATCTATGAAGGAAGAGATGGAGTCGATGGACAAAAATTTTGTTTTTGAGAAAA

AACCGCTGCCTGAAGGTGCAAATGTTGTTGGATGCAAATGGGTCTTCAAGACCAAAAGAGATTCTGAAGGGAAGATAGAA

AGGCACAAGGCTCGGCTTGTTGCGAAAGGATATAATCAGAAGTTTGGAATTGATTATTCCGACACCTTTTCTCCAGTTTC

TAAGAAAGATTCTCTCAGAATTGTGTTGTCTCTAGTAGCGCATTTTAAATTGGAACTCCATTAAATGGATGTCAAAACTG

CATTCTTAAATGGAGATCTCCAAGAGGATATATATATGGAGCAACCGGAAGGTTTCAAAGTAAAGGGAGAAGAACATTTA

GTTTGGAAACTAAAGAAGTCGATTTATGGATTGAAGCAAGCATCTAGGCAGTGGTATTTGAAATTCAATAAAGTGGTTGT

AAGATTTGGCTTCAAGGAAAGTCTTATGGACAGATGTATATATCACAAAGTCAGTGGGAGTCGTATCATATTTCTAGTAT

TGTATGTATATATTGTTAGCCAGCAATGATATTCATCTCTTACTTGATACCAAGCGTTTTTTGTCTGGAAATTTTGATAT

GAAGGACATGGGAGATGCTTCATATGTTTTGGGAATTAAAATTAAGAGAGACCGAAAAGCAGGAGTTCTTGGTTAATCTC

AAAGTGCCTACATTGATAAAGTTCTCAAAATATTTAACATGCAAAACTGTGCACCTGGAGATACACCCGTTGTGAAAGGA

GATGTCTTTAGCAAAGATCAGTGTCCAAAAAATGAGATTGAGAAGAGGCAGGTGAACTCCACTGAGTATGCATCAGCTGT

AGGAAGTTTAATGTACTTGATGATTTGTACCAGACCAGATATTGCGTACATTGTTGGAATGCTTGGAAGATTTCAACAAA

ATCCTGGTATTGCTCATTGGAGAGCTGTTAAGAAAGTTTTAAGGTACCTACAGAAAATTAAAGATTACATGCTTGTGTAT

AGAAGTTCCGATTCTTTGGAAGTCATTGGATAATCTGATTTAGATTTTGCTGGTTGTAGAGATAGTTTTAAATCTACTTG

AGGCTACATTTTCATGCTTGCTAGTGGTGCTATATCATGGAGAAGCGGCAAGCAATCGTTAGTGACATCCTCAACCATGG

AGGAGGAATACGTTGCTTGTTTCGAAGCTACACAACATGCTATTTGGTTGAAGAATTTCATTACTGAATTGTCGGTTGTG

GATTCTATAAATCACCCAATCAAGATATTTTGTGACAATCAAGCTGTTGTCAAATATGTGAGGAATGATACCACCTCTAG

TGGGAACAAGTACATTATGATCAAGTATATGAGTGTTAGAGAAAAGTACCGGCTGGGACTTATTATAGTTTCTCATCTCT

AAACTTATTTCATGATTGCTGACCCCTTTAACTAAAGGACTTCCTCCTAGGGTGTTCAAGGAACATGTGTCCCATATGGG

AATTCTAGAGTCTTTGGATACAGTTTTGTAGTGGGAGTTGTACCTTCATATTTTGCTTATGAATTTTAGTTTTGTTTGCT

GGATGGTATTTTTCGAATCTGTTAATGTAACTTTGTGGTGGAAGTTTTTTTATCTTCCTTGTTATGTTTCAGAATAAATT

TTAAAGAAGTAATTTTGTTACTATTTATTACAGACCTAAAGTAGCCTGATAAAGGTTTGCAAAAGTTTTAGTTTTTGCTT

GGGCTAAATATGAATAGGCATCATGTTTGTTAAGCCTTTATCTATGATAATTGGTATGTAACGGATTGTTACGCGCAGAA

GCAAATAATATATCTAGATAACATGCACGTTTACTGCATGAAGCATGGACAACTAAGAATAACACATGATTTAGCAACAC

CAAAGTTGAGATCATGGAATGAACTGTACATCGTAGTGAATGCACTGTGTTTTCATGATCCAATTTTAGTGGTGTTATTT

CACCTAGGTGAAAGTTATGGATATGTCTCAGTAAAGTACCCAGTGTGACTGTTTTCAAACATAAAGTTCTTAAAAGATTG

ATCCAATACTGGTTGTTTTCATAAAATTCATCTTAATGTTTTGTGTTAACCAAGTGGGAGAATGTAAGTTTTATTTAGCC

TTATTGTTTGTTCAAATAAAACTGACCCAAAACAAGGACTAGAATCGAAAATAGGTACAGTCCGAAAGCTCAAATTCTAG

TACTTCCACTTCCGGATCTTCAAGTTGGATGATCCTCTAACGTCGCGGTGTTCTACTCTAGATTCATTTATGGACTCTGG

TTTGGTGGTGAAATAGGTAGAAAGTAGATAGGGTGTTGGTTTCTGCTGCCTAATTCTTCTTTTTGATGCTGTATGCCAGC

GGAACCACTCCTCTGTTAAACACCTCGACGTACATTGCACCACCAGCAACAATCAACTGCAAACAAATGCCAATTCATGC

TTGTTCAAAAATAGAACTAAAAAGCACAGCTTCAATATGAACAAGCAGGGGGACAAAGAGTAGAATTTTAAGTCTTTTGA

TAGTGCAACAAAGTAGTGTACTATGCAATACTTCCAATGTGTTTGCGTGACTGCTCATAGGTTGAAGCTACATTCTTAAA

CTAGAAGCGATTTTCGTTTCAGATTTCGCAATACGCAGGACAAATGGTACAATGAAGCTCCAAGTACTAAGGAAAAAAGA

ATGCAACACCAAAAAGTTATCTAAGATCTTCGGAGGAAACTTTCGAAACTGGTTAAGCTTAAAGAACAAATTCCTTCATA

AAGCTAAGAAATCGACACAACACTTCTTAGAGAGCAATACATCTACTACTTCTTACACTCAATTGCACAATCGCATATAG

CGGCGTAGCTAGTTTTGTAACTGCCCCAAACTATTTTCCCATTGCAAGCACATTGAGAGCTCCCACCAATTTGACTTGTT

ACAAGAAGGAAAAGACACACATCTCCACCAAACTGTTCTAATTGTTCTTGTCAATGCAAATCTCTAAGCTTTCAACAAAC

ACCCAAACTGCATATTCATGGTGGAATCTTACAAAACAAATTTGAGAAAAACTGAAAAAAAAATGAGGGTTTCATCTAAT

TTACCTCTTCATACTTCTGTGTAACAGTAAGACATTCATCTTCAGTCATATCAGGTCATAAAACAGCCATATTTTCGTAC

CGTCTCTAACCAGGTGGACATTGAGGTTCTTCTTTCTCTACAGTGGCTCCTAAAATACCACCAGAAGAAGTAGAAGGAGA

AGAAGTGGAATCTAATGAATATAAACGTCCCATCAAAGACAAATCCCGCTCCAATTGAAGAAACCCAAGACTTTTACCTC

TATCAAACTTAACAACCTATAACATGTGTTATCAAACAGAGAGACGAGATCCTTAAATACAAGGTGGTTATGTGAAGTGA

ATCCTAAATGTAGAATCTCATGCCTCTGCTACCATACACTAGTGGAAACATGGTTCAAAAATGCGCTTTAAACTTTCACA

TCCGCGTGTAGCAGATATGAATCAGACATTTTGTCAACCAAAGATGGGCCTTGGAAGAGACATATATACCAAGTGAGATT

GCAGTTTCCGTTCATGGACTACCTAGAAAGAAAAATAAAGAAAAAATTTCAGTGACTACGATATGTGTATGTGTACAGTT

TTAATATAGCTGGTATTGGAAGAATATGAAGAAATTTGTTAGTATAAAGGAAGTAAACAGGAGATGAGAAACTCACAGGG

GTTTGAATTTACATCTAGCCCATCAATGGGGTGCTAATTAAAGACAGCAATACGAACCTAGAAAATGCAAACCACTCATC

AGATCAGAGTATAGTAGATAACTATCAAGTTATGCATCCAGATAAGTATATGAAACAGTTATAGCAAAATAATCATTTAC

CAACTCAGGAAAACACAAGTCAATAAACTCAAATAATCGAGCAAGCCTAAAGGTTTTGTAAAAGCAGAATGCACCCAACA

GAGCTAAGAATTCACACAACATTGTTTGAAGTTCGTATCCTCTTAGAGAGCAATATATATACTAGTTCTTACAGTCAATT

CAACTGTTGCCTACAACAGCTTCTGAGCTTATGACGTTGGTTAGCTACTTTCGAAACAGGCTGTTTATGTATAAACAGGT

GTGGAAGTTGTATTCAGTATTCCTAAACTAGATATCCTTCTCCTTCAAGATCTGGCGAAACATAGAACAGACGCTGCAAG

TACTAATCAAAAATATGCAACACCGAAAATGTTACTTTTGCTCTTTTGGAGAAAAATTACAAGACGCAAAAGATTAAAAA

ACCTTTGCAGAATCCCAAAGCTAAGGAAAATCAATCTCATTTACCACTTCCAAATTCACAGAATTTGCAGCATCTGAACA

GAAAACATGCTCAAAAAACTAATTTGGGGGGAAATTGCAAACACAAATTTTATCATTTTCAGAGACAAACTTAAATCAAC

AAAGACCAATTGATTATCAGGCCAATTGCTACTCAATCTTAAACTAGATATCCTTCTTCATCCATATATATCTAAGAACA

TTCAACAAACACCTGAATTGCATTTTCTTGGTGGAACCTTACAAACAAATTCAGGAGCAACTAAAAAACATAAGGGTTTA

ATTAAATTTACCTCTTTACTAAGAAGATATGATATCAGGTCTTAAAACAGACATAGTTTCATATTGTCACAAACCAGGTG

TACATTGAGGTTCTTCTTTCTGTTCTTCAATAGCACCTAAAATACCACCAAATGACGGTGTTTCAAAATCATCTTCTACT

CTTCTCCAACACCTCCTTCAAAGAATAAACCTGAAAAATCTAAAGTTAATGCTTTAACTGATAAACTTTCATGGGTTTTT

TCTCAGATTCTCCATTTTGTAATTACAACATTTGTTTCGAACGAAAACTTGAGGTGTGAATGAAACAAGGGATGAAGAAG

GAGTGGAGAGAAGTTTGGGTGATTAGAAGGAAGTGAAAGCTAATGAAATTGATGATGTTGCCATTGTTGGGAAGAGGAGG

AGGAGGAATGAGTCAAAAGGGTTTTCACTGGTCACTTCTTTTCAAGCGAGAAAGTGAAGAAGCAGAAGACCAGAAAGTCG

TACCCATGGGTGGATGATGCCATTAAACCCAACTTTTGTTTTTGCTTACACCTCACTTACACCGGAAAGAGAAAGTCAAT

TTGTCGAAAATATTTACTCACGGTAAAAGTACATTTTCTGAGATCGTGATCAGTTAGTTGCATGGTTGACACAATTACCA

CCTTCGAGGACGGAAGCACAGGTAGGCCTGTCAATGGAAGAATATCCATCGGATATTTAGAAATCCGATATCCGATAGGT

TAAGCACTACCGGATATCTGATATCCGATAATATCCTATAGGATATCAAAATCAGATATCGATTCCGATAGGCTAAGCTG

TCGGATATCCGATAAATATCCGGTTATACCAAAAAAGCTTAAAAACCCTTGTAAAACATTTTCATAATTGACACTTATCG

GATATTTATCCGACAATATCCGATACTATACTCGAAATTAGGATAAATTAAAAATAAGTTCCTATAATATCGGATATTAA

CATTTCGGATCCGTTTTCGATATGTTAAGGAAAAATATCCGATAAAATATCTGATTCGATATCCGATATTCGATAAAACG

GATAAAATCCTATAGGATCTCGAATATTCGGAAACGAATTCCGGATATCGGACAGATATTGACAGGCCTAAGCACAGGTT

GACTAAACCTGGCTGTAGCCTCCCTTTTTTCTGGAAAATTTCACCAAGTTAGAATTTCACCGTTATCCAAAATAACAAAT

TTAGTTTAGTCCACCAAGTTCATCGATTTAGTCCACCAAGCATAACAGTTAGTCCACCGAGAAGAATAAATAAATTAAAA

AAAATGGAAAGTTAAATGCTTAGTGGAAATTCAATACAATTATTGTTCACAAGATCAATGAAGAATGAAATTTTCTTCCA

TGCTTCTTATTATACCTAGGTATTATTTCGTTTCATGATACTCAATTAGGTTTTGCCGATTTGAAAGGAGTTAATGATGT

GAATTTTTTTTCCCAATGGTTTTGATTCTTCTTTAGGTTCCGTTTAGCTTTTTTTATCAATGAATTTAGGGGGTTTTGTT

AAAATTGTTTCTAAATAGCTGAGGTTATCGCAGTTATGTCTACTAGTTGTCAAAACCCTAAACGAGGAATAACTATTATT

GGAAACATATTTCATACCTATACAATTCCTGAAATTCTAGTTTCTAAACATATTTCTAACTTCTTATTACTAGCATGTAG

AACTCTACGTCCTCATAGGTCTTTGGATGCTTAGGACACTGGAACGCTTTGGAGTTGGAGTAGGTTTTGTACGTATACCT

CTCGTAAGCCCTCAAGAGACTATAACTCGTCCACTAGGGACACCTAAGGGTTTAAAGGCTTGTTGCACATGCTAAGTGCA

CCCGGGTCCTCAACAAAAGGGAATTGTTGAATTTTAGGTTAGATAGTTTGCTCGAGGACTAGCAAAAGCCAAGTGTGGAG

GAATATGATAAGTGCATAATTCACATGATTTTAGTGTCAATTCCATGCTAGTAATTGTTTAAATTCTTGCAAATTATAGT

GTATTACGTTTATTCTCCCTTATTTGTGTTTTATTAGGAGAATCATCCAAAAGAAGTGAATTTGCACTTAATTGTGTAAT

AAAAGAAGCTAGCTACAAATGGAGAAGGAGCCAAAGACCAAGAGAAACTCGTTCAAACCCAGGAGTTCTTGTTATCACGT

TGAAGAGGACGAAAATACAAAGTTAACGGCGAAAGAACATAGTCATTCTGAGTTCGTATAAAGAAGTTAGTCCGTGCCAC

ATATGGCTTATGTTTTTTTTTCAAAATATACTTGTTAAAATTAGATAAACATCAAATCAACCATAGATTTGCAGTATATG

CTTGCATATTTGCTGATCCAAAATAGTCTGTGCCACATATGGCTTATGTTTTTTTTCTTGAGAAAATTTTGTAATTGTGT

TTAGAAAAAATTACTTACGACTGTTCATATAATAGGCAACAAAACTTTATATCATAGATTTATTGGTTTTGAAATCCTCT

TCTTCATACAGATACGAATTTTGGATTTAATCTTGTCAGATTGCATTCTTAATCTATGAATTTGTAAATATCATGTTCAT

GCTAAAAATATCCCTTATTATTTTGTTAAGAATTAAACTGGTGAAAAAAGATTCGAGTTCTTCCTTCTTTAAAACAACAT

AATGGGTATAAATTACCATTATTGAACTCAACTCATTTCCATTATGGAATATGTATAAGCAAATCGAAAAAAATATTTAA

AAGGGCAGATAAATGTAAAAGTAAAAAGTCTAAGACAAGGAGTAATTATTGCTTTAACGTAAGGTATAAATGGTTACACA

CAGAGGGTATAAAGTAAGAAATAAAAACCAACCACCATAAAACTGACACGTGTCCTTTATCAGTATGGTTTTGTAGTGTT

TTGAATCTTATCTCTTTGACTTGTGGTCATTGAGAAAAAAGGAGGAAGAGGAGAAAAATTGAAAGAGAAATAAGTAGCGT

GTGAAATATTGAAATTTTGCCTGTCTTTAAGAATGAAGGTAAAACAATTTACCTTCTTCATTTATTGTTGTCATGCCTTA

AAAAATTTCATTGATGCCTTTAATGTCACTATTGTTGTGGGTTTAATTCATGTTTCATAGATAAATTTCCTAAAATTTTG

TTGTTGTTCCTTAAAAGATTAGTTAAAAGAAGAGTGTAAAATAAAATTACTCAGATATGACGATTTTGTTGACAACAAAC

TCTCTCACACCTTTGTTTTTTTTCAGAGAGATAATGGTAGCAGTTTTCAACTTCTTAAAGAGAGTGAGGTGGTTAAAAGT

ATTGCTGACAAACTTCCTATTCAATCTGTGATTGATTTATCCTGTATGTGTTGGCGGTTAAACTGGTCTCTCCAAAAATA

TGTACGAGATAAAAAAAGCAATTACACGAGGTAACTTTATTATTTTTCTTCGGTAAATTTCTGTGTGTTTCAATTTGGTT

GCTATCAGATAGACAAATTTCCTTGTTTCATAGATTGATTTGGAGTTTTGAAATCATCTTCTTCATACAGAGGTGATTTT

TGGATATAATTTTGTATCAGCACATCCTAAATCAAAGATAATGAACTCCTCTTCTTCATTCATCTTTTGATGAATATATT

GTAAACTAAGGATGAATGACTTTGTAAATTCCCTCCTCTTTCCTCTTATGCTAAAACCAGTTGAACATTTTTTTTTAAAG

CTGACATAAAAATCATTTTCTTAATTGGGTAATGTTAGGAACTAAATTGTTAAAAATTGAGTTGAGTATTCTTACCACTA

ATTGAAATCCACTTCTCTTTTACTTTTAGCAAGACTATTATACTAGCTTTTTACAAATTTTTGAAGACTATAAGGAGGAT

AAAATCATTACCGAGAAGCTTCCGACTTAATTCATCATTCATTTATCATGCGTGAGTTTGCATTTGAAGTATTCTCTCGG

AAACTGTCTACAACACATAAAACGATTACATGAGGTAATCTTTATTATTTTTCTGTCATAAATTTCAATGTGTTTCAATT

TATGTTTTTATCAAACAACTTTCCTTGTTTTAGGAGAAAGTACTTTTTTATCTATGAAAAAAATATAACATTCCCATATT

GTACAGGTGAGGTTACATAGTGGAGGAGGAACGACGTGAATAATATATACAAAAGAGAAAGAGGAATACTAACTGGGAGG

CAGAGATGAATAAGGAGAAAGAGATACACTTTTTAACAACATTTTCATGTATACCGAAGTGGCTAGACACATTATGGACA

AGATTTATGTTCAAGTTGTTGCAAATCTACAAGTTGTATCTAAGAAAATGAAGCAAAGGCTTACTATTCGAAAATATCTA

GAGGTAATTTTTTTCCTGGAGTTTTTCATTATTGTTGGTAATTATGAGGTATTTTCTTTCTCTGTGAATTGAATATAGTT

GGTGGTTTTCTCCCTTTACTCTATGAAAGTAGATACATTTGGTATCTTCCTTGACTAGCATATACAGTACTTATTCTTGG

TAATTACGAGGGACACAATGTTTATTTCCTTAAAAAGTATATATTATTTGAAATTATAAATATTCATTATAAAATTGGAA

GAGATATACATATATTTGGTATTTTTTTCACTTCAATACCATACTTATTCTTGGTAATTAAGAATGACGAAATTTATTAT

TTTTTCCTGCATTTTACAGGATGAGAGGAGGAAACTTAAAATCCTTGAACTAAGTAATCACATACTCAGTCATATTTCTT

CTAACGCTCTTCTCACATTCAAATGTATAGATTACTATATTTCATCTAACGCTCTTCTCACATTCAAATGTGTTTCTAGA

ACTTGGAATTCTGTTATTTCTGATTCTTCATTCTTAGAGTTACATCTTTGTCAAGGAAAATCACAACCTTAAAATATCAT

TGTTGTTTCTGGTTATAGTTTAACCAATGTCTTTTCACCAAAATTTTGCTTCGAAAGTGGTGGTGAAGCTCTTCTTAAAG

ATTTAATTAGATGGGATGAAGGCGATATGCTAGAGCCTTTGAATGGCTTGTTCTGCTTTCTTGATACAACGAATTACAAA

ACTCGTATATATAACATGGGAACAGGACGAATCATTCCATCTATTTTCACTTCTCTTCCACTAAAATCCAGTGGTACTAT

AATCTTAAAAATATCTAATAAGCTTGCTTCCTCTCTCTTTTTCTTTATACCTCTTTGGAAATAAAATCAGTAAACCAATT

AAATACTTTTTAGTAGTTATGTGTTTTTAATACGTTTATGTCAGGATATATGTACATCCAATTATGAGAAACCAATGTTT

ATTTAAAATGGTCCAAAGTATGTTATAAAATTCTATTTCTTTCGTTATATGTGTAAGAACTAGGTTTAGTTTGATATAAT

TCATATAATCATCAATATTGGTTCTTTATTATAATGTAAATTTATTCTTTTACGTAGGTAGATTCATTAAAATGGACGTA

AAAGCACTACATATTCCAATATAAAGTTTGTTTGTTATGCCTTTGTTTTTTTTAGGAATATCCGATACTTATGTTAATTC

ATCCATTTTTGAGCTACGCATGGTGTTGTTGGTAGCTCCTTTCTTCACGATCCATGAAATTTATTATGAGTGAGTCTTAT

GTTCTAGATGTATGCTTTGTCCTTAAGTTTTTAACGATCATATTTTAGGAAAATAAAAAACTGAAAAGCATCATAATTAA

AGATGACACATGTTGCGGTCTTATCCCTTGGATTTCATATCTAAGGAGAGAGATGAATTAAATAACTAGATCCGAAGAAG

TGAGAAACAAGGGAATAATGGCTGCAAACGCTTTATAAATGCTAGAGAGAGAGAGATATAAGAAAAAGAAGAAAAAGAAC

AAGAAAAAGTAAAACTTTGTAGAAGTAAGCAACAACAAACAGAGGTGAAAAAAAAATGTAAAAGATTTTATTTCCTTATT

TATAGTAACAACAATTACTACCCATAATCAATTTTGTATATATATTTCAGGTTTTCATTCCCTTAAATAATCTTTTGGGT

GATGCATGCCTTAAATTTATCTCATTCTATTTAAGAAAAAAAAAAAAAACAAACGATTTATTCCTTGTTTTTAGAGAGGG

ATTTTTTTTTGTATAATATTTTAGTGCTGGTAGTACAATTAGGTTTAATTGGTTTACTAACTTTCTTTCGGTGAAATCTG

ATGCAGTTTCTTTTTTCTTTTTAAAATGGGGGTATAACAACCATACCCAATAACAACCTGTGATAGAATTGAAATATCAC

AGGTTGCGCATAGTTCAATCAATTGGTAAATCCAACCTTGATTGTTCGATAGGAATTGATTGACACTTGGGCATTGGTCT

TTAGTATCGCCCAAGTTATTTCTCATAACAACCGGGCTCACAGGTTTTTCTCTATTCGATTAGAGACTGTATTGATAAAT

TGAGATATAACTCTTTGATGTCTTTCTTGGTTGAGCTCGCTTTATAAGTTGGTGCTCTCAGAATTATATTGGAGTTATTC

CATACAGATTGCCGAACGAATTATTGGTTATGGTTGTTATACCCCCATTTTTTCAGAAAAAATAAACTGATCCATAAATG

GTTTCACCTGTTTCTTTTGATTCATACGGTTTAAAAACGAATGGGTTTCACCGTTTCACCTATCCAAAGCTCAAGTTCAA

CAAGAAAAATGATTAAATAAGCATATTATTTTTCGAAAAAAAAGTGGTGATAACTAAACTTACTTTCTGTACTGAGGAAT

AACTAGCAAGAATCATCTCCTGTCAAAACATAGGCAAGATATCAGCATAAAAAATGGCTCTTCTTAAACATAGCCCACGC

TTTTATTATTGATAGCCCACATGACCTAGTCAATGAAACTGACCAACAAATCTAATTTGACCATTTATCGTTATTATTTA

TAGCCCACGATATTTAACTGGGATAGATGGAAGGCTGATCCAACACGTTCTTCTTCCCTTGAACCATATAAGTATTTCCA

GATTTTTCTATCTCCCTATTTATTATGATTAACACTACATGTTTCTTTTTTATTCCTTTTCAAAAAATCCAATAAAGTCG

TTGGCTTAGTTGTTCTCTTTACAAAGGATTCATGCAAACCAATAAAAGAGATTTCATGAATTTCTATCTTTTCATTCAAC

AGATTGAGAATCAGGAATGGTGTTATATCACCATTTCCAACACTTCTTAATTCTTTTTCACAAGATGTTTATGTAAACGA

TAGTCGATGGGGATGGTTTATAAACAGAATCATGGACAGGTTGCAAAAATCTCGCTAATAACATAGTTAACATTGCCACA

AACACATTTAAATCTTCGAGAAAATGGATTAGTATGAAAAAACCTCATCAAATGGAGGAAACCCATTAAACTTCTCAACG

AATTGAAAGGGACAGACAGAGTCACGGACTGCAAATCAATGGATGCAAAAATGGGAAAAACTCTATTAGATGAGGACATG

GTAGATTGAGTTAAGTTTCTGATAACAAAATCCAAACTCTCGCAGCTAGCTAGATCAATTAAACTCCTTTAGTTCCATAA

TTTACTTTTACAAAAAAAGAAAAAGAAAAGTATTGCTATGGTTTGATATCAATCAAAATAGGATAATAATAACAACAGAA

AAATAATGATGAAGAACGAGTTGGATCGGCCTTCGCTTAACTTCGTGGGCTATGGATAATAACCTAAAAAGGTCAAATTT

GATTTATAGGTCAACTTTGTTGACTTAGTTTCATGGGTTATCAATAATAAAAGTGTGGCCTATATTTAAGAAAAGCCAAA

AAAATTTATGATTGAAGTAAACTTTTTCTTACATACACTGCAACCCAAATCAACACTTCAGCTTCTTCATAACAGAATTA

GAAAAAATGAACCCTAAATATCCATCCATACTTCAATCACGAGAAAACCCCTCATAGATTATTCAATTTATAAATCAATT

GAGAAACCCTAAATCAACTTATCATTCTGAGTTAATTTGAGAAACCCTAAATCAACTTACCGTTTATGCTTAGTTGAATC

AATACTTGATTTAACATCACTACTATGAATCACCGGAGTTCTCAACATATCCATCATCTTTTCATGATTTCTTCACCTTA

ATAATAAACCCTAACCCTAAATCTCTCCCTGATTCATACACAGAACCTAAACTCCTACTTGTTCTCCTCCGTACATCAAC

AATTATAGCATCACTCTCATCATCTATACAAATTCATTCTCTATTTTCTTATAATCAACTTTCACTTCCCAGAATTGCCA

TAGAGAAGAACAAAAAGAGAGAAGAAGATAAATGATTGAATGGTCGGGAGAGGAGAGGAAATACCGAAGAATAAGGAGGG

GGAGGAAACATCCTCTAGTTCATTTTTTTGAGGGTGGAAAAATCTGTCATGCATGTGAATGGCACATGCATAAATAACAG

GTGTTACGTCCCTAGTGTGACCAAGCGTGTGTGTAGTTGCAGGAATTCCCACACTACACCCTATTCAAGAATTTATCAAT

ATTTATGACACAATTTGAAGAATAAGGATCTTTTTTATTAATATGATCTTAAAAGAATTATAGAAATCAAGAACACAAAA

CACTTAATAAGATTTTCTCTCTCCTAAAAACTTTCTTAGCACTCAAAAATTCATCTTCTTTCCTATGGAACTATTCATTT

TCTTTTTATAGGTATTTACATAGTGGATTACAGCTCATGACAGCTCATTTACAGCTAAGATTCCCTTTAATTTCGGATCT

AGGTTCTTGCGATAAATCGCAGTTAGCACCGCCTTATTCATTCGCAAGTCTTATTGGTTCGTCGCAGAAATGTCACACTT

TACTCATTTATCCGTCGTTTTCATCTGATATGTCATTCCTTACGTCACTCTGATACATGTCGCAGAATATCGTTGATGTG

TCGCAGAACTGTTGTTTATTTCACAAGAATATTGTTGTGACAATATTCTGATCCTACATCTTGCCTCTTCTCATATCTTT

TCTGAGAAGGAGATAAAGATCATGAGAATTTACTATTCTTGATTTGCCGCAACTCCTTACTTTTTACCCGACAAAGTTCC

TTATTTTACGTTTTACCGTTGGCACGTGTCAATCTTTACTTTTTTACCTATCAACTCGGTTACCCATCCTTCATCTATAA

ATAGATGTCCCACTCGAGGAAAGATTTTCTTTTATTCTCTCTTTTTTCTTCAACTTCTTTCTTTTTCGTCTGCAATTTTT

TCTTTCCTAGTATCCTTCCTGCTGCTGCTCTAATGGAGACTCAAAGGTATGTTCTTATCTTGAACCTTATTTTGTTCATG

TTTCATCATTTCTATCTCCTGATTTTCTTTGTAATCCCCATATTCTATATTATCCTTTTCAGTAACACTACTTCTCAGAT

TAAGATTACTGTCATAAACCCTAACCCTAAAGTTTCTCAAAAAAGGCGTGTTGGGAAGGCGAAAAGATCTGAGAATGATA

AGGTAGTAAGAAAAACTTATTTCTTTGAAATAACAATATTGTTGCCTCTGTTTCTTATCTACATTATTATTCGCAGGGTG

AAGAAAATATTTCTTGACCTCTTAAGCGTTCTCGTCAAATTAAAGCTCTTCCAGATGAAGTGGATATTCTTGATTTCTTG

CCTGCAGACCATGTTCCGCCGCAAGATAAATCTTCTCCTTCTCCTCGCGTAAATTCTAATATTAACACCACTACTACTTC

TTTATCAATTCTAAATGAACCTATTGATGCTGATATTTCCATCAATATTTCAAGAGTTGCAATGGATGAATCTTTAATAG

CTAAAACTAAAGTGAATGAATCAGCAGCCAATAATCTTGCGATTAATGAAACTGTGGTGAATGAAACCATGGTAAAAACT

GCGGCGAATGAAGCCGTGATGAATGAAGTTGCGATGAAGGATGTTACGGTGAATGAAGCCGTGATGAATGAAGTTGCGAT

GAAGGATGCTGCGGTGAATGAGACTGTAACTGCGACTAACCAAGTTCCTTCTTCCACCCCTACTACCGCGAGTAATTCTA

ATCTTCAAGTTGTTGTCACCAAAGCTTCTTCCTCTACTTCTCTTCCTTCTTCTCAAGAAATAGCAACAGGGGTTGAAGCT

GCTTTTCAAATATTATCTGAAGTTTTGAATGATGCTATGAAATCTTCTACAGACCCTGCGAGATAAGATATTTGTATGAT

TCTAAGAAAGCATCTTGGATCCTGCGGTTCTTCGTCTAAGCTAACTTCTTTAGAAAGTGAAATTGAAACCTTGAAGAAGA

ATCTTCAAGCAAAAACTTTAGAATGTGAACTTCTCGATCATAAGTTTCGCAATCAATTAGATGAAAATCTTCGCAGAGAA

ATGAAATGATGACCTGAAAGGTACAACTTTCTTGCCTCTGTTTTTGCCTCAATAATTCTTTGTTTTCTATATACTCAATT

TTTCTTTATTTATTTTTTGACGCATATTTAGCTTCAAAACATAACCAATTATTGAAGATACTTGAATGGGAAACAATCGC

TGATGAGGCCCGCATTGGGAAAGAGTCTTGGAAGAATCAATTTGATAGATTGAATACCATTCACACATATTCAATTGACG

AGATAAATAATCTTAAGAATGAGAATAATTAGCTAATTTGTGAACGAGATCGCCTAAGTAATGAAGCCACTCAACTTTTA

TATTCGCTAAATCAGACTAATTTGAGGGTGGAATCTTTATTAGATGAAAACAAAACCTTATCTCAATAGAAACGTAGCTT

TGTGAGTAAAACAATGGTTTATTCGCAGTAATTGAGTAATCTTCAAAAGCTGTGTGACAACCAAAAGCAATCTCTTCGTG

TATTGAAAGATGAGCTTAAAAAGATAACCGAATCATCTATTGCCAACAAGAGAAACTTATAAAAGGTAGATTAGCTCGAG

TGTAAAAGTCAACAGTCTCAAAGATGACTATGCTCTCTTTGAAGAACTCCCATTCCTCCTTAAAAGAAGAAAATACTCAT

CTTCTTTCTAAAGAAAAAAGTATAAAATCTCGTTTGAAAGGTAAAGTTGGTGATACTTTCGATGATACCATGAATCAATG

GAAGAATTGCTTCCTTAACATAATTCAAAATTTATCCTCGCAAAAGGAAGGTAGTCATTTTGACTGCCTCGTCTATTTAA

TTATTTCATTGTATTTTCCCATCTTATTTATTGCGAAAATTCCATCTAATATGTGATTTTTCTTCTGCAGAATCTGAGAA

GCATCTTCAAGCTTCAATCTCTGATCTGTCTATGAGGTTGGCCAAACTTCGCAAAAAGAACACTGCGCTTTCTAACTCTC

GCAACAGAGGTTTAAGAAGATTTGACTATCTTGAATATTTCAAAGATATAGAGATCAGAAACCGGGAAAAGAATTGCCTC

CGTAAGGCGCATTTAGGCGTTAAGGTTCTCATAAACAATGTTCTATTCTCTCATTCTTTACCTCCTATGAAGATCAAACC

CTTGGAGATAGATAATAACGAAGTTCCTCGTCTCATAGATGAAGATCTAAATTATTCAAGTAATGAAGAAGATTTTTCTG

AGGATGAACAAAATCCACCTAAAGAAATAGAAGAAGAAGAAAATCCTCTCAATGATGGAGAATATCAAGAATATCCTCAT

AAAGAATCAGAAGAACATAATTCTCCTCACAAAACAATCAATGGTGATGGAGAAACCCAAACTGAAAAAGGTGTCGCTGA

TGAAGATAATAGTAGCGAACAAGAATGATAAATTTAAGGAAGAATATTCTCTGGAGTTATATATTCTTGCGAAATATCTT

GTTCTTTTCCTTATCCTGCGAATGACTGTAATTTCTGAATCTCTTTTGTGAAACATTGCAATTTTTGTAACCCCATCATC

TTCTTCTAATTATATATTTTTATTAATCTGAATCCAATCTTCATAAATTCTTGTAACTTAATTATCTTCTTTTTTATTAT

TTTAATTCTTCTCAAGGAAGATAATGAAATATTGTTTTACAATCCCCATATTTGCCCCTCATAATCTTCCTTGTGAAGTA

TTGAATTTGTTAGCCTTGTAAGAAATCCCATACTTGCCTCTTCTTATCAATCTGCAATATGATAAATGAATTATCGCAAT

CAATGCTCCGGAAAATTTGTTATATGCTTTGCAGATATAACCTCTCTTGTGGTCTTATTTCGCCTTCCTAGTTAAAGGTC

TTATTATGCCACTTTTATGCGACAAGATCGCAGAGATTCTTACTTTTACATTGATCTTGTAATAATCCATTGATCTTGCC

TTATCATTTAATTTTAATCTCTGCGACAATATGATAAATCTAAGCCTTCTTGTGAGATTAATCTCTCTTTGCTTTTTAAG

ACTTATGACTTTCTCGTCTTTTCGATGAAATCGCAGGACGTCTTTGAGATTTATATCCTCTTATTATTGATTTCATTTAT

CTCGTCTTTTGATCTTATCGCAAAAACCCATTGATCTCATATTAACATTTCTCTTTAGGCTTTTTGGCCTGTCACGCGGC

AATATGACGGGCCTAAATATTCTTGCGAATATAAAGTCCCATGACATTGTTTATGCGTGTCACGCGACAATATGACGGGC

CTAAATATTCTTGCGATTATAAAGTCCCATGACATTACCGTCTTTTTTGATACACACTCCCTAATGGAGGGTATCGCCCT

ATCTCTCCCCGGTTGCCCCTTCAAGGATTATACATCGGCCATACAAGACTAGCACTTTCCCATCCGGTTCTTCCAACAAC

CAGTTGTTTTTGCAGCCTTGTCCTTTATGCCTTATATGGCTTATAGTTACAAGACTGCACCCTAAGTGGGGTTTTCTTTG

GGCCGAACACATCGTAAGCCAAGACTTGCCGTGGATGGCCAGGGGTAGGTTCATCCATCCCGGAACCCACAAACTCAACC

GTGTACCTCGGCGCCTTGATCATACTTTGCATTCCTTAGGAGAAACTCTCGGGCGCCCAACCGGGTATAACCTTATATAG

TTGGGAATCCAAGTTATCTCTCCCGGATGGATTTTCTATAATCCCAAATCTCCATGACTAGGTTCCAGCAGCGGTATAGC

CTTTCCCATAGCCATGCAAAGAGATACCCCCCTTATGACGCACTTTAAGGACATTGCCTCTTACGGAATAGAATCCGCGA

ACTAAGGTGTACGCCATAAAATTTCGTCCCATTCTTATATGTTTTTAGGCAATTTTCGCCCATCTTATGCATTTGCATTA

AAATTCTTAATTATTTTGCAAATTATTGCGGAAAATAGATTTTATATTTAGCCACCTTTGTTCTTGCAAAACATCTCTTT

TATTCTCTCTCAATTTTTTGAGTGTACTCCATTCATTCTAGTCTTTCAAACCTTTATTGATAATATTTTCTTAGATATTG

TCTATTCCATGGTCTGCCAAGTCTTTGACCCATGTATCTTCCTTCTGGATCCATTAATTTATATGCTCCAGTTCCAACTA

TTTCTTTGATGATATATGGCCCGTCCCATTTTTTCGCTAATTTTCCTCCATTTTCTCGTTGATATTCTGGTGTTTCTCGC

AGAACTAGATATCCAACTTGGAATTCACGCTTCTTTACTCGTTTATTATATTCACGAGCTAATCTGCGATGGTAGTTTTC

CATATGTTGTAATGCCTTTTCTCTCTTATCTTCTACCTCATCAAGTTTGTCTAGAATCATACCAGCACTAAGATTCTTTT

CCCAAGCTTCTCTTTTTGTTGTAGGAATAGCAACTTCTGTTGGTATCTTTGCCTCAACCCCATATGTTAAACAAAAAGGA

GATTCCCAAATTGCTTCTCTTCGTGTTGTATTATAAGCCCAAACTGCATTATGAACTTGTTGACACCACGTTCTATGATT

CCCTTCTAACTTCTTTTTTAGAATATCTGCGATTGTTTTGTTAGTTGCTTCAGCTTGACCATTGGCCTGTGGGTATAAAG

GAGTCGATTTTCCGCACTTTATCTTGAATGCATTAAGTAACATCTCTATATTTTGTCCTTCAAATTGTTTTCCATTATCT

GACACCAACTGCGCTGGAATTCCGAATATGCAAATGATATTTTCAAATATAAATGTGAAAATGTCCTTATCACGAATATG

CTGTACTGCTTTAACTTCCGCCCATTTTGTGAAGTAATCTGTTGCGACAATTAAGTATCTCTTTTGCCCTGTTCCCGGCA

TAAATGGTCCCATAATATCTAAACCCCACTTTCCAAATGGCTAAACATTTTTTGAGGATGTTAATGTTTCACCTGGTGCA

TGTATTCTTTTTGAAAATCTCTGACATTCCTCGCAACTTCTTGATAATTCTTTGCATCTTCATGCATATATGGCCAATAA

TAACCTTGCATTTTAGTTCTGTATGCTAAAGATCTTCCGCCGCTATGATTACCAACATTCCCACTATGTAACATTTTTAA

TATTTGCTTTCCTTCTTCTTGTGTCAAACATCTGAGGGAAGGTCCATTTAAAGATTTTCGATATAACACCCCATCTCTTA

ACTCATAATTTGTTGCTCTACTTTTTAACTTATGTGCTTCTAATATGTTCCTTGGAATCTCCCCCCTTGCTAGATATGCA

TGAATTTACGTACGCCAGTCTGCGATAATTCGACTTGATTCTTCGCCATTATTTTCTACTACCATTGCCTCAGCTTCCTC

TTCTTCTTCTTTATTAATTGATGGTGCTATGAGTGTTTGTATTTTTATATTTCTTGAAGTCGGATCTACCATCATACTTG

ATATAAACGCAAAAGCATCTGCAAGCCTGTTATCTCTTCTTGAAATATGCCTCCATTTGAGTTTTGGAATTTTCACAGAT

AATTCTTCAACAAGTTTCTTATATTTTTTCAAAGATATATCGTTAGTTGCATATTCCCCTTCAATTTGGCGGATAATCAA

CTGTGAATCACTGGTAACTCTTGCATTCTCCACCTTCATTTCTATCGCAAATCTTAGAGCATGTACTATAGCCTCATATT

CTATCTCATTATTGGTAGATTCAAATTCCATTCGAAATGCAAATGCCAGTTTTGCGCCTTCTGGCGAAATAAAAACGACT

CCCACTCCATTTCCTTCTCCATTGGATGATCCATCTACTAATATCTCCCACCTATTTGGTTATGTTAAAATATCTCTAGG

ATCCCCATATTCTTCTTCTATATCCATCATGTCTTCTACAGCTTCATCATCCTCCAAAGGAAATTCTGCGAGAAATTCTG

CGACAATTTGAGATTTTGGCGAAGATAGAATTGCATATTGGATCTCAAAATGCCCCACTTGTGCGTTCCATCTTTCGACT

CTGCCTGATCTCTTTGAATTTTTCATTACTGACTCAATTGGTACTTTTGTTAGTACTTTGATTTTATTCTCTTGGAAATA

TATGTGAAGCTTAGAAGACGCATACACTAATGCCAAAATTAGCTTTTCAATCTTCGAATAATTTTTCTCCGCAGCATTAA

ATGTTTTGCTGATGTAGTAAATAGGTTTTTCAACTCCTGCGTCATTTCTCAAAAGTACTGCGCTTAAAGCATGTGGAGTT

GTCGCAAGATATATCAGCAATTCTTCTCCTGGTTCAGCCTTCTGTAAAATGGATATATTCATCAAATATTCCTTAATTCC

TTGAAAATCTTTCTCACATTCATCTGTCCATTTAAATTTTGTTCCCTTCTTTAGTATGTTGAAGAAATGCTTGCATTTAT

CTGATGATCGCATGAAATAAATCTTCCTAGTGAGGCTAAAAGTCCATTCAATTTTTGCACATCTTTTATTGTTCTTTGGA

GTTGGCATATCTCGAATCGCTTGCACCTTTTCAGGATCAACTTGTATTCCTTCTTTTGATACAATATATCCCAAGAATTT

CCCAGATGATACTCCGAAAGTACATTTCTCAGGATTTACTTTGATATTAAAAAGCCGCATCTGATCAAAAATTTCCCTTA

AGTCCTGCACATGATTTTTAGCTTCTTTGCTTTTTACCAACATATCATCCACATATACCTCTAAAGTTTTATGTATCCAT

TTCGCGAACACCTTTTCCACCATCCTTTGATAAGTTGCTCCTGCATTTCGTAACCCAAACGGCATCTTTGTGTAGCAATA

TAGTCCTTTGGGAGCAAAGAATGCAGTATGTTCTTGATCTTCTTCCGCAAGTGGGATTTGATTATAGCCCTTGTATCCAT

CTAACGAAGATATTCGATCATTTCTGCTGCATTCCACCAATTGGAATATCTGGCAAGGTAAAACTATCTTTAGGACATGC

TTTATTCAGATCAGTAAAATCAATGCAAATTCTTATCCCCTTATTTTTCTTGGGAACTACTACCATGTTGGCGATCCATT

CTGGATATTTTGCTTGCCTTATAATTCATGCGTCTAGCATCTTCTGCAATTCTTCTTCTATCTTGGCATGATAAGTAGTC

TCAATCTTTATTATTCTTTGTTTGAATGGTATGAATTTTTTGTTAATCTCCAATATATGGCACGCAACAGATGGATCTAT

TCTAGGAATCTCGTCCATATTCCACTCAAAAATGTCTTTGTATTCTCGCAGAATATTTACAGTTTGCTTTTCTTCCTCCA

CATCCATCTTAGTTCCAATTTTCAGCATTTTTGGGTCTTCTTCACTGCCAACATTTACTTCTTTTGTGGGTTCTGCTGCA

ATATAATTGGCTTCATGTTCTCCCATTGGTGTTGGTGTTTTTATTACCTCCGCTGGTTCATCACTTTCTTGCGATACATC

GTTAAATATTCCTTTTCCTTCTTCTGATCTTATCATGTAAATCTTGAATTCTTCTTCTTTCTTTTCTTCTTTCCTCATTT

TTCTACGCTGTTTTCGCAACTGAATTTGATTTTCACAATTCTTTACTTCATTCTGGAAACACAACGCTGCACAATCTGTA

TCTCCTCTAATCTCGCTAATTCCGTTAGGCATAGGAAATTTTATGCATTGATGTAGAGTTGATGATACAGCCTTGATTGC

ATGTATCCATGGTCTTCCCAACAACATATTATATGGTGATTTCACGTCCACTACACACAAAGTTATTTGCGTCTATATTT

CCCCCAGCGGAATTCGCACTATAATTTCACCTTTAGGTTTTGTTAGAGATTTTCCAAATCCATGAATATGATATGTTGGA

CTAAGCATTTCTTCATCTTTGTATCCCATTTTCTTGAATGTGTGGTAAAACATTATATCCACATAACTTCATGCGTCCAC

GTAGTCCTATTCACAAAAATTCACTCTTCTGTAGATCTTATTGATTTCTCTTTTCTTGTTGGTACAACAGACATTGTGAT

CACTAATGGAGATGTATGACTTAAATTTTCTTTCGGAATTTCTGCTGCCGTGAATGTGATCTTCTGTTTTTCCCAATCTT

TTAGCGGTGATACCTCCTCTTCCGACATGATTTTCTTTCCTTCAAAATATCGCTTGTGTATCCTTCCTCTAATATCTTCA

TGGAACTTATCTATCATCATCTTTGTTATTACATCACACTCCACTTTATTTTGATTATCTTTGACTTTAACCATCTTTTC

TTCCATCAATTTTCTTGATATTCTTCTAACTCTTGAATAATCTCTTTCTTTTACCTAACTTTTACTATGATTTTCAACTA

TTCATCCTCGTTTGGGCGCCAAAATGTAGTTGCAGGAATTCCCACACTACACCCTATTCAAGAATTTATCAATATTTATG

ACACAATTTGAAGATTAAGGATCTTTTTTATTAATATGATCTTAAAAGAATTATAGAAATCAAGAACATAGAACACTTAA

TAAGATTTTCTCTCTCCTAAAAACTTTCTTAGCACTCAAAAATTCATCTTCTTTCCTATGGAACTATTCATTTTCTTTTT

ATAGGTATTTACATAGTGGATTACAAATTTCATTTTACACTAAGATTCCCTTTAATTTCGGATCTAGGTTCTTGCGATAA

ATCGATTAGCACCGCCTTATTCATTCGCAAGTCTTATTGGTTCGTCGCAGAAATGTCACACTTTACTCATTTATCCGTCG

TTTTCATCTGATATGTCATTCCTTACGTCACTCTGATACATGTCGCAGAATATCGTTGATGTGTCGCAGAACTGTTGTTT

ATTTCACAAGAATATTGTTGTGACAATATTCTGATCCTACATCTTGCCTCTTCTAATATCTTTTCTGAGAAGGAGATAAA

GATCATGAGAATTTACTATTCTTGATTTGCCGCAACTCCTTACTTTTTACCCGACAAAGTTCCTTATTTTACGTTTTACC

GTTGGCACGTGTCAATCTTTACTTTTTTACCAATCAACCCGGTTACCCATCCTTCATCTATAAATAGATGTCCCACTCGA

GGAAAGATTTTCTTTTATTCTCTCTTTTTTCTTCAACTTCTTTCTTTTTCGTCTGCAATTTTTTCTTTCCTAGTATCCTT

CCTTCTTGCTTGCTCTAATGGAGACTCAAAGGTATGTTCTTATCTTGAACCTTATTTTGTTCATGTTTCATCATTTCTAT

CTCCTGATTTTCTTTGTAATCCCCATATTCTATATTATCCTTTTCAGTAACAATACTTCTCAGATTAAGATTACTGTCAT

AAACCCTAACCCTAAAGTTTCTCAAAAAAGGCGTGTTGGGAAGGCGAAAAGATCTGAGAATGATAAGGTAGTAAGAAAAA

CTTATTTCTTTGAAATAACAATATTGTTGCCTCTGTTTCTTATCTACATTATTATTCGGAGGGTGAAGAAAATATTTCTA

GACCTCTTAAGCGTTCTCGTCAAATTAAAGCTCTTCCAGATGAAGTGGATAGGTATTTACATAGTGGATTACAGCTCATG

ACAGCTCATTTACAGCTAAGATTCCCTTTAATTTCGGATCTAGGTTCTTGCGATAAATCGCAGTTATCACCGCCTTATTC

ATTCACAAGTCTTATTGGTTCGTCGCAGAAATGTCACACTTTACTCATGTTTCTGTCGTTTTCATCTGATATGTCATTCC

TTTCGTCACTCTGAGACATGTCGCAGAATATCATTGATGTGTCGCAGAACTGTTGTTTATTTCGCAAGAATATTGTTGTG

ACAATATTCTGATCCTACAGTGTGAACGGCACTCCTAAAAAAAATTGTAACGTCTTAGCTCTGTTATGAATTTTTAAAAA

ATTAGAAACGGCTTCAGCCTGTTACGAATATTTTGTAACAGGAATTTTGTAATGGCCACAAAGCCGTTACGAATCCGGGA

ATTTGGTGTAGTGAGAGGTACAAAACTCGTTGAAAAGCTTAACCTTAGAAGCTTGAGCATTTCTTTGGGTCATCTATGCA

ACTTTAGCAGCATCTATCTCCATCTTTCTCAAATCCTCGTTCAAATACTGGATATCTTCCTTGAGCTTGGAAACCTCATT

GATAAGACCGTCTTTCTCACGAGTTACGTGAATAATTTGGTTCTTGAAGTGTTGCTCAGAATCTAAGAAGACAAAACAAA

TGAGATCATAAGCTTGATTAATAAAGCATAGTATAAACGAAAAATTACCTACAACATCTAAAACTACCTTCATTTTGACA

GATAGTAGAGCCCAAAGTCTTTTGAATATTAGCGTTAAAATGGTCAAGCCTAGAAATTTCAAGGTTCGCAGCTTCAAGTT

GAATACAGAGCTCGTGATAAGAGTTGTCCCATCCTTTACCATGACCATATTCTGAACCATCTTCGCCAGAACGAATGACA

ATATGTCCCATATTCTTAAAAAGCTCTATTGGTTTTGTCTTTGAAGGAGGGTCAACATGTTTTTTACATTGAAGTTTCTC

CAAGGCGAGATAGAAGAAAGGATACTTCGTCCTTCAGATGACCGTTCTCAATTTTCAGACCGTTGGCTTCGTTATGAGAT

TTGCGTAGATCAGCTCCAGACTTATCTTTTCTCTTGATTATACGAGTTTTATCTTCTTCTAACTATTTAATTAAAGTTAA

AGATGAAGAATGAAGATTCCTTGAAGTCGTGAGGAGATTCCCAACCCGTATTAACTCGATTTTCAAGTCCTCCTTCTCTT

TATGGAGATGAAAATTTAAACCTTCATTATTCGCACATTCAACAGTCAAATAATCCACTTGCGAGAAAAGGGTCTCTACT

TTCTCATTCAAAATTTCATTCTCAGTCACATAGCTATCAAGGTGAGTTACAAAATTAAAATTCTCTGCACGAGTATCTTC

AAGCAATTCTTTCGCCTCGTTAGCATCCATTAGACATTTTGTACATTGAGCTTGACGAAAAGTAAAATAAGAATATAACA

AAAGCAAGCCAAAGACAAGAATCAAGCACATGAAAGAAAGAAACTTACTGCGAAGTTCACCAGCTTTCTTCCATAGCTTG

TCTGCAAATTTATTTTCATCATAAATTTCCTTTTCCATATCTTTAATACAAGCAACAACCTATCTCCAATATTTCATGGG

ACATCCTAGTCTCGACGAGGTTATCCATATGATGATTGAATTCCTGCGAAGAATCAAGTCGAAGTAAGAGAAAGATTCGT

GAAATATTGCAAATTAAGAAACTAAGACTCACCAATCCCATTAAGGAAACATGTTGTTGAAGAGACAAAGCCAGTGAAGT

GTTTTCCAAGAAAGATTGATCTTCAACAAGGAAGGTATCAGAACAAAAAGAATCCAAGGATTCGAGGGCAAGGGATCTAA

GAGACGGATTCTGCCGAGAGGAAATGTAGATATTCTTCATAAAATCAAGGTCAGGAACATAAACAGGATCGTCCACTTCA

TACATCTTCTTCTTGCCTTTGTATGCTTCCTTCTTGTCCTTGGGAAGTTTATCAGGAGAATCTTGCACTTCAATGACTGA

AGGAGTAGGAAGAGGAAGATGGGAATTCAAATCCTTTGAAGGCTTGGAAGGGATAAGTTCTTGGCAAGAACCAGTTGGTT

GATGCTTAGGAAGAGCAGTATATTGACTCGAAGCAAGCTTAGAATCCTTTACTCGGGTCAGTGCAGCCAGATTCAAGACC

CTAAGATTACGCTTCGTGTGAACATCAAAAATTCAAGCATTGGTGGCATCATTCTGCAAGATGCGAAGTTAGAAAATAAT

TCGAAGAAAAAACCCGCGAAATACAGAAAACGAAAATACCTGATCAGTGATGGGAGATACTTCAGAACGATCACGTTTTT

TATGACCTCTAGTTCCAAAATTCGATTACATCGTCGTCGCAATTGAGGAGTCCAAAGACGTCGAAACCATGACCGTGGAT

GAGCTCATGGGATCGCTACAAGCCCATGAAGAAAGAATGAAAAGAGAAGCAAAGAGGGAGCGAGTAGAACAAGTTCTACA

AGCAAAGCTCACTTTGAAAGACAAAGGAGAGAGTTATGGTAGTAATGCTAGAGGAAGAGAGACGGCTACTATGGAGGAAG

AGGAAGAGGCAAGCGAGAATGACGATGATCAAAGGGGCCAGAGGTCAAGTTCCACAAGAGGTCGTGGCATGAGGAGATAA

TTCATGGCGCAACAATAATAGGCCAAGGAATAACAAATCTCAAGTTGAATGTTATAATTGTCATAAGCGTGGTCATTATG

CTCGAGATTGTTGGTTTTCATCGGATAACGTCGAAGAAAAAGCTAATTATGCTAATAAAGAAAAGAAGGAGACCAGACTT

TGTTGGTGGCATGCAAAGGAGGTGACCACAGTGAGAATTGTACATGGGTGCTTGATACATGAGCAAGCAATCATATGTGC

AGCACTAAGGAGAAGTTCGTGGAGCTTGATGAATCAGTTTCCGACAACGTAACATTTGGAAATGGAGCGAAAACTCCGGT

GAGTGGTCTTGGCAAAATTTTAATTAAGTTGAAAAATGGAAATCATGAGTTTATTTCCGATGTTTATTACGTCCCTAACG

TAAAAATGAATATATTAAGTTTGGGACAACTCTTAGAAAAAGGCTATAGGATGAATATGGAAAACCTTAGTCTTTTCATA

AGAGATAGGAGAGGAAAATTGATTGCCAAGGTGCCAATGACGGGGAATAGGATGTTCTATCTCGAACATTCAAAATGACA

TCGCGAGATGTATGACGCTTGTATTAAAGACTCGTCATGGTTATGGCATCTTAGGTATGGACACCTTAATTTTGGAGGGT

TAAAATTATTGTCCAAACAAAAGATGGTTAGAGGTTTGCCAAGAATTGACCACCGGATCGGTTGTGTGAAGGATGTTTAC

TTGGAAAACATTTGAGAAAAAGTTTTCCCAAGGAAACATTCTCAAGAGCCACAAAAACTGCGAGTTGGTACACACGGATG

TTTGTGGACCAATAAAGCCAAGCTCGTATGGTGGAAATAACTATTTCCTCATATTTGTCGATGAATTTAGTCGAAAAACA

TGGGTTTATTTCTTGAAACAAAAATCGGAGGTTTTCTCCAATTTCAAGAAATTTAAATCCCTTGTGGAGAAAGAAAGTGG

CTTTCAAATTGTGGCTATGAGATCGAGATAGAGGAGGAGAATTTACTTGAAGGAGTTCGAGGCATCTTGTGAAGACAATG

GAATCCGTCGGCCTCTTACGGTTCCATACACACCTCAACAAAACGGAGTTGCCGAAAGGAGAAAACAGAACAATTCTCAA

CATGGCCCGGAGTATGCGAAGACAAAGAGAGTGCCCAGGGAGTTTTGGGCGAAGCGGTTGATTGTGCGGTTTACGATAAA

CCGAAGAAGTAGAGTTTTTAAAGAGTTAGTCTTGCTTCCTAGGAGCTCTAGCAACACAACTCAACCTAGAAGCAACGTGA

GAACCAGCAGATTCATCATCCTGCGAGATTCGAATCAGGAAATGGTTTCATGACCAAGACAGCGATGTACAAGAGACTAA

AAAATACTTGTTCCGATGCAGGAGACTCAGTAGTAGGATCCCGATTTTTACCATCCTTAAGTAAACCAGCAGGAGGAGTA

GGGTACTGAAACAAGGAGGATACAGTTAAAAATGCCACTGATGAGAAGTAAACGAATTAGAAAAAGAGCAAGACCCTCCC

CTAATGGCATCTCAGACCATTGAAACCTCCCAGGCTCATAAGCTGCGAAGTGACTATGCTTATGAAGAGGACCAGAATGA

ACACTGCCATCAATATTCAAACCCCAGATCAAGGGGCCCTGCACGACCAAGGGAAACATCATCCAATCTTCATCATTAGA

CCAACGAAGATGATCGTTCTTTGTACGACCAAACGGGTCCACATCTTGAAGGAGAGCTTTGGAAGAGTCTTCAAGAGTAT

TTCGAGTAAGCTTGACACCCCACTCTTGATACTTCCTCGTGGATATTGATTTACCAGAGTAGTTCACAACAACAAAATCA

ATATTATAATCCTTCGTGAATCTCGGAATTTGAGAGGAGCCGCCTTTTGAACGAATCAAAAACTCGTTAAATATACGAAT

GGCATTACCGCTGAGCTGAAATACACCTCGTTGGATCTTGGTGAGAGCCTCGTAAAAGAAAGGATTCCCCGGATCATATA

AAGGGCAGGGTAGACCATCCCTTAACTGGCCAGCAATGATGATGACCTTGTTGGAATTCCATATATCCGACTGAATCCAT

CAATAGTTTAACTCCATGTTCTTGGCAGACGGGATATGAGGAGTGACAGATGAATCGACCGCTGGGGAGAGAGATAGACC

TATATTTCTAAATTCGACTTGTAAATTCATCGAAATTCTTAGATGATGTTAGAGACGGGGTCTTCTTTGGAGCCATGGAT

CTATAAATTAAAGGATTCTGTAGATTTAGAATAAGGATAAAGTTGCAGTTGAAGAGCCGGTAAAGTCTGAAAGAAAGGTA

ATACCAACAGTAGCAGCAGCGGAGGAAAAGAAGAAGAAGATTGAAGGAAAATAAGAGAAGTGAAGAGAGGCTGAGATATT

TATTGTGAGGTCGAAGGTAACCGGATAAGGCAGGTCATAGAGACGGTGGATAAAGTGGAACCGGTAATAGGCGGTTATGA

ATGAACATGGTGACGTGGATGTGAGAATGACGGAGATGTGGCGGTTACAAATATTTCCATAAGAATCTCTTTCCATTCTT

AGATCCTGCGAATGGAACGAGAAAAGACAAAATGTAGGTACATAAAACCAATTCTCCACGTGGTCGGACGAAGTGGCCCG

AAATGAACGAGAAAATCTCAAACAAAAGATCTTTCATTAACCCTGAGGATGAATGTCAGTGACAATTCAAATCAAGTGTC

AGCGTCACTTTTATGAAAATTGGATCTGTGAAGTATGAATTCTTCCCCGAAATGAGAGATCTGCGAGATTCACAACAAAA

GAGATAGCCAATAGGCGAGATAAAGCGACTGACTGAAGAGATAGTTAACAAAGAAGAAGTTATGAAGCAAAGTTAATAAT

GGCTCCAAACGAAGATGAATTGTCGAACAAGAAATTTGGACAGATGTACAGACAAGTTGCCGAAGATATCGGTGAATGAA

TTGGGAGAACTCTGTTGAAAATGGCTTGGGCGAAGTATAAAGTAGAATCACGTGAAAAATGAGCTATACTCCACTATTTA

GTTTGCCTATAAATAGAGACCGTAGATCATTGTAAAGGGGGTTGGATAATTTGTATTGATAGAGGTTATGTTTAGAGAAG

GAAAGTTTTCTTGTGAGTCTTCATACTTGTAATATCATCATCATAAAATGATCAATAAAAGAGAACTTTGATTTTATTCT

TCTAAACATGTCTTAGAACTATATAAATTACTACTGGATTTTCAGTAGTTACACTACTGACACGAAACGACATGCCAAAA

AGTCCAAAGAAAATTGCTTTTGATTAAATTTGCAGAGAACACTCCAATTGAGAGAGAATATATACCTCACTGGAACCTTT

TGTACGATGGTTTCCGGTTCCTTTTTTGACCAGAAATCCTGGATATTGAAGGATTCAATATAAGGAAGGAACCCACGTAT

GTGCATGCATGACGGGTCATAGGTGATTCTCTAATTAGCCTAATAAGTTTCTCAAACATCATTAACAAAATGAGAGCTTT

GTATTGCGAATTATCTAAGAAAATCTAACTAAACATAACTCAAACTACAAGGCTCCACCCCAAGATTCCCAACAGTTTGT

GAGGTGTCTCCTTCCATCTTTGAACCAGTACTGGTGTTGCCTGCAGGTTGATGGAGGTTCTTTTCTGCATTTTCAATAGG

TAGAAGACTTTCATGATAAATAAACACGCCGTACCAGCCACTATATGCAGGGGTCAACAACCGTGATATACCAGTGATTT

CAATCTCCCGGAAAGTCTTCATACTCCGATTGTAAATATACAGACTTACAATTCCTCTGGGCTGCCATTCAGTATACGAG

ATATGTGATTTTATTACTATTTCGTCTGTTCCTTGGACTGGATTAAGATCAAGACTGAATCCTCTTTGAAACGAGGGAGG

AAGCTTCATGGTTTCCTCGGTCCAATTTGTGGTCATCCTCTTCTCCATGTAACTATCATCATCAGATATCCATAACTTAA

CAACGTAGTCGTCTACTCTATTCACTACAGCCATGTGCCCATCTATCTGTAATAGAAAAGTATTTAAAGTCCGGGGAGAA

GGAGCGATGAAGTCTGGGATCTCAATGACTCTAAACTTTTCGGTTCCAACATCAAATACCGCAATCAACTCAGCATCTCG

AGGTGCGAAAAAACTAATCATACTGTCATTCCTCCAGTAAATCGAACCGTTTGCATAAACACCGGTGTCTCCAAATAATC

TAACTGGTGGCACCTCATCTATCTTCCTCCACTTATTTTCTCCTCCCCCCACTCTAAAAACTTCGCAACTATGTTCGACT

CCTTTTGGATTCTTCAAAGAAAGAGACGTGCTAGAATATGTCTCCCAAACACATGCTACTTTGTAGTTCTTAGACAAAAG

ATCAAATCCAAATCCATAACTCGGCTTGTGAACAACAGAAGCGCCAATCTTTGCTGGTATGGATGTCTTTGCCCATGGAG

TTACCTGCCCAGTGCCGAGGTTGTATATGCGAGTACCTCTAACAGACCTGCTAAATAGACAGAACAACCCATGGATGGGG

TTTAACATAACAGCACGAAACCATGGGATTTTCACCTTATGAAGAGCCGCTCCGCCTTTGAACCCATCCTCCGGTGAAGA

GACCACTACGGAATCAGTGTCTTTAAACGTGTTGAGAAGTTGAGGGTGTGCTTTTGATCGACTTAGATGCGACTCTATGA

AAGATGGATCCTCAATGAATGACTGCCAACTTTTGGACACACACTTAAAACGCATAAGAGACTTGACAGGAAGGTTGACA

AGTATATCTTCTGCTGATTTTTCATCAAGACGGTTTGAAAGTGATTTCCTTTTTGATCGGTTTCCTCCCTCCATCTTTTT

GATCGGAACCGCACCTACGCAAAAACTCCAGAGAAAGTTACTGTTCCTTTATATGTAGTTGTGAATTTATTTCTTTCCCA

ATTCCTTTTTTAACTTGAAGTCCACGATATATATAGAAGAAGTGAATTTAAGGAAATCATGCATGCATGACTGGTCATAG

ATGAGTTATTTCCTAATTAGCCTAGGTTTCTAAAAGGTTCTCCCAATCTGAGTTGCTCAGTTTCACCCATTGCACGTAAA

TAAGGTTTACTAGCATAAATATACTGTCCAGGTGAGGAAAAAAAAATGGCTGCCATATCTAGATACCCATAGATATTGAA

AAGTAGGAAATACCATTTCGTCCTAGAAATAATATATTAGAGAGGGTCTAGTCCAGTTGACCCAGTCAACTGTCTTTTTG

GTCCTATTTGGTAATATAAATTATCATTTGGTCCTAGATATTATCATTTGGTCCTACGCTGATGTCATGGATGATGTAAA

TGTCACCCTGTAGTGGCAGAAATACCCTTTTGGATAAGGGCCATATATAGTCAAAAGGTGAGAGAGAAAAATCATTTCTC

AGATCTACTTTCTCTTTCCTCCCGGTTCTCTCTTTCTTCTCCGCCGCTGCAATTTTTCTAGGGTTTTTAATTTTTTTCTG

CTGCTGCAATTGATTACAGAGATCGAGTTACGGAGGTTGATTACAAAGATTTGATTTACAATGATGTTGAAGATCAATTT

ACAGAGATTATGAAGATAAATTCGAAGATTTACAAAGAGATTATGACGGAATCTGCTGATGTTGTTGAAGAGATTATGAC

GAAATCTCTTGCTGCTGTTGAAGATTCATCTGTTGCTAGAGTTAATGAAGAAGATGATGGATATAATGAAGACGTTGTTG

TTGTTGATTTACAGAATTCATTCTATTTGATGATGTTCTTGATTCTTGATTTTTCTGCTGTTGTTTGGTTACGATCTTGT

TTTTGAATGATTTTGTCTCTGTTGATGATGATGATGAAGATGAAGATGAAGATCTGCTGCTGTTATGAGCGCTGCTGATG

ATGAAGAAGATGAAGACGAAGGAGTTCATTTTCCGGAATGAACTGGTTTGAAATGAATATTTGTGATTTTTTTCGTCTTT

TTTGTTATTGTTGATGATATAGACGATGAAGATGAGTGATGCTGATGAAGAAGATGAAGATGAGGGAGTTCATTTTCTGG

AATGAACTGTTTTTGTATGGAGTTCATTTCTCGGAATGAACTATGTTTTTTTAGGTTTTTGTATGGAGTTCATTTCACGG

AATGAACTCTGTTTTTTAGGTTTTTATATGGAGTTCATTTCACGGAATGAACTCTGTTTTTTTAGGTTTTTATACGGAGT

TCATTTCACGGAATGAACTCTGTTTTTTTAGGTTTTAATACGGAGTTCATTTCACGGAATTATTTATAGGAGTTCATTTC

TCGGAATGAACTCTGTTTTTAGGTTTTTATGTGGAGTTCATTTCTGGAATGAACTCGTTTTTAGGTTTTTATGTGGAGTT

CATTTCACGGAATGAACTCTGTTTTTTTAGGTTTTTATGTGGAGTTAATTTCACGGAATGAACTCTGTTTTTTAGGTTTT

TATATGGAGTTCATTTCACGGAATGAACTCTGTTTTTTTTAGGTTTTATATGGAGTTCATTTCACGGAATGAACTCTGTT

TTTTTTTTAGATTTTTATATGGAGTTCATTTCACGGAATGAACTCTGTTTTTTTATGTTTTTCTATGGAGTTCATTCCTG

GGAAAATTGTCGGGGGAAAGAAAATGGTATTTTTGTCCATTTAAGTATTTTTATAAATTATGGACCAAACAGTAAAGGCG

TTTGACCAAGTGACTAAATAGACATGGGCCCACCTAAAAAAGGACCAAACGATAATTTTCCCTATTGAAAAAGCTCAACT

AGCATAAAAATACTGTCAAAAAAGATGCACGTAAAACATGGAGTAATGAAGTTTTGGAGCTAAAAAAACAATATACATGG

TGTCCGATTAGTAATCTAAGGATTTCTAACTAAACATAATCATCCAATACTAGTCCATAAACTAAAAGGCGGTCAGTTCC

GTGGGTTGATCAAGTTTCTTTTCTGCATCTTGTTGAATAGGTACAAGACTTTCATGAACAATGCGTACGTCGTAACAATA

TGGATAGCTCAATAACAGTGATATGGTTTCCTCGGTCCAATCTGTTGTCTTCTTCTTCTCTTTGTTAGAACTGTCTGTTG

GAAATAAGTTACAAATTTCGTATGCAACTGCGAAATAATACTTTGATTACACTGATAATAGTAACAAACAACTCAAGAAA

CATGAGAAAAATACTTAACACTCACTTGTTATTCTTTATACTTGAGATGCAGATGATAGTATCAAAAACTTAGTAGAGGC

ACGTATGAGATACGCTTAAAGGTAATATTGATGCAGGTGATGAAGGAAGACGCTTCTCGAGGCCGCGGATATCCAAAACC

GAGAAGGCAAAGAAATAAAATTCTTTGCATGTGTGGAGAAGCGTGTGGAAAAATTCAATTTTAAACCCACACCCTCACCC

CCACCCTCACAATCAAAGGGCCATACCCTAAGGCTAAACCAATGCTATTTAAGTCTAAAACTCTAAGTGTTTTAGTCTAT

ATGGGACTAATGTTTTATTCATTCACATGAAAAAACATCTAGGGAGTAATAGGTCTTGGGATCAAAACATGGTCCATGCA

TTTTTTGATTAAAACAAAACATTGTTTTTTTCTAACAATCCCCCACATGAATAATAAAACATTTCGATAAAACACGAGAA

AACATAAGGCAAAATATAGGCTAAGGTGTTCCTAACTTAGTGAGAGGTTACTCTTTGTTGATACGTCAATTGCAAGTATA

CCGCTTAATCTACTCACGTAGTAAGCAATATCAGGTCTTGTACCATCAAATACATAAGACTACCAATAATGCGAGAATAC

TCAAGTTGATCAATACCCTTACCCTTGTTCTTTTTCAGTTTACAAGTGGGATCAAAAGGTAGATGCAGGTTTAGCATTTT

CATAATTAAATCTCTTTAATACGGTTTCAATAATGAGATTGACTAAGACTATATCCATCGACGTCTTTTCTGATTTTCAT

CCCTAAGATTACATCAGTAGGGCCTAAGTCTTTCATGTCAAAATTATCTCTAAGCATGCTTTTAATGTTATTAATAATAT

CAAGGTTATCGCCTAATATGAGCATATCATCAACATATAGGCACACAATAACATGAGAATCATCCACGGATTTAATGTAA

ACACATTTATCCATTAACCTTGAAACCATTAGTTATCATAACATGATCAAATTTTTCATGCCATTGTTTAGGTGCTTGTT

TTAAACCGTAAAGAGATTTGTTCAACTTGCATACTTTGTTTTCACAACCTTTTACAATAAAGCCTTCAGTTGATCCATAT

AAATTTCTTCATTCAATTCACCATATAAAAATGCGTTTTAACATCCATCCGATGTATATGCAAATCATAAATTGAAGCTA

TAGCAATCAACATCCTAATCGAGGTGATTCTAGACTGGTGAATAAGTATCGAAGAAATCTAATCCTTCCTGTTGTCTATA

GCCTTTAGCTACCAACCTATGGATTTACTACCAGGAGGTAAATCTACAAGCTCCCAAGTTTCATTTTGCTGAATGGACTC

CCATTCATTTAATGAAGCTTCATCGAAATCTCTTCGTAGTCTGGTTATAAATATACAAACTTATAGGTTCTCTTGGAAGC

CTTTCATGATTTGTGATCTGAATATGTGGTTTCACGACAAATTTATCTGTTCCTTGAACTGGATAAAGATGAAGATCATA

CCCGCTTTGAAACGAACAAGGAGGTGATATGGTTTCCTCGGTCCAATCTGTTGTCTTCTTCTTCTCTTTGTTAGAACTGT

CATCATCAGATATCCATAATTTTACGACATGCATATCAAAACTCGATCTATTAAAGCCATGTGCCCATCTATCTGTTGGA

AAATTGGCACTAAAAGTGGGTGAAGAAAAGAATGTTGTATTAGGGTTCAAGGCTCTTATGAAAAGATTCTTTTCGACAAT

TATTCGACCCTTTCCCAAATAGGAAGAAGTGTATCTATCTCGTTGGAAAAAGTGGGTAAAGAAAAGAATGTTGTAAGGTC

AAAGAATATCTCCGTTTCCCAAATAGGAAGAAGTGTACTTATCTCCGTCAGAAGCCCCACAACAAATGTTGGATTCAAAG

CTTGTACAACCTTGATCCAACCAGGAACACACTGGTTTTGAAAACTGACGATGCTTGTTTAAAAGGAAAACAAAGAAAGT

TGAGAGATGTGTTTTGAATGAAAACAAGTCTGATATTTAAAGAGAAATCATACCCTTTGAAGATGACTTTTCATCTTTAC

AGGAATCCTTTCAGAGTGAACAGGCGTGTCTCTTCCGTCAATGCGTCCATTGGAGAAACAATGTTTCTGTTCAGGTTTTC

TAATTGTCATTAGAACCCCGACAACATGGTAAATGACTGCATTCAAAGGGTTCTTTGAAAATATCCAACATTCTCCCCTT

ATTTTCAAAAAACCAAAATAAGTGATGAAAAATTTAAAACTGGAGTGCCTACATCACCAAAGTGACTGCCTACGTACCCG

AGTGGGATCAAGCCAACGTAGTTCAAGCTAAGTTGAGACAAAGCATCATCGTCGAGACCAATACATAAATGATAGGTACC

TTCTGGGTTTGAACCTCCACTTAGTGTAAGATATTCAAAGCTTCATCGAGGTCCTATGGTGAATTTAATCTTGAACCAAG

TACCCCTTGTGATAAAGCCGGAACCTCCACACATATTGATTTCATGGGAATAGTGTGTTCCATAGCTCGCACATTAATGG

CCCTGAGCTTTCCCATGAAATTCATGAGTCCTTTTCTGAGAACGGTCTTATTCTCATAGGAAACGGCCTGATTTCCAAAC

ACTCATATAGGTAAGTCTCAAAGGTGTTTCTGCGAGACACCTTGTGATAAAAGTTAGACTTATTAAGGATTTTCATCCAT

CCCGACAACAGGGTAAATGACTGCATTCAAAGGGTTCTTTGAAAATATCCAACACTATCTATGTTATTGTGAACTATTCT

GCCTGTAAACCGCCGGATTTTTTCAAAGATCCAACATCATATGCTGCATTTGATAAGTGAGTATTCATAATTCAATATTG

ACTAAGCCATGAAGTTTGATGATTTTGTATCTAGCATTCTTCTAAAATGTTGGATTCTTTTTTGCTTTTACTATAAATTT

TTGTTATTAGTCAACCACTAATTTGCAAAAAAAAAATTATCATCTTGCAAAAATAAATGCACAGATGCATTATTTCAATT

TCGAAAGACCTAGCACTAGTGTGAGAGAAAAATTATACAGGAAAGGGTTAACTGCATATCCCTCAAAACTTTTGCTATAC

TGACCTGACTGCGTTTTATGGGAGAAAATGAAGTCTATTGTGACTCAAACCATTGGCGTGAGTTAGTTGATGCGTTTCTT

TTCTGCATCTTCAATATATACCTAGAGGACTTTCATGAAAAATGTGCACATCATAATCATACGGATAGCTCAATAACGGT

GTTATACCAGTGATGTCAATCATTCGAATACTCTTGTTTGTCCGATTATAGATATTGAGACTTAGGGATCCTATCGGACT

TCTCGTGTCGTCCCGAACAAGTGTTTGTATGACTATCTCATCTGTCCCTTGGACTGGATGAACATAGAGATTCTTCTCTT

CTTTGAAGGAAGGAGGAAGTAATATGGTTTCCTCGCTCCAATTTGTTGCTGTAATCTTCTTCTCCTTGTTAGAACTGTCA

TCATCAGATATCCATAACTTAAGGATACAGGCGTCTTCTCTGTCAATTAAAGCCATCTGTCCATCTATCGATGTTAATGT

GAACTTATCATCCCGTTGTTTTTCATCTTTTAAAATCTCGGGAGGACCAACAATGAAGTCTGGGATCTGAATGACTCTAA

ACTTTTCAGTTCCAACATCAAATGCAACAATTAATTCCTTGTTCCTCCAGTAAATTGAACCGTTTGCATAAACGCCAACA

GGTTCATACAACCGAACTGGTGGCACCTCATCTATCTTCCTCCATTGATTTTCTCCTCCCACTGTAAGAACTTGGCAAAT

ATGGTGGAGTTCTACTTCCAAATTTGGGAAACGGATCCTAAATATATTCCAGACACATACTAGTTTATAGGTTTTGGTCA

GAGGATCAAAACCAAACCCATAAGTTGGCGTCCGAATAATATCAGGACCGAACTCTAATGGTACGGTTGTTTGTGCCCAC

GGAGTAATTTGTCGAGTCGCAAGGTTGTAGATGCAAGTAGCAGCACTATGTACGGGAGAGGAGGGGTCAATAAAACAGAA

CAAGCCATCTATAGACGATGGTGTTTTCTTTAATTCCATGCACCTTGGAACTGTCACTTTATGAATGGCCACTCCACCTT

GAAAACAAGCCTTTGGTGAATAGACTATTATGTGATTACCGCACTGATGGGAGATGAGAAGTTGAGGAGGATGTGGTTTT

GATCGATACATCTCTATGAAGGATGGATCCTGAATGAATGACTGCCAACTTTTGGATACACACTTGAAACGCAAAAGAGA

CTTGATAGGAAGGTCGACTAGTATATCTTCTACTGATTTTTCATCAAGACGGTTTGAAAGTGTTGTTCTTTGTAATTTCT

CTCTTCTTCTGTCCATCTTCTCTTTTACTACTGATCGAAAACACACCAACGCAAAAACTCCAAAGAAAATTGCTTTTGAT

TTACGCCATACTGTTCTTTTATTTATTTAAGGGAATTTTAAAAAAGTGTCCATGTGAGGAAATGCATTTAATAAAATACC

CACGTTTTTTTCAAATTTTATTAAATGCCCATACCGCTGGGTTTTCCGTCTGAGCCCACCAAATAAGTAAAATATCGTGT

ACTTAGTCAATTTCTCATCTGACGAGATGAGATTTTTATGGGCACAATTACAAATCCGCCCATTACTCTAGCAGATGAGA

ATATCATCTTTGACCGTCTACCCGTTTTCCCATCGACGGCAGAAGGTTAAGGAAACCAAATTCTTACACGTGTTTAAAAA

ATGCGCATCTCAACATTTAACAAATAACTAGGATCTTATTTGGTCTCTCCGCCTCCTCCTCTTTACCATTTCTCGCTGCA

TCGTCTTTATCGGCTACACCTAGGGTTCTTCCTTTTCTTTGCTATATCTTGTTCATCGGCAACGCTGTCTCCATCACCAG

CTATATCTAACCAACACCAAGTATTTTTTGCTTGTTTGAAATGATTAGTTAGATCGAATTGAACTTTCAGTTTGGGGAAT

CAATGGCATTTATGTTGGGTTTTCAGTTGGGGGTTTGAATTGGGTTTCTAGTGGTGGTGGTATGAAGGTGGTGAAAGGGT

TAATGGTGTTTAAATTGGGATAGGGTGTTCTTGCAGTAGATAGTAATGGCGGTTGTTTTGAAATTGGTCATGTATTTTGA

GGTTAGAGAATTCTGAACTTTTCAATTGTAAATAAGGTGTTTGATTAATGTGCAATCAGGAACTAGGATACCGAAATTTT

ACGTTTCAGTTTGGGGTTCAATTGGTCAATTGCTTGTTTCAATCTGTGAATGAATTAAGAAGAGTTTGCAACATTTTGTT

TAATTATTCAATAGCAATTCCGGTGGATTTGATTGTTAACTCGCTAACATACCTATACGTAAAGAGACATAGTAGCTAAG

TGCTTCAGAGATTACCCTAATTTTTGAATTAACAACATTCTATCATGTTTCATACTGTTTTATCTAAATAATATATATGC

TTCTCTAGAAGTTAGCAACTTAGGAATTGTGTTAAATTGGGGTTATTTTCATTTGAAATTTTAGTATCAACTGGGACTTT

TTGGTTTGATGCAGCAAGAGATAATGGAGGTGATTTTTCCTAATTAATTCCATTTTTATTTAATTGAAGAACTGTGAGTT

GTAATTGTATGAGTTATGTAAATAACTGTTGGGGAAGAAGATGAAGTACTTTTTCTCTTACAGTTACAGCAGCAGTGACT

CGAGTCAATGCAACTCGGTACTGACCCATTCGTTTTGGCTCGGGTCTATTCAAGCTATTCAGGGGCTAGAATGTCTTTTT

AGAAGGCCGATTTATTGCCACCTGTCTACTAACGGCTGTTAGCCGTTTTTTCAATAAAACGGGACGGAAATGGTCAAGGG

ATGGGCATTTAATAAATTTTGAAAAAAACGTGGGCATTTTCTTAAATGCATTTCCCCACATGGTCATTTTATTAAAAACC

CATTTATTTAATGGCTGTCCAAATTCCTTTTTTAACTAGAAGTCCAATATAGAAGGACTGAATATAAGGAAGGCATGCAT

GCACCCACAGATGTGCATGCATGACTGGTCTTAGGTGAGTGATTTCCTAATTAGCCTAAGTTTCTAAAAGGTGCGCAGAT

TTCCCAATCTGAGTTGCTCAGTTTCACCCATCGCATATAAATAAGGATTCAACTCTGGTAGATGTTTGAGATTTCAAACG

CAGGAGGTGAGTTTGGTTCCATATTAATCGGTTTCTCTTTTTCAGACCAGTAGAAAGAACATAATGAATATGGAGGAAAC

AAACCTTTCAAACCTTCTTAATAACGAATCAATGCATCAAATACTTAGCAACCTTCCTGTGAAATCTCTTATGAAGTTTA

CATGTGTATCCAAGAGCTGGGAATCACTTACTCATGATCCATCTTTCATTAAGCTACATCGATCAAAAGCACAACTTCAG

CTTCTCATCATGATTTACAAGGTTGATTCCATAATAGTGTATACACCGAAATATGGGTTCGAATGCGGTGAAGCTCTTTA

TAAAGCGACGATCCCATGGAGTGGAGGAGTTATACCCAAGCCTGTAAACGGGTTGTTCTGTTTTCTTGATTCCAGACGTT

CTTTGTCCTGTGTTTACAACCTTGCCACTCAACAGAGTACTCCATGGGTTTCTGTACCAGTTCCATCGGTGACAAAGCCT

GGATATGAAATTATTATGGATTCGGTTTCGATCCTTTAACTGGCCAACACAAGGTATTGTGCACTTGGGTTATATCTCAG

CACGGTGCAATGACATTTGGGAAAGCTGACCATATTTGTGAAGTTTTTACCCTAGGAGAAAATCAATGGAGGACAATAGA

TGAAGTGCCACCGGTTAGGCCAGATGGAGTCTCTGTTTATGCAAACGGTTCGATTTACAGAAGAAACCGCGATGATTACT

TTTTCAAAGAGCCAGAGGTCGAAGTCATCGTGGCATTTGACGTCGGAACCGAAAAGTTCAGAGTGATTCCAATCCCAGAC

TTTGTCATTGCTTCTTCTGAGACTCCGGAAGACTCATGTCGCCACTTTGAGCAACTCTTACTAGAAGTAGATGGACACAT

AGCTCTGATTAACAGGATAAGCGAGAGTATGGTGAAGTTGTGGATTTCTGATGATGACTACATACAGAAGAAGGCCGATG

TGAAATGGGTTGAAGAAACCATCATACTTCCTTTTCAGTGGCCAAGGGGAGAACGTCTGCATTTCCGTGCGGTTGAAGGA

ACAAAACAAATGATCATTAACAAAGCACCTGAAGCGAGAGGTTTTGTGAATATGTACATCTATGACCGGGTTATGAAGAG

ATTTAGAGATGTCAAAATCGATGGTTTACCACGACTCCCCAGCCCATTTCGATACCGCATGTTCGCAACCCATGAGAGTC

TTCTGCCTATTCAAAATAAGGAAAAGAATGGACACTAAAATCACATGTAGTGATCTGAGTTTGAAACTCGTAAGCGCTAA

ATTCTTTAACAATAAAAATAAACCCACACATGGAATCATGCTCATAGATTGATAGCTTTTCAGTTGATCAGTGTTTGCTA

ACCGTTATTCGGTGGTGGAGGGAATTAAGGACAAGTAGGTGCAACTGCTCCTCCCGCTTGACAACGAGCTCATTCAAGAT

TTTTATACTTTTATTTACACAAAACTTTATAAACTCCTTATTATTCATATTTTTCCCGCATATAGCAATAATGTTCATTA

TTTTACCCTTATACTTCAAAAGGGTACAAGGGTATTCGTGACACAAAAACTACTTAGGTTGAGTGCAGTGGAGTAAAAGA

AGGGAATGATAGCAACATATGGGCGTGGCTAAAGCCTTTATTCAAAACCCTTCCCTTTCTCTTTGCTTGAAAGCAGGGTG

GTTAGCGCGGATTTCAAAATAGAATCCTCGCCCAAAAAAAGTGAAAAGCGAGGGGACAATCCTCGTTTTGTCTTTTATTA

TAATAAAAAACCTAATATATATTTTTTTGATTTTCAGATTGTTTATGTAGCGCTCCCTAAGCTAGCAGCTGATTAATCCA

ACAAATTAACTAAAACATGTACTTAAGCATTCAATTAAAAAATGCGATACAAAACTTAACCCAAAGCTAACCCCGCTCAG

AAATATATATACAAGATCTTCTCTCAAATGATACATATATAATAGTCATTTCGTTTTTAAATAAAATATACAACTTAATA

AATATAATAGAAGTTAATCAACTAGCGAAGTCAACCCTTCAAGCTTTCGCTGCGCAACTCTGATCTGTCACAGAAACTGA

AGTGGGAATAATTAACATTTAAACTTCTAAGTAATCATAGGCAAGATTTGGAAAACAATACATGTTTTTCCGAAACATTA

AACAGTGACTAAGTCACTGAAATAATCATACATGTATATGGACAAACTCCTTCTTCAAACAATGTATAATATGGTTGTAA

ATCCCTACCCGGCAATTCACACCTAATAGTTATGGATTCCACCAAGACCATGGTGACCAATACTCTAAACAATAAAAGCT

GAATTAAAGTAGGTATAAATGTGAAGGAGCGTATCCTATATACCACAAGTGGAAATTCTCATTTCCCATAACAATTCATA

TTGACAACGAAACAATACAAGTCCTTTCCAAATCGTGGAAAGATTCGCAGAATCAAAAGGATTATCATAATGCATTTTAA

ACAAAGCATGGAATGCACGGACATACAATGCATGAGTTTGTTAATCATAAATTTTTATAAAAGCAACAATAATGATTACA

AAAGAGTCAAATGCATTCCCACACCTTTTGATGACAAGCCACGCCTACCTCGCGATCCACGATCCACTGGTTCATCCTTT

GAATCGATCGTTGTTAATAATTAACTGTTAATCACACAAGAATTCTAAGATTCTAGCCAAAATGGCTCACACAAGAATTA

TACAAATCTAACTAATAGTCCTAAGCCCATTTACGTTTTTATACCCTTTTATTATCTATCTTAAGCCCATCTAAGGTTCT

TCACATCTTCATAATCCATCTCTTGCATATCTAAAGATTTACGAATTCATTGGGTTGGTTCTCTAGTCCAATAGCTAAGT

CTTATGAGTATCTATAAGAGTGTAGAAGAAATTAAAGGCAAATCCATAACATAAGGGATCCAACATTCATAATAGAATTA

CTAGGTCTAAGCCCAAGTACACTAGACTCGGCCCATTAGTACAAGGTTTGACCCAACTGGGTCAACTAACAGTCACTGAC

GGTCAAATGGTCAACTAACAGTCCACGTCCGTCCAAAGTCAAGTCCAGGTCAACTCAGTCAAGGTTCATTGAGTCAGCAA

GGTTCAACTCATGTTATCAGGGTCAACTCGACTCAGTCAGATTAACTGAGTCAGCTAAACAAAATTGAATCAGACTGAAT

AATCTCAGTCAGACAAGCCAAGGATTTTTCTCTCACATCTGTAACTCTAACTCTCTAACGCTACAACTCTAACTTTACAA

TTCTAACTCAAAGCATCAATTCATATCATTTAAATTATAACTCAAAATTGATTTACAAGTTAACAAATTCTACCTGCAAA

TGCAGTTTCCAAGCCTCACACACCACTACTATCTTTAGCTACCATTGGTCTGCAACTGCAATTTATTATCTTACAATCAA

CACAAAACCATAACAGAATTAGCTTAAGTCTCAACACTTCAAACATCCAACCATACACACCTGCAATTCATATCACATCT

CCACCAGACCAAACTCAGATCAACTATTATCCTCAGCCACAATAATCCAATCCCCATATTACCAATAATCAGTTCAACTC

AGAACTATATCATTAACATCTCTAGGATTCATTTGAACAGTTCACTCAAACCACCATAACAATCTAACCATACATCCTGC

ATAACCAACATCTTTATCTCCAACTCCTTGTATTGATCAAGGTATAGTACCTCCACTTCTAACACACACACAACCAGACC

TTCATTCCAGCAAAGATGAATCTACAACTCCACTAGAATTACAACTAACACCAGCATCCAAACACATAACAAAACATCTA

AACACAGTTCCCCTGCAACTCCCTACCAACGCCATTAGTTTAATTCAAACAACATCTTCAATAACTCAATCTAATTACCA

TTCACTATGTTTTTACTGCAACTGAAATAACATATCAACAACATCAATTATTCATATACCCTAACTGAGATAGTAACTTC

ATGACATCTCATAGTTCATAATTCAGTTCACTAGTAATTTAATTCCAACATACAGACAACACAATTCCTCATAACTGTAA

CTTCAAAAGAAACACCATAACCATAACTCATCTCATACATTAGCGTCAAAACAATCACAGTTCCATCACAAACCAACCCT

GTCCATCTCAATCTTAATCAGTAGCTACTCATAATCTCAATGATAGAAACAAATATCTTTATTAGGTTTACTGAGAAAGA

CAGTTACCTCAAATTAGTTTTGTTGAAGAAATTGAATGCAAACTCTTCTTCACCATTTTCTCCCGAGTCTTGAATCAACA

CCAACAATAACTCCATCACCTTCTAATTGACAGAATCGACTCCAAGTCTTCATCACACAATCCCTATCTCAAATACTTAT

GCTCACAAAATCCTAAAAGGTTTGACTCGGTCTCCATCTTGTCTCTCACAAGTTTTCTGTTTCGAATTCATATATAATTC

TAACTCCTCAATTCAACCAAACCCATCTCAAATCCCCTTCTTAAAACATCAAATTCATCTTATAATCGAACTTCTACACA

ATCTTCTGAAAAACCCTAACTTCATACTCAATTTCCTACAAAAACTACGATTAAACACAATACATCTATTTATTCTCCAA

TTCAGAACAAACCCAACATTCATTCAATCCTCTAAGCCATTAACTTCATCTTCAGACCCTCCTCTGAAATTTTATTATGA

ACCCTAGCTCTTGAATCAATCCTAGAATCAACTTCTAAACAACCTCACAAATTTCATCAAACACCAATTAAAACGAACCC

AACCTCAGTTTCATCTTAATCATCACTAATTTGAAACCCCAATTCATAATCACAGAAAACCCCGACTTGGTTCTCTGAAA

CATATTTTCTTTTTCATCAACAACAACCTCGACTCCTCCATCATCTATACAAGTAATAACCTGATTCTTATCAACCAATT

CTCTAGATTTTCTCTCAAAAATTCGCAGAAGAAGAAGAACAGATGAGAATAAAAGAAAGAAGAAGAAGAAATGAATTCCT

CTCGGGCGGTCTGGGGATAAGGCCGACCCAATCTTCTGAGTCACCCCCTTTCTTTAACACGTGCAGATACAAGGATTACT

ATCCAAAATTACTATTTTACCCTTCTTGCAATCTAAGTGATATCTTCTTTGCCCAGTGTCCGAATGACACGCTCAAGTAG

CCCATTCCGCGTAACTTTTCAAGATCTATCTAACGATACTAGTTTCATATCTAAATCATTGTTAGATTAATTTATATTAA

TTAACATTCGTGTTAAACTAATCGAGTTATTATCATCTAATTCGTTTCTTGACTGGTTTAATATCGTTGACGAGCTTGAG

GGTCCTTACAGTTTATTTCATCCTCCCGAAATGCTACAATGTGTAAACATAGTGTTCTCACCACCACCATAATCTCGATG

GGTTAGAGTGGAGTGGAGAGTTTTTAATGACGGCTTTTGATCTCTATCTATTTTCTCACGTCGTTCTCTACTCGTGAATT

TGATAGTTCCTTTTTAGTTAGCTTTTCTGTTTTATTAATTTGCTACCACTCAACTGTTCTGATTCCCTCACCATGAATAC

AACTTGTAATTTGGTGGCTTCAACGGCGGATTTCGGTTTTTTAGTTGGTGCTTCAGCTGAAAGCACCTAGCATGTTTGTT

TTTATGCCTCAATGACTGCCGCAGGTAAAGGTGTTAGGCTATGTCCCTTTAAGGATTTTCATGGGTGCCCGGATGGGGGT

ATTGGGGCTAAGGGGTTTGCATGTTTGATACCCCATTGGAATTGACTACACTTCTACAATTTGGAACGAAAAAATGCACT

TCAATAGGCCATAAGTCGGGACTTGGATATTTTTATTAGTTTAGAGGAAACTTTAAGGTCTTTGAAGCAGTGGATGTGTT

TTGAATGTATGAGCATCCTAGCATGGAGTAGGGAATGCAATCATGGAGGGGATGATGATATTAGGCAGGTATTTGATGAT

TCGGGGTATATAGTTAAAACAATCATAGGGGTATCTAGGCCAGTGATGCAAAAGGTTTCTTCTCAATTTGGGGGTAATGA

TTTTGAACTGAATGAAGAGACAGTAGATGCGATTTTCAGAGCTCCCATATGCACTGTAAAGAGTATTCCAAGAGGATGTC

GGCTCGCATTTTCTCATACCCTTAAGGTAGCTTTCCAGAAAGTTACAACAAAACCTTGTTCCATCAGGCATTGGGTGAGA

TTATTGGTACTTCCTCGCTGTAAACTCCGAGTCTTTCGACCAAAGGGTAGGGTTGAATGTAAGTCATGTAATCGCAAAGT

AATGTAGCAGAAACAAATCCTTCAAGCGCTTGCCATTTGGGGTGAACCTGGTGGGGTTCAAAAGCTTGTTTGAGGGTTTT

TGAAAGCACCATTACCAGGGTGTGTTGTGTTATTAAGGCCATCAGCTCCTCAGGCGTGGCGCCACACAATGAGACTAATA

GGAAAGCGTTAAAGGAAAAATATCTGACCATGCACCCCCCCCCCCCGGAAACCAACTATTCCTCTGCGTGAGACATCTTT

GGTGGCTGAGGAAGTAACCGTTCTCAATTGAATTAAATCGTTTCCTAAGGGGACTACGTGTGGTAGGGATGGTTTGCGAG

AACAACATTTGTTGGATGCTTTTTGTGGGGAAGGTTCCGTTGTTTCTGCTGATTTGGTCACATCAATTACAAGGGTTGTC

AATTTGTGGCTTGGAGGGAGATGCCCACTTAATCTGGCGGAGTTTTTAGCATCGACACCCTTGACACCTCTCCTCAAACC

GGACAAGAGCATTCTTCACATTGCTGTTGGGACGGTTTGGCGTCGCCTTGTGTCCAAAATAGCAATGAAAGGTGTTGGCA

AGGACATGGAAGCGTACCTTCAAGATTTCCAGTTTGGAGTTGGGGTCTCAGGAGGGGCGGAGGCAATATTTCACGCTGTG

AACAGGTTTGTCAGTAAACATCATACACATGTTAGCCTTACCATGTTAACTGTAGACTTCACCAATGCCTTTAACCTAGT

TGACTAAACTGATATGCTTCTTGAGGTACGTAAGCTTTGTCCTTCTATATCGCCGTGGGTGGAATATTTGTATGGCCAAC

CTGCAAGATTGTATGTGGGGGAATCCCATATCTTTGCTTCTGCAGGAGTGCAACAAGGTGACCGTCTGGGGCCATTGCTA

TTTGCTTTAGTACTTCATCATCTTGTAGCAAAGATAAACGAGAATTGCATCTTATACCTACAGGCTTGGTATCTTGATGA

TGGCACTATTATTGGTGACACTGCAGAAGTGGAAAAGGAACTTACTGTTGTCCAGGCAGAAGGGCCTGCTTTGGGGCTTA

TATTGAATATTAAGAAAACATATATTTTTTGGCCCTCTTGCGATGGACGGAGGTTAAGTGATGGGATGTTTCCCGATGAA

ATCTCAAGACCAGGTAATGGAGTTAAACTTTTAGGAGGTGCGGTTAGTCTTGATGATGGGTTTGTGAAGGGACTTGCCAT

TAAGGGAGCAGAAAAGACGGTGGATTTGAAGCAGGTTTTATCTAAGCTCCGCGACCCTCAGAGTGAATTGTTGCTTCTAC

GTGCATGCATGGGCATCTCCAAGATACTTTTTGGACTTCGGGTTTGCTTCTCGGAATACATTGTGGAGGCTATAGATATA

TTTGATAGAGCCTTGAGAGAGGTGATAGAGGGAATTGTGGTTTGTGGAGGTATGTTCTTTGGGGAACTTCAGTGGCGGCT

ATCCACTTTACCTATTAGATATGGAGGTCTTGGGTTTTACACTACTAAGGATGTAATGACCCGTCCCTCCACCGATATTG

TCCCCACTTAGCCACTGCGGTCATCCGGCAGTGTGGGGTTTTCAACGGGCATCGCTGCGGTAATCCAGCAGAAACTTCCC

GGGAGGTCACCCATCCCTGGATTGCTCCCGTACGAGAACGCTTAACTGCAGAGTTTTCTTCCAACTCTGGATCCAATTGT

GCTGAAAAGGCCTCGGTGTTAGGAGAGGACAAACCATTACCTATATTCCATTCCATCAGCGAACCACTGCCAAAATCGGG

GTATTACAATACCACCACCTTAAATTGGAGCCGTCCTCGGATCCGGAATATATTTTCAAGCCCAGATCTCCGACGTTCCC

TGCCCCTCATGAGAGAAGTACCTGACATATCCCGCCCAGCGTACTCTCTCACCCTCGGAAGGTGAACTGTCTTCAGCTCT

GATACCATTTGTAATGACCCGTCCCTCCACCGATATTGTCCCCACTTAGCCACTGCGGTCATCCGGCAGTGTGGGGTTTT

CAACGGGCATCGCTGCGGTAATCCAGCAGAAACTTCTCGGGAGGTCACCCATCCCTGGATTGCTCCCGCCCGAGCACGCT

TAACTGCAGAGTTTTCTTCCAACTCTGGAGCCAATTGTGCTGAAAAGGCCTCGATGTTAGGAGAGGACAAACCATTACCT

ATATTCCATTTCATCGGCGAACCACTGCCGAAATCGGGGTATTACAAAGAAAGCATCACAATATGCTTTCCTTGTTTCAC

GTGTGCAATCGTGGGGGTTACAAAATCACATAATGAGGAGTAGTGGGGTGGAAGGCATGGATGACAACTTTAATAGGGCT

TTGAAGACACTACAAGTAACTCTTCCTTATTTTGACTTTAGCAGTTTTACCAATAAAGATACCTCCCCCCTAAAGCACAA

AGTACTTTGGCGAGTGCTCTTTTTGGTAAAGTGGTTAAAGAAATGGATAGAGTTTTTGAGCTTTCACATAGACAGAAAGC

GGTTTTTGGTTGCCTACAAGCAACGCATGCCCAGGATTTTCTCCTTGCTATTCCAATTGAGGGACTTGGACAGAAGATGA

CACCCTTGGAGTACCGATCAATTCTCAAATACAGGATAATGATCCCTCTTTACCGTTTAGATTCACAATGTCCCGCTTGT

TTTACAGGATGCTTGGACTTGTATGAGGACCATGCAGTTCATTGTAAGGTGGACCTTGGGTTTAAATATAGACATGATCA

TGTGAGGGATACTTTATATGATGTGCTATGGAGAGCCGGCATTTCAGCAAATAAAGAAGTCGCTGTCAATTTTTTGACAG

ACCCACTCGAGGGGAGATCAACACTCAGGCCAGCGAATGTTCTTATTTTTGGGTGGGTTAATGGGAAACACGCTTGTGTT

GACCTTACAGGAGTCTCGCCGTTAGTAGGTATTGGACATGGTACTTTCACGGTTGGACAAGCAGCACTAAGGGCTGCCTC

GGGTAAGATATCAAAACATGAGAAAGCGTGTATTGACAATGGACATGCTTTTATTCCCTTTGATTTCGATACTTTTGGTT

ACCTTGCTCCAGAGGCGGTTGGACTTCTCAAAAGAGTTCAGAAAGTCATGAACAGTAATGTCATGACTCCTGGCTCGAGA

GAGTTTGTTTTTAAGAGGATAGGTTTTGCAATTCAAAAAGGGCTTGCGGCGCAGCTTGTTGCTCGCTTGACTGCTGTGTA

AATTCTGAAGTTTTTCGGACCCTAGGTTTATTTATTATGATAAGATATAAATAAAACATATTAATATCAGATTTTGGTCC

CACCTTAAGAGTTTTAAATAAATAAAACACTCTTGGGCGAGGATTGCTTCCTCGCCCAAATCTAAATTTCGTCCCCCTCC

ACTACGCTCAGGCCTATCTTCAACGTGATGTGGCGTCACTGTGCAACCCCTACCTTCCCTCCTCAACTACACTCACCGAT

AAATCATGATACGGTAAAAGTGTGTCAATTCATGACTCATGAGTGATAGTAAAAAGGTGATTGTGATTTCTTAACTAAAA

TAAAATAAGATAAGATAAAAGATCATTCATGGCCTACTTATGATCTTTCTCCTTGTGCTGTGCCTTAATTGTATATATCT

TGGAAGATCAATAAGTAGGGACTAAAAAGATTGAGCACTTACTTTGTCAAGTGTGCATCCTTAATCATTTTACTTTGTCA

AGTGTGCATCCTTAATCATTTGAAGAGAAGTGAGAGTGAAAGATCCAAGTATATCCAACACCAACATCATATGCTGCATT

CTTCCATGTTGCATTTGACAAGTGAGTATTCATAATTCCATATTGACTAAGCCACGAAAGTCGATGATGGGCACTATGAT

TTTATAGCTAGCATTCTTCTAAAATCTTGCATTTTTTGCTTTTATCATAAATATTTGTTATTGGTGAACCTCTTGTGAGT

GTGCAAGGCTATAAACGTTAAATGGAGGCGTTGCCACTATTTTGCAAAAGAAGACATATACATCATCTTGCAAATACCAG

AAAATCGGTCAAAACGGGAATAAATGCATGGTTATATTGTGGTAGTATATTGCTAATCCCACATCACCCAATACCTCCAT

CACCTTGTAAGCATGCTTATATTGCATAATGAACAAAAGTAAATCATGATAGGGTTTTTTTATTCAACAGATAACTTGCA

CTACAAACTCATTAGAATTTGGTATTAAGTTTGTAGTTAGCTTACAACTACTAATTAATTAAGTTGTCTAAAAAACCAAA

GGATATCTATGTTTAATATTGAGATGGACATAAGAATGTAAAGTAAAATAGAAAACTCTGAAAAGATGATATAAGTTGGG

TACAAAATCAAAGATCATCATCATCTTATATTAAAACTATAAATATACTTCCAATATGCTCATTCATCTTCATAACTTCT

ACTCATGAATTGCCGCCTACCGACAGTTACCATCAAATGTTGACACTATCGTTTATCTTATTACTTCTTATCACTTCCAA

TGATTTTTATTTTTTTTTGATGAAAATCCCTTGATGAAATGAAATCACACCTTTGCAGATCCTTCCTAAATTTGGGCCTA

GCAGTGATTGAGACTTTTCATTCCTCTCGAGCGATTTGTTTATTGTAGAGGCTTATAGTTTAATTGGTTGACTGGTAATA

CCACTTAAGGGAGGATAGACCGTCCACCCAGTACCACTACCTACTTCGACTAAGGCTGAGCTTAATAGGAGCAAGAGAAC

TTAATTGGGGCCTATAACTACATGACAAAATAGAGTCACTAGGAAGTAATTTGTGGGAACCCCTTATCTGTTATTTATAT

TAATGCCTAATATACCCTTATTAAATTAGTGATGATTAATTAAGTTAATGTTTATTAAATTAATAAGTTTTGAGTAGGAT

TATAATGTTTGGATTAGAGTAGAAGATCATTTTTCTTTTTAAGGTTGGGAGGGAAAGAAGTAGAGGGAAAAAAGGAAAAA

CTAGGGTTTGTGATTTTTTTAGCAAAAATGAAACTTTATAGTGATGATGAAGACTCTAGTAGTGATGAAGAAGGGGGTGC

ATCATATTATAATGATTTCGATCCTACTGAATTGGAAAATTTGTTGAATGAAGAAGAGAAAGTGAACCAAAAAATGCTTA

AGGATATTATTGAAGCACAAGAACACTTTCACCATGAGGAAGAAGATGATAATGCTTCCAATAAACATGTATGAGTTGTA

ATGATCTACAAACATGTATTTGCATGTCCCAATTGCCAAATAAGGAAAAAAATTCCAAACTTTAAACCAGAATCGGTTTA

TGCCATGCTAGCGAATAAACCGATTCTAGGTAAAATCTCCAAATTAGGTACAGAAATCGGTCTTTGTTCATCACCAAATA

AACCGATTGTAGTAAATTTTTGGCGCGTTGGGTACGTGACGAGAATCGGTTTTTGTTGGTGTTGTGCATAATCCGATTCC

TTGACTAGAATAATCACGTAGAATACACAAAATTGACATTCGGTTTATTTTCGTGAATATGTAATCCGGTTTTGTAGAAA

AAAAGATCCAGGAGAAATGTGAACAATAATCGGATTATATGACACGATACGTGAACCGATTGTATTCTATGTATTTTTTT

CGCCAATACTTTAATCGGTCTTTATTATTGCACATATAAACCGATTACTTTAATCGGGTTAATCGACTGTAACATATAAA

CCGATTTTGAATCATCAAAGTTTTGGTATTTGTTGTAGATTGTTGTGGCGTTTGATGAAGATCAACCCAAGACAAAGAAG

TGATTTTTGAAAAAATGGTTGCTAGAGTACAAGGGCTAAAGGGGAATGGACCAATTACCCAAGAGCATAGGATGCAGATG

CCTCTATGTGGGCATATCTTTGCGGAAACGTGGAGTTGCATTGTTCATCTATTTGGTAAGGGATCATGTTATACATACGC

ACCAAAATACACCATTTGGGATGAATCGCTTAAGGATCGGAGAATTGTTTTATTCAACCATAACAACATACATTATATCG

GTCTTAGAGTACGTGATGATTGTCCATTACCACAGGTAGATGGGTTTCATGAGGTCACGGAGGAGTGGCTAAAAACATAC

GATACCAACATGAGGCGATGGGAGGAATTGATCGTGCCGGATCAATTCGATGGTCTCCATTTGTTGGAATGAGTATTTTC

CTTATGTTAAAAACTTGATCTATCGGATGCTTATATTTTGTCGCTTTCTTATGGAATGAGTATCTTAATGAATGAAAGTG

GTTTTTTTTTTTTTTTTTGGTAACTAGGAATCATGAATTTTGCTTAAGAATGGACTTAAAATACGTTTAAGAAGTCCGAT

TCTGAAGATAGTTTGTTGCATCTTTTTAAGAATCGGTTTATATGTACGACAATGTAAACTGATTGTTGATACAGAGATTT

TAAATTACAAAATCGGATTACATATGCTGCAAAATTATCCGAACATGAAACCAGAATTGGTTTATGTGTGTACTAATGTA

ATCCGATTATAGTTGATGTTCGTCAAATCCACCCAGAATCGGGTAATGTAGGTTCACATTAAAAACCGATTATGGCCATA

ATCGGATTATTATGGTGCACATATAGACCGATTCTGGATGGATTTTCAGAAAATTTGATGAGTGGATTTTAAAGATTTTG

AGGTAAATCATTGTAGAGTATCCCTTAGAAGGAAAGGTTTAAAGGGATTCAACTCAACACTTGAATTTTTTGTTTTTTGT

TTTAATTTTGATGAAAAATAAATTGAGTTTTAATTGTTGTTTGTGATTGATTAGGATTTAGTTTTAATTTTGATGAAAAA

TAAAAACTATTTTTGATTAATGATTTTGGTTTTTGTTAATTAAGTTTTAATTTTGTATTTGTATTTGTGTTAGAATAATT

AAGGTTTCTTTAAGGTTAATTTAGTAATTTTACAATATTTTGGACACCCCTTATCATTTTCTATTGGGGGATGAAGAAAT

CTATAGGCCCCAATTAAGTTGGCTAGGCCCCAATTTAGGAAGGTTTGCAGATGGGTTATGTATATAAATATTGAGAGATT

AACAAACTTTGAGATTGAATTCTATTATAAATTTTACTGTGTTGCCTCTTTAATGGGTTAGAAGATTTTTTTAAGCTTAT

GGCCAAACTCACCTTTGTAATGGGTTAAATATTTTATTAAAATTTATGGGGCTAACTCACCTTGAAAACCCGTTTTACAA

GGTTACAGTTGATCAAAACATGCAGATACACTTCACAAATTAAGATAAGCTTAATAGCTCCTTAATAGTGGGATAGAAAT

CTAGAGATTGTGGTAATTACCATGAATGGAAGGTTAAAGTGAGTTAAACAATCCCTTGAAGGGTTAAACAATGTCCACAC

ATCACCTACCCAAGCTCAAACTCCATACTAAAAGTCCATGAACCTAATTCATGTTGGAAATCGACTTACAAGGTTAAGAT

TTTTAAAAGACATACAAACTTCACAAATCCACGAGATTGTCCACTATTTAAACCATATTTATACTCCAGCTTTAACACAA

ATTTAGAATATCAATTTACTAGGAGGGTGCCAAAAATCATTTAAGAGTATCATATATGGGTTTCAGCACCTCCCAATAGC

GGTGTTGATCAGTTTTTTTCCTGCATGAATCATTTACGAAAGTGGTAATAGTAGTGGAGCAAAAGGAACCAATGAGGCAA

TGAGTTTAAAAGAGGGGTTAAGATGAATGGGTTAAAAAGTTTAAGCATTTTGAGGGCCATATCATAGTGCTTTAAGAAGT

TATGGTGGGCTATTGTCCCATCATGCAAAAAAGAAAAATGTCTATCAACACACTTAACAATAATTTGTCTTGTTGATGAA

TTTTTTGACACATAAAAAAAGTATATCTATTTTGTATCATTATTAAAGGTTCAATATACTTATCAAAAACACTTGCACAT

TTCTTTTCATGGAGGGGGGGTCGACAAACACTTTCTCTTGATAAGATATTTCTCATGTGCCTACTATACCTATTCTTGCA

TCTTGTTCAACTTAAACATGAAACATTACATTATCAAAAAAATCCCAACTTTTTCTTCTCTTTAAAGCTCGCTCACTCCT

TAAGAATCCCACTTACTTAGTCTTGGGAGTTGGATGCATATATATATATATATATAAGAGTGATGTACCCTCAAAATCAC

AAAGTATATAGTTCAAACTTCAAAGATTATTGAGTTGCCTAATTTTAGAAGAAAAGGAAGAGATTGTTTATAGTAAAAAG

AGATTAATGGGTGTTCCTAATCAAAATCAAAACCAAAATGATCATTTCTATCCTCAAGAACTTCAATTAAAACTGTACCA

AGCATTCATATTTTCAGTACCAATTCTTTTCTCTATAATTCTTCTCCTCTTGTTTTATCTCTTTTACCTTAAAAGAACTA

GAGATCAAGCAAGCAATCACCCCACTTCTCTTCCAACAAGTGCTCCAAGAAGTACTAGTTTGAATAATCAACCCACTTCG

ATCCTCCCATTCATGGTACGTACTTACCATAATCTCTTAATTTCTCTTCAACTATTGAGTGTTTTTTTTTTATAGATTTT

TCTTGTACTAATCTACTTTTTTGTTGTTTGTAGTCGATTGATGTTGGTTTGATCAAGGAAGAACTAAAGGAGAAGCTTCC

AATAGTCACATTTGATGAAAGCTTAAGAGACAAAGATTCATTGTGAGTACATACACATCCATCTAAACTTGGTGAAATTT

GTTACTTTTTATTTATTTTATATCTGAGTTTCGAATTCAGGTAACTGATATCATCATCTAGTGACTTGACTTCACCACTG

TGATAATCTTACGAGATTTGTGGGCATTGTTTTCTTGCATCCAACTTTGAAAACATGCATAACAATGTTGCATATGGCAC

TTGAGTGATCAAGAAATTGAAGGCATGAATCTTGCAGTAAATTTTATCCAGGACATGATAGTTCCATTTACAGTATTTTT

TTATTTTATTTTTATGGTAAAAAGTGTTGGTAGGTCAAATAGCTGGATGGTTAAGATCTATGATACTAAAGTGATGGTGT

ATATAAGTGGCAGTGTCATCATTCCAAGTGCTTTGTGATTTTCAGTGTCACTAACTTTTCATGTAGAATCTCTTATACCA

TCCATTGTTTTCTTTTGTCATTGTCAAGAAAAAAAAAAAAAAAAAAAAAAGTAGAATGTGATGAATAGTAGTATACAAAC

TCAATGGTTCATACAAATTCAGCATCTTCAACATATAAAACCACTAGTCATATTGCTAAAGGCATCATATGGGTCAGAGA

TGCAAATTCTCCTAGTGAAATTTAGCTTTCGAAAGAAAAAAATGTCAACAATAAACAAAAATGTTCATTCATTAAAAAAA

AAAAAATCATTTTTTAGAATTTGGTAACTCTTTTGTAACAGCCTTGATATTAAGGAAAGACCGTTATGAATTTTTTTTTA

TAATTCGTGGAAAAAGAATCCTCGAAAACCACCGTTTGATCAGAAACGGTGAGTCCTATCATAAAACCACCAAAAAAAAA

AAAAATTTACTTTCCACGGTTTGTTTAGAATTTCTCAAAAGATTTAACTGTCTAAAGGAGAATCAAACACTTTCCTCCAT

CTGACAGTGAGAAAGTGTTGATAAGAATCTCCATTGTACTAGCTACCCAGTGGGTTCAAAATTCAAATAAATAAATAATT

ACTGGTTAATAGTATCGTACCATGGCCATGGTGGTCTCAAAAGCCTCCTCCTCCTCCAATATGCGCCAACCTATGTGCGG

AGGTAGACGTACCCATGCATTCAAAAATCTCCATCAATCAGAGATATACAGTGTCGTGCAGTGCTTAGTGCTCAAAAAGA

TTCTCTTCTTCATGGTGGTACCAAAATCGTAGTACGGTTTTTTGTTTTTTGTTTTAAAGAAAAAAAAAGGAGATGATGAT

ATTTACTTGCACCAAATTCCCTAGTGGTATATGTTTGGTTTGTGATGCTGATATTTCGTCTATTTTTCATGTATTTTTGC

AGATGCTGCGTATGTCTGGGTGATTTCGAGCTCAGGGAGCAGTTGCAGCAGATTCCGTCGTGCAAACATGTGTTTCACGT

CGAATGCATTCATCACTGGCTCCGCACCAACTCGACGTGTCCGCTGTGTCGGTGCTTCATTCTTGCGGTTGGGAAGGCGG

ATCCTGCAGTAGTAGACCCTCCACTTCCACCGGTTTGATCGTCGGAGCTCAGTTTTCATGGCTACGGAAAAAAAAAACAG

ACAAAAAGAAAGCAAATTCTTGCACCCTTGGAACCAGAAGCAATTGCTGGGTGGTTTTACAAGGTGGCCGGAAAAGTGCG

CATCTCCTGATATGTTTTGCCCGAGGTGGCAACCAAATGAGCAGTCTATACACACATGCAGCATATACTTAGATAAAGCC

TAATGTTTCATTTCGATATCGTTTTAAGTTTTCTGCGACCGGAAATCTCAATTAACACCGCCAACCAGTGGCTGAGGCGG

AGCTAGGTAATAGGACTTGTAATCAATTTGTCCATGTTATGACATTTTTCCCCTTTAATCAATTTCAATTCTCCACCACA

ACAACCAAATGTAACATATTTTCCCGGTTACCGGAACTAGGACTGTAATCAACTTGTCAGGTCATGTTATGACAGTAAAT

TATTTCATCATTCGAATTGCTGCCAAGCATCCAGTTAAATTGCAGGCACCACCACCACCACGAAAGAGATCAATCATACG

TCTGTCCAATGTCTCAATTCAAATAATTGAGAGATAGAGATTGAGAGAGATAGGTCATACATTTGTGTCTCTCTTAAATA

TGCACATCTCTATCTCTCTTTTACACTTAAAAGGTCCTTTTCCTTGTCCGATTTCCAATGGGATTCTTCTTGTGTTTCAA

CTTTCAAGTGGAAAAATTCATCTTCTTCTTCTCCACCCAAACACAACAGAGATAATTGAATTCATAGACCTCGTACAAAC

TTTGGCCCACTGATTCAACCAACATTCTACCCCATAATCACAGTTGAGAGATCAAACATCTGGATTGACCGCTCCGTTAC

GGAACGCGCGTGCACACCCAACGTGTTTAGGACTCTAGTTCCGACTCCAACAGTTGCTTCTAACGAGACGAGATTCGGTT

AAAATACTCGAAAAAAATCGATAAGGAAATAAACTGGCTTTGGCGAAACCAAACTTGCATATATACAGGAAGTGTTTACA

AAATATATTAAGAAAAGCAGCAAGTACAGATTCATCTACCTCGCATGGCTACGGTGACACGCCTATATTCACACCTTCAA

AAGGTTAACCCTGCTCTACTGCCAATACTACTATTACTACTACTACTACGTGCTAGCTAATATATATACCAAACAAAGAG

TTATCGTTGTTACTACTTTGTGATCACAGATCCATCATTTCATTCTCTTTTGAATTGCAAGTCTTTTGAGCCTATTCACT

TCTCCAACAAACCATTACCAGGAAGCTCTAAAAAAACCGAGCTGCGCAAAATTAGTTACACTTTTTGAACTTTTGAATTC

GCAGAGCTGAGTTCAAAATATTTTCCTCCTCAAAATATGTCATTGCTTTCAAAAGAGAACCGACCTCCCAAGAAAAGATA

ATGGAACGACCTACGGCCTTTTCATTTATCTGTTTTTAAATTCGAGTCTAAGCTTACAAGAGTACCTTCGCCACCACTTA

AAGACGATAATAACTTGCATCCTGGACTCATCCTTCTGACTTCTTGGCTCGAATAGATGAAAATCTTCTTCACCATCTTG

CAGAATTCCCTGCAAGTACAAAAGGATAGCGACGACTCTCGGTCAGATTCGAAATGATGCAGTAGAAAAAATTGAAATGA

AATGCAACTGTAGACAGGGAATTGATAGATTATGAAGTGTTAGGAAGACGCTTACAGCCATGGATCATCACCAACGAGCA

TCATATCCCCTTCGTCATCGGTAAATACAACTTCCCATTTGTCACGAGGGGATAACTCTCCTCTAATCTCAAACATCTCC

TCCAACTCATTTATAAGCTGATCGTACCCATCCAAAGTGGTCAAATCCACTGCGCGACCAACTGCTATCCCTTGCATGTG

TACCTATATTAACCAAATGAAGAATCAGAGTTAATTCTCTAGTGGATTAAAAACAAAATCAATCGAGGACATAAAAAGTT

TATTCACTAGACTGCCCAGGTTCAATACCTTAGTCCGACTTCTGGTAGCTGTCGAGAAGCTCTTTGGCTGTGCTTGCAAT

AGTACTTGTTGCTGTTCTTTGGATGCATTCGAAAGAGCAGATTGTGGATCTGACTCGGACCCGGATAAGGCTGAGGGGGC

AGTTATTTCAGCAACCACACTGGAACAAGTGATGTCATTTACTGGTTCCTTTACTACATTGGCTGGGGTATTTGTATTGT

TAATCAGCTCAATCCCAAATAGACGGTAGCCGCTTGGAGTCTCAGATTTCTTTTCCGATTCTACTTGGTCCTGCATCTGA

TCGTTCTTTGTTATTGGTGGATAGTCGGCGTTTGAATAGCCCAAGTGCCCTGACCACATCGAAGCTGTTCTATCATCTTG

AAACAGATCAAGGGAAGCATTCCCAAGGGACTCTCTCTGGATCCTCATGATGCCGCCGTTGTTGTTGACAAAATTTCCCA

TGTTTTCATTTTGTTTGGGAAGCCAGATGCTTTGACTTTCGCTACTTTGAATGTCCATGCCACTCGATGCAGTTACTTCA

TTGGATTGCGTCGCACCAGGATACCAGAGTGGTGAAGCTACAGAACTAGCAGTAGTTCCTCCTACTGACGAATGAATAAA

AGAACAGGATCAGCACATATTCCGACTACCAAATGAAATCATATGCAGGCAGAAACTGAAAGATCGGTATCAGATTTCAT

ACCATGAATTGGGAGATCTGTGGAGATTCGAAGCCTTTTATTCTTTGATACCAAAGGTTGAATCGGCTTTGGAGGAGCTG

GAGAAGCAACAAAAGGCTCAATCTCCCATGGAGAAACTCTGTCAGGTCTAGGTGTGGAGCAAGGTTCATCCCATTGAACC

TATAAAATAAACATATAGTTTGAGACAGCTGACACGGAAACAAGATAATAAACCACTGAACTACATAAATGTTATATCGA

AGATCTACCTTCAATGATCTCCACTTGGAATCTGACCACTGGGGAGAAACATCATTCGTTCCAACTATCGTACCTGAAAA

CCTGACAAAATCATAAACATCATTCAAATTAGATGGATAATCCCGGATAACAAATGATTCAAGAAACCACCGGTAGAATT

AATCTTATACCTTCTCTCAGGGGAATCTTCCCCTTCAAATCTCATCTTAAAACGCATTCCCACAGAAAATCCATTTTTCA

AAGACTCTAAATACTTGTTCAGGCCAATTATAAATTGACTTGTCCTGCAAAAACAGAAGGACAAATTACGTTAGCCAAAG

GAAGGGAATCCAAGAAACTTTCGTTAGAGGCGGTGGAATAGTTCAAACCTTGGCTTGTTGTAAACTATGAAGAGGGTTTT

CGTAGATACAGCATGAGATGCGGTGGCAAGCACTCCAAGATGCATGCTCTGGCTGGATATCACAGATGAGGGCATATTGC

TTTGTTGCCTAGCATGACGCCTTACTCCAACACGCAGATTCCCATTCTCCCCTCTATATATACATGAAATAAACAAGAGA

TCAGAAACAATCCTAAATAGAGAGATCGAAAGAGTTGGGAATTTAAAGCTGTGGTTACCTGAGAAAAACAAAAGTGTCCC

CTGCAACCAATCTCTTAGAAGTGACAAAGGTACTCCAACCAGTTGTAAGTAAATGTCTTCGAGGTTGACCTGTCCATGGA

ATAATCATCAGTAGTCAATTACAAGGACATGGATCTACACTAAATAATTGCTAGATCTACATCTTGTTCTTGGTATCGGC

TAATGTTATGATCATTAAAATTTGCAATAAACTACTACTGTTACATACCTCTAAAAATATGCTTGAATCGCCACTCGTAG

CCGTGAAGATCCTTGGCAATCAATTCCTGAGTTGGTGTTGCTTGGGTCATATCCTAAAAACAAACCTCACAGAATTAGAT

CCAGGCTAAACAAAAACAAGAAACATCAGCCTGAAAAGAAATGAAACAAGTCTTTACCAACGGAGGTAAACTCTCATTAG

CATGTTTCCGAAGAACGGAGAACCCTCCATGAGTACTAGTATCAGAAGCGGTCAAAATCTTGCAGAAAGATTGAACACCC

TCTCTTGGTGGTCCTTGTACACATGGATCAGGACTATTAGGTTCATTCGGCTAAATACAAAAACAAAAGCTTATAAGTAG

ATGTAAATGTAAATCAAACAATTGATTTCGACAAAAGAATAGACTCCAATAAAGAGATTCCTTACATCATCTTCTGGTTG

CAAAGTAACTCTTGCATAAACCTCATCGGTTTCTTGTTCAGCCTAAAGACATTCCAACTCCAATTAGCATCATAACATAA

AGATCTAACCATAGAGAAGATGAATACATACCAGCAATTCGATATGAACAACTCGACAAAGTATCTTTGATGGTAGATTA

AACAAAGGAATTCTCTGAACTAATTCTTCATTTGTTGATGCTTCTAACTACAAAAACAACCCCAAATAATGTACCAAATT

TAGCTCAAAATTTCACCAGAAAAACAATGGATGAAGAACAAGGGAGAATGAAAATCAGTTGAGGGATGATGATTCTTACT

TGCTCCATATGACCTTGAGGGAAATAGTAAACCCTTTCACCAATTTTTGGAACATCAACAAGAGGACCAGCACAAGCTCT

CCATAACTCTGAATATAGATCATCACATCCATAAACTACACAAAATCAAAGTCAACCAGAGTCAAAATAGACACAAAAAT

CAAACTTAGAAGCTCTGAATCAAAACCCAGAATTTTTTTTTTAAGAAAACCCCATTATCAAACACAGAAAAGGGAGCTTC

AATTTCTAATTTCTGAATCAAAACCCAGAAAAATTAACCCCCATTATCATGAAAACCATTAAAAAAAGAATCTCACCTGT

ATTATTTTGTACTTCCCCAGGAAAACCTGAGCTATTATCAAACATAAACGCCATTAAAACCAGCAAACTACCACCACCTT

CTCCTTCCTCTGTCTTCTTCTCTCAAACACCAAAACCCAAACCTCAAAAATCACCCCAACTTCAATCAACACCTTCTTTC

TACATATAAACTCCCAAAACCGTTTCTCCACAATCAATAAACCCACCACCACCACACTATAAATGCTCTTTAATCACCAG

CACCAGGCAGCAGCAACAGTTTTCTTCCGTTACAAAACACCCAAAAACGTTTCTTCTCCTTCTTCTATCTCCTCTGAGAA

AAAAAGAAAAGAAACTGATCCACAAAGCTTGAGCTTTAATAATAAACTTTGAACAAACAAAAAGAAAACAAACCCACTTT

TTAAGTTATTTTCTTCTCTCTCTCTCTCCCTCTCTCTCTCAGTAGAAAAGACTCTCTCAGACTGCAGACCTGCAAGTACC

TCGTTTTAACATGAAAGAGAGCTCTTTATCACTTTTTTTTCTTTTTGCTCTGTTTTTACTTATTATTTACTAACCGGAAA

ACGACGTTACCAAATCCCCAAACCAAACTTACTTCTGTTTTTCTCTCTCCGTCCATCTCTCGGATTATTGATTAAAGTTC

AGGGTTCAGATCACAGTTTTCTAGTAAGAAAAACAGAGCAAACTGGTAAACGGAAAAGTTGGGTTTCTGATGCCGTCTGG

TAACGGCGGCGCCTTTTTAATGTCTCTTTATCTTTTTCTGCCTTTACTCACTGACAATCTCTCTGCTCTGTACACATTTA

TGCGTTAGGGTTAGTAAACGGTAACGGTTTTCTTTATGTTAATGCCTTAACGATGTTTGCATAAATAGTTTTCGGTTATG

GGATACAAAGTCTATCATCGCATACTCGAGTGCATAATGAATATACATATTTTGGCACGTTAATTTATTTATGTGCATCA

GTACGATAGGAAGTCACATTTGTGGACTCGTGGTATTATTATTATTATTTACAAAAATATATAACAACATAATGAATATG

TAAGGTCAGGGGCGGAGCCAGGAAGTTTTGGATGGGAGGATAAATTTATATGGGATGAAGGATGATATCGTCGCACTATA

TTGCGGCCGCAAAAAAAAAAAAATTAGACGTATTGGTGGTGATCCATTATTACTACCATCTCGATCAAGGAGGTATTTAG

ATAGGTTTTGGATTGGTGAAGAAGCTTTGTAAAAGGTAACTTTCAGAAAGTGTATCTGCTAACTGTAAACCAGAAAAACT

AGGGGGGACAAATATTGCAACTTGACCAAATCATCTCGACTGATTTTGTGTTTACCGGAAACTAGGGAGGACAACTGTCC

CCCCTAGTCCCAATGTGGCTCCACCCCTGTGTAAGGTGATGGTAAAAACGTTACTTCAGCAATTTCAAGAGAAACAAGAG

GCATTCTCTGGATTATACGCTCGGTGTTTCCGGAGATTCGAAATGAAGTTACTTCTTTTTGCAGATACAAAATTTCAGCA

TATATTGAAAACTGTTAATCATCTGGCACAGTCGTTATGCCAATTTTTTCAATGCATGATGATGTAAATTTCAAGTAGAA

TGTTAATTCTCCACTAAGTTGTATCTCTTCAGACTTTTGTTTCAATTTGTTAGACTTTTTTAACATGACCCCTAATAAGG

GGTACCAGTTTACTATGAAGGGCGCCAAAACTTATATAAGACAAACTAGAATTTTTCTGTTTTATATAAATTTAGGTACC

CACAAGTAATTTGGCATCCTCATTAGCAATGTTGCTTCTTTAACCCATGTTTCCTTCCAAAATTAAAAGAAAAATAAATA

TGCACATGGCTTTTATTAATGTACAACTCCGTTTTAGCAATTCAGATAGCTCAGAGAGTTGAGCTTGTTATGCACACATC

TGCGTGATAGCAGTAGTAGATTTAAGCAAAATTACTTGAAACATACGAAAACACGCAAATACTTGCAATTATTAACTATC

ACTAGGGAGTAGAAAATGACCGATACATAGACATCCAGGAAATGAGATCCTAAAACTAAATTTATTCGTGCATAACTGGA

TTTTTGCTCTTACAATCTCCACAAAGATTATTTGTCCCGAACTCACTCCATTTCTTTTAATTTTTCATCGACTTCTTTTT

GTTGATAACAAATCTCGAACTCAAGTCCTTAGTATCATCAGAGTCACCCGGCAACTTAACCACCGTACGAGGAGCGACAT

CGGCCCTAAAAAATTGACCTGCACCGCGAATCTGGACACGACATGGTGGTGTTGTAAACTGTGGGTACATTTTGGTTGAA

TGTCAAGTTTTCAACGGACCTGCCTGACATATTCAGGTGGGGTGTGCTCTTAACCAGAGAGAGGAGGTACATGTATCACG

AGCTTACATAATTTCCCGTGAAACTTTCCGTCACCGTCACCGTTAAAAAAGAACCCCATTTTTGTCTGCTTCAAAGCAGG

CAGTCCTGCTCCTGCAATCAGAACTCCTCTTTCTTTCCCATCAAAGTCGACAAAGAAAATTTGTGGGACAGGACCTACAC

GTGGCGTTATCATAATGGTTTCAACTTGGTTCGGTTACACGTCCGTTAATACCGAGACCTTTTGGATTATGGGTTTCGGA

TGAGATTGAAATGTCACGAAATCACTCTTTGAAACAGGCGTTCATCAAGATCAAGATCAGTAAAAACTATTTTACACTTA

TGTTGATTTTTCTTGGGATTTATTTCAAAAAGAGATCCAACGGTAAAAGAATAACCCTTCTAACATTGAGATACATGATG

GTTAAAATGGGTCCCAATCGTTTTCCCAGTAGAGATTTTTTTCAAGATGATTCTCGAAATATGCAATATTAGAGATACGG

TAATATCGATTTTCACTAGTAATGCGGACAGAATCGAGTGAACTGTGAATATACACTTAACTAAACCAAAACGGATAGCA

TCTAAAACCACACGGCAAGATGGTTTATGTTGTGGACCATTTTCAACTTTTATTATGCACGTATGGTGAAAGCTATTTGA

GAGTCTTATGACATCATCACTTTTGTTTATTTAAGTACCGAAAATGACTTAAAAGTAAAATAAATTAGAGTTCCAATAAT

ATGCAATTTAGGGCTTTAAATAAATATGAAAAAGACTGGTTAGTTTGACATGTATGTAGCCCCCATGGGTATCACTTTTC

ACCATTTGTGTCAAAAAGGATTCTTATTTAGGAATTGCAGCCAATCTATGATGTTGATTCATTATTCGGTGTTGTATAAC

ACAAATTACACAGTTTTATTTATGTAAACTAGCCCTTAACCCATGCCGTTAATATATCGTATAATTTGTGTCCCGAATAC

TTATCAACTCCTTATGATTTAATATGTGTTTGTCATAAAAATAGTTGGGATTTTCCTCTCTTTTTTTCTTTGTTTTTTCC

TTTAATATTTTACCGCCGGAGATTTGCTCAAACGAAAGAAATATAATCTTAACTTCCAGACACTTTTTCATTTAGGTTCA

TTTCTTATAATGAGATAATGCTCTACATGAACATGGAAGTTTAAATATTTTTTGGAAATATGTCGTCAGCTAGCAGCATC

TCATCTCGGAAATCAACTAATCAATATCGCCTATAAAAGTCTCCACATGCTCCTATTTCGCTCCAGCTGACTCATCTATC

ATGGTCATATTGTCGATCCTTCAAATAGGCAGAGAATAGTATTTTTTGCATCGAACGAATAATACAAACTATCCAATGTG

ACATGATTTTCTTGTAATTTTTCAACACCTATTAGATGACAATGTTTAAAATGTGTACCCATTACCTTTTCAAAGGGATT

GTATATATGCGAACCTCTGCCATTAAATCTTCAACAAAGTATTGTTCCTAGCAATTCGTCTTAACGAAGGCTTTGGTAAC

TTCAAAAGTCTTGGACATTTTTTCACTTCTGTATGTTCCTACCTTCACTTGGAATACAATATTCCGAAACATTACCATAT

ACAGAACCTTGAACACATATTGTACTCCCTGATCCTATTTCAAATTAATAGGTTGTCAATAGATCCTTTACCAAAAGTTG

ACATATTCTACGTTGAATATTTAAAAAAAAAATTGGTATTGTGGACCGTTGCTTTCTAAGATTGTAGTTTTCAAAATTGG

CCTCTTCAAGTGCCACGGCTTTCGAAGTTTATATTTCTTGTTTGGTTTTTTTAATATTTTAGTTAGACATATTGTGTTTG

TTACTAATTTTAAACTTGATGTCATCTTCTACTGTGAACTTTATGTGATTTCAGAATCAAAATAGTTATATCTCTTTCAT

ACGTATCAAGTAATTCGTTGTATATCCCAACAATTCTTCTACAACAATTCCTTTCTTTTGTCGTTTCCGGACCATCAGTA

GCCATTATTGTCAAACGCAATTTGTTTAATATTTGTTGATGTTATGTATTTTCAATGTTCATTTTTCTTAAATAAATAGA

TTGTGGAAAAACTAAAAAAAAAAGAAAAAAAGAATATAAGTATTACATTGGGTCTAAGTTACTATATCAAAAATACTCCG

GACACTGCTTCTGGTGCCTTGTTTCCTCAAATTTCTCGTTGCATTCTAGACAGTACATGCAACACCAATCATTTGCGAAG

TGAGCTATTTGCGCAAGTGTGGTCAATGTAGCCCTTGCCTGTAATAATAATAACAACACGTCAATTTTTTTTCAAGGCAT

CCAGTTGTAAATGGTATGAATCTAATTGATTTGAATTATACCTCAATCTTAGATATCACCTTCTTCTCACATAACATAAC

AATTGAGGAGGGACAAATCTTGACCACTCTTTACATAGGCAGAGTTAGCCAAACTATGTTTTGCAATTGTCTTACGTGAA

AATTATTATTAATCCATATTTTAGTCAGCTCCTATCGAAGCCGATCTAATTAGTCCTATAGAAAGTTGATCTTAAATATT

TGACCTTATCTCTTGACCAAATTATTGAAGTATTCTTAGTCATGTTCTGCTTTGTGTTATCTACCCATCTGTGTCTACTC

TACCTTCTAACCCAGTGAATGAAACCTATACATATTGATCCGTAATTTGATATTGTTGATTTGTCATCAAGAGGTCACAT

GATTTGAGAATGTAAAATATAAAAAATATTCTATAAACATCGGAGGTGAAATTTAGATTTATGATTTAGAGTTTTTTGTG

ATGATTTTGATGAACATACATAATTTTCAAACAAATATTTTAGAATGCAAATACATTTTCCCGGTCAATAAGGTTTTATA

TTGGGAATTGAGTGAAAGTTGTTGTCATTGTAGGTAATTTTCGCGAAACAACTCGACATTGATGGTCTTTCAGGAAAAAG

AATGTAATAGTACTTGAGAGAGCGATAATGTTCTCTTACACGGGTACACCAATTGCCTTTATATAATACGGGTGATCGAA

CAGTCATGATCATATTGTCTTAGATGTTCCGGTATCATCCTCGAAATATTTGGTTTTTGTCCTTAACAATCGTGCTAAAA

ACAACCAATTACATCTTTCACAATACAATGTTGCATGTTTTGTGAAGATTATTTTTTTACTTGCTCAATAAAACTCACTA

AAAAGAAAAAATTATTATTATACACTACTATTACCAACACCCAACACCATAAAAACCATCCACCGCCACTATTTCTTCCA

CCAATAGTAATGTTACCGCCTCCACCATTCACAAACACCCGCCACCGCTTATACTAACCCTACTTCCACTACCATCATTA

GAATTAATATAGTTTTTAGTAAAATTATTTAGTAATAAAATTATGTATTTATGTTAGATTATTAAAGAGACATTTATAAT

AGAAATTTAATTAATCATTATGAACAATCAATGAGACATTAATTAATAAAACGTTTAAAAATGATTAAGTTGAACAAACC

CCAGCGATAAATTTATCTTTACTGAGGACAAATCCTGACCACTATTTACATAAAAAAATCAATAATAAATAATTTCATAT

GTTCTGATCTTGATTTGTGTCGTTCGTTTAGTTATTACATAACATGGCAATTGATTGCACCAATTTTTTTAAAATGGATA

TGTGTTTTATCGAGTTTATTTTAACATTAAAATGATCAACCACGTTTTATACTTTATAATATAACGTTACACATTTTTTG

GCGTTACGAAATTAGCATAATATTTTTTTGTTCACCAATACCCACCACCACCATAAAACCATATGAATCCACTTTTTTAT

CCACCACCACCTAGTGGCGGAGCCAAGGGTTGACTAACCTGGCTCTAACCCCTTAAATTTTTAGTAACTACATAATTTTT

TTAGTTTATTTTATAAATAGTACTTTGTCCATATATATTTATTGTTAAGTCCACCAAAGTTTACATGTTTCTTTTTAAAA

AATAATAATAGACCTCCTCAATGTCAATTTTTGACTCCACCACCGCCACCATCACTGCCGCCCATCCCATGTAGGTTGTT

GGACGTAATTCTAATTTTCTATCACAAATGCCATGTATGAGATTTATATTGAACTAAATTTCGCCAATTGATTAGTTTCT

TTATTTTTTTCCGTGTATATATTCTTTTTTGTATTGCGATCAATTTTATTTGAACATTAAAAAAATCATGATTTATACTA

TCTAGTACCAAAGATAATATGGGTACCATTTGGGTGATCCAACTACGATCATGAAAATATTGTCTTGTATAATATCATCC

CTGAAATATATGGTACCTTCCCTTATCAAAACTGTTAAAAACAACCAACCACATCTTCCACACTACGATGTTGTAATGAC

AACATAAAATTAATATTGTCCATCACTACTCACCATCATTCACCGTCACTACTTATTCCACCACAAACCAGGGGTAAAGC

CATGATTTTGAGGAGAGAGGGACAAAAAATTCTTGGCAAGTTGGATATATGTGAAAAATGGACTTTGTATCCCATTTTAC

GATAAAAGAACAAAAATAAATATAATTGTGCTTGGCTTCGTCCTTAACCACCACTAATCTTTCGGCCCCCACCTCTTCTA

CTACTCATTGTGGCAGAAAATGTTTATTGATATTATTTATCAAAATTACTTATTAATAGAAATAAATATTATAATTAATT

AAGAAAACATTTATAATTAAAAAATGATTAGTCATTTATTATGAGCAATTAGTGAAACACTAATTAATAAAACACTTTAT

AAGGAATATATTAATTATTATGTGTAATTAGTGAAATACTAATTAATAAAGCAATTATGGTATTTAAGTTGAAAAACCCA

CCACTGTTGATTTATCTTTTCTATCAATTCCAAGCAACGTGGAACATTGGTTAGAGGCCACGCTTATGACACAATAGGTC

ATGAGTTCGAACCTAACCTTTGATTTATCTTTCCCACGTATCCTCTTATAAATGAATTCCTAACATGTTTTATGTATGTA

TCCTTACTTATAATTGATTTTTTTTCACTTTTCACTATTTATGTCGAAATGAATTCCTAAGTTACGTATCCTCTTATAAT

TGATGCCGTATTACACAAATTACACTGTATTAAGAATATGTAAATATCAAGAGCACGGAAATAACCTGGATGAGACGTGT

TATAAACGTACGAACAAATCTAGTTTAAAACAAGTTGATTCGCTCTACTGTTGAATCATGGCAGTAGGGTTGATGGCATG

ATTGTTTATTTGGTTAATCTTGTGATTACAAAGGACAGGTTAATTTTAGTATTAGATTAAACTGTCTGTCAAACTCCCTA

GATTTAGGACCACAAAACCCTTACCTGTTAACACTCAAACTAATCACCAATTATTAACCTTAACATTAACATTGTATCCC

TTTCTTTTCTCATGATGTCATTTCAACCTATAATTAGCGCATAAAAATTAAAAACTATCTAATAAAGTATGAATTTAGTC

GAATTAGTCGAAACTTACGACATCGATCGACAAATTATCGTCCGCTTATTGGTTAACTTGGATAAGAATCTTATTTAGTG

TGTGTGTGATTAGTATTTAATCCAATAATTTACAAAAAAAAGAAAGAAAAATATTTTGTTTAATTAAGAACATGGATTAC

AAGCAAGGACAAACCAATCAAGTTGTTAATACGTGTGTGGGATTGTCCTTATAATCAGGTTTAGGGTCTGCTTTAAAAAA

AATTTACAATTATTATACTTAATTAATACAATATATTTATTAAGGGAAGTGAAGTACAACAATTTTGATATTGCCTCATG

AATGAAACTAAGAGGACGAAAATGACTCATTCTTATTATCAAGGGTCATTGAATTTGGTTAAGGGTCAAAATACTTTAAT

TTCCTCATCACAACTTTAAACCCTAAAATATTGTTTTGGTATATAAATGGGAGAAGCACATTATTGATTATTCTTCCTCT

TCTCTAGCTAGCATTCAACTAGCAATTACAAAGTTTTTTTTATTTTTTATTTTTTTTTATCTATTTTAAAATTAAATTAC

ATATTTTTTAGTTGGTAACCTAGCGACAAGCTGGGCTCCAACACCTTTTTGGATTGCAATACCTATTCTATGAAAAATAA

AACTACCAACTCCACTACTAGCATCATTATTAACTAAACAATTCTTCAGACGCTTGAAAAAACATAAAGTATCATCTCCC

AATTCCCCTAAGGTAGAGAAAGCTAGAACACCAAGTCCATACCCAAGTGAAGCACATTTGTCCCAGTACTTAGTGCGTTT

CCATGCGACAGCATTGGAAATAGCTTTGCCAGGAACGAAAGAGCGAATACCATCACCAGTGAATGGAGAAACCACTGTGA

CATCCATGCACACATTTTTTTTTATTAGTTCGTAAAACGGTTAAAATTGCACACATTTGTGTTGGTGAATATTTTTTGAT

GGCGTATTTGGTTGGAGGAAATGGGATCAGTTTGCTAGGAATATAATTTCATTTTCATGTTTGATAAACAGTAACAGAGG

GGGAATAGAAATTCAATTTCCAAAAAAAAAAGAGAATTCCAAAATCTATATTTTATTTTTTTTTGATCATCAAACCACAA

GTATTTCATTCCATAAGAAACGTTTACATGATGGTTTTAAAGAGAAACAAATACATCATAACATGAAAAAAATACAAAGA

AAGATACAAAACGTAAATAAATCGCGAAGAGAAAGTGCGATCTACCACGATAGCACCCAAATCTTGTACTATGAGATTGG

AGAAGGAATGGTATACGAAATCAAAATCTCTATGGATGATCAGATAAGTTCTTGGGAGTGGGATTCCCTTTCAACCATCC

AAACACGCCACTAAAGCGAATTGGAGAATTTAATTTGTGTTTGGGGTTTGATTCAGAAGTAAATTATCAGGGTTTTTAGT

ATTTTCATATTCTCTTTTGAATATATACAAGTATAGCAACAATTACCATCTAGAGTATAAAGCACAGTTTGGCTAAGTTA

GCCTATATGAGTGGTGGTCAGGATTTATCCTTGGCAAAGATAAATCAAAGGCTGTGGTTCTCCGTGCCACACCCTACAAG

TCGCACATGTGGTGTTCATTCCAATATTATGAATTAAAAATGAATACTCAAGTCAACATATGAAATTTAAAAAATTATAA

ATTTTTTATTCGAACTCATGACATATTGTGTCATAAGAGTGGCCTCTAACCATTGTTCCACCTTGGTTGGAATTGATAGA

AAAGATAAATCAACAGTGGTTGTTTGTTCAACTTAAATATTAATTAGTATAAGTGTTTTATTAATTAGTATAAGTGATTT

GTCTAGAGAGAAAATAAATCTATGGCTACGGTTTGTTCAACTTAAATATTAATTAGTATAAGTGTTTTATTAATTGATGT

CTCATTAATTTCTCATGATAAATGATTTATTAATTTTACATCATAAATATCCCTTAATTAATATTAATACATATCTCCAT

CAATAAATAATAATTTTGACAAAAATTATTTCAATAGATATTAGTGGTGGTGGTGGTAGTAGAGGAAGAAGTAAGTGGTG

GAAGTGGTGGAAGGTGTTACTAAGTGATTGAGGCGGTAACATTAGTTTTGGTGGAAGGAATAGTGGCGTTGGATGGTTTT

AATGGCGGTGGTATTGGTTTTTTCTTTTTGAATCAAAAATATTAATTTTTTTTACATGACGGATAACATTATATTGTGAA

AAGTATAGTTGGTTGTTTTTGACACGATCAAAAAAGGCGGATAACAAATACTTTTGGGATGATACCAAATCATATAAGAC

AAAATAGTTGTTTTTGATACCACTGCTATCTTTGGTAACCGTTCGTACGCATGATTTTCAGTATTAACAAAAAATATATA

GTTGAATAAATTAATTGACGACCTAAGTTTGTTAAATCTAAACTTTATACATGACGTGTTGGATTAAAAGTTATAAGTGT

GCCCAATATTCTACGAGGGATGAAAAATGGTGGTCGTGGAGAAAGAAGTGATATCTGGTGGTTTGTGGTACTGGCCGGTT

TTGATGAGTAGTAATTGACTATATTAATTATGTGCTGCTACAAAAGGATAACACTATAAATTAGAAATGGGGTTGATCGT

CTTAATGCTAATATAAACTCGGTTAAAATACATGAAAATTAAGAAGAAGAAAAAAACATTTGGATGTCACGTTATATAAT

AAATAACACTAAAAACTACATATTTAGTATCAATATTTCTCCATCCAATAGATAAATCTAAGGTTGTATTGAGATCCCGT

AAGCAGAGGGCGAGTTGGTCTTGGATTTCTGAATGATAATGAATATAATAAAACGTTCCCTCTCGGTTTCTCCAATGAAA

ACCGAAACCTTCTAAAAAATAATCAATCAAACAACACTTAGATGGAACGCATCACCACCAGTGTAAATTTCTCTGAATTG

CTACATATCCAGTGAAATTTCTACAGATTAATCTGGTGATTTCGTCCTACCGTAGTTTTTAACGAGCCATGCAATTAAAT

ATCTAATAGTGATTTTAAAGGTATTTAATAATTACAAAATTCTCTTATAAGTTTTGTTTATGCAAGTATTGATTTAACTT

GGGTTCTTTTATATTTTTGTGTTAAGTTGATGACTTTTAAATTTGAATACATGTTTGGTTAACAAACTTTGATACATATA

GGGTATTTTCTTTGTAGAACTGATGTTAGTCTCTCATTTTTCCCGTTAGCACTTTGTTTCGATCCCGTATTTGTAGACAT

TATTTGACTAATGTTGGGGGTATTTGTTTAATCTCAAGGCCGATTAAACCCAATGTGTGTTATATACATTGAAGGCGACA

TAAGTTTTTGTCGGATCTACGTTGAAACTAATAATTTTTCTTCCGAAGGTTAGTTGAATTTATGTAGATACAAATATCAA

TATGACTATGCTATTTAAAACTCCAAATTGCTTGAGTAACACTAGATGTGAGAACGTCATTCATGCACTACTTAGCATAG

TTGGTGTGAAATCTTTTTATCCACGAATTTCTTCAATTTTATGTTATCAATATGATTGAGAAAGAATCTTACACATGTTT

TGATTTACTATTATAAGTTTATTTTTTAAAATCACATGTGTATATCTCAGCCTTCCAATAGCCAAGGGTTCTTCATTTTT

TCAACTCACATGGGTTTTTGCTTCTCTCCTTAGAGCCTGAGTTATCAATAATTTCGGAACAAATGTATTCATAAACATAT

GGAGTTGGATTGGCAGTTTAGGTATATGTAATGTTAGATGAAATATTCTTTTTGAAGTTTGAAATTATGGTTTGATGGAA

CAATAAAGGGGGAATTTTTTTTATTTACTGTATTTTTTATTTTATTTTCATGTTGGCACAACATTATTACTCCGTCCAAA

TAATAAATTGGAATAACATGCTTAAGTAAAAAATAAAAAAAATCAGGTAACCCATTTTGTTTCCCAAAACCTATTTGAAT

ATCAAAATAATTACTCACGAATAAAATGAAGCAAGAGGAATGTGAACAATATCGTATTAGAGACAAAACATCCCGTGCCG

TTACGGCACGGGTTAAGTTCTAGTGGTGATATATAAAATAAATAATAACCTACGTAAAATCATAAACTATCTGGTTTTGC

AATGTCCTTTAGTAAGGTAGTAAAATTGTCAAGTAATGATAATGATGATCAGAATTCAAGACTCATCAGCATCGTATTTC

TATTAGCATCATATTTCTATCGACATATCGAGAGAAAAAAATGTCATTTTTTTTTTTCCACCAACAATACTATATAAATC

AATCTCATAAATTTTTGCTCATGTGCTCCTCGAGTTTTTTTTTTAATTAATAATATCACAATTATCTGTTGCAATATGTT

CGGAGAAAAACCTTTTCGGTTCCAGAAAAAAAATTCCCATCACATCTTTGAATTTTTAGATGACGAGATATTTATACGAG

GAACAAAGTGTAACATTAACAATAGGTAAAATATAGGGAATGTCAAAAACTTTGAATTATACTAAGAATCTTGGAACTTT

CATCCTTTTTTACCTTCCTTCGGATTGTTTACACAAGAACTTTCTCAAATGGAACATGCTAAAGTTTGAAAAGGGGGTGT

GATGAGGAAATCCTTACATAAAATAATAAAAATGGATCAATAGGAAAATAGTTTTGAAAAGTGAGTTTTCTCCTATCTCA

ATAGAAATGACATCCATCTACAAGAATGAGGGCATCCTTCCTTTACTTTATTTTTTGATGATAGGGAGTGACCTATTGCT

AGCTAGGAGGAGAAGGAGTAAGTGATGGTGTAATGTACACCCCATGTGGAGACCAATTTGGTTGGTGTTCCTTGGTTGCC

CTTAATCTTGACCAATCTCATTCTTCTCTTTATACCCACCTACAACATTTTTTTAAGTACTAACTTTATTTTTAATTGAT

GTCATTGGGGGTTCCACAAAGTCACCCCATAAGTTTGAACACTCTTTTGTGATTAGACCAATAATTATATGATTTCATTT

GCTTAATTAGCAGGATAATTAGCATTAACAAATTATTAATAGGTACATTTAGGATCTTTTTTTAATTCATGTCTGCTCCC

GCAAGAACTCAATTGGGGTTTGGAATCTTATAAGTCATTGAAACATGTGGAGAGATCATTATAGGCAAATTAATGAAGTT

TTGTAAACCTTTGATTTATTGGAATCCTCATGTATATACAATAGTAAAAGCTACCATTAGAAAAGATAATGGAATCAAAT

TGTGTGTTTAGTGTAGCTAGTTAAGGTAGGTTAGATATTAGGATCTTGACTTTGGTCGATCATATTTGACCTAAATGATA

GACCTTGGTCATCATCAAACAGTTGGAATCAAATCCAACCAATTAGGGTTGTTCATTTGACTAACACTAATTGGTTCAAC

AAATCTTTTCTGCCTTCCAAACATTTAATTTTCCTTTTTATTCAATTTATTAAGGTTGGTAAGATTCAATAATGTGACCT

AATTTTTGCTATTTCTATGCCAATAAAACTCTCAATTGGTCCTCTGATTATATAGTGTACAGATTGGCTTTAATATAAGG

TTAGGACCAATTAAGTTAGCAGCCTTTTGCTAAGACTATTCCTTAATAATGCATGATTGCTATTTGTGTAAGGAGAAATT

TAGGTCATGTTTTACCCGGCATTTTTAGGGGCGACCCAAAAGAATTAGGGACGACATAAAAGTACAAAAAAGGTCACCCA

AGCGTAAATTTAAGCTATCCCTTATCTCCGATTTTTTGAAATGGCTAATATGCCCTTTTATAATCGGTAGCCAACACTAC

AATCGGTAGGTTAGAAGACAATCGATAGGAAATATAGAATCAGTAGTTACCGATTAGTGTGTTAATATTTCGGTTAATAG

GCCCTTTTACAATCGGTAGCCAAAACAATAATCGGTAGTTAAAAAAAAATTCGGAAGCAAACTCTTGTAATCGGTAGTCT

GGGATCTCTATATTAGTGTTTGGAATTTGTTTTTGAGTCATTGTGAGATATGTCCCTAACTTTCGATTGAGTGAGAGAAG

AAGAAGGAAAAAAAAAGTGGGTATATGTTGCCAATTTTCGAAAACATGGATGGGTACGAAGAAGAACGTGAACTTGTTGA

AGATCAAGGTTTGCTACCATTTAGAGAAGACGATTTGGGGTATATGTTGAATAATCCAAACTTTTGTGGAGATGAAACCC

TAGACGACGCTTTCATGGAAGACAAATTAGAATCACCCAAGATCGAAGCTAACGATGCATCAAACATACATGTATTGTGT

TAACAAACTCTAATACCCATGTTATGTGCGTTTGTGTGTGTTTTTAAAAGAGAAAATTCGATTTATGCATGCATTGAATG

CCATGCACGACACATAATCTGTAGCTAATGGAGTACCATATCTACCGATTACAAGAAACTGCTTCCAAAATATCTACAAT

CGGACCTTTTTTGATGTATAAAACTACCGACATTGAAAAAAAACCCAAATTGAAAGTTGTAATTTTTCATAATCGGTAGT

TTTATACATTTACCTTACTACCGATTTTCAAAAGTCGCCCCAAATACGAATTTCTCCAAAGATGAAAAAATTGTAATTCT

TGATATTTCATAATCGGTTGTTTATGAATGCATTTATCTTCCGATTATGGTAGTTGTCCCAAAATCCAAATAATAATTTC

CCTCCATCCTGCGGTAGCCCAAGTATTACACCTTCAGGAATATCCTTCACCTTCAGTTCTTTTGGGACAAGGTGTTTCGC

TTCCTTTCGAGGAGGATTTTTCTTTTCCGCTTCTCCTTCTTCTTGCGAACTGCTAGGTTTTCCTACTTCTAGTACTCTAG

CTTCTGGTTGAATTTCATCTTGTGATTGTGGAGCACTTGGTTCTACAGCTTCTTGAACAGCAGTTGGTTGAACACCTTCT

TAAACTGGTCCTGATTGTGGGGCAGTTGGTTGAACACCTTCTTTACTTCCTTGTTGAGTTCCATCTTGAGCACTTGAAGT

GGATTTAGCAGTCCTAGATCTCCTTGCACTAGCTGTTAAGTTCTTACGCCTTTCCATCCTTGCACTATCTGAGACAATTG

CATCTTCCGGTTTTTTACATCTCCTATCCCTAAACAACAAAATAAAGTACAATAGGTTCGTAAACATATGTTACCAAACT

ATACAGTATATACAATTGGAAGTTTTTCTACCGAATTTGTTTATCCAATCTAAAATATCTATCATAATCGGAAGACATAT

TTCAACACTCTCTTCTGAATACACATAATCGAAAGACAAGTTAAAGTCTATCTTGCCGAATAAACAACAATCGGAAACCA

TACAAAACAAATGTTTACCGAATATGCATCATTTTGTGTCCAGAAAATACTTAACTTGTTGAGTCTAAGAAATCGGAAGC

TAATTACAAAACATATCATCTGATTACTGCGTCTCTATAACTTTTAGTTCCCTAATTTATTCGGTAGGCAAAATCACATT

TAACTTCCGATTATGGTCAGATTAGGGCTCAGTAATAAACAAGGTTACACAATCGGTAGGAAACCTAAAAATTTATAAAC

CGAATATGACCATCGTTCTTCTGAAATGAAGAATTCGACCACAATCGGAAGTTAAAAACAATTACTAACCTACCGATTCT

GTGCAATAACCGAAAACCCTAAAAAAAATATTCATGGATTCGACGAATTGAAGCAATAGAAACCAAAAAAATGGTAGAAT

CTTACCTAATGGGGGCCATTTGAAGTTGCTTGAGAGTTTGGCTATCAGAATCATCTACATTAGGGACTTCATCTTCTTCA

GTGAGAATCTCGTTATTACGGGCATGCAGTCCAATGTTTGGATCAACACCACGAACAACGTTTTTGGTTCAAGATTTCTT

AACGATAATTCGATTCGACGATTAACGATTAACGATGTATTGAAGCAAAGTTGAACGAGGGAAAAAAGTTTTCCTCTCTG

CAATCAGAGGATAAGATGAGAGAAGAGGAGTTGAAGTTGAAAAGAAAAGAAAAGAAAAGAAAAGAAATGGAATTAGGTTT

ACATTGGCTTTGTCGGTCAAAGGATATAATTGGTTGTTAATAGCCATCTCATTGCTACCAATTTTTACATTAGCCCCAAA

TTTTAAATAACCAAAATTATAAAGTTTTCGAATTTCTCGACTTTCCTGATCGGTAGTTTATACATTTTATTTTCTTCTGA

TTTTATAATAGCCCCAAATTTCATAATAGCAAAATTCTAAAGATTTCGAATTTCTATACTTTAACAATCAGTAGTTTATT

GAGGTTCATGTTTCTACCGATTAGGATAGAAGCCAGATACACACTTATATATTATTCGGTAGTATGTTTGCGTTACTTAA

CTCCCGATTTAGTATTAACAATCAGACACAAAATAACTTCCGATTATGAAAGGGCATTATCGTATTCTACTGAACTTTAG

GACATCCCTTAACGTTGTATATGGGTTGGAGATAAAAAAATCGCCCCTAATTCTTTTGGGGTCGCCCCTAAAAATGCTCG

GTTTTTTTGGTCACCTAAAATTGAAAAAAAAAACTTTTGCATATTCAAAATGGTTTAGTTACTTTAGTTCTCGCTGAAAT

TCTAGCCAGGTACTAAATATCCATAACAAATTAAAAACACCTCATAAAATTGCTTTTAACTCCCAAAACCTCTCGAGTAT

TTTAGGCTTTTGAGGATAATGTTTGTTCTTGGGGAGCCTTGGTGCTAAAACTTCCCAGGCAAGGGAGGAGCCAAGGATTG

ACTAACCCTGATTGTAGCCCTCTCTAAAATACCAAAAATGATGTAAAAGCCTTTGAGTTAGAAATTTTTTTAGCGTTTGT

TTATAATAATTCCTGGTTTCGCCACTGTCCATAAGCCTCAATAATCTGGACATGTAAAAAGATGGTTATTCTTCCTTGAC

AATGCAGCAACAAACTTACCCATTGTTTTCAAACAAGATACTTGACCAAACTCAAAAAATCAAATCCTAAACTAATTAAA

CATGTAGTGACCCAATGCAATGCAGGAACTTTAAACAACTAAGTTACATGCATACATACATAAACTTAGTTACTAGCTGC

TGCTAGTAGGAATATGTTCCAATCATGGAGTTAACTTGTTTTGATGGGATTTGGGACCACTGTACCACTTGTCTAAACAA

GCTGTTTGTTTAAGTTTTTAGGTCATTTTGTTTTTGCTTTTGGCTTTGGGAATGGTGGGTAGTGAAGCTAGGGTTACTAT

CATTGGAGATCAAGTCCCCTTAATTTTAATTAGGTATTCATTTATTCAAAGACAAGGTACCAATAATATAACCCTGATTT

ATTTTCTTAGTTAATCAATAGTTAGATGTGGTACCACCATTGTAAGGCTCAAAGTATATTCTTTAGTATCCACCATTATG

AAATGTTTAATCAAGGAGATAATAACAATCATGTAGGATAATCATACTCGGGAAATCCGGTTTATAATGTTTTATTTGTT

GGGTGTCAACATAACTAATTAATAGGCTAATTTTGATTCTCCTGCTCGAAAAAATAAAATAAATAACAAAATCCATATAT

TTTGGAAATGTGATATTTTTGTATCGAAATTGTGATTTTGCTTGAAGTAGTAAAAGATTTTCAGGTTGGAAGTTTGGAAA

AAAATGAAAGAGTCATTGAATGGTATTGAGCAAATGACAAACAATAAGACCTAATTTTTTTGTTATATATTATGTTTAAA

ATTTCTTTACCTAATTAATTAATTAAACAAACACTTTCAATTTAGAAAATTATATTAAAAGATAAAAATTACATACTTCT

TCTTCTTCTTCAATCCTATTAAAAATAAAATCTCGTGAACCAGGGGTCATTAATTAATTAATTAAATTTTTTTTTTTTTG

CTTCTTCTTCTTCTTCTTCCAATTGGGTAAGTGGGTGAAAACAATAGGAAACTTCCAACTTTTGAATTTTGGTCCCAAAT

TATATGTTTGTAAATAATCAGACAATGACCAAATTTTGGGTACTAGACTTGTTCAATGTGAGTTCTTTTAATTTTCTTAT

CAATTTGGAAAAAGTAGGCACTATGAATTACACTAATACAATAGTAGTACTCATTCCTTTTTGGGCATTGCCCATTTGAT

GATGCAAAAGATGCAATATCCTCCCACCAAGTGTTTGGAAATTTTCAATAGGAGTTGGGCTTATCATATACATGTATAGA

GATTAGTTTGGTTCTTGTCTAGCTAACCTCTATGTTCAAACCTTTTTATCTATCTAACCTAGTCTACCACAGTTTTAATT

TAATAATGTTTGAAGAATCAAAGAAGCACCAGTTTGAGGGTTGTGGGGATGATCGATAACGTCATGGTTGAAGAGTTAGG

CACGATAATAAGCTTAATCTAAATGCTTCTGCGCGGATGTGGTAAAGTTGTTAGCCAATGATATTCATGAGTTACAAGTT

AAATTACAGATTTATTTTTTTCTTTTGGTAAATTTATGGTGGCTCTTCCTAAACAGCCGTCTTTGTCATATTAGTAGAAC

GTCATCAAGTGGTTCTATGGTCTAGCGGTTAGGACATCAGACTCTGAATCTGGTAACCCGAGTTCAATTCTCGGTAGAAC

CTTTTTTTTTTTTTTTTGTTTGTTGGGTTCTATTGTTCTGATCAGGGTTTGATTTTCTTTTAGAGTATGTTTTAATGGGT

ATGCATCATTTTATAATTGTTGCAGAAGCCGTGCCATACGACATCTAGAAATGTAAGACAGGTAGTTCAAGAACGATTCG

GCTTATGCATTTAAAATTAACACATGATTACTGTGTTTATATGAAAGACGATCATACATGTAAACTTACATTCAGTTCTT

CGATTGCTTTTCTTGATTGCTTTTTTCCTCACATCCTCATGCAAGTCTTATCTAGTTGATTTTGTTTTTCAAATGGAAGT

ACCTTGTCATTTCACAAAATAGAACTTCAGCTATTTTACACACACAGCTATTTATTTGCAAGCGCTTCCAATGTCACCTT

AATAAAGGTGTGATTCTGCTTCCTTTTCTCATTTCGTCTTGCCATGGAAAAAATGTGAGTTTTTGGTATCTGAAACCAAG

TTCTAAATTAAAAAAAAACACAAGGACGATCACAGTTTTATTTTATGCAAACATCAACAACATAATCGGGAATACGAGGT

AGAAGTAATACTCTACTCACAAGCTCATCAAAATCATTTTCAGTTTCCCCCAATATCTCATTATGTTTTCCAAGTTAAAT

GGTCTCAAATGAAAGTATATAAGTGCCCTTCTGAGTCGCCTTAAAAAGAAAAAAAGTTTGCTCTAGCAGCTGGTGAATCC

AACGGTGGTGGTTCCCAGTTCAGTTTTTTACTCCAAGGGCAAAAATGAAATTTCCATCATAAACCACCACCACCCTTAGA

TTCACCCATCCAAGGGCTCTAGCAGCTGCTAGAGCAAAATCGCCTCCCGTGTGATTCTACTTCATTATTTCATTTGAATG

CATTTTTGACTTACCATGGAAAAAAAAAAAAAAGTGATAATAGTTCTGTTTTAATGGAAGTCGAAAGACTAGAAGAAACC

AATTGGAAATCAATATCACAATTAGATACATGGAGAAATACTGTAACTATACAGCTGCAGTATAAAATAATTTCCAAAAC

GCAGGGATGAACGTCTAGCCAGCTATGGCAGTGCAAAGCTCACAAGCTCTTTGCACAATCAAATCTTAGACACCCCACAG

TAGATTCGTTGAATCTTTTGCAAGTCCCAATCCAACCAGCCCCAAAGATGGGAAGCAAGGTTGTTGCAATTCAAGTACTA

AACTCAAGTCATGATGAAAAATAAAAAAAGATGAAATGAGTAACAACAACTACTAAAAAACAACAGTCTAGTAGACTAAG

AAGATGTATGAATGATTTTCCATATGAAGTGCCCAGTACTATATGAGGTGACCGACTCCGACAACTTTTCCCCAAAGAAA

AGTGCCTCAAAAACTGACCGACTCCCATCAACAACACTATCTATACTCTCGACGACAACTTTTCCCCAAAGAAAAAGTGA

CTCTATTTAACTGACTCATCAACAACACTATACTTTGGGAGGGACACACTTTTTTCCTCTTTCTCCCTTTTTCCCATTAC

AAAAGACACCAAGTAAAAAAAGACAGTGCTATGCGCATATTTGAACCCCTCCAAATATGAAACTGAGAAAAAGTTAGTAG

GAAAAAGACAACAAAATAAATGCAAAATGACATGATGACCCGAATCTTCAATATCCGTCAGGCATAGGAGACACAAATGA

CACAAACCAGTTCACTGCAATCTCCAGTGTCTAAGCAACCAACCCCAGCTCACAAATGCATTGTAGGATTCACCAGACCA

AAAAAAAAGAAAGAAAGAATACCCTTTTGTTCTTTCTCCTGTAACCTACTATTCCTGCAAAATTAACATAGAACATGGCT

ACAAATTAGCTTCTTTCGCTTAATGCAGATATTAAGAAAAAGCTGAATTAAGAAACCTTCACAAGAAATGAACAGACGAA

CAAGACTACAGGCTTAAATTACATGTTGAAGCTAAGGAAGCAGCAACATCTTTTGAACTTTAAGCTTAGATCCGTTGAAT

TGGTTATATATGGCTTCATTATGCAACAGCACAAGGGTTCACAGATTACGTGAAATTGTCAAGTTTCATTGTTTTGGAGA

CCATGCAAGTAAGTCCCAAGGTTACTACTCCCAAACCCAAATCAACCACAAAAGTCTGCCAAATTAGTTCAAGCACCTTA

TTCCTATACAAACTTGAACAGAAAGGTTCCCTTTCGGTCCATAACCACAATTATCAATGCATGTAAATGTATCCAAGTAT

TAGTTCTATCGATCATTCAATGCCAAGATTGTGCTATGGAAGTGTTGCAAGGTTTGCACAATTCATATTCTGCCATTGGG

AGCCAGATCCCAGTAATGGAGCCATTTCCATCCACCACAAATATTTATTCCTTGATTCGCCAAGAGGAAAAACAGCAAAA

GATCAATACAACTCTTATTTCTAAAAAGTTAGTATTGGTCTCTTGTTGTCGAACAGACGAATAAATCTTTGTGGTAGATG

CGTAAATGGTTCCATTAGTATGATCGGACTCCTAATGACAAAAAAGTGACTCTGTAGACCTTGCAAGAATTTCATAGCAC

AGTCTTGTTGCAAACATTCCTTCATAGATTTTCTGTGTCCACATGAAACTAAGTTGGCTGAAAAAAAACCTTCCTACGTT

CAAGTTCTGCTATCTATTAATTTGGACTTTAGCATACACCAGGAAAAAAGAAAAAAAATCCTGTGAAATGCCTGAGTAGT

GGTCTGTGAGCAGATTCACAAGCAGAGAAATAGAACGCATGAAATGTGAATTCTATTTAACAAACAATTTCATACTTATA

TGCAGATTCTGTTGAGATTTGTGACATACCCTTTTTAAGAGAAAATTGGATATCTAAACAACAAAATCAAGGAGCAAAAA

GGCACTACAAAAATGATAAATGATGCCATTGGTCACTATGGTTAACCTAAATAGACAAAAGATGATTTATGGTAAAAAAG

GCACATTCAAGTTTAGAAAACTTGTTTCAGACAGAATTGATTTCTAAACATAACAGCACCTAATTCTAACAAACCAACAA

GAGAGACCTAACAAGCAAGGCCACACTAGGCCCTCGATAATAGATGTAAGAGCTAAGTGAAAAGTAAACCATAAAAAGTA

CATATCACAAAAGGAAAAAAGAAGCAAAAAGCGCACAACAATAATAAGAAAAAAACAGCTTACGTTAAAACAACTTACTC

AGACTATGGTGCAATTGTATGTGCTGCAGAGCTGGAGCAATGCTGTTTGATTGGCTGGATCCAAAGACTGTTGAATGATT

GTTGGATACCTTCCAGGACTCATAGCTGAGAGCCTTGCCAATGACCTTACCAAGAACTCCTTCGGGTCCCTTATATCACT

AACAGGATCTTCATCTTTCTTTCCAGCATTGTGGAGACGCACAAAGCTAGCACTGTAACCCACATTCTCCCCAATTTCTG

GCACTTCTGGGTCTTCATCAACCCTTTCTTCTTCTGGTCGAGACAGAAGGGTAACAATGCTGTTCAACATTTTCCCCCAA

GTAGCAGCAGCTGCAGGATCCAAGAGGACTGTAGATTCACATATTAATCTGGTCGAAGCAACAGCAGTTAACTTCAACTC

GATTGTCCCAATTATCGATTTCAGATGAGGTATCCAAAACTGTTCGATTATACTACCAAGTAACTTTCCATCGTTCTGGA

TGGCGTTGATCGAGTCAGTGAGTGCTGGAGGTCCATACTTGACCAAAAATAGTGACATATATATCACCATACACTTGATG

AATTTAACCTTGGGACTCTGCATGAGCTTTGTAAAGAGCGCGGTCCAGATGTGAGTAACAAAAGGTGCGATCACATCATA

CCTGAGGTTCTCTATAATGGTGTTCAGCACATAAAACCCTAGCTCTTCTGTGCTTGGTACGTCGACGAGCTTTTGAAATA

TCCCGAGCACTTGGCCAAGCCTACCCCCTTGGCTCAGTTCATCAGGAGCCTTCTTCAGATAGGCCCGAAGAAGCTGAACA

AGAGCAGGTGTGTTAGCGTGTCCCTTCCATGTCTCAGTTGCAAGGAGAAGCTCGAAGATCGACATATAACTAGCTGGAAG

AGGTGGCTTATTCAGATTGACGAGCTGTGCTAATAGCTGGAAAGCATAAGGCCAAAATTCAGTGACATTCTGGCCCAAAA

TAAATTGGAGTATTGGAAATATGCTAGCTTCAAAGGTTGGTATCAAAGAAGGATCCTTCTCACACACTCGCCCAACCAAA

CCCGCTACTGCCTCAAACAGATAGTGATTGTACACAGGGTTCTTCGGATTTTTGCAAACCTCGCTGAGAAGATGTGTCAA

ACCAGTAATACAATGTCCAGCAACTTCTTGAGAAAAATCAGCAACTCCCAGAACCCGCATTATACACTTCATCACATAAG

AATTCTCTTCTGATTCAGGAAACTTAAGGGCATTGAAAAGGTTAGTCATGAGACTCGGAAGGAAGGGGTTGATGTCAGTG

GAATTAAACCTCCTCTGCCCTCCATCCTTGATTAACAAGAGCTTTTCAATACAATTTGCAGCATAAGAGTGAACGACATT

TGACTCTGATGCGAGGAACCGAACAACATCACCCATCAATCCAACAGCAGTCTGCTTCGGTATCTCATTCCTGAAAACTG

TAAAGAACTTCAAAGCACCAGCTTTCAACATTGCAGAACCGGACTGGCTTTGAATCTCTGGGATAATAACTGATCCAAAA

AATTCACCGACATCGACAAGATCAGTTGAGGCAGAAGTTCCTCCAGCCTTCCTTGTTGCAAGGGAAACAACCAAGTAAAT

AGCACAATCCTTATCCCGCCAATTCTCAGCTGGATTTGCATTAAAAACAGCCAACATATTTTTAATTTGTTCTACAACAA

TGGATTGAACTTCTGTTTTATAGTTGGTGGCAATTCCTTTGAGTAGCTCACATGCTATCCTCCTCCTAGTGTCAACATCA

CTCCCCTCAATATCCCTCCGAACAAACTCCACATAATTGAACTCAAACAACTCCTCATCCTCTTCCCTTAATCTTGCATT

CGGAATCACAATAGACTGACAAATCTGCTGCAAGACATCAGCTCCAGCAAACAAGTTGTGGTGTACACTGGTACTAATAG

TAGTTAAAAACTTAATCGCAGTAATAGTTAACCTGTCACGACCAGTCGCCGATGAAACACCAACCAAAAGTGTCCAAACT

GCAAGTGCGAAATCCTTCAAGTAACCTGCAAACTCCTCCTCATACCTCTCCAAATATAAACTAATATTCTCACATACAGC

CGCTCTAAGATCATCCACAAGCGCCAAACCATCACCACCAGCACCTTCCAGAACCGGATAAGTACTAGTCAAATACTTCT

TAAACTCCCCCATCCACTCCTTCATATGATCTTCGAAAAACTCAGGTATTTCCATAAAATTCAAACTAAAGAAAATCCTA

CAACAAAGCCTCTGACTCTCCATAAGTGGTTTCAGGTTCGCTGCGGTAACAGCACCACCACCAACCATACTATCAATCAC

ATTGGAAGTCCTCATAAAAACCTCAAGCAATGGTGCAGCAAAACCATCAAGACAATACTTCAAATCAAGCAACAAATCAT

TCGTCTTATACTGATACCTAAACTTCTTAAAAATGGAATTCGCAGTACCTAATATCCCATTGACAGTAACATAATCATTA

TTCTCCATACCTTGTCTCAAACTGGAAACAATCTGAGGTAACAACGCAGGCCAAGCTTTAGGAAAATCATGTTTCCCAAT

CACAACCAACGCTTCACTCAACTGACTCTGTATCCTCGGCGAAGCCGACAACATCAATGGTAAAATCAACTGTTTAATCT

GTTCCTTCTCAGTATCATCAATTGGTGACAAAACAGGAGCATCAGGTTGATTATTAGTACCACCGCGATGATTCTTGTTC

GTTTCTGCTGAAATCCATCTCGATTTAAGATGATTTTTGAAATTAACAGCAGCAGAAATACGGATCTGATCATCAATGTT

GGTCTCAGCAACTAAACGAAGAACGGCGAGACCGAAATTAGGTTTATCAGATGCTTCTGATAAGTATGATTCAGCTTGTC

TACGAGGTTCATGATTTGGTGATAGTGTGTGTAGGAAACATTCTGATAAGATTTTTAGGGTTTCTGCATTCCACTCCATG

ACGATGGATTTTCTAGGGTTCAACGATGAAGAAAGAAATCAAAATCTGTAAACTAGGGTTTAAGATGATGATGATGAAGA

GATTTAGGGTTTTACCTTTGAAGAAGATGAAGGACTGAACTGAGGAGAGAGAAACTTCTTCGGCTTTCGTCTGTCTCTGA

TTTGAAAAATGAAGGAACTGACTGAGTGATATGACGAAACAGGGTTGAATCGGGGTTTATATAGAGGAGGTCATTTTATT

TTAGGTCCTTTGATTTGTTGCTTTATTTGTGATACCCCTTTGATAAATGTTATTTTTGGTAATTACTTGACGGTAAATGC

ACTTATGAAAAAGGATATTCTGTTTTTGAGGTTTGGTTTTGACGTTGGGGATGGATCGGTAACGGGGAGTGTTTCAAAAC

CCTCCATTTACCGCATTCATCTCAACGAATTTATCATGCGCCCCATAGGACAAAATGTATTCCATGGTTTGTTCTCGGAT

TGCTCTTTATAGAATTTGAAGAGCAAACCAGTGTGTTGTGATTCAATTAGAGTTGATAAACAATTTTCTGTGAAAAGTTT

GTATGCTAAATTGATGGAACCTAAGACTGCAAATGTTGATACTAAAAGCTTTTGGAACTTTTGATATATCTCAAAGGATA

AAACGTTTTCTTTGGAAATGTTTAAATGACAACAAGGCAAAAATTAGCTTTTATTCTTAGGGGAAGGGGGTTTGAGATCT

CTTGTCCTTTGTGTGATTTATTCTTTGAGTTCTCTTATTCCAAATCTGTTTTGTCTCTTTCCGAATGTTTCTAATGTTTA

TAGTAGTGCTTCCTTCCTAGATCATTATGTTGAATGGGTTAGAGATGACCCGGATGATGAGTCTTTAAAAACAGCAGCAA

CTAAAGCGTGGTTTATATGGAAAGAGATGTGTTCACTAATAAATCGAGTACTAGTATTAGTTTGTCCATAAAAATTCAGA

GATACATTCATTTTTGGACTCCTTCTAGGACTCTGCCGAAAACGTTAGTGCATCTACGATTATTCCAAGGACTATATTGC

CTTTACCTAATAAACATCGGACTAGACCCGAAGGTGATACTACAAAGATAAATGTGGATGGGGCTTGGATTTCTAGGGAT

ACAAAAAGCAGGTTATGGTTTGGTTATTCGTAATGCTGCAGGTTATCAAGGAGCACCCTGAAGTGGCTGAATCATTAGCC

GTTTTCGAAGCTGTAAAATGGGCGAAAGATACGGCCTTAACAAATTTTTTTCTAGAAGGAGACTGTAAAAATGTAATGGA

TTATTTGAATGGCAAGGATTCTTCTATCTCTTGGCAGGTACAGAGAATTCTAGATGAAGTTAAAAAGATGATGGCAGAGT

TTACAAGCTTTTTGGGTTTCCAATTCGTACACAGAACAGGGAATACTTCGATCGGCAAGTACTTCTCAACAAACATGGAG

TCTCATTCCAGAAGTTATTAGTCTGTCTCTTTTAGTTGAACAATCTAATGCTATTTCAAGTTCATATGTTCCTATATCTT

TAAATTTGCTTCGTGAATACTCTTGTAATCAAAACATTTTGGTTTAGGCTTTTGTGAATGAAAGTTTGTTTTTCAAAAAA

AAAAAAAAAAACAATTTTATTCAAATTTCAAAGGAAGTTCAATACAAGACTTAAGGCTGACCTCATCAGGTTAGTAAAGA

GAATAATGCTAAGAACCATAAAACAAAAAATTGCAGGACAGGAGGACCAAGTGTGCCTGCTGCGCAATATGTAATGCAGC

ATAGCTAGTTGCGAAAGAAACATGTTGTCTGCATTAGGGAAATAAAAAACATGCATCGCCGCTGGGTTTCTATCTGACAT

TCTGAAGGAGTGAAGAGGAGGAGGAGGATGTTTCTTTTAATAATTTAAGGCTTGAAAGGAGCAAAGACAACAACTGAATT

GTGCCCGCCGAACCCGAATGAGTTAGATATACCTGCAGGTAATTTGATAACTGGTGTAAACTGTTCGAACTGGAAATTTG

AGCAAAGGAAAGCTAACTACTTGGGAATTTTTCTGCAGGATTAAGAATCATGCAATAGGCATTTCGATGATCTTAATGTA

TTTAGAAGAGAATACATACCAACATGAACTTCATGCTGCTTCTTCATATTTGGGACGGTGTCAATTGTAACGTCAGACTC

CAAATTCTGCGTCGGATTCAAAACAGCAAATATCGTGAATTTCAGTCAGATTATGAAGATATATATCGTTCAGTTAAAGA

AAATTAGTCTTAATGGAGCTATACATACATCTTGGTTAATGGTTGGATGTAACCATCCAGTGGTGATAGCTTTGATTGTA

GCTATGGCTTCTAAACCACCAGCTGCACCGAGCCCGTGTCCAATCATCGACTGAAAACGAAAGAAGACATAAAAAAAAAA

AAAAACCTCAAAAAGTCGCCAGGCTAAAATGTTGCTGCACTGATAAAAGAATGGCTAGAGCTTATCACTGTCAGAAAGTA

GAAACAGCGTGTCAGAGAATGAGCAATCTTGCCTTAGTTCCGTTCATTTTGATCTCCGATGTGTTCTTAAACACCTTCTT

GATGGCATTAACCTCTGCTAAGTCACCCGCAAGTGTTGATGTTGCATGAGCATTGATGTAGTTCACCTAAAAATTTCAGA

TATCTATCAATTTCCGGTGAAATGTAGGAAACTGTACTAAGACTAGAATGTAAAATAAACGGGCATCGTTTATTCTCACC

TCTTCAGGAGAAACTCCAGCATCTTGCAAGCTCTTGTTTATGCAGGATGAAACTCCAAGTCCGTCTGAACGAGGATCTGT

CATATGATGAGCATCACAGGTAACAGCGCCTCCCAAGTACTCCGCTATTATGTTTGCTCCTCTTTTTGTTGCACTCTCTA

AGCTCTCCATAACCTACTCAACAGGAAAAAAAAAAAAAAAAACAACAACAACAGCCCCAACAACTCAGTTATAACAGCAT

TCACTGCCTCCTCGTGCGCCTTTCACAACATAGGCTCAGAGAGGTAAAATGGGGCAATTACCAGCACACCAGAACCTTCA

CCCATAACAAATCCGTCACGATCTCTATCCCACGGCCTCGAAGCTTTTTGGGGTTCGCCGTTTCTCTGAGACAAAGCTCT

GCAAGCAATGAAACCACCAACACCAATTGGGATGATAGCAGCCTCAGTCCCACCAACTACCATGATATCAGCTTCACCTC

TTCTGATATGATTTGCCGCAGCATAAAAGCAGTAATTTGCAGTGGCGCACGCAGTGGAAATGGAGTAGTTTGGACCCATT

AGACCTGTATCAATGGCAAGTAATGCTGATCCCATGTTCGTGATGGAGTACGGAATGAAGAACGGAGTGATCTTCTTGTA

TCCTTTCTGGAGAAGAGATTCAACTCCACTGCTGAATGCAGTTAATCCTCCCATTCCTGTCCCAACTAAAACTCCAATTT

TCGACCGATCCATCTGTTGCAAAATCCAGTAAGCCAAACAGTATATGCAAACACAAACTACGTTACAGATTGTTGCAATA

AGACTTACGGATTCAAGAGTTTCGGTTCCCAATTTGGCATCTTCAAGAGCTCTTTTACCAGCAACTAAGCAGTACCTCCA

ACAATCATCCAGACGCCGGTCATTTTTTCCGTCAATGTAACCTTCGGATGAAAAGTTGCGAATCTGACCAGCAAACCTGA

CAGAAAAGGGTGAAGCATCAAACCTATCTATAAGACTAATACCACTCTCTCCTTCAAGTAGTCTATTGTAGAAGGTATCG

ACATCATTCCCAAACACCGAAACAAGTCCCATTCCTGTTATAACAATTCTTTTCTTCGGATCGCTCTCTCGTTTCGGAGC

TGATGGGGTTCGCGAAGCCATAGCTCTGACTTGCCTACTCCTCAAAACTGAACGAAGGCAGGAAACGCACAAAACCATAA

TCAATTCAAAGATGCATTTACATTACACGTACCGCGGCCGGAAGAAGCTGAAATAAGCCTAAATCTCGTCCTAGGAATCG

ACTCGAACACTTAATGTGTAACATGACAATTACAATGCGCATCTAAAAAAAAAAACACAAAAATCGTACCTGATTTCGCC

CGCACAGAACCAATCTGCTTGGTTTCAAAAGGTCTAATAAGACCATTTGAATGAACCCCAGATGCTCCTAATTCATTTCT

GAGTAAATTTTTGTTTCCGGAGAGAAGAGTACTAGTAGAACAAGAACTACCAGTAATCGCTGACATTGTTACACGAACTG

ATCAAGAATCGGCACAATCGATATAACCGAAATTGTGGAAAAACAAATGATGGGGCGATCAAGAAAAGGATTAGAGTATT

AAAAAGAGAATTTTTTTAATGAAGAGATGATGATTTAGGTTTGATATTCATTGGCAAGTCCGTCACCTAACTTGTTATCA

CCTCCATTCGACGATGTTTACAATTTTCATGAAATAATTATGGTTCTTATTATTTGCATCACAATGACAACCCACACACA

AATTTCATTTTGTTTAACAACTGTAGTCTGTAACTACCCCTTGTATGGGGTAATTATCATATACTTTTTCGTACTATTTT

GACAAATAGTGTCCCGATAATCGTAGTCGTTAATCTTGTTCGGATTTATTTTTATTTACATGCATTTTCAAGTAGTATTC

CGTCTGTGGACATAATTAGGAAAAGGCTGGTGGCAACTGGCAATGAAGCCATTGTCTTGATTGAAAGATTGATTGGCGTA

CCTAGTAAGACAGACAAGATTTTTAGCGTATGGGAGATGTGATATGTCCATGATGCACCCCTTGCAACCGGCCTAGTAAA

AGTGTCTCTTTAATTTTCCAATGGCACACACTGTGCGTTGGTATCGGGACCTGTTAGGGGTAAGTGAGGTTAGTCAATTC

TAGGATTCTTCCAACTAAACGAGGTGTCAGTAGTTATGGAATACGAGATGAGATTCTTAGAATGTACGGATGTGTGGCAT

GTTTCAAACCAAAATGATTCGCATACATCCCAACGGTTTTACTGTTCAAGGATGATTAGGGTGTCCAAAATACGTCTGCA

ACCTTATGCACAAATTGTATTCATGGAATTTTATTTAATTCTCAACTCGGGTGACCTTATAGAATTTGAATCAGTTTTTT

GGTCTATTAGTCCTTGGATTTGAATTCCTGGGCGTAAAAGACTTGTAAAGCATGTTTGTCGCTCAAAAAGACCAAATCGT

AAGGATAAACCTGATACCCTCAGATGGCTCCTCACCAAATCTGGTATCTTCTCGGTGAAATCCATGTATAAGAAGTTAAC

TGAGAGAACAGTGAATCACTACAACTTAGGACAACCAGAATCCTTTTGGAATAGATTATGGAACCTTAATGTATCATGAA

TTGTCGAGAATACTTAGAAATTCATTTCGGAAGATCTAACCGTTGATTTAACCTAAATTTTTATGACTCTACATGCTGGA

CCGCTAAGCGGCTGCCTACGTACCCAAAAGGATCAAGCACGACCGTAGTTCATCAGACGATGCACCAAATACAAAATAAA

CAATTTTAAGATGATCAAATCATTAAGGTTAAACTTACCCAAGACACTTGTTCGACCCATTTAAAATATTGCTCCTCCTC

TGGGAGCCCATATCAAATGAAGAAAAGAGAACCAAATTAATCAACTGCTCGCCCCAGGGAGCCAACACATGGAAACACCA

CATCCATAAACAATAATTAAATTCAATCAATCTACTCTCCCCAGGGAGCCAACATATGGAAACACCACATCCATAAACAA

TAATAAATTCAATCAATCCGCTCGCCCCAGGGAGGCAACATATGGAAACACCAAATCCATAAACAATAATAAATTCAATC

AATCCGCTCGCCCCAGGGAGCCCTACGTGGGTTTAGTAAACAAAACCGTTCCCAAGGATATCTCGTATCCCATCAAAAAA

TATAATACAACAATAAATTCTATCAATTCGCTCACCCCAGGGTGCCCCACGTGGGTTTAGGAAACAAAACCGTTTCCAAG

GATATCTCGTATCCCGTCATAAATAATTTTTAAGAATTACAATCTATGGACCGCAGCCCAACATTCATAACACTACACCA

AAACGGCCATTTAGGGACGTTTATAAACGTTGCGCATAATTTGTGCAACGGAAATAAACGTTGGCAAAACGGCGTTGCGC

ATGGGTGTGCAACACATATTATCCGTTGCACAAATTAATTTGTACAACGCTTATAAACGTTGCACAACCGTTGCACCTTT

TTCTATTTGTGCAACATATAGGTTAATTTGTGCAATGTATATAATTTATAGAATAACACTTCTATGGGTCCCATTTTGAT

TTTGTCAACGTTTTATTTATTTGGTTACAGTTACATCCGTTGCTCATTTCCGGTAAATTGTGCAACGTATAATTGTACAA

CGTTTATATCTGTTGCAAAATAATTTTCAGGTAAAAAAAAACCGCCGGCAAAAGAATTCGGCAGATTTCTGCTCCTGAAT

ATATACTGATTTGCACATGTATAATAGATGAACAATCATAAACCCATACAAATGATTGGTATACACTTTCATACTCTATA

TCATTCATGTACAACAATAAAATAAAAATGATAGGAACATTAAACTTCCATTTCCCTTAGACATCAATATGGGGCATTTG

AAAAGTTCCTTCTTTAGCTTCTGATGGGATGGACTTACTTGTTGAGTATGAAATATCTCGATGTAAGGATATTATTTTTG

CTCTGTGATGATACAGAATCCAGGATACTTGAGATATTAAACCTGAATGACCCAATAGTTAAAGAAAGTACACCAATAAT

AGATTTAAAAATTTCATGCCAGCAAAAAATGCTTACAAGAAAGGTGATCTCTTAGTGTTCCATTTGGCATGAACTCATAA

ACCAACATCTGCATTAAGACATTTAATTATAACATCTGCTAAATAAAATCCACATATGAAAAAATGTAGCAGTTGAGTAA

GAGTAAGTTACAAATAAGTACTTGTTCGCTTGCTTCATCGCAGTATCTAATCAAAGAAACAAGATTTCGATGATGAACCC

TTGATAAAAGCTCAATTTCATTAAAAAATTCCTTGTCCCCTTGAAATGACCCCTCTTGTGCACGCTTAATGGCCACAAAT

AATCCATCAGATAAAACACCTTTATATACCTTACCAGAGCCTCCTTGGACAACCTGAGTAGAGATGTCAAAATTGTAAAC

CCTTTGACACCATTAGCTTCAAAAGTAAACCCATTTCTTCAAAAGTAAACCCTTTGACACCATCAACTTAGATTGGGATC

CTCTTCGACGCTGCGCATATACATACGAGGTGTTAATTTGGATTACACCATCTAATTCAATATCAAACAATCAGACAAAG

CTTTGATTAATGATATACTTACATTGATGTTTTTTAGATACTGCTCGAGAATAGTCTTTCATATGTCTTCTCGCAATAAC

AATTGAAACAAGTGCAGTTACCATAATTGCTCCAGCAATAACCCCTAGTATAATTCCCACAAATGCACCCTTGTTTAACC

CTGTGTTTTGTGATTCTAGGTCCCTTGCACACAAAAGTAGACAATAAACATTCCTTACTTGCATTGTTTCATATTAAGCA

TCTTAATGACCATCACTCTCATTACACGTGATATACATATAAAGATATCACGTGTGGAGACATGGGTTCAAATTCAACTT

ACCCGAAAGTTCCCATTACACGTGTGGAGACATGGGTGTTGGTATCAGACGCGAGTTTCGCCAGTCCAAAATCCGCAAAC

TGAAGAAGAAACCAACTTCTTTTTAGCAATACTATAAAATGTAAAAGATTATATTTAGTATAGTAAAGATAAATCAAAAA

TCATTAGTTAGGAACTTGTGACTGTATATTATAGACTCTATATCTGAGCCTCAAATTTAAAATCAATAAGAATATTAGCA

TCCTTTATATCACGATGAATTATTTTAGGATGGCCTAAAAAAAGAAATCAGCCAAATCAAATTCAGGTATAGATGCCCTA

AATCAGGAAAAATCTAAGAAATTTACAATTGAAGTATAACATGGGAAATGAAAACTCACAATCGTGTAAATATGCTAGTC

CTTTTGCAGATCCCAGGGCAATTTTCAGCCTTGTACTCCAGTCCATGGTTGGTCTACCCTTTCCTGAAATCATTCTAAAA

AATTTCAAATCAGATATTAATAAACAACGAAAAGGACCAGAAACATCACGAATATAGAGTTGATGCACAATGGAATCTTA

TGCATTTCAAGTTTTTGTTGGATCATACGACTTTAAATAGAACTAACTCTCACCACAGACCCACTAATTTCACCCAATTT

TTCGAAGCAAGTTGTTGTTGCCATGTAAGTGGAACTCCAAGGTGTTGTTGGGCACAAACTCATAAACCAACAACACAGAG

TGGAAGCTAGACTTGCCATGTAAGTGGAACTCCAAGGTGTTGCTGGGAAAAAACTCATAAACCAGCATTCTTTTGTCTCC

AACCATGCAGTAACCGACCAGCGAAACTAAGTGTCTATGATGAACTCGGCTAATAATTTCGACCTCAGCCTAAAATTCAC

GTTCCTCTTGACCACTTCCAGCTTTAAGTGACTAAACTACGACTTCTGTACCATTTGGAAGGACTCCTTTGTGGGCATAC

CCAAACCCACCTTTCCCAGAAGGTTGGCATCTGAAATCCATTCGTTGCCATTTTTAATTCTTCATACGTGAATGTGCTCT

CTGAGAATCCTAGTGACATGCCAGGAGATGGTGATGGATATGGATTTGTCGACCCTGAATAATTTGACCCAGAACCTCCA

CTGCTGCTCATAAAAGGTTGAGGTGGCATCGAGGGAGGCGGAAAACGACCCAGAACTTGCAGTTTTCAATGTTCATACGT

ATGCTACCTGTAAGACTGTTATTTCTTACATCACTGCGAATCTTCGATAGGTAAAACACGATGAAACCAATAAACAAAGA

TCTTCCCTAGAATGTGTAGAGAAATAGACATTAAATATTGTATAAAGCATATATAAACTTACAAAAACCACAATCATGTT

AATTGGTAGATATCGGAAAGAGTTATAACCTGACATATATAAATTTGTTCTCACGCAATCCACTGCATTACAAGACAATC

CTGAAACAGTTATACCCTTGCCTGCAAACAATTGCACGAAAAATAATGGCAACAAGCAAAGGATTTAATTATAACTCAAA

AGTGAAATGAACACTCAAATAATCCGCACTCATATCTAAACTTACAGATATTCTAGAAATTCATTCAGTAGATAAGTCTA

GGTATCTCACCACTAAGTTTATTTTGAGCCAAGTCCCTACAACAAAAGAATTTCAATGTTAGATAAAATCGCAGAACAAT

AGAATACAAATTAGGACCACTCAATGGCAGTAAAAGCTTACAACATTTCTAATTCTTTCAACTTTGATGCTGAAAATGAG

TATGTCACCCATAAATTTGATGCGTATCTCATTGAAGGATAAATAACTGGCATGAAAAATTGATAGTCAGTTAATACAAG

GAAGAAAATGCGAGACCAAGCAGTAAAATATCAGGGTTTGCAGGGATGAACATCCGACATTTGTCCCGAAAGATGGTTCC

AACTCAAAGAATCCATAAGCATGACAATCAAAAGAGCTCCAACTCAAAGAAAGCATTCTGTAGTATTAGTATGAACAATC

AAAAAAAAAAAAAGAATAGGTACATACATAGAAACTAGTCTTTTCAGATTACCAATGCTAGATTCATATCAACCTCAAAC

CCATACAGATATGAAAGGTATGTATGAACAAGTTTCATATCAACTTCAAATCTACACAAAACTAAAACTGAAATAAGAAA

CAAACCCATGCTTTGAAAGTTAGAAACTGCAACCAAGTACAAATCACATACAGAAAATAAGAAAATCTTTACAAAATAAC

ATCCAAAGAGCTGGATCTGAATAAACTACTGAAATAAAAAATAAACCCATGCTTTGAAACCAAGACAGAACATCTACCAA

ATGAAGTAAAATCACCAATAATTGCAAACACTCCATATTAATTGCTAAGAATCTTAAGCTTAATCATTTTGCCAAATGCT

AGCATCAAAAATAAATAAAGCATTTACTCAAGCATCTGAAATCTACCTCAAGCATCTGAAATCTACCTAAATGAAAAATG

AAGAAAAAAAAGCAAAACCCATATTTTATTTTCTTTCATACAAGAGTTATAACAAAAAAATAAATACTTGAATCTAGTTT

ACAGAACCTCTTCCAAATGAAGTAAAATCACTACAAACACCTTGGAAAGCATTAGATGACTTCAAAGACTCTACACAGAA

ACAATCTCTGCACAAAAAAGACCCAAATAAACAACAGATGACTCAAAAACCAAAAGCATCAGATGACTCAAAAACCAAAT

AAAACTTCATCTTTAACTGAATCAAATCATACCCAATACTTCAAATCTACATATTACAACAAAAAAAAAATCAAAAAAAA

AAAACATTACTCTTTCAGATACTAGTTAGGTCAAAATCAAAAACCCTAAATCTATCAAATTATTTCAGATAAAAATAAAA

TTAACAACAACCTAGTTAATACCTGATGATAATCCACCAAAATCATAGGTTGAATGACTTGATTTTTTATTTCAGGGAAA

AATTAGGTTTAGAAAAAAAAGTATAAATTTGATTTCAGATGGGTTTTGATAAGCTTGCAGATTTGTTGAAACAAGGGTTT

TGATTTCATGGAACGAGGTAAAATATACGAGATCTGTTGAAGAAGAAGAAGAAGAAGAAGAAGAAGAAGAGAAAAATTTC

TCTCGATTAAATATAAAGCTAGAGACCCAGAGAAGAGTTCTTCTTCCTCGATGAAAACTATGAAGTGAGAGAGAAATAAG

GTGAGTGAAATAAAATTTTAATCTTAAAAAAAAAGCAGAAAATTAATAAATAAAAAAATAACTACCATTCGACATATTAC

AAAAAAGCCATGTGTTGTACAACGGATGTACGCGTTGACCATCCGTTGCACCAGTAATTTATTTTAGTAATTGAAATATA

AATGATTTGTGCAACGGATAATCTGGTGTGCAACGGATACTAACGTTGTTAATTGTGCAACGTCTATATCCGTTGCACAC

CTCCAAAAATGAAGCGTTGCTAAAACGAGGATTTGGTGTAGTGTAATCATGTTCACAACATCGATTATCATTCATAAAAA

CATTATCATTACATAATTTTTTTTTTCTTTTTAATATACTAATCAAAAGATAATCAGATAAACTTTTTATTTTCAAAAAT

TTGTGTAGACTTCATTACACAACGAAAATGAGTGATTCTTATCTCATTTTAAAGAAGAATAAATAAGGTAAATTATTATC

AAAAACAAACAGTTGAAGAACAGGTCCAAACCACGCTTGAAAATTCTAACAGTAAACTGTCCACAGAAAACTAGCAGTTT

TGGTAGTCTTTGCTAGGAAATTCACGTAAGCTTCATTATTTATCGGAATCTATTGATTCTTGGTTCTTTTTAAAGAAAAG

AAGATAACCTCTAATATAAAAATTATAGGTAAAAACAATAAATTCAGAAACCAAGTTGAAATCTCATCAGGAAGTCCGAC

ACTAAAAACGTCACAGAGAAACTACAGTTTTGGGAGTTCCAACTTAAAAATTCACACAAGATTCATTACTCAACCAAATT

TATTGGTTCTTGGCTCATTTTAAAGAACAAAACATAAGGTTTCATATAAAAATTATATGTAAAATCAATGAGTTTTATAT

CTATCTCAAAGTGTTTCACAAAAACAGAACAGAGATCACGTTTCTCAAAAATTGCAGTTTCGACAGTGTACATTCAAAAA

TTCGTATAAAAATCATTACACAATGAAATTTAGCAAACTTAACGCACAATTAAAGATAAAACATAGATATACAACTTTTA

TTAAGACCCAAAATCATTTTGATATCATCAAGTTTGCCGAAACACATCCAAGAACAGACAAAGAGAAACTGTACAGAAAT

TTCAGAAGTCAATATCCAAATTTAAACAAACATTCAATGGTGAATCTTTGGTGAAATATCCTCAATTTTTAATTAGTCTT

TCCCAAACATGTTTTCTATCAATATTACAAGGCTGATCATCAATTGGATGCATATACATACACGAATAAATTTCAGAAAC

CTAACTATGTAAAATTAACGAAATCAAAATCAAAACACAAAGAGAAATAGAATTTAGCATTCTACCTCTTAATTTGCTAC

AAACACCTCCTGATTTTCTTTATCTTTCTTCTCCAAAAAACCTCCAGATCCATCTTCCTCCCTTCCCTTTCTATCTAGTT

TTTTTTTTTTTATGTTTTATAACTCTTTCCTTTTAATTGTCCCTTCCTTTTCTTTTAATAAAAATAATTAATAAGAATTG

TAATTAAGTAACTATTTCCACCCTAACATTTTACTAAGTTGATAAGAGAAGTTCAAGCAACCCATATAGGCTAAGGCAAT

GCAAACAACAACCTAACTAACTCGGTTAAAACCCGACCCGAAACGATAACTAAAATTACAGGGTATTACAAACACTTCTC

TCCCTATTGAAATCATGCTGACAAAGGCTTGGTTCATTTGGAAGGAAAGGTGCAATAGAGCCTTTGAACAAAAACAACAA

TCTAGTGTTCAAATAAGTTTAGAAATACAAATATTTCTGGACTCTCGGTATAAGGACAATATCCCAACTGGAAATCTACC

TTTGGTTGTGTCTAAATGGAATCCTCCTAGGACATTTTAGTCTAAACTTAATATAGATGCAACTTGGGTTTCTTCGAATT

TGCCTACAGGTTTCTCTTTAATCTTTAGGACTGATACATGTGAGTTTGCAAGCGGAAAGGCCGGACCTATCTCAGCAACT

ACCTCTGAAGAAGCGGAAGCCTTAGGACTACTCCAAGGAGCACAATGGGCAGTGGAGGAAGGACTGACAGATTTTACAGT

AGAAGGCGACTGCAAGAACTTTTTTGATTACTTGAACGGGTAATCTTCTCAATTGGAATGGCAGAATCAATCCATTTTAG

ACAAAGTAAAAAGAACCTTTAATCTTTATAATAATTTTTTGGGTTTTCACTTTGTTCCTAGAACTGCAAACAATGTGGCA

GATACTTTGGCAAAGGAAGCCAAGACTTTTAGGGATATCCTCACTTGGAGAAACAATCCACCTTTTTGTATTATCAATAC

TTTAGAAGTGGATAAATCAAATGTTAGGGTTGTAATTAACCCCCGACATTAGATGGATCTACTCTAACCAATGTAAGGGT

TACTAACTCCCAAAGTTAGTTGGTTGGAAATGCACGTAACTTTCAGGAAAAAAAAAAAAAAGGGCAGTTCACAAGTTAAT

GAATAATAAGCTACGTTTAGATCACTCCTATCCACCATGCCGCAAAAAAAAAAATTAGATTGTTGATTAGTGAGTCTAGA

AGACTAGGTACAGGTCACATGTATAATTCTACATATCATTCAGGTTACTATTTGGCGCATTTAACTATTTTGCAAATGTC

AGATTTTAGGTTACAAAAAAGATTTTGGAAGCATATTTAAGACCCCGATTCTAAGTTTTGGTCCCTTGGTCCTTGGCAAA

TTTTGGGTTACAAATAATTCTGAGCCACAATGCAGATTTTATTCAAAGAAAAATGTGCTTGTGCATTGATATGCTTCGAG

AACTGAAAGAAAAATATAGAGTTATTTTGTAAAAGGGTCGTATGTTTATCATCTAATTAGGGTCTGGGTCGTAGAATATG

CAGTTGATGCATTAGGGTCGTTGACTAACCAAAGACTAGGTTGACCATCTGAAATTCTAACCAAAATACCCATACGAAGT

AGTAAAATGCCATTTTATCCAACTCCGTTTCACATCCAAATAATTTCGGATATTTTATTTAAACCATCGATCTTAAGCAC

TAGTGGACGGCTGAGGATAAACTTGTGCTGATTAATTGATCTTTCTCCGTTGACGATGGAATGGGTCTTCTCTCACCTCG

TCTTTTTATGCTATTTGTTAGAATAATGGCGTCCGATGATCTTCTTTTGTCTTCCTCAGAGAATAATCATCAATTCTAAC

AATAATAATCTCATTAGTAATGAGTCATTACTTGGACGTCTCTGATTACTCACTTCTCACATACCCATAAATTTAATGAA

GTATTAAACCTTTTTAATACTTTCTTAGCTGTTGTATTACCAGTTTCTGGTAAAAAAACTAGTGTCCTACATGGGAAACA

GATACATTGTTTGGTAGTGAAATATAGGTTTGATTATGATGTGTTTGTTGAGAGTGGTTTGCTCGATATGTATGGGAAGT

GTTATGGACTGAGTTAGGCGGAGAAATTGTTCGATGAAATGCCTAAAAGAAATTTAGACCCATTTTAGCGAGGGATGGAG

ATGAAGGAGTGCTCATGTTGCGGCATTAGCACAGGTTCATCTTGTTCATCATTTATGGCTTATCTTGATTATACTTCCAT

TGATATTAGATGGTTTTGTTTCTGTATTGGCTCCAAACAATGAAACATTGGTTTTCTATTATATAGTCTCCATAGATTAA

ACATTGGTTTTCATCGTATTATCACAGGAAATTTCCGGTCAGAATCGTTCTCTAACATAAGCCTAGCTTAATTTTCGGTA

CCGGAAGAGACAGACGTATTGTCGGTCCAATTAACAGCCGTTTATTTTAATGGGTTCGGATGAGATGATTGGTTGGATAA

CAAAAAAATTTATTTAGGGGCAGATAGAAAACTCATTTTATATTAGTCAAAACAGTTAACGTCAGTCAACCCATAGTCAG

GGGGTTAAACAAAACAAGAAAAGTCAACTGACAGTTTTATAACGGTCCTTGCTAGTCTACGACCCTGATACATCAATTGC

ATGTTCTACGACCCAGACCCTAATTAGATGTTAAATATAAAACCCTTTTACAAAATAACTCAAAAAGATATAAATGTTTT

CAGGTAGCTACACATTTGAATCCTTCGCAAATGAATAACCATGCCAAGAACAAAATACAACAAATAACCAAATCTACCAA

CAAAACTATTCTAACATTTTCTCTCCATCTCTTCCGAGCTTCTTGCTGCAATCTTACTTCATCCATGTAGCTACATTCTC

TGCTCGACGTGCTTCGATTCGATGCATTCACTCCTTGTAATACGCCGTGGACGAATTCCTCTTCTTTATGATCAGTACTT

TTCTTATTCTTCTTCTTGTTCTTTATACCTATATTAATCTTATTCTCCGTTCTTCCTAGCAACGGTGTCGAACAAATTTT

ACTTCCTGTAGACGATCTCGAGTCAAAGCTTTCGTTTCGACTAGCATTTGAAAAAGCATCCGGTGACGATACTACCGATA

ATTCATCAGTAAAAGATACTAATAGCTCTCGAAATATCGGTACAAATTCTTCGTTATAGCAACCGGGAGCTATGTCAATC

AGCGGTTGTTCATTGTTGCAGAAGTTTGCAACGAACGCGTCTTTAACAAGCGCAAGAAAGTCAAGGCCGCGAGACAATTC

CCGTTCGCTGCCATCCACATTAGGAATAGTGTCTTTCTTAAAGATTGCGAAATATGTAAAAAGACCGTCTATCAAGAAAC

TATACGTTCTGGTTGAAGTTGTCTGAGAGAATTTTTGATGAAGTTTTGGAGTTTTTTCAAGATATTGTGAAGCGAGTTTT

TCGATATCGGTATCACCGGCAATGTTGAGTTCGGCTAAGATCATCGTTCCTTTTGAAACACAAGCGTAAAGGAATTGATT

ATTCGGCTCCGCCGGAATCATTTAGTTTTTTGTAATGCAAAAAATAGAAAATTAACGGACAAGCAGAAGAAGACTTATCT

CATGCATGCATGGTTTCTCCGGCAAGCATGCATTAACCGGAAAAGCTTCCGAGAAGAATGAACGAACATTGAAGAGATTC

AAGAGAACACATGAAATGAATTGGGGTGAATATGATATTCAATGGGGTTGTTCTATTTATAGAATATTGTTGAGATCTGA

AATTTAAATCCACTGGGATTTAAAAAATGCATCATAATTTGAGAAACGACGGAATTTTCATTTTCATTTTGATTTGCTTA

AAAATCTTCAGATTTGTGGAACTTTGGGAGGGGTAATTTAATTGACAAGAAATCCCATGAGAATTGAAAAAACCAATGAA

AATTGGCGAGTCTTTTCATAAATGTTGCTTGGGTTATGAAGAAAATTGGCGAGTCTTTTCAAGAAAAAAACTCTCCTAAT

GAGATGAGAGATAATTTTGTGAGTTGGAAAATATTTTGGTAAAACAATACCCTAAAGCCTTGACACCAATACATCTTGTA

GCGTCCTAGACTTGGGATGCACCAGGACTAGCCGAGAAGGCTTCCATGAATTCAGCATCATTCTATATAGCCTTAAAAAC

CTCAACTGGAAGGCAAATTTCAATTTCAACACTTGCTTCAACGGTCCTGGATTCACAGTAGTTATTCCATAAGATTTACC

GTTTGCACCGGTTTTTACAGCAATAGATCGTCCCCAACCGAAATCGTTACCATACATGTTGAACCACTAGGAACTCCTCG

CCATAAATATATTGCTGAAAATTCCTGTATTTGCATCACCAGGCCCGGCATAAAAGGTTTCACCATCCATGATTCTACAA

AGTTTCTACTCATTTCAAAATTGTTGGAGTTAACCATCTCCTTCAATAACTAGGCTAGGAAACCAAATCCCCTCTTGACC

AGTTCACCCTCTTTCAGACTTACCAATCCCCATGCTAATGAGTTGCCAAAGTACGTTTCCGACAGCGGTGGGATCAACTT

AGTTCTGTTATTCATCAATAGTGAAACTACAAGTACTCGGCTATCATCATAATTATCATTGAAACAGCTCCTACAGCGTA

TTACAGCTATCCAAACATGAGCTATTAGAGCCTGTAATGATGATATTACCATATTTTGTTTTGTTTCAGAAATGATCTCT

GAATTAGCCCTCGATTTGAGTTCTGCAATGTTTGTGTTGAAGATTAGAAAAGGAAGAAACTTAACACAGTAATTACGTGG

TTCGGCTATAGCCTACGTCCACGGGAGAAGAAAGTGATTATATTTTTCTAATTAGCATCAATGGGGTTACAAGGATTATA

TAGATTACACTATATCTATCAATAGTAAGAGTACTGCGGTATATACTCCATACAGTAAGTACTAAACCCTATATACGGAA

GTATTCAACACTCCCCCTCAAGTTGGTGCGAAGATATCGATAGAGTCCAACTTGGACAGTATCCCTTGAAATTGCGTAGC

AGGTAAACCTTTAGTGAAAACATCCGCAAGCTGTTGATGGCTACGAACAAAAGGAGTGGAAATTACTTTGGCTAACATCT

TCTCTCTAACGAAGTGACAATCTACTTCAATATGCTTCGTACGCTCATGAAAGACAGGATTAGAAGCAATATGTATAGCA

TCCTGATTATCGCAAAACATCTTGGTTGGCTGAGATGAGAGGAATCCCAAGTCTTTCAATAATGCCCGAAGCCAAACAAT

CTCGCAAGTAGTTGATGCCATGTCCCTATACTCGACTTCGGCACTAGATCGAGAAACAACATTTTGTTTCTTACTTTTCC

ATGTGACCAGGTTACCACCAAAGAAAATGCAGTACCATGTTGTTGATTTGCGATCAATTGGACATCCGGCATAGTCAGCA

TCGGAGTATGCTGTTATACAGTAACTAGAAACTGAATCATGTCCACTTTTACTCATCAGAACTCCTCTTCCCGGAGACCC

TTTTAAGTATTGTAATATCCTAGTTACCGCATTCATGTGTTTAACCTTTGGAGAATGCATATATCGACTAACAAGGCTTA

CGGCATATGATATATCGGGTCTAGTAATGGTAAGATAAATTAACTTTCCAACTAATCTTTGGTATGAACCTGAATCTTCT

AGCAAATCACCATCGTTATCAAGCTTATTACCACTTTCCACAGGAGTATCACACGGTTTTACTCCCATTTTTCCTGTTTC

CCGTAACAAGTCCAGAACATATTTTCTTTGATTAAGAAAAATTCCCTTCTTTGAATAAGCCACTTCTAGACCCAAGAAGT

ACTTCAGTATCCCTAAATCCTTGATGGCGAACTTGGAATGAAGCATTGCTTTAAGATTCCGAATTTCTTGTTCATTATCT

CCTGTTATGACAAGATCGTCGACATAAACTAAAACAATAGTGGTTCCAAGGTTACCTTTCTTTATAAATAAAGAGGGATC

AGCAGAGCTTCTCTTAAGCCCACATTTAACAAGAACTGATCTCAACTTTGCGTACCACGCACGCGGTGATTGCTTTAGCC

CGTATATCGCCTTCTTTAGTTTACAAACCAAATTGATGTTATTTTCTTGTGGATGCCCAGGTGGAATATCCATATACACT

TCTTCTTCAAGTTCCCCATGTAAGAAAGCATTTTTCACATCCATTTGAAATAACTTCCAATCTTTATTTACTGCAAGAGA

CACAAGAACACGGAAGGTATTCATCTTTGCAACTGGTGCAAATGTTTCCGTATAGTCTTCTCCATATGTTTGAGTGAATC

CTCTAGCTACCAATCTCTCCTTATAACGTTCCACAGTTCCGTCACTTTTGTACTTGATTTTAAACACCCACCGACATCCG

ACAGTCTTCTTTCCTGGTGGCAACTTGACAATACTGAATGTATCATTTTCGTCTAAGGCTTGTAGTTCTTTTTTAATTGA

TGTGTTCCATACCGGATTCAATTTGGCCTCATTAAAAATTTTAGGCTCTTGAGTACTGGATATGGCACTTAAAAATGCAG

AATGGGAAGAACCTATATGATCATAGGTTAAGTGGTTCTGCATAGGATAAGCAGCGGAATGATATGTGACAAAATCCTTA

AATTTTTCAGGAGCTTTCTTTTGCCGATTTGAAGTCCTCACTGGTGGTCCCACGGTATCTTCTATTACTATAGGTTCAGG

TTGTTGCGGCACATTTTCCGCAACCAAATCATGATCGACAACATTATTATCTGAAGCCAAATTATGAATTACTTCATCTT

GTGGAGCCACAAATTCCTCAACTGCAATGTTATCTCTTTGGATACCTTTTAAAGTAGATAACTCATTACCGACAATTGGA

AGTGGTGACAAATCATTGAAAAACTCCCCTTGGGCACGACTTCCTGATTCCCCTTGAAAATATGTTGTAGTTTCATCAAA

CACAACATCACGAGAAATTATAATTTTTCGAGTAGCTGGATTGTAACAGACGTACCCTTTTTTAGTCGAAGAGTATCCAA

AGAACACACACTTAACAGCTCGTGAGTCAAGTTTGTCACGATCTTTTGCCTGTAAATGCACATAACAGACGCATCCAAAT

ATTCGAAGATGAGAAATATCTGGCGTGTGATTTTTTAATGCCTCTAACGGAGACTGAAAATCCAGAACACGACTAGGAAG

GTGATTGATTAAGTATGTGGCTGTCAAGGACCCATGCGACCAATATTTTTTTGGGACATTCATTTGAATCATAATAGAAC

GAGTTGTTTCCAGAATGTGACGGAGTTTGCGTTCCGCAACACCATTTTGCTGAGATGTACCAACACAACTTGTTTGATGA

ACAATGCCATGACTGCGAAGATAACCTGGTAAAAAACCCTTGACAAACTCGGTACCATTATCTGATCTAAAAGTTTTGAC

TTTAGCATCAAATTGATTTCTTATCATAAGATGAAAATCTTGAAAAACAGATGATACTTCGTTTTTAGATTTGAGTAAAT

ATATCCATGTAGCACGAGACCAATCATCTATGAATATGACAAAATATTTGTATCCATCATACGAATCAATGGGAGCAGGA

CCCCATAAGTCTGAATGAATTAATTCAAAAGCCATTGTAGAACGAGACACAGAATTAGAAAATGGTAAGCGAGATTGTTT

AGCAAATTGACAAGCATCACAAACTTGCGACTGAACAGAAATCGAAGGAAAATCCTAGACAAAATACGACTGGAAGGATG

ACCAAGTCGTTTATGAAAAATGTAACTTTGATGCGAGCTTTTGGTGAGACACGCCTTGTCACTGGAAGTTAACAGGTAGA

GTCCATGAGAGTATATGCCTTTACCAATCGTCTTCTTCGTCTTTCGATCCTGAAAGATCACCGAAGTAGGAGTAAATGTA

ACATCACAATTCAGAGAATTAGTGATTTTCCCGACAGATAGTAACCGAGTAGGAAAAGATGGAACAACTAGCGCATCAGA

AGGTGGACAATGAGGAAAAAAGTTAACTTGCCCACTCCCTAGCACAGGAACAGTAGTTCCATTAGCAACAGAAACATCTT

TATGAGAACTAGGAATAAAATGATGAACTGACGAAGAATCATTAGCCATATGATCTGTTGCACTTGAGTCGATAATCCAA

GTACTGAATCTGGAAATAGTCGAGGTGAGACAAGCGAGTAAATTACCTGCATCATGAGTGGACTTTGGATCCGAGTTATT

TCCCAATTGGTGGAAAAATTCCTTCAATTGATTCATGGTAAATGAAGTACTATTAGCTACCGTAGCTCGGGAAGCCTTAT

TCTTTTCGAAATTAGCCTTAAGATGAGGATAAATATCCCAACATCTATCTTTGATGTGCCCAGATTTGTTACAATGATCA

CAATGAAAAACTTCACGCTTACCCTTACCACGAAAGTTCTTTCCTTTCTCTACAGCAAACGCCTTTTTTTATCATCTTTC

TGGGTTAAAAGAGCACTGGATTCAGCTACATCCAGATCTGATGTCACTTTAGAAATAGGATTCATGACTTTCTTTCGTGT

TTCTTCATGCTGCACAGTTGCACAAACACTAGACAAGCTTGGTAATTCAGCACTCATGAGAATATTGCTTCTAAGATATT

CATATTCTGGTTTGAGGCTGCTAAGTACATCAAAGATTTTATCTTCTTCAACCCTTTTTAAAAGGATTTTTGCATCAGTT

GTGTGAGGACGATAGGAATCGAGTTCATCCCACTTTTTATTTAAGTACCCATAATGTTCAGTAAAGCTTTTCTCACCTTG

TTCAGCATCAGCAATTTCACGTTTCAATTCAAACACTCGAGCAGCATTATTTTGTAGACCATAGAGGCCAACAACAGAAC

TCCACAGATCTTTCGCGGATTCAGAGAAAGAAAAAATTTCTGCTATATGTGGTTCCATTGAGTGTAGAAGCCATGTTCTA

ACAAGTTGATCATCACCAACCCAGTCATCATATTTTGGATCTTTAACTTCTGGACATACGCTTTTCTCACCAATATATTT

GGATTTTCTTTTACCCCCAAGAGCAAGAGCAGCAGCCCTTGACCAAGTTAAACAATTAACATCGTTTAAGAGAACATATG

TTAACCTTAAACTAGTATTGTTATCATATTGTGCCATGATAAAGAAGAAAGAAAAACAAATCAAGAGACTAATACTTGAT

TGTAAAATAATGGCCAGGATCTTTAGAGATGCTCTGATACCATGTTGAAGATTAGAAAAGGAAGAAACTTAACACAGTAA

TTACGTGGTTCGGCTATAGCCTACGTCCACGGGAGAAGAAAGTGATTATATTTTTCTAATTAGCATCAATGGGGTTATAG

GGATTATATAGATTACACTATATCTATCAATAGTAAGAGTACTGCGGTATATACTCCATACAGTAAGTACTAAACCCTAT

ATACGGAAGTATTCAACAGTTTGCTTTAGTGAAATGGAAACATCTCTCTATAAGACCTTCAAATGGTGAAACTTCAGTAG

TAGTAGTAGTAGTAACCAGCGAAAATTTATCAGCGAAAGAGAACGGAAGACGGATGGGGCAATCGGTTTCCTTGATGAAG

TACCGCTTAAAAATTGGAGGAGGACATGAAGTATGATTGTCGAAAGATCTCGCAATCTCAGACCACGAATTAACGAAATG

CCAAAATGATGTACCATCACATACCGAATGGTTTGCGCTGCATCCTATAAAAATAGCACAGTGGCGCAGCTCAGTTACCT

GGATAGAAAGTAAAGGATGTGACCGACCTTGGTAGTTTGGTACCCCAGTATGGGTAAATAGTGGATTAATTAAGCTCTGA

GGAACATAGGTTGGAGAGACGATGTCTTGGATGGATATGTCTGCAGTGGCATGGATGAACTCTACACCCTCAAAGTTGCA

GTCAATATAGACGGAGATGGTGTTATCATCTTCATGTTTTTAAATAGCAAGACGACCGGCTATAGGAGAGAAATGATCAA

GTGTACGTGACAAAGAAGTTTTTTTGTTGGACGCCTACTCTGCAGGAACAAAGAAAAATAAAAAAAACAAGCTACATAAA

GACTAGCAGACAAAATCAACGTCTTGCTAAAGAATAGAAGAGGGGATTTCGGAAAATGTCCACCAAGTTTGGACATTATG

GGACTTAGCATATTTTGCAAGAATATGAGCCAAGGTGTTGACTCTCTTTGGAATGAATCGGAACTCCACAAAGTTCATAG

ATATGGCCTTAGACAAAGAAGTTTTAAGATGACTTATCTTGTGGTTGATAATCCCATAATTTGTTACATCTTCTTGCTGC

TTGGTGAAGAGAAAACCCCTTTGCAAATTCAGAAATTTAGCGTGCATAAGAAATTTAAGAATGGTCCTGATAAGGTGGCT

CCGTCTAGCCGAGAAAATTCTCATAACAGGTTTCAGGGTCTTGAAGATGAAGAAGAAGTTGCTACATCTATTTCAGCTAC

ATATGAGGCCGTGGTCAGTTACTCTAAGCCTCTTAAACCTTCTGTTTCTGAACCAAATCTTGCGCAAATTAGAAGAGCGA

ACTCGAAAGCCAAAGCTTCGGGTTTATTTCCGACTGGATCGCTGAACACAAATTATAAAGGTTCGTCTAAGTCTCTAAAT

GATCCTACTAAAATTTCTGCTCCTGCCCTCATGGAATCTCTATCAGCCATAGGAATGAGCTTTCCTACGTCTAAGAAGGA

TCGTGAATCTATGTTAGATAATCTTGCAGACGATCATAAACACCTGGACATTGAGCAGATTTAATTTGTTCCTTTCCACT

CTTCTGGGGGATTGGGGTGGTTTACGTTTCGTTTTAGTGTTCTGGCAGCGTTCTCCCTTTTTTTTTCTCTTTTTGAAGTT

ATTGGGATGCCTTTTCTTTCTCTTTATTTTAGTTTGTTCTTCTTAGGTTGTCTTTGCTGTGGGCTTTCAGCACTTCTTTC

GTTGTAATCGTCTGCACACGGCTTGTGTGTTTTTCAATTTATAATCTTTGCCGAGTAAAAAAAAAAAAGAGAAAACCCCT

TTGCATATATGTACTTCTTAGAAATAATAGATCCCACGGATTCAAATCAATCCGTCTATGATTGTCGGTTTTTTCATCGA

TATAACTGGCTGGTCGGACCGTGGTTGTAGAGACATGGCAAACTTTTTTTGGCAGAACTCATTCTTTGATCCCGCTCAAA

TCTTTGCTAAAACAGCTCCAATGTCTTATATAGTTACATAAAGTTGGTATTCTGATATCAAACCATAAATGCCTAGACCT

AAAATACATGTACTAGGCAAGTAATATTTGATAAACTTAAATTTCAACCACCAAGAATAGTTTTTCATTTCGGTGCCCAC

TGTTGTGTGAAGCCAATGTCAAAATTAAACAACGTAAGTACATATTTTATTCGAGTAGATATGAATTTGTAGTTGCAGTA

AACCCTACAAGTTACACCCAAGTGATATTATCAAATAAATCTAAGACATGATGGAAGAATCTCAAAGTTATTTTATTAAG

AACATGAACAAATACAAGTATGATATGCGAAAACCCTAACTTGGAGAACAAATAACTTTCTCTCTCCTAAATTCCTCACA

TAAACTTTTAAAAGGGTCTCTCTCTTTTACAATGGCCTCTGGTTCCTTTTATAGGAGGTTACGTAGTGGAGGACAGCTAA

TAAAGCTCTTTAATTCGGATATGAGAGAACGATATTATTGCGCATGTGTTATCATCACGTTGGACTTCACATATTAAGTG

TGATTCTTGTGACGTCATTGCGCTCGTCAGTAGTGACGTAGTGGACTTGTTACTAATTCGTCTTTGAGTAATGATCTTCT

CACGTTCTTATTGGTTGAGCGGTATTTTGTATACACATTTTGCCTCTTCTCACATTGATCTTTTGGAAGAAAGGGAGATA

TGAGAAGTCTCAATCTATCGCATCAACCCTGTCTTTTCATATTCTGTAGATCTACTAACACTATTCCGGGTTTTCAGCAT

GACCCTCTACATGTGCTAACTGTCACTCCTTTAAATCGACACGTCTTCTAATTTGACCGGTGATTTCTTTTGACGGTTTT

CTTACCTTAACAACTGTGGAGGAGAGAGACTTATTTTCATCTCTGCAAACTTCCTTCATCTTCCAAAGGTTTTCTTCAAC

ATCGATCTTCGCTCAAAACAATGGACCCATCATCCTGGTTAGTAGTTCTTATGTATGCTATTATTATCTCTGATTTTCTT

TACTTTGTTTTTGATTTTAAATCTTTTCCTGTTGTTGCTGCTGCTACTACTCTTCATTTTTTTGCTACTGTTACTCATAT

TCCCATACTTTTATTATCCCCACCAACACAATTAAAGCATCTATCCACAAGAAACCCTCTTATGATACCCATGATATTCC

AGGTAAGAGTAAACTTGTTCTTATAATCTTAGGGTTTCTCCCAAGTCTTCCTCTTCCTTAGCGGATCAAAAGAACCACGA

GTCTAGCAAAAGGAACCTTCCCACAACAAAGGTGAAATAAATTCTTTATTATTCAATTTTTTCGCTTTAAGAAAAATAAT

ATTGTTGCCTCTGTTTCTTATCTGCTTTGTTGTATGATGGGTGACGAAGGCGAAGATAATCTCGAGAAGCGAAGACTCAA

GAAATCCCATCATATCGCTAAAATTGACTCTTCTGTTAATATGCCTGAAGTTTCTTTTACATTTGGTGATGAGGTTCTGT

TGCCTCATGCATCTGCCACGCCTACTGTTTCATCTACTGAAACTCTCTCATCGAATCCAAAGGATATCATAGCCTCTCCT

TCAAAGAATATTATCGCTCTTTCTTTGTCTATTGTTATAGTTGCACTGTCCACTCAGTTGGTTAACAAGACTTCTCTTAC

TAGTACAAGCGTTTTGGTGGTGTTAATGGTGGTTTTTAGAAAAAGCGTAAAATCGTAAAACCCCGAATAACCTAATGTCA

CCGGTGACACATTTATTAGAACATTTAATGCATGTAGTCAGTAATTAATAGGGGGAATTTTTTCAAATTAGGGTTACAAC

GAGCAAAACCCTAATTCCTTTCCGTGCCAATGGAAAATCATGCCAGCCACACCTAGCCCGAGATAGCCGCTCCATGCCAA

ACACACTCTTCCTTGGCCCTAACTATACTAGTTACACCAAACGCGCCAATGCATGCCAACCATGCTAGCCGCATCCCTGT

GCCAGCGCCAAAGCATGTTAGCCACACCAAACACGCCAAAGCATGCCAACTATGCTAGTCGCATCCCTGTGCCAACAACA

AATCATGCCAGCCACACCAAACACGCCAAAGCATGCCAACCATGCTAGCCGCACCTCCGTGCCAATGGAAAATCATGCCA

GACACACCTAGCCCAAGCTAGCCGCACCATGTCAAACACACTCTGCCTTGGACCTAGCTATGCTAGTCACACCAGACGCG

CCAAAGCATGCCAACCATGCTAGCCGCAGCTCCGTGCCAATGGAAAATCATGCCATCCATACCTAGCCCAAGCTAGCCGC

GCCATGCCAAACGCACTCTGCCTTGGACCTAGCTATGCTAGTCACACAAAACGCGCCAAAGCATGCCAACCATGCTAGAT

GCACCTCCGTGCCAATGTCAAATCATGCCAGCCACACCTATCCCAAGCTAGCCGCGCCATGCCAAACGCATTCTGCCTGG

ACCTAGCTATGCTAGTCACACCAAACGCGCCAAAGGATGCCAACCATGCTAGCCGCACCTCCGTGCCAACGGAAAATCAT

GCCAGCCACACCTAGCCCAAGCTAACCACGTCATGCCAAACACACTCTGCCTGTCCCTATCTATACATAGTCACACCAAA

CGCGCCAAAGCATGCCAACCATGCTAGCCGCACCTCCGTGCCAATGAAAAATCATGCCAGCCACACCTAGCCCAAGCTAG

ACGCGCCATTCCATACGCACTCTTCCTTGGCCCTAGCTATGCTAGCCACTTTCCTTACCAAGACGCTATTCCACTGGAAT

ATGCCAAAGTAGCGCCCGCGCCCTAGGCTTGCAAACCGTGCAACCTAGGATAATATGGTTGTGCGTGCCATTGGCACATG

CCAACACGCACCGCTAGCCACTTTCAAGGCCACTATTCCATTGGCACATGCCAAAGTAGCGCCCGCGCCCTAGGCCTGCA

AACCGTGCAACCTAGGCCAATATGGACGTGCGTGCCATTGACATATGCCAACACGCACATCTAGCCACTTTCCAAGCCGT

TATGCCATTGGCACATGCCAAAGTAGAGCCTGCGCCCTAGGCTTGCAAACCGCGCAACCTAGGCCCATATGGACGTACGT

GCCATTGGCACATGCCAACACGCACCACTAGCAGCTTTCCTTACCAAGTCGCACATGCCATTGGCACATGACACATGCTC

CCCTAGCAACTTTGGCATTGCCGCTTGCACTCCAAGTGCAACCCGCGACCAAGGCATTCCGCACATGCCACACACTCCCC

TAGCAGCTTTGGCATTGCCACTTGCACTCCAAGTGCAACCCGCGACCAAGAAATTCCGCACATGTTATTGGCACATGCCA

CACGCTCCCCTAGCAGCTTTGGCATTGCCACTTGCACTCCAAGTGCAACCCGCGACCAAGGCATTCCGCATATGCAACAC

GCTCCCCTAGCAGCTTTGGCTTTGCACCTGCACTCAAAGTGCAACCCGCGACCAAGGCATTCCGCACATGCCACACGCTC

CCCTAGCATCTTTGGCATTGCCACTTACATTCAAGGTGTAAATTGCGGCCAAGGAATTCTACACATGCCATTGGTACATG

CCTCACGCTCCCCTAGCAGCTTGGCATTTCCACTTGAACCCAAAGTGCAACCAACTCTCGAGACATCAATACAAAATTGT

GGGATGCTTTCTTTTACACTTATTTTTGTTATTTTTTAATTTTTTAATTTTTGCTTTTGCTTTAAAACTTAGAAACCCTA

ACTACCTCTTTTGAGCTCAGCGGTCTTGTTAGGCTGTTGTAAAAAAGGAAGCAAAGAAGTTTTTTATTTTAAATTAAAAA

AAGGGTTTTGGTTTGGCGGTCACAATATGAAGCGTACACTACGCTCGTATTAATAAAGAGTGAAAAGGAATAAAAGTGTT

CAATTAAATGGGGAAATAAAAGTGAAAAACGTTTTCACTATTAAACACGCCGGTTACCACCTCCTCCCATTTAACTACCT

TTTATATCCTTCCGTAATGAGGCTAGGGTAGAGCATGATACTTATTTGGGTCTTGATGGATTGTTACATGAATGTGAGCA

TCTTATCTAAACTTTTTTAAGAGTACCTAATCTAGGATTGGAAATATTTTGCGATTAAATATAATACCTTTGATGAAATT

CACTCATGTCTCTCCATCCATCCCTTTCCACTTTGAACTAGATCTTGTGCATGGTGCACCCCTGGTCTCACCTATTGCTG

TCTGTTATAATTATGTAAGGTCATTGATGTTTCCGCACTACGTCTTTTGGGTTGATCCTCAACTTTTAGATTGTTGGTAC

ATGGTGAGTCATCATTGTATAATCTAGTAGGTAAAATTTTGAATCTTGTTGATATATTTGTGTGCTAACCAAACTTTTTG

TCTTGCGCTCATATTATTTGCTTTAAAAGCATTCATTCAATTATGGTTAGCACCACCCTTTATGCATCGGGTAATCTCTT

TCCTTTAAATCTTCTCTACCTTGTTCTCTTGATATGTGTTCTTATCATCTTTTGAAACATTGAGGACAATGTTAGATTTA

GGTTTGGGGTGTGTTTTGCATATAGAGTTTTTATTCTTTTATTAAAAAAATCAAATTAGTCTTTTATTAAAAAAAAAAAA

AAAAAAAAAATCAAAATTTTTCGATGACCCAATTTGTGTGTAATTTTTTTTTAGTCTTTTGCATTTTATTTGTTGTTGTT

TTGGATGACCAGAGCTCTCTTTTGTATGTTTTTTTTTCCTTTGTAGTCTTTTGCATTTTTTTTATGACCAGCTGTGTAGC

GGGTCTTTTGTAGGATTTTTTTTTTGTAGTGTTTTGCATATAGATTTTTTTAGTCTTTTATTAAAAAATAAATAAATCAA

AAATCAAATTTTTTGTATGATGACCATCTGTGTAGCGGGTCTTGTGTTTGGTATCGATGATCGACATATCGGTAAAAAAA

AATGATGATCGAGTGTGTAGCGGGTCACTAAAAAAATATGATGATCGTGTAAGCGGGTCTTAAAAAAAATGGTTACAAAT

TATCAAGGTTCAACACCATTCTTTGGTAAGCCCTCACGAGACTACAAAAAGCATCTCTTTGTTTTGATGAGTGCCAAATA

TTGCATATTCATACTTCATTGGCACTTAACTCATCTTTCGCTCTTCAAAGTCATATATTATCCCAAATTCGGCATTTTTA

CCATTTTCAAGAATAAATATTTGTATTAATAATTTTGTGTTTTTAAGTACTAAATAAAGCTTGGAAGAATTGAGAAGACA

AAAGAGAAGAAAGTTGGAAGCGTTGCAACCCGCTCGCGCATCGGTGGCATATCAAACCCAAGACCCAAATCCATTTCCAC

CTCACCCCATTGGATTCTTTCCAAAACATCTACCTCACCCACATTCTCTCACTCAATCACTCCCAATTTGTTTCCCACAA

CCAAATATATTCCATCAATTAAGAAAACATGTTTTGTTGAGTGGATGAGTGCTTGTGTATAAATAGAGAGGTTGTTGTGT

TGCAAAACCATCCAAGAATAGCTTGTTCATGTTAGTTTCTTAAATTTCAGTTTTGTTATAATGTTCTTCATGTTCTAGTT

CATTTGCTAGGGTTTTAATGAAGCCATAACTCATTGGTATGTATTCATATCTTGTATCTTGTTCTTGAAAAATTGTTTAA

AGTATAATTTAATCCAAGTTTGTCAACTCTTGCTCTCACAATGTGCCATGTATCCATTGTAGTTAGAATATCTCCATGTT

GATTATGCTACAACTCATGTATTGATCCTTAGATTATTTGTGCTTTAATAAGACATCTTTGGAGAAATACTTTACCTTAT

AAGGTAGACTTGAATCTACTATAAGGACAAGAATAATATTACAAGTTGAATGCATATCATAGGTTATTGGTGGATTTGAA

TACCAGTATCCATATTCTCATTGTTTACATATCTTGTTATTTGCATCTTTTATTATTTCAAACTTGAAAATCATATCGTG

ATTCAAACCCCTTTTGGTTACTCTTTGAATCGGGATTTAGAATCAACTTCACACTCTCCAAATCTCGTTTTGTACTTCTC

AATATGTGGAGGAAAATCTCAGAAGCTTCCGACAAAATAATGCTAGGAGTGTATTGCATGATAAACCATATAACATTGAT

TATGAATTTCCCTCTGTTGAAGACTCTCCTTCCATTGAACAACTTAATGCATTTTTAAAAATGAATTTGATTCATTTACA

AAATGTAGGAAAGGATGAATTTCAGGATGAAATATTTGTTTGTCCTAATGAAATAAAATGTGAAAATAGTATGCCTAGTC

TTACTTGTGCAGCTGATATTGCATATGAACCTATTTTAAATGAAGGGACTAGAAATACTACGATTTTAAATAAGGAAATG

CATTTCTATGATGATGATGATGATGATGTTATGTTAGAAGAATATGTTTTACTGAAAATATTGTCGAACCGATTGTGAAC

TGGGGCTTATTGATTTGTTTTATGAGCATGAGCATGATGTTGTACTTGTTTCAATCTAAACTTATTTTTTTTATGACATA

TAAATTAAAAAATAAATAAATCAAAAATCAAATTTTTTGTATGATGACCATCTGTGTAGATGATCGAGTGTGTAGCGGGT

CACTTCTCACCTATTTTCTAGAGACCAAAGTGAATAGTTAGATTTCGGATTTGTTTCACTTGAGGACAAGTAAAATTCAG

GTTTGGGGGTACAACAAGTCATTGGCACTTAAATCTTTCGCTCTTGAAGTATTTGTATTAATTAATTTTGTGTTTTTAAG

TACTAAATAAAGCTTGGATTCAAAAATTTCCACCAACACCATATATTCAAAAAAGCATCTTTGCTAGGGTTTTAATGAAG

CTTGTTCTTGAAAGTTTAGAATTGTTTAAAGTATAATTTAATCAAGCTTCCATTGTAGTTAGAATATCTAATGCTACAAC

TCATGTCTTCCTTAGATTATTTGTGCTTTAATAAGACAATTAATTCTTTGGAGAAATACTTTATAAATTATCCATTGGAG

AAACACCTTTTTGTTTACATATCTTGTTATTTGCATCTTTTATTATTTCATTCACTTGAAAATCATATCGTCATTATCGA

CAAAATCAAACCCCTTTTGGTTACTCTTTGAATCGGGATTTAGAATCAACTTCTACACTTCTATGTGGTTCGATACACAA

TCTAAACTTATACAAGTCGAGACACTACCCAACCAAAAACTCTCTGGTCTCATCACCTTTCTTCGATATCCTCCCCAAAT

AAACCCTTCTCCTCCACTTTGTGACCGAAGAAAGTCTGGAACGGCCATTTCTTGGTTTAGGCCGGACTTGTATAGATTGA

TCTCTCGAATCTAGAAAGTGCTCACGTGCAGCACACTTTTTTAGGGTTTAAGCCAGTTTCTCACTCACCTACCGAGTTTA

CCAAAATCGGCAGAAACGATTTTTACCCATAAACAATTGGCGACCACAGCGGGAGGATTGATCTATCGATTACAATACCA

ACTCTGATTCCTGATTTCGTTTACCGAACGCCAATCCCAACATGGTGGAACTCAGGTGTGGAGCAGCTTCCGAGAATACC

AGCACAGAGCATGTACCTGATGTTAACGTCTCTCCCAATGACGTTGTAACATCACCTGCTGACTACAGTAACTCTGAATG

CACTCCCTCGAGCGCAACACCGCCAATCACCAGGCCAGTGCCGCTACCTCCTCTAGTGTCTCTGCACAACCCTCTAATCA

CGGCAACGGAAATACCTCTTTGGGTGTCACGCCACCGGTCACCAGGGCCAGAGCCGCAGCTGTGGACCCTAATGATTCAT

CTAGCATGGTACCTCCGACCACAACTAAGGTTCCAACTACCCCACCGACGGGACGAGAGACCGGAGGGACCAGAGGAAAC

CGTCCTTATGTCACCATCGCTAATTTGATGGAACGACATGAGATGCTCGCCAGATCCCAGACATATATAGTTTCAACACA

GAAAGAGGTACTTGTTTTTCTCAAGACCATGACAGAATGACTACCACGTGAGTCGCATCCTCCAGAGCCGCTTAGGAACG

TCTCCAACAACCCCGACACCAGCACCTCGGCGGAGCACGCCTTTGCGGACGAGGAAACTCGCGCTAGAATCTCGTTGGAA

AGAAATTAGTTGTCTAATTTCATCACACGAGAAGATCTGGAACAACTTCTCCACAACCGAATGAAGAATGATGAATTGGA

CATACATCATCACCAACCTCCTTACCCATCCTAAATGCAGAGAATCCCTCTTCCAAGGGGGTACTCCTCTCCTCAATTTA

CCCTGTACGATGGTACCGGGAATGCACATGAGTATGTCTTTCGTTTCCTGGAATCCCTATGTGAACATGAATATAATCAC

GTTCTTCATTTAAGGGAATTCTCAAAATCACTTACCAGAAGGGCGTATACATGGTACAACATCATCCCGCCAAACATCAT

TCCCAACTAGATTAGCATTTTCACTGCGTTTTATAAGAAATACTTCTTCGTGTCTGAGCAAGTTACTTTATCGGACCTCG

GAAGAATGTATCAGTGGGATAACGAACATCCGAATGATTTCTTAAAAAGGTTCAGGGTTCAAGCACTCGACTGTCATGAC

CCAAACGTGACTGAGCAACAGCTGGTGAACGCATGCATCAATGGTATGGTTCTTATTTATCGCGCCTTGCTAGAGAACTT

ACGCTTCCAAACTTTCTCCGAACTCCATGAATCCTCCAGACGATCTGCAACGACAGCACCGGCATTACTGGAAAAAACTA

GGCCAACTCGAACCGAAGACACGCATGAAAATCGAGGGAATAGATGCCTCATCAACAAGCAATTTAATGTCGGCTTATCC

TCCGTGAACATAATAAACGAAAGGGGAAAAAGGAAGTCCCCAACTATGGGCCACCAGCCAAAACGTGAATCGTCGGCGCA

ACGGGAACCCCCGCCACGGAAAACCGTCATTAAAGCTCCCATGCAACATCATGAGACCGAAGGAAATACCATTGATTTCT

CATTTCCCATCGAGGAGGTGATTGACCTCCTAGAAGCCTGGGTTCAAGATGATGTTATAAGATTTCCACCTCCCAGATGT

CCACCAACTGAGGCTGAGATGGAGAACCCCAAGTACTGCCATTACCACAGGTTCGTTCACCACCCAACCAGCGATTGCAA

CAGACTAAAGCATATTTTCAGAGAAAGGATAGAAGCGGGAGAACTCCAACTAGGATACGAGAGAGTGCATCGAGACCCAC

TCCAAATAAGAAACTGCGGGGTCCTCTGGGGACTCCGTGCACGAGACTGTACGATCCATGATGCAGCATATATGCGAAAT

TCTCTTTTTTTCCAAGGTGCAGTGCCAAGATATTTTCGCAGCTCTCAATCGCGTTGTATCCGGCAAACGACTTTTTTTCT

CCTCGGAAACCACGTCAGGGGCCCAATATTTCCAAACATGCGGAACCAAAAGGGAGAACTGGGGACTGCTGACCACGTCC

CGTTTAAGAGGTAACGAATTTGACAACACGCTCATCGACGCGGCCTCTGACCTCAACATCATTACTTTCAAGGCACTAAA

GGCAGTAGGATTTACAAAGCAAGACGGTACCTATTACCCGTTCGTGATTAAGAACTCGAGAGGAGAAGCTATGAACGCGC

ATGGCTACATCAACCGAGAGGTCAGGGTGGGAGATGCTATAACGTATAGCAAATTCCACATTGTCAAAGAACTGGGTCAT

GACATGATATTGGGACACTTATGGGTGCTCGACAACAAAGCGAAATTAGATGTGCACCCTATGCCGATGCCCACGCCCGA

AGGAACTTCCAACAAATCATCCCACTAGTTATCGCATTGACGAACTATCAAAGGAACCCAAATATCCGGAGAGGACCCGC

TGGCGCTATCTCATGGTTTTCAAATGCAAGTAAGTCTGACCGAGCAACCTTTTCTGCAACCAGTCATTCCTCCCCCCATT

CCGTCATGGGGACTCGGCGCAATTATTGGCACAGACGCTGTCAAGAAATGTGAATGGGGCGCCAGACCATTTGAGCACTT

TTCAAAAAGCTTGGAAACCTCAATTGAAGCCAACGAAGCCAAGACTCAAACGTTGGCACGATGTCTGAACCCGTTTTCAA

TCGGAGATGTAGTGGTTAAGGAAAGATCGTTTCAAGTTGGGAGGATGACACTAAAAATGGCTATTCAGACTGATTCGCTC

AATGGAAGAGAAGACGAGCTGTATCTGGTGGCGGACACCCTTTTAGGAGGATATTGCAAGCTCATCAACGCAGACACACC

TGAAGGCGTGATAATTCACTAGCGATGGATCAAGGCCTCCAATCCCTAAAATGTTGTACTTCCCAGTCACTCCCCCTTTT

AGTATTTCCTTTTACAATTCCTCTTAGTAGTTGTATTCCCCCAAAGATGAATACACCGTTTGTATTTTTCCTTGGGGATT

CCTTTTCCCGTCCAAAAAAAAAAAAGAGAGCATAAATACATCTCTTTATAAATAAGTTCTCCAGATATCATAGAAATACA

TCAGATTAACCCAGAAACACCATTACAGAGCCATCATACCACAACCTCTTTCAATTTTTTTTTTATATATATAAAGCAAA

ATGTTGACTGATGAAGAAGAAGGTAAAGATGATTTTACGAAAAAAAAAAAAAAAAAGACATCAGGGGAACTAAGCAGCTA

ACAAAAACAACAAAAAGGAGGAACTAGCCCTCTAAATATGGAGGACAAGTCCCCTAGGTCAAGGCTATCTGCAAGCACTT

CCCCTAGCAATCCCTTAGAAAACAAAGGTTCCATTCAACGGTGTTAGCAGCAGCAACCTCAACATCCCATCCCGAAGCAA

CCACAGCAGCAGCAATGTCCGCCCCGACGACCGGTCCATCATACACCACCATGACCCCTCCGGTACCGCCAACGACACCT

CTAGCGTCAACTTCGAAGGCTACTTCATTTCTGTCAGATGCTCCTTTGAAGGTTTTCTCGTTGACACTTCGGAGAAAATC

TTTTTCATCCGCATCAAAATCAAGAACCATGGCACTCTCCTCCAAGAGATTTACATGTTCGTTTTTTCGTCGTTCGATCT

GCCCCATACCTGGGCCGAGGAAAATCTTTACCGCACTGGAACTTCTCAATATGCCAAGGTTCTTCCCCGTTTTCTCGCCC

CTCCTCCATACTCCCTGAACCCGACAAAGTCCGAGCACGCGCATGTACTTCATGGGGATAGATGGAAGGGAGGTCCTTAA

TGATTTCGCTAGCTTCATGGAACATAGATCCAATCCCAGCCAATGCCTCCTCAAGAGAAGAAGGTCGATCTCCGCCAATT

CTGAAGTCATTCTCCGAAGAGTTAAGGGCGTGATCATCCATGGAGGATCCCATCATACACTGCAAAAAATAGGGAAAATG

ATCAAGTTTCATCCTATCTATACACATATAAAAGAGGATAACATGAAAACAGAAGAAGAAGAAAACACACCAGTATGATC

TTTTTATCCCAAACAACCTTAGTCGGGATGCTTTTGCAAGAGTCATACCTCGCGCCTTCACCTGGGGAATCTGGCCGGTT

AAAGGATGATTCGTCTCTGTGCGGCATGACAAAGAGAGAAAGTCAACAAGAAGAACCTTTTCTCAAACAAGGGAGACGGT

ACAATTGCGCTGGGTATGAATACGGAGCCTTAAAAAAGGAAAGGAAACAATGAGAGACATAACGCTTCCATATTTTTAGC

ATGGATGATGCCCTGATTGAAGCCGCGTGTGAATAAAGGAAAAATAAGGAAACATTGGTATCTGTCGAAATACCTACTCC

CTCCTTGTTTGCACCGGTCCCACCAAGCTCGCGCCCAGCTTCCAAGTATCCCAATTGCTCCGGTCCCACCAATTGCACCA

ACTTCCCAATGACCGCCCAAGTCGCTCCAAGCTAGCACGCTCAGTTTTCCCACCTTGCGTCTATCCCACCAGCTAGCACG

CTCAGTTTCCCAACTCGCGTCAACTTCTTAAGTCCGCTCAGCTTCCTGTCTCTCAATGATGCTGACTTCCCAATCGTGCC

AGCCTCCCGATTTGTACAAAACTGCCAGAGGCAAGGATTCCAAATCCTATCTATAAAAAGGTCCGACAAGCCTCCTCAAT

TCTCATCACAATTTACTGCGGATAGTTTGTCACCATCTTATGCTTACCTCGCAACAAATCTCCTGTTACTTTGTAAAACT

CCGAATAACCTAATGTCACCGGTGACACATTTATTAGAATATTTAACGCATGTAGTCAGTAATTAATAGGGGGTATTTTT

TTCAAATTAGGGTTACAAAGAGAAAAACCCTAATTCCCCTCCGTGCCAATGGAAAATCATGCCAGCTACACCTAGCCCGA

GCTAGCCGAGCCATGCCAAACGCACTCTGCCTTGGCCCTAGCTATGCTAGTTACACCAAACGCGCCAAAGCATGCCAACC

ATGCTAGACGCATCCCTGTTCCAACGGCAAATCATGCCAGCCACACCAAACGCGCAAAAGAATGCCAACCATGCTAGACG

CACCTCCGTGCCAATGGCAAATCATGCTAGCCACACCTAGCCCAAGCTAACCGCGCCATGCCAAACGTACTCTGCCTTGG

CCCTTGCTATGTTAGTCACACCAAATTCTCCAAAGCATGCCAACCACGCTAGCCGCACCTCCGTGCCAATGGAAAATCAT

GCCAGCCACACCTAGCTCAAGCTAGTCGCGCCATGCCAAACACACTCTGTCTTGGCCCTAGCTATGCTAGTCACACCAAA

CGCGCTAAAGCATGCCAACCTTGCTAGCCGCACCTCAGTGCCAATGGAAAATCATGACATCCATACCTATCCCAAGTTAT

CCGCGCCATGCCAAACACACTCTTCCCGGCCCTAGCTATGCTAGTCACACCAAACGGGCCAAAGCATGCCAACCATACTA

TCCACACCTCCTTGACAATAGCAAATCATGCCAGCCACACCTAGCCCAAGCTAGCCGCGCCATGCCGAACGCACTCTGCC

TGGCCCTAGCTATACTAGTCACACCAAATGCGCCAAAGCATGCCAACCATGCTAGCCGCACCTCCGTGCCAGTGAAAAAT

CATGCCAGCCACACCTAGCCCAAGCTAGCCGCGCCATGCCAAACACACTCTGCCTTGGCCCTAGCTATGCTAGCCACTTT

CCTTACCAAGCCGTTATGCCATTGGCACATGCCAAAGTAACGTCCATGCCCTAGGCTTGCAAACCGTGCAACCTAGGCTA

ATATGGCCGTGCGTGCCATTGGCACATGCCAACACGCACCGCTAGCCACTTTCCAATCCGCTATGCCATTGGCACATGCC

AAAGTAGTGCCCACTCCCTAGGCTTACAAACCGTGCAACCTAGGCTAATTTGGCAGTCTGTGTCATTGGCACATGCCAAC

ACGCACCGCTAGCCACTTTCCAAGCCGATATGCCATTGGCACATGCCAAAGAAGCGCCCTATCCCTAGGCTTGCAATCCG

CGGAACCTAGGCCCAAGTGGCCGTGCGTGCCATTGGCACATGCCAACATGCACCAATAACAGCTTTCCTTACCAAGTCGC

ACATGCCATTGGCACATGCCACATGCTCCCTTAGCAGCTTTGGCATTGCCGCTTTCACTCCAAGTGAAACCCGCGACTAA

GGCATTCCGCACATGCCATTGGCACATGCCACACGCTCCCCTAGCATCTTTGGCATTGCCACTTGCACTCCAAGTGCAAC

CCGCGACCAAGGAATTCCGCACATGCCACACGCTCCCCTAGCAGCTTTGGAATTTCCACTTGCACTCCAAGTGAAACCCG

CGACCAACGCATTCCGCACATGCCACACGCTCCCCTAGCAGCTTTGACATTGCCACTTGCACTCAAAGTGCAACCCGCGA

CCAAGGCATTCCGCACATGCCACACGCTCCCTAGCAGCTTTGGCATTGCCACTTACATTCAAGGTGTAACTTGCGGTCAA

GGAATTCCACACATGCCATTGGCACATGCCACATGCCATTGGCACATGCCACATGCTCCCCTAGTAGCTTGGCATTTCCA

CTTGAACCCAAAGTGCAACCAACTCTCGAGACATCAACACAAAACTGGGGGATGCTTTCTCAGGGTTTTGGTTTGGCGAT

CACAGCATGCAGCGTACACTACGCTCGTATTAAGAAAGCGTGAAATCGAATAAAAGCGGTCAATTAAATGGGGAAATAAA

AGTGAAAAACGTTTTCACTCTTAAACATGCCGGTTACCACCTCCTCCCATTTAACTACCTTTTATTTCCTTCCGTAATGA

GGAAAGGGTACGTTTCAATCTCAACTTGTATAAATAGGTCTCACCTATTTTCTACTAGACAACAAGTCTTGGTCAGGATA

AAAATACAACACCATCAAAAAGCATCTCTTGCAAGCTTCCTAAGCTTCTACTTTTTCTGATACAAGTCGAGACACTACCC

AACCAAAAACTCTCTGGTCTCATCACCTTTCTTCGCTATCCTCCCCAAATCAACCCTTCTCCTCCACTTTGTGACCGAAG

CAAGTCTGGAATGACCATTTCTTGGTTATGGCCGGACTTGTATAGATTGATCTCTAGAATCTAGAAAGTGCTCACATACA

TCACATTTGTTTAGGGTTTAAACCAGTTTCTCACTCACCTACCGAATTTACCAAAATCGGCAGAAACGATTTTTACCCAT

AAACAAGTGGATGCTATTTCCGGGGTTGACTCAATGTTGAAGATTTTTGAGGCCGTTGGTTCGTCTAGCATTGATCCTTA

TCTGTCTGAGGTTTGCGGTATTCTCACAAAGTTGATGTGTGATTCCCCACCCGACGAGTCATCCAAGGCGCAAAATTTTC

TCCAATTTAGGCCATTAATGTTGGAAATTTGGGCTAAACGGAGGATGAAAAACAATACAAAACCGAAAAGAATGTTGTTT

CACAAGCTTCAATGCCTTACGTTTCTTTTCAACAATTATTTCGACCATTCCCAAATAATTGGCGAGTACAACCGATCAGC

GAAATATTACCTTCAGGATACAATGAAGCCCAAACAACGTTGTTGAATTCAAGAGTTGTACTCCCTTGACCCAACCAAGA

ACACATAATATTTAGAATCTGATGATGCTTATAAGAGAAAAGAAATCAGAGATTTTTATTGTGTTTTTCCATTTGTTTGA

ATGAAAACAAATCATCACTACTTATAGGGAATTCATGCCCATTGGAGATGCCTTTTCACTTCCAAAAGACACCAATAATG

TCTAATGGGAATGGATGTGTCTCTTGTGAACCATCACACCCTTTGGAGATGCCTTTTCATCTCCAACTAACATCCATTGC

ATCTATTAGGCTCAGGACGCTCCCCATCGTAGTGTCTAACCAAAAATGAAAACCATTTTATTTTCTATATTATTGAAAAT

ATCCAACAATCTCCCCCTATTTTCAATAACCAAAAACCATGAAAATAAAACAAAGTAAAAACTTACAAAACTGGAGTGGC

TGCATCACCGAATTGACTGCCTACGTACCCGAGTGGGATCAAGCCAACGTAGTTAAAGCACAATTGGGAAATAAACATCA

TCGTTGAAATCAATACATAAATGATATGTATCTTTTGGATTTGAACCTTCACTTAGTGTAAGAAATTCAAATCCTTATCG

GGGTTCAGTGGTGAAACTAGTCTTGAACCAAGTACCCCATGTGATAAAACCGGAACCTCAACACATACTGATTTCAGGGA

TTGTGTGTTCTGTAGCCATCACATTAATGGCCATGTGCTTATCCTGTATTCATAAGTCTCTTAGAGAACAGACTTTTTCT

CTTAGGAAACGGCCTCATTTCCATGCTCATATAGGTAAGTCTCGAAGGTGTTTCTGCGAGACACCTTGTGATAAAAGTTA

GACTTATTAAGGATTTTCATCCATCCCTTTTCTTCTTGCAGAGAACCTGCACTAAACAAAAATAAGAATTTTTACTTAGT

GCACCATAATCACTTCCATAAATTGTTGGTACCACATTGAACCTTGTCTATCTTTTTCAAAACATAGGTTGGGTTTCTTC

TATGGGTGATCGAACTTCATTCAAAGACCTTAGTCTCATTTATGTATACTTCATTGAAAACAATTCCTTTGGGAGACTCT

TCATGATAAGTCTGCCAAGTTTTTTTCTCCGTTTCAAAGGCTAACTGCAATTACTCCTTTCTTGAGTAATTGACTTACCG

TTCAATGTACAAGACTTGTATGTCTTGAATTTTCATTATAAGTCTTTGTTAGAGATGTTATAGATCATAGCCTGATTCAC

AATTAATCAAGACCTCTTGATATGGCTTCTTCCAAAAAATGGGATCTCTACAAGTTAGGTTCTAAGCCATTCCTCCTCAT

TAATTGCATCTACTCGTGCAATCAATTCTAAAAACATTATTAAAAATGTTCCTATTTGATTTTACAAAATAATTGTAACT

TCTCTAGTGATCCTAATTAACATTAACGGTTAATTCACAAGAAGTCATTACAGACTTGCATACACTAGAGGCATAGAAAA

TTTGTAGGCAACTTACTTGCTATTTAAACAAGCACATTATTACGATGAGTAACTCCTCTACTCATGCCCAATCCACTACA

AATTTATACCCAAAAGACATAGACAATGTCTATTTAACATTTTACATTATGTAATGTAAATGTTCATAATTTATGCATCG

GTAATATCATAACACCTAATTGTTGGATATTTGGTCGCGCAATCTTCCAAATCGTTACATATTAGCCGGTATTCCACCCC

TTTCGATGTCATTTAAGACATGCCAATTTCCAAATTTGGAACCATATAATTCTAAATGCAAGAATTATGTTCACATCTAA

ATGTTTCTCAACATTTATTGATGATCTCAGACTTCTAAATTTATAATTCCTACCGCAAGAATTATATCCGTACCAAAAAT

TGTAGTCTCAATAATCATTGGTGATCTCTATTGGTTCTCTAATCTTAGAGCATTCATGACAGTACTGCTCCAATGTCATG

CGACCAAAACTGTGTCATTTAACACTTCTTCGAACTCATGTCAAGTACCGTACATGGGACATAATTCCTGAATGGAAATC

TTTAAATCCATACCGGTACGGGCACTATTACTATGGATTTTGGTTTTCTTGCAGATATTCCAATTGATGCCTGCACGTAT

CCAATTAATACATTTTTTTATTCATGGATCAAAGATTCTAGTCTCATGCACCCAAAACAAAAAAACCCTCAGCACATCAA

AGAGATAATCATAACACTGCTCACAATGACCACAGAATTCCTTAATAGAGAAAAGGAACCTTCAAATCCAATATTACGTG

CCGACAAAACTCTTACAGCTTTGGTATTGGATTCGGCTCTCTTTGAGCATTATGATGTGGCTGTCGGAGAAATTGGCTTT

TTCCTGCTCAACCGTACTCATACGTTCACCAGTATGTAGTTATGAATTAAAATGTGTCAACCATAAGAACTCAATTAAAA

TAAATAAATAAATAATAGCCCCACAGCGCTATACAACATTTCTTAAGAAACATAACATGGATGACCATTCAAACGATACA

CTTTATGGAAGGAACCTCAATCATGTACTGTTCACAACAAAATCCTCTTAATTCATATGAAACACCTCAACTAGTAATAT

AAAAAAAAATGGTCCAGCCAACTATCTCTTGTTTCTTTATTCACTAATCTAAAACAGTGAATAGATCTGGTTGCTCGTGC

AACATGAAACCCATATAATGAATTCAAAATAATCAACTATATGTCACCTTGAATCTACACGGTACATTTCAAAATGAAAC

ATGCTAAATATGTAATCATAAGATCAACGTCCCGTACATGCTCAACATTATCAATTAAGACACGATCCAAGTAAAAACAA

TTTCAGTGCACGATTATTGGTCATAAACTGGCCTTTTATGCGACTCCCTGCTATCTTACTTTTAAAATAAAAAGGGTTGC

TATTTCACGAAATAGGTCAGGATCTTAGAACAAGTAAGAGGTGTGACTACCAGAAAATATATACTAACAACACGGTAATC

TGGTCAATTTATTATCGTCATTGAAGAATAAATTCTCAAATGATCTTATTAATCTAAGTGATTCATTGTATATATATATA

TTCACTAGTTACGGCACATACATGTATCACATTTATACCAGAAAAGGAAGCAAAAGGAAGAAGAAAACGAGATTTATAAT

ACTTATCTCAACCTTGAAGAGAGAATGATGTTCCATCATCGCCTTTTGTCTTTACCTCTTCATTAGATTTTACTGGTGCA

TCAATCATGCTTGTTTCTCCTTCAGAAGAAGTATACTTCAAAACGATCAGAAAACCTAATCTACCTATCCCGTGTATGGA

AGAAAAAAAAAAAATCCAACTTCTTGATATATGTTAAAGCCCAATATTGTGGCCCGGTTAGACTTTACATGCAACATATA

TAATACTTCTCCCAAAAACAACAGCCATGACGGAAACCTTACGAACCAGGAACATTCTTAAGAAATTCAACCATACCAGA

ACCAGTTTCATCCTCTATTGTCAAAATACAATTAATTGCTGAAAACTGTTGGAAATTTGGGCTAAACGGAGGATGAAAAA

CAATACAAAACCGAAAAGAATGTTGTTTCACAAGCTTCAAGGCCTTACGTTTCTTTTCAACAATTATTTCGACCATTCCC

AAATAATTGGCGAGTACAACCGGTCAGCGAAATATATTGCCTTCAGGATACAATGAATCCCAAACAACGTTGTTGAATTC

AAGAGTTGTACTCCCTCGACCCAACCAAGAACACACGGTATTTAGAATCTGATGATGCTTATAAGAGAAAAGAAATCAGA

GATTTTTATTGTGTTTTTCCATTTGTTTGAATGAAAACAAATCATCACTACTTATAGGGAATTCATGCCCGTTGAAGATG

CCTTTTCACTTCCAAAAGACACCAATAGTGTCTAATGGGAATGTATGTGTCTCTTGTGAACCATCACACCCTTTAGAGAT

GCCTTTTCATCTCCAACTAACATCCATTGCATCTATTAGGCTTAGAACGCTCCCCATCATAGTGGCTAACCAAAAATGAA

AATCATTTTATTTTTTATATTATTGAAAATATCCAACAATTAAATGGGCATGTGACAGCTAATAGATAAGTGTGTAGTGC

ATGTATATGATCTTATATTTCTATAACATCTAAATTTAAAAACTAAACTATAAATTACAGGTAGCCATAAATTATGTACT

CTAAGGCACTATAAAATTGAAAATGTTTTAGATACCGACGGAAAATCCCGACGATTAGCCCACAAGACAGATTGATCGGG

GGTCCCACTTTACCATTTATACCCTAGATGTTAGAACGGCTGCCCATCTCCACACGATCCCCTGTCGGGATGATTCACCG

GGTTGTTTTGTCGGCCAGTGTAGCATTCCTCTCTATAAAAAAATATCAATCAATTAAAGCCACAAAAAATGGTACAACCG

ATTACACCATCACAACCATCTCCCAGCTAGTATTGGTGTTTCTTTGGAGAAAATTAGAGTACCCCTGTTTTACCTTTTGG

TCTTTAGGTTTATCTGTTAGATTTTCTCATTTTTATTGTTTTATCTTTTAACTCTTGTGAAAGGATTAATTCACTATTTA

ATGTTGAGATTATATTAGTCCCACATTATCTGGATAATTTAAGATTTTTCGGTTTATAATCTTTTGGGCCTCTCCACTCA

TTGCCAATTTATTTTGAGTTAGATCCCCACATTCTAACATTTAACTCTTGTAAGGTCATTGATAACAATCTCCTATCTCC

TCACGAAAAGGCTAATGGTGTACCAAATTTCATAGTTCAATTGTGAACCAATGAAATCCAACCACTAGATTCAATTAAGC

AGGCCCCACTTGCATCTAAAAACCTCACCCCTCCACCGTCAAGGAAATAGGTCCACCATTGATACACCTTTTTGGTCCGC

CCCTGGACGGGTAGATCTCCTCATGTGACTCTTGTAAGGTCACTGATGATAATCTGAAGTACTTTCTCTTATCAACAGCT

TCTAACACATCAACACCGACAATTTCATCTATTTAAGTCTCTCATGTTACTCTTGTAAGGTCACTGATGATAATCTCAAT

TATATGAAGTACTTTCTCTTATCAACAGCTTCTAACACATAAACACCGACAATTTCATTAATCTTAATCAGGAAACATCG

GCCATTTAATCCATAACTTAAAATTTGGAACAATACGTTGCCAAATTTCAAAAATTCTGAATTTCCTAGCATGTTCTGCT

CGGCTTATTACAACTTCTACTAATGTGCTGGAAAATCCTGCTATTAAAATTACTAGATAAAATCTAATGCATTCAATAAT

CTTTTGATTTTTCCAAGATAAGTTAGCAAGAGATCCATTAACAGCAATAGCTACATTCTTTGAGTTTGTCTCCAGTTTAG

GTATTTTATTCAATAGGGTTGTGTATTTATAGAATTCCGTTGATATATGAAAATTTGTATCGATTGGGATTTTAGAAAAT

GCATTATAATTCGAGGGAGAAAGGATTTAAATAAATGCAGTATAATTTCGGGTCACGCTTTGGCTCAATAATTTTGCATT

TTTGTGGTCGAATAATTCATTAGATTGTCGTCAAAGTCTCTCATGTGATAAAATAGTATTTGCAAAGACCAATTATAATT

CCTCTTTTTTGTTTTTTTTAGGAAAAACTTAGGCCTAGAAAGTCACTAAGAGCAACCACAGTCTTCGACTATAACTAAAA

ATTTAGTCGAAACAGCAGGCGCTACGGGGCGGAGTAAAGACCAAATCTTAGACTAAAACTAAAAACCGGAGTATATTTGG

TCTTTGGTTTAGTTATAGTCAGGCGTAATTCACGATTACGCCTGATTGGGACGGTACTGCAGGTTACGCCTCACCTTGAG

GCATAATTAATATTTGCGCTGGGTTGGGGCGGGAGTGCATGTTACGCCCAATGGGGCGGAATTTCAGGTTGCGCCTGATG

CCAGGCGGCTTCATAATGTTACGCCCGATTCAGGCGTAATTATAGGTTACGCCCCATCAGGCGCAATTATCAATCCCGCC

GATCGATGGCGTAATCATAAATTACGCCTGATGGTGAATGGAGCGCAACTTGGTACTCCGCCTGTATTTTCATGGGGCGT

AATCATATTTTCATGGGGCGTAGCTCCCAACTTGCATGTAATGCCTGATTCATGCACCAAAAATATGAGAACATGACATG

CAGACGTGGAAATGAACCACTTAGGTATGCATGGTAAAGAGAAGATAATCTTACAAGCGTTTCGTCCACTGTTTTGGAGT

CCAAAATTAATTTGTAGCCTCTACAAATGAGATCCTCTTCGTCTGACGATTTTTCGTCTTCAATAATCTCCGAATCAAGC

TCAACTGGGAGTCTGTTTTTTTTATGGGCGTAACTTATGTTTTATGTGTGGTGGGGCGGAGGTCTGAGTTAGTCCCACGA

AGCGGCGAGCGATTTTTCACAGTTGCGCCCCATCATGCGTAATCAGTTTTCCGTTTAGAGTTACGCCCCATCGGGCGTAA

TCAGTTCTGCATGTTTTGGGTTGCGCCCCATCAGGCGTAATTTCAAATTACGCCTCTCCATCGGGCGGTTTTTAGATATT

ACGTCCCACGACAAACGCAATCGCAGGTTACGCCCGACAACAGGCATAACTTATAGTTACGCTCCAACCAAGCGTAATTT

ATTACTACGCCCGTATCAGGCGAACTCTTATACATCCGCTCCACCAAATTTACTCCTGCACGGTAGCGTCGTTCGACGGA

CTAAACCAAATTATCTTAGTTTTTTAGTCTTTAGTCTTTGGTTTATAATCGTATGACTGTAGTTGCTCTAAGGGTAGTAC

ATAGGAGTTGTGTTGGCAGTGACTGTGGATGCACCACTCTGCCTGCCAGCCTCGTTCTTACAAGAGAGAGCTAAATTGAA

AGTCTCCATATTTTGGTTACGATGGACAATACTAAGATTAAGTAAAACAAAAAATATGGGGTTTATGACCATATTTGTAC

TGTCAAGAAGAAGACAGTTTCTTCTTTCATACTAGTGAAGGTGAAATCTTACATGGACTGCTTCATTGTAAAGGAATTTG

AAACAATCAATGTTGGTGTGCTGGGGAATGTCTATTAATTACCCCGAAGTCAAAGTGATGTTATCATTCCAGCGACTTTT

TGGAGTCTTGCAATTATATTTTCTTGGCTTCTTTTGTTCTTTGTACTAGAGTATGTGCTTGTTAGAGACCAAGACAAGAC

CAAAAACAAAACAAACATCAAAGTTTATATAATCCGAGATACATTGCATAAAGAGAAAAATATATAGAGGTGAATGAAAT

GATCGCCCGGCCGGGTGCAAGAATTTCTTCTGTTCTACTAGGAAATTAAGCAGAGACAAGTACATAAAATAAAAATTCGG

ATTTAGGAATGTGAATGACGGACGACAAGCATAAACCCCAGAGAAGCAACCAGCAATTATTGATATAATAAGGCGCAGAA

ATGCTGGTTGAATATGTGCCTAGACCACTCATTCATTACTTCTTCTTGAAAAGCAGTTGTGCGTTCATCCATTCATCTCC

ATCGTTGAATGAGAATTGTACATACCCTCCCAAACACAATGTCCTGATGTGAGTAATGACGCCATTTACTGCTTCATGGT

TCCCACCAAGTCCTGGTAATAGAATTTCTTTTACCGCTTCCATATTTTTATCTAGCTTCTTTCTCCACGCATCCCTGCAG

TCAAGATGAATTTTAATCACTCTCAAAAGAGTTTGGTCTGTTGTTCCATTAAGTAGGATTTATTTTTTGTGTCGAGATTA

ATTATAAAAATCCTTACATTGTACGAGCCATGTGCATCCCGTTCACAACCCAATGATCCACAACGGAAACATCGTCCTGC

AAATAGATAATACTATTACAAACTCCACAAATATATATTATGAAGATTGAACATAATAGGTAATGACTTACTCAAGCTTT

ACCTGGAAATAGAGGAGACTATCAGCAGATAGAACTGTAAGACAACCCGGGGGAAAGATGTAACCAGAGTACCAGTCATC

CTCGTCGATTGCCTGCACAAGCCAAGCCAAGCTTTAAATTAAATACACATACGTAAATTAAAATTGCCCCAACTACAACT

CGTGAAATTCTGACCTCAAATAAGTGACTAAATGTTTTATGGCAGAGATGATCCACAAAAAGAAAGCTATCTTCGGTCAT

CCAGGTTGATAGTTTCTTCATAAACAGTTGTATGTTTTTCATGTGCTACAAAACATGAACAGAGAAAGGTATTCAGCTTC

GGCTTGCGCCGTTCTACTTTTCTCTCTATAGATTGATAATTGGATACAAGTGCTTATGAGAATTATTTATTCATGAGTGT

ACCTCTACAGCTCCAATTACAAGTATTCGATCGTATGTCTTGTCAGATTCGTATTGAGTAACATCTGCCAGTAGGACGTC

CACGTTTTTCAGCCCAAGCTTCCTATATATGGAAAGATATGAAATGCTTAGTGTAATACTACTATAGTCGTCACCATATT

GTCGTTTACTCATAGAAATTAACTTACTCTGTTTGTTTGATTATATAATTCACTTGTTCCTTTGAATTAGTGAGCCCTGT

AACATGACAGTTCTTATATTTCTGAGCAACGTATAGAACTAGACCACCTTGACCGCAACCGATATCAAGAACAGTTTGTC

CATCTTTGATTTGTGCTCGCTCGCAGTATAATTTATGTGCAGCTTCTTCTGCTTCGTCTATTGTTGATGATTCATGTTCG

AAGTAACATCCACTGCGTTGATTTTTTTAAGATAATGATCAGAAACTATATGATAGAACACATATTGCATTGCTGATTTT

GTTGCTCGAGTTGTACCTTAGTTTCATACTTTTTCCATAAGCTGCTTCTAAAAATTCTGTTGGTAATTCATAGACTTCGT

TTGTGAATGCCTCGACCTCTCCTGACATATCCATCTTTTTTATAGCTACATAGGAAATGAGTTTGTCGGATTATTATGTC

AATAGTATATATCGAACTTAAAACAAATAAGTTGTATTACAAGTTCAAAACTTGTCAACACCAAAATCTTTACAAATGAA

AAGAAATTTGTACTATAATTTAAGGGGAAGGGAGAAATTTACATACATTTAATGAATTCAAGGTTGAAGGAGAGTTGTTC

TTCATGAGATGATTTATAACCCCATTGAAGACGTTTCTCGAATTGAAACTTAATTAATTTCTTCAATTCCTTTTCATTAA

TCTCTCCTCTTAACAATCTCCCTAGAGTTTCCTCTGCTGATTCCTTCTTCACTTCTTGTATTGAACCCATGATGACACAA

TCCAGGCTAAACTAGGGAGAGAAACAAAATGAAATACTTGCGAAATATTGAGAGAGGTGGTGCGTTATGAGCCAGGCTTA

TTACACAGCTTATATAGAGATCAACATCGGGTTTATGGCACAACATTACGCACTAACAGGTCGGCACAAGAAAACATATC

TCAAAATTTGCATGCCAACTTGGTTCTAGAAAGTACATGAATTCCGCGTGTTTTGTTATGATTTACCAAAAAAAGATTTA

AAAAAAATCGAATACTTGAACCCTAAACCCTAAAAAAGGACAAGCAGATGTGAAATTATCATACATAATTCAGAAAAACG

TGTGGACAAGTCTGAACACCTAGAAACTAGATATGGATTCCTCTAGATAGCATCCACAACTAGTTATCATTTTGATTTAG

TATACCGCAGAAGATGATAAAGGTTACCAACCCTACCACAACTGTGCAGTTAAAAAAATTTAATGCTGCTATGAGAGTTA

GTAATTGTACAATTTTTGGGTCACTCGATAGACTCTATTATTCCATTGTGCAGTTAAAAAAAACTTATTGCTGCTGTGAA

TGGACTATTATTGCCTTTACGGATTTACAGGAGTGGTCCTGCCATTTTTCATGGCTTGAAGCTGCTCAAGAAAATAATTT

TTGTTTCTTTGCTGATATCATTTCAAATACCCCTTTCACGAGATAAGCAATTCAATTTGCACCGATAAAATCTATTCAAT

TGGACCAGACTAGTGTGAGTCTAGCAGCTATCCTCTTGCATTCTGCAGATCCGAAACCAAGAGTTTTGTAAGAAGTGGAT

TTGGTAGAATCGAAGAGCTTAGAAAAGCATCCTGAACAGCCATTACAGGGTACCCAAAGCTGCTTCACTACTAGCATCCA

TGGGCCATGGCCATCAGCATTGTTTGAGCTGGTGTGCCAATGTTAACTCGAACAATGTAAGTAGGACTCTGGATAATCTG

TCTGGCAGATGCAATCAGTACAAGGCTAGAAAGATATCGAAGACGATTTTCGCCTTTACGAGCCATATCAAACAACAATT

TTTTCTTCCCATAAAAGTGATTTTGATGGTTTAAATGGTGAGCATTTGCCGTAGATATGCATGACCTTTAGGGTAGTATT

TAAAGTAGTGCAGGTTGAAGGTGGTTTAAATGCTGATTTGGAATGAGTCTTGGAAGTTTTAAATTTTATAAGGGTAAAAT

GTGAAATATACAAATTTAATATTAGTGGTTAAATAAACTTAGATAGTGGTTAAATGAACATAAATTTGAAAGGCTCTAGG

TACTGGATTTATATAGAAACATAGAAATAGATATGCAGGTAGAGGAGAAAACAAACATGTTAGTAATATTATTATAATGG

TGGTGCTGCAGCTGCTAATGCAGGAAAGAGGTCTGAACATTCAGTAGAAGACCAGAAGGTAGTTCAGCTCAAATTCGAAC

AAGTGTAGCTTATTTGTTTTACTGCTGAACTAAATTTTAATAGGTTTTCGAATCTTAAACAAATACTAAAGTTGGTATTA

CACATGGTGAAATCAATCAATAATTAATCTAAAACAAATACTAAAGTTTGTATTTCTTGCAAAGTGTTTGCCTGACAGTT

GCAGGCAAATTCTAGAAATGTTCATGATTATGAACATGTGTAAATCTTTTTTCCTTAAGAAATCTGAATCCATGTAATCA

CATAGTGGTGTTGTTTCGTCGGGTTCTTCAGTCAGCCCTTCCACCATTGGCAACGGCTGTACTAGTAATAATCTACTACT

TACTTTGTAGTTTAGCTTCACAACTACTGCACCAAAGATCTTTTTTCCAACTCTGTTATGCCAATCATATTGAATGTCAG

TATCACCTGTGTACAAAAAGAGTTCAAAAGCAGTTACCATTCAGGAGTTCACATGGGCAAGAGATAGGTTGGTAATACAT

AAAACCAAGTCAATGGTTGGTATTTACATGCATAGAACTATTAGTGGAAAGGAAAAAACAATGAACCCACCCACATCAAA

ATATTACACCAAATGACATAAATTATTTAGATATATTATATAATGGTGCAAACATGATAATACATCATAACTATTTTCGA

AATAGTCGTACAGTGGATATGGTACTAAACATGATAAGTAAAATGTAGAAAACCTGAAACACAGATAACATTAACTAAGT

ACTCAGAATGTCTTATTCAATCTAATAATGATCAAGTGGTAAGCCAGAACTCATGTTCATCATGTTATACCACTTGGTCG

GCGTGCCTTCTTAGTCGCGCTTTCTTCTTCAGTCTCTGACCTCCCTTCAACTTTGAAAGGGGCAAAATGCCTTTCTTTTC

ATGCAGTAGAAACTTGGAGGGCATGTAGGTCAAAGAAGGAGATGGTGTACCTGGTTGTAACATTCTTTTACATGTTGAAG

AAATTGTGAAAGTAATTTGAAATCCACGGCCTAATGCATGATCCGTGAGAATCTCAAAAGTAAATGGTAAAATCCCAGCC

ATCAACACTACTACTTCCATCTTAGCCACTAAATCAGAAAGAACGCCTTGCCCTAAAGGCCTTGAACATAGACCCCCCAC

AGTTGAAGAGTTTAATTGGATGATTGGAGTTGAATGGACTGCACAAAAAAATGGAGCGGAAGTTAATCAATGGGAAAACT

CAACAAGTTAGTTACCCTTGGCCAAAGCAAAAAGTTAGTTACCCTTGAAGTGTCACTAACCCAAGAAGTTAGTTACCCTT

GGCCAAAGCAAAAAGTTGATGCCTAGAATCATGTTTGAAAAAATGAAAATTTTAACAATAAAGTGTCATTGTTAAGAGTC

TAAAAAACCGGTTTATAACCATTTTCCATGGACCACTTGCACGATGCAGTCAAATGGAACAAAGGAAGAGTAAATTAATA

AATAAATATAATATGCCAATTACCATCTCCTATCCTTGAGTTCAAATTACGCCTTCCGTTAGACAACCAAATTAAGATTT

CATAATACGTTTCAAGTACATAAACTATCTACCTGCTCGAAGATCACTTATAGTCATACTATCGAATGCAAGGCAAGGAA

AACATCGCTATCATATTGGCATCTAAAAGAGGATTAATTTATCTCGCAATAACTGTTGTCATTTTTGGGTGCGTCCAAAG

CAATCTCTAATCTCTAAGTAAGATTCATTTAACTGCTTTGATATGCACCAATTTTATGCCTATAATTATGAACACTATAG

AATATGATTTCTAAACATATAATATTACAAATTCTTCTAAATGTGAGCAAATGTTTTTGTACCTGGAATGTCGTCTGATG

GCGGATATTACCCTTTCACATCTGAGAATAAGGGTTTGGTAAGTCGAAAATATTCGCATCTAAATCAGAACCCAAACCAC

ATTTTCCTTGGTGTGAATCACCTCAAACATGACCTAGCATAATGCAACCTGGGGAACTCATTCAAATTGGTATATGATGT

GTAATTCTTAAACATATGCCGGGAAATATCTAGTCAAACAGAAAAATTATTGTTAGGCTCCAATAGAAAGGCAGTACATT

CCGAGAAATCCTATTGCATTGGGATCGATATCCTAAATTAGCTAGCAGCAAGAAAATCAAACACTAAACTTAATGACATT

GCATCGAATCAATACCCTAACTTTAATAATTCTTAACCCCTGCCGTCCACTGTATGCCGCATGTCTAAAGGATGATTATC

TTAATTTTAGCAGTAAAAGGTACATAGGAAAGGGAAGAAGTATAAGAACACACCTTGATTAGCTGATGTTGAAAGACGAT

CGAATAGGAAGAAGCGCAAGAAAAAACTAACAAAGAAGCAAAAATTTAGGGGTTGCCAGTTCTATAAGAACCTTCTTCTC

CTTGTGTCTACAACCTTTCTACCCAAGGGGAATACGGTTTTTGTACGATCCAGCTGACATTGTCTTCTCAATTTATTCCA

AAAGCAGTTCTCGTGGTTCTAGTTAGTTGCAATTGGCTAGTAACCTATAAGAAACTTCTTCTCCTCATCTATAAAAACCC

TTCTACCCCAAGAGGAAGACTGTTCTTATAAGATACCGACCAATAGCACAAACTGGATACCGATGCCTAAAACTCGAAAC

CTAGTGATAAAATTGTCACGTGAAGTCTGGGCATGTAAGATCGAGCCAAGAAATTCAAAACATCACTGGAACTTCTTTCT

GAACCGAGAAACTATTGCAAAATAACTAAAGAATTTTTAAATCAACATCCGTCTAAACCAACAGCGCATACATACTAATC

TTGAAACATATATAGGAGCAACACTAAGGCATCAAATATGATGCAGGCGCAAAATATCAACTGAAAATATGGCAAACACA

GATGCAACACAATAACAATTAACAATCTATTTGTGGGATTTCCACAGAAGAAAAGAAAATGGAAGAAAATATGATATAAA

AAGAGAACTCTATTGCGAGTGTAATCTTTGAAAATCTACATCACCCTAATTGTCTTCATCTCTACGAAAACTTCAGACTA

ACTTTTGACATGGAAACTATATAGATAACAAAATCAAGACCAAAAACATAAAATGTAGTATGAAAACAAAATTGTTGACA

ACATGCCATGTTTCAAAATTTAAAAATTGAGTACACATTGCACCTTGGTCCTTCCTTAGACAATATAGCCGTCCGAGTAA

ATGCGAAATGGATCATATATGTTATCGAATCGGAATGCGAAACCAAATGGGCATAGAAAGTTAGACATACAGTTACCTTG

TGTAAACCATTATAGGAGATTTTGAGATCGGGCAACGAAATCGACCTGTATTTAAGTATAATTTAAATTCAAACTTTCTC

CACAGTAGGCCAGTAATTAATAGCACATAAGACCAATAACACATTTATCACCACCATACATTTTTTGCATTAATCTTATT

TACTGATGCCCAAAAGGCCAAAACATTCATATTCGTCTCATAAACGATTCAGATTATAACAGCGACCGGCATCATAGCCG

AAATAAAAATGAGAAAACAATTATACTCACCATAAATCTGTGTCACTTGATCAATAATTTTATTTCTTTCTCTATTTTTA

CCTGTTGTTTGGATTTTATTGTTTGTGTTTGGCTGGGGAGCTTCCTATTCCCCTGTCACAGTATCTTCTGCTACAAACTT

CACTAGATCAATCATTCCTTGCAAATCAAACTGCAAAATAAAATACCCATAAGAATAAATCAAGGTAAAATATCGTCATT

CAAAGTGGATTCCAAGAGGAAATCAAGTAAATGAGAATAAAGTAATGTTAAACCTGAAAATTGTTCTGCATTCCTTAACT

GACCTGATCTTGCGAGAACTTTGCACATTCAGACTGAAATTTCTCGTATAATGCCTCTATTATATCAGAACTAATCTGCT

CATCCTAGGAAATCAAAATTAGGGTTTTGAACGAAACCTCAAAATTTCAAATTTGGTGCTTTGAAAATTAACAAATCCAG

GGTTACCTGGTATCCCTGTAAAGACTCATTTTGTAAGTCTATCAAACTCATTTTTTTAAAGAAGTAAATAAAAACGAAGA

ATCAATCAAGTTAATACGCACCTGATCAGGTAGAGAAAATAATATCCTAGAATTGCCGAGAGGAACAGGGAGAGGTTGAT

TTAATCATGGACGAAGACTGGCTGCCACTTATTTTGATTTTTATTGTTATTATGTTAATATCTTGGTGAACACTTCTCTC

TTAATCTCCAATAATTTTCTGAATCTCGTTGAAAAGTTTTTCGCATGATATTCTTCGTGGGAGATTAAAAGTTTGTTGGT

GAACTAATGGGAGTCCTTGGAATTTAAGAGAGGTGGGGCATGTCCAATGAACTTGGGTTAATATAATCACGACCGTTGGA

TTTAAAGTATAAGATCAAATCTTGGAGGGGGAGGAAAGATCATGGCCTCTAGTTGGAAAAATGAATATATCTCAGCCGCT

GCGTGACACACCAAACCCTCCTGCTGTATAAGCATTGATAGATCTTGCAGAAATTTACACAATTCAGACAGATAGAATTG

AGCATTACCCTGCTGGCCCATGTATGCACAAAAAATTGATAAGCTTGCAAGTTATACCTTAAGAAGATCCTGCAGCAACA

CACGCAAAGCACTCGTGACACTACATAAAAGAAACAAGGAAAAAACATCAAAAAACTCGAAACACTACAAAGATTCAAGA

GCTAGACAGATAAACAACTAGGTAAATCATAAAGAAAGTTAAAAAGAAGAGCATACGGATAGGACTGGAATAACCTTCCT

TTGGATTGGTGTTGGCTCTTTAAATTGAGGTAATAACAAAATATTGAGATATTAGCTCAACTATCGCCAAGTCAAAACTG

CTTACAAAACAGTAAGTGCCTGCATCAGACTAACAACAAATAAAACTTTCCCCGTTAGTCTGTTTCCTTGAATAGAAAAC

CTGGAGTGCAAATAAATCAGTTCATAAACGATCATGCCAAAACACTAAGATCAACACAAGTAAAGCGCATAACGCAAGAG

AACCAACCAACCAGTAGTATTACTCATCCTAAATTTATTGATGTAAACCTCAAAGCGTGAAAGATATGCCTAACTGAAGA

AAGACAAGGATGTTGGGAGAATTAAGTGCATGAAATGCCTGAATTACAAAGAAGTTCCAGCAATTCACATCATGATCCAG

TAAGCAGCTCATTTAGTTAAAGAGTTAATAAAAAAAGCATATAAAGTCAAATAAAACTCACCTGAAACTAAACATAATCT

ATAAGAACACAGAAGTACAGTACATTTAACACACGAACTACTTCCTAAGAACAAAACATTAAAAAGGAAAATGAATCTAA

AAGCTCAAAACCAATTTAATAGAAACCATTAAACTATCAGATTACCCCCATTCGAGAAAACAAAACTAAGCCAAAGAATA

GTACACAAAACAACCCATTACACCCATTTATAGATTCATAGTAGTGTATAACTATGTAAGGTCTCCCAATGTTCAAAGGT

CTCCCAAACCTAACTCTCTGCATACAACCACTAAAAAGAACAACCAAATTCAGTAAAATTGAAGGGGGGTCACTACTCTC

AATGCACAATTCAGTGAAAATTGAGTACTGAGAATCCGAAGGATGAGAGACATTATACCTAAAAAAACCATTACTTTATG

CACAACTCACTTCTCTCAGTGAACAAATTAAGGGAATCCTTCATTTGAATTATCAGCTTTGGATTCAATGAAATGAGATT

GTGCTTCATAACCATCGGAACCACTCCTTGAAAAACTACCAGAATCTGAATCTGAATCTGCAATCAGTACCAATTACATC

ACGACTTGAACAATAAACAATTGTAGAACAAGAACAAGATTGATACAGGGGAAACAAAATTATATAGCCCATACCAATAT

CAGAAGATGAATCACAAGGAACTTATTAGAACCACTACCAGTTTTGTCACTTGCATAACCCGCATCTTTCTCAATCTCAA

CTGGAGGGGAATTACTAGTAGGAATATCCTCACCAATTTCAGTATTCCATTCATACCAATATTTTCAGCTTCCCTGTTCT

TAATAGCTGCAACTTCAGGTGCATCTCCACAGCAGTATGAGACTACATCGTTCAAAACAAAACCATTAATTAGTAAAACT

TCATTGCAACAGAGCTACCAAAACAAAAACTAAGGTCTTTAGTTGTTTTACCTTGAAGCAATCTCAACCCCATGAAGCAT

ATACATCATTCCCTTCTGCATCTTTAAAGATACAAATCCAGCTGAAAAATCAGTATTCATTCACAGATTCATCCCAAAAT

CAAAATCAAAAACAAGTAATAAATTTAGAGTTAACCAGATGTTTCTTCTCAGAAATTACAAAATAAACATTTGATTCTTC

TTAAGAGTCTAAAAATCTAATCTCTAGGGTTTGCGAAATCAGGTTATGCAAAGATGGATTTGATTTCTGTAAAATGGGTT

TGCAGATTTAAGATTAGGTTATTGAATTCTCGCAGGTTTAGGTTTTATTTTTGGGAAGTTTAGGGGGTTTTTCTGGTTCA

TCAATTGGAGGAGAAAGAGATGAGGGAGAAGGAGAAAGAGGAAGAGGAAGAGAGGCGGAGAAGAGAAAGAAGTGAGTGAG

AAGGATAATGTTTTTCTCTCTCCTTTGGGAGTTTATTTTTGCTGTAGTTTCTCGTAAGTTGTTTTTTTTTTTTGGTGGAT

ACATTGTCCTTCCCGTGTAACATATGTCTTCTATTAATAAAAATCCTTATTACCGATCAAAAAATAATAATAATGTTTTT

CTCTCTCCTTTGGTGTAAAGATAGGTTTGGGGCGAGATATTTGATTCAATCATTTAGATTAAATTTTAACTGTAGAAGGT

GGAATATCCTACCAAGGAAACTAGACTTTTCAATTGTCGTTTTTAAGTCAACAAAAGTCCTAACACAACGACATTTTTAT

ATGTAGTGAGTCTACGATATATGGTTTGTGTCGTTTCATTGTGTCTCAGAAAACCTATTTTCCACTAGTGGTAGGGATAA

GAGTTATCAAATTTGGATCAATTTCAGGCTTTAAAGTAATAGCAATATCATGGAAATGTTTAGTTATACAATCTGTTTCG

TATTTACCAAACATTATCGTCATTTTTTATTATCTCTATTTTATTTCTTTTATACCTTTTTTAGCTTTACTATGAAAAAA

CATTGTGTTCCTATCTCCTTGGGAAATATTCTGGTCTCTACTCTTGTTTTTCCAAAATCTTCCTTTAATAGCATACCAAT

TAGAGAGGTCATTTCTCAGGTAAATTAAATCATGCATCTTGCACTAGTGGAAAAGACGAATTCCGCGACTGCCAGGACAC

GTCATATACTCGAGTTAAACGACCGGTTATGGGGTTCTCTGAATGGGTCGTTTATAAAGCGTCTTTTAATTGCACGACCC

TTATTGCATGGGTCTCTGAACAGAGACCGATTCTTACAGTCACTAGATGGTGACCCGCGATACTGGGCGTAGATTTGAAT

TATAATAAAAAAAAATCAGATTCAGATCAGCTGAAATGTCTGAATGTGTGAATCTGAATTAATCTTACTGGTATTTAAGT

CAAAAAGTCAAGTCAATTCAAGTCTACTTAAGTCAATTAGTCAAGTTTAGTCTAGTCAAAAAGTCAAGTCTAGTCAAGTG

AAGTCAAACTGAAATTCAAATGCAACCTCAGATTGAACTGAAATTCAAATGTAACCTAAAATTCAAATGCAACTCAAATT

AACTGAAATTCAAGTGCAACTTCAAATTTAGAGAATTTTAACATTCATTCCATTGGATGTAAAACTACAAAAAAAAAATG

AAATACTGAGTTCAGTTAATACATGTCCAAGATTGCTAAGTTCTGACAACAACATATGCTAAACTAATAAACTGATAAAA

AAACAAAATGTAAATTCAGCTCCTCCTATGACAATACTTCACAAATGATATCAAGCAGAGCATCAATGTTCCCTGTCCTT

GGTTCACATGCGACTTCCTGCAAAGCAAATCAATAACAATTCAGTTCACAATACAAGGTAGAACAAAATAAGTTACTAAA

TAGGTCATTTACAACAGTGCGAGGTAATTCGACGGACATTCTAGCAATGGGTAGTAGTAGCTGAAAGCACATTTGTAAGG

TAATTTCACATATAGGTCATGTACGTAGATAATAGTTATTGGCTCATTAATTGCAGGGATCCATAATCAAGAACACAACT

ATAATTCCCATCATTTGAACAACTTATACAATCCAAGCAAAACTCACTGTAATATACAATCATAAACTTTAATATATGAT

AGAAAACTTTAACTATTAGTGTTGCAACCTTCACAAGTCATCAAATAGTGAACATAAGGAAACAGATGCTCAAACTTGAA

TAACCTATTAACCCATCACATAAAACATAGGAAAATTAACCAACTGCAAACCAAAAGGAAGAGCAAAGAAAAATATATGG

GTAACATCCATAAAACATGATTCTTTTCAACAACACAAAACCTAAAACATGATCCACAGACCCAATCAACAATATAACTT

AACTAACCATAATACATGATCAACTATTATCACCCAAATCGATAAAATATGCTATACCTGATGAGAATCTGAACTGCAAA

GCTGTTGAGATGAACCTGAAATCAACCAACAAAACAATAATTATTAAACCCCATGACCAAAATCGAACAGTAAAAGAAAA

AGAAAACTTCAATCCAAAATCCTAACTGTACCTGTTGAAGAATTACAACTAGATCGCTATTGAGATTTACCTGAAATCCA

CTAATAAAACAGTTATTGTTATACCCCTTGAAGAAAATCGACAGTAAAAGAAAAAAAAAAGGGAAAAGAAAACTTCAATC

AAAAACCCTAACTTTACTAGTACCTGATGATTTTGAGAATTAAGCCTAACTGCTGCACAATTGAGAGTTGTAATAGGTTA

ATATTAGTAAAGATTTTGTGAAATAAAGGCTTAATTAGTTTTAATGGTGGAGTTCTGGGAGAGGAGGAGAAATTTTAATG

GTGTCTGGTTTTTTTCTCTGCAAATGAACTATTGGAGAGAGAGGGGCGATATGAACGAGATAAATACAATCGTTACTAAC

GAGGTACATAGAAGTGGAGGGAAACGAGTGAGACTTCCCGCCTTAGAAAATATCACTGGATCAACTAGGTGGAGCCGTCT

CGGTCTGGGTTACTGAATTAGGATATGTTTGGAATTAAAAAATTAATTCCACGACCGTTTATGAGGCTCCGGGATATGCT

CCGGAATCTTACAGTATCCGCGACGGTCAATTTTGGTCAACAGTACGTTGACCGGTAATGAGGGTCTCTGATGTGGGTGG

TAAAACACCCATTTTCCACTAGTGTTGTTTGGGTAATTGACCAAAGAATTGGTTCTTTTTCCGGTTTTTTCTTTATATTT

ACTATTTATATTTCCAAATCTAGTTCTATTCCATAATTTCAGGATGTGTTTCATTGTGTCTAGTTTCTTGTTTACTTTAT

AGGCTGAGGACCCTTTGACATTTGTGTTCCAATTGTCCTGAACTAATAATTTACAGTTCTCATGATTTGTCCACTCTCCA

AAGATTTTAAAAGGAGTTGGACCATTATTTTACAGTCCCCATGCTGCGTTCGTTGTCCAAATAATTTAAAAGGTGTTGGT

CCATTATTTCAATGGTTATACATAGTCACTTTTATAGGCATATGATCAGAAGAGTTGGGGTACTTCCTATGCCATTTTGA

AAATACGAGGGTACCCAAATACACCACAATCTTTTATTTATCAACTTATAACTCCTCTACCGAATGTGATCGTCTATGGA

AAGAGTCAAGACAATATGACAATTTAGTTCACACTTCGTATGATCGTCTACAGATACGAGATAGAGACAATACAACAAAA

AGATCACTTGTGTGATTGACTATGGATACAAGGTCAAGACGATACGAAAACAAAGTATGTACTTGATAATAGGTTCGGTA

ATAACAGAAACACTATAGGATCAATATCAAGTATTACAGAGTTAACATACTTGTGTATTTTACTTAACTATAATAACAAT

TATAATACAGAAAACACATAACAAAATTTTTTTAACGAGGAAACCGCAAATGCAGAAAACCCCCGGGACCTAATCCAGTT

TTGAATACTCTCAGGATTAAGCCACTATACAAAATTAAATACCAACTTCGTATAGTTGAGACCAGGCATACTACCCCTAG

CTACTTATTTCCATAGAATCCATGCGCCTTCGACATCTACAGTCACACGCGTGAATAGAAAGTCCTTTGGATCGTATTAC

AAACAACAAATGAAGAATCTGTTTTGTAACCACATTAATCAATATTGCTAGAGGATAAGATGAGTCGTTTAAAAAGGCTC

TTATGTTTCATCTAATAAAATCCTTTGTCTGGTTAGATCAATCTATCCAATAACTACCGAAATATTCAAGTTTAGATTCG

CAGTCAAACGATATAGATCTCAAATAAAAACTATAGAAAATGCCAATCTCACTAAACTAATCAATCATATCTATCAAATA

TAAATTGATTCTAGTTGGATCCCAGCCGATTAAGGTTGTGTTAGAGCATTGCTCGGTTGAACCCACCAAGCGTTGGTATG

TCAAGTTTGGTTGTCATATTTTAGTGAGCCAAAACTCATTTAAGAAGTCACTTGATTTAAATGTACTAGAGTCAACTTTG

TATAGGTTAGCTTGAAAGTATTAGGGTGAGACATTACAAGTATTGCGAAGACTTGAAGAAGTACGGAGCTACAACGACAA

CATCATCATTCCACTTGAGGTTAGTGATATTTAACTTGAATTGTTTCATTCCCTAACGTATCTTTCAAGTCGTGCATATT

AAAAACATAAAACTGCGAAGCATGAACACTCTAGATAGACATAGTATTAAGGAATATAATATGAAGTTTCTTGCTTAACC

ATTAAACTTTGTATATAAGACATTGACATAATCGTATGAATGCTATTGTGATTATGTATGGGTATGAGGTGATGATTTCA

TCCTAGGGAACAATGTTTACATGTGTTCTAAGGAAGTAAGTTCATGATATGTGTTGTGAACCGAAAAGGAAATCGCCAGG

CGTTATTGGGTATGTTCTTCATTGCATATCTTATGAACGACCAATATGTGTGATTAGTATAACCGCTCATGACTTGTTTG

TGTTCTTGGTAAAACTATTCACATGAGGCCTGACTTATGAATTGGTATGACTTTTATTAGTGAAACCGATCTTAAGTAAT

CACCTGGGGTATGATCGGGTTGTGTATGTCTGAACTTATTACTGGGATAAGGGGAACCGATCCTTCTAAGAGGTGCAGTG

CATCACAAGGGGAACCGATCCTTGTATGAGGTGCAACAAGGTTTATGGAAGAAAGGGGAACCTATCCTATGAACATGTGC

AACACGTTTTTGGAAAGGGGGAACCGATCCTATGGACATGTGCTGTGCATGCAAGTTAGATACCAAATATATGTGGGGAA

CCGATCCTAGTACCTAGTCAACCGTATCTTTGGTAAGATAGTATAACTATGCACAATACTCATATGGAGGTAGAACCGAA

ACTTGTTTTGGTAGAACCGTAAACCCATAAGTTGTGATTGAGTGGTACTTGATCAATCACATAGTTCTTGAAAGTCATAT

GAACCAATTATAAACTTGTTTGGAAGCGTGGTAAATCGGTTTCAAGTGAGCAAGCATGGAAAAGACTTGCAAGAGTAAGA

TCTCGTCACTCTTTGAACACGGTTTTTGATGGTGGATTTTAGTTCAGGGCTAAAGTTGTAAAACCAAATTTTGTGTAACC

ATCCTGCAGAGGATGAGACTATCTTATAAGTGATGAATACTTCTTTGCTTTATTAATTTACCTAGAATGGAGCATGTAGT

AACAATCCCCTGAATATTCTGTGAATTTTAGCATTATTGATTAATACATAAAACGCCTTGTATTTCCTGTGATGTTCCTA

ACATAACTATCATTGTTGGTATGGCATGATGACTGATTGTTACTCACAGTAGAGTAACATGAATATCCAAGTACCAATAG

TGAGGAATGCACGAAATGCCCGTATAGGCGGTAACTCTCGGAGTGTTATACTAGACCGATTGAGCCTACATCTCTCGGAG

TGTTATACTATCCCGATTGAGCATACATCTCTCGGAGTGTTATACTAGCCAGATTGAGTATACATATCTCAGAGTATTAT

ACTAGCCTGATTGAGCATCCAGACTGTCCATAGCAGTCTAGGAATGATCGTGATCATCCAACATCACCAGCGTGATTGGA

TGGCCCAGATCGAATCCATGACCCCAAGCAGGTATTAAAATTTTCACTGCCGACCCAACGCGGGTCCCGCATGGTCGGCC

TCCCAAAAAGGGGTGGCATAGCTTTCCAAACCGTCCAACCCAAGCAGAATGGACAAGAAACCCTAAATTAGGGTTTGCGG

CACATTACATGTGTCGCAAAATGATCTCAACCATCCATCTTCGCAGCCATAGATGGAGGGTGTAGATGTAGATTCAAAGG

GCCCAGAGTATGTCAACATAATCACCGTAGGCCCGTCTACACACATGAGCAGTCGTTCAAGACCAACTATGGTTGGTTGC

CCTAATTCGTGCGCCCAAAACAAGGCATGACGCACGGCCAAACCCTAATTAGGGTTTGTATTGATCACGAGCATCCGTTT

TCTCCTCATCGAATGAAGTTTTTAAGAGATAATTGAATAGGTCCAGTGAACATCAACATGATCACCGTGGATCCATCTAT

CTCCTCACCCGACCGACCAAGGCATACCATGCCTGGTTTTTCCAAATGGCACACCCAAACTAGCATGGTCACACTTCCCA

GATCATAAACTGATCTCGACCACTCAACTTCGCCGGACGAATTAAGTGGTTTAGATCACGTTTCCATGGGACAGTCAGGT

ATAGGCGTAAATAACAGTCTACCACCTACACGCCGTACCAAACGACTGAGACAGACCAAGCTTAGTTGTCTCGAAGCGCG

CGCCTTGAAGGTGGCATGCCGCGCACCTTTAGGAATCCTTTGCGACATAGGATTTTTTTCACAAATCGATTTCAGCCACT

CAACTTCACCGGTAAATTTGAGTGGCCTAAATCAAACCTTCGTGGGACAGTGAAGCATGGACATGGATATCAACATGCCA

TCTATACGCATGCCTGATCGGTCCTCCATGAAACGTCTCCCAAGCTTGGATTCTACCAAGATGAAACAGTGCACTCGCGC

ACCTCCTAAACCCTAATCTGATGGTCACAGATGTGGGTAGCAAGAATATCCAAGTACCAACAGTGAGGCATGCACAAAAT

TCTCGTACAGGCTGTAAATCTCGGAGTGTTATACTTGCCCGATTGAGCCTACATATCTCGGAGTGTTATACTAGCCCGAT

TGAGCATACATATCTCGGAGTGTTATACTAGCCCGATTGAGCATCCAGACTGTCCATAGCAGTCTAGGAATGATCGCGAT

CATACAACTTCACTAGCATGATTGGATGGACCAGATCAAATCCATGACCCCAAGCAGGTATTGAAAATTTCACTGCCGAC

CCAACACGGGTCCCGCATGGTCGGCCTCCCCAAAAGGGGTGGTATAGCTTTCCAAACCGTCCAACCCAAGCAGAATGGAC

AAGAAACCCTAAATTAGGGTTTGCGGCACATTACATGTGTCGCAAAACGATCTCAACCATCCATCTTCGCATGCATAGAT

GGAGGGTGTAGATATAGATTCAAAAGGCACATAGCATGTCAACACAATCACCGTAGGTCCGTCTACACACATGACCAGTC

GGTCAAGACCAACTATGGTTGTTTGTCCCAATTCATTCTCCAAAAACAAGGCATGATGCACGGCCAAAACCTAATTAGGG

TTTGTATTGATCACGAGCATATTTTTTCTCCTGATCGAATGAAGGGTTTAGGAGACAATTGAATAGGTCCAGTGAACATC

AACATGATCACCGTGGATCCATCTATCTCCTCACCCGACCGACCAAGGCATACCATGCCTGGTTTTTCCAAATCGCACAC

CCAAACTAGCGTGATTACACTTCCCATATCATAAACTGATCTCGACCACTCAACTTCGCCGGAAGAATTGAGTAGCTTAG

ATCACGTGTCGATGGGACAGTCAGGTATAGGCATTAATACCAGCCTACCACCTACATGCCGGAACAAACGGCCAAGACAT

ACCAAGCTTGGTCGTCTCGAAGCACGCGCCTTGAAGGTGGCATGTCGCTCCCCTTTAGGAATCCTTTGCGACATAGGATT

TCTGTCACAAATCGATCTCAGCCACTCAACTTCACCGGTAAATTTGAGTGGCCTAGATCAAACCTTCGTGGGACAGTGAA

GCATGGACATGGACATCAGCATGCCGTATACACGCATGCCTGATCGGTCCTCCATGAAACGTCTCCCAGTCTTGGATTCG

ACAAAGCTGAAAGAGTGCACTCGCGCACCTCCTAAACCCTAATCCGATGGTCGCATATGTGGGTCGCAAGTCATCTTGTC

CATCCAACTTCGCCAGCCTGGACGGACGGCCTAGTATAGGAGTTAGGAGACCAGTGAGGTATCACCGTGATCAGCGTCTT

CCACTCTACCACACCATGTGGTCTTCCAAGGGAGGGATTGCTAGTGCAACCTCCAAACCATCACATGAAGTAGGTAGGGC

ATAATGCTCAACAACGATGCTTCATATATGTAGAATACATTCGATGCATAACATGCATAGAATACATTCGATGCATAACA

TGCATAGAATACACACGATGCTTTACAAGTATCGACTTCCTAAATAACTTGGTTATTTAACTAGCATTACATGTTACCTT

CTGTGGGTACCACGATCCACACACTTGAGACATCTATCATGTCACAAACTGGGGGATACATACTAGGGTATTGGTCTGGC

AGTCTACAGCGTGCAACGCGCCATCACAAGAACGTGTCAGGAAGACATGGAAGGTTAGTGATGATGAGAGATGTGGGAAA

GACTGATCGTGTGACGCTAGAACACAACTCCACTACTTCATAACTCCACTTCCCACGAGAGACGTGAATAAGGCTCAAAT

GACTTGTATCAATAGGTTCTCCTCCCTATTTCCACAAGAGAACACAACAAATAAACCAAGTGTTAATACAAAAACGTATT

CAAACACCCAGAGCCTTCTTGCGTTAATTGTAAGCCTTCTCACCTTTCTGTTACAAGTCATAAACAAGCCTACACCTTCA

CAATCTCAACACCTTCTTCGCTTCCCTCCCTAAGATCAACCCCATCTCTTTCACTTTGTGACCGAAACAAGTCTGGAACG

GCCATTTCTTGGTTTAGTCTAGAATTGTATAGATTGATCTCTCGAATCAAAAGCACTCCCATGCAGTGCATTTGTTTAGG

GTTAAGATTCACTTCTCATCCACACACCCACAAATTACCAAAACCAGCAGAAGTAGTTTTTACCCATAAACAATTGGCGA

CCACAGTGGGAGGTTAATCTCTCGGTTGTAAAGTCATTTCATCAATACCCAATATCATTGATTGATCTCAATATGGTTGG

TCTTAGGACTCACCAAACAGTTTCATATTCCGGTCGAAGTTCCTCCAACGAAGATGCTCCTCCTCCCAATGCCGTGTCTG

ATGGTGTAACCGGAAGAATCCCAACATTGATTTAGTTAGCACGGGTGCAGAGGATTCACACGAAGAACATGGACCTGATG

AAAGAAACGATAAACGGCATACTCGATCTCCTCATCAAGATGACATCAGAAATGAGCGACTCCCATGTCATAGAGATTTC

TATACTAAGGACTGATCCTTACAATGATCCTCCTCAAGTCAACAGTTTCACTATGAATCATAGGATGGTGGACCAGGAAG

CGGCTTCCCTAGTGGAAATATTGTGTGAACTATTACACCTCTCGAAGAATCAGCGGGCGGAAACGTTCGCTGCAATCAAC

CAGATAGCTTTGGACGTGCAACTAACTCCTTTACGTCCAGAAACCCTCAAAACAACAGATATATTTGTGAATAATGTTAC

ATTTACAGAAGAAGACATGATGGCGGAAGAAGGTCACAACCGGGCCCTACATGTCACTGCGCGTATCAAGGATTCAGAAT

TCAAGAGGGCCCTTGTCGACACAGGAGCATCTTCGAATATGATTACCCTCAAGAAACCCAGGATGTCCAAGGTTAGCCCA

AGCAAGATCGTGCAGCAGCCTACCACGATGACATATTTTGAAGGAAACCAAACCAATACGTACGGGTACGTCAATCTGAA

TTTATCTGTGGACTCGTCTAGTCTAAAGTGAAATTCGAAGTCATAAAAAAGGAGCCGGACTACCACATGATATTGGGGAG

ACCGTGGATTCATGACAATATAATTGTTCCTTCAACATATCACCAGTGTTTGAAGACTATGCTGAATAAGGAAGTCGTTC

GCATCCCTGCATCGTTCTCTCCTTACGCACCGATCTGTGGCGTAGAGTTGTTTGAGCCGAAAGAAACTTTGTGGAAAGAA

TCAGGTAAAGTTCATTACATCCCACTCCCCACGTGGGCATCAATCCAGGAGGAAGATCGACTTGCATCATTGAAGGCTAA

GCCCCATGATATGTCAGAACATCCTTCTGAATCTACGACTCATATTCACACTAAAAGAGAGCGAACTTTTGTGGCAAGCC

GTGATCAGAAGGGAAAGACTGTGTACCGATGTCGTTGTTAGAAATTAGCAGGGGAGGCTACTACCCAGCCTCTCTGCCCA

ATGGAATGTCGCAACAACGAAAAGCCGCAGGAGGAGTTATTGGATGCTCCACGATAGCTGCAGGACGGCACCAACTCCAC

GACCGACGAAATCGAAGTTGTTAATATCGGGAGTGAGGAATTCCCCAGGCCTATCTTCATTAGCTCAACCTTGTCCACAA

ATGAACGGACAAAACTTGTGCAACTTCTCAAGGAATACCAAGACATATTCGCTTGGACGTATGAAGAAATGCCCGGTTTA

GATGAAAAATTGGTCACCCATCATCTGCATGTCATACCTGGATCCAAGCCGGTCAAACAATCTCCGAGACAATTCAGGCA

CGAAGTGGAAAAATAGATCAAGACTGAGATACAGAAGCTTCTAGAAGCTGGCTTCATTAAACCCATTCATCACCCAACTT

GGTTGGCTAATGTGGTTCCAGTAAGAAAAAAATTGCCAGATCAGATGCTGCGTGGACTTCAGAGATTTGAACAGATGTTG

CCCCAAAGATGATTTTCTTTTACCTAACATCGACATGTTAGTGGATGCAACCAGTGGGAATGGCATGTTCTCCTTCATGG

ACGGATACAATGGGTACAATCAGATAAGAATGTACGAGCATGATGCAAGTAAGACATCTTTTCAAACTCCTCTCGGAAAT

TTTTATTATACAGTAATGCCATTTGGTTTGAAGAACGCCGGTGCGACCTATCAATGCGCTATGATGGAAATCTTCCACGA

CATGATGCACAAGAAAGTTGAAGACTACGTGGATGATATAGTAGTCAAATCAAGGGCTCGGTCGGATCACACAGGAGTCC

TACCACAAGTGTTCGAAAGGTGCCATGAGTACAAATTGAAGATGAACCCATTGAAATGCGCCTTCGGTGTTTCTTAAGGC

AAATTTCTGGGATTCCAGGTAACCGCTAAGGGAATCGAGGTAGACCCATCCAAGACTCAAGACATCCTCATGATGCCACC

GCCACAAACTGTGAAGGAACTCCAAAGCTTCATGGGAAAGGTGAACTACATTCAGATTTCATACCCGGCTTGGCCTAGCT

CGTTGCTTCATTCACTTCCTTGCTAAATAAGGGAGCAATATTTATCTGGATCGCACTACAACATGAAGCGTTTCGGAAGA

TTCAACGGATGTTGTCGTTTCCAGTGGTCATGAAGTCCCCTGTCCAAGGGAGAACGTTGTATCTCTATACAGCCTTCAGC

GATACTGCCGTAGGGGCTTTGCTCGCATTTGAGGATGATGAAGGAGTCGAACATCCCATATACTATTTCAGTCGAATACT

AAGAGATGTTGAGCTCAGATATCCCAAAGCGGAGAAGGCGTGCCTAGCCCTGATTCATTCTATTCAGAGGTTCAGACATT

ACCTGATGTCCAACAAGGTAGTACTCATATCTAAATTCGAACCTGCAAAGCTTTTACTTTCAAATGCGGTCTTGATGGGA

AGGGCGGCCAAGTGGATTCTCTAAATGTAGGAGCTAGACATAACGTGCACGTCACTGAGAGCTATCAAAGGACAAGATGT

TGATGATTTACTTGCAACATTCTAGAGAAGGCAATACTGCATTACACGAGGATCTTCCTAGAGAATTTCCAGAGATCTCC

ATTGTCGAAAGAGAGGCATGGCTACTGTATTTCGATGGATCCTCCACTCCTAGAAATAACACTGGAGGAGCAGGCGTGGT

CCTAGTATATCCAACTGGCGAAATCTTTTCACACTCTTTTAAGTTGGACTTTCAATTCACCAACAACTTATCGGAGTATG

AATCCTTTCTCATAGGACTGTCAATAGACAAGCAAGCAGGGGACACTCGCCTAGAGATAAGAGGTAACTCTAAATTACTG

GTTAATCAGATGAACGGAGTCTATGCATTGAAGGAAGCAACACTGGCCCCATACCGATCAGAAGTGCAAAGGCTGTTGAA

CTACTTTGCAGATGCAACCATAACCCATGTTGGTCGCAGCAACAACATACATGTCGATTGTTTAGCCACACTGGTGTCCA

AATTGCAGTTCGAAGGTTTGGAAGAGACTCTGATGGTAAGAAGGAGGACAGTGGAATCAACATGGCTCTCACAACGCGAA

TACATTGGGACCTGTGATTGGAGAACTCCAATCATTCAAGAACTCAGCAGCTCACTTTCTCATGGAAAGGTTAGTCTCAA

AACTCTGCAAAGCGATTTCATGCTTCATGGTATGCTATATCATCGAAATCCGGATGGTTCCATATCGAGATGTCTCTGAG

ATGAAGAGGCGCAGTTAAAGCTTAACCGAGTTCATACTGAAACCTGTGGACAAACGTTGGTGGTAATGCTCTATCGTCGT

CTTCAACGCCTGGGCTATTATTGGCCATAAATGGAAGCTCAATATCGTTTACTCCAGAAGTCTTGCACCAACTTCCAAGC

GCAGCCACACCAACTAGAGGTCCTTAATGTAAGCCATGCTGGGGACTAGATAGAACCGTACATTAGTTATCTCTGCGATG

GAGTAACTCCTGCCAACCAAAGTGATGCTGTCAAGATGAAACAAAAGGCGAAGAGATTTGTATTTCATGAGGGGATCCTA

TATCATAAAAGTTTTGGAGGCGATCTATTACGGTGTTTGGCTAAGGACGATGTTTCAACTATGTTGAAAGGAATGCACGA

AGGGGAACATCAAGGGAAGAGGAAACTATTTATACAAATACATGAGAAGTACTATTGGCCGACCATGGAGGATGATGCAG

CTTCATTCGTGCAACACTTTCACGAGTGTCAAGTTCACAACTTCACATGAACCTCGTTCACGCACCCTCCACTCCTTTAC

ACTCGATAAGCAGTCTGTGGCCCTTTTATAGCTGGGGACTTGACATTATTGGGAGGATTAATCTGTCATCGTCGAAGAAG

CACGAGTATATCATAACTGTGACGGAGTATTTTACTAAGTGGGTTGAATCCATCCCTCTACGGGGATCTACAGGAGAAAC

AATTGCCGCCTTCATCAAAGAATACATCATTTGTCAGTTTGGTGTACCCAAGCACATCATCACCGACAATGGCACTCCCT

TTGCAAATAAACATGTAAGGGAATTGCTTGAAGAATATGGAATTAAACAGGTCTTCTCCACCATCTACTATCCTCAAGGA

AATAGACAAGCTGAGAGCACCAACAAGAAAATGATCCGGATTCTCAGTCGAATAGTGCATAACCCTAGGAACTGGCACGA

ACAGGAACAACATTCACTATAGTAACAACATTCTTTGAACCGCCTGCTCCACCTTCTTTGATTCCGCAGCCAGAGCCTTT

TCCTTCTCCAATATAGCTTTTGTAGCAGGATCAACATTCACGAAGATACCAGCTCGGTCAACCTTGATGATATCAAGACG

CTGCCGCAGCCAGCCCATGTTGAACTCCATGGATTCGCAGGTAGATACCATGTCCTCCCACTTTTCGATATTTGATTCAG

CGACCTTGCGAACGAAAATATTCTCCATCTCAGCAATCATGGGAAGAAGGCAGTTTACTGTAGTAACCAGAGCAGATGAG

TAGTGCCGAACTACATTCCTAGTTGCGATGTGGCCATACTTTTTCCACTCTTTTTTATACAGAGGCACAAACTCCTTCCG

CACTAAGAATCCACCAACATCTCGGTAGCTGTAAGACAGGCATTTCATGTGTACATCGAATGCAAGACCCGCATTGGGGT

ATTCGTAAGAAGAGATACCCAGATCAGCATCTTCAAGGTTTGCAGCGTCAGGCGTGTTCGAGTTTGAAGGAAGAGGACTG

GTTAAGTCATCATCATCGATAACCGTTACACATGTTCCAGTGGGGTATCTCTGCAAAAAGGAAAAGATCATTCGTCTGAT

CGTCATGAATAACAAGAAACTGTGCGGAGAATAAAAGGAAAAGAGAGGCAAACCGGTATGACCTTTTTGACTCGGAGAAC

ACGAGCATGGGACTCATCATGAAGCCCGGACGCAGATCGTGTACGCTTAATATAAGATCCTGGGTGGTATGAAAAAGATG

ATCCACTGTTGTTCGACATGATGAGGAAACAGAGTTTCTTCCAGAGAACCCTATTGTAAAATTATAAAAAGTGGTTCAAA

ATCTCCACGCATGCATATATACTGAGGAAACCCTAAAGGAAGAGTAAAGTGGCCGCAGCGAAGCCTACCCGATATATGAG

GATTCAACACTGACCAAGTCATGCTTTCTTTCAAACCCTAATCAAAGATTGAGAGAACAACTACATATGAACGATTTGGC

CAGAGCAGTGTGTAAACACCCACGCCGGGGAATCCCAGTAGAAGAAGACGCCTAGGGGAGGAAGCCGGTCCCACTAAGGC

CCGTGGTCACGCATGCATGGAGGATTTCCCACGATTTGTCAGAGTCCCTCCAGGTAGACTTCCCTCCTCTCATCCTCTTA

TGTTATATATGTTACCATCTCATGCCTACCCCACGATAGTGTAACCATCATGTGCTAGGCAGGGGACTTAATGTTGATGG

TGGATTTTAGTTCAGGGCTAAAGTTGTAAAACCAATTTTTGTGTAACCATCCTGCAAAGGATGAGACTATTTTATAAGTG

ATGAATACTTCTTCGCTTTATTAATTCACCTAGAATGGAGCATGTAGCAACAATCCCCCGAATATTCTGTGTATTTTAGC

ATTATTGATTAATGCATAAAACACCTTGTATTTCTTGTGTTGTTCCTAGCATAACTCTCATTGTTGGTGCGGCATGATGA

CTGATTGTTACTCGCATCAGAGTAGCATGAATATCCAAGTACCAACAGTGAGGCATGCACGAAATGCCCGTACAGGCTGT

AAATCTCGGAGTGTTATACTAGCCCGATTGAGCATCCAGTTTGTCCATAACAATCTAGGAATGATCGCGATCATCCAACT

TCACTAGCATGATTGGATGTCCCAGATCGATTCCATGACCCCAAGCAGGTATTGGAAATTTCATTACCGACCTAACGTGG

GTCCCGCATGGTTGGCCTCCCCAAAAGGGGTGGCATAGCTTGCCAAACCGTCCAACCCAAGTAGAATGGACAAGAAACCC

TAAATTAGGGTTTGCAGCACATTACATGTGTCGCAAAACGATCTCAACCATCCATCTTCGCAGGCATAGATGGAGGGTGT

AGATGTAGATTCAAAGGGCCCAGAGCATGTCAATACAATAACCGTAGGCCCGTCAACACACATGACCAGTCGGTCAAGAC

TAACTATGGTTGGTTGTCCCAATTCATGCGCCCAAACAAGGCATGGCGCACGCCCAAACCCTAATTAGGGTTTGTATTGA

TCACGAGCATCCGTTTTCTCCTCATCGAATGAAGGGTTTAGGAGACAATTAAATAGGTCCAGTAAACATCAACATGATCA

CCGTGGATCCATCTATTTCCTCACCCAAACGACCAAGGCATACCATGCCTGGTTTTTCCAAATCGCACGCCCAAACTAGC

ATGGTCACACTTCCCAGATCATAAACTGATCTCGACCACTCAACTTCGCCAGACGAATTGAGTGGATTAGATCACGTTTC

CATAGGACAGTCAGGTATAGGCGTGAATACCAGCCAACCACCTACACACCGGACCAAGCGGCCGAGACATACCAAGCTTG

GTCGTCTCGAAGTGCGCGCCTTAAAGGTGGTATGCCGCGCCCCTTTAGGAATCCTTTGCAACATAGGATTTCTGTCACAA

ATCGATCTTACCCACTCAAACCTTATATCAAACCTTCATGGGACAGTGAAGCATGGACATGGACATCAGCATGTCGTCTA

CACGCATGCTTGACCGGTCCTCCATGAAACGTCTCCCAGGCTTGGATTCGACCAAGCTTAAAGAGTGCACTCGCGCACCT

CCTAAGCCCTAATCCGATGGTCACAGATATGGGTCGCAAGTCATATTGTCCGTCCAACTTCGCCATCCTGGACGGACGGC

CTAGTATAGGAGTTAGGCGACCAGTGAGGTATCACCGTGATCACCACATGCTTTACAAGTATCGACTTCCTAAATAACTT

AGTTATTTAACTAGCATTACATGATACCTTCTGTGGGTCTCATGATCCACACAATCGAGACATCAATCATGTCACAAACT

GTGGGATACATACTGGGGTATTGGTCTGGCGGTTTACAACGTGCAGCGTGCAACGCGCCATCACAAGAACGTGTCAGGAA

GACACAGACGGTTAGTGATGATGGGAGAAGTGGGCAAGACTGATCGTGTGACGCTAGCACACAACTCTAATACTTCATCA

CTCCACTTCCTCTACTTCCCATAAGAGACGTGAATAAGGCTCAAATGACTTGTATAAATAGGTTCTCCTCCCTATTTCCA

CAAGAGAACACAACACATAAACCAAGTGTTAATACAAAAACATATTAAAACACCCAGATCCTGCTTGCGTTAATTGTAAG

CCTGCTCACATTTCTGTTACATGTTATAAACAAGCCTACACCTTCATAATCTCAACAACTTCTTCGCTTCCCTCCCTAAG

ATCAACCCCATCTCCTTCACTTTGTGACCGAAACAAGTCTGGAATGACCATTTCTTGGTTTAGGCCAGAATTGTATAGAT

TGATCTCTCGAATCAAAAGCACTCTCGTGCAGTGCATTTGTTTAGGGTTTAGATTCACTTCTCATCCACACACCCCAAAT

TACCAAAACCAGCAGAAGTAGTTTTTACCCATAAACAACGGTGCATTAATGCTTATTCTATTTTGTTCAAAGATATTCCT

TAAAGGCTAAGAAGGAAGTAAATCCCAAAGATCGAATTAAATAAGTTAAGAATCTTTTTAATTAAGGGTTTTTAATTATA

TATGGAAAATAAGAATTAGTAATGTGCATTTGCTAATTAAAGATTTTCCAAGGAGATTTCGATCGTTATTTTTGAATAGT

GCATTCAGAGGAATTATGAAAATCGAATTTGGTGGAGTAATAAAAAATATCTTGATAATATTTTCGGTTTTGGAAATTCC

TTGGTGTCCAAACTTCCTTGTTTATAAATACTTTGAGTTTGCATTTCTATCAAACTAATCCTGCGTGACATAGACTTCCT

CTTTTGTCTTTGTTACTGGTGAAGCCACCTATCGGAGAGGAGAGTAATCTAATTAGGTGAAATCTCTTACGGCCGCTAAA

TTTAAAGTCTTCTTTGGGATTGAGAATCTATAGCGCGACCCGTTGGTGGGAAACTAGATATTGCGGTTTATATTTAGTTT

TCGTATTGATTTGATTGACTAACGGTGGTTGAACTTTGATTGCACCTAGTTTGTTTATGCTTGAGAATCTTCTCTTCTGA

TATAAGATTCACTCAAACTAGTTCAAGGTTTCAACATGGATCTTTAAACTGTTGTTAGATCTAAAGACGATCTTGTGATA

ATCCATTGTTAACAGACTCCATTCTGTATGTGATTCATCACAAGAGATTCAAGTAATTGTGTGTAGGTGTTTATTGAAGA

TTTAAGAAGATATTGAAGATTTCTGACTTAGTTTTTACAATCTTTGGTGTGCACAATACTTGTTTGGGAAAAATAAGATC

TAATTAATAATCGGTTTATCCTTGTGTTAGATTGATTAATTGAGGAAGATCGGCATCAATACGTTTACTTGAGATTGAAG

GTATTGATTGGCTTAATATTAACGATTACCTTTGGGTAATTGAACATAAGATAGATCTTGGAACCTAGCAAAGGAGTTTA

TGTTAAGATAAACAAGGGAACCTTTGCACAACTCATATCACTTGGTTGAAGAGAGTTGATACCAAACAGATTTGTTGTTC

CTTAACTTTTTGGAATACGATCCAAAGGAATTGGTCCAAGTACGTGACTTGTTTATAGGTTGGAGGCGTGGGAATATAGA

AGGAACTAGGTGAATTATAGGTTTAGTTGCTTTGTATCAACTATACGAAGTTGGTATTATTTTGTATAGCGGATTAATCC

TGAGAGTATTCAATTCTGGACTAGGTCCCGGGTTTTTCTATATTTTCGGTTTCCTCGTTAACAAAATCTTGTTGTGTCAT

TTACTTTATATTTCCGCATTATAATTGTTTTATTATAATTAAAGTAAATTGCACAAACGTTAATTCCTATTTACTTGATG

CGTGAATCCTATTGAGTTTGATTAAGTCTGAACCTTTGTATCAAGTAAACACACTTTGTTGTTGTATTGTCTCGATCTCG

TATCCATAGACGATCACACAACGTGCGAACCGATTTTTTGTATTGTCTCGACTCAGTCCATAGATAATCACTTTCGGGGA

AAGGACTTATAGGAAGGAAAAGTAATATCTTGAGGTATATTTAGGTACCCCCTCCTTTTCAGGTTTGTGCACTAAATCCA

CAAGATACGAAAACTAATAAGAATTCTTCTTCGTCTTCAAATATGCTATTTCCTTCAAATACCTGCACAAAAACACTTGA

ATACTCATTTGGAGTCTGTTAACAATGGATTATCACAAAATGTATTTAGATCTAACAACAGTTCTAAAGATCCAGTCGAT

ACTTTGATCTAGTCTGAGTGAATATTATATGAGAAGAGAAGATTCTCAAGAATAAACAAACTAGGTGCAATTAAAGTTTC

AACAACTTTTAGTCAATCAAATAAATAATCAAAAACTAATAACACTGCAATTATCTAGTTTCTCACCAACAGTACTCTTA

GAGCTTCTTGATCCCACACATGTCTTTAAACGAGCGGTCTTAAGAGATTTCACATAATTAAGATACTTTTGTCTCTAAAT

AGACGGCTCTACTAGAAACAATAAGAAATTAAGTTTTTCTGGATCTTAGAATAGTTTGATAGAAATGAAAAATTAGGTAT

TTATAAACCAAGGTTGTTTGGACAAGGCTTCGAGTCTCCCTCAAGGTTTTGAGTTTCTCTCTAGCATATTTTCGGCAAAT

TAGAGACAAAACAGTTAGTTGTGAATCCGAGCAACAGTTCCCAGGGTCGAATCTCCCTTCACGCAATATTTTTTTGACTT

TTAAAAAAAAAAAATGGACACCAAGGAATTTCCAAAACCGAAAATATTCTTAATATATGCAATGAATGCCCAAATTCTGT

TTTTCTAATTCCAATAAATGCAGTCCACTATTTTCCGAAATATCTCATAGAAAATCTCCAATTAGTAAATGCAAATTACT

AAGTTTTATTCTCTAGAGATATGCATTAATTGTTGTAATTAAAGCATAAACCCCAAAAACCTTAATTAAAAGATTCTTAA

TTATTTCGGTTACAGGATCACCTTGAGTGCCAAGGAATATCTATGAACAATAAATGATAAGAGTTAATGCTCATGTTCAA

ATTATGTTGACATCTTTACACAAAAAACCTAAATCCGAAACTACACAAGTTCATGAAGAAAATATGTAATTCATGGAAAA

TCGGATTTGCATCTTGGAATCAGTTCCACCACACTTCCAAACAAGTTTAGAATTGGTTCTACCAGAATTCCAAGACTACT

ATGTGATCAGTCATATCAAGTCATAATGGTTAGTTGTGATCGGTCCTACCAAGTCACATACATGGTTCGGATCGGTTATA

CCAATCACAACCTGTGTATTGTTAGGTTAATTAGGGTATTAAGATTCATGTGTATTGTTTGTTGTTTTTCAGTAGGTAAG

TATGATTATTGATTTATGCTAAGATATACCTTTTTTTTTTACATTGATGCATGTTTGATGTTTGCTTCAATTTGAACCTC

AACATCTACTAAATGGGTTCTATGTAAAACCGATGTTGTATTCTTACACATTTCAAACTCTATGAAATGCAAAAAATGAT

CTTGTCTTCTTTTCTTTGTTGATAGATTATAATTCATGCTGATGAGAGTTTCTTAGCCTTATTAGAATTATGTCTCATCT

GAACTTGTACTTCATACTATTAATTAGAGATTTATGTGGCGGATGGTATTTTTTCTTTGCCTGATTTGATCTATTTATAA

GACAATTCTTGCAAGGTTGCAGTTGTTATTCTTTATTTTAAGGTGTAATTTTTCTTCTTGTTCGTGTTTTAGTCTATATG

AGTATAAAAAATAACATATATAGATTTATTAACAGTGTACCCTGTGAATTTTGACAAAATGCACATTAAGTGTTTGTATG

ATTTACTTTAAAGATATGGGTTTTCTTAAATACTTGCGTTGTTATTGTCTTCCAGTTGCATAATACTTGAGTTTTCAGAA

CCGAAATTTATTAGGTCTATTTGAACCCTGTATGACATGGTTTTCAGCCAATATTGAAGCAAATCAAATCATATTCAAGT

AACCTGTTTGTAAGGTAAAATTGTTTCTACCCGAAACACAGAAATTTGTATGATGTTGTTGTTGGGAATCGATCGCTGCC

TCGATCTTCAGTTGAATTATGGAGAAAGGGGGGTACCTACAAAAAACCCTCCGATGCTAAAGGCAGTATAGGATTTTTTT

CACAAGAGTTTTTGGTATGGAAGATTGCTTACCTTTTTGGTTGTAAAACTCTCCATTTATATGTGGTGAATTTCCGTTTT

ATTTCTTTGATTTTATGGCAGATATCAATTGCCATAACTGGCCCCTGCATGCCTCCCTCCAATAATAGTCCCTGGAATAA

TTCACCTGCATGCTTGCATGACTGCATGGTTGCATGCCCAGGCTTCCCCTGCATGGTTCCCAGAATATTCCAGAAACTTC

CATTATGGTTAAGGAATTATAATGGGGTGTGGATAACCAAAGGTGTAAGAGAAATGGGTGTAAGCAAACGGGGCGTAAAT

ATTCTTAGAGTAATAGGACTGGGGCGTAAATATCCAAGGGTTACGGGATTAAGGCACATCGAGCAATACCAGATCAGTCC

CATCAATTAATCGCCACCGATTAATCGCCACCGATTAATCAGCGATTAAACGCCAAGAACGCTTAATCCACGATTAGTCG

CCGATTCATGGCCGCCGGCTTCGGCCATCGTCCATACTCCCGTTCAGAATGACGTCATGCTGACATCGCAAATAGAATAT

TCCGCCTCCACTATGAATATGCTGACGCTGTTAGTGACGGCGTTAATTTTAACGAAGATGGTGACACCATTAGCATTTAA

CAGAATATTCTCGGGATATGCTGACGCTGTTAATGATGTCGTTAAACTTAATGGTATTACTGATGTGGCGTCAATTTCAC

CAATCTGGCCCTATTTGCTGACGTGTCACACACCACGTCAGCACCTTTGCCACGCCATCCTTCCTGGTGAAGTGGCGCGT

GGTCTGCTGACATGGCACCTACCACGTCATCACCTTCCGCTGACGTGGACACACTTTGCTGAGTGGCAGATGCCACATCA

TCACCTACTGATGATGTGGCCACACTTTGCTGATGTGGCTTATGACATGTCATCGTCCTTGATGACGTGTCAAATGCTCT

GCTGATGTGGCACTTGTCTAGCCTTAATTTGTGTATTTTGCATATGGGCTTGCATATGGGTGTTTGTTTGTCCTAGTTAG

CCATACTTGGGTAACCAAGGGAAGCACCTTAGGCCCAATGATCCCTCCTTTGCTAATAGTAGGAGTTAGGTTATCCCATT

TATTTGTGTAGAATATAAGGGGTACAAACATAACCCAATTTTATGTGCAGGTACTATTAAACATTTCTATGGAAAATGCA

GCACCAATTGGTAAGAAACGTCAAGTTAGACCTACAATGTGGGATCACTTTGATATGGAAGAGGCGCCTTCAAGATATGC

AAGGTGTCATTATTGTGATGCAAGATTTGCTTCTGATCCTATTGTTTATGGGTAAAAATCGTTTCTTCCGATTTTGGTAA

ATTCGGTAGGTGAGTGAGAAACTGGTTTAAACCCTAAATAAATGTGTTGCACGTGAGCACTTTCTAGATTCAAGAGATCA

ATCTGTACAAGTCTTGCCTAAACCAAGAAATGGCCGTTCTAGACTTGCTTCGGTCACAAAGTGGAGGAGAAGGGTTGATT

TGGGGAGGGAAGCGAAGAAATGTGTTGAGACCAGAATAATTATGGAAGAGTAGTGTCTCGACTTGTATTAGAAAGTGAAA

GCTTCGGAAGCTTGCAAGAGATGAAAGCTTTTTGGAGAGCTTGCAAAGTTTCCTTTTATGAGAGTGTTGTGTTTTTCTAC

TGAACGAGATTTATTATCTGGTGGAAATAGGTAGGACCTATTTATACAAGTCGAGATTGAAACGTACCATGGCCTCATGA

AGAAAGAGAGTAAAGGGTGAAATAATGGGAGGAGGTGGTAACCGCATGGTGTTTATGGGTGAAAACGTTTTCACCATTAT

TTCCTCATTTATTTGACCTCTTTATTCATATTCACACTTTCTTAAAGTGAGCGCAGTGTACGCTGCATGCATTGACCGCC

AAACCAAAACCCTAGAAAGCATCCCCCAGTTTGTGTTGATGTCTCGAGTATTGGTTGCACCTTGAGTGCAAGTGGAAATG

CCAGGCTACAAGTGGAGCGTATGGCATGTGCGGAATGCCTTTGATGTGGGTTGCACCTTGAGTGCAAGTGGTAATGCCAA

AGCTGCTAGTGGCGCATGTGGCATGTGCGGAATACCTTGGTTGTAGGTTGCACGTTGAGTGCAAGTGGCCATGACAAAGC

TGCCAGTGGCGCGTGTGGCATGTACCAATGGCATGTACGGAATGCATTGGTGGTAGGTTGCACGGCTTGCAAGTTAGGCT

ACCAGCGTGCGTGTTTGGCATGTGTCAATGGCACACACGTCCATTAGGGCCACGACTAGGTAATGAAAGCTACCAGTGAG

CGCGTTGGCATGTGCCAATGGCACGGTCGGACATTAGGGCCACGGCTTGGTAAGAAAAGCTGCCAGTGAGCGCGTTGGCA

TGTACCAATAGCACGCTCGGCTATTAGGGCCACAGCTAGGTAATGAAAGCTACCAATGAGCGCGTTGGCATGTGCCAATG

GCATGCTCGGCCATTTGGGCCACGACTTGGTAAGAAAAGCTGCCAGTGAGCGTGTTGGCATGTGGCAATGGCATGCTCGG

CCATTAGAGCATGGCTTGGTAAGAAAAGCTGCCAGTGAGCGTGTTGGCATGTGCCAATGTCATGCGTGGCCAATTTGGAC

TAGGTTGCATGGTTAGGACCAATGTAAGGGTGTTTGGCTTGGCGTGGTGTGACTAGCATAGCTAGGTCCAAGGAAAAGTG

TGTTTAGCATGACGTGGCTAGCTTGGGCTAGGTGCGGTTGGCATGATTTCCCATTGGCATAGAGGTGCGGCTAGCATGGT

TGGCATGCTTTGGCGTGTTTGGTGTGGCTGGCATAGCTAGGGCCAAGGCAGAGTGCGTTTAGCATGGCGCGGCTAGCTTG

GGCCAGGTGTGGCTGGCATGATTTGACATTGGCATAGAGGTGCAGCTAGCATGGTTGGCATGCTTTGGCGCGTTTGGTGT

GGCTGGCATGATTTGCCATTGGCACAAGGGTACAGCTAGAATGATTGACATGCTTTGGCGCGGTTGGTGTGACTAGCATA

ACTAGGGCCAAGGCAGAGTGCGTTTGGCATGGCGCAGCTAGCTTGGGCTAGGTGTGGCTGACATAATTTGCCATTGGCAC

ATAGGTGCGGCTAGCATGGTTTGCATGCTTTGACGTGTTTGGTGTGGCTGGCGTAGCTAGGGCCAAGGAAGGGTGTGTTT

GGCATGGCGCGGCTAGCTTGGGCCAGGTGTGGCTGGCATGATTTTCCATTGGCATATAGGTGCGGCTAGCATGGTTGGCA

TGCTTTGGCGCGGTTGGTGTGACTAGCATAGCTAGGGCCAAGGAAGAGTGCGTTAGGCATGGCGCGGTTAGCTTGGGCTA

GGTGTGGCTGGCATGATTTGCCATTGGCACAGAGGGAAATTAGGGTTCGCTCGTTGTAACCCTAATTTGAAAAAATAGCC

CCCATTAATTACTGACTACATGCATTAAATGTTCTAATAAATGTGTCACCGGTGACATTAGGTTATTCGGGGTTTTACAA

TTTTACCCTTTGTCTAAAAACCACCATCAACACCTATGAAGAATGGTACCTCTGGGTTGATTGCTCATCTAAAGAGATAT

AGACTAATAGTTAATCAGATAAGAAATTGATTGAAGTTTACTTTGTATTAGTATAAGTACATATTTGAAATCTTAATGCC

TCAAATATTATGTTTGTATCAGGTTAAATTCTATCAGTTTCATGTTACTAGAAATTTTCATTGAAAATTTTACTTGTCGG

ATATGGAAAACTGACTAAAATGAAAGTCTGTTTATAAATATGTGGTTAACCAAAACCTACCAATTTAAATCCGAAACCGA

GAGGTTCAAAACTGAATGGAACCGAATAGGCTAAATGGTTTCATTATTGGAAAACCGAGAACTTTGGTTTTGCCAATTAC

CGATGAATACCGAACCATGTGCGGCCCTAATCACAAGGATCGGTTCTACCTAAATATGGTACCTACTTGTGATCGGTCAT

ACCAGTCACTAGGGTCGGTCATACCAATTACAAGGATCGGTCACACCAATTACCAGAATTGGTCACAACAATTATAAGGA

TCGATCATACCATCTCATGGTGATTACTTGGGATCGTTTACACCAATTACCAGGACCGGTCATACCAAATCATAAGTCTA

GGTATTGTGATTGGTTATACCAAGATACATAATCTGAGTTAATATTGGTTCTACCAACTCACACATATTGGCCATCCAAA

GAATATGCAATGAATAGCTGGACCAGTATACCTACTGATTTCCATTTCAATTCATGAAACAAGTTCACAAACTTACTTCC

TTTAAAAATATGTAAAACATTGTTTACTAGGATGAAATTTTCACTATACACCCATACACATAATCATAAAACTATGTACA

AGATTATGTCGATGTCATATCTACGAAAAAGATAAGTGTTATACTTCATAATCTAATTCCTTACTACTATGACCATACTA

TTATGATCATGTCTATCCACTACAATATTATATACAGTATATACAGCTTCGCATGTTATGTTTTAAATATAGCACGACTT

GAAAGATACGTTAGGAATTGAAACAAGTTCAAGTCAATATTACTAACTTCAAGTGGAAGGATGATGTCCTCGTTTTAGTC

GTTACTTCTTCTAATTCTTTAGGTCTTCGGAGTAATAATTGTATATCTCACCATTCCTAGACTTTCTAGTCTAACCTAAA

CGAAGTTGACTCTAGTATTTAATCAAGCGACTCTAGATGAGTTTCGATACTAAAATATGACAACCAAAATTGACATACCA

ACTCTTGGTGAGTTCAACCGAGCTATGCTCTAATACATTTATCGTTCACCAAAGCTCTGTCTAATCTTTATTCAATATTT

TGATAATTTTGCCTATGGTTATTCCAGGTGAAAGTGCAGCCTGTAAATCCTAGATCCATAAGATTGCACTTGTTGATTAA

CTTGGCCACATTTTTCTTAAATGGATAACGGCTTTGATTTTCTTCTTCATGAAAAACAACATTTAAATCCCCTATGATTA

TCCAAGGTTTTTTTATTGCAGCAGCCATTTATTTTATGGTTTTCCAAGATTTTAGTTTCAAATTTATGTATAGACTCCCA

TAAAAACAGGTGAGAGTCCACTGTCATTTATATGATTCGTAATAGTAGCATTTAGTTAAAAGCATGGTCCTCTATTTTTG

TGTTAACATTATTTTTCCAAATGAGACAAAGACCTCTGACACTACCTCTAGGAGGAACTAAAAAATAGTTATGAACATTC

ACAATCTCATTATATTTCCCATCTCCTTATTTTGTTGTTTAGTTTCTGATATGAATATGATATCTGGGTTTTCCTTGTTT

AAAGTATCTAGGAGAATGTTTCTGGTGTTTAATTTCCCTAGTCCTTGACAGTTCAGGATAGCTAAATTATACTAGAAGTA

TGATATTTCAGGATAGCTAAAATATACTAGCACAATACAATTTACTATGTCATGAGTACTACTTAAATAATTTTAAAATT

TTCTAAACCCTAATAGCTTGAACAGTAGTTTCTGCTCTTGAAATATCAATCAGTATACATACAAAACCAAAAAAATAAAA

TAAAATGCAATTATGTGCGCGCACGAGAGATGGAATCATTAAACAAGCAAACATCCATTTTACAATAAGTAAAGCCTCAA

TACTTGTACGTAGGCTACAGTATTTCATAACATATTGGATTAATGCTGGTGAAAAGCGAATAGATTGATTATTAATGATC

TTAATAAGCGATGATATTTGTTATTTTGTTATTTTTTTTGGTTCAGCTAGGTTTTCTTTTTTAAACGAAGAAATAAACAT

TGATATGATTGATTCTTCGGAGGTTAAGGATCGAGGATATTAACGACAGGTGGTTTTCGAGAATATTTACTTGGTATTTG

TTGTATTTTACTTTGACTGCATGTTGACTAGAATTCGGTCATAATTATTCCCTCTTCATATCCATTATATTATTTCCTAC

CTAATAACATGACTCATGATTTTGTTTACTCATTATAAATTGAACTTGATAATTAAATTTTGAGTTCTAGTTAAAAAAAT

CAAAGTTGTAATGAAAACCCCAAAAAAACCAAAAAAACGGTGAGACGGTTTTGGTTTCTCATATTATAAACCGAGCATAG

TCTAGTTGGTTTTTGGTTTTTCAGAAGAGACATTGGTTTGTGCCAAACCCAAACCCAAGAAAAACCAAACCACGGGAAAC

CCTAGTCAAGACACTTAGAATGTTGCTTCAGGTAGTCTTCTTTGGTGATTGTACACTTACATATGCATATTTATAACCAA

AGTCTGATTATTGAAATTTAATCATTATAAGATAAATTTTATTATCCCATATCAGGACTCGAAAGAGCAATTTTTTTTGT

CACATATCAGGACTTGGAAGAGAAAAAATTGTGCAGAAATATAGATATGCAAAGACATTAAATATAGCAGTAATTTCCAA

GGAAAGTTGATTCAGCAGCCAGAACGAGTTCTGGACAAGGTTGCGAGATGTATACCAGATATGCGCATCTTAGTTTAATG

ATGCATCCCGAAAGTTCTGGAAATGAAATTGGCATGAATGTTTTTTTAGAACTTGTCAAATTAAACAACTCATTTTACTG

GTTGCTTGCAGTAAGACATCCAATTTAGGCGCAAGGTTTTGAAGGAAAGTGGCGAAACACATTGCTCCAGGGGACATATC

AATCTGTTTTAAGATTAATATCATGCCAATGCAAATTGGGAAATATGGATTCTGCTACAGAGCCAGACGTTCCTTGACGG

GATAAATTAGTTGAGTATTCCTACTGAGACATCTCATTATGGGGTTTTATAATTTTTACAAGCCCCGAGAAAATGATATA

GCCATTGTGTTTGTGGACAAAGTATTGAGAGAGCGATCCTCTTGAACCCAATAACCCAAAAAATGATCCGAGTATCTTTA

AAACTATGGGCTCATGGGAAAGCAGAAACATTTTTCCGACAATTACTGAAAAAAGATATCTAGGAACCTATAGTGTTCAA

TTCTCTGGAATGATGAACTTGAGCCTGCGCAGCATGGGCACACTACACCATATTTTGCGTTTAGTAACTTCCTATTTCAG

TTACTAAATGGGGTTGTAACACAAATAAGCCGTTACAAAAGGGTGTTAACAAATTTTCGTAACGAAGATTATCCGTTACT

AATCTTGAGTCACAACTGATTTCGTAATGGAAACTCGGCCGTTACTAATATTTTATTTCGTATCAGACTCCCAGCCGTCA

CTAAATTTTTACTATAGTTGCAAAATCTTGACTAGGTCAAAGAAGACCGACTCCTGTTTCCTTGGTAAACATTAGTAACA

TATTTTTGCCGTTACTAAATTTTTGCAACTATTAAAAAAAGGAGGATGTCGGTCTTCATTGACCTAGTCAAAATCAGTTT

GCCCCCAAATATTTTTCCCTGTAACTACTCAATTCACCCAGATTTATCCAATTCTAGAAACATAAACAGAAAAAGAAATG

GTAGCATTCCAAGTTATAAAAAAGTTTCATATTACACAGTAAGAAAAACACTACAAGAATTATGAGACTACGAAAATCGT

CATTTGGATAATGTCTGTACACTCAATAGATACATAATGTTTAAACAATAATAAAGTCTAAAATTTTACGTCTTCAATAG

AAAAATGAAATCTGAAAGACAAATACATGAAGTACCTGTCCATCCATCGATACCAAACTCTCAGCTGGTTCCTGCATCAT

CAGATTTAGTCAATCAGTAAAGGCACCTTTAACCAGACTAAACAAGCGAAATAACGAATGAAATCTTGGAGTGTAATTCA

GTTTTGAACTGATAGTGCATTACCATGGTTTTATCCCCTCTTTCGAGAATTGCGATTTGTAGGTGGTAGAGATCTTGACA

CCAGTACTTCATGCTCAGGCTCTTTTCTAGGCTCTGCAGGACGCCCGGCACTTGTATTCTGATATGTTGGCTCTGCATGA

CGAATGAAACCTGTATTCTGATAACAAACGTGTTTTAATTTAATAGTCAAATGCTTACATAATAAACACACACTTAACCA

CATCTTATGACTATTAAAGGCGAATCAAGCACATCAACACAACATACCTATTTCTTCTTGTCCCTTCTCCTCTCTACTTC

AAGAAATGAATCTGTGCAGCATACGGTACAGTGAAGTAAGAAACAAAATTATGCAGAAATTAGCTAGCACCAATATTTCT

TAAAGAACTTATACACGTAAGAAGGTAAAACATTATAGATTTTTCTATCCCCAATTCTGTATATATACAATACTGAGATG

CTGCAAAAGAGGCAAAGGAGTATAAAATATTTGTCACAGTGACACTAGAATGAACGTTAACGAGATTCAACTCATATAAT

AGGCAATTGATAAACGATGTCCTTAAATAATCACTGACTGTGTCAAACTATTATAGTAGCATCCAAGTAATGGTAGACTT

ACCAGGTAAGCTAGACAACTTAGATTTTTTCGGTGTATTGTGAACCAATGATATTTTATTTACATCCTTGAAGTTCAACT

TGTCTTCAATCTTCTCTTCAACTTTCAATCTTGTCTTCAACTTGAAGTTCAACTGCAACCTCCTTTCCTACACTTAACCT

GCTACCATCGGCTTCTCCGTCATCTGGAGACTCTCCATCAGATTCACTAACCGAATTTCCATCTTTGGTAGAATCACCAC

TTTCGAGTATGGAGCTCTGGAAGTTGACAAGCTTGTGTACCAACTCAGGGTGGATAGCTAATGTAAGTGATATGAAATGG

GAGTAATGAAGAAATGGGTTGTTAACTGCCTGCAATTAGAAGGAATAAGATAGTGCAATTAATTACAAACATTAACGAAT

AAAACGGTGGTACACACGCTAAAGCGAGCATAGAAAACAATTAATACGCAATGCGCTTGGTTTTGTAGTTATGCTAGTTG

AGCAAGAAGCAACTCAGGATCAAGCACATAAAAAGAGAGTTGCTTAAAATGCAGTAATGCAAATTAGGCTAGTCAATTAT

GTTGTTGTAGAGTTGGGCCTTGTAACACAAAGTAGGACCTCGTTTTCTATTCAGCATTTCGAGAATCTAATGGAGTTAAT

AAGTCATCAAGATAAACAAAGAGATTCACCATGCACATTCCATGGAGTCGCCCCCAATAACCGCAAAAAGATACATGAGT

CAGACATTTCCTTTGTCATGCAGTACTATAGAGGATGTTGTATATGCGGTAGAAATATCCACACGTTATATCTTACTACA

CACTCAAAAAGAGTGAACAATATCTGAAAAGTACACAAACTTAGAACTTGCAAGCAAAAACAATGCATGGATATCATCAA

TGAAAACTATGAATACTGACCACCATGAACAAAAATGTACAGACAACATTGGACATGTAAATCCAACAACATATACTCAG

GACTTTTTGCAATTAATTGCCAACCACCTATGCAACTTCTAGTCGTTGCCACACTGTGGCACTGAGGTCAAATAAAACAT

TGTAGAGTTTAATCATAGGTTACAGAAAATAAATCTAACAAAATGTGGACAGTAATCATTTTGGCCCCAGAAACTTACCA

ATGGAATCCTCCTCCTTAAAAGAGGAAAAACTATTTTTACACCAATCCCTCCCTCAATTTCTCTCTGGAGAGACCCTCTG

CAACAAAGTTATGCGTTAGAAAAGAACCACATGCATAAAGCCAATAACATGTCTACTACTAATAGACACAAGAATTAAAC

AGTCAAATTTAGGGGGGGGGCAAAAAGAAACTCAAGATAAAAGTAGTACCCTTATCCTTTGAAAAAAGGCCTCAGGGAAG

CAGGTACCTAGGAAAAAGCCCCAAAAAATGAGCGTCATACATTTTAATAACATACTTGAGAACCGATGGTACAGTTAAAC

CACATTATACGGAATGCTAAAACACTATTTATTAGGATATTATGTCAATTTAAAATCTAACATTTAAGTAAACATTGAAA

CAGATTCATGAGTTATTACATAGAAGGCAAGCGAGCAACAAGCTGCGCCGCAAGCCCTTTTTGAATGGCAAACCCTATCC

TCTTAAACACAAACTCTCTAGAATCGGGAGTCATGACATTACTATCCTCTTAAACACAAACTCTCTAGAATCGGGAGTCA

TGACATTACTGTGCATGACTCTTTGAACTCTCGCTAGGAGTCATGACATTTAAGATAAAAAAAATACATATTAAGATGAA

ATTACCTTTGTCGATATTGAGCATTTTGCGCCCACATTGACAAAATGAACACCTTCCCCCTCTACATTTGCCGAAACTAT

CGACCCCACTTCTCCAATACCTGATCCATCAGAACTTACTAATCTGACCGACTCGGGTACAGTTTTACCATTATCCTTAG

AAGTATTAGTACCAACTACACATCAAGTACAGGTACAAAACTTCAGGAGTTCAAATAATCCAAAAATGCTTAGTTGACCT

GTGTCTGTGGAATACAAAAAACTTAACAACCAATACAAAGTAAACAAACCAAAATAATCAAAATCCGAAACAAAAATATG

GAAAACTAAGAATTGCCTCAATGAAATTAGATTCGAAAATAAATAAATAAAAGAGCTAAAAATATGGAAATCACAAAGTA

GATTCGAAGAATTGCCTCAATGAAATCATACCCTTCGGTAATTTTTGAGCTGGTCCACCTGCTAATGACCATCAAACAAC

AAAAACAAAGTTAGGGTTCTTAACAATCTTCAAAATTTAGTTGAAGTTAACTCAAGAAATCTAATTTTTATTAGGGTTCA

GATCGATTGTTCAGATCATCCCAATGGAACAACTGAACTTCGAATCTGTAAGAAAAATTAAAACAAAGCACGTAACATCA

CCTGATTAAGAACCTCATGTAGATCAATCGAAATGATCTGCTACATTTTTTATTCAATTCGCATCCTCTTATAGATTAAT

TGGAGTAATCTCTTGTTCATCTCTAATTAAAACCCTAGATAAGAAATAGAAATCATGGATCTAGGGTTTCTAGTGTAGAG

ATTACACAAGAGGGTTATAAGAAAATTTGATTTAAGAATAAGAAGAAGGTAGATAACGGCGCTGGTATCAGTGAGATGGT

GCTAGAGCAGAAAGAAAAGAAGAAAAGAAAGAGAAGAAGACATCGTCTGCTCTTCTCTGCTCGGTGAGGGAAGAGAAGAG

GAGAGGAGAAAAAAAGAAGATAATGTGGTTGGGCTAAGGATAAGAGATAAGCATAGGGTAGAATTATCTTTCGACTTGAA

AACAGTTACTAATATTTGGGCGGAAATAGGAGGTCTTGTGTTACCATGCACACAGACTCTTGGCACGATATTTTCCTAGC

TGTTGCTAATAGTAACATACACAAAGCCGTTCTAATCCATTCTAAATTCGTAAACGACAATGGGCCGTTACTAAATTTTA

GTTCCTAATTTGCATAAATGGTGTTGTGGCATACTCCTAGTATATAGGAATATGTAAGTGATGTAAGAGTTTGGTGGCAA

AGCTGGTTTTGTAATGGCTTTAGAAACTTAATTTGGGGTGTTGGGCTCTACTTATACAACGGGCATCGTTCATGGGGCTA

TACTATAAGGTTTAACATATGTGTAGAGAACCAATTGACTTTGATTTGGACGCTTGGGTCCTAATATAATCGTAATTCCA

GGTTGCAAATGATCAATCCGGCATGTGTGTTTTCTTCCTTTTCTGGTCTTACAATCGAATCAGCAGTGCCAAACACGGTG

TTGCATTTTTTTGCTCCCTTGCAGTTCTGCCTGTTGCGAACCAGTTCGTAACAGCTTAAAACTTTGACTAACTTTTGTCG

AACAGATGTTCTTAATTTGCAACATATATCGCAACCCGAAAAAAAATCGAAAATGATTTAGCTATTCCGTAAAAAATATG

GTTCATAAATCGCCATGGCTATATATATATATATTTTTGTGGACGGACAAAATGCACTACCTACTGGCACTATTATTCAA

ATCATCTAGAACTAGAAGGTTTTCATCTTTTTGGAGATGAATAATACCACAGCGTAATAATCACATGGTTCCAATTTCCG

CCCATGTACCCCTATGTTTTAACTTTTAAGTAGAAATATAATGTAAATGATGTGTAAATCGGACGTAGATATGATGTAAA

TTGTTAATTGCTAAACCTCCGGAGTTAAGTTTATTGACACTTGTATCGGAAGGGCTACACTTCCCCCACCATCTTCCCCC

ACTCCCTTCTCCCGTTAGGTGTTATCGTTCTTAGCCCTTGATCCTTGGATTGAGATTAGCAGGGGGTCCCATTATTATTG

GATTTAACCAACAAATACTTGATACATAACGGGGGAAATGAGTGGGGGAAGGGAGTGTTGGAAAAAACTGGTTGCAATAG

AAGATGAAATAACACAGGAAGAGATTGTTGTACTCCAAACTTTCAAGGCTCATAAGATTTCTTTTAGACAATTATTTCTG

CCCTCCCAATAAATTGGCGATTACAACAGGTATGTGAAATAATGCCTTCAAGATACAATGAAAGCCCTACAACAAAAATT

GAATTCAAGGTTTGTACCCCTTGATCCAATCAAGAACACATGTAAACAAAATTTCTAATGATGCTTTAGAAAGAGAGAGA

ACAGAAAAGGTTTTTTCGATATATTTTAAACGAGTTTTGGGAGATGCTATTTATAGAGGTTTTCGCATCTGTTGGGAATA

AATATCTTTTCACCAACAAGTTTCTCTTCACGAAACGATGAGTCTCTTCATCATCAGACATCAATGGCGTCTATTGGATT

CGAAGATTCTGGGTATATTTGTAACTGTCGAAACAGTTACGTACCACAAAAATATTCCAGGACACCCCTTGGATAGCGTA

GTTGAAAATATTCCAACATTCTCCCCCTATGCATATCATTTTCGCACTTGTATTGGAGCCTAGTAACGCTGCCTTCTTTC

TCTTTTTTATTCTACAAAAAAACAGTCGCTTGTAGGTTGTAGCTTTCGTGTGGCATTAACCAATAACGAAGAATTCTCCT

ATCTTATCTTATTCTTATACTGATATAAAGAAAAGAGGTATTTTGGTTGGTCAGGAAAAGACAAAATAACTTAGTGTAAC

CCTAATATTACTAATATGTCCTTAGATATCAATCACCATTATAAAGGAGGTATTTTGGTTGGTCACAAAAGGACAAAATA

GCTTGGTGTACCATCTGAATATCTAATATGACTTTAAATATCAACCACCATTATACATATATATTCAAGGGTAAAAATGT

ACATTAAATCAATATTTATATATTTACAGGAATGCCATTATTTTAAAATAGATCACAAAATTTAAAAACTTGATATCTTT

TAAATTACACATCGGAATTTCGCAAAATTAGTATATTTGGAAAACTCTTTGAATTATCTACGTAATGAGTGCAATTAAGA

ATATTAAATTTTTCATTTTAAGATTTTTTTTCATTATTATTAAGAATCAAATTCAAGCATGTGCATCGTACATGTGGTCG

TATTAGTATATATAGGAAATTCCCCAAATTCTCTGAACCTTCACCGGGCAAAATTTTTCTGGCCCGGATAAAAAACTAGT

ATACTGATGGACCTGTAAGGAGCTAACCTGGGGTTTAGAAGATTACATGAACCAAAAACTTAGAGCCTTTTTGGTACAAT

CTCTGTCGTTATGTGAGTTTGGCACCTATACTATTCCACCTCCAAGTTGCTCTTGGACATTTCACGAGCTGGAGCGTTCC

AAAGAATTTGGGTCATGTCAGAAACTATATAAAACTTGTGCAAAAATCTTGTTCGCATCTATGTATAATTATTGACGTAC

GCCTATTGTCTTCATAGCCAGGGGCGGAGCCAAGGGTTGACTAACCCTGGCTCTAGCCCCCCTTAAAATCACCAAAATAA

AAAAATAGCCCTTGATTTTATAATACATTTTACATTTAGTCCACCTAATCTCTAAATTTAGCCCACTAATGTATGCATAA

CTTTTTCTTAGTCCCCTCATTTTCAAAGTCTGGCTCCGCCACTGTTCATAGCTGTAGCAATCTAGGTGTTCTAATTTTCA

ACCAATTCTCCTTCTCTTTTATTGCAATTTGATTGTACATCGCCGTAGTCAGTACATTTCTGATAAGCATGAAGCCGATA

TCTCCACTTTTAATATATTTACACATCAAGAAAAGGTACTGATTTAGAACTTTATTGTGTAAATTTTCAGTATATACCTC

TTAGAACATTTTTTTTAAAAACAGTTTCCTTTTCACCGGCATTCTTGGATAGTTTAGCCACACCCAACAATAATTCCCGA

GTCAGTACAAACGCCTGCATGGGCGACCTTAATTAATAATTATTCCCACGAGGTTCTTGCATTACTCTCGTGGGCTGATA

TGCGACAGTGCTCCACCCCCACAATCAGACACCTTTATAAAAAGCATCGGCATATTACATCAATCCTTTATAAAAGATAA

TTTACATATATAAGGGCTGAGATGGGACACTGCTTCAACACACAATCAAACACCATTATAGAAAGCATTAACATATTACA

TCGATTCTTTATAAAAGATCATCTTTATATATGCATGGGCTGATACACGTGACACTGCTCCACCACATAATCAACAAATC

AGTCCTTGATAAAACATTGAATTTTGATAGGGTGACCGTCGATTCAGTCGCGTATATTAGTAAGGTAGCCCCGTTTTAAA

TTGCTTTTTTCAACCAAAATCACTTTATAACAACTAATAAACATTTTCCAAAAATTATGTTTTGTAAAATTATATTACTA

TTAAAATGTTTTTTTGGAGAAAAGCAGAAGTTGTGGATTAAAATCCAAGCTTAAGTTACCACGAGAAGCACCGAAATAAA

ACAATCCTCCTATTCATGTGCTACGTGTCTGGAATAACTTGATTTGCTAAAGGCATATGTAATGATATCAGCGGGGCGGG

TACCATTTCAAATAAGACACAGAACTTGCAACTAGTGGCTCAACCATTTCCATTACGATTAGAGTTATCAGATATCAAAT

AATTTTTCAATAATTTTCGTTATGTAACAGAAATCATACAAACTTGTTTTTTATCCAAAATTCAGTTAAAACAGTTCAAA

ATATCAAATCCGGTCCAACCGTTTTGTAAAAGATCTGTTGGGTAGGAAGGTCCAGCTGTTTTATAAAATAAAATGTCGGG

TAAAAAGATCCTGACACTTTCAGGGGCGACCAATTTGTAATTAGGAGCGACATAAAACGACAAAATAACAAATAATCGGT

AGGTTATTTCACAATCGGAAGGTTATTTCCGAGGCGTTTTTTTTAATGGCCAAACTACCCTTTACAATCGGTAGATAACT

ATACAATCGGTAGGTAATCAAAAATCGGGAGTAAATTTAGTTTGTATGACTTCCGATTATACAATGATTCTCTTTGCAAA

TATCCTACCATAATCGGTAGTCAAGAAAGTTCATATAACTACCGAATATAGAGTAGTCCCAAATTCAGATTTTGCAAAAA

TCCTTTTTTTTTTGAATTTTGGTTGAACCTATAATCGGTAAGATATGAAGAGTTATCATGACTTCCGATTATTCTGGCTC

TCTTTGCAAATATCCAACCATAATCGTTAGTCAAGATAGTTTATATAACTACCGAATATAGAGTAGTCCCAAATTCAGAT

TTTGCAAAAATCCTCTATTTTCTGAATTTTGGTTGAGCCTATAATCGGTAAGATATGAAGAGTTATCATGACTTCCGATT

ATACAGTGGATCTCTTTGCACATATCCTACCATATTCGGTAGTCAAGATAGTTCATATGACTACCGAATATAGAGTTGCC

CCAAAATCAAATTTTGCAAAAATCCTCTTTTTTTCTCAATTTTGGTTGAGCCTATAATCGGTAGTAAATGTAGTGTTACC

TGACTTCCGATTATCTCGGCTCTCTTTGCAAATATCCAACCAATGGTTTCTCTTGAAGAAACCCTATGGGAACACAGCCG

GGAGGGGCTACGCCACCCGGATCCCCCGGATAGCGGCAAACAATCTTGAAAATATCCAACACATATAACCACCGAATATA

CAGTTGCCCCAAAATCCTCTATTTTTCGAATTTGGTTGAGCCTATAATCGGTAGTAAATGAAGTTACCATGACTTCCGAT

TATACAGTATAATCTTTGGCGAAATCTACCATAATCGGTAGTCAAAATAGTTCATATAACTACCGAATATAGAGTTGCCC

CAAACTACCGATTATGGATTAACCTCCCGATTATGAAGGGGCATATAAGTATTTTCAATCTATTTTGTCTTTTGTGTCGC

TCCTAATTACAAATTGGTCGCCCTAAAAGTGTCAGGAAAAAGATCCAACCATTTTATAAAAGAAACTAGCGGGTAAGAAG

GTCCGCCCACTTTATAAAAGATATGGCCGAGTAAAAAAAAAAAATCCTTATTTTTCTATATCTGATTCACTGATTCGTAA

GTGATAAAGAGAAGATATTAGGAGAGTGTGACTGAAACAAATTATATTGAATAAGAAGAATTTAGAAGAAAAAGATATTT

TGCTCAGCTATAAAACAATAAAGGATTAATAACAATGTGTATATATTATCGGTATTAATGTAACGGAAACTGTTTCCTCC

GGTTCCGTTACTTTAAGCCAAAAAGTTCTAATAACTTCCCGATTCCGACGCTAACTGCTGTGTCGGACAAAAATCCTATC

AGATCTTGAATTATCTGTAACGGCTATAGGATTTCGGCATCGCATTGACAGCCCTGCTTACAACTGATTGGGACCGTCGG

ATCACTCAAATAATACCAGAAACGACGTTAATGCCAGTGGTGCCTCTAACAATCATGTGGAATGTGTTTTTTTGTTTTTT

TTTTCTATCAACTTAGTGGGACTCACTATACCTGATAGTGGAAGGTGACAAGGTGTGGGGAAATGTGCGTGGGAGGAAAC

TCACGCTCATGGCCAGAAGTGTATACTCTTTCTCTTGTTTGTAATGGATAGCAGCTAGTCATTTATTTGTATTTATTGTG

CGAGGAAATGAGGATTAGATATAGCTGGGTTAGTAGCGGGTTGGGTATGGGAAATAGTTCCCCTCCTCACACGTGATAAA

CCTGGAGATTGGACCCACGAAAGCTTTATGACTAGTGCTTATAAAAAAGAATTAAAGATCTTTAGCACGCTAGCTATATA

TATACATTTGAGTTCACCATAGATATATCATATCATTAAAATCTATCATCCTACATACTGTACTGGGTTGAATCATGGAG

CTCCAATATTTTTCTTATTTTCAACCAACTTCCTCCGTTGTTGCTCTTCTACTTGCTCTTGTATCCATCTTATTCAGTGT

CGTTGTTTTGAGGAAGACATTTTCGAATAACTACTCATCACCTGCATCATCAACAGAAACAGCGGTACTTTGTCATCAGC

GGCAGCAGTCGTGTGCATTGCCAATTTCCGGTCTTCTTCATGTTTTCATGAATAAAAACGGCTTAATTCATGTAACTCTT

GGAAATATGGCGGATAAATACGGACCGATTTTCAGTTTCCCAACAGGTAGCCATAGAACTCTCGTTGTGAGCAGTTGGGA

GATGGTAAAAGAGTGTTTTACAGGCAACAATGACACTGCTTTCTCAAACCGTCCTATCCCGTTAGCTTTTCAGACTATAT

TCTATGCATGCGGTGGCATAGACTCATACGGTCTTTCGAGTGTACCTTATGGAAAATATTGGAGGGAGCTACGAAAGGTC

TGTGTGCATAACCTCCTCTCTAATCAACAACTACTCAAGTTCAGACACTTGATAATTTCTCAAGTCGATACGTCTTTCAA

TAAGCTGTATGAGTTATGCAAAAACTCTGAAGACAACCAGGGAATGGTGAGAATGGATGATTGGCTCGCCCAATTATCGT

TCAATGTGATAGGAAGAATAGTCTGCGGATTCCAATCAGACCCAAAGACAGGTGAGCATTTGCCTAGAGTTTCTGTAGAA

AATGATGGATTTTATACTATTAGATCATCAAAGACTATTCTGTATATCTTAATCAAGCATACATATCTTCTAACAGTGCA

CATGTATAAACAGGTGCTCCTAGCAGGGTGGAACAATTTAAAGAAGTAATTAATGAAGCATCTTATTTTATGTCGACATC

TCCAGTGTCAGATAACGTTCCAATGCTAGGGTGGATTGACCAATTGACTGGTCTTACGAAAAATATGAAGCACTGCGGAA

AGAAATTAGACTTGGTGGTAGAGAGCATAATTAAAGATCATCGTCAAAAGAGACGATTCTCTAGAACTAAAGGAGGAGAT

GAGAAGGACGATGAACAAGATGACTTCATCGACATTTGCTTGTCAATTATGGAGCAACCACAGCTTCCTGGCAACAATAG

CCCTCCTCAGATACCTATCAAATCTATTGTCCTGGTATGCAATATTGATCCGTAGTATGATGACAAAACAAAAGGCTGGA

GCATTAATTGTATTCCAGCGATTAATTGGTTGCGGTAGCTCATAGAGTGCAGAAGCATGATTTGGTTTTGATTGGTATTA

TTGTATAAATGCAGGACATGATAGGTGGGGGCACTGACACCACAAAACTGACCACCATCTGGACCCTTTCCTTGCTGCTG

AACAACCCCCATGTGTTGGACAAGGCAAAACAAGAAGTGGATGCACACTTTCGAAAGAAAAGGAGATCAACAGATGATGC

AGCAGCAGCCGTGGTGGATTTTGATGATATTCGTAACCTTGTCTACATCCAGGCAATCATCAAAGAATCAATGCGGTTGT

ATCCAGCCAGCCCCGTGGTGGAGCGACTGAGCGGCGAGGATTGTGTGGTCGGTGGGTTTCATGTACCAGCAGGGACGAGA

TTATGGGCTAACGTATGGAAGATTCAACGAGACCCTAAAGTATGGGATGATCCATTGGTGTTTCGACCAGAGAGATTTTT

GAGCGATGAACAGAAGATGGTTGATGTAAGGGGTCAAAATTATGAGCTGTTACCATTTGGAGCCGGTCGACGTATATGTC

CAGGTGTTTCCTTCTCTTTGGATCTAATGCAACTGGTACTGACTCGTCTTATTCTTGAGTTTGAAATGAAGTCCCCTAGC

GGGAAAGTGGACATGACAGCAACACCAGGATTAATGAGTTACAAGGTGGTCCCTCTTGACATTCTGCTCACCCATCGGCG

CATAAAGTCGTGTGTGCAGTTAGCATCCTCTGAGAGAGACATGGAGAGTAGTGGTGTACCAGTAATCACTCTGAGCTCGG

GCAAGGTGATGCCTGTTTTAGGCATGGGAACATTTGAGAAAGTTGGTAAAGGGTCTGAAAGAGAGAGGTTGGCGATTTTA

AAAGCCATAGAGGTGGGTTACAGATACTTCGACACAGCTGCTGCATACGAAACTGAAGAGGTTCTTGGAGAAGCTATTGC

TGAAGCACTTCAAATTGGCCTAATCAAATCTCGAGATGAACTTTTCATCAGTTCCATGCTCTGGTGCACTGATGCTCACC

CTGATCGTGTCCTCCTCGCTCTTCAGAATTCTCTGAGGCAAGTCTTTTTGATGCAAATTAGGCTTATATATATTTGCACA

TACCAGCAGGTACACTTGAACATCTATTTTCAGATAAATGAATTTGTTTTGTGTGATATGTATAGGAATCTTAAATTGGA

GTATCTGGATCTATATATGTTACCCTTCCCGGCAAGCTTGAAGCCAGGGAAGATAACGATGGACATACCAGAGGAAGATA

TTTGTCGAATGGACTACAGGTCTGTATGGTCAGCCATGGAAGAGTGTCAGAACCTTGGCTTCACTAAATCAATCGGTGTT

AGCAATTTCTCCTGCAAAAAGCTTCAGGAATTGATGGCGACCGCCAACATCCCTCCAGCCGTGAATCAAGTGAGTACTCT

TACTTGAAAGTTTTGATAACACATCGGAAGATTTCTAATGCTTATTAAATTTCCAACATTATGCATAAAACTACTACTCT

TTATAATGTTGATTAAGGTGGAGATGAGCCCGGCTTTCCAACAAAAGAAGCTGAGAGAGTATTGCAACGCAAATAATATA

TTAGTCAGTGCAGTCTCTATACTGGGATCAAACGGAACCCCATGGGGCTCCAATGCAGTTTTGGGTTCTGAGGTGCTTAA

GCAAATTGCTATGGCCAAAGGAAAATCTGTTGCTCAGGTTGGTCAAATTCTCTGCATATGCCATAGTGTGGTCTCTGTTC

TTTGGCCATCGTATTTGCATCACATTGTATTCCTTCCTTATACGGAATGTAGCTTACATTTGTATTACGTACTGATCAGG

TTAGTATGAGATGGGTTTACGAGCAAGGCGCGAGTCTTGTGGTAAAAAGTTTCAGTGAAGAGAGATTGAGGGAAAACTTG

AACATATTCGACTGGGAACTCACTAAGGAAGACAATGAAAAGATCGGTGAGATTCCACAGTGCAGAATCTTGACTGCTTA

TTTTTTAGTCTCGCCTAATGGACCTTTCAAATCTCAAGAAGAGTTGTGGGATGACAAAGCTTGAATATCGATCATTTAAC

TCTCTACATGTGCATTTATAAGAGAAGCTTCCCCTGCAGTGTGCTCAATCTTGATTTATTTTATCAAATTACATCTTACT

ATATGGGAGTCACAGATCTCTTTCCCTTCCTATATGTTTGAATACATTTGTATTGAGTGATGTAGCTAAAGAAATAAATG

TAGTAAGTTTCTTTCCTAAGCTTGTTTTCCTTTACAAAAACTAGATGAGAGTCAGAGGAAAATAATGGTGAGCTCGTGTT

CATCAGCAACAATAGTGCCATTAGGCACAATGTATATACCTTAGAAGAAAACACTATTCCAGACACAAAGAAATGTACGA

ATGCGTAAAGAGTATTCTGCCATGTATATACAATAGCTGATGCGGAAAAAAAAAATACAAAACCTTCTGTGAAACCAGAC

GTATTATTGTCCATGTGAGAGTAAAATATCTAATCGCCATGTGTAAGCATCACATGAGGTGATCATAAGATAAGACGGTA

AAACATATTCTCTGAATAAAGATGACTGAACAATTGTATAAGCTCTCGTACCCTAATAAGGTGGTAGAGGCATGTATTTA

TAATGAGCTTAATGGACATATTACAATGGACTCTTTACATAACACCGACTTAGATATATGGACTCTATATGGACAGTACA

TATGCATTGGATTATATTCTTCACCCTCCCTCAAGCTGTACACGGTGGAAGACAACGGGAAAGCTTGCTACAGAGATAAC

GAAATCTGGCAGTCGAAAAACCCTTCGTGAACAGATCAGCAAGTTGATGCGTAGTAGCAATATAACATAAATGGAGATGC

CCTGATGATATCCGTTCACGGACAAAATGATAATGAATATGAATATGTTTCGATCTTGAATGTTGAATAGGATTAATAGC

AAGATAAGTGGCACTTATATTATCACAATATAGAATCGGTGGTTGCTTTAAGGATAATCCAAGATCAGACAATAAGTATT

GAATCCATGTAATCTCAGCAGATGTAGAAGATAAAGCACGATATTCAGCTTCCGTACTACTGGTAGAGACTGTGGGCTGC

TTTTTAGAACCCCATGAGATGAGATTCTTACCTAGAAAAATACATTGACCTGTAGTAGAACGAGAAGTATCGGGATCACC

ACCCCAGTCAGCATCAGAAAAGGAAGTGAGTTGCATAGGATTGTTACGCCAGAAAGAAATACCATGGCCAAGTGTGCCTT

TAAGATACCGGAGGATACGCTTCACAGCTAGAAAATGAGTTTGAGTTGGTCGATGCATGAATTGACTGACCTGATTAAAT

GCATAGGTGATATCGGGGAGAGTCATGTTGAGGTATTGCAGGCCACCAACTAGACTGCGATAGAGTGTTATGTCAGAAAA

CAACTCAGCGTCATTAGAAGATAATTTAGATCGAGGCGCAACAGGTGTAGAAACTGGCTTTGAATTAGTGAGATCAACCC

GTTCCATAAGATCCATCGTATATCTGTTTTGAGTGAGAACCAAGCCATCCTTTGTACGTGTAGCTTCCAAACCAAGGAAA

TAGTGCAAATCCCCTAAATCTGTCATGGAGAATTCCTTTTGAAGTGTAGGCAAGAGATTGTGAAGACCAATCGTATCACT

TCCAGGAGAATTATATCATCAACATAAAGTAATAAGATAAGAATACCAGCACTGGAATGTTTTATAAACATAGAGTGATC

AGAGTGGGATGATTTGAACCCAATATGCAGTAGAAAGGCAGCAAAGAGATGATACCAGGCACGAGGAGCCTGTTTCAAGC

CCTAAAGAGCTTTGTGCAATAAGCACACGTGATCAGGATAACGAGGATCAACGAACCCTTGAGGTTGTTCCATATACACT

TCTTGATCAAGTGAACCATGGAGAAACGCATTTTTAACGTCAAGTTGTTTAATAGGCCAAGAAGAAGAGACTGCAATCTT

TGAGAACCAAACGTCACCAGAATCTCTTTACAACAGGACTAAATGTCTCATCAAAGTCAATACCATACTGCTGTGAATAA

CCCTTAGCAACGAGACGAGCTTTTCGACGATCAAGAGAGCCATCTGGTTTTTGTTTCACACGATATACCCACTTGGAACC

AATGACATTCATCCAAACTTGACGCGGAACAAGAGACCAAGTGCGATTCCGCAAAAGAGCATTAAATTCCTCTAGCATGG

CTGCCCACCATTCCTTGAACTTGGATGCAGTAGTAAAGCAGGTGGGTTCAGAAGGCAGCAATTGAAACTGGGCTGCAAAA

CTGTGTAAAACAGGATACCGAGAAGAAAAACAACCTTTAAAACTGCGAGGCTGGATGGTATTATATTTAAGTCGAGTAAC

CATGTGATGAGTATTACCAGTCGACTGAGGGAGCTCCTGTACACTGGTCAAGTCCGCTGGCAATGTAGAACTGGACGAAG

CAGATGACGGAGATGATGAAGCGGGAGATGGTGGAACCAAGGGTGATGATGTAGAAGGAGAAGGAGAGGAATGTGTTGGA

GGAGTAGGAGATTGAGAAACAACATCGTCAGAAAAAGATAGGCGTGTAGGAGACAGATATGAAGAAGAGCGCACCGGAGA

AGACAAAACTGGGTTAGAAGCTGTAGAGGAGAAAGGAGATAAATGCACCGGAGGCAAAGAAATTGGAGTAGCCGGGTCTG

GAGAAGAAACAGGTGAGGAAGAGTGTAGATGAGTGGTAGCAATAGGATTGGGCCAAGCCAAAAGAGTATTGGTAGACAGA

TTATATTTTAAGAGAGGAATGGAGGTACCAGTATCTGTAGGAAATGGAAAAATAGACTCATCGAAAACAACATGACGAGA

GATATAAACCTTATTACTCTTAGCTTCAAGACACTTATAACCTTTGTGGCGATCACTATAACCAATAAAAACACAAGGTA

GTGTTTTGGGTTGTAATTTATTTTGACGAAAGTGATTGAGAAGTGGATAACAGCGACAACCAAACACTTTCAAAAACGAG

TAATCAGGAGCACGATTATGAAGTATAGAGTAGGGAGATTTATTGGATAAAGTAAGAGTTGGTAGACGGTTTATTAAGTA

TACAGCTGTTGCATAAAACTCAGGCCAAAAGGTTTCAGGAAAACCACTTTGAAGCATCAAAGCTCGGCCAGTCTCAGTAA

TATGGCGAATTTTTCTTTCAGCAAGACCATTTTGTTGTGGTGTGCTAGGACAAGAGGATCGACTGAGAATACCATGCTTG

GATAAAAAAGATTTTAATGATTCAGCATATTCAGTGCCACCATCAGTTTGTAGAATTTGTATGGTGGTCGAAAATTGAGT

TTCAATCATAGCTTTAAAATTAAGGAAGGAACGAAATACTTGGGTTTTGTCTGTCAAGAAATATATCCAAGAAAAGCGAG

AGCATTCATCAACAAACAAGAGAAAATAGCGAAAACCACAATTTGAAAAGAAAGGTGCAGGACCCCATAAATCAGAATGT

ATAATAGAAACAATTGTATTTGAGTAAGTACTTCGAGATTAAAAGGGTAAACGATGATGCTTGGCACCAACACAACTAGA

ACAAAGAGAGCTAGATAAAGAAGAACTAGTAGAATCCAAAAGTAAGCTAGACTTAAGAGATTGAAAGACTGCATTGTTGG

GATGACCTAAACGCATATGCCATATAGTTGGAGTAGCACGAATACCAATGGTAGCTGCGGAAAAGGCTGATAACTGCGGT

GATGGAGATGGAGAGGAAACACTAGAAAATATAGGGTACATGCCAGCGTCATTCTTGCGCTGGTACAGAAGAGCTCCCGT

TAATCGATCCTGTATACAAAAACCATGACGATCAAAGTGAAAAACACAGTCAGATTCTCGTGTAAGTTGACCAACAGAGA

GTAAATTTTTAGCAAGAGAGGGAACATGAAGAGTATTCTTCAAAGTCAACTGTGATGATGGTGACGAAAGAGTTACATCA

CCAATTCCTGTTATAGGTAGGAGTGTACCATCACCAACGAGAACTTTAGCATTACCATTGTAATCATGGCGATTGATCAT

CTGGGAAGAATCAGACGTCATATGGGCGGAAGCAGCAGTATCGGAAAACCAATGATTCTCATACAACATGGGCTGAATTT

GCATTGCAGAAAAAGCTTGCGGAAGATCATAGTTGGAAGCAGAGGCAAGGCGGTTGCAAGTTTGTGATGTATGTCCTCTC

GTATTGCACCACTGACAAACAATAGCATGTGTTTGAGGTGGAGGACCAAGAACCGAGCCAACTTGAGTATTATGATTATA

CGAGCGTCCATAAGATTGGTTTTGATAACCACCAGAGGGTTGAAAACCGTTGTTATTACCACGACCAGTGCCTGCAGTCC

AGTGTTGATTATTCCTGTTATTATTCCAACCAGCATTCCAAGTAGATCGACCTCCAAAATTGTTATGACCACCACGATTA

GAATTGTTATAGGGACCACGACCAGAGCCTCCACCACGACCACGATATGGAGTACTGTTCGAACCACGACCGCGACTAGT

ACAATGATAGGAATTTGATGCAGGATGCGAACTACCACGACCAGGAGGAGCACCAGAAAAGAAGGATACCGGACTTTGAA

TATCAGTAGCGTGCGTCTGTAGGTTTTGCAGATATTGAAGGTGAGATTCATGATGTAAACAACAAAGCATGAAGCTCGGT

AAAGGTGGGAAGAGTGGTACGGGAAATGACTGCTGTGTAAATGTTCGACCAATCAACACCCAAGCCTCGTAGTGTTGTAG

AGACAAGCTCAAGATCAGATACGGGTTGAAGGACAGCAGCTAGAGAATCAGATATTTCCTTGATTTTCAAAATATATTCT

GAGATGGACGAGTTACCCTTTTGAATAGTTGACAAGTTTGCTTTCAATTGTAGAATACGAGCAGGATTGATTGTTGCATA

GGTGGAACGAAGATAGTTCCATACGTCACGAGAGTTTCCACGACGTGCACATTCAGATTTCATAGTTTCAGTAAGAGAAC

AATTTATCCATGATATGATCGTTTGATCAATAACTTCCCATTCCTCATAAGCCGGGTTGGGAGTTCTAGCTTGAGTTTCT

TCATCAACAAGCTCTTGTGCTGGTATTTAATAAGAGTCGGCAAGATACTTAAACAGCCTCTGTGATTTAAGAATAGGAAG

AATCTGTGATTCCCAGAGAAGGTAATTGGTTCTATCTAAGAGAACTATCTTGGTTGAAGAAGGCATGAAAGGGGATTTTT

CAAATATAGAGGAACAAGAGTTAAACATAAGAGAACCTATGCTTTGCGAGATAGAAGCAGGGATAGTTGGAGAACCAAAG

CTTTGATGAGAAGCCATTGTGCTAGCAGTAAGAGAAGCAGATGAAGATGATTGAACTGGGTAAGTCATGAAGAAGGGAAG

GTAAAAGAGGAGCAGGCTAGGGTTTTGAAAAAAAAAAAATTTGAGGCTAGGGTTTTAGAGAGATCGTTCTGGCTCTTGGT

ACCATGAACAATTGTATAAGCTCTCGTACCCTAATAAGGTGGTAGAGGCATGTATTTATAATGAGCTTAATGGACATATT

ACAATGGACTCTTTACATAACACCGACTTAGATATATGAAATCTATATGGACAGTACATATGCATTGGATTATATTCTTC

AATGACAACAAAACAAGATTATCTACAGCTTCTCCTCATGAAGAACGCGTCTAATCTCTAGTAAAATCATACTGACGGTC

AAGTTTAGAAGAAAGCAGTGCATTTAATGAGAATCGGAAGAGGTGCGATACAGCTGTAGAAGGAGGTACATCCCTCCCGT

GAATTCGGACGAAGGGCGAGGTACTGCTACCCGAAGAAGGCGCCACACGATATGACCCAAAGGGCGATATATAATATCAT

TCAAAGAAGACGACACACGAGGTAAGTCGAAGGGAGCAATGACGAGCCAGTCCGAAGCGAAGAGGCGTATTGTATCGTCA

TACAACCAGGAGCACCATATCGGACCTGAAGGAGTTTACTTTGCCTATAAATACCAGTCCATGTTAAGGGAGATGGACAT

GTTATGTAATATTAGATGGTCTTTCTACAGAGAATTCTAGAGAGAGATTTAGAGAAGGACAGAGTGGGAAGTCTAAGTGA

TATTTGTAATTCATTCAAGGGTAAGACCTTAATCAATAAAGAAATACTTTCTTTCTCGTGGATGTAGGTCAATACGGCCG

AACCACGTTATATGCATGTGTTCATCTTGATTTTCATGCTCTTTACATCTTGTTTATCTTTGAATCTTGAATGTTAGTAG

CAATATAGTCCTTAGATTTACTTGGGGTGTAATAGTCAGAAAAGATACAACTACATTTTGGCGCTAGAAACATGAAGGTA

TGAAGATTTAGTCTAACTCCCTTGCAAAGTTTTCAAAGTTCTTTTCATAATCTCTTAAAAAGAGTTATTTTGTATATCGT

TGTATAGATCTCTATAAGACTTTACGAGTTGATCTCGGAAACAACATGATCTACACAATCTCAAAGTCTTGTAGACTCTC

AAAGAGCATCAAATCTATAATTGTATCATTAGTTTTCCCTTAGATTCGTATTTTCTACTGTTTTTTATTGCCTTCCTAAA

AGAGACTCAGGAGAGTAGATGGTTGGAACGTAAGTGGAGTCATAAAACTTCAAACTAAAGTTCAAACAACCTCCTCCATG

AACGAGTTGGGAAGATATCTGAGAAAGTAAAATTCGAAGAAAGAAAGTACACTACGATAGTTGAACAAACTCAAAAGAAG

GACGGTTTGAAATGTTCGTCATCTTCTAAATCACTATATAGCCTAAAATCTTCTTCTTAACTTAAATCTAAGCAAAGGGG

AAAGGTTAATGAGTACACAATAATTCATCAAAAGTAAGTGTTTCAAAATCATGGGTTTAAGACATTGAGAAATATTTCCT

AATATGGGAAAACATACTCAATCATCATAACTTGGGTTTTACCCATTAAAGGTATGACGTGCAAGAGGAGGTAATTTCCT

CGAAAAGAGAAAAAAGAAAAATACCTTTTCATGTCAGCAAAACTCTATCCATGTGGGGAATGTTTTAACATCATCACATG

TCTCCAAAATTTAATGAAAAGTCTTCATGCACTAAAACTGCATAGTTTGAACTATGACTTAGTAATTGCACATGAGTTGC

AAACTTAGGGTCCCACCTTTCTGAAAAAAGAGAAAAGTTAAGCATTATTGAACATGTCACAGGTAAGGTGATGGTGACGA

TATGATCTGATCGTACAAGCCAAGGAGAAACTGGTGGTGCATTACCCGAAAGACTTGACGCGATGGAGCCAGGAGAGGCT

GTGATCAATGGTGTGCTCCCATTCAAGGGACATCCCGATAATCGATATCAAGAAGGGTTAACAGATAACTTTAGCTCCGA

CGAAGAGGAAAATCCCTTCGAGGAACAAGATCAACCAGGAGGGTTGAATCATTTTATAAAAAAGACGACAAACATTGGGG

CTAAGAAGGATGACTCCCCTATACTTGATTTATAGTCTACTTATAGATATGTCTCGATCTTCAAGCTAGATTAGAGAAAG

GAGGCTCCCCGGATCCATTTCTTCAAGATTAGGATAGAAGTGTGTAGCTGAGATTTAGACTTCATGTCGTTCACCAATTG

AAGAAGATCTATTGAATAACTTGGAGGAATTAATAGTGGAAATGGGTATTATACAACCCACATCTGAGACCCTCATTGAC

CGTACTAGGACCGTAGACCAAATATAACAGTCCCTGATATGGTAAAAAAACGGACAAAATCCGATAGTATATTTAGCCAT

CTCTGAATAGGAAAATATATTTACCATTCTGCCATCCAAAACAGAATCCTTTAAACTGCGATTCAGAACTGGTAGTTCCG

TGTTAGAATTTTTTGGGCCTACGCAGTGTATTGCCACGATTGTGCCATCTGAAGTGAACGGCTGGATGAAGATTAAGATA

CCTTATCTCTTTGTTTTTATCACTAAACCAAATACCCTTCGTACTCTGCCTCACATGTCTCCCTCTCCCTCTAAACCTAT

CTTCTCTTCCCGTTTAGTTTTACAGAAACTAAAAGTAAAACCTCGTTGTTTCTCCTCAAGAACAAATGGGTTGAATTAAT

CGAAGAAAAAATCTCCCACCATCACTAAATGTAAGTAGTTGTCGGAAGTAATTTCTTTCCATCTCAGAAAATCAGATAAT

TGATTTTTCGGAATTGAAGAGATGGATAAAAATGGAGGAGGGTATAATCTAAGAACAATAGAAGAAGTATCTAGAGACTT

TAAAGGTAGAAGAGATGGAATGATTAAAGCTCTTACTTCTGGTTAGTATTCAATTTAATTCTTCTACTCATCTACAATCT

TATTTAACTTTGAGTTTTTTTTTTTTTTTTGTGTGTTTATTTGTGGATTGTACATATGGAAGAATTTTTCACGCAGTGCG

ATCCTGGTGAGGATATTTCTTCTCCTTCAATCTTATCTCTATTTACACCATTTTTGTTAATTAGCGACTATTTTTTCATG

TTTTTTGGGATTTTTTATCGAGTTTATTTATCAATGTTGGTATGATTAGTAGATTAGGGTTGATTTGAGGGGTTTTGCTT

GTTAGGGATTGTGTTTTCTTGATTATTTGATGTACTGATTTGACCGAATTAGAGTATTTGATTCGATTTAGATTAAATAA

TTGTTATTGTTTGAAGAGGATTTTAGTGTGTTTTAAGAGGATTTTAATGTGGGGTTTCAAGATGGGTGACTTTATGTTGT

TCGGTATTGTTGCTCAAATTGAGTTATAGTAGTCAGCCTCTGGGACTGATCCTAAAATTGGGTTTTTAATTGATCATGTT

AAGCCTTGGTTTGGAATTGTTGGCGATGTAACTTGTGTGCTTTTATGGTTTACAATGGACAAAAACAATGCAGGTTATGA

AGCCTAGCTATGATATGTTTCAGCTGCTCAAGATCCTAACATGTATTCTAACCAAGGCTATGGCGGTCCTGGATATGGTA

ACATGTATTCGGCCAGCTACGGGGTCTATATTTATACTTCCACAGGATATCTTATAAATGCATTTTGTTTTATATAAATT

GGAATTAATTAGCTTATGGGATTCATGTATAGGCAACGACTAGAAGACTAGAATTTGAGAATGATACCGTGCCAGATTGA

ACGTGGTATGGCTAACTAGTCCAAACTGGTAGCTAGGATTATCTTTTAAATTTTTATATGGTAGATTGATTAGTCCTATT

TTGGCTAACTATTGTAATGATATTCAACTGGCGTAGGCTAGGAGTATCGTAGCTTGTGGATTGGTTCCTTGTTTGTCTTT

TAAATGCGGAATACTGCGAATGTCTCCATAGAATATGATGGCTGTTACGGCTTGGGATGTATGGATTTTATTTCAAATAC

AAAATAAATTCCATTGGGTACTTATGATAAGCTAGGAATTCTTCATGTTAAATGGGTTTTCCCCCAAATTAATTTAAAAT

GTCAGGGTGTCTCTTTTGGTTGTGTGTACTCTATTTAACTTGTAAATCTCATTATCTAGATTTAGACCATCCTCATCTAA

ACATACATTTTTAATATCAGGGATTTGTTATACTGTAGGTGTAGATTATCCACAGGGCTACGTAGAAACATCTCAAAGCT

ATGATGTACCCAAACATGAAGAGACGAGAAGCACAGGATAGAGGCTGGCAGGCGCGAGATTTAAGGAGCCACGTGGATCG

GGCATGATCACTCTACAGTTGTCGGACAAGACTCGACCCCGGGCCTCTGACAAAGCTGACAGACGAGACGTATTGGGTCA

AACGATAAGCGGTAGTCAAAGGGATAAGAAGATACGTGTCATATCGGCCAACTTCATGGGCGACAGACTGGAAGGGCGAC

ATCGCACCGAGATAATACCTGAATGGATCGGCTCGGACGAGAGACTACAAAGGAATCAAAGAACATCGGCCATAATTACA

GAAGAAGGGACGACGAAAGCAGGCCCGACGACAGGACTTCAATTTAGATATGTTGTGATCAACACCCTAGTTTGGTTATA

GGTAGCCTTTCCTATAAATAGAAGAACTGGTATCTAAGAGAGGGAGTGGTAATTTCGGAGAGAGAGAGAGAGACTTGAGA

TCACATTAAGGAAGGAGAGAATTGAGAACTATCCCATCTTGTAAGTGAGTTCAACTTGGAGAGCAACACCATTGTAACCC

TTGAAATAATAAGATTCATCCTTTCACCCCCATGGACGTAGGTAATCATACCGAACCACGTATATCCTTGTGTCTATTAT

CATTGTTCTTACTTTCACTTGCATGAGTAAATGGTTTTTATCCTTTGGATGTAGTTTGTAGTAAAATTCTTCTACATTCT

GGCGCCGACTAAGGGGATCCGACTAAGATCTCCGGATACCTTTCCTAGGAGGCATTCAGAGACAATCCAAAGCTCCTGTA

TACACGTTCAATTGTTGTAGTTGAAGCTATGATAAACTCTGATTGCCTAAATTGTTGTTATTTTGAGTCATTTAGTTTAC

ATGAAAAGTTACAGCAGTAGCAGCGTCTTCACAAGGTGGGGTAAAGTTCTCGTTTCTTCGCCCACCAGCATTTTGATAAG

CAGAAAGTCTTCATAAGCTCACCGGCGATTAACAACAAAGGTTCGGAGCTAGGTTGAAAGTATGCTGATATGATGACTGC

TTGTTTTTGATATTTGATTGAGTTTTAGCGGTAGATAAGCTGTTTTCTTGCGAGATTACTTTACTAAATGGGTTGAGGCA

TCTCCACTAAGGCACATCCGAGACAAAGATGTTTTCAAGTTTATTTAGGAGCATGTCATTTGCCGCTTCGGGGTCCCAGC

AGCTATCGTCTCCGATAATGGGAAACAGCTCCAAGGGGAAAACATTGACCTTTTGGTCAATACGTACAACATCAGGAAAA

GTAAGGCTACTCCCATTTACCCCCAGAGTAATGGGCAGGCCGAGATCACTAACAAAATAATCTCCAACAACATGAAGAAG

AAGTTGGATAGGGAATATGGTAACTGGTGCGAGGAGCTACACAACGTGTTATGGGCATACAGGACCACACGGAGGGAAGC

TACCGGGTTCTTACCGTTCATGCTTACTTACGGAACGGAGGCGATTTTACCGACGGAAGGAATGAACCCCACTACTAGGA

CAGAAGCTTGGAGGAGAAACGTCTCGTCCGATATGATTTTGGAAAAACTAGACGATTTGGAGGAGACACGAGAGATGGCG

CTACAAAAAATGGAAAACTATCAAAGGAGGTTACAACGAGAATATAACAAACGTGTTCGCCCAAGAGAATTCCAACCAGG

AGAGCAAGTGATGAAGTTTCTGGCCGAATGGGAGCGCGACAAAAAGGGTGGCAAGCTAGCACCAAAGTGGGGCATGCCGT

ATATAATTGCATCTAAGGCAGGCGAGGGATCGTATCGGCTACTGAGACTCGATGGAACAAAAGAGCCCAAACCCTGGAAC

GTCAAATACCTAAAACTATACCACCCATGAATGGAGGTGCGATGAAAAGGTGAGCATACTTGTCTTCTATTTCTATGAAA

CCGTGACGGGTATGAAATAGGGCATGAGCGACTAATGGTCATTCGTACTCGGGCAAAGCCAGTGTGGAAGTGCCGAGGTG

CCTTACAGTTGCCTGGCTATTCGTACTCGGGGAAGTGGCAGTGTGCAAGTACCGAGGTAGCCTAAGGGTTCTGGTGGTTG

GAAACCAGTTCCACAGGTTGTTAAGACACGAGGTCTTCTCAGCTAGGTGGCCTACTTACATTTGAAAGGCCTCCCCCATC

GCAAGTGGCTGTGACCGCTTGCCCGAGATTGGTTGATCGATAACCGCTCGGGTTAATCGAGCCTAGAATTAGGTTCACTG

GTACCCTTCCACTGAATCCAAGAGATAGTGATGAGATACCCCAATCCGGACATGGCATTGGGCCCGATGACCCTTCTTGT

TGGGTGTGTTCGACAGAAAAGGTTAAAATAATAACTTACATAAATGACAATTGCAGGAATGACATAAGAAACAAAAGTTA

ATTCGATAGCGTCATGCGTCCATTAAGAAAGTGCAGAAAGTGTCATGCAACCATACGAAACGAAAAAGTTTAACGGAAGT

AAAACAAAGAGCATCGGCGATGCAGCACCCCCATTTAAACAAACAAGTAGTGTATCAGAAGTAAAAATGGGAGAACTGTT

CACAGCGATTCACAACCCATGGCAAAGGGCACTTTAGTGACGTAGCACTACAAAAGGTGGATTAAAGCCTTGGGAGGAAT

AGTGTCGGCCCTCTTTTGCGCGACCTCGCGCCGCTTATTCTCCACCCACTCTTCAATACCCGACCGAACCTTGTCGTCTA

TATCCTCTGCATGGGCGACGCGTTCATCCTCAAGTTTCTGGGTAAGATGAGAAGTAGTTTCCTCAGCATCGGACAACTTC

GAACCTAACTCAGCCTCGACCTGTTTCAACCGTCGGACCTCCAGATCCAATTCGTGAGAACGTTTCTGCTCGCCTACAAG

ATCACAACAACGTGGGATTATTCATTGAATAAGGTTTAGCAATTTCACAACTACAACAAATAATCTAGACCTAAGGAACT

GCCTCAACACGTCGCTAACAAGACCTGAAAGGGCTGGTAACACTGACTGAAGCGACTCTACTGACTTGGACAACTCATCC

TCGGCCTTGGACAATTTCATGGTTAGCTTGTTATAGCATTGTTCATTGAATTTATTACTGGGGAAGTACTTCTTAAGTTC

ACTCCACTCCTCCATTAAACGATTGCGTGATACTTGAAGCACTAAAAGTTGCGACCTACTCCATTTTATCTCACCCACCA

TCTTTTCAAACTCGCCCACTTTGCCCAGTAGATCCCTATTGTCTCGATCCACCACATCTTTATCCCTCAATATTCGTCGG

CGATCTTCGTGAAGGTCTCGATTCTGCAATTGAAGAGCGTCTACCCGTTGCTCTAAATTCGTGGCCAGAGCAGTTTTCTC

AGCAGCAAGGCTTTGGGAAGAGCCCAAGTGAGCCCGTAAATTTTCTATCTCTACATGGTCATACTCAAACTGACGATATC

GTTCTAGTTCACCTTCGAGTTCTTCGATCTCCCCAGATTGGCGGTCTATTTTTCCATCTTTGTTTTCCATTCGCCGACGG

AGATTCTCCACTAAATCCTTTTCAATATCCCACTTCCTCTTCATGATCCTAAACTTCGAAGAAGCATCTAAACCTATACT

ATCTATCTCGGCTAGAAGAAGGATACCCGACTCAGGATACTGGAACGATAAACTCAATCGACTTAAGTTAAGGTTTATAG

AACTATAACATACCTTGAGCATCGGCGAGTTTGATTTCCAGCTCGCGCGCACGACTCTCATCGTTTTGCAGTTGAAGTTG

AAGAGTTTCCACCAACTTCCTCTCCCGTTGATGATTAGCGGCATCCAACCGTACTCGATTAAGGATCTGACCCAGTAAGA

AAGTTAGAATGGCAGGTAACGATCAAGAAGAAACTAAGGAGCGGGACATCGCCCAAACATGCAGAGCCACTCACCACTGT

GTCAAGAGTCTGAGAAATATCACCATAGAGGGGAACCGCATCAAGCATTTTTTCAGCAGTATACGCTCCAAAGTCGGGGG

TAATAAGCGCTCCAAGAGAGTTCTTGATAGGGCCGGAGACAGTGAAGGAAGTTTCAGCACTAGACGACGATCCATCAGTA

ATCCGGGGCACACGAGGTAGTCCAACAGTAACAATCTTCTCTGAGTCGGACGTCTTCAGGGGTTCTTGTTTGTCTCCCAT

CTCTTTCCCATCGCCCTTGTCATCCGAGACATCCTCGCCGAATATGTCATCTTCATCGGGAATATCGATGATTGGTTTCT

GCCTCTGCTACGGTTTAGTAACAACATGCTTGCCCCTAAGAGGATTACGCCTAAGAGGCTGAGGATCATACTCTGGCACA

TCGGCTTCATCATCATCGTACCTAGGAAATTTCTGCACAAGGCTTCTCGTCAGAACATGTGAAGGCAAAAAATAGGGGAA

AGTACATGTATTTATAAGAAAATAATTGTGTTTGTTACCTTCCCATGGTCTTCCTCAACCCTTGTGCCAATACCCATTTT

TCTCGACGAAGCGGCGACTTTGGTAGAGTCGCCCTGGGGCTTCAATAACAACAAACTGGCGATTGGCGTCAAAACTAACA

AGATAGAAATTGAGGAGAGTAGAAGGAAGCACATACCTTATCATCCATATCGTCGTTAACCACCCAGAAAGGATAGGGCT

CGGCGGGAAGTGGAGGCGACGGGAAGCTACCATGTAAAATGGGACCTCGGACAAGAAGTGGAACTCGGGGATAAATTGGG

TCATTAGTATGGCGGGGTCGCCTATTAGAACCTTGGCCCATGGGATCAGCATCCAGCATCATCCTCTCAACATTCTTGGG

CATAGTTCTTTGACGATGCGGACTGGCATAAAAACCAGCAAGATCACCGTTGGTTGAGGTACCGACTCGATAGTTGACAA

GGAAATTGGCAACAATATACTCTGGAGCGTGCTCGGGCTTGTAGGTCGACCACTCGACGGTAGAACCATGACCATATCGG

CGGGCTCTCCATTCATCCATAATGCGAAAAGTGTTGCCAGTCCACTGAGCCACGGCATGATAAGGCTTGAGCTGGGAGAG

AACCTCATAATAAAAAGGGCTATCTGGGTTGTACAAAGGGAGACGAAGTCCTAACTCCAGCTGTCCCTGAGTAATGACCA

CACACTTAGCCGACTGATCGTACTTATCCATGTCTGACTTCTTCAGAATACTATTTTGGCCCGAGATCAGGGTAATTTTG

AAGATCGTACTTCTCCTTCAACTGCTCAATGAATTCTTCATCAGAGAGATCGTTCCCCCTTTTCTTGGACCTGCAAGAAA

TAGAAGGAGAATCATCAGGAAAGATTGATTTCATGGCGCGGACATCGTACTTGGAGGGACTATTAAATGAAAACATTGAA

TAAAGGGGAAAAGCATCAATCTCGAAAAAAACATCATCAACACACCTTTTCTCGGTCATAGAGGGAGAAGAAGCCATTGA

CGGGAATAGTGCAGAGCTTGGAAATCGGTGAAGAAGGTTGGGACGACGAGAACGTAACCACAACTACTGTAGAAATCGGT

GATGAAGAGGTGAGGGAAGAAAATCAAGAAAACAAAGGATTATAGAGGAAGAAAGGTAATGGAGAGTCTCGAAGAGAAAG

TTGCTGTGTCTGGAAACTAAAAAAGAAATGGTAAAGAAGAAGAAAGAGAACAGGAAATGGGATATTTAAAGAAGAGAAGT

TGTAACCGACGCCGACGGTTACAGGTAATAAAGTGGAGTTAATGGGGGACTTCATGGCGGAACTCGCGGCATGAACGAGG

ACGTGGGAAGCTGGGACGACTTTCTTCCACTCGTGTCTACCACACGAGCGAAAGAAGGGGCATAAATGTAGGTGTAGATT

ATCCACAAGGTTAGGTGGAAACATCTCAAAGCTATGATGTACCCAAACATGAAGAGACGAGAAGCACAGGATAGAGGCTG

GCTGGCGCGAGATTTAAGGAGCCACGTGGATCGGGCATGATCACTCTACAGTTGTCGGACAAGACTCGACCCCAGGCCTC

TGACAAAGCTGACAGACGAGACGTATTGGGTCAAACGAGGAGCGGTAGTCAAAGGGATAAGAAGATACGTGTCATATTGG

CCAACTTCATGGGCGACAGACTGGGAGGGCGACATCGCACCGAGAGAATACCCGAATGGATCGGATCGGACGAGAGACTA

CAAAGGAATCAAAGAACATCAGCCATAATTACAGAAGAAGGGACGACGAAAGCAGGCCCGACGACAGGACTTCAAGTTAG

ATATGTTGTGATCAACACCCTAGTTTGGTTATAGGTAGCCTCTCCTATAAATAGAAGAACTGGTATCTAAGAGAGGGATT

GGTAATTTCGGAGAGAGAGAGACTTGAGATCACATTAAGGAAGGAGAGAATTGATAACTAGCCCATCTTGTAAGTGAGTT

CAACTTGGAGAGCAACACCATTGTAACCCTTGAAATAATAAGATTCATCCTTTCACCCCCGTGGACGTAGGTAATCATAC

AGAACCACATATATCCTTATGTCTATTATCATTGTTCTTACTTTCACTTGCATGAATAGATGGTTTCTATCCTTTGGATG

TAGTTTGTAGTAAAATGCTTCTACAGTGACCTAGGTTGCAATTTATAAAGATGGTAAATGCGAAAACATGAACTGTCTAT

TTTGTATGTATTACAATTAATGCCTTTTTCCAGTGATATTTGCTCGATCTAATACTTCTCTGATGGTTTTCAGTAGTAGA

CACTAGATGCATTTTTGTGTTCGCACTATCGGTTCTCATCTGTTCACTCATCTAGAGTTCAGAAACATCATGATCCTAGG

TGTCATGATGAGCTTGGACTAAACTTCTGTGCTAAAATATTTATCTGGCCAAGACTTGCATAACATTACTTTTGTCTTGT

TAATGACATATTTTTAACTGCCAAGGAAGAGGTTGGTCCGTGTGCGGTGATGTATATGGGTGGAGTTACATCTAACAGTG

GGTTGATCTATTGAATGATCACGTGGATAGAAATTTCTTGTTTTCATTTGTAACTGTTGATTTTCTTAAACATGCCGCAC

ATGGGTTGTATATTCAAGCAGTACTTGGCTGATGTTTATTTTCCCTTTTTGCTAAGGTAGTAACAACTTACGGTAAAACT

TTTTAAAAATAATACTCAATAATATAATAATTTCGCTAAAATAATAAAATTTTCCAGTCCAGACTCGGGCCATTAGTGGT

AAAATAATAAATTCAATAAAATTATAATATAGTTCATTTCAAAATTTCCCCTTAAATATTAATAGGCCCCAAATATTATA

TAAAATAATAATTTCATTAGAAAATAAAAAATAATTTTCATACACATTAAATTTTCATACACCAATTCCATTAGAAGACA

TCCTAATGATTTAGAAGCTTGTATTGTTAGTGACTAAATTGAATTGTATGTTAAATACAAAAATTTTCTGTATAAAGTTT

TTTTTCTCATGGAAATTAATAAGGTTTATCACTTTAAAAATGCATGCCGTTCTAAAAATATTTCTGTTAATGTTTTTTAA

TCTTTTTCTATCCATTTTTAAAAATCTATTTCGGTATTTATTCTAAGATACTTTATTTTTATTTTATTTTTAATATGTAA

ATTCTTGTTTTTTCCAAAATAGAAAAAGAAAAGAATCTCAAATTCAAAATCAAGCTGCTCTCTACATATTTTTTTTTTTG

ATAAAAAGATTGGAAATGACTTTTAATTATATAAAGTTAAAAATTACATACTAGGCAAGCGAGCGACAAGCTGTGCCGCT

AGCCCTTTTTGAATAGCAAAACCTATCATATTAAAAATAAAATCTCGTGAGCCAGGGGTCATTACATTGGTGCGCATGAC

TTTCTGGACCCTCTTGAGTAGTATTAAATTATTATTTTATCGATTAAGTAATATCGCTTTAAAATAATAGGATTTGGGTG

GTCTCAAGCCTATTATTTTATAGATGTTTTACCGTACTCTGAACTCATCATTTCCAATTTCTTGTTTTTCGTTTCTATGA

AATTTGGTATAACATAGAATGTACAAGCGGCCTAGAAATAGAAATTCTAGGTGGCAGCACCTGATAGGGTTTTGTCGGTA

AATTTAGGAGCCCTTAGGATATGAATCAGTCCAGGGAGTCATCTAAAAAAATTAGGTGGACTTCTCTAGTTTCCAAGGAG

ATATCAGATACTTATTTTTGTCGAGTATGAATCATGTAGGAATTTCTGAACACTGCTGGTCCAAAATAAAAAATTTAGGG

TGGAAGTTCCGTTATATATTTTCTTGGCGATATTATTCTTTGTTCTCGAATCATTTAGAGTTTAAACTAAACTTGAGTGT

CTTGTAATGAAATATAGTCCTTGGGATGACATTTTCGGATTCATACTAGGGGACTCATGCCCAATTATGGTTACGTATAT

ATCCTGGAACACTCATTTTCTTTGAGTGGAGTCCTTCTTGTGTCCTTTTTAAAAGGTCGGCAGATACCTCAACTTTAACA

TATCTGTCGACTTGTAGACAGTAAATGTAAGTTTGCATGGTCATTAGACTATGATTCCTACTTATTTGGACCTCTTTATT

AGTCCATGGTTACGTTATTAGAATGTGATTTCCTACATATTGAGGACCCTTTTATTAGACAATGGTTTGGTCACTAATAA

AGATATTTTTCATTTAATTTGATTCCATTGTCCTTTACCCTGCAGCCCAATTGTTTAGTTTTAGCAATTTCACCTATATT

ATCATGCACGTTTGTTGTGGAACAAAGCAAAGGGACAATCAGATGATTTTCTTTTCTTGTGTTGTCTGGATGTTCCTAAA

ATTATACTTGGTTATCTATGAGCTTTGGTTTATTTCATACTCAAATTGATGAATAATGGTCGATTTTGCAGCACTTGGAA

ACTATAATCAAGTCTCGCATCTCAGGCCTCCAGTCGCAAATTAATAAAAGTGTTGCTGAACTAGAGGCGGAATTGAGTCG

TTTTGGCAAGCCTGTTGCTTCTGATGCTGGAGTAAGTCTATCTTAAATATCAGTCTTCAATTTAGAGCCTATTGTCTTTA

GTCTTCAATATGCACTTATGAGCAAACATGCACTCAGTTATGGAGATTTGCCGTATTTTGGGCTAATTCACTAATTGTGG

TTTTTGCAAGGTAAATTGTACTCAATTATGGAGATTTGTCGTCTTTATGATGGAATTTACAAAGAAAATCTTGACGGCAT

GTATGTAGCTAGTACATTGGAATGTAGCTGGTGTTACTTTCTGGTATATTGGAATGTAGGTGCCCCTGCGCACACACGTG

AGTTGACAATAAAACTGGAACCTCCTTTTGTGATTTGATTTGTGAGTTTTATTTGTTGCCTTATAAGGATCTATTTGTTC

CCCTGTGATGTTGTTACTTAAATATTAGTCTTCAATTCAGAGCCTATTGTCTTTAGTCATCAATATGCACTTATGAGCAA

ACATGCACTCAGTTATGGAGATTTGCCGTTTTTTGGCCTAATTCACTAATTGTGGTTTTTTGCAGGGTAAATTGTACTCA

ATTATGGAGATTTTTCGTCTTTATGATGGAATTTACAAAGAACATCTTGACGGCATGTATGTAGATGGTAGATTGGAATG

TAGCTGGTGTTACTTTCTGGTAGATTGGAATGTAGGTGCCCTTGCGCACACATGTGAGTTGACAACAAAACTTGAACCTC

CTTTTGTGATTTGATTTTGAGTTTTATATGTTGCCTTAGAATGATCTATTTGTTCCCCCGTGATGTTGTTAATTACAACA

AAAAAAACATGAGTTTCCAAAGGATTCAACCCATTTCTTTGTATCTGCCTTCGATTTCCCTGTCAGCTACAAATATCAAT

TTTGACTGATGAGCCAACTGATGTTTACGTGAGCCTTATACCTTAAACATTACATTTATGGTATTGGTGGTTGGTGTATG

TTTAGGTGATTTTATCAATAGAATATTGTTTTCCCTGTAGTGTCTTAGGATGAGTTTGTAGTCTTATATATATGTAACAA

GATGTTGGGGACATAAGTTCAGTAAATTTGCTATAAGATTGATGCTAGTTATAATACAGTCATGTAGATGTTATGCAATG

AACCAATATGTATTATATTTGTTTGCCATTTTAGGAAAATGATTGTGATATGTTGAAGTTGATCTGGTTTTGCCATATGA

TTAGTGTTACAAACGCGGCTGTTACAAAGAACGATAGATAAAATCTGTTTCTGTACAACTACGACCCTAAACCAACTGTC

GCTTCTACTGTAGTATTAACGATTGCTTTTGGGGGTCAATGTTCTTCGTTCTCCTCCTCATGCAGCAGCAGAAAAGAATT

TTCTGCAGCTTTTTTTTCTGCTCTACAGCGTCCCCGAACTCTCCATCCTTCTCATAACCCTCAGAACCGTTTATATACTC

GACAGGGGCAAAATATCACGCAGTAAAGTCTCCAAATTCTCTCACAGAAAAGCCATTACTCGGGAATAATATATTTTCAT

ATTTACGGGATTCTTTCCTTCTTTATCTCTTTCTTGCGTTGCATGTAGGCTAACACACTCCTAACAGACCACAACATGTA

CTAGAAACGCTGTAGACTGAGTAACATTCACTGCAATCACACGAAATCCCGCGTATTTCTCCTCCCGCAACCGTGCTGCT

CTGGTTTTCTTCACTCGGAAATTCCAGCCAATTCTAACCGATTCCAAAGCCCATACCAAGCCTGTTAAGATCAATCAAGT

ATAACCATGAAGTTTCAGCCATTGAATCTCACTGAAACGCCTTCAAAATACAAACCCTAAATCTGTTTTCTGAAGAACTC

CTTTTTCCCGCCAAAACTGAATTTGAATGGTAGGGATGGTGTGCCCCGATCCAAAAGTCGGGTGCCTTTATCAACTTGGG

GTGCCTGGGGTGCCCTTATCCAAAGCGAGGGTGCCTTTATCACTTTTCGAGCAACTTTTTCTACACAAGTGTATTCCTCC

AAAAACACCTAAACAAACATAAAAACACCAAAATTAGTACAAAAATCGAGACTAACAATAGAGACACTGAGGACAATTCA

GACACAAAAATGTGTCTATCAAATACCCCCAAACTTATTATTTGCTAGTCCCGAGCAAATCTAGTCTAGAAAATAAAATG

ACTACACTTAGCACGTGCAGCAAGCCTTTAAACCGCTAGGTGGCCCTAGTGGCGGAGTGTCGTCTCCGGAGGGTTTACCA

GAGGTGTACCCACAAAACCTGTACTTCTAATTGGTCACCAGTATCCAAAGAGCTATGAGGACATACAAATTCTCAACCTA

TCTCCAAGTGACTAGAATGCCAGAGAAATTAAAGGTGTCAACTCTAAAGCTAACTGAAGAAAAGGGGAGACACATACGCA

ACACTGCTAGATAAAGAGATATCCGCTACACAGCTAGATAAACATTGTAAGATGCGTCCGCTGATTTACAGCTGGATAGG

ATTAGGAGAGAGATAAAAATGAGAGGGACATCTGCTACACAGCTGGACTAATTACGTGTGATGAGTTAAACCAGTGCTAG

AAAGATCTTGTGCCAGATTGAAAGCGGACTAACAAAGAAACCAAATGCATCTCTCTTCGACTCTCTCATAGTGCTCAGTA

GAAACAACGTCTTCTTCGGTCTTCAACTGTTGATGATAAATTCTTGAAACTCGTAGAACCTTGACAATTAACTCTTCTCT

TTGATTTCTCACTTGAAATTTCCTTATAGATTTCTATGGCCTTATTGAGCAAAATTTAAACTTCTTTCTTTCTTTCTTTC

TTTCATTTTTTTTTTTTTTGACATTATTCCTCTTTTTTTTTCATTTTTTTTAAAGAATCAACAAGACAAAAATTTACATG

GCCATCATAGAAGGACTTTAACACTTGGATCTTTGCAACTTGCAATGTCTTAGTATCATGAACTTCAACAAATTATATCG

TTTTTCTTGCTTTTATAACTTCTTGTAACCCTTGATTTTTTAATTCTGATTTTCTATTGTTGCTTCTAAACCTTAAACCT

CTTCAACTTTCTTCATAGATTTTGATGTCGCTCCGTTTGTATATGATGATAAGTTTCCACTGAGAGAGAGAGTCATATTC

CAGTAACTAAGACTACATTGTGAGGTTGCTTTGTCTTTCTGGCATTTCCTGACCTACTTGCCTTTCCATCATGGATGGTT

AGGTCCATCACGGTTACCCTCTAAAAGGAACAAGTTCTCTCCTGAATTTCAAGGATCTCAATGTCTTTTTCTCTAATGTC

TCAATAGGTTGTTATCCCTAGCATTCCAATTTCTATCTTTTCGGTGTGAAACAGTATGTAAACTTAGCTAACCGGATACC

ATGTGACGCTAGAAGTTTCAAAAGTGCAACTAAAAAGTTCTTCCTCACCCCCAAACTTAAACCTAACATTGTCCTCAATG

TTCTAAAGATGAAATTAAAAGCATGAACAAGGAGAAACTGTTACCACTTGAATAAAAAGAGTAAGGAAAGATATTACCGT

GTTGCAAGAATATTGGGTTACCTCCCAAGAAGTGCTAAGTTTAAAGTCTTCATCCAGACATCAGAAGGAATTAGTCACCT

CATAGAATCAAAAAGTAATAGCCGGAATAACTGCGGATCACGAAAACCAAAAAGGGCTATCACAAGGAATCTACAACATC

CCAAGAAAATTAGCAAATAAAGCGTACCCTTGTCTAGTTTCCTGTTTACGACAACTACAAATAACTGTGGTTCAGGTTCA

GGTTCTATAACTGGGTCTAGATAACATATTTTCATAGGTTCCAGTTCCTCAAAAGTTAGATCCGAATGTTCAGGTCCTAG

AGTCTGGAAAAACTCGAGTAAGAACTTAGAAGCACATAACATTAACCGGAATAACTGGGGATCCTCTAAGTCAATAAGAT

TTGACTCACACATCTGACCACAGTGGTGGTCATCCTTAAGAAAATGTGTCGACCCTAATCTCCTAAAGTAATTAGGTTTA

TTCACAAAACTTAATAACTTACACATCTGAAATTCATATGTTCCCACAACTGGAAGGAAGGTTTTTGGTGGGAAAACAAA

GTCAATCGTGGTATCATATCCTAGGTTAACCACATCAACCAGGGGATGGGTTTCTAACAACTGAACTTCTTCCTGGACAT

TATTAGGTTCGGGAGAGCTAGCATGAAGGTAATCTGGCATAATAGTTGAGGCACATATATCAAGTCCCAAGTGAGGGATC

TCTGTAAGAGCTAAAGGTAGGGCACAAGGAGAATGATAATCACCCCCAAACTTAGAATTTTGGGTGGCTCTAGATAGACT

AGCTACAATTTCCCTAAGTTCTAGATCATCAGAATCCTGAAAATGGTCAATTGCTTCTTGTAGGTTATACTCAGGTGATG

CATCCTCACTCAGATTTAAAAGCTCTACGAGTTCATCATCTTTTTCTAAGTCTGATATTTCTAAATCACTAGACTCTAAG

ACAAAGTTCTCATAATAAACTCGTTCCTCAAAACTATCATCGGCTTCGTAAACAGGAAATACTACATCGTCTAAAACAGT

ATCCCTAGTCAAATCCTCGTCCTTTCAAATAGGTGAATAATTGTTAAAATTATTTGGATTTGTACTAGAAACAACACTAT

CATTATAAAGATCAATCGGGGTAACAAAATCCTGATCACTATGCCTACACATTTCATGTTCCTCATCAATACTATCCTCA

TCAAAACAAGTAGTGTTACCAATTCTAACCTCGTCATCTTGATTATGTGAATAAAAACTATCTTTAATTTCAAGGGTGGT

TTTAGGAACACTATAATGGAAATTAAGACAATCTTGAGCAAGTTTTTCCACAATCTTATTTGTACTCTCAGTAAATTGCT

TGAGGGACTCTTTTAGAGACATCTTAGTTGACTGCTCCATGGATTCTTCTAGGGGAAGAATATTATAAGTCCCTTTCATA

AATAAGTGAAACTTCCTTGTTGATTCATTGAAACTCTTGAGAGACTCTTCTAGATTAATAGAAGGATTGTTTTGCTTGTA

TGACCTCTGGGTATGTGGATAGTAATTTGGATCACAATGGTATGGACCATAACCTTCAAAAGTTTCGCGTTCCCCATCAC

TATTCACATTATGGTCATAAAAAGGATAATGTCCAAGCTGCTCAGTCGGATACTCATTGTTTTGGCTATTATAATACCAG

TTCGACATCCTAACTGCAAGGGAATTATAAACAAGCACAAACAAGGATGACTTGACCACAACAAACCTACTGAATTTTAG

CAAACAAAAAACATGATGGCTCCACTTAGATTGTTTCTAGACCAGCTTCTAATACTTCGAAAGGGAATTCGTTACAATCT

GAGCAAACCCCTCTGGAATCAATCTGCGTTTAAGTAAGTTGAATAGAGGCGAGGGAAGCTGATGGAGCTTTGATACCCAA

GGCCTCACCGCATTACAGGGCGGCGCAGTCACGCATTCAACTCACAGAAACTTTCATGAACTTCGAAGAATGCTCAAAAG

AGTAACCAATATTTTTCGAACGAGTTTCCAATTAAGCTCGTTACCCTATCGGTCTCGTTCTATTCCAAATTTTAAGGATT

AGGTTCGCGTTAAGTATCGTTTTCCTAAGGCGGGCAAGAAGAGAACGGTGATGAAATCCGAACCCTTATCTTGTATGGCC

AGTCCTTGCCGTTTACTAGGAAATTAAAGCAGCCTTTTTCAGTTCCTCGACATATATGCAAACGAAATCATCCAGTAAAC

CCGCTGACAGGGGATTCGCGGGTGTTTATAATTCTTACCTCCCGTTCCAGAAGGGGGATGAACCGTTGAAGTCGACTCGG

GCCACGACCCCTATGTCATGTACGAACCCGAGGGGCCGAGGTGATATCGTAATCACCGTCCTTCCCTGTGCAGTTTATAT

TTAAAACTACCCTTCCGTAGGGTTTAAAAAAATAAAATAATGTCCCAAATGTCCAAGTCCAAAGTCCAAAATAAAAATAA

AAAAATATTAAAGTGCAAAAGAAAATTACAAGAATAAAGAAATCCTAATACAATCTAAAATAAATAAATAAATATAAACT

ATTTTTTTTTCTTTTTATAATAATAAATAAATAAAAAATCTTCTTCTTTCACTCCTTTCGCTTTGTCTTTATTTCCAAAT

CTTTCAAGTATTCACCAAAATCATTGGCAACTTTTCTTTTTGCTCCAAGTTCCAAATTCTGTAAGGAAAAGACAAAAAGC

AAAAACGTAAAGAAGAACAAATAAAATAAAAACCTAAAAAAAAGAATAAAACTCTAAAAACTCTAGCCTAAAGACAAGCT

CCGTGTCGACGGCGCCAAAAATTGATGGTATTTATCAATGATGTTGTAGTAGTAAAAGGTTCGTGAGACTTGTGAAAACA

AAAATTTTTAGATTTAATAAAACTAAGATAAGATAAAAATATATATAAAAAATTGTCACAAGATGAAGAGATAACCGAGA

CTCAGAATTCCACCAGACTTCAATTTCATGCGGTTCAAATACCAATTCTAAACAATAATATCTCATAATATAAAGTTGCA

TTGACTCTCATTCATTGCCTAAAGTAGATTTCGAGAGTAATGGATGTAAATCTAAAGCATGGGATATCAAAACATTTAAG

CAAGCATAACCCATCAATTGAAATGACAACCATTCAAAGATAATCATAATTCATTTAAATAATTGTGCAATTAGTCATAA

AAAGAATTAATATAATTACCGCATTAGTGAAATTCGGCTTCCTTCATCGCCCCAGAGGATGGGTCTAGCACATCATGTTG

GAAACACGCTCAAAAATCATTATTATTGCTCAAAGGAGGTGTACAAATGATGGAATGAAGAGAAAATGATGAAAACCGGG

TTTGTAACAGTTATACGTGTTACAAACGCGACTGTTACAAAGAACGATAGATAAAATCTTTTTCTGTACAACTACGACCC

TAAACCAACTGTCGTTTCTACTGTAGTATTAACGACTTCTTTTGGGGGTCAATGTTCTTCGTTCTCCTCCTCATGCAGCA

GCAGAAAAGAATTTTCTGCAGCTTTTTTTTCTGCTCTACAGCATCCCCGAACTCTCCATTCTTCTCATAACCCTCAGCAC

CGTTTATATACTCGACAGGGGCAAAATATCACGCAGTAAAGTCTCCAAATTCTCTCACAGAAAAGCCATTACTCGGGAAT

AATATATTTTCATATTTACGGGATTCTTTCCTTCTTTATCTCTTTCTTGCATTGCATGTAGGTTAACACACTCCTAACAG

ACCACAACACGTACTAGAAACGCTGTATACTGAGTAACATTCACTGCAATCACACGAAATCCCGCGTATTTCTCCTCCCG

CAACCGTGCTGCTCTGATTTTCTTCACTCGGAAATTCCAGCCAATTCTAACCGATTCCAAAGCCCATACCAAGCCTGTTA

AGATCAATCAAGTCTAACCATGAAATTTCAGCCATTGAATCTCACTGAAACGCCTTCAAAATACAAACCCTAAATCTGTT

TTCTGAAGAACTCCTTTTTCCCGCCAAAACTGAATTTGAATGGTAGGGATGGTGTTCCCCGATCCAAAAGTCGGGTGCCT

TTATCAACTTGGGGTTCCTGGGGTGCCCTTATCCAAAGCGAGGGTGCCTTTATCACTTTTCTCCCGGGGGTCCAAAAAAA

ACACTTTTCGAGCAACTTTTTCTACACAAGTGTATTCCTCCAAAAACACCTAAACAAACATAAAAACACCAAAATTAGTA

CAAAAATCGAGACTAACAATAGAGACACTGAGGAAAATTCAGACACAAAAATGTGTCTATTAATTAGTAATACTTGCTGC

ATAGGTTTTTGATTGTTCCATATTAATGGTTTATTATATTCACTAGTGTCCTAGTATTGTGTCACCTGTAAGTCTTGCTC

ATTTTTAGTTTCCTACATTAGGATCCTGCAATCATCAATTTACTTAGGTCTTTGGTGTAGAATTGTTACTTTCATTTTGG

ATTTAAAAGTGTGGATGTCTATCTTGAAGTTGTTGCTCTGCAGCAAATACTACTTTCAGTTCAGGATTTTATAAATTTAT

TGGTGTTTTAGCATTGTGTCAAAGTTTTGGTTCAGTTTCAGTTTCTTATATCGGGATCGTGCGGCCATCAATTTACTTAG

TTTCTTATATTAGGATCATGCGGCCATTAATTTACTTAGGTCTTAAGTTATATATGACCGTTATGTTAATGTATGCAACT

CTGCGCTTATCTTCATACTGCTTTTTTAATTATTCACTTCGTTTGTGACTTAACCTTATGTTAATGAGAGAATATGTTCT

CTACCATGTATTATGAACTGAACTAAGTCATTGTTTATTTGCTTTGCAGGTACGATATTCTTTGTCATATGTGAAGTAGG

GACACAACACATGGATGCTATGCTTGTTATGAAGTATTGGCTAAGTTCATCGTCATTGGAGGAGCTTATCGTTGTTTAAT

CAATTGTCTATTAGTTTAACATGTTTTTGTTAGAACTTACTAGCTAGCTTGGAAATGAACTCAGAATTTCATCCTTTTTT

TTTGTAATTTTCTAAACCCAATGAATAAAATGTTGAATTCTCAATTTGAGTTAAATTTGAATTTGAATTTGAGTTTGAGT

TTGGGTTCCATTTGACTTTAACTTGAGTTCCATTTGACTTTGAATTTAATTGGGTTTCATTTGACTTCAATTTGAGTTGC

ATTTGATGCACTCAAGATTGATTCAGATTCTACATTTCAGCAAGTTTAAATCTATTTTATATATTATATTTTAAACCTAC

GACCGATTTAATGGGTCTGCATCTCGTTACGGTTAGAATCGGTCGTAATTATGAAACTCACGGCTAAGGGTTGTTCAATT

AAAAACGCTTTATCAGCGACCCCATCAGAGACTGTTAAAACAGGTCGTTTAATTCGTATATATGACCTGTCCTGGCGGTC

GCGGAACTTCTATTTTCCACTAGTGTATAAACTTCATCAATAAAATGTATGTGGATACTAAAACTTATATATCCCTTAGA

TGTATATTTTATTCTATCTTCTAAAGAGACTAAATCGTACAGCTATATAGACTTTTACATTATACACTATTGATTGATTA

ATCACTTGAAAATTACTTGAAGCTAGTGGTATGTGTAAGATTACCATTGTTGTCTTCTAAGGATGTTTCAATTATTAAAC

ACGAGTTTAGAACGACTACCAATGACTAGATATGACATAGCATGTATACTTTGTATGTGCATTTTGAAGCTCTAGTTCAA

GTCCGGAAATCTAGTATGCGAACTATGTTTGCGAACAGGTTTACTTGTGAAAAGGTCCAGAGCTGTTGTTCGCGTACTCT

ATTTGCGAACGAGTAGACAAGGTTGAGTCCGGAACTATTGTTAGCGTATTATGTTTGCGAACAACACAGTAAGGACAATG

TACGGATCCGTGGTTCGCGTACTCTGTTTGGGAACATAGTGGTTAGGTTCTAAATTCTGTAAGTATAAAATTCATACTCA

TGAACTCAGGTTCGTTTATGAACTAAAGTCACTGGAACTTTAATGAAATAAGGTATGTGAACTAGTTTTACAAACCGTGG

CATTATGTTTATGAATTGGTTATTGTATAAGTACTTTGCACAATTACAAACCAAATCGATTTTGTTTCAAATGGTTCATA

TATATTTCTATGAGATAGTGAACATTTGAACAACTCTCTTGAAACAAATTTAGATTCATTTGAGATTATCTATCAGGATT

GATTGATCATCATATTTAATCTAGAAGTATTAGATGAATGTGGCTAAACTATAAGTGTTCATGTGGCTAACTTCGGTTAA

CTATTATTGAGCCAACAATATATACACGTTTGGGTACTGTTCCATATCTAAATATAAGTATATTTTATTGTGTGTATCAA

GCGAATACCATCTAACGGTGGAGATTGATTGCTTAATTTCTAAGCAGACTTATCTTGAATCGTACATTAGGATCTCATCT

AACGGTGAATATCAATTAATTTGTTACTAAGCCATCTTATCTTTGATTGTAAGCAACCCTGATTTGAAAGACTACATAAG

GGAGAACTCTAGCATCTTGGAAACCTAATACCTGTTAGAGCATTGCTCAGTCTAACTCGCATGCATTGCTATCTCAAGCA

TGTTTGTCAATGTTAGTGATCAAAACTATAAGTCTTGATTTCTAGTCTACTGTTAGGGTTCTAAACCAAAACAACGGTTA

AGAACACAAACTAAACAGAACCCTAAAGATAAAAATCCTTTTTACTAGGGATTAGCCATCTCTACTGTGAAGAACACAAA

TAAGAACTTGGTTTATATTAATTGATATCTATTACAGAACTAAATAGGAGTATTTATACTCACCAATACTTGGCTAACAA

GGAAAGAATCTCACGAGTTTAATTAATAGAACTCTACCTAAAACAGGAAACTACTCAATTTATCTAAGCATGAATAATTA

AGGAAACTAGTAAGTAAAGGAGTTGGCATCGCAACATCCCACCCCGCTGGAAAAATTTCTTGTCCTCAAGCACCAAAATG

TGGAAAACGTAAGAGTAGTGCATCAGCATCTTCCCAAGTTGCTTCCTCTAAAGGATGATCTTTCCACTGCACTAACCATT

TAGTACCTGCAGTATTACCGCGCTTGAACATCTTTCGATCCAAAACTTTATACGGTTCCCACTTGTCATAATCCATAGTA

ATGGGTAAATTCGCATCTACCATGTCTGTCGATCCAAGTTTTAGTTTCAGCTGTGAAACATGAAACACCGGGTGAATACG

ACTTGCAGTAGGTAAGTCCAGTTTATAAGCCACATGACCAATCTTTTCTAGGACTCGAAAAGGTCAATAAAATTTAGGAG

AAAGCTTAGAATAAGCTTGAGTGGCCACTGTGTGCTGTCTATAAGGATGTAAGCGGAGAAAAACCAAATCGTGGACTGCA

AAGTCGCGTTCAGTACGCTTGGTATCAGCATAAGTCTTCATCCTTAATTGATCAGCTTGCAAATGTGATTTCAAAATCTT

TAAGGTATGATCACGGGCTCGTAAAAGAGAATCAACAGCATTGACAGAGGTAGAACCGAGCAAATAAGATGCAATTGTTG

GTGGTGCACGACTATAAACTTCTTGGTATGGAGACATCTTGATTGCAGAATGAAAACTTGTGTTATACTACCATTCTGCC

CACGGTAACCATTTCGACCATTCACTAGGTTTTGTTCCATCAAAACACCGTAAATAACATTCCAGTGTTCGATTAGTAAC

TTCTGTTTGCCCATCCGACTGTGGATGATATGCCGAACTTCGACATAATTTTGTATCTTGCAAAGAAAAATATTCCTCCC

AAAAATTGCTCATGAATATGCGATCACGATCGTTGACTATACTTCTTGGCATGCCATGGAGTCGAACAATTTCTCGTACA

AAGATTTCAGCAACAGAGCTGGAAGAATACGGATGGGAAAGGGCAATAAAATGTGCATATTTAGATAGTCTGTCAATGAC

TACCAGAATAGAAGTCTTCCTATTTGAGGGTGGAAAACCATTAACAAAATCCATTGAAATGTCTAACCAAATATCAGCCG

GAATTGGTAAAGGATTTAACAATCCGGGAGGTGCTACTGCCTCAGATTTGTTACGTTGGCAAATATCATAATGGGAGATA

AAAAATTTAACTCGTTTTTTCATACCTTTCCAATAAAAGCTTTGTTGGAGACGCTTGTAGGTACGAAGAAATCCAGAATG

TCCAGGGTTTGAATGAAACTCGTTTAGGATTTTATTGCACCATATTGAGGATGATGGAACAACAACCCGCTTCTTATAAC

GTAAAACTCCATTCTCATAAGAATAATGTGGTTTACAAGAAGCATATGTTCGTAGTTGTTGAATAAGAGTACTCAATTCC

GGGTTATCATGGCACTCTTTAATGATATCAGCAAGGCCAGAAAAAATTGGCGCAAAAATGAGAGAAAAAGATCCATGAAG

ACGTGATAGAGCATCAGCAGCAACATTCTCTTTACCATGTCGAAAAATTATTTCATAATCATATCCCAATAATTTGGATA

ACCACTTCTGTTGCTCCATAGAAGATAGCTTTTGATCCAAAAAATACTTTAAGCTTCGATGATCCGTATAAATCTTGAAA

TGTCTTCCAAGTAAATAGGGACGCCATTTTTGAACAGCGGAAACAACAAACAACATCTCTTTATCATAGATAGAAAGATT

CAAATTCTTTCTAGATAAAGGTTTACTGTAATATGCAATTGGTCTATTAGATTGCATCAAAACATCACCTAATCCATTCC

CAGAAGCATCACATTCCAACACAAACTCTTTGGAGAAATATGGAAGTATTAAAACCGGTGTTGTAGTAAGAGCTTGTTGT

AGTTTTTTGAAGGTTGTATTAGCTTTGTCATTCCAAACAAAAGCATCCTTCTTGAGTAGCTGAGTTAATGGAGCACTTAT

TTTTCCAAAATCCTTAACAAACTTCCTATAATACCCAGCCAAACCCAAAAAACCGCGTAATGCCTTCACTGTGGTTGGAA

TCGGCCATGAGATAATGCACTGTATCTTATCACTCTCAACTGACACACCTTCAGAAGAAATCACATGTCCCAAATACCCA

ATTGAACTTTGAGCAAATGCACACTTTGAGTCTTTTACAAACAACTGATTAGTTCGAATAATATCAAACACTAAGGACAA

ATGCTTCAAATGATTAGCAAGATTCTTACTGTATATCAAAATATCATCAAAAAAACAAGCACAAACCTGCGTAAGTATGG

CCTGAAAATATGATGCATGAAACTCTGAAAGGTGGTCGGTGCATTGGAAAGACCAAAAGGAATGACAATAATTTCTTAAT

GACCATCGTGAGTTCGAAAAGCTGTCTTAGGAATGTCAGATGCGTGAACTCTGAGCTGGTGATAACCAAAACGAAAATCT

ATCTTAGAAAAAACCGTTGCACCATGTAATTCATCCAACAACTCATCCACCATGGGTATAGGAAATCTATCTTTGATAGT

CAATTTATTTAAGGCGCGATAATCCACACACATTCTCCACGATCCATCTTTTTTACGAACCATTAAAATTGGAGAAGAAT

AGGGACTATTGCTGGGACGTATGAAACCCGCTTGTTGCAATTCTGATACAATTTTTTCTATCTCATCCTTTTGAAAATGT

GGATATCGATATGGTCTCACATTAACTGGAGCAGAACCATGAACTAACGGAATATGATGATCATGCAAACGTTCAGGTGG

AAGAATATTTGGAATCTTGAAAATATCTGCATATTCAGAAATCAATTGCTGAATTGAAGCGGGCACTATTGAACTAGGAG

TAAGTGCATTGCTTTCCAAAGATAGTTGAAAAACAAAACCATAAGTTTCTTTGCCTAATAAACGATTCATGGGAGCACCA

TCCATAAGCATGACTGATGAAGTATTGTTGCCACGGAAAATGACATTTTGACCTCCAATGGTGAACTGCATCATTAAATT

TGTAAAATCCCAAGATATTTGACCCAAGGTGCGTAACCATTGAACACCCAGAATTGCATCACAACCACTGATTTCTAAAA

GATGAAAATCGACCAGAAAAGTAAAATCTTGTAACTTAACTAGTACGTGACACAATACCCATGAGTATGTAATTGACCTC

CATCGCTTACTGTAACAGACAAGGAAGAGTTGGATTCAATATTAAAACCACATTGTTTGGCAATAGTTGGATGAATAAAA

TTATGTGTAGCTCTAGAATTCACCAGAAGGGTAAGAGATTTTGCCTTAATATAGCCTTGAAGTCGCATTGTATTTGGAAA

AGAAGAAACCAATAAAGAATGTAAGGAAATTTTGAGTTCTTGTTGCTCTTCGACAATATTAGTATCAGGAGGCACAGGTT

GAATTTAGTCATCAAAATTTTGATCATTGTCATAACATTCCGGAGCGACATCTAAAATCAACAATCTCGGTTTTGCACAT

AAGTGACCAGGATTGTAATATTCATCGCAATTGAAACAAAGACCCTTCTCTCTCTTCTCTTTTTGTTCTTCCCAAGATAA

TCTTTTAGCGCCAGGAGGCAAAGTGATTTTTCGGGTTTGAGAGTTTATAGCTCCTGGTGATCTGAACTGAGTTGAACGAG

AAGACTTTGAAATAGAAGAAGCCGCCAGTATAGTATCTTCCTGTTGGATAGCCTTTTTGAAAACCTCAGGTAAAGATGTT

GGCTCAAACAAACTTACACTAGTGCCAATATCCACTCTTAATGCAACAATAAATAAATCAACTAGGTATTGCGTCGGAAG

ATCATGGACAAAATTTAATAATTTTTCGAAGTCAGTAATGAGCTCTCGAACAGTACCTGTTTGTGTTAATTTTGTCAAAG

CAATATGAGGATTTATAAACTTGTGATCAGAAAATATTTTACGTACTAAGGAACAAAAAGTTTCCCATGTTGGTGTAGTA

AGTGTTTGCTTCGTCCAACGATACCAAGAATTTGCATCGCCTTTAAGATGAGCAGCAGCAATTGGAATTTTGAAAGCATT

AATGACGCCGTGTAGGCTGAAATACTGATCTTCCTGGAAAATCCAACCTTCAGGATCAGTTCCCTCAAATGTTGGAAATT

TGAGGTTAATATCAGTTGTACGTATGGTAGTTTCAGTTCCACCATTACCACCATTACCATTATTAAGTGTATGATTACCA

GAAATAAAAACATCTTCAGGTTTCTTCGTACTAAAAGCATCAGCTATAGCCTCTTTCATGGAAGATGCGAGTGCATGGGC

TAATGATGTCGTCAAGGCCGTAGAAAATGATGTTGATAAAGCAGTGATATCATCTTTTCGATTGGCTCTCATAGTGCCTT

CATTAGCCTCAGCCTTATGAATCTCATCAAGACGATGTTGTGCTAAACCATCTATCAAGTTTTGAATTTGGTCCTTCAAT

CCAACTTCATTTGTTAACTCAACGATTGTCATACTTAAAAAAAATATTTTTGCCCAATTTGATGATATAATAACCTAGCT

CTAAACCAGATGTTAGGGTTCTCAACCAAAACAACTGTTAAGAACACAAACTAAACAGAACCCTAAAGATAAAAATCCTT

TTTACTGGGGATTAGCCATCTCTACTGTGAAGAACACAAATAAGAACTTGGTTTATATTAATTGATTACAGAACTAAAGA

GGAGTATTTATACTCATCAATACTTGGCTAACAAGGAAAGAATCTCACGAGTTTAATTAATAGAACTCTACCTAAAACAG

GAAACTACTCAATTTAGCTAAGCATGAATAATTAAGGAAACTAGTAAGTAAAGGAGTTGGCATCGAAACATATACTATAG

CTTAGGTATCGGACTAGGATAGAAAGTGTAGTTGAGCTCAAGAACTCCATGACAATCATTATACAAGACGAAGGACTGCT

CAAGGAACTGGTGGATCTTTATCGACTAAAAGGTATGTGGAGACTTGAACTTATCTATCACTCAAAAGTTTATTTATCTC

CTATCCTGAGACAAAAGTCGTATTACTATATAGACTAAGATATATTATGCACATTTGGTATTTCGAGCTGAGTATATCTC

GCCTATCTATTTCTCTAAATATGTGTTGGTAAAGCTTTCGCTTTAGCCAAGTTCATCTTTACCTAGTGACGAAAGTCATG

TTAAGTTTCAATCACTTTGAAAATTGCTCTGACGAAAAATGGTTTCTGAATAACAACTATATAACATCGTCTGAGAATGT

TTCAATGATTGAAATGAGAGTTTAGATAATATAACCAATGGAGGATATAATCATTGTTGTGGAAACACATATGTGTATAA

GTCCTTATTCCTTGAACCAAAGTTTGCAAACTTTGTTGATCAAGAGAACCGGAGAAGTGCGTGAGCTTCAGTCCGCGAAC

TGGCAAAGTTCTCAAACCCGAGAATTTCTGCTGGATTTTGTGAACTCCATCCGGGAACTTAATTCCGCGAAATTGAGTAG

GTTATGTCTAAAAACGATGTTTGTGAACTCATCTCTATAAACTAAGGAATGCTATTGCAAACCGTGGATATATATTTCAT

GAACCGATTCGAGTGAATCAAATCATTTTTGGTTCAATTGTATCTTGTGTAATACATAAAGATATAAGCAATTGAACAAG

TCATTTGAACTAGTTATGGTGAAGAAGAATATGGTTGATATGAAAGTTATCATATGGCTAACCATTTGGTTAACTATTGT

TGAACCAACTAATGTACAAGTTTGGGTACGGTCTTCACAAGCCTACAAATGTGCATTTCATGTGTGTATAACAAGCTATG

TTTTCGATCTAACGGTTGATAAATATTAGCTTAAATCTAATCATGTTTTCACCTAACGGTGAATATTGAATGCTTTGTTA

CTAAGATAACATTGATTGCAAACCCTGATTTGAAAGACTATATAAGGGTGAACTCTAGCAACTGGGAAACCTAATCCCCA

TACCTTACGTGTGATACTAGTTGCATAGCTAGAGTCGATTCTCCTTTAACCTTTGGTATCTTCTTCTAAACCAGGTTAAC

GACTTAAAAACTTCATTGGGATTGTGAAGCCAGACCAGTACTACTTTTATCGTAGTTGTGTGATCTGATCTTGTTGTTTC

TATTGTACGAGTACAATTATAATAATTGGCTTGAGATTTCCATCTGATCTTGGTGATTTTATTGTACGAGTAAAATTGTA

ATAATTGGCTTGAGATTTCCATCTCCGATACGCAAGATATAAAGTAGTCACAAACATCTTCGTCTCATCGTTTGTGATTC

CACAATATCTTTTTTCGCTGCGTCGACTAAGACTATTATGAGGTGATTGATAAAGCTGAGTTGTTCTTCGGGAATATAAG

TCTGGTTTTATTAATTGGTTCATGTTCACCTTGATTTATCAAAAGACGGAACAAAACTCGTAGGTATATTCGTGGGAGAT

GGATTTATCTATTACCGTAGACTTTTATGTGTGATACAAATTGGTTTATTAAAGTCTTCGACTTTGGGTCGTAGCAACTC

TTAGTTGTGGGTGAGATCATCTAAGGGAATCAAGTATGTAGTATCCTGCTGGGATGAGATACGTAGGAGCATAACTGTAC

CTTAGATCAGTGTGACATTGATTGGGGTTCAACTAAAGTCCATACCGAAGTTAGCTTGGAGTAGGCTAGTGTCTATAGAG

GCTTAATACAGTGTATGTTTAATCTGTACTAGGTCCCAGGTTTGCATTTGCGGTTTCCTCGTTAACAAAATTCTGATGTT

TGTGTTATTTCATTTTCCGCATTATATTGTTTATCTTTATAATTGAAATAATACAGGTTGTGCGTTGAAGATCAATCAAT

TAGAATATCCAACCTTGGGTTGTTGATTTACATTGATTGACACTTGGATATTGGTCTTTGGTACCATCCAAGTTATTCCT

TGTATTTGATTAAAGACTCGATGATTTATATAAGCTTGAGTAAATCAAAACAAGAGAGAGATAATAACTCCTTGAGATAT

TTCTATCTTGATTGAGTCTGACTATCTAGTTGATTCTCTAGAACGTAACTCGGAGTTAGTCCATACAGATTGCTAATTCG

AAATATTGGGTGTGGTTGTTAGACCCTCGCTTTTTCAATATTGACACTCTTGTGTGTCCTAGTTGCAAACTAAAGTCGAA

TCTCCTTTAACTTGGGTTTTCCAAAACCATTATTAGGTTAACGACTTGAAGACTTAATTTGGGATTCGTGAAGCCAGATC

CAACTATTTTATTTGTAGTTTCATATTCTGATCTAACTATTTCTAACATATTGAGTTTTATCTTCTCTAAGATTTACTCG

ATATTTATCTCCGATAAGTGAGATATAAAAAGTAATCACAACAGTTCTTCGTCTCAGACTCATGTGATTCTGCAATAACT

TGTTATGTTAATCAGTTAGGTTATTATGAGGTGATTAATATTTCTAGGCTACTCTTCGGGAGTATAAGACCGAATTATTA

GTTGAGTTTCCTGTTTAATTACCTTGATTTATCAAAATACGAAAACAAAAACAGAATAAACATATATGTAGGAGAAATAT

TTGTTTATACGTCTTCGACTTTGGGTCATAGCAACTCTTAGTTGTGGGAATCAAATCTTAATCGATGTGAGACTTGGTTA

GGATCAACTACATTCTAGTTCGAAGTTAACTTGTAGTAGGCTAGAGTCTGTAGCGGATTAATACAATGTAGTGTTCAAAT

CTGGACTAGGTCTCGGGGTTTTTCTGCATTTGAGGTTTCCTCGTTAACAAAATTTATAGTGTTTGTGTTATTTCTTTTCC

GTATTATATTTGTTTATATAATTAAAGTATCACCGGTTGTGCGTAGTTCAATCACAGATGATATATCTAACCTTGTTTGT

TGGATCGTAGTAGATTGAAACTCGAACATTGGTCCTTGGTATCATCCGAGTTATTCTCATATCAATCAGTCTCACAGATT

TCTTTCGAGCTGATTTACTGATTGTATTGAGAAAGAGATAAAGCTCTTTGATATATTTCTTTATTGAGTCTCACTTTTAA

GTTGGTGCTCTCAGAATTATATTGGAGATTAGTCTATACATATAGCCGAACGAAATATTAGGTGTGGTTGTACCCCGCGT

TTTCACAAGCACCTAGGGGTAAATCTTTTTTTTTCCTGGCCAAATTGGGGTATACCAATTTTAAACCCATCCAAGCTAGT

GTGTTAGGAGGTGTCTACAAGATGCTAAAAACTATCAAAACAATCTTCAAAATGTCTTTAAAAATCTACATGATGAAATC

AAAAAATAAAAATCAAAAAATAATAAATACGGGTTTTGAGATTGGGTACAAAATCATACCCAAAAGTCAGCAGGGATCTT

CTTTTTTTTCTGAACTTGCCGACTAGTATTAGTCGGCAGGGTATTTTTTTGTCGACCATGCCGACTAGTATTAGCCGGCA

GGGTAATGTTTTGTCGAACTTTCCGACTAGTATTAGTCGTTATCTTCTTAGGTTGAATATCGTAATGACTTTTCATCTGT

GAAATTAGCATTTACAGGGTCGTCAAGGTATTCATCCTTATACTTTGCTGACTATAGTTTAGTCGACAGGTTATGAAATT

AATGCCTTGCCGACTAAACATGCATTCTTAACACCTTTCTCTGTATCACTAGTACAAAAATGTCTTTTAGTCACGGTAAT

TCACCGTGTCCAAAGCTCTATTGTCTCGGTATTTCATTTCGGAAAAACAGTGACTAAAGCCTGCGAGGCCAAAAGTGCAC

CATTTGGTCACGGTACATTTTGGGGCTAATTCTTTTAATCACTGTGACCAAAATTTCAATAGTCACGGGTTTTTATGGGT

CGCCGTGACAAATTTGCTCATTAGTCTCGGAATGCTGAAGTACACCGTGACAAAATTTGTTCAATGGCCACGAATTCCAT

ACTAATACCGTGAACTTTGTGTGTTGGTCATGTATCCCATAATTCACGTTGTCCAACCCATCAAACATATAATACTTTTT

TGGGTACCGTTTTGAGCCACAGTAACAAATAATCTTAATACAATCACCAAACATTTTATTGAATCTGATAATATCGTTGA

AAATCAACATAGTTCACATTATCTTGATATATATATATATAAATTTTATTACAGTTGATAACAAGGACATCAATATAATT

AGTTAGTATCAGAATATATGTAAACACGCTCGCATAAAAGAATAGGAAGGGGAGCCTCCAACGAAGTTTCTGAAGGACAA

CCACTTATTGCGTTCCTACTCGAGGTCTTGTCTACTGAAGAATCCTGTGCAGTCACCCTGCAAGAGTTACTGCACATTCT

TGCACTACGATGTCGCTTAGCTGCAAACAAGAGAACAATAAAACAGAATGTGCAAGTCAAGTTACGATGCAAGATGTTAT

ACTGCACAAGTGCGTAGTAATTCATCAAACACTGAATATACTCTTTCATCCGGGAGAGCATATGCTAGTTTGTAAGTCAC

TGCTAGCAATCTTTATCATCATTAACCTTACAAGCAATTCATCAGATTACAGTACCATTTCGAATCTACCATGTATAAAT

CCATTGGCAAGTGGAAAAAATCATGTCAAATATGATGAAGAAGCTCAGTAAGAATGTTGAGATGCATAATGAACAAACCA

CTCTCTTCATATAAGTTCGGTTAAAAACATAAATCCAGCTTGAGTGGAAAATTTACCTTATCAACCTTGTTAATAACCTG

AAGCTTTTTGGACGGTACTTTATGCAAGGGAATGGTCGCAGACGACTGCACAGTGAATTTGCTTATGCTCCCTAGAAAGC

AAACTATAAAACTTTGCCAAGTGGAAACACCTGATGTTGTATATCAAAGCAACTTCAGTTGAAACCTGATGTTCGTTCCA

GGTGTCACCTAACTCACAATTCTTAGCATAATTAGTGAAGAGAAAATAAGACCCCTGTCTAAAACAGTACATTCCTTACA

TGTAGAGCTTTTAGAAAATGCGGCTTTTCCTGGAGTTCCGTTTCTCACAGAGGCTATAGGAGTCGGAAAGATCAGTTGTA

GTTTGCCAATAATAAACAGTGAGATACCCAAACCTTTGAATTAAACATGATAATATCATTACTGAAACACCATTTTACAG

CTAATTCTAAGGCTCCAAATTAACCCCCATTGCAGATTGACAAACCCTAAAGGCTAAAAGATAGATGAAAAGATGAAGAG

TAGATTATGGCTTCCAAACCTAAATCAATTGGAGGTAACATACAAACATGGAACAGATCAAACCAGTAACTTAACTAGAT

ACCAAGTGAACAAAAAGAAGGCAGAAGTTGAATCTTTTAAGTATATGCGCGCCTTCACATATCCAAAACCTGGCAGGAAA

AGTAAACATTAATTCATTCAGTTACCTTCCCAAGTCAAAAATGCACTTAGAAGAACATACAATAACTTGCTACACTCATC

ATTAGTTTTTAACGGTGTAATCTCCATTTTTTTCTGACTAATGGGAAATTACAAGCTCGTCAAATATTTGAAGTCAGTCT

TATAGCTTTAGGAATAAAAGGATCTAAAACACTATGTTTGAAGGAATATAATCGCCCCGATCTTAAAGAAGTTGTATCCA

ACAGCCCAAAAAATTGACAAAAAGGAAAAACTCAAGATCCTTTAATTAAATCATACACACCCAATACTGTTAAATTTACA

GCAGCAGAACGTGCAAAAGAGAATCAATTTAGTAGCTTATAACCAGCATAGTCTCCCTAACTTAATTTCGTGACATTTAG

TATTCAATTTCAAGAACTGTAAAATCAAAATACCAAAGAGATCTCAAATGAACATAAAGGAACATACCCTTAGTAAAGGA

GGAGATTGGCCCAGGAAATCCACTTAGTGCATGAATTTCCTAGCTTACTTTTATGTCCCAATAACTGGGCAAATAAATGT

CACAAATTACTAAACTTCTCAACTTTCTTATATATTGTATCTTCCAACGGATATAATCAATGTACATAAAAACAAGGAAT

TGTGATAATTAATGTAACAAGTTACCTGAGGTGGCACAAACACAATCTACACAGACCACATTTAGTAATCCTTCTGTACT

GATGATAATAATCAATTTATAGGAATGGATTTTAGTTGCATCTCCCATGATATTACACTGATTGAACACATATTTGATCT

TTAACAATAGATTGAATCCATTGATTGGAAGGCCATCATTGTCCTAATAAAGATGCAAGATTCACTATCAACTTCAGAGA

TCACTTATCTGAACCAAGCCGGCACCATTATGGCTAAATGGTGGATAGATTTCGCTACATTAAAACGTATTCATATCAGG

CAAACATAATAATAGCAATTGTAAATCTGGAACATAAAAAGAATGGATTTTTGCTCTTAACTGTTTCTCTATCATGGTAA

TAACGACAATGGAACATCTCATTACAGCAAGGACAAAGGGCCTCGGATTTCACATCAGCATTCTTACATACTATACTAAA

AATACACATATCAATCCCTGATAATTCTGAAAACAACCAAATCGATAAATAAATCATCATCTTCCACATAATTGGGTGGG

TGAAACAGAAATTGTGGTTCTCTTCAAAAATTGAGGAAAAAACTAACTTACCTTTCACATAAAGAGAAAAGGCACGGATC

TCCCACACTCTAGCAATGCTTATTTTAGGGTACCTAATTTCAAATTCAAACACAATCAAAATTGTAAAAAATAATGTGCC

ATGATTTCAAATAAGAAAACCCTAACATACAAAGAATTCACTAATTAACACAAATTATTGCCTCTATTTCAGTCCAAAAA

TTAACGCCTTCATAATCTTACCAGAAAAAAACCAAAGTTTAATGGAACATCAGTTTCACCGTGAAATAGATAAACGCGTT

GATTACTGCGTAAACCTCTAATTGTAATAATAAAAATTCCCGAACTATCTGATCTAGAGACTAGAGAGACAAAACATTGA

CTAACTGATTCGAAACTTCCGATTGATTACGCATAAGAACATGAAATCAGATCGAACAGTCTTGATATAAGAGGGTTTTT

ATTTTGGATTGAATAAAAGAGAGACGAAGAAGCAGATAGGAATGATTACCCTAATACTTGAGGAGGTGGGCGGAGGTGTA

GGAGATCGTGGGAGACGGCAGTGTAGAAGTTAAGTTAAGAAAGAAGTAAGAACCAAAGCAGTTCTAACTTTTGATCGTGT

TTAAAATTGTGGATTGATTTACAGTTAATGTACATTAGACACGCGTCTAATGCACCCGTAGCCAAAGAGATGTGAAATGG

TCCCGGAAATACATTTACCCGGGACCAACAAAGCTTTAGTCCCGTTTATAGATACAGCCGTGACAGGGTCAGCCGTGACC

AAAGACCATAATTGTACTAGTGTATGTCATAATTCCTATAATCGGCATGGTATTTTTTTGTCGACCTTTCATACTTTCAG

AATTGAAAGTGTTGATTTCTTCTGTAATTTTGAGTATAAAAGCACTCAGTAGCTTTGCAACCCCTTTGTAGAAATATTTG

GCTAGTGTGGTTTTATTTCCGTTACAAAACAAAATCTCATCAAAAAAGTCCTCAAAAATCATAACACATAATCCATGAGT

TCAATTCAAATAATACCTAATTTCATAATTTTAATTCAACTAATTAACTAACTAAATTAATTAACCAAAACACATTATTA

GTGTTAATTAAGCAAGGGTAGGCATTTTGAAAATACATGGTTAAGGGTTGTCCTATTTTACTACAAAATGACCCGGATTT

TTTTGTCTCATTAGGTATACCCCAATTAATCTGGGTATACGCTAATCTCGCCAAGTTTTTTTTTGCTGAGAACCTAGGTG

TAAATCAATAATGACATTTGAGCGATCTCTTGTGTTAGATGAAATGCCATCTGTTTTATGGTTTATGTAACCCATAGTTT

ACTTTATTATTCGATACTCCCTATATTTCAAAAAGACCGTCCTCTATTCCATTTTCGTCATTTTCAAAATTATGTCCAGC

TTCTGTTTATAATCGTATTTTTCCAATGTACTCTCTAATATAACCTTAATTTTTTTGTATTTATAAAAGCTCAATCATGT

GAGACTTAGATCACTAAGAGAAAGCTCTATGGTGCATAAGATATCTGAATAATCAGTATCAGAAAGCAAAATATAAGTTG

CATTACTACAATAATGATTATACAAAGTGATATAACACGTAATCATTGTATGCTACCTTGCAAGAGTGTTTTGAAGCTGT

GAAATTATGGGATTTGAGTATTGAGATCCCATAAATGCACCGAACCATCACCATTGTGTTCCATCAGGGATGGACACAAT

GATAATACCAACACCATTATGATATGCAGGTCAAGAGTACATGAAAAACAAACGTAATCAACCTCCACAAAATTACTTCC

TCGTCGAGAATCAAAATCACAACAGGAATAAATAATGTAAGATGACACCTCAACGGCTGATTGGAAACATGAATTAACTG

CATAGAGACCCAATCATGTTGTCTCCCTCAATTAAGAGATAGGAACAAGGCAATTTTGTCTGGGTTGTACACAACCTTCA

AGTAAGATGCCACCTCCTTAACAATTTTATTGTATCTAGCCTTCGAGTACTTGGGGGTGGTGGCATCCAGAAAGTTATTA

CAAAATAAAAGTACTATTAGTTGGTATCATATAAAAATGTAGATAAAGATATTTACAAACTAGCTTATATAAACAAGATG

GTATTTGAAACTAGTTATACCTTGTTACAGCAGTAGATCATCTGCTTAACACAAAATGTGAAAGTAAGAAGGGCGTGCTC

AGGAGTCTGGCCATCCTTGGAAATACCAGCCTCAAAACCTCCTATGGTAGAGTCAATAAGGACGACACATTTGGCCTGAG

AGGTACTAGTGATCATGTTCTTAATAAAGTCACGATGTCCAGGAGAAGTACTACTTGGTGGTCTCAAACTTCCACAAGGA

AATATCAATGGTAATACCTGATAGGTGTCTAAAACACCTATATATTTATCTATTGTATATGCTTTCTTTGCTTAGTTTTC

TTGTAAAATATGTATCTTATGTTTTTCTTTTGAATATTTAGGTACTTTGGAGTCACTTGGAATAAAAGGAGAAGAAACGG

CGCAAAAATCTCATCTCATGTGCAAATGCTTTTGGAGGCAGATTATTTCTCACAATTTACTTTTATTTCTTCATCTCTGT

CTGAAAAGTATGTCGGCTTTAGTGATGCAAATTTTCTGGTTCAAAAGCACGACAACTTTAGTGATGACTCTCATTGAAGA

GAAGAAGTAGATTTGAGAAGTAAAAGACGGTTACTATTGACGGAAAGAATTGAAGGGAGCAAAAGCGCCAAAATGGATTG

CATCCGTGCACGTGGCTAAAAGCTCATTGGAGGCCAGGGACACTGTAAGTGAAGCTGGAGCTAGAAGAGAATTTAATTCT

CTTTGTTGAGGTCTGAGAAATAAACGGACATGGACTAGAAATGGAGCTTGGGTGCTGGTAGTATGAGCCGTAGGTGTTGA

GCTGGCTTCTGAGACAAATTATATGATGATGGCTTAACATTTGCTTGGACTATAATTGTCCAGTGGGCTTCATCTCAGGG

TCGTACACATTTGTACCATAAGAGAAAACAATTAGGTTTTCATAGAAACTCAATTTCAATAGTAGAGGAGCCTAGTGTTA

ATCATTGAGCTGCTACTAATTTTATTTCCGAAACCAGTTTGAGAACCAAGTTCTTGCCACCGGTTACCACAGGTTCGATG

GGTGCCTGCAAAGAAGAAGATCGAGGCCAAGGGTTCATACAAGAAGGAAATGGGCAGATACAGATTTGGTCGATCTGTTT

TGGAACCAAGAATACAGGGAAGAAACTTAGCTGCTATTGTTGATTTTTGATGATAAACGGAGGAGATTAAGAAGCTTATC

AGGGTACATGAAATGTCAGAAAGAATGGGTCAACATCAAAATCAGATTGCAATCAGACATGGACAGGAATTTGGCCGTGA

AATTCAGATGAAGATATGGTTTTGTCGCCGTACTATTGATGCTGTTGTTGTTTAACTCACGAATGGACGAGATTTCCAAT

CGATTTTGGAGTCTTGAATGAATGTGGGTTGTGTTTGAATGCGAATGTGCAGGGAAGTAGTTGTGGGATGGTCTTCGAAT

CAAAAACAGTTTATCTCTGAAGCACCAGTAGACTGCACATTATATGTTCGAGAAAATGTCTGACTGAGTAGTTCTTCACA

CTTCAGATTCACCTGCGACCAGGTTGCTGTACCAGGTTTGCCCTGGACTTTTGTCATTCTTTGTGAATGGTTAGTACAGG

ATTTTGAGTTTTAGGACAACACAGATTGGGTACAAATTGTGATCGGTTTTGGGTAAGTTGTTTGAGCTCAGATTCAGCTG

AAGTATTGAGTTGCTTCCATGGTTCAAGGCTTATATGCAAGGGTTTTGTGGGGATTAAATGGGACATATTTGGAATGGTT

GAGACACTGATTTGTGCTGTGACTTTTGTGGCTTGTACCAAGGGTTAGAAGGAGTTCAGTATTGGCCGAGAAGTCAGGTT

ATAGGAGATAGTTATGAGCCAGAAAAACAGGGAAGATCCAAAGCTTTCTGCCGTTGGTTTACGGAGCAATGGAAGCTGCT

GAGTTGAGTTGCTAAGTAAATTGCAGGGAGCATGAGCTGGGGGCGCTAGGAAGAAGTGCAGCTGTTGGATTACAGAGCTA

CAGGGGAGTGTTGATGTTGCAGCTGCTCAACTATAATATCTCAATTGATGTGCTGCTGGATTGGTGCTGTAGTGAGGCCA

GAGAACCAGTGCTGTTGATTGCTGCGAACAAGTAGCTAGTGCTAGTGATATTGGAATGCAGCTGGTGGTGAACTGAAGCT

GCTATAGATGGTTGTGTTGATGGATCCATGTGAGGAGGAATGAAGAGTTTAGCAGCAATGCCTGGTGGTAATGGAATTAC

TGTGCATTTGGAGAGCTGGCAGGTGAACTTGAGAAGGTATTGGATTGCTTGGCTAAAAAATGAATCAGATGCAGTGGAAC

TGGTGACTGAATGTAACTGGTTAGCTGTGACTGACGGTTAACTGGAGACGTAAAGAAGAGTTATTGATGTCGTAGTCTCC

TATTGTTGTGAAGCGAGTTTGCCGCCGATCAAGCCATGAGTTCACCAGATGATGCTAAGCAACACAATGGCAGAGAAGAT

GGGTTTGCAAAGTGGTTGAAATTGCAGGCAAAGTTGGAATAGATGGTTGTTGCAGGATAGTGTGCAGCTGCAGTCGCAAT

AGGAATGAACTGAGTATCGCAGGCAAGGAAGGAACTGAAGCAAGTGGAGAAGTGTAGCTCAGATGGGGTTGTACAAAGGA

TATGTTGCAGATGGTGCTTCTCAGATGGAGTAGCAGAACTATGAACTGAAATGAGAAGCTGTTGAACTGAAGTACAGGTG

TAATTCAAGGGAAGAAGCAAGGATTAAGTAAAGCAGCTGGAGTTGTGTTTGTGAAGCTACAGGCAACATGAATTTAAGGT

TACTTACTCATCAACCAGGACTCAGTGATTAACTGAGTTGGTTGACCGGCCGGCGACTTTTCCGGCGCAATTTCGGCCAA

GTTCCGGCAACTTTCTGGGCTCTCCGGTGACTACTTTCTGAGCATATTTTCTATGTGTGTTTTCTCCTATGTGGGCTTGG

TGGATATACTGGACTTGGACTTTTGGGGGTATGACCTAGTTTGAAGAGAAAAGGCTACAAAAGGGAAAGTTTTCTACAAC

TTTAATATCACGTATTTTCTATTCTACTGGGAGTTTTGCAGAGAGCTGGAACTAGGGTTTGCTTCCTTGTTATTTTCATC

TTTATTATTTCATTTATTATATTTGATGATGAACTCCAATATTATGAGTATCTAAAACTCTCTTTGATTAAGGGCGATTT

CGAAGCCCGAAATATGAATCTTCAATTGATATGAAATAACTTGTTCTCTCTTGCATTCATTTATGTTATTGATTGATTTT

CTACCATGCTATTATTGTCTATGGATTATTTGTTAAGGGTTTAATTTCGAAATTGAATCATTAATTCTTAATCTATTTGC

TAGTTTAGATTTTCTTCGACCCGTAATTGTCTAGAGAAGATAAACACAAGTAGCAATATGTGGGTACTGATGATGGGACT

TTGTTAAGTATCCATATGTAGAAATCAACACTAGTAGGTGATTCATGCTTTTGAGTTCATCACTAGGAGCAACCTTATCT

CACGAACTTTAGAGCATTTGTTTAAGTCAAACCTAGTAAGCGTACCTCTAGGAAACTTGGCAAGTAGGAACCGGTAGTCG

AGCGCTTCCGTATCCGGTGAGCTAGGAGATAATAACGGAATAGTTGAGCATACTAATTATTCTAGGGTTGTTAGTAACAT

GATAATGCAACGAACTTGGCTATATATCAATTTGGTTTGAATTTCGTGGTCTAGAATGAACCTTTAATCATTTCATCTCA

TATTTGAATACATTTTAAAATTTTCAGATTTTCTACAACACAATCATCTTTTGTAAGAGTATTTGCATAATAACTTAGAC

TTCATTCCTCCCTTGGGACGATCCTGCTTACATTATACTACTTGATAGTGTAAAAGTAGAGAAATATTATTTTTGCGAGA

TTCGACACCTCGTCAATACCTCGCTCATGTTCATCCCTAAGTTTGTCAAGAACCCATGTGTACTTGAATGACCTCTTGTT

CATCTCAGCAGCCTCTTTCTAGAACCTCTCGCTCACACGCTTGTCAATACCTCCAAGATTGTAGATCAAGTGACCAGTGA

TGGTCAATTTCTAGGGTCAATACGTCCAGTGAAAATAATGTTGATGTGAACCTTCCACTTGACCATATTGAGTTATTCAA

GAAACCTCAAGCATCCCATCAGAAGAACCATATAATAAGTGACATCTCTAAGAATCAAAAACAACATAAAATTTCTAAGA

AAGAAAAACTGAAACAACAAGCAATTATAACTCATAAAGGTGTCAATGATTTATATTGGTGAATGGTTCCACATAACATC

ACAGGTTTTTCTTAATATCATGGTAAGTTGGAAGTAAATAAAGAAAACTGAACAGACAAAAAAACTTAGTACCAACAAGA

TGCAGAACAGTTATAGTGTGTGACATTTTTACAAAAACCGCATGATCTCGGTGGCATGCTAAATAATTTCAGTAAGTATG

ATTACTATCCACATTGAACCAACTGAACCAAGCACGAATTAATAATATTATGAAACCAGACCAGCACTGACGATCAAATT

AGGCATAATCCATAGATGCAAATATAAAGAAAACATATAAATCTATGAGAACTGATTAACGAAATAGAATAAATCAACCA

CCAATATGGTATCACTGATTTATAGGATATGTTCTTCATCTCATCTCTTCTCTTCTTTTCTTTTCTGATCCAATTTCTTG

AAATCCGTTGGGATAATTGTGTTTCTAGGACTCCATCTCTTTACAGCATACCTTGAATCCCATCTCCTCCTGTAATAAAA

TCAAATTCAATTTGAATTTCAAAACTCTAATCTTATCTTGACATATTTATTGAAATTTCAATTACACAATCAACATATAT

CAAGTAAACAAACAAACCCTTTTCTACTAACCCTAGTTTCGATGACGACCATTAGGTTTTGGGATTGGGGGATCTTCGGA

CAGAAAAAGATTTCTATGGTTTCGAAAGAAAAAAGGGAAAACAATTTGGGGGTTTATGACGTTGATGATAACGATGGTGC

TGGTAGTAGAATGGGGGGAAGAGAACGAGAGACGATCAGAGAATTAGAAACAAGAACGTGAGAGGAAGGGAGAAGAGAAT

GAAAGGTAGAGGAAGAGAAGATAAGAAAAAAAAAGGTGTTCTCGATACAGATAAAGCACGAGTCAGTTTTGAAGCTGACT

CCGAGCATCTGAATAGATAGAAACCGAGTCATATTGAATGTGTGGGAAACACACACACGAGTCGCACGCGAATCACAACA

TAAGAAAACAGTTGCGAGTTTCGTAACTGTTCCAGGATGTTGCGAATTTTTCACAACTGAAATTAGTAGCAGACAATTCG

CAACATCTTTTAAGTTGTTGCGAATCTGAGTTGCTAAACCCCATATTTGGTGTAGAGAACACCATTTTTTTTTTTCAATT

TTCATGATAATCAAAATCTAAAACATAATCACATAACAACTCAATTCTCAATTGTTTTAGGTTTTCAAACAATATTTCTC

GGCGATAAACAATTCGAGTGATTATGTTCAAATCCTTTCAAGACAAACCTTTTAAGAGGGATTCATCAAAGATTTAACTA

TGAATCTCTGAAATTTCTCATCGAAATTTTCTTAAACTCTACTAGAATTGGTTGATGTAGACATGTAACATAAACCAATT

GTTATTGATGTTCCCCGAAATCGGTATCTGTTTTACACCCACATAAACCGATTCTCAGTTGATGTTCTTCGTGTTTGATG

AACATCAACCAAAATCGTTTTATGTTAATATTTACATCAACCGATTCATTAACCATGTTTGTATTTTTTTCACAAAACCT

GAAATTTCTATGTTTATATTATAGAGCATGAAAATACGAATTGAACGCAAAAATAATGAGGTCATTCCGACATCGGAAGT

TACGAGCAAAATTGTCGAAGGAATGCCCAGATTTACAATCGGTTTATGTGGGCATTAACATAAACCGATTCTAGCGAAAA

AAATGTTCAGAAAATCATAAACTGGGACGGTCTGAGGCTTATTTATTTTTGCATTTAAAAAAAAAAAAGGCTCTTAGAAT

CTGTTTATGAGAGAGCGCATGAACATCCACCGATTTTAGGGTGATTTTTCTCAAAAAAAAAAAATTCATATTTTAATATT

GATTTTAAGATTAATTAAATTCTAATTTAATTTGATTAAACATCAAATAATCACCAAATTAATTTGGAATAAATTAGTCA

TTATATAAAAAGTTGGATAAGAAGTCTTCCACTTTACTTCTAAATGTCCCATTTTTATCTCATAGTGGTAAAAGGCCCCA

AATAAAATGGGCCCAAACAAGGAAGGTTTGGATTCTATATTTGTTTTGGGTTATGAATCGATTAACCAACTCCATCCCAC

GTGCATCAACGAGTAAGCATCCAACCAACTCCATCCCACGTGCATCACGAGTAACCATCCCACTTCCTATATCTAAAGTG

TAAATCCTATTCCTAAGATTCCCAACACACTCAAAATACACAAACACGGGCTAGAGACTAAGATTTCCAGAGAAGATCAT

GGATATCAAATTAGAAGATGAAGAACAAGAAATAAAATACCAAGGTCATGTTTGGAACATGATCTGCGGTAGTGTTAATT

CATCCGTACTTAAGTGTACGATTGTTTTAGGTCTTTTCGATACAATTCATAATTCATCCAAACCCATGACCCTATCCTCC

CTATCATCAGCATCACCTTCTTTATCTCCTCTCAAAACCGAAAACTTATATAGACTATTAAGATACTTATCTCATATGAA

TCTCATCGCCATCAATTCAGTTCAAGGAAGTGATGATACCTTTTCATTAACAGATACATCTAAGTTACTACTTAGTAATC

AAGAAAGAAGTTTGAAGGATTGGGTTCTCGGTATCGACGATCCACTCGCAATTAACGGGTGGCATGAGTTAACTGATTAT

TGTTTATCCCCCGCTGATGCGCCGGTTCCTTTTGTAAAGATGCACGGGAAAACCATGTGGGAGTTTGCTGCAGAAATTCC

CGAAGTTAATGAGCTATTCAATAACTCCATGGCTTGTGATACTAAACTAGTAATGCCTGCATTTTTGCAAGGTTGCCATA

GTATCTTGAATAAAGGTACTGTAAAAAAGATGGTTGACATCGGTGGTGGTACAGGTGCTGCTATGAGTTATGTTGTTAAG

GCTTTTCCCGACATAAAATGTATGGTTTTTGATTTGCCTCATGTTGTTGCTAAAGCACCGGAGATTCAAGGTGTGGAATT

GGTTGGTGGTGATATGTTTGAATTTATCCCTCCTGCTGATGCTCTTTTGTTGAAGGTACGTACAAATTTTATGCACACTC

CATCAATAAGTTTACATTGGTTAGGTACTTTGATGGGTGGTTTTGTTGATGGTTAATTTTTTTCCTCCGCGTGTGTGTTT

TAATTAATAGTTCATTCTCCATAACTGGGATGATGTCGGATGCATAAAGCTACTAAAGAAGTGTAAGGAGACAATTTCAG

CTGATCATGGGAAAGTTATCATAATAGAAATGGTTTTGGAAGAAGACGACCATGACAATGATGCTTTGACACAAGCAAGA

TTGAGTCTTGATTTAGATATGATGGTATGTTCAGGAGGAAGAGAAAGGACTAAAGATGAATGGACAACTTTAGTGGAAGA

GGCTGGTTTCAGCAAGATTGAATTCTTTCCTATTTTTGCCATTCAATCAGTTATTGTTGCATATCCGTAAAACGATTTAA

CCTAAAAACTCTTCAACTAGCCTCCACCACTTTGCTCCGACCTACTTCAATTGACTTTTGGACTAGATATCAGCAGGAAT

TCCACTGTATTACATGGTTGCAAGTTTTTTAGCTTAGGTAAATGGGCATGCGTGGGATATTCATCACGTATGTGTACCTG

TCTCTTCCAGCCAAGTGTGTCAATAACGGTTTCGTATGTATAACGTCTGTTACTGCGACTGTCTGTTCGCTTTGTTTCAG

TTGTTGTTTTTAATCATGCTAGCTATATATGCCTCCTATCAATTAGCTGTGATTATGGTGTCTTGTAATTTTTTCCCCCG

CTTTTTAAAAAGAACGCGCAATTGGGGATATGGGTTTTAATTGAGGGATATTGCTACAAGACAAAATAGGGATTCATATA

TCAATTGGGGGTCACCCCTTATGTAATTATATTTTAAATTCCTAATCTAGCCCCCACTAATCAGGATTAGTGATTAATTA

TAATTAGTAAAATGTTAAGTTTTTGGGGGATAAATTGATGTGTATTTATTTTTATTTGAATTTGTGTGGGTTAGTTGAGA

GTGAAGGAGTGAAAAATATTTTTGAGTGGATTTTTTTTTTGAAAATGGAAGATGATTGTGATGAGAGACTTGCAGCTGCT

GAATCATTAGCTCAACTACAACAATAATCATATCTTCAATCTTCGGTTAATGATAACAACTCACAATTGGAGTTATATGA

TGAACCAACACCCTTGGATCATGATTTTCGTGAGCATGAAGTGTTTTGTGAAGAACCAACCCAAGAAAGTAAACATGTTC

AGCATACAAGCAACCCCAACACACAGGTAATGCTTTAAAGAGGTCGAAAATTTGATTTTTCTTTTTCAAACCACCATTTT

TTTCTCTGAAAAAGCTTAGTGCCGGCAAACCAAAAATCCATGCCGGTAACCATTTACACTTTTTACCTGCCACCTAATAT

TCCAAGGAATATCCATACCGACATGCTTCACTCTTAACTAATTATGTAGGTACCACCATAAAAAACATACAGGAGTTTTG

GATTGTCGAACTTAAATACCGGCTTTCTATTTAATTTTTCGTGCACGCTGGCACTTGGTACCAGCGTGGTTTATATATTA

TGGGACAAGCCGGCAGTCAGTACCGGATTGAAACCTTACATATATACCATGCCGGTTTTCCAAGAAAAACATACATGAGT

TAATGATATTTTGAACTTCAGTACCGGCGTCGAATGTAAATTATCGTGCACGCCGGCACTCGATCCCGGCGCGAGTTTTA

ACTTTTCGACAATGCCGGTATGAGCAATCCCGACATTGCAGTTCAATAACATACAATGCCGGAACGTTATTATGTGGAAC

TAATTCAATTTTCTTTTCATTCATGCCTTAGATTGTGACATACATCGATCCAAATTCCAAACCTCTCTTTCGGGATACGT

CCGAATACTATAAAATTCCTCGAGTATGTTACCAAACAATACTTTCACCTAGTTTTTACAATTTTATGTTGGTATTGCTT

GTAATTAACCTTATAATCTCCCATTGTTGTCCTTATATAGGGAATGTAATCTCTCCAGAGGAAGCAATTGCATGGGCAAA

AGAGACGGCTCTTAAGAACATGTGTGTCTTAGTGAGGAATAGCCAGCGTTCAGATTCGTTTTGAGATGGTTTGTGAGAGA

AACGGAAAATACAAGGAGAAGAATAGCCATAATAGAAAGGGTTATGTATATGAAAAGAAGACTAATAGGGTATACAAGAC

TCATACGAGGAAGGATGGCTGCCCCTTTAAGCTTGTATTCTATCTAAGAGACAAGAAGAAGAAGAAGAAGAAGAAGAAGA

AGAAGAAGAAGAAGAAGAAGAAGAAGAAGAAGAAGAAGAATGGGATTGTACGGTGTGGGTTGGGCGTCATAACCATCGGG

ATCCAAAAGATTTTGTTGGTCACTCCCTAGTTACGAAGCTAAAACCTCATCAATTCGAGGATGTAAAGATAATGACCAAA

GCATGTATCAAACCAAGAGTGATTCTCAGAGGATTCAAGGAAAAGGATGAGACGAACGTTTCTTCACTAAGTACAATTTA

TAGCGCACAAGCAACCATTAGAAGGTTGGAATGGGAAGGGAGGACCGTTATGCAAGAATTTGAGAAGAGATTTTGGGATC

ACAACTACACGCGTATCATTAAAAGAGGACCGGACAACAAACCCCTTTAAATATTCATTGTACATCCTTTGCTTTCATAA

TTGGCTCATACGTGTTGTGATGTTCTTATGATGGACTACACCTACAAGACAAACAAATACAACATGTCGTTGTTCAACAT

CGTGGGGCAAACATCGAACAAGGTAACATTCACATTGGCTTGGTGTTTAATAGAAAACGAGAGGGATTACAATTATCATT

GGGCATTGCTACAATTGAAATTATTGTTCCCGAAAAATCAATTTCCGAGGGTCATCATAACTGATCAATTTGATGCACTA

ATGAATGCCATATCCGACGTATATCCGGATGCACAAAATTTCCTATGTACATATCATATAGGTGAAAATGTGAAAAAACA

TTGTCATGCTTTGTTTGAACCCTCTAAGGCGGATATTAAGAAGAGAATGGTTGCTCAACTTGAGGAAGATGTTCGCAAAG

AGAAACTATCACCCGAAGAAGAAGAAAATGAAAGAGGAAAGATTAAAGAGCGAGTCGACAAAGAACACGATGAAAATCAT

AAAAGGTGGTTGGAATTTCTAAAGGATTGGGAGAAAGTTTATTGGTCTCTTACCGAGGACATGTATGAAAAGAGATTGAA

CATGTTTATTGCCAAATATAACGAAGTGTATCCAAGTGCCGTCTCATATTGTCGGGATAATTGGTTGGATAAGTTCAAGG

AAAAATTTGTTTGTGCGTGGAAAAATGGTTATATACATTACGGGAATGAAGCAACTAGTATAGCGGGGTCCTCTCGTGGG

AGATTTAAAGATCTCATCCAGTCCATCCAAGGAAATGTGGTTACGGTTACCGAGGCAATGGAGCAACACTTTAAGGCTGA

TATCGATAGGATAAAGAAGGCTTTTGATAAAAGCTCAATGTAAAGGATGAATCAATTTCTTTAATATACTAAGTTGCTCC

AAGGAATAGAGTTCAACGTTTCCCATTGGGCAATAAAACATATGATGAAACAAATGTAATATGAGAAAAATTATGGAAAA

CCGGGTGATGTGTGTATTTGTCCCAACATGTCTTCATTGGGGCTTCCGTGTCGTCATATGCTTGTGAAGTATGAATAAGT

AATACCTATTGATGTTATTGATCCGTTTTGGAAGCAACTATCTTTCGACCTTCCCCCTACGGAAGCTCCCTGGAAATCAC

CATGGGATACAAATGAGGCCAAGGAATTCTATGAAGCTTACTCACGTGGCAACTCGGCTAGCCGACAAGTATTGTTGAGC

CAACTAAGGCTAATTACACGCCCATGGATAGCACCAAGTGAAGAGCCAACAAAGGGAGATCCGCCCGGTAGACCAAATAG

TACTTTCGGTTTCAGAGCTTCACCCCAAATTTTACCACGTTCAAGGTCGGGTATCATGTCATCAATGTGAGGAAGGTGAC

AACCTTTAACTTTGGGTTTCCCCTTGCCCTTGCCCTTGCCTTGTATTGGTTGTTCACTCGGCCCACCAACTTCGCTTTAG

CTTGTTAATTGACTTGGGTGAACCTCATCATCTTCCTCCTCACTTTCCCCTTCATCATGCTCATCGTCCTCAACTGTTCC

AGTACTTTCACGGATTACAAGTGCACTAAGGTCACCACCACTCTCCCAAATTTCCCCTCTTTTTTCAAACCCCAAATGCT

TCTTGCGAGCTTCATCTTTCCCTCGTGGAGGCAAAGACTGAGGAGTATCAAGACCGTCCTTCCTCTTTCTCTTATTGACA

ACATCTTTATTTTTTATTTTGTTGGATTCCTTTGCTTTTGACCTATAACATAAGCACACGAAACAAAAATAAGACAAGTC

ATATTGGAAGTTTAAGTCAAAGTCTTAATAAATTATGATTTAACTTTAGTTCATTGAGTCTTGTTAATGGGTTAATTTGG

ATAATCTATTTAACCTAGCTAAAAAACTAAAAAAAAATAATACCCTTTCTCAGATGATCAAAGAGAATCCTCAATTATGA

AATCCCGAATCTATGATAACAACTACCTGAGATATCAATTGAAGACATCAAATTTCGAAGATTTCCAAGTCGATCAACAT

TAAGCAGATAAAAAGCACCATTGTTTCTCTATTTTGTTCTCGTTCTAGAAGCAAACATAAATTGCACGAGACTATCATCA

ACACACATACTGTTTCTGCAGTAACATACATATCCAAGAAAATGCTCCATTAATGTTACAAATAAGAAGACTGGGTATCT

CACACTAGAATTGCAAAATGGATTGTGCATAAGGTAAAGTGGGAATTGTTTTCGAGAGCTGGGGGGTTTTCATGAATTTC

AACATTAGCCACCTTCCTAGAATTGTACATCGTGTATTGTTGTTTTATATCTGTTGATTTGGCCTTATAGTTACAGGTAG

GTATACGAAATAAAATTTGAACATGTAAAACGAAAACTCAATTCATTCCTATCTGATTCATGCAAAAAAACTAAACAATT

TTATTCCTTTCCAATTTCTACATTAACCAATTTATCAAACTAACCAAAATCTTTGGATCTTATTAGATAATATTTTGGTC

TCGAGAAATCAATTTTCTGATTAACCAGACATAAGCTTCATCAAAAAATAGAATCAAAGATGAAAAGAGATTTGAAACTT

ACTTTAATTTGATATCTTCTTCTTCTTCGATCTCCTCTTAGATTTTTCTAGCAAAATAAAGAGACAAAGGTTTGAAGAAG

TGAATTCAGAAACCATTTTCATGCAATCAAACTAATCAAAATCACTCCTTCCTAATTTCCCATTTATAAAATCAAAATCT

TCGCATATACAATTTTGGTCTCCAAAATCAATTTTTTAGGGACCAGATAGAAACTTGAGAGCAAAACTCAAATACAAAGA

TGAAAAGAATTTGAAACTTACTCGTCTTCTTCTTCTTCCTCTTTTGAGTTAGAGTCTCCTCTAGGGTAGTTAGGTTCCCG

GAGCGAAAATAACAGACGGCGGGGTGAAGGTGAAAAATGAACTCAGCTCTTTTGGTTTTGGGGTTCGTTAGATCTGAACC

AATATGGCTCTATATAAAAGATGTATGGCAAGTACGCTGGTGGCTAACCACGCTAACCATTTCAGATGCATTTTGCTAAG

CCGGTGGCTAGCCATTTTTGAGCACATAATAAAAGAGTTATGGTGGGCATATAAAGACTAAAATATATGATGGATTTATT

TTGTAATTTAATGTTCTATCATTAAATCACACACCTACTAAATTACCATGTTTTTAAATGACAGTTTTTTTACCTTAAAT

GACGGTATTTTTTTTGTAATACCTTAATCACGTAATTAATTTGTCATTTAAAAACGTCATTTGTAGCCCTTTCTGGACTA

GTGCTCCATCATCAGGAAAGCTCATCCAGTACCACACTAATTCGTCCGCAAAGAAACCTAATTTATGAACTCCTCCTAAC

ACTCTTCTTCTTCCCCTCATTTTCTCTAAAGTCCCAACTTCAAACAGAGTTTTTTGTTCAAAGACTTACGTACTCATGTG

GCGGCGCTCGTAATCGGTGCTCTGTTTCTTCCTTTCCTCTTACAGAAAGTGAGGTTAATTATCAAGCCTCCTCCTCATCT

CCAGAGGTCAATTGAACCACACAGAACATAAATACTAAGACCCTTTCAGTTCCCGTTAGGGTCGGCTTATTGCTTATCCA

ATATAATTGATTTGATGGGATTAATGACAGAGAATCTAGCAGAAGGCACTGGTTGAAGAAGAATTCGTTTAAACGTTTTT

AAATTACGACTAATGGAAGTTAACACGGAAAGTAACGGTTTGACCATTTGACCGTTTTGATTAACGGGTCAACGACCCAA

ACACTATAAAGAACAAATTTATGACTCAAACACTAAAGGACACCTAGGATCCAAACACTAATTAACTCTCATTTTTACTC

TTTTTTGTTTAATCTCCCCATTAGTCTCTTCTCTCTTAAGGACTTCAATGTATGTTTCTCCCTCAAAGGAATATGAATTT

TATTGTTTAAACACTTCTGGTTTTATCGGGAGGTTGTTAATGGAGGCAACTATAGGAAGATGATGATGATGGTAACTGGC

TTTTTTCGTTTTCATTTTCTTAAAGTTCCACTTAAGAAAACAATAGTCACAAGATGAAAACAACTAATATCTGTTAGATT

TTAAATTTTTCTGATGATTAGATTGAGAAACTAATTTTTCTATATTTGAAAGAAAATTGGTTGAAAATTTACGCGGGACA

CCAAACAAACAATTTTACTAAATACCAGATCGCAAAATGATCTAATGATGAAACACAAAAGATACATAAAACAAATTATT

ACTGAATACCAGATCGCAAAATGATCTAATGATGAAACACAAAAGATACATAAAAAAAATATCAAGGCTCACAATACTCC

AATTTCCAACTTAATTAATTAAGTGAACCACGTCGACCGTCAAGATCTCCACAAGTTATCCCGAGACTGCTACATAAGTA

ATACCTTAGGACGAATACATACCGCCATCGGTGCCCTTGTCATTAAGGTTACGCAGTCAAATCCTGCACAACAAGGGTTA

TTTCCACAGATTTCACCGCCTATGGAACAAGCTGATGAGTGAACAGAATGAATGAAAACGGAAGAAACCATTACCAATAC

AAAGAAACAATTCAGAAGATGCTTCCTTTCCATCTTGGAGTTCAGTAACTTCAGTCGGTGTTTTGGCGATAGGAGTGCTA

CGATACATTTATATATAGAGAGAGAGTTGGGAACTCATTATTTTCTAAATTTGAGATTATTACTTTTTCTGGGACCAGCC

ATACAGTAGTTTCCACTAAATAATTACTTTTTATTATGGGACACATAATAAAAGCTTTACAAACTTTATATGGTACTATC

AATTATTTGTCTACTTTTCATTTTTCTATTTCTTTTTGTGTTGGGTGAGAAGATGGTGACCGAACTTGATGTGAACACAA

CATTTTTTTTTTCTTTTTTTTTTTTTTTTGTCGAACTTGATCACAAATCACACTCACAACAGTAGTTTCCCCTTTGAAAA

ACTTCAGCGTTTTTCGCGTTTTGCTATAAAATATTACGTTTACTTATAAATATTAAGGGTAGTGGAGGAGGCAAGGTGGT

ACTTGAATAGTGCGCCAAGTCACGTAGGATCAGTTTTAGTATCTACCGACGAGGGAGGTAATTAGGACATTGATAAAGAA

ATAATTCTCACCCAAAATATATTTTATTTTATTTTAATGAGTTTTATTATTTTTAAATAATTATTTTTCTTTTTTATTTG

AATAGTGGTTCATACGGTTTTCTTTTTAATAAATTGGGTGTCACTATTCAAAAGGGAGCCGACGTCTAGCTTGTTGCTCT

GTTACCAACCAAAGCTGTGTATGAATCCTAATCCTCAGATATAAAGTAATAATTAATAAGATATTTATTTTTATAAATTT

TATTGTTTACAAATATGTAAAGTAGGTCCACTATCAGATTCTGTTCGTTTTTCTCTTATTCTTGGCGTGGATTTCTAAAT

CCTCTCTTACCTTACCTACTTTCAGATAGAGAGAGCGATAACTTAATTTTTCTGGTTTTAGACCATGATTTTTGGTCAAC

ATCATCATACCCAATTTCTCCACTAGACTCAGCCTAACGAATGTTCGTATATATACCCCAGAAACACTCTGCATACTTGA

TAAGCTAATCCTTTTACTCCCATCAAAAACCACGATTTATAAAGGTACTCGTACTAATTGTGTCCCGGAAACTTATCAAC

TCCTGTTGTAGCGGTTTTTGTCTTCTAAATATGATGTGGCGCTTAACCCATGTCGAAACGGCATGGGGATAATTTTTATC

TGGATATTAAATGAGGGAGGATCCCTTCACACTCTTAGATTCACACTATCTCACATTTTGGAAGATATTTTTCTTAATAA

AACTTAAATGAGAGAGTATTTGAAGTTAATATTCTGGGAATATGCCGTCTCGCAAAGCTCTACATATCCACAAAAAATGA

GCATATTCTGAAATACTAAACCCCACCACCATCTTCCGATCTTAATTTTAACCGTTGAAAATTCGTTGATTTTCGATGGT

TGATTTTCAAAGTAGTAGATTACGGAGATATACATTTCAGAATTAACATAGTCTATTATTACTCACCAATACGCGCCACC

ACCACAAAACCACATGATACCATTTCTTTCACCACAAAACTACCCAGTAAATCCTTAATTTTATTTTATAAATAATACTT

ATCACATCTATATGTAAGGTTTAGTCCACCAAAGTTTACATGTTTATTTTAAAAATAATAATAATAGCCCTCTCAGTGTC

AAGTTCTAACTCCGCCGCTGTTACCACCACTGCCTCCTACCCCCTGTATAGGTTGTGGGGAGTAATTCTAATTTTCAATC

ACAACCACCATATCCGAGGTTTAGATAGAACAAAATTAAGCCGTCAATTAGTTTGTCCCATATTTGTTTTTTAGTCCTCT

TTTGTATTGCAATCAATTTTATTTTAACATTAAAAACCATGATTTATACGATATGGTACCAAAGATAGCATTGGTGCCAT

TTGGTTGATCCAGCTATCATGAACAATTTTTTATATATGATTTGATATCATCCCCGAAATATATGGTACATGCCTTTTTC

AATCTTGTTAAAACAACCAACCGCATATTTCACAATACAATGTTATCCGACTATCTGTGTTGTAATGACACAATACTAAT

ATTGTCCACCATCACTCACCACCCACCGCCACTAAGTCTTCCAACACCACCACCAGGGGTAGAGCCAGGATTTTGAGGAA

GGGGGATAAAAAATTCTTGAAATTGGATACGGGGGCAAAATACATTTAATAACCCATTTTATGATTTAAAAAAATCAGAA

AAAATCAAAATTGGGATAGGCTACGCCCCTGACCACCACTAATGTATATGCTTCCAACACTTTTTAAGAGCGCCACTGCT

TCCACCACCTATTACCGCCACCAGTGTCTATTGATATCATTTTTATTTTTACAAAAATTATTTATTAATGGAAATACATA

TTTATTAAATTAATTAAGGGAACATTTATAGTGAAAAAAATTTAATAGATCATTTATTATGAGCAATTAGTGAAAATCTA

ATTGGTAGAACACTTTATAATGAAAAAAAAATTAATATATCATTTATTATGAGCAATTAGTGAAACAATAATTAATAGAT

CATCTATTATGGGACGGGTGTTATATAAATTACTTATAATTATTATATTAAATTATTTTGTGAAAATTGTTTCTAATCAA

ATTCATCTAGCCTTTTGAATAGAAGCCCGGAAAACGGAGTATTCTATATACATATGTAAATGCCGGTCGATAGTCAAATT

ATAATGTGCAATAATTTTAGTAGTTTCTCGCTAAGTGTATCTATCAAGATAAATCCAAAATGCTACTTCCACTAATTAAT

GGTAAGGAAGTAGATAATGCGATGGCTGAATTGGGGAGGTTACTTATCTATGAAGCGTCAAAGGACCTAGTACGTTGGCT

TATCTATGAGGTTACTTAACAGGTCAGAGCCTGTGTGACTACTAAACACTGTTGATGAGCAATACGATTGGGAGTCCCAA

GCTGGTGGATCATGCACTGTTACCAGAGATGTAGATGGTTAGCAGTTGAGTAGAGGAGTCAGAATCAGAATCACTCTCTT

CCTCGAAGAAGGTAAGGTAAGAAAAAAAACTTCGGTGATTATTGACTGAGTCTTACCAGGTCAATGTTGACTAGCAGTAA

TTTTTTTACTGTTACAAAAAGTTCGTAATGGCTTAGGGCTGTTACGATGTGTTACAAAATAATTTTGTAGCGGTTGTTAC

GAAATTTCCACGTAGTAATAAATAGCGGCCGTTACAACAGACGTTGTAAAATATATTCGTAACGGCTGGTAGGCCATTGC

AAAAAAAATTATAGTACCCCTTTTCATTACAATGGAAAACCTTTATAACCCAGATTCGTAACGCTAATAAACCGTTACGA

ATTGCGAAAAATGTTGTAGTGTTAATTAGTTGCCCCAAAGTTGTTCCAAAATCTTTAAACAGTTTAACTAAACTGTGGCA

AGTTTTTTCGTTCGCTTTACAGAAACCTAAACAATCTTCATCAAGAAGTAAATGGCTTAAAGAAGGATCCTTGTGAACTA

GTTTGATACCTTTGATTATACATGAATTTTCAGCACGTAAAATGGTTCGAGAAAGCGTTAATTCCATGCAAAAAAGAAAG

AGATAAGGGGAGATTGGGTCTCCCTGCCTCCAACCCCATGTTGGTTTAAAGAACTTTGTGGGAGACCCATTTAGAAAAAC

AAATTAAGGTTGTTGTCGAAATACATTGGTGTACTGGTGAGATTTTCTTGATGTTGTAGTAAGATAATTCATTCAAGACT

TGTGAAAGCAAAACATTTTAGACTTAATAAAACTTAAAAAAAAAAAATTGATTCAAAATTTTATTTCAAGATATTAAAGA

TAACCAAAACATTGGATTTCACTATTACACATGAATGAATGATGGTAAATGATGGTTATATTCGTCCAAAGATAGAAAAT

TCAGAAACTGATTTTAAACACTTAGTAGAGTTTTCGAACGAAGAAAAGGCTCTTGCAAAGCAGAATTCAGATGGTTTGAA

TGTTATAATACATGCGGTAAGTCATGATCTACAACATCATGTATCAACATTTCAAACTTCTAAAGAAACTTAATTTAATC

TTCAGATCATATTCGAAGGAAATTCATCAAAGAAAGAAGCTAGACTTCAAACTCTCATTTCCGATTGGGAAAACCTTCGA

ATGGATGACAACGACACTTTTGAAGAGATTCATATTAAACTCTATGAGATAATTAATGCCTCTTTGTCGTTAGGAAAGAC

TATTTCTGAAAAACATATAATATGCAAGATTCTTAGATCGTTCCCATCTAGATACGATTCTAAGAAGCATGCAATCAAAG

AAGCAAATGATTTTTCAACTCTATCACTTAGCACACTCGTTGGAAAGTTACAGATCTTTGATCATGAATCACAGTCTATT

CAAAGTAAAGGAATCGCTTTTAAAGCTGCAAAAATTCTAAAAGATCAAATAATAGAAAGGAATAGACAGATGGATGATTC

AGATGGATGATTCAGATGAACAGATTACAGAAAATTTTGATGATAAAATTGAACAATCATTGTCATTAATTACTACACAG

TTTCGATATCTCTTAAAGAAAAGAAATAAGAGATTCGTTAAGGATACTTATATATCTTCTCGCCCACATGATCGAGTTCC

TGTCAAAAAAGATCATGAAGATGATGATGAATATCATCCGCAGTGTTTCAAGTGTAAAGGGTTTGGACATATTGCTAAAG

ACTGTCCCAATCTTAAGAAATACACTGGAAGCAAGGCATTGGTTGTGACTCTAGATTTGACCTCTGATTCATACGATAAA

GAGAAAATAACTAATATTGCATTAGTTGTCAATCTTATTTCTTCCTCCTCCATTAGTGATTTACCCATTTTAGAATGGAC

ATGTCTTCCGATGTTATTAAATCACCTAATGGTAAAGGGAAAAGTAAGATAAGATGCAAAAACTGTCTCAAAAACAGAAT

TGCTAAGAGTTGCACATGCAATTCTAGTCCATCCCAAGATACTTCGTTAGTTTGAGGTAACAAAAGGAGACTTTCTCATA

CTATACTAGATCAAGAACACTCTGGTTATTCAAAATTCCATAATGAGAGAAAATCTCATAACAATGCCACTTGATCGGTA

TTAGAATTCTTTCCTCCCCATGTGTGGCAAATATTTGAGTAAGGAAGAAGAAAAATCAAGGGACCTTTGTGTAGATTATG

GAAATAATATCTAGTATTCGAAGATATTTGATATATTCATATTTTTAGGAATATTATATTTTCGGTAGTATGAAAAATAT

AAAAAGATTTTTCTCCTATTTTGATCATTCCTTGTCCGAACTATGGGTCTGGATAAAATGGTTTCTCTAGAGAAGATACA

AGATCAGAAAGATCATCTTAAGACTAAGTCTGGATTGTCTCATGATATTAAGGGTGAAATTAGAAAGGGAGATTGGCTTA

GCAAGAAGGAAAGAGAGAATACTCTGAAGAAAGATCAGTGTATTGAGGAATTGACACACTTTTGTGTTCAATCAAAGACA

GAATTACATGCTCTTGCAGAATCCTTCATAAAAATTTCTCATCTGCAAGAGACTCTGGTCAAGCAACAAGGTTTTCTACT

TGAAGAAATTCAAAATCTGAAAAGTAGTAATCCACCTTCATTCAGCTCTGATATGAAGGACTAGGTTGCAAGAATAATGG

ACATCAATTTTTGATTTCTTTCATTAATTGTATTGGTATATTAATAAATAAAACTATTATGAATAAAATTATTTGATTGG

TAATTTTTCTTGTGATGGTTTTCGGTTTACATAGATTGTGCTTGAATGCTTTTATCTTATTGCTATGTATGTTTATGGGA

TGTTTGATTTAGGTTTTACTACCTTAATATTCGATCTCATATATTGTGAACTGTTATGGTTTGGGTGACTTTTTTATTTA

GCCGGATTAAGTTCTATCCCAAGTTGGTAGGCTTTATCAAAGAATCAGGAGGGGTTCTAGTAGAAACAATATGTGAAAAA

GTCACAATATGTTGAATCGGTTTTGGTTTAGCATAAATGCATATGTGTTTCGACGGTTTTCAACTCTTGCTTGCAATTGT

TAAAACCGGTTTTCAACCTTTTTCTGGTACAGGTTGAGGGTTGTTGTTATTTTTTTGTTTTGGAATAAGGATGACAAAAT

GATGATATTTCGATATCAATTCTCCCTGGACGAAAATGCAATCATATTGGAGAAGTGGATGTGAAATTTGAGGTAACAAT

CTTGTTAGTTAATTGTTTTCCTATTTAAGAAAAGCCTGAGTTTATATTATATATATTTATTGCTTTTGCTTAACAAAATT

TCAGTTCAATTGATGTTTTATTCCATGATGTGTGTGGATGAGTGTGATTCAGTTGGTTCCGGTTAAGAAAAACTATTGTT

ATATTGCTTTATTGTGTGATATCAATTGTTTAGGCTTACAAAATTTCGGTGCAATTGCGGTGCTTTTAATGTCTATGTTG

TTTCAATTGCTTCCGGTTGATGTGAATAATTATGCAAGTTGATTTATGTTGTTTTGTATGAATTATTTATGTCTTAACTA

AATAAAAGGTGTACGGGATGAGTTGTTTAGTCCAATTCGATTCCGGATAAGAGAAACTAAGTTAATTCTGATCTTAGTTA

GATTTATCGAAGTGAGGTTTCGGTTATTCAAGTCTAATCGAATGCCTTAACAAGAAATGCTAGTTAACTAACCTAGTACT

TGTCTTGTTTAAAATTAAAAGGTCTAAGTTTATGGATAATTAGAACCTAATGAAAAAAAATAGAGTTATCTTTATTTTCG

TCTTATCGAATTAAGAAGTTGTTTTTGAACTATTGATTTAGCAAAGGGACTAGGGTGATTAGTTGATTTTTGTTTTGCTA

AACTAAATTGGAAAAATTGTTTCAGACAATTGTTTCCTTGGTAACATATCAATATAAAACTACTTGTAGTTTCGGTTTTG

ATATTGGTTATCATAACATGATGTGTGGAACCCTCGTGCCCAACTCTACAGGTTGCAAGTCTATTTTGAGATTTGTAAGA

TTCTTTTCGTGTTGTCCTTTATTTCTTCGTTTCTTTGTCGTTTTTGTGACAAAAAGGGGGAGAAATATATGGAGTAAACA

ATTGATACTGGCATTGATTTTATATTGATTGGTATCGCTAAGTAAAAGAACATTGGTGCTTGAACATTTATCTAACGAAA

GAGTGAAATCACGGACTAAGGGGGAGTAACATGTCATATAAATTGGTATAACAAAGACGAGCGGATTGAATATCTACCTA

TCTTCCTTAGGGGGAGTATTAGCTTTGTTATTATAATGTCAACAACGACATTTTCAAAAATTGAATGTAAGCAGTGTATT

ATGTTGTTGAATTCGGGAATCAAGCGTATGTGTAATGAATTCTTGTAATTTGTTTATCCATATGATTTAAGAGTTTTTTC

ACAAAAATTAACAAAGGGGGAGATTGTTAGAGAATGGGTCGGTTGAACCCACTAAGCATTGGTATGTCAAGTTTGGTTGT

CATATTTTAGTGAATCAAAACTCATGTTAAGAGTCGCTTGATTATGTACTAGACTCAACTTCGTATAGGTTAGCTTGAAA

GTATTAGGATATGAGACATTACAAGTATTTCGAAGTCTTGAAGATATGAAGAAGCAAGGAGCTACAACGCCAAAAATCAC

CCTTCCACTTGAGGTTAGTGATATTTGACTTGAACTGTTTCATTCCCTAATGTATTTTTCAAGTCGTGCATATTGAAAAC

AAAACTGCGAAGTATATTTGAACTTTATATAAACATAGTATTAAGGAATACAATACGAGGTTTATTTCTTCACCATTAAA

CTTTGTAGATAAGACATCGCTATAATCATTCGAATGCTATTGTGATTATGTATGGGTATAAGGTGAGGACTTCATCCTAG

GAAACAATGTTTTACATGTGTTCTAAGGAAGTAAGTTCATAAACTTGTTTGTGAACCGAAAAGGAAATTACCAGGTGTTA

TTGGTTTTGTTATTCATTGCATATCTTATGAACAACCAGTATGTGTGATTGAGTATAACCGCTCACAACTTGTTTGTGTT

CTTGGTAAAACTATTCACAAAGGCCTGACTTATGTATTGGTATGACTTTTACTAGTGAAACCGATCTTAAGTAATCACCT

GAGATGGTATGATCGGGTTTGTGATTTTATGTCTAGACAAATATGGGAAAGGGGAACCGATCTTAGTAAGAGGTGCAGTG

CATTACAAAGGGGAACCGATCCTTGTACAGGGTGCAACAAGGTTTATAGTAGAAAGGGGAACTGATCCTATGTACATTTG

CAACACGTTTTTAGGCAAAGGGGAACCGATCCTATGGACATGTGGAACACATATAAGTTAGATACCATATATATGTGGGG

AACCGATCTTAGTACCTAGTCAACCAAATTTCGGAAAGCTAGCGTGAGTATGCACAATACTCACATGGAAGGTAGAACCG

AAACTTGTTTTGGTAGAACCGTTAAACCCATGATTGTGATTGAATATTTGTTTGATCAATTGCATAGTTCTTGAAAGTCA

GATGAACTAATTCTAAACTTGTTTGGAAGTGTGGCAAATCGGTTTCAAGGTTGTAAGTGTGAAAGCGAACTTACAAAGTA

AGTATGTCGACATACGTTGAACACGTGCTGTGAATGTTTATTTCTTTAATTGTTCAAAGTTATTCCTTAATAGCTAAGGG

AAGAGAATCACAGGATCGAAACATAAGTAAGTTAAGAATCTTTTAATTAAGGTAATTAACTTCATTTTATAGGGAAATAT

AAGAATTAGTAATGTGCATTTACTAATTATATTTTCCAAGAGATTTCGATCATTATTTTTGGACAGAGCATTTCCATGAA

TTATGGAAACCGAATTTGTGCTTTAATGAATATCTTGAGAATATTTTATGTTTTGGAAATTCCTTGGTGTCAAACTTCCT

TGTCTATAAATACTTGAAGTTTGTCTTCTAGCAAACTAATCCTTCGTAAAAACAGACTTCCTCTTTTCTTGTTGTTACTG

GTGTAGCCGCCTATTCAGAGAGGAGAGTAACCTAATTAGGCGAAATCTCTTACGGCCCCTCAGTTTAAAGTCTTCTTTGG

GATTGAGAAGCTCTAGCATGTACTATTGGTGGGAAACTAGATAATTGCGGTTTATCTTTTGTTTTTATTGATTTGATTGA

CCAACGGTGGTTGAAATCTGATTGCAACTAGTTTGTTTATGCTTGAGAATATTCTCTTCTGATATAAGATTCACTCAAAC

TAGTTCAGAGTTTCGACAGGGATCTTTAGACTGTTGTTAGTTCTAAAGATGATCTTGTGATAATCCATTGTTAACAAACT

CCGTTCTGTGCGTGATTGATCACAAGAGATTCAAGTGATTGTGTGCAGGTATTTATTGAAGATTTAAGAAGATTTGAAGA

AAAAAAAAGATATTGAAGATCTGACTTGGGTTTATAATCTTTGGTGTGCACATACTTCTTTCAGTAAAAGAGGATCCAAT

TAATAATTAGTTTATCCTTGTGGTAGATTCGATTGATTAATTGAGTAGATCGGCATCAACACAATTCCTTGGATTAAGAG

TGTTGTTGGCTTAATCTTAAACGATTACTTCGGTAATTGAACATAAGATAGATCTAAGGACCTGACGAAGGAGTTTATGT

TAAGATAAACGGAAGAGCCTTTGTCCGACTCATATCACTTGGTTGAATAGAGTTGATACCAAACAATTTTTTTGTTCCTT

TACTGTTTGGAATACGAACCAAAGTAATTGTTTCGAGTACGTGACTTATTTATAAGTTGGAGGCGTGGGAATACAAACAG

AACTATGTGAACTGTAGGTTTAGTTACTTGATCTCAACTATACAAAGTTAGGTGTAATTTTTTTTAGCGGCTTAATCTTA

AGAGTATTCAATTCTGGACAAGGTCCCGGGATTTTTCTGCATTTGCGGTTTCCTCGTTAACAAAATCTTGTTGTGTCATT

TACTTTATATTTCCACATTATAATTGTTTTATTATAATTAAAGTAAATCACACAAACGTTAATTCCTATTTACTTGATAA

GCAAACCTATTGTGTTTTATTAAGTCCGAACCTTTTTATCAAGTAACATAATTCGTTGTTGTATTGTCTCAATTTCGTAT

CCATTGACGATCACACGAAGTGTGAACCGATTAGTTATATTGTCTCGACTCAATCCATAGACAATCACTTTCGGAGAAAG

GACTTATAGGTAGGAAAAGTTTAGCTTGAGGTATATTTGGGTACCCTCGCCTTTTCAGTTATTACATAATTAGACCTATA

AATTATAACAATTACCAGATTTGTCCTAACGTCAAGTTACAATATGCGTGTGTCTCAGGATATCTCTCACTGCTTATATT

AAAAACCATTTCTTTCTCTCTTTCTGTTCTGTTAGTTGTCGAGAAAAAAAGAAGAATTGTTGATACCATCAGAATTGTGG

TTAAGTTTGAATGTCCTAAGATGGAATGGAGCTGGTGATAGATAAGAAAGTGAGGATGTAGTAGTTGTTGTTGCTTCCCT

TGATTTTTATAGGTTTCGAAAGTTGCTAAGAAGACAAGAAATGGGTTTGATGGATGTGGTGACAAAAAGAAGAGATGGGT

TTCCATTGGATGTGTAATTGGCAGAGAAATGGGTTAGTGGTTTGAGACTGCAAAGAACATAGAAGTAACCGGGAAGGAAT

AGTTGGAAGATTTATGAGCAGTGAAGTGGAAGATGAAGTTACTGCTAGGTTTTTTGATGTTGAATGATTAAGAGAAATTG

ATGATTTTGGATGATTAAAATGAATGGGTTTATCTTGCATTCGTGATTAATTACAGGGAACAAGATGCAATTGGATTATA

ATCAACAACACAGATATCTCTTGATTACTAAATTCACAGACTTCATGTGTTTATAAGAATTCCTAAGAGACGAGTACTTT

TTATAAACTATGGTTCTGATGGGTTGGTTCGACTAGGAAGAGTTGTAAAATTTCAAGCAAGGAATGAGTCAGTTTTTTCC

GTGACAGTGCTGAAATAATGAAGTTGAGATGGTTAAGCATAGATGGAGTCTGGTTTTCACAGTAAAAACTCGGTGATGTT

GCTATTTTCTGTGGTGAAGTGGTAGTTTCAAAGAATTAGAGCAATGATCATGATGAAGATGAGATGGTGCTATGAGTCTG

GGGGTGCATTGGTTAGGTCTAAGGATTCAGGGATGTATTTACAATGATGGAGATGAATGAAAGTGAAGCAATTGTAGTTT

TCTCGAACAGCCGAGGTTGATGCCATACTTAAGGCTCAGGTGGAGCACCTCCTTACACTCTTGGCTCCCCAAGAAAGTTA

TGACAGCCAATGGAAAGCTTTCGCACTTAAACGTGAGAGTAAACATGACCTGCGCACCTACACCAATTTCAATAAAAGAT

GAAGAAGTACCAGAGGTGCTAGATGAAGAAACACATATCATAATTACAAGTTCAATAAAAGAAGCCAGGATATTGGCCGA

CACCAAAAATCTAGTGCAGGCTTGGCCAAAGTTGATTGCCACCGGTAAATCACTTATAGAGGTTGATGCTACATGAGGCT

CAGGTGGAGCACCTCCTTAGTGATCGAAACAATGTTTGAATCTAAATAGTGTTTCAATGTCAGATGAACGTACTTCTAAC

CATCCGACCAAATCACTCATATCCCAATTTCTTTTAATAAGTCAAATGTTGAAGATCAGTTTATGCCCGTTAATTATTTA

TCATTTATTATTTATTTTAATTATTTAGTTAACAGTGTGTTAAGTCAACTGTTTGACCAGATAGTCAATAACAGGTCAAG

GTACCAACCCTTAATATTTGAAATGTTGGGTACCAAAGCTAGGTGTTGGACATTTGCTGGGTACCAACCGTTAAATAACC

CAATAATTATTTATGCAAAATATTTGTAACTATTTAACTTATGAATTTTTATTATAAAAACAGATCAAACACGGCTTTTG

GTTGGTAGCCGAGCAACAAGCTAAGCTCCAACCCCTCTCTGAATAGTAATTCCTAATCTATGAAAAATGAAACTACCAAA

ACCACTACTAGCATCATTATTAACTAGGCAATTCTTTAGACGCTTAAAGAAACGCAAATTGTCATCGCCAAGCTCACCTA

AAGTAGAGAAGGCCAAAACGCCCAAACCAAAACCCAAAGCAGAACATTTGTCCAAATATTTAGTGCGCTTCCTTGAAAAT

GCATTAGAGATAGCCTTCCCTGGAACAAAAGATCGAACACCATCACCGGTGAAGGGAGAAACAACAGTGACATCCATACA

AACATCTTACCCATTTTCCCAATTGAGAACAAGAATATCTACAAGCCTCAAATCCTTATCATCATCTGACTGGATGCCAA

GAGATACTTCCTTACGCGTCGGTGCATTAGCCTTATAGCAAATGTCAAGTGCAACATCTCAAACAAGGTCATGGATAAAC

TTTCTCCCAATATATTTAGCACAGTGAAGCGCATGGTCTCCAAAAATGTCCATAGATCGGTTGCAACTTGAGTATAAGCC

ATTCTCAACAAACAAAGGAATAACAAGTCTATAACAGATCACAACCCGAAACTGCCAAGGTCCAATACACTGATTAAGCC

CACTAATAGGCCCCGCTAATAAATAATCATGAGCATGCTTGACACGGTTACACTGCCATAAAATAGAGTCTCTTGCAGAC

ATGTGAATTTGGTATGGGATTCGCTTATTGACTGCATCAAAATAAGTGACTTCCAGGGAGTGCATGGAGGAGGGGGGGGG

GGAAGTATCATAGACACAATAAGAAGGGGACATTCCACATACCTGGATAAAGTTTTGAAGATCCCACTACAAGGTAGAAC

CTTTGTTTTCTGAGAATAAATCACCAAGTTTAATCTTCTGTACTGAAGCAGTATGACTTTGAGAAGCAAGGTAGCAATAA

GCACGAGTGTCATCCATGGTGTAGATGCCAAGGCCACCCTCCTTGATAGGAAAGGTGGCTAATATCTGCTGCAGAGGACC

AAAACCAGCACCATCTCTAGTGATGAGTAATCGTAAATAATTAAGTAAATGGTCATCAAAGAGATCTGTGGATGGTTGCA

AGGAAGCTGGGTTGGTAGTTCGCATTGCAAAATATAAATGAGAAACACCAGTTTTATTCACTCTGTGGGTCCTTGTGTTT

TTTGATCGCTGCCATTAGTTGAACAGTTTTATTCACTCTGGTCAACATCATGTCACTAATAAAGTTCAAGTCTAAACTCA

CCGGACCACCCAAAAGTTTAACCCCATTCGAGGGTATACCAATATTAGGAGGAAAAACACCATTATCCATACTCCTGGTA

TCAATAGAATGCCAGAAAACTTCTGTCTTCTTCACATTAAGATGAAGCCCCCTACTTGGCCCTTTAGTCTTTATTATGTT

CAAAGCCTTAGCCACCTCCAAAGTGTCACCAATAATGGTTCCATCATCAAGGTACCAAGCGTGTAAATTAAGTGTGCATC

GAGAAGCAATGGACTTCACTAGGGGGTGAAGTGTCAACGCAAATAATAAGGGACCAAGGGGGTCCCCTTGTTGAACTCCA

AGTGCAGGCGATAGAATATACTGATCATAATATAAATTACCTGGACGCAAGTAAAAAAATTCAACCTAGCGAGAAATACC

TGGGAAATGTAGGCGAACCTCCTTGATAAGATGAGATCTACTAACCATGTTAAAGGCATTAGAAAAAGGAAAAGACTCTC

ACAATTAGGAGAGTCATGAAAAAGGAAAATAGTCAAAACCCTAGCTTAACTTTCTAGATAACTTATTCACTATACATAAT

TATAACGTGAATGATAAACTTTTTTTATCCCTTTTGGAAAATTGGTGTTAGCTTTATAAGGTTGTTTTCCATGTCTTCTG

TTCTTTGTGAACAAGAGTGAGATAAAAGTCTTTGTATTGGGGATCACAAAGCCGAGTTGGATTTAATCTTCGTGATTAAT

TATAAAACTTGTAATATAGGTTTTTCACCTCTTGTCTATTCTAAGGTTTTCTCTTCTGATAATAAACCATTCAAGAGTTG

TTGATCATAAACCCTAGGTTTTGATAGATTTCAATTTTTGATCGTACACCAACACTTGTTAGTCAAAATTGGAAGATAGA

AAGAAAACTTCGATTAAGGTATCTCGTAAAAATCCTTAAGAAGTATTAAACAAGATATCTCTAGATTTATTCTAGTTGTT

CTTGATGTAGAATCATTAGGTTTCTCTTCATATTTTTGAAGAGTATAATAATCCCTAATCTACAAGTTTGTTTTGATATT

ACTTTTAGGGTTATTTGGTTTTTGGTACTTACTTTTTGACCTAAACCTGGGTTTGGTCTAACATTTGTCCAGGCTTTGCG

CTTGATCTAAAGACCAACGTTGACCCGTGTTGACTCGGTTATACGATGACATGTCTAACTTTTGACCTAATTAATTTTTT

TGTGGCCCTTCGAATTGTAACTTATTTTCTTATTCCACCCCCACCCCAAATCACTCTTCAACTGTAACATATAAAACGAC

CATTAACATGATGAAGTTTATGTGACCATTAACAGATCAACGACCTATAAATAGGTCAAACGACCATTAACATATCACTC

TTCAATCCATGCAATTTCAAATTAGAGACAAACCCACTATCCAAAGCATTAAAAAAAAATCAAAATACATGCTCAATCAC

CCATTATTCATTGTCAATCATATCAATCTCTTTCTTTCCTAGAATTCTCAGTGGGACTATACCAAAAACATTATACATCT

CAAACGGCCAATAGTTATTTCCCATATGTTTGCAGCATCATCTCTATAAAACAACATACCCTGTTGATGTAAACCAACTC

GAATTATCTATCATCAATTATCACCATCATCACTGTGACATCTTATAACTCAATCTCTCAATCATTCGTCCTTGACAACA

TCAGCCATTCTTCTTTTTCTTTGTTCTTCGGACTCGGTCCCTGAATCACCAGCAACTATAACTCATTAAATTTATCTTCA

GAGTTATCTTCATCATCCGAAATTTCCTTGCTGACTCTGTTCTAATCATAGACTAAATCAACTTTCATTAATCTGCTTGA

GTTCTTCTCCACCAGTCACCGATCCAAAAATGGCAGCACCATCACTTGCATTTCAAACTCCATTTCGAACGCGACCCAAT

ATTCACCACCTTGTCCATCATCTCTTAAGTTCTGATTTAACATGATTTCCATCATTGAAAAACCATTGATCCCACCAGTA

CTATCACCACCAACAGCAGTACCACTACATCATCCCTATCATTTTCACCTTATTTATCCCAATAACCCGATTCAATTCAA

ACATTTAACCAAAACTCAATTAAACATAGTCAATCTTCATTATCATTCAATTAATTTCAATTATAAACCCTTATATATTT

AAAGAACAGAAAACCCCCCAAATTAGCTTCAATTATAAACCCTAATTTAACAATCGAGTCAAATCAAAAATGAAATACCA

ATTGTAATCATCAAAACATTATGCTTCATTTAATTTGGGGAAAAAGAAGCTCCGGGTTTGGTTACTTACTTGAATTCACC

TTAGATTTGAATCGATTTTGAAATCTCTGATTGTTTCTACTTCTAATCGTCTTCCTTTATCTGCCCGTGCCAAAAAAAAC

CTAACTAACCATTACAAAAGATAGAAAGTGATTTTGGTATATTATCTTCCACAAGAGTTTGTGGAGATATTGAACTCTTA

TTTATACTTGAACGAAGTATAATATTCAAAGCATAGAAAAAAAGGAAATACAAGTTGGATCCTAGTATATAGTCGGAGTA

ATCCGTTGGAGACCAGACTAAGTACTTGGTTATTGAATAGCTTACTGAATTGATGCAGTTGATGTAGACCCAGTTGTTGG

CTCTTGATGAATTTTGCTGACACCCTCCCTCAAGTTTAACTGGAGTTGGCGAATGTTCAACTTGTCTCTTATGTGCTGAA

AACGCAGTGAAAACAAACCCTTCATAAAGATGTCAGCTAGCTGGTCCTTGGTGCATATAAACTTGACATGAAGCTTCTTA

CTTTCCACTCTCTCACGTACAAAGTGGTAGTCAATATCTATATGTTTTGTGCGCGCATAAAAAATAGGATTTGCTGTGAG

ATAGGTTGCGCCTAAGTTATCACACCAGATGAAAGGGGTAGAGACATGAATACCCAATTCTTTTAGCAAGGATTGTATCC

AAATGATCTCTGAAGTTGCGATTGCCACCCCACGATATTCTGCTTCTGTTGATGACTTCGAAACAATTTTCTGCTTTCGG

GCACTCCATGAGATTAGATTCCCACCAAAGTAAATACAGTAGTCGCTCGTAGACCTCTGATCATCCAAGTTTCCAGCCCA

ATTATCATCGGAGTAGGCATGAAGCGAGTAGTCCTTTTGAGGCTTGATGAACAGTCCATAGTCAATTGTATTTTTCAGAT

ACTTTAGAATGCGTTTGACAACCTCCCAGTGTTCTACATAGGGTTCATGCATACACTGGCACGCTTTATTTACCGCAACT

GAGATATCCGGCCTGGTGATATGTAAGTACTGCAAAGAACCTACTATACTTCTGTAATGTGTAGGATATGAAAACTTCTT

TGTGAGAGCACTAAATTGGGTCCTTGACGCAGCACTTCAATGCCGAAGAAAAATGACAGTTCGCTCATATCCTTAACCGC

AAAGGTAGCACTCAAATATGAAACAAGTTTGTTAATCACCTTTGAATCAGAGCCTGTTATGATTATGTCATCGACATAGA

TTAAGACATAGAAAATGGACTTGCTAGTAATCTGCATAAACAATGAAGAATCAGCAATGCAAGCGGTAAAGACAAGTTGT

ATAAGATGGTTTCTTAGCCTCACAAACCAAGCACGAGGTTCCTGTTTGAGTCCATATAAGTACTTGTTCACTCTGCACAC

ATGATTAGGGAAATTAGGATCTACGTAGAAAGCTGGTTGCTTCATGTAGACCTCTTCTTCCAAAACCCTATGCAAAAACA

CATTTTCGACATCTAGTTGACGTAAATTCCATCCATTCATAACAGCAATGGAGAGTATGAGCCTGATAGTGCTGGGATTT

ATCACTGGAGAAAATGTTTCATCATAATCAATGCCCTCTTGTTGGTTAAAACCCTTTGCTACAAGAAAAGCCTTGTGCTT

CTCTATACTTCCATCATGTCTTCTTTTTATTCAAAAAACCCATTTTGAACCTACCACGTTCATCCCATCTTCATACTTCA

CCAAAGAGTAGGTACCATTAATTATTAGCGCATTAATTTGAACATCCATTGCGTCTCTCAATTTAGGAATCTTACAAGCC

TGTGAATAGAAAGTAGGTTCCATGAAGTCCCCATCTTGTAACGGATGTTTAGTTGTGGTTAGACACAACTTCTGGATTGG

TTTTGTTATGCCATTTCATGCACGAGTAACCATGGTATGTCTTGGAAACAGATCAGGTTGATGTGGTTGAACTTGCATAT

CACTCGACGGTGACAAGGAAACGGCAGGAATTACTGTGGATGAACTGGAAGTTTGCTTCTATGAATATGAATGAGTCGTT

GCAGGTGGATCAAGCTGACCAACATTATCAGGTTGTTGTTGTATGTCAAAACTTGCTGCTACAGAAAGATGTTGTGGCTA

TATTGATACTGGTGCTTGCTTTCTGAAAACCGGAGAACCAAGAATATCCATACTAGAGCAAGTTATTTGCTAACAAAGTT

CAGGTGCAGGATGCTGAGGAGGAGAGAATGGGTACGAAGATTATTAAAACCTGACATGACGAGAAATGTAATCCCTATTT

GAAGGATGATGCAAGCACACATAACCTTTGTGTGCTGGGCTGTACCTTATAAAAACACATGGTAGAGATCTGGGTTCTAG

CTTGTTGAAATTGTATGGATGCAGACATGGATATGCTAGACAACCGAAGACACGCAAAAACTTTTAATCTGTATTCTTAT

TGAATAAGAGTTCAAGAGGAGTTTTAGGCTCAGTGGAAGACTTTGGAATTCTGTTAATGAGATATGACACAGTTGTAAAA

GCCTCAAACCAAAATTATTTAGGCATACGTGCCTGGAATAGAAGGACTAAGCCACTTTCATTGACATTCCTTATTTTACG

TTCTGCTAGACCATTCTGCGCAGATGTATGTGGACGGGAAAAGCGATGCTGAACACCATAATTGTTTAAGTGAGATGTAA

ACTTTTTATATTCAAGAGCATTATCCTACTGGAAAACTTGGATTTTATGACCTGTTAGATTTTCTACTTGTTTTTGAAAC

TGGATAAAAGTAGCCAAAGCATCAGAACGATTTACTAAAGGGAATACCCATGTATAATGTGAAAAAACATCCATAAATAA

TAAATAATAACGAAAACCATGTCGAGATAGTTGAGGAGAAACCCAGATATCACTAACAACTAAACTCAGAGGTTTATCAA

ACACTGTTGAACTGACATAAAAGGGAAGTCTCTTGCTCTTGCTTATGTGACAAGAGTGAAAAAAATCAAATTTCTAATTC

AAAACATAAAGGGAAAAACTATTGATTATTTTGGAGACTGTGGATAGCATTGGATGCCCTAAGCGTTGGTGCCATATAGG

TAAACTAACAGATAGAAAAGACTTTGGAGATTTGGTTTGATTGATGCCACCAACCCCATCGAAGATATAAAGCCCATTAT

CATTCCTCCCCTTTAGCAGTATTGCTCCCGTGAACTTGTCCTTCACAACAAAAACGTTTGTGTGAAACTCAAAGCAAACA

TTGTTATCATTGCAAAATTGATAAATGAATAGTAGATTTGTTTTGATTTCAGGCACATGATATATATTATTCAAGAGGAA

ACACCGTGTATTATGAGTAAAGGTAGCGTTACCAATATTAGCAATCTGCAGGCCATTACCATTACCAACCCTAACTTGGT

CAGTTCCTTCATACTTATATGGAATGCAAAAGATTTTGAATATCAGGAGTCAAATAGTGAGTTGCACCAGTGTCATTCAC

CCATTGTTCTCTCCTGTTTTCTCTCAATAGAATTTTTTTCTCTGAAGCCGCTTCGAATGAATAAGACGATGGATATGGTT

AGTTTGCACGTGTGGTTTAGGATGAAATAACAAAACGAAATATAATTTGAGGGCCACAGAAAAATTAATTAAGTCAACAG

TTAAACAAGTCACCACTCAAGGGAGTCAACACGAGTCAACGTTGGTATTTATACGAAATGCAAAGCCTGGAAAAATGTTA

TACCAAACCCAAGTTTAAGTCAAAAAGTATGTACCAAAAACCAAATAACCCTTACTTTTACCTAGTTATTTTTTTCTCAA

AACAAACACAAGATTTCCAAAAACTTGAATTACATCTAAAGGGAATCGATTCGGTGGATTGTGAAAACAAAAGATCGAAA

AGTATCAACCGTGTTGTTGTATCTTCTGAAGATAACTTTGTTCTGCCAGACGATTTTCGGGAATAAATTTTAAAGAGAAT

TTGTGTTCATCTAGAAGTTATACGAGAAGAATTCCTAAAGAGAATCGGTGTTCTGCTAGGAGTTCTATAAGTGCTTGAAT

ATAGCTGAAGCAGGTATTACATCTAGTCCGAATAGGTAGTAGGAAATTGGTGTAATAACTTATAATAAGTGTGTGTTTAT

TTTGGACTAGGTCCCTGGGTTTTTCTTCATTTGCGGTTAACAATTTTTTTGGTGTCTGTGTTATTTCTTTTCTGCATTAT

ATTGTTTATCTTTATAATTGAAATATCACAGGTTGTGCGTTAATCAATCATTATGGATAAATCCGACCTTGTTTGTCAAA

TAGGATTTTTATTGATCTTGGATATTGGTCTTTGATACCATCCAAGTATTTCTCACATCAATCAGGTTCATAGATTTCTA

TCTGTTTGATTTGCCGATTGATTTGAGAAAAAAAGATAAACTCTTTAGTGTTATTCTTTGATTAACTTTCACTCTCGAAA

TATTATGAGGTTTAGTCCATACAGATTTCCAAACGAAATATTGGGTGTGGTTGTTAGGCCCCCGCTTTTTCAATAATGGC

TCTGCAGTTGTTCTGATTATCCTTAGAGAATAGGTGTCTATCTTGCAAATGACAAAGAATGGAACCCCTGGCATAACCTC

TGCCATTCACTCTGTCACGGAAGCCATCAAATTTTTTGAATGGGAAATGAACGTAACCATTAAGCGGAACATCATCTGCG

AGATATGGAATCTTCTGTTTTTTGTTAAAATTAATATCACATTGAATTTCATCTCTAATTTTCCTTATCATCACAAGAAT

TTCTAGTGTTGTTTCTCATAACACAGCGAGAACTTATTAATGTGATTATCACTTTAACAACCTCATTTTGTGAATGGGGA

TCTATTATAGAACTTTCATCATATTTTTTTTCCATCTTTAACAGTTCCCATAATGCTCTCAAATATTTCTTTATCAGTCA

ATAATTTTGTAGCTTCAATTTTTTCCATATATTTAAGATATCTTGACATTCACTAAATTTCTATAACCCATCTTTTCAAT

CAAATTATGAAAGTTTTGAGTGATATCACCATCTAACTATTTATCTGAACTGATTAAATTCATGTTATCTGTTGATCAAA

GTTAAATTTATGTATAAGTTAATAAATATTAATTTATAAATTTACGTATAAGTTTGTTTGTAAGTGTTTATAGGTGAAAA

TGGTTTCTGCTGATTTTGGTAATTGCAGGTGTGAAAGTGAGAAACAATTTAAACCCTAAACAATGTACTCCAAGGGAGTA

CTCTTAGATTCGAGAGATAAATCTATACAAGTCCGTCCTAAACCAAGAAATGGTCGTTCCATACTTGCTTCAGTCATAAA

ATGGAGGAGAAGGGTTGGTTTAGGGGGGGAAGAGAAGAGAGTGTTGAGGACAGAATAATTCTGGAAGTGCAGTGCTTGAC

TTGTATCAGAAGGTGAGAGCTTTGAGAAAAGATAACAAGAGTGTTGGAATATTTTCAACCGCGCTACGAAGGGGGGTCCT

GGGGGGCGTAGCCCCCCAGGCTGGTTCTGAACACTGGAAAGCGGTTTCCAGAGTTATGGCTTACTTGAAAGGAACAATTG

ATTTTGGTTTGCATTACCAAGGCTATCCCGCTGTATTGGAAGGTTACAAACGATGCTAACTGAAACAATATAGAATCCAA

GTCCAAGTCGACATCCTGGATGGATTTTTACTTTAGGGGTGTCTTTCTTGTGTCTTGGAGTTCCAAGAAACAGACTTGTA

TATCTGATTCAACAATGTTGTCGAATTGATAGCTTTAACGAATGCATGTAAAGAGGCGAGAATGGCTTAGGGACCTTCTT

ATGGAAATTCCACTTTGGAAGAAGCCATCTCCGGGAGTTTTGATTAATTGTGATAATCAAGCTACAATCTATAATGTCTC

TAATAAGACTTATAATGGAAAGTGTAGACATGCGAGTCTCGTGACACCAATTGGTTAGGCAACTACTCAAGAGGGGAGTG

GTTACGGTTAACTTTATTGAATCAAAAAGGAATTTGGCAGACCCATTCACAAAGAGTCTACCATATTCAGTTGCTTCAAT

AAAATGTAATGAAATGGGACTAAGGTCTCATGAAGGAAGGTTGATCTCCCATAGCAGAAACCCAACCTATGTTTTGAAAA

GGATAAACAAGGTTCAATGTGGTACCAACAAGTTATGGTCAGAGATTATGGAGCACTAGCATAAAACTTCCCATTTCAAG

TTAGTGCTGGTTTTTCTGCAAGAAAATAGGATGGATGTAAATCCTTAATGAGTCTCAGATTAGTAAAGGGTGTCTCGCAG

AAAACACCTTCGAGACTTACCTATATGAGTATGGAAATGTGGGCCGTTTCCTAAGAGAAGAAGACCGTTCTCTAAAGAAG

ACTCATGAATAAGGATATAAGCACAAGGCCTTTATAGTGCTTGGCTACAGAACACATGTCTTATGGAATCAATATGTGTG

GAGACTCCGGCTTTGTCATAAGGGATTCTTGGTTCAAGACTAGTTTCACCATAGAATATCGATAAAGCTTTGAGTTTCTT

ACACTAAGTGAAGGTTCAAACCCAGAAGGTACCTATCATTTATGTATTGGTTTCAACGATGATGCTTAGTCTCAATTGTG

CTAGAACTACGTTGGATTGATCCCACTCGGGTACGTAGGCAGTCACTTCGGTGATGCAGCCGCTCCGGTTTTAAAAGTTT

TTAACTTTGTTTCATGGTTTTGAAAATAAGGGGAGAATGTTGGAATATTTTCAACCGCGCTACGAAGGGGGGTCCTGGGG

GGGGTAGCCGGTGTTGTGTGATTGTTTCCATATTACGTAGGGGTGAGTGTTGATGAAGAGACGCATCTCTTTGTGAAGAG

AAACCTGTTGGTGAAAAGACATGTCTTTTCCCAACAGATGCAAAAACTCCTATAAGTAGCATCTCCCAAACCCATTTCGA

AATCATCAAAAAAAAAAAAACTATCTTCTGTTCTTCTTTCTTACAAAGCATCATCAGAAACTTATTTACATGTGTTCTTG

ATTGGGTCAAGGGTGCAAACCTTGAATCCAATTTTCGGTTGTAGGGCTTTCACTGTATCCTGAAGGCAGTATTTCACATA

CCTGTTGCAATCGCCAATTCGTTGGGAGGGTAGAAATATTTGTCTAAAAGAAATCACCAGAGCCTTGGAAGTTTTGAGTA

CAACCACAAAAGATTTCTGTCATTCATCTCCATTGCAGCCCGGTAACTTTTTTCCAACAAAGAGTTTCCTTTCTGAGGAT

GGTATTTTCCCTGACCAAGAACTCGTGTCTCTGGTGGAAATAGGTAAGCCTACTTATACAAGTCCAAAAAGAAACGTATT

CTGGCCTCTTAAGAAAAGAAAACGGATGAGTTAAATGGGAAGAGGTGGTAACGCCTGGTATTGATGGGATTTGATGGAAT

TAATGTCCCATAATGAATGAGCATTTCACCGTTGCTTCTTGCATTTACTAACCGCCTTAATCTTATTACACTATCTTGAA

ACGGGCATTTGTGTATGTCGCACGCTATGTACCACCAAACCAAAACCCTAATGAGCATCCCCCAGTTTGTGTTGATGTCT

CGAATATTTTTGGAAAACATGGAGCATGTTGTTGTGGCTTGGAAAGTCAAGCTAGGGAAACTTGCTATCCTGGTGACCTT

CGACGGTCGATATTTTGCATCTTTAGAGGAGGGGAAACCGTTGATCGTTGCGCGCCTTTGTTTTTGGTAGCCGCGTAGTG

GTGGCGCGACTAGGTTGGCATGCCGTTTGCACGGCATGGTTTGGCATGGCCAAACTTAGGTTTTCGGGCCAAAGGTTACC

ATGTGTCGTCTGGTAACCTTGGATGGCTAGGATTTTCTCCTGGGATTGAAGGGAAACCGTTGATTGTTGCACGCCTTCGT

TTTGGTAGCCGCGTAGTGAGGGCCAACAGGGTGGCGCGGATAGGGTGTGTTTATGCTTGGCATATGGGGCCAAGGGTTCT

CCTTGGGCGTGGCTAGCTTTGGCTTGGCATGTGGGGACAAGGGTTTGCCATGGGCGTATGGCGTGGCCAAGTCTAAGGCA

AGGGCGCTTGGATTGGCATGACGTGACTAGCATAGCTAGGGACAAGGAAGGGTGCATTTGGCATGGTGCAGATAGCTTGG

CTAGGTTTGGTTTGGCTTGGCATGTGGGTCCAAGGGTTTTCCATGGACGTATGTCATGACTAGGGCCAAGGTAAGGGCGT

TTTGGCATGGCGCGGCTAGTTTGGCTAGGTGGGTGACCAAGCTAGGGTTTTGGGCTAAAGGTCGTGAAGTGACCTTAGAC

GGCCAAGATTTTCTCCTGGGATTGAAGGGAAACCGTTGATTATTGCACGCCTTCACTTTTGTAGCCGCGTGGGGAGGGAC

AGCTTGCTGGTGTGGCATGAGGGGCCACGGCGAGATGTGTTTGGCATGGCGCGGGTAGCTTGGCTAAGTGCGTAACCCAT

CTAGGGTGTTTGGGAAAAAGGTCGTGAGGTGACCTTAGACTGTCAGGATTTGCTCCTGGGATTAAAGGGAAACCGTTGAT

TGTTGTACGCCTTTACTTTGGTATCCGCGTGGGGAAGGCTAGCATGGTGGCGCGGCATGAGGGGCTATGGAGAGATGTGT

TTGGCATGAGGCGGTCGGTGTGGCTGTCATGATTTTCCATTGGCACAGAGGTGCGGCTAGCATGTTTGTCATGCTTTGGC

GCGGTCGGTGTGGATGGCATGATTTTCCATTGGCAAAGAGGTGCGGCTAGCATTGTTGGCATGCTTTGGCGCGGTTGGTG

TGGCTGGCATGATTTTCCATTGGCACAGAGGTGTGGCTAGCATGGTTGGAATGCCTTGGCACGGTTGGTGTGGCTTTCAT

GATTTGCCATTGGCACAGAAGTGCGGCTAGCATGGTTGGCATGCTTTGGCGCGGTTGGTGTTGCTGGCATGATTTGCCAT

TGGCACAGAGGTGCGGCTATCATGGTTGGCATTCTTTAGCGCAGTCGGTGTGGCTTGCATGATTTGCCATTGGCACATAG

GTGCAGATAGCATGGTTGGCATGCTTTGGCGCAGTTGGTGTGGCTTGTATGATTTGCCATTGGCACAGAGGTGCGGATAA

CATGGTTGGCATGCTTTGGAGCGGTCGGTGTGGCTGGCATGATTTGCCATTGGCACAGAGGTGCGGCTAGCATAGTTGGT

ATGCTTTGGCGCAGTGGGTGTGGCTAGCATAATTTTCCATTGGCATAGAGGTGCGGCTAGCATGGTTGGCATGCTTTGGA

GCGGTCGGTGTGGGTTGCATGATTTGGCATTGGCACAGAGGTGGGGCTATCATGGTTGGCATGCTTTGGCGCGGTCGGTG

TGGGTGGCATGATTTGCCATTGGCACAGAGGTGCGGCTAGCATGGTTGGCATGATTTGGGGCGGTCGGTGTGGCTTGCAT

GATTTTCCATTGGCACAGATGTGCGGCTAGCATGGTTGGCATGCTTTGGCGCGGTCGGTGTGGCTTGCATGCTTTTCCAT

TGGCACAGAGGTGCGGCTAGCATGGTTGGCATGCTTTGGCGCGTTCGGTGTGGCTGGCATGATTTGCCATTGGCACAGAG

GTGCGGATAGCATGGTTGGCATGCTTTGGCGCAGTCGGTGTGGCTTGCATGATTTGCCATTGGCACATAGGTGCGGCTAG

CATGGTTGGCATGCTTTGGCGCAGTTGGTGTGGCTTGTATGATTTTCCATTGGCACAGAGGTGCGGATAACATAGTTGGC

ATGCTTTGGCGCGGTCGGTGTGGCTGGCATGATTTTCCATTGGCACAGAGGTGCGGCTAGCATAGTTGGTATGCTTTGGC

GCAGTGGGTGTGGCTAGCATATTTTTCCATTGGCACAAAGGTGCGGCTAGCATGGTTGGCATGCTTTGGAGCGGTCGGTG

TGGGTTGCATGGTTTAGCATTGGCACAGAGGTGGGGCTATCATGGTTGGCATGCTTTGGCGCGGTCGGTGTGGGTGGCAT

GATTTTCCATTGGCACATAGGTGCGGCTAGCATGGTTGGCATGCTTTGGCGCGGTCGGTGTGGCTTGCATGATTTTCCAT

TGGCACAGATGTGCGGCTAGCATGGTTGGCATGCTTTGGCGCGGTCGGTGTGGTTTGCATGATTTTCCATTGGCACAGAG

GTGCGGCTAGCATGGTTGGCATTATTTGGCGTGGTCGGTGTGGCTAGCATGATTTTCCATTGGCACAGAGGTGCGGCTAG

CATGGTTGGCATGCTTTGGCACAATAGGTGTGGCTGGCATGATTTGCCATTGGCATAGAGGTGCGGCTAGCATGGTTGGC

ATGCTTTGGCGCGGTAGGTGTGGGTGGAAGGATTTGCCATTGGCACAGAGGTGCAGCTAGCATGGTTGGCATGATTTGGC

GCAGTTGGTGTGGCTGGCATGATTTGTCATTGGCACACGTGGGGAATAGGGTTTGGCTTGTCGTAACTCTAATTTGAAAA

ATATACCCTGGTAACTATTTTAGAGACACATTAAAGGTTTCAGTAAATGTGTAAAATGCCAAGCCCATGAAGATTAATTC

ACTTAAGCCCATAATGGGGCGTATGAATACGCAATCCTAACAACATGGGGATGAAAAATCCATCAAGCTATTGGCTTGAC

TTGGCTAATCAACTCAGCTGGTTGAGTTGCCTCAGTGGGGCAAAAATATCCTTTAGTGTAAAGAAATGGGGCGCAAATAT

CCCGAGGGTACGAGAATAAATTACTTCGTGATGCACTGGAATTCTCTTGTCGTTGCCGAACAGAATTCAGTCCAGCTGCC

GGACACAGACGCATGACCTCGTTTCCTGACGCCGTCGCCGGACACAGACGCATGACCTTGTCGCCTGATACCGCCGTCGG

GCATGGATACCGACCAACTACTGCTGCTACCGTTGCCGTAGAAGATGTCGTCGTCGACCAACTGTTGTTGTTGCCGTTGC

CGGAGAAGACTATGATGACGTCATGCTGACATATGCTAACGACGTTACTGACATTGACGGAATATATTAACGGCATTAGT

CTCTGTTGGCGTGAACTGCTTATTGTGCTTGTGATGCTGATGTGGCAGTCTTTGATTGGCCCACACTGGTGACGTGGTTC

CTTCTTGCCTTGACTTGGATATCTGCATATGAGCTTCTATGTATAGTGCTTCTGAGGCCTAGTTAGCCATACTTGGCTAA

CAAGAGCAGCATATTAGGCCCAATGAACCATACTTAGCTAATAAGAGGAGCTTTTTAGGCCTAAGTAAGCCCGTTTTTGG

TGTAAAATATTAGGGTAGAAACAAAGTTAATAAGATATTAATTTATCGGTTAATTAATATCTCGCTTAATTAATAAATTT

TCGTAGTCCCAACAACATTAATTTATAGAGTTTCTGCTATACTTAGCAAACAGATGCCCTCTCCACCAGTTGCTTATCAA

ATAATCACATGGTTCTAATTTAAGCACCTTAATGCTGAGGATAATAGTCCCACACTGTTAATGTCCCCTTAATAAACTTA

AAATATAATATGGAAGGGCATCTCCACTCATTGCCAATTGATTTTGAGTTGGATGCACGCAGACTTTAATTTGGTATCAA

AGTTAGGCCCGTGTGTGAATAGGCCCAACGGTCACCACACAACGTAGGTCCACGTGTTAGACCCAACGAGGCTTCACATG

AGGGAGCGTGTTAAGGATAATAGTCCCACACAGTTAATGTCTCCTTGATAAACTTAAAATATAATATTGAAGGACCTTTC

CACTCATTGCCAATTGGTTTTGGGTTGGATGACCGCAAACTTTAATTATCCTATGTATTTGTTTCTCACCCCTTATCCCC

TCTATTTATATATGCTTGTTTAAATTTGAGTAGTACATAGCTTTTTTAGTCGTACTCTTCATAAGCTGTGTTTGTTATTT

CTTTACTTTTAGTGTCCTTTTGACAATCGAGAGGGAAAGTCTCTACCGTTTTGTGAGTTTGGCACCTATATTATTCCACC

TCCAAGTTGCTCTTGGACGTTTCACGAGCTGGAGTGTTGTTCCAAAGAACATGGGTCACGGCAGAAACTATATAAAACTT

GTGCAAATATCTTGTTGCCGTCTATGTATACTTGTTGTTAGTCCCGTTTCTACGTACGCCTATTGTCTTTATAGCTGTAG

GAATCTAGCCGTTCCAATTTTCAACCAATTCTCCTGTTCTTTTAATGCAATTTGAATGCACATCGCTGGAGTCAGTACAT

CTATGATAAGCATGAAGCAAATATCTCCACTTTTAGTACATTTACGCATCAAAAGAAGGTACTGCTTTAGAACGTTATTG

GGCAAATTTTTAATATATACCTCTTACAACATTCTTTTAAACAGGTTCCTTTTTACCAGCTTTCTTGGATGGTTTAGCCC

CACCAAACAATAACTCATTAACTCCCGACACAGTACAAACGCCTTCTATAAAAGCCACATATAGCGTGGGTGACCTTAAT

TATCCCCACGAGGTTCTTGCATTACTCTCGTGGGCTGATATGGGACACTGCTCCACCCCCACAATCAGAAACCTTTATAG

AAAGCATCAGCGACAGCGTATTACATCAATCCTTTGTAAATCTATCTATATATAAGGGATGATATGGGACACTTTATAAA

AGATAATTGATTGATATATATAAGTGATGATATGGGACCCTGCTCCAACACACAATCAAGCACCATTATAAAAAAAATAA

TCAAACACCATTATAGAAAGCATTAACATATTACATCCTGATATGTGACACTGCTACACCACATAATCAACAAATCAATC

ATTTATAAAATCCGGCTAATTTTGGGTCTATTGATCAGGCCTATGAATTTCTTCAACCAACCTAAAGAGAAGGATATGAG

GTATCTAAAAGATGGTAAAAATACTATATTAACCTTTAAAGAAACCGCAATTATTCTAATATAAATACATAATTATAATC

ACTTAATTAACACATACAAAAATAATTAACCAAAACTCACTTTTTTTTTCAATTTCCATAAAAATCAATCACAAACAACA

TTTTTTTTTAAAAAGAATACAAAAGGAAAGACCATTAAAAAATCAATTTCAAAACAATCACGAGTTATTACATAGTAGCC

AAGCGAACAACAATTGAAATATAATTTCTTTTTGGGTTTTGAATAAAAAATCCTGAAAACAAACAATTCGAGTGATTGAG

TTAAATCCCTTTATCCATTCCTTCTAAGAGTAACTCAACAATGATTTACCTCTAAATCTTTGAAATTCACTCATCAAATT

TTTTGAAAATTCACTCAGAATCGGTTTATGTGTGCACTGTGTTAGTCCGATTGTAGTTGATGTTACAAGCAATCGGTTTA

TATTGTACACTTACATAAACTGATTAACATCAATCACAATCGGTTTACACTAATGCACACATAAATCGATTCTGGGTTCA

AGTTGGGATATGTATAGAAATAAAAAAAAACGGATTATTCGTGCATCTATATAATCCGACTGTAACTGTTAATGGCTTTT

TTTTTTTTTTTGAAATTATAGTCGTTGGTCCTTAGCGTATATAAAGAGGATAACCTATTCTCTATCATATACTGCCTAAT

TCAAAGAAATCTTTTTCCATTGTTAATTTTTTTCATTTGAAGATGTCAAGTCCATCATCATCAACTACTAGTTCAATCGA

ACACATACTTAGAAGATGCAAGCGAACAACCAAATTAATTATACGGTAATAAAGAAAAGATACGCAAAAAAATAAAAAAA

TTATGATACTAATAGCTTGTGTTCGAAACTCGCGGATGGGGGGGGGGGGGGGGGGGGGGGGCAAACTCGCCAATTAGAAC

GGGCCCTAGCTCAGCTGGTCATCCCCGTTTCCCAAAGTTTCATGCGTTGCAGGAGGTCACAGGTTCAAGCCCCCAATGGC

GTAATATTGTAGGAATTAACATGAAAAATAGTGTTAGTTTGCCCCCTTGTGACACAATCTCAGCCCCCATCCGCTTCGGC

TCCCTTGTCTAGGTTACCCTTGAGGCTCGCTGGTAGGTTAACATAGCAACCTGTTCAATTAATGTAGTAGTCTTAGCTTA

TTAGGCTTCCAACTAGTTAATGTAATACTCTTTATCCAATAATATTAATTATCAAGAAAATAAATAAAAAAATGTTAATG

TTAATAATTAATGAGTGATTCTTAATGGCCTGTTCAAAAATCCGGCCGTTTTTTAGTTGTGATTGATTTTAGGACAAGGT

GTGCCCCCTCTAAACTATGGTTTTGGATTCTCCACTGCACGAGTTTCTCGTGCATGATGGTTTGTTATACCGATGTCCTC

TTTCCCTTATAAATGTAGTTCTCGGTGCTTGAATACCTAACACATTTGGGTTTTGATTCAATCTCTATTAATTAATAGTC

AAATTCAATAGAGTAATTCCCTAATCTTTGCAAACTGTCTTATATTTCTGTAATTCATGGCTCCGATTAATATAGAGGAG

AATGATTTTTGGATGATAGCATGCACTGTAATAATAGTATTTGCATTGATGAAGTTCATGGTTTCTTTTTATCAATCTGC

AAATACAACGGAATGGCCAGAAGGTCCAAAAACACTACCCATAATTGGAAACCTTCATCAGTTGGGAGGAGGTGTTCCCT

TACAGGTTGCTTTGGCAAACTTAGCCAAAGTTTATGGAGGTGCATTTACAATTTGGATTGGAAGCTGGGTTCCAATGATC

GTCATAAGCGATATCGACAACGCTCGGGAAGTTCTTGTTAATAAATCTGCTGATTATTCCGCTAGAGATGTACCTGATAT

TCTTAAAATCATCACAGCAAATGGGAAGAATATTGCTGATTGTGATTCTGGTCCATTTTGGCATCATTTAAAGAAAGGTC

TTCAAAGTTGTATAAATCCATCAAATGTTATGTCTCTATCTCGTTTACAGGAAAAAGACATGCAAAATCTCATCAAATCC

ATGCAAGAAAGAGCGTCACAGCAGAATGGAATTTTAAAACCTCTTGATCATGCCAAAGAAGCGTCTATTCGATTACTGAG

TAGAGTTATATTTGGTCAAGACTTTTCAAATGAGGATCTCGTTATTGGTGTGAAAGACGCCCTCGATGAGATGGTACGCA

TAAGTGGGTTGGCAAGTTTAGCTGATGCTTTTAAAATTGCTAAATATTTACCAAGCCAGAAAAAAAATATTCGGGATATG

TACGCCACAAGAGACAGAGTATATAATTTGATTCAACCACATATCGTCTCTAATCTTCCTGCAAATTCTTTCTTACATTT

TCTTACATCTCAAGATTACAGTGATGAAATTATTTACTCAATGGTACTTGAAATTTTTGGTTTGGGAGTAGATAGTACTG

CAGCAACGGCAGTTTGGGCACTCTCCTTTTTAGTCGGCGAGCAGGAAATTCAAGAAAAACTCTACCGCGAAATCAACAAC

CTGACGGGTGGGCAAAGACCAGTGAAAGTGGTAGATTTGAAAGAGCTGCCATATCTACAAGCCGTGATGAAAGAAACATT

GAGGATGAAACCCATCGCACCACTAGCGGTCCCTCATGTAGCAGCAAAAGATACTACATTCAAGGGGCGGAGAATCGTTA

AAGGTACAAAAGTAATGGTGAATCTGTATGCTATTCATCACGACCCTAACGTTTTCCCTGCACCGTATAAATTCATGCCA

GAGAGATTCTTAAAGGATGTTAACAGTGATGGACGTTATGGTGATATCAACACAATGGAAAGTTCATTGATACCATTTGG

TGCTGGTATGAGAATTTGTGGAGGTGTAGAATTAGCCAAGCAGATGGTAGGTTTTGCTCTTGCAAGTATGGTCAACGAAT

TCAAATGGGATTGTGTTTCTGAGGGGAATTTGCCTGATCTTAGTGAAGCTATTAGCTTCATTCTCTACATGAAAAACCCA

CTTGAAGCCAAAGTTACCCCTCGTACAAAACCGTTCGACAGTAGATAAAATCTTATTTTTATCGAACTGCTAGTGTATGT

TTATGTAAAATCAAGGGTGAAGATGCTTGATCGTAGTACACCAAAATGGTGGTGGCTATGTAGGAGTTAGATTAGCTCTT

GATTTAGCAAATGTTTTTCTTTGCTAGAGACTGTCTCAAGTGTGTGTGTTTTAAGCGTGTGTTGTGTGTGTAATTACTTC

CACCTGACAATTAAAGTGTACTTTAGTCGTTGGCTGTATCTGTCACATCATTCTTGTGTTTTTAAATATCTTTATAAATG

GTTTTCCCAATTTACTGCCCTACATTTCCTATTTCCCATCCCTTGAAGAGTTATTAGGCCTATTCCTATGTACATGCCAA

AACATGATATTTGATATATATGCACCCATTATGGATATGCCAAACTGCCAAAACTCATATGCCAATATGTTATTCCTGGC

ATATAGGCATATACCACAGAGTTTCCAGCGAGCGATAGAGTCACGGTCGTTCGATGGTCAACAAATCGCTCGATGGTCAT

TAAATCGACCGCTAGTAAGATCAACGCTTGTCATTCCCGTATCTAACGGCTACAATTCCCCCCTCTATATAAATGCATAA

TTTCAAACTCATATCAACTCACATCGGAAATTCTAAATCTCTCAATCTTTTTCAAATCTAAAACTATCACACCTCCTCTA

CAATTATATATTTTCTTGTAAAATGGATTCACTAGCACAAACTCAAGGAAAAGGTAAGAAGAACCGAGTCCGTGGTGCAA

AATACAGCATGATAGAAGATGAATGTATTTGTCGTAATCATGTTTTTTTTACCCAAGATTGTATCGGTGATGCACAACAA

CATGGTAACACCATGTTGGATAATATATTTCAAAAATATGAAGAACAAACCATGAACATCAATAGTCGTGATGCAAAAGG

ATTGTCAGAACACTTCATTGTAATCAACCGGGAAGTAGCCGCTTATGTTGCATTGATAATCCAAGCTAAGAGAAAAGGCA

AGGATAGTGGTCAGAGTGATGTTGATTTTGAAAGAGAGGTTCGTACAGATTGGCAACGAAAACATGGTAAACAGTTCACT

TTCGAAAGTTGTTATCATATTTTGAAGGGTTTGAACAAATATAATCTAGATTACTTAGCAAATAATCAAATAGTACCTGA

AAAGTCACCATATAATTCTTCTCCCTCAACCCCAAATTCTTCACCATTTTCACAAGGTCCATCAAATTCTTCACCAGATA

CCCCTACAATCTCAAATTCCAACGTTGTTTTCAATAATTATGTTGATGGTAATAATAAGGGAAAACTTCCAGGGAGGAAT

CGGGCAAAACTGGCTAGAAAATTAGCTCAAGAAGGAGGGAGTTCCGGAGGATTTAACATGGAGGAGTTTATGGAAAATCA

AAAGACCGTCGAGAAGCAAAGATCCATTGAAAGGAAATTTCAAAACCGGGAGAGTAAGAAAAGTCGTCTTTCTCAAGAAA

AATTTGGGTTTGACTATGACAAGCATAACATTCTTCAAGCTAACACCTCAATTATGAACGCCCAACAAAATCATGTATGG

CAAATCGAATTGGATAGGATTCAAGCACAAATTGAAAAGCAAGCTGGATTTACTAACAACCATGATGAAGATGATGATGA

TGATAATGATGATGGAATTGATGATGAGTTTGATGAAATAATTCCTCTAAATGATTAATGTACTAGTTTAGTTTTAGTTT

AATGTAGCAGTTTAAGTTTTAATGTTTTAGTTTAAGTTTAAATTTGATATGTTGTTATGCATTCCAATTAATATTAATGT

GAAATGAAATGTTGAAGTTTATTAATTAGATAAATAACAGTACATAACGTAACTTAAAATTAAGCTAAAACATTAAAAAT

AATAACATAACTTAAACTGAAGCTAAAACATTAAAAATAAATAACTTAACTTAAACTGAAGTTAAAACATTAAGAAAAAT

ACATAAAATCATCTTTCATGCTCTGCCCAAAGATTTAATCTCAGATCATTTCTTAACTGGTCATACAAATTACGGTTCTG

AATGTAGTTAGTCATTTGACCATAATTCCTTGCAGGAGGCCTCTTTCAGGTCGAATCTTCGGCCTCAAATCTTCATCATC

ATAGTTAATCCAATCTAAATCACGCCTAGTTTCCTTAATTACCATGTTATGGAGAATTATGCAAGTGAGCATAGTCTTGT

GCATTTCACGAGGACTCAACCCACGATATGGGCCACAAATGATTGCGAACTTCCTCTTCAAAATTCCAAAAGTGCGTTCC

ACATCCTTCCTCAGTTCCATTTGGGCATCGTTAAAATATCGGTATGAAAGGCCTAATGGACTAGCAGGAGGCTGACGGTA

GCATTGAACCAAGGTAGACCATTCTGGGTAGATACCATCCTCAAGATAATAGCCATGAGTGTACTGATGATGGCCGTTGA

TCATGAAACGAACAGATGGAGAAATTCCATACTTAAAATCTTCAAACAAAGGAGACTTGTGCCAAACATTAATATCATTT

TGTGAACCCGGAAGACCAAAAAAAGCGTTCCATATCCAACAATCATAAGAAGCGGCAGCTTCAAGGATAACAGTTGGTCT

TGGATAATGACCTTTATATTGACCATCCCAATAACCAGGGCATCCATGCCATACCCAATGCATACAATCTAGATTACCTA

GCATTCCTGGGATCCTATTTCCGCATTCTCCCTTAATATTTGTCTAACATCTTCGTCAGTTGGTTTTCGTAAATATGTTG

GACCAATATGATTAATCATTACTTCACAGAACAATGAAAGTTAAGTGTAAGAAGTTGTTTTACCCATACGAAGGTACTCA

TCGTTCTCATCTGCTGGAGTGCCATAACCTAGAATCCTTAAAGCCGAATAATTTTTTGTTCAGGACTATCACCTCTAATA

TTCAGTGCATCAAACTGATAATTGAATTGAGGTTCTACTCGACAAAGCTCTTCAATAATCTTTTTCACTAGATGACGAGG

CATGCGGAATCGACCTTGAAAATCTCGATCAGAGTACACACAATTGGGAAGAAAATAATCGTGGATGAGCTTATGATGGT

AAAATTCCCTCCCTCGATACGTATATCTTCTTGATAATACTTCTTTGGGCTCTGGAACTCTTGGTATTTGACCCGTATGT

ACAAGCATCAAAATTTCGATTAGTTTATTATCTTCAGCTTCCTCTTCGTCAATTTCTTACAACCTCATACGATCCTTTTT

TTGTTGTAACCTGAACCGATTCATAATAATATCTCTTTCTTCATTGGATCTCGACATTGAGATGATTAATGAAACTTAAA

ATTGAGAATATAGGTGAAATATTTGTAGAAGCAGGTGTGAGTTATATAGAGAGGTAAAAGGGCGTATATATAGGAATCAT

AGGGGGGAAATAGCCGTTGCAGAATAACTACTGGGCGATAACCATTGTATCGCCATCGTTATCATTTAAAGTGAGCGATT

TTAACAATATCGGTAGCGTTAAAGTCTTTAACGCCAGCGATATTGAGAATATCGCTCGATAGCAAGTCAGTCGCTAATTG

GTTTCCCCAAAGAGGCACAGGTGGTTTGCCATGCCACCTGGCATGCCAAACACCATTTTTTTGACATGTGCATAGGAATA

GGCCTTAAAACCAATTCACTTTTCAATTGTTATTTTTCCTATTTACCACCTATAAGAGTCAAATCCATTTTGACTATAAT

TATTACTGTTTTTATAAGTCTTCTTCTTCATTAATCTTCTTTTTTCATAAACAAAACCATCAAACATCAACTTAACAAGC

CTGGGTTAGATTGCGTTAAGTTTTTCCTACATTTTCAAAGGTGTATGCCGCATTTGTGTTTTCCCTTACAGTCCAATGAT

CTAATTAAGATTCAAAAACAAAATCCAAAAATTGATTGAAATGTATCACTAGCCGGACAAACATAGTAGACGATACCTCG

ACTACAGTTCAGCCATTAGACGTTTATAGATGACCGACTAGTTCTTATAGCATCAAAGAATCAACATTCAGTCATCCATG

CTCGGCCCCCAGTAGTTCCCAACCCCGCTAACCCAAACTCTTCTCCACATGTTCATTAATTCAGCTGTAATCTTCGCACC

GACCGGAATTAAGAGCAATCCAACTAATACAATCAAATATTATTCAACGATTAGTTCTTTGTACAGAAAAAAATATCGGA

CTTAAGCAACAATTTGATTATACAATCAAACTTGGATTAAACGACATACTAGGTTTAGATTTAACTTTTTCTGTGGAGAT

GATTTTGGTTGAACGACCACTTGTTTTCTTGGAGGCATCTATGGGTGATCAACAATGAGAAAAAGAAATAAGACTTGAAT

ATGGCATCGGCCATAGAAGAAGAAGTTAAAGACGATTGTGGTGGGAATAAAGAAGAAGAAAGAGGATTTGTGTTCTTAGG

GTTTTTAGGAGTGGGGTGAAATAAAATTACGTACAGTTTAAAAAACAGAGTCAAAATAATTGAAGGTGGTATTTAGGATC

TTTTAAAATACAGATGGATGGTACTTGATCTAGTTTTTATGTTGTTCTAAAGTGGTAGTAAAGATTTGTTCAACTCGGAC

TTGATGAGATTGATTTCAGATTTAAAAATAGAAAATATATACAAAAGTTTGTCACAATATCGAGAGGTACTGAGACTCGG

GATTCCACCATAAATTCATTCATGCGATTCATATATTTATATCAAACAATTATAGCTCAAATAATATTGACTGCAATTCT

TTGCCAAGGTAGATTTTTAATGACTATAAATCACTAGCATGACGCATCAAAAGCACTTAGACCAAGCATACATCATCATA

CGACATCACAATCAATTAAGAAAAATCATAAACCGATTGATAATAGTGCAAGTAGTCATAAAAGAATTAAATATAATTAC

CACATGCATGAAATATGGCTTTCTCCGTCGTCCCAGTATTAGGTTTTAGCTCATTGGGCTCAAAGCACGTTAAAAAAAAA

ATTCCATTGCTCAAAAAGGTGTTCTATTGGTGAAAATATAATAGACAGAGATTCGCAATGGCTTATAGAGGTTACAGAAC

ACACTGGATGACTGTTATAGAGAAAAGATGACATGGCTGTTATATGTTGTGATCTATTATTCGTCTTCTTCGTCGCCTGG

AACTGCATGAATTGAGTTCTGCTACTTCTTTTTCTCCGACCTCAGCTCCTGTTGCTTCTCTTCTATCCTCCGGTTCTACC

CCCAAAGTCTAGACACCCTTGTTTATGACCCCAACACCCTTTTTAATACTCCCACCGGTCGAACAAAGCCCAAAAATATC

TCAAATATCTTTTCTTTCTTTCCGCGGCAAGTTACGTTGATTTCTCCTCCTCTTTCTATCTCTTATACGCGTCTCCCCTG

TGTCAATCCATTCTCTATTGATAAAATCAAATCTTAAAGGAAGATATTCACTATCCTATTGCTCACGTACTGCCCAATAT

AACTCACAAAATTCCCGATATAATATCAGGGGCGGAGCCAAGATTTCAAATTTGGGGTGCAGAAATTTGTTTTGGGGAGA

TAAATTTTTTTTAGGGGGTGCAAAAAGTAAACTTAGGCTTATAGGTAATTAAAAAAAATATGCTTTGGGATCAAATTAGG

CTTTGGAGCCAGCTGGGCAGGGGGTGCAAGTGCACACCTTCTTTCCCTGATTCATCGAGAACTGATTATCCAGTCCACTT

TAATCCGTTAAATCTCTTATACCATCCATGTTTAGTCGAGTATAGTCCAACGCAAAAAAGCTCCGTGATTAAATCTCTCC

CAGACACGTTCAAAAACCGAGTCCAACTCTGCCAGTAAAACACGCAAGAGAAAAAAAAATTCAGCCTTTTTCAAACTTTA

AAAGTAGGGGATGGAGTTCCCTGTTTCAAAAACGAGGTGCCTTTAACACATTGGTGCCTGGGGTGTAAACATCAGGTGGA

CGCCCCTTATCAATTAAAGTGCCCCTTATCCAAATGAGGGGTGTCTTTATTAATTTTCTCCCACGAGTGCAAATTCCACT

TTTCGAGATAATTTCAACACAACTGCATTTCTCCAAAAACACCTACAAAGACATAAAAAACCATAATAAATACAAAATCG

AGCACTAACAATATACACAATTGAGAAAAATAGACACATAAATGCGTCTATCAGGACTTATGATGACTGGGGATGTGATT

TTATATGGATGGGAAATAGGAAAAACGAGGATAGGAAATAGGGAAAACCTTGTAAATCTATCCTAAGTCTGTAACAGAAA

ACATCAGATCGAAAATCAATGAAAATCACTGAAAGAATTGTCAAACGACATTCATCTTTTTACATTCTTCATTTAGTCAC

GATTTATTCCGATTTCTTGTTGAATTGCCAAAATTACTCCTCGAGTTGCTCGATCGTAGTGAAATGTTACTTGGATGCAA

CTGTGTATATCTGTAACTATGTGAATTACCTGTACGTGTGTGTGTTGCTCGTGCTACTTCCTCTTTTAATTTAGACTCTC

AGGTGTGTCTGAGAAACTTCAACGCTCTTATTATTTTCCTTTTGTGAGATTTACTTTCTCTCTACTGTTCTGAATATTAT

TTCACTTTACATTGAAGGTTGTTGTTATGTTTACTAGATCGTACAAAACCTTTCGACAGTAGATAAAATCTTATTTTTAT

TTATTTTTTTGATGGAAGAGAGAATTATATTAGATAAAGAGAAAAGAAAGAAAACGGAAGAACTAAGAAAATTAAGAAAA

AACAGTTTCCTAGTTGTCGAGTGTGTAAAGAAGTTGATGTGCACTCTCTTCCCAAACACAGCTGCCCAAGATAGAACAAG

ACTTTTTGCGTCATTTGCCAAATCTTTGTCAGTCTTAAAACTATAATTTTCTTAAAACGTACGTCAATTCTTTTCTCTCC

ATAGAACCCAAATAACAGTCGCAGGGATTCGTTTCCAAACTATTTGACCCGAAGCTTAGAGATAATTGCAAGGCCACGAA

TGAGCAAATGAATTCATTGTCTCCGAAAAAGCCCAATCATGAAGCTTTGCAGGTGAAAAATCAAACTGCATGTGTATGTT

TATGTAAAATCAAGGGTGAAGATGCTTGATCTTAGTGCACCAAAATGGTGGTGTTTATGTAGGTGTTAGCTTAGATCTTG

ATTTAGTAAATTTTTTTCTTCATTTAGTTAATGTTTTCTTGATTTATTCTGATTTCTTGTTCAAGTATAGTTGTATTGGT

GTGCTTTTCATGTTTTTTTGAAATAAATTCTCTTCTTTTCCTCTTAGTTTTTTTTTTGATCGGAAAAGAGTAACAATATA

TTAAGGGCCCTACCATAGGTAAGCAAAACAAGCTCAAGAGATACAACACAAGCAAGAGACAAGTCTCTAGCAATCCGAAC

CCAGAAACAGGAAAACAAATAGCTAGAAAATAACTAAGTTATGAAATACATTCCCCAATTATCCAGGATGGTATCAAGAG

TAAAACCATTTAACGGGCACAGTTTAAGACTAGAGTCTGGACAAGAGTAAAACTAGTTCCTCTTAGTTATTGAATGATTA

TGAGTAACCACTAGTTTGGTTTGCAGCATTCAGTTGTGACTACAGTAATCAAGTGAGTTGTTATATCTTTGTGGAAGATA

ATTAATTACGAGTTATTAGATTTATCAATTATAAAACCTTGCAGCATCCTTGGCACAATAGAACTATTTGAGATCAATGT

AAAGTTCATGTTTTCTAGCGAAACATGCAATTCGTTGCAAATTCTAAAAATTCGGTGATTTGAAATCGGTCCGAGTTGTG

CTCTGAGGATCGGAGATTGACTCGGCGACTTTTTTATTCTCAGATCTGTTTTCTGTGTCTCAAATTCATCAATTCAGATG

TTGTTTTTTGTTAATTCTTTCTCTTATTGTGGTTTACAATGATTGTTGTTCTTTTATTCAAGTAATTTATTTGGATCTTG

ATCCTAAAATCACTTTGGTCGATCAAAAATAGTTCGATGTGGCTAGTTTCAGTAAAAATGAAATTTCAAATTCTATTCCA

TCGATTCAGTTACATGATGATTTATTTGGCATGGTCCTGTGGAAGCTTGATCTTGACAAGCATATTATCCACATATACCT

CTAGGGTTTTATGTATCCAACAGTCAAACATTTTTTCTACCATTCTCTGATATGTTGCAGCGGCGTTTTTGAGTCCGAAT

GGCATCTTGGTATAACAGTAAAGACCTCGTGGGGTAAAAAATGCAGTGTGCTCTTGATCCTCCTCTTCTAGTGGGATTTG

ATTAAATCCCTAGTATCAATCCATCATCGACAGCAATCCGTATCCGGATGTTGCTTCCACGAGCTGATCGATGTTCGGGA

GTGGAAAGTTGTCTTTTGGATACGCTTTATTAAGATCGCTAAAATCAATGCATATTCTCACTCCCCAATTTTTCTTCAGC

ACGATCACCATGTTGGATATCCATATCGTATACTTCACTTCTCGGATGACACCCGAGGCTTCCATCTTTTTGAGATCGCT

TTGGATAGCCTCATGATAACCGGGTGCCACTTTTCGCATCATTTTCCTTAAGGGCTTGTGCTTTGAATTAATTTTAAAAT

GATGGAAACAAACTTCTGGGTCTATCCTTGGCATGTCATCCATGGGCCATGAGAAGACATCATAATATTCTTTTAGTACT

TGTACAAATTTCTCCTCTTTCTCTATATCGAGGAGGGTTCCAATTTTGACCATCCCTGGATTTTTAGGTGACCTGATGTT

CACCTTCTACGTCGGTTACGAGGCCGAGAAATTTAATCTTAGTTCTCCGATATCGGAAAGATTCTGTAATTGCTATAAGG

GATGTTTAGGTTTTCCCGAAATAGAAATTGGTAAATCGTTTGTACATATAGCTTCATTTGTTATCGCTACGTAGGCGTCA

ATTTCCTCCCGGTTCTTAGCATCTTTAGCTCTACGCTTTCTTGCTTTTGGGGAATTGATTTTATTCTCACCATTTTGTAC

ATCCATAATGTAACACTACTTTGCTTCGCCTGAATCTCCTATGATCTTTGGGATCCCGTCTGGTGCTGGAAACCTTAGTC

TTTGGTGATAAGTTGATGTGACTCCCTTGATTCCATCAACATAGGATCGCCCAAAAATAGCGGCATAAGGTGAAACAACA

TCCACTACACAGAAAGTAGTTAAGTTCAGCATTTTTCCCACACAAATTTCCAGGTTTACCTCTCCCCTTAGAAGGGTAGA

TGAACCGTTGAAGCCAAAGATATTATATGTGATGGTATTAAGTACTCATCTTTTAGGATCATTTGTTTGAACGTCTCATA

TAATAAGATCTCAACAGAACTTCCTCTATCAATTAGTATCTTGGGAATTAATTTCCCATGGTAGTGCCTTCGGCTTCTCC

ACTTCATCTTCATGTCCTGGGAGTACGATTGCCATCATGAACACTAGAGGATTGTTATGATTCCTTCTGTCATCTGGTGT

TTCAGTGGATGAAAAGATGATAGGATGTTCATCCAACCCTCCATTAGAAGTTCTTTTGCCACACTAAAAACCTCGTTTCC

TTCAAAATTTCTCTAATGGATTCGTCCACTGATGTTTCCCTCCAATATAGGTGTTGTAATAGCTGAATGGGAAATCGTGT

TGGATTCCAGGTATTGTTCTCCGCGTGGGATTTCCAATGGTGGTCCAGTCACATGTGGGGCTGTCGGCTATGCAATGTAT

ACTTAGAGCTTACCATCATCAATCATGTTTTGCATCATCTTCATTAGATTCCTACATGAATCTGTCGGATGGCTGTAAAA

TTGATGGAACTCACATAAATCTTTTCTGTTCTTAGCATGTTCAGGATGCATCCCTTTTTTCCATGGGAAGTTAAACTTGT

ACTTTTCGTCAACCTTCTTCAAGATTTCAAAGATAATTATGTTTAACTTAGTGTAGACGGGGTCTAAGAAATTTCCCTTC

TTCTGTTGGGCCTGGTCTCTCCCTTACCATCCTCCACTTATGTGTTTATCGAATCATGCCGATTCATTATGAGGTCGCTC

AGGACCTTGAGTTTTCACTTGGGGGTTTTATGGTTCTACTACATTTTCACCTCTATTATTTCCTCAGACTAGCTTATCAT

ATGTCCCATCCTGGAGTTCTTCGAGATCTATGTAGTCCTTTTGTATTTCTTTTAGTTTCCTTAGATAGGGTGGAATAGTC

TCATACATTATGAAAAATATGGGTTATGTCTTTCGCAAACTGTTTTTGAAACCTAAGATCGCATAGGCCTTAGGGAATCC

TCCGATTTCAACACATAATTTCCTCCAGCGAGTGACCAGATATCAGAGGGACTCATTTGGTCCTCTAATTAATTGGAATA

ACATATAAATTCCTGGTTTAACTATGCTATTGTGCATATAGGCCTCTAAGAATAGCTCGGACAAGTGAGAAAAAGACTTA

ACTGTCTCTCTCATCAGGTTGTTGAACCACATCAGGGCTTCCTCATTTAGGCTGTATGGGAAGAATTTACACATCATGAC

TTCGTACTGGCTCCATTGTGTTAACGTCGTGCTTTAGGATTAGAGGTGTTCAATTGTGTCGCTAGCCCATTGAGGGGAGT

TGCAAAAGTCGAGAGAAAACATTTCCTTGTAAATGGTTTGGCTAGAAGACTCACAGCTAGGGGCGATTGTCCTGCTTCCT

GAATGACTTCTGCTAATTTCTGGCCTCATTTCCTTTGTACATCGCTTCAAGTTGGCAGAGTCTGACCTACAATTGATCAT

CGATATGCCTCGTTTGGACCTTCTTTCTTCGGATTTGTTCTCTTTAAAGAGAGAAGATAAATCCTGGATGGAGATACCTT

GATCGCCTTGAATCCTTATCGGATTGCTCAACATATGATATTTGTTTGGAATGAGATGACTTTTCTTTTAAAGTCTCCCC

TTGCTTTGGATTTTATCCCGCCGGGTTCTTGGGAAGGCCTTGGCTTTAGTTTCTTCCTCATTTTTATTATTTTCTCGCTT

ATGCTTTGGATTTACATCAGATGACCCTTCCGAGGGACTATCCTCATCGTTTAAGACTGCCTTACTTCCTTTTTGGAACC

GATCTTATGTCTTCTCGTTCGAGGTATCTGGTATGTCCATATGTTCGTCGTCTTTTAAGTTCTCAATAGGATTTTCCTCC

TCGGGGGTCTCCTTTGGGATGGTAAGTGGAACCTTAATCCTCGTCCTTCATTTCAATCTCCAATTTTTCTTTTCCAATCT

TTTGTTTTGTTTTTCTAGATTTCTCATTCATTCCTCCTATACCTTTTGTGCTTTCATTATAGCTGCAAGGGCAGAGGTGT

TGTTATCGAGGCTAGAAGAGTCATCCTTCTTTGCCTCATGGTTTTCCATTTGTTTTGGCAAGTGATTCAACCCTCTTGGT

TGATTCAAGTGAATGTCATCCTGTCATTCGAAGGAATTTTCCTCTTCATCGGAGCTAGAATTATTTGTTAACCCTTCCCG

GTATTGGTTATCGGGATGTCCCTCACTTGGGAGCACACTATTGATCTGAGCCTCTCCTGGATCCATTGTGTCAAGTCCTT

TGGATAAATATATACCAATTTCTCCTTGGCTAGTGCGAGTGGATCGTCTCTTCACCATCGCCTTACCTGCGATATATGTA

ATAATGCTTAACTTGTCTTTTCTTTCAGAAAAGAGGTATTAGAGCATTATTCGGTCGAACTCGCAAGCGTTGCTATGTCA

AGCTTGTTTGTCAATATTAGTGATCAAAACTATAAGTCTTGATTTCTAATCTACTTATAGTAATGTCTTGGACCAGGACA

GATTGTGTAGTTGAGCATTAGACTTCACGGCGGTCATTAATTGAAAACGAAAAACTATTAAGGAGAGCTTGTGGAACTTC

ATCAACAAAAGGTATGTGGAGACTGAAACTCATCTATCACTTGAAAAGTCTAATTCTACTCTATCTCCAACATTGAGACA

AAAGTCGTATTACTATATAGTTGTCGGTTATACACATTTGAGATTTCGAGCTGAGTTTATCTTTGTTACATATTTCTCGA

AATATGTGTCGTTAAGCCTTTGTTTCAACCAAGTTCATCTTATATTCTTGACGAAATTCAAAACATGATCATGTGAAAAT

CGTCGAGTAACATCTTACATGGTTTGTGTGATATAATCATTTGGTGCAAACTTGGACTGTTTTGTTATGATTATTTCAAT

AGCTTGAAAATTGCTTTGATGCTAATAGTGTGTGAAAACGGCTATTGCCATCCTCTAAGAAAGTTTTAATAATTGAAATA

AGAGTTTATAATACTTAACCATCATTGGATTATACACATATTGTGCATTCTTGCATATATGTGATCCATGACCGGAACTA

AAGTATGCATACCCGTATGCGTATTTGTAGTTGTGAAATTCCGTTTACCAGGTACATATACTGGTACGCATACTTGCGTA

AGGTATAGGTCCGGGAATTTCTGTTGTGTTTTGGAAGTGTACTAAGTATGCGTACATGTTTGCATACTGGAGAACCCAAA

CTCAGACCGGCTACTTAAATATGTGTACCCGTTTGCATACTTGAGTGGTTAAAGTTCTAAAATCGGTTGGCTCATGAACT

AATACATTTATATATTAAGGAATGCATTCTTTGCAAACAGTGGCTATATTTCTCATGAATTTATTTGAGTAAATCAAACC

GATTTTGCTTCAATTGTGTTCTTGTATACTTCTATGAGAATATAGCAATTGAACAACTCTTTAACTAGTTTCATTTGAGT

CATTTGAACTAGTTATGGTTAATAAGAATAAGGTTGATATGAGAGTGTTCATATAGCTAACCTCGGTTAAATACGGTTGA

GCCAACATGGTGTACACGTTTAGGTACGTTTACTAAGCCTAAATGAGGGTACATTTCATTTTTGTATAACAAGCTAAGTT

CGATCTAACGGTTAAAAGATATTAGATTGAAGAACATAATTGTAATAATCCAAGCTTTTGATAATAACAAAACTTCAATT

AGTTAACAAATATTTATTTAGATTTGGAAGAATAGAATTGTAATCCAAGCTTTTGATAGTAACAAAACTTTATTTTATTT

TTAACAAAGTCTACTTTACATGAACACTTCAAAAGAACAAAAGAATTTTGCTTGCATCGCTATGCAAATAACTTAATAGT

CCTACAAGGGCTATTAGCTAAAAACAACTAAATCCTTAAAGAAGTAACTTCAGGGCCTCTTGCAATGCAAACAAGCTAAA

AGCATCTAAATCTTAAAGAAGTAACTGCCGGACATCTTTCTATGCAAATAACTTAATAGTCCTACAAGGGATATTATCTA

AAACATCAAAATCTTATAGAAGTAACTGCAGGGCCTCTTGCTATGCAAACAAGCTAAAAACATCTAAATCTTAAAGAAGT

AACTGTCTCTTGCTATGCAAACAAGCTAAAAACATCTAAATCTTAAAGAAGTAACTGCCGGCCTCTTGCTATGCAAACAA

CTTAATAGTCCTACAAGGGCTATTAGCTAAAAACATCTAAATCTAAAAGAAGAAACTGTCGGGCCTCTTTATTCGCAGCT

GATTGGTTTTTACATTTTGATTACTTTGCATTTTGGTCACTTATAACTTTAAAGGTCATTCGGAAAATGGGTCATATGAC

CACAATTCATAAAAGTGCGAGTGTAATGGACTTGTAAAATATAGGTCTGGGTGAAAAAGACACAACAAAACTTAGTCATA

TATACCCTTTACTTTAAACTTTATTCTTTATCACATCCTTTTCCCTCTCTTATTTATTTTCTTTTTCCAATTTAAAATTA

ATCGCGAGAAGGGTCTTAACTTCTGAAATATAAGTCGGATTTTAATAAAATTTACTCCCTCCGTCCCAGTATTAAGTGAT

CTAGTTTAAGTTAGCACAATTTTTAACGCAAGCAAGGAAAATAGTACTTTTAAGCATTTTTTACAATTATACCCTTATGG

ATAATAACTAGTGAAATTTAGAAATCATTTATCTCTCAAATACTCCACGGTTGTTCACAAACTTCATACCATTGAAAAGC

ATTTTAAAACACCTACATAAAGAATATAAACATGCCTATCAAATTATGCATATTTCTTATATTTTTTATAATTAACTAAA

AGGATAATTTTAGAAATATCTCCTTGTTTAGTGATGTAGGTCACTTATTAGTGGTACAAAATCTAAAATCAAATAGGTCA

CTTAATAGTGGGACGGAGGGGGTATATTTTTGGAAAGTGCTCGAAAATCTCTACAAAATAAGATCCATTGTTGATATGAA

ATTCGAAAAAAAATAATATTTTATTTAAAGCCATGTGCAACCAGGCAAAAATTTATTTCCTATAAAAATATGCATTATGC

ACCCAGATGCAACTTGGCAAAAATTATTACATGTAAAAATATGCATTATGCACCTAGGTGCAACCTGACCAAAATTTATT

TTCTATAAAAATAAGCATTATACACCTAAGTGCAACCTGGCCAAAACATTTTCCTATAAAAAATGCATTATGCACCTGGT

CAAAACATATTTCCAATAAGAATATGCATATACAATATACACTCGGGTGCAACCAGACCAAAACCTATGTCCCTCCAGAA

TATACCTTATGCACTCAGGTGCATTTTGTGTTTTTCATGGATAAATTTAGGTTTATCCTACTAATCAACTACAATATATT

ATGTGCAATCAAATCACTAGTTCATGATCTCAAAAAAAAAAAAAAAAATCCAGCACCATTACCATCACATTCACCATTAA

AACATCAACTAAATATAAGTACCATATAAGAAAAATAAGACGAACAACAAAATATTTAAAAATAAATAAATAAATATATA

TATTATGCACTTGGGTGCATTTAGACCAAAACCTATTTTTCTGACAGATTATACACTAAACATTCAGGTGCATCTTATGC

ATTTCATGGATAAATCTAAGTTTAATCTAATAATCACCTGCAATTTATTATACAAAACTAAAATCACTAATCCAATATCT

CAAATAAAATAATCCACCGCCATTATCATCACATTCACCATAATAAAAAGGTCACCAAACTTGTAAGAAAAATCGTAGCC

GGGATCAATGTTCCCATGTATATATGCCAGTGGGATATTCAGAGATGATATATCCCCAATACGTAATGACTGAAGGTGTT

TCATACTAATTCCAAAAGCAACATGCCAATAGCCTTCCACTGCCACATAAAAAAGACAAAAGTAAGAAGTACTCACCACT

AAATAGATAGATCAAAATAAGTACATCATGAGGTGCAACAATTAAAAACCAGTTGCAGGCTTTCATCACATGGTCTAAAA

TCCAGAAGTATTAGTGTCTCCATACCGATATGAGTGGACTAACTAATCTTTCTCGATATAAATTTCAGTCCCTCGGAGCA

AATTAATAATTAAAATTCAAACTATTAACCCATGACTTTTTGCGACAATGTCACCACTTATTACTACTGAGGTGTACACA

CAACAAACGATGAAGTATTGTTGACTTAAAAGTATATGTATGCAAGTATGAGTATGCAACTATGTGGATTCTTCCAGGGG

AAATTTCACAAACATCATACATGCATAACCAAATTCTCAAGCTAACTTAAGCAGGCAGAAATTCAAACACAAACCCAATA

TAAGAATTAGAGTTTACAATTAAAACTCATCTTAAAAACACATCTTAGCCAGAAAATATTCATGCACAAAGAAAACTAAC

CAAACGAATCCAAATGCAGCTATCACCAGAAATATTAAAACCTTAATAGAGGAGTTTACCTGATGGATTGGTTCTTCAAT

TCATTAAAACCAAGTGAATCCCATCTCCTTACTACCCACGTGCCATTATATTTCAACATGTTTTAGCCTAACCACCTGCA

GGACTTTTTCTAAACCACTAATAAGTACGTAGTTTGCAAAATAGTAAAAAAGAACAAAATACTTATGACTAATCATCTCG

TTTTGCTAAGCCACACTATGGTTATAAATTAGACACTCTCAAAGTTGGCAAAAAACCAGATGCAAACACAACCTTATTTA

AATGTTATTTCAATTCCCTAAGATAAACAGGTTCATCCAACAAGGGATGTTATACACCCCATCTCTGCTAAAACATATTA

TAAAAAAGAAGAACTAAATAACATATTATTAGGGAAGACAATTATTTGACGTTAGAACTTGGAGAAAAGAAGAAGTGACT

CTACTTCTCAAATCCCTAGCTAGACTCTTGTAACCTGCTAAGTGTTCAGGTCACCTTTCCGATGTGCGATGAATTAATAT

TACTCTTGGTTCTTAAAATACCCTATCGCCCTTCAATCAGTACGGGTAGGGACTGGAGTAGGAGCAGTAAATGTTGAGGG

CGTTGTGGAGACATCTATGGGTGGAACTATTGCAGTTGATGAGAGGGTGGAACGACGGTGAGTTGGAGTGTATATATGTA

GTGCAATAGGTTGGAGAGTAGAGGTTTGGTTGACAGTAGGAGCAGTAGATGTTGACGGGTAAGAGAGATTACTAGTGTAT

AAGGAATGAATGAGTGAAGGAGACGAATCAATATCAGAAAAATTATTTGGAGGAGGGTATGTTAAGGGAAATACTTTTTC

ATCAAAAATCACGTGACGAGAAAAAATGATCTTGTTTGTGGCAAGATCTAGGCAGCGGTATCCACGGTGGTTGGGTGGAA

AAACAAGAAAGACACACCTAGTGGAACGAGGGGCAAGTTTGTGAAGAGTTGTGGAGAAGAGATTTGGATTGCAAAGGAAA

CCAAATGTACGAAGATGGTCATACGTAGGTTGACGTTGATAAAAAAATAGACACATGCGAGTGAAAATTACGGATTTTGG

TGGGTAGAGGGTATGGAGATAGACAACCATATGAAGTGAGTAAACCCAATATTTGGGGGGAACGGAAGCATGAGACATAA

GGGTGCGAGTGATGTCATTGACATGGCGTATCATGCGTTCAAAATTGCCATTTTGGGAGGAAGTTTGGGTGCATGAGAAG

TGAAAACAAAGACCATTATCACAAGCAAAGTTATGGAAGGAGGAGTTGTCAAATTCACGACCCATGTCGCATTGAAAACA

TTTAATTTTACGTTCAAATTGAGTTTGAACAAAAGAACGAAATTCCAAGAACTTGTCATAAACTCCGGATTTAAACTTCA

AAGGATAAACCCAAAGATTGTTGGTGAAATCATCTAGCAAGATAAGATAATAACGAAAACCAGTATCAATATGTAATGGA

GAGGTCCATAAATCGTTGTGAATGATATCAAATGGTGTAAAAGTAATGGAATTTAATTCATAAAATGGCAATCGAACATG

TTTACTAACTTGACATGAATGAAAAAGTTTAGAATACTGATTCTTATTATAGTGAATAGAATTATTGGTACGTAAGACAT

CTAAAATAGCCTTGCGGGGTGGCCAAGGAGCCTATGCCAAATGTCTGGCGAACAGACAACAAGAGAAATGAGGGAAAACT

GGGACGATATGGGAGGTGGCACGACATAGTTGGCGAACAGACAACAAGTGTCATTGTTGATCTTAAATATGTGTTGATTA

TCATGAATTCTTGTGAGGGATGTATTGTGGGATTTCATGCAACTACATAATGACGCTAGAAACAGGGACTGAAGGTTGAA

AATTGTGGAAAAACAAAATAAAAGATTGTTATTAAGATCGCTTAAATCAAAAAAATGATGATACCAACCGGTAAAACCAG

TTAAGGGAAAAATAATCAAAATCAATGAAGGTACCAGAAAGTTGGAGTGTAGTTTAATCACAAAAATAATGATTAATGAA

TTTCATGAAAGCATTAGAGGAAGAATACATAAACAAGATTTTGAAGGAAATAAAATTATGACGGTAGAGAAGGTAGCAAT

GCTGGGAAGGCAATGGATACACGCAATTAAAGACGTAGCGTCGACATTGCATCAATGCGTCAAATTTCCTATGCCGAATG

AGATTGGTGAAATTAGAGGAGATGTTGATAAAGAGAAATTATGCCATCAAATTGAAGTGAAGCATTATGAAGGACGTGCA

AAGAAGCGAAAAAATCGCAGAAAAATGGCGAAAGAGGCAAAGAAAGAAGAAGAATTTAGAATTTACATGATAAGAGTAGA

AGAAGGAAAAGGAATACCCATCGACATTTAAGAAGAAAGAAACTAACTTGCGAAGGTGATAAAAGAACCAACACCCATGG

GAGAACCTAAAGCCATTTACACCGCGGCGGAACCAACAAAGAAAGTGAATGTTGGAACTAAAGAAGAACCCAAAATGCTA

AGGATTGGGACCAAGATGGAGGAAGATGAAGAATAAAGAACTATTAATCTTTTGCTTGAATACCAAGATATTTTCGCATG

GAACATGGATGAGATGCTTGGACTAGATCCGCCTGTGGCGTGTAATAAGAAAAATCGCAACAACATATCATCCAAAAATA

TAAGAAGAGCTGCAGAAGATGCTTGATGCAGGAAATATAAGATAAGCCAAATATCCGGAGTGGATAGAAAACATGGTTGT

GGTCCCAAAAAATAATAAAGGGATAAGAATTTGTATAGACTTTACTGATCTAAACAAAGCTTATCTGAAAGACAGTTTTC

CATTACCAGATATTCCACAAATGGTTAAATCAGCGGCGGGAAATGATAGAATATCTTCTCTAGATGGGTACAAAGGTTAC

AATCAAATATCTCTCGCTGAAGAAGATCAAGAGAATATTGCTTTTTTCGCTCCAAGAGGATTGTACTGTTATTCAAAAAT

ACCATTTGGTTTGCGAAATGCTGGAGCAACATACCAAAGGATGGTAGAGAAGGTGTTCACGAACTGGATACACAAAACAT

TAGAGGTGTATGTAGATGACATGTTAATAAAAAGCAAAGAAGCAAAAGACCATGTGCAAGATCTGAAGGAAATTTTGAAC

AGATGCGGAAATTTTGAACAGATGCGGAAATGTATCTTTGGGGTTGCGTCAGGAAAAAAATTAGGCTACATTGTATCGAA

AGAAGGAATACAAGTTGATCCAGATAAAGTGCAAGCAATTCGTGACATGCCAACACCAGCGACAGTAAAGGACGTTCAAA

ATTTAAACGGACTCTTAGCTTCACTAGAAAGGTTCATCCCGCGATCATCAGACAAATATAAACATTTTTTCAACATACTC

AGAAAGGGGACAAAGTTTAGATGGACAGATGAATGTGAGAAGGCTATTTAAGGAATAAAAGAACATCTGATAAATATATC

TATTTTGCAAAAGGCAGAGCCAGGAAAAGAATTATTAATCTATCTTGCGACAACTTCGCAGGCATTAAGTGCTGTGTTAT

TAAGAATAGATGCAGGAGTTAAAAGACCAATCTACTACATCAATAAAACCTTCAATGCTGCAGAGAAAAATTATTCAAAA

ATTGAAAAACTAATTCTGGCATTAGTTTATGCAACGTTTAAACTTCGCATATATTTTCAGGAACATAAAATAAGGGTGTT

GACAAAGGTACCAGTTGAATCAGTGATGAAGAATTCAAAAAGGTCAGGCAGGGTTGAGAGATGGAATGCACAAGTAGGAC

ATTTCGAAATCAAGTATGAAATATTATCTTCACCAAAGTCACAAGTTATCGCAGACTTCTTAGAAGAATTTCCTATAGAA

GATGATGAAGCTGTGGAAGAAATAATGGATGTGGAAGAAGAACATGGAAAACCAAGAGATCTATTAACATAGACCAATAG

ATGGGAAATATTGGTGGATGGATCGTCAAATGGGGAAGGTAATGGAGTTGGAGTTGTTTTGATTTCGCCAGAAGGAACCA

AAATGACTTTTTCTTTTAGATTGGAATTCACATCCACTAATAATGAAATAGAATATGAAGTTGTGATACATTCCCTAAAA

TTCCCAATAGAAATGAAATTGGAAAATGTAAGAATCACTAGTGATTCGCATCTAGTCATTCGTCAGATAAAAGGAGAATA

CACTACAAATGAACCATCCTTAAAGAAATACAAGAAACTTGTTGAAGAATTATCAGCACAAATCCCAAAAATAAAATGAA

AACATATATCAAGGAAAGATAATAGACTTGCAGATGCCTTTGCTTTCATCTCAAGTATGATGACAGACCCAGTTGCGAGA

TGCATAAAGATACATACACTTCTAGCACCATCAATCAATAAATAAGAAGAGGAAGAGGTGGACGTAATGATAATATAAGG

TGGAGAAGAAGAACAAGTGCCCAATATCGGAGATTGGAGAGCAGAACTTCATGCATATTTGGGGAAATGAGAAATACCAA

GAAATAGATTGGAGGCACACAAACTAAAGATATGTGCGAAAAATTATGAACTAAGAGATTGGGTATTATATTGAAGATCC

TTTAATGGACCATCACTCAGATGTTTGACAAGAAAAGAGGGAGAGGAAATATTAAAGGTGTTACATTGTGGGGACGCTGG

CAACCACAATGGAGGAAGATCTTTGGCGTACAGAGAGAAAATGCAAGGTTATTACTGGCCATATATACATGAGGATGCGA

AACAAATATCAAGGAATTACGAAGATTGTCAGCGGCACGGAAAGAAAATACATGCATCTGGAGCATCATTAACATCTTCA

ACAAGTGTATGGCATTTTGGAAAGTGGGGGTTAGATATTGTGGGACCAGTTTTGCCAGGATCTGGACAAAGAAGATATTT

TATTGTCATTGCAGATTATTTCACAAAATGGGCAGAAGTAAAGGAAGTACAACATATTCGCGACAAAGACATTTTTACAT

TCATATTTGAAAATATCATTTGTAGATTTGGGATCCCTTCACAATTGGTACATGATAATGGGAAATAGTTTGAAGAGCAT

AACATAGAAATGTTACTCAATGAATTTAAGATAAATTGTGGGAAATCAACACCTCTATATTCGCAGGCCAATGGACAAGC

AGAAGCAACAAATAAGACAATCACATATATATTGAAGAATAAATTAGAAGGAAATAATAGGGCATGGTACGAACAGGTGC

ATAATGCAGTATGGGCTTATAACACAACTAGGCGAGAAGCTACTGGAATGTCACCTTTCTGTTTAACATATAAAGTTGAG

GATGTGCTACCAACAGAAGTTATTATTCCCACAACAAAGAGTGAGGCTTGGGAGAAAAACCTTATTGCGGGTATGATTTT

AACCAAGCTTGATGAATTAGAAGAAAAGATAGAAAAGGCTCTGCAGTATATGGAAAATTATCAGCGAAGGTTAGCCCGAG

AATATAATAAACGTGTCAAAACACGTGAGTTTCAACCAGGAGATTTGGTGCTGCGAGAAACACCAATATATCAGCGAGAA

CATGGTGGAAAACTAGCGAGAAAATGGGATGGGCCATACATTATTAAAGAGATAGTCGGTACAGGGGCATATAGATTGAT

GCATCCTGAGGGTAAAGATGTGGGTCATAGACTTGATAGACCATGAAACAAATTATACTTAAAGAGATACTACCCATAAG

AGGTTCTGGAAAAGAGTATATTGTGAAATGATATCTGATTGAAACAAAATGGATGTATGAAAAAATCTTATGAAAGAGGA

ATTGCGAAATTATAAATATCCAGAAAGAAATGAAAATACGCGAAAATTTTGCAGAAATAATGATATTACCAAGAGATAAT

CAAAGATAAATGATATACTAGAAAGGGGCGAACCTACTATGTCATACACCCTAGTTCGAAGGATCAAAATTTCGCAAGAG

GCAAATGTAATACCTAAAGTACGTCATAGAAGGGAGTACCTGTTGCATGACTCAGGGAAAAGGTTACACCGCCCCTGGAA

CCCGAGTCATGGAGATTTGGGGTCAATAAAGTCCATCCAGAATAGGTACTTTAGATTTTCAACCTAAAGAGATAGTCGGT

ACAGGGGCATATAGATTGATGCATACTGAGGGTAAAGATGTGGGTCATAAACTTGATAGACCATGAAACAGATTATACTT

AAAGAGATACTACTCATAAGAGGTTCTGGAAAAGAGTAGATTGTGAAATGACATCTGATTGAAAAAAAATGGATGTATGA

GAAAATCTTATGAAAGAGTAATTGAGAAATTATACATATCCATAAACAAATGAAAATACTCAAAATTTTTGCAGAAATAA

TGATATTACCTAGAGATAATTAAAGATAAATGATATACTATAAAGGGGCGAACCTACTATGGCATACACCCTTATTCGCG

GGATTAAAATTTCTCAAGAGGCAAATGTATGACCTAAAGTACGTCATAGAAGGGGGTACATGTTGCATGACTCATGGAAA

AGGTTACACCACCCATGGAACCTGAGTCATGGAGATTTAGGTTCAATAAAGCCCATCCAGAAGAGGTACCTTGGATTTTC

AACCTAAGATTATGACTTAGATTGGAAGGCTGAGGGTCTCTTCTAAGGAGTGAAAAATGTGATCAATGCGCCGAGGGGCA

CGTTTGAGCTAAGAGTGCCTGGGACGTCTGGAACGTACCTGGCCATTCTTGTAAAGTCTTGACCATGATGTACTCGGCCT

GAAGAAAACCCCACTTACGGTGCAGTCTCATAACCATAAGCCATATAGGCAAAATGGATAAGAGGCTGCGAAAACAACTG

GTTGTTGTTAAGACCGGATTTGAGGATCCAACCTTGTATGGTAGAGGTACGTCTCCTTGAAGGGGAAACCAGGGGGAAGA

TAATGGCGGCACCTTCCATTAGGGAGATGATAAGTGTCTTAAGACGCAAATGTCATATGAGGCTTTATATTCGCAAGAAT

ATTTAGGCTTGTCATATTTGTGCAATAATCCTGAAGAGGCAATAATACAAAACAGAAAGATATGACGAGATCAATGGGTC

AGTACATGAAAGTGTAAGGAATTCGCAGGACGACAATATCATGAGCCTTTATTTTCGCAAGAATATTTAGACCTGTCATA

TTGTCGTGACAAAAATGATAAAAGAAAGAAGTTAAGGCAAGATCAATGGGTTTCTTCATGAAAATGCAAAGACGTCCTGC

GATTTTGCCGCAATGACAAGAGGTGGAAAAATAAGACCACAATTATTTTAACATTAACTATTATTTATGCAGATCATTCA

TCGCAAGAAAGATAATGAGAGGCAAGTATGGGAATTAATAAGCAAGAGAAATTATCTTTCTCGCCAACAAGAACAATACT

AAACGAGTAAATGAATCTCAAAGTTGACTGAAGTATATCATTCACAAAAAGTAGGAAAGATGAAACAACTTAAGAAACTA

AAATTACACAAGAAGTTCAGATTTCAACATTAATTTCTTCATCAACGTTTTCGCTAGTTTCTCCACTTTTGCTATCTTTT

TCATTTTCGCCAGTCTTTTCTTTGTTCTGATCTACTTCAACATCAATTCTCTCTTCATCTTGGTTTACTCTAGCTTCTTC

TTCATTTGGCTTATCATCAACAGTTGCTTCCTTAAGAGGGATATCTTCCTCATTTGCAAAATATTCTTGAGAGATAGTCT

CTGCACCACTTTCGTAATCATAGATAGTCACTATCAGCAGCACGAGGAACTTCATCATCATCTATTTCCAAATGATTGAT

CGATGTAGGAGGAAGAGATTTGGATAGTAAAATTTCATTCACAAGAACTTCAGCCTGGAGATGAGCTTGGCGAAGGAATG

ATTTCTCACGGTTTATAATTTCCATATATTTGAAATAAGAAAGATAATCATATCTTCTTTCTACTCGATCGCGAGAACAA

GGAAGGGTAGAGACAAGTAATGCATTCTCTTTGCAAATATTGGCAAGTTGAGAATCCAAAATAGAAATGGTGGAATGAAG

TTTTTTCTCAGATTTTGCGAGAAAGAAGAACAGAGAAAATTATTCAGGATTGTGCTTCGCAAAGAAAATATCACTAAAAG

ATTTAACAAAGAAAGGGTGATTGAGGCAGTCAAATTTTTATGACTACCTTCCTTTTGCGAAATAAGATATTGCGTTAAAG

CCAAATAACTATTCTCCCAGTAATCCATTGCATCATCAAACTTATCTCCAACTAAACCTTTCAGGCGAGATTGAAGATTA

TTTTCTTTAGAAATGAGTAAGGTGTTCTCCTTCGCAAGAGAAGTGTATGATTCTTCTAGTTTGGAGTATTAATCTTTGAG

TTGACTTTCCTTTATAGTTAAAGTTATTCTACCCTGGATAAGGTTTTCTCTCTCAGCGGTAAATGATTCTGTTATTTCCT

TAAGTTTATCCCTCAATGAGTGAAGAGAATCTTCCTGATCATCACATACTTTTTGGCAAATGATAAGTTGCTGTGAATAG

GTCTCCACCTTGTTTAAGAGAATTTGCTTCTCCTGAGATAGGGTTTTATTCCCATCTGATAAAGTTTCCAACTCCAAGTC

AACTTTTGTCAAAGAATAAGAGAGGCGAGATATTTCACTACTTAATAAGTTTTTCTTTTGAACTAATTGAGTATTCTCCT

TGGTAATATTATCTATTTGATCGACAGAGTATGTATGAAGGTTATCCAATTAGTTATATTGATCCATCCAAATATCTCTA

TCCACACGGGCTTCTTCAGCGACAAATTCGACTCTCCATTATTCGTTTCTGGTTTAAAACTTAAAGGGCGAACAAAGGAT

CTTAAGAAAATTGAATAAGGAAAGGGAAAAACTATTAAAAAGGGTAAATCAAAGGCAAAAATAAGGAAATTATACCTTTT

AATTCTTTGTTCTTCTTGCAAAGACCATCACAGTCTAGCGAAATGTTTTGAAGCTTTTGGTTCTCAAAATGGAGAGTGCC

ACATTCCAAGGTCATGGATTGGAGTTGCTTCTTTAAAGATTCAACTTCACTTTCTAAAGATGTTTGCGAAATCATTCTAT

CAGAAACAAAGTACTTACTCAAAATTTCGTAGACACCAAATTTAATAGGATCAACGAAAGATTTCTTTAGAACGTCCATC

ACATCAGACAAAACCTTAAAATAAATATCTATCCCATCTAAAGTTTCTTTAGAGAGAGGGAGAGATTCATCCAAAGAGGA

TGTGATAATAGGAAGTTGGGGTACTGAATCAATCTTTGGTTTAGATGGTTGAGGAAAAGTTTGAGGAAGAATAATGATCG

AGGTTGAGGTGGAGTCAAGTTGAGGGGTATTGGTAGTTGGAGAAGAAAAGGTTTCATTCGAAACAAGATTAATGAAGGGA

GTCTCATTCATAATAAGATCAGTCGAAGGGGTAGAAGCAGTAAAGATATCTTCGCAGACAGGAGGTAAGGATTCAACCTG

CGAAGGTATTACGGGATACATTGAAGGGTTAAATATAGAAAATGGGACAAAAGTTGCAATAGATTTAAGATGGAGAGATC

TCTTGGGGGGTTTATTGGTATCTCTATCACCCTACGAATCACAATCCAGATAAGAAACATGCAACAATATTGTTATTTAA

AAAAATAGTGTTTCTTACTACCTTGGTATTCGCAGGCTTTCTTTTACTAGCAGCAACTTTGAGCTGCGAACCCTCGGGAA

CATTAGGGTTTCAGACTGTAAATTCGGTGTTGAAGGTGCTTTTGCTGAAACAGAAAAAGTATGGGAATTATATAAAGTTG

AGCAAAATTAAATAAGGAGATTAGTTCCAGCAAAATCAGAAGAAGAAAAGAGGAAACTAACCTTGATGACTCCATGAAAA

CAGTAGCAGAAAGATGAATCCGTAAAAATTAGAGAAAAAAGTTACAGACAATGAAGATTTACAGAGAGCATGGAGGTGAG

GAAGATGAAGTAAAAGAAGAAAATTATCTCTCTGCTAAATCTCTTCTTAAATAGACAAAAAAATGGTGACCGGATAACAT

AGAACGTTCCGGTGAGCAAGGGAATATGTGAGATGTGTAGGCGGGCTTAATTCATGGAGTCATGTCGGTAGAAAATTTTG

CAGAAGATAAACAGCTGCGGCATTTTGAAGTATAATTTATCATGACATTCTCTCATTCGCAGAGAAGACATGATAAGAGG

CAAGATGTAGGATCAGAATATCGCAATAAGATTACTCTTGCGAACTCGTCAACAACAATACACCTGCAAATTTCGCAAGA

TGATTACAAAAGTGACACAAAAGATGACGTTAGCATAAACACGAGAAAGATGTGAAGATTTTGCGAAATCTAAGAATGTT

GTGAATAAAATATATTACGCTTGCGATACCAGATCCGAAAATAAAGGACAACTTAGCTGTCATCCACTATGTAATATCCT

ATATAAAGAGTAGAATAGGCTGAAGAAAGTGGGAGATCAATTTTGGAGTGTTGAGAAAGTTTAGAGGAGAGAGAAAGTCC

AAAACAGTGTTGTATTCTTGATTTCATTTCCTTCTTGTAAAATTTCTAGATTCAATTAATAAAAAGTGTCATTGTTGATC

TTAAATATGTGTTGATTATCATGAATTCTTGTGAGGGATGTATTGTGGGATTTCCTAAAACTACAAAACCCGTGAGGAGT

ATTCATCTTGTTTGATATCTAAGAGAACATGGATTAGGGTTTTGATGTTGGTGACGACAAAAGCCGGGTGAAGAGATTTC

TTCTCACCTGCCATTGTTCAACGATTTAGAAAAAGAGGGATCGAAGGGTGGCTGATACCATCTTTGATTGGTGATAACTT

CCTTAGGGACTAAAGATCTCCAGGGTTCAGTATTATATAAAATCAATAATATTTTTTGGGAATACAACAGTTATTGTCGG

AGATAATCTCCTTATCATAAATAAGGTAAACATGATAATATAACCCCTACATTTGTTTGCGGAAAATCGTTATATATATA

GTAGATGGCTAAATGAGTATCCAAATTTAAGATAATGATTAGGGAAACATGGTGCTCATATTGTACCAACGACCCGGCTT

GTACTGTCGCTAGTTTGGTTCCCGCCAAACATCCTTGCCAATCCAAAGTCTAAAATTCTTGGGACCATATCTTAATCTAA

TAAAATGTTACTCACTTTGATATCTCTATGAATAACTCTTAGTCATGAATCACAATGAAGGTAAAGATGGCCACGAGAAA

TCCCTCCAATAATATTGAACCGTTTACCACAATCTAGGCATGCTTTGTTCCTTGGACCTGCAACAAATTTCAAAATGTTT

AACATGCAAGGCCAGCCTTCATGAAGGTTTCGGTATTTTATGTTCATAGACGAAGGATTATTTGTGATAATAAAATACAA

GAAAAGAAACAAACCAAATATAAATACATCTAGTATTTTGTTGGGCATGTACTCATAAATTAAGATGTTCTCTCCTCCTT

GGATTCAGCATCCAAGAAGCTTCACAAGGTTACTGGGTTGAAGTTTGGAAATCAATATAACCTCATTTTTGAACTCTTTA

ATACCTTGTCCAAAGCTTTTCAACAAGATTTTCACTGCTATTTGCTTCCCGTTTTGTAATTTACCCTATAAACCATCCCC

CAATCAGTGAGCATATAGCGATTATAGTAAATAGATTTATGCAAAAAAAATAAAAAATCATCATAATTAATACAATAGTC

AAACAAACCTTGAAAACTGGCCCAAACCCCCATTCTCCCAATTTATTTTCCAAGGAGAAATTGTTTGTAGCAGTGGCTAA

GCAAAAAAGATCAAATATTTGCAATTCATGTGTTTCTCCTTCCGGTCGGTAAGCATGTTTGTATTCTGAGTGTTGTTGTA

GGTAGCTTTGGTTATCAACAAATCAGTCAACACCCCCTGGAGTTCTATTAAACTTCCTGGGAAGCCATATGAACATGAGA

AGGAATTGAAGAGATGATTAGTCAAAGTAACTCATAAATACAACTAGCCTGAATTCAAGGGTGAGCATCACCAAAATAGT

TCTCCTTGTATTAGTCCATAATAAGATTAATTGTTTTTTATGATGTAAAGCAATGTACCCAGCCAATATAAAGGAAAATG

TATTTACCTCTTTTATTAGCCTTATGTCTATTGCACCAATACATGTACCCTAATACGCCCATAATAACTGCCACCAAAGA

GATTATTACAAGTATCATGATCCTCCAAATTTTTAGCTTTCTCTTTCTAACTGGATTTCCGACCGAAACTTCTAAAGAAT

TAAAATAAGAATGCAGATCCATGTTAGCGCTAATTGATGTCAGTGTTTTATGATGTTTTATTTTCAGTTTAATAAGTTAC

AGTCAGTTATGTGGTTATGCACTTGTCACATAAGATTGCAAGTCTTAGATGCATAATTTCGGGAAAAAGAATATTCGACT

ACTTACCTTTAGGGTGATTTTCATGGAGAATTTCTGAAGATGCAAATCTGAGATAGAGATTCCTTGCTTTTCCATCTGAT

TTCTGTTTGATACCCTGTATATTGTTCGTATATACTTTGGCAAGATAATTTGCATTCTTCCGTGCTATTGACTGGTTGAG

GTTGAGAAATATCATGAAAACTCGAGGTTGGAATTCGTTTAAAAATATCTTTATTCTTACATTGAAAAGGCACCTTTCTC

TGACTCCCACCAGTTAAATCCCGAAGATTCCAATCTGAAAGATAATTTTTAACGAAACCAGAGTTATGCCAAGTTTCCTC

ACCATAGCTTCTACATTATAAAATGCATATATCGGTCAATAGAAAGCGAGGGTTTTATCACCATTCAATATGAGAGCTTA

TTTTGTCAAGGCTTTTAAGGGAGAAGATAGGCGTCCGTAGAAACGGGACTAACAATAATTATACATAGATGCGAACAAGA

TTTTTGCACAAGTTTTATATAGTTTCTGACATGACCCAAGTTCTGTGGAGCTCTCCAGCTCACATAATATACCATATTGG

GTGGTAGTGTATATGGTGATATGATATGTTTACCACATTTTGTATATATAATTGGAGGACAATTTAAAGAACTAGGGTCA

AAAGTAGAACATAGCTAGAATTCTAGAGCTTTTCGCCCCACTTAAGAGAAGTGCCAGATCAAAATATTTTCATGTTTATA

CGGAATCTCAGTGGTGTAAGAACATGGATTTGGCTAAAGCCAAAAGCACTAGAATGGGGGCTTCGAATGATGACTAACCA

AAGTCACCATTGATGATAACATGACAAGCTGAGGTTGCCAGCAACGATGACATGGCACAGACTGGATGCCACCAAGAGAT

GATTAGTGATTACTTAAATAACCCCAACTGTAATAGAACACGTGTACGGATAATGACAGAAGACTTACTAGAAAAGGTAT

TAAGGTACACGTGTAAGGTATTTCCATATAAGGATTTGCCTATAAAAGCATAGCAATCCTCCATAGAGAGGCAGAGCTTT

TCAGAGAACTGTACTGTGTATTGGGATTAGTCTCCTCATAACCTTATAATAATAATAGGGTGTTTACATTTAGCGACTTC

ACTGGGGGCTAATCCTCAATACCACTATAAAATATAAAACCATGGCACCGGGTGCATCACACGACCGTGCATCCAAAACC

AAGACGAACCTATTAATCAACCACCTAATCACTGGATTGAATCAGATAGCTTTCCTGGAACACCAACGAAAAGAGAAAGA

ATCGGACCATATGTACACCGAAGTAGAAACTGAAATAGATTCCCCATTTTGAGTCGAAGAAAAAGGAAATCCGGAGATAG

AGCTTACATGAGGAAAATGGGAAAATCTTCTCTAAGTTCATTCGGGAAATCCACTTCACAAAACATCAACAAGAAATGGC

TTGTATGCATATACTAAACATTTTTTCTAGGAAAACCCACACAAGTATATTGAAAAGTACTTCCCAAAGCAAATAACCTT

TTCGGAAGGAGATAGAATGGTTGAAATCGGGCATTTACGTCCCCTTTATATTACGGTGTCGGTGGAAGAAAATACCATGA

ATAGAGCTTTCATAGATACGGGAGCTTCTTTGAACCTAATCTCTCAAGATATCATTCATCGAGTACGCATCCCCCGATGG

AAATCAGAGGTTTCTATACCAGAATACGAGACTTCAGTCGGAAGAGGCAGCGAACAATTGGAACAATATAATTAGAGATC

ACTCTCGGACCTATCCAAGCCATCGAACTTTTCATGTCACAGAGGTAAACCTAACTTGTCATATCATATTAGGGCAATCA

TGGCTACAAAGACATGGAATAATACCATCAACGTACACCAATGCATCAAATTCTTACATGGATGTAAAGAATGGCGCATC

ACAGCCACGAGAAACCCATTCACGAAAGCAGAAAAAGATCTTTTCCCCGAATCAGCCTTCTATGAACAACAAAGTGGAAT

AATTTTCGAGGACAATTCTCGAAACAACCAACAAGGTAGAGGAAAGACAAGCAACAGCAAAAAACACTTTCAAAGCCATA

CTAAAATTGATGGGAGAAAAATGTACTGCTATGTCGATAAAGAAAACACAGAAGTCGAAGAGGAGAAAATAATGCACGCG

ATGGAGGATCTAGCTATCACTGAACCCGAACACGCATGGGAGTGTCCAACCATTGTTTCCAGCGAAAGTGCCGAACCAAC

GATTCCTCTAAATATAGGGACTTCAGACAATCCCAAAATACTCCAGATAAGCAATAACTTAACAAGGGAAGAACGCAAAG

AATTGGAAAGATTACTAATGGAATTCCAAGATGTTTTCGCCTGGACATATGAAGACATGCCTGGGTTAATTCCAACTCTC

GTTCGGCATGAACTCAAAATTTGCCATCCTTCAAACCTGTAAAACAGTCGTGTATAAAGTTCTCCCATACATTCGAAGCA

CAAATTAAAGAGGAAGTCTCCAAATTACTCGAAGCTGGGTACATCAAGCCCATACACCATCCGACATGGTTAGCCAACAT

CGTCCCTGTAGAGAAGAAGAACGGAAAAATCTGATGTTGTGTGGATTTCTGACATCTCAACCGAGCATGACCCAAAGACG

ATTTCCTTCTTCCTCTAGTCGATCTTTTGGTCGACTCAACCGACGACCACGAAATGTACTCTTCATGGACGGGTATAACG

GCTAAAACCAAATTAGAATGAAAGAATCTGATGCATTCAAAACCGTGTTCTCCACACCGGTGGGAAACTTTTACTATATC

GTCATGCCTTTCGGTCTGAAAAAAGATGGCGCTACGTACCAACGAGCCATGAACTTAATCTTTCACGATCTAAAACACAG

AGAAATGGATGTATATGTAGATGACATAATTATCAAAGCTAAGAAGAAGTCCGAGCATGTACCGAACCTCCGGCATGTAC

TCTCCCGATGCCTAAGTTATGGATTAAAAATGAATCCAGCAAAATGCACGTTTGGGGTAACTTCGGGAAAAATCCTCGGC

TTCAAGGTTACCAAGTCCGGAATTGAATTGGACGAAGATAAAGCGAAAGCTATTTTAACAATGACGGCACCACGAAGTAA

AGACGAGCTACAAGCATTAATAGGAAAAATCTCATACGTGAGGCGTTTCATCCCTGTCTTGGGTACTAGTATGTAAAACC

TCCTACCGCTACTCAATAAGGATGCTAAATGGACATGGGGCGAAAAGCAACAATCAACACTCAACAAAATAAATGAATGC

TTAACTCACCCACATGGGGCGAAAAGCAATAATCCCCATGGGAACCTTTATACTTGTATGTGGAAAATACAGAGAAATCC

ATAGGAGCAGTCTTAGTACTAGATATGGGAAATGACAAGTTATCTCTAATATACTATATTAGCAGAGTACTGAGGGAAGC

GAAACTCCGATACTCCCTGGAAAAAAAGAATTGTCTCCCACTGGTACTGAGAGCACAAAAGGTGAGACATTATAGTCGCA

TGTCGCCTCGGCCAACCCAATTGCTTACTTTTCCACAACAACATTCCCATCAGGTCGGATGGCTCGGTTGTCTGTGCAAC

TACTCGAATTTTCCCTTCATCCATCCAAGCCAAGAGGAATCCGAGGATAGTCTATCGCAGACTTTATAGTGGATTTTCCG

GGATGCGACGTATCTCCACTCCACACAGATATACCATGGGAAATAGCATTAGCAGAAGAAACAAAACCATGGCTCTTATA

TTTCGATGGTTCCTCACACAGATCTAATGGTGGATCATGGATAGTCATGGTTACCCCGACTGATGAATTGATATCTAAAG

CATTCAAGCTTGAAGTAAAAGGTGATTCACGTCTACTCATACAACAAATGCTTGGATAATTCAGTGTCAAGGAACCCGCT

CTTGCTGTATACCGAGATAAATATCAAAAAACAAATAAAGGATTTCTATTCAGTGTCACTTGCACACATGGGAAGAACCT

CAAATAGGCACGCTGAAGCTCTAGATACCTTAGCATCTAGTCTACAAATGTTGCATTCAAATGAAGAAACCATCATCGTA

ACCAAAAACTCGATGCCGTCAACATGGTTTGAAGACATCATCTTTGAAGACAAGAATGACTAGAGGCGTCCAATTCTGCA

AGACCTTAAGCAACAACCCGAAGACCGTCAACTGTCATGGAAGCTTTTAAGAAAATATAAAATTATTGATGGAAGTCTCT

ATTTTATCACCACGACTGGCATTTTATGCAAGTGCATCCGAACTGAACAGGCAAAGAAACTACTACACGAAATTCATGAA

CAAACCTGTGGTTTCGGACTCGAAATACCCCTACGACGACGAATTCAACGAGCAGGGTTCTATAGTCCATACATGATACA

CAAGCCCAAACGTGACAAGACAACTTCATAATCTGCCAAGGAGCCCCGCTACATGAAGAAGCTTGTTGCGTCACCACAGA

AGAAGATTGGTGTTGATAGGGTAAAAGTGGGAAGGAGAAGCAAGTCCAAGCCACAACAATCCCACCCCCTTAGAGATGAA

GAGAGATAATGGATCAGTTAAGAATAATTTTCTAAATATCCTTTCTTTCCTTTTATAATAATTTCTCCTCCTCTTCTTAC

GAAGAAGCACTCATAAGATCAATATATTATAAAACTACCATTTCATTTCCTTTAAATTATTAGTAAACCCTAAGAAGCTT

TCTTTCTTCCTTGAACCAAAGCCCAACTAAACTAACATGCTCTTTTATGGGCTAGAGCTATCCCAACCTTGTAGGTTAGG

TCTAGTCCCCAAGTCTCTCTATCCTAAAAGGAATCTCTAATAGGTTAAGAGGACTCATTTTCTAAGGATAATTATTTTCA

TGTATCAACATTAGTCCCCTGACTGTTGACTGGTCAGAGGCCATGTCAACAGTCACACGTTGATTCGTCAGTTTATGATT

GCAACTGATGGGATTTTAGATAGACACTTCCGTGCATGTTGAACTATCATGTGTTCCCTTCATCCTCATCTGATGGTCGG

ATTATCCGTCCCAGGCGGATGTGTCCTCGATAGCCTCTTTAGAGGAGGTACCTCGCCTAGAATCTTATAGGGAGACTGTA

TCTTCTCGACAATAATTAATTGCACTTCGCTCGTCCACTAAATCTCCACCCGCTTGGGAGCCTTGGGTGGACTACATATA

CTCCAACCCATATTATAGGGAAATACTCCGCTATTTAGGCATACATGACGCCATAGCATCCTCTTGGTATAGAACTGTCC

AGTGAGATCATGTCAATTTGGTGCGCTTATTTGCTCGTTGGATGATTGATCCTTACATGTTTAATTTTCTTGGGGGAGGC

GACTTCAGTGCTGGAAGACATGGTCGCGCCTAATGGTTTGCCTATTCACGGAAACACTTGCTATTCTGAGTCTTGTATAT

TCAAAGCTATGAATAACACCGACAGGGATCTTCAAGATCATTCGATTCTAACCAAGGAAAATTCGATAATCTTCCAGTTT

TGGACGAAGTTAAAGTGGAGGCTCCGGAGGAAGGGGAGGATAAATAGGAGTTGGGAGCTTCTTCCTGATGATGAATCTCC

TCGGAAGGTTATTAGAAGGGGCAGTTCTTTACTCCATTTGTCCGTCTCAAAGAATAGGAGGCGTCCTTCCGTGGTGAAAC

AGTTGGCAAGATTCATGGGAGTTGTTAAAACACGATGAATAGTTTATATCCGCGATTCTAAGTCCCCAATATTATCATTC

AAATCAATGTTTATTGTCTTCTAATAATAATCCAATAAGATTATTGTATGTCAGTAATAGGTCCCCGGGTGCCAAAGTGA

ACAGATTCTAGGCCATGATGAGTATATTATATTTAATGTGCTTGCATATGCGGCTATGCCTCATGACGCGCGAGTATGTC

GTAGTTATGATGTATGACTACGCGTCTTTCAGAGAGGCATTTCAGCTATAGCAGTCACATGGTCATAATCAGCTAGCTCG

AGCGGATACTCTTGGAGAAAATATGTATGGGCATGTGACAATGGTTGAAAGAGCTTTTGGATGTTGTTAAGATAAATAAA

CAATGTGCGTAATCTTCTTCCAAAAGAGATTGCTAACCCGGAAAAAAATCCTGCTCGCCCGCTAAATGTGGGAAGCCAAG

AAGGGAGTATATGATCAGCCAACATCGGGCATAGGAAATGCTATTGCAGAGCATGAAAGGAATGGGTTGGCTTTTCTTGA

CGGGTGGCATCCGTTTACGAAGTTGTTTTATTTCACTCATTCCCTCATTCTTCTGTTGTTTGCGTTATCTGGTTTAATTG

TGTGTTGTAACATTTGAAAGTTACTCCTTTGTTTAAGTGTTGCAGTTTTATGAATTCGTTGCTTACACCCTATCTCTATT

TTTTTTTAGTGTTCAAGTGGAGGGTGGATGAGTGCAACGTGCGCGCCTCCTAAAATCTTATTCAAGTGTTTAAGTGGAGG

GAGGATGAGTGCGACTTGCGCGCCTCCTATAATTTTATTTAAGTGTCCAAGTGGAGGGCGGATGAGTGCAACCTGCACGC

CTCCTAGAATCTTATTCAAGTGTCCATGCGGAGGGCAGATGAATGCAACCTGTGCGCCTCCTAGAATTTTATTCAAGTGT

TCAAGTGGAGGGCGGATGAGTGCAACTTGCGCGCCTCCTAGAATTTTATTCAAGTGTTCAAGTGGAGGGCGGACGAGTGT

AACCTGCGTTCCTCCTAGAATTTTATTCAAGTGTTCAAGTGGAGGGCGGATGAGTGCAACCTGCGTGACTCCTAAAATCT

TATTTAAGTTTTCAAGTGGAGGGCGGATGAGTGCAACTTGCGCGCCTCCTAAAATCTTATTTAAGTGTTCAAGTGGATGG

CGGATGTGTGCAACCTGCGCGCCTCCTAGAATTTTATTTAAGTGTTCAAGTGGAAGGCGGATGAGTGCAACCTGCGCGCC

TCCTAGAATTTTCTTTAAGCGTTCAAGTGGAGAGTGGATGAGTGCAACCTGTGCGCCTCCTAAAATCTTATTTAAGTGTT

CAAGTGGAGGGGATGAGTGTAACTTGCGCGCCTCCTAAGCAACATGATTTTGGGTTTTATATATGCCTCATTGTTTATGC

GCCTCTGATGCGTATGCTTTCTAGATTTATTATTTAAGTGTAAATCAACCTGCAAATCTTCAATGTGTATCGGTTTTGTA

AACTACTTAATTAAAGCGAAAGTTTGACTATTGGGTGTTCATGCTGCATAAGTATAACCCCCTAGATGTTAGTTATGCGT

GCCTCCATGGTAGGCGGTGTTTGGAGCTTAACTGATTTGACAATTCATGCATATTATGTAAGTATAATCTTCTAATAATC

GGTAAGAGCATTTAATTTATATTTTTTATATATATTTATTTTTTTGTATATGTCAACCACATGGTAAGTATTGTTCTTTG

ATAAATTACTTCATTTCTTTATTATGGCTCGAGTTTGCTTGTTTTTTGAGATGAAGCACCTCAATCATCTAATGAACGTA

CTCCGCCGCATGTGATCATTTTGATAGTGAAAATACACTTTGATGTTTCCGACTAATAAATGTCGGTCGGCTTGAGATGG

AAGCTCATAGGAGTGGTGTACTACCTTCTTTTTTGCTTCTTAACGACATCGTGAGCAGCAGCTACTTACAAATTTTCACC

GTTCGAGCATATGTAGGCCGACACGGTCGTTTAATCCAGCGTCAAATGATGTGCGCTTAGCATGCTTGCCGTTAGAGCTT

CTTCAATGGTGAATGTGGATATTTCATATGTGGCATCACTTTTAAGACATCTTTTAGTATAAAATCCTAAATATTGGGAA

ATATAAAAATTTATCTCCAACCCATAAAGTTTTCTTTATTTTTTTACTTATTTGTTGACATACAATTGGTTAAAATTCTT

TACATATTCATTTTTCAATGATTTTTATTAATAAAAGTGTGACTAGGATGGGAAGATATCCTCTAAGTGAATGTCTAAAA

CTAGAGGTTCACCTAGAGGATGTCTACCTTTTTTTTTTTGCCACATCACCTTCACATCAATTGGTTGGAGATAATTTTTT

ATAAAAATAGTTATGGATCCTATGTGGATTAAGACTTTTTTCTTCACACACATTCCATTGGAGAGGCTCTTAGTTGAGGA

GGTTTTCATCTATGCACAGTTATTGGAATCATTTGGCGATGCCGATAAAGATGGGTGATCATAAGCATGAAATGACATTT

GCAATTGTCCGCTTAGTGGGTGAAAGCGCGTCAGTGGCTTTGCTCTCAATGTACTTGAAGTGTGTTAATTTAAAGAACGA

GGCGGACGAGTGCTTGGTGTTCTACTTGTGTGCACGAGTCTGTATAGTCACTTGGTATGCGGAATTATTAGACGCGCTTG

AATAGGTGAAGTTGGCGGACGAGTGTATGGCATTTCACTTGCTCGCCTCCCGTGCAAGGGTTCAAGGAGTGATGATATAT

TTCTGGAGATAAGTCTCCTCATACCACTTTAAATACGCTCGAGACGTGGTGGATAGCTATGATCATGAATTAGCGATTGC

GTCTTTTCATTCATACTAAGTTGTGGGAGCTTTGTCATAATACTTTACTAATAATCTCATTCACACTTGATTATATCTCA

TATCTGAAATGATTTTCGAATATTGTGGCCGCCCTGCACTTTTTTTTTTCATCATCATCATCTCATAGCCGGATTGTGTC

ATCGTTCAGCGCTGACTTAATTCATGGCGCATTCGAGCTATCTATCTATTTTATTTTTCACTTCTCATAGTACATTCTGG

ACCACCCTGTCGTGGCTATTTTCAGTTCATGGTGCACTTCATCTTATCTCACCGCTCGGCTATTTTCATTTTATCGGTCG

TACTCGCCGCCTAGTCATACAGCTTTACTCATTTGTCATGCTCGAGGCTCTATTCAATCTAGCCATACTCGGGGCTTTTA

ATCAATCTAGCCATACTCGGGGCTATAGTGTATTTCATTGGCCATGCCCAAATCCTAGCCATACTCGGGGCTACTGGTGC

ACTTCATTGGTCATGCACGAGATCTAGCTAGAATCGGGGCTATTGGTGCACTTCATTTGGTCATGCACGGAGCCTATCCA

GACTCGGGGCTATTGGTGCAATTCATTTGGTCATGCACGGAACCTAGCCATACTTGGGACTATTGGTGCACTTCATTTGG

TCATGCACTGAACCTAGCCACATTCGGGACTATTGGTGCACTTCATTGGTCATGCACGGAACCTAGCCAGACTTGGGGCT

ATTGATGCACTTCATTGGCCATGCACGGAACCTAGCCATACTCGGAGCTATTACTCATTTCATGGGTTGTACTCGTAGCC

TTTTTCATTTCTCATGGTGCATTTGAGGCCACCTTTTCTCATTACTCGGCTGTACTCATGGCTATTCTCAATTCATGGTG

TTATCACATTTAAATTTATATAGTTCAATGCAATATGACGCGGATATTGTACACTTAGTGAAATTAGTAAGTTGGTAGTT

TACACATGCGATTGGATGATATACGCGCGTTTCATGGAACTCAAATATTGATACATAAATAGGCATTATTTTGGACACCT

CTTGGTTTAATACACTAAATATATGAGGTGGGTGCATGTATTCATAATGGTGAGAACAATTGTAAATCTGTAAAAATGAA

ATCTGAGATAATGCAATCATATTTTAATCGCCACATGTTATTGACTTTTACTTAATTAAAATAAAGCGATATAGAGAAGT

AACACAGGAAAAAATCTGATGTTAAAACTTTCAAACTAAGTGGTGCAAGTATGTACTTATATCCTAATGACGAGGTGACA

ACATACTCACATGAGATGAAGCTCATACGAGCCGATGCATGGCAAGATTGATTAAATCGGTAAGATGACAAATCTACGTA

TGAGTCATGCTTTCTGAATGTTTTACTAGGCATCAGAGTTCCACATAAAGCTAACTTATTCATTTGTCAATTCAAGAGAC

ACAGAACATCCTGTCGGGAGGCTTACGAAATCCGAGCCGCTATGCGCAATACGAAACCTACACAAGCAATGATATACAAA

TGCGGAATAACAACAAACTACGCAATATGGAAACAATCACACAACACCGGCAATTACGTGGAAAAACCCCTCCAAGGTGA

AGGGTAAAAAACCACGGGAAAATCTTCCACTATAATCAAACAATGGGTTACACACAATTCTCCCAAATTTGGGGATAAAC

TCTTCTTCAAGGTGCCTATAGCACTTGGATATCAATTTGCCTAATTACACTCACCCCTAAGGATGGATCAACAATTCATT

CATATATAATATGAGATTCACTATCAATTTGAGAATTAGAAGTGATTTGAGATACTTACTACTAGGGTTTGACAAACCCT

AGATTTTTCCTTCTTTGAAGAGATGTGGAAATTGATAGCTCAATCTTCACCACAACAACCAAAGATATCCTTAGCTAAGA

ACCCTTTCTTCTTCTCTTCAAGATGATACCAATCTCACCAAAAATGGTGGCTTAATTCTCTCTCACTTTCTTTCTCTTCT

TCCTCTCTTTCTTTCTTCTTCAAAAATGAAGAGATAGAACCCGTAACACAAAAGAGTGTCTCATGATGCCTTTTGTACAC

AAAACCAAAATAACAGGCATCCACTAACTTCCTCCCTTGAATGGTTAAGGGGAATCCACATGTTTGGGCCAAAAGGAGTA

AAACCCAACAATCTCTCCCTTACGACTTATGGCAAGGTACCCGTCACCACCAAACCTGCCCTGCTGCAGAAATACACATG

CTTGTATTTTGGTAGCGACTTTGTCATCATATCAGAACCGTTGTCATCTGTATGGATGTTCTCAATATACAAGACCTTCT

CCTCGAGAACTTCTTGTATCCACTGATACCTAATATTGATGTACTTGGACCTCGAGTGAAAACTCGAATTCTTGCTGAGA

TGGATGGCACTCTGGCTGTCACAGTAAACGATGAACCTCTGTTTCTTCACACCGAGTTCTCTAACAAACCTCTTCATCCA

TATCATCTCCTTGCAACTCTCTGCAGCTGCAATGTACTCTGCCTCTGAAGTAGACAATGCCACACATTTTTGTAACTTGG

ATTGCCAAGACACAGCTCCCCCTGCATAGGTAAATCAGTAACCCGATGTAGACTTGAGACTGTCTTGAGAACCTGCCAAA

TCTGAATATGTGTAACCAACTAACTCTGGCTTGTCACCTCCATAGCACAAACACATATTAGAAGTACCCCTTAAATACCG

TAGTATCCATATTTCACTGCTTCCCAATCCTCCATAGCAGGATTTGATAGAAATCTGCTTACCACTACAACTGCATGAGA

TATATCCGGCCTTGTACACACCATTGCATACATCAAACTACCAACTGCGAACGCATAAGGAATTTTCTCCATCTTCCGTT

TCTCAACATCATTCACAGGACATTGATCACGAGTTAACTTAAAATGGTTTGCAAGTGGACAACTTACCGGCTCAGCCTTG

TCCATGTTGAATCTTTGCAACACCTTATCAATGTAGCTTTCTTGTGATAGCCACAACTTCCTCGCTTTCCTATCACAAGT

GATCTCCATGCCAAGAATACGCTTTGCTGCACCCAAGTCTTTCATAGCAAAAGACTTGCTCAGATCTGACTTCAACATGT

CAATCTTTTTCTTATCTTTGCCAACAATAAGCATGTCATCAACATATAGCAAAAGAGTAAGAAAATCACCATCAGCAAAT

TTTTGAATGAATACACAATGATCTGAAGCTATCCTCTTGTACCCATGTTCCTTCATGAACGATTCAAACTTCTTGTACCA

TCTTGTTGGAGCTTGCTTCAAACCATATAGACTCTTCTTTATCTTACACACCATGTGCTCTTTGCCTTTGACTTCAAAAC

CCTCGGGTTGTGCCATATACACATCTTCCTCCAACTCACCGTGAAGAAATGCTGTCTTGACATCTAGTTGTTCTATCTCA

AGATCCAAACTAGAGTCTATTTCCAAGAAAACCCGAATAGAATAAAATCTCACCACCGGTGAGAAAATCTCGTCAAAGTC

AACACCCTTTCTCTGCTCATAACCCTTCACAACTAAGCGTGCTTTGTACCTTGGTTGTGAGTTACCATCTTCAGTCTTGA

TTCTGTACACCCACTTGTTCTTCAGAATTTTCTAGTTCTCAGGCTTGTCAACCAAATCAAAAGTGTCATTCTCATATAAA

GAGACAACCTCTTATTTCATGGATTTCAATCACTTCTGACTTTCAGGAACACTAAGATCTTCTTCATAAGACTCTGGTTC

ACCTGCATCTGTGAGCATCACATACTCATGTGGTGTATACTTTGTTGAAGGTAATCTGGTCCTGCTGGATCTTCTCAACT

CAGATTCTGGTTGTTGTTGTTGCACTTCTTGCTCAACATCACCCACTGGAACATCTTCATGAGCTTCATCTTCATTGCCA

TTACCATAAACATCATCATCATGTTGTTCAAGAGTCGGAATAACTGGATTTGGATCCAAGGTCATATCATCTTGAAGCTT

TGGCATCTCAATCTTCTCAAAGTCTTCAATAGTCTGATCTTCAAAGAATACTTCATCCCTACTTCTGATAACCTTCTTCT

CAACTGGATCCCAAAATCTGTAACCAAACTTCTCTATTCCATAGCCTAGGAACACACACTGCTTTTCTTTGTCATCAAGC

TTTGATCTCTGATCTTTTGGGATATGAACAAATGACCTGCAACCAAACACTTTCAAGTGATCATAAGTAGGCTCTTTGTC

TCTCCATACCTTTTCTACTACTTCTCCATCTAGTGGTCTTGAAGGAGATCGATTGATCAAATATAAAGTAGTATTCATTG

CTTCACCCCATAAATGCTTCTTCAACTTGGAATGAGATAACATGCACCTGATTCTCTCAAGAATAGTGTGGTTCATCCTC

TCTGCAACACCATTATGTTGTGGAGTCTTCGGTACTGATTGTTCATGACGTATACCCATGCTCTTGCAATACACTTCAAG

TGGCCCAATGTACTCTCCCCCATTTTCAGAACGAAGGCACTTCAAAATTCTACCCGTTTCTCTTTCCACCATGCTGTGAA

ACATCTTGAACACAAGAACAACATCATCTTTGGTCTTGATTGCATATGACCAAACCTTCCTTGAAGCATCATCTATGAAA

GTAACAAAATAAACAGCACCACCATATGTTTTGACCCTTAAAGGTCCACATACATCAGAATGAACAAGATCAATAATACC

AGAATTGCGCTTGGGTAAAGCTTTATGAAAGGAAACTCTAGTTTGTTTACCAACTAAACAATGAGTACACTTACCCAATT

GCATACCTTTGTCATTTGGCAAAACTTGTCTTTTGGTGAGGATTTGAATTCCCTTCTCACTAAGATGACCAAGTCTTATA

TGCCACAGATCGGATATTGAATCTTTGTCAACATCATTCAACTCACATTTCAACACCTTCAATTGTGTCTTGTACAATGT

GCCACACTTCTTACCTCTTGCAATAACCATGGATACCTTGCTTAGCTTCCATTTTCCATCACCCAAGTGATTGTTATACC

CTTCATCATCTAACACTCCAGGTGACATTAGATTCAAACGAAGATCCGGTACATGTCGCACATTTTTGAGAACCAACGTA

CAGCCAACACTGGTCTCAACCTTCACATCACCCATTCCTACAATTTTCGCAACACCACTGTTACCCATTCTGACTGACCC

AAAGTCACTAGCCTTGTAGGAGGTAAATACATCTTCATGAGGAGTAGCATGATAAGATTCACCTAAATCAACTATCCATT

CTGAATCAGAACCTGCAAAACTCACACAAAAATCATTACAATCTGAGAATACAATGATATCATCATTTGTCACAGTAGCT

GCAACACCATTATCTTGGTTATCATTTCCTTTCCCCTTCTTGTGTTCTTGCTTCGGCTTTCTGCATTCATTTGAATAATG

ACCTCTCGTATGACAGTTCCAACATTCAACTTCCTTCCTGGACTTTGATTTACCCCTTGACTTGCTACGACCAACTTGTC

TTCTGTCTTTTCTTCTACCCCGATTTTCTGTCACAAGAGCTTCTGAATGAGATGATCCTAATGACATTCTTCTAACTTCT

TCACTCAGAAAACTGCTAGTTGCAACAGTCATGTTCAACTTCTTGTCTGGAGAAGAATAATTGAGACAATTCCTCAATGG

CTCCCAACTGTCTGGTAAAGTATTTAACAACAAATAAGCTTGAACCTCTTCAGGAATTTTGAGATCCATAGATGTAAGCT

GATTCATGATGCCCTTAATTTCATTTAAATGTTGCGCCATTGAACTCCCTTCCTTGTAACACAGCTTTCCAAGCCTTTTG

AATAGAAGACCTTTTTCACAAGGAGACTGCTGATCATATATCTCCTATAACTTGGTCCACAAATAATATGCTAGTGTCTC

ACTAGCCACATGATGAAAAATACCATCATCAATCCACTGACGAATAAAACTAACTGCTTTTCGATTAGTTACCTTCCAAT

TGCCTTCACTTGTCGTCTCAGGCTTTACTCCCTTAGTCTCTATTGGACCATAGAGATCTTTACAATATAACCAATCTTCT

ATCTTCACCTTCCATATCACAAAGTTCTCACCGGTCAAATTGACCATTCTATTACTAGTGCTAGCTTCCGCCATTGCACA

TGATTAGCTCAACCAAAAAGCTCTGATACCACTTGTTGGGGGAGGCTTACGAAATCTGAGCCATTATGCGCAATACAAAA

CCTACACAAGAAATGATATACAAATGCGGAATAACAACAAACTACGCAATATGGAAACAATCACACAACACCGGAAATTA

CGTGGAAAAACCCCTCCAAGGTGAAGGGTAAAAAACCACGGGAAAATCTTCCACTATAATCAAACAATGGGTTACACACA

ATTCTCCCAAATTTGGTGATAAACTCTTCTTCAAGGTGCCTATAGCACGTGGATATCAATTTGCCTAATTACACTCACCC

CTAAGGATGGATCAACAATTCATTCATATATAATATGAGATTCACTATCAATTTGAGAATTAGAAGTGATTTGAGATACT

TACTACTAGGGTTTGACAAACCCTAGATTTTTCCTCCTTTGAAGAGATGTGGAAATTGATAGCTCAATCTTCACCACAAC

AACCAAAGAGATCCTTAGCTAAGAACCCCTTCTTCTTCTCTTCAAGATGACACCAATCTCACCAAAAATGGTGGCTTAAT

TCTCTCTCACTCTCTTTCTCTTCTTCCTCTCTTTCTTTCTTCTTCAAAAATGAAGAGATAGAACCCGTAATGTTACGTGT

TTAAGTAAGCTCAAAATGCACAGGTTTGTATTCTCAAACAACCAAGGATCATAAGATAAATGATGAGTGAGAATAAGCAA

ATAAAATAGGTACTTATCTGAAGCGGTTTGAAATTTTCCTTATTCTTGGAGAGGGGTGGTCTTGTAGAGATGAGAAGCGT

TGGTAGTGTCCCCGCTAGCTGAGATGGTTGCGACTAAGGATGACGGATACGGGTGCTGTTACTGACAGTGGAGGGATGGC

GCTTGGTAAGTAAAGCAATGCCGATGAGGTCAGGTGCGAAATTATGGGGTAGTCATTCTGCACTAATCAACGAATTTTCT

CTCCATTGCTTGCAGGTTGAGGACTGTATCCTCAATGCAATAGCGCGTAGCATACACCTAATTGTTGCAGTAATTAATAG

TTTTCAGACGACGGCGTTTGGAACTTCTCCACCTAGTGAAACCATGGAGGTGCCTTAACCGATGCTGATGAGCTTTACGT

TAGTTGCTACTCGCTTCTAGTTTTGTTTGGGATGGTGAGATAAACTTGGTCAAGTACCTGCGCCTTTTATATCTCTATCT

CTCGATTTAAATATTACTCAAGCAAACTGCGAGTCTCAATTTAAATATAAGAGAGATAAACTTGGATGATACCAAAGACC

AATATCCAAGTGTCAATCAATTTAAATAAACAACCAAAAGGTCGGATATTCTAATTGATTGAACAAGGCACAACCTGTAT

TATTTTAATTATATAATAACAAATATAATGCGAAAAATAAATAACACAGACACCAGAATTTTGTTAACGATAAAACCGCA

AATGCAGAAAAACCACGGGACGTAGTCCAGATTAAACATACACTGTATTAAGCCGCTATAGACACTAACCTACTCCAAAC

TAACTTCGGTCTGGACTGTAGTTGAACCCCAATCAATCTCACACTGATCCAAGGTACATTTATGCTCCTACGTCTCTGAT

CCCAGCAGGATACTATGTACTTGATTCCCTTAGCTGATCTCACCCACAACTAAGAGTTTCTACGACCCAAATTCGAAGAC

TTTAATAAACAAATCTGTATCACACAGAAAAGTCTACGATAATAGATAAATCCATCTCCCACTAATATACCTACGAGTTT

TGTTCCGTCTTTTGATAAATCTAGGTGAACAAGAACCAATTGATAAACCAGACTTATATTCCGAAGAACAACTCAGTATT

ATCAATCACCTCACAATAGTCTTAATCGACGCAGCGAAAAAAGATATTGTGGAATGACAAACGATGAGACGAAGATGTTT

TTGACTACTTTTTATCTTGCACTATAAGAGATATAATCTCAAGCCAATTATTACAATTGTACTCGTGCGATAGAAAATGG

CAAGATCATATCACACAACTACAAGAAAAGTAGTATAGGTCTGGCTTCACAATCCCAATGAAGTCTTTAACCTGGTTTAG

AAGAAGAAACCAAAGGTTAAAGGAGAATCGACTCTTCCTTAGCACAACTAGTATCACATAGAATGTGTGGAGATTAGGTT

TCCCAGTTGCTATAGTTATCCCTTATATAGTCTTTCAAATCAGGGTTTGCAATCAATGTTAGCTTGGTAACAAAGCATTC

AATATTCACCGTGAGATGAAAACCTGATTAGATTCAAGATAATATTTATCAACCGTCAGATCGAAAACATATCTTGTTAT

ACACACATGAAATGCACGTTTGTAGGCTTTTGTAACCGTACCCAAACTTGTACATTAGTTAGTTCAACAATAGTTAACTA

AATGGTTAGCCATATGATAAATTTCATATCAACCATAACTAGTTCAAATGACTCAAATGAACTAGTTAGAGAGTTGTTCA

ATTGCAAGGAAATCTCATGTACTACACAAGACACAATTGAAGAAAAAACGATTTGATTTAGTCAAATCGGTTCATGAACT

ATATATCCACGGTTTGCAACTGCATTCCTTAGTTTATAAAGATAAGTTCACAAACATCGTTTTTAGACATAACCTACTCA

AGTTTGCGGACTGGGTTCGCGGACTTAAGTTCCCGGATGGAGTTCACAAAATCCAGCAGAAATTCTTGGGTCGAGAACTT

CCGCCAGTTCGCGGACTTAGATCACGCACTTATCCGGTTCTCTTGATCAACAAAGTTCGCAAACTTTGGTTCAAGGAATA

AGGAATTATACATATATGTGTTTCCACAACAATGCTTATATCCTCCAATGGTTATATAGTCTAAACTCTCATTTCAATCA

GTGAAACATTCTCAGAGGACGTTATATTGTTATTATTCACAAACCATTTTTCGTCAGAGTAATTTTCAAAGTGATTGAAA

CTTATCATGACTTTCGTCACTAGGTAAAGATGAACTTGTCTAAAGATAAAGCTTTACCAACATATATTTCGAGAAATATA

TAGGCGAGGTAAACTCGGCTCAAAATACCAAATGTGTATAATCTAAGTCTATATAGCAAAACGACTTTTGTCTCAAGATA

GGAGATAAATAGATTTTTGAGTGATAGATAAGTTCAAGTCTCCACATACCTTTTAGTCGATGAAGATCCACCAGTTCCTT

GAGTATTCCTTCGTCTTGTATGATGATTTCCATGGAGTTATTAAGCTCAACTACACTTTTTATCCTAGTCTGAGACCTTA

GCTATAGTAGACTAGAAATCAAGACTTGTAGTTTTGATTACTAATATTGACAAGCATGCTTGAGATAGCAACGCATGCGA

GTTCGACCAAGCAATGCTCTAACACTATGCTTTTAAGATACATGGCTAGATTAATCATTAGGAAATAATCTCTTTTGAGC

TCAATTCTTCCACAGGTGCTGCTGACAGGTGTAGATGTTCTATGAGATTTCGACTCATTAAAACATCAGTCCGTAGCGCT

CATTAGGAAGCAACTGCTTCATCCTCCCTAGATATTCTGTATAAGATCTGCTATTACCATAAGGATAACAACAATGTTTT

CAATTAGATATCACATTTCCATGGAAGATATCGGATCATATTTTCTATGAACAACCCATCAAAATTTGATTTTATATCAG

ATAATTAATCTAGAATAGATATACGCTCAGTAACATATCAGAAAATTGAAATTTCATGAAAACGGAATCAATAAAAAATG

GGGAGGTAAAGATTCGTGTACGCTGATTTTGAATCATATATATGGGCAACGGGTCTCTAAGATTGAATATCGACAAGTTT

ATTCAACAACAAGATATTGTAAATGGAGGTAGAAGTAAATATCAAAGGGACTTTCCCCAAAAAAATTTTTGTGAGATTCT

AAATATTTTTCCCTAAGATGATAATCACAAATTGGATCTTAGGTTTCAAATCAGATCGTAAAATATAAATAAATCTCGAA

TTTGAGTTATGAAAACCCTAAAATATTTTCCCGAAAACCAGAAATTAATTAATTATTACGGTAAATGACAACAAGTAGAT

TGAAATTATCTGTTTGATGGATTTTTGTTGATGAATCTATGATTTGGTTGTTGGTATTGTTCCTCAGAAACGGCAGAAAG

GAGTAATGGTTGATCTGTTAGGAAGGAAGAATAAGTCGCCTTGTTGTAGAGGACGACGATGATGTAGTTCGGCAGCGCGG

ATGTGGTAAAGCGGCGGTGGTTGGCGCAGAGATTTACTGTATCAAGTTAACCGATCGGATGTTATTCCTCGGAGTTGATG

GTAGTCGATCTTTATCTGGATCTAGTTGGTGCTCGATTTAGTGTAGTTCAATGTAACCGGAGAAAGATGAAAAGTCTGGT

AGTCTGACTCGGCGTAGGTGAGCGGAAAGAACAAAGTTTCTCGGCAGTTTAGAAGATAAGTCGAGTAAATAAGCACCGAT

ATAGAAGATAAAAGAATATAATGTCTCTTGATGCTAGAAAAACGAATGAGATACGGAGGAAAAGAGGCGCTGATCATGTT

GAGTAGCACATAAATACTTCCTAAGCGAAAAATAAGTGGAGAGATGGTGTAAGAGAAGAGAGGCGTGCAAGAGAAATAAG

AAATGTTTGTGATAATTTAACGGATGTGTCTTGCTGGGCGAGAAACTAAACGTCTTCCACTGTGGTCGCAATGTTGATAG

GATAAAAGTATGATAAAGAAGCAAGTCCAAGCCACGTCAAGCCCACCCCCTTAGAGATAAAGAGAGAGAATGGATCAGTT

AAGAATAATTTTCTAAATATCCTTTATTTCCTTTTATAATAATCTCTCCTCATCCTCTCATGAAGAAGCACTCATAAGAT

CAATATATTACAAAACTACAATTTCCTTTCCTTTAAGTTATTAGTAAACCCTAAGAGGCTTTCTTTCTTCCTTGGACCAA

AGCCCAACTAAACTAACATGCTATTTTATGGGCTAGAGCTATCCCAACCTTGTGGGTTAAGTCTAGTCCCCAAGTCTCTC

TATCCTAAAAGGAATCTCTAATATGTTAAGGGGACTCCTTTTCTAAGGATAAGTATTTTCATTTATCAAAAATTGGCGAA

CCCCGTACATCGATTACCTAATAATGGGAACATTGCCAGACGATAAGAAGTTAGCAAAACAACTAGAAAAAAGGGCTAAA

CGTTTCTTCGTCATCAATGCCCAACTCTTCCGAAGATGCTTCAACCAAAAAACTCTCAAATGCATCGACAAAGAAAACAC

CACCAGGGTAATAGAAATAAGTCATGATGGGGAACACCAAGGAAAAGCAAGACTCTTTATCCATATTCACCATGTATGAT

ATTATTGGCCTACAATAGAAAGTGACATCACTGAATATGTTCATAAGTGTGAACAATGTCAAATTCATGGTAACTTAATC

CATGCACCACACACCATGCTCCAGAGAGTCTGTACTCATTGTCCTATTCATAGTCGGGGACTAGACATTATCAGACCCAT

CAACCCGAAATCTTCAAAGCAACATCGTTACATCAACAGAGTTCTTCACCAAATGGGTGGAAGCCATACCATTAAAGGGT

ACAACTGGGAAAGTAATATCAACATTCATCAAGGAGCATATCATCTATCGTTTCGGAATTCCAATGTATATCATATCCGA

CAATGGTACACCTTTCGTGAATAATCATGTGAAGGACCTCCTAGAATCATTCTGAATCATACACAAGACTTCGACTCTTT

ACTATCCATAAGGAAACGGTCAGGCTGAATCTACAAACAAAACGTTACTCAAAAAACCTGAGCAGGAGCCTATATGACAA

CAAACACTTATGGTATGAGGAACTTACATTAGCACTGATGGCATACCGGATTGCTAAGAGAACATCAACTGGGGCGTCAC

CATATAGCCTAGTCTATGGCGAGGATGTCATCATACCAGCAAAAGTAGTTGTGTCTTCGGCTAGGATATCATCTGCAAGT

GGGGTTGACCTAGACAGTCAGTGCTTCTGACCACTTAACATGATAGACGAACGCCGAGACACAGCTGAAAAACGATCTCA

ACAATGTCGAGATCGGATAGCAAGATCTTACAATTAACACATTAAGGAAAGACAATTTTAAAAGTGAGACCTGGTACTCG

TTGTAACGCCCCACGTGCAAAGAGAACACAACACTAGAAAATTTGCTCCAAACTGGGATGGACCTTTCCAAAATCGCAAA

GTAGGGGTACACGCTGATGGCACTAAAATCAAAGGGAATTTGTTTTCCAAAATCAACGGAATCTGGTTGAAAAAGTTTCA

TTCCTAGGAAAAACTTACCAGCATCGTGCATGTATCCAACGTCCATCACAAGGATAATGTTTAAAATACTCTTCTACTAC

AGCAACTCTTGTTTTCTGAATAAAACAGTGACACTTATGATGACATGAGTCATTCTTAACTTCGATTCATACCGAAACAA

CCACCATTCGCTGCAAACACATGTTCAGAATCAGCTACTCACAAAAGGTACTGGGATTCCATCCCAAAAAGTCAATTGAG

AGCTCTTTCTAAAAAAGAACATGACCCCACAAAAGGTAGCCAAAGACCTACCGGATGGGAGGATTATAACTCATAAAGCG

CAAGCTTAGAAAACTAACATTGCAAAAATGCGTGTATATTTTTCTAATCACTTCTTCTTATACCATAATAGGTGGTATAT

TTGGGTTTCCCCCGCTAAAGACACAAATTAATCTTTTAAATATTCTTACACAGTCTGCTGAATGTCATATCTAACGATAA

AACTAAAAGCAAAGGGCCAACGTAGTACATTTCAAGGGACCCCAGCAGTGGATAAGGGGCACCTCCTCCTTCCACACGTT

TAAAACAGATTAGGCGGGAAGAATAACCATACCAGTATGAGTTTCTGGATTTGATTTGGGATGAGTTTAAACGGATTTTC

TCACATATATGCCTCGTTTTGGGCCTGTTAACATAAACCAGGGTTGGTAATAGAATTGGTTAACGAAAAAGGACAAGAAA

ATCATTGTTCACGATAAAACAGGGAAGAAAAGAGTTTCTCGGGTTCTTCTTGGTGGTATATTTGGAAGATACGTGCAAAT

ACAGGGAATGTATACGATGGGATTCAATTACTTTCCATCATTGAGAGATTCTTACAACCCACAGACGCGTAAGAGAGAGA

GACAAGAGGGAAGAAATCCCGTGGAGTGGCATGAAAGAAAAGGGAGTATTTATGAATATTTCGTGGAGATACAATCACCT

TGCCGTGTATATAAAGAGTGGGGAGAAGTCATAGAGGGATTATCAAGAGTTTGGGGAAGTTGCAGGGAGACTAAATCACA

AGTCGCGGGATTCTGTTCCTGCTGCTGCTGAAGAACGAAGAACAGACACCCTAAAACAGTCGTTTATCAATAGTGGTTGC

AATAGTCTGACATCAGTCTTATACGCTACAGTTACAACGACAATTTATTTATATCGATCTCTGTAACAGTGGACTTGCAA

CACCATGAAAGTGTTACAGTTTCTTGGTTTATTCAATTTTTCTCCCTTTATCATTTGTAAACACTTTTGAGCAATGAAAT

CAACTTTTGAGCGTGTTTCCATGATGAAGAGCTAAATCCCAATACTGGGATGAGGGAGGAAGGAAATTCTTATCATTGTG

GTAACTTTATTTAATTCTTTTTACGACTTTTGCATTAATATAATTTGAATTATGATTTTAAATTAATTACTTGTCATTTA

GTTTGATGGGTTATGCTTAGCTTATGTGTTTTGATAATCCTTGCTTACCATTTACAATTAATGTTTTAAAAAATCTACCT

TGGCAAAGATTTAGAGTCAATAGTTATTATTTTTGGAGCTATAATTGTCAAAGAATTAATAACTGAACCATAAGAATATG

AGTTTGTTGGAATCCTGATTCCCAGTAATCTCTCAATATTGCTATCATCTTTTGCATATTTTTGCTAGTTTTAATATTTT

ATTAAGTCTAAAATCAATATCTTCCCAAGTCTGAGTATCAAACCCTTTCTTACAACAATTAAAACTACATCAGTTTTTGG

CGCCGTCGACGCGGACTTGTATTTAGGTTAGGTTTTAGGTTTTGCATTTTTTTTAGGTTTTGTGTATATATTTTTTTTCT

ATTATTATTATTTTTTAGATTATTTTTTTGTTCTATTTTACGTCTTTGGTATTTTTGTTTTTGTTGTGCAGGATTGGCTT

GAAGCGAAACGTGGAAAGTTGAACTTGCCCATAAAAGGGGGCGTTACTCCTTGTTCTCATGTTACGTTTCCGGTTTTGCT

TCCGAATCCCAGTGCTATCAGCATAAAAGCAAAACGCTAGATTGTGTGTCTCAGACACGTATTTCCATATTTTCTCACCG

AACCCTAATTATTTCGCACAACCAGGCATGAGACTTTCCCTTACATTAGCTGGCAGAATAGGGTTTAATTAAGACCGGAT

ATCATGCCTTAATTCACGGCTAGTAGAAATCTTTCCTTCTCGGGACTGCATCGTTCTTCTAAATATAACAGATGATCTTG

AATCCTTCATTCGCACAGTTCGTCCTGAAATCGTTCCCATCCTCTTTACATGTATCGTCTCGTGTTTTCTGGTTTCCATA

TTCTTTGTTATGTATGTTGCTGGTGTGGATAGGGTGAAAAGCTTGCCTTCTTTATCTGGTTTTGTAGTGCATAATGGGAT

CTTCTAGAGGTAGATGTACCTCTAACTCTTCGGAGAAAAACTTTGGAATCGGTGAAAACCAGTTTTTTTCTCATGAGGAG

GCATTGGATAGAATTGAATCTCTGTTCCATAAAGTTTGCAGAATTATTCACGGCCCACCTTCCATCCGTCTAGGTGAAGT

GCGGGTGCGTATTCATACCTTCTTAGGTTCGGTGAACATGGAATAGGAATGGGAAAATGACGAAGCGCCCCGGTGCGTTG

AGAGGCTTCAGTGCAGTAAAACTTCTTCAAGGCTCAGTGCTAGGGCGGAAAAGCCCGCTTTTAGGTTGATACGACGGAAA

ACTGATTTTGTGAAACTTTTAGGGAAAAGCCAACGTAACCTCTTTGTTCAGGACGGCGTCGTAATTCTCGACTCTGATGC

GGATGAAACAGGAAATTCTCCTCGAAAGGTCGCTGAGAAAACCTTTGAGGAAGGCTCTGCGTTTAATGGGACTGAATTAG

CCTTCGGAGTATGCGTCGAAACTGTTGTCGGAGATATTTGGCGGGCCATGGTGGCATATGCTAGATCGGTTTTTGAAGCG

GGTGTCGGGAAGGAGACGGGTGTTGTGGCTTCTAAGGAGGCTTCGGGGTGGGATGTTGAGGTTACTACTACTGACGCGGC

CGTTGGATAGAACATTTTTTTCCCGAGTGATTGTTAGGGAAAGTGCTTGAAAAGAGTTATGACCGAGGCAACTCACGCTC

CACCTTTCGATGACTAACTTCTTCCTTTTATCGTTTCTATTTGTTTTGTAGTCGTCCCAATGTTTTGTAGACTCCTCTTT

ACCTTCTTTGTATACGTGGCGAAGTTAGGGTTTCGCCAATTAAGATCCCGTTTGCGATTTTTCATTTTTGAGTAATTGAG

TAAGGTATTTGTGACACGGTTTTGGCTTTTTGGTTGTGATATGATGGTTAATGGATTCTCAATGATTTTGGATTAGTTTG

GTGTGTTTTTTTGAGATTTAGAAAACCCATTCGGAAAGAAATGTATGTTTTTTTTTTTTTGAATGAAAATGAAACTCCGA

AGAAGAATACAAACGGTGTACTTATCTCGGGGAAATAAAAATATCAAGATAAACTACTTAAGGAAATACTAAATGGAGAG

TGGCTGGAAAGGTATAGCATTTTAGGGATTGGAAAGCTTGAGCCATCGCGATTGAACTATCACACCTCCCGTCTTGTCTG

TGTTGATGAGCTTGCAGTATCCTCCTAAAAGAACGTCTTCCACCAGATAAGGCTCGGCTTTTTTTTCCTTAGGCAAACCA

ATCCAAACATGCCTTTATCGTAATTTCCCAACGTGAAACGAGTTCACTTTGACCATTATATCTCCGATTGAAAACGGGTT

CAGGTGTTGTGCAACCGTTTGAATCTTGGTCTCGTTAGTTTCGATGGCAGCTTCCAAATTCTTTGAAAAGCATTTGAATG

GACCATTCACATCTCTTGACAGCGTCCGTGTCAGTAATCGCGATGAGTCCTCATGACAGAATAGGGGAAGAAATGACTGG

TTGTAGAGAAGATCGTTCGGCCAGACTTACTTGCGTTTGGAATCCATGAGTTAGCGCCAACAGGTCTTTTCCAGGCGTCC

GGGTTCCATCGGTAACTCGTCGATGCGGCAGGAAGTGGGATGGTTTATTGGAGATTCTTTCGGTCGTTGACATTGGTAGA

GGGTATGTACCCAATTTTTCCTTGTTTTCGAGTGCCCACAAGTGCCCCAATATCATGTCGTAACCTAGTTCTTTGATAAT

ATGGAATTTACCATGCATTATAACGTCTCCTATATTGACATCGAGGTTGATGTAGTCGTGCGCGTCCAAGGCTTCCCCTT

CAGGGTTCTCAATCACGGATGGGGATCGTGTAGCCTCTTGTTTCATAAACCTCGATGCTTTTAGAGTCTTGAAAGTAATA

ATATTGAAGTCAGAGGCCGTGTCGATGAGCGTGTTGTCGAATTCAATACCTCTTAAATAGGACGTGGTCAGGAGTCCCCA

GTTGTCCTTTTCGATTCCGCTCGCTGGGAGAGCTTGGGCTCTTGGCGTGGCTTCCATGGAGGGAAAAAGCCGTTTTCCCG

ACACAACGCGATTGAGAGCCGCGAAAATATCTTGGAGCTGCACCTTGGAAAGGTAGAGAATTTCACATACGTGTTGCATC

ATGGATTTCAAAGTCTCATGCACGGAGTCCCCAGAGACCCCACAATTCCTCATTGGAAGTGGGTCTCGATGTACTCCCTC

GTTTCCTAGTTTTAGTTCCCCTACTTGTATCTTTTCTTTAAAGATATGCTTTAGTCTATTACAATCATTGGTTGGGTGGT

GACCAAATATGTGGTTGTGGAAGTATTTGGGGTTTTCCATCTCGGCCTCAGTCGGTGGGCGTCTGGGAGATGGTAGTCTG

ATAACATCATATTGAATCCATGCTTCTAAGAGATTGATCACCTTCTCAACGGGAAACATTAAATCCAGGGTATTTCCATC

AGTTTCGTGATGACGCATGGGAGCTTTAGCGGTGGCCTTCCGTGGCGGAGTTCCCTATTTTGCTGGTGCTTCAAGTTTAG

GATGTTGGTCTGCAGCTGGGGCCTTCCTTTTCCCTCCCTCATTTACTACCCTCACATAGGAGGGGCTAGTGTTAAATTTC

TTGCTGATGAGATGTTTATTCCTGCGAGTTTAGCGCATGTCTTCGGTTCTGGTCGGTCTAGATCTTTCCAATAATGCTGG

TGCATTCGTTGCGGATTGTCTAGCAGCTTCATGGAGTTCGGAAAATGTCTGGAAATACAAGTTCTCTAGCAAGGTGCAGT

AAATGAGAGTCATACCGTTGATGCATGTGTTCACCAGTTGTTTCTCGGTCACGTTTGGGTCGTGACAGTATAGTGCTTGA

ACCCAGAATCTCTTAACGAAATCATTCAGATGTTCGTTGTCATGCTGGTACATTCTTCCTAGGTCCGAGAAGGTAACGTG

TTCATACACGAAGAAGTACTTCTTATAAAATGGGGCAACCATGCTACCCCAGTTGGGGATGCTGTTTGGAGCGATGTTTT

TGTACCACGTGTACGCTCTCCCGGTAAGAGATTTCGAAAACTCTCTCAGACGGAGAACGTGATTGTATTCATGTTCGCCT

AAGGATTCGAGGAAGCGAGAGATGTGTTCACGTGCATTTCCAGTGCCATCATACAGGGTGAATTTAGGTGAGGAGTATCC

TCTTGGGAGAGGAATTCTTTGTAATTCAGATGGGTAAGGAGGTTGATGATGATGCATGTCTATTTCATCATTCTTGACTC

GATTCCGGAGAAGTTCCTCCATATCCTATCGCGTGACAAAATTGTACGGCTGACCTCTTTCCAATGGGGCTCTGGCATGA

ATCCCCTCATTCGCAGAGACGTGTCCTGCCGAGGTGCTGACATCGGGGGTGTTGGAGGCGTTTCTAAACGACTATGGAGG

GAGTGATTCACGCGGAAGATGCTTGATAAGTGCATATTTCACATATATTTGGTGTTAATTCCATACTAGTATTTGATGGT

TTTCTTGCAGATTATTGTGTGTTACGCGTGTTTTCTCTTATTTGTGTGTTGGAAGGTGAATAATCCAAAAAGAAGTGAAT

TCGAGCTTAGTTGTGCAGTGAAAGAAGCTAGCTAAGAGTGAGGAAGAGCCAAAGACCAATTGTAACTCGTTCAAATACAA

GATTTCTACTTTTCATGGGGTAGAGGACGAGAAGGCGAAGCCAACGGCGAAAGAACGGAGTCATTCGAAGTTCGTATGAA

GAAGTTATGGGCAAAATAAGAATGAACTTGGAATTAAACTGTCAGTCTTCCAGGACTGACAGTTGAATGGGCATCTCTTG

TTAAGGGGCACCCTCACTGTACGTTAAGGGACACCTTCATGTTAAGGGGCAGACTAATTCAAGGGAACTGCTAAAGGCAG

TCAGTCTCTGTGCATAAGGGGCTGCCACCAATTTGAATGTTGTTAAAGGCGCCCGGCTAATTCATCCGGGAACCCTTCTT

CTTCTTCACGAGTAAAGAAAAGTGGCGGGAAACCTTTCAATTCAAATATCTACACCTTCATTTCTTCTGGGATGATTTTG

GTCGTGGATTAGAGAGATTCAAGAGCAGATATGAGTTGGATTAAGCCTGTTTCGATTAAACAAAGATGGTAAGTCGATTG

GTTTTAAGTATAACAAAGGTATGAGCTCGACGATGAGAAGAAAGGTTGCTGCCATTAATGGTGATGCTCGGGCAGAGAGC

ATAGCATGCATGGGTGATCGGTAACTGATGGAGTAGCTCGAGATTAAAAGCAATGGAGAAGGTGATGTTCGGTGTTTGGT

TGGGATGACAAACATGTCATGAGAGCTCGAGTTTACGCTGCATGCATGGCCGGAGGAAAAACATAGCTGCTGTTCGAAGC

AGACCTCAACAATGGGTTGTCTTCCTGAGATGGCAGTGAACGTGGGATAAACTCGAGCAGCTGAACGTGGCAGACATGAG

AACGTGAGAGCATGGCAGACAACAAAACAAAGCGTGAAGGCATGGGAGAAACAGAACACCGTGGTCCATAGAATGCTCAT

GATGGGCTATTGGCATAAAAACAAGAGAAAGATTTGTTGCAGTCGAGAGCATGGGTTGTCTTACTGAGATGGCAGTGAAC

GTGGGATAAACTCGAGCAGCTAAACGTGGCAGACAAGCAGAACTCGAGAATGCCGGAGTGTTACTGTCGAAGCTATTGAA

CATACGGTGTAATGGTTCATGAAGGGTGCTGTTCAAAATGAAACAAAGAGACAACATGGATGTACGTGAAGAAATGGTGG

CTTGCACGTGAAGAGCATAAGAGTATGGTCTGTTGGCATGGCAGAGAACGGATATGGCAAGGAGTCGAGCAAGTTCAAGG

GACTTTCCGTGCACGTGAAGCAGACAACATCATCTGCAGGCAGTAAATAAAGAAAAGGATAAAGTACTACTCGCGTGCTA

TGGCTGTGAAGGAAGGTGAACTCGAGTTTGGCAGAGAGAACGTGATGATGTTGGAGCTAATGGAAGAAAGATGGAGGAGC

TCGAGCATGGGTCAACAATGCGCATGGGAAAGCAACACATGGAGCTGTTGGCAGTCATGATGATACCCATCGAACATAGC

ATGCGCATGGCCTGAAAGCAGAAAATTGTTGGTTATTTTGCCCTATAAATAGAACTCGAATTCATCCATTCAGGGGGAGC

CGGCTGGAGACCTCTTCTGCATATTCTTCTTTCATTTCTTCTTCTGTTTTTAATTTTTCTTTCATCTGTTGCGAGAAACA

GTGAAAGCTCCTGGCTTTCACTCTAGGCTTCCATTTTTAGTTTTATTATAAATTCCTTCTTATTTATTTATTTATTTGTT

AGTACTTTTAATTATGAGTAGCTAAATTTTTATCTAGGATTAGGGAGGAAATCCTACTATGTGTGCTATAATGTTTTGAT

TATTTTCTATAATAATTCTCTTGGTCCTTATTCTTGATTCGTAACGGTAATAATTTTCTTCATGTGTCCTATGATGCTTG

ATAATTGATTTATATATTCTATGTTAGAAACTAGCATTTTATAGTGAAGAAGATTGTTAATTCTTTGAGAAAATTAATTG

CTTGAAAAGATTTATTTTCCGAGACGTATTATTTGTCCTAAATCGTCTTGAGTAATTTTTGGAGAGTATTATAGGGAATA

GTTATAATCATTTAGTATGTGTAGTAGTGATGGAATCTCGACATCCTAGACTCTTTCAATTGAACACAAAAATCATTTTT

AGTATTTCCATTATTTTATAAGTTTTATAAAAATCACCATCTTTTTGTCTAAAAACCAAATCTTTAAAAAAAGTCCGAGA

ATCAAACCCCTATACTTACCACTTGTAAAACTGCATCAGGGTTTTTAGGAAGACGAGTACTTCTTTCTGTGTCAAAACCA

TATCCGTCTGGGCTCTAGCGAGATCCTCCTGGCATTCCATCAAATCGGCGATGGTAATATAGGGTCATCTTCCTATGGTT

CCTCCAGTTTCTCGTCCCGTCGATGGGGTAGTTAGAGTGTTAGTGGTGGTTGGAGGTGCCATGCTTGACGAAGCATTAGG

AGCGGCAACTCTGGCTCTGGTAACTGGTGGCGTGATGCCCGAAGGAGTGTTTTCAGCACTACCATGACTAGCAGGTTGTC

CAGGTATATTAGAAGCGGCAGCAGCTCTGGCCCTGGTGATTGGAGGTGTTACACTCGAAGGAGTGCTTTCGGTGTTACTG

TGGTCAGTAGGTGACACAGTGACGTTGTTGTAAGTAACGTTAACACCAGGGATGGGTTCCGCGCTGGTGTTCTCGGCAGA

TGATCCGGATCTGAGTTCTACCATGGTGGAGTTATAAGTGAAAGGTGATATTGGAAATTTGGGAATCGATATTGCAACCG

AGAGATTTATTTCCCACTGTTCTCGCCAATTGTTTGAGGGTGGAAATGGTTTATGCTGATTTCGGTAATTTTGGGTATAT

GGGTGAGAAACGAATTTAAACTCTAAAAAATGTACTGCACGGGAGTACTTTATATTCGAGAGATCAATCTGTACAAGTCC

GGCCTAAACCAAGAAATGGCCGTTCCGGACTTGCCTCAGTCACAAATTGAATGAGAATGGGTTGGTTTTGGGGAGGGAAG

CAAGGAAAGTGATGAGACGAGAATAGTTGATTTCTGGAAGAGTAGTTATTTTCTATGACTTGTATCAAAAAGTAGGAAGC

AAACAAATGGAAATCTAGAAAATGTTTTCTGAGTGTTGTATGCTCCTGACTGAACTTGTTGTTTGGTGGAAATAGGTAAG

ACCTATTTATACAAGTCGAAGTGAAACGTACTTTGGTCTCATTAAGGAATGGAAAAATGGGTGAGTAAATGGGAGGAGGT

GATAACCATTAACACGTGGAATTTATGTTCCATAAAAAGAGCGTTTCACCATTAACTCCCTATATTTACTAACCGCATCA

TCCTTATGACACTTCCTTATGACGGGCGTAGTGTACGCTGCACGCAGTAAACCACCAAACCAATACCCAAAGAGGATCCT

CCAGTTTTTGATATGCGTTGATGTCTCGGATGTTTTTGTGAAAATCAGGTAGCATATTGTTGTTGCCTGGCAAGTCGAGC

TTGGGAGACTTGCTAGCTTGGTGACCTTCGACTGTCGAGATTTTGCATCTTTAAGAAGAAAGGTAGTCGTTGATCATCGC

ACGCTTTCGTTTTTGGTGGCCGCATAGGGAATGACAATATGGTGGCATGGCTAGGTAGACATGCCTTGGCGCCATTTGGC

GTGGCCAAATTAGGGTTTCTGGTCAAAGGTTACCATGTGTTGTCTGGAAACCTTGGACAGCTTGGATTTGCACCTGGGAT

TGAAGGGAGACCGTTGATTTTCGCACGCCTTCGTTTTGGTAGCCGCATAGGGAAGGCCAGCATGGTGGCACCGCTAGGTC

GGCATGCCATGGCACGGTTTGGCGTAGCCAAACTAGGGTTTCGGGTCAAAGGTTATCATGTGTTGTCTAGTAACCTTGGA

CGGATAGGATTTGCTCCTAGGATTGAAGGGCGACCGTTGATTGTCGCATGCCTTCGTTTTGGTAACCGCATAGGGAAGGA

AAACATGGTGGATTGGCTAGGTCGGCATGTCATTGGCATGGCATGCCTTGGCGCGGTTTGGCTTGGCCAAACTAGGGTTT

CGGGTCAAAGGTTACCATGTGTTGTCTGGTAACCTTGGACGGCTAGGATTTTCTCCTGGGATTCAAGGGCGACCGTTGAT

TGTCGCACGCCTTCGTTTTGGTAGTCGCATATGGAAGGCCAGCATGGTGGCGCGTCTATGTCGGCATGCCATTGGCATGT

CATGCCTTGGCGCATCTTGGCGTGGCCAAACTAGGGTTTCTGTTTAAAGGTTACCATGTGTTGTCTGTTAACCTTGGACG

ACTAGGATATGCTTCAGGGATTGAAGGGCTGTAGAAATAGAGGACTATGCACGGGTCGAGGGCCTGTCTATTCTTTGTCC

ATAATTGTTATGGGGCGAGGTCCATGTGCTCATCCGTTCGTGGATTTTTATCACACAGGGTCTGGCAGAGATAATTTCTT

ACTAGTTTATTAGCAAGAGGGTCAATTCTGCAAAGAGTAAAAGAGAGGTAGATTGGAGTCGCCACCTGAATAAGGGGAGT

CAGGACCCTTTTAACCAGTTTGAATTATTTCGGGAAAACAAAACCAGTTATGACTTACATGTTTATGGTCTGCAATCAGA

GATTCTAGGTAAGGAGATTCTAGTACAGGATGGGAAGGTGTTAGGCACCCATCAACCGCCCATTCGTAAGATTGGCCTCT

ACTTCATGATTAAAGGGGAAAAATGTTTATAATTTATTGAGAATAGATTTTCATGCATGAGCTATTTAGAAAAGAGACAT

AGTTAGAAGTTGGGTGGAACTTCGACTATGGAGATGTGGTTTCTTTCTTTTTATGGTGGTTTGGGTGGAAACCTTGGTTT

TTTTTATCCTTTTGTTGTGGTTTGGGTGGAAACCTTGTTTTGTGGTTTGGGTAAAAAACTTTGTTTTGTGGTGGTTTTGG

GTGGAAAACCTTGTTTTGTGGTGGTTTGGATGGAAACCTTGTTTTGTGGTGGTTTTGGGTGGAAAATATTGTTTTTGTTG

TGGTTTGGGTGGAAACCTTATTTTATGGGGGTTTTGGGTGGAAAACCTTATTTTGTGGTGTTTTGGGTGGAAACCTTGTT

TTATGAGAAGAAATTTGGATGAAATTTCAAAATTAAACAATTAACCTGCAAATTAGCTCAATAAAAAAATAGAATTAATG

CAAAGCTTATTATTATTTCTTTTTGACAATGGGTCTAATGTGGGAGGGAATTCCGCTTTATACCCACACTTTGATTCATC

CCACTTGGAAAATTAATAACAAAACACAATTTTTTATGTAATGCAAAGAGTAGTCTCGGTGTTACTACGAAATGGAAAAC

CATTCCGTTCCCGTATGACTACATAATTTACCGAAGAGATATGCAATTAATATACTTGAAAATATGATATTTAAATTTGA

AAGGATTCTTACGATGATTGCTTGCTTCCGTTAAGTATGATGAAAGTAGTTAAAGTACTAGCAATTTAATGCTAATCTTT

TAAAAACTAAAAGACAATACTTGATTTAATCTTCTTAGTTTTTGAGTTTTCTCAAAATTAACTATAAGTGCACAAGACAA

TGATAACTATTGTGTAGGAAGAAAATTTGATTTCTCTTTGATTAAATGACACAATACATGGATACTTGAAACATGTGAGA

AGAAATATTTTATATTTGAGAAACAATAATTCTTCTTGGTTTAGGTGTAAACCTAGATTTTTGGATTGTGGTTTGGGTGG

AAACCTTGTGTTGTGTGGTTTGGGTGGAAACCTTGTGTTGTTGTATGGTTTGGGTAGAAACCTTGTTTTGTGTGGTTTGG

GTGGAAACCTTTGTTTTTGTGTGGTTTGGGTGGAAACCTTGTTTTGTGTGAAGAAGTAAAGGAAGAAGTCTCCCTCCTTT

TCTCCCTCTTTCTCAAGTTCCTCTTCTCAGTTTTTTTTGTGGTTCGAAGAAGTCCTTTCTCTCTCTCACTCGATCTCTAT

TTATACATAAAATTAACCAACCATTTCCTCTAATTCCTCCGCCAATTCCGGGCTGGAAATGTTGAACGAAAATTAAAAGG

CGGCTTCTGCGTCGTCCTTTTCTTCTTCGCATGCATTTTTTTGAAATTTGCATCTGTCCTTTTGTTCGTCGTCCTTTATT

CTTCTTTTTCTTCCTTTTTCTTCAATTTTTTCAATTGCATGCGGCAGTTTAACAAACATGATTATGACCGTGGGTCCCCC

GCACTGTTATCGTCCACAGTACTGTTACTGTTTATGTTGATGTTACTGTTCACACTGATGTTACTGTTTACTGCGTCAAA

AACATGATTGATGTGAAATCAGGCTCTTACCCAAGTTCCCTATCGTCCCCAGGTGCATGGTGCATCTAGAATGGCTAGGG

TCAAGGCAAGGTGCGGTTTGGCTTGGCATGGTGGGGCTAAGGGTTTGGCGTGACCAAGGTAGGATCATGACCAAGTTAGG

ATCGTGACCAAACTAGGGTTTTGGCATAGTTTAGGCGCGGCCAAAAATTAGGGTTTGACGTTGGGCCAAAGGTTACCACG

CGTTAGCTGACAATCTTGGACAGCTAAGATTTGCATCTTAGATGGAAAGATAGCCGTTGATTGTTGCAGTCTTTTGTTTT

GGGCAGCCAGCGTTGTGCATTGTGTGTTGGGACGGCAAGCATGATTGGTATTTTCTAGCGTGGAGATGTAGCTGGCACGA

CTTACCATTGGCACAGGGGTGCGGCTAGCATGGTTGGCATGCCTTGGCGCGGTCGGTGTGGCTGGCACGGCTTTCCATTG

TCATAAGGGTGCGGCTAGCATGGTTGGAATGCCTTGGCGCGGTCGGTGTGGCTGTCACGGATTTCCATTGGCACAGGGGT

ACGGCCAGCATGGTTGGCATGCCTTGGCGCGGTGGTTGTAGCTGGCGCGTCTTTCTATTGGCACAGGGGTGTTGCTAGCA

TGGTTGGCATGCCTTGGCACGGTGGGTGTGGATGGCGCGGCTTGTCATTGGCATAGGGGTGTGGCTATCATGGTTGGCAT

TCCTTGGCACGGTAGGTGTGGCTGGCATGGCTTGTTATTGGCACAGGGGTGTGGCTAGCTTGGTTGGAATGCCTTGGCGT

GGTGGGCGTGGCTTGCATGGCTTGCCATTGGCACATGGGTGCGTCTAGCATGGTTTTCATGCCTTGGCGTGGTGGGCGTG

GCTGGCACGGTTTGTCATTGGCACAGGGGGTGCGGCTAGCATGGTTGGCATGCCTTGGCGTGGTAGGTGTGGCTGGCACG

ACTTGTCATTGGAACAGGGGTGCGGATAACATGGTTGGCATGCCTTGGTGTGGTAGGTGTGGCTGGCATGGCTTGCCATT

GGCATAGGGGTGTGGAAATTAGGGTTTGGCTTATTGAAACCCTAATTAGTATTTTAAAAAAAAATATACCCTGGTAATTT

TCTCGGAGACACATTAAAAGGGCTCGATAAATGTGCCTAGTAGAGACATTATTACTTAAATTTTGTGACGTCATTGTCTG

ACTATGGTTTTACGATTTTAACCCTAAGCTAAAAACCACCATCAACACAAACATATACACATTTAAACTAAAAATTCATT

CTTAAAACCAACAAGGGTGCTGAAATATTTTAATTGGGTGGCCTAGTTTCAAAGAGAAGTTGTGAACCTGCATTCTTGTA

CAAGAAAGAATCCCTAGAGTCTCCCTTTTTTAGCTAACGAACACGATAAAGAACACATGCATAGAGTTGAATATTTATAA

AACTGATAAACTACATGTTCTTTAATATCGGCGTAATTCTTCTTGCTTCAAACCTCTCTTGAACCTACACATAAAGTGAC

CAATTAAATTGCAATCCCTTTCGTACCAAATCAACCTGCTCTGTCAAACCTGCAAAGAGATACTACTAAGTTATATTGAG

CTCGTACAGTACTACAATGTCGTATAAACAAAGACCAATTTAAGTACAAACTTAAAGGAATGAAGAAGAAACAAGAGCTC

AAAAACAAATCTTGTGTAAGTAAGTATCAGCACAATCAGGCATATCATAATTGATCACAATATTAACGCGTTCAATGTCT

ATTCCTGTATTAACCAAATCCTTGAAACATTTATATCGAGTCAACCTATGAAATGTAAAAAAAAATCCAACTTCAGGGTA

TGCCTTCGCTTTTCTTAGTGATTTAATTTTCAGCAGTAAAGGCAGTATTATGATGACTTAACCATAAGATTATTTTAAAA

TAAATTACAGTAGTATACTTTTACATTAACCAAACCACTAGAAGCTTCACTAGGCACTTGCACCTTCATGCATACACATC

AATGTAACAAGACCGAAAGTAATTAGATAAAATCTAAGAAAAAAGAATGTATACCTTTCTTCCCGGGACATTCCAGAATG

AGTGCAAATTGATGGGAAGTTGCACTGAACATGAGGCACTCTTTTTTCAATAATTCCTTGGGGGTCCTGATGCTAAAACT

ACCATTGAACACAACAACATTAGTATCAGGCGGGGTAAGTGTTGAACCTCTCAAACTCTTGGAAGATCTCAACTTTAGAT

ATAAGCATTAACATAAAATGATCAAAATAAAGTTTAGTTGCAAACCTTGTAAGACAACCTGGCGTTGGATCACAAAATTA

GAACAACTTATCCATCCACATGCTCAATTTTCTACAGTGTCAATAGGCCAAATACAATAGTTTTTCTCATCCCATATTTT

CCTTGACAGATAACATCCATAACCAAAAATAGCTTCATTCATGTTGAACTGATTGAAAACCAAAAGTTCCAAAGGACTAG

GCTCTTCCATAAAGAAAAATACAATTACTAAATAAGTCCAAGTTTCTAACTAACAAAAACTATCTATGGGACAACACAAA

ACAAGTCTCGGTATTACTACTAAACTGCAAACTATCTATAGTGGAAGCGTTCCTTTGTATAGAACATAAAATCACACAGC

AAACGGAAAAAAAAATGGTCACTATAACTAGAAAGATACGTGGAGTCATGGAGTACGCCTTATTTAGATCTAAACACACT

AACAAGGAAGCGAATAAAAGTTGAACTTCTGAGGAAGTTATCTAATACGAACTGCCTTTAATTATCCAGTTAAGAGATAT

AGACAATTTAATTTAGATGCACATAACACCAAGATAAATAAAAAAGACGAAAAACCATATTTAGATGAGAGAAAACCTCA

AGGGGAGTTGAAATGACTAAAATATATCTGATAGCTTGTTCCTCACCACTGTATTTTGGACATTTACCATCCATATGAAA

ATATAGTGAAGCAAAAATATAAAAGGTTGATTACTAAAGATCCCAGAAGTCTTCATTACTCTTTAATTTACATTCAATCA

CAAAAATGCCTTCAGATCTTGTGGCGTGAACTATATGGGAACAAAAATATTATATGATTAAAGCAACTCTCTGTCCCACA

ACATGAAGACTTATATCTTCTCAGATAGTACCTAGTTTTGATATTCTTATGAACCAGACGAAATCTTTCATACTATGAAT

TTTTCACCAACACAAAATAAAAATTCATAGATTCATATACTAGCTTCAAGTGTCCAACTACCCACACACATGAACGAAAC

AACACACAACATGATACAACTATAAATCTGTACACACAATACCTTTAAGTGATAAAACCCTTGAAAATAAAGCGAATAAA

AATCAATTGGATATATGTTGAATCCCAAATCGACCACTGCTCTTAGAAGCTCCGGATTTTAATAAAAATATCTTAGTCCA

TAACTGTACATTCCAACATACCCCATGTACCGTTACAAAAGATTCACATTAGCCTCAAACTTTAAAATTTAAAGTTCCAT

ACATACTACCACAAATCGAAAAATAATTCCAATTTCTAAGGATTTAGGAAAAAACCAAATCAATCTCATAAATCCAAAAT

CATATCAATTTCACTCGCAAGATGTAATTAACTTTCAAAAATCCAAAATCAATATCAATTTCATAAAAAATTATTTCTAC

ATTCCAGCAAAATCAATCAAAATACAAAAGTTACATCAATTTCGTTAGAAAAATTTCTCAAATCTAGGGTTTTGAAAAAT

ACTAAAACAAGCTCAGTCTCAAAAGACCATTCAAAAAACAAAATCTAAGAACCAAAAATAGAAACTAAATAAAACTCATA

AGTAAAATCTATCAATTATAGATAACTAACCTCACGAAACAGAGAAATCTTTCTAAGAACCCTAGATTTCTTCCCACAAG

ATACATATGAGAGTTTCATTTGTGATGGATCTCTTTGAAGAATAAGAAGAGGAAGGGGAAGAAAAACCCAATATTGATTC

ATATGCGATTTATTGTTATGATTGAGAAAGAAAATCGACTGAAACATAGAATTCTAAGAAAGGGTTAAAGGAAGAAGAAG

GAAGGAGGCCGAAGGAGCAAGACCGAGATAGAGATAAGATAAGAGGGAGAAGATTATAGAAGTTTCTTTCTCCCACAGAC

GTTTCTCAACTCCAAATAGAGATATGAATTTTCACAAATATACCCCCTAAGAAAACTACTGTATCAGAGAGAAGTTTTAG

TAAAACGATTGTGGCTCACAGTACCTATAGTTACAGCTGCACGGAAACATTGGTTTTCGTGAAGACCAGAAAGTGGTTAA

AAATGCATTTTACACTAGTGGAAGAACCCGAGAAAGCCTTTCAATATGAGATATTTCCAAACTTAAACGAGCGCGAGACA

TCATGAAAAGGAAAGCAGAAGAACATGCCAAAATTGAAACAGCTAAACTTCCTTTTATAAGGAAAGAAATCTGGAGATAA

TTAATTATAGCCATTAATGGACCTAATTATGGCTGATAACACTGAAAACAAGACAAACAAAATGGGAGCACCCATAAGCA

ATTTATATCAGCACTCTGCAACTCAGCTTCTCGACATCGGATGATAGCTTGATCCATTCGGCCACCTCCTCCATAAAAGC

ACGGGCTTCAACCAACGAACGACATGCCGTCACATATTCGACATCTACATCTAGGAAAAGTTATGGAGTACGCTCCAACG

CTTCCAGAGAAACCTTTTATTGAAGCGTCATCCCAGACGTCTCTGATTATTCTACCCATTTGGCACAAAGGGGAGTGCGC

ACAGCTTGACGATGGGCCAATAAGCATTCCACTTCCTCTCGTACCAATTGTAAATGGTGAATCACTTTCAAAGAACTACT

CCTAATTACCAAGCCGAACGATACTCCTAGGAAAACCCGAATGACCTTCTCGATGACAAAACAACCATGGGGCTAATGTA

AGCACATGGATTTGGCTAAAGACGAAAGCACTAGAATGGGGGCTTCGGATGATGACTAACCGAAGCATGACAAGCCGAGG

TCGCCAGCAACGATAACGTGTCATGCCGAGGCTTCCAGCAACGATGACATGGCACAGACGGGCTGCCCCGAAGAGATGAT

TAGTGATTGCCTAAATAATCCAAATTGTAATAGTACACGTGTACGGCTAATGATAGAAGACTTATTAGAAAAGGTATTAA

GGTACACGTGTACGGTATTTCCATATCAGGCTTTGCCTATAAAACAAAACAATTTTCCAGAAAGAGGCAGAGCTTTTCAG

AGAATTATATTGTATATTGCGATTAGTCGCCTCATAACCTTATATAATAATAAGTAGGAATTACCTATAGGTACTAGTTT

AGTGATACTAGTTAGTGGAAGGTCCACCCACTAAAGCCCAGTAACTAGAGGTCCATAATAAAAAGAAAACATAAAAAATG

ACCCAACTTTTTATCACTTGGTCCTATACACGTGCGAATCTAGGAAGATATTTTCGGTAAATGTCTAGTTTACCCTTCAA

ACATAATAATGATAAGACTAAATCATTCACAGACATTTCATTTTTATTTTCATCTTCTATTTCCACGACTCGAGCTCTGC

GTTAAAGATCGAAGAAACTCCATTTGAAAGCATCACCGACAAGTTTTATTGAGAATCATCAATAAAATCCAACAAATAAG

ATAAGTTATTGGTTAGATCGGTCATTTCTGTTTTTTTTTATGAATCTTGTTTGTATTTATTCAGTTATTGGTTAGATTTA

TTAAGTTTTTATCTTATTTTAGCTTTCGATTGGATTTTTTTTCTGATTTCTTTTCAAAATCAGGTTTGTTTGCACTTTCA

GATCGTATTATCCAGCAGTACATATATGACATACTCATTTTTTCCACTACAGTTTTTGAATCATGTTTAAGTTTTTAGCT

TATTTTAGCTTTCGATTGGATTTTTTTATGTTTTCTTTTCAAAATCAGGTTTGTTTGCACTTTCAGATCGTATTATCCAG

CAGTAAATATATGACATACTCATTTTGCACTTTCAGATCGTATTATCCAGCAGTACATATATGAAATACTCATTTTTTCT

TCTATAGTTTTTGAATCATGTTTAAGATTTCAATTTTGGATCATGGAGATCGGTCGTTTCTGTTTATTTTTATGAATCTT

GTTCGTATTTATTCAGTTATTGGTTAGATTATTAAGCTTTTAGCTTATTTTAGCTTTCGATTGGATAGTTTTTCTATTTT

CTTTTCAAAATCAGGTTTGTTTGCACTTTCATATCGTATTATCCAGCAGTACATATATGACATACTCATTTTGCACTTTC

ATATCGTATTATCCATCAGTACATATATGAAATACTCATAAAAACTGCTCTTAGATTATGTTTATTTCATAGTTTTCTCC

CTTTTTTTAGTTTGATCTATCAGTACGTATATGAAATACCATATTATGTGCTTCGATTCTGCTGTTTCTCCTAGATTCTT

AATTATTCCTATCATGCAGTTGATTTGTAGGTCATGAATCTGCAGTACATATATGGCATACTCATAGAAACTACTCTTAG

ATTCTGTTTCTTTCATAGTCTTCTCACTTATTTTAGTTTGATCCATCAGTACATATGTGAAATACTGTGTTTTAATACTG

TTACATCTTTGTCACATTTGCAGGTTCAAAACATGTTTTGTGGGGTCCAACCGGATTTTATTGGTTTATGAATCCGCAGT

ACGTATATGTCGTACTGATAAAAACTTCTTTTGATTATGTTTTATTCATAGTTTTTGCCACTTTATTCTAGTTTTGTTGT

TTTGTTTGAATATTGTTATATTCTGGTCACATTTACAGGTTCAATATGTCCAACGAATACATACTCTATATGTAAGCAGT

GCAGAAGATATATACTCAGATTTGTAAGCAATGTAGCATATATGTACTCGAATTTATAAGCAGTGTAGCAGATATGTACT

CGAATTTATAATCAGTGTAATAGATATGTACTCCATATAATGAACGTGTAAACACACACGTTTGGTAACAATGGGTATTA

TGGTCATTTCCATGTTTTAATTAATTTCGGACCTCAAGATAAAAAAAAAATCCGGTTGGACCCTACTATAAATAAATTTG

TGGGCTTAGGACCAAACAAGAAATTTTCCTAATAATAATATAGGGTGTTTACAAGTGGCACATTTTTTCGCAAGAAACCT

CTAACAAATTTCGGTTAAATCAGCTTAGATATGTCCCTCCAACGGAATCAACCCAAAAAAAAAAACCTGGCTCCATCATC

GTCGGTAGACCAAATCGTGTATATTATGGACCAAATCATTATTTCCCTAAAAATAAATGACAATAGGGATAAAATCGGAA

GAAAAATCTCATGCAAATCTCAAGTTGATTTATTATTTGTCATCCAAAAAAAGATTCGGTACATTAGAAATGGATCGATC

CTCTGGACATTCGTCACACAAAACAAGCTAAAAACAGAAGCAAAGTGATAAAGCTATCAGTAGATAGGACAGTACTAGCA

AACAACTGATGCCATTTTTATTCATTTGGCCGCTGAATCGATATAAAATCAAAATGCAGATACTTCTAAACAATCATAGA

AAAAACCAGAAGGTCCATCGTCGGGGAAAAGAGCTATTCTGACTACATGTTTAGCACCTTCGTCGGCAGTATAATTTCCA

ATGCCGTAGCACATTTCTGTTTTAACCGAACCAGGACAAACACAATTGACCTGAAATTTGGGAATTTTCTTTGCTAACAC

CCTTGTGTAAGCATTCAAACATGCTTTTGATGTTGTGTATGCAGCTCCGAAACTAGGCCACCGATTTGTTTCGATCAAAT

TTTCCTTAAAATCCTTAAGAAGCATATTCCCTACCATGTCAATTCTCTCTTCTGTTAATGCATCACCATCTCCAAGTATC

TCTAGAGCTGTTTCATTGGATACATACTGAAACCAAACCAGCGTAAACAAGGTGTAAGAAAAACATTTTCGGATTTCAAG

TTGATAGTTGCAATAATAATTACTACCTTGAGGCTTCCCCTGGATGATGAAACATTAACAATTCTTGGTGAATCAGATAG

TCGAAGTAAAGGAATTAGAACTTCGGTTACCGATTTAACACCGTAGTAATTTATTTTCAGACATTCTTCTGCTAATGCAT

ATGTCTCTGTCATTAATTCTTGGGCTTCTGGTTTTTCGTAGATCTCCACGATCTCCTCTGAATCCTGGAAAAAGTGCGAG

TAACTAATTCATAATAGGAAAAGCATGCAACTATAATAACTTTCGATTTAGCCATTTAACTCTTTAGATTTCAAGAATTG

ATATGGCGCCGATGTCACTATAGTATAGTTACAGTTGTGAAGTCTTTGGCCTAAGTTTGAAACTCATCGGCATCAAATTT

TTATTACCTCTCCAATGTCACTTATCATTGCCTTAAAACGATCAGCATCAACTGAAAAACCTGCAACCCCAGCATTGTTT

ACCTACGCATAGTAAAACACAAATTTTTGAATGAACTTGTGGGAAAGATTGGAATATAGCTAGTTCAATAGGGAGTTTAA

AGAAAGAATTTGGATGCATATATACCAAGATATCAAGCTTTCCGAAGCGTGCTTTGATGAAATCCGCTAAAGAAGACATA

GTAGTAATTGGATCCGTAACATCAAGTTGATGAAAAACCACATTCTCATGATTAGAATTTTTGAGATTTTCAACAGCTTC

AAGACCTCTAGTTACATCTCTACAAGTTAAAACAACCATGATTCCACTAGAAGATAATTGCTTACAAATCTCAAATCCAA

TACCCTTGTTTCCGCCCGTAACAACTGCACACCTGAAAATAAAATCCAGAAAGAAAAGTTTCCATTGATTAGAACGACCC

TTAATCAGCTTGCTAGCACTAGATATATATCCAAATCCAAATTATGTAACCATTTTTTTTGTTAGTACCTCATCTTTGTA

ACAGTATTTGAACATGTTTCAGCCATACTTGTTGATGATTACTGGATTGGATTAACAACTATAAAAAGAATAAACGAAGA

CTCAAGCTATGAAAATCAACGTAAGATTCTTTCCGACTGATTTTGTTAAGAGAACTTTCTATTGATTATAATTTCTCATG

ACCTGAACAGAATAACAAGGCTCTCTTTATATATAATATATAGGAACTGATGTTTAGTCCCGAAAGAGTTGGACTTGTTA

CTCTCTGGTCCTGTCCTTTCCGAATTTTAGATTTCTTTTGGTGGTTTAATCCAAATATTTACGATCTGTCTAACGGGACA

CATGATAACGTCAGTGATTTACAGGTTAAAACCATAGCTAAAGTGTTTGGCACTCTCTTCACTTTTTTTATATTCGAATA

AAAAGGATTATATAAATAAAAATATTTGATTATTGTAAGAGCTAGTTATAGCTCCATCCATGTCATCATCAATAGTCACT

AAATCTGTCATGTGTTCAGTGACAGCCCGCTGTCTAACTCGCTTATTATATAATGACCGAAGTCTTTTAGTGGGAAACCT

TTGTTTTGTAGTTGAATAGCAGTTGAAGACCATATTATATATATATCGATGTAATAGAAAGCCGTCTGATTGTGGGGGTG

GATCAGTGTTCCATATCAGCCCACCAGAGAAATGCAACCTCGTGGGAACAATTAAGGTCACCCATGCCATATGTGGCTTC

TATAGAAGGCGTTTGTACCGTGTCGGGAGTTACTGTTTGATGGGGCTAAACCATCCAAGAAAGGTGGTGAAATTATAAAA

AGGCTGTAAGAGTATATATTAAAAATTAGGAATTTGAAAAAAAATGTCCATGTGGAAAAATACATTTAAAAAAACCTGGC

ATATTTAGGGGCCACTCAGTCGCAATTAGGGGCGACACTAAAAGACAAAAAAGATCACCCAAATGTAAAACTGAGCTATC

CCTTATCCGCGATTTTTCAGAATGGCAACTATATCCTTTTATAATCGGTAGCCAAAACAACAATCGGTAGGTTAGTTGTT

AATCGGTAGGTTAGTTGTTAATCGGTAGGTTAAATTGAAACTATACCTACCGAATATTGTGTAGATACTTCGGCAAATAA

TCCTTTGTTTATTTGGTAGCTTATCCAAATTTTAAGGCTACCGATTATCAGGAGACCCTTATATAACCTTGGCCAAATCC

CGGAGGTTCTGAGGGGTGCAGAGCCAACCATAACCACTTGGCTTGAGTTGTGGATTTATCGATAATCGGACGGATAATAT

GTTAACACCCTACCGATTATAAGTTGCCCCAAAATGATATTATTTCTAAAATCCTTCACGTAGAGAATTTTGAAATCTTC

GATAATCGGTAAGTTATACAACTACAATATCTGCCGATTATATCGAGCCCCAGATCAAACCTTTGCCATATTTCAAAGGT

TTTGAGTGTACACAACGAGCCATAACCACTTGGTGCGTGTTTTGGGTGCAGCTGTAATCAGAAGTTTTTTATATTACAAA

ACCATCCGATCATAAGTAGCCCCAAAATGATTATTTTCGAAAATCCTTCATGCATTGAATTTTGAAATCTGACATAATCG

GTAGGTTATCTAACTTAAACAACTTACCGATTATAAGGAGCCCCAGATCGAACCATTGACAAATTCCATAGGTTTTGAGT

GTTCAGAGCCACGGATAACCACTTGGTTCGTGTTTTGAATGTAGACATATTCGTAAGGAAAATTGGTTTCACAACCTACC

GATTATTGGTTGCCCTAAATTCAAATTTTGTAAGCTACTATATTCGGTAGGTATTATTAATCAAATTACTACCGATTATT

GGTTTCTCCACTTTCAACCTTCGTCAACCTTCGTCAAATTAGTCGTTGCATGTTGAACCAAATTTCATAAGTATCCGTTG

GAGTTTTTGGGAAAAGGAAAACAACCGTTGGATCCTAAGTCTATATAAAGAAACTAATATCTATCAGCCTAATTGCACCA

AACTCTTTTCTCTATTCAGTTCTAATTTAGAAAACCTAAAACATGGATATTGCTTTTGCAGAAGAAGAAGATTGTGCTAT

TTACAGAGCGTGGGTTTTGGTAACTGTTCATGATATTGTTCACAAGCTCAACACACAAGAATCCGAAAAGCAGTTATGGA

ACCGTATTTTAATGGTATTTGGGGCGTTAGATGGTAATCGAATTCGAAGATCATCCAATGATATTAAGATGAGAATTAAG

GCGATCGATAAGGACATTGACAAGCTTGAAGATGAAGAACGATTTGTTAGGGATGCTTTTCCTCAATGGACGTATCAAAA

ACGAGTAAGTATGTTTCATACTCCGGCTCACTTGTTACCCATTCGTTAATACTAACTTGGTCATTCGTAATTTCATAGAA

ATCTTGTTGATTATCGGTATATGGACCTCTACGGGAAACGATTTAAACATGAAGGATGTTACAAAATCAAGAGGGACAAT

TTCTGCGGTGTCTGGACGTTATGATCTTTCGTAATACTTTTATAGTCAATAAGGAGTATGGATTTAGGTGCTTTTGACGA

AATTGTAAGGTTGTGGTCTTTTGAGGTTTATGTAATGCACTTAATCGGATGAATATTAATCTAAAAAAGTTTCTGATTAT

TCATATTCGGTAGAATCTCTCTCCAACTAAGTTCCCGATTAACATATTCGGTAAATTTGTGAGTCCACATACTTTCCGAT

TATGTTGTTATTTGGTTCCAGGAAACCAAGTTTTTTCACACAGAATAATCTGATATTAAAAAAAACATATATCCTACCGA

ATATAACTATTCGGATGTCAGGGTGAACATTTACCTTCCGATTGTTGGGTTGTTTGGCTACAGAAAATAAAGTTTTTGGG

CACAGAATAATCGTAAGGAATTCATAAAAAAAAACTACCGTTTAATATAATCGTTAACCAGTGAAGGAATATAGATTCTC

TATTAGGGTGTTATTTGGTTCCAGGAAACAAATTGTTTTCACACATAATAATCAGAAATTAACCATTAAAAATCCTACCG

AATATTTATATTCGGGAGGATTGTTTAAAAATAACCCTCCGATTGTGGTGTTGTTTCAGTACAGAAGACAAGTTTTCTTT

ACACCTAATAATCGAAAGGCAGATTAAATTATTAACCTTCCGATTCTGACTAAATGTTTTGCAAAATCTCCAAAATCGGT

TGGTTAAAAATATATAAAGCTTCCGATTCTGACAAAAAAAATACAAAAACCCCAGAATCGGTAGGTTCTTATATAAAAAA

TACAGTTATAGGTTCTTATATAACTGTTCCAGAATTCAACAGTTACATGTCTTATCGTGTGATTCGCTTTCGAGTCCTCG

TTAAATGCATAAACATAGCAATATAGTCATTCATAATCCATAAAAGGAAGAATTCACAATCAAAAAACATAACGTACCAA

CTAATAATCCGACACAAACTTTTAAAACGACATAATCCGACATAAACTTTGAAAACCATAAAGTTAACTAAAAACAGCGA

GAGTTTACAACATCGAAAGTTTATTTCTTCTTTTTTCCCATGACTTTGCGACCTTTACTGCCTCTACCACTTGGACCAAC

ATCATCACCGGTTTCATCAACTTGAGTGCTCGATGACGCCCGAGTTCTTTTTTTCACAACCTTTTTAGGTTGCTTGCCTT

TATCCCCAAACCGGGATGCATAATCTTCATTGTCAATGTTATCAATCATCTCTTGGTTCTCTTTTATCTTGTCAAGTGGC

ACATGCTCTCCGGCTTTTATGCAATTAGTGCACCATTTGGACAAACCTCTTAACCTGTTCATCTGTTTTAAAACATAGCA

TACCATCAAGGACGGTTCTAGTCTGATTTTGACCCATAAAATAAAACTAAATTAAGAAACTAATACTTACCAATCTATCA

AAACCCGGAGGTTTATCTTTGAGGATACTCTTATACGGTGCGGGCTTTGCTTTAAGATCACGGATTACGCGATAATGAGA

AATATCCAGATACCAATCCATATAGTTGGGAGCATTTTCAAAACCGTTGTTGGCTTGCCTTCCACCGTTGATCAAGTAAT

CCATTATTTTATCCCAATGGTGTTTGCAGGATGGGTTACCCTTGTAAACCACTGTGATAGAAGCTTTATCTGAATTACAT

TTATCCACATCCAAAAAAAAATTCCCATTGATTTGCCACTTTGGTAAAGCTTGTACATAACCGTTTTGTCGCATCACCCT

CGAGGCATTGTACATCACGTATCATGTTGGGTGCCACAATGGCCCAAAATAAAGGGCCAAATCGGACCGACCTGCGATAT

GTCCATTATCTCGGTCTTCCTTGTATGGATCAAAGCATATGTCTTGGGCTTTCAGTGAGTCCAAAACCTCCCTCAAACGA

ATTAGATGCTTCTCCTTTGTTCTAGAATGGTTGTCATCGAAGACGTACTTCTTTCCTCTAGGATGAACTGGCTGTGATTG

TGGGGCACTTGGTTGAACACCTTCTTCAAGTGGGTCTACTTATGGGGCAGGTTGTCGAGTTCCATCTTTAGCACTTGAAG

TGGATTTAGTTACCCTATCTCTCATGGCACTAGCTGTCAAGTTCTTACGCCTTTCCATCCTTGCACTATCCGAGACAACT

GCGTCTTCGGGTTTTTTTACTCGCCTATCCCTAAATAGCATGAAAAGGTAAAATATGTTTATAATCAGAAGTATAAGTTA

AGCATAAGCTTCCGATTATAAATAATCGGAAGCCTAATTTCAACATACTCCAGCGAATACACATAATCGGTTGCCAAAAA

AATATTATACTTTCCGAATATGCAGCACTTTGTGTCCAGGAGATACATAAGCCTATTATAATCGGAAGGAAACCTATGAT

TTTAGCAACCGAATATGATAAATGTTCTTTAACAGTGAGAATTGGAGCAATAATCGGACGTTTACAAAAATATATAACCT

CCCGAATATGTGTAATAACCGAAAACCCTAAGAAAACATTTCGTGGTTTCTTCGAATTAAAGTAATACAAACCCAAATCT

AACGGTAGAATGTTACCTAATTGTGGCCATTTGAAGTTTCCTGAGAGTTTGGTTATCAGAATCATCTACATCAGGGACTT

GTTCTTCATCACCTTGTTCGCGTTCTTCGTGTTCTTCGGTAAGAACCTTTTTATTACGGGTAGGCACTCCAATGTTTGAA

TCACCCCCACGAACCACATTTTGGTTTCGACCCGATTTACGTACTCTATATTTCATCTTCTTAGAGAATATTCGACGATT

ATTTGATTCGACGATTAATGATAACGATGTATTGTAGCGAAGTTAACGAGGGGGGGGAGTTTTGCTCTCTGTATTCGGAG

AGCAAGAATAGGATGGGAGAGTTGAAGTTTAAAAGAAACGAAATGAAAAAAGAATTTAGGTTGACATCGGTTTTATCTTT

CAAAGGTACTATAATCGGAAGGGTATTATACTAAATTAACTTTCGATTATAGGTTGCCCCAAAAATGAGTTTTTCCAAAA

ACACTTCAATTCCAAAATTTTGAAATTAAATATAATCGGTAGAAAATTTTGCTACCGATTATATGTTTCCCCTAACCACT

TGGTGCGTGTTTGCGACTTATCATAATCGGGATGTTTCTATACAAAATTACTTTCGATTATAGGTTGCCCCAAAAAGGAG

TTTTTCCAAAAACCCTTCTATTCCAAAATTTGGAAATTAAATATAATCGGTAGACAATTTGGATACCGATTATATGTTTC

CCCTAACCACTTGCTGTGTGTTTGCGACGTATCATAATCGGAAGGGTATTATACTAAATTAACTTCCGATTATAGGTTTC

CCCAAAAATGAGTTTTTCCAAAAACCCTTCAATTCCAAAATTTTGAAATTACATATAATCGGTAGACTATTTGGCTACCG

ATTATATGTTTCCCCTTAACGAACCTTTGCCAAAATTGCACAGGTTTTAGGTTAATGAGCTAGCACTAACCACTTGGTAT

GTGTTTGCGACGTAGCATAATCGGGGGGTTATTATACTAAATTAACTTCCGATTATAATAATCAGACTCTTTCACATGCA

ATCGGTAGGAAAGGTACTATATTATGGTGTTGGGAATTCGTTTTGGAGTAATAGTCCCAATGACTTCCGATTGAGTGAGA

GAAGAAGAAGAAAAAAAGGAGAAGTGGGTATATTTTTCCACTACTTCCGATTAAACATGAATGGGTACGAAGAAGTACGT

GAAGTTGTTGAAGTTCAAGATTCATTACCGTTTAAAGATGACGATTTGGGGTATTTATTGAATGATCCACACTTTGATGG

AGATGAAACCCTAGAGGACGATTTCATGGAAGAATATGGACAATCACCCGAAATCAAAGGTAACAATGCGTCAAACAATC

AGGTATGTTGTTAACAAACTTTAATAGCAATGTTATGTGTGCTTGTGTGTGATTTTGAACGTGAAAATTTGTCACATGTT

ATGCATTGAATGACATTAACGACTATTAATCGGTAGTCATATTAGTATAATTATCTACCGATTATTGAGAATCGGGAGTA

TTTTATGTATTAACTTACCGAATACAATAAAAATCCCAAATTGAGATGTTCAATTTGTATTGATCAGAAGTTTTATACCT

AACAATAATCGGTAGAATATCTCAGAACTAAACTTCGATTATCAATAATCGGTAATGTGTTTTAGTATTTGATTACCGAT

TGTTATACAATCGATTGAACAATCTTTAAACTTTTATCCGATTATTAATGCGTAACTTGTTGTTGTGGTGCGTAGGCTGT

TGATGACGAATATTATTTAAGGCCTGATACTACCGTACACTATGAAAACGATTTGGTATGTTCATGAATTATCTTTTCTT

TTTCGAATCATGTATGTTATATGGTTATGCTTATTTTGTTTGTTTTGTTTTATGCGCATTTAGATATTTTAAAGTAAGAA

GGAGGCCAAGGCTTGTCTAATGAACAATGCAAAAGAGAACATGTGCGTCGTAGTTCAAAACCCTTGTGTGAAGAATATTC

GGTTTGAAATGACTTGTGAGCGATGTGGGGTGAAGAATAGTCACGAGGGAAAGAATAGTAAGTATGTAAGAAAGATGGAG

AGAAAGAAATAACGAACTCATACAAAGAAAATTGGATGCCCATTTAAGATTGTCTTCTATAAGTCCAATGGTAATGATAG

TGTAGAGTGAAAAATGTCTAAAGTTGCTAATGGTTGTCATAACCATGATGATCCGGATACTCTTGTTGGACATGTAATGG

TTGCGAAGTTAAAACCACATGAAATGGAGATGGTGCGGGAAATGCGGATCCAAAAGGCGAGTGCGATCTTAAGTAAAATT

AAGGCGGACGACCCCAACAACTTGTCTTCTTTGTCTACTATTCAGTCGGCGCTAGCTACGATCAAAAGAATGGAATGGGA

TGGTAGAACGGTCATGCAACAATCTCAGTGGTTAGCTGCATTACACGACTACACATTGAGAACGGAAGAAAAGGATGGTA

AAGTGGTACATATTTTGTGGGTTTCACGTCGCTACGGAACAAATTGGTTACTTTGGACAAGTCGTGGATGAAGATGTCAC

CGAATGGAAACATTTAAGAACTAAGATAGCGGTACAACTAATAAAGGACAAGGATTTCTATATGGGGATGGTGATTGGTG

AAGCAAGGAGATAGACACGACAAAGAGCTAGAGTTCAAAAACTTAGTTGCACGTGTAAAGTGCCGAAAGGGATTGAAGAC

TATCGGATCCAAGTATTGGATGAGAATTCCTTCATGTGGATATCTCTTAGCGGAAGTTTTGAATTGCGTGGTACATTTCT

TTGTTCCACCTAAGTATGGACATTCCTTTACGTTTGCACCTACAAGGACGGTTTGCGATGATTCGGTTAAAGATAGAAGA

ATTGTGATGGCATTTGTGAATGAGATGCATTTTATCGGTTTAAGAGTAAAGAGAGATTGCCCATTACCTCTGTTGGATGG

TGAGACGGATTACTTCCCAACGAATCATTGTAAGGATTGTTTGAAACAATATGAGACCAACATGTTCGATTGGGATCGCG

TCATGAAAATCAACTTTCCGGCAGGCATACTCGAATTGGTTCCCTCAGACGATGAAGATGAATGATCGTTAATTGCGGAA

TTTGTATTTTTTGGAATTTTTTGAAATGCTCTGTGAAGTATAAACCTATAATTGGTAGTTAAATTATAAATATGTCAACC

GATTATAAATGTTTTTGTAAACAGACTCAATCAAATTTATAATCGGTCATTAAAACTTCTAACTTACCCTCCGATTGCAA

GTCTAACAAAATTACTGAAATTTATTCCCCAGAATCGGTAAATAACATCTTACATTAACATCCGATTGTCACCAAACTCT

GGAAAAAAATTGAAAAAGTTTCCAAGAATCGGTAGACAAAACAGAACATATACGTCCGATTATTACCAAACTCTGACATA

AAAAGATGAAAATCTCCCCAGAATCGGTAGAAAAGATATAACAATAACTTCCGATTGTAACAATCAATCAATATTTATAC

TGTGGTCATGGACTTCAGAAATTCATAACTCCCATCATAAGTGAATTCTTCGCCCTTTTCCACCAAGTACTGTCTTTTTG

CAAGTGCCTCCTACATAAACAAACAAAAAATAATTCCAAGTTGGTTTTAGTAAGAAATATTAGACACCCGGTGTTAAAAA

TAAAGAATTTAAACCATACTTACAAATTGTTGGATTGACAAGCTTAAGTGGCTAGTATTAAACATCCTCTTTTGTTTCGT

TATATAATCATTTACCCTTTTTTGGACAAGTCGGAAACGATCATGAACTTGTTGGAGTGATCTTTGCTTTGGATTTCCAT

AATCTCGCACAAATTGGCTATATACCCTCGACCAAAAAACTTCGACTGTTAATGTTCCCGCTATGACATCTTCACGTCTT

GTTTCATGATACCAGGTTTTCACAATGACTAAATCTTCAGCTGCGTCAAACGTTGCCATGCTAATAAGTTGAGATATATA

AGGAATGAACTAGAGTAGACGATGGAAACGATATTTGTAGAATACACAATGTTATGTATGTGGTTTTTGCTGCATAAATA

AGAATAGTTTACCCACCTTATATAGATGACACTCCCAATGTCTCTTTTTTTTGGAAAATATGGCCGTTGATGCTACTTTA

CAATACTGTACTGAAGTTGCGTACAATTATTACACATAATCGGTAGGCAATTATGTAACATAAGTAACCGATTATGTTAA

CTCGGAACCTTAGGATACTTGTGAAACTTCCGATTATAGACTGTTTTGTTAACAGGGTAGGTAGAAAGTTTCATTAGAGA

GGTATAATCGGAGGTTTATCAATATGTTTTCGCTACCGATTTTGTGTGGTTTTCGAGGCTCGGCTTCAAAAATATTCGTT

GGAGTATAATCGGTAGTTAAATAAATAACATACACTTCCGATTTTCATAATTTTTGGGAAATTCATTACAATATCACCTC

AACATCATCATAATTTTACGTTGAACTTCCACAAATTCTATTAAAATGTATTGTGATTAGCGTCGTGGATGTGTTGGACG

AGTCCTAAATGGATTAAATCTTTTGATCTCCATAAGGATTCTAAAATGAGCTTCATAACGGAAGATTGATATTTTCCATC

TCCGCCATTCCCCGAGAGATTGTTCTATAACCTCATCATTGGTTGCACCCCTCAATCGATATTGATTCACAAGATAAGTT

AAATCGACAAATTCTCCGACGCGTCCGTTTATTGAACCAATTCGGCAGAAAAGATCAGCAGAGTGACGGTTGTTTGGGTT

AATTACCGGTGTCGGCGCAGAATTTTTCGTGAATTCTCCCCAAATAACTCATACTCATTATGCCTCCTTACCTCCGTTTT

GCTCCCCGTCTCAGATAGAATTGAACGTAGCGCCTACATAAAGATAAATCTTCTTCAATCGTGAACTCCGTTGGATTAAT

GAATGGTTGGGCCATTATGTTGAAGATGAAGTTGGAGAGAGTTAAAAGGTATTTGTGTTCCATAGAGGGTACAAGGATAC

AATCCTTATATAGGGTTTAGAGTTCCAACGACTCTTTTTTTGTTGGGAAACTGTATAATCTGTAGCCAAATAAGGTATTT

AGCTACCGACTATACAGTTGCCCCAAATTTTGTTTTGGGAGAAATCTCAAATTTACGAATTTGAGAACTATAATAATCGG

TAGTTTATATGACTACAGATTATAAGTTGCCCCAAATTTTGTTTTGGGAGAAATCTCAAATCATACGAATTTGAGAACTA

TAATAATCGGTAGTTTATATGACTACCGATTATAAGTTGCCCCAAATCTCTTCTTTGGGAGATACTATAATCGGTAGTAT

AGTGAGGTTATTATTACTACCGATTGTTATAGGACCAAGATTCACGTTTGGCACGAAATCTATATTAATCGGTAGTCAAA

TATGATTACTAACCTTCCGATTATATAAGGACATTAACGTAATCTCCACGTTTTTCGACACCCCTTAACGTGGCCTCTGG

GTTGGGAAAGAAAAAGTCGCTCCTAATTGCGACTGAGTCGCCCTTAAATGTGCCAGGTAAAAAAATGCCTACATTTTTTT

CATTATATATTTAATGGGGGAACTTATTTCCCATGTCGTTTGCATTTCCGTCTAGGATCGTCAAACCAGTAAAATCTCGT

GTACTCGGTCAATTTCTCATTTGATGAGATGAGATTTTGGTGGGTGTTACTTAACTGGGATACAAACAGGTACGGGTGGA

GACGTGTCCCTGGATGTCGTCTTGTTGTTTATCACGACAAAGTGTATATTTGTTCTTCCGATTCCAACAACAACAACGAC

GATCTCTAAACACGAATTTAAGTTGGATTTAAGAGATTTTCGATTGTTATAGACTGCAGCAATTAAGTCTCTCGATTTGG

ATTAATTGGGGGTAATCGATTAAGTTCCGGGGTTGATTAAAACAAGAGTTTTTTAAGATTTTGATGGTTGGGTTCATATT

CGATTTGAAAAAAACGAGTTATGGGGGTCGAATCCCAGGACCAGCAACACCGCCAAAACACAAGGCTTTTATTAATTCGT

TAAGTTAAGATTGCTGTGGTATTAAGTTGATAAGAACAAATTGAAGACGAATTGTAAGAATAATCGATCGAAACTGTAAG

TACTGAAACCCATTTCATCTGTTTGCTGAAAATTTTCATTTGAAATTTGTTTGGATGATTAACTTTTTTTTTAGGGTTTA

TGCAATTTGGAGTTTTTATGTTTGAACATTTACCAAGTGTTAACACGTTTTACTGATTATGGTTGTTTTATGTGGTGTAA

GTGAACATGTTTCTGTTAATATGTTTGAATATTAAATCCCCTAATTTTCATTTTGGTGCTAACTTTGATTATGTAAACAG

TCAGTTTAAGGTTTTTAAGACATGGGCTTGCTAATTAGGAGTAGGGTTTTACCTTTTCCTGATTGAGTGAATCTTTGGAT

ATTTTGATGTAGTGTTAGGCTTTTGAACAGTTGGGTTATCCAAAAAGTTGATCATGATGGGTTAAAGCTAAATGAGTAAG

AATTGGTAGGAATTAATCTTTGTGTTTGTCTCAAATAGGTACTGAAGAACATAGATTGGCCAACCTGTTAGATGAGTATG

GGTTATCCAACTTATTCTAGGTAAACAAAGGATTGCAGATTATTTATCTACCTAAATCCCCAATTTGAGCTCCAACTTAT

GCGGCATGCTCAGTATAAATGACTGGTTATGAAACATAGGCATTGTAGGAAAGATGCTGTGCTGCTTATCAAAATCGCAA

AACAAGAGTTAATGCAGCAGGTTGTAAGCTTACAATGCTCATCCCATAAGTAAGGATTTAACAACATAATAATAAGAAGA

AGGAGCTTGGAGGTTTGATAGATATTCTTCCTCCTGTCGAATTCTGTTGCTCGTATGGATCGTCTCTCCTTCCTAACAAC

AATGACAAGGTTTCTTATCTTGTGCTTGCCTTCCTTTTGACTACCACTTTTTTGTCTATTCTCGTTCTTGGGTGTTTTTT

TACATAGCTAAAAATTGTTTTCCTGTCTTTTCAGAGTACAATGGTATACTCAATTATAGGATTTTCGGATCCCTTGCAGT

GGCATGAACAGGTAAATTTTGGAAATTAACTGTTTCAGAATAAGAAAATTGAAAGGGCCCCAACTGACCATGTTGTGGAT

TCTAATTGAGACTGGTTTTGAAAATGAACAGAAATCATTATGCCTCATGGATGATGCAATTTGGCGTTGCAGAAGTGGTA

TTCTTTTTTCCCATTTATCATCTGTGTATCGTCAAATTTTCTATCCAAAGCTCTTTACTCATCCCAACCTTCACAAAACG

TATGTAATTTATCTGTTCTTTATCTTTGATCGGTGTTCTTGGCCTTGTTAGATTATGGGAGATGCTGAGAGCATTGGAGT

CGGAGTACACTTTAACCCATTTGTCACTTGGGAAGACGAGGTACTAATCTTTCAACTCATTGGCATGAGAAGTTTGTTCC

ATGTATGGAAATGTAACAAAGTTTACAGCGTTAATCCTTATTCTTGGTCAAGTATGGTGTTGTTCGCATAACTTGGTACA

AGATATTTTAAATTGGGAAAGCTTCTATATAAGTGGGGGATTGCAGAAACCGGTATGTCTACTTATGCATAATGAGCATT

TGGAAGAGTCTTTCTGATGCTAGATATTCTTCCTGTCGTTGCAGGTTCTTCCCTGTTTTTATTCTCAAAATCATCATATC

ATTACTAAGCGAAAAGTACATTTTGGATTTCCCTTCGTCGTTGCAGCTTCAGTACTAGTTCTTGCAACTACCATCTTATT

CATGTTCTTATCTATATCCTCCTCACCATTGGATCCCATTGTTGGGGTTAGTTTCGATAGTCTTTACCACATATTTTCAT

CTCCACGTGGGCCTTTTTGTACATCAAACAGTAGCATTTCTGGGCTCCAAGCAATGCTTAGTGCATATAGCATCTTGCCT

GTTAGTAATGCAAAGTGGAGAAATCTACTGTTATTATTTCTAATGGCAGTTGGCTATAAAATTATGTCTTTTATGCATTT

TTGATGTGAATCCTATTTCGGTATAGTTAGATTGTGTTTTTCGTGAATAGTATGCAGGGATGGAGTTGTGTTTAGGATCT

AGAAATCTAGCTAGCTTATCAATTTTCATTTGGGGTCATGCTTAGCAGGTGTTTGACAGAATTCCTGAACCAGTCATGTA

GTAAAACTGTGCAAAAGGGAGGAGAAATAGTTAGATTGTGTTTTTCGTGAATAGTATGTGTTTTGAGTCACGTTTTCGGT

TAAGTTTCCGCCAGAATTGTATTCTACATCTCTGAACATTTTTCAGCTGATTTCAATTTAACTCTTCAACTATATACAAT

TTTAGAAATTACTTTACATCTTTTTTACAAAGAAATGTGATATCTTGTTTGAACATCTCTTTGCTTTGCAATTTCGTTTT

ACAAATTTACAAGTCCGATTCCATGCCATTATGGAGCAACTAAATTATGAGTACAAAATTTCATCTTGCGTCAATTTTTT

TGGTCGGCCGACCTATCGTGGCAGCGGTCATCTAGTTAGTTTTGTTTTTTAAAGAAATTTAGGGATTGTATCGTATGTTA

ATTTCTTTGATTTTTGTCAAAGGAAGAGCAAAAATGGATTCGAGGAGAGCGAAAACTGTATCTGACTCACTAACCTGACT

GATTCCTATAATGAGTTTTGACTCGGACAAGTTTGCCCTTCACACGCTCCCCTCACATGCTTGCATTTCTTAACTAAATT

AAACTCGCTAGCCAAGCTCCTAACCTGTCAAACCGCCTCCTTCTCCTTACTTCGTCTATACACGGTGGAAATCTATATGT

AACCATAAGCCAAATCCATGTTATGTGGTGATCATTAAAAACCAACATCCAATAAGTGGTTTAATAAGTGACGACGGATT

ATGTATCTCAGGAAGTTGAAAGTGAACATCCAAAGCTAGTTATGCAGTGATAATCAATCGACGGACGTCGAAGCCAAAAA

TTGGTAATATCTGCAACCATTACTACACAGAGAAATGTACATTATCCATCAAAAAGTTGGAAATATTATAAGTTGACTTC

AAGAAAAGAAAAAAAAAAAAAACTAGTAACTTGTGCAAATTGATGTGCACACATTGCTGCTACATCAGAATTGGTATCAC

AGAGCTACAGTTAGAATATATTGGCCAACACAGAACGTTTTGTTTTGATATTCAAAAAGCATACCTGAAAAATATAATGT

TGATACACTATCTTTGCTTCTTATTCCTCTTTTCTTTTCTCAACTTGAATAAGAGAAAATTGGAATAACTACTTTATTAA

ACAATGGTTTTTAAGACCACTATTCATTCACGCCTCTACGATCAGTTCATGGAAGCTTGAATTTCTTCTAGCGTTCGGCC

TTTTGTTTCTTGTACTATCTTTGCGACGAATAAAATTGCTAATAGATTAACCGCGACATAGAGTAGAAATGTAGCTAACA

ATGGCGTAGAGTATGACATACATTGATTGATTGAATAACATTAAGAAAGGAAAAAAAGAATTGCCGACAACATTTCAAGA

TGCAAATCTACATAAGGATTTGAGATTTAGGGTGACAATGAAAATTCTGATTTTAAGACTCCTGTCAAAATTGCAATCCA

TTCAATTTGGAAAATGGGTCATTTGTCCAAATATTTTTAAAACATGATTTTAATGGACAAATAAAAATTAGTATGGGCAA

AATGGACACCAAAAAAATAGCAAGGATGAAACTGGATTCATCATGGCTTAAACTTAAAAAATATCGAGGATGAAACTGGA

TGCATCCTAATATAAATTAAAAATAAGAAAAAGTATTTGAAAATGGGTAGGATGAAACTGTTTACATCCCGGCTATTTTT

ACATTTTTGTCCATTTAAACAGCATCAAAATCGATTTTTTTTTTACACCCAGAAATTGTTGATTTTGATCTTTTTAACCA

ATTTTATGAAGTTAAATCTACCGAGCCAAAGGAGAAGTACTACTGCCTAGTTCAAATCTTCATTAATTAGTTGGGCCACG

TCAAAACGAGTTTCAAACCCAAGTTAAAAATATCCTAAAACAGATTGCATTGCCTTGAAAATCTACATAGGTTAGGTTAG

TAATCACGGTTAGGGTAAATGGAATTTTGGGCAAAAATAAAAAAAAAAATTAACTAAGTACACAAATCGATCTAGTTAAT

GAGTCAATCACGATTTTCGCTCTCCTTGAATTTATTTTTGCTCTTCCTTTAACAAAAATCATTTCTTTTTCTTATTCATT

AAATGGGCTGTTGTAATCTTGAATGAATTGATGATTTATCGTTTATGATTTTCAATAGATGCAATTGGATTGTTACCGGT

GATTTCTAGACCGGAGAAGGTACCGGGGCTAGAAAAGAAAGAACACATACTGCAATCGTGGACCCCCTCGAACAAGTGGG

TGACCTTAACCATCCATTATTTGAACGGGTTTACACCCATGCCATTTAACTAGACAGTGACTCGGGCACTTAACATGGTT

CAACCAGGTCGTTATCACATGACTCAGGGACCTGAATGTCTTTTCCCAAGCCTGAATTATTGCCACGTATACAGTAACGT

CCGTTTGCCCGTTTTTTTTTTTTGAAAAAACGGGACGAAAAAAGTCAAAGTATGAGCATTAAATATATATTGAAAAAAAT

GTGGGCATTTTCTTAAATACATTTCCCCACATGGGAATTTTATTAAAAACCATAATAAAGTTCTAAAGCATTACCTTTTT

TTGATGTGTAATTGTACTATGCTTCATTCTTATCAAAATGTACCGACTACGGCGGTGTACAATCAAATTGCATTAAAAGA

AATATACTGACTATGGCAGTGTAATTGTGCTATGCTTTATTTAACAAAAATCATTTCTTTTTCTTATTCATTAAATGATT

ATTTTCCCTTCATAATACCTATTTGTTGTATTGTCTCGACTCAGTGCATAAACAATCACTTTCATTAAGAGGACTTATAG

GTGGAAAAGTTTTAGATTGAGGTATATTTGGGTACCCTCGTCTTTTCAAACTTGTTTTACTACCATAAAGTTAAGTTATT

GTGAGGTGATTGATATTTCTAGGATGTTCTTCGGATATAAGACCGGATTATCAATTGGTTCCTGTTCACCTTGATTTATC

AAAAGATGGAACAAAACTTCATAGGTATTTCTATGGGAGACAAATTTAATTATCTATCTGAATAGACCTTTCTGTGGGAG

ATAGATTGGTTTATAAGTCTTCAACTTTGGGACGTAGCTGTAACAACCCTCAAGTTCGTCAACTCAATTAGACCAGTCAA

GAGACGACTAAGACGGTAATGACTCGATTAATTAGAGATGAACCTTAATTAATAGAAATTAATCTAACAACGGTCTAGAA

TCGAAACTAGTACCTCTAGATAGATCTAGGAAAGTTACGCGGAATGGACTACCCGAACGTGTCATTCAGATACCGGACGA

AGAAAATATCACTGGATTATTATGAAGGGAAAATAATCATTTTTTACATCTAAACCGCCCGGCTGCCCTTATCTATTTTT

GGGCGCCTTAATAACTATTCTTGCCTTATCATCACCAAAACGTGTCGAGATAAACTCATTTCTATTTTCTTCTTTCTCTT

TCTACTTTTTTCTTCTCTGTTCTTCTTTTTCCCGATCTTTCTCTTTCTCTTCCTTGTTGATCTCAGTTCAGAAGAAATAG

AAAATTAGTTAGCTTGATGAATCTGAAGAAGATGAAGTTAGAACCGAGGTTCACTACAAGAAAATGGGGATTTAGTGTCA

AAAAAAATTGTCAGGAAATGCTATGTTTTTGTCTCGAAATATTTTCTGAGATGTATTTTTTTTTGTCACGAAGTTGTCTC

AAATACTCCGTCTGGGAAAGTATTTCGGGACAAAATTATGAACGGTCACGAGAAAAATTAATTAACAACTAAAAATTCAT

TGTTAACAAAAGTCTCTTTCACCAACGAAATAAAATTGTCTGCGAAAGTTATTTTAATGACCAAAAAATTAGCCACCAAA

AGGCACTTTTTTGCAGGCAAGTTTTTTGTCGCAAAAATGTAAATTTAAGGGAAAAATAATTTGTCTTTAATGATAACTAT

TTAAGACCAATAAAATTGTCTCAAAAACGGAAGTTTTGTGTCAATTTAATTTGTCTTTAATGATAACTATTTAAGACCAA

ATACATGAAGACTTGGTGACATATATTCATGCTCAGTTGACAAGGATACCACCAAGACTTAGTGATGTCTAAGTGACACC

TAGAAAACAACAGTATAACTTATTTGTATATAAGCATAACTCACGAATCTAAACTTCTTCGTAATCTTAAAACAACAATC

AAATTTCAGGCACATGACATATTCCATTCCATAGCCTTAATAACAGCTATCACTTCAGCTGGCAAGGTGAGGAGGCATGG

GGAGCATCTTTCCTGAGGATTCATAACTAATTGGACCCAAAGTCTGATACAAAAAAATAGTTCCTAGGATAAAAGAAGCT

CCTTGGAATATAATGCATCTCAAACATGATCAAGTCAAGCCTTTGAAATTTGATTGTTGTTGGATCCCAGGATAATGAAA

AAGACTACTATGGATCTCTGCAAACCAATGCAAGCATGGTTCTGAACAAAACATGACAACATACAAAAATCAAACCACCT

ATGAACTGAGATTAAACTGGTGGTTATACAACAAACTCACCTCTACCATCTTTACAATAAATCTAATGTCTAGAACCACA

GCACAAGGAGAAAAAACAAACAAAAAATTGTTGTTTTTGTTTTTTACTAGAAAAGATAAAGCGAATTGATAATCAAGATA

TTTATAAGTCATCCATGATAACTCAACTTAACCAATTTGGTGTTAGGAATTAGGTCAAATCGCAACGGAATTCCAATTTT

AAATTTTAACCAAAACACAACATGCAACTACATAGATAATAATATCAATTAGGGGATTTTTACCTTAGCTCGCTAATCCA

AATCTCCAAGAACACCACACGAATTTATTCTCTTGAAGTTGTATCTTCAAACACCAACAATTGTTGCTTATGAATGGAAG

AAGGAAAAACAGCTACTATTGCTTTGTTTATGGTTACCCAACCTTTAACTTTTCACCAGCTACTATTGATTACTTCAATT

CATAAGCAATAATTGGTGAATAAGAACTTATGTGATTCCTTTATAAGCTGATCCATCACCGACTGCAAACTTGAGAAGAG

TCTAAATATAATTTTTCTTCTATTCCCAAGTATAGTCTATTTCGAGAATGTCTCTGCTTAAATTACTTTATCGCCCTCAT

GAATATCAAACCATATGCCCGGTCCTCAAGGTTGGTTTCGTCATCCACTGTTATAAAACAAATCTTACCTAATTAATTTT

TTGTTTCTTTTAATTTTCCATATCCGGGGTGTTGTCTATTTTCCAAGAGAATGTCTCCAATGAAAATACATTAGCGCCCC

CGAAAATATGGGTACGTATACCTTGTCCTTAGGGGTGGTTTTGTCATCTAACTGATATAAAACCAATACTGACTAATTAA

TTTACTTTTTATTTTATTTATAAAAAAAAAATTATAATTATTAAGAAAAATACAAAATAATTTACACAGGAAGCCTAGTA

TAATAATTATCACTATTAATAATTATTTTATTATTGGATAAAGAGTAACTAGTTTCTTTTTTTAGGAGGTTTCAATCCAG

AGAGAAGCAATATCAACTGGGAGTTGAGTACTTGTCCACATAGCTTGGGCTTTGTGAGATAAGGCGTAATCCGCCAGGGA

GTGGGATGCAGTGTTGGCCGATTCCGTGACATATGATTGACTGTGCTTGAGCTAAAACTATCTGTTAGACACTTCACTAC

GCCTGATAGTCTCCTTCAATGATGATATGCTGTGGACTAACAAGCTAACCTCCAACAACGTACTATTACATTAGTTGAAC

AAGTTGTCTTGCTAGATAGCCTACTAGCGAGCTCCAACAACGTAATATTAGTTGAACAGGTTGTCATGCTAGTCTACTAG

CGATCTAACTTTACCTTCCATGTATAATTTTAATAAATTCTCCTTTTTAGCCAATAAAAAAAAAAAATTAATACAAATTT

TATTTTTATCTTTTATTAAAATAATAATATATTAATAAAACTGGAGTTTGAGGACTAAATCAGTCAGTTTCCGAGAAAAT

TCAGTGTCGTCCTCCACTATAGATTGGGCATCCCGTTTTTCGCTAAGGAGAGCTTGTGTTCTATCTGTAAACGAGGTTGG

TCTCAAATTCCGACATGATATGGTGAGAGACGTTATTGCGGGTATGTTCGCATGTTTCTTTATACACACCTCCTTTTTGC

TTTTTGCAATATGCCTACAACACATCTAGCAATTATTCCATTAGCAACTGAAATTGAATGCTCATGTATAAAAGAAGCAT

CGAAGTTGATTTTCATAGCCAATTAGCAGTTGGAGGTTTCCAAACTGGAATATGCATATTGTCAGTAGATATAGCACACT

CTCAATCAAATATAAACTGCGTATCAAACAAAATGAAATGGTATGTTTAAATTGTGCATCGTCAAAACAAATCGAACATA

TATACCTCCAGATGAGGCATAGAACATAACAAATAAATCTAATATTTTCTCTGAGGTAAAGCCAACTAAATTTTCATGTC

TTTGGATAAACTTGGGACTCAATTGATGTCCAAAAACTTCGAAACTCATTTAATATTTCTCAAAAATAATTTTTGGATTT

GGAAATCCAATTTTGTTTGTTCCAACACGAAATTTAGACAGGAATGTAGCAGCATTCAAAATTTACGTATCCAAATTTTC

AAACTTACAAAAGTGTGTATCATTTTTTTTTTTAAATGATTTCGGATCAGACTTATTTGTTTAACACCAGTAGATGAACT

TCAAATTATCGAGAACATAAGTAACTTACAAAATGTTACAAAGAACTCTTTCCTTCTTTTGGAAAGATGTTGCTTGCAGG

GTCTTACAACATTGGCTCGTTTTCTCCCTGGTTGTTCAAAGGGTAGTTGTTATTTTTTGGACCTCATCCTTTAAACAGTG

GTAATACCTCGATATCCTTCACTCCAATATCAGTGGTTGCAAGAGAGTATCAATTAGACCATTTGTAAAATCGAGATGTC

ACTTAAAATCTATCACTTTATATTGTGGTTCAAAGTTTGTCGTCGTAATCCAAAAAATGTATAGAGAATAAAAATTTATT

ATATAACATGGAACTTTCACATGACCCATAATCTAAAACTTGACCAATAATTCCAGAATTTTTCATTATTTTGATGGAAA

GCTTCCATTTTCCCTTAATAATCTCAAATAAAGGTTACATGAGTAGCTTTACTGATTCTTTGGTACTGTGATGCTGGTAA

CTAAATGTCTCCCGATCTCATGAGTAGATTTATCTTCTTGGATCAGAGCTGAAACACAGAAGTTTCTTCATTAGCTTCAT

GTCTTAGGAGACACTAATTTTAATGTATCTAAATTTTCAGTATGGGGGTAGATGCAGTTAATGCTGCCATCGAGCATGGG

AATCTGGATCATTTAAGGAGATGTGTGACATCTGACTTAGCCATTGTAGGTGCAGAGGGGTGGAAACCTTTAGCATTTAT

GGAGTTGGCTCAATCTTATGTACGCAGGAAGCAGATAAGGGAGGGTAAAGTGGTGCCAGGAGAATTAAACACATCAATTT

CCAAGAACCTCTCTTGTGCATATGGAAGCATGGCCACGCAAGTGGCCTGGCTATTGCACAGGTTGGAATTTAGAATATTA

ATAGCCATTTTCTCATTGTGGTTTGTCATTGATGAAAGGACACCATCATTCCAAGATTGGGACGTAACCCATCCCTTGAA

TGGTGATGTTTTTATTATTTTTATTAACTTATGTGGTGGTAAGAATTACAATACAAGCATAGTTCACTGATGTTTTTTTT

TGTTAAGGAGCCATTCTTTGAACTCCTTCATCCCACAATTGTTTCATGACGCTGCTCCACAAGCACATTTAGATTGTATA

CATTTATGAACCAGCTCTCTCATTGGATACAAGAGGAAGATCTTGGGAAAAGGATTTCCCATGGTTACCCTTATCTAGAG

GCTGAAATCCGTTACTGTGCCAGGAATGAGTACGGTGTGACTGCCATCGATTTCATTACCAGAAGAACGAGGCTGGCCTT

TTTGGATGTTAATGCTCAGTAATGTCACTCGTGGAATGTGTGTTGAGTCTTAGTAATGTCACTCAAAATTTTGTTGCAGT

TTTTTTTTTGATCGATTATTGTAGTATGAACAGACTATATCACTAGCAGTCACCGAATATCGGGATTTGCAAACTACAGT

TTTTCTTGTCGCAACAAAATTTTAAAGACAAAATTTCGGCTTTGCTTCGTATTCATTTGGATGACGAATCAGTTTTTGTC

AGCATAAATAGAGACTGTAACACCCTGAGTTCCGGACCAGGGTCAAACCCATATGGAGATCCGAATTCGAAGCGTTAAGG

GTCGGGAAGAGACTTATGCATATATATTTTTCTTAGTTAATCTTTTTTTATTACAACATATATAATTTAGTCTTTTTAAT

ATGAATTTAAACATATGAATTCTCTAATCATTTGTAACAACATTTCCAACAAACGTATTATTTTAAATTACATCTCAAGC

AATGCAATAAAACCATTCAACCAATAAGCTACTCAATCCTAGATTCCGTTGCGCAACTCTAATAAATCTGCAAGGATGTC

AATTGGGGTGAGATGACATCTCAGTAGAATGCATTTCCAATTCTCTAAAGATCCGCAAACATAATCATTTTGTCAAAGCA

AAACATATAACAATAATAGCGTACAACTTCAACGAATCAATTCACAACAACAATATTATTTTGAAAAATTCATCCATTTG

AATTTCACAAATGAGTTCATAACACCAATTATATTTCCAAAAACTCATTTAATTTAGATTCCGAACACATTATTCCATAT

TACATTGTATGTTCCATAGTAATAATTTTATCTGATCATCCCTAAATTTTACAGGAAAATTCTTGACACATTCTTATATT

ACATATCTAAATTTCACACCAAACCAACGGTTCAAACAAAAGATGTTGAATTTATAATGACAACCTAAAAACTGGGCAGA

TTACATGTTGTTACCAAACAATTCATGATAAATTCATAACAATCTGAAATTAACCCAATCCAAAGCACACTGAAAGATAA

TATGTCTATCTAATTTTTTTATTAAGACCTCATCAAATCCTTTTAATTCTATTAAGACTTCCTAAGAACATCAAAATCAC

AGCAAGGAGAAACTATCCAGAAATTCAGAAACTATCACCCAATTTCAAACAAACACTCAACGGTGAATTTACATTGGAAA

TTTTCCAAATTTTAACAAGAGGTACTCAAACATGTTTAATATCAATTAGAGAAGCCTGATCATCAACAAGATGCACATAT

TTATTCTAATAAATTACTGAATCATTACAATACAAAATCTTATAAAGTAAACACTAAATAAAAACACCTAAAGCGAAAGG

TTTTCACATTCTACCACCCTTAAAATAAATTTCGTCCCGAAATTAAGAACATGCTTATAACCTAATACAAAAAAAACATG

ACAATTCAATTCAACTCAGCACACAATTAACTATCATGCGATTTCCGACAAAACAAAATATCCATTTGCAAAATGTTAAG

AAACATCACCAGCAAGCATAACAAACAATCATGCACATCAATTCATACGATTTTTTTTTAACGTCTCATGTAAGATCCTG

GAGCTCTGATACCAACTTGTAACACCCCGAGTTCCGGACCAGGGTCAAACCCATATGGAGATCTGAATTCGAAGCGTTAC

GGGTCGGCAAGAGACTTATGCATATATATTTATCTTAGTTAATCTTTTTTTTATTACAACATATATAATTTAGTATTTTT

AATATGAATTTAAACATATGAATTCTCTAATCATTTGTAACAACATTTCCAACAAACGTATTATTTTAAATTACATCTCA

AGCAATGCAATAAAACCATTCAACCAATAAGCTACTCAATCCTAGCTTCCGCTGCGCAACTCTAATAAATCTGCAAGGAT

GTCAATTGGGGTGAGATGACAGCTCAGTAGAATGCATTTCCAATTCTCTAAAGATGCGCAAACATAATCATTTTGTCAAA

GCAAAACATATAACAATAACAGCGTACAACTTCAACGAATCAATTCACAACAACAATATTATTTTCAACAATTCATCCAT

TTGAATTTCAAAAATGAGTTCATAACACCAATTATGTTTCCAAAAACTCATTTAATTTAGATTCCGAACACATTATTCAC

ATATCTATAAATCACCTCAAAACATGAGTTCACGGTACGAAAACCTTGGTTCGTACATCACATATCGTTTATCGGCACGG

ACAGCCTCCATCGATGGTCAGGAACAAAAGTTCCTATGGGGATCATACCCATTTTCTCTCTGGAATCATTACCCTTTTCA

TTAACAGTACAAAAACCTTGGTTCGTACACCACCTATCGTTTACCGGCACGGACAGCCGCCATCGATGGTCGGGAAACAC

AAGTTCCCATGGGGATCATTACCCATCTAACTCTCGGGGATCATTACCCGCCGTTAGAATGAGAATAAATTCCATCTAAC

GGAAAAACATGAACTCATTTCCTTTTATAAATCATTTTCACAAACAACACAAGCAACCATCGGTTATCAACATGAATTTC

TTTCAAGCATTTATGCAAACAAGTTAACTGCAGCCATATTACATTGTATGTTCCATGGTAATAACTTTATCTGATCATCC

CCAAATTTTACAGGAAAATTCTTGACACATTCATATATTACATATCTAAATTTCACACCAAACCAACGGTTCAAACAAAA

GATGTTGAATTTATAATGACAACCTAAAAACTGGGCAGATTACATGTTGTTACCAAACAATTCATTATAAATTCATAACA

ATCTGAAATTAACCCAATCCAAAGCACACTGAAAGATAATATGTCTATCTACAAATTTTATTAAGACCTCAAATCCTTTT

AATGCTATTAAGACTTCCTAAGAACATCAAAATCACAGCAAGGCGAAACTGTCCAGAAATTCAGAAACTATCACCCAATT

TCAAACAAACATTCAACGGTGAATTTACATTGGAAATTTTACAAATTTTAACAAGAGATACTCAAACATGTTTTATATCA

ATTAGAGAAGCCTGGTTATCAACAAGATGCACATATTTATTCTAATAAATTACTGAATCCTTACAATACAAAATCATATA

AAGTAAACACCAAAGAAAAACAACTAAAGAGAAAGGGTTTCACAGAGACATAGGGACAAGAGTTATAATTGTCAGCAATA

GTATTTTCCGAGACAAATTTGTCTTTCGTCCATAAAAAACAACCCTTTTGTGACTATAATTTTTTCGAATAGTAATAGTA

CTAATGGCAAAATCCTCATTTTCCATGCAATCATTTCGGCAACAAACTAGTTTTGTCAGCATAAGATACATTCAGGGACC

AAAGTTTCGAATTGTTGGCAATAAGATATTTGGATAAGTTATTTTACCTTTGTCGCTAGATTCCAATTCCGTGACCGAAA

TAATTGGTCACAAAGAAATATTTTGAGATGGAGACAGAATTTGTCACTATATTTGGTCATAAAAAAATATATTCTGAGAC

GGTGACAGGTTTGGTCACAATATTTGGTCACAAAAATTTACAGTGACAACTTTAAAGACTAAATTTACCGTTTGTCGCGA

AAAAAAAAAACGCTGACAAAATAAAAATCCCTCCCTAAAAACTAGATTTGCAAACAAAATTTTATTGGTCGTATAATTTT

TTTTCACTAAACGTGTGTTTTGTTGTAGTGGTTGTTTGAGCTCCTGGAGAAGAAGAATATGATTTTGATGTTGATGTTGA

ATTGAAGTTAGGGTTTCTAAAATTAGGGATGAAGAATCGAGGGGTTGTTATTGGATTGACAAGTTGATATTTTTCTGATG

ATTAAACGAATAAAAAAGGTAAATCAGAGTTTCAATTAGGGTTTCTATTTTGATTTTTCTTTATGGAATTGATGTTATTT

GATTGAATTATTACTTGGGTTTGTTCTTAATCGAGTGAAGATAGTTGTATAGAGATTTAATTTTAATTCTTTAGTAATTT

AATTTGAGAATTGGGGTTTGATGAGTTTAAAATGAACCTGTTTGATTTGAGTTACATGGAAAAGAGTTAGATGATTATAA

AGATGTAAATGATGTTGGTTGTTTGAATGAGTTGAGGTACAAGTTGAGATGATGTTGAGTCATGAGATGCTTTTACAGGT

GGTTGTAATTGAGTATGTTGGTGGTTCTTTTGGAATCTGGTGGTGATGGAATTGTTGTTGGTGGTGGTGGTTTAGTAAAA

AACAGGTTTATGGAGATGGTGTTGAGACCAAATTGGCTACTGGTGCAGGACATGGGAGAGTATTTGATGTTAAACTGCTA

ATGACGATGCTGGTGGTGTTGAGGTTACATGGAGTTTTGCGAATTATGAAATGGTTGTGTGGATTGTGTTGATTAGAAGG

TGTGGTGGCTATCTGAAGGCTGAGGTGGTAGAGATGGAATTGCAAGTCATGTAAGGTAGATTGGATGGTAATGAAAGTGA

AGTTAAAACCCTGGTAGTTGTGTTGCAAATTAGTGATAGTAGTAGCTAGGGATGGTGGAGTTATGTGTGGTATTGTGATG

TAAGGAAGAGGCTTGAGATATATGCAGTTGAGCTGAAATAAATCAAGAATAGAGATGGATCTACAGAGTTGAGTTGTATA

TACAAATGGTAGTAAATGAAGTGTTTCAAGACTTCGGTGATGATGCCGAGGCTAACCTAAGTATGGTGTTGCTTTGTTGG

ATTTGATTGAAGTGATTACAATTCCGGAGAACTGATAGTAGTGATGTTACAGTGATGGTGAAAGTAAACTGTTAGGATGT

ATCTTGACCTAAGATGAAGATATTGATAGTTGTAATGAGGTTTAGATGTATGGTGTGAAGTTGATGTTTATGTGTAAGCT

AGATGAATGAAAGTAGGAGTTGTTTTGTGTTTGGATTTTTTTGTATTGTATCTTTTAATCTTGGAAGGTTAGGTGTTGCA

CCTTGTTGGTGCTCGTATAATAAGTTACTGGTAAGGTACATATCGTTTCTGACTTCGAGGACGGAAAACCTCACGGCGCT

TTAATGATCATTGGAGATTCCACAACATTCATGCCTTGCTACTCCGAAGTAAGACGTCGCTTGAAAAAGCGTGGGGGTTT

AACAACCACACCCCATATTTCGATTAGCAATCTGTATGGACAACTCCGAGTTACTTTCTAGAGAATAAACTAGACTTAAT

CAATAGAAAATATCTCAAGAAGTTATTATTTCAATCTCTCAATCTGATTTACTCAAGAAAATACAATCTGCGAGTCTTAA

TCAAAGAGAGATAACTTGGATGGTACCAAAGACCAATATCCAAGTGTCAATCAATTTAAATAAAATACCAATAAGGTCGG

ATATTCTAATTGATTGAACAACGCACAACCTATATTATTCAATTATATAAAAAAATATAATACGGAAAAGAAATAACACA

GACACCAGAATATTGTTAACGAGGAAACTGCAAATGCAAAAAAAACCCGGGACGTATTCCAGATTAAATGATGCAGGACA

TCCTACTCACAAGACCTATTTCCTAATACTAGACTATTTCTTAATTTAGGATTTCTTTCATAATTCTAGTTTGTTTCTTT

GTTTAGTTTGATTTCCATTTCAAGTCCTAGTTATATTTAGAATCCTTGAAGCCTATATAAGAAGGCCTAGACATAGTAGA

GAGACACAATTCTACAACCAATATTATTATTTTATTAATATCAAATTTATCTTTATTTTCTTATCACTTTTGTGAATCTA

TTGATCTCACACTAGTTTTTATCTTTTATCGGATCTTTGATCCAAAACATACACTTCCGCTACCTCAAGTGGCATCAGAG

CCACGGTTGAGATTCTCTTTTGAGTATCTTTAATTTGTGTTCTATCCACGGTTAAGTGCCTCCATTGAGTGCCTTGACTT

TCAAGTTCTTCGTTATGGAAAAGGAATTTCGTACTAGATTCAGTGTAGATGGAATGATCACTGACTTTATTATTGATCCA

TCAAGCCCGAAAAACTATATATCTATTGGAGCAGCAAATTACTTCAGAGAAAGGAATTGTGGTGAATGGGATTTATTTGG

CGGTTTTCTTGATTTTAGCTTCGACGGAGGTAATTATTTCAGTTTTGCTTATTGTGAACTGTCTTCTATGAATAATTGTC

ATGTGTTGTTGGGCGAACCCTGGCTATCGAGTTTAGGTGCTTTGTATAATAAAAATTTGCACAGTTTTGATTTTCATCAA

GGGAACAAAATACATTGCTTAATGGGATTACAGGAACCAACACCATTACAACGTTGGAAAAAAAAAGAAAAGTCAAATTT

TTTGAAAGAATTCCATGAGATACTTGCCAGTTTCCGAGAGAAAATATATGTTGAAGATCCTACACCAGCGCTGGAATCCC

AAGAGCCTATCTTAAATAAAGAAGAAGAAAACAACGACAATGAGGAGGAACCTAGTGTTGATGAAATCAAGCACCCACAA

TCACTTGATCTTACTGTTCCTCAAGAGAAATCTGATACTTCCTATAAGAATCCACTTCCACCATGTCTTGAGATTGCTGA

TGTTTCTCGTACTTCGGTTGATCTAGTAATTCCAAAGCCAAACTTCAATGACATCAGTAGTTATGTCAAGTTGATGGATT

TTCCTATTTTATGGTATAAGCAACTGAAGATACAAAAACAGTCAAAGTATATTGTTCGCTGGGATGGGCGCTCGCAATAC

TTACTAGACTTGAAGGCGAGTCCGTCAAAAGATGGCGGACATGTTCCTCTAAATGAATCACTTGAACTGATTCCAAAACA

AAAGTATATCTCCAAGATGATTTCGCAGGTTTCCATAGCTGTGGAACAAGAGTCAGGCACACTTACTTCGAGAGATCAAC

ATGTTACTATATGCAAAAAATCGCAGAAACACCTACTGATACAAAGGAATTTTCCTACTAGGATAGAGGTGTTGAGATCT

TTCATTAAGCAATCTCGTTCTCACAGCTCTCCACGGGTCGCCCATGATATTTCGGCAAGGGTACTAGAGATTTACACTTC

ACTTCCCAAATATTTAGGTGATGATGTTGAATTTTTAGGCATCGACAATACGAGAAGAAAATTAATTAGGAAAGACCTAG

AGGATAACAAATTGATCAGGAGAAAGAAAGTTAAGAAAGGGTTTCAAATAGAAGATGATAACAATTTCACTGACGGTGCT

CCTCCTCCAACAACGGATTTTGGTGTTCCGAAGAAGCATGTTCCTGATTATTCGTGCTTCTCGGAAATTTTAACTATCTG

CGTGAGTAAGGATTCTATATTTTTCGATTCTATGAACAAGCTTATACAAGTAAAACTCCTTGAAGATGGATACTTGCACT

GGATTACTGACTCGAATTCAAGTTCGCGGTTGAATTCTTCAAAATGGATTAGTAACTATGAAGCTTCATTCAACAAATGT

GACGACGCTTTATCAAATCCTCATGGTGTTCATGAGATTCCTAGAAATCCTCCTTGGCTGGTTGTACTTTTGCGGTGTAA

AAGAAGAAAAAGAAGTAGGGATACTGTTCCATCCCAGACAGCAACAATTACGGCAGAATTCTTCGCTCAAAAGGAGGAAG

ATACAGGCAAGCTGTTAACGGCAAATGCACACAGCGGAAATTTTAATTTCCAGATGGAGAGATTCAAAGGAGTAGGTGAT

GGCATGTATTTTATAAGACCTGGAAAAACACCGGAGAAGTTTACAATGACAGCTAGATCTGTTGTTTCCATAGAGCACCC

ACATGCCGTTATTGTGCAGTCTGGTAGTCCATATTGGCTGGGGACTGTAATGAGAATTTCTTCCTATATTAATGCTCATC

TACCAGAAGGAAGACATGCCCAGGAAATGTTCACTAATCAGATGCCTGATCCGCATGTTACCGATGCAAATATTGACTGC

CAACCAATCAAGGTAGCTGCTCTTAAGAGCCATCTGATAGATGAATCAGGTCAGTTTCTCTCGAGACTACCATTTTCAAA

GAATTTCTTGAAGACATATCTTCTACCCACTATTCATCAGCAGCTTAGAACTGTGCTAAAGGATTATGAGATTCCACGAA

AACCTCTTTGGTTTATTCGTGTTGGCTTTCGAAATCCATATCTAAATCGTGCAGGAGTTCTAAAACCTAATGGTTCATTT

ACTATGGAAAATTCATGGAATGAGGTGTCCTTATCTACCTCTTCTGTATCAAAGATTACAGTTAACAAACGCTACGAGGA

AGATACGCAACATAGTTTTGTTGTTTATACAATATTAGAGATCGGGTTTAGGTCTTCTGATGTTAATATTTCATTTGGCA

ATATAACAGTTTGTATAGCTCATGGTTTACTGAAAGACCTTGGGCATAAGGCTGTTAGTGAGACCGTGGAGATAAAGAAG

TACCCTACTCTCGGCATAGGGGTCATAAAAGAAATTAGATTGCTCTTGCAACATACGGCTCAAGTTTTCGGAGTTAAGAT

AGTGTATGCTTTGAAAGTTGAGGATCAAACTCCTCATATTATTTTTCAAGAAGCAACGAATTACTTAACTACAGCTCATA

CGGGCGCCTCCACTGTCTCGAAGGATATGGATTATTATAATCAATTTATTTGTATGAATAGAGCTGCAGATACTAAGCTG

AATAGATGGGAATATGATGAAGTGACAGTTAAATTTTGGATGGTGCACTTCTGGCCATTTTAATCTGAAGATTTTAAGAA

TGTTATCTGCAAGGACACCCATCTCATCATTCACGTTCTGAAATCACATTCAGCATTGCTTCACATACTTATTTACCATG

GAGACTATTTGACAACTGTACAAGAGAACTATAATCCAAAAGAGATTTTTGAATTTTTAACTCAAAGTTTGAGTTGGTCC

GAATATCAGTTCGATGCCTTGGAAGAGAAAGAAAATGATGAATTGTCAACTGACATAAGAACAACACAATGGGGCCGTTA

TGAATTCTTGGACCCTCATTATAAGTTAGCTATCATTCGGGAGACAGTTTGTTGTGCTCTTGCAACTAATATAATTGGGG

ACCAGTTGGATGGATCTACTGAACAACAACAGGTTCTTGCGTCCAATCAAATGGGAGTTGCGCTGGGGGAAGGTATAATG

AGGAGGAATGATAGACGTTCCTGGAAAGGTGTTTCACATTTTCAATTATTCACACAAAAGTTTTCAGAAGTGGACAATGA

GAAGTATTTGATATCTAATGATCTTCGTCGACGTTTTAGACCTTATGATCCTGGCGTCGACAGTGTCTGCCATTTCATAC

TGAAACTGAGGTATTTGATCGACGACGATCTTCGCGGTCATTTTAGGCCTTATGATCCTGGCGTCGACGGTGTTTGTCAT

TTCTTACCGAACATGAGGAATTTGATCGATGACAATCTTCGCCGTCGTTTCCGACCATATGATCCTGGCAAGGACGTTTG

TATCCCTACAGTAAACTCGTGGTCGAGTTCTTTTCAAGAAGGGGGGCCTGATGTAGGACATCCTACTCACAAGACCTATT

TCCTAATACTAGACTATTTCTTAATTTAGGATTTCTTTCATAATTCTAGTTTGTTTCTTTGTTTACTTTGATTTCCATTT

CAAGTCCTAGTTATATTTAGAATCCTTGAAGCCTATATAAGAAGGCCTAGAAATAGTAGAGAGACACAATTCTACAACCA

ATATTATTATTTGATTAATATCAAATTTATCTTTTGTTTTTTTATCTTTATTTTCTTATCACTTTTGTGAATCTATTGAT

CTCACACTAGTTTTTATCTTTTATCGGATCTTTGATCCAAAACATACGCTTCCGCTACGTCATTAAACATACACTGTATT

AAGCCGCTACAGACAATAGCCTATTCCAAATTAACTTCGGTCTGGAATGTAGTTGAACCGTAACCAATCTCGCACTGATT

CAAGGTACAATTATGCTCATACGCCTCTAATCCCAGCTGGATACTGCATACTTGATTCCCTTAGCTGATCTCACCCACAA

CTAAGAGTTGCTACGACCCAAAGTGGAAGACTTTAATAAACAAATCTGTATCACACAGAAAAGTCTACGATAATAGATAA

ATCCATCTCCCACTAATATACCCTATGAGTTTTGTTCCGTATTTTGATAAATCTAGGTGAACAAGAACCAATTGATAAAC

CAGACTTATATTCCGAAGAACAGCTCAGTATTATCAATCACCTCACAATAGTCTTAATCGATGCAGCGAAAAAAGATATT

GTGGAATGAAAAACGATGAGACGAAGATGTTTTTGACTACTTTTTATCTTTCCTTATCGGAGATATAATCTCAAGCCAAT

TATTACAATTGTACTCGTACGATATAAAATGGCAAGATCATATCACATAACTACAAGAAAAGTAGTATCGGTCTGGCTTT

ACAATCCCAGTGAAGTTTTTAACCTAGTTTAGAAGAAGAAACCAAAGGTTAAAGGAGAATCAACTCTTACTTAGCACAAC

TAGTATCACACAGAATGTGTGGAGATTAGGTTTCCCAGTTGTTAGAGTTCTCCCTTATATAGTCTTTTAAATCAGGGTTT

GCAATCAATGTTATCTTGGTAACAAAGCATTAAATATTCACCGTGAGATGAAAACCTTATTAGATACAAGCTAATATTTA

TCAACCGTCAGATCAAAAACATAGCTTGTTATACACACATGAAATGCACGTTTGTAGGCTTGTGTAACCGTACCCAAACT

TGTACATTAGTTGGTTCAACAATAGTTAACCAAATGGTTAGCCATATGATAAATTTCATATCAACCATATTCTTCTTCAC

CATAACTAGTTCAAATGACTCAAATGAACTAGTTAGAGAGTTTTTCAATTGCAAGGAAATCTCATGTACTACATAAGACA

CAATTGAAGAAAAAACGATTTGATGTACTCAAATAGGTTCATGATATATCCACGGTTTGCAACTGCATTCTTTAGTTTAT

AAAGATAAGTTCACAAACATCGTTTTTAGACATAACCTACTCAAGTTTACGGACTGGGTTCGCGTACTTAAGTTCCCGGA

TGGAGTTCACAAACTCCAGCAGAAATTCTCGGGTCGAGAACTTTCGCTAGTTCACGGACTTAGATCACGCACTTATCCGG

TTCTCTTGATCAAAAAAGTTCACAAACTTTGGTTCAAGTAATAAATAATTATACATATATGTGTTACCAAAATAATGCTT

ATATCCTCCAATGGTTATCTTGTCTAAACTCTCATTTCAATCATTGAAACATTCTCAGAGGACGTTATATAGTTGTTATT

CACAAACTATCTCTCGTCAGAACGATTTCCAAAATGATTGAAACATAACATGACTTTCGTCACTAGGTAAATATGAACTT

GGATAAAGCGAAAGCTTTACCAACATATATTTCGAGAAATAGATAAGCGAGATATACTCAGCTCGAAATATCATATGTGT

ATAATCTTAGTCTATATAGTAATACGACTTTTTTCTCAAGATAGGAGATAGAGTAGATAGACTTTTGAGTGATAGATAAG

TTCAAGTCTCCACATAACTTTTAGTCGATGAAGATCCACTGGTTCCTTGAGTAGTCCTTTGTCTTGTATGATGATCGCCA

TAGAGTTCTTGAGCTCAACTACACTTTCTATCCTAGTCTGAGACCTTAGCTATAGTAGACTAGAAATTAATACTTTATAG

TTTTGATCACTAACATTGAAAAACATGCTTGAGATAGCAACGCATGCGAGTTCGACCGAGCAATGCTCTAACAGTCTCCA

CCTTTGTCAATTTTAGTGGCAAAACTATTAATACATATGAAATATAAAAACAATATAAATAAACTTGGTAGCTCCTATTC

CACATGTCTTATCTTCAACATTACTCAAAATCTTCGTCACTTCCAAGTAGTCCAATGATCCCAAAGGTTGTAAGTTTAAC

ATCATCGTTGTTGAAAATCCATCGCTAGAACAATGAGAAAAGCAGTGGTTCTCAATCATTGTTATAAAATGTCATATTAT

TATTACACAGCGTCAAAGTCCAATTGCATCACAACTTTAACAACAATACTATGGTGATACTTCACTCCCCCTTAGTCAAT

AATCCATCTCACATGGAAACCACTCCCCCTTACATAATGATCCGAAAACCATATGTATTTGTAGTATGAACTACATATTA

ATTCTCCCCCCTTTTTGTCAATAAAATTGGCAAAGGTACAAGAACGGGATCCTAATAAAATTTATGTAAGAGACATTTCA

TGACCAAAGGTAAATGTATATCATCTTGTTTAGATGCAATCATAAAGCCGAAGCTAAATTCATTCATCAAGGAGTTTATA

AAGATACAAGATAATCCCTCTAATATTCCACATCCATACTCCCCACTTCAAACTTGAGAACAAAATGGACGTGTTGTGTA

TAAGGCCTATTTTTTCTAAATTCAATATATTCATGATATATGTAAGACTATAGCTAACATCCAAGTGGAGATGAAATTGT

TTGCTTCACTACTGATGGTGGCTGCAAAGATGAGATGACTCTGCAACTTGATCTTCAGCTGGTATACTACTTTCAGTAAT

CTTTCATTGAGTAAAACCTCATGATTCTACTTCCCTGGTTTGTCTTATCTCCTCGATCATCCTTTCAATTTGCATAAACT

TAGTTTTTCTTTCTTATACCCTATTGGAAACCGGTAGACACTCTCCTGTTGCACAATACTTGTTCGTGAAATTGTGATAA

TGAATTTTTTAATCTTTCCTAATGCCTCCGTCGACTGCTAATGACAAACATCCGTAATCTTGTCTTGCATCTCTTCCCCC

TTCATTGAGATCTTTCTCCCCTCTTTTTTTTTTGTTTTGCATCTTTTATAACTATGCATTACTTTGTTTAAACTGTTGTT

TTCGGTTGTTGTATCTTAGCTGAGTTTAATTTTTCAATTGGTTTTGGTCTCATTCCTTTTTTCTTAATCATGAGTGAGTA

ACCACCTGCAGATCTCTGATCCTAATTCTATGGGTAACCAGTGGTGCTTTTTAAGCATGAACAAAGCAGTTGAGCTAAGA

ACAGTGGTGCGGCCGGCAGTTGTGCTAACCATCTTCTTTCTGATGATGGTCCATATTAGTAACTCGGAAGATTCTCTTCA

TTCAAACATCAACCATACAAAATCAAATCGTTCATTACCTTCATCCCTATTTCAAACAATAAGTTGTCTTAAATCAAATT

CAAATGTTAATCTAACCTTTTGCGAGGTTATACATGGTACGCTGGAATTGGGTTCTTTTCAAAGTTCTTGGCCTCCAATA

ATTCGGACATCACTGCCGATTCATCACCGGTGCTTCATCCATCATTACCTATCACACCTACTGCCAGTAAACTTGAGTCT

GCTTTACCCTGCTAAACATAATGATGGTATTTTTATTAATTTTCTAAAAGCCTCTTTTGGAAAACAACATTTTAAAGATT

CCAACTCGAAGGTGTTAAAGAAAACCTTGGGTGGAAAGTGGATCTGGAAATACTTTTTCCAAAAAAAAGCTCTTTGGATC

ATGCCCATCAATCAATTTCTCTCCAATTTGTTAGTCTGGACAACGGTCACCACAGACTGTCAAACATATTAAACTCTTAC

TTGAATATTCTAAAACTTTCAAACCTGGACAAGCAATATATATTATTCACAAAGACTCAAGTATTCTCCCTCATCATTTA

TCCGTTACCTTCAATTCGTCCTCCTACCCAATCTAAGTCCAAAATCAAAATGTCTGCTTCTCAGAATGATCGCATCAACA

ATATTACTCAGCAACTTCGTCATACAAACATTGATTTATCTGCTTTCACTAGGATAACTGTTGGGTCGTCCCAAACCATT

AATCAGGCAGCGAATGTGTGGATTTTTGGAATTTTGGGGAAACTAATGAAAAAGGAAAAAATAAAGATGCAGATTTTTTC

AGCTGAAATTAATTCACTTTGGGGAAAATTCCGTAACAAAGAAATAAGAGTAATGGGGCATAATCTTATGCTTGTCAAGC

TTAACAGTGAATATGAAAGAGATGAAGTGATTGCTGAAGGTCCTTGGTTAATTTATGGAGTGCTCTTCCATGTACAGAAA

TACTACTCAAACATTCCTATTATCGACTACACCTTCGACAAACAAATCTTTACTATACACTTCATATATCTCAGGCTTGA

GCATTTGAATGCAGTGGTTATTGATGAAGCAATAGGTTTTATGGGAACCAAGATATCCACTAATCCTAGGAATTGCAGAC

CTCGCGCAGGAAACACGGTTAAAGCAAGAATAGAGGTTGATCTTATAAAACCTCTGCATAGAGGAGGATGGTGGAAGACA

ATTACAGATGATGATGTGTGGATCCGTTACCACTGGGAACTTCAGAAAAAGAATATATGTCAAAAATGCTATGTTATTGA

TCATGTGGATGAGAGTTGTGATCACACTGCTGGTCTTCTTTATCTGGACAGTCTTTCAGATTCAGACTATGAATCTTGGC

TCAGCGAATCTGTCGTTGAACAACATGAGGAGCAAATCCAAAAGGAAAACAATACTATTGTGGAGGAAGTAGAAATGGTG

CCTACACCTAGTGATGCTCCTAGAAATGTCAAAAGAATGAGAAACTCACCTCTGGATCAACAACCAATGATGAACCCACA

TAACATTCATATTCATTCTCCATGCAATGACAATTTTGAGGTTACTAATATGGACCAACAATCATCTTCGGAGATGGATA

CGGAAAACTCAGCTGCTACTGTTCATGCTCCCATGGATGATGGCGTCGGAATTCTAATTCAGGTTTACCATCAACTTCTT

TCTCGCATTTACGTTACCCATTCACGTCAGTTCAGTTTTGCTTTAAATATGGACATTGTTAGTAAAAACTGTGTCAATAT

GAAAATTGTTGCTTGGAATTTCCAGGGATTTAATGATGTTAAAACTAGGAACCATATTAGAAATATCTCTAAAACTCATA

ACCCCGACATTATATTCCTGCAAGAAACTAAAATTTGTTCGGACAAAATTCTTAAGTTGGCCAAAAACCTAGATTTTCCA

AATAAAGTATTTCAACCTTCGATTGGCCTCTCTAGAGGTATTCTTTTTCTTTGGAAAGACGGTTTTCATCTAGATATTGT

GTTCTCTTCAAAAAACATGTTTCATTGCTTGGTTAATCATGATCCTTCTAAGCCGGAGTGGCTGCTTTCCTGTGTGTATG

GCTCTTGTAAACCAATGGAAAAAAAAAGAGCAGTGGAATTTTATAAGGGAAATAGATCTAGAAATAAATCAGCCGTGGGC

GCTGATAGGAGATTTAAACATTATTCTTGATCCTAGTGATAAAGTCAATAGTAAAAACCAATCTAGTTCTACCTCCCCTT

CAATTATTCAATGCATTGAGGAAGCAGGTCTCACTGATTTAAGCTATAATGGTACTCCATTTACTTGTTCGAATAACACA

CACAGCACTGGGAGTATCAAAAGTCGGTTAGATAGAGCCCTAACCAATGATGACTGGCTTATTTCTTTCCCGGAATCTAA

TTTGTTCCATTTAACTCATCTTTCTTCTGATCATACTCCAATTTGTCTAAACTTATATAAGCACGATAATTATAAGGGCA

AAACGTGGAAGTTTTTTGGTGTTTGGTTAAAAGACAGCTCGTGTGAGGATCAAATTAAACTCGCATGGAAATCAAGTATA

ACAGGCAGTGCTGCGCACAAGTTCAATAACAAGCTGAAGATCACAAGAAGGTTTCTGAGTGTATGGAATAGAAATACTTT

TGGCAACATCGACACTAATATCAGAGAGCTCTATAATCAATTGGACAACATGCGCAATTCGGTTGCTCTATCGAATTCTT

CTGATATGATCATTGGAGTGGAAAAGAAATCGAGAAATGGGAGGATTTCAAAAAAGAATTCTGGATGCAGAAAAACAAGG

TTCTTTTCTTCAAAGAGGTTGACTACAACACGAGATTCCATCATGCTAATGCAAACAGGCGCAGATCGATAAATAGGATT

GATGCTCTGCGAGATAGTTCTAGGAATTGGGTTACGGACAAAAACGATTTATCTACTCTACTAACTAAGCATTTCCATAG

TATCAGCTCCACCTCATCTCCAACTATTGATCTAAACCTGCTCAGCCATCTTCAGTTAGTTATCACTGCCAACGATAATG

ATAAACTAATAGCAATCCCCTCGGAGAAAGAAATCCATGATACTCTTAAAACAATGGCTCCATGGAATGCTCCCGGAGCT

AACGGTTTCCCACTGGGCTTTTACTTATCCCACTGGGAAACTGTAAAAGAAGACGTTGTTCTAACCGTACAAAATTTCTT

CAGCAGTGGTTTTATATCCAAAGAACTGAACCAAACAAGACTGATACTTGTGCCGAAAGTCAAGACTCCAGGTTCTCCAG

CTGACTATCGTCCGATAGCTCTTTGTAATACTTCTTATAAACTTATATCTAAACTGATGGCAATAAGAATGAAGGTGGTT

CTCACAAAAGTTATCTCTCCCCTACAATCAGCTTATGTCTCAGGCAGAATTATCTCGGATAACATATCTCTAGCACATGA

AATTATACATACTCTTAAAAGAAAGAAGAAGGGTGATAGTTTTCTTGCGCTAAAGCTGGATATGTCGAAGGCCTTCGACA

AATTAGAGTGGTATTTTATCATGGAGGTTATGAAGCGTATGGGGTTTTGTGAGGAATGGTGCTCACTAATATTTCAGTGT

ATCTCAACCACTGAAATTTCAGTGCTTTTAAATGGCTCGCCATGTATCCCATTCAGACCTACTTGTGGCATTCGCCAAGG

AGACCCAATATCCCCGTACTTGTTCATAATCTCTATGGAGGCATTCTCAAGGGCAATGGTGGCGGCAGAGTTCTCAAAAG

ATATAAGCGGGATTAAAATCGCAAGAAGAGCACCTCCGATAAGTCATTTGTTGTTCGCAGACGACTGCCTTATCTTTGTT

AAGGCTAATCTGAATAACATCAACAACCTGCAACATATTCTTAACTCTTTTAGCACAAGCTCAGGTCAGGTGCTCAATTA

TCAAAAATCTAGTATCAGATTTAGCAAGAAAATTGCTGCAGAGGCAGCTACTAATATAACAAGATTCCTGGACGTAAAGA

AAATGGGCCCTGAAGAGAAATACTTGGGTATCCCGTTATTTATCAACAAAAAGAAGAGAATATCGTTCAACAATTTGGTA

GTTAACATGGAAACAAGACTATCCAAATGGAGAGGTACAACTCAATCACAAGCTGGAAAAATCATAATGGTGCGTAACGT

TTTAAGCACCATTCCTTTACATCAAATGTATTCTTTTAAGTTGCCTGAGGAAACTATCAAGAAGATGAACTCAGTTCAGT

TAAGATTCTGGTGGAATAAGGCAGATAAAAAAGGCTTGTTTCTTACCTCCTTTGAAAGTTTGTGCAAGACAACAGATGAC

GGCGGTATGAACTTCAGGGACTTACGGACATTTAATCGTGCTCTCTTGGCAAAAACAGCTTGGAGAATTTGTTCTACTGA

AGACTCTCTCCTGGATCGTTCTCTAAAAGAAAAATACTTCAAAAATAATCATCCTCTTCATGCCCCTAGAAAGGAAGACT

CAACATGGTCTTGGAAGAGCGTAAGTTCAGAGTTTGATTTCATTAAGAGATTCAGTTGTTGGTCAATCGGTAGCGGCAAA

AAAATTCAAATATGGAATGACAAATGGGTCGCAGAACAAGACTCTCCTCTTATCCCGTACAACAATGCAACAAATCCGTT

ATCTTATACATTTGTCTCATAACTGATTGATGAGAACACTAAAAGTTGGAAGTTACAATTGATCAATAATCTTTTTGATC

CAAACAATGCTCAGAAAATCCTATCGTTAAGAATTCCGCATCACTCGGAAGACAAATTGTTTTGGTCACTAACCAAGAAT

GGGAAATTTAAAGTCAAGTCGGCATACTATGACCTATACAATATCAAGTATCATAATGGTATGGCAAATTATAAACCTTC

GCCATCTAACACATGGAGGAAAGTTTGGAAAATGGATGTTATCCCAAGGGTGAAACTTTTTATCTGGAAATGTATAAAGG

ATATTGTTCCTTCTAAGGAGAGACTGGGCAGAAGAGTTTGCCAAGATACAGTTCGATGCAGCAGATGCGAAACATCTGTT

GAATCTACCATGCATATTTTAACTGATTGTGTTTATGCAAGGGAAGTTTGGTTTGCATTTGGTTACAGGCTGGATGTCGA

CCAGTGGGAAAACCAATTCGACTGTATAGCCTCCTGGTTCTCTATACATGACTCCTCAACAATCAGTAAGATGGCTACTA

TTTGTTGGTTCTTATGGAAAGACAGGTGTTGTCTGGTTTTTGACAATAAAACTCAAACACCGCAACAGCTAACGCTTTCA

ATAAACAGGTTCCTTAACGGTAGGGGCTCAGTCCAACATAAAATGGCTAATTCGGCAGAGTGCTTGTCAACCCATAAATG

GAAACCACCACCCAACAATTTCATTAAATTAAATACGGATGCATCGTATTGTGACACTACTAAAAGAGGCGATTATGGAC

TAATATTTCGTGATTCTGCAGGTACTCATCTAGCTTCGAAATGCAGCAGTTTCAGGGGGGTTTCTGATGAAAACTATCTA

GAATCTGTGTCCATTATGGAAGGCATGCGTTGGGGGCTGAAGTTGGGCCTAAAAAAGATCCTGATAGAGAGCGACTACAA

AAGTCTAATACATGCTATCCAAGGAGAGGAGTCGGACTTACCATGGGAACTGCGCGGCATCATTGAAGATACTATTAATT

TACTTAAGCAATTTGATTTTTATTCTTGTAAGCACATAAACAGACTGGCTAATACATGTGCGGACAAATTAGCTAAGCAT

GCCCGTAAGAAGGGTAATTGTAATAGTTGGGAAGGGGTAGCTCCTTATTTCCTGTCAAACTCTATTATGTATGACTAATC

CTGATCAATGAAATTTTTCTTTTTGTAGCAAAAAACAACAACAAAGATTTGGAAATCAAGCACAAATTCAAAAGAACTCT

CCCCCATAAAATGTCATTCCCAAGGGAACAACAAAAGCGACCTTAATTTCAAAAGAAAATAAGGATTTCTTTAGACTCAA

CAAATTACATACAAGTATGAATTTGGAATCCAACAAGAAATATTCAATCATATTAACCTAAGCGAACCAAGATTAATTTG

ACTAAATATGCTCAACACAAGTAAACTTATGGAGAATTAATAACTTGATTAGATTATTCACAAGAGAACTCATAATTAAT

CTAACCGGAATACACAATTAAACTAGTCACAAAAGTAATCAATTTAATCGTTTATACTCACATAAGAAAATCTATGGAGC

GATAACTAAGTTAATCATAAAAATAACCAACCCAGTCAATCATGCTCAACATGAGAAAAACTTACGGAGTCATAACTAAA

TAATCACACAAGATGATTGATTTAGTTCAACATGTTCAACATAAAGTACCTTATGGAACAAAAACTAAGCTAATCACCAA

AGCAATCAACTTTTGTTGTTTTAGTGCTCAACATAAGACACTTAATGGAACAACACATTATTTACACAAAAATTGTGGAT

CGGAGAATCAACCTTATACCGTGGAGTAAGCAAGTATTCATTCCATTTCCATCACATTTGCACAATGACATACAATAGCC

ATAATACTTGAATCACAAAAGATTTTAACCTATCTTACATCAATAATTGACAATATAGGCTTAACTTTTGTGTTTTGCCA

AAAGTCTATTCATTCTTTTATCAATATTTGCATATCAATTCACGAACGACTTTACTTTTGACAAAATATGGGACAATCAA

GTTCACGGACGTAAACACACATATCCCATCAAAATATTGCAATATATAAAACTGTAAAGATTAATACTGCAACCATCATC

TTCCAAACAAGTTTAGAATTTAACACAATAAATCTAAAAACAAGAAGATGAAAACTTTGGACATACCTACGTGTAATCAC

AATAATTCCTATTCTAAAAACACTAGTTATTCTTCTAAACAAAAGAGAAAAATATAAGATTTACTAGGCAAACAGAAATC

AGAGTTCATTAAGAACCTCGTATCTTTCAATGAAACCATCAAAGAATGTATCGCTTATTTCCTTTGTTCTCTTGACCGTA

GACACACCATGTTCGTAAGTTAGAACACTAACTTTGCGATCAACCACCCGAACAAGTTGCCTAGCTTTGGCATAGTCCAT

GATAAGATTCTTTTGATTCCGTATCAGAATGTTCTGATTAGCAAGAATTCTGGACTGACCATCTAGAAGTTGGTTTATTT

GCAATTGAAGATTTGCAAACTCGACCCTTGAATCGTTCTAAAAACGAGTAAAATCCTTGACAGAATCATCAATCCAGGAG

TTTTGATCCTTTATTTCAAGCATCTTCTTTAGAACCAGCTCCTGAATCGACTTACGTTCCTAATTGTCTTCCTCTATATC

AAGACGCACAATCTCTTGAGATTTTGCATAGTTCTCCAAAGAGTTTTAGGGAAGGGAGAACACAATATATAACAAGTATA

TATGCATTCGTCAACAAGACTAGAAAACTCTGGTTTTGAAAACCCTAGAAACTCACAAGAGGAGGACGCGGACGTTTGTG

GACTACCAACAAAAGGACTCAAACAGAAATAATATACAACATACTTGAGTATTCCCCAATTATTTTCCTTGCATCTCTCA

AGTTAGAAACAAGGAAACAAATACCTGGAATCAGGTAGTAGAGTGGAAATTAATTCCATTATGATCATCGAAAAAGGAGT

TGTATATATCATATGACTGTTTGATCTTTGTTTTATTCGCATTTCTTCGGTTAATTTTCATCCCTGAAGGATAATACATG

GCGTACTTAACAAATCCATCAGATGGATTTCTCTAAGGACTAAGTCTAGGACCTCTTACAATTGGTTTAAGAGGATCAGA

ATAGATAGGAATCTATTTTCTAGACTCAAATTTACCAGAAACATTCTTATAAGTGAAGGGAATACTTTCCTTAGCAACAC

AAGAATTTTTGCCAGAATGCTCTTGAAAACCTGTAATGATTCCTAGAGTAGAGATCATTTTTATAAGGTTTCTGGTTTTC

TGTGGGCATGTGAATCACTGCAGTAGTCGTCCGACTGTTTGTTCCATTGACAGTGAAGAGAAGCAAATTATGAAGCTTAG

AAATTTGTTTCCTTCTAGCAAAACAAAGGATAGAATAGTGATTCTTTTCCCCACAAAAAGTACAGGTTCGTTGAGGAGAA

GCAACAATTGTCACCTATCCTAATTTCATAGCCATATGAGAGGATTATAAGTTATGTAAACTTTTCTTGACACAATCTTC

TACTGGAGAACACTTCTTCATATGCGGTAATTCATGAGAATTATTTATAGAAGAATTCTCCTTAAGACAGAGTTTTCCTT

TTCCAAGGATTTACACTGTTCATGGGCTAGAGTAAGATATGCACTTAGTTTTTCTTTTTCAACACGAAGTCTATTCAGAT

CTGATGAATGAGTCTTGAGCAAACATTTTTCACTAATTGAGTCCTCCTTAAAAATATCCTCAAGAGTACTAATCTTTTCA

CGTTGAAGAGTAGTTTCTTGGAGAAGTTTCTCTATATTCTTAGAGAAGGACTTAATGATGTCATAGAGATCATGTTCTCG

ATCAAGAGACTTTTCAAATTCATTAATGAGATGATCATGATCATCAAAGTCAACAATCTCAGCTGTATTTTTTGAAATTC

TGGTGAGCGCCTTTTTATCTTTTGCCAGAGTGTCACATGCCTCATGAGAGGAAGAAATGTCATTATCAAATTGAACATTC

TTTCCACCTCGAGATTCGGAAGAATCTAATAAGGAATAATCCTTTGAGGTGTTTCTAAGACACTTGACATAATCAAAAGG

TTTAGGAGACATTTTGGTATACGCAAAGGAGATCTTGTCCTCAAAATCAGATTGCCACAAACACAGACTTGTAAGGTCTT

GAACGTGTTTGCCTGATCTGATACCAATTGAAAAAGCGGGGGTCTAACAACCACACCCAATATTTTGATTAGAAATCTGT

ATGGACAACTCCGAGTTACTTTCTAGAGAATCAAGTAGACTCAATCAATATAAAATATCTCAAGGAGTTATTATCTCAAT

CTCTCAATTTGATTTACTCAAGCAAATACAATCTGCGAGTCTTAATCAAAGAGAGATAACTTGGATGGTACCAAAGACCA

ATATCCAAGTGTCAATCAATTTAAATCAACAACCAATAAGGTCGGATATTATAATTGATTGAACAACGCACAACCTATAT

TATTTCAATTATATAACAAAATATAATGCGGAAAAGAAATAACACAGACACTAGAACTTTGTTAACGAGGAAACCAAAAA

TGCATAAAAAACCCCGGACCTAGTCTAGATTAAACACACACTGTATTAAGACGCTACAGATACTATCCTACTCCAAATTA

ACTTCGGTCTGGACTGTATTTGAACCCCAACCAATCTCATACTAATCCAAGCTACAGTTATGCTCCCACGCCTCTGATCC

CAGCAGGATACTGCGCACTTGATTCCCTTAGATGATCTCACCACAACCAAGAGTTTCTATGACCCAAAGTCGAAGACTTT

AATAAACAAATCTGTCTCAGACAGAAAAGTCTACAAGATAGATAAATTCGTCTCCAACGAATAAACCTATGAGTTTTGTT

CCGTCTTTTGATAAATCAAGGTGAACATGAACCAATCAATAAAACCAGGCTTATATTCCCGAAGAACAACTATGTTTTAT

CAATCACCTCACAATAGTCTTAATCGACACATCGAAAAAAGATATTGTGGAATCATAAACGATGAGACGAAGATGTTTGT

GATTACTTTTATATCTTGCCTATCGGAGATAGAAATCTCAAGACAATTATTACAATTGTACTCGTACAATAGAATATGCA

AGATCAACTCACACAACTGCGATAAAAGTAGTATCGGTCTGGCTTCGCAATCCCAATGAAGACTTTAAGTCTTTAACCTG

GTTTTAGAAGAAGAAAACCAAAGGTTAAAGGAGAATCGACTCTAGAGTAGCAAACTAGTTTCACATAGAATGTGTGGGGA

TTAGGTTTCCCAGTTGATAGAGTTCTCCCTTATATAGCCTTCAAATCAGGGTTTGCAATCAATGTTATCTTAGTAACAAA

GCATTCAATATTCACCGTTAAATGAAAACCTAATTAGATTCAAGCTAATACTTATCAACCTTTAGATCGAAAACTTATCT

TGTCACACACACATGAAATGCACGTTTGTAGGCTTGTGAAAACCGTACCCAAACTTGTACATTAGTTGGTTCAACACTAG

TTAACCAAATGGTTAGCCAAACACTAGTACAAAATGGGCTTTGGTGACAGTTCAAATTTCAAAACCGTGACAGTTCACGA

TTTGAATGTTGCAAAAGAATTGCTTTCCAGTCATAGGTAACAGTTTCTTACCAAAACTCACACCAATAGTGCTAAGAAAT

AGTGACAGTTTTTGAACTGTTACTGAACACTTGTGTAATGACAGTTTGTGTAGAAAAATGTTATTGTACAATGATATATA

ATAATAGTTATTAACTGTTACCATTCTTAAGCTAATGTAACCGTTTAATTTGAAAACTGTTATCGAAGTGTAAGCTAATA

GCTAATAGTTTTTCTAATAACCGCCACTTCCATTTATGGTGACAGTTCTGATACAGAACAACTGCTATTCATATAATTTA

CCATAAAGCATTCTTAAATGCATGATGTTGTCAGTTTGACTAGTAGTGCCATGGTAGGTCGTTTTTGTAACTTATGATAG

TGGGTCTCTCCACACCTTCGCCTTTCATTTTTGTTGGAAACCCAAAAGGAAGAAGCTTAAACACAATTTCTCAATACTAT

CAACCATATTTCTCTATTTCCATCGAACGATAAACCCAATCTATTTTAGGAATTTCTTATTTATGTACAAGCAGATATCA

TATCTCTCTTTACATGGTTTTTTATCTTGGAAAAATCTGGGATCTCATCTTTTACTTAAATCTAGGGTTTCAACCTTTGC

TTCAATCCTTATCCCTCGATCACTCACTATTTATCTCGCTTGCTCTAAAAAAATTTAATTTCGCTTTTGTGTGCAGAAAT

TGAGCTTCAATGCTCTCTCGTTATTTCCATAAAATTATGGTCTTGAGTTACTTGATTTGGTTCTCGATGGTGCTCGCGAA

GATTCTAAATATGTAATTTCCTACAAGGAACAATTTTTCGTTTTTTAGTGAAATCCTTAGTTTCTGATTTGGGTTTATCT

TATTTTCTCTATTTAGGTTGTTTTTCTGATTTATATTTATTTGTATGAATCCCTAGCTTTTAATAGAAACATTATTTTTT

AGTTCTTTTCTTGGTTGATTTCGTAGGATTTCAAGCTTGTCATTTGTATGAATCCCTAGATTTATATTTATTTCTTGGCT

GTCATTCACCCCGGAGAAAGAAAAATACGAACGAAACATGGAGGGAGGAGGAGCAGGAGAAGGATTGATGGAGATGACAC

ATACTAACAATTATACTATATCGCTAATAGCTCAACTTCAAAATGGTGTTGAAACTGTTGTCGTCACTACCACCTGTACT

TTGCGAGGATTTGCTTATGGTGTTTTTGTTGGAAGCATTTTGAAGATTATTCCTCCTTATCAGGTGATCAATGATTGTAT

TAAGCTGAATTCATCAAACCTAGATTTTAATTTTTTTCGAAACTTTAGTTGCATTCACGTGATTGAGTAACTGACTCCTT

GTATTTTACAGGGTATAGTTTGTTTGCAAGGAAACTTAGCTCGTGAATTTTCTGTTTGCAATAGGAAGGTATGATTTGTG

CGTTGAAAACAATTAGAGGGAAAGATGATACCAAGGCTAGGTAATTACCTAATTTAAAAACGTACCTCTCATACAATCAT

TTACTGCTTCTAATTCTTTGCTAATTTTCAAAAAATGCATCCCAAATATGGTAGATAAACATAGAAATAGACATCATTGA

GCTTGTTGCTTCTGCTCGTTAAAATCATGGATGCTTTATGGATTGATAAACAATTTGTTGTTGAACTTGGTTCACTACCT

TTTTTTCTTGAAATATGAATGAGTCGGGTAATCATTTTTGGACCTAATTGAGGGAAACGTTTAGTTTTGCGGAGGGGTGG

CAAGTTTTACTTCTGGAAAATGTTAACTTCGGTCGGGAATACTTCCCCTAATGGGGCCTTAGAGGCAAACGTAACATGTC

TCATATTTGTGCTTTGTTTTAGTGTTCTGCATAAGGTAACTTTCGTCTTCCACTTTCTTAATTGTGGAAAAACAATCATT

TAAAATTTTGTCGTTATCGGTCTTGATTGTGTTCTGCTTCAAGGTAGTTTCGTAAGTTAATTATCCTTGGTTCAAAATTT

TAAAGTGTGTAATTTGGTTAATGATAAGTGAGATTGGAACTTGAGTTTGTTGGTAAACAAATTATTTGTTTCTGGACTAC

GGTTTTAGTTGTAGCTGGTATGTGATGTGACTTCTTTATGGTTGTAGTCTTCTATTCAAGGAAGTTGATGAGTAATAAGA

GATCAGTTATGGGTCTGCGCAACCACGTACAGAGGAGGAAATACTTTGGTAATCGTTTTCCATGTTATTGGCTTTTGGTA

CATACATAGTTTTCTCTCTACCAACCTTGTGTGGTGATCATCTCTCTTTCTTGTTATTCTGAATTCATACTAGGTAATTT

AATTTTCATATTTGAATTTAAGTTAAGGTCTGTGATATTTATTTGATTCCCAGTAGAGGTTATTTTTGCAGAAAAGAACA

CTAGTTCTCACTTTTTTGTAGAAGACTTCACTCATCTGCCTAAGAAAGAGCTTTCATTTGAATAGTTCAGTTAAATTAGT

TATTTTCTCAAATACATTTGTTGCTTCAGGATTTAAACGTTGATCTAGTTGCTTGGTAGGAAGAAAAAAATGTCGCAGAG

AATCCCGTTTCCATGCTTAGGCATAACAAACTCATGGTTCTAACGTGTTTTCCTGTCCTGTGTAGATTCGTTGTTTAGAT

GCACCATGTTTGTGCTAATAAAAGAATGTACAATCTATATATGTTTACTACTAGCCTAACTTAGGTTTTATATGCACATT

GCTTTTCTAAAGTTCCTATTGGTGATATGAAGAATTCTACCGGGTGGACTAGATATGAAGATATTTATGGAAGGACTTCC

TGTTAGACTAACTCCCAACATGTTACATTGTTGTTTTTGTTTCTTCAGCTTTCCTTTGAATTGTTGGTTGCAGAATTTAA

ATTCAAATGTATGACATCTACTTCTGCTGGTATAAAGATTAGAGAGAGATGAACAACACATCCTTTTTCATCAGAAAACG

GGAGAACATAACACTGCAAAATTAATAAAAGTGAATTATACTCCTGAAGACTCCAACTGTTAGGAATCTAAACCGGCTTC

TGGAGCAACAAGTGAATTTTTGCTAATAAACAATACACATTCAAGTATAAATTGAGAAAATATATTATTTTATCTGGATC

TCCCTTTGAATTTGGTGTTTGTGATGGCTCTTGTAGGAACTGAGACATGTTACGTATTACAAAAGCGGGAGTAAAGTTTA

TGAAAATGAAGTACGACGAGGATAAGCATGAGGTAATGCCAGATGAAAATTGTAATACTGAAACTAGCAAATGCTGCTCT

AAAAAAAAAGGTTCTGGAATCTCGACAAAGTCATCAGGTAATATACAAATTCAATCTTCAAAATATGTAAGTTTCCATTT

GGATATTTCTTGAATAAATAACACATAAGATATCATCTTGAGCCTCCTGTATTGAGTATCGATTTTCTTTTAAAAATTTG

CATCTTGCCCTAATCTATGGCAGACAACTATATCTGAATCGTCAAGAAGTCCCAAAACTAAATAGGTAATGTTACCACGT

AACTGTTACTATTATGTTGCGACTCTTTCCTTTCTTTCTTTTGCTACTCAAATTTTAAATCTTGTTGTGATCAAACTTAA

ATTTCTTTTCCAAAAATTTGTTACTGCTCTAATACAAGGAGTTCTAAACTTAAGAAGGTAAGTCTTGTAACATTTTTTTT

TGTCCATATGTGAAACGAACTGTGTGTATATATTTGAGTTGTTTTTGGTTATCTATCAGTTTGATGAAACTGTTTAAATC

ATCTCACTAGTGATTTATTTCATGTCTTTTAAGTATAAAAATTCATGTTTTCAGACAACGGCTATTTCGTTACTAAAAGA

AAATGTCCCTGAGAGGAAGAGATCCGGAAATGATATGTTGCTGAAGTCTTACCATAAGAAGGTAACCGAATATCCATTAC

CTAATATAGTTGTATTCACTAGCATACCATAATAGTTGATTAATGAATCTATTGATGCTATCTCATAATCAATATTCTGA

AACGGCTGTAAATACCTTGCTCAAATATCATTGGTAATTTTGACATATCCCATGTTTGAGTTTGAGTTTTAGTATTATAT

GTTTGTATTGTTTTTAATTATTTCGTACAAGCTAATTGGTTAACAACACATAGAACAAACCACAAGTTACTGATAGGCGA

TTAAACTCTGGTGAGGTGGCAACTCAAAATGTTGGGAGCACCAGCGACAAATTGGTACATGATGTTCCCTTGGGTGAAGA

TAATTTCCATGTTCAGATATGCTAAAGATCCCAAAGCATTAGTCCCAATAATGTTGAAGATGAGATATTCACAGTCAGAG

AGACTATAGGGTCATATTTGGCGTGGCACAAAAAACTCATTATTATTCTTTTTGTAGGGGAAGTCAAAGTATGATGGACA

TATCATTGAAGTAGAATGGATAAGCTGGGAATTGAAACATGAACATAGTCGGATGATCAACTTTACGAGAAATGGATATG

ATTTCCGATTCCATAGTTAACTTTTTTTTTTCTTTCCTATACAGTACCAAGGAATGTTTTTATATGAAATCTGTTGAACT

TTTTATGTCCTGTGCTAACCTGTAATATAACTTCGGCTCAGAATTTGTGGTTGCGTACTATATTTATTAAATCAATTGAA

TTCTATAAATTATATTTTTTTTTGATGTTAATTATCTCACTGTGCATATTACCAAACCTGTGAGGTGAACTCACTGCAAA

CTAGGGTGTAAGACGACAGTTTTATCTTAGAAATGTTACCTACTATGACGGCTTCATGAGTATAACTGCTACCGTAGTAC

AAAAAGATCAAGTGTTCGATGACAGTTTTATTTAGAAGATGTTACCGTAACTTGTCAATGGTGACAGGTTAATGAAGAAA

CTGTTACGTTATTATAAGAAGTTAAATTTCTGGTGACAGTTTTATGTAGAAAATGTTGCCGCAAGTTGGTAGGCTCGGAA

ATATACCTACTTCTATGTATAAACTGCAAGTGCACAGTGTCGACAAGCAACACAGGGCAAGCACAGGTCGAACCACAGAG

ACTTGGAGAGTGTATAAGTTGATTCTAAATCCTGATTCAGAGAGTAACCAAAAGGGGTTTGATTTTGTCGATACGACGAT

ATTATTTTCAAATGACTGAAATAATAAAAGTAAAAGATGCAAATAACAAGATATGTAAACAATGATAATATGGATACTAG

GGTATTCAAATCCACCAATAACCTATGATATGCATTCAACTTGTAATATTATTCTTGTCCTTATAGTAGATTCAAGTCTA

CCGTATAACTAAGGTGTTTCTCCAATGGATAATTTAATCACAACTAAAGTATTTCTCCAAAGCATTAATTGTCTAATTAA

GCACAACTAATCTAAAGATCAATATTTGGTTAAGCATGAGTTGTAGCATAATCAACATAGAGATATTCTAACTACAATGG

ATACATGGCATACATTGTGAGAGCAAGATGTTGAAAAACTTGGATTAAATTATATCTTTAAACAATTCTAAACTTTCAAG

AACAAGATACAAGATATGAATACATACCAATGAGATTAGGGCTTCATCTAAACCCTAGTAAATGAAATTAGAACATGAAG

AACATTACAACAAAATTGAAATTTGATAAACTAACATGAACAAGCTATTCTTGGATGAGTTTTCAACACAACAACCCTCT

CTATTTATACACAAAAATTCTCTCACACTTGGGATCCGAATTTCACTGTAGCAACCACTCAACAATTAAACAAAACATGT

TTTCTTAATTGATGGATAGAATGGAATACAATGGGGGTGAGGTGGATGTTTTGGAAAGAATCCAATGGGTGTGAGGTGGA

TGTTTTGGAAAGAATCCAATGGGGGTGAGGTGGAAATGGATTTGGGTCTTGGGTTTTGGATCATGGGTTGTTTAATCCCA

TGAATTTGTAAACAGGTTGCCACTGATGCGCGAGCGGGTTGCAACGCTTCCAACTTTCTTCTCTTTTGTCTTCTCAATTC

TTCCACGCTTTATTTAGTACTTAAAAACACAAAATTAATTAATACAAATATTTATTCTTGAAAATGGTAAAAATGCAGAA

TTTGGGATAATATATGACTTTGAAGCGTGAAAGATGAGTTAAGTGGCAATGAAAAGGTGTATGAATATGCAATATTTGGC

ACTCATCACAAGTTGATGACAATGACAGGTTTACAGAGAAACTGTTGCCATATATTTATGATTCTACGATAGTTTCTTAT

AGACCTTTCGTAACAAGAGCTTTGGTGACAATGCAAAAAAATTTGAAAACTGTCACATAAATATTTTAGTGACAGTTTTT

GGCTGTCACCTAAGCCCATTTTTCTACTAGTGAAATGATAACTTTCATATCAACCATATTCTTCTTCACCATAACTAGTC

CGAATGACTCCAATGAACTAGTTAGAGAGTTGTTCAATTGCTTAGATCTTTATGTACTACACAAGACATAATCGAAGAAA

AAACAATGTGATTCACTCGAGTCGATTCATGAACTATAAAGCCACGATTTTCAATAGGATTCCTTAGTTTATTATGATAA

GTTCACAAACATCGCTTTTAAACATAACCTACTCAAGTTGGCGGACTGGGTTCGCGGACTTAAGTTCCCGGATGGAGTTC

ACAAACCCCGGCAGAAATTATCAGGTTCGAGAACTACGCCATTTCGCAGACTGGGTTCAAGGACTGGGCTTGTATGTTGA

GTTCGCGGACTAAGCTCACACACTACTCCGGTTTCACTTGATCAACAAAGTTCACAAACTTTGGTTCAAGCAATAAGGAC

ATATACATATATGTGTTACCACAACAATGCTTATATCCTCCAATGGTTGTATTGTCTAAAACATCATTTCAATCATTGAA

ACATTATCATAGGACGTTATATAGTTGTTATTCACAAACTATCTCTCGTCAGAGCGATTTCCAAAATGATTGAAATATAA

CATGACTTTCATCACTAGGTAAAGATGAACTTGGGTAAAGCGAAATCTTTACCAACACATATTTTGAGAAATAGATAAGC

GAGATATACTTAGCTCGAAATATCATATGTGTATAATATTAGTCTATATAGTAATACGACTTTTGTCTCAAGATAGGAGA

TAGAGTAGATAGACTTTTGAGTGATAAATAAGTTCAAGTCTCCATATACCTTTTAGTCGATGAAGATCCACTAGTCCCTT

TAGTATTCCTTCGTCTTATATGAGGATCGCCATGGAGTTCTTGAGATCAACTACACTTTCTATCCTAGTCTGAGACCTTA

GCTATAGTAGACTAGAAATCAAGACTTTATATTTTTGATCACTAACATTGACAAACATGATTGAGATAGCAATGCATGTG

AGTTCGACCGAGCAATGCTCTAACATCACTCACTATTATTGCTGTTGTGAATGTTTGCCGAATGATTGGTTGGTAGAACT

TTATATTTCCGTACAACTTATCATTATATTATGATTTATCTGTTTTTCCGTGTTTTATCGGTTTTTAGTAAAACCTGCAT

TGTCTTCAATTTATTTTAGTGATTACTTTAAAGTTAGTTATTATTTCTACTGGGCACCATTTTTAGGGATGATGTGCTGA

CCCATTGCCACTTTCTGTTTCAGAGACTGGTCAAGATGCGTACGTGGCGATAGAAGATTCGAGGAGTTGGTTTTGTTAGT

TGGTTAGCTTCCGCTATGTTTATTTTTGTGTTTATATTCTATTTGTAAACCAACTCATTATTAAATATGTATCATTTGAG

AGGATTTCTTGTATTTATTCTATAGAAGGTTTTGTTTCGGGTTAATCTTTGTTTAGAACTTCCTTCTCAGTGTTTCAAAT

AGATGTTTTAGATCGTTAGTGCCTCATTTTATAGTTAATCTCTTGGATTAGTCTGCTACTAGCTTTGGGGGCGCCATAGT

AGCAACTTTTAGTTGTGGGTGAGATGATCTAAGGGAATCAAGTGCATAGAATCCTGCTGGGATTTAGAGACGCAAGGAAC

GTGACTGTACCTTGATCAGTGGGAGATTGGTTAGGGCTCAACTACATTCTAGTCCGAAGTTAATCTTTTAGTAGGCTAGT

GTCTGTAGAGGCTTAATACAATATGGTGTTCAATCTAGACTAGGTCGCGGGGTTTTATGCATTTACGGTTTCCTCGTTAA

CAAAATTTCTGGTGTCTGTGTTATTTCTTTTCCGCATTATATTTCTTTATACAATTGAAATATCATAGGTTGTGTGTAGT

TCAATTAATTGGTAAATTCGACCTTGATTGTTGGATAGGAATTGATTGACAATTGGATATTGGTCTTTGGTACCACCAAA

GTTATTTCTCATTTCGACCAGGCTAACAGATTCCAATCTGTTCTATTACAGATTGAATTGAGATATAACTCTTGGATACT

TTTATGTGATTGAGTCTGACTGTCTAGTTTATTCTCTAGGAATTGTATTTGAGTTGTCCATACATATTGTCGAACGGAAT

ATTTGGTATGGTTGTTAGACACACTCTTTTTCACTAACATACTTTCAACTCTGCTAACTGCAAACTCATCATCACAATCG

TTTCAATCCTAGGATTTGAATCAGTTTGTGAGTGTGCTAATTGAACCTCTTCGCTGTCCAAGGACGGATATTCGATCCTT

CCCCTCGGGGACGAGATTTATTGATTAGTGGAGATTTGATTTCGCTGTGAGTTTTTTCCTTTTTGTTTAACTTTATAGTT

CTATGTTTACTGAAAGATCTAAAGAAAGAAAAGACCGTATGATGAAGGGGTCAGATAAATATCAAACGACGGCGAGCTAT

TGGAAAATAGATACATGATCTAGTCTTGGTAATGGAGAAGGTAGCACTAATCCTTCGAGAAGATTCATTCAACTGGATAG

AAAATCTTTTACCATTGAGGAAGATAATATTGGGAATACAAAGGTCATCAAGTTCACTGAAAGAGGCAGAGGAGGATTCA

TATCTTTGTTGGCGATTGATAAAAAGATGATTTCATGGATGATTCATATATTGGTGTCTACATCGGATGGCGGATATGCA

AAGGATAAATGGTCTGAAAGATTAAGTGAGGATGTTTACCTGATGCAACTTCAAAACAATGAAGCTGGTGAATAGCTAAG

ACTATCAAGATTTAGAAAAGATGGAGAAAGTAAGAAATTCAGTGTTTGTTTTCCTGGTGGTGATGGTAGAGAGTATTGGT

CTATTTTAGCTGAATTGATGGAGTCTTTTTTGAATAAGAAGAAACATAACTCAGTGGGAGCTTTATATTCTGATTCAAAC

TTCCCGAAGTTACAATCAATGGAATCTGTACAAAAGATATGGGGTCTGGTAAAAATTTCGCACAAAACCTCATGGAGGAA

GATCGCTGGGATTATCAAAAGGAAGATTAAGTCTTCTAAATTCATTGAGTTGAAAGAGATTAATTTACTCTCTGCTGTTT

TCGAAATGGAAGAAGAGGAATGGAAAGAAACTATGGAAGCCAATAATATGGACTCAGAACGATATTATCATCAAAGTCTG

TAAATGGTTCGGGAGTGAGGACAAGGACACTGGTATTATTTTCAAAAATAAAAACCATGTAAAGGTTTCAAAAGCTCAAG

TTTTGAAAGGGAATACTGAAATTCAGCAAACTATATGTATTGATAATTTCTATAAACAAGTCCAATGGTATAAAATTAAG

GGTTTACTATCCAGATTATGGAGTTCAAAATCAATAGCCTTGCTTGGTAACAAATTGGAGATATGCACAGAATTTAAATG

GAAACTTTGCAAATTAAAGACAAGAGAGGTTGTCATTTCTTTAGGATATCTGTCAATGGGGGTTGCCAAGGAGTTCCAGC

GGTGTTACTAATGTAGATAGAAGGGGAAAAGACGGTGTTTCAAATTAATTGGGATCCTGAATATTTGCCGAAGGATCAGT

CATCTGATTTGGAGTATGCAAGGAAAGTTCAAATATTCGAAGAGTTCTTTCAGGTAACAAACTTGGAAAGTTCAAAATTT

TCTTCTCCAACACGAAATTCACGCAGCAAGATGGACGGTGAAGATGGATCAAATCCAAAGGTGGTGAGTATGGAGACTAT

ACAAGGTTACATTAAAGAGGGTGTGGGATATATTGTTAGTGAAGATCATGATGGCAGTTGGATTTTCGTTTCTAAACCTA

GAATGAGTACTTCATTGGCTACCTCAAACAATAATAATTCCATCAGAGCATCTCCAATAGTGAGTGCTAAAAATCTTATG

TGGAATTTTTATTTAGATCCTTATTTATGGGGTCATATGTATGTGGCAAAAATAATAATGTTTTGCTCATGAACCATCTC

CAATAGTATAAGGTCTTATTTTAAAAATTTGTTGTTTTGGATAATGATAAAATTGATTCCCTCTTTCCTATTGGTTCGTA

ACAATTTATTTAAAAATTTAAAAATTTAAAAATAGTTAAAATGATGATGTGGAGGCCATTCTCAAGGTCTAAATAAGACC

TTCTCCAAGACCTTCATATTTATTTTAACACCCTTCCTATTTTCTATGACCCACTATTGGAGATGAATTTAATTTAAATG

AGGGTCTAAAATCCTATGTGTCATGAAAAAAATAAAACCTCCACCCTCTCTTGGAGATGCTCTCAATGCGTCAGGGTCTC

GTTTTAGATACTTACGTGACGAAGAAGACTTTGTAAACGGAAGTTTGGAGAAATCTGTAAACGAAAAAAAAAGAAGACAG

AATCGATGACTGAAGAAAGTGCTGGAAAGGAGATGGAAGACGAAGTTGTTCAAAATCTTAACATGACAAAGAATTCGTCA

GAAAGGAAGGAGGAGAACATAGACAAGACTGTGAATTCTTTTGTTAATTTAGCTTGCAACTATGGTCATATTAGCAATGA

GAATATAATGTCTCTAAAAACGAACATGACACGACTTGTTCTTAACTACAAAGGAAGGTTTCAAAGTCCGAGAAAGGAGA

ATGCAAGCAAGGGTACGGAGGAAATTAATAAGGATAATGAGACTTTAATTCAGCAGTCTCAGGAGGAGGAGTTTTTTGGA

CTCACTGAAGTACAAGGGAGAAGCATCACTGATGGATTTTAAAATTATAAGTTACAATATAAGGGGACTTAATAATTTTG

ATAAGATTACCTCTCTTAAAAATATCATCAACAATCAAAGATGTGATGTTTGCCTCATTCAAGAAACTAAAATTATGAAG

TTCTCAAACTGGTTTATATAAAACAGTTATGGTACGATGATTTTTTGAACGGTCATTCATTCTATCAGTGGGTGCTAGTG

GTGGTGTTTTATCCATTTGGGACAGTAACAGATTCACTAAGGAGGATGAGAGAAAAGGATACAATAATTTATCTACCTTA

TTTGTATATAAAGCTAATGGTTTTAAGTGTGATGTAACTAACATCTACAGTCCATGTGATTATCATGCTAGAGCTGAATT

CTGGGAAACTTTGGAGGAGGTTAGACACTGGTGGGCTGGACCAATTTACTTTGCAGGTGATTTCAATGCAGTCAGGTCTG

ATGATGAAAGAAATAGGGGTGAAGCATATAGAAGGAACTCTGGATTTCTGAACAGTTTTATTCTTCAGCAAGAATTGGTA

GACCAAACTTTGGTAGGAGGCTCTTAAACCTGGTCTAATAATCAAACTGACCCTCTGCTTTGTAGGGTTTATAGGTTTTT

ATTCAGTGTTGATTTTGATGAGGCATTTACTAATGATTTTCAAATTGCACTGACTAGAACAATGTCTGATCATAACCCTC

TTATGGTTATTACAAGCCACTCTATAAGAAGTAAACCTTACTTTAAACTGGAGAAAGGTTGGGTTGAACATAAAGACTTT

GAAAAGAAGGTTGAAATTTGGTGGAATTCTATGCAATACAGTGGAAGCTCTAGTACAATTGTTTTCTTAAAACTTCAGAA

CTTTAAACACTTTGTTAAGCCGTGGGGTTTGTCTGAGATTGGTGCTGTAGGGAATGAGAAAAAGCTGTTGACTGAGAATA

TTGGTGAAATTAATCTTTTAGAAGAGAGTGGTGCATTACTTCCTGCTAAGTTTGAAGAAAGATTACAATACAAGATTAAG

TTGAAGACTGTCGAAAACTCTGAAGCCAGGAAATGGCATCTTAGAGCTAAATAAAATGGTTTTAAGTGGGGGTGATGCTA

ACACTGGATACTTTCACAGGATTGCTAGCTCCAGGAAGAAGAGAAACACCATTTCTAAGCTGCAAATAGATGGAGTAGAC

TGTTTTAATCAAGATGCAATTAAAGAAGAATTAAGAAACTACTATATTAACCTATTTTCAAATCATAACAGTGTATATTC

TACACTAAATTCTTTAAGTTTTCCTAGTGTGGGTGAAGAGAAAAAGATCTGGCTGGAAAGGAATTTCACTGAGGAAGAAG

TTTGGGCTGTAATAAAAAACATGGGAGCTAAAAAATCTCCAGGTCCCGATGGTTTCTCAGCTGAGTTCTATAAGAAGTGT

TGGAAGACTAATAAAGGTGATTTTATGAAAATGCTAGATGATTTTATCAGGTTTGGTTCTCTAGACTGGAGACTAAATTG

TTCTTTTATTACTCTAATTCCATAAAAAGAAGACTCTTGCACTCCTAAAGATTTCAGGCCTCTAATCTTAATTGGAAGTG

CTTACAAAATCATCTCTAAGCTTCTTGCTGAGAGGTTAAAAACTGTAATGCCTTCTCTTATTTCAGATTTTCAGGGAGCT

TTCATTAATGATAGGAAAATTCTGGATGGTGTGCTTATTGCCAATGAATGTGTGGATAGCAGGTTGAAGTCTAAGATGCC

AGGAGTTTTGTGCAAAGTAGATATGGAAAAGGATTTTGATAACATTAATTGGCAAGACCTTTTTTGTATTCTTCATAAGC

ATGGTTTTGGTGAGAAGTGGATTGGTTGGATTAAATGGTGTGTTTCTTCAACTCATCTTTCTGTTCTTGTGAATGGAAAC

TCAACTGAGAAGTTCAAACCAGGTAGAGGTCTTAGACAGAGAGATTCACTGTCTCCTTATCTCTTCCTTCTTGTGGTAGA

GATTATTTCAAAACTAATGAATGATGCAGTTGAGAGAGGACATATTAGTGGATTTAAAGTGGCTGATACTGGGACAATCA

TTTCTCATCTCCAGTTTGCTGATGATACTCTCATTTTTTTGAATGATGACCCTACTGAAGTTACAAGACTTTTCATTATT

GTTTCAGTCTTTGAAGCTCTTACAGGTATGAGACTAAATCTATAAAAAAAGTACCATGGTTAGTGTGGGTGCAGACAATG

TCATTGAAGTTTTTGCAAAAGAACTTGGTTGCAAGACAGAAAAGCTGCCTATAAAATACCCTGGTCTCCTTATTGGTGTA

AACTCAAGAAGTACTTTAATTTGGACTTAGTTTTAGAGAAGATGGAGTTGAAGATGGCTACATGGAAGAGGAAGTACTTA

AACAAAGCAGGAAGACTAGTACTTATCAAGCATTGGGGGGGGGGGGGGGGGGGGGGGGAATCAAGATATAAGAAAGATGG

TTTGGTTGGGATGGAAAAAGATTTGCAGGCCAAATCATTATGGTGGCTTAGGTATAAAAAGTTTGAGGGTGACTAATAAG

TCTTTATTGGTTAAATGGGTGGTTAATGGTTGTCTTAAAGATCTTTTCCCAGTTATTTTTAATGCTTGTCGTAATAAAAA

AGCCTCAATAGTAGACATGATTGTGGAAGGAAGATGGTCTGGGATGTTTAGACATCAATTGAATGCAAATGAACAACTCG

AATGGGACCTGTTGAGAAGAGATTTGGGAGCAGTACCTTTCTTGGTTGATGGAGAAGATGAGGTTCAACTTTTTGATGGT

AATTTCTCAACAAAATCTTGTTATGATGTCTTGGCAGGAACTATGGAGACATGTAACTTTAATAAATTCTTGTGGAGGAG

TATCATTCCTTCAAAAGTTAGATTTATTCTGTGGGATGCTTTCAACAACTCTCTGCCTATAAGAGATGTGTTGATTAATA

GAGGGGTGGATACAGAAAGTGTTGTTTGTGTAATGTGTAACATGGTGGATGAAACAACGGATCATTTATTCATTCATTGT

CCTACAACTTTCAAAGTCTGGGATTACTTTATAAAGGCGTTTCATATTGACTGGCCTCTTCCTGGTACAATCCTGAATCT

TTTTGAAGCTTGGGAATGGAATGCTCTAACTGGCAGGTGTAAAGATTTGTGGAAAATCATGCATTTTGCGCTGATCTGGA

AAGTTTGGGAGGAAAGAAATAGTAGAGTATTTGGAGGGAGGCATAAATTTTTTATGAGATAATCATGCACACTAAACAGA

CTCTCATTCTCTGGTGTTGCGAGAAAGATACTTTTCGTTTTTTGGACAGTACGCATATTATGCGAAACTGGGAAACTGTT

TTTCATATGCAACCTAGTCTGCTGTGACTAATTTTGTACTTGGCTGATTTTCTTTTTAATATAACCTTTTCAATTTCGAA

AAAAATATATTTAACTAGTGAAGTATGGATATGTCAGGTACGTTTTGAAACTTGTATTTAATATTTTGGTGTAGCAAAAT

TGAATAATCCTAGATCATGTTATTCATGCCAGAATTGTTTCAAAGGCACGATGAAAATTCAATATGATCCCTACCATATC

TGCACAAATTGTTGATAATTGTGATAACTCTTTTTTGAAGATTGTCTATTGTTTGACATGTATTGAGGTATGCAGTCCAT

ATAAAAAAAAAACATAACTTAGAAGGAACCAAGTTAATCCACTTTTTCTTTTTCTTTTTTTTGAAGGAAAGAAGAGGATA

CACCTCTTTAGATAACTTAGAAGGAACCATTATCGTAACCAATTCCAGTAGCCACCGATCTAGCTTGATCGAATCTTGAT

CATCGTCACACCGAAATTGATTGCCTTCCAGATATACTTGAATATATCATCGGAATCCAATTGAAGCCCCATAAGGCTAT

AGGAAATCACACTAGTGACGCATATAAATATAAGTATATGATATGAAATAGATATCACCCGTAAAATAAGAAAAACAGAC

GTAAAACCTACAATCACAAATAATGTGATTTTTTTTTTTTTTTTTTTGAAATGAAAGGTTATATTAGAAAAGAGCTCATG

CATAAGCAATGAGAGAATTACAATACAGAAAGTCTGCACACGAAGAGAAAGAGGAAATGGTTTTTTTTTTAATTAAAATC

ATGAAAAACTAATTTCAAAACTCAAATCTATTTGGAAAATCTGATATCAGATCCGAGTAGATTTAGATTTGAAGATGAAG

AATACATAAGAACGGAGTTTTTAGATAATTTTTTGGAAAATCGATTATGCAGATAGGGAAGAGAGGGAATTTACCGCGAC

GGAAGAAATTTCCTAGTTAATCCAATTTGCATGCATAGGATATAATGCTTTCATCACCATCATTTATGAATTTTGCATAC

ATACGTTTGACCGTGTACGTGCCACTTGGATTAAAGCCCCACTCTCGTGAATCTTTCTTATTAACTTGGATATAAGTACC

AACAACATTTATTTATTTTTTGATTGGTAAAAGTAGATATATTAAGAAGTAGGAAACAAGCTCAAGAGATAGATACAACA

AAACACCATGACAATAGAGAAATAATTGTTATACAAACCCAAGAGAAAAGAGCTAAACAAGAGAAGCTATCAAATAAAAA

TAACTACCCCAGTTAACCATAACCAACAACATATTAAAAGAACAAATACCTAACAAGCAAATCAAATGCAATGAAGGAGG

CAAGGCAATAACATCTACTGACGAGGCGTGACCATAACTTCAAGAGGTGCTACCTGAGCGCATACAAAGCCTGGTATTTC

GGTCATATCCATTTTCCCATCTGCACTTACTGCTTTCATATCAAACCCGTGAATAAGACGGGGCACCACTAGATGCATCA

CCTGTAATGCCGAAAGCATACCAGCGCATATCCTCCGACCTATCCCAAATGGTAATAATTCAGATTTTTGACCTTTGAAA

TCCATGTTCACCTTGTTATTATTATCATCATTACTCAAAAATCTCTCTGGTTTGAACTCCAGAGGATTCTGCCACACGCT

CGGATCTCGTTGGATCTTCCAAAGGTTCAGGATTATGCGTCCCCCAGCTGGTACGTGGTATCCAGCAACTTCACACTCTT

CCGACGTGTACCGATCAATCAATGACAAATTTGGATACAATCGCATCGTTTCTTTGATAATTGCATTAATATAAACGAGC

CTAGGAATATCCGAGTCTTCTACGTGTCTATCGCGTCCAACGTGCATATCTATTTCCTCTTTTGCCGTCTTTAACACTTG

TGGGTGGTTCAATAATAAAGACAGGGTCCAACTTAAGGTTTTTGACACGGTATCGACCCCAGCTGATACGGTATCCTGCG

CAGCGCATTCATATGCACGAGGTAAAAATTTAAGCAAGTGCATACTCAAAATTCTATAGACATATATAAGCATGCATTTT

CTTTACTTTTTCTTAATTATAATATCTTGATTACACTAACTAGATCGATAACATTACCAGGATCATAGTCTTAATAACCA

AATCAGGGTCATCTCCAGGAAGCTGTTGTTCAGCAATCGACAAGAGAACATTAATGAAGTCCTGTTCATCACCAACTCCT

TTATCACTTTTAGATATGGATTGCGATCTCTTCCGGATATGCTCTTCAATAAGACACCCTGCAATTGAATTCAAATCATC

TCTAAAGTGTTTTGAGTCCCTTATGAGACCTCTTGATCTATCTACCCACTCAAGACATGGAAATACGTCTGAAATAGCAA

AGCTTGACAATAGAAAAGCAAAAGCTTCTTGGATCCTCGTCTTGTATTTCTCCCGGTTGGATGCCGTATCACCTAATAAG

AACGTTAAGTTAGTAAAGAAGCAAAAACCATGCGATTGGGATGGAGGATAAATGCTACTTCTGTGGTATGAACAGAGAGA

CCATCTCCCATCTTATTCTAAACAAATTTGGAAGCACTTTAATCCTAAATTGCTGCCGCCTAACGGGACAATGCCTAACT

GTATCCATTCAATCCTCAAGCACTGGGAAAGAAGGTCTTTGTCGTCTGCGGTAAACAACGGTGGGCTAGAATTCCAACAG

TGCTTGCTTGGACTGTTTAGAAAGCCCGTAAAGATATATTGTAATAAAGTGGCCACGGTGCATCATCCATGTGAAACTGC

AAGCTTTCTATTGGTGTATAAATTATAAAAGAGAAATCTGTGAAGGGTTTTTGGTGTATAAATTATAGGGAAAATGGGTA

ATATGCCCCAAAATGGGCTGCAATCTTAGGGATAAGCCCCAGGATTCCATTTCTCGGGAATATGCCCCGCACTAACAGTT

TCCATCCAAGGACGTTAAAAAGTCAATTTCCCGTACATTGACCGGTCAATATCTCAATACATGAGATCTCACATTATAAT

TGAAATGACATATTTGACCTTATACAGGTGTATACATCCTCTGGTGCCCACGTGTTTTCATCACGTCGAGTTTTAATCTC

GAGAATTACTAACGGCGTTTGATCAACCTAATCAAATCTAGGAATCCTATGTCTCACGGGTTTCTTTTCTATTCTTCTCT

TATCTTCATCTTCCTTTAGAAAAAAAAAAAGAAGAAAAAAAACAGACCACAGTTGAAGAACGGTCGAGAGATTCAGAGGC

TTTTGATGTGAGAATATAGGATTGGTTTGATTAGTTGGAGTAGTTATGGTAAACCTAGAATCTATCACCATTTTTGCTTT

TCCTTTCGGTCCGTATAGTTGAAGAACAGTTGAAGTTAAAGGGAGAAGAAGAAGAAGGAAAATTACCAATTGAAGCGGCA

TGGGTCTTCAAAAAGAGTAAGTAATAGAAACCCAACAAAGATTTATACTTGTTTTCGTTATATTTTGTGGTTAATTGTGT

TAAATTGATAAAAAGTGAAATTAGGGTTCTTAATTTTTTTTTCCTGAGTTAGGGTAATTTTGTGGAAAAATCTTAAGTAA

ATGAGATTAAAAGTGTACTGATCTTCGTTTTTGTTGTATTGTGTTGTTAATTGTGATACTCTGATAAAAAAAGTGAATTT

AGGGATTTGTTGTTAATTTTTTTTTCTGATTTAGTTAATTTGGGAAAAATGGATACATTATTAGGGTTATTATTAGTGAA

TTTGTGATACATTATTAATTTGGGAATTGTTAAACAGTACATAGATTATTATTATTTTTTGTTGTGAATTTGTGATGAAA

TAAGACTCTAACAAGGCGGTTGATATAGAGGAGGTAGGTTAATATCATGGGCATGTTTCTCTGTGTATGTAATAGTAGGA

GTACCCCCATTCTCATCGCTTGACACATTTAGCTTTAGGAGGCCAATGAGTACTTGAATTCAGCTAAGTCAATCACTTGA

ACGCTGCCAATACATTTTTTTTACTTACATCTAAAACTAAATTTGATTACTTGAATTCAGACCATAGTTTAGCTAAAACT

AAATTTGCCTTAGGGTACTTGAATTCATCTAAGACCATAGTTTTAGCTTTAGGAGGCCAATGAGTACTTGAATTCAGCTA

AAACTAAAAATGTCGGTCAGTGGCTAAAGCATTATTTCTTGAATTATATGGCTTCTTTAAGAATGTTGAAAATGCCGGTC

AGTTTTTTTTATGCTTGTTTGTTTGCGAAATTTCCTTTCATTGTTTATCTGAAAACATATGAGTGCCAATATTTAACATT

TTTTTAAATTTTATGAGTGCCAATTAGAATGCTGTGTATACATGATATATTTCAGGAGGGTTAAGAAAAGGAATTTTATT

ATATCTATAAACAATGGAACTTTGAAGGGTCATAAAATGTATTTTTGTGAAGTGAAAGGTGATTCCGTTAGTTTTGATCA

TTTTAAAGATATGGTAAAGGAAAGGTTGAAATGTTTGGGTGCGAAGCCGGATTGTGGTGTGGAAATATATTGGGATTCAG

ATTTGATTGGTTGTGAATTAGAATTTTTTGACATGTGGAGAGATGTAAAGCCTGATCAATGGGTGTTGGAAATTATGACA

CGGAACCATCATCAGAGATGAGAGTGTATGAAGTGACTAACAAAACTAGCAAGGCAGTTTTTACAATTGACATTGTGAAG

CAAGCATGCACTTGCGTGCAATGGAAGCTCAGGGGATTTATGTGTCAGCATGATGTGTGTGTATTGGACCCCTATAACCC

AGATTGGTCTGAGTAAGTAATACTTTCATTTACTATTATAGTGTGGCTGTCAATAATGATCCCAATAATGAATAAATGAG

CTGTTGGTTGAACTGAATTTGCAGGTATTGTAGTAAATATTACACAGTCAAATCATATAGGACCACTTACTCATGGCCAG

TAACTCCACTGGGTCACATTGATGACTGGCTAGAGGTTAGTGGACACATGTACATCCTTCATTTAATATTATGTTTTTGT

CCTTTCACCATTGTTTTACTTTGTCGACATTCTACATCTAAACCCTAGTTGTATATAAAGAATACTAACCCTATCTTTCT

TCTTCATATCTTTTCTTGCACTTGCAGGTCACATCTCCAAAGATTATCCATCCGGAGCTTCATAGATCATCAAGAAGACC

TTGTGTTAACAGGAAGAAGTCTTATATGGAGAAGAAGAAAATCATAAAGTGTAGTACTTGTGGAGAGTCTGATCATAACA

GCAGAACTTGTAAAGGGGGGGCAGTTGGATCCAACTCAAAGAAGAAAAGAATCAGAACAGAGTACCATCTTGAGGGTGAA

ACATTTGTTGAGAGGATTGCAACTCCAAAGAAGCAGAGGAAAAGCATTGGAAACACTGCAAGTACATCTTCAAGAAAGAA

ATCTGCACCAATTGCAACTGCATCTTCAACAAAGAAATCTGCACCAAATGCAACTACATCTTCAAAGGCTACAACTACTT

CTACTTCTTGTGCACTTGGTGGTAGAAAAGCTCCAGCTGCTAGTTCAAAGGTATCAAATTCCCTATCAAATACTTTCAAA

GGTAAAAACAAGAACAATGCTATCCACCAAACAATCAACAACAATGAACCTGCATCAAAGAGGCAAAGAAAGCCTAATTC

TAAATATGATGTGTAAAGTTATATTGGTTGTTGTGCTGTTTTTTTTATTTTACCTTTTCAAAGACAATACTTAATGTGTG

TACCCTGTTTTTTAGGGAGTTCACTTCCCCAGTTCATCTAGATTTTTGGTTTAATTAATCTTTTAAAGACCACTATTTAT

GTTATTGTACTTACCATCATGAGCTAGCTTCCATTTCTTTTAATTCATGTGGATATGGGGAAGAAAATGACCGTTGTATT

ACAACAGTCATGAAAATGTACATCTTCTGCCGTCCAACCAAATAAAAATACACGTGGAGATAATCTAAACCAAGTGAATA

TAAGCTTAATAATTATCTAAAAATAAAAGGTAAATTAGTAATTGGGGACAAATGTGAAATCTCGGCCGTTAGATGATATA

TGACGTGTCGTTACCGGATGAGATCTGGTTGTATTGTGAGTGTGTGATTCCCACAAGTGGTGAAGGGTACATCTGTCTTT

CATTTCGCACATGTAATTCCACATGTGTGCCAAAATTGACTAATTAACGTCCTTGGATGGAAATGTTGACGGTCGGGGCA

TATTCCCGAGAAATGGAATCCTGGGGCATATCCCCAAGATTGCAGTCCATTTTGGGGCATATCCCCAATTTTCCCATTAT

TATAAAAGAGAAAGTTGTGAAGGGTTTCAAGTCGGATCAAATTGTAGTAAACTGGGATACTATTTTAGCTTCCCGGTAGC

TTTTCTTCGGGTCTAGTTGCTTGTACCTTTTTGGGTCTTTTATGGTTTTTTTGGGTTGGATCTTTGGAATCTCTACTTCT

TGTAACCATTTGCCATTTCTGATGGCGTTGCATTAATCTATATAATTTTGCCTATCAAAAAAAAAAAAAAAAAACAAGCA

AAATATAATAATCACTTAAGTCGATACAGAGGAGTGTCATGCATATTACCTGTTATTGTCGCCTTTCCTCCATCTTGGTA

CCCAAAGAACATGCTCATGAACACGTTAAATATCAAATCGACAAGCCAGTTGTCCATCCTAATTGGGGTAGTTTTTCCAG

TACCGTTGTTGCTGCATAACTCGTTAAGTCTACTAAATGAAGCATTTATTTCTTTGATTTTCGAATGTTTCCACGCCTCT

AAACATTGTTTAGAGAGTAGATGTTGCACCGTTATCTTTCGCACTTCTCTCCAATATGTTCCAGATGGTGTAAACGTGAC

AGAAATTGGAACGTTAAATATATATTTCAGAACTAATGGAGTTGTCCGATTCGAAAATAACTTGTCATTTTGGATACTAA

ACCATTTCTTTACCATCTCCGAATCACTCAGAACTAGGATTTCTTGGTTACCTAAACGAATGTTGAATGCAGGTCCATAC

TTATCAGCCATGGTTCCTAGCTTTTTATATACTGTTTCCTGATCGTTCATGAATAAACGATAGAAACTATTAATGGCCAT

GCCCCCGATGCCTTTGGAGCCTTCTTTATCGTGTTCTTGTGAGTAACCAAGAAGAAGACATTGTATAGAACAACGATAGA

AACTATAACAACAACAATACTAACAGTAGCAATCGGATCAAGTAAGAATAACAACTCCCTCAAAATCTCCATTGATGAGA

TAATTTGGTAATATTTGGTGCACGACCCAAAGATACTTGATGCAGGAAACCAAGATCCTTTATAGATTAGTTGGTCTTAG

AAGTACACACTTATTATTAGTTGGTAATTATATATAGAGAAAGATTAGTTGGTATTTCATTCTTTCTTTCTTTCTTCTGG

TGTTTCTAGATTCGTTATAGGAAAACTCTAGAATTAGGACAACTGTTATTTTTCCAGATATGGGACAATGAATCTGAACT

TTTAATCATTCCCCTTTTCACCCCACTACTAATCCCGATGCTAGAATCATTCCCCTCTCAACCATTAAGATTGTGCCACC

TCATGTCCAAGTTTTGGACTATCCCTGTGTTGCGCTAAAACCTTGCGGAAGCGATTTGCACAGCTACAAACGGTACTCCA

TAATATTGCCAAGAACACAAACTAAAGAAGAAAAGAGAACAACAATTTACGTGGTTCGGCAAGTTGCCTACATCCACGGG

AAGAACAAGAGATTCATATATTATCACAAGGGAGAATACATATAAACCTTAAGAGAGAAAGAAGAATTTTTTGGTGATGT

TCATTTGCTTTCTCTAATGGCCTATTTATACTAGTTAGCTAAGCCTCCAAGTGACCAACTCCAAATTGAGTTCTACTTCC

TTAGCCTACAAACTAGAACTAGCCTAAAATTAGACTTCCTTTGCTTTCAAACTAGGACTCAAAGACCTACCTTGTTTGAA

TGCATGAATTTACAAGTGTGAAAGGGAAAGCTATGGAACTCTTCCACTAACCAATATTTTTCACATGGTTTTATTCCCAC

CTACCTACTAGCCACCATCCAAACATGGTTATTAGAGTTCCAAACAAAATTGGACTCTCAACAGCCCCTTATTCTAGAAG

TAAATTAGGCACCGTCCAACAATTCTCCACCCCGGATCATGCATTCATGCATGAGATGATCAATACAAGAAATCTCCTTC

GTCTTCCAACGTCGATTGCGCCATAGGCTGCCTACGTATCCTCTGCAAGGAATCAAACCGACCGTAGTTCACTCAATGGC

CTTGCAAGAAAGGCCTCCACCAGGTTCCTAAGAACCTCTACCCCAAAGGGTATCACCAAACTGTAAGAATGTGGATCAAT

TTCGAGCAATGACGAAACTTGACCCCAGATACTACTTTCGTTAACATATCAGCTGGATTGTCTTTGGTATCGATCTTCAC

AAGGAGAATGTTTTCTTCTTCCAGAATTTCTCTAACAAAGTGAAATCGAACATCTATATGTTTGGTTCTGGCATGATGAA

CTTGATTCTTAGCCAATTAAATTGCGCTAAGACTATCACAATGTGCATGCAGCTGTTCCTGCACAACACCCAAGTCATAT

AGCAAACCTTGTAACCATATAGCCTCCTTAAATGCCTCCGTCACTGCCATATACTCCGCTTCAGTAGTTGAGAGAGCCAT

AGTAGACTGCAAAATCGATCTCCAACTAACCGGTGCTCCATCCAAGGTAAATACATACCTTGTAGTTGAACGCCTTTTGT

CCAAATCACCAGCATAGTCAGAATCCACATAACCAACACACAGCTGATTGTTCCCATCTTCCTTCATAAACTCCAAGCCA

ACATCAACAGTACCATAAAGATACCTCAAAATCCATTTCACAGCTTGCCAATGTCCTTTACCAGGATTATGCATATAACG

GATGATCATACTTACTGCATATGGGACTTGGGCCATATACTCACGCTCTTCTACAGTAGTAGGAGACATATCAGAACTCA

ACTTAAAATGAGGACCAAGGGAGTACTTACAGATTTAGTTCCTTTGGAGATACCAAATTTCTGTAACACTTTCTTCAAGT

ATTCCTTCTGAGACAAACGAACTTTACCCTTCTCTCTGTCACGTTGAATCTCCATGACAAGAATCTTCTTAGATTCTCCT

AGATCTTTCATCTCGAACTCAGTTGACAATTTCTGCTTCAAATTATCAATCTCTTTCTTGTTATTCGATGCTATCAGCAT

AGCATTGACATACAAGAGAAAATATATGAAATACCCATCACGTAGCTTCTTGAAGTATACACAATGGTCATAGTGACTTC

TTGTGTATGTCTGGCCAATCATAAACTGATCAAATCGCTTGTACCACTGTCTTGGAGACTTCTTCAGTCCATACAACGAT

TTTTTCAGCTTGCATACACAATATTCTTTTCCAGCAACTTTGAACCCACATGGCTGAGTCATATAGATCTTCTCTTCTAG

ATCACCATGTAAGAACGCCGTCTTGACATCTAGCTGAACTAGGTCTAAAACATATTGTGCTACCAAGGCCAACAAAATTC

GGATTGATGAGTGTTTCACAACTGGAGAGAACACCTCATTGTAGTCAATCCCTTCTCTTTGGCCACAACCTTTCGCAACT

AACCTTGCCTTGTAGCGTACTTGATTCACCTGAGATCCTTCTTTCTTAGCATATACCCATATGCACCCAATGGCCTTCTT

ACCGTTCGGAAGCTTCGTAAGAACCCATGTGCCATTCTTGTAAAGAGACCCCATCTCGTCCTGCATTGCACCTTCCCATT

CTTTACACTCTGCACTACGTACAGCTTATAAATATTAAGTAGGAGTACCACCTTCTATAACTGGTAATGCATAATAAATG

TAATCATTCATCCAACCAGGCTTCCTGGTTAATCTTCTTGGTTTGCTGGTTGCTATGGTCTCAGTGACCGTAGCTTCTGT

ATCTTCCACTGCTTCATGTTCCTCTTCATCCAGATCATCACTATCATTTGGAACTTCAGTATTTTCCTCCCTTGTAGACA

CCACGCCTTAGGAAGTATTCGTAACAGATACAGACTTAGCTAGAATTAGTGGAGTTGCATCAATCTCCACCTGCTGCAAA

GGTTCACTGGTACCGGTGCTTCTGTTCTCTACCTGCAGAGAACCCCTAGACTTCACCATAGACGCTTCATCAAATGTAAC

GTCTCTACTCATGACTATCTTCTTCTCAACTGGATTCCAAATTTTGAACCCTTTAACTCCTTTCTTCACTCCCATAAAGA

TCCCTTTAACAGCACGGCTATCAATTTTGTTTTCACTAACATGATACCAAGCAGGGCAACCAAACATATGAATTGAGTCA

TAATCATAAGCTGGTTTACCATACACTAGTGGAAAATACAAAAAAAAAGTCTGCTAGAAAGACACTTAAAATCGCAAAAA

AGGACTGCTTTTGGGAGCCCTAACATTGTAGTCTTATTTTCCAAAAGGACTGCGCCTGGGAGCCCTATAATTGTTAAAAT

TACGTCTTCCTTCACCAGTTTAAATATTACTACTCATAGACACAGTCCTTTTGCTGGATCATATTTATAAAAAAAATGAA

ATTGATTCAGCTGAATACTAAACTGATTCAGCTGAATACTAAACTGACTCAGCTGAATCCAAAACCAGAATAAAATAAAT

TGCAACGAACAAGAAACTAAAATTCAATTTATATTCTGAATTCTTCAGTTGAAAACATCCCCCATTCAATTAATTTCATA

ATTCATTATACTACAACATAACTGATCATAACTGATTTCAATTGATACTTGGATTCAGAATCACCAAAAAAAAATTAAAC

AAAACCTGAACTAGGACAAGTACAAGCTCTAAGTTTTCTTAGAGATAAGCGCAAACTAACAAAATGATGATCGATGATGA

ACTAACTGTCTAATGAAGAACTGAGCCGCACAACTTAGAATCACATATACCTGCTGCAAAAATAATATTGGGCATTAGTT

TACAAACTAGTTTAAACACGACAAGAGTTAACTATTCAGAGATGATATAGAAAACAATGTGCAACTCATTCGCCATACAT

ATCTTGACCGAAAGAAGTGTGGTATACAAAACAATACCTGCTTATGTTGTTATTCGATATGCGAGTGAACAGTTTGTCTA

AGAATCCCACAAGAACATGCAACGACAACATCAGCTTGAACCTGCATCAATATATTTGAATAAAATCCAAGCTTATATAT

CAGTGTAACCTTATCCTCTAATACATAGAACAATATTCTGCAGCTGCAAGTTAATAAATAATCATCGCTTCAGCTCACAA

ATACCATAACCAGAAAACAATAAATGATAAAGAATCTTAAATAATCTCGGCAACCAACAAATACTGGAATCAAATAAGCA

CACAACACATAACAAAAACGAATATAGCAACCGACACAAGCAAAACAGTTTGTACTTCCTAACTGTCCCTAGTATCAACT

GATGTTCCACTTACCTGATCTTCCTATAATTTTTACAGGGAACTCAAAATGACCTGGAGTGAAACCTATAAAAATTTCAT

GGAATTCTAACGGTGAAATCTTGAGATACCAGGCATTCTTCGATTCTGTCCTAAGGAAAAAATCTGTAGAGGCAGTCAAT

GTTTCGAGGGAAAATGGGGTTCCTACCCATTTACGATCCTTAAAAAATTTGTACAGGATATTAATAAGGTTAGAAATAGA

TGCATACCAAAAATTTAGAAAAATCTAATGGTTACATTCATCAGATCAAGAGCTACAGATATCCATTTCAAAATTAATGT

TCTGCAGAAAACAGTCAATTCCATTGATTGTTACAGGGGCTTACCAGAGTCCGATGAACCTAAAAATTGAACCAAAGGTA

CCTGACATACTCAACTTCAACACAACCAAAATTCATAGGAAGACAACGGTTGGAAAATTATACACAGGCATTACAGTAAT

CATTAACCACGACAGTTTAGCATTTCTAACATGTGACGTTTCATTGATTACCTTAAGAAAAACCTCCTCAAGCTCTCTTG

CACCCAGTTGTTCACCCATAGGCACGTTCATCTATAGGCACCCAATCATCTTCAAGTCGCACCTCGTCATCAATCTGAAT

ACCTCAGCAAAGAAAGGACAAAGGGATATTAACAAGATGGAATTTGATTACTAAATTCTAATTTGATTAACAACCAGAGA

TGATTTCTAACATACACCTAACTAATACCAAATTGAAATTCTAGATAAGGAATCATCAAGTTATGGTTTGTCAGCAGTGA

CAAATAGAAATAACTCTTACTAGTTAAGTAAAAAAAAAGTAGTAAGAAAACACCTTGAGGAACACATCATGGTCTTTGCT

CCTCTACAAAGGGTAACGTCATATGTCCTGTCACAGAGATCTTGCAGGAATAGTTCCAATGCTTTAGCTGCAGAGTTAGA

ATTCGAAGTGGTTAATGTACCAAGAGAGGTAAGGTTAAACTTAATGCGATATGAACAAATAATGTCGGTAAATGTTTCCA

GCCCCTTTTATAGCTAATTGTCCCCTGCAAACATGAATCACATACAGACACAACAATTAGAAGCACATACACAAAGTATT

CAGTTCACATTTTGAAAATGCAAAGAGAAGTGACCAACCTGGTTTTAATTGTTCATATAGGCCTTCTTCGGATTCAATTT

TTCAGACATTCCAACTTTACTTCACCCCTGTAGAGTTAACTTCACCCCTGAGACCGCACAATCCATCGTCTATATTGGAG

GTACCCAATACAAGATAAAATCCAAGCTTGATGTGAGCCAAAGGAAAGAGAGATGCTAACATAAATTCCAGCACCATTTT

AATACGAGCTTGGATAACCTTGAGTTTCTCAGCATTTTATCTCCCATCAACCGGTACACCAAATTGTAACACCAATGACT

GAAAGTGTATGCAGTACATATTACAAAAATAAGTAGAATGAGTCTGCTTTTCAGTCTTGCCTAAATGATCCACACATTTC

CAATTGGCACGAATGAACCATGAATTAAGATGCAAACACAAAATAGAATTAGAAAAGGATAACAGAAATTGTACGGGTGA

TAAAAATGATGGCAAAAGATGTAACATTGCAAATTCCGCAGCACACTTAGACAGTTCTTTCATCTGTTTAACATCCTCTA

CCATCTTAAATGGAATGCTTTCTGGTGGCAGGGGTTTTTTGGGTTCACAATCACCAATATTTTTATCAACAAAATCTAAT

ACAGAGATTTTTGTGAACGGAAAATTACTCTTTTATTTTTTGTACTTCATAAAAATCAAAAAAATTGCCAAAATCAAATC

AAATTACACCAAAGGGGGAGATAGAGACTACCTTGCATTCAGATTTCTGGTGTGTATAGAAATCCAACCCATTCCATAAT

TATCAAATGCAGAGACCTCAATTTCATTTCTTACTCTGTTGAAATCATCACTAAACTAAATGAATCGTTTCTAATGATGT

TCTTCTCATCATTTGGTGTCCTAAGTAAGACCCTTAATTTGAGGTCCTCAAGAATCATGTGTTCCACCTGATCCTGTGTT

CATTAATCAATCAAACTGGTCAAAACTCACCATTAAAACACCAATAAAAGCACGAATTTGATAAAAATGAATCAAATTTA

TTTATTATTAAAAGAAATCAGAAAATTAGGGTGTTCATTTTGTGTTTTCTTACCCTGCTAGCTGCTTCCTAGATGTATTA

CTTAGAAATAAAAGGGGTTCGATTTCTTCTTCGGTATAAACAGAAATAAACCCATCAATAATTCTAACCAAAATATCTTC

TCCGAAATAAAAGGGGGTCAATTTCTTCTTCCGTATAACAGATGAAAGCAAAAAAAAAAAAAATTACAGTTAAAATTCAA

GAAGAGAATGGTTTACAATTGAGATTAAAGTACAAATCGAAGAAATTGATTACAGATCTAAGAATCTTGAGATCCAATCA

TGGGTTCTGAGAAATGGCTAGGTTTTGCTTTTGTTTTCAGAAAACGAAAATAAAATAAATAGAATACATCGATAAGAACG

AGGAGAAGTAAAGGGATCGATTAACGATGATGAACATAATTCAGAAGATAGGTTTTCGTTTTGTAGATGAGAAAGAAGCG

ATTGCGTTTTGTAAAGGAGAAGAAGAAAGGGGTTTGGGGACAGGATTACAAAGAGCAAATTTCTCTTGTTTATTTTTTAA

TTTCACCATTATTGTTGAATAGGGGGTTCAATTTCTTGGTGATATGAAGATGGAGAATATATGCAGACAATAAAGAAACA

GATGAGTGAGGTGAGTTCTGGATAAGTGAGAGAGAAGTGATGAAAGAGGCAGGTTTGTTTCTTGGATTATGGAAACTAGT

TTCTGGCACAGTCTTAATTAAAGTGGGATCACCGAAACAATTTTTTCTGCCAGTTTTAACCGAAATCCACGTGGAAATGT

TTTTTATTTCCAAAACTACCCCTAATATTAGGGCTGCTAAGAGAACTTGATTTCCTTTTCCATGTATTTGTTATAAAAAG

GACTGTTTTGGGAAGCTGAATATAAAAAGAAAGCCACCAGAAGTCTTATTTATGAATCATAATCAGACTTAAAAAACCCA

TTTTCCACTAGTGATACCACTTTTCCATCGGAGTTTTACCTTCTAATGCAGCTGATGGCAACTTATTAATGAGGAAGAAC

GCATATGTAACTACCTCAGCCCAAAACGCTTTACCTAACCCAGCATTAGATAACATACATCGTACCTTCTCCAGCAAAGT

ACGATTCATGCGTTCATCTACCACATTTTTTTGCGGTGTCTTCTTAACTGTGAAGTGCCTCTGTATTCCCTCATCCTGAC

ATACTTGCAAGAATGGATCACTTTTGTACTCTCCACCATTGTCCGAACGTAGTACCTTGATATTTCTGCCAGTTTGAGTC

TCAGTTGCCTTTTTCCACCTCAGAAAGACTTCTAGGACTTCATTCTTATGCCTCATGGTGTACACCCATACGCGCCTAGA

ATGATCATCAATAAAAGACACGAACCAATGTTTCCCTCCCAATGATGCATTCTTGGAGGGACCCCAAACATCTGAGTGAA

CATAATCAAGAACTCCACCAGTGTTGTGGATTGCTGTGCCAAAGCTTGTTCTTGTCTTCTTTCCTTTCACACAATGTTTA

CAAAACTCTAATTTGGAGGCAATAGCACCTTTCAATAATTATGGACGAATTAAAGAATGTAACGACTTATCACCAGGATA

TGCAAGTCGCATATGCCATAACTTCGTGGTCTCCATAGCTGCAGTCTCGATGTGTTCTGAAATAGACACATCTCCCATTA

TTGTACTCCTAGTTAAGTAGTACAAGTTATGATGACTTGTGCCCTTCATGATTATCAATGCTCCAGCGATCACCTTTAGA

ATGCCATTTTCAGCGACTAACTTGTAGCCCTTAGCTTCCAAATTTCCGAGCAAAATCAGATTCTTCTTCACAGCAGGCAT

GAACCTTACTTCCATCAGCTCCCGAACCATACCATCATGCATCTTGAATCGAATACTACCAATCCCAATCACCCTACATG

TGTGATTATTCCCCATACGCACTTCTCCATCAAACTCTTCAAAGTCTGAGAACTAGTCCCGATGAGGACAAATATGATAA

GTAGCACCAGAATCTAATACCCACGTATCGTCAGTTATGATAAAAGTTGATGGTATAACAGTCAGCAAGAAATATGAAGG

TTCATCTGAATCTTCACTTGCCTTAGCAATGTTCACCTCGGTGTTGTCCCTTTCTCGATCCTTACGCTTGGTACAATCCT

TAGACCAATGGCCAAAGTCATGACACCATGCACATTCATCCTTCTACTGACGCGCTCTTGACTTTGAACGTCCTTTAACT

TTTCCACGATCATCATTCTTCTTTCTCATGTTTGTTGAACGACCTCTTGCAAGAAGCAAGTCACTGTTAGCTTCACTTGA

CAGGTCTTCACGGTCCATATTCCTGAATTCCTCACTGCGCAATGTTGTAGTGACCTCGGCGTATGTCATCTTATCCTTTC

CGTGCATTAAAGCTTTAATCATGGGTTCATACCTTTCAAGAAGGGTGTTTATCAGACACAGAGCTTGTTCCTCGTCGTTG

ATCTTCTCATCGTAGTTAACCAGCTTAGCAAGAAGTTTATTATATGAATCTAGGTGCTCAGTTAGAGAGGCACCCTTCTT

CATGTTATAGCGATACAAATTTCTCTTAAGATGTATCCTATTAGCCACGTTCTTCACAAGATATTCCTTCTCTAGCTTTT

CCCAGAGATCCTTAGCTGAAGTCTCGTGCTGATAATTAACACTTACTGATGTTGCTAAACACCCTCGAATCGAACCAAGA

CATAATTTATTCATCTTGTTCCACTGCTTGTCAGACATCTCCGCTGGCCGACCTTCTAGAGCTTCTTCTAGGTCAAGATG

AACAAGAGCATCCATTACGTCTGTTCTCCATAAACCAAAGTTATTGGTTCCGGTGAACTTTTCAACCTTCAGTTGTGATC

TATGAGAATGAAGTATTTCTTCTTTTGTTTCTTTATCTTTATTTGTGGACTCCGAATGTTCTCCTTCAGGATCGTTCGGA

TCAACTCCAAATTGAGTTCTACTTCCTTAGCCTACAAACTAGAACTAGCCTACAATTAGACTTCCTTTGCTTTCAAACAA

GGACTCAAAGACCTACCTTGTTTGAATGCATGAATTTACAAGTGTGAAAGGGAAAGCTATGGAACTCTTCCACCAACCAA

TATTTTTCACATGGTTTTATTCCCACCTACCTACTAGCCACCATCCAAACATGGTTATTAGAGTTCCAAACAAAATTGGA

CTCTCAACAACCCCTTATTTTAGAAGTAAATTAGGCACCGTCTAACACCATGTTGTCCCAGCTGTAGACGCTCCCTTAGT

TATATGCAAATGGTAAGCTAGACGGTATCTTTGAGTATAATATTAAAGGGAGGACCGACACGTATGTGCACGAATAATAT

AAAACGAAATTTGGTCCTTGAAAGTGGCAAACCAACCGCTCTTTTGTATGCACTGTATATTGAGTTTGTGGAATATGTTT

ATCGTTTGGATCGTTTAGGATAATATAGAAGGGAAAACGGGTCCTGATACTCGTTGGGATAATATAGAAAGGAAAACGGG

ACCTGAAACGTGCAAATGGTAAGCTAGACGCTATCTTTCAGTATAATATTGAAGGGAGGCACGACAGGCATGTGCACGAA

TAATACACAATGAAATTTGGTCCTTGTGACAAACCAACCACTCTCACCCACTCTTTTGTACTTCATATTTAGCTTGTGGA

ATGTGTTTCTCGTTTGGGATAATATAGAAAGCAAAACGGACCTAATACTCGTCAGGATAATATAGAAAGCAAAACGTGGA

CGCTATCTTCATGTAATATTGAAGGGAGAAACGACAGGCATGTGCACGAATAATATACAACGAAATTTGGTCCTTGTGGC

AAACCAACCACTCTCACCCACTCTTTTGTATGCACTTCATATTTAGTTTATGGTATGTGTTCATCTTTTGGGATAATATA

GATCGGATAACGGTTCCTTTTGATTTTAATTCCCAGCAATTAAATGCATATCTCTGGAAAATTAAAAATTGTTAATGTGA

ATTTTACTAATTGGAGATTTTCTAGCGAGATTTCGGAAATATTGGACAAACATTTCCAAGAATTATGAAAACTGATTTGG

GCATTTATTGCATATCTTGAGAATATTCGGTTTTGGAAATTCCTTGTTGTTCAAACATCCTTGGTTTATGAATACCCAAT

TTTTTATTTCGAGCAAACTATCCTAAGAGCCAACAAAACTATTTCTTTGTGTTGTTACTAGTGGATCCGTCTATTCGGAG

AGGAAAGTACCCTAATTAGGCGAAATCTCTTACGACCGCTCGTTTAAAGACTTCTGTAGGATCAAGAAGCTCCACGACTA

CCGTTGGTGGGAAACTAGATAATTGCAGTTTATTTTAGTTTCGATTGATTTGATCGACTAAGGGTTGTTTATCTTTGATT

GCACCTAGTTTTTTTATTCTTGAAAATCTTCTCTTCTGATATAAGACTCACTCAAATAAGATCGAAGTTTCGACGGGATC

TTTAGAAATGTTTGTAGATCTAAAGACGTCTTGTGATAATACATTATTAACAGACTCATTTCTGTGTGTGATTGATCACA

AGAGATTCAACTGGTATTGTGCATGTGTTTATTGAAGATTAAAGAAGATTTGAAGACAAAAAAAAAATTCTGATTGGGTT

TTTATATCTTTGGTGTGCACAAACCTTGATCGACTGGGATCCAACTAGAATCGGATTTATTTGATTGATTAGTTGTGTAA

GATCGGCATCTCCTTATAGTTTCTTGTTTGATTTTATATTGATTGATTGCATATCTAAACTCTGCTACTTTGGTAGTTGT

TGGATATATTGATCTGACCAGGCAAAGGAGTTTATTTGTTAAACAGAAGAGCCTTTGCTTATGACTTGAGATATCTTCAT

CTGAAAGAATCAAGAGAGTGGTTACCAAACAGATTTTTTGTTCCCTTCCTGTTTGGAATACGATCAAAATGAATTGTTCC

AGTGCGTGCACTTATTAAATATCGGAAGCGCAAGGATACTGAAGAAACTATGTGAACTATAGGTTTAGTTGTTTGGTCTC

AACTATACGAAGTTTGTTAGGTTTTGTATAATGGATTAATTCTGAGAGTATTCAATTCTGGACTAGGTCCCGAGATTTTT

CTGCATTTGAGGTTTCCTCGTTAACAAAATCTTGCTGTGTTATTTACTTTTATTTTGCGCAATTATAATTGTTTTTATTA

TAATTTACAGCAAATTACACAAATGTTAACTCCGCATTACTTGATAGTGATCCTATAGAGTTTGGTTAAGTCCGAACCTA

TTATCAAGTAATCACACTTTGATTGCTGTATTGTCTCGATCTCGTGTCCATAGACGATCACACAAAGTGTGAATACCTAT

TTTTAGTATTGTCTCGATTTTGTCCATAGACAATTACTTTCGGTAAGAGGACTTATAGGTTGATAAATTAAAGATTATGG

TGTATTTGGGTACCCTCGTCTTTTCAACTAGGATTTGGACAAACCTTAGAAACCCTTTGAAGAGATGTAGGATGTGGTGG

ATAAATCATCACCAACCAGACTGGTTCTTCGCTAGAACATCCTTATTATTCAAGCTCATATAATCTTCACCCTTCTCCTC

GACTGGTGTCTGCCTTCTTTTTTATTCATCTTGTTTCATCACGGAAGCAAAAACCTCTCGGTTTTTCTCGCCCCCTTCGG

TTTCCACGAACAAGAGATTTATACCAAAATTGTTTCCTTTTCTAACTTCACCCCAACTTCTTTATTTAGCTCCCCTTTAT

TTGGTATGGGATTCCTGACTGCTGGGACATAAGGCGTGAAAACCCAACAATCTCCCTCTCACGACTTATGGCGAGGTACT

CGTCACCACCGCATCCGTCATGCTGCAGAAATGTACATGATTGTACTTTGGTAGTGACTTTGTCATCATATCAGAATCAT

TTTAATCTGTATGGATTTTCTCAACTAACTCCCCCTTCTTGTAACTGCATATTTTCTGATGGCTACACCCAAGTAGATCT

AAAGGCTATAAAACTACTAACATTCAAGATTCAAATATGAATAAGATGTAAAGAACATGAAAGTAAATATTAACACAAGC

ATATAACGTGGTTCGGCCATGAAGACCTACGTCCACGGGAAAGAAGTTGTAATATTTATTGATTAAGGATTTACCCTTGA

AAGAGTTACAAATATCACTTAGACTTCTCACTCTTTCTTTCTCTAAGTCTCTCTCTAGAATTTTCTGTAGAAAAACACCA

TCTAAGATTACATGCCCATATCCTTCTACATGGACTGGTATTTATAGACAGATAAAAATTGTGAAGGTATGGCCCGATCT

CGCCGAGTCGATGAATCTATCGTACATCTTGTTATTACTTCGGACGAGATTTAAACTATCTATCGTGATGACCTACTCGA

TTCTCCTTCCGATTTGTCTCGTCTTGTGTGTAGTCGCCTCGCTCTTCTTTGGAGAATCTCGTGTTGCATCATTCTCGAGT

TATAGTATAGATCATATGAGTCTTCTGAAGTGATGTAGATTCACCTATGCCTGTTCTGATCCTTCATTGAAGGCATCCCT

TCTTTTTATACTTGACCGTATGAGTTGATTAGACCACTCCACACACGCGTCATTCATGTAGTGTTGTAGCAGGTGCCTTG

ATTCAGACAAAATCACATTTTAGAGTATGATTCGATACTTTTATATCGTCACCTCATCATGAAACCATCCCTTGATATGT

TGACACGTGGTGATCAGGTATTTTACCCTCACAATTTGATCTTTTTCTTTGCATCTTGCCAAGGGCACTATGGGAAGAAA

ACTTCGTAACCGCCCCATGTTTTTCGCCACGTCTTTTCTTTTAATTCTTTTTCACCTGTTAACTGGGGACACCGGTTCGC

TATCTAAGAAGTTATATCTCCTATATAATCTTCCTAGGATAGAGAAAACATGTTTTTATTTTCCTCTCCCTTTTTCACAG

AGTTCAGAACTTTTGTTCTCTTTCTTCTTTCTCTTTTGTTGTTCCTTCTTAACTCTTTGCCTGCTATAAAAAAAAGTCAT

CATGTCTAGGAAGGATTCTCCATCTCATGCTTTAATAAGATAAGGGCTTTCTTCTCCTTCTTTTTTCTTTTTCAGGTTAC

TTTAATCAATGGTCCTCACCATTGTTTTTCAACTGTAAAGTTCCTTGGGAATCGCAATTGTCGTCTCCGTCCTTCTCTTG

ATTTTTTCGCCGGTTTTTCGTCAGTGTAAGTTCTGATTCTTTTCTTCTGTCTACACTTTATAGTGCTAAGAAGAGACCTG

CGAGAAATCTGAGTACCAGTCGTTCTTACGGCATGTATGATGCTACATATTTAGAGATGGTGAAGGCTAAGCATCGTCTC

TAGAGATTTCAAGTCACTTTTGCTAGCCTGGAGGGTCAAGATGTTGTTTTGACCAAGGATATTCTTGAGAATAACCCTTT

TGGTCCCAATGAGATAATCGTGATAGTGGGACAACTAGATACTGGGCTTCGTCTCCCTTTGTACTTTCCGCTGATTTCTT

TCTAGTACAAAGTACTCTGCCTGCTCGATCCTTAGTGAGCCATCTTTCAACCAAATGGTAAATTTTACGGATCGTCAGAG

AAAGTGATCGGCGATATAAAGGTAAGGTTACTGATCATGAATTGAATCAGTTTGTTCCTTCTCATAGGGATTTGTATTCT

CCCAAGAGATTCTTGGAGGACTACTGGTGTCAGCAGTCCGATAAGTTTGATGGCTTCGGGGTGGATATCTCCAGGATTCT

CTGAACTCTGAAGGGTCCCTTTCTCATGTCCGATTTAGATCCTTCTAATCCCAGAAATTTCACCCTTCATAAGCACTACA

CCATTTTTTCCGATTTGAAACGGTTTTTGAGAATTACGAATCTGGGTTGTAACGGTTCCTGACCGTTACGAAAAGGGGTG

CTATAATTTTTTAGTAACGTCCCACTAGCCGTTACGAATTTTTTTCGTAACGACCCTTGTAACAATTAGGAGCCGTTACT

ACGTGGCACTATCGTAACATGCTCGGGTCATTACAAATTTATTTAGTAACAAGACATTGTAACAAGTTCAAGCCGTTATG

AATTTGTTTATAACTGTAAAAAAATTACCGCCGGTCAATCTTGACTGGGTCAAAATTAGTCAGCAATGACCAATTTTGAC

CGGGTCAAGGTTTACCGGCAGCTCATTCTTGGACGGAATATGCCGGAATTCCAAATAACCACCTGCATACATATTTCATT

CTACCAACCATCCATATCTAACAATAGTTTCAATCAATTCATACCATACTCAATCAATTCATAGCTACACATTCATATAA

AAAAACACAAATTTTTCTAAGTACCAATTTTTTTTTGTTAAGTATCAAATTTTGAGGGAACACATAGATAGAAATTTACT

AAGTTCCAAAGTTCAGTTCATAGAAACTTTAAAAACTTTCTAAGTTCCTAACTTATATCTAACTTTTCCCATGTCACGGG

CTAACAGAAACAGAAACACGAATCACATAAAAGCCTTCTCATGCCGCAGACTGCTACATCTGAAGCAGGATAGGCAAAGC

TGATGGAATCCAACGCGACAGCAACTATGCATCTTCAATACAATGGCCATTTCTCTTACAGGAGCTCTCTACCAAGAAAT

AAATAGCTACGCAGATGACACAATGATCCTTTATTTCATTTCATGAACGACATATAGTTTTCTAATCGTACTGTATTATT

CAACCCGAATGATCAATGATTCTGATCTAGTTTCATTAGTAAAATTAATCGACTGAGAAGGAGTAATAGTAATCAGATGA

CACAGGGATTACTAGGAGATATGTACCACTAAATTCACACTTGTTTGTCGTCTTGTTGCTGAGTCCAACTATGATTGAGA

TGAGATTTAGGAGGGAGGAGTAAATTGATGGAGGACAATTTACTAAAATGAGTGTCTTAACTTGGACTCAGCGTGCCTTT

CTCAACCATATTCTGTGTTAGGTGCCTTTCCATCCCTTCCAATTGTTCATTGGGGTCTCTTCTCATCCAACTGCTAATAA

CATCAGAGATACAAAAAAATGGAAGGAAGAGCTTCCCACTAAAAAATCAATTTAAAATTTGGATTGCTAGCTGAGAGTAT

GTCCACTTTTTTCTTTAGTAGCATGTCATCTGGTGAGATTTTGGCATATAAAAAAGAAATCTGAAAGACCAAAACATAAA

ATTGAAGATGGTCAGATCCCACTATAGTGTATGAACTTTGATATGGTTGACACTTAGTGGTTACAATGACAAAATAGCCA

TAAAAATTACATTTAAAACAGTCTATGCTAGTATGTGTATATGACTTTCTTTCTAGTTCTCAGTTAACTTCCAGTATGGA

GCCTGGGCAACATAAAATACTACATGTTTGTGCTGCGTTGTATATCTCGGCATCCTGAAATGCCTTGATAAAATGATTGT

GGTCTAGATAATCAATTTCATAGTCTTGAAATGCCTTGATAGAATACTACATGTTTGTGTTGCGTTGGTTTATAAATTAC

AGACTCTGGTCTCTGCAAGAAAAACAAAAAAGAAGACAATACAAATCATTCTTAAGCAGATAAATAACCAATAGAGTGGA

TAGAAACAAACATGAGGGTTATCTAAAAGAAAATGCTTGAGTTTACATCTATCTAATAATTAAAATCCTCAAAATCCAAA

CACATCGACCCATCTAAGTATGTTCAGGTTGGAAAAATCCATCTATACTTGATAGACTTTGTCATTATACATTAGGTAAC

AAACCCCTCTCCGAAGTATGTTCATGTAGAGATAAATCATATTCTCAGGTTAAATTGTTTAAGAACGAAAATTATGATCT

TCAGAAAAATACGCAAATAGGCTCACTGACAAAGAAAACTAAACAAATGATGAAAAATGTTACTAAAGGAGACCCACAAG

TCTTACCTCGGAGCAAAATTTCATTTCACATTAGGTTATCCATTCAAAACCAGTCAATCCACCATCCCACAATCTGAAGA

CACAAAACAACACACTATAATATAATATAATTTGATATTCAGGATATCTTTAGAACTAAAGCCTTGCTCTGAAGCTTCAG

ACCCCTTATGTAGTACAGCTTTAACTTTCTGCAAATTTCAAAAATCATAAATAAGCACTCTCTGCTTAGGACTGACTTCC

TTAACAATACAAATTTACACAGCTAAGATAAAATTGGCAACCCAAAACAAAATCTCAAAAGAATTAGGGATGACCTGCAA

TATGTCGATGTAACGCTGTGAGTTTTTCAATTTAGAGACACTATCCAGATCCTCAAATTTTATAACTTCTATATCAGCAA

TGTCACCACTATCATCATCCATATCGCTCTCACGTTAATGTGGAAGTTAAACTTTTAACTTATTCATCAAAAAACAAGAA

AAAGTATTACTCACCAAAACACTAGCTTCATTATCTGATGAAATGCCAAGAACTTCGAAGAAAGCATCAGCAAGTGTAGC

CATAATTTAAATTTAATTGAAAGTTCAAAATTGGAACAACATGCCACCCAAGAACATTAACCTTTGCAAGTAAAGATTGA

ATAAAAACTTTGGTATCTGCATAATGAGCATCACAAATCCATATTTCTTTGACAAATGAACTCTTCCTTGCAAGCTGGTA

ATGCAAATGGGCATTAGCTGATAATATTTTTCAGTTAAGAGTGTAACCACTTAATATATTAGTAATCATCAAAACATACA

AGCAGTAATGATAGTAATGATATATTTGAATTGTACAATTACAGATCCGAACAATAAAGTGACAATCATGCTATGTGCGA

TACCTGAAATCAGAGAGATAAAGGTCAGATAAAACCTGCAGACCAGAAAACTTCCATCATTAGGAAAAACAAGTAGAGTG

AAAAGGACATACAAGTTAAAGGTTGGGTGGGTGAGATAAGAGCTTTACGTAACAAGTCCTTCTGGGATCGCTTTCTGAAC

CTCCGGTGCGAGTGACACAGAAGTAAATGGCACCCGTGAGTTTTATGAAAAAGGTCATTCAAAGACATTCATTAAGCTCT

ACCACAATAAGCTCTGAAAAATGCAGCAACTTAACCCTTGTCTTAAGATGATCAAACCACCCCTCTTCAATCAACCACAA

ACCTCTGCAACCAAATATAATCATGCACAATCAAAAATCTACTGAAGTAAAACCCACAGAACTCAACATAAGGAAACATG

AACATACCTTGATTTTGGAAAGAATGACCTTAAACCTCTCGAAATCATTAAGGGAAGATAACCTTGCTATCCCATGAACT

ACTCCCCCACTTATTCTTCACATATATAAAATCGAGACCAAAAAACATCATTGTATAAAAAAACACAAATAAATACGTAC

CAAAATTGAATGGAAACAAAACCATTGATATATAACTGCAACTTCCATGGAAGCAAGGAGTTCTTTCTTGTTAGGTAACC

ATATCAGGTGCATCAACAAGTGAATCATCTTATTTTTACTCACCTGTCAAGACGTCTAAGCATGGTACCATTCATAAGAT

TAAACATGTATTTTTATTGCATAGAAATCAATTATCCCAAAGAGTAATTTTTTGTCTCTATCCTTAAGAGACATAACCTA

AATTAAATCTTAAAATTCCCCTAACATAGAGAAAATTAAATTTTCCTTAAACTCCACAGTACGCAACCTAAATCTAAACA

AATCTAAAATCAATTAAATCTAAAAAATATCAAAAAAAATCAGAAAAAAATAGAATTTGGGATTTGTTTTTATACCAAGA

ACATTTTGAATACCAGTTTCTCTGATAATGCTTTCAGTTCTCAAATGAGTAAACCCTAACTTCAATACCAAAGTAACCAT

ATTTAGCGAGCTCTCTAGTCAATCACCTCGGAAAAAATTACAACTTTAGCAACAAACTACAAATCAAATCGAAACAATTG

ATTTCAGCGTCAAAATAATCACACAAAAATCAGAAATAAATGATACAGAAAGAGGGTTTGAAGGAGGGTTTAACTTTCGT

TTCTTAATCACATGTGTTGCTAAGATTAATCAATACTAAAGCCTAACTTGAAAACTTTGAAAACCCAATAGAAAACCTAG

ATCTAGAACTCTAGGGTTACAAAACCTTGTTTTAGGAAATAAGGTACGAAATATAATCTCTAGTTTAGTGATTGTTAAAG

GGATTTGGGTGTTTTTATGTTGCTGATGATGGTGGTGGTAGTACAGAATCGATTGTTAGATGGATCGGCCTAGGTTCTTC

AAGGAACAGAAGAAGAGAATAAAGAATGAAGAAAGAAGAAAGAGGCAAAGAAAAAGATAAGGGTGGGAATGAGGTCGAGC

GGTAGATAGATAACCATGGGAAGAAATATCCTCAAGTTTGATCAAAGTGGGGTGGAAAAAAATGGCACCCATGTGAATGA

CACACGCACAATGAAAACATGTTACCAGCTCTAGTGTGACCAAACGTGTGAGTAGCACGCACAAAAACGTAGTCTGTTAC

AATTCGTAACGGTGGTACCGTGTTACGAATACTAAAAAAAATTCGTAACGGCTTCCATGGGTTACAACAGGGTTGTAACG

GCTTTTTGTAACAGGAAATTTCGTAACGGCTACTGGGCCGTTACGAATTTTTGTTGCGAATATGGAAAAATGGTGTAGTG

AAGACACACGACCCAAAGTGGTTCTTAGTTCCTCTTCGTGTTGTGGGTTATTATATTTATGGCTGGGACGAGCTTGACAA

TGTTGTTTTGGAGCTTCTCGAGATGCACAACTATATGCCGCCTTATAGACCATGGGAGGTTAGGTGGGTCATTCCTCGTA

AGCGTCAATCGTGAAACCACTCGAGCTATTGAGGATGATAAGGTAAAAAATATTGTCTTAATATTTTAACGTATATATAC

CTTGCCCCTGTTTTATGCCTTAGCATTTGTTGTCATTTTTTAACAGAGGATGAGGGTTGCTGATAGTTTTGGCCCATCTA

CTGCTCATGTGGAAGTTGCAGAACATGCTTTGGATGTTGATATTGGTGCTATTTTTGGTGAAAACGAGATGATGGATCTT

GGTGTGGGATTTTTCGACGATGTGTATCAAGCTTTGTTGCATGAACTTGTGAATCTCGATGGAACTGTTACCATATCTTA

GATGGGGGTTCCCCAGGCGAGTAATCTCACTGATATGGGTGTTCCCTCCGATGGAGGTGCTCAGATTGGAGGCAGTGGTG

TTGGTGTTGTTCAAACTGGTGGTAGTGTTAGTACTGGTGTACCAAGTGCTTTTACTGGTGGTTTTCCTTTTTCTCCGACA

GTTCCTTCGAGTTCGTCTCGCACCATTTTTGGTGGAGGATCCGAGGCGATGGCAATGTTATTCTCGGATCAATATGCTCG

GCTGAGCGAGGACAAGAAAGCTCATATCTTGAATTCCTTGGACAGATCTGGTCTGCTCAGCATCAGCTTGTGCAGGCGAT

ATGTATTAATACTTTACTATATTTTAATAGTATGTACTTGTTAGTGCCACAGTCTAACTGGTGATGTTGTGTGATTTTCT

AGAGCAAAAATATGAGGGTTGGTATGTTATCCGAGACTCTTGTTTCTAGGAGGCAAGCCAGAAATGTGGAGCGCGAGAAC

CAGCAATTGTGTCTCGCTATTGAGATGGTTATGAAAGAGGATTTTTCCTTTCGCCAGGATACGAAGGAGTTGGAAGGTAT

TTATTCTTACCTTTGTCTAAGTTTTTCAATTTTGTAGTAGGTTCTAGAGTTATGAATTAGTCCTTCTATTGGTATCTACA

GTCAAACGTTTGGAGGATAAGATAGATAATAGTGGTCATAGGACTGCTAGATAACTTCAAAGCCTAAAGCTCGATGTGAT

CTCTCGTCAGTGTAAAATTGATTCCTTTGATAATGAGGTTACACAACTTCAGACTGATGTGGATGTTCACCGAGATCGCA

TTGTTGAGATTTGTGCGATGTTAGATGAAGCTAAGCAGGTGGATGTAGCTCGCTCGGATGAGATCAAGCTGCTCCGAGCT

AAGAAGGATTGGCTTATTGAGTGGCAGTCCGATCATTTGCCTCAGCTAGCGCAAGCAACGAATGACAGAGAAGATCTTGC

TCGTCTTAAGAGAGAATTTGTCCAACTCGATAATGAGCGTGATGGGCATATGTTATCATGCCCGGCTGGTGATGCTACTA

TAACGTCCATTGAAGTACATATTAATACTCTAGACAATCAGAAAGACCAAGTGGAACAGACCTTAGATTCTATTTTAGTT

GAGTATGATGGTTGCCTATTAGCATATTTTGATTGTATATATTTCATTCCTTGTAAAATTTCCTTTGGATTAACTGTCTC

CATTTGTTATATTGTAGAGGCTCGTAAGTGCAAAGCAGAGTTTGAGATACAGTTGAAGGATGTGAAGGAAAAAAAGAAAA

ATATTGATGACGAGCTTAATGATTTGGTGACCCGCCATGAGGAGGAATTAGAGGTCGAACGTGCTGACAAGCAAAAGAAG

ATGAGCAATCTTCATGCTAAGTACAAGGTCAAGATGACCAATGCATGTAAGGCTGCTATTTTGCGTGAGCATCAAAGGTT

TTCTGCAATTGATACACCCAATGTTGTGGTTCCGCCTGAACCTGCTGAGTCCGAGTCCGATCATGTTGCTCATGATACTG

CCGAGGGAGAGGATCAGATGGATACCGAGCAGGATGCTCCTGGTGATGGTCAAGCTTATATAAGTTCATAAATTTTCTTG

CTCGTGTAGGCATTACTTTTTAAGAACTCTTTATTCTCTTCTCTTTTTGTGGTTAAGCCTCTTGTTGGTTTCCGAGGTGG

ATTCTAAAAAATATTGCTTTGATTGGTAAGTCCTTATGCTCTTTCCGATGAGATGTTTAAATGGTTTGATTAGTTGGTAT

GTTAACTCCTTTTTTTTTCGGAGGCATTATTTTGCTAATCATTTTTTGGTAAGTCATCATATGTGGGAGGCAACTTATAT

TCTTTACTAAAGTACGTAGTAATATGCACGATAACTTCTGTTGTTTAAGTGTACCATGTATTGTTCTTATTTTTATTGAC

TCTGAGGGTAGCTATTGTGCATACTAGCGGTCCAACCATGGGTGACAAAGCCAAGGAGTGATCAATTCCTTAATTGGATA

GGGCATCTCTTACGGTGTTGGGCTTTAAGTATAGCACACTTACCTTCTAGTCTTAGTTTTGTGCCTTTAAGGACTTTTCT

TTTTGATAGTGTCCAGTTTAGGTTACTTTCAAAAAAAAAAAAAAAAAAAAAAAAAAAAAAAAAGTGGCAAAGGCACAGTG

GGTAGCAAATTTTATTAAAAATCAGAAAAAGAAAGAAAGAGGGGGACTAGTCCTTACCTCCCACAAGCAAGAGAAGAAAT

GAAAAAAAAAAAAAACTAAGAGCAAACAAATTACATATTACGAAAATGGTAAAGCGGAAATACATAATTGTAGATGCATT

TTACTACAAACTACATCCAAAGGATAGAAATGTATGACTCTTCTCAAACGAAAGTAAACACTCAACCATGAACACAAGAT

ATACGTGGTTCGGTATGATTACCTACGTCCACGGGGGTGAAATGATGAATCTTATTACTTCACGGTGTTACAAAGGTGTT

GCTCTTAGTGGGATTTCCTTACAAGATGAACTTACTTCACTTCTTTCTAATGCTAACTCGGGTTTCTCTCACTTTTCTCT

CCCAAAAATTACAACCTACCTCTCTAGAATACAAGTCCCGCTATTTATAAGAGAAAGTTTATACAACGAAGTTAGAATGT

TGACCTTGTTACATCTAATTAGATGTTCTGTGATCGGGCCGACTCACGTCGTCTTGTGCGGTCGATGCATCTTGGATCCT

CGCCTGTATTTCATTCGAGCCAATCTCTTATGGTCCTCTGTTGACGCGATGTCGGCTGTCTCGATAGTCGCCCTTATGAT

CAGCCGATGTGTTCCACACTCTCTTTCCTGTTTGACTGTAGCTCCTCGTTTGTCCCATCCCGCCTTATCTTGTTAGCTTT

ATCAGAAGTCTGGGTCAAGTCTTGTCCGACATTATTACGCCCGATCCACGTGGCTTCCTTAAATATCGCGCTCTTTAATC

TTTGCTCTATACTTCGTGTTTCTTTGTATTTGGGTGCACTGGGTCTAGGATGTTGCCACCTGGATATGTAGATAATTTAC

ACCTACATTTATGCCCCTTCTTCCACCCGTGTGGGAGGCACGAGTGGAAGAACATCGTCTTAACCGCCCATGTCTTCGCT

CATGCCGCTAATCCCGCCATGCAGCGCCCCACTAATTTCCACTTTATGACTGGTAACCGTCAGCGTCGGTTACAAAGTTT

CTTCTTTAAATATCCCGCTCCTTTCTTCCTTCTTCTTCTCTGCCCTTCTGTTCTGCTGCTGTGGTTTTCTTCCAAGTTTC

TTCATATCTTACGCTTCTCAAAAAACCCTAGCTTTCTTTATTTTCTTTCGTCCTTTTCTCCTCATCGATCGTTACAGTAG

CTGTTGTCGCCTCATCTTTCTTTGTTGAAGACCGAGTTCTGCATTATCTTCGTTAATGGCTTCTTCTCCTTCCATGGCTG

AAAATAGGTCTGTTGATGATGTTTTTTCCGAGTTTTATGTTCCCCCCCTTGCTTCTATGTTTTTGCTTGAGAGACCTTCT

AGGTATGATGATCGCGCGATGAGATCTATTTTTGCTGATAGCTCTTCTTCTATTTCGTGCAGGTCCGAGAAAAGGGTGAA

AGATCTTTCCGATGAAGAGTTCATTGAGCAGTTGAAGGAGACGTACGATCTTCAAAATTACGAAGTGACCCTTCTCTCGG

GCCGAAACGGTAGTTTGAAGAAAACCTATATGAAGAAGCACGAGCAATCGGCTAATTGTGTGGTCATCACCTTGGGTCAG

TTGGAGCTAGGCCTTCGTATTCCCTTATACAACCCTGATAGCCCATTTTATTACGAGATTTTGTCCCAGATTCGGCCTTC

TCGTGCTTTAGCTCAATGGACTGGCAATACTTTCCGAATTATGAACGAGTGGAGGGATCGCCGCGATGGTCATTGTTCTA

CTGCCGAGTGGTCGACCTACAAGCCCGAGGACGCTCCTGAGTATACCGTTGCCAATTTCCTTGTCAACTATCGAGCTGGT

ACTTCGACCTTGGGCGATCTTTCTGGGTTTGCTGCAAGTCCTCATCGGGTCAAAACTATGCCCGAGGGTGTCGAAATAAT

GATGCTGGATGCTGATCCTATCGGCCATGGTTCCAACAAGAGAGCCTGACATTCTAATGATCCGACCTATCCCCGAGTTC

CGCTTCTTGTCCGACGTCCTATCCTTCATGGTGACTTCCTGTCGCCTACTTTTCCCACCGAGCCCTATTCTTTCTAGGTG

GTTAATGATGACAAGGCCGATAAGGTACGCGCTTTTCTTGATTTCTATCTTGTTAGCTTCGGGCGTCAATTAGTCATTTG

TTGTTGTTTGAAAATATCAGGTCAGTTCCACAAAAACTATCGGGTCTAGGAAAAGAGGTGCTGGCACAAGGGTTGAGGAA

GACCAGGGGAAGGTAACAAACACAATGATTTCCTTATAAGTACATGTACTTGCCCCTGTTTTTTTCCTTCACATGTTCTG

ACGCGACACTTGTGCAGAAACACCCGAGGCGCGATGATGATGACGTTCACGATGTCGACCCTACTCCGCTTAGGCGTAAT

CTTGGTAGGGGTAAAGAAATTACTAAGACACCGCCAAAAACGGGCCCCCTCATTGATCTTTCCGATGAGGATGAGTTGTT

TGGAGAGACTGTGCCAGGTGACAAAGTCGGTGAACAAGGTGTGGGCGACCAGCAGGAGCCCTCGGAGACGCCCGCCGTAG

AGAAGGTTGTTACTAGTGTTCCGGTAGAGATTCCATGTGTGCCCCGGTTTACTGAGGGATCGTCTTCTAGCGCCGAGACT

TGCTTCACTGTTTCTGGACCGATCATGAATTCCTTGGGTGCGCTTACTTCCTCCGACTTTGGGAAGTATACTGGTGAAGA

AATGTTGAATGATGTACCCTGTTATGGCGACATCGCCTTAACACTATGGTGAGCGACTCGGTTCTGTATAGTAATGCCCC

ACTTCTGATTTCTGTCCTTAGTGTCGCTTATGATGCAAACTCGTTTTTTTGTGTTAGATCATGAACCATGTACGCTTGGA

TGCCTCCAATCGCCTTCGAGATAAACAGTTGATAGAGTCTCTCTAACTTCAACTTCAGAATGAGAAGGGTCGCGCGAACG

ATCTGGAGATCAAACTTGCCGATGCTCAAGGTATATGATTCTTCTTGACTTGTGTTGATTGAGTTTATCGTGCCAGTATC

CTGAGTCGGGTATCCTTATTCTTAGCCGAGGTATCTAGTATAGGTATAGACGCTACTTCGGAGTTTAGGATTCTGAAGAG

AAAATGGCATATTGAGAAAGACCTTGTGGAAAATCTTCGTCAACGAATGGCAAACAAAGACGACAAAATAGACCGTCAAA

CTGAGGAGATTGCAAGTCTTGAGGACGAGTTAGAACGACATCGCCGGTTTGATTATGATCCTTCCGAGATGGACAATTTA

CGGGCTCATTTGGGATCTTCTCAGCGTCTTGCTGCCGAAAAAACTGCTTTAGCCACGAGTTTGGAGCAAAGGGTAAACGC

TCTTCAATTACAAAATCGAGATCTTCACGAGGACAGCCGACGAATATTGAGAGATAAGAATGTGGTGGAGCGGGCTAACA

GCGATTTACAGGGCCGAGTGCGTGAATTTGATAAAGTTATGGGCGATTTAGAGTGGAGTAGATCGCAGCTTGTGGTTCTC

CAAGCGTCGCGTAATCGGTTAATGAACGATTGGGCCGAGCATAAGAAGTATTTCCCTAGTAACGAGTTAAACGAGCAACA

TTATAACGAATTAATAATGAAGTTGTCCAGGACCGAGGATGAGTTGTCCAAGTCGGTAGAGTCGCTTCAGTCGGTGTTAC

CCGCCATGTCAGGTCTCTTAGCCGATGTGTTGAGGCCATTTTTTAGATCCAGATCGCGCTTTATTCTTGCAAAAATGTTA

AGTTGTATCCGATGAATAACATCCTATATTTGTGGTTTTGTAGCCACGCAAAAACGTTCGCAGGAGCTTGATCTGGAGGT

TCGCCGATTGAAACAGGTCGAGGTCGATTTGGGATTCAAGTTAGCTGAGACTGAGGAAAACTCTTCCCAGATTACCCAAC

AACTCGAGGATGAACGGGTCGCCCATGCGGAGGAGTTAGATCAGAAAGTTCAGGCTGGCATTATGGACTGGGTGGAGAAG

AAGCGGCTCGAGGTCGCGCAGAGGAAGGCCGGCGCGGCTCCTCCCAAGGTTTAAGCTTTCTTTTTGTGGTGCTGCGTCAC

TATGTTTCCCCTTCCATCGGGTTGTAAATCGCTCTAAACACTTCTCCCCTTTTTAATTTCCGCTACCTTGCTTGTTCGTT

TTAATGGGGGTGCTGCGTCACCGATGTTGTTTGTTTCATTTCCGCTAAACTTTTCATTTCTTATGTCCGCGCGATACTTT

CCGTACTTTGTTATCAGACGCTTGGTGTTATCGATTTAGCTTTTGTTTCTTATGTTATTCTGGCGATTGTTGATTGATGT

ACCATGCGATCGATGTGGTTTTCGTTTTGTATGCTTCTCCTCTGGGTGTTGGTTGTTACACGACACGATCGATTTGTTTC

TTGTTCCGCATGTTTCCCCGACCGTTTGTTGATTAACACACGATGTTATTTGTTTCGCTTTTGTTCCTGGTGTTATGCCC

TACAATTGTTGCTTTTATATGTTAGAACTCTAACCATTTCTGTCGAACACACCCAACAAGGAGGGTCATCGGGCCCTATG

CCATGTCCTGATCGGGGTATCTCATCACTATCTCTTTGGATTCAGTGTAAGGGTATCAGTCATCCTATTTCTAGGCCCGA

TGGTCCCGAGTGGTTATCGATAAACCAATCTCGGGCAAGTGGACACGCCCACTTGCGATGGTGGTGACCTTTCAGATGTA

AGTAGGCCACCTAGCTGAGTAGACCTCGCGTCTAAACAATCTTTGGAACTGGTTTCCAACCACTAGAACCCTTAGGCTAC

CTCGGCACTTGCACACTGCCACTGCCCCGAGTTCGAATAGCCAGGCGACTGTAAGGCACTTCGGCACTTCCACACTGGCA

TTGCCCCGAGTACGAATGCTCATTCGAGTGTAAGTCACCTCGGCACTTCCACACTGGAATTTCCCCGAGTATGAATGACC

ATTAGTCGCTTGTGCCCTATCCCATACCCCTCACGGTTTGACAGAGATAGAAGACACGTTTGCTTACCGTTTTTTTTTTT

TTTTGCATTTGCCCGTCAAGGATGATAGAGTTTCAGGAATCGGATGTTCCATGGTTTGAGCTCTCTTGTTCCGTCGAGTC

GCAGTATTCGATACGATCCCTCGCCCGCCTTAGATTCGATTATATATGGCCCGCCCCACTTTGGTGCTAGTTTTCCGCCT

TTTTTGTCGTGCTCCCATTCAGCTAGAAATTTCAATACCTACTGCCCGGGATTAAACTATCTTGGTCGAACGCGCTTGTT

GTACTCCCATTGTAACCTCCTTTGATAATTTTCCATTTTTTGTAGCGCCATTTCTCTTGTTTCTTCCAATTCGTCGAGTT

TAGCCAAGATCAGATCGGACGACACGTTTCTCCTCCATGCTTCAGTTCTTGTGGTAGGTATCATTGCTTCTGTTGGTAGA

ATCGCCTCGGTTCCGTAGGTAAGCATAAAAGGGGAGAACCTCGTAGCTTCCCTCCATGTGGTTCTGTATGCCCATAGGAC

GTTATATAATTCCTCGCACTAGTTGCCATATTCCCCGTCCAGTTTCTTCTTCATGTTATCAGCGATGGTCTTATTAGTGA

TCTCGGCCTGCCCATTGCACTGGGGTTAAATCGGGGTAGCTTTGCTCTTCCTGATGTTGTAAGTGTTAAATAGAAGATCA

ATGTTCTTGCCCTGTAGTTGTTTTCCATTATCAGAGACGATGGCAGCCGGGACCCCGAAGTGACAAATGATATGTTCCCA

AATGAACTTAAAAACATCTTTGTCTCGAATATGCCTCAGTGGTGCCGCCTCGACCCACTTGGTAAAGTAATCCGTAGCCA

CGATCAAATACTTCCTTTTCCCCGTTCCTACATGTAACGGCCCGACAATGTCCACTCCCCATTTAGAAAAAGGCCACGGA

CTTATGATCGAGTTTAACGGTACGCCGGGATCATGGATTTTCTTCCCGAAACGCTGACATTCTTCACAAGCTCTTGCCAT

TTGCTTTGCGTCCTCGTTCATATACATCCAAAAGTAGCCCATTGTTTTTTCCTTAGCCGACAAGCTTCTTCCCGAGCTAT

GATTTCCAGCCGCCCCATAGTGTAGCTCATGTAGGATTATATGGCCTTCTTCCCTGCTTAAACATCGTAGGAGTGGCCCT

AGGTAGGAATTCCGGTATAAGATCCCCCTCCTTAACTCGTAGGCGCCCGATTTACTTTTGATCTTGTTCATCTCGGGCCG

ACCCTTAGGTAGCTCACCCGTCTCTAAGTATTGATGTATCTTCTTTCTCCAGTCGCCTTATTTGTATCTCTCGGCTGTGT

TAACGGCCACCTCGATCCGTGCTTCTGGTGTTTCAGGGATAGATGGGTTAAGTAGCCTCATAACGCGTATCCCTTCAATG

CTCGGGTCTACTAACTGTGAGGCGATATATGATAGAGCATCCGAGTGTCTGTTATCCTTTCGGCAGATGTGGCGGAATTT

AATATCTGGGATTTGCTTGATGTAAAATTGGGCGAACTCCCAATATTTTTGCAGCACCGGATCGTGTGTGGCGTATTGGC

CCGTAATTTTCCGGATCACTAGTTGAGAGTCACTAGTCAGTCGAACCTCATGCAGCCCCATTTCCACAATTATCCGTAAG

GCATGGATCACCGCCTCATATTCGGTGACGTTGTTCGTTGCCTTGAATTCTAACCTGAACGAATGCACAATATTCATCCC

CTTTGGCGTGGTGAACACCAACCCGAGACCCGATCCGTCTTTGTTAGATTCCCCATCAACAAATACTTCCCATCGTGTTG

GGCTACTAGCGGCTAGCAGGTCCGCCGGCCTCTCTTGTTCTTCTGTCATTCCCGGATATCTTCTACCTCGTCCTCATCAT

TGAGGGGGAAGTCTGCCAAGAAATCGGCCACTACTTGTCCTTTGTTCGCGGTTCTTGTCTCGTATATCACATCATATTGT

TTGATTATGTCTCCCCACTTGGAGATTCTCCCCGATCGCCCAGCACTATCCAAGACCGCTTCGATAGGCGATTTGGTGAG

CACCCGTATTCGATGAGCTTGAAAATATGTTCGTATTTTAGAGGCAGCAAAGGCTAGCGTGAGCACCATCTGCTCCATTC

TAGTGTAGTTCTTCTCGGCCGGACTTAAAGTTTTGCTGGAAAGTATACCGGCTTTTCCACGCCCATCTCTGTTTTAACCA

CTACCGCACTTATGGCATAGGGTGTCGTGCCTAAGTATAAAGTGAGAATTTCTAACGGCTCAGGCCTTTGAAGGACCGGG

AGGCTAGATAGATGTAGCTTGATACTTTGAAAATCTTCTTCTCTTGCGTCCGTCCATCGGAATTTTTCTCCTTTCTTCAA

GATGTCGTAAAAAGCCTTGCATTTGTCCGATGAACGTGACACAAATCGTCCCAATGAGGCTATGCAACCATTTAGTTTCT

GCACGTCTTTCACCTTCGCGGGAGATGGCATGTCAAGGACGGCTCGTACCTTCTCGGGATCCGCTTCTATTCCCCTGGGA

GTGATGATGTAGCCTAAGAATTTCCCCGATGTGACCCAGAAGTCACACTTGGCAGGATTTAATCTTCATCTTGTGTTTCC

TCATTGTTTGAAAGATCTCTCGAAGGTCTCGGACGTGGTTTTTTGATTTCCGACTCTTAACTAACATGTCGTCCATGTAA

ACTTCTATGGTATCGTGGATCTTGTCTTCGAACATATCGCCTACCAATCTCTGGTACGTAGCCCCAGCATTCCTAAGTCC

GAATGGCATCTTGGTGTAACAATACAATCCTCTTGGCGTGAAGAAGGATGTGTGTTCCTGGTCCTCGGGCGCCAGTGGTA

TTTGGTTGTATCCCATGTATCCGTCCATTAAAGTCATCGCCTCGTATCCCACTATTGATTCTACTAATTGATCTATACTG

GGAAGGGGGAAGCTGTCTTTCGGGCAAGCCTTGTTCAAGTCACTGAAATTAATACATATTCGGACTCCCCCTTTATTTTT

TGGTACCACGACCATGTTAGATATCCACTGTGGGTAGTGGGACTTCCTTATTATTCTGGATTCTTGTAACTTCTTCATCT

CGACCTCGACCGCCGCTTGCAATTCTGGCGCGATACGCCTCATTTTTTGTCTAACCGGTTTGAACTTCGGATCAATTTTT

AAGTGATGACAGCAGACCTTTGGGTCAATACCCTCCAAATCATGCACTTCCCAAGCGAATGCCTCCAGGTTTTCTTTTAA

TAGCGTCGTGAGTCTACTGGTTTCTTCGTCCGATAGTAGTGCCCCGATCATCACTACTTTAGGTTCCTCTATCGTTCCCA

AGTTCACTTCTTTAGTCGGCTAGACTGTCGAGAAGTTTGATTTGGTCTCCCTGACTGGGGATGCTTCCTTTAATTGCTAT

AAAGGGTCTTGTTCTCGCGTTTCGTGTGCCAGGACCGCTTGAGAAATCGCCTCCTCCAATTCTTTTGCTGCCTTCACTTC

CTTTTCTTTGTTCTTTTCTTGCCTTTTCCAAGCTCGCCGTCGCTCATTTATTTATATATCAACTTTCATGCATTGTCGTG

CCGCATGGGAGTCGCCCATAACTTCCTCTATTCCCTTGCAAGTCGGGAACCTTAACCTTTGATGGTAGGCCGATGCTACG

CCCTTTATCCCCGCAACCCATGGGCGCCCGATAATGGCCTCATAAGGTGAAGCGACGTCGACTATACATAAAGTGGTTAG

TGTGTCGAGGGAGCCACCACCATCTGAAACTCTTAAAGTTACTTCGCCCTTCGGGATGGTTACTGTGTCGTTGAAGCCGT

AGATACGGTATGTGGATGGGACGAGCATATCGTCTGAGAGTTTCATTCTTCTGAACGTGTCGTAGAACAGCACCTCGACT

GAACTTCCGCTATCCACAAGAATCCTTCTTACTCCCCACTTTTCAATCAACAGTGTGATGACCAGTGGATCGCTGTGTGC

CTGTCCGTTTGCTGGGACATCCTTAGCCGAGAACGATATGGGCACCGTCATCCAGGGTTCCATGGGAAGAGACTTGGCTA

CGATGAATATCTCTTTACCGTCACGATCCCTTTTATGGATCCTAGTGGTGATTTCTGGGACGAGAGTATAGTTTTTCATA

GCCGAATGTGAAATGGTGTTTACGAATTGGTTCGCCCTATCGATGCGGACTTGATGAACTGGCGCATCGGGTGTTGCTTC

CGTCCGAGGGGGGTGGCGGACATACTTCCTCAAATATCCATCGCGGATGAGGTGTTGAACAATATCTTTCACCTCTCGAC

AGTTATTGGTGGAATGGCCCCTATACTGATGGTAGTGACAGTATTCGGGATGATCTCGGGTCTCTTGGAACTGTACCCCC

CTCGATGCTGGGTAGTTGATGTCGCTCCTCTTCTCAACTTCCTTGAACACCTCTTCCAAAGAAGCGTTTAGCGGGGTATA

CGATCGGGCTTCCTTCTTTGGCTTCTTCCCTTTGGCCATCCATTCCCTCTTCTCACTGTTCCTCGGTGGAGGGTCCGATC

TTTTTTCTGATCGCCTGATATGGTCGTCCGTGACTGTTTTAGTTTCGATCGACTTCAAAGACACTGGCTTCTTGTACCCC

TCTATCTCTTGATTCGGCCTGAATTTCTTCCAAGGCGATGTAGTGTTCTTGCATTCTTCTCATCTCCTTTAAGGTTTTAG

GTTTTTCAGAAAACATGGAAATACAAATGGGGTCTGATCTCCCCAACGCGTTTTCAAACCCAAAGATGGCTTGGTCCACC

GGTACCTTCCCAATTTCAGTACACAGTTTCCTCCATCTGTCGGTCAAGGTTCTCAATGTCTCTCTGAATCTTCGGGCGAG

GGTAAACAGCTTGTTCATTCTTGGCCGAGGTCGACTGTTATGCATATAGGTCTCAAGGAAGGCCTCGACTAGCGCATCGT

ACCCGTTGATCGAACCAGGTGCGAGCGAATAAAACCAATCTCGCGCCTCGCCCTCCAAACTAGCTGGAAAGAATTTACAC

ATAACTACCGGGTTACTTCTCCATTGGATCAGTGACATGGTATACATCTTAAGGTGTTCGGTCGCATCGCCGGTTCCATC

AAATCTAGAGGGGAATGAAGGTAAAGTACACTTAGGTGGGAACGGCATTGTTCCAATTCTTGAGAGAAAGGCGTTCTCTC

GGCTTCACTCATAATCTCTGCCAGCTTATCCTCTTCGCTTCCGCCCGAAAGTTCCTTGATTTTTGCTTCCAACTCCTTAA

GCATGGCTTTAGTTGTCTGATCTTGTCGCCCGTCCCTTCCTCGCCTTTGTTCATCGGTGGACGAGTCCTCGGGTGGTCCA

TATCTCTGCGTTGGGGGAACCGGAGGTAATGGCGGCTGACCTGACTGATTGGGCTGATTAATGTTACTCTGCACCATCAT

GGGTGGCAAGTGATCGGATCGGCCACGATAACCTGATGTTCCTCGGCTTGACGACCCTCTTGATCTGGTCCGTTGGTTCC

TCAGTTGGTAGTTCACATACTCCAAGGCTAAATTCCGCTGCTGGAGTTCCCTCAGTGCCGCTTCTTGTCTCCTCTGACTC

TCCAAAATGGCCAACAACGTCGGGTCCGTACTCTCATGACTGGAATGTCCATTGGGTTGGTGTGGCCGAGGTTGTAATGG

TGCTTATGTGGGTGGTGGGGGCGGTATTGGATCGGTTATATGATTGACAGCTTGAACGGTTGTCGGGCCGATGTTACTTG

CTATTGCTCGCCCCAACTGAATCTGCCTAAGACGCTCTTGTTCCCTTCCCAAAGCTTCCTGGTGTTCAATTTGTCGCCTC

GTCGATTGCCTCTGATCTCGCGCTGCAAGTGCTTCCTCATCGTCATCGGTGTCTGAAGCTGTGAGTCCATCGACTTGATC

GACTCTTATCTGAGTTAGTATAATCTGGTGCAGAGGTAAGGTGTTATCGTCCCGCCCCAAATCTACTTCCTCGTCTACAA

CAGGCATCCCTCGCGCAGCTTCATGAGATTCCTCGGGCGGTGGTTGTCGACTGTTCCCTTGCCTCGGCCCCATTATCCCT

GCGTGCATGCTGCTGCTAGTGACCGTGCGATTCCTCAACATCCTTATCGTCCCTGAAGTGCTATCCATTAGTTACGTCGC

GTGATAAGCCGAGTATCGCCCGTCCCTACGTCAGCAAATACGAAAAAGAACCTCTTACTTCGACCTGATTTTTCTAACAA

AATGCACAGACAAAAAAGGCGAAACGTTCGACTCTACGTCTATTGTCACTATGCAGGGTCAACTTTGGTTATCAAGTGGG

TTCCGTTTATCTACCCTATTTTGGTCTTATGCTGAATCTGTGGCTGTACTCACTTCCCCGTCACTATCACTCCGTACGAC

GTTGCTCTCTGTACTCGACCTATGCATTTAAGTTGGTTTGTCCTTTGTTCTACATGCTCCAAATCATTGATATTATGCAC

GTGATGCTTAGAAAGTACGGCAAGTAACCCCTAGAATTAATTTGAACTTGTACCAACTCCACTTTCTCGAGATTTAATTA

TGACCTTTCCAACGACTTACGTTATACCGTCGGCTAAAAATTATACAAGGTCGACAGTTCATTACACGATGAAACATCTA

AAACGCTAACTTTTGGTAACTCCGAGCACCCAACTTCTTACTGCAGCTAAAAACAACTCACTTAACGCTAACAAACACAC

AATTACCAAGAACAAACAGTCGTCCCTTCAACAGATTCTTCACGTAGCTTCTAACCCGAACTGTTAGTCGCTGGTGAGCT

CGTGAAGTTTCTCTGTCGATCAAACTGTTGAAGATGCTAAGAAGACGCTGCTACTAATATGACTTTAAATGCTTAGTAAA

TGACTCAAAACAACCATTACTTCGGCCAACATAGTTCATCGTAGCTTCTACTCCAACACATTAATGTGTATTGAGGAGAT

GTTGATCGTCTCTGAAGACTTTAATGGTAAGGTATCCGGATATCTTAGTCGTGTCCCCTTAGTCGGCGCCAGAATGTAGA

TGCATTTTACTACAAACTACATCCAAAGGATAGAAATGTATGACTCTTCTCAAACGAAAGTAAACACTCAACCATGAACA

CAAGATATACGTGGTTCGGTATGATTACCTACGTCCACGGGGGTGAAATGATGAATCTTATTACTTCACGGTATTACAAA

GGTGTTGCTCTTAGTGGGATTTCCTTACAAGATGAACTTACTTCACTTCTTTCTAATGCTAACTCGGGTTTCTCTCTCTT

TTCTCTCCCAAAAATTACAACCTACCTCTCTAGAATACAAGTCCCGCTATTTATAAGAGAAAGTTTATACAACGAAGTTA

GAATGTTGACCCTGTTACATCTAATTAGCTGTTATGTGATCGGGCCGACTCACGTCGTCTTGTGCGGTCGATGCATCTTG

GCTCCTCGCTTGTATTTCGTTCGAGCCAATCTCTTATGGTCCTCTGTTGACGCGATGTCGGCTGTCTCGATAGTCGCCCT

TATGATCAGCCGATGTGTTCCACACTCTCTTGCCTGTTTGACTGTAGCTCCTCGTTTGTCCCGTCCCGCCTTATCTTTTC

AGCTTTATCAGAAGTCTGGATCAAGTCTTGTCCGACATTATTACGCCCGATCCACGTGGCTTCCTTAAATCTCGCGCTCT

TTAATCTTTGCTCTATACTTCGTGTTTCTTTGTATTTGGGTGCACTGGGTCTAGGATGTTGCCACCTGGATCTGTAGATA

ATTTACACCTACATTATCTTAATATTGTCTTAATATTTTAACGTATATATACCTTGCCCCTGTTTTATGCCTTAGCATTT

GTTGCCATTTTTTAACAGAGGATGAGGGTTGCTGATAGTTTTGGCCCATCTACTGCTCATGTGGAAGTTGCATAACATGC

TTTGGATGTTGATATTGGTGCTATTTTTGGTGAAAACGAGATGATGGATCTTGGTGTGGGATTTTTCGACGATGTGGATC

AAGCTTTGTTGCATGAACTTGTGAATCTCGATGGAACTGTTACCATATCTTAGATGGGGGTTCCCCAGGCGAGTAATCTC

ACTGATATGGGTGTTCCCTCCGATGGAGGTGCTCAGATTGGAGGCAGTGGTGTTGGTGTTGTTCAAACTGGTGGTAGTGT

TAGTACTGGCGTACCAAGTGCTTTTACTGGTGGTTTTCCTTTTTCTCCGACAGTTCCTTCGAGTTCGTCTCGCACCATTT

TTGGTGGAGGATCCGAGGCGATGGCAATGTTATTCTCGGATCAATATGCTCGGCTGAGCGAGGACAAGAAAGCTCATATC

TTGAATTCCTTGGACAGGTCTGGTCTTCTCAGCATCATCTTGTGCAGGCGATATGTATTAATACTTTACTATATTTTAAT

AGTATGTACTTGTTAGTGCCACAGTCTAACTGGTGATGTTGTGTGATTTTTTAGAGCAAAAATCTGAGGGTTGGTATGTT

ATCCGAGACTCGTGTTTCTAGGAGGCGAGCCAGAAATGTGGAGCGCGAGAACCAGCAATTGTGTCTCGCTGTTGAGATGG

TTATGAAAGAGGATTTTTCCTTTCGCCAGGATACGATGGAGTTGGAAGGTATTTATTCTTACCTTTGTCTAAGTTTTTCA

ATTTTGTAGTAAGTTCTAGAGTTATGAATTAGTCCTTCTATTGGTATCTACAGTCAAACGTTTGGAGGATAAGATAGATA

ATAGTGGTCATAGGACTGCTAGAGAACTTCAAAGCCTAAAGCTCGATGTGATCTCTCGTCAGTGTAAAATTGATTCCTTT

GATAATGAGGTTACACAACTTCAGACTGATGTGGATGTTCACCGAGATCGCATTGTTGAGCTTTGTGCGATGTTAGATGA

AGCTAAGCAGGTGGATGTAGCTCGCTCGGATGAGATCAAGCTGCTCCGAGATAAGAAGGATTGGCTTATTGAGTGGCAGT

CCGATCATTTTCCACAGCTAGCGCAAGCAACGAATGACAGAGAAGATCTTGCTCGTCTTAAGAGAGAATTTGTCCAACTC

GATAATGAGCGCGATGGGCATATGTTATCATGCCCGGCTGGTGATGCTACTATAACATCCATTGAAGTACATATTAATAC

TCTAGATAATCAGAAAGACCAAGTGGAACATACCTTAGATTCTATTTTAGTTGAGTATGATGGTTGCCTATTAGCATATT

TTGATTGTATATATTTCATTCCTTGTAAAATTTGCTTTGGATTAACTGTCTCCATTTGTTATCTTGTAGAGGCTCGTAAG

TGCAAAGCAGAGTTTGAGATACAGTTGAAGGATGTGAAGGAAAAAAAGAAAAATATTGATGAAGAGCTTAATGATTTGGT

GACCCGCCATGAGGAGGAATTAGAGGTCGAACGTGCTGACAAGCAAAAGAAGATGATCAATCTTCATGCTGAGTACAAGG

TCAAGATGACCAATGCATGTAAGGCTGCTATTTTGCGTGAGCATCAAAGGTTTGCTGCAATTGATACACCCAATGTTGTG

GTTCCGCCTGAACCTGCTGAGTCCGAGTCCGATCATGTTGCTCATGATACTGCCGAGGGAGAGGATCAGATGGATACCGA

GCAGGATGCTCCTGGTGATGGTCAAGCTTATATAAGTTCATAAATTTTCTTGCTCGTGTAGGCATTACTTTTTAAGAACT

CTTTATTCTCTTCTCTTTTTGTGGTTAAGCCTCTTGTTGGTTTCCGAGGTGGATTCTAAAAAATATTGCTTTGATTGGTA

AGTCCTTATGCTCTTTCCGATGAGATGTTTAAATGGTTTGATTAGTTGGTATGTTAACTCCTTTTTTTTTCGGAGGAATT

ATTTTGCTAATCATTTTTTGGTAAGTCATCATATGTGGGAGGCAACTTATATTCTTTACTAAAGTACGTAGTAATATGCA

CGATAACTTCTGTTGTTTAAGTGTACCATGTATTGTTCTTATTTTTATTGACTCTGAGGGTATCTATTGTGCATACTAGC

GGTCCAACCATGGGTGACAAAGCCAAGGAGTGATCAATTCCTTAATTGGATTGGGCATCTCTTACGGTGTTGGGCTTTAA

GTATAGCACACTTACCTTCTAGTCTTAGTTTTGTGCCTTTTAAGGACTTTTCTTTTTGATAGTGTGCAGTTTAGGTTACT

TTAAAAAAAAAAAAAAAAAAAAAAAAAAAAAAAAAAAAACACTATTGGCAAAGGCACAGTGGGTAGCAAATTTTATTAAA

AATCAGAAAAAGAAAGAAAGAGGGGGACTAGTCCTTACCTCCCACAAGCAAGAGAAGAAATGAAAAAAAAAATAACTAAG

AGCAAACAAATTACATATTACGAAAACGGTAAAGCGGAAATACATAATCATCAATTAGAAGATTCTGCCTTGTGAAATAT

GGAGGCTCAATCCACCAGCGAGTTTTAACCAGCGCACCATGCTTGGCTAATAGGTCCGGCACCGAGTTTCCTTCACGGTA

TATATGAGTACACACAAACTGAATAGAAGAAACAAATTTTAAGCAATTTTTCCAGTGAGTAGAAAGAGTCCAAGGGACCA

AATTTTAATAGCTTGTAAGATAAAAGTAGAGTCATATTCAAGCCAGGGACATTGTATTCCCTTTGTACAGGCTATACGAA

GAGCATGAACAGCTCCCCAAATCTCGGCACAAGAAGAATTGTTGAAACCCAAGTTCTGAGCCAAGCAACCTTTAGCCAAA

CCATTGGAGTCTCGAAAAGTCACCCCACAACCATAAGGACCTGGATTACCCATAGAACATTCATCGATGTTAGCTTTAAT

CTAATCAGGAAGAGGAGGCTTCCATTTAACCATGATGGCTTTTGGCATCTTAGGAGGCAGTCTTTTAATGCCCAGGAAAT

CAAGAATGATCTTGTCTGCTGAAATGTAAAAGGAGAACCCCCCATTGTGTCTACTCAACTCAACAATTAAGGTTTTCAGG

TGAGTAATAGCTTGATGGAGGAAGTAAGGCAATTCTCAAACCTTAGTCTATTACGACAATACCAAATGAGCCAAAAACAG

ATATCCAGAAAAGATTTTTTGTTTGAGAACAACCAATGAAGCTTCTAATAGAAGTAAACAGCGAGCTTCCATTTGCACAG

TCCAACTGATGCCTAAACACCTGCGTATTCCAGCTCCATTGAGAATTAGAAAAAACACAGTGAAATAAAATATGAGATTG

AGTTTCATTACAGCAAGGATAGGGGCATAAGTTTCAGCAGTTCGTTGCATCCACACCTTGCACTTTCACGTTATCATCAG

TGGCGATTCAATTATGCAGAATCTTCCAGGCAATACAAGCACGTCTAAGATGGACTTTTACAGCCCATAGATCTTTGAAG

TCTTTATTTTCCGAATCTCTAGGCCGATGATAGTTATAGGAGTCTTTCATAGTGATGCATCCCGACTTACTAACAGTCCA

AACCTGTTTATCATTTTTGTGAGAAGCAAAAGGGGTTTCCCTTATAAACTGTTAAACCTGACCTTATAGTTCAGGCATGT

ACGAGGGTATCTGCCATTTCCCTTCAATGATAAAGTCTTCCACAGTTTTTGAAAATTCGAGTTATCAAGATCCAAGTAAT

CTGAAATGGAAGAACCCATATTGACACACTAAGTATCATTCCAGAAATCGATATTGCCACCATGACCAACTATCCAGATC

AAGTTCTCTTGAATGATGCTCCACTGTTCTTTGAACACATGCCACACAGAAGATTTTTTGTAAGTAGCGCCTCTGAAACG

AGATTTAATAAAATCACCAAAATTTGAGATTTCAGCCATTATCTTCCATGTCATTTTCATAAGGGCAGCGTCATTCAGGC

TTCTCAAAGATTTTATGCCAAGACCACCTTCGTCTACCGGCAGACATACATTGTCCCAAGACACAAAGTTGTGATGAATC

ATGTTGATATCTCCAGACCACACAAAATTACGAATTCACTTCTCAGGAGTTGTGATGCATGAAATAAGCCATTTGTAAAC

ATGAAAGCTATATAGGGAAGAGCTAGCTATTACAGTTTTAACAAGTTCAACACGCCCCATCATGGATAAATATTTACCCT

TCCAAGCGGCAAATTTGAGCTTGATTATGTCCATAATCTCATTGAAATGAGAATTCCTTGGTTTACGTTAGAAAATTGGA

ACTCCAAGATAACGAAAATGGAGAGTACCCATCGGGATATTATAGGAGTCAGCAATAATAGAGACTCTATAATAAATGTG

CTTACTTGAATAGATCGTGCATTTAGCCGCACTAATAAACTGCCCAGAGTTCTGACCACATTCTTCAAAAATATTTATAA

GAGATTGCACGTCTCGTTTAGTCCCTCTACAAAAAACCATGATGCCATCTGCATAAAGGACATGAGAAGGAGCATTCGTA

CATCTTGGAGAGGAGATGTGATGCACTAGACCTTGATTAGAAGAATTAGTAATAGATTTTAGGTTACACTTGGACCCTCG

GTTCATGTGAGGGTAAATTACACAATGGACATATGTCAACATCTCAAGGGATGATATCATGATAAGATGATAAGGTGAAG

ACATCGAATCATAATCTATAATGCTGATATCGTCTAAATCGAGACACCTACTACAACTGTTACATAAATGACGCGTGTGA

GGAGTGGTTGAATCGACTCAAACGGTCAAGTATAAAAAGAAGAACTGCATTTAATGAAGGATAAAAACAGACGTGGACGA

ATCTACATCAGAGCATAAGGATCAAGCGATCTATACTATGACCCAGAGGTGATATAACACGAGGTTCTCCAAGGAGGAGT

TATACGATGACGAACGAAGAGAGACGACTCGGGAGGAGAACCAAGCAAAGCACTATGAGGGATAGTATAAAACTCGTCCA

AAGTAATGAGGAATATGTACGATAGTTATTGCTTCAAAAATAATTTCTTCAAAATAAATAAATAAAATGTGCGATAGTTC

CACCAGCTCGGCGAGACCATGACTTCACGAATCTAACTTGCCTATAAATACCGGTCCGTGTAGAAGGAGATGGATAACTT

ATGTATTATTAGATGGTCTTTCTACAGAGAATTCCTAAGAGAGATCTAGATAAAGAGAGGGTGGGAAATCTAAGTGATAT

TTGTAGCTCTTTCAAGGGTAAGACCCTAACTAATAAAGAAAATATCTTATTTCCCATGGGTGTAGGTCTTCATGGCCGAA

CCACGTTATATGCTTGTGTTAATCTTTACTTTCATGTTCTTTACATTATGTTTGTCTTTGAATCTTGAATGTTAAAGTAA

TTTATAGCCTTTAGATTCACTTGGATGTAGCCATCACAAAAGATGCAACTACATTTTGGTGTTAGAAACAGGGAGTTATG

AAGATCTAATCTACATGTGAGGGTAAAATACCCGGTGGCCACGTGTCAACATCTCAAGGGATGATCCCATGATGAGATGA

TAAGTTAGGAGCACCGAATCATACTCTATAAATCTAATCTCATATGAATCGAGACGCCTTCCACGAACGTCACATGAACG

ACGCATGTGAAGGGTAGTCTGATCGATTCAGTCAGTCAAGTCTAAAAAGTATCACCGCATTTAATGAAGGATCAGAAGAG

GCGAAGGCGAGTCTATGTCACAGCAGAAGTCTCGGAAGATCTATATAATAACTCATAGGTAAGGCAGCACGAGGTTCTCC

AAGAAGACGATGACGAGCATGGCGAGACAACCGGGAGGGGAACCGAGCATACCACCACAAGGAATAGTAAAGAATATGTC

TTAAGCTATGAATACGATACATGATAGCTCCACAAGCTCGGTGAGGCCAGATCTTTACGAGTTTATTTTGTTTATAAATA

CCATTCCATGTATCAGGAGATAGACAAGTTAGCTAATGTATTATTAGATGGTCTTTCTACAGAGAATTCCTGAGAGAGAT

CTAGATAGAGATAATAAGGTTTCTAGGTGATATATGTAACTCTTTCAAGGGTAAGACCTTATGATCAATAAATAAAACAT

CTTCTTTCCCGTGGACGTAGGTCTTCAAGGACGAACCACGTTATATGCTTGTGTTAATCTTTAATTTCATGTTCTTTATA

TCTTGTTTATCTTTGAATCTTGAATGTTAAATAGATTATAGCCTTTAGATTCACATGGATGTAGTCAACAGAAAAGATGC

AACTAAATTTTGGCCCTAGAAATAGGGAGGTATGAAGATATAATCTACCTCTCTAATAATAGCTCTATATGGTTTTCATA

ACATCTCTGATAGATAATTCCTTTCTCCGAAAGTGTTTGTCTATGGAATGAGTCGAGACAATACAACTAATCGGTTCACA

CTTCGTGTGATCGTCTATGGATACCAGATCGAGACAGTACGACAATAAGATAACTTGTGTGATTGATTATGGATACAAGA

TCGAGACAATACAACAATGAAGTATGTTTACTTGATAATATGTTCGGACTTAACCAAACACAAAAGGATTTCTATCAAGT

AAATAGGAATTTGTGTATTTTACTTCTAATTATAATAAAACAATTATAATTGCGGAAATAGAAAAGTAAAATACACATCA

AGATTTTCTTAACGAGGAAACCGCAAATGCAGAAAAACCCCGAGACCTTGTCCAGAATTGAATACTCTCAGAATTAAGCC

GCTATACAAAATCTAAACCAACTTCGTATAGTTGAGACCAAGCAACTAAACCTATAGTTCAACCTGTTCCCTCAGTATCC

CTGCGCCTCCAACTTGTAATAAGTCACTCAATTAGAACAATTCCTTTGGTTCATATTTCGAAAGTAAAATAACAACAAAT

TTGTTTAGTAAAAAATCTTTTCACACAACATGATATGAGTTTGACAAAAGGCTCTTCCGTTTATCCCAATAAACTCCTTT

CTCAGGTTCTTAGATCTATCTATTAAACAACTACCAAAGTAATTATTTAGACTATGCAATCAATACTATTAATCACAAAG

AATTGTGTTGAGGCCGATCTACTCAACTAATCAATCAAGGTTATCACAAAGATAAACCGAGTATAGTTGGATCCCCAGCC

GATCAATTTTTGTGCAAACCAAAGATTATGAACTCAAATAAGAAATCTTCTTTGTCTTCAAATCTTCTTAGATCTTCAAT

AAACACCTGCACACAAACAACTTGAATCTCTTATGACCAATCACACATAGAATAGAGTCTATTAACAATGGATTATCACA

AGAAGTCTTTAGATCTACAAACAGTTCTAAAGATCCTCGTCTACACTTCGATCTAGTTTGAGTGAATCTTATATCAGAAG

AGAGGATTATCAAGCATAAACAAACCAGGTGCAATCAAAGTTCAACAACTTTTAGTCAATCAAATCAATCGAAAACTAAT

AATAAACTGCAATTATCTAGTTTCCCAACAACGGTACTCGTAGAGCTTCCCAATCCCAAAGAAGTCTTCAAAATGAGCGG

TCGTAAGAGATTTCGCCTAATTAAGGTACTTTCCTCTCCGAATAGACAGCTCCACCAGTAACAACACAACTAGGTAGTTT

TGATGGATTTGAGTATTAGTTTGCTTGAAATGCAAACTTCAATATTTATAGACCAAGGAAGTTTGGACACCAAGGAATTT

CCAAAACCGAATATTCTCAAAGTTATGCAATAAAGTCCAAATCGGTTTTCATAATTCCTGGAAATGCTCTGTCCAAATAA

TGACCGAAATTTCAGTAGAAAATCTCCAATTAGTAAATGCATATTACCAATTTTTATTTTCTAAAGATATGCAATTGATG

GAAATTAAAAGCTTATAAAAACCAAAAACCTTAATTAAAAGATTCTCAATTTATTTCGATCAGGGATTCTCCTTTAGTTA

TTAAGGAATATCTTTTGACATTAAAAGATAAGAATTACTGCACATGTTCAAAGTATGTTGACATCTTTACTTTGTAAATC

CTTTTTCATATTTACAATCTTGGAACCGATTTTCCACACTTCCAAACAAGTTTAGAATTGGTTCATATGACTTCCAAGAA

CTATGTGATTGATCAACCTATCAAATCACCAATCATGGGTTCAACAGTTCTACCAAACAAGTTTTGGTTCTACCTCCATG

TTGGTACTAGGATCAATCACGTTAGTTATGAAAATTTGATTGACTAGGTACTAGGATCGGTTACCACATATTTATGGTAT

TTAACTTTTATTCGGTTGCACAAGTCATAGGATCAGTTACCAATTACTAAGACTTGTTGCACCTCTTAATAGATTCCACA

TACTTGTGAACGGTTGCACCTCTTACTAGGATCGGTTCCCCAATGTCTAGAGTTGGTCATACCAATTACAATATATCGGT

CATACCATCTCAGGTGATTACTTAAGATAGGTTTCACTAATAAAAGTCATATATACCAACACAAAAGTCAGGCATTGTGA

ATAGTTTTACCAAGATACATAAACAAGTTATGATCGGTTATACTTAACACACATATTGGTAATCCAAAGATTTGCAATGA

ATAACAATACCAATAAGCCTAGTGATTTCCCTTTCGATTCATAAAACAAGTTTATGAATTGTACTTCCTTTAAACGAATG

TAAAATATTGTTTCCTAGGACGAAATCTTCACCCATACCCATACATAATCACAATAGCATTCTTCGTAGTTATGTTTTCA

ATATGCACGAATTGAAAGATACGTTAGGAATGAAACAGTACAAGTCAAATATTACTAACCTCAAGAGGAAGGATGATGTC

GTCGTTGTAGCTCTTTACTTCTTCACATTCTTCAAGTCTTCACGTAATACTTGTAAGTCTCATATCCTAGTAACTTTCTA

GCTAACCTATACGAATTTGACTTTACTAAATAATCAAGAGACTCTTTAAATGAGTTTTGATTCACTAAAATATGACAACC

AAACTTGACATAACAACGCTTAGTGGGTTCAACCGAGATATATATGCTCTAACAATATTACATGTCATCATCATCGATGT

TTATATTCTTGATAGTTATCTCACTGTCAATCCATACATCATGCTATATATGGAATTGGACTTATTTCCAAAATTGTTTT

CTTATTCCAAATATATATACTTTTGTATAGCAATCAATGTTTCTTCTCCGAAATGTTTTTTGCTTCGTGTGGAAGTATGT

CGTCATTTGTTTGTAACTTTGATGAAAAATATTTTCCTTCCCAAATGATCATTATCTTTATGATAATATTTCCATTTTCA

ATTTAATTCCGAAAATATATCCATTTTCGAAATTCTTGCCAAACTCTTGTAATCAATTCATTTATTTTGGAATTTTTTCC

CTATTTTCCATCTTTTTCCTACATATCCATCTTTTGTTTCTGAATATTATTCTGAAAATATAATATCTTTTCAGATTTGG

TAACATCATTTTTGTTACAATGTTCTCTCTTCTTTTTATGTTTTTGGAAAACATAAGTTTCGTACTGTGAAACTCATCGA

TAATATCTCTTGGGCTTCTTCACAAGCCAGCACGTTGAACTATCAAAGATTGGTATAGTTTTTATTTATTTATTTATTTT

TCTTTTATATATTGGAAATTCGCGGTAGAAATTACTTTTCTATCCATTCATGGTGTACCACCATCGATCAGCAATGGTTA

GCGTCTACAGCCACTGCAATAATCATTATTGGTCAATGAAAGATTGTGTTCTCATTGAAAAATCAGTCTTTATCTAACTG

ATTAAGGAATAATACTCTTCGGAGTTTTTCATATATCCACACAGGTTAGATTAAACCATGTGTATTCCTCTCAGTATCTC

ATCACATTTGTATGCCTCTCAATATCTTATGACCTCATTTGTTAACCTTAATGTTTTGTGATTTTGTGTTTCCTTATATT

CATACCTTATTAAGGTTTAACAGAAACATTGGTATGAAAGACTCATTCGCGTAACTGTATCAACCTTTGTCATCAACAAG

GTTTATTCATGTATGATTTTGTATCGTTTTCACGCAGTTATATCCATCATCATACATATTTTGCTGATAGCAAATTCTAA

ATTTTCCTGCCATTTTTGTAGTCTATTTTACATCATATATTTTTATATCTCCGAAAATGATGATTTTTTCTCCATATCAT

TTCCAAGTAGTAGTTTTCATAAATGGAAATTTTCTTAGACAGCCATCTTCTTTTCAAAGTTGTTGGTTTGTCTATCATGT

CTTGTCTTGGTTAGTAATCAACAAACCAAGTCTATATTACTCTTCTAAACCTTCTCCAAAGCCTAATATCCAATCAAGTC

CATTTGCAACCACAAGATCCCATATATCTCAACAGTTATGTGAAGATCATTTTGCTGATTGTTGTTATATTTCACTATTG

TTCATGATCGAAGGTTAAGCGACCTTGGTGCTGAAGTCCGTGTTACAAGATTTTACTTGGAGTTAAAGCAACAAGATTCT

GCACTTTTACTTAAGATTTTTCAAGTTATACACTTTTACTTGGGGGGGAGAGAATTTTGTGTGCACGTACAATTGTGAAG

ATAACGTCAGAGATATTTAGAAGACAAAGTCATTCTAATCTAACTAGTAAATCTTTCTGTTCTAGTTTGATTACCTGGTC

AAGATATGTTAAGTTTGTTACTATCTTTTAGGGTTTAGGGTTTACTAATTATCATATCATTCCTATAAATAGGAATACTA

ATGTAAATGTAAATTATCTCGGAAGTTTTGATGATCAATCTATTTCACCCTACGTGATTTCTTGGTAAACGTAGGTCAAG

TGACCAAACCACTTTAATATTCGTCTTCTTTATTTCTGTTTGATAATCGATGATCCGTATCGATACCCATAATTTATTAA

TGACTTCTCCCAAACTGGAGTTTCACACAACTTCAAGATGCATATAGCACGCTCTTTATCTTTACTCACCGCTAACGATG

GATAACCAAAATTCTTTACCAAAATAATTAGGTAAGATCAACCAATAACAAAACAAGGAAGATGATCACTGCTAGGATTT

GAACAAACCCTAAAAACCCTTTGAAGAGATGTAGGATGTGGTGGATAATTCATCACCAACCAGATCGGTTCTTCTCTAGA

ACACCCTTCTTTTTCAAGCTCACATAATATTCACCCTTCTCCTCGAATGGTGGCTTCCTTCTTTTCTCTTCATCTTGTTT

CCTCACGGCAGCAAAATCCTCTCAGTTTTGATCTCCCCATCCCCGTTTCCACGAACAAGAGATTTATACCAAAACGGTTT

CCTTATCTAACTTAACCCCAACTTCTTTATTTAGCTCCCATTTATTTGGTAAGGGATGCTTGACTGCTAGGCCATAAGGC

GTGAAAACCCAACAATGCCTTGTCCAATATTTTCCTAAATTTCTCGATAGAAAATCCGCCCTTCTAAACAACCCAAAAAC

AATATTCTCTTTTCCTCTCACATTTTTTTCCGGTGGGTTCCAGTGGGGCCACAGAGGACAGCAGTGGGTCCCGCCATTGT

TTTTGGTATGTGTCTCGTTGTTTTACCCTTGTTTGCTTTAGGCGGTACGCCAATCATTTTTCATAGAAAGTCTTCAATTA

GCAAATGCATCAAATCAATTCCAGGATTTCACTTCATCCCCTAATATTTGAAACTCTTTTCTGATGGGAATTTATGGATG

AATATGGAAATGTCAAAGCATCGAGCAGTACAAGTTTTTTAATTTTCCTTTATTTCGTTCTTTGTAACCAACTACGGAAA

GGGAAATGGAGAAAATCATGATTACGGAACACATGTAGATAATGCAATATCAAATCAATTCAAGGATTTCACTTAGGTGA

AGCTCGAACTTAGCCATATCATCCTCAAGAAAGCTCGCCCAAACTTCTATTCCTTGACCACATTTCGTATCATTCATGAA

AAGACAATTCTTATTCGGCTTGATATTTGGATCAATTGTAACCCAAATTGGCTTTCCCCAACCAAAATCAATCTCGTATA

ATCCCATCCTGCACCAGCTACTTATCCAAAAGTTTATCACCTTATCTTCAACATCATTGATTTTCTTTATTGAAGCATCA

TAACCGGAAACAAAATTTTGAATGACTTTCTCCAAGCTACCTTTATCACCCTTGACCTTATCTATTTCACTCCTTAGCTG

AGCTATAAAATCATGTAACTCATGAAGTACCACTTGCGATTCACTACTCGTACTATTGATTGTCTTGTTGACCGCATTCG

TCGTTGTCGTTGTTGTTGCTGGTAAGAACGCCGAAACAGTTACAGATAGATTTCCGAATGACGCATCTTGTAATGGTGGG

TCCATTTTCTTTCTAAGGTTCACGGCATGATCTACGACTGGTAAAAAACCGGCTGGAGTGGATTTCATCACGGACTTCCA

TATTAGAGAAGTAACCACCTCAACCCTTGTTGGCCTGCGGCTTTTGTAGTTATCATGCATCAATACTTGCAGCTTTTCGC

TAACAGATGTTATCTTTGCAAAATCGAACACAAATCTCTTGCTGACAGTTTTATCATCTTCTGTATCCTCCGGGATACTG

GAAAATGGTATCTCTGCCATCCCTGATGGAGATGCCAATCGTTCACTAGGTGGGAATAGAGATGCCGAGTCGAAAGAAGG

AATCAATTTCTGATCTGTGATAACGGCAGCAGTTGCACCCCCGCTACGAGATGTTTTGGTGGTGCTTGCCCAACTACGAA

TGAACGTACCCATTGTAGCTACATCGGAAATCTTATGTGATATACTCGAACAAATGGCTGTTCCACCACAGTCAAACATG

TTCACTTGAACGATCACCAGTGCTTCCTTAGGGACATTCGCGGAAACAACTTCAGATGGAAGAAGCTGGCTTAGAGGCAC

ATCCGGTTGCGACATGAAGTCACACATTTTACCTCTAATTTTTACTTTGTAAAAGTCAATCCCTTGGTCATGACAGTCGA

CCAGGATATTGTCTATCATTCTACCAGCCATTGGATAAAAGTGAACTAGTGTTTTGGAAAGAGAACTCTTAAGCAAGTCA

AGATCATCATGATGGCTACTGCTACCGGTACTATTAGCGACGGTGTCCGGGTAGAAAAGAATGATTGGAACATAATAATA

TAAAGGAAAACATTGATCGAGAAGTGACAAATTGAAGTTTTTAAGTTGAGATGGGGTTGGAGTTGTGGGTTTAATGGTTT

CCTTAGAGATCACTTCAACAGCAGCACTATACATTGTTGCCTGTTTAACAGGATTGATAATGGTAGTTGAATAAGATAGT

TAGTTCGCAAGTGCTGGAGAAGCTCGTGGTCTCCCTCTCTTATATAATGACCAAGTCTTTTGGTGGGAAACTTTTGTTTT

GTAGTTGAATAGCAGTTGAAGACCAAAGGTTATATATATATATATATATATATATAATAGTATTATCTTTTATATTGTAA

AGGATCGATGTAATATGCTGATACTTTCTATAAAGGTATCTGATTATATATATATATATATATAATAGTATTATCTTTTA

TACTGTAAGGGATCGATGTAATATGCTGATGCTTTCTATAAAGGTATCTGATTATATATATAATAGTATTATTTTTTATA

CTGTAAAGGATCGATGTAATATGCTGATGCTTTTTATAAAGGTATCTGATTGTGGGGGGTGGATCAGTGTCCCATATCAG

CCCACGAGAGTAATGCAAGAACCTCGTGGGGATAATTAAAGGTCACCCACGCTATATGTGGTGGCTTTTATAGAAGGCGT

TTGTACATGTGGCTTTCATAGAAGGCGTTTGTACCGTGTCAGGAGTTATTGTTTGGTGGGGCTACATCATCCAAGAAAGC

TGGTGAAAAGCAAACTGTTTATAAAAATGTTGTAAGAGGTATATATTAAAAATCAAAAATACTAGGCGTACGGCCTAAAG

CAAAAAAGAATAAAACAACTTGACATATACCAAAAATAATGGCGGGACTCCTGCTGTCTTCCGCGGCCCAACCGGGACGC

ACCGAAAAAAATGTGAGAAGAAAAGGGAATATTTTTTTGAGGTTGTTTAGAAGGGCAATACGTTTAGAAGGGCTGATTAA

AAATTTACACAATAAAGCAGTACCTTTTCTTGATGCGTAAATGTACTAATAGTGGAGATGTTTGCTTCATGCTTATATGA

AATGTACTGACTACGGCGATGTACAATCAATTTGCGACAAAAGAGCAAATAAGAGACAATAGGCGTACCTAGAAACGGGA

AGAACAATAAATTTACATAGATGCGAACATGATTTTTGCACAAATTTTATATAGTTTCTGGCATGACCCAAGTTCTTTGG

AACACTCCAGCTCGTGAAATGTCCAAGAGCAACTTGGAGGTGGAATAGTATAGGTGCCAAACTCACATAACGGCAGAGAC

TTTCCCTCTCCATTGACAAAAAGACACTAAAAGTAAAGAAAAAACACCGCTTATGAAGAGTATGGCTAAAAATGCTTAAG

TGCTATTCAAATTTAACAAGCATCAAACAACCTAAAAACTCAATCCCATCTCTCACTCCAGAATAAAAGTAGCACTGGTG

GAGCTTATCCTCCCTCTTATTGGCTAGTGAGCGTAACTGAACGCTTTTATTGGCTCGGAGAAAAACCATTCGTATTCGTC

AGATGAAATTTCGTGAAATATGTTACGTAGATCTGACAACGTGAGGTGTAAATTGCGTAGGTGGTTAGTACACACCCTAC

GGACGGTCCTCATTATTTGATATATCTAATCGTTGTATAGCCAAGATTTTTGATTTGACGACCTAAAATTATTGCATATA

TAGTGACACCGAGTTTACATGATTTTTAAAAATCATGACTGTCGAATTTTTCTAAAAGAATTAATATATTAATTTCGTGT

ATTTATGATAAAAATCATGAATATAAAAAAAAAACTAGTATTATATTAATTTCTAACATTTATGATAAAAATCCAAAAAA

TTGAGTAAACATATATATTGGGTAAAAAAATCATATTTATATTTTTTGGATCGTCTCTAGGGTAAAAAGTCTAAAAGTGA

CTATATTAATTTCAAGTATTAATGATAAAAAACTGAAATAAATAAAAAGAATATTATATTAATTTCGGATATTTATTTAA

AAATCCAAAATATTCAAGCATCTTTATGGGGTAAGAAAAAATCAAATTTTATTTTGTGGATTGAAGATAAAAAATTTAAA

AATTACTATATTAATTTTGAGTATTTGCAAAAAAAAAAAAAAAAAAAAAAACTATATTAATTTCGGATAATTATGAAAAA

AATTTCAAAACATTGAGTAAACAAATTTACAGGGAAAGCTTGAGAAAACGTGAGAATGGACATGCTTTTATTCCCTTTGC

TTTTGATACTTTTGGTTACCTAGCTCCAGAGGCGTTTGGATTTCTCAAAAGAGTTCAGAAAGTCATGCACAGTAATGTCA

TGACTTCCGTTTGAAAGAGTTTGTTTTTAAGAGGATAGGTTTTGCAATTCAAAAAGGGTTTGCGGACATCTTGTTGCTCG

CTTGCTTTTTATGTAATTATCTTCATTTGGTAATAATCAATTAAAAACTGTTATTTATTAAATTAATAATAAGCAAACGA

AAAAATAATTACCGGATACAAAATTAGATTTATATTTTTTGGATCGTTTGTAAGGTAAAAAATATAAAAATTACTATATT

AATTACTACCCGAAATTTCGGATAATTATGATAAAAAATCCAAAAAGTGAGTAAACATATTTACAGGGTAAAAAATCATA

TTTATATTTTTTGGATCGTTTGTAGGGCAAACAATCTAAAAACTACTCTATTAATTACTACCCGAAATCAATATTTATAT

TTTAGAATATCTAAAAGAAATCAATATCTAAATTAATCTTTATATAATCTTTAAAATCTAAAAGGATTAATAATCCAAAA

AAAATCGAAATAATTTAAGGTAAAAGGATCTTTGCAGGTTTTCTTAGGATCTGTAAGGTGAGTTGTATTTGAAGAGACGA

TTCAAGAAGTAAAAGCAAAATACGTTGTATTGATTAATGAGTTGTTATTATAGAGTAAAACCTCTTTATAAAAAAAGAAA

GAGAAAAGATACGGGTGTAGATTGAAAAAGCGGGGGTCTAACAACCAAACTCAATATTTCGATTATCAATCTGTATGGAC

AACTCCGAGTTGTTTTCTAGAGAATCAACTAGACTCAATCAGTAGAAAATATCTCAAGGAGTTATTATCTCAATCTCTTA

ATTTGATTTACTCAAACAAATACAATCTGCGAGTCTTTATCAAAGAGAGTTAACTTGGATGGTACCAAAGACCAATATCC

AAGTGTCAATCAATTCAAATCAACAACCAATAAGGTCGAATATTCTAATTGATTGAACTATGCACAACCTGTATTATTTC

AATTATATAAAAATATAATGCGGAAAAGAAATAATACAGACACCAGAATTTCGTTAACGAGGAAACCGCAAATGCAGAGA

AACCCCGGGACCTAGTCCAGATTAAACATACACTGTATGAAGCCGCTACAGACACTAGCCCACTCCAAATTAACTTCGGT

CTGGACTGTAGTTGAACCCCAACCAATCTCACACTGATCCAAGGTACAATTATGCTCTTACGCCTCTGATCCCAGCAGGA

TACTGCGCACTTGATTCCCTAAGCTGATCTCACCCATAACCAAGAGTTGCTACGACCCAAAGTCTAAGACTTTAATAAAC

AAATCTGTCTCACACAGAAAAATCTACAAGATAGATAAATTCGTCTCCCACGAATAAACCTAAGAGTTTTGTTCCGTCTT

TTGATAAATCAAGGTGAACAGGAACCAATCAGTAAAACCAGACTTATATTCCCGAAGAACAGCTCAGTTTTATCAATCAC

CTCACAATAGTCTTAATCGACGCAGCGAAAAAAGATATTGTGGAAATCACAAACGATGAGACGAAAATGTTTGTGATTAC

TTTTATATCTTGCCTATCGGAGATAGAAATCTCAAGCCAATTATTACAATTGTACTCGTACGATAGAAACAGCAAGATCA

GATCACGCAACTACAAGAAAAGTAGTATCGGTTTGGATTCACAATCACAATGAAGTCTTTAAGTCGTTAACCTGGTTTTA

GAAGAAGAAAATCAAAGGTTAAAGGAGAATCAACTCTAGCTTAGCAAACTAGTATCACACAGAATGTGTGGGTTTAGGTT

TCCCAGTTTTTATGGTTCTCCCTTATATAGTCTTTCAAATCATGGTTTGCAATCAATGTTAGCTTAGTAACAAAGCATTC

AATATTCACCGTTAAATGAAAACCTGATTAGATACAAACTAATATTTATCAACCGTTAGATCGAAAACATAGCTTGTCAT

ACACACATGTACTTGACATTTTTGGGTTTGTGTAACCGTACCCAAACGTGTACATTAGTTGGTTCAACACTAGTTAACCA

AATGGTTATCCAAATGATAACTTTCATATCAACCATATTCTTCTTCACCATAACTAGTCCAAATGACTCAAATGAACTAG

TTAGAGAGTTGTTCAATTGCTATGAGATCCCATGTACTACACAAGACACAATTAAAGTAAAAACGATGTGATTCACTCGA

ATCGATTCATGAACTATAAATCCACGGTTTGCAATAGCATTCCTTAGTTTATTAAGATAAGTTCACACACATCGTTTTTA

GACATAACCTACTCAAGTTCGCGGTCTTGGTTCGCGAACCCAAGTTAACCGGTTGGAGTTCACAAACTCCAATAGAAATT

CTCGGGTCGATAACTTTCGTCAGTTCGCGGACTGAAACTGAGCTACAACTCCAGCAGAAATTCTCGGGTTCGAGAACTAC

ACCAGTTCGCGGAATGAGTTGGCTACTCACGTTCCGGTTTCACTTGATCAACAAAGTTCGCAAACTTCGGTTCAAGCTCT

AAGGACTTATGCATATATGTGTTACCACAACAATGTTTATATCCTGCAATGGTTATATTGTCTAAACTCTCATTTCAATC

ATTGAAACATTCTCAGAGGGCGTTATATAGTTGTTATTCACAAACTATTTTTCGTCAGAGCGATTTCCAAAATGATTGAA

AAGTAACAAGACTTTCGTCACTAAGTAAAGATGAACTTAGCTAAAGCGAAAGCTTTACCAACACATATTTCGAGAAATAG

ATAAGCGAGATATACTCAGCTTGAAATATCACATGTATATATTCTTAATCAATATAGTAATACGACTTCAAGATAGGAGA

TAGAGTAAATAGACTTTTGAGTGATAGATAAGTTCAAGTCTCCATATACCTTTTAGTCGATGAAGATCCACTGGTTCCTT

GAGTAGTCCTTCGTCTTGTATGATGATTGCCATGGAGTTCTTGAGCTCTAATACACTTTATATCCTAGTCCGAGACCTTA

GCTATAGTATACTAGAAATCAATACTTTATAGTTTTGATCACTAATATTGACAAACATGCTTGAGATAGCAACGCATGAG

AGTTTGACCGAGCAATGCTCTGACACAGATCCTCTGTCCCGGAATCCCCTCTGTGCCCCTGTCCCAACTAATCCTAAAAT

GCCACATGGATATCATTTTAACAGCATCCAAACCCGTTTTCTTTTATATCTTTCTTCATTCTTCCTAATCCTATCTTCTT

TTATTATTCGAAATCTCAATTTGAACTCTGCAAATATAATTCAGTTTTTGCATTACCACTAAGACACAACACCGTTGGTG

ATTTATATTGAGCTCAGAAAGGTGACAGAAAAATTAATTTTAGCTAGGATTTCTAGGAGGCAGGCACCTAGCAATCAAGA

TATGAAATTGGGTTTGATCTGTATTTAAAGGTGGTGAAATTTGAGCTGATCCATGTCTTTGATATTGTATATGATAATGA

ATTTCCAAGTAACTCCCTATTAATTCCCATAACACAATGGCTTATATGATTTGGTATGTAATCATCACAAAGGCAAAGAG

AATTTTAAGGAATGAAAATTTATCCGTTTTTTATTTAAACCATCAAGGTTATAAAAATCAAAGATAATCATCGCAAAGGC

AAAATCAATTGGTTGCTTCTTTGATTTAAGATATCTATTTATTAAATCCCAATGATTCATCACAGTATATGATATGATTT

GGGTTACACAAAATCAATTTTCCTCACCACACCAAATTTCTAGAAATTTGAAACAAAACTAAGCCATCACAAAAAAGGGT

TGTAACGGTTTTTACCTGTTGCAAAAAGTGGTATTACAAATTTTTTGTAACAACAATGTAGCCGTTACGAATTTTGGGTT

GTAACGGCGTTTCTAACAGTCTCGAGCCGTTACTACCTGACGAGTCGTAACAGACGTGATCCGTTACGAATATTTTTGTA

ACAAGAATGTCGTAACAGGCTACGGGCGTTACTACGTGTCATATTTGTAACAAGCATTAGCCGTTACTACGTGTCTGGTT

GAAACAGTTGACAATCGTTACGAATTTTTCTGTAACATCAAAAAAATTACTGCTAGTAAAAATTGACCTGGTCAACATTA

GTCAATACTGACCAATTTTGACTAGGTCAATATTGACTCATAATTCATTTTTAGCCTGAAATAACTTGAAATTCAAAAAT

AATTACCGAAAATACACCGGAATGTAATTCCTTTTTCATTCAACCCAATATTCAATCACATGTTCAGCATAATGAAATCT

TACACATTATGTTAAGGAAAATGAAAATCTTACATATTGTCCTAAGTCAACTTATTTGAAAGACTAGATAAATTCATGTC

CTAAGGGAACACACATTATTCATATGTCCTAAGGGAATACAAATTATGTCTAGTACAAAGATTAAACATAATCTTACCAA

CGAAATTGTATACCTAATGGCAGTTATAACTTTGCGACATAACACTTCCAAACCTAACTTCTTCATGCAAACAACGATAA

CTTGCATTAGAAAGTAATCAAAGAAGTGTAAACGTGCATAAGAAGGATAAAAACCATGGAATACAAGGAACTGAAGAGAA

TTGAAGACATCAAATAGGATATGCAAAGATATTAAAGAGTCTAAAGTTAGAATAATTATACAAGGATCTAATTCAATCAA

GAGCAACATATTGTCGATAAGTCTAACTCAACCCATGAAACTAACTGATACTTATAACAACATAATACAAGGATCTTTCC

CACTAACAGTCTGAATCTCAATTACGCAATTTTAACTTAATTAGTATAGATGATCACCGATTTTAGGTAACTACAATTAG

TAAAGGAACTTCAGAGAGAAAAAAAGGTGAAGTTGAGAACTTCACATGTCCAAAGTAGTTCCTTGGATTCACATCTGCTC

CTTACTCAAGCAATAATCTAATAAGAACATCAAAATGGAATTAGAGATGGTATGTGCCAAGATCAAGATTTAGCAGAACA

ATCACAGTGAAATCTTCCAACATAAATAACGGGGTTAATTACTTCATTATGTCCACTAGCAGCTGCAAAATGAAGTGGTG

AATTAAGACCACCAAAAGTATAATATTTAGATAATCCAGGATTGAGATAAGTACCTGTTCAGAGAGCCATCTCAGCCTCT

AGTACAGCCCAACGCATCTGCCTCAGTTTTGTTCTTTTGAGGCGCAAGTTGTGTTGGTTTCTTTCTTGCCAATAAATATG

TTGTAGCTGCAAGAGTTTGTGTTGCAGGTTGTATGAGTTTTTCTTATAACTCTGCTCCTTCGAACTACTCAATTTCAGCT

TCTCATCCTGCAACCAACAAAGATAATTAACTATCACTCTCTGGTTAAAGTTAATTTAACTCTGAACAAATACAAATAAA

ATGTAAAAGGTTACCTCATCAAAATCGGCAGCGGCTCCAATGGGTTTTGACTTCATTACAGCAGCTTTATTTCTTAACGC

ATCAACATTCTCTGTTGTAAGCCATCGAAGATGGGATACTGGACGCATCAGGAGTTAAAAAGAAGCAATGATCAGGATAA

ACATATGCAGGTTCTTGTGGCACCATAGTGTATTCATATGGTGAAGTCTTAGGTTTCATCTTCCACATCATAAATGTATA

AAAAACACAAATTAATTGATAAAAACTGTGAGAAGATTAGAAACACTTGAGAAAAATCATAGTATTAAGAAAGTAGTACC

TTAAATATGGGATACTTACAAATACCAGAGACCGGTCAATTACACGTATCAAAAGATAGAGTACCAGTTGGAGAATTCCC

TCTCAGGTCTCTGTATCAAATCAGATCCACGTGGTAAGAACATAACGAAGTTGTAACCCACTGAAAATTTGATGCATCAA

ACAAGATTATTCTAAGGAAGGAATGTACTCACAAATACTGCAGTACTTCATCCCAATATATTAACCTAGGAATGTCACCA

GTTAACTGGTTCTGAACCAGATCCCTGATACAGAACCAATTAGCAACTTGACAAAAAAATAAATAATAAATTTGTAAATA

CAATTAGCACTTCAACAGGGTAAAGCTAATGGGATTGTAAACATGGTGTATCCTACAATACTCAATATGCTTCTTAAACC

ACCAGTCACATTCCTCTCAAAGTACTCAAGTTATCAATAGTAAAACTTACAAAGTATTCCGAATAAGGAGTTGTGATAGC

GACAAAGGAATATGACCAGTTAACTTTTTGTTCTTCAATGATCTGCATAGCAACCCAACGCAAACTTAATAAAATTAAAA

CAAGTTCTATTGTTCAAGAACTATGGTGATCAGCATGTGGTGCTCTACCAGTTTCAGCAGTTTTCACACAATCTCAGGTG

GTAAATTCAAAAAATTCCTCACAAACTCAACGGATTACCCAAGAACTATCTTTAAAAACAATCAGAAAACAACATGACAA

TCTCCACTAACACTACCAAATGCAACACAAACAATCTTGCAAAAATACAAATAGTATAACACATACAAACTCTTAACACT

TACAAATCCTATTAGAGATAACATATCAGTGAGAAACAAGACAATATGCATGAGAAATTCAGGCATTACATATCAATGAC

CAACAACACAATATGCAGGAACAAATTCCATCTCTTATTCAGATATAATAAAGCAATAAACAGTTCATTTATAACTAAAG

GGACTAGATTTAAACCTTAAACTATGGCCAATACTCTGTTGGAGAAACCACCTTTATCTCCTCCAAACTCCCCTAGTGCT

CTCCCTCTTGGACCACCACGGTATCCATCTCGGTCTGTCACCACCTTCATAATTGCGCCTAATTACATAACAACATCAAC

ATCAAGCTCCAAAACCATACACAAATCAGAAATCAAATGCTACAAACACATCAAATCTTACCTAGGTCTGTCACCACCTT

CATAACAACATCAACATCATAGGGCGATTACCCTATGGTGCGCCTAATTACATAACAACATCAACATCAAGCTCCAAAAC

CATACACAAATCAGAAATCAAATGCTACAAACACATCAAATCTTACCTAGGGCGATTACCCTATGGTGCTCTACTAGTTT

CATAAGATTTCTTCTAAGTATTAGGCACAATCTCAGATGGTATATTCAAATAAGTCCTCAAAAACTCAACACCATCATTA

GTCAAAAACCAATAGTGCTCCCTCTGCAAACCAAAAACAAATCGACCATTAACTAAAAAGAAAAATCCTTGTCCAAATCA

AACCCTAACTTAAGTAGCGATTTGAGAAAAATAAAATTGCAATTACATGTTAAAAGAAAATCAAACCCTAATTAAATGAA

TTCGTATGCACATAATCAAACCCAACCCAAAAAATTTCAACACCGAATCAAAAGTAAAATCAATGAAATTTTTACAAACT

CAATTTAGTAAAACCCTAACAAAAACCCCCAATCAAATCCTCCAAATCAAAACCTAATATCATCAACGGAGTAGCAGTTC

TAGAAAGGTAGGGTTTTCAGTGGAGAAAGAAAGCAGATATGGGAGAAAGTTAGAGACTGAAAATCAAAACGAGTAACCAC

AAACCCTAAAATCGATTGAAATACTGAAACCCTAATATCTGATTAACAAACATAAATACTACAGATGATGAAATACTTAC

ACAGATCTGTGAAATCGAAAACTTGAACTCGATTAACATATGATTGAAGATTAGTTCTTCAAAGATCATCTGAATAATCT

GCAGTTGAGAAGAAGAGAAGAGAGGCGGCAGAAGAAAAGAGAGAAAAAAGTAGAGATAAGGGAGAAAAAAATCTTAGGCA

CGGGAGGAAATATCCGTTAGTATAACCAATCGTGGGTGGGAAAAAAAATGTCGTACATGTGAATTACACACGCGCAAATA

CACATGTTACTACCCTAGTGTGACCAAGCGTGTGAATGGCACGCCTAAAAAAATATTCTTAACAGCTGCCCCATGTTACG

AATTATTAAAAGATTCGTAATGGCTACATCCTGTTACAAAAGTTTCGTAACGGCCATTTGTAAAGGAAAATTCGTAACGG

CTCATAAACTATTACTAATCTGGAAATTTGGTGTAGTGCCTAACCATAATTCAAAAATTGAAATAAAGATCTATAAAAAG

AACTTACTTGAGTGATCAACATAAAATTGGATTCTATGATTAGGACAAACAAAAAATAGTGTACTTAGATACGAAGAAGG

AGAAAAACTGGTTTCCTTTGATTTTTTCGATTTAATCTAAATATTAAAAGAAGTAACGGCGTTTGGATCTGCATCCAAAA

GACACTTATACTCTTATAGTATCATTACAGGACTGGACATCTAATAATGACAGACTGTTAAAAACAGTTAGACAACGAGC

TGTCACTGAACACATGACGGATAATCAGTGAATGATGATGATTACATGAATGGAGCTATAACTAGCTCTTACAATAATCA

AATATGATCTTCCTTATTCGAATAGGAAAGAGTGAAGAGAGGGCCAAACACTTAAGTTATGGTTTTAACCTATAAATCAC

TAACGTTATCATCTTATGTTGGTAGACCTTGCTTTCTTCATTCTGTTGAAGGGCTTGGATCTACTTATTATCGTTTTAAA

TATCTTTTTGGGCCCATTGTAGATACTCTTTGTAATTTCTTCCCATTAGACATATCGTAAATATTTGAATTAGGTAGTGA

CTAAATACAACCCATAATAAAAATAAATACGACCCTTATCTCAAAAGTTTAAATTTTGTCAACCTAGCTAATGTGGGTAA

TCAAAACTCACCTTAACTTTACCCTTTTTTTACCTCCACAAAGCACCGAAAAACAAAACTACTGAAAAACAGAAACCAAC

CATAAACCAAGAAAGTATGTCGATCATTGATAAAATTTAATCATATTTAGAAATTAAACTTAAACCATTTAGATCAAGCA

ATTAAAAATAATTAGTTAATGATAACTATAAAATAGGTTTAGTCTTCCTTTTCCTTTGATTCTACATACTTCCTAGCATA

ATTAATGTGATTCGGATAGATGAAAAACTATTACAGTTGGTAGCTAGATCAGAAGATGACAACAAAGGTGAGTTTTAACT

ACCTACATCCATTGAGGTTGATAATATTTAGATTTGGAAATACAGATTGTATTTAACCAAACGAGGGTTGTATTTAGTCA

CTGTCTTGAAATCTAAAATTCGGAAAGGACAGGACCAGAGAGTAACAAGTCCAACCCTTTCGGGACTAAACATCAGTTCC

TACTTATATATAAAGAGAGCCTTGTTATTCTGTTCCGGTCATGAGAAATTATAATCAATAGAAACCTCTCTTTACAAAAT

CAGTCAGAAAGAATCTTATGTTGATTTTCATTGCTTGAGTCTTCGCTTTTTCTTTTTATAGTTGTTAATCCAATCCAGTA

ATCATCAACAAGTATGGCTGAAACATGTCCAAATACTGTTACAAAGATGAGGTACTAACAAAAAAAAATGGTTACATAAT

TTGGATATATATCTAGTGCTAGCAAGCTGATTAAGGGTCGTTCTAATCAATGGAAACTTTTCTTTCTGGATTTTATTTTC

AGGTGTGCAGTTGTTACTGGCGGAAACAAGGGTATTGGATTTGAGATTTGTAAGCAATTATCTTCTAGTGGAATCATGGT

TGTTTTAACTTGTAGAGATGTAACTAGAGGTCTTGAAGCTGTTGAAAAACTCAAAAATTCTAATCATGAGAATGTGGTTT

TTCATCAACTTGATGTTACGGATCCAATTACTACTATGTCTTCTTTAGCGGATTTCATCAAAGCACGCTTCGGAAAGCTT

GATATCTTGGTATATATGCATCCAAATTCTTTCTTTAAACTCCCTATTGAACTAGCTATATTCCAATCTTTCCCACAAGT

TCATTCAAAAATTTGTGTTTTACTATGCGTAGGTAAACAATGCTGGGGTTGCAGGTTTTTCAGTTGATGCTGATCGTTTT

AAGGCAATGATAAGTGACATTGGAGAGGTAATAAAAATTTGATGCCGATGAGTTTCAAACTTAGGCCAAAGACTTCACAA

CTATACTATAGTGACATCGGCGCCATATCAATTCTTGAAATCTAGAGAGTTAAATGGCTAAATCGAAAGTTATTATAGTT

GTATGCTTTTCCTATTATGAATTAGTTACTCGCACTTTTTCCAGGATTCAGAGGAGGTCGTGAAGATCTACGAAAAACCA

GAAGCCCAAGAATTAATGTCAGAGACATATGAATTAGCAGAAGAATGTCTCAAAATAAATTACTACGGTGTTAAATCAGT

AACCGAAGTTCTACTTCCTTTACTTCAACTATCTGATTCACCAAGAATTGTTAATGTTTCATCATCCACTGGAAGCCTCA

AGGTAGTACTTAATTATTATTGCAACTATCAACTTGAAATCCAAAAATGTTTTTCTTACACTTTGTTTACGCTGGTTTGA

TTTCAGTATGTATCCAATGAAACAGCTCTAGAGATACTTGGAGATGGTGATGCATTAACAGAAGAGAGAATTGACATGGT

AGTGAATATGCTTCTTAAGGATTTTAAGGAAAATTTGATCGAAACAAATGGGTGGCCTAGTTTCGGAGCTGCATACACAA

CATCAAAAGCATGTTTGAATGCTTACACAAGGGTGTTAGCAAAGAAAATTCCCAAATTTCAGGTGAATTGTGTTTGTCCT

GGTTTGGTTAAAACAGAAATGAACTACGGCATTGGAAATTATACTGCCGACGAAGGTGCTAAACATGTAGTCAGAATAGC

TCTTTTCCCCGACGATGGACCTTCTGGTTTTTTCTATGATTGTTCTGAACTATCTGCATTTTGAGTTATATCGATTCAGC

GTATCGTGCAGCAGCCAAATAAATAAAAATGGCATCAGTTGTTTGCTAGCACTGTCCTATCTACTGATAACTTTATCCCT

TTGCTTCTGTTTTTAGCTTGTGTTGTGTGAATGTCCAGAGAGGATCGATCCATTTCTAATGTATTGAATCTTTATTTGGA

TGACAAATAATAAATCAGCTTGAGATTTTTCTTCCGATTTTATCCCTATTGTCATTTATTTTTAGGGAAATAATGATTTG

GTCCATAAAATACACGATTTGGTGTACCGACCGTGATGCAGCCAAGTTTTTTTTTGGGTTGGACTCCGTTGGAGGGACAT

ATCTAAGCTGATTTAACCGAAATTTGCTAGCCGTTTCTTACGACAAAAACGTTCCACTGAGATACCATACAAACAGGAAA

ATATTTGATCTGGCACTTCTCTTAAGTGGGCTTAAAGCTCTAAAATTTTAGCTTTGTTAAACACTTCTTTCAATTGGCCT

CAAATATATATACAAAACAAAATGTGGTAAATATATACCATATCACCATATCCACTGCCACCCAATATGGTATATTATGC

GAATCCTCTAATAAATCCGAACAATATCTTGGCTCATCTGGAAGTACCGATGCTCAGTGGTCTTTGACAAAATAAACCCA

AACCCAAGTTCATTTATTGAACAAATAAACAAATACATGCTACAATATCCACAAGAGAATATCAAGAAAAGCATAACGAT

TCATCCAAACACGGATTACAAGGACAAATGGTCACGTTTATCTTCAGAATGGATATTATTCATAGACGCGGCATTCAAAA

AGGAAACCTCAACTATGGTGTATGCCTTTATATTATACTCAGCAGACAACAAAACTTTGATGCACAATGAAGAAGGAGCA

GATAGGGCCTTATCATCCTTCCATGCAGAGTCGAAAGCCTTATTAAAAGCGACTTCTTGGTCACTTTAGCTACGGATTGC

AAAATGTTAGCGGAAACAATCAGAAAGGAGGCGAAGATCCCAGATTGGAGCGCGGATAACACAATAAGGAAGACTCAAAA

TATTTTGCAAGCACTTCCACAGTCAAAGGTGAAATATATAAATAGAAGGTTTAATTCAGCAGCAGATCACATAGCAAAAG

AAGCCCGTAAAAAGAATATGCAGAATATATCTACCAAAATCCAACTAAGGATATCGAAATCCAGTGTGATGTCCAATATC

TTTGAAAAGAAAGACATTGTAAATTTGATGTACTTTATTGCTTAATTAATATATTTAATCTTTCCATCAAAAAAAAAAAA

CCCGAACAATGATAAAAGAAGAAACGCTAATTATTAATGTAAATAAATAATACAGTGGAGTAGTGGACCATGGCCCGCAT

CACGTATGGCCCTCATTATCTGAGTTTGGCACCTATACTTAGAGGTGTAACATCACGCTAAAATTTAAGCACGGCCCAGC

CTAGCCCAGCACGTGATTTATCCCAGCACGGCACAACACGTTAGTTAAATGGAACGTGATGGGTTTGATTAATTGGCACA

ACAACACATCACAATACGCGGCACGGATAAGTACGTGGCCCAGCCAAGCACAACACGTTAAGAAAGCACGTCACAACATG

TACAAATCTAGAATACAAAACAGAGCTAGGCTAAGTTTGCATGTGTTAGTGGTGGTCAGGATTTGTCCTTGGCAAAGATA

AATCAAGGTCGTGGTTCTCCTGGCCACTCCTATAACTCGCACAAGTGATATTAATTCCAATATTGTGAATCAAAAATGAA

TACGCAAGTCAAGATATGGAATTTCAAAAATTACAACGGGGAGGTTCGAACTCATGACCTATTATGTCATAAGAGTGGCC

TCTAACCATTGTTCCACCTTGGTTGGAATTGATATAAAAGATAAATCAACAGTGGTGGTTTGTTCAACTTAAATACCATA

ATTGGTCCATTAATTAGTGTTTCACTAATTGGACATAATAAATAATCTATTCATTATAAAGTGTTCTATTAATTAGTGTT

TCACTAATTGCTCATAATAAATGACTAATCATTTTTTTAATTACAAATGTTTTCTTAATTAATTATAATATTTATTTCTA

TTAATAAGTAATTTTGATAAATAATATCAATAGACATTTTCTGCAACAATGAGTATTAGAAGAGGTGGAGGCCGAAAGAT

TAGTGGTGGTTAAGGGCGGAGCCAAGCACAATTATATTTATTTTTGTTCTTTTATCGTAAAATGGGATACAAAGTTCATT

TTTCACATGTTTCCATCTTGCCAAGAATTTTTTCTCCCTCTCTCCTCAAAATCCTGGCTATGCCCCTGGTTTGTGGTGGA

AGAAGTAGTGACGGTGAATGCTGGTGAGTAGTGATGAACAATATTAATTTTATGTTGTCGTTACAACATTGATAACATCG

TAGTGTGGAAGATGTGGTTGGTTGTTTTTAACACTTTTGATAAAGGAAGGTACCATATATTTCAGGGATTATATCATACA

AGACAACATTTTCATGATCGTAGTTGGATCACCACAAATGGTACCACTGTTATTTTTGGTACTATATCGTATAAATCATG

ATTTTTTAATGTTAAAATAAAATTGATTGCAATACAAAAAAATATATACACGGAAAAAAATTAATGAAAGTAATCAATTG

ACAAAATTTTGTTCAATCTAAACCTCATAATGGCATTTGTGATTGAAAATTAGAATTGCGTCCAACAACCTACATGGGAT

GGGCGGAAGTGATGGTGGCAGTGGTGGAGTCAAAAATTGACATTGAGGAGGCCTATTTTTTTTTTAAAGAAACATGTAAA

CTTTGGTAGACTAAACCATAAATATATATGGACAAAGTACTATTTATAAAATAAACTAAAAAAATTATGTAGTTATTCAA

AATTTATGGGGTTAGAGCCATGTTAGTCAACCCTTGGCTCCTCCACTAGGTGGTGGTGGATAAAAAAAGTGGATTCATAT

GCTTTTATGGTGGTGGCGTGTATTGGTGAACAAAAACATATTATGCTAATTTCGTAACGCCACAAAATGTGTAACGTTAT

ATTATAAAGTATAAAATGTGGTTGATCATTCTAATGTTAAAATAAACTCGATAAAACACATGTCCATTTTAAAAAATTGG

TGCAATCAATTGCCATGTTATGTAATAACTAAACGAACGACACGAATCAAGATCAGAACATATAAAATTACTTATTATCG

ATTTATTTATTTAAAGAGTGGTCAAGATTTGTCCTTGGCAAAGATAAATTTACCGTTGGGGTTTGTTCAACTTAACCATT

TTTAAACGTTTTATTAATTAGTGTTTCATTGATTGTTCATAATGATTAATTAAATTTCTATTATAAATGTCCCTTTAATA

ATCTAACATAAATACATAATTTTATTAATAAATAATTTTACTAAAAACTATATTAATTCTAATGGTGATAGTGGGAGTAG

GGTTAGTATAAGCGGTGGTGGGTGTTTGTGAATGGTAGAGGCGGTAACATTACTATTGGTGGAAGAAATAGTGGCGGTGG

ATGGTTTTTATGGTGGTAGGAGTTGGTAATAGTAGTGTATAATAATAGTTTTTGCTTTTTAGTGAGTTTTATTGACCAAG

TAAAAAAATAATCTTCACAAAACATGCAACATTGTATTATGAAAGATGTAATTGGTTGTTTTTAGCACGATTGATAAGGA

CAAAAAACCAAATATTTCGAGGATGATACCGGAACATATAAGACATTATGATCATGATTGTTCGATCACCCGTATTATAT

AAAGGCAATTGGTGTACCAACGTAAGAGAACATTACCGCTCTCTCAAGTACTATTACATTCTTTTTCCTAAAAGACCATC

AATGTCGAGTTGTTTCGCGACAACTACCTACAAAGACAACAACTTTCACTCAATTCCCGAAATAAAACCTTATTGACCGG

GAAAATGTATTTGCATTCTAAAAATTTTGTTTGAAAATTATGTTTGTTCATCAAAATCCTCATAAAAAACTCTAAATCAT

AAATTTAAATTTCGCCTCCGATCTTTATAGAATATTTTTATATTTTACATTCTCAAATCATGCGACCTCTTGATGACAAA

TCAACAATATCAAATTATAGATCAATATGTATAGGTTTCATTCACTGGGTTAGAAGGTAGAGTAGACACAGATGGGTAAA

TCACACAAAACAGAACATGACTGGGAATACTTCAATAATTTGGTCAAGAGATAAGGTCAAAAATTTAAGATTAACTTTCT

ATATGACTAATTAGATCGGCTTCGATAGGAGCTGACTAAAATATGGATTAATAATAATTTTCATGCAAGACAATTGCAAA

ACATAGTTTGGCTAACTCTGTCTATGTAAAGAGTGGTCAAAATTTGTCCTTCCTCAATTGCTATGTTATGTGAGAAGAAG

GTGATATCTAAGATTGAGGTATAATTCAAATCTACTAGATTCATACCATTTAGCCTTGAAAAAAAATTGACGTGTTGTTA

TTATTATTACAGGCAAGGGATACATTGACCACACTTGCGCAAATAGCTCACTTCGAAAATGATTGGTGTTGCATGTACTT

TCTAGAATGCAACGAGGAATTTGAGGAAATAAGGCACCAAAAGCAATGTCCAGGGTATTTTGGATATAGTAACTTAGACC

CGACGTAATACTTATGTTTTTTTTTCCACAATCTATTTATTTAAGAAAAATGAACATTGCAAATACATAACATCAAAAAA

AATTAGACAAATTGCGTTTGACAATAATGGCTACTGATGGTCCGGAAACGACAAAAGAAAGGAATTATTGTAGAAGAATT

GTTTGGATATACAACAGAGTACTTAATACGAATGAAATAGGTATAACTATTTCGATTCTGAAATCACATAAAGTTCACAG

TAGAAGATGACATCAAGTTTAAAGTTAGTCAAAAATATAATATGGCTAACTAAAATGGTGGATCAACATCATCTACGTAG

GTTAGTTAGTAGATTAGTTTGGTTTTTATTATGTTTTCTACTTATCTACACCATTTTGGTGGTTTTATCTACTATTTTGA

GGTTTATCTTCGCTTTAATGGCGATTCTTGGTTATTGGGTTGAGATTTAAGATCAATAGTGATGCTCCATGCTCTCCAAC

AACGAAACCTTCATCTTGGTTGTCTCCGACGACGAAATCTACATCGGAATCTTCATCATCAACTTATCCGGCGACGGAAC

ATCCATATGTGGTCTTTTTGGCAATGGAAGCTTCATCAAAAAAATACTATTCTCTACCTATTTGGAGGATCGACAACATG

ACCATGATAGATGAGTCAGCTGGAGAGAAATATGAGCATGTGGAGACTTTTATGGACGATATTGATTAGTTGATTTCACT

GAGATGAGCGCTGCTGACGACATATTTCCAAAAAAAATTTAAATTTCCATGTTCATGTAGAGCATTATCTTATTATAAGC

AATGAACCTAAATAAAAAAGTGTCTGGAAGTTAACATTATATTTCTTTCGTTTGAGCAAATCTCTGGCGATAAAATATTA

AAGGAAAAAACAAAGAAAAAAAGAGAGGAAAATCCCAACTATTCTTATGACAAACACATATTAAATCATAAGGAGTTGAT

AATTATTCGGGACACAAATTATACGATATATTAAAAATGAGTCAACATCATGGTCCGTGCCGTTACGGCACGGGTTAAGT

GCTAGTACACCATATAACAAGGCATAATACATCACATTTTTTGTATATTTCACAAAAGAGTCACTATTCTAGCTAGTTTT

TAACATTTTATGCTGGTTTATTTATAACAACAAATATAATACCTAAATATTTGGAATCATATTTCTATCGTTGTTATAAT

GTCGTCACCTATATTTAACTTGAACATTAATTTACATGACAATAATTCAATTTTATATTTGATGGGCTATTTTCATTGAA

TATAAACTTGTTAATTTTACTTGAAATAATTTCAAATGATATGTTGTCTAATTAAATTCTTGTGATAATGTCCGACTTAA

TGTATACATTAAGTGATTTCAAATTGTTTATAGCACGTGTGCCAGCACGGCACAACACGACACGTTTATTCATATGTTAT

GCCCGTGGGCTGTGATGTGCCTAGCTTTTGACTAGTAAAGCACATCACGGCACAACACGTTTAAATCGTTGGCACAACAC

AACATGACACATAGCACGCGACTTCGTGTGCCTGGATGGCACGTTTTACACCTCTACCTATACTATTCCACCTCCAAATT

TCTCTTGGACATTTCACGAGCTGGAGTGTTGTTCCTACTTGGGTCATGTCAGAAACTATCTAAAACTTGTGCAAAAATCT

TGTTAGCCTTTGTGTATACTTATGGTTAGTCTCGTTTCTTCGTACGTCTGTCTTCTCCCTTAAAGGCCTTGAAAAAATAA

GCTCTCACATCGAACGGTGATGAAACCCTCACTTTCTATTGACCGAGATATGCCTTTTATAATGTAGAACCTATGGTGAG

GATACTTAGCATCACTCAATTGGTTGAGATATTAAAAATTGAGTTATATTTTGTTAAGATGAAATATTTAACCGTTTGTT

AGTAACGGCGTCGACAAAGAAGTAAGTGAACAACACTTCCGTCAAAGATTTAACCTAATTAATTTTGTGGAGATGATAGA

AAAACCAAAACTAAGAAAACCCACCATTTATTTTTACTTTTGATTGATTTAAAAATTCTATTCGATTAATTTGAGAAACA

AAAAAGTTTCTGAAACGGTATAACGGCTGGGAGTTCACACTTTCCCAACCGTGAACCCTGTATTAAAACAAGCAATTTGA

TTCCCGTTTTTTTTTCTCTTCTTCTTCTTCTTCTCTTCCCTCTAATTTTCTTTTTGTTTTGATTTTCATTTTCATCACTT

CACTTAAAGAGATGAATTAGGCTTAATTGATTTACTAATCATGATTAATAATGTTCATCAAGTTTATTATCTAAAATCAT

ATGGGGTCATATTAGTCTTTATATGAATATATGATGAGGAGTGTCCCAATTGATATTTGGTGACCCTAGAAAGCCCCTAA

AAAAACTTGACTTTCTAAGCCTAAAATTAGGATTCTAAAAGAAGCATCACTAATTATTAATGTAAATAATACGGTGGAGT

AGTGGACCGTGGCCCGCATCACGTATGGTCCTAATTACGTGAGTTTGGCACCTATACTATTCTACCTCCAAATTGCTCTT

GGACATTTCACGAGCTGGAGTTTTGTTCCAAAGAATTTGGGTCATGTAAACTATATAAAACCTGTGCAATCTTGTTCGCC

TCTATATATACTTATTGTTAGTCTAGTTTCTTCGTACGCATGTCTTGCCCCTTAAAGGTCTTGGAAAAATTAATTCTCAC

ATCGAATGGTGATGAAACCCTGGCTTTCTATTGAATGAAATATACCTTTTATAATGTAGAAGCTATGGTGAGGACACTTA

GCATAACTCTATTGGTTGAGATATTAAAAACTGAGTTATACTTTGTTTACATGAAGTATTTAACCGTTCGTTAGTAATTG

CGTCTCTAAAAAATTAAGTGAACATCACTTCCGACAAAAATTTAACATAATTAATTTTGTTACATCGTGGAACGAAAAAT

ATATTAGGAATCACAAAACTAAATTTAGAATTTTGTGCAAAAAAGAATTCAATTTGGAAGGCCTTGACAGCCAATTGTTT

TAAAACAATTAGGAAAAAAAGCAAAAATTCGGAGAAAAAAAAGGAAACCACAAGTTTATTTACGATAGACAAAGGAAATT

CCTGAAGTCACTTTAGCCGCTCAGCTCAGAAGAGGGTTACTCAAATAGTTTGGCGTAAGCGAGAGTTATGCTATTAGACT

ATGAGATACTAAAACTTTATATATATTCTTATATTAAATTAAACTTTATATGTGGAGACATATGTCTGGAATCTGGATTG

ACTATTGGTGGATCGCCCAATTGGCTGGCAACCACCCATGACTGAGATCAAAGTGCATATCAATAAAATTGAACTATCAC

ATGGAATTTTCGCATTATCTCTGTTACTCAATCTTTTTATGCATAAAACGGGCAGCTTAAGAGATGAAACTTGATCAAGG

GTTCTGTAATTACATAAGTTAAGTATACTGCTTAACTTTTTTTGGATAACCTTTCATGAATGTTTGTGTAGTGAATCTTT

TTGTATAGTCATTGTGCTTTTAATTTTTGGTGCCATGCATTTTACCAATCTAGCTGTTCCAATTTTGAACTAATTTTCCT

GTTTTTTTCTTGCAATTTGAATGTACATCGTGGTAGTCGGTACATTTCTGATAAGCATGGAGCAAATATCTCCACTTTTA

GTACAATCACACATCAAGAAAAGGTATTGCTTTAGAACTTTATTGTGTAAGTTTTCAATATATACCTCTTACAAACAGTT

TCCTTTTCGCCAATGTTTTTCGATAGTTTAGCACTTTAGCTCCGCCCAACAATAATTCCCGAGCCAGTCTATACCCCAGC

AGTACAAACGCCTTCTATAAAAGCCAGTAAGCCACATATAGCATGGTGACCTTTAATTATTATATTCCCACGAGGTTCTT

GCATTACTCTCGTGGGCTGATATGGGACACTGCTCCACCCCGACAGTCAGACAGCTTTATAGAAAGCATCAGCATATTAC

ATCAATCCTTTATAAAAGAAAAAATGATAGATGAACCGAAGTATTACACTTCGTTATACACCGTGCTCTAATAACAAAAT

TTCGGTCCATCTAGCACTGCTGAAATAAAAGATAGTGCGTATATATATAAGGGCTGACATGGGACACTGCTCCAACACAC

AATCAAACACCATTATAGAAAGCATTAACATATTATTAATAAAAGACCATCTTTATATACGTATGGGCTCATATGTGACA

CTGCTTCACCGCATAATCAACAAATCAATCTTTTATAAAACACTGAACTTCATACATTATTTATATAAGTTATGATAGGT

TATCTGCGGATTTAGTATTAGTAGCAGCCCCGTTGTAAATTGCCTTTTTCAACCAATATCACTTTATAACAAACTTGTTA

CAAACTTTTACTAAAAAAAATTGTCATTGGCCAATATACCATGACCTGAATTCGAAATACGTCATCACCAAATTCTTTAC

CAATTAAAAAAAAAAAAAACAAAACATAGTTTGCTACCAACTAACTATTTGTATGATCCCAACAATAATTAAGTAATTAG

AAGATCAGCTTCAATTCTTATAATTTTATTATAAAAAATTATTATAGAAAAAAAGAAAAAAGTTTTTTATAAATTGAACA

CGTTTTTTTTTCTTCTTTTCTTTCATTATAAATAAAAATATAAGCTAATTTTTTTCAAATTTACGTGTAGGTTCAAAAAG

TGATCTCCTACATCGTAGATTCTAAAAGTGTCTGATTCCAGGTTTGGCTTCCTGTCAAACAGTTAAACTATTTTGACCTC

CTTTCATATTTGAAAACTAATTACAATAGCATCATGTAATATTTGCATTTTTGTTCGTTACTACCCTCGGCAGTGTATGA

TGGTTCCAAAGTTACAAACTCAAAGGCAAAACTTAATTTATCGTGTTTGGAATTTTATGGAAGTCTGAATTCAATTCGGA

AATCGAACACCTATCATATTAGGGCCCATACTTCCAAAATAATACATAAAGGCGGGAAAAGAACTAAAAAAAATAATATT

TCACACGTATTTTCGGCGAAAATAAAATGTATAAATAAATAAAATCCATACAGGCTTCATATCTTTCTTCGCCTATTCAA

TGCATTCGTCCCGCCATACCACATCAAAAATGCATCGCCTACGGGGCAAAGCTCCGCGCATCCCAAATCAAAAATGTACC

GCTACGGGGCGAGACGAAGCCCCACAAACGGCAAGCTAAAAGAAAAAAAATTCTCAATGCGTAGTATGGGCACAAACCTA

GATTAGTGCACGTGCTACACACCGGGCGTTTTTCTCTATAACCATTTTGTTTAGTGTTGTCTCTCCCGTATCGATGCATT

AAGTATGTGTAAAAAATATACGAAATATTTGCGGATTTTATTTGTTTTTTTCATTTATTTTGGTAGAAAATATGTGAGTT

TCTTTTTCTTTTTTATGTATTATTTTGCAAGCGCGGGTTTTCTCACCGTCTCAGTAACTTGGTGTGCCCGTGCTACGCAA

GTGGCCATTTTCTTTCGTGGAAATTGAATTATTGTCCTTACTTTTGATTGGACTTCGCAAATATCCCTGCGCACTATATT

TATATCCCTATCAAAAAAATCAGAAACTGATTTTAAAATTAATCCCACAAATACCCTTCATGTATACCAAGAGTCATCGT

TCATCTCTTATTCTTCTTTCATTTCAGATCTAACAAAGTTCTATGTCTCTTCTCTCTTTCTAGCCGCCACCGCCATCACC

ACCACAACAACCGTTACCACCTCTGCGGTGACCATCATCTGGTTAATCGATTTTGTTGTCGATGTTGATAGTGATCGACA

GAAAAACTAGGTCGTGATGTTGGTTTGATGAAGTCCGAGATGGTGAGGCAACAACGATGGGAATTAATTAGTGGTTTGAA

ACTGAATCCATATGTGAAGAATTTCTCAAGCTAGTTTCCATACTTATCCTTCCTCTTTTAGGTTTCGATTCACTTCCTCT

TTCGATTTTTCTTTGTTTCTCTCTCTCTCGGTTCCTTTCAAACACCAACAACACCTCTACAACGACCACCAACAGCTCCG

CCGCCGCCACCAGTTCGCTCCATCACAAGCACAACAAACCACCACCGTTACGACGATCACCACTTCCTTTTTCCGAATTA

AATGATAAATTATGATCAAACAGAAATCGAGGTTATGATTTAGAGAATTGAAATAACAAATCAAATCAAAAACTACTATG

GTTTTTCAGATCTAGAATTTTAGGTTTCAATTTTTGATTTTTTTTCGAATTGATGTTTATTTTGATTCGAATATCTTTAG

ATTCGATTCGTCTTTGAAATAGGTATGTTTCTGCTTTCTCTTAGGTTTATTAATTTTATTTTGGTATATAGTTTTGATTT

ATTGCAGTAACTCAAGGTCACTGAATCATGCAAATTGCAACCGGTTTAAGGAAAAACAAAAGATGTGTATGATGTTTTCA

ATTGTACATATACAATGGCTTTATTATTGTTGACGAAATTTTTGAACCATTGTTGTGTTTTAAGCAGTGGTGGTGGTAGT

GGATTTTGTAGTGGAGGTTGTGGTAGTTATTTCAGTGGCGGCAATTGCTGCAATACTGGTGTTTGTGGTTGGTGGTAGTG

ATACTATTGTAATTGTATTTGTATTGGTGGTAGCGGATGTTTTTGGTGGTGGTGGAAGTGGTGGTGTTTGTTGTGTTTGC

AGTGGTGTGGTGGTTGATGGTTTTGTTGGTGGATCTGAAACTGCAGCTGGTGAAATGGTAGCATTTTTCCATCAGCATGG

ATAAACATCCATTGTTGATCCAACTGCAATGGATAAACTTCTACTGTTGATCCAACTACAATGGATAAACATCCATTGTT

GATCCAACTGAAATGTATGAGTTGGTGGTAATGTTGGTATGAGTTGTTGTTACAGAGATTGAAGCTTGTTGCTGAATGAA

ATATGGTGGTTATCTGGGTTGAAGCAGGCAGTGATATTGGATCAACTATGGTTGTTGATCTATCTATATAGGATCAACTA

CTGTTGTTGATCCTTTGAACTCATGCTTTGTAGTTTAAATCATGTAAATTTGTTCCAATGTTAAACTTGTGGTGCAATGG

TAGCATGACTGACTCCATGTCAGATGATACGTGTTCGACTCACGCCAAGTTCACTCCATTTTACATGGATCAACTTTCAT

TGTTGATCCAATTTTAGTGGATCAACAATGGAAGTTTATCCACTAAATTGGATCAACTTCTACTGTTGATTCTTTTTTTG

ATCAAATACTTTTGGTTCTATTTTTATTTGGATCAACTACTGTTGTTGATCCAAAAATATCTAAACCAACGACGATGGTG

GCGGTGGTGGCGGCGGCGACGGACTTGTGGTGGAGGAGGACTGTTGGTGGTGTAATGGTGGTGGTGAAGGAGTGGTGGTG

GAATATTAGTGGTGGTGGCAGTTGGTAGGTGGTGGTTGTTGAACGGTGGTGGTGGAATGGTAGTTGAATGTTGGTGGTGG

TGGTGCTAGTGGTGGTGGTGAAGTGGTAGAGGAAGAAATTTGTTGTATTTTGAAATAAAGAAAAATATTATATGGGTGAG

GGTATTAATGTAATCTAATATTAAATGGATAAGGTTATAAAAAAAGTATGGACATATTTATTGTTGTATAAGGACATATT

TATTGGGTGTCAAAAGTAGGGACATATAAATAATGTACCTTTTTCTTTCTTTTAACTTGGTATTTGTGGAGTTTCACCAT

TCCCCATAACGATGCATTTTTAATTTGGCATGCGCCCCCGTAGCAATACATTATTGATTTGGTATGACAGGACGAATACG

TTAAATAGGCGGTACAAAGATAGGAAATATGTGCCGCAAATATGTGCCATGCATTTTACCAGTCTAGCTGTTCCAATTTT

GAACCAATTTTCCTGCTTTTTTGTTGCAATTGGAATGCCCATCGCGGTAGTCAGTACATTTCTGATAAGCATGAAGCAAA

TATCTCCACTTTTAGTACAATTACACATCAAGAAAAGGTACTGCTTTAGAACTTTATGTGTTAGTCTTACAAACAGTTTC

CTTTTCGCCAATGTTCTTCGATAGGTTTAGCTCCACCCAACAATAATTTCCGAGCCCGTCTATACCCCAGCTGTACAAAC

GCCTTCTACCACCCAACAATAATTTCCGAATCCGTATATACCCCAGCAGGGCCGTCGCTGGCTATAGGCCAGCTAGGAAA

CAGCCTAGGGCCCCCAAAATCTTGGGGCCCCAAATTCTAATAATTGTTGGTAGTCAATACGTTTGAAACTCTAGAAAAAA

TAATGGTTTCCTTATGTGTGAATCAGCAGTGACCACAAAATCCACGGAAAATTAAATTTCTTTTTTGATAACTGCATATC

AGAAGAAATTGAGCTTTGGGTATATTGCAATTTGTGAACAACTGATTTTTTATGTATTGTTGAATGCCTATTCATTTCAA

GGTGTATATTTTTTTTGCTTTTCCAATTCTTATTTAGTTATCTTTCTTTAGGCCTTTATCTTTTAAGATTTACTAACTGG

TTCAAGCTTTATAAGTTTTTACTCTTTAGACTTAGTTTATGTTGTGTTGTCACTGTTTCCAAATGGTGATTATTAAATTT

TAATAACTGACATGGACATATAAATTTGGGATGTTTGTGTACAGGTTTTGTATATTTTCATCCTTAAAGTGACCAAATAT

CAATTCCAATCATGGGTCCTAGAAAGAAAAAATGCAAGAGTGATGTTTCTAAAATAAAGAAGAACATAAGAGTCGAAAAA

CTTAAGAGTTCTCTGCGAGGAGCAATGAAAAAATATTTAGTTTCAAATGATGACAACCATGAAAAACAATGAACGAAAGA

GGTGAACAGATGGCAAATTTTGCTAATGAGATGGAGACAAATGGTGATCTTTTGGACAATGATGAGAATGCATTTAGGAA

AAATAAAGAACCAAGATGGCAAGAATTGTCACCACCAGATACAAATGAGATTGGTCAAGATGAAGAGCAAGTGAGGAAAC

AATCGTCACTATCTAATACAAATGAAATTGAACAAGATGAAGTTGAACATGAATAATATTGACCAATGAATATTTATGAT

TCGGGAAATTGGGGTAACCATATTAACCAAAGTTTAACAGATATTTGGGTAAAGAAAGATCCAATAAGAAAAAAATATAA

GAATTTTCGGTTTCCTTTTTATAAATATATGATAGATGCTTCTATCATGATCACTATATATGGAGGATGGAGAATGGGAA

ATTCGGGACATGAAATGGCAACAAAGAAAATGTATTGTGCTTATAGGCCTCCGTTTCAGTAAATTCTCCTAGGGCCTCTA

AAAACATTGCTACGGCCATGTACCCCAGCTGTCCAATCGCCTTCTATAAAAGCCAGTAAGCCACATATAGCGTGGGTGAC

CTTAATTATCCCCACGAGGTTCTTGCATTACTCTCGTGGGCTGATATGGGACACTGATCCGCCCCCACAATCAGAGACAC

ATTTATAGAAAGCATCAGCATATTACATCATCAAACCTTTATAAAAAGATAATTTACATATACAAGGGGTGACATGGGAC

ACTGCACCAACAAACAATCAAACACGCATTAACATATTACATGAATCCTTTATAAAAGAAAAAGATCATCTTTATATATG

TATGGGCTCATATGTGACACTGCTTCACCACATAATCATCAAATCAATCCTTTTTTTTCTTTGGAAAGGACATCAATCCT

TTAAAAAACACTGAACTTTATATATATTTATATAAGTTATGACAGGTTGGCCGCAGAACAGTCCCCTTGTAAAATTGCCT

TTTCAACCAAACCACTTTATTGTTGGAAAGTACTGGGCTGCAATGGAGATGAAAACAGAAACAGATTGTGGTTGTACTCT

AAAACTTTCAAGGCTCTGGATGATTTCTTTTAGACAAATATTTCTACCCTCCCAACGAATTGACGATTGCAACAGGTATG

TGAAATACTGACTTCAGGATACAATGAAAGCCCTACAACCGGAAATTGGATGCAAGGTTTGCACCCCTTGACCCAATCAA

GAGCACATGTAAAACAGATTCTGATGATGCTTTGATTAAGAAAGAAAAACAGAAGAGATTTTCGATTGATTTGAAAACGA

CTCTAGGAGATGCTACTTATAGGGTTTTTCACGTCTGTTGGAAAAAGACAAGTTTCTCTTCACGAAACGTCAGGTCCCTT

CATCAACAGACATTAATGGTGTCTATTGGATTTAGAATTTTCTGGAAACGTTTCTAACGGTCCAAACAGTTACGTATCAC

CCATGACCCGAATATTCCAACATTTATAACAAATTTTTACTAAAAATTGTCACCGGCGATATACCATGACCCGAATTCCA

AATATGTCAGCACCAATTTATTTCAGTTTTACTAATTAAAAAAAAATATAAAAAAATAAAAGGTTTCCGAAGTGTGGGAC

TTGTTTTGGTGGGAGAGACAGTAGATGCGAATCTACCTGGATTTCACTAGAATGGATTTTAATTGATAGCAACATGCCAA

AGTTAGACACGTACAACTCATAATGCCACCTTGAACAGTTAATCCACCTTCTTACTTTTCTATATAAACTCGTTCAACTG

ACATGGATTTCTATTCTTCCATATACTTCTGTGCATATTTTAGGGAACTAATCATCAAATTGTTGATGGCTCCTCGTGGT

GTTTCAGGTCTAGTCGGGAAACTTTCAACTGAATTAGATGTCAATTGCGATGCTGAAAAATATTATAACATGTATAAGAA

CGGAGAAGATGTACAAAAGGCAGTTCCTCATCTTTGCATGGACGTCAAAGTTATCAGTGGAGATGCAACCCGTTCAGGTT

GTATCAAGGAATGGAATGTTAACATTGGTAAAGTACAAACACACTATTGCAATTGATCAATTTACTTCACATACATTTTG

GTTCTCCGTGTATCATAATATACTCACAATTGCAGTGATATATTTCACGCTTAATTAGATTTGATTTGGGTTTCACAACT

TTGCGGTAAAATGTTTGTCTTCACGCATGCAGATGGTAAGACGATTCGCTCAGTAGAGGAAACAACACACAATGATGAAA

CGAAAACGTTACGTCACCGTGTATTTGAAGGAGACATGATGAAGGATTACAAGAAGTTTGATACGATAATGGAAGTCAAT

CCAAAGCCGGATGGAAATGGATGTGTTGTGACTCGGTCAATTGAGTATGAGAAAGTCAACGAGAATTCTCCAACTCCCTT

TGATTATCTACAATTTGGCCATCAGGCCATGGAAGACATGAACAAGTACTTACGCGATTCTGAATAACTGATCGATGCCA

TTTATACATGAACCCATTCGGTGTTGGTGTACGTGTACCTATGCACGTACATATTTTATATTTATCAGTTGTGACTTGTG

TGTATTTTATTGTTATAGTACATGGTAATACTCGTCTCAGTGACTCCATATTATGTCAGTGACCCAGTGTATGGGTGTGG

TTTGATTTGTCATGTATGTTTCTACTAGAATAAATAAATTTGGGATAAGTTGTTTCGGAAATGATTTATCCACCAAGGGA

GAGATATATCAATAATTCTTCATTCGGCTCTGGCAACATTGACAAGTTCGCCGGATAATCTTTTTTCCTATAGAGGCAGG

GCCGGCCCTCCCAGTAAAGCAAGGAAAGTCCTGCTTGAGGCCACCAATGGGAGGAGGCCATAAAATCTAAATTAGGTTAA

TTAAAAAAGAAAAAAATGGAAGGAAAAAAAAATAAACTGCGGAAGGAAAAGTGAAAATGAAACGTACATGAAGAGATTCA

TAATGATCTCATTAATACTATTGATGCGCACGGTAAATTTTAATCGTCTCCTTTCATAATATTTCTCTGTTTGAAGCACT

ACGGGAAAAAACAACGTTTACAACACAATCTAAGAGTTGTAGTACCCCTACGACAACTCCATTTCATGGGTTATGGAAGG

GGGTATTGTAGAAAGTCCAAGCTTTTCTACAATACTTGACTAAGAGTTGTAAAGGGTGATGTGATTCATGGTTCGACAAT

ATTTTTCCGTAGTTGTGATTCTCTTTCGATTTTTTTTGAAAGCCTAACACCACTCTTGCTTGTGTTGTAGGACTATCTTG

AACGACACAAGCGGGATATCGTGAAAATATATGTAACAACAATTACTAGTATTATGAATAAGATTTGCACGACTCTTGTA

TGAGATGTCGAACTATCTTGGACAACATATAGTTATGTTGTAAAGAAACTATTTCACAATATTTGGCTTGTGTAGTTGAA

ATCTGGAAGACAATATCTATTAGTATTGTGAATACTGCTGTCACAACACTTCATTGAGTTGTATAACTGTCATGAACAAC

TCCTGATGATGTATCAAGATTTTATTTTACAATGATTGTTGTGGGTAGTAGAACTTCTTAACACAACACTTATTAGAAGT

TGTGTGCATAAAATTATTTTGAAAAAAAAAATTTATAGCTGAAACCGAATTGCCGAAAGCATTTCACATTCATAATTTCA

TATTAACCTGCCAGTATAACATATTTACATTCATTCAAGACCTTCCACTCAAATTACATTACCTATCTAACTTCAAATCA

CATTGAATGAAAGTGTGTTGCGCAACGAAAGAAAAGAAAATGATAGCATCAGGACCACAACGAACTGAAACTAAGTCTTG

GCAAAAATGCCACAGACCAACCCATCTTGCCATGACTTGTATCAATAATCTTTTCCAATTCCTGCAACCCATACCCATGG

CAGAGCACTTATCTTCTTCTTTCAATTCAGAATCAAACCAGTTTCTTGTCCTTTCTAAACTGATCATGACTTATCATTTT

GCATCCTCCACATTCATTTGCCTAAGGGTACATTTCAGCTACAATTTCAGCAGCTCAGGCCATTTAGCATTTTCATTCCA

AACTATAGCTCTAGCACCATGATCCATTCCCTGCATATCAGCACCATTCTTCTATTTTATTGCAAGGTTCGTTCACAACA

CCCAACTTGACCTATAACCACCGTAGACACTACTTCCTACTTTGTAACTTAAGTACGGGCAGCCGCAGGAGGAGCTAGCT

TCACCAGGAGTGGCACCAGCAGCACCACCAGTACCGGTACGTGCGCCTTGTCCTGGATTAGCTGCATCGTCATGTACATA

AAAAAGAAATCAAATAAATAACAAATCAACAACAATAATTAAAATATCAACAATTCAACTTAACTGAATCAAAGATATAA

TCTCACCAATAACTAATTAAAGTCTTCCCAGTCTCAACACAACTTCGATTTAGACCCAATTCATAGTTAAGGTTACCATT

TATTTTTTTATTTTTCAGACAGAATCAAATTCATTATCTACTACCATTTGTAATTACGATTTTGTTCCTTTTCGCTTAAC

TATTGATGATCAACAATAAAACGAATACGAAAAAAAATCCCTTTACATCTTTAATAATGGGACCAAAACTTACTAATTGA

GTAATGGGGTTAGATCGGCTGAAGATAAGAAAAAAAAATGGTAACCCTTGCTATCTAGCTAGTAAGGAAACTGAAATATT

GATAATGGGAACGTTGGGACAATCCTCATTAAGAGGTCATTCCTTATACTTCATAATGGGACCAAAAAATGAACTTATTA

GTTACATTTTGCTGTATGCTAAAAATAAGTTATTAACAAAAAATAACAATATTGAATATTCTAAAGAACTCCATAAGTTC

TAATGATTTGGAAAGGAATTGCGGAATGACAATAAATCAAACTCTAGCCTAATAATTTATTGACTATTTACAATTACCTT

TAGTGATTCCTGCAGATTCCAGCACTAAAGATTGTGAGAAGGACGAGAATATCGAGAGCCATAACTTGGGAATTCTGGGA

TTGAGTAATAATGGTGGCTGTGGTGATCAAATCCACCATTGCCAACTGAATTTCTCCCTGGGTGAGACTAAGTGAAGCTG

AAATCAGCACAGATGATAATGATTTAAGTTACCAATAAGCAAATTCTATCCTACATGAGCTCAATTCAGAAGCTAACAAT

ATTGCACGACATTGTAAATCACAGAATGTACTAGTGATCTAACACAATGTACGAATTACCAAAATATGTAGCACTAATCC

TGTATTTGTCCGTCTGTAGTCACACCATCTGCAATGATCATCTGTAGGATTGGGTTTTAAATTTTCAAAAACCTCTTTAT

TCCTACATTTCCAAATAGCGAGCCCAGGCATAAGCCATTTGTGCACCACTGAAGATGATTTTTCCCATAGTGCAGTATCA

TCTTCTGTAGATTCACAACCACCTTGGACATGGAAACAACTCACGGAGAACCACATAAACTACCTGCAACTTGGATTACA

CAGCACCTTCAGTTCATAGTGGAAAAACTGGGCAGAAATGTCGCAAAACTGACGTCTCAAACAGCCTGCGTTGTCGAGGC

TCCACAGACTTCATTATAATTCCAAATTCTGAGCTTCACCCCAATTCTAAAACTAGAAAGACCCATCTTTCAAATTCACC

TGAAAAATCCAGATTTGAGGTTCGTTTGGTCTGTGAATAATATGCGCAAAAATGGTCACAGAACTGCGAAGCTACAGATT

TCAGTTCTGTCAAGCTGTGTTTCATGGGACAGATCTCCTAAATGATGAATCCAAAATCTAAGCCCTTTTAACTCATTATA

GTGGAGACATATATGAACATTTTATGTCAAGGAAGTACTATCTAAATCCCTTCCTATTGGCGGCAGTTTCGCAGATAAAA

GAGCACAGAGAACAAGTGACATAAAAACATGGTCTTCAACATTCATTTTACATCTAACTCTCAAAGACAAATCACCATTT

AATAAATCCTTTAACGTGTATAAAATAAACTTATAGGATAGTAATTCACATGTCTAAGTATTCCACATTGATTCCTTGAA

AGAAAAACACCAAAATACATACTTAGCCTTGAAATCAAAGAAAAACACCATGGTACATATGACGCCAAGAGCAAATACTT

GTATATGGAGTACTTAGCCTTGAAATCAAAATCCGATAAACATAGCATACTAATTGGAAAGCTAGTAGTGTGAAACATTA

CCTTTGAACAGCTGAAGAATTTTCCTTCGCTAGCTTCTTTCCGTTCTCGGTTTCCTCGTGTGCAAACTTCATACCAGAAC

GGCCGAATTCCATGAGCACCAACTCCTGAATCACAAAATCAAACACAACAGTAAAAAAAAAAATCGGTTCATAATGTATC

AAGGCTAGCTAATTTATATCATAATTACTAAGGTTCACTTGCTGAATGACAACCCCATCTTTTCCTGTATCAAATCAACA

CACAACAACAACAAAAAAAAATCAAAATCTAATCCCATTACCTAAACCAACACAGAACATTGTTTTCAAATTAAAAAAAT

CAAAATCTACATGATAACTCAAATCCAATCCAAACCCAGCTTCTAAATCTTGTTGTAGAACCCGATTCTCCATTTTTCGA

TTCAAACCATCTCACATCAGTAACATAAAGAAACCCAATCTATGATCATTGCAACACCAACTTCACAAATTATCCCTTGA

TTTCCTCTCGAGGTTGAACATAAACACAAATCTATCTCTGTATTTCTTGAAACCCTAACTTCTCTGCTTCAATTTCTTCA

TCCATCAAACCCATCTAAGAAACCTTCGGAGAAACCACCATCACTTCCTCATCTTCATCTAGTTCTCTGGATTAGGTTAA

TGCTTATCGAGATTTAGGCTTTGAAGGTCGAATCGGTCGATTTTTTATATTCGATTTAGGTTTTTCAAATTGATGTGAAT

TGAATCGAGTATTTCCAATGCAGATTTGGGTTTCTGAAACGCAGATGAGAAGAATAAAAAATTCCCAAATATCCCTGAGT

CTGTGTTCTTCCCTGAGTCTATGTTCTTCCCGACCAAGACCTAACCTAACCTAGGATAACCATAGAAAGTAAATGGTTTG

ATTTTATGTAAAATAATTAGGGTTTTTGAAAAGATTTGGTGAAGCAGGTGAAGAGAGGGAGAGGATATGGCGATGAAGAG

TAGAGATGGGTTGAGTAGTAATGCCGCCGGTCATCTGTGGTGTGTGGAGGAGACGAAGAGTAGAGGAGATGAAGATGGAG

CAAAACAGATTGAGCTCGATTGGTTGAGGGAGGTTTTGGTAGGTGAATAAAAAATATGGACGGACCAGATTTACTTTGAT

CCCCTCATACGGATTGGGAAATTAATCTAACGGTTGATATTCTATTTTTTATAGTCATACGGATTGAGAACCATTTGTTT

TTCTATTTGTGCTAATACGATCTCATGTATCCGATTTATTATCAACCTTCGACATGAAGAAAAAAATCCAAAAAAATATT

AATAAATTTAGGACATTGTTATCGACATTCTGATATAAAAAAAATCCAAAAAATGTATAAGAATGAATCGGGAAATCAAA

AATGAATTCGATAATTGGTGTATTTCTTTGTTCTTCAGATATGGTATAACGGCATACATACTATGAACCAAGAAGCTCCG

GGAGGGTTCTGACTTCAAACTAAGCATGATACATCATCGAAATCGAAAAATATGAAGTTTATGGTTGATTCGAACTTCAA

ATGTGGTATAACGACATACAAACTATGAACCAAGAAGCTCTGGGAGGGTTCATACTTCAAACTTATCAGGATACATCATC

CAAATCAATAAATATGAAGCTCAGTGTTGATTCGAACTTCAAATTTGATATAACGACATTCAAACTATGAACCAATAGTC

TCCGGGAGGGTTCTGACTTCAAACTAAGCATGATACATCATCGAAATCGATAAATATGAAGCTTAGCGTTGATTCGAACT

TCAGATATGGTATAACGACATACATAATATGAACCAAGAAGATCCAGGAGGGTTCTGACTTCAAACTAAGCATGATACAT

CATCGAAATCGATAAATATGAAGCTTAGGGTTGATTCAAACTTCAAATGTGGTATAACGACATACAAACTATGAACCTAG

AAGCTCCGGGAGGGTTATGACTTCAAACTTATCATGGTACAACATCGAAATCGATAAATATGAAGCTCAGTGTTGATTAG

AACAAATATGATATAACGGCGTACAAAGTATTATCCACAAAGCTGCGAGAGGGTTATGACTTCAAACTTAGCATGTTACA

CCATCAAAATCGATAAATATGAAGCTCAGTATTGATTCGAACTTCAATATGCGAGAACGAACCCTATTCTGGCATATTGT

ATGCATACTCTGGATCCGAGATCGAAACATGGTCGTGAGTGGACCATTATAAGCTGAAAACCTTATAATTAAGAACAATA

AATCCATCCATTTACAGATATTATTCCCTCGAAATATTCCCCAAAACTTTAAACAAACCCTATTCTGGCATATTATATGC

ATAACCTAGCTCCGATGTCGAAACCTAGCAATGAGTGGACCACTATAAGCCGAAAACTTGGTAAGAAAGATGAATCCATC

CATTTATGGGTAAGATTCCTTCGGAATATTCCCCAAAACTTTAAACAAACCCTATTCTGGCATATTGCATGCATAACCAA

ACTTTAAATTTGGTTAAAGGGCATACGAACCATGAAACATCAGTGTACGAAGTGGTTCTGACTCCAAAATTTGCAGGATA

TATCATAGAAACTGTGATGCCACGATTTTCGTGTCCCGGATCCCACGCCCGAAGACCCGAAGTGCTACTTGGAAACCATA

GCCGAAGTTCATTCCCACCAATTAACAAATAAAATCACTTAATTATCAAAACACCATCCTAAGGTAAAAGACTATTTCAC

CCTTTCTAGTAATAAATACCTATGTATTTACTATGCCAAGTGACATCTTCTCTATGCATTTACTATGCCAGGTGATATCT

TCCCTATGTATTTAGTATGCCAGGTGACATCTTCTCTATGCATTTACTATGCCAGGTGACATCTTTTCTATGCATTTACT

ATGCCAGGTTACATCTTGTCTATGCATTTACTATGCCAGGTGACATCTTCCCTATGTATTTACTATGCCAGGTGACATCT

TCTCTATGTATTTACTATGCCAGGTGACATCTTGTCTATGTATTTACTATGCCAGGTGATATCTTTTCTATGCATTTACT

ATGCCAAGTGACATCTTCTCTGGAATATTCCCAGAAACTTTAAGCAAACCCTGTTTTGGCATATTGTATGCATAACTTGG

CTCTGAGATCGAATCTTGGCCGTGAGTGGACCACTATAAGCCGAAAACTTGGTAAGAACGATGAATCCATCCATTTACAA

GTAAGATTACTCTGGAATATTCCCCGAAACTTTAAACAAACCCTGTTTTGGCATATTGTATGCATAACCTGGCTCTGAGA

TCGAAAACTGGCCGGGAGTGGACTACTATAAGTCGGAAACTTGGTAAGAACGATCAATCCATTATGGAGACTGTGAAAAA

TGACCAGGCTGTGCTTATCGAAAATAGGTAAAGTAAATATCACAGAATATGAAAATGACCATATTACCCTTAATGGAAAT

AAGTGCTATAAATATAAAGGAGACTGTGAAAATGACCAGATTACCCTTATTGGAAATAGGTAAAGTAAATATCATAAAAT

GTGAAAAATGACCATATTACCCTTAATGGAAATAAGTGCAATAATTATTATGGAGACTGTGAAAAATGACAAGACTACCC

TTATCGGAAATAGGTAAAGTACATATTTCATAATATGAAAATGACCATATTACCCTTGATGGAAATAAGTGCAATAAACA

TTATGGAGACTGTGAAAATGACCAGACTACCCTTATCGGAAATAAGTTAAGTAAATATTATAAAATGTTAAAATGACCAT

ATTACCCTTAATGGAAATAAGTGCAACAATTTTTATGGAGACTGTGAAAAATGACAAGACTACCCTTATCGGAAATAGGT

AAAGTAAATATTATAGAATATGAAAATGACCATATTACCCTTGATGGAAATAAGTGCAACAAACATTATGGAGACTGTGA

AAATGACCAGACTACTCTTATCGGAAATAAGTAATTTTCAGAGAAGGCTTCGCGCTGTTTGCGGCTCAGTCACTGAGCAT

TATTTTATTGGCATTACATTTTGTTTTACAACACTTATGAAGGGTTGTCATAAAGTTAGTGTATTAAAGAACCACAAACT

TGACAGAACCTGTGATAGTGTAGGATGGTTAGTGGTGTATATTTCCATGCCCGAGGCTCAGGATCGAATCCCCCTAACCT

CATTTTTTCCAACATTATATATTTTTACTTCAACAACACTTATAAGTGTTGTTATAGGTTTCTATTTAAAAAAATGAAAA

AAAATCAAAACTTGTGAACTGAAAGGATGGTTGATGGGCTAGACTTCCATGCAGTAGGCTCAGGATCGAATCCCCCCAAC

CTCATTTTTTCCAACATTATATATTTTTACTTCAACAACACTTATAAGTGTTGTTATAGGTTTCTATTTAAAAAAAATGA

AAAAAACCAAAACTGGTGAAGTGAAAGGATGGTTGACGAGCTAGACTTCCATGCAGCAGGTTCATGTTCGAATCTCCCCT

CTAACATATTTTCATTATCATATTTTTTTGTTCTTCAACAACATTTGTGAGAGTTGTGATAGGATAAGTCACTACACTAA

TAAAACAGAGTTGTTATAGCTAAATGTCGTCACAACCTTTTTTCCTAAAGAGTTGTCGTTTGTTTTCTTAGCGCAACCCC

TTGGTCCTAAGTGTTGGTAGTGATGATATACGACAACGCTTCCTTTGAAATCAGTTGTAAAAGATAGGACTTTCTACAAT

GTTTAGAAAAGTGTTGTAAAACCCCAGTTGTAGATGTGTATTTTTCCCGTAGTGAAGCATCGTAATTTATCAATTTTTCT

CATGACTTACCTATTTTTGTGCATTTCCTTTGAAATTTGTGAAGTACGAGATATATTTAGAACTTAGGGAGTTAGGTGCA

TCTCACATATTCATTGATCTACACATGAACTCAAACAAAATCCGAATACATTGCAATACTTAACTGTACATTATGAACTG

TAACTGTGAATTTCCAGTTCTGAGTTATCATAAATTAGGATACTGAGTGAGTGTTACAAACATTCATCGATTAATTTGTA

GCTTTAATGAGCATTTCGATAGTGAACACGGGTTTTTAAATGATGAATTTGTGGTTTTCAACAATGTCTTACATGTCTGC

CGTTTGATCACATTTTAGTTAATTTTTTGCACCCGTATAATCTTTACACCTAGACTTAGTTTGATTGTTGTTTAGTCGTT

GCATTAGCTATTTGTATATTAAAGTATTTAGATATAGACTTAAAGTGGTGACAATAATCTAAGTGATTCAAGTCAGGTCC

GCTATCTTATTCAAGTACTCGTGTATGATCAAAGCTTGCTATCTTGATGTTTTTTATATTGACAAAATTGATATGAAAAA

AGGGCCACCAAAAAAAAATCTTGCATTAGGCCACAAAAAACCCAGGACTGGCCCTGGTTGTATAATACCCATTTTCCACT

AGTGATTGGATGAAGTATCCCCAGATCATTGAGGGTCCCATACTGTGAGGTCATTGTTCCGATGGTCTTCCTCGCATTGT

TCCTAAAGAATCGCATGATTCCCAAGAGTATTTCCCTTGGGTTTTCAAATAGCCTGAAAGGGCTGGAGAGGTGCTTATTC

ATGTTTTCACGTTCTTAGTAATTCGCTTCGCCTTTCTACTAGTTTAACTAGATGCGTGCTTGTCTCCTAACTATTGTTAA

CTTCAGTTGGAGAAGGAAAAGGAAACTCGTAAGCGGAAGTCCTCGCAATCCAAGGAGGTAGGAAATATTTATGTTGTCGT

GCTTTTCCAATATTGTTTCCTCTATTTTTTACTCTTCTTGGCTTGTGCGGGACGGATCTGAGGTGGAGAAATCTGACAAA

GATGTGGATGATGGAGGATCTTCCATTTCTCCAAGGCCCAAAAAGGAGAAAACTAGTAATGTCTCCAAGAGCAAGCCCAA

GATCCAAATACAAAGTCCTATGATTGTTGTTAATTTGCCTCATCCAGAACAGCCTGTCAATAAACCTGCCTCAGCTGGTG

GTGTTTCTTCCTCCAAGCTTCCTCCTGCTAAAGGGTATACTCAAGCTAAGTGGTTTGCTGGTACCGATGGTGCGCAAAAT

TCAGTTGTCTTTCCAACTGACCCAATGCCTTTTGTTAGCTCGTCCGTCAAGTCTTATGTCTTGTCATCTATTTTGCATGA

GCAACCGCCTTCCCGTAAAGACAAACAACCTTTGATGTCTGGGACTCTTGCCATTTAAGATGTTGACACAGGGTTCGCAT

CCCAGATTTTCATTGCTTCTCAGTCATCCATCTTTGACCCCAATCTTTTGAAAATCTGTGGTGCGATGATGAAGCTATGC

CTTCGATCTTCTCTTGATGAACAAACCAAGTATGATGTTGCCTCAGCTTTTAGTCTGGAGAAAAATATCGCGATCGAATA

TTTGGTGAGTTATTCTTTTGTATTCTTTTTTCAAATATATCGCACCGCCCTTCCTGAAAATTTCTTTCCTAATTTTAGAA

CTTTCGTCGCAAGCTGACTCAAGTTGAGGACTACAGCAAGGCTATGAACTTGACTCGCTCTCAAGGTGAATAAATTTCCC

ATCTTCAAGCTGAGATTTCCAAGTTACAGGGTGATATCTCCAAGCTTCGAAAGGAATATAACAAACTTGAAGGGGAGTCA

TTTTCCAATGCCCATTCTGAGATCTCGCATTTCAGAACACTCAGCTTCTTGGTAAACATTAATGATATTTAATGATATTA

TTCAACACATGACTACAATATAATGTTAAAAAAATAATTACTAAAAAATGATTTTTAAGGTTTTAATAAAATTATAATTT

ATTATTTTATTTTATAAATGAAAAACTCATTAGTCCCACATTTTGGAGTTTCCACTTTGTAGTTGTTTTAATAGACTATA

TAAGTTTTTTAGTCCCACATTGGGGAGTTCATTTTCTTAACAAATTCAATACTTTTATAAAATCTATGGGAAAAGGTTGC

TCTATATTTAGAGAGACCCCCAAGGGAAAAAATATTTTATATTGTTTTCTCAAGCGTTCGCCATTTTCCTTTATGGTTTT

TTTGGATTTTCCAAGCTCAAGTTGAGCATCTACTACATATGCTAGTAGTAGGTGTATTAGGGTGTTTTATCCTGGAGATA

TTCATCCTGTGAGGGCTATAGCATCACTCTTGAGTGTAGTCGGGCGTTAATTTCTTAAGGGCAACGTGTTGAACACGTGA

CTCACTCTGTTTTTCAATGTTTTGCCTTGTTGCTGTTGTGGAGATACGGGAAGCTCGCTCGTTTCGTCAAATTCATCACT

TCCATTACAAAGGCGTTAAGTATCAATAACTTTTGCTTATTTGATTTTTCTTTGTTTTGATTATTGCACCCAACAATCTT

AAGACATTAAAATTTGTAATAATCACGGATGTTAATTATGGAGTGAAAAAAAAAAACGAATTTTTGGTCAGCTGTGGAGT

TTCGAATTTATGTCTCAACCTAGAAGGAATTTCGATGAACCATTTTGACACAACGTAGTGGACATCCTGATAGTTACCCA

CGAAAATTTCAGAATATTTGGAGTTGTAAAAGTATTTTTTTGATATTTTTCTGAACAGAGAAACGTTCCTGAAAAATTCT

GACAGGTAGAAATTTGTTGTTAACTAAAGTAGTTTTTACGGGTAACCATGAGTTTTTTGATATGGTGGTTCAAACGAAGT

TTGTAAATCTAGATATTATCTTCAAAAATCATATTTTATCTTCCGTTTCGGAACTTAGGTTTGTGAGATGTGGTGTATTA

GGTGATTGGAAAATTGCCGTGTCCGTGGTCAGAGTATAAACGAAGATTGTGATATGAAATTGAAAAGGAACATGGCTACC

AGGCGCATGTGGATGAACACAAGTCTTACAAAGATGACAAAGGCGAGAACATCAAACACTACTCTAAGCGTAGTCTTCAT

AAGAAAGGTATGTTTCGTAAAACTGAATCCGACATTACTTTAATTAAGGGTGATTGTTATGTTTGTAAAATTCCTGGCCA

TACGGCAGTAAAGTGTAGACAACATAAAACCTTAATAAGTAGAGAGTTAATGCTCATTTAGTTGAAACAAATTAGAACGA

GTTCATGTGAGAGACTGGTGGGTGGACACTGGAGCCACTAATTATGTTTGTTGAAATAGAGACATGTTCACCTCTTATCA

GAGGATAAGGGATGTCGATGTAGTACCAGAATATCGCTACAACATTATTCCCGCGATATTAACAACATTTCTGCGAGCGT

CACTAACAAATCTTCAACGACATTTCACGAGCGTTACGGAACAGATGAAAGAAATTGACGATGGAAGAAATAAAAGAAGA

ACATGATTGATCCTGCGATGGTCATCATGTTTGCGATAAAGAAATATTGTGATGATCGTGATAAGATCGCAGAGTAGATC

CGAATTAATGGGAAATCTTAGCTATCTTCCATTATATATTTTCCTATATAAAGAGCACGATAGATCCAGAAAAGGATGAT

GCATTTTTGAGAGCTGAGAGAATTTTGAGGAGAGAGAAAATCCAGAAAGAAAGTTGTACTTCTCAGTTTTAATATTTCTT

GTAAACCTTCCTGAAAGTGAATTAATAAAAGTGTTGTTGTTTATTCACAATTATGACTTAGTTTTCATAGGTTCTTGACA

AGGGTGTAGTGTAGGAAATCCTGCAACTACATTTTGGCGCCCGAAACAGGGATGAATTAACGAAAATAGAATAAAAAGTG

AAGAAGAAGATAAAAATTGTTCAAGATCAGGGAAATCATAAAGTAAATCGGTGAGAAAATTGAAGGAAGAATGATCAGAA

TCAAAAAGGATTGTAGAAAATTGAAAGGTGATGTAATAACAAAAGAAATGATTGGTAAATTTCATGAAAATATCAAGGGA

AGGATACATAAACGGGATTTTGAAGGAAGGAAGATTATGACAGTGGAGAAGATATCATCATTGAAAGATTGGGAAAAACA

AGAAATCACGTTTATAGCAGCAGAAATACCAGAAGGAGATTCAAGTCATACATCTCCATTGGTGGTCACAGTAACCACTA

TACCAATGAGAAAGGAGAAATCGATAAAATCTGAAGAAGAATGGGATATGAATAGAACTTTAGTAGACACAGGAAGTTCT

GTACACATGATATTTTATCATACCTTCAAGAAGATGGGATATAAGGATGAAGAAATGTCTCATCCAACGTATTATGTTCA

TGGATTTGGAAAGTCTATAACAAAACCTAAAGGCGAGATAATGATACGAATTCCACTCGGAGAAAGTGAAACACAAATAT

CATTATGTGTGGTAGATATGGAATCGCCGTACAATATGATACTGGGAAGATCGTGGATACACGCAATCAAGGTTGTGGCT

TCAACATTACATCAATGTGTTAAATTTCCTATGCCTAGTGGAATTGGAGAAATTAGAGGAGATGTGAATTTCGCGAGATT

GTGCTACCAAAAATTGAAATAAGAAATTGTGAAGAACGTCCAAATATGCGAAAAGATCGCAGAAAATTGAACAGAGAAGT

AAAGAAAGAAGATGAATTTAAAATTTACATGATAAGGGCAAAAGAAGGAAAAGGGATACCAAGTGAAGTTTCGCGAGAAG

GAGATGAGTTCGCGAGAGTTATGAAAACGCCAACACCAATGGGAGAACTTGAAGCCGGTTACACTTCTGCTGAACCAACA

AAAGAAATAAATATTGGAAGTAAAGAACCTAAAATGCTGAAAATTGGGACTAGAATGGAGGCAGATGAGGAAGAAAGAAC

TATAAATCTTTTGCGAGAATATGCAGATATCTTTGCATGGAACATGGATGAGATGCCAGGGATAGACCCGTCAATAACAT

GTCATAAATTAGAAATCAACAAAAACATAAGACCATTTAAGAAGAGAATAAGGAAGATCGCAACAACATATCACTCTCAA

ATAGAATAAGAATTACAGAAGATGCTGGACGCAGGAATTATAAGACAAGCAAAATATCCAGAATGGATTGCAAACATGGT

GGTGGTCCCAAAGAAGAACAAAGGGATAAGAATTTGCATATATTTTACGGATTTAAACAAGGCATGTCCAAAAGATAGTT

TTCAGTTGTCAGATATTCCACAAATGGTCGAATCAGCAGCAGGAAACGCAATAATATCATCATTGGATGGGTATAAAGGA

TATAATCAAATCCCTCTCGCTGAGGAAGATCAAGAATATACTGCATTTTTTGCTCCAAGAGGATTATATTGTTATACAAA

AATGCCATTTGTCTTACGAAATGCTGGTGCAACGTACCAAAGAATGGTGGAGAAGGTGTTTGCAAAGTGGATCCACAAAA

CATTAGAAGTGTATGTAGATGATATGCTGATCAAAAGCAAAGAAGCTAAGAATCATGTGCAAGATTTGAGAGAGATTTTC

GATCAAATGCGGCAATTCAATATTAAATTGAATCCTAATAAGTGTACCTTTGGAGTAACTTCAGGGAAGTTTTTGGGGTA

TATTGTGTCAAAAGAGAGAATACAAGTTGATCCAGAAAAGGTACAAGCAATTCGCGATATGCCAACTCCTGCGATGGTAA

AAGATGTTTAAAAACTAAATGGACTCTTAGCCTCGCTAGGAAGGTTCATTTCGCGATCTTCAGACAAGTGTAAACATTTT

TTCGATATACTCAAGAAGGGAACAAAATTCAAGTGGATAGATGAATGCGAAAAAGCTTTTCAAGGAATAAAAGAACATTT

TGCGAGTATATCTATATTACAAAAGGCTGAACCGGGAGAAGAATTATTGATTTATCTCGCAACAACTCCATATGCGTTAA

GTGTTGTGTTATTGCGAGTAGACGCAGGGGTAGAAAAACAAATTTATTATATCAATAAAACATTTAATTCATCAGAGAGA

AATTATTCAAAAATTGAGAAACTAATTCTAGCATTGGTACACGCAACCTTTAAGCTTCGAATTTACTTCCAAGCGCATAA

GATCAAAGTATTAACGAGAGTTCCAATTGAATCGGTGATGAAAAATTCAAAAAGATCAGGCAGAGTTGAAAGATGGAATG

CGCAAGTAGGGCATTTTGATATTCAATATGAAATTCTATCTTCGCCAAAATCACAAGTTATCGCAGATTTCTTAGCTGAA

TTTCCTTTAGAAGATGATGAAGTTGTAGAAGAGATGATGGATATAGAAGAAGAACATGGAGACCCAAAAGAGCTATTAAC

AGAACCTGACAGATGGGAAATATTATTGGATGGATCGTCAAATGGAGAAGGAAATGGGATTGGAGTTGTCTTCATTTCAC

CAGAAGGAGAAAAAACATCTTTTTCATTTAGATTGGAATTTGCATCCACTAATAATGAGACAGAATAAGAAGATGTAATC

CATGCATTGAAGTTTGCGATAGAAATGAAGTTGGAAAATGATAGAATCACCAGCGATTCACAATTGGTTATTCGCCAGAT

AAAGGGAGAGTATACCACAAATGAACCATCTTTGAAAAAGTACAAAAATCTTTTTGAAGAATTGTCTGCGAGAATTCCAA

AAATAAAATGGAGACACATATCAAGGAAAGATAATAGGCTTGCAGATGCTTTTGCTTTCATATCAAGTATGATGACAAAC

CCACTTGCGAGATGCATAAAAATTCAAACACTTTTGCAACCGGCGATAAACAAAGAAGAAGAAGAGGTAGATGTAATGAT

GGTAGAATCTGGCGAGGAAGAAATAGGGAATAATGCTACAGATTGGCAAGCAGATCTTCATGCTTATTTGGCGAGAGGAG

AAATTCCCAGAAATCGATTAGAAGCACATAAATTAAAAAGTAGAGCAACCAATTATGAACTAAGGGATGAGGTATTATAT

CGCAGATCATTTAATGGACCATCGCTAAGATGTTTGACGAGAGAAGAAGGACAAAAGGTGTTAAAAATGTTGCATAGTGG

TGATGCTGGTAATCAAAGCGGGGGTAGATCTTTGGCATATAAAACAAAGATGCAGGGTTATTATTGGCCATATATGCATG

AGGATGCAAAGAAACTTTCAAGGCGTTGTGAAGATTGTCAGCGATTTGGGAAGAAGATACATGCACCAGGAGCACCATTA

TCATCTTCGTCAAGAGTATGGCCTTTTGGAAAATGGGGATTGGATATTGTGGGACCATTTGTGCCTGGTTCTGGACAAAG

AAGATATTTAATCGTTGCTACAGATTATTTCACAAAATGGGGAGAAGTTAAGGCAGTACAGCACATCCGCGATAAGGACA

TTTTTACGTTCATATTTGAAAATATAATTTGCAGATTTGGGATCCCAAACACGTTGATGTCTGATAATGGAAAGCAATTT

GAAGAGCAGAATATCGAATTGCTTCTTAATGCATTTAAGATAAAATGTGGAAAGTCAACTCCTTTATATCCACAAGCAAA

TGGTCAAGCTGAGGCAACTAATAAGACAATCGCAGACATCTTGAAAAAGAAGTTAGAAGGTTATCACAAATCATGGTGCG

AGCAAGTGCATAATGCAATATGGGCATATAATACAACTAGACGAGAGGAAACTGGAATGTCACCTTTCTGTTTAACATAT

GGAGTTGAAGCTGTCCTACCAACAGAGGTTGCTATCCCAACAACAAAAAGAGAGGCATGGGAGAAAAATCTAAGTGCTGG

CATGAATTTACATGAGTTAGATGAACTAGAAGAAAAGATAGAAAAAACTCTACAACATATGGAAAATTACCAGCGAAGAT

TAGCTCGCGAGTACAACAAACGAGTCAAGAAACGTCAATTCCAGTCTGGCGATCTAGTTTTGCGAGAAACACCAAAATCA

GCAAGAAAGTGGTGGAAAATTAGCGAGAAAATGGGATGGACCATACATCATTAAAGAAATAATTGGAAATGGAGCATACA

AATTAATGGATCCAGAGGGAAGAGATGTTGGGCAAAGACTTCATAGACCATGGAATAGATTGTTTCTGAAAAGGTATTAT

CAGTAGCAGTTTTTGCGAGGTAAGAAATATAAATGTAAGATGAAACAATTAAAAAATTCTTCTTTCTCTTTTATAATTTA

TGCGAGCATCATCGCAAGAATAGCGAATAAAATTTCGCAGAGATTAAAAGAATGGGACAAACCTTTATGGCGTGCACCCT

AGTTCGCGAATAAAATTTCGTAAGAGGAAAATGTAAGACCTAATAGCGCGTCATATGAGGGGGTATCTCTTGACATGACT

CAGCGATAAGGTTATACCGCTACTGGAACCTGAGTCATGCAGACGATGGGGTTAAAGAAAACCTATCCGGGAGAGAGATC

TTGGATTTCCAACTATATAAGGTTATACCCTGGTTGGACGACCGAGAATCTCTTCCAAGAAGTGCAGAGTATGACCAAGG

CGCCGAGGTACACGGTTGAGCCAAGGGTGCCCGGAGCGTACGAAACTCTCTTGACATCCTTGGTAAGTCTTGGATACAAT

GCACTTGGCCTGAAGAAAACCCCACTTAGGGTGCAGTCTTGTAACTATAAGCCATATAGGCATAAAGGACAAGAGGCTGC

GAACACAACTGGTTGTTGGCAAATCCGGATGGGGGAGCTAGTCTTGTATGGCCGATGTGCCCTCCTTGAAGGGGCAACCA

GGGGGAAGATAAGGGCGGCACCCTCCATTAGGGAGCTGATAAGTGTCTTAAGACGGCAATGTCATGGGGCTTTATGATCG

TAAGAACATTTAGGCCTGTCATATTGTCGCGTGACGTAAATAATGCAGAGGCAACAAGCCATGAGAGCAATTTTAAGATG

AGATCAATGGGTTTTTACATAAAAGTGTAAAAACGTCCTGCGATTTCATCGCAAGACGGCAATATCATAAGGGTAAATTC

TTGCGAGAGAATGTATGCTTATCATATTGTCGCAAAGATTTTGAAAGAAACATGGATAAGACAAGATCAAAGGGTCTTTA

CATGCAGATGTTAAGATATCCTGCGATTTTGTCGCAGACAGAAAAAGTTTGGCAAAATAAGACCTTTAACTAGGAAGGAA

AAATAAGACCAAAAGAATAATCTTACCAATTAACTATTCTGCTTTAAGATTGCAGGAAACATTCACTCATCGCAGAATTA

TATATAAGGAGGCAAATTTTTAACATCACAGATTCCCTCAGAATTTTTCAGAAAAGTGCAACAGTATTACATATGTCGCA

GATGTATTCATATTTTCGCGAAAATGCAGCAACACTATTTCTTAAAGAAGATAATGAATATTCACCATTTCAAGCAAGAT

AATGAAGGGAAAGAAGGGGAACAGAGAATTAAGATTACATTATCTTACTTGAAAATATTTAAAGTAAGCTAAATCGAAAG

TTAAAATTCTATAAGTCTCGAAGATTACATCGCTCGCAAGAAAGAAGAAGGAAAAGAAAACAAAAGAAAAATTCAAGCCT

CGCCAGATTTATTGTTCTTGTCAGATTCATCAACATCATCAGCATCCTTTTCACCGCGAGCTTCTTCTTCTCTGGATTCT

TCATCAGCGCTTACTTCTTTTGGAGGACTTTTTTCTTCTTCTTCACCATCCACTTGATCTTCTTGAGTGGTATTTTCTTC

ATCGTTAGTAAACTCTAAATCTTTATCCATGGGAAGAGAAATTTCATCATCGTCAACCTCTAGAGGAGGGACAGAAACAG

GATAAAGAGAACGGGGAAGCAGAATATTGTTCACAAGAACTTCATCCCTAAGATGGACTTGGCAAAAAAGAGCTCTTTCA

CGATTCCTAATTTCTATATTCTTAAAGTATTCAAGATAATCAAGTCTTCTTTCTGCTTTATCGTGAGCACTAAGGCAATT

AGAAGCAAGAGATTTATTTTCTTTGCGAACTTTATCCAATGTAGGGGTCAGATCAGTGATGGTAGAATGAAGATTTCTTT

CGGATTCTGCGAGCAACAAAATAAAAAATCAGTTAGACAAGAATTTCACAAGGAATATAATATAAGATTTTATGAGAAGA

TCAACAAGAAAGGCAGTCAATTGACTACCTTCTTTTTGCGAGGTTAAGTTTTGGATTAAAGCAAATAAACAGTTCTCCCA

CTGATTCATGGCATCATCGAATTTATCGCCAACCATACCCTTCAGACGAGATTTCATATTTTCTTCTTTGTTGCGAAGAT

ATGAATTTTCCTTTGCTAATGAAGAATGAGATTCCTTTAACTGAGAGTATTGGTCTTGAAGTTCGCTGGCTCTTAAACTA

AGAGTTAATATACCCTTGATAAGTTTATCCTTTTCGGCGGCAGATGATTCATTTAACTCCTTTAGCTCCTTCTTAAGAGA

ATGAAGAGATTGCTCTTGACTATTACATGTATTTTGAAGATTACTTATTTGCTGCGAGGAAATCATCATCAAGCTTGAAA

ATCTATGTTTTTCTTGGGACAGGGTTTTATTCTCATCTGATAAAGATTACATCGCCAAATTAGCCTTAAAAAACAAATAT

GAGAGGCGATCAAGTTCGCCACTCAAAATTTTTTGCCTTTGCAATAATTGATTGTTCTCATTAGTAAGATTATTTATGCG

ATCAATTGAATAAGTGTACAGATTTTCTAGTCTAGTATGTTGATCTATCCAAGAATCTCTTTCAACGCGAACTTCTTCAG

CAATATATTCAGATTTGAGTCTTTCTATGAAACGAGGCCTATTTGAAACTAATATATGAATCAAGAATTTAAGAAGGAGG

TCTAAGTACAAAAAGTATGTATATGAAAAAAAAAAAAAAAAAAAAAAAGAGAGCAGAGGCAAGAAAAATATACCTTTCAG

CTCGTCATTTTTTATGCGAAGATTCTCGCAATCTAGAGAAGAATTACGGAGCATGTGATTCAAGGTATCACACTCAGCAG

TTTTAGCTTGAAGAGTTTTCCTCAGAACTTCAATATCATTCTCTAAAGCCACTCGTGAATCATTACAACCAGAGCCAAAA

TACTTAAACATAGTCTCGCAAGCTTCTTGTTGGAGAGGGTCAATGGAAGACTTTCTAGCATTGTCTAAATCATTAGACAA

GACTTTAAAAGCAATATCAACACCTGCGAAAGTTTCGCTTGATGAAGGAAAGGATTTAGAAGGTGAAATAGTAGTGAGAG

TAAGTTGGCGATCATTTGTGGTGGTTGAGTTATCAGGAGGAGTATGTGGAAAAACAACAGTTGAAGGGCACCGTAAAGGA

CTATAAGGAACATCAGTCACAGCAGCTTTAGAAGCTTGGTTTGTCAAAGTGTCAGAAGGGCTAGAGGTATGAACATCGAT

ATTCTCTCGGATAGGAGAAGTAAGATTTGGTTGCGATGGTGTGGGAGTAATCTTCACAGATGTTAATGGGGGCATAATGA

GATTTTCCCCGAGTGTCTTAAAATGGAGAGACCTTTTGGGAAGTTTATCAGGATTCACGTCATTCTGCGAATAATAAACC

AGATAAGAAACAGAGGCAACAATATTGTTATTAAAAAATAAAGGGTGTTTCTTACTACCTTATCATTCGCAGAACTTTTC

TTCCTCTTCTCAGTCTGTCTTTTGCGAGGATTAGGGTTGTGTGTAGTAGATTGTGTGTTAGAATTCTTTCCGCTAAAAAG

AAGACATAACAAGGAGATAAGAAAGATTGGAGAATGAAAGCAAAAACTGACTTAAAATAAACAACATAAATCAAGAAACA

AACCTTGAGGACTCCATTAGAACAGTAGCAGAAGAATATGAAGAAGAGGCAAAGTTGCAGACGGAAGATGAAGAATATGA

AGAAACAGAAAAGTAAAATTAATTTTGAAGATTAGTTCAACCTTTTATAGGCAAAGGGGAGTGGACCTATGGAGTAGGAA

CGTGTCAGTAGATGAAGGGATTCACAACACGTGTAAGAAAGGATGGATTAACGGGATCTTGTCGAGGAGAAAGTATAAAA

GGGGAAGAAGTTGCGAAGAATAATAAAGAATTTTAGATTCTCATATCTTCTCTTCTTCGCAGAAAAGACATGAGAAGAGG

CAAGATGTAGTACCAGAATATCGCTACAACATTATTCCCGCGATATTAACAACATTTCTGCGAGCGTCACTAACAACTCT

TCAACGACATTTCACGAGCGTTACGGAACAGCTGAAAGAAATTGACGATGGAAGAAATAAAAGAAGAACATGATTGATCC

TGCGATGGTCATCATGTTTGCGATAAAGAAATATTGTGATGATCGCGATAAGATCGCAGAGTAGATCCGAATTAAGGGGA

AATCTTAGCTGTCTTCCACTATTTATTTTCCTATATAAAGAGCACGATAGATCCAGAAAAGGATGATGCATTTTTGAGAG

CTGAGAGAATTTTGAGGAGAGAGAAACTCCAGAAAGAAAGTTGTACTTATGAGTTTTAATATTTCTTGTAAACCTTCCTG

AAAGTGAATTAATAAAAGTGTTGTTTTTTATTCACAATTATGACTTAGTTTTCATAGATTCTTGACAAGGGTGTAGTGTA

GGAAATCCTGCAACTACAGTCGAGAAACTCAATATGAGTAACTCATCTGCGACAGAGGTTGCATAAAAGGGAAAGGTCGA

GCATAAGCTCATATATGTAATTTTCTCACACTGAATGAAGTTTTCATGTTCCGAGCATATGCAAAAATCTTGTATCTAGT

TCTCTTGAAGATGGTAAAAGATTGAAGATCTTAATTAAATGTGGAAAACTTGTTATAAATAAGGGCAGTGATTTTTTAGG

CAACAGTTATAGGACTTAGGGTCTATATAAGCTTAAAGGAAAATCTGATGAAGTGAACATAGTTGATTCTTGTGCTTTCT

TTTGTGTGTCTTTGAATGTTTTTCATGGTAGACTTGGAACCGTCAAATTATAAGTCAATGCATAAACTTCCTAGCATAGG

CTGCGTGCCCAAATTTAGTTTGGATTTTGAACACAAAAGTGAAATCAATATGTTATTAAACCTTTTAGAAAAAATGTTCA

GAGTAATTCTAATCCCTTAGAATTAATTCGGTTAGGCCTAGTTGAAATGAGTACAACCCAAACTATGGTGGTAAAAGGTG

GTTTGTAACTTTCATAGATGATTGTTAGAGGTACTCTCTTGTATAGGGGTAAGGATGATTCCTTAGAAGCCTTAATAATA

TAAACTTTAAATTGAAAGCGAAGTATATGTTACCACTAAATTGGTTAGGTCTAAACATGATGGAAAGTATGAAATTCATA

TAGGAGAACTTTACTTGTTCAATGACATTATTCATGAGATCGTACCTCCTTATTCACCATAGTCTATTGGCATAGTTGAA

CGTAAGAACAAAATTATTAAGGAGATGATTTATGCCATGTTGTTCAAAGGATAACCGCAAACTTGTGTGTGTGTGTGTGT

GTGGGGGGGGGGGGGGGGTCGTTATCTAATTTTGTTATATCCTGAATAGAGTAACCTTTAAAGGATCAGATAAAAATTCA

TATGATTATGGAAAGATAGACAACCTTTTTATGCATACGTCAAAGTGTGGGGGTGTTTAGATAAGGTTTCTTGAGTACCT

TACAACATATAGATTTTTGGTTGTGTGTTCTGATTTTTCAGACTTTGGTGTGATTTTTCTGACATTGGTATGAAGATTAT

TACGGAATCTAGGGATGCCGAGTTCAAATAAACATGTACCTTATTAGAGATGTGTTGTTGATCCCCTAACTTTATTTTCA

AATAGTCAGAACTTACTTAAAAGGAAGATGGAGTAGAGTTGATTCTAGGATAATTAAAAGGATTATAACTGAGACTTCTT

ATGGACCTGATAACATTCCTACCTTAGTCTGAGGCCCAGACTTGTAAAGAAGCCTTGTCATCTACTGAAACCCCATTCTG

GTAAGAACCTTCATTTAGTAAAATGGACTCAGTCCGTCGGAACCAGACTTGGGAGCTTGCTAGTTTACCTCAAGGGAGTA

AGACCATGGGTTGTAAATGAGTCTTTAATAGGAAACGTAAGGTAGATGGAATTGTGGAAAATATAAGGCTAAGTTGATAG

ATAAAGGCTATAAATTAAAAGAAGGTGTAGATTTCCTTGATTCGTATTCACTTGTGACGAGAATTATTTCTTTTGAGATG

TTAATTGCTATTAATTGTTGTCATAAACAACTTAGAGATACATCAGATGGATGTTAAGATAGCTTTTCTAAAACCGTGAA

TTAGATAAAGAAATTTACATATACCAACCTGAGGACTTTGCAGTGAAAGGTTATGGAGACAAAGTTTGTAAGTTGAACAA

ATCTTTGTATGGTTTTCAAATAAAAACGTAAACAGTGACATGGAAATTTTGATCATGTGATAATGAGTAGTGGATTTAAG

TCAATGAATCTGACAAGTATATTTATAAGTAACTTGTTAAGGATGCCTGTGTGATTGTATGCTTGTATGTTGATGATATG

CTTATAGTTGATACAAACATAGATGTGATTAATTCCACTAAAAACATGAGCCGAATGAGAACGTTGACTTGAAAGACTTA

GGCCCTATTGATGTAATCTTAGGGATGATGATTAGAAGATGATCTAATATTTATAGTATTAGTCGTTCTCATTATGTTGA

ATTTGTTCTTAAGAGATTCAATCAGTATGATTGTAAGCTTGCTTGTACTCCGTACGATTCTTCTTGTAACTCAAGAAAAA

TAAGGGTAATAGAGTATCTTAACTTGAATACTCAAGAGTTGTAGGATGTCTAATGAATTTAATGAACTGTAAGAGTCCAG

ACATTGCCTATATTGTGAGTAAGTTAAGTAGAAATACTTGTAGTCCAAAGCAAGAGCATTGGGATGCACTTAGTAGAGTA

TTACGGTACCTAAAATACTCTATTACCTTTTGTTTGATTTATGAAAGGTATCTTGCTTTCCTTGTGGGACTTTGTGATGC

AAATTGGATAGTTGACTACGAGTGAATATGTTTTCACTCTAGCAGGAGGGTTTGTTTTTGGAAGATTTCCAAACAGACAT

ATATTGCTCAATTAATTATGGAATCTGATAATATTGCGTTAGATAAAGAACGAGAGGGGGCCGAGTGCCTAAGATAATTT

TGAGAAGACATTCCTCTCTGGCATAGGCATGTGCCAGCTATATTTATACATTGTGTTAGCCAAGATATAATATCTAAAGC

TAAAAGTAACTAATCTCAATCGGCGTTATTTCCATTGATTGGATAAAGTCCAAGGAGAATATCACGCAACCTTTGACGAA

AGGTTTATCCAAAGAGATAGTTAGAAATGCATCGAGGGGGATGTGTCTTAAGCTCATAAATTAAACTTTCCATGAAGGAT

ACTCAACCTTGCTGACTGGAGATCCCAAGATCGAGGTTTCGAATGAGACAACTAATTTGTGGTGGGTAAAGGTAAACACT

ATCAGATAATTTTATTCTCTGTCCCTTCCCTATGGTGTAGATGTGATAGTGTGACTGCATGTGAAGGATGACTTTTAAGA

ATTAATGAGTTCTATAGTTTCAATTTAAGATTGAAGTGGGGTGTAGCAGTAACACTCTTTATGGAAATTCACCTATCTGA

ATGAGGAAGTGGGGCCGTTTCCTATGAGAATATGAGCTTTGATTCTCTAAAGCATTTGAGAAACATGATATGTCTAGGGC

CAAATTGGACAAAACAGCACGAGTTTGGCAGCACCCTTGGAGATATCATCTGTGGTCGTTATCGCGAATTAAATCAAATG

CTAGCAGTTCAAGACATAGTTCACTGTATCTAGCAAGTAATTACGGTAATATCTCGCTGAGCAAAGGTTTAAGACCTCAT

GGACACCTCTGCCAAAAATGGTATTTCCTGTGTTTTTTATGTGAATTTTCGTTTTGATGGTTTTGGTATTATTGTAACTT

AGGACCTAAGAGTCACTAAGGGTTCACTAGTTCATGATTTTTCACTTTGGTGAAAGAACCTTTAGTCTCACTTATGAGGA

ACTAAAGATGAAAGCTCTCTATACTATATGATTTTTCGTATGACCCTGGGGTCAACACACTGTTGTGTGAGCGAGATGAC

GTTGGAATGGCTAGTATGAATTCAACATGCACGATTTGTATGTTGTTCAGTCATTTCGGGAGATTGGGTTGTTGATTTAC

CAAAATCCATCCGTGTTTTCGTGTTTTATTGAGTTTTCATTCATGTGGGGGATTGTTGGAAAAAAATTACTAAAAAATGA

TTTTTAAGGTTTTAATAAAATTATAATTTATTGTTTTATTTTATGAATGAAAAACTTATTAGTCCCACATTGTGGAGTTC

CCACTTTTTAGTTGTTTTAAGAGACTATATAAGATTTTTAGTCCCACATCGGGGAGTTCTTCTTCTTAAGTTGTTTTGTT

TCATTATATAAACAAATTCACTACTTTTGTAAACTGTATGGGAAAGGGGTTGCTCTATATTTAGAGGGACCCCCAAGGGA

AAACATATTTAATTTATATTGTTTTCTCAAGCGTTTGGGATTTTCCTTTATAGTTTTTTTCGGAGTTGCCAAGATCAAGT

TGAGCGTCTACTACATATGCTAGTAGTAGGTGTATTGGGGTGTTTTATCCTGGAGATATGCATCATGTGAGGGCTATAGC

ATCATTCTTGAGTGTAGCCGGGCGCTAATGTCTTAAGGGAAATGTGTTGAACACGTGACTCACTCTGTTTTTCAAAGTTT

TGCCTTGTTGCTGTTGTGGAGATACGGGAAGCTCGCTTGTTTCGTCAAATTCATTACTTCCATTACAAATGAGATAAGTA

TCAATAACTTTTGCTAATTTGATTTTTCTTTGTTTTGATTATTGCACCCAACAATCTTAAGACATTAGGATTTTTAATAA

TCATGGATGTTAATTATGGAGTGAAAAACAAAAAACTTATTTTTGGTCAGCTGTAGATTTTCGAATTTATGTCTCAACCT

AGAAGAAATTTCGATGAACCCTATTGACACAACGTAGTGGCATCCTGATAGTTACCCACGTAAAATTTCAGAATTTTTGG

AGTTGTAAAAGTATTTTTTTGATATTTTTCTGGACAGAGAAATGTTCCTGAAAAATTCTGACAGGTAGAAATTTGTTGTT

AACTAAAGTAGTTTTTAAGAGGTAACCATGAGTTTTTTGATATGGTGGTTCAAACGAAGTTTGTAAATCTAGATATTATC

TTCAAAACTCATATTTTATCTTCCGTTTCAGAACTAAGGTTTGTGAGATGTGGTGTATTAGGTGATTGGAAAATTGTCAT

GTCTGTGGTCAGAGTATAGACGAAGATTGTGATATGAAATTGAAAAGTAACATGGATACCAGGCGCATGTGGATGAACAC

AAGTCTTCCAAAGATGACAAAGTCGAGAACATCAAACACTACTCTAAGCGTAGTCTTCATAAGAAAGGTATGTTTCGTAA

TACTGAATCCGATATTACTTTAAATAAGGGTGATTGTTATGTTTGTAAAATTCCTGGCCATACGGAAGTAAAGTTTAGAC

AACATAAAACCTTAATAAGTATAAAGTTAATGCTCATTTAGTTGAAACAAATTAGAACAAGTTCATTGACATGATGTCGG

AAGTTATTTTAATAACTAATGTGAGAGACTGGTGGGTGGACTCTGGAGCCACTAAGTATGTTTGTTGAAACAGAGACATG

TTCACCTCTTATCAGAGGATAAGGTATGTCGAGAAACTCAATATGAGTAAATCATCTGCGACAGAGGTTGCATAAAAGGG

AAAGGTCGAGCATAAGCTCATATCTGTAATATTCTCATACTGAATGAAGTTTTCATGTTCCGAGCAAATGCAAAAATATT

GTATCTAGTTCTCTTGAAGATGGTAAAAGATTGAAGATCTTAATTAAATGTGGAAATCTTGTTATAACTTAGGGCAGTGA

TTTTTTAGGCAACAGTTATAGGACTTAGGTCTATATAAGCTTAAAGGAAAATCTGATGAAGTGAACATAGTTGATTCTTG

TGCTTTCTTTTGTGTGTCTTTGAATGTTTTGCATGGTAGACTTGGAACCGTCAAATTATAAGTCAATGCATAAACTGCCT

AGCATAGGCTCCGTACCCAAATTTAGTTTGGATTTACCAAGATCTATCTGTGTTTTTGTGTTTTATTGAGTTTTCATTCA

TGTGGGGATTGTTGGAAAACAATTACTAAAGAATGATTTTTAAGGTTTTAATAAAATTATAATTTATTGTTTTATTTTAA

GAATGAAAAACATATTAGTCCCACATTGTGGAGTTTCCACTTTTTAGTTGTTTTAAGAGACTATATAAGCTTTTTAGTCC

CACATCGGAGAGTTCCTCTTCTTAAGTTTTTTTATTCCATTATATAAAAAAATTCACTACTTTTGTAAAATCTATGGGAA

AGGGGTTGCTCTATATTTAGAGGGACCCCAAAGAGATAAAATATTTTATATTGTTTTCTCAAGCGTTCGCGATTTTCCTT

TATGGTTTTTTAGGAGTTGCCAAGCTCAAGTTGAGCATCTACTACATATGCTAGTAGTAGGTGTAGTAGGGTGTTTTATC

CCGGAGATATCCTTCCTGTGAGGGCTATAGCATCACTCTTGAGTGTAGCCGGGCACTAATGTCTTAAGTGAAACGTGTTG

AACACGTGACTCACTCTGTTTTTCAAAGTTTTGCCTTGTTGGTGTTGTGGAGATACGGGAAGCTTGCTCGTTTCGTCAAA

TTCATCAATTCCTTTACAAAGGAGCTAAGTATCAATAACTTTTGCTTATTTGATTTTTCTTTGTTTTGATTATTGCACCC

AACAATCTTAAGACATTAGAATTTGTAATAATCATGGATGTTAATTATGGAGTGAAAAACAAAAAATGAATTTTTGGTCA

GCTGTAGAGTTTTGAATTTATGTCTCAACCTAGAAGGAATTTCGATGAACCCTACTGACACAACGTAATGGACATCCTGA

TAGTTACCCACGTAAAATTTCAGAATTTTTGGAGTTCCAAAAGTATTTTTTTAATATTTTACAAAACAGAGAAACGTTCC

TTAAAAATTCTGACGGGCAGAAATTTGTTGTTAACTAAAGTAGTTTTTATGGGGTAACCATCAGTTTTTTGATATGGTGG

ATCAAACGAAGTTTGTAAATCTAGATATTATCTTCAAAACTCATATTTTATCTTCCGTTTCGAAACTAAGGTTTGTGAGG

TGTGGTGTATTAGGTGATTGGGAAATTATTGTGTCCATGGTCAGAGTATAAACGAAGATTGTGATATGAAATTGAAAAGG

AACATGGCTACCAGGCGCATGTGGATGAACACAAGTCTTCCATAGATGACAAAGGAGAGAACATCAAACACTACTCTAAG

CGTAGTCTTCATAAGAAAGGTATGTTTCGTAAAACTAAATCAGGCATTACTTTAATTAAGGGTGATTGTTATGTTTGTAA

AATTCCTGGCCATACGGCAGTAAAGTGTAGACAACACAAAACCTTAATAAGTAGAAATTTAATGCTAATTTAGTTGAAAC

AAATTAGAATGAGTTCATTGACATGATGTCGGAAGTTATTTTAAAAACCAATGTGAGAGACTGGTGGTTGGACTCTGGAG

CCACTAAGTATGTTTGTTGAAACAGAGACATGTTCACCTCTTATCAGAGGATAAGGGATGTCGAGAAACTCAATATGAGT

AACTCATCTACGATAGAGGTTGCATAAAAGGGAAAGGTCGAGCATAAACTCATATTTGTAATATTCTCACACTGAATGAA

GTTTCCATGTTCCGAGCAAATGCAAAAATCTTGTATCTAGTTCTCTTGAAGATTATAAAAGATTGAAGATCTTAATTAAA

TGTGGAAAACTTTTTATAACTAAGGGCATTGATTTTTTAGGCAACAGTTATAGGACTTATGGTCTATATAAGCTTAAAGG

AAAATCTGATGAAGTGAACATAGTTGACTCTTGTGCTTTCTTTTGTGTGTCTTTGAATGTTTTGCATGGTAGACTTGGAA

CCGTCAAATTATAAGTCAATGCATAAACTGTCTTGCATAGGCTGCGTACCCAAATTTAGTTTGTATTTACCAAGATCTAT

CTGTGTATTCGTGTTTTATTGAGTTTTCATTCATGTGGGGGATTGTTGGAAAACAATTACTAAAAAATGATTTTTAAGGT

TTTAATAAAATTATAATTTATTGGTTTATTTTATGAATGAAAAACCTAATAGTCTTACATTGTGGAGTTTCCACTTTTTA

GTTGTTTTAAGAGACTATATAAGCCCACATCGGGGAGTTCATCTTTTTAAGTTGTTTTCTTCCATTATATAAACAAATTC

ACTACTTTTGTAAAATCTATGGGAAAGGGGTTGCTCTATATTTAGAGGGACTCCGAAGGGAAAAAATATTTTATATTGTT

GTCGCAAGAGTTCGCGATTTTCCTTTATGGTTTTTTCGGAGTTGTCAATCTCAAGTTGAGCATCTACTACATATGCTAGT

GGTAGGTGTAGTATGGTGTTTTATCATGGATATATCCGTCCCGTGAGGGCTATAACATCACTCCTGAGTGTAGCCGGGCG

CTAATGTCTTAAGGGAAACGTGTTGAACACGTGACTCACTCTGTTTTTCAAAGTTTTGCCTTGTTGCTGTTGTGGATATA

TGGGAAGCTCGCTCGTTTCGTCAAATTCATCACTTCCATTACAAAGGAGCTAAGTATCAATAACTTTTGCTTATTTGATT

TTTCTTTGTTTTGATTATTGCACCCAACATATAAGCCCTGCTCAACAATCTTTATCAAATACTTCTACTCACCGAGAATC

AAGAACACCGTTGTTATTTGATAAAGGAATAAGAACATATGGAAGACTAAGTTCTTTACAAACCTTAATATATATAACTC

ACTATTTTTTCCACTGCTAACTCTCTCAATTTCTCTATAAAAGGTCTGTCCCTCATTTACAAAAATCTCAAGGACTTTAT

ATCAGCATTTTTCATAGTGGAAGACATCCAATTCTCACATTCAGGACCACATAGTGGAAGACATCCAATTCCCACATTCA

GGACCACCATTGACAGATTATGTTCTTAATTTTTTTTTACTTGCCATACTTTTCAGAAGTTGTCTCCTATTATCGCACTG

TCATTCGGTTCTTCTCTATTCCTTCACGAGTTTGGTCTTCGCACTGCGATCATCGCATAACTGCTATTCGCCTTCTTCCT

TGTTAAAATTATGTCTTCATGCTCGGGGATTATTCTTACCAAGGCGAAGTAATATTTATAAGTATGCTTTGCCTTGTGCG

ATATTTCGCCCCTACAAGAATTGAAGAGTTTCCAGCTGATCTAGAGGTGGTTGAGAAAGCTCAAGACACTCGCTTTGGGG

CTGTGGAGTAAGCTGAAAAGAAATTAAAGATTCATCTCAAGGTTTTGAAGGAAGTTGTGAAGATCAAAGGAACGAAGACA

AAAATTTAACAGGAGAAGATTTGGATAACAAGAAATATAACGTTATACCTAATAGGATGTTGGAGTCTTTCTCTGAAAAT

GTGGAGGCTCTTCATAAGTGTTTTGCATTGTTGGGAGCTCCTAGAGGGGGTCGAAAGAGGAAAGATTGAGTATGTTCTAT

GAGTTAGCTCACAACTACCTACAAAGAACGAGAAAGGAGACTAAGAAAGACAGGGAGCTTGAAAAGCTGGGTCCGAAGGC

TGAACTGATCAAGGAGTTACTTGGCAGAGGACATTGGAAATCAGCAAAGAATAAATTTTGATGTAACCAGTGGTTCTTTG

GTTTTTTGGCGCGTTGGTTAGTTTACCTTTTTATTTGTTGCTTGGCTTTTCTTCTTGTATGATCCTAGCTCTAGTTAGAA

TCTTGTTTGCTCTTTCATGAGTTTTTTTTTTTTTTTTTTTTTTTTTCCTTTTTCTTTTCTTTGTTCATTATCTTTCAGTT

CAATGAATTTTATCTTTCAAGTCAAAAAAGAAAAGAAAAAAGATTTGTATCTATACGTGAGATGTATCGTGACCGAATGT

TACCTGCACACCCGTAACCACACGTACGTAGTATGATATGTACGTGGGGTGCACCGTGACCTAATATCACCTACACGCCC

GTGACCACACGTACATAGTTGCACGTGTTAGGCCGAATGATTGGCCCTACACATGAGGGGGATTGTGAATTGACAATAGT

CCCACATCACTAGATTCCACATAATTAACTTAAATATAATATTAAAGGGTTGTTCCACTAATTGTTAATTGGTTTTTAGT

TGGATGACCGCATACTTTAATAGGATTAGATATGGATCATTCTAAGTCGGGGTTGTCTCACTACTAATAATATTATAATA

CTTTCCTATATAGTTCGAACAGTCTAGGTCGAAGATGTCTCACTGCTAATAATATATAATACTTGCCTATATAGTTCAAT

GCAGAGGAACGAGAGAGAAAGGAAACCTTGAGACTCTCTTTGGCATCGTCACCATTTATTTGGAATACTTCTTTTGCCAA

ACCAAAATGCTTGAACTCCAACCAATACAGACATAACAATTTCAATGAACTAGAACCTAAATACTACTGTATTTCGAAAA

TCTACTATCTCCATACATACCTGTATTTCGCATCCGGCCAGTAACTAGTTCATCTTAGGATCCCCTTGTACATGAACAAA

ATGCAATCCGGATTGCAGGAATCACTTCCCAAAAGGAACAATATCTACAATTAAGATCACGATAGACAATTTTATCATAA

AATAACAAATGCATTTATACATATTCAGTTATCCCACTCATCAATGGCTCCTGAAGTCTATTTTATATACATTTTTACAT

TTTTTTTAAATTTAAAATCTTAATTAATTTCTAAGACAACTTTAAAAATCGAATTTTAATAATTCAAATTACAGAGTTTT

TGGTTGGCATCCGAGCAACAAGCTGAGCACCAACCTCTTTCTGAATAACAACACCTAGTCTATGAAAAATAAAACTACTC

GAACCACTGCCAGCGTCATTGCTAATTAAACAATTCTTCAGGAATTTGAAAAAACAAATGAAATCTTCACTAAGCTCCCC

TAAAGTAGTAAAAGCTAAGACGCCCAACCGTTATCACGTTTTGGCGGTATGTGCCTCACTTTTTATCTCAATCCAAACAA

TCTAACATAAACGCGTATATTGGCCAAATTATTTAACACAAAATTGGTTAAAAATAATGAGAATCTACTCACCGTTAAGG

AGAATACAGGGAGTAATGAGGATCAACTGACTGTTGAGGAAAATACTCCAGATCCATAGAAGAAGGACACCAATTAATCC

TAGAATTTCTCCACAGGTTTATGTTTAGAAAGATAGTTATATTGATTCCATATATATTGTACATACTTATCTTTGGAAAG

TTATTCACATCCTCTATAAAATGCTATGTAATGCTTGATGTAATTCAAGAGAGTGTAATAGTATTCTTTGTTATTCAAAC

TTCAATTCTATACATGGTATCAGAGCACTCTATTTGCGACGAGTCATTTGACCGCGCCTCTACTCCGCGTCACCCGATTT

ATTATTGTCCATGTGTTAGACCCTACGAGGCTACATGTGAGGGGGCGTCTTGAGATTATTAGTCCCACATTGTTGGGTTA

TATAAGATCCTGCAGTTTGTAATCTCTTAGGGCCTTTCCACTCATTGCCAATTGGTTTTCAGTTGGATGCCCATATTATA

ACATGGTATCAGAGCTATATTGCTCAAACGAGTTAAGACAATCCCGTGTAACAGAAACATTACATCAAAGATATGACGCC

ACAAAACAAAGAGTCATCTTTTTCTTTTTCCGCTGTAACTCTGCCGCACCTAATAAACATTGTGTCATTGAAACTCAATT

CTAAGAACTATTTATTATGGAAGATGCAGAACCAATCGTTGATCCAGGGTTTTAAACTCATATATTATATCAACAAGGAT

GCTCCTCCGAAGATGATTACAGAGGAAGGTGTTGAAAATCCAAAACAAGCCTTTGAAGAATGGCAAGAAACTGATGTGTT

ATTGAAAAGCTGGATTGCTGGTACACTTTCAGAAGATGCCTTAAGTCATGTCATTGGTCATACTACCTCAAGAGAGGTAT

GGGTAGCCCTTGAAGATGCATACCAATATGCCAACAAAGATCGCGAACTTCAACTTAAACAACAATTGCATGCTCTTAAG

AAAGGTAACAACAATATGACTGATTTTCTAAAGAGTTTTAAGACTATTTGTGATGATCTTGCAGCAATTAAAAAAACATG

TCCCTGATGATGATAAGGTTCTGCAACTTTCACAAGCTCTTGGAGATAAATATGATACTTTTGTGACAGCAATGTTATCA

AAACCTCCATTTCCCACTTTTAGTCAGTTTGTCACAGCCTGGAATGGATATGAATTGAGGCATCCAAATGCTACTGAAAA

TAATGATCAAGACGTTGATCATAATTTAGCATTTTCCGCTCAGCGATCTGGACGTGGTAAAACTTACCTTGGAGGACGAG

GTAGTAATAATCATCATCATTTTAGTGGACGCGGAAGAGGGTACCCACCATATAATGGTTCAAAATCATCTCCAAGAAGC

CCAACCTATCAAGGAAATTCTTCCGACGGTAATAAATTCCGCTCACAGTATAATCCCTCTCGTGATTCTAACAACCGTGG

GTTTCATGATTCAGAAATATGTCAACTGTGTTGTCGTTTTGAACATGATGCAATACGTTGTTGGTACCGTTTTGATTACT

CTTTTGAAAATGACGCTAATATGAATGCACTACCTGCTGCAACTAATCCATCTTAATCTGATCCACGCTGGTACGCTAAT

TCAAGTGCGATATCTCATATGATATCTGATACTGGTAATCTTCAATCTTAAATGCCATATACCAGTAATGATAAAATATT

TGTTGGAAATGGATCTGGACTATATATCTCTCATATTGGATCAAAATATATAGCATATTTTTTAGGACATAACATTAGAT

TGAATAATGTTTTAGTAGTGCCTAAGATAACCAAAAATTTACTATCGATTAGACAATTTACTCGTGATTATTTATGCTCT

TTTGACTTTTCTGATTCTGATTTCAAGATAAAGGACCGACGAACGGGGGCAATATTAGCAACCAGAAATAGATATGGTAG

TTTATATGCCTTAGATACAGGTGGAGGAAGTGATGCAGCTCTCTATGCAATAAAAAGTGGAACAGCAGATTGTGATTTAT

GGAACCAAAGATTAGGCCATCCACATTCCAAATGCTTAAACAAATTAGATTCTAATGATTTAATCTCTGTTTCATGGAAG

AAACATTTTACTCTTTGTTCTAGCTTCCAAATGGGAAAACGTTATCGTCTTTCATTTTCTCATTTGAACAACAGATCTGA

AACTCCGCTGCATAAAATTATCATATCCCTCCTTAGTCATATATTCTGGGTGCCCTTATCCTTTCTATGACTCTATTTAG

TTGAGTTCATGTTGAGTACCCTTATCTTTTCTATGAATGTATTTAGTTGAGTTCCTGTTGAGAGTATTTGGTACGTGTGA

GCCAAAACTATCTTTTCCTTTTATTGGTTTGTATCCTCCATGGAGGGGGTGTGAAGATTTTTATCCTGAAAATTCATTTT

TCGCCTCTCTGAGGTTCTTTATCAATATCACAGTGTAGATCCTGTGACTTGATCCCCAATTAGAACTAATTGTAGTGTAG

ATCACTACAAGTGGTATCAGATTAGGTCTATCCTTGGCAATTATCATGGCAGATGGTACTAGGTTGCAGAAATTGGATGA

ATCTCAGAAGACTATGGGTTTGGAATTGAATGAAATCCGTGAAGAGATGCAGTCGTTTCATCAGGAGATGCAATCGTTTC

GATCTGTTAACACCGATGTACAATCATTGAAAACTACTCTTACAGATCAAGCTAACGTTCTTCAATCGATTCTCCATAAA

CTGAAAATCTCTCATCGAAATTTCCAAGATGACAATCTTAATCATCCTTGGGAAGATGGTGAACCTGTTACCCCCAACAC

CTTTACGGGTCATCAACGAATTCCACAACAACGTCTACTTCATCATCATCAGTTGGGTGAAGGTCAGTTTTCTAGTTATA

GACCTAAAATCGATTTTCCTCGTTTTAATGGCACGAATCCTAGAGCTTGGGTCTGTAAATGCAAGAAATAGTTCTTCCTA

CACCAAATGAATGATGAAGAGAATGTGAATCTAGCAACCCTGTACTTGGAAGGAAAAGCATATGTTTGGTTCCAGGATTA

CCAAATTGGGAAAGGTATTTTATATTGGGAAGATTTCATTATTGATGTTTGCACTAGGTTTCAATAAATAGGCCATGATA

ATGTTGTAGGTGAGTTCAATAAATTAGAACAAACTCATTCCGTTTTGGACTATCAAGAAAAATTTGAAGAACTCAAAGCC

TTAATGTTAGAAAAAAATCCTTCATTAACTAAAGATTACTTCATTCGTAGTTTTATCAGTGGTCTAAAGGAGGATATCAA

ACTAGTGGTAGAAATTTTTTCCCTTAATACCCTACAACAAACTATCTTTTTAGCCAGAAGGCAAGAATGTATCCTAGAGA

AAACATCTAAGGCTCCACCCAAATACTTAAGAAAGTTTTCTTCTGGTAGCAAACCTTTCAACATTCCTATAAAACCTCCT

AATCTCACTCCTCTAGTCACTAGCCCCATTGTTACTAGTCCAACAACAAGTCCTAAATTGCAGTCTGTGAAGAAACTCAC

TTATGCTGAAATGCGTGCAAAGCGAGAGAAGGGCCTTTTTTATGACTGTGATGAGACTTTTTCTAGAGGACATAAGTGAA

ACAAACAACAATTATATCTGATTGTAGCTGATGATGAAGATACCTCACAGCCTGCAGAAACAAGTGACAAATCAACTCCA

ACTATTGAAGAATTACAAGAGGACATGGATATATCTGTACATGCCTTGGATGGATCCATTTCACACACAAACATTAAAAT

AAAGGGCTTAGTCAAGTAACAGGATCTCACCATCTTAATTGATAGTGGAAACACTCATAGTTTCTTGGATACCAAAGCTG

CAAGTAAAAGTGGATGTATGATCAGTTCTACACAACCCTTGCAAGTGGCAGTAGTTGATGGAAACAAATTGTTAAGTCAA

GCACAATGTTCTGCATTCACTTGGAGTATGCAAGGTCATCAATTTTCTCATGACTTGCACATTCTAGAACTAGGGGGTAT

GATATGGTACTTGGTGTAGACTGGATGCGAGACCTCATTTCTATGATCTTCGACTTCAAGAAATTACTTATTAGGTTTCA

ACACCAAGGTCAAGAAATTGAACTGGTAGGTCTACCAGAAGCAGCCACCTTATCTCAAATCTCAGGTTCCCTGCTGAAGC

GACTACTACAAAAAGGTACCCAAGGTCTCATTGGTTAATTTTTTTTTCTTCAAGCCAACACTACCCCTGAAGTCCTGGTA

ACACCCATTGCTGAAGTTCTTGAACAATTTCCTACCATTTTTCAGACACCTACCACACTACCTCCTGAAAGAACTCATGA

CCACCACATCAATATGGTTCCCAATGCTACACCTCCAAATCTTCGCCCTTACAGAGTCCCATATGTTCAAAAGGAGGTGA

TTGAGAAACTTGTGCAATAAATGCTTGAAGCATAAGTGATACAGCCCAGCCATAGCCCTTTTGCATCACCAGTTATATTA

GTCAAGAAGAAATATGGAGGATGGCGGTTCTGTGTCGACTATCGCAAGATGAATGCCATCACTATTAAAGATAAGTACCC

TATACCAGTTATAGAGGAATTACTAGCTGAATTAGCAGGAGCTACTATATTCACCAAACTTGATCTTCTAGCTGGTTATC

ATCAGATTAGGGTTTACTTAGATGACATAGCTAAAACTACTTTTCGTACTCACCACGGTCATTTCGAATTTCGAGTGATG

CCATTTGGGCTTACAAATGGTCCTGCTACCTTTCAGTCATTGATGAATGACATTTTTCATCCATACCTCAGAAAATTCAT

CTTAGTCTTTTTCGATGATATACTCATTTATAGTAAAGACATGGCTACTCACTGTAAACATCTGCATATTGTTTTGTCTA

TTTTGCACCAACACTCATTGTTTGTTAAGCGAAGTAAATGTTCTTTCGGTGAACCACAAATAGAGTATCTAGGCCACATA

ATCAGTAGAGAGGGTGTAGCAGCTGACCATGCGAAAATCGAATTTATGTTAAATTGGCCTCGACCAACTACTATTAAAGG

GTTACGAGGATTTCTGGGCTTGACAGGATACAACCGCAGATTCATTAAGGGTTATGGTTTAATTTGTAGACCATTTACAG

AGCTACTTAAAAAAGGAAACTTTGCTTGGAATGATACTGCAGAGTTAGCCTTTGAGCAACTGAAGCGAGTAGTTACTTCT

ACACCTATTCTCATTCTGCCATACTTCACTAAGGAGTTTGAACTTGAAATTGATGCTAGTAATATAGGAGTGGGAGCCGT

ATTAATGCATAGTGGTAGACCAATTGCCTTCTTCAGCAAAGGTATGGGAATCCAATATCAAGCCTTATCCACTTACGAAA

AAGAGTTGCTTGCAATTGTCTTATCTGTCACCAAATGGAGATCATATTTATTAGGGTCTCATTTTACAATATTTACGGAT

CATCAAAGCCTCAGATTTTTTATGGATCAGCAACTGCATACTGTTCTTCAACAGAAATGGATGACTAAACTATTTGGATA

TGACTACACTCTTAAATACAAGAAGGGTTCATAAAATAGGGTGGAAGATGCTCTTTCTAGAGCCAATTGTCATTCTCTCT

CAGCAGCAACACCAAAATGGCTTGAACACATTCTTTTTAGCTATAAAGATGACCCCATAGCAGTTTATTGGATTCCTAAA

CTGACTATCTCTTCCATGAATGATACATAATATACTCTTACTAATGGCATTCTGAAGTTTCAGGGTAGAATTTACATTGG

TAAAGCTGGGACTGCTAGGCAACAAATTCTGGCAACTGTCCACTCCTCAGCCATTGGAGGCCACTCAAGCATGCAAGCCA

GCTATCAACGTGCCAAGACTTACTTTCATTGGCATGGAATGAAGCGTGATTTAATACAATTTTTCCAACACTGTGATGTC

TGCCAACAACATAAAGTGGAACACAAGCTACCTGCGGGTCTTCTTCAGCCTCTCCCCGTTCCTGACCAAGCCTGGAAGCA

CATCAGCATGGATTTTATCTTTGGGTTTCCGAAATCCGAAGGTCGATATGCCATTATGGTAATTGTTGATAGTTTCACTA

AGTAATCTCATTTCATTGCTTTAAGTCATCCCTACACAACACAAACTGTATCCAAACTTTTCATCAACACAGTCCTTAAT

CTCCATGGTACTCCTGAAACCATTGTATCAGATAGAGATGCCATTTTTCTTAGTGGGTTTTGGCAAGCCTTATTTCAGGA

TGTTGGTACTTCTCTTCACATGAGCACATCATATCATCCACAAACAGACGGCCAAACGGAGCGTGTCAATGCTTTTCTAT

AGTCTTATTTACGTTGTATATGTAGCTACAGGCCAGTTAAGTGGGTTTCTTGGATTTCTTTGGCAGAGTGGTGTTTCAAC

ACCTCTTACCACACTAGTTTGAAACTCACTCCTTTTCAAGCTTTGAACGGCTACTCTCCTCTACAGTTTGGACTACATCA

GACTGGTGGCATTCGTCACTCTATAGTCGATGATTATCTACACCAAAGGGGAATAATGACAACATGTATTGCGCCAGAAT

CTTCTGCACTCTCAACACCGAATTAAGCAACAAGATGACAAGAGTAGAATGGAGAGAGTTTTTGAAGTTGGGGATTTGAT

GTATTTACTCCTTCAGCCGTATCGTCAAACTTCTGTCCTACTACGCAAGAACCTCAAGCTGGCAGCCAAGTTTTTTGGAC

CATATAAAGTTTTAGAGAAAATTGGAGCGGTGGATTATCGACTTCAACTACCTGCTTCTTCTCGAATCCATCCCATTTTT

CACGTTTCACTGCTAAAGAAGAAGCTTGGACAAGGATTAATTCCACACACATCATTACCCAACATTGACAAGGAAGGGTT

GTTTAAGGTAAAGCCTGTCAAGTTTATCAACTATCGCACCATTTTCAAAGGCCGTAAACTTATTCCAAAGGTGAAAATAC

AATGGTCTCATACATCTGAGGGTGACAGTTCCTGGGAATACGTAGCTGCTATCAAATCCTCTTTTCCCAAGTTCATCCTT

AAGGACAACGATGCTCTGTAGGAGCCGGCATTTGTCATATCCCTCCTTAGTCATATAGTCTGAGTGCCCTTATCCTTTCT

ATGACTATATTTAGTTGAGTTCATGTTGAGTACCCTTATCTTTTCTATGACTGTATTTAGTTGAGTTCATGTTGCGAGTA

TTTGGCGCGTGTGAGCCAAAACTATCTTTTCCTTTTATTGATCTGTATCCTCCATGGAGGGGGTGTGAAGATTTTTATCT

TGAAAATTCATTTTTCGCCTCTCTGAGGTTCTTTCTCAATATCACAGTGTAGATCCTGTGAATTGATCCCTAATTAGAAC

TAATTATAGTGTAGATCACTACAAAAATTCATTGTGATTTACGGGTCCGGCCCCTGTTAAATCATATCAAAATTTTAGAT

ATTATGTTGTTTTTGTCGATGATTTTTCTCGCTATACTTGGGTTTATCCTCTGAAAAAGAAATCAGATTTTTATTTTACT

TTCATTACATTTCATAAAATGGTTGAAAAACAATTTGATAATCGAGTTTAAAAATTTCAATGTGATGGAGGGGGAGAGTT

CAGTTATATTCATTTCTTTGCTTATTTACAAGATAATGGCATTATTCGTTTTGTTTCGTGTCCTCACACCCCTAAACAAA

ATGGTGTGGCGGAATGAAAACACCGACACATTGTTGAGACTGGTCTTACACTAATGTTTCATGCAAATGTTCCTCTCCAA

TATTGTACGGATGCCTTTCTGATGGCTACGTACCTTATTAACCGTCTTCCAAATTCGAATCTTAAAATGGAGACACCATT

TTTCACATTATATAATACTAACCTCAGTTATGAGATGCTTAAAGTTTTTGGATGCATGTGTTATCCTTATCTCCGGGACT

ATGCCAAAAAAAAGTTTGGCCCTCGATCATATCTGTGTATGTTTTTAGGTTATAGTCTCATCCACAAGGGCTAGGTGTTT

ACAACCATCAACTAAACGCGTGTTTATTTCCTGACATGTCGTCTTCGATGAATGTGTTTTTCCTTTTAAATAAACCATTT

GCAAAAGTTCTCCATCTAGTAACTTGAGAATTACATCCTATGATGAGTGGACTACAACCACAGAACCTTTAAAATCATCT

TTATCTAGCTTAGAAATTGAACCTTCATTGGATTCACCAACTCCTACTGTTATTCCTATTGCTCATACAACTCCACCTCC

GGCTACAAGGATTACGGACGTACCCAACATCCACCACATCTAAATATCCAATTCCACTTGATCCTATAACACCAAACCAA

AGTGAAACTGATAGTACGACATTAGTTCCACTTTCTCAAACTCTTGTCTTTACTGATACCACGTCACCGGCACCTATAGA

ACCTAATCTTAATAAAATTAATTACATTTCATATCTACCCTCTCAAACTTCAGAAATAATCAACATCCCTATGGGTTCAT

CATCCCAAAAAAATCTCCCAACACAGCCTTTGCTCAACCATTGTCAACCCCAACCATCTACGGTTTCTAATGCTCCAGTG

CATTATATGAGAACCAGATTTAAGGATGGAATATTCAAACCAAACACCAAGAACAAGAACTTCTTAGCCATTTCATCTAC

TTCCCACGCTGATCTTCCATTATTGCAGTCCACTGATCCTAGTGGGCCCAAAACCATTAAAACTTCTCTTAACAATCCTC

ATTGGGTTTCTCCATGCTTGATGAACTACGAGATCTTCTTGAAAATGGTAATTGGGTACTAGTTCCATGGAAATCTAGCT

ACAATGTTGCTGGTTCTCGTTGGGTTTTAAAAACCAAACTCAAATCTGATGGTTCGATTGAACGATTTAAGGCTCGACTC

GTTGCTAAAGGTTATAATCAAGTTGAAGGAATCGACTTCGATGAGACATTTAGTCCTGTTGTGCAACACGCTACTCTGCG

GATAGTCCTTTCAGTTGTTCTTGTACGTGGTTGGTCACTTGCCAACTGGATGTAAAGAATGCATTTCTACATGGCCATCT

TAAAGAATCCGTCTATATGGAACAACCACCTGGATTTGTCGAACCTAACAAGCCGGATCATGTCTTCTACCTTTGTAAGG

CGATTTATGGACTTAAACAATCCCCGAGGGCCTGGTTTGAACGTTTCAGTAACTATCTTATTTCTATTGGTTTTAATTAT

TCACGAGTGGATTCTTCACTTTTTACTTGGCATTGTGATACATGTATTATATACTTGATTCTGTACGAGACGACATCATT

GTCACGGGATCTTGCGATCAACTTCTGCAACGTCTTATCACCAATCTCAGTTCCGAGTTCGCAATGAAAGATTTAGGTCC

TTTACACTACTTCTTGGGTATTGAGGTTATCAAATTTTATGGTGGGTTGTTTTTGTCACAACAGAAGTATATGAAGGATA

TTCTTTGTAGGACATCCATGACAAAGAGAAAACCTATTGCCACACCACTCTGTCTCAAACATGATCTTCACCAAGACAGT

AGCACACTTGTTGCTACTCAGTACCGTCAAATTGTTGGCTCATTACAATATCTTACTCTTACTAGGCCGGACATTTCTCA

TGCGGTTAATCTCGTTAGCCAGTTCATGCATCGACCCACTATTATAATCTCACAGTAGTAAAAAGAATCTTAAGATACCT

GAATGGTACCTTGGATTATGGGATTCGACTCCTCTCTAAGTCCACACTTAACCTTTATGCCTTATGTGATGCAGATTGGG

CTGGTTGCTTGGAAACACGACGCTCCACTACTGGTTATAGTATCTGTATTGGTGCCAATTGCATCTCTTGGTCTTCAAAG

AAACAACCAACCGTGTCTCGCTCGAGTACAGAAGCTGAATACAAATCCATGGCCGTCAGTACTGCTGATTTCACTTGGGT

AACCTATCTGCTACGAGAACTTGGTATTAGTCTTGACAAACCTCCTGTGCTATTCTACGACAACATTAGTGCTCTTTCTC

TAGCTAAAAATCCCAAGTTTCATGCTCGGACGAAGCACATTGAGTTGGATTATCATTTTATACGGGAGAAAGTTGTTAAT

GGTTCGCTTATTACTCGATATGTGTCCACCACCTCTCAACTAGCTGACTTGCTAACTAAGCCATTGCCGAAATATACATT

TCAGTCTTTACGCAACAAGCTCGGTGTACTGTCAATACCCACTCGCTTGAGGAGGAGTAATGAGAATCTACTGACCGTTG

AGGAGAATACAGGGAGTAATGAGGATCAACTGACTGTTGAGGAAAATACTCCAGATTCGTAGAAGAAGGACACCAATTAA

TCCTAGAATTTCTCCACAAGTTTATGTTTAGAAAGATACTTTATTGATTCCATATATACTGTACATACTTATCTTTGAAA

AGATATTCACATCCTCTATAAAATTCTATGTAATGTTCGATGTAATTCAAGAGAGTGTAATAGTATTCTTTGTTAAGCAA

ACTTCAATTCTCTACAAAAAAAGACCAAAATTAACAATTCCTGGATGAAAATGACATTTAGATTTTGATACTGTTTAAAT

GGACAAAAATGTAAAAGTAGCCAGGATGTAAATAGTTTCATCCTACCCATTTTCATATATTTTTTCTTATTTTTAATTTA

CATCAGGATGCATCCAGTTTCATCATTGCTATTTTTTAAGTTTAAGTCAGGATGAATCCAGTTTCATCTTTGCTATTTTT

TTGGTGTCGTTTCACCTATACTAATTTTTACTCGTCCATCTGAACAATGTTTTAAAAATATTTGGACAAATAACCCATTT

TCCGATTATTTAACATTTAAGAGAAAATCTTGGGAAACATTTGTTCATACAGTAATTTATGATGCTTCTTGATCCTCTAA

TCTTTTGGGATCGCTTTCCACTTTCTCGATTTTCTCATGTGTTTCATTATCACCCACTTTATTCATCATCTTGTATTCTC

CGTAGAAATAAGAAGAGAAACCCCAAAGACATAATGCCAATGCCATTCCTTTTAATCCCGTAAACCTCTCATGGTAAAAT

ATGACGGCTGCAACTTCTGTGAATGGAACGAGGACTGATACGTATATACCGTTGAAAAGAGCATTTCTGTACAAAATTAG

GCCTACGAGTCCAATTCCTGATAATTGCCATGTCAATGCAGTCAGGCATATTATCATATAATACTTGCCTTTCCCAAGCT

CGAAGTTTTCTGCTTCTCTGGGTATAGCCTGTGAGACAAATCAAATTTCATATTTATTTGACTCCTAAGAAAAACAAATT

TGATACAAACTTCAACAGAGTTTTATTTATGCGGATGAATTAAAAAAAAAAAAAAAAGGCTGTAAGCCCCCTATAAAAGT

TAGCATACCTGAAAATCTTTATTTATTAGCATCCCAATGACGTTCAGGCAGGTTCCAAACAGGCCAAGAGTGATTTGAAA

CTGCAACAAAGTAGAATAAGTAATATTTCTGGTTGCTTTAAAGAAAGCAAGCTCAATGAAAGGCGTCCCGAGGCCAGTTA

GAGCAGCAGCAGCCAGCGTCATAAGAAACCCTATCAAGTACTGAGCTTTTGATACCCCAATTGGTCTATCACCATTGGTA

TTAATACCCAATAAGATCGATCCTAGTGTCATCACAATCACTGCATTTATAATAAATGCTGTAAATTTTTGTTTCACAAT

CAACCATGCAAAAAATGCTGTGAAACACAGCTGAGTTGCAAAGAGAATTGTCGCAGTTGATACTGGAAGATAAGATAAAC

CCAATGCGTACATGAAATTAGAAACACCAAACACAACTCCTATTACGGCACTCCAAAGAAACAACTTTGGTTCCATAAAA

AAGGAAGAACTTTGATCGTTGTTTTGTGTTGGCGAATTCGATTTAATATAGAGAAAACTGAGAGGAAATATGAGAACTGG

GAAACCAGCAGTTTGCAGAAAACTGCTAAGCCATTTACGACTACCACCATGAAGATAGTACAGTCTCATTAAGAGAGGAC

CACCTATAACACCACAGAAAACTAAGGCACAGTTAATAATGATTAACGGCCAGTTTCTGGCTTTGATTGGTTGCTCAGTA

TGGGTTCCTGAATTGCCATTCTGATTTGGTCCCAGGATATCTAGTGTTTCAATGATCATCGTGGAAGTAGGCACTTGAAA

TGATGATCGGTTAGAGAGAAATTGAGAGATTTAAAGTGAAAGTTGTGAAGTAGTTCCAGGTGAAGGTGTTTCTTATATAT

TGCATGATGACCAACAGAGTCGTGCTAAGCTTTGATGCTCCTTGCATCGCTGTTGTGTAAAAATCTTCGTGAATCTAACA

GTCTAATATATCTAGATAGAGAGAGTTACAATCGAGGATATATATGAAATGTATTGCAATGTAAAGGTGTTCTCTTCTGC

TTGATATCAAAGTTTATGGCGACTGGTAGCTCATCTATCATGATTGAGAAAAGTGGAAAACAACCCCCACTCCCCCAGTT

TTTGTACATATTTCTCCCTGTTTCTTGTTTGAAATTTAAAAAGAACACGCACACAAAAAACAAAATAAAATAAAAAAAGG

TATTTGTCCACATGATACTTTCTATCGAAGCATTAAAAAAATGTTTTGATTCCCTCCCACTTTCAGTCCTGATTCTGCCT

TTGGATCGTTGTTATGGAACTATTCCATGAAGTTATATGTTTGCGTTTGACTACGATAATAAGATTCTACCCTATATTGA

CCTAGACTAGTTGTTCTGAACAATTACAAGCTAGGGCCAATCAGTACACCAAGTATTCAAATTTTGCAAATGAGAGGCAG

TTTCGGCCGCCCCATCAATTAAGTTAGTAAAAACATTATACACAAAATTGGTTAAAAAAACAAAAATCAATAATTTTTGA

GTAAAAAAGACTTGTAAAATTTGATATTATTTAAATTGACGAGAATGTAAAAATATCCAGGATGTAAACAATATCATCCT

ACCAATTTTCAAATACTTTTTTTATTTTTAATTTACACCAGGATGCATCAAGTTTCATCCTCGCTCTTTTTTAAGTTCAA

GACAAGATGAATCCAGTTTTGTTTTTGCTATTTTTTGGTGTTCATTTCACCCATACTAATTTTTACTCGTCTATTAGAAT

CATATTTTAAAAATATTTGGACAAATGACCAATTTTCCGAACATTATACTATTCCTTTCTGCACAAAAAGAGTTATTATC

ATTTTTTTTCATTTTAATAAAAATAGGTAAAAGAAGATGCAGAATTTGAAATAACTTACAAGCGGAGCTAAATTAAAGCG

CACACACCTGTGTAAATTCATGGTTCATTTTTGAGAGAAAATCAAGATGAGTAAAAAAGAAAAACCTATTCTTGTATTGA

AAAAAAGGGAGAATGGGTATGCCCAAAAATATCTAAAAACAATCAACAATACATGAATAAAGACAGGAAAACACACGCTC

CAAGGAATATCTGAATAAGCACAAAAATTTGCGTCTCTAAGAGCAAGGGCTATGGAGTGAGGATCCTCATTCAATATGAA

TGAGCCTCACTCGTTTTGGTCTAATTTGATGCTGCTATGGGCAGTGACTGAGTCCACTTTCCCCCAAAATCATTCATACA

CACATTGAGACGAGTGAGTGTTGGAAAGTCTTCACTAGGCGGCTGAAAATTAGCCGTTTGGGAAAAAAAACACAAAGAGC

GGCTGAAAATATGCCTCTTACAAATATTTTCAGAGTTTTATTAAATTAAAACTCTTTAAGATGGGCCCTTTACAAGTTTT

AATATTAAAATAACCTAAAAAAAAAGGGGGAAACAACTAATGTTTAGCCGTTTGTGTTTTTTTAATAAATAAATAAAAAG

AAACAGCTGAAAATCAGCCGCTTAGACCCATTAAATTTGTTCCCAATGGTCACAAAGTCCCTCTTTCACCTAGTCTCTCA

TTGATTTGGGTAGTGACTGAGTGTTAAAAACACTCATTCCATAGCCCTTGCTCAAAGTGAAAAATCTTTAACCTCCTAAA

CCGATTTTCAAATCCCAATATTTCCCATTCTTTTGAATCGTCAATACTTAGTACGAGTGGTAACAACTAACAACTAGTCG

TATGAAAGCTAACACTGAGCATGACAGGCCCTTTCTAGCTTGAACTAACATTTAAAAAGTATATAAACCATGTCTACCCA

TTTAATTGGAACCAGCTTGCTTACGAACTGGACATAGGAGTGTCAAAGGAATCAGCAAAGTGTGATAAAATCTGTATGTA

TAACTTGTGGACTACCAGCCGCTACGTGTCTGCAGCTGGAACCTTACGATTGACATCCCCTGTACCCCTAGATGTAAGTT

CACAAAGCTAAACCTTTTAAGGAAGAGCAGTAATGTTATCGACAACAACATCCAGGGCCGTTCCTGAATATATTTTTGCC

CCGAAGCGAGTAATTAAATGTTTTTCCCTGATAAAGAAAAAATTAGCACAAACTTAACTACATTCGTATCAGTTGCTACT

AAGATACATTATCCTTATCGGCTTTTTTTTTTTAAAGTGGAGGAAGCTAGAATTTAAGTATACAAGGAATAAATATTTGA

ATACAATGTGATTAACGGGGGTTAGTCAAATAAGATTTCAGATGACTTTCAGAGAGTTCTCAAATCCAATGTTAGTCCAT

TGAGACTTTTAGTGAGTCTCGAATGTCTCTTGTTATTGGTTAGAGACCTTTTTCAAGTCATTTAAATTCTCTGAAACTCC

ATGTTATTGGATTAGGATTTTATAAGAGTCACTTCAACATCCCTGTTATTCTAATAAGAATTGAAAGTCATTGATTTATT

GTTTTCAGAGATTTGAAGAGACTTTTTTTAGGAAAATAGGCATGGTAGAAATGTCTTCTCAAGAGGTGATATTTCTAGAG

ACTTGGAAACTTGGGGAATATTTGTTTTTGAGAGTAAGACCAATTTTTTTTTTGTTTGAGCTGATGGCCAAGTAACCGGC

CATTTGCGAAGTAAACTAGTGGCGAGATTGCATAGAAAATAAAATGAATCTGGGATGAGTGAAATAAAATAAAATAAAAA

ACATAGGCATAAGGCAACAACGAGATTTATTTTGACAATGAACACGTATTTTACCAGGTATTCATGGTATTTTTATTTAG

ATTATAACGTGCATAATAAAGTAACTAGTAGATGTTTTGAATTTTTGATTAGATATTGTACATTTTTTTGTCTCTACAAA

TCATAGAGACTCTTTACGAATATCAATTTTTTGTCTCTACATGTCACAAAGACTCTCTAATATACTCTCATAGAAACACC

AGAAATCTCTATATTTTTAGAGAATCTCCGGGAGTCACTCAAATAAAAAGTCTATATAAATCTCAGTTGACTACCTCCTT

GTAAATCTTCTTGGTGATCTTATGCTACACAATATATATATATAACCTAAATTAAAAAAACTCGTTAAAACATTATGTAA

AAGTACATTTAATGATATATTTTTTTCTTGATAAATATATGATGACATGGTTGTAATTTTCTAAACAGATTCAATTCAGT

TATTAGTAGATTAAAACCTTGATTCCCTAAAATCAATCAACCATATAAAATTTAACATCAATCAAGGGTTTATATTTTAC

AAAAAAATCAATAATAATTAGATAATAATAATAGGTAAAGAAAACTGAAGAATACAAACAAAAGATGCAAGCAAATTAGG

ATTGAGAGAGTATAACAAGAAAGCTAAAAATTATTTGATCCAAATTAATCTCCTTAAACTTCTGCCTTTAGATTTTACAT

TAATTATATGTAGGTCCCTAATAATTATTTATTATAAGAAAATACCCAAATGTGAAATACTTATTTGGTTACCCCCCATC

CCTTGTTGCCCCAAAGTGGATCTTGTCCTGCTTACCCTTCCGTACCGGCCCTGGAAACATCCCGTGTTTCATCTCCCTGC

TCGAGTCGTGGTTGTGGTTTTATGAAGAAAAGACCATGTTGCATATATGGTGTAGGAGCGAAATATCGCACAAGATCTTC

TTCCCAGATAAGATAGCCCTAAACACGAAATTACTACGATAATCAAAGATCAAGGCGAAGAATATCATCGCGACAATAAA

GAAATAATGTGAATGAAACTATCATGCGATGAAGATAGAGCGAAGATCGGTTCAAAAGGAGTTGTGCGAAAGACAAAGCC

AACAGCGTGAAAGGCAGCGACATCTTCCCAAATCCTGGAGAATAAGACAAACTGAAATAAGATCAGTCAACAGAGATTTT

GCATGTGAACCTTAGTTGTCTTCTACTAATATATTTTCCTATATAAGGGCTTATAAATGATAGAGAGCGGGAGAGATCGG

TGAAGAGAGAAGTGCTAAGGAAATTATCAATAGAATCAAGAAGTACCATAAGAACCTAGCACTCATCTTATATTTCTCCA

TTGTTGTATGCAAACTCATCTCGTGATCATTGTAAGAGTTCCATACATTCAATACAAAGTGTTCAAGAAGATCATACTAG

TGTCAAAAGTGGTGAATAATATCATTAATGTTTGAGTACGTTGGTCCTTATGACTTAAGCTTTGTGTTATTATCATTACA

ATGGGTGTAGTTGTGGGATTTTCTGCAAATACATTTTGGCTCTAGAAACAGGGAGGATTTATCCTCACCTGCTAGTTACC

AGTTAGAAATCATCCGCATTCAAATTTTATTTGTTATTCTCATTACCATCATTTTTCAGTCCACCATGAAAACCTGATAG

TAGAGCGTTTGAAGTCATCATTGCTTTAAAGACCTACATTACTTTTGAGTCTAACGTGAAGTTTTGGTAAAAACATCACC

GCTGTATTATTTGGAAAAGTTTATGACGAAAAAACCTTATACCTTTCAAAGTTCAGAGCAACACTACATAGGATTTCTTT

CTTAAGATCCTTTCTCGGTAACTGAATACCAAACCAAAATTTATTCACCAACACTCTTCACTAGAATAATGGTGCGAACC

ACCTCAAAAGCAGGGCATGTCATCACTACACGGAGAAGCAACAAAGTGGTAAACAGAAGAGGAGCAAGAGATGAAGCTGA

AACTTCAGCTGCTGGAGAACGAACAGGTTTAGAGCGACATAACCGAAATCGCAGCCTACTCTCCATCATATAAACAAATC

AACAGAACAACAATGTTGTCGCCCAAAATCCAGTACAACAAAGTAACACCGAAGAGAGAGTCATCTACGAAGAAGGTGTG

GTACACACAACTGACACTGAAACTATGGAGGAAGAGGACCAGGCGGTACATGGAAGCAGTGAAAACCCTAAAAATGAGTA

CAACTTGACCATTCAGCAGCTAAGACAGCGTCTCACATAAGATATGCGAATTGAAAGATAAATGAGGCAAGGACTCGAAC

AGAGCAGAGCAAGAGATGCGGAGCTAGAAGAAGAACGAGCAAACTTGGCATGACAGAACGCCCTGTTGACAAAAGAAAAT

AACCAGCTGAATAAGGATATACAAGCGTGAACCAATATTATAGAAACAGAAACTCAGACTAGTAGGAATTCGACAAGACG

AAGAGAAGAACATCACATTGGAAATAGCGAAGTGCAAAATAGCCGTGTGGAATATGAACGAGAAGTGCGTAAAAGAGAAC

GAGAAATGGAAAACGCGCGCGACATAGAAAGAAGACTAAAAGATCGAATAGAAATGAGAAGGTATCGAGAAAGAATAGAA

AGTCCCGACCGCAAAATTGATATACAACTATATGATGGAAGGAGAGGTAGCAGTTCTGAGAATGAAATGGAGAAATGTAA

TCGCATGAGAGAAAGAAGTTTTGAATGAGATACCAGAAGGGCTATAAGAAGAAGTTTGAGCAAACAATGAGGGTTGGAAG

CAGTAGTGGAACCAGAACGAGCAGAACAAACTAATGCGAATCAAGCTATACTCCAAAATCTTGAAGAACTGAAGGAATCA

ATAGAAAATAATAATAGAAGATGAGGGGGAAAATGAAATTAGCAGAGGCTATAGAGGATACATAACGATCACCATTTTCA

CAAGAAATATTACAAGCACCCATACCCGCCAAATATGTTTTTCCCACATTCTCCAACATCTTTAATGGAACATGAAACAC

AGTACAACACATGAAAGACTACAATCTTACACTAATTCCCGGGGCCCAGCATCAAGCGGTGCTATGCAAGTATTTTCCTG

CTAGTCTAACATGGGAAGATTCACTATGGTTCGACAACTTGGGCGAAGGATCAGTAAATTCATTCGCGCAATTGCAAAAA

TTGTTCCTGTGAATAACTGGCAATGTCGAAAGTACCCACAACATCGGACATGGAAAGTCAGCAAAAGAAAGTACCGTGAA

AGAAGTATCTCTATCAATCAAGGATGAATAACGAATTCAGAGGGAATATGGGGATAGGCTTCACACATATAAACAATCAA

AGATGTTAGATAATTGCTCGGTTGAACTCGCAAGTGTTGTTATCTCAGACTTGTTTGTCAAATTTAGTTGTCAAAACTAT

ATGTCTTGATTTCTAGTCTACTTATAGCTAAGTCTCGGACTAAGATAGTTAAGTGTAGTTGAGCTACATACTCCACGACG

ATCATCTTATGAAGACGAAGAACTACTCAAGGAACCAGTGGAACTTCATCCGACTAAAAGGTATGTGGAGACTTGAACTT

ATCTATCGCTCAAAAGTCTATGTACTCTATCTCATACTCTTGAGACAAAAGTCATATAAGTATGATAGTTTTCATACATA

CACATTTGCTATTTCTAGCCGAGTTTACTCGCCTATCTATTTCTCGAAATATGTGTTGGTAAGTTTTCGCTTTAACCATT

TTTCATCTTTACCCATGATGACATTTCAATCTTGAAAAATAGCTTTGATGACGATAGTTGTGAATAACGACTGTTATAAA

ATTATAGACGAATGTTTCAATGATTGAAATGTAGAGTTGAGATTACGTAACCATCTATGGATATAAGCATATATAGTGTG

TTCGCACATTAGTGTATAAATCCATGTGCCGGAAACCAAGTGTGTGCATATGTGTGCATGCAGTATTGGTGAAGGAGACA

GGTTGGGTATGCGTACTGGCAGAACTTTTCCTCACGGAATATTCTGCCGAGTTTGGAGTTTATGAACTTATAAACAAGTC

ACCTTAGGTACGCATACTGGCGAAACTTTTCTACCCAAAAATTTCTGCTGAGTTTGGAAACTCAAATCGGGTTGCTAAGG

TATGCATACCCGTACGCGTACTGGTAGGTTGTGAAACCCACAATAAAATACTTCAAGTGCACAGTGTCTAACTAGTAGAA

CAATGGCAAGTACAGGTCGTTCCCACGAGGAGTGTGAAATACCACAACTAATTGATACCAAGAATAACTAATCGACAAGA

TGAAGTTTTAACGATTTTTTAGAAACTTGGAACAAAATGAAATAATGTAAAATTCTAACAATAAACAAAATTGGGAATGT

GTTATTAGGATTTTCCGTTTCCACCAATTAATCATGCAAACTCTACTAAACATTAAATATATATTCCATGCATTTTCAAT

TAGTGTTTCTCCGCTGTAATTTTCTTCGCTTTACGGGTAGTGATTATAAAGCATTACCTATCTATCCTTGCATACCACGT

CTAGGGATTTAACCGAAGCACAAAATATTAATTGAAAAGCATAAATAATCTAGAAAATCACATGAGCAATTAGATACAAG

CTTTTATGAAGAGAAATTACTAATCAATCAACTCATTATTAAGCATATAGGTGTAGAGTTTTCACAATAAAATTGGGTTC

TACCTAAACCCTAACGTTGCTCTAGCCAGCCATGGCTTTCTCAAAAGCCATAAAAATATTAAAATCAAAACTAGCTAAGC

AAAAACCAAATTGAGAGAATAAAACTTCAAACCCTCGTCTTCTCCCCAGCTGGCTTTGTAGCCGCGCCTCCGGCCTCTTT

TTCCCAGTTGCGGCTGTCTGTCGCTTCCTCCCCCCTCAACTGGCGTCGACCTCTCCCCCTTTTTCTTCCCTTTTGTGTTG

CGAGAATAAACCTTAAGACCTAGAAAATCATTTGCTCAAGGCCTGCTTCAATCATCAAAATTCATTTCCAAATTTAGGCC

CAAGTAGTAGCTGAACGATGGCCCAGTCCACTTATACTCTTTTAACGTCCAGGCAAAGCAATTGCCAGCTCCCCCTCTCC

ATTCTGCGCGTTCAAATTCTTCTTAGGCTGTGTTTGATAACTGAGAAAGGAAACAATTCCATGGAAAGTGTTTCCATGGA

ATCAGTTCCCAGGGGAGGCAAAGAAAGACTTTCTATGGAATTCCACGGAATTGAGTTTAAAACATACTTTAGGCCACGTT

TTTTTTGTAATTCCACGGAAACATATTTTTGACCAAACGACAGAAAACACGCTATTTCCATGGAAAGATGGCTAATTCCA

TGGAATTAACTCCTTTCAAACACAACCTTATTATTCTCCATCTACAGACGTGCATCACCGCCAATTCTCGGCATCAAGAC

CAACACCTTGTTTGTTTGCAGCTGACTCGGCATCTGGATCTGAGTCAATTCCTGACTCGGGCCGAGTCAGCAGTCAGACC

GTTTATTTTGCATTTTGAGTCAGATCTGACTCGACCCCTGACTCAGACGTAACCCCTGACTCGGACCCATTTGAGTCAGG

TAACAAAATACCCCTGACTCATGGAACCAAACCACTGACTCATATAATTTTGAGTCAGATGAGTCAGACCTGACTCAAAA

CAAACATATCGAGTCAGATCCAGATGAGTCAGGTGATTTCACTCAGATCTAGATGACTGAGATCCAGACGACCCAGATGA

GTCAGGAGTAAACAAACATGTTGCAAGTCCGGCTGCAGACTCAACATTAGTTCTCAGCTTCTTGCCCGTCTGGTCATAGT

AGAGACCTTTCTCCGAACTCCCAGTTCGATCACAGCAAGAAACCCCCGCCTGTGTTCAATGCAGATAGTCTTCTTGTTAT

TCTCTTGGTACCGACAGACCCAGCTGTTCAACCATCGCCACAGACAACTGTTGATGAAAACCAACACCAAAATCGACCAC

CATCTGCAAATACAAAGTTGCTTACTTCATGAAACTCAAATACCAATTCCAGCCAAATACACCTCCACTTTGCTGCAAAT

TCAGAGTCCTCCCATCTTTAGTATCTTCACGGCAACCTGACATAGATAATCTAGACAACTACCAGTGCAGCAAACCATTC

GTTCTCATGCCTTCAACACCTTCAGTTTCCGTCTCCCTGACCAACTGAACACCTATCGGCTGCCATAGCGAGCACAAATT

CTCGAAACTGGATTAATCTGATTCTGCCACAAACCCAAATTTCAGTCTTATTCACGGCTACATTTGATTGCTGCCATCTA

CCCATCACCACTGGTTTCCTTGGTCCGTGCAAGCTTCTTATTCCTTGTAAGCACCTTCCTCCAAACTTTGGTCATCTCCT

TGTATTGAAGTTCACAACCAATACCAAGGAAACACCACCAGTTCGAGCTCATACTTTTGCTACTGTTTCTGTTGTTTTAT

CTAAGTTACGGCCACCATTTCTCCACTGAGAATCTATTCAGCCTTGCACCTTATCCGGTTGCAAACGATCACCAATCATC

TTCATGCAATCTGAGCTTTATCGACACATCACCACTGCTGGTTATTGTTTCAGCCAACCCATAGCTCGTCCTTCATCCAA

GACAAACTCCTAACTTCATCCATTACTTAGAAAGTTTCCCATTGTTGCTACACTGACGAAAAACTAGAGAGTAAACTCAT

GCCCATTATCTCAAGAACTTCAACTAACCCCTACTTCAGTCGACATCCAAGACCCCACAAAAGAAATCATATAGCACCAA

CAGTAAACATCACATTTCTTCACAATCCAAAACGTTAATCACTCCTACAATTCCACCGTTTCTGCTGCTGCTCGAACTAA

AGACAGATGAGGCCTTCCCTTGTAGTTCACACGAAAATATGTGGAGTCATGGTACCATGCCATGTCACTCATCCTTTGAG

CCCTAGTGGAGTTTAGGTCATCCTCCCTTTGGAATCTTTCTTCACCTCTTGTCTGTAGGATGTAGGCTAACTGCTGCCAA

CTGGTGCAAGTGGAATATAGCCTCCTCAGTTGATTCTTGGAGATGCGGCTTAGTAGTTTTTGTCATCAAACGCCACTTTA

GCTTTAAATTAAGCTGTTTATTCTCCTGTGATCGGAATCGCTCGTACTTTCTACAAAAGCACTGAAATTGTGTTATTAGC

AAAATATGGAGTCCTAGATAATATAAATATGGGTATAAAATGTAGCACATACGTGCTTATCACGTACCCAAGCTGGTTAT

TTCTCAAATCGGTAGTTCATGAACTTAAAACAATAAATTATAAGGAATGAAATTTATGCCAACCGTGACTATATTGTTCA

TGAATTGATTCAAATGAATCAAACCGATTTTGTTTCAATTGTATCTATGCATAAAGACCTAAGCAATTGAACAACTCTTT

AACTAGTTCTTTTGAAGTCATTTGAACTATATGAGAAAGATGAATACGGTTAATATGAAAGTGCTCATATGGCTAACCAT

TGGTTAACTATTTGTGAACCAACTAAGTGTACACGTTTAGGTACGGTTACTCAAACCTAAATGAAATACATTTCATTTGT

ATGTGAAAAGCTAAGTTTCGATCTAACGGTTGAAAGGTATAAGCTTGGTTGAATCAGTTTTTTCATCTAACAGTGAATAT

TGAATGCTTTGTTACTAAGCTAACATTGATTGCAAACCCTGATTTGAAAACTATATAAAGGAGAACTCTAGCAACTGGAA

AAACTAATCCCCATACCTCATGTGTGATACTAGTTGGTTTTGCTAGATTCGATTCTCCTTTAACATTAGGTTTCTTCTCG

AGACCCTGTAGGTTAACGACTTAAAGACTTAGTTGGGATTGTGAAGCTAGACGAAACTACTTCTCTTGTAGTTGAGCGAT

CTAATCTTGCCATTTTCTATCGTACGAGTTTAATTGAAAGATTTGCTTGAGATTATATCTACGATAGGGCAAGATAAAAA

GAAAGCACAAATATCTTCGTCTCATCGTTTGTGATTCCGCAATATCTTGTTTCGCTACCATATGATTAAGATTATTATAA

GGTGATTGATAATTCTAGGCTATTCTTCGGGAATATAAGTCCGGGTTATCGATTGGTTCCTGTTCACCTTGATTTCTATC

AAAAGACGGAACAAACTCGGAGGTTTATCTGTGGGAGACAGGTTTGTTTATTCAATACGGAACAAACTCGTAGGTTATCG

ATTGGTTCCTGTTCACCTTGATTTCTATCAAAAGACGGAACAAACTCATAGGTTTATCTGTGGGAGACATCGAAAAGCTT

GATCTCTTCATCGAAAAGCTTGATTTCTACAGGGTTCGTGGGAGACATCGAAAAGCTTCATCTCCTTTAACCTTAGGTTT

ATTTGTGGGAGGCATCGAAAAGATTTATATTTTACGTAGGTTTATCTTTCTCCCATTAATTTTACCAGCTTCGACACTAA

ATCGTCGAAGCTGGTAAAATTAATGGGAGAAAGAGATGAAGCTTTTCAATGTAGAGATAGGGCATTAGAATATATGCGGT

TAATTCAAGATGTTACCCGTCAGCCTATAATATCGGGTGACCTTTGTGTAATTGTTTCGCAGAAAGAAGGTAGAGGTAGT

CGTGAAGGCTGTTGTAATTGTTGTCTCAGTTTTATATATTTATTTAAGCTGTGTCGTGCTGACTTGTCTTTCTTTCTTAT

TTGGCAGAATCGTTCCAACAGGAAGTTGTCGCGCTAAAGCAAAAGTTGGAAGATTCTATCAAGGCGCGGGATTCTTCTTC

TCAATAATTTTATATAGTTTCGCGCATGGTGTCTGATTTTACGCATGAACGAGATAACTTGCGGAAGTCAAAGGAGAAGT

TGGAGTAGGAATATCAGTCTTTCGTGGATGGGGCCAGCAGGAAATATTCGAATGCGACTCAACGTCGTGTGAACGAAGAG

ACCCAACTCGCGGTTAACTGGGTTTGCGATAAGCATGGACTTCGCAGCGTGAATTATCCTCATGTTCCTGTCAGCGATGA

TGAAGAAGATGAAGATGTGAGTGCGAACATCTCTGAGGGCGAGCATGTTGATGATGAAGGGTATGGTCCTGTCAATGTTG

ATGATGTTGAGAAGGAGGGAGATCGTTCCCAAACCAAGGATGAAGGTTTTATAAGCGAGCAAGATGGTGATGCTTTGGTG

CAAACTATCCAGTCTGTTCCTGAAGTTACTGATGTTGTTAATTCTTAGATTTCACTCTTTTGTTTTTCCCTCTTTTTTCC

TTTTTTTCATGTTGACGTCCTTATCTTCTTTTGAGTTTATCTGGGTTCTCCCAAGATTGGGGGGGCGTAGCTTGACTTTT

GAATACTCCTTGTTTTCCTGTTTTTTAATATTTAGACAAGTAATTCTAATTCTATCGTTGTTGTGTACTTAAAATTCGAA

ATCCATCTTAGTGTGTTTCTAGGCATTTGACTGAGGAGGTGGAAAGCATATGAATTTTAGCATTCCTGTACTTTCCTATG

CTTTCCACCTTCGAAGGAAGACTAGTGTTCTATCATGTTATACGAATATAGTCTATGCAAATGAATCTATTTCAGTGGTT

GAGGGCGAATATTTGTTTATGACCCATGCGAACGTTAAATTTAGGTTTTGGCGAATTCTATGGTAATACTAGTAGTAATT

TAGGGATAAAATTAATTTTATATAGTAATTTAGGGATAAAATTAATTTTATATCTATTTCTTATCATGCCATTCATGCAT

ATGAGGTTATGTCGCATCTTTGGGGTTTGATGTGTGTGCGAAGCAAGTTTTTAGTGTGATGTTATGATTGGATATAGTGT

ATGTCGCATGGTATTGGGATAATGGTGACATAGGCATTTATGCAGATACTCTCTAACATGTTATAAATATATCAAAATAT

ATTTTTGAATGATACTTAGCTCCTTGGGATCGCATTTCCTCATTGCTTCTTCGTGCCTCAGCATGATCTTTTGGTAGCCT

CAGTTTCTTTCGCGCTCCTTTGTTGTTTTTATTCGGATGATCTTTATAGGTCTTATTTTCCTCCTAGATAAGGTCTTATT

TCCTTCCCCCACATAGAGAACTTAAAGGACTTATGGTCGCCTTGGGTAGGGTTTTCATCATTGGGCGACGAAATCTCAGT

TCCATGTGCTCTTGCGAGTCATTTTTCATCGATCTCTCCTTAACTTTGTTAGTATTTTTACCTCCCATGTCATATTTGTG

ACAATATACCAGGTCTTAGATACTCCCTTAAGTAAAAATCCCCAACACATTTCTATCTTTGACACAATTCATCTCCCTAA

TGGAGGGTTCCGTCCATTTATATCCCCCCCTGAATTTCCCTTTCAAGGAGGTCACCCCTAACCATTTCAGTTGGGCTCCT

CCCATCCAGTTATTCAACAACCAGTGTTGTTGTGACCTCGTACCTTTCGCCTATGTGGCTTATGGGTACAAGATCACACC

CTAAGTGGGGTTTCTTTGGACCGAGTGTATCCAAGCCAGGAACTTCTTCTACTCTACTGGTCATGTTTCTACACCCCATG

ACGGAGAACTTATTCGCTGAGTGGTTCTTTCTCAGTGGCTTCCAGCGGATAGGCTTTTTTTTTTGCCCTAATATAAGTGA

GTTTTCCAAGATTCTCCCAACATGACGCGCGTTAGGGATTATTGACGCCTCTTACGTGATCGTGCTAAACGATGTCGCAC

GACCATGCGAACTAAGTTGACGCGCAAAAATTGGGTACCCACCCTCCCAAATTAGAGGTTTTAAATAGTGACTTGCTTCC

GCCTATTGAGGTCTTATGAGATGATTACGTAGACTTTGTTTCGCCTTTGGCTTGCTCGCGTAGGTTTATCTTTTCGATGT

CTCCTTTAACCTTAGGTTTCTTCTCGAGACCCTGTAGAAATCAAGGTGAACAGATAAACCTACGAGTTTGTTCACCTTTA

GATACCTCAAGAGCTCTGGCAAGCTGACCAGATATGGAGCAATGTTGATCCGACCGAGCACGCCAACCAGAATTGTCAAT

CTGTAGCCCCACAACTTGCTCGTGTAAACGCAAACGACCATTCTCCAATTGAGTAATTTGAGCCTTCAGTGCAACACACT

CATCTCTAGCATCTTTTGCGGCGAATTCGAATTCATATCGTTCGGACGTATGTTACCGGTTTAAAGCTTTAAAGCTTATG

CACGAAAGGCCTAAAAATCAAGAAGGGCGACGCAAAATCAAAACATGCACGAAAGACACAAAATTTTATAGAATGTCATC

TTTAAGTTCCTCATTCTTCGCACGAATCTTAGTAACATCATTTGCTCCAGCAAATAGCTCAGATACTTGCTTTTTCAAAT

CATCAATAATTTTTTAGACTTGCAGGGTTCATCAATCCGAACCAAACGATGGCGAAATTCCTGCAAATACCAAAACTTCA

GAGCTCCGATAAACAAGGAAAGGAGTAAAAACCAATAAGATAAAAACTTACCTCCGATTCAATGGCGATAAGTTTATCGC

GCTTAGAAAGTTGGGACGACTTAGAATCAGTCTTGGGAGGCGACATACCCAGCGCGATAGAGATTAATCCGGTCCACTGC

GAAGCTTTTGCAGCATCATTGGCCACAGCAGTACCAGAGGCCTGCTCATCCTTTGCGATAGCAACATCAAGGGAAGAGCA

TACTATTGTCGCAGTAGAACCTTCATAAGCACGACCAGGAGCAAATGTAGAAAGAGGTCGAGGAGAAAGACTCGAGGACA

TCTCTCCCTTTTCCTGCACGACAGATTTTGGCGGCATTAGCGTCAAGGGACTCACAACATGCACGAGAGCATTTCTATGC

GAATCCATGTCCTTTTGGAAAGAACCCTCTTTACCCGGCAAATGTTTCCGTTATCGAGATATTTTTTTCGTAACTGTTTT

CCGAACACCAGCGGGATTCACCATAGGTTTGGATAGAATACGTCTGAGGCAGCAGAAATAGTACGGGAATACTAAAATCG

TACAAAATACCAAGAACACTAACAACATCAGAAATAAGAAAAACCCAAACAACTCAATCAATTGTTAGAAACTCAAGCAA

AATATAGATATTTACTGGGAGGATAATTCATTAGAAGAAGGGATAGTGGATTGGCTCGACATAGCGGTAAATCTCTCAAA

TCAGACGAACTTTATGGAAGTAGGTTTTATCGAAGCAAAGAGGAAGAAGAAGAATCTCTATCTTCGAAAGTATTTATTGA

AACAGACGGACCGGCTGAAGCAGTAGCCGGTCAAGAATTGGAACCTTGATGTGTCAATCGAGAAACGGTGAAGTTACTGC

ACGTGGAAATAACCATACGGAAAAACCGGAGAAATGTCGGGAAAAAGAAGGCAAAAAGTCACAAACCCAAAAATGAGATT

CTCGTATTATTCCACTTACACAAGAACAACACGAGAAGAGGCAAAATGAAGGGGCAAAATATCGCACAAGACATAACTTA

ACTTCGCCCCAGATAAGATAGCCCTAATCAAAGATCAATGCGAAGAATATCGGCGCGACAATAAAGAAAGAATGCGAAGG

AAATAATCATGCGATGAAGACAAAGCGAAGATCGGTTGAAAAGGAGCTGTTTGAAAGACAAATCCAATAGCGCGAAAGAC

AGCCACATCTTCCCAAAGGCTGACGAATAAGACAAACTGAAATAAGATCAGTCAACAAAGATTTTGCATATGAACCTTAG

TTGTCTTCTACTGATATATTTGCCTATATAAGGGTTCAAAAATGATAGAAAGAGGGAGAGATCGGTGAAGAGAGAAGTGC

TAAGGAAAATATCAAGAGAAACAAGAAGTAGCCATTAGGTATTAGCCTTCATCTTATATTTCTCCATTGTTGTATACAAA

TTCATCATGTGATCATTGTAAGAGTTCCATACATTCAATACAAATTGTTCAAGAAGATCATACTAGTGTCAAAGTGGTGA

ATAATATCATTAATGTTTGAGTTGGTCCTTATGACTTAAGCTTTGTATTATTATCATTACAACAGGTGTAGTTGTGGGAT

TTTCCGCAACTACACATGGAATTCCAAGATGTATTAGATCCCATGAATTATGATCCATGCGTTCACAACTATCAGAATTT

TTGTCAATGTAACTCGCTGGTTGGACCTTGGTTGTAGATACACGACGAACATTGGGTAAACTCAATCTCTGATTCCTAAC

ATCTTGTGCTTACTGCAATTGCACACTAAAACATCTCCAGTGTTTGCATCTTATATAGTATTTGAATTCTGAAACTGAGC

TACAAAAGTACAACTTATCTTGTGCCACCACAACCGCCCAAGAATTTTGAATCCCGCAAGAAAAGTTGTCCATCCACCAG

TGCTGGCTCGAAACGTTAATGATGTTATACGTAGGCTCGTGTCACATGATATCAAATCTAAACGACAATATTATGTATAT

GTACGAGCAGTTTCATAAACCGGACATTTTAACTCATGAAAACTCTCTCTCTCTCTCGCATATTTCCACTCTAGACACTA

AACATCAATAAATGTTTGAAAGTTTGAACAATCAAAGAGCTTTTAGGGTGTCTTCATTAAATTTTAGGAAGCTGGCACAA

GGAGTAGAGACTATCGGCAATGGGGTCGATAACAATATGGGAACCACTACTATTTCCTGCAAAGTGCAAGTGTTTAGGTG

CTTTTGCTCTATCGAAATAAAACCCGCCAGATGCAAGTTTCTCCTTAGGCTGCAATGCTAACCATACTATTGTGTCTGCA

CCCTCGTCACTTGTTCGAAGCATCCATGACATGCTGCAACACCAAATAGAAATTAACTAATCATCGCACACACAACAAAA

TAAGATATCAAGAGAATTTATTTTCCTTGCCTTTTACAAAGCTCAGGTAAGCCTTTGACAACACCTGGTGTCATAGCCCA

ACCAGGATGCATAGAATAGAATTCAGTTCCTTTCTCGCCGTATAGTTTTGCCCATCTTTCTGTTAATGCCACCTTGAAAA

GAAAAGTATTTTACCATCTGGTGAGCCGGGTTTTTCATCGAATGTTTGGTTCTATTTATATGAGGCTGAAAATATATATC

AAGAATTCAAGCGCATGTATGAGCAAACCTGAACTCTTTTGGTTCTAGCATACTGCAGATCCCCGTCAAAATTTCCTTCA

CTGAACTATAATAAGAGAATGAAATTAGAATCCAATAAAACAGAAAAGGAATGAAGTCGCAGTTTGTTATGTAACCAAAT

ATTTGTATGTTACCTGCAAATCTTCAGTCAAAGGAGCTGTGTACATCCCATCAGATGAGACTGTAATGACTCGAGCATCA

GGTGCAGCTTTCTCCAATAACGGCAACATCAACTCTGTCAAGGAAAACGTTCCTGCTACGTTTACGGCAAAGTTCAACTC

TAATCTGGTAAATATAAACAGAAGACGAAAATGCATCAGTTCTTGTGCAACTAGAATTCTAACATGTACGTAATAATTCC

GTTGCTCAAGATAGAAGACTACGCACCCTTCTGGAGTAGTAATTCGGTTTTTCTCAAGCAAGCCAGCATTGTTAACCTAT

GATTACCAAATTTCATCAGCAATGAATTGAGCACCAGCCGTTATGTAATTTGCACAGTCACTGATATTAAATTCTAAACA

AAAGAAGACATACCAAGACATGAAGTGGTTGTTTCCTGGAAATAAATTTGGATGCCCATGATTTGATATGACTGAGAGAG

GATAGGTCGCATAGCTACAAGTAAAATGCGACTTTAAATGGAAGGGTAATGATCACCAATAAACCAAAGGAGAAAATGGA

AAGCGAGTATACGTACCTCCAAGTACACATTCTTGTTGCCAGTTACAGACTGAACTTTCGAAACAGCAATTTCCCCTCGT

TCCTTATTTCGACACACCATATACACATTTCCCCCACTGAATTTTAACAAGAAAATTAATAGGTTCAACATATAAGAAAA

AAATTAAATTCGGAAGTAAGTACAAACCGGAAAGCAAGAGCTTCGGCAGCAGCATAACCAATGCCTGAATTAGCTCCGGT

CACTACACAATTCTTTCCATCGAATTGAATTTTCATATCTTCTTCATTGAAATTTTTCGAGTGTTCACTATAAAATTTAC

CCCAAATGTTGTGGGGAGATTAAATCATATGAACAACATTAATGAAACTAGATTATGCAATAGACGTACGTACATGAAAC

CAGATTTGGTAAAGTTGAGGTGTCCATGAACACCAAAAGCAGCTGCTTTCCATGCTTTGTGAGCAGAAGACATATTTGAT

TATGTGAAATCAATTCCAATTTGAATTTTTTTACTAGATTATTAGTAGACATACGCCATTACACAATAATGGATGACCAA

TAAAAAAACTGTCGTTGGCGTACGTGGTACGTCTGTCATCATAAATTGTATCATTTTATGCTGTCGTCGACCCACGGACT

ACTCGTGCATCAAGTGGTATTAGTACTCGTTTCCAAAAGCGGACCGCCGCCGGCGGCGGCCAAAAGCTGATAAAGCGTTG

CTTTTGTCTTTTATCGTAACAACAACTACTAGATGACACCAAGATGTTATTAATTTTTTTTATTATGGAAACATCTCCTT

TAGACATTTACACTAGTACGGTGTATATGTGCACTTCCCATGAGGACAATAATATATCAGATATCAAATAGCACGCCTTA

TACACAGAACGATTGAGTAACTAAATGGAACTTCTCAATCTCCTCCACTTCCAACCAATTTCTCTAATAATAGCACCTCT

TTTACTTGGTTTCGTTGTGCTATCTTGGAACAACTATGGAAGGAAATCCAACAATAATCAAAAACAAGAAGCACCAGAGG

CACCAGGAGGAAGACCAATAATGGGTCACCTTCATCTTTTCGATGAAATGGATTCGATGCATCGAAAACTCGGTTCCATG

GCGGATGAATATGGTCCGGCTTTCACTATCCGACTCGGTTCTCACAAAACTCTAGTTGTGAGTGACTCTGAATTGGTCAA

GGATTGTTTTACTAGAAATGATATATTCTTCTCAAATCATCCTGCTATCCTATCTGTTAAACTCTTGTTTTACGACATCG

TCTCGGTTGGTTATGCTCCATATGGAGCTTATTGGAGAGAACTCCGAAAATTTTCAGCTATAAAACTTCTTTCGAATTAT

CGTCTTGAAACGTTGAAGGGTTTGAGAGCCTCTGAAGTCGACTCGAGCTTTAAATCTTTGTATAGTCAATGCAAGGGAAA

CGACTTAAAGTTTGCTTCGGTGAGGTTGGATAGTTGGTTGGGGGATCTGGCATTTAACGTGATAGCGAGAATTGTTATCG

GAAAGAAGAATTTCGCAACAAATGGAGCGGAGAGATATAAAGCAGCTATTCAGGAAGCTATGAGACTAGTGACAGTTTTC

GCATTTGCAGATGTGTTTCCGGCATTAAGTTGGTTAGACAATTTTACAGGTTTAACTAGAGATTTGAGAAAATGTGCATA

TGAAATTGACCAAATACTTTCCGGTTGGGTTGAAGAACACCGTTCGAAAAGAACTGACGGGAATAAATCATCGGAACTTG

AACAAGATTTTATTGATGCTTGTTTGGATAATCTTCAAGGGATTTCTTCGTTGCCTGGTGTTGATCCTGATATTGTCATC

AAATCTACTTGTCTGGTATGCACGTACAATTATTTTGCTAGCATTTTTCGTATATATATGTTACAATAAAAAAATTAACC

ACCTTCAAAACTTTGGTCTCTGCAGGATACGATAATGAACGGGAGCGACACTCTAGCCTTAACACTGACATGGGCAATTT

CCTTACTCTTGAACCATCCGGACGCGCTAAAGAAAGCACAGGAGGAGTTAGATGAACGCGTGGGAAAATATCGAAAAGTG

GAAGAATCCGATATCCACAATCTTGTCTACATCCCGGCGATCATCAAAGAGACGATGAGGTTGTATCCAGTCGGACCATT

GTCTCGGAGTACATTCGAGGACTGTGAAGTAGGTGGGTACCATGTTCCGGCCGGCACGCGTTTGCTAGTCAATCTTTGGA

AGATGCAAAGAGATGGGAACGTATACAGAGATGATCCGTTAGAGTTTAGACCTGAGAGGTTCTTGACGAGTAATGCAGAT

GTGGATCTTAAGGGGAAACATTATGAATTGCTGCCGTTTGGAGCGGGTCGAAGGATATGCCCCGGTGTGTCGATGTCTGT

TCAACTGTTGCATTTGATTCTTGCTAGAGTAATTCATGAGTTTGAAATTACGACAGAAGGAAAAGTTGATATGAGTGAAA

GAATGGGATTTTTGTTTTATAAGAAAATGCCATTAGAAGTACTCATCAGACCTCGACTCACTGGAGTTGAGAAAAACTGA

TTAAATATGTAAGGAGCATCTTAATTGCGGCAAAATCAATCAGAGTCAATAAACTATGGATCATTTTATTCTGTTTTACT

GGGTTGAGCACCTTTAGGCTAAGTCCTTCGGATAGATTGAGCTGCTAGATTAGCAGAAAAGTGTTGCAGTTAAAAAAAAA

ATGTGTTGTCTTTTTATTAGCACTAGTTCTCTCTCCAAGCATTTGCTCGCACTTGAAATAGTTGCGCTGAATGGTTCAGC

GCAACAAAAATCACCTTTTCGCTGAATGGTTCAGCGCTGGAAGAATATTCCAGGGAATCTGCGCTGAACCATTCAGCGCC

CCAACTCTGGTGTTGTTCTATTCAGCGCTTGACCTTTTTTCTTTTCTTTTTCTTACACCTCCTTTACCCCACCAACGGCT

ATTTTTCTCAAATTTTGAATTTTTAAATGTTCCATTCTTTCCTACACCTAGTTACACCACCAACGGCAATGTTTTCAAAT

TTTGAATTGTGAAATGTTTGACAATTTGTTCCAATGGTTATTTGATTCACTTTAATCTCGTTCCTCCATTTTTTTGTAGA

GGGGTTTCTCTCAATCCGCTACTTCATCCAAATTTTGGCGTAGGATCTTCAAGTCACTTTCTCTTAGTTGGGTTTGTGCC

AAGATTGTGGAGTTTTCCATATACATCTTAAGGTACGCTTCCTCTGTTTCATATTTTGAAACTCTTTATGCTTGCGATTT

TTCAAGAAGTTCGATGAGCTTATCCATTTTGTCATATGCTCTCTGCGACGCTTTAGCCTCACGAGCCCTTTTTTTGGAAC

TATCTCACCCTGCACCTCTAACAATATCTGTTTTTTCGAAATAACGCCTTCTACGAGATATGTCTGAGGTTTCATAACCA

GGATTAGATGATTAGATACCAAACATTGCCATATCTTCCAAATCTTCATCCATAGTTACTGTAGCGGAAGTACCATCTCC

ATTGTCTTGAGTATGTTGACTGTCTTGAGTAGGTTGAGCGAAAGTACCAGCTTCATCATCATTATTCACCTGATAGGTCT

TCAGGATCTGGAAAACCGATTCATGTTTCCAAGCTCTTTTAAACTCTGATTCATAAAAAAAAAATCCATGATAATACTAG

GGGGGGAGTGGTGCATTAGAGAAAGAATGAAAGAAAAGATTTAGAGAAAGAAAGAAAGAGACCATAATGGAAAAACAACA

TCAAAACAAATTCAGGTACTTACATAATCATGGACTTTCAATCCCATAGGTTTGTTTCGGAAGTAGGTCCTAATGTGTGA

AATATATCTTCTCACTTCGGATCTCAACGTGTTACACCTTGATTGCAATGAATTTTTGTCCCTCTCTCCACCATTAGGAA

ATCTTTTAAGGTACTTTTCAATAATCTTCGCCCAAAATTGTGAACCTTTTTTTTTTTAAATCTTTTTCTCTAATCGGATC

GTTGCTTACTATAACCCAACCAAAAATTAGTGCCTCATCCTATTGTAATAAAGAAAAGGAAGAAAATCAATGAAGAAGAG

AATCAATTGATTTTGGTGTGAGGGATGAAGGAGAAGTAGTTCTCTATTTATAGGGTAGACTGAAAATATAGCTATTGATA

TTTCAAACATCCAGTAGATCATATTTTTGATATGATGGATGAAATTATACAACGTTGAATGGTACAACGCATCCCTTTCT

TCCATTGAAAATAACGGCGATTTTTTAATCTTGCGGCTATTAGGCAAAGCGATGTGGCATTGTTTGGTTCGGCGCTCACT

GTTGTACCATTCAGCTTGTCTAACGGATAAATTTTCACCTCAGAATCAGATCATTTCTTCCCTCTATTTAAACTCAACAC

ATGAATTCAACATCCCCACCCTAAACTTTTGAATTTTCACTCTTTTCTTTTTCTCCTAAATTCTCAATTCAAGCATGGCC

GGATTGGTTGGATTCTATATTGCCGAATACAATGACTTGGTGAGAAGGTTTTGCAAAATTTGCGACACAAATCTTTCGAA

GACATTGTTTGAAACACGCAACAATTTTTGGGAACAAGTTCTTGAAGAGTTTTTGATAATCCATGGTAATGCTAATGGAA

CAAGATGAGGAATCCTACAATGTCGGTTAGCAAGCATAAAGCGTTATTGTAAGATTTTTGTAGCTTGATTAAGGGTATCA

AATAGTAATCGTGAAGATGCTATGGTTTTATACCAGGCGGATCATGGAATGTTGTTCATTCGTACCACCGCTTTGGATAT

ATTTGGTGGAGCTCGAAGATGATTTGAAGTATATGCATGAAGAAATATTATGAAGTTCTAGTTAAATATGTAATGTACTT

GTAAACGTACTTAAATGAAAAAAAGAAGAAGAGTATTATGGAGTTATTACTAGAAATAATTAAAATGTCTACTTAATGAA

AACATGATCGAATGGAAAAAGGAGTATTATGGTTAAATAAAGTGGTGAATATGAAATTTACCCTCCATTCTTATAAAATG

CATCATCTCCTTCATGGTCATATTGGTCTTCATCGGTATCCATGTCGGTTATACCCTCATGAACTTTCGGCAAGGCATCA

TCCATCACCATGTTTGGAATATCACCATCTCTGAAACCTTATCCATGTCGTTTCCAAATGTGTGTAGCGAGTTCCAAGTG

AAGTTACGCCTAACGCGAAGGGTTGTAAAATATATGAGGAGCTAGCCTTATGTTTCCTTGAATTTGCGGTTGAGGTTCAT

CTCCCGTGATCCTCCCCAGTCCTCATTGTGATATTCCATCTCAACACACATATTGTGCATTATTAAGCACCCCATCATAA

TTGCCTTCATGTCCTATTTTTTATAATAACGACATGGTTTCAATATAATGCCAAACTTACCTTTTAGAGCACCAAATGCA

CGTTCTATGTCCTTCCTTTTGGCTTCTTGGTATTGATTGAAAATATATTCCCTATTATTAGCTGGTGGTTTGTATGCCTG

CATAATGCAACCATACATAGGATATCTACCATCTCCTAGGTAGTACCCCTTGTCGTAGTTCCTGTCATTAATTGATAATG

ACAAGGAGGTATTATACCATTAAGTTGTCTATCGAACAAACCCGAAGCATGCAAGACATTAAGGTCATTGTTCTGCCCAC

CCAACCCAAAATAGGAATGACAGATCTATCTATCATATGAAGCTACCGCCTGCAAAACAATTGAGGGATATGACTTATAG

CCGCAGTGGGATCCTTCCTGACCCATTGGACATGCTCTGCATGCCCAGTGAAAACAACCCACACTACCAAGCATCCAAGG

AAATCCTCGAGCTTCATTTTCTTCCAAAATCCTTTTTGTATCTTGAGCGGTAGGCTGACGCATGTATCGACCATTATAAA

TCCAAATAATTGCATTCATAAACTTCATAGCATAATAGTATATAGTGGTTGCTCCCATTTGGGTGTAATCATCTAAGGAG

TCGGGTGCCACGCCTTTGGAAAAATATTTCATGACAGCATCCATTTTCATATCTGGAGTATGACCAGGAATACCGCAAGC

ATCCATTTTTTGGCGAAAATCATGGTCATAATCACAAACTTGCTCTAAAATTCTCAGAAAAATGGGACGAGGCAAAGAAA

ACCGACATTTGAACTTCTTTTACCCATACATACAATTGAAGTTGAAGTATTGTTGCATCATATGAGCATCTGCATGTACA

CATTTTCTGTGCAGCCTCTAAAGAATCTTGACTTTATTCGGGATGGGTTGATGTATTCCTTGCCTCCTCTGTGATTCCAA

AAGCGTAGCCACTCCTAGAACATTCTTCTCAGTATCATCTTCATCATCCGGAAGGCCGAACAAAATTCTTCTCAAATATC

CTTGCATAAAGTACCTAATAAAACAAAAAGTTTTTATAACACTCTATAAAAATAAAATATTTAAAAACAATCTATAAAAA

TAAAACACTCCATTAAATAAAAAGAACTTATAACACGCCATAAAAAAATAACACTTAAAACACACCATAAAAATATAACA

CTCTATAAAAAAGAATTATCACTCCATATGTTTTCATAAACAGTACAAGTGGTTTCACAAACACTGCAAGTACTTTCATT

CAATCAAAAGAAATGACGGGGTTGTTGGGTGTATTGAGTATCAAAAACAATGTTAGTCTCGCTTATCGCCAACAACGCAT

GCTTTGGTGATGTTGTTGTTGATCTTGTTATGTAGTGGCTCCATCTTCACGCCTAGGATGGTAATAATCTTATGGATTAT

TTGAAGGACAATGTGCATATGTACCACCCAATGAAAGATCTTTTGGAATATTTGGTTGTTGGGAAAAGGTAGATGGTATT

GAAGAAGAAGAAGATCTTCCAGATGAGGAACATGAAGTTGTAGCTCTTCGGTATGTCTCATTCTATACGACTTGACGAGC

TCCAAGATGGAATGAATATTGTCTTGCCATATACTTGTAGCAATCCTCCTATAATTATATATAAAGTAAAGAGATTTAGT

TAAATATTCAAGAAACTTAAAATGAAAATGCAAAAAAATGTAAATCTACTTAATTACCAATGTCAAAGCCATCTGGGAAA

AGTCAAACGGACAAGGTACTTCTGGTTCTGGTATTTGCTCTTCTACGCCGGCTTTATTAGAAAATAGTAATATGAAATCT

ATATCCGGCAGTTGTAACAGGGGCTTATTATGCATGCCATAAGTATATGTCTGCGACATTATGTTAGGAGGCGGATTACT

AGGAAACAAAGAGCGAGGAACAGCTGAGCTACCCAACGCTATCTCATCGATGTTCCGTGCATGGTATGACTCAGTGAGAT

AAGGTCCAACAGTTTTATTCTTCCACCAAGTGTTATATTGATCCTTTGTGTGGTGCCAAGCTACATCTACTCTCGCTTTG

TCCTCAATTAGTTGCAAATTTGGAAGATTTGTTGGGACTACAATCTTACCTGATATCTTCCTATGGAATCTTTCAACCAA

GTAACAACTAATTCTTGTGTTTTTATCCAGGTAAAAAAAAACACAACTCTTCTTCTGGATAACTGTAGTGCGTAGTCATA

ATGGTCTCCCACAAAAAAAAAAAAAAAACAAACTATCTACGCATGTTTGTCAAATACCTGATGCACTTGTTCTAGTGTCA

ATCTGTGTTCTTGCGCTTTTGACTGAGTATTTCCTAATATCCTTTTGTCCAACTCCATGTCTTCTCTTCCTTGATATAAT

GTTCGTTTTCTTGAAGAGACAAATCTTTGGGAAAACATCACAATGTTCTCCAGATGTCTGACAAAGCAAGGGTTGCATCT

TGCGGAAGTAATAATAGTACCAATATTCAACTACCATAATCATGCCAAATATATCCTTACCCTTGGTTGACCATTCGCCA

AGAAATTTATAGGTATTACCTAACGAAGCTATACCCCAATCATATTCCCCAACTCTATCTAAATAAGCCAACAATATATC

CCAACGATTATCACTGATCTTCCATTAGAATCCGCAAGAAGAAAATGACCAAAACACCAAAGAAAGAATGCTCGTGTCAA

TGTCTCTTCACATGCAATATTGTCCTGGTTTTCTCTACTTTTTATGTAGTTTTTCAAGATAGCCAAGGATATTGAATTTT

TCTTAGAGTCCCATGCACCATGATGATTTTTGATATCCAAAAAATATAAATCATCCAAAGCGGGATATGTTAGGTCACCC

AACTGCTGCATAACTACTGGTCATTTACTAGGGTCGCCAGTGGGAATTCCCGTTAACATATACCAATCTAAACGAAGGAT

GCCAATTTTCAAATCCTGAAAATAAAAGCTCTTCGTTTTTTCCCACCATCTCTCAGCTATGGCTTCAACTAAGCCTCTAT

CCCCATTGGCGAAATCGAACTCAAATACCGAAGAAAACCCACAATTATCACAATATCGTTGTGCTAGCGGATTATTTTGC

ACTATAATTTTATAAAACGCTTCAACATCTTTCAAATTACTTCTAAAGTTTACATTTCGAGGATCTTTCGTATCTTGACA

TACATGTTTATCAATATTCTTTGACGATACTATATCCAAACGATCTAGCAATTCATCCATACTCATACTACAATCAATAA

CAAACATGAAAAAAAAAAAAATCAATCACTAAATTTAGCTACAAATTTTCATTACAAATGTAACTACAAACTAATAATAC

AAATTATTATTTCATCACAAATAATCAAAATTTCATTTTTTTTCTTCATCACAAATAATCAAATTAAATCCCCAAAAAAA

TTCATCACAAAAAATCAAATTAAATCATACAAGACTGACACATATCATCTAAATCATCTACATCATCTAAATAACTAATA

TCAATCATCAATTTCATCAAAATATCATGATAATCAAAAATTATAAAACCTAGAACTTAAATTTCTTACCCCAATTGATT

TCGTGGACGGAATCGAACCAAACCACCCTCAAACGATCACAAATTTGTTACTGAACAAACCCATGCGGAAGGAATTGAAG

AAAAAAAATGTAGAAATGTTGAATGAGAACATGCATTTTCGGGAGAGTTACGGAGAGAAATGAAAAATGGGGGAAGAAAT

GAGGAAGATCAATCTTCTGTTGAAACTATCCATTTTCTGAACGTTGTTTGGTTCGGCGCTGGGAAATATTTGGTGTCGCG

CGGTCCACCTAGTTGACTAAGTCAAGCGTTGTTTTATTCGGCGTTTCATATTGCGTTGTATTATTCAGCGCCAAGCGCTG

TTTTATTCAGCAACTAGCATGCTAGATTAGTACTACCATAGGATTTAGGAGGTTGCCCTAGTTGACGTGGACTGCTAATC

TAGCTACTAGATTAGCACACCATAGGACTAAGCCTAAAGTATCGCTTTTCAGTGAATTTTCTTGCCAACTGATAAAAAAT

AAAAAAATAAAAAAGTTGAAATTTTCCATAGAATACTCCTTCCGTCCCAAATTAGATGAGGTAGTTGTAGTTTGCACAAT

TCTTAAGGCAAGGAGAAGAGGGAAGTATGTTTAAGTATTTTTTTACAATTATACCCTTACGAATAATAACTAGTAAAACT

TATAAATGATTTATCTCTTAAACTATACCACGGTTTTTCGTAAACTTTATACTATTCAAAAACATTTTAAAACATCTATG

TGACAAATATAAAAATGGCTATCAAATTATATATATTTCTTATAGTAACAATAATCAATTAAAATAATAGTTTTATAAAT

ATCCCCTTAATTAAAGAGACAACTCATCTATTTATGAGACAAACTTTAAAACCAACTAACTCATCTAATATGTTAACTAA

CTTTAAAAAATCGGCTTATAAGGATAAAAATGTCTAACAACATGTTTGTTTTTTCCTCTATATAATTAGAGGGTTCTACA

TAGAGGTTTACCTCTATCTCTATATAACAAGAAGAATTAATTGGGGGATATGGAAACTAATTGGGGGATATAGCTACAGG

ACAAAAACGGTGTCCCAATAGCAATTCTGGGTGACCCCCTATGTAATTTTTTTTTAATTCCTAATCTAGCCCTCACTAAT

CATGTTTAGTAGTTAATAATAATTAGTATATGTTTAAGTATTTGGGGGATAGTTTAGTGTGTGTTTTGATTTTAATTTGT

GTGAGTGAGGAGTAGAAGGAGAGGAAAAAAAAATAAGTGGAATTTTTTTTTTTTGAAATGGAAGATGATTGTGATGAGAG

AATTGCTGCTAAGAGGTTGAAAATTTGATTTGTTTCCTTTGAATTCACCATTGTTGCAGTTGAAAAAGCTCGGTGCCGGC

ATGGTATTCAACCAAAAAACCATGCCGGCATTTGTCGGAAATTTTCAAAAATTGTTTGCCACTATCCAGGGGGGGTTCGG

AGGGCGTAGCCCATGGTCTTCATATCCCATGGTCTTCATATCCAGGGGGGGTTCGGGGGGGGCGTAGCCCATGGGCGTAG

CCCAGTTGATATGAAGACCATGCCGGTATTGTTTTCCGGCATAGAAAATAAAGCAACATGCTATGCCGGAAAGCAATACC

GGCATGGTCTTCATATCAACTGACCATGCCGGCAGTGAAGTGATTAAAACCAGAAAAAAAATCAAAAAGGAAGTAATGAA

AACCATGCCGGAATTGATGAAGACCATATGTCGGAACATGATACCGGCATGCTCGAGAATCAATTCTGGCATGGTCATCA

TCACTTTCAGCATGATCTTCGTCACTTCCGAATAAAATAAAAATTCGTTTCCAAAGTGAAGAGAAGAGAATTTGATGTAG

AGTGGGGAGAGTTTAGAATTTTAAAGGGAATGGGGAATATTTAGGTTTCAAAGCTCTACCCCTAAATAAGATTAATAAGA

AGTGGAGATGGAAATGGGGGATATGATTTTGTTTTTTGATTTTAAGTTTTTAATTTTAATTTTGTGGGAAAGTTATTTTA

GTATTTTCATAACCAAAAATAAGCCATTAACACATACCATGGGTTGGGGGAAGTAATTCATATCCCCCAATTAATTGTTC

TATAACAACCACTATATAGCATCTTTAAAACTCATGTTTGTTTTACCACTATCTAATACCTCTATATTTATAATCAATAT

TTTCAATTTCTCATTTCACCCATATGAGAATTTACAAATTCCTCGCAGATCCATTCCATCCTATTAACTTATTAAGATGA

AATACAAAAGGTCTAAAATTCAATTACATACATTTTTATTTTTTTTAGAATTCATACAGTTCAAATTTGAAATTTAAATG

TTAACGATAATGTTTTAATTTGGCTCTAACAGCACCAAAAGCTCTCTTACAAAACCTTTACCCGAGCAGATCATCCATAC

ACCAACATAAAACCCATTATTTGTTCGCCGCCATCAGCAGCTTCCAGAGTCACCAGTTTATCACTGGTTGGCATCCATTA

CCATTACCACCGGTCAGTCGTTTCCACCACTAGTCCAACATCAGCTCATCATTCACGACCAATCACCAGCAGTACACCAC

CCCATATCCATTCATTCAAAAGAAAAAATTAAAAAAAAAAAAAAAAAAACTGGCTTATTTTTGGTATAGTGATGAGTTAT

CCACTCATTCAAACTGGCATATTATTGTACAATCACTACTTATGAACATCGCAGATACATACAGATTTAGATTGTGGATT

CAAATTTATGAAATTTAAGAAGTATGACACTCACCTTAAGATCTCTGAAATCAAGTGCAAAAGATCTTACGTTGTTCGTC

GCTATCAAGTGCAAAAGATCCTTCAACGCTGTTGACTAAACTAACAATACCAGTAGCCTTCAATGTATCGATCCACACAA

CTGAAGATGCAACTACTTTTGCTAATATTGGTTTGCTTCCAACCCAGGGAATTAAACTTGCATTCGTGCAATATGAAATG

AACAAAATGTACTCACAGCAATGGCAGGTGCAGCAAAACTGGCTCTTTCCCTTGGGTCCTTGCGATAATAATTCTCGAAG

TAGAGTATCGATGCGTAGTACCACATTAGTGTAGTGGTATCACAAATCCAAGTAGGAAGCTGGAACACAAAAGTAAAAAT

TTATATTGATAATGATCAATACGAGAAGGATAAAATGAGAGAGAAAACTTAAAAAAACCATTTTATCTACAACCAAAGCA

CGGAAGCAGTTATTTATACAATCTTGGAGGGTGGTCATGCCAGTCTTTAATTTGGGTACATCTACCTTTTTCACAGTCAC

TAAAATCGTTAGCATCGGCTGCATTCTATATGATACACATTGCTTAGGCATCTAACACCAAACTTTAGCATCTGCTGCAT

TTTATAAGAACAAAATTTACAGCTCTCGAATTTCAGTCTAGACTAGATAACCCATTGACAAACAAGTAATAGAATCAGAA

GCATTTGGAGTAGCGAGTGTAGCTTAAATCTTATCATGCAGTGAGCAAAGATGCACGGGAAAATGAATAAACAACTGGTA

CTTGTTCTTCTACACATACATTCGATAGGATTAACACAAAAATCTAGAACCATGAGCCCGTGATATAGCATGAGGGAAAT

CATAATTGACAGCTACTTGTACATGGTACCCAAATTGCAAGTGAAATTAACTTGGACATATTAGGTGGGAATGAAATGAG

ACTACACATACGACAAGCAATTGATTGGGGTGCTTCAGCAACAAAAGATAACACTATTGGTTGGGCGGAGCTTTTTTTAA

TTAAATCACAAATACTTGTATATCTCTATTGGATTAGTTACGTGTGGCTGTTTGCAATTAGTATTCACATGTGATTATCC

AATTATGGCTGCTCTAGTACTAATTAATTTTCGATAATTAAGATAATACCCACACTGAAATTTTATCAAATACTTCATAT

ACAATAGATTTAACCTCCAGATTGTACATAAGTAGTAACACTCCCAAGTCTCAAATAAACTAAACTTAACCGAAACTTTC

GTGTTTAACCTTTTGGAGTATGGAGTTTGAACTTCGCTGAAATAGAAAAATTAGGGAAAATGAAGCTCATGCAGCAACCA

AACGAAAAGATAACACCAGGATCGGCAAAATGGAAATTACCTGAAAATTATTACGATCGATATTGAAACTAATTCCAATC

AACCCTAATCTTCTGATTTGAATATACCAAACCTATCTCTCAATGAGATTTAATTGCTCGAGAGAAAGCAAACCCTTAGA

TCTTAATGAAATCTTAAACCTAACCTAGAAATCAATTGAATCCCTCTATGAATTTGATGAGGTTCCAGAGAGAAATAGAG

AAATGAGAGAGACGAGCACCACTGATTGTCCGATGATTTTTTTATTTTTATTTTTTTATCGAGGTTAAAGCTAAAAATTT

TAAATTAGTCTGTGGGTATATCGGACATTTCACGAGTAAATAGAGGTTTCCCCGGAAGATATCACCTATTTCATTTTCTC

TATCTGATAGAGGTTTAGCCTCTATTTTAATAGAGGTAGTGGAAAAAACAAATAACTAAAAAAGTGGGGGCCAACTCTAC

ACATAGCAAAAAAACAAACATGTCGGTAAAGCTAGCATCTTACATCTTCGTAGTGGTTTTACTGTATCAGCAAGTGACAT

ATACCTACTCCTACGACAACAAAACAGATACAGTAGCTCAAATAATTTTATACAAGATAAATAATGTGATACACAATGAC

AAATCTGATTTTGGAGTTTGTCACTTTCTCGAGTATTTTATTTCGTGGTCATGCTAAGAGCCATGTTCAGTGAACGTCTT

TTGGATTCCAAATGTCTTTTCTTTATCTTTTTATTTATATTTAGAGTAGAGGGGATAAATTAATATTGATTTTTCTTTTT

CTTTTTCTTTTTATGATAAACAATGAAATTTTTCATTGACAGAAATCTGGGGTCACAATGAGTTTAAGATCGAAAGGAAG

ATAATATAAAGTAACTAGAACATACAGTAAAAGTACAATACCGAGAAATCCGATAATTACCGGAGTAAGACAAATACAAG

TATGAATAAGATCCGATTAAATCGGAGGAAGAAGGGTTTTTCTCGGAATTAAAATCCAACTATTTAGTTTGTTTTTCTTC

TTTAGAAAGATATGCTCCCGGAATAAAAGTGATGCGCTCAAATCTCTTGTAAATAGTGATGAAGCTTTCGTCGTCAAAGA

AATCACCGTTAGAGATTTCGTCGCCAGATAAGTTGTTGATGAAGATTTCGTCGTTGGATAGCATCATTATTGATCTTAAA

ACCCAACTCAATAACCAATAATCGTCATTAAAGCGAAGATAAACCTCAAAAGAGTAGATAAAACCATCGTAATGTTGTAG

ATTAGGTAAAGCATTATAATAGCTAAGTTAAAACTAGTAACACGCATAGAATAGTGAAGAAATCTTCTCAGATCAGGTCC

AAAATGAACAAGATCTGAGCAGAGAAGGGTTGTAGGCGATGGTTTTGTTAAAGAAGAAGAATGAGTGAGAGAGAGAGAGA

GGAGAGAGAATATAAAGAATAAAGAAAAGAGATAATTTTGTTTGATCAAAACTAATGTTGATAAACCATCGATCCGTAGT

AATTATGAAGTTGTTGTTGATGAATTTTGAAATCAGACCACATCGTGTGTATTAACACCAATAAAATAAAATGGTAATAA

TGAGTTTAGTGTTCTTGCCTTAACGAATCTGGGAGAACTTGGTGCAGACACATGCGTATTTTTGAAAAGATTGAAGAACT

GTCTTGCAAGTCAAAACACAAACCGCAAGATATGAGGTTCTTTATTTCATAGGTTAGGTATTATTAGGAGTGGGTGCCCA

GCTTGTTGCTCGGTTACCTTCCACCGACGTGTAAGATTCTTCTTATAAAATATGTGAATTTTTTTGAGGAACCCGAGTTT

GAGCTCTTTCTCTTCTATAAACCCATCAATAAGATGATGTATATCTATAGACCAAAACGTAAAATCTACGGTTAAAATTA

CTAGTTTAGGTTGAATTTAGATTTTTAGATTTTTAACAAGACAAAACTAACATTTATCCATTAAGCATTACTAATAAATT

TATCTACTTATTTTTATGGCCAATAAATTGAATGAATTATGTATGGCAACATATTTACGGCTAGTTAATAGATTTCAGTT

TTTATCCTTTATATTCTTCTTTCTCCTTCTTAACATCTGTTTAATCAAATGCTAGCTCTTGATTGATTTGTTATTGGTTT

GATTTCGTTAATTTCTTTGTTTATAATTTCAATGTGAGATAGTAATAGTTCCATCCAGGAACACCTTTCGTAAATAAAAT

TAAAATCCATAAACTCACCGTTTGGGGACTAAGGTTCTAAGAGCATCCGTAGCGGGCGATCAAATACCAGAGTAAAGATC

AAACACGAGACGCAGTGGAGAAGGAGTAAAGATCAAAGACTAGATTAAAACTAAAAATCATACCAAATTTGGTCTTTGGT

TTGGTTTTAGTCGGGCGTAATTTAAAGTTGCGCCTGATGGGGCGTAGGGAAGAATTATGCTTGATATAAAAAAAAGAAAA

AAATTAGAGTGGGGCGCAACTTAAAGTTGCGCTTAATTCCACCCCATATATATATAAAAAAAAAAGAGTGGGGCGCAACT

TTAAGTTGCGCCCCACCAAATTTGCTCCTCCCACCATAACGTTCAAACACATACTAAATCCAAAATTTAGTCTTTTTTTT

GCTCTTTGATCTTTGGGTTTGATCGCACCACTGCAGTCGCTCTAATAGGTGCTTGGGAATGTAAACTGATATGCTAAGGT

TTACACTTATGTGATAAGATTTTTAATTTTTATTATATATTAAATCTGACTATCAGATGTATATATATTTTTTTTTTCTA

CCTATCTACTAGGTAAATCCGATTTTTCACTTATGTTTTATATAAGTGAAAATTTGGGTTTTCATCGGGAATTGAATTTT

CACCCTTAATAAAAGGCTCTACCTATTTGTCATCCCTACTTTTCCTATTTGCCACTCTCATTAATTACTACCCCCATTTA

TCCTATTTTCCACCCCTATACTTTTTCATATATCCTAAATACTACATCTTTGACTTTTCGTTTACACCTAGAAATGGAAC

CACCCCAACTAATAAACTTAAAAAAACCAAAAATGATATTACGGCAATCAGGGATTGCTATGTAGAAGCTGCAACATCAA

TTGAGAGAAGAAGTAGACGTAAAACTCCACCCCTAACAAAAACTACACCGTGGCTTGTAGTTCCATATGGAAAAGGTAAG

AAAAACCAAGCTTTGTATAATATCTATGAACCTAATAACATAGCTATTATAAAATGAATAACAGAATTGAATGAAAAAAA

GTTGTGGAAAAAACCTTCTCATAAAGGATGGTTAATTATTATATGCCATTATAAAAGTTATGATTACACACCAAAATAGA

ATTATGGGGATCGTTTCCTTTGGAATCCGTCATCGTTTGAGACAGTGCAGTTACCTAGTCTATTACATAGGATTGATAAA

CCCAAGCAATATGAATTGGTTGATTGTGTTTTGTCCTCACCTCCTACACTCACTACTAATACTCAGAAAAATTGTTATGA

TTCAATGGTTATGCCTGAAATATACAATTGGTATGGTGAAACTTCTAGATATATCTTCTTATATTGCAACCCTGAATATC

AACAATGGAGGTCGGAGGAAGTGTCTCAAGATAGTCCCAGGAGCCCCTTATCACTCCATTACCTTAGAGATAAGCTATAT

GCAATGGCGAATATGCATTCATGCTTAGAGATCGAAATACAAAATAAAAATGACAACAATGTTAACCTATGTGTAAGGCG

ATTTGAACCAAGGGAAAGTATTGATTTTCCATGTGAACGAGGATCTTCGTACACTTGTCTTTTTCAGATACATTTTGGTG

AGAGTGAGCAGCCAACCCATGTGATATATGTAGTTGTCCAGAAACGATCTTGTGTCAAAGTATATAGTCGTCCAGTTATA

GCCTACTATTTCTTTACAAAGTCGTCCTTGTTCAGACGGCTGAAGTACTTTGTGGTCGTCCAGAAAGTAGAGACCAGTTC

AACGAAAATTGGAAAAACAAAAAACGGAAAAAAAAAACACTAATCAAACGAAAATTAATCAGTTACTTATGTACCTCCAA

CTCTTCTTTAACGACTGTATTTTGAGCTAAATTAACAAATTCTCGTTGGATGTTTCATTCGTTTAAGGTTTTGATATCTT

GCCCTAAATTAAATTGTTGATTTCTATTAGTTAAATGATAAGAGTTAAAGAAATAATAAACGTAGGGTTAATATTTTTTA

ATGGATTTTCAGTAAAAATAGAGATTGATGGAGAAGAAGACAAATTAGAAGTGGTGATAGTTATAGTCAAATTGAATTTG

ACTTTTGTAGGTGGTAAATAGGAAAAAAATAGAATTGAGAAGTAGAATTACTTTAGAAATTCTTATGAGAGTGGCAAATA

GGAAAAGTGAGGGTGGCAAATAGGTGGAGCCTAATAAAATGGAATTTTCGCCAACTTTGCAAAATTACAAGTGAAAATCG

GAATTTTCATCAACAACTGCAATTAGCGATAGTTGCGATTTTTATTAAAAGTAATAAGTAGAAGTTAAAAATTGCGATTA

TCACTGATCCCTAACTCATGAAAACTGGAATTTTCACAAGGCAATGAAAATCAGAATTATCATAAAATTCGTGAAAATTT

GAAAATTACCAAATTCTCATGCATGATAATTTGAGTTTTTTTTTGTTGGAATCTACCGAATAATTATTGAGAAATTTATT

CTTATGACCCTGGAAATTATGGGTCTTACTCTTTTGGCCCCGGGGTTATTTAATAAAATTTTCAAATTATTCTAGTGGCC

TTGGAATATTATAACCGGTGTGCATATTGTATGCACATAAAATTAGTGTGCATATTGTCTGCACACTATAAAATATGAAG

AAAATTTAAAAAATTATTTACAAAAATCAGTGTGCATATTTTATGCACATAAAATCAGTGTGCATATTGTCTGCACACTA

TAAAATATGAAGAAAAATTAAAAAATTGTTTACAAAAATCAGTGTGCATATTTTCTGCACTCTATAGAATATGAAGAAAA

TTTTAAAAATTACTTAAAGAGTTAAAAAAAATTATATGAAATTAAAATTTGATAGTTGTAGTTACGCTCGTTATGTAGAT

CTTTCAAACAACTTTCCAAATATATAAAATTTGTCGAATTCTTACGTATGGTTCGTGAGACTTGTTATGTTTAAGTTGGC

ATGACTATTATACCCTTAAAGGATATAGTTTATTTTTTTTATACTATCGTCAACCATATTTATTCAAATGACCCTCCTTC

TTTTTTGAGGGTCATTTCAATAAAATGGGTTAATTTTTTTTAAAATTAAGTTTTAAACCAAAGGCCATTTGAGTAAGAGA

GAGTGAGGAGAGGGTCAAACACTTAATTCTGACATAATTATTATTGGATAATTATTAGAGAATTATTATAGGAAAATTAT

TAGGGGAGTTATTTCATGGGAGTTTATTTATTTTAGGAGAGATTTTCTTGGAGAGTAATTTTAGGATTTATTATTTGATT

TTTATAACTAAAGGGTATTTTAGTAATAAATAAATTTATTTAATTTCCTAAAGTAGAAAGGGTATTTTTGTAATCATAAA

TTATTATAACTAATTTTGGGCTCTTCATAATATAATCTTAATGATGGGTTTTAGATACACAATTTTTTTTATTTAGGCTA

GAAATAAATTTTTCCAATAAAATATACATTTTTTTACCGATCCAAAGCTTTGTTTAATTTTTTAATAAAATATAATCGTT

TGTGCATAAAACGATAATGAATTATTTATTCAACAAACAGAATCACAAGGATTCATACACAGCTTTAGTTAGTAACCGAG

GGACAAATTGGACAAACTTACAAAGAATCTTAAGTTAAAATTTACTTATAAAAAAAGTAAAAAATGTACCCACTCCACGA

AGAAGTTAGTAACCTGGTCGTGAGCTAATCGAGCAAGACATGTCACGAGCATCCCGTGGTCGGAAGACAGTAGCTGCCTG

ATTATATATAATGGTCAAAGTCACCAAAGCGAAATCACCTAATAGTCAAAGTCAAAAATGGTGAACCAAGAAATATTACA

CTCAAAACGTCACTTCACGGTATACAAAAACGAGATAAATTAATTCTCGATTTATGATCAGTTTCACTTCACCAATCAGT

CAATCAAAAGAAACATGGATTTTCTTGATCTAATTCTTCTCCGGTCCCAACCGATTTCTATTGCACTTCTTGTGTTAACA

CTAGCTTTGGCATTTTGGAACTATGGAAGAAGAAAATCGGCGAAAAAACTAGCACCTGAAGCCTCAGGTGGGAGACCTAT

AATGGGTCATCTCCATCTTTTCAACGACGGTGAGTTAACTCATCGGAAACTCGGGGTCATGGCGGATGCTTACGGTCCTG

TTTTCAATATCCGATTCGGTTCGCATAAAACGTTAGTTGTAAGTGATTGGGAAATCGTTAAAGAATGTTTTACTACCAAC

GATAAGCTCTTCTCGAATCGTCCGGGTACGTTAGGTATCAAACTCATGTTTTACGACGCCGATTCGGTGGGTTATGCGCC

TTACGGTGCTTACTGGAGAGATTTACGAAAGATTTCGACCTTAAAACTTCTTTCGAATCATCGTATCGATACGATAAAAC

ATCTGAGGAGTTCTGAAGTTGAGTCGTGTTTCGAGTCGCTTTACAATCAGTGGAAAAACGGAGAAAAGCGTGGCCAAAAG

TCAACGTTCACTCCGGTGAGAATGGATAGTTGGTTAGGCGATTTGACGTTTAACGTCGTGGCGAGAATCGTTGCCGGGAA

AAAGAATTTCTCGGCAAACGGTGACGTTGGAGCTCAGAGATATAAAGCAGCAATGGATGAAGCTATGAGATTAATGAGAT

TTTTCGCATTTTCCGATGTAATTCCGTCGCTAAGCTGGTTAGACAATTTAAGAGGTTTGGTTCGAGATATGAAAAAATGT

GCATCTGAAATCGATTCAATTATGGCTTCTTGGGTTGAAGAACATCGTGTTAAAAGAATTTCAGGAGATAACTCTCAGCT

TGAACATGATTTTATTGACGTTTGCTTGGATATTATGGAGCATTCTTCTTTACCTGGGGACGATCCTGATCTCGTCGTCA

AATCTACGTGTCTGGTATGTATTTTTTAAATTTATTAGGACGTGCTAATTTGTTTCAGAAAAAATGATACTTTTTATATT

TTTTTAATTTTATTTAGAAAAACATCACTTTTTTCTGAAACGAAAAAAAATATCATTCTCACCATTTAATGATTTAAATG

GAATTGAAGGACATGATACTGGGTGGGAGTGACACAACAACAGTTACCCTAACATGGGCAATGTCTTTACTCTTAAACCA

TCCCCAAGTGCTACAAAAAGCAAAAGAAGAACTAGACACACAAGTAGGAAAGAACCGGCAGGTCGAAGACTCGGACATCC

CGAATCTTCATTTCATCCAAGCGATCATCAAAGAGACGATGAGGTTGTACCCTGCCGGACCGTTAATCGAACGGAGGACG

ATGGAGGATTGCGAAGTAGCGGGGTACCAAATTCCGGCCGGCACGCGCTTAATAGTTAACGTCTGGAAGATGCAAAGAGA

TGGGAACGTATACAAAGGCGATCCGTTGGAGTTTAGACCCGACAGGTTTTTAACTAGTAATGCGGATGTGGATCTAAAGG

GTCAGCATTACGAATTGATACCATTTGGAGCGGGTCGACGGATATGCCCGGGAGTTTCGTTTGCGGTTCAGTTGATGCAT

TTGGTTCTTGCTCGATTACTTCATGAGTTCGAGATCACGACGGTGGAGCAAGAAGCCAAGGTGGATATGGCTGAAAGTGG

TGGTTTGCTTTGTTATAAGATAATGCCGTTGGAAGTACTGATCAAACCTCGACTCGAGATATGATCAAAGGAGGAGTCCA

ACATATAAACAAGAATGCGAATCGCCGTATCTCAGGTCCAACATAGTATAGTTATGTATGATACTTGTTTTCTCTTTTGT

CTTGATTTTATGGGCTTTCTAAAGTTGTAAGAGCAGTACAATAAAATTAAAATTACGGGGACTTTAGCCCTACATATATA

TATATATACTGGCCTAGCATCAGCTGCAACAGTACTTGCAGAAAGGTGCTCCAGTTAGCAACGCCAACACCGCCTTTCAT

GGTTTGCCAGATGAAGTTGACAACATATTCATCCGCGAGCTTGCCTTTTCAGCCTTCAAATTCCAGGATAGATAAAACGT

TAGAGAATGATGTAAGGAACTAGAAACGGTATTGTTATGACCAATGGCTCTTCCAAGCACAATCAATAGAGAATTGAAAT

AGAAATGAAATCAAAAGTAAAACATGCACACACACACAACACAAGGATTTACAGTGGTTCGATCAAGATGATCTACATCC

ACTTTGGAAGACGACGATAACGTTGCACTATGATTTGAACAAGTTGGTAGCATCATGTATGCTATGGTTTGTACAAGGCC

GGATATCGCACAAGCTATGAGTGTGGTAAGCAGATACATGGGAAGCCCCGGGAAGGCTCATTGGCAAGCTGTAAAGTGGA

TTTTGAAGTATCTAAGGGGAAGTACAGATGTTGGTTTAGTGTTTGATAAACACTATGGTGATTCGACGAATGTTGTGGGT

TATGTTGATTCTGATTATGCAGGTGATCTTGATAGAAGAAGGTCATTGACAGGTTATACTTTTACATTGTGTGGTAGTAC

TATCAGTTGGAGGTCGGTTTTACAAAGCACAATTGCATTATCTACAACTGAGGCAGAGTACATGGCAGCCGCGGAAGCAG

TGAAGGAAGGTATATGGTTGATGGGTTTGGTCGGTGATTTGGGTTTCAAGCAACATAATGCAACTTTGTTCTGTGACAGT

CAAAGTGCCATTCACTTGACCAAGAACCAGATGTATCATGAGAGAACGAAGCACATCGATGTCAGGTATCATTTTATCAA

GGATGCAGTTGAGGAAGGTAGGATAGCGATGCTAAAGGTTCCTACAATGGATAACCCTGCGGACATGTTGACGAAGCCGA

TTCCAAGTATCAAGTTCAAGCACTGCTTGGACTTGGTTGGTGTCAACAGGTGTTGAGAGCCCTTGAGGGGCTGAGAGAAG

ACGATGATGATGAGTGGTGATGGTATTCAAGTCAAGGTGGATATTTGTAATATATGTCTTGAATATACTAATCACTCAAG

TGGAGCTAGCTTTCGGTGGAGTGGATCTAGCCTTTCGGTGGACAGTGGAGCACTCAAGTGGAGATAGCCTTTCGTCTTAG

TTGGCACTCAAGTGGAGATAGTCTTTCGGCTTAGTGGGCACTCAAGTGGAGATAGTCTTTCGGCTTAGTGGGCACTCAAG

TGGAGCACATAAGATGGTGATATATGTTGACATTTAACACCTATTCTAATGTGCTCCACTTGAGTGCCCACTAAGCCGAA

AGACTATCTCCACTTGAGTGCCCACTAAGCCGAAAGGCTATCTCCACTTGAGTGCTCCACTGTCCCCCGAAAGGCTAGCT

CCACTCCACCGAAAGCTAGCTCCACTTGAGTGATTAGTATATTCAAGACATATACTACAAGTATAATTTTGAAGGAGAAT

TCACCTGCATATCCCAATCACATCTAATGGTGATTATGATTAGTATTAACGTTTGAACGGCGACACCAGATATCATCCCA

CTCCATAGACCCTGCAAGTTTTCAATTTTGGTTGAGCAGATGTACTTTCGATTTCCGTTTGGAAGAAAAATTCTTAACAA

AAAATGGGCATTGCAACATTGGAATACGTACTCCAACACCAAGATGCAAAACCCATCCAAGGGCAGCCCCAAGTGGAAGT

CCGATGATATAGTAGCAACCGAGGTTAACGTATGCTACGAAGGCTTGCCATCCTAAGCCAATGGCTACCCCTGATAAAGC

ATACCACTATTTAACTAATGTTTGTAAAGATGGGGTACAAAATTGAAGTGGTGTGTACCCGAGAGAACTTGTTGAATGCT

GTTGAGGAGAATGGTGAAAGCCAAAAGGAACGCAAGGTTGTCAACTGCGAGGAGGACAACATTGCTCGAGGTGAAAACAA

TCGCCAACTTGTCATGAAAAATCCCTATCAATGTACAAATAAATATCCCTATTATCAAAGATGTAATTGTTGAGATTTCG

GCAGCAAATTTTGCAGCTTTTCCATTCCCTGCTCCAAGCTCATTTGCGACCCTTATTCTGATGCATGGATAAAAAACACA

AGGCGCAAAAATTAAAAGCATGTAGGACAATATTGACTATATCTCTTTACTACCATGTTGCAAATTCATCTGATTCAAAA

TAGTCAAATAATTACCCAGCTGAAGCGAAGAAACCAAGTGAAATCATCGACTCCAATCCATTTATGCTCATACTGTGAAC

CATCATATATATCATATAAGTACAAATCAAACTCTAATATCAATGGCTATCGACTTCAGGGCACCCTAAGATGTTACAAC

TTTTATTACATATCGGGTCCTTAGAAGCCAAACAAAAGAAGAATACACACCAGACAGTTAGAGCTGAGACCGCAACCGTA

GCGTTTTTCAAGTTCCCAGCCATCAAAACAAGTATATTGTAGTACCAATTCTCCAAGCTGTATATTCCCCGTGATTAGCT

ACATATAATATTTAGCCATAAAAAAATCAGTGAGAGAGAAAGATATACCATAACATGACACCAGAAGCCACGGAGAGTTT

GAAAAACTCCCAAAGGCCAGAAAAAGCTTTCATAGAAAAACCAGTCCAAGTTTCAGGGCAACCACCACCAACAATATAAG

AAAATAATACCAGATCTATAATCCACCACGAGACATTCGACGTAGCAGCAACTCCAATAACACCAAAACCAAGTATGTGA

ACAGCTAACCAGTTGAGGAATATATGAAGGAGTAGGGCCCCGAAACATACCCAAGCAATTATAGGTGTCTTAAGTTGACA

CTGAAGAGATATTTGTAAAGGAAATTGTAAAGCTAAGGCAAAATGAAGTGGAATCAACCAGATTGTTATTGATCCCGATA

GCTCGGCTAGATCATCTGGTTGTCCCAAGAGCTTAAGAATTGGTGTAGTGAATACGAAAAGCGGTAGAAGTAGAATTGAA

CAGAGTGTCAAAATAACCAATGACCGCTGCGTGTAAACTCCTATCATATGGTATTGTCTAGCTCCGTAAGCTTGACCGCA

TAATGTCTCTAATGCACTTGAAATTCCCATCTGAATGAAAGCATTTTAAATAAAAACCTAAATGAAAAGCTAGAGTATAA

AACAGAGTAAGTGAAAGAGATGTGGAATATTGTTTACCAATACGCCGAAGCTAAAACCAATAACAACCGTAAAGGAAAGG

GCGATTGCAGCAAGCTCAAGGTTACCAAAATGACCAGCAAAAGCTTGGGTGACAATGATCATAGAAAAAGAAGCTATACT

ACTGAAGATAGATGGACCAGCAATGTACCATAACTTCTTTGATTCAACCCAAACTCTAGTCGGTAAATCGTCTTGTATTT

CATTATGATCATGATTCAGTTGCAGTGTAGTGGCCGGCGATAAATGTTCCTCTAATAACGGAACGTTTATTGGTTCCTTT

CTTGCTCCATCTCCACTTTTGCCGTCTTCCATTATACTTGGTCTAGCTAGTAGTTCAATCAGTCACCAGTTGAACTACTA

GTAGAGGAAATATGTATACTCTGATCTATTTCATGTGAACAAATATGTATGTATATATACATATATAGACTCATAGTTGA

GATTTCCAAGACTTCTTGTTGAGATATTCTTAATTAGGTAAGTAAAGAGCACAATATCCAAAAAGAATGAAGAAGATCAT

ATAACTTTGAAGGTGTAGAACAGTAAAATGAAGCAGAAAGAGAGAACTCTGGGTTTGCTGCGAGGGAAGAGAACATTGAA

TGAAGGCCAATGAAGCAGCTAAAAGTTGATGGTGATTTTTATTTTATTTTTCGGAAAGTCTAGATTCACCGTGAAAAATT

CACGGTGATATCACCGACCCACACAAAAACAGTGGAGCCCACCGTCAATCAAAACCCCAACCTAGTTAGCAATTCGGGGT

ACGGAGAGGAGTTCGGGACGCCATACGTTCGAGAATTATACGGATTTGGATAGGTCGGATTCGGTCAAAAAATCGGTCAA

CGGTTCGTTGTTCGGAGTGTAGTTCGGAACGGATACGTGCGTGTATAAATCGTTTTAGAAATATGAGTTCGGTACGGAAA

ATTCGGCTTACATCAAATTAATAGAAATTATATTTTAAAGTTTGTATAGAGTTGAAAGAATGATTTAAACCAATTTTATT

GTATTATATAGATAATACATTAAGAATAGACACAAACAAGCAAATAGCTCAATGGTTTGAGCCTTAGTTAGCAAGTAAGA

GGTCATGGGTTCAATCCCCACCAACAACTTCTTTTTAATAACAAAACTAGTGACAAAACCCCTAGGTGAACAAACTACTG

TAAGGTGGATCCTGGTTTCAAATATAGACATGATCATGTGAGAGATATCTTATATGATGTATTATGGAGGGCTGGTATTT

CGGCAAAGAAAGAGGCGTCGGTCAACTTTTTGACGGATCCTCTTGAAGGCAGATCAACACTCAGGCCAGCGGATGTTCTT

ATTTTTGGGTGGGCCAACGGAAAACATACGTGTGTTGATCTTACTTTGGGGTTTCTCCATTAGTAGGCATCGGGCAAGGC

TCTTTCACAGTTGGACAAGCAGCTTTGAAAGCTGCCTTGGGTAAGATAACGAAGCATGAAAAGGCTTGTATTGATAATGG

ACATGCTTTTATTCCCTTTGCCTTTGACACTTTTGGGTATTTAGCTCCGGAGGCGGTTGGACTCCTAGCGAGAGTTCAAA

GAGTCATGCACAGTAATGTCATGACTCCCGATTCTAGAGAGTTTGTGTTTAAGAGGATAGGGTTTGCCATTCAAAAAGGG

CTTGCGGCGCAGCTTGTTGCTCGCTTGCCTTCTATGTAATAACTCATGAATCTGTTTCAATGTTTATTATTGATTTAACA

AATAAAAACCGTTTCAAATAAAATTGGCACAAAAACTCCACTTCAAACAAAATTTTTAAAATTTAGTTTTTATTTTATTT

CTAAATTCCAAAATTAACCTTCCCTATATAAAACCACTTTCAAAGGAAAGTAGTTTGGCGGAAACAGGAAACTAAAAACA

AATTCGGAGCAATTTTGATTCTTCATTCGTAGTTTTTCTTCGTTTTTTTCCAGATCTCGTTTCAAAGATTCGTCATCGTA

TTCTCGTTTCGATTCAAAGATTGAGAGAGCAATTTTAGGTTTGATTGAAGAACCAGAAGAGAAGCTTACAAGTTTTGAGA

TCCGTAGAAGAATCCAATTCCGTTGTTCGAAGAAGAGAAGATAATCGTAGCTGTTTGAAGAAGAGAAGAATCCAATAAAA

ACCCCGAGATCTGGTGGCATTCTGTTTAGCTGAAATCATATTTCCGCTTTTCAAGTTGAAATTGAATCTCGAATTGAAGA

ATCCAGCTGGACAATCAATCAATTTCTTGCTTCTCATCTTCGTCTTCATATTAATCTGTGTAGTATTCGGATTATTCATC

AAGGAATCAATCAATTTTTCAGGTATTCAATTCTCTTTTTTGTGTTGATTTTTGATTTGTTTTCGTTAATTTTTCTAGTT

ACTCTAGACGAATAAACTGAATTTCTGTTAGCATATGATTTGTTTTATGTTTGTGTTGATGTTCTTATGAACGATTTCAT

TTGTTTTGGAATGACTGGTTTTCAATTTTTGTTGTTGTTGATATTCACATCTAGTGAAACTGTTTTTGGTTGCATCATTT

TGTTCTGTAGATTATGGATTCTGTTCTCGCAGTTGTTTGTCACAAAGGTGATTGTTTAAATTTTCGTATTAGCCTCCAAG

AGTCTTTTGGTAATTTCAAGACTAGTATATGCTCAAGTTGGGATGATTTCTCTCCTCCTTCTATGGAAATTTTTCATATG

GTTGATGATCAGCAAAAGGTCATTGATTCTGATTTTGATCTTCGAGCATTGGCTTTATTTAGTCTGCATAATAAAGAAGG

AATTATGGAGTTACACGTACGTCATAAAGCTGAGAGTTCTGGTTCAGTAAGTTCAGGAATTATAGCTTTTGATGACCAGC

CACGGAGTTATGGAATTATACCTTTTGAGAAGCCACGAATTGATTATGATGTTCCAGATAAAATTAAGGAACCGTTGAAG

TCTGCTCACTGGTTACATCTTTTTCAAGGAGTTGAACAATTGTTTCCGGGAGGCGTTCAACAAGTTCGTTGTGATTTACA

GAAGTATCATGCAGCAGCTGGTTATGCATTCAAGATATACAGGAATGAGAAGTTGAGGATTGGTGCATGTTGTGCTAATA

AGAAAGAGGAGAAATGTGATTGGCATTTATATGCTGTGTATGTTGATGGTGTTGTAGGAAGTCCTTTTATTATCAAGGAA

CTAAATAATGAGCATACTTGTGTTGGTGGATATGTTAACACACCACAGATATCGAAGAAACTAGTTAGTAGCTTGATTAT

TGATGAAATTAAACAGGATCCTAATAAGAAAACTAAGCATATTATGGAACAGTTTACGAGAGAATACGGCTTTGACATTA

GTCGTTATTATGCATACTCAGGGAAGAAGCATGCGCTGAATACTACATGGGGGGAGAACGAGAAATATTTTATGTTCTTG

AACTGGTATAAAAACAAGTTGATTGAAACCAATCCTGGATCTCACGTGGTTTTGGAAGTCGAAGAAGGGACTCAAAAATT

TCAGAGGATGTTTATCTTTTTTGAAGCTTGTAGTCATGGATTCAACTTCTGTCGTCCTGTATTATTTGTTGATGCGGCAC

ACTTGAAAAGCAAGTACCTTGGTCATATGATGGCAGCAACGGGTCAAACTGGAAATGATGGTAAGGTTTTCATAATTTTT

TTATGTTGGTTTTATCATTTTTTCATGTTGGTTTCATCATTTTTTCATGTTGGTTTCATCATTTTTTATGTTGGTTTCAT

CATTTTCTATTTGTAGTTTTCATCGCATCTTATGCTTGTGTTATGTTTTTTTGTTCATGTTACAGAAATATACCCTCTTG

CTTATGATGTGGTATCATCGGAGAATGAAGAGAACTGGAAATGGTTTTTGGATAATTTGAAGCAAGTCCTTTCTCCTTTG

CGACCAAAGCTTACTTTTGTGAGTGATCGAGGTATTGGATTGGTAACTCAAATTCATGTTGTATTTCCTGAGGCTTTTCA

TGGTTGGTGCTACTGGCATATGGAATGCAATATCAACACATGCTTACCAAAGAATTGCAAGGTTTATAACAAATATTTAT

TGGGACTGTTTAAGAAATGTGCATATGCTTCTACTCATAAAGATTTTACAGAGAGTTGGGATAAGTTGAGGAAGGTTGAA

AATCATAAATTGCAAGACTATCTAAGTAGAGCTCATGTTGATAAATGGGCAAGTTTTTTCTTCAAGGGTCGACGATATGG

TAGGTTGTGTTCAAATATTGCTGAGAGTTTCAATGGTTCGGTCGTTGAAGAAAGAGAGAAGCCTATAGCCGTCATGGTTG

ATGCGATCAGGGTGAAGCTTATGGACCAAATGTGTACACGTCGCGATGATTGTGCTAACTTAATGAATTGGCGTCAAACT

CTATGTCCAGATTATGAAGTTATGCTCCACGAAAACAAGATTCATGGCTGTAACTACCCAGTCACCAAGTCGTCGGAATA

TGTGTTTGAAGTTTATGCTTCATTAACACAGACAGTCGATTTGAAAATAAAAACATGTTCTTGCAATCGATGGAGTATTG

ACGGTTTCCCTTGTGCCCATGCTGTCCGTTGTAACATGGTGAAGGGAGGAGACCCTTACAAGTATGTTGAACACTACTTT

ACAGTAAAGTTATATCGAGAATCATATAAACATGCAATCAATCCTGTTCCTACAGTTGAAAAGCCTGTGACAGTCTCCAA

AATCAATGGTTTTTGGTCCGAATAATGGACCACAGCCAAGACGTCCACCGAAGAAGAAGAGGATTCCTAATACAGGTTCA

GTTCCCAATAAGAAGATGAGAACATGTGGTTCTTGCAATGTTTTGGCCACCCATAATAGGATGACATGCCCTTCTCGTCA

GATTTCATAAGAACTTGTAGTCATTTACTATTTCGTGCGACTACAAGTCATTGAATTTCGTTTTTAGACTACAATATGTG

TTTCATTTACTATTTGAGATTTCTGTTTTAGAAGTTAATGTGATGCAGGTGTTCTTATTTTATAATTTGTCTATGAATAA

GTAATTTTGATCAAGTTTGTCAGACTTTTTGTTTGATTAAGTTTGACAGAATTTCTGTATGCCTTTAGAGGTTGGTGTAA

CCGAATTAGGTTTCATCATTTTTTTGATGGAACCAGAATATGTTTCATCATATTTTGGTGGAACCCTTTTTGGTTTCACC

ATTTTTTTACTGTCACTTTTTCTTCTATATTGTTTGACAACAGAGTAGATGATGTTTTGTGTACTGTTTATTGGTTCCCC

AAATATATGATACATCATTTATGGATACTGTTTTATAAAAATGAGACATCATATTTCTGTAGACAAGCTTTTAAAGGAAC

ACAACTGAAACCGCACGAAATCGTTTTTTATATAAACTGTGACCATTTCACAAGTACTTTAAACTTCAACTAAAATTTGA

ATCTGCAAGAAGAACTATAGTATCATCATTACAAACTACATTTCAAAAGTAATTTAAACTTCAACTAAAACTGAACGTTC

AGGAGGATGGGCACAACATGAGTATTATTCTATATTGTCAAGTAGTATCTTCAACGCCAACTTTGGTCTCATATTATGCA

CCTTTTCTCGTATATCATCTTCACGAGACATGTTTCCTTTCAAGAGGCTCTTCATGTAATAGAAAACATGTAGTCCACAA

TCGTTCCTGAAAAACCAAAACAATGTTTCAACTGTTATAAAGTGACTAGCTGAGAATATTATGATATTAGTAGCTATATG

TAAAGCATTCGGTTTTTAACTATTTACAAGTGTTGTAATGATTTCAAGCATATTCATTTATGAACTAATAAAATTCTAAA

CATAGCTTACCCAATTTTTTGACAACATGAAGGGGTGTGCAGAGTGTATGGTGTTGGTTGACTAGTCTGCTCGTCACGCG

GGAGCTTTGCGTCGTGCTGTGCAAATGCTGCTATTATTCGCTGTCTCATGCTTTCAACACTTTTTCCGCATTCTTCTTCG

GTGGAAACCAAGGAGTTGTAGTGATAAAATTCTCCAGTTTCAAAATCAAAAGCAAGCAGCGTCCAATTGTTTACATCTCC

AATTGATGTCAGCAACAGAACGAACAAAAAACGCACGCTATAATCCATGTTGTCGATGAAACCTTCTATTACAGCACCGC

ATTTCTTTTCTAATGAGTGACATGTCTGAAACAGAATACCAGATAACAGCTCCTTAATGAGTTACAAACATGCAAGTTTG

ATGAAACCAAACTTGGTTCCATCAAAATTAACCATATAAACAACATCATTTTTTGCTTTCCACAACTATTTTTTCTGCCA

AAAAAATTACATTAAAGAATTTTAGAGTAAACTTACATATGCAAATGTGCTTAGAAACACAGCTCTCTGGTACTTTGGAG

AACCATCTTGTTTCAACTCATTTTTTTTTATGTTGTTTTGCAGCTTCAGAATATAGAACTCAATTATGTCTCATGCGACG

TCACCTTTTCTTATTAAACTATCCATATTCTTCCCCGATATATGCAAATGAAGATTTAAATGCTCCCACGCCGTTGAGCT

GTTAATTTTGTTTTGAAAAAGCTAGTCAATGATGGATCAAAAAAGGAGAAACAATGCATAACTTGGAACAAAATTTTGAT

GAAATCAGTCTTGGTTTCATCAGTTATTTCCCACTTACCTTTCACTTTTCTTTCTGAAAAAAGTGGTCACGAGGGGTTTC

TCATTTCTGTCGAGTACCCTCAAGATCTTCAAGTTATCAACTGGTTTCAACTTCACCCCCTGTACTTCCTGAACCTCCTT

TTTATCCTTGCACTTACTAGCTACAGCCTTTCTTCTTTCTTTGCACTGATTTTGTAATCTTGCATCTTAATAGGTGGTCC

CAAATTCCTTCTATCTCTTCTATCTTCCCTCATTCTTGTAACCATGTTACTCATACTCTGTTCAGCACTGAGTTCTCCCT

GTCCTTGCTCTTGATTTTCATGCTGCGACATACCTAAATCAAATCCAGGTTGTTCATTTTCTAACTCGAGTAAATCATTC

GCCATTTTTACGGCTGAGTAGTAGTATACGCTTCCAACATTTGGCATTGATGACGAACAACTTTTCTTATTGTATTCGTC

TTCCATTTTCCGAATTTCTTCTTCATCAATTACAAATTGAGTTTCATTGTCTTCTTGCTCAATATTTATCTTCAACGAAG

TTGCTTTCTCAATTTAAGATTCGACCCGTTCTGCCACCTTGATATCCTCGATAGTGTTCGGAGTCAACTCAGTACCATCT

CCTTCTTGTTCTTGCTCGGTTACTATCTTCGTCGAGGTAGCTTTATCCATTTCAGATTCCACCCGCTCTGCCGCCTTGAT

ATCCTCAATAGTATTCGGAGTCAACTCAGTACCATCTCCTTCTTATTCTTGCTCGGTGACTGGCTTCGTCGAGGGAGATA

TGACCTGCTCTTTTTCTGCTACTCCTATTTGGTTTAACAATTTCGAAAGGTCCACAGTTGAAAAGACACTATCTAGTGTA

ACGGGATCTTCACCTTTTGAAGATTCTTGTTGATGTTCCACATCTTGTGGTGTTTCTCCTTGGGTTGAATCCTTTTCAGG

TTGCATCAGAAAATCTTCAAGAGTTTGAGCAGAATCTTCCCATGTTTTGAAACTTGAAACCTTTTCTCCAGGAAGCAAAC

CCCCACCTTGTTCCATCACAGCATTGTCATCATGTGGTAGCCTTTCTTCAGACAGTGAAACTTTTTATTGTTCTATCATG

TTAACATCCCTTTTTATCACCTGCAATATTAAGATATCTAAGTTATTTTTTTCTTCATCTTCTTATGTTTGAAAACCCAA

TGCTGTAAAATCTAAATTTCTGAAAGAAAAGAAAGAACTGGCCAATTCGTCATTAACTTGAAATGATGGAACCAACTCTG

GTTTCATCAAAAAAATGATGGAACCAACTCTGGTTTCATCAAAAAAATGAATTAAAACAAACATTAACTTGAACTGATGA

AATCAACTCTGGCGACAACAAACCAAATATGACAGAAAGGGTACATAATACCACACAAGTTTATGAAACACTACCAATAA

ACTGAAATTTAAGAAAGAAGTTAGAAATTTAAACTTTCTCCTCAATTAAACTCATACAAGTTTCTTTAATCTTAGAGTCT

ACAATGTTATTCATAGGCAATGAAACCATTTATGTTTCCATCAAAAAAGGAATTTCATAGCAAACAATGCACAAAGCAAA

CTAATAAAACTAGATCTAACATATTACCTCAACATCTTCTTCACGGTTCGAAACCGGTTCAGGTACTATTGGAACATCCG

TTTGTTTTCTTTTACGTCTCAAATAAGTGCTCTGTACCGGTTGTTGTTCCCTGTCCTCGGCATCATATTTCGACAGCAGT

GATTCTTCTTCTTCGGAAAGAACATTTTTAGCATTGGTCCTAAACTTTTCAATCAAGGCTCGTGCAAATTGGTATTTGGA

TTTGTAGGAGTTAGCTTCACTGCTAGAACCAATATTTTCTTCAACTGGAACTTCATTTTCTTCAACTGGCACTTGATTTT

CATTATCACCAATCGTCACTTCATGTTGAACATCATCTTCAGCAATAATTTCCACCAAGCGCATTTCTATTTCGTCATAT

GGAAGCAAGAATGATCCCTTTCGGTGACCAATATTCTCCTATAGTACACAAGAAAATTTTCATAAAAAAAAAATGATGAA

TCCAAAACTGGTTTCATCAAAAAAGTAAATCAAAAACTGGTGGAACATGGCATGATATACAATTGAGACATGGCATGATA

TCTAAAACACGGCATGATATACAATTTAGGCATGGATGACATCTAAAACATAGAAGATTTCGAATATTGTCTAACAAACA

TAAAAGATTTCGAACATGGCATGATATACTAACCAAATTAAACATATAAGAACCTTGAAGTTATCTAAGAAACAAAAATT

CAGCTAGAATTTATGATGAAACCAAAATTGGTTTCATCAGGAAAGTAACTATTTACCTCAGACCAAAATTCTTTTTTATC

GACAAGATTATCAAAGTTAGACAATACTTGGTGACAGATGTCGCTAAGGTTCCATCTAGCAAACCTCAAATACTTTTCGA

GCCCATCCTTTCTTTGTGCAATCTTTGTCATCTCCGCCAACCAATACTGGAAAAATAAAAATAAATATTCAAATGAATCA

TCAAAAAAAAGTCGCTTATTAATCAACAAACCAATACGGAAAAAAAAAAAAGTTAACCTATGAAGAAAAAAAAAAAGAAA

AACTCTTAACCTACCAAAGGATATATGACGCAACCAACTGCGTTTTCTACTTCTTTTCTGTTTTTCATTATCGAATCAAG

TAGATAGTCATGGAAGACTACCGGCCAACACACCCTGTTCATATCAAACACCATGTACAAGTACTTTGCTGCCAGAGTTT

TCCCACTTTTGTTGGGAAAAAATACTGAAACCAAAAGAAACATCACGAAAAACTTGACAAATTCATACGCTCTGGCTTCA

TCAGCTGTGGATTCCATCATACAATCTTCAATAATTTTTTTACTCGTCTTTTTATTTCCTTGAAAATACGTACCCACGAA

GACGTTCACCTGTTTGAGCAGCTCTTCGTCAATGTTATCTACTTCTACATAATCCACCATTTGGAGACCAAAAGTCACAA

CAAAATGTTCGGGAATGGTCCGCAGAGTCATTTCCCTTGATCTTGTTACTTTGAAACAATACGGGCGCTTACCATCCTGT

CTATTTTCTATTGAACTTAAAAACATAATTACCGCTTTCGTCTTTTTTATGAGCTCTTCTTCGTTCATTACACCATCATA

GAACAGCTCAAAGAAACGCCAGAATGGACATGATTGAAGCGCTTTCAAATGCATTTGCTTGTTTTCTATCAACGCCTTAA

TTTTCTCTGCTAGCTTGCACAAATGATGCAATGAACACCTGTATGGTTCTGTTTGTCTTTTTTTACCTATTGTGAAACCA

GAAGTAAACCATAAGAGAATAATAAACAAATAATAATGATGAAACCACAAGATACACACTTGAAAAATGACGAAACCATT

AGCACGAACATAGTGATGAAACTGTGCATAATTGAAAAGTGGTGAAACCACAAGATACACACGTGAAAAATGACGAAACC

ATAAGCACGAACAAAGTGATGAAACTGTGCAAAATTGAAAAGTGGTGAAACCACACGATATACACTTGAAAAATGATGAA

ACCATAAGCATAATTAGGTCCACAAACAATAACTCTAGATGTAAACCAACAACTAAAAAATGATGAAACCAATTTTGGTT

TCATCAACAAACAGAATCCATAATCTACAGAACAAAATGATGCAACCAAAAACAGGTAAAAACAACAACAAAAAATGAAA

ATGAAATGTGGTTTCATCATTTATCTGTTTACTCCAACAACAATAGAAAATTGTTTTTCACCATTAATTGAACATGATGT

AACTCGATTTTGGTTTCATCAAACAGAATACACAAGAAAAAAAACACCCAAAATTGAAATTAGTTTTCAACTGAAACCTA

AAGCTAACAAATGAAATCTATAACTAATAGATGAAACCTAAAATTAAACTACAAAACCCAGAAAAATGACGAAACCTAAA

ATCAAACTGAAAACCCAAAAAAAAGAAAAAAACGAAACCTAAACCAAAAAAATCTGGTTACATCAAAAAATTGAAGAATA

ATGATGATATTTCTTACCTTTTGTTGCTGGTTTTGCAGTTTTCCTAAGCTTATCATCTTTATTTTTCTCCTGTTAATTTT

TTTTCTTCGCCATTGTTGCTGTTCTTAGACAATTTATGACAGATTTTGATGAAACCGTTGAAAAACGAAAGCATCGACAA

AGATAACAAGTTGAAATCGAATATCAGTTGACTAATTTCGCTACAGGATGATGATTTTGTTTCAGATCTGAGAATTTTGT

GGGTAGTTTTGAAGGAGATGCGGAAGAGAGGGGAAGTTTTTGTTTCTTTTGTTTCTTATGAAACGTTATAAAGAAATAAA

TGCAGGCGGTTGTTTGGTCTTTCTTTTTTTTGGTGGGGTAGTTTAGACAATTTAGGATATTTGGGGTTTTTGTGTCATAT

TTGTTCGGACGGGTTTTTTGTAGATGGGTTATAACCGCGCCGTTTTAATAACTCTAAAAAAAACACTTAAATGGGTTCGA

ATCGCCGAGTCACTGTTCGAATCGCCGAGTCACCTGTCGAATCGCCTAGTCACCTGTCCGAATTGATTCGCGGAAAATTC

GGATAACCCATATAATGCGCGAACCGGGTCGGTGCTAGAGTAATTCGGCCGAATACATCGGATTCGGTTGATTCGTACGA

ATTCGCGAATCATTCGCCGAATTACTAACTAGGAACCCACCCCTAATTAGTAATTCGGCGAATCATACACGTATGTATGA

TTCGCGAATTCTTGCGTATTATTCGGGACAGAATTATACGGCCGTCCAACTGAATCACCGTATCTAATACGCGCGTTTTG

TGGAGTTTTCGAATAATCCGCGAATTATACGGACATTCGGTTCGCCAGCTGGATTTATTTTCGCAAAAATGGGAGTGGGG

AGAGTCGAACCTACGACCTGTGGCCTGCTAAACCAACCTTCAACCACTGCACCAGGTGCGTGTTGATGTTATATTTGGGT

ATATTAAATATATATTGCATTTTTTTATCACAAATCATGCCCCTCAACACTTAAAACATATTATTTTATTAATTAATGTT

TGCCGTATTTTCCATACCGAACTCAGACTAGTAAAACGAATTATACACGTACGTATCACGTTCCGTACTACTGCCGAATC

ACGAATCGTTGACCGGATTTTTGACCGAATTTAATTTTTACGAATCCGTATAATACTCGTACGTTTGCCGTTCCGTACGT

ATGCCGTATCCCGAATTGCTAACTAGGCCCCACCCCCTAAAAAACTCGTGGTGGGTCCCAGGGCATTCTCTCACCCGGTG

ATTTTTGCGGTGAGAAAACCCGATGTGTGTAGCACATCCGTTTATTTTTTCATTTCCTTTTTCGGTATATTGAATTTTTT

TAATAAAATGCTTTATGCACCGTGCAGTGTACACAGCGGTGAGAGAGTGAGAGAAAAAAACATGTGGGTCCCACATATAG

CCCTCACCCCTGCTAACTCCTCCCCAGGACGCCCCTAACAATCTTTGGATTTTCCCGATATGTGCATCACGGTACACACC

TTTGGTGTGTAGCAAAGCTGAGGAAAAAAGGCTCATGACAACACAGAAAGACCGACAAAAAATAAAATAACAAAAGGCAT

AAGACAACCAAAAAGAAGAAAAGCAAAATGAACTTACAATTGGATTTGAAGTTACCCACATGAATTCCTTTCCATATGAT

GCAACCATAAGCATGCTTCACTAACATGTTTTTCTTCTTTCTTTTTTTTTCTTTTTTTGTTAAAAAGGTTATGCTATTAT

TGAAAAGAAAAGGAAACATACGGAAAAGAAAGCACCTCACTAGTGCAAACAAAAGCACCAAAGGAGAAAGGCTAGTTCAA

ACACAGAGCCACACACAAAGCCCAGCCGGAGCAAAACAGCCTTACCGGCCTAAAACAAGAAAAAGACTAAGAAAGTATAG

CCAAATGGAAGAGCAGCTACTCGGGAATTCAAATCCCAAGGGGTTATTCAACCAAAAAAAGTGTTTCTTCTGACTAAGCT

ATTGGCCCTTGTTATCAGGAGATGTGCATATTTGTACATATATTCTATATATGGTTAAGAAATATGTTAGTTATGTTTAT

ATTATTTTTGGTGTAAATTTGTTTTTGGGTCTATATATATTTAGTCTACAGAGAAACATATTGTAATCTTAATCTTTTGA

TAATACACATCGAAATGTCTCAAGGAGCTGATGGACTATTTCTATCTCAACGAAAATACACATTGGATATTCTGTCTGAG

ACTGGACTTTTTGGAGCCAAACCAGCTACTTCACCCATCGATCAACATCATCGTTTAGCTCTTGATGATGGACCTCTGTA

TTCTGACTCATCCCAGTATCGTCGCATGATTGGACGTTTGATTTATTTGACCGTTACACGTCCTGAATTGTGTTACTCGG

TGCATGTGCTGGCTCAATTTATGCAGACTCCTCGGGTATCTCATTGGAAAGTTGCTGTCTGAGTTCTTCGTTATCTAAAG

GGTCATCCTGGTCAAGGAATTGTTTTACGCAAGGATAGTATTCTTCAACTTACTGCTTATTGTGACTCTGATTGGGCCAG

GGCCGGCTCTGAAGCCCGTCAACTTGTGCAGCCGCACAAGGCAGGAAAAAAATGTAAGCTACCGTGGGCTAATATTTTAT

CGCCCAATATTTTTGGGTGGTTCACAAAGAAACACGGTCAAGAAAGTAGTTGACGGAAACTACAAATTTATCTATTTATC

TTCTCCTGTAACTTCTCTCAACCCTAAAGGATTTGGATTCCAAACTGCAATTGAATTTGAAGAAGCAAGCAGGGGATAGT

GAGAATGGAAAAATTTAGTAAATCCCATACGGTATTATTATTCATGGTAGTTAGAATTTGTGTTTACAAATTCCATTCAG

TACATATACTTATGTTCAATTTTTGTCTCTACCTAACCTAGTATGTTTGACTATTTCATAGAATCACAATCAAAGATTCA

CATTCACAATTGACAATTAATGTTTTGATTATCCATTTAGTACATACACCATACTGATAGTTTCCGTTTGATATTGTTAT

TATCGAATCAATCTTTGTTACTTGTTAGTCGTGATTTGCAAACAGTGAACTGATTACAAGTTGCAACATTTTTTTATAGG

TGACGTTGTTGGACTTTGGATTTTCTTTGATTTTTTCAAGTTGCAAGATTTAAACGGAAAGAAAAAAGCGATGCATTTTG

TAATGTCTCAATCAGGAGCCATGGATAAGTTTGTTGTTAAAATGCCACGAAATTTTGTTGAAAGCTCGTCAGCCGCGAAC

AAAGGTGTTCATAATGTAATTCGTGAGCATGTTACTTAAACATGTAGACCACAACATAGTCATGAATCGTGGTTCTACAA

CTCTATAAATACAGGGTTCGTATTTGGAATTGAAAAGCACAAACAAAACTTATCAGTTGCTAAACAGTAACAACATATTC

GGAAGCATGGCAAAAAGTTGTGTCATTTTCTCTCCTTTCTTTTATGGTTTACTTCTCCTAGTTTTGATTTCATTTCTGAG

CAAGTACTCATTCTACTACTTCTGTAATGTACATGGCTTTGAAAGAACGCGCAATTTGGGGATACTTTTATTAATTGGAG

GATACGAATTACTTCCCCCAACCAGGGGTGGCTGATAAGGGGTAACTTTTGATTATGAAAATACCAAAAAACCCTTTCCC

TAATTAAAAACCTTTATGAAATGACTGTTATGCCCTTTCCCCTAAAACAATAAAATCAAAAACCAAAACAATTTATCCCC

CATTTCAATTTTGTACTGGCATAATTAAGTTAGGGTTCTGTTTTTTTTTCTTCATCTTCTTCATCGTTCTTCATCGACGA

TTAATCGTTAAATCAAAAGTTTCTTCGTCGATTAACCTACCTACCAGAATCATAACCATGGCTCCACGAAAGAAAGTAAA

AGCCAAACCGAAAACCAAATCGATTTCTCAACAAAAACACGAACAAAGAATTAGTTCGATTGCTGAAGAACAAGAAGAGG

TCCAGCCGGCGAACTCGGACGATCAACCACTCGAAGTCGTGGCTTCTTTGAGAAGGTTAGTGATTTTGAACTCATTTATG

ATTGTTTTAATCGATTTATAGCAATGGGTTTAGGTTAGAATCGCCAAACAAAGATGAAACAGTCGAGGTACACAGAGTAC

CGGCATTCAATCTTGGATGAATACCACGCCGTTTTCCAGTACCGGCATTCAATATGGGATGAATACCACGCCGGTTTCAA

TCCCTGAATGTGAAATTGCAGTACAGGCATACAAATTGGGTTGAATACCATACCGGTATAATGTACCGGCATTGCGTAAT

AGTAATTCGTGTATGCCGGTAGTATATTTAAAACACACAGAAACATTATGAGGTTTGTCACCAGTACCGGCATTGAATTT

AGGTTTCTTGTATGCCAGTATAGAAGAAACGGCATAGATTTGATTAAATTCGTGTATGCCGGCAGTAAACTTTAAACATA

CAGGAACATATATGATTTGTTGATCAATACCGGCAAGCTCTTTCGCTTTCATTCCTTGCCGGTACACTGTTACTGGCGCT

GCATTGATTAACTTTTAAAGTGCCGGTAGTTGTCTTTTATTTACCGGCATTCTCGATAGATAAGTTTGAATGCCGGTAGA

GCTCAACCGGCTTCGTCTGTTTAAGAGTCTTAATGCCGGTACTTAAACTTGCAATATGACTTGTCTTATTTTTTTTTCGT

GTGCTTATGTTATAGGTCAAAAGCCAAGGAAGGTAAAAAAGGAAATAATAAAGATGCTCTCACTAAGAGAAAGAGGAAGG

ACGGTCTTGATACTCCTCAGTCTCTGCCTCCACGAGGGAAAGATGGGGGTCGCAAGAAGCGTTTGGGGTTTGAAACAAGA

GGGGCAATTTGGAAGAGTGGTGGTGACCGTAGTGCACTTGTAATCCGTGAACTTACTGGAACAGTTGAGGAGGATAAGCA

TGATGAAGGTGAAAGTGAGGAGGAAGATGTTGAGGTTCCCCCAAGTCAATTAGCAAGCCAAAGCGAAGTTGGTGGGCCGA

GTGAACAACTAACACCAGGCAAGGGAAAGGGCAAGGGCAAACACAAAGTTAAAGGTTGTCACCTTCCTCACATTGATGAC

ATGATACCCGACCTTGAACGTGGTAAAATTTGGGGTGAAGCTCCGGAACTGAAAGTATTATTTGGATACAAGCACTCTTG

GGCTCGTACTATCTATCAAACCTATGTAAGCATTTTTTATTTTGTGCATATGTATTGTTTTGGATTTATGCGAAATCTCT

TAATCGTATTATCTTCTAATTATAGGATCATACTAGGGCATTCCCCGACACTAATGGTAATAGGGTAAGTGCAAACTTTC

TTCAGTACCTGGACCCGTTTCAAGATGTTTATATCTATTCTTGGGGCATCGCAGTAATGGCCCATCTGATGGCGGAGATG

AGAAGGGCCTCTTTGGATAAAACAAACCAATTTGTCGGGAATTTCACTCTACTCCAGGTAAAAATATTTTCTTAACTTAT

TTTCTTATGTATATTGTTCACATGTATATATTCATCATGAACTTAGAATAAAAATTTACTTCTTCACATTGAAATAGGTG

TGGTGTTATGAACACTTCCCAACCTTGTTCAAGAATTTCCCTTTCTTGAAGACCATTGACCACCCCGAACCCGATGCTCC

TAGAGCTAAGAAGTTCGAGTATAACAGCATTCCAAAGCCCGGTAACAATGGTCGACTGATCGAGATGAGATTGGCTCTAG

AAGAAATGAGTACATAAGACATTGTGTTTGACCCATATAAGGAGGATAGAGAAAACCGTCAACTTGTGAAGTTTGGTAAC

GTTGCATTTTACAATGGGCTGTTGTTCCACCCAAAAGGATATATCATGGCTGATCCATGTCTTGTGATGCGCTAACATGG

ATATAAGCAACTATTACGTTTTGAGTCGGCCTCTAATGATAATTATCTTCTTCACCTTTCTCTGTGAAATTCAACTCCCA

ACAAGTTGCATGTAGTGTATGATCCAGATCCGGAACCAATACACTGGAGGGATCGGGAGCCACGTCATTTTGTAGATATT

AGTAAGTGGGATATTGCGGAAAATGGTGATGAAGCTGATAACGACTACATGGAAGATTACCTGGAATAGTCATATATGTA

TATCGTGCGCCCTCCCTCCATCGAAGTTCCACCCCAGAAAAAACCTCATGTTGAAACTCCTACACCTCCACCACCTACGA

CGGCCCAGCTTCATAAGACTATTGGATTGATAGTAAGTTTTATTAATTTCTTAGTTGTTTAATGTCAACATAATGTTATA

TATTAGTATATATTAATTATGTGTTTATGTATTAAACTAGCGTCGTCGGCTTCGCAACTTGAAGAAGATGTTTGGTTGCA

AGTACGATGAAGGAATGGTGCTATCCCTTGAAGAAATCAAGGAAGTCGAGCAAAATCTTGATAGAATTGAAGAAACCGAC

CCTAGTGCAAAAGCTTTGTTTGGAGAAACGAGGAAGGTGCCCGCCAAAAAGAAACTAAAGACCAAGTGAACAATGGGTGG

TTATTCTAATTGTGTTGAACATGTCTCGATTAAACTTGTTTTCGTTTGAACAATGGGTGGTTATTTTAATGTGTTAACTC

TTTAGTTATTTGGTTTGCTTTTGGTTTATGCTACCTGGATTAAAAACACTTAAGCAGAATCCAGTTGTCTGGTTTATGTT

GAACATGTCTGGATTTCTAGTGAAAGATTAAAAACACTTATACCAGCGTCCTATAAATTTATAATTCAATGCCGGCTTTA

AAGTGTCGGCATTGAAATTGTCTCTAATTCTATTCCGGAACAAGAAGTTAAAAATTTTCATGTTCCTGTAAAGATTTTGA

CAACTACCGGCATGGTTATTTTTTAAAATTCAACGTCGGTATTATAATTTTACAATTTCCTACATTGTTGACAGAAAACC

GGCATAGGTATGTCGATTCAATCAATGCAGGTACACTTCTAAAAATTTGTTTCAAATATTCTATGATTTCATTCCTAAGT

AATCTATTACATTGGATTGGATACTTTAAATTAAAACATGATAAAAAAACATTCAATGGATTGCACTTTGTTAGAGATAA

TCTATTTCCATCCCTTTTCAACAATTGCGGAACCTTTGAGCCTTTCCCGAATCTGTTCGAAGACATCGTCCGGGATTGGG

GTGTCAAGGATCATCTTCACTTGTTCTTCTTATTTTTCAACATATTCCTTTCCCTTCATATAACACCCATCATACCCATC

GATAAGCTTGCATTGGCCCTTATGGTTTTCTCTTGACTCTGCTAGTTCAACAGCTTTGTCAAACCATTGAAACTCATCGC

ACTTTGGGTGATTCAAACACCTTAGAAACATTCTTCCATTTTCAGCATCATCAGTTTTTGCAATCCAAAACCGGCGTGTC

CCATCACAGCTTCTACACTTTGAATAAATAAAAGGGCAGTCCATTGCTAGATGAGATGGTGACTTACAAATTTGACACTT

GCAATGATGACTCTGTAACAGAAAATTATATTAGTTATGAGTCATGGTAGTGAAGATTGTATTGTTAGTAATGATTAATA

TACTTACTTTGCAAATAGAGCTAAATGATGAATCTCCCATCGCTTGTGAAGCTTTAACCGTGCTTGATTTGTTGAGAGAA

GGATTTTTTTTTATTTTTTGTGTGGTTTGTATAGATTTTCGCCAAGGAATGGGGGTATTATATACAAATAATTCTCCAAC

GGTCTTATTTTCAAAAAAACTAGCCCTTTTTTTTTAATTTCACCAATACCGGCATGTTCGATATTGAAGATTCAATGCCG

GAATTTCGTACCGGTATAGTTGTTCACAATGACATCAATTTCGGTTCTGGTTATTTTCAAAAAAAACTAGTCGTTGAGGT

TTCTCACTATATAAAGAGAGCTAATACTTGGAACCAAAAGTCCAAAACCGTTGTGTTCTTATATATCTCCTTTAGCCTCT

TAACTTAGAAACCATCCACCCTAGAATAATCACCTATAAATAACTTAGTATAGTACCTAACCCCACATTCAACAAATGGC

ATCGTTTGACTCGGAGGAAGATTTAGCAATCACCCAAGCATGGTATACCGAAACTCGTCCTTCAATTATAGATAACAGCA

TGCGGGATTCCATATGGTTGAATATTTTTAACAAATTTGTTCACGACTATGGTAACCCAAAAGGAAGAAGTGCTAAACAA

ATTGAGGAATGACACGACATCATCCTAAATTCGGTGGTTGAGTATCTAAAATTGAAACACTGGATTCATCAAGCTAACCA

CAATAGATTGTCTCCTCAACAACTTGTAAGTATTTTTATGATTAATTCGACATAATTCACTATTTTGGAATTCGTATGTA

ACAACCATAATGTGTGCATTGTTTTTGTATCATGAAGCCGTGAAAATGCGTTAATTACAAGAAAATGGAGAACCATTCTT

GTTTGAAGCCAACGTTAACCATATTGTAACCAAAGTCACGGAGTTCCACCAGCTGGGTGAATGGGAGACGGATTCTTCTT

AAGAATGTTGATAGTTTAGTGGTAATTTCTTAGGTTTTTAGGGTATTTTATTAGGTTTTTTAGTGGGTAGGTATCTATGT

AAACCCGTAATACCCTGCCGGCGTAGACTTAATATATATTTGAATGCCGGTAGTAGGTGCCTAATTTCTGTATGTTTCTG

TGTACAATTTATGTATTTCCGGCACACACTTTTGATACTATACATTGCCGGTTCAGAGTGTCGGCATCCACGTAAACCAT

AGGTCAGTGCCGATACTTAAGTTTCAATTTCCAACGGCTATTTTATCAAACTATAAACCTGTAAAGTAAATTTCCGGCGT

AGATTTGAGGTTACCGTCAATGCCAATACCATGAAATGGAAAAAAGCCGTTGTGTAGTCTTATATTTAAAGTGGCTACAC

CTATTGCAGTTGATACATTTATATGATGGAAAAAAACTCACCCACTACCGATTTCAGAGAATATTTGTATGAAAAAAAAA

ACTCAACTCCGGTAGGGAATCAATCATCATCCACCACCAACTCAATTCCCATGATACTAGAAAAACTTAGAAGGAAGGAC

CAAACCACCAAGTCAATTTCCGCAATGGAAGAAGGGATACTTATAAGGAAACGAGAAGAAGAAGAGGCAGCAAAGGAAGA

TTACTATCGCGAAGAAACAATGAGAATATACGAACGTGTAAAACAAGCGAAGATTGAAGCGGATGTTGAGGAAGAAACCA

AATGGATCAAAAATTACTACATAGTATCGGATTTCCCGGAAGATCACCGCCCTCCAAACCTTTTTAGGGGTCTTGTTGCG

GATGGTCCTTATATGAGGGAGTTCGTGGACGATAAGTGCAAAAAACATAAAATAGTTTACGGGGATGAGTTTGTATCGCA

TCAGACTACTTCCCTCATAAAATTATGTCGCAAATACAACGAGTTTCTCGCAAGATACCGAGATAATAGGTTTCAAGCTC

AGGCTGCATACACCAAATGGAGAAAACAACCTTTTAGGCATTTCGAAGCCACTTTGTTTGTTGAATCTCGTACCAAGAAA

TGATGTTAGTTGCAATTATCTTGAGATCTTTCATTTTATGTGTAAGGTTGTTCAAGTGTTTAATGATTTTCAGTATATTT

CCTTATGATTAATATGAAATTTCTTTGGTTTGTGGTTAATGTTGTTTATTTCATCAAGTTTCCTTCACATACCGGCATTG

ATAAATACACCAAATTATTTTTAAGTGCAGGCACCGTCAATGAACGGTGCACCAAATATTAAGCCGGCATCATATTGACA

ATTTAAACATATTGTCCGAATGTCGGAATTGAATTGTTTCATGTATCCATGCCGGCATTATGACGGTTACAATTTAAAAC

TCATTCATTCCACTTGTTTGACGGTTACAAACCACTTTAACCCATTTATTGGACGGTTACAAAATCCTTAATGCGTCGGT

TACAAACTTATCATTCTATTTAAGTGACGGTTTTGTCTACGAAATTATTAAATCTTCCTTTGTATTCAATATCCATATTA

TTTCATCTTCCATACCAAAGACAACGGGTCCTCGCAAATCTCGTGAAAAACCTCGTCCACAAATACATGTAGATGAACCG

CAAGAAAGTCCTGAAGAGTCTGTGGTGAGGCGAGCATAAAAAATTTGATGCGGTAAGTGAAGTATTTGTTGAATTGAATT

ATTTTTAGGGTTTATCCCATAATATATCCCACAATTATAATTTGAAATTGCAATCTAGTCAATTGAAAATCTGTTCTGTT

CGCGGTAGAATTCAATTTGAAAATCATGCCGGATTTTATGTCTTCAATCCCGATTGTATTGCATGAGATTGCCGGCATTA

GAAATTAGTCAAATTGTATGCCGGTTACAAGTCCCGGCATTATATATTTGATCAATACCATGCCGTATTTATTTTCCTGC

CTTGTTAAAAACTCTGTTCAAAAATCACACAGTTCCGGCATGCTCATAAATATGTCTGACAATGCCGGCATTTGTTTCGA

CATTCTCGATAGTTTTAAAAACTATGCCGGTTTGGTTTCCTTAGACTTATTATTCCCTGTATAGAATTCAATAAAACCGG

CATGCAATTTATAATTTCCTACTATGCCGGTTCGAGTACCGGCACTCATATTTGGTTTCATTTATATGTCGGCGTCCAGT

GTATAACTCAGTTTAACGTATTGGAACATAGGTCCAAAGCATCAAAAAAGGACAAAGGAAAAGGTAAAGCTATTTTACAA

CCAAAAAGTAAGTCTTCTCGACCTATCCGTCCTGATCCGGCTACTTCTAAGGTCACAATTCGGGATCAACCTATAAGGGA

CTCAACCAGTAGTGGTGATGATGACGATGACTGTGAAACTGCGCATGTACCTGAATGAGCTGGTGGCTCATCCTCACAAG

CAGTTGATGTCGGAGTTGAATCCTCTAAGGAAGATGAAAAGGCACCCCCAGCCCACCTGCCTGATAAAGATGACATCATT

CCAGACCTCGTTGATGGAAAACATTGGGGTCAACCGAAAAACCCGAAACTCTTATTTTCCTACAAAGACTCTTGGGATCG

TTGGATCTATCAAAATTATGTAAGACGTTTCCATCTTGTTATTAAAATATGTATCCATCGATCAAGTGAATTCAACTAAG

TTTGGTGTGTTTTTATAGGACCACAAGAAAGCCTTCCGTGTTAGCCGCCATCAACAATCGTATGTATGGACTTTCAAGGA

TCAGTGTGCGACTGTTAAGGCAGTAGTGAAAGCATCTGGGTTGTAACCTGCGATAAATAATTATGTGGACACTGATACCA

TCATGGAAAACTTCTTTTTGGAGAGGTTTTATGCTGAAACTAACACCATGCATTTTCCTTTTGGCGAGATGACCATTATC

CCTGAAGATGTCCAAAGGATAATAGGTTTAGAGGTTGTTGGGAAAGCCGTTAAGGTAGGAGATGACTCTAAGGACCTAGA

ATGGGACAATATCTATGAGCTCATTAAAAGGTTGTTTGGATGGAACAAGGAAACCACTTTGGAAAATTTGTACAATACAA

GGACTTCCCAGACAAAGACTATCACATTCGTCAAACTCTTTGAATTGTTTGGGGATATGAAGAACAAAGTATTGACTCCC

AAACAAGTTCAGCAAAGTGTCGCTGCTTATATGTTGTTCATCTTGGGCTCCGCTATATTCCCTGACGCAAATGGTAATAG

GGTACATATGAACCTCCTACAATACTTGGATCCGTTGGAAAAGGTTCATGAGTATTCTTGGGGCACTGCTTTCTTGGCAC

ATTTTTTGGCAGAAATGCGTCGAGCTTCTAAGGCAAAAGCCCGTCAATTTAATGGGAATTTCACTGTGATTCAGGTATAG

AGAACTACCTATTTTGTGATCGTTATCATTAAATTTATTAGTTTCTGTTTTTTTTTTCTTGCACATTTCATTGAAACGAT

TTTTCATGGGCACAGGTGTGGGCTTATGAACATTAGAAAAATGACTTTTGGTAACAGTTCAAGTTTTCAAAAGGTGACAG

TTTAGAAAATACGTGTTACTAAATATTGAGTTTTTGAGTAATAGCAACAGATTCATGTAAGAACTGTTACGAATAGTTTC

AACATATAAATATTATACTGTAATGGTTTCAAAGCGTCATCTAAGTTCCCTACTGTAACAGTTGTATAACAACTGTGACT

GTATACCACTTATACTATAATAGATTTAATCTGACACCGTTGTTGACATTATCGTGACAGTTTCTTAAACAAGCTGTGTC

TAATATCCGAGTTACTGTGTCAATGGGCCTCCACATTTAGTTATCAGGGCATCCACTTTAGAAAACTTCTGAAGCTGGCG

GGGATTATAATCTGGAGTGTGGACCAACACGTCACAGACTCACAGTACACTTCTTTGGGGTTATTTATGCCGTACTAGGT

TGGTCATCACCGTGTAGCCTTGGCTCGTATGGCTGGTTACTTAGGTTTAAGAGAGATTCACTATTTCAAGATATAAGAAG

ACCCGTCAAACTAACCCTCATCTTCGACCTACGGAGTACCATTCTTCTCTGCAACCACCTCGGATTGAGTTCGGTCATAT

GATAAGAAATTTGAGCAGTGGTTCTCTAAGTGGTCACTGGGATTCCTTGTGAGTTTTTTTCCTTTTTATTTCATAATCCT

CCTTCAATTTCAGAATTATATATCTGCTCTAAGCTCAAAGAATCCCCTGCTCAATCGATAATTTTCATCTGTGAACCCAA

AATGGAAGCTCAAAGAATCCCCTTTCGAATAATAAGTATTTTATGTTAATGAATTGAGTATGTTAAGTGTTCGCTAAAAA

AAATTCTTCAAATGTCATCGTGTTTACGAACTAGGTTGCATTTCTTATATTTGGATGTCGTATTTTGGGATCGTATCCAC

TGGTTTGTTACCCTTTAAGTATATACCAAATTTCTGGATTAAGAAATGCATACAAAACTTGTCCAGTAGAAATTGTTAGA

TAATTGATGAGGATCAGTTTCTGAACCGGGTTTGAGTTCCTGCAAATTTTGTTTCTTAGTCCATTGTGGGTTGTTAAAAT

ATTCCATGGTGATACATTGGTGCTTAGTAATTCGTAAGTGCTATTTCATTTTTGCTTATGTTATAAAGCATTAGACCTGT

GCTATCACTATTAGGAGGTTTGAGGTGTTGATTTCGTATATAAGCGCTGACGATAGTTCCCCACATAAATAATATGAACA

ATTATTTAAAATTATCCCACCAGTACAACATATATTGGCCTCGTTTTAACGGTCATCATAAGGTAAAGTTGCACCTGGAA

ACCTGATTGTTTGGGATCGTATAATCGCATCTTGAGGTACCCTATCACAAGCAAAAGCCACATGCATCTTAGTTATTTAC

CTAATTTCAACTAACTGGTGTTACTGGATTCCAAAATTTGTTCTTTTCCCTATTGTTTAGTTCCCATATTGTTTGTAATT

AACATTCTTTATTGTCAATATTCAATTGTTCGGTCTATATCCTACTATTCATTGCCATGAATTCAGTTGTTTCAGAAAAA

TTAGTTCATATTGTGGTCTCATCGGATTGATTCCCTTAGTGGTTGTAATTCTTTAGTGGGGAAAAGCGAACAATAGAAAG

AAGTGGACATTTAAGTGGGATTTACTATTTTGGTCTCATCAGATTGATTCCCTTAGTGGTTCCCTTAAAAGAACATAATG

TTGAATTACAATTTTACTGTTAGTCTGCTATCTGTTTCAGTTGGTAATTATTATGTGGACATTGCAGGACAGAAGCTGAT

TCTTGCAAAACAAGTTGTAGCATTGTTTAAGAAGGAAAAATATATTAAAGACCTACAAAATCTTTGTGCAGAGGACTTAA

TGGAATTTCTCGGTCAAGGTATTTTTGTTGGGCCAGTCACAACTTATGGTCTAACATAGGTGTAACCCCTTAACCATTTA

AGGGATCTAACTAACAAGTGTCCTATGCTATATGTTTTCATTCAAAAACATCCAGGAGAATATCTTCTTCATTTAGTCAT

GAAGAAAATAAAGAGAAAAAAAGAGAGGAAGAAGAAAAAAAAGAGTGAGTGAGGTAGAGAGATTAAGCCACCAATTTTTG

GTGAGATTGAAGCATCTTGAAGAGGAGAAGAAAGGGTTCTTGTCAAAGGATATTTTGGTTGTTGTGGTGAAGATTGATTG

ATCAATTTCCACATCTCTTCAAAAGGGGAAAAATCTAGGGTTTGTTCAAACTCTAGTAGTAAGTATCTCAAATCACTTCT

ATTCTCAAATTGATAGTGAATCTCATATTATATGTGAATGAATTGTTGATCCATCCTTAGGGGTGAGTGTAATTAGGAAA

ATTGATATCCAAGTGCTATAGGCACCTTGAAGAAGAGTTTATCCCCAAATTTGGGAGAATTGTGTAACCCATTATTGTTA

TAGTGGAAGATTTGCCCGTGGTTTTTTACCCTTCACCTTGGAGGGGTTTTTCCACGTAATTGTCGGTGTTGTGTGATTGC

CTCTATATTGCATAGTTTGTTATTTCCGCATTTGTATGTCATTGCTTGTGTATGTTTCGTATTGCGCATAGCGGCTCGGA

CTTCGTAAGCCTCCCCCAACAAGTGGTATCAGAGCTTTTGGGTTGAGCTAATCCTGTGCAATGGAGGAAGCTAGCACTAG

TAACAGAATGGTCAATTTGACCGGTGAGAACTTTGTGATATGGAAGGCGAAGATGGAAGACTGGTTATATTGTAAAGATC

TCTATGGTCCAATAGATACTAAAGGAGTAAAGCCTGAGACGACAAGTGAATACGCTTGGAAGGTTACTAATCGAAAAGCA

GTCGGTTTTATTCGTCAATGGATTGATGATGGTATTTTGAATCATGTGGCTAGTGAGACACTAGCATATTCTTTGTGGAC

CAAGTTAGAGGAGATATATGATCAGCAGTCTCCTTGTGCAAAATATCTTCTATTCAAAAAGCTTGGAAAGCTGTGTTACA

AGGAAGGGAGTTCAATGGCGGAACATTTAAATGAAATGAAGGGTATTATGAATCAGCTTACATCTATGGATCTGAAAATT

CCTGAAGAGGTTCAAGCTTATTTGTTGCTGAATACTTTACCAGACAGTTGGGAGCCATTGAGGAATTGTCTCAGTTATTC

TTCTCCATACAAGAAGTTGAACATGACTGTTGCAACTAGTAGTTTGCTGAATGAAGAAGTTAGAAGAAGGTCATCAGACT

CATCTCATTCAGAATCTCTTGTGACAGAAAATCGAGGTAGAAGAAAAGACAGAAGACAAGTTGGACGTAGCAAGTCAAGG

GGTAAATCAAAGTCCAGAAAGGAAGTTGAATGTTGGAATTGTCATACGAGAGGTCATTATTCAAATGAATGCAGAAAGCC

GAAGCAAGAACACAAGAAGGGGAAAGGAAATGATAACCAATACAATGGTGTTACAGCTACTGTGACAAATGATGATATCA

TTGTATTCTCAGATTGTAATGATGTTTGTGTGAGTTTTTCAAGTTCTGATTCAGAATGGATAGTTGATTCAGGTGCATCT

TATCATGCTACTCCTCATGAAGATGTATTTACCTCCTACAAGGCTGGTGACTTTGGGTCAGTCAGAATGGGAAACAGTGG

TGTTGCGAAAATTGTAGGAATGGGTGATGTGAAGTTTGAGACCAGTGTTGGCTGTACGTTGGTTCTCAGAAATGTGCGAC

ATGTACCAGATCTTCGTTTGAATCTAATGTCACCAGGAGTGTTAGATGATGAAGGGTACAACAATAACTTGGGTGATGGA

AAATGGAAGCTAAGCAAGGGCTCCATGGTTATTGCAAGAGGCAAGAAGTGTGGCACATTATACAAAACACAAGTGAAGGT

GTTGAAAGGTGAGTTGAATGTTGTTGACAAATATTCAACCTCCGAGTTGTGGCATAGAAGACTTGGTCATCTTAGTGAGA

ATGGAATTCAAATCCTCACCAAATGACAAGTTTTTCCAAATGCCAAAGGTATGCAATTGGGTAAGTGTACTCATTGTTTA

GTTGGTAAACAAACTAGAGTCTCTTTTCATAAAGCTTTACCCAAGCGCAATTCTGGTATTATTGATCTTGTTCATTCTGA

TGTATGTGGACCTTTAAGGGTCAAAACACTTGGTGGTGCTGTTTATTTTGTTACTTTCATAGATGATGCTTCAAGGAAGG

TTTGGTCATATGCAATCAAGACCAAAGATGATGTTGTTCATGTGTTCAAGATGTTTCACAGCATGGTGGAAAGAGAAACA

GGTAGAACCTTGAAGTTCCTTCGTTCTGACAACGGGGGGGAGTACATCGGACCACTTGAAGTGTATTGCAAGAGCATGGG

TATACTTCATGAACAATCAGTGCCGAAGACTCCACAACATAATGGTGTTGTAGAGAGGATGACCGCACTATTCTTGAGAG

AATCAGGTGCATGTTATCTCATTCCAAGTTGCAGAAGCATTTCTGGGGTGAAGCAATGAATACTGATGTGTATTTGATCA

ATCGATCTCCTTCAAGACCTCTAGATGGAGAAGTAGAAGAAAAGGTATGGAGAGAAAAAGAGCCTACTTATGATCACTTG

AAAGTGTTTGGTTGCAGGGAATTTGTTCATATCCCAAAAGATCAGAGATCAAAGCTTGATGACAAAGAAAAGCAGTGTGT

GTTCCTAGGCTATGGAACAAAGAAGTTTGGTTACATATTTTGGGATCCAGTTGAGAAGAAGGTTATCAGAAGTAGGGATG

CAGTATTCCTTGAAGATCAGACTATTGAAGACTTTGAGAAGATTGAGATGCCAAAGCTTCAAGATGACATGAACTTGGAT

CCAAATCCAATTGTTCCTACTTTTGAACAACATGATGATGATGTTGATGGTAATGGAAATGAAGATGAAGCTCATGAAGA

TGTTCCAGTGGGTGATGTTGAGCAAGAAGTATAGCAACAACAACAACCAGAAGCTGAGTTGAGAAGATCCAGCAGGACCA

GATTACCTTCAACAAAGTATACACCACATGAGTACGTGATTCTCACAGATACAGGTGAACCAGAGTCTTATGAAGAAGCT

CTTAGTGTTCCTGAAAGTCAGAAGTGGCTGAAATCCATGAAAGAAGAGGTTGGCTCTTTAGATGAGAATGACACTTTTGA

TTTGGTTGACAAGCCTGAGAACCAGAAAATTCTGAAGAACAAGTGGGTGTATAGAATCAAGACTGAAGATGGTAACTCAC

AACCAAGGTACAAAGCACGCTTAGTTGTGAAGGGTTATGAGCATAGAAAGGGTGTTGACTTTGACGAGATTTTCTCACCG

GTGGTGAGATTTTCTTCTATTCGGGTTGTCTTGGCAATAGCCGCTAGTTTGGATCTTGAGATATAACAACTAGATGTCAA

GACAACATTTCTTCACGGTGAGTTGGAGGAATATGTGTATATGGCACAACCCGAGGGTTTTGAAGTCAAATGCAAAGAGC

ACATGGTGTGCAAGCTAAAGAAGAGTATATATGGTTTGAAGCAAGCTCCAAGAAGATGGTACAATAAGTTTGAATCGTTC

ATGAAGGAACATGGGTACAAGAGGACATCTTCAGATCATTGTGTATTCATTCAAAAATTTGATGATGGTGATTTTCTTAC

TCTTTTGCTATATGTTGATGACATGCTTATTGTTGGCAAAGATAAGAAAACGATTGACAGGTTGAAGTCCAGATATGAGC

AAGTCTTTTGCTATGAAATACTTGGGTGTAGCAAAGCGTATTCTTGGCATGGAGATCACTCGTGATAGGAAAGAGATAAA

GTTGTGGCTATCACAAGAAAGCTACATTGAGAAGGTGTTACAAAGATTCAACATGGACAAGCACTACACCAAAAAATCCA

AATTTACCACGTTTATATTCGTTGCTAATAATTTAAGCAACGCTTATGTACGTTGCCCACACGGCGTCACAAATATTTGC

CAACGTTTGTAAACGTTGCTTAAATTAATTAAAGCAACGTTTCATGTGTTGCTTTATCCGTTGCTTTAATTAGACTTAGT

GCAACGTATAGTATAATTTAAACAACATATATATCCATTGCAAAATATATTACCAGAATTAAAAAAAAATACAAAATCTC

TATCAATAATTTAAAATTCCAGTATCACAAAAAAATATAGAACGTTGACCTGATAACACAATCACTCATCCTTCCAACAT

TTTACAACATAATAATGAACTTAGTTTGAATGTACACCATCAAATTCCATTCCGAAACATAAAGCCACTGTTACTAGATG

TTCAATAACCATGTGTGATGTAGTGACCTGTAAAATGCATAACGATGAGGATCAAGATACAACAACGATAGTTGATAATC

ATGGCAAGTGCCAATATGTGTCTCTGGTAAGTCAAAAAGTAACCTAAAACTAGAGTATCCTAAACCTGGAATAGCCTTAG

AATAAATTACACTTCTCTGAATGATTATCAGAAGCTAGAAAAAAAAAACCTGAACAACCACCATCTCACATAACCATGTT

TTAATAAGAGAGAGTTCATTACCAAAAAGAGATGAGCATATATCTGCTGAAAAGAAAGCATAAAACATATAGGTAAAAGA

TGCATTTTAAAATTAACAGAAAGATGATGGTATAAAAATCCTAACTGCCTAGCCATAATATTTTCAAACATTAAAATTTA

TATGAGTAAATGGTAGGAAAAGTGTGTGGTTATAAGATCTAATTCACCTGCATTACTGAGCAAAACAAAAGTGGACTGCA

TTTCATGATGTAGAAAGAATTGGTTAGTTTCCCCTCACGGTCGATTTAGATTCGGTGCATCTTATATCCCAAGAGCAATG

AAAAATTAAACCACTCTGAAATGTTCAATAACTTTTTTTTTGGGGTTGATCATTCAGTTTGTATAACTGTCTAATTGACT

CACCTGATTTTGGTATTCTGTCACATGCCAAGCAAAACATTTCCATCTGCCAGAAGTGCTATAATTTCATATTCTGATTG

GTATTCTTCGACACCCGTCATCATGGTATCCTGAAACAATTCAAATAAATATCATGTAACTTTTGACTGTACATGGAAGT

AGTTGCAAAAAATTCACACAAAAAGGTTTTTATTCTAGCAGTCCCTGTTCTGTGTAAATTTCTGAAGTACTAAGGTAAGC

CTATTAATTTGTGGCACCACATCCTAATTGAATTGATTAGGATTTAAAATTTTCAATATGTATCCTTGCTTAGGGCCTAC

GAAATACAGGAAGAGTAATACATTACTTGAAACTTATGTCAGCTGTCAGCATACCTGTAACTCAATATTATATTCTAAAT

GATAGTGGTCACCCACTACAGAATTCTGCACACTACATTCTTGTAAAAAGCATCCGTGGAAAATTATTGAATTCAGGACC

TGCAAGAGATAGCAAAAAGAACATGAGAAGTGAAAGATAAAAGAATAAAAGAATAAAAGAATTTTGATTATATGCGAACC

CATATACCCAGCATTTTTGAACTTTGGTTGGCGATAAGAAGCGTGGAGATGTAAAAAAAAGGTGTCTTTGGATCAGAAAA

ATAAAAACTTGAAGAATGCAGAAAGAGATTAAAATAATCAATTGACTGAAAGTTTAATAGCATAGCAAGGTAACATGTAA

TCACATTGGTGTCTCGTGCATCAATAATTTTCCAGTGAAATGTTTGGTGGCTGAACAAAGTTAGCAAGAACGAAAACTAA

CTTGTGCAGTGAGCGCCAATAAGTATTTAATTGTTCCAATCCCAAAATTCATTGAATAAATATGCATGTTCAGTTTACCA

TTCAACAAACTCAACTGCAAAATTTTACATATTATACCTTTATTCACTGACTTCTTTTAAATGAAATAAAAAAGCCAGAA

AGGTGCAAGATGCAATTGCAGACTGTAGAGTAGTTGAGAATTTAACCCATACTCATTGTGGTCCTTAACAGCTGATGGAA

TGATTGCACCCAAGAATACCTTTTTGATGCCAGTGACCCAAAGCTCTGTTCCTAAGAACACCTTTACTTATTCATCCAAC

ATATCAGCAACCTTATACGTGACTTAACCCAAATGATCCTCCATATTTTTACTAGCATTCCTCATCATTTATCAAACTAA

AAAGAAAAAATCACAAAAAAATAAATTCATCAGAAGAATGCCCTTTTAGACCCTTCGAACCCAAATGAATTGAATAAAAC

AAAATTATATAGAAATTGAAATAATGCTACCTACATTTGTTTTTGGTATATGATACTCAATGAAGTAATTATCATGTTCA

AGTTATCCTCCCTCTCAAGCCTTTCTTAAAATATAGAAAGTTGGTCAACTTAAGGTAGATAGTCTATAAACTACTAGTAG

AGTGCTGCCACCTCTACCTATCCCCATGATTTTTCTCATCTTCCTTAATAATCATAACATTTCTTTGTTGTTCACCTATG

GCCTATCATTGCTTTACCTTATCTTTCCAAAATAGTTAAATAAGCAAAACTATTAAAATAAGCTAACGAAACAATATCTA

AGTCTTGAATAAAAATTAACACCGACCACGTATATGTCTTCACTTAACTATGCAGCGGGATTTAGAACAAGTAGTAATAT

ATTCCAACAAAAATCTGCACTTTGTGGTCCCTGGATCGAATAATTAGACGACAAAAAGAAACATAAACTAAAAGAAAGTA

CTGCAAGGACATAAGTCTAGTCTCCATTTACCAAATAGCCACACTTATGACCAATTTCTTGCAGCACTCCAACAGGATTC

AGGAAGGGAAGACTTTAAACTTCAATTTCCCTAGTGTTGGTAGATGCAAAGAAGAACCAGTAACTAAGTCAACTTCAAGT

AACAATATCAGTCTAGCATTGAAGGTTACCAAAAACCACAAAAATAATTTAGCATAAAGTTAATTTCTTAATTGGTAGAT

AAATTTTAATCTCCACCAACTGTAACAAACCAATAGTAATGAGAAATCTCCAACAACTACTGCCAATCCCCAACTTTTAC

CTGCATCACCTCCACAACCACCCTTTCTGATAGCAGATCCAGAGTTTTTTTCACAAGCACTAACAAAATAAGGCACCACC

ACCAACTGCAGCACTAACCACGCCACCATACCACACCACCAACACCTGCTACCACCATACCACCATAAAAATCATCAACA

TCGTAACTACAAAACTGAAATCAGAAGGAAACCCTACTTTTAAAATCCATCAAATACAGTAAACAAAAACAAAAATTGCA

AAATCGATCTGTACAAAATATTACCTACCAAAAGATGTACGTAGTTCATAGAATTAGGTCAAAATAAATCAATAATAGCA

AGAACAGGTAAGAAGAATATGGTTATACCCACCAAGATCTATTAGCTCATCATTATAGAAATATTTGACACCAAGCACAC

CTTACAGAATCATTTGATTTTGGAGTATAGAAACGATTTCTAGGGTTTGAGTGATTGAAGATTTGATTTGATTTGATTTG

ATTTCCGAGATGAGATTTTATTTATAAGATCGGGGTTTCGGAGAAATGAGTTTTGATTTCTGAAAACTTGGGTTTGATTT

CACAAATCTAAGTTCAGGGTTAGAAGTTAGAAGAGAAGAAAAATGGGAGGAGGAGGAGGAGAACTCGAGGAGATGGATAA

GAAATAATAAGGGAAAAGAAAATAGAATGGAAAAAAATGATAATAAAATATCGAAAGACATTTAAGACCTCAACTATATT

CATAAAGGCTAGCGTGTCCTAGGTTCTCTTTTTTCACTGACAATTAAGATTTAATGCAACGAATCTTTGATATTAGCAAC

GGATAAATCCCCGCAACATATATATCCGTTGCAAACCCAAAAACTCAAACGTTGCAAAAACCTAGATTTGGTGTAGTGAA

GGCTGAGACGGTAAGTTTTCCACTTGCAAACCATTTTAAGTTATCTCGTGATCAATGTCCTGTGATTGATGTTGAGAAAC

GGAAGATGGAGAAAATTCTTTATGCGTCAGTAGTTGGTAGTTTGATGCATGCAATGGTGTGTACAAGGCCGGATATAGCT

CATGCAGTTGGAGTGGTAAGCAGATTTCTATCAAATCCTGCTATGGAGCATTGGGAAGCAGTGAAATGGATACTAAGGTA

TTTAAGGGGTACTTCTAATATGTGTTTGTGCTATGGAGGTGACAAGCCAGAGTTAGTTGGTTACACATATTCATATTTTG

CAGGTGCTCAAGACAGTCTCAAGTCTACATCGGGTTACTTATTTACCTATGCAGGGGGAGCTGTACGTCTTGGAAATCCA

AGTTACAAAAATGTGTAGCATTGTCTACTACAGAGGAAGAGTACATTTCAATCTACAGAGAGTTGCAAGGAGATGATATG

GATGAAGAGGTTTGTTAGAGAACTCGGTGTGAAGCAACAGAAGTTCATCGTTTACTGTGACAGCCAGAGTGTCATCCATC

TCAGCAAGAATTCGAGTTTTCACTCGAGGTCCAAGCACATCAATATTAGGTATCAGTGGATACAATAAGTTCTCGAGGAG

AAGGTATTGTATATTGAGAAGATCTATACAGATGATAATGGTTCTGATATGATGACAAAGTCGCTACCAAAATACAAGCA

TGTGTATTTCTGCAGCAGGGTAGGTTTGGTGGTGACGGGTACCTCGCCATAAGTCGTGAGGGGGAGATTGTTGGGCCATT

CAAAACTTATGGTCTAACATAGGTGTAACCCCTTAACCATTTAAGGGATCTAAGTAACAAGTGTCTTATGCTATATGTTT

TCATTCGAAAACATCCAGGAGAGTATCTTCTTCATTTAGTCGTGAAGAAAAGAAAGAGAAAGAAAGAGAGGAAGAAGAAA

AAAAAGAGTGAGTGAGGTAGAGAGATTAAGCCACCAATTTTTGGTGAGATTGAAGCATCTTGAAGAGGATAAGAAATGGT

TCTTGTCAAAGGATCTTTTGGTTGTTGTGGTGAAGATTGATTGATCAATTTCCACATCTCTTCAAAAGGGGAGTAGTAAG

TATATCAGATCACTTCTATTCTCAAATTGATAGTGAATCTCATATTATATGTGAATCAATTGTTGATCCATCCTTAGGGG

TGAGTGTAATTAGGAAAATTGATATCCAAGTGCTATAGGCACCTTGAAGAAGAGTTTATCCCCAAATTTGGTCGTATTTT

GTAACCCATTATTGTTATAGTGAAAGATTTTCCTGTGGTTTTTTACCCTTCACCTTTGAGGAGTTTTTCCACGTAATTGC

CGGTGTTGTGTGATTGCCTCTATATTGCATAGTTTGTTATTTCCGCATTTGTATGTCATTGCTTGTGTAGGTTTTGTTGC

GCATAGCGACTCGGACTTCGTAAGCCTCCCCCAACATTTTTTAGTCAAAATCTATATTTCCTATCGCACTTTCAGTAAAG

ATTCCTTATTCCTACTGTTTTAAGAATACTACACTCAAATATTTTTCTCGGTTCAGGCTAGAACACGAACCTCAACTACT

AATACACAACTTGAAATCTTATACTTAATCCATGTCTTTAACCAAAATACGGTAGATACTAATTCTGACTTGCTTCCTTT

GTAGAGGTGGACCTTCAGCATTGTAGTAAAGCTACAGTAACTTAGAAGTGGTTAAACTTGTAGTGTGGAGTGGACACCTC

CAAGGTCTGGTAAGTTACACCTAAATTTCATTCTAACTCAGAAAATGAATTGGTCAAGTATTATCTAATCCAGCATTTCA

GAATTTCAGTTGTTAGGGAGACTGGAAGGTGTAAAAGAGACTGATACAGATTAAAACAAAACAATGGAGCTGCACATATG

GGTTGATTACTTACAAGATGTTGTTATTTGAAGCATATTGTTTCATGTTTTTTTGTTTTTGCTTTTTTAAAATAATTGTA

ATTCTTTTCTTTGGATTTTTCATTTAACTTTTTTTATATGATTAAAAATATTTAAGACATCATTAATACATTTATTAAGT

ATATTTGTGAAATAGTTTACTTCTGTCTACGTAGGAATATTGTAGTAATACTACTGTAGGCCTAAACCTGCTTTGCCGGA

TCTTAACTATAGGGCCTCCCCCCGTGCAGAGTTCGCCCAATCCAACATATGCTCGTTTGGCCCACCTCGCTAAGGTGACA

TATGATAACTCTTTATAGAATTATACGTCACTTTAACCCAACATTTAGTGACAGATTTTTTTACAAATTTCACTGTAACA

CTTTACAGTAACAGTTTTATCATATTTATACCATGTTCAAGTCGAGTCTGTTCACAGTTTGTGAAAAATCTGTTATAGTT

ACTCTAACAACAACAACAGTTTCTGTAAAAACAGTTACCATAATAATTATTCAGTAACATAAATTCATATAAAGTGTTGC

TGAAAGCAACTAACGGTGACAGTTTTCAAACTAACGCAACATTTCTTAAGAACCATTCTGTAATAGTTCTTATAAACATT

TCGTAATAGGGGATTTAGTAACACAGTGGAAAAAAATAAAAATGTTACAGAAACAATATAGTAATAGTTTGTAATTGTCA

CCTAATGTCATTTTTCTACTAGTGACTTCCGAAGTTTGTTCGTAGGGTATAAAGGATTGAAGACTGTCCGTGGATATGAA

CATAATAAGGCTCTAACTAATAAATACTACTTCGACAATACCCCGAGGCCCAATAACAAGAAACGGTTAATTGCAATGAG

GATATATTTGGACTCGGTGGCTGTGGAAGACGTTGTGTTTGACCCATATAGGAATGCCAGAGGAGTTCACTTACAAAGGC

GTGACAATGTTTCATTTTACAACGGACCTTTATTTCACCCTAAAGGATTTTTTATGCACAACCCTCGCCGTGTAATGCGA

CAACTTGGGTACAAGAAGAAATATCTCACGGATACATCATATCAGAAATTCTCTCATAACTTTCCTCAATGTCATTCAGA

TAAGAACCATACTCTCATAGTTTATGACCCTAAACTTGTTCTTGCACATTGGACTGAAAGGGAAGAAAAGGGACTTGTGT

TGAATATGTCTGATTGGGAATTGGCTGAGAATGGTTGTCAGAGTGATCCAACCTTCCTAAGAGGCCATCGTAAGTACTCG

CATCCTTATGTTGTGCGCCCGGATACAGTACAAGTTCCTTCACAACCATCACGTCCCTCCACAACTCCGTCAACTGTGCC

CGACCTTCTACATCTCGTTACGTTGTTGGTACGTACGTCTTGTGAATTACTATTCTTTAAATCTATATGTATGAACTTAT

ATAATTGTACATACTAAATAAATGTTGAAAAATCCCTTAACAGCGTCAGAGGATTGGAAACTTGTCAAGGTTCTTTAAGA

AGAAGACCAAGAAAGGCGAGTTGATGACCCAGAAAGAAATGGAGAACTTCGTCCAAAAGATAGACCAAATTGAAGATCCT

GAGGCAAAATACTTGTGGGGAGAAAGGAAGAAGAGGGTACACAAGAGGCCAAGAACCGAGTGATCGTATGTTTCTTTTAT

AGTATTATTTTGGTTTAAACAATTAGAGTTACTTTATGTTTGTTTGTGTGTCTTGTTAAACTTTGAACATCTTTCTTTCA

CTTGTTTTAGGTTTGGAACTTATTAACTTTACATTTGGTTTAGTAAAACTTTGCTTATGTTATTTTATGGTATTAGGTCT

TTGCTTATGTTATTTTATGTGTCTTGCAATTTACAATTTATAACTTTTTATTTTCCGAAATAGGCACAGTCCATATCGTG

TGTACTATGCCGTAATTCTGTTTAGTTCCAGAATGAAAAAAAAATAGAGTTCCGGCATGGTCGTTAGCGCCACAACCATG

CCGAAAATGGAAAAATCATATATGTGTACTTATGGTGTTATTCCGGCACAAAAGAGAACATATATTGCAACGTCGTACGA

GTACCGGCATAGTAAAGTTTGAATCAAATCACACTGGCACATACATGGCATATGGTTTAATGATTAAAATTCAAACTTCC

AAAAAAACAAAAGGTGGCGTATTAGTAAATCACATATATCCAAATTACTCATCATCGCCTCCATACGTATGAAGTTTCAA

CTCCTCCTCTTCAAGAGGTTTCTTTGATAAGGCCAAAGCATCCCATAATTGGTGATTCTCCTCATACAATGCCATCCATC

CCTTTCATATATTTGGATCTAGTTCCTTGTTTTTAACCATCCCACAAACAGGTGGCAATGGACAATTGTCCTTGAGTTTT

AGAACAACGAAATGATTTCCGTTCACGTACGCCAAAACAACTCTTCTCGTCTTAAGGGATCCCTTGCATTTTTTCCACTT

TGGTGTAAATGTTGTGACTACGGTGGGAGTAAAATAATGAACAAAATAATTAAATGTATCCGCTACTAGATGCCCACAAA

GTGGCATCCTCATCCAATATTGAGAAGGAAGAAATACCATTTCCTGCTCGGGTCCTAACACTTGAGCACACATATCATCA

AACACCTCGTCGTGACCAACCAATTGCTCATAAAATCTCCTCTTCTTCACAAGTTGATCGGCCATCCTCATTCTAGCATA

TCGGCATGGTGTCATTCCTTTACTAACATCCGTCTCGAAGATTCCTAATTGTTCGTTCACGGCATGATAGCCACAATTTC

CATCTCCATCGACATCATCCGTGTACATAATAAACTCGCGAATAACCTCAGGAAGTTGGTCTATGTAATACTCTATCTTC

GTATGATAAGTTCTAATGGTCCTACCTGTTCTGGTATCCTTATACATCTTTATATTACCAACTTTTAGCTTAATTATATC

CAAACCATCATCTGCAATCCCTACAATCCAAATGTCTTTGGTATTAGACGAGAAAATATTCTTCCTTGGCCTACCCCTAC

CCATTTTCCTTTGAACCGCGGATACATCGACTATGGGTATAGCTTCAGAATGCTCACCTTGTTGTCCTTGGTATCTTAAT

ATTGGTACCTTCAGCGGTCTTTCCTTTTTCCTTTGAACCTCGGATGTAATAACCTGTAATTTTTTTATATACTGTTTCGG

GTCGGGTTATACCTAGATTGGTTAACCTTGTGTTTAGATTGTTATTCCTTAAGACCTAACCTTAAACTTAACCCTAACCT

TAAACTTAACCTTAAATGGGTGGATTGAACAAGAAAGGCTCCAAGTTAGGGTGGAAATAGAGCCACCCAAATTCATTGTT

TGTTTTTCCTTAATAATGTTATTAAGGTTTGATAACTAAAACTTTAGAAATTTTAGAGAGAGAAACTAAAAACTAGGGAG

AAGAAGGAGATGAAGTTAGAGTTTATTTTAGAGGGGGAAAACCATGGAAAATCAGTGGAGAATTAAGAGGAGTTTTTTTT

TAAGCTATTGAGGTAGTATGCTAGATTCCCTTTCTCTTTACGTTTTTTTTTATACTTGATTTTGAATTTGTATGATTAGA

TCTAAGTGATTTATTGATTAAAAACTATGAATCTAATTGATGATCGGCATTTAATATTGATAGATAACATGCTTAGGTAT

CTCTAGTTAAAATTTGAGAATATTTCATCACATATTCACCATAAAATGTTTGTTTGTACTTGGATGTTGATTTCTAAATT

GTTTTGTACATTTTCGCTTTGTATGTTCTTGACTGTACTTATGACATCTTGATGATATCAAAACGATTTTTGATCTTGAT

AAAAGTTGTAGATCTATGTCTTATCTTTTTATTAGTGGGACAAAGAATCTTTTGATTTTTTTGAAATTAGTTTTGATTAA

TTATTTTCTTTTTCTTTAGATTAAAGAATTCTTAGTGGGACAAAGAATCTTTTGTTTTTTTTGGAATTAATGGTTATTAA

TGATGAAAGTTAAGGAATAATAGGTATTAATAACTTATTATATTCAAATTTGAATTCTTTTTGTTTTTTTGGTGGAATCC

TAGGAAACAACTTTTCTTTATAGTACTGTTTGTTTTAATAAAGAATAAAAATGAAGAATTATGTTAGAGAGAGAAATAAA

TTAAAAGAAGGTGGTAGTGGTAGACATAAATAATTAGCTATCACTTTGAAGACATAATTATTAGATTCTTATCTTGACCA

AACTTTTAAACTTGAGGTTAGGTGGAACTTGTTTTAGGTTGTGAGTATATATATATGTATATATCTTATACTAAGAAGAA

AAGAGTAACCATACTAAGAAGAAAAGAGTAAACATACTAAGAAGAAAAGAGTAAACAAAAATCTAAATTAATAGAAATAA

CGAGAGATTCTTTTAGTTCATATCAAAGACAATTTCTTCGGTGATAAAGCTTTTCCTTCTATTATTTATAGTTTTTGGAT

TGCGATTCGAGGTAGGCGCAACACAATCTAAAACTCTTCTTTTATTTAAGTTTCTTTTAATTAAATTTATGTTTGCAAGT

TTCATTAATATATGGATGATAATCGATGTTGTGAGCATGATTATGAATGTTGGGCTGCGGTCCATAGAATGTGATTCTTA

AAGATTATTTATGACGGGATACGAGATATCCTTGGAAACGGTATTTGTTTCCTAAACCCACGTGGGAGCTCCCTTGGGCG

AGCGGATTGATTGAATTTATTGTTGTATAATGTTGGGATACGAGATATCCTTGGGGACGGTTTTGTTTCCTAAACCCACG

TGGGGGGCCTCCTTAGGCAGCGGATTGATTGAATTTATTAATGTATATGGATGTGGTGTTTCCATATGTTTGTGGTGGTG

TTAACCATGCAACTTGTTTTGAAAAGAAATTGAAAAATATTGCATAAAATGTTTTGTTGCTTTTCCTACTGAGTGTCATA

ACTCACGTCGTATTATGACAAAAATTCCAGATTTTATGACACCACAAGGAGCTAATGGCGGAAAAGCATAGAGATCGGTT

TTGGTGTTTTAACTAGTAGTTGCATATTATGATTGGAAATTTGTAATAAAATATCATCAAACTCTCTTTTGGATGATTTA

TTGAGTGCTTAATGAGTATTTATATTTTGGAGATTATTGGGTTAATGTATAATTAATAGAACAATGTGTTATTAAGACAA

TTTTATTTGGTTAGATAAATGAAAATTTCTGAGTCATGTCAGTTTGATCAAGTAATGATAATTATTTTACAAGATATTTA

TGACTCAATATTTTATGGCACGCGCAAAACTCACGGTTTATGAAGTTGTTAAAATATTAGAATTTTCTGAGTCCCGGAAG

TATTTTTAAAACAATTGTTCCTCACTGTAAGAAATTTTAACGTTTTCTGACAGATTCAACTATATTGAAGTATTTTTGTT

TTAAATTTATTCTACAACATGTTTTAGGATTTGGTGTCCTCTGTAAACTTGAAGAGCTTTCTTTTTAGTTTCCAAAATGG

TAAACTAAACCTTATTTGGAGTTGTGAGTGATTTTATAAAATGGACTATCTCTGTTTTTCTTTTCCGCAGTTCTGGTCAA

CTACGTATTTGACTGTGGTCAAAAGGTCTTTAAGATAATCTAAAGTTTTTGTAAAATTTATGAATGGAACTATACTCAAT

TAGGATTCAGAAAGTATAATAATAATAATATTTGGATTTGAAATAGGTTCGGTAAAATTCGTGGAGTTAATTAATGTCTG

TTTTTAATTTATAACTCTTCCTTAAATATGCGATTCTTTTGAATTAAATCTGGAACGTTTAACGATGAATTTTTTTTTTA

TGGTAAACTATAGTTCTCTATTGTTCATGTGAGTTAACAATAACGACAAGTTAACGGTTAGATGTTCCCAAAATAATTTT

TATGTATCCGCTGCGACAAAATTTCGGAATTGAACATAAATAAAGAATAGACAAAGAATCGGTTGTTTGGACTTGGGCCA

AGTGTAGAAGTCTAGGATTGTTATTAAATGTAACATTTCGGATTTTGGGTTTGGATTTGGGTCGAAATCCAGGCCCGGAA

TTTGGGGTGTCACAGTTTGGTATCAGAACTATTTTCTGACATAAGAAATTCAGATATTACTATGTTTTCTGATATAACAG

TTTGGGGACGTAATTATTTTTAAAACAATTGTTTATTACTATAAGAAATTCAGATATTTTCTGACAGATTCAACTATGTT

GAAGTTTTCCTGTTTTTAAACTATTCTATGGCGTGTTTTAGTCTTTGGTGTCTTCTGTAAACTTGAAGAGATTTCTTTTT

AGATTCCAAAATGGTAAACTAAACCTGATTTGGAGTTGTGAGTGATTTTATATGATTTTATAAAATGGACTATCTCTGTT

TTTCTTTTCCGCAGTTCTGGTCAACTACGTATTTGACTGTGGTCAAAAGGTCTTTAAGATAATCTAATTTTTTTTATAAA

ATTTATGAATGGAACTATACTCAATTAGGATTTAGAACATATAATAATAATAATATTTGGATTTGAAATAGGTTCGGTAA

AATTCGTGGAGTTAATTGATGTCTGTTTTTAATTTATAACTCTTCCTTAAATATGCGATTCTTTTGAATTAAATCTGGAA

CGTTTAACGATGAAATAAATTTTTATGGTAAACTATAGTTCTTTATTGTTCATGTGAGTTAAAAATAATGACAAGTTAAC

GGTTAGATGTTCCCAAAATAATTTTTATGTATCCGCTGCGACAAAATTTCTGAATTGAACATAAATAAAGAATAGACAAA

GAATTGGTTGTTTGGACTTGGGCCAAGTGTGGAAGTCTAGGATTGTTATTAAAAAGTAACATTTCGGATTTTGGGTTTGG

ATTTGGGTCGAAATCCTGGCCCGGAATTCGGGGTGTCACATGATTTGGTATCGAGCCATCAGTCCTTTGGGAAGACTAGT

TTCTGAATGAGTCAATGTGTGAATATGCTTGTATGTGAATTGATTGCAGCAATTGTTGATTGTATTTTGGTAGAATGCAT

AATAGATTGCACTCTTTTTTTTTGCATAGAGAATTCTCTCTGTGATACTTATGATTTAATGTCCCTCATTCAGGACATGT

CTCGTCGTAATAATGGTTCAACCGGCCCACAAGATGGGGGTTTTGGTGATGTTGTTCGTGCTTTGAATGTTGTTGCTCAA

GCAATGCAGCAACAAGTTACTCTTAATCAGAATGCACCTCCCCCTCCCCCACCACCTAATCAACGGGCTTTGTTAGTTAG

AAGATTTAGTGAACAACACCCCGATTCCTTCAAGGGTAGTCCTGATCCATTAGTTGCTGAAGATTGGATAGATAAGATTG

AGAAAATATTCACCCTATTAGGCGTGAATGATGAGAGTAAGCTGGATCTTGCTGTTTATAAGCTTGAGGGTGAGGCCACT

CGTTGGTGGGAATTGACTCGTCGTTCTAGGAATGATGGTTTGTTTACTTGGATGGAATTTCGCCTCGCCTTCCTAAATAA

GTACTTTCCACAAACTGCACGAAACCAGCGCATGATCGAATTTATGCAGTTGACCCAGAGGAACATGACAGAGCACAAAG

ATGCAAGCCAAGTTTGAAGAACTCTCTCGTTTTGCTACTAACCTGGTTGATAATGAAGAATTTAAGACTTTTAAGTTTCA

GGAGGGACTTAGGCCTTCAATCAAGAGTAGGTTGTCTATTCTAAAGATCACCTCTTATAATGAGATAGTGGAAAGAGCAA

TGATTGCTGAGAGAGACATTGATGAGGCAAATAGAATTCGAGAACGTCATAATGGGGATAAGAACAAGAACAAAAATAAC

AACACTCCATACCAGAATAAAGGAAATGGACCGAAGATGGAGCCTCCCGCACCTTCATGTTATCACTGCAAACAACCTGG

ACATTTTAAGAGGAACTGTCCCAAGACTTAATTCTTAGAGGAACCAACAACCATTTCCACCTCATAATCAGCGTCAAGGG

TACCAACAGAACCGTCCTCAGAACAATCCTCCACAATATCAAGCCCGACAACCCCTCAAAGAACCTAAGGCCTTCAATAT

AGCTCATGTTGGACAAGATCCTGATAACTCGTTGTTGAAGGTACATTCTTAGTTTACAATTCTTGGGCTAAGATATTGTT

TGATACTGGGTGCTACTCATTCGTTTATTGCTTCTTCGTTTGTCTTTTCCTTGGGTTTGGAAACCGAATTACTTGATAGA

ATTCTCGGCGTAGCTAGTCTTTGTAGGAGGTAGTGCACGTATTGATCGCGTGGCACGAAAGTGTATAGTACGTATATCTA

GGAATGAGTTTTTGATTGACCTTTTCGTGATGAAAATGTCTAGTTATGACATAATCCTGGGTATGGATTGGTTATCCGCT

CATCATGCTGTTTTTGATTGTTTTCAAAAACGTGTGAATCTGCACTCGGGTGAAGGTTGTTGTCCACTTTACCTAACTCG

TAGTTTTGCAGCTCCTTATCTTGTCGGAAAGACATTTTCGACAGACTCATTCTAGAGGTTTTCTTTATCACTTAGAGATG

ATAGAAGATGAGCATCAAGATTCCGTCTCTATTATGGGTATTCCAGTGGTAGAAGAGTTTGAAGATGTTTTCCCTGAGGA

ATTACCAGGTTTGCCCCCTAGAAGAGAAATGGACTTCACTATTGAAGTTCCACTGGGTACTTCGCCAATTTCGATGCCTC

CTTATCGAATGGAACCAATAGAGTTGAAAGAACTGGACAAACAACTACTCGAGTTGGAAGCTAAGGGTTTTATTCGAGGT

AGTACGTCACCTTGGGGTGCACCAGTCCTTTTTGTGAAGAAGAAGGATAATTCACTAAGATTATGCATCGATTATAGAAA

ACTCAATGAAGTTACAGTGAAAAATAAGTATCCCTTACCTAGGATTGAGGATTTGTTTGATCAGTTGAAGGATTCCCAAT

TTTTCTCGAAGATTGATATGCGATCAGCTTATCATCAACTGCGAATCAAGGATGAAGATATTCCGAAAACCGCTTTTCGA

ACTCGATTTGGACATTTTGAGTTTGTAGTGATGCCATTTGGACTCACAAATGCTCCAGCAGCATTCATGGATCTTATGCA

TCGAATTTTTCGACCGTATTTAGATAAATTCGTAGTGGTATTTGTAGATGACATTCTTATTTATTCGCGGTCACGAGGAG

CACCTGAGATTAGTGTTGCAATTGTTGAGATCTCATCAGTTGTATGCTAAGTTGAGTAAATGTGACTTTTGGCTTTCTGA

AGTCCGGTTTTTGGGACACGTAGTTTGTAGAGAAGGAATAAAGGTGGACCCATCTAAGGTTGAAGCAGTTACTAAATGGG

AAAGGCCAAAGAATGTGTTTGAAGTTCGTAGCTTTCTGGGCTTATCGGATTACTATAGACGATTCATTCAAGATTTCTCA

AGGCTTGCTTCACCATTGACGAGGTTGACTCGAAAGGATTCCAAGTTCGAGTGGAATGATGATTGTGAGACGGCTTTCTC

CATTCTGAAGGAGAGATTAACTACTGCACCAGTTTTGACGACACCAGTAAGTGGAATTGGTTATGTGATCTTTTATGATG

CGTTGTTGAAGGGTTTAGGATGTGTTCTTATGCAAGAAGGTAAAGTGGTAGCTTATGCCTCAAAACAATTAAAGACACAT

GAGTTGAATTACGCAACTCATGACCTCGAGTTGGCAGCTATTGTGTTCGCTCTGAAGTTATGGAGACATTATCTGTATGG

TGAGAAGATAGACATATTTTGTGACCATAAGAGCTTGCATGTATATATTCTACCCAACGAGATTTGAACCTAAGACAGAG

GCGTTGGGTGGAATTTCTTAAGGATTACACGTATACCTTCCATTACCACCCGGGAAAGGCAAATGTTGTAGCAGATGCTC

TTAGCCGAAAATTAAGGGTCGGTGTAGCAAGTATGACTCTTCATGCTTGGGATTTGTGCAGACTTATGGAGGAATGCGAC

TTGAGCCCTGAAGAGATGATTTACAATACTGTAGTTGTTCCAGAGTTACAACAACATATCGTAGAAGCTCAACATACTGA

TAGAGAGATGATTTCGATTCTTGATCAAATAGAGAAAGGTCAAACACCAGGTGGTTGGAATCAAACTGATGGTGTACTTC

GTTATATGGGGAAACTTTTTTTGCCAGCAAAATTGAGGATGGAAGTACTAGAACAAGCTCACAATTCTAAGTTTATCGTT

CATCCGGGAGGATCGAAGATGTACCATGACATGAAAAGACAGTATTGGTGGAAAGGGATGAAGAAAGATGTTTCCTTGTA

TGTGTCGAGATGTCTTACTTGCCAGCAAGTGAAGATTGATCATCGGAAACCAGGAGGGTTACTTCAACCATTACCCATAT

CAGAATGGAAATGGGAACACATCACTATGGATTTTGTCATGGGATTACCCAAAACTGCACATGGCAAGGATGCTATATGG

GTTATCGTTGATAGACTTACCAAGTCAGCGTACTTCTTAGCAATCAAGAAGACTGATACGTTAGCCACGTTAAGTCATAT

GTATGTCCATGAGATAATAAGGCTGCACGGTACGCCGGTGTCAATTGTATCTGACAGAGATCCTCGTTTTACTTCTCAGT

TCTGGAAAAGTTTACAGAAGGCATTGGGTACAGATGTCTCTCTTAGTACTGACTTTCATCCCCAGACCGACGGACAGTCT

GAACGGGTGATTCAGATTGTGGAAGATATGCTACGATCTTGTGCCTTGGATTTCAAAGGAAGTTGGAGTCAGCATTTGCC

ATTAGTGGAGTTTGCTTATAATAATAGCCATCAGACTAGCATTGGTATGGCACCTTTTGAAGCATTGTATGGACGACCTT

GTAGATCTCCAGCGTGTTGGACCGAGATCGGCGAACGAAGCATTTTTGGACCTGATTTGGTACGAGAGACGACAGAGAAG

GTTCATCTAATCCGGCAGAGGTTAATGACAGCTCAGAGTCGTCAAAAGAGATATGCAGACAAGAGACGTCGACCTTTAGA

GTTTACTGAGGGTGACTTGGTGTTACTCAAAGTGAGCCCAAAGAGAGGTGTTAAGCGTTTTGGCAAGAAAGGAAAGTTAT

CAGCACGTTACATTGGACCTTTCCAGATTATGCGGCGTGTTGGTGCAGCATCTTATCAGTTGGAGTTACCTCAGCGAATT

GGGCATGTTCATGACACTTTCCACGTGTCAATGTTACGGAAGTACGAGAGAGATCCTTCATATGTTATTGACCCTTCACA

TATTATTGCTTGGGAAGATTTGGAGATTCAAGAGGATGCTACGTACGTGGAGAGACCGGTCCGAATTTTGGACTTTAAGG

AACATGTGCTTCGCACAAAGACCATCCCTATGGTTAAGGTGCAGTGGCGGCGTCGCCAAGTCAAGGAAGCAACGTGGGAA

TTAGAGTCCGAGATGCGGAGCAAATACCCTGAGATTTTCCAAGGTAACTTCTAAATTTCGTGACGAAATTTCTTTTAAGG

GTGGTAGAATGTAATAACCTGTAATTTTTTTATATATCTTTTTGGTCGGGTTATACCTAGATTGGTTAACCTTGTGTTTA

GATTGTTATTCCTTAAGACCTAACCTTAAACTTAACCCTAACCTTAAACTTAACCTTAAATGGGTGGATTGAACAAGAAA

GGCTCCAAGTTAGGGTGGAAATAGAGCCACCCAAATTCATTGTTTGTTTTTCCTTAATAATGTTATTAAGGTTTGATAAC

TAAAACTTTAGAAATTTTAGAGAGAGAAACTAAAAACTAGGGAGAAGAAGGAGATGAAGTTAGAGTTTATTTTAGAGGGG

AAAACCATGGAAAATCAGTGGAGAATTAAGAGGAGTTTTTTTTAAGCTATTGAGGTAGTATGCTAGATTCCATTTCTCTT

TTTTTTTTTATACTTGATTTTGAATTTGTATGATTAGATCTAAGTGATTTATTGATTAAAAACTATGAATCTAATTGATG

ATCGGCATTTAATATTGATAGATAACATGCTTAGGTATCTCTAGTTAAAATTTGAGAATATTTCATCACATATTCACCAT

AAAATGTTTGTTTGTACTTGGATGTTGATTTCTAAATTGTTTTGTACATTTTCGCTTTGTATGTTCTTGACTGTACTTAG

GACATCTTGATGATATCAAAACGATTTTTGATCTTGATAAAAGTTGTAGATCTATGTCTTATCTTTTTATTAGTGGGACA

AAGAATCTTTTGATTTTTTTGAAATTAGTTTTGATTAATTATTTTCTTTTTCTTTAGATTAAAGAATTCTTAGTGGGACA

AAGAATCTTTTGCTTTTTTTGGAATTAATGGTTATTAATGATGAAAGTTAAGGAATAATAGGTATTAATAACTTATTATA

TTCAAATTTGAATTCTTTTTGTTTTTTTGGTGGAATCCTAGGAAACAACTTTTCTTTATAATACTGTTTGTTTTAATAAA

GAATAAAAATGAAGAATTATGTTAGAGAGAGAAATAAATTAAAAGAAGGTGGTAGTGGTAGACATAAATAATTAGCTATC

ACTTTGAAGACATAATTATTAGATTCTTATCTTGACCAAACTTTTAAACTTGAGGTTAGGTGGAACTTGTTTTAGGTTGT

GAGTATATATATATGTATATATCTTATACTAAGAAGAAAAGAGTAACCATACTAAGAAGAAAAGAGTAAACATACTAAGA

AGAAAAGAGTAAACAAAAATCTAAATTAATAGAAATAACGAGAGATTCTTTTAGTTCATATCAAAGACAATTTCTTCGGT

GATAAAGCTTTTCCTTCTATTATTTATAGTTTTTGGATTGCGATTCGAGGTAGGCGCAACACAATCTAAAACTCTTCTTT

TATTTAAGTTTCTTTTAATTAAATTTATGTTTGCAAGTTTCCATTAATATATGGATGATAATCGATGTTGTGAGCATGAT

TATGAATGTTGGGATGCGGTCCATAGAATGTGATTCTTAAAGATTATTTATGACGGGATACGAGATATCCTTGGAAACGG

TATTTGTTTCCTAAACCCACGTGGGAGCTCCCTTGGGCGAGCGGATTGATTGAATTTATTGTTGTATAATGTTGGGATAC

GAGATATCCTTGGGGACGGTTTTGTTTCCTAAACCCACGTGGGGGGGCCTCCCTTGGGCGAGCGGATTGATTGAATTTAT

TAATGTATATGGATGTGGTGTTTCCATATGTTTGTGGTGGTGTTAACCATGCAACTTGTTTTGAAAAGAAATTGAAAAAT

ATTGCATAAAATGTTTTGTTGCTTTTCCTACTGAGTGTCACAACTCACGTCGTATTATGACAAAAATTCCAGATTTTATG

ACACCACAAGGAGCTAATGGCGGAAAAGCATAGAGATCGGTTTTGGTGTTTTAACTAGTAGTTGCATATTATGATTGGAA

ATTTGTAATAAAATATCATCAAACTCTCTTTTGGATGATTTATTGAGTTCTTATTGAGTATTTATATTTTGGAGATTATT

GGGTTAATGTATAATTAATAGAACAATGTGTTATTAAGACAATTTTATTTGGTTAGATAAATGAAAATTTCTGAGTCATG

TCAGTTTGACTGTTAATGATAATTATTTTACAAGATATTTATGACTGAATATTTTATGGCACGCGTAAAACTCACGTTTT

ATGAAGTTGTTAAAATATTAGAATTTTATGAGTCCCGGAATTATTTTTAAAACAATTGTTCCTCACTGTAAGAAATTTTA

ACGTTTTATGACAGATTCAACTATATTGAAGTATTTTTGTTTTATATTTATTCTAAAACATGTTTTAGGATTTGGTGTCC

TCTGTAAACTTGAAGAGATTTATTTTTAGTTTCCAAAATGGTAAACTAAACCTGATTTGGAGTTGTGAGTGATTTTATAT

GATTTTATAAAATGGACTATCTCTGTTGTTCTTTTCCGCAGTTCTGGTCAACTACGTATTTGACTGTGGTCAAAAGGTCT

TTAAGATAATCTAAATTTTTTGTAAAATTTATGAATGGAACTATGCTCAATTAGGATTCAGAAAGTATAATAATAATAAT

ATTTGGATTTGAAATAGGTTCGGTAAAATTCGTGGAGTTAATTAATGTCTGTTTTTAATTTATAACTCTTCCTTAAATAT

GCGATTCTTTTGAATTAAATCTGGAACGTTTAACGATGAATTTTTTTTTTATGGTAAACTATAGTTCTCTATTGTTCATG

TGAGTTAACAATAACGACAAGTTAACGGTTAGATGTTCCCAAAATAATTTTTATGTATCCGCTGCGACAAAATTTCGGAA

TTGAACATAAATAAAGAATAGACAAAGAATCGGTTGTTTGGACTTGGGCCAAGTGTAGAAGTCTAGGATTGTTATTAAAT

GTAACATTTCGGATTTTGGGTTTGGATTTGGGTCGAAATCCAGGCCCGAAATTCGGGGTGTCACATCGGATACATCGACT

ATGGTTGTAGCCTCGGAATGCTCACCTTGTTGTCCTTGGTAGCTTAATGTTGGTACCTTCGGCGGCCTTCCCCTTTTCCT

TTGAACCTCGGATACATAGACTATGGGTGTAGCCTCGGAATGCTCACCTTGTTGTCCTTGGTAGCTTAATGTTGGTACCT

TCGGCGGCCTTCCCCTTTTCCTTTGAACCTCGGATACATCGATTATGGGTGTAGCTTCGGAATGCTCACCTTGTTGTCCT

TGGTAGCTTAATGTGGGTACCTTCAGCGGCCTTCCCCTTTTCCTTTGAACCTCGGATACATCGACTATGGGTGTATCCTC

GGAATGCTCACCTTGTTTTCCTTGGTAGCTTAATGTTGGTACCTTCGGCGGCCTTCCCCTTTTCCTTTAAACCTCGGATA

CATCGACTATGGGTGTAGCTTCGGAACCCGTCGTTTGATGTTGACTTGATATCAGTTCCTTCTTTGGCCTACCCCATTTT

TTTGGAACCGGCGCTTCCGCGAATCTGGCTTCCGAAATCTCACTTGCGGATAGATCTCGTCTCTTAGTTTCCGCGAGTTC

TTCTTCACGTCTAGTCATTTCTTTCAATTCTTTCCTTTGTTTTCTCCTTGGAATTTTAGTTTGTGGTCTACCGGGCGGAT

CTCCCTTTCTTGGCTCTTCACTTGGTGTTATCCATGGCCGTGTAATCATCCTTAGTTGGTTCAACAACACTTGTTAGGTA

TCCGAGTTGGCTCGTGAGTAAGCTTCATAGAATTCCTTGGACTCCTTTGCAACCCATGGTGATTTTCCGGGATCTTCCGT

AGGGGGAAGGTCAAAAGTTAGTTGCTTCCAAAACGGATCAATAACATCAATAGGTATCACTTCTTCATACTTCACAAGCA

TATGACAACACGGAAGCCCCAATGAAGACATTTCGGGACAAATACACAAATCACCCGGTTTTCCGTAATTTTTCTCATAT

TGCATTTGTATCATCATATGTTTTATTGCCCAATGGGAGACGTTGAACTCTACTCCTTGATCAACTTAGTATATTTAAGA

AATTGAGTCATCCTTTCGATTGATATTTTCTCAAAAGCCTTCTTGATCCTATCGATATCAGCCTTAAAGTGTTTCTCCAT

TGCCTCGGTGACTGTAACCACGTTTCCTTGCCCGGACTGGATGAGATCTTTAAATCTCCCATGAGAGGACTCCGCTATAC

TAGTTGCTTCATTCTCGTAATGTCTATAACGATTTATACACGCACGCACAAATTTTTCCTTGAACTTATCCAACCAATTA

TCCCGACAATATGAGACGGCACTTGGATACACTTCGTTATATTTGGAAATAAACATATTCAATCTTTTTTCATACATGTC

CACGGTAAGAGACCAATAAACTTTCTCCCAATCCTTTAGAAATTCCAACTACCTTTTGTGATTTTCACCGTGTTCTTTGT

CAACTCGCTCTTTAATCTTTCCTCTTTTATTTTCTTCTTCTACGGGTGATAGTTTCTCTTTGCGAACAACTTCATCTAGT

TGAGTAACCATTCTATTCTTAATATCCTCCTTAGAGGGTTCAAACAAAGCATGACAATGTTTTTTCACATTTTCACCTAT

ATGATATGTACATAGGAAATTTTGTGCATCCGGATATACGTCGGATATGGAATTCATTAATGCGTCAAATTGATCGGTTA

TGATGACCTTCGGAAACCGATTTTCCGGGAAAAATAATTTCAATTGTCGCAATGCCCAATGATAATTGTAATCCCTCTCG

TTTTCCATTAAACACCAGGCCAATGTGAATGTTACCTTGTCCGATGTTTTCCCCACGATGTTGAACAACGGCATGTTGTA

TTTGTTTGTCTTGTAGGTGCAGTCCATCATAAAAACATCACAACACGTATGAGCCAATTGTGAAAGCAAAGGATGCGAAA

TGAATATTTGAAGGGGATTCTCGTCCGGACCTCTTTTAATGATTCGCGTGTAGTTGTGATCCCAAACTATCTTCTCAAAT

TCTTCCATAACGGTCCTCCCTTCCCATTCCAACCTTCTAATGGTTGCTTGTGCGCTATAAATTGTACTTAGGGTAGAAAC

GTTTGTCTCATCCTTTTTCTTGAAGCCTCTGAGAATCACTCTTGGTTTGATACATGCTTTGCTCATTATCTTTACATCCT

CAAATTGATGAGGTTTTAGCTTCGCAACTAGGGAGTGACCAACAAAATCTTTCGGATCCCAATGGTTATGACGCCCAAAA

TAAACCGTATAATCCGATTCTTCTTCTTCTTTTCCTTTGTCTCTTAGATGGAATACAAGCTTAAAAGGGCACTCATGCTT

CCTCGTTTTAGTCTTGTATACCCTATTATTCTTCTTTTCATATACATAACCCTTTCTCTTGTGGATATTCTCCTTTTATT

TTCCGTGTCGTTCACAAACCATCTCAAAACGAACATCTGAACCTTGGCTATTCCTCACTAAGACACACATGTTCTTAAGA

GCCGTCTCTTTTGCCCAACCAATTGCTTCAGCTGGAGATTTTATTCCCTATATAAGGACAAAAAGGGAAGATTATAAGGT

TAGTTACAAGAAATCCCAACATAAAATTGTAAAAACTAGGTGAAAGTATTGTTTGGTAACATACCCGAGGCATTTTATAG

TATTCGTACGTATCCCGAAAGAGAGGTTTGGAATTTGGAGCGATGTATGTCACAACCTAAGGCATAAATGAAAACAAAAT

TGAATTAGTTCTACATAATAATGTTCCGGCAATGTATGTTTTTGAACAGCAGTGCCGGGATAAGGATGTCGGCATTGTAG

AAAAGGTAAAATCCTCGCCGGTACCTAGTGCCGGCATCCACGATAATTTACATTAGATGCCGGTACTGAGGTTAAAAATA

TCATTAACTCATGTATGCTTTCCATGTAAAACCGGCATAATATATATGTAAGGTTCCCTGCCGGTACATAGTACCGGCTT

TCACGAAAAATTCCAAACTACGCCGGTACTAAGTGCCGACGTTCACGAAAAATAAAATAGAAATACGATACTTAAGATCG

AAAAACAAAAACTCATGTATGTTTTACTTGTGGTACCGGCATAGTAAGTTTGGAGTAAACCATGCCGGTATGGATATTCC

ATGGAATATTCGGTGGTAGGTAAAAAGTGTAAATGTATACCGGCATGGATATTTGGTTTGAATACCACGCCGGCACTGTT

TCAAAAATGGGGGATATACTAAACCGGCATGGAAAGGAGTATGAATACCACGCCGGTACCACTAGTTCCGGCGTAGTATT

CATACCACGGGTGTGCCGGCACTAAGCTTTTTCAGAGAAAAAATGGCGGTATGAAAAGAATAATCAAATTTTCGACCTCT

TTGAAGCATTACCTGTGTGTTGGGGTTGCTTGTATGCTGGACATGTTTACTTTCTTGGGTTGGTTCTTCACAAAACACTT

CATGCTCACGAAAATCATGATCCAAGGGTATTGTTTCATCATATAACTCCAATTGTGAGTTGTTATCATTAACCAAAGAT

TAATGATATGATTGTTGTTGTAGTTGATCTAAAGATTCTGCAGCTGCAAGTCTCTCATCACAATCATCTTCCATTTTCAA

AAAAAAAAAAAAATTCACTCAAAATTTTTTTCCCTCCTTCTACTCTCAACTAACCCACACAAATAAAAATAAAACTAAAA

ACACACTAATTTATCCCCCAAATACTTAACATTTTACTAATTATAATTATCACTAATCCTAATTAGTGGGGGCTAGATTA

GGAATTAAAAAAATAATTGCATAGGGGGTCACCCAGAATTGTTATTGGGACACCATTTTTGTCCTGTAGCTATATCCCCC

AATTAATAAAAGTATCCCCCAATAGCGCGTTCCTTTGAAATTGTGAAATTAATAGGAGTAACATTTTATGTTGTCTTGAA

TATAATTCATATTATTGTTTCTCTAACAGATTTTTATTTTTCCTTTGGGTATTATTTTTGAAGGCTGTTTTTAAGCATAC

ATAAAAATTTCTGCGCATACAAAATTATTTTCCCATCTAGGATGACTTTATAAGGGGTAACTAATGGTAGTGCAATTACT

TATATGCCCTAAAACAATTTAAATTACTAATATATCCTTACCCTTAAAAAAAACTAAATTCAAAATCAATAAAATCAAAA

TCAGTTTCCTATCATATACATCTCTTCTTCTTCCTTCTTCCTTTTGCTCAACCCTCGTGAACCGTCGAAAATTTATTTTT

CCATCGTTATAATTAATCATCGATTGGTAAAATTGATCATGGATTAAATTAAATTATATAATGACTTGCACAATGCAAAA

AGTAGGAAATAGGCGTTCTGAATCCTTTTCTAATCCATCAATCGAAGAACAAATAGAAGTTGAATGTGCAGAATCTAAAA

TTGAACCAAAAATTCAACTGATTGAAAAAACAACTCCTAAAATGGAATTAGAAGGTATTTATCACAAACCCATCTCTTAT

TTTGTTATTTATGATTGATTGAATAGGTTTAATTCGGAACAAAGTTGGTTTTTAACGATCACTGATTGATGTTCGGATGA

TAATTTGTCTACGTTATCAGCCGAATTGTTCATTATTTTTCACCTTGAAAGTGTATATGAATAATCCAGCTGATAATCTG

TCTATGTTATCAGCTGAACTACTCTCTGTGTCAGCAACACAATGAAAGTTACCTTAGTTCGGCACATATTTTTTTATTTC

GGGACAGCCGAACATTTTTAGATACTACAAAGAAAAATGAATGGTTAGGTTCGGCTTGTAAGATAATCATAATATGTGCC

GAACCTAGTGCATTTATAAGGCTCGGCTACTACGATAATCACAATATGGGACGAATCATTAACAAATTATGAATGGTATG

CTTGGCATGTGTACTACTTAGGTAACAAGACGAACCAAATATTTCAAACAGGTTCGTCTCATACGATTTTCATTAAATAA

GCCGAACTAGTAACATATTTTCTTCCAAAACTCTCAGGAATAGGTTCGGCTTGTTATTAGACTTTAATATAAGCCGAACT

AGGATCTTGCAAATAGGTTGGAAGTTGGAAAATGAAGGTTCGGCGTATTACGTTAACAAATTCAGCTAGCCGAACTGTTC

ATCAATTTAGTGGGTTTGGCTTATATTTTTAAGCCGAACGTAGTCCTGGTTCGACGAACCATGATTAAAAATACAATCTT

TTGATTATTTCAATCTACAAAAGTGAGATTAAACATTAAATAGAAGTGTACTTGTGTGTTAGAAGCACTGGGTTCCTCAT

CTATGTATTCATCATCTGGATTTGGCTCATAAAAACTCGATTTTTTTTCCTCCATTGAATCAAGAATTATTTTTTTCTCA

CTTTCTCCTTTTCCTTCTCTCCTTCTCAATAAAAACTTGATTTATTTTTTCCCCAAATTTTTTCATCTACACAAAACTTT

ATAATTCTGAAAAATTATCTTAATCACTAAACAAAAGTTTTAATCACTAATTCAGATTACTAACACTAATACATAAATGG

AAGATTTGCCATTATAAAAAAATTTGGTTAAGGGAGTTTATGATTTTGTTATTTGACACCCTTTTTTGTCTTTATTGAGT

ATGCCTTGAAAGATTTAGGTATGCCTAAAAACAGTCTTCTTATTTTTACAAAAGGAGGAAGTGTGAGAGGATAATAAACC

AAGAATGAATCCACTTATGGGCTTTAAACCATGAATGAATCCATTTATGGGCTTTCTAAAGAATAGTTATCTGCTCTGTC

TTGATTTCATGGGCTTTCTAAATTTTTCTGTAGTTGAATTTCCAACAATACTACTATTAACCAACAATTCAATACTGTAC

TATTAAATGATCATCGAAGAAAATTCAAATTCGTCAAACTGAGTAAATGGATTAAGACTGCTGCTGCTCGAGCAACCAGG

TATCCCGTAGCGGTGGACGACCACCGGACCCGTTTCGGTTCACTTTAGCATTTCAGGGGTCCATAAACCCAGCCTCAATT

AAAGAAACAAGTGCTTCTGTTGAGATTTGAAGGCATGATGGTACCTGGGGTCAGAAAACAGGTGGTTGTAGCGACTTGGA

CTACTAACAAGTTATTTCAGAATCTCCCGTCTCATCTGAATAACTGAGAATTAAGCCAGCCAGCCAACCACCTACTAACC

TCTAACAAAGTTGATTAATTGTCCCTAAATGTGCATGATTGTTAAAGGTTGATGTCAAATCTTTCGGGGATACCATTACA

TATAAGACAAAAAGGTCAAAATGATCCTGACCGTTGGATCATTCAAACGGTATCCCTGAAAGATTTGGCATCAGCCTTTA

GTAATCATGCAAATCTGGCCCGTGTTAGTTGTCTCGGCTTAAAAGTTTTTCCAAAAAACTCAACTGGGACGTGTGCTCCT

AGTTAGTAATTCGGCGTATGATTCGCGAATCATACGCGTACTTTTCCTTATAATTCGTTACAGAATGATACGGCCGCATT

TCTCTACTACCGAACCTAGTACGCGTGCTTTCTGGGCTTAGCGAATAATTCGCGCTTTATACGGATCCGCGAATCATTCG

TTCATCGGATTCGCCATCTGGGCTGTTACTTTAGCCTAAGCATATCTCACAAGCCCAACCGTATAAACCTTCGACATATT

GTTTTCTAAACTGACATTTAACCACTACACCAGCCATCTGTTGGTGACATATATTGGTATATTAAATATATATAGCATTT

TATATCACGAATCATGCCTCCTACCTCCTGAAACATATTATTTTATTTATTAATGTTTGCCGTATTTTCCATACCGAACT

CAACCTAGTAAAACGAATAATACACGTACGTATCCCGTTCCGTACTACTGCCGAATCACGAACCGTTGACCGGATTTTTG

ACCGAATTTGACTTTTACGAATCCGTATAATACTCGTATGTATGTCGTTCCGTACGTATGCCGTATCCCGAATTGCTAAC

TAGGCGTGTGCTTCGTCTTAAATAATTGTCATATTTTTTTGAGTGTCTGACTGGTGGTCTGACTCAGTTCAAGCATCTAA

CAATATTTGAGTCACATATAACTTTATACTTGATTCGTCGAGCACGCAGTAGACTAGATTCTGGAAAAAAAGAAAACAAT

AATCCTAAAATTTAATCGCATTCAAATATAACGAGGCACATTCAGATGACTTGGAAGAAAAAAAAAATATCTAGAAATGG

AAATCGTGAGTTATTTACGTCTAAATCTTGAACTCTGAACCGTATAAAAAACGTGTGTGTATATATATATATATATATAT

ATATTTACTATTATAAAAAGAGCATAGAAATGGAAACAAAATGTTAGTGAAGGTGGGTAGGAAAAAGAAAAAAAAAAGGT

TGAAAACTAGGAATATTCTGTGGTGTTGAGTGACTAATGGGTACATAATTGGTGGTGGTGGGAAGATAAGATAAAAGAGT

TTAAAAATGTAAGCTTAACACCTAAGCAAAGGAGGAAAGAGAGATGAAAGCTATCTATCGAAATCAAATCAATGACTTCA

TGACTGACACATTGACCACTCAAGTAATTCCAAATGTTTTTTCTAATTTCCTTCTAGAGAAACGCGGTTTGGTTGGTGGG

TGGTGTCATGTGACTCAAAATCACATCTTACTCAAACACACGCAATTCTCACCTACAAACCAAAATTAAAGATATTATTA

ATGTTACCACCGTTAGGAGTCTTTTTATTTCTCAGTTACCCCAATTTACTTTTTACAGAATTGGTACCACCTACCTACGT

TTCAGTCGAAAGTATCATTTTTTTTTTTTTTTTCTGAAACAAAAGGATAGCGTGTTTGACAACTTGTATCAGAAATAGCT

TTTGGATGTTAAAAGCAATTTATTACTCCCTCTTTCCTAAATTATTAGTCCGCTTTGACTAAGTCAAGATATTAAGAAAA

CCGGAAGAATGTACAAAACTATCCCTATTAAATGCTATGTTATTACTAAAATTTAAAGGAGTTAATGGAGCATATAAGAA

AAATACATATTTAATAATTTTATTACTTATAGAAATGTTTAAATAGAAGATAAAAATAAAAATATTTATGGATTAATTTA

GTAATTTTGTTTCCTATTGAGGAAGATATCATTTAGGATGTTTAAATAGAAAATAAAAATAAAAATATTTAAGGCTTGAT

TTAGTAGATTTGTTTCTTATTGAGAAAGATATCATTTAAGATTTTTGTTTGGTGATATTTAAAAATTATTTTCTAATTAT

AAAAATTGAAGAATATATTTGGGAGCTTGACTAAAAAAATATGATTATAAAGAACAATTAATTGGGGGATATGGGAATTA

ATTGGGGGATATGAATTACTTCCCCCAACCCATGGTATGTGTTAAGGGGTTATTTTTGGTTATGAAAATACTAAAATACC

CTTCCCACAAAATTAAAATCAAAAACTTAAAATCAAAAAACAAAATCATATCCCCCATTTCCATCTCCACTTCTTATTAA

TCTTATTTAGGGGTAGAGCTTTGAAACCTAAATATTCCTCATTCCCTTTAAAATTCTAAACTCTCCCCACTCTACATCAA

ATTTTCTTCTCTTCACTTTGGAAACGAATTTTTATTTTATTCTAAGTGATGAAGATCATCTTTGAAAGTGATGATGACCA

TGCAGAATTGATTCTCGAGCATGCCGGTATCATGTTCCGGATATGGTCTTCATCAATTAAAACATGGTTTTCATCACTTC

CTTTTTGAATTTTTTCGGTTTTAATCACTTCACTCTTAAGATGGTCAGTTGATATGAAGACCATGCCGGTATTTCTTTCC

GCATAGCATGTTGCTTTATTTTCTATGCTTGGGAAAACAATCCAAGATGGTCTTTATATCAACCGACCATGCCGGAGTGG

AGTGACTAAAACGAAAAATTCAAAACGTAAGTGATGAAAACCATGCCTAATTCATGAAGACCATATGCCTGAACATGATA

CCGGCATTCTCGAGAATTTCCTATCAATGCCGGTATTCAAGTGAAAAAAAAATGAACATTCTGGATACTTTTCATCCTGT

TCCGGCATAGAAATTTAAGCAAAATATTATGCCGGAACATGATACCGGCATGCTCGATAATTTCTTGTCAATGCCGGTAG

TCAAGTGAAGAAAAAAATCAACATTCTAGGTACTTTTCATCTTGTTCCTAGATAGAAATGTAAGCAAAATACTATGCTTA

ACGGAGCAGTGCCGCATGCTCGAAAACTAGATTACCATGCCGGTACCTGTTCAAACCCAGAAAAAAATCAACATTATGAA

GATCAAAACTATAAAAAAATCGTAACGAGAGGGCGTAGCTCCCTCGTGTGGGCTAAGCCCCCCCGAACCCCCCTGGATAG

TGGCAAACAATTTTTGAAAATTTCCGACAAATGCCGGCATGGTTTTTTGGTTGAATACCATGCCGGCACCGAGCTTTTTC

AACTGCAACAATAGTGAATTCAAAGGAAACAAATCAAATTTTCAACCTCTTAGCAAAGAATTCTCTCATCACAATCATCT

TCCATTTTCAAAAAAAAAATTCCACTTATTTTTTTTCCTCTCCTTCTACTCCTCACTCACACAAATTAAAATCAAAACAC

ACACTAAACTATCCCCCAAATACTTAAGCATATACTAATTATTATTAACTACTAAACATGATTAGTGAGGGCTAGATTAG

GAATTAAAAAAACAATTACATATGGGGTCACCCAGAATTGCTATTGAGACACCGTTTTTGTCCTGTAGCTATATCCCCCA

ATTAGTTTCCATATCCCCCAATTAATTCTTCATTATAAACAGAAGCTGACCAGCATTTTAGGACAAACAAAAATAGAATA

GAGAACAAAAATTTTAAGACAGAGGGAGTATTATTATTTTTTTATATTTCAAGAAGTCATGCTAAAATTTAATATTGATT

CCGGATCAATATTTAACTTTTTCCGCACGAAAAGCTACTTCTTTAGATGGTATACAAACAACCACAAAATCTAACTCACC

CAACTGAAAACAAAACGAACAAGCTGATTTTAAAAAAAAGTTTTTGACTGATAACAAGTAAATAGAGAGATGTAAAAGAC

TTAACTTTTTGTTGAAGAGAGATAATATTTTTACCGAGTGACCTTTGACTCGATCATTCGTATACATCAAGTCAACCTAA

AGGACCAAAATAGTCAATTTCACTTTTAACCATCAAGGGACCCATTTCGAATATGCTACACTAGAAGGGATCTAAGCTTT

GAAAAAGCCAACATTGTAACACACACGGCTTCGCGAGCACTTTCCTTGACATTTCCTCCTCCTTTTTGCTTCCTTATAAA

TTCTTCTCACTCAGAAAATCAAACCAACTCCATCCCCCATTCTCTTGCTTCTTCTCTGCAACTAAGAAAAATCAAAACAA

GAAATCAAGACCAAGAAGAAATGGAAAGCAATCAAGAATCTTTCTGGCAATTCAGTGATCAACTAAGACTGCACAACTCG

AGCTTTACGAATCTCAGCATCAATGATTCGATATGGGGATCATCTCTCAATTCAAAGAAGCCTGAAGAGAGGAGGAACTT

CGATATCAGAGTTGGTGGCAATCGAAATTCATCACCTTCTTTCCCATCTCCTACCGTCAACAACAACAACAAGAACATTG

ATTACAGTAATAATACTAAACAAGTGCAGCAGCAGAAAAATGGGGCGGGGTCTGATTATCTGAATGGATTCAATGATGCT

TGGAAGATTGCGAGTAAGTTTGGTGGAATTGGCGATGGAGGAACATCGTTGATGCAGAAGAATATGAATTTTGATGTTAA

CGGAGGGTTTAACAAAGGTGTTTACTCCAAGACTGACAACTTCAAACAGAGTAACACGAAAACTGTTAACAATAATAATA

ACAGCAATCACAAGAAGAATGAAGAAGTTTATGAAGGAAAATTCGGGAAGAAGAACAATAACAACAACAACAATAATAAT

AACAAGAACAGCGTCGATAAGAGGTTCAAGACACTTCCTGCAGCAGAATCTCTTCCTAGGAACGAAACGGTTGGTGGGTA

TATCTTTGTCTGCAACAATGATACCATGCAAGAGAATTTGCAAAGGCAGCTTTTCGGTATGTCCTCTAAATCTCTAGATT

TTGAACCTTTCATCAATTTACATTCATGGGTTTTTCTTTAAAATTTCAGGTCTTCCCCCAAGATATCGAGACTCGGTCAG

AGCAATCACACCTGGACTGCCTCTGTTTCTTTACAACTACTCTACCCACCAACTCCATGGAATTTTCGAGGTATGTACAC

ATAAAAGTCAAATAACATTTCATATTCCTAGTTCGGTCTCGGTCTGGATTAAGAAATATTCAATATCTGGTTCAGGCTTC

AAGTTTCGGAGGAACAAACATTGATCCAACAGCCTGGGAAGACAAAAAATGTCAGGGAGAATCTCGTTTCCCTGCTCAGG

TAATAATCCACTCGATCTCTAAATCTTAAAAACATATGTTTCGCTAGGAAATTCTTAATATAAACATTACTTGATTATTT

CAGGTCCGAGTTCTTCTAAGGAAAATATGTGAACCGCTAGAGGAAGACTCATTCAGGCCAGTCCTTCACCATTACGACGG

ACCCAAATTCCGTCTTGAACTCAACATACCAGAGGCATTAGCTCTCTTGGACATATTTTCCGAGAACAATCCTTGAATGT

CAGAGAAGATTTACTATACAAGTATAAAAACTAGATTAGATGATGAACTACAGAAGCACTGCGTCTTCCCTTTTTTGAAT

CTCTTGATGCGATATTAGTAAATTTTAAATTACCAAGATGAAAATCTCCCTATGTAAATTATTTGGAAGGTTTATATATA

TTTGAGATTTCAAAATATTACTGAATGAGGTTGTCAAAATCGAAACGTTATGAGCAAAATGATTGTAATCACATAAAAGA

GATTTTCAGTTGAACAACATATCTGACTTTATTCATTATTTTATTTTGTTCGACATATCTCTAAGTAATATAATAATAAA

ATTGCAGCAACTTATGTGACTATAAACAAGTTTACCAAAATTCATCTACGCGCTATAATAACATTAATGGTGTTAGTACC

TCATTTAACATAAGGAGGATCCATGAAAGCCCATTTAAAAACTTGTTCGTACTGAGCAGCAGCTTTGTCCCATGTAAAGT

CCTGTTCCATTCCTCTCTTCATCAATCCCTTCCATGAATCTTTATGTTTCCTATACGTGTCCATGGAAATTCTCAACACC

TGTTTTCAATAACGATAACATAACACAGTAAATTACTGAACAACCAAAATAAGAACAAGTTAAAAAAAAAAAAAAATACC

AGTGTAGCAAAGATGAAAAGGACAAGTTTGTATTTTTGAAACTTGACTGTATCTAGCAATTCCTTAAAGACACGATGAAT

GGGTTAGAGGCCTTGAGTCATTATCACTTCTACGATTTTTCAAGCTAAAGCAAGAAAAAAGAGGTCGTGCTGCACATTAT

TGATATCATATTCTAGAATGCATACCCTTTTATCATTGACTGGCGACTGATACGTGTTTCAGTTGAACAAAGTATTAGGT

AGCTGGAACTACTAATAAATGAAGAGAAAATCAAACCAAAAGGTTCACAAGTTTTATTCACGTTATAAAACCACGACCCC

ACAACCACGAGAAAAGTGAGGTTTCTTTTCTTAATAGGCACCTGAATGAGTCAAGATTCCATAATATTCTACACCTACTG

GTACTTGGAATTTCTATGAGTTGGCTGTTTTGGGCTCTTCCTGAAATTTCTATGAGTTATATGTCAACACTTCCTGTTTG

CTATCTACCGAATTTGATATTAAAAACCTACAAGATTTACTTGAGGAATACGATTTACATAAGATACTTGAGAAGAACTT

ACCCCCAACATAGGGTCCTTCGCTAATGGTGAAAAGGTCCATCTGTCAAAAATACCACCAAAACATTCATGAGTTTTTTT

TTACCTTACTGCATTTCTGTTTGGTCTGTAAAATTTATTCTAAACAAGATAAAAAATTGATTGCATACCCTGTTCCTTCA

CCGCTGCCTTCATTAGCAAAAGGATCAAAGTTTTCCACCGTGTCCTGCAGCAAGGCACCAATTTGTCAAAACAGATAGCC

AACTAGATAAGCATAAAAGTACAAAGACACTGTAGTTACTTGAACTCACCCTTAGTCCTCCAGTGCAATGCACTACAGGT

ATAGTTCCGTATCTCATAGCATACAACTGGTTTAGACCACAGGGTTCGAATCTCGATGGCATCAACAGTATGTCAGCTCT

GTTTGGTTCAAAAAGATGACGGACCAATTAGTTTCATAAAGTTGTGAAAAATATTTTATTTATGCAGCTAAGTTCAGATA

TATAAACCCACCCTGCAGTTATTCTGTGAGAAATAGGTACATTGAATCCAACCCAACCCCGGTATTTGTCCTTATAAGCT

GCCTCTGTTGCCTTCATCCAGTCTTCCATACGTGCATTACCAGACCCAAGCATAATCTTGAATAAAGAGGCATCACACAC

AAATAAATAATTTATGCAGAATATTTTTCATAAGACTAATTATTGTGACCCAAAAGAAAAATCATAACGGTTCAATTAAA

CAAAATTCGCCAAGTGCTTTTATTTGATAGAAAATTGAAATATCAAGCTTACAAACTGAACATCGTCCATCATAAGCTCT

GGCATTGCTGCTTCAATCATATCTATCCCTTTCTGGTAGTCCAGTCTCCCGATGAAACCAATCTAAGAGTCAGAAAAAAA

AATGAATTACGAAACAGGATAACGTTATAAAATTCTAACAAGATAATGTGAAGGGATGTAAATCTTACCAACGGGACATC

AGGCCTGACTGGCAACCCTAGTTCCTTTTGTAAAGCCGCCTTGCATTCTTGCTGTGTAACAAGTCCTATTTGATGTAAGT

GATAGACCTTTGAAGTGATTTGAGCAAAAATGGAACATCAAACACACATGGACGTACCTTCCCAGATAAGTCATCAATTG

AGTAATGGAAAGGAATGTGTACATCTGAAGATGGATCCCACTCATCAACATCAATCCCATTAGTGATTCCTGTCAGAAAA

ATAAATTAGATTAATATAAAAAAATCATCTAACAGTTGCACAACAAAGTAATCTGCCCAAAAGATAAGAGGGATATTCAT

TCGTAAATAAATACCATCCAAGACACTCTCTCGACTGCTTATGAGTCCGTCAAGACCATATCCACCTTCAACAGTTGTTA

TTTCCCATGCATAACCCTGGCACAAGGAGGTGGAAAAACAATAAGCTCCGCAAAAGCAGAAAATATATGTAACCTTCAAC

AACTGCAGTCGAAATACTTTTCAACGAATTGAAAACACCTCAATCAACACTCCTGGTCCATCTCATTCTCTCAGTTGTAA

AAGAATGATGATTCGGTAAATCAGCTAGTGGCTTCATCCAAGAAAAAGAAAGAGGTGCAAAGATGAAGAGAGAACCGATT

ATGACCTTGCTCACTGTTAGAATCCGGTCGGCTGTTACAATTGCACCCTTCATAATATTGACAGCTTCCCCTGTGTCAAG

AGCATGAGTTCTTGCCCAAGTAGGAAATACCCATTCTAATGCTCCATGCCACTCAGCTGGTAACCCAAAATTCTTGTAAG

TTGCAGCTGGTTCCACACCCTTGAACAAATGTACATATAACAATAATCAGCAACAATTTCCAGAAACTACACTTGCAGAT

TAGTACTCCTCAGAATAGTATTCCAACAGGTTGAAATGTTCAATCATACAAAGTGAATCGATGCATGCATAGTTCATATA

TGTAGAGCAAATTAAAAGAAGAATGTAAAAAGAGTTAACCTGATGTGCAAGGTTATGTATTGCAACAATACTGCGAGCAT

CCTTATAAACTCCATATGGGCGATATTTAGATGCCAAGAGGCTGAAAATACATGGCATTCGGTTACTTCATCAGATTCCA

GCGAAATACTAGTTATGCAAGTCTAAGCTGATTTAGTAAATGCTAGAAAATGATTGCTAGCCAATAAAAACACTAAATGG

CCTTTCAGAAACTGGTATAATTGGATTTTTTTGCTACAAAACTCGGTATGAAAATGAGTTCCACAAGCAACCCATTTATA

AAGGAGAGTTTCAAGAACTGATTTTCAAGTAGAAGAGTTAAAAGAAAAGTCCAGAATATCGTACACTGGTACAAGACTTG

CATGCCAATCATTAACGAGGAACATACACTTTTCCCCGTATGTGAATCCTCCCAGTGGAAGTACTAATGGAGCCTCACAT

GCTGCATGGCTCAGTAGCGTGAACCTAAACTGTTGGGTTAAAAGATAGAATGTCATGACATTGACAATATAAACAAGTAA

TAGTTCGTCAACCTCTTAACCGTATACCGTATGAAGTAGAGCCAGTACCTGATTATCACCATAAGCACCAAATTGATCCC

CATAAGGATTTCCAGGCCTATGATAAGACGGGTGATCCACGAACACCTGTGCGCGATTAAGTAAGACATTAAATAAAAAG

ACTATCAAAGCAATCATATGTACGAGCAAATAAACGCTACCAAGTTAGAATTTGTATTACCCAATCAACACCCGCCCTGT

ACTCGTGGAAGAATGCAACCTCATGCTCAGCTTCAAAGCACGGAATCTTAATCCTACATTGCGCATCAAGCGCACCTTTA

TATATCTTATCGGAAGCACTACCATTCACATATCTTGGAGACACTACCATGACTCGATGTCCCCGAGCAGCAAGCGCGAT

AGGTAATGAACCACAAACATCTCCTAAACCTCCTGTTTTAGAATACGGCACCGCCTCCGATGATACAACAACAATGCTGT

ATGTCTCTCCTGCTGCTTTGGGTTTCACCACTTCTTCAGCAGCTTCACCAATTTCAGCCACAGACTCATCAATCACTTCA

CCTGCAATGAATAAACAATAACAACTAGTACTAATTCAGAAACTAAAATTGAGACCCAGAGACACAATTTCTCTTCTGAA

TCTTCAAAACCCAGATTGAATAATCAAAAACGGAAACAAACCCAACATCAAAATCACAATCTTAAGCAAATCAAATGAAG

AAAAATAATCAAGAAATGTAGAACTAACCAATTTTAGAAACCCCGGAATCAGTTTTTTTCTTGTAAGTCATAAAAATGCC

GTCTTTATCTTTCCCATCAATTTCTTCTGCGTCAGAAATTGAAATCCCATCTCGATAACATCGAAATCTGATTTCTCTGT

CTGTAATTGAGGTTTTCAGTGAGAGTAAAATCCCATTACTAGTACTAAAAATGTTTGATCTTTTTAATGAAGTAGAAGTT

CTGGGTTTGAATGTTGTTGTTGTTGGTGATGATTTACAAGTAACAACAGTACAAGAGATTTGCAGAGCTTCCATTAGAGA

TAGAGAGAGAAATTGTTAGGCAAAATTAGTGAAGAATGAATAAAAAATTGGTGATATTTTTGTTCGAAACCGTTAAGAGT

TTCGCGGCTAGAATTTGACTGGCTCCCACCTGAGTAGCGACGTGTCAGTTAATTGTACACGTACAGTTGAGGTCGGTCAT

AATTTGTGCCACACTCGTGAGGTACTTACTTGGGATCTCAAAGGGTACCTTTGTCCTAAACCCAACTTTTGTCTTGGGGG

TGCGGATTTGCGGTTTGCCTCAATCCTACTTGGTTGGTTGGGGGTGTGTGTGTGGGGAGGGCCAAGAGCAATGAGTTTAG

ATTAGCAAATCGCCAAAACTCGGTTTAAACTCGGATTGATTTAGAATCCCGTTGAGTATATAATATCAAAATTAAGAAAA

TAAGTTACTCAGAATACGGGAAAATTGGACTTAAAATAGTAAAATCGGATTTGGAACTCGCTAAAATTCGGAATCGGTGA

GGAATTGTAGATCTCTTCAGCAAAACTAGCGAATAGAGAGTATTCGTCTGGAAATTCTCCTTGGAAAACGATTGTAGGTG

TAAATTATCCCCAGGGCTAGGTGGCAACATCTCAAGCTATGAGGTACCCAAACATGAAGAGACGAGAAGCACGGGATAGA

GGCTGGAAGGCGCGAAATTTAAGGAGCCACGTGGATCGGGCGTGATCACTCTACAGCTGTCGGACAAGACTCGACCTTAG

GCCTCTGACAAAGCTGGCAGACGAGACGTATTGGGTCAACCAAGGAGTGGTAGTCAAAGGGATAAAAAGATACGTGTCAT

ATCGGCCAGATGCAAGGGCGACAGGTTGGAAGGGCGACATCGCACCAAGAGAACACCCGAAAGGATCGCCTCGGACAAGG

GACTACATAGGAATCAAAGTACATCGGCCATAGTGACGGAAGAAGGGACGACGCAGGCGGGCCCGATGACAGGACTTCAA

GTTAGATATGTTGTGACCAACACCCTAGTTTGGTTGTATGTAGCCTCCCCTATAAATAGAAGAGCTTGTATCCAAAAGAA

AGGAGGAGAGAGAGACATGTAATCAGAAAAGAAATAAGTAAAGTTGAGAGCTAGTTCACTTTGTAAGTGAGATCACTTGA

GAGAGCAACACCATTGTAACACCTTGAAGTAATAAGATTCATCCTTTCACCCCTGTGGACGTAGGTAATCATACCGAACC

ACGTATATCCTTGTGTCTATCGTCATGTCCTTACTTTCACTTGCATGAGTAGAAGTTTTCTATCCTTTGGATGTAGGTTG

TAGTAAAATGCATCTACATTCTGGCTCCGACTAAGGGGAACCGACTAAGATCTCCGGATACCTTGACTTGGAGACCTTCA

GAGACGATCCAAAGCTCCTGTATACACATTCAAGTGTTGTAGTTGAAGCTAGGATGAACTCTGATGGCCGAAATCATTGT

TATGTCGAGTCATTTAGTTTACATGCATAGTTACAGCAGTAGCAGCGTCCTCGAAAGGCGTGGTAAAGTTCTCATTTCTT

CGCCCACCAACAGTTTGATCAGAAGAAAGTCTTCATAAGCTCACCGGCGACTAACATCATAGGTTCGGAGCTAGGTTGAA

AGTTTGTTGATATGATGACTACTTGTTTTTGATGATTGATCGAGTTTTAGCGGTAGATAAGCTGTTTTCTTGCGATGCTA

TAGAGCTTGAGTGTTTGGAGTGACAAAAGGTCATAGCTAAGGTGTCTTTTTGCATAACAAGATGCCAATCTTGAGATGTC

TTGCCCGGCGATGTTAACTAGTCATCGGAAAGTTCTTGGAGAAGATTTAGTTAAGACGGCCATAATAAGATCTTAGGAGA

ATAACAACAATAGGAATTTAACTTAAGTTTTGGATTTATGCGATGTACTGTTCAATCATCATGTGCATGAGTTTAATGAT

TTGGAGCATATAGAAGTAAGGACAAACTAGCTTAAATACGGAGGTCAAGTACTGAAGGCGACGTCGTACGGAGTGATAGC

GACGTAGCGTATCGAAGTCAGCATAAAGTCAAAATAGGGTAGATAGGCAGAGCCCATTTGATAACCGGAGTAGGCTTTAT

ATCTGCGAGCTGTGGCGTAATGACAGTTTTTGTGTCAGTCGAATTTGTTTCACCTTTGTGAATGTTTATTTCTAAAATAT

TAGGTAAAAGTAAGGTGTTCTTTGTATTTGCTGACGTAGGTAGGGACGACACTCGGCTCACCGCGCGCCACACCCGATAG

CTAGCACATCAGGGACAAGAAGGACGTTAAGGAATCGCATAGTGACTAGCAGCAACATGAAGACAGGGGTAACAGGGCCG

AGGCAAGAAGGCAGTCGACAACCACCACTCGAGGAATCTAGGGCAGCACCACGAGGCATGCCAACTGTTAATGAGGGAGT

GGATCTCGGGCGAGATGATGACACCCTACCTTTGGACCAGGTCATCTTGACCCAGAGGAGAGTTGACCAAATTGATGGGC

TCACAGCTTCCGATACAGAGGACGATGAGGAAGCACTTTAAGCCCAGGCCCAGAGACAAAGAACAAGGCAACAAGCAGAG

TACCAGGAAGCCTTAGGAAGAGAGCAGGAGTGTCTCAGGAAGATTCAGCTGGGCCGAGCAGCAGCGAGCAACGTCGGCCC

TACCCCCGGCCATGCGACTGGTCCGATACTACCCCCACCGCCAATGTCGATACCATTACAGACTCGGCCACACCACCCTA

ATGGACACTCTAGCCATGAAAGCACAGACCCGACACTCTTAGCCATTTTAGAAAGTCAAAGAAGGCAAGAAGCAGCATTA

AGGGAGCTCCACCAGCGGAATCTCGCCATAGAGTTCGAAAATTATCAGTTGAGGAATCATCAAACCAGATCAAGAGGTTC

GTCCAGCCGATAAACGTCGAGTCATCATGGCCGATCAGGTCAATCACCGCCTATGATGGGGCCAAGCAACCCTAATCAGT

TCGAACGATCAGGTCAACCAACACCACCTCCCGTACCCCCCCCCCCCCCCACAAAGATATGGACCACCCGAGGACTCGTC

CACTGATGAGGAAAGGCGAGGAAGGGGCGGACGACAAAATAGGGAAACAGAAGCTAAGCTCAAAGAGTTGGAAGCAAAGA

TCAAGGAGCTATCGGGAAGTAGCGAAGAGAACAAATTGGCAGAGATTATGAGCCAGGCCGAGAAGACGCCTTTCTCCCAA

GAACTCGAACAAATGTCGTTCCCACCAAAGTGTACTTTTCCTACATTTCCTTCTAGATTCGACGGAACAAGAGATGCGGT

CGAACATCTAAAAATGTATACCATGTCATTAATTCAATGGAAAAGTAACACGGTAGTCATGTATAAATTCTTTCCAGCCA

GTTTGGAGGGATTGGTTCTATTCGCTTCCACCCAGATCAATCGACGATTACGACGCTCTTATTGAGGCCTTTCTTGAAAC

ATACATGCACAATAGCCGACCTCGGACAAGAATGAACAAACTATTCACCCTCGCCCGGAGGCTTAGGGAGCCCTTGAGAA

CCTTGACCGATAGGTGGAGGAAGCTTTGTACTGAAATCGGAAAAGTTCCGATAGATCAAGCTATCTTTGGATTTGAAAAT

GCACTGGGAAGATCGGACCCCATCTGGATAGCTATGTTCTCTGAAAAGCCAAAAACCCTAAAGGAGATGCGAAGAATGCA

AGAACATTTCATCGCCTTAGAAGAAATCCATGCCGAGTCGAGAGATAGGGGAATACAAGAAGCTATTACTGAAGAGGGTG

CCGAGACTAAGACGGTTATGGAAAACCCAGCTAGGAGACCCGAGAATAGATCAGACCCTCCACCAAGAGGTAATGAGAAG

AAATAATGGGTGGCCAAAGGGAAGAGGCCGAGGCAGGAATCCCGATCGTACACCCCTTTAAACACTCCTTTGGAAGAAGT

ATTCAAGGAAGTCGAGAAGAGGAACGATATCAATTACCCAGCATCAAGGGAGTCCAGTTTCAAGAAACCAAAGGCTACCC

AGACTACTGCCAGTACCATCAGTATATATAGGGGCCACTCCACTAATAATTGTCGAGAAGTGAAGGATATCATCCAACAC

CTCATCCGAGACGGGTACCTGAGACAGTTTGTCCGACATCCATCCCAGTTGACAGCAGCACCCGATATGCCGGTCCATGA

AGTCCACATTGATAGGTCAACTCAATTTGTGAATACCATTTCACACTCCACTATGCAAGAATACACTTTCGGACCAATTA

TCACTTCAAGGATTCACAAGAGGGACCGCAATGGCAAATAAATATCCAGTGTAGCTAAGACCTTACCAATGGAACCTTGG

ATGATGCAACCTATTTTTTCCTCGGAAAAGGATGTCCCAGCAAATGGGCAAGCACACAGTGATCCTTTGGTCATCACGTT

ATTAACTGAAAGATGGGTAGTAAGAAGGATTCTCATAGATAGTGGAAGTTCAGTCGAAGTGTTGTTCAACGACACGTTTA

AAAGAATGGATCTATCGGACGACATCCTCGTCCCCTCAACATATCACATCTACGGTTTTAACGGCACAGTTACCATCCCT

AAGGGCGAGGTAACACTGAGGGTTTCTGACGGTGGGGGCTATCTTGACACACTAACCACCTTTTGTGTGGTCGACGTCAT

CTCGCCTTATGAAGCAATTATCGGGCGTCCATGGATCACTGGGATTAAAGGGGTGGCATCGACATATCATCAGAGATAAA

GGTTCCCTACCTACAAGGGAATAGCTGAAGTAGTGGGCGACTCCCAAGCGGAGCGACAATGCATGCAGGTTGATACCGAA

AATTATAAAGATCGGCGAGCTCTACAAAGACGAGAGAAGAATAAATCCAGAGAGGTTAAAGCTGCAGAAGAGTTGGAGGA

AACTATTTCACGAGCGGTCAAGGCGTACGAGACCCAATAGCAAGACCATTTATAGCAATTAAAGGAAACGACTCCATTCG

AGGGACCTAAATCCAATTTCTCAACAGTTGAGCCGACTCGAGAGGTGAATTTGTGGATGATGGAGGAGCCAAAGATTGTG

AGGATCGGGACGTTACTCCCGGATGAACAAGCAAGTCAACTCATGACGGTGTTAAAGGAAAACTTAGACGCGTTCACTTG

GGAAGTACACGATCTAGAGGGAATTGAACCGGAAGTCTGTTGCCACCATTTGAAGATTGATCCAAGGTTCAGACCCGTTA

GACAAAAAATAAGACGAATCGCACCAGAGTTACAGGCAGCGGTCGAGGTCGAGCTAAAAAAGTTGTGGGAATCCGGGATA

ATCAGGAAATCCCATTACCCGCAATGGATATCGAATATGGTCGTGGTACCAAAGAAAAAAATGAGGAGTTAGGATATGTA

TTGATTTAAGTGACTTGAACAAGGCTTGCCCGAAGGACATCTTTCCTCTTCCTAGCATAGACTAATTAATAGAATCCATA

GTCGGGTACGAGGCGATGACATTGATGGATGGATACATGGGCTACAACCAAATACCGTTAGCGCCCGAGGATCAAGAGCA

TACCTCGTTTTTCACACCAAGAGGGTTGTATTGTTATACTAAGATGCCATTTGGACTTAAGAACACTGGTGCTACTTACC

AGAGATTAGTGGGATACATGTTCGAAAACAAGATCCACGACACCATAGAAGTTTACGTAGACGATATGTTAGTCAAGAGT

CTCCGAGCTAAGAATCACGTCCGAGACCTTCGAGAGATTTTTCAGACGATAAGGAAACACAAGATGAAGATTAACCCCGC

TAAGTGCAACTTCTGGGTTACATCAGGAAAATTTCTGGGATATATCATCACCCCCAGAGGGATAGAGGCAAACCCCGAGA

AGGTCAGGGCTGTCTTAGAAATTCCTTCTCCCGCGACGGTAAAGGACTCGCAAAAGCTAAATGGCTGCATAGCAGCATTG

GGGCGATTCGTGTCACGGTCGTCGGACAAATGTAAAACCTTTTACGACATCCTGAAGAAAGGAAAACAATTTCAATGGAC

GGATGCAAGTGAAGAGGCCTTTCAAAACATCAAGTTACACTTAGCCAACCTCCCGGTTTTGCAGAGGCCCGAGCCGTCAG

AGATCCTCACCCTATACTTGGGAACGACACCATATGCCATTAGTGCGGTATTGGTCAAGACGGTGGAGAATGTGGAAAAA

CCGGTATATTTCACTAGCAAAACACTAAGCCCGACCGAGAAAAACTATACCAAAATGGAGCAGATGGTTCTGGCACTAGG

TTTTGCCACTTCTAAACTCCGAACGTATTTTCAAGCTCATCGCATACGGGTGCTCACCAAATCGCCTATTGAATCAGTCT

TGGATAATGCAGGGCGATCGGGAAGAATCTCTATATGGGGAGCCTTAATCAAACAGTATGATGTGGTATACGAAACAAGG

ACGGCGATCAAGGGGCAAGTAGTGGCCGATTTCTTAGCTGACTTCCCACTTAGTGATGAAGACGAGGTGGAAGATATACC

GGGGGTGATGGAAGAACGAGAAGAACCTGCTGACCTATTAGCAGCCAATAGTCATACAAGATGGGAAGTTTTCGTCGATG

GGGCGTCAAACAAAGAGGGATCAAGCCTTGGGTTAGTGTTCACCACGCCGAAAAGAAGATAGATAGTGCATTCTTTCAGA

CTAGAATTCAAGGCAACAAACAATGTCACCGAGTATGAAGCGGTGATCCATGCCTTACGGATAATTGTAGAAATGGGGTT

GCAAGAAGTTCGCTTAACTAGTGACTCACAATTGGATGATTAGAAGTTTTCTATCCTTTGGATGTAGGTTGTAGTAAAAT

GCATCTACAACTATTTTAACAACTCTGATTTGGATGTGGAAAAATAAATACCTCAGAGCTGGATGTATGAGCCGAGCCAG

ATCTCGAGTCAGATATCTGAAAAAATGAAAAAACATATGATTTTATATTTGTCTCGGATGAAGAAAAACATATATGAAAT

TCATACTTCTACAGAAAATAGATATGCTTGTTTCGAGTATTCGTACCTTATACTTTTACGGAGAGAGATAGCTAGCCAAA

TTATGTGTCACAAGAAGAGATATTTTTGTTTCAGTTATTTTTCTAGATTTTTTTTTAAAATAAATATAAAAGTATAAAAT

CATTAATTTGCACTCCGATGTTTGTATAGAGAACCGGATAAAGTAATTCGAGATAAGGTTTAGGATGTGAGGTACGAACA

ATCCTCGCCTAAAACAAGATAACAGTCATTTGAAAGATGGCTCGGTCTCATGAAGGGAAGCAGGAAAAGAGAGATTATGA

TTCTAAGTTGCAGAAATAAATCTCTAAGAGAGAAGTTATAGTTAATGTTGTTGTGTTTAGAATGGTTGATTTATGGCCTA

TTTATAGATAGTGAACTCTTATCTCATCAAGTCGGTGGGATAAGGATGGAGGATAAGATAAGCTCCTCAAATGTGATTAC

AGGAATAACTATAGATATAGTTGGATATATCGAGTTTATCCCTTGATTTGTGTAACCTCCACCGACGACTCTTACGCGGT

GAGAAATACAGCTGGGTATCCTCATTTTTTTCCTTACACCACCAGACCAAAATCCTAATTCAGATTCTTCGTTTGTGCGA

CTGAATAAGCCAACGTGTGTGATGTGTTCGCAAGGAAGAAATACTTTCCATATGTGAGGTGGATGAGTATTCCTTAATTT

GTGATTATGATTAATATGAGTCATGTGGAGAGCATGAGATAGCATGCTACAACATGACCATCACACATTAGAGGACTTGA

TGGTGACTATCAACCTTTGGGGTCTGCCATGCATGGCTTATCGACTTGTAAGTCCCTAAGGGCAATTCCTATGGCAATCA

ACAAAAAAACAAAATGTTGAGCCACGTGGATGAACAACCCGTGTTTTCCACTATGGAAAACTAGAGCAAAATGAAAAAAT

TGGGTTTCCCAGGGCCCCACTAATATTTCTTTTTAATATAAAATGCTTTATAGGATAAAGTTAAAATATAAAATATAATA

TTTTGGTGAGATAAAATAGGATGAAGTGAGAAAACACAGGATAAAAGTGAGATAAAATATGATAAAGTGAGATAAATCTT

AGTAAAATAACAGAATTTGAAGTGGGCGATTGTGGTAGAAACGCCGGAGTTTGTGGTAGGAACGCCCGCGGTTGTAGTAG

AAACGCCCGCGTGTTTACTTGGCACGCTAGAATTACCTAGGCTTGCGCGACCCTCCATTAAACGCGCGCTACATGTTCCA

TCTCAATATTCAAACCAACATATTTGTTGGTTTGAACATCCATTTTTCATGTTTGTTGAGTCCATAGTGGGAGCTAGGAA

CTTCTCCAAGATGAGTTGTGAAGCTCTAAGGTGTCTTGAACACTTTTTATTAGATGTTCGAACGAATTTTTATGGCTCAT

ATCTATAATATTAGTCATGAACATGCTTATGAAACTGAGGATGAGGCTTCTAGTACACTCGAGTCTCGTGTTTGTATGAA

ATTAATATGAGATCTTGCCTCTCGACGAGTGATCATAGCATGCCTAGATGTGACTAATGGACCTAGTGGAGGACATGATT

AATAGCTAATCGCGAGGGTATGTGAGACTCATACGGTCACTACAAGAAACTTAATATATTGCAGCACCCAATGGTTATGC

CATAAATCCCCACTAGTGTCGCGATAACCTATGTAGAAAGCAGTCTATTGCTGCACCCAATCTTTATTGCTGCAGCAAGA

GATTATACTTTTTTGCCACACTATTTTCATATTGTTGCTACAGTTGTGTGGCAAAAATTTCATTTTGCAGCAGTAAATTA

TATAACTTTTACTGCAACTGAAAAGTATTGTGCCTAGAAATTTTCTTGTAGTGGGTGTTTGGTGCCCTTGATTCACAAAG

TTAGGCTTAGAAGCCTAGCTTTACCTTATACAACAGAGTGTGAGTTGTAATCATGAGGTTCAGGCTAGGTTCAATGTTAG

CATGCAGGCGTGGTAATATGGGTTACGAGTTTTGCTTTTGGAGAACAATATGATTTATATGATATTCTTACTTAGATGAT

ATTTATCAGCACCCAACAATTTTATCATTCTAACAGAAAGATTTTTTATTGTGAAAGAAAAATCTTTAACAAGTCACAAG

CCAAGGATTTGTGGAAGATTGAATACAACCCAAACCCCAGTGTTCCAGTAACACAACAAATTACGAAGATTTTGTTGGAA

ACGAAAAATTATTTGATTAACCGAATCCACCGCTTCAATTAGGCTAAGCCAGGGCTACCTTACTGCTTTAATAGGCTTTC

CCATCTTCAACAACCGCATCATCTTTTGACTACTTGTTGGGCTTGGGCTATTAATTGGACCTAGTTGGTTCATTGTCCTC

ACTTCTGTTCCAGTTCCGATTCCGATTCCATCAGTACTTGAACTCATTGACAAATCATTCAAAAATATCCTCTTAGTTGA

TGATGAAGAGCTTAACAAAGAATTCTGTGCCGATTTTGCCTTTGCTTTTGCAGACTGAGTCGTAGCCATGTAACTCGGTG

TATTAACATTTTCTGTTTCCACTGAAACCATAACTCTTTGTTTCCGCTGCCGTATCTTAACCGAGCCATCCGAGTTTAAA

CCGAGCTTTGCCAGGTTAGCAATTGGGTTAAACCGGAGCTGATAACTTGGTGATAACGATACCAAGTCCGTATCTGCCGT

TCTTTCAGATGCATTGTCATCGTCGGTATTGTAGGTGCGGGAGGTGTATGAGCTCTCTGGTATTGCTGCGACATGTTTAT

CACGATGCCATGGTTGCGAATCAATCCAACGATCCAACAAGTTCCACCCACCACCTTGGTGTGTTTTTTCATCCGAGTAT

GAATCGACCGAGTTCTGACGGTTTGACACCGAGTCCTTTAAAAGAAAAAAGAAAAAAACTTACTGATTGCACAAATGCAC

AGCAACTGCAACATCATGCACAAAGTAGAACAACCAACCTGGTAGGGATAAGCATAAGCAAGAGCTCTTTCATTTTTAAT

CGTCATAGAATTACTCGAAAGATAATGTTCCTGTTTGTTGATCTTTTCAGTCGATGACCTTGCGTATTCTTTCTCATAGC

TAAAACTATTGCGCTGCCTTCCCCTACTATTATCCCACGTCTCTGTCGCTCTCTTTTTACCTTTGTTTTTCTCATTGTCG

TGTGAAGTTGCGCCTTGTAAGCGGAGGCGGTCAGCTTTTGTTCTTGCCTGCACGCGTACAATTGCTTGCATACATCGCAT

TGTCATCTCAGCTTGCTTCCTGACGTTATGTCCTCTAACTAGTGCTTGTAGCTTCACTAGTCCTCTCAGTGCCCTCAAAG

CTCTTCTTGCCTAATTTGAGATTTTAGATTATTAATCCGTCGATTAATGAATATCAAAAAGATTAGGATGAATTAGCAGT

CCTTACCAAGTATCCTTTGTAATGGGATTGTATTTTGGTGGCAGCTCTTTCTTCTTTAGAGTAACCATAAAACCTGCGAC

GATTATAACTAGAAAACCTAGACACCTTTGTAGCCTCATGAGCAGCTAAAACCGCCGCTTCTGCAGCCACAACCGCCATT

GCAATCGCATGGTTACTACGTCTACGATCTTCATCTCTCATCTCTAGTACAGCTTCAGCATTTTCATCTATCGTAGTCGT

CAGAGACGAAGTATCGCAAATCGGATAATGCTCAAGCGACACAATCTCCGGTATCTCGTGCTCCTCCTCTATTATCTATA

ATTATTAGCACAAGTAATCAAAAATTAGCGCAATTGATCAACTAATAATAGGAGCATTGCCACAAAGGGGGACCAATATG

TTGATGTTGAGACTACAAAGTGAATAATGTGAAGATGCAACCCATCTCACCAACATTTACTTGTGGATCATTTTATAAAG

CGGCAATCGCATATGCTCAAGGGACCATCAATTTTTTTGTATTTTTTTCATAATGTGTTAAAAGTCTCCTCCACTGTTGA

TGAAAATTGCAGCAACTTCGAACTTTGGATAGTGCAAGCAAGTTTTTTAGTGCTCTAAAGTCAGAGAGAAAGAAAAAAAA

ACAAGAAAGTTTGTTGACTAGAGTTGACTAAGAACAGAAGTTATGGCTATTAATTTGGGGTATCCTGTTCCAGAGGGTAC

TCTCATTATCACAATCACCATCACTTACCCCCCCGGTATCCCCATGACATGATATTAATATCTTTTGTCCAAGTTGGTAT

GTGTATTGTCACTGTCTCTGTAGGGCTGTGCCCATCTGTAAATTTATCAGCTCAGATTGCCCATTTTCTCCGGGCTACCA

CTGCAAAAATCTCCTTGTATAGCAAGAATCTAAATCTCATTTCTCTGAATTTTATACAGTACAAGGCTCGGTTTGGAGGT

TTCATTTGGGAGTGGCCGCCTTAAATCAAAAGCTCAACAAGTGTTTTTTCCGGTGAGAAAATTTAAAAAAATATGCTAAC

ATAACATAATATGTAGTTAAGTGAGTTTAAGTTGTCATCAACAACTGAATTGATAAAATGTTATACTTGCAGAAATATAA

GGCTTACAAAGCATATATGGTTGTACAAACTACGCTTACCTTATTGGCTAGTGAGTAATCCTTGGATGGAGAAGATGATG

ATATGGGTGATGAGGACTTGAATGCATTCTTGACGGAGGTGAACCAGTTCTTCGCCGTCTTCTTTTTCTTCGACATTCCC

ATCATTTCTAATGCAATCACAAAATCATGACCAACTTCAGATGGATATGAAATTCGTATTTGAATGGAAAAAGATGATCA

AGTATGAGAGGAGGAAAGCAAAGAGAAGGAGTTAGAAATCTAGCATTTATGCTTACCCTTTCGTGAGGGGCTTTATATAG

TGGGAAAGCTGCCTTTAATCATCTCACGGGGATTTTAACGATTTGTTGCAGTATGTAGTTGTACTTTATACACAGTTTGT

ATTTGTTTGGTTATAATTGAATTTTGTATGACACGTGTCGACTGGGTAATAGACGTGCCAAGGCCCTGTGCTTGCTTGCC

TATTATGCAGGAAGCCCGTTGGTAAAAAGTCCACATAGGTAAAATTAAAAGAAACCAATAATTAGGGTCCTAGGATTTTC

GTTAAGGGCTACATTAGGATTAAAATGTCCTAAACGAGGATAGGGGACCTAATATGTATAACAAATGTCAAAATTACTCT

TTAATTGGATAAACCTAAATCTATCGTAAACCAAAGAAAATAAAATAAAATAATATATATATTCTTCTTCTTCTCTTCTG

CATGCATGCATTTCAACTTATCCTTCTTTCTTATTTTTTTTTTTCAAACAACGATTCACCAAAATGATTTATGTTAGGTA

AGAATCGGAAAAAACCCACTATTTTTCTTTGCTTTGGATTGTTTAAATAAGTTATTCGAAATTCAATTTTAGATTCGGTT

TAGGAAACACCAATAGCTGGGAATTGATGAACTACCGGTCGTAAACAACAAATTACGGAAATCGATGAGCCCCAACCGTA

GCTTAGATAGTTATATAATTATTGGTGGAAAATGAAGAACTCCTAGTCGTAGGTTAGATTTAAAAAAAAATTCTCTAGCC

GCAAGTTAGATTTAATGTTGTTATTATAATAGGTTTAATGATTCCGGCCGTAATGAAAAATATTAAATCTAACTTATAGG

TAGGAGTTCATCAATTCTCATGCGTAAGTAGTTCTTGAGATTGAAACTTGAAATTGAATTTCAAAAAATCGAACCTATTT

AAGCAAGAAAAAACAAGAAAAAATGATATGTTGGAAGGCACACCTTTAAGGTTTAGGTTTTTGGATAAGCCTTAATTTCG

AGATAAGTTATGCCTCATGTAGCAAATCATCTATAATGTTTCAGAACTTTCCTCTATGATGACTTGTCCGTCAAATCTTT

CATAATGTTTCATAACCTTTCTCTATAGTGACATAGCCAACAAGTACTCCCGAAACTTCTGGAACCTTTTTTTTTTTAGG

ATTTTGGAACTCAGGGGACGAAAATTTTTAGTGGGAGTGGATTGGCCAAAGTGGACCGACAAAATGGCCGGCGAAGTGGG

TTTGCGAAAATGAGTTGTGAAGATGCAATTGCAGAATAGGCTGAAGGAATGTGCATACTTTTCCGGCGGACTCTTCCTAC

GAACATTTCCAGCGGACTTTACCAGCTAAAGTTTTCGGGGAATCTTGACCGGAAGGAGTGGGATGGGTAGGCAGCGGATA

CCTACAAAAACCCTCCAACGTTCAAATTAACATAGTGATTTCACATGAGTTTTTAGGGTTGGAAGGAATACCTTTTAGGT

TTTAGGTCCCCTACTATATAAAGTTTATCCCTTGGAATAAACCCTTATTTCAAGATGAGTTCTTTCTAATGGTGACTTAT

CCTGCAAATTTTCCAAAATGTTTAAGAACACTCCTTTGTGGTGACTTAGCCAACAAGTACTCCATTAACTTTTAGAACTT

TCTTCTAAGGTTTTTGAAATGGGACAACGGAAAAAACAGTGGAGGCGGACCGGCCAAAATGGACCGAAGTAGCAAGGCTT

CGTGCTTGCAAAGTAACAAGTATATGGAGCTCGCTGCTAGGGAAGGATAACATATCACTATTTTCAAATACGAGGCTTTG

GACCAAAAACGAATGTTGGGATCACGATTGAGACGACGTTTAATTTGTGCTTGTTATGCTGCAAGCATTTATACTCGCGT

GTAAGTTGAAATTTTATGCTTGAAATTAGAGCAAGGATTAAATGCTAACATGTAAGGCGAGGCTCTAGGAGCTTGCATCA

TGAAAGAATAAAGGAATTTAAGAAATCCTCAACTGCACTATTTGCACATATTGTTTAATTGAAAGTGTCACATTTAGTAC

CTCTAAGCAAAGTTGTTGCAGTTGAGGTGATATGTGACTTACCCCTCGGTATATGTTTTCTAATGCTTGTGTATTTGTAT

ATGGAAGAAGGTCCAAATTTTTATCTAGTGGACCTACTGGCGAACTTTATATGCCTCTAGGCATGTTAGCGATATTAGTT

TGTTACTGTACCGGCAACATTAACATGAAATTGGTGCCTACAATTGTTGCCGAAAACTGGCCTGGCAGCAGCTTTGCACC

AAAGATGTTTGGGAAAAAACATATTTTCTCCTCAATAGAACATCGATGATGATTACCATGATGATGTTTATGACCATATT

GATGACCTCACCTCATGATGAATAAATTATAGAAGGGGGTGGCGGGTTTCTACATAAACTCTATGATTCTCAAGTTAGCA

TAAAGTTTTCACACGAGTTATTAGAGTTGGAAGGCATGCTTTTTAGGGTTATGTCCCCTATATTTATAGGCTTTAACCTC

AGGGATAAAACCTAATTTCGAAATAAGTTCTGCCTCATGATGACTTATCCAAGAAATCCTCCACAATGTTCTATGTAGTG

ACCTAAGCAACAAGTACTCCAGAAACTTATAGAACGACCTTCTTTTAAGGTTTGGAATTAGATCGGCGAAAAACGTCAGT

GGAGGTGGGACGACCTAAGTGGACCGGTGATATGGGATGGCTAAGTAGGTTGGCCAAAATGAGTTATGGAATGAGTCAAC

GTAATGCCAAAAATGAGTCAGAGGTGTTAGAGCACTACTCGGTCGAACTCGCAAGTGTTGCTATCTCAAGATTGTTTGTC

AAGTTTAGTTGCTAAAACTACAAGTCTTGATTTCTATTCTACTTATCGGATTAGGATAGAATGTGTAGTTGAGCTTTAGA

CTTCACGACGCTCATCGATTGAAGAAGAAGAACTACTAAGGGGAGATTGTGGAACTTCATCAACAAATGGTATGCGGAGA

CTTGAACACATCTATCACTTAAAAGTCTATTTATACTCTATCTCCTATTTGAGACAAAAGTCGTATAGCTATATATACTT

TACATTATACACATTTGATATTTCGAGCAGAATTTAACTCGCTTAGATGTTTCTTGAAATATGTGTTGGAAAGCTTTCGC

TTTAACCAAGTTTATCTTATATTCATAACGTAAGTCAAAAGATGATCATGTGAAAATCTCCTGGTAACATCTTATATGAT

TTGTATGAGACATTCATTTGGTGTAGACTCAGAATGTTTCGTATTGATCATTTGATCACTTGAAAATTTCTTTAAAGCTA

ATAGTTTATGTGAGACAGCTATTCCCGTCTTCTAAGAATGTTTCAATGATTGAAATGGGAGTTTAGAACAATTAACCATT

GATTGGATATAGAGCAATATGCATACTTGTATGCTAACTGTTGCAAACTATTCCAAGTCCGAGAACCATAGTATGTATTC

CCGTATGCGTACTAGTTTGATAGTTGAAGTCTGGGAACTTAGTATGCATACCATATGCATACCGGCGTAACAGTTCAAGT

CTGGGAATTCAAGTGAGTTTGGTTGCACGATAGTATGTGTACCCATTCGCATACTGGCGAACCTAGACCAAGTCTGACTA

CTTAGGTATGCATACCCATTTGCATACTTGAGTGGGTTAAGTTCTAAAATCGGTTTGTTCATGAACTAATACATTTATAA

AATAAGGAATGCAATGTTTTGAAAACCGTGGCTATAATGTTCATGAATTGATTCGAGTGAATAAAAATCGATTTTTCTTC

AATTGTGTCTTGTATACTTCTATGAGAATATTAACAATTGAATAACTTTATAACTAGTTTCATTTGAGTCATTTGAACTA

GTTGTGATTGAGATGAACAAGGTTGGTATAAAATTGTTCATATGGCTAACTTCGGTTAACTATTGTTGAGCCAACAATGT

GTACACGTTTAGGTACGGTTACCCATATCTAAATGAAGTCACTTTTCACTTGTGTATAACAATCTAAGTTCGATCTAATG

GTTGAAAGATATTAGATTGAATCTAATTAGAATTTCATCTAACGGAGAATATAGAATGCTTTGTTACCAAGGTAGCATTG

ATTGCAAACCCTGATTTGAAGACTATATATGGGAGAACTCTACCTACTGTGTGATACTAGTTGCGACTAGAGCCGATTCT

CCTTTATCCTAGGTTTTTCTAAAACGATTATCTGTTAATGACTTAAAGACTTCATTGGGATTCTGAAGCCAAACCAAACT

ATTTTCCCTGCATTTATGTGTTCTGATCTTGCTTTGTTCTATCGTATTGAGTATTATCTTCTCTTAGATTTTCTCGAGAT

TTAATATCCGATAGGCAAGATAAAAAGTAGTCACAAACATCTTCGTCTCATCGTTTGTGATTCCACAATATCTTGTTTCG

CTACCATACGATTAATATTATTGTGATGTGATTTATATTACTAGGTTGTTCTTCGGGAATATAAGCCTGGTGTATTAATT

GGTTCATGTTAACATTGATTTATCAAAAGACGAAACAAAAACTCATAGGTATTTCTGTGGGAGATATATTTATCTATTCA

ATATACTTTTCTGTGTGAGACAGATTTGTTTATCAAGTCTTCGACTTTGGGTCGTAGCAACTCTTAGTTGTGGGTGAGAT

CAGATAAGGGAATCAAGTGCGTAGAGTCCTGCTGGGATTCAGATGCGTAAGGAACGCGACTGTACCTTAGTCAGTGTGAG

AATGGTTAGGGCTCAACTATATTCCAGTCCGAAGTTAACTTGTAGTATGATAGAGTCTGTAGCGTATCAATATAGTGTGG

TGTTCAAATCTGGACTAGGTCCCGGGGTTTTTCCTGCATTTGCGGTTTCCTCGTTAACAAAATTTCTGGTGTCTATGTTA

TTTTTTTTCCTCATTATATTTTTATATGATTGAAATATCACAGGTTGTGCGTTGTTCAATCAATTAGATAATCCAACCTT

TGGTTGCTGATATAAATTGATTGGCACTTGAACATTTGGTCTTTGGTAACGTTCAAGTTGTTTCTGATAGTAATCAGGCT

CGCAGATTTCTATCTGTTTGATTTGTTGATTACATTGAGAAACAGAGATATAACTCTTGGATGTATTTTCCTTGATTGAG

TCTGAGTATCTAGTTGATTCTCTTGGAAGTATATTGGAGTTAGTCCATACAGATTGTCAAAACGAAATATTAGGTGTGGT

TGTAAGACCACCGCTTTTTCAAGAGGAATGTGCGAAATCTTCTACGAACATTCCAACGGACTTTACCGACAGCTGTTTCC

GAAAAGACTGGTCCGACGACGATTTTTCAGTGAAACTGGACCGACAACGGTTTCCTGAAAAACTGAACTGATAATGGTTT

CTGGTGAATATGGGTTGGAAGACTGGGCCGACAGACTATTAACTACACGATTTTAAGTTTCTCATACAATATTGGACTAC

TCTCAACATGCTTCCTCACGTGTAGACATGTTAGGTCTAACACATGGACAGTTAACCGGGTGCGGCGTAGTATGCGAAAT

CCGTGGGCCTAATATCCTCGGAAACCAGCTTAAAAGGTTAGGGTTTTCCAACCCCGTTAGATTTGACACATGGACATTTA

GTCAGGTGCAGACTCATCACATTAATCAGGTGTGACGAACTAGGTGAAGTTCATGTGCCTAACTATTCTTAAAAACCGGT

TTACATGGTTAAGGTTTTCCAGTCCCGTTTGATCTAACACATGGGCATTTAATCGCTTGTAGTCAAAAATGGGATCTAAC

ATAACCTCATCGGATCTAACATGTGGATATTTAATTGGGTGAAATCAAAGAACTCATCACAAATATAAGGACTCCTCACA

AACAAATTAACCCCCACTAATACCATGTTGAAGTCTATTGGCCCAGCTAATCTTGAAAATCGACTTTAAAAAATAGGATT

GTCCAAAATTTCTTAGCCGCATGATCCCATATTTCCGGTATAATATGGGAGTAATCTCAACATGTTCAGATTATAACAGT

TTTCGTAGGGCAGATGCGCTGATGATAGTATCAAAAAACTTAGTAGAGGCACGTATGAGATACTACGAGCTAAAGGTAAT

ATTGCTCGCTACTCCTCCGAGATGCTATAACAATTAACAAATTACCTCCAGGATACAACTATCGTTAGTTCTCCTCTATG

TGTAGCATACACAGGAGAGAACCCTTAGAACTTGGTAAAAATAATGAAAACAAAAACATAAATCGAAACTACAACCAGTT

TAGAAATGTAAAAAATAATTTATGTTTCTAAACTAACTATGTTTTTGTAAACAGAGAAGTAGTATTACAAAATTATGTTT

CGTAAACTACAACTCGTTTTTTATGTTTCGTAAACTAAGAACTCTATTTAATGTTTCTATAAACTTAAATAAAATTTCGA

AAGAAAGAAGAAGAATTTTCGAGAGAAGAGTTCTGAAGGTATGGTTTCAAATGAAGAATTCATCCGGTATTTATAGCAAA

AATTATTTGAACGTTTTGACTTAGAATCCAATGTGCGATGACTGTCTCGTGATTCACGCGTTTGCTTTTGCTCTGACGAG

TAAAAGCATCCGCTAAAAATACAAGAAAATTCTTTTTGTCGGACATTTAAAATTCTCGAGATTCTTCATGCAGGCACAGT

GTCAAATGCACCGTGTGACACGCGTGAAGGAAGATGCTTCAGAGGCATTGACATCCAAATCGAGAAGGCAAAGAATTAAA

ATTCTTTGCATGTGTGGGAGAAGCGTGTGGAAAATTTCAAGAAAAAGATAGGGTCGGTAAACCCACACCCACACCCACAC

CCACACCCACACCCTCTCCTCGCCCTCTCCTCGCCTCTCCCTCGCCTCGCCCACGCTGAGCTTTGCCGAGCCGAAGCCGA

AAGGTGCTTCATGCATAAGCACCGCGCGAACTAGAAAGGCAATAAGAAATGCAATAAGGTTTAGGCAACCTTGCCATAGT

CTAGGGCTTTCCCCATACACAAAGGTCTAAAGACCCAAAGGGCCATACCCTTAAGGCTAGGGAGTAATCTCAACATGTTC

AGATTATAACAGTTTTCGTAGGGCACACACCACTATTCTCTAACTCTGAAGCAGTGGCATTGCCACATCAAATTTGTTGT

CAACCAGCAATTTTAGGGTGGTATACTAGCGTGTCAAACAACATTAAATTAATCATAAATTACAATGAAACTAACAAATT

TTATTCTTGGCACAATGATCATGCAATGAACTTGTAAAAAGTTTAAATTCATCAAAATTTGAACTTTGAATCATTTTAAA

ATACAACAATGGCAGGAATCTCCAGAGACCCAAATGGATATGCATTGAAAAACGTTGTCTAAGTATGTCGTATTCCCCCA

CCTTCATATTTCTTTATGCTTATATGTCAGTTAATAATATCAATCTATATTATCATGTCCCTCCCCATTATTTTTAGATC

ATTGGATACTTCCCAACGGTCTAATTTAAATCGAAATATCAAAATTTTGAACAGCATTTTTTTGTCTCCTACGTACCTCG

TGATACCTCATTGCCAACTTAATATAAATATATGCATTTCATTTAGTGTTTCTTCATTCATCTTAAGAAAAGAATCGAAT

TCTAACCCCAAAAAAAATAATATCTAGCTAGCTACAATGGGGAACTGTTCTCTTAAAGGAATTGCAGGAAATATTAGTGA

TCCAATTCGTATAATGACGGATTCCGGTGGAACTGTTGAATTCCGAGGTCCAACACTAGTGTGCGACGTTGTTAGTGAGT

ATCCGGGGTATGGTGTTTTCCGGGAAGGTTGTCTATCATCACCGCCGTTGTTTGGCAATGAGCAACTTTTTAGTAGTCAT

TTATATTATCTTCTTCCGGTCACTGAAGAAGAAGCAAAGAAAGAAGATAATGTGATTGACAAGGATGAGAAGGAAAAGGA

GGAAGTGGTGGTGGTGGATAAGAGGAGGAAATCGGATGCGGAATTGTCCGGGAACCAAGCAGTAAGACCATCATTCAGTA

GAGTTTCGTCTGCCACGGTGACGGATCTTGGAGGTAAGAATTTGACGATGAAACCGGCACTAGAAGTTTTGCCGTCGTTA

GGGAATGGTGTTTGGAGAGTTAAGATGGTGATTGATACGAAACAACTGGAAGAAATATTGTCAGAACAAGTAAATATTGG

CGAGTTGATTGAGAAAATGAGAGAAGTTGCCGAAGTTAGTAGAAATAGTGATAGTGACAGTAATGCGAGCACGCCAAGGA

GAAGAAGCTGGAGGCCTAGTTTTTCTAGTGTTTTTAAAGCATCGTCAACTGCAGGACCACCACCAGCACCGCCAGTAGTG

GCTGATGGCAAGTAAATTTTGTTATAGGCTTGGATCAAGATCCATAATTTCTTTGTTTTTTGATTGTTGGGGGAGTTTTA

ATTAGATCTAGCTAGATAAAACGAATCAAAGAGTTCATGATTCAGTTTTTGTTTGTCTTTTTAGTTTTTTTGGGTCGACT

CATCAGACACTTCTCTGTACATATGATGTTGCCTTATCTTTCTGCCATGAACAGTGAGTGTCAAGAGAAGAGTCTGTGAA

TTTTTATGGTCATAGGAATGGCCGGCCGGATGGGTTATATATTATCTATGCTTGGTATACATTTTCTTGGGGGGACGCTG

TTAAAATAAATCGTGATAGTAATTGATAGTATTGTTTAACGACCTTTAAGTAATTCATTTTTCCCTTAAAAACACGTGCA

ATGTAACGTATCAGTAACAGTTGGGGGACAAATTTTCGGCCCTTGAGAAAGAGAAACCAATAGGACACCTGGTTCGGGCT

CATTTGTCAGCACCTATGGCCTGGAACTTGTTAGGCAATGAAGTCGGAGTACAACAGTCAGTAGTACAACTGTACCTAAT

GAGTTATGTGCCATTAAGGTGCATCTATAATGAACCAAAATCTTGCCTCATTTAAGTTGTTGACTTATTGGCACACTTCC

AATTGGAAAAGTTTGTCTTAAATTCTCTCCTGGATCTGATTCTATTATCACAGATTCACAATCAAGCAAGGACTCCAAAA

TTGTTCAAGCAATTCATTCACTTGATGAAAAGCTAAATCACAATTTTAATGAAGGGTATTAAACAAAAATGTCCCTCTCA

TAATATGTTAATTACAAAAGTGTCATTGTTTGAAACAGAATTATAAAAATGTCCCTCACCTAGTGTTTTCCATCCAAACG

GTGTTAACTCGTCGTTAACTGATTTTGGTCCCACCAAAAAACATTAAATAAACCCAATCAATTACCAATTTATCCCTACT

AAAATTAATCTGAAATCTAAACTCGAGTTAATCTGAAATCTAACACTGATTCGATCTCGAGGATGATCGTACGTCAATAA

GAAGGACACGTGTAACAAACCCCAAGAATGATCTAACGGCTAATAGATGATACATGTGTGATCTTACTGCAGAGGCTCTT

GAGATATAAAGCACGCGAAGAAACCCTAAAATCACAACCTACCCACTTTCCCTCAGTTCCGCCTCTCTCTTATCTTCTTC

CCCTTCTTCTTCTCTTCAAGGTTTCTGTTGATGAAGAACTCATCCTATATAACCGTGGTAAGTGAAGATGGGTGGCGATG

AAAGTGATTCGAGCAATGGAAGGTAATAAACTGTTCTTGTTATTTTGATTCTCGAAAGGATGCTGTAATTTTTGAGTTTA

TGTTTATCTCGAGAACTGTTTCTGTAATTTTAGGGTCTATGTTTATCTCTCGATATGTTTCTGCAGTTTTAGGGTTTATT

TGTCGCTCAAATCACTGATAAAAGTGTAGTTTTAGGGTTCTTTTGATTCTCGAACTGATTTTGTTGTTTTAGGGTTTATG

ATTTTTTCGATTTGATCGATGGAATGAATATGATGAAATCCATTTACCCTTCTTTGATTTTTGATTTTCATTTTTCAAAC

CCCAATTTTGAAATTTGGGGATTTTTTCATTTCTCGATTCATGGATGTTATTGACCGGTAAAAATGCTCCATATGTATGT

TAGATTAGTAACTGTTTGTATGATTTTTATTTATGTATGTATGTTAGTATAGATTTTGAATTGTTTGTATGGTTTTGTCC

TTGTTGTATGTGCAGAAGATTTGATGTAGGAGGACAGGTAGATGGGTGATTTAGAGTCTGAGAAGTAAGAGAAGAACGGA

ACTGTGGAGGTATTGAAGCCTAGAAAATGGAGGAAATAGTTTGTTGTTGTTGATTTGTTGGTTCATGGTGGTTAGATTTT

GGAAAGAGATTAGAAGTTGGTGGTGCTGAGATACGAAGATCGAAACGCAGTAGTTGTTATTGGTGGTATTGAGAAGTCAG

GGATGGAGGTGATGTTGCAGAGAAGTTGGAGTTTGTAACCATGGTTTGTCTCTGAATCGGTGAGGATATTAATAGGGATT

TTGAGAAGTTGAGATATAATCAGTGGTGCTGTGGTTCATGGGTTTGGATGTGGTGATGTTTTGAAATTGCGGTTCGAGTT

AGTTGCAGATCGAGAACTCAATGAAAGAGTTGTTGTCGATGTTGAGCAATTGAGATGTTATTGTTCCGTGAACTGGTGGT

GGGTATAATAATTGGTGATGGTTGATGATGTTTACATTGGTCTTAAAGGAGCTGCTACTGGTGATTACTCTTTTTTATAG

GTGCAAAATGGAGAAATATCTGATGTTGTTGCTTACAGTAGCGAGTTGATGGAGTTTGAGATGGTGCTGCTGCTGATGGG

AATGTGGTTCTTCATTTGAGAATTGGTAGTGATGCATAATGGATATCCAAGTTGTGCGATTGAAGCAAATGGAAAGGAAA

TTATGGAGCTGATACTATTGGTGATAAATTGATATGGAGTCTGTGTTGATGTTGAGCTCAAGATGAGCTGGTGGTTGGAC

TAGTCAAATGGGAGCTGCAGGATTGAGAAGTTGATGATGTTGGTTAAGTTGATGTGTTGTGCAGATGTGCAGGTGGTGTT

AGCTCTATGCATGGACTGATATTGCAGGTGAACTTGAAATGTTGCGGGTTTGGTGTCTTGAGAGATGAAGAAATGGTGTG

CTTGCACTTGCAAGAAATGATACAGTGCAGAGATGACCTGTACAAGTCGCCAGGTGTAAATGATTCCACGTCAACTTGTA

CTGTTAACTACCTAGAGTTAAATGTTGAGGGCAGGTTTGGCCAATATAGGGTGTAAATATGTGTGACGTGGCACCGTTTC

TTGCTTTTGTAAATTTCGGATGGAAAAACTTAACAAAGGGATACTTTTCTAATTCTGTTTGAAAGAGTGACATATTAGTA

ATTACAATTATAAAGAGGGACATATTGTCAAAAAACCCTTTAATGAATGAGTGTTTATTTGTTTTCTACTTTCACCATTT

GACATTTGTATTTTCTGCTTTTTTTTTTCTTGAAGTATGAAATTTACTAAGCTACTAATGGACTCGTTGCAAGGAATGTT

ATTAACGAGCAGGGGAAGAGTTAGGAATTACGAGATGGGTACAAAAATTTGCTAGGGGTGAAAAAAATATATTTTTAGGG

GTGCAAAAAAGCGAAAATAGGCTTGTATATGAAAAAGAAAATCTTATTAGGCCTAAATTAGGCTTTGAAACCAACTGGGC

AGGGGTGCAATTGCGGCCCCATACAATGGTTTGGCTCCGCCCCTGTCAATGAGACCTTTTGGCATTTGTATTTTCTGCTA

GCCACATTTCGAGGATTTTTTTTTTCTTGACGTATGAAATTTACTAAGCTACTAATGGACTCGTTGCAAGGAATGTTATC

AATGAGCAGAGGCAGAGTTAGGAATTATGAGATGGGGTACAAAAATTTTCTAGGGGTGCAAAATAATATTTTTAAGGGTA

TAAAAAAGCAAAAATAGGCTTGTATATGAAAAAGAAAATCTTATTGGGCCTAAATTAGGCTTTGGAGCCAGTTGGGCAGG

GGGTGCAATTGCAGCTCCATGCAATGGGCTGGCTCTGCCCCTGTCAATGAGATCTTTTTTTCTTCTTTTTTCTTTTGGCC

CCCGGCCGGTAATCAATGAAAATAACATCTAATTGAGATCAACGGTATCAGCATATTTCTATACCAAATTTAATAACAGT

TCGATCACCCGACACTTTTGAGAATATAATTCACAAAACATATATTCGACTTTTTAGTAAATATGAATTTTATTTTTCCT

TCTACAGAAACATTAAAAAAGTTAAAAACTTGGTGCATTCTATCAATTTAAATTTCAACGGTAGTTATCAATAATTAAAT

TTGAGAGTCAACTTAAAAATTGATAGGTAGTCAACGAGAATTCTATAAACTATCATAGATTTTTTATTCGAGTGACTTCC

GGAGATTTTCTGAAATCTTAGGAACTCTAGTGATTTAATGAGAATATTGCAGAGAGTTTTTGTGCCTCGTAGATACAAAC

TTAATGGATTTTTAAAGAGTTTTTGTGATTTGCTGAGAATAAAATAAAATAATATCTAATCAATAATTCAAAATATTTAA

CTACTTTTTTTATTATGCAAGTTATGTCAAAAAAATATTAGTATATGAATCACGCCATCTAAGTTATTCTTTTCTTATTT

TTCTTCGTTTATTTTCTATTAATATTTTTTTCTCTTGGCAATGACAAAGTACTATCATGCGACATAAGTGAAACATAATC

AGTATACACTGAATGTACAAAAGCTAGTCTAGCTACTAGAATATTTCAAAATAAATAAAAAGAAGGTTTTTATTGAGGGG

AAGGACCCGATGACATGATTAAATTGACATTATCATTACATGGTAACGCCATTTGTTCTATAAAACCCAAATAGTGATGA

ATTTGAAGCTAAGTTTTTAGTGACATGTTTCTCTTATAAAGCTTTAGACTAAGTCCTATGGGCAAGATTTGGCTGCTACA

TTAGACAAAAAACGTGGTAGATTACACAAAAAGAGTTGCCTATTCTTCAACACTACTTCTCTTTCTAGATATATCTCGCA

TTCCAACTAGTAGTGAGATAGTTTCGCTGAATGGTTCAGCGCAGAAAAAATCAACTTTTCGCTATTTGGTTCATCGCAGA

TTGATTTATTTTTTGATCCGAGGATTGATATGGTTGCGCTATTCCATTCAGCGCGACCACTGGGAATAGTTCAGCGCATC

TAGATGCTAGATTAGAACTGCCAAAGGACTTAGGAGGTTATTCTAGCTGACATGGACTTCCAATCCAGATACTAGATTAG

AATACCATAGGACTTAGCCTTACACCATGTCATCTAAATTTAATGCTTATGAGAAAATAACCCAAACTCCTAACTCTCTA

GAATTCTCACATATTTAGGAGAGATTTTTGTCATGCCTATTTTCTTATAAAAGTCCACCAAAGTCTCTAAAATCTACAAA

TCCATGACTTATAAATATAAATCTAATACCAAGGGTATTTGAGAGACTCGCACTTGCAAGTGATATAGGTCTTAAATTTT

GTCAAGAGCTCGTACTTGACACTTTAGTGGATGCTTGTTTCAGAGATGGGGTCCCTCCAAAGAAAGAAAGACTGCCGGTT

TAGCTCTTGATGTTACTAGTGTCTCTCCTTTTTCTCCGCGGTAGGTGCTTATGCTTTTATGCCCGACCATGCTATTTTCA

ATACTGTAGTCTCTCGCAAGCGTATTAAATACTTTGGTGTCATATATTGCTTTCTATACGCTCGATGAACTCGATGATGA

CATGGAGGCCCATCTAAAGAGATTGAAGAATTGTATTGCGGTTAAAATTGGCAATTACTTTTTCATAGAATAGTATGGTC

ATTAAGAAAGGGGTTGGAGCCCACTTCTTAGGAGAATACATCGTCGCACCCCTCACACCCCTTCTTAAACCGATGGGGGT

AGGTTGGTTCGGAATTGAGACCTGCCGATATCATTGTTTACAACTGGAACAGTGGTTAAACATCGATCTTGATGTTACTA

GTGTCTCTCCTTTTTCTGCGGTAGGTGCTTATGCTTTTATGCCCGACCATGCTATTTTCAATACTGTAGTCTCTCGCAAG

CGTATTAAATACTTTGGTGTCATATATTGCTTTCTATACGCTCGATGAACTCGATGATGACATGGAGGCCCATCTAAAGA

GATTGAAGAATTGTATTGCGTGTCGTGATTTTTATGTTAAAATTGGCAATTAGTTTTTCATAGAATAGTATGGTCATTAA

GAAAGGGGTTGGAGCCCAGCTTATTGTTAGGCTCCCCTCCCCCCTGCTAACTCTTTGTAATTCACTTTCTCTCATGTCTT

AGGAGAATACATCGTCAGCGCACCCCTCACACCCCTTCTTAAACCAGATGGGGGTATCAGGCCTATCGCCATTGGCACCA

TTTGGAGAAGACTCTTCTCCAAGATTACTGCCAGTTCTGTAAGTGCTGTAATGTCTGCGTATTTAGGTGACCACCAGTAC

GGTGTTATCATCCCATGCGATACTGAGGGGATTATGCATTCTGCCAATAACTCCTTGAACTCAATGGTACACAGAACTGT

CTATCTATGTTGCTCATCGATTTTAGGAATGCATTCAATTTGGTCAACCAAACTGCACTCATTAGAGAAGTGAGAAGTAA

GTGTCCAAGCATTTCAAAGTGGGTAAAATTTTGCTACGCAAGTCCTGCAAGATTATACTATGGTGAATCAACTATGTCCT

CAGCTCAGGGTGTGCAACAAGGCGACCCACTTGGTCCCTTGATGTTCACATTGACCCTTCACCCACTTGTATGCAAAATA

TCATCTCAATGCTCGTTGGACCTTCATGCCTGGTACTTAAACGACAGAACCATAATTGGGGATACTTTAGAGGTTTCTAA

GGAACTCAATATAACCCAGCAGGAGGGCGGATCCAGTGGTTTACATTTAAATATCCACAAGACGGATCTATTTAATTTGG

CCATATCCTAATCCTAGGAGTTTGGAAGATGGTGTATTCCCCCGTAACATTGGATGACCTAAGCCTGGCGTTAAGTTGTT

GGGAGGTCCAATCATCCTCGACTCACAGTTCTGTAGGTACCTTGTGTTGTCTAGGGTTTTGTTAACCATATTCTTTATGG

ACTGTGTAAGGAAATTACATGACCACCAATGCGAGCTTCTTAATTTGCACAAATGTACAGGCATATCCAACCTATACTTC

ACCCTACGCATTACCAAACCACAGGTGCTCGACGCTGCAATTAATCCATATGATTCCCATCTTTTTCAGTATTTGCAGCA

CCTTATTACATGCGAAGCGGCTGTCTTTGGACCTCTACAGTACATGCTAGCCACTTTACCCATCAAAGATGGAGGACTTG

GCGTATACACTATGGCCAATACAAGCCAATATTATTATCTTTCCTCATGCATTCAAATTGATGCTTTACAGACTACCATT

CTCCGCAACATAGACGTATCTCCTTCGTGCACCAATTTCCAATCAGCTCTCAACAACTACATGCAAACCTGTGGACTTTC

TTCTTCTTTATTCGACATCAAAGACACTGCCCCACAACCTATGAAAGCTTTAGCAGTTCGTTACTTTGTTGTTGTCAAGA

GGAAATTACCAAATATGTACATTTGCATTGTCGTCTAGAGAGGTTGTTCTTTGGCAAAGTAATCAGATGGCTCAGGCAAT

GGACTTTGTTAAGGAAATACCTATCGCTGGCCTTAATCAGAACCTTGGGCCTCACTAGTAAAGGGACATCATCCAATACC

GTCTAGGAGTGCCGATTTTTGTTGAATACAATGTGTGCCCTTGCTGAAATCGTGTCATGTATATTTTTGGCGACTACGCA

ATGCACTGTGCTAGCGAAGTTGGCCTTAAATTTATACACGACCTAGTTCATGATGTGGATGGACATATGCTACAAAGCAG

GTGTCGCCGCCAGGAAGGAAGTCTCATTGGCATTTTTAGATGATGGCAAAAGTTTAAAACATGATGATATTATGGTGTAT

AACTGGGAGAATGGTCAAGATACATGCATTGATGTCACTGGAGTCTCACCATTTGCAGGCGATGGTCTTCATATCTTCGT

ACTCGGGCGTGCTATTGCTGCTGCTATTACTCATAAACGCACTAAATATTTAGAAAAATGCACGTCTCATGGTTACAACT

TTGGTGTTTTAGCCTTTTCTATATTAGGTGGATTAGGACATGATTTTGTTATTTTTCTTAATAGATTGACGAACTGTCTA

GCCAATTATGACGCTAACTATAAGATATGCAGTTCGCTCTTTCAGCGAATTGGAATAACCATCCAAAAAGATGTTGGGGA

GCAACTTGTTTCTCGGCTCCCAACCCTTCCAACCAGCTCTGTGTAAATATTTCTCTTATTGGTCAAAAATATAATGAATC

TTTGAATATTCCGCAGTATTGATCTTTCCTTGATCCACTTCTATGTGAAAGTATTGAATGGCTCCGTAAGTTCTCTTATG

TTGAGCATTTCCGATTAAATTAATCATGGGTTCTATTGTGGTTAATTTAATTGAGTTTTCTGGATAAAAATTCATGTTTC

ATGTAATTTTTCAATGTCCAGAGAAATCCTTATTTTCTTGTAAAAGTAAGGTCGTTGTTGTTGTTCTTTCGGGAATGACA

TTTTATGGGGGAGAGTTCTTAATTGAACTTGTGCTTAATTGCCAAATCTTTGTGGCGAATGCGGTTGTGGAATATTGTAT

GAGTTTTATTGTATCTTTATAAACTCCTTGATGAATACACTAAGTTTCGACTATGTGAAATCATCAAAAAAAACTTGATA

TGTTCTCTTTTAGTCATGAAGTATCTCTATGAGAATTTCATTATGATCCCGCTAGTTTTCGTACCTTTGCCAATTTATAT

TGACCAAAAGGGGAAGAATTATTGTGTAATCACACTACAAATACATATGGTTTATGGAGCATTATGCAAGGGGGAGTGGT

TTTCATTGTGAGATGAAGTATTGACTAAGGGGGAGTGATACATATCATCGTAGTATTATTTTCAAAGTTGTGATACAATT

GAACTTTGATGTTGTGTAATAATACTCTGACACTGTATAACAATAACAAAACATCTTGTTTTTATATATTGTTATGGATA

CGGATCTTCAACAACTATGATGCTGAGTTGAAAACGTTCAGAATTTCTAGAGTACTTGGAAGTGACGAAGATTTCGAGTA

ATGGTGAAGCACCAAGGAAAACAAGTATTTGGATGAGAAGCTACAAAGTTTATTTATTTTGTAATCCATATGTATTGATA

GTTCTCAAGCTTGTTTGTCAAGTTTAGTTGTCAAAACTATAAGTCTTGATTTCTAGTCTACTTATAGCTATGTCTCGGAT

TAAGATAGAATATGTAGTTGAGATTTAGACTTCACGACGTTCATCGATTGAAGACGGAGAACTACCAAGGGGAGCTTGTG

GAACTTCATCAACAAAAGGTATGTGGAGACTTGCACTCATCTATCACTCAGAAGTCTATTTCTATTCTATCTCCTATTGA

GACAAAAGTCGTATAGATTTATAAACTTTATTTTATACACATTTGATAGAGCTGTGTTTAACTTGCTTACATATTTCTCG

AAATATGTGTTGGTAAGCTTTTGCTTAAATAAAGTTCATCTTTATTCTTGACGAAAGTCAAAATATGATCATGTGAAAAT

CGCCTGGTAATATCTTACATGATTTGTGTGAGACAGACATTTGATGTAAAATCAGAATGTTTCGTAATTGATCTATTTAT

CACATAAAAATTGCTTTGAAGCTAATAGTTTGTGTTAGATAGCTATTCCCGTCTTCTACGAATGTTTCAATGATTAAGAT

GGAGTTTAGAACAATTAACTTTTGATTGGATATATCACATTATGCGTACTTGTATGCTAACTATTGCAATTTATTCCAAG

TCCGGGAACCATAGTATGCAAACTTGTATGCGTACTGGTTTAACAGTTAAAGTCCGGGAACTTAGTATACATACCTGTAT

GCATACCGGCTTAACATTTCAAGTCCGAGAATTCAACTGAGTTTGGTCGTGAACGACAGTATGCTTACCCGTTCGCATAC

TGGCGGACAGACCAACTCCGGAACTCTAAGTATGTGTACATGTTTGCTTACTTGAGTGGGCTAAGTTCTAAAATCGGTTT

GTTTGTGAACAAATATATTTATATATAACAAGGAATGCAATCTTTTGCAAACCATGTCTATAATAACCATGAATTGATTT

TGCTTTAATTGTGTTTTGTATACTTCTATGAGAATATAAACAATTGAACAACTCTATAACTAGTTTCATTTGAGTCATTT

GAACTAGTTGTGATTAAGATGAACAAGGCTGATATGAAAGTGTGCATATAGATAACTTAGGTTAACTATTTTTGAGCCAA

CCAAGTGTACACGTTTAGGTATGGTACCATATCTAAATGAAGGAACATTTCATTTGTGTGTAACAAGATAAGTTCGATCT

AATGGTTGAAAGATATTAGCTTGAATCTAATCAGGTTTTCATCTAATGGTGAATATTGAATGCTTTGTTACCAAGGTAAC

ATTGCTTGCAAACCCTGATTTGAAGACTATATAAGGGAGAACTCTAGCAACTGGGAAACTTAATCCCGTCACCTCTTGTG

TGATACTAGTTGTGACTAGAGTCGAATCTCCTTTAAACTACGTTTTTCTAAAACCATTATAGGTTAACAACATAAAGACT

TCATTGGGATTATGAAGCCAGACCCTTAGTTATGTGTTCTGATTTTACTTTGTTTTATCGTATTGAATACTATCTTCTCT

AAGATTTTCTCGAGATTCAATCTCTGATAGGTAAGATAAAAATTAATCACAAACATATTCATCCCATCTTTTGTGATTCC

ACAATATATTGTTCCACTTCCATGTTGTTAAGATCATTGTGAGGTAATTGATATTATTAGGTTGTTCTTTGAAAATATAA

GTCTGGTGTATCAATTGGTTCATGTTCACCTTGATTTATCAAAAGACGGAATAAAAACTCGTATGAATTTCTTGTGGGAG

ACAGATTTATCTATTCAATAGACATTTCTGTGTGAGAGAAATTTATTTATCAAGTCTTCGACTTTGGGTCGTAGCAACTC

TTAGTTGTGGGTGAGATCAGCTAAGGGAATCAAGTGTGTAGAGTCCTGTTGGGATTCAGAGGCGTAAGGAACGCGACTGT

ACCTTAATCATTATGAGATTGGTTAGGGCTCAACTACATTCTAGTATGAAATTAACTTATAGTAGGCTAGAGTCTGTAGC

GGCTTAATACAGTGTGGTGTTCAAATCTGGACTAGGTCCTAGGGTTTTTATGCATTTGTAGTTTCCTCATTAACAAAATT

TCTGGTATCTTTGTAATTTCTTTTCCGCATTATATTTTATATAATTGAAATATTACAAGTTGTGTGTAGTTCAATTAATT

GGTAAATCTAACATTTGTTTGTTGATTGAAATTGATTGACACTTGAACATTGGTCTTTGGTATAGTTCAAGTTGTTTCTC

ATAATAATCAGGCTCGCGGATTTTTGTCTGTTTGATATGCTGATTACATTGAGAAACAAAGATGTAACTCTTGGATGTAT

TTTCTTTGATTGAGTTTGACTGTCTAGTTGCTCTTGGAAATATATTGGATTTAGTCTATAAATATTTCCTAAACAAAATA

TTGAGTGTGGTTGTTAGACCCCCGCTTTTTTAGAAAGGTACATCTATTTCCATGTCCATGATACTCATTGACTTCACTAA

TGCTTTCAACTTGGTTGACATATTAACTCTCATTCAAGAAGTTAGAATCGAGTGCCCATCAATTTCTAAGTAGGTGGAGT

TTTACTATGCAAAACTAGCTAAATTATACTATGATAACCTCGTTTTGTCCTTATCCCAAGGTGTCCAACAAGGAGACCCT

CTTGGTCCCCTATTATTCGCATTAACCTTACATCCATTGGTGCAGAAAATTGCTACTCAATGTACCTTAGATTTGCATGC

CTGGTACCTCTTGAAGGCATTATTATTGGTGATACTCTTGAAGTTTCTTGTGCGTTAAGTATTATACAGTTGGAGGGTTC

AAGTAGAGGGTTATATCTTAATGTCAGTAAGACCGAACTTTTTTGGCCAACTGAAGATCCAAGGAGGCTAACTTATGGCA

TTTTTCCGAAGAATATTAGTAGGTCTACTGTTGGCGTGAAACTATTAGGGGGTCATGTGAGCCTTGAGTTACAATTTTGT

AGAAATATGGTGTCTAGTAGAGTTGATAAGATAATTCAATTGATAACGGCTATTAAGAAACTCAAGGATCCGTAAAGTGA

AATGTTGCTTCTTTGTAACTGCGCAAGAATCTCTAAACTGTACTTCACAATACACACTACAAACCCATCAGCACTCCAAC

ATGTTACATCTCATTTCGATGGTTACCTAGTATTTGCGACACCTAATTACATGGCTTGGGTACTTATATTATGTCTGATA

CTGTTATTTAGCCTCCCAAAGTCAGACGTGTTTTACCCAGTCTACCATACTCAATTTCCCCCCTCAAGAGTTCATGGACA

CACTTTTGATCAAGCTCTTCAAACCTTCCGCCAGGTATGTGGATTACCTTTTTCCTACCGCATCAACGTTACTCCCCCCA

GTCTATTCACACCTTGGAAGTTGTAAACTTTGATGCCGTTAAGAAACAACTTCCCATCCAATTTCATATGTCTAAAAGGG

ACGCTGTTGTGGCAATGCAATAGAATTAATCACTCGCAAGATTATCTTTTATCCATACCATTTGCGGGGACCTAGACAAT

TTCGGGATGTGTTAGTTATCGGCTGAGAATCGCCTTCTTTGGTACAATCAACTTATGCTCCGTTTGCAATACAGAAATGG

GAATTTTCAGAGACCACGCTCTACATTGCGCAAGTGATGTTGGCCTCAAGTTCCGACATGGCCTTGTCCATGATTTCCTT

TTTGATATTTGTTATAAAGCTGGTGTGCATTCTAGAAAAGGGGTTGCGTTGGGCTTCTTGTCAGATAATAAAAAATAGTT

GAGACCTGCAGATATCCGCGTATAAAATTGGGAAATTGACCAGGATGTTTGTTTCGATGTGGATGGTGTTTCACCTTTCA

TCGACAGCAGCATTCGTACCTTTAGACATGGGCAAGCCATCTCCAATGCTATTCTCCGTAAACACAACGAATACTTGGAC

AAATGCTTAGCACATGGATATCGTTTTAGTATTTTGGCCTTCTCAACCCTGGATGAACTTAGTGAAGATACTATCGTCTT

TAAGAGGTTTAGGAATTCTCAGGCTAACAATGATGCTAATAATGGTTTAAGAATTTTTCTTTTTCATAAAGTAGGTGTGG

CTATTCAAAAGGTTGTTGGGGCTTAACTTTTCGCTCGGCTTCCAATTAACTTTGTGTAATAACTCTTTATATTTTGTACT

CTCTCTGTTCCATTTTATTTGATGTTTTAGGGTTTATTTTATGTTCCAAAATAGATGAGAGCTTTAGTATTTTCCCCAAT

TTCTTTCAGATTTACCTTTGCATTGGAATCAAATTAAAACATAAACATAAATTCTCATCGTAATCAACACATAAACAATT

ACTCAAATTTTTAAATATTTCTCATGGAGGATATACTCTTAATCTATAAAAATTTGGTACATCCCAAGTAAATTACATTG

GGATTGGTAACTTTCCCTAGAATTTAGAAACTAAAAACTAGGTCATTTAATTTATTAAAAAGGTATCAATATGGGTAGAA

TTGGAAAAATTTAACTCTCTCTGATGTTCCTTAATCTGCGTGAAAAACTCTACAACATCAAGTAAAATGGAACGAAGGGA

ATACGTTATTATTTAAAAGAGAGAGAGAGAGAGAAAAAAAGAGTCTCCAGCGCCATGGCAATTCCGTGAAGTTTAACAGA

AGGAGGATGGGTTTGTTTGATCAACCCTCATCTTCTGTTGTTTTCTTCTCATGTTTTGCTTCTTCAAACTTTCCTTTTCT

TCTTTATCCCTATTATGTTGCACCAACCTTAACCCTAGTCATGTATTGCTTCCACGTTGCTTCACAATATAGTGAGTTTT

AGGGTTTCAATGCCTAACGTCGAATCTTCATGCAGTTCTTCGTTGGATTCACAAGGAGATCCTTTCGTCCATTGTCCTTT

CCGTGGATGTGATAGGTGTAAAGATGGAGTTGGAGGAAGGGGTTTGGATAGATATTCGTTGCTTAACCATTTAAATGACA

AATTTTTTGTGTCAATGGTGTTGAAAACGAACGTTGCAGGCAAAGGTTAGCTAACATCCAGGGAATTTACAAAGCATGGG

AGGAGGCTTTACATATCTTCAAATGTGGTTGTGTACTGAATGTATGTAACTCCATTTGTGGAAGAGATCATGCAGGAAAT

ACAGTCATGTAGGGGGTTTCTATCGAGTCCTTTCAATGGTACGTACAGACGCCAATTTCCTTGTTCATGGCATCGTTTTC

CCTCCATCTCCTGATGAGATCCCTGTAGCTGTTAACGATGAAGGTATGTATTATATTCGTCTGCCATGCCTCAATGTTGA

ACTGTTGGACAAGGTTTTTCGGAAACATATACAGACAGTAAAGAATATTCCTTCCAAGTGTAAGGTTCAATTCTCTAGGG

TTTTGAAGTTCTCTTTGGATAAGGTTATTGCATGACCTAGGGATATTTATGGTTTGATACAACTATTATTACAGCGTGCC

TACGTTCTTCACATTTACGTTCCCAAAAACAACAATGAGGAGAAATCTGGGGTTCACAATAAACTACAAATTTCCTCCAT

TAATCAGTCCATTTCTCAGTGGTATGAGCCTGATGGATGCATTGGTTTAATTTATGAGCGATTGAATGACAGTCAATAGT

CGGGTGACAAGATTGCAGACATCCTAAACACCAAGAGGCTCTTAATTTGAGGACTTGTATGAAGAAGATTCGCAATGGGC

TTTTCACGGCTGCTATCAAAGTCCTTACATCTAGTGGACTGGCACCATCAGGAGAAGATACTCTCAACTGTTTGGTGGAT

AAGTATCCTTACGTCCCTCCCCCAATTATTCCTACCAGCCCTATCATTACAACCAGCGTAATTTCTACCACGAAATCTTT

CTTGGGGTGAATCAAGAGCTTTCCTAAAGGCACGTCGTGTGGAAGGTATGGTTTGAGGGTTCAACACCTTCTTGATGCGC

TCAATGGGCCAGATGCTGTTGTGGCTGATGACTTGATGTCATCTATTAATAGCGTGGTTAATTTGTGGTTAGATGGTTTA

TTCCCAGCGCTGTTAGGATAATTCGTGGAGAGTGAACCTCTCACACCTACTTAAGCCGGATGAGGGTATTCGACCAATTG

CATTTTCCACGGAGTGGATGCGACTCGTTTCTAAGGTTGCTGCTCATTCCATTGGTAAGGATATGTATGCATATCTTGAA

AACTTCCAGTTTGGGGTGGGTGTAACGTGTGGTGATGAAGATATCCTTCATGTTGTCAATATGATTATCCGAGATAAAGG

TGACTCCCAAACCATGACTATGATGCATGTAAATTTTTAGAATGCCTTCAACTTGGTTAATATGACTTCTATGCTTCAAG

AGGTAAGAAATCATTGTCCTTCTATATCCAAATGAGTGGAGTTTTGTTACTTTAACCCTCCAAGGCTATACTATAATGAA

TATGTCCTGTCGTCTTCTAAAGGAGTTCAACAAGGAGATCCTCTTTGTCCTCTCCTGTTTGCATTAACCCTAAACCCTTT

AGTTAAATTGACTGCGAAATAGTGCCAGTTAGATATGCAGGCATTGTATTTGGACGACTGTACCATCATTGGAGACACAT

TAATTGTCTCTTAATCTCTCAACATTATCCAGTCTGAGGGTGGTAAGTTTGGTCTTCACCTTAATATCACCAATACTGAG

ATTTATTGGCCCACAATAGATGTTAGATTTAGTGATGCTTTTCTTCCTCAGATTGGTCACCCGTCGAAAGGGGTTAAATT

GTTAGGGGGGCCTGTGCGTCTCGACTATGGTTATTGTAGTAACCTAGTTAAGTCCATGGTTTTTAAAACAACTGATCTTA

TGGATGCAGTTAGAAAGTTGGATGATCCACAGTGTGAGTTGCTCTTGCTGAGAAGTTGTGTGGTGGTGTTGAAGATATAC

TTCTCCATGAGAACACCTATGCCATAGTACCTTACTGAGGGGTAATATGTTTTCGACTCACATTTAACACAACATGTACG

TCATTCAATCACTAATGATGGCCCGGATTTGGGATTGCTACAACTCCAATTAGCTTTTGTACCCCTTCGACATGGTGGTT

TGGGTATATATTTTATGGTTGATGCGGTTCATTATTGTTATCTTGCATCATGTATTCAAACTCAGCCCCTTCAGGCCATC

ATTTTGCGTAATACTCCCATGCTTGGTCCAGATTTAGGTTTGCAGCGCGCCATGGATTTATTTTTCAGTGTATGCCCTAA

TGCCAATGTTGATTTATCATCCCCCCATCCCATGAAGTCTTTGGAAGTTAAATACTTTGACGTTGTCATGAATGAGATGC

CGGAAAAAATTTATATGTCCACACGTGACTATGCTCTTTGGAAGTGTAACAAGTCTGAACATGCTCATAACTTCCTCAAA

GATTTACCAATTGAGAGCCTCAATCATCAAGTGGGTGCACGATAATTTAGTGTTGTCTTGTGTTATCATTTGGGAATACC

ATTATTTTCTGACGACATCCAATGTGCTTTCTGTGATCGAAACATGGATTGTTATGGGGATCATGCCTTACATTGTTCAC

ACGAGGTGGGGACCAAATTTTGGTATGATTTTGTGCGTGACATCATAGTGGACATGTGTTTTCGAGCTGGTGTTCCAACT

AGGAACGAAGTTTCCTTGGGGATTACGGATAATGATGGTAGCGCCTTACGTCCTACGGATATTTTTGTCTAGAACTGGGA

AAGTGGTGGGGATATGTGCTTGGGGTCTCGCCTTTTACTGGTGGGGGGGTTCGGATCTTTCGGCCCAGCTTAGCTTTGAG

CAATGCTATTACTCGCAAAAAGAAAAAAAAAAAATACTTGGATTAGTGTTTGGCTCAAGGTTATGGCTTTGGTACCCCGA

CTTTCATGACGTTTGGCGAGCTCGATGAGGATACAATTGGTTTACTTAATAGGTTGCATAATCATCTGTCTAGGTTTGAT

GTTGGTGCAAAGAAATGTAATTTCCTTTTTCACATATTAGGGATTGCGATCTAGAAAGGTGTTGGAGCCCAGCTCGTTGC

TAGGATTCCAACTAATTTCTTGTCGTCTGATGATGTACTTGATATTTAAATTTCATTTTATTAGTGAATTTTTTTATTTA

AAAAATAAAAAATAATGATAAATAGAAATAGAAATTTAAAATAATATTATTATTTTTTGGGTAGTTTTATTTTTTATAGG

CCAGGTATTTCTATTCAAAAAGGTGTTGGAGCTCAGCTTGTTGCTAGGCTACCAACCAAAGTTCGGTATTAAAGTTACTA

AAGAAATTTAAAATGATAAAAATATAAATAAAAACACTATTACTATTCATATTTATTTAAACAATATTCAAAATAAAAAA

TATATTTAATGAATGTCCTCTCATTAGCTACATCATGAACACTTCAACATACCAACTCATCAAACAGAATTTCCAGCTTG

TTAGTTCAACAACACTTTACATATTATAGTCTACCAAACGTTCAACAACTACTTTATTTTCCTCGGAAGTGTTTGACCCA

CTGAAGTTTCCTCCCGAAAATAATCTTGATATATGAATGGACGGTCACGATTCGTCGTTCTAGCATTTGGATTTGTGCAA

TGTGAATTGGATCCCCTTGTCCCTCAGTTGGGATATTCATCGTTCTCGATAATACCAGCATTTTCATATGTTTATGATAC

CGCACAAACAACATAACAAAAAAGTGTTTTTCTAAAACTCGCTGAAATACCCAAACTATCCTTTAATAGCAGTCGAAAAT

TTGTATCCCTGACCAACTCAACTGTTTGACACTTAGAGCAGTTCCTATGGCAATGGCCAAACACAAATATTTGTTCAACC

ACGTGGATGGTAAAATGAGTTTCTCCACTATGGTAAAGCAGGTCAAAATGAACATATTTGGTTTTCTAGGGCCCCACTAC

TATATCTTTTTAATATAAAATGATTTATATGATAAAGTTGAAATATAAAATATAATATTTTGGTGAGATAAAATAGGATG

GAGTGAGATAAAATAGGATAAAGTGAGAAAAATATTAGTAAAAAAATCTGGATTTGTAGTGGGCGTCAGTAGGGATGTCG

CCCGCGCTCGGGCTAGTGTCACCCGCGCTCATGCTAGTGACGCCCGTGTTTTTTCTGGGCTCGCGCGTCAGTAGGTATGT

CACGCGCCATTTCACCAGACACGCGTCTGGTAGAGTGTCTCGATGTTCAAATGAACAGATCTGTTCATTTAAATATCCAT

TTTCTTGGTTTGTTCATCCCATAATGGGATCAAAATGAACAAACTCTAAATTTGTTCATCTCATAGGAATTGCCCTTACA

CACCATAACAAAACCCTTAGTGGTGGGTCCTCCTCCCAACTTCTAGAAAAATTATAAATCCATATTTTTCTGCTTTAACA

AAAAGTCATATGACGACTTCTCCGAACTGACTGACTGCGCACGAGAGAGAGAGAGAAAATCACTTCGTCTTCTTCAGTCA

GAATCTACTCGACTCGACTCACCGAGTCGAATCGATTCTGGGAGGTTGTTCAACCATGTATACACCTAATCTTCACCATC

ATCCGGTTCGTTTCCTTAAACCCTAAATACCAATCCCCCCATTCTCTTTCAATTTTAATTGAATTGTCAATTGCTTTTAA

AAAAAAAAAATCTAGGGTTTTTAATGAATTCTACTTGCATGTCTGAATTACTTGTTTAAATCACTCATAAATGTTTGTTT

TTGGGGTTTTAATCATTGTTGGTCTTGATTTGGTACATTTAGATTGGGTTTTCTTCACTCAATTTGGGAATGTGAGGTTA

GGGAGTGATGATTTTGTACTGCATTTCAATCAAAATGGTGTGGGTGTGTGTGTTTTGAGATGGGTTTTTGGAGTTTGTCA

AGTTTTTGTGATTAGATGAGATTTTTCGGCGTTTCCGATATTATTGGTGTAATGGGTGTTTGAATGGTGATTTTGTGTTT

TTGTATTGTGTAGATTCATGTTCACGTAGCACCATTACAAGTTCAAGACCCGTCCTCGATGGATTACAATAACAGTATGA

GTCCTGGTGGAGGAGTTTATATGCAAAATGCAAGTCCTGGAGTTGCAGGAGCTGCTTTTATGCAGAATGTGACTCCTGGA

GTCGGAGGAGTTCATAGTCCTGGAATTGGAGGAGCTGCTTATACGCAAAATGCAAGTCCCGGGGTTGGAGGAGCTGCTTA

TATGCAAAATGCAAGTCCTGGAGCTGCTTACATGCAAAATGGAAATTTCAGTAAACATGGAACTGTTGTGGACATTACTA

AGGTAGCTAGTCTTATACTCCATATGAATGCTAGTTAGTCTGACTTGTTAGTTTGATGATGGTTATGGTATATGTACAAA

GAGTGTTTGTGTGGCCGGATACATTCCCATTTAAGATGTGTTAGTTCGGGTTTTGACTTTGTGAATTGATGCACTTGTTG

ATGGTGGCAGAGATTACAAGATGAACTGACGAAGATAGGGCTTAAAATCAAAGCTCACGAGGAGAATCTGAGATTCCTTA

AGAAAGAGATGGACAATGTAGATGAATCTATTCTTGATTTGCAAGGTATCTTTTTATATACGCTCATACTTGTTATGTTA

TTACTACTTTAGGTAGTTTACAAGGTTTAGAGTTGGATGTTTTTAGTTAATAAAAGGAGTTGACCAATTGATGGTCTAGT

TTGCACAATAGGATGATCTTAATTTCTAATGCGTATTCTGCCCCTATCATGTTCGAGTGACAGCTAAATATATCAACTGT

TTCCTTTTTTCTCATCTTCTGGATTTTCTGCAGCTACACTCGGAAAGCATCGGTCTGAGAATGAGGGTACCGCAGCAAAT

GATGACTCAACCCATGTTCAGACTGAGGAAAACACGGTTCAGCAGATATTGAAGCAGGAAAAGTCTGCAGCTGGTGTGAT

ATGTCAGCTGAAAACTCACCATGCAACACAGGCATCGAATCTACCCTTAACCAAGGATGTGCTGGGTATTGTAGCTACTC

TCGGCCATGTGGATGATGATAACCTTAGCAGGTAATGTTTGTTATCTTATACTGGGTCATAGAACAATACCACCTAGATT

ACATTTACTCCTCAGTTGGGCTTGACATGATACTTTTTCTGTGTAACTTTCTGTGCGCAGACTTTTATCAGAGTTCTTAG

GGTTAGAAAAGATAATGGCAATTGTATGCATGACCTATGAGGGTGTTAAATGTATCGAATCATATGAAAAGGAAGGAAAA

ATCAGTAAGGGTTCTGGTATACACGGTCTTGGAGCATCAATTGCAAAACCCGTGGATGGAAGATTTGAAGTCATCTGTCT

TGAAGGTTTAAGGCAAGTCTGTTACTTTCACAGCCTGGGCAAATATATCTGAGAGCCGCCCTAAAAATGTTTTTCTTTGC

TTGTGCAGACCCTATGTCGGTGATATAGTTTCTGATGACCCACAACGGAAGCTTGCCCTTATGAAACCAAGACTACCAAA

TGGGGAGTGTCCACCTGGATTTCTCGGCTTTGCTGTAAATATGATCAATTTGGACCATTCGTACTTGTGTTATGTAACAG

CCAGCGGTCATGGTCTTCGAGAGACCTTATTCTATAGCCTTTTCTCTCGCCTCCAAGTGTATAGAACTCGAGCAGAAATG

TTACTTGCCCTTCCTTTTATATGTGATGGAGCCATGTCTCTGGATGGTGGAATGGTTAGGAGTGCTGGTATGTTGGTCCT

TGGTGATAGGTACGCATCAAGTCCATTTAGCAATGAATAAAGAGTCTGTTTTTGCGGATAAAGCCAGTACAGGTTTTGGT

TTGGTATTGTTGGGGTAGTTTTATTCCGTAAATTCCTTGAGTCTATTGTTAACTGAGCCATTTTCAATCTGGTTCCAGGA

AAGACGTGGAGGTTAGATTTCCACTTACTTCTCCAACGAAGAAAATGGCTTTAAACTACTTGGATACTGAAGAAAAGTTG

AAGCTGAAGAAATGGGAAAAGGAAAGGATTTTGGAGGATGTTCTAAGAGAACAACAGTTATTGATCCACGTGAAACAAAG

TTTTGAATCTAAACGGCAAGAATTTGTTAAGAATCTTGCTGAAAGCTCATCCCGCATAACTCAGGTATGGCATTTCGATC

CCCTCTTTGACTTTTTTAATTGTAGAAATGAAACTGTTTTTGTTCCCTCTTTGTTGGGTCGACTAGAAAGTGGTTTTTTA

GGCTAAATCTTTTTAAACGTCAACAGTGCGACCCCAAATTAATTTTAGTGAGGGAGGTTCTGTTTACATGCTTAAATGAC

ATTCTCCATAGCGCTCAAATGGTGAAGACCCGATGGCTGATTGTTTGTGGATGATGATCTACCCCTTGGGTGGAATATGT

TCACCAAAATAATCTTGTAATTCCATCATCTCGGGATAATGGTTGTAATTACACAAGTCGAATATTGTAATCAAGGTTCC

GAATTTGAATGGACTAAGTTTTGGCGAGAATGTCCTTCGGGACATACATCATCGTCTTCGTAAAATCAGCCATCCGGTGA

CGGTCTTCACTGTTTTAGCGCTATGGAGAATGTTATTTTAAGCATGTGAACGGGACCTTCCTCATTAACTCTTTATGTGG

TTAAGATATATGAACCGAAAAGCTGATCTTTATATTTCTTTGCGTTCTGTGATTGGCTCGTTACAAATGACAGCAGCAAT

TCCAGCCAGGAAGTAGAAGCAGCACCCCAAGATGAGCCGTGTACAAACTGAATCAACTTTACTCTTTTAACAATGAGTTG

ATAAAGATCTGTAAATCCACCTCAACTCCTATCATGGACGAAGATTTTTATTTCGTGTCAGTTGAAAGAATGCATCTGAG

AAGTTAACTGCCTTATGCAGTGCAGAACAGTTGGTGGAATTTTCCCCCAACCCTCCAAAGTTTTGCATATATCAGACCTT

CCATGTGCAAGTAATGAATGAATGTCTGTCACCTGTGTAACTGAAAGAACTTTAATGTAGCGGTAGTCTTTCGTTGTATC

GGTTTCGGTCATATTCTCTGTCTTCGGCCTGGTAGCAGTTGTCTGTTTTCTTTTCTTCATCCTTGATGATTTTTTGTGTG

CAAGTTGTCCTGATTAACATTGTTACTTGTTTCTGATGAATTACTGCACAGCAGTAACCCATAGAAAACTACTTCTATTT

GAATTTCAATTTGATTCGTAACTCCGTTTTATCACGCGGAGACGGGATTAACCCATACGGAACCAACCTGCCAAATGATG

GAATCTGGATTTGCACTGCCCCATGACTTTTTGCAGTGTTTTTTTATACTGTAGGTTATACGAGCAAGCAATTCAGATGT

TGATTCTGAAGATTCTCTTGAACAAATGTTCCGTAAGCGGAATGTGGAAGATGAGGAGGATGAGAAAATGGAGGCAATTG

CATTTCAGTGGCACCAGTCGGCTGCAATAGCGATTCTAAAATCAGAACAAAACAAGCTTAAACACAGGCAATTCACCGAC

AGGCAAAGGAAGGGCACACCACAAGGTAAAGCGCAGCAACAGGAGAAAGGTCCTTCCTGATGCCTGAAGTGGATTTTTGG

CGCATATTTAACCTTCAATTGCACTGTTTGTGCCATCGACTACAACCCAATAGCGTCCACCAGCTGTTGTCTCCAAATAT

TGATTTAATATCTTGAAATTTTTGATCAATTATAGTCTGATTCATTTTGCTAAAGTTGGCAACATTTTTCCTACAGAAAT

CTGATATGTGCTGAATCCAACCATGTCGTGCAGCAGGTCAGTTTTACAGCAAACCTTAATATCACAAGAACTACTCCAAA

AAAATAACAGCCCAAAAAATAGGAAAAAAACACGACAGCCGAAGAACATTACAGTCCTGTCCAAAAACTTTGCAAGTTGC

AGAAAATTTTAGTCCTGTTCTTCACGTGCAAATTGAAACTGATTAAATAAACAAAATAAGTCTTCCTTGAGTTAAACAAC

CAGCAGTGGATACCTCTACTCTATTTTCTACCGAGACAAATGGCTTGCTCAAATTAATATAACTCAAAATCAATTGGTAG

AACTTATAAACTCTTAGATTACCCAGAAAATGTGGGATTAATAGTCTCAACAGATGCAGGAAACCCTATCTTATCCGATA

AAATGACGTAATTCACCTAAAAATGTTGTCTTTCCATTATTTTTACCAAAAGGTTCTTTCTCTTGTATTTTTCATTACCT

CATGTCTCCAAATATGAACTATGAAACTAGATGGCCGCTGCCACAGTGGGAGGGCCGACCGAAGAAAAAAGGAAACACTG

CTGCAATTGGACTGTAAAAGAAAATTTATGCCTAAAGTGAAAGTATCATTTTTCTTGAAACGGGAAGAGTACATCGTTTT

ATTAGTTGCAAAGGAAATTTATTGATGAATAAACCAAAATATGGTTACATAAGGTATTATGTATCCTTTGTGCATAATGG

ATCCACATTTATTTTCAAAATTGTGCCTAAAATAATAAATACCTCCGAATTTTTCGTATACACTCTAAATTTTATACCGT

TGTTTGTGCTCGTTGGATAGGTTTTTTTTTATTAGCTTTCCAACGAGATAAAATTTGTAAAATTCCAAGGCACGGATTTT

TAGATATGTTATGTTAAAATTTGTTTTCCAATTATAACCCTAAAGAAATAGGATACATAAGTAATTTTAATAAAATTTGG

CTTTTATATCTAGTTGCCCTTCTTCTTTGGTTTATTTACGAAAATGTCTAATATCTTCCTGCCCTTCCTCTTTGGTTTAT

TTACGAAAATGTCTAATACCTTACTGCCCTTCCTCTTGGGTTTATTTACGAAAATGTCTAAAGGAAAAAATGAAACTTTT

TATTTAGAGGGGCAGTGTCTTCCTTTCATCGTTGAACCAGATTCAATCTTTTTTATGAAATTGGAAAAAATTGTCACGCT

TCACATTGACACAAAGTCACCATATATAGAGTCCTCCCTCGCTTCGCTCGGTCGGCCTAATAAACACATACTGTAGGGTG

TGCACAAAGGGTTCTGATAAAGGAAGAATCCAAAGCCCGTGATCTGTGTGTGTATGGATAAGCAGGATACATGCATATAG

AGTAATTTGGACGCTGATTTTAACAGATAACCCATCCGTAATTGTATCTGTTATGCTCAAGGTAAACTGAGTTGCTTCAC

TAAGTTATTTCTTTAAAAACTTTGAACCTGCCATGTAACTTTGGTGACACTAATGCTTAATGGATTTATATGTTACAGGA

AATACAAGACTGTTACCTCATCGACTGACGCAGAGATCAGGAACAATCCATGAATGAAAGATCTTGGTGGACATAGATGC

GGTGACGATACTAACTAATATGCTACTTATATATAACTCTGGCAACTCAAGTGCTTGTTTTTTTGAATACAAGGGAATGG

GATGGTCTCTCTCTCTCTTTTTTTTTATGCCATCAGGTGTGTTCCTATCATGTCCCAAACTTTCCACAGGCATTTTGGAT

GTTGATAATCTTAGTTACCAGCTTTTTGAACTCTCTGGGTCTTGTTCTATTGCCCAGATTTATGTTTTTTGAACAGTTAA

ATATCTTACTGTCGGATATTTATGGTGCATCAATGTAAAACTTTGTTATGAATGGTATAAATTTCTGGTTCAGTATGAAT

GATGTGAGTTTGTGGTTCAATATGAATGGTATAAGCTTTTTAATTTTTGTGTGAATGCTTAAGAGTTCTGAGAATTTTGT

AACTCTGGGAGTCTAAAAATCTGGCGTTGAAGAACAGTAACGATGACCGGTATCGATGTCCGGCAAACATCGGCATAAGA

CTGGCGCCAAATAACTGACGGAGGCAGGAAAATACTTTATGACGGTTTGACCAAATGCCACAACTAAAGTAACTATTCGT

GACAGTTGACTAACTGTCACAAATGATCTCAAACCGTCACAAAAAACATGTGTGACCCCAGTAGTAACGACGGATTGTCC

ACCTCACAAACGTGTTTTGTGACGGTCTTTTTAATATTTGTGACGGTTTGTAAACCGTCACAAAGGCCATCTGTTTTGTA

GTGTATCCAAATATTAGCGATTTACAGTTCACTAGACTAATTGGCTCATTATTTTCTCTAAAATGTGAGTCCTTTCACAT

TCATTAAAGGAAATTTATAGATGCATTCCCATTCTTACAATGAATTTTTGTACGATAATCGTTCGCACAGATTTCGTAAG

AACTTGACAAAGTGTGATTATCTCTTAGTTCTTACAGAGCTTATGTAACTTTTAATCGTTATACCAATTAGGCAACAAAT

ATTGAGCATTCCTTCGCTCGGCTATTACATGTAATTATCCAATCATCACATAGTATTATATAAGGAAACAATTCAACAAG

CAGTTTAATATATTTCCTTAATTAAGAGCTTCGTGTAATAAGTGGTGTGGCCTTATTACGTAACAAAAGAAAATTTATCT

CATTTAATTTAGAGTGTTATAGTTCATGAAGATGATATACTTTTCATCATAATGTACGAATCATCCTTTTAACTCAAGTA

CTTTTGAGTATATCTCAAGTGCTTTAGAGTATTTAAACCTCATGTAAATGATATACTATTCATTCTGTAAGAACATACAA

TGGATATAATTGTAACAAAGTTATTTAGACTCCTGATTAAAGTAGACATTTGACAAAAATGCTACCATAATCGGTAGGGA

ATTTAGGTTATTTAACTCTCGAATATAGAGTAGCCCCAAAATCAGATTTTGCAAAAATACTCTTTTTTTGAATTTTGGTT

TAGCCTATATAATCGGTAAGAAAGGAAGAGTTATCATGACTTCCGATTATACAGTAGACATATTTGCCAAAATCTTACCA

TATTCAGTAGTCAAGATAGTTCATATAACTACCGAATATAAAGTAGCCACAAAATCAGATTTTGCAAAAAATAATAATTC

TGAATTTTGGTTGACCCTATAATTGTTAGAAATAAAGAGTTATCATGACTTCTGATTATACAGTAGACCTCTTTACCGAA

ATGTTACCATATTCGGTAGTTAAGATAGTTTATATAACTACTAAATGTAGAGTAGCCCCAAAATCAGATTTTGCAAAAAA

AAAATTTAAATTTTGGTTGAGCCTATAATCGGTAAGAAAGGAAGAGTAATTGTGACTTCCGATTATCCTGGCCCTCTTTG

CAAATATCCAACCAATAGTGTCTCTTGAAGAAACCCTATGAGAACATACACGTTTTGTGAAAATACCCAACACATATAAC

TACCGAATATATAATTGCCCCAAAATGAGATTTTGCAAAAATCATCTATTTTTCTAATTTGGTTGAGCCTATAATCGGTA

GTCAAAATAGTTTTTTTTTTGTTTTTTTTTTTTGATGGAAAAAGGATATGTATTAAAGAAAAAAATGTATGTACACAGAG

CACATAAGGCTTCTATCCCAGCTTAAAATTACAGAGAAAATCTTTCACAAATCCATGATTTCTTCACATTTGAAAAGCAC
